# Lysosome Lipid Signaling from the Periphery to Neurons Regulates Longevity

**DOI:** 10.1101/2021.06.10.447794

**Authors:** Marzia Savini, Jonathon D. Duffy, Andrew Folick, Yi-Tang Lee, Pei-Wen Hu, Isaiah A. Neve, Feng Jin, Qinghao Zhang, Matthew Tillman, Youqiong Ye, William B. Mair, Jin Wang, Leng Han, Eric A. Ortlund, Meng C. Wang

## Abstract

Lysosomes are key cellular organelles that metabolize extra- and intracellular substrates. Alterations in lysosomal metabolism are implicated in aging-associated metabolic and neurodegenerative diseases. However, how lysosomal metabolism actively coordinates the metabolic and nervous systems to regulate aging remains unclear. Here, we report a fat-to-neuron lipid signaling pathway induced by lysosomal metabolism and its longevity promoting role in *Caenorhabditis elegans*. We discovered that lysosomal lipolysis in peripheral fat storage tissue up-regulates the neuropeptide signaling pathway in the nervous system to promote longevity. This cell-non-autonomous regulation requires the secretion from the fat storage tissue of a lipid chaperone protein LBP-3 and polyunsaturated fatty acids (PUFAs). LBP-3 binds to specific PUFAs, and acts through a nuclear hormone receptor NHR-49 and neuropeptide NLP-11 in neurons to extend lifespan. Together, these results reveal lysosomes as a signaling hub to coordinate metabolism and aging, and a lysosomal signaling mechanism that mediates intertissue communication to promote longevity.

---

Aging is a progressively declining process occurring at the levels of cells, tissues, and the whole organism. Mechanisms that govern the crosstalk across different organelles within a cell and among different tissues contribute to the regulation of longevity^1,2^. In particular, lipids play important signaling roles in mediating organelle crosstalk and tissue interactions^3^. Lysosomes actively participate in lipid metabolism and signal transduction, and lipid breakdown by lysosomal acid lipases releases free fatty acids from triacylglycerides and cholesteryl esters^4^. Here, we demonstrate how lysosomal lipid signals actively mediate the communication between peripheral metabolic tissues and the nervous system to systemically regulate longevity.

LIPL-4 is a lysosomal acid lipase, specifically expressed in the intestine, the peripheral fat storage tissue of *C. elegans*. Intestine-only constitutive expression of *lipl-4 (lipl-4 Tg)* prolongs lifespan and increases lipolysis (Supplementary Fig. 1a) ^5,6^. Through transcriptome profiling analysis, we identified a series of genes that are differentially expressed in the *lipl-4 Tg* worms compared to wild type (WT) worms (Fold Change > 1.5, *p*<0.05, Supplementary Table 1). DAVID functional annotation of the genes up-regulated by *lipl-4 Tg* revealed the enrichment of distinct biological processes including innate immune response, defense response and neuropeptide signaling pathway (Fig. 1a). While “immune response” and “defense response” are gene ontology (GO) categories commonly associated with longevity regulation, the enrichment of “neuropeptide signaling pathway” is unexpected. This GO category consists of genes that encode neuropeptide processing enzymes and neuropeptides. We first confirmed the induction of neuropeptide processing gene expression by qRT-PCR (Fig. 1b), including *egl-3, egl-21, pgal-1/pghm-1*, and *sbt-1* that encode proprotein convertase subtilisin/kexin type 2, carboxypeptidase E, peptidylglycine alpha-amidating monooxygenase, and neuroendocrine chaperone 7B2 ortholog, respectively (Fig. 1c). Among them, *egl-21, pgal-1* and *pghm-1* are specifically expressed in the nervous system^7,8^. Their induction suggests an interesting cell-non-autonomous regulation of neuronal genes by intestinal lysosomal lipolysis. These enzymes catalyze different steps in the processing and maturation of various neuropeptides (Fig. 1c), including insulin-like peptides (ILPs), FMRFamide-related peptides (FLPs) and non-insulin, non-flp peptides (NLPs). The transcriptome profile revealed that 3 ILP genes, 12 FLP genes and 19 NLP genes are transcriptionally up-regulated in the *lipl-4 Tg* worms (Fig. 1d, Supplementary Table 1).

**Figure 1.**
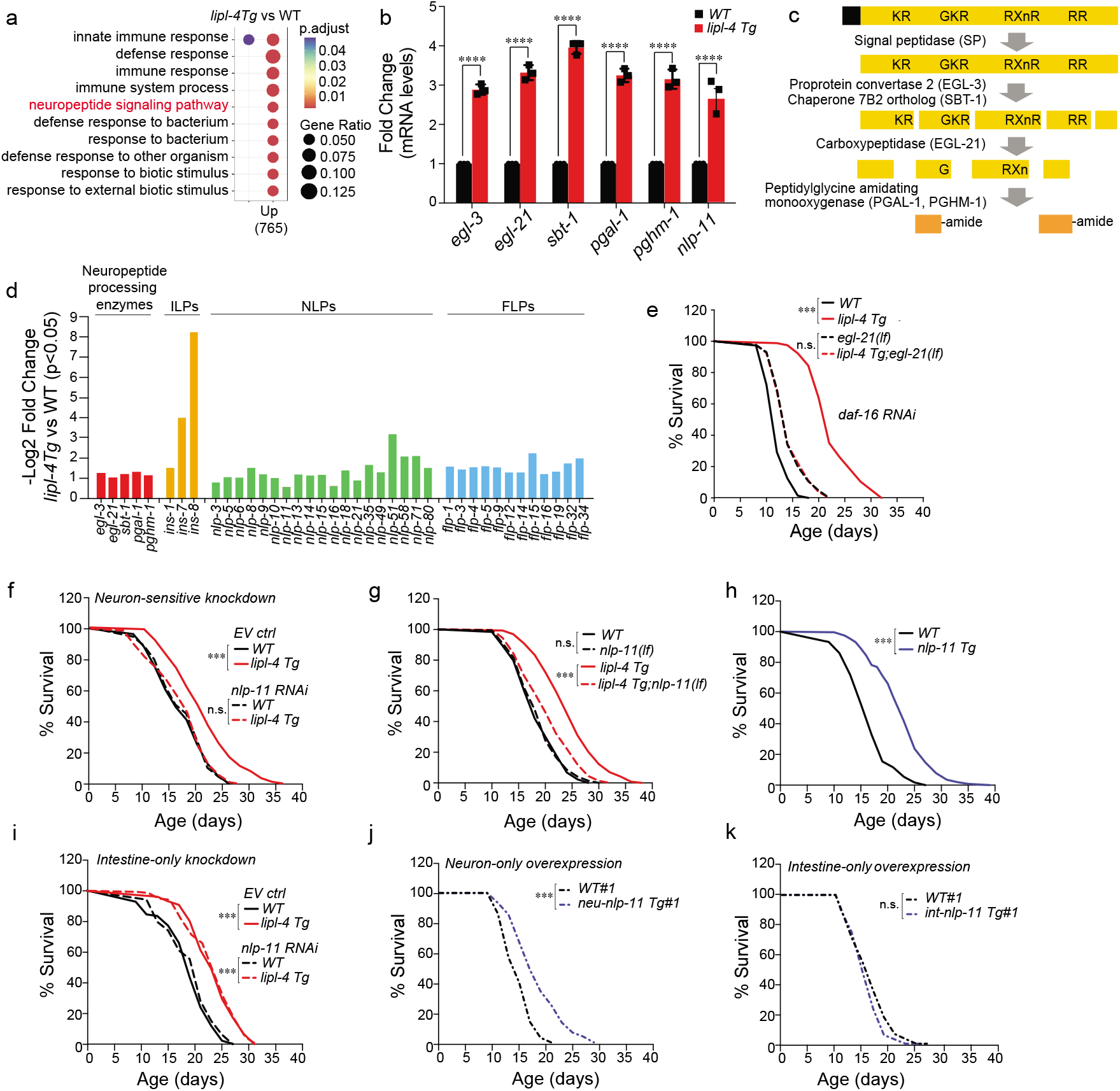
Peripheral lysosomal lipolysis induces neuropeptides to promote longevity. **a)** Gene Ontology of the up-regulated genes in the *lipl-4 Tg* worms compared to WT. **b)** Genes encoding neuropeptide processing enzymes, *egl-3, egl-21, sbt-1, pgal-1* and *pghm-1* and the neuropeptide *nlp-11* are transcriptionally up-regulated by *lipl-4 Tg*. **c)** Schematic diagram of the neuropeptide processing and maturation pathway. **d)** List of the up-regulated neuropeptide genes by *lipl-4 Tg* shown in RNA-seq profiling. **e)** The loss-of-function mutation of *egl-21(n476)* suppresses *lipl-4 Tg* longevity in the background of *daf-16* RNAi knockdown. **f)** Knockdown of *nlp-11* in a neuronal RNAi sensitive background suppresses *lipl-4 Tg* longevity **g)** The loss-of-function mutation of *nlp-11(rax51)* suppresses *lipl-4 Tg* longevity. **h)** Constitutive expression of *nlp-11* driven by its endogenous promoter extends lifespan. **i)** RNAi knockdown of *nlp-11* selectively in the intestine shows no suppression of *lipl-4 Tg* longevity. **j** and **k)** Neuron-specific overexpression of *nlp-11* prolongs lifespan (**j**), but intestine-specific overexpression has no such effect (**k**). Error bars represent mean ± s.e.m. *** *\*p*<0.0001, two-way ANOVA with Holm-Sidak correction (b), log-rank test (e-k).

Next, we examined the role of the neuropeptide signaling pathway in longevity regulation. Down-regulation of neuropeptide processing genes decreases the levels of mature ILPs, which can lead to reduced insulin/IGF-1 signaling that is known to prolong lifespan through the FOXO transcription factor^9^. Consistently, we found that the loss-of-function mutant of *egl-21* has extended lifespan, and the lifespan extension was suppressed by RNA inference (RNAi) inactivation of *daf-16*, the FOXO transcription factor in *C. elegans* (Supplementary Fig. 1b, Supplementary Table 4). On the other hand, the lifespan extension in the *lipl-4 Tg* worms is not suppressed by *daf-16* RNAi inactivation (Supplementary Fig. 1c, Supplementary Table 4). We then crossed the *lipl-4 Tg* worms with the loss-of-function mutant of *egl-21*, and further employed *daf-16* RNAi. We found that the inactivation of *egl-21* fully abrogates the lifespan extension conferred by *lipl-4 Tg* (Fig. 1e, Supplementary Table 4). These results suggest that the up-regulation of neuropeptide genes is required for intestinal lysosomal lipolysis to promote longevity, which is likely associated with NLPs or FLPs but not ILPs.

To identify specifically involved neuropeptides, we performed a RNAi-based screen in a neuronal RNAi-sensitive background to search for neuropeptide-encoding genes whose inactivation suppresses the lifespan extension in the *lipl-4 Tg* worms. We discovered that RNAi inactivation of *nlp-11* specifically suppresses *lipl-4 Tg* longevity without affecting WT lifespan (Fig. 1f, Supplementary Table 4). We also generated a CRISPR deletion mutant for *nlp-11* (Supplementary Fig. 1d) and crossed it with the *lipl-4 Tg* worms. We found that the loss-of-function mutant of *nlp-11* suppresses the lifespan extension in the *lipl-4 Tg* worms but has no effects on WT lifespan (Fig. 1g, Supplementary Table 3). Moreover, *nlp-11* is transcriptionally up-regulated in the *lipl-4 Tg* worms (Fig. 1b, d). We then made a transgenic strain to overexpress *nlp-11* driven by its endogenous promoter, and we found that these transgenic worms live longer than WT controls (Fig. 1h, Supplementary Fig. 1e, Supplementary Table 3), suggesting that *nlp-11* is sufficient to prolong lifespan. *nlp-11* expresses in both neurons and the intestine (Supplementary Fig. 1f). To further test whether *nlp-11* functions in neurons to regulate longevity, we knocked down *nlp-11* selectively in the intestine and found its intestine-only inactivation does not affect the lifespan extension of the *lipl-4 Tg* worms (Fig. 1i, Supplementary Table 4). We also overexpressed *nlp-11* in either neurons or the intestine using tissue-specific promoters and found that only neuron-specific overexpression of *nlp-11* is sufficient to prolong lifespan (Fig. 1j, k, Supplementary Fig. 1g, h and Supplementary Table 3). Together, these results demonstrate that neuronal *nlp-11* is specifically responsible for the longevity effect conferred by intestinal lysosomal lipolysis.

Lysosomal acid lipase is known to release free fatty acids from hydrolyzing triacylglycerides and/or cholesteryl esters ^4^. Through lipidomic profiles of free fatty acids, we found that the levels of polyunsaturated fatty acids (PUFAs) are significantly higher in the *lipl-4 Tg* worms than those in WT worms (Fig. 2a). We hypothesize that these PUFAs serve as cell-non-autonomous signals to regulate neuropeptide genes. To test this hypothesis, we utilized the loss-of-function mutants of *fat-1* and *fat-3* that encode ω-3 fatty acid desaturases and Δ6-desaturase, respectively (Fig. 2b). These mutants lack PUFAs ^10,11^, and we found that they fully suppress the transcriptional up-regulation of neuropeptide genes (Fig. 2c), as well as the lifespan extension in the *lipl-4 Tg* worms (Supplementary Fig. 2a, b and Supplementary Table 3). Both FAT-1 and FAT-3 express in the intestine and neurons and catalyze PUFA biosynthesis locally ^12,13^. We then selectively reduced intestinal PUFAs by knocking down *fat-1* and *fat-3* only in the intestine using RNAi. We found that intestine-only inactivation of either *fat-1* or *fat-3* is sufficient to fully abrogate the lifespan extension in the *lipl-4 Tg* worms (Fig. 2d, e and Supplementary Table 4). Together, these results suggest that intestinal PUFAs mediate both the up-regulation of neuropeptide genes and the longevity effect induced by lysosomal lipolysis.

**Figure 2.**
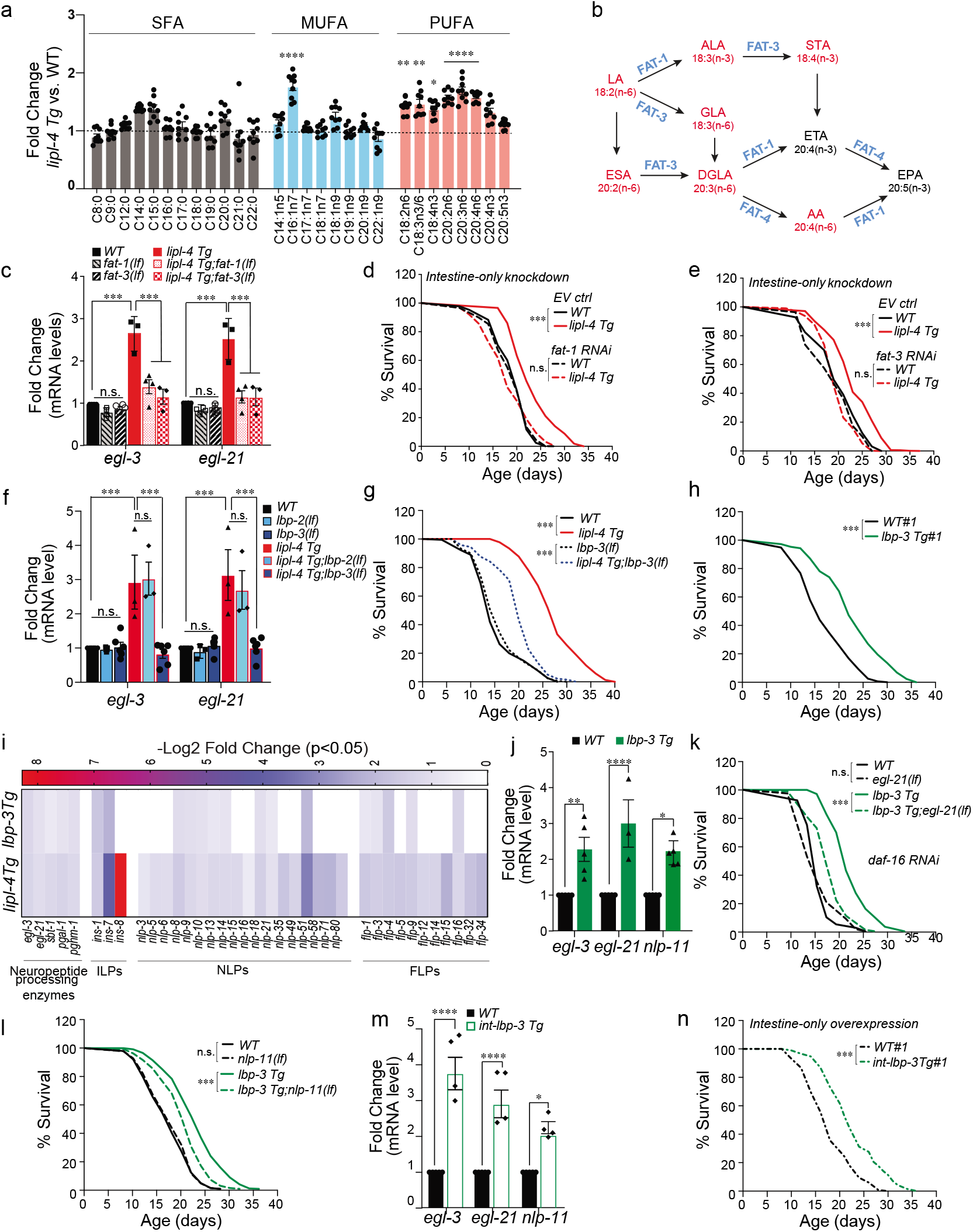
Peripheral lipid signals mediate neuropeptide induction and longevity. **a)** Relative levels of free fatty acids quantified by LC/MS in *lipl-4 Tg* vs. WT. **b)** Schematic diagram of polyunsaturated fatty acid (PUFA) biosynthesis in *C. elegans*. PUFAs enriched in *lipl-4 Tg* are highlighted in red, while desaturase enzymes are highlighted in blue. **c)** The loss-of-function mutation of *fat-1(wa9)* or *fat-3(wa22)* desaturase suppresses the transcriptional up-regulation of *egl-3* and *egl-21* neuropeptide genes by *lipl-4 Tg*. **d** and **e)** RNAi knockdown of either *fat-1* (**d**) or *fat-3* (**e**) selectively in the intestine suppresses *lipl-4 Tg* longevity. **f**) The loss-of-function mutation of *lbp-3(rax60)* but not *lbp-2 (rax63)* lipid chaperone suppresses the transcriptional up-regulation of *egl-3* and *egl-21* by *lipl-4 Tg*. **g)** The loss-of-function mutation of *lbp-3(rax60)* suppresses *lipl-4 Tg* longevity. **h**) Constitutive expression of *lbp-3 (lbp-3 Tg)* driven by its own endogenous promoter prolongs lifespan. **i)** Out 39 neuropeptide genes induced by *lipl-4 Tg*, 22 are also induced by *lbp-3 Tg*. **j)** The transcriptional levels of *egl-3* and *egl-21* are induced in the *lbp-3 Tg* worms. **k**) The loss-of-function mutation of *egl-21(n476)* suppresses *lbp-3 Tg* longevity in the background of *daf-16* RNAi knockdown. **l**) The loss-of-function mutation of *nlp-11(rax51)* suppresses *lbp-3 Tg* longevity. **m)** Intestine-specific *lbp-3* overexpression up-regulates the transcriptional level of *egl-3, egl-21* and *nlp-11*. **n)** Overexpression of *lbp-3* selectively in the intestine extends lifespan. Error bars represent mean ± s.e.m. n.s.*p*>0.05, \**p*<0.05, \*\**p*<0.01, \*\*\**p*<0.001, *\*\*\*\*p<0.0001*, two-way ANOVA with Holm-Sidak correction (a, c, f, j, m), log-rank test (d, e, g, h, k, l, n).

Fatty acids have low aqueous solubility and must be bound to proteins in order to diffuse through the lipophobic environment. A family of proteins termed fatty acid binding proteins (FABPs) can function as lipid chaperones, which reversibly bind fatty acids and fatty acid derivatives to mediate their trafficking and signaling effects ^14,15^. To test whether specific FABPs facilitate the action of intestinal PUFAs on neurons, we focused on three FABPs, LBP-1, −2 and −3 that carry putative secretory signals. We found that RNAi inactivation of *lbp-2* or *lbp-3* but not *lbp-1* specifically suppresses the induction of neuropeptide genes caused by *lipl-4 Tg* (Supplementary Fig. 2c, d). To confirm the results of *lbp-2* and *lbp-3* RNAi knockdown, we generated their CRISPR deletion mutants (Supplementary Fig. 2e) and crossed them with the *lipl-4 Tg* worms. We found that only the *lbp-3* but not *lbp-2* deletion suppresses the induction of neuropeptide genes in the *lipl-4 Tg* worms (Fig. 2f). Deletion of *lbp-3* also suppresses the *lipl-4 Tg* longevity without affecting WT lifespan (Fig. 2g, Supplementary Table 3).

Next, to examine whether *lbp-3* is sufficient to promote longevity and induce neuropeptide genes, we made transgenic strains that constitutively expresses *lbp-3*. We found that these transgenic strains live longer than WT worms (Fig. 2h, Supplementary Fig. 2f and Supplementary Table 3). The transcriptome analysis showed that among 39 neuropeptide genes that are up-regulated in the *lipl-4 Tg* worms, all five neuropeptide processing genes and 16 neuropeptide genes are also up-regulated in the *lbp-3* transgenic strains (Fig. 2i and Supplementary Table 2). The up-regulation of *egl-3, egl-21* and *nlp-11* was also confirmed using qRT-PCR (Fig. 2j). Moreover, we found that inactivation of either *egl-21* or *nlp-11* suppresses the lifespan extension in the *lbp-3* transgenic strains (Fig. 2k, l, Supplementary Fig. 2g and Supplementary Table 3, 4), which supports that specific neuropeptides act downstream of the LIPL-4-LBP-3 signaling cascade to regulate longevity. Next, we made transgenic strains to selectively overexpress *lbp-3* in the intestine, and we found that intestine-only overexpression of *lbp-3* is sufficient to up-regulate neuropeptide genes (Fig. 2m) and prolong lifespan (Fig. 2n, Supplementary Fig. 2h and Supplementary Table 3). Together, these results support that the specific lipid chaperone, LBP-3 mediates the cell-non-autonomous communication from the intestine to neurons to regulate neuropeptide genes and longevity.

To further understand the lipid chaperone function of LBP-3 in this endocrine regulation, we examined whether LBP-3 can be secreted from the intestine. In *C. elegans*, coelomocytes are scavenger cells that take up secreted materials from the body cavity and serve as a monitor of secreted proteins ^16^. We generated a transgenic strain expressing an intestine-specific polycistronic transcript encoding both LBP-3::RFP fusion and GFP, such that GFP indicates cells expressing *lbp-3* and RFP directly labels LBP-3 protein. Without tagging with any proteins, GFP was detected ubiquitously within intestinal cells (Fig. 3a). LBP-3::RFP fusion, on the other hand, was detected within intestinal cells and also in coelomocytes (Fig. 3a), which shows the secretion of LBP-3 protein from the intestine into the body cavity. We also discovered that in the *lipl-4 Tg* worms, the level of LBP-3::RFP in coelomocytes is elevated (Fig. 3b, c), revealing increased secretion of LBP-3. Within intestinal cells, the LBP-3 protein is detected in the cytosol and also at lysosomes that are marked by LMP-1 and stained with LysoTracker (Supplementary Fig. 3a, b). Moreover, we generated transgenic strains that constitutively express LBP-3 without its N-terminal secretory signal only in the intestine and found these transgenic strains do not exhibit lifespan extension (Fig. 3d, Supplementary Fig. 3c and Supplementary Table 3) and lack the up-regulation of neuropeptide genes (Fig. 3e), in contrast to the transgenic strains that do express secretable LBP-3 (Fig. 2m, n, Supplementary Fig. 2h and Supplementary Table 3).

**Figure 3.**
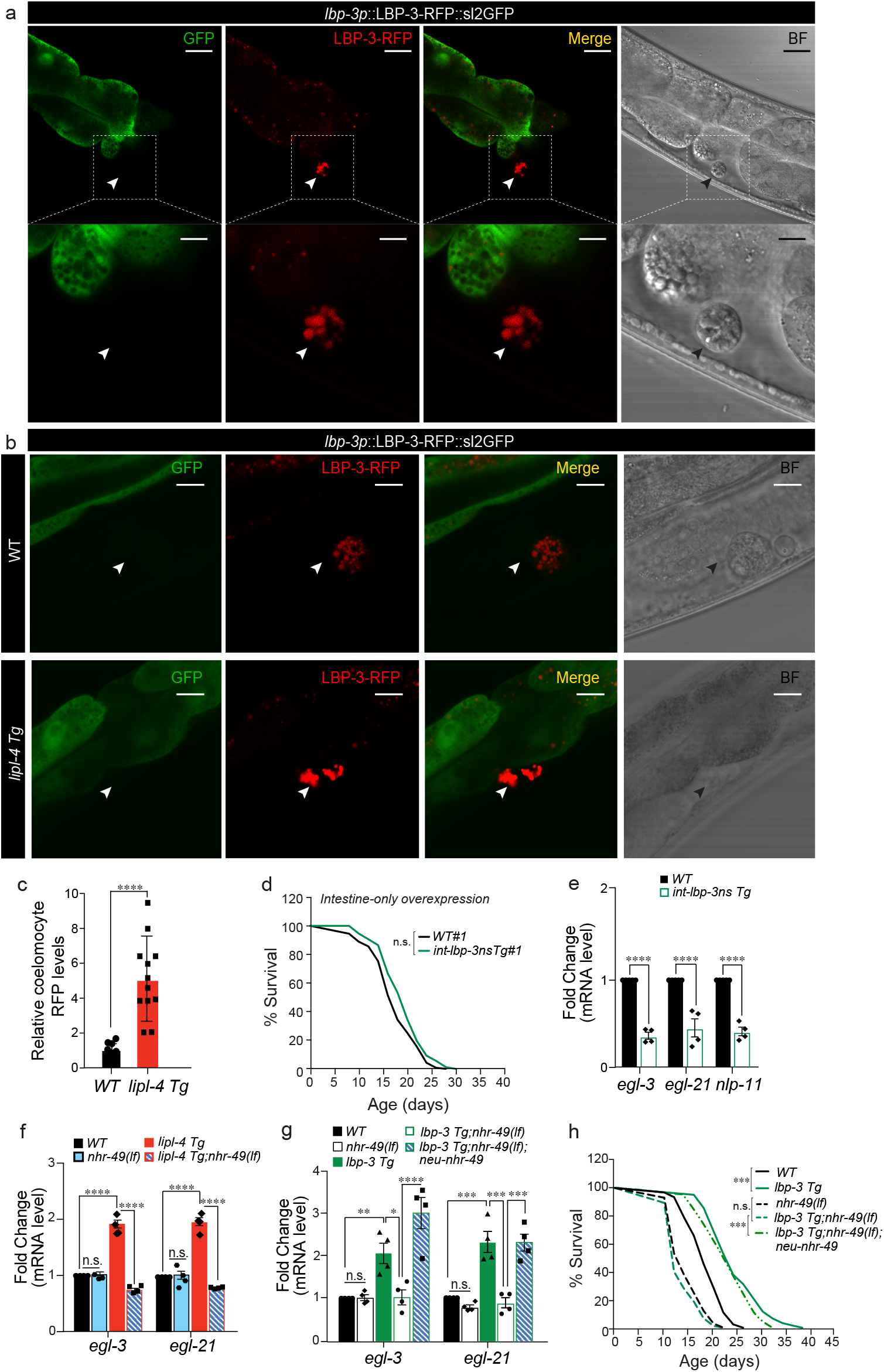
Secreted lipid chaperones act through neuronal nuclear receptor to regulate neuropeptide and longevity. **a)** The gene expression of *lbp-3* is indicated by polycistronic GFP, while the LBP-3 protein is visualized by its RFP fusion. Secreted LBP-3::RFP fusion is detected in coelomocytes marked with arrowheads. Scale bar 30μm and 10μm in the inset. **b, c)** The level of secreted LBP-3::RFP fusion is increased by *lipl-4 Tg*. Secreted LBP-3::RFP fusion proteins in coelomocytes are marked by arrowheads **b**), and their levels are quantified in *lipl-4 Tg* vs. WT **c**). Scale bar 10μm. **d)** Intestine-specific overexpression of *lbp-3* lacking its secretory signal (*lbp-3ns*) fails to extend lifespan. **e)** Intestine-specific overexpression of *lbp-3ns* decreases the expression of neuropeptide genes. **f)** The loss-of-function mutation of the nuclear hormone receptor *nhr-49(nr2041)* fully suppresses the induction of neuropeptide genes by *lipl-4 Tg*. **g)** The loss-of-function mutation of *nhr-49(nr2041)* fully suppresses the induction of neuropeptide genes in the *lbp-3 Tg* worms, and the neuronal restoration of *nhr-49* fully rescues their inductions. **h**) The loss-of-function mutation of *nhr-49(nr2041)* fully suppresses the lifespan extension in the *lbp-3 Tg* worms, and the neuronal restoration of *nhr-49* fully rescues this lifespan extension. Error bars represent mean ± s.e.m. n.s. *p*>0.05, * *p*<0.05, \*\**p*<0.01, \*\*\**p*<0.001, \*\*\**\*p*<0.0001, two-way ANOVA with Holm-Sidak correction (c, e, f, g), log-rank test (d, h).

In searching for factors mediating the transcriptional up-regulation of neuropeptide genes, we discovered that the nuclear hormone receptor NHR-49 acts in neurons downstream of LIPL-4-LBP-3 signaling. Its loss-of-function mutation fully suppresses the up-regulation of neuropeptide genes caused by the constitutive expression of LIPL-4 (Fig. 3f) and secretable LBP-3 (Fig. 3g). Importantly, in the *nhr-49* mutant background, neuron-specific restoration of *nhr-49* fully rescues the up-regulation of neuropeptide genes (Fig. 3g) and the lifespan extension (Fig. 3h, Supplementary Table 3) conferred by the LBP-3 constitutive expression. Together, these results suggest LBP-3 proteins are secreted from the intestine, which is triggered by LIPL-4-induced lysosomal lipolysis, and secreted LBP-3 acts through neuronal NHR-49 to up-regulate neuropeptide genes and promote longevity systemically.

Now, we have found that both intestinal PUFAs and LBP-3 mediate the neuropeptide-inducing and lifespan-extending effects downstream of lysosomal lipolysis. To test whether they coordinate with each other, we first examined the effect of PUFAs on LBP-3 secretion. Using the RNAi inactivation of *fat-3*, we reduced PUFA biosynthesis in peripheral tissues. We found that the *fat-3* inactivation suppresses the increased level of LBP-3::RFP in coelomocytes by *lipl-4* Tg, without affecting the level in WT (Fig. 4a, b). Thus, the PUFA induction by *lipl-4 Tg* promotes LBP-3 secretion from the intestine. It is known that the *fat-3* mutant lacks 20-carbon PUFAs ^10^, including dihommogamma-linolenic acid (DGLA) and arachidonic acid (AA) that are induced by *lipl-4 Tg* (Fig. 2a). We then tested whether LBP-3 binds to DGLA and/or AA. Using purified LBP-3 proteins, we performed a competitive fluorescence-based binding assay. In this assay, when bound to LBP-3, the amphipathic molecule 1-anilinonaphthalene-8-sulfonic acid (1,8-ANS) shows enhanced fluorescence that is quenched once out competed by free fatty acids. Based on this competition assay, we found that both DGLA (Kd=10.96μM) and AA (Kd=2.9μM) bind to LBP-3 (Fig. 4c). As controls, we also examined eicosapentaenoic acid (EPA) and eicosatetraenoic acid (ETA), the two 20-carbon PUFAs that are not induced by *lipl-4 Tg*, and found that EPA (Kd=4.76μM) but not ETA bind to LBP-3 (Fig. 4c). Next, we supplemented DGLA, AA or EPA to worms and measured the expression levels of neuropeptide genes. We found that in the *fat-3* mutant, the supplementation of DGLA is able to restore the upregulation of neuropeptide genes caused by *lipl-4 Tg* (Fig. 4d). Neither AA nor EPA supplementation shows such an ability (Fig. 4d). These results suggest that the induction of PUFAs by lysosomal lipolysis promotes the secretion of the LBP-3 lipid chaperone from the intestine, which binds to DGLA and mediates its long-range signaling effect on the neuropeptide pathway.

**Figure 4.**
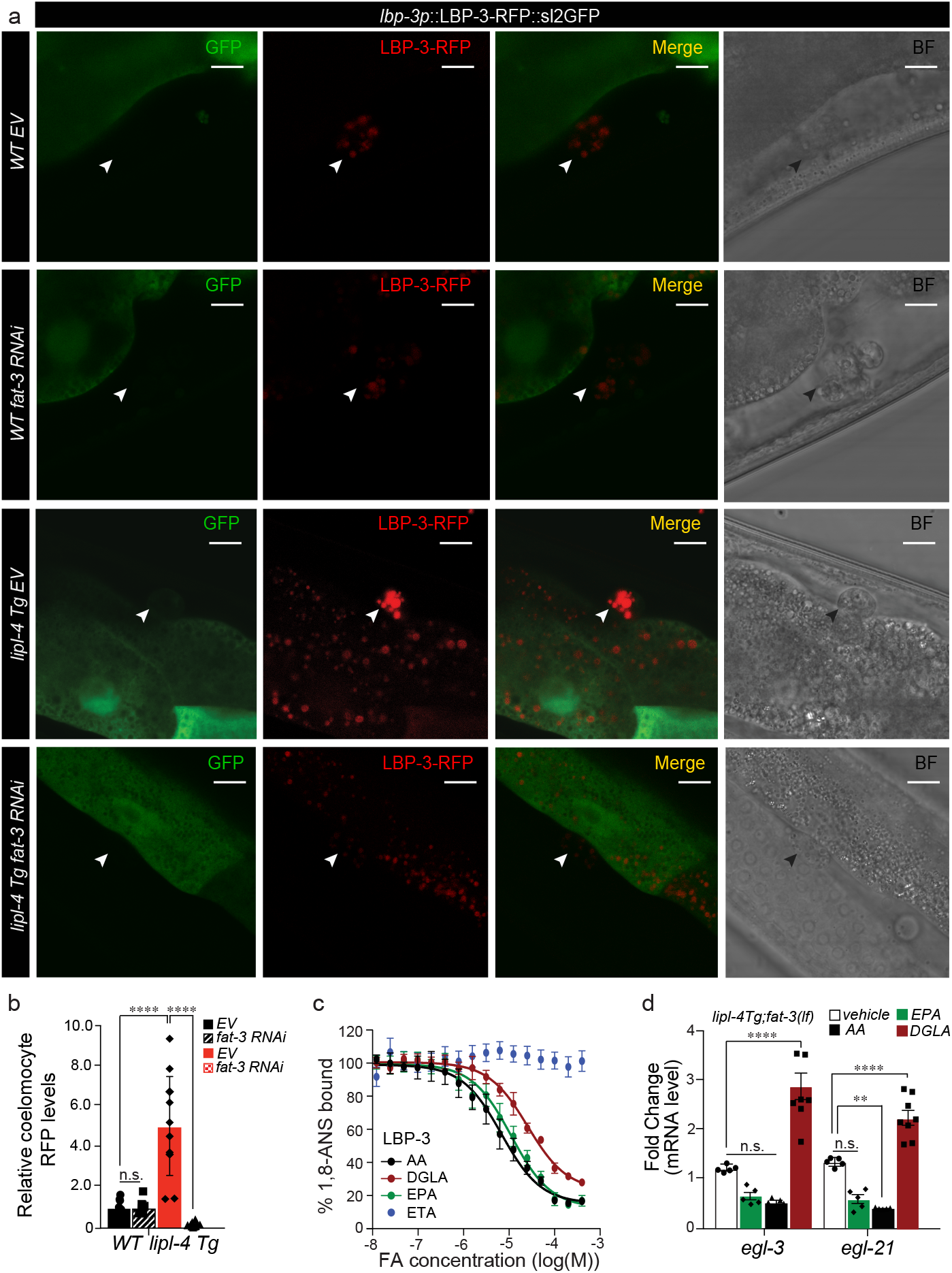
Lipid signals induce the secretion of lipid chaperones to regulate neuropeptide and longevity. **a, b**) In WT conditions, LBP-3 secretion is not affected upon *fat-3* RNAi knockdown. However, the increased secretion of LBP-3::RFP fusion by *lipl-4 Tg* is suppressed by RNAi inactivation of *fat-3*. Secreted LBP-3::RFP fusion proteins in coelomocytes are marked by arrowheads **a**), and their levels are quantified **b**). Scale bar 10μm. **c)** Fluorescence signals derived from 1-anilinonaphthalene-8-sulfonic acid (1,8-ANS) bound to LBP-3 are decreased, when 1,8-ANS is competed off by the increasing amount of arachidonic acid (AA), dihommogamma-linolenic acid (DGLA), and eicosapentaenoic acid (EPA), but not eicosatetraenoic acid (ETA). **d)** DGLA supplementation restores the transcriptional induction of *egl-3* and *egl-21* by *lipl-4 Tg* in the *fat-3(wa22)* mutant. DMSO serves as the vehicle control. Error bars represent mean ± s.e.m. n.s. *p*>0.05, * *p*<0.05, \*\**p*<0.01, \*\*\**p*<0.001, \*\*\**\*p*<0.0001, two-way ANOVA with Holm-Sidak correction (B, D).

In summary, we discovered that LIPL-4-induced lysosomal lipolysis in the peripheral fat storage tissue up-regulates DGLA, which binds to a secreted lipid chaperone protein, LBP-3 and cell-non-autonomously induces the neuropeptide pathway to promote longevity. Downstream of LIPL-4-LBP-3 lipid signaling, the nuclear hormone receptor NHR-49, a *C. elegans* homolog of PPARa, specifically acts in neurons and mediates the transcriptional induction of the neuropeptide genes to promote longevity. Interestingly, mammalian FABP4 secreted from adipocytes has been implicated in the hormonal control of metabolism^17^, and FABP5 at the brain blood barrier contributes to the brain uptake of docosahexaenoic acid, a PUFA essential for cognitive function ^18^. Therefore, FABP secretion may function as an evolutionarily conserved mechanism to facilitate the transportation of lipid signals from peripheral metabolic tissues to the central nervous system. In response to lipid signals, nuclear hormone receptors are best known mediators of transcriptional responses. In particular, PPARa has high binding affinity for fatty acids and plays a crucial role in metabolic tissues to regulate lipid catabolism ^19^. PPARa also expresses at a high level in the nervous system, however its neuronal functions and regulation remain poorly understood. Our studies reveal that in the nervous system, NHR-49 regulates neuroendocrine gene expression in response to peripheral lipid signals. Previous studies also show that neuronal NHR-49 mediates the longevity effect conferred by AMPK activation in neurons ^20^. Thus, lipids may be key endocrine signals that link peripheral metabolic status with neuronal transcription via nuclear hormone receptors like PPARa. Distinct to transient neurotransmitter and neuropeptide release, transcriptional up-regulation of neuroendocrine signals enables long-term adaptation in the nervous system, which will be consequential for the long-lasting aging process. Interestingly, lysosomal lipid metabolism is able to fine-tune this endocrine signaling mechanism to actively regulate longevity. This study supports an emerging paradigm that lysosomes are the critical signaling hub for longevity. Strikingly, lysosome-derived signals are crucial not only for organellar crosstalk in the cell ^21^, but also tissue coordination in the organism, making them exciting targets for optimal pro-longevity intervention at the systemic level.

## Supporting information

Supplementary Table 1 - Transcriptome analysis of WT and lipl-4 Tg animals

Supplementary Table 2 - Transcriptome analysis of WT and lbp-3 Tg animals

## Acknowledgments

We thank A. Dervisefendic and P. Svay for maintenance support, J. Mello (Harvard Medical School, USA) for providing strains JM45, S. Mutlu, P. Rohs, S.M. Gao, X. Ma, L. Ding, C. Herman, H. Dierick, and B. Arenkiel for critical reading of the manuscript. Some strains were obtained from the Caenorhabditis Genetics Center (CGC), which is funded by NIH Office of Research Infrastructure Programs (P40 OD010440).

## Author Contributions

M.S.., A.K.F., and M.C.W. conceived the project and designed the experiments. M.S., A.K.F., J.D.D., L.Y., P.H., I. N., J.F., Z. Q., M.C.T., performed experiments and Y.Y conducted the bioinformatic transcriptome analysis. M.S. and M.C.W. wrote the manuscript. M.S., W.B.M., E.A.O., L.H., J.W. and M.C.W. edited the manuscript.

## Competing interests

The authors declare no competing interests.

## Funding

This work was supported by NIH grants R01AG045183 (M.C.W.), R01AT009050 (M.C.W.), R01AG062257 (M.C.W.), DP1DK113644 (M.C.W.), March of Dimes Foundation (M.C.W.), Welch Foundation (M.C.W.), and by HHMI investigator (M.C.W.), NIH T32 ES027801 pre-doctoral student fellow (M.S.). We thank WormBase.

## Methods

### *Caenorhabditis elegans* strains maintenance

*C. elegans* strains obtained from Caenorhabditis Genome Center (CGC) or generated in this study are listed in the Supplementary Table 5. Strains were maintained on standard nematode growth medium (NGM) agar plates seeded with corresponding bacteria at 20°C and analyzed for imaging and qRT-PCR analysis at day-1 adulthood.

### Molecular cloning and generating transgenics

Tissue-specific promoters driving *lbp-3* or *nlp-11* vectors were generated using Multisite Gateway System (Invitrogen). Promoters were cloned to 1’ position Donor Vector pDONR 221/P1-P4; cDNAs were cloned to 2’ position pDONR 221/P4r-P3r; and *sl2-RFP::unc-54 3’UTR* or *GFP::unc-54 3’UTR* was cloned to 3’ position pDONR 221/P3-P2 using Gateway BP reaction. Entry vectors were then recombined into the destination vector, pCMP1 (pCFJ151 modified to contain Gateway Pro LR recombination sites) using Gateway LR reaction. For expressing LBP-3::RFP::sl2-GFP in the intestine, LBP-3 was cloned into 2’ position pDONR 221/P3-P2 and *RFP::unc-54 3’UTR* without SL2 sequence to 3’ position pDONR 221/P3-P2. Then, upon LR reaction and sequence verification, LBP-3::RFP was amplified and cloned into 2’ position pDONR 221/P4r-P3r followed by Gateway BP and LR reactions as described above.

For generating transgenic lines driven by endogenous promoters, the whole genomic region of *nlp-11*, or *lbp-3* including the promoter, 5’UTR, coding sequence and 3’UTR was first PCR amplified and then fused together with *sl2-GFP::unc-54 3’UTR* using fusion PCR. 2.2kbp or 1.1kbp of the upstream promoter region was used for *nlp-11* or *lbp-3* respectively.

Transgenic strains were generated by injecting a DNA mixture consisting of 10 ng/μL transgenic construct, 10 ng/μL co-injection marker and 80 ng/μL salmon sperm DNA into the gonads of young adult worms using the standard *C. elegans* microinjection protocol. For generating integrated strains, late L4 worms containing extrachromosomal arrays were exposed to 4000 rads of gamma irradiation for 5.9 min, and integrated strains were backcrossed to N2 at least five times.

### CRISPR methodology for generating deletion mutants

For *nlp-11*, mutations were generated using saturated sgRNA targeting throughout the *nlp-11* locus. sgRNAs were identified using the http://crispr.mit.edu/website. Possible sgRNAs were then screened for predicted efficacy using http://crispr.wustl.edu/. Following the protocol suggested by Dickinson *et al.* ^22^ and Ward et al. ^23^, candidate sgRNAs for *nlp-11* were cloned into pDD162 using the Q5 site directed mutagenesis kit and universal primer. 60 animals were injected with the following injection mix: two *nlp-11* specific sgRNAs created from pDD162 (active concentration 25ng/μL each), *dpy-10(cn64)* repair oligo (500nM), pJA58 targeting *dpy-10* (active concentration 25ng/μL) and *myo-2p::mCherry* (active concentration 25ng/μL).

For *lbp-3* deletion mutations, following the protocol suggested by Paix *et. al.* ^24^ and Arribere *et al.*^25^, specific crRNAs for *lbp-3* were identified using http://crispor.tefor.net/ and predicted efficacy using http://crispr.wustl.edu/. Together with *dpy-10* crRNA and tracrRNA, *lbp-3* specific crRNAs were purchased from Dhamarcon, while Cas9 protein from PNABIO. All the sequences are provided in Supplementary Table 7. CRISPR injection mix were added in the following order: 0.8μl *dpy-10* crRNA (4.0μg/μL), 1.0μl gene specific crRNAs (8.0μg/μL) or 1.0μl if using two gene specific crRNAs (4.0μg/μL), 5.0μl tracrRNA (4μg/μL), and RNAse free water up to 15μL. The mix was heated at 95°C for 2 minutes and cool down to RT. 1μL of purified Cas9 (1.0μg/μL) was added at the end to minimize pipetting. After spun for two minutes at 13000rpm and heated at 37°C for 15 minutes, the mixes were injected into the gonads of 60 animals as described above.

In both strategies, F1 worms were screened for the co-crispr injection marker *dpy-10* that was introduced through either pJA58 or *dpy-10* crRNA co-injections. Candidate F1 worms were individually picked, allowed to lay progeny, and then lysed to get genomic DNA. Genotyping PCR was performed using *nlp-11* and *lbp-3* spanning primers listed in Supplementary Table 6. Candidate worms with notable band shifting were saved and back crossed at least four times with N2.

### Genetic crosses

Genetic crosses were performed using standard methods. In brief, *lipl-4 Tg* or *lbp-3 Tg* transgenic males were generated by crossing N2 males to the transgenic strains that carry fluorescence markers. The transgenic males with the fluorescence markers were then crossed to the desired strain. Full list of *C. elegans* strains is provided in Supplementary Table 5.

### Preparation of synchronized worms

Gravid adults were collected in a 15 mL conical tube with 10 mL of M9 buffer (22mM KH_2_PO_4_, 22mM Na_2_HPO_4_, 85mM NaCl, 2mM MgSO_4_). A bleach solution (60% NaOCl, 1.6M NaOH solution) was freshly prepared. Worms were spun down in a centrifuge for 30 seconds at 3000rpm, and M9 was aspirated. 2 mL of bleach was added to each 4mL aliquot of adult worms. Worm bodies were dissolved in the conical tube by shaking for 1 minute and repeating this step once more. After immediate spinning and aspiration of the bleach solution, collective embryos were rinsed four times in 10 mL of M9. Embryos were allowed to hatch in M9 on a rotator at 20°C.

### Lifespan assays

Worms were synchronized by bleach-based egg isolation followed by starvation in M9 buffer at the L1 stage for 24 hours. For all experiments, every genotype and condition were performed in parallel. Synchronized L1 worms were grown to the first day of adulthood, and Day 1 was determined by the onset of egg-laying. During adulthood, worms were transferred to new plates every two days. 60-120 animals were assayed for each condition/genotype with 30-40 animals per 6cm plate. Death was indicated by total cessation of movement in response to gentle mechanical stimulation. All lifespan assays were carried at 20°C. For integrated transgenic strains and newly isolated CRISPR deletion mutants, the strains were backcrossed at least five times prior to lifespan analysis (Supplementary Table 3, 4).

For lifespan assays involving strains containing mutation of *egl-21* or *fat-3*, FUDR at the final concentration of 100μM was added at L4 stage to prevent aging-irrelevant lethality due to internal eggs’ hatching. All the other lifespan assays did not use FUDR.

### Bacteria culture

For preparing worm plates with OP50, a single colony of *E. coli* OP50 was cultured in lysogeny broth (LB) at 37°C shaking overnight with 100 μg/mL streptomycin, and 300 μL of OP50 culture was seeded onto NGM plates and dried. RNAi HT115 colonies were selected by carbenicillin (50 μg/mL) and tetracycline (50 μg/mL) and verified by Sanger sequencing. For preparing RNAi feeding plates, RNAi clones were cultured in lysogeny broth (LB) with carbenicillin (50 μg/mL) at 37°C shaking for 14 hours, and 300 μL of RNAi culture was seeded onto IPTG-containing RNAi plates that were dried in a laminar flow hood and incubated at room temperature for 24 hours. Bacteria transformed with the L4440 vector (used to construct all the RNAi clones) were used as empty vector controls for RNAi experiments.

### Quantitative RT-PCR

Total RNA was isolated from more than 2000 age-synchronized young adult worms for each genotype in three or four independent biological replicates by Trizol® extraction followed by column purification (Qiagen). Synthesis of cDNA was performed using the amfiRivert Platinum cDNA Synthesis Master Mix (GenDEPOT). Quantitative PCR was performed using Kapa SYBR fast PCR kit (Kapa Biosystems) in a Realplex 4 PCR machine (Eppendorf), and values were normalized to *rpl-32* as an internal control. All data shown represent three-four biologically independent samples. Primers used in this study are listed in Supplementary Table 6.

### Fluorescent microscopy

Day 1 adult worms were mounted on 2% agarose pads containing 0.5% NaN3 as anesthetic on glass microscope slides. Fluorescent images were taken using confocal FV3000 (Olympus). Polygon selection tool was used to select coelomocytes’ area to be quantified and average pixel intensity was calculated with the “analyze-measure” command. After subtracting the background intensity, all measurements were averaged to obtain mean and standard deviation. In each imaging session, around 15-20 animals were analyzed. LBP-3 fluorescence intensity in *lipl-4Tg* were normalized to wild-type levels (Fig. 3), while LBP-3 fluorescence intensity in *lipl-4Tg* animals fed with *fat-3* RNAi conditions were normalized to empty vector WT animals (Fig. 4).

### Lysotracker staining

LysoTracker Red DND-99 (Molecular Probes) was diluted in ddH_2_O to 1000μM, and 6 μL were added to each 3.5 cm standard NGM plate (containing 3mL of agar) seeded with OP50. The plates were kept in the dark for 24 hours to allow the lysotracker solution to diffuse evenly throughout the plate. 10~20 worms were added to each plate at the L4 stage and kept in the dark for 1 day at 20°C before imaging with a confocal microscope.

### Free fatty acid profiling

For each sample, 40,000 age-synchronized worms were grown on NGM plates seed with OP50 bacteria and collected as young adults. The worms were washed three times in M9 buffer and returned to empty NGM plates for 30 minutes for gut clearance. Following intestinal clearance, worms were washed twice more in M9 buffer, pelleted in a minimal volume of M9 and flash frozen in liquid nitrogen. Proprietary recovery standards were added to each sample prior to extraction, and the samples were extracted using the methanol, chloroform, and water. The extracted samples in chloroform were dried and resuspended in methanol and isopropanol (50:50, vol/vol). The samples were analyzed using a Vanquish UPLC and an LTQ-FT mass spectrometer with a linear ion-trap (LIT) front end and a Fourier transform ion cyclotron resonance (FT-ICR) back end (Thermo Fisher Scientific Inc.). The mobile phase A was 5 mM ammonium acetate with pH 5 and mobile phase B was 2-propanol and acetonitrile (20:80, vol/vol). The free fatty acids were first identified using lipidsearch software, and then validated using standards.

### Lipid feeding

Age synchronized worms were grown on NGM plates seeded with OP50 bacteria to the day-1 adulthood. Arachidonic acid (AA), dihomo-γ-linolenic acid (DGLA) and eicosapentaenoic acid (EPA) (Nu-Check Prep) was dissolved in DMSO and diluted into OP50 bacterial food to a final concentration of 1mM. 300 μL of each mixture was added to standard 6 cm NGM plates that were dried in a laminar flow hood under dark conditions. Worms were collected after 12 hours of lipid feeding under dark conditions followed by RNA extraction and qRT-PCR.

### Competitive Fluorescence-based binding assay

LBP-3 cDNA was subcloned into the pMCSG7 plasmid containing a N-terminal His_6_ tag followed by a TEV protease cleavage site to assist in purification. Recombinant expression of His_6_-LBP3 in *E. coli* BL21 (DE3) was induced with 0.5 mM isopropyl-1-thio-b-D-galactopyranoside (IPTG) for 18 hours at 16°C. LBP-3 was purified using nickel affinity chromatography, followed by tag removal and size exclusion chromatography. Quantification of ligand binding was conducted via competition of the probe 1-anilinonaphthalene-8-sulfonic acid (1,8-ANS), a small molecule whose fluorescence increases drastically when surrounded by a hydrophobic environment and which has been shown to bind an array of intracellular lipid binding proteins (iLBPs) with varying affinity. Briefly, binding of 1,8-ANS was carried out in a buffer containing 20 mM Tris HCl pH 7.4, 150 mM NaCl, 5 % glycerol, and 0.5 mM TCEP with a constant amount of 1,8-ANS (500 nM) and increasing amounts of pure LBP-3 (40 nM – 400 μM). Blank measurements containing LBP-3 alone were subtracted from each protein concentration tested, and the resulting values were fit with a one site binding curve to determine the binding constant, K_d_. Competition assays were then carried out in the same buffer using a constant concentration of 500 nM protein and 10 μM 1,8-ANS, with ligand added via 50X ethanol stocks to maintain an ethanol concentration of 2%. Following a one-hour incubation at 37 °C, data were collected on a BioTek Synergy NEO plate reader using an excitation wavelength of 360 nm and an emission wavelength of 525 nm. Blank wells containing only ligand and 1,8-ANS were subtracted from wells with protein at each ligand concentration tested. Background subtracted values were fit with a one site (Fit Ki) curve to calculate the Ki in GraphPad Prism 8. All curves are the average of three independent experiments.

### RNA-seq preparation and analysis

Total RNA was extracted from around 3000 worms for each genotype in three different biological replicates using Trizol extraction combined with column purification (Qiagen). Sequencing libraries were prepared using the TruSeq Stranded mRNA Sample Preparation kit (Illumina) following the manufacturer’s instructions. Libraries were pooled together and sequenced using Illumina NextSeq 500 system. RNA-seq reads were aligned to the *C. elegans* reference genome using hisat2 with the default setting. HTSeq was used to count the read numbers mapped to each gene. The DESeq2 was used to normalize the raw counts and identify differentially expressed genes (|fold change| ≥ 1.5; FDR < 0.05).

### Statistical information

For survival analysis, statistical analyses were performed with SPSS software (IBM Software) using Kaplan-Meier survival analysis and log-rank test. Details on samples size, number of biological replicates and statistics for each experiment are provided in the lifespan tables (Supplementary Table 3 and 4). For Free Fatty Acid profiling, statistical analysis was performed using two-way ANOVA test with Holm-Sidak correction to identify compounds that differed significantly between *lipl-4 Tg* (n=9 biological replicates) vs. WT (n=6 biological replicates) or between *fat-3(wa22)* (n=5 biological replicates) vs. WT (n=5 biological replicates). For qRT-PCR, all comparisons of means were accomplished using two-way ANOVA and multiple t-test using Holm-Sidak correction as indicated in the corresponding figure legends. For all figure legends, asterisks indicate statistical significance as follows: n.s. = not significant p>0.05; * p<0.05, **p<0.01, ***p<0.001, **** p<0.0001. Data were obtained by performing independently at least three biological replicates. Figures and graphs were constructed using GraphPad Prism 7 (GraphPad Software) and Illustrator (CC 2019; Adobe).

## Notes

### Competing Interest Statement

The authors have declared no competing interest.

