## Supplementary Materials for "Lysosome Lipid Signaling from the Periphery to Neurons Regulates Longevity"

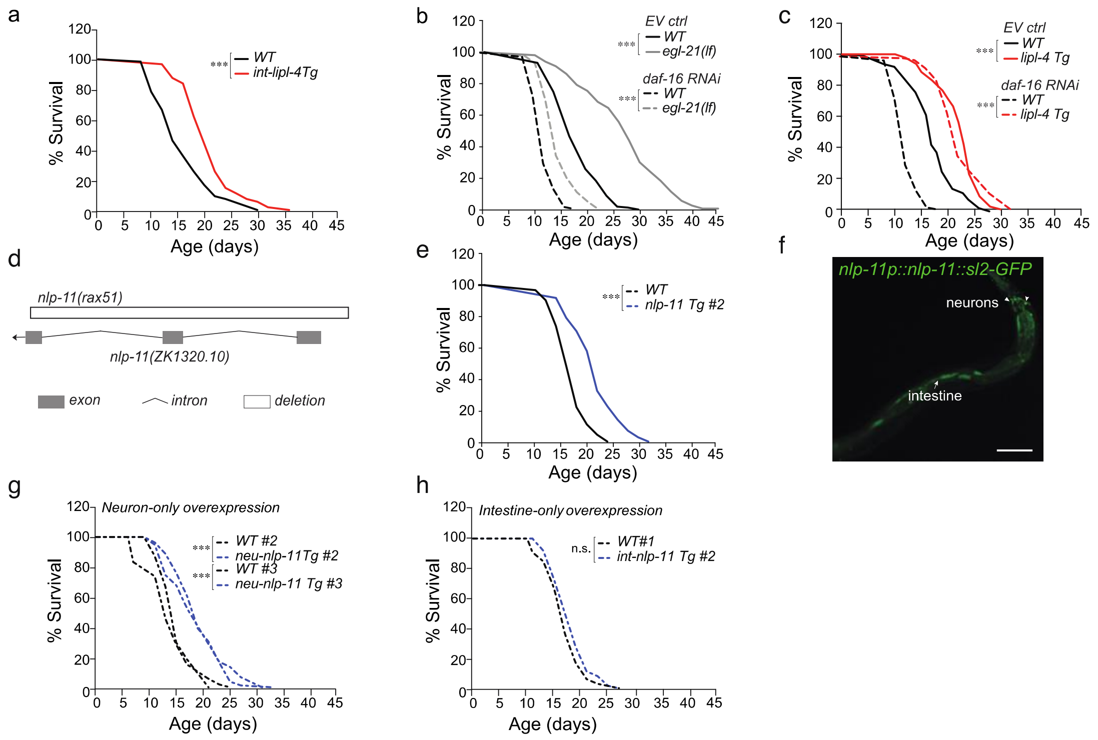


**Supplementary Figure 1: Peripheral lysosomal lipolysis induces neuropeptide to promote longevity (related to Figure 1)**

**a**) Overexpression of *lipl-4* selectively in the intestine extends lifespan.

**b**) In the background of *daf-16* RNAi inactivation, the longevity effect of the *egl-21(n476)* loss-of-function mutants is suppressed.

**c**) Inactivation of *daf-16* does not affect the longevity effect of *lipl-4 Tg*.

**d**) A schematic representation of the *nlp-11(rax51)* mutation. White boxes represent the *nlp-11(rax51)* deletion, black lines represent introns. The mutants lack the three exons and the transcriptional start site.

**e**) Constitutive expression of *nlp-11* driven by its endogenous promoter extends lifespan.

**f**) The transgenic strains expressing *nlp-11* under its endogenous promoter *(nlp-11::sl2-GFP)* reveals the expression of *nlp-11* in intestinal and neuronal cells. Scale bar 100µm.

**g** and **h)** Neuron-specific overexpression of *nlp-11* prolongs lifespan (**g**), but intestine-specific overexpression has no such effect (**h**).

Lifespan analyses were performed using Kaplan-Meier survival analysis and log-rank test (a, b, c, e, g, h).

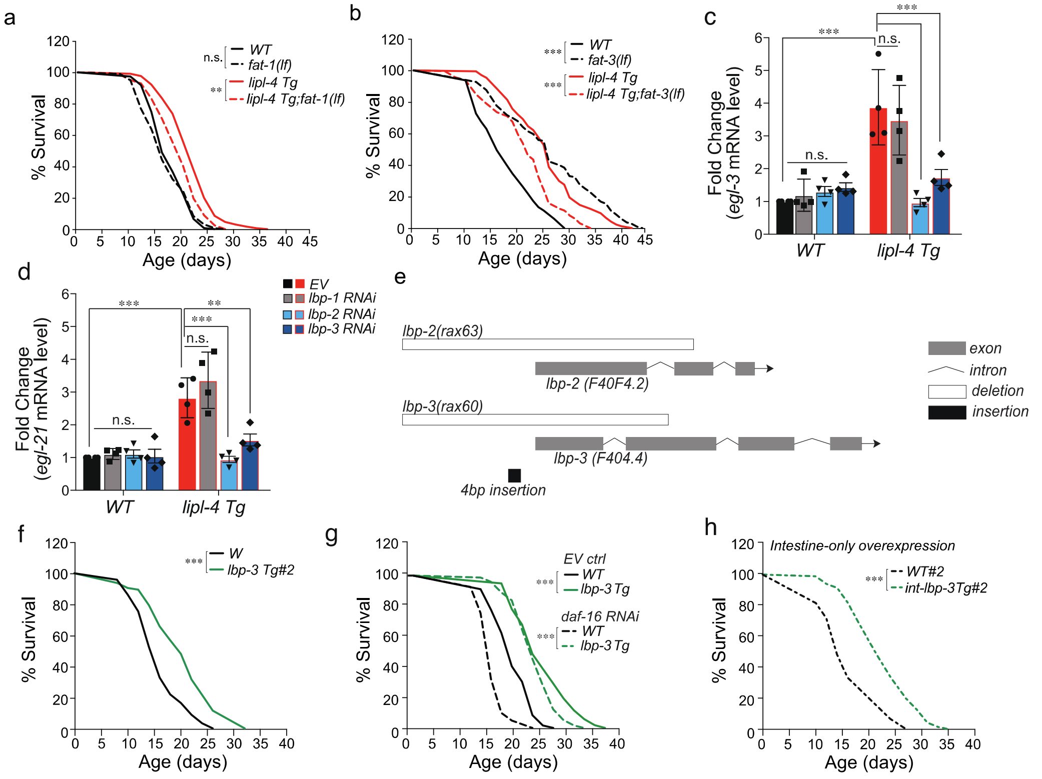


**Supplementary Figure 2: Peripheral lipid signals mediate neuropeptide induction and longevity (related to Figure 2)**

**a)** The loss-of-function mutant *fat-1(wa9)* suppresses *lipl-4 Tg* longevity.

**b)** In the *fat-3(wa22)* loss-of-function mutant background, the lifespan-extending effect of *lipl-4 Tg* is shortened.

**c** and **d)** The induction of *egl-3* **(c)** and *egl-21* **(d)** in the *lipl-4 Tg* worms is suppressed by RNAi inactivation of *lbp-2* and *lbp-3*, but not by *lbp-1*.

**e**) Schematic representation of *lbp-2* and *lbp-3* loss-of-function mutants. White boxes represent the *lbp-2(rax63)* and *lbp-3(rax60)* deletions, black boxes represent insertions, while black lines represent introns. The mutants lack the entire first exon and transcriptional start site.

**f**) Constitutive expression of *lbp-3* (*lbp-3 Tg*) driven by its own endogenous promoter prolongs lifespan.

**g**) Inactivation of *daf-16* does not affect the longevity effect of *lbp-3 Tg*.

**h**) Overexpression of *lbp-3* selectively in the intestine extends lifespan.

Error bars represent mean ± s.e.m. n.s. $p>0.05$, **$p<0.01$, ***$p<0.001$, two-way ANOVA with Holm-Sidak correction (c,d), log-rank test (a, b, f, g, h).

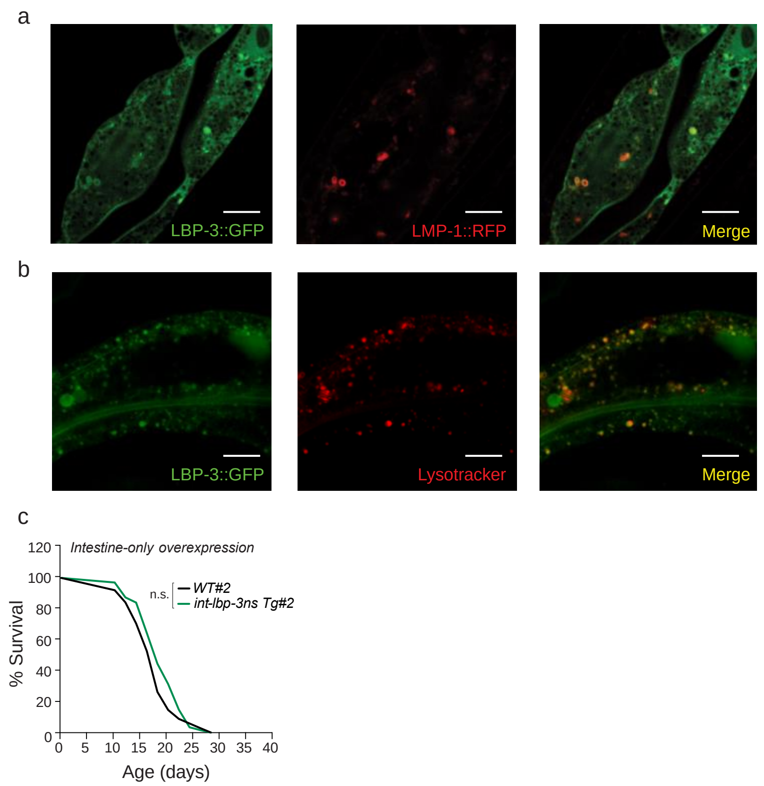


**Supplementary Figure 3: Secreted lipid chaperones act through neuronal nuclear receptor to regulate neuropeptide and longevity (related to Figure 3).**

**a)** LBP-3 and the lysosomal membrane protein LMP-1 are visualized by their GFP and RFP fusions, respectively. LBP-3::GFP colocalizes with LMP-1::RFP at lysosomes and is also detected in the cytosol. Scale bar 10µm.

**b**) LBP-3::GFP colocalizes with Lysotracker Red that marks lysosomes, and it is also detected in the cytosol. Scale bar 10 µm.

**c**) Intestine-specific overexpression of *lbp-3* lacking its secretory signal (*lbp-3ns*) fails to extend lifespan.

Lifespan analyses were performed using Kaplan-Meier survival analysis and log-rank test (c).

**Supplementary Table 1. (separate excel file)**

Transcriptome analysis of WT and *lipl-4 Tg* animals.

**Supplementary Table 2. (separate excel file)**

Transcriptome analysis of WT and *lbp-3 Tg* animals.

**Supplementary Table 3: Summary of longitudinal survival analyses using mutants (related to Fig.1, Fig. 2, Fig. 3, Supplementary Fig. 1, Supplementary Fig. 2, Supplementary Fig. 3).**

| Group | Genotype | Replicate | Lifespan (Mean ± s.e.) | *p*-value 1# | *p*-value 2& | Number Total (censor) | Lab Code Strain Number‡ | Figure Number |
|---|---|---|---|---|---|---|---|---|
| A | *WT* | #1 | 18.788 ± 0.607 | | | 88 (20) | | Supplementary Figure 1a |
| | | #2 | 15.972 ± 0.733 | | | 89 (34) | | |
| | | #3 | 17.162 ± 0.685 | | | 92 (17) | | |
| | *lipl-4 Tg* | #1 | 21.169 ± 0.660 | 0.023 | | 91 (27) | MCW1404 | |
| | | #2 | 20.837 ± 0.664 | < 0.001 | | 82 (27) | MCW1404 | |
| | | #3 | 21.309 ± 0.644 | < 0.001 | | 95 (26) | MCW1404 | |
| B | *WT** | #1 | 17.440 ± 0.426 | | | 70 (2) | | Supplementary Figure 2b |
| | | #2 | 17.432 ± 0.523 | | | 65 (2) | | |
| | | #3 | 17.751 ± 0.742 | | | 89(12) | | |
| | *lipl-4 Tg** | #1 | 24.338 ± 0.699 | < 0.001 | | 70(3) | MCW14 | |
| | | #2 | 25.312 ± 0.592 | < 0.001 | | 65 (3) | MCW14 | |
| | | #3 | 26.416 ± 0.793 | < 0.001 | | 89 (12) | MCW14 | |
| | *fat-3(wa22)** | #1 | 23.983 ± 0.984 | | < 0.001 | 65 (9) | BX30 | |
| | | #2 | 24.132 ± 0.672 | | < 0.001 | 70 (10) | BX30 | |
| | | #3 | 26.356 ± 0.935 | | < 0.001 | 93 (3) | BX30 | |
| | *fat-3(wa22);lipl-4 Tg** | #1 | 19.423 ± 0.495 | < 0.001 | < 0.001 | 65 (3) | MCW1117 | |
| | | #2 | 19.932 ± 0.823 | < 0.001 | < 0.001 | 70 (4) | MCW1117 | |
| | | #3 | 21.187 ± 0.675 | < 0.001 | < 0.001 | 99 (3) | MCW1117 | |
| C | *WT* | #1 | 17.762 ± 0.692 | | | 80 (11) | | Supplementary Figure 2a |
| | | #2 | 17.143 ± 0.163 | | | 80 (12) | | |
| | | #3 | 17.360 ± 0.452 | | | 80 (15) | | |
| | *lipl-4 Tg* | #1 | 20.835 ± 0.628 | < 0.001 | | 83 (19) | MCW14 | |
| | | #2 | 21.143 ± 0.218 | < 0.001 | | 83 (18) | MCW14 | |
| | | #3 | 20.550 ± 0.457 | < 0.001 | | 84 (26) | MCW14 | |
| | *fat-1(wa9)* | #1 | 16.985 ± 0.438 | | 0.951 | 80 (8) | BX24 | |
| | | #2 | 17.185 ± 0.262 | | 0.983 | 80 (9) | BX24 | |
| | | #3 | 17.346 ± 0.525 | | 0.612 | 80 (11) | BX24 | |
| | *fat-1(wa9);lipl-4 Tg* | #1 | 18.214 ± 0.459 | 0.103 | 0.002 | 70 (10) | MCW643 | |
| | | #2 | 18.134 ± 0.528 | 0.382 | 0.002 | 77 (11) | MCW643 | |
| | | #3 | 18.176 ± 0.517 | 0.419 | 0.003 | 75 (14) | MCW643 | |
| | *WT* | #1 | 16.132 ± 0.671 | | | 89 (15) | | Figure 2g |
| | | #2 | 14.927 ± 0.409 | | | 92 (10) | | |
| | | #3 | 15.372 ± 0.538 | | | 90 (4) | | |

| | | | | | | | | |
|---|---|---|---|---|---|---|---|---|
| D | *lipl-4 Tg* | #1 | 26.051 ± 0.676 | < 0.001 | | 106 (26) | MCW14 | |
| | | #2 | 28.961 ± 0.506 | < 0.001 | | 90 (13) | MCW14 | |
| | | #3 | 27.423 ± 0.385 | < 0.001 | | 90 (14) | MCW14 | |
| | *lbp-3(rax60)* | #1 | 16.706 ± 0.588 | | 0.756 | 95 (16) | MCW1118 | |
| | | #2 | 15.167 ± 0.400 | | 0.619 | 88 (4) | MCW1118 | |
| | | #3 | 15.183 ± 0.325 | | 0.728 | 90 (3) | MCW1118 | |
| | *lbp-3(rax60);lipl-4 Tg* | #1 | 20.017 ± 0.628 | < 0.001 | < 0.001 | 91 (15) | MCW1143 | |
| | | #2 | 19.894 ± 0.508 | < 0.001 | < 0.001 | 91 (28) | MCW1143 | |
| | | #3 | 19.422 ± 0.723 | < 0.001 | < 0.001 | 90 (12) | MCW1143 | |
| E | *WT* | #1 | 20.223 ± 0.597 | | | 80 (13) | | Figure 2h |
| | | #2 | 19.526 ± 0.823 | | | 80 (13) | | |
| | | #3 | 15.871 ± 0.461 | | | 101 (9) | | |
| | *lbp-3 Tg #1* | #1 | 25.631 ± 0.712 | < 0.001 | | 80 (26) | MCW966 | |
| | | #2 | 24.824 ± 0.834 | < 0.001 | | 80 (26) | MCW966 | |
| | | #3 | 20.002 ± 0.653 | < 0.001 | | 100 (18) | MCW966 | |
| | *WT* | #1 | 17.435 ± 0.534 | | | 90 (3) | | Supplementary Figure 2f |
| | | #2 | 17.578 ± 0.476 | | | 90 (7) | | |
| | | #3 | 17.693 ± 0.638 | | | 89 (7) | | |
| | *lbp-3 Tg #2* | #1 | 22.643 ± 0.427 | < 0.001 | | 90 (12) | MCW1163 | |
| | | #2 | 22.528 ± 0.523 | < 0.001 | | 90 (15) | MCW1163 | |
| | | #3 | 22.394 ± 0.732 | < 0.001 | | 90 (16) | MCW1163 | |
| F | *WT* | #1 | 19.231 ± 0.432 | | | 70 (7) | | Figure 2n |
| | | #2 | 18.672 ± 0.342 | | | 70 (7) | | |
| | | #3 | 16.200 ± 0.668 | | | 90 (30) | | Supplementary Figure 2g |
| | *ges-1p::lbp3 nonTg* | #1 | 18.923 ± 0.723 | 0.982 | | 70 (10) | MCW1129 | Figure 2n |
| | | #2 | 18.346 ± 0.234 | 0.927 | | 70 (9) | MCW1129 | |
| | | #3 | 15.999 ± 0.699 | 0.938 | | 95 (23) | MCW1130 | Supplementary Figure 2g |
| | *ges-1p::lbp3 Tg* | #1 | 22.823 ± 0.817 | < 0.001 | | 70 (15) | MCW1129 | Figure 2n |
| | | #2 | 22.623 ± 0.349 | < 0.001 | | 68 (16) | MCW1129 | |
| | | #3 | 22.723 ± 0.446 | < 0.001 | | 95 (23) | MCW1130 | Supplementary Figure 2g |
| G | *ges-1p::nslbp3 nonTg* | #1 | 17.161 ± 0.491 | | | 97 (71) | MCW1381 | Figure 3d |
| | | #2 | 17.662 ± 0.547 | | | 81 (17) | MCW1381 | |
| | | #3 | 18.599 ± 0.500 | | | 100 (30) | MCW1381 | |
| | *ges-1p::nslbp3 Tg* | #1 | 18.571 ± 0.518 | 0.085 | | 75 (14) | MCW1381 | |
| | | #2 | 19.764 ± 0.582 | 0.005 | | 90 (21) | MCW1381 | |
| | | #3 | 19.360 ± 0.571 | 0.382 | | 72 (20) | MCW1381 | |
| H | *WT* | #1 | 17.650 ± 0.520 | | | 90 (14) | | Figure 1g |
| | | #2 | 19.448 ± 0.469 | | | 89 (17) | | |
| | | #3 | 17.438 ± 0.284 | | | 90 (5) | | |
| | *lipl-4 Tg* | #1 | 24.003 ± 0.552 | < 0.001 | | 135 (27) | MCW14 | |
| | | #2 | 27.127 ± 0.601 | < 0.001 | | 90 (22) | MCW14 | |
| | | #3 | 24.735 ± 0.328 | < 0.001 | | 90 4) | MCW14 | |
| | | #1 | 17.628 ± 0.513 | | 0.940 | 93 (12) | MCW630 | |

| | | | | | | | | |
|---|---|---|---|---|---|---|---|---|
| | *nlp-11(rax51)* | #2 | 19.986 ± 0.477 | | 0.951 | 90 (13) | MCW630 | |
| | | #3 | 17.826 ± 0.835 | | 0.978 | 90 (3) | MCW630 | |
| | *nlp-11(rax51);lipl-4 Tg* | #1 | 21.140 ± 0.577 | <0.001 | 0.001 | 119 (26) | MCW629 | |
| | | #2 | 20.282 ± 0.552 | 0.551 | <0.001 | 102 (35) | MCW629 | |
| | | #3 | 20.736 ± 0.382 | <0.001 | <0.001 | 90 (24) | MCW629 | |
| I | *WT* | #1 | 18.595 ± 0.439 | | | 105 (91) | | Figure 1h |
| | | #2 | 17.086 ± 0.458 | | | 90 (20) | | |
| | | #3 | 16.967 ± 0.387 | | | 75 (34) | | |
| | *nlp-11 Tg #1* | #1 | 22.892 ± 0.549 | < 0.001 | | 104 (83) | MCW231 | |
| | | #2 | 21.568 ± 0.551 | < 0.001 | | 109 (19) | MCW231 | |
| | | #3 | 22.407 ± 0.551 | < 0.001 | | 104 (18) | MCW231 | |
| | *WT* | #1 | 18.375 ± 0.402 | | | 105 (14) | | Supplementary Figure 1e |
| | | #2 | 15.762 ± 0.351 | | | 114 (39) | | |
| | | #3 | 14.455 ± 0.549 | | | 90 (14) | | |
| | *nlp-11 Tg #2* | #1 | 20.636 ± 0.528 | < 0.001 | | 109 (27) | MCW644 | |
| | | #2 | 24.224 ± 0.529 | < 0.001 | | 120 (20) | MCW644 | |
| | | #3 | 22.474 ± 0.599 | < 0.001 | | 90 (14) | MCW644 | |
| J | *ges-1p::nlp-11 nonTg* | #1 | 17.121 ± 0.412 | | | 76 (10) | MCW289 | Figure 1k |
| | | #2 | 15.912 ± 0.349 | | | 77 (9) | MCW289 | |
| | | #3 | 16.683 ± 0.395 | | | 80 (5) | MCW290 | Supplementary Figure 1h |
| | *ges-1p::nlp-11 Tg* | #1 | 18.419 ± 0.470 | 0.048 | | 78 (16) | MCW289 | Figure 1k |
| | | #2 | 16.608 ± 0.454 | 0.110 | | 75 (13) | MCW289 | |
| | | #3 | 17.153 ± 0.438 | 0.328 | | 80 (8) | MCW290 | Supplementary Figure 1h |
| K | *rab-3p::nlp-11 nonTg* | #1 | 15.000 ± 0.337 | | | 72 (7) | MCW291 | Figure 1j |
| | | #2 | 13.742 ± 0.530 | | | 66 (11) | MCW292 | Supplementary Figure 1g |
| | | #3 | 15.234± 0.389 | | | 82 (28) | MCW293 | Supplementary Figure 1g |
| | *rab-3p::nlp-11 Tg* | #1 | 19.007 ± 0.700 | < 0.001 | | 86 (27) | MCW291 | Figure 1j |
| | | #2 | 18.553 ± 0.527 | < 0.001 | | 84 (8) | MCW292 | Supplementary Figure 1g |
| | | #3 | 19.320 ± 0.490 | < 0.001 | | 94 (15) | MCW293 | Supplementary Figure 1g |
| | *WT* | #1 | 16.935 ± 0.538 | | | 100 (14) | | Figure 2l |
| | | #2 | 17.103 ± 0.525 | | | 100 (13) | | |
| | | #3 | 18.072 ± 0.467 | | | 90 (7) | | |
| | | #4 | 17.671 ± 0.454 | | | 90 (6) | | |
| | *lbp-3 Tg* | #1 | 21.650 ± 0.643 | < 0.001 | | 100 (15) | MCW966 | Figure 2l |
| | | #2 | 21.281 ± 0.633 | < 0.001 | | 101 (12) | MCW966 | |
| | | #3 | 24.940 ± 0.482 | < 0.001 | | 93 (12) | MCW966 | |

| | | | | | | | | |
|---|---|---|---|---|---|---|---|---|
| L | | #4 | 24.448 ± 0.470 | < 0.001 | | 100 (9) | MCW966 | |
| | *nlp-11(rax51)* | #1 | 17.260 ± 0.553 | | 0.837 | 100 (14) | MCW630 | Figure 2l |
| | | #2 | 17.281 ± 0.518 | | 0.581 | 100 (11) | MCW630 | |
| | | #3 | 18.154 ± 0.440 | | 0.887 | 100 (10) | MCW630 | |
| | | #4 | 18.044 ± 0.424 | | 0.642 | 100 (9) | MCW630 | |
| | *nlp-11(rax51);lbp-3 Tg* | #1 | 19.839 ± 0.536 | 0.002 | 0.007 | 100 (14) | MCW1368 | Figure 2l |
| | | #2 | 19.215 ± 0.568 | 0.003 | 0.009 | 101 (12) | MCW1368 | |
| | | #3 | 21.191 ± 0.466 | < 0.001 | < 0.001 | 81 (9) | MCW1368 | |
| | | #4 | 21.442 ± 0.488 | < 0.001 | < 0.001 | 101 (20) | MCW1368 | |
| M | *WT* | #1 | 18.449 ± 0.407 | | | 100 (16) | | Figure 3h |
| | | #2 | 18.674 ± 0.303 | | | 100 (8) | | |
| | | #3 | 18.582 ± 0.565 | | | 100 (5) | | |
| | *lbp-3 Tg* | #1 | 24.333 ± 0.671 | < 0.001 | | 80 (19) | MCW966 | |
| | | #2 | 23.399 ± 0.697 | < 0.001 | | 101 (18) | MCW966 | |
| | | #3 | 23.932 ± 0.628 | < 0.001 | | 100 (12) | MCW966 | |
| | *nhr-49(nr2041)* | #1 | 14.480 ± 0.322 | | < 0.001 | 100 (0) | STE68 | |
| | | #2 | 13.811 ± 0.303 | | < 0.001 | 100 (16) | STE68 | |
| | | #3 | 14.238 ± 0.397 | | < 0.001 | 100 (5) | STE68 | |
| | *lbp-3 Tg; nhr-49(nr2041)* | #1 | 13.680 ± 0.342 | 0.105 | < 0.001 | 99 (24) | MCW1387 | |
| | | #2 | 13.316 ± 0.312 | 0.290 | < 0.001 | 100 (24) | MCW1387 | |
| | | #3 | 13.542 ± 0.368 | 0.096 | < 0.001 | 90 (12) | MCW1387 | |
| | *lbp-3 Tg; nhr-49(nr2041); rab-3p::nhr-49* | #1 | 22.947 ± 0.599 | < 0.001 | 0.111 | 95 (25) | MCW1389 | |
| | | #2 | 22.250 ± 0.516 | < 0.001 | 0.032 | 101 (29) | MCW1389 | |
| | | #3 | 22.853 ± 0.542 | < 0.001 | 0.052 | 90 (16) | MCW1389 | |

Note:
### *p*-value 1: compare *WT vs. Tg* or *Mutant vs. Mutant;Tg* in the same group with the same replicate number using a log-rank test.
& *p*-value 2: compare *WT vs. Mutant* or *Tg vs. Mutant;Tg* in the same group with the same replicate number using a log-rank test.
‡ Details on the strain genotype is provided in Supplementary Table 5.
* FUDR is added to inhibit reproduction in the experiments where certain mutants cause internal hatching of progeny.
Combined data from multiple replicates of one line are used to generate lifespan curves shown in figures.

**Supplementary Table 4: Summary of longitudinal survival analyses using RNAi treatments (related to Fig. 1, Fig.2, Supplementary Fig. 1).**

| Group | Genotype | RNAi | Replicate | Lifespan (Mean ± s.e.) | *p*-value 1# | *p*-value 2& | Number Total (censor) | Lab Code Strain Name‡ | Figure Number |
|---|---|---|---|---|---|---|---|---|---|
| A | *WT** | *vector* | #1 | 16.324 ± 0.832 | | | 90 (4) | | Figure 1e |
| | | | #2 | 16.271 ± 0.842 | | | 90 (8) | | |
| | | | #3 | 16.438 ± 0.857 | | | 90 (6) | | |
| | *lipl-4 Tg** | *vector* | #1 | 24.278 ± 0.259 | <0.001 | | 90 (15) | MCW14 | |
| | | | #2 | 25.216 ± 0.395 | <0.001 | | 90 (18) | MCW14 | |
| | | | #3 | 24.538 ± 0.428 | <0.001 | | 90 (8) | MCW14 | |
| | *WT** | *daf-16* | #1 | 13.681 ± 0.975 | | <0.001 | 100 (2) | | |
| | | | #2 | 12.253 ± 0.791 | | <0.001 | 89 (17) | | |
| | | | #3 | 13.147 ± 0.825 | | <0.001 | 90 (4) | | |
| | *lipl-4 Tg** | *daf-16* | #1 | 21.162 ± 0.432 | <0.001 | <0.001 | 100 (6) | MCW14 | |
| | | | #2 | 22.630 ± 0.494 | <0.001 | <0.001 | 91 (14) | MCW14 | |
| | | | #3 | 21.864± 0.468 | <0.001 | <0.001 | 90 (7) | MCW14 | |
| | *egl-21(n476) ** | *vector* | #1 | 25.245 ± 0.264 | | | 90 (10) | KP2018 | |
| | | | #2 | 25.935 ± 0.238 | | | 90 (11) | KP2018 | |
| | | | #3 | 25.487 ± 0.275 | | | 90 (12) | KP2018 | |
| | *egl-21(n476);lipl-4 Tg** | *vector* | #1 | 23.613 ± 0.672 | 0.032 | | 90 (12) | MCW1365 | |
| | | | #2 | 23.840 ± 0.836 | 0.029 | | 90 (13) | MCW1365 | |
| | | | #3 | 23.538 ± 0.742 | 0.035 | | 90 (14) | MCW1365 | |
| | *egl-21(n476)** | *daf-16* | #1 | 16.452 ± 0.313 | | <0.001 | 98 (14) | KP2018 | |
| | | | #2 | 14.496 ± 0.312 | | <0.001 | 91 (10) | KP2018 | |
| | | | #3 | 15.352 ± 0.334 | | <0.001 | 90 (16) | KP2018 | |
| | *egl-21(n476);lipl-4 Tg** | *daf-16* | #1 | 16.241 ± 0.291 | 0.731 | <0.001 | 96 (11) | MCW1365 | |
| | | | #2 | 14.597 ± 0.346 | 0.845 | <0.001 | 90 (9) | MCW1365 | |
| | | | #3 | 15.352 ± 0.548 | 0.684 | <0.001 | 90 (8) | MCW1365 | |
| | *Intestinal RNAi* | | | | | | | | |
| B | *WT* | *vector* | #1 | 18.510 ± 0.481 | | | 102 (4) | | Figure 1g |
| | | | #2 | 18.850 ± 0.447 | | | 110 (3) | | |
| | | | #3 | 18.634 ± 0.456 | | | 90 (5) | | |
| | *lipl-4 Tg* | *vector* | #1 | 23.375 ± 0.551 | <0.001 | | 65 (1) | MCW1056 | |
| | | | #2 | 23.618 ± 0.548 | <0.001 | | 72 (4) | MCW1056 | |
| | | | #3 | 23.543 ± 0.586 | <0.001 | | 90 (4) | MCW1056 | |
| | *WT* | *nlp-11* | #1 | 19.591 ± 0.476 | | 0.087 | 96 (8) | | |
| | | | #2 | 19.000 ± 0.488 | | 0.502 | 95 (5) | | |
| | | | #3 | 19.234 ± 0.473 | | 0.094 | 90 (7) | | |
| | *lipl-4 Tg* | *nlp-11* | #1 | 23.602 ± 0.579 | <0.001 | 0.702 | 70 (8) | MCW1056 | |
| | | | #2 | 22.622 ± 0.592 | <0.001 | 0.277 | 80 (11) | MCW1056 | |
| | | | #3 | 23.426 ± 0.584 | <0.001 | 0.754 | 90 (12) | MCW1056 | |
| C | | | #1 | 17.051 ± 0.761 | | <0.001 | 70 (5) | | Figure 2d |

| | | | | | | | | | |
|---|---|---|---|---|---|---|---|---|---|
| | *WT* | *vector* | #2 | 17.252 ± 0.674 | | <0.001 | 70 (7) | | |
| | | | #3 | 17.148± 0.694 | | <0.001 | 70 (4) | | |
| | *lipl-4 Tg* | *vector* | #1 | 23.154 ± 0.692 | <0.001 | | 70 (10) | MCW1056 | |
| | | | #2 | 23.253 ± 0.463 | <0.001 | | 70 (13) | MCW1056 | |
| | | | #3 | 23.148 ± 0.527 | <0.001 | | 70 (14) | MCW1056 | |
| | *WT* | *fat-1* | #1 | 17.801 ± 0.764 | | 0.614 | 68 (9) | | |
| | | | #2 | 16.702 ± 0.771 | | 0.865 | 64 (7) | | |
| | | | #3 | 17.426 ± 0.753 | | 0.753 | 70 (8) | | |
| | *lipl-4 Tg* | *fat-1* | #1 | 17.744 ± 0.442 | 0.551 | <0.001 | 71 (15) | MCW1056 | |
| | | | #2 | 18.286 ± 0.622 | 0.839 | <0.001 | 64 (8) | MCW1056 | |
| | | | #3 | 17.932 ± 0.548 | 0.762 | <0.001 | 70 (12) | MCW1056 | |
| D | *WT* | *vector* | #1 | 17.412 ± 0.491 | | | 80 (9) | | Figure 2e |
| | | | #2 | 18.711 ± 0.554 | | | 80 (8) | | |
| | | | #3 | 17.842 ± 0.523 | | | 80 (5) | | |
| | *lipl-4 Tg* | *vector* | #1 | 22.403 ± 0.762 | <0.001 | | 74 (12) | MCW1056 | |
| | | | #2 | 21.832 ± 0.832 | 0.001 | | 84 (10) | MCW1056 | |
| | | | #3 | 22.623 ± 0.649 | <0.001 | | 80 (13) | MCW1056 | |
| | *WT* | *fat-3* | #1 | 15.621 ± 0.710 | | 0.927 | 60 (24) | | |
| | | | #2 | 17.423 ± 0.701 | | 0.087 | 60 (27) | | |
| | | | #3 | 18.862 ± 0.732 | | 0.822 | 60 (17) | | |
| | *lipl-4 Tg* | *fat-3* | #1 | 15.180 ± 0.944 | 0.055 | <0.001 | 66 (27) | MCW1056 | |
| | | | #2 | 15.984 ± 0.713 | 0.351 | <0.001 | 67 (24) | MCW1056 | |
| | | | #3 | 16.521 ± 0.772 | 0.072 | <0.001 | 65 (38) | MCW1056 | |
| | *Neuronal sensitive RNAi* | | | | | | | | |
| E | *WT* | *vector* | #1 | 17.103 ± 0.525 | | | 100 (13) | | Figure 1f |
| | | | #2 | 16.935 ± 0.538 | | | 94 (14) | | |
| | | | #3 | 16.845 ± 0.582 | | | 90 (12) | | |
| | *lipl-4 Tg* | *vector* | #1 | 21.281 ± 0.633 | 0.001 | | 101 (12) | MCW84 | |
| | | | #2 | 21.650 ± 0.643 | <0.001 | | 80 (13) | MCW84 | |
| | | | #3 | 21.492 ± 0.627 | <0.001 | | 90 (11) | MCW84 | |
| | *WT* | *nlp-11* | #1 | 17.281 ± 0.518 | | 0.045 | 100 (11) | | |
| | | | #2 | 17.260 ± 0.643 | | 0.084 | 90 (12) | | |
| | | | #3 | 17.245 ± 0.735 | | 0.098 | 90 (4) | | |
| | *lipl-4 Tg* | *nlp-11* | #1 | 17.215 ± 0.548 | <0.001 | <0.001 | 101 (12) | MCW84 | |
| | | | #2 | 17.839 ± 0.536 | <0.001 | <0.001 | 71 (17) | MCW84 | |
| | | | #3 | 17.462 ± 0.586 | <0.001 | <0.001 | 70 (8) | MCW84 | |
| F | *WT** | *vector* | #1 | 19.002 ± 0.455 | | | 95 (33) | | Figure 2k, Supplementary Figure 2g |
| | | | #2 | 19.238 ± 0.471 | | | 95 (35) | | |
| | | | #3 | 19.148 ± 0.436 | | | 90 (12) | | |
| | *lbp-3 Tg** | *vector* | #1 | 23.635 ± 0.724 | <0.001 | | 90 (33) | MCW966 | |
| | | | #2 | 24.920 ± 0.701 | <0.001 | | 92 (38) | MCW966 | |
| | | | #3 | 24.436 ± 0.635 | <0.001 | | 90 (22) | MCW966 | |
| | *lbp-3 Tg; egl-21(n476)** | *vector* | #1 | 22.376 ± 0.636 | | | 105 (23) | MCW1391 | Figure 2k |
| | | | #2 | 24.679 ± 0.608 | | | 100 (18) | MCW1391 | |
| | | | #3 | 23.269 ± 0.582 | | | 90 (15) | MCW1391 | |
| | | | #1 | 17.344 ± 0.423 | | <0.001 | 92 (14) | | |

| | | | | | | | | | |
|---|---|---|---|---|---|---|---|---|---|
| | *WT** | *daf-16* | #2 | 15.519 ± 0.333 | | <0.001 | 100 (19) | | Figure 2k, Supplementary Figure 2g |
| | | | #3 | 16.472 ± 0.548 | | <0.001 | 90 (9) | | |
| | *lbp-3 Tg** | *daf-16* | #1 | 22.004 ± 0.499 | <0.001 | <0.001 | 80 (17) | MCW966 | |
| | | | #2 | 23.479 ± 0.531 | <0.001 | 0.048 | 80 (22) | MCW966 | |
| | | | #3 | 22.831 ± 0.472 | <0.001 | <0.001 | 90 (15) | MCW966 | |
| | *lbp-3 Tg; egl-21(n476)** | *daf-16* | #1 | 19.600 ± 0.377 | <0.001 | <0.001 | 100 (0) | MCW1391 | Figure 2k |
| | | | #2 | 17.140 ± 0.505 | <0.001 | <0.001 | 100 (0) | MCW1391 | |
| | | | #3 | 18.852 ± 0.438 | <0.001 | <0.001 | 90 (4) | MCW1391 | |

Note:
[#] *p*-value 1: *Tg* compared to *WT* controls under same RNAi conditions in the same group with the same replicate number by a log-rank test.
[&] *p*-value 2: RNAi treated animals compared to empty vector controls with the same genotype in the same group with the same replicate number by a log-rank test.
‡ Details on the strain genotype is provided in Supplementary Table 5.
[*] FUDR is added to inhibit reproduction in the experiments where certain mutants cause internal hatching of progeny.
Combined data from multiple replicates of one line are used to generate lifespan curves shown in figures.

**Supplementary Table 5. List of mutant and transgenic *C. elegans* strains generated in this study**

| ***Strain*** | ***Source*** | ***Lab Code*** |
|---|---|---|
| *WT* | CGC | |
| *raxEx547[ges-1p::lipl-4::3xFLAG::sl2-mRFP; myo-* | This Study | MCW1404 |
| *raxIs3[ges-1p::lipl-4::sl2-GFP; myo-2p::mCherry]* | Folick *et al.* [6] | MCW14 |
| *egl-21(n476)* | CGC | KP2018 |
| *raxIs3[ges-1p::lipl-4::sl2-GFP; myo-2p::mCherry];egl-21(n476)* | This Study | MCW1365 |
| *fat-3(wa22)* | CGC | BX30 |
| *raxIs3[ges-1p::lipl-4::sl2-GFP; myo-2p::mCherry];fat-3(wa22)* | This Study | MCW1117 |
| *fat-1(wa9)* | CGC | BX24 |
| *raxIs3[ges-1p::lipl-4::sl2-GFP; myo-2p::mCherry];fat-1(wa9)* | This Study | MCW643 |
| *lbp-3(rax60)* | This Study | MCW1118 |
| *raxIs3[ges-1p::lipl-4::sl2-GFP; myo-2p::mCherry];lbp-3(rax60)* | This Study | MCW1143 |
| *raxIs141[plbp-3::lbp-3::sl2-GFP; myo-2p::mCherry]* | This Study | MCW966 |
| *raxIs119 [lbp-3p::lbp-3::gfp, myo-2p::mCherry]* | This Study | MCW1163 |
| *Ex440[ges-1p::lbp-3::sl2-GFP;myo-2p::mCherry]*<br>*Ex441[ges-1p::lbp-3::sl2-GFP;myo-2p::mCherry]* | This Study | MCW1129<br>MCW1130 |
| *Ex545[ges-1p::nslbp3 Tg::sl2-GFP; myo-2p::mCherry]* | This Study | MCW1381 |
| *raxIs22[nlp-11Tg::sl2-GFP;myo-2p::mCherry]* | This Study | MCW644 |
| *nlp-11(rax51)* | This Study | MCW630 |
| *raxIs3[ges-1p::lipl-4::sl2-GFP; myo-2p::mCherry];nlp-* | This Study | MCW629 |
| *raxEx70[ges-1p::nlp-11::sl2-GFP; myo-2p::mCherry]*<br>*raxEx71[ges-1p::nlp-11::sl2-GFP; myo-2p::mCherry]* | This Study | MCW289<br>MCW290 |
| *raxEx72[rab-3p::nlp-11; myo-2p::mCherry]*<br>*raxEx73[rab-3p::nlp-11; myo-2p::mCherry]*<br>*raxEx74[rab-3p::nlp-11; myo-2p::mCherry]* | This Study | MCW291<br>MCW292<br>MCW293 |
| *raxIs119 [lbp-3p::lbp-3::gfp, myo-2p::mCherry];nlp-11(rax51)* | This Study | MCW1368 |
| *raxIs86[lbp-8p::lbp-8::3xflag::sl2-RFP; myo-2p::GFP]* | Ramachandran *et al.* [21] | MCW897 |
| *raxIs119 [lbp-3p::lbp-3::gfp, myo-2p::mCherry];*<br>*raxIs114[sur-5p:lmp-1::RFP-3XHA;unc-76(+)]* | This Study | MCW1318 |
| *raxEx509[ges-1p::lbp-3::RFP::sl2-GFP]* | This Study | MCW1264 |
| *raxIs3[ges-1p::lipl-4::sl2-GFP; myo-2p::mCherry];*<br>*nre-1(hd20); lin-15b(hd126)* | This Study | MCW84 |
| *nre-1(hd20); lin-15b(hd126)* | CGC | VH624 |
| *raxIs119 [lbp-3p::lbp-3::gfp, myo-2p::mCherry];egl-21(n476)* | This Study | MCW1391 |
| *Is[ges-1p::RDE-1::unc54 3'UTR, myo-2p::RFP3];rde-1 (ne219)* | J. Mello | JM45 |
| *nhr-49(nr2014)* | CGC | STE68 |
| *nhr-80(tm1011)* | CGC | BX165 |
| *raxIs119 [lbp-3p::lbp-3::gfp, myo-2p::mCherry];nhr-49(nr2041)* | This Study | MCW1377 |
| *raxIs3[ges-1p::lipl-4::sl2-GFP; myo-2p::mCherry];nhr-49(nr2041)* | This Study | MCW1387 |
| *raxIs3[ges-1p::lipl-4::sl2-GFP; myo-2p::mCherry];nhr-* | This Study | MCW1388 |
| *rab-3p::nhr-49::unc-54 3'UTR, myo-3p::mCherry* | CGC[20] | WBM179 |

| | | |
|---|---|---|
| *raxIs119 [lbp-3p::lbp-3::gfp, myo-2p::mCherry];nhr-49(nr2041); rab-3p::nhr-49::unc-54 3'UTR, myo-3p::mCherry* | This Study | MCW1389 |
| *raxEx402[ges-1p::lipl-4::sl2-GFP];rde-1 (ne219); Is[ges-1p::RDE-1::unc54 3'UTR, myo2p::RFP3]* | This Study | MCW1056 |

**Supplementary Table 6. List of primer sequences**

| *Primer name* | *Primer sequence* | *Source* | *Experimental assay* |
|---|---|---|---|
| *rpl-32 - rev* | AGGGAATTGATAACCGTGTCCGCA | This Study | qRT-PCR |
| *rpl-32 - rev* | TGTAGGACTGCATGAGGAGCATGT | This Study | qRT-PCR |
| *nlp-11 - fwd* | AGTGCTCCAATGGCAAGCGACTAT | This Study | qRT-PCR |
| *nlp-11 - rev* | AAGCTGGAGAAATTGCAGGGCTTC | This Study | qRT-PCR |
| *egl-3 - fwd* | TGCTGGACTCATTGACACTCCACA | This Study | qRT-PCR |
| *egl-3 - rev* | TTGAGATCCGGCACATCCATCAGT | This Study | qRT-PCR |
| *egl-21 - fwd* | ACTTCCTCGATTCTGGGAGGACAA | This Study | qRT-PCR |
| *egl-21 - rev* | ACCATTCCCTTAACTCCGCTGTGA | This Study | qRT-PCR |
| *sbt-1 - fwd* | ACACGGATGCTTGGAGGAGTTTGA | This Study | qRT-PCR |
| *sbt-1 - rev* | TGAACATATGCTCCTGGTCGCAGA | This Study | qRT-PCR |
| *pgal-1 - fwd* | TTCCCGGAGAGGATCAAGCTCATT | This Study | qRT-PCR |
| *pgal-1 - rev* | TGTTACAGTACCCATCGGCCACAA | This Study | qRT-PCR |
| *pghm-1 - fwd* | TGGATGCACCACCACTTGAATTGC | This Study | qRT-PCR |
| *pghm-1 - rev* | TGCATGACAAGGTGACGGATGTTG | This Study | qRT-PCR |
| *lbp-3 - fwd* | *TCATTGTCCATCTGCTCTTG* | This Study | Genotyping |
| *lbp-3 - rev* | *ACCTTGGCTAATCTGTTCTATG* | This Study | Genotyping |
| *nlp-11 - fwd* | *GTGAGTTGTTGGTGCAAAGAG* | This Study | Genotyping |
| *nlp-11 - rev* | *AGGTGGAGAGGGATGTATGA* | This Study | Genotyping |
| *egl-21 - fwd* | *CTGTTGAGATCGACTCCGTTGATG* | This Study | Genotyping |
| *egl-21 - rev* | *TTCCTCGTTTCTTTCACGTCCGCT* | This Study | Genotyping |
| *fat-1 - fwd* | *GAAAGAGATCTCGTTAAATCAATC* | This Study | Genotyping |
| *fat-1 - rev* | *TTGTCGTTAAAACATTATTTGTCGC* | This Study | Genotyping |
| *fat-3 - fwd* | *GATATCAATGTATCAGCATATGATG* | This Study | Genotyping |
| *fat-3 - rev* | *CGTGATGAGTGTTATGCTGTAAA* | This Study | Genotyping |
| *nhr-49(nr2041)ext - fwd* | *CGTCCGTGAAATGAGATCGG* | This Study | Genotyping |
| *nhr-49(nr2041)ext - rev* | *CTCTCAAGGCTCTGACTCAG* | This Study | Genotyping |
| *nhr-49(nr2041)int - fwd* | *GTCATTCAGGTCCATGTTCTATC* | This Study | Genotyping |
| *nhr-49(nr2041)int - rev* | *GCACAACCGAGGTAGTGAGT* | This Study | Genotyping |
| *lbp-3p_genomic - fwd* | *GCTGTGTAAGATAGTGTAAG* | This Study | Cloning |
| *lbp-3p_genomic - rev* | *TTTTTGTGAGATCATGCAATTTTTG* | This Study | Cloning |
| *lbp-3_cDNA - fwd* | *ATG AAT CTG TAC TTA ACT TTA TTC* | This Study | Cloning |
| *lbp-3_cDNA - rev* | *CTACTTCTTTCCGGTCGAG* | This Study | Cloning |
| *lbp-3_ns_cDNA - fwd* | *ATG GCT GAG GCG GCT TCT GA* | This Study | Cloning |
| *lbp-3_ns_cDNA - rev* | *CTACTTCTTTCCGGTCGATCG* | This Study | Cloning |
| *nlp-11p_genomic - fwd* | *TCAATTCCACTGCTGTCTTGTTG* | This Study | Cloning |
| *nlp-11p_genomic - rev* | *GTCTAACAAAACACTTTATTCTCGTC* | This Study | Cloning |
| *nlp-11_cDNA - fwd* | *ATGATGAGCACATTGGCAC* | This Study | Cloning |
| *nlp-11_cDNA - rev* | *TTAACGACGTCCGGCTCG* | This Study | Cloning |

**Supplementary Table 7. List of sgRNAs, crRNAs and tracrRNAs**

| ***crRNA name*** | ***Sequence*** | ***Source*** |
|---|---|---|
| *lbp-3 - fwd* | *AAGAAUACACUCUACCACGUUUUAGA* | This Study |
| *lbp-3 - rev* | *UUGGUUCAAGCGAUCGACGUUUUAG* | This Study |
| *nlp-11 - fwd* | *CCTCCCAAGACGTAATGATGACG* | This Study |
| *nlp-11 - rev* | *ACATCAACAAATAAAATACCATACCAACGA* | This Study |
| *dpy-10* | *GCUACCAUAGGCACCACGAG* | Arribere *et al.* [25] |
| *tracrRNA* | *AACAGCAUAGCAAGUUAAAAUAAGGCUAGU CCGUUAUCAACUUGAAAAAGUGGCACCGAGU CGGUGCUUUUUUU* | Deltcheva *et al.* [26] |
