## Supplementary Table 1 - Transcriptome analysis of WT and lipl-4 Tg animals for "Lysosome Lipid Signaling from the Periphery to Neurons Regulates Longevity"

| ID | log2FoldChange | pvalue | padj | SYMBOL |
|---|---|---|---|---|
| WBGene00000001 | -0.111103144 | 0.45819651 | 0.64973678 | aap-1 |
| WBGene00000002 | 0.537332727 | 0.03070026 | 0.10513307 | aat-1 |
| WBGene00000003 | 0.36405257 | 0.10830359 | 0.2588431 | aat-2 |
| WBGene00000004 | -0.211912368 | 0.20696034 | 0.39624772 | aat-3 |
| WBGene00000005 | -0.48123759 | 0.0315158 | 0.10703384 | aat-4 |
| WBGene00000006 | 1.016342444 | 0.00032943 | 0.00272493 | aat-5 |
| WBGene00000007 | 0.352248155 | 0.10995749 | 0.26143027 | aat-6 |
| WBGene00000008 | 0.91146593 | 0.01834064 | 0.07220277 | aat-7 |
| WBGene00000009 | 3.725402978 | 0.00062595 | 0.00472157 | aat-8 |
| WBGene00000010 | -0.026961486 | 0.81075954 | 0.89531472 | aat-9 |
| WBGene00000012 | 0.087691801 | 0.54697536 | 0.71995249 | abf-1 |
| WBGene00000013 | 0.395285716 | 0.04906873 | 0.14821855 | abf-2 |
| WBGene00000014 | NA | NA | NA | abf-3 |
| WBGene00000015 | 0.181789487 | 0.04369581 | NA | abf-4 |
| WBGene00000016 | 2.636103463 | 3.47E-10 | 9.27E-09 | abf-5 |
| WBGene00000017 | 1.969442175 | 1.16E-11 | 3.80E-10 | abf-6 |
| WBGene00000018 | -0.260683046 | 0.06232879 | 0.17654399 | abl-1 |
| WBGene00000019 | 0.324117495 | 0.06253191 | 0.17694814 | abt-1 |
| WBGene00000020 | 0.56169293 | 0.0027764 | 0.01639744 | abt-2 |
| WBGene00000021 | NA | NA | NA | NA |
| WBGene00000022 | 0.90671473 | 6.32E-11 | 1.88E-09 | abt-4 |
| WBGene00000023 | 0.064408474 | 0.682214 | 0.8161756 | abt-5 |
| WBGene00000024 | NA | NA | NA | abu-1 |
| WBGene00000025 | 0.020925399 | 0.84643798 | NA | abu-2 |
| WBGene00000026 | -0.040794878 | 0.53203165 | NA | abu-3 |
| WBGene00000027 | -0.023056499 | 0.78121253 | NA | abu-4 |
| WBGene00000028 | NA | NA | NA | abu-5 |
| WBGene00000029 | 0.056188364 | 0.42092875 | NA | abu-6 |
| WBGene00000030 | -0.006738828 | 0.92480778 | NA | abu-7 |
| WBGene00000031 | 1.868022228 | 0.01174921 | 0.05069853 | abu-8 |
| WBGene00000032 | NA | NA | NA | abu-9 |
| WBGene00000032 | NA | NA | NA | pqn-57 |
| WBGene00000033 | 0.347750535 | 0.02239347 | 0.08348636 | abu-10 |
| WBGene00000034 | 0.277853395 | 0.11043232 | 0.26217604 | abu-11 |
| WBGene00000035 | 0.648751472 | 0.00843755 | 0.03939255 | ace-1 |
| WBGene00000036 | 0.54603302 | 0.00886065 | 0.04094972 | ace-2 |
| WBGene00000037 | 2.779607582 | 3.70E-05 | 0.00039844 | ace-3 |
| WBGene00000038 | 0.19087931 | 0.19337723 | 0.37998067 | ace-4 |
| WBGene00000039 | 0.194902854 | 0.37125884 | 0.57063899 | acn-1 |
| WBGene00000040 | 0.488750507 | 2.32E-14 | 1.03E-12 | aco-1 |
| WBGene00000041 | -0.150374443 | 0.00080572 | 0.00585147 | aco-2 |
| WBGene00000042 | -0.066492402 | 0.62362851 | 0.77505748 | acr-2 |

| | | | | |
|---|---|---|---|---|
| WBGene00000043 | 0.148021412 | 0.07455439 | NA | acr-3 |
| WBGene00000044 | 0.131416208 | 0.34385832 | 0.5436749 | acr-5 |
| WBGene00000045 | 0.063223962 | 0.33019567 | NA | acr-6 |
| WBGene00000046 | 0.102918357 | 0.35599754 | 0.55590444 | acr-7 |
| WBGene00000047 | 0.203092151 | 0.33352895 | 0.53304196 | acr-8 |
| WBGene00000048 | 0.105444527 | 0.45916567 | 0.65047359 | acr-9 |
| WBGene00000049 | 0.155591171 | 0.1197606 | 0.27797684 | acr-10 |
| WBGene00000050 | 0.28247476 | 0.17498295 | 0.35564444 | acr-11 |
| WBGene00000051 | 0.053479713 | 0.76520707 | 0.86670277 | acr-12 |
| WBGene00000053 | 0.031584814 | 0.69986754 | NA | acr-14 |
| WBGene00000054 | 0.124614128 | 0.5456247 | 0.71910085 | acr-15 |
| WBGene00000055 | -0.050930351 | 0.79978974 | 0.88817669 | acr-16 |
| WBGene00000056 | 0.489926227 | 0.05535898 | 0.16213458 | acr-17 |
| WBGene00000057 | -0.095355236 | 0.6300744 | 0.7799881 | acr-18 |
| WBGene00000058 | 0.394980552 | 0.06665893 | 0.18469974 | acr-19 |
| WBGene00000059 | 0.167314572 | 0.29899879 | 0.49955966 | acr-20 |
| WBGene00000060 | 0.042796001 | 0.78430392 | 0.87847449 | acr-21 |
| WBGene00000061 | 0.329650472 | 0.00669025 | 0.03289183 | lgc-11 |
| WBGene00000062 | 0.06579847 | 0.52371461 | 0.70133592 | acr-23 |
| WBGene00000063 | -0.213378988 | 0.0184933 | 0.07258991 | act-3 |
| WBGene00000063 | -0.213378988 | 0.0184933 | 0.07258991 | act-1 |
| WBGene00000064 | -0.269927171 | 9.36E-11 | 2.70E-09 | act-2 |
| WBGene00000065 | 0.121358883 | 0.46344277 | 0.65337175 | act-3 |
| WBGene00000065 | 0.121358883 | 0.46344277 | 0.65337175 | act-1 |
| WBGene00000066 | -0.301710079 | 2.22E-07 | 3.93E-06 | act-4 |
| WBGene00000067 | 0.37562872 | 3.70E-07 | 6.28E-06 | act-5 |
| WBGene00000068 | 0.539171991 | 0.02538529 | 0.09170148 | acy-1 |
| WBGene00000069 | 0.182853462 | 0.31858007 | 0.51889317 | acy-2 |
| WBGene00000070 | 0.344506376 | 0.09741101 | 0.2417018 | acy-3 |
| WBGene00000071 | 0.553825586 | 0.01344848 | 0.05671058 | acy-4 |
| WBGene00000072 | 0.101140002 | 0.60840394 | 0.76399837 | add-1 |
| WBGene00000073 | -0.039988925 | 0.79081702 | 0.88222736 | add-2 |
| WBGene00000074 | 0.063854503 | 0.64814211 | 0.79160333 | adm-2 |
| WBGene00000075 | -0.285605344 | 0.04396954 | 0.13708648 | adm-4 |
| WBGene00000079 | -0.295351616 | 0.00382295 | 0.02112265 | adr-1 |
| WBGene00000080 | -0.163678188 | 0.09977678 | 0.2450649 | adr-2 |
| WBGene00000081 | 0.138571276 | 0.0559319 | 0.16348551 | ads-1 |
| WBGene00000082 | -0.026364718 | 0.88146107 | 0.93716888 | adt-1 |
| WBGene00000083 | -0.115850843 | 0.38030063 | 0.5797656 | adt-2 |
| WBGene00000084 | 0.272696771 | 0.16767894 | 0.34527078 | aex-1 |
| WBGene00000085 | 0.032970343 | 0.82908045 | 0.90653972 | aex-2 |
| WBGene00000086 | 0.902277388 | 0.0009025 | 0.00645508 | aex-3 |
| WBGene00000088 | 0.797727291 | 0.00031779 | 0.00263758 | aex-5 |

| | | | | |
|---|---|---|---|---|
| WBGene00000089 | 0.40687909 | 0.10250282 | 0.24950022 | aex-6 |
| WBGene00000090 | -0.026609022 | 0.75396098 | 0.86072068 | age-1 |
| WBGene00000092 | -0.0586382 | 0.68128004 | 0.81558635 | ags-3 |
| WBGene00000093 | 0.17605851 | 0.31832049 | 0.51858577 | agt-1 |
| WBGene00000094 | -0.47128106 | 0.03072019 | 0.10516117 | agt-2 |
| WBGene00000095 | 0.158742664 | 0.35849237 | 0.55878705 | aha-1 |
| WBGene00000096 | 0.104463479 | 0.4851049 | 0.66991162 | ahr-1 |
| WBGene00000097 | -0.402303772 | 0.00261536 | 0.01558463 | aip-1 |
| WBGene00000098 | -0.156868027 | 0.00817239 | 0.03843341 | air-1 |
| WBGene00000099 | -0.028051853 | 0.772989 | 0.87123004 | air-2 |
| WBGene00000100 | 0.431540492 | 1.53E-05 | 0.00017927 | ajm-1 |
| WBGene00000101 | -0.040584921 | 0.50770117 | 0.68813672 | aka-1 |
| WBGene00000102 | -0.086162606 | 0.17453959 | 0.35515336 | akt-1 |
| WBGene00000103 | 0.104007693 | 0.59586086 | 0.75477619 | akt-2 |
| WBGene00000105 | -0.052570729 | 0.60703913 | 0.76297727 | alg-1 |
| WBGene00000106 | -0.18767841 | 0.03739687 | 0.12180813 | alg-2 |
| WBGene00000107 | 0.087507445 | 0.16866157 | 0.34677349 | alh-1 |
| WBGene00000108 | NA | NA | NA | alh-2 |
| WBGene00000109 | 0.449654801 | 6.26E-07 | 1.03E-05 | alh-3 |
| WBGene00000110 | -0.034498351 | 0.6765214 | 0.81229431 | alh-4 |
| WBGene00000111 | 2.761471729 | 2.59E-25 | 2.45E-23 | alh-5 |
| WBGene00000112 | -0.453321276 | 1.92E-05 | 0.0002203 | alh-6 |
| WBGene00000113 | 0.21305348 | 0.12749527 | 0.28874635 | alh-7 |
| WBGene00000114 | 0.076588778 | 0.15996558 | 0.33471329 | alh-8 |
| WBGene00000115 | -1.168807254 | 3.25E-41 | 6.18E-39 | alh-9 |
| WBGene00000116 | -0.741336485 | 3.60E-06 | 4.87E-05 | alh-10 |
| WBGene00000117 | -0.05296878 | 0.69675766 | 0.82701491 | alh-11 |
| WBGene00000118 | 0.370675311 | 0.00056116 | 0.00427754 | alh-12 |
| WBGene00000120 | -0.356962485 | 0.01828648 | 0.07209301 | aly-1 |
| WBGene00000121 | -0.163580659 | 0.0158681 | 0.0647505 | aly-2 |
| WBGene00000122 | -0.386721622 | 3.10E-08 | 6.38E-07 | aly-3 |
| WBGene00000123 | -0.034514716 | 0.61723642 | 0.77061127 | ama-1 |
| WBGene00000133 | -1.516773 | 0.00310626 | 0.01794031 | amt-1 |
| WBGene00000134 | 0.122833479 | 0.25726816 | 0.45540601 | amt-2 |
| WBGene00000135 | 0.038588586 | 0.83888728 | 0.9127791 | amt-3 |
| WBGene00000136 | 0.314463316 | 0.0224116 | 0.0834932 | amt-4 |
| WBGene00000137 | -0.211745213 | 0.17455663 | 0.35515336 | amx-1 |
| WBGene00000138 | 0.716796923 | 0.00205405 | 0.01276458 | amx-2 |
| WBGene00000139 | -0.293817295 | 0.20259174 | 0.39131209 | amx-3 |
| WBGene00000140 | 0.402593981 | 0.00284682 | 0.01674577 | anc-1 |
| WBGene00000142 | -0.126100512 | 0.12748456 | 0.28874635 | aos-1 |
| WBGene00000143 | -0.276558296 | 0.00322338 | 0.01848929 | apc-2 |
| WBGene00000144 | -0.201459307 | 0.13613173 | 0.3010903 | apc-10 |

| | | | | |
|---|---|---|---|---|
| WBGene00000145 | -0.383832951 | 0.0041114 | 0.02238567 | apc-11 |
| WBGene00000146 | -0.102802132 | 0.11029572 | 0.26197918 | ape-1 |
| WBGene00000147 | -0.01532002 | 0.89970904 | 0.94665211 | aph-1 |
| WBGene00000148 | -0.213273896 | 0.03965574 | 0.12725972 | aph-2 |
| WBGene00000149 | 0.692677901 | 2.96E-17 | 1.78E-15 | apl-1 |
| WBGene00000150 | 0.158704534 | 0.31729049 | 0.51740497 | apm-1 |
| WBGene00000151 | -0.061976211 | 0.52831932 | 0.70537326 | apn-1 |
| WBGene00000155 | -0.108322724 | 0.05571169 | 0.16301149 | app-1 |
| WBGene00000156 | -0.007958481 | 0.91681315 | 0.95361334 | 1-Apr |
| WBGene00000157 | -0.160259956 | 0.33289187 | 0.53260517 | aps-2 |
| WBGene00000158 | -0.172988409 | 0.04841627 | 0.14670169 | apg-1 |
| WBGene00000159 | -0.169074483 | 0.13326303 | 0.29688131 | aps-1 |
| WBGene00000160 | -0.090958049 | 0.31518346 | 0.51565238 | apb-1 |
| WBGene00000161 | -0.151295694 | 0.14510992 | 0.31448819 | apa-2 |
| WBGene00000162 | 0.155718955 | 0.11851338 | 0.27590307 | apd-3 |
| WBGene00000163 | 0.07724031 | 0.31680936 | 0.51698682 | apb-3 |
| WBGene00000164 | -0.226191651 | 0.15204048 | 0.32451635 | apm-3 |
| WBGene00000165 | -0.143271906 | 0.22988501 | 0.42425231 | aps-3 |
| WBGene00000166 | -0.007763717 | 0.95806043 | 0.97694406 | apt-9 |
| WBGene00000168 | 0.030261142 | 0.73209939 | 0.85011851 | apx-1 |
| WBGene00000169 | 1.160275828 | 5.34E-16 | 2.78E-14 | aqp-1 |
| WBGene00000170 | 0.455074092 | 0.00070406 | 0.00519825 | aqp-2 |
| WBGene00000171 | 0.290915872 | 0.13666924 | 0.30194271 | aqp-3 |
| WBGene00000172 | 0.55790528 | 0.00946062 | 0.04309672 | aqp-4 |
| WBGene00000173 | 0.150041977 | 0.24238754 | 0.44051588 | aqp-5 |
| WBGene00000174 | -0.060016243 | 0.56110759 | 0.73159138 | aqp-6 |
| WBGene00000175 | 0.385178555 | 2.88E-06 | 3.98E-05 | aqp-7 |
| WBGene00000176 | 0.055880425 | 0.68873618 | 0.82075423 | aqp-8 |
| WBGene00000177 | 0.049462832 | 0.8117297 | 0.89587496 | aqp-9 |
| WBGene00000178 | -0.286340652 | 0.0326296 | 0.10999998 | aqp-10 |
| WBGene00000179 | -0.264475237 | 0.05122368 | 0.15314802 | aqp-11 |
| WBGene00000180 | 0.140795375 | 0.42560259 | 0.6215184 | arc-1 |
| WBGene00000181 | -0.049572622 | 0.52461102 | 0.70205458 | ard-1 |
| WBGene00000182 | -0.189710233 | 0.00038232 | 0.00309085 | arf-1.2 |
| WBGene00000183 | -0.193625627 | 0.00038175 | 0.00308792 | arf-3 |
| WBGene00000184 | -0.12070737 | 0.31288375 | 0.51333563 | arf-6 |
| WBGene00000185 | 0.020232312 | 0.84643798 | NA | arg-1 |
| WBGene00000186 | 0.101876401 | 0.40894135 | 0.60680149 | ark-1 |
| WBGene00000187 | -0.046009088 | 0.68453184 | 0.81794097 | arl-1 |
| WBGene00000188 | -0.15552484 | 0.34312094 | 0.54313812 | arl-3 |
| WBGene00000189 | 0.051336287 | 0.75836418 | 0.86329917 | arl-5 |
| WBGene00000190 | 1.271066455 | 0.01034369 | 0.04607299 | arf-1.1 |
| WBGene00000191 | 0.081350485 | 0.62328646 | 0.77472935 | NA |

| | | | | |
|---|---|---|---|---|
| WBGene00000192 | 0.023992197 | 0.74280198 | 0.85444405 | arl-8 |
| WBGene00000193 | -0.03147554 | 0.70637395 | NA | arl-6 |
| WBGene00000195 | 0.026914696 | 0.86900753 | 0.93000745 | arr-1 |
| WBGene00000196 | -0.193385271 | 0.13255753 | 0.29580404 | aars-1 |
| WBGene00000197 | -0.355942182 | 6.44E-11 | 1.91E-09 | aars-2 |
| WBGene00000198 | -0.219453656 | 0.00290462 | 0.01701059 | art-1 |
| WBGene00000199 | -0.088625859 | 0.29967196 | 0.50010521 | arx-1 |
| WBGene00000200 | -0.147449028 | 0.07620879 | 0.2038686 | arx-2 |
| WBGene00000201 | -0.127753458 | 0.22241619 | 0.41509184 | arx-3 |
| WBGene00000202 | -0.073579249 | 0.52776886 | 0.70495975 | exos-4.2 |
| WBGene00000203 | -0.224507735 | 0.03110948 | 0.10597732 | arx-5 |
| WBGene00000204 | -0.108888749 | 0.29653463 | 0.49751505 | arx-6 |
| WBGene00000205 | -0.11969361 | 0.42617757 | 0.62179031 | arx-7 |
| WBGene00000206 | -0.358180818 | 9.87E-09 | 2.21E-07 | asb-1 |
| WBGene00000207 | 0.065707585 | 0.37647024 | 0.57565543 | asb-2 |
| WBGene00000208 | 0.349248608 | 0.00536232 | 0.027654 | trip-4 |
| WBGene00000209 | -0.145648018 | 0.03118264 | 0.10614858 | asg-1 |
| WBGene00000210 | 0.034485225 | 0.70886136 | 0.8357577 | asg-2 |
| WBGene00000211 | -0.079451453 | 0.66294181 | 0.80196075 | asm-1 |
| WBGene00000212 | 0.42028004 | 0.05225038 | 0.15558222 | asm-2 |
| WBGene00000213 | -2.435442983 | 6.83E-07 | 1.11E-05 | asm-3 |
| WBGene00000214 | 0.994207086 | 1.39E-73 | 4.53E-71 | asp-1 |
| WBGene00000215 | 0.623745021 | 1.56E-25 | 1.49E-23 | T18H9.11 |
| WBGene00000215 | 0.623745021 | 1.56E-25 | 1.49E-23 | asp-2 |
| WBGene00000216 | 0.455418941 | 8.54E-12 | 2.88E-10 | asp-3 |
| WBGene00000217 | -0.556383325 | 8.70E-19 | 5.93E-17 | asp-4 |
| WBGene00000218 | 1.194288656 | 2.55E-96 | 1.38E-93 | asp-5 |
| WBGene00000219 | 1.705404327 | 3.59E-243 | 5.84E-240 | asp-6 |
| WBGene00000220 | 0.13337948 | 0.4881119 | 0.67216387 | atf-2 |
| WBGene00000221 | 0.224087682 | 0.08005941 | 0.21113383 | atf-5 |
| WBGene00000222 | 0.021427493 | 0.89966641 | 0.94665211 | atf-6 |
| WBGene00000223 | 0.022227628 | 0.74445731 | 0.85556581 | atf-7 |
| WBGene00000224 | 0.844506126 | 7.33E-08 | 1.41E-06 | atgp-1 |
| WBGene00000225 | 0.265309172 | 0.140909 | 0.30860862 | atgp-2 |
| WBGene00000226 | 0.017891473 | 0.83243448 | 0.90857859 | atl-1 |
| WBGene00000227 | 0.162206679 | 0.11508371 | 0.27038196 | atm-1 |
| WBGene00000228 | 0.238232271 | 0.00906828 | 0.04161332 | atn-1 |
| WBGene00000229 | -0.287730619 | 6.90E-12 | 2.35E-10 | atp-2 |
| WBGene00000230 | -0.141245539 | 0.0084403 | 0.03939255 | atp-3 |
| WBGene00000231 | -0.048219261 | 0.44331554 | 0.6368667 | atx-2 |
| WBGene00000232 | 0.361841087 | 0.13726601 | 0.30289549 | avr-14 |
| WBGene00000233 | 0.196976151 | 0.30506234 | 0.50565347 | avr-15 |
| WBGene00000235 | -0.145120031 | 0.0898094 | 0.2277675 | baf-1 |

| | | | | |
|---|---|---|---|---|
| WBGene00000236 | -0.000576901 | 0.9947977 | 0.99684027 | bag-1 |
| WBGene00000237 | 0.283034886 | 0.13531864 | 0.30000003 | bam-2 |
| WBGene00000238 | 0.442089232 | 0.03054552 | 0.10475917 | bar-1 |
| WBGene00000239 | -0.430075322 | 0.01111228 | 0.04859465 | bas-1 |
| WBGene00000240 | -0.912556297 | 1.37E-15 | 6.90E-14 | pah-1 |
| WBGene00000241 | 0.128826435 | 0.24887962 | 0.44726236 | bbs-1 |
| WBGene00000242 | 0.027847144 | 0.79386641 | 0.88399982 | bbs-2 |
| WBGene00000244 | 0.127859194 | 0.40676216 | 0.60473241 | bbs-8 |
| WBGene00000245 | 0.12441713 | 0.35930362 | 0.55945595 | bca-1 |
| WBGene00000246 | -0.228070979 | 0.29568451 | 0.4967871 | bcc-1 |
| WBGene00000247 | -0.080239225 | 0.54007373 | 0.71451837 | bec-1 |
| WBGene00000248 | 0.137192049 | 0.36379052 | 0.56356544 | ben-1 |
| WBGene00000249 | -0.31267506 | 0.00917306 | 0.04199585 | bir-1 |
| WBGene00000250 | 0.368641683 | 0.01486728 | 0.06146019 | bir-2 |
| WBGene00000251 | 3.047723582 | 0.00263838 | 0.01569629 | bli-1 |
| WBGene00000252 | 0.152457348 | 0.27357147 | 0.47383885 | bli-2 |
| WBGene00000253 | 0.272088817 | 0.07591318 | 0.20338625 | bli-3 |
| WBGene00000254 | 0.001367228 | 0.98276149 | 0.98999663 | bli-4 |
| WBGene00000255 | -0.111313683 | 0.42275399 | 0.61864479 | bli-5 |
| WBGene00000256 | -0.039173014 | 0.84707735 | 0.91761389 | bli-6 |
| WBGene00000257 | -0.178104594 | 0.03530806 | 0.11674465 | bmk-1 |
| WBGene00000259 | -0.08205383 | 0.4899768 | 0.67384467 | bpl-1 |
| WBGene00000262 | -0.073656108 | 0.69425967 | 0.82485238 | bra-1 |
| WBGene00000263 | -0.084228261 | 0.25146123 | 0.44991306 | F23H11.5 |
| WBGene00000264 | -0.222687719 | 0.08437062 | 0.21872063 | brc-1 |
| WBGene00000265 | -0.132963139 | 0.21869363 | 0.41049916 | brd-1 |
| WBGene00000266 | 0.500230153 | 0.0045424 | 0.02431638 | bre-1 |
| WBGene00000267 | 0.097857957 | 0.63647107 | 0.78445295 | bre-2 |
| WBGene00000268 | -0.086084585 | 0.323416 | 0.52329307 | bre-3 |
| WBGene00000269 | 0.054163898 | 0.75145593 | 0.85953643 | bre-4 |
| WBGene00000270 | -0.10838697 | 0.49665751 | 0.67888737 | bre-5 |
| WBGene00000271 | -0.312904234 | 0.00363865 | 0.02034117 | brf-1 |
| WBGene00000272 | 0.399255004 | 0.00431015 | 0.02329456 | bro-1 |
| WBGene00000273 | -0.049639742 | 0.4709122 | 0.65895302 | brp-1 |
| WBGene00000274 | 0.01578434 | 0.8650975 | 0.92790421 | btf-1 |
| WBGene00000275 | -0.112649462 | 0.08477617 | 0.21933343 | bub-1 |
| WBGene00000276 | -0.205478172 | 0.01628142 | 0.0660505 | byn-1 |
| WBGene00000277 | 0.82484017 | 9.22E-14 | 3.87E-12 | cab-1 |
| WBGene00000279 | -0.108941949 | 0.59133583 | 0.75215398 | cah-1 |
| WBGene00000280 | 0.124023863 | 0.48422359 | 0.66945397 | cah-2 |
| WBGene00000281 | 0.414666513 | 0.00148748 | 0.00975287 | cah-3 |
| WBGene00000282 | 0.216229327 | 0.00248398 | 0.01494161 | cah-4 |
| WBGene00000283 | 0.200887615 | 0.13851639 | 0.3048735 | cah-5 |

| | | | | |
|---|---|---|---|---|
| WBGene00000284 | 0.169669264 | 0.33825322 | 0.53789273 | cah-6 |
| WBGene00000285 | 0.523947766 | 0.0469182 | 0.14367902 | cal-1 |
| WBGene00000286 | 1.092099046 | 0.00020717 | 0.00182282 | cal-2 |
| WBGene00000287 | -0.024138639 | 0.89986165 | 0.94665211 | cal-3 |
| WBGene00000288 | 0.996257927 | 0.00639705 | 0.03175966 | cal-4 |
| WBGene00000289 | 0.147507134 | 0.10501292 | 0.25337741 | cam-1 |
| WBGene00000292 | -0.092411754 | 0.2333422 | 0.42875992 | cap-1 |
| WBGene00000293 | -0.076154213 | 0.33051611 | 0.53007847 | cap-2 |
| WBGene00000294 | 0.109999612 | 0.25143548 | 0.44991306 | cas-1 |
| WBGene00000295 | 0.182169067 | 0.29851577 | 0.49933345 | cat-1 |
| WBGene00000296 | 0.094682347 | 0.43905953 | 0.63370437 | cat-2 |
| WBGene00000298 | 0.429792847 | 9.59E-05 | 0.00093425 | cat-4 |
| WBGene00000301 | -1.440629514 | 2.37E-45 | 4.88E-43 | cav-1 |
| WBGene00000302 | 1.134131105 | 0.00977213 | 0.04410661 | cav-2 |
| WBGene00000308 | NA | NA | NA | NA |
| WBGene00000310 | NA | NA | NA | NA |
| WBGene00000311 | NA | NA | NA | NA |
| WBGene00000366 | 0.078053923 | 0.25595509 | 0.45436804 | cbp-1 |
| WBGene00000367 | 0.666548305 | 0.00387872 | 0.02135701 | cca-1 |
| WBGene00000368 | 0.522035101 | 0.02124461 | 0.08028973 | ccb-1 |
| WBGene00000369 | -0.189602505 | 0.04513197 | 0.13961065 | ccf-1 |
| WBGene00000370 | 0.248074036 | 0.04919382 | 0.14853688 | ccg-1 |
| WBGene00000371 | -0.181099099 | 0.00141507 | 0.00936622 | cox-5B |
| WBGene00000372 | 4.032096834 | 0.00029382 | 0.00246807 | cyp-13A7 |
| WBGene00000373 | -0.710755411 | 0.00221386 | 0.01352238 | cyp-14A5 |
| WBGene00000374 | 0.670428762 | 0.0011777 | 0.00804984 | NA |
| WBGene00000375 | -0.125467311 | 0.28124991 | 0.48075477 | cyp-44A1 |
| WBGene00000376 | -0.20321248 | 0.00031253 | 0.00260281 | ccr-4 |
| WBGene00000377 | -0.221188547 | 6.52E-05 | 0.0006657 | cct-1 |
| WBGene00000378 | -0.313660916 | 1.21E-07 | 2.25E-06 | cct-2 |
| WBGene00000379 | -0.471145965 | 1.18E-16 | 6.80E-15 | cct-4 |
| WBGene00000380 | -0.294518098 | 1.12E-10 | 3.20E-09 | cct-5 |
| WBGene00000381 | -0.327071622 | 1.75E-08 | 3.75E-07 | cct-6 |
| WBGene00000382 | -0.183208181 | 0.02669964 | 0.09536704 | cdc-6 |
| WBGene00000383 | -0.257042925 | 0.00103311 | 0.00722338 | cdc-14 |
| WBGene00000386 | -0.256481102 | 2.59E-05 | 0.00028884 | cdc-25.1 |
| WBGene00000387 | -0.268768832 | 0.01033374 | 0.0460577 | cdc-25.2 |
| WBGene00000388 | -0.133497545 | 0.28205208 | 0.48180195 | cdc-25.3 |
| WBGene00000389 | NA | NA | NA | cdc-25.4 |
| WBGene00000390 | -0.112738266 | 0.08386662 | 0.21778844 | cdc-42 |
| WBGene00000391 | 0.189375188 | 0.38443012 | 0.58374211 | cdd-1 |
| WBGene00000392 | -0.312958034 | 0.01803022 | 0.07120946 | cdd-2 |
| WBGene00000393 | -0.318538713 | 0.03029264 | 0.10408694 | cdf-1 |

| | | | | |
|---|---|---|---|---|
| WBGene00000395 | -0.134911326 | 0.48389028 | 0.66930917 | cdh-3 |
| WBGene00000396 | -0.274313264 | 0.0104835 | 0.04648337 | cdh-4 |
| WBGene00000397 | 0.045511931 | 0.80927933 | 0.8944409 | cdh-5 |
| WBGene00000399 | -0.060782953 | 0.55960335 | 0.73003231 | cdh-7 |
| WBGene00000401 | 0.279625901 | 0.20730338 | 0.39645783 | cdh-9 |
| WBGene00000402 | -0.065476614 | 0.65402167 | 0.7949984 | cdh-10 |
| WBGene00000403 | 1.009823053 | 1.16E-13 | 4.79E-12 | casy-1 |
| WBGene00000405 | -0.131820399 | 0.026334 | 0.09445274 | cdk-1 |
| WBGene00000406 | -0.043558763 | 0.80694778 | 0.89296858 | cdk-4 |
| WBGene00000407 | -0.045893917 | 0.75216967 | 0.85970214 | cdk-5 |
| WBGene00000408 | -0.084392369 | 0.40374306 | 0.60207804 | cdk-7 |
| WBGene00000409 | -0.026597393 | 0.77640077 | 0.87335186 | cdk-8 |
| WBGene00000410 | -0.080355728 | 0.56537836 | 0.73494669 | cdk-9 |
| WBGene00000411 | -0.281340401 | 7.49E-05 | 0.00075082 | cdl-1 |
| WBGene00000412 | -0.067584835 | 0.66069738 | 0.80010508 | cdr-1 |
| WBGene00000413 | -0.37130976 | 0.00019124 | 0.00170104 | cdt-1 |
| WBGene00000414 | -0.084686563 | 0.53242765 | 0.70853237 | cec-1 |
| WBGene00000415 | 0.147163758 | 0.07819318 | 0.20762176 | ced-1 |
| WBGene00000416 | -0.19524145 | 0.03894671 | 0.12548828 | ced-2 |
| WBGene00000417 | -0.027913682 | 0.66749388 | 0.80567007 | ced-3 |
| WBGene00000418 | -0.290329069 | 0.02811291 | 0.09877637 | ced-4 |
| WBGene00000419 | -0.214321584 | 0.00628666 | 0.0312908 | ced-5 |
| WBGene00000420 | 0.145054134 | 0.18077173 | 0.36295651 | ced-6 |
| WBGene00000421 | 0.006237696 | 0.92882479 | 0.95998218 | ced-7 |
| WBGene00000422 | -0.363919149 | 0.1165877 | 0.27280814 | ced-8 |
| WBGene00000423 | 0.244850122 | 0.01699465 | 0.06809486 | ced-9 |
| WBGene00000424 | -0.216858157 | 0.04698729 | 0.14380037 | ced-10 |
| WBGene00000425 | -0.241321269 | 0.26809174 | 0.46699955 | ced-11 |
| WBGene00000426 | 0.020286514 | 0.84509026 | 0.91692355 | ced-12 |
| WBGene00000427 | NA | NA | NA | ced-13 |
| WBGene00000428 | 0.051038635 | 0.61191785 | 0.76645472 | ceh-1 |
| WBGene00000429 | 0.009183187 | 0.93015602 | 0.96068422 | ceh-2 |
| WBGene00000430 | 0.1368789 | 0.30034392 | 0.50066452 | ceh-5 |
| WBGene00000431 | 0.168482975 | 0.38764764 | 0.58619593 | ceh-6 |
| WBGene00000432 | NA | NA | NA | ceh-7 |
| WBGene00000433 | -0.023449708 | 0.69192352 | NA | ceh-8 |
| WBGene00000434 | -0.008807557 | 0.92153097 | NA | ceh-9 |
| WBGene00000435 | -0.086827908 | 0.5160284 | 0.69567629 | ceh-10 |
| WBGene00000436 | NA | NA | NA | ceh-12 |
| WBGene00000437 | -0.194767053 | 0.32077876 | 0.52079418 | ceh-13 |
| WBGene00000438 | 0.760729481 | 0.02644033 | 0.09469492 | ceh-14 |
| WBGene00000439 | -0.047290119 | 0.74202131 | 0.85412964 | ceh-16 |
| WBGene00000440 | -0.002914971 | 0.95718083 | NA | ceh-17 |

| | | | | |
|---|---|---|---|---|
| WBGene00000441 | 0.548285625 | 0.0076135 | 0.03633287 | ceh-18 |
| WBGene00000442 | 0.25049378 | 0.17309515 | 0.35340762 | ceh-19 |
| WBGene00000443 | -0.028110505 | 0.87704489 | 0.9345835 | ceh-20 |
| WBGene00000444 | -0.003731801 | 0.9805167 | 0.98875577 | ceh-21 |
| WBGene00000445 | 0.380860625 | 0.09343778 | 0.23380732 | ceh-22 |
| WBGene00000446 | 0.148148177 | 0.24764536 | 0.44607264 | ceh-23 |
| WBGene00000447 | 0.100625191 | 0.38388795 | 0.58327725 | ceh-24 |
| WBGene00000448 | 0.386766201 | 0.11466549 | 0.26968955 | pros-1 |
| WBGene00000449 | -0.367011322 | 0.13210814 | 0.29529699 | ceh-27 |
| WBGene00000450 | NA | NA | NA | ceh-28 |
| WBGene00000451 | 0.0372604 | 0.77625488 | 0.87330165 | ceh-30 |
| WBGene00000452 | -0.128511654 | 0.35456308 | 0.55451895 | ceh-31 |
| WBGene00000453 | -0.067847168 | 0.73441803 | 0.85122027 | ceh-32 |
| WBGene00000454 | 0.047655942 | 0.68860137 | 0.82066819 | ceh-33 |
| WBGene00000455 | 0.136916711 | 0.49831286 | 0.68033156 | ceh-34 |
| WBGene00000457 | -0.837756444 | 0.01630319 | 0.06612052 | ceh-36 |
| WBGene00000458 | 0.739559433 | 0.00990975 | 0.04456039 | ceh-37 |
| WBGene00000459 | -0.220984676 | 0.00218913 | 0.01341047 | ceh-38 |
| WBGene00000460 | -0.327829119 | 0.0169227 | 0.06794401 | ceh-39 |
| WBGene00000461 | -0.660778623 | 0.00537696 | 0.02770023 | ceh-40 |
| WBGene00000462 | 0.001079472 | 0.99622786 | 0.99769349 | ceh-41 |
| WBGene00000463 | -0.792905987 | 0.00104141 | 0.00726409 | ceh-43 |
| WBGene00000464 | 0.041862124 | 0.61525207 | 0.76921987 | ceh-44 |
| WBGene00000465 | 0.08434803 | 0.08198827 | 0.21450864 | cpg-1 |
| WBGene00000466 | -0.198224672 | 0.0417184 | 0.13180818 | cel-1 |
| WBGene00000467 | -0.30888194 | 0.00332801 | 0.01896669 | cep-1 |
| WBGene00000468 | -0.06730507 | 0.31931151 | NA | ces-1 |
| WBGene00000469 | -0.06139947 | 0.74730724 | 0.85713477 | ces-2 |
| WBGene00000472 | -0.10509165 | 0.09426215 | 0.23530983 | cey-1 |
| WBGene00000473 | -0.192038182 | 9.69E-07 | 1.54E-05 | cey-2 |
| WBGene00000474 | -0.125511752 | 0.00651202 | 0.03218362 | cey-3 |
| WBGene00000475 | -0.164940802 | 0.00229934 | 0.01396296 | cey-4 |
| WBGene00000476 | 0.576500533 | 0.04817878 | 0.1463202 | cfi-1 |
| WBGene00000477 | 0.265950427 | 0.23382267 | 0.42932 | cft-1 |
| WBGene00000478 | -0.04440845 | 0.82846896 | 0.9062566 | cfz-2 |
| WBGene00000479 | -0.213426921 | 7.40E-06 | 9.36E-05 | cgh-1 |
| WBGene00000480 | 0.049826184 | 0.5326738 | 0.70866671 | cgp-1 |
| WBGene00000481 | 0.788035214 | 0.02464344 | 0.08988653 | cha-1 |
| WBGene00000482 | -0.305748128 | 0.00177304 | 0.01131116 | chd-3 |
| WBGene00000483 | -0.002914971 | 0.95718083 | NA | che-1 |
| WBGene00000484 | 0.119979054 | 0.31074161 | 0.51164732 | che-2 |
| WBGene00000485 | 2.158167297 | 2.39E-05 | 0.00026732 | che-3 |
| WBGene00000487 | -0.032552334 | 0.73270008 | NA | che-6 |

| | | | | |
|---|---|---|---|---|
| WBGene00000488 | 0.131030251 | 0.38239642 | 0.58180047 | che-7 |
| WBGene00000490 | 0.266265332 | 0.10835642 | 0.2589271 | che-11 |
| WBGene00000491 | 0.26627317 | 0.13238463 | 0.2955991 | che-12 |
| WBGene00000492 | 0.14149427 | 0.36095613 | 0.56083865 | che-13 |
| WBGene00000493 | -0.058151087 | 0.73635097 | 0.85197681 | che-14 |
| WBGene00000496 | -0.092974691 | 0.06167083 | 0.17508681 | chs-1 |
| WBGene00000497 | 0.587417211 | 0.01776526 | 0.07052636 | chs-2 |
| WBGene00000498 | -0.116241154 | 0.24271353 | 0.44083966 | chk-1 |
| WBGene00000499 | -0.098898959 | 0.58324658 | 0.74703117 | chk-2 |
| WBGene00000500 | -0.247024771 | 0.01119549 | 0.04887096 | chn-1 |
| WBGene00000501 | 0.5070485 | 0.06298853 | 0.17782783 | cho-1 |
| WBGene00000502 | -0.327317629 | 0.00143892 | 0.00948123 | chp-1 |
| WBGene00000503 | -1.521660943 | 2.76E-35 | 4.25E-33 | cht-1 |
| WBGene00000506 | -0.150751056 | 0.29585095 | 0.49696578 | cic-1 |
| WBGene00000507 | -0.186594168 | 0.15661142 | 0.33058647 | cit-1.1 |
| WBGene00000508 | 0.095404518 | 0.44442 | 0.63794923 | cit-1.2 |
| WBGene00000509 | -0.070912703 | 0.49011501 | 0.67393674 | cka-1 |
| WBGene00000510 | 0.332909898 | 0.15889008 | 0.33321705 | cka-2 |
| WBGene00000511 | 0.254480747 | 0.25935307 | 0.45741564 | ckb-1 |
| WBGene00000512 | -0.202937968 | 0.20783988 | 0.39677283 | ckb-2 |
| WBGene00000513 | NA | NA | NA | ckb-3 |
| WBGene00000514 | -0.276204744 | 0.16788196 | 0.34551024 | ckb-4 |
| WBGene00000515 | -0.775343832 | 6.24E-05 | 0.00064047 | ckc-1 |
| WBGene00000516 | 0.223422531 | 0.31161754 | 0.51262836 | cki-1 |
| WBGene00000517 | 0.024534635 | 0.82730782 | 0.90561599 | cki-2 |
| WBGene00000518 | 0.505626588 | 0.06581663 | 0.18318234 | ckk-1 |
| WBGene00000519 | 0.11189687 | 0.35156701 | 0.55166504 | cku-70 |
| WBGene00000520 | -0.188349459 | 0.05887326 | 0.16924474 | cku-80 |
| WBGene00000521 | 0.020317245 | 0.84643798 | NA | cky-1 |
| WBGene00000522 | 1.040720238 | 3.89E-08 | 7.88E-07 | clc-1 |
| WBGene00000523 | -0.112973217 | 0.53343462 | 0.70899686 | clc-2 |
| WBGene00000524 | 0.222282638 | 0.31357151 | 0.51405222 | clc-3 |
| WBGene00000525 | -0.131786469 | 0.22320503 | 0.41587488 | clc-4 |
| WBGene00000526 | -0.085776245 | 0.65154751 | 0.79364585 | clc-5 |
| WBGene00000527 | 0.097320983 | 0.4258715 | 0.62160822 | cle-1 |
| WBGene00000528 | 0.390449713 | 0.0013442 | 0.00895784 | clh-1 |
| WBGene00000529 | 0.970150833 | 0.00994101 | 0.04467353 | clh-2 |
| WBGene00000530 | 0.243862294 | 0.09905169 | 0.24407452 | clh-3 |
| WBGene00000531 | 0.580303738 | 0.04766784 | 0.14524554 | clh-4 |
| WBGene00000532 | -0.218260973 | 0.00904244 | 0.04156684 | clh-5 |
| WBGene00000533 | 0.220667177 | 0.05302445 | 0.15721567 | clh-6 |
| WBGene00000534 | 0.056482527 | 0.4397848 | 0.63430988 | cpi-2 |
| WBGene00000535 | -1.221491107 | 7.96E-25 | 7.37E-23 | cpi-1 |

| | | | | |
|---|---|---|---|---|
| WBGene00000536 | 0.074608392 | 0.59584864 | 0.75477619 | clk-1 |
| WBGene00000537 | 0.067048874 | 0.56941772 | 0.7382268 | clk-2 |
| WBGene00000539 | 0.854382834 | 0.00065534 | 0.00489283 | cln-3.1 |
| WBGene00000540 | 0.294668466 | 0.07742353 | 0.20598907 | cln-3.2 |
| WBGene00000541 | 0.336789102 | 0.15881056 | 0.33319818 | cln-3.3 |
| WBGene00000542 | 0.179486126 | 0.00661157 | 0.03257066 | clp-1 |
| WBGene00000543 | -0.075704901 | 0.52916296 | 0.70598459 | clp-2 |
| WBGene00000544 | 0.012518106 | 0.92490827 | 0.95767907 | clp-3 |
| WBGene00000545 | 0.008856027 | 0.95642321 | 0.97575028 | clp-4 |
| WBGene00000546 | 0.460008677 | 0.05767896 | 0.16709094 | clp-6 |
| WBGene00000547 | 0.232545313 | 0.00652651 | 0.03222751 | clp-7 |
| WBGene00000548 | -0.028864907 | 0.73377035 | 0.85087373 | clr-1 |
| WBGene00000549 | -0.274152842 | 0.00343301 | 0.01945905 | cls-2 |
| WBGene00000550 | -0.301728874 | 1.21E-06 | 1.88E-05 | clu-1 |
| WBGene00000551 | NA | NA | NA | clx-1 |
| WBGene00000552 | 0.058065218 | 0.31731297 | 0.51740497 | cmd-1 |
| WBGene00000553 | 0.823058023 | 0.02125676 | 0.08031491 | cmk-1 |
| WBGene00000554 | 0.43946682 | 0.00298176 | 0.01739975 | cnb-1 |
| WBGene00000555 | NA | NA | NA | cnc-1 |
| WBGene00000556 | 0.186513185 | 0.35158709 | 0.55166504 | cnc-2 |
| WBGene00000557 | -0.038454169 | 0.84999349 | 0.91926207 | cnc-3 |
| WBGene00000558 | -0.577286264 | 0.04367086 | 0.13647493 | cnc-4 |
| WBGene00000559 | NA | NA | NA | cnc-5 |
| WBGene00000560 | 0.020925399 | 0.84643798 | NA | cnc-6 |
| WBGene00000561 | -0.366784956 | 0.10651318 | 0.25602685 | cnd-1 |
| WBGene00000562 | 0.034293201 | 0.84822185 | 0.9180823 | cng-1 |
| WBGene00000563 | 0.010413888 | 0.90992859 | NA | cng-3 |
| WBGene00000564 | -0.113200064 | 0.24251241 | 0.44063343 | cnk-1 |
| WBGene00000565 | 0.12361199 | 0.29601209 | 0.49712239 | cnt-1 |
| WBGene00000566 | 0.014377311 | 0.88266777 | 0.93795463 | cnt-2 |
| WBGene00000567 | -0.198161306 | 0.00105938 | 0.00736835 | cnx-1 |
| WBGene00000583 | 0.145836221 | 0.47335878 | 0.66130209 | cof-2 |
| WBGene00000584 | -0.028047269 | 0.84994578 | 0.91926207 | cog-1 |
| WBGene00000585 | -0.258264166 | 0.01033408 | 0.0460577 | cogc-2 |
| WBGene00000591 | -0.142928475 | 0.32565871 | 0.52475997 | coh-1 |
| WBGene00000592 | -0.018532808 | 0.88675049 | 0.94033043 | coh-3 |
| WBGene00000593 | 0.040107227 | 0.58693163 | NA | col-2 |
| WBGene00000594 | 0.079401127 | 0.41122513 | 0.60883276 | col-117 |
| WBGene00000594 | 0.079401127 | 0.41122513 | 0.60883276 | col-3 |
| WBGene00000596 | 0.159247053 | 0.27978227 | 0.47949107 | col-7 |
| WBGene00000597 | -1.412580896 | 4.78E-172 | 5.00E-169 | col-8 |
| WBGene00000598 | NA | NA | NA | col-9 |
| WBGene00000599 | 0.020925399 | 0.84643798 | NA | col-10 |

| | | | | |
|---|---|---|---|---|
| WBGene00000601 | 0.597876708 | 0.03358843 | 0.11248666 | col-12 |
| WBGene00000602 | NA | NA | NA | col-13 |
| WBGene00000603 | 4.095341621 | 8.47E-08 | 1.61E-06 | col-14 |
| WBGene00000606 | 5.291220306 | 8.54E-05 | 0.0008411 | col-17 |
| WBGene00000608 | -0.788293165 | 1.59E-62 | 4.46E-60 | col-19 |
| WBGene00000609 | -1.043215835 | 1.03E-110 | 6.54E-108 | col-20 |
| WBGene00000610 | 0.020317245 | 0.84643798 | NA | col-33 |
| WBGene00000611 | 0.239823054 | 0.27944661 | 0.47925241 | col-34 |
| WBGene00000612 | NA | NA | NA | col-35 |
| WBGene00000613 | 0.038125098 | 0.61119482 | NA | col-36 |
| WBGene00000614 | NA | NA | NA | col-37 |
| WBGene00000615 | 1.700520658 | 0.00042844 | 0.00340192 | col-38 |
| WBGene00000616 | 2.336662393 | 1.56E-06 | 2.34E-05 | col-39 |
| WBGene00000617 | 0.020925399 | 0.84643798 | NA | col-40 |
| WBGene00000618 | 0.204737186 | 0.03089539 | 0.105465 | col-41 |
| WBGene00000620 | -3.063069601 | 3.13E-05 | 0.0003415 | col-43 |
| WBGene00000621 | 0.020317245 | 0.84643798 | NA | col-44 |
| WBGene00000622 | 0.011538334 | 0.91199233 | 0.95112162 | col-45 |
| WBGene00000623 | 0.020925399 | 0.84643798 | NA | col-46 |
| WBGene00000624 | NA | NA | NA | col-47 |
| WBGene00000625 | 1.084605631 | 0.0126882 | 0.05409667 | col-48 |
| WBGene00000626 | 2.23211202 | 0.00763056 | 0.03640244 | col-49 |
| WBGene00000627 | NA | NA | NA | col-50 |
| WBGene00000628 | NA | NA | NA | col-51 |
| WBGene00000629 | -0.023109762 | 0.78121253 | NA | col-52 |
| WBGene00000630 | 0.07163168 | 0.68023541 | 0.81488454 | col-53 |
| WBGene00000631 | 0.109591702 | 0.21566448 | NA | col-54 |
| WBGene00000632 | NA | NA | NA | col-55 |
| WBGene00000633 | NA | NA | NA | col-56 |
| WBGene00000634 | 0.442817743 | 0.00351924 | 0.01984026 | col-58 |
| WBGene00000636 | 0.195066035 | 0.22409937 | 0.41711331 | col-60 |
| WBGene00000637 | -0.030550281 | 0.70468744 | NA | col-61 |
| WBGene00000638 | 0.249583013 | 0.22122898 | 0.41371995 | col-62 |
| WBGene00000639 | 1.120970442 | 0.01380685 | 0.05787387 | col-63 |
| WBGene00000640 | 0.107377299 | 0.31369484 | 0.5141086 | col-64 |
| WBGene00000641 | 4.655134044 | 4.06E-05 | 0.00043377 | col-65 |
| WBGene00000642 | NA | NA | NA | NA |
| WBGene00000644 | -0.138130672 | 0.42190059 | 0.6184468 | col-68 |
| WBGene00000645 | -0.004864426 | 0.94835335 | NA | col-69 |
| WBGene00000646 | -0.078514061 | 0.31122914 | NA | H17B01.2 |
| WBGene00000647 | 1.090859586 | 0.01042338 | 0.04632929 | col-71 |
| WBGene00000648 | -0.023056499 | 0.78121253 | NA | col-72 |
| WBGene00000649 | NA | NA | NA | col-73 |

| | | | | |
|---|---|---|---|---|
| WBGene00000650 | 3.757150698 | 3.04E-09 | 7.23E-08 | col-74 |
| WBGene00000651 | 0.020317245 | 0.84643798 | NA | col-75 |
| WBGene00000652 | 0.091280098 | 0.64544288 | 0.79001558 | col-76 |
| WBGene00000653 | 3.215474259 | 1.60E-08 | 3.46E-07 | col-77 |
| WBGene00000654 | NA | NA | NA | col-78 |
| WBGene00000655 | -0.008969483 | 0.9173367 | NA | col-79 |
| WBGene00000656 | -1.322674055 | 1.14E-27 | 1.24E-25 | col-80 |
| WBGene00000657 | -2.132545251 | 9.54E-34 | 1.36E-31 | col-81 |
| WBGene00000658 | -0.053591705 | 0.45278076 | NA | col-83 |
| WBGene00000659 | 0.056304773 | 0.44260385 | NA | col-84 |
| WBGene00000660 | -0.027535543 | 0.68063395 | NA | col-85 |
| WBGene00000661 | -0.05458785 | 0.5305143 | NA | col-86 |
| WBGene00000662 | -0.049845644 | 0.48082527 | NA | col-87 |
| WBGene00000663 | 2.454420015 | 0.00618393 | 0.03086907 | col-88 |
| WBGene00000664 | 0.177616853 | 0.0463229 | NA | col-89 |
| WBGene00000665 | 0.619137483 | 0.02179981 | 0.08183873 | col-90 |
| WBGene00000666 | 4.219594161 | 0.00105412 | 0.00734226 | col-91 |
| WBGene00000667 | NA | NA | NA | col-92 |
| WBGene00000668 | NA | NA | NA | col-93 |
| WBGene00000669 | -0.003108065 | 0.95728506 | NA | col-94 |
| WBGene00000670 | -0.541068744 | 7.13E-13 | 2.72E-11 | col-95 |
| WBGene00000671 | -0.301901498 | 0.17893368 | 0.36025413 | col-96 |
| WBGene00000672 | 1.299475884 | 0.00853067 | 0.03970042 | col-97 |
| WBGene00000673 | -2.228773084 | 3.75E-171 | 3.66E-168 | col-98 |
| WBGene00000674 | 0.056830639 | 0.78560642 | 0.87882516 | col-99 |
| WBGene00000675 | -1.389590948 | 2.90E-168 | 2.65E-165 | col-101 |
| WBGene00000676 | NA | NA | NA | col-102 |
| WBGene00000677 | -2.118875455 | 3.01E-148 | 2.59E-145 | col-103 |
| WBGene00000678 | 1.950229742 | 0.00042846 | 0.00340192 | col-104 |
| WBGene00000679 | 0.048485426 | 0.45975815 | NA | col-105 |
| WBGene00000680 | -0.957995573 | 2.10E-91 | 1.02E-88 | col-106 |
| WBGene00000681 | 2.765411897 | 4.25E-05 | 0.00045313 | col-107 |
| WBGene00000682 | 0.020317245 | 0.84643798 | NA | col-108 |
| WBGene00000683 | 3.911776767 | 0.00040206 | 0.00322901 | col-109 |
| WBGene00000684 | -0.632992939 | 0.02783954 | 0.09802757 | col-110 |
| WBGene00000685 | -0.101826549 | 0.59136588 | 0.75215398 | col-111 |
| WBGene00000687 | 0.198228458 | 0.30310341 | 0.50377307 | col-113 |
| WBGene00000688 | 0.085537126 | 0.24698522 | NA | col-114 |
| WBGene00000689 | 2.244282888 | 0.00739426 | 0.03550653 | col-115 |
| WBGene00000690 | 0.071910655 | 0.70740537 | 0.83484782 | col-116 |
| WBGene00000691 | 0.07892265 | 0.25795693 | NA | col-117 |
| WBGene00000691 | 0.07892265 | 0.25795693 | NA | col-3 |
| WBGene00000692 | 0.501644189 | 0.00327884 | 0.01873751 | col-118 |

| | | | | | |
|---|---|---|---|---|---|
| WBGene00000693 | | -0.681172176 | 1.91E-60 | 5.09E-58 | col-119 |
| WBGene00000694 | | 1.816926908 | 0.00607175 | 0.03046488 | col-120 |
| WBGene00000695 | | 0.100557393 | 0.51441888 | 0.69441331 | col-121 |
| WBGene00000696 | | -1.078120167 | 3.13E-91 | 1.48E-88 | col-122 |
| WBGene00000697 | | -0.071306693 | 0.34233497 | NA | col-123 |
| WBGene00000698 | | -1.017992074 | 3.59E-92 | 1.81E-89 | col-124 |
| WBGene00000699 | | 3.158085696 | 3.04E-08 | 6.27E-07 | col-125 |
| WBGene00000700 | NA | | NA | NA | col-126 |
| WBGene00000700 | NA | | NA | NA | col-127 |
| WBGene00000701 | NA | | NA | NA | col-126 |
| WBGene00000701 | NA | | NA | NA | col-127 |
| WBGene00000702 | | -0.001993815 | 0.95612996 | NA | col-128 |
| WBGene00000703 | | -2.128614455 | 1.27E-67 | 3.88E-65 | col-129 |
| WBGene00000704 | | 3.689824182 | 0.00066452 | 0.00495375 | col-130 |
| WBGene00000705 | | 0.05787416 | 0.40594086 | NA | col-131 |
| WBGene00000706 | | 0.075726029 | 0.27693214 | NA | col-132 |
| WBGene00000707 | | 0.297684806 | 0.16334772 | 0.33887458 | col-133 |
| WBGene00000708 | | 0.529662139 | 3.30E-06 | 4.48E-05 | col-135 |
| WBGene00000709 | | -0.003421551 | 0.95689871 | NA | NA |
| WBGene00000710 | | 0.131317611 | 0.10121567 | NA | col-137 |
| WBGene00000711 | | 2.045869635 | 0.00363552 | 0.02034117 | col-138 |
| WBGene00000712 | | -1.922547997 | 1.33E-34 | 1.96E-32 | col-139 |
| WBGene00000713 | | -0.50586129 | 7.39E-31 | 9.58E-29 | col-140 |
| WBGene00000714 | | -0.392423623 | 0.06614824 | 0.18362328 | col-141 |
| WBGene00000715 | | -0.771640218 | 6.42E-08 | 1.25E-06 | col-142 |
| WBGene00000716 | | -0.553618305 | 6.34E-20 | 4.73E-18 | col-143 |
| WBGene00000717 | | 0.013097374 | 0.89046473 | NA | col-144 |
| WBGene00000718 | | 0.020925399 | 0.84643798 | NA | col-145 |
| WBGene00000719 | | 3.82298918 | 0.00196293 | 0.01231059 | col-146 |
| WBGene00000720 | | 1.633894563 | 0.00067851 | 0.00504405 | col-147 |
| WBGene00000721 | | 0.020232312 | 0.84643798 | NA | col-148 |
| WBGene00000722 | | -1.894111753 | 0.00015756 | 0.00143284 | col-149 |
| WBGene00000723 | | -1.536373824 | 9.26E-05 | 0.00090711 | col-150 |
| WBGene00000724 | | 0.060105893 | 0.41982798 | NA | col-151 |
| WBGene00000725 | | -0.010487137 | 0.91553956 | 0.95298026 | col-152 |
| WBGene00000726 | | 0.546321923 | 0.0339251 | 0.11329777 | col-153 |
| WBGene00000727 | | 0.041956259 | 0.74147635 | 0.85407613 | col-154 |
| WBGene00000728 | | -0.139144284 | 0.40504217 | 0.60312812 | col-155 |
| WBGene00000729 | | 0.031770744 | 0.6954204 | NA | col-156 |
| WBGene00000730 | | 0.099038939 | 0.43298947 | 0.62797258 | col-157 |
| WBGene00000731 | | -1.995884756 | 6.42E-07 | 1.05E-05 | col-158 |
| WBGene00000732 | | -2.386450107 | 4.67E-26 | 4.62E-24 | col-159 |
| WBGene00000733 | | -0.447567161 | 1.08E-08 | 2.40E-07 | col-160 |

| | | | | |
|---|---|---|---|---|
| WBGene00000734 | 2.397473736 | 0.00590131 | 0.02978792 | col-161 |
| WBGene00000735 | 0.105013672 | 0.47963689 | 0.66640541 | col-162 |
| WBGene00000736 | -0.059246987 | 0.34871219 | NA | col-163 |
| WBGene00000737 | 0.020317245 | 0.84643798 | NA | col-164 |
| WBGene00000738 | 3.905125307 | 2.61E-10 | 7.13E-09 | col-165 |
| WBGene00000739 | 0.507308641 | 0.00851441 | 0.03964995 | col-166 |
| WBGene00000740 | 2.015008437 | 0.00277591 | 0.01639744 | col-167 |
| WBGene00000741 | 0.204978633 | 0.03076956 | 0.10520716 | col-168 |
| WBGene00000742 | 0.126299345 | 0.16983897 | NA | col-169 |
| WBGene00000743 | 0.090103632 | 0.20323472 | NA | col-170 |
| WBGene00000744 | NA | NA | NA | col-171 |
| WBGene00000745 | 0.042065281 | 0.558937 | NA | col-172 |
| WBGene00000746 | 0.181472875 | 0.20122728 | 0.38993842 | col-173 |
| WBGene00000747 | 0.086430603 | 0.29509282 | NA | col-174 |
| WBGene00000748 | 1.713408533 | 0.00973597 | 0.04399515 | col-175 |
| WBGene00000749 | -0.131840768 | 0.29639449 | 0.49751505 | col-176 |
| WBGene00000750 | 0.12158807 | 0.31166878 | 0.51265506 | col-177 |
| WBGene00000751 | -0.77742396 | 2.68E-52 | 6.55E-50 | col-178 |
| WBGene00000752 | 0.059690168 | 0.38915366 | 0.58756303 | col-179 |
| WBGene00000753 | 0.294412269 | 0.07698791 | 0.20527774 | col-180 |
| WBGene00000754 | -1.159426002 | 4.47E-131 | 3.27E-128 | col-181 |
| WBGene00000755 | 0.110545227 | 0.1354043 | NA | col-182 |
| WBGene00000756 | NA | NA | NA | col-183 |
| WBGene00000757 | -0.97479147 | 3.47E-86 | 1.54E-83 | col-184 |
| WBGene00000758 | NA | NA | NA | col-185 |
| WBGene00000759 | -0.010139963 | 0.94751096 | 0.97044477 | col-186 |
| WBGene00000760 | -0.060526604 | 0.59267979 | 0.75314295 | col-187 |
| WBGene00000761 | -0.213403071 | 0.08873916 | 0.22611347 | coq-1 |
| WBGene00000762 | 0.228193472 | 0.10402737 | 0.25210364 | coq-2 |
| WBGene00000763 | 0.191023791 | 0.25635072 | 0.45460486 | coq-3 |
| WBGene00000764 | 0.171669807 | 0.29997272 | 0.50044446 | coq-4 |
| WBGene00000765 | -0.351500863 | 0.0040033 | 0.02190295 | coq-5 |
| WBGene00000766 | 0.169930852 | 0.29156635 | 0.49225357 | coq-6 |
| WBGene00000767 | -0.201587346 | 0.03673459 | 0.1202203 | coq-8 |
| WBGene00000768 | 0.12876783 | 0.49750745 | 0.67973185 | cor-1 |
| WBGene00000770 | 0.039873266 | 0.72817781 | 0.84850209 | cpb-1 |
| WBGene00000771 | 0.079061785 | 0.41293424 | 0.61056051 | cpb-2 |
| WBGene00000772 | -0.220777274 | 0.00132781 | 0.00887289 | cpb-3 |
| WBGene00000773 | -0.21286515 | 0.01728504 | 0.06899409 | cpf-1 |
| WBGene00000774 | -0.127582916 | 0.30466527 | 0.50545453 | cpf-2 |
| WBGene00000775 | -0.306270066 | 0.17061001 | 0.34931872 | cpin-1 |
| WBGene00000776 | 0.38611621 | 4.91E-13 | 1.91E-11 | cpl-1 |
| WBGene00000777 | -0.193766811 | 0.00289522 | 0.01697944 | cpn-1 |

| | | | | |
|---|---|---|---|---|
| WBGene00000778 | 0.115920614 | 0.54689273 | 0.71995249 | cpn-2 |
| WBGene00000779 | 0.097795575 | 0.17655463 | 0.35738765 | cpn-3 |
| WBGene00000780 | 0.636616903 | 3.96E-05 | 0.00042419 | cpn-4 |
| WBGene00000781 | 2.573677456 | 0 | 0 | cpr-1 |
| WBGene00000782 | 0.260521751 | 0.01485574 | 0.06144149 | cpr-2 |
| WBGene00000783 | 0.536056035 | 0.02326026 | 0.08573853 | cpr-3 |
| WBGene00000784 | 2.884576857 | 0 | 0 | cpr-4 |
| WBGene00000785 | 0.217122733 | 0.00124438 | 0.0084191 | cpr-5 |
| WBGene00000786 | -0.219019002 | 1.14E-06 | 1.78E-05 | cpr-6 |
| WBGene00000787 | 0.157852612 | 0.21349081 | 0.40425749 | cps-6 |
| WBGene00000788 | -0.650588323 | 2.57E-28 | 2.92E-26 | cpz-1 |
| WBGene00000789 | 0.430277117 | 0.00109288 | 0.00757619 | cpz-2 |
| WBGene00000792 | 0.148327832 | 0.42931852 | 0.6241339 | crb-1 |
| WBGene00000793 | 0.127935569 | 0.33946847 | 0.53917963 | crh-1 |
| WBGene00000794 | -0.051352941 | 0.50151293 | 0.68305968 | crn-1 |
| WBGene00000795 | -0.096726633 | 0.61573035 | 0.76945272 | crn-2 |
| WBGene00000796 | -0.119532562 | 0.10459398 | 0.25284464 | crn-3 |
| WBGene00000797 | -0.146752495 | 0.40557913 | 0.60370923 | crn-4 |
| WBGene00000798 | 0.02049827 | 0.89139637 | 0.94346376 | crn-5 |
| WBGene00000799 | 1.047294911 | 5.68E-08 | 1.12E-06 | crn-6 |
| WBGene00000800 | -0.104969835 | 0.14407824 | 0.31302114 | cars-1 |
| WBGene00000801 | 0.591686466 | 0.02653962 | 0.09500297 | NA |
| WBGene00000802 | -0.097806193 | 0.02700751 | 0.095975 | crt-1 |
| WBGene00000803 | -0.145669849 | 0.33649204 | 0.53590558 | csb-1 |
| WBGene00000804 | 0.014256345 | 0.88078 | 0.93662842 | csc-1 |
| WBGene00000812 | -0.018819768 | 0.86245441 | 0.92672958 | csk-1 |
| WBGene00000813 | -0.000352174 | 0.99633061 | 0.99769349 | csn-1 |
| WBGene00000814 | -0.204563154 | 0.00901463 | 0.04147808 | csn-2 |
| WBGene00000815 | -0.229144032 | 0.00652477 | 0.03222751 | csn-3 |
| WBGene00000816 | 0.009888398 | 0.91365619 | 0.95181729 | csn-4 |
| WBGene00000817 | -0.001167951 | 0.98925796 | 0.99398816 | csn-5 |
| WBGene00000818 | -0.018349996 | 0.86552908 | 0.92809516 | csn-6 |
| WBGene00000819 | 0.04225313 | 0.75108794 | 0.85938701 | csp-1 |
| WBGene00000820 | 1.922044112 | 0.00312823 | 0.01802454 | csp-2 |
| WBGene00000821 | 0.018860755 | 0.77262519 | NA | csp-3 |
| WBGene00000822 | -0.364554728 | 0.00013019 | 0.00121665 | csq-1 |
| WBGene00000829 | -0.359210521 | 0.0068541 | 0.0334828 | CYTB |
| WBGene00000830 | 0.204768966 | 0.34925108 | 0.54917678 | ctl-1 |
| WBGene00000831 | 0.356474796 | 0.00165772 | 0.01070606 | ctl-2 |
| WBGene00000832 | 0.120478114 | 0.44900195 | 0.64173691 | ctn-1 |
| WBGene00000833 | -0.039606128 | 0.41524399 | 0.6126146 | cts-1 |
| WBGene00000834 | -0.057734923 | 0.49526189 | 0.67792907 | cua-1 |
| WBGene00000835 | -0.389578162 | 0.00142052 | 0.00939805 | cuc-1 |

| | | | | |
|---|---|---|---|---|
| WBGene00000836 | -0.276264442 | 4.73E-05 | 0.00050016 | cul-1 |
| WBGene00000837 | -0.134743268 | 0.04361468 | 0.13632845 | cul-2 |
| WBGene00000838 | -0.210486369 | 0.00034966 | 0.00285677 | cul-3 |
| WBGene00000839 | -0.149166235 | 0.10715778 | 0.25715409 | cul-4 |
| WBGene00000840 | -0.223089003 | 0.01302465 | 0.05528962 | cul-5 |
| WBGene00000841 | 0.470739735 | 0.07121271 | 0.1939142 | cul-6 |
| WBGene00000843 | -0.052209449 | 0.53473919 | 0.71000989 | cup-2 |
| WBGene00000845 | -0.140197523 | 0.49145067 | 0.6750473 | cup-4 |
| WBGene00000846 | -0.011741509 | 0.8966934 | 0.94573597 | cup-5 |
| WBGene00000851 | NA | NA | NA | cut-1 |
| WBGene00000853 | 0.072473389 | 0.54345983 | 0.71696212 | cut-6 |
| WBGene00000854 | -0.00371026 | 0.97135453 | 0.98427495 | cux-7 |
| WBGene00000857 | -0.042265277 | 0.83227404 | 0.9084712 | cwn-1 |
| WBGene00000858 | 0.132030522 | 0.53373064 | 0.70904186 | cwn-2 |
| WBGene00000859 | NA | NA | NA | cwp-1 |
| WBGene00000860 | 0.033463284 | 0.6365037 | NA | cwp-2 |
| WBGene00000861 | NA | NA | NA | cwp-3 |
| WBGene00000862 | 0.477883897 | 0.05086545 | 0.15231452 | cwp-4 |
| WBGene00000863 | -0.206410188 | 0.01642017 | 0.06646604 | cya-1 |
| WBGene00000864 | 0.110863018 | 0.50253208 | 0.68391651 | cya-2 |
| WBGene00000865 | -0.186419994 | 0.00184144 | 0.01168132 | cyb-1 |
| WBGene00000866 | -0.235027128 | 0.00118539 | 0.0080948 | cyb-2.1 |
| WBGene00000867 | -0.291863751 | 1.47E-07 | 2.68E-06 | cyb-2.2 |
| WBGene00000868 | -0.323098239 | 2.85E-11 | 8.83E-10 | cyb-3 |
| WBGene00000869 | -0.177632035 | 0.00188657 | 0.01192116 | cyc-1 |
| WBGene00000870 | -0.57078654 | 0.00614218 | 0.03071169 | cyd-1 |
| WBGene00000871 | -0.18620852 | 0.02706083 | 0.09609451 | cye-1 |
| WBGene00000872 | 0.219843047 | 0.03889799 | 0.1253589 | cyk-1 |
| WBGene00000874 | -0.21317688 | 0.01130742 | 0.04924465 | cyk-3 |
| WBGene00000875 | -0.03866653 | 0.59940372 | 0.75732394 | cyk-4 |
| WBGene00000876 | -0.262014493 | 0.00409776 | 0.02231969 | cyl-1 |
| WBGene00000877 | -0.208024157 | 0.00538812 | 0.02773477 | cyn-1 |
| WBGene00000878 | 0.513797947 | 0.06238157 | 0.17662512 | cyn-2 |
| WBGene00000879 | -0.194713454 | 0.00305117 | 0.01771796 | cyn-3 |
| WBGene00000880 | -0.073712853 | 0.46256769 | 0.65252509 | cyn-4 |
| WBGene00000881 | -0.317008172 | 2.60E-06 | 3.65E-05 | cyn-5 |
| WBGene00000882 | -0.415804022 | 0.00759414 | 0.03627593 | cyn-6 |
| WBGene00000883 | -0.396268006 | 1.79E-20 | 1.39E-18 | cyn-7 |
| WBGene00000884 | -0.221569811 | 0.10899638 | 0.25977796 | cyn-8 |
| WBGene00000885 | -0.046300744 | 0.67681841 | 0.81237075 | cyn-9 |
| WBGene00000886 | -0.226787797 | 0.07738738 | 0.20593032 | cyn-10 |
| WBGene00000887 | 0.138843741 | 0.42731688 | 0.62252204 | cyn-11 |
| WBGene00000888 | -0.290463844 | 0.01140571 | 0.04954906 | cyn-12 |

| | | | | |
|---|---|---|---|---|
| WBGene00000889 | 0.008134217 | 0.93253928 | 0.9620984 | cyn-13 |
| WBGene00000890 | -0.18161381 | 0.07464272 | 0.20089046 | sig-7 |
| WBGene00000891 | -0.118822306 | 0.22312926 | 0.41583474 | cyn-15 |
| WBGene00000892 | -0.146601371 | 0.21273968 | 0.40320754 | cyn-16 |
| WBGene00000893 | 0.122532655 | 0.26103761 | 0.45919656 | cyn-17 |
| WBGene00000894 | -0.197918381 | 0.00102432 | 0.0071722 | dab-1 |
| WBGene00000895 | 0.194492696 | 0.29480073 | 0.49577045 | dac-1 |
| WBGene00000896 | -0.262379648 | 0.00070181 | 0.00518689 | dad-1 |
| WBGene00000897 | -0.03494785 | 0.76814802 | 0.86844708 | daf-1 |
| WBGene00000898 | 0.452547976 | 2.99E-09 | 7.12E-08 | daf-2 |
| WBGene00000899 | 0.086385664 | 0.56866291 | 0.73757917 | daf-3 |
| WBGene00000900 | 0.144974506 | 0.19078997 | 0.37682446 | daf-4 |
| WBGene00000901 | 0.498775379 | 0.00080539 | 0.00585147 | daf-5 |
| WBGene00000902 | 0.056421825 | 0.7149855 | 0.84013666 | daf-6 |
| WBGene00000903 | 0.881693332 | 0.00278617 | 0.0164485 | daf-7 |
| WBGene00000904 | -0.088968694 | 0.64368722 | 0.78910028 | daf-8 |
| WBGene00000905 | 0.041986586 | 0.81725374 | 0.8995874 | daf-9 |
| WBGene00000906 | 0.209293393 | 0.16134213 | 0.33592293 | daf-10 |
| WBGene00000907 | 0.316766166 | 0.10030322 | 0.24594532 | daf-11 |
| WBGene00000908 | 0.277678555 | 0.12183428 | 0.28082112 | daf-12 |
| WBGene00000910 | 0.767990836 | 0.00192244 | 0.01209618 | daf-14 |
| WBGene00000911 | 0.063212053 | 0.37441374 | 0.57328922 | daf-15 |
| WBGene00000912 | 0.637674248 | 4.63E-16 | 2.49E-14 | daf-16 |
| WBGene00000913 | 0.01657197 | 0.74447841 | 0.85556581 | daf-18 |
| WBGene00000914 | 0.250381338 | 0.10875678 | 0.25941805 | daf-19 |
| WBGene00000915 | -0.428791022 | 5.53E-27 | 5.74E-25 | hsp-90 |
| WBGene00000917 | -0.059562128 | 0.65867844 | 0.79882238 | daf-25 |
| WBGene00000920 | 0.105934325 | 0.58110847 | 0.74581663 | daf-28 |
| WBGene00000923 | -0.277198588 | 0.00565101 | 0.02882742 | daf-31 |
| WBGene00000928 | -0.313662488 | 0.17508948 | 0.35564444 | dao-2 |
| WBGene00000929 | -0.357978104 | 0.02055046 | 0.07816408 | dao-3 |
| WBGene00000930 | -0.003614866 | 0.95700117 | NA | dao-4 |
| WBGene00000931 | -0.229453525 | 0.00189115 | 0.01194495 | dao-5 |
| WBGene00000932 | 0.042199095 | 0.55922995 | NA | dao-6 |
| WBGene00000933 | -0.271019956 | 0.01208834 | 0.05200863 | dap-3 |
| WBGene00000934 | 0.711556634 | 0.02907855 | 0.10136642 | dat-1 |
| WBGene00000935 | -0.041440012 | 0.50861285 | 0.68894903 | daz-1 |
| WBGene00000936 | 0.602939348 | 0.01361713 | 0.05724882 | dbl-1 |
| WBGene00000937 | -0.112804771 | 0.31609564 | 0.51674366 | dbr-1 |
| WBGene00000938 | -0.225703955 | 0.00595323 | 0.02996259 | dcp-66 |
| WBGene00000939 | -0.15557298 | 0.02740414 | 0.0969709 | dcr-1 |
| WBGene00000940 | -0.165696273 | 0.20612339 | 0.3952138 | dcs-1 |
| WBGene00000941 | -0.308822891 | 0.00436638 | 0.02353911 | ddp-1 |

| | | | | |
|---|---|---|---|---|
| WBGene00000942 | 0.272656928 | 0.0145513 | 0.06038706 | deb-1 |
| WBGene00000950 | 0.211657878 | 0.32304817 | 0.52302147 | deg-1 |
| WBGene00000951 | 0.257958166 | 0.25053319 | 0.44909133 | deg-3 |
| WBGene00000952 | 0.000818013 | 0.99632799 | 0.99769349 | del-1 |
| WBGene00000953 | 0.269943228 | 0.01368153 | 0.05746165 | del-2 |
| WBGene00000955 | 0.432457186 | 0.09332365 | 0.23360431 | des-2 |
| WBGene00000958 | 0.623852001 | 0.02504502 | 0.09094035 | dgk-1 |
| WBGene00000959 | 0.147971866 | 0.3265475 | 0.52572927 | dgk-2 |
| WBGene00000960 | 0.275147046 | 0.06584168 | 0.18319803 | dgk-3 |
| WBGene00000961 | -0.597922639 | 0.00033238 | 0.00274137 | dgn-1 |
| WBGene00000962 | 0.166654605 | 0.00287809 | 0.01692291 | dhc-1 |
| WBGene00000963 | 0.64822135 | 0.00023751 | 0.0020419 | dhp-1 |
| WBGene00000964 | 1.135661618 | 2.48E-07 | 4.34E-06 | dhp-2 |
| WBGene00000965 | -0.113614325 | 0.25034771 | 0.44895131 | dhs-1 |
| WBGene00000966 | -0.266320998 | 0.23902278 | 0.43606953 | dhs-2 |
| WBGene00000967 | -0.922879891 | 5.01E-13 | 1.94E-11 | dhs-3 |
| WBGene00000968 | 0.407770416 | 0.10485411 | 0.25324464 | dhs-4 |
| WBGene00000969 | 0.175787115 | 0.41536316 | 0.61272866 | dhs-5 |
| WBGene00000970 | -0.128988967 | 0.12072821 | 0.27950375 | dhs-6 |
| WBGene00000971 | 0.095220825 | 0.64897822 | 0.79237696 | dhs-7 |
| WBGene00000972 | 0.440840504 | 0.07927902 | 0.20970625 | dhs-8 |
| WBGene00000973 | 0.066951238 | 0.66189177 | 0.80128637 | dhs-9 |
| WBGene00000974 | -0.027296217 | 0.84816368 | 0.9180823 | dhs-11 |
| WBGene00000975 | -0.19406327 | 0.0833747 | 0.21707176 | dhs-12 |
| WBGene00000976 | -0.222380078 | 0.02025592 | 0.07735185 | dhs-13 |
| WBGene00000977 | -1.048097443 | 2.39E-07 | 4.19E-06 | dhs-14 |
| WBGene00000978 | 0.126375685 | 0.45925758 | 0.65054085 | dhs-15 |
| WBGene00000979 | 0.107533449 | 0.21907273 | NA | dhs-16 |
| WBGene00000980 | 0.119649246 | 0.47450572 | 0.66189389 | dhs-17 |
| WBGene00000981 | 0.3014245 | 0.07591734 | 0.20338625 | dhs-18 |
| WBGene00000982 | -0.091691593 | 0.54004188 | 0.71451837 | dhs-19 |
| WBGene00000983 | 0.111755757 | 0.59312229 | 0.7533533 | dhs-20 |
| WBGene00000984 | 0.765142358 | 7.27E-05 | 0.00073171 | dhs-21 |
| WBGene00000985 | -0.089445697 | 0.25651205 | 0.45472731 | dhs-22 |
| WBGene00000986 | 2.466857454 | 4.04E-07 | 6.83E-06 | dhs-23 |
| WBGene00000987 | -0.216520555 | 0.04803868 | 0.14604119 | dhs-24 |
| WBGene00000988 | -1.818498416 | 2.86E-21 | 2.33E-19 | dhs-25 |
| WBGene00000989 | -0.153215506 | 0.24067512 | 0.43821967 | dhs-26 |
| WBGene00000990 | -0.175610801 | 0.37505247 | 0.57390712 | dhs-27 |
| WBGene00000991 | 0.180300383 | 0.05361798 | 0.15852603 | dhs-28 |
| WBGene00000992 | -0.195475521 | 0.35187178 | 0.55181487 | dhs-29 |
| WBGene00000993 | 0.096178499 | 0.48270928 | 0.66843342 | dhs-30 |
| WBGene00000994 | -0.174853825 | 0.09909426 | 0.24412027 | dic-1 |

| | | | | |
|---|---|---|---|---|
| WBGene00000995 | -0.467999037 | 6.18E-06 | 7.90E-05 | die-1 |
| WBGene00000996 | -0.324847104 | 0.00048973 | 0.00381797 | dif-1 |
| WBGene00000998 | 0.64133603 | 2.76E-06 | 3.84E-05 | dig-1 |
| WBGene00001000 | -0.364332162 | 2.45E-05 | 0.00027327 | dim-1 |
| WBGene00001001 | -0.214625084 | 0.00961705 | 0.04363284 | dis-3 |
| WBGene00001002 | -0.256398715 | 0.00445969 | 0.02395243 | div-1 |
| WBGene00001005 | -0.28054795 | 8.63E-08 | 1.64E-06 | dlc-1 |
| WBGene00001006 | 0.303534884 | 0.01802737 | 0.07120946 | dlg-1 |
| WBGene00001007 | -0.151910527 | 0.12783673 | 0.28914839 | dli-1 |
| WBGene00001008 | 1.043173747 | 6.45E-15 | 3.06E-13 | dlk-1 |
| WBGene00001016 | 0.061435053 | 0.53472928 | 0.71000989 | dna-2 |
| WBGene00001017 | -0.154108657 | 0.05446861 | 0.16015662 | dnc-1 |
| WBGene00001018 | -0.026673927 | 0.80220418 | 0.88992108 | dnc-2 |
| WBGene00001019 | -0.081550937 | 0.38192844 | 0.58114886 | dnj-1 |
| WBGene00001020 | -0.252847672 | 0.02712334 | 0.0961998 | dnj-2 |
| WBGene00001021 | -0.428650676 | 0.04508493 | 0.1394946 | dnj-3 |
| WBGene00001022 | 0.206416458 | 0.16404883 | 0.33991494 | dnj-4 |
| WBGene00001023 | -0.17737998 | 0.00094657 | 0.0067243 | dnj-5 |
| WBGene00001025 | -0.19154209 | 0.09352998 | 0.23396077 | dnj-7 |
| WBGene00001026 | -0.052389819 | 0.54828553 | 0.72092678 | dnj-8 |
| WBGene00001027 | -0.301365785 | 0.00706137 | 0.03423361 | dnj-9 |
| WBGene00001028 | -0.158225033 | 0.12131212 | 0.28041217 | dnj-10 |
| WBGene00001029 | -0.367819053 | 4.82E-09 | 1.13E-07 | dnj-11 |
| WBGene00001030 | -0.273979296 | 2.17E-06 | 3.13E-05 | dnj-12 |
| WBGene00001031 | -0.177344958 | 0.00816727 | 0.03843341 | dnj-13 |
| WBGene00001032 | -0.029804588 | 0.82201739 | 0.90246087 | dnj-14 |
| WBGene00001033 | -0.157840177 | 0.24133831 | 0.43915414 | dnj-15 |
| WBGene00001034 | -0.187430402 | 0.08766103 | 0.22426713 | dnj-16 |
| WBGene00001035 | -0.026603428 | 0.80135341 | 0.88934884 | dnj-17 |
| WBGene00001036 | -0.017789925 | 0.90006361 | 0.94668216 | dnj-18 |
| WBGene00001037 | -0.2997451 | 2.08E-07 | 3.70E-06 | dnj-19 |
| WBGene00001038 | 0.018860976 | 0.77582124 | 0.87314926 | dnj-20 |
| WBGene00001039 | -0.17440914 | 0.28636408 | 0.48638707 | dnj-21 |
| WBGene00001040 | -0.084055657 | 0.3875121 | 0.58611204 | dnj-22 |
| WBGene00001041 | -0.125123925 | 0.15729456 | 0.33126434 | dnj-23 |
| WBGene00001042 | 0.083079398 | 0.6744517 | 0.81099272 | dnj-24 |
| WBGene00001043 | -0.079280833 | 0.41522034 | 0.6126146 | dnj-25 |
| WBGene00001044 | -0.302187711 | 0.06841123 | 0.18834313 | dnj-26 |
| WBGene00001045 | -0.487379642 | 0.00128286 | 0.00863156 | dnj-27 |
| WBGene00001046 | 0.023146486 | 0.84567192 | 0.9169413 | dnj-28 |
| WBGene00001047 | -0.234382398 | 0.00035311 | 0.00288015 | dnj-29 |
| WBGene00001048 | -0.027590688 | 0.81906084 | 0.90049334 | dnj-30 |
| WBGene00001049 | -0.040026882 | 0.70783193 | 0.83521658 | dog-1 |

| | | | | |
|---|---|---|---|---|
| WBGene00001050 | 0.194333974 | 0.23958815 | 0.43670695 | dom-3 |
| WBGene00001051 | NA | NA | NA | cks-1 |
| WBGene00001052 | 0.641977306 | 0.0380435 | 0.12333809 | dop-1 |
| WBGene00001053 | 0.391154618 | 0.11696567 | 0.27352701 | dop-2 |
| WBGene00001054 | 0.196952641 | 0.36591709 | 0.56563063 | dpf-1 |
| WBGene00001055 | 0.793740594 | 5.18E-09 | 1.20E-07 | dpf-2 |
| WBGene00001056 | -0.133599823 | 0.03490933 | 0.11581859 | dpf-3 |
| WBGene00001057 | -0.076657424 | 0.4122175 | 0.60976496 | dpf-4 |
| WBGene00001058 | -0.082583711 | 0.20059335 | 0.38919788 | dpf-5 |
| WBGene00001059 | 0.585421899 | 0.04270054 | 0.13418731 | dpf-6 |
| WBGene00001060 | -0.05363159 | 0.61791493 | 0.77099816 | dpf-7 |
| WBGene00001061 | -0.323875523 | 7.92E-05 | 0.00078548 | dpl-1 |
| WBGene00001062 | NA | NA | NA | dpr-1 |
| WBGene00001063 | 0.406853111 | 0.06830314 | 0.18815169 | dpy-1 |
| WBGene00001064 | 0.092162487 | 0.62051257 | 0.77298771 | dpy-2 |
| WBGene00001065 | 0.406436438 | 0.0408407 | 0.12993235 | dpy-3 |
| WBGene00001066 | 3.287469535 | 0.00013608 | 0.00126101 | dpy-4 |
| WBGene00001067 | 5.925572357 | 1.16E-05 | 0.0001399 | dpy-5 |
| WBGene00001068 | -0.32596013 | 0.14675392 | 0.31699972 | dpy-6 |
| WBGene00001069 | 0.18130048 | 0.35469367 | 0.55451895 | dpy-7 |
| WBGene00001070 | 0.637401202 | 0.01978608 | 0.07599371 | dpy-8 |
| WBGene00001071 | 0.083790205 | 0.6425279 | 0.78833475 | dpy-9 |
| WBGene00001072 | 0.312899209 | 0.17061527 | 0.34931872 | T14B4.19 |
| WBGene00001072 | 0.312899209 | 0.17061527 | 0.34931872 | dpy-10 |
| WBGene00001073 | -0.156937408 | 0.04352969 | 0.13610879 | dpy-11 |
| WBGene00001074 | 1.623137671 | 0.00326316 | 0.01866732 | dpy-13 |
| WBGene00001075 | 0.425086608 | 0.0376386 | 0.12240486 | dpy-14 |
| WBGene00001076 | 0.285799621 | 0.09133431 | 0.22992189 | dpy-17 |
| WBGene00001077 | -0.080669369 | 0.58616665 | 0.74887137 | dpy-18 |
| WBGene00001078 | -0.091825261 | 0.63207466 | 0.78140716 | dpy-19 |
| WBGene00001079 | 0.048485426 | 0.45975815 | NA | dpy-20 |
| WBGene00001080 | 0.045485847 | 0.67817233 | 0.81306265 | dpy-21 |
| WBGene00001081 | -0.074293122 | 0.40965466 | 0.60757234 | dpy-22 |
| WBGene00001082 | -0.168194877 | 0.14113753 | 0.3087858 | dpy-23 |
| WBGene00001085 | -0.094214373 | 0.20757365 | 0.3966957 | dpy-26 |
| WBGene00001086 | -0.372647323 | 0.00048316 | 0.00378082 | dpy-27 |
| WBGene00001087 | 0.048206346 | 0.54974743 | 0.72232362 | dpy-28 |
| WBGene00001088 | -0.072527183 | 0.42442662 | 0.62029117 | dpy-30 |
| WBGene00001089 | 0.674149526 | 8.20E-06 | 0.00010266 | dre-1 |
| WBGene00001090 | -0.11789994 | 0.25377189 | 0.45176267 | drh-1 |
| WBGene00001091 | 1.056217619 | 8.08E-15 | 3.79E-13 | NA |
| WBGene00001092 | -0.197993508 | 0.08485111 | 0.21938866 | dro-1 |
| WBGene00001093 | -0.159037915 | 0.01914613 | 0.07448196 | drp-1 |

| | | | | |
|---|---|---|---|---|
| WBGene00001094 | -0.214544207 | 0.00039776 | 0.00320155 | dars-1 |
| WBGene00001095 | -0.13029259 | 0.3143452 | 0.51491699 | dars-2 |
| WBGene00001096 | 0.086430603 | 0.29509282 | NA | dsc-1 |
| WBGene00001099 | 0.523099248 | 0.00016569 | 0.00150118 | dsc-4 |
| WBGene00001101 | -0.007418762 | 0.95863292 | 0.97725557 | dsh-1 |
| WBGene00001102 | -0.314741467 | 7.74E-05 | 0.00077209 | dsh-2 |
| WBGene00001103 | -0.30768695 | 0.18659013 | 0.37097584 | dsl-1 |
| WBGene00001104 | -0.756086092 | 0.00012893 | 0.00120694 | dsl-2 |
| WBGene00001105 | -1.427492254 | 0.00325743 | 0.01864426 | dsl-3 |
| WBGene00001106 | NA | NA | NA | dsl-4 |
| WBGene00001107 | 0.040107227 | 0.58693163 | NA | dsl-5 |
| WBGene00001108 | 0.144224636 | 0.20464571 | 0.39387641 | dsl-6 |
| WBGene00001109 | 0.054696727 | 0.53979411 | NA | dsl-7 |
| WBGene00001110 | -0.162257876 | 0.19018552 | 0.37608132 | duo-1 |
| WBGene00001111 | -0.03323923 | 0.69122143 | NA | NA |
| WBGene00001112 | -0.127017712 | 0.1147806 | 0.26991691 | duo-3 |
| WBGene00001113 | 1.57593109 | 2.87E-08 | 5.94E-07 | dur-1 |
| WBGene00001115 | 0.586754816 | 0.02539714 | 0.09172163 | dyb-1 |
| WBGene00001116 | 0.314763051 | 0.17531255 | 0.35594084 | dyc-1 |
| WBGene00001117 | 0.121755954 | 0.46127726 | 0.65201394 | dyf-1 |
| WBGene00001118 | 0.255838998 | 0.24665556 | 0.44495002 | dyf-2 |
| WBGene00001119 | 0.239046478 | 0.27459648 | 0.47493999 | dyf-3 |
| WBGene00001121 | 0.086426234 | 0.42213848 | 0.61846519 | dyf-5 |
| WBGene00001122 | 0.27014264 | 0.17824346 | 0.35975496 | dyf-6 |
| WBGene00001123 | -0.019582799 | 0.91044144 | 0.95112162 | dyf-7 |
| WBGene00001127 | -0.024258867 | 0.86159488 | 0.92665912 | dyf-11 |
| WBGene00001129 | -0.002641891 | 0.95121282 | NA | dyf-13 |
| WBGene00001130 | -0.091065412 | 0.32779933 | 0.52710708 | dyn-1 |
| WBGene00001131 | 0.470276616 | 6.20E-05 | 0.00063618 | dys-1 |
| WBGene00001132 | -0.053289965 | 0.64291214 | 0.78861233 | alp-1 |
| WBGene00001133 | 1.58382233 | 0.00754213 | 0.03613363 | eat-2 |
| WBGene00001134 | -0.226734072 | 0.00776411 | 0.03690724 | eat-3 |
| WBGene00001135 | 0.524424217 | 0.05794286 | 0.1674909 | eat-4 |
| WBGene00001136 | 0.02311099 | 0.86906769 | 0.93000745 | eat-5 |
| WBGene00001137 | 0.096200164 | 0.0599227 | 0.17138664 | eat-6 |
| WBGene00001145 | 0.004502798 | 0.97928366 | 0.98812489 | eat-16 |
| WBGene00001147 | 0.016453512 | 0.83636998 | NA | eat-18 |
| WBGene00001148 | -0.358630537 | 0.01083461 | 0.04756089 | eat-20 |
| WBGene00001149 | 0.365550355 | 1.01E-05 | 0.0001248 | bcat-1 |
| WBGene00001150 | -0.920536218 | 0.01360535 | 0.05724882 | ech-1.1 |
| WBGene00001151 | -0.226820397 | 0.08810609 | 0.22512414 | ech-2 |
| WBGene00001152 | 0.107598524 | 0.5866126 | 0.74937572 | ech-3 |
| WBGene00001153 | -0.176728732 | 0.06269154 | 0.17733131 | ech-4 |

| | | | | |
|---|---|---|---|---|
| WBGene00001154 | -0.04873763 | 0.68979828 | 0.82175237 | ech-5 |
| WBGene00001155 | -0.524396834 | 9.67E-18 | 6.05E-16 | ech-6 |
| WBGene00001156 | 0.179702442 | 0.12220373 | 0.28134075 | ech-7 |
| WBGene00001157 | 0.347090506 | 0.06694694 | 0.1853925 | ech-8 |
| WBGene00001158 | 4.745427822 | 2.53E-06 | 3.58E-05 | ech-9 |
| WBGene00001159 | 0.402433476 | 0.10698756 | 0.25682978 | eff-1 |
| WBGene00001160 | -0.052368668 | 0.54487455 | 0.71824149 | efk-1 |
| WBGene00001161 | -0.109020144 | 0.24665049 | 0.44495002 | efl-1 |
| WBGene00001162 | -0.001555572 | 0.99010801 | 0.99446943 | efl-2 |
| WBGene00001163 | -0.284032139 | 0.21539106 | 0.40643645 | efn-2 |
| WBGene00001164 | -1.217185702 | 0.00055042 | 0.00421258 | efn-3 |
| WBGene00001165 | 0.133018217 | 0.47770771 | 0.66470097 | efn-4 |
| WBGene00001166 | -0.12930193 | 0.05494891 | 0.16132082 | eftu-2 |
| WBGene00001167 | -0.286956781 | 3.07E-12 | 1.08E-10 | eef-2 |
| WBGene00001168 | -0.285083714 | 1.52E-15 | 7.60E-14 | eef-1A.1 |
| WBGene00001169 | 0.20845362 | 0.00194877 | 0.01224546 | eef-1A.2 |
| WBGene00001170 | 0.026266105 | 0.81604796 | 0.89891744 | egl-1 |
| WBGene00001171 | 0.359083241 | 0.13999871 | 0.30737534 | egl-2 |
| WBGene00001172 | 1.287477008 | 7.10E-25 | 6.62E-23 | egl-3 |
| WBGene00001173 | 0.164362395 | 0.00394017 | 0.0216222 | egl-4 |
| WBGene00001174 | 0.20222564 | 0.30536221 | 0.50592194 | egl-5 |
| WBGene00001175 | -0.225900623 | 0.29402253 | 0.49520117 | egl-6 |
| WBGene00001177 | 0.629185589 | 6.19E-08 | 1.21E-06 | egl-8 |
| WBGene00001178 | -0.034665216 | 0.81421976 | 0.89752986 | egl-9 |
| WBGene00001179 | 0.116921475 | 0.44326408 | 0.6368667 | egl-10 |
| WBGene00001182 | 0.754737536 | 0.00044607 | 0.00352447 | egl-13 |
| WBGene00001184 | 1.243811121 | 2.26E-07 | 3.99E-06 | egl-15 |
| WBGene00001185 | -0.007811648 | 0.92797367 | NA | egl-17 |
| WBGene00001186 | -0.214576201 | 0.09691767 | 0.24083022 | egl-18 |
| WBGene00001187 | 0.608184703 | 0.00124199 | 0.00841458 | egl-19 |
| WBGene00001188 | 0.567157948 | 0.03824882 | 0.12378446 | egl-20 |
| WBGene00001189 | 1.063273376 | 7.11E-19 | 4.89E-17 | egl-21 |
| WBGene00001190 | 0.069909355 | 0.72239068 | 0.84436547 | egl-23 |
| WBGene00001193 | 0.003019084 | 0.98687256 | 0.99236271 | egl-26 |
| WBGene00001194 | -0.080301286 | 0.23332281 | 0.42875992 | egl-27 |
| WBGene00001196 | 0.332890488 | 0.0006692 | 0.00498356 | egl-30 |
| WBGene00001202 | 0.268002792 | 0.2372419 | 0.43353077 | egl-36 |
| WBGene00001204 | 0.057058552 | 0.69072914 | 0.82235004 | egl-38 |
| WBGene00001207 | 0.577983193 | 0.04043153 | 0.12905124 | egl-43 |
| WBGene00001208 | 0.975857125 | 1.76E-08 | 3.77E-07 | egl-44 |
| WBGene00001209 | -0.259067657 | 1.13E-05 | 0.00013788 | egl-45 |
| WBGene00001210 | -0.108312475 | 0.5373712 | 0.71232305 | egl-46 |
| WBGene00001211 | 0.101383319 | 0.3983416 | 0.5965142 | egl-47 |

| | | | | |
|---|---|---|---|---|
| WBGene00001214 | -0.121737394 | 0.13563818 | 0.30048096 | ego-1 |
| WBGene00001215 | 0.047459423 | 0.60313867 | 0.75988622 | ego-2 |
| WBGene00001223 | 0.020232312 | 0.84643798 | NA | ehn-3 |
| WBGene00001224 | 0.009339062 | 0.93524177 | 0.96340496 | ehs-1 |
| WBGene00001225 | -0.041578669 | 0.54581497 | 0.71922212 | eif-3.B |
| WBGene00001226 | -0.269514274 | 1.18E-06 | 1.84E-05 | eif-3.C |
| WBGene00001227 | -0.014791567 | 0.83616348 | 0.91095093 | eif-3.D |
| WBGene00001228 | -0.320325779 | 7.17E-06 | 9.10E-05 | eif-3.E |
| WBGene00001229 | -0.419797889 | 3.17E-08 | 6.51E-07 | eif-3.F |
| WBGene00001230 | -0.164701559 | 0.05857894 | 0.16856412 | eif-3.G |
| WBGene00001231 | -0.209385246 | 0.00685619 | 0.0334828 | eif-3.H |
| WBGene00001232 | -0.151624587 | 0.03902848 | 0.12564115 | eif-3.I |
| WBGene00001233 | -0.233460325 | 0.00399354 | 0.02185771 | eif-3.K |
| WBGene00001234 | -0.322128657 | 0.00185175 | 0.01172639 | eif-6 |
| WBGene00001235 | -0.157390951 | 0.23919639 | 0.43606953 | elb-1 |
| WBGene00001236 | 0.012532565 | 0.91995842 | 0.95544619 | elc-1 |
| WBGene00001237 | 0.015194928 | 0.8086441 | NA | elc-2 |
| WBGene00001239 | -0.198474266 | 0.00103419 | 0.00722745 | elo-1 |
| WBGene00001240 | -0.309166158 | 0.00071155 | 0.00524559 | elo-2 |
| WBGene00001241 | -0.066697368 | 0.42679093 | 0.6223751 | elo-3 |
| WBGene00001242 | -0.202771096 | 0.287706 | 0.48798697 | elo-4 |
| WBGene00001243 | -0.038724105 | 0.79957777 | 0.88800866 | elo-5 |
| WBGene00001244 | -0.256107625 | 0.12824506 | 0.28971392 | elo-6 |
| WBGene00001245 | 0.033463284 | 0.6365037 | NA | elo-7 |
| WBGene00001246 | 0.116929069 | 0.32003852 | 0.52005372 | elo-8 |
| WBGene00001247 | -0.529522803 | 0.02616074 | 0.09396943 | elo-9 |
| WBGene00001248 | 0.151355807 | 0.45377759 | 0.64589961 | elp-1 |
| WBGene00001249 | -0.426687475 | 0.0237366 | 0.08716516 | elt-1 |
| WBGene00001250 | 0.521257269 | 0.00832996 | 0.03900191 | elt-2 |
| WBGene00001251 | 0.380460782 | 0.02115838 | 0.08004648 | elt-3 |
| WBGene00001252 | 0.108796787 | 0.30818879 | 0.50908094 | elt-4 |
| WBGene00001253 | 0.151002787 | 0.46475101 | 0.65458581 | elt-6 |
| WBGene00001258 | -0.231491565 | 0.00094087 | 0.00669029 | emb-4 |
| WBGene00001259 | -0.196584358 | 0.00020565 | 0.00181384 | emb-5 |
| WBGene00001262 | -0.162926866 | 0.02755291 | 0.09734607 | emb-8 |
| WBGene00001263 | 0.445607146 | 6.56E-11 | 1.94E-09 | emb-9 |
| WBGene00001281 | -0.218097196 | 0.01902449 | 0.07413829 | emb-27 |
| WBGene00001284 | -0.036148416 | 0.67272679 | 0.80965005 | emb-30 |
| WBGene00001303 | -0.274236245 | 0.00046478 | 0.00365261 | emo-1 |
| WBGene00001309 | 0.043057503 | 0.65297604 | 0.79443429 | emr-1 |
| WBGene00001310 | -0.663736398 | 0.03223728 | 0.10905407 | end-1 |
| WBGene00001311 | -0.154951773 | 0.28272083 | 0.48260647 | end-3 |
| WBGene00001320 | -0.111632691 | 0.15565703 | 0.32914134 | ent-1 |

| | | | | |
|---|---|---|---|---|
| WBGene00001324 | 0.053001016 | 0.57608012 | 0.74318345 | eor-1 |
| WBGene00001325 | 0.116700395 | 0.44311444 | 0.63685467 | eor-2 |
| WBGene00001328 | -0.02404981 | 0.76865279 | 0.86844708 | epi-1 |
| WBGene00001329 | 0.304648647 | 0.00012821 | 0.00120252 | epn-1 |
| WBGene00001330 | 0.41949992 | 0.00132147 | 0.00884258 | eps-8 |
| WBGene00001331 | -0.207594826 | 0.06661009 | 0.18459934 | erd-2.1 |
| WBGene00001332 | 0.125114696 | 0.26033099 | 0.45840351 | eri-1 |
| WBGene00001333 | 0.024945367 | 0.65616348 | 0.79672329 | erm-1 |
| WBGene00001334 | -0.009790598 | 0.9175243 | 0.95412315 | ero-1 |
| WBGene00001335 | -0.177642455 | 0.29289428 | 0.49375534 | F35A5.8 |
| WBGene00001336 | -0.254126071 | 0.00123115 | 0.00834887 | qars-1 |
| WBGene00001337 | -0.309985214 | 4.79E-06 | 6.30E-05 | ears-1 |
| WBGene00001338 | 0.08071839 | 0.57697835 | 0.74401444 | ears-2 |
| WBGene00001340 | 0.026367319 | 0.7938261 | 0.88399982 | etr-1 |
| WBGene00001345 | 0.503283097 | 0.00800042 | 0.0378669 | fos-1 |
| WBGene00001352 | -0.051341891 | 0.48499471 | 0.6698982 | evl-14 |
| WBGene00001358 | -0.061579906 | 0.71912378 | 0.84237564 | evl-20 |
| WBGene00001362 | 0.185255125 | 0.27487206 | 0.47516409 | exc-1 |
| WBGene00001365 | -0.305556288 | 0.01378376 | 0.05782463 | exc-4 |
| WBGene00001366 | 0.465802524 | 0.01610376 | 0.06543857 | exc-5 |
| WBGene00001368 | 0.217624169 | 0.27749995 | 0.47750252 | exc-7 |
| WBGene00001371 | 0.170177823 | 0.33382637 | 0.53328444 | exl-1 |
| WBGene00001372 | -0.128231904 | 0.15248398 | 0.32515555 | exo-3 |
| WBGene00001373 | 0.187598338 | 0.30202001 | 0.50260002 | exp-1 |
| WBGene00001374 | 1.605825253 | 5.12E-05 | 0.00053541 | exp-2 |
| WBGene00001377 | -0.942109619 | 0.00060458 | 0.00457926 | eya-1 |
| WBGene00001385 | 0.06478847 | 0.25989373 | 0.4579683 | far-1 |
| WBGene00001386 | 0.287254786 | 3.80E-08 | 7.71E-07 | far-2 |
| WBGene00001387 | -1.995836385 | 2.31E-06 | 3.31E-05 | far-3 |
| WBGene00001388 | -0.053348766 | 0.53925327 | NA | far-4 |
| WBGene00001389 | -0.419525607 | 0.08539359 | 0.22030613 | far-5 |
| WBGene00001390 | 0.877905108 | 6.47E-18 | 4.12E-16 | far-6 |
| WBGene00001391 | 2.572376693 | 8.44E-08 | 1.61E-06 | far-7 |
| WBGene00001392 | 0.993442437 | 3.67E-24 | 3.26E-22 | far-8 |
| WBGene00001393 | 0.149639952 | 0.00665701 | 0.03275042 | fat-1 |
| WBGene00001394 | 1.007313685 | 4.06E-93 | 2.12E-90 | fat-2 |
| WBGene00001395 | 0.874733462 | 3.23E-27 | 3.43E-25 | fat-3 |
| WBGene00001396 | 0.486040011 | 7.55E-16 | 3.86E-14 | fat-4 |
| WBGene00001397 | 0.259747098 | 0.16298179 | 0.3384224 | fat-5 |
| WBGene00001398 | -0.035520453 | 0.58102245 | 0.74581663 | fat-6 |
| WBGene00001399 | 0.027405468 | 0.8120935 | 0.8958773 | fat-7 |
| WBGene00001400 | 0.060728101 | 0.52119783 | NA | fax-1 |
| WBGene00001401 | -0.066370115 | 0.69149023 | 0.82289754 | fbf-1 |

| | | | | |
|---|---|---|---|---|
| WBGene00001402 | -0.0697718 | 0.6842227 | 0.8176383 | fbf-2 |
| WBGene00001403 | 0.268470816 | 0.08208749 | 0.21469149 | fbl-1 |
| WBGene00001404 | -0.096600368 | 0.21101131 | 0.40096257 | fbp-1 |
| WBGene00001405 | -0.074906018 | 0.40820953 | 0.6061102 | fce-1 |
| WBGene00001406 | -0.116811495 | 0.44518336 | 0.63844021 | fce-2 |
| WBGene00001410 | 1.542468742 | 2.34E-05 | 0.00026265 | feh-1 |
| WBGene00001411 | -0.123905302 | 0.20485384 | 0.39397451 | fem-1 |
| WBGene00001412 | -0.027454844 | 0.72578106 | 0.84643624 | fem-2 |
| WBGene00001413 | -0.172168376 | 0.10495424 | 0.25334286 | fem-3 |
| WBGene00001414 | 0.064127358 | 0.62384507 | 0.77522625 | fer-1 |
| WBGene00001423 | -0.566951054 | 7.86E-28 | 8.65E-26 | fib-1 |
| WBGene00001424 | 0.106997704 | 0.16248985 | NA | fis-1 |
| WBGene00001425 | 0.079636984 | 0.60240532 | 0.75960867 | fis-2 |
| WBGene00001426 | 0.068965645 | 0.42365755 | 0.61965737 | fkb-1 |
| WBGene00001427 | -0.251457282 | 1.31E-05 | 0.00015578 | fkb-2 |
| WBGene00001428 | -0.355037176 | 0.05194502 | 0.1547989 | fkb-3 |
| WBGene00001429 | 0.283959964 | 0.14126011 | 0.30896166 | fkb-4 |
| WBGene00001430 | 0.047357331 | 0.77191179 | 0.87031066 | fkb-5 |
| WBGene00001431 | -0.38284922 | 1.17E-10 | 3.32E-09 | fkb-6 |
| WBGene00001432 | -0.078018371 | 0.6937834 | 0.82442032 | fkb-7 |
| WBGene00001433 | 0.363685622 | 0.13158563 | 0.29439872 | fkb-8 |
| WBGene00001434 | -0.476729972 | 0.06942816 | 0.19031974 | fkh-2 |
| WBGene00001435 | -0.473987973 | 0.05363364 | 0.15854031 | fkh-3 |
| WBGene00001436 | -0.450319076 | 0.06361897 | 0.17908966 | fkh-4 |
| WBGene00001437 | -0.108216615 | 0.58458684 | 0.74789723 | fkh-5 |
| WBGene00001438 | 0.165326373 | 0.41635681 | 0.61345276 | fkh-6 |
| WBGene00001439 | 0.247385599 | 0.14130613 | 0.3089954 | fkh-7 |
| WBGene00001440 | 0.020392401 | 0.90289016 | 0.94829375 | fkh-8 |
| WBGene00001441 | 1.218790799 | 4.90E-16 | 2.58E-14 | fkh-9 |
| WBGene00001442 | 0.009099132 | 0.93427495 | 0.96290283 | fkh-10 |
| WBGene00001443 | -0.218296162 | 0.00818889 | 0.03847675 | fli-1 |
| WBGene00001444 | 1.62156655 | 6.43E-14 | 2.75E-12 | flp-1 |
| WBGene00001445 | 0.698999298 | 0.0250438 | 0.09094035 | flp-2 |
| WBGene00001446 | 1.475819581 | 7.04E-09 | 1.61E-07 | flp-3 |
| WBGene00001447 | 1.586285872 | 0.00017635 | 0.0015821 | flp-4 |
| WBGene00001448 | 1.636224703 | 5.36E-07 | 8.92E-06 | flp-5 |
| WBGene00001449 | 0.014496092 | 0.93017277 | 0.96068422 | flp-6 |
| WBGene00001450 | 0.404810032 | 0.09042453 | 0.22876518 | flp-7 |
| WBGene00001451 | 0.857957319 | 0.00116131 | 0.00796384 | flp-8 |
| WBGene00001452 | 1.577641917 | 4.63E-14 | 2.01E-12 | flp-9 |
| WBGene00001453 | 0.62650143 | 0.00597845 | 0.03004642 | flp-10 |
| WBGene00001454 | 1.091002555 | 4.31E-05 | 0.00045853 | flp-11 |
| WBGene00001455 | 1.328166895 | 4.29E-07 | 7.23E-06 | flp-12 |

| | | | | |
|---|---|---|---|---|
| WBGene00001456 | 1.205623495 | 2.81E-05 | 0.00030953 | flp-13 |
| WBGene00001457 | 1.329797552 | 6.27E-12 | 2.16E-10 | flp-14 |
| WBGene00001458 | 2.27950857 | 2.75E-07 | 4.77E-06 | flp-15 |
| WBGene00001459 | 1.254565959 | 1.76E-05 | 0.00020268 | flp-16 |
| WBGene00001460 | 0.870385645 | 0.0008586 | 0.0061861 | flp-17 |
| WBGene00001461 | 1.217850522 | 7.25E-07 | 1.18E-05 | flp-18 |
| WBGene00001462 | 1.370214857 | 2.60E-06 | 3.65E-05 | flp-19 |
| WBGene00001463 | 0.77689094 | 0.02399274 | 0.08792932 | flp-20 |
| WBGene00001464 | 1.226715992 | 0.00434031 | 0.02344889 | flp-21 |
| WBGene00001465 | 0.165917182 | 0.43880236 | 0.6335804 | flr-1 |
| WBGene00001466 | 0.789700723 | 0.02295071 | 0.08507051 | flr-2 |
| WBGene00001468 | 0.705739177 | 0.0278702 | 0.09808835 | flr-4 |
| WBGene00001470 | -0.171435698 | 0.1207038 | 0.27949144 | baz-2 |
| WBGene00001475 | 0.340312774 | 0.15161882 | 0.32414722 | fmi-1 |
| WBGene00001476 | -0.003161919 | 0.98621342 | 0.99217691 | fmo-1 |
| WBGene00001477 | 7.263981033 | 5.67E-08 | 1.12E-06 | fmo-2 |
| WBGene00001478 | -1.227061227 | 6.12E-06 | 7.83E-05 | fmo-3 |
| WBGene00001479 | -0.171168271 | 0.28124878 | 0.48075477 | fmo-4 |
| WBGene00001480 | 0.095518549 | 0.61784244 | 0.77099816 | fmo-5 |
| WBGene00001481 | -0.283665622 | 0.07696482 | 0.20527774 | fog-1 |
| WBGene00001482 | -0.281504645 | 0.03433472 | 0.11435273 | fog-2 |
| WBGene00001483 | 1.794241509 | 0.00475042 | 0.02511767 | fog-3 |
| WBGene00001484 | -0.233875141 | 0.14492443 | 0.3142999 | fox-1 |
| WBGene00001485 | 0.074029858 | 0.52032915 | 0.69916842 | fre-1 |
| WBGene00001486 | -0.070873679 | 0.69343405 | 0.82420587 | frh-1 |
| WBGene00001487 | 0.321093834 | 0.02494705 | 0.0907002 | frk-1 |
| WBGene00001488 | 0.084043604 | 0.2466863 | 0.44495002 | frm-1 |
| WBGene00001489 | 0.34239387 | 0.04325623 | 0.13538145 | frm-2 |
| WBGene00001490 | 0.849767808 | 0.00114431 | 0.00786566 | frm-3 |
| WBGene00001491 | -0.01351312 | 0.8752884 | 0.93344293 | frm-4 |
| WBGene00001492 | -0.185053029 | 0.3711827 | 0.57063899 | frm-5.1 |
| WBGene00001493 | 0.224872549 | 0.03730977 | 0.12160559 | frm-7 |
| WBGene00001494 | 0.472823551 | 0.04947868 | 0.14908775 | frm-8 |
| WBGene00001495 | 0.091692696 | 0.62365632 | 0.77505748 | frm-9 |
| WBGene00001496 | 0.233172658 | 0.01596948 | 0.06501923 | frm-10 |
| WBGene00001497 | -0.473959538 | 1.40E-10 | 3.90E-09 | fars-1 |
| WBGene00001498 | -0.444980995 | 1.29E-07 | 2.39E-06 | fars-3 |
| WBGene00001499 | -0.203969576 | 0.02154495 | 0.0811316 | fsn-1 |
| WBGene00001500 | 1.272284029 | 0.00768925 | 0.03663661 | ftn-1 |
| WBGene00001501 | 0.171416232 | 0.03946581 | 0.12682593 | ftn-2 |
| WBGene00001502 | -0.026877396 | 0.6882574 | 0.82045382 | ftt-2 |
| WBGene00001503 | -0.022332029 | 0.71967665 | 0.84280801 | fum-1 |
| WBGene00001505 | 0.480479638 | 0.03465458 | 0.11520837 | fut-1 |

| | | | | |
|---|---|---|---|---|
| WBGene00001506 | 0.015597454 | 0.8447034 | NA | fut-2 |
| WBGene00001507 | 0.030256826 | 0.86758512 | 0.92947386 | fut-4 |
| WBGene00001508 | 0.443013143 | 0.03589002 | 0.11808221 | fut-8 |
| WBGene00001509 | 0.019185411 | 0.84479973 | 0.91675274 | fzo-1 |
| WBGene00001510 | 0.040449508 | 0.69175069 | 0.82310107 | fzr-1 |
| WBGene00001511 | -0.286953853 | 3.79E-06 | 5.10E-05 | fzy-1 |
| WBGene00001512 | 0.346878709 | 0.08083831 | 0.21252536 | gab-1 |
| WBGene00001513 | 0.051525345 | 0.50552152 | 0.68651707 | gad-1 |
| WBGene00001514 | -0.190015096 | 0.02055404 | 0.07816408 | xnd-1 |
| WBGene00001515 | 1.239679686 | 0.00119114 | 0.00811893 | gap-1 |
| WBGene00001516 | 0.032355126 | 0.77523061 | 0.87262026 | gap-2 |
| WBGene00001517 | 0.90446606 | 0.01787733 | 0.07077935 | gar-1 |
| WBGene00001518 | 0.448695594 | 0.08392414 | 0.21789915 | gar-2 |
| WBGene00001519 | 0.219051125 | 0.31972718 | 0.51987478 | gar-3 |
| WBGene00001520 | -0.113053756 | 0.17844494 | 0.36000072 | gas-1 |
| WBGene00001522 | -0.606552259 | 0.00292429 | 0.01709845 | gbh-1 |
| WBGene00001523 | 0.433723537 | 0.04821427 | 0.14638841 | gbh-2 |
| WBGene00001526 | 0.147285857 | 0.14021076 | 0.30763161 | gck-1 |
| WBGene00001527 | -0.103621011 | 0.21013883 | 0.39971971 | gcs-1 |
| WBGene00001528 | 0.020317245 | 0.84643798 | NA | gcy-1 |
| WBGene00001529 | NA | NA | NA | gcy-2 |
| WBGene00001530 | 0.036821884 | 0.59344795 | NA | gcy-3 |
| WBGene00001531 | NA | NA | NA | gcy-4 |
| WBGene00001532 | 0.048978668 | 0.62587391 | 0.7769414 | gcy-5 |
| WBGene00001533 | 0.056411843 | 0.41891328 | NA | gcy-6 |
| WBGene00001534 | 0.131827335 | 0.16097389 | NA | gcy-7 |
| WBGene00001535 | 0.304452338 | 0.00960023 | 0.04358355 | gcy-8 |
| WBGene00001536 | -0.137290216 | 0.37240155 | 0.57164302 | gcy-9 |
| WBGene00001537 | 0.763994036 | 0.01044865 | 0.04641344 | gcy-11 |
| WBGene00001538 | 0.303670944 | 0.04967207 | 0.1495166 | gcy-12 |
| WBGene00001539 | -0.027728334 | 0.67598605 | NA | gcy-13 |
| WBGene00001540 | NA | NA | NA | gcy-14 |
| WBGene00001541 | 1.480403338 | 0.00130841 | 0.00877932 | gcy-15 |
| WBGene00001542 | -0.02791392 | 0.6763823 | NA | gcy-17 |
| WBGene00001543 | 0.313734909 | 0.17789011 | 0.35919032 | gcy-18 |
| WBGene00001544 | -0.020099257 | 0.88530004 | 0.93966057 | gcy-19 |
| WBGene00001545 | -0.027350315 | 0.68023343 | NA | gcy-20 |
| WBGene00001546 | 0.213832099 | 0.32326576 | 0.52320738 | gcy-21 |
| WBGene00001547 | 1.875028321 | 0.00780254 | 0.03706098 | gcy-22 |
| WBGene00001548 | 0.143170539 | 0.35876873 | 0.55903927 | gcy-23 |
| WBGene00001549 | 0.020925399 | 0.84643798 | NA | gcy-25 |
| WBGene00001550 | -0.003421551 | 0.95689871 | NA | gcy-27 |
| WBGene00001551 | -0.040033582 | 0.75258826 | 0.85995822 | gcy-31 |

| | | | | |
|---|---|---|---|---|
| WBGene00001552 | 0.209122417 | 0.0663397 | 0.18402418 | gcy-32 |
| WBGene00001553 | 0.020220349 | 0.88150994 | 0.93716888 | gcy-33 |
| WBGene00001554 | 0.015896709 | 0.80422001 | NA | gcy-34 |
| WBGene00001555 | 0.352335288 | 0.10066076 | 0.24661549 | gcy-35 |
| WBGene00001556 | 0.083594176 | 0.47969591 | 0.66640541 | gcy-36 |
| WBGene00001557 | 0.694708375 | 0.02131268 | 0.08046387 | gcy-37 |
| WBGene00001558 | -0.197760802 | 0.00010089 | 0.00097953 | gdi-1 |
| WBGene00001559 | 0.741437898 | 0.00017193 | 0.00155094 | gei-1 |
| WBGene00001560 | 0.727181835 | 6.81E-08 | 1.32E-06 | nhr-27 |
| WBGene00001561 | -0.237294011 | 0.00556363 | 0.02849588 | gei-4 |
| WBGene00001562 | -0.202655685 | 0.01620437 | 0.06577438 | lin-66 |
| WBGene00001563 | -0.198676868 | 0.02884386 | 0.10074021 | gei-6 |
| WBGene00001564 | 1.356376092 | 5.80E-110 | 3.54E-107 | icl-1 |
| WBGene00001565 | -0.082610383 | 0.42813505 | 0.62303203 | gei-8 |
| WBGene00001566 | -0.401398795 | 0.00027215 | 0.00230318 | acdh-13 |
| WBGene00001568 | -0.223776209 | 0.00829424 | 0.03885949 | snpc-4 |
| WBGene00001569 | -0.295121956 | 0.00068532 | 0.00508017 | meg-3 |
| WBGene00001570 | 0.366722773 | 0.13349227 | 0.29716592 | gei-13 |
| WBGene00001571 | -0.226536194 | 0.01593342 | 0.06492656 | gei-14 |
| WBGene00001572 | 0.266395785 | 0.18259984 | 0.3654243 | gei-15 |
| WBGene00001574 | -0.066479414 | 0.37807163 | 0.57726006 | gei-17 |
| WBGene00001575 | -0.109207217 | 0.47694675 | 0.66409676 | gei-18 |
| WBGene00001577 | 0.136983648 | 0.50200308 | 0.68359084 | gem-4 |
| WBGene00001578 | 0.636620278 | 0.00061653 | 0.00466009 | ges-1 |
| WBGene00001579 | -0.039539914 | 0.67729803 | 0.81260807 | gex-2 |
| WBGene00001580 | -0.202194098 | 0.02018307 | 0.07717428 | gex-3 |
| WBGene00001581 | 0.570823077 | 0.00824759 | 0.03869045 | gfi-1 |
| WBGene00001582 | -0.02640316 | 0.72026054 | 0.84328945 | gfi-2 |
| WBGene00001583 | -0.034406426 | 0.80841635 | 0.89382448 | gfi-3 |
| WBGene00001584 | -0.217974375 | 0.01049737 | 0.04650288 | nrde-4 |
| WBGene00001585 | -0.161203398 | 0.18210588 | 0.36488466 | gfl-1 |
| WBGene00001586 | 0.303137914 | 0.10661819 | 0.25615308 | ggr-1 |
| WBGene00001587 | 0.257294578 | 0.144862 | 0.31425759 | ggr-2 |
| WBGene00001588 | 0.423484858 | 0.08277025 | 0.21582177 | ggr-3 |
| WBGene00001589 | -0.346208108 | 0.00055401 | 0.00423122 | gip-1 |
| WBGene00001590 | -0.320845879 | 0.0482406 | 0.14641182 | C45G3.3 |
| WBGene00001591 | 1.665332413 | 1.84E-19 | 1.34E-17 | glc-1 |
| WBGene00001592 | 0.110160217 | 0.58938925 | 0.75111006 | glc-2 |
| WBGene00001593 | 0.092396429 | 0.45519267 | 0.64717418 | glc-3 |
| WBGene00001594 | 1.70674362 | 7.10E-06 | 9.02E-05 | glc-4 |
| WBGene00001595 | -0.154277486 | 0.00083105 | 0.00601154 | gld-1 |
| WBGene00001596 | 0.066707568 | 0.27213582 | 0.47224612 | gld-2 |
| WBGene00001597 | -0.10591358 | 0.0419322 | 0.13228385 | gld-3 |

| | | | | | |
|---|---|---|---|---|---|
| WBGene00001598 | | -0.471383657 | 2.89E-19 | 2.05E-17 | glh-1 |
| WBGene00001599 | | -0.100538098 | 0.09929561 | 0.24443111 | glh-2 |
| WBGene00001600 | | -0.09470077 | 0.36099607 | 0.56083865 | glh-3 |
| WBGene00001601 | | -0.276890507 | 0.00014709 | 0.00134678 | glh-4 |
| WBGene00001602 | | 0.119245721 | 0.46609902 | 0.65560147 | gln-1 |
| WBGene00001603 | | 0.937986876 | 0.01733153 | 0.06908548 | gln-2 |
| WBGene00001604 | | 0.200297589 | 0.34668994 | 0.54639908 | gln-3 |
| WBGene00001605 | | 0.038125098 | 0.61119482 | NA | NA |
| WBGene00001606 | | -0.033333676 | 0.56933207 | 0.73818551 | gln-5 |
| WBGene00001607 | | 0.046711516 | 0.29039952 | 0.49092715 | gln-6 |
| WBGene00001608 | | 0.242887728 | 0.08338641 | 0.21707176 | R07B1.8 |
| WBGene00001609 | | 0.065154531 | 0.32450391 | 0.52364838 | glp-1 |
| WBGene00001612 | | 0.710279518 | 0.02305707 | 0.0852902 | glr-1 |
| WBGene00001613 | | 0.227027029 | 0.20356761 | 0.39247213 | glr-2 |
| WBGene00001614 | | 0.1161173 | 0.12124304 | NA | glr-3 |
| WBGene00001615 | | 0.388059922 | 0.11020119 | 0.2618821 | glr-4 |
| WBGene00001616 | | 0.285439481 | 0.20006266 | 0.38868331 | glr-5 |
| WBGene00001617 | | 0.247888572 | 0.26366648 | 0.46181851 | glr-6 |
| WBGene00001618 | | 0.012453785 | 0.89141911 | NA | glr-7 |
| WBGene00001619 | | 0.053579067 | 0.64433586 | 0.78947268 | glr-8 |
| WBGene00001620 | | -0.115359819 | 0.23908057 | 0.43606953 | glt-1 |
| WBGene00001621 | | 0.855935693 | 0.00226954 | 0.0138221 | glt-3 |
| WBGene00001622 | | 0.517739961 | 0.01638136 | 0.06636401 | glt-4 |
| WBGene00001623 | | 1.756183416 | 3.83E-06 | 5.14E-05 | glt-5 |
| WBGene00001624 | | 1.762819573 | 0.00028782 | 0.00242252 | glt-6 |
| WBGene00001625 | | -0.011008483 | 0.94240101 | 0.96728563 | glt-7 |
| WBGene00001626 | | -0.003421551 | 0.95689871 | NA | gly-1 |
| WBGene00001627 | | 0.28804431 | 0.16333051 | 0.33887458 | gly-2 |
| WBGene00001628 | | -0.163428986 | 0.06712065 | 0.18562778 | gly-3 |
| WBGene00001629 | | -0.060793526 | 0.6325378 | 0.78177194 | gly-4 |
| WBGene00001630 | | -0.023215658 | 0.79552341 | 0.88521082 | gly-5 |
| WBGene00001631 | | 0.315559956 | 0.04087328 | 0.1299513 | gly-6 |
| WBGene00001632 | | -0.145693659 | 0.08021982 | 0.211507 | gly-7 |
| WBGene00001633 | | -1.114778276 | 1.18E-06 | 1.84E-05 | gly-8 |
| WBGene00001635 | | -0.113389253 | 0.22056966 | 0.41295062 | gly-10 |
| WBGene00001636 | | -0.315641206 | 0.15608864 | 0.32975175 | gly-11 |
| WBGene00001637 | | -0.188762703 | 0.36580076 | 0.56554265 | gly-12 |
| WBGene00001638 | | -0.215858133 | 0.14798394 | 0.31876649 | gly-13 |
| WBGene00001639 | | 0.372673725 | 0.12593706 | 0.28697495 | gly-14 |
| WBGene00001640 | NA | | NA | NA | gly-15 |
| WBGene00001641 | NA | | NA | NA | gly-16 |
| WBGene00001642 | NA | | NA | NA | gly-17 |
| WBGene00001643 | | 0.104538598 | 0.39370335 | 0.59205811 | gly-18 |

| | | | | |
|---|---|---|---|---|
| WBGene00001644 | 0.294541757 | 0.20186697 | 0.39045849 | gly-19 |
| WBGene00001645 | -0.066190863 | 0.39538796 | 0.59330482 | gly-20 |
| WBGene00001646 | -0.087535652 | 0.66976653 | 0.80768073 | gna-1 |
| WBGene00001647 | -0.162446313 | 0.02532129 | 0.09156065 | gna-2 |
| WBGene00001648 | -0.081523619 | 0.23891346 | 0.43598803 | goa-1 |
| WBGene00001649 | 0.196153105 | 0.10306585 | 0.25058216 | gob-1 |
| WBGene00001650 | 0.451831271 | 0.00489146 | 0.02571488 | gon-1 |
| WBGene00001651 | 0.264773482 | 0.01511035 | 0.06226588 | gon-2 |
| WBGene00001653 | -0.154827471 | 0.0567399 | 0.16518093 | gon-4 |
| WBGene00001660 | -0.039646216 | 0.76605956 | 0.86713124 | gop-1 |
| WBGene00001661 | -0.140290488 | 0.16505795 | 0.34132958 | gop-2 |
| WBGene00001662 | -0.118637291 | 0.22386242 | 0.4168345 | gop-3 |
| WBGene00001663 | -0.137374051 | 0.46293235 | 0.65296652 | gpa-1 |
| WBGene00001664 | 0.109245085 | 0.46855612 | 0.65741545 | gpa-2 |
| WBGene00001665 | 0.224761343 | 0.21010051 | 0.39969875 | gpa-3 |
| WBGene00001666 | -0.269703718 | 0.15959642 | 0.33423705 | gpa-4 |
| WBGene00001667 | -0.050042403 | 0.47734215 | NA | gpa-5 |
| WBGene00001668 | 0.074958105 | 0.3349622 | NA | gpa-6 |
| WBGene00001669 | 0.544472181 | 0.04471125 | 0.13851405 | gpa-7 |
| WBGene00001670 | 0.14657853 | 0.12995995 | 0.29224835 | gpa-8 |
| WBGene00001671 | 0.896779432 | 0.01970913 | 0.07578168 | gpa-9 |
| WBGene00001672 | 0.12265071 | 0.31029352 | 0.51113945 | gpa-10 |
| WBGene00001673 | -0.079918275 | 0.48767415 | 0.67181381 | gpa-11 |
| WBGene00001674 | 0.457433918 | 0.06013608 | 0.17176135 | gpa-12 |
| WBGene00001675 | 0.048205879 | 0.71100571 | 0.83727456 | gpa-13 |
| WBGene00001676 | 0.200947052 | 0.23969224 | 0.43670695 | gpa-14 |
| WBGene00001677 | 0.074558528 | 0.49307975 | 0.67607985 | gpa-15 |
| WBGene00001678 | -0.122277363 | 0.2040396 | 0.39307402 | gpa-16 |
| WBGene00001679 | 0.033559617 | 0.62127581 | 0.7735436 | gpb-1 |
| WBGene00001680 | 0.451487032 | 0.0235982 | 0.08674398 | gpb-2 |
| WBGene00001681 | 0.289233967 | 0.12524666 | 0.28607432 | gpc-1 |
| WBGene00001682 | -0.073877166 | 0.50591042 | 0.68674779 | gpc-2 |
| WBGene00001683 | -0.125277073 | 0.02903399 | 0.10125934 | gpd-1 |
| WBGene00001684 | -0.190632018 | 0.01231401 | 0.05276248 | gpd-2 |
| WBGene00001685 | -0.191311141 | 0.00424819 | 0.02301064 | gpd-3 |
| WBGene00001686 | -0.290209704 | 7.31E-05 | 0.00073439 | gpd-4 |
| WBGene00001687 | 0.122801286 | 0.55653509 | 0.72797554 | gpn-1 |
| WBGene00001688 | -0.140584851 | 0.35284359 | 0.55286632 | gpr-1 |
| WBGene00001689 | -0.069054256 | 0.59554728 | 0.75477619 | gpr-2 |
| WBGene00001690 | 0.452338819 | 0.08470931 | 0.21927671 | grd-1 |
| WBGene00001691 | 0.132031485 | 0.16048086 | NA | grd-2 |
| WBGene00001692 | -2.848855322 | 4.67E-74 | 1.63E-71 | grd-3 |
| WBGene00001693 | -2.030916532 | 0.00014083 | 0.00129841 | grd-4 |

| | | | | |
|---|---|---|---|---|
| WBGene00001694 | -2.437557273 | 1.39E-68 | 4.42E-66 | grd-5 |
| WBGene00001695 | 0.403882535 | 0.00454825 | 0.02433879 | grd-6 |
| WBGene00001696 | 0.020317245 | 0.84643798 | NA | grd-7 |
| WBGene00001697 | 0.020925399 | 0.84643798 | NA | grd-8 |
| WBGene00001698 | 0.040019663 | 0.56356856 | NA | grd-9 |
| WBGene00001699 | -0.53335283 | 0.04643862 | 0.14255181 | grd-10 |
| WBGene00001700 | -0.078146297 | 0.41725748 | 0.61426198 | grd-11 |
| WBGene00001701 | 0.038125098 | 0.61119482 | NA | grd-12 |
| WBGene00001702 | -2.044309829 | 8.75E-06 | 0.00010915 | grd-13 |
| WBGene00001703 | -1.81125417 | 3.10E-07 | 5.35E-06 | grd-14 |
| WBGene00001704 | 0.020317245 | 0.84643798 | NA | grd-15 |
| WBGene00001705 | 0.038595855 | 0.57520533 | NA | grd-16 |
| WBGene00001706 | NA | NA | NA | NA |
| WBGene00001707 | 0.858867498 | 0.00776087 | 0.03690382 | grh-1 |
| WBGene00001708 | 0.066868572 | 0.62177879 | 0.77393862 | grk-1 |
| WBGene00001709 | 0.146093456 | 0.44090592 | 0.635052 | grk-2 |
| WBGene00001710 | 0.115914333 | 0.27363721 | 0.47384061 | grl-1 |
| WBGene00001711 | -0.00346345 | 0.95708614 | NA | grl-2 |
| WBGene00001712 | -0.056943998 | 0.52719684 | NA | grl-3 |
| WBGene00001713 | -0.603102107 | 0.00596113 | 0.02998416 | grl-4 |
| WBGene00001714 | 0.402554336 | 0.00385393 | 0.02125584 | grl-5 |
| WBGene00001715 | 0.038325924 | 0.57925535 | NA | grl-6 |
| WBGene00001716 | 4.967536988 | 0.00021803 | 0.001899 | grl-7 |
| WBGene00001717 | 0.077895643 | 0.60366105 | 0.76024502 | grl-8 |
| WBGene00001718 | NA | NA | NA | grl-9 |
| WBGene00001719 | 0.075601728 | 0.63376697 | 0.7826402 | grl-10 |
| WBGene00001720 | -0.006239126 | 0.93041881 | NA | grl-11 |
| WBGene00001721 | 0.033034883 | 0.64219503 | NA | grl-12 |
| WBGene00001722 | 0.065014541 | 0.37399213 | NA | grl-13 |
| WBGene00001723 | 0.233276312 | 0.13336063 | 0.29700836 | grl-14 |
| WBGene00001724 | 0.076689517 | 0.45471893 | 0.64678155 | grl-15 |
| WBGene00001725 | 4.121375576 | 1.87E-06 | 2.74E-05 | grl-16 |
| WBGene00001726 | -0.151227801 | 0.0649288 | NA | grl-17 |
| WBGene00001727 | 0.033815842 | 0.73608433 | 0.85184122 | grl-18 |
| WBGene00001728 | NA | NA | NA | grl-19 |
| WBGene00001729 | -0.054932793 | 0.45568274 | NA | grl-20 |
| WBGene00001730 | 0.603041471 | 0.00975241 | 0.04405211 | grl-21 |
| WBGene00001731 | -0.196272191 | 0.28567072 | 0.48577294 | grl-22 |
| WBGene00001732 | -0.023056499 | 0.78121253 | NA | grl-23 |
| WBGene00001733 | NA | NA | NA | grl-24 |
| WBGene00001734 | -0.013545023 | 0.89941055 | 0.94665211 | grl-25 |
| WBGene00001735 | -0.002950686 | 0.96771748 | NA | grl-26 |
| WBGene00001736 | -0.027535543 | 0.68063395 | NA | grl-27 |

| | | | | |
|---|---|---|---|---|
| WBGene00001737 | NA | NA | NA | grl-28 |
| WBGene00001738 | NA | NA | NA | grl-29 |
| WBGene00001739 | NA | NA | NA | grl-30 |
| WBGene00001740 | -0.157559106 | 0.286783 | 0.48692913 | gro-1 |
| WBGene00001743 | -0.095081404 | 0.27633016 | 0.47632079 | grp-1 |
| WBGene00001744 | -0.47355264 | 2.24E-11 | 7.08E-10 | gars-1 |
| WBGene00001745 | 0.100115414 | 0.61997079 | 0.77264865 | gsa-1 |
| WBGene00001746 | -0.130666739 | 0.19886012 | 0.38722051 | gsk-3 |
| WBGene00001747 | -0.236701589 | 0.00022335 | 0.00193607 | gsp-1 |
| WBGene00001748 | -0.050869713 | 0.37174137 | 0.57098881 | gsp-2 |
| WBGene00001749 | -0.041621669 | 0.54126447 | 0.71522926 | gst-1 |
| WBGene00001750 | -0.150326718 | 0.19957203 | 0.38803905 | gst-2 |
| WBGene00001751 | 0.292471947 | 0.15896594 | 0.33321705 | gst-3 |
| WBGene00001752 | -0.518872725 | 0.0058075 | 0.02946209 | gst-4 |
| WBGene00001753 | 0.791896262 | 0.00014412 | 0.00132212 | gst-5 |
| WBGene00001754 | 0.885255598 | 0.00201867 | 0.01256025 | gst-6 |
| WBGene00001755 | 0.136270188 | 0.04163817 | 0.1315831 | gst-7 |
| WBGene00001756 | 0.075726029 | 0.27693214 | NA | gst-8 |
| WBGene00001757 | -0.27745601 | 0.12586304 | 0.28690032 | gst-9 |
| WBGene00001758 | -1.671144717 | 1.58E-13 | 6.44E-12 | gst-10 |
| WBGene00001759 | 0.164164867 | 0.28102378 | 0.48055 | gst-11 |
| WBGene00001760 | 0.342305779 | 0.14992581 | 0.32125686 | gst-12 |
| WBGene00001761 | -0.628587808 | 3.41E-05 | 0.00037011 | gst-13 |
| WBGene00001762 | 0.032011836 | 0.76544766 | 0.86673775 | gst-14 |
| WBGene00001763 | 0.290925722 | 0.18666713 | 0.37103756 | gst-15 |
| WBGene00001764 | -0.238410948 | 0.27536449 | 0.47572289 | gst-16 |
| WBGene00001765 | -0.022930767 | 0.78121253 | NA | gst-17 |
| WBGene00001766 | -0.023109762 | 0.78121253 | NA | gst-18 |
| WBGene00001767 | 0.148471211 | 0.22741443 | 0.42103879 | gst-19 |
| WBGene00001768 | 0.433706004 | 0.03112317 | 0.10599542 | gst-20 |
| WBGene00001769 | 0.187275372 | 0.25608823 | 0.45448602 | gst-21 |
| WBGene00001770 | 0.15288008 | 0.47445744 | 0.6618896 | gst-22 |
| WBGene00001771 | 0.17643325 | 0.15385691 | 0.32713026 | gst-23 |
| WBGene00001772 | 1.592054871 | 1.06E-05 | 0.00013009 | gst-24 |
| WBGene00001773 | 0.087076318 | 0.29487856 | NA | gst-25 |
| WBGene00001774 | 0.238708933 | 0.12471471 | 0.28518106 | gst-26 |
| WBGene00001775 | 0.288014885 | 0.04335268 | 0.13565432 | gst-27 |
| WBGene00001776 | 0.441319238 | 0.02489759 | 0.09058787 | gst-28 |
| WBGene00001777 | -0.01743761 | 0.89082172 | 0.94312826 | gst-29 |
| WBGene00001778 | -0.050231235 | 0.75999663 | 0.86410737 | gst-30 |
| WBGene00001779 | 0.132442484 | 0.10147609 | NA | gst-31 |
| WBGene00001780 | 0.063417569 | 0.59071659 | 0.75179777 | gst-32 |
| WBGene00001781 | 0.559315176 | 0.05181968 | 0.15448829 | gst-33 |

| | | | | |
|---|---|---|---|---|
| WBGene00001782 | -0.006711909 | 0.92834175 | NA | gst-34 |
| WBGene00001783 | 0.486692682 | 0.06297539 | 0.17782783 | gst-35 |
| WBGene00001784 | 0.064593612 | 0.65702764 | 0.79757414 | gst-36 |
| WBGene00001785 | NA | NA | NA | gst-37 |
| WBGene00001786 | -0.004877215 | 0.97240924 | 0.98478548 | gst-38 |
| WBGene00001787 | 0.065308889 | 0.75090466 | 0.85937584 | gst-39 |
| WBGene00001788 | 0.105444527 | 0.45916567 | 0.65047359 | gst-40 |
| WBGene00001789 | 0.263004036 | 0.24250141 | 0.44063343 | gst-41 |
| WBGene00001790 | -0.67772231 | 0.00016104 | 0.00146087 | gst-42 |
| WBGene00001791 | 0.11471102 | 0.57744011 | 0.74411462 | gst-43 |
| WBGene00001792 | 0.221557973 | 0.31194315 | 0.51266736 | gst-44 |
| WBGene00001793 | -0.280244496 | 2.29E-05 | 0.00025865 | gsy-1 |
| WBGene00001794 | -0.274313074 | 3.28E-06 | 4.46E-05 | gta-1 |
| WBGene00001795 | 1.012797885 | 9.72E-15 | 4.53E-13 | gtl-1 |
| WBGene00001796 | 0.222431265 | 0.31525547 | 0.51565807 | gtl-2 |
| WBGene00001803 | 0.521197876 | 0.06066518 | 0.17290225 | lite-1 |
| WBGene00001804 | 0.0398788 | 0.58971221 | NA | gur-3 |
| WBGene00001805 | -0.116329124 | 0.45732666 | 0.64918009 | gur-4 |
| WBGene00001806 | 0.089126141 | 0.49432499 | 0.67727983 | gur-5 |
| WBGene00001808 | -0.34715077 | 0.00906979 | 0.04161332 | gut-2 |
| WBGene00001811 | -0.244379032 | 0.2476905 | 0.44607264 | haf-1 |
| WBGene00001812 | 0.073890912 | 0.48326135 | 0.66879098 | haf-2 |
| WBGene00001813 | -0.147462546 | 0.13313494 | 0.29668625 | haf-3 |
| WBGene00001814 | 0.302688154 | 0.04277746 | 0.13437132 | haf-4 |
| WBGene00001815 | -0.084616297 | 0.43348677 | 0.6285086 | hmt-1 |
| WBGene00001816 | -0.117366365 | 0.37743401 | 0.57652718 | haf-6 |
| WBGene00001817 | 0.242590808 | 0.2048166 | 0.39397451 | haf-7 |
| WBGene00001818 | 0.020232312 | 0.84643798 | NA | haf-8 |
| WBGene00001819 | -0.218571073 | 0.15291065 | 0.32587552 | haf-9 |
| WBGene00001820 | -1.26103263 | 1.08E-05 | 0.00013203 | ham-1 |
| WBGene00001821 | -0.003741765 | 0.98368684 | 0.99058801 | ham-2 |
| WBGene00001823 | 0.068065991 | 0.67520376 | 0.81135495 | hap-1 |
| WBGene00001824 | 0.696907285 | 0.00356896 | 0.02006648 | hbl-1 |
| WBGene00001827 | -0.16903554 | 0.00932052 | 0.0425645 | hcf-1 |
| WBGene00001828 | -0.399478432 | 0.10878506 | 0.25944323 | hch-1 |
| WBGene00001829 | -0.105866869 | 0.07516047 | 0.20165374 | hcp-1 |
| WBGene00001830 | -0.023557265 | 0.72077297 | 0.84351221 | hcp-2 |
| WBGene00001831 | -0.333264704 | 0.00306135 | 0.01775799 | hcp-3 |
| WBGene00001832 | 0.005653874 | 0.94540439 | 0.9693722 | hcp-4 |
| WBGene00001833 | -0.193259554 | 0.00948593 | 0.04318518 | hcp-6 |
| WBGene00001834 | -0.182142181 | 0.0059268 | 0.02988097 | hda-1 |
| WBGene00001835 | -0.003996148 | 0.97154858 | 0.9843905 | hda-2 |
| WBGene00001836 | -0.066991969 | 0.48623193 | 0.67071054 | hda-3 |

| | | | | |
|---|---|---|---|---|
| WBGene00001837 | 0.272686574 | 0.05584088 | 0.16328465 | hda-4 |
| WBGene00001838 | 0.267854599 | 0.03796153 | 0.12312687 | hda-10 |
| WBGene00001839 | 0.173763119 | 0.0833979 | 0.21707176 | hdl-1 |
| WBGene00001840 | -0.242127245 | 4.23E-07 | 7.15E-06 | hel-1 |
| WBGene00001841 | -0.032289844 | 0.72411808 | NA | hen-1 |
| WBGene00001842 | -0.204537979 | 0.24580396 | 0.44402415 | her-1 |
| WBGene00001843 | -0.179133191 | 0.07117665 | 0.1939142 | hgo-1 |
| WBGene00001844 | 0.001510439 | 0.99341344 | 0.99586212 | hid-1 |
| WBGene00001851 | 0.378540638 | 3.05E-06 | 4.19E-05 | hif-1 |
| WBGene00001852 | 1.387347124 | 6.00E-06 | 7.71E-05 | hil-1 |
| WBGene00001853 | -0.678033654 | 1.21E-09 | 3.04E-08 | hil-2 |
| WBGene00001854 | -1.027493959 | 2.28E-09 | 5.52E-08 | hil-3 |
| WBGene00001855 | -0.104002788 | 0.11489627 | 0.27014553 | hil-4 |
| WBGene00001856 | -0.10949052 | 0.11735552 | 0.27380421 | hil-5 |
| WBGene00001857 | -0.123061034 | 0.28173111 | 0.48136599 | hil-6 |
| WBGene00001858 | 1.174647887 | 0.00593408 | 0.02989207 | hil-7 |
| WBGene00001859 | 0.035512071 | 0.72830584 | 0.84850209 | hil-8 |
| WBGene00001860 | -0.231706615 | 0.00012739 | 0.00119944 | him-1 |
| WBGene00001862 | -0.181768424 | 0.0495266 | 0.14920144 | him-3 |
| WBGene00001863 | 0.232690182 | 0.04738149 | 0.14455352 | him-4 |
| WBGene00001864 | -0.098559223 | 0.54750922 | 0.72018982 | him-5 |
| WBGene00001865 | -0.056800207 | 0.61974626 | 0.77262475 | him-6 |
| WBGene00001867 | -0.208023962 | 0.08006386 | 0.21113383 | him-8 |
| WBGene00001869 | -0.156989224 | 0.08380485 | 0.21770525 | him-10 |
| WBGene00001872 | 0.115478401 | 0.41908152 | 0.61579411 | him-14 |
| WBGene00001874 | -0.141159217 | 0.05902813 | 0.16959004 | him-17 |
| WBGene00001875 | 0.147839603 | 0.11747252 | 0.27391546 | his-18 |
| WBGene00001875 | 0.147839603 | 0.11747252 | 0.27391546 | his-37 |
| WBGene00001875 | 0.147839603 | 0.11747252 | 0.27391546 | his-67 |
| WBGene00001875 | 0.147839603 | 0.11747252 | 0.27391546 | his-1 |
| WBGene00001876 | -0.073771486 | 0.46841036 | 0.65738945 | his-2 |
| WBGene00001876 | -0.073771486 | 0.46841036 | 0.65738945 | his-59 |
| WBGene00001877 | -0.179348401 | 0.19024188 | 0.37613324 | his-3 |
| WBGene00001878 | 0.048485426 | 0.45975815 | NA | his-4 |
| WBGene00001879 | -0.054524006 | 0.53093834 | NA | his-5 |
| WBGene00001879 | -0.054524006 | 0.53093834 | NA | his-18 |
| WBGene00001879 | -0.054524006 | 0.53093834 | NA | his-37 |
| WBGene00001879 | -0.054524006 | 0.53093834 | NA | his-67 |
| WBGene00001880 | -0.153834358 | 0.2676011 | 0.46647787 | his-59 |
| WBGene00001880 | -0.153834358 | 0.2676011 | 0.46647787 | his-6 |
| WBGene00001881 | -0.061534917 | 0.5872152 | 0.74981371 | his-7 |
| WBGene00001882 | 0.032306427 | 0.76663584 | 0.86741232 | his-8 |
| WBGene00001883 | NA | NA | NA | his-9 |

| | | | | |
|---|---|---|---|---|
| WBGene00001883 | NA | NA | NA | his-25 |
| WBGene00001883 | NA | NA | NA | his-59 |
| WBGene00001883 | NA | NA | NA | his-13 |
| WBGene00001884 | NA | NA | NA | his-14 |
| WBGene00001884 | NA | NA | NA | his-26 |
| WBGene00001884 | NA | NA | NA | his-18 |
| WBGene00001884 | NA | NA | NA | his-37 |
| WBGene00001884 | NA | NA | NA | his-67 |
| WBGene00001884 | NA | NA | NA | his-10 |
| WBGene00001885 | NA | NA | NA | his-11 |
| WBGene00001886 | NA | NA | NA | his-12 |
| WBGene00001886 | NA | NA | NA | his-43 |
| WBGene00001887 | NA | NA | NA | his-9 |
| WBGene00001887 | NA | NA | NA | his-25 |
| WBGene00001887 | NA | NA | NA | his-59 |
| WBGene00001887 | NA | NA | NA | his-13 |
| WBGene00001888 | NA | NA | NA | his-14 |
| WBGene00001888 | NA | NA | NA | his-26 |
| WBGene00001888 | NA | NA | NA | his-18 |
| WBGene00001888 | NA | NA | NA | his-37 |
| WBGene00001888 | NA | NA | NA | his-67 |
| WBGene00001888 | NA | NA | NA | his-10 |
| WBGene00001889 | NA | NA | NA | his-15 |
| WBGene00001889 | NA | NA | NA | his-11 |
| WBGene00001889 | NA | NA | NA | his-44 |
| WBGene00001890 | NA | NA | NA | his-16 |
| WBGene00001891 | NA | NA | NA | his-59 |
| WBGene00001891 | NA | NA | NA | his-17 |
| WBGene00001892 | -0.027728334 | 0.67598605 | NA | his-18 |
| WBGene00001892 | -0.027728334 | 0.67598605 | NA | his-37 |
| WBGene00001892 | -0.027728334 | 0.67598605 | NA | his-67 |
| WBGene00001893 | NA | NA | NA | his-51 |
| WBGene00001893 | NA | NA | NA | his-21 |
| WBGene00001893 | NA | NA | NA | his-53 |
| WBGene00001893 | NA | NA | NA | his-19 |
| WBGene00001894 | NA | NA | NA | his-52 |
| WBGene00001894 | NA | NA | NA | his-22 |
| WBGene00001894 | NA | NA | NA | his-20 |
| WBGene00001895 | NA | NA | NA | his-51 |
| WBGene00001895 | NA | NA | NA | his-21 |
| WBGene00001895 | NA | NA | NA | his-53 |
| WBGene00001895 | NA | NA | NA | his-19 |
| WBGene00001896 | NA | NA | NA | his-52 |

| | | | | |
|---|---|---|---|---|
| WBGene00001896 | NA | NA | NA | his-22 |
| WBGene00001896 | NA | NA | NA | his-20 |
| WBGene00001898 | -1.581510406 | 9.13E-30 | 1.14E-27 | his-24 |
| WBGene00001899 | 0.020317245 | 0.84643798 | NA | his-25 |
| WBGene00001899 | 0.020317245 | 0.84643798 | NA | his-59 |
| WBGene00001900 | NA | NA | NA | his-14 |
| WBGene00001900 | NA | NA | NA | his-26 |
| WBGene00001900 | NA | NA | NA | his-18 |
| WBGene00001900 | NA | NA | NA | his-37 |
| WBGene00001900 | NA | NA | NA | his-67 |
| WBGene00001900 | NA | NA | NA | his-10 |
| WBGene00001901 | NA | NA | NA | his-49 |
| WBGene00001901 | NA | NA | NA | his-59 |
| WBGene00001901 | NA | NA | NA | his-27 |
| WBGene00001902 | NA | NA | NA | his-50 |
| WBGene00001902 | NA | NA | NA | his-28 |
| WBGene00001902 | NA | NA | NA | his-18 |
| WBGene00001902 | NA | NA | NA | his-37 |
| WBGene00001902 | NA | NA | NA | his-67 |
| WBGene00001903 | NA | NA | NA | his-29 |
| WBGene00001903 | NA | NA | NA | his-34 |
| WBGene00001904 | NA | NA | NA | his-30 |
| WBGene00001904 | NA | NA | NA | his-33 |
| WBGene00001905 | -0.218692005 | 0.13989908 | 0.30727009 | his-18 |
| WBGene00001905 | -0.218692005 | 0.13989908 | 0.30727009 | his-37 |
| WBGene00001905 | -0.218692005 | 0.13989908 | 0.30727009 | his-67 |
| WBGene00001905 | -0.218692005 | 0.13989908 | 0.30727009 | his-31 |
| WBGene00001906 | 0.015448378 | 0.93943902 | 0.96589373 | his-32 |
| WBGene00001906 | 0.015448378 | 0.93943902 | 0.96589373 | his-59 |
| WBGene00001907 | NA | NA | NA | his-30 |
| WBGene00001907 | NA | NA | NA | his-33 |
| WBGene00001908 | NA | NA | NA | his-29 |
| WBGene00001908 | NA | NA | NA | his-34 |
| WBGene00001909 | 0.123953249 | 0.34873937 | 0.54884373 | his-35 |
| WBGene00001910 | -0.14795143 | 0.27640563 | 0.47632473 | C50F4.6 |
| WBGene00001911 | 0.761406754 | 1.37E-08 | 2.99E-07 | his-18 |
| WBGene00001911 | 0.761406754 | 1.37E-08 | 2.99E-07 | his-37 |
| WBGene00001911 | 0.761406754 | 1.37E-08 | 2.99E-07 | his-67 |
| WBGene00001912 | -1.59989181 | 0.00101155 | 0.00711342 | his-18 |
| WBGene00001912 | -1.59989181 | 0.00101155 | 0.00711342 | his-37 |
| WBGene00001912 | -1.59989181 | 0.00101155 | 0.00711342 | his-67 |
| WBGene00001912 | -1.59989181 | 0.00101155 | 0.00711342 | his-38 |
| WBGene00001913 | NA | NA | NA | his-39 |

| | | | | | |
|---|---|---|---|---|---|
| WBGene00001914 | | -1.840837317 | 0.00072932 | 0.005347 | his-59 |
| WBGene00001914 | | -1.840837317 | 0.00072932 | 0.005347 | his-40 |
| WBGene00001915 | | -0.052581163 | 0.73347975 | 0.85087141 | his-41 |
| WBGene00001916 | NA | | NA | NA | his-42 |
| WBGene00001916 | NA | | NA | NA | his-59 |
| WBGene00001917 | NA | | NA | NA | his-12 |
| WBGene00001917 | NA | | NA | NA | his-43 |
| WBGene00001918 | NA | | NA | NA | his-15 |
| WBGene00001918 | NA | | NA | NA | his-11 |
| WBGene00001918 | NA | | NA | NA | his-44 |
| WBGene00001919 | | 0.036894305 | 0.59120501 | NA | his-45 |
| WBGene00001919 | | 0.036894305 | 0.59120501 | NA | his-59 |
| WBGene00001920 | NA | | NA | NA | his-46 |
| WBGene00001920 | NA | | NA | NA | his-56 |
| WBGene00001920 | NA | | NA | NA | his-18 |
| WBGene00001920 | NA | | NA | NA | his-37 |
| WBGene00001920 | NA | | NA | NA | his-67 |
| WBGene00001921 | | -0.045959069 | 0.50707733 | NA | his-47 |
| WBGene00001922 | | -0.394664181 | 0.08763561 | 0.22426713 | his-48 |
| WBGene00001922 | | -0.394664181 | 0.08763561 | 0.22426713 | his-62 |
| WBGene00001923 | NA | | NA | NA | his-49 |
| WBGene00001923 | NA | | NA | NA | his-59 |
| WBGene00001923 | NA | | NA | NA | his-27 |
| WBGene00001924 | NA | | NA | NA | his-50 |
| WBGene00001924 | NA | | NA | NA | his-28 |
| WBGene00001924 | NA | | NA | NA | his-18 |
| WBGene00001924 | NA | | NA | NA | his-37 |
| WBGene00001924 | NA | | NA | NA | his-67 |
| WBGene00001925 | NA | | NA | NA | his-51 |
| WBGene00001925 | NA | | NA | NA | his-21 |
| WBGene00001925 | NA | | NA | NA | his-53 |
| WBGene00001925 | NA | | NA | NA | his-19 |
| WBGene00001926 | NA | | NA | NA | his-52 |
| WBGene00001926 | NA | | NA | NA | his-22 |
| WBGene00001926 | NA | | NA | NA | his-20 |
| WBGene00001927 | NA | | NA | NA | his-51 |
| WBGene00001927 | NA | | NA | NA | his-21 |
| WBGene00001927 | NA | | NA | NA | his-53 |
| WBGene00001927 | NA | | NA | NA | his-19 |
| WBGene00001928 | NA | | NA | NA | his-54 |
| WBGene00001929 | | -0.02791392 | 0.6763823 | NA | his-45 |
| WBGene00001929 | | -0.02791392 | 0.6763823 | NA | his-55 |
| WBGene00001929 | | -0.02791392 | 0.6763823 | NA | his-59 |

| | | | | |
|---|---|---|---|---|
| WBGene00001930 | NA | NA | NA | his-46 |
| WBGene00001930 | NA | NA | NA | his-56 |
| WBGene00001930 | NA | NA | NA | his-18 |
| WBGene00001930 | NA | NA | NA | his-37 |
| WBGene00001930 | NA | NA | NA | his-67 |
| WBGene00001931 | 0.017708609 | 0.82438962 | NA | his-47 |
| WBGene00001931 | 0.017708609 | 0.82438962 | NA | his-57 |
| WBGene00001931 | 0.017708609 | 0.82438962 | NA | his-65 |
| WBGene00001932 | NA | NA | NA | his-58 |
| WBGene00001932 | NA | NA | NA | his-62 |
| WBGene00001932 | NA | NA | NA | his-66 |
| WBGene00001933 | 0.211509152 | 0.33483145 | 0.53413188 | his-59 |
| WBGene00001934 | -0.053052915 | 0.79294726 | 0.88379574 | his-60 |
| WBGene00001934 | -0.053052915 | 0.79294726 | 0.88379574 | his-18 |
| WBGene00001934 | -0.053052915 | 0.79294726 | 0.88379574 | his-37 |
| WBGene00001934 | -0.053052915 | 0.79294726 | 0.88379574 | his-67 |
| WBGene00001935 | 0.156595543 | 0.46509401 | 0.65481694 | his-61 |
| WBGene00001936 | -0.158810246 | 0.42702761 | 0.62241028 | his-62 |
| WBGene00001937 | -0.109486302 | 0.5808272 | 0.7457591 | his-63 |
| WBGene00001937 | -0.109486302 | 0.5808272 | 0.7457591 | his-59 |
| WBGene00001938 | -0.017711638 | 0.91515088 | 0.95281497 | his-64 |
| WBGene00001938 | -0.017711638 | 0.91515088 | 0.95281497 | his-18 |
| WBGene00001938 | -0.017711638 | 0.91515088 | 0.95281497 | his-37 |
| WBGene00001938 | -0.017711638 | 0.91515088 | 0.95281497 | his-67 |
| WBGene00001939 | 0.020232312 | 0.84643798 | NA | his-47 |
| WBGene00001939 | 0.020232312 | 0.84643798 | NA | his-57 |
| WBGene00001939 | 0.020232312 | 0.84643798 | NA | his-65 |
| WBGene00001940 | -0.072377862 | 0.54072055 | 0.71476071 | his-58 |
| WBGene00001940 | -0.072377862 | 0.54072055 | 0.71476071 | his-62 |
| WBGene00001940 | -0.072377862 | 0.54072055 | 0.71476071 | his-66 |
| WBGene00001941 | -0.2153108 | 0.32072431 | 0.52076352 | his-18 |
| WBGene00001941 | -0.2153108 | 0.32072431 | 0.52076352 | his-37 |
| WBGene00001941 | -0.2153108 | 0.32072431 | 0.52076352 | his-67 |
| WBGene00001942 | 0.10903011 | 0.55947098 | 0.73003231 | his-68 |
| WBGene00001943 | -0.003421551 | 0.95689871 | NA | his-69 |
| WBGene00001944 | 0.0398788 | 0.58971221 | NA | his-70 |
| WBGene00001945 | 0.803584343 | 1.35E-05 | 0.00016041 | his-71 |
| WBGene00001946 | -0.353818549 | 1.88E-06 | 2.75E-05 | his-72 |
| WBGene00001947 | 0.178740886 | 0.25410934 | 0.45227508 | his-73 |
| WBGene00001948 | -0.161682098 | 0.44780959 | 0.64068438 | hlh-1 |
| WBGene00001949 | -0.350554484 | 0.00902308 | 0.04150389 | hlh-2 |
| WBGene00001950 | -0.561630008 | 0.0435351 | 0.13610879 | hlh-3 |
| WBGene00001951 | -0.001842784 | 0.95621055 | NA | hlh-4 |

| | | | | |
|---|---|---|---|---|
| WBGene00001952 | 0.039745363 | 0.76861705 | 0.86844708 | hlh-6 |
| WBGene00001953 | -0.026730413 | 0.85362474 | 0.92153958 | hlh-8 |
| WBGene00001954 | -0.055120925 | 0.45606622 | NA | hlh-10 |
| WBGene00001955 | 0.702856916 | 1.08E-08 | 2.40E-07 | hlh-11 |
| WBGene00001956 | 0.050147126 | 0.73549364 | 0.85166527 | hlh-12 |
| WBGene00001957 | 0.038125098 | 0.61119482 | NA | hlh-13 |
| WBGene00001958 | -0.27316824 | 0.20283203 | 0.3914648 | hlh-14 |
| WBGene00001959 | NA | NA | NA | hlh-15 |
| WBGene00001960 | -0.07937498 | 0.55943786 | 0.73003231 | hlh-16 |
| WBGene00001961 | -0.025848508 | 0.67901638 | NA | hlh-17 |
| WBGene00001962 | NA | NA | NA | hlh-19 |
| WBGene00001963 | 0.185194765 | 0.37356399 | 0.572527 | btb-1 |
| WBGene00001964 | -0.046650166 | 0.5657231 | NA | hlh-25 |
| WBGene00001965 | -0.049845644 | 0.48082527 | NA | hlh-26 |
| WBGene00001966 | -0.133906003 | 0.29805136 | 0.49896857 | hlh-27 |
| WBGene00001967 | -0.022930767 | 0.78121253 | NA | hlh-28 |
| WBGene00001968 | -0.102692111 | 0.21667182 | NA | hlh-29 |
| WBGene00001971 | -0.329394068 | 7.89E-08 | 1.51E-06 | hmg-1.1 |
| WBGene00001972 | -0.15089161 | 0.0699296 | 0.19109846 | hmg-1.2 |
| WBGene00001973 | -0.14497467 | 0.03078647 | 0.10524042 | hmg-3 |
| WBGene00001974 | -0.064990267 | 0.38023283 | 0.5797656 | hmg-4 |
| WBGene00001975 | -0.154960193 | 0.12064075 | 0.27938963 | hmg-5 |
| WBGene00001976 | -0.532424513 | 0.00191652 | 0.01207913 | hmg-11 |
| WBGene00001977 | 0.102981825 | 0.20001142 | 0.38863533 | hmg-12 |
| WBGene00001978 | -0.202690717 | 0.00960589 | 0.04359573 | hmp-1 |
| WBGene00001979 | 0.015656079 | 0.86391196 | 0.92738279 | hmp-2 |
| WBGene00001980 | 0.089962256 | 0.18401544 | 0.36740352 | hmr-1 |
| WBGene00001981 | -0.416689378 | 0.09681055 | 0.24060486 | hnd-1 |
| WBGene00001983 | -0.016701808 | 0.83211379 | 0.9084712 | hoe-1 |
| WBGene00001984 | -0.071310397 | 0.6403696 | 0.78675996 | hog-1 |
| WBGene00001985 | 0.103716601 | 0.41646821 | 0.61354691 | hop-1 |
| WBGene00001986 | 0.186456149 | 0.38642925 | 0.5851299 | hot-1 |
| WBGene00001987 | 0.670607741 | 0.02674502 | 0.09546083 | hot-2 |
| WBGene00001988 | 0.009813971 | 0.92662957 | 0.9586234 | hot-3 |
| WBGene00001989 | 0.165903372 | 0.31535892 | 0.51576966 | hot-4 |
| WBGene00001990 | 0.167035936 | 0.27736811 | 0.47747657 | hot-5 |
| WBGene00001991 | 0.253787473 | 0.24549486 | 0.44365517 | hot-6 |
| WBGene00001992 | -0.039434656 | 0.74266589 | 0.85444405 | hot-7 |
| WBGene00001993 | -0.394923643 | 4.41E-08 | 8.81E-07 | hpd-1 |
| WBGene00001994 | 0.076731122 | 0.55044263 | 0.72291268 | hpk-1 |
| WBGene00001995 | 0.260533789 | 0.24630836 | 0.44469671 | hpl-1 |
| WBGene00001996 | -0.170887983 | 0.08028045 | 0.21156663 | hpl-2 |
| WBGene00001997 | -0.31427559 | 0.05786663 | 0.16740276 | hpr-9 |

| | | | | |
|---|---|---|---|---|
| WBGene00001998 | 0.134518052 | 0.29070579 | 0.49119717 | hpr-17 |
| WBGene00001999 | -0.222553996 | 1.66E-05 | 0.00019288 | hrp-1 |
| WBGene00002000 | -0.361564593 | 1.23E-12 | 4.56E-11 | hrp-2 |
| WBGene00002001 | -0.396489841 | 1.01E-06 | 1.59E-05 | hars-1 |
| WBGene00002002 | -0.192918616 | 0.07276389 | 0.19673797 | hsb-1 |
| WBGene00002003 | -0.108720765 | 0.26153474 | 0.45962431 | hse-5 |
| WBGene00002004 | -0.048737404 | 0.55101826 | 0.72360376 | hsf-1 |
| WBGene00002005 | -0.339265613 | 4.25E-15 | 2.06E-13 | hsp-1 |
| WBGene00002006 | -0.162537021 | 0.37090628 | 0.57036433 | NA |
| WBGene00002007 | -0.239923119 | 1.74E-06 | 2.57E-05 | hsp-3 |
| WBGene00002008 | -0.25775543 | 2.85E-06 | 3.95E-05 | hsp-4 |
| WBGene00002010 | -0.353697745 | 2.80E-13 | 1.12E-11 | hsp-6 |
| WBGene00002011 | 0.31307975 | 0.05138901 | 0.15344072 | hsp-12.2 |
| WBGene00002012 | 2.639568813 | 3.68E-11 | 1.14E-09 | hsp-12.3 |
| WBGene00002013 | 0.820747244 | 0.02323325 | 0.08568211 | hsp-12.6 |
| WBGene00002015 | NA | NA | NA | hsp-16.1 |
| WBGene00002015 | NA | NA | NA | hsp-16.11 |
| WBGene00002016 | 0.032192384 | 0.61853721 | NA | hsp-16.2 |
| WBGene00002017 | NA | NA | NA | hsp-16.1 |
| WBGene00002017 | NA | NA | NA | hsp-16.11 |
| WBGene00002018 | -0.006738828 | 0.92480778 | NA | hsp-16.41 |
| WBGene00002019 | NA | NA | NA | hsp-16.48 |
| WBGene00002019 | NA | NA | NA | hsp-16.49 |
| WBGene00002020 | NA | NA | NA | hsp-16.48 |
| WBGene00002020 | NA | NA | NA | hsp-16.49 |
| WBGene00002021 | 1.455661791 | 8.02E-17 | 4.70E-15 | hsp-17 |
| WBGene00002023 | 0.086815991 | 0.40625808 | 0.60428981 | hsp-25 |
| WBGene00002024 | -0.387683503 | 0.00166895 | 0.01075962 | hsp-43 |
| WBGene00002025 | -0.760806698 | 8.63E-42 | 1.66E-39 | hsp-60 |
| WBGene00002026 | -0.157989692 | 0.31673294 | 0.51698682 | hsp-70 |
| WBGene00002027 | -0.289851707 | 1.32E-08 | 2.90E-07 | hsr-9 |
| WBGene00002028 | -0.03748224 | 0.70371937 | 0.8318388 | hst-1 |
| WBGene00002029 | -0.279812725 | 0.15377114 | 0.3270785 | hst-2 |
| WBGene00002030 | 0.13437921 | 0.39515501 | 0.59325928 | hst-3.1 |
| WBGene00002031 | -0.071183622 | 0.68710793 | 0.81974798 | hst-6 |
| WBGene00002032 | -0.166630259 | 0.08448574 | 0.21889147 | htp-1 |
| WBGene00002033 | -0.079428297 | 0.46174193 | 0.65228105 | htp-2 |
| WBGene00002034 | -0.092768726 | 0.3100011 | 0.51077268 | htp-3 |
| WBGene00002035 | 0.042763542 | 0.62274569 | 0.77445168 | hum-1 |
| WBGene00002036 | 0.027177377 | 0.70411366 | 0.83203617 | hum-2 |
| WBGene00002037 | 0.470042766 | 0.00162109 | 0.01050194 | hum-4 |
| WBGene00002038 | 0.09507207 | 0.38664046 | 0.58527739 | hum-5 |
| WBGene00002039 | 0.889532653 | 0.00031688 | 0.00263307 | hum-6 |

| | | | | |
|---|---|---|---|---|
| WBGene00002040 | 0.251071502 | 0.1578024 | 0.33195186 | hum-7 |
| WBGene00002041 | 0.649130534 | 3.26E-06 | 4.45E-05 | hum-8 |
| WBGene00002042 | -0.018771326 | 0.9023368 | 0.94802873 | hus-1 |
| WBGene00002043 | -0.139987751 | 0.20346124 | 0.39242209 | hyl-1 |
| WBGene00002044 | -0.105449424 | 0.3497414 | 0.54959362 | hyl-2 |
| WBGene00002045 | -0.377734233 | 4.05E-16 | 2.20E-14 | icd-1 |
| WBGene00002046 | -0.388190559 | 0.00010961 | 0.00105302 | icln-1 |
| WBGene00002047 | -0.225324218 | 0.01016878 | 0.04543213 | icp-1 |
| WBGene00002048 | 1.374340777 | 1.94E-12 | 6.98E-11 | ida-1 |
| WBGene00002050 | 0.820579755 | 5.75E-08 | 1.13E-06 | ifa-1 |
| WBGene00002051 | 0.046672806 | 0.77458639 | 0.87229593 | ifa-3 |
| WBGene00002052 | -0.274267389 | 0.02646461 | 0.0947587 | ifa-4 |
| WBGene00002053 | -0.199274886 | 0.02528296 | 0.09144462 | ifb-1 |
| WBGene00002054 | 0.999504496 | 1.34E-22 | 1.15E-20 | ifb-2 |
| WBGene00002055 | 1.034186512 | 8.40E-10 | 2.14E-08 | ifc-1 |
| WBGene00002056 | 0.829122096 | 4.75E-20 | 3.60E-18 | ifc-2 |
| WBGene00002057 | 0.205878414 | 0.31696158 | 0.51716884 | ifd-1 |
| WBGene00002058 | 0.573265519 | 0.00294407 | 0.01720722 | ifd-2 |
| WBGene00002059 | -0.343965021 | 6.67E-08 | 1.29E-06 | ife-1 |
| WBGene00002060 | -0.242005872 | 0.06327035 | 0.17846903 | ife-2 |
| WBGene00002061 | -0.197162945 | 0.00081245 | 0.00589447 | ife-3 |
| WBGene00002062 | -0.275064031 | 0.19264498 | 0.3791015 | ife-4 |
| WBGene00002063 | 0.308981036 | 0.12409821 | 0.28442955 | ife-5 |
| WBGene00002064 | -0.38603159 | 8.47E-12 | 2.86E-10 | iff-1 |
| WBGene00002065 | 0.500161192 | 2.23E-08 | 4.70E-07 | iff-2 |
| WBGene00002066 | -0.112080835 | 0.02475573 | 0.09019242 | ifg-1 |
| WBGene00002067 | 1.120708421 | 0.00042977 | 0.0034078 | ifp-1 |
| WBGene00002068 | -0.199330778 | 0.00551476 | 0.02832047 | ify-1 |
| WBGene00002069 | 1.565130665 | 1.39E-06 | 2.11E-05 | ikb-1 |
| WBGene00002070 | -0.262845698 | 0.00013524 | 0.00125561 | ile-1 |
| WBGene00002071 | -0.165751311 | 0.14685065 | 0.31716187 | ile-2 |
| WBGene00002072 | -0.08934199 | 0.23496994 | 0.43077821 | ima-1 |
| WBGene00002073 | -0.085945002 | 0.09997819 | 0.24539492 | ima-2 |
| WBGene00002074 | -0.288099862 | 4.33E-06 | 5.75E-05 | ima-3 |
| WBGene00002075 | -0.422883349 | 1.69E-08 | 3.64E-07 | imb-1 |
| WBGene00002076 | -0.176678285 | 0.00354229 | 0.01996256 | imb-2 |
| WBGene00002077 | -0.102180562 | 0.06298514 | 0.17782783 | imb-3 |
| WBGene00002078 | -0.229084903 | 6.65E-06 | 8.46E-05 | xpo-1 |
| WBGene00002079 | -0.125133912 | 0.02690411 | 0.09577028 | xpo-2 |
| WBGene00002080 | -0.164582071 | 0.07024106 | 0.19172247 | xpo-3 |
| WBGene00002081 | -0.036913677 | 0.76032845 | 0.8642162 | ina-1 |
| WBGene00002083 | -0.485266796 | 2.03E-19 | 1.47E-17 | inf-1 |
| WBGene00002084 | 1.557438181 | 0.00405186 | 0.02211263 | ins-1 |

| | | | | |
|---|---|---|---|---|
| WBGene00002085 | -0.125872228 | 0.10485217 | NA | ins-2 |
| WBGene00002086 | 0.075194005 | 0.67295024 | 0.8097193 | ins-3 |
| WBGene00002087 | 1.109621397 | 0.00952603 | 0.04335425 | ins-4 |
| WBGene00002088 | 0.071801919 | 0.64975907 | 0.79273174 | ins-5 |
| WBGene00002089 | 0.222315711 | 0.28209647 | 0.48182156 | ins-6 |
| WBGene00002090 | 4.030539961 | 3.74E-21 | 3.01E-19 | ins-7 |
| WBGene00002091 | 8.279822373 | 4.32E-10 | 1.14E-08 | ins-8 |
| WBGene00002092 | -0.00936818 | 0.91368063 | NA | ins-9 |
| WBGene00002093 | 0.127918313 | 0.10854122 | NA | ins-10 |
| WBGene00002094 | 0.175586955 | 0.04633883 | NA | ins-11 |
| WBGene00002095 | -0.37275143 | 0.04121082 | 0.13071222 | ins-12 |
| WBGene00002096 | NA | NA | NA | ins-13 |
| WBGene00002097 | -0.035877804 | 0.75669453 | 0.86226195 | ins-14 |
| WBGene00002098 | -0.001488154 | 0.95639912 | NA | ins-15 |
| WBGene00002099 | NA | NA | NA | ins-16 |
| WBGene00002100 | 0.60117034 | 0.02023019 | 0.07731405 | ins-17 |
| WBGene00002101 | 0.236785857 | 0.23998958 | 0.43708017 | ins-18 |
| WBGene00002102 | 0.155122056 | 0.06782466 | NA | ins-19 |
| WBGene00002103 | 0.020232312 | 0.84643798 | NA | ins-20 |
| WBGene00002104 | NA | NA | NA | ins-21 |
| WBGene00002105 | 0.161433691 | 0.35885446 | 0.55911335 | ins-22 |
| WBGene00002106 | 0.332745373 | 0.077705 | 0.20658778 | ins-23 |
| WBGene00002107 | 0.198171302 | 0.26450532 | 0.46290011 | ins-24 |
| WBGene00002108 | 0.038125098 | 0.61119482 | NA | ins-25 |
| WBGene00002109 | 0.381793314 | 0.12169341 | 0.28070284 | ins-26 |
| WBGene00002110 | 0.196354134 | 0.34521375 | 0.54493526 | ins-27 |
| WBGene00002111 | 0.792285519 | 0.02035336 | 0.0776227 | ins-28 |
| WBGene00002112 | -0.020817452 | 0.87542065 | 0.93344293 | ins-29 |
| WBGene00002113 | 0.251459 | 0.08065705 | 0.21212499 | ins-30 |
| WBGene00002114 | 0.038195847 | 0.57893903 | NA | ins-31 |
| WBGene00002115 | -0.027728334 | 0.67598605 | NA | ins-32 |
| WBGene00002116 | -0.10505942 | 0.55285126 | 0.72508974 | ins-33 |
| WBGene00002117 | -0.433991386 | 0.07633692 | 0.2040995 | ins-34 |
| WBGene00002118 | 0.100264625 | 0.38646629 | 0.5851299 | ins-35 |
| WBGene00002119 | -0.003270206 | 0.95698316 | NA | ins-36 |
| WBGene00002120 | 0.3355211 | 0.06959554 | 0.19056449 | ins-37 |
| WBGene00002121 | NA | NA | NA | ins-38 |
| WBGene00002123 | 0.477272455 | 0.05688471 | 0.16542055 | inx-1 |
| WBGene00002124 | -0.959387608 | 6.32E-13 | 2.42E-11 | inx-2 |
| WBGene00002125 | -0.587106511 | 6.99E-11 | 2.05E-09 | inx-3 |
| WBGene00002127 | -0.131338382 | 0.47807639 | 0.66503719 | inx-5 |
| WBGene00002128 | 1.553993716 | 0.00072568 | 0.00532829 | inx-6 |
| WBGene00002129 | 0.252594144 | 0.1852781 | 0.36906893 | inx-7 |

| | | | | |
|---|---|---|---|---|
| WBGene00002130 | 0.149712101 | 0.05735994 | 0.16643023 | inx-8 |
| WBGene00002131 | -0.043437277 | 0.7807524 | 0.87640849 | inx-9 |
| WBGene00002132 | 0.382681019 | 0.08847656 | 0.22573322 | inx-10 |
| WBGene00002133 | -0.069068952 | 0.70481903 | 0.83266537 | inx-11 |
| WBGene00002134 | 0.27041271 | 0.06464758 | 0.18118633 | inx-12 |
| WBGene00002135 | -0.083014747 | 0.53764105 | 0.71255568 | inx-13 |
| WBGene00002136 | -0.081710431 | 0.26262219 | 0.46064466 | inx-14 |
| WBGene00002137 | 0.540102969 | 0.00090243 | 0.00645508 | inx-15 |
| WBGene00002138 | 0.243981706 | 0.10653136 | 0.25602851 | inx-16 |
| WBGene00002139 | 0.322200151 | 0.10240254 | 0.24938049 | inx-17 |
| WBGene00002140 | 0.135027272 | 0.41288012 | 0.61056051 | inx-18 |
| WBGene00002141 | 0.054220244 | 0.72892579 | 0.84861661 | inx-19 |
| WBGene00002142 | 0.329054399 | 0.08246776 | 0.21537824 | inx-20 |
| WBGene00002143 | 0.130372404 | 0.3073367 | 0.50792602 | inx-21 |
| WBGene00002144 | 0.106265437 | 0.24014255 | 0.43730436 | inx-22 |
| WBGene00002146 | 0.077804045 | 0.667817 | 0.80599363 | ipp-5 |
| WBGene00002147 | 0.047494982 | 0.51970491 | 0.69871438 | ire-1 |
| WBGene00002148 | -0.270512759 | 0.00021074 | 0.00184865 | gon-14 |
| WBGene00002149 | 0.533884554 | 0.03897176 | 0.12554138 | irk-1 |
| WBGene00002150 | 0.102236848 | 0.50583357 | 0.68674779 | irk-2 |
| WBGene00002151 | -0.045030099 | 0.82621865 | 0.90477236 | irk-3 |
| WBGene00002152 | -0.199123554 | 0.0011554 | 0.00793443 | iars-1 |
| WBGene00002153 | -0.041471328 | 0.63977238 | 0.78634212 | iars-2 |
| WBGene00002162 | -0.137391209 | 0.0194167 | 0.07514669 | isp-1 |
| WBGene00002163 | 0.318961289 | 0.16310096 | 0.33862183 | ist-1 |
| WBGene00002169 | -0.251543487 | 7.51E-05 | 0.00075211 | isw-1 |
| WBGene00002173 | 0.516566272 | 2.13E-06 | 3.09E-05 | itr-1 |
| WBGene00002174 | 0.761450041 | 9.88E-07 | 1.56E-05 | itx-1 |
| WBGene00002175 | -0.037718747 | 0.79247649 | 0.88340554 | jac-1 |
| WBGene00002176 | 0.289855768 | 0.20929485 | 0.39876248 | jip-1 |
| WBGene00002177 | 0.169950175 | 0.19619667 | 0.38376959 | jkk-1 |
| WBGene00002178 | 0.431662109 | 0.09124299 | 0.22981054 | jnk-1 |
| WBGene00002179 | 0.408170103 | 0.01134208 | 0.04934899 | jph-1 |
| WBGene00002180 | 0.010379891 | 0.94395306 | 0.96829095 | jtr-1 |
| WBGene00002181 | 0.28594844 | 0.17977333 | 0.36139795 | kal-1 |
| WBGene00002182 | 0.270554237 | 0.23369519 | 0.42913976 | kap-1 |
| WBGene00002183 | -0.396080552 | 7.63E-09 | 1.73E-07 | kat-1 |
| WBGene00002184 | 0.870226182 | 0.011569 | 0.05009809 | kel-1 |
| WBGene00002185 | 0.056069258 | 0.6454762 | 0.79001558 | kel-3 |
| WBGene00002186 | 0.843963211 | 0.01711871 | 0.06849822 | kel-10 |
| WBGene00002187 | 0.370253472 | 0.01491874 | 0.06159767 | kgb-1 |
| WBGene00002188 | 1.464801126 | 0.00368952 | 0.02057229 | kgb-2 |
| WBGene00002189 | 0.733032051 | 2.41E-09 | 5.78E-08 | kin-1 |

| | | | | |
|---|---|---|---|---|
| WBGene00002190 | 0.364375142 | 0.00074472 | 0.00545171 | kin-2 |
| WBGene00002191 | -0.27865617 | 3.27E-06 | 4.45E-05 | kin-3 |
| WBGene00002192 | -0.096848112 | 0.12408226 | 0.28442955 | kin-4 |
| WBGene00002193 | 0.347750535 | 0.02239347 | 0.08348636 | kin-5 |
| WBGene00002194 | NA | NA | NA | kin-6 |
| WBGene00002195 | 1.002441968 | 0.01428819 | 0.05946374 | kin-9 |
| WBGene00002196 | -0.31077058 | 2.31E-05 | 0.00026027 | kin-10 |
| WBGene00002198 | 0.124608589 | 0.55487166 | 0.72686037 | kin-14 |
| WBGene00002199 | -0.992507315 | 0.00099543 | 0.00701353 | kin-15 |
| WBGene00002200 | 0.126157756 | 0.5465854 | 0.71978385 | kin-16 |
| WBGene00002201 | -0.124126255 | 0.08644096 | 0.22214886 | kin-18 |
| WBGene00002202 | -0.223035423 | 0.00013596 | 0.00126063 | kin-19 |
| WBGene00002203 | 0.331540279 | 0.01857206 | 0.07282098 | kin-20 |
| WBGene00002204 | 0.042516415 | 0.55991373 | NA | kin-21 |
| WBGene00002205 | 0.05366068 | 0.40968658 | NA | kin-23 |
| WBGene00002206 | 0.018795205 | 0.77532208 | NA | kin-24 |
| WBGene00002207 | -0.00378212 | 0.97305449 | 0.98528046 | sid-3 |
| WBGene00002208 | 0.148467664 | 0.07389511 | NA | kin-26 |
| WBGene00002210 | 0.543333135 | 0.00200767 | 0.01250823 | kin-29 |
| WBGene00002211 | 0.428975613 | 0.07604212 | 0.20357153 | kin-30 |
| WBGene00002212 | 0.2155296 | 0.12850355 | 0.29016354 | NA |
| WBGene00002213 | -0.105823821 | 0.56402996 | 0.73404113 | kin-32 |
| WBGene00002214 | -0.117014947 | 0.1119779 | 0.26464382 | klc-1 |
| WBGene00002215 | 0.121080676 | 0.33095031 | 0.53051972 | klc-2 |
| WBGene00002216 | -0.152045712 | 0.40064058 | 0.59897669 | klp-3 |
| WBGene00002217 | 0.184670987 | 0.19488798 | 0.38182188 | klp-4 |
| WBGene00002218 | 0.058399858 | 0.51469406 | NA | klp-6 |
| WBGene00002219 | 0.006787781 | 0.93589216 | 0.96380369 | klp-7 |
| WBGene00002220 | 0.153226613 | 0.40853968 | 0.60645133 | gcc-1 |
| WBGene00002221 | -0.315465308 | 0.00404979 | 0.02211263 | NA |
| WBGene00002222 | 0.335567379 | 0.16033433 | 0.33484732 | klp-11 |
| WBGene00002223 | -0.025158135 | 0.74024649 | 0.85342581 | klp-12 |
| WBGene00002224 | -0.305929438 | 0.12902971 | 0.29103743 | klp-13 |
| WBGene00002225 | -0.085214933 | 0.24365212 | 0.44163915 | klp-15 |
| WBGene00002226 | -0.213286005 | 0.00189649 | 0.01197347 | klp-16 |
| WBGene00002227 | 0.183370727 | 0.08738565 | 0.22379103 | klp-17 |
| WBGene00002228 | -0.168668928 | 0.02722148 | 0.09650112 | klp-18 |
| WBGene00002229 | -0.08383267 | 0.17133089 | 0.35029402 | klp-19 |
| WBGene00002230 | -0.037177345 | 0.82900949 | 0.90652983 | klp-20 |
| WBGene00002231 | -0.157793598 | 0.01310201 | 0.0554893 | knl-1 |
| WBGene00002232 | 0.412554975 | 0.08709442 | 0.22324045 | kpc-1 |
| WBGene00002233 | -0.089350781 | 0.66270188 | 0.80173676 | kqt-1 |
| WBGene00002234 | 0.172063314 | 0.40169372 | 0.60010592 | kqt-2 |

| | | | | |
|---|---|---|---|---|
| WBGene00002235 | -0.082669676 | 0.6840385 | 0.8176383 | kqt-3 |
| WBGene00002238 | -0.416147863 | 7.43E-16 | 3.82E-14 | kars-1 |
| WBGene00002239 | -0.375099889 | 0.03463239 | 0.11516075 | ksr-1 |
| WBGene00002240 | -0.118891673 | 0.25730189 | 0.45541065 | ksr-2 |
| WBGene00002241 | -0.050150385 | 0.58445773 | 0.7477974 | kup-1 |
| WBGene00002242 | 0.684061633 | 0.03341074 | 0.11204001 | kvs-1 |
| WBGene00002243 | 0.037929215 | 0.85141695 | 0.92034307 | lad-2 |
| WBGene00002244 | -0.272967441 | 7.56E-05 | 0.00075624 | laf-1 |
| WBGene00002245 | -0.052801944 | 0.51932994 | 0.69848084 | lag-1 |
| WBGene00002246 | -0.840689807 | 0.02168159 | 0.08145757 | lag-2 |
| WBGene00002247 | -0.311093866 | 0.0012189 | 0.00828116 | lam-1 |
| WBGene00002248 | -0.371799186 | 0.02238351 | 0.08348636 | lam-3 |
| WBGene00002249 | -0.249283557 | 0.00067566 | 0.00502658 | lap-1 |
| WBGene00002250 | -0.136990754 | 0.01355907 | 0.05712759 | lap-2 |
| WBGene00002251 | -0.058796325 | 0.38902261 | 0.58748633 | lat-1 |
| WBGene00002252 | 0.503132213 | 0.06611445 | 0.18362328 | lat-2 |
| WBGene00002253 | 0.327083855 | 0.0003325 | 0.00274137 | lbp-1 |
| WBGene00002254 | 0.33407833 | 2.08E-08 | 4.40E-07 | lbp-2 |
| WBGene00002255 | 0.367127294 | 0.00134823 | 0.00897653 | lbp-3 |
| WBGene00002256 | -0.247682476 | 0.0412095 | 0.13071222 | lbp-4 |
| WBGene00002257 | 0.76784492 | 0.00024454 | 0.00209256 | lbp-5 |
| WBGene00002258 | 0.201916289 | 0.00694624 | 0.03377717 | lbp-6 |
| WBGene00002259 | 0.423175586 | 0.05715249 | 0.16595987 | lbp-7 |
| WBGene00002260 | 0.009857261 | 0.96139758 | 0.9791195 | lbp-8 |
| WBGene00002261 | -0.181999429 | 0.18700941 | 0.37155717 | ldb-1 |
| WBGene00002262 | -0.360524452 | 0.00021499 | 0.00188115 | ldh-1 |
| WBGene00002263 | 0.53410837 | 1.80E-16 | 1.01E-14 | lea-1 |
| WBGene00002264 | -0.010350579 | 0.86476903 | 0.92782379 | lec-1 |
| WBGene00002265 | 1.002406926 | 1.13E-47 | 2.55E-45 | lec-2 |
| WBGene00002266 | 0.684799063 | 0.00024218 | 0.00207838 | lec-3 |
| WBGene00002267 | -0.289220318 | 0.0126869 | 0.05409667 | lec-4 |
| WBGene00002268 | -0.12974045 | 0.10821531 | 0.25875883 | lec-5 |
| WBGene00002269 | 0.467071673 | 5.81E-10 | 1.51E-08 | lec-6 |
| WBGene00002270 | 2.874297173 | 0.00018726 | 0.00166769 | lec-7 |
| WBGene00002271 | 0.61380756 | 6.46E-09 | 1.49E-07 | lec-8 |
| WBGene00002272 | -0.035020737 | 0.71540817 | 0.84039298 | lec-9 |
| WBGene00002273 | 0.466218028 | 1.18E-06 | 1.83E-05 | lec-10 |
| WBGene00002274 | 0.109291993 | 0.53781183 | 0.71258851 | lec-11 |
| WBGene00002275 | 0.186719935 | 0.06733813 | 0.18608862 | lem-2 |
| WBGene00002276 | -0.207715053 | 0.02201648 | 0.08244075 | lem-3 |
| WBGene00002278 | 0.011305423 | 0.93994973 | 0.96628311 | lep-2 |
| WBGene00002280 | 0.41388202 | 0.00050176 | 0.00389518 | let-2 |
| WBGene00002282 | -0.132552917 | 0.45741014 | 0.64918009 | let-4 |

| | | | | |
|---|---|---|---|---|
| WBGene00002285 | NA | NA | NA | NA |
| WBGene00002295 | -0.037445855 | 0.69792847 | 0.82793475 | let-19 |
| WBGene00002297 | -0.081134787 | 0.29266616 | 0.49354126 | ect-2 |
| WBGene00002299 | 0.746822575 | 3.58E-05 | 0.00038751 | let-23 |
| WBGene00002324 | -0.109828638 | 0.44909749 | 0.64173691 | let-49 |
| WBGene00002335 | -0.104298591 | 0.23157807 | 0.42654268 | let-60 |
| WBGene00002344 | -0.166058362 | 0.00053603 | 0.00411737 | let-70 |
| WBGene00002348 | 0.632284063 | 5.02E-08 | 9.95E-07 | myo-1 |
| WBGene00002363 | -0.17862372 | 0.000524 | 0.00403995 | let-92 |
| WBGene00002368 | -0.312348837 | 0.00072273 | 0.00531311 | let-99 |
| WBGene00002393 | 0.100603117 | 0.4683547 | 0.65738945 | lpr-1 |
| WBGene00002497 | 0.150627626 | 0.38940774 | 0.58773772 | let-268 |
| WBGene00002583 | 0.044752124 | 0.50354565 | 0.68467923 | let-363 |
| WBGene00002601 | 0.09945472 | 0.5327719 | 0.70870744 | let-381 |
| WBGene00002632 | 0.090839788 | 0.47470313 | 0.66192665 | let-413 |
| WBGene00002637 | -0.156682637 | 0.00502762 | 0.02623283 | let-418 |
| WBGene00002694 | -0.147296821 | 0.02390172 | 0.08768355 | let-502 |
| WBGene00002696 | -0.379749343 | 1.62E-06 | 2.42E-05 | let-504 |
| WBGene00002717 | 0.061589466 | 0.32459708 | 0.523741 | let-526 |
| WBGene00002783 | -0.126374208 | 0.10764486 | 0.25791858 | let-607 |
| WBGene00002804 | -0.101332606 | 0.30820938 | 0.50908094 | let-630 |
| WBGene00002827 | -0.479980258 | 0.05664758 | 0.16499724 | let-653 |
| WBGene00002845 | -0.086273394 | 0.05674203 | 0.16518093 | let-711 |
| WBGene00002850 | -0.402821172 | 9.44E-09 | 2.12E-07 | let-716 |
| WBGene00002855 | -0.257595473 | 0.00010986 | 0.00105473 | let-721 |
| WBGene00002879 | -0.25649669 | 0.00050158 | 0.00389518 | let-754 |
| WBGene00002881 | 0.540580668 | 0.03580101 | 0.11794837 | let-756 |
| WBGene00002889 | 0.096394593 | 0.15163691 | 0.32414722 | let-765 |
| WBGene00002891 | -0.212495129 | 0.00128105 | 0.00862335 | let-767 |
| WBGene00002915 | 0.343853559 | 0.00023702 | 0.00203891 | let-805 |
| WBGene00002957 | 0.039851484 | 0.62231398 | 0.77415962 | let-858 |
| WBGene00002974 | 0.080283248 | 0.70126908 | 0.83028308 | lev-1 |
| WBGene00002975 | 0.083048408 | 0.63270196 | 0.78177194 | lev-8 |
| WBGene00002976 | 0.752338881 | 0.01852931 | 0.07271176 | lev-9 |
| WBGene00002977 | 0.0328853 | 0.6130758 | NA | lev-10 |
| WBGene00002978 | -0.030787171 | 0.64951008 | 0.79273174 | lev-11 |
| WBGene00002979 | 0.616562453 | 6.47E-05 | 0.00066198 | lfe-2 |
| WBGene00002980 | 0.138724131 | 0.03146555 | 0.1069376 | lgg-1 |
| WBGene00002981 | 0.466422446 | 1.33E-06 | 2.03E-05 | lgg-2 |
| WBGene00002982 | -0.195095741 | 0.23213124 | 0.42717867 | lgg-3 |
| WBGene00002983 | 1.535021855 | 0.00218894 | 0.01341047 | lgx-1 |
| WBGene00002985 | -0.119305841 | 0.10729131 | 0.25730584 | lig-1 |
| WBGene00002986 | 0.146145137 | 0.27627716 | 0.47632079 | lig-4 |

| | | | | |
|---|---|---|---|---|
| WBGene00002987 | 0.07356829 | 0.43672709 | NA | lim-4 |
| WBGene00002988 | 0.303485633 | 0.19275332 | 0.37926373 | lim-6 |
| WBGene00002989 | 0.615897062 | 0.02123176 | 0.08026661 | lim-7 |
| WBGene00002990 | 0.883466103 | 0.00031301 | 0.00260535 | lin-1 |
| WBGene00002991 | 0.489189164 | 0.02406308 | 0.08816507 | lin-2 |
| WBGene00002992 | -0.151480272 | 0.14241271 | 0.31058875 | lin-3 |
| WBGene00002993 | NA | NA | NA | NA |
| WBGene00002994 | -0.171684678 | 0.01492684 | 0.06161373 | lin-5 |
| WBGene00002996 | -0.145708056 | 0.35754252 | 0.55772214 | lin-7 |
| WBGene00002997 | 0.195059965 | 0.30837479 | 0.5092967 | lin-8 |
| WBGene00002998 | -0.119016425 | 0.22888733 | 0.4226967 | lin-9 |
| WBGene00002999 | 0.21349769 | 0.09056959 | 0.22888093 | lin-10 |
| WBGene00003000 | 0.491495357 | 0.06801387 | 0.18759283 | lin-11 |
| WBGene00003001 | -0.069780706 | 0.71228157 | 0.83815617 | lin-12 |
| WBGene00003002 | -0.096957953 | 0.1546409 | 0.32770262 | lin-13 |
| WBGene00003003 | 0.768347863 | 9.87E-06 | 0.00012202 | lin-14 |
| WBGene00003006 | -0.280572261 | 0.19658144 | 0.3841045 | lin-17 |
| WBGene00003007 | -0.121984532 | 0.52377763 | 0.70135617 | lin-18 |
| WBGene00003008 | 0.071810825 | 0.50121915 | 0.6828513 | lin-22 |
| WBGene00003009 | -0.109720179 | 0.11449256 | 0.26936936 | lin-23 |
| WBGene00003010 | -0.047925514 | 0.66409203 | 0.80296004 | lin-24 |
| WBGene00003011 | -0.30553624 | 0.05386222 | 0.15899127 | lin-25 |
| WBGene00003012 | -0.254818767 | 0.00646716 | 0.03201533 | lin-26 |
| WBGene00003014 | -0.068644307 | 0.52695815 | 0.7041981 | lin-28 |
| WBGene00003015 | -0.071816875 | 0.72493429 | 0.84572464 | lin-29 |
| WBGene00003017 | 0.111543706 | 0.48431136 | 0.6695121 | lin-31 |
| WBGene00003018 | -0.085611125 | 0.53226715 | 0.70853237 | lin-32 |
| WBGene00003019 | 0.040412264 | 0.84074392 | 0.9139688 | lin-33 |
| WBGene00003020 | -0.188953825 | 0.0130894 | 0.05548382 | lin-35 |
| WBGene00003021 | -0.078231436 | 0.30534026 | 0.50592194 | lin-36 |
| WBGene00003022 | -0.226728842 | 0.03250323 | 0.10970026 | lin-37 |
| WBGene00003024 | 0.357890632 | 0.1227963 | 0.28238651 | lin-39 |
| WBGene00003025 | 0.079131407 | 0.31245388 | 0.51325449 | lin-40 |
| WBGene00003026 | -0.090743664 | 0.11837792 | 0.2757031 | lin-41 |
| WBGene00003029 | 0.31466939 | 0.11751013 | 0.27395953 | lin-44 |
| WBGene00003030 | -0.142900969 | 0.06708412 | 0.18562158 | lin-45 |
| WBGene00003031 | 0.234549636 | 0.2252617 | 0.4182697 | lin-46 |
| WBGene00003033 | 0.015899792 | 0.91775838 | 0.95418978 | lin-48 |
| WBGene00003034 | -0.177345599 | 0.06844287 | 0.18839482 | lin-49 |
| WBGene00003035 | 0.071507263 | 0.60710916 | 0.76297727 | lin-52 |
| WBGene00003036 | -0.223527293 | 0.00037588 | 0.00304891 | lin-53 |
| WBGene00003037 | -0.153211086 | 0.08367706 | 0.21752767 | lin-54 |
| WBGene00003038 | -0.229387234 | 0.1322994 | 0.29556762 | lin-56 |

| | | | | |
|---|---|---|---|---|
| WBGene00003039 | NA | NA | NA | NA |
| WBGene00003040 | 0.186740994 | 0.07716597 | 0.20560272 | lin-59 |
| WBGene00003041 | -0.149631654 | 0.13149369 | 0.29428297 | lin-61 |
| WBGene00003043 | -0.097192729 | 0.28553556 | 0.48576878 | lip-1 |
| WBGene00003044 | 0.094965391 | 0.34445499 | 0.54420692 | lir-1 |
| WBGene00003045 | -0.162249151 | 0.2546325 | 0.45298595 | lir-2 |
| WBGene00003046 | 0.125651198 | 0.54897241 | 0.7216939 | lir-3 |
| WBGene00003047 | -0.09788306 | 0.34594989 | 0.54556789 | lis-1 |
| WBGene00003048 | 0.002041422 | 0.98429441 | 0.99099536 | lit-1 |
| WBGene00003052 | -0.136715119 | 0.00759075 | 0.03627376 | lmn-1 |
| WBGene00003053 | -0.120084104 | 0.12432542 | 0.2845917 | lmp-1 |
| WBGene00003055 | -0.401275777 | 0.00160365 | 0.01041201 | lon-1 |
| WBGene00003056 | -0.086120913 | 0.59713292 | 0.75563078 | lon-2 |
| WBGene00003057 | 0.466694148 | 0.03777975 | 0.12267318 | lon-3 |
| WBGene00003058 | NA | NA | NA | lov-1 |
| WBGene00003059 | -0.324769772 | 0.00469266 | 0.02488418 | lpd-2 |
| WBGene00003060 | 0.10620712 | 0.17851275 | 0.36000072 | lpd-3 |
| WBGene00003061 | -0.267547306 | 0.0009229 | 0.00657525 | lpd-5 |
| WBGene00003062 | -0.24697051 | 0.00209118 | 0.01295113 | lpd-6 |
| WBGene00003063 | -0.124807447 | 0.08470912 | 0.21927671 | lpd-7 |
| WBGene00003064 | -0.050320837 | 0.60002881 | 0.75785213 | lpd-8 |
| WBGene00003065 | -0.07258498 | 0.46943613 | 0.65798188 | lpd-9 |
| WBGene00003066 | -0.092711833 | 0.51795455 | 0.69719339 | lpl-1 |
| WBGene00003067 | 0.459772726 | 0.01662759 | 0.06708308 | NA |
| WBGene00003068 | -0.046177127 | 0.70267831 | 0.83114502 | lrk-1 |
| WBGene00003071 | 0.30252065 | 0.00493573 | 0.02591493 | lrp-1 |
| WBGene00003072 | 0.032038934 | 0.75601193 | 0.8618244 | lrp-2 |
| WBGene00003073 | -0.256907243 | 2.65E-05 | 0.00029431 | lars-1 |
| WBGene00003074 | -0.035656227 | 0.75035204 | 0.85901198 | lars-2 |
| WBGene00003075 | 0.102923266 | 0.39605932 | 0.59403606 | lrx-1 |
| WBGene00003076 | -0.20924323 | 0.04401524 | 0.13709778 | lsm-1 |
| WBGene00003077 | -0.257019762 | 0.04405972 | 0.13713401 | lsm-3 |
| WBGene00003078 | -0.221713883 | 0.12869025 | 0.29040597 | lsm-4 |
| WBGene00003079 | -0.331227897 | 0.00881324 | 0.04076926 | lsm-5 |
| WBGene00003080 | 0.239866096 | 0.09436927 | 0.23549692 | lsm-6 |
| WBGene00003081 | -0.021190771 | 0.84676305 | 0.91761389 | lsm-7 |
| WBGene00003082 | -0.478160361 | 1.37E-05 | 0.00016175 | lsm-8 |
| WBGene00003083 | -0.157911104 | 0.06592391 | 0.18332232 | lst-1 |
| WBGene00003084 | -0.04598034 | 0.74577372 | 0.85611361 | lst-2 |
| WBGene00003085 | -0.054370343 | 0.33468665 | 0.53413188 | ccar-1 |
| WBGene00003086 | -0.127331429 | 0.1405729 | 0.30822025 | lst-4 |
| WBGene00003087 | -0.208785033 | 0.1636022 | 0.33922955 | lsy-2 |
| WBGene00003088 | NA | NA | NA | NA |

| | | | | |
|---|---|---|---|---|
| WBGene00003089 | 0.074629982 | 0.7217395 | 0.84380644 | ltd-1 |
| WBGene00003090 | 0.423172697 | 2.85E-08 | 5.91E-07 | lys-1 |
| WBGene00003091 | -0.558745875 | 0.00022856 | 0.00197421 | lys-2 |
| WBGene00003092 | 0.158529162 | 0.27740442 | 0.47747657 | lys-3 |
| WBGene00003093 | 3.277165419 | 0 | 0 | lys-4 |
| WBGene00003094 | 3.162427553 | 1.14E-05 | 0.00013853 | lys-5 |
| WBGene00003095 | 2.222988561 | 8.37E-08 | 1.60E-06 | lys-6 |
| WBGene00003096 | 3.324303352 | 0 | 0 | lys-7 |
| WBGene00003097 | 0.922818134 | 8.12E-23 | 7.00E-21 | lys-8 |
| WBGene00003098 | 6.989180843 | 2.01E-07 | 3.58E-06 | lys-9 |
| WBGene00003099 | 5.021818051 | 1.11E-24 | 1.02E-22 | lys-10 |
| WBGene00003100 | 0.006757176 | 0.95198564 | 0.97244274 | mab-3 |
| WBGene00003102 | 0.320075661 | 0.14878616 | 0.31960586 | mab-5 |
| WBGene00003104 | 0.156379156 | 0.38440829 | 0.58374211 | mab-7 |
| WBGene00003106 | 0.024510693 | 0.84702992 | 0.91761389 | mab-9 |
| WBGene00003107 | 0.119982374 | 0.51519669 | 0.69495068 | mab-10 |
| WBGene00003111 | -0.413969947 | 0.00723456 | 0.03489988 | mab-20 |
| WBGene00003112 | 0.451659717 | 0.0852108 | 0.22003021 | mab-21 |
| WBGene00003114 | -0.039636691 | 0.54579661 | NA | mab-23 |
| WBGene00003119 | -0.263158627 | 0.01514412 | 0.06231733 | mac-1 |
| WBGene00003123 | 0.044446078 | 0.59280342 | 0.75314295 | mag-1 |
| WBGene00003124 | 0.075861705 | 0.71491437 | 0.8401205 | mai-1 |
| WBGene00003129 | -0.097380074 | 0.46766454 | 0.65691994 | map-1 |
| WBGene00003130 | -0.360526363 | 1.32E-05 | 0.00015721 | map-2 |
| WBGene00003132 | -0.09348746 | 0.15988953 | 0.33461158 | mat-1 |
| WBGene00003133 | -0.137536136 | 0.07080801 | 0.19298214 | mat-2 |
| WBGene00003134 | -0.133247177 | 0.20125424 | 0.38993842 | mat-3 |
| WBGene00003136 | -0.07133453 | 0.50089678 | 0.68269142 | mau-2 |
| WBGene00003142 | -0.077876164 | 0.55626369 | 0.72794301 | mau-8 |
| WBGene00003143 | 0.374280332 | 0.12126378 | 0.2803447 | max-1 |
| WBGene00003144 | -0.027815964 | 0.70339997 | 0.83172985 | max-2 |
| WBGene00003148 | -0.293904653 | 0.00094079 | 0.00669029 | mbf-1 |
| WBGene00003149 | 0.160262254 | 0.22076189 | 0.4132159 | mbk-1 |
| WBGene00003150 | -0.087393778 | 0.08980956 | 0.2277675 | mbk-2 |
| WBGene00003151 | 0.307380074 | 9.86E-05 | 0.00095996 | mca-1 |
| WBGene00003153 | -0.075957956 | 0.19921081 | 0.3876456 | mca-3 |
| WBGene00003154 | -0.150827421 | 0.00483161 | 0.02547339 | mcm-2 |
| WBGene00003155 | -0.040953178 | 0.46251826 | 0.65252509 | mcm-3 |
| WBGene00003156 | -0.141911483 | 0.0164658 | 0.06656443 | mcm-4 |
| WBGene00003157 | -0.148885038 | 0.00495004 | 0.02595754 | mcm-5 |
| WBGene00003158 | -0.119060663 | 0.04822289 | 0.14638841 | mcm-6 |
| WBGene00003159 | -0.311368451 | 5.18E-09 | 1.20E-07 | mcm-7 |
| WBGene00003160 | -0.132867018 | 0.10150459 | 0.24793605 | mdf-1 |

| | | | | |
|---|---|---|---|---|
| WBGene00003161 | -0.203192867 | 0.01782178 | 0.07065493 | mdf-2 |
| WBGene00003162 | -0.424207253 | 2.97E-15 | 1.45E-13 | mdh-2 |
| WBGene00003163 | 0.117755861 | 0.52047682 | 0.69923849 | mdl-1 |
| WBGene00003164 | -0.152048282 | 0.1173687 | 0.27380421 | mdt-6 |
| WBGene00003165 | 0.678012209 | 0.02284955 | 0.08480109 | mec-1 |
| WBGene00003166 | 0.441515004 | 0.08417205 | 0.21842663 | mec-2 |
| WBGene00003167 | 0.035818975 | 0.78840203 | 0.88033682 | mec-3 |
| WBGene00003168 | 0.208035801 | 0.06528848 | 0.18214342 | mec-4 |
| WBGene00003169 | 0.103120334 | 0.58393056 | 0.74771098 | mec-5 |
| WBGene00003170 | 0.052935385 | 0.75333827 | 0.86027811 | mec-6 |
| WBGene00003171 | 0.4747059 | 0.03582681 | 0.11798028 | mec-7 |
| WBGene00003172 | -0.210584174 | 0.17931493 | 0.36082247 | mec-8 |
| WBGene00003173 | 0.4621886 | 0.07995387 | 0.21103381 | mec-9 |
| WBGene00003174 | 1.123055209 | 0.01453759 | 0.06034729 | mec-10 |
| WBGene00003175 | 0.555667221 | 0.00739395 | 0.03550653 | mec-12 |
| WBGene00003176 | 0.015507174 | 0.80923808 | NA | mec-14 |
| WBGene00003177 | -0.067775908 | 0.43814864 | 0.63294862 | mec-15 |
| WBGene00003178 | 0.196804934 | 0.30635623 | 0.50701475 | mec-17 |
| WBGene00003179 | 0.082290045 | 0.65336343 | 0.79469207 | mec-18 |
| WBGene00003180 | -0.05445452 | 0.53143225 | NA | med-1 |
| WBGene00003181 | 0.040529223 | 0.56245259 | NA | med-2 |
| WBGene00003182 | 0.923641048 | 0.00017416 | 0.00156725 | mef-2 |
| WBGene00003183 | -0.130409496 | 0.13369582 | 0.29743895 | mei-1 |
| WBGene00003184 | 0.030311209 | 0.73343708 | 0.85087141 | mei-2 |
| WBGene00003185 | 0.167118002 | 0.02874098 | 0.10050078 | mek-1 |
| WBGene00003186 | -0.081028116 | 0.47236862 | 0.66042293 | mek-2 |
| WBGene00003196 | 0.041344076 | 0.58889758 | 0.75085338 | mel-11 |
| WBGene00003209 | -0.191167335 | 0.00157206 | 0.01022046 | mel-26 |
| WBGene00003210 | -0.110694639 | 0.08823993 | 0.22534813 | mel-28 |
| WBGene00003214 | 0.065567954 | 0.17609562 | 0.35699473 | mel-32 |
| WBGene00003216 | -0.370283435 | 0.00021662 | 0.00188891 | memi-1 |
| WBGene00003218 | -0.290031518 | 2.63E-06 | 3.68E-05 | mep-1 |
| WBGene00003219 | -0.15862601 | 0.2316731 | 0.42654268 | mes-1 |
| WBGene00003220 | -0.020320937 | 0.85310977 | 0.92125536 | mes-2 |
| WBGene00003221 | 0.119389803 | 0.20843632 | 0.39762802 | mes-3 |
| WBGene00003222 | -0.017244957 | 0.84480759 | 0.91675274 | mes-4 |
| WBGene00003224 | -0.030299127 | 0.7769987 | 0.87386987 | mes-6 |
| WBGene00003225 | -0.150709628 | 0.12743595 | 0.28874635 | mev-1 |
| WBGene00003228 | -0.079417487 | 0.20569091 | 0.39466399 | mex-1 |
| WBGene00003229 | -0.09665562 | 0.05637354 | 0.16451366 | mex-3 |
| WBGene00003230 | -0.081471602 | 0.07936162 | 0.20988682 | mex-5 |
| WBGene00003231 | -0.003354617 | 0.9460003 | 0.96977947 | mex-6 |
| WBGene00003232 | 0.114405274 | 0.47407494 | 0.66173431 | mgl-1 |

| | | | | |
|---|---|---|---|---|
| WBGene00003233 | 0.287457599 | 0.20710976 | 0.39643012 | mgl-2 |
| WBGene00003234 | 0.504244082 | 0.00528024 | 0.02730007 | mif-1 |
| WBGene00003235 | -0.028677092 | 0.82816547 | 0.90617864 | mif-2 |
| WBGene00003236 | -0.135831114 | 0.37588726 | 0.57500725 | mif-3 |
| WBGene00003237 | 0.093276871 | 0.45181988 | 0.64411829 | mif-4 |
| WBGene00003238 | 0.129205478 | 0.50093258 | 0.68269142 | mig-1 |
| WBGene00003239 | -0.158482014 | 0.3014551 | 0.50211651 | mig-2 |
| WBGene00003241 | -0.104256959 | 0.17535463 | 0.35594084 | mig-5 |
| WBGene00003242 | 0.454114288 | 1.20E-09 | 3.01E-08 | mig-6 |
| WBGene00003243 | 0.130883583 | 0.42230677 | 0.61854675 | mig-10 |
| WBGene00003245 | -0.019960757 | 0.92074003 | 0.9557288 | mig-13 |
| WBGene00003246 | -0.113136067 | 0.34303034 | 0.54313812 | mig-14 |
| WBGene00003247 | -0.16424985 | 0.22288748 | 0.41556803 | mig-15 |
| WBGene00003248 | 0.563903607 | 0.02885917 | 0.10076964 | mig-17 |
| WBGene00003249 | 1.16357479 | 0.00029508 | 0.00247725 | mig-18 |
| WBGene00003252 | -0.08541265 | 0.56136662 | 0.73181094 | mig-21 |
| WBGene00003253 | -0.288907635 | 0.00025198 | 0.00215244 | mig-22 |
| WBGene00003254 | -0.048347854 | 0.75798606 | 0.86309488 | mig-23 |
| WBGene00003258 | -0.00621531 | 0.95036944 | 0.97182989 | cogc-1 |
| WBGene00003260 | NA | NA | NA | NA |
| WBGene00003261 | NA | NA | NA | NA |
| WBGene00003262 | NA | NA | NA | NA |
| WBGene00003263 | NA | NA | NA | NA |
| WBGene00003264 | NA | NA | NA | NA |
| WBGene00003265 | NA | NA | NA | NA |
| WBGene00003266 | NA | NA | NA | NA |
| WBGene00003267 | NA | NA | NA | NA |
| WBGene00003268 | NA | NA | NA | NA |
| WBGene00003269 | NA | NA | NA | NA |
| WBGene00003270 | NA | NA | NA | NA |
| WBGene00003271 | NA | NA | NA | NA |
| WBGene00003272 | NA | NA | NA | NA |
| WBGene00003273 | 0.020925399 | 0.84643798 | NA | NA |
| WBGene00003274 | NA | NA | NA | NA |
| WBGene00003275 | NA | NA | NA | NA |
| WBGene00003277 | NA | NA | NA | NA |
| WBGene00003278 | NA | NA | NA | NA |
| WBGene00003279 | NA | NA | NA | NA |
| WBGene00003280 | NA | NA | NA | NA |
| WBGene00003281 | NA | NA | NA | NA |
| WBGene00003282 | NA | NA | NA | NA |
| WBGene00003283 | NA | NA | NA | NA |
| WBGene00003284 | NA | NA | NA | NA |

| | | | | |
|---|---|---|---|---|
| WBGene00003285 | NA | NA | NA | NA |
| WBGene00003286 | NA | NA | NA | NA |
| WBGene00003287 | NA | NA | NA | NA |
| WBGene00003288 | NA | NA | NA | NA |
| WBGene00003289 | NA | NA | NA | NA |
| WBGene00003290 | NA | NA | NA | NA |
| WBGene00003291 | NA | NA | NA | NA |
| WBGene00003292 | NA | NA | NA | NA |
| WBGene00003293 | NA | NA | NA | NA |
| WBGene00003294 | NA | NA | NA | NA |
| WBGene00003295 | NA | NA | NA | NA |
| WBGene00003296 | NA | NA | NA | NA |
| WBGene00003297 | NA | NA | NA | NA |
| WBGene00003298 | -0.023056499 | 0.78121253 | NA | NA |
| WBGene00003299 | NA | NA | NA | NA |
| WBGene00003300 | NA | NA | NA | NA |
| WBGene00003301 | NA | NA | NA | NA |
| WBGene00003302 | NA | NA | NA | NA |
| WBGene00003303 | NA | NA | NA | NA |
| WBGene00003304 | NA | NA | NA | NA |
| WBGene00003305 | NA | NA | NA | NA |
| WBGene00003306 | NA | NA | NA | NA |
| WBGene00003307 | NA | NA | NA | NA |
| WBGene00003308 | NA | NA | NA | NA |
| WBGene00003309 | NA | NA | NA | NA |
| WBGene00003310 | NA | NA | NA | NA |
| WBGene00003311 | NA | NA | NA | NA |
| WBGene00003312 | NA | NA | NA | NA |
| WBGene00003313 | NA | NA | NA | NA |
| WBGene00003314 | -0.001993815 | 0.95612996 | NA | NA |
| WBGene00003315 | NA | NA | NA | NA |
| WBGene00003316 | NA | NA | NA | NA |
| WBGene00003317 | NA | NA | NA | NA |
| WBGene00003318 | NA | NA | NA | NA |
| WBGene00003319 | NA | NA | NA | NA |
| WBGene00003321 | NA | NA | NA | NA |
| WBGene00003322 | NA | NA | NA | NA |
| WBGene00003323 | NA | NA | NA | NA |
| WBGene00003324 | NA | NA | NA | NA |
| WBGene00003325 | NA | NA | NA | NA |
| WBGene00003326 | NA | NA | NA | NA |
| WBGene00003327 | NA | NA | NA | NA |
| WBGene00003328 | NA | NA | NA | NA |

| | | | | |
|---|---|---|---|---|
| WBGene00003329 | NA | NA | NA | NA |
| WBGene00003330 | NA | NA | NA | NA |
| WBGene00003331 | 0.079798406 | 0.30849364 | NA | NA |
| WBGene00003332 | NA | NA | NA | NA |
| WBGene00003333 | NA | NA | NA | NA |
| WBGene00003334 | NA | NA | NA | NA |
| WBGene00003335 | NA | NA | NA | NA |
| WBGene00003336 | NA | NA | NA | NA |
| WBGene00003337 | 0.020925399 | 0.84643798 | NA | NA |
| WBGene00003338 | NA | NA | NA | NA |
| WBGene00003339 | NA | NA | NA | NA |
| WBGene00003340 | NA | NA | NA | NA |
| WBGene00003341 | NA | NA | NA | NA |
| WBGene00003342 | NA | NA | NA | NA |
| WBGene00003343 | NA | NA | NA | NA |
| WBGene00003344 | NA | NA | NA | NA |
| WBGene00003345 | NA | NA | NA | NA |
| WBGene00003346 | NA | NA | NA | NA |
| WBGene00003347 | NA | NA | NA | NA |
| WBGene00003348 | NA | NA | NA | NA |
| WBGene00003349 | NA | NA | NA | NA |
| WBGene00003350 | NA | NA | NA | NA |
| WBGene00003351 | NA | NA | NA | NA |
| WBGene00003352 | NA | NA | NA | NA |
| WBGene00003353 | NA | NA | NA | NA |
| WBGene00003354 | NA | NA | NA | NA |
| WBGene00003355 | NA | NA | NA | NA |
| WBGene00003356 | NA | NA | NA | NA |
| WBGene00003357 | NA | NA | NA | NA |
| WBGene00003358 | NA | NA | NA | NA |
| WBGene00003359 | NA | NA | NA | NA |
| WBGene00003360 | NA | NA | NA | NA |
| WBGene00003361 | NA | NA | NA | NA |
| WBGene00003362 | 0.020925399 | 0.84643798 | NA | NA |
| WBGene00003363 | 0.0398788 | 0.58971221 | NA | NA |
| WBGene00003364 | NA | NA | NA | NA |
| WBGene00003365 | NA | NA | NA | NA |
| WBGene00003366 | NA | NA | NA | NA |
| WBGene00003367 | -0.113988731 | 0.13607168 | 0.30108582 | mix-1 |
| WBGene00003368 | 0.230245817 | 0.24488877 | 0.4429796 | mkk-4 |
| WBGene00003369 | 0.020990356 | 0.73672278 | 0.85204655 | mlc-1 |
| WBGene00003370 | -0.118899972 | 0.02740798 | 0.0969709 | mlc-2 |
| WBGene00003371 | 0.032017395 | 0.63229067 | 0.78160821 | mlc-3 |

| | | | | |
|---|---|---|---|---|
| WBGene00003372 | 0.009456487 | 0.89069793 | 0.9430654 | mlc-4 |
| WBGene00003373 | 0.203754443 | 0.07015894 | 0.19160549 | mlh-1 |
| WBGene00003374 | 0.495801445 | 0.00467724 | 0.02481138 | mlk-1 |
| WBGene00003375 | 0.07476891 | 0.58229271 | 0.74624349 | mlp-1 |
| WBGene00003376 | -0.042082175 | 0.76848598 | 0.86844708 | mls-1 |
| WBGene00003377 | -2.141650254 | 0.00042793 | 0.00340192 | mls-2 |
| WBGene00003378 | 0.03576583 | 0.78969734 | 0.88144841 | mml-1 |
| WBGene00003380 | -0.127971537 | 0.48573247 | 0.67027418 | mnm-2 |
| WBGene00003384 | -0.310468346 | 0.10813873 | 0.25869594 | moc-1 |
| WBGene00003385 | 0.140257012 | 0.49657831 | 0.67886649 | moc-2 |
| WBGene00003386 | -0.025669709 | 0.88496961 | 0.93940914 | mod-1 |
| WBGene00003387 | 0.911535013 | 0.01436544 | 0.05976824 | mod-5 |
| WBGene00003388 | 0.884091925 | 0.00276532 | 0.01635178 | moe-3 |
| WBGene00003389 | -0.147025223 | 0.03580008 | 0.11794837 | mog-1 |
| WBGene00003390 | -0.306598873 | 0.00104089 | 0.00726409 | mog-2 |
| WBGene00003391 | -0.198213994 | 0.0390003 | 0.12560568 | mog-3 |
| WBGene00003392 | -0.102872972 | 0.28887859 | 0.48912587 | mog-4 |
| WBGene00003393 | -0.105076862 | 0.14933191 | 0.32058188 | mog-5 |
| WBGene00003394 | -0.311386071 | 0.14208996 | 0.31012806 | mom-1 |
| WBGene00003395 | -0.086558722 | 0.32959961 | 0.52913689 | mom-2 |
| WBGene00003396 | -0.000321201 | 0.99674256 | 0.9978865 | mom-4 |
| WBGene00003397 | -0.038285022 | 0.53355149 | 0.70899686 | mom-5 |
| WBGene00003400 | 0.47058602 | 0.00155974 | 0.01015846 | dapk-1 |
| WBGene00003401 | -0.216114193 | 0.00201945 | 0.01256025 | mpk-1 |
| WBGene00003402 | 1.360557568 | 4.71E-07 | 7.91E-06 | mpk-2 |
| WBGene00003403 | 0.298029921 | 0.0302759 | 0.10405388 | mps-1 |
| WBGene00003404 | 0.877661204 | 0.00010958 | 0.00105302 | mpz-1 |
| WBGene00003405 | 0.070370399 | 0.38474239 | 0.58409511 | mre-11 |
| WBGene00003406 | -0.226057717 | 0.0029222 | 0.01709307 | mrg-1 |
| WBGene00003407 | 0.157264148 | 0.18424659 | 0.36766449 | mrp-1 |
| WBGene00003408 | 0.40851331 | 0.0339686 | 0.11334213 | mrp-2 |
| WBGene00003409 | 0.568521093 | 0.00176255 | 0.01124914 | mrp-3 |
| WBGene00003410 | -0.194641398 | 0.00254993 | 0.01525688 | mrp-4 |
| WBGene00003411 | -0.006568245 | 0.93800266 | 0.96517113 | mrp-5 |
| WBGene00003412 | 0.071316993 | 0.71152824 | 0.8376204 | mrp-6 |
| WBGene00003413 | 0.014553989 | 0.85261688 | 0.92106277 | mrp-7 |
| WBGene00003414 | -0.034244275 | 0.8572364 | 0.92325428 | mrp-8 |
| WBGene00003415 | -0.249086015 | 6.17E-05 | 0.00063431 | mars-1 |
| WBGene00003417 | -0.127644622 | 0.42470099 | 0.62055982 | mrt-2 |
| WBGene00003418 | -0.100921082 | 0.16998105 | 0.34842782 | msh-2 |
| WBGene00003420 | -0.10638375 | 0.60827689 | 0.76399837 | NA |
| WBGene00003421 | -0.052343703 | 0.65398569 | 0.7949984 | msh-5 |
| WBGene00003422 | -0.027444344 | 0.69313219 | 0.82398087 | msh-6 |

| | | | | |
|---|---|---|---|---|
| WBGene00003423 | 0.184788527 | 0.13446031 | 0.29873041 | msi-1 |
| WBGene00003424 | 1.474238239 | 0.00621614 | 0.0309981 | msp-3 |
| WBGene00003425 | 0.090723942 | 0.20045316 | NA | msp-10 |
| WBGene00003426 | 2.43659394 | 5.40E-05 | 0.0005606 | msp-19 |
| WBGene00003429 | 1.094977804 | 0.00261125 | 0.01557283 | msp-31 |
| WBGene00003430 | NA | NA | NA | msp-32 |
| WBGene00003431 | 1.633621355 | 0.00013909 | 0.00128402 | msp-33 |
| WBGene00003432 | 1.040046445 | 0.00649474 | 0.03212483 | msp-36 |
| WBGene00003433 | NA | NA | NA | NA |
| WBGene00003434 | 1.302589926 | 0.01068896 | 0.0470668 | msp-38 |
| WBGene00003435 | 1.549243455 | 9.54E-05 | 0.0009309 | msp-40 |
| WBGene00003437 | 0.05366068 | 0.40968658 | NA | NA |
| WBGene00003438 | 0.909163932 | 0.0171561 | 0.06861036 | msp-45 |
| WBGene00003442 | 0.203343447 | 0.30974307 | 0.5105386 | msp-49 |
| WBGene00003443 | 0.804598225 | 0.0168001 | 0.06756644 | msp-50 |
| WBGene00003444 | 1.887074084 | 2.43E-05 | 0.00027152 | msp-51 |
| WBGene00003445 | NA | NA | NA | NA |
| WBGene00003446 | 1.188988541 | 0.00719976 | 0.03477784 | msp-53 |
| WBGene00003448 | 2.489100008 | 1.19E-05 | 0.0001434 | msp-55 |
| WBGene00003449 | 0.677360714 | 0.02841269 | 0.09963837 | msp-56 |
| WBGene00003450 | 1.652327508 | 0.00048069 | 0.0037655 | msp-57 |
| WBGene00003451 | NA | NA | NA | msp-58 |
| WBGene00003452 | 1.825015478 | 1.33E-05 | 0.00015781 | msp-59 |
| WBGene00003456 | NA | NA | NA | msp-63 |
| WBGene00003457 | 0.517037422 | 0.05714433 | 0.16595987 | msp-64 |
| WBGene00003458 | 0.471157867 | 0.04457432 | 0.13827441 | msp-65 |
| WBGene00003460 | 0.039034462 | 0.57625038 | NA | NA |
| WBGene00003462 | 0.0398788 | 0.58971221 | NA | msp-74 |
| WBGene00003462 | 0.0398788 | 0.58971221 | NA | F58A6.9 |
| WBGene00003463 | 0.377299739 | 0.12648466 | 0.28757277 | msp-76 |
| WBGene00003464 | 1.088720643 | 0.00841188 | 0.03931975 | msp-77 |
| WBGene00003465 | 0.705777002 | 0.03132307 | 0.10657465 | msp-78 |
| WBGene00003466 | 0.253787807 | 0.10508675 | 0.25347201 | msp-79 |
| WBGene00003467 | 1.365316442 | 0.00071063 | 0.00524147 | msp-81 |
| WBGene00003468 | 0.314514949 | 0.06554172 | 0.18274583 | msp-113 |
| WBGene00003469 | 1.903163542 | 0.00335883 | 0.01909031 | msp-142 |
| WBGene00003470 | 0.566370648 | 0.04870002 | 0.14746418 | msp-152 |
| WBGene00003471 | 0.41149006 | 0.07951354 | 0.21021268 | mtd-1 |
| WBGene00003472 | -0.081545347 | 0.38808462 | 0.58667495 | mtk-1 |
| WBGene00003473 | 1.775451189 | 0.00099434 | 0.00700924 | mtl-1 |
| WBGene00003474 | -0.106021429 | 0.23609936 | 0.43192937 | mtl-2 |
| WBGene00003475 | 0.084003939 | 0.56819143 | 0.73716355 | mtm-1 |
| WBGene00003476 | 0.073210882 | 0.19706668 | 0.38459787 | mtm-3 |

| | | | | |
|---|---|---|---|---|
| WBGene00003477 | -0.007218274 | 0.9580291 | 0.97694406 | mtm-5 |
| WBGene00003478 | -0.008695566 | 0.93458207 | 0.96313199 | mtm-6 |
| WBGene00003479 | 0.306366401 | 0.02096264 | 0.07945118 | mtm-9 |
| WBGene00003480 | 0.572193967 | 0.0299712 | 0.10343186 | klf-3 |
| WBGene00003482 | -0.710208689 | 1.02E-06 | 1.61E-05 | mua-3 |
| WBGene00003485 | -0.793006645 | 3.69E-10 | 9.81E-09 | mua-6 |
| WBGene00003495 | -0.196850829 | 0.02435436 | 0.08900953 | mup-2 |
| WBGene00003497 | -0.193112256 | 0.04201293 | 0.13248144 | mup-4 |
| WBGene00003499 | -0.078912835 | 0.43235811 | 0.62736252 | mut-2 |
| WBGene00003504 | -0.14570742 | 0.0920932 | 0.23147407 | mut-7 |
| WBGene00003507 | -0.233558714 | 0.05557331 | 0.16266469 | mut-14 |
| WBGene00003508 | -0.023387671 | 0.7782205 | 0.87443793 | mut-16 |
| WBGene00003509 | -0.020180288 | 0.86552528 | 0.92809516 | mxl-1 |
| WBGene00003510 | 0.134722404 | 0.35491882 | 0.5548117 | mxl-2 |
| WBGene00003511 | -0.407270534 | 0.00305202 | 0.01771796 | mxl-3 |
| WBGene00003514 | 1.326214567 | 1.07E-25 | 1.04E-23 | myo-2 |
| WBGene00003515 | 0.550719412 | 4.85E-06 | 6.36E-05 | myo-3 |
| WBGene00003516 | 0.281973638 | 0.17383926 | 0.35433393 | nab-1 |
| WBGene00003517 | 0.262755085 | 0.19437305 | 0.38124084 | nac-1 |
| WBGene00003518 | 0.296476743 | 0.15610336 | 0.32975175 | nac-2 |
| WBGene00003519 | 0.290545846 | 0.10146868 | 0.24788969 | nac-3 |
| WBGene00003520 | 0.017708609 | 0.82438962 | NA | nas-1 |
| WBGene00003521 | NA | NA | NA | nas-2 |
| WBGene00003522 | -0.380635059 | 0.12061439 | 0.27938963 | nas-3 |
| WBGene00003523 | -0.087499584 | 0.55415905 | 0.72629511 | nas-4 |
| WBGene00003524 | 0.007316719 | 0.96985704 | 0.98346632 | nas-5 |
| WBGene00003525 | 0.12905577 | 0.23139092 | 0.42640584 | nas-6 |
| WBGene00003526 | -0.054169556 | 0.78804557 | 0.88005428 | nas-7 |
| WBGene00003527 | 0.067954055 | 0.58910847 | 0.751016 | nas-8 |
| WBGene00003528 | -0.15386976 | 0.4427073 | 0.63670704 | nas-9 |
| WBGene00003529 | -0.001842784 | 0.95621055 | NA | nas-10 |
| WBGene00003530 | -0.486683307 | 0.00116207 | 0.00796529 | nas-11 |
| WBGene00003531 | 0.457127889 | 0.04319937 | 0.13526131 | nas-12 |
| WBGene00003532 | -0.00856524 | 0.92861797 | 0.95995875 | nas-13 |
| WBGene00003533 | 0.077469432 | 0.56952839 | 0.73824393 | nas-14 |
| WBGene00003534 | 0.095053003 | 0.34474636 | 0.54449098 | nas-15 |
| WBGene00003535 | NA | NA | NA | nas-16 |
| WBGene00003536 | 0.020232312 | 0.84643798 | NA | nas-17 |
| WBGene00003537 | -0.040794878 | 0.53203165 | NA | nas-18 |
| WBGene00003538 | 0.033463284 | 0.6365037 | NA | nas-19 |
| WBGene00003539 | -0.256749589 | 0.17376249 | 0.35422677 | nas-20 |
| WBGene00003540 | 0.065014541 | 0.37399213 | NA | nas-21 |
| WBGene00003541 | 0.227044512 | 0.12471924 | 0.28518106 | nas-22 |

| | | | | |
|---|---|---|---|---|
| WBGene00003542 | 0.020317245 | 0.84643798 | NA | nas-23 |
| WBGene00003543 | 0.020232312 | 0.84643798 | NA | nas-24 |
| WBGene00003544 | 3.253281937 | 0.00083624 | 0.00604316 | nas-25 |
| WBGene00003545 | -0.004047623 | 0.95527667 | NA | nas-27 |
| WBGene00003546 | 0.089530144 | 0.45239285 | 0.64474678 | nas-28 |
| WBGene00003547 | 0.155329439 | 0.28170043 | 0.48136599 | nas-29 |
| WBGene00003548 | 1.187156544 | 0.01228964 | 0.0526889 | nas-30 |
| WBGene00003549 | 0.772179285 | 0.01552179 | 0.0636211 | nas-31 |
| WBGene00003550 | 1.710903588 | 0.00133472 | 0.00891498 | nas-32 |
| WBGene00003551 | 1.098262874 | 0.01184883 | 0.05109039 | nas-33 |
| WBGene00003552 | 0.06931055 | 0.68374007 | 0.81761922 | nas-36 |
| WBGene00003553 | 0.929799502 | 1.09E-08 | 2.40E-07 | nas-37 |
| WBGene00003554 | 0.156265885 | 0.25060284 | 0.44909133 | nas-38 |
| WBGene00003555 | 0.693990989 | 0.0056129 | 0.02866357 | nas-39 |
| WBGene00003556 | NA | NA | NA | NA |
| WBGene00003558 | -0.16384618 | 0.43760265 | 0.63253434 | nca-2 |
| WBGene00003559 | -0.055095778 | 0.60330014 | 0.75988622 | ncl-1 |
| WBGene00003561 | 0.907493667 | 1.50E-06 | 2.25E-05 | ncr-1 |
| WBGene00003562 | 0.927312503 | 0.01227057 | 0.05263796 | ncr-2 |
| WBGene00003563 | 0.201909316 | 0.21417922 | 0.40507948 | ncs-1 |
| WBGene00003564 | 0.708363304 | 5.43E-14 | 2.34E-12 | ncs-2 |
| WBGene00003565 | 0.188592179 | 0.38495078 | 0.58429334 | ncs-3 |
| WBGene00003566 | 0.693419518 | 1.86E-06 | 2.74E-05 | ncx-1 |
| WBGene00003567 | 0.764063341 | 1.55E-09 | 3.84E-08 | ncx-2 |
| WBGene00003568 | 0.077607035 | 0.65313352 | 0.79455985 | ncx-3 |
| WBGene00003569 | 1.778784156 | 0.00011364 | 0.00108672 | ncx-4 |
| WBGene00003570 | 0.033463284 | 0.6365037 | NA | ncx-5 |
| WBGene00003571 | 0.038053842 | 0.85487739 | 0.92241579 | ncx-6 |
| WBGene00003572 | 0.109173499 | 0.53902239 | 0.71373539 | ncx-7 |
| WBGene00003573 | 0.676674984 | 0.02666894 | 0.09532712 | ncx-8 |
| WBGene00003574 | 0.159347697 | 0.28801881 | 0.48834782 | ncx-9 |
| WBGene00003575 | 2.057923594 | 0.00050607 | 0.00392441 | ncx-10 |
| WBGene00003576 | -0.123166283 | 0.12659894 | 0.287682 | ndc-80 |
| WBGene00003577 | -0.116022591 | 0.36272909 | 0.56256996 | ndg-4 |
| WBGene00003578 | 0.487207285 | 0.06880932 | 0.18897717 | ndx-1 |
| WBGene00003579 | 0.30283852 | 0.1928688 | 0.37932398 | ndx-2 |
| WBGene00003580 | 0.223254556 | 0.25549571 | 0.45380659 | ndx-3 |
| WBGene00003581 | -0.115637871 | 0.47620638 | 0.66344444 | ndx-4 |
| WBGene00003582 | 0.07305012 | 0.48913093 | 0.67323737 | dcap-2 |
| WBGene00003583 | 0.13911493 | 0.41629673 | 0.61345276 | ndx-6 |
| WBGene00003584 | 0.005769425 | 0.9735366 | 0.98538191 | ndx-7 |
| WBGene00003585 | -0.191907759 | 0.36445403 | 0.56412276 | ndx-8 |
| WBGene00003586 | -0.147170745 | 0.19436788 | 0.38124084 | ndx-9 |

| | | | | |
|---|---|---|---|---|
| WBGene00003587 | -0.096741234 | 0.32422494 | 0.52363682 | ned-8 |
| WBGene00003588 | -0.07656372 | 0.31033395 | 0.51114853 | nex-1 |
| WBGene00003589 | 0.186621377 | 0.17535696 | 0.35594084 | nex-2 |
| WBGene00003590 | -0.296947019 | 0.10342078 | 0.25103223 | nex-3 |
| WBGene00003591 | 0.115720028 | 0.50348527 | 0.68467923 | nex-4 |
| WBGene00003592 | -0.267236996 | 0.00242605 | 0.0146402 | nfi-1 |
| WBGene00003593 | 0.369992511 | 0.03144986 | 0.10690908 | nfm-1 |
| WBGene00003594 | 0.058592733 | 0.64354287 | 0.78898938 | nft-1 |
| WBGene00003595 | -0.153554953 | 0.44021996 | 0.63450679 | ngn-1 |
| WBGene00003596 | -0.296680407 | 0.00059832 | 0.00453888 | ngp-1 |
| WBGene00003597 | 0.785853911 | 0.00063002 | 0.00474247 | nhl-1 |
| WBGene00003598 | -0.096920387 | 0.12287551 | 0.28250947 | nhl-2 |
| WBGene00003599 | 0.303775155 | 0.07263715 | 0.19646786 | nhl-3 |
| WBGene00003600 | 0.068737463 | 0.65842922 | 0.79870748 | nhr-1 |
| WBGene00003601 | -0.744397875 | 2.42E-06 | 3.43E-05 | nhr-2 |
| WBGene00003602 | 1.238043055 | 5.42E-08 | 1.07E-06 | nhr-3 |
| WBGene00003603 | 0.260937175 | 0.12968376 | 0.29201626 | nhr-4 |
| WBGene00003604 | 0.16555761 | 0.4150607 | 0.6125911 | nhr-5 |
| WBGene00003605 | 1.09436818 | 0.00464597 | 0.02471217 | nhr-6 |
| WBGene00003606 | 0.219336325 | 0.32139908 | 0.52145434 | nhr-7 |
| WBGene00003607 | 0.429929504 | 0.02017356 | 0.07717428 | nhr-8 |
| WBGene00003608 | 0.020317245 | 0.84643798 | NA | nhr-9 |
| WBGene00003609 | -0.032492526 | 0.81692296 | 0.89935852 | nhr-10 |
| WBGene00003610 | 0.031916165 | 0.86816794 | 0.9299032 | nhr-11 |
| WBGene00003611 | -0.226588572 | 0.22392322 | 0.41689469 | nhr-12 |
| WBGene00003612 | -0.012056221 | 0.93971131 | 0.96610584 | nhr-13 |
| WBGene00003613 | 0.167621354 | 0.30107503 | 0.50165465 | nhr-14 |
| WBGene00003614 | -0.411240808 | 0.09757152 | 0.24196217 | nhr-15 |
| WBGene00003615 | 0.204652087 | 0.2694927 | 0.46849235 | nhr-16 |
| WBGene00003616 | -0.066866983 | 0.71419903 | 0.83968426 | nhr-17 |
| WBGene00003617 | 0.869889303 | 0.01079013 | 0.04741951 | nhr-18 |
| WBGene00003618 | 1.566935313 | 1.41E-07 | 2.59E-06 | nhr-19 |
| WBGene00003619 | -0.143181877 | 0.34336588 | 0.54313812 | nhr-20 |
| WBGene00003620 | 1.086824273 | 1.10E-05 | 0.00013461 | nhr-21 |
| WBGene00003621 | 0.173658441 | 0.31984733 | 0.51991615 | nhr-22 |
| WBGene00003622 | -0.342099176 | 0.00190432 | 0.01201213 | nhr-23 |
| WBGene00003623 | -0.802091988 | 0.0140745 | 0.05870789 | nhr-25 |
| WBGene00003624 | 0.483528326 | 0.01577079 | 0.06446124 | nhr-28 |
| WBGene00003625 | 0.346199349 | 0.02203114 | 0.08247453 | nhr-31 |
| WBGene00003626 | 0.594920576 | 0.0086141 | 0.03999752 | nhr-32 |
| WBGene00003627 | 0.350961223 | 0.032684 | 0.11015803 | nhr-34 |
| WBGene00003628 | 0.829661523 | 3.66E-05 | 0.00039449 | nhr-35 |
| WBGene00003629 | -0.023056499 | 0.78121253 | NA | nhr-38 |

| | | | | |
|---|---|---|---|---|
| WBGene00003630 | 0.190753834 | 0.25155504 | 0.44992318 | nhr-40 |
| WBGene00003632 | 1.040368551 | 0.00038067 | 0.00308262 | nhr-42 |
| WBGene00003633 | -0.53362653 | 0.05613054 | 0.16396791 | nhr-43 |
| WBGene00003634 | 0.180255238 | 0.34990488 | 0.54967354 | nhr-44 |
| WBGene00003635 | 0.521944092 | 0.01640736 | 0.06644002 | nhr-45 |
| WBGene00003636 | 0.143580863 | 0.21552096 | 0.40653717 | nhr-46 |
| WBGene00003637 | -0.230580352 | 0.18285367 | 0.36573231 | nhr-47 |
| WBGene00003638 | 0.173174983 | 0.24947224 | 0.44781778 | nhr-48 |
| WBGene00003639 | 0.091865001 | 0.27939491 | 0.47921987 | nhr-49 |
| WBGene00003640 | 0.996860698 | 0.00726035 | 0.03500126 | nhr-50 |
| WBGene00003641 | 0.033463284 | 0.6365037 | NA | nhr-51 |
| WBGene00003642 | 0.017604133 | 0.79437959 | NA | nhr-52 |
| WBGene00003643 | 0.114125284 | 0.58184008 | 0.74607818 | nhr-53 |
| WBGene00003644 | 0.249810745 | 0.16682448 | 0.34396243 | nhr-54 |
| WBGene00003645 | 0.71536461 | 0.00613262 | 0.03068613 | nhr-55 |
| WBGene00003646 | 0.380983033 | 0.12295964 | 0.28261415 | nhr-56 |
| WBGene00003647 | 0.441918732 | 0.08176482 | 0.21403875 | nhr-57 |
| WBGene00003648 | 0.524196244 | 0.05917312 | 0.16990659 | nhr-58 |
| WBGene00003649 | 0.318764139 | 0.10252135 | 0.24950298 | nhr-59 |
| WBGene00003650 | 0.239800877 | 0.12774577 | 0.28904364 | nhr-60 |
| WBGene00003651 | 0.291985181 | 0.15871229 | 0.3330875 | nhr-61 |
| WBGene00003652 | 0.378562414 | 0.10767013 | 0.25791858 | nhr-62 |
| WBGene00003653 | 0.127178381 | 0.43788417 | 0.63281632 | nhr-63 |
| WBGene00003654 | 0.222095786 | 0.28458867 | 0.48455201 | nhr-64 |
| WBGene00003655 | -0.178152768 | 0.39421852 | 0.59240002 | nhr-65 |
| WBGene00003656 | -0.037053778 | 0.68416875 | 0.8176383 | nhr-66 |
| WBGene00003657 | -0.203450515 | 0.27038176 | 0.4696116 | nhr-67 |
| WBGene00003658 | 0.430227021 | 0.00041901 | 0.00334318 | nhr-68 |
| WBGene00003659 | 0.516808532 | 0.00582788 | 0.02953099 | nhr-69 |
| WBGene00003660 | 0.033412729 | 0.86426233 | 0.92755718 | nhr-70 |
| WBGene00003661 | -0.0310046 | 0.81002262 | 0.89497909 | nhr-71 |
| WBGene00003662 | 0.090723942 | 0.20045316 | NA | nhr-72 |
| WBGene00003663 | -0.003108065 | 0.95728506 | NA | nhr-73 |
| WBGene00003664 | -0.257833644 | 0.24266549 | 0.44083966 | nhr-74 |
| WBGene00003665 | NA | NA | NA | nhr-75 |
| WBGene00003667 | NA | NA | NA | nhr-77 |
| WBGene00003668 | 0.032225462 | 0.87721322 | 0.93466842 | nhr-78 |
| WBGene00003669 | 0.401250858 | 0.09439021 | 0.23550903 | nhr-79 |
| WBGene00003670 | 0.650580914 | 0.01023874 | 0.04570284 | nhr-80 |
| WBGene00003671 | 0.037818748 | 0.65938735 | NA | nhr-81 |
| WBGene00003672 | NA | NA | NA | nhr-82 |
| WBGene00003673 | 1.642310676 | 0.00501408 | 0.02619953 | NA |
| WBGene00003674 | 0.305911926 | 0.16517578 | 0.34141236 | nhr-84 |

| | | | | |
|---|---|---|---|---|
| WBGene00003675 | 0.758015292 | 0.02668748 | 0.09534686 | nhr-85 |
| WBGene00003676 | 0.023373059 | 0.86588927 | 0.92841339 | nhr-86 |
| WBGene00003677 | 0.236822873 | 0.27802928 | 0.47777309 | nhr-87 |
| WBGene00003678 | 1.079407291 | 2.03E-07 | 3.63E-06 | nhr-88 |
| WBGene00003679 | NA | NA | NA | nhr-89 |
| WBGene00003680 | 0.600294587 | 0.02157737 | 0.08119255 | nhr-90 |
| WBGene00003681 | 0.114569223 | 0.57816473 | 0.74429877 | nhr-91 |
| WBGene00003682 | 0.30295721 | 0.1892258 | 0.37468961 | nhr-92 |
| WBGene00003683 | 0.007620206 | 0.94711745 | 0.97031324 | nhr-93 |
| WBGene00003684 | 0.154613467 | 0.45545345 | 0.6472815 | nhr-94 |
| WBGene00003685 | -0.211967807 | 0.28850169 | 0.48865725 | nhr-95 |
| WBGene00003686 | 0.493227197 | 0.06837335 | 0.18830968 | nhr-96 |
| WBGene00003687 | 0.4060995 | 0.00010066 | 0.00097794 | nhr-97 |
| WBGene00003688 | 0.218153311 | 0.2551965 | 0.45365856 | nhr-98 |
| WBGene00003689 | 2.206503937 | 2.36E-06 | 3.36E-05 | nhr-99 |
| WBGene00003690 | 0.391859332 | 0.03399918 | 0.11336414 | nhr-100 |
| WBGene00003691 | 0.517437965 | 0.01007947 | 0.0451295 | nhr-101 |
| WBGene00003692 | 0.149237782 | 0.48184594 | 0.6678599 | nhr-102 |
| WBGene00003693 | 1.181766871 | 0.00505898 | 0.02637765 | nhr-103 |
| WBGene00003694 | 0.009262836 | 0.96540224 | 0.980746 | nhr-104 |
| WBGene00003695 | 0.097553359 | 0.60605635 | 0.76212732 | nhr-105 |
| WBGene00003696 | 0.200712002 | 0.33538092 | 0.53454301 | nhr-106 |
| WBGene00003697 | 1.08248636 | 0.00081522 | 0.00591168 | nhr-107 |
| WBGene00003698 | 0.183795136 | 0.2621 | 0.45995993 | nhr-108 |
| WBGene00003699 | 0.48031343 | 0.07048925 | 0.19218598 | nhr-109 |
| WBGene00003700 | 0.352470862 | 0.12710904 | 0.28843823 | nhr-110 |
| WBGene00003701 | NA | NA | NA | nhr-111 |
| WBGene00003702 | 0.991167622 | 0.0099653 | 0.04475518 | nhr-112 |
| WBGene00003703 | 0.020317245 | 0.84643798 | NA | nhr-113 |
| WBGene00003704 | 0.585151145 | 0.00300108 | 0.01747072 | nhr-114 |
| WBGene00003705 | 0.337446225 | 0.15422295 | 0.32715947 | nhr-115 |
| WBGene00003706 | 0.210404655 | 0.29277961 | 0.49361888 | nhr-116 |
| WBGene00003707 | 0.403718554 | 0.10904919 | 0.25986152 | nhr-117 |
| WBGene00003708 | NA | NA | NA | nhr-118 |
| WBGene00003709 | 0.246205491 | 0.2003206 | 0.38887481 | nhr-119 |
| WBGene00003710 | 0.675226832 | 0.01951716 | 0.07529662 | nhr-120 |
| WBGene00003711 | 0.596019912 | 0.01404083 | 0.05861757 | nhr-121 |
| WBGene00003712 | 0.336793202 | 0.11741669 | 0.27387251 | nhr-122 |
| WBGene00003713 | 0.016239541 | 0.90653234 | 0.94941364 | nhr-123 |
| WBGene00003714 | 0.015516711 | 0.87392635 | NA | nhr-124 |
| WBGene00003715 | 0.15128992 | 0.39008425 | 0.58793736 | nhr-125 |
| WBGene00003716 | 0.415497869 | 0.10199911 | 0.24872901 | nhr-126 |
| WBGene00003717 | NA | NA | NA | nhr-127 |

| | | | | |
|---|---|---|---|---|
| WBGene00003718 | 0.16996842 | 0.38980255 | 0.58786842 | nhr-128 |
| WBGene00003719 | 0.283528955 | 0.17848019 | 0.36000072 | nhr-129 |
| WBGene00003720 | 1.842336816 | 1.75E-08 | 3.75E-07 | nhr-130 |
| WBGene00003721 | 0.251137784 | 0.26120952 | 0.45924511 | nhr-131 |
| WBGene00003722 | 0.329020104 | 0.16661096 | 0.34361897 | nhr-132 |
| WBGene00003723 | 0.223777296 | 0.31104402 | 0.51198812 | nhr-133 |
| WBGene00003724 | 0.389765444 | 0.11106474 | 0.26303767 | nhr-134 |
| WBGene00003725 | 0.295320207 | 0.20015426 | 0.3887619 | nhr-135 |
| WBGene00003726 | 0.712121233 | 0.02516965 | 0.09123765 | nhr-136 |
| WBGene00003727 | 0.151647038 | 0.4619563 | 0.65228105 | nhr-137 |
| WBGene00003728 | 0.562540863 | 0.00245025 | 0.01475696 | nhr-138 |
| WBGene00003729 | -0.291528274 | 0.20396486 | 0.393031 | nhx-1 |
| WBGene00003730 | 0.607606445 | 6.65E-07 | 1.09E-05 | nhx-2 |
| WBGene00003731 | -0.064689646 | 0.75384769 | 0.86065846 | nhx-3 |
| WBGene00003732 | 0.160369478 | 0.43380759 | 0.62886402 | nhx-4 |
| WBGene00003733 | 0.238819485 | 0.14208604 | 0.31012806 | nhx-5 |
| WBGene00003734 | -0.006015451 | 0.92944803 | NA | nhx-6 |
| WBGene00003735 | 0.377534637 | 0.02779286 | 0.09799649 | nhx-8 |
| WBGene00003736 | 0.64841079 | 0.01314802 | 0.05566807 | nhx-9 |
| WBGene00003738 | -0.012660505 | 0.89545938 | 0.94517796 | nid-1 |
| WBGene00003739 | 0.548953481 | 0.04370596 | 0.13655546 | nlp-1 |
| WBGene00003740 | 0.261304788 | 0.10935678 | 0.26039175 | nlp-2 |
| WBGene00003741 | 0.837332622 | 0.00201559 | 0.01255221 | nlp-3 |
| WBGene00003742 | -0.060366122 | 0.6178919 | 0.77099816 | nlp-4 |
| WBGene00003743 | 1.096287834 | 0.01083691 | 0.04756089 | nlp-5 |
| WBGene00003744 | 1.073398393 | 1.72E-05 | 0.00019913 | nlp-6 |
| WBGene00003745 | 0.566594685 | 0.036196 | 0.1188752 | nlp-7 |
| WBGene00003746 | 1.562750512 | 1.31E-07 | 2.43E-06 | nlp-8 |
| WBGene00003747 | 1.235145478 | 3.56E-05 | 0.00038578 | nlp-9 |
| WBGene00003748 | 1.056229184 | 0.00034504 | 0.00283009 | nlp-10 |
| WBGene00003749 | 0.617361555 | 0.02118924 | 0.0801425 | nlp-11 |
| WBGene00003750 | -0.197025345 | 0.33010572 | 0.52954078 | nlp-12 |
| WBGene00003751 | 1.228235058 | 7.26E-07 | 1.18E-05 | nlp-13 |
| WBGene00003752 | 1.179013761 | 2.60E-06 | 3.65E-05 | nlp-14 |
| WBGene00003753 | 1.215148036 | 5.43E-05 | 0.00056269 | nlp-15 |
| WBGene00003754 | 0.658865784 | 0.00645921 | 0.03199229 | nlp-16 |
| WBGene00003755 | 0.139177552 | 0.44452961 | 0.63794923 | nlp-17 |
| WBGene00003756 | 1.432642473 | 5.70E-06 | 7.36E-05 | nlp-18 |
| WBGene00003757 | 0.211994539 | 0.21355948 | 0.40428294 | nlp-19 |
| WBGene00003758 | 0.145988643 | 0.4372761 | 0.63224959 | nlp-20 |
| WBGene00003759 | 0.936136572 | 0.00014979 | 0.0013681 | nlp-21 |
| WBGene00003760 | NA | NA | NA | nlp-22 |
| WBGene00003761 | NA | NA | NA | nlp-23 |

| | | | | |
|---|---|---|---|---|
| WBGene00003762 | -0.974906509 | 3.84E-15 | 1.87E-13 | nlp-24 |
| WBGene00003763 | 0.017749406 | 0.90460118 | 0.94913088 | nlp-25 |
| WBGene00003764 | -0.266628515 | 0.20970763 | 0.3991069 | nlp-26 |
| WBGene00003765 | -0.660080537 | 8.62E-11 | 2.50E-09 | nlp-27 |
| WBGene00003766 | 0.068716678 | 0.67326109 | 0.80998475 | nlp-28 |
| WBGene00003767 | 0.128498034 | 0.51660833 | 0.69612149 | nlp-29 |
| WBGene00003768 | 0.148271134 | 0.38860779 | 0.58710213 | nlp-30 |
| WBGene00003769 | -0.149591013 | 0.38955924 | 0.58786842 | nlp-31 |
| WBGene00003770 | 0.020232312 | 0.84643798 | NA | nlp-32 |
| WBGene00003771 | 0.172538917 | 0.22753619 | 0.42116549 | nlp-33 |
| WBGene00003772 | -0.012931117 | 0.94802233 | 0.97044737 | nlr-1 |
| WBGene00003773 | 0.395674298 | 0.00363677 | 0.02034117 | nlt-1 |
| WBGene00003774 | 0.068351763 | 0.55915919 | 0.73003231 | nmr-1 |
| WBGene00003775 | 2.675099675 | 0.00360039 | 0.02021606 | nmr-2 |
| WBGene00003776 | 0.296186788 | 3.10E-06 | 4.25E-05 | nmy-1 |
| WBGene00003777 | -0.01361163 | 0.77731898 | 0.87398508 | nmy-2 |
| WBGene00003778 | 0.458919282 | 0.08072454 | 0.21226437 | nnt-1 |
| WBGene00003779 | -0.055003348 | 0.76879623 | 0.86844708 | nob-1 |
| WBGene00003783 | -0.110553221 | 0.48582839 | 0.67034337 | nos-1 |
| WBGene00003784 | -0.057152 | 0.51571155 | 0.69532487 | nos-2 |
| WBGene00003785 | -0.048169062 | 0.48873388 | 0.67283053 | nos-3 |
| WBGene00003786 | 0.47525409 | 1.88E-13 | 7.65E-12 | npa-1 |
| WBGene00003787 | -0.339648712 | 9.84E-06 | 0.00012186 | npp-1 |
| WBGene00003788 | -0.187448742 | 0.00844013 | 0.03939255 | npp-2 |
| WBGene00003789 | -0.140463408 | 0.03001794 | 0.10353184 | npp-3 |
| WBGene00003790 | -0.025223352 | 0.80772639 | 0.89346646 | npp-4 |
| WBGene00003791 | -0.162412841 | 0.03300315 | 0.11092725 | npp-5 |
| WBGene00003792 | -0.071605112 | 0.31091964 | 0.51188287 | npp-6 |
| WBGene00003793 | -0.140489737 | 0.00924589 | 0.04228961 | npp-7 |
| WBGene00003794 | -0.070545288 | 0.33760748 | 0.53731692 | npp-8 |
| WBGene00003795 | -0.126799689 | 0.00831156 | 0.0389282 | npp-9 |
| WBGene00003796 | -0.00889197 | 0.87067387 | 0.93091263 | npp-10 |
| WBGene00003797 | -0.033059951 | 0.68824301 | 0.82045382 | npp-11 |
| WBGene00003798 | -0.179106997 | 0.00179948 | 0.01143497 | npp-12 |
| WBGene00003799 | -0.281792667 | 0.00052184 | 0.00402543 | npp-13 |
| WBGene00003800 | -0.068443677 | 0.23366807 | 0.42913976 | npp-14 |
| WBGene00003801 | -0.142635171 | 0.06528194 | 0.18214342 | npp-15 |
| WBGene00003802 | 0.027361742 | 0.74878633 | 0.8582013 | npp-16 |
| WBGene00003803 | -0.305745651 | 0.0002212 | 0.00192193 | rae-1 |
| WBGene00003804 | -0.166363208 | 0.04428276 | 0.13759422 | npp-18 |
| WBGene00003805 | -0.236521867 | 0.00379373 | 0.02099964 | npp-19 |
| WBGene00003806 | -0.17523215 | 0.06137298 | 0.17451191 | npp-20 |
| WBGene00003807 | 0.047071372 | 0.71852369 | 0.84199658 | npr-1 |

| | | | | |
|---|---|---|---|---|
| WBGene00003808 | 0.466678764 | 0.06464841 | 0.18118633 | npr-2 |
| WBGene00003812 | 0.379324722 | 0.00184869 | 0.01171212 | nrf-5 |
| WBGene00003813 | 0.063885982 | 0.49936096 | 0.68137407 | nrf-6 |
| WBGene00003815 | -0.376565895 | 9.27E-10 | 2.35E-08 | nars-1 |
| WBGene00003816 | 0.011608898 | 0.95039003 | 0.97182989 | asns-1 |
| WBGene00003818 | -0.049542156 | 0.60578679 | 0.76196945 | nsf-1 |
| WBGene00003821 | -0.27000113 | 0.0001485 | 0.001358 | nst-1 |
| WBGene00003822 | 0.265640613 | 0.02418092 | 0.08850822 | nsy-1 |
| WBGene00003825 | 0.055097317 | 0.52837684 | 0.70538573 | ntl-2 |
| WBGene00003826 | -0.182791886 | 0.00680363 | 0.03333735 | ntl-3 |
| WBGene00003827 | -0.129053057 | 0.07456943 | 0.20076701 | ntl-4 |
| WBGene00003828 | 0.87628629 | 5.44E-27 | 5.69E-25 | nuc-1 |
| WBGene00003829 | -0.246552535 | 0.00126844 | 0.00855422 | nud-1 |
| WBGene00003830 | -0.216757274 | 0.00975459 | 0.04405211 | num-1 |
| WBGene00003831 | -0.323895691 | 2.09E-09 | 5.07E-08 | nuo-1 |
| WBGene00003834 | -0.05923242 | 0.4392246 | 0.63387751 | nxf-1 |
| WBGene00003835 | -0.168269555 | 0.15969052 | 0.33433848 | nxf-2 |
| WBGene00003836 | -0.138970933 | 0.28075267 | 0.48036693 | nxt-1 |
| WBGene00003837 | -0.319714202 | 0.13781659 | 0.30365277 | oat-1 |
| WBGene00003838 | -0.032895608 | 0.74022276 | 0.85342581 | ocr-1 |
| WBGene00003839 | 0.058277345 | 0.69673206 | 0.82701491 | ocr-2 |
| WBGene00003840 | 0.132152599 | 0.16128998 | NA | ocr-3 |
| WBGene00003841 | 0.073458642 | 0.29111983 | NA | ocr-4 |
| WBGene00003842 | 1.115146968 | 0.00109059 | 0.00756386 | 1-Oct |
| WBGene00003843 | 0.343488175 | 0.12734411 | 0.28871852 | 2-Oct |
| WBGene00003844 | -0.886972073 | 3.61E-07 | 6.15E-06 | odc-1 |
| WBGene00003845 | -0.138064904 | 0.4566809 | 0.64869855 | odd-1 |
| WBGene00003846 | -0.023656001 | 0.9070591 | 0.94967137 | odd-2 |
| WBGene00003847 | -0.687843996 | 0.0043703 | 0.02355009 | blmp-1 |
| WBGene00003848 | 0.131827335 | 0.16097389 | NA | odr-1 |
| WBGene00003849 | 0.075314074 | 0.69877267 | 0.82846224 | odr-2 |
| WBGene00003850 | 0.111471905 | 0.5256772 | 0.70293542 | odr-3 |
| WBGene00003851 | 0.042384297 | 0.70013322 | 0.82952793 | odr-4 |
| WBGene00003854 | 0.0398788 | 0.58971221 | NA | odr-7 |
| WBGene00003856 | 0.230235782 | 0.2407222 | 0.4382509 | odr-10 |
| WBGene00003858 | -0.129463437 | 0.10582167 | 0.25483506 | ogt-1 |
| WBGene00003859 | 0.46274199 | 0.05839735 | 0.16833044 | oig-1 |
| WBGene00003860 | -0.866797785 | 4.92E-05 | 0.00051798 | oig-2 |
| WBGene00003861 | 0.402058395 | 0.0697968 | 0.19082013 | oig-3 |
| WBGene00003862 | 0.018333177 | 0.89402846 | 0.94488347 | old-1 |
| WBGene00003863 | NA | NA | NA | old-2 |
| WBGene00003864 | -0.199432082 | 0.00017893 | 0.00160325 | oma-1 |
| WBGene00003865 | -0.065141192 | 0.18805327 | 0.37307424 | oma-2 |

| | | | | |
|---|---|---|---|---|
| WBGene00003868 | 0.05745888 | 0.5482904 | 0.72092678 | ooc-3 |
| WBGene00003870 | -0.262412585 | 0.001021 | 0.00715578 | ooc-5 |
| WBGene00003876 | 0.370308633 | 0.02002434 | 0.07668751 | pept-2 |
| WBGene00003877 | 0.721222293 | 1.03E-14 | 4.77E-13 | pept-1 |
| WBGene00003878 | 0.225501606 | 0.2617219 | 0.45968092 | pept-3 |
| WBGene00003879 | -0.004944805 | 0.94514249 | NA | ora-1 |
| WBGene00003882 | -0.138923633 | 0.1853663 | 0.36919439 | orc-2 |
| WBGene00003883 | 0.166454039 | 0.37271647 | 0.57191473 | osm-1 |
| WBGene00003884 | 0.157819677 | 0.45470049 | 0.64678155 | osm-3 |
| WBGene00003885 | -0.006380379 | 0.9626889 | 0.97924853 | osm-5 |
| WBGene00003886 | 0.626061651 | 0.03098623 | 0.10567653 | osm-6 |
| WBGene00003887 | 0.935209283 | 0.00031755 | 0.00263716 | osm-7 |
| WBGene00003888 | 0.0627133 | 0.50876942 | NA | osm-8 |
| WBGene00003889 | 0.068577315 | 0.58771728 | 0.75003656 | osm-9 |
| WBGene00003890 | 0.43068162 | 0.08680629 | 0.22278071 | osm-10 |
| WBGene00003891 | 0.295649228 | 4.78E-05 | 0.00050485 | osm-11 |
| WBGene00003892 | 0.300785675 | 0.02950964 | 0.10243197 | osm-12 |
| WBGene00003893 | 0.261494899 | 0.00023613 | 0.00203246 | ost-1 |
| WBGene00003898 | 0.021301282 | 0.82418206 | 0.9034778 | oxi-1 |
| WBGene00003901 | -0.304144549 | 7.06E-09 | 1.61E-07 | paa-1 |
| WBGene00003902 | -0.120584321 | 0.00308449 | 0.01785688 | pab-1 |
| WBGene00003903 | -0.252736279 | 0.02455597 | 0.0896345 | pab-2 |
| WBGene00003904 | -0.444103629 | 1.72E-07 | 3.12E-06 | pabp-2 |
| WBGene00003905 | -0.014639625 | 0.86745658 | 0.92947386 | pad-1 |
| WBGene00003906 | 0.065014541 | 0.37399213 | NA | paf-1 |
| WBGene00003907 | -0.055927997 | 0.78092663 | 0.87653686 | paf-2 |
| WBGene00003909 | 0.307610173 | 0.02505037 | 0.09094035 | pag-3 |
| WBGene00003911 | -0.154787012 | 0.24861005 | 0.44694251 | pak-1 |
| WBGene00003912 | -0.343324804 | 1.56E-05 | 0.00018202 | pal-1 |
| WBGene00003914 | -0.445537161 | 4.10E-18 | 2.64E-16 | pam-1 |
| WBGene00003915 | -0.021762178 | 0.88154429 | 0.93716888 | pan-1 |
| WBGene00003916 | 0.094666381 | 0.19695943 | 0.38449106 | par-1 |
| WBGene00003917 | 0.013893704 | 0.90337007 | 0.94859364 | par-2 |
| WBGene00003918 | -0.113734957 | 0.05875267 | 0.16896442 | par-3 |
| WBGene00003919 | -0.181871024 | 0.0352864 | 0.11671402 | par-4 |
| WBGene00003920 | -0.210318869 | 1.38E-05 | 0.00016336 | par-5 |
| WBGene00003921 | -0.18383948 | 0.07984337 | 0.21081818 | par-6 |
| WBGene00003922 | -0.204658875 | 0.00156426 | 0.01017879 | pas-1 |
| WBGene00003923 | -0.305110455 | 1.61E-07 | 2.93E-06 | pas-2 |
| WBGene00003924 | -0.227408539 | 3.29E-05 | 0.00035766 | pas-3 |
| WBGene00003925 | -0.319479028 | 2.41E-06 | 3.43E-05 | pas-4 |
| WBGene00003926 | -0.206032142 | 0.00035225 | 0.00287475 | pas-5 |
| WBGene00003927 | -0.338310241 | 3.31E-08 | 6.78E-07 | pas-6 |

| | | | | |
|---|---|---|---|---|
| WBGene00003928 | -0.226163944 | 0.0002274 | 0.0019665 | pas-7 |
| WBGene00003929 | 0.009204821 | 0.94740371 | 0.97044477 | pat-2 |
| WBGene00003930 | -0.069910811 | 0.33127219 | 0.53082588 | pat-3 |
| WBGene00003931 | -0.054470752 | 0.72871563 | 0.84857437 | pat-4 |
| WBGene00003932 | -0.036119978 | 0.79340925 | 0.88399982 | pat-6 |
| WBGene00003933 | -0.099153914 | 0.6337213 | 0.7826402 | pat-9 |
| WBGene00003934 | -0.217871149 | 8.09E-05 | 0.00079945 | pat-10 |
| WBGene00003936 | -0.204191067 | 0.02915423 | 0.1015094 | pat-12 |
| WBGene00003937 | -0.00346345 | 0.95708614 | NA | pax-1 |
| WBGene00003938 | NA | NA | NA | pax-2 |
| WBGene00003939 | -0.440359949 | 0.0778349 | 0.20682048 | pax-3 |
| WBGene00003941 | 0.284657957 | 0.13026534 | 0.29250015 | pbo-1 |
| WBGene00003943 | 0.056362788 | 0.76265526 | 0.8652488 | pbo-4 |
| WBGene00003947 | -0.25635908 | 0.00064114 | 0.00481379 | pbs-1 |
| WBGene00003948 | -0.334228485 | 5.08E-06 | 6.64E-05 | pbs-2 |
| WBGene00003949 | -0.193775497 | 0.00220076 | 0.01346481 | pbs-3 |
| WBGene00003950 | -0.241598242 | 0.00013908 | 0.00128402 | pbs-4 |
| WBGene00003951 | -0.277995682 | 7.47E-06 | 9.43E-05 | pbs-5 |
| WBGene00003952 | -0.314156192 | 1.87E-06 | 2.75E-05 | pbs-6 |
| WBGene00003953 | -0.257850216 | 1.96E-05 | 0.00022411 | pbs-7 |
| WBGene00003954 | 0.09617032 | 0.61892683 | 0.77186608 | pcm-1 |
| WBGene00003955 | -0.171806165 | 0.00097354 | 0.00688913 | pcn-1 |
| WBGene00003956 | 0.461419173 | 7.40E-05 | 0.00074246 | pcp-1 |
| WBGene00003957 | -1.429218091 | 7.31E-10 | 1.87E-08 | pcp-2 |
| WBGene00003958 | 0.279392618 | 0.07325896 | 0.19782081 | pcp-3 |
| WBGene00003959 | 0.17573832 | 0.38544512 | 0.58460167 | pcp-4 |
| WBGene00003960 | 0.241875046 | 0.15050529 | 0.32214315 | pcs-1 |
| WBGene00003961 | -0.088610049 | 0.26800063 | 0.46689639 | pct-1 |
| WBGene00003962 | -0.1804811 | 0.00074364 | 0.00544657 | pdi-1 |
| WBGene00003963 | -0.230048019 | 2.33E-05 | 0.0002616 | pdi-2 |
| WBGene00003964 | -0.157831316 | 0.00049245 | 0.00383509 | pdi-3 |
| WBGene00003965 | 0.044798912 | 0.79177166 | 0.88268703 | pdk-1 |
| WBGene00003966 | 0.209065011 | 0.23529697 | 0.43115338 | pdl-1 |
| WBGene00003967 | 0.119203602 | 0.29532887 | 0.49643054 | pdr-1 |
| WBGene00003968 | 1.779237865 | 0.00012846 | 0.00120405 | peb-1 |
| WBGene00003969 | 0.30267643 | 0.18667484 | 0.37103756 | pef-1 |
| WBGene00003970 | 0.299062343 | 0.00668243 | 0.03286442 | pek-1 |
| WBGene00003975 | -0.117698306 | 0.447571 | 0.64049331 | pen-2 |
| WBGene00003976 | -0.455608778 | 0.08226645 | 0.2150827 | pes-1 |
| WBGene00003977 | -0.046394786 | 0.50231372 | NA | pes-2.1 |
| WBGene00003977 | -0.046394786 | 0.50231372 | NA | pes-2.2 |
| WBGene00003978 | 0.699021591 | 0.00062765 | 0.00472947 | pes-4 |
| WBGene00003979 | -0.503151522 | 0.00266419 | 0.01581126 | pes-5 |

| | | | | |
|---|---|---|---|---|
| WBGene00003980 | 0.781793597 | 1.16E-06 | 1.80E-05 | pes-7 |
| WBGene00003981 | -0.466008169 | 0.0720359 | 0.19541923 | pes-8 |
| WBGene00003982 | -0.283819302 | 2.08E-05 | 0.00023624 | pes-9 |
| WBGene00003983 | -0.724641084 | 0.00104129 | 0.00726409 | pes-10 |
| WBGene00003986 | 0.158228792 | 0.1914529 | 0.37761847 | mct-4 |
| WBGene00003987 | 0.308794361 | 0.14506093 | 0.31445617 | pes-23 |
| WBGene00003989 | -0.247925958 | 4.69E-05 | 0.00049661 | pfn-1 |
| WBGene00003990 | -0.105507723 | 0.41201678 | 0.60964575 | pfn-2 |
| WBGene00003991 | -0.376754272 | 0.09712689 | 0.24122307 | pfn-3 |
| WBGene00003992 | -0.197548711 | 0.00030885 | 0.00258207 | pgl-1 |
| WBGene00003993 | -0.270568881 | 0.01719442 | 0.06872605 | pgl-2 |
| WBGene00003994 | -0.128270259 | 0.04157501 | 0.13149833 | pgl-3 |
| WBGene00003995 | 3.429736514 | 6.69E-16 | 3.46E-14 | pgp-1 |
| WBGene00003996 | -0.040695314 | 0.72492814 | 0.84572464 | pgp-2 |
| WBGene00003997 | 0.692660723 | 0.00091006 | 0.0064996 | pgp-3 |
| WBGene00003998 | 0.101495568 | 0.59930617 | 0.75732394 | pgp-4 |
| WBGene00003999 | 0.161703509 | 0.41158818 | 0.60930865 | pgp-5 |
| WBGene00004000 | 0.498474332 | 0.05794272 | 0.1674909 | pgp-6 |
| WBGene00004001 | 0.040087849 | 0.56144502 | NA | pgp-7 |
| WBGene00004002 | -0.034198367 | 0.83298858 | 0.90884461 | pgp-8 |
| WBGene00004003 | 0.573325005 | 0.00173954 | 0.01113144 | pgp-9 |
| WBGene00004004 | 0.696272775 | 0.01950775 | 0.07529356 | pgp-10 |
| WBGene00004005 | 1.227692244 | 8.52E-09 | 1.93E-07 | pgp-11 |
| WBGene00004006 | 0.446760733 | 0.08599344 | 0.22146524 | pgp-12 |
| WBGene00004007 | 0.032926043 | 0.87422821 | 0.93298863 | pgp-13 |
| WBGene00004008 | 2.083992287 | 7.01E-21 | 5.58E-19 | pgp-14 |
| WBGene00004009 | 0.23038633 | 0.14850036 | 0.31935867 | NA |
| WBGene00004010 | -0.23042027 | 0.24548763 | 0.44365517 | pha-1 |
| WBGene00004011 | 0.10719576 | 0.31431546 | 0.51491699 | pha-2 |
| WBGene00004013 | -0.040666746 | 0.82970863 | 0.90708661 | pha-4 |
| WBGene00004014 | -0.538904021 | 1.31E-13 | 5.38E-12 | phb-1 |
| WBGene00004015 | -0.346016243 | 2.74E-07 | 4.77E-06 | phb-2 |
| WBGene00004016 | -0.32693823 | 0.01200918 | 0.05168322 | phf-5 |
| WBGene00004017 | 0.054048556 | 0.63849631 | 0.78556509 | phg-1 |
| WBGene00004020 | -0.52523721 | 0.00024315 | 0.00208554 | pho-1 |
| WBGene00004023 | 1.17376539 | 3.02E-07 | 5.21E-06 | pho-4 |
| WBGene00004024 | -0.018788525 | 0.92376059 | 0.95682907 | php-3 |
| WBGene00004025 | -0.301239947 | 0.10290106 | 0.25026154 | phy-2 |
| WBGene00004026 | -0.031590944 | 0.84136385 | 0.91430328 | phy-3 |
| WBGene00004027 | 0.062758514 | 0.54620472 | 0.71941196 | pie-1 |
| WBGene00004028 | -0.123155511 | 0.31562217 | 0.51614254 | pif-1 |
| WBGene00004029 | -0.224795896 | 0.14579502 | 0.31553362 | pik-1 |
| WBGene00004030 | 0.25235028 | 0.26146992 | 0.45956556 | pin-2 |

| | | | | |
|---|---|---|---|---|
| WBGene00004031 | -0.232990644 | 0.03285317 | 0.1106146 | pis-1 |
| WBGene00004032 | 0.135598845 | 0.11549908 | 0.27117095 | pkc-1 |
| WBGene00004033 | 0.328071442 | 0.00905244 | 0.04159974 | pkc-2 |
| WBGene00004034 | -0.058674493 | 0.45820178 | 0.64973678 | pkc-3 |
| WBGene00004035 | NA | NA | NA | pkd-2 |
| WBGene00004036 | 0.432607076 | 0.01496322 | 0.06171168 | plc-1 |
| WBGene00004037 | 0.308292585 | 0.14024827 | 0.30766781 | plc-2 |
| WBGene00004038 | 0.474550394 | 0.00698732 | 0.03393078 | plc-3 |
| WBGene00004039 | -0.150555939 | 0.16227474 | 0.33739454 | plc-4 |
| WBGene00004040 | -0.129388994 | 0.13769873 | 0.30355699 | pld-1 |
| WBGene00004041 | -0.102429853 | 0.61871842 | 0.77167189 | NA |
| WBGene00004042 | -0.27428265 | 1.52E-06 | 2.28E-05 | plk-1 |
| WBGene00004043 | -0.23756692 | 0.01231334 | 0.05276248 | plk-2 |
| WBGene00004044 | -0.212082785 | 0.00051043 | 0.00395406 | plk-3 |
| WBGene00004045 | 0.81527829 | 0.00718798 | 0.03474386 | pll-1 |
| WBGene00004046 | -0.037192668 | 0.55691384 | 0.72827577 | plp-1 |
| WBGene00004047 | 0.058164076 | 0.56278985 | 0.73275288 | plx-1 |
| WBGene00004048 | 0.352052819 | 0.14185142 | 0.30983837 | plx-2 |
| WBGene00004049 | 0.142402543 | 0.12056924 | 0.27938963 | parp-1 |
| WBGene00004050 | -0.13878928 | 0.11837014 | 0.2757031 | parp-2 |
| WBGene00004051 | -0.220427686 | 0.01532314 | 0.06293019 | parg-1 |
| WBGene00004052 | 0.074923135 | 0.59286406 | 0.75314295 | parg-2 |
| WBGene00004053 | 0.56369921 | 1.90E-06 | 2.77E-05 | tank-1 |
| WBGene00004054 | 0.750211011 | 0.01871326 | 0.07317863 | NA |
| WBGene00004055 | -0.017178114 | 0.91227213 | 0.95126775 | pmk-1 |
| WBGene00004056 | 0.21908167 | 0.27928877 | 0.4791501 | pmk-2 |
| WBGene00004057 | 0.481668752 | 0.02585092 | 0.09303916 | pmk-3 |
| WBGene00004058 | 0.258189951 | 0.07406921 | 0.19956705 | pmp-1 |
| WBGene00004059 | -0.12547876 | 0.35954381 | 0.55958116 | pmp-2 |
| WBGene00004060 | -0.065576294 | 0.4321236 | 0.62727758 | pmp-3 |
| WBGene00004061 | -0.218835762 | 0.26774078 | 0.4665971 | pmp-4 |
| WBGene00004062 | 0.601209521 | 1.34E-16 | 7.65E-15 | pmp-5 |
| WBGene00004063 | 0.022318318 | 0.81748481 | 0.89973842 | pmr-1 |
| WBGene00004064 | -0.004117263 | 0.97004231 | 0.98346632 | pms-2 |
| WBGene00004068 | 0.111753265 | 0.20650252 | 0.39568163 | pnk-1 |
| WBGene00004075 | 0.14178681 | 0.0054302 | 0.0279254 | pod-1 |
| WBGene00004076 | 0.280376267 | 0.00013897 | 0.00128402 | pod-2 |
| WBGene00004077 | -0.412026625 | 0.00105731 | 0.00736092 | pop-1 |
| WBGene00004078 | -0.106300189 | 0.02308932 | 0.08534299 | pos-1 |
| WBGene00004083 | 0.623650102 | 0.00204653 | 0.01272327 | pph-1 |
| WBGene00004084 | NA | NA | NA | pph-2 |
| WBGene00004085 | -0.180065149 | 0.17638926 | 0.35724376 | pph-4.1 |
| WBGene00004086 | -0.109546182 | 0.17993224 | 0.36162163 | pph-4.2 |

| | | | | |
|---|---|---|---|---|
| WBGene00004087 | -0.089489916 | 0.33986895 | 0.53958158 | ppk-1 |
| WBGene00004088 | 0.184421588 | 0.18973211 | 0.37544235 | ppk-2 |
| WBGene00004089 | 0.046069296 | 0.65133892 | 0.79364585 | ppk-3 |
| WBGene00004090 | -0.175943383 | 0.11426132 | 0.26895499 | eif-2Bgamma |
| WBGene00004091 | 0.117704162 | 0.20427722 | 0.39337403 | pps-1 |
| WBGene00004092 | 0.157816706 | 0.46107831 | 0.65192158 | ppt-1 |
| WBGene00004093 | -0.211321178 | 0.01887472 | 0.07371161 | ppw-1 |
| WBGene00004094 | -0.174223045 | 0.08980315 | 0.2277675 | ppw-2 |
| WBGene00004095 | 0.09389081 | 0.17337822 | 0.35364036 | pqe-1 |
| WBGene00004096 | 1.810217687 | 1.29E-24 | 1.18E-22 | pqm-1 |
| WBGene00004097 | NA | NA | NA | pqm-96 |
| WBGene00004098 | 0.036687355 | 0.59489568 | NA | pqn-2 |
| WBGene00004099 | -0.00346345 | 0.95708614 | NA | abu-15 |
| WBGene00004100 | 0.330880642 | 0.15179843 | 0.32424969 | pqn-8 |
| WBGene00004101 | -0.049937455 | 0.53200047 | 0.70833968 | hgrs-1 |
| WBGene00004102 | -0.104347724 | 0.47004851 | 0.65837369 | pqn-10 |
| WBGene00004103 | -0.132192812 | 0.13071783 | 0.29308418 | pqn-11 |
| WBGene00004104 | 1.628987482 | 0.00857532 | 0.03985833 | pqn-13 |
| WBGene00004105 | -0.146293083 | 0.11616532 | 0.27208069 | szy-20 |
| WBGene00004106 | -0.211053577 | 0.05059318 | 0.15169666 | pqn-15 |
| WBGene00004106 | -0.211053577 | 0.05059318 | 0.15169666 | C52B9.8 |
| WBGene00004107 | -0.040794878 | 0.53203165 | NA | pqn-16 |
| WBGene00004108 | 0.831227897 | 0.00012794 | 0.00120149 | pqn-18 |
| WBGene00004109 | 0.01753203 | 0.8383115 | 0.91254414 | stam-1 |
| WBGene00004110 | -0.138892252 | 0.03007408 | 0.10365222 | pqn-20 |
| WBGene00004111 | -0.309566781 | 0.00895091 | 0.04128866 | pqn-21 |
| WBGene00004112 | 0.047474584 | 0.62214753 | 0.77413738 | pqn-22 |
| WBGene00004113 | 0.307406657 | 0.06330436 | 0.17847856 | pqn-24 |
| WBGene00004114 | 1.083284436 | 0.00131789 | 0.00882511 | pqn-25 |
| WBGene00004115 | 0.144712595 | 0.33946496 | 0.53917963 | pqn-26 |
| WBGene00004116 | -0.082117073 | 0.37661457 | 0.57581566 | pqn-27 |
| WBGene00004117 | -0.129401933 | 0.04190151 | 0.13221552 | sin-3 |
| WBGene00004118 | 0.105619007 | 0.22574074 | NA | pqn-29 |
| WBGene00004119 | 2.170010461 | 3.16E-27 | 3.38E-25 | pqn-31 |
| WBGene00004120 | 2.217799952 | 0.00434302 | 0.02345488 | pqn-32 |
| WBGene00004121 | 0.28239313 | 0.04291847 | 0.13464974 | ptrn-1 |
| WBGene00004122 | 1.709641733 | 1.10E-08 | 2.44E-07 | pqn-35 |
| WBGene00004123 | 1.778287701 | 2.68E-06 | 3.74E-05 | pqn-36 |
| WBGene00004124 | -0.142949598 | 0.48539382 | 0.66999896 | pqn-37 |
| WBGene00004125 | -0.087455585 | 0.40525039 | 0.60332664 | mdt-30 |
| WBGene00004126 | 0.035994104 | 0.79451348 | 0.8844641 | pqn-39 |
| WBGene00004127 | 0.106568874 | 0.42433422 | 0.62027529 | abu-12 |
| WBGene00004128 | 0.2533201 | 0.01948804 | 0.07529356 | pqn-41 |

| | | | | |
|---|---|---|---|---|
| WBGene00004129 | 0.50906337 | 0.06481434 | 0.18144297 | pqn-42 |
| WBGene00004130 | 0.175346088 | 0.22884295 | 0.42266805 | ketn-1 |
| WBGene00004131 | 0.825117874 | 3.99E-13 | 1.57E-11 | pqn-44 |
| WBGene00004132 | -0.083710439 | 0.03995947 | 0.12787904 | ifet-1 |
| WBGene00004133 | 1.377955579 | 0.01643564 | 0.06651029 | abu-13 |
| WBGene00004134 | 0.056495059 | 0.72598499 | 0.84660663 | myrf-1 |
| WBGene00004135 | 0.484548655 | 0.04388833 | 0.13691215 | pqn-48 |
| WBGene00004136 | 0.012091632 | 0.87493366 | 0.93332632 | pqn-51 |
| WBGene00004138 | 0.102798134 | 0.17130105 | 0.35028193 | pqn-53 |
| WBGene00004139 | 2.800200744 | 0.00392185 | 0.02154588 | pqn-54 |
| WBGene00004140 | -0.302804604 | 0.00737189 | 0.03545724 | ebax-1 |
| WBGene00004141 | NA | NA | NA | abu-9 |
| WBGene00004141 | NA | NA | NA | pqn-57 |
| WBGene00004143 | 0.057079991 | 0.28527695 | 0.48544163 | pqn-59 |
| WBGene00004144 | 1.420832488 | 0.00048849 | 0.00381072 | pqn-60 |
| WBGene00004145 | 0.931046973 | 0.00186825 | 0.01182154 | pqn-62 |
| WBGene00004146 | 0.065014541 | 0.37399213 | NA | pqn-63 |
| WBGene00004147 | 0.001244805 | 0.98470453 | 0.99127193 | larp-5 |
| WBGene00004148 | 0.105036931 | 0.18096036 | 0.36323562 | pqn-65 |
| WBGene00004149 | -0.05616772 | 0.63153035 | 0.78099644 | trpl-5 |
| WBGene00004150 | 0.787424706 | 0.02467421 | 0.08997636 | pqn-67 |
| WBGene00004151 | -0.088205342 | 0.6519613 | 0.79392543 | pqn-68 |
| WBGene00004152 | -0.202033163 | 0.00360635 | 0.02023351 | pqn-70 |
| WBGene00004153 | 0.143932909 | 0.06890664 | NA | pqn-71 |
| WBGene00004154 | 1.560486522 | 0.00013019 | 0.00121665 | pqn-72 |
| WBGene00004155 | 0.570016393 | 0.03066173 | 0.10503472 | pqn-73 |
| WBGene00004156 | -0.03502557 | 0.86246274 | 0.92672958 | pqn-74 |
| WBGene00004157 | 1.983810152 | 5.36E-13 | 2.07E-11 | pqn-75 |
| WBGene00004158 | NA | NA | NA | pqn-76 |
| WBGene00004159 | NA | NA | NA | pqn-78 |
| WBGene00004160 | NA | NA | NA | pqn-79 |
| WBGene00004161 | -0.03149798 | 0.82746547 | 0.90562762 | pqn-80 |
| WBGene00004163 | -0.045660216 | 0.77338605 | 0.87148095 | pqn-82 |
| WBGene00004164 | -0.072348501 | 0.41400551 | 0.61154897 | pqn-83 |
| WBGene00004165 | 0.038125098 | 0.61119482 | NA | pqn-84 |
| WBGene00004166 | 0.024285804 | 0.70991581 | 0.83666404 | pqn-85 |
| WBGene00004167 | 0.038544558 | 0.44491051 | 0.63824562 | pqn-87 |
| WBGene00004168 | 1.446698384 | 5.61E-05 | 0.00058005 | pqn-88 |
| WBGene00004169 | 0.619509464 | 0.00465749 | 0.02474248 | pqn-89 |
| WBGene00004170 | NA | NA | NA | pqn-90 |
| WBGene00004171 | NA | NA | NA | pqn-91 |
| WBGene00004172 | 0.955372226 | 7.63E-08 | 1.47E-06 | pqn-92 |
| WBGene00004173 | 1.881665037 | 2.89E-10 | 7.85E-09 | pqn-94 |

| | | | | |
|---|---|---|---|---|
| WBGene00004174 | 3.486128409 | 1.47E-07 | 2.68E-06 | abu-14 |
| WBGene00004175 | -0.030867233 | 0.81855423 | 0.90020675 | pqn-96 |
| WBGene00004176 | 0.157098662 | 0.06528165 | NA | pqn-97 |
| WBGene00004177 | 0.148021412 | 0.07455439 | NA | pqn-98 |
| WBGene00004178 | -0.084189943 | 0.22281793 | 0.41556803 | prg-1 |
| WBGene00004179 | 0.002104113 | 0.97526792 | 0.98597553 | prg-2 |
| WBGene00004180 | -0.024830601 | 0.83875275 | 0.9127791 | pri-1 |
| WBGene00004181 | 0.009184269 | 0.9267385 | 0.9586234 | pri-2 |
| WBGene00004182 | 0.725572429 | 2.37E-05 | 0.00026497 | prk-1 |
| WBGene00004183 | -0.363004756 | 0.11325348 | 0.26692598 | prk-2 |
| WBGene00004184 | 0.500555621 | 0.01426169 | 0.05938718 | T19D2.2 |
| WBGene00004185 | -0.173163881 | 0.23439808 | 0.43000863 | pro-1 |
| WBGene00004186 | -0.245506156 | 0.00047027 | 0.00369378 | prpf-4 |
| WBGene00004187 | -0.078246715 | 0.11614432 | 0.27207504 | prp-8 |
| WBGene00004188 | -0.266345759 | 0.00809076 | 0.03821187 | prp-21 |
| WBGene00004189 | -0.383789901 | 3.25E-10 | 8.71E-09 | pars-1 |
| WBGene00004190 | -0.276556391 | 0.05524979 | 0.16191195 | pars-2 |
| WBGene00004191 | -0.104498929 | 0.60724 | 0.76307361 | prx-1 |
| WBGene00004192 | -0.059846762 | 0.73282416 | 0.85051752 | prx-2 |
| WBGene00004193 | -0.12403607 | 0.49725939 | 0.67958319 | prx-3 |
| WBGene00004194 | 0.047842986 | 0.7848823 | 0.87855213 | prx-5 |
| WBGene00004195 | 0.122731185 | 0.55177408 | 0.72414165 | prx-6 |
| WBGene00004196 | 0.091373564 | 0.63822876 | 0.78552676 | prx-11 |
| WBGene00004197 | 0.086240475 | 0.67178149 | 0.80894366 | prx-12 |
| WBGene00004198 | 0.247244962 | 0.21310331 | 0.40368036 | prx-13 |
| WBGene00004199 | -0.150009128 | 0.14271672 | 0.31103237 | prx-14 |
| WBGene00004201 | 0.012267791 | 0.90554022 | 0.94916953 | prx-19 |
| WBGene00004202 | 0.259263938 | 0.19095004 | 0.37698214 | pry-1 |
| WBGene00004203 | -0.092168801 | 0.24969951 | 0.44813074 | swsn-1 |
| WBGene00004204 | -0.134484792 | 0.0147441 | 0.06103149 | swsn-4 |
| WBGene00004205 | -0.274653547 | 0.00819253 | 0.03848149 | psr-1 |
| WBGene00004206 | -0.129195486 | 0.26684942 | 0.46561105 | pst-1 |
| WBGene00004207 | 0.457023154 | 0.03249582 | 0.10970026 | ptb-1 |
| WBGene00004208 | -0.05228798 | 0.24801804 | 0.44637149 | ptc-1 |
| WBGene00004209 | -0.023109762 | 0.78121253 | NA | ptc-2 |
| WBGene00004210 | 0.505792447 | 0.00216137 | 0.01329049 | ptc-3 |
| WBGene00004211 | 0.016219169 | 0.9155536 | 0.95298026 | ptd-2 |
| WBGene00004212 | 0.573294887 | 0.02967212 | 0.10282354 | ptl-1 |
| WBGene00004213 | 0.755554897 | 0.00272395 | 0.0161267 | ptp-1 |
| WBGene00004214 | -0.109053242 | 0.21842503 | 0.41031093 | ptp-2 |
| WBGene00004215 | 0.406834024 | 0.00010466 | 0.00101145 | ptp-3 |
| WBGene00004216 | -0.245718585 | 0.26174889 | 0.45968092 | ptr-1 |
| WBGene00004217 | -0.097843189 | 0.11235484 | 0.26542401 | ptr-2 |

| | | | | |
|---|---|---|---|---|
| WBGene00004218 | 1.678255132 | 0.00124397 | 0.0084191 | ptr-3 |
| WBGene00004219 | 0.793342677 | 0.02213769 | 0.08278873 | ptr-4 |
| WBGene00004220 | 0.655524377 | 0.014374 | 0.05978683 | ptr-5 |
| WBGene00004221 | 0.086891199 | 0.67796896 | 0.81292048 | ptr-6 |
| WBGene00004222 | 0.201162332 | 0.29277566 | 0.49361888 | ptr-8 |
| WBGene00004223 | NA | NA | NA | ptr-9 |
| WBGene00004224 | -0.060782953 | 0.55960335 | 0.73003231 | ptr-10 |
| WBGene00004225 | 0.106972975 | 0.61021173 | 0.76529981 | ptr-11 |
| WBGene00004226 | 0.351745319 | 0.14094992 | 0.30863628 | ptr-12 |
| WBGene00004227 | -0.124878984 | 0.41746238 | 0.61433981 | ptr-13 |
| WBGene00004228 | -0.384462055 | 0.05346645 | 0.15817383 | ptr-14 |
| WBGene00004229 | 0.071133416 | 0.54701751 | 0.71995249 | ptr-15 |
| WBGene00004230 | -0.082490559 | 0.68180101 | 0.81594316 | ptr-16 |
| WBGene00004231 | 0.178768928 | 0.09068156 | 0.22888093 | ptr-17 |
| WBGene00004232 | 0.274783857 | 0.21679669 | 0.40829951 | ptr-18 |
| WBGene00004233 | 0.012622814 | 0.93254946 | 0.9620984 | ptr-19 |
| WBGene00004234 | 0.489348476 | 0.02264625 | 0.08415323 | ptr-20 |
| WBGene00004235 | 0.26864235 | 0.02311215 | 0.08534299 | ptr-21 |
| WBGene00004236 | 1.718799813 | 0.00447282 | 0.02399655 | ptr-22 |
| WBGene00004237 | 0.706466863 | 0.00012339 | 0.00116628 | ptr-23 |
| WBGene00004238 | -0.18490572 | 0.11396012 | 0.26828914 | ptr-24 |
| WBGene00004239 | -0.17175946 | 0.00065156 | 0.00486708 | puf-3 |
| WBGene00004240 | -0.034296832 | 0.71935516 | NA | NA |
| WBGene00004241 | -0.085028107 | 0.04701237 | 0.14383988 | puf-5 |
| WBGene00004242 | -0.123814357 | 0.12973238 | 0.29203748 | puf-6 |
| WBGene00004243 | -0.124925987 | 0.12636386 | 0.28746011 | puf-7 |
| WBGene00004244 | -0.131065115 | 0.15911155 | 0.33341238 | puf-8 |
| WBGene00004245 | -0.095989415 | 0.40993613 | 0.60772325 | puf-9 |
| WBGene00004246 | -0.152998966 | 0.17319472 | 0.35347679 | NA |
| WBGene00004248 | -0.103977582 | 0.39547059 | 0.593368 | pus-1 |
| WBGene00004249 | 1.36325651 | 1.78E-09 | 4.35E-08 | pvf-1 |
| WBGene00004254 | -0.021669399 | 0.78977988 | 0.88147334 | pxf-1 |
| WBGene00004255 | -0.008152874 | 0.92103705 | NA | epac-1 |
| WBGene00004256 | 0.029941736 | 0.84523262 | 0.9169413 | pxn-1 |
| WBGene00004257 | -0.050515361 | 0.80562893 | 0.89208355 | pxn-2 |
| WBGene00004258 | 0.019097776 | 0.7430679 | 0.85461565 | pyc-1 |
| WBGene00004259 | -0.225852496 | 0.00209203 | 0.01295113 | pyr-1 |
| WBGene00004264 | 2.112908999 | 7.64E-06 | 9.63E-05 | qua-1 |
| WBGene00004265 | 1.186813503 | 0.00264377 | 0.01572195 | qui-1 |
| WBGene00004266 | -0.111136357 | 0.04990457 | 0.14996978 | rab-1 |
| WBGene00004267 | 0.998759999 | 0.00110962 | 0.00767407 | rab-3 |
| WBGene00004268 | -0.096246342 | 0.22767714 | 0.42120558 | rab-5 |
| WBGene00004269 | -0.151543777 | 0.08945802 | 0.22726891 | rab-6.1 |

| | | | | |
|---|---|---|---|---|
| WBGene00004270 | -0.254582931 | 0.08487779 | 0.2194025 | rab-6.2 |
| WBGene00004271 | -0.167815165 | 0.08029113 | 0.21156663 | rab-7 |
| WBGene00004272 | -0.070297853 | 0.45411627 | 0.64625937 | rab-8 |
| WBGene00004273 | -0.129435458 | 0.10511637 | 0.25350169 | rab-10 |
| WBGene00004274 | -0.088897208 | 0.06708406 | 0.18562158 | rab-11.1 |
| WBGene00004275 | -0.004047623 | 0.95527667 | NA | rab-11.2 |
| WBGene00004276 | 0.001945512 | 0.98836412 | 0.99352139 | rab-14 |
| WBGene00004277 | -0.206266481 | 0.04621507 | 0.14214188 | rab-18 |
| WBGene00004278 | -0.002320249 | 0.9873952 | 0.9927519 | rab-19 |
| WBGene00004279 | 0.110902271 | 0.40461697 | 0.60270598 | rab-21 |
| WBGene00004281 | 0.097567122 | 0.4396195 | 0.63425986 | rab-28 |
| WBGene00004282 | -0.207336766 | 0.21737776 | 0.40904904 | rab-30 |
| WBGene00004283 | 0.085017855 | 0.5025789 | 0.68391651 | rab-33 |
| WBGene00004284 | 0.110358687 | 0.38976434 | 0.58786842 | rab-35 |
| WBGene00004285 | 0.095876171 | 0.57713135 | 0.74411462 | rab-37 |
| WBGene00004286 | 0.199205201 | 0.18230466 | 0.36517643 | rab-39 |
| WBGene00004287 | -0.023056499 | 0.78121253 | NA | rac-2 |
| WBGene00004296 | -0.097137414 | 0.18826595 | 0.37339499 | rad-50 |
| WBGene00004297 | -0.332633399 | 0.00013654 | 0.00126446 | rad-51 |
| WBGene00004298 | -0.004062505 | 0.97284141 | 0.98515501 | rad-54 |
| WBGene00004300 | 2.11387872 | 1.06E-07 | 2.00E-06 | ram-2 |
| WBGene00004301 | 0.063290468 | 0.41068021 | NA | ram-5 |
| WBGene00004302 | -0.198751423 | 6.14E-05 | 0.00063104 | ran-1 |
| WBGene00004303 | -0.043421272 | 0.55888227 | 0.72974184 | ran-2 |
| WBGene00004304 | -0.065883558 | 0.3019807 | 0.50260002 | ran-3 |
| WBGene00004305 | -0.241120268 | 0.00066496 | 0.00495451 | ran-4 |
| WBGene00004306 | -0.018138434 | 0.84496238 | 0.91685276 | ran-5 |
| WBGene00004307 | -0.150966929 | 0.04229096 | 0.13312876 | rap-1 |
| WBGene00004308 | -0.166070897 | 0.15673736 | 0.33070929 | rap-2 |
| WBGene00004309 | -0.139350704 | 0.44253514 | 0.63652669 | rap-3 |
| WBGene00004310 | 0.72867561 | 0.00087431 | 0.00628413 | ras-1 |
| WBGene00004311 | 0.131975982 | 0.41202409 | 0.60964575 | ras-2 |
| WBGene00004312 | -0.177698038 | 0.02486219 | 0.09048156 | rba-1 |
| WBGene00004313 | 0.139386149 | 0.11094523 | 0.26296689 | rbc-1 |
| WBGene00004314 | -0.165366748 | 0.14609298 | 0.31589857 | rbc-2 |
| WBGene00004315 | -0.317968916 | 0.00165749 | 0.01070606 | rbd-1 |
| WBGene00004316 | 1.69270151 | 2.32E-08 | 4.86E-07 | rbf-1 |
| WBGene00004317 | -0.002796951 | 0.98482364 | 0.99132368 | rbg-1 |
| WBGene00004318 | 0.005256114 | 0.95952011 | 0.97781959 | rbg-2 |
| WBGene00004319 | -0.13773306 | 0.12019978 | 0.27876523 | rbr-2 |
| WBGene00004320 | -0.373422048 | 1.46E-06 | 2.20E-05 | rbx-1 |
| WBGene00004321 | 0.772969232 | 0.00079635 | 0.00579204 | rcan-1 |
| WBGene00004322 | 0.015836909 | 0.89643558 | 0.94558453 | rcq-5 |

| | | | | | |
|---|---|---|---|---|---|
| WBGene00004323 | | -0.021222315 | 0.81192188 | 0.89587496 | rde-1 |
| WBGene00004324 | | -0.012270513 | 0.90335004 | 0.94859364 | rde-2 |
| WBGene00004326 | | 0.067203044 | 0.47828652 | 0.66520309 | rde-4 |
| WBGene00004328 | | 0.079162379 | 0.57764612 | 0.74411462 | rdy-2 |
| WBGene00004332 | | 0.270245021 | 0.12221435 | 0.28134075 | rec-1 |
| WBGene00004333 | | -0.09330044 | 0.40398996 | 0.60226219 | rec-8 |
| WBGene00004334 | | 0.069587419 | 0.73096178 | 0.84986592 | ref-1 |
| WBGene00004335 | | -1.686350207 | 0.00036261 | 0.00294944 | ref-2 |
| WBGene00004336 | | -0.185040335 | 0.00024344 | 0.00208674 | ret-1 |
| WBGene00004337 | | 0.030864927 | 0.64485002 | 0.78968348 | rfc-1 |
| WBGene00004338 | | -0.155187671 | 0.09991965 | 0.2453335 | rfc-2 |
| WBGene00004339 | | -0.194551913 | 0.01193447 | 0.05137855 | rfc-3 |
| WBGene00004340 | | -0.20265505 | 0.00565483 | 0.02882742 | rfc-4 |
| WBGene00004341 | | -0.068209612 | 0.50857687 | 0.68894903 | rfl-1 |
| WBGene00004342 | | 0.039554613 | 0.84792325 | 0.9180823 | rfs-1 |
| WBGene00004343 | | -0.149245447 | 0.06083319 | 0.17334737 | rgr-1 |
| WBGene00004344 | | 0.179240871 | 0.39938059 | 0.5976422 | rgs-1 |
| WBGene00004345 | | 0.687881173 | 0.01683875 | 0.06765535 | rgs-2 |
| WBGene00004346 | | 0.034864219 | 0.84694878 | 0.91761389 | rgs-3 |
| WBGene00004347 | | 0.920635955 | 0.00225012 | 0.01372096 | rgs-4 |
| WBGene00004348 | | -0.243950519 | 0.1753514 | 0.35594084 | rgs-5 |
| WBGene00004349 | | 0.83469605 | 0.00780657 | 0.03706098 | rgs-6 |
| WBGene00004350 | | -0.052369249 | 0.67207297 | 0.80906269 | rgs-7 |
| WBGene00004351 | NA | | NA | NA | rgs-8.1 |
| WBGene00004351 | NA | | NA | NA | rgs-8.2 |
| WBGene00004352 | | -0.412035269 | 0.00957552 | 0.04351372 | rgs-9 |
| WBGene00004353 | | 0.060243949 | 0.76204002 | 0.86495294 | rgs-10 |
| WBGene00004354 | | -0.159225728 | 0.32645038 | 0.5256622 | rgs-11 |
| WBGene00004355 | | -0.136104464 | 0.05455325 | 0.16031998 | rha-1 |
| WBGene00004356 | | 0.149135474 | 0.24915242 | 0.44753289 | rhi-1 |
| WBGene00004357 | | -0.20301998 | 0.0090123 | 0.04147808 | rho-1 |
| WBGene00004358 | | 0.134356967 | 0.18890286 | 0.37415135 | rhr-1 |
| WBGene00004359 | | -0.244576257 | 0.26585362 | 0.46448245 | rhr-2 |
| WBGene00004360 | | 0.067406638 | 0.57050971 | 0.73912332 | rib-1 |
| WBGene00004361 | | -0.353762455 | 0.00688334 | 0.03354476 | rib-2 |
| WBGene00004363 | | 0.269153866 | 0.23127085 | 0.42626706 | ric-3 |
| WBGene00004364 | | 0.663441826 | 0.00150177 | 0.00982992 | ric-4 |
| WBGene00004367 | | -0.175647931 | 0.1769944 | 0.35797417 | ric-8 |
| WBGene00004368 | | 0.269068032 | 0.23562486 | 0.43143867 | ric-19 |
| WBGene00004369 | | 0.307136141 | 0.13444215 | 0.29873041 | rig-1 |
| WBGene00004370 | | 0.504247056 | 0.01694676 | 0.06799868 | rig-3 |
| WBGene00004371 | | 0.092297042 | 0.26169581 | 0.45968092 | rig-4 |
| WBGene00004372 | | 0.010664262 | 0.94547534 | 0.96937707 | rig-5 |

| | | | | |
|---|---|---|---|---|
| WBGene00004373 | -0.19921641 | 0.00267189 | 0.01585052 | rme-1 |
| WBGene00004374 | -0.17248655 | 0.00018504 | 0.0016509 | rme-2 |
| WBGene00004375 | 0.074702271 | 0.57607816 | 0.74318345 | rme-4 |
| WBGene00004377 | 0.124677829 | 0.35936692 | 0.55949501 | rme-6 |
| WBGene00004378 | -0.128636489 | 0.06944139 | 0.19032037 | rme-8 |
| WBGene00004380 | -0.152719649 | 0.20671712 | 0.39598918 | rnf-1 |
| WBGene00004381 | -0.119788305 | 0.2979159 | 0.49893568 | rnf-5 |
| WBGene00004382 | -0.231634486 | 0.14204463 | 0.3101216 | rnh-1.0 |
| WBGene00004383 | -0.095272642 | 0.35867149 | 0.55894723 | rnh-2 |
| WBGene00004384 | -0.1742733 | 0.06155141 | 0.1748834 | rnp-1 |
| WBGene00004385 | -0.096360638 | 0.43588137 | 0.63104313 | rnp-2 |
| WBGene00004386 | -0.136593628 | 0.31132887 | 0.5122686 | rnp-3 |
| WBGene00004387 | -0.248947417 | 0.00067911 | 0.0050446 | rnp-4 |
| WBGene00004388 | -0.092262716 | 0.40020868 | 0.5984532 | rnp-5 |
| WBGene00004389 | -0.12406382 | 0.16223001 | 0.33738773 | rnp-6 |
| WBGene00004390 | -0.051905568 | 0.54051171 | 0.71476071 | rnp-7 |
| WBGene00004391 | -0.203480903 | 0.00013185 | 0.00123036 | rnr-1 |
| WBGene00004392 | 0.007680531 | 0.86447974 | 0.92771735 | rnr-2 |
| WBGene00004393 | 1.348584315 | 0.00747822 | 0.03586262 | rnt-1 |
| WBGene00004394 | 0.439081064 | 0.02989525 | 0.10337655 | rol-1 |
| WBGene00004395 | 0.205838466 | 0.32558757 | 0.52475997 | rol-3 |
| WBGene00004397 | 0.402401601 | 0.00424517 | 0.02300279 | rol-6 |
| WBGene00004398 | 0.180638168 | 0.29840979 | 0.49925925 | rol-8 |
| WBGene00004400 | -0.23895416 | 0.22205074 | 0.41483283 | rom-1 |
| WBGene00004401 | 0.034188945 | 0.81293386 | 0.89638233 | rom-2 |
| WBGene00004402 | 0.131317611 | 0.10121567 | NA | rom-3 |
| WBGene00004403 | -0.01274253 | 0.90217434 | 0.94794994 | rom-4 |
| WBGene00004404 | 0.011241552 | 0.93939821 | 0.96589373 | rom-5 |
| WBGene00004405 | 0.204223247 | 0.00640121 | 0.03176953 | rop-1 |
| WBGene00004408 | -0.268987072 | 2.19E-10 | 6.03E-09 | rla-0 |
| WBGene00004409 | -0.386432036 | 1.36E-11 | 4.42E-10 | rla-1 |
| WBGene00004410 | -0.266805464 | 1.52E-05 | 0.00017848 | rla-2 |
| WBGene00004411 | -0.323124906 | 0.00019279 | 0.00171249 | rpc-1 |
| WBGene00004412 | -0.162604874 | 0.00218288 | 0.01338338 | rpl-1 |
| WBGene00004413 | -0.318381433 | 3.38E-12 | 1.18E-10 | rpl-2 |
| WBGene00004414 | -0.311793023 | 1.95E-12 | 7.00E-11 | rpl-3 |
| WBGene00004415 | -0.16696229 | 0.00021646 | 0.00188867 | rpl-4 |
| WBGene00004416 | -0.18637667 | 2.32E-05 | 0.00026149 | rpl-5 |
| WBGene00004417 | -0.300084487 | 3.47E-10 | 9.27E-09 | rpl-6 |
| WBGene00004418 | -0.335533978 | 6.46E-19 | 4.46E-17 | rpl-7 |
| WBGene00004419 | -0.163489202 | 0.00089378 | 0.0064052 | rpl-7A |
| WBGene00004420 | -0.256374707 | 2.80E-06 | 3.89E-05 | rpl-9 |
| WBGene00004421 | -0.260661046 | 2.29E-08 | 4.81E-07 | rpl-10 |

| | | | | |
|---|---|---|---|---|
| WBGene00004422 | -0.405952384 | 6.34E-19 | 4.40E-17 | rpl-11.1 |
| WBGene00004423 | 0.234795507 | 3.92E-05 | 0.00041995 | rpl-11.2 |
| WBGene00004424 | -0.309736445 | 4.07E-08 | 8.20E-07 | rpl-12 |
| WBGene00004425 | -0.235417602 | 7.28E-08 | 1.40E-06 | rpl-13 |
| WBGene00004426 | -0.368771057 | 3.86E-16 | 2.11E-14 | rpl-14 |
| WBGene00004427 | -0.443180564 | 1.09E-20 | 8.50E-19 | rpl-15 |
| WBGene00004428 | -0.2636053 | 8.69E-09 | 1.97E-07 | rpl-16 |
| WBGene00004429 | -0.305492114 | 6.19E-13 | 2.37E-11 | rpl-17 |
| WBGene00004430 | -0.253220647 | 3.61E-08 | 7.36E-07 | rpl-18 |
| WBGene00004431 | -0.055158038 | 0.25761317 | 0.45573917 | rpl-19 |
| WBGene00004432 | -0.389785339 | 1.46E-14 | 6.63E-13 | rpl-20 |
| WBGene00004433 | -0.244632812 | 9.08E-07 | 1.45E-05 | rpl-21 |
| WBGene00004434 | -0.254918851 | 3.94E-06 | 5.28E-05 | rpl-22 |
| WBGene00004435 | -0.206283336 | 3.73E-06 | 5.02E-05 | rpl-23 |
| WBGene00004436 | -0.365342011 | 9.03E-14 | 3.80E-12 | rpl-24.1 |
| WBGene00004437 | -0.357705483 | 0.00131032 | 0.00878811 | rpl-24.2 |
| WBGene00004438 | 0.402465299 | 7.91E-05 | 0.00078448 | rpl-25.1 |
| WBGene00004439 | -0.364718311 | 9.22E-11 | 2.67E-09 | rpl-25.2 |
| WBGene00004440 | -0.353748532 | 4.95E-11 | 1.49E-09 | rpl-26 |
| WBGene00004441 | -0.328638545 | 1.36E-12 | 5.02E-11 | rpl-27 |
| WBGene00004442 | -0.270232735 | 3.18E-06 | 4.35E-05 | rpl-28 |
| WBGene00004443 | -0.318105766 | 7.66E-07 | 1.24E-05 | rpl-29 |
| WBGene00004444 | -0.107749024 | 0.05293604 | 0.1570598 | rpl-30 |
| WBGene00004445 | -0.19790798 | 0.00011596 | 0.00110676 | rpl-31 |
| WBGene00004446 | -0.247804834 | 1.70E-06 | 2.52E-05 | rpl-32 |
| WBGene00004447 | -0.277585397 | 2.47E-06 | 3.50E-05 | rpl-33 |
| WBGene00004448 | -0.348839143 | 1.12E-07 | 2.08E-06 | rpl-34 |
| WBGene00004449 | -0.090626729 | 0.06456241 | 0.18104292 | rpl-35 |
| WBGene00004450 | -0.397576156 | 9.73E-13 | 3.68E-11 | rpl-36 |
| WBGene00004451 | -0.040568855 | 0.75130535 | 0.85949848 | rpl-37 |
| WBGene00004452 | -0.366629986 | 1.89E-08 | 4.03E-07 | rpl-38 |
| WBGene00004453 | -0.181056449 | 0.10424625 | 0.25244404 | rpl-39 |
| WBGene00004454 | -0.249919282 | 1.27E-05 | 0.00015153 | rpl-36.A |
| WBGene00004455 | 0.059762547 | 0.38965825 | NA | rpl-42 |
| WBGene00004456 | -0.176661983 | 0.00251381 | 0.01508728 | rpl-43 |
| WBGene00004457 | 0.568485588 | 0.00780534 | 0.03706098 | rpm-1 |
| WBGene00004458 | -0.16330897 | 0.00115385 | 0.00792751 | rpn-1 |
| WBGene00004459 | -0.200806936 | 0.00016857 | 0.00152256 | rpn-2 |
| WBGene00004460 | -0.36243868 | 5.16E-11 | 1.55E-09 | rpn-3 |
| WBGene00004461 | -0.339462424 | 4.52E-09 | 1.06E-07 | rpn-5 |
| WBGene00004462 | -0.268255545 | 4.91E-05 | 0.0005165 | rpn-6.1 |
| WBGene00004463 | -0.286068952 | 1.27E-06 | 1.95E-05 | rpn-7 |
| WBGene00004464 | -0.280765967 | 7.92E-07 | 1.28E-05 | rpn-8 |

| | | | | |
|---|---|---|---|---|
| WBGene00004465 | -0.357371369 | 1.72E-09 | 4.23E-08 | rpn-9 |
| WBGene00004466 | -0.216932407 | 0.00150123 | 0.00982992 | rpn-10 |
| WBGene00004467 | -0.16600905 | 0.011502 | 0.049867 | rpn-11 |
| WBGene00004468 | -0.269216878 | 0.0003756 | 0.00304833 | rpn-12 |
| WBGene00004469 | -0.307071751 | 9.42E-12 | 3.16E-10 | rps-0 |
| WBGene00004470 | -0.297990256 | 1.33E-09 | 3.31E-08 | rps-1 |
| WBGene00004471 | 0.13172006 | 0.02630649 | 0.09437719 | rps-2 |
| WBGene00004472 | -0.181841936 | 9.57E-05 | 0.00093259 | rps-3 |
| WBGene00004473 | -0.222194827 | 1.18E-06 | 1.84E-05 | rps-4 |
| WBGene00004474 | -0.391535901 | 1.64E-15 | 8.21E-14 | rps-5 |
| WBGene00004475 | -0.230090565 | 2.77E-08 | 5.74E-07 | rps-6 |
| WBGene00004476 | -0.421118239 | 3.35E-18 | 2.18E-16 | rps-7 |
| WBGene00004477 | -0.31186562 | 5.71E-14 | 2.45E-12 | rps-8 |
| WBGene00004478 | -0.280089631 | 1.87E-11 | 5.92E-10 | rps-9 |
| WBGene00004479 | -0.265744613 | 3.94E-09 | 9.31E-08 | rps-10 |
| WBGene00004480 | -0.312633292 | 4.75E-10 | 1.24E-08 | rps-11 |
| WBGene00004481 | -0.292490015 | 1.37E-09 | 3.40E-08 | rps-12 |
| WBGene00004482 | -0.307814402 | 1.62E-10 | 4.49E-09 | rps-13 |
| WBGene00004483 | -0.225384137 | 9.70E-08 | 1.83E-06 | rps-14 |
| WBGene00004484 | -0.365379931 | 7.48E-12 | 2.54E-10 | rps-15 |
| WBGene00004485 | -0.37303973 | 5.60E-10 | 1.46E-08 | rps-16 |
| WBGene00004486 | -0.201729735 | 1.15E-05 | 0.00013986 | rps-17 |
| WBGene00004487 | -0.327263018 | 3.70E-12 | 1.29E-10 | rps-18 |
| WBGene00004488 | -0.214500152 | 0.00019709 | 0.00174782 | rps-19 |
| WBGene00004489 | -0.29751211 | 3.94E-12 | 1.37E-10 | rps-20 |
| WBGene00004490 | -0.275316875 | 3.65E-06 | 4.93E-05 | rps-21 |
| WBGene00004491 | -0.178550338 | 0.00065143 | 0.00486708 | rps-22 |
| WBGene00004492 | -0.392668668 | 2.25E-16 | 1.26E-14 | rps-23 |
| WBGene00004493 | -0.341682336 | 6.11E-14 | 2.62E-12 | rps-24 |
| WBGene00004494 | -0.229412603 | 1.30E-06 | 2.00E-05 | rps-25 |
| WBGene00004495 | -0.323048551 | 2.01E-08 | 4.27E-07 | rps-26 |
| WBGene00004496 | -0.317277688 | 5.14E-08 | 1.02E-06 | rps-27 |
| WBGene00004497 | -0.32675492 | 8.40E-07 | 1.35E-05 | rps-28 |
| WBGene00004498 | -0.116663945 | 0.08730688 | 0.22366755 | rps-29 |
| WBGene00004499 | -0.095950861 | 0.03961827 | 0.12717629 | rps-30 |
| WBGene00004501 | -0.211569203 | 0.00014973 | 0.0013681 | rpt-1 |
| WBGene00004502 | -0.265784674 | 8.44E-07 | 1.36E-05 | rpt-2 |
| WBGene00004503 | -0.22984675 | 9.49E-05 | 0.00092747 | rpt-3 |
| WBGene00004504 | -0.203607342 | 0.0003419 | 0.00281066 | rpt-4 |
| WBGene00004505 | -0.280386832 | 1.64E-06 | 2.45E-05 | rpt-5 |
| WBGene00004506 | -0.05911684 | 0.41596094 | 0.61311629 | rpt-6 |
| WBGene00004507 | 0.258626477 | 0.213779 | 0.40459388 | rpy-1 |
| WBGene00004508 | -0.060855491 | 0.5043342 | 0.68554053 | rrf-1 |

| | | | | |
|---|---|---|---|---|
| WBGene00004509 | 0.289991948 | 0.14012582 | 0.30749132 | rrf-2 |
| WBGene00004510 | -0.122467003 | 0.16098776 | 0.33556248 | rrf-3 |
| WBGene00004512 | NA | NA | NA | NA |
| WBGene00004513 | NA | NA | NA | NA |
| WBGene00004567 | -0.472451419 | 0.00590188 | 0.02978792 | NA |
| WBGene00004622 | 0.104336625 | 0.61330219 | 0.76759765 | NA |
| WBGene00004677 | -0.190626379 | 0.37460954 | 0.57340912 | NA |
| WBGene00004679 | -0.230004022 | 0.00121058 | 0.00823993 | rars-1 |
| WBGene00004680 | 0.058678657 | 0.60104516 | 0.75861226 | rars-2 |
| WBGene00004681 | -0.064745599 | 0.36018836 | 0.55995869 | rsd-2 |
| WBGene00004682 | 0.11696191 | 0.3649465 | 0.56453912 | rsd-3 |
| WBGene00004684 | -0.210841909 | 0.00442249 | 0.02378753 | rsd-6 |
| WBGene00004698 | -0.276036444 | 7.67E-05 | 0.00076534 | rsp-1 |
| WBGene00004699 | -0.155821554 | 0.04373479 | 0.13655812 | rsp-2 |
| WBGene00004700 | -0.276388369 | 2.30E-06 | 3.30E-05 | rsp-3 |
| WBGene00004701 | -0.202306218 | 0.00592503 | 0.02988097 | rsp-4 |
| WBGene00004702 | -0.373011314 | 6.77E-05 | 0.00068742 | rsp-5 |
| WBGene00004703 | -0.235923719 | 0.00146236 | 0.00961399 | rsp-6 |
| WBGene00004704 | -0.239513175 | 0.00143759 | 0.00947671 | rsp-7 |
| WBGene00004705 | -0.147664473 | 0.0563506 | 0.16447948 | rsp-8 |
| WBGene00004706 | 0.100137019 | 0.40217381 | 0.60068853 | rsr-1 |
| WBGene00004708 | NA | NA | NA | NA |
| WBGene00004716 | NA | NA | NA | NA |
| WBGene00004717 | NA | NA | NA | NA |
| WBGene00004719 | 1.319940971 | 0.00083358 | 0.00602687 | sad-1 |
| WBGene00004721 | -0.130122665 | 0.21220262 | 0.40249495 | san-1 |
| WBGene00004723 | 0.108210053 | 0.30157322 | 0.50214187 | sap-49 |
| WBGene00004726 | 0.102615283 | 0.37127976 | 0.57063899 | sas-4 |
| WBGene00004727 | 0.284059666 | 0.07159804 | 0.19466424 | sax-1 |
| WBGene00004728 | -0.122880365 | 0.0735364 | 0.19846018 | sax-2 |
| WBGene00004729 | 0.075653842 | 0.49187471 | 0.67512065 | sax-3 |
| WBGene00004732 | -0.049668455 | 0.41292111 | 0.61056051 | sax-7 |
| WBGene00004735 | 0.520560129 | 0.00069715 | 0.00515711 | sbp-1 |
| WBGene00004736 | 0.2806934 | 4.89E-08 | 9.74E-07 | sca-1 |
| WBGene00004737 | -0.241002149 | 0.01590277 | 0.06481972 | scc-1 |
| WBGene00004738 | 0.140309861 | 0.06849911 | 0.18847613 | scc-3 |
| WBGene00004739 | 0.507280811 | 6.30E-05 | 0.00064486 | scd-1 |
| WBGene00004740 | 0.847094862 | 0.00703348 | 0.03414364 | scd-2 |
| WBGene00004742 | NA | NA | NA | scl-1 |
| WBGene00004743 | 0.013184085 | 0.9110805 | 0.95112162 | scm-1 |
| WBGene00004744 | 0.126041632 | 0.22786992 | 0.42139368 | scp-1 |
| WBGene00004745 | -0.168232971 | 0.21252136 | 0.4029951 | sdc-1 |
| WBGene00004746 | -0.258669948 | 0.09362148 | 0.23406969 | sdc-2 |

| | | | | |
|---|---|---|---|---|
| WBGene00004747 | -0.154068119 | 0.27700788 | 0.47702569 | sdc-3 |
| WBGene00004748 | NA | NA | NA | sdf-9 |
| WBGene00004749 | -0.243922959 | 0.04672788 | 0.14324601 | sdn-1 |
| WBGene00004750 | -0.549350229 | 0.00022601 | 0.00195569 | sea-1 |
| WBGene00004751 | 0.175724692 | 0.11847163 | 0.27584972 | sea-2 |
| WBGene00004752 | -0.066012132 | 0.4994101 | 0.68137763 | sec-5 |
| WBGene00004753 | -0.133547309 | 0.20472721 | 0.39397451 | sec-8 |
| WBGene00004754 | -0.129424311 | 0.1047268 | 0.25297889 | sec-23 |
| WBGene00004755 | -0.208932033 | 0.00294923 | 0.01723054 | sec-24.1 |
| WBGene00004756 | -0.109430739 | 0.25637583 | 0.45460486 | sec-24.2 |
| WBGene00004758 | 0.025053134 | 0.85638853 | 0.922958 | sek-1 |
| WBGene00004759 | -0.085510587 | 0.22434184 | 0.41720746 | sel-1 |
| WBGene00004760 | -0.012395326 | 0.8718909 | 0.9314787 | sel-2 |
| WBGene00004762 | -0.056714148 | 0.48610895 | 0.67066727 | sel-5 |
| WBGene00004764 | 0.459359789 | 0.03851579 | 0.12440099 | sel-7 |
| WBGene00004765 | 0.140952252 | 0.29461854 | 0.49569181 | sel-8 |
| WBGene00004766 | -0.147766431 | 0.05580664 | 0.16321715 | sel-9 |
| WBGene00004767 | -0.197391158 | 0.00509312 | 0.02651793 | sel-10 |
| WBGene00004768 | -0.009768822 | 0.8968491 | 0.9458163 | sel-11 |
| WBGene00004769 | -0.110783068 | 0.49995773 | 0.68187063 | sel-12 |
| WBGene00004771 | -0.091898006 | 0.58438981 | 0.74778226 | sem-2 |
| WBGene00004773 | 0.46531547 | 0.02984468 | 0.10325045 | sem-4 |
| WBGene00004774 | -0.265667353 | 0.02912895 | 0.10149284 | sem-5 |
| WBGene00004775 | -0.142915851 | 0.04585701 | 0.14128628 | 1-Sep |
| WBGene00004776 | 0.140290652 | 0.48779757 | 0.67192062 | ser-1 |
| WBGene00004777 | 0.117178412 | 0.51500267 | 0.69482643 | ser-2 |
| WBGene00004778 | 0.323856537 | 0.14966307 | 0.3210104 | ser-3 |
| WBGene00004779 | 1.000328276 | 0.01127664 | 0.04913727 | ser-4 |
| WBGene00004780 | -0.330042791 | 0.16112383 | 0.33570713 | ser-7 |
| WBGene00004781 | -0.315989553 | 0.00127528 | 0.0085942 | set-1 |
| WBGene00004782 | -0.165041422 | 0.03524441 | 0.1166132 | set-2 |
| WBGene00004783 | -0.077862976 | 0.36822688 | 0.56797406 | seu-1 |
| WBGene00004786 | -0.623385047 | 1.20E-05 | 0.00014452 | sex-1 |
| WBGene00004787 | 0.053798968 | 0.76272917 | 0.8652656 | sft-1 |
| WBGene00004788 | -0.231514491 | 0.00570128 | 0.02905409 | sft-4 |
| WBGene00004789 | 0.795019506 | 7.81E-07 | 1.26E-05 | sgk-1 |
| WBGene00004790 | -0.239088917 | 0.28289423 | 0.48274758 | sgn-1 |
| WBGene00004793 | 0.241052512 | 0.17503685 | 0.35564444 | shw-3 |
| WBGene00004795 | 0.138182087 | 0.14520361 | 0.31460967 | sid-1 |
| WBGene00004796 | 0.039472344 | 0.84045304 | 0.91385614 | sid-2 |
| WBGene00004798 | 0.188573285 | 9.51E-05 | 0.00092874 | sip-1 |
| WBGene00004800 | -0.1006466 | 0.27553711 | 0.47572289 | sir-2.1 |
| WBGene00004801 | 0.570551826 | 0.00714286 | 0.03453719 | sir-2.2 |

| | | | | | |
|---|---|---|---|---|---|
| WBGene00004802 | | 0.8451863 | 0.00708355 | 0.03431807 | sir-2.3 |
| WBGene00004803 | | 0.152077799 | 0.34272392 | 0.54300333 | sir-2.4 |
| WBGene00004804 | | -0.057430049 | 0.38567174 | 0.58471782 | skn-1 |
| WBGene00004806 | | -0.061269439 | 0.40281546 | 0.60131712 | skp-1 |
| WBGene00004807 | | 0.039610342 | 0.5306252 | 0.70725139 | skr-1 |
| WBGene00004808 | | -0.344489261 | 2.47E-06 | 3.50E-05 | skr-2 |
| WBGene00004809 | | 0.505041484 | 0.00126371 | 0.00853317 | skr-3 |
| WBGene00004810 | | 0.950641413 | 2.92E-06 | 4.03E-05 | skr-4 |
| WBGene00004811 | | 0.207056196 | 0.02971382 | 0.10287067 | skr-5 |
| WBGene00004812 | NA | | NA | NA | skr-6 |
| WBGene00004813 | | -0.892049045 | 5.93E-13 | 2.28E-11 | skr-7 |
| WBGene00004814 | | -1.111804619 | 2.98E-07 | 5.15E-06 | skr-8 |
| WBGene00004815 | | -0.879403925 | 0.00032934 | 0.00272493 | skr-9 |
| WBGene00004816 | | -0.968309319 | 3.78E-07 | 6.41E-06 | skr-10 |
| WBGene00004817 | | 0.241402123 | 0.26187391 | 0.45972373 | NA |
| WBGene00004818 | | -0.850159784 | 0.00019226 | 0.00170907 | skr-12 |
| WBGene00004819 | | -0.664491403 | 0.00414781 | 0.02254018 | skr-13 |
| WBGene00004820 | | -0.752625256 | 6.05E-06 | 7.76E-05 | skr-14 |
| WBGene00004821 | | -0.176130249 | 0.33953815 | 0.53923181 | skr-15 |
| WBGene00004822 | | -0.238998135 | 0.23518278 | 0.43100652 | skr-16 |
| WBGene00004823 | | -0.246657799 | 0.02999559 | 0.10347913 | skr-17 |
| WBGene00004824 | | -0.053130525 | 0.64337344 | 0.78886575 | F56B3.4 |
| WBGene00004825 | | 0.381646171 | 0.025452 | 0.09185178 | skr-19 |
| WBGene00004826 | | 0.149382861 | 0.45232654 | 0.64471503 | skr-20 |
| WBGene00004827 | | -1.058897314 | 0.00066105 | 0.0049304 | skr-21 |
| WBGene00004829 | | -0.191027923 | 0.16041384 | 0.33484732 | sli-1 |
| WBGene00004830 | | 0.462486824 | 0.06710573 | 0.18562158 | slo-1 |
| WBGene00004831 | | 0.490625899 | 0.00771307 | 0.0367123 | slo-2 |
| WBGene00004833 | NA | | NA | NA | NA |
| WBGene00004834 | NA | | NA | NA | NA |
| WBGene00004835 | NA | | NA | NA | NA |
| WBGene00004836 | NA | | NA | NA | NA |
| WBGene00004837 | | -0.022930767 | 0.78121253 | NA | NA |
| WBGene00004838 | NA | | NA | NA | NA |
| WBGene00004839 | NA | | NA | NA | NA |
| WBGene00004840 | NA | | NA | NA | NA |
| WBGene00004841 | NA | | NA | NA | NA |
| WBGene00004842 | NA | | NA | NA | NA |
| WBGene00004843 | NA | | NA | NA | NA |
| WBGene00004844 | NA | | NA | NA | NA |
| WBGene00004845 | NA | | NA | NA | NA |
| WBGene00004846 | NA | | NA | NA | NA |
| WBGene00004847 | | -0.023109762 | 0.78121253 | NA | NA |

| | | | | |
|---|---|---|---|---|
| WBGene00004848 | NA | NA | NA | NA |
| WBGene00004849 | NA | NA | NA | NA |
| WBGene00004850 | NA | NA | NA | NA |
| WBGene00004851 | NA | NA | NA | NA |
| WBGene00004854 | 0.538186219 | 0.04905598 | 0.1482106 | slt-1 |
| WBGene00004855 | 0.287356395 | 7.47E-05 | 0.00074904 | sma-1 |
| WBGene00004856 | 0.367356756 | 0.12432206 | 0.2845917 | sma-2 |
| WBGene00004857 | 0.029027146 | 0.88912349 | 0.94194334 | sma-3 |
| WBGene00004858 | 0.790534617 | 8.36E-05 | 0.00082347 | sma-4 |
| WBGene00004859 | 1.086800809 | 3.70E-07 | 6.28E-06 | sma-5 |
| WBGene00004860 | 0.236705795 | 0.208478 | 0.39764544 | sma-6 |
| WBGene00004862 | 0.277641258 | 0.00213688 | 0.01317864 | sma-9 |
| WBGene00004873 | -0.151469981 | 0.03495666 | 0.11589685 | smc-3 |
| WBGene00004874 | -0.027975419 | 0.64640467 | 0.79064418 | smc-4 |
| WBGene00004875 | 0.089670205 | 0.41050471 | 0.60813512 | smd-1 |
| WBGene00004876 | 0.15823344 | 0.38496351 | 0.58429334 | smf-1 |
| WBGene00004877 | 0.020925399 | 0.84643798 | NA | smf-2 |
| WBGene00004878 | 0.051503522 | 0.8035671 | 0.89074999 | smf-3 |
| WBGene00004879 | -0.138513629 | 0.0779362 | 0.20705205 | smg-1 |
| WBGene00004880 | 0.089511146 | 0.28928391 | 0.4895758 | smg-2 |
| WBGene00004881 | 0.070827506 | 0.45909514 | 0.65047359 | smg-3 |
| WBGene00004882 | -0.163390171 | 0.08438992 | 0.21872063 | smg-4 |
| WBGene00004883 | 0.025806292 | 0.84637462 | 0.91736533 | smg-5 |
| WBGene00004884 | 0.222981331 | 0.01218212 | 0.05233522 | smg-6 |
| WBGene00004885 | 0.015446705 | 0.9108091 | 0.95112162 | smg-7 |
| WBGene00004886 | -0.089961696 | 0.5346062 | 0.71000989 | smi-1 |
| WBGene00004887 | -0.136922271 | 0.20285797 | 0.3914648 | smn-1 |
| WBGene00004888 | -0.249398606 | 1.42E-05 | 0.00016746 | smo-1 |
| WBGene00004889 | 0.223339753 | 0.05222086 | 0.15552596 | smp-1 |
| WBGene00004890 | 0.180647463 | 0.26385202 | 0.46203293 | smp-2 |
| WBGene00004891 | 0.023727569 | 0.81772076 | 0.89976324 | smr-1 |
| WBGene00004892 | -0.097988201 | 0.25634977 | 0.45460486 | sms-1 |
| WBGene00004893 | -0.380162509 | 0.03097319 | 0.10565669 | sms-2 |
| WBGene00004894 | 0.093320383 | 0.62806085 | 0.77868057 | sms-3 |
| WBGene00004895 | -0.299581521 | 0.00169734 | 0.01091384 | smu-1 |
| WBGene00004896 | -0.107367151 | 0.22540425 | 0.4184813 | smu-2 |
| WBGene00004897 | -0.015883067 | 0.84232383 | 0.91500691 | snb-1 |
| WBGene00004898 | 0.853676451 | 0.00011633 | 0.00110809 | snb-2 |
| WBGene00004899 | 0.078633594 | 0.26310689 | NA | snb-5 |
| WBGene00004900 | 0.151038699 | 0.21562399 | 0.40661397 | snf-1 |
| WBGene00004901 | 0.236124961 | 0.01963238 | 0.07562583 | snf-2 |
| WBGene00004902 | 0.08991834 | 0.57484496 | 0.74227635 | snf-3 |
| WBGene00004903 | NA | NA | NA | NA |

| | | | | |
|---|---|---|---|---|
| WBGene00004904 | 1.208031987 | 0.00633965 | 0.03150674 | snf-5 |
| WBGene00004905 | 0.156410867 | 0.16968107 | 0.34803874 | snf-6 |
| WBGene00004906 | -0.054282928 | 0.52914905 | NA | snf-7 |
| WBGene00004907 | -0.00346345 | 0.95708614 | NA | NA |
| WBGene00004908 | 0.349919523 | 0.07196399 | 0.19526033 | snf-9 |
| WBGene00004909 | 0.065942193 | 0.61473329 | 0.7687973 | snf-10 |
| WBGene00004910 | 0.913269496 | 0.00376964 | 0.02090579 | snf-11 |
| WBGene00004911 | 0.199035135 | 0.32991617 | 0.52934824 | snf-12 |
| WBGene00004912 | 0.395377121 | 0.10868366 | 0.25932815 | sng-1 |
| WBGene00004913 | 0.156262314 | 0.44461636 | 0.63801118 | snn-1 |
| WBGene00004914 | -0.206934467 | 0.0022694 | 0.0138221 | snr-1 |
| WBGene00004915 | -0.401223389 | 2.64E-09 | 6.30E-08 | snr-2 |
| WBGene00004916 | -0.38720166 | 1.52E-06 | 2.28E-05 | snr-3 |
| WBGene00004917 | -0.311836224 | 5.03E-06 | 6.58E-05 | snr-4 |
| WBGene00004918 | -0.31932102 | 0.00037627 | 0.00305036 | snr-5 |
| WBGene00004919 | -0.166627198 | 0.02660382 | 0.09514082 | snr-6 |
| WBGene00004920 | -0.36289318 | 7.94E-05 | 0.00078683 | snr-7 |
| WBGene00004921 | 0.760398729 | 0.00412403 | 0.02244606 | snt-1 |
| WBGene00004922 | -0.278801076 | 0.13114615 | 0.29368472 | snt-2 |
| WBGene00004923 | 0.794234736 | 0.02114088 | 0.08000094 | snt-3 |
| WBGene00004924 | 1.27069336 | 6.41E-12 | 2.19E-10 | snt-4 |
| WBGene00004925 | 0.565154093 | 0.01731261 | 0.06902885 | snt-5 |
| WBGene00004926 | 0.305487525 | 0.19109273 | 0.37703623 | snt-6 |
| WBGene00004927 | 0.139637461 | 0.16439008 | 0.34038115 | snx-1 |
| WBGene00004928 | 0.090176229 | 0.64983517 | 0.79273174 | soc-1 |
| WBGene00004929 | 0.374297605 | 0.02341186 | 0.08621053 | soc-2 |
| WBGene00004930 | -0.20968345 | 0.00766275 | 0.03653219 | sod-1 |
| WBGene00004931 | -0.190030282 | 0.02193171 | 0.08220176 | sod-2 |
| WBGene00004932 | -0.221730227 | 0.30592869 | 0.50666281 | sod-3 |
| WBGene00004933 | 1.025937809 | 2.79E-06 | 3.89E-05 | sod-4 |
| WBGene00004944 | 0.143537461 | 0.42032533 | 0.61700252 | sol-1 |
| WBGene00004945 | 0.147244706 | 0.32424046 | 0.52363682 | sop-2 |
| WBGene00004946 | 0.03098877 | 0.73139539 | 0.84995249 | sop-3 |
| WBGene00004947 | -0.028017725 | 0.78734914 | 0.87949869 | sos-1 |
| WBGene00004949 | 0.410909481 | 0.04614588 | 0.14196717 | sox-2 |
| WBGene00004950 | 0.124862738 | 0.50092297 | 0.68269142 | sox-3 |
| WBGene00004951 | 0.341069897 | 1.29E-05 | 0.00015377 | spc-1 |
| WBGene00004952 | -0.18763903 | 0.10102008 | 0.24712364 | spd-1 |
| WBGene00004953 | -0.060903668 | 0.41868762 | 0.61546239 | spd-2 |
| WBGene00004954 | -0.015061731 | 0.92302042 | 0.95651506 | spd-3 |
| WBGene00004955 | -0.115922947 | 0.04453728 | 0.13820905 | spd-5 |
| WBGene00004958 | 0.420807683 | 0.05548922 | 0.16245104 | spe-4 |
| WBGene00004959 | 0.375024342 | 0.05646606 | 0.1646196 | spe-5 |

| | | | | | |
|---|---|---|---|---|---|
| WBGene00004960 | | 0.091364597 | 0.66300037 | 0.80196534 | spe-6 |
| WBGene00004962 | | 0.086632749 | 0.29414331 | NA | spe-8 |
| WBGene00004963 | | 0.063527295 | 0.41294485 | NA | spe-9 |
| WBGene00004964 | | 0.078992087 | 0.41352528 | 0.61119077 | spe-10 |
| WBGene00004965 | | 2.878136408 | 0.00344266 | 0.01948897 | spe-11 |
| WBGene00004966 | | 0.040405337 | 0.64936961 | NA | spe-12 |
| WBGene00004969 | | 0.658935976 | 0.00300567 | 0.01749048 | spe-15 |
| WBGene00004971 | | 0.040107227 | 0.58693163 | NA | spe-17 |
| WBGene00004972 | | 1.045165693 | 0.0169719 | 0.06804097 | spe-26 |
| WBGene00004973 | | -0.022930767 | 0.78121253 | NA | spe-27 |
| WBGene00004974 | NA | | NA | NA | spe-29 |
| WBGene00004975 | | -0.224529475 | 0.01950143 | 0.07529356 | spe-39 |
| WBGene00004976 | | 0.705074924 | 0.01024823 | 0.04573128 | spe-41 |
| WBGene00004978 | | -0.235726857 | 0.01047606 | 0.04648337 | spg-7 |
| WBGene00004979 | | -0.114853183 | 0.5623502 | 0.732441 | sph-1 |
| WBGene00004980 | | -0.029002804 | 0.60897459 | 0.76433751 | spk-1 |
| WBGene00004981 | | 0.050791204 | 0.55335933 | 0.7254418 | spl-1 |
| WBGene00004984 | | -0.160800882 | 0.00042758 | 0.00340192 | spn-4 |
| WBGene00004985 | | -0.203346209 | 0.07911317 | 0.20946513 | spo-11 |
| WBGene00004986 | | 0.916184552 | 4.99E-18 | 3.20E-16 | spp-1 |
| WBGene00004987 | | 3.509877856 | 5.55E-104 | 3.13E-101 | spp-2 |
| WBGene00004988 | | 0.414733898 | 5.83E-06 | 7.52E-05 | spp-3 |
| WBGene00004989 | | 0.153381374 | 0.23978639 | 0.43681877 | spp-4 |
| WBGene00004990 | | 0.43862072 | 4.70E-13 | 1.83E-11 | spp-5 |
| WBGene00004991 | | 0.108538718 | 0.15420469 | NA | spp-6 |
| WBGene00004992 | | 0.033034883 | 0.64219503 | NA | spp-7 |
| WBGene00004993 | | 1.965510834 | 1.59E-66 | 4.76E-64 | spp-8 |
| WBGene00004994 | NA | | NA | NA | spp-9 |
| WBGene00004995 | | 0.475471537 | 4.27E-11 | 1.30E-09 | spp-10 |
| WBGene00004996 | | -0.023109762 | 0.78121253 | NA | spp-11 |
| WBGene00004997 | | -0.004864426 | 0.94835335 | NA | spp-12 |
| WBGene00004998 | | 0.034505823 | 0.68373529 | NA | spp-13 |
| WBGene00004999 | | 0.026328299 | 0.65063809 | 0.79317171 | spp-14 |
| WBGene00005000 | | 1.186099217 | 9.66E-09 | 2.17E-07 | spp-15 |
| WBGene00005001 | | 0.794271048 | 0.00151878 | 0.00993143 | spp-16 |
| WBGene00005002 | | -2.457651069 | 2.40E-61 | 6.62E-59 | spp-17 |
| WBGene00005003 | | 2.834295898 | 4.36E-74 | 1.56E-71 | spp-18 |
| WBGene00005004 | | 0.107582918 | 0.22104628 | NA | spp-19 |
| WBGene00005005 | NA | | NA | NA | K04A8.21 |
| WBGene00005005 | NA | | NA | NA | spp-20 |
| WBGene00005006 | | -0.191159055 | 0.0803991 | 0.21171511 | spr-1 |
| WBGene00005007 | | -0.126236009 | 0.06707414 | 0.18562158 | spr-2 |
| WBGene00005008 | | -0.40089455 | 0.01155212 | 0.05005181 | spr-3 |

| | | | | |
|---|---|---|---|---|
| WBGene00005009 | -0.32612212 | 0.00024854 | 0.00212423 | spr-4 |
| WBGene00005010 | -0.188493391 | 0.08394876 | 0.21792444 | spr-5 |
| WBGene00005011 | -0.089721092 | 0.59794635 | 0.75608389 | F26F4.8 |
| WBGene00005012 | 0.134146994 | 0.43982747 | 0.63430988 | F26F4.2 |
| WBGene00005013 | 0.135879867 | 0.08173087 | NA | jmjd-1.1 |
| WBGene00005014 | -0.04713086 | 0.67936585 | 0.81416022 | spt-4 |
| WBGene00005015 | -0.06634908 | 0.26997577 | 0.46922071 | spt-5 |
| WBGene00005016 | 4.537499 | 0.00055635 | 0.00424685 | sqt-1 |
| WBGene00005017 | 0.31499014 | 0.06178312 | 0.17533759 | sqt-2 |
| WBGene00005018 | 1.160611303 | 1.99E-07 | 3.55E-06 | sqt-3 |
| WBGene00005019 | -0.166687867 | 0.02702151 | 0.09600143 | sqv-1 |
| WBGene00005020 | 0.004140927 | 0.97680073 | 0.98670757 | sqv-2 |
| WBGene00005021 | -0.263718053 | 0.02929205 | 0.10184395 | sqv-3 |
| WBGene00005022 | -0.178071638 | 0.0049933 | 0.02613761 | sqv-4 |
| WBGene00005023 | -0.231701603 | 0.00021579 | 0.00188505 | sqv-5 |
| WBGene00005024 | -0.150044771 | 0.17294508 | 0.35315048 | sqv-6 |
| WBGene00005025 | -0.132960356 | 0.14610908 | 0.31589857 | sqv-7 |
| WBGene00005026 | -0.145685273 | 0.18341994 | 0.36639736 | sqv-8 |
| WBGene00005027 | -0.022930767 | 0.78121253 | NA | sra-1 |
| WBGene00005028 | NA | NA | NA | sra-2 |
| WBGene00005029 | 0.020232312 | 0.84643798 | NA | sra-3 |
| WBGene00005030 | NA | NA | NA | sra-4 |
| WBGene00005031 | NA | NA | NA | NA |
| WBGene00005032 | 0.020317245 | 0.84643798 | NA | sra-6 |
| WBGene00005033 | 0.020925399 | 0.84643798 | NA | sra-7 |
| WBGene00005034 | NA | NA | NA | sra-8 |
| WBGene00005035 | NA | NA | NA | sra-9 |
| WBGene00005036 | -0.053348766 | 0.53925327 | NA | sra-10 |
| WBGene00005037 | 0.040107227 | 0.58693163 | NA | sra-11 |
| WBGene00005038 | 0.020925399 | 0.84643798 | NA | sra-12 |
| WBGene00005039 | NA | NA | NA | sra-13 |
| WBGene00005040 | 0.00667944 | 0.97210319 | 0.98476102 | sra-14 |
| WBGene00005041 | NA | NA | NA | NA |
| WBGene00005042 | NA | NA | NA | NA |
| WBGene00005043 | NA | NA | NA | sra-17 |
| WBGene00005044 | NA | NA | NA | sra-18 |
| WBGene00005045 | NA | NA | NA | NA |
| WBGene00005046 | NA | NA | NA | sra-20 |
| WBGene00005047 | 0.0398788 | 0.58971221 | NA | sra-21 |
| WBGene00005048 | NA | NA | NA | sra-22 |
| WBGene00005049 | NA | NA | NA | sra-23 |
| WBGene00005050 | NA | NA | NA | sra-24 |
| WBGene00005051 | 0.020232312 | 0.84643798 | NA | sra-25 |

| | | | | |
|---|---|---|---|---|
| WBGene00005052 | NA | NA | NA | sra-26 |
| WBGene00005053 | NA | NA | NA | sra-27 |
| WBGene00005054 | -0.023056499 | 0.78121253 | NA | sra-28 |
| WBGene00005055 | NA | NA | NA | sra-29 |
| WBGene00005056 | NA | NA | NA | sra-30 |
| WBGene00005057 | 0.017141582 | 0.79355725 | NA | sra-31 |
| WBGene00005058 | -0.166965874 | 0.22152487 | 0.41411461 | sra-32 |
| WBGene00005059 | 0.169967924 | 0.15950102 | 0.33408505 | sra-33 |
| WBGene00005060 | 0.020925399 | 0.84643798 | NA | sra-34 |
| WBGene00005061 | 0.006744913 | 0.95285511 | 0.97312721 | sra-35 |
| WBGene00005062 | 0.0398788 | 0.58971221 | NA | sra-36 |
| WBGene00005063 | 0.076452416 | 0.42707996 | NA | sra-37 |
| WBGene00005064 | NA | NA | NA | sra-38 |
| WBGene00005065 | -0.003108065 | 0.95728506 | NA | sra-39 |
| WBGene00005066 | NA | NA | NA | srb-1 |
| WBGene00005067 | NA | NA | NA | srb-2 |
| WBGene00005068 | -0.023056499 | 0.78121253 | NA | srb-3 |
| WBGene00005069 | NA | NA | NA | NA |
| WBGene00005070 | NA | NA | NA | srb-5 |
| WBGene00005071 | NA | NA | NA | srb-6 |
| WBGene00005072 | 0.020925399 | 0.84643798 | NA | srb-7 |
| WBGene00005073 | NA | NA | NA | srb-8 |
| WBGene00005073 | NA | NA | NA | srb-9 |
| WBGene00005074 | NA | NA | NA | srb-8 |
| WBGene00005074 | NA | NA | NA | srb-9 |
| WBGene00005075 | NA | NA | NA | NA |
| WBGene00005076 | NA | NA | NA | srb-11 |
| WBGene00005077 | -0.000848723 | 0.99109796 | 0.9949664 | src-1 |
| WBGene00005078 | -0.009207842 | 0.96261908 | 0.97924853 | src-2 |
| WBGene00005079 | NA | NA | NA | srd-1 |
| WBGene00005080 | NA | NA | NA | srd-2 |
| WBGene00005081 | 0.056411843 | 0.41891328 | NA | srd-3 |
| WBGene00005082 | NA | NA | NA | srd-4 |
| WBGene00005083 | 0.180007984 | 0.0451335 | NA | srd-5 |
| WBGene00005084 | NA | NA | NA | NA |
| WBGene00005085 | -0.002914971 | 0.95718083 | NA | srd-7 |
| WBGene00005086 | NA | NA | NA | srd-8 |
| WBGene00005087 | NA | NA | NA | srd-9 |
| WBGene00005088 | 0.020925399 | 0.84643798 | NA | srd-10 |
| WBGene00005089 | 0.020317245 | 0.84643798 | NA | srd-11 |
| WBGene00005090 | 0.020232312 | 0.84643798 | NA | srd-12 |
| WBGene00005091 | NA | NA | NA | srd-13 |
| WBGene00005092 | 1.037280985 | 0.00362297 | 0.02030772 | NA |

| | | | | |
|---|---|---|---|---|
| WBGene00005093 | -0.045959069 | 0.50707733 | NA | srd-15 |
| WBGene00005094 | NA | NA | NA | srd-16 |
| WBGene00005095 | -0.023109762 | 0.78121253 | NA | srd-17 |
| WBGene00005096 | 0.087256674 | 0.23800185 | NA | srd-18 |
| WBGene00005097 | NA | NA | NA | srd-19 |
| WBGene00005098 | -0.022930767 | 0.78121253 | NA | srd-20 |
| WBGene00005099 | NA | NA | NA | srd-21 |
| WBGene00005100 | NA | NA | NA | NA |
| WBGene00005101 | 0.128253632 | 0.25054479 | 0.44909133 | srd-23 |
| WBGene00005102 | NA | NA | NA | NA |
| WBGene00005103 | NA | NA | NA | srd-25 |
| WBGene00005104 | 0.020317245 | 0.84643798 | NA | srd-26 |
| WBGene00005105 | -0.003421551 | 0.95689871 | NA | srd-27 |
| WBGene00005106 | 0.020232312 | 0.84643798 | NA | srd-28 |
| WBGene00005107 | NA | NA | NA | srd-29 |
| WBGene00005108 | NA | NA | NA | srd-30 |
| WBGene00005109 | NA | NA | NA | srd-31 |
| WBGene00005110 | 0.037992904 | 0.65794732 | NA | srd-32 |
| WBGene00005111 | -0.055120925 | 0.45606622 | NA | srd-33 |
| WBGene00005112 | 0.038125098 | 0.61119482 | NA | srd-34 |
| WBGene00005113 | 0.042516415 | 0.55991373 | NA | srd-35 |
| WBGene00005114 | NA | NA | NA | srd-36 |
| WBGene00005115 | -0.023056499 | 0.78121253 | NA | NA |
| WBGene00005116 | NA | NA | NA | srd-38 |
| WBGene00005117 | NA | NA | NA | srd-39 |
| WBGene00005118 | NA | NA | NA | srd-40 |
| WBGene00005119 | 0.033463284 | 0.6365037 | NA | srd-41 |
| WBGene00005120 | -0.003270206 | 0.95698316 | NA | srd-42 |
| WBGene00005121 | NA | NA | NA | srd-43 |
| WBGene00005122 | NA | NA | NA | srd-44 |
| WBGene00005123 | NA | NA | NA | srd-45 |
| WBGene00005124 | NA | NA | NA | srd-46 |
| WBGene00005125 | NA | NA | NA | srd-47 |
| WBGene00005126 | 0.020317245 | 0.84643798 | NA | srd-48 |
| WBGene00005127 | 0.056304773 | 0.44260385 | NA | srd-49 |
| WBGene00005128 | 0.020925399 | 0.84643798 | NA | srd-50 |
| WBGene00005129 | 0.020317245 | 0.84643798 | NA | srd-51 |
| WBGene00005130 | -0.025848508 | 0.67901638 | NA | NA |
| WBGene00005131 | 0.174185285 | 0.32632395 | 0.52559615 | srd-53 |
| WBGene00005132 | -0.067064794 | 0.32104386 | NA | srd-54 |
| WBGene00005133 | 0.06726311 | 0.45792378 | NA | srd-55 |
| WBGene00005134 | NA | NA | NA | srd-56 |
| WBGene00005135 | NA | NA | NA | srd-58 |

| | | | | | |
|---|---|---|---|---|---|
| WBGene00005136 | | 0.036687355 | 0.59489568 | NA | srd-59 |
| WBGene00005137 | NA | | NA | NA | srd-73 |
| WBGene00005137 | NA | | NA | NA | srd-60 |
| WBGene00005138 | NA | | NA | NA | srd-61 |
| WBGene00005138 | NA | | NA | NA | srd-75 |
| WBGene00005139 | NA | | NA | NA | NA |
| WBGene00005140 | | -0.001842784 | 0.95621055 | NA | srd-63 |
| WBGene00005141 | | -0.066863888 | 0.6400607 | 0.78663047 | srd-64 |
| WBGene00005142 | | 0.020317245 | 0.84643798 | NA | srd-65 |
| WBGene00005143 | NA | | NA | NA | srd-66 |
| WBGene00005144 | NA | | NA | NA | srd-67 |
| WBGene00005145 | NA | | NA | NA | srd-68 |
| WBGene00005146 | NA | | NA | NA | srd-69 |
| WBGene00005147 | | -0.070877205 | 0.33324932 | NA | srd-70 |
| WBGene00005148 | NA | | NA | NA | srd-71 |
| WBGene00005148 | NA | | NA | NA | srd-72 |
| WBGene00005149 | | 0.019325396 | 0.77340931 | NA | sre-1 |
| WBGene00005150 | NA | | NA | NA | sre-2 |
| WBGene00005153 | | 0.261780902 | 0.12423495 | 0.28458189 | srf-3 |
| WBGene00005159 | | 0.020925399 | 0.84643798 | NA | srg-1 |
| WBGene00005160 | | 0.019675302 | 0.61254092 | NA | srg-2 |
| WBGene00005161 | | 0.036687355 | 0.59489568 | NA | srg-3 |
| WBGene00005162 | | 0.020232312 | 0.84643798 | NA | srg-4 |
| WBGene00005163 | | -0.003270206 | 0.95698316 | NA | srg-5 |
| WBGene00005164 | | 0.065014541 | 0.37399213 | NA | srg-6 |
| WBGene00005165 | | 0.020925399 | 0.84643798 | NA | srg-7 |
| WBGene00005166 | | 0.020317245 | 0.84643798 | NA | srg-8 |
| WBGene00005167 | NA | | NA | NA | srg-9 |
| WBGene00005168 | | -0.045959069 | 0.50707733 | NA | srg-10 |
| WBGene00005169 | | 0.020925399 | 0.84643798 | NA | srg-11 |
| WBGene00005170 | | 0.017279211 | 0.79380376 | NA | srg-13 |
| WBGene00005171 | NA | | NA | NA | srg-14 |
| WBGene00005172 | | -0.001993815 | 0.95612996 | NA | srg-15 |
| WBGene00005173 | NA | | NA | NA | srg-16 |
| WBGene00005174 | NA | | NA | NA | srg-17 |
| WBGene00005175 | | -0.022930767 | 0.78121253 | NA | NA |
| WBGene00005176 | NA | | NA | NA | NA |
| WBGene00005177 | NA | | NA | NA | srg-20 |
| WBGene00005178 | NA | | NA | NA | srg-21 |
| WBGene00005179 | NA | | NA | NA | srg-22 |
| WBGene00005180 | | 0.038125098 | 0.61119482 | NA | srg-23 |
| WBGene00005181 | NA | | NA | NA | srg-24 |
| WBGene00005182 | | 0.056188364 | 0.42092875 | NA | srg-25 |

| | | | | |
|---|---|---|---|---|
| WBGene00005183 | NA | NA | NA | srg-26 |
| WBGene00005184 | 0.0398788 | 0.58971221 | NA | srg-27 |
| WBGene00005185 | NA | NA | NA | srg-28 |
| WBGene00005186 | NA | NA | NA | srg-29 |
| WBGene00005187 | 0.040107227 | 0.58693163 | NA | srg-30 |
| WBGene00005188 | NA | NA | NA | srg-31 |
| WBGene00005189 | 0.074865281 | 0.33025729 | NA | srg-32 |
| WBGene00005190 | NA | NA | NA | srg-33 |
| WBGene00005191 | 0.190220605 | 0.2399004 | 0.43697209 | srg-34 |
| WBGene00005192 | NA | NA | NA | NA |
| WBGene00005193 | NA | NA | NA | srg-36 |
| WBGene00005194 | 0.038125098 | 0.61119482 | NA | srg-37 |
| WBGene00005195 | 0.020925399 | 0.84643798 | NA | srg-38 |
| WBGene00005196 | NA | NA | NA | srg-39 |
| WBGene00005197 | NA | NA | NA | srg-40 |
| WBGene00005198 | 0.038125098 | 0.61119482 | NA | srg-41 |
| WBGene00005199 | NA | NA | NA | srg-42 |
| WBGene00005200 | -0.003614866 | 0.95700117 | NA | NA |
| WBGene00005201 | -0.003421551 | 0.95689871 | NA | srg-44 |
| WBGene00005202 | NA | NA | NA | srg-45 |
| WBGene00005203 | NA | NA | NA | srg-46 |
| WBGene00005204 | NA | NA | NA | srg-47 |
| WBGene00005205 | 0.01912147 | 0.77587372 | NA | srg-48 |
| WBGene00005206 | 0.065014541 | 0.37399213 | NA | NA |
| WBGene00005207 | NA | NA | NA | srg-50 |
| WBGene00005208 | NA | NA | NA | srg-51 |
| WBGene00005210 | 0.020232312 | 0.84643798 | NA | srg-53 |
| WBGene00005211 | NA | NA | NA | srg-54 |
| WBGene00005212 | NA | NA | NA | srg-55 |
| WBGene00005213 | 0.020317245 | 0.84643798 | NA | srg-56 |
| WBGene00005214 | NA | NA | NA | srg-57 |
| WBGene00005215 | NA | NA | NA | srg-58 |
| WBGene00005216 | NA | NA | NA | srg-59 |
| WBGene00005217 | NA | NA | NA | srg-60 |
| WBGene00005218 | NA | NA | NA | srg-61 |
| WBGene00005219 | 0.038125098 | 0.61119482 | NA | srg-62 |
| WBGene00005220 | NA | NA | NA | NA |
| WBGene00005221 | NA | NA | NA | srg-64 |
| WBGene00005222 | 0.138293571 | 0.13257743 | 0.29580404 | srg-65 |
| WBGene00005223 | 0.020232312 | 0.84643798 | NA | srg-66 |
| WBGene00005224 | NA | NA | NA | srg-67 |
| WBGene00005225 | NA | NA | NA | srg-68 |
| WBGene00005226 | 0.038633797 | 0.57999349 | NA | srg-69 |

| | | | | |
|---|---|---|---|---|
| WBGene00005227 | 0.033463284 | 0.6365037 | NA | srh-1 |
| WBGene00005228 | 1.281019106 | 0.01365314 | 0.05737531 | srh-2 |
| WBGene00005229 | 0.020232312 | 0.84643798 | NA | srh-3 |
| WBGene00005230 | NA | NA | NA | srh-4 |
| WBGene00005231 | NA | NA | NA | srh-5 |
| WBGene00005232 | NA | NA | NA | NA |
| WBGene00005233 | NA | NA | NA | srh-7 |
| WBGene00005234 | -0.023056499 | 0.78121253 | NA | srh-8 |
| WBGene00005235 | 0.056304773 | 0.44260385 | NA | NA |
| WBGene00005236 | NA | NA | NA | srh-10 |
| WBGene00005237 | NA | NA | NA | srh-11 |
| WBGene00005238 | NA | NA | NA | NA |
| WBGene00005239 | NA | NA | NA | NA |
| WBGene00005240 | 0.020925399 | 0.84643798 | NA | srh-15 |
| WBGene00005241 | -0.066883576 | 0.32235182 | NA | srh-16 |
| WBGene00005242 | -0.005581345 | 0.94060131 | NA | srh-17 |
| WBGene00005243 | 0.038125098 | 0.61119482 | NA | srh-18 |
| WBGene00005244 | NA | NA | NA | srh-19 |
| WBGene00005245 | NA | NA | NA | srh-20 |
| WBGene00005246 | 0.020317245 | 0.84643798 | NA | srh-21 |
| WBGene00005247 | NA | NA | NA | srh-22 |
| WBGene00005248 | NA | NA | NA | srh-23 |
| WBGene00005249 | 0.020232312 | 0.84643798 | NA | srh-25 |
| WBGene00005250 | NA | NA | NA | srh-27 |
| WBGene00005251 | -0.024740619 | 0.69507748 | NA | srh-28 |
| WBGene00005252 | NA | NA | NA | NA |
| WBGene00005253 | 0.036687355 | 0.59489568 | NA | srh-30 |
| WBGene00005254 | NA | NA | NA | srh-31 |
| WBGene00005255 | -0.023056499 | 0.78121253 | NA | srh-32 |
| WBGene00005256 | -0.00346345 | 0.95708614 | NA | srh-33 |
| WBGene00005257 | -0.027373974 | 0.68245543 | NA | srh-34 |
| WBGene00005258 | 0.016985298 | 0.837043 | NA | srh-35 |
| WBGene00005259 | NA | NA | NA | srh-36 |
| WBGene00005260 | NA | NA | NA | srh-37 |
| WBGene00005261 | 0.020232312 | 0.84643798 | NA | srh-38 |
| WBGene00005262 | 0.020317245 | 0.84643798 | NA | srh-39 |
| WBGene00005263 | -0.023109762 | 0.78121253 | NA | srh-40 |
| WBGene00005264 | NA | NA | NA | srh-41 |
| WBGene00005265 | NA | NA | NA | srh-42 |
| WBGene00005266 | NA | NA | NA | NA |
| WBGene00005267 | NA | NA | NA | srh-44 |
| WBGene00005268 | NA | NA | NA | srh-45 |
| WBGene00005269 | 0.085309563 | 0.24820057 | NA | srh-46 |

| | | | | |
|---|---|---|---|---|
| WBGene00005270 | NA | NA | NA | srh-47 |
| WBGene00005271 | -0.096217903 | 0.56649263 | 0.73548095 | srh-48 |
| WBGene00005272 | 0.020925399 | 0.84643798 | NA | srh-49 |
| WBGene00005273 | NA | NA | NA | srh-50 |
| WBGene00005274 | NA | NA | NA | srh-51 |
| WBGene00005275 | -0.023056499 | 0.78121253 | NA | srh-52 |
| WBGene00005276 | NA | NA | NA | NA |
| WBGene00005277 | NA | NA | NA | NA |
| WBGene00005278 | NA | NA | NA | srh-55 |
| WBGene00005279 | NA | NA | NA | srh-56 |
| WBGene00005280 | NA | NA | NA | srh-57 |
| WBGene00005280 | NA | NA | NA | srh-305 |
| WBGene00005281 | 0.020317245 | 0.84643798 | NA | srh-59 |
| WBGene00005282 | NA | NA | NA | srh-60 |
| WBGene00005283 | NA | NA | NA | srh-61 |
| WBGene00005284 | NA | NA | NA | srh-62 |
| WBGene00005285 | NA | NA | NA | srh-63 |
| WBGene00005286 | NA | NA | NA | NA |
| WBGene00005287 | NA | NA | NA | NA |
| WBGene00005288 | NA | NA | NA | srh-67 |
| WBGene00005289 | 0.020925399 | 0.84643798 | NA | srh-68 |
| WBGene00005290 | -0.003108065 | 0.95728506 | NA | srh-69 |
| WBGene00005291 | 0.172497267 | 0.31711677 | 0.51727506 | srh-70 |
| WBGene00005292 | 0.052360174 | 0.60300068 | 0.75988622 | srh-71 |
| WBGene00005293 | -0.027350315 | 0.68023343 | NA | srh-72 |
| WBGene00005294 | NA | NA | NA | srh-73 |
| WBGene00005295 | 0.036692668 | 0.59311964 | NA | srh-74 |
| WBGene00005296 | 0.056188364 | 0.42092875 | NA | srh-75 |
| WBGene00005297 | 0.020317245 | 0.84643798 | NA | srh-76 |
| WBGene00005298 | 0.020317245 | 0.84643798 | NA | NA |
| WBGene00005299 | NA | NA | NA | srh-78 |
| WBGene00005300 | NA | NA | NA | srh-79 |
| WBGene00005301 | NA | NA | NA | srh-80 |
| WBGene00005302 | NA | NA | NA | NA |
| WBGene00005303 | 0.020317245 | 0.84643798 | NA | srh-82 |
| WBGene00005304 | NA | NA | NA | srh-83 |
| WBGene00005305 | NA | NA | NA | NA |
| WBGene00005306 | NA | NA | NA | srh-85 |
| WBGene00005307 | 0.020317245 | 0.84643798 | NA | NA |
| WBGene00005308 | -0.026384613 | 0.67299636 | NA | srh-87 |
| WBGene00005309 | NA | NA | NA | srh-88 |
| WBGene00005310 | NA | NA | NA | srh-89 |
| WBGene00005311 | 0.020925399 | 0.84643798 | NA | NA |

| | | | | |
|---|---|---|---|---|
| WBGene00005312 | NA | NA | NA | NA |
| WBGene00005313 | -0.046394786 | 0.50231372 | NA | srh-92 |
| WBGene00005314 | NA | NA | NA | NA |
| WBGene00005315 | -0.023056499 | 0.78121253 | NA | srh-95 |
| WBGene00005316 | NA | NA | NA | srh-97 |
| WBGene00005317 | NA | NA | NA | srh-98 |
| WBGene00005318 | -0.023056499 | 0.78121253 | NA | srh-99 |
| WBGene00005319 | NA | NA | NA | srh-100 |
| WBGene00005320 | NA | NA | NA | NA |
| WBGene00005321 | NA | NA | NA | srh-102 |
| WBGene00005322 | NA | NA | NA | NA |
| WBGene00005323 | 0.038125098 | 0.61119482 | NA | srh-104 |
| WBGene00005324 | 0.020925399 | 0.84643798 | NA | T27A1.1 |
| WBGene00005325 | -0.022930767 | 0.78121253 | NA | NA |
| WBGene00005326 | NA | NA | NA | NA |
| WBGene00005327 | NA | NA | NA | NA |
| WBGene00005328 | NA | NA | NA | srh-109 |
| WBGene00005329 | NA | NA | NA | NA |
| WBGene00005330 | NA | NA | NA | srh-111 |
| WBGene00005331 | 0.040107227 | 0.58693163 | NA | srh-112 |
| WBGene00005332 | NA | NA | NA | srh-113 |
| WBGene00005333 | NA | NA | NA | NA |
| WBGene00005334 | NA | NA | NA | srh-115 |
| WBGene00005335 | NA | NA | NA | srh-116 |
| WBGene00005336 | NA | NA | NA | NA |
| WBGene00005337 | NA | NA | NA | srh-118 |
| WBGene00005338 | NA | NA | NA | srh-119 |
| WBGene00005339 | NA | NA | NA | srh-120 |
| WBGene00005340 | NA | NA | NA | srh-122 |
| WBGene00005341 | 0.040107227 | 0.58693163 | NA | srh-123 |
| WBGene00005342 | 0.0398788 | 0.58971221 | NA | srh-125 |
| WBGene00005343 | NA | NA | NA | NA |
| WBGene00005344 | 0.020317245 | 0.84643798 | NA | srh-127 |
| WBGene00005345 | NA | NA | NA | srh-128 |
| WBGene00005346 | 0.018795205 | 0.77532208 | NA | srh-129 |
| WBGene00005347 | NA | NA | NA | srh-130 |
| WBGene00005348 | 0.020925399 | 0.84643798 | NA | srh-131 |
| WBGene00005349 | 0.108942554 | 0.15594412 | NA | srh-132 |
| WBGene00005350 | 0.020925399 | 0.84643798 | NA | srh-133 |
| WBGene00005351 | NA | NA | NA | srh-134 |
| WBGene00005352 | NA | NA | NA | srh-135 |
| WBGene00005353 | NA | NA | NA | srh-136 |
| WBGene00005354 | NA | NA | NA | srh-137 |

| | | | | |
|---|---|---|---|---|
| WBGene00005355 | NA | NA | NA | srh-138 |
| WBGene00005356 | NA | NA | NA | srh-139 |
| WBGene00005357 | NA | NA | NA | srh-140 |
| WBGene00005358 | NA | NA | NA | srh-141 |
| WBGene00005359 | NA | NA | NA | srh-142 |
| WBGene00005360 | NA | NA | NA | NA |
| WBGene00005361 | NA | NA | NA | srh-145 |
| WBGene00005362 | -0.040794878 | 0.53203165 | NA | srh-146 |
| WBGene00005363 | -0.023056499 | 0.78121253 | NA | srh-147 |
| WBGene00005364 | 0.048485426 | 0.45975815 | NA | srh-148 |
| WBGene00005365 | NA | NA | NA | srh-149 |
| WBGene00005366 | NA | NA | NA | NA |
| WBGene00005367 | NA | NA | NA | NA |
| WBGene00005368 | NA | NA | NA | NA |
| WBGene00005369 | NA | NA | NA | NA |
| WBGene00005370 | NA | NA | NA | srh-154 |
| WBGene00005371 | NA | NA | NA | NA |
| WBGene00005372 | NA | NA | NA | NA |
| WBGene00005373 | NA | NA | NA | NA |
| WBGene00005374 | 0.020317245 | 0.84643798 | NA | NA |
| WBGene00005375 | NA | NA | NA | srh-159 |
| WBGene00005376 | NA | NA | NA | NA |
| WBGene00005377 | 0.040107227 | 0.58693163 | NA | NA |
| WBGene00005378 | NA | NA | NA | NA |
| WBGene00005379 | NA | NA | NA | NA |
| WBGene00005380 | NA | NA | NA | NA |
| WBGene00005381 | NA | NA | NA | srh-165 |
| WBGene00005382 | NA | NA | NA | srh-166 |
| WBGene00005383 | NA | NA | NA | srh-167 |
| WBGene00005384 | NA | NA | NA | NA |
| WBGene00005385 | 0.020317245 | 0.84643798 | NA | srh-169 |
| WBGene00005386 | NA | NA | NA | srh-170 |
| WBGene00005387 | 0.038125098 | 0.61119482 | NA | srh-171 |
| WBGene00005388 | NA | NA | NA | srh-172 |
| WBGene00005389 | -0.023109762 | 0.78121253 | NA | srh-173 |
| WBGene00005390 | NA | NA | NA | srh-174 |
| WBGene00005391 | NA | NA | NA | NA |
| WBGene00005392 | NA | NA | NA | srh-177 |
| WBGene00005393 | NA | NA | NA | srh-178 |
| WBGene00005394 | NA | NA | NA | srh-179 |
| WBGene00005395 | NA | NA | NA | srh-180 |
| WBGene00005396 | NA | NA | NA | srh-181 |
| WBGene00005397 | 0.042065281 | 0.558937 | NA | srh-182 |

| | | | | | |
|---|---|---|---|---|---|
| WBGene00005398 | | 0.020317245 | 0.84643798 | NA | srh-183 |
| WBGene00005399 | NA | | NA | NA | srh-184 |
| WBGene00005400 | | -0.002914971 | 0.95718083 | NA | srh-185 |
| WBGene00005401 | NA | | NA | NA | srh-186 |
| WBGene00005402 | NA | | NA | NA | srh-187 |
| WBGene00005403 | | -0.022930767 | 0.78121253 | NA | NA |
| WBGene00005404 | NA | | NA | NA | srh-190 |
| WBGene00005405 | NA | | NA | NA | srh-192 |
| WBGene00005406 | | 0.038125098 | 0.61119482 | NA | srh-193 |
| WBGene00005407 | NA | | NA | NA | srh-194 |
| WBGene00005408 | | -0.003270206 | 0.95698316 | NA | srh-195 |
| WBGene00005409 | NA | | NA | NA | NA |
| WBGene00005410 | | 0.020232312 | 0.84643798 | NA | srh-199 |
| WBGene00005411 | NA | | NA | NA | srh-200 |
| WBGene00005412 | NA | | NA | NA | srh-201 |
| WBGene00005413 | NA | | NA | NA | NA |
| WBGene00005414 | NA | | NA | NA | srh-203 |
| WBGene00005415 | NA | | NA | NA | srh-204 |
| WBGene00005416 | NA | | NA | NA | srh-206 |
| WBGene00005417 | | 0.020317245 | 0.84643798 | NA | srh-207 |
| WBGene00005418 | NA | | NA | NA | srh-208 |
| WBGene00005419 | | 0.020925399 | 0.84643798 | NA | srh-209 |
| WBGene00005420 | | 0.040218942 | 0.56174727 | NA | srh-210 |
| WBGene00005421 | NA | | NA | NA | srh-211 |
| WBGene00005422 | | 0.090327171 | 0.34257245 | 0.54287296 | srh-212 |
| WBGene00005423 | | 0.07892265 | 0.25795693 | NA | srh-213 |
| WBGene00005424 | | 0.020317245 | 0.84643798 | NA | srh-214 |
| WBGene00005425 | NA | | NA | NA | srh-215 |
| WBGene00005426 | NA | | NA | NA | srh-216 |
| WBGene00005427 | | -0.006293139 | 0.94680499 | 0.97026471 | srh-217 |
| WBGene00005428 | NA | | NA | NA | srh-218 |
| WBGene00005429 | NA | | NA | NA | srh-219 |
| WBGene00005430 | | 0.020925399 | 0.84643798 | NA | srh-220 |
| WBGene00005431 | | -0.023056499 | 0.78121253 | NA | NA |
| WBGene00005432 | NA | | NA | NA | srh-222 |
| WBGene00005433 | NA | | NA | NA | srh-223 |
| WBGene00005434 | NA | | NA | NA | NA |
| WBGene00005435 | NA | | NA | NA | srh-228 |
| WBGene00005435 | NA | | NA | NA | srh-227 |
| WBGene00005436 | NA | | NA | NA | srh-228 |
| WBGene00005436 | NA | | NA | NA | srh-227 |
| WBGene00005437 | NA | | NA | NA | srh-229 |
| WBGene00005438 | | 0.048485426 | 0.45975815 | NA | NA |

| | | | | |
|---|---|---|---|---|
| WBGene00005439 | NA | NA | NA | srh-231 |
| WBGene00005440 | NA | NA | NA | srh-233 |
| WBGene00005441 | 0.020232312 | 0.84643798 | NA | srh-234 |
| WBGene00005442 | NA | NA | NA | srh-235 |
| WBGene00005443 | NA | NA | NA | srh-236 |
| WBGene00005444 | 1.110681035 | 1.46E-06 | 2.20E-05 | srh-237 |
| WBGene00005445 | NA | NA | NA | NA |
| WBGene00005446 | NA | NA | NA | srh-239 |
| WBGene00005447 | -0.023109762 | 0.78121253 | NA | srh-240 |
| WBGene00005448 | NA | NA | NA | srh-241 |
| WBGene00005449 | NA | NA | NA | srh-243 |
| WBGene00005450 | NA | NA | NA | srh-244 |
| WBGene00005451 | 0.020317245 | 0.84643798 | NA | srh-245 |
| WBGene00005452 | NA | NA | NA | srh-246 |
| WBGene00005453 | NA | NA | NA | srh-247 |
| WBGene00005454 | NA | NA | NA | srh-248 |
| WBGene00005455 | NA | NA | NA | NA |
| WBGene00005456 | -0.045959069 | 0.50707733 | NA | srh-250 |
| WBGene00005457 | NA | NA | NA | srh-251 |
| WBGene00005458 | NA | NA | NA | srh-252 |
| WBGene00005459 | NA | NA | NA | NA |
| WBGene00005460 | NA | NA | NA | NA |
| WBGene00005461 | NA | NA | NA | srh-255 |
| WBGene00005462 | NA | NA | NA | NA |
| WBGene00005463 | 0.020232312 | 0.84643798 | NA | srh-257 |
| WBGene00005464 | NA | NA | NA | srh-258 |
| WBGene00005465 | NA | NA | NA | NA |
| WBGene00005466 | NA | NA | NA | NA |
| WBGene00005467 | NA | NA | NA | srh-261 |
| WBGene00005468 | -0.006738828 | 0.92480778 | NA | NA |
| WBGene00005469 | NA | NA | NA | NA |
| WBGene00005470 | NA | NA | NA | srh-264 |
| WBGene00005471 | NA | NA | NA | srh-265 |
| WBGene00005472 | NA | NA | NA | srh-266 |
| WBGene00005473 | NA | NA | NA | NA |
| WBGene00005474 | 0.020232312 | 0.84643798 | NA | srh-268 |
| WBGene00005475 | 0.127758703 | 0.16886741 | NA | srh-269 |
| WBGene00005476 | NA | NA | NA | srh-270 |
| WBGene00005477 | 0.085537126 | 0.24698522 | NA | srh-271 |
| WBGene00005478 | 0.020317245 | 0.84643798 | NA | srh-272 |
| WBGene00005479 | NA | NA | NA | srh-274 |
| WBGene00005480 | NA | NA | NA | srh-275 |
| WBGene00005481 | 0.020317245 | 0.84643798 | NA | srh-276 |

| | | | | |
|---|---|---|---|---|
| WBGene00005482 | 0.020232312 | 0.84643798 | NA | srh-277 |
| WBGene00005483 | NA | NA | NA | srh-278 |
| WBGene00005484 | NA | NA | NA | srh-279 |
| WBGene00005485 | NA | NA | NA | NA |
| WBGene00005486 | 0.020925399 | 0.84643798 | NA | srh-281 |
| WBGene00005487 | 0.042199095 | 0.55922995 | NA | srh-282 |
| WBGene00005488 | -0.00346345 | 0.95708614 | NA | srh-283 |
| WBGene00005489 | NA | NA | NA | srh-284 |
| WBGene00005490 | 0.020317245 | 0.84643798 | NA | srh-286 |
| WBGene00005491 | NA | NA | NA | srh-287 |
| WBGene00005492 | NA | NA | NA | srh-288 |
| WBGene00005493 | NA | NA | NA | srh-289 |
| WBGene00005494 | NA | NA | NA | srh-290 |
| WBGene00005495 | NA | NA | NA | srh-291 |
| WBGene00005496 | NA | NA | NA | srh-292 |
| WBGene00005497 | NA | NA | NA | srh-293 |
| WBGene00005498 | NA | NA | NA | NA |
| WBGene00005499 | NA | NA | NA | srh-295 |
| WBGene00005500 | 0.020925399 | 0.84643798 | NA | srh-296 |
| WBGene00005501 | -0.022930767 | 0.78121253 | NA | srh-297 |
| WBGene00005502 | -0.003108065 | 0.95728506 | NA | srh-298 |
| WBGene00005503 | NA | NA | NA | srh-299 |
| WBGene00005504 | 0.020317245 | 0.84643798 | NA | srh-300 |
| WBGene00005505 | NA | NA | NA | srh-301 |
| WBGene00005506 | NA | NA | NA | NA |
| WBGene00005507 | NA | NA | NA | srh-303 |
| WBGene00005508 | NA | NA | NA | srh-304 |
| WBGene00005509 | NA | NA | NA | srh-57 |
| WBGene00005509 | NA | NA | NA | srh-305 |
| WBGene00005510 | NA | NA | NA | NA |
| WBGene00005511 | NA | NA | NA | NA |
| WBGene00005512 | 0.020232312 | 0.84643798 | NA | srh-105 |
| WBGene00005513 | 0.031735686 | 0.7706112 | 0.86948778 | sri-1 |
| WBGene00005514 | NA | NA | NA | sri-2 |
| WBGene00005515 | NA | NA | NA | NA |
| WBGene00005516 | NA | NA | NA | sri-4 |
| WBGene00005517 | -0.079857863 | 0.31359805 | NA | sri-5 |
| WBGene00005518 | NA | NA | NA | sri-6 |
| WBGene00005519 | NA | NA | NA | sri-7 |
| WBGene00005520 | -0.052235464 | 0.65137345 | 0.79364585 | NA |
| WBGene00005521 | NA | NA | NA | sri-9 |
| WBGene00005522 | 0.058314668 | 0.40208118 | NA | sri-10 |
| WBGene00005523 | -0.045959069 | 0.50707733 | NA | sri-11 |

| | | | | |
|---|---|---|---|---|
| WBGene00005524 | -0.045771477 | 0.50913401 | NA | sri-12 |
| WBGene00005525 | NA | NA | NA | sri-13 |
| WBGene00005526 | NA | NA | NA | sri-14 |
| WBGene00005527 | NA | NA | NA | sri-15 |
| WBGene00005528 | -0.0260089 | 0.6772113 | NA | sri-16 |
| WBGene00005529 | 0.020232312 | 0.84643798 | NA | sri-17 |
| WBGene00005530 | NA | NA | NA | sri-18 |
| WBGene00005531 | NA | NA | NA | sri-19 |
| WBGene00005532 | -0.046394786 | 0.50231372 | NA | sri-20 |
| WBGene00005533 | 0.020232312 | 0.84643798 | NA | sri-21 |
| WBGene00005534 | 0.065014541 | 0.37399213 | NA | NA |
| WBGene00005535 | 0.116737817 | 0.11962812 | NA | NA |
| WBGene00005536 | NA | NA | NA | sri-24 |
| WBGene00005537 | NA | NA | NA | sri-25 |
| WBGene00005538 | NA | NA | NA | sri-26 |
| WBGene00005539 | NA | NA | NA | sri-27 |
| WBGene00005540 | -0.00346345 | 0.95708614 | NA | sri-28 |
| WBGene00005541 | 0.015451648 | 0.80337623 | NA | sri-29 |
| WBGene00005542 | 0.033034883 | 0.64219503 | NA | sri-30 |
| WBGene00005543 | 0.020232312 | 0.84643798 | NA | sri-31 |
| WBGene00005544 | 0.020317245 | 0.84643798 | NA | sri-32 |
| WBGene00005545 | NA | NA | NA | sri-33 |
| WBGene00005546 | 0.020232312 | 0.84643798 | NA | sri-34 |
| WBGene00005547 | 0.038125098 | 0.61119482 | NA | NA |
| WBGene00005548 | NA | NA | NA | sri-36 |
| WBGene00005549 | NA | NA | NA | sri-37 |
| WBGene00005550 | -0.022930767 | 0.78121253 | NA | sri-38 |
| WBGene00005551 | NA | NA | NA | sri-39 |
| WBGene00005552 | -0.921954802 | 3.28E-12 | 1.15E-10 | sri-40 |
| WBGene00005553 | NA | NA | NA | NA |
| WBGene00005554 | 0.033463284 | 0.6365037 | NA | sri-42 |
| WBGene00005555 | NA | NA | NA | sri-43 |
| WBGene00005556 | -0.022930767 | 0.78121253 | NA | NA |
| WBGene00005557 | -0.066883576 | 0.32235182 | NA | sri-45 |
| WBGene00005558 | NA | NA | NA | sri-46 |
| WBGene00005559 | 0.020232312 | 0.84643798 | NA | sri-47 |
| WBGene00005560 | NA | NA | NA | sri-48 |
| WBGene00005561 | NA | NA | NA | NA |
| WBGene00005562 | -0.023109762 | 0.78121253 | NA | sri-50 |
| WBGene00005563 | NA | NA | NA | sri-51 |
| WBGene00005564 | NA | NA | NA | NA |
| WBGene00005565 | 0.020317245 | 0.84643798 | NA | sri-53 |
| WBGene00005566 | NA | NA | NA | sri-54 |

| | | | | |
|---|---|---|---|---|
| WBGene00005567 | -0.11213261 | 0.12265889 | NA | NA |
| WBGene00005568 | 0.018998992 | 0.77286019 | NA | NA |
| WBGene00005569 | -0.121342735 | 0.24147567 | 0.43933988 | sri-57 |
| WBGene00005570 | 0.020232312 | 0.84643798 | NA | NA |
| WBGene00005571 | -0.031265512 | 0.69836883 | NA | NA |
| WBGene00005572 | NA | NA | NA | sri-60 |
| WBGene00005573 | NA | NA | NA | sri-61 |
| WBGene00005574 | NA | NA | NA | sri-62 |
| WBGene00005575 | NA | NA | NA | sri-63 |
| WBGene00005576 | NA | NA | NA | NA |
| WBGene00005577 | 0.020232312 | 0.84643798 | NA | sri-65 |
| WBGene00005578 | 0.020925399 | 0.84643798 | NA | sri-66 |
| WBGene00005579 | 0.020925399 | 0.84643798 | NA | sri-67 |
| WBGene00005580 | NA | NA | NA | NA |
| WBGene00005581 | NA | NA | NA | sri-69 |
| WBGene00005582 | 0.020925399 | 0.84643798 | NA | sri-70 |
| WBGene00005583 | NA | NA | NA | sri-71 |
| WBGene00005584 | 0.020232312 | 0.84643798 | NA | sri-72 |
| WBGene00005585 | NA | NA | NA | sri-73 |
| WBGene00005586 | 0.020317245 | 0.84643798 | NA | sri-74 |
| WBGene00005587 | NA | NA | NA | NA |
| WBGene00005588 | NA | NA | NA | NA |
| WBGene00005589 | 0.020925399 | 0.84643798 | NA | sri-77 |
| WBGene00005590 | 0.020317245 | 0.84643798 | NA | sri-78 |
| WBGene00005591 | NA | NA | NA | srj-1 |
| WBGene00005592 | 0.038125098 | 0.61119482 | NA | srj-4 |
| WBGene00005593 | NA | NA | NA | srj-5 |
| WBGene00005594 | 0.020925399 | 0.84643798 | NA | srj-6 |
| WBGene00005595 | 0.020925399 | 0.84643798 | NA | srj-7 |
| WBGene00005596 | NA | NA | NA | srj-8 |
| WBGene00005597 | 0.016985298 | 0.837043 | NA | srj-9 |
| WBGene00005598 | 0.020232312 | 0.84643798 | NA | srj-10 |
| WBGene00005599 | -0.003270206 | 0.95698316 | NA | srj-11 |
| WBGene00005600 | NA | NA | NA | NA |
| WBGene00005601 | NA | NA | NA | srj-13 |
| WBGene00005602 | 1.276645707 | 0.00031071 | 0.00259062 | srj-14 |
| WBGene00005603 | NA | NA | NA | srj-15 |
| WBGene00005604 | NA | NA | NA | srj-16 |
| WBGene00005605 | NA | NA | NA | NA |
| WBGene00005606 | 0.038125098 | 0.61119482 | NA | srj-18 |
| WBGene00005607 | NA | NA | NA | srj-19 |
| WBGene00005608 | NA | NA | NA | srj-20 |
| WBGene00005609 | NA | NA | NA | srj-21 |

| | | | | |
|---|---|---|---|---|
| WBGene00005610 | NA | NA | NA | srj-22 |
| WBGene00005611 | 0.042516415 | 0.55991373 | NA | srj-23 |
| WBGene00005612 | NA | NA | NA | srj-24 |
| WBGene00005613 | NA | NA | NA | srj-25 |
| WBGene00005614 | 0.015070939 | 0.84337138 | NA | srj-26 |
| WBGene00005615 | NA | NA | NA | srj-27 |
| WBGene00005616 | 0.179129814 | 0.09050034 | 0.2288455 | srj-29 |
| WBGene00005617 | -0.003421551 | 0.95689871 | NA | srj-32 |
| WBGene00005618 | NA | NA | NA | srj-33 |
| WBGene00005619 | NA | NA | NA | NA |
| WBGene00005620 | NA | NA | NA | NA |
| WBGene00005621 | NA | NA | NA | srj-37 |
| WBGene00005622 | NA | NA | NA | srj-38 |
| WBGene00005623 | NA | NA | NA | srj-39 |
| WBGene00005624 | NA | NA | NA | srj-40 |
| WBGene00005625 | NA | NA | NA | srj-42 |
| WBGene00005626 | NA | NA | NA | srj-44 |
| WBGene00005627 | NA | NA | NA | srj-45 |
| WBGene00005628 | NA | NA | NA | srj-49 |
| WBGene00005629 | NA | NA | NA | srj-50 |
| WBGene00005630 | NA | NA | NA | NA |
| WBGene00005631 | 0.098261066 | 0.23575476 | NA | srj-52 |
| WBGene00005632 | -0.027535543 | 0.68063395 | NA | srj-53 |
| WBGene00005633 | NA | NA | NA | srj-54 |
| WBGene00005634 | NA | NA | NA | srj-55 |
| WBGene00005635 | NA | NA | NA | srj-57 |
| WBGene00005638 | -0.049904855 | 0.54559715 | NA | srm-1 |
| WBGene00005639 | 0.020317245 | 0.84643798 | NA | srm-2 |
| WBGene00005640 | -0.023109762 | 0.78121253 | NA | srn-1 |
| WBGene00005641 | 0.298682843 | 0.14916714 | 0.32036908 | sro-1 |
| WBGene00005642 | 0.672234173 | 0.00064914 | 0.00485892 | srp-1 |
| WBGene00005643 | 0.105695152 | 0.54619347 | 0.71941196 | srp-2 |
| WBGene00005644 | 0.120889669 | 0.5300545 | 0.70678761 | srp-3 |
| WBGene00005646 | NA | NA | NA | NA |
| WBGene00005647 | 0.496960255 | 0.02662035 | 0.09517668 | srp-6 |
| WBGene00005648 | 0.032347395 | 0.67047539 | 0.80824686 | srp-7 |
| WBGene00005649 | 0.642367725 | 0.03566522 | 0.11763336 | srp-8 |
| WBGene00005650 | 0.020317245 | 0.84643798 | NA | NA |
| WBGene00005651 | NA | NA | NA | NA |
| WBGene00005652 | 0.280182341 | 0.14298466 | 0.31138456 | srr-1 |
| WBGene00005653 | 3.77427933 | 0.00216702 | 0.01330289 | srr-2 |
| WBGene00005654 | 0.168867312 | 0.29229789 | 0.4930906 | srr-3 |
| WBGene00005655 | 1.768123444 | 8.52E-21 | 6.74E-19 | srr-4 |

| | | | | |
|---|---|---|---|---|
| WBGene00005656 | 0.018476367 | 0.76699386 | NA | srr-5 |
| WBGene00005657 | 2.43039302 | 0.00111548 | 0.00769794 | srr-6 |
| WBGene00005658 | NA | NA | NA | srr-7 |
| WBGene00005659 | 0.038125098 | 0.61119482 | NA | srr-8 |
| WBGene00005660 | NA | NA | NA | srr-9 |
| WBGene00005661 | NA | NA | NA | srr-10 |
| WBGene00005662 | -0.218415564 | 0.05034853 | 0.1510559 | sars-2 |
| WBGene00005663 | -0.302848363 | 1.91E-05 | 0.00021922 | sars-1 |
| WBGene00005664 | 0.01912147 | 0.77587372 | NA | sru-1 |
| WBGene00005665 | 0.056188364 | 0.42092875 | NA | sru-2 |
| WBGene00005666 | -0.023056499 | 0.78121253 | NA | sru-3 |
| WBGene00005667 | NA | NA | NA | sru-4 |
| WBGene00005668 | NA | NA | NA | NA |
| WBGene00005669 | NA | NA | NA | sru-6 |
| WBGene00005670 | 0.0398788 | 0.58971221 | NA | sru-7 |
| WBGene00005671 | 0.033463284 | 0.6365037 | NA | sru-8 |
| WBGene00005672 | 0.020232312 | 0.84643798 | NA | sru-9 |
| WBGene00005673 | NA | NA | NA | NA |
| WBGene00005674 | NA | NA | NA | sru-11 |
| WBGene00005675 | NA | NA | NA | sru-12 |
| WBGene00005676 | NA | NA | NA | sru-13 |
| WBGene00005677 | -0.023056499 | 0.78121253 | NA | NA |
| WBGene00005678 | NA | NA | NA | sru-15 |
| WBGene00005679 | NA | NA | NA | sru-16 |
| WBGene00005680 | NA | NA | NA | sru-17 |
| WBGene00005681 | 0.020317245 | 0.84643798 | NA | sru-18 |
| WBGene00005682 | NA | NA | NA | sru-19 |
| WBGene00005683 | NA | NA | NA | sru-20 |
| WBGene00005684 | NA | NA | NA | sru-21 |
| WBGene00005685 | 0.020232312 | 0.84643798 | NA | sru-22 |
| WBGene00005686 | 0.060105893 | 0.41982798 | NA | sru-23 |
| WBGene00005687 | 0.020317245 | 0.84643798 | NA | sru-24 |
| WBGene00005688 | -0.125229132 | 0.27784332 | 0.47767779 | sru-25 |
| WBGene00005689 | 0.020232312 | 0.84643798 | NA | sru-26 |
| WBGene00005690 | -0.022930767 | 0.78121253 | NA | sru-27 |
| WBGene00005691 | 0.020925399 | 0.84643798 | NA | sru-28 |
| WBGene00005692 | NA | NA | NA | sru-29 |
| WBGene00005693 | NA | NA | NA | sru-30 |
| WBGene00005694 | NA | NA | NA | sru-31 |
| WBGene00005695 | NA | NA | NA | NA |
| WBGene00005696 | NA | NA | NA | sru-33 |
| WBGene00005697 | NA | NA | NA | NA |
| WBGene00005698 | NA | NA | NA | sru-35 |

| | | | | | |
|---|---|---|---|---|---|
| WBGene00005699 | | -0.023056499 | 0.78121253 | NA | sru-36 |
| WBGene00005700 | | -0.003421551 | 0.95689871 | NA | sru-37 |
| WBGene00005701 | NA | | NA | NA | sru-38 |
| WBGene00005702 | | -0.001993815 | 0.95612996 | NA | sru-39 |
| WBGene00005703 | | 0.065014541 | 0.37399213 | NA | sru-40 |
| WBGene00005704 | | 0.020925399 | 0.84643798 | NA | NA |
| WBGene00005705 | NA | | NA | NA | sru-42 |
| WBGene00005706 | NA | | NA | NA | NA |
| WBGene00005707 | | 0.020317245 | 0.84643798 | NA | sru-44 |
| WBGene00005708 | NA | | NA | NA | sru-45 |
| WBGene00005709 | NA | | NA | NA | NA |
| WBGene00005710 | NA | | NA | NA | sru-47 |
| WBGene00005711 | | 0.020317245 | 0.84643798 | NA | sru-48 |
| WBGene00005712 | | 0.244463787 | 0.20729251 | 0.39645783 | srv-1 |
| WBGene00005713 | | 0.040218942 | 0.56174727 | NA | srv-2 |
| WBGene00005714 | NA | | NA | NA | srv-3 |
| WBGene00005715 | | 0.020232312 | 0.84643798 | NA | srv-4 |
| WBGene00005716 | | 0.020925399 | 0.84643798 | NA | srv-5 |
| WBGene00005717 | | 0.0398788 | 0.58971221 | NA | srv-6 |
| WBGene00005718 | | 0.049146204 | 0.78220925 | 0.87737414 | srv-7 |
| WBGene00005719 | | 0.020925399 | 0.84643798 | NA | srv-8 |
| WBGene00005720 | | 0.038125098 | 0.61119482 | NA | srv-9 |
| WBGene00005721 | NA | | NA | NA | srv-10 |
| WBGene00005722 | NA | | NA | NA | srv-11 |
| WBGene00005723 | | -0.023109762 | 0.78121253 | NA | srv-12 |
| WBGene00005724 | | 0.020232312 | 0.84643798 | NA | srv-13 |
| WBGene00005725 | | 0.020317245 | 0.84643798 | NA | srv-14 |
| WBGene00005726 | | 0.093320698 | 0.44732956 | 0.64038659 | srv-15 |
| WBGene00005727 | NA | | NA | NA | srv-16 |
| WBGene00005728 | NA | | NA | NA | srv-17 |
| WBGene00005729 | NA | | NA | NA | NA |
| WBGene00005730 | NA | | NA | NA | srv-19 |
| WBGene00005731 | NA | | NA | NA | NA |
| WBGene00005732 | | -0.023056499 | 0.78121253 | NA | srv-21 |
| WBGene00005733 | NA | | NA | NA | srv-22 |
| WBGene00005734 | NA | | NA | NA | NA |
| WBGene00005735 | | -0.071706397 | 0.33859663 | NA | srv-24 |
| WBGene00005736 | NA | | NA | NA | srv-25 |
| WBGene00005737 | NA | | NA | NA | srv-26 |
| WBGene00005738 | | 0.020317245 | 0.84643798 | NA | srv-27 |
| WBGene00005739 | | 0.038125098 | 0.61119482 | NA | srv-28 |
| WBGene00005740 | NA | | NA | NA | srv-29 |
| WBGene00005741 | NA | | NA | NA | srv-30 |

| | | | | |
|---|---|---|---|---|
| WBGene00005742 | NA | NA | NA | srv-31 |
| WBGene00005743 | 0.077351407 | 0.40869513 | 0.60654062 | srv-32 |
| WBGene00005744 | NA | NA | NA | ops-1 |
| WBGene00005745 | 0.040107227 | 0.58693163 | NA | srv-34 |
| WBGene00005746 | NA | NA | NA | NA |
| WBGene00005747 | NA | NA | NA | NA |
| WBGene00005748 | NA | NA | NA | srw-1 |
| WBGene00005749 | NA | NA | NA | srw-2 |
| WBGene00005750 | NA | NA | NA | srw-3 |
| WBGene00005751 | NA | NA | NA | srw-4 |
| WBGene00005752 | NA | NA | NA | NA |
| WBGene00005753 | NA | NA | NA | srw-6 |
| WBGene00005754 | 0.017604133 | 0.79437959 | NA | srw-7 |
| WBGene00005755 | 0.033034883 | 0.64219503 | NA | srw-8 |
| WBGene00005756 | NA | NA | NA | srw-9 |
| WBGene00005757 | 0.020317245 | 0.84643798 | NA | srw-10 |
| WBGene00005758 | 0.058314668 | 0.40208118 | NA | srw-11 |
| WBGene00005759 | 0.020232312 | 0.84643798 | NA | srw-12 |
| WBGene00005760 | NA | NA | NA | srw-13 |
| WBGene00005761 | NA | NA | NA | srw-14 |
| WBGene00005762 | 0.039888668 | 0.56326497 | NA | srw-15 |
| WBGene00005763 | NA | NA | NA | srw-16 |
| WBGene00005764 | -0.003421551 | 0.95689871 | NA | srw-17 |
| WBGene00005765 | NA | NA | NA | NA |
| WBGene00005766 | 0.038125098 | 0.61119482 | NA | srw-19 |
| WBGene00005767 | NA | NA | NA | srw-20 |
| WBGene00005768 | NA | NA | NA | srw-21 |
| WBGene00005769 | NA | NA | NA | srw-22 |
| WBGene00005770 | 0.020925399 | 0.84643798 | NA | srw-23 |
| WBGene00005771 | NA | NA | NA | srw-24 |
| WBGene00005772 | -0.027373974 | 0.68245543 | NA | NA |
| WBGene00005773 | NA | NA | NA | srw-26 |
| WBGene00005774 | NA | NA | NA | NA |
| WBGene00005775 | NA | NA | NA | NA |
| WBGene00005776 | -0.023056499 | 0.78121253 | NA | srw-29 |
| WBGene00005777 | NA | NA | NA | srw-30 |
| WBGene00005778 | NA | NA | NA | srw-31 |
| WBGene00005779 | NA | NA | NA | srw-32 |
| WBGene00005780 | NA | NA | NA | srw-33 |
| WBGene00005781 | 0.020232312 | 0.84643798 | NA | srw-34 |
| WBGene00005782 | 0.063731418 | 0.41159439 | NA | srw-35 |
| WBGene00005783 | NA | NA | NA | srw-36 |
| WBGene00005784 | NA | NA | NA | NA |

| | | | | |
|---|---|---|---|---|
| WBGene00005785 | NA | NA | NA | srw-38 |
| WBGene00005786 | NA | NA | NA | srw-39 |
| WBGene00005787 | -0.022930767 | 0.78121253 | NA | srw-40 |
| WBGene00005788 | NA | NA | NA | srw-41 |
| WBGene00005789 | NA | NA | NA | srw-42 |
| WBGene00005790 | NA | NA | NA | srw-43 |
| WBGene00005791 | 0.020232312 | 0.84643798 | NA | srw-44 |
| WBGene00005792 | NA | NA | NA | NA |
| WBGene00005793 | 0.033463284 | 0.6365037 | NA | NA |
| WBGene00005794 | NA | NA | NA | srw-47 |
| WBGene00005795 | -0.022930767 | 0.78121253 | NA | srw-48 |
| WBGene00005796 | NA | NA | NA | NA |
| WBGene00005797 | NA | NA | NA | NA |
| WBGene00005798 | NA | NA | NA | srw-51 |
| WBGene00005799 | NA | NA | NA | NA |
| WBGene00005800 | NA | NA | NA | srw-53 |
| WBGene00005801 | NA | NA | NA | srw-54 |
| WBGene00005802 | NA | NA | NA | srw-55 |
| WBGene00005803 | NA | NA | NA | srw-56 |
| WBGene00005804 | NA | NA | NA | srw-57 |
| WBGene00005805 | NA | NA | NA | srw-58 |
| WBGene00005806 | NA | NA | NA | srw-59 |
| WBGene00005807 | NA | NA | NA | srw-60 |
| WBGene00005808 | NA | NA | NA | srw-61 |
| WBGene00005809 | NA | NA | NA | srw-62 |
| WBGene00005810 | NA | NA | NA | srw-63 |
| WBGene00005811 | -0.001842784 | 0.95621055 | NA | NA |
| WBGene00005812 | NA | NA | NA | srw-65 |
| WBGene00005813 | 0.020317245 | 0.84643798 | NA | srw-66 |
| WBGene00005814 | -0.002914971 | 0.95718083 | NA | srw-67 |
| WBGene00005815 | -0.023056499 | 0.78121253 | NA | srw-68 |
| WBGene00005816 | NA | NA | NA | srw-69 |
| WBGene00005817 | NA | NA | NA | NA |
| WBGene00005818 | 0.040107227 | 0.58693163 | NA | srw-71 |
| WBGene00005819 | NA | NA | NA | srw-72 |
| WBGene00005820 | NA | NA | NA | srw-73 |
| WBGene00005821 | NA | NA | NA | NA |
| WBGene00005822 | NA | NA | NA | srw-75 |
| WBGene00005823 | NA | NA | NA | srw-76 |
| WBGene00005824 | NA | NA | NA | srw-77 |
| WBGene00005825 | 0.052893861 | 0.5912442 | 0.75215398 | NA |
| WBGene00005826 | NA | NA | NA | srw-79 |
| WBGene00005827 | NA | NA | NA | NA |

| | | | | |
|---|---|---|---|---|
| WBGene00005828 | NA | NA | NA | NA |
| WBGene00005829 | NA | NA | NA | srw-82 |
| WBGene00005830 | NA | NA | NA | srw-83 |
| WBGene00005831 | 0.020232312 | 0.84643798 | NA | srw-84 |
| WBGene00005832 | 3.926329297 | 0.00038172 | 0.00308792 | srw-85 |
| WBGene00005833 | 0.267466646 | 0.01389687 | 0.05816582 | srw-86 |
| WBGene00005834 | 0.033463284 | 0.6365037 | NA | srw-87 |
| WBGene00005835 | -0.023056499 | 0.78121253 | NA | srw-88 |
| WBGene00005836 | 0.009723132 | 0.92667978 | 0.9586234 | srw-89 |
| WBGene00005837 | 0.020232312 | 0.84643798 | NA | srw-90 |
| WBGene00005838 | NA | NA | NA | srw-91 |
| WBGene00005839 | 0.020317245 | 0.84643798 | NA | NA |
| WBGene00005840 | 0.020925399 | 0.84643798 | NA | NA |
| WBGene00005841 | NA | NA | NA | srw-94 |
| WBGene00005842 | NA | NA | NA | srw-95 |
| WBGene00005843 | 0.073887753 | 0.28837975 | NA | NA |
| WBGene00005844 | -0.046394786 | 0.50231372 | NA | srw-97 |
| WBGene00005845 | -0.029029491 | 0.71849588 | NA | srw-98 |
| WBGene00005846 | 0.040107227 | 0.58693163 | NA | srw-99 |
| WBGene00005847 | NA | NA | NA | srw-100 |
| WBGene00005848 | 0.020232312 | 0.84643798 | NA | srw-101 |
| WBGene00005849 | 0.038125098 | 0.61119482 | NA | srw-102 |
| WBGene00005850 | NA | NA | NA | srw-103 |
| WBGene00005851 | NA | NA | NA | NA |
| WBGene00005852 | NA | NA | NA | NA |
| WBGene00005853 | NA | NA | NA | NA |
| WBGene00005854 | NA | NA | NA | srw-107 |
| WBGene00005855 | NA | NA | NA | srw-108 |
| WBGene00005856 | NA | NA | NA | srw-109 |
| WBGene00005857 | NA | NA | NA | NA |
| WBGene00005858 | NA | NA | NA | srw-111 |
| WBGene00005859 | NA | NA | NA | srw-112 |
| WBGene00005860 | NA | NA | NA | srw-113 |
| WBGene00005861 | NA | NA | NA | NA |
| WBGene00005862 | -0.022930767 | 0.78121253 | NA | srw-115 |
| WBGene00005863 | -0.046394786 | 0.50231372 | NA | NA |
| WBGene00005864 | NA | NA | NA | srw-117 |
| WBGene00005865 | NA | NA | NA | srw-118 |
| WBGene00005866 | NA | NA | NA | srw-119 |
| WBGene00005867 | NA | NA | NA | srw-120 |
| WBGene00005868 | 0.042065281 | 0.558937 | NA | srw-121 |
| WBGene00005869 | NA | NA | NA | srw-122 |
| WBGene00005870 | NA | NA | NA | srw-123 |

| | | | | |
|---|---|---|---|---|
| WBGene00005871 | NA | NA | NA | srw-124 |
| WBGene00005872 | NA | NA | NA | NA |
| WBGene00005873 | NA | NA | NA | NA |
| WBGene00005874 | NA | NA | NA | srw-127 |
| WBGene00005875 | NA | NA | NA | NA |
| WBGene00005876 | NA | NA | NA | srw-129 |
| WBGene00005877 | NA | NA | NA | srw-130 |
| WBGene00005878 | NA | NA | NA | NA |
| WBGene00005879 | NA | NA | NA | srw-132 |
| WBGene00005880 | NA | NA | NA | srw-133 |
| WBGene00005881 | NA | NA | NA | srw-134 |
| WBGene00005882 | NA | NA | NA | srw-135 |
| WBGene00005883 | 0.020232312 | 0.84643798 | NA | srw-136 |
| WBGene00005884 | NA | NA | NA | srw-137 |
| WBGene00005885 | NA | NA | NA | srw-138 |
| WBGene00005886 | 0.020317245 | 0.84643798 | NA | srw-139 |
| WBGene00005887 | NA | NA | NA | srw-140 |
| WBGene00005888 | NA | NA | NA | srw-141 |
| WBGene00005889 | NA | NA | NA | srw-142 |
| WBGene00005890 | NA | NA | NA | srw-143 |
| WBGene00005891 | NA | NA | NA | srw-144 |
| WBGene00005892 | 0.04909525 | 0.45360729 | NA | srx-1 |
| WBGene00005893 | 0.098475636 | 0.33611531 | 0.53553861 | srx-2 |
| WBGene00005894 | NA | NA | NA | srx-3 |
| WBGene00005895 | -0.023056499 | 0.78121253 | NA | srx-4 |
| WBGene00005896 | NA | NA | NA | srx-5 |
| WBGene00005897 | NA | NA | NA | srx-6 |
| WBGene00005898 | NA | NA | NA | srx-7 |
| WBGene00005899 | NA | NA | NA | srx-8 |
| WBGene00005900 | NA | NA | NA | srx-9 |
| WBGene00005901 | NA | NA | NA | srx-10 |
| WBGene00005902 | NA | NA | NA | NA |
| WBGene00005903 | 0.040937706 | 0.64530356 | NA | srx-12 |
| WBGene00005904 | 0.040107227 | 0.58693163 | NA | srx-13 |
| WBGene00005905 | NA | NA | NA | srx-14 |
| WBGene00005906 | 0.020317245 | 0.84643798 | NA | NA |
| WBGene00005907 | NA | NA | NA | srx-16 |
| WBGene00005908 | -0.023109762 | 0.78121253 | NA | srx-17 |
| WBGene00005909 | 0.020232312 | 0.84643798 | NA | srx-18 |
| WBGene00005910 | -0.024016858 | 0.7036901 | NA | srx-19 |
| WBGene00005911 | NA | NA | NA | srx-20 |
| WBGene00005912 | -0.023056499 | 0.78121253 | NA | srx-21 |
| WBGene00005913 | NA | NA | NA | srx-22 |

| | | | | |
|---|---|---|---|---|
| WBGene00005914 | NA | NA | NA | srx-23 |
| WBGene00005915 | NA | NA | NA | srx-24 |
| WBGene00005916 | NA | NA | NA | NA |
| WBGene00005917 | NA | NA | NA | srx-26 |
| WBGene00005918 | NA | NA | NA | NA |
| WBGene00005919 | NA | NA | NA | srx-28 |
| WBGene00005920 | -0.022930767 | 0.78121253 | NA | srx-29 |
| WBGene00005921 | NA | NA | NA | NA |
| WBGene00005922 | NA | NA | NA | srx-31 |
| WBGene00005923 | NA | NA | NA | srx-32 |
| WBGene00005924 | NA | NA | NA | srx-33 |
| WBGene00005925 | -0.022930767 | 0.78121253 | NA | srx-34 |
| WBGene00005926 | 0.020232312 | 0.84643798 | NA | srx-35 |
| WBGene00005927 | NA | NA | NA | srx-36 |
| WBGene00005928 | NA | NA | NA | NA |
| WBGene00005929 | 0.018342935 | 0.76675379 | NA | srx-38 |
| WBGene00005930 | 0.020317245 | 0.84643798 | NA | srx-39 |
| WBGene00005931 | -0.013911416 | 0.89431427 | 0.94489559 | fbxa-199 |
| WBGene00005932 | 0.033180193 | 0.61386379 | NA | srx-41 |
| WBGene00005933 | NA | NA | NA | srx-42 |
| WBGene00005934 | 0.020232312 | 0.84643798 | NA | srx-43 |
| WBGene00005935 | 0.020317245 | 0.84643798 | NA | srx-44 |
| WBGene00005936 | 0.393460285 | 0.05804579 | 0.1677222 | srx-45 |
| WBGene00005937 | NA | NA | NA | srx-46 |
| WBGene00005938 | -0.023056499 | 0.78121253 | NA | srx-47 |
| WBGene00005939 | NA | NA | NA | srx-48 |
| WBGene00005940 | NA | NA | NA | NA |
| WBGene00005941 | -0.002914971 | 0.95718083 | NA | srx-50 |
| WBGene00005942 | 0.130070456 | 0.24730079 | 0.44573814 | srx-51 |
| WBGene00005943 | NA | NA | NA | NA |
| WBGene00005944 | NA | NA | NA | srx-53 |
| WBGene00005945 | NA | NA | NA | srx-54 |
| WBGene00005946 | NA | NA | NA | srx-55 |
| WBGene00005947 | NA | NA | NA | srx-56 |
| WBGene00005948 | NA | NA | NA | NA |
| WBGene00005949 | -0.054151501 | 0.53354431 | NA | srx-58 |
| WBGene00005950 | NA | NA | NA | srx-59 |
| WBGene00005951 | NA | NA | NA | srx-60 |
| WBGene00005952 | -0.023056499 | 0.78121253 | NA | NA |
| WBGene00005953 | NA | NA | NA | srx-62 |
| WBGene00005954 | NA | NA | NA | srx-63 |
| WBGene00005955 | NA | NA | NA | srx-64 |
| WBGene00005956 | NA | NA | NA | srx-65 |

| | | | | |
|---|---|---|---|---|
| WBGene00005957 | NA | NA | NA | srx-66 |
| WBGene00005958 | -0.003614866 | 0.95700117 | NA | srx-67 |
| WBGene00005959 | 0.018657025 | 0.77508599 | NA | srx-68 |
| WBGene00005960 | NA | NA | NA | NA |
| WBGene00005961 | NA | NA | NA | NA |
| WBGene00005962 | NA | NA | NA | NA |
| WBGene00005963 | NA | NA | NA | NA |
| WBGene00005964 | NA | NA | NA | srx-73 |
| WBGene00005965 | 0.015896709 | 0.80422001 | NA | srx-74 |
| WBGene00005966 | NA | NA | NA | NA |
| WBGene00005967 | NA | NA | NA | srx-76 |
| WBGene00005968 | -0.040794878 | 0.53203165 | NA | srx-77 |
| WBGene00005969 | NA | NA | NA | srx-78 |
| WBGene00005970 | NA | NA | NA | NA |
| WBGene00005971 | 0.020317245 | 0.84643798 | NA | srx-80 |
| WBGene00005972 | NA | NA | NA | NA |
| WBGene00005973 | NA | NA | NA | srx-82 |
| WBGene00005974 | NA | NA | NA | NA |
| WBGene00005975 | NA | NA | NA | srx-84 |
| WBGene00005976 | 0.020232312 | 0.84643798 | NA | srx-85 |
| WBGene00005977 | 0.020925399 | 0.84643798 | NA | srx-86 |
| WBGene00005978 | NA | NA | NA | srx-87 |
| WBGene00005979 | NA | NA | NA | srx-88 |
| WBGene00005980 | NA | NA | NA | srx-89 |
| WBGene00005981 | NA | NA | NA | srx-90 |
| WBGene00005982 | NA | NA | NA | NA |
| WBGene00005983 | -0.039636691 | 0.54579661 | NA | srx-92 |
| WBGene00005984 | NA | NA | NA | srx-93 |
| WBGene00005985 | NA | NA | NA | NA |
| WBGene00005986 | NA | NA | NA | srx-95 |
| WBGene00005987 | 0.020925399 | 0.84643798 | NA | srx-96 |
| WBGene00005988 | -0.02791392 | 0.6763823 | NA | srx-97 |
| WBGene00005989 | 0.007709255 | 0.93892705 | 0.96577427 | srx-98 |
| WBGene00005990 | NA | NA | NA | NA |
| WBGene00005991 | NA | NA | NA | NA |
| WBGene00005992 | NA | NA | NA | srx-101 |
| WBGene00005993 | NA | NA | NA | srx-102 |
| WBGene00005994 | NA | NA | NA | NA |
| WBGene00005995 | 0.040107227 | 0.58693163 | NA | srx-104 |
| WBGene00005996 | 0.020317245 | 0.84643798 | NA | srx-105 |
| WBGene00005997 | NA | NA | NA | NA |
| WBGene00005998 | NA | NA | NA | NA |
| WBGene00005999 | NA | NA | NA | srx-108 |

| | | | | |
|---|---|---|---|---|
| WBGene00006000 | NA | NA | NA | NA |
| WBGene00006001 | NA | NA | NA | srx-110 |
| WBGene00006002 | 0.065014541 | 0.37399213 | NA | srx-111 |
| WBGene00006003 | 0.020925399 | 0.84643798 | NA | srx-112 |
| WBGene00006004 | 0.020232312 | 0.84643798 | NA | srx-113 |
| WBGene00006005 | -0.006392486 | 0.92872129 | NA | srx-114 |
| WBGene00006006 | 0.017279211 | 0.79380376 | NA | srx-115 |
| WBGene00006007 | NA | NA | NA | srx-116 |
| WBGene00006008 | 0.020925399 | 0.84643798 | NA | srx-117 |
| WBGene00006009 | -0.050042403 | 0.47734215 | NA | srx-118 |
| WBGene00006010 | 0.020232312 | 0.84643798 | NA | srx-119 |
| WBGene00006011 | NA | NA | NA | srx-120 |
| WBGene00006012 | NA | NA | NA | srx-121 |
| WBGene00006013 | NA | NA | NA | srx-122 |
| WBGene00006014 | 0.020317245 | 0.84643798 | NA | NA |
| WBGene00006015 | NA | NA | NA | NA |
| WBGene00006016 | 0.342749564 | 0.09457239 | 0.23584302 | srx-125 |
| WBGene00006017 | -0.023109762 | 0.78121253 | NA | NA |
| WBGene00006018 | NA | NA | NA | NA |
| WBGene00006019 | NA | NA | NA | srx-128 |
| WBGene00006020 | 0.020232312 | 0.84643798 | NA | srx-129 |
| WBGene00006021 | 0.095967469 | 0.18297344 | NA | srx-130 |
| WBGene00006022 | NA | NA | NA | srx-131 |
| WBGene00006023 | NA | NA | NA | NA |
| WBGene00006024 | -0.023056499 | 0.78121253 | NA | srx-133 |
| WBGene00006025 | NA | NA | NA | srx-134 |
| WBGene00006026 | NA | NA | NA | srx-135 |
| WBGene00006027 | NA | NA | NA | srx-136 |
| WBGene00006028 | 0.020232312 | 0.84643798 | NA | srx-137 |
| WBGene00006029 | NA | NA | NA | NA |
| WBGene00006038 | NA | NA | NA | ssp-9 |
| WBGene00006038 | NA | NA | NA | ssp-11 |
| WBGene00006039 | 0.404622589 | 0.10215069 | 0.24897424 | ssp-10 |
| WBGene00006040 | 0.059980168 | 0.38780842 | NA | ssp-9 |
| WBGene00006040 | 0.059980168 | 0.38780842 | NA | ssp-11 |
| WBGene00006044 | 0.147476436 | 0.20068817 | 0.38927869 | ssp-16 |
| WBGene00006047 | -0.072889732 | 0.33672315 | NA | ssp-19 |
| WBGene00006048 | 0.333719708 | 0.02383526 | 0.08746165 | ssp-31 |
| WBGene00006049 | 0.020232312 | 0.84643798 | NA | ssp-32 |
| WBGene00006050 | 1.482787299 | 0.00011362 | 0.00108672 | ssq-1 |
| WBGene00006051 | 1.522965569 | 7.63E-05 | 0.00076269 | ssq-2 |
| WBGene00006052 | 1.913371364 | 0.00021533 | 0.00188213 | ssq-3 |
| WBGene00006053 | 2.202469387 | 1.67E-10 | 4.62E-09 | ssq-4 |

| | | | | | |
|---|---|---|---|---|---|
| WBGene00006055 | | 0.026559602 | 0.87116557 | 0.93126126 | ssr-2 |
| WBGene00006056 | | 2.207357061 | 2.51E-06 | 3.54E-05 | sss-1 |
| WBGene00006057 | NA | | NA | NA | NA |
| WBGene00006058 | | 0.118140836 | 0.36445852 | 0.56412276 | sst-20 |
| WBGene00006059 | | -0.18495273 | 0.06754981 | 0.18653278 | stc-1 |
| WBGene00006060 | | -0.097013169 | 0.62665138 | 0.77732804 | sth-1 |
| WBGene00006061 | | -0.019798051 | 0.82895086 | 0.90652983 | stl-1 |
| WBGene00006062 | | 0.054081886 | 0.79413369 | 0.88410854 | stn-1 |
| WBGene00006063 | | -0.5302371 | 0.00765563 | 0.0365101 | sto-1 |
| WBGene00006064 | | 0.037616261 | 0.78155736 | 0.87704309 | sto-2 |
| WBGene00006065 | | -0.024655542 | 0.73435809 | NA | sto-3 |
| WBGene00006066 | | -0.044663729 | 0.77768428 | 0.8741977 | sto-4 |
| WBGene00006067 | | 0.429195461 | 0.09103852 | 0.22953245 | sto-5 |
| WBGene00006068 | | 0.342157234 | 0.05159542 | 0.1539451 | sto-6 |
| WBGene00006069 | NA | | NA | NA | str-1 |
| WBGene00006070 | | 0.020232312 | 0.84643798 | NA | str-2 |
| WBGene00006071 | NA | | NA | NA | str-3 |
| WBGene00006072 | | 0.020925399 | 0.84643798 | NA | str-4 |
| WBGene00006073 | NA | | NA | NA | str-5 |
| WBGene00006074 | NA | | NA | NA | str-6 |
| WBGene00006075 | | 0.039135871 | 0.65930174 | NA | str-7 |
| WBGene00006076 | NA | | NA | NA | str-8 |
| WBGene00006077 | | 0.020317245 | 0.84643798 | NA | str-9 |
| WBGene00006078 | NA | | NA | NA | str-10 |
| WBGene00006079 | NA | | NA | NA | NA |
| WBGene00006080 | | -0.023056499 | 0.78121253 | NA | str-12 |
| WBGene00006081 | NA | | NA | NA | str-13 |
| WBGene00006082 | NA | | NA | NA | str-14 |
| WBGene00006083 | NA | | NA | NA | str-15 |
| WBGene00006084 | | 0.033042231 | 0.73176645 | 0.85001776 | str-16 |
| WBGene00006085 | NA | | NA | NA | str-18 |
| WBGene00006086 | NA | | NA | NA | str-19 |
| WBGene00006087 | NA | | NA | NA | str-20 |
| WBGene00006088 | NA | | NA | NA | NA |
| WBGene00006089 | NA | | NA | NA | NA |
| WBGene00006090 | NA | | NA | NA | str-23 |
| WBGene00006091 | NA | | NA | NA | NA |
| WBGene00006092 | NA | | NA | NA | str-25 |
| WBGene00006093 | | -0.023109762 | 0.78121253 | NA | str-27 |
| WBGene00006094 | NA | | NA | NA | str-28 |
| WBGene00006095 | NA | | NA | NA | NA |
| WBGene00006096 | | 0.020232312 | 0.84643798 | NA | str-30 |
| WBGene00006097 | NA | | NA | NA | str-31 |

| | | | | |
|---|---|---|---|---|
| WBGene00006098 | NA | NA | NA | str-32 |
| WBGene00006099 | NA | NA | NA | str-33 |
| WBGene00006100 | NA | NA | NA | NA |
| WBGene00006101 | NA | NA | NA | str-35 |
| WBGene00006102 | NA | NA | NA | str-37 |
| WBGene00006103 | 0.020232312 | 0.84643798 | NA | str-38 |
| WBGene00006104 | NA | NA | NA | str-39 |
| WBGene00006105 | NA | NA | NA | str-40 |
| WBGene00006106 | -0.023056499 | 0.78121253 | NA | str-41 |
| WBGene00006107 | NA | NA | NA | NA |
| WBGene00006108 | NA | NA | NA | str-43 |
| WBGene00006109 | 0.070327578 | 0.44827578 | NA | str-44 |
| WBGene00006110 | 0.020317245 | 0.84643798 | NA | str-45 |
| WBGene00006111 | NA | NA | NA | str-46 |
| WBGene00006112 | 0.020925399 | 0.84643798 | NA | str-47 |
| WBGene00006113 | NA | NA | NA | str-48 |
| WBGene00006114 | -0.025848508 | 0.67901638 | NA | NA |
| WBGene00006115 | NA | NA | NA | NA |
| WBGene00006116 | NA | NA | NA | NA |
| WBGene00006117 | NA | NA | NA | str-52 |
| WBGene00006118 | NA | NA | NA | NA |
| WBGene00006119 | NA | NA | NA | NA |
| WBGene00006120 | 0.015062379 | 0.80838968 | NA | str-55 |
| WBGene00006121 | NA | NA | NA | str-56 |
| WBGene00006122 | 0.020232312 | 0.84643798 | NA | str-57 |
| WBGene00006123 | NA | NA | NA | NA |
| WBGene00006124 | NA | NA | NA | str-60 |
| WBGene00006125 | NA | NA | NA | str-61 |
| WBGene00006126 | 0.020925399 | 0.84643798 | NA | str-63 |
| WBGene00006127 | 0.020232312 | 0.84643798 | NA | str-64 |
| WBGene00006128 | NA | NA | NA | str-66 |
| WBGene00006129 | NA | NA | NA | str-67 |
| WBGene00006130 | -0.022930767 | 0.78121253 | NA | str-68 |
| WBGene00006131 | NA | NA | NA | str-69 |
| WBGene00006132 | NA | NA | NA | NA |
| WBGene00006133 | NA | NA | NA | str-71 |
| WBGene00006134 | NA | NA | NA | str-73 |
| WBGene00006135 | -0.04044783 | 0.53612608 | NA | str-74 |
| WBGene00006136 | NA | NA | NA | str-265 |
| WBGene00006136 | NA | NA | NA | str-76 |
| WBGene00006137 | NA | NA | NA | str-77 |
| WBGene00006138 | -0.023109762 | 0.78121253 | NA | str-78 |
| WBGene00006139 | 0.020317245 | 0.84643798 | NA | str-79 |

| | | | | | |
|---|---|---|---|---|---|
| WBGene00006140 | | -0.023056499 | 0.78121253 | NA | str-81 |
| WBGene00006141 | NA | | NA | NA | str-82 |
| WBGene00006142 | NA | | NA | NA | str-83 |
| WBGene00006143 | | 0.017279211 | 0.79380376 | NA | str-84 |
| WBGene00006144 | NA | | NA | NA | str-85 |
| WBGene00006145 | NA | | NA | NA | str-86 |
| WBGene00006146 | | 0.038633797 | 0.57999349 | NA | str-87 |
| WBGene00006147 | NA | | NA | NA | str-88 |
| WBGene00006148 | NA | | NA | NA | str-89 |
| WBGene00006149 | | -0.023109762 | 0.78121253 | NA | str-90 |
| WBGene00006150 | NA | | NA | NA | str-92 |
| WBGene00006151 | NA | | NA | NA | str-93 |
| WBGene00006152 | | 0.059980168 | 0.38780842 | NA | str-94 |
| WBGene00006153 | NA | | NA | NA | str-96 |
| WBGene00006154 | NA | | NA | NA | str-97 |
| WBGene00006155 | | 0.020317245 | 0.84643798 | NA | str-99 |
| WBGene00006156 | NA | | NA | NA | str-101 |
| WBGene00006157 | | 0.040107227 | 0.58693163 | NA | str-102 |
| WBGene00006158 | NA | | NA | NA | str-103 |
| WBGene00006159 | NA | | NA | NA | str-106 |
| WBGene00006160 | NA | | NA | NA | str-108 |
| WBGene00006161 | NA | | NA | NA | str-109 |
| WBGene00006162 | | -0.022930767 | 0.78121253 | NA | str-111 |
| WBGene00006163 | | 0.058439243 | 0.52349407 | NA | str-112 |
| WBGene00006164 | NA | | NA | NA | str-113 |
| WBGene00006165 | | 0.020232312 | 0.84643798 | NA | str-114 |
| WBGene00006166 | NA | | NA | NA | str-115 |
| WBGene00006167 | | 0.0398788 | 0.58971221 | NA | str-116 |
| WBGene00006168 | NA | | NA | NA | NA |
| WBGene00006169 | | -0.001993815 | 0.95612996 | NA | str-118 |
| WBGene00006170 | NA | | NA | NA | str-119 |
| WBGene00006171 | NA | | NA | NA | Y45G12C.6 |
| WBGene00006171 | NA | | NA | NA | str-120 |
| WBGene00006172 | | -0.002914971 | 0.95718083 | NA | str-121 |
| WBGene00006173 | | -0.027728334 | 0.67598605 | NA | str-122 |
| WBGene00006174 | NA | | NA | NA | str-123 |
| WBGene00006175 | NA | | NA | NA | str-124 |
| WBGene00006176 | | 0.370692785 | 0.01663903 | 0.06709228 | str-125 |
| WBGene00006177 | NA | | NA | NA | NA |
| WBGene00006178 | | 0.020317245 | 0.84643798 | NA | str-129 |
| WBGene00006179 | | 0.033034883 | 0.64219503 | NA | str-130 |
| WBGene00006180 | | -0.022930767 | 0.78121253 | NA | str-131 |
| WBGene00006181 | | 0.020232312 | 0.84643798 | NA | NA |

| | | | | |
|---|---|---|---|---|
| WBGene00006182 | NA | NA | NA | str-134 |
| WBGene00006183 | 0.018998992 | 0.77286019 | NA | str-135 |
| WBGene00006184 | NA | NA | NA | str-136 |
| WBGene00006185 | NA | NA | NA | NA |
| WBGene00006186 | NA | NA | NA | str-139 |
| WBGene00006187 | NA | NA | NA | str-138 |
| WBGene00006188 | NA | NA | NA | str-140 |
| WBGene00006189 | 0.020317245 | 0.84643798 | NA | str-141 |
| WBGene00006190 | -0.001488154 | 0.95639912 | NA | str-143 |
| WBGene00006191 | -0.045308129 | 0.49733342 | NA | str-144 |
| WBGene00006192 | 0.015896709 | 0.80422001 | NA | str-145 |
| WBGene00006193 | 0.09279984 | 0.48565054 | 0.67027418 | str-146 |
| WBGene00006194 | 0.020925399 | 0.84643798 | NA | NA |
| WBGene00006195 | 0.004135627 | 0.97469894 | 0.98580873 | str-148 |
| WBGene00006196 | NA | NA | NA | str-149 |
| WBGene00006197 | NA | NA | NA | str-150 |
| WBGene00006198 | -0.023109762 | 0.78121253 | NA | str-151 |
| WBGene00006199 | NA | NA | NA | str-153 |
| WBGene00006200 | NA | NA | NA | str-267 |
| WBGene00006200 | NA | NA | NA | str-154 |
| WBGene00006201 | 0.017604133 | 0.79437959 | NA | str-155 |
| WBGene00006202 | NA | NA | NA | str-156 |
| WBGene00006203 | 0.020925399 | 0.84643798 | NA | str-158 |
| WBGene00006204 | NA | NA | NA | str-159 |
| WBGene00006205 | 0.020232312 | 0.84643798 | NA | str-160 |
| WBGene00006206 | NA | NA | NA | str-161 |
| WBGene00006207 | -0.004047623 | 0.95527667 | NA | str-162 |
| WBGene00006208 | 0.121783946 | 0.26340991 | 0.46153476 | str-163 |
| WBGene00006209 | 0.038125098 | 0.61119482 | NA | str-164 |
| WBGene00006210 | -0.023056499 | 0.78121253 | NA | str-165 |
| WBGene00006211 | 0.020317245 | 0.84643798 | NA | str-166 |
| WBGene00006212 | 0.131491284 | 0.24291251 | 0.44097731 | str-168 |
| WBGene00006213 | 0.0398788 | 0.58971221 | NA | str-169 |
| WBGene00006214 | NA | NA | NA | str-170 |
| WBGene00006215 | -0.003270206 | 0.95698316 | NA | str-171 |
| WBGene00006216 | 0.056411843 | 0.41891328 | NA | str-172 |
| WBGene00006217 | 0.020925399 | 0.84643798 | NA | str-173 |
| WBGene00006218 | NA | NA | NA | str-174 |
| WBGene00006219 | NA | NA | NA | str-175 |
| WBGene00006220 | -0.081503648 | 0.62463619 | 0.77568265 | str-176 |
| WBGene00006221 | NA | NA | NA | str-177 |
| WBGene00006222 | NA | NA | NA | str-178 |
| WBGene00006223 | 0.003823937 | 0.97327774 | 0.98532835 | NA |

| | | | | | |
|---|---|---|---|---|---|
| WBGene00006224 | | 0.014294003 | 0.85724546 | NA | str-180 |
| WBGene00006225 | NA | | NA | NA | str-181 |
| WBGene00006226 | | -0.003614866 | 0.95700117 | NA | str-182 |
| WBGene00006227 | | 0.011769853 | 0.86975693 | NA | str-183 |
| WBGene00006228 | NA | | NA | NA | str-185 |
| WBGene00006229 | | 0.020317245 | 0.84643798 | NA | str-187 |
| WBGene00006230 | NA | | NA | NA | str-188 |
| WBGene00006231 | NA | | NA | NA | str-190 |
| WBGene00006232 | NA | | NA | NA | str-193 |
| WBGene00006233 | NA | | NA | NA | str-196 |
| WBGene00006234 | NA | | NA | NA | str-198 |
| WBGene00006235 | NA | | NA | NA | str-199 |
| WBGene00006236 | NA | | NA | NA | str-200 |
| WBGene00006237 | | 0.036687355 | 0.59489568 | NA | str-204 |
| WBGene00006238 | NA | | NA | NA | str-205 |
| WBGene00006239 | NA | | NA | NA | str-206 |
| WBGene00006240 | NA | | NA | NA | str-207 |
| WBGene00006241 | NA | | NA | NA | str-208 |
| WBGene00006242 | NA | | NA | NA | str-209 |
| WBGene00006243 | | 0.020925399 | 0.84643798 | NA | NA |
| WBGene00006244 | | 0.020925399 | 0.84643798 | NA | str-211 |
| WBGene00006245 | NA | | NA | NA | NA |
| WBGene00006246 | NA | | NA | NA | NA |
| WBGene00006247 | NA | | NA | NA | str-214 |
| WBGene00006248 | NA | | NA | NA | NA |
| WBGene00006249 | NA | | NA | NA | str-217 |
| WBGene00006250 | NA | | NA | NA | NA |
| WBGene00006251 | NA | | NA | NA | str-220 |
| WBGene00006252 | NA | | NA | NA | str-221 |
| WBGene00006253 | NA | | NA | NA | str-222 |
| WBGene00006254 | NA | | NA | NA | str-223 |
| WBGene00006255 | | -0.075452738 | 0.28904131 | NA | str-224 |
| WBGene00006256 | | -0.003108065 | 0.95728506 | NA | str-225 |
| WBGene00006257 | | 0.020925399 | 0.84643798 | NA | str-226 |
| WBGene00006258 | | -0.031690166 | 0.70451177 | NA | str-227 |
| WBGene00006259 | NA | | NA | NA | str-228 |
| WBGene00006260 | NA | | NA | NA | str-229 |
| WBGene00006261 | | 0.063527295 | 0.41294485 | NA | str-230 |
| WBGene00006262 | | -0.023056499 | 0.78121253 | NA | str-231 |
| WBGene00006263 | NA | | NA | NA | str-232 |
| WBGene00006264 | NA | | NA | NA | str-233 |
| WBGene00006265 | NA | | NA | NA | str-236 |
| WBGene00006266 | NA | | NA | NA | str-238 |

| | | | | |
|---|---|---|---|---|
| WBGene00006267 | -0.003421551 | 0.95689871 | NA | NA |
| WBGene00006268 | NA | NA | NA | str-240 |
| WBGene00006269 | NA | NA | NA | NA |
| WBGene00006270 | 0.020317245 | 0.84643798 | NA | str-243 |
| WBGene00006271 | 0.020317245 | 0.84643798 | NA | str-244 |
| WBGene00006272 | -0.08416487 | 0.38570875 | NA | str-245 |
| WBGene00006273 | NA | NA | NA | str-246 |
| WBGene00006274 | 0.020317245 | 0.84643798 | NA | str-247 |
| WBGene00006275 | NA | NA | NA | str-248 |
| WBGene00006276 | NA | NA | NA | str-249 |
| WBGene00006277 | NA | NA | NA | str-250 |
| WBGene00006278 | NA | NA | NA | str-252 |
| WBGene00006279 | NA | NA | NA | str-253 |
| WBGene00006280 | NA | NA | NA | str-254 |
| WBGene00006281 | NA | NA | NA | str-255 |
| WBGene00006282 | NA | NA | NA | str-256 |
| WBGene00006283 | NA | NA | NA | str-257 |
| WBGene00006283 | NA | NA | NA | Y19D10A.1 |
| WBGene00006284 | NA | NA | NA | str-258 |
| WBGene00006284 | NA | NA | NA | Y19D10A.2 |
| WBGene00006285 | -0.003614866 | 0.95700117 | NA | NA |
| WBGene00006286 | NA | NA | NA | str-260 |
| WBGene00006287 | 0.018998992 | 0.77286019 | NA | str-261 |
| WBGene00006288 | 0.018860755 | 0.77262519 | NA | str-262 |
| WBGene00006289 | NA | NA | NA | str-263 |
| WBGene00006306 | 1.590325711 | 0.00026888 | 0.00228084 | sue-1 |
| WBGene00006307 | -0.15207431 | 0.05528496 | 0.16198262 | suf-1 |
| WBGene00006308 | 0.045791898 | 0.81669118 | 0.89923859 | sul-1 |
| WBGene00006309 | 1.740064412 | 0.00044754 | 0.00353423 | sul-2 |
| WBGene00006310 | 0.35861953 | 0.13596485 | 0.30093142 | sul-3 |
| WBGene00006311 | -0.250434264 | 0.00164165 | 0.010621 | sun-1 |
| WBGene00006314 | NA | NA | NA | NA |
| WBGene00006315 | -0.003270206 | 0.95698316 | NA | NA |
| WBGene00006316 | NA | NA | NA | NA |
| WBGene00006318 | -0.160137543 | 0.32921999 | 0.52869474 | sup-9 |
| WBGene00006319 | 0.030138851 | 0.88064812 | 0.93655619 | sup-10 |
| WBGene00006321 | 0.52765888 | 0.02994521 | 0.10341402 | sup-12 |
| WBGene00006324 | -0.065839682 | 0.27211626 | 0.47224612 | sup-17 |
| WBGene00006327 | NA | NA | NA | NA |
| WBGene00006330 | NA | NA | NA | NA |
| WBGene00006331 | -0.295320751 | 0.00191914 | 0.01208522 | sup-26 |
| WBGene00006333 | NA | NA | NA | NA |
| WBGene00006334 | NA | NA | NA | NA |

| | | | | |
|---|---|---|---|---|
| WBGene00006338 | NA | NA | NA | NA |
| WBGene00006342 | 0.433538994 | 0.00309614 | 0.01791015 | sup-37 |
| WBGene00006344 | NA | NA | NA | NA |
| WBGene00006349 | -0.016705184 | 0.8590082 | 0.92455628 | sur-2 |
| WBGene00006351 | -1.035962804 | 6.28E-08 | 1.22E-06 | sur-5 |
| WBGene00006352 | -0.118072459 | 0.05843329 | 0.16836992 | sur-6 |
| WBGene00006353 | -0.025552851 | 0.85721103 | 0.92325428 | sur-7 |
| WBGene00006359 | -0.148749829 | 0.18316964 | 0.36616603 | swp-1 |
| WBGene00006363 | 0.725377383 | 0.02864456 | 0.10028336 | syd-1 |
| WBGene00006364 | -0.043363917 | 0.75000275 | 0.8587464 | syd-2 |
| WBGene00006365 | 0.234734225 | 0.28963707 | 0.49001345 | syg-1 |
| WBGene00006366 | -0.960965215 | 0.00028911 | 0.00243126 | sym-1 |
| WBGene00006367 | 0.26710097 | 0.23169824 | 0.42654268 | sym-2 |
| WBGene00006368 | -0.096339072 | 0.5856008 | 0.74847501 | sym-3 |
| WBGene00006369 | -0.025806695 | 0.83415525 | 0.90957526 | sym-4 |
| WBGene00006371 | -0.117838734 | 0.54922426 | 0.72192916 | syx-3 |
| WBGene00006372 | 1.018784642 | 0.00117892 | 0.00805438 | syx-2 |
| WBGene00006373 | -0.249275329 | 0.00491329 | 0.02582035 | syx-5 |
| WBGene00006374 | -0.163232711 | 0.22719137 | 0.42089568 | syx-4 |
| WBGene00006375 | -0.069390178 | 0.55642158 | 0.72795714 | syp-1 |
| WBGene00006376 | -0.25385294 | 0.02946109 | 0.10233448 | syp-2 |
| WBGene00006377 | -0.230094403 | 0.10495388 | 0.25334286 | syp-3 |
| WBGene00006379 | -0.276795762 | 0.04478232 | 0.13870486 | sys-1 |
| WBGene00006380 | -0.010834285 | 0.9091334 | 0.95068624 | tab-1 |
| WBGene00006381 | 0.125703716 | 0.255968 | 0.45436804 | tac-1 |
| WBGene00006382 | 0.01866965 | 0.82510365 | 0.90394661 | taf-1 |
| WBGene00006383 | 0.099954569 | 0.40428845 | 0.60246181 | taf-2 |
| WBGene00006384 | 0.076639898 | 0.54134806 | 0.71522926 | taf-3 |
| WBGene00006385 | 0.062738968 | 0.58483211 | 0.74805087 | taf-4 |
| WBGene00006386 | 0.023113164 | 0.79144531 | 0.88250563 | taf-5 |
| WBGene00006387 | 0.043463677 | 0.75584141 | 0.86179224 | taf-6.1 |
| WBGene00006388 | -0.146377378 | 0.48413789 | 0.66945397 | taf-7.1 |
| WBGene00006389 | -0.067081698 | 0.61714153 | 0.77055851 | taf-7.2 |
| WBGene00006390 | -0.166295913 | 0.1320365 | 0.29522699 | taf-8 |
| WBGene00006391 | -0.34858305 | 0.00988493 | 0.04447611 | taf-9 |
| WBGene00006392 | -0.051312041 | 0.66393678 | 0.80289902 | taf-10 |
| WBGene00006393 | -0.37005817 | 0.05945806 | 0.17062133 | taf-11.1 |
| WBGene00006394 | -0.522288272 | 1.17E-05 | 0.00014082 | taf-11.2 |
| WBGene00006395 | -0.311498286 | 0.17317117 | 0.35347679 | taf-11.3 |
| WBGene00006396 | -0.218859658 | 0.05292903 | 0.1570598 | taf-12 |
| WBGene00006397 | 0.117276038 | 0.26018683 | 0.45840351 | taf-13 |
| WBGene00006399 | NA | NA | NA | tag-4 |
| WBGene00006402 | -0.023584372 | 0.79851454 | 0.88715885 | fut-3 |

| | | | | |
|---|---|---|---|---|
| WBGene00006404 | -0.368152304 | 0.00473731 | 0.02506648 | tag-10 |
| WBGene00006405 | -0.133799875 | 0.08514986 | 0.21991164 | itsn-1 |
| WBGene00006406 | -0.11011633 | 0.08139743 | 0.21348074 | srgp-1 |
| WBGene00006407 | 0.293804261 | 0.00689296 | 0.03356165 | lim-9 |
| WBGene00006408 | -0.367658442 | 0.00014539 | 0.00133277 | tag-18 |
| WBGene00006409 | 0.124133031 | 0.25369644 | 0.45170492 | hdl-2 |
| WBGene00006410 | 0.043940581 | 0.71557707 | 0.84042707 | nck-1 |
| WBGene00006411 | 0.365712755 | 0.1329909 | 0.29656413 | octr-1 |
| WBGene00006412 | 0.110745697 | 0.5904227 | 0.75162641 | nlg-1 |
| WBGene00006414 | -0.090097375 | 0.27104026 | 0.47056806 | raga-1 |
| WBGene00006415 | -0.356473695 | 0.00052673 | 0.00405792 | tag-30 |
| WBGene00006416 | 0.044613311 | 0.82851203 | 0.9062566 | sams-5 |
| WBGene00006418 | -0.233787367 | 0.28387978 | 0.48373881 | spl-2 |
| WBGene00006419 | NA | NA | NA | oac-39 |
| WBGene00006423 | 0.211242455 | 0.19689823 | 0.38442285 | asd-2 |
| WBGene00006424 | 0.303222307 | 0.19395918 | 0.3808686 | ctbp-1 |
| WBGene00006428 | 0.221807513 | 0.23539452 | 0.43118853 | tkr-3 |
| WBGene00006429 | -0.001488154 | 0.95639912 | NA | arrd-3 |
| WBGene00006430 | -0.321810159 | 0.07860083 | 0.20851508 | tag-51 |
| WBGene00006431 | 0.225771797 | 0.1111459 | 0.2631162 | tag-52 |
| WBGene00006432 | -0.286238775 | 0.01049525 | 0.04650288 | atrn-1 |
| WBGene00006433 | -0.010069009 | 0.90197416 | 0.94780763 | sdhb-1 |
| WBGene00006434 | -0.13765395 | 0.00629593 | 0.03130005 | prdx-2 |
| WBGene00006435 | 0.078633594 | 0.26310689 | NA | stdh-4 |
| WBGene00006436 | 0.790537383 | 2.50E-09 | 5.99E-08 | ttn-1 |
| WBGene00006437 | -0.107514544 | 0.14284988 | 0.31113924 | mrck-1 |
| WBGene00006438 | 0.325152937 | 0.00197247 | 0.01233326 | nrfl-1 |
| WBGene00006439 | -0.131290662 | 0.00061975 | 0.00467957 | ant-1.1 |
| WBGene00006440 | -0.321144929 | 0.00216643 | 0.01330289 | tag-63 |
| WBGene00006441 | 0.971907873 | 0.0156706 | 0.06408748 | cpna-4 |
| WBGene00006442 | -0.19293421 | 0.05501345 | 0.16141321 | tag-65 |
| WBGene00006443 | -0.203677955 | 0.02931408 | 0.10189634 | pak-2 |
| WBGene00006444 | 0.796491127 | 0.0013812 | 0.0091752 | shn-1 |
| WBGene00006445 | 0.258841263 | 0.25192038 | 0.44993917 | tag-68 |
| WBGene00006446 | -0.172376308 | 0.11100219 | 0.26297459 | atx-3 |
| WBGene00006447 | -0.103142386 | 0.42391491 | 0.61990993 | tag-72 |
| WBGene00006448 | 0.146615046 | 0.22777107 | 0.4212729 | glod-4 |
| WBGene00006449 | 5.922702358 | 1.19E-05 | 0.0001434 | alg-4 |
| WBGene00006450 | 0.017197748 | 0.86843458 | 0.92992376 | tag-77 |
| WBGene00006452 | 0.116894041 | 0.21402963 | 0.40486495 | heh-1 |
| WBGene00006454 | 0.184772721 | 0.39461038 | 0.59262392 | aex-4 |
| WBGene00006459 | 0.552687267 | 0.0483524 | 0.14662955 | tag-89 |
| WBGene00006460 | -0.098873845 | 0.08627315 | 0.22186041 | ppm-1.A |

| | | | | |
|---|---|---|---|---|
| WBGene00006461 | -0.247288887 | 0.24834972 | 0.44674877 | tag-96 |
| WBGene00006462 | 0.657733588 | 0.02281995 | 0.08473949 | svh-5 |
| WBGene00006463 | -0.391691602 | 5.84E-07 | 9.66E-06 | nduf-2.2 |
| WBGene00006464 | 2.819578946 | 0.00074743 | 0.00546711 | bath-47 |
| WBGene00006465 | -0.295433066 | 0.00221377 | 0.01352238 | fntb-1 |
| WBGene00006466 | -0.157363263 | 0.18163765 | 0.36434537 | tag-115 |
| WBGene00006467 | 0.308045569 | 0.16293987 | 0.33838335 | magu-2 |
| WBGene00006468 | 0.48833265 | 0.04383585 | 0.13678616 | rhgf-1 |
| WBGene00006470 | 1.186113091 | 1.01E-07 | 1.90E-06 | tag-120 |
| WBGene00006471 | NA | NA | NA | nhr-233 |
| WBGene00006472 | 0.045159023 | 0.62235221 | 0.77415962 | tag-123 |
| WBGene00006473 | -0.45135326 | 0.00558557 | 0.02857382 | tag-124 |
| WBGene00006474 | -0.208121742 | 0.01904236 | 0.07417063 | wdr-5.1 |
| WBGene00006475 | 0.042365862 | 0.82187469 | 0.90243492 | tyra-3 |
| WBGene00006476 | 0.117154981 | 0.55360454 | 0.72569828 | rhgf-2 |
| WBGene00006477 | -0.088091208 | 0.59400488 | 0.7536897 | chup-1 |
| WBGene00006478 | -0.054710214 | 0.65699185 | 0.79757414 | tag-131 |
| WBGene00006479 | 0.096658668 | 0.38966056 | 0.58786842 | tmbi-4 |
| WBGene00006481 | -0.282389097 | 0.00034237 | 0.00281294 | plrg-1 |
| WBGene00006483 | 0.096746302 | 0.19483272 | 0.38181426 | dgk-4 |
| WBGene00006484 | 0.020232312 | 0.84643798 | NA | tag-138 |
| WBGene00006486 | 0.047779262 | 0.8062476 | 0.89246487 | zipt-16 |
| WBGene00006487 | -0.005377232 | 0.94281242 | NA | zipt-17 |
| WBGene00006488 | NA | NA | NA | F56D6.6 |
| WBGene00006489 | -0.339193141 | 0.00473566 | 0.02506648 | zhit-3 |
| WBGene00006490 | -0.108690201 | 0.18484769 | 0.36856171 | kdin-1 |
| WBGene00006491 | -0.234047864 | 0.064902 | 0.18163274 | acl-3 |
| WBGene00006492 | -0.066567576 | 0.53370226 | 0.70904186 | let-391 |
| WBGene00006493 | -0.250686477 | 0.00458103 | 0.02448735 | ivns-1 |
| WBGene00006494 | 0.010087808 | 0.9127204 | 0.95126775 | zipt-7.2 |
| WBGene00006495 | 0.450055265 | 0.00044985 | 0.00354861 | cpna-1 |
| WBGene00006496 | 0.017397961 | 0.89970295 | 0.94665211 | cgef-1 |
| WBGene00006497 | -0.269989937 | 0.00148268 | 0.00972579 | tag-151 |
| WBGene00006498 | 0.254938219 | 0.02191415 | 0.0821939 | ten-1 |
| WBGene00006499 | -0.198133293 | 0.00722301 | 0.03486716 | tag-153 |
| WBGene00006503 | -0.100008952 | 0.30439886 | 0.50529521 | snx-3 |
| WBGene00006504 | 0.197168868 | 0.01867497 | 0.07305758 | kcc-1 |
| WBGene00006505 | 1.107724034 | 0.01066408 | 0.04698549 | lagr-1 |
| WBGene00006508 | 0.03645328 | 0.78808768 | 0.88005428 | tns-1 |
| WBGene00006509 | 0.099238659 | 0.51697095 | 0.69633202 | tag-164 |
| WBGene00006510 | 0.577769001 | 2.67E-13 | 1.07E-11 | mtrr-1 |
| WBGene00006512 | -0.422985318 | 1.22E-09 | 3.06E-08 | abcf-1 |
| WBGene00006513 | 0.411033009 | 0.07695629 | 0.20527774 | rimb-1 |

| | | | | |
|---|---|---|---|---|
| WBGene00006514 | -0.195398307 | 0.01513796 | 0.06231733 | tdp-1 |
| WBGene00006515 | -0.202665481 | 0.02746831 | 0.09711749 | txdc-9 |
| WBGene00006516 | 0.051543615 | 0.62558914 | 0.77666841 | vps-16 |
| WBGene00006517 | -0.018152987 | 0.87069992 | 0.93091263 | madd-3 |
| WBGene00006518 | -0.307035174 | 5.56E-05 | 0.0005758 | bckd-1B |
| WBGene00006519 | -0.002006133 | 0.97773484 | 0.98731056 | cox-6A |
| WBGene00006520 | -0.015134202 | 0.92961969 | 0.96051954 | tag-175 |
| WBGene00006522 | 0.203433334 | 0.15165688 | 0.32414722 | abcx-1 |
| WBGene00006523 | -0.201121421 | 0.00730585 | 0.03515114 | tam-1 |
| WBGene00006524 | -0.074647607 | 0.68647996 | 0.8193326 | tap-1 |
| WBGene00006525 | 0.179490365 | 0.09068203 | 0.22888093 | tax-2 |
| WBGene00006526 | 0.195455661 | 0.06397992 | 0.17976012 | tax-4 |
| WBGene00006527 | 0.548271141 | 1.45E-06 | 2.20E-05 | tax-6 |
| WBGene00006528 | -0.323716466 | 1.05E-12 | 3.97E-11 | tba-1 |
| WBGene00006529 | -0.179225262 | 1.64E-05 | 0.00019029 | tba-2 |
| WBGene00006530 | -0.225648171 | 0.04047616 | 0.12911897 | tba-4 |
| WBGene00006531 | 0.259772513 | 0.153179 | 0.32633809 | tba-5 |
| WBGene00006532 | 0.133396135 | 0.15669766 | 0.33067317 | tba-6 |
| WBGene00006533 | 0.420135484 | 0.0774962 | 0.2061075 | tba-7 |
| WBGene00006534 | -0.022930767 | 0.78121253 | NA | tba-8 |
| WBGene00006535 | 0.531349267 | 0.04029492 | 0.12869937 | tba-9 |
| WBGene00006536 | -0.245478089 | 1.76E-07 | 3.18E-06 | tbb-1 |
| WBGene00006537 | -0.13866717 | 0.00027411 | 0.00231576 | tbb-2 |
| WBGene00006538 | 0.120319138 | 0.543046 | 0.71679918 | tbb-4 |
| WBGene00006539 | 0.173206748 | 0.40396065 | 0.60226219 | tbb-6 |
| WBGene00006540 | -0.129090536 | 0.12415165 | 0.2845053 | tbg-1 |
| WBGene00006541 | -0.448145886 | 1.21E-12 | 4.51E-11 | tbh-1 |
| WBGene00006542 | 0.064628812 | 0.51781413 | 0.69706847 | tbp-1 |
| WBGene00006543 | 0.43863462 | 0.03966691 | 0.12725972 | tbx-2 |
| WBGene00006544 | -0.207435123 | 0.02535182 | 0.09164841 | tbx-7 |
| WBGene00006545 | -0.389282839 | 0.08885863 | 0.22627625 | tbx-8 |
| WBGene00006546 | -0.164212787 | 0.30133123 | 0.50196729 | tbx-9 |
| WBGene00006547 | -1.135442753 | 0.00030129 | 0.00252583 | tbx-11 |
| WBGene00006549 | NA | NA | NA | tbx-30 |
| WBGene00006549 | NA | NA | NA | tbx-42 |
| WBGene00006550 | 0.007561332 | 0.94067768 | 0.96669207 | tbx-31 |
| WBGene00006551 | -0.103351434 | 0.41907513 | 0.61579411 | tbx-32 |
| WBGene00006552 | 0.066488561 | 0.72828735 | 0.84850209 | tbx-33 |
| WBGene00006553 | 0.177616853 | 0.0463229 | NA | tbx-34 |
| WBGene00006554 | 0.061275255 | 0.51782541 | NA | tbx-35 |
| WBGene00006555 | 0.04370938 | 0.78072183 | 0.87640849 | tbx-36 |
| WBGene00006556 | -0.235501215 | 0.16234611 | 0.33739454 | tbx-37 |
| WBGene00006557 | -0.019655792 | 0.91517253 | 0.95281497 | tbx-38 |

| | | | | |
|---|---|---|---|---|
| WBGene00006558 | -0.107872345 | 0.46171755 | 0.65228105 | tbx-39 |
| WBGene00006559 | -0.064434069 | 0.62078057 | 0.77318999 | tbx-40 |
| WBGene00006560 | -0.023109762 | 0.78121253 | NA | tbx-41 |
| WBGene00006561 | -0.153926712 | 0.42695825 | 0.62241028 | tcl-2 |
| WBGene00006562 | -0.079577354 | 0.24204839 | 0.44005348 | tdc-1 |
| WBGene00006563 | 0.149644982 | 0.42839667 | 0.62335079 | teg-1 |
| WBGene00006565 | -0.062527048 | 0.49153216 | 0.67506222 | tfg-1 |
| WBGene00006566 | -0.095824244 | 0.4895051 | 0.67351228 | tgt-1 |
| WBGene00006567 | -0.183537367 | 0.14281183 | 0.31113924 | tgt-2 |
| WBGene00006568 | -0.083611292 | 0.42649786 | 0.62219562 | thk-1 |
| WBGene00006570 | 0.063544337 | 0.7554453 | 0.86140769 | tig-2 |
| WBGene00006571 | -0.036299102 | 0.58039794 | 0.74546945 | tim-1 |
| WBGene00006572 | -0.046130246 | 0.73798393 | 0.85272061 | tin-9.1 |
| WBGene00006573 | -0.194535678 | 0.11652488 | 0.27270472 | tin-10 |
| WBGene00006574 | -0.436192711 | 0.00048068 | 0.0037655 | tin-13 |
| WBGene00006575 | 0.509536266 | 0.00367047 | 0.02047992 | tir-1 |
| WBGene00006576 | 0.151264273 | 0.34796755 | 0.54786459 | tkr-1 |
| WBGene00006577 | -0.159249456 | 0.08401681 | 0.21806242 | tlf-1 |
| WBGene00006578 | -0.202955618 | 0.24312361 | 0.44114175 | tli-1 |
| WBGene00006579 | -0.115425868 | 0.04217695 | 0.132827 | tlk-1 |
| WBGene00006580 | 0.130604551 | 0.49470777 | 0.67742391 | tlp-1 |
| WBGene00006582 | 0.73463236 | 0.00166194 | 0.01072387 | tmd-2 |
| WBGene00006583 | 0.43188763 | 0.02560505 | 0.09229039 | tnc-2 |
| WBGene00006584 | 0.478705697 | 0.02314824 | 0.08541165 | tni-1 |
| WBGene00006585 | 0.501335889 | 0.00318507 | 0.01832322 | tni-3 |
| WBGene00006586 | 0.464270364 | 0.00042894 | 0.00340385 | tni-4 |
| WBGene00006587 | -0.448736955 | 7.57E-10 | 1.93E-08 | tnt-2 |
| WBGene00006588 | 0.306545877 | 0.00760211 | 0.03629034 | tnt-3 |
| WBGene00006589 | 0.707035917 | 0.00054155 | 0.00415341 | tnt-4 |
| WBGene00006590 | -0.258315203 | 0.00362735 | 0.02031679 | toc-1 |
| WBGene00006591 | -0.364978631 | 0.07693529 | 0.20527774 | toh-1 |
| WBGene00006592 | -0.036674142 | 0.81676266 | 0.89924966 | dpy-31 |
| WBGene00006593 | 0.004128933 | 0.97412688 | 0.98568926 | tol-1 |
| WBGene00006594 | 0.573568155 | 0.00118661 | 0.00809561 | tom-1 |
| WBGene00006595 | -0.268025366 | 1.64E-05 | 0.00019066 | top-1 |
| WBGene00006596 | 0.134154656 | 0.27972431 | 0.47944787 | top-3 |
| WBGene00006597 | 0.124661631 | 0.3079132 | 0.50870652 | tor-1 |
| WBGene00006598 | 0.038726127 | 0.57551883 | NA | tor-2 |
| WBGene00006599 | 0.527434245 | 0.00175438 | 0.01121654 | tpa-1 |
| WBGene00006600 | 0.352568115 | 0.05953574 | 0.17068 | tph-1 |
| WBGene00006601 | -0.097176927 | 0.12583808 | 0.28690032 | tpi-1 |
| WBGene00006602 | 0.425667906 | 0.00372405 | 0.02070785 | tps-1 |
| WBGene00006603 | -0.073988118 | 0.57162777 | 0.73987023 | tps-2 |

| | | | | |
|---|---|---|---|---|
| WBGene00006604 | 0.24931176 | 0.1211435 | 0.28014596 | tra-1 |
| WBGene00006605 | -0.289763701 | 0.00428616 | 0.02319056 | tra-2 |
| WBGene00006606 | -0.569215971 | 1.51E-05 | 0.00017757 | tra-3 |
| WBGene00006607 | -0.009187805 | 0.8988147 | 0.94659373 | tre-1 |
| WBGene00006608 | 0.206029707 | 0.05575189 | 0.16308959 | tre-2 |
| WBGene00006609 | 1.338520331 | 4.73E-19 | 3.33E-17 | tre-3 |
| WBGene00006610 | 0.377981568 | 0.12148638 | 0.28051644 | tre-4 |
| WBGene00006611 | 2.883411414 | 2.42E-11 | 7.62E-10 | tre-5 |
| WBGene00006612 | 0.020317245 | 0.84643798 | NA | trf-1 |
| WBGene00006613 | -0.160572818 | 0.10936775 | 0.26039175 | trm-1 |
| WBGene00006614 | 0.619697311 | 0.03651927 | 0.11961492 | trp-1 |
| WBGene00006615 | 0.290825117 | 0.20553279 | 0.39464797 | trp-2 |
| WBGene00006616 | 0.167953207 | 0.42523866 | 0.62114248 | trp-4 |
| WBGene00006617 | -0.409518743 | 2.46E-11 | 7.70E-10 | tars-1 |
| WBGene00006618 | 0.143911888 | 0.22093365 | 0.41337886 | trt-1 |
| WBGene00006619 | 0.078524143 | 0.46849269 | 0.65738945 | try-1 |
| WBGene00006620 | 0.375350319 | 0.0922736 | 0.23180814 | svh-1 |
| WBGene00006621 | 0.291123488 | 0.18640311 | 0.37075504 | try-3 |
| WBGene00006622 | 0.013986715 | 0.92884636 | 0.95998218 | try-4 |
| WBGene00006623 | 0.030735895 | 0.78211024 | 0.87737414 | try-5 |
| WBGene00006624 | 0.157258146 | 0.17499775 | 0.35564444 | try-6 |
| WBGene00006625 | 0.078633594 | 0.26310689 | NA | try-7 |
| WBGene00006626 | -0.067757217 | 0.17148705 | 0.3505154 | tsn-1 |
| WBGene00006627 | 0.675129217 | 0.03184664 | 0.10795708 | tsp-1 |
| WBGene00006628 | 0.389575059 | 0.08105649 | 0.21286962 | tsp-2 |
| WBGene00006629 | 2.483930356 | 0.00040873 | 0.00327901 | tsp-3 |
| WBGene00006630 | NA | NA | NA | tsp-4 |
| WBGene00006631 | NA | NA | NA | tsp-5 |
| WBGene00006632 | 0.30120157 | 0.14628341 | 0.31612332 | tsp-6 |
| WBGene00006633 | 0.240068295 | 0.24083313 | 0.43839833 | tsp-7 |
| WBGene00006634 | 0.093090225 | 0.61111199 | 0.76604108 | tsp-8 |
| WBGene00006635 | -0.118123305 | 0.52692712 | 0.7041981 | tsp-9 |
| WBGene00006636 | 0.301686245 | 0.17429125 | 0.35495871 | tsp-10 |
| WBGene00006637 | 0.290085049 | 0.20507421 | 0.3942347 | tsp-11 |
| WBGene00006638 | -0.129338074 | 0.13628859 | 0.3013746 | tsp-12 |
| WBGene00006639 | 0.098687168 | 0.57920468 | 0.7449645 | tsp-13 |
| WBGene00006640 | -0.167624619 | 0.43127015 | 0.62628707 | tsp-14 |
| WBGene00006641 | 0.310270394 | 0.17681009 | 0.35765081 | tsp-15 |
| WBGene00006642 | 0.106447325 | 0.51431869 | 0.69434209 | tsp-16 |
| WBGene00006643 | 0.106253557 | 0.37088808 | 0.57036433 | tsp-17 |
| WBGene00006644 | NA | NA | NA | tsp-18 |
| WBGene00006645 | 0.058314668 | 0.40208118 | NA | tsp-19 |
| WBGene00006646 | -0.135281809 | 0.51607467 | 0.69567629 | tsp-20 |

| | | | | |
|---|---|---|---|---|
| WBGene00006647 | -0.082924118 | 0.5741326 | 0.7419131 | tsr-1 |
| WBGene00006648 | -0.139178508 | 0.1576819 | 0.33182531 | ttb-1 |
| WBGene00006649 | 0.401892864 | 0.00484616 | 0.02553173 | tth-1 |
| WBGene00006650 | 4.288195901 | 0 | 0 | NA |
| WBGene00006651 | 4.240593477 | 2.26E-09 | 5.46E-08 | NA |
| WBGene00006652 | 0.784435721 | 0.02160843 | 0.08122441 | ttx-1 |
| WBGene00006654 | -0.050223056 | 0.47776088 | NA | ttx-3 |
| WBGene00006655 | 0.201477748 | 0.18624237 | 0.37058637 | tub-1 |
| WBGene00006656 | 0.169615621 | 0.39761222 | 0.59578758 | twk-1 |
| WBGene00006657 | 0.358169905 | 0.1385034 | 0.3048735 | twk-2 |
| WBGene00006658 | -0.039636691 | 0.54579661 | NA | twk-3 |
| WBGene00006659 | 0.033463284 | 0.6365037 | NA | twk-4 |
| WBGene00006660 | 0.06634931 | 0.73256348 | 0.85034979 | twk-5 |
| WBGene00006661 | -0.022930767 | 0.78121253 | NA | twk-6 |
| WBGene00006662 | 1.556634053 | 6.89E-05 | 0.00069749 | twk-7 |
| WBGene00006663 | 0.356126482 | 0.10172222 | 0.24832686 | twk-8 |
| WBGene00006664 | 0.083775379 | 0.51663521 | 0.69612149 | twk-9 |
| WBGene00006665 | 2.964175219 | 0.00213431 | 0.01316835 | twk-10 |
| WBGene00006666 | 0.922971522 | 0.01934575 | 0.07499104 | twk-11 |
| WBGene00006667 | 0.146548948 | 0.46309777 | 0.65301113 | twk-12 |
| WBGene00006668 | 0.381368912 | 0.11905175 | 0.27684826 | twk-13 |
| WBGene00006669 | 0.131264594 | 0.37406234 | 0.57293092 | twk-14 |
| WBGene00006670 | -0.253519815 | 0.25879416 | 0.45678183 | twk-16 |
| WBGene00006671 | 0.099297564 | 0.63308918 | 0.78213304 | twk-17 |
| WBGene00006672 | -0.050106206 | 0.77726524 | 0.87398508 | twk-18 |
| WBGene00006673 | 0.038125098 | 0.61119482 | NA | twk-20 |
| WBGene00006674 | 0.231946026 | 0.08873442 | 0.22611347 | twk-21 |
| WBGene00006675 | -0.608450551 | 0.01372845 | 0.05764217 | twk-22 |
| WBGene00006676 | 0.036785715 | 0.7118385 | 0.83785091 | twk-23 |
| WBGene00006677 | 0.460771294 | 0.06266137 | 0.1772802 | twk-24 |
| WBGene00006678 | 0.520681286 | 0.03595037 | 0.11822761 | twk-25 |
| WBGene00006679 | 1.305232836 | 9.86E-06 | 0.00012202 | twk-26 |
| WBGene00006680 | 0.117155162 | 0.42110452 | 0.61783659 | twk-28 |
| WBGene00006681 | 0.937008405 | 0.01395669 | 0.05834947 | twk-29 |
| WBGene00006682 | 0.018825944 | 0.92607427 | 0.95834417 | twk-30 |
| WBGene00006683 | 0.45750872 | 0.07855411 | 0.20842891 | twk-31 |
| WBGene00006684 | 0.114183493 | 0.37036365 | 0.57006877 | twk-32 |
| WBGene00006685 | 0.364198566 | 0.1170257 | 0.27352701 | twk-33 |
| WBGene00006686 | 0.08843159 | 0.66420661 | 0.80296004 | twk-34 |
| WBGene00006687 | -0.002988003 | 0.98190257 | 0.98949726 | twk-35 |
| WBGene00006688 | 0.403394549 | 0.0331567 | 0.11136139 | twk-36 |
| WBGene00006689 | NA | NA | NA | twk-37 |
| WBGene00006690 | 0.001427146 | 0.99554794 | 0.99731901 | twk-39 |

| | | | | |
|---|---|---|---|---|
| WBGene00006691 | 0.425190033 | 0.05306663 | 0.15730886 | twk-40 |
| WBGene00006692 | -0.074872918 | 0.71759817 | 0.84158561 | twk-42 |
| WBGene00006693 | -0.070617324 | 0.73234428 | 0.85023927 | twk-43 |
| WBGene00006694 | 0.059980168 | 0.38780842 | NA | twk-44 |
| WBGene00006695 | 0.037825269 | 0.8293267 | 0.90674126 | twk-45 |
| WBGene00006696 | 0.2554909 | 0.25294978 | 0.4510337 | twk-46 |
| WBGene00006697 | 0.025381702 | 0.72951108 | 0.84916296 | uaf-1 |
| WBGene00006698 | -0.324030147 | 0.00017255 | 0.00155531 | uaf-2 |
| WBGene00006699 | -0.191630661 | 1.17E-05 | 0.0001411 | uba-1 |
| WBGene00006700 | -0.123862784 | 0.12970289 | 0.29201626 | uba-2 |
| WBGene00006701 | 0.209757525 | 0.0954404 | 0.23757788 | ubc-1 |
| WBGene00006702 | 0.0028947 | 0.97503166 | 0.98591235 | ubc-3 |
| WBGene00006703 | -0.079600464 | 0.48932113 | 0.67332243 | ubc-6 |
| WBGene00006704 | -0.218506534 | 0.00014297 | 0.00131321 | ubc-7 |
| WBGene00006705 | 0.91045447 | 1.16E-05 | 0.00014003 | ubc-8 |
| WBGene00006706 | -0.235199209 | 2.72E-05 | 0.00030179 | ubc-9 |
| WBGene00006707 | -0.183634387 | 0.05544594 | 0.16235681 | ubc-12 |
| WBGene00006708 | -0.273550943 | 0.00103526 | 0.00723151 | ubc-13 |
| WBGene00006709 | -0.057802876 | 0.65743812 | 0.79794011 | ubc-14 |
| WBGene00006710 | -0.193795532 | 0.08986907 | 0.22785184 | ubc-15 |
| WBGene00006711 | 0.111016533 | 0.36184147 | 0.56165797 | ubc-16 |
| WBGene00006712 | 0.222005758 | 0.25886847 | 0.45685792 | ubc-17 |
| WBGene00006713 | -0.204834937 | 0.00166383 | 0.01073135 | ubc-18 |
| WBGene00006714 | 0.07114573 | 0.73279447 | 0.85051752 | ubc-19 |
| WBGene00006715 | -0.226666692 | 0.00520252 | 0.0269619 | ubc-20 |
| WBGene00006716 | 0.056411843 | 0.41891328 | NA | ubc-21 |
| WBGene00006717 | -0.08769705 | 0.42790504 | 0.62288304 | ubc-22 |
| WBGene00006718 | -0.220993213 | 0.30656674 | 0.50711327 | ubc-23 |
| WBGene00006719 | 0.011913685 | 0.89608828 | NA | ubc-24 |
| WBGene00006720 | 0.104051877 | 0.10584332 | 0.25483506 | ubc-25 |
| WBGene00006721 | 0.357716 | 0.05739249 | 0.16643091 | ubh-1 |
| WBGene00006722 | -0.124668814 | 0.39858374 | 0.59675473 | ubh-2 |
| WBGene00006723 | -0.267351193 | 0.01407364 | 0.05870789 | ubh-3 |
| WBGene00006724 | -0.208726453 | 0.01859752 | 0.07286407 | ubh-4 |
| WBGene00006725 | -0.078387371 | 0.10438217 | 0.25259961 | ubl-1 |
| WBGene00006726 | -0.382621185 | 0.01677864 | 0.0675436 | ubl-5 |
| WBGene00006727 | 0.094131529 | 0.03227495 | 0.10910588 | ubq-1 |
| WBGene00006728 | -0.330551234 | 1.00E-10 | 2.87E-09 | ubq-2 |
| WBGene00006729 | 0.186285149 | 0.26845512 | 0.46735451 | ucp-4 |
| WBGene00006730 | -0.185753822 | 0.02356279 | 0.08663555 | uev-1 |
| WBGene00006731 | 0.051292135 | 0.60829666 | 0.76399837 | uev-2 |
| WBGene00006732 | -0.163066101 | 0.25140085 | 0.44991306 | uev-3 |
| WBGene00006733 | 0.024990761 | 0.77259991 | 0.87092973 | ufd-1 |

| | | | | |
|---|---|---|---|---|
| WBGene00006734 | -0.088234933 | 0.20352728 | 0.39247213 | ufd-2 |
| WBGene00006735 | -0.049405241 | 0.64816206 | 0.79160333 | ula-1 |
| WBGene00006736 | -0.026277354 | 0.73058646 | 0.84986592 | ulp-1 |
| WBGene00006737 | -0.255835947 | 0.00296519 | 0.01730994 | ulp-2 |
| WBGene00006738 | 0.039292282 | 0.80704469 | 0.89298227 | ulp-3 |
| WBGene00006739 | -0.0266255 | 0.76240127 | 0.86502767 | ulp-4 |
| WBGene00006740 | -0.159500215 | 0.32509311 | 0.52436798 | ulp-5 |
| WBGene00006741 | 1.118016382 | 0.00044136 | 0.00349104 | unc-1 |
| WBGene00006742 | 0.507628525 | 0.01752 | 0.06972284 | unc-2 |
| WBGene00006743 | 0.230509619 | 0.21353516 | 0.40428294 | unc-3 |
| WBGene00006744 | 0.134010361 | 0.30738503 | 0.50794855 | unc-4 |
| WBGene00006745 | -0.118599363 | 0.52242978 | 0.70031995 | unc-5 |
| WBGene00006746 | 0.210106523 | 0.32248352 | 0.52270521 | unc-6 |
| WBGene00006747 | 0.998106285 | 0.0070731 | 0.03427914 | unc-7 |
| WBGene00006748 | 0.065150006 | 0.64051355 | 0.78685676 | unc-8 |
| WBGene00006749 | 0.653347608 | 0.0225718 | 0.08400197 | unc-9 |
| WBGene00006750 | 0.137113705 | 0.50854556 | 0.68894903 | unc-10 |
| WBGene00006751 | -0.050597463 | 0.60044071 | 0.75824154 | unc-11 |
| WBGene00006752 | 1.017393586 | 7.26E-08 | 1.40E-06 | unc-13 |
| WBGene00006753 | 0.528486008 | 7.14E-05 | 0.00071976 | unc-14 |
| WBGene00006754 | 0.107265205 | 0.38735794 | 0.58593942 | unc-15 |
| WBGene00006755 | 0.047825316 | 0.55801937 | 0.72887515 | unc-16 |
| WBGene00006756 | 0.802040865 | 0.01923444 | 0.07473764 | unc-17 |
| WBGene00006757 | -0.022452628 | 0.83168831 | 0.90832691 | unc-18 |
| WBGene00006759 | 0.652461697 | 8.87E-15 | 4.15E-13 | unc-22 |
| WBGene00006760 | -0.547231724 | 0.00131825 | 0.00882511 | unc-23 |
| WBGene00006761 | 0.216671859 | 0.21534348 | 0.40639906 | unc-24 |
| WBGene00006762 | 0.541187785 | 0.0194822 | 0.07529356 | unc-25 |
| WBGene00006763 | 0.039105031 | 0.74839794 | 0.85798248 | unc-26 |
| WBGene00006764 | -0.518468559 | 1.84E-15 | 9.16E-14 | unc-27 |
| WBGene00006765 | 0.010454042 | 0.9562196 | 0.97561053 | unc-29 |
| WBGene00006766 | -0.026643337 | 0.87914011 | 0.93529214 | unc-30 |
| WBGene00006767 | 0.294964609 | 0.19687978 | 0.38442285 | unc-31 |
| WBGene00006768 | -0.121799215 | 0.06668723 | 0.18474319 | unc-32 |
| WBGene00006769 | -0.142467764 | 0.20729008 | 0.39645783 | unc-33 |
| WBGene00006770 | 0.000880956 | 0.99507352 | 0.99691196 | unc-34 |
| WBGene00006771 | 0.194042818 | 0.03085704 | 0.10543241 | tln-1 |
| WBGene00006772 | 0.592097029 | 0.01046042 | 0.04645164 | unc-36 |
| WBGene00006773 | -0.372970671 | 6.49E-06 | 8.28E-05 | unc-37 |
| WBGene00006774 | 0.143731544 | 0.45312852 | 0.64528841 | unc-38 |
| WBGene00006775 | -0.220908261 | 0.21420158 | 0.40507948 | unc-39 |
| WBGene00006776 | -0.176697919 | 0.01341085 | 0.05658451 | unc-40 |
| WBGene00006777 | 0.55192331 | 0.04678724 | 0.14339795 | unc-41 |

| | | | | |
|---|---|---|---|---|
| WBGene00006778 | 0.337018486 | 0.08041443 | 0.21171511 | unc-42 |
| WBGene00006779 | 0.281976588 | 0.00826385 | 0.03874194 | unc-43 |
| WBGene00006780 | 0.238335259 | 0.00118604 | 0.00809551 | unc-44 |
| WBGene00006781 | 0.14839704 | 0.10631111 | 0.25568726 | unc-45 |
| WBGene00006782 | 0.166470253 | 0.1986263 | 0.38697108 | unc-46 |
| WBGene00006783 | 0.168439554 | 0.29442145 | 0.49565557 | unc-47 |
| WBGene00006784 | 0.973694352 | 1.95E-05 | 0.0002228 | unc-49 |
| WBGene00006785 | -0.012262229 | 0.9098251 | 0.95107449 | unc-50 |
| WBGene00006786 | 0.614892627 | 1.99E-06 | 2.89E-05 | unc-51 |
| WBGene00006787 | 0.317927827 | 6.38E-05 | 0.00065271 | unc-52 |
| WBGene00006788 | 0.414629568 | 0.04160427 | 0.13153275 | unc-53 |
| WBGene00006789 | 0.488544961 | 5.10E-05 | 0.00053389 | unc-54 |
| WBGene00006790 | 0.077898987 | 0.48089182 | 0.66716666 | unc-55 |
| WBGene00006791 | -0.183893232 | 0.02300023 | 0.08520911 | unc-57 |
| WBGene00006792 | 0.212875304 | 0.32361866 | 0.52343138 | unc-58 |
| WBGene00006793 | -0.197062081 | 0.00645478 | 0.03199202 | unc-59 |
| WBGene00006794 | 0.0317104 | 0.82998671 | 0.9071919 | unc-60 |
| WBGene00006795 | -0.141439889 | 0.07723858 | 0.20566135 | unc-61 |
| WBGene00006796 | -0.026916295 | 0.83809898 | 0.91238064 | unc-62 |
| WBGene00006797 | 0.005534914 | 0.98025153 | 0.98860925 | unc-63 |
| WBGene00006798 | 0.498309382 | 0.00463971 | 0.02469359 | unc-64 |
| WBGene00006801 | 0.482919158 | 9.03E-07 | 1.44E-05 | unc-68 |
| WBGene00006802 | 0.113267171 | 0.44334031 | 0.6368667 | unc-69 |
| WBGene00006803 | 0.189710626 | 0.00757863 | 0.03624917 | unc-70 |
| WBGene00006804 | 0.151050615 | 0.04729672 | 0.1444454 | unc-71 |
| WBGene00006805 | 0.323684284 | 0.00508141 | 0.02646633 | unc-73 |
| WBGene00006807 | 0.150735692 | 0.31239047 | 0.51320791 | unc-75 |
| WBGene00006808 | -0.119418572 | 0.19412798 | 0.38110708 | unc-76 |
| WBGene00006809 | -0.110822646 | 0.59624812 | 0.75503103 | unc-77 |
| WBGene00006810 | 0.012583656 | 0.92207733 | 0.95620205 | unc-78 |
| WBGene00006811 | -0.074119168 | 0.71805012 | 0.84184591 | unc-79 |
| WBGene00006812 | 0.816863983 | 8.55E-05 | 0.0008411 | unc-80 |
| WBGene00006814 | 0.292890488 | 0.07624754 | 0.20393501 | unc-82 |
| WBGene00006815 | 0.603239431 | 9.77E-07 | 1.55E-05 | unc-83 |
| WBGene00006816 | -0.145947233 | 0.14611469 | 0.31589857 | unc-84 |
| WBGene00006817 | -0.09596184 | 0.24717376 | 0.4456592 | unc-85 |
| WBGene00006818 | 0.198018606 | 0.30348564 | 0.50423664 | unc-86 |
| WBGene00006819 | -0.122735085 | 0.07280037 | 0.19680025 | unc-87 |
| WBGene00006820 | 0.512235552 | 1.74E-06 | 2.57E-05 | unc-89 |
| WBGene00006822 | -0.107071468 | 0.57461326 | 0.74227217 | unc-93 |
| WBGene00006823 | 0.028031489 | 0.76153182 | 0.86480288 | unc-94 |
| WBGene00006824 | 0.165116575 | 0.39874872 | 0.59687966 | unc-95 |
| WBGene00006825 | 0.482977289 | 0.02497499 | 0.09075671 | unc-96 |

| | | | | |
|---|---|---|---|---|
| WBGene00006826 | -0.201408549 | 0.1273277 | 0.28871852 | unc-97 |
| WBGene00006827 | 0.372212328 | 0.10194525 | 0.2486805 | unc-98 |
| WBGene00006829 | -0.228486134 | 0.03678788 | 0.12036008 | unc-101 |
| WBGene00006830 | 1.002464788 | 0.00049483 | 0.00385159 | unc-103 |
| WBGene00006831 | 0.477476998 | 0.00811474 | 0.03827205 | unc-104 |
| WBGene00006832 | 0.190326758 | 0.38186141 | 0.58110726 | unc-105 |
| WBGene00006833 | 0.22242272 | 0.00419764 | 0.02277898 | unc-108 |
| WBGene00006836 | -0.004234074 | 0.97426492 | 0.98568926 | unc-112 |
| WBGene00006839 | 0.081578731 | 0.46826654 | 0.65738945 | unc-115 |
| WBGene00006840 | -0.123589033 | 0.06499703 | 0.1817459 | unc-116 |
| WBGene00006843 | -0.287880416 | 0.10308599 | 0.25058216 | unc-119 |
| WBGene00006844 | 0.00608244 | 0.97660908 | 0.98670757 | unc-120 |
| WBGene00006845 | 0.01958786 | 0.91951415 | 0.95531294 | unc-122 |
| WBGene00006852 | 0.623788536 | 0.02871033 | 0.10044665 | unc-129 |
| WBGene00006853 | -0.045987583 | 0.80234445 | 0.88993372 | unc-130 |
| WBGene00006856 | -0.082165204 | 0.24821811 | 0.44662178 | usp-14 |
| WBGene00006861 | 0.126928501 | 0.37221569 | 0.57146318 | cal-5 |
| WBGene00006862 | -0.337496055 | 0.03295225 | 0.11080706 | pnk-4 |
| WBGene00006863 | 0.467536398 | 0.00044812 | 0.00353691 | gyg-1 |
| WBGene00006864 | 0.264810676 | 0.17832259 | 0.35986506 | npr-16 |
| WBGene00006865 | 0.646348392 | 0.03582413 | 0.11798028 | slcf-2 |
| WBGene00006868 | 0.113365395 | 0.14478463 | 0.31413631 | vab-1 |
| WBGene00006869 | -0.322416575 | 0.17030404 | 0.34882784 | vab-2 |
| WBGene00006870 | 0.549561132 | 0.03774018 | 0.12259907 | vab-3 |
| WBGene00006873 | -0.000282921 | 0.99680781 | 0.9978865 | vab-7 |
| WBGene00006874 | 0.110236333 | 0.48526345 | 0.66999896 | vab-8 |
| WBGene00006875 | 0.797267084 | 0.01266744 | 0.05405536 | vab-9 |
| WBGene00006876 | 0.419758145 | 6.96E-12 | 2.37E-10 | vab-10 |
| WBGene00006881 | -0.09334204 | 0.60891015 | 0.76432215 | vab-15 |
| WBGene00006882 | 0.386118066 | 0.06085353 | 0.17337158 | vab-19 |
| WBGene00006886 | 1.372403416 | 2.68E-05 | 0.00029689 | vap-1 |
| WBGene00006887 | 0.07086081 | 0.6474868 | 0.79141821 | vav-1 |
| WBGene00006888 | -0.344692763 | 9.75E-12 | 3.25E-10 | vbh-1 |
| WBGene00006889 | -0.399015253 | 5.27E-05 | 0.00054848 | pfd-3 |
| WBGene00006890 | 0.247877908 | 0.06139455 | 0.17453933 | vem-1 |
| WBGene00006893 | 0.465686024 | 0.00365042 | 0.02039137 | spon-1 |
| WBGene00006894 | -0.05198345 | 0.74305912 | 0.85461565 | ver-1 |
| WBGene00006895 | -0.04044783 | 0.53612608 | NA | ver-2 |
| WBGene00006896 | -0.291154704 | 0.04067866 | 0.12961399 | ver-3 |
| WBGene00006897 | -1.566342834 | 0.00373859 | 0.02074932 | ver-4 |
| WBGene00006902 | -1.213887705 | 4.99E-16 | 2.62E-14 | vet-6 |
| WBGene00006910 | 0.07117469 | 0.2728813 | 0.47297917 | vha-1 |
| WBGene00006911 | 0.028095765 | 0.56758236 | 0.73663447 | vha-3 |

| | | | | |
|---|---|---|---|---|
| WBGene00006911 | 0.028095765 | 0.56758236 | 0.73663447 | vha-2 |
| WBGene00006912 | NA | NA | NA | vha-3 |
| WBGene00006912 | NA | NA | NA | vha-2 |
| WBGene00006913 | -0.022323327 | 0.76123879 | 0.86464679 | vha-4 |
| WBGene00006914 | 0.380634755 | 1.05E-06 | 1.64E-05 | vha-5 |
| WBGene00006915 | 0.403726812 | 5.52E-10 | 1.44E-08 | vha-6 |
| WBGene00006916 | -0.033467788 | 0.65882348 | 0.79889303 | vha-7 |
| WBGene00006917 | -0.096052714 | 0.0993184 | 0.24443111 | vha-8 |
| WBGene00006918 | 0.11566154 | 0.14505677 | 0.31445617 | vha-9 |
| WBGene00006919 | 0.05969815 | 0.40665549 | 0.60471122 | vha-10 |
| WBGene00006920 | -0.027659584 | 0.66827567 | 0.80641425 | vha-11 |
| WBGene00006921 | 0.144679877 | 0.01122846 | 0.04900026 | vha-12 |
| WBGene00006922 | 0.054349241 | 0.79068174 | 0.88218122 | vhl-1 |
| WBGene00006923 | 0.593979869 | 3.97E-05 | 0.00042541 | vhp-1 |
| WBGene00006924 | -0.348800479 | 1.31E-16 | 7.50E-15 | vig-1 |
| WBGene00006925 | -2.579623174 | 1.46E-46 | 3.19E-44 | vit-1 |
| WBGene00006926 | 0.574457551 | 0.00039137 | 0.00315529 | vit-2 |
| WBGene00006927 | -2.879227638 | 6.14E-74 | 2.09E-71 | vit-3 |
| WBGene00006928 | -1.518362892 | 4.39E-16 | 2.36E-14 | vit-4 |
| WBGene00006929 | -0.970459165 | 3.47E-13 | 1.37E-11 | vit-5 |
| WBGene00006930 | 0.621838482 | 0.00033529 | 0.00276255 | vit-6 |
| WBGene00006931 | -0.138501862 | 0.12464525 | 0.28511378 | vps-26 |
| WBGene00006932 | -0.036088501 | 0.73478201 | 0.85123781 | vps-34 |
| WBGene00006933 | -0.313812967 | 2.59E-05 | 0.00028813 | vps-35 |
| WBGene00006934 | -0.128285305 | 0.15368418 | 0.32702528 | vps-54 |
| WBGene00006935 | -0.071742676 | 0.52005133 | 0.6989234 | vars-1 |
| WBGene00006936 | -0.326125497 | 1.48E-08 | 3.22E-07 | glp-4 |
| WBGene00006937 | -0.0572153 | 0.50460176 | 0.68577689 | wah-1 |
| WBGene00006938 | -0.239842731 | 0.26675493 | 0.46550166 | wee-1.1 |
| WBGene00006939 | -0.051723619 | 0.80156661 | 0.88947527 | NA |
| WBGene00006940 | -0.195053646 | 0.00686306 | 0.03348286 | wee-1.3 |
| WBGene00006941 | 0.055874377 | 0.35167848 | 0.55169019 | wnk-1 |
| WBGene00006942 | -0.384917686 | 0.01040549 | 0.04630624 | wrk-1 |
| WBGene00006943 | -0.260927317 | 0.00421853 | 0.02288382 | wrm-1 |
| WBGene00006944 | -0.042318367 | 0.67706091 | 0.81252859 | wrn-1 |
| WBGene00006945 | -0.260223207 | 0.00969458 | 0.04387585 | wars-1 |
| WBGene00006946 | 0.072022637 | 0.55289269 | 0.72508974 | prx-10 |
| WBGene00006947 | 2.317574075 | 0.00040079 | 0.00322063 | wrt-1 |
| WBGene00006948 | 2.144987794 | 0.00801554 | 0.0379178 | wrt-2 |
| WBGene00006949 | -0.200120721 | 0.34704425 | 0.54665893 | wrt-3 |
| WBGene00006950 | 3.919336683 | 0.00167014 | 0.01076257 | wrt-4 |
| WBGene00006951 | -0.092566344 | 0.63682666 | 0.78460587 | wrt-5 |
| WBGene00006952 | 0.179263092 | 0.15013324 | 0.32159485 | wrt-6 |

| | | | | |
|---|---|---|---|---|
| WBGene00006953 | 0.020317245 | 0.84643798 | NA | wrt-7 |
| WBGene00006954 | 0.219871264 | 0.02304865 | 0.0852806 | wrt-8 |
| WBGene00006955 | -0.016478531 | 0.89116397 | 0.94335418 | wrt-9 |
| WBGene00006956 | -0.39856987 | 0.11104244 | 0.2630274 | wrt-10 |
| WBGene00006957 | -0.038935754 | 0.6835323 | 0.81748051 | wsp-1 |
| WBGene00006958 | -0.026361955 | 0.82394906 | 0.9034778 | wve-1 |
| WBGene00006959 | 0.008839097 | 0.8959808 | 0.94530914 | xbp-1 |
| WBGene00006960 | 0.026732797 | 0.82414184 | 0.9034778 | xbx-1 |
| WBGene00006961 | -0.189413276 | 0.01164414 | 0.05036393 | xnp-1 |
| WBGene00006962 | -1.008120969 | 0.01048579 | 0.04648337 | xol-1 |
| WBGene00006963 | -0.083038255 | 0.50508408 | 0.68624127 | xpa-1 |
| WBGene00006964 | -0.204306832 | 0.01472217 | 0.06095794 | xrn-2 |
| WBGene00006965 | 0.135980078 | 0.33822471 | 0.53789273 | xtr-1 |
| WBGene00006966 | 0.015507174 | 0.80923808 | NA | xtr-2 |
| WBGene00006967 | NA | NA | NA | NA |
| WBGene00006968 | 0.045043347 | 0.70033654 | 0.82964862 | yars-2 |
| WBGene00006970 | 0.856506229 | 0.01685048 | 0.06768394 | zag-1 |
| WBGene00006971 | 0.145017422 | 0.48848939 | 0.67255719 | zak-1 |
| WBGene00006974 | -0.278949881 | 2.51E-05 | 0.00027962 | zen-4 |
| WBGene00006975 | 0.065669793 | 0.37178715 | 0.57099924 | zfp-1 |
| WBGene00006976 | -0.252885788 | 0.02668539 | 0.09534686 | zhp-3 |
| WBGene00006977 | -0.246260714 | 0.00213371 | 0.01316835 | zif-1 |
| WBGene00006978 | 0.240681347 | 0.25193643 | 0.44993917 | zig-1 |
| WBGene00006979 | 0.281959211 | 0.21858175 | 0.41042776 | zig-2 |
| WBGene00006980 | 0.526159305 | 0.05817685 | 0.16790209 | zig-3 |
| WBGene00006981 | 0.657586619 | 0.0359835 | 0.11830998 | zig-4 |
| WBGene00006982 | 0.8072994 | 0.01692451 | 0.06794401 | zig-5 |
| WBGene00006983 | 0.15485085 | 0.46674554 | 0.6563217 | zig-6 |
| WBGene00006984 | 0.894678571 | 1.11E-07 | 2.08E-06 | zig-7 |
| WBGene00006985 | 0.361963013 | 0.11929945 | 0.27724812 | zig-8 |
| WBGene00006986 | 0.84981423 | 0.00021062 | 0.00184865 | zip-1 |
| WBGene00006987 | 0.045301335 | 0.72652391 | 0.84710015 | zmp-1 |
| WBGene00006988 | -0.194174217 | 0.06376618 | 0.17929721 | zyg-1 |
| WBGene00006993 | 0.093052699 | 0.44971225 | 0.64204323 | zyg-8 |
| WBGene00006994 | -0.022568058 | 0.70814168 | 0.8352455 | zyg-9 |
| WBGene00006996 | 0.171314313 | 0.00626445 | 0.03120712 | zyg-11 |
| WBGene00006997 | -0.101656925 | 0.15081044 | 0.32252639 | zyg-12 |
| WBGene00006999 | 0.118764634 | 0.20188312 | 0.39045849 | zyx-1 |
| WBGene00007000 | -0.440171083 | 1.62E-11 | 5.23E-10 | tufm-1 |
| WBGene00007001 | -0.159172924 | 0.09714656 | 0.24122307 | tufm-2 |
| WBGene00007006 | 0.207682688 | 0.26473363 | 0.4632443 | npr-3 |
| WBGene00007007 | 0.249056892 | 0.15829888 | 0.33261394 | mak-2 |
| WBGene00007008 | -0.146471378 | 0.04943757 | 0.1490232 | rfp-1 |

| | | | | |
|---|---|---|---|---|
| WBGene00007009 | 0.143995409 | 0.02521041 | 0.0913601 | wwp-1 |
| WBGene00007010 | 0.225378723 | 0.01139614 | 0.04952533 | alx-1 |
| WBGene00007011 | -0.231498269 | 0.18343642 | 0.36639736 | mdt-1.2 |
| WBGene00007012 | 0.230363855 | 0.11130728 | 0.26339905 | mdt-4 |
| WBGene00007013 | -0.262361827 | 0.01459436 | 0.06051727 | mdt-8 |
| WBGene00007014 | -0.169814125 | 0.22670416 | 0.42036164 | mdt-10 |
| WBGene00007015 | -0.204999443 | 0.12347081 | 0.28356644 | mdt-11 |
| WBGene00007016 | 0.164671981 | 0.02300707 | 0.08521119 | mdt-15 |
| WBGene00007017 | -0.094014007 | 0.23885294 | 0.43593192 | mdt-17 |
| WBGene00007018 | -0.226279129 | 0.09904157 | 0.24407452 | mdt-18 |
| WBGene00007019 | -0.126800551 | 0.22649164 | 0.42005699 | mdt-19 |
| WBGene00007020 | -0.009060185 | 0.94952762 | 0.97122452 | mdt-20 |
| WBGene00007021 | -0.041021377 | 0.8322558 | 0.9084712 | immp-1 |
| WBGene00007022 | -0.316864453 | 0.01834265 | 0.07220277 | mdt-22 |
| WBGene00007023 | -0.195021763 | 0.07748953 | 0.2061075 | mdt-27 |
| WBGene00007024 | -0.074382708 | 0.17399004 | 0.35444388 | plin-1 |
| WBGene00007025 | -0.063367491 | 0.61107793 | 0.76604108 | mdt-29 |
| WBGene00007026 | -0.221503992 | 0.21898033 | 0.41087928 | mdt-31 |
| WBGene00007027 | 0.008837428 | 0.89806967 | 0.94632825 | ssl-1 |
| WBGene00007028 | -0.013131992 | 0.86509412 | 0.92790421 | trr-1 |
| WBGene00007029 | -0.156848853 | 0.09832255 | 0.2432018 | mys-1 |
| WBGene00007030 | 0.108108499 | 0.21207727 | 0.40241359 | epc-1 |
| WBGene00007036 | -0.071706397 | 0.33859663 | NA | sod-5 |
| WBGene00007041 | 1.451998465 | 7.88E-05 | 0.00078263 | tag-180 |
| WBGene00007042 | -0.065204537 | 0.32290699 | 0.52291574 | pbrm-1 |
| WBGene00007043 | 0.011819147 | 0.92159182 | 0.95613845 | algn-10 |
| WBGene00007044 | 1.543735163 | 9.67E-06 | 0.00011997 | cpna-5 |
| WBGene00007045 | -0.107347131 | 0.37489732 | 0.57372966 | pgap-2 |
| WBGene00007047 | -0.04449142 | 0.76022142 | 0.86416163 | wts-1 |
| WBGene00007048 | -0.124186405 | 0.05236684 | 0.15586559 | nfx-1 |
| WBGene00007049 | 0.248915664 | 0.12356696 | 0.28365379 | tag-191 |
| WBGene00007050 | -0.235296575 | 0.01832857 | 0.07219493 | utp-20 |
| WBGene00007053 | -0.117007802 | 0.16782831 | 0.34544838 | chd-7 |
| WBGene00007054 | 0.158591534 | 0.08695621 | 0.2229252 | scpl-1 |
| WBGene00007055 | 0.828175172 | 3.99E-16 | 2.17E-14 | tag-196 |
| WBGene00007056 | 0.093383738 | 0.62831798 | 0.77880152 | crn-7 |
| WBGene00007057 | 0.106267305 | 0.22564025 | NA | ant-1.2 |
| WBGene00007058 | 0.516488014 | 0.00176013 | 0.01124346 | dmd-6 |
| WBGene00007059 | 0.072625232 | 0.67491599 | 0.8111554 | vps-52 |
| WBGene00007060 | 0.344154528 | 0.1261413 | 0.28708764 | wht-6 |
| WBGene00007062 | NA | NA | NA | ebp-3 |
| WBGene00007063 | 0.007457729 | 0.96970069 | 0.98346632 | 2L52.1 |
| WBGene00007064 | 0.079869272 | 0.62911119 | 0.77958671 | rga-9 |

| | | | | | |
|---|---|---|---|---|---|
| WBGene00007065 | | 0.008966268 | 0.96208711 | 0.97924853 | pot-3 |
| WBGene00007066 | | 0.020317245 | 0.84643798 | NA | NA |
| WBGene00007067 | | 0.287148381 | 0.15790668 | 0.3320758 | 4R79.2 |
| WBGene00007068 | | -0.369027424 | 0.01785643 | 0.07073539 | cTel55X.1 |
| WBGene00007070 | | 0.152257071 | 0.32875031 | 0.52822125 | ugt-49 |
| WBGene00007071 | | -0.272937775 | 0.00860911 | 0.03998922 | AC3.5 |
| WBGene00007072 | | -0.322123625 | 0.09905681 | 0.24407452 | ugt-1 |
| WBGene00007073 | | 1.059328826 | 2.05E-05 | 0.00023367 | ugt-2 |
| WBGene00007074 | NA | | NA | NA | NA |
| WBGene00007075 | NA | | NA | NA | AC8.3 |
| WBGene00007075 | NA | | NA | NA | AC8.10 |
| WBGene00007076 | NA | | NA | NA | AC8.4 |
| WBGene00007076 | NA | | NA | NA | AC8.11 |
| WBGene00007077 | | 0.007946235 | 0.93790307 | 0.96517113 | NA |
| WBGene00007078 | NA | | NA | NA | NA |
| WBGene00007079 | NA | | NA | NA | AC8.7 |
| WBGene00007079 | NA | | NA | NA | AC8.12 |
| WBGene00007080 | | 0.095576857 | 0.24390154 | NA | sfxn-1.1 |
| WBGene00007081 | | 0.0398788 | 0.58971221 | NA | AH6.3 |
| WBGene00007082 | | 0.318421974 | 0.09277703 | 0.23259392 | acs-10 |
| WBGene00007083 | | 0.018795205 | 0.77532208 | NA | AH10.2 |
| WBGene00007085 | | 0.038125098 | 0.61119482 | NA | AH10.4 |
| WBGene00007087 | | -0.08309422 | 0.25628753 | 0.45460486 | B0001.2 |
| WBGene00007088 | | -0.067678533 | 0.51531539 | 0.69504677 | B0001.3 |
| WBGene00007089 | | -0.23255598 | 0.01245818 | 0.05328667 | B0001.4 |
| WBGene00007090 | | 0.204463444 | 0.05950441 | 0.17068 | B0001.5 |
| WBGene00007091 | | 0.230164338 | 0.05979025 | 0.1711081 | eri-12 |
| WBGene00007092 | | -0.241564071 | 0.00723275 | 0.03489988 | B0001.7 |
| WBGene00007093 | | 0.038728301 | 0.71622538 | 0.84075838 | B0001.8 |
| WBGene00007094 | | -0.057716226 | 0.64627322 | 0.79060565 | B0019.2 |
| WBGene00007096 | | 0.133262947 | 0.42750712 | 0.62265977 | B0024.3 |
| WBGene00007097 | | 2.39378132 | 4.76E-05 | 0.00050264 | B0024.4 |
| WBGene00007099 | | 0.039291823 | 0.76155342 | 0.86480288 | trx-2 |
| WBGene00007100 | | -0.054792455 | 0.55489132 | 0.72686037 | asps-1 |
| WBGene00007101 | | -0.073752774 | 0.45148483 | 0.64382872 | B0024.11 |
| WBGene00007102 | | -0.139244584 | 0.21049249 | 0.40018446 | B0024.13 |
| WBGene00007103 | | 0.707735518 | 4.22E-05 | 0.00045046 | crm-1 |
| WBGene00007104 | | 0.127903362 | 0.3216407 | 0.52161514 | B0024.15 |
| WBGene00007105 | | -0.085632415 | 0.25864553 | 0.45662959 | znf-207 |
| WBGene00007106 | | -0.280401084 | 0.00817442 | 0.03843341 | B0035.3 |
| WBGene00007107 | | -0.215865469 | 0.0301157 | 0.10372113 | pfd-4 |
| WBGene00007108 | | -0.117431776 | 0.16900788 | 0.34728789 | gspd-1 |
| WBGene00007109 | | -0.125100027 | 0.17609505 | 0.35699473 | edg-1 |

| | | | | |
|---|---|---|---|---|
| WBGene00007110 | -0.070628033 | 0.36721045 | 0.56679025 | leo-1 |
| WBGene00007111 | -0.276196067 | 0.00015683 | 0.0014271 | sart-3 |
| WBGene00007112 | 1.437231014 | 2.05E-33 | 2.84E-31 | B0035.13 |
| WBGene00007113 | -0.201190053 | 0.08382994 | 0.2177318 | B0035.15 |
| WBGene00007114 | 0.356174321 | 0.09635808 | 0.23968376 | mttu-1 |
| WBGene00007115 | -0.243976055 | 0.2518442 | 0.44993917 | B0198.2 |
| WBGene00007116 | 0.173374378 | 0.41434615 | 0.61181736 | B0198.3 |
| WBGene00007117 | -0.006738828 | 0.92480778 | NA | spe-42 |
| WBGene00007118 | -0.140159735 | 0.03887233 | 0.12530379 | npp-22 |
| WBGene00007119 | 0.010455302 | 0.94746535 | 0.97044477 | calf-1 |
| WBGene00007120 | -0.18684577 | 0.30263049 | 0.50315842 | NA |
| WBGene00007121 | 0.072482412 | 0.72856673 | 0.84853595 | NA |
| WBGene00007122 | -0.448795841 | 1.79E-06 | 2.63E-05 | B0250.5 |
| WBGene00007123 | NA | NA | NA | srbc-79 |
| WBGene00007124 | -0.005218063 | 0.97342649 | 0.98538191 | B0250.7 |
| WBGene00007125 | -0.025226573 | 0.69184128 | NA | B0250.8 |
| WBGene00007126 | 0.028137767 | 0.88085897 | 0.93664436 | dhcr-7 |
| WBGene00007127 | 0.020317245 | 0.84643798 | NA | srbc-78 |
| WBGene00007128 | -0.299494278 | 0.10882162 | 0.25948817 | memb-1 |
| WBGene00007129 | -0.364072995 | 0.00049944 | 0.00388537 | B0272.3 |
| WBGene00007130 | -0.221545533 | 0.27991005 | 0.47959777 | B0272.4 |
| WBGene00007131 | 0.43976969 | 0.0144199 | 0.0599437 | pals-26 |
| WBGene00007132 | -0.026384613 | 0.67299636 | NA | pals-27 |
| WBGene00007133 | -0.219160321 | 0.29891486 | 0.49947643 | B0284.3 |
| WBGene00007134 | 0.106997704 | 0.16248985 | NA | pals-28 |
| WBGene00007135 | -0.173169981 | 0.07969733 | 0.21058447 | cdk-12 |
| WBGene00007136 | 0.132412756 | 0.33281008 | 0.5325325 | B0285.3 |
| WBGene00007137 | -0.256898956 | 0.01490322 | 0.06155091 | lcmt-1 |
| WBGene00007138 | 0.743130919 | 0.02395822 | 0.08782481 | B0285.6 |
| WBGene00007139 | -1.06395387 | 1.00E-05 | 0.00012412 | mnp-1 |
| WBGene00007140 | -0.106562042 | 0.57724032 | 0.74411462 | cyp-29A4 |
| WBGene00007141 | 0.020317245 | 0.84643798 | NA | pitr-4 |
| WBGene00007142 | -0.721075301 | 4.98E-06 | 6.53E-05 | ttr-18 |
| WBGene00007143 | 0.183839878 | 0.02006361 | 0.0768178 | B0334.3 |
| WBGene00007144 | -0.1561337 | 0.10325427 | 0.2507873 | B0334.4 |
| WBGene00007145 | -0.260194559 | 0.08114208 | 0.21290521 | B0334.5 |
| WBGene00007146 | 0.203422726 | 0.13459523 | 0.29882541 | B0334.6 |
| WBGene00007147 | 0.098399041 | 0.57908067 | 0.74488842 | NA |
| WBGene00007148 | 0.020925399 | 0.84643798 | NA | B0334.10 |
| WBGene00007149 | -0.034050838 | 0.7451853 | 0.85590582 | B0334.13 |
| WBGene00007150 | -0.079748267 | 0.15178046 | 0.32424969 | acly-2 |
| WBGene00007151 | -0.003270206 | 0.95698316 | NA | NA |
| WBGene00007152 | 0.274746247 | 0.18487507 | 0.36856171 | clec-225 |

| | | | | |
|---|---|---|---|---|
| WBGene00007153 | 0.949636217 | 6.28E-16 | 3.26E-14 | clec-41 |
| WBGene00007154 | 1.908166758 | 0.01132117 | 0.04928731 | dhc-3 |
| WBGene00007155 | -0.17332514 | 0.29459947 | 0.49569181 | B0379.1 |
| WBGene00007156 | 0.099958276 | 0.38721674 | 0.58584689 | B0379.2 |
| WBGene00007158 | NA | NA | NA | B0379.6 |
| WBGene00007159 | 0.506746507 | 0.0378061 | 0.12271091 | B0379.7 |
| WBGene00007160 | NA | NA | NA | NA |
| WBGene00007161 | NA | NA | NA | B0391.3 |
| WBGene00007162 | -0.084286698 | 0.67735847 | 0.81260807 | fbxa-153 |
| WBGene00007163 | 0.056188364 | 0.42092875 | NA | fbxa-154 |
| WBGene00007164 | 0.020317245 | 0.84643798 | NA | fbxa-139 |
| WBGene00007165 | -1.093540082 | 5.43E-05 | 0.00056309 | NA |
| WBGene00007166 | 0.036687355 | 0.59489568 | NA | fbxa-152 |
| WBGene00007167 | -0.147268321 | 0.02792129 | 0.09822094 | rbg-3 |
| WBGene00007168 | -0.193059161 | 0.00838363 | 0.03921557 | B0393.3 |
| WBGene00007169 | 0.110039823 | 0.21624696 | NA | B0393.4 |
| WBGene00007170 | 0.148936023 | 0.44943816 | 0.64203572 | B0393.5 |
| WBGene00007171 | -0.057839417 | 0.55767797 | 0.72877494 | B0393.6 |
| WBGene00007172 | -0.008807557 | 0.92153097 | NA | B0393.7 |
| WBGene00007173 | 0.040107227 | 0.58693163 | NA | B0393.8 |
| WBGene00007173 | 0.040107227 | 0.58693163 | NA | VB0393L.2 |
| WBGene00007174 | 0.081768833 | 0.41230943 | NA | mboa-1 |
| WBGene00007175 | -0.423489369 | 0.00115801 | 0.00794863 | B0395.3 |
| WBGene00007176 | 1.417970892 | 0.0010857 | 0.00753353 | kcnl-1 |
| WBGene00007177 | 0.015584271 | 0.80362925 | NA | oac-1 |
| WBGene00007178 | 0.322013203 | 0.1172858 | 0.27380421 | B0457.2 |
| WBGene00007179 | NA | NA | NA | B0457.4 |
| WBGene00007180 | -0.446469254 | 0.05531534 | 0.16203919 | B0457.6 |
| WBGene00007181 | 0.266841745 | 0.01361552 | 0.05724882 | NA |
| WBGene00007182 | -0.475458767 | 0.07366265 | 0.19869102 | fbxb-119 |
| WBGene00007183 | -0.007516521 | 0.93068503 | NA | B0462.4 |
| WBGene00007184 | -0.149211319 | 0.04871818 | 0.14746418 | ctr-9 |
| WBGene00007185 | 0.065041938 | 0.5707784 | 0.73934057 | nlp-36 |
| WBGene00007186 | -0.144200814 | 0.03123952 | 0.1063175 | B0464.6 |
| WBGene00007187 | -0.152470254 | 0.15685065 | 0.33080529 | tag-342 |
| WBGene00007188 | 0.048013381 | 0.74554155 | 0.85598121 | B0464.9 |
| WBGene00007189 | -0.277378707 | 0.04214132 | 0.13274334 | pigm-1 |
| WBGene00007190 | 1.58892726 | 0.00560117 | 0.02863365 | rmd-3 |
| WBGene00007191 | 0.191116214 | 0.1930171 | 0.3794258 | lgc-20 |
| WBGene00007192 | -0.19954323 | 0.00653946 | 0.03228058 | B0491.5 |
| WBGene00007193 | -0.076996849 | 0.5523774 | 0.7248035 | B0491.6 |
| WBGene00007194 | -0.388694591 | 0.03057005 | 0.10481876 | B0491.7 |
| WBGene00007195 | -0.275078828 | 0.00686008 | 0.03348286 | thoc-7 |

| | | | | | |
|---|---|---|---|---|---|
| WBGene00007196 | | 0.218561535 | 0.01885661 | 0.07368018 | B0513.4 |
| WBGene00007197 | | -0.144189062 | 0.32442133 | 0.52363682 | prdh-1 |
| WBGene00007198 | | -0.050545243 | 0.47515706 | NA | NA |
| WBGene00007199 | NA | | NA | NA | NA |
| WBGene00007200 | | 0.399653561 | 0.11055808 | 0.26229761 | vamp-8 |
| WBGene00007201 | NA | | NA | NA | exos-4.1 |
| WBGene00007202 | | -0.101400155 | 0.54600048 | 0.71940182 | B0564.2 |
| WBGene00007203 | | 3.345219378 | 1.31E-11 | 4.26E-10 | best-1 |
| WBGene00007204 | | 0.020232312 | 0.84643798 | NA | best-2 |
| WBGene00007206 | NA | | NA | NA | B0564.9 |
| WBGene00007206 | NA | | NA | NA | B0564.6 |
| WBGene00007207 | | 0.005419693 | 0.96652456 | 0.9812694 | B0564.7 |
| WBGene00007208 | NA | | NA | NA | B0564.9 |
| WBGene00007208 | NA | | NA | NA | B0564.6 |
| WBGene00007209 | | 0.108942554 | 0.15594412 | NA | BE10.1 |
| WBGene00007210 | | 4.937824456 | 0.00021594 | 0.00188529 | agmo-1 |
| WBGene00007211 | | -0.429891627 | 0.09246809 | 0.2320976 | BE10.3 |
| WBGene00007212 | | 0.107540866 | 0.5096789 | 0.68965343 | BE10.4 |
| WBGene00007213 | | -0.293030427 | 0.11980751 | 0.27803165 | C01A2.1 |
| WBGene00007214 | | -0.165725505 | 0.43157499 | 0.62660546 | C01A2.2 |
| WBGene00007215 | | -0.278874402 | 0.00845954 | 0.03946545 | oxa-1 |
| WBGene00007216 | | 0.119928773 | 0.4111633 | 0.60881041 | C01A2.4 |
| WBGene00007217 | | -0.186036068 | 0.08204101 | 0.21460826 | tads-1 |
| WBGene00007218 | NA | | NA | NA | C01A2.6 |
| WBGene00007219 | | 0.743077851 | 0.02092519 | 0.07937594 | nlp-38 |
| WBGene00007220 | | 1.104426806 | 0.00126065 | 0.00851739 | C01B9.1 |
| WBGene00007221 | | -0.197510676 | 0.02523296 | 0.09139553 | cpna-3 |
| WBGene00007222 | | 1.510106694 | 0.00396492 | 0.02174172 | C01F6.2 |
| WBGene00007223 | | -0.19741632 | 0.05296501 | 0.15707123 | C01F6.9 |
| WBGene00007224 | | 0.082768056 | 0.46000565 | 0.65119895 | C01G6.2 |
| WBGene00007225 | | -0.515776197 | 3.55E-10 | 9.47E-09 | C01G6.3 |
| WBGene00007226 | | 0.167977176 | 0.15797654 | 0.332175 | C01G6.4 |
| WBGene00007227 | | -0.056021242 | 0.36991627 | 0.56985944 | C01G6.5 |
| WBGene00007228 | | -0.320259608 | 0.08759529 | 0.22421025 | acs-7 |
| WBGene00007229 | | 0.151572367 | 0.28642748 | 0.48643832 | C01G6.9 |
| WBGene00007230 | | 0.058314668 | 0.40208118 | NA | C01G10.1 |
| WBGene00007231 | | 7.621993853 | 1.18E-08 | 2.59E-07 | C01G10.4 |
| WBGene00007232 | | 7.47754613 | 2.15E-08 | 4.54E-07 | C01G10.5 |
| WBGene00007233 | | 0.06560776 | 0.524929 | 0.70219146 | C01G10.6 |
| WBGene00007234 | | -0.107206559 | 0.44599542 | 0.63930108 | C01G10.7 |
| WBGene00007235 | | -0.354044353 | 3.73E-08 | 7.59E-07 | ahsa-1 |
| WBGene00007236 | | -0.306113453 | 0.15506413 | 0.32821944 | C01G10.9 |
| WBGene00007237 | | -0.104521563 | 0.54684719 | 0.71995249 | C01G10.10 |

| | | | | |
|---|---|---|---|---|
| WBGene00007238 | NA | NA | NA | NA |
| WBGene00007239 | -0.026384613 | 0.67299636 | NA | C01G10.14 |
| WBGene00007240 | 0.096462808 | 0.24109807 | NA | C01G10.15 |
| WBGene00007241 | 0.013560238 | 0.88662252 | NA | C01G10.16 |
| WBGene00007242 | 0.181445976 | 0.37157038 | 0.57085779 | madf-5 |
| WBGene00007243 | NA | NA | NA | nspb-7 |
| WBGene00007244 | -0.173238805 | 0.4178498 | 0.61452833 | C01G12.3 |
| WBGene00007245 | 0.020317245 | 0.84643798 | NA | C01G12.5 |
| WBGene00007246 | 0.038125098 | 0.61119482 | NA | nspb-10 |
| WBGene00007247 | 0.111481314 | 0.38115484 | 0.58051472 | C01G12.7 |
| WBGene00007248 | 0.424806404 | 0.09272365 | 0.23249991 | catp-4 |
| WBGene00007249 | 0.232344911 | 0.0475789 | 0.1450349 | C01G12.9 |
| WBGene00007250 | 0.020317245 | 0.84643798 | NA | nspb-8 |
| WBGene00007251 | NA | NA | NA | nspb-9 |
| WBGene00007252 | 0.030356701 | 0.88205579 | 0.93750845 | mlt-2 |
| WBGene00007253 | 0.048399149 | 0.71683338 | 0.84118707 | C01H6.3 |
| WBGene00007254 | -0.077456848 | 0.69597504 | 0.826488 | C01H6.4 |
| WBGene00007255 | 0.162069652 | 0.37284071 | 0.57195734 | cnnm-4 |
| WBGene00007256 | -0.263188336 | 0.00075123 | 0.00549082 | swsn-9 |
| WBGene00007257 | -0.364186546 | 0.1334771 | 0.29716592 | C01H6.8 |
| WBGene00007258 | -0.220012966 | 0.07706853 | 0.20538048 | hasp-1 |
| WBGene00007259 | NA | NA | NA | C02B4.3 |
| WBGene00007260 | 0.160300205 | 0.44961877 | 0.64204323 | olrn-1 |
| WBGene00007261 | -0.308782438 | 0.09310123 | 0.23320702 | lron-3 |
| WBGene00007262 | 0.038125098 | 0.61119482 | NA | jud-4 |
| WBGene00007264 | 0.314968583 | 0.02581753 | 0.09294182 | C02F4.4 |
| WBGene00007265 | 0.020317245 | 0.84643798 | NA | C02F4.5 |
| WBGene00007266 | 0.226879798 | 0.25948694 | 0.45750822 | C03A3.1 |
| WBGene00007267 | 0.034471635 | 0.74358343 | 0.85511642 | C03A3.2 |
| WBGene00007268 | 0.036444334 | 0.71682131 | 0.84118707 | C03A3.3 |
| WBGene00007269 | 0.059409224 | 0.42187437 | NA | C03C10.2 |
| WBGene00007270 | -0.209520065 | 0.03279627 | 0.11051096 | rei-1 |
| WBGene00007271 | -0.002950686 | 0.96771748 | NA | C03C10.5 |
| WBGene00007273 | 0.038125098 | 0.61119482 | NA | C03C10.7 |
| WBGene00007274 | 0.30089425 | 0.02992993 | 0.10341402 | C03C11.1 |
| WBGene00007275 | 0.210595152 | 0.31633614 | 0.51694731 | C03D6.1 |
| WBGene00007277 | -0.19255248 | 0.05013371 | 0.15056568 | asfl-1 |
| WBGene00007278 | 0.070987467 | 0.58973619 | 0.75112054 | lab-1 |
| WBGene00007279 | NA | NA | NA | C03E10.1 |
| WBGene00007281 | 0.444141632 | 0.05898827 | 0.16950879 | C03E10.3 |
| WBGene00007282 | 0.875276551 | 4.17E-13 | 1.64E-11 | clec-223 |
| WBGene00007283 | 0.506673777 | 0.00135906 | 0.00904451 | clec-222 |
| WBGene00007284 | -0.188580883 | 0.23544855 | 0.4312237 | C03H12.1 |

| | | | | |
|---|---|---|---|---|
| WBGene00007285 | 0.322714853 | 0.10365803 | 0.2515176 | C04A11.1 |
| WBGene00007286 | -0.156191794 | 0.45381936 | 0.64589961 | C04A11.2 |
| WBGene00007287 | -0.196221141 | 0.09899586 | 0.24407452 | gck-4 |
| WBGene00007288 | 0.139678833 | 0.29835322 | 0.49925925 | C04A11.5 |
| WBGene00007289 | 0.038125098 | 0.61119482 | NA | C04B4.1 |
| WBGene00007290 | -0.294749555 | 0.00299521 | 0.01745733 | C04B4.2 |
| WBGene00007291 | 0.040107227 | 0.58693163 | NA | lips-2 |
| WBGene00007292 | 0.020925399 | 0.84643798 | NA | C04B4.4 |
| WBGene00007293 | NA | NA | NA | C04B4.5 |
| WBGene00007294 | NA | NA | NA | C04B4.6 |
| WBGene00007295 | -0.047890151 | 0.80936258 | 0.89446539 | C04C11.1 |
| WBGene00007296 | 0.223061881 | 0.11241111 | 0.26542401 | arrd-25 |
| WBGene00007297 | -0.118114191 | 0.26945808 | 0.46848779 | C04F12.1 |
| WBGene00007299 | -0.008276754 | 0.96229974 | 0.97924853 | C04F12.5 |
| WBGene00007300 | 0.038633797 | 0.57999349 | NA | NA |
| WBGene00007301 | 2.048910977 | 2.12E-06 | 3.07E-05 | C04F12.7 |
| WBGene00007302 | -0.084077668 | 0.6853454 | 0.81837876 | C04F12.8 |
| WBGene00007303 | 0.21063059 | 0.33773825 | 0.53742264 | rnh-1.3 |
| WBGene00007304 | 0.438192793 | 0.08890975 | 0.22633561 | ttr-39 |
| WBGene00007305 | 0.124120287 | 0.30981572 | 0.51058217 | ttbk-5 |
| WBGene00007306 | 0.017604133 | 0.79437959 | NA | C04G2.5 |
| WBGene00007307 | 0.929477403 | 0.01745839 | 0.06951546 | spch-1 |
| WBGene00007308 | 2.884284223 | 9.98E-08 | 1.88E-06 | C04G2.9 |
| WBGene00007309 | -0.340751625 | 0.15405075 | 0.32715947 | C04G2.10 |
| WBGene00007310 | -0.040794878 | 0.53203165 | NA | irld-21 |
| WBGene00007311 | NA | NA | NA | C04H4.1 |
| WBGene00007312 | -0.231422827 | 0.13165465 | 0.29450812 | C04H5.1 |
| WBGene00007313 | 0.727724969 | 0.00914728 | 0.04189095 | clec-147 |
| WBGene00007314 | 0.025750003 | 0.84187105 | 0.91465079 | gcy-29 |
| WBGene00007315 | 0.243073277 | 0.12692401 | 0.28824805 | C04H5.7 |
| WBGene00007316 | 0.893588719 | 0.01657271 | 0.06689854 | nlp-41 |
| WBGene00007317 | -0.105114169 | 0.61531406 | 0.76921987 | C05A9.2 |
| WBGene00007319 | 0.014072415 | 0.87856141 | NA | C05B5.1 |
| WBGene00007320 | 1.171520288 | 0.01537401 | 0.06310077 | C05B5.2 |
| WBGene00007321 | -0.016179431 | 0.92039271 | 0.95550377 | C05B5.4 |
| WBGene00007322 | -0.089522706 | 0.58922385 | 0.75107317 | C05B5.5 |
| WBGene00007323 | 0.033463284 | 0.6365037 | NA | fbxa-155 |
| WBGene00007324 | -0.014237991 | 0.94333715 | 0.96789535 | C05B5.8 |
| WBGene00007325 | 0.084908438 | 0.30232871 | NA | C05C9.1 |
| WBGene00007326 | 0.020232312 | 0.84643798 | NA | C05C9.2 |
| WBGene00007327 | -0.215153248 | 0.32646994 | 0.5256622 | C05C9.3 |
| WBGene00007328 | -0.083781919 | 0.43771677 | 0.63263684 | pho-10 |
| WBGene00007329 | -0.278278293 | 0.00014795 | 0.00135381 | C05C10.2 |

| | | | | |
|---|---|---|---|---|
| WBGene00007330 | -0.585415582 | 5.14E-15 | 2.45E-13 | C05C10.3 |
| WBGene00007331 | -3.3181832 | 1.16E-114 | 7.69E-112 | pho-11 |
| WBGene00007332 | -0.022125417 | 0.74554892 | 0.85598121 | C05C10.5 |
| WBGene00007333 | -0.206104089 | 0.00064914 | 0.00485892 | ufd-3 |
| WBGene00007334 | -0.170270017 | 0.33118079 | 0.53077152 | C05C10.7 |
| WBGene00007335 | 0.017557163 | 0.82585836 | NA | C05C12.1 |
| WBGene00007336 | 0.551197198 | 0.00251438 | 0.01508728 | C05C12.4 |
| WBGene00007337 | 0.383950972 | 0.00435982 | 0.02352825 | C05C12.5 |
| WBGene00007338 | -0.043785352 | 0.7393023 | 0.85323387 | C05C12.6 |
| WBGene00007339 | 0.052218507 | 0.79347627 | 0.88399982 | C05D12.1 |
| WBGene00007340 | 0.636962537 | 0.01393207 | 0.05827984 | C05D12.2 |
| WBGene00007341 | NA | NA | NA | C05D12.4 |
| WBGene00007342 | -0.292626391 | 0.20314032 | 0.39190636 | C05D12.5 |
| WBGene00007343 | 0.327886392 | 0.13448962 | 0.29875019 | C05E7.1 |
| WBGene00007344 | -0.037348399 | 0.82178826 | 0.9024077 | C05E7.2 |
| WBGene00007345 | 3.903384765 | 0.00176178 | 0.01124912 | C05E7.3 |
| WBGene00007346 | 0.122975561 | 0.30952627 | 0.51040962 | frpr-2 |
| WBGene00007347 | 0.830567058 | 0.00121364 | 0.00825689 | C05G5.1 |
| WBGene00007348 | -0.294813804 | 0.07905416 | 0.20943503 | C05G5.2 |
| WBGene00007349 | -0.1920289 | 0.37087414 | 0.57036433 | C05G5.3 |
| WBGene00007350 | -0.071517081 | 0.4478601 | 0.64068438 | sucl-1 |
| WBGene00007351 | -0.09784065 | 0.46924691 | 0.65783171 | C05G5.5 |
| WBGene00007352 | -0.277329337 | 1.39E-09 | 3.44E-08 | cdc-48.1 |
| WBGene00007353 | 0.149067622 | 0.47205028 | 0.66010393 | C06A1.2 |
| WBGene00007354 | 0.250401798 | 0.13369618 | 0.29743895 | C06A1.3 |
| WBGene00007355 | NA | NA | NA | rpb-6 |
| WBGene00007356 | 0.015062379 | 0.80838968 | NA | C06A1.6 |
| WBGene00007357 | 0.254589039 | 0.21430744 | 0.40517501 | C06A12.3 |
| WBGene00007358 | 0.038595855 | 0.57520533 | NA | lact-6 |
| WBGene00007359 | NA | NA | NA | C06A12.8 |
| WBGene00007360 | NA | NA | NA | C06B3.1 |
| WBGene00007361 | -0.195857192 | 0.15963003 | 0.33425963 | oac-3 |
| WBGene00007362 | -0.060177734 | 0.69789547 | 0.82793475 | cyp-35C1 |
| WBGene00007363 | -0.114286167 | 0.50590939 | 0.68674779 | stdh-1 |
| WBGene00007364 | 0.038125098 | 0.61119482 | NA | stdh-3 |
| WBGene00007365 | 0.362791775 | 0.13299912 | 0.29656413 | C06B3.6 |
| WBGene00007366 | 0.258149735 | 0.1910756 | 0.37703623 | C06B3.7 |
| WBGene00007367 | -0.05123092 | 0.55432822 | NA | nhr-150 |
| WBGene00007368 | 3.442121567 | 1.18E-12 | 4.42E-11 | C06B8.2 |
| WBGene00007369 | NA | NA | NA | C06B8.3 |
| WBGene00007370 | NA | NA | NA | str-264 |
| WBGene00007372 | -0.000551732 | 0.99621162 | 0.99769349 | C06B8.7 |
| WBGene00007373 | NA | NA | NA | C06B8.11 |

| | | | | |
|---|---|---|---|---|
| WBGene00007374 | 0.01476227 | 0.84668595 | NA | C06C3.3 |
| WBGene00007375 | -0.223840576 | 0.15421654 | 0.32715947 | C06C3.4 |
| WBGene00007376 | -0.023056499 | 0.78121253 | NA | NA |
| WBGene00007377 | -0.058723914 | 0.68053439 | 0.81496025 | C06C3.7 |
| WBGene00007378 | 0.129926662 | 0.40947056 | 0.60746362 | lido-17 |
| WBGene00007379 | NA | NA | NA | C06C6.1 |
| WBGene00007380 | NA | NA | NA | C06C6.6 |
| WBGene00007381 | 0.086430603 | 0.29509282 | NA | C06C6.7 |
| WBGene00007382 | NA | NA | NA | C06C6.8 |
| WBGene00007383 | 0.040107227 | 0.58693163 | NA | C06C6.9 |
| WBGene00007384 | -0.523060535 | 0.02498491 | 0.09077023 | swt-3 |
| WBGene00007385 | -0.153821833 | 0.00289698 | 0.01697944 | atp-5 |
| WBGene00007386 | -0.294716057 | 0.02240406 | 0.08348636 | C06H2.2 |
| WBGene00007387 | 0.018412518 | 0.89777611 | 0.94623621 | jmjd-5 |
| WBGene00007388 | -0.071256495 | 0.72878862 | 0.84859187 | folt-1 |
| WBGene00007389 | 0.738690385 | 0.02782404 | 0.0980051 | glb-3 |
| WBGene00007390 | -0.15559607 | 0.20899998 | 0.39838156 | lmtr-3 |
| WBGene00007391 | 0.104469399 | 0.44831319 | 0.64099154 | C06H2.7 |
| WBGene00007392 | 0.620400825 | 0.02914254 | 0.10149284 | fbxa-156 |
| WBGene00007393 | 2.314412315 | 0.00156647 | 0.01018865 | fbxa-157 |
| WBGene00007394 | -0.063760714 | 0.70807385 | 0.83523278 | C06H5.6 |
| WBGene00007395 | 0.438249677 | 0.00900561 | 0.04147188 | dcar-1 |
| WBGene00007396 | 0.240445456 | 0.28297133 | 0.48275265 | tiar-3 |
| WBGene00007397 | 0.069945147 | 0.66909237 | 0.80700069 | C07A4.2 |
| WBGene00007398 | 0.175344803 | 0.27472965 | 0.47510119 | C07A4.3 |
| WBGene00007399 | 0.056411843 | 0.41891328 | NA | clec-162 |
| WBGene00007400 | -0.095063606 | 0.35924903 | 0.55945595 | bud-31 |
| WBGene00007401 | 0.020925399 | 0.84643798 | NA | C07A9.5 |
| WBGene00007402 | -0.878714939 | 0.0019982 | 0.01245979 | ugt-60 |
| WBGene00007403 | -0.210010496 | 0.0842503 | 0.21851349 | set-3 |
| WBGene00007404 | 0.311293881 | 0.15181556 | 0.32424969 | best-5 |
| WBGene00007405 | -0.14220109 | 0.47363752 | 0.66152502 | C07A9.9 |
| WBGene00007406 | NA | NA | NA | C07A9.10 |
| WBGene00007407 | -0.045771477 | 0.50913401 | NA | C07B5.2 |
| WBGene00007408 | 0.415165222 | 0.00405094 | 0.02211263 | C07B5.3 |
| WBGene00007409 | 0.953900865 | 0.00396795 | 0.02174231 | C07B5.4 |
| WBGene00007410 | 0.032835971 | 0.73334935 | 0.85087141 | C07B5.6 |
| WBGene00007411 | 0.0398788 | 0.58971221 | NA | C07C7.1 |
| WBGene00007412 | -0.126764421 | 0.17135981 | 0.35030424 | stip-1 |
| WBGene00007413 | -0.500734816 | 1.36E-07 | 2.50E-06 | pro-2 |
| WBGene00007414 | 2.451218772 | 8.40E-10 | 2.14E-08 | C07E3.3 |
| WBGene00007415 | 0.0398788 | 0.58971221 | NA | C07E3.4 |
| WBGene00007416 | 0.037023624 | 0.59153183 | NA | ceh-57 |

| | | | | |
|---|---|---|---|---|
| WBGene00007417 | 0.079423729 | 0.6585068 | 0.79870748 | ceh-58 |
| WBGene00007418 | -0.079048478 | 0.39823903 | NA | C07E3.8 |
| WBGene00007419 | -0.370325304 | 0.04922037 | 0.14858462 | C07E3.9 |
| WBGene00007420 | 0.072653761 | 0.34256154 | NA | C07E3.10 |
| WBGene00007421 | 0.090398591 | 0.66658643 | 0.80470744 | C07H4.1 |
| WBGene00007422 | 0.602356761 | 4.69E-05 | 0.00049661 | ugt-17 |
| WBGene00007423 | 0.84607382 | 0.02157881 | 0.08119255 | C08B6.2 |
| WBGene00007424 | 0.357152912 | 0.13633073 | 0.30142226 | C08B6.3 |
| WBGene00007425 | 3.991795671 | 0.00030898 | 0.00258207 | C08B6.4 |
| WBGene00007426 | 0.020232312 | 0.84643798 | NA | C08B6.5 |
| WBGene00007427 | -0.022930767 | 0.78121253 | NA | C08B6.6 |
| WBGene00007428 | -0.082361816 | 0.24594222 | 0.4442191 | wdr-20 |
| WBGene00007429 | -0.134805472 | 0.266473 | 0.465176 | C08B6.8 |
| WBGene00007430 | 0.181461235 | 0.39598866 | 0.59402356 | C08B6.10 |
| WBGene00007431 | -0.001103568 | 0.99421343 | 0.99639119 | C08B6.11 |
| WBGene00007432 | NA | NA | NA | srxa-18 |
| WBGene00007433 | -0.200382796 | 0.00129041 | 0.00867841 | swsn-7 |
| WBGene00007434 | -0.007731098 | 0.9381401 | 0.96523607 | arp-6 |
| WBGene00007435 | -0.151876831 | 0.22127063 | 0.41374499 | algn-6 |
| WBGene00007436 | -0.222573404 | 0.08859701 | 0.22594476 | C08B11.9 |
| WBGene00007437 | 0.108717649 | 0.15669083 | NA | C08E8.1 |
| WBGene00007438 | NA | NA | NA | C08E8.2 |
| WBGene00007439 | -0.004944805 | 0.94514249 | NA | C08E8.3 |
| WBGene00007440 | 3.615475355 | 0.00028962 | 0.00243413 | C08E8.4 |
| WBGene00007441 | NA | NA | NA | C08E8.5 |
| WBGene00007442 | -0.023109762 | 0.78121253 | NA | C08E8.6 |
| WBGene00007443 | -0.203765184 | 0.03507174 | 0.1161206 | pfd-1 |
| WBGene00007444 | -0.169508042 | 0.11316615 | 0.2668275 | C08F8.2 |
| WBGene00007445 | -0.100766838 | 0.25747794 | 0.45566717 | C08F8.3 |
| WBGene00007446 | 0.138080903 | 0.42172288 | 0.61831011 | mboa-4 |
| WBGene00007447 | -0.213050898 | 0.0241216 | 0.08835736 | fbxb-9 |
| WBGene00007448 | 0.23719449 | 0.16029246 | 0.33484732 | C08F8.6 |
| WBGene00007449 | -0.163109464 | 0.43524263 | 0.63045722 | C08F8.9 |
| WBGene00007450 | 0.047003838 | 0.777279 | 0.87398508 | C08F11.1 |
| WBGene00007451 | NA | NA | NA | NA |
| WBGene00007452 | 0.159232204 | 0.24509745 | 0.44329485 | C08F11.3 |
| WBGene00007453 | 0.134367017 | 0.48678088 | 0.67134127 | fbxa-98 |
| WBGene00007454 | 1.904698733 | 0.00738833 | 0.03550653 | C08F11.7 |
| WBGene00007455 | -0.763321959 | 2.34E-13 | 9.45E-12 | ugt-22 |
| WBGene00007456 | NA | NA | NA | srz-29 |
| WBGene00007457 | 0.200099185 | 0.12439168 | 0.28469886 | C08F11.10 |
| WBGene00007458 | 0.122009567 | 0.05769088 | 0.16709241 | ule-4 |
| WBGene00007459 | -0.034981743 | 0.85444711 | 0.92208742 | C08F11.12 |

| | | | | |
|---|---|---|---|---|
| WBGene00007460 | 0.860190461 | 0.00092146 | 0.00656821 | C08F11.13 |
| WBGene00007461 | 0.037023624 | 0.59153183 | NA | NA |
| WBGene00007462 | 0.24605251 | 0.18965494 | 0.37543779 | ctsa-3.2 |
| WBGene00007463 | -0.052909704 | 0.24637204 | 0.44469671 | vgln-1 |
| WBGene00007464 | -0.005638522 | 0.96246237 | 0.97924853 | algn-8 |
| WBGene00007465 | 0.090416984 | 0.26156585 | NA | chil-1 |
| WBGene00007466 | 0.020317245 | 0.84643798 | NA | chil-2 |
| WBGene00007467 | 0.193189808 | 0.16019835 | 0.33477934 | chil-3 |
| WBGene00007468 | NA | NA | NA | NA |
| WBGene00007469 | -0.001842784 | 0.95621055 | NA | chil-4 |
| WBGene00007470 | 0.055099827 | 0.63661445 | 0.78445295 | chil-5 |
| WBGene00007471 | 0.122730824 | 0.25820345 | 0.45633859 | chil-6 |
| WBGene00007472 | 0.085038347 | 0.30255919 | NA | chil-7 |
| WBGene00007473 | 0.02686966 | 0.86426713 | 0.92755718 | chil-8 |
| WBGene00007474 | -0.420812263 | 0.06188561 | 0.17559101 | C08H9.15 |
| WBGene00007475 | NA | NA | NA | NA |
| WBGene00007476 | 0.172087679 | 0.29739389 | 0.49843299 | C09E9.1 |
| WBGene00007478 | -0.002914971 | 0.95718083 | NA | C09F9.1 |
| WBGene00007479 | -0.762426112 | 0.0063857 | 0.03171404 | C09F9.2 |
| WBGene00007480 | 0.396347884 | 0.08530766 | 0.22020265 | glna-1 |
| WBGene00007481 | NA | NA | NA | NA |
| WBGene00007482 | 2.487875949 | 3.48E-07 | 5.94E-06 | C09F12.2 |
| WBGene00007483 | 0.055503542 | 0.44925503 | NA | C09F12.3 |
| WBGene00007484 | -0.003099287 | 0.9660382 | NA | C09G1.2 |
| WBGene00007485 | -0.087037154 | 0.46256578 | 0.65252509 | C09G1.3 |
| WBGene00007486 | 0.086749523 | 0.67412449 | 0.81066584 | C09G1.4 |
| WBGene00007487 | 0.124807572 | 0.5325552 | 0.7086377 | C09G1.5 |
| WBGene00007488 | -0.368407429 | 0.01856665 | 0.07281926 | dph-2 |
| WBGene00007489 | 0.15585044 | 0.2779532 | 0.47769842 | C09G5.7 |
| WBGene00007490 | -0.09033157 | 0.50124996 | 0.6828513 | mks-5 |
| WBGene00007492 | -0.066890143 | 0.44451395 | 0.63794923 | C09G9.1 |
| WBGene00007493 | -0.159149123 | 0.08307217 | 0.21637781 | npp-23 |
| WBGene00007494 | NA | NA | NA | C09G9.3 |
| WBGene00007495 | 0.167884487 | 0.29696692 | 0.49801628 | C09G9.5 |
| WBGene00007496 | 0.059806133 | 0.74882187 | 0.8582013 | npax-4 |
| WBGene00007497 | -0.067044232 | 0.44588628 | NA | C09G9.8 |
| WBGene00007499 | 0.056251223 | 0.61136997 | 0.76616175 | C09H10.5 |
| WBGene00007500 | -0.084684872 | 0.2108162 | 0.40069583 | nasp-1 |
| WBGene00007501 | -0.214073501 | 0.14105847 | 0.30869378 | apc-17 |
| WBGene00007502 | 0.130647715 | 0.31336723 | 0.5138324 | glb-4 |
| WBGene00007503 | 0.132442484 | 0.10147609 | NA | C09H10.9 |
| WBGene00007504 | 0.179143136 | 0.32707593 | 0.52623283 | nbs-1 |
| WBGene00007505 | 0.10739896 | 0.16005247 | 0.33476117 | pezo-1 |

| | | | | |
|---|---|---|---|---|
| WBGene00007506 | 1.101979817 | 0.00935319 | 0.04268706 | C10C5.2 |
| WBGene00007507 | 0.334130016 | 0.11540193 | 0.27098631 | C10C5.3 |
| WBGene00007508 | 0.429767774 | 0.02933816 | 0.10195584 | C10C5.4 |
| WBGene00007509 | 0.18078169 | 0.40080217 | 0.59903477 | C10C5.5 |
| WBGene00007511 | 0.212226111 | 0.25925491 | 0.45731942 | C10C5.7 |
| WBGene00007512 | 0.020925399 | 0.84643798 | NA | C10C6.3 |
| WBGene00007513 | -0.200091837 | 0.04652888 | 0.14276439 | wht-2 |
| WBGene00007514 | -0.113484473 | 0.13126305 | 0.29383159 | catp-8 |
| WBGene00007515 | -0.227747056 | 0.27972113 | 0.47944787 | C10C6.7 |
| WBGene00007516 | -0.517462497 | 2.80E-05 | 0.0003091 | gpx-5 |
| WBGene00007517 | 2.227365218 | 3.05E-10 | 8.20E-09 | gpx-3 |
| WBGene00007518 | 0.017279211 | 0.79380376 | NA | del-8 |
| WBGene00007520 | 0.355477465 | 0.10078463 | 0.24671128 | C11E4.6 |
| WBGene00007521 | -0.56626126 | 0.00815103 | 0.03837274 | C11E4.7 |
| WBGene00007522 | 0.141490413 | 0.25763691 | 0.45573917 | C11E4.8 |
| WBGene00007523 | 0.03555743 | 0.72324271 | 0.84472009 | C11G6.2 |
| WBGene00007525 | 0.132109332 | 0.53289804 | 0.70870744 | C11G10.1 |
| WBGene00007528 | 0.056643884 | 0.77512562 | 0.87262026 | C11H1.2 |
| WBGene00007529 | -0.053212409 | 0.73405596 | 0.85093534 | C11H1.3 |
| WBGene00007530 | -0.004864426 | 0.94835335 | NA | C11H1.5 |
| WBGene00007531 | -0.094481878 | 0.55491467 | 0.72686037 | C11H1.7 |
| WBGene00007533 | -0.584352231 | 4.31E-06 | 5.72E-05 | cbl-1 |
| WBGene00007534 | -0.066737029 | 0.3022623 | 0.50276842 | fubl-1 |
| WBGene00007535 | NA | NA | NA | ttr-19 |
| WBGene00007536 | 0.027458049 | 0.87687756 | 0.9345148 | daf-36 |
| WBGene00007537 | 0.038125098 | 0.61119482 | NA | fipr-8 |
| WBGene00007538 | 0.020925399 | 0.84643798 | NA | C12D8.9 |
| WBGene00007539 | 0.05787416 | 0.40594086 | NA | NA |
| WBGene00007540 | 0.074892148 | 0.28555128 | NA | C12D8.13 |
| WBGene00007541 | 0.060105893 | 0.41982798 | NA | fipr-4 |
| WBGene00007542 | -0.004747605 | 0.9473792 | NA | C12D8.15 |
| WBGene00007543 | 0.088571513 | 0.35419363 | 0.55415141 | fipr-7 |
| WBGene00007544 | 0.038125098 | 0.61119482 | NA | fipr-6 |
| WBGene00007545 | -0.221525717 | 0.02480847 | 0.09030851 | C13B4.1 |
| WBGene00007546 | 0.506084877 | 0.06470274 | 0.18126919 | nhr-153 |
| WBGene00007547 | -0.12486132 | 0.54359246 | 0.71696212 | nhr-154 |
| WBGene00007548 | -0.13934359 | 0.50105882 | 0.6827999 | C13C4.4 |
| WBGene00007549 | 0.522183219 | 0.00289015 | 0.01697341 | spin-1 |
| WBGene00007550 | 0.158609602 | 0.4418409 | 0.63595638 | C13C4.6 |
| WBGene00007551 | 0.981033968 | 0.01542332 | 0.06326234 | C13C4.7 |
| WBGene00007552 | 0.039292341 | 0.65808492 | NA | C13C12.2 |
| WBGene00007553 | 0.310966738 | 0.1744832 | 0.35510266 | C13G3.1 |
| WBGene00007554 | -0.171125979 | 0.03471319 | 0.11525777 | pptr-2 |

| | | | | |
|---|---|---|---|---|
| WBGene00007555 | -0.228501771 | 0.06375346 | 0.1792959 | dohh-1 |
| WBGene00007556 | -0.156537337 | 0.09640056 | 0.23974871 | algn-9 |
| WBGene00007557 | -0.038514639 | 0.84444283 | 0.91656071 | C14A4.6 |
| WBGene00007558 | 0.429941554 | 0.09110474 | 0.22960929 | C14A4.7 |
| WBGene00007559 | 0.065014541 | 0.37399213 | NA | C14A4.8 |
| WBGene00007560 | 0.344448119 | 0.00614181 | 0.03071169 | C14A4.9 |
| WBGene00007561 | -0.051419423 | 0.66373135 | 0.80271689 | ccm-3 |
| WBGene00007562 | 1.013755163 | 0.00139135 | 0.00923843 | C14A4.12 |
| WBGene00007563 | 0.156889814 | 0.11568537 | 0.27143421 | C14A4.13 |
| WBGene00007564 | -0.298639301 | 0.00793509 | 0.03759793 | mrps-22 |
| WBGene00007565 | 1.191275763 | 4.53E-17 | 2.68E-15 | clec-48 |
| WBGene00007567 | 0.036687355 | 0.59489568 | NA | C14A6.3 |
| WBGene00007569 | 0.036687355 | 0.59489568 | NA | C14A6.5 |
| WBGene00007570 | 0.054314019 | 0.54747109 | NA | C14A6.6 |
| WBGene00007571 | NA | NA | NA | C14A6.7 |
| WBGene00007572 | NA | NA | NA | C14A6.8 |
| WBGene00007573 | -0.29034257 | 0.00260097 | 0.01553683 | C14B1.2 |
| WBGene00007574 | 0.272508533 | 0.12653198 | 0.28761911 | C14B1.3 |
| WBGene00007576 | -0.056140949 | 0.68987335 | 0.82175538 | dph-1 |
| WBGene00007577 | 0.034735823 | 0.73428098 | 0.85112879 | nrde-1 |
| WBGene00007578 | -0.300993796 | 0.00210271 | 0.01301173 | C14B1.7 |
| WBGene00007579 | -0.128052866 | 0.32279434 | 0.52291098 | C14B1.8 |
| WBGene00007580 | -0.180086091 | 0.0085644 | 0.03983209 | C14B1.9 |
| WBGene00007581 | -0.036904602 | 0.75930423 | 0.86379292 | alkb-8 |
| WBGene00007583 | -0.230536615 | 0.21851714 | 0.41037833 | C14B4.2 |
| WBGene00007584 | 1.094683893 | 0.01379825 | 0.05786887 | C14C10.1 |
| WBGene00007585 | -0.059942505 | 0.75105413 | 0.85938701 | C14C10.2 |
| WBGene00007586 | -0.066549638 | 0.38883043 | 0.58725668 | ril-2 |
| WBGene00007587 | -0.032539802 | 0.69226818 | 0.82342176 | mma-1 |
| WBGene00007588 | -0.05410175 | 0.35961336 | 0.55958116 | C14C10.5 |
| WBGene00007589 | -0.039272894 | 0.83970119 | 0.91324209 | erg-28 |
| WBGene00007590 | 0.260854026 | 0.16596567 | 0.34248109 | ttr-43 |
| WBGene00007591 | -0.808029036 | 2.83E-06 | 3.92E-05 | zipt-13 |
| WBGene00007592 | -0.072204569 | 0.65917388 | 0.79896991 | C14H10.2 |
| WBGene00007593 | 0.53043967 | 0.00327736 | 0.01873637 | C14H10.3 |
| WBGene00007594 | 0.059980168 | 0.38780842 | NA | lgc-23 |
| WBGene00007595 | 0.11411957 | 0.12533078 | NA | C15A7.2 |
| WBGene00007597 | 0.038550418 | 0.65345073 | NA | C15A11.2 |
| WBGene00007598 | -0.040764755 | 0.81896083 | 0.900451 | C15A11.4 |
| WBGene00007599 | 0.956966609 | 1.82E-05 | 0.00020968 | C15A11.7 |
| WBGene00007600 | 0.249858065 | 0.25312326 | 0.4511782 | C15C6.1 |
| WBGene00007601 | 1.923320382 | 0.00019748 | 0.00175022 | C15C6.2 |
| WBGene00007602 | 0.02361128 | 0.82121395 | 0.90200331 | C15C6.3 |

| | | | | |
|---|---|---|---|---|
| WBGene00007603 | -0.056163564 | 0.73078226 | 0.84986592 | C15C6.4 |
| WBGene00007604 | 0.603178825 | 0.04402263 | 0.13709778 | xbx-9 |
| WBGene00007605 | -1.179945425 | 8.83E-14 | 3.74E-12 | hrg-7 |
| WBGene00007606 | -0.046421737 | 0.63476178 | 0.78338999 | C15C8.4 |
| WBGene00007607 | 0.014937555 | 0.87003897 | NA | C15C8.5 |
| WBGene00007608 | 0.057890378 | 0.58124336 | 0.74581663 | C15C8.6 |
| WBGene00007609 | -0.023077974 | 0.86914958 | 0.93000745 | C15C8.7 |
| WBGene00007610 | 0.114325509 | 0.27126915 | 0.4709096 | C15H7.3 |
| WBGene00007611 | 0.020925399 | 0.84643798 | NA | C15H7.4 |
| WBGene00007613 | 0.199461643 | 0.06869713 | 0.18884617 | C15H11.1 |
| WBGene00007614 | 0.0990293 | 0.33366152 | 0.53319563 | gnrr-2 |
| WBGene00007615 | -0.328329277 | 0.01106713 | 0.04844063 | set-31 |
| WBGene00007616 | -0.252299794 | 0.11066819 | 0.26243813 | rpoa-12 |
| WBGene00007617 | -0.359431528 | 0.00012012 | 0.00113982 | rrbs-1 |
| WBGene00007618 | -0.022930767 | 0.78121253 | NA | C15H11.10 |
| WBGene00007619 | 0.071340939 | 0.62723862 | 0.77799057 | C15H11.11 |
| WBGene00007620 | -0.176667807 | 0.06288235 | 0.17772319 | ocrl-1 |
| WBGene00007621 | 0.024247822 | 0.78198534 | 0.87735392 | C16C2.4 |
| WBGene00007622 | -0.003722263 | 0.97507075 | 0.98591235 | C16C10.1 |
| WBGene00007623 | -0.141618411 | 0.19026315 | 0.37613324 | C16C10.2 |
| WBGene00007624 | -0.178489377 | 0.00033196 | 0.00274137 | hrde-1 |
| WBGene00007625 | -0.136571652 | 0.14524647 | 0.31460967 | C16C10.4 |
| WBGene00007626 | -0.071620033 | 0.51774635 | 0.69704132 | rnf-121 |
| WBGene00007627 | -0.061986096 | 0.50253557 | 0.68391651 | ccdc-55 |
| WBGene00007628 | -0.369890051 | 0.01674994 | 0.06744659 | C16C10.8 |
| WBGene00007629 | -0.005017292 | 0.94670066 | NA | C16C10.9 |
| WBGene00007630 | -0.30708705 | 0.00170183 | 0.01093789 | har-1 |
| WBGene00007631 | 0.035402431 | 0.72436751 | 0.84556796 | wht-3 |
| WBGene00007632 | 0.055503542 | 0.44925503 | NA | C16C10.13 |
| WBGene00007633 | -0.043884293 | 0.49359635 | NA | C16D2.1 |
| WBGene00007635 | -0.026388072 | 0.88227547 | 0.93762317 | npr-4 |
| WBGene00007637 | -0.057843018 | 0.62591515 | 0.7769414 | dhhc-7 |
| WBGene00007638 | NA | NA | NA | C17D12.3 |
| WBGene00007638 | NA | NA | NA | Y47H9A.1 |
| WBGene00007639 | 0.018998992 | 0.77286019 | NA | C17D12.5 |
| WBGene00007640 | -0.110810047 | 0.39419965 | 0.59240002 | C17D12.7 |
| WBGene00007641 | 0.139881928 | 0.21242661 | 0.40286762 | C17E4.1 |
| WBGene00007642 | -0.985779716 | 0.00183257 | 0.01163514 | C17E4.2 |
| WBGene00007643 | -0.122674968 | 0.08895558 | 0.22638599 | 3-Mar |
| WBGene00007644 | -0.01493851 | 0.93622802 | 0.96394616 | C17E4.4 |
| WBGene00007645 | -0.21914595 | 0.02240199 | 0.08348636 | C17E4.6 |
| WBGene00007646 | -0.266977494 | 5.61E-05 | 0.00058016 | nkb-1 |
| WBGene00007647 | -0.117174896 | 0.2051575 | 0.39429128 | C17E4.10 |

| | | | | |
|---|---|---|---|---|
| WBGene00007648 | 0.020232312 | 0.84643798 | NA | folr-1 |
| WBGene00007649 | 0.070062154 | 0.57805956 | 0.74423743 | C17G1.2 |
| WBGene00007650 | 0.045924666 | 0.77943513 | 0.87526536 | ugt-23 |
| WBGene00007651 | -0.045654628 | 0.69100591 | 0.8224095 | nra-3 |
| WBGene00007652 | -0.046909917 | 0.81810604 | 0.90005189 | C17G1.5 |
| WBGene00007653 | 0.012889188 | 0.93353504 | 0.96267218 | cysl-1 |
| WBGene00007654 | 0.020232312 | 0.84643798 | NA | NA |
| WBGene00007655 | 0.020232312 | 0.84643798 | NA | C17H1.2 |
| WBGene00007656 | 0.085309563 | 0.24820057 | NA | pals-2 |
| WBGene00007657 | 0.05787416 | 0.40594086 | NA | pals-3 |
| WBGene00007658 | 0.842383882 | 0.02262033 | 0.08412097 | pals-4 |
| WBGene00007659 | 0.15038247 | 0.07114191 | NA | pals-5 |
| WBGene00007660 | 2.332132879 | 0.00681987 | 0.03339732 | pals-6 |
| WBGene00007661 | 0.058314668 | 0.40208118 | NA | pals-7 |
| WBGene00007662 | 0.095576857 | 0.24390154 | NA | pals-8 |
| WBGene00007664 | 0.257431432 | 0.23328978 | 0.42875992 | seb-3 |
| WBGene00007666 | 0.258410283 | 0.04488958 | 0.13900768 | C18B12.4 |
| WBGene00007667 | -0.048018882 | 0.81108218 | 0.89548708 | fbxa-97 |
| WBGene00007668 | -0.22025505 | 0.15810533 | 0.33235034 | C18B12.6 |
| WBGene00007669 | -0.001993815 | 0.95612996 | NA | C18D1.2 |
| WBGene00007670 | 0.03732969 | 0.59229443 | NA | stg-1 |
| WBGene00007672 | 6.543583514 | 1.17E-06 | 1.82E-05 | fbxa-136 |
| WBGene00007673 | -0.001842784 | 0.95621055 | NA | C18D4.3 |
| WBGene00007674 | 0.040218942 | 0.56174727 | NA | C18D4.4 |
| WBGene00007675 | 0.696959613 | 0.01859798 | 0.07286407 | C18D4.6 |
| WBGene00007676 | NA | NA | NA | NA |
| WBGene00007677 | 1.197948428 | 0.00099237 | 0.0069987 | NA |
| WBGene00007678 | NA | NA | NA | srz-90 |
| WBGene00007679 | -0.025309808 | 0.89381336 | 0.94472433 | C18D11.1 |
| WBGene00007680 | -0.068065257 | 0.74002797 | 0.85342581 | maa-1 |
| WBGene00007681 | -0.020531542 | 0.82378282 | 0.9034778 | C18D11.3 |
| WBGene00007682 | NA | NA | NA | C18D11.6 |
| WBGene00007683 | -0.194786234 | 0.02245541 | 0.08361638 | C18E9.2 |
| WBGene00007684 | -0.328525104 | 0.00171093 | 0.01098193 | C18E9.4 |
| WBGene00007685 | 0.120513063 | 0.40137638 | 0.59970932 | C18E9.5 |
| WBGene00007686 | -0.384975253 | 2.17E-07 | 3.85E-06 | tomm-40 |
| WBGene00007687 | 0.154324145 | 0.3991502 | 0.59738086 | C18E9.7 |
| WBGene00007688 | 0.057303378 | 0.52744004 | NA | C18E9.8 |
| WBGene00007689 | 0.501050483 | 0.02033661 | 0.0775895 | C18E9.9 |
| WBGene00007690 | -0.142966719 | 0.25173694 | 0.44993917 | sftd-3 |
| WBGene00007691 | 0.079262432 | 0.46762717 | 0.65691994 | C23H4.2 |
| WBGene00007692 | -0.068939076 | 0.7318224 | 0.85001776 | C23H4.3 |
| WBGene00007693 | -2.243996619 | 0.00416256 | 0.02260533 | C23H4.4 |

| | | | | |
|---|---|---|---|---|
| WBGene00007694 | -0.05102092 | 0.80113435 | 0.88919778 | C23H4.6 |
| WBGene00007695 | -0.120798003 | 0.53775583 | 0.71258851 | C23H4.7 |
| WBGene00007696 | -0.165440218 | 0.0136161 | 0.05724882 | tram-1 |
| WBGene00007697 | 0.34282616 | 0.04731655 | 0.14447582 | C24F3.2 |
| WBGene00007698 | 0.010876823 | 0.90560372 | 0.94916953 | qns-1 |
| WBGene00007699 | NA | NA | NA | C24H11.1 |
| WBGene00007700 | NA | NA | NA | C24H11.2 |
| WBGene00007701 | 0.05787416 | 0.40594086 | NA | srd-74 |
| WBGene00007702 | 0.025609219 | 0.90110594 | 0.94730323 | C24H11.5 |
| WBGene00007703 | 0.086663531 | 0.298168 | 0.49908194 | gbf-1 |
| WBGene00007704 | -0.447075511 | 0.04092748 | 0.13008348 | mdt-21 |
| WBGene00007705 | -0.073006112 | 0.47199556 | 0.66010393 | C25A1.1 |
| WBGene00007707 | -0.028819537 | 0.73020785 | 0.84963627 | fath-1 |
| WBGene00007708 | NA | NA | NA | nola-3 |
| WBGene00007709 | -0.042896353 | 0.311984 | 0.51266736 | clec-87 |
| WBGene00007710 | -0.092077506 | 0.26030986 | 0.45840351 | rsa-1 |
| WBGene00007711 | -0.096388029 | 0.28295979 | 0.48275265 | lid-1 |
| WBGene00007712 | -0.116341896 | 0.38392479 | 0.58327725 | mrpl-34 |
| WBGene00007713 | NA | NA | NA | C25A1.15 |
| WBGene00007714 | 0.185427242 | 0.12556502 | 0.28666356 | C25D7.1 |
| WBGene00007715 | -0.027373974 | 0.68245543 | NA | NA |
| WBGene00007716 | 0.834413394 | 0.02282138 | 0.08473949 | fbxa-82 |
| WBGene00007717 | 0.441943058 | 0.08950431 | 0.22726891 | C25D7.5 |
| WBGene00007718 | -0.293621604 | 0.16907812 | 0.34728855 | otub-1 |
| WBGene00007719 | NA | NA | NA | C25D7.9 |
| WBGene00007720 | -0.304690603 | 0.0544923 | 0.16017302 | C25D7.10 |
| WBGene00007721 | NA | NA | NA | NA |
| WBGene00007722 | 0.175409541 | 0.31304137 | 0.51343714 | C25D7.15 |
| WBGene00007723 | -0.001488154 | 0.95639912 | NA | C25F9.2 |
| WBGene00007724 | 0.627314275 | 0.02875304 | 0.10051891 | C25F9.4 |
| WBGene00007725 | 0.638349014 | 0.01800983 | 0.07118842 | C25F9.5 |
| WBGene00007726 | 0.034618355 | 0.68540017 | NA | C25F9.6 |
| WBGene00007727 | NA | NA | NA | C25F9.8 |
| WBGene00007728 | 0.399981711 | 0.06192633 | 0.17564188 | C25F9.9 |
| WBGene00007729 | 0.178978873 | 0.09042234 | 0.22876518 | clec-185 |
| WBGene00007730 | 0.242512155 | 0.08767878 | 0.22426713 | C25G4.2 |
| WBGene00007731 | -0.328584517 | 0.06878451 | 0.18895467 | C25G4.3 |
| WBGene00007732 | 0.010822832 | 0.95078325 | 0.97201103 | spe-44 |
| WBGene00007733 | 0.608405791 | 0.04207066 | 0.13262743 | smz-1 |
| WBGene00007734 | 0.098747568 | 0.38520852 | 0.58437861 | C25G4.7 |
| WBGene00007735 | 0.231871756 | 0.02096847 | 0.07945118 | C25G4.8 |
| WBGene00007736 | -0.063278102 | 0.63357643 | 0.78260005 | igdb-2 |
| WBGene00007739 | 0.048730085 | 0.71626113 | 0.84075838 | C26C6.4 |

| | | | | |
|---|---|---|---|---|
| WBGene00007740 | -0.003421551 | 0.95689871 | NA | C26C6.6 |
| WBGene00007741 | 0.138287108 | 0.38576097 | 0.58472269 | glb-8 |
| WBGene00007742 | -0.040794878 | 0.53203165 | NA | C26C6.8 |
| WBGene00007743 | 0.06460902 | 0.74031954 | 0.85342581 | C26C6.9 |
| WBGene00007744 | 0.157419081 | 0.46021925 | 0.65136993 | C26D10.3 |
| WBGene00007745 | 0.454245912 | 0.08230504 | 0.21510675 | C26D10.4 |
| WBGene00007746 | 0.204020537 | 0.30975441 | 0.5105386 | C26D10.6 |
| WBGene00007747 | 0.020925399 | 0.84643798 | NA | C26E1.1 |
| WBGene00007748 | 0.801485233 | 0.00929036 | 0.04245323 | C26E1.2 |
| WBGene00007749 | 0.586072177 | 0.0320665 | 0.10860181 | ceh-79 |
| WBGene00007750 | 0.403299021 | 0.08653408 | 0.22223216 | syg-2 |
| WBGene00007751 | 0.59086659 | 0.00206251 | 0.0128063 | C26G2.2 |
| WBGene00007752 | -0.238820589 | 0.01867724 | 0.07305758 | wdfy-3 |
| WBGene00007753 | 0.586314653 | 0.00216102 | 0.01329049 | C27A7.1 |
| WBGene00007754 | -0.023109762 | 0.78121253 | NA | NA |
| WBGene00007755 | 0.568394684 | 0.04897184 | 0.14801747 | C27A7.3 |
| WBGene00007756 | -0.133908809 | 0.27353689 | 0.47383502 | C27A7.5 |
| WBGene00007757 | -0.149538505 | 0.19293452 | 0.37936534 | C27A7.6 |
| WBGene00007758 | -0.410664167 | 0.09859426 | 0.24347628 | C27A7.8 |
| WBGene00007759 | 0.020317245 | 0.84643798 | NA | C27A7.9 |
| WBGene00007760 | 0.122478977 | 0.45816665 | 0.64973678 | C27B7.2 |
| WBGene00007761 | -0.061097042 | 0.46509233 | 0.65481694 | rad-26 |
| WBGene00007762 | -0.320433966 | 0.00092602 | 0.00659426 | C27B7.5 |
| WBGene00007763 | 0.095967469 | 0.18297344 | NA | C27B7.6 |
| WBGene00007764 | 1.363838271 | 4.84E-06 | 6.35E-05 | C27B7.7 |
| WBGene00007765 | -0.661791653 | 0.00053629 | 0.00411737 | C27B7.9 |
| WBGene00007766 | 0.867458542 | 1.49E-14 | 6.76E-13 | C27C7.1 |
| WBGene00007767 | NA | NA | NA | NA |
| WBGene00007768 | 0.033034883 | 0.64219503 | NA | C27C7.5 |
| WBGene00007769 | -0.15455794 | 0.33049966 | 0.53007847 | C27C7.7 |
| WBGene00007770 | NA | NA | NA | nhr-259 |
| WBGene00007771 | -0.258408893 | 0.13302384 | 0.29657409 | C27C12.1 |
| WBGene00007772 | -0.011495065 | 0.93513414 | 0.96336188 | egrh-1 |
| WBGene00007773 | -0.347683351 | 0.0217391 | 0.08163173 | C27C12.3 |
| WBGene00007774 | 0.143785159 | 0.36098152 | 0.56083865 | C27C12.4 |
| WBGene00007775 | 0.248368687 | 0.26114503 | 0.45922927 | acd-3 |
| WBGene00007776 | 0.042199095 | 0.55922995 | NA | dmd-4 |
| WBGene00007777 | 0.394063692 | 0.04304605 | 0.13486781 | C27D8.1 |
| WBGene00007778 | 1.95681835 | 0.00289345 | 0.01697944 | C27D8.2 |
| WBGene00007779 | -0.135495808 | 0.3896885 | 0.58786842 | C27D8.3 |
| WBGene00007780 | -0.10040632 | 0.42906735 | 0.62389265 | C27D8.4 |
| WBGene00007784 | -0.363789518 | 6.09E-05 | 0.00062683 | ruvb-1 |
| WBGene00007785 | 0.021687266 | 0.835387 | 0.91048532 | tofu-1 |

| | | | | |
|---|---|---|---|---|
| WBGene00007786 | -0.083166887 | 0.26221249 | 0.46009745 | rmd-2 |
| WBGene00007787 | -0.310944226 | 0.00020748 | 0.00182401 | C27H6.8 |
| WBGene00007788 | -0.117184814 | 0.46548762 | 0.65511912 | C28A5.1 |
| WBGene00007789 | -0.156557636 | 0.3457039 | 0.54529744 | C28A5.2 |
| WBGene00007791 | 0.801914465 | 0.01800651 | 0.07118842 | C28A5.6 |
| WBGene00007792 | 0.073887753 | 0.28837975 | NA | C28D4.4 |
| WBGene00007793 | 0.127435179 | 0.16853418 | NA | C28D4.5 |
| WBGene00007794 | 0.380182381 | 0.01864761 | 0.07299992 | C28D4.7 |
| WBGene00007795 | 0.020317245 | 0.84643798 | NA | C28D4.8 |
| WBGene00007796 | -0.013029346 | 0.89540716 | 0.94517796 | C28D4.10 |
| WBGene00007797 | -0.217625018 | 0.03402844 | 0.11341006 | C29A12.1 |
| WBGene00007798 | -0.27928162 | 0.17505314 | 0.35564444 | NA |
| WBGene00007799 | 0.67694437 | 0.02193581 | 0.08220176 | nrx-1 |
| WBGene00007800 | 0.06111288 | 0.51882185 | NA | C29A12.6 |
| WBGene00007801 | 0.032894921 | 0.87456777 | 0.93307197 | trpa-1 |
| WBGene00007802 | NA | NA | NA | cutl-27 |
| WBGene00007804 | 0.020317245 | 0.84643798 | NA | C29F3.3 |
| WBGene00007805 | NA | NA | NA | clec-231 |
| WBGene00007806 | 0.208247896 | 0.06511123 | 0.18184975 | clec-230 |
| WBGene00007807 | -0.381622026 | 0.02158959 | 0.08119255 | C29F3.7 |
| WBGene00007808 | 1.398914599 | 1.03E-08 | 2.29E-07 | best-7 |
| WBGene00007809 | 0.057961644 | 0.59023878 | 0.7515163 | C29F4.3 |
| WBGene00007810 | 0.360343126 | 0.1242352 | 0.28458189 | C29F7.1 |
| WBGene00007811 | 1.246132516 | 2.75E-14 | 1.21E-12 | C29F7.2 |
| WBGene00007812 | 0.060793721 | 0.59077164 | 0.75180249 | C29F7.3 |
| WBGene00007813 | -0.092753003 | 0.64116126 | 0.78738819 | jmjd-3.3 |
| WBGene00007816 | 0.117061468 | 0.17857748 | 0.36004827 | C30F2.2 |
| WBGene00007817 | 2.593425666 | 0.00011439 | 0.0010932 | C30F2.3 |
| WBGene00007818 | 0.729937731 | 0.01813433 | 0.07156462 | C30F2.4 |
| WBGene00007819 | 0.41155827 | 0.10457748 | 0.25284464 | C30F2.5 |
| WBGene00007820 | 0.043242326 | 0.81539758 | 0.89862521 | clec-199 |
| WBGene00007821 | NA | NA | NA | NA |
| WBGene00007822 | 0.020317245 | 0.84643798 | NA | clec-202 |
| WBGene00007823 | 0.085537126 | 0.24698522 | NA | C30H6.5 |
| WBGene00007824 | -0.107951647 | 0.24973801 | 0.4481449 | dlat-2 |
| WBGene00007825 | 0.160694484 | 0.2906272 | 0.49114651 | C30H6.8 |
| WBGene00007826 | -0.238889556 | 0.15369585 | 0.32702528 | C30H6.9 |
| WBGene00007827 | 2.736717918 | 5.19E-08 | 1.03E-06 | trh-1 |
| WBGene00007829 | 0.149903014 | 0.19597885 | 0.38361181 | oac-5 |
| WBGene00007831 | NA | NA | NA | srbc-83 |
| WBGene00007832 | 0.032192384 | 0.61853721 | NA | srbc-82 |
| WBGene00007833 | -0.094721951 | 0.644972 | 0.78968348 | oac-6 |
| WBGene00007834 | NA | NA | NA | srxa-7 |

| | | | | |
|---|---|---|---|---|
| WBGene00007835 | -0.11213261 | 0.12265889 | NA | oac-7 |
| WBGene00007836 | -0.3071994 | 0.00789174 | 0.03741674 | C31C9.2 |
| WBGene00007837 | -0.30499923 | 0.18256845 | 0.36541631 | fbxa-167 |
| WBGene00007838 | -0.167164262 | 0.42581989 | 0.62160822 | fbxa-168 |
| WBGene00007840 | 1.080497417 | 0.00134794 | 0.00897653 | C31C9.6 |
| WBGene00007841 | -0.233608035 | 0.29170644 | 0.4924333 | C31C9.7 |
| WBGene00007842 | NA | NA | NA | C31E10.1 |
| WBGene00007844 | 0.063731418 | 0.41159439 | NA | irld-24 |
| WBGene00007846 | -0.172250153 | 0.20683648 | 0.39611418 | C31E10.5 |
| WBGene00007847 | 0.14914245 | 0.44626263 | 0.63961334 | C31E10.6 |
| WBGene00007848 | 0.093014446 | 0.41695492 | 0.61396329 | cytb-5.1 |
| WBGene00007849 | 0.236434363 | 0.24377458 | 0.44166608 | tbc-19 |
| WBGene00007850 | -0.142354525 | 0.49269486 | 0.67574197 | C31G12.1 |
| WBGene00007851 | 0.736765825 | 0.02870871 | 0.10044665 | clec-245 |
| WBGene00007852 | NA | NA | NA | C31G12.3 |
| WBGene00007853 | 0.035512071 | 0.72830584 | 0.84850209 | C31G12.4 |
| WBGene00007854 | 0.038125098 | 0.61119482 | NA | C31H5.1 |
| WBGene00007855 | -0.175680562 | 0.27457649 | 0.47493999 | C31H5.4 |
| WBGene00007856 | 0.080564285 | 0.68520558 | 0.81834529 | C31H5.5 |
| WBGene00007857 | 0.157075628 | 0.30365055 | 0.50437151 | C31H5.6 |
| WBGene00007858 | 0.047717713 | 0.62950231 | 0.77968456 | C31H5.7 |
| WBGene00007859 | -0.557765901 | 0.00063646 | 0.00478786 | mrps-31 |
| WBGene00007860 | -0.004216897 | 0.96585826 | 0.98082345 | rilp-1 |
| WBGene00007861 | 0.015451648 | 0.80337623 | NA | C32A9.1 |
| WBGene00007862 | 0.319315205 | 0.11697469 | 0.27352701 | kvs-2 |
| WBGene00007863 | 0.064938666 | 0.46494484 | NA | C32C4.3 |
| WBGene00007864 | 0.048790507 | 0.24485145 | 0.4429796 | irg-6 |
| WBGene00007865 | NA | NA | NA | srz-15 |
| WBGene00007866 | -0.097617052 | 0.40394701 | 0.60226219 | C32H11.3 |
| WBGene00007867 | -0.095887013 | 0.64551304 | 0.79001558 | C32H11.4 |
| WBGene00007868 | 0.182023446 | 0.04351057 | NA | C32H11.5 |
| WBGene00007869 | NA | NA | NA | C32H11.6 |
| WBGene00007870 | -0.060485643 | 0.76734646 | 0.86792389 | C32H11.7 |
| WBGene00007871 | NA | NA | NA | C32H11.8 |
| WBGene00007871 | NA | NA | NA | C32H11.11 |
| WBGene00007872 | NA | NA | NA | C32H11.9 |
| WBGene00007873 | 0.020232312 | 0.84643798 | NA | dod-21 |
| WBGene00007874 | NA | NA | NA | C32H11.8 |
| WBGene00007874 | NA | NA | NA | C32H11.11 |
| WBGene00007875 | 3.564629883 | 1.24E-211 | 1.65E-208 | dod-24 |
| WBGene00007876 | 7.146600193 | 3.98E-10 | 1.05E-08 | dct-19 |
| WBGene00007877 | 0.791399433 | 1.22E-05 | 0.00014679 | nfki-1 |
| WBGene00007878 | 0.434252226 | 0.05438313 | 0.1599806 | C33A11.2 |

| | | | | | |
|---|---|---|---|---|---|
| WBGene00007880 | | -0.087781471 | 0.32718293 | 0.52634714 | C33A12.1 |
| WBGene00007881 | | 1.700558855 | 0.00251564 | 0.01508868 | nlp-35 |
| WBGene00007882 | | -0.11554544 | 0.28079058 | 0.48037565 | C33A12.3 |
| WBGene00007883 | | 1.25864869 | 0.00085898 | 0.0061861 | C33A12.4 |
| WBGene00007885 | | 1.486048688 | 1.98E-06 | 2.89E-05 | ugt-21 |
| WBGene00007886 | | -0.272153189 | 0.19134147 | 0.37744954 | ethe-1 |
| WBGene00007887 | | -0.372743049 | 0.10753099 | 0.25779617 | gadr-2 |
| WBGene00007888 | NA | | NA | NA | ttr-9 |
| WBGene00007889 | NA | | NA | NA | NA |
| WBGene00007890 | | -1.077527377 | 7.86E-05 | 0.00078105 | C33A12.19 |
| WBGene00007891 | | -0.079824684 | 0.4472736 | 0.64038659 | C33B4.2 |
| WBGene00007892 | | -0.198522429 | 0.34052195 | 0.54032533 | C33B4.4 |
| WBGene00007893 | NA | | NA | NA | C33B4.5 |
| WBGene00007894 | | 0.047071372 | 0.71852369 | 0.84199658 | C33D3.3 |
| WBGene00007895 | | 0.020317245 | 0.84643798 | NA | C33D3.4 |
| WBGene00007896 | | 0.01912147 | 0.77587372 | NA | C33D3.5 |
| WBGene00007897 | | -0.002914971 | 0.95718083 | NA | clec-182 |
| WBGene00007898 | | 0.555230217 | 0.02017995 | 0.07717428 | C33D9.3 |
| WBGene00007899 | | -0.003948282 | 0.98301664 | 0.99018553 | NA |
| WBGene00007900 | | 0.07867533 | 0.70475097 | 0.83265486 | C33D9.6 |
| WBGene00007901 | | 0.137795582 | 0.33794157 | 0.5376293 | C33D9.8 |
| WBGene00007902 | | -0.001488154 | 0.95639912 | NA | NA |
| WBGene00007903 | | 0.178316987 | 0.25943338 | 0.45746887 | lgc-21 |
| WBGene00007904 | | -0.601335421 | 0.00346477 | 0.019586 | C33G3.4 |
| WBGene00007905 | NA | | NA | NA | C33G3.5 |
| WBGene00007906 | | -0.129753198 | 0.49568278 | 0.67825155 | C33G3.6 |
| WBGene00007907 | | 0.080838789 | 0.57771675 | 0.74411462 | C34B4.2 |
| WBGene00007908 | | 0.524675659 | 0.04239868 | 0.1334105 | C34B4.3 |
| WBGene00007909 | | 0.015874474 | 0.83570347 | NA | C34B4.4 |
| WBGene00007910 | | -0.079472169 | 0.3156132 | NA | C34B4.5 |
| WBGene00007911 | | -0.110909677 | 0.57841506 | 0.7444901 | C34B7.1 |
| WBGene00007912 | | 0.005650989 | 0.95131864 | 0.97203268 | C34B7.2 |
| WBGene00007913 | | 1.646784412 | 1.90E-10 | 5.26E-09 | cyp-36A1 |
| WBGene00007914 | | 0.090355288 | 0.49584912 | 0.67841575 | mys-4 |
| WBGene00007916 | | 0.227252075 | 0.27597365 | 0.47597246 | C34C6.3 |
| WBGene00007917 | | 0.254959198 | 0.07465682 | 0.20089148 | C34C6.4 |
| WBGene00007918 | | -0.078957004 | 0.56190387 | 0.73225052 | sphk-1 |
| WBGene00007919 | | 2.071153445 | 5.52E-10 | 1.44E-08 | cup-16 |
| WBGene00007920 | | 0.018657025 | 0.77508599 | NA | C34C12.1 |
| WBGene00007921 | | -0.241018297 | 0.03995596 | 0.12787904 | C34C12.2 |
| WBGene00007922 | | -0.16753253 | 0.1208297 | 0.27965132 | pph-6 |
| WBGene00007923 | | -0.008873276 | 0.94832425 | 0.97044737 | C34C12.4 |
| WBGene00007924 | | -0.035328897 | 0.77767978 | 0.8741977 | rsu-1 |

| | | | | |
|---|---|---|---|---|
| WBGene00007925 | -0.390312033 | 0.11448429 | 0.26936936 | C34C12.6 |
| WBGene00007926 | 0.607745118 | 0.02640504 | 0.09461484 | C34C12.7 |
| WBGene00007927 | -0.232954943 | 0.033245 | 0.11158645 | C34C12.8 |
| WBGene00007928 | -0.138803899 | 0.5111981 | 0.69127657 | C34C12.9 |
| WBGene00007929 | 0.144346207 | 0.24938843 | 0.447792 | dmd-10 |
| WBGene00007930 | 0.020232312 | 0.84643798 | NA | dmd-11 |
| WBGene00007931 | 0.391107402 | 0.09643158 | 0.23978513 | C34D1.4 |
| WBGene00007932 | -0.012402851 | 0.94965338 | 0.97122452 | zip-5 |
| WBGene00007933 | 0.031160622 | 0.79719548 | 0.88643875 | C34E7.3 |
| WBGene00007934 | -0.604112008 | 0.00212855 | 0.01314941 | C34E7.4 |
| WBGene00007935 | 0.009733915 | 0.95098181 | 0.97202769 | C34E11.2 |
| WBGene00007937 | -0.011288461 | 0.94292348 | 0.96758451 | C34E11.4 |
| WBGene00007938 | 0.031516717 | 0.79377878 | 0.88399982 | C34F6.1 |
| WBGene00007939 | -0.332586051 | 0.16131123 | 0.33590637 | C34F6.5 |
| WBGene00007941 | -0.197145834 | 0.26033913 | 0.45840351 | C34F6.7 |
| WBGene00007942 | -0.003521102 | 0.97618075 | 0.98662587 | idh-2 |
| WBGene00007943 | -0.307738526 | 0.09369213 | 0.23420634 | C34F6.9 |
| WBGene00007944 | 0.40370096 | 0.0080022 | 0.0378669 | C34F6.10 |
| WBGene00007945 | 0.084157082 | 0.68758815 | 0.81998681 | C34F6.11 |
| WBGene00007946 | 0.148021412 | 0.07455439 | NA | ugt-33 |
| WBGene00007947 | -1.753092646 | 8.59E-26 | 8.44E-24 | C35A5.3 |
| WBGene00007948 | 0.0398788 | 0.58971221 | NA | C35A5.4 |
| WBGene00007949 | 0.020232312 | 0.84643798 | NA | C35A5.5 |
| WBGene00007950 | 0.70734224 | 0.00642246 | 0.03186417 | C35A5.6 |
| WBGene00007951 | 0.325075468 | 0.13544142 | 0.30022678 | dmsr-8 |
| WBGene00007952 | -0.079906469 | 0.17813252 | 0.35958061 | C35A5.8 |
| WBGene00007953 | 0.168739287 | 0.33472413 | 0.53413188 | hda-11 |
| WBGene00007954 | -0.054524006 | 0.53093834 | NA | gcp-2.2 |
| WBGene00007955 | -0.118280439 | 0.29001963 | 0.49054729 | selt-1.1 |
| WBGene00007956 | -0.191520697 | 0.15863278 | 0.332981 | trpp-9 |
| WBGene00007957 | 2.45280887 | 1.25E-13 | 5.15E-12 | C35C5.8 |
| WBGene00007958 | 0.983964834 | 0.00243196 | 0.01466487 | C35C5.9 |
| WBGene00007959 | -0.380144411 | 0.02741703 | 0.0969709 | C35C5.10 |
| WBGene00007960 | -0.151795781 | 0.20931175 | 0.39876248 | C35D6.3 |
| WBGene00007961 | -0.016367563 | 0.92882067 | 0.95998218 | C35D6.4 |
| WBGene00007962 | 0.179490365 | 0.09068203 | 0.22888093 | NA |
| WBGene00007963 | -2.773774459 | 1.07E-13 | 4.47E-12 | cyp-25A1 |
| WBGene00007964 | -0.781507331 | 0.00018059 | 0.00161513 | cyp-25A2 |
| WBGene00007965 | -0.223550717 | 0.00056124 | 0.00427754 | C36A4.4 |
| WBGene00007966 | -0.115483399 | 0.17943918 | 0.36082531 | maph-1.3 |
| WBGene00007967 | 0.792206191 | 0.01272564 | 0.05422471 | cyp-25A4 |
| WBGene00007969 | -0.012339374 | 0.85189791 | 0.92048984 | acs-19 |
| WBGene00007970 | -0.088550894 | 0.19967951 | NA | C36A4.10 |

| | | | | |
|---|---|---|---|---|
| WBGene00007971 | -0.229631543 | 0.00498462 | 0.02610149 | rpb-3 |
| WBGene00007972 | -0.162320387 | 0.16903036 | 0.34728789 | prp-4 |
| WBGene00007973 | 1.977740953 | 2.66E-05 | 0.00029547 | C36B1.6 |
| WBGene00007974 | -0.280913798 | 0.00612616 | 0.03066429 | dhfr-1 |
| WBGene00007975 | -0.241349313 | 0.00072708 | 0.00533596 | gls-1 |
| WBGene00007976 | 0.026849612 | 0.8542816 | 0.92206349 | C36B1.9 |
| WBGene00007977 | 1.525463611 | 0.00676767 | 0.03318345 | gska-3 |
| WBGene00007978 | -0.240937587 | 0.00576133 | 0.02928877 | C36B1.11 |
| WBGene00007979 | -0.056085972 | 0.32960374 | 0.52913689 | imp-1 |
| WBGene00007980 | -0.531789754 | 2.53E-15 | 1.24E-13 | C36E8.1 |
| WBGene00007981 | 0.323520777 | 0.15601301 | 0.3297036 | glb-11 |
| WBGene00007982 | 0.429795636 | 0.02987014 | 0.10331412 | pxd-1 |
| WBGene00007983 | -0.008467941 | 0.96573764 | 0.98082345 | C36E8.4 |
| WBGene00007984 | -0.169209786 | 0.4223556 | 0.6185564 | irx-1 |
| WBGene00007985 | 0.163407262 | 0.21718735 | 0.4087973 | swah-1 |
| WBGene00007986 | 0.021251951 | 0.85464241 | 0.92223021 | C36F7.5 |
| WBGene00007987 | 0.614168298 | 0.02096354 | 0.07945118 | C36H8.1 |
| WBGene00007988 | 0.077410951 | 0.65150643 | 0.79364585 | best-8 |
| WBGene00007989 | -0.147668177 | 0.29131438 | 0.49199836 | fipr-22 |
| WBGene00007990 | 0.020925399 | 0.84643798 | NA | fipr-23 |
| WBGene00007991 | 0.680180859 | 0.00064513 | 0.00483383 | C37A5.7 |
| WBGene00007992 | -0.027373974 | 0.68245543 | NA | fipr-24 |
| WBGene00007993 | -0.025246577 | 0.77685804 | 0.8737788 | idhb-1 |
| WBGene00007994 | -0.065736487 | 0.6228444 | 0.77450866 | C37E2.2 |
| WBGene00007995 | NA | NA | NA | C37E2.3 |
| WBGene00007996 | 0.065014541 | 0.37399213 | NA | C38C6.3 |
| WBGene00007997 | -0.039195877 | 0.73588415 | 0.85181646 | sre-13 |
| WBGene00007998 | NA | NA | NA | C38C6.5 |
| WBGene00007999 | 3.197658617 | 0.00193618 | 0.01217683 | tag-297 |
| WBGene00008000 | 0.0697467 | 0.26988866 | 0.469125 | slc-17.2 |
| WBGene00008001 | 0.090103632 | 0.20323472 | NA | C38C10.3 |
| WBGene00008002 | 0.295126297 | 0.03005615 | 0.10361481 | nlp-48 |
| WBGene00008003 | -0.046999685 | 0.7597616 | 0.86405055 | enu-3.2 |
| WBGene00008005 | -0.028809633 | 0.74986757 | 0.8587464 | C38D4.4 |
| WBGene00008006 | 0.112404177 | 0.33378167 | 0.53327122 | tag-325 |
| WBGene00008007 | 0.061829618 | 0.74882328 | 0.8582013 | C38D4.7 |
| WBGene00008008 | 0.021168473 | 0.91565096 | 0.95301384 | C38D4.9 |
| WBGene00008009 | NA | NA | NA | fbxa-171 |
| WBGene00008010 | 0.383998465 | 0.05089362 | 0.15234787 | C38D9.2 |
| WBGene00008012 | 0.020232312 | 0.84643798 | NA | fbxa-133 |
| WBGene00008013 | 0.020232312 | 0.84643798 | NA | NA |
| WBGene00008014 | 0.44296708 | 0.04323403 | 0.13534092 | fbxa-172 |
| WBGene00008015 | 0.036988272 | 0.57084766 | NA | fbxa-174 |

| | | | | |
|---|---|---|---|---|
| WBGene00008016 | -0.73200463 | 0.0278652 | 0.09808835 | C38D9.8 |
| WBGene00008017 | 0.212921434 | 0.15703493 | 0.33108565 | fbxa-176 |
| WBGene00008018 | -0.025402267 | 0.90037226 | 0.94680387 | tbc-8 |
| WBGene00008019 | 0.543975264 | 0.00354659 | 0.0199714 | C38H2.2 |
| WBGene00008020 | 0.019126951 | 0.92149285 | 0.95613845 | C38H2.3 |
| WBGene00008021 | 1.059428339 | 0.00769278 | 0.03663954 | C39B10.1 |
| WBGene00008022 | 0.033144996 | 0.68360425 | NA | lgc-41 |
| WBGene00008023 | 0.03557932 | 0.72309875 | 0.84472009 | C39B10.3 |
| WBGene00008024 | NA | NA | NA | C39B10.4 |
| WBGene00008025 | 0.042199095 | 0.55922995 | NA | C39B10.5 |
| WBGene00008026 | NA | NA | NA | NA |
| WBGene00008027 | 2.626814338 | 8.87E-19 | 6.01E-17 | scl-5 |
| WBGene00008028 | -0.099568767 | 0.62928503 | 0.77968456 | scl-6 |
| WBGene00008029 | 0.020317245 | 0.84643798 | NA | scl-7 |
| WBGene00008030 | 0.040107227 | 0.58693163 | NA | scl-8 |
| WBGene00008031 | 0.533539759 | 0.05441752 | 0.16004959 | C39E9.7 |
| WBGene00008032 | -0.690421323 | 2.03E-09 | 4.95E-08 | C39E9.8 |
| WBGene00008033 | 1.713956805 | 5.20E-05 | 0.00054255 | spin-2 |
| WBGene00008034 | -0.014369773 | 0.87353891 | 0.93276403 | C39E9.11 |
| WBGene00008035 | -0.064984858 | 0.51130475 | 0.69129309 | C39E9.12 |
| WBGene00008036 | 0.412855205 | 0.04660331 | 0.14292399 | C40C9.3 |
| WBGene00008037 | 0.020317245 | 0.84643798 | NA | C40C9.4 |
| WBGene00008038 | 1.323073039 | 0.00018699 | 0.0016663 | C40H1.2 |
| WBGene00008039 | 0.034618355 | 0.68540017 | NA | C40H1.3 |
| WBGene00008040 | 0.440044726 | 0.012264 | 0.05263796 | ttr-5 |
| WBGene00008041 | -0.22976374 | 0.10828064 | 0.2588431 | ufc-1 |
| WBGene00008042 | 1.665352178 | 1.75E-09 | 4.30E-08 | C40H1.7 |
| WBGene00008043 | 4.336524313 | 3.88E-10 | 1.03E-08 | C40H1.8 |
| WBGene00008044 | 0.202987723 | 0.14001999 | 0.30737534 | C40H1.9 |
| WBGene00008045 | 0.257971364 | 0.18341612 | 0.36639736 | C40H5.2 |
| WBGene00008047 | -0.051558841 | 0.74040125 | 0.85342581 | lact-7 |
| WBGene00008048 | 0.0829707 | 0.5403346 | 0.71470223 | NA |
| WBGene00008050 | -0.008099706 | 0.95265362 | 0.9729893 | chst-1 |
| WBGene00008051 | 0.78430307 | 0.02524988 | 0.09139553 | lron-4 |
| WBGene00008052 | -0.016184904 | 0.84893773 | 0.91871515 | ctns-1 |
| WBGene00008053 | -0.113169303 | 0.0368094 | 0.12040358 | cdc-48.2 |
| WBGene00008054 | NA | NA | NA | C41C4.9 |
| WBGene00008055 | -0.003270206 | 0.95698316 | NA | NA |
| WBGene00008056 | 0.020925399 | 0.84643798 | NA | nhr-164 |
| WBGene00008057 | 0.020232312 | 0.84643798 | NA | irld-25 |
| WBGene00008058 | NA | NA | NA | C41G6.12 |
| WBGene00008059 | NA | NA | NA | Y75B12B.8 |
| WBGene00008059 | NA | NA | NA | C41G6.13 |

| | | | | |
|---|---|---|---|---|
| WBGene00008060 | NA | NA | NA | C41G6.16 |
| WBGene00008061 | -0.356954491 | 0.00042827 | 0.00340192 | C41G7.3 |
| WBGene00008062 | -0.238750287 | 0.18338921 | 0.36639736 | set-32 |
| WBGene00008063 | 0.603272793 | 0.0098741 | 0.04444106 | C41G7.6 |
| WBGene00008064 | 0.056188364 | 0.42092875 | NA | C41G7.7 |
| WBGene00008065 | 0.208868193 | 0.13613946 | 0.3010903 | npr-18 |
| WBGene00008066 | -1.502182383 | 0.00876686 | 0.04060599 | fbxb-65 |
| WBGene00008067 | 0.087256674 | 0.23800185 | NA | C43D7.4 |
| WBGene00008068 | -0.100648358 | 0.39264273 | 0.59094184 | sdz-6 |
| WBGene00008069 | -0.053591705 | 0.45278076 | NA | C43D7.7 |
| WBGene00008070 | -0.139326529 | 0.36813022 | 0.56788479 | C43D7.8 |
| WBGene00008071 | -0.107047092 | 0.29651565 | 0.49751505 | fbxb-64 |
| WBGene00008072 | -0.003270206 | 0.95698316 | NA | C43F9.4 |
| WBGene00008073 | 1.129619778 | 0.01452857 | 0.06032693 | C43F9.5 |
| WBGene00008074 | 0.299059939 | 0.00899991 | 0.04146247 | nkb-2 |
| WBGene00008075 | 0.030724443 | 0.74618255 | 0.85644863 | C43F9.7 |
| WBGene00008076 | 0.304049889 | 0.19288758 | 0.37932398 | lgc-43 |
| WBGene00008077 | 0.151799006 | 0.34704858 | 0.54665893 | oac-8 |
| WBGene00008078 | -0.002355698 | 0.98798123 | 0.99323079 | vps-50 |
| WBGene00008079 | 0.325619382 | 0.08475854 | 0.21933343 | C44B9.2 |
| WBGene00008080 | -0.146766548 | 0.2834866 | 0.4834064 | C44B9.3 |
| WBGene00008081 | -0.141557471 | 0.12585278 | 0.28690032 | athp-1 |
| WBGene00008082 | 0.032221943 | 0.78722749 | 0.87949869 | com-1 |
| WBGene00008083 | 0.020925399 | 0.84643798 | NA | NA |
| WBGene00008084 | 0.080843675 | 0.69582511 | 0.82637698 | C44C10.3 |
| WBGene00008085 | -0.144386298 | 0.44253843 | 0.63652669 | smcl-1 |
| WBGene00008086 | -0.51933501 | 0.06029219 | 0.17200661 | C44C10.5 |
| WBGene00008087 | NA | NA | NA | C44C10.6 |
| WBGene00008088 | NA | NA | NA | C44C10.7 |
| WBGene00008089 | -0.125885159 | 0.47026909 | 0.65840969 | C44C10.9 |
| WBGene00008090 | NA | NA | NA | C44C10.10 |
| WBGene00008091 | -0.482723683 | 0.06732324 | 0.18608862 | C44F1.1 |
| WBGene00008092 | NA | NA | NA | gmeb-3 |
| WBGene00008093 | 0.040107227 | 0.58693163 | NA | lron-1 |
| WBGene00008094 | -0.289716319 | 0.06556487 | 0.18277517 | C44H4.4 |
| WBGene00008095 | 0.105469621 | 0.456805 | 0.64869855 | C44H4.6 |
| WBGene00008096 | 0.06854046 | 0.61533386 | 0.76921987 | C44H4.8 |
| WBGene00008097 | 0.022102615 | 0.86129549 | 0.92654132 | ugt-15 |
| WBGene00008098 | 0.038125098 | 0.61119482 | NA | NA |
| WBGene00008099 | 0.274065804 | 0.20980955 | 0.39924897 | C44H9.4 |
| WBGene00008100 | 0.626125474 | 0.00964519 | 0.04371986 | C44H9.5 |
| WBGene00008101 | 0.257670354 | 0.21803292 | 0.40988956 | C44H9.6 |
| WBGene00008102 | 4.084581475 | 0.00022132 | 0.00192193 | C44H9.7 |

| | | | | |
|---|---|---|---|---|
| WBGene00008104 | NA | NA | NA | NA |
| WBGene00008105 | 0.109591702 | 0.21566448 | NA | C45B11.2 |
| WBGene00008106 | NA | NA | NA | C45B11.5 |
| WBGene00008107 | -0.200744795 | 0.01594775 | 0.06496689 | aspm-1 |
| WBGene00008110 | 0.135197171 | 0.22644905 | 0.42004821 | C46C2.2 |
| WBGene00008111 | 0.282530242 | 0.13106376 | 0.29364202 | C46C2.3 |
| WBGene00008112 | NA | NA | NA | NA |
| WBGene00008113 | -0.627958926 | 0.00494369 | 0.02594283 | C46C2.5 |
| WBGene00008114 | 0.177674915 | 0.20420638 | 0.39328935 | C46C2.6 |
| WBGene00008117 | -0.089235642 | 0.16357754 | 0.33922644 | gsr-1 |
| WBGene00008118 | -0.001004325 | 0.99340367 | 0.99586212 | madf-8 |
| WBGene00008119 | 0.026890254 | 0.77733992 | 0.87398508 | C46F11.4 |
| WBGene00008120 | 0.120522839 | 0.3713215 | 0.57064323 | C46F11.5 |
| WBGene00008121 | 0.376198954 | 0.10536631 | 0.25397895 | C46F11.6 |
| WBGene00008122 | 0.017279211 | 0.79380376 | NA | C47A4.1 |
| WBGene00008123 | 0.097131274 | 0.58568574 | 0.74851824 | ent-4 |
| WBGene00008124 | 0.078203897 | 0.26561294 | NA | C47A4.3 |
| WBGene00008125 | 0.040107227 | 0.58693163 | NA | C47A4.5 |
| WBGene00008126 | NA | NA | NA | sre-21 |
| WBGene00008127 | 1.263864752 | 0.00010899 | 0.00104845 | ddo-1 |
| WBGene00008128 | 0.05376816 | 0.53681718 | NA | srab-12 |
| WBGene00008129 | 0.018476367 | 0.76699386 | NA | C47A10.12 |
| WBGene00008130 | 0.10836943 | 0.60169617 | 0.75908213 | fbxa-140 |
| WBGene00008131 | -0.187716875 | 0.06809018 | 0.18770633 | C47B2.2 |
| WBGene00008132 | 0.274660229 | 0.00382317 | 0.02112265 | gale-1 |
| WBGene00008133 | 0.348423358 | 0.04115117 | 0.13057958 | efsc-1 |
| WBGene00008134 | -0.034328978 | 0.83362568 | 0.90929864 | C47B2.9 |
| WBGene00008135 | NA | NA | NA | NA |
| WBGene00008136 | -0.143714099 | 0.06650772 | 0.18442035 | C47D12.2 |
| WBGene00008137 | 0.276631457 | 0.16061599 | 0.33512594 | sfxn-1.4 |
| WBGene00008138 | -0.514033949 | 0.0619767 | 0.17572849 | C47D12.4 |
| WBGene00008139 | 0.544342038 | 0.00536958 | 0.02767559 | C47D12.5 |
| WBGene00008140 | -0.029074526 | 0.78325949 | 0.87801104 | xpf-1 |
| WBGene00008141 | 0.033463284 | 0.6365037 | NA | C47E8.1 |
| WBGene00008142 | 0.164250871 | 0.25533163 | 0.45376287 | C47E8.3 |
| WBGene00008143 | -0.221460092 | 0.0720959 | 0.19550956 | C47E8.4 |
| WBGene00008144 | -0.050921541 | 0.47213035 | NA | gasr-8 |
| WBGene00008145 | -0.148327412 | 0.23964202 | 0.43670695 | set-5 |
| WBGene00008146 | 0.063421086 | 0.41095452 | NA | C47E8.9 |
| WBGene00008147 | -0.061701501 | 0.55114744 | 0.7237085 | C47E12.2 |
| WBGene00008148 | -0.1355145 | 0.17324974 | 0.35347679 | C47E12.3 |
| WBGene00008149 | -0.02488294 | 0.7285553 | 0.84853595 | pyp-1 |
| WBGene00008150 | 0.146240597 | 0.12970301 | 0.29201626 | mam-2 |

| | | | | | |
|---|---|---|---|---|---|
| WBGene00008151 | | -0.319066171 | 0.01247121 | 0.05332682 | rrp-1 |
| WBGene00008152 | | 2.034391004 | 0.0013957 | 0.0092589 | C47E12.9 |
| WBGene00008153 | | -0.046394786 | 0.50231372 | NA | C47E12.10 |
| WBGene00008154 | | 0.03732969 | 0.59229443 | NA | C47E12.11 |
| WBGene00008155 | | -0.023056499 | 0.78121253 | NA | C47E12.12 |
| WBGene00008156 | NA | | NA | NA | C47E12.13 |
| WBGene00008157 | | 0.188293707 | 0.38352381 | 0.58321273 | C47F8.1 |
| WBGene00008158 | | -0.023056499 | 0.78121253 | NA | nhr-165 |
| WBGene00008159 | NA | | NA | NA | C47F8.3 |
| WBGene00008160 | | 0.020925399 | 0.84643798 | NA | glct-4 |
| WBGene00008161 | NA | | NA | NA | C47F8.5 |
| WBGene00008162 | NA | | NA | NA | C47F8.6 |
| WBGene00008163 | | 0.015584271 | 0.80362925 | NA | C47F8.7 |
| WBGene00008164 | | -0.041814953 | 0.75472048 | 0.86091644 | C47G2.3 |
| WBGene00008165 | | -0.073217518 | 0.41586617 | 0.61310005 | C47G2.4 |
| WBGene00008166 | | -0.161699531 | 0.01896454 | 0.07398343 | saps-1 |
| WBGene00008167 | | 1.478798706 | 9.60E-19 | 6.48E-17 | acox-1.5 |
| WBGene00008168 | | 0.148402091 | 0.11646869 | 0.27261681 | C48B4.3 |
| WBGene00008169 | | -0.169524664 | 0.25915965 | 0.45720647 | C48B4.6 |
| WBGene00008170 | | 0.144461262 | 0.32881691 | 0.52822125 | C48B4.7 |
| WBGene00008171 | | -0.16566984 | 0.35834215 | 0.55876045 | C48B4.8 |
| WBGene00008172 | | -0.196498702 | 0.17936673 | 0.36082247 | C48B4.9 |
| WBGene00008173 | | -0.22777277 | 0.10626663 | 0.25568726 | C48B4.10 |
| WBGene00008174 | | -0.224681853 | 0.03261423 | 0.10999882 | C48B4.11 |
| WBGene00008175 | | 0.0398788 | 0.58971221 | NA | C48B4.12 |
| WBGene00008176 | | -0.039590353 | 0.73555761 | 0.85166527 | C48B4.13 |
| WBGene00008177 | | -0.159599823 | 0.44685286 | 0.64010124 | C48D1.1 |
| WBGene00008178 | | 0.398022376 | 0.10854619 | 0.25911216 | C48D1.5 |
| WBGene00008179 | NA | | NA | NA | NA |
| WBGene00008180 | | 0.05787416 | 0.40594086 | NA | C48D5.3 |
| WBGene00008183 | | -0.001013025 | 0.9887283 | 0.99380144 | rin-1 |
| WBGene00008184 | | -0.003270206 | 0.95698316 | NA | C49A1.1 |
| WBGene00008185 | NA | | NA | NA | best-10 |
| WBGene00008186 | | -0.006388227 | 0.92874308 | NA | best-11 |
| WBGene00008187 | NA | | NA | NA | C49A1.5 |
| WBGene00008188 | NA | | NA | NA | clec-115 |
| WBGene00008190 | NA | | NA | NA | clec-114 |
| WBGene00008191 | | 0.520019792 | 0.02527439 | 0.09143621 | C49A1.10 |
| WBGene00008192 | | 0.140933312 | 0.36270577 | 0.56256996 | NA |
| WBGene00008193 | | -0.023056499 | 0.78121253 | NA | NA |
| WBGene00008194 | | 1.101765853 | 3.28E-37 | 5.39E-35 | C49C3.4 |
| WBGene00008195 | | 0.208921254 | 0.33010851 | 0.52954078 | ceh-88 |
| WBGene00008196 | | -0.066780621 | 0.55518475 | 0.72703012 | C49C3.6 |

| | | | | |
|---|---|---|---|---|
| WBGene00008197 | -0.190159937 | 0.08648309 | 0.22217016 | C49C3.7 |
| WBGene00008198 | -0.097160411 | 0.63898839 | 0.78584032 | C49C3.8 |
| WBGene00008199 | 0.133440854 | 0.4766439 | 0.66386447 | C49C3.9 |
| WBGene00008200 | 0.198145291 | 0.33868005 | 0.53821932 | C49C3.10 |
| WBGene00008201 | -0.040794878 | 0.53203165 | NA | C49C3.11 |
| WBGene00008202 | 0.059980168 | 0.38780842 | NA | clec-197 |
| WBGene00008203 | -0.053284153 | 0.73611741 | 0.85184122 | clec-198 |
| WBGene00008204 | 0.192897351 | 0.37273481 | 0.57191473 | C49C3.15 |
| WBGene00008205 | 0.420653567 | 1.16E-11 | 3.81E-10 | sams-1 |
| WBGene00008206 | -0.073073925 | 0.69837003 | 0.82825724 | set-6 |
| WBGene00008207 | -0.080409231 | 0.57648655 | 0.74357674 | C49F5.3 |
| WBGene00008208 | 0.107186719 | 0.31444014 | 0.51495727 | nhr-167 |
| WBGene00008209 | -0.022930767 | 0.78121253 | NA | C49F5.5 |
| WBGene00008210 | -0.029651409 | 0.80139193 | 0.88934884 | C49F5.6 |
| WBGene00008211 | -0.020866305 | 0.86764013 | 0.92947386 | C49F5.7 |
| WBGene00008212 | -0.081511571 | 0.51230035 | 0.69212785 | C49F5.8 |
| WBGene00008213 | 0.975301422 | 0.00577132 | 0.02932934 | C49F8.1 |
| WBGene00008214 | 0.077422651 | 0.69480389 | 0.82536501 | gem-1 |
| WBGene00008215 | 0.298978162 | 0.16664126 | 0.34363304 | C49F8.3 |
| WBGene00008216 | 0.059403107 | 0.68419169 | 0.8176383 | mec-19 |
| WBGene00008217 | NA | NA | NA | C50B6.1 |
| WBGene00008218 | -0.133335693 | 0.00064184 | 0.0048166 | nasp-2 |
| WBGene00008219 | -0.191553644 | 0.01598114 | 0.06504861 | C50B6.3 |
| WBGene00008220 | 1.305067406 | 7.41E-13 | 2.81E-11 | C50B6.7 |
| WBGene00008221 | 1.899320987 | 1.09E-05 | 0.00013374 | nhr-168 |
| WBGene00008222 | -0.02166915 | 0.89990465 | 0.94665211 | C50B6.9 |
| WBGene00008223 | NA | NA | NA | lgc-48 |
| WBGene00008224 | 0.160513097 | 0.10261288 | 0.24964361 | slrp-1 |
| WBGene00008225 | 0.084959629 | 0.56114823 | 0.73159138 | nuaf-1 |
| WBGene00008226 | 0.220794385 | 0.22984858 | 0.42425231 | C50B8.4 |
| WBGene00008227 | 0.283514174 | 0.21528948 | 0.40639906 | C50B8.5 |
| WBGene00008228 | 0.225018284 | 0.29460993 | 0.49569181 | C50B8.6 |
| WBGene00008229 | 0.744252995 | 5.65E-07 | 9.39E-06 | C50F4.1 |
| WBGene00008230 | 0.107005324 | 0.31506195 | 0.51557662 | pfk-1.2 |
| WBGene00008231 | 0.042065281 | 0.558937 | NA | tag-329 |
| WBGene00008232 | 0.101917188 | 0.62009555 | 0.77270574 | C50F4.4 |
| WBGene00008233 | 0.61687689 | 0.00102075 | 0.00715578 | C50F4.8 |
| WBGene00008234 | 1.541766687 | 0.00459869 | 0.02454589 | C50F4.9 |
| WBGene00008235 | 0.142225333 | 0.07970925 | NA | C50F4.10 |
| WBGene00008236 | -0.048930905 | 0.69651673 | 0.82690507 | C50F4.12 |
| WBGene00008237 | -0.158080053 | 0.08543712 | 0.22030613 | nstp-10 |
| WBGene00008238 | 0.068198802 | 0.54355536 | 0.71696212 | C50F4.16 |
| WBGene00008239 | 0.250881604 | 0.10489728 | 0.25330711 | fshr-1 |

| | | | | |
|---|---|---|---|---|
| WBGene00008240 | 0.018795205 | 0.77532208 | NA | C50H2.4 |
| WBGene00008241 | -0.023109762 | 0.78121253 | NA | C50H2.5 |
| WBGene00008242 | NA | NA | NA | ceh-75 |
| WBGene00008243 | -0.027373974 | 0.68245543 | NA | C50H2.7 |
| WBGene00008244 | -0.105667918 | 0.21716877 | NA | fipr-11 |
| WBGene00008245 | -0.064768995 | 0.62851992 | 0.77898587 | fipr-10 |
| WBGene00008246 | 1.17801777 | 0.00380736 | 0.02106708 | C50H2.13 |
| WBGene00008247 | -0.003614866 | 0.95700117 | NA | srsx-26 |
| WBGene00008248 | -0.03323923 | 0.69122143 | NA | srsx-27 |
| WBGene00008249 | 0.020925399 | 0.84643798 | NA | srsx-28 |
| WBGene00008250 | NA | NA | NA | srsx-29 |
| WBGene00008251 | -0.022729726 | 0.70048408 | NA | srsx-30 |
| WBGene00008252 | 0.080142712 | 0.68511628 | 0.81830567 | C51E3.6 |
| WBGene00008253 | NA | NA | NA | NA |
| WBGene00008254 | 0.202729016 | 0.26742106 | 0.46635384 | C51E3.9 |
| WBGene00008255 | 0.020512425 | 0.92038737 | 0.95550377 | C51E3.10 |
| WBGene00008256 | 0.119757623 | 0.26107051 | 0.45919656 | glb-12 |
| WBGene00008257 | NA | NA | NA | C52A11.3 |
| WBGene00008258 | 0.015020554 | 0.91075495 | 0.95112162 | mans-2 |
| WBGene00008259 | 0.077524279 | 0.40780354 | 0.60572706 | C52E4.7 |
| WBGene00008260 | -0.311935354 | 0.14878994 | 0.31960586 | C52G5.2 |
| WBGene00008261 | NA | NA | NA | C52G5.3 |
| WBGene00008262 | -0.160980792 | 0.00511631 | 0.02661027 | ril-1 |
| WBGene00008263 | -0.003306275 | 0.98001596 | 0.98845506 | C53A5.2 |
| WBGene00008265 | 1.046643714 | 0.01461163 | 0.06056877 | kcnl-4 |
| WBGene00008266 | -0.228555629 | 0.00966942 | 0.04378903 | C53A5.6 |
| WBGene00008267 | 0.020232312 | 0.84643798 | NA | C53A5.9 |
| WBGene00008268 | NA | NA | NA | srt-32 |
| WBGene00008269 | 0.033034883 | 0.64219503 | NA | C53A5.11 |
| WBGene00008270 | -0.031282667 | 0.86861465 | 0.93000745 | C53A5.13 |
| WBGene00008271 | 0.13169099 | 0.16083826 | NA | C53B4.1 |
| WBGene00008272 | 0.03555743 | 0.72324271 | 0.84472009 | C53B4.2 |
| WBGene00008273 | -0.277803534 | 0.175711 | 0.35646179 | C53B4.3 |
| WBGene00008274 | 0.00337407 | 0.96524165 | 0.98071499 | C53B4.4 |
| WBGene00008275 | -0.021615443 | 0.89363476 | 0.94468695 | nstp-1 |
| WBGene00008277 | 0.188290841 | 0.22180762 | 0.41448442 | mltn-12 |
| WBGene00008278 | 0.006679547 | 0.97309989 | 0.98528046 | npr-10 |
| WBGene00008279 | -0.131188188 | 0.08273944 | NA | C53C7.3 |
| WBGene00008280 | 0.078236818 | 0.47921183 | 0.66615262 | acc-2 |
| WBGene00008281 | -0.290013409 | 0.0230959 | 0.08534299 | C53D6.4 |
| WBGene00008282 | 0.033034883 | 0.64219503 | NA | NA |
| WBGene00008283 | -0.245593878 | 0.22761065 | 0.42118901 | C53D6.6 |
| WBGene00008284 | 0.027601861 | 0.88457137 | 0.93922761 | C53D6.7 |

| | | | | |
|---|---|---|---|---|
| WBGene00008285 | -0.023056499 | 0.78121253 | NA | C53D6.8 |
| WBGene00008286 | NA | NA | NA | C54C6.4 |
| WBGene00008287 | 0.125676313 | 0.47365391 | 0.66152502 | C54C6.5 |
| WBGene00008288 | 0.031298586 | 0.84592235 | 0.9170788 | C54C6.6 |
| WBGene00008289 | -0.027188909 | 0.68205301 | NA | nhr-169 |
| WBGene00008290 | NA | NA | NA | bgnt-1.8 |
| WBGene00008291 | NA | NA | NA | C54C8.3 |
| WBGene00008292 | 0.020925399 | 0.84643798 | NA | C54C8.4 |
| WBGene00008293 | NA | NA | NA | glct-5 |
| WBGene00008294 | NA | NA | NA | clec-11 |
| WBGene00008295 | 0.108942554 | 0.15594412 | NA | nlp-39 |
| WBGene00008296 | 1.118563376 | 3.72E-06 | 5.02E-05 | cdr-2 |
| WBGene00008297 | NA | NA | NA | cdr-3 |
| WBGene00008298 | 1.46912814 | 4.04E-37 | 6.57E-35 | C54D10.3 |
| WBGene00008299 | 0.168170932 | 0.05190965 | NA | C54D10.4 |
| WBGene00008300 | 0.130176683 | 0.3191215 | 0.51931286 | C54D10.5 |
| WBGene00008301 | 4.737761555 | 2.85E-05 | 0.00031312 | pals-39 |
| WBGene00008302 | 0.202375559 | 0.0697073 | 0.19079913 | pals-38 |
| WBGene00008303 | 0.131026573 | 0.24406054 | 0.44207477 | C54D10.9 |
| WBGene00008304 | 0.293599443 | 0.15384862 | 0.32713026 | C54D10.10 |
| WBGene00008306 | 0.020232312 | 0.84643798 | NA | C54E10.1 |
| WBGene00008307 | 0.191996912 | 0.15872901 | 0.3330875 | ncs-5 |
| WBGene00008308 | NA | NA | NA | C54E10.3 |
| WBGene00008309 | 0.52100251 | 0.03751083 | 0.12209782 | nhr-170 |
| WBGene00008310 | 0.175617814 | 0.36098166 | 0.56083865 | C54E10.6 |
| WBGene00008311 | 0.598667519 | 6.32E-07 | 1.04E-05 | rskn-2 |
| WBGene00008312 | 0.350818909 | 0.00635585 | 0.03157649 | C54G4.2 |
| WBGene00008313 | 0.127792695 | 0.25173677 | 0.44993917 | C54G4.3 |
| WBGene00008314 | 2.65592656 | 0.0047823 | 0.0252541 | C54G4.4 |
| WBGene00008315 | 0.034173114 | 0.81821162 | 0.90007974 | C54G4.5 |
| WBGene00008316 | -0.068861987 | 0.73840563 | 0.85304428 | dod-18 |
| WBGene00008317 | 0.030961904 | 0.8248584 | 0.90387067 | C54G4.7 |
| WBGene00008318 | -0.036970036 | 0.78436245 | 0.87847449 | C54G4.9 |
| WBGene00008319 | 0.019325396 | 0.77340931 | NA | C54G10.1 |
| WBGene00008320 | 0.660074484 | 0.01650886 | 0.06671438 | slc-25A29 |
| WBGene00008321 | NA | NA | NA | C55A1.1 |
| WBGene00008322 | NA | NA | NA | C55A1.4 |
| WBGene00008322 | NA | NA | NA | C55A1.7 |
| WBGene00008323 | 0.09250164 | 0.53370556 | 0.70904186 | C55A1.6 |
| WBGene00008324 | NA | NA | NA | C55A1.4 |
| WBGene00008324 | NA | NA | NA | C55A1.7 |
| WBGene00008325 | NA | NA | NA | C55A1.9 |
| WBGene00008326 | NA | NA | NA | C55A1.11 |

| | | | | |
|---|---|---|---|---|
| WBGene00008327 | NA | NA | NA | srz-44 |
| WBGene00008328 | 0.065014541 | 0.37399213 | NA | srz-43 |
| WBGene00008330 | -0.194511086 | 0.03161986 | 0.10728769 | C55A6.1 |
| WBGene00008331 | 0.055721228 | 0.53110943 | 0.70766758 | ttll-5 |
| WBGene00008332 | -0.085184386 | 0.65180762 | 0.79387034 | C55A6.3 |
| WBGene00008333 | -0.09389486 | 0.64106421 | 0.78733505 | C55A6.4 |
| WBGene00008334 | -0.010101906 | 0.95349624 | 0.97344247 | sdz-8 |
| WBGene00008335 | 0.077228947 | 0.70797955 | 0.83523278 | C55A6.6 |
| WBGene00008336 | 0.238568084 | 0.28578773 | 0.48591548 | C55A6.7 |
| WBGene00008337 | NA | NA | NA | NA |
| WBGene00008338 | -0.247235941 | 0.00139741 | 0.00926605 | pafo-1 |
| WBGene00008339 | -0.314134324 | 0.01190933 | 0.05129876 | C55A6.10 |
| WBGene00008340 | 0.041444886 | 0.72659604 | 0.84711678 | C55A6.11 |
| WBGene00008341 | 0.145269483 | 0.42063663 | 0.61727382 | ttr-44 |
| WBGene00008342 | 0.026083459 | 0.89731819 | 0.9459022 | frpr-5 |
| WBGene00008343 | -0.093454557 | 0.47687421 | 0.66409108 | C56A3.4 |
| WBGene00008344 | -0.2051202 | 0.08168091 | 0.21393385 | C56A3.5 |
| WBGene00008345 | -0.102502977 | 0.43598371 | 0.63111939 | C56A3.6 |
| WBGene00008346 | -0.116126776 | 0.5144935 | 0.69445002 | C56A3.8 |
| WBGene00008348 | 0.180587198 | 0.39114691 | 0.58911449 | C56G7.3 |
| WBGene00008349 | 0.027818693 | 0.89027981 | 0.94282725 | CE7X_3.1 |
| WBGene00008350 | 0.327739215 | 0.12318076 | 0.28303351 | NA |
| WBGene00008351 | -0.002914971 | 0.95718083 | NA | NA |
| WBGene00008352 | 0.453224483 | 0.06524028 | 0.18207833 | fbxa-6 |
| WBGene00008353 | -0.078002411 | 0.69982621 | 0.8293796 | D1025.1 |
| WBGene00008354 | -0.786389276 | 0.01337307 | 0.05645763 | gcsh-1 |
| WBGene00008355 | 0.04002455 | 0.78340394 | 0.87807405 | fbxa-141 |
| WBGene00008356 | 1.352192169 | 2.83E-12 | 1.00E-10 | nspc-20 |
| WBGene00008357 | 3.050274791 | 2.45E-11 | 7.69E-10 | nspc-16 |
| WBGene00008358 | 1.548986532 | 9.59E-07 | 1.52E-05 | nspc-18 |
| WBGene00008359 | 1.876323279 | 1.40E-08 | 3.05E-07 | nspc-17 |
| WBGene00008360 | 1.521462816 | 1.65E-05 | 0.00019215 | nspc-19 |
| WBGene00008361 | 0.185177904 | 0.03841264 | 0.1241575 | D1043.1 |
| WBGene00008362 | -0.219084612 | 0.01928085 | 0.07487821 | cfim-2 |
| WBGene00008363 | -0.163708763 | 0.13512871 | 0.29966971 | D1046.2 |
| WBGene00008364 | 0.052802611 | 0.71685616 | 0.84118707 | slc-25A26 |
| WBGene00008365 | 0.277006836 | 0.20693441 | 0.39624772 | D1046.4 |
| WBGene00008366 | 1.395628188 | 6.19E-08 | 1.21E-06 | tpra-1 |
| WBGene00008367 | NA | NA | NA | D1053.2 |
| WBGene00008368 | 0.636466558 | 0.03873788 | 0.12500801 | D1053.3 |
| WBGene00008369 | 0.146879174 | 0.44448072 | 0.63794923 | D1053.4 |
| WBGene00008370 | 0.184813232 | 0.29869431 | 0.49942984 | D1054.1 |
| WBGene00008371 | -0.213516121 | 0.01308149 | 0.05546656 | D1054.3 |

| | | | | |
|---|---|---|---|---|
| WBGene00008372 | 0.089092193 | 0.67056721 | 0.80824686 | D1054.5 |
| WBGene00008373 | -0.067064794 | 0.32104386 | NA | NA |
| WBGene00008374 | 0.108717649 | 0.15669083 | NA | NA |
| WBGene00008375 | 0.972373503 | 3.75E-10 | 9.94E-09 | D1054.8 |
| WBGene00008376 | 2.341117405 | 0.00683087 | 0.03341488 | D1054.9 |
| WBGene00008377 | -0.027898317 | 0.60285535 | 0.75988622 | D1054.10 |
| WBGene00008378 | 0.098324659 | 0.06512111 | 0.18184975 | ule-3 |
| WBGene00008379 | 0.062571732 | 0.6745808 | 0.81103194 | secs-1 |
| WBGene00008380 | -0.046689447 | 0.65936643 | 0.79896991 | prp-38 |
| WBGene00008381 | 0.020232312 | 0.84643798 | NA | D1081.3 |
| WBGene00008382 | -0.008303499 | 0.92600956 | NA | D1081.4 |
| WBGene00008383 | 0.260273078 | 0.03986972 | 0.12764763 | D1081.5 |
| WBGene00008384 | -0.339993807 | 0.1032378 | 0.2507873 | D1081.6 |
| WBGene00008385 | -0.109566927 | 0.05509872 | 0.16159863 | D1081.7 |
| WBGene00008386 | -0.161287793 | 0.00772368 | 0.03675087 | cdc-5L |
| WBGene00008387 | -0.156679598 | 0.33346945 | 0.53304196 | zhp-2 |
| WBGene00008388 | 0.495111003 | 0.02084964 | 0.07913473 | D1086.1 |
| WBGene00008389 | 1.135537187 | 0.00820172 | 0.0385123 | D1086.2 |
| WBGene00008390 | -2.350838769 | 3.53E-24 | 3.15E-22 | D1086.3 |
| WBGene00008392 | 0.549230066 | 0.04037616 | 0.1289026 | D1086.5 |
| WBGene00008393 | -5.311377312 | 1.89E-17 | 1.16E-15 | D1086.6 |
| WBGene00008394 | 0.382722902 | 0.0026562 | 0.01577663 | D1086.7 |
| WBGene00008395 | 0.026926593 | 0.89753125 | 0.94605868 | D1086.8 |
| WBGene00008396 | NA | NA | NA | D1086.9 |
| WBGene00008397 | NA | NA | NA | D2005.1 |
| WBGene00008398 | -0.072517511 | 0.52133779 | 0.69975307 | D2005.3 |
| WBGene00008399 | -0.151107047 | 0.16572877 | 0.34228169 | D2005.4 |
| WBGene00008400 | -0.164942221 | 0.03812494 | 0.12349277 | drh-3 |
| WBGene00008401 | 0.203966906 | 0.31597618 | 0.51660605 | D2005.6 |
| WBGene00008402 | -0.111304747 | 0.24746234 | 0.44591953 | wdfy-2 |
| WBGene00008403 | -0.391907345 | 0.10631952 | 0.25568726 | csa-1 |
| WBGene00008404 | -0.109199063 | 0.48195778 | 0.66787768 | D2013.6 |
| WBGene00008405 | -0.144298875 | 0.10439107 | 0.25259961 | ttll-12 |
| WBGene00008407 | 0.230019907 | 0.15395198 | 0.32715947 | D2023.3 |
| WBGene00008408 | -0.115817907 | 0.40482829 | 0.60295941 | D2023.4 |
| WBGene00008409 | -0.291813531 | 0.02027575 | 0.07740737 | mpst-1 |
| WBGene00008410 | -0.038081347 | 0.69203147 | 0.82320708 | D2023.6 |
| WBGene00008411 | -0.11244717 | 0.1648719 | 0.34108936 | mans-1 |
| WBGene00008412 | 0.515251189 | 0.0066405 | 0.0326912 | D2030.2 |
| WBGene00008413 | -0.118857223 | 0.2790214 | 0.47882175 | D2030.3 |
| WBGene00008414 | -0.36468745 | 7.16E-05 | 0.00072181 | D2030.4 |
| WBGene00008415 | -0.417587315 | 0.00083799 | 0.00605279 | mce-1 |
| WBGene00008417 | -0.209039868 | 0.02311172 | 0.08534299 | D2030.7 |

| | | | | |
|---|---|---|---|---|
| WBGene00008418 | -0.177679229 | 0.05853063 | 0.16849851 | D2030.8 |
| WBGene00008419 | 0.252302264 | 0.01104358 | 0.04835201 | wdr-23 |
| WBGene00008420 | -0.301670012 | 0.09768409 | 0.24215929 | D2030.11 |
| WBGene00008421 | 0.164565518 | 0.36702797 | 0.56666208 | D2030.12 |
| WBGene00008422 | -0.218625181 | 0.00446133 | 0.02395247 | D2045.2 |
| WBGene00008423 | 0.12249296 | 0.3088748 | 0.50968174 | D2045.5 |
| WBGene00008424 | 0.150668559 | 0.4800516 | 0.66658152 | D2045.7 |
| WBGene00008425 | 1.518152857 | 7.34E-06 | 9.30E-05 | D2045.8 |
| WBGene00008426 | -0.265029708 | 0.20678761 | 0.3960724 | D2045.9 |
| WBGene00008427 | 0.063293301 | 0.75358454 | 0.86049222 | atg-10 |
| WBGene00008428 | -0.108039008 | 0.3427289 | 0.54300333 | eif-2Bepsilon |
| WBGene00008429 | -0.07436562 | 0.39956607 | 0.59782443 | etc-1 |
| WBGene00008430 | 0.180386436 | 0.03237708 | 0.1094006 | hgap-2 |
| WBGene00008431 | 0.001277975 | 0.99334577 | 0.99586212 | piga-1 |
| WBGene00008432 | 0.177775596 | 0.40350856 | 0.60185094 | D2085.7 |
| WBGene00008433 | 0.098822211 | 0.38506743 | 0.58434621 | 2-Mar |
| WBGene00008434 | 0.464754542 | 0.033969 | 0.11334213 | D2089.3 |
| WBGene00008435 | 0.020303396 | 0.89698291 | 0.94582116 | glna-2 |
| WBGene00008436 | 0.582458474 | 2.87E-06 | 3.97E-05 | DH11.2 |
| WBGene00008438 | 0.98539769 | 8.86E-05 | 0.00086953 | DH11.5 |
| WBGene00008439 | 0.678520688 | 0.00503866 | 0.02628107 | mfb-1 |
| WBGene00008440 | -0.09249276 | 0.24363431 | 0.44163915 | DY3.8 |
| WBGene00008441 | -0.037837143 | 0.74160819 | 0.85407737 | usp-50 |
| WBGene00008442 | -0.194106061 | 0.22830149 | 0.42198739 | E01B7.2 |
| WBGene00008443 | 0.491998442 | 0.02367071 | 0.08694502 | pde-3 |
| WBGene00008444 | 0.182661045 | 0.06648114 | 0.18438158 | tbc-14 |
| WBGene00008446 | -0.012322166 | 0.92604724 | 0.95834417 | E01G4.3 |
| WBGene00008447 | -0.839922396 | 0.00180476 | 0.01146353 | E01G4.5 |
| WBGene00008448 | 2.077477567 | 0.00070402 | 0.00519825 | E01G4.6 |
| WBGene00008449 | 4.294249843 | 0.00088614 | 0.0063598 | E01G6.1 |
| WBGene00008450 | 0.391345918 | 0.11528983 | 0.27076651 | E01G6.2 |
| WBGene00008451 | 0.240703186 | 0.28127487 | 0.48075477 | E01G6.3 |
| WBGene00008452 | -0.012787727 | 0.89963939 | 0.94665211 | mrps-5 |
| WBGene00008453 | 0.764439148 | 0.02228006 | 0.08315891 | E02A10.3 |
| WBGene00008454 | 0.117823748 | 0.50219193 | 0.68364407 | E02A10.4 |
| WBGene00008455 | -0.12360107 | 0.44371749 | 0.63728347 | E02H1.1 |
| WBGene00008456 | -0.204652167 | 0.09110915 | 0.22960929 | eral-1 |
| WBGene00008457 | -0.056167815 | 0.68605126 | 0.81895447 | E02H1.5 |
| WBGene00008458 | -0.225985876 | 0.15411446 | 0.32715947 | E02H1.6 |
| WBGene00008459 | -0.228234 | 0.11723201 | 0.27380421 | mrpl-53 |
| WBGene00008460 | -0.098797583 | 0.56506429 | 0.73479938 | E02H4.2 |
| WBGene00008462 | 0.212034572 | 0.14700907 | 0.31731679 | E02H4.4 |
| WBGene00008463 | 0.037127706 | 0.59421403 | NA | E02H4.5 |

| | | | | |
|---|---|---|---|---|
| WBGene00008464 | -0.064792683 | 0.64434793 | 0.78947268 | E02H4.6 |
| WBGene00008465 | -0.027535543 | 0.68063395 | NA | NA |
| WBGene00008468 | 0.020925399 | 0.84643798 | NA | E03G2.1 |
| WBGene00008470 | NA | NA | NA | E03H4.2 |
| WBGene00008471 | NA | NA | NA | E03H4.3 |
| WBGene00008472 | NA | NA | NA | E03H4.4 |
| WBGene00008473 | NA | NA | NA | E03H4.5 |
| WBGene00008474 | 0.033463284 | 0.6365037 | NA | nhr-174 |
| WBGene00008475 | 0.020232312 | 0.84643798 | NA | oac-13 |
| WBGene00008476 | 1.311861484 | 5.90E-06 | 7.59E-05 | E03H4.8 |
| WBGene00008477 | 5.461186839 | 5.08E-05 | 0.00053315 | clec-17 |
| WBGene00008478 | -0.023056499 | 0.78121253 | NA | E03H4.11 |
| WBGene00008479 | NA | NA | NA | glct-2 |
| WBGene00008480 | -0.22295977 | 1.11E-05 | 0.00013529 | eif-2A |
| WBGene00008481 | 0.089449525 | 0.57695706 | 0.74401444 | E04D5.2 |
| WBGene00008482 | -0.021481305 | 0.89990179 | 0.94665211 | cut-4 |
| WBGene00008483 | 0.187499109 | 0.30497476 | 0.50562059 | E04D5.4 |
| WBGene00008484 | 0.526819408 | 0.05503702 | 0.16145 | E04D5.5 |
| WBGene00008485 | 0.250620912 | 0.24777883 | 0.44611982 | ugt-43 |
| WBGene00008486 | -0.89715428 | 8.02E-06 | 0.00010067 | ugt-44 |
| WBGene00008487 | 0.015451648 | 0.80337623 | NA | NA |
| WBGene00008488 | -0.271456168 | 0.03811 | 0.12347169 | F01D4.5 |
| WBGene00008489 | 0.015908114 | 0.8416225 | NA | srb-17 |
| WBGene00008490 | 0.008755578 | 0.93314632 | 0.96246532 | F01D4.8 |
| WBGene00008491 | -0.053591705 | 0.45278076 | NA | bgnt-1.1 |
| WBGene00008492 | 2.900968046 | 5.07E-16 | 2.65E-14 | F01D5.1 |
| WBGene00008493 | -0.205014568 | 0.27934978 | 0.47919861 | F01D5.2 |
| WBGene00008494 | -0.419020518 | 0.09455838 | 0.23584302 | F01D5.3 |
| WBGene00008495 | 2.913213889 | 2.55E-13 | 1.03E-11 | F01D5.5 |
| WBGene00008496 | 0.308076585 | 0.05276115 | 0.15675243 | F01D5.6 |
| WBGene00008497 | NA | NA | NA | F01D5.7 |
| WBGene00008498 | 0.415830633 | 0.09862395 | 0.24347628 | F01D5.8 |
| WBGene00008499 | 0.347016811 | 0.08510865 | 0.21988278 | cyp-37A1 |
| WBGene00008500 | 0.059711191 | 0.42255368 | NA | F01D5.10 |
| WBGene00008501 | NA | NA | NA | NA |
| WBGene00008502 | -0.182267769 | 0.07949023 | 0.21018899 | skih-2 |
| WBGene00008503 | -0.183097144 | 0.07671036 | 0.20493339 | F01G4.4 |
| WBGene00008504 | 0.198828581 | 0.23847979 | 0.43546802 | pigq-1 |
| WBGene00008505 | -0.125793411 | 0.00359154 | 0.02017794 | F01G4.6 |
| WBGene00008506 | -0.059474529 | 0.34484344 | 0.54458556 | tkt-1 |
| WBGene00008507 | 0.020925399 | 0.84643798 | NA | F01G10.4 |
| WBGene00008508 | 0.207958431 | 0.06563442 | 0.18286028 | F01G10.5 |
| WBGene00008509 | 0.084162846 | 0.39428309 | 0.59243625 | F01G10.6 |

| | | | | |
|---|---|---|---|---|
| WBGene00008510 | 0.289320669 | 0.09237503 | 0.23194354 | lipl-7 |
| WBGene00008511 | -0.951029042 | 0.01343657 | 0.05667672 | F01G10.9 |
| WBGene00008512 | 0.236561341 | 0.25037139 | 0.44895131 | F01G10.10 |
| WBGene00008513 | 0.234869228 | 0.15504528 | 0.32821944 | F02A9.1 |
| WBGene00008514 | -0.267237085 | 0.02913882 | 0.10149284 | F02A9.10 |
| WBGene00008515 | 0.000186237 | 0.99950191 | 0.99981268 | F02C12.1 |
| WBGene00008516 | 0.056677809 | 0.43780437 | NA | F02C12.2 |
| WBGene00008517 | -0.035554943 | 0.7427605 | 0.85444405 | F02C12.3 |
| WBGene00008518 | 0.033144996 | 0.68360425 | NA | irld-2 |
| WBGene00008519 | -0.357547651 | 0.0861209 | 0.22169907 | cyp-13B1 |
| WBGene00008520 | 0.020232312 | 0.84643798 | NA | F02D8.1 |
| WBGene00008521 | 0.565292851 | 0.00011036 | 0.00105882 | F02D8.4 |
| WBGene00008522 | 0.65370715 | 0.03606499 | 0.11849809 | nlp-59 |
| WBGene00008523 | NA | NA | NA | F02D10.2 |
| WBGene00008524 | 0.020232312 | 0.84643798 | NA | F02D10.3 |
| WBGene00008526 | -0.471956106 | 0.0540317 | 0.15928466 | F02D10.6 |
| WBGene00008527 | NA | NA | NA | set-8 |
| WBGene00008528 | 0.037441967 | 0.80781311 | 0.89349488 | F02E9.1 |
| WBGene00008529 | 0.072492309 | 0.64402285 | 0.78931424 | F02E9.3 |
| WBGene00008530 | 0.194729581 | 0.20952453 | 0.39896587 | F02E9.5 |
| WBGene00008531 | -0.112252052 | 0.32346064 | 0.52329307 | F02E9.7 |
| WBGene00008532 | -0.282115897 | 0.00011498 | 0.00109808 | dpt-1 |
| WBGene00008533 | -0.223854335 | 0.03288187 | 0.11064664 | F02E9.10 |
| WBGene00008534 | -0.089727974 | 0.65928852 | 0.79896991 | NA |
| WBGene00008535 | -0.121099487 | 0.3287093 | 0.52822125 | F02H6.2 |
| WBGene00008536 | 0.127413407 | 0.31669161 | 0.51698682 | F02H6.3 |
| WBGene00008537 | 0.024446197 | 0.86904224 | 0.93000745 | NA |
| WBGene00008538 | -0.762476894 | 0.01264956 | 0.05399482 | sqrd-1 |
| WBGene00008539 | NA | NA | NA | F02H6.6 |
| WBGene00008540 | NA | NA | NA | F02H6.7 |
| WBGene00008541 | 2.213184099 | 0.0001952 | 0.00173211 | F07A5.2 |
| WBGene00008542 | 0.164245343 | 0.38304071 | 0.58260911 | scav-6 |
| WBGene00008543 | 0.206424592 | 0.15348991 | 0.32677064 | F07A5.4 |
| WBGene00008545 | -0.012132735 | 0.93874975 | 0.96565974 | F07A11.1 |
| WBGene00008546 | 0.004799949 | 0.93216378 | 0.96185847 | gfat-1 |
| WBGene00008547 | 0.055388715 | 0.74722821 | 0.85713477 | F07A11.4 |
| WBGene00008548 | -0.110804459 | 0.53700181 | 0.71191877 | F07A11.5 |
| WBGene00008549 | 0.188237745 | 0.05832346 | 0.16822305 | din-1 |
| WBGene00008550 | 0.071566689 | 0.49302248 | 0.67606464 | F07B10.4 |
| WBGene00008551 | NA | NA | NA | lgc-36 |
| WBGene00008552 | -0.022930767 | 0.78121253 | NA | srsx-31 |
| WBGene00008553 | 1.524991234 | 0.00054125 | 0.00415327 | F07C6.2 |
| WBGene00008554 | 0.821440686 | 0.00080437 | 0.00584742 | F07C6.3 |

| | | | | |
|---|---|---|---|---|
| WBGene00008555 | -0.130307766 | 0.17937547 | 0.36082247 | F07C6.4 |
| WBGene00008556 | -0.023056499 | 0.78121253 | NA | F07D3.3 |
| WBGene00008557 | -0.027728334 | 0.67598605 | NA | F07H5.6 |
| WBGene00008558 | 0.092421167 | 0.47364008 | 0.66152502 | F07H5.7 |
| WBGene00008559 | 0.422347129 | 0.03522458 | 0.11657394 | F07H5.8 |
| WBGene00008560 | 1.114372323 | 7.75E-12 | 2.63E-10 | pho-13 |
| WBGene00008561 | 0.056032022 | 0.60328287 | 0.75988622 | F07H5.10 |
| WBGene00008563 | -0.743444744 | 0.01013573 | 0.04532598 | F07H5.13 |
| WBGene00008564 | -0.255586479 | 0.03868131 | 0.12485296 | acox-1.1 |
| WBGene00008565 | -1.058726654 | 1.04E-06 | 1.63E-05 | acox-1.2 |
| WBGene00008566 | -0.766905132 | 0.00196788 | 0.01231909 | acox-1.3 |
| WBGene00008567 | -0.365132896 | 0.03189025 | 0.10805489 | acox-1.4 |
| WBGene00008568 | 0.123334683 | 0.41402045 | 0.61154897 | F08A8.5 |
| WBGene00008569 | -0.193320072 | 0.14802683 | 0.31876649 | fbxa-101 |
| WBGene00008570 | 0.427706633 | 0.06947958 | 0.19038938 | kcnl-2 |
| WBGene00008571 | 0.329615274 | 0.11730832 | 0.27380421 | prmn-1 |
| WBGene00008572 | 1.688241401 | 4.34E-39 | 7.75E-37 | F08B12.4 |
| WBGene00008574 | NA | NA | NA | srbc-61 |
| WBGene00008575 | -0.176644877 | 0.04116724 | NA | scl-24 |
| WBGene00008576 | 0.158296246 | 0.42478318 | 0.62055982 | F08G2.4 |
| WBGene00008577 | 2.952974302 | 6.45E-11 | 1.91E-09 | F08G2.5 |
| WBGene00008578 | -0.058726493 | 0.77371111 | 0.87164546 | F08G2.7 |
| WBGene00008579 | 0.037234531 | 0.80221149 | 0.88992108 | F08G2.8 |
| WBGene00008580 | -0.158120515 | 0.14764326 | 0.31835714 | dsb-1 |
| WBGene00008581 | 0.06111288 | 0.51882185 | NA | acl-13 |
| WBGene00008582 | 0.548846762 | 0.04623179 | 0.14214188 | F08G5.3 |
| WBGene00008583 | -2.518071407 | 2.73E-06 | 3.80E-05 | ugt-65 |
| WBGene00008584 | -1.193442624 | 0.00164533 | 0.01064015 | irg-4 |
| WBGene00008585 | -0.077466134 | 0.6809835 | 0.81536469 | F08G12.1 |
| WBGene00008586 | -0.139614001 | 0.21532777 | 0.40639906 | snrp-40.1 |
| WBGene00008587 | 0.085831482 | 0.68177019 | 0.81594316 | F08G12.3 |
| WBGene00008588 | 0.057857293 | 0.7688825 | 0.86844708 | F08G12.5 |
| WBGene00008589 | 0.362921102 | 0.08257494 | 0.21546804 | nlp-57 |
| WBGene00008590 | 0.373838858 | 0.04869925 | 0.14746418 | F08H9.2 |
| WBGene00008591 | -0.05972283 | 0.59752871 | 0.75586814 | F08H9.3 |
| WBGene00008592 | -0.040794878 | 0.53203165 | NA | F08H9.4 |
| WBGene00008593 | 2.266980628 | 3.18E-07 | 5.45E-06 | clec-227 |
| WBGene00008594 | 1.136968764 | 3.27E-06 | 4.45E-05 | clec-57 |
| WBGene00008595 | 0.760165374 | 1.27E-06 | 1.94E-05 | clec-56 |
| WBGene00008596 | 1.612549647 | 2.61E-11 | 8.15E-10 | clec-54 |
| WBGene00008597 | 3.992470253 | 0.00187156 | 0.01183652 | clec-55 |
| WBGene00008598 | -0.480314582 | 0.06957975 | 0.19055688 | spin-3 |
| WBGene00008599 | 0.7808907 | 0.01914829 | 0.07448196 | F09A5.2 |

| | | | | |
|---|---|---|---|---|
| WBGene00008600 | -0.032616144 | 0.7322195 | NA | F09A5.3 |
| WBGene00008601 | 0.290933728 | 0.02043137 | 0.07787962 | mob-2 |
| WBGene00008602 | 1.941239749 | 2.16E-05 | 0.0002447 | oac-14 |
| WBGene00008603 | 0.158703727 | 0.38154331 | 0.58074595 | F09B9.4 |
| WBGene00008604 | 0.081937479 | 0.62445232 | 0.77558589 | F09B9.5 |
| WBGene00008605 | 2.344310858 | 0.00320926 | 0.0184334 | mlt-9 |
| WBGene00008606 | 0.040945262 | 0.81794648 | 0.89994397 | dhhc-1 |
| WBGene00008607 | -1.809501141 | 5.72E-25 | 5.37E-23 | F09B12.3 |
| WBGene00008609 | -0.00346345 | 0.95708614 | NA | F09B12.5 |
| WBGene00008610 | 1.57150029 | 2.26E-06 | 3.25E-05 | F09C3.2 |
| WBGene00008611 | -0.931018686 | 0.01858773 | 0.07286292 | fbxb-68 |
| WBGene00008612 | -0.119703227 | 0.37889403 | 0.57827452 | fbxa-103 |
| WBGene00008613 | -0.178262568 | 0.13368605 | 0.29743895 | fbxb-69 |
| WBGene00008615 | 2.315230679 | 0.00308423 | 0.01785688 | F09C6.1 |
| WBGene00008616 | 0.395470409 | 0.01759848 | 0.06997809 | fbxa-44 |
| WBGene00008617 | 0.091075881 | 0.25872286 | NA | F09C6.3 |
| WBGene00008618 | -0.023056499 | 0.78121253 | NA | fbxa-45 |
| WBGene00008619 | NA | NA | NA | nhr-262 |
| WBGene00008620 | 0.019325396 | 0.77340931 | NA | F09C6.10 |
| WBGene00008621 | 2.972071338 | 1.82E-74 | 6.65E-72 | F09C8.1 |
| WBGene00008622 | -0.292488367 | 0.0292109 | 0.10161007 | F09C8.2 |
| WBGene00008623 | -0.024917643 | 0.69549488 | NA | spe-43 |
| WBGene00008624 | -0.266770427 | 0.03060884 | 0.10492717 | crld-1 |
| WBGene00008625 | 6.319624928 | 3.01E-08 | 6.20E-07 | scl-14 |
| WBGene00008626 | 1.021715802 | 0.01360111 | 0.05724882 | nlp-79 |
| WBGene00008627 | NA | NA | NA | hpx-2 |
| WBGene00008628 | 1.730101373 | 5.86E-09 | 1.35E-07 | ttr-21 |
| WBGene00008629 | -1.017977105 | 2.37E-07 | 4.16E-06 | cpt-5 |
| WBGene00008630 | NA | NA | NA | nhr-175 |
| WBGene00008631 | NA | NA | NA | F10A3.1 |
| WBGene00008632 | NA | NA | NA | fbxa-88 |
| WBGene00008633 | 0.065014541 | 0.37399213 | NA | fbxa-18 |
| WBGene00008634 | 0.403642836 | 0.10932212 | 0.26039175 | F10A3.4 |
| WBGene00008635 | 0.085537126 | 0.24698522 | NA | NA |
| WBGene00008636 | NA | NA | NA | F10A3.11 |
| WBGene00008637 | NA | NA | NA | F10A3.12 |
| WBGene00008638 | NA | NA | NA | F10A3.16 |
| WBGene00008639 | 0.044704654 | 0.72047409 | 0.84342114 | F10B5.2 |
| WBGene00008640 | -0.022930767 | 0.78121253 | NA | F10B5.3 |
| WBGene00008641 | 0.188761939 | 0.07821727 | 0.20764806 | pch-2 |
| WBGene00008642 | -0.152354742 | 0.08228635 | 0.21509631 | F10B5.8 |
| WBGene00008643 | 0.033463284 | 0.6365037 | NA | tmem-138 |
| WBGene00008644 | -0.166631934 | 0.42259605 | 0.6186446 | F10C2.3 |

| | | | | |
|---|---|---|---|---|
| WBGene00008645 | -0.118786448 | 0.09232684 | 0.23187677 | F10C2.4 |
| WBGene00008646 | -0.144799475 | 0.23197533 | 0.42694542 | F10C2.5 |
| WBGene00008647 | 0.059762547 | 0.38965825 | NA | F10C2.7 |
| WBGene00008648 | -0.243347137 | 0.03687202 | 0.12052764 | F10D11.2 |
| WBGene00008649 | 0.012696942 | 0.89378933 | NA | F10D11.3 |
| WBGene00008650 | 0.078203897 | 0.26561294 | NA | F10D11.4 |
| WBGene00008651 | 0.081964019 | 0.5169817 | 0.69633202 | F10D11.5 |
| WBGene00008652 | 0.551464804 | 0.0194097 | 0.07514433 | F10D11.6 |
| WBGene00008654 | -0.391079386 | 0.00479352 | 0.0252908 | pfas-1 |
| WBGene00008655 | 0.312323905 | 0.07402882 | 0.19953164 | lips-3 |
| WBGene00008656 | 2.584574549 | 1.63E-08 | 3.52E-07 | lron-5 |
| WBGene00008657 | 0.106997704 | 0.16248985 | NA | NA |
| WBGene00008658 | NA | NA | NA | NA |
| WBGene00008659 | 1.528663773 | 0.01494865 | 0.06166899 | clec-151 |
| WBGene00008660 | 0.036687355 | 0.59489568 | NA | clec-153 |
| WBGene00008661 | 0.061275255 | 0.51782541 | NA | F10G8.1 |
| WBGene00008662 | 0.106997704 | 0.16248985 | NA | F10G8.2 |
| WBGene00008663 | 0.020317245 | 0.84643798 | NA | eak-6 |
| WBGene00008664 | 0.00781199 | 0.94033717 | 0.96654567 | nubp-1 |
| WBGene00008665 | 0.031887463 | 0.77806068 | 0.87435671 | ercc-1 |
| WBGene00008666 | 0.363375848 | 0.0472313 | 0.14433594 | F10G8.8 |
| WBGene00008667 | 0.280545323 | 0.09388106 | 0.23455846 | F10G8.9 |
| WBGene00008669 | 0.529960658 | 2.12E-07 | 3.77E-06 | acs-14 |
| WBGene00008670 | -0.195221687 | 0.03950559 | 0.12689805 | eif-2Bdelta |
| WBGene00008671 | 0.056188364 | 0.42092875 | NA | F11A5.3 |
| WBGene00008672 | 0.020317245 | 0.84643798 | NA | F11A5.4 |
| WBGene00008673 | NA | NA | NA | F11A5.5 |
| WBGene00008675 | 0.020925399 | 0.84643798 | NA | irld-26 |
| WBGene00008676 | NA | NA | NA | oac-15 |
| WBGene00008677 | 0.598475235 | 9.12E-06 | 0.00011335 | F11A5.9 |
| WBGene00008678 | 2.289178664 | 0.00346063 | 0.01957011 | stdh-2 |
| WBGene00008679 | 0.020925399 | 0.84643798 | NA | F11A5.13 |
| WBGene00008680 | -0.110336878 | 0.53985923 | 0.71445958 | F11A5.15 |
| WBGene00008681 | 0.366427169 | 0.13413076 | 0.29820707 | scrm-4 |
| WBGene00008682 | -0.228243976 | 0.00030724 | 0.00257043 | lex-1 |
| WBGene00008683 | -0.2582088 | 0.01457735 | 0.06047802 | repo-1 |
| WBGene00008684 | -0.068640741 | 0.45662566 | 0.64869555 | mig-32 |
| WBGene00008685 | -0.08302904 | 0.34144278 | 0.5414344 | mon-2 |
| WBGene00008686 | -0.155975935 | 0.05012599 | 0.15056568 | F11A10.5 |
| WBGene00008687 | 0.030059498 | 0.83544034 | 0.91048532 | F11A10.6 |
| WBGene00008688 | -0.574706712 | 1.23E-06 | 1.89E-05 | rbm-34 |
| WBGene00008690 | -0.255590967 | 0.2471329 | 0.44565497 | F11C1.1 |
| WBGene00008691 | -0.123133551 | 0.53628736 | 0.71133513 | mff-1 |

| | | | | |
|---|---|---|---|---|
| WBGene00008692 | -0.215428034 | 0.30369961 | 0.50437151 | scav-4 |
| WBGene00008693 | 0.025242718 | 0.83914562 | 0.9128413 | F11C1.4 |
| WBGene00008694 | -0.8695473 | 1.10E-25 | 1.06E-23 | vwa-8 |
| WBGene00008695 | 0.077249732 | 0.71016283 | 0.83679989 | F11C3.1 |
| WBGene00008696 | NA | NA | NA | clec-255 |
| WBGene00008698 | -0.003614866 | 0.95700117 | NA | F11D11.3 |
| WBGene00008699 | NA | NA | NA | F11D11.22 |
| WBGene00008699 | NA | NA | NA | F11D11.4 |
| WBGene00008700 | NA | NA | NA | clec-254 |
| WBGene00008701 | NA | NA | NA | F11D11.6 |
| WBGene00008702 | -0.045771477 | 0.50913401 | NA | clec-252 |
| WBGene00008703 | 0.0398788 | 0.58971221 | NA | clec-251 |
| WBGene00008706 | 0.654223787 | 0.00373088 | 0.02073008 | gba-3 |
| WBGene00008707 | -0.707051891 | 6.72E-20 | 4.99E-18 | F11E6.3 |
| WBGene00008708 | -0.068113617 | 0.65186392 | 0.79387287 | F11E6.4 |
| WBGene00008709 | 0.25675632 | 0.14098377 | 0.30863628 | F11E6.6 |
| WBGene00008710 | -0.183831606 | 0.00814662 | 0.03836851 | F11E6.7 |
| WBGene00008711 | 0.699943244 | 0.02101504 | 0.07956585 | F11E6.8 |
| WBGene00008712 | -0.003614866 | 0.95700117 | NA | F11E6.9 |
| WBGene00008713 | -0.008603946 | 0.9233394 | NA | F11E6.10 |
| WBGene00008714 | -0.130384213 | 0.52171014 | 0.70003975 | F11F1.1 |
| WBGene00008716 | NA | NA | NA | F11F1.4 |
| WBGene00008717 | NA | NA | NA | F11F1.5 |
| WBGene00008719 | -0.069906374 | 0.4226639 | NA | ttr-52 |
| WBGene00008720 | -0.006210079 | 0.92646948 | 0.9586234 | F12F6.1 |
| WBGene00008722 | -0.052815515 | 0.55636106 | 0.72794301 | F12F6.7 |
| WBGene00008723 | -0.401925114 | 0.10374961 | 0.25169811 | NA |
| WBGene00008724 | 0.423743218 | 0.04790023 | 0.14583223 | F13A7.1 |
| WBGene00008725 | 0.033463284 | 0.6365037 | NA | F13A7.7 |
| WBGene00008726 | 0.206706027 | 0.33616156 | 0.53555401 | F13A7.11 |
| WBGene00008727 | 0.307058696 | 0.00931656 | 0.04255968 | F13A7.12 |
| WBGene00008728 | -0.108290402 | 0.33530964 | 0.53448758 | sre-36 |
| WBGene00008729 | -0.104413569 | 0.12595458 | 0.28697495 | F13B12.1 |
| WBGene00008730 | 0.106637186 | 0.61214421 | 0.76660707 | F13B12.2 |
| WBGene00008731 | -0.322320328 | 0.17112813 | 0.35007502 | F13B12.3 |
| WBGene00008732 | 0.077484302 | 0.42130829 | 0.61801169 | F13B12.4 |
| WBGene00008733 | -0.05545338 | 0.53701461 | 0.71191877 | F13B12.6 |
| WBGene00008734 | NA | NA | NA | F13B12.7 |
| WBGene00008735 | 0.090235139 | 0.64496076 | 0.78968348 | F13D2.1 |
| WBGene00008736 | 0.570019388 | 0.01051141 | 0.04652743 | gnrr-6 |
| WBGene00008737 | 0.022452728 | 0.84543921 | 0.9169413 | gnrr-7 |
| WBGene00008739 | -0.220323939 | 0.25140824 | 0.44991306 | F13D12.3 |
| WBGene00008740 | -0.395561342 | 0.00058953 | 0.00447912 | F13D12.5 |

| | | | | |
|---|---|---|---|---|
| WBGene00008741 | 0.949923268 | 5.79E-29 | 6.94E-27 | ctsa-1.2 |
| WBGene00008742 | 0.104059064 | 0.60529619 | 0.76161409 | F13D12.8 |
| WBGene00008743 | 1.883260948 | 0.0001358 | 0.00125993 | F13D12.9 |
| WBGene00008744 | 0.084662079 | 0.60709632 | 0.76297727 | F13D12.10 |
| WBGene00008745 | -0.021622691 | 0.85331798 | 0.92134429 | F13E6.1 |
| WBGene00008746 | 0.005320816 | 0.9755371 | 0.98617957 | F13E6.2 |
| WBGene00008747 | -0.128260469 | 0.54289386 | 0.71666297 | phf-31 |
| WBGene00008748 | 0.718225442 | 5.24E-06 | 6.83E-05 | yap-1 |
| WBGene00008749 | -0.234572944 | 0.16385074 | 0.3396006 | plpp-1.3 |
| WBGene00008750 | -0.037524927 | 0.70307653 | 0.83144159 | F13E9.1 |
| WBGene00008751 | NA | NA | NA | fipr-14 |
| WBGene00008752 | 0.020925399 | 0.84643798 | NA | fipr-15 |
| WBGene00008753 | NA | NA | NA | F13E9.4 |
| WBGene00008754 | 0.015908114 | 0.8416225 | NA | F13E9.5 |
| WBGene00008755 | -0.243962187 | 0.19953893 | 0.38803905 | sdz-13 |
| WBGene00008757 | 0.020232312 | 0.84643798 | NA | F13E9.8 |
| WBGene00008758 | 0.020232312 | 0.84643798 | NA | F13E9.9 |
| WBGene00008759 | NA | NA | NA | F13E9.10 |
| WBGene00008760 | 0.286926868 | 0.20888264 | 0.39824923 | F13E9.11 |
| WBGene00008761 | 0.073766056 | 0.53310666 | 0.70892048 | F13E9.12 |
| WBGene00008762 | 0.249888121 | 0.17504325 | 0.35564444 | ztf-2 |
| WBGene00008763 | 0.070265188 | 0.50058947 | 0.68260505 | F13G3.3 |
| WBGene00008764 | -0.295678262 | 0.00216416 | 0.01330204 | dylt-1 |
| WBGene00008765 | -0.03401703 | 0.72812014 | 0.84850209 | ttx-7 |
| WBGene00008766 | 0.019369829 | 0.89670834 | 0.94573597 | F13G3.6 |
| WBGene00008767 | -0.201351907 | 0.122202 | 0.28134075 | F13G3.7 |
| WBGene00008768 | -0.210051155 | 0.05953275 | 0.17068 | F13G3.10 |
| WBGene00008769 | -0.272217713 | 0.05370219 | 0.15868983 | mrpl-13 |
| WBGene00008770 | 0.320529627 | 0.17446373 | 0.35510266 | F13G3.12 |
| WBGene00008772 | 0.040107227 | 0.58693163 | NA | irld-4 |
| WBGene00008773 | -0.008969483 | 0.9173367 | NA | F13H10.1 |
| WBGene00008774 | -0.114277898 | 0.22957906 | 0.42386722 | F13H10.3 |
| WBGene00008775 | -0.05134125 | 0.46662191 | 0.65621087 | mogs-1 |
| WBGene00008776 | 0.147118377 | 0.35595659 | 0.55590444 | F13H10.5 |
| WBGene00008777 | 0.092554273 | 0.61272466 | 0.7670458 | F13H10.6 |
| WBGene00008778 | NA | NA | NA | nhr-264 |
| WBGene00008779 | -0.253442863 | 0.25227491 | 0.45017504 | F14B4.1 |
| WBGene00008780 | -0.097113016 | 0.08223045 | 0.21502698 | hxk-1 |
| WBGene00008781 | -0.41831918 | 1.49E-05 | 0.00017559 | rpoa-2 |
| WBGene00008782 | -0.748424147 | 0.00995978 | 0.04474416 | sdz-14 |
| WBGene00008783 | 0.33675251 | 0.15524445 | 0.32850614 | F14B6.2 |
| WBGene00008784 | -0.763400709 | 0.00130111 | 0.00873831 | F14B6.3 |
| WBGene00008785 | -0.061342743 | 0.61398195 | 0.76825141 | F14B6.4 |

| | | | | |
|---|---|---|---|---|
| WBGene00008786 | NA | NA | NA | oac-16 |
| WBGene00008787 | 0.231902338 | 0.29120268 | 0.49186645 | F14B6.6 |
| WBGene00008788 | NA | NA | NA | F14D7.1 |
| WBGene00008789 | -0.354945184 | 0.00013261 | 0.00123504 | F14D7.2 |
| WBGene00008790 | 0.020317245 | 0.84643798 | NA | F14D7.3 |
| WBGene00008791 | 0.076452416 | 0.42707996 | NA | NA |
| WBGene00008792 | -0.087157727 | 0.2060916 | NA | F14D7.5 |
| WBGene00008793 | -0.085919754 | 0.67134049 | 0.80891252 | F14D7.6 |
| WBGene00008794 | -0.101752954 | 0.3872665 | 0.58586162 | F14D7.7 |
| WBGene00008795 | -0.009295168 | 0.91004451 | NA | F14D7.8 |
| WBGene00008796 | NA | NA | NA | F14D7.9 |
| WBGene00008797 | -0.009664907 | 0.91096914 | NA | F14D7.10 |
| WBGene00008798 | -0.002914971 | 0.95718083 | NA | F14D7.11 |
| WBGene00008799 | -0.163860865 | 0.41652345 | 0.61354691 | F14E5.1 |
| WBGene00008800 | -0.105029569 | 0.11570971 | 0.27144783 | F14E5.2 |
| WBGene00008801 | 0.038125098 | 0.61119482 | NA | acp-3 |
| WBGene00008802 | -0.120770987 | 0.56219463 | 0.73230632 | acp-2 |
| WBGene00008803 | -0.017082785 | 0.92996037 | 0.96066816 | lips-10 |
| WBGene00008804 | -0.027728334 | 0.67598605 | NA | acp-4 |
| WBGene00008805 | -0.144615881 | 0.34956095 | 0.549546 | git-1 |
| WBGene00008806 | -0.020798714 | 0.89584272 | 0.94523157 | mboa-7 |
| WBGene00008807 | 0.020232312 | 0.84643798 | NA | F14F3.4 |
| WBGene00008808 | -0.035738516 | 0.75158125 | 0.85956565 | ntr-2 |
| WBGene00008809 | 0.038726127 | 0.57551883 | NA | cyp-13A11 |
| WBGene00008810 | 2.371026096 | 0.0006787 | 0.00504405 | cyp-13A12 |
| WBGene00008811 | -0.013029346 | 0.89540716 | 0.94517796 | F14F7.4 |
| WBGene00008812 | -0.067905328 | 0.31502275 | NA | F14F7.5 |
| WBGene00008813 | 0.020317245 | 0.84643798 | NA | NA |
| WBGene00008814 | NA | NA | NA | srz-1 |
| WBGene00008815 | NA | NA | NA | srz-103 |
| WBGene00008816 | 0.108538718 | 0.15420469 | NA | F14F8.8 |
| WBGene00008817 | NA | NA | NA | F14F8.9 |
| WBGene00008818 | NA | NA | NA | srz-101 |
| WBGene00008819 | 0.058295004 | 0.7631702 | 0.86563177 | shw-1 |
| WBGene00008820 | 0.747768821 | 0.0085236 | 0.03968013 | nlp-55 |
| WBGene00008821 | 0.0398788 | 0.58971221 | NA | best-12 |
| WBGene00008822 | -0.214439919 | 0.16646232 | 0.34340916 | F14H3.3 |
| WBGene00008823 | -0.164434465 | 0.16336157 | 0.33887458 | F14H3.4 |
| WBGene00008824 | 0.103061315 | 0.53805686 | 0.71275968 | F14H3.5 |
| WBGene00008825 | -0.070594061 | 0.34407619 | 0.54381673 | F14H3.6 |
| WBGene00008826 | NA | NA | NA | fbxa-100 |
| WBGene00008827 | -0.066883576 | 0.32235182 | NA | NA |
| WBGene00008828 | NA | NA | NA | F14H3.9 |

| | | | | |
|---|---|---|---|---|
| WBGene00008829 | -0.348029024 | 0.14625733 | 0.3161136 | cyp-35D1 |
| WBGene00008830 | 0.23271795 | 0.29592735 | 0.49703708 | nhr-176 |
| WBGene00008831 | 1.816140217 | 2.78E-05 | 0.00030695 | F14H3.12 |
| WBGene00008832 | 0.2910781 | 0.10934217 | 0.26039175 | obr-2 |
| WBGene00008833 | 0.065014541 | 0.37399213 | NA | F14H8.2 |
| WBGene00008834 | 0.017631418 | 0.81668032 | NA | F14H8.4 |
| WBGene00008835 | NA | NA | NA | F14H8.5 |
| WBGene00008837 | -0.003614866 | 0.95700117 | NA | sre-38 |
| WBGene00008838 | -0.758809859 | 0.01327838 | 0.05609023 | F15A4.2 |
| WBGene00008839 | -0.022930767 | 0.78121253 | NA | sre-37 |
| WBGene00008840 | 0.863042115 | 0.01844438 | 0.07249509 | F15A4.5 |
| WBGene00008841 | 1.917493373 | 2.40E-39 | 4.34E-37 | F15A4.6 |
| WBGene00008842 | 5.377138836 | 9.03E-05 | 0.0008863 | NA |
| WBGene00008843 | 0.098839497 | 0.52078461 | 0.69928849 | arrd-9 |
| WBGene00008844 | 0.137968469 | 0.44975038 | 0.64204323 | F15A4.10 |
| WBGene00008847 | -0.62655943 | 0.01963348 | 0.07562583 | fbxb-102 |
| WBGene00008848 | 0.190504854 | 0.03562366 | 0.11754924 | inft-2 |
| WBGene00008849 | -0.115944686 | 0.58032438 | 0.74546945 | try-10 |
| WBGene00008850 | 0.114359445 | 0.36876764 | 0.56868833 | F15B9.6 |
| WBGene00008851 | 0.414966701 | 0.03285062 | 0.1106146 | F15B9.8 |
| WBGene00008852 | -0.208523614 | 0.00405219 | 0.02211263 | ubql-1 |
| WBGene00008853 | -0.023056499 | 0.78121253 | NA | clec-101 |
| WBGene00008854 | 0.463727327 | 0.07072702 | 0.1927973 | F15D3.4 |
| WBGene00008855 | 0.020925399 | 0.84643798 | NA | F15D3.5 |
| WBGene00008856 | 0.063425274 | 0.51934925 | 0.69848084 | F15D3.6 |
| WBGene00008857 | 0.014828872 | 0.8692621 | 0.93000745 | timm-23 |
| WBGene00008858 | 0.423404817 | 0.08357061 | 0.21732811 | pals-1 |
| WBGene00008859 | 0.195182461 | 0.16698836 | 0.34425184 | F15D4.2 |
| WBGene00008860 | -0.110505862 | 0.39967011 | 0.59789211 | romo-1 |
| WBGene00008861 | 1.243455406 | 0.00212439 | 0.01313477 | F15D4.4 |
| WBGene00008862 | 0.322877339 | 0.15599633 | 0.3297036 | F15D4.5 |
| WBGene00008863 | 0.0745366 | 0.58019557 | 0.745406 | F15D4.6 |
| WBGene00008865 | 0.3038583 | 0.00807691 | 0.03815877 | F15G9.1 |
| WBGene00008866 | 0.020232312 | 0.84643798 | NA | F15G9.2 |
| WBGene00008868 | 0.402973515 | 0.10853879 | 0.25911216 | F15G9.5 |
| WBGene00008869 | -0.030755733 | 0.70286946 | NA | F15G9.6 |
| WBGene00008870 | 0.040107227 | 0.58693163 | NA | F15H9.1 |
| WBGene00008871 | 0.108942554 | 0.15594412 | NA | tag-314 |
| WBGene00008872 | 0.020232312 | 0.84643798 | NA | F15H10.5 |
| WBGene00008873 | NA | NA | NA | NA |
| WBGene00008874 | 0.016036363 | 0.86497314 | NA | F15H10.7 |
| WBGene00008875 | 0.040107227 | 0.58693163 | NA | F15H10.8 |
| WBGene00008876 | 0.175957945 | 0.06218625 | 0.17617432 | F16A11.1 |

| | | | | | |
|---|---|---|---|---|---|
| WBGene00008877 | | -0.258658217 | 0.03316073 | 0.11136139 | rtcb-1 |
| WBGene00008878 | | -0.079729804 | 0.29658571 | 0.49751505 | ppfr-1 |
| WBGene00008879 | | 0.116640199 | 0.28845567 | 0.48863582 | F16B12.1 |
| WBGene00008880 | | 0.067643143 | 0.71818546 | 0.84186977 | F16B12.4 |
| WBGene00008881 | NA | | NA | NA | F16B12.5 |
| WBGene00008882 | | 0.102600159 | 0.33517116 | 0.53432502 | F16B12.6 |
| WBGene00008883 | | -0.023109762 | 0.78121253 | NA | F16B12.7 |
| WBGene00008884 | | 0.020925399 | 0.84643798 | NA | nhr-281 |
| WBGene00008885 | | 0.141500222 | 0.47311027 | 0.66116737 | F16C3.1 |
| WBGene00008886 | | 1.552245304 | 4.16E-06 | 5.56E-05 | F16C3.2 |
| WBGene00008887 | | -0.246650621 | 0.00165238 | 0.01068099 | tbcd-1 |
| WBGene00008889 | | -0.023056499 | 0.78121253 | NA | F16D3.6 |
| WBGene00008890 | | 0.179812924 | 0.37766823 | 0.57676443 | ser-5 |
| WBGene00008891 | | 6.973393115 | 2.30E-07 | 4.04E-06 | clec-42 |
| WBGene00008892 | | 0.136478325 | 0.42526687 | 0.62114248 | clec-246 |
| WBGene00008893 | NA | | NA | NA | F16H6.3 |
| WBGene00008894 | | 0.094421322 | 0.54753316 | 0.72018982 | F16H6.4 |
| WBGene00008897 | | 0.625229402 | 0.03512612 | 0.11627437 | F16H6.7 |
| WBGene00008898 | | -0.022930767 | 0.78121253 | NA | F16H6.8 |
| WBGene00008899 | | -0.006558791 | 0.92999162 | NA | F16H6.9 |
| WBGene00008900 | | 0.147932085 | 0.47383094 | 0.66163899 | F16H6.10 |
| WBGene00008901 | NA | | NA | NA | nhr-27 |
| WBGene00008902 | | 0.020317245 | 0.84643798 | NA | phf-32 |
| WBGene00008903 | NA | | NA | NA | F17A2.4 |
| WBGene00008904 | | -0.214122856 | 0.26609547 | 0.46473861 | F17A2.13 |
| WBGene00008905 | | 0.356202027 | 0.00628979 | 0.0312908 | F17B5.1 |
| WBGene00008906 | NA | | NA | NA | oac-17 |
| WBGene00008907 | NA | | NA | NA | clec-109 |
| WBGene00008908 | NA | | NA | NA | F17B5.4 |
| WBGene00008909 | NA | | NA | NA | clec-110 |
| WBGene00008910 | | 0.3981696 | 0.11244814 | 0.26542401 | F17C8.3 |
| WBGene00008911 | NA | | NA | NA | F17C8.6 |
| WBGene00008912 | | 0.570518452 | 0.03214957 | 0.10880816 | F17C8.7 |
| WBGene00008914 | | 1.268806107 | 4.37E-06 | 5.78E-05 | F17C11.2 |
| WBGene00008915 | | 1.136684668 | 1.64E-08 | 3.54E-07 | F17C11.4 |
| WBGene00008916 | | 0.260297341 | 0.06408155 | 0.17987307 | clec-221 |
| WBGene00008917 | | -0.038531578 | 0.8506271 | 0.9197276 | F17C11.6 |
| WBGene00008918 | | -0.059184168 | 0.55725161 | 0.72847563 | pigt-1 |
| WBGene00008919 | | 0.207802909 | 0.11438828 | 0.26921055 | vps-36 |
| WBGene00008920 | | -0.164627577 | 6.94E-05 | 0.00070189 | eef-1G |
| WBGene00008921 | | -0.098141884 | 0.14254101 | 0.31069569 | ctf-4 |
| WBGene00008922 | | -0.20508903 | 0.18562602 | 0.36961112 | F17C11.11 |
| WBGene00008923 | | 0.110364334 | 0.52803497 | 0.70521324 | F17C11.12 |

| | | | | |
|---|---|---|---|---|
| WBGene00008924 | -0.218330939 | 0.22479589 | 0.41761662 | F17E5.2 |
| WBGene00008925 | 0.47055737 | 5.51E-06 | 7.15E-05 | F17H10.1 |
| WBGene00008926 | 0.356343404 | 0.0291907 | 0.10156392 | F17H10.2 |
| WBGene00008927 | -0.223652687 | 0.25536762 | 0.45376287 | snx-17 |
| WBGene00008928 | NA | NA | NA | F17H10.4 |
| WBGene00008929 | 0.192987265 | 0.19871159 | 0.38708573 | F18A11.2 |
| WBGene00008930 | 0.080447133 | 0.70074259 | 0.82979394 | F18A11.3 |
| WBGene00008932 | 0.349359955 | 0.14816085 | 0.31884712 | F18A11.5 |
| WBGene00008933 | -0.234578609 | 0.28248581 | 0.48236717 | NA |
| WBGene00008934 | -0.298581698 | 0.01487607 | 0.06147346 | F18C12.3 |
| WBGene00008935 | 0.05787416 | 0.40594086 | NA | F18C12.4 |
| WBGene00008936 | 0.60691919 | 3.53E-09 | 8.35E-08 | papl-1 |
| WBGene00008937 | -0.049826614 | 0.47909173 | NA | srt-43 |
| WBGene00008939 | -0.089984714 | 0.66245335 | 0.80166643 | F18H3.1 |
| WBGene00008940 | -0.169323195 | 0.3023092 | 0.50276842 | F18H3.4 |
| WBGene00008941 | -0.00346345 | 0.95708614 | NA | NA |
| WBGene00008942 | NA | NA | NA | NA |
| WBGene00008943 | 0.020925399 | 0.84643798 | NA | NA |
| WBGene00008944 | 0.473583659 | 0.00467524 | 0.02480978 | F19B2.5 |
| WBGene00008945 | 0.074955426 | 0.70566654 | 0.83334029 | F19B2.6 |
| WBGene00008946 | NA | NA | NA | F19B2.7 |
| WBGene00008947 | 0.0398788 | 0.58971221 | NA | srz-16 |
| WBGene00008948 | -0.338825428 | 2.21E-07 | 3.90E-06 | F19B6.1 |
| WBGene00008949 | -0.024917643 | 0.69549488 | NA | F19B6.3 |
| WBGene00008950 | 0.124326451 | 0.11293212 | NA | wht-5 |
| WBGene00008951 | 0.055493009 | 0.74728691 | 0.85713477 | F19C6.2 |
| WBGene00008952 | 0.23843122 | 0.25415462 | 0.4523007 | F19C6.3 |
| WBGene00008953 | -0.299574825 | 0.17619435 | 0.35699717 | F19C6.4 |
| WBGene00008954 | -0.072738541 | 0.7102597 | 0.83679989 | F19C6.5 |
| WBGene00008955 | 0.314240239 | 0.12748786 | 0.28874635 | F19D8.2 |
| WBGene00008956 | 0.155943271 | 0.44630079 | 0.63961334 | nekl-3 |
| WBGene00008957 | 0.253370554 | 0.22475999 | 0.41761662 | glb-13 |
| WBGene00008958 | 0.067254953 | 0.5627682 | 0.73275288 | F19H6.3 |
| WBGene00008959 | -0.160232337 | 0.26771414 | 0.4665971 | F19H6.4 |
| WBGene00008960 | -0.009428459 | 0.92392378 | 0.9569304 | F19H6.5 |
| WBGene00008961 | 0.020232312 | 0.84643798 | NA | F19H6.6 |
| WBGene00008963 | -0.163379263 | 0.44192475 | 0.63595638 | F19H8.2 |
| WBGene00008964 | -0.119189894 | 0.57013428 | 0.7387023 | mltn-9 |
| WBGene00008965 | -0.042331728 | 0.82750402 | 0.90562762 | mltn-10 |
| WBGene00008966 | NA | NA | NA | NA |
| WBGene00008967 | 0.235245962 | 0.05065786 | 0.1518284 | F20B10.3 |
| WBGene00008968 | -0.210037017 | 0.02308722 | 0.08534299 | elli-1 |
| WBGene00008969 | 0.269665239 | 0.21706716 | 0.40870373 | F20C5.4 |

| | | | | |
|---|---|---|---|---|
| WBGene00008970 | 0.023145529 | 0.89228001 | 0.94374205 | F20C5.5 |
| WBGene00008971 | 0.036859857 | 0.8558369 | 0.92263556 | F20C5.6 |
| WBGene00008972 | NA | NA | NA | F20C5.7 |
| WBGene00008973 | 0.054178339 | 0.63813541 | 0.78552676 | F20D1.1 |
| WBGene00008974 | 0.220515141 | 0.10204945 | 0.2487689 | tbc-1 |
| WBGene00008975 | 0.077715507 | 0.46688862 | 0.65645984 | F20D1.3 |
| WBGene00008976 | -0.104273348 | 0.2145641 | NA | plp-2 |
| WBGene00008977 | 0.192532818 | 0.2472795 | 0.44573814 | iglr-1 |
| WBGene00008978 | 0.083086907 | 0.61072761 | 0.7658156 | cutl-3 |
| WBGene00008979 | -0.307228751 | 0.0280165 | 0.09848953 | slc-25A18.1 |
| WBGene00008980 | 0.516050255 | 0.0158385 | 0.06466576 | emre-1 |
| WBGene00008981 | NA | NA | NA | srz-48 |
| WBGene00008982 | NA | NA | NA | srsx-2 |
| WBGene00008983 | 0.040107227 | 0.58693163 | NA | F20E11.5 |
| WBGene00008984 | 0.020317245 | 0.84643798 | NA | F20E11.7 |
| WBGene00008985 | 2.975427788 | 2.72E-11 | 8.47E-10 | F20G2.1 |
| WBGene00008986 | -0.256229856 | 0.10636026 | 0.25574324 | F20G2.2 |
| WBGene00008987 | 0.213412237 | 0.32414965 | 0.52363682 | F20G2.3 |
| WBGene00008988 | 0.733034749 | 0.0183059 | 0.07212506 | F20G2.5 |
| WBGene00008989 | 0.155122056 | 0.06782466 | NA | F20G2.6 |
| WBGene00008990 | -0.077166851 | 0.3372358 | 0.53685652 | smgl-1 |
| WBGene00008991 | 0.077615635 | 0.67491941 | 0.8111554 | F20G4.2 |
| WBGene00008992 | 0.063731418 | 0.41159439 | NA | F21A3.2 |
| WBGene00008993 | -0.865242211 | 0.00444512 | 0.02389169 | F21A3.3 |
| WBGene00008994 | -0.00586362 | 0.94588716 | NA | F21A3.4 |
| WBGene00008995 | -0.283231828 | 0.09180473 | 0.23089202 | prde-1 |
| WBGene00008996 | -0.006485285 | 0.96842911 | 0.98272599 | glb-14 |
| WBGene00008997 | 0.20801616 | 0.31653063 | 0.51698682 | lgc-31 |
| WBGene00008998 | -0.351342276 | 0.12541767 | 0.28637557 | ncs-4 |
| WBGene00008999 | -0.001940141 | 0.9839043 | 0.99073885 | myrf-2 |
| WBGene00009000 | 0.081768833 | 0.41230943 | NA | arrd-28 |
| WBGene00009001 | -0.031417647 | 0.69702864 | NA | tyr-3 |
| WBGene00009002 | 0.018005318 | 0.87156759 | 0.93143219 | hint-1 |
| WBGene00009004 | -0.270922347 | 0.00194656 | 0.01223685 | pfd-6 |
| WBGene00009005 | -0.008099777 | 0.92782023 | NA | F21C3.6 |
| WBGene00009006 | -0.451049638 | 1.36E-12 | 5.02E-11 | F21D5.1 |
| WBGene00009007 | -0.220876735 | 0.00196603 | 0.01231692 | otub-3 |
| WBGene00009008 | 0.083785507 | 0.60012403 | 0.757907 | F21D5.3 |
| WBGene00009009 | -0.248573012 | 0.25090206 | 0.44951751 | F21D5.4 |
| WBGene00009010 | 0.089189699 | 0.50128309 | 0.6828513 | F21D5.5 |
| WBGene00009011 | -0.1269827 | 0.21492513 | 0.40587776 | F21D5.6 |
| WBGene00009012 | -0.063349747 | 0.37635836 | 0.57554447 | F21D5.7 |
| WBGene00009013 | -0.305365225 | 0.03767371 | 0.12249185 | mrps-33 |

| | | | | |
|---|---|---|---|---|
| WBGene00009014 | 0.043757326 | 0.68560617 | 0.81848988 | F21D5.9 |
| WBGene00009015 | -0.203614548 | 0.21786233 | 0.40967407 | F21D9.1 |
| WBGene00009016 | 0.207958431 | 0.06563442 | 0.18286028 | F21D9.2 |
| WBGene00009017 | NA | NA | NA | F21D9.3 |
| WBGene00009018 | 0.094368351 | 0.43654405 | 0.63161511 | F21D9.4 |
| WBGene00009019 | -0.14270574 | 0.44522058 | 0.63844021 | NA |
| WBGene00009020 | -0.218564149 | 0.22710731 | 0.42089568 | NA |
| WBGene00009021 | NA | NA | NA | F21D9.8 |
| WBGene00009023 | 0.512283669 | 0.04721295 | 0.14430999 | F21G4.1 |
| WBGene00009024 | NA | NA | NA | F21G4.3 |
| WBGene00009025 | -0.011120603 | 0.95485264 | 0.97448749 | phf-34 |
| WBGene00009026 | -0.097924613 | 0.5716421 | 0.73987023 | F21G4.5 |
| WBGene00009027 | 0.103240765 | 0.54049898 | 0.71476071 | F21G4.6 |
| WBGene00009028 | 0.224895389 | 0.04935798 | 0.14881595 | F21H7.2 |
| WBGene00009029 | 0.124133031 | 0.25369644 | 0.45170492 | F21H7.3 |
| WBGene00009030 | 0.010111268 | 0.92440225 | 0.95729051 | clec-233 |
| WBGene00009031 | 1.646620711 | 0.00179035 | 0.01139673 | F21H7.5 |
| WBGene00009032 | 0.056411843 | 0.41891328 | NA | bgnt-1.3 |
| WBGene00009033 | 0.063421086 | 0.41095452 | NA | F21H7.12 |
| WBGene00009035 | -0.232196915 | 8.78E-07 | 1.40E-05 | gfat-2 |
| WBGene00009036 | NA | NA | NA | F22B3.5 |
| WBGene00009038 | NA | NA | NA | F22B3.7 |
| WBGene00009039 | 0.0398788 | 0.58971221 | NA | F22B3.8 |
| WBGene00009041 | 4.513169208 | 1.12E-10 | 3.20E-09 | cut-3 |
| WBGene00009042 | 0.336504015 | 0.0068872 | 0.03354476 | F22B5.4 |
| WBGene00009043 | 0.229784742 | 0.15977447 | 0.33441858 | F22B5.5 |
| WBGene00009044 | -0.009115904 | 0.96019793 | 0.97823797 | F22B5.6 |
| WBGene00009045 | -0.259575153 | 0.00385454 | 0.02125584 | F22B5.10 |
| WBGene00009046 | NA | NA | NA | F22B8.3 |
| WBGene00009047 | -0.003614866 | 0.95700117 | NA | NA |
| WBGene00009048 | -0.179350782 | 0.32402278 | 0.52363682 | cth-1 |
| WBGene00009049 | 0.265190237 | 0.17446107 | 0.35510266 | F22B8.7 |
| WBGene00009050 | 0.263741919 | 0.00063738 | 0.00478831 | F22D6.2 |
| WBGene00009051 | -0.111735531 | 0.26506135 | 0.46365025 | nduf-6 |
| WBGene00009052 | -0.133437978 | 0.16511715 | 0.34135557 | ekl-1 |
| WBGene00009054 | 0.020826809 | 0.90990254 | 0.95108753 | F22D6.9 |
| WBGene00009055 | 0.142639656 | 0.48418831 | 0.66945397 | NA |
| WBGene00009056 | 0.095576857 | 0.24390154 | NA | F22D6.15 |
| WBGene00009057 | -0.066043635 | 0.73298841 | 0.85064071 | cept-1 |
| WBGene00009058 | 0.600538498 | 0.00030095 | 0.00252507 | F22E12.1 |
| WBGene00009059 | -0.098169737 | 0.63522187 | 0.78364369 | chw-1 |
| WBGene00009060 | 0.010402222 | 0.94780625 | 0.97044737 | F22E12.3 |
| WBGene00009061 | 0.162923794 | 0.209657 | 0.39906243 | pals-14 |

| | | | | |
|---|---|---|---|---|
| WBGene00009063 | -0.217744403 | 0.17769093 | 0.35893665 | F22G12.3 |
| WBGene00009064 | 0.184501321 | 0.08273134 | 0.21576282 | F22G12.4 |
| WBGene00009065 | 0.191669026 | 0.01155489 | 0.05005181 | F22G12.5 |
| WBGene00009067 | -0.295987864 | 0.14365571 | 0.3123813 | F23A7.1 |
| WBGene00009068 | 0.216561804 | 0.31408505 | 0.514606 | F23A7.3 |
| WBGene00009069 | -0.974287508 | 4.06E-19 | 2.87E-17 | F23A7.4 |
| WBGene00009070 | 0.542045936 | 0.02677413 | 0.09549848 | F23A7.5 |
| WBGene00009071 | -0.308320588 | 0.17654794 | 0.35738765 | NA |
| WBGene00009073 | -0.001842784 | 0.95621055 | NA | delm-1 |
| WBGene00009075 | 0.10726575 | 0.22063221 | NA | F23B2.7 |
| WBGene00009076 | NA | NA | NA | NA |
| WBGene00009077 | 0.063527295 | 0.41294485 | NA | F23B2.10 |
| WBGene00009078 | -0.078518244 | 0.63593246 | 0.78416454 | rpb-12 |
| WBGene00009079 | 0.078633594 | 0.26310689 | NA | F23B12.1 |
| WBGene00009081 | 0.24276111 | 0.18852918 | 0.37361339 | F23B12.4 |
| WBGene00009082 | -0.173512851 | 0.00610135 | 0.03056103 | dlat-1 |
| WBGene00009084 | 0.032249759 | 0.7225239 | 0.84438636 | F23B12.7 |
| WBGene00009085 | -0.05458785 | 0.5305143 | NA | F23D12.1 |
| WBGene00009086 | -0.375986657 | 0.02910446 | 0.10143261 | F23D12.2 |
| WBGene00009087 | 0.703544117 | 0.02363235 | 0.0868259 | F23D12.3 |
| WBGene00009088 | -0.698265168 | 0.03017752 | 0.10386204 | F23D12.4 |
| WBGene00009089 | -1.282925358 | 0.00050067 | 0.00389084 | jmjd-3.2 |
| WBGene00009090 | 0.565200087 | 0.04924591 | 0.14860045 | fipr-3 |
| WBGene00009091 | 0.201713015 | 0.34315102 | 0.54313812 | F23D12.7 |
| WBGene00009092 | -0.161050812 | 0.06915741 | 0.18968408 | tomm-20 |
| WBGene00009093 | 0.52932531 | 0.04818043 | 0.1463202 | F23H12.3 |
| WBGene00009094 | 0.440978174 | 0.07254611 | 0.19625787 | F23H12.5 |
| WBGene00009095 | 0.07847479 | 0.31620173 | NA | F23H12.7 |
| WBGene00009096 | 0.032222623 | 0.87364502 | 0.93276403 | fipr-1 |
| WBGene00009097 | 0.899315172 | 1.66E-06 | 2.46E-05 | fipr-2 |
| WBGene00009098 | -0.054783063 | 0.49082934 | 0.67457359 | ehbp-1 |
| WBGene00009099 | 0.063290468 | 0.41068021 | NA | F25B3.2 |
| WBGene00009100 | 0.667634431 | 0.03021355 | 0.1039616 | rgef-1 |
| WBGene00009101 | -0.006377843 | 0.96517001 | 0.98071026 | F25B3.4 |
| WBGene00009102 | 0.431940922 | 0.08987428 | 0.22785184 | F25B3.5 |
| WBGene00009103 | -0.201591331 | 0.0032139 | 0.0184528 | rtfo-1 |
| WBGene00009104 | -5.486112184 | 6.61E-05 | 0.00067452 | NA |
| WBGene00009106 | -0.203156708 | 0.30339077 | 0.50413623 | acs-12 |
| WBGene00009107 | 0.033034883 | 0.64219503 | NA | F25D1.2 |
| WBGene00009108 | -0.075452738 | 0.28904131 | NA | F25D1.3 |
| WBGene00009109 | 1.698707874 | 0.00555705 | 0.02847768 | degt-1 |
| WBGene00009110 | 1.627030941 | 3.28E-08 | 6.73E-07 | F25D1.5 |
| WBGene00009112 | -0.092928885 | 0.31256578 | 0.51332311 | tag-353 |

| | | | | |
|---|---|---|---|---|
| WBGene00009113 | 0.036404791 | 0.61199497 | 0.76648575 | maph-1.2 |
| WBGene00009114 | 0.020925399 | 0.84643798 | NA | F25D7.5 |
| WBGene00009115 | 0.228653863 | 0.21150075 | 0.40161457 | F25F2.1 |
| WBGene00009116 | 0.130091658 | 0.38821851 | 0.58676004 | snx-27 |
| WBGene00009117 | -0.001842784 | 0.95621055 | NA | F25H2.3 |
| WBGene00009118 | -0.135240819 | 0.23532181 | 0.43115338 | F25H2.4 |
| WBGene00009119 | -0.281116793 | 5.16E-09 | 1.20E-07 | ndk-1 |
| WBGene00009120 | -0.028939499 | 0.74167599 | 0.85408827 | F25H2.6 |
| WBGene00009121 | -0.022378739 | 0.88570706 | 0.93975194 | F25H2.7 |
| WBGene00009122 | -0.277906555 | 9.30E-09 | 2.09E-07 | tct-1 |
| WBGene00009123 | 0.070244245 | 0.64786115 | 0.79156598 | F25H2.12 |
| WBGene00009124 | -0.087157826 | 0.41554195 | 0.61275434 | rtel-1 |
| WBGene00009125 | -0.053640783 | 0.74022999 | 0.85342581 | F25H5.2 |
| WBGene00009126 | -0.370678495 | 1.97E-12 | 7.04E-11 | pyk-1 |
| WBGene00009127 | -0.249261643 | 0.00989775 | 0.04452009 | clsp-1 |
| WBGene00009128 | -0.202879041 | 0.20653968 | 0.39570105 | mrpl-54 |
| WBGene00009129 | 0.077155384 | 0.58856603 | 0.75058302 | F25H5.7 |
| WBGene00009130 | 0.201255314 | 0.2553354 | 0.45376287 | F25H5.8 |
| WBGene00009131 | -0.373252547 | 0.00332449 | 0.01895399 | F25H8.1 |
| WBGene00009132 | -0.026147963 | 0.81887463 | 0.90042384 | F25H8.2 |
| WBGene00009133 | 0.328419772 | 0.1670255 | 0.34427992 | bed-3 |
| WBGene00009134 | 0.017279211 | 0.79380376 | NA | lurp-2 |
| WBGene00009135 | -0.001320437 | 0.993333 | 0.99586212 | F25H9.2 |
| WBGene00009136 | 0.040107227 | 0.58693163 | NA | lurp-3 |
| WBGene00009137 | 0.085121253 | 0.68041294 | 0.81494812 | bath-39 |
| WBGene00009138 | -0.06409498 | 0.64802845 | 0.79160333 | F25H9.6 |
| WBGene00009139 | 0.031120515 | 0.85429897 | 0.92206349 | F25H9.7 |
| WBGene00009140 | 0.107539899 | 0.46987361 | 0.65831766 | F26A3.1 |
| WBGene00009141 | -0.216235903 | 0.02578904 | 0.0928621 | ncbp-2 |
| WBGene00009142 | 0.831839451 | 4.26E-07 | 7.19E-06 | F26A3.4 |
| WBGene00009143 | 0.201839397 | 0.15427286 | 0.32715947 | F26A3.5 |
| WBGene00009144 | 0.129401398 | 0.10528579 | NA | del-3 |
| WBGene00009145 | -0.212442983 | 0.10120562 | 0.24737325 | F26A3.7 |
| WBGene00009146 | 0.108651045 | 0.59451235 | 0.75407219 | F26C11.1 |
| WBGene00009147 | NA | NA | NA | F26C11.3 |
| WBGene00009148 | -0.013403255 | 0.91262295 | 0.95126775 | F26D2.3 |
| WBGene00009149 | 0.129578469 | 0.09326699 | NA | F26D2.10 |
| WBGene00009150 | NA | NA | NA | clec-234 |
| WBGene00009151 | NA | NA | NA | F26D2.13 |
| WBGene00009152 | NA | NA | NA | F26D2.14 |
| WBGene00009153 | 0.010372999 | 0.90944755 | NA | F26D2.15 |
| WBGene00009154 | -0.010973364 | 0.90310817 | 0.94845469 | F26D2.16 |
| WBGene00009155 | -0.029683071 | 0.71268666 | NA | lhfp-4 |

| | | | | |
|---|---|---|---|---|
| WBGene00009156 | -1.08109316 | 0.00019882 | 0.00175991 | clec-196 |
| WBGene00009157 | -0.233278329 | 0.29179354 | 0.49252355 | F26E4.2 |
| WBGene00009158 | -0.398006833 | 0.04069976 | 0.12962481 | F26E4.3 |
| WBGene00009159 | -0.147255782 | 0.17076996 | 0.34953767 | F26E4.4 |
| WBGene00009160 | 0.04919219 | 0.62443867 | 0.77558589 | F26E4.5 |
| WBGene00009161 | 0.019971943 | 0.80465841 | 0.89140028 | cox-7C |
| WBGene00009162 | 0.591858654 | 0.00696192 | 0.03382991 | F26E4.7 |
| WBGene00009163 | 0.212261577 | 0.01724312 | 0.06888309 | drsh-1 |
| WBGene00009164 | 0.017035776 | 0.81263172 | 0.89618417 | hrdl-1 |
| WBGene00009165 | 0.008447884 | 0.96233872 | 0.97924853 | gpx-1 |
| WBGene00009166 | 1.970580281 | 0.01041897 | 0.04632376 | F26F2.1 |
| WBGene00009167 | NA | NA | NA | F26F2.2 |
| WBGene00009167 | NA | NA | NA | F26F2.9 |
| WBGene00009168 | 0.176849354 | 0.04526585 | NA | F26F2.3 |
| WBGene00009169 | 0.102280207 | 0.23256935 | NA | F26F2.4 |
| WBGene00009170 | 0.076434456 | 0.48279798 | 0.66845238 | F26F2.5 |
| WBGene00009171 | -0.001993815 | 0.95612996 | NA | clec-263 |
| WBGene00009172 | 0.144878022 | 0.20392389 | 0.39300376 | F26F2.7 |
| WBGene00009173 | -0.042421404 | 0.68947313 | 0.82143185 | prom-1 |
| WBGene00009174 | -0.131281253 | 0.20958534 | 0.3989779 | F26H9.2 |
| WBGene00009176 | -0.121157728 | 0.49558228 | 0.67825155 | nmat-2 |
| WBGene00009177 | -0.380432378 | 0.0005189 | 0.00400878 | F26H9.5 |
| WBGene00009178 | 0.072023211 | 0.52867162 | 0.70564594 | uggt-2 |
| WBGene00009179 | -0.123408053 | 0.2463883 | 0.44469671 | kbp-3 |
| WBGene00009180 | -0.066798251 | 0.25101909 | 0.44956215 | nurf-1 |
| WBGene00009182 | -0.25423303 | 0.24335973 | 0.44140607 | F26H11.4 |
| WBGene00009183 | 0.7189324 | 0.02507339 | 0.09100138 | F27C8.2 |
| WBGene00009184 | NA | NA | NA | F27C8.3 |
| WBGene00009185 | 0.341809448 | 0.0221126 | 0.08271637 | F27C8.5 |
| WBGene00009186 | -0.015781122 | 0.75439639 | 0.86089996 | trcs-1 |
| WBGene00009187 | -0.193423418 | 0.00305923 | 0.01775273 | F27D4.1 |
| WBGene00009188 | -0.380983041 | 8.30E-05 | 0.00081794 | lsy-22 |
| WBGene00009189 | -0.379407542 | 1.73E-07 | 3.14E-06 | F27D4.4 |
| WBGene00009190 | 0.261803809 | 0.21154105 | 0.40161457 | F27D4.6 |
| WBGene00009191 | -0.312319812 | 0.02589855 | 0.09315154 | F27D4.7 |
| WBGene00009192 | 1.674945012 | 8.35E-27 | 8.49E-25 | asah-2 |
| WBGene00009193 | 0.044519488 | 0.23878159 | 0.43591038 | F27E5.3 |
| WBGene00009194 | 0.039292341 | 0.65808492 | NA | F27E5.5 |
| WBGene00009195 | 0.020317245 | 0.84643798 | NA | hpo-33 |
| WBGene00009196 | 0.020232312 | 0.84643798 | NA | F28B1.2 |
| WBGene00009197 | NA | NA | NA | NA |
| WBGene00009200 | -0.100435711 | 0.27580404 | 0.47593777 | F28C1.1 |
| WBGene00009201 | -0.125218796 | 0.19472852 | 0.38171613 | F28C1.3 |

| | | | | | |
|---|---|---|---|---|---|
| WBGene00009202 | | -0.910381679 | 0.00010426 | 0.00100891 | aptf-4 |
| WBGene00009203 | | -0.130666989 | 0.2475633 | 0.44604655 | aptf-3 |
| WBGene00009204 | | -0.054902042 | 0.66884955 | 0.80684075 | pigg-1 |
| WBGene00009205 | | 0.063880945 | 0.60606846 | 0.76212732 | F28C6.5 |
| WBGene00009206 | NA | | NA | NA | NA |
| WBGene00009207 | | -0.222699784 | 0.060456 | 0.17233961 | F28C6.8 |
| WBGene00009208 | | 0.04909525 | 0.45360729 | NA | F28C6.9 |
| WBGene00009209 | | 0.159164655 | 0.33316144 | 0.53291999 | F28C6.10 |
| WBGene00009210 | | 0.033463284 | 0.6365037 | NA | NA |
| WBGene00009211 | | -0.360489689 | 7.58E-05 | 0.00075767 | wdr-46 |
| WBGene00009212 | | -1.066650935 | 8.51E-07 | 1.37E-05 | atz-1 |
| WBGene00009213 | | -0.326828435 | 0.15705411 | 0.33108565 | thn-1 |
| WBGene00009214 | NA | | NA | NA | thn-3 |
| WBGene00009215 | | 2.167210649 | 4.91E-86 | 2.12E-83 | thn-2 |
| WBGene00009216 | | 0.216145844 | 0.3031554 | 0.50380229 | F28D1.6 |
| WBGene00009217 | | 0.020232312 | 0.84643798 | NA | spe-45 |
| WBGene00009218 | | 0.022097517 | 0.91290573 | 0.95130625 | acs-20 |
| WBGene00009219 | | -0.186350921 | 0.18010839 | 0.36182313 | dpm-3 |
| WBGene00009220 | | 0.020875279 | 0.92084638 | 0.95577144 | F28D9.4 |
| WBGene00009221 | | 0.651373556 | 9.39E-07 | 1.49E-05 | acs-2 |
| WBGene00009222 | | 0.007578588 | 0.97118485 | 0.98427495 | fbxa-95 |
| WBGene00009223 | | -0.118170542 | 0.35960086 | 0.55958116 | mdt-28 |
| WBGene00009224 | | -0.476836033 | 0.03030229 | 0.10409568 | F28F8.7 |
| WBGene00009225 | | 0.081587519 | 0.47549961 | 0.66264895 | fbxa-94 |
| WBGene00009226 | | 5.464750824 | 1.77E-06 | 2.61E-05 | cyp-37B1 |
| WBGene00009227 | | -0.029327418 | 0.71586712 | NA | F28G4.2 |
| WBGene00009228 | | 0.078810597 | 0.30833291 | NA | F28G4.3 |
| WBGene00009229 | | 0.130463311 | 0.16095178 | 0.33556248 | F28G4.4 |
| WBGene00009230 | | 5.812673904 | 2.06E-05 | 0.00023408 | NA |
| WBGene00009231 | | 0.049337506 | 0.79569771 | 0.88526377 | ceh-89 |
| WBGene00009232 | | -0.034792449 | 0.7155449 | NA | nkat-1 |
| WBGene00009233 | | -0.107941078 | 0.42245287 | 0.6185751 | F28H6.4 |
| WBGene00009234 | | -0.185983006 | 0.27836532 | 0.47818218 | F28H6.6 |
| WBGene00009235 | NA | | NA | NA | F28H6.7 |
| WBGene00009236 | | 0.195360286 | 0.11504103 | 0.27035566 | F28H7.2 |
| WBGene00009237 | | 0.432235118 | 8.15E-06 | 0.00010216 | F28H7.3 |
| WBGene00009238 | | 0.09059338 | 0.49208023 | 0.67515197 | selt-1.2 |
| WBGene00009239 | | 0.091012931 | 0.55770667 | 0.72877494 | irld-6 |
| WBGene00009240 | | -0.057467219 | 0.59968046 | 0.75754284 | F28H7.7 |
| WBGene00009241 | | -0.004557919 | 0.98136089 | 0.98926637 | F28H7.8 |
| WBGene00009242 | | 0.157098662 | 0.06528165 | NA | sre-6 |
| WBGene00009243 | | 0.094831453 | 0.49615494 | 0.67870733 | F29C6.1 |
| WBGene00009245 | | 0.294888035 | 0.00062141 | 0.00468975 | rict-1 |

| | | | | |
|---|---|---|---|---|
| WBGene00009246 | -0.44586206 | 2.14E-05 | 0.00024308 | gfm-1 |
| WBGene00009247 | 0.213019417 | 0.23554242 | 0.43134167 | bath-45 |
| WBGene00009248 | NA | NA | NA | F29C12.6 |
| WBGene00009249 | NA | NA | NA | NA |
| WBGene00009250 | 0.063290468 | 0.41068021 | NA | F29D10.1 |
| WBGene00009251 | -0.003614866 | 0.95700117 | NA | F29D10.2 |
| WBGene00009252 | -0.005691347 | 0.93660638 | NA | F29D10.3 |
| WBGene00009254 | -0.209220766 | 0.00738962 | 0.03550653 | capg-1 |
| WBGene00009255 | 0.130951132 | 0.15809096 | 0.33235034 | ugt-34 |
| WBGene00009256 | -0.018830712 | 0.88947972 | 0.94225256 | tut-2 |
| WBGene00009257 | 0.647192249 | 0.01999016 | 0.07657663 | F29G6.1 |
| WBGene00009258 | 0.034973496 | 0.81609252 | 0.89891744 | ccdc-149 |
| WBGene00009259 | 0.468051417 | 1.06E-10 | 3.04E-09 | hpo-34 |
| WBGene00009260 | -0.19856852 | 0.35663871 | 0.55666817 | calm-1 |
| WBGene00009261 | 0.127693253 | 0.10904601 | NA | F30A10.2 |
| WBGene00009262 | -0.201650145 | 0.05156506 | 0.15388586 | F30A10.3 |
| WBGene00009263 | NA | NA | NA | F30A10.4 |
| WBGene00009264 | 0.01371866 | 0.85621024 | 0.92283379 | sac-1 |
| WBGene00009266 | -0.159968819 | 0.28601073 | 0.48612528 | F30A10.9 |
| WBGene00009267 | -0.165131318 | 0.0204597 | 0.07796732 | usp-48 |
| WBGene00009268 | -0.054146032 | 0.57401669 | 0.74187918 | F30A10.11 |
| WBGene00009269 | 0.181588631 | 0.04307337 | NA | F30A10.12 |
| WBGene00009270 | -0.164186345 | 0.04195712 | 0.13233396 | F30F8.1 |
| WBGene00009271 | 0.299319838 | 0.19534347 | 0.38258628 | glna-3 |
| WBGene00009272 | -0.079700925 | 0.37105221 | 0.57052882 | gras-1 |
| WBGene00009274 | -0.133361268 | 0.52274225 | 0.70044791 | F30F8.5 |
| WBGene00009276 | -0.152677984 | 0.26640877 | 0.46511935 | F30F8.9 |
| WBGene00009277 | -0.029567434 | 0.7718522 | 0.87031066 | F30F8.10 |
| WBGene00009278 | 0.132442484 | 0.10147609 | NA | npr-33 |
| WBGene00009279 | NA | NA | NA | F31B9.2 |
| WBGene00009280 | 0.075052726 | 0.65045748 | 0.79307985 | F31B9.3 |
| WBGene00009281 | NA | NA | NA | F31B9.4 |
| WBGene00009283 | -0.03891762 | 0.73758869 | 0.85246644 | F31B12.4 |
| WBGene00009284 | -0.076759493 | 0.44135096 | 0.63556795 | F31C3.2 |
| WBGene00009285 | 0.085433255 | 0.22879552 | 0.42263376 | F31C3.3 |
| WBGene00009286 | -0.05089336 | 0.55038735 | 0.72290492 | zipt-1 |
| WBGene00009287 | -0.096621727 | 0.3856495 | 0.58471782 | psf-2 |
| WBGene00009288 | 0.358031888 | 0.12868909 | 0.29040597 | F31C3.6 |
| WBGene00009289 | -0.312199114 | 0.02508961 | 0.09103766 | exos-7 |
| WBGene00009290 | 0.023176941 | 0.78531877 | 0.87870478 | F31D4.2 |
| WBGene00009291 | 1.407521092 | 4.61E-05 | 0.00048889 | clec-264 |
| WBGene00009292 | 0.314195978 | 0.12385529 | 0.28418199 | F31D4.5 |
| WBGene00009293 | -0.607344923 | 0.04108671 | 0.13048818 | F31D4.8 |

| | | | | | |
|---|---|---|---|---|---|
| WBGene00009294 | | -0.189466198 | 0.07175274 | 0.19494002 | F31D4.9 |
| WBGene00009295 | | 0.469999013 | 0.07677952 | 0.20505819 | NA |
| WBGene00009296 | | 0.084341568 | 0.56143575 | 0.73183589 | F31E9.3 |
| WBGene00009297 | | -0.687877233 | 0.01834535 | 0.07220277 | sdz-16 |
| WBGene00009298 | NA | | NA | NA | srz-58 |
| WBGene00009299 | | 0.44098214 | 0.07130834 | 0.19412893 | F31E9.6 |
| WBGene00009300 | | -0.250115512 | 0.1702864 | 0.34882784 | F31F6.1 |
| WBGene00009301 | | -0.410551334 | 0.05232809 | 0.15578195 | F31F6.2 |
| WBGene00009302 | NA | | NA | NA | F31F6.3 |
| WBGene00009303 | | -0.023109762 | 0.78121253 | NA | lips-8 |
| WBGene00009304 | | 0.71994462 | 0.02709006 | 0.09617495 | eva-1 |
| WBGene00009305 | | -0.053529976 | 0.58978213 | 0.75112054 | F32A7.4 |
| WBGene00009306 | | 0.275392213 | 0.05087207 | 0.15231452 | maph-1.1 |
| WBGene00009307 | | -0.178907806 | 0.04451692 | 0.13817515 | F32A11.1 |
| WBGene00009308 | | 1.29472066 | 0.0051079 | 0.02657597 | F32A11.3 |
| WBGene00009309 | | -0.03147554 | 0.70637395 | NA | NA |
| WBGene00009310 | | -0.093274527 | 0.63876628 | 0.78569916 | NA |
| WBGene00009311 | | 1.46907069 | 0.00011629 | 0.00110809 | mltn-1 |
| WBGene00009312 | | 0.056188364 | 0.42092875 | NA | F32B4.1 |
| WBGene00009313 | | 0.076416372 | 0.53138767 | 0.70779174 | F32B4.2 |
| WBGene00009314 | | 0.067307265 | 0.55934526 | 0.73003231 | F32B4.4 |
| WBGene00009315 | | 0.461972099 | 0.07220396 | 0.19569384 | F32B4.5 |
| WBGene00009316 | NA | | NA | NA | abhd-11.1 |
| WBGene00009318 | | -0.041805562 | 0.73885983 | 0.85307041 | F32B4.8 |
| WBGene00009319 | | -0.248773187 | 0.00068697 | 0.00508744 | mccc-1 |
| WBGene00009320 | | -0.129106306 | 0.33936606 | 0.53913394 | F32B6.3 |
| WBGene00009321 | | 0.322195002 | 0.04997076 | 0.15013789 | F32B6.4 |
| WBGene00009322 | | -0.037323959 | 0.67078907 | 0.80831463 | tbc-3 |
| WBGene00009323 | | 0.355786794 | 0.10725333 | 0.25725689 | best-13 |
| WBGene00009324 | | 0.065014541 | 0.37399213 | NA | ttbk-3 |
| WBGene00009325 | | 0.066591731 | 0.69246788 | 0.82359239 | F32D8.15 |
| WBGene00009326 | | -0.054932793 | 0.45568274 | NA | F32D8.1 |
| WBGene00009327 | | 0.108717649 | 0.15669083 | NA | F32D8.2 |
| WBGene00009328 | | -0.023410703 | 0.87406097 | 0.93293917 | F32D8.3 |
| WBGene00009329 | | -0.213029647 | 0.14960362 | 0.3209605 | F32D8.4 |
| WBGene00009330 | | 0.193347797 | 0.1866972 | 0.37103756 | F32D8.5 |
| WBGene00009331 | | 0.184021517 | 0.22693418 | 0.42062835 | F32D8.7 |
| WBGene00009332 | | -0.046394786 | 0.50231372 | NA | F32D8.8 |
| WBGene00009333 | | 0.529607116 | 0.0456382 | 0.1407397 | F32D8.10 |
| WBGene00009334 | | 0.712688173 | 4.90E-16 | 2.58E-14 | F32D8.12 |
| WBGene00009335 | | -0.478991791 | 0.07311566 | 0.19754317 | F32D8.13 |
| WBGene00009336 | | -0.256373695 | 0.0104446 | 0.04640954 | F32D8.14 |
| WBGene00009337 | | 0.645720721 | 0.00585087 | 0.02957962 | uig-1 |

| | | | | |
|---|---|---|---|---|
| WBGene00009338 | NA | NA | NA | F32G8.2 |
| WBGene00009339 | 0.0398788 | 0.58971221 | NA | F32G8.3 |
| WBGene00009340 | -0.028631491 | 0.83703848 | 0.91149724 | best-14 |
| WBGene00009341 | -0.181366946 | 0.10426399 | 0.25244404 | thoc-3 |
| WBGene00009342 | -0.326083616 | 4.35E-06 | 5.76E-05 | fasn-1 |
| WBGene00009343 | -0.057692394 | 0.65964437 | 0.79909426 | F32H2.6 |
| WBGene00009344 | 0.907252242 | 0.01982959 | 0.07608766 | F32H2.7 |
| WBGene00009345 | NA | NA | NA | F32H2.8 |
| WBGene00009346 | -0.096639288 | 0.49659587 | 0.67886649 | F32H2.10 |
| WBGene00009347 | 0.466938262 | 0.00174557 | 0.0111651 | F32H5.1 |
| WBGene00009349 | 0.235649035 | 0.28434582 | 0.48430749 | F32H5.3 |
| WBGene00009350 | 0.271777027 | 0.21953984 | 0.41166531 | F32H5.4 |
| WBGene00009352 | -0.157066786 | 0.13402622 | 0.29808261 | F33A8.4 |
| WBGene00009353 | -0.089413012 | 0.2756015 | 0.47572289 | sdhd-1 |
| WBGene00009354 | -0.122663876 | 0.5470898 | 0.71995249 | ilkp-1 |
| WBGene00009355 | 0.148399287 | 0.12769527 | 0.28900702 | F33A8.7 |
| WBGene00009356 | 0.079571953 | 0.41776011 | 0.61446914 | F33A8.10 |
| WBGene00009357 | 0.063421086 | 0.41095452 | NA | F33C8.2 |
| WBGene00009358 | -0.377742414 | 0.10958461 | 0.26079784 | F33C8.4 |
| WBGene00009359 | -0.023056499 | 0.78121253 | NA | clec-102 |
| WBGene00009360 | 0.094654016 | 0.59112677 | 0.75212367 | F33E2.4 |
| WBGene00009361 | -0.237610703 | 0.20716309 | 0.39645769 | F33E2.5 |
| WBGene00009362 | NA | NA | NA | F33E2.6 |
| WBGene00009363 | 0.020925399 | 0.84643798 | NA | F33E2.7 |
| WBGene00009364 | 0.086388148 | 0.35298026 | 0.55296212 | F33H1.3 |
| WBGene00009365 | -0.014516444 | 0.82467434 | 0.90381444 | F33H1.4 |
| WBGene00009366 | -0.037334976 | 0.68751781 | 0.81998681 | F33H2.2 |
| WBGene00009367 | -0.207679889 | 0.03771742 | 0.12255944 | F33H2.3 |
| WBGene00009368 | 0.060433214 | 0.36948733 | 0.56931846 | pole-1 |
| WBGene00009369 | 0.000178136 | 0.99974581 | 0.99983667 | F33H2.6 |
| WBGene00009370 | -0.237806839 | 0.23005678 | 0.42442809 | set-10 |
| WBGene00009371 | -0.014383545 | 0.90655029 | 0.94941364 | F33H2.8 |
| WBGene00009372 | -0.013820684 | 0.87623155 | 0.9339623 | evl-18 |
| WBGene00009373 | -0.011376986 | 0.94818772 | 0.97044737 | F34D10.3 |
| WBGene00009374 | -0.149487602 | 0.15606749 | 0.32975175 | F34D10.4 |
| WBGene00009375 | -0.095227159 | 0.63162476 | 0.78099644 | F34D10.6 |
| WBGene00009377 | 0.40417804 | 0.04773799 | 0.14542902 | F34D10.8 |
| WBGene00009378 | NA | NA | NA | F34H10.1 |
| WBGene00009380 | 0.06589064 | 0.68729076 | 0.81989929 | F34H10.3 |
| WBGene00009381 | 0.038726127 | 0.57551883 | NA | F34H10.4 |
| WBGene00009382 | 0.208713848 | 0.12787766 | 0.28919628 | F34H10.5 |
| WBGene00009383 | 1.926325067 | 2.74E-18 | 1.79E-16 | F35B12.3 |
| WBGene00009384 | 1.212026321 | 2.07E-05 | 0.00023555 | piit-1 |

| | | | | | |
|---|---|---|---|---|---|
| WBGene00009385 | | -0.117206734 | 0.2014809 | 0.39024763 | sas-5 |
| WBGene00009386 | | 1.487772707 | 1.34E-07 | 2.48E-06 | tag-290 |
| WBGene00009387 | | 0.411505905 | 0.04184989 | 0.13210959 | glb-15 |
| WBGene00009388 | | 0.744897632 | 0.00050331 | 0.00390513 | F35B12.9 |
| WBGene00009389 | | 0.518369938 | 0.03111062 | 0.10597732 | F35B12.10 |
| WBGene00009390 | | 0.027743969 | 0.79793973 | 0.88691671 | F35C5.1 |
| WBGene00009391 | | -0.003108065 | 0.95728506 | NA | F35C5.3 |
| WBGene00009392 | NA | | NA | NA | F35C5.4 |
| WBGene00009393 | | 2.35737673 | 3.51E-42 | 6.94E-40 | clec-62 |
| WBGene00009394 | | 1.738122943 | 5.76E-175 | 6.49E-172 | clec-63 |
| WBGene00009395 | | 0.223708871 | 0.0534604 | 0.15817383 | clec-64 |
| WBGene00009396 | | 0.383674243 | 0.00133952 | 0.00893489 | clec-65 |
| WBGene00009397 | | 0.34307551 | 0.08369427 | 0.21753379 | clec-66 |
| WBGene00009398 | | 0.056411843 | 0.41891328 | NA | nspb-11 |
| WBGene00009399 | | 0.21032908 | 0.13642314 | 0.30153552 | ddn-2 |
| WBGene00009400 | | -0.178088452 | 0.3771443 | 0.57623349 | F35C5.12 |
| WBGene00009401 | | 0.21099373 | 0.06480333 | 0.18144297 | F35C11.2 |
| WBGene00009402 | NA | | NA | NA | F35C11.3 |
| WBGene00009403 | | -0.187129284 | 0.38703679 | 0.58569567 | F35C11.4 |
| WBGene00009404 | | -0.134269055 | 0.06939591 | 0.19026694 | F35C11.5 |
| WBGene00009405 | | 0.036274395 | 0.85437562 | 0.92207824 | F35C11.6 |
| WBGene00009406 | | 0.135118223 | 0.39488747 | 0.59297923 | F35C11.7 |
| WBGene00009408 | | -0.023056499 | 0.78121253 | NA | F35C12.3 |
| WBGene00009409 | NA | | NA | NA | F35E2.1 |
| WBGene00009410 | | -0.138634502 | 0.12394756 | 0.28431306 | F35E2.2 |
| WBGene00009411 | | 0.020232312 | 0.84643798 | NA | NA |
| WBGene00009412 | | 0.032241129 | 0.76934983 | 0.86867054 | F35E2.5 |
| WBGene00009413 | | 0.037023624 | 0.59153183 | NA | oac-19 |
| WBGene00009414 | | -0.023449708 | 0.69192352 | NA | NA |
| WBGene00009415 | | -0.259334167 | 0.19649358 | 0.38404251 | F35E2.8 |
| WBGene00009416 | | 0.157226365 | 0.28832687 | 0.48858723 | F35E2.9 |
| WBGene00009417 | NA | | NA | NA | F35E8.1 |
| WBGene00009418 | NA | | NA | NA | F35E8.2 |
| WBGene00009419 | NA | | NA | NA | F35E8.4 |
| WBGene00009420 | NA | | NA | NA | F35E8.6 |
| WBGene00009421 | NA | | NA | NA | F35E8.7 |
| WBGene00009422 | NA | | NA | NA | F35E8.9 |
| WBGene00009423 | | 0.038125098 | 0.61119482 | NA | F35E8.10 |
| WBGene00009423 | | 0.038125098 | 0.61119482 | NA | F35E8.13 |
| WBGene00009424 | NA | | NA | NA | F35E8.13 |
| WBGene00009425 | NA | | NA | NA | NA |
| WBGene00009426 | | -0.673738114 | 0.0077117 | 0.0367123 | F35E12.2 |
| WBGene00009427 | | 0.440885387 | 0.08246494 | 0.21537824 | fbxa-181 |

| | | | | |
|---|---|---|---|---|
| WBGene00009428 | 2.519773408 | 0.00183798 | 0.01166442 | NA |
| WBGene00009429 | -0.905606193 | 0.00558231 | 0.02857021 | irg-5 |
| WBGene00009430 | 1.005535806 | 9.41E-18 | 5.91E-16 | F35E12.6 |
| WBGene00009431 | 4.31714481 | 4.47E-36 | 6.97E-34 | dct-17 |
| WBGene00009432 | 0.113925348 | 0.58615578 | 0.74887137 | cld-9 |
| WBGene00009433 | 0.336601057 | 0.11635981 | 0.27240549 | F35E12.9 |
| WBGene00009434 | -0.996295472 | 8.09E-11 | 2.35E-09 | F35E12.10 |
| WBGene00009435 | 0.259022985 | 0.00011851 | 0.00112667 | F35G2.1 |
| WBGene00009436 | 0.270230256 | 0.01939317 | 0.07511518 | marb-1 |
| WBGene00009437 | -0.036289591 | 0.82494219 | 0.90387067 | F35G2.3 |
| WBGene00009438 | 0.085309563 | 0.24820057 | NA | F35G2.5 |
| WBGene00009439 | -0.074124546 | 0.53046477 | 0.70714146 | mlcd-1 |
| WBGene00009440 | -0.153841521 | 0.01745693 | 0.06951546 | idhg-1 |
| WBGene00009441 | 0.010830986 | 0.90889258 | 0.95068624 | wdr-48 |
| WBGene00009442 | -0.062404083 | 0.63929981 | 0.78615728 | F35G12.5 |
| WBGene00009443 | -0.261669827 | 0.19914777 | 0.38757444 | F35G12.7 |
| WBGene00009444 | -0.277512179 | 0.01209423 | 0.05201867 | erh-2 |
| WBGene00009445 | -0.23938933 | 0.00728955 | 0.03511618 | F35G12.12 |
| WBGene00009446 | 0.038125098 | 0.61119482 | NA | F35H8.1 |
| WBGene00009447 | -0.047495966 | 0.81716958 | 0.89956239 | F35H8.2 |
| WBGene00009448 | -0.010982753 | 0.90866078 | 0.95060397 | zfp-2 |
| WBGene00009449 | 0.111319389 | 0.1344031 | NA | F35H8.4 |
| WBGene00009450 | 0.342971983 | 0.00335671 | 0.01908564 | ugt-58 |
| WBGene00009451 | -0.344068976 | 3.62E-05 | 0.00039134 | cids-2 |
| WBGene00009452 | -0.09304927 | 0.42868233 | 0.62364249 | F36A2.2 |
| WBGene00009453 | 0.390717402 | 0.00067958 | 0.0050455 | F36A2.3 |
| WBGene00009454 | -0.258224636 | 8.53E-06 | 0.00010643 | F36A2.7 |
| WBGene00009455 | -0.053161513 | 0.70946543 | 0.83640239 | phip-1 |
| WBGene00009456 | 0.045782929 | 0.60311436 | 0.75988622 | F36A2.9 |
| WBGene00009457 | 1.994831824 | 0.00622821 | 0.03104773 | F36A2.10 |
| WBGene00009458 | 0.019975058 | 0.61254052 | NA | F36A2.11 |
| WBGene00009459 | 0.191024347 | 0.12986891 | 0.29221004 | F36A2.12 |
| WBGene00009460 | 0.097946545 | 0.06386466 | 0.17951954 | ubr-5 |
| WBGene00009461 | 0.145027479 | 0.48569063 | 0.67027418 | madf-2 |
| WBGene00009462 | 0.020317245 | 0.84643798 | NA | sre-22 |
| WBGene00009463 | -0.050580107 | 0.47300881 | NA | F36D1.4 |
| WBGene00009464 | NA | NA | NA | F36D1.5 |
| WBGene00009465 | 0.033034883 | 0.64219503 | NA | F36D1.6 |
| WBGene00009466 | 2.997137648 | 6.42E-11 | 1.91E-09 | F36D1.7 |
| WBGene00009467 | 0.020232312 | 0.84643798 | NA | F36D1.8 |
| WBGene00009468 | -0.035163513 | 0.86342212 | 0.92712602 | F36D1.9 |
| WBGene00009469 | -0.123729638 | 0.44086717 | 0.635052 | F36D3.1 |
| WBGene00009470 | 0.360086084 | 0.01998507 | 0.07657663 | F36D3.4 |

| | | | | |
|---|---|---|---|---|
| WBGene00009471 | 0.108942554 | 0.15594412 | NA | F36D3.5 |
| WBGene00009472 | NA | NA | NA | NA |
| WBGene00009473 | 0.0398788 | 0.58971221 | NA | F36D3.8 |
| WBGene00009474 | NA | NA | NA | NA |
| WBGene00009475 | -0.133237609 | 0.52387747 | 0.70139111 | F36F2.1 |
| WBGene00009476 | 0.551068605 | 0.0096982 | 0.04387865 | F36F2.2 |
| WBGene00009477 | -0.052147364 | 0.43103881 | 0.62607533 | rbpl-1 |
| WBGene00009478 | -0.025899098 | 0.82239373 | 0.90266637 | syx-7 |
| WBGene00009479 | -0.03347758 | 0.69098622 | 0.8224095 | fcp-1 |
| WBGene00009480 | -0.081021825 | 0.51186203 | 0.69179101 | F36F2.7 |
| WBGene00009481 | 0.015899775 | 0.84174438 | NA | F36F2.8 |
| WBGene00009482 | 0.09864678 | 0.53808707 | 0.71275968 | F36G3.1 |
| WBGene00009483 | -0.195599083 | 0.34670087 | 0.54639908 | F36G3.2 |
| WBGene00009484 | 0.085309563 | 0.24820057 | NA | F36G3.3 |
| WBGene00009485 | -0.263690675 | 0.23969517 | 0.43670695 | F36G9.3 |
| WBGene00009486 | NA | NA | NA | F36G9.7 |
| WBGene00009487 | 0.016059264 | 0.84014476 | NA | clec-232 |
| WBGene00009488 | -0.190664255 | 0.35379481 | 0.55364577 | oac-20 |
| WBGene00009489 | NA | NA | NA | F36G9.13 |
| WBGene00009490 | -0.011535809 | 0.91042331 | 0.95112162 | fbxa-99 |
| WBGene00009491 | NA | NA | NA | F36G9.15 |
| WBGene00009492 | 0.083209383 | 0.30786773 | NA | F36H1.3 |
| WBGene00009493 | -0.309530891 | 0.18188599 | 0.36461879 | hrg-4 |
| WBGene00009494 | 0.019325396 | 0.77340931 | NA | hrg-5 |
| WBGene00009495 | -0.219754665 | 0.27877929 | 0.47871443 | hrg-6 |
| WBGene00009496 | 0.049346887 | 0.47367826 | NA | F36H1.11 |
| WBGene00009497 | 0.215827431 | 0.05807304 | 0.16776784 | F36H1.12 |
| WBGene00009498 | -0.051966473 | 0.46861441 | 0.65743423 | tat-5 |
| WBGene00009499 | -0.211412637 | 0.02554454 | 0.09216305 | ent-6 |
| WBGene00009500 | 0.92112808 | 0.00590127 | 0.02978792 | F36H2.3 |
| WBGene00009501 | -0.027188909 | 0.68205301 | NA | F37A8.1 |
| WBGene00009502 | 0.063527295 | 0.41294485 | NA | F37A8.2 |
| WBGene00009503 | 2.084188692 | 1.30E-06 | 1.99E-05 | F37A8.5 |
| WBGene00009504 | 0.121409754 | 0.56480804 | 0.73464869 | F37B12.1 |
| WBGene00009505 | -0.150332895 | 0.12425422 | 0.28458189 | F37B12.3 |
| WBGene00009507 | -0.144019481 | 0.08485745 | 0.21938866 | mus-101 |
| WBGene00009508 | 0.079027406 | 0.31969357 | 0.51987478 | row-1 |
| WBGene00009509 | -0.012374601 | 0.9071207 | 0.94967137 | F37D6.3 |
| WBGene00009510 | -0.178024231 | 0.21573034 | 0.40673326 | F37D6.4 |
| WBGene00009511 | 0.340527163 | 0.15407939 | 0.32715947 | F37H8.2 |
| WBGene00009512 | 1.883290269 | 1.37E-07 | 2.51E-06 | bigr-1 |
| WBGene00009513 | 0.07616368 | 0.48795915 | 0.67201673 | sfxn-1.2 |
| WBGene00009514 | 0.162972778 | 0.09413508 | 0.23503269 | F37H8.5 |

| | | | | |
|---|---|---|---|---|
| WBGene00009515 | 1.970395009 | 1.05E-10 | 3.02E-09 | clec-170 |
| WBGene00009517 | 0.13389452 | 0.15713149 | 0.33111142 | clec-167 |
| WBGene00009518 | 3.018041677 | 1.19E-37 | 2.02E-35 | clec-166 |
| WBGene00009519 | NA | NA | NA | F38A1.6 |
| WBGene00009520 | 0.03732969 | 0.59229443 | NA | clec-168 |
| WBGene00009521 | -0.124481061 | 0.19912866 | 0.38757444 | F38A1.8 |
| WBGene00009522 | 0.312751762 | 0.16653938 | 0.34351974 | F38A1.9 |
| WBGene00009523 | 0.435098307 | 0.09181431 | 0.23089202 | clec-165 |
| WBGene00009524 | 0.059980168 | 0.38780842 | NA | F38A1.11 |
| WBGene00009525 | 0.020317245 | 0.84643798 | NA | F38A1.13 |
| WBGene00009526 | 6.767089945 | 2.12E-28 | 2.46E-26 | clec-169 |
| WBGene00009527 | NA | NA | NA | NA |
| WBGene00009528 | 0.094575086 | 0.53333796 | 0.70899686 | F38A6.4 |
| WBGene00009529 | 0.582690906 | 0.03492975 | 0.11583886 | F38B2.2 |
| WBGene00009530 | 0.052920013 | 0.6423149 | 0.78820992 | F38B2.3 |
| WBGene00009531 | -0.309916743 | 0.05840575 | 0.16833044 | F38B2.4 |
| WBGene00009532 | 1.051990281 | 1.54E-13 | 6.32E-12 | ccch-1 |
| WBGene00009533 | -0.569688053 | 0.04403196 | 0.13709778 | F38B7.2 |
| WBGene00009534 | 0.187679696 | 0.28934495 | 0.4895758 | F38B7.3 |
| WBGene00009535 | NA | NA | NA | F38C2.1 |
| WBGene00009536 | -0.061567612 | 0.58709361 | 0.74981371 | F38C2.4 |
| WBGene00009537 | -0.131006211 | 0.51674401 | 0.69614457 | ccch-2 |
| WBGene00009538 | NA | NA | NA | clec-191 |
| WBGene00009539 | -0.485557035 | 0.06994718 | 0.19109846 | F38C2.7 |
| WBGene00009540 | NA | NA | NA | hlh-31 |
| WBGene00009541 | 0.016453512 | 0.83636998 | NA | cutl-17 |
| WBGene00009542 | -0.128812664 | 0.03980601 | 0.12750514 | copb-2 |
| WBGene00009543 | 0.091768519 | 0.52068454 | 0.6992609 | F38E11.6 |
| WBGene00009544 | 0.313414083 | 0.18248083 | 0.36537689 | F38E11.9 |
| WBGene00009546 | -0.507247617 | 0.06172739 | 0.17521339 | fbxb-3 |
| WBGene00009547 | 1.151746977 | 0.00438577 | 0.02360737 | F38H4.3 |
| WBGene00009548 | 0.036988272 | 0.57084766 | NA | F38H4.4 |
| WBGene00009549 | 0.137938171 | 0.3348676 | 0.53413188 | F38H4.5 |
| WBGene00009550 | 0.036692668 | 0.59311964 | NA | F38H4.6 |
| WBGene00009551 | 0.029470505 | 0.83413044 | 0.90957526 | mob-1 |
| WBGene00009552 | 0.143776183 | 0.2284551 | 0.42211145 | piki-1 |
| WBGene00009553 | -0.16815007 | 0.02777616 | 0.09799295 | hinf-1 |
| WBGene00009554 | -0.387027072 | 0.00585378 | 0.02958412 | F39B2.3 |
| WBGene00009555 | NA | NA | NA | NA |
| WBGene00009556 | -0.026469052 | 0.87305776 | 0.93245617 | F39B2.5 |
| WBGene00009557 | 0.10196281 | 0.56517955 | 0.73488399 | mtcu-1 |
| WBGene00009558 | 0.790089559 | 0.01553725 | 0.0636488 | F39B2.8 |
| WBGene00009559 | 0.252441188 | 0.0529397 | 0.1570598 | mtx-1 |

| | | | | | |
|---|---|---|---|---|---|
| WBGene00009560 | | 0.035894636 | 0.83253626 | 0.90860486 | psa-3 |
| WBGene00009561 | | 0.163672294 | 0.16750002 | 0.3451014 | F39D8.3 |
| WBGene00009562 | | 1.312989324 | 6.70E-09 | 1.54E-07 | flp-22 |
| WBGene00009563 | | -0.246161652 | 0.03501268 | 0.11599279 | F39H2.3 |
| WBGene00009565 | | -0.138565284 | 0.20701701 | 0.39630441 | F39H11.1 |
| WBGene00009566 | NA | | NA | NA | srsx-1 |
| WBGene00009567 | NA | | NA | NA | srbc-25 |
| WBGene00009568 | NA | | NA | NA | srbc-26 |
| WBGene00009569 | NA | | NA | NA | srz-66 |
| WBGene00009570 | NA | | NA | NA | NA |
| WBGene00009571 | | -0.023056499 | 0.78121253 | NA | F40D4.12 |
| WBGene00009572 | | 0.146769602 | 0.20207537 | 0.39052078 | F40D4.13 |
| WBGene00009573 | | 0.141741219 | 0.25587374 | 0.45436596 | F40E10.5 |
| WBGene00009574 | | -0.163273513 | 0.12702556 | 0.28833817 | F40E10.6 |
| WBGene00009575 | | -0.043801175 | 0.63953481 | 0.78619899 | F40F8.1 |
| WBGene00009576 | | -0.179316231 | 0.38170818 | 0.58093445 | F40F8.3 |
| WBGene00009577 | | 0.366462654 | 0.05447578 | 0.16015662 | F40F8.4 |
| WBGene00009578 | | 0.5223825 | 6.07E-05 | 0.00062551 | F40F8.5 |
| WBGene00009579 | | 0.576365472 | 0.03821811 | 0.12373979 | arrd-6 |
| WBGene00009580 | | -0.00779573 | 0.9478099 | 0.97044737 | xbx-6 |
| WBGene00009581 | | -0.013476838 | 0.89898666 | 0.94663864 | F40F9.3 |
| WBGene00009582 | | 0.36879829 | 0.04468841 | 0.13851405 | F40F9.5 |
| WBGene00009583 | | -0.126071935 | 0.05426351 | 0.15978923 | aagr-3 |
| WBGene00009584 | | -0.031219438 | 0.72327103 | 0.84472009 | drap-1 |
| WBGene00009585 | | 0.036687355 | 0.59489568 | NA | cal-7 |
| WBGene00009586 | | 0.193160093 | 0.16040287 | 0.33484732 | F40F9.10 |
| WBGene00009587 | | 0.073194838 | 0.40265737 | 0.60119382 | mig-38 |
| WBGene00009588 | | -0.037267312 | 0.76798611 | 0.86834482 | F40F11.3 |
| WBGene00009589 | | -0.053591705 | 0.45278076 | NA | F40F11.4 |
| WBGene00009590 | NA | | NA | NA | NA |
| WBGene00009592 | | 0.176461773 | 0.09037499 | 0.22872606 | F40F12.3 |
| WBGene00009593 | | -0.07885766 | 0.48761998 | 0.6718024 | mam-7 |
| WBGene00009594 | | 0.102008175 | 0.14706415 | 0.31738888 | cyld-1 |
| WBGene00009595 | | 1.484599903 | 4.90E-14 | 2.12E-12 | cbp-3 |
| WBGene00009596 | NA | | NA | NA | sre-8 |
| WBGene00009597 | | -0.049665371 | 0.48040244 | NA | F40G12.2 |
| WBGene00009598 | NA | | NA | NA | NA |
| WBGene00009599 | | 0.033034883 | 0.64219503 | NA | F40G12.4 |
| WBGene00009600 | | -0.001993815 | 0.95612996 | NA | F40G12.5 |
| WBGene00009601 | | 0.059281883 | 0.42158309 | NA | F40G12.6 |
| WBGene00009602 | | 0.060721284 | 0.5731562 | 0.74127107 | F40G12.7 |
| WBGene00009603 | | -0.023109762 | 0.78121253 | NA | srt-33 |
| WBGene00009604 | NA | | NA | NA | F40G12.9 |

| | | | | | |
|---|---|---|---|---|---|
| WBGene00009605 | | 0.139235459 | 0.25624634 | 0.45460486 | F40G12.10 |
| WBGene00009606 | | -0.094009315 | 0.43526136 | 0.63045722 | F40G12.11 |
| WBGene00009607 | NA | | NA | NA | oac-26 |
| WBGene00009608 | NA | | NA | NA | nhr-265 |
| WBGene00009609 | | 0.020232312 | 0.84643798 | NA | oac-27 |
| WBGene00009610 | NA | | NA | NA | oac-28 |
| WBGene00009611 | NA | | NA | NA | F41D3.6 |
| WBGene00009612 | NA | | NA | NA | F41D3.7 |
| WBGene00009613 | NA | | NA | NA | F41D3.8 |
| WBGene00009614 | | 0.057855329 | 0.58501057 | 0.74805087 | F41D3.9 |
| WBGene00009615 | NA | | NA | NA | oac-25 |
| WBGene00009616 | | 0.020232312 | 0.84643798 | NA | F41D3.11 |
| WBGene00009617 | | -0.348208027 | 0.12911542 | 0.29114663 | F41E7.1 |
| WBGene00009618 | | -0.086038083 | 0.67332111 | 0.80998475 | F41E7.2 |
| WBGene00009619 | | 0.302050668 | 0.13451358 | 0.2987581 | npr-6 |
| WBGene00009620 | | -0.622250668 | 0.03456417 | 0.11503843 | fip-5 |
| WBGene00009621 | | -0.811652368 | 2.86E-10 | 7.78E-09 | fipr-21 |
| WBGene00009622 | | -0.189578268 | 0.15329235 | 0.32640391 | F41E7.6 |
| WBGene00009623 | | 0.227134235 | 0.28541561 | 0.48562115 | F41E7.7 |
| WBGene00009624 | | -0.705555629 | 0.00015677 | 0.0014271 | cnc-8 |
| WBGene00009625 | NA | | NA | NA | F41E7.9 |
| WBGene00009626 | | -0.992267773 | 2.97E-13 | 1.18E-11 | F42A8.1 |
| WBGene00009627 | | -0.09256952 | 0.41393723 | 0.61154897 | cup-15 |
| WBGene00009628 | | 0.96413697 | 2.37E-28 | 2.73E-26 | tatn-1 |
| WBGene00009629 | | -0.601477574 | 0.04282095 | 0.1344791 | sprr-2 |
| WBGene00009630 | NA | | NA | NA | F42E8.1 |
| WBGene00009631 | NA | | NA | NA | NA |
| WBGene00009632 | | -0.020061502 | 0.86251778 | 0.92672958 | ttyh-1 |
| WBGene00009633 | | -0.02419332 | 0.70411432 | NA | F42E11.3 |
| WBGene00009634 | | 1.360150618 | 4.82E-16 | 2.56E-14 | nspc-9 |
| WBGene00009635 | | -0.508372874 | 0.0289117 | 0.10090492 | F42F12.3 |
| WBGene00009636 | | -0.372480983 | 0.08423453 | 0.21851129 | F42F12.4 |
| WBGene00009637 | NA | | NA | NA | nspc-13 |
| WBGene00009638 | | 1.507404216 | 4.61E-28 | 5.16E-26 | nspc-14 |
| WBGene00009639 | | 0.065014541 | 0.37399213 | NA | nspc-12 |
| WBGene00009639 | | 0.065014541 | 0.37399213 | NA | nspc-15 |
| WBGene00009640 | | 1.56895295 | 4.63E-21 | 3.71E-19 | nspc-10 |
| WBGene00009641 | | 0.096406411 | 0.47317203 | 0.66116737 | F42G4.2 |
| WBGene00009642 | | 1.443685067 | 9.35E-11 | 2.70E-09 | F42G4.5 |
| WBGene00009643 | | 4.317362955 | 0.00013078 | 0.00122116 | F42G4.6 |
| WBGene00009644 | | 0.489508263 | 0.07018385 | 0.19163776 | F42G4.7 |
| WBGene00009645 | | -0.035290889 | 0.81639365 | 0.89911384 | F42G10.1 |
| WBGene00009646 | NA | | NA | NA | F42H11.1 |

| | | | | | |
|---|---|---|---|---|---|
| WBGene00009647 | | -0.043433535 | 0.75109051 | 0.85938701 | phlp-2 |
| WBGene00009648 | | -0.070790404 | 0.65222076 | 0.79404326 | zhit-2 |
| WBGene00009649 | | -0.232801689 | 0.28144087 | 0.48098235 | F43C1.5 |
| WBGene00009650 | | 0.070751443 | 0.47104617 | 0.65901452 | ccnk-1 |
| WBGene00009651 | | -0.013175483 | 0.91108067 | 0.95112162 | NA |
| WBGene00009652 | | 0.027912924 | 0.82840275 | 0.9062566 | F43D2.3 |
| WBGene00009653 | | 0.238609166 | 0.26509942 | 0.46365025 | F43D9.1 |
| WBGene00009654 | | -0.241274588 | 0.02800225 | 0.0984821 | sly-1 |
| WBGene00009656 | | -0.209760591 | 0.10517896 | 0.2535691 | NA |
| WBGene00009657 | | 0.080803767 | 0.69904422 | 0.82858699 | F43G6.4 |
| WBGene00009658 | | -0.362921671 | 0.12083015 | 0.27965132 | F43G6.5 |
| WBGene00009659 | | -0.171806121 | 0.32671868 | 0.52577359 | F43G6.7 |
| WBGene00009660 | | 0.781505775 | 0.00196418 | 0.01231059 | F43G6.8 |
| WBGene00009661 | | -0.156300165 | 0.01114705 | 0.04870306 | patr-1 |
| WBGene00009662 | | 0.067243289 | 0.72050208 | 0.84342114 | F43G6.10 |
| WBGene00009663 | | 0.107290688 | 0.60729005 | 0.76307361 | hda-5 |
| WBGene00009664 | | -0.098400208 | 0.17020458 | 0.34872169 | idha-1 |
| WBGene00009665 | | -0.187548012 | 0.20169176 | 0.39039503 | lmd-1 |
| WBGene00009666 | | 0.06295621 | 0.67489597 | 0.8111554 | slc-25A42 |
| WBGene00009667 | | -0.091888212 | 0.40442971 | 0.6025966 | F43G9.4 |
| WBGene00009668 | | -0.297893752 | 0.01522601 | 0.06259108 | cfim-1 |
| WBGene00009670 | | 0.039888668 | 0.56326497 | NA | F43G9.8 |
| WBGene00009671 | | -0.238347379 | 0.0019496 | 0.01224546 | mfap-1 |
| WBGene00009672 | | -0.182754319 | 0.0577254 | 0.16712627 | F43G9.12 |
| WBGene00009673 | | -0.111140528 | 0.49764778 | 0.67976712 | F43G9.13 |
| WBGene00009674 | | -0.32586141 | 0.00768966 | 0.03663661 | nucb-1 |
| WBGene00009675 | | -0.049278988 | 0.46988128 | NA | F44A6.3 |
| WBGene00009676 | | -0.089811309 | 0.654037 | 0.7949984 | F44A6.4 |
| WBGene00009677 | | 1.239399318 | 1.42E-06 | 2.14E-05 | F44A6.5 |
| WBGene00009678 | | 0.525926707 | 0.00493549 | 0.02591493 | magu-4 |
| WBGene00009679 | | 0.509484996 | 0.02450144 | 0.08948005 | F44D12.2 |
| WBGene00009680 | | 0.065014541 | 0.37399213 | NA | msd-1 |
| WBGene00009680 | | 0.065014541 | 0.37399213 | NA | msd-3 |
| WBGene00009681 | | 1.843915057 | 3.74E-08 | 7.60E-07 | gipc-2 |
| WBGene00009682 | NA | | NA | NA | msd-2 |
| WBGene00009683 | | 0.033180193 | 0.61386379 | NA | F44D12.6 |
| WBGene00009684 | NA | | NA | NA | msd-1 |
| WBGene00009684 | NA | | NA | NA | msd-3 |
| WBGene00009685 | | 0.058314668 | 0.40208118 | NA | F44D12.8 |
| WBGene00009686 | | 0.966532717 | 1.25E-06 | 1.92E-05 | ent-7 |
| WBGene00009687 | NA | | NA | NA | F44D12.10 |
| WBGene00009688 | | -0.256353788 | 0.00090613 | 0.0064747 | F44E5.1 |
| WBGene00009689 | | -0.139293192 | 0.37206208 | 0.57136154 | F44E5.2 |

| | | | | |
|---|---|---|---|---|
| WBGene00009690 | -0.058259153 | 0.59476533 | 0.75426238 | F44E5.3 |
| WBGene00009691 | NA | NA | NA | F44E5.4 |
| WBGene00009691 | NA | NA | NA | F44E5.5 |
| WBGene00009692 | NA | NA | NA | F44E5.4 |
| WBGene00009692 | NA | NA | NA | F44E5.5 |
| WBGene00009693 | NA | NA | NA | NA |
| WBGene00009695 | 0.129401398 | 0.10528579 | NA | clp-8 |
| WBGene00009696 | -0.32869945 | 0.14693838 | 0.31725775 | F44F1.4 |
| WBGene00009697 | -0.750007311 | 0.01841446 | 0.07241112 | NA |
| WBGene00009698 | -0.763167104 | 0.00899762 | 0.04146247 | F44F1.6 |
| WBGene00009700 | 0.187261134 | 0.28093787 | 0.48048327 | lurp-1 |
| WBGene00009701 | 0.0385743 | 0.56208407 | 0.73230632 | egg-3 |
| WBGene00009702 | 0.003490488 | 0.97787291 | 0.98738188 | F44F4.3 |
| WBGene00009704 | 0.033463284 | 0.6365037 | NA | F44F4.9 |
| WBGene00009705 | 0.256731711 | 0.04143003 | 0.13119448 | F44F4.10 |
| WBGene00009706 | -0.088856577 | 0.52677931 | 0.70408762 | argk-1 |
| WBGene00009707 | NA | NA | NA | srxa-16 |
| WBGene00009708 | 0.301778588 | 0.05789082 | 0.16740658 | F44G3.7 |
| WBGene00009709 | 0.097476395 | 0.3367938 | 0.53627204 | fbxa-144 |
| WBGene00009710 | 0.036894305 | 0.59120501 | NA | F44G3.10 |
| WBGene00009711 | -0.28785996 | 0.00227472 | 0.0138421 | F44G4.1 |
| WBGene00009712 | -0.200363037 | 0.00891124 | 0.04114337 | F44G4.2 |
| WBGene00009713 | -0.457573104 | 0.01227637 | 0.05264745 | F44G4.3 |
| WBGene00009714 | 0.155122056 | 0.06782466 | NA | F44G4.5 |
| WBGene00009715 | 0.308590227 | 0.12054388 | 0.27938624 | F44G4.6 |
| WBGene00009716 | 0.275387966 | 0.07722155 | 0.20566135 | F44G4.7 |
| WBGene00009717 | 0.395418952 | 0.05697644 | 0.16561287 | dep-1 |
| WBGene00009720 | 0.020232312 | 0.84643798 | NA | F45B8.3 |
| WBGene00009721 | -0.243897968 | 0.18123618 | 0.36368954 | F45D3.1 |
| WBGene00009722 | 0.241171385 | 0.26524355 | 0.4637486 | F45D3.2 |
| WBGene00009723 | 0.48840823 | 0.00089909 | 0.00643698 | F45D3.3 |
| WBGene00009724 | 1.922510539 | 3.58E-06 | 4.85E-05 | F45D3.4 |
| WBGene00009725 | 0.020925399 | 0.84643798 | NA | F45E6.1 |
| WBGene00009726 | 0.222257845 | 0.21065747 | 0.40044612 | tbc-13 |
| WBGene00009727 | 0.102252662 | 0.5094917 | 0.6895487 | lips-1 |
| WBGene00009728 | -0.105120254 | 0.34515761 | 0.54490539 | sdz-19 |
| WBGene00009729 | 0.367389939 | 0.1124058 | 0.26542401 | F45E10.2 |
| WBGene00009730 | 0.478711211 | 0.05137983 | 0.15344072 | myo-6 |
| WBGene00009731 | -0.148156826 | 0.14187393 | 0.30984133 | exo-1 |
| WBGene00009732 | -0.144521794 | 0.15882727 | 0.33319818 | cope-1 |
| WBGene00009733 | -0.085788228 | 0.63662796 | 0.78445295 | F45G2.7 |
| WBGene00009734 | -0.405759057 | 0.00608504 | 0.03050652 | F45G2.8 |
| WBGene00009735 | -0.003275043 | 0.98544889 | 0.9917485 | F45G2.9 |

| | | | | |
|---|---|---|---|---|
| WBGene00009736 | -0.068679803 | 0.61712417 | 0.77055851 | F45G2.10 |
| WBGene00009738 | 0.245486254 | 0.25718607 | 0.45539154 | hecw-1 |
| WBGene00009739 | -0.026005211 | 0.76166133 | 0.86485835 | F45H10.2 |
| WBGene00009740 | -0.085227797 | 0.35354818 | 0.55343729 | F45H10.3 |
| WBGene00009741 | 0.0332672 | 0.8669311 | 0.92912219 | drr-1 |
| WBGene00009742 | 0.077811957 | 0.61646427 | 0.77004124 | F45H10.5 |
| WBGene00009743 | -0.576919219 | 0.04023593 | 0.12853901 | sptf-1 |
| WBGene00009744 | -0.073005503 | 0.50911731 | 0.68943312 | hpo-35 |
| WBGene00009745 | 0.080151898 | 0.62455781 | 0.77565111 | cox-19 |
| WBGene00009746 | NA | NA | NA | F46A8.3 |
| WBGene00009747 | NA | NA | NA | F46A8.4 |
| WBGene00009748 | 0.038125098 | 0.61119482 | NA | F46A8.5 |
| WBGene00009749 | NA | NA | NA | F46A8.6 |
| WBGene00009750 | NA | NA | NA | F46A8.7 |
| WBGene00009751 | 0.038125098 | 0.61119482 | NA | F46A8.8 |
| WBGene00009752 | 0.020232312 | 0.84643798 | NA | NA |
| WBGene00009753 | 0.181789487 | 0.04369581 | NA | scrm-6 |
| WBGene00009754 | 0.086426234 | 0.42213848 | 0.61846519 | F46A9.1 |
| WBGene00009755 | 0.156925762 | 0.10796859 | 0.25837987 | F46A9.2 |
| WBGene00009756 | 0.038125098 | 0.61119482 | NA | F46B3.1 |
| WBGene00009757 | -0.003614866 | 0.95700117 | NA | NA |
| WBGene00009758 | NA | NA | NA | ttr-11 |
| WBGene00009759 | 0.078203897 | 0.26561294 | NA | ttr-12 |
| WBGene00009760 | 0.038125098 | 0.61119482 | NA | F46B3.7 |
| WBGene00009761 | NA | NA | NA | F46B3.8 |
| WBGene00009762 | NA | NA | NA | F46B3.9 |
| WBGene00009763 | NA | NA | NA | srz-18 |
| WBGene00009764 | 0.020232312 | 0.84643798 | NA | NA |
| WBGene00009765 | NA | NA | NA | F46B3.14 |
| WBGene00009766 | NA | NA | NA | F46B3.15 |
| WBGene00009767 | NA | NA | NA | F46B3.16 |
| WBGene00009768 | 0.074892148 | 0.28555128 | NA | F46B6.2 |
| WBGene00009769 | -0.052259064 | 0.70233128 | 0.83093594 | F46B6.4 |
| WBGene00009770 | -0.166666474 | 0.01909037 | 0.07429614 | F46B6.5 |
| WBGene00009771 | -0.135116698 | 0.16927473 | 0.34754612 | F46B6.6 |
| WBGene00009772 | 0.105767006 | 0.05338268 | 0.15810973 | ztf-7 |
| WBGene00009773 | 2.131167119 | 1.40E-06 | 2.13E-05 | lipl-2 |
| WBGene00009774 | 0.092764015 | 0.53664614 | 0.71162359 | F46B6.9 |
| WBGene00009775 | NA | NA | NA | F46B6.10 |
| WBGene00009776 | 0.062540986 | 0.59869436 | 0.75675421 | F46B6.12 |
| WBGene00009777 | 0.850929601 | 0.01922387 | 0.07471639 | F46C3.2 |
| WBGene00009778 | 6.003036825 | 1.14E-12 | 4.29E-11 | F46C5.1 |
| WBGene00009779 | 0.22798035 | 0.21553177 | 0.40653717 | F46C5.2 |

| | | | | |
|---|---|---|---|---|
| WBGene00009780 | 0.109547845 | 0.44213794 | 0.63611514 | F46C5.4 |
| WBGene00009781 | 0.224413057 | 0.30514983 | 0.50573904 | F46C5.6 |
| WBGene00009782 | 0.020232312 | 0.84643798 | NA | F46C5.7 |
| WBGene00009783 | -0.079880606 | 0.43407065 | 0.62916824 | rer-1 |
| WBGene00009784 | -0.134128125 | 0.1155802 | 0.27123262 | F46C5.9 |
| WBGene00009785 | -0.530028065 | 0.04587785 | 0.14129104 | F46C5.10 |
| WBGene00009786 | -0.022979331 | 0.86926095 | 0.93000745 | NA |
| WBGene00009787 | -2.168678689 | 4.87E-20 | 3.67E-18 | F46F2.3 |
| WBGene00009788 | -0.001488154 | 0.95639912 | NA | F46F2.4 |
| WBGene00009789 | -0.968392401 | 0.00310178 | 0.01792843 | F46F2.5 |
| WBGene00009790 | 0.099313616 | 0.63656345 | 0.78445295 | lgc-44 |
| WBGene00009791 | 0.122730824 | 0.25820345 | 0.45633859 | F46F3.3 |
| WBGene00009793 | 0.08329139 | 0.42467645 | 0.62055982 | pkn-1 |
| WBGene00009796 | 1.115093381 | 4.20E-16 | 2.27E-14 | F46G10.1 |
| WBGene00009797 | 1.05790834 | 1.01E-05 | 0.00012428 | F46G10.2 |
| WBGene00009798 | 0.282052547 | 0.19878982 | 0.38713512 | F46G10.4 |
| WBGene00009799 | 0.069615514 | 0.72247988 | 0.84438636 | lgc-47 |
| WBGene00009800 | 0.150305082 | 0.48188457 | 0.6678599 | rrc-1 |
| WBGene00009801 | 0.37985092 | 0.01667637 | 0.06720581 | ipla-2 |
| WBGene00009802 | 0.020317245 | 0.84643798 | NA | F47B8.1 |
| WBGene00009803 | 1.467899273 | 1.06E-06 | 1.66E-05 | F47B8.2 |
| WBGene00009804 | 0.356019086 | 0.14005197 | 0.30737534 | F47B8.3 |
| WBGene00009805 | 0.075450426 | 0.33451267 | NA | F47B8.4 |
| WBGene00009806 | -0.003270206 | 0.95698316 | NA | F47B8.5 |
| WBGene00009807 | 0.03363967 | 0.7669638 | 0.86758224 | pepm-1 |
| WBGene00009808 | 0.065014541 | 0.37399213 | NA | srm-4 |
| WBGene00009809 | 0.427171866 | 0.06623951 | 0.18378107 | F47B8.8 |
| WBGene00009810 | 0.009497318 | 0.9341572 | 0.96290283 | srm-6 |
| WBGene00009811 | -0.023056499 | 0.78121253 | NA | F47B8.10 |
| WBGene00009812 | 0.191665183 | 0.01076225 | 0.04734678 | suca-1 |
| WBGene00009813 | 0.305986753 | 0.00476461 | 0.02518365 | haly-1 |
| WBGene00009814 | 0.011724584 | 0.89772193 | NA | F47B10.3 |
| WBGene00009815 | -0.023056499 | 0.78121253 | NA | F47B10.4 |
| WBGene00009816 | -0.159415604 | 0.45374823 | 0.64589961 | F47B10.5 |
| WBGene00009817 | NA | NA | NA | F47B10.6 |
| WBGene00009818 | 0.498661943 | 0.01603514 | 0.06520911 | acbp-3 |
| WBGene00009819 | 0.124329259 | 0.25873913 | 0.45673976 | F47B10.8 |
| WBGene00009820 | -0.041549293 | 0.83037401 | 0.90741199 | F47B10.9 |
| WBGene00009822 | 0.020925399 | 0.84643798 | NA | clec-113 |
| WBGene00009823 | -0.267480833 | 0.11058668 | 0.26229761 | gadr-6 |
| WBGene00009824 | -0.429291312 | 0.0923326 | 0.23187677 | gpdh-1 |
| WBGene00009825 | 0.14492256 | 0.43893935 | 0.63365323 | F47G4.4 |
| WBGene00009826 | -0.02419332 | 0.70411432 | NA | F47G4.5 |

| | | | | | |
|---|---|---|---|---|---|
| WBGene00009827 | | -0.050740777 | 0.47171914 | NA | hmg-6 |
| WBGene00009828 | NA | | NA | NA | NA |
| WBGene00009829 | | -0.228776746 | 0.0011647 | 0.00797957 | tmed-10 |
| WBGene00009830 | | -2.460358209 | 5.51E-09 | 1.28E-07 | cutl-18 |
| WBGene00009831 | | 0.243557727 | 0.26626027 | 0.46491551 | F47G9.4 |
| WBGene00009833 | NA | | NA | NA | F47G9.6 |
| WBGene00009834 | | 0.126566078 | 0.35259604 | 0.55253758 | lsy-27 |
| WBGene00009835 | | 3.471462494 | 1.13E-06 | 1.77E-05 | F47H4.2 |
| WBGene00009836 | | 0.020317245 | 0.84643798 | NA | fbxa-185 |
| WBGene00009837 | NA | | NA | NA | NA |
| WBGene00009838 | | 0.455338906 | 0.05616864 | 0.1640465 | NA |
| WBGene00009839 | | 0.763296716 | 0.02678745 | 0.09550554 | fbxa-188 |
| WBGene00009840 | | 0.997994185 | 0.00537029 | 0.02767559 | fbxa-189 |
| WBGene00009841 | | 0.057263786 | 0.43360027 | NA | fbxa-134 |
| WBGene00009842 | | 1.170725461 | 0.0096753 | 0.04380212 | igeg-2 |
| WBGene00009843 | | 1.200994673 | 0.01057914 | 0.04668147 | F48C5.2 |
| WBGene00009844 | | -0.006968242 | 0.87034079 | NA | cwp-5 |
| WBGene00009846 | | 0.550915356 | 0.04403872 | 0.13709778 | NA |
| WBGene00009848 | | 0.078748631 | 0.61178436 | 0.76638272 | F48F5.6 |
| WBGene00009849 | | 0.075076776 | 0.28090944 | NA | F48F7.3 |
| WBGene00009850 | | 0.620820155 | 0.03700552 | 0.12082912 | F48F7.5 |
| WBGene00009851 | | 0.009078889 | 0.95929513 | 0.97767605 | F48F7.6 |
| WBGene00009852 | | 0.17283442 | 0.18424019 | 0.36766449 | arrd-24 |
| WBGene00009853 | | 0.056188364 | 0.42092875 | NA | arrd-27 |
| WBGene00009854 | | 0.717902537 | 0.02951015 | 0.10243197 | clec-31 |
| WBGene00009855 | | -0.084044353 | 0.52599177 | 0.70322761 | clec-22 |
| WBGene00009856 | | 5.252097012 | 3.80E-06 | 5.11E-05 | clec-24 |
| WBGene00009857 | | 4.309452436 | 1.87E-05 | 0.00021487 | clec-28 |
| WBGene00009858 | NA | | NA | NA | thn-4 |
| WBGene00009859 | | 6.896908352 | 2.95E-07 | 5.10E-06 | clec-33 |
| WBGene00009860 | | 0.189177615 | 0.03619054 | 0.1188752 | clec-32 |
| WBGene00009861 | | -0.169369425 | 0.26621571 | 0.46489315 | fbxb-8 |
| WBGene00009862 | | 0.062748575 | 0.75466622 | 0.86091644 | fbxb-67 |
| WBGene00009863 | | -0.153246843 | 0.10890683 | 0.2596319 | F49B2.3 |
| WBGene00009864 | | 0.020317245 | 0.84643798 | NA | NA |
| WBGene00009865 | | 1.451512141 | 0.00068782 | 0.00509116 | F49B2.6 |
| WBGene00009866 | NA | | NA | NA | F49C5.3 |
| WBGene00009867 | | -0.004864426 | 0.94835335 | NA | F49C5.4 |
| WBGene00009868 | NA | | NA | NA | F49C5.5 |
| WBGene00009869 | | 0.020232312 | 0.84643798 | NA | F49C5.7 |
| WBGene00009871 | NA | | NA | NA | F49C12.1 |
| WBGene00009872 | | 0.059675786 | 0.4188661 | NA | F49C12.2 |
| WBGene00009873 | | 0.127266368 | 0.1668541 | NA | F49C12.3 |

| | | | | |
|---|---|---|---|---|
| WBGene00009874 | 0.05787416 | 0.40594086 | NA | F49C12.4 |
| WBGene00009875 | NA | NA | NA | F49C12.5 |
| WBGene00009876 | 0.457821521 | 0.07231475 | 0.19588535 | F49C12.6 |
| WBGene00009877 | 1.877709773 | 1.91E-15 | 9.47E-14 | F49C12.7 |
| WBGene00009878 | 0.006546452 | 0.95358014 | 0.97346024 | F49C12.9 |
| WBGene00009879 | 0.04600504 | 0.24849803 | 0.44690575 | F49C12.10 |
| WBGene00009880 | -0.271633126 | 8.08E-05 | 0.00079877 | F49C12.11 |
| WBGene00009881 | -0.047070829 | 0.60294747 | 0.75988622 | F49C12.12 |
| WBGene00009882 | 0.094873612 | 0.32612342 | 0.52545097 | vha-17 |
| WBGene00009883 | 1.130234729 | 2.94E-05 | 0.00032253 | F49C12.14 |
| WBGene00009884 | 2.080544454 | 6.94E-08 | 1.34E-06 | F49C12.15 |
| WBGene00009885 | -0.01819312 | 0.92250711 | 0.9563426 | F49E2.1 |
| WBGene00009886 | 0.302284329 | 0.04342995 | 0.13586708 | F49E2.2 |
| WBGene00009887 | 0.142365279 | 0.42442981 | 0.62029117 | glb-17 |
| WBGene00009888 | 0.711950605 | 2.01E-12 | 7.19E-11 | F49E2.5 |
| WBGene00009889 | -0.110907368 | 0.51486705 | 0.69476207 | F49E11.2 |
| WBGene00009890 | NA | NA | NA | scl-9 |
| WBGene00009891 | NA | NA | NA | scl-10 |
| WBGene00009892 | NA | NA | NA | scl-11 |
| WBGene00009893 | 0.075029222 | 0.33362495 | NA | F49E11.7 |
| WBGene00009894 | NA | NA | NA | F49E11.8 |
| WBGene00009895 | 3.850771699 | 1.86E-135 | 1.43E-132 | scl-2 |
| WBGene00009896 | 1.800014457 | 2.63E-10 | 7.18E-09 | scl-3 |
| WBGene00009897 | -0.129751329 | 0.02871179 | 0.10044665 | skpo-1 |
| WBGene00009898 | 3.705859877 | 0 | 0 | dod-23 |
| WBGene00009899 | -0.443897309 | 0.03333225 | 0.11182802 | efl-3 |
| WBGene00009900 | NA | NA | NA | F49E12.7 |
| WBGene00009901 | 0.009065483 | 0.93103325 | 0.96103059 | F49E12.8 |
| WBGene00009902 | -0.605252607 | 0.0001503 | 0.00137192 | drd-1 |
| WBGene00009903 | 0.020798779 | 0.90552276 | 0.94916953 | F49E12.10 |
| WBGene00009904 | 0.151587434 | 0.12223267 | 0.28134075 | F49E12.12 |
| WBGene00009905 | NA | NA | NA | clec-240 |
| WBGene00009906 | NA | NA | NA | clec-241 |
| WBGene00009907 | -0.003614866 | 0.95700117 | NA | F49H6.3 |
| WBGene00009908 | 4.658929831 | 5.37E-06 | 6.98E-05 | F49H6.5 |
| WBGene00009909 | -0.023109762 | 0.78121253 | NA | F49H6.8 |
| WBGene00009910 | NA | NA | NA | NA |
| WBGene00009911 | NA | NA | NA | NA |
| WBGene00009912 | NA | NA | NA | srz-94 |
| WBGene00009913 | NA | NA | NA | F49H6.12 |
| WBGene00009914 | 0.033034883 | 0.64219503 | NA | F49H6.13 |
| WBGene00009915 | -0.162081518 | 0.03215716 | 0.10880816 | F52A8.1 |
| WBGene00009916 | 0.104079018 | 0.46027675 | 0.65136993 | F52A8.3 |

| | | | | | |
|---|---|---|---|---|---|
| WBGene00009917 | | 0.565630747 | 0.04736758 | 0.14454121 | glb-18 |
| WBGene00009918 | | -0.128595467 | 0.25687772 | 0.45510003 | gcsh-2 |
| WBGene00009919 | | -0.208323675 | 0.05853312 | 0.16849851 | kbrl-1 |
| WBGene00009920 | | 0.137513283 | 0.10568306 | 0.25461671 | abts-1 |
| WBGene00009921 | | -0.219387153 | 0.00593389 | 0.02989207 | F52B5.2 |
| WBGene00009922 | | -0.176025772 | 0.01945282 | 0.0752467 | F52B5.3 |
| WBGene00009923 | | -0.05884644 | 0.75447083 | 0.86089996 | F52B5.7 |
| WBGene00009924 | | 0.005825262 | 0.94906362 | 0.97101611 | cfp-1 |
| WBGene00009925 | | -0.154849057 | 0.22837469 | 0.42206941 | F52B11.2 |
| WBGene00009926 | | -0.635057158 | 0.00275043 | 0.01627032 | noah-2 |
| WBGene00009927 | | 0.083342601 | 0.65014923 | 0.79283982 | F52B11.5 |
| WBGene00009929 | | 0.411283265 | 0.10459942 | 0.25284464 | abts-2 |
| WBGene00009930 | | -0.118543239 | 0.42603122 | 0.62170069 | F52D10.2 |
| WBGene00009931 | | 0.155748879 | 0.28072787 | 0.48036693 | F52D10.4 |
| WBGene00009933 | | 0.020317245 | 0.84643798 | NA | F52E10.2 |
| WBGene00009935 | | 0.467050933 | 0.06997854 | 0.19114847 | F52E10.4 |
| WBGene00009936 | NA | | NA | NA | NA |
| WBGene00009937 | | -0.017207453 | 0.87125008 | 0.93126126 | lsl-1 |
| WBGene00009938 | | 0.056931656 | 0.59060466 | 0.75172067 | F52F12.5 |
| WBGene00009939 | | -0.692959646 | 3.31E-05 | 0.0003593 | ztf-11 |
| WBGene00009940 | | 0.112883824 | 0.26904506 | 0.46815887 | strl-1 |
| WBGene00009941 | | 0.034276484 | 0.68560721 | NA | F52F12.8 |
| WBGene00009942 | | 0.075254164 | 0.33558625 | NA | F52F12.9 |
| WBGene00009943 | | 0.479522888 | 0.03391073 | 0.11327561 | F52G2.3 |
| WBGene00009944 | | -0.105714685 | 0.34144172 | 0.5414344 | mtcu-2 |
| WBGene00009945 | | 0.342678676 | 0.0477886 | 0.14555292 | bath-38 |
| WBGene00009946 | | 0.051554731 | 0.7231683 | 0.84472009 | ogr-2 |
| WBGene00009947 | | 0.096533914 | 0.62061732 | 0.77305243 | ttc-36 |
| WBGene00009948 | | 2.286709754 | 0.0008735 | 0.00628141 | F52H3.6 |
| WBGene00009949 | | 0.126750219 | 0.44922115 | 0.64185097 | F53A2.1 |
| WBGene00009950 | | 0.012433524 | 0.90600628 | 0.94916953 | sdz-20 |
| WBGene00009951 | | -0.685558772 | 0.00909252 | 0.04169231 | F53A2.3 |
| WBGene00009952 | | -0.158665355 | 0.01309622 | 0.05548382 | acaa-2 |
| WBGene00009953 | | -0.111072172 | 0.59091811 | 0.75192352 | F53A2.9 |
| WBGene00009954 | NA | | NA | NA | F53B2.1 |
| WBGene00009955 | NA | | NA | NA | eak-4 |
| WBGene00009956 | | 0.177616853 | 0.0463229 | NA | F53B2.5 |
| WBGene00009957 | | 1.37665891 | 7.53E-06 | 9.51E-05 | F53B2.8 |
| WBGene00009958 | | 1.449295208 | 4.68E-06 | 6.16E-05 | madd-4 |
| WBGene00009959 | | 0.742900058 | 0.0218619 | 0.08202975 | F53B6.4 |
| WBGene00009960 | | 0.037967508 | 0.71286512 | 0.83851998 | F53B6.5 |
| WBGene00009961 | | -0.023449708 | 0.69192352 | NA | cutl-7 |
| WBGene00009962 | | 0.018735388 | 0.89931317 | 0.94665211 | F53B6.7 |

| | | | | |
|---|---|---|---|---|
| WBGene00009963 | -0.027728334 | 0.67598605 | NA | fipr-26 |
| WBGene00009964 | -0.113029026 | 0.4495312 | 0.64204323 | fip-6 |
| WBGene00009965 | 0.179238063 | 0.31723746 | 0.51739709 | frpr-9 |
| WBGene00009966 | -0.246482327 | 0.00659119 | 0.03251129 | isy-1 |
| WBGene00009967 | 0.059980168 | 0.38780842 | NA | F53B7.4 |
| WBGene00009968 | 2.293554059 | 1.14E-19 | 8.34E-18 | fig-1 |
| WBGene00009969 | 0.358657202 | 0.06878822 | 0.18895467 | F53B7.7 |
| WBGene00009971 | -0.398612317 | 0.04157542 | 0.13149833 | F53C11.1 |
| WBGene00009972 | -0.012087302 | 0.90618753 | 0.94916953 | NA |
| WBGene00009973 | 0.193160044 | 0.03841323 | 0.1241575 | F53C11.3 |
| WBGene00009974 | -0.111500971 | 0.20220752 | 0.39072461 | F53C11.4 |
| WBGene00009975 | -0.109955363 | 0.15378789 | 0.3270785 | F53C11.5 |
| WBGene00009976 | -0.16691278 | 0.1674716 | 0.3451014 | swan-2 |
| WBGene00009977 | -0.411543922 | 0.00145398 | 0.00957181 | swan-1 |
| WBGene00009978 | 0.031199019 | 0.75594329 | 0.8618244 | F53E4.1 |
| WBGene00009979 | 0.038125098 | 0.61119482 | NA | cutl-1 |
| WBGene00009980 | -0.228314477 | 0.18944933 | 0.37508149 | F53F1.2 |
| WBGene00009981 | 0.002808438 | 0.98800718 | 0.99323079 | F53F1.3 |
| WBGene00009982 | -0.594705793 | 0.003968 | 0.02174231 | F53F1.4 |
| WBGene00009983 | 4.367811751 | 3.57E-07 | 6.08E-06 | cut-2 |
| WBGene00009984 | 3.307254764 | 1.37E-12 | 5.05E-11 | F53F1.6 |
| WBGene00009985 | 0.063731418 | 0.41159439 | NA | F53F4.1 |
| WBGene00009986 | -0.025848508 | 0.67901638 | NA | F53F4.2 |
| WBGene00009987 | -0.096995922 | 0.30862203 | 0.5095326 | tbcb-1 |
| WBGene00009988 | 0.054658285 | 0.74968057 | 0.85873723 | F53F4.4 |
| WBGene00009990 | NA | NA | NA | F53F4.7 |
| WBGene00009991 | 0.834856947 | 0.0222711 | 0.08315891 | NA |
| WBGene00009992 | -0.176553028 | 0.02516585 | 0.09123765 | F53F4.10 |
| WBGene00009993 | -0.393653573 | 3.09E-06 | 4.24E-05 | F53F4.11 |
| WBGene00009994 | -0.103092133 | 0.46307937 | 0.65301113 | F53F4.12 |
| WBGene00009995 | 0.937181001 | 1.10E-14 | 5.05E-13 | F53F4.13 |
| WBGene00009996 | 0.000916292 | 0.9930053 | 0.99586209 | F53F4.14 |
| WBGene00009997 | NA | NA | NA | F53F4.15 |
| WBGene00009998 | -0.059168106 | 0.58774465 | 0.75003656 | klf-2 |
| WBGene00009999 | -0.02791392 | 0.6763823 | NA | srsx-35 |
| WBGene00010000 | 0.072536146 | 0.52394745 | 0.70139111 | F53F8.3 |
| WBGene00010001 | 0.81071837 | 0.0007567 | 0.00552009 | F53F8.4 |
| WBGene00010002 | -0.067588272 | 0.50592594 | 0.68674779 | F53F8.5 |
| WBGene00010003 | 0.085537126 | 0.24698522 | NA | F53F8.6 |
| WBGene00010004 | 3.638831938 | 0.00074266 | 0.00544209 | F53H2.1 |
| WBGene00010005 | -0.035932958 | 0.73828688 | 0.85300333 | cnc-7 |
| WBGene00010006 | -0.163199773 | 0.1777257 | 0.35895738 | F53H2.3 |
| WBGene00010007 | -1.266352039 | 2.91E-32 | 3.94E-30 | F53H4.2 |

| | | | | |
|---|---|---|---|---|
| WBGene00010008 | -0.083864977 | 0.57893383 | 0.74483042 | F53H4.3 |
| WBGene00010009 | 0.22390727 | 0.28014394 | 0.47984024 | F53H4.4 |
| WBGene00010010 | 0.082104143 | 0.61348935 | 0.76776627 | gmeb-2 |
| WBGene00010011 | 0.175690791 | 0.32779337 | 0.52710708 | F53H4.6 |
| WBGene00010012 | 0.601204196 | 1.87E-06 | 2.74E-05 | saeg-1 |
| WBGene00010013 | -0.001520545 | 0.98239868 | 0.98983546 | F54B3.1 |
| WBGene00010014 | 0.017604133 | 0.79437959 | NA | F54B3.2 |
| WBGene00010015 | -0.50285149 | 1.01E-11 | 3.33E-10 | atad-3 |
| WBGene00010016 | 0.040107227 | 0.58693163 | NA | F54B8.1 |
| WBGene00010017 | 0.086764008 | 0.294364 | NA | nhr-282 |
| WBGene00010018 | 0.547826982 | 0.03546854 | 0.11714298 | fbxa-69 |
| WBGene00010019 | 2.767549593 | 3.66E-05 | 0.00039449 | F54B8.4 |
| WBGene00010020 | NA | NA | NA | srbc-48 |
| WBGene00010021 | 0.020925399 | 0.84643798 | NA | srbc-49 |
| WBGene00010022 | NA | NA | NA | srbc-50 |
| WBGene00010023 | NA | NA | NA | srbc-51 |
| WBGene00010024 | NA | NA | NA | NA |
| WBGene00010025 | NA | NA | NA | srbc-54 |
| WBGene00010026 | NA | NA | NA | srbc-56 |
| WBGene00010027 | NA | NA | NA | F54B11.4 |
| WBGene00010028 | 0.223622045 | 0.22109688 | 0.41356214 | F54B11.5 |
| WBGene00010030 | 0.020232312 | 0.84643798 | NA | F54B11.7 |
| WBGene00010031 | 0.036443458 | 0.71434249 | 0.83978548 | F54B11.8 |
| WBGene00010032 | 0.086470338 | 0.29130368 | NA | F54B11.9 |
| WBGene00010033 | 0.040529223 | 0.56245259 | NA | F54B11.10 |
| WBGene00010034 | 2.301048843 | 2.08E-11 | 6.58E-10 | F54B11.11 |
| WBGene00010035 | 1.325439815 | 0.01305979 | 0.05540666 | F54C8.1 |
| WBGene00010036 | -0.19021296 | 0.0715715 | 0.19463354 | cpar-1 |
| WBGene00010037 | 0.014306702 | 0.84644058 | 0.9173689 | F54C8.4 |
| WBGene00010038 | -0.133455639 | 0.24334764 | 0.44140607 | rheb-1 |
| WBGene00010039 | 0.022725499 | 0.90345129 | 0.9486109 | F54C8.6 |
| WBGene00010040 | -0.100222219 | 0.36298691 | 0.56280507 | F54C8.7 |
| WBGene00010041 | 1.099979156 | 2.61E-14 | 1.16E-12 | F54C9.3 |
| WBGene00010042 | -0.007540051 | 0.94617673 | 0.96982452 | bcs-1 |
| WBGene00010043 | 0.345591504 | 0.07344654 | 0.19829076 | F54C9.7 |
| WBGene00010044 | -0.197696622 | 0.02896523 | 0.10104359 | F54C9.9 |
| WBGene00010045 | 0.327947589 | 0.16771845 | 0.34527078 | F54C9.11 |
| WBGene00010046 | 0.145211821 | 0.20736794 | 0.39645783 | F54D1.1 |
| WBGene00010047 | 1.159775684 | 5.32E-11 | 1.59E-09 | F54D1.6 |
| WBGene00010048 | -0.292850432 | 0.03015897 | 0.10382257 | F54D5.2 |
| WBGene00010049 | 1.540039451 | 4.28E-75 | 1.61E-72 | F54D5.3 |
| WBGene00010050 | -0.880936396 | 4.09E-11 | 1.25E-09 | F54D5.4 |
| WBGene00010051 | -0.049219378 | 0.61768545 | 0.77097465 | F54D5.5 |

| | | | | |
|---|---|---|---|---|
| WBGene00010052 | 0.109684322 | 0.20590813 | 0.39495113 | F54D5.7 |
| WBGene00010053 | -0.029601804 | 0.77808849 | 0.87435671 | F54D5.9 |
| WBGene00010054 | 0.138972973 | 0.17762187 | 0.35884667 | gtf-2E2 |
| WBGene00010055 | -0.774437067 | 1.79E-27 | 1.94E-25 | F54D5.12 |
| WBGene00010056 | -0.02480213 | 0.70519572 | 0.83297867 | smc-6 |
| WBGene00010057 | 0.524675981 | 0.06013384 | 0.17176135 | F54D5.15 |
| WBGene00010058 | -3.161866347 | 2.61E-07 | 4.55E-06 | F54E4.2 |
| WBGene00010059 | 0.281854334 | 0.18194896 | 0.36467005 | F54E4.3 |
| WBGene00010060 | 0.053293819 | 0.59664761 | 0.75522171 | F54E4.4 |
| WBGene00010061 | -0.081389718 | 0.15069948 | 0.32243039 | F54E12.2 |
| WBGene00010062 | 5.884818056 | 2.98E-22 | 2.52E-20 | lipl-1 |
| WBGene00010063 | 0.770849862 | 0.00034546 | 0.00283192 | dhrs-4 |
| WBGene00010064 | -0.327556624 | 0.04657152 | 0.14285641 | F54F7.2 |
| WBGene00010065 | 0.136850805 | 0.38373256 | 0.58327725 | F54F7.3 |
| WBGene00010066 | -0.573595271 | 0.00487837 | 0.02567368 | F54F7.6 |
| WBGene00010067 | -0.079660709 | 0.31592878 | NA | dot-1.2 |
| WBGene00010068 | -0.010141451 | 0.90973981 | NA | F54F7.8 |
| WBGene00010069 | -0.03147554 | 0.70637395 | NA | F54F11.1 |
| WBGene00010070 | 0.312884959 | 0.00010505 | 0.0010145 | nep-17 |
| WBGene00010071 | NA | NA | NA | sre-45 |
| WBGene00010072 | 0.603358383 | 0.01041575 | 0.04632376 | F54F12.1 |
| WBGene00010073 | -0.056642891 | 0.73140847 | 0.84995249 | F54F12.2 |
| WBGene00010074 | -0.009579513 | 0.92340349 | 0.95666222 | irld-34 |
| WBGene00010075 | -0.247738308 | 0.00153123 | 0.01000389 | F55A11.1 |
| WBGene00010077 | -0.120448512 | 0.29741744 | 0.49843299 | F55A11.4 |
| WBGene00010079 | 0.031916307 | 0.87154872 | 0.93143219 | F55A11.6 |
| WBGene00010080 | -0.076148198 | 0.47078443 | 0.6588409 | F55A11.7 |
| WBGene00010081 | -0.043770977 | 0.75143208 | 0.85953643 | F55A11.8 |
| WBGene00010082 | 0.170696496 | 0.28561078 | 0.48577294 | F55A11.11 |
| WBGene00010083 | 0.708178724 | 0.00022297 | 0.00193395 | xdh-1 |
| WBGene00010084 | -0.624256099 | 6.97E-11 | 2.05E-09 | F55B11.2 |
| WBGene00010085 | -0.527997539 | 3.90E-05 | 0.0004191 | F55B11.3 |
| WBGene00010086 | -0.418718128 | 0.03976044 | 0.12746499 | F55B11.4 |
| WBGene00010087 | 0.079118509 | 0.40655535 | 0.60467054 | F55B11.5 |
| WBGene00010088 | 0.020925399 | 0.84643798 | NA | F55B12.2 |
| WBGene00010089 | 0.018294352 | 0.88314482 | 0.9382283 | hpo-31 |
| WBGene00010091 | 0.235539276 | 0.20717832 | 0.39645769 | ssp-35 |
| WBGene00010092 | 0.13169099 | 0.16083826 | NA | NA |
| WBGene00010093 | -0.154600262 | 0.04372618 | 0.13655812 | capg-2 |
| WBGene00010094 | -0.127787993 | 0.30031432 | 0.50066452 | tsfm-1 |
| WBGene00010095 | -0.136315634 | 0.21674097 | 0.40824708 | NA |
| WBGene00010096 | -0.065989529 | 0.4695315 | 0.65798188 | rskd-1 |
| WBGene00010097 | -0.151422807 | 0.04161632 | 0.13154244 | srpa-68 |

| | | | | |
|---|---|---|---|---|
| WBGene00010098 | 0.014958179 | 0.93418845 | 0.96290283 | F55C5.10 |
| WBGene00010099 | -0.060470622 | 0.63814343 | 0.78552676 | fbxb-117 |
| WBGene00010100 | -0.567212215 | 0.02296379 | 0.08509563 | F55C9.3 |
| WBGene00010101 | -1.617361246 | 0.00355578 | 0.02001063 | fbxb-57 |
| WBGene00010102 | 0.009830039 | 0.91298006 | NA | F55C9.5 |
| WBGene00010103 | -0.004944805 | 0.94514249 | NA | F55C9.6 |
| WBGene00010104 | -0.127871019 | 0.35295937 | 0.55296212 | fbxb-60 |
| WBGene00010105 | -0.2016414 | 0.3343994 | 0.53378105 | fbxb-62 |
| WBGene00010106 | -0.215106154 | 0.28616972 | 0.48616975 | fbxb-61 |
| WBGene00010107 | -0.067905328 | 0.31502275 | NA | F55C9.11 |
| WBGene00010108 | -0.023109762 | 0.78121253 | NA | F55C10.4 |
| WBGene00010109 | 0.054740883 | 0.7900762 | 0.88173685 | F55C10.5 |
| WBGene00010110 | -0.220454158 | 0.27317993 | 0.4733847 | F55D12.1 |
| WBGene00010111 | -0.342192374 | 0.0804968 | 0.21185578 | F55D12.2 |
| WBGene00010112 | NA | NA | NA | NA |
| WBGene00010113 | 0.012426581 | 0.91137086 | 0.95112162 | F55D12.5 |
| WBGene00010114 | 0.207056196 | 0.02971382 | 0.10287067 | F55D12.6 |
| WBGene00010115 | 0.099194164 | 0.59713463 | 0.75563078 | aakb-1 |
| WBGene00010116 | 0.683706761 | 0.00260849 | 0.01556274 | F55F3.2 |
| WBGene00010117 | 0.313981521 | 0.00326401 | 0.01866732 | nkb-3 |
| WBGene00010118 | 0.090903669 | 0.45631587 | 0.64831836 | F55F3.4 |
| WBGene00010119 | -0.148264266 | 0.36944297 | 0.56931003 | F55G7.1 |
| WBGene00010120 | -0.286775958 | 0.2110687 | 0.40101958 | dot-1.4 |
| WBGene00010121 | 0.05787416 | 0.40594086 | NA | F55G7.3 |
| WBGene00010122 | 0.14077579 | 0.25824838 | 0.45633859 | F55G11.1 |
| WBGene00010123 | 0.336201334 | 0.15895872 | 0.33321705 | F55G11.2 |
| WBGene00010124 | 4.425275436 | 7.86E-122 | 5.48E-119 | F55G11.4 |
| WBGene00010125 | 3.530317315 | 0.000375 | 0.00304513 | dod-22 |
| WBGene00010126 | -0.022930767 | 0.78121253 | NA | F55G11.6 |
| WBGene00010127 | 0.056411843 | 0.41891328 | NA | F55G11.7 |
| WBGene00010128 | -0.831759657 | 0.01631972 | 0.0661509 | F55G11.8 |
| WBGene00010129 | NA | NA | NA | F55G11.10 |
| WBGene00010130 | 0.130794961 | 0.03758562 | 0.12231408 | vha-14 |
| WBGene00010131 | 0.090245062 | 0.50205944 | 0.68359084 | F55H2.5 |
| WBGene00010132 | -0.241735594 | 0.01908018 | 0.07427624 | F55H2.7 |
| WBGene00010133 | 0.780031516 | 0.00017986 | 0.00160963 | F55H12.2 |
| WBGene00010134 | 0.431728807 | 0.016217 | 0.06580739 | F55H12.3 |
| WBGene00010135 | -0.901043934 | 4.87E-11 | 1.47E-09 | F55H12.4 |
| WBGene00010136 | 0.234240661 | 0.14253228 | 0.31069569 | F55H12.5 |
| WBGene00010137 | 0.203237968 | 0.32986958 | 0.52933278 | ztf-26 |
| WBGene00010138 | 0.267367022 | 0.01351721 | 0.05696759 | anoh-1 |
| WBGene00010139 | -0.405796701 | 2.33E-06 | 3.33E-05 | F56A8.3 |
| WBGene00010140 | -0.049589902 | 0.71322826 | 0.83883106 | F56A8.4 |

| | | | | |
|---|---|---|---|---|
| WBGene00010141 | -0.039285302 | 0.77898015 | 0.87495577 | F56A8.5 |
| WBGene00010142 | -0.776179585 | 0.00170938 | 0.01097677 | F56A8.8 |
| WBGene00010143 | 0.09339839 | 0.43604446 | 0.63111939 | F56A12.2 |
| WBGene00010144 | 0.157384033 | 0.1160067 | 0.27192405 | F56C4.1 |
| WBGene00010145 | NA | NA | NA | F56C4.2 |
| WBGene00010146 | 0.185555864 | 0.33792351 | 0.5376293 | glb-19 |
| WBGene00010147 | 0.0398788 | 0.58971221 | NA | F56D5.2 |
| WBGene00010148 | 0.138627158 | 0.51198103 | 0.69188797 | F56D5.3 |
| WBGene00010149 | -0.436858675 | 0.04641526 | 0.14255181 | F56D5.5 |
| WBGene00010150 | 0.544879866 | 0.00370536 | 0.02062743 | F56D5.6 |
| WBGene00010152 | 0.065014541 | 0.37399213 | NA | F56D5.9 |
| WBGene00010153 | 0.020232312 | 0.84643798 | NA | srxa-2 |
| WBGene00010155 | 0.175549375 | 0.30667279 | 0.50722959 | F56F3.4 |
| WBGene00010157 | 0.079571953 | 0.41776011 | 0.61446914 | oac-34 |
| WBGene00010158 | -0.046394786 | 0.50231372 | NA | pes-2.1 |
| WBGene00010158 | -0.046394786 | 0.50231372 | NA | pes-2.2 |
| WBGene00010159 | 0.104599499 | 0.32523041 | 0.52447389 | F56G4.4 |
| WBGene00010160 | 0.098324675 | 0.43500261 | 0.63033187 | png-1 |
| WBGene00010161 | -0.11482837 | 0.46874491 | 0.65749215 | F56G4.6 |
| WBGene00010162 | NA | NA | NA | F56H6.1 |
| WBGene00010163 | 0.052061634 | 0.60828039 | 0.76399837 | F56H6.2 |
| WBGene00010164 | NA | NA | NA | F56H6.3 |
| WBGene00010165 | NA | NA | NA | F56H6.4 |
| WBGene00010166 | NA | NA | NA | gmd-2 |
| WBGene00010167 | NA | NA | NA | bgnt-1.6 |
| WBGene00010168 | NA | NA | NA | F56H6.7 |
| WBGene00010169 | NA | NA | NA | clec-18 |
| WBGene00010170 | NA | NA | NA | F56H6.9 |
| WBGene00010171 | 0.020317245 | 0.84643798 | NA | oac-35 |
| WBGene00010172 | 0.020232312 | 0.84643798 | NA | oac-36 |
| WBGene00010173 | NA | NA | NA | F56H6.13 |
| WBGene00010174 | 0.618774187 | 4.86E-11 | 1.47E-09 | F56H9.2 |
| WBGene00010175 | 0.583322644 | 0.04708441 | 0.14400729 | F56H11.2 |
| WBGene00010176 | NA | NA | NA | F56H11.6 |
| WBGene00010177 | 0.662882915 | 0.00179689 | 0.01142842 | F57A8.1 |
| WBGene00010178 | -0.134908324 | 0.18850564 | 0.37361339 | yif-1 |
| WBGene00010179 | 0.154431772 | 0.2838513 | 0.48373881 | F57A8.4 |
| WBGene00010180 | -0.022930767 | 0.78121253 | NA | nhr-192 |
| WBGene00010181 | 0.037127706 | 0.59421403 | NA | F57A8.6 |
| WBGene00010182 | 0.040107227 | 0.58693163 | NA | F57A8.7 |
| WBGene00010183 | -2.010146432 | 6.13E-06 | 7.84E-05 | fipr-13 |
| WBGene00010184 | 0.086430603 | 0.29509282 | NA | F57A10.2 |
| WBGene00010185 | 0.0528687 | 0.78888268 | 0.88074053 | F57A10.4 |

| | | | | |
|---|---|---|---|---|
| WBGene00010186 | 0.020925399 | 0.84643798 | NA | nhr-283 |
| WBGene00010187 | -0.015701823 | 0.9001272 | 0.94668216 | F57B1.1 |
| WBGene00010188 | -0.026376047 | 0.89630669 | 0.9455167 | F57B1.5 |
| WBGene00010189 | 0.135418156 | 0.41116845 | 0.60881041 | F57B1.6 |
| WBGene00010191 | 0.571147359 | 0.03096016 | 0.10565399 | dmsr-1 |
| WBGene00010192 | 0.18428029 | 0.38869743 | 0.58712904 | F57B7.2 |
| WBGene00010193 | -0.596828598 | 0.00365352 | 0.02040093 | btb-20 |
| WBGene00010194 | -0.143133219 | 0.27005071 | 0.46923955 | btb-19 |
| WBGene00010195 | -0.036881335 | 0.78712022 | 0.87949869 | pot-2 |
| WBGene00010196 | 0.744954275 | 1.30E-09 | 3.25E-08 | F57C2.4 |
| WBGene00010197 | 0.067932351 | 0.42045095 | 0.61706319 | F57C2.5 |
| WBGene00010198 | -0.094338649 | 0.31975086 | 0.51987478 | spat-1 |
| WBGene00010199 | -0.280471084 | 0.00356151 | 0.02003232 | bet-2 |
| WBGene00010200 | 0.18479106 | 0.23908902 | 0.43606953 | F57C7.4 |
| WBGene00010202 | 0.116182835 | 0.30040637 | 0.50071148 | F57E7.1 |
| WBGene00010203 | NA | NA | NA | F57E7.2 |
| WBGene00010204 | 0.407458756 | 6.94E-10 | 1.79E-08 | F57F5.1 |
| WBGene00010205 | 0.035512071 | 0.72830584 | 0.84850209 | F57F5.3 |
| WBGene00010206 | 0.149813019 | 0.07294962 | NA | F57G4.1 |
| WBGene00010209 | 0.001977299 | 0.99368578 | 0.99606692 | NA |
| WBGene00010212 | -0.182282708 | 0.37255716 | 0.57176928 | fbxa-192 |
| WBGene00010214 | 0.002412426 | 0.99049902 | 0.9947106 | F57G8.5 |
| WBGene00010215 | 1.935271221 | 1.65E-05 | 0.0001912 | nhr-193 |
| WBGene00010216 | NA | NA | NA | F57G8.7 |
| WBGene00010217 | -0.022930767 | 0.78121253 | NA | sre-31 |
| WBGene00010218 | NA | NA | NA | sre-30 |
| WBGene00010219 | -0.288942354 | 0.16588433 | 0.34245805 | F57G9.3 |
| WBGene00010220 | NA | NA | NA | sre-29 |
| WBGene00010221 | -0.069481533 | 0.57220958 | 0.74047379 | F57G12.1 |
| WBGene00010222 | 0.066203175 | 0.5655613 | 0.73505397 | F57G12.2 |
| WBGene00010223 | 0.34412342 | 0.13702299 | 0.30247674 | F58A3.3 |
| WBGene00010224 | 0.056667729 | 0.70679941 | 0.83433445 | F58A3.4 |
| WBGene00010225 | -0.543366073 | 0.01731207 | 0.06902885 | ttr-31 |
| WBGene00010226 | 0.012892043 | 0.89216955 | NA | F58A4.1 |
| WBGene00010227 | -0.209068812 | 0.10405145 | 0.25210364 | F58A4.2 |
| WBGene00010228 | -0.046394786 | 0.50231372 | NA | clec-161 |
| WBGene00010229 | -0.19213664 | 0.27883865 | 0.47871443 | F58A4.6 |
| WBGene00010230 | -0.390341771 | 0.00099652 | 0.00701781 | rpac-19 |
| WBGene00010231 | -0.358607701 | 0.00017737 | 0.00159022 | F58B3.4 |
| WBGene00010232 | -0.088236401 | 0.28650826 | 0.48651907 | F58B3.6 |
| WBGene00010233 | -0.19010392 | 0.1424943 | 0.31068638 | rbm-17 |
| WBGene00010234 | -1.134114558 | 7.38E-29 | 8.72E-27 | ttr-50 |
| WBGene00010235 | 0.020317245 | 0.84643798 | NA | F58B4.2 |

| | | | | |
|---|---|---|---|---|
| WBGene00010236 | 1.183687522 | 0.00484277 | 0.02552302 | F58B4.3 |
| WBGene00010237 | NA | NA | NA | scvp-1 |
| WBGene00010238 | 0.980083618 | 1.54E-08 | 3.35E-07 | F58B4.5 |
| WBGene00010239 | 0.354781717 | 0.13807008 | 0.304074 | F58B4.6 |
| WBGene00010240 | -0.211985409 | 0.24431553 | 0.44248191 | NA |
| WBGene00010241 | 0.277430491 | 0.15168386 | 0.32415756 | F58D2.2 |
| WBGene00010242 | 0.078718519 | 0.70639226 | 0.83398831 | F58D5.2 |
| WBGene00010243 | 0.065730147 | 0.58222898 | 0.74624349 | F58D5.3 |
| WBGene00010244 | -0.083010659 | 0.52276445 | 0.70044791 | F58D5.5 |
| WBGene00010245 | 0.020925399 | 0.84643798 | NA | F58D5.6 |
| WBGene00010246 | 0.24308816 | 0.08345371 | 0.21713111 | F58D5.7 |
| WBGene00010247 | 0.056411843 | 0.41891328 | NA | F58D5.8 |
| WBGene00010248 | -0.143041203 | 0.49925319 | 0.68129052 | F58D5.9 |
| WBGene00010249 | NA | NA | NA | F58D12.1 |
| WBGene00010250 | 0.220957478 | 0.12603725 | 0.28699798 | F58D12.3 |
| WBGene00010251 | 0.175574315 | 0.39697161 | 0.59488857 | sta-2 |
| WBGene00010253 | NA | NA | NA | F58E6.4 |
| WBGene00010254 | 1.929626959 | 0.00459428 | 0.02454589 | F58E6.5 |
| WBGene00010255 | 0.018860755 | 0.77262519 | NA | F58E6.6 |
| WBGene00010256 | 0.065014541 | 0.37399213 | NA | hrg-3 |
| WBGene00010257 | 0.018476367 | 0.76699386 | NA | F58E6.8 |
| WBGene00010258 | 0.056449805 | 0.68221822 | 0.8161756 | F58E6.11 |
| WBGene00010259 | 0.124143176 | 0.15026652 | 0.32179087 | ric-7 |
| WBGene00010260 | -0.310569576 | 4.97E-11 | 1.50E-09 | ddx-17 |
| WBGene00010261 | NA | NA | NA | srh-308 |
| WBGene00010262 | 0.678725354 | 0.01218759 | 0.05234335 | droe-4 |
| WBGene00010263 | -0.016602498 | 0.83892023 | 0.9127791 | wago-4 |
| WBGene00010264 | -0.188690489 | 0.10447541 | 0.25274694 | F58G1.2 |
| WBGene00010265 | 0.186331422 | 0.12800753 | 0.2894006 | F58G1.3 |
| WBGene00010266 | -0.031361287 | 0.70060749 | 0.82976818 | dct-18 |
| WBGene00010267 | 0.22417086 | 0.02255296 | 0.08395572 | lips-9 |
| WBGene00010268 | -0.131010918 | 0.53123996 | 0.70772377 | arrd-10 |
| WBGene00010269 | 0.316393375 | 0.17668427 | 0.35749508 | F58G1.7 |
| WBGene00010270 | NA | NA | NA | F58G1.8 |
| WBGene00010271 | -0.005581345 | 0.94060131 | NA | NA |
| WBGene00010272 | -0.08664549 | 0.49656416 | 0.67886649 | amph-1 |
| WBGene00010273 | 0.078948492 | 0.40741673 | 0.60539819 | srm-3 |
| WBGene00010274 | -0.210481353 | 0.28095193 | 0.48048327 | F58G6.3 |
| WBGene00010275 | 0.062747866 | 0.70613753 | 0.83382205 | acc-1 |
| WBGene00010276 | NA | NA | NA | NA |
| WBGene00010277 | -0.435428418 | 0.01185049 | 0.05109039 | F58G6.7 |
| WBGene00010278 | 0.097867894 | 0.33847824 | 0.53801541 | F58G6.8 |
| WBGene00010279 | -0.129230945 | 0.08844589 | 0.22571664 | letm-1 |

| | | | | |
|---|---|---|---|---|
| WBGene00010280 | 0.087932816 | 0.14058431 | 0.30822025 | rde-12 |
| WBGene00010281 | -0.249601573 | 0.00091473 | 0.00652979 | F58G11.3 |
| WBGene00010282 | 0.238529326 | 0.18189848 | 0.36461879 | F58G11.4 |
| WBGene00010283 | 0.003694171 | 0.97500353 | 0.98591235 | ccz-1 |
| WBGene00010284 | 0.050513669 | 0.50985637 | 0.689717 | aman-2 |
| WBGene00010285 | 0.210596279 | 0.06467643 | 0.18123016 | F58H1.2 |
| WBGene00010286 | -0.127994657 | 0.34406393 | 0.54381673 | F58H1.3 |
| WBGene00010288 | 0.15015282 | 0.48253854 | 0.66839748 | F58H1.5 |
| WBGene00010289 | 0.026222354 | 0.87542551 | 0.93344293 | F58H1.6 |
| WBGene00010290 | 1.075767301 | 0.00113767 | 0.00783472 | F58H1.7 |
| WBGene00010291 | 0.171262732 | 0.20570186 | 0.39466399 | F58H10.1 |
| WBGene00010292 | NA | NA | NA | F59A1.6 |
| WBGene00010293 | -0.010305273 | 0.95717577 | 0.97638197 | fbxa-108 |
| WBGene00010294 | -0.115125237 | 0.57956769 | 0.74518754 | NA |
| WBGene00010295 | 0.319370148 | 0.11045622 | 0.26219025 | NA |
| WBGene00010296 | -0.361471658 | 0.13488767 | 0.29927116 | dgat-2 |
| WBGene00010297 | -0.157894121 | 0.16923951 | 0.34752253 | F59A1.11 |
| WBGene00010298 | 0.020232312 | 0.84643798 | NA | F59A1.12 |
| WBGene00010299 | -0.083839105 | 0.57887007 | 0.74481384 | F59A1.13 |
| WBGene00010300 | 0.056411843 | 0.41891328 | NA | F59A1.15 |
| WBGene00010301 | NA | NA | NA | F59A1.16 |
| WBGene00010303 | -0.407949522 | 5.12E-08 | 1.01E-06 | cri-3 |
| WBGene00010304 | -0.205091111 | 0.10498129 | 0.25334286 | clpf-1 |
| WBGene00010305 | -0.129453337 | 0.41111484 | 0.60881041 | F59A2.5 |
| WBGene00010306 | -0.059252437 | 0.53602956 | 0.71119246 | golg-4 |
| WBGene00010307 | -0.18073696 | 0.06758224 | 0.18658713 | skat-1 |
| WBGene00010308 | -0.004183378 | 0.96877162 | 0.98300543 | F59B2.3 |
| WBGene00010309 | -0.15029089 | 0.33324934 | 0.53300235 | rpn-6.2 |
| WBGene00010310 | -0.13212655 | 0.22235714 | 0.41503454 | F59B2.8 |
| WBGene00010311 | 0.082852087 | 0.56105431 | 0.73159138 | F59B2.9 |
| WBGene00010313 | 0.037992904 | 0.65794732 | NA | NA |
| WBGene00010314 | 0.206575708 | 0.02995553 | 0.10341402 | F59B2.12 |
| WBGene00010315 | 0.181561614 | 0.39213174 | 0.59023345 | F59B2.13 |
| WBGene00010316 | 0.042731542 | 0.75881232 | 0.86363774 | F59B8.1 |
| WBGene00010317 | 0.133675298 | 0.25686462 | 0.45510003 | idh-1 |
| WBGene00010318 | 0.065014541 | 0.37399213 | NA | F59B10.2 |
| WBGene00010319 | 0.396233284 | 0.05780879 | 0.16730153 | F59B10.3 |
| WBGene00010320 | 0.156359065 | 0.42210579 | 0.61846519 | F59B10.4 |
| WBGene00010321 | 0.10275069 | 0.48705737 | 0.67155416 | F59B10.5 |
| WBGene00010322 | -0.002914971 | 0.95718083 | NA | F59B10.6 |
| WBGene00010323 | 0.062171771 | 0.51210308 | NA | dhhc-12 |
| WBGene00010324 | 0.018795205 | 0.77532208 | NA | F59C6.3 |
| WBGene00010325 | -0.11054882 | 0.322564 | 0.5227072 | exos-3 |

| | | | | |
|---|---|---|---|---|
| WBGene00010326 | -0.129926004 | 0.05092469 | 0.15237134 | F59C6.5 |
| WBGene00010327 | -0.041916026 | 0.74653544 | 0.85672656 | F59C6.8 |
| WBGene00010328 | -0.047237011 | 0.79400142 | 0.8840285 | F59C6.11 |
| WBGene00010329 | -0.030935094 | 0.87218963 | 0.93168892 | F59D12.1 |
| WBGene00010330 | -0.067694303 | 0.63507762 | 0.78353176 | F59D12.2 |
| WBGene00010331 | 0.096619834 | 0.33829084 | 0.53789273 | F59D12.3 |
| WBGene00010332 | -0.004200584 | 0.98158551 | 0.98942468 | F59D12.5 |
| WBGene00010333 | -0.267503455 | 0.00306443 | 0.01776885 | copz-1 |
| WBGene00010334 | 1.823903912 | 6.65E-06 | 8.46E-05 | del-5 |
| WBGene00010335 | 0.055752568 | 0.75069211 | 0.85919975 | F59F3.6 |
| WBGene00010336 | 0.094305516 | 0.61291144 | 0.76717418 | acox-1.6 |
| WBGene00010337 | 0.147417483 | 0.09840486 | 0.24324591 | F59F4.2 |
| WBGene00010338 | -0.214011538 | 0.26210271 | 0.45995993 | F59F4.3 |
| WBGene00010339 | -0.627503339 | 2.90E-05 | 0.00031826 | acl-1 |
| WBGene00010340 | 0.252903965 | 0.2256483 | 0.41882819 | slcf-1 |
| WBGene00010341 | 1.371031257 | 5.66E-06 | 7.32E-05 | glo-3 |
| WBGene00010342 | 0.259809973 | 0.17119518 | 0.35016327 | F59F5.3 |
| WBGene00010343 | NA | NA | NA | F59F5.4 |
| WBGene00010344 | 0.088295023 | 0.52652898 | 0.70381729 | F59F5.5 |
| WBGene00010345 | -0.024929993 | 0.8570745 | 0.92325428 | F59F5.7 |
| WBGene00010348 | -0.236206719 | 0.26778967 | 0.4665971 | H01G02.1 |
| WBGene00010349 | -0.045231251 | 0.58189544 | NA | dyf-18 |
| WBGene00010350 | 0.178719022 | 0.34237605 | 0.54273795 | H01G02.3 |
| WBGene00010351 | 0.070353994 | 0.12714685 | 0.28847931 | cbd-1 |
| WBGene00010352 | -0.026384613 | 0.67299636 | NA | fbxb-76 |
| WBGene00010353 | -0.305039341 | 0.00012392 | 0.00117049 | memi-3 |
| WBGene00010354 | -0.217563066 | 0.00688628 | 0.03354476 | cyp-31A2 |
| WBGene00010355 | 0.0398788 | 0.58971221 | NA | clec-229 |
| WBGene00010356 | -0.010304572 | 0.94829229 | 0.97044737 | famk-1 |
| WBGene00010357 | 0.201045816 | 0.34705244 | 0.54665893 | H03A11.2 |
| WBGene00010358 | NA | NA | NA | H03G16.1 |
| WBGene00010359 | -0.067064794 | 0.32104386 | NA | mam-6 |
| WBGene00010360 | -0.0059938 | 0.93322519 | NA | fbxa-96 |
| WBGene00010361 | -0.01155814 | 0.9041121 | 0.94889643 | H03G16.5 |
| WBGene00010362 | 0.534237684 | 0.00538563 | 0.02773477 | enu-3.1 |
| WBGene00010363 | -0.33625134 | 0.01249322 | 0.05338972 | H04D03.2 |
| WBGene00010364 | 0.13201922 | 0.51174373 | 0.69169498 | H04D03.3 |
| WBGene00010365 | -0.963854498 | 0.00296133 | 0.01729429 | H04D03.4 |
| WBGene00010366 | 0.211979021 | 0.10766797 | 0.25791858 | H05L14.1 |
| WBGene00010367 | -0.150571792 | 0.14961788 | 0.3209605 | H05L14.2 |
| WBGene00010368 | -0.024112202 | 0.90516509 | 0.94916953 | H06A10.1 |
| WBGene00010369 | 0.106247313 | 0.30389595 | 0.50463203 | chd-1 |
| WBGene00010370 | 0.132738128 | 0.30484091 | 0.50557043 | ctg-1 |

| | | | | |
|---|---|---|---|---|
| WBGene00010371 | -0.012630797 | 0.89855366 | 0.94658742 | NA |
| WBGene00010372 | NA | NA | NA | H08J19.1 |
| WBGene00010373 | 0.110039823 | 0.21624696 | NA | H08M01.1 |
| WBGene00010374 | 0.527024948 | 0.0318354 | 0.10794396 | rga-5 |
| WBGene00010375 | 0.239907666 | 0.16468116 | 0.34084094 | H09F14.1 |
| WBGene00010376 | NA | NA | NA | NA |
| WBGene00010377 | NA | NA | NA | nspa-1 |
| WBGene00010377 | NA | NA | NA | nspa-3 |
| WBGene00010378 | -0.022930767 | 0.78121253 | NA | H12D21.2 |
| WBGene00010378 | -0.022930767 | 0.78121253 | NA | ZC412.9 |
| WBGene00010379 | NA | NA | NA | H12D21.3 |
| WBGene00010379 | NA | NA | NA | ZC412.8 |
| WBGene00010380 | 0.925121645 | 0.01791564 | 0.07089266 | mpst-2 |
| WBGene00010381 | 0.056188364 | 0.42092875 | NA | H12D21.5 |
| WBGene00010382 | NA | NA | NA | H12D21.6 |
| WBGene00010383 | 0.449598702 | 0.00857549 | 0.03985833 | mpst-3 |
| WBGene00010385 | 0.256823182 | 0.18314759 | 0.36616603 | H12D21.9 |
| WBGene00010386 | -0.119260977 | 0.56747601 | 0.73659312 | H12D21.10 |
| WBGene00010387 | NA | NA | NA | H12D21.11 |
| WBGene00010388 | NA | NA | NA | srz-42 |
| WBGene00010389 | NA | NA | NA | srz-31 |
| WBGene00010390 | NA | NA | NA | NA |
| WBGene00010391 | -0.099452571 | 0.17595618 | NA | oac-37 |
| WBGene00010392 | NA | NA | NA | oac-38 |
| WBGene00010394 | NA | NA | NA | srz-59 |
| WBGene00010395 | NA | NA | NA | H12I19.8 |
| WBGene00010396 | -0.020363922 | 0.9059895 | 0.94916953 | H13N06.2 |
| WBGene00010397 | -0.099365208 | 0.4803376 | 0.66672571 | suox-1 |
| WBGene00010399 | NA | NA | NA | clec-15 |
| WBGene00010399 | NA | NA | NA | clec-13 |
| WBGene00010400 | NA | NA | NA | NA |
| WBGene00010401 | -0.075452738 | 0.28904131 | NA | H16D19.3 |
| WBGene00010402 | 0.056753339 | 0.53102759 | NA | NA |
| WBGene00010403 | 0.055897183 | 0.44636755 | NA | otpl-8 |
| WBGene00010404 | NA | NA | NA | H19J13.2 |
| WBGene00010405 | -0.340684353 | 3.98E-08 | 8.03E-07 | erfa-3 |
| WBGene00010406 | -0.159937171 | 0.00775482 | 0.03688706 | math-33 |
| WBGene00010407 | 0.583814665 | 0.00148094 | 0.00971951 | H19N07.3 |
| WBGene00010408 | 0.197197315 | 0.04959822 | 0.149325 | mboa-2 |
| WBGene00010409 | -0.061824624 | 0.53232762 | 0.70853237 | H21P03.2 |
| WBGene00010410 | NA | NA | NA | nhr-267 |
| WBGene00010411 | 1.465734912 | 0.00084433 | 0.00609555 | H25K10.1 |
| WBGene00010412 | NA | NA | NA | H25K10.2 |

| | | | | |
|---|---|---|---|---|
| WBGene00010413 | 0.1298478 | 0.10432033 | NA | H25K10.4 |
| WBGene00010414 | -0.002914971 | 0.95718083 | NA | hpa-1 |
| WBGene00010415 | 0.033034883 | 0.64219503 | NA | irld-39 |
| WBGene00010416 | -0.140150341 | 0.1659313 | 0.34245844 | hxk-2 |
| WBGene00010417 | NA | NA | NA | clpr-3 |
| WBGene00010417 | NA | NA | NA | clpr-4 |
| WBGene00010418 | -0.054661807 | 0.64993879 | 0.79273174 | H27A22.1 |
| WBGene00010419 | -0.144039373 | 0.0006173 | 0.00466354 | atp-1 |
| WBGene00010420 | -0.20691591 | 0.08452746 | 0.21892209 | mcrs-1 |
| WBGene00010421 | 0.144119331 | 0.49863448 | 0.68063653 | slc-36.4 |
| WBGene00010422 | 0.036372075 | 0.84772163 | 0.9180823 | H32K16.2 |
| WBGene00010423 | 0.075029222 | 0.33362495 | NA | zmp-5 |
| WBGene00010424 | 0.010481661 | 0.93466856 | 0.96315332 | lep-5 |
| WBGene00010425 | 0.544683075 | 1.91E-14 | 8.56E-13 | lpin-1 |
| WBGene00010426 | 0.137567171 | 0.46248068 | 0.65252509 | H37A05.2 |
| WBGene00010427 | -0.176019728 | 0.01681421 | 0.06759387 | hpo-11 |
| WBGene00010428 | -0.319214958 | 0.00053089 | 0.00408665 | dcn-1 |
| WBGene00010429 | NA | NA | NA | H38K22.4 |
| WBGene00010430 | NA | NA | NA | NA |
| WBGene00010431 | NA | NA | NA | NA |
| WBGene00010432 | NA | NA | NA | H39E23.2 |
| WBGene00010433 | -0.199949101 | 0.12364702 | 0.28379306 | H40L08.1 |
| WBGene00010434 | -0.029818428 | 0.88197382 | 0.93748937 | H40L08.3 |
| WBGene00010435 | -0.361967742 | 4.39E-05 | 0.00046597 | JC8.2 |
| WBGene00010436 | -0.071843769 | 0.7026684 | 0.83114502 | JC8.4 |
| WBGene00010437 | -0.153948533 | 0.27864984 | 0.47855862 | cox-11 |
| WBGene00010439 | -0.14026735 | 0.27262922 | 0.4726499 | JC8.7 |
| WBGene00010440 | 0.023517137 | 0.77498365 | 0.87260906 | ttr-51 |
| WBGene00010442 | 0.12622451 | 0.17121794 | NA | bus-12 |
| WBGene00010444 | 0.186204552 | 0.10650546 | 0.25602685 | magi-1 |
| WBGene00010446 | 0.036821884 | 0.59344795 | NA | K01A6.4 |
| WBGene00010447 | 0.036687355 | 0.59489568 | NA | K01A6.5 |
| WBGene00010448 | 0.098606405 | 0.56849591 | 0.73742789 | K01A6.6 |
| WBGene00010449 | 0.11698322 | 0.57262597 | 0.74081619 | K01A11.1 |
| WBGene00010450 | -0.150979162 | 0.37378721 | 0.57275766 | K01A11.2 |
| WBGene00010451 | 0.025972078 | 0.877309 | 0.9347017 | K01A11.3 |
| WBGene00010453 | 2.354189988 | 1.73E-12 | 6.26E-11 | fozi-1 |
| WBGene00010454 | 0.281690468 | 0.20820315 | 0.39732825 | K01B6.3 |
| WBGene00010455 | -0.00346345 | 0.95708614 | NA | K01B6.4 |
| WBGene00010456 | -0.034264787 | 0.69931429 | 0.82884 | K01C8.1 |
| WBGene00010457 | 0.033034883 | 0.64219503 | NA | K01C8.2 |
| WBGene00010458 | -0.282846854 | 0.02041761 | 0.07784747 | mrpl-10 |
| WBGene00010459 | -0.064178613 | 0.70229294 | 0.83093594 | slc-25A32 |

| | | | | |
|---|---|---|---|---|
| WBGene00010460 | 0.025265704 | 0.84471115 | 0.91675274 | clec-142 |
| WBGene00010461 | 0.229957869 | 0.02159441 | 0.08119255 | K01D12.1 |
| WBGene00010463 | 0.099958276 | 0.38721674 | 0.58584689 | NA |
| WBGene00010464 | -0.003270206 | 0.95698316 | NA | K01D12.5 |
| WBGene00010465 | -0.274873143 | 0.00572014 | 0.0291198 | K01D12.6 |
| WBGene00010466 | -0.023109762 | 0.78121253 | NA | K01D12.7 |
| WBGene00010467 | 0.056411843 | 0.41891328 | NA | K01D12.8 |
| WBGene00010468 | 3.503686929 | 0.00102679 | 0.00718607 | K01D12.9 |
| WBGene00010469 | 0.020317245 | 0.84643798 | NA | K01D12.10 |
| WBGene00010470 | 0.768058757 | 4.54E-05 | 0.00048165 | cdr-4 |
| WBGene00010471 | 0.02390295 | 0.86186203 | 0.92668569 | cdr-6 |
| WBGene00010472 | 0.002530616 | 0.99173161 | 0.99519825 | cdr-7 |
| WBGene00010473 | 0.274907766 | 0.20785791 | 0.39677283 | hrg-2 |
| WBGene00010474 | NA | NA | NA | K01D12.15 |
| WBGene00010475 | 0.552719489 | 0.04727318 | 0.14440365 | K01F9.2 |
| WBGene00010476 | -0.066219661 | 0.54667748 | 0.71984037 | rnf-113 |
| WBGene00010477 | 0.003912659 | 0.98644986 | 0.99221025 | enu-3.6 |
| WBGene00010478 | -0.524377988 | 1.20E-14 | 5.51E-13 | K01G5.5 |
| WBGene00010479 | -0.065859367 | 0.56429376 | 0.7342539 | K01G5.8 |
| WBGene00010480 | 0.022429291 | 0.83312136 | 0.90892174 | tasp-1 |
| WBGene00010481 | 0.120239366 | 0.56926135 | 0.73815919 | K01G5.10 |
| WBGene00010483 | -0.581797506 | 0.02674546 | 0.09546083 | K01G12.3 |
| WBGene00010484 | 0.033695602 | 0.85844766 | 0.92422474 | dph-3 |
| WBGene00010485 | 0.357924427 | 0.11466539 | 0.26968955 | ant-1.3 |
| WBGene00010486 | 0.35603113 | 0.05833402 | 0.16822305 | K01H12.4 |
| WBGene00010487 | 0.012740904 | 0.89342361 | NA | K02A11.2 |
| WBGene00010488 | 0.611588997 | 0.00126415 | 0.00853317 | endu-1 |
| WBGene00010491 | -0.103979822 | 0.52174263 | 0.70003975 | K02B7.3 |
| WBGene00010492 | -0.133188927 | 0.04517216 | 0.13964644 | meg-1 |
| WBGene00010493 | -0.152612449 | 0.1906206 | 0.37668729 | meg-2 |
| WBGene00010494 | NA | NA | NA | K02B9.3 |
| WBGene00010495 | -1.00277526 | 0.0001887 | 0.00167946 | K02B12.2 |
| WBGene00010496 | 0.024350277 | 0.82407436 | 0.9034778 | sec-12 |
| WBGene00010497 | 0.00046475 | 0.99992729 | 0.99992729 | axl-1 |
| WBGene00010498 | 0.018946333 | 0.85517732 | 0.92241889 | K02B12.5 |
| WBGene00010499 | -0.045771477 | 0.50913401 | NA | K02B12.6 |
| WBGene00010500 | 0.002607315 | 0.9796348 | 0.98820677 | K02B12.7 |
| WBGene00010501 | -0.00695846 | 0.94125224 | 0.96684409 | K02C4.2 |
| WBGene00010502 | 0.010187677 | 0.87063671 | 0.93091263 | K02C4.3 |
| WBGene00010503 | 0.643041761 | 0.00153347 | 0.0100096 | K02C4.5 |
| WBGene00010504 | -0.22920724 | 0.24332173 | 0.44140607 | K02D3.1 |
| WBGene00010505 | 0.0603226 | 0.52039473 | NA | K02D3.2 |
| WBGene00010506 | -0.024917643 | 0.69549488 | NA | K02E2.1 |

| | | | | |
|---|---|---|---|---|
| WBGene00010507 | 0.219651494 | 0.32022772 | 0.52007255 | K02E2.6 |
| WBGene00010508 | 0.131317611 | 0.10121567 | NA | K02E2.7 |
| WBGene00010509 | 0.034981685 | 0.68236555 | NA | K02E2.8 |
| WBGene00010510 | 0.196167709 | 0.18617917 | 0.37052463 | ent-3 |
| WBGene00010511 | NA | NA | NA | sre-7 |
| WBGene00010512 | 0.159147731 | 0.45766458 | 0.6494153 | K02E11.3 |
| WBGene00010513 | 0.669781424 | 0.02557441 | 0.09222534 | K02E11.4 |
| WBGene00010514 | 0.155756667 | 0.37034275 | 0.57006877 | K02E11.5 |
| WBGene00010515 | 0.521320676 | 0.0564631 | 0.1646196 | K02E11.6 |
| WBGene00010516 | 0.074300855 | 0.72082132 | 0.84351221 | K02E11.7 |
| WBGene00010517 | NA | NA | NA | NA |
| WBGene00010518 | -0.003270206 | 0.95698316 | NA | K02E11.9 |
| WBGene00010519 | -0.425030854 | 0.02775907 | 0.09795628 | K03A11.1 |
| WBGene00010520 | 0.346401743 | 0.15076432 | 0.32252198 | cpx-2 |
| WBGene00010521 | -0.029234136 | 0.7166623 | NA | K03A11.4 |
| WBGene00010522 | 0.318225599 | 0.17721503 | 0.3583214 | K03A11.5 |
| WBGene00010523 | NA | NA | NA | K03B8.4 |
| WBGene00010524 | 0.197026926 | 0.3611429 | 0.56098439 | K03B8.6 |
| WBGene00010525 | -0.322115326 | 0.17126083 | 0.35024861 | K03B8.8 |
| WBGene00010526 | -0.039636691 | 0.54579661 | NA | K03B8.11 |
| WBGene00010527 | -0.022930767 | 0.78121253 | NA | srz-74 |
| WBGene00010528 | 0.108538718 | 0.15420469 | NA | K03D3.2 |
| WBGene00010529 | NA | NA | NA | NA |
| WBGene00010530 | NA | NA | NA | srz-61 |
| WBGene00010531 | 0.036687355 | 0.59489568 | NA | K03D3.5 |
| WBGene00010534 | NA | NA | NA | fbxa-102 |
| WBGene00010535 | NA | NA | NA | K03D7.8 |
| WBGene00010536 | NA | NA | NA | K03D7.9 |
| WBGene00010537 | -0.213079559 | 0.12143963 | 0.28048552 | mys-2 |
| WBGene00010538 | -0.180028577 | 0.18636407 | 0.37072776 | ttr-3 |
| WBGene00010539 | -0.016317615 | 0.83794484 | 0.91228067 | ttr-2 |
| WBGene00010540 | 0.675281333 | 0.00178333 | 0.01135697 | K03H1.5 |
| WBGene00010541 | 0.277700717 | 0.20043824 | 0.38900004 | ttr-1 |
| WBGene00010542 | -0.034993013 | 0.80805926 | 0.89369963 | K03H1.7 |
| WBGene00010543 | -0.003108065 | 0.95728506 | NA | NA |
| WBGene00010544 | 0.020232312 | 0.84643798 | NA | K03H1.9 |
| WBGene00010545 | 1.444580792 | 7.55E-09 | 1.72E-07 | cbp-2 |
| WBGene00010546 | 0.011333516 | 0.94404285 | 0.96831521 | K03H1.11 |
| WBGene00010547 | 0.132031485 | 0.16048086 | NA | K03H1.12 |
| WBGene00010550 | 0.158887578 | 0.32861474 | 0.52812825 | K04B12.2 |
| WBGene00010551 | -0.0126715 | 0.9127647 | 0.95126775 | smg-8 |
| WBGene00010552 | -0.235338541 | 0.14790623 | 0.31876649 | K04C1.2 |
| WBGene00010553 | 0.035565884 | 0.86327265 | 0.92710151 | K04C1.3 |

| | | | | |
|---|---|---|---|---|
| WBGene00010554 | 0.046484472 | 0.80004831 | 0.88826172 | mlc-6 |
| WBGene00010555 | -0.0908635 | 0.36940572 | 0.56931003 | K04C1.5 |
| WBGene00010556 | -0.384415842 | 1.04E-14 | 4.82E-13 | rack-1 |
| WBGene00010557 | 0.04887786 | 0.63803857 | 0.78552676 | mspn-1 |
| WBGene00010558 | 0.246870688 | 0.24523731 | 0.44340768 | ptp-4 |
| WBGene00010559 | 0.00911803 | 0.93063136 | 0.96081261 | K04D7.6 |
| WBGene00010560 | -0.433355935 | 1.58E-09 | 3.89E-08 | eif-2beta |
| WBGene00010562 | 0.009910614 | 0.93690118 | 0.96443579 | cdc-48.3 |
| WBGene00010563 | 0.199358631 | 0.20784657 | 0.39677283 | K04G2.4 |
| WBGene00010564 | -0.131943516 | 0.11902158 | 0.27682208 | ath-1 |
| WBGene00010565 | -0.13520614 | 0.17715294 | 0.35824533 | vacl-14 |
| WBGene00010566 | -0.154338002 | 0.17364298 | 0.35408173 | K04G2.7 |
| WBGene00010567 | -0.283479034 | 0.13236926 | 0.2955991 | K04G2.9 |
| WBGene00010568 | -0.633290544 | 0.00064947 | 0.00485892 | K04G2.10 |
| WBGene00010569 | -0.322051103 | 0.10462818 | 0.25285927 | scbp-2 |
| WBGene00010570 | 0.020925399 | 0.84643798 | NA | K04G11.1 |
| WBGene00010571 | -0.10444141 | 0.59365715 | 0.75344508 | K04G11.3 |
| WBGene00010572 | -0.060512403 | 0.76645974 | 0.86728009 | wdr-5.2 |
| WBGene00010573 | 3.102298088 | 1.66E-06 | 2.46E-05 | K04H4.2 |
| WBGene00010574 | 0.163629632 | 0.33820027 | 0.53789273 | K04H4.5 |
| WBGene00010575 | 1.771732606 | 0.00052016 | 0.00401461 | flp-25 |
| WBGene00010576 | NA | NA | NA | clec-116 |
| WBGene00010577 | NA | NA | NA | K04H8.2 |
| WBGene00010578 | -0.106222287 | 0.21522989 | NA | K04H8.3 |
| WBGene00010579 | -0.298918423 | 0.00072078 | 0.00530301 | K05C4.2 |
| WBGene00010580 | 0.016059264 | 0.84014476 | NA | K05C4.3 |
| WBGene00010581 | -0.075681975 | 0.69320603 | 0.82400174 | K05C4.4 |
| WBGene00010582 | -0.177334195 | 0.04839947 | 0.14670169 | K05C4.5 |
| WBGene00010583 | -0.202633152 | 0.03335556 | 0.11188059 | K05C4.7 |
| WBGene00010584 | -0.031867744 | 0.72748876 | NA | K05C4.8 |
| WBGene00010585 | -0.377960389 | 0.12089469 | 0.27975647 | K05C4.9 |
| WBGene00010586 | -0.023056499 | 0.78121253 | NA | K05C4.10 |
| WBGene00010587 | -0.033790569 | 0.83631304 | 0.91104607 | sol-2 |
| WBGene00010588 | 0.036687355 | 0.59489568 | NA | srz-14 |
| WBGene00010589 | 0.296448629 | 0.0602594 | 0.17198008 | cyp-33D1 |
| WBGene00010590 | NA | NA | NA | srz-13 |
| WBGene00010591 | -0.023056499 | 0.78121253 | NA | K05D4.9 |
| WBGene00010592 | 0.072338251 | 0.61456191 | 0.76874831 | K06A4.2 |
| WBGene00010593 | -0.343578478 | 0.02733894 | 0.09679887 | gsnl-1 |
| WBGene00010594 | 0.092938702 | 0.19548342 | NA | swt-5 |
| WBGene00010595 | -0.52458889 | 0.00081906 | 0.00593472 | haao-1 |
| WBGene00010596 | NA | NA | NA | K06A4.6 |
| WBGene00010597 | 0.264644917 | 0.01596586 | 0.06501923 | K06A4.7 |

| | | | | |
|---|---|---|---|---|
| WBGene00010598 | NA | NA | NA | K06B4.3 |
| WBGene00010599 | NA | NA | NA | K06B4.4 |
| WBGene00010600 | 0.020925399 | 0.84643798 | NA | nhr-196 |
| WBGene00010601 | NA | NA | NA | nhr-268 |
| WBGene00010602 | 0.020232312 | 0.84643798 | NA | nhr-197 |
| WBGene00010603 | 0.016864519 | 0.82560414 | NA | nhr-198 |
| WBGene00010604 | 0.033034883 | 0.64219503 | NA | nhr-199 |
| WBGene00010605 | 1.603355857 | 4.28E-90 | 1.96E-87 | K06G5.1 |
| WBGene00010606 | -0.699537886 | 0.00984231 | 0.04433344 | cyp-13B2 |
| WBGene00010607 | 0.0833815 | 0.40410587 | 0.60231233 | K06G5.3 |
| WBGene00010608 | -0.496547927 | 5.24E-05 | 0.00054592 | K07A1.1 |
| WBGene00010609 | 0.020853195 | 0.71619741 | 0.84075838 | dut-1 |
| WBGene00010610 | 0.143230984 | 0.50080756 | 0.68269142 | K07A1.3 |
| WBGene00010611 | 0.040107227 | 0.58693163 | NA | K07A1.4 |
| WBGene00010612 | 0.14714586 | 0.12797728 | 0.28937689 | K07A1.5 |
| WBGene00010613 | -0.160434353 | 0.33425063 | 0.53367105 | K07A1.6 |
| WBGene00010614 | 0.095885866 | 0.50359237 | 0.68467923 | cri-1 |
| WBGene00010615 | -0.069570063 | 0.50661195 | 0.68739475 | ate-1 |
| WBGene00010616 | -0.373062692 | 0.00268738 | 0.01593596 | K07A1.10 |
| WBGene00010617 | 0.140491874 | 0.45676334 | 0.64869855 | K07A1.13 |
| WBGene00010618 | NA | NA | NA | NA |
| WBGene00010619 | 0.003011386 | 0.98943948 | 0.99398816 | K07A1.15 |
| WBGene00010620 | -0.099139937 | 0.2181809 | 0.41011511 | K07A12.1 |
| WBGene00010621 | -0.121163563 | 0.03795129 | 0.12312096 | egg-6 |
| WBGene00010622 | -0.150769391 | 0.21306387 | 0.40365788 | K07A12.4 |
| WBGene00010623 | 0.109304433 | 0.5967082 | 0.75522171 | K07A12.5 |
| WBGene00010624 | -0.372778013 | 0.00197923 | 0.01236787 | mrps-15 |
| WBGene00010625 | -0.201751848 | 0.14993157 | 0.32125686 | K07C5.2 |
| WBGene00010626 | 0.118830936 | 0.2408747 | 0.4384195 | K07C5.3 |
| WBGene00010627 | -0.647107321 | 8.63E-19 | 5.91E-17 | nol-56 |
| WBGene00010628 | -0.117529707 | 0.53515585 | 0.7103551 | ceeh-2 |
| WBGene00010629 | -0.113989752 | 0.09719729 | 0.24127934 | K07C5.6 |
| WBGene00010630 | 0.153901279 | 0.47078707 | 0.6588409 | ttll-15 |
| WBGene00010631 | -0.067203119 | 0.24091581 | 0.43843981 | cash-1 |
| WBGene00010632 | -0.000917122 | 0.99403465 | 0.99634841 | K07C5.9 |
| WBGene00010633 | 1.904519217 | 1.60E-05 | 0.00018655 | nspd-2 |
| WBGene00010634 | -0.003108065 | 0.95728506 | NA | K07F5.6 |
| WBGene00010635 | NA | NA | NA | K07F5.7 |
| WBGene00010636 | -0.019342581 | 0.87789644 | 0.93480719 | K07F5.8 |
| WBGene00010637 | -0.427544411 | 0.0075912 | 0.03627376 | K07F5.12 |
| WBGene00010638 | -0.284745274 | 0.00817356 | 0.03843341 | K07F5.14 |
| WBGene00010639 | NA | NA | NA | K07F5.22 |
| WBGene00010639 | NA | NA | NA | K07F5.15 |

| | | | | |
|---|---|---|---|---|
| WBGene00010640 | 0.113268997 | 0.58292211 | 0.74668089 | K07F5.16 |
| WBGene00010641 | -0.05663522 | 0.41049453 | 0.60813512 | crml-1 |
| WBGene00010642 | 0.302740095 | 0.13252988 | 0.29580404 | mks-6 |
| WBGene00010643 | 0.017604133 | 0.79437959 | NA | K07G5.4 |
| WBGene00010644 | 0.073685737 | 0.72431109 | 0.84556796 | K07G5.5 |
| WBGene00010645 | -0.008660674 | 0.92230225 | 0.95626565 | fecl-1 |
| WBGene00010646 | -0.223867593 | 0.24477212 | 0.44292812 | K08C7.1 |
| WBGene00010647 | 0.029393842 | 0.81314698 | 0.89654981 | K08C7.4 |
| WBGene00010648 | -0.00226322 | 0.98931146 | 0.99398816 | K08C7.6 |
| WBGene00010649 | 0.091645381 | 0.61615668 | 0.76979828 | K08C7.7 |
| WBGene00010650 | 0.100806964 | 0.4256516 | 0.6215184 | K08C9.1 |
| WBGene00010651 | 2.515950454 | 0.00039531 | 0.00318354 | K08C9.2 |
| WBGene00010652 | 0.020925399 | 0.84643798 | NA | K08C9.5 |
| WBGene00010653 | NA | NA | NA | K08C9.6 |
| WBGene00010654 | 0.095418869 | 0.34312661 | 0.54313812 | K08C9.7 |
| WBGene00010655 | 0.12568073 | 0.22469281 | 0.41759166 | K08D8.1 |
| WBGene00010656 | 0.020232312 | 0.84643798 | NA | K08D8.2 |
| WBGene00010657 | 1.190827864 | 1.70E-12 | 6.17E-11 | K08D8.3 |
| WBGene00010658 | 0.356596473 | 0.0828817 | 0.21603542 | K08D8.4 |
| WBGene00010659 | 0.23282587 | 0.27810686 | 0.47785031 | K08D8.5 |
| WBGene00010660 | -0.451832972 | 0.0008776 | 0.00630156 | K08D8.6 |
| WBGene00010661 | 1.529861639 | 1.83E-11 | 5.85E-10 | tyr-2 |
| WBGene00010662 | 0.035719489 | 0.66203875 | NA | K08E3.2 |
| WBGene00010663 | 0.049684211 | 0.63033441 | 0.78011209 | toca-2 |
| WBGene00010664 | 0.011019026 | 0.8901482 | 0.94282427 | dbn-1 |
| WBGene00010665 | -0.221956612 | 2.38E-06 | 3.38E-05 | K08E3.5 |
| WBGene00010666 | 0.023498578 | 0.88852519 | 0.94151388 | K08E4.2 |
| WBGene00010667 | 0.294435791 | 0.08710999 | 0.22324127 | K08E4.3 |
| WBGene00010669 | 0.256261337 | 0.04134953 | 0.13101029 | bath-37 |
| WBGene00010670 | -0.177591471 | 0.06363897 | 0.17910023 | tmed-1 |
| WBGene00010671 | -0.071772833 | 0.42753901 | 0.62265977 | eak-7 |
| WBGene00010672 | NA | NA | NA | K08E7.4 |
| WBGene00010673 | 0.202103033 | 0.18451654 | 0.36805269 | K08E7.5 |
| WBGene00010674 | 0.187911372 | 0.13101477 | 0.29361506 | K08E7.6 |
| WBGene00010675 | -0.040610057 | 0.81343564 | 0.89680054 | K08E7.8 |
| WBGene00010676 | -0.075756105 | 0.39679353 | 0.59468258 | ctf-18 |
| WBGene00010677 | -0.17402225 | 0.00623231 | 0.03105762 | gtbp-1 |
| WBGene00010678 | 0.092652419 | 0.27556924 | 0.47572289 | K08F4.3 |
| WBGene00010679 | 0.166964812 | 0.20281346 | 0.3914648 | K08F4.5 |
| WBGene00010680 | 0.074572661 | 0.49475847 | 0.67742997 | K08F4.12 |
| WBGene00010681 | 0.648938423 | 9.99E-15 | 4.64E-13 | mak-1 |
| WBGene00010682 | -0.063457996 | 0.62322412 | 0.77471763 | K08F8.5 |
| WBGene00010683 | NA | NA | NA | K08F8.7 |

| | | | | |
|---|---|---|---|---|
| WBGene00010684 | 2.296132339 | 0.00087669 | 0.00629816 | K08F9.1 |
| WBGene00010685 | -0.18472139 | 0.00298807 | 0.01742964 | aipl-1 |
| WBGene00010686 | 0.271279547 | 0.01086327 | 0.04766232 | K08F9.3 |
| WBGene00010687 | -0.221535621 | 0.02095527 | 0.07945118 | K08F9.4 |
| WBGene00010688 | NA | NA | NA | srz-12 |
| WBGene00010689 | -0.174424065 | 0.40693937 | 0.60487303 | K08H2.2 |
| WBGene00010690 | -0.336684274 | 0.04933675 | 0.14878257 | K08H2.3 |
| WBGene00010691 | -0.02841942 | 0.84920192 | 0.91889144 | K08H2.4 |
| WBGene00010692 | 0.020925399 | 0.84643798 | NA | K08H2.5 |
| WBGene00010693 | -0.250502796 | 0.10725159 | 0.25725689 | K08H2.7 |
| WBGene00010694 | 0.016748122 | 0.90589365 | 0.94916953 | bgnt-1.5 |
| WBGene00010697 | -0.0530764 | 0.47060713 | 0.658715 | uda-1 |
| WBGene00010698 | 0.288872802 | 0.09063428 | 0.22888093 | K08H10.6 |
| WBGene00010699 | -0.124498492 | 0.35862572 | 0.55893541 | trpp-6 |
| WBGene00010700 | 0.150064203 | 0.223544 | 0.41645346 | nipi-3 |
| WBGene00010701 | -0.13196012 | 0.27781517 | 0.47767779 | ent-2 |
| WBGene00010702 | -0.137387504 | 0.3433061 | 0.54313812 | usp-33 |
| WBGene00010703 | 0.376952502 | 0.04641282 | 0.14255181 | K09A9.6 |
| WBGene00010704 | -0.247782801 | 0.21789117 | 0.4096757 | K09A11.1 |
| WBGene00010705 | -0.03731551 | 0.85569879 | 0.92262213 | cyp-14A1 |
| WBGene00010706 | -0.037517955 | 0.85000819 | 0.91926207 | cyp-14A2 |
| WBGene00010707 | 0.0398788 | 0.58971221 | NA | cyp-14A3 |
| WBGene00010708 | -0.080472653 | 0.40010183 | NA | phf-33 |
| WBGene00010709 | 0.098823724 | 0.49771946 | 0.67976712 | nol-9 |
| WBGene00010711 | -0.161903541 | 0.29307208 | 0.49399819 | K09B11.4 |
| WBGene00010712 | 0.083479586 | 0.30566331 | NA | K09B11.5 |
| WBGene00010713 | -0.170324061 | 0.11500059 | 0.270304 | uso-1 |
| WBGene00010714 | 0.046470303 | 0.62612299 | 0.77700176 | mam-3 |
| WBGene00010715 | 0.020317245 | 0.84643798 | NA | K09C8.2 |
| WBGene00010716 | 0.08378722 | 0.40219586 | 0.60068853 | lge-1 |
| WBGene00010717 | 0.132031485 | 0.16048086 | NA | K09C8.7 |
| WBGene00010718 | 0.038125098 | 0.61119482 | NA | K09C8.8 |
| WBGene00010719 | 0.226872498 | 0.20279214 | 0.3914648 | K09E4.1 |
| WBGene00010720 | 0.103677019 | 0.36310785 | 0.56290195 | algn-3 |
| WBGene00010721 | 0.163516566 | 0.2826487 | 0.4825396 | K09E4.3 |
| WBGene00010722 | 1.419588883 | 0.00154785 | 0.01008545 | K09E4.4 |
| WBGene00010723 | 1.514048868 | 0.00012781 | 0.00120134 | cpg-7 |
| WBGene00010724 | 0.510787995 | 0.01974784 | 0.07588664 | K09E9.1 |
| WBGene00010725 | -0.018005773 | 0.90860925 | 0.95060397 | erv-46 |
| WBGene00010726 | 0.049754783 | 0.74404312 | 0.85546846 | K09E9.3 |
| WBGene00010727 | -0.2880204 | 0.00460363 | 0.02455786 | K09G1.1 |
| WBGene00010728 | 0.105820647 | 0.22502924 | NA | K09G1.2 |
| WBGene00010730 | -0.19432015 | 0.00344673 | 0.01949905 | ensa-1 |

| | | | | | |
|---|---|---|---|---|---|
| WBGene00010731 | | -0.301361051 | 0.00026421 | 0.00224901 | K10C3.4 |
| WBGene00010732 | | -0.234138097 | 0.04076275 | 0.12976894 | K10C3.5 |
| WBGene00010733 | NA | | NA | NA | NA |
| WBGene00010734 | | -0.170796565 | 0.2947074 | 0.49577045 | parn-1 |
| WBGene00010735 | | 0.003696665 | 0.97674232 | 0.98670757 | frpr-15 |
| WBGene00010736 | | 0.009862039 | 0.9105854 | 0.95112162 | istr-1 |
| WBGene00010737 | | 0.011470707 | 0.89993902 | NA | K10C8.4 |
| WBGene00010738 | | 0.537932463 | 0.04880578 | 0.14761753 | K10D3.4 |
| WBGene00010740 | | 1.924038998 | 6.00E-08 | 1.17E-06 | K10D3.6 |
| WBGene00010741 | | 0.057774287 | 0.59140746 | 0.75215398 | lgc-49 |
| WBGene00010742 | | 0.82487665 | 0.01226728 | 0.05263796 | K10D6.2 |
| WBGene00010743 | | 0.020317245 | 0.84643798 | NA | K10D6.3 |
| WBGene00010744 | | 0.110525273 | 0.57953041 | 0.74518754 | K10D6.4 |
| WBGene00010745 | | 1.130960765 | 0.00223622 | 0.01364754 | dod-17 |
| WBGene00010746 | | 0.020232312 | 0.84643798 | NA | K10D11.2 |
| WBGene00010747 | | 1.438293042 | 0.00132357 | 0.00885265 | K10D11.3 |
| WBGene00010748 | NA | | NA | NA | K10D11.4 |
| WBGene00010749 | | 0.160933435 | 0.41654643 | 0.61354691 | K10D11.5 |
| WBGene00010750 | | 1.120370764 | 0.00097855 | 0.00691456 | K10D11.6 |
| WBGene00010752 | | 0.939339739 | 0.01894457 | 0.07392961 | K10G4.3 |
| WBGene00010753 | NA | | NA | NA | NA |
| WBGene00010754 | | 3.773467764 | 0.00053505 | 0.00411266 | K10G4.5 |
| WBGene00010755 | | 0.853609915 | 5.31E-05 | 0.00055207 | vglu-2 |
| WBGene00010756 | | 0.008984052 | 0.96680679 | 0.98141983 | K10G9.2 |
| WBGene00010757 | | 0.116702699 | 0.57440591 | 0.74213528 | pad-2 |
| WBGene00010758 | | -0.200878013 | 0.12994103 | 0.29224835 | vnut-1 |
| WBGene00010759 | | 0.048676793 | 0.73344544 | 0.85087141 | cysl-2 |
| WBGene00010760 | | 0.077387215 | 0.23878538 | NA | K10H10.4 |
| WBGene00010761 | | -0.003108065 | 0.95728506 | NA | K10H10.5 |
| WBGene00010762 | NA | | NA | NA | K10H10.6 |
| WBGene00010763 | | 0.060484201 | 0.41382805 | NA | K10H10.7 |
| WBGene00010764 | | 0.020232312 | 0.84643798 | NA | K10H10.9 |
| WBGene00010765 | | 0.059980168 | 0.38780842 | NA | K10H10.10 |
| WBGene00010766 | | -0.431640715 | 4.97E-05 | 0.00052159 | mrps-27 |
| WBGene00010767 | | -0.107760073 | 0.48505263 | 0.6699043 | blos-9 |
| WBGene00010768 | | 0.468603541 | 0.07842263 | 0.20815549 | K11D2.1 |
| WBGene00010769 | | 2.446502439 | 5.22E-78 | 2.07E-75 | asah-1 |
| WBGene00010770 | | 0.429361516 | 0.0127339 | 0.0542321 | K11D2.4 |
| WBGene00010771 | | 0.057068005 | 0.75120677 | 0.85945287 | K11D2.5 |
| WBGene00010772 | | 0.396296884 | 0.02895327 | 0.10102592 | K11D9.3 |
| WBGene00010773 | | 0.184474566 | 0.34534139 | 0.54496048 | K11E4.1 |
| WBGene00010774 | | 0.169100567 | 0.37743413 | 0.57652718 | K11E4.2 |
| WBGene00010775 | NA | | NA | NA | poml-1 |

| | | | | | |
|---|---|---|---|---|---|
| WBGene00010776 | | -0.375325714 | 0.01646691 | 0.06656443 | pix-1 |
| WBGene00010777 | NA | | NA | NA | NA |
| WBGene00010778 | | -0.146427233 | 0.03073698 | 0.10518593 | gpdh-2 |
| WBGene00010779 | | 0.145611791 | 0.20114132 | 0.38993141 | K11H3.2 |
| WBGene00010780 | | 0.009091723 | 0.89455347 | 0.94489559 | K11H3.3 |
| WBGene00010781 | | -0.259048699 | 0.06702658 | 0.18557794 | K11H3.4 |
| WBGene00010782 | | 0.220374423 | 0.23710446 | 0.43347869 | K11H3.5 |
| WBGene00010783 | | -0.090659618 | 0.55725079 | 0.72847563 | mrpl-36 |
| WBGene00010784 | | 0.196796168 | 0.30174622 | 0.50237281 | twk-48 |
| WBGene00010785 | | -0.071639878 | 0.14770894 | 0.31845186 | top-2 |
| WBGene00010786 | | 0.057474242 | 0.58731396 | 0.74981371 | K12D12.4 |
| WBGene00010787 | | 0.05324373 | 0.65211096 | 0.7939756 | K12D12.5 |
| WBGene00010788 | | 0.173656847 | 0.41899774 | 0.61579411 | sulp-4 |
| WBGene00010789 | | 0.416002568 | 0.08374291 | 0.21762158 | sulp-5 |
| WBGene00010790 | | 3.630254538 | 5.77E-300 | 1.06E-296 | sodh-1 |
| WBGene00010791 | | 4.045110343 | 0.00139381 | 0.00925059 | sodh-2 |
| WBGene00010792 | | 0.007229767 | 0.95127221 | 0.97203268 | K12G11.6 |
| WBGene00010793 | | 0.490500526 | 4.42E-06 | 5.84E-05 | LLC1.2 |
| WBGene00010794 | | -0.283342371 | 3.13E-06 | 4.29E-05 | dld-1 |
| WBGene00010795 | | 2.220739011 | 4.83E-07 | 8.10E-06 | M01A8.1 |
| WBGene00010796 | | 0.324227278 | 0.05674886 | 0.16518093 | clip-1 |
| WBGene00010797 | NA | | NA | NA | srbc-75 |
| WBGene00010798 | NA | | NA | NA | srbc-76 |
| WBGene00010799 | | 1.043186704 | 0.01288215 | 0.0547482 | chil-12 |
| WBGene00010800 | | 0.020317245 | 0.84643798 | NA | srsx-37 |
| WBGene00010801 | | 0.04909525 | 0.45360729 | NA | M01B2.8 |
| WBGene00010802 | NA | | NA | NA | srsx-39 |
| WBGene00010803 | | 0.06298612 | 0.76307196 | 0.86558739 | M01B2.10 |
| WBGene00010804 | | 0.020317245 | 0.84643798 | NA | M01E5.1 |
| WBGene00010805 | | 0.160749997 | 0.39192567 | 0.58998393 | M01E5.2 |
| WBGene00010806 | | -0.011035719 | 0.87847588 | 0.93499312 | M01E5.3 |
| WBGene00010807 | | -0.072501383 | 0.50940107 | 0.6895487 | M01E5.4 |
| WBGene00010808 | | -0.980429302 | 9.14E-09 | 2.06E-07 | sepa-1 |
| WBGene00010809 | | 0.260694674 | 0.01079174 | 0.04741951 | M01F1.3 |
| WBGene00010810 | | 0.129579726 | 0.2415004 | 0.43933988 | M01F1.4 |
| WBGene00010811 | | 0.29531893 | 0.07196003 | 0.19526033 | hmit-1.3 |
| WBGene00010812 | | -0.602046929 | 0.00091568 | 0.00653019 | mrpl-35 |
| WBGene00010813 | | 0.286241358 | 0.08296784 | 0.21618298 | pitp-1 |
| WBGene00010814 | | -0.122254935 | 0.38050537 | 0.57992875 | M01F1.8 |
| WBGene00010815 | | 0.130768457 | 0.23750204 | 0.43384497 | M01F1.9 |
| WBGene00010817 | NA | | NA | NA | M01G12.2 |
| WBGene00010818 | NA | | NA | NA | NA |
| WBGene00010819 | NA | | NA | NA | M01G12.5 |

| | | | | |
|---|---|---|---|---|
| WBGene00010820 | NA | NA | NA | M01G12.6 |
| WBGene00010821 | NA | NA | NA | M01G12.7 |
| WBGene00010822 | 0.762617208 | 0.01678365 | 0.06754519 | M01G12.9 |
| WBGene00010823 | NA | NA | NA | M01G12.10 |
| WBGene00010824 | NA | NA | NA | NA |
| WBGene00010825 | 0.321093834 | 0.02494705 | 0.0907002 | M01G12.14 |
| WBGene00010827 | 0.177257625 | 0.38589756 | 0.58481794 | M02B1.2 |
| WBGene00010828 | 0.138875547 | 0.05694982 | 0.16556837 | M02B1.3 |
| WBGene00010829 | -0.007949484 | 0.92915667 | NA | M02B1.4 |
| WBGene00010830 | 0.078236818 | 0.47921183 | 0.66615262 | M02G9.1 |
| WBGene00010831 | 0.105191732 | 0.16565281 | NA | M02G9.2 |
| WBGene00010832 | -0.022930767 | 0.78121253 | NA | M02G9.3 |
| WBGene00010833 | -0.360120916 | 0.13745537 | 0.30322195 | M03B6.1 |
| WBGene00010834 | 0.054862227 | 0.62112156 | 0.7734831 | mct-3 |
| WBGene00010835 | 0.219593233 | 0.12775093 | 0.28904364 | M03B6.3 |
| WBGene00010836 | -0.327580852 | 0.12689863 | 0.28824805 | M03B6.4 |
| WBGene00010837 | 0.017905199 | 0.82252206 | NA | M03B6.5 |
| WBGene00010838 | 0.278419119 | 0.22376856 | 0.41676573 | M03C11.1 |
| WBGene00010839 | 0.050105441 | 0.65089822 | 0.79329067 | chl-1 |
| WBGene00010840 | -0.128459831 | 0.15484861 | 0.32795291 | M03C11.3 |
| WBGene00010842 | -0.022339239 | 0.83570913 | 0.91065923 | ymel-1 |
| WBGene00010843 | 0.014880509 | 0.94058802 | 0.96666778 | M03C11.6 |
| WBGene00010844 | -0.123156542 | 0.12868093 | 0.29040597 | prp-3 |
| WBGene00010845 | -0.170287804 | 0.02046935 | 0.07798381 | M03C11.8 |
| WBGene00010846 | 0.147614548 | 0.34532893 | 0.54496048 | M04B2.2 |
| WBGene00010847 | -0.199278299 | 0.07507178 | 0.20153163 | M04B2.4 |
| WBGene00010848 | 0.559530073 | 0.04874332 | 0.1475095 | M04B2.6 |
| WBGene00010849 | 0.107916049 | 0.3351699 | 0.53432502 | M04B2.7 |
| WBGene00010850 | 1.246662485 | 4.00E-14 | 1.75E-12 | M04C3.1 |
| WBGene00010851 | 2.173044944 | 5.83E-06 | 7.52E-05 | M04C3.2 |
| WBGene00010854 | 0.056411843 | 0.41891328 | NA | M04C7.3 |
| WBGene00010855 | 0.017279211 | 0.79380376 | NA | M04C7.4 |
| WBGene00010856 | 0.225264848 | 0.29644982 | 0.49751505 | M04C9.1 |
| WBGene00010857 | 0.020925399 | 0.84643798 | NA | M04C9.2 |
| WBGene00010858 | -0.278616847 | 0.20311419 | 0.39190636 | M04C9.3 |
| WBGene00010859 | 0.264804674 | 0.22456601 | 0.41754834 | M04C9.4 |
| WBGene00010862 | NA | NA | NA | M04D8.4 |
| WBGene00010863 | NA | NA | NA | M04D8.5 |
| WBGene00010864 | 0.113207936 | 0.37685042 | 0.5758759 | xbx-3 |
| WBGene00010865 | 0.614487138 | 0.04177028 | 0.13194362 | M04D8.7 |
| WBGene00010866 | NA | NA | NA | M04D8.8 |
| WBGene00010867 | 0.244122446 | 0.1694671 | 0.34787245 | tag-260 |
| WBGene00010868 | 0.680863445 | 0.00012574 | 0.00118539 | somi-1 |

| | | | | |
|---|---|---|---|---|
| WBGene00010869 | 0.155122056 | 0.06782466 | NA | M05B5.1 |
| WBGene00010870 | 0.317008034 | 0.14815937 | 0.31884712 | let-522 |
| WBGene00010871 | 0.262652301 | 0.03980871 | 0.12750514 | M05B5.3 |
| WBGene00010872 | -0.038811706 | 0.77446739 | 0.87222901 | M05B5.4 |
| WBGene00010873 | 1.101201188 | 0.00484899 | 0.02553743 | trpa-2 |
| WBGene00010874 | -0.079660709 | 0.31592878 | NA | M05D6.1 |
| WBGene00010875 | -0.147652082 | 0.05775372 | 0.16717521 | M05D6.2 |
| WBGene00010876 | 0.085309563 | 0.24820057 | NA | M05D6.3 |
| WBGene00010877 | 0.441900395 | 0.05061968 | 0.15174504 | lact-4 |
| WBGene00010878 | -0.08425699 | 0.57859176 | 0.74452118 | M05D6.5 |
| WBGene00010879 | -0.277336531 | 0.02621726 | 0.09411792 | M05D6.6 |
| WBGene00010880 | 0.038125098 | 0.61119482 | NA | M05D6.8 |
| WBGene00010881 | -0.103096652 | 0.38511821 | 0.58436271 | M05D6.9 |
| WBGene00010882 | 0.011210793 | 0.92371581 | 0.95682907 | atg-7 |
| WBGene00010883 | 0.09249737 | 0.19271823 | NA | M7.7 |
| WBGene00010884 | 0.513133013 | 0.06142566 | 0.17459389 | M7.8 |
| WBGene00010885 | 0.033034883 | 0.64219503 | NA | M7.9 |
| WBGene00010886 | 0.020232312 | 0.84643798 | NA | M7.10 |
| WBGene00010887 | -0.034740775 | 0.86215059 | 0.92672958 | M7.12 |
| WBGene00010888 | 0.628068586 | 0.00812342 | 0.038292 | dlc-2 |
| WBGene00010889 | 0.090983851 | 0.49169868 | 0.67507131 | M18.3 |
| WBGene00010890 | -0.000864165 | 0.99061634 | 0.9947106 | ddb-1 |
| WBGene00010891 | -0.002955458 | 0.98259848 | 0.98995618 | M18.6 |
| WBGene00010892 | -0.138326949 | 0.11619913 | 0.27211634 | dhhc-6 |
| WBGene00010893 | 0.003808313 | 0.97608652 | 0.98659874 | cutl-9 |
| WBGene00010894 | 0.059762547 | 0.38965825 | NA | M28.2 |
| WBGene00010895 | 0.058314668 | 0.40208118 | NA | M28.4 |
| WBGene00010896 | -0.610566958 | 3.06E-17 | 1.83E-15 | M28.5 |
| WBGene00010897 | 0.423377611 | 0.00324969 | 0.01861451 | lact-3 |
| WBGene00010898 | 0.103103655 | 0.39535886 | 0.59330482 | nphp-1 |
| WBGene00010899 | 1.217600071 | 4.45E-07 | 7.48E-06 | M28.8 |
| WBGene00010900 | 0.218324784 | 0.15583216 | 0.32941649 | M28.9 |
| WBGene00010901 | 0.948042945 | 2.44E-12 | 8.66E-11 | M28.10 |
| WBGene00010902 | 0.259161079 | 0.10689939 | 0.25674441 | M79.2 |
| WBGene00010904 | -0.25014561 | 0.02712034 | 0.0961998 | ugt-62 |
| WBGene00010905 | -0.332507922 | 0.00834305 | 0.03905072 | mrps-34 |
| WBGene00010906 | 0.045333246 | 0.785933 | 0.87892197 | M88.3 |
| WBGene00010907 | -0.115324768 | 0.51227183 | 0.69212785 | M88.4 |
| WBGene00010908 | -0.088276211 | 0.23041062 | 0.42486674 | imph-1 |
| WBGene00010909 | -0.228190012 | 0.06595888 | 0.1833499 | cisd-3.1 |
| WBGene00010910 | -0.072149435 | 0.46080458 | 0.65170165 | M106.2 |
| WBGene00010911 | -0.08478135 | 0.30232724 | 0.50276842 | M106.3 |
| WBGene00010912 | -0.132323565 | 0.09093244 | 0.22938351 | gmps-1 |

| | | | | | |
|---|---|---|---|---|---|
| WBGene00010913 | | -0.134813851 | 0.45881569 | 0.65022945 | M110.3 |
| WBGene00010915 | | 0.056411843 | 0.41891328 | NA | M110.7 |
| WBGene00010916 | NA | | NA | NA | M110.8 |
| WBGene00010917 | | 0.020317245 | 0.84643798 | NA | M110.9 |
| WBGene00010918 | | 0.275746086 | 0.12710509 | 0.28843823 | M117.1 |
| WBGene00010919 | | 0.21011606 | 0.13481343 | 0.29915178 | M117.3 |
| WBGene00010920 | | 0.019892966 | 0.89292827 | 0.9441978 | M117.4 |
| WBGene00010921 | | -0.242647152 | 0.1800774 | 0.36181052 | NA |
| WBGene00010922 | | -0.193338936 | 0.15854313 | 0.33293603 | M142.5 |
| WBGene00010923 | | -0.096305865 | 0.24656498 | 0.44495002 | rle-1 |
| WBGene00010924 | | -0.273088698 | 0.19163264 | 0.3778203 | M153.1 |
| WBGene00010925 | | -0.097859337 | 0.63871846 | 0.78569916 | M153.2 |
| WBGene00010926 | NA | | NA | NA | M153.3 |
| WBGene00010927 | NA | | NA | NA | clec-259 |
| WBGene00010928 | | -0.027373974 | 0.68245543 | NA | clec-258 |
| WBGene00010929 | NA | | NA | NA | srt-45 |
| WBGene00010930 | NA | | NA | NA | NA |
| WBGene00010931 | | -0.05264144 | 0.73374706 | 0.85087373 | M162.5 |
| WBGene00010932 | NA | | NA | NA | NA |
| WBGene00010933 | | 0.046709512 | 0.72096605 | 0.8435758 | M162.7 |
| WBGene00010934 | | -0.023449708 | 0.69192352 | NA | fbxa-118 |
| WBGene00010935 | | 0.868645754 | 1.01E-05 | 0.00012419 | M163.1 |
| WBGene00010936 | | 0.108148892 | 0.38647493 | 0.5851299 | ztf-14 |
| WBGene00010937 | | 0.017141582 | 0.79355725 | NA | M163.5 |
| WBGene00010938 | NA | | NA | NA | M163.6 |
| WBGene00010939 | NA | | NA | NA | M163.7 |
| WBGene00010940 | | 1.427744547 | 1.99E-05 | 0.00022734 | M163.8 |
| WBGene00010941 | | -0.178129167 | 0.07386458 | 0.19912564 | gss-1 |
| WBGene00010942 | | -0.068242037 | 0.54361017 | 0.71696212 | chch-3 |
| WBGene00010943 | | 0.84236356 | 3.72E-06 | 5.02E-05 | M176.4 |
| WBGene00010944 | | 0.351265893 | 0.14622966 | 0.31610047 | M176.5 |
| WBGene00010945 | | 0.072088865 | 0.73007522 | 0.84956474 | chil-13 |
| WBGene00010946 | NA | | NA | NA | M176.10 |
| WBGene00010947 | | 0.327611687 | 0.14526066 | 0.31460967 | M176.11 |
| WBGene00010948 | | 0.014875112 | 0.87573887 | NA | M195.2 |
| WBGene00010949 | NA | | NA | NA | M195.4 |
| WBGene00010950 | NA | | NA | NA | srt-44 |
| WBGene00010951 | | 0.0573358 | 0.75291107 | 0.86007127 | M199.2 |
| WBGene00010952 | NA | | NA | NA | NA |
| WBGene00010953 | | -0.788096732 | 0.00478208 | 0.0252541 | clec-190 |
| WBGene00010954 | NA | | NA | NA | clec-189 |
| WBGene00010955 | NA | | NA | NA | M199.7 |
| WBGene00010956 | | 0.020317245 | 0.84643798 | NA | M199.8 |

| | | | | |
|---|---|---|---|---|
| WBGene00010957 | -0.453115404 | 0.08150088 | 0.21367925 | ND6 |
| WBGene00010958 | -0.131827039 | 0.26703536 | 0.46587997 | ND4L |
| WBGene00010959 | -0.255765629 | 0.00042711 | 0.00340036 | ND1 |
| WBGene00010960 | -0.463251469 | 0.01663233 | 0.06708372 | ATP6 |
| WBGene00010961 | 0.061014849 | 0.74488064 | 0.85585245 | ND2 |
| WBGene00010962 | 0.118917513 | 0.46998744 | 0.65835115 | COX3 |
| WBGene00010963 | -0.358278181 | 0.04002166 | 0.12803875 | ND4 |
| WBGene00010964 | -0.023132716 | 0.8220767 | 0.90246087 | COX1 |
| WBGene00010965 | -0.283570857 | 0.07997734 | 0.21104039 | COX2 |
| WBGene00010966 | -0.353678031 | 0.13696424 | 0.30241193 | ND3 |
| WBGene00010967 | -0.307887137 | 0.00012214 | 0.00115669 | ND5 |
| WBGene00010968 | 0.020232312 | 0.84643798 | NA | R01E6.2 |
| WBGene00010969 | -0.023056499 | 0.78121253 | NA | R01E6.5 |
| WBGene00010970 | 0.597289611 | 0.04180208 | 0.13201558 | glb-20 |
| WBGene00010971 | 0.274548772 | 0.10288087 | 0.250254 | R01E6.7 |
| WBGene00010972 | 0.355536702 | 0.12895887 | 0.2909673 | R01H10.4 |
| WBGene00010973 | 0.144174961 | 0.40738419 | 0.60539819 | rip-1 |
| WBGene00010974 | 0.09989433 | 0.59644153 | 0.75521062 | bbs-5 |
| WBGene00010975 | 0.062537451 | 0.50348383 | 0.68467923 | R01H10.7 |
| WBGene00010976 | -0.23421244 | 0.28335613 | 0.48329649 | R02D5.1 |
| WBGene00010977 | 0.568531411 | 0.04433412 | 0.13772456 | R02D5.3 |
| WBGene00010978 | NA | NA | NA | R02D5.4 |
| WBGene00010979 | -0.079064367 | 0.48997006 | 0.67384467 | R02D5.6 |
| WBGene00010980 | 0.206817716 | 0.23620561 | 0.43206975 | R02D5.7 |
| WBGene00010981 | -0.186066412 | 0.38499268 | 0.58429334 | blos-8 |
| WBGene00010982 | 1.779284437 | 3.08E-05 | 0.00033617 | flp-32 |
| WBGene00010983 | 0.198710882 | 0.34278863 | 0.5430392 | mocs-1 |
| WBGene00010984 | -0.073609946 | 0.6022709 | 0.75950459 | nkat-3 |
| WBGene00010985 | 0.537851231 | 0.00369264 | 0.02057229 | R03A10.5 |
| WBGene00010986 | 0.173125878 | 0.32979816 | 0.52933278 | sprr-1 |
| WBGene00010987 | 0.599800232 | 0.00253192 | 0.01517391 | R03C1.1 |
| WBGene00010988 | 0.945561035 | 3.98E-06 | 5.33E-05 | metr-1 |
| WBGene00010989 | -0.458195223 | 0.0049746 | 0.02605834 | R03D7.2 |
| WBGene00010990 | -0.24494491 | 0.00253095 | 0.01517391 | tceb-3 |
| WBGene00010991 | 0.042065281 | 0.558937 | NA | R03D7.5 |
| WBGene00010992 | 0.085309563 | 0.24820057 | NA | R03D7.8 |
| WBGene00010993 | -0.010091081 | 0.90568704 | 0.94916953 | vha-20 |
| WBGene00010994 | 0.212584816 | 0.31671643 | 0.51698682 | lgc-25 |
| WBGene00010995 | NA | NA | NA | ceh-90 |
| WBGene00010996 | NA | NA | NA | R03G8.1 |
| WBGene00010998 | 1.249072741 | 0.00184631 | 0.01170209 | R03G8.3 |
| WBGene00010999 | -0.523874846 | 0.04012531 | 0.12821359 | R03G8.4 |
| WBGene00011000 | -0.824940008 | 1.30E-07 | 2.40E-06 | R03G8.6 |

| | | | | | |
|---|---|---|---|---|---|
| WBGene00011001 | | 0.020317245 | 0.84643798 | NA | R04B5.1 |
| WBGene00011002 | | -0.134602599 | 0.33991989 | 0.53960394 | nhr-205 |
| WBGene00011003 | | 0.794711255 | 4.17E-05 | 0.00044516 | R04B5.5 |
| WBGene00011004 | | -0.064193791 | 0.62965899 | 0.77968456 | R04B5.6 |
| WBGene00011005 | NA | | NA | NA | R04B5.7 |
| WBGene00011006 | | -0.007401379 | 0.95091419 | 0.97202643 | ugt-47 |
| WBGene00011007 | NA | | NA | NA | NA |
| WBGene00011008 | | 0.020925399 | 0.84643798 | NA | R04B5.11 |
| WBGene00011009 | | -0.010723471 | 0.91920705 | 0.95522113 | cyp-14A4 |
| WBGene00011010 | | 0.0261468 | 0.80815736 | 0.8937406 | R04D3.2 |
| WBGene00011011 | | -0.087129087 | 0.19089941 | 0.37693301 | R04D3.3 |
| WBGene00011012 | | -0.228078364 | 0.1656013 | 0.34206669 | R04D3.4 |
| WBGene00011013 | | 0.020232312 | 0.84643798 | NA | srxa-8 |
| WBGene00011014 | | 0.084488565 | 0.56208123 | 0.73230632 | hpo-30 |
| WBGene00011015 | | -0.112068782 | 0.0602719 | 0.17198224 | R04F11.2 |
| WBGene00011016 | | -0.171197283 | 0.0222491 | 0.08312045 | R04F11.3 |
| WBGene00011017 | | -0.427639961 | 0.00460428 | 0.02455786 | R04F11.5 |
| WBGene00011018 | | 0.014602567 | 0.85416114 | NA | R05A10.1 |
| WBGene00011019 | | 0.345538187 | 0.14242831 | 0.31058875 | R05A10.2 |
| WBGene00011020 | | 0.133066977 | 0.45084493 | 0.64322016 | R05A10.3 |
| WBGene00011021 | | -0.028668027 | 0.7217843 | NA | R05A10.4 |
| WBGene00011022 | NA | | NA | NA | NA |
| WBGene00011023 | | 0.064941241 | 0.4010185 | NA | R05A10.6 |
| WBGene00011024 | NA | | NA | NA | R05A10.7 |
| WBGene00011025 | NA | | NA | NA | R05A10.8 |
| WBGene00011026 | | 0.222564976 | 0.30227993 | 0.50276842 | R05D7.1 |
| WBGene00011027 | | 0.015194928 | 0.8086441 | NA | R05D7.2 |
| WBGene00011028 | | 0.252822732 | 0.18734361 | 0.37201923 | R05D7.3 |
| WBGene00011029 | | -0.019754817 | 0.88443286 | 0.93914865 | abhd-11.2 |
| WBGene00011030 | | -0.18079384 | 0.3289782 | 0.52842236 | R05D7.5 |
| WBGene00011032 | | -0.320278369 | 0.00370744 | 0.02063117 | ddx-52 |
| WBGene00011033 | | -0.414463882 | 0.00361459 | 0.02026856 | R05D11.5 |
| WBGene00011034 | | -0.297817668 | 0.00683563 | 0.03341852 | paxt-1 |
| WBGene00011035 | | -0.147086383 | 0.16353888 | 0.33919432 | snrp-27 |
| WBGene00011036 | | -0.165759996 | 0.01452427 | 0.06032621 | edc-3 |
| WBGene00011037 | | -0.313761768 | 0.00333335 | 0.01898974 | R05D11.9 |
| WBGene00011038 | | -0.139145096 | 0.00849256 | 0.03957339 | R05H5.3 |
| WBGene00011039 | | -0.141413989 | 0.45165936 | 0.64401487 | R05H5.4 |
| WBGene00011040 | | 0.142420174 | 0.34088021 | 0.54077658 | arv-1 |
| WBGene00011041 | | 0.038595855 | 0.57520533 | NA | R05H5.7 |
| WBGene00011042 | | 0.319580325 | 0.06490694 | 0.18163274 | R05H10.1 |
| WBGene00011043 | | -0.436341645 | 2.96E-05 | 0.00032386 | rbm-28 |
| WBGene00011044 | | 0.003852909 | 0.97647517 | 0.98670757 | R05H10.3 |

| | | | | |
|---|---|---|---|---|
| WBGene00011045 | 0.024651663 | 0.79510989 | 0.88499346 | gpx-2 |
| WBGene00011046 | 0.156062369 | 0.10430569 | 0.25250325 | R05H10.7 |
| WBGene00011047 | 0.000643514 | 0.99683891 | 0.9978865 | R06A4.2 |
| WBGene00011049 | -0.948176623 | 0.00310297 | 0.01792843 | sdz-26 |
| WBGene00011050 | -0.090620439 | 0.36467192 | 0.56433374 | agl-1 |
| WBGene00011051 | -0.109160841 | 0.24550879 | 0.44365517 | pfs-2 |
| WBGene00011052 | 0.127693253 | 0.10904601 | NA | arrd-11 |
| WBGene00011053 | 0.109291922 | 0.21520653 | NA | arrd-12 |
| WBGene00011054 | 0.095133228 | 0.46123519 | 0.65201394 | arrd-13 |
| WBGene00011055 | 0.176457681 | 0.3701135 | 0.57005643 | arrd-14 |
| WBGene00011056 | 0.114508231 | 0.54402831 | 0.71744897 | R06B9.5 |
| WBGene00011058 | -0.170579827 | 0.13311388 | 0.29668448 | fdps-1 |
| WBGene00011059 | 0.787227954 | 3.33E-22 | 2.79E-20 | R06C1.4 |
| WBGene00011060 | 0.636250314 | 0.0265523 | 0.09500297 | R06C1.6 |
| WBGene00011061 | -0.270409811 | 1.38E-09 | 3.41E-08 | wago-1 |
| WBGene00011062 | -0.16228029 | 0.39079828 | 0.58871053 | R06C7.2 |
| WBGene00011063 | -0.065675808 | 0.13287488 | 0.29637737 | cpg-3 |
| WBGene00011064 | -0.083876354 | 0.40305436 | 0.60141856 | adsl-1 |
| WBGene00011065 | 0.106825636 | 0.60325746 | 0.75988622 | R06C7.6 |
| WBGene00011066 | 0.031614647 | 0.75444769 | 0.86089996 | ztf-15 |
| WBGene00011067 | 0.03128167 | 0.76446794 | 0.86623548 | vps-11 |
| WBGene00011068 | 0.071981184 | 0.57228154 | 0.74050146 | set-14 |
| WBGene00011069 | -0.011727068 | 0.94093741 | 0.96675541 | ceh-62 |
| WBGene00011070 | 2.687928735 | 1.66E-17 | 1.02E-15 | msrp-7 |
| WBGene00011071 | -0.185740296 | 0.04409125 | 0.13720298 | R06F6.8 |
| WBGene00011072 | 1.223521743 | 2.03E-09 | 4.95E-08 | tag-209 |
| WBGene00011073 | 0.607217805 | 0.03586993 | 0.11805091 | R07A4.2 |
| WBGene00011074 | -0.031727027 | 0.72861504 | NA | R07A4.3 |
| WBGene00011075 | 0.103598762 | 0.32446098 | 0.52363682 | R07A4.4 |
| WBGene00011076 | 0.376993952 | 0.07503199 | 0.20149364 | scav-5 |
| WBGene00011077 | 0.020232312 | 0.84643798 | NA | R07B1.5 |
| WBGene00011078 | 0.020317245 | 0.84643798 | NA | R07B1.6 |
| WBGene00011079 | NA | NA | NA | R07B1.7 |
| WBGene00011081 | 2.47612733 | 0.00017262 | 0.00155531 | R07B1.9 |
| WBGene00011082 | 0.099908963 | 0.57351485 | 0.74143178 | R07B1.11 |
| WBGene00011083 | -0.281728229 | 0.21770035 | 0.40942207 | glo-1 |
| WBGene00011084 | 0.020317245 | 0.84643798 | NA | srsx-21 |
| WBGene00011087 | -0.392243121 | 0.00045902 | 0.00361318 | R07B7.2 |
| WBGene00011088 | 0.047426939 | 0.77277393 | 0.87105883 | kmo-2 |
| WBGene00011089 | -0.219866855 | 0.23020136 | 0.42455339 | kmo-1 |
| WBGene00011090 | 1.033897793 | 0.00016774 | 0.00151691 | R07B7.6 |
| WBGene00011092 | -0.086798893 | 0.64871993 | 0.79221858 | R07B7.8 |
| WBGene00011093 | 0.244864006 | 0.27488832 | 0.47516409 | R07B7.9 |

| | | | | |
|---|---|---|---|---|
| WBGene00011094 | -0.10046772 | 0.60745921 | 0.76322065 | R07B7.10 |
| WBGene00011095 | 0.44527466 | 0.02510062 | 0.09105506 | gana-1 |
| WBGene00011096 | -0.001488154 | 0.95639912 | NA | R07B7.12 |
| WBGene00011097 | 0.475130848 | 0.03777644 | 0.12267318 | nhr-206 |
| WBGene00011098 | -0.03147554 | 0.70637395 | NA | nhr-207 |
| WBGene00011099 | 1.513078933 | 0.01015361 | 0.04539205 | nhr-208 |
| WBGene00011100 | 0.126401354 | 0.52188546 | 0.70005196 | nhr-209 |
| WBGene00011101 | -0.04607539 | 0.79645138 | 0.88581317 | R07D5.2 |
| WBGene00011102 | -0.255839142 | 0.06890385 | 0.18909489 | R07E3.1 |
| WBGene00011103 | 0.230192667 | 0.09497363 | 0.23664209 | R07E3.2 |
| WBGene00011104 | 0.061275255 | 0.51782541 | NA | cut-5 |
| WBGene00011105 | -0.052432218 | 0.78428345 | 0.87847449 | R07E3.4 |
| WBGene00011106 | 0.074651699 | 0.58767632 | 0.75003656 | acl-5 |
| WBGene00011107 | -0.333685691 | 0.14171795 | 0.30966242 | R07E3.6 |
| WBGene00011108 | 0.935863023 | 0.01720706 | 0.06875778 | R07E3.7 |
| WBGene00011109 | -0.159500859 | 0.09026278 | 0.22852106 | R07E5.1 |
| WBGene00011110 | -0.06221424 | 0.35843486 | 0.55876045 | prdx-3 |
| WBGene00011111 | -0.308263577 | 0.00089226 | 0.00639741 | snfc-5 |
| WBGene00011112 | -1.732122521 | 1.26E-10 | 3.54E-09 | R07E5.4 |
| WBGene00011113 | -0.236319464 | 0.11604272 | 0.27192405 | R07E5.5 |
| WBGene00011114 | 0.251474207 | 0.0953713 | 0.23751169 | R07E5.6 |
| WBGene00011115 | -0.135625721 | 0.1479573 | 0.31876649 | R07E5.7 |
| WBGene00011116 | -0.389419621 | 0.00577382 | 0.02933186 | pdcd-2 |
| WBGene00011117 | -0.445945076 | 0.01051708 | 0.04652743 | R07E5.11 |
| WBGene00011119 | -0.114452567 | 0.23128649 | 0.42626706 | mpc-1 |
| WBGene00011120 | 0.271818331 | 0.03743535 | 0.12190635 | R07E5.15 |
| WBGene00011121 | -0.005691347 | 0.93660638 | NA | R07E5.17 |
| WBGene00011122 | -0.161396014 | 0.02973383 | 0.10291561 | cpt-2 |
| WBGene00011123 | -0.070290364 | 0.66261001 | 0.80169186 | nuaf-3 |
| WBGene00011124 | 0.006965276 | 0.95463038 | 0.97432857 | sdz-27 |
| WBGene00011127 | NA | NA | NA | fbxb-79 |
| WBGene00011128 | 0.059022371 | 0.31884652 | 0.51898076 | R07H5.8 |
| WBGene00011129 | 0.128762944 | 0.4075003 | 0.60546092 | R07H5.9 |
| WBGene00011130 | -0.023056499 | 0.78121253 | NA | zip-6 |
| WBGene00011131 | -0.205585557 | 0.29338044 | 0.49444046 | R07H5.11 |
| WBGene00011132 | 0.157815695 | 0.10232055 | 0.24922229 | R08A2.1 |
| WBGene00011133 | 0.048710032 | 0.62265298 | 0.77440216 | R08A2.2 |
| WBGene00011134 | 0.040107227 | 0.58693163 | NA | ssp-33 |
| WBGene00011135 | -0.023056499 | 0.78121253 | NA | fbxa-147 |
| WBGene00011136 | 0.074892148 | 0.28555128 | NA | R08A2.5 |
| WBGene00011137 | 0.222505058 | 0.16278343 | 0.3381544 | R08A2.7 |
| WBGene00011139 | -0.442437446 | 0.04666584 | 0.14308578 | R08B4.3 |
| WBGene00011140 | 0.020317245 | 0.84643798 | NA | R08B4.4 |

| | | | | | |
|---|---|---|---|---|---|
| WBGene00011141 | | 0.198206308 | 0.17524152 | 0.35590388 | nlp-66 |
| WBGene00011142 | | -0.116127622 | 0.25666408 | 0.45488293 | R08D7.1 |
| WBGene00011143 | | 0.089371298 | 0.44013558 | 0.63450679 | rpap-2 |
| WBGene00011144 | | 0.078754295 | 0.4907608 | 0.6745427 | R08D7.4 |
| WBGene00011145 | | 0.072934931 | 0.62244635 | 0.77421095 | R08D7.5 |
| WBGene00011146 | | 0.085745462 | 0.29515785 | 0.49625701 | pde-2 |
| WBGene00011147 | | -0.213708898 | 0.31781303 | 0.51804728 | R08D7.7 |
| WBGene00011149 | | -0.023056499 | 0.78121253 | NA | NA |
| WBGene00011150 | NA | | NA | NA | nhr-269 |
| WBGene00011151 | NA | | NA | NA | R08H2.10 |
| WBGene00011152 | | -0.367636421 | 0.07608588 | 0.20365144 | R09A8.1 |
| WBGene00011153 | | -0.328116051 | 0.01959715 | 0.07554551 | R09A8.2 |
| WBGene00011154 | | 0.02551193 | 0.8947678 | 0.94498272 | R09A8.5 |
| WBGene00011155 | | -0.204226534 | 8.02E-05 | 0.00079347 | rbm-3.1 |
| WBGene00011156 | | -0.26206016 | 1.97E-05 | 0.00022518 | rbm-3.2 |
| WBGene00011157 | NA | | NA | NA | chil-15 |
| WBGene00011158 | NA | | NA | NA | chil-16 |
| WBGene00011159 | NA | | NA | NA | chil-17 |
| WBGene00011160 | | 0.05787416 | 0.40594086 | NA | NA |
| WBGene00011161 | | 0.207056196 | 0.02971382 | 0.10287067 | chil-18 |
| WBGene00011162 | | 1.657725685 | 0.00810566 | 0.0382699 | chil-19 |
| WBGene00011163 | | 0.057307995 | 0.52728879 | NA | chil-20 |
| WBGene00011164 | | 0.020925399 | 0.84643798 | NA | chil-21 |
| WBGene00011165 | | 1.041308488 | 0.01694742 | 0.06799868 | NA |
| WBGene00011166 | | 1.623269334 | 4.61E-06 | 6.08E-05 | chil-22 |
| WBGene00011167 | | 0.522961012 | 0.04522228 | 0.13974237 | chil-23 |
| WBGene00011168 | | 0.35500987 | 0.06401187 | 0.17977214 | R09D1.12 |
| WBGene00011169 | NA | | NA | NA | R09D1.13 |
| WBGene00011170 | NA | | NA | NA | R09D1.14 |
| WBGene00011171 | | 0.336504015 | 0.0068872 | 0.03354476 | R09E10.1 |
| WBGene00011172 | | 0.036821884 | 0.59344795 | NA | R09E10.2 |
| WBGene00011173 | | 2.036984774 | 0.00978889 | 0.04415254 | acs-18 |
| WBGene00011174 | | 0.020925399 | 0.84643798 | NA | NA |
| WBGene00011175 | | 0.180085223 | 0.39172567 | 0.58975942 | R09E10.5 |
| WBGene00011176 | | 2.659405354 | 1.76E-11 | 5.64E-10 | R09E10.6 |
| WBGene00011177 | NA | | NA | NA | R09E10.8 |
| WBGene00011178 | NA | | NA | NA | NA |
| WBGene00011179 | | 0.010950708 | 0.87860515 | NA | R09H10.1 |
| WBGene00011180 | | 0.092092532 | 0.6199449 | 0.77264865 | R09H10.2 |
| WBGene00011181 | | 0.540404175 | 0.00096408 | 0.00682547 | R09H10.3 |
| WBGene00011182 | | 1.130709541 | 2.23E-37 | 3.75E-35 | R09H10.5 |
| WBGene00011183 | NA | | NA | NA | R09H10.6 |
| WBGene00011184 | | 2.750241562 | 5.19E-07 | 8.66E-06 | R09H10.7 |

| | | | | |
|---|---|---|---|---|
| WBGene00011185 | 0.151084584 | 0.44787956 | 0.64068438 | R10D12.1 |
| WBGene00011186 | NA | NA | NA | clec-226 |
| WBGene00011187 | NA | NA | NA | R10D12.6 |
| WBGene00011188 | 0.034981685 | 0.68236555 | NA | R10D12.7 |
| WBGene00011189 | 0.088546802 | 0.65922094 | 0.79896991 | R10D12.8 |
| WBGene00011190 | -1.247745525 | 0.00264902 | 0.01574678 | swt-6 |
| WBGene00011191 | 0.131317611 | 0.10121567 | NA | R10D12.10 |
| WBGene00011192 | 0.020925399 | 0.84643798 | NA | srh-24 |
| WBGene00011193 | 0.028086816 | 0.85534517 | 0.92251261 | algn-13 |
| WBGene00011194 | 0.049396293 | 0.71743714 | 0.84158547 | R10D12.13 |
| WBGene00011195 | -0.37850283 | 5.86E-10 | 1.51E-08 | sao-1 |
| WBGene00011196 | NA | NA | NA | R10D12.15 |
| WBGene00011196 | NA | NA | NA | T26F2.1 |
| WBGene00011197 | -0.003270206 | 0.95698316 | NA | srw-145 |
| WBGene00011198 | -0.294492946 | 0.03971977 | 0.12739039 | R10E4.1 |
| WBGene00011200 | 0.03886122 | 0.78639071 | 0.87903088 | R10E4.3 |
| WBGene00011201 | 0.319039568 | 0.05153377 | 0.15382384 | nth-1 |
| WBGene00011202 | 0.121282048 | 0.56419247 | 0.73418736 | R10E4.6 |
| WBGene00011203 | 0.320795808 | 0.14172856 | 0.30966242 | R10E4.7 |
| WBGene00011205 | -0.016522694 | 0.93058486 | 0.96081261 | R10E4.9 |
| WBGene00011206 | 0.008461754 | 0.94353337 | 0.96799608 | R10E4.11 |
| WBGene00011207 | 1.008298657 | 0.00031137 | 0.00259461 | R10E8.1 |
| WBGene00011208 | NA | NA | NA | R10E8.2 |
| WBGene00011209 | 0.136259019 | 0.33496088 | 0.53416428 | R10E8.3 |
| WBGene00011210 | 0.059980168 | 0.38780842 | NA | NA |
| WBGene00011211 | NA | NA | NA | R10E8.5 |
| WBGene00011212 | 1.374432642 | 2.35E-05 | 0.00026321 | R10E8.6 |
| WBGene00011213 | -0.007799522 | 0.96724569 | 0.98172923 | R10E8.8 |
| WBGene00011214 | 1.771121175 | 0.00035805 | 0.00291559 | R10E9.2 |
| WBGene00011215 | -0.030960139 | 0.87428843 | 0.93298863 | R10E9.3 |
| WBGene00011216 | -0.123979471 | 0.2327701 | 0.42792404 | usp-46 |
| WBGene00011217 | 0.520323699 | 0.06106861 | 0.17381522 | R10E11.5 |
| WBGene00011218 | -0.031808302 | 0.85259687 | 0.92106277 | R10E11.6 |
| WBGene00011219 | 0.038125098 | 0.61119482 | NA | srxa-10 |
| WBGene00011220 | 0.063731418 | 0.41159439 | NA | R10E11.9 |
| WBGene00011221 | -0.036273396 | 0.84822753 | 0.9180823 | R10E12.2 |
| WBGene00011222 | -0.087912802 | 0.42701374 | 0.62241028 | R10H10.3 |
| WBGene00011223 | 0.035884754 | 0.71848022 | 0.84199658 | R10H10.4 |
| WBGene00011224 | -0.090002255 | 0.63613207 | 0.78430397 | R10H10.6 |
| WBGene00011225 | 0.161637976 | 0.1571718 | 0.3311487 | R10H10.7 |
| WBGene00011226 | -0.461399879 | 0.00896263 | 0.04132973 | slc-25A21 |
| WBGene00011227 | 0.502930647 | 0.03846407 | 0.12428877 | nlp-67 |
| WBGene00011228 | 0.175603548 | 0.39639903 | 0.59439556 | R11.3 |

| | | | | |
|---|---|---|---|---|
| WBGene00011229 | -0.295316675 | 0.19914237 | 0.38757444 | R11.4 |
| WBGene00011230 | -0.157026042 | 0.06859548 | 0.18865836 | nud-2 |
| WBGene00011231 | -0.966836713 | 0.00177895 | 0.01134392 | R11A5.3 |
| WBGene00011232 | 0.024091696 | 0.59589222 | 0.75477619 | pck-2 |
| WBGene00011234 | 0.367300822 | 0.04030902 | 0.12871633 | R11A5.6 |
| WBGene00011235 | 0.26255263 | 0.23506252 | 0.43087597 | suro-1 |
| WBGene00011236 | 0.272340615 | 0.04595812 | 0.1415085 | R11A8.1 |
| WBGene00011237 | 0.010507988 | 0.9039836 | 0.94889643 | gkow-1 |
| WBGene00011238 | 0.055621344 | 0.63350346 | 0.78257882 | ugt-59 |
| WBGene00011239 | -0.283376296 | 0.00136487 | 0.00907496 | pges-2 |
| WBGene00011240 | -0.0435481 | 0.465817 | 0.65551967 | mask-1 |
| WBGene00011241 | 0.061275255 | 0.51782541 | NA | mpz-5 |
| WBGene00011242 | 0.284312599 | 0.01048662 | 0.04648337 | R11D1.1 |
| WBGene00011243 | -0.003108065 | 0.95728506 | NA | R11D1.2 |
| WBGene00011244 | 0.053245733 | 0.78502643 | 0.87864628 | R11D1.3 |
| WBGene00011245 | NA | NA | NA | R11D1.4 |
| WBGene00011246 | 0.059762547 | 0.38965825 | NA | R11D1.7 |
| WBGene00011247 | 0.001104499 | 0.99593537 | 0.99763888 | mrpl-49 |
| WBGene00011248 | 0.159559545 | 0.29461346 | 0.49569181 | R11D1.10 |
| WBGene00011249 | 0.031134574 | 0.70125581 | NA | R11G10.2 |
| WBGene00011250 | -0.160837541 | 0.0670972 | 0.18562158 | R11H6.2 |
| WBGene00011251 | 0.106072193 | 0.45533375 | 0.64723703 | glb-22 |
| WBGene00011252 | 0.227938685 | 0.28024063 | 0.47984024 | R11H6.4 |
| WBGene00011253 | -0.187479065 | 0.03622178 | 0.11893317 | R11H6.5 |
| WBGene00011254 | 0.015584271 | 0.80362925 | NA | R12G8.1 |
| WBGene00011255 | 0.065014541 | 0.37399213 | NA | R12H7.4 |
| WBGene00011257 | 0.056790603 | 0.58474897 | 0.74803929 | npax-3 |
| WBGene00011258 | -0.140781284 | 0.50313898 | 0.68442421 | best-15 |
| WBGene00011259 | 0.502061798 | 2.98E-06 | 4.11E-05 | miz-1 |
| WBGene00011260 | 0.064225876 | 0.59856812 | 0.75672531 | cnnm-5 |
| WBGene00011261 | 0.040988747 | 0.77770815 | 0.8741977 | nphp-4 |
| WBGene00011262 | 0.297115257 | 0.20074395 | 0.3893353 | pho-8 |
| WBGene00011263 | 0.87092632 | 0.00430869 | 0.02329456 | R13H4.5 |
| WBGene00011264 | 0.020232312 | 0.84643798 | NA | R13H4.6 |
| WBGene00011265 | 0.166645955 | 0.20118447 | 0.38993141 | R13H4.7 |
| WBGene00011266 | 0.075726029 | 0.27693214 | NA | R13H4.8 |
| WBGene00011267 | -0.330751129 | 0.13721655 | 0.302832 | fbxb-85 |
| WBGene00011268 | -0.100415684 | 0.37398144 | 0.57286695 | pde-12 |
| WBGene00011269 | 0.172959314 | 0.40487741 | 0.60297123 | R17.3 |
| WBGene00011270 | 0.111644313 | 0.21837734 | 0.41031093 | R31.2 |
| WBGene00011271 | 0.031508535 | 0.77600558 | 0.87322248 | flad-1 |
| WBGene00011272 | -0.043513286 | 0.70703727 | 0.83454795 | dtmk-1 |
| WBGene00011273 | -0.128137807 | 0.00643712 | 0.0319158 | R53.4 |

| | | | | |
|---|---|---|---|---|
| WBGene00011274 | 0.364473469 | 0.00600089 | 0.03013 | R53.5 |
| WBGene00011275 | -0.129413822 | 0.20957361 | 0.3989779 | psf-1 |
| WBGene00011276 | -0.022398963 | 0.83413076 | 0.90957526 | aakg-5 |
| WBGene00011277 | -0.003421551 | 0.95689871 | NA | R53.8 |
| WBGene00011278 | -0.0702384 | 0.66560667 | 0.80398872 | R74.2 |
| WBGene00011279 | -0.278478414 | 0.00383513 | 0.02116476 | asd-1 |
| WBGene00011280 | -0.054206087 | 0.74087103 | 0.85369847 | pelo-1 |
| WBGene00011281 | -0.043349444 | 0.64936977 | 0.79268158 | R74.7 |
| WBGene00011282 | -0.125339984 | 0.06479508 | 0.18144297 | R74.8 |
| WBGene00011283 | 0.398585633 | 0.01916772 | 0.07453775 | ttbk-7 |
| WBGene00011284 | 0.412965334 | 0.05371673 | 0.15868983 | ttr-27 |
| WBGene00011285 | 0.381684327 | 0.07619449 | 0.2038686 | ttr-28 |
| WBGene00011286 | 1.039685533 | 0.00550881 | 0.0282998 | ttr-29 |
| WBGene00011287 | 0.257671759 | 0.1788156 | 0.36021452 | glb-24 |
| WBGene00011288 | 0.127299467 | 0.16839118 | NA | R102.1 |
| WBGene00011289 | 0.896189809 | 1.99E-07 | 3.56E-06 | R102.2 |
| WBGene00011290 | 0.087256674 | 0.23800185 | NA | R102.3 |
| WBGene00011291 | -0.276458285 | 0.05428224 | 0.15981225 | R102.4 |
| WBGene00011292 | 0.039790421 | 0.58936429 | 0.75111006 | allo-1 |
| WBGene00011293 | -0.403750337 | 0.08059478 | 0.21199931 | R102.6 |
| WBGene00011294 | -0.039636691 | 0.54579661 | NA | NA |
| WBGene00011295 | NA | NA | NA | R102.8 |
| WBGene00011296 | 0.059980168 | 0.38780842 | NA | glb-21 |
| WBGene00011297 | 0.011113023 | 0.90389313 | NA | R102.10 |
| WBGene00011298 | 0.026978567 | 0.86673202 | 0.92904485 | R107.2 |
| WBGene00011299 | -0.07964973 | 0.29876476 | 0.49942984 | ikke-1 |
| WBGene00011300 | -0.348087619 | 1.79E-05 | 0.00020599 | R107.5 |
| WBGene00011302 | -0.235702362 | 0.00921067 | 0.04214169 | R166.2 |
| WBGene00011303 | 0.040728402 | 0.81007191 | 0.89497909 | R166.3 |
| WBGene00011304 | 0.52899403 | 2.16E-13 | 8.75E-12 | mnk-1 |
| WBGene00011305 | 1.139664395 | 0.00906697 | 0.04161332 | R186.1 |
| WBGene00011306 | -0.20984167 | 0.05245101 | 0.15605269 | R186.3 |
| WBGene00011307 | -0.076672822 | 0.6973571 | 0.82745808 | mpst-7 |
| WBGene00011308 | 0.082794429 | 0.37329231 | 0.57229033 | pnn-1 |
| WBGene00011309 | -0.424132952 | 0.01603226 | 0.06520911 | R186.8 |
| WBGene00011310 | 0.353772349 | 0.08821531 | 0.22532455 | T01B4.3 |
| WBGene00011311 | -0.03159829 | 0.75058836 | 0.85919975 | T01B7.5 |
| WBGene00011312 | 0.038683759 | 0.62950079 | 0.77968456 | trcs-2 |
| WBGene00011313 | 0.038595855 | 0.57520533 | NA | T01B7.8 |
| WBGene00011314 | 0.253722974 | 0.13032446 | 0.29256062 | T01B7.9 |
| WBGene00011315 | 0.213274481 | 0.31817218 | 0.51845953 | mbr-1 |
| WBGene00011317 | 0.093157605 | 0.54569872 | 0.71913367 | T01C1.4 |
| WBGene00011318 | -0.029843055 | 0.75372592 | 0.86058654 | cdt-2 |

| | | | | |
|---|---|---|---|---|
| WBGene00011319 | -0.201564009 | 0.06977814 | 0.19082013 | T01C3.2 |
| WBGene00011320 | -0.28926717 | 0.00021414 | 0.00187514 | T01C3.3 |
| WBGene00011321 | 2.140339919 | 0.00021149 | 0.00185414 | fil-1 |
| WBGene00011322 | 0.110039823 | 0.21624696 | NA | irld-14 |
| WBGene00011323 | 0.110513173 | 0.36622096 | 0.5658338 | mut-15 |
| WBGene00011324 | -0.171069061 | 0.16586894 | 0.34245805 | hrde-2 |
| WBGene00011325 | -0.055120925 | 0.45606622 | NA | T01C3.11 |
| WBGene00011326 | 0.942155066 | 0.00041463 | 0.00331728 | T01D3.1 |
| WBGene00011327 | 0.038125098 | 0.61119482 | NA | hlh-34 |
| WBGene00011328 | 0.632553616 | 0.02782573 | 0.0980051 | T01D3.3 |
| WBGene00011329 | 0.23340408 | 0.03682649 | 0.12043258 | zipt-9 |
| WBGene00011330 | 0.880821474 | 1.17E-16 | 6.74E-15 | T01D3.6 |
| WBGene00011331 | 0.882685376 | 3.80E-05 | 0.00040874 | T01E8.1 |
| WBGene00011333 | -0.263791087 | 0.00198937 | 0.01241451 | nrde-2 |
| WBGene00011334 | -0.105675152 | 0.41045843 | 0.60813512 | mrps-14 |
| WBGene00011335 | 0.157872234 | 0.45811064 | 0.64973678 | T01E8.8 |
| WBGene00011336 | 0.394644168 | 0.04296367 | 0.13469288 | ubxn-5 |
| WBGene00011337 | 0.622930116 | 0.02801782 | 0.09848953 | T01G1.2 |
| WBGene00011338 | -0.262356423 | 0.00048478 | 0.00378741 | sec-31 |
| WBGene00011339 | 0.010312842 | 0.95931249 | 0.97767605 | T01G5.1 |
| WBGene00011340 | -0.086998533 | 0.67194171 | 0.80897713 | ugt-30 |
| WBGene00011341 | NA | NA | NA | T01G5.6 |
| WBGene00011342 | -0.22681531 | 0.20493614 | 0.39402101 | T01G5.7 |
| WBGene00011344 | -0.036434084 | 0.72486034 | 0.84572464 | T01G9.2 |
| WBGene00011345 | 0.229012442 | 0.27353231 | 0.47383502 | dma-1 |
| WBGene00011346 | NA | NA | NA | NA |
| WBGene00011348 | 0.297101963 | 0.00643722 | 0.0319158 | T01H3.2 |
| WBGene00011349 | -0.195293078 | 0.02325203 | 0.08572978 | T01H3.3 |
| WBGene00011350 | -0.268990104 | 0.00094635 | 0.0067243 | perm-1 |
| WBGene00011351 | -0.087726894 | 0.48696193 | 0.67152771 | T01H3.5 |
| WBGene00011352 | -0.1854082 | 0.00184542 | 0.01170151 | rskn-1 |
| WBGene00011353 | -0.260262966 | 0.18611072 | 0.37047546 | T01H8.2 |
| WBGene00011354 | 0.098073944 | 0.39294661 | 0.59115611 | lgc-13 |
| WBGene00011355 | 0.020232312 | 0.84643798 | NA | lgc-14 |
| WBGene00011356 | NA | NA | NA | lgc-15 |
| WBGene00011357 | 0.020925399 | 0.84643798 | NA | NA |
| WBGene00011358 | NA | NA | NA | lgc-16 |
| WBGene00011359 | NA | NA | NA | lgc-17 |
| WBGene00011360 | NA | NA | NA | lgc-18 |
| WBGene00011361 | 0.071312951 | 0.69097788 | 0.8224095 | lyst-1 |
| WBGene00011362 | 0.179773052 | 0.2342723 | 0.42987602 | cest-1 |
| WBGene00011363 | NA | NA | NA | ocam-1 |
| WBGene00011364 | -0.058057218 | 0.77615247 | 0.87325354 | T02B5.3 |

| | | | | | |
|---|---|---|---|---|---|
| WBGene00011365 | | -0.190281678 | 0.32687042 | 0.52595997 | T02C1.1 |
| WBGene00011366 | | -0.119810962 | 0.56551697 | 0.73505397 | T02C1.2 |
| WBGene00011367 | | 0.029850734 | 0.80214297 | 0.88992108 | snpc-3.4 |
| WBGene00011368 | | -0.108475546 | 0.29734198 | 0.49843299 | tftc-5 |
| WBGene00011369 | | -0.003108065 | 0.95728506 | NA | NA |
| WBGene00011371 | | 0.085152163 | 0.50532795 | 0.68637729 | T02D1.4 |
| WBGene00011372 | | 0.179660049 | 0.30031061 | 0.50066452 | npr-26 |
| WBGene00011373 | NA | | NA | NA | T02D1.7 |
| WBGene00011374 | | 0.150242878 | 0.35218153 | 0.55200619 | T02D1.8 |
| WBGene00011375 | | -0.093028413 | 0.4060441 | 0.60415566 | T02E1.2 |
| WBGene00011376 | | -0.340971434 | 1.77E-05 | 0.00020366 | gla-3 |
| WBGene00011378 | | 0.396207378 | 0.03373301 | 0.11281065 | T02E1.6 |
| WBGene00011379 | | 0.404694189 | 0.00463985 | 0.02469359 | T02E1.7 |
| WBGene00011380 | | 0.146027321 | 0.38762807 | 0.58619593 | T02E1.8 |
| WBGene00011381 | | 0.430364478 | 0.03975218 | 0.12746499 | npr-25 |
| WBGene00011382 | | 0.299720638 | 0.16234851 | 0.33739454 | dop-5 |
| WBGene00011383 | | 0.227402214 | 0.28616521 | 0.48616975 | T02E9.5 |
| WBGene00011386 | | -0.023109762 | 0.78121253 | NA | T02G6.3 |
| WBGene00011387 | | -0.046394786 | 0.50231372 | NA | T02G6.4 |
| WBGene00011388 | | -0.158304081 | 0.36658225 | 0.56621276 | T02G6.5 |
| WBGene00011389 | NA | | NA | NA | T02G6.6 |
| WBGene00011390 | | 0.200240708 | 0.03227172 | 0.10910588 | T02G6.7 |
| WBGene00011391 | | -0.053663731 | 0.70409169 | 0.83203617 | mrps-12 |
| WBGene00011392 | | 1.234204635 | 2.94E-16 | 1.62E-14 | sbt-1 |
| WBGene00011393 | | -0.108051607 | 0.52763251 | 0.70484193 | T03D8.6 |
| WBGene00011394 | | 0.020232312 | 0.84643798 | NA | T03D8.7 |
| WBGene00011395 | NA | | NA | NA | T03E6.2 |
| WBGene00011396 | | 0.020317245 | 0.84643798 | NA | nhr-271 |
| WBGene00011397 | | 2.012277912 | 1.06E-08 | 2.35E-07 | T03E6.8 |
| WBGene00011398 | | -0.497231615 | 9.79E-06 | 0.00012142 | qdpr-1 |
| WBGene00011399 | | 0.056004863 | 0.48460219 | 0.66978766 | T03F6.3 |
| WBGene00011400 | | 0.169025208 | 0.35317946 | 0.55300783 | T03F6.4 |
| WBGene00011401 | | 0.145703839 | 0.20637314 | 0.39553725 | T03F6.6 |
| WBGene00011402 | | 0.033034883 | 0.64219503 | NA | T03F7.5 |
| WBGene00011403 | NA | | NA | NA | T03F7.6 |
| WBGene00011404 | | 0.152947873 | 0.43713605 | 0.63217196 | T03F7.7 |
| WBGene00011405 | | 0.033034883 | 0.64219503 | NA | NA |
| WBGene00011407 | | -0.148828696 | 0.39594816 | 0.59402356 | ppat-1 |
| WBGene00011408 | | -0.441204401 | 2.23E-05 | 0.00025186 | nifk-1 |
| WBGene00011409 | | -0.310378479 | 2.80E-05 | 0.00030921 | T04A8.7 |
| WBGene00011410 | | -0.041489298 | 0.58501349 | 0.74805087 | T04A8.8 |
| WBGene00011411 | | 0.015115802 | 0.9074762 | 0.94995226 | sel-13 |
| WBGene00011412 | | -0.33291432 | 0.00891442 | 0.04114337 | mrpl-16 |

| | | | | |
|---|---|---|---|---|
| WBGene00011414 | 0.017141582 | 0.79355725 | NA | T04A8.13 |
| WBGene00011415 | -0.012788558 | 0.89222731 | 0.94374205 | him-18 |
| WBGene00011416 | NA | NA | NA | T04A11.1 |
| WBGene00011416 | NA | NA | NA | T04A11.4 |
| WBGene00011417 | NA | NA | NA | T04A11.5 |
| WBGene00011417 | NA | NA | NA | T04A11.2 |
| WBGene00011418 | NA | NA | NA | igdb-1 |
| WBGene00011419 | NA | NA | NA | T04A11.1 |
| WBGene00011419 | NA | NA | NA | T04A11.4 |
| WBGene00011420 | NA | NA | NA | T04A11.5 |
| WBGene00011420 | NA | NA | NA | T04A11.2 |
| WBGene00011421 | NA | NA | NA | NA |
| WBGene00011422 | -0.38067666 | 0.11918457 | 0.27702513 | T04B2.3 |
| WBGene00011423 | 0.212873843 | 0.1355554 | 0.30038847 | ipla-7 |
| WBGene00011424 | 0.272074949 | 0.21217384 | 0.4024925 | dhs-31 |
| WBGene00011425 | 0.214956244 | 0.10395465 | 0.25207022 | T04B2.7 |
| WBGene00011426 | 0.020232312 | 0.84643798 | NA | T04C10.3 |
| WBGene00011427 | 0.127693253 | 0.10904601 | NA | T04C12.1 |
| WBGene00011428 | 1.550202667 | 8.97E-07 | 1.43E-05 | nlp-80 |
| WBGene00011429 | 0.114829426 | 0.57490293 | 0.74227635 | T04C12.7 |
| WBGene00011430 | 0.031406188 | 0.86391461 | 0.92738279 | T04C12.8 |
| WBGene00011431 | -0.236681556 | 0.0875114 | 0.2240347 | T04D3.1 |
| WBGene00011432 | -0.982835918 | 3.66E-08 | 7.45E-07 | sdz-30 |
| WBGene00011433 | 0.157334117 | 0.46306517 | 0.65301113 | pde-1 |
| WBGene00011434 | -0.071275941 | 0.5049367 | 0.68610471 | T04D3.5 |
| WBGene00011435 | 0.215416267 | 0.2904182 | 0.49092715 | T04D3.8 |
| WBGene00011436 | 0.358592341 | 0.00151082 | 0.00988377 | T04F3.1 |
| WBGene00011437 | 0.148405894 | 0.43146636 | 0.62650987 | smoc-1 |
| WBGene00011438 | 0.070134426 | 0.57769863 | 0.74411462 | T04F3.3 |
| WBGene00011439 | -0.030987878 | 0.87864626 | 0.93503852 | T04F3.4 |
| WBGene00011440 | -0.013208622 | 0.91168718 | 0.95112162 | sfxn-1.5 |
| WBGene00011441 | -0.168395483 | 0.1464623 | 0.31646318 | T04F8.2 |
| WBGene00011442 | 0.042972894 | 0.75722972 | 0.86256908 | T04F8.3 |
| WBGene00011443 | -0.013272875 | 0.90577115 | 0.94916953 | cutl-11 |
| WBGene00011444 | 0.268648143 | 0.1197649 | 0.27797684 | T04F8.6 |
| WBGene00011445 | 1.83002828 | 7.80E-07 | 1.26E-05 | T04F8.7 |
| WBGene00011446 | -0.322843748 | 0.00021884 | 0.00190373 | T04F8.8 |
| WBGene00011447 | 0.00184829 | 0.98963069 | 0.99411203 | T04F8.9 |
| WBGene00011448 | 0.0398788 | 0.58971221 | NA | NA |
| WBGene00011449 | 0.047703711 | 0.4525119 | 0.64479092 | T04H1.2 |
| WBGene00011450 | 1.440563823 | 0.00478474 | 0.0252541 | ttr-22 |
| WBGene00011451 | 0.102914618 | 0.42696479 | 0.62241028 | T04H1.5 |
| WBGene00011452 | 0.081671655 | 0.58796884 | 0.75019188 | ugt-55 |

| | | | | |
|---|---|---|---|---|
| WBGene00011453 | 0.167641235 | 0.16100856 | 0.33556248 | ugt-56 |
| WBGene00011454 | 0.049073037 | 0.79874473 | 0.8872854 | lron-14 |
| WBGene00011456 | 0.352054099 | 0.07907635 | 0.20943503 | T05A1.5 |
| WBGene00011458 | 0.020317245 | 0.84643798 | NA | irld-47 |
| WBGene00011459 | 0.020317245 | 0.84643798 | NA | NA |
| WBGene00011460 | 0.16222405 | 0.37413046 | 0.57297532 | ttr-14 |
| WBGene00011462 | 1.143855523 | 0.00372762 | 0.02071987 | vap-2 |
| WBGene00011463 | 0.227156829 | 0.16497622 | 0.34123996 | T05A10.6 |
| WBGene00011464 | -0.262383961 | 0.00012132 | 0.00114962 | T05B9.1 |
| WBGene00011465 | -0.027535543 | 0.68063395 | NA | T05B9.2 |
| WBGene00011466 | 0.700094256 | 0.0196395 | 0.07562913 | T05C12.1 |
| WBGene00011467 | 0.133396135 | 0.15669766 | 0.33067317 | decr-1.3 |
| WBGene00011468 | -0.028533941 | 0.70959323 | NA | T05C12.4 |
| WBGene00011469 | 0.020925399 | 0.84643798 | NA | dylt-3 |
| WBGene00011470 | NA | NA | NA | NA |
| WBGene00011471 | 0.204632929 | 0.1672016 | 0.34459441 | T05C12.8 |
| WBGene00011472 | 0.020317245 | 0.84643798 | NA | T05C12.9 |
| WBGene00011473 | -0.059528357 | 0.55776396 | 0.72877494 | T05C12.11 |
| WBGene00011474 | -0.0328867 | 0.75525002 | 0.86125209 | aldo-1 |
| WBGene00011475 | -0.795929476 | 0.0011403 | 0.00784914 | T05D4.2 |
| WBGene00011476 | 0.133771856 | 0.47462407 | 0.66192665 | T05D4.3 |
| WBGene00011478 | 0.122470862 | 0.32795351 | 0.5272971 | T05D4.5 |
| WBGene00011479 | 0.135182857 | 0.4990426 | 0.68106662 | T05E11.2 |
| WBGene00011480 | -0.152495501 | 0.0012433 | 0.0084191 | enpl-1 |
| WBGene00011481 | -0.242327746 | 6.63E-05 | 0.000676 | imp-2 |
| WBGene00011482 | 0.085549112 | 0.49831813 | 0.68033156 | pigk-1 |
| WBGene00011483 | NA | NA | NA | NA |
| WBGene00011484 | 0.308533252 | 0.17542914 | 0.35603799 | T05E11.8 |
| WBGene00011485 | NA | NA | NA | srbc-14 |
| WBGene00011486 | 5.067244658 | 1.02E-135 | 8.28E-133 | T05E12.3 |
| WBGene00011487 | 0.364370278 | 6.99E-05 | 0.00070602 | T05E12.6 |
| WBGene00011488 | -0.295024991 | 7.23E-06 | 9.16E-05 | nra-2 |
| WBGene00011489 | -0.015426678 | 0.78437291 | 0.87847449 | T05F1.2 |
| WBGene00011490 | 0.08937311 | 0.45474826 | 0.64678155 | T05F1.4 |
| WBGene00011491 | 0.217055999 | 0.10104807 | 0.24715081 | T05F1.5 |
| WBGene00011492 | 0.084852855 | 0.40078218 | 0.59903477 | T05F1.7 |
| WBGene00011493 | 0.279185448 | 0.09949151 | 0.24461044 | T05F1.8 |
| WBGene00011494 | 0.263580702 | 0.12498246 | 0.28566356 | T05F1.9 |
| WBGene00011495 | 1.609772408 | 0.00343532 | 0.01946462 | T05F1.11 |
| WBGene00011497 | -0.15233579 | 0.45281052 | 0.64509087 | T05F1.13 |
| WBGene00011498 | 0.068809443 | 0.71054524 | 0.83693426 | T05G5.1 |
| WBGene00011499 | 0.19451062 | 0.2828008 | 0.48268671 | T05G5.4 |
| WBGene00011500 | 0.210697605 | 0.10838775 | 0.25895969 | T05G5.5 |

| | | | | |
|---|---|---|---|---|
| WBGene00011501 | -0.218411351 | 4.84E-05 | 0.00051095 | rmd-1 |
| WBGene00011502 | -0.074580577 | 0.53266555 | 0.70866671 | vps-53 |
| WBGene00011503 | -0.096215913 | 0.30020739 | 0.50066452 | gcc-2 |
| WBGene00011505 | 0.116586079 | 0.50537171 | 0.68637729 | pzf-1 |
| WBGene00011506 | NA | NA | NA | srbc-77 |
| WBGene00011507 | -0.111201318 | 0.0698062 | 0.19082013 | T05H10.1 |
| WBGene00011508 | 2.049540252 | 0.00048854 | 0.00381072 | T05H10.3 |
| WBGene00011509 | -0.441967905 | 3.25E-06 | 4.44E-05 | T05H10.4 |
| WBGene00011510 | -0.069596499 | 0.19612521 | 0.38373234 | pdha-1 |
| WBGene00011511 | -0.095953969 | 0.28048099 | 0.48012652 | gpcp-2 |
| WBGene00011512 | -0.039548418 | 0.83842757 | 0.91260264 | T05H10.8 |
| WBGene00011513 | -0.303102656 | 0.16325753 | 0.3388508 | fbxa-197 |
| WBGene00011514 | 0.178520741 | 0.40793641 | 0.60581891 | NA |
| WBGene00011515 | 0.020232312 | 0.84643798 | NA | oac-42 |
| WBGene00011516 | 0.020925399 | 0.84643798 | NA | T06C12.9 |
| WBGene00011517 | 2.49908377 | 0.00020625 | 0.00181659 | cgt-1 |
| WBGene00011518 | NA | NA | NA | T06C12.11 |
| WBGene00011519 | 0.020925399 | 0.84643798 | NA | T06C12.12 |
| WBGene00011520 | 0.34957088 | 0.0891826 | 0.22684545 | nhr-213 |
| WBGene00011521 | 0.085537126 | 0.24698522 | NA | T06C12.14 |
| WBGene00011522 | 0.857319774 | 3.25E-07 | 5.56E-06 | srap-1 |
| WBGene00011523 | 0.111319389 | 0.1344031 | NA | T06D8.2 |
| WBGene00011524 | 1.937225902 | 0.00312176 | 0.01800857 | plpr-1 |
| WBGene00011526 | -0.077584784 | 0.44637365 | 0.63965515 | cox-15 |
| WBGene00011527 | -0.290814752 | 7.67E-05 | 0.00076534 | cchl-1 |
| WBGene00011528 | -0.028608029 | 0.80916175 | 0.89437847 | fndc-1 |
| WBGene00011529 | 0.01664353 | 0.84367724 | 0.91613725 | T06D8.9 |
| WBGene00011530 | 0.318890724 | 0.02475734 | 0.09019242 | T06D8.10 |
| WBGene00011531 | 0.509402219 | 0.0229512 | 0.08507051 | rsbp-1 |
| WBGene00011532 | -0.05830432 | 0.48179093 | 0.66785656 | chaf-1 |
| WBGene00011533 | NA | NA | NA | T06E4.5 |
| WBGene00011534 | -0.00346345 | 0.95708614 | NA | T06E4.7 |
| WBGene00011535 | 0.063217264 | 0.41230173 | NA | T06E4.8 |
| WBGene00011536 | 0.065014541 | 0.37399213 | NA | T06E4.9 |
| WBGene00011537 | 0.020232312 | 0.84643798 | NA | T06E4.10 |
| WBGene00011538 | -0.49087694 | 0.00748341 | 0.03587575 | T06E6.1 |
| WBGene00011539 | -0.006015451 | 0.92944803 | NA | fbxa-135 |
| WBGene00011540 | 0.690898463 | 0.00299834 | 0.01746169 | T06E6.10 |
| WBGene00011541 | NA | NA | NA | srbc-63 |
| WBGene00011542 | -0.227152092 | 0.1338554 | 0.29774794 | fbxa-104 |
| WBGene00011543 | 0.344814567 | 0.00848568 | 0.03955391 | acl-2 |
| WBGene00011544 | NA | NA | NA | T06E8.2 |
| WBGene00011545 | 0.059938442 | 0.52286458 | NA | T06G6.3 |

| | | | | | |
|---|---|---|---|---|---|
| WBGene00011546 | | 0.009588798 | 0.94958645 | 0.97122452 | T06G6.4 |
| WBGene00011547 | | -0.023109762 | 0.78121253 | NA | T06G6.5 |
| WBGene00011548 | | 0.033034883 | 0.64219503 | NA | T06G6.6 |
| WBGene00011549 | NA | | NA | NA | NA |
| WBGene00011550 | | 0.081500299 | 0.51938809 | 0.69848084 | NA |
| WBGene00011551 | | 0.063731418 | 0.41159439 | NA | T06G6.12 |
| WBGene00011554 | | -1.09210249 | 0.01188216 | 0.05119681 | T07A5.1 |
| WBGene00011556 | | -0.315920355 | 0.17665951 | 0.35749508 | vglu-3 |
| WBGene00011557 | NA | | NA | NA | ostf-4 |
| WBGene00011558 | | -0.205486348 | 0.16015854 | 0.33477934 | T07A5.5 |
| WBGene00011559 | | -0.214193873 | 0.02457873 | 0.08967287 | umps-1 |
| WBGene00011560 | | -0.112782873 | 0.08505196 | 0.21977511 | T07C4.3 |
| WBGene00011561 | | -0.249802009 | 1.50E-05 | 0.00017611 | ttr-15 |
| WBGene00011562 | | -0.143645975 | 0.17880038 | 0.36021452 | sas-7 |
| WBGene00011563 | | 0.055761315 | 0.7892456 | 0.88107852 | jmjd-4 |
| WBGene00011564 | | 0.69215028 | 0.00072436 | 0.00532127 | ugt-50 |
| WBGene00011565 | | 0.020925399 | 0.84643798 | NA | nhr-272 |
| WBGene00011566 | NA | | NA | NA | nhr-214 |
| WBGene00011567 | NA | | NA | NA | nhr-215 |
| WBGene00011568 | | 0.0398788 | 0.58971221 | NA | nhr-26 |
| WBGene00011570 | NA | | NA | NA | sre-28 |
| WBGene00011571 | | -0.346390872 | 0.0002879 | 0.00242252 | ttr-46 |
| WBGene00011572 | | 0.073458642 | 0.29111983 | NA | mam-5 |
| WBGene00011573 | | -0.153403712 | 0.35597453 | 0.55590444 | anmt-3 |
| WBGene00011574 | | 0.020317245 | 0.84643798 | NA | T07C12.10 |
| WBGene00011575 | | 0.102593339 | 0.57201423 | 0.74028643 | madf-4 |
| WBGene00011576 | | -0.091569875 | 0.43627516 | 0.63130111 | rmh-2 |
| WBGene00011577 | NA | | NA | NA | fipr-12 |
| WBGene00011578 | | 0.395771979 | 0.10777969 | 0.25813879 | npr-20 |
| WBGene00011579 | | 0.266593637 | 0.21399865 | 0.40486495 | T07D4.2 |
| WBGene00011580 | | -0.203443792 | 0.00618386 | 0.03086907 | ddx-19 |
| WBGene00011581 | | 0.020925399 | 0.84643798 | NA | T07D10.1 |
| WBGene00011582 | | 0.030033944 | 0.78465586 | 0.87854069 | ntr-1 |
| WBGene00011583 | | -0.413135483 | 0.09547965 | 0.23761986 | T07D10.3 |
| WBGene00011584 | NA | | NA | NA | clec-15 |
| WBGene00011584 | NA | | NA | NA | clec-13 |
| WBGene00011585 | NA | | NA | NA | NA |
| WBGene00011586 | | 1.024687293 | 0.00423283 | 0.02294439 | flp-33 |
| WBGene00011587 | | -0.112898375 | 0.37355235 | 0.572527 | anp-1 |
| WBGene00011589 | | -0.029603436 | 0.77304484 | 0.87123004 | T07F10.3 |
| WBGene00011590 | | 0.915996155 | 0.01399735 | 0.05850278 | bus-19 |
| WBGene00011591 | | 0.26192996 | 0.24828805 | 0.44669273 | T07F10.5 |
| WBGene00011592 | | 0.159279986 | 0.25767278 | 0.45573917 | T07F10.6 |

| | | | | |
|---|---|---|---|---|
| WBGene00011593 | -0.080242334 | 0.51565996 | 0.69531934 | T07G12.2 |
| WBGene00011594 | 0.153236558 | 0.06820274 | NA | T07G12.3 |
| WBGene00011595 | NA | NA | NA | T07G12.4 |
| WBGene00011596 | 4.569113185 | 5.88E-05 | 0.00060655 | T07G12.5 |
| WBGene00011597 | -0.055827494 | 0.59348543 | 0.75344508 | zim-1 |
| WBGene00011599 | -0.60377428 | 0.02789039 | 0.09813585 | NA |
| WBGene00011600 | 0.001435189 | 0.99153799 | 0.99507216 | zim-2 |
| WBGene00011601 | -0.204083998 | 0.10972984 | 0.26097378 | zim-3 |
| WBGene00011604 | -0.17982977 | 0.04444821 | 0.1380204 | T08A11.1 |
| WBGene00011605 | -0.147238831 | 0.04055766 | 0.12931288 | sftb-1 |
| WBGene00011606 | -0.190435441 | 0.33481368 | 0.53413188 | tmed-12 |
| WBGene00011607 | 0.221096184 | 0.31290281 | 0.51333563 | NA |
| WBGene00011608 | 0.741764702 | 0.00084962 | 0.00613071 | NA |
| WBGene00011609 | -0.00346345 | 0.95708614 | NA | T08D2.4 |
| WBGene00011610 | 1.3983588 | 0.00069743 | 0.00515711 | T08D2.5 |
| WBGene00011611 | 0.314429421 | 0.13027754 | 0.29250015 | T08D2.6 |
| WBGene00011612 | -0.09020305 | 0.66016617 | 0.79959405 | T08D2.7 |
| WBGene00011613 | 0.156813681 | 0.45110802 | 0.64335404 | T08D2.8 |
| WBGene00011614 | -0.148369825 | 0.33339665 | 0.53304196 | nfya-1 |
| WBGene00011615 | -0.303474142 | 0.06406616 | 0.17986437 | lsd-1 |
| WBGene00011616 | -0.161269848 | 0.35622706 | 0.5561442 | T08D10.3 |
| WBGene00011617 | 0.090723942 | 0.20045316 | NA | NA |
| WBGene00011618 | NA | NA | NA | T08G3.6 |
| WBGene00011619 | 0.033034883 | 0.64219503 | NA | T08G3.7 |
| WBGene00011620 | 0.020232312 | 0.84643798 | NA | T08G3.11 |
| WBGene00011621 | 0.020232312 | 0.84643798 | NA | T08G3.13 |
| WBGene00011622 | 0.110536695 | 0.41659247 | 0.613553 | T08G5.1 |
| WBGene00011623 | NA | NA | NA | T08G5.2 |
| WBGene00011624 | -0.105991329 | 0.4496938 | 0.64204323 | T08G5.3 |
| WBGene00011625 | 0.057804631 | 0.47927768 | 0.66615262 | vps-39 |
| WBGene00011626 | NA | NA | NA | T08G5.7 |
| WBGene00011626 | NA | NA | NA | F58D12.5 |
| WBGene00011627 | NA | NA | NA | T08G5.8 |
| WBGene00011628 | NA | NA | NA | T08G5.9 |
| WBGene00011629 | 0.176678887 | 0.00195988 | 0.01230472 | T08G11.1 |
| WBGene00011630 | 0.09351994 | 0.44866466 | 0.6413954 | T08G11.3 |
| WBGene00011631 | -0.027580792 | 0.78526825 | 0.87870478 | T08G11.4 |
| WBGene00011633 | -0.078669441 | 0.70371288 | 0.8318388 | T09A5.4 |
| WBGene00011634 | -0.119850806 | 0.30957158 | 0.51040962 | T09A5.5 |
| WBGene00011635 | -0.206113764 | 0.10563551 | 0.25454403 | mdmh-35 |
| WBGene00011636 | -0.120100923 | 0.17427191 | 0.35495871 | cec-3 |
| WBGene00011637 | 0.015558064 | 0.87802598 | 0.93485407 | sds-22 |
| WBGene00011638 | -0.154179566 | 0.02880439 | 0.10067023 | ostb-1 |

| | | | | |
|---|---|---|---|---|
| WBGene00011639 | -0.038254363 | 0.69360061 | 0.82427001 | ztf-17 |
| WBGene00011640 | NA | NA | NA | T09A5.14 |
| WBGene00011641 | -0.191470026 | 0.31492906 | 0.51557662 | T09A5.15 |
| WBGene00011642 | -0.084431632 | 0.55131422 | 0.72373276 | T09B9.1 |
| WBGene00011643 | 0.122328087 | 0.29039128 | 0.49092715 | slc-17.9 |
| WBGene00011644 | 0.206536681 | 0.16403155 | 0.33991494 | T09B9.3 |
| WBGene00011645 | 0.463328629 | 0.00020125 | 0.00177819 | T09B9.4 |
| WBGene00011646 | 0.020232312 | 0.84643798 | NA | T09B9.5 |
| WBGene00011647 | 0.04427749 | 0.45757056 | 0.64934483 | noca-1 |
| WBGene00011648 | -0.203416013 | 0.05416719 | 0.15960189 | cni-1 |
| WBGene00011649 | -0.023109762 | 0.78121253 | NA | T09E8.4 |
| WBGene00011650 | NA | NA | NA | glct-1 |
| WBGene00011651 | NA | NA | NA | nhr-217 |
| WBGene00011652 | NA | NA | NA | T09E11.3 |
| WBGene00011653 | NA | NA | NA | oac-43 |
| WBGene00011654 | NA | NA | NA | oac-44 |
| WBGene00011655 | NA | NA | NA | T09E11.6 |
| WBGene00011656 | NA | NA | NA | oac-45 |
| WBGene00011657 | NA | NA | NA | T09E11.8 |
| WBGene00011658 | NA | NA | NA | T09E11.9 |
| WBGene00011659 | NA | NA | NA | T09E11.10 |
| WBGene00011660 | 2.470672142 | 4.14E-06 | 5.53E-05 | NA |
| WBGene00011661 | 0.196402257 | 0.31506482 | 0.51557662 | ztf-27 |
| WBGene00011662 | -0.141263058 | 0.06322965 | 0.17843972 | T09F3.2 |
| WBGene00011663 | -0.07774694 | 0.6663533 | 0.80462505 | T09F3.4 |
| WBGene00011664 | 0.258183644 | 0.07152163 | 0.19456489 | T09F3.5 |
| WBGene00011665 | -0.002641891 | 0.95121282 | NA | T09F5.1 |
| WBGene00011666 | NA | NA | NA | T09F5.2 |
| WBGene00011668 | 1.864125375 | 2.64E-50 | 6.24E-48 | clec-47 |
| WBGene00011669 | 0.017279211 | 0.79380376 | NA | T09F5.10 |
| WBGene00011670 | 0.342353232 | 0.13954153 | 0.30694525 | fbxa-84 |
| WBGene00011671 | 2.271122233 | 1.78E-07 | 3.22E-06 | cyp-13A4 |
| WBGene00011672 | 4.585702084 | 1.20E-73 | 4.01E-71 | cyp-13A5 |
| WBGene00011673 | 2.307279539 | 0.00235344 | 0.01426189 | cyp-13A6 |
| WBGene00011674 | -0.007565754 | 0.93022802 | NA | cyp-13A8 |
| WBGene00011675 | 0.290763925 | 0.03132964 | 0.10657465 | cyp-13A3 |
| WBGene00011676 | 0.651705239 | 0.00405906 | 0.02214184 | cyp-13A2 |
| WBGene00011677 | 0.20280353 | 0.06932421 | 0.19010598 | cyp-13A1 |
| WBGene00011678 | 0.211929806 | 0.1840473 | 0.36741703 | T10B9.9 |
| WBGene00011679 | -0.114661823 | 0.33340149 | 0.53304196 | ucr-2.2 |
| WBGene00011680 | 0.01915291 | 0.86719908 | 0.92929001 | T10B10.3 |
| WBGene00011681 | 0.497851431 | 0.00322661 | 0.01849678 | T10B10.4 |
| WBGene00011682 | 0.09508162 | 0.44320405 | 0.6368667 | snt-7 |

| | | | | | |
|---|---|---|---|---|---|
| WBGene00011683 | | 1.96097454 | 8.88E-17 | 5.16E-15 | phat-6 |
| WBGene00011684 | | -0.015681218 | 0.93718215 | 0.96465719 | T10B10.8 |
| WBGene00011685 | | 0.048485426 | 0.45975815 | NA | T10B10.9 |
| WBGene00011686 | NA | | NA | NA | T10C6.2 |
| WBGene00011687 | | 0.093284585 | 0.18803017 | 0.37307424 | cwc-15 |
| WBGene00011688 | | 0.027074586 | 0.7852733 | 0.87870478 | T10C6.6 |
| WBGene00011689 | | -0.071023375 | 0.62020169 | 0.77273193 | T10C6.7 |
| WBGene00011690 | | -0.001993815 | 0.95612996 | NA | NA |
| WBGene00011691 | | -0.13094664 | 0.41771556 | 0.61446914 | T10C6.10 |
| WBGene00011692 | | 1.023544363 | 0.00390245 | 0.02146348 | T10G3.1 |
| WBGene00011693 | | 0.016453512 | 0.83636998 | NA | T10G3.2 |
| WBGene00011694 | | -1.051389921 | 0.00010812 | 0.00104148 | T10G3.3 |
| WBGene00011695 | | -0.002914971 | 0.95718083 | NA | T10G3.4 |
| WBGene00011696 | | -0.132146919 | 0.18317055 | 0.36616603 | eea-1 |
| WBGene00011697 | | 0.129401398 | 0.10528579 | NA | T10H4.4 |
| WBGene00011698 | | 0.020925399 | 0.84643798 | NA | cyp-34A1 |
| WBGene00011699 | | 2.544630932 | 1.65E-06 | 2.46E-05 | cyp-34A2 |
| WBGene00011701 | NA | | NA | NA | srab-17 |
| WBGene00011702 | NA | | NA | NA | srab-18 |
| WBGene00011703 | NA | | NA | NA | NA |
| WBGene00011704 | | -0.053243802 | 0.73405283 | 0.85093534 | sdz-31 |
| WBGene00011705 | | 0.454752294 | 0.07979863 | 0.21076336 | clp-9 |
| WBGene00011706 | | -0.965321702 | 7.12E-05 | 0.000718 | T11B7.1 |
| WBGene00011707 | | 0.06162375 | 0.74206324 | 0.85412964 | T11B7.2 |
| WBGene00011708 | | 1.511123258 | 4.09E-08 | 8.23E-07 | T11B7.5 |
| WBGene00011709 | | -0.045787961 | 0.74188097 | 0.85412964 | T11F9.1 |
| WBGene00011710 | | 0.020317245 | 0.84643798 | NA | T11F9.7 |
| WBGene00011712 | | 0.097858538 | 0.42588293 | 0.62160822 | T11F9.10 |
| WBGene00011713 | | 0.44546082 | 0.00291317 | 0.01705318 | T11F9.12 |
| WBGene00011714 | NA | | NA | NA | T11F9.13 |
| WBGene00011715 | NA | | NA | NA | T11F9.14 |
| WBGene00011716 | | -0.003108065 | 0.95728506 | NA | NA |
| WBGene00011717 | | -0.170134895 | 0.26664649 | 0.4653679 | T11G6.2 |
| WBGene00011718 | | 0.364129216 | 0.07157346 | 0.19463354 | T11G6.3 |
| WBGene00011719 | | 0.106403258 | 0.58138141 | 0.74581663 | T11G6.4 |
| WBGene00011720 | | -0.227243024 | 0.00539124 | 0.02773477 | T11G6.5 |
| WBGene00011721 | | -0.001842784 | 0.95621055 | NA | NA |
| WBGene00011722 | | -0.158567839 | 0.0430359 | 0.13486485 | rbm-22 |
| WBGene00011723 | | 0.695309262 | 0.00628525 | 0.0312908 | T12A7.2 |
| WBGene00011724 | NA | | NA | NA | scl-18 |
| WBGene00011725 | | 0.295517055 | 0.19804895 | 0.38605175 | lips-5 |
| WBGene00011726 | | 1.375621774 | 0.00027245 | 0.00230439 | lpr-2 |
| WBGene00011727 | | 1.870006269 | 6.01E-06 | 7.72E-05 | T12A7.6 |

| | | | | |
|---|---|---|---|---|
| WBGene00011728 | -0.026619844 | 0.72800589 | NA | T12A7.7 |
| WBGene00011729 | 0.024455292 | 0.77402369 | 0.87186343 | set-16 |
| WBGene00011730 | -0.16167251 | 0.05952362 | 0.17068 | drr-2 |
| WBGene00011731 | 0.039866295 | 0.74217863 | 0.85412964 | acbp-5 |
| WBGene00011732 | 0.370817571 | 0.09266894 | 0.23240253 | arrd-17 |
| WBGene00011733 | 0.678996079 | 0.00214743 | 0.01323256 | T12D8.5 |
| WBGene00011734 | -0.242069565 | 0.00221087 | 0.01351539 | mlc-5 |
| WBGene00011735 | -0.114031761 | 0.04639471 | 0.14255181 | hip-1 |
| WBGene00011736 | 0.406473988 | 0.00340927 | 0.01933619 | T12D8.9 |
| WBGene00011737 | 1.00726706 | 6.08E-15 | 2.89E-13 | sqst-1 |
| WBGene00011738 | 0.354269631 | 0.00032889 | 0.00272358 | T12G3.2 |
| WBGene00011739 | -0.278589478 | 0.03634674 | 0.11918312 | T12G3.4 |
| WBGene00011740 | -0.097561011 | 0.34010792 | 0.5398439 | mrpl-51 |
| WBGene00011741 | -0.175415428 | 0.02620181 | 0.09409385 | T12G3.6 |
| WBGene00011742 | -0.132943891 | 0.20868277 | 0.39798416 | tgn-38 |
| WBGene00011743 | -0.188017351 | 0.03744795 | 0.12192028 | T13F2.2 |
| WBGene00011744 | -0.046394786 | 0.50231372 | NA | T13F2.4 |
| WBGene00011745 | 0.056253204 | 0.53433155 | NA | sre-4 |
| WBGene00011746 | 0.014067514 | 0.84286955 | 0.91545974 | T13F2.6 |
| WBGene00011747 | -0.113466876 | 0.13871519 | 0.30526516 | sna-2 |
| WBGene00011748 | 3.539648607 | 6.26E-07 | 1.03E-05 | NA |
| WBGene00011749 | 0.056188364 | 0.42092875 | NA | ssp-36 |
| WBGene00011750 | 0.033463284 | 0.6365037 | NA | nhr-218 |
| WBGene00011751 | NA | NA | NA | T13F3.4 |
| WBGene00011752 | 0.330524286 | 0.14477124 | 0.31413631 | fbxa-2 |
| WBGene00011753 | NA | NA | NA | T13F3.6 |
| WBGene00011754 | 0.074833405 | 0.3346945 | NA | T13F3.7 |
| WBGene00011755 | 0.339529924 | 0.07877065 | 0.2088899 | T13H5.1 |
| WBGene00011756 | 0.59483881 | 0.02341926 | 0.08621609 | ctg-2 |
| WBGene00011757 | 0.011279172 | 0.94160445 | 0.96696575 | cht-2 |
| WBGene00011758 | -0.238266262 | 0.00811657 | 0.03827205 | T13H5.4 |
| WBGene00011759 | -0.305752705 | 0.00610911 | 0.03058943 | mrps-18B |
| WBGene00011760 | -0.195237233 | 0.35217078 | 0.55200619 | T13H5.6 |
| WBGene00011761 | -0.08736785 | 0.40446122 | 0.6025966 | T13H5.8 |
| WBGene00011762 | 0.108396595 | 0.38624234 | 0.5851299 | T13H10.2 |
| WBGene00011763 | 0.46594366 | 0.05336896 | 0.15810907 | T14B1.1 |
| WBGene00011765 | 0.229048801 | 0.2586147 | 0.45662959 | frpr-17 |
| WBGene00011767 | -0.13298593 | 0.3530577 | 0.55297888 | agxt-1 |
| WBGene00011768 | 0.305252993 | 0.11558092 | 0.27123262 | oac-46 |
| WBGene00011770 | -0.14301707 | 0.39829379 | 0.59650361 | T14G8.2 |
| WBGene00011771 | -0.146638508 | 0.12900655 | 0.29103002 | T14G8.3 |
| WBGene00011772 | 0.394644168 | 0.04296367 | 0.13469288 | T14G8.4 |
| WBGene00011773 | -0.270055514 | 0.01904292 | 0.07417063 | ttr-53 |

| | | | | |
|---|---|---|---|---|
| WBGene00011774 | 0.367285639 | 0.11526268 | 0.27074616 | ttr-54 |
| WBGene00011775 | -0.121753576 | 0.05974603 | 0.17105736 | copg-1 |
| WBGene00011776 | -0.000897784 | 0.99152574 | 0.99507216 | pigs-1 |
| WBGene00011777 | -0.134570978 | 0.52352163 | 0.70116211 | T14G10.8 |
| WBGene00011778 | NA | NA | NA | T15D6.1 |
| WBGene00011779 | NA | NA | NA | bgnt-1.7 |
| WBGene00011780 | 0.020317245 | 0.84643798 | NA | T15D6.5 |
| WBGene00011781 | NA | NA | NA | glct-3 |
| WBGene00011782 | NA | NA | NA | T15D6.8 |
| WBGene00011783 | NA | NA | NA | T15D6.9 |
| WBGene00011784 | -0.001842784 | 0.95621055 | NA | T15D6.10 |
| WBGene00011785 | 0.038125098 | 0.61119482 | NA | T15D6.11 |
| WBGene00011786 | 0.105191732 | 0.16565281 | NA | T15D6.12 |
| WBGene00011788 | -0.344206514 | 0.01680277 | 0.06756644 | hal-3 |
| WBGene00011789 | -0.004968869 | 0.94188684 | NA | T15H9.4 |
| WBGene00011790 | 0.080388946 | 0.40624029 | 0.60428981 | T15H9.5 |
| WBGene00011791 | -0.728541017 | 0.01173605 | 0.05067164 | T15H9.6 |
| WBGene00011793 | 0.19447955 | 0.35891264 | 0.55914449 | T16A9.3 |
| WBGene00011794 | 0.623768511 | 0.01603293 | 0.06520911 | nep-21 |
| WBGene00011795 | 1.841992899 | 0.00559427 | 0.02860833 | T16A9.5 |
| WBGene00011796 | 0.121439489 | 0.26182825 | 0.45969869 | T16G1.2 |
| WBGene00011797 | NA | NA | NA | T16G1.3 |
| WBGene00011798 | 4.656144098 | 3.19E-28 | 3.59E-26 | T16G1.4 |
| WBGene00011799 | 2.721938824 | 0.00012289 | 0.00116226 | T16G1.5 |
| WBGene00011800 | 1.074305848 | 0.00038615 | 0.00312008 | T16G1.6 |
| WBGene00011801 | 0.638454761 | 0.00426117 | 0.02306389 | T16G1.7 |
| WBGene00011802 | -0.223401284 | 0.05032266 | 0.15100924 | T16G1.9 |
| WBGene00011803 | -0.237547109 | 0.07992392 | 0.21099282 | T16G12.1 |
| WBGene00011804 | -0.021771339 | 0.85217661 | 0.92072303 | T16G12.3 |
| WBGene00011805 | -0.058435902 | 0.68771466 | 0.82007088 | T16G12.4 |
| WBGene00011806 | -0.21012933 | 0.02226427 | 0.08315592 | ekl-6 |
| WBGene00011807 | -0.102585299 | 0.2962418 | 0.49737336 | T16G12.6 |
| WBGene00011808 | 0.042065281 | 0.558937 | NA | T16G12.7 |
| WBGene00011809 | -0.562221059 | 0.00117587 | 0.0080448 | T16G12.8 |
| WBGene00011810 | 0.024917767 | 0.88174923 | 0.9373187 | T16G12.9 |
| WBGene00011811 | -0.193720498 | 0.36563276 | 0.56534261 | T16H12.1 |
| WBGene00011812 | -0.200732673 | 0.35574457 | 0.55568722 | T16H12.2 |
| WBGene00011813 | 0.122901392 | 0.36722712 | 0.56679025 | T16H12.3 |
| WBGene00011814 | 0.091344972 | 0.44696939 | 0.64013292 | gtf-2H2C |
| WBGene00011815 | 0.007760381 | 0.942322 | 0.96728563 | bath-43 |
| WBGene00011816 | NA | NA | NA | srt-55 |
| WBGene00011817 | 0.538959441 | 0.04636942 | 0.14250776 | T16H12.9 |
| WBGene00011818 | NA | NA | NA | NA |

| | | | | |
|---|---|---|---|---|
| WBGene00011819 | -0.191695658 | 0.14349514 | 0.31207848 | hal-2 |
| WBGene00011820 | -0.267843318 | 0.06605963 | 0.18353384 | T18D3.1 |
| WBGene00011821 | 2.133242778 | 1.47E-10 | 4.10E-09 | cdf-2 |
| WBGene00011822 | -0.094751992 | 0.44171829 | 0.63591328 | T18D3.5 |
| WBGene00011823 | 0.332885519 | 0.14697996 | 0.31730073 | T18D3.6 |
| WBGene00011824 | 0.979492028 | 0.00065607 | 0.00489575 | T18D3.7 |
| WBGene00011826 | -0.028603107 | 0.85278667 | 0.92117822 | T18D3.9 |
| WBGene00011827 | -0.126066122 | 0.22276988 | 0.41554004 | T19A6.1 |
| WBGene00011828 | -0.242690308 | 0.03142747 | 0.10688259 | nepr-1 |
| WBGene00011829 | 1.737787249 | 9.07E-06 | 0.00011289 | T19A6.4 |
| WBGene00011830 | -0.067053692 | 0.72440126 | 0.84556796 | cyp-29A2 |
| WBGene00011831 | -0.227119719 | 0.11316871 | 0.2668275 | T19B10.2 |
| WBGene00011832 | 0.083117083 | 0.61797858 | 0.77101187 | bgal-1 |
| WBGene00011833 | -0.406949289 | 0.06370209 | 0.17922028 | T19B10.5 |
| WBGene00011834 | -0.19393935 | 0.01948877 | 0.07529356 | dvc-1 |
| WBGene00011835 | -0.212112642 | 0.00140128 | 0.00928752 | pgap-1 |
| WBGene00011836 | 0.130209587 | 0.35116109 | 0.55117384 | mam-4 |
| WBGene00011837 | NA | NA | NA | srsx-33 |
| WBGene00011838 | 0.181129468 | 0.31670494 | 0.51698682 | T19C4.1 |
| WBGene00011839 | 1.444302502 | 5.14E-05 | 0.00053703 | T19C4.5 |
| WBGene00011840 | NA | NA | NA | srbc-62 |
| WBGene00011841 | NA | NA | NA | scl-25 |
| WBGene00011842 | NA | NA | NA | NA |
| WBGene00011844 | 2.712104889 | 0.00104717 | 0.00729726 | T19C9.8 |
| WBGene00011846 | 0.350672571 | 0.09758847 | 0.24196323 | chil-25 |
| WBGene00011847 | NA | NA | NA | chil-26 |
| WBGene00011848 | 0.19432963 | 0.12218946 | 0.28134075 | chil-27 |
| WBGene00011849 | 0.313021002 | 0.01585867 | 0.06473007 | T19H5.4 |
| WBGene00011850 | 0.193399567 | 0.36495887 | 0.56453912 | T20B3.1 |
| WBGene00011851 | 0.175825429 | 0.08377095 | 0.21765581 | NA |
| WBGene00011852 | 3.239929911 | 1.74E-17 | 1.07E-15 | clec-26 |
| WBGene00011853 | 0.287941555 | 0.07688186 | 0.20521919 | clec-40 |
| WBGene00011854 | 0.181759116 | 0.31195882 | 0.51266736 | T20B3.14 |
| WBGene00011855 | NA | NA | NA | clec-183 |
| WBGene00011856 | 0.7667694 | 1.66E-22 | 1.42E-20 | T20D3.2 |
| WBGene00011857 | -0.387900471 | 0.00408189 | 0.02224153 | coa-4 |
| WBGene00011858 | 0.000419631 | 0.99829788 | 0.99898019 | T20D3.5 |
| WBGene00011859 | 0.022484833 | 0.87175586 | 0.9314787 | T20D3.6 |
| WBGene00011860 | -0.210753608 | 0.07799826 | 0.20714176 | pigc-1 |
| WBGene00011864 | -0.048121682 | 0.70726868 | 0.83475378 | T20F10.2 |
| WBGene00011867 | -0.051704114 | 0.33188413 | 0.53151558 | chc-1 |
| WBGene00011868 | 0.590343579 | 0.0281935 | 0.09903576 | best-18 |
| WBGene00011869 | 1.912838471 | 9.34E-37 | 1.47E-34 | dod-6 |

| | | | | |
|---|---|---|---|---|
| WBGene00011870 | 2.074963719 | 4.27E-35 | 6.44E-33 | T20G5.8 |
| WBGene00011871 | -0.006946851 | 0.96425125 | 0.98045715 | T20G5.9 |
| WBGene00011872 | -0.05479708 | 0.73033695 | 0.84971895 | blos-1 |
| WBGene00011873 | -0.023056499 | 0.78121253 | NA | T20G5.12 |
| WBGene00011874 | -0.022930767 | 0.78121253 | NA | T20G5.13 |
| WBGene00011875 | -0.10988786 | 0.45316955 | 0.64528841 | cox-14 |
| WBGene00011876 | NA | NA | NA | srz-6 |
| WBGene00011877 | NA | NA | NA | T21B4.3 |
| WBGene00011878 | 0.163085767 | 0.23914921 | 0.43606953 | ador-1 |
| WBGene00011879 | -0.338091722 | 0.13250043 | 0.29580404 | pho-7 |
| WBGene00011880 | 1.134435198 | 6.62E-11 | 1.95E-09 | T21B6.3 |
| WBGene00011882 | -0.007169147 | 0.93387772 | NA | nstp-9 |
| WBGene00011883 | -0.142178095 | 0.21862756 | 0.41042776 | mrpl-50 |
| WBGene00011884 | -0.220302853 | 5.29E-06 | 6.89E-05 | enol-1 |
| WBGene00011885 | -0.207545602 | 0.00104539 | 0.00728836 | T21B10.3 |
| WBGene00011886 | -0.345894832 | 0.00017906 | 0.00160345 | T21B10.4 |
| WBGene00011887 | -0.25576307 | 0.03385567 | 0.1131433 | set-17 |
| WBGene00011888 | 1.12799459 | 0.00598014 | 0.03004642 | cutl-15 |
| WBGene00011890 | -0.032181285 | 0.73463552 | 0.85123061 | mics-1 |
| WBGene00011891 | -0.03222324 | 0.7515989 | 0.85956565 | del-6 |
| WBGene00011892 | -0.244635235 | 0.0168012 | 0.06756644 | erh-1 |
| WBGene00011893 | 0.384686464 | 0.01950106 | 0.07529356 | T21C9.6 |
| WBGene00011894 | 3.646819624 | 2.62E-06 | 3.67E-05 | ttr-23 |
| WBGene00011895 | 0.180007984 | 0.0451335 | NA | T21C9.9 |
| WBGene00011896 | 0.239907249 | 0.01930642 | 0.07487821 | T21C9.11 |
| WBGene00011897 | -0.228167951 | 0.01284668 | 0.05464503 | scpl-4 |
| WBGene00011898 | -0.037413403 | 0.63537329 | 0.78369842 | T21C9.13 |
| WBGene00011899 | 0.756273926 | 0.02682003 | 0.09558717 | nlp-68 |
| WBGene00011900 | -0.001301201 | 0.99238388 | 0.9955752 | T21C12.4 |
| WBGene00011902 | -0.001993815 | 0.95612996 | NA | T21E8.4 |
| WBGene00011903 | -0.024369794 | 0.78114619 | NA | T21E8.5 |
| WBGene00011904 | 0.237947605 | 0.09484637 | 0.23636523 | hlb-1 |
| WBGene00011906 | -0.09732218 | 0.5645457 | 0.73446933 | hsp-12.1 |
| WBGene00011908 | -0.254623971 | 0.0057364 | 0.02919242 | pash-1 |
| WBGene00011909 | 0.0398788 | 0.58971221 | NA | T22A3.6 |
| WBGene00011910 | 3.352094416 | 0.00062947 | 0.00474074 | alg-3 |
| WBGene00011911 | 2.273251207 | 0.00711437 | 0.03443356 | T22B3.3 |
| WBGene00011912 | -0.080382598 | 0.36377206 | 0.56356544 | T22C1.1 |
| WBGene00011913 | -0.023056499 | 0.78121253 | NA | glb-26 |
| WBGene00011914 | -0.148989851 | 0.1550074 | 0.32819427 | pigu-1 |
| WBGene00011915 | 0.201774869 | 0.32304975 | 0.52302147 | ctf-8 |
| WBGene00011916 | 0.020094752 | 0.86646362 | 0.92889316 | T22C1.5 |
| WBGene00011917 | -0.121065775 | 0.19622355 | 0.38377091 | T22C1.6 |

| | | | | |
|---|---|---|---|---|
| WBGene00011918 | 0.193244597 | 0.06508246 | 0.18184587 | T22C1.8 |
| WBGene00011919 | -0.028098241 | 0.71579894 | NA | T22C1.9 |
| WBGene00011920 | -0.226881161 | 0.19244085 | 0.37885255 | T22C1.11 |
| WBGene00011921 | NA | NA | NA | T22C1.12 |
| WBGene00011922 | 0.065014541 | 0.37399213 | NA | T22C8.1 |
| WBGene00011923 | -0.360899659 | 0.08253914 | 0.21546315 | chhy-1 |
| WBGene00011924 | 0.099185832 | 0.63241539 | 0.78169639 | T22C8.3 |
| WBGene00011925 | -0.410436398 | 0.09944382 | 0.24461044 | T22C8.4 |
| WBGene00011926 | 0.19257742 | 0.25447561 | 0.45281689 | sptf-2 |
| WBGene00011927 | 0.288371975 | 0.15458709 | 0.32763602 | T22C8.6 |
| WBGene00011928 | 0.51302692 | 0.03011582 | 0.10372113 | cutl-12 |
| WBGene00011929 | NA | NA | NA | T22G5.1 |
| WBGene00011930 | -0.378745687 | 0.09401242 | 0.23480653 | T22G5.3 |
| WBGene00011931 | -0.027350315 | 0.68023343 | NA | srsx-19 |
| WBGene00011932 | -0.53410998 | 4.83E-06 | 6.34E-05 | sptl-3 |
| WBGene00011933 | 0.020232312 | 0.84643798 | NA | NA |
| WBGene00011934 | NA | NA | NA | T22H2.4 |
| WBGene00011935 | 0.125175132 | 0.48489716 | 0.66987915 | scrm-1 |
| WBGene00011936 | -0.024664683 | 0.87116708 | 0.93126126 | pgrn-1 |
| WBGene00011937 | 0.09249737 | 0.19271823 | NA | ilc-17.1 |
| WBGene00011938 | -0.617580457 | 0.00021871 | 0.00190373 | alh-13 |
| WBGene00011939 | -0.245235731 | 0.00116057 | 0.00796249 | prmt-9 |
| WBGene00011940 | -0.065055963 | 0.55013399 | 0.72270181 | T23B5.3 |
| WBGene00011941 | 0.029542753 | 0.86423871 | 0.92755718 | T23B5.4 |
| WBGene00011942 | NA | NA | NA | T23D5.3 |
| WBGene00011943 | NA | NA | NA | T23D5.8 |
| WBGene00011944 | -0.15728331 | 0.05587089 | 0.16333979 | T23D8.3 |
| WBGene00011945 | -0.175732703 | 0.04901892 | 0.1481292 | alg-5 |
| WBGene00011947 | NA | NA | NA | T23F1.3 |
| WBGene00011948 | 0.337435172 | 0.02344883 | 0.08628156 | rocf-1 |
| WBGene00011949 | 0.1390172 | 0.29820154 | 0.49908194 | T23F6.1 |
| WBGene00011950 | NA | NA | NA | T23F6.2 |
| WBGene00011951 | 0.136454535 | 0.33441356 | 0.53378105 | T23F6.3 |
| WBGene00011952 | 0.017279211 | 0.79380376 | NA | T23F6.5 |
| WBGene00011953 | 0.467615098 | 1.75E-06 | 2.59E-05 | ppm-2 |
| WBGene00011954 | 0.226089023 | 0.09799313 | 0.24259678 | T23F11.2 |
| WBGene00011955 | 0.604589472 | 0.03105184 | 0.10587565 | cdka-1 |
| WBGene00011956 | -0.188836312 | 0.27909345 | 0.47887111 | T23F11.4 |
| WBGene00011957 | 0.223678308 | 0.20844171 | 0.39762802 | T23F11.6 |
| WBGene00011958 | 0.048485426 | 0.45975815 | NA | T23G4.2 |
| WBGene00011959 | 0.057065547 | 0.69747957 | 0.82753634 | nyn-1 |
| WBGene00011960 | 0.08924158 | 0.26408462 | NA | oac-48 |
| WBGene00011961 | NA | NA | NA | T23G4.5 |

| | | | | |
|---|---|---|---|---|
| WBGene00011962 | -0.007066805 | 0.92974065 | 0.96057673 | T23G5.2 |
| WBGene00011963 | -0.093876942 | 0.57545839 | 0.74264312 | T23G5.3 |
| WBGene00011964 | 0.183414735 | 0.3806998 | 0.58012342 | saeg-2 |
| WBGene00011965 | -0.20948466 | 0.1225439 | 0.28192414 | T23G7.2 |
| WBGene00011966 | 0.001502597 | 0.9865346 | 0.99222732 | T23G7.3 |
| WBGene00011967 | -0.086601686 | 0.45578593 | 0.64769114 | pir-1 |
| WBGene00011968 | 0.657749329 | 0.03627776 | 0.1190227 | T23G11.1 |
| WBGene00011969 | -0.00133527 | 0.99137502 | 0.99504501 | T23G11.4 |
| WBGene00011970 | 0.002411193 | 0.98603339 | 0.99213215 | rlbp-1 |
| WBGene00011971 | -0.1413552 | 0.46083377 | 0.65170165 | lron-9 |
| WBGene00011972 | 0.040378655 | 0.66626547 | 0.80458535 | T23G11.7 |
| WBGene00011973 | 0.00822867 | 0.84531204 | NA | T23G11.10 |
| WBGene00011974 | 1.025143014 | 0.00855433 | 0.0397979 | xbx-5 |
| WBGene00011975 | -0.072570238 | 0.48281428 | 0.66845238 | T24B1.1 |
| WBGene00011976 | -0.075664074 | 0.4431065 | 0.63685467 | T24B8.2 |
| WBGene00011977 | -0.585298329 | 1.87E-05 | 0.0002148 | T24B8.3 |
| WBGene00011978 | 0.906634582 | 2.24E-08 | 4.71E-07 | T24B8.4 |
| WBGene00011979 | -1.124842824 | 0.00010745 | 0.00103563 | T24B8.5 |
| WBGene00011980 | -0.029414425 | 0.63002647 | 0.7799881 | T24B8.7 |
| WBGene00011981 | -0.431915177 | 0.05712444 | 0.16595987 | T24C2.2 |
| WBGene00011982 | 0.018795205 | 0.77532208 | NA | T24C2.3 |
| WBGene00011983 | NA | NA | NA | fbxa-81 |
| WBGene00011984 | 0.185759006 | 0.39002214 | 0.58790427 | T24C2.5 |
| WBGene00011985 | -0.162396705 | 0.04069234 | 0.12962481 | T24D1.2 |
| WBGene00011986 | -0.095190324 | 0.31667599 | 0.51698682 | T24D1.3 |
| WBGene00011988 | -0.111773572 | 0.2478439 | 0.44616778 | har-2 |
| WBGene00011989 | -0.06730507 | 0.31931151 | NA | T24D5.1 |
| WBGene00011990 | -0.048123126 | 0.80434289 | 0.89116459 | T24D5.2 |
| WBGene00011991 | -0.023109762 | 0.78121253 | NA | T24D5.3 |
| WBGene00011992 | NA | NA | NA | T24D5.4 |
| WBGene00011993 | -0.047052602 | 0.79776891 | 0.88680697 | T24D5.5 |
| WBGene00011994 | -0.207438714 | 0.00694882 | 0.03377745 | samp-1 |
| WBGene00011995 | 0.270034502 | 0.14432937 | 0.31342723 | rsf-1 |
| WBGene00011996 | 0.056430134 | 0.44291702 | NA | T24F1.4 |
| WBGene00011997 | 3.222319248 | 1.38E-38 | 2.44E-36 | nlp-51 |
| WBGene00011999 | 0.015597454 | 0.8447034 | NA | T24F1.7 |
| WBGene00012000 | -0.383271007 | 4.20E-06 | 5.59E-05 | T24H10.1 |
| WBGene00012002 | -0.024462508 | 0.83365314 | 0.90929864 | T24H10.4 |
| WBGene00012003 | -0.126597633 | 0.24719612 | 0.4456592 | T24H10.5 |
| WBGene00012004 | -0.106496331 | 0.39404197 | 0.59231709 | dyrb-1 |
| WBGene00012005 | 0.497523258 | 0.01112717 | 0.04864523 | 1-Jun |
| WBGene00012007 | -0.694629808 | 0.00127554 | 0.0085942 | T25B9.1 |
| WBGene00012008 | 0.040107227 | 0.58693163 | NA | T25B9.2 |

| | | | | |
|---|---|---|---|---|
| WBGene00012009 | 0.0038468 | 0.97645698 | 0.98670757 | T25B9.3 |
| WBGene00012010 | 0.132442484 | 0.10147609 | NA | T25B9.4 |
| WBGene00012011 | 0.193036759 | 0.0748227 | 0.20111604 | T25B9.5 |
| WBGene00012012 | 0.249238568 | 0.04148586 | 0.13130013 | T25B9.6 |
| WBGene00012013 | 1.851903656 | 5.83E-07 | 9.66E-06 | ugt-54 |
| WBGene00012014 | -0.107926224 | 0.51500966 | 0.69482643 | T25B9.8 |
| WBGene00012015 | 0.022122142 | 0.700178 | 0.82952793 | T25B9.9 |
| WBGene00012016 | 0.327510969 | 0.16608163 | 0.3426721 | inpp-1 |
| WBGene00012017 | 0.160557691 | 0.27751307 | 0.47750252 | T25C8.1 |
| WBGene00012018 | 0.903115788 | 5.12E-41 | 9.62E-39 | T25C12.3 |
| WBGene00012019 | 0.094725114 | 0.36145079 | 0.56130685 | dkf-2 |
| WBGene00012020 | 0.007104532 | 0.89831685 | 0.94647791 | gyg-2 |
| WBGene00012021 | -0.822213941 | 0.00041842 | 0.00334026 | T25E12.6 |
| WBGene00012022 | -0.027373974 | 0.68245543 | NA | clec-39 |
| WBGene00012023 | 0.149717874 | 0.0720891 | NA | clec-34 |
| WBGene00012024 | -0.050223056 | 0.47776088 | NA | clec-29 |
| WBGene00012025 | 0.0398788 | 0.58971221 | NA | clec-38 |
| WBGene00012026 | 0.020232312 | 0.84643798 | NA | srz-100 |
| WBGene00012027 | -0.009101738 | 0.94808436 | 0.97044737 | fbxa-127 |
| WBGene00012028 | 0.25072866 | 0.07969035 | 0.21058447 | srz-99 |
| WBGene00012029 | -0.207377379 | 0.14100872 | 0.30864234 | T25G3.1 |
| WBGene00012030 | -0.168871591 | 0.03807813 | 0.12340873 | T25G3.3 |
| WBGene00012031 | -0.218499866 | 0.00273744 | 0.01620003 | T25G3.4 |
| WBGene00012032 | 1.975810627 | 0.01058413 | 0.04668942 | T26C5.2 |
| WBGene00012033 | -0.045313328 | 0.61963154 | 0.77258029 | T26C5.3 |
| WBGene00012034 | 0.573664094 | 0.04871449 | 0.14746418 | T26C5.4 |
| WBGene00012035 | 0.266617628 | 0.17655269 | 0.35738765 | T26C5.5 |
| WBGene00012036 | 0.036692668 | 0.59311964 | NA | clec-103 |
| WBGene00012037 | -0.345758784 | 0.00172842 | 0.01107474 | T26E3.4 |
| WBGene00012038 | -0.649643335 | 0.03564201 | 0.1175833 | T26E3.5 |
| WBGene00012039 | NA | NA | NA | T26E3.6 |
| WBGene00012040 | -0.041190003 | 0.73550288 | 0.85166527 | T26E3.7 |
| WBGene00012041 | -0.954425982 | 0.00882051 | 0.04078997 | T26E3.8 |
| WBGene00012042 | 0.049950653 | 0.56395944 | NA | T26E3.10 |
| WBGene00012043 | 0.020317245 | 0.84643798 | NA | irld-52 |
| WBGene00012044 | 0.038125098 | 0.61119482 | NA | T26E4.2 |
| WBGene00012045 | 0.065014541 | 0.37399213 | NA | T26E4.3 |
| WBGene00012046 | -0.04044783 | 0.53612608 | NA | T26E4.4 |
| WBGene00012047 | NA | NA | NA | NA |
| WBGene00012048 | NA | NA | NA | NA |
| WBGene00012049 | NA | NA | NA | T26E4.7 |
| WBGene00012050 | 0.020925399 | 0.84643798 | NA | nhr-223 |
| WBGene00012051 | NA | NA | NA | T26E4.9 |

| | | | | |
|---|---|---|---|---|
| WBGene00012052 | NA | NA | NA | T26E4.10 |
| WBGene00012053 | NA | NA | NA | NA |
| WBGene00012054 | NA | NA | NA | srsx-38 |
| WBGene00012055 | 0.056550688 | 0.43750329 | NA | srsx-36 |
| WBGene00012056 | NA | NA | NA | nhr-285 |
| WBGene00012057 | NA | NA | NA | R10D12.15 |
| WBGene00012057 | NA | NA | NA | T26F2.1 |
| WBGene00012058 | 0.31795584 | 0.17728798 | 0.3584194 | NA |
| WBGene00012059 | -0.480143148 | 4.17E-06 | 5.56E-05 | T26G10.1 |
| WBGene00012060 | NA | NA | NA | T26G10.3 |
| WBGene00012061 | NA | NA | NA | T26G10.4 |
| WBGene00012062 | 0.301072797 | 0.10080096 | 0.24671128 | T26G10.5 |
| WBGene00012063 | -0.40392996 | 0.06770345 | 0.18688655 | fbxb-1 |
| WBGene00012064 | -0.156258503 | 0.10784881 | 0.25826208 | fbxb-115 |
| WBGene00012065 | -0.137060469 | 0.42381237 | 0.61982188 | fbxb-2 |
| WBGene00012066 | -0.217188769 | 0.2198824 | 0.41199107 | fbxb-116 |
| WBGene00012067 | 6.188870896 | 4.64E-06 | 6.12E-05 | sqst-3 |
| WBGene00012068 | -0.005221332 | 0.94448901 | NA | oac-49 |
| WBGene00012069 | 0.124326451 | 0.11293212 | NA | T26H5.4 |
| WBGene00012070 | 2.9179136 | 0.00322404 | 0.01848929 | T26H5.8 |
| WBGene00012072 | NA | NA | NA | T26H8.4 |
| WBGene00012073 | 0.165364897 | 0.42138454 | 0.61806162 | cpd-2 |
| WBGene00012074 | 0.094929623 | 0.63655453 | 0.78445295 | T27A8.2 |
| WBGene00012075 | -0.023056499 | 0.78121253 | NA | T27A8.3 |
| WBGene00012077 | NA | NA | NA | T27A8.5 |
| WBGene00012078 | NA | NA | NA | NA |
| WBGene00012079 | 0.020317245 | 0.84643798 | NA | NA |
| WBGene00012080 | NA | NA | NA | clec-244 |
| WBGene00012081 | 0.038125098 | 0.61119482 | NA | T27C5.8 |
| WBGene00012083 | NA | NA | NA | T27C5.10 |
| WBGene00012084 | 0.502188169 | 0.04379408 | 0.13668496 | npr-15 |
| WBGene00012085 | 0.384162653 | 0.01736018 | 0.06918085 | T27D12.1 |
| WBGene00012086 | -0.023109762 | 0.78121253 | NA | clec-144 |
| WBGene00012087 | 1.874390846 | 0.00116618 | 0.00798596 | T27E7.1 |
| WBGene00012088 | NA | NA | NA | T27E7.3 |
| WBGene00012089 | 0.020317245 | 0.84643798 | NA | T27E7.4 |
| WBGene00012090 | NA | NA | NA | T27E7.5 |
| WBGene00012091 | 0.09249737 | 0.19271823 | NA | pals-29 |
| WBGene00012093 | NA | NA | NA | T27E7.9 |
| WBGene00012094 | -0.230634475 | 0.00502475 | 0.02623283 | T27E9.2 |
| WBGene00012095 | -0.218579895 | 0.04750359 | 0.14486567 | pssy-2 |
| WBGene00012096 | -0.006194943 | 0.97226224 | 0.98476102 | T27E9.6 |
| WBGene00012097 | -0.338710126 | 1.21E-10 | 3.43E-09 | abcf-2 |

| | | | | |
|---|---|---|---|---|
| WBGene00012098 | -0.141488534 | 0.42747759 | 0.62265977 | fbxb-86 |
| WBGene00012099 | 0.034358027 | 0.8568667 | 0.92320152 | acc-4 |
| WBGene00012100 | 0.748679124 | 0.00034709 | 0.00284054 | sipa-1 |
| WBGene00012101 | -0.685394536 | 0.02219149 | 0.08296873 | zip-10 |
| WBGene00012102 | 0.084908438 | 0.30232871 | NA | T27F6.1 |
| WBGene00012103 | -0.303230532 | 0.19205323 | 0.37824204 | clec-12 |
| WBGene00012104 | 0.036781244 | 0.69999977 | 0.82951814 | sygl-1 |
| WBGene00012105 | 0.15828231 | 0.2208271 | 0.41323231 | T27F6.6 |
| WBGene00012106 | -0.030057668 | 0.85151064 | 0.92034307 | T27F6.7 |
| WBGene00012107 | 1.127302443 | 0.00075366 | 0.00550337 | T27F6.8 |
| WBGene00012108 | 0.020317245 | 0.84643798 | NA | T28A8.2 |
| WBGene00012109 | -0.189051051 | 0.09882603 | 0.24383415 | T28A8.3 |
| WBGene00012110 | -0.13662215 | 0.22851774 | 0.42217391 | T28A8.4 |
| WBGene00012111 | -0.262062672 | 0.11991863 | 0.27820134 | T28A8.5 |
| WBGene00012112 | -0.410083451 | 0.00143118 | 0.00944717 | T28A8.6 |
| WBGene00012113 | 0.381404439 | 0.0212807 | 0.08038461 | T28B8.1 |
| WBGene00012114 | 0.070933025 | 0.58136842 | 0.74581663 | T28B8.3 |
| WBGene00012115 | 0.032688266 | 0.77146716 | 0.87012177 | T28B8.4 |
| WBGene00012116 | 0.065883259 | 0.59562036 | 0.75477619 | del-4 |
| WBGene00012117 | 0.176494364 | 0.02204888 | 0.08251985 | T28B11.1 |
| WBGene00012118 | -0.040350788 | 0.81771909 | 0.89976324 | grsp-2 |
| WBGene00012119 | -0.310824804 | 0.17060197 | 0.34931872 | T28C6.3 |
| WBGene00012120 | 0.001487708 | 0.99294772 | 0.99586209 | T28C6.5 |
| WBGene00012121 | 0.023711838 | 0.78541717 | 0.87874773 | T28C6.7 |
| WBGene00012122 | -0.549228858 | 0.01890715 | 0.07381856 | T28C6.8 |
| WBGene00012123 | 0.683693294 | 2.65E-06 | 3.70E-05 | T28D6.3 |
| WBGene00012124 | 0.171384484 | 0.42715661 | 0.62247436 | T28D6.4 |
| WBGene00012125 | 0.338952195 | 0.09260529 | 0.23228269 | T28D6.5 |
| WBGene00012126 | -0.33325306 | 0.00022776 | 0.00196851 | T28D6.6 |
| WBGene00012127 | 0.080189124 | 0.68157134 | 0.81586837 | T28D6.7 |
| WBGene00012128 | -0.030543688 | 0.74609836 | 0.85641914 | nra-1 |
| WBGene00012130 | -0.493764233 | 0.01004232 | 0.04500449 | slc-17.4 |
| WBGene00012131 | 1.551857698 | 0.0007275 | 0.0053363 | T28F3.5 |
| WBGene00012132 | 0.017753724 | 0.82398962 | NA | ifta-2 |
| WBGene00012133 | NA | NA | NA | NA |
| WBGene00012134 | 0.976070404 | 1.65E-07 | 3.00E-06 | T28F3.8 |
| WBGene00012135 | 0.059790696 | 0.66903154 | 0.8069938 | T28F3.9 |
| WBGene00012136 | 0.15214336 | 0.36299217 | 0.56280507 | T28F4.1 |
| WBGene00012137 | 0.089549418 | 0.49189669 | 0.67512065 | asic-2 |
| WBGene00012138 | 0.039332645 | 0.65776843 | NA | T28F4.3 |
| WBGene00012139 | 0.187474708 | 0.32437845 | 0.52363682 | T28F4.4 |
| WBGene00012140 | 0.322535561 | 0.0589721 | 0.16949559 | T28F4.5 |
| WBGene00012142 | 0.228286631 | 0.30370461 | 0.50437151 | T28H10.1 |

| | | | | | |
|---|---|---|---|---|---|
| WBGene00012143 | | 0.836050191 | 0.0028965 | 0.01697944 | T28H10.2 |
| WBGene00012144 | | 1.605156903 | 1.39E-39 | 2.54E-37 | T28H10.3 |
| WBGene00012145 | | 0.010018959 | 0.92686789 | 0.9586234 | nlp-46 |
| WBGene00012146 | NA | | NA | NA | B0393.8 |
| WBGene00012146 | NA | | NA | NA | VB0393L.2 |
| WBGene00012147 | NA | | NA | NA | VC27A7L.1 |
| WBGene00012148 | | -0.077072416 | 0.32401144 | 0.52363682 | inos-1 |
| WBGene00012149 | | -0.075615066 | 0.29019054 | 0.49077968 | VF13D12L.3 |
| WBGene00012150 | | -0.031366256 | 0.81556039 | 0.898737 | syx-17 |
| WBGene00012151 | | 0.158950036 | 0.45852095 | 0.64987466 | VH15N14R.1 |
| WBGene00012152 | | 0.501905692 | 0.05160733 | 0.15394924 | cnc-10 |
| WBGene00012155 | | -0.035136393 | 0.72019208 | NA | VW02B12L.2 |
| WBGene00012156 | | 0.045374461 | 0.65928287 | 0.79896991 | ebp-2 |
| WBGene00012157 | | -0.105136914 | 0.414623 | 0.61208134 | adbp-1 |
| WBGene00012158 | | 0.204142994 | 0.02565522 | 0.09244844 | ucr-2.1 |
| WBGene00012159 | NA | | NA | NA | VY10G11R.1 |
| WBGene00012162 | | -0.050717078 | 0.78221192 | 0.87737414 | sek-6 |
| WBGene00012163 | | 0.005084406 | 0.97001007 | 0.98346632 | VZK822L.2 |
| WBGene00012164 | | -0.28178806 | 0.19641533 | 0.38396663 | W01A8.2 |
| WBGene00012165 | | -0.023109762 | 0.78121253 | NA | cutl-6 |
| WBGene00012166 | | -0.242976572 | 0.00235918 | 0.01427895 | nuo-6 |
| WBGene00012167 | | -0.092536991 | 0.33257841 | 0.53237822 | tofu-5 |
| WBGene00012168 | | -0.12691662 | 0.52874724 | 0.70564594 | W01A8.6 |
| WBGene00012169 | | 0.097743371 | 0.43891278 | 0.63365323 | ttbk-4 |
| WBGene00012170 | | 0.004174637 | 0.97387262 | 0.98551763 | W01B6.3 |
| WBGene00012171 | NA | | NA | NA | W01B6.4 |
| WBGene00012172 | | 0.109291922 | 0.21520653 | NA | W01B6.5 |
| WBGene00012173 | | 0.087256674 | 0.23800185 | NA | W01B6.6 |
| WBGene00012174 | | -0.003421551 | 0.95689871 | NA | W01B6.8 |
| WBGene00012175 | | 0.246800427 | 0.08140585 | 0.21348074 | W01C9.1 |
| WBGene00012176 | | -0.073067159 | 0.42051181 | NA | W01C9.2 |
| WBGene00012177 | | 0.074107881 | 0.33444485 | NA | decr-1.2 |
| WBGene00012178 | | 0.046423204 | 0.72607783 | 0.84664746 | glb-27 |
| WBGene00012179 | | -0.371829413 | 4.63E-11 | 1.41E-09 | W01D2.1 |
| WBGene00012180 | | -0.023056499 | 0.78121253 | NA | W01D2.3 |
| WBGene00012181 | NA | | NA | NA | srt-11 |
| WBGene00012182 | | 0.384395348 | 0.00339443 | 0.01927022 | osta-3 |
| WBGene00012183 | | 0.204110432 | 0.06872563 | 0.18885359 | W01D2.6 |
| WBGene00012184 | | -0.06373433 | 0.66420528 | 0.80296004 | mnr-1 |
| WBGene00012185 | | -0.207985967 | 0.10095606 | 0.2470083 | W01F3.2 |
| WBGene00012186 | | -0.146023147 | 0.48725632 | 0.67155416 | mlt-11 |
| WBGene00012187 | | -0.283145812 | 0.02696993 | 0.09588801 | rpb-11 |
| WBGene00012188 | | 0.019289499 | 0.86305144 | 0.9270525 | W01G7.4 |

| | | | | |
|---|---|---|---|---|
| WBGene00012189 | 0.071420261 | 0.54955527 | 0.72221805 | NA |
| WBGene00012190 | -0.025848508 | 0.67901638 | NA | W02A2.5 |
| WBGene00012191 | 0.037992904 | 0.65794732 | NA | W02A2.8 |
| WBGene00012192 | -0.079539434 | 0.64976 | 0.79273174 | W02A11.1 |
| WBGene00012193 | -0.0160491 | 0.90827873 | 0.95041108 | vps-25 |
| WBGene00012194 | 0.207516078 | 0.20262408 | 0.39132292 | toe-4 |
| WBGene00012195 | 0.28170457 | 0.20526833 | 0.39439004 | bath-34 |
| WBGene00012196 | 0.020129102 | 0.92204354 | 0.95620205 | bath-35 |
| WBGene00012197 | 0.318260149 | 0.17617765 | 0.35699717 | W02B8.1 |
| WBGene00012198 | -0.267579746 | 0.03158487 | 0.1071938 | W02B8.2 |
| WBGene00012199 | 0.019541525 | 0.85882582 | 0.92455628 | mltn-3 |
| WBGene00012200 | 0.0398788 | 0.58971221 | NA | mltn-4 |
| WBGene00012201 | -0.573238994 | 0.00500578 | 0.02619357 | W02B12.1 |
| WBGene00012202 | 0.180179092 | 0.39503641 | 0.59314204 | W02B12.4 |
| WBGene00012203 | 0.333751488 | 0.01109049 | 0.04852835 | rga-1 |
| WBGene00012204 | -0.17809687 | 0.14424506 | 0.31333708 | mfn-1 |
| WBGene00012205 | -0.20252913 | 0.0442237 | 0.13745399 | W02B12.10 |
| WBGene00012206 | -0.164549662 | 0.10514235 | 0.2535226 | W02B12.11 |
| WBGene00012207 | 0.993334613 | 0.01770508 | 0.07034467 | W02B12.12 |
| WBGene00012208 | 0.057984889 | 0.57115649 | 0.73951536 | W02D9.2 |
| WBGene00012209 | 0.05636627 | 0.70952241 | 0.83640239 | hmg-20 |
| WBGene00012210 | 0.052686356 | 0.78117771 | 0.87672852 | W02D9.4 |
| WBGene00012211 | 0.248202004 | 0.25908382 | 0.45718286 | ssp-37 |
| WBGene00012212 | 0.550252384 | 0.00021508 | 0.00188115 | W02D9.6 |
| WBGene00012213 | 0.44715559 | 5.03E-06 | 6.58E-05 | W02D9.7 |
| WBGene00012214 | NA | NA | NA | W02D9.8 |
| WBGene00012215 | NA | NA | NA | W02D9.9 |
| WBGene00012216 | 0.294081964 | 0.19592267 | 0.38359238 | W02D9.10 |
| WBGene00012217 | 0.350740374 | 0.09453392 | 0.23582741 | W02H3.1 |
| WBGene00012218 | NA | NA | NA | NA |
| WBGene00012219 | -0.026851213 | 0.72588365 | NA | W03C9.1 |
| WBGene00012220 | -0.246499133 | 0.00927045 | 0.04237549 | W03C9.2 |
| WBGene00012221 | -0.132181413 | 0.36436667 | 0.56409986 | W03C9.5 |
| WBGene00012222 | 0.190877283 | 0.38032999 | 0.5797656 | W03C9.6 |
| WBGene00012223 | 0.033034883 | 0.64219503 | NA | W03C9.8 |
| WBGene00012224 | 0.623686732 | 0.0395251 | 0.12693289 | W03G11.2 |
| WBGene00012225 | 0.21877629 | 0.29362628 | 0.49476146 | W03G11.3 |
| WBGene00012226 | 0.312657162 | 0.13989464 | 0.30727009 | W03G11.4 |
| WBGene00012227 | NA | NA | NA | W03H9.1 |
| WBGene00012228 | -0.199079511 | 0.29227193 | 0.4930906 | W03H9.2 |
| WBGene00012230 | -0.037152512 | 0.64807883 | 0.79160333 | cacn-1 |
| WBGene00012231 | -1.625274856 | 4.40E-12 | 1.52E-10 | W04A4.2 |
| WBGene00012232 | NA | NA | NA | W04A4.3 |

| | | | | |
|---|---|---|---|---|
| WBGene00012233 | 0.020232312 | 0.84643798 | NA | clpr-1 |
| WBGene00012234 | -0.044586499 | 0.61242433 | 0.7668267 | W04A4.5 |
| WBGene00012235 | 0.174227498 | 0.41480847 | 0.61228055 | W04A4.6 |
| WBGene00012236 | -0.183057986 | 0.21464808 | 0.40560951 | W04A8.1 |
| WBGene00012237 | 0.172290497 | 0.3543172 | 0.55428551 | W04A8.2 |
| WBGene00012238 | -0.19498541 | 0.33097241 | 0.53051972 | W04A8.3 |
| WBGene00012239 | -0.185008545 | 0.33215678 | 0.53189406 | W04A8.4 |
| WBGene00012240 | -0.900539643 | 0.00299645 | 0.01745763 | W04A8.5 |
| WBGene00012241 | -0.095608645 | 0.27736153 | 0.47747657 | W04A8.6 |
| WBGene00012242 | 0.058314668 | 0.40208118 | NA | srt-39 |
| WBGene00012243 | -0.224021436 | 0.00593717 | 0.02989207 | W04D2.4 |
| WBGene00012244 | -0.048855396 | 0.65357129 | 0.79476224 | mrps-11 |
| WBGene00012245 | -0.137365754 | 0.06745437 | 0.18633952 | rbm-25 |
| WBGene00012246 | NA | NA | NA | fbxa-131 |
| WBGene00012247 | -0.138463962 | 0.51268729 | 0.69245891 | W04E12.2 |
| WBGene00012248 | NA | NA | NA | W04E12.3 |
| WBGene00012249 | 0.108942554 | 0.15594412 | NA | W04E12.4 |
| WBGene00012250 | -0.00346345 | 0.95708614 | NA | W04E12.5 |
| WBGene00012251 | 0.52635183 | 0.00786304 | 0.03730483 | clec-49 |
| WBGene00012252 | -0.039805974 | 0.84571084 | 0.9169413 | W04E12.7 |
| WBGene00012253 | 0.587542401 | 1.50E-15 | 7.55E-14 | clec-50 |
| WBGene00012254 | 0.040107227 | 0.58693163 | NA | W04E12.9 |
| WBGene00012255 | 0.180783597 | 0.29803258 | 0.49896857 | lpr-6 |
| WBGene00012256 | 0.342483738 | 0.1034237 | 0.25103223 | lpr-5 |
| WBGene00012257 | 3.505742447 | 0.0004142 | 0.00331562 | lpr-4 |
| WBGene00012258 | -0.011919178 | 0.89455137 | 0.94489559 | W04G3.5 |
| WBGene00012259 | 0.318543798 | 0.12924094 | 0.2913792 | sulp-7 |
| WBGene00012260 | -0.03323923 | 0.69122143 | NA | W04G3.7 |
| WBGene00012261 | 0.256185881 | 0.2559557 | 0.45436804 | lpr-3 |
| WBGene00012264 | 0.020925399 | 0.84643798 | NA | W04G5.3 |
| WBGene00012265 | 0.058088375 | 0.58008525 | 0.745406 | W04G5.4 |
| WBGene00012266 | 0.0398788 | 0.58971221 | NA | W04G5.5 |
| WBGene00012267 | 0.020317245 | 0.84643798 | NA | W04G5.7 |
| WBGene00012268 | 0.13389452 | 0.15713149 | 0.33111142 | W04G5.8 |
| WBGene00012269 | 0.086920836 | 0.29201443 | NA | W04G5.9 |
| WBGene00012270 | 0.106231916 | 0.14359628 | NA | W04G5.10 |
| WBGene00012271 | -0.050364259 | 0.47474184 | NA | W05B2.2 |
| WBGene00012272 | 0.064923556 | 0.7345456 | 0.85123061 | dhc-4 |
| WBGene00012274 | 0.311868051 | 0.09861391 | 0.24347628 | W05B5.1 |
| WBGene00012275 | 0.015507174 | 0.80923808 | NA | npr-14 |
| WBGene00012276 | -0.269783225 | 0.00013838 | 0.00127982 | his-74 |
| WBGene00012277 | 0.30147821 | 0.01515038 | 0.06232215 | ccch-3 |
| WBGene00012278 | -0.023756719 | 0.6882956 | NA | W05B10.3 |

| | | | | |
|---|---|---|---|---|
| WBGene00012279 | 0.39443695 | 0.0898895 | 0.22785184 | W05B10.4 |
| WBGene00012280 | 0.216467107 | 0.32370299 | 0.52345213 | W05E10.1 |
| WBGene00012281 | 0.017279211 | 0.79380376 | NA | W05E10.2 |
| WBGene00012282 | 0.287104693 | 0.07464071 | 0.20089046 | W05E10.5 |
| WBGene00012283 | 0.020925399 | 0.84643798 | NA | W05H5.1 |
| WBGene00012284 | -0.039636691 | 0.54579661 | NA | W05H5.2 |
| WBGene00012285 | -0.003108065 | 0.95728506 | NA | pitr-6 |
| WBGene00012286 | NA | NA | NA | sre-35 |
| WBGene00012287 | NA | NA | NA | sre-33 |
| WBGene00012288 | -0.023056499 | 0.78121253 | NA | sre-32 |
| WBGene00012289 | 1.535316168 | 5.32E-05 | 0.00055271 | W05H12.1 |
| WBGene00012290 | 0.020925399 | 0.84643798 | NA | W05H12.2 |
| WBGene00012291 | -0.456267062 | 0.03603617 | 0.11845657 | W06A7.2 |
| WBGene00012292 | NA | NA | NA | NA |
| WBGene00012293 | -0.344556231 | 0.13414328 | 0.29820707 | W06A7.4 |
| WBGene00012294 | 0.137638672 | 0.33139382 | 0.53090458 | nspa-8 |
| WBGene00012295 | -0.836393735 | 0.01174281 | 0.05068587 | nmat-1 |
| WBGene00012296 | 0.119894552 | 0.45941438 | 0.6507 | spe-46 |
| WBGene00012297 | 0.035371374 | 0.66749535 | NA | W06D4.3 |
| WBGene00012298 | -0.083071386 | 0.47923819 | 0.66615262 | prmt-7 |
| WBGene00012299 | -0.071494245 | 0.65081774 | 0.79325861 | W06D11.1 |
| WBGene00012300 | -0.146640163 | 0.44816689 | 0.64088487 | W06D11.2 |
| WBGene00012301 | -0.131742958 | 0.39395391 | 0.59230634 | W06D11.3 |
| WBGene00012302 | 0.020925399 | 0.84643798 | NA | dot-1.3 |
| WBGene00012303 | NA | NA | NA | W06D11.5 |
| WBGene00012304 | NA | NA | NA | cutl-4 |
| WBGene00012305 | NA | NA | NA | W06D12.6 |
| WBGene00012306 | 1.249272181 | 2.40E-08 | 5.03E-07 | W06F12.2 |
| WBGene00012307 | 0.149813019 | 0.07294962 | NA | W06F12.3 |
| WBGene00012308 | 0.272767029 | 0.14044067 | 0.30799757 | oac-53 |
| WBGene00012309 | NA | NA | NA | W06G6.2 |
| WBGene00012310 | NA | NA | NA | mltn-11 |
| WBGene00012311 | 0.015062379 | 0.80838968 | NA | W06G6.9 |
| WBGene00012312 | 0.059762547 | 0.38965825 | NA | W06G6.10 |
| WBGene00012313 | NA | NA | NA | W06G6.11 |
| WBGene00012314 | NA | NA | NA | W06G6.12 |
| WBGene00012314 | NA | NA | NA | F14F8.15 |
| WBGene00012315 | -0.244574928 | 0.00196949 | 0.01232276 | immt-2 |
| WBGene00012316 | -0.398442182 | 8.23E-08 | 1.58E-06 | ctps-1 |
| WBGene00012317 | -0.2487141 | 0.0585661 | 0.1685603 | ztf-6 |
| WBGene00012318 | 0.056188364 | 0.42092875 | NA | srxa-1 |
| WBGene00012319 | 0.111327775 | 0.21261922 | 0.40312846 | ipla-3 |
| WBGene00012320 | 0.057565879 | 0.43429308 | NA | NA |

| | | | | |
|---|---|---|---|---|
| WBGene00012321 | -0.05076076 | 0.47342176 | NA | srxa-6 |
| WBGene00012322 | 0.205664089 | 0.31810908 | 0.5184144 | W07A12.4 |
| WBGene00012323 | 4.161000565 | 6.93E-23 | 6.00E-21 | oac-54 |
| WBGene00012324 | 0.198222795 | 0.20194456 | 0.39049949 | rhy-1 |
| WBGene00012325 | -0.081550477 | 0.65791265 | 0.79831751 | W07A12.8 |
| WBGene00012326 | -0.20929558 | 0.00266173 | 0.0158031 | W07E11.1 |
| WBGene00012328 | 0.020232312 | 0.84643798 | NA | W07G1.1 |
| WBGene00012329 | -0.008807557 | 0.92153097 | NA | sre-44 |
| WBGene00012330 | -0.375164379 | 0.01973887 | 0.0758721 | zip-3 |
| WBGene00012332 | 0.118807528 | 0.30848382 | 0.50936185 | rpi-1 |
| WBGene00012333 | 0.012730745 | 0.86589641 | NA | sre-43 |
| WBGene00012334 | NA | NA | NA | W07G1.7 |
| WBGene00012335 | 0.073887753 | 0.28837975 | NA | W07G4.1 |
| WBGene00012336 | 0.056411843 | 0.41891328 | NA | W07G4.2 |
| WBGene00012337 | -0.025583939 | 0.76370457 | 0.86603661 | W07G4.3 |
| WBGene00012339 | 0.329983318 | 0.04285197 | 0.13454766 | W07G4.5 |
| WBGene00012340 | 0.098607935 | 0.48732029 | 0.67157909 | dct-15 |
| WBGene00012341 | -0.22849795 | 0.0028809 | 0.01693265 | catp-6 |
| WBGene00012342 | -0.144050997 | 0.06726541 | 0.18599298 | mtr-4 |
| WBGene00012343 | -0.05197483 | 0.64138649 | 0.78746664 | W08E3.2 |
| WBGene00012344 | -0.387461927 | 3.32E-11 | 1.02E-09 | ola-1 |
| WBGene00012345 | -0.063178667 | 0.61562873 | 0.7693914 | W08E3.4 |
| WBGene00012346 | 0.108538718 | 0.15420469 | NA | W08G11.1 |
| WBGene00012347 | 0.207533253 | 0.07048963 | 0.19218598 | W08G11.3 |
| WBGene00012348 | -0.399629893 | 0.00043265 | 0.00342774 | pptr-1 |
| WBGene00012349 | NA | NA | NA | W08G11.5 |
| WBGene00012350 | NA | NA | NA | NA |
| WBGene00012351 | -0.20844196 | 0.01054968 | 0.04661203 | W09C5.1 |
| WBGene00012352 | -0.16854481 | 0.33120201 | 0.53077152 | dkf-1 |
| WBGene00012353 | -0.241769838 | 0.01489845 | 0.06154859 | W09C5.7 |
| WBGene00012354 | -0.010219926 | 0.87017721 | 0.93075432 | cox-4 |
| WBGene00012355 | NA | NA | NA | NA |
| WBGene00012356 | 0.346114179 | 0.03462769 | 0.11516075 | W09D6.1 |
| WBGene00012357 | 0.239118525 | 0.04923912 | 0.14860045 | W09D6.4 |
| WBGene00012358 | 0.495776127 | 0.00033002 | 0.00272831 | W09D6.5 |
| WBGene00012359 | 0.014173249 | 0.89158114 | 0.9434794 | W09D10.1 |
| WBGene00012360 | -0.002726698 | 0.97049096 | 0.98379549 | tat-3 |
| WBGene00012361 | -0.33355089 | 0.00325446 | 0.01863454 | mrpl-12 |
| WBGene00012362 | -0.071451018 | 0.4801528 | 0.66665106 | W09D10.4 |
| WBGene00012363 | -0.060400417 | 0.75903065 | 0.86374691 | W09D10.5 |
| WBGene00012364 | -0.04572804 | 0.72493984 | 0.84572464 | zmp-4 |
| WBGene00012365 | 0.002313469 | 0.98924526 | 0.99398816 | W09G3.1 |
| WBGene00012366 | -0.072296885 | 0.66357415 | 0.80259308 | maea-1 |

| | | | | |
|---|---|---|---|---|
| WBGene00012368 | 0.170625272 | 0.42673943 | 0.62236199 | NA |
| WBGene00012369 | 0.102911854 | 0.25701621 | 0.45523523 | W09G3.6 |
| WBGene00012370 | 0.018917614 | 0.91530403 | 0.95285597 | W09G3.7 |
| WBGene00012371 | -0.014643403 | 0.93656299 | 0.96422324 | W09G3.8 |
| WBGene00012372 | -0.015946848 | 0.89462076 | 0.94489559 | W09H1.1 |
| WBGene00012373 | 0.063731418 | 0.41159439 | NA | W09H1.3 |
| WBGene00012374 | 0.30372333 | 0.08284245 | 0.21597158 | W09H1.4 |
| WBGene00012375 | 0.007478274 | 0.94083169 | 0.96674008 | mecr-1 |
| WBGene00012376 | -0.283517839 | 0.00023181 | 0.002 | nduf-7 |
| WBGene00012377 | NA | NA | NA | NA |
| WBGene00012378 | 0.065014541 | 0.37399213 | NA | W10G6.1 |
| WBGene00012379 | 0.193624399 | 0.07476626 | 0.2010662 | Y1A5A.1 |
| WBGene00012380 | NA | NA | NA | Y1A5A.2 |
| WBGene00012381 | 0.260007756 | 0.22725805 | 0.42094567 | Y2H9A.4 |
| WBGene00012382 | 0.205038666 | 0.00142393 | 0.00940917 | ttr-16 |
| WBGene00012383 | 0.12432891 | 0.37030346 | 0.57006877 | ttr-17 |
| WBGene00012384 | NA | NA | NA | NA |
| WBGene00012385 | -0.061905481 | 0.59600294 | 0.75485113 | Y5F2A.4 |
| WBGene00012386 | -0.182852437 | 0.02826873 | 0.09920481 | agef-1 |
| WBGene00012387 | 0.534948611 | 0.05625562 | 0.16426777 | Y6B3B.1 |
| WBGene00012388 | NA | NA | NA | Y6B3B.3 |
| WBGene00012389 | 0.065774112 | 0.49467689 | 0.67742391 | Y6B3B.4 |
| WBGene00012390 | 0.207843373 | 0.28820697 | 0.48849576 | gpap-1 |
| WBGene00012391 | 0.148742837 | 0.12742652 | 0.28874635 | Y6B3B.7 |
| WBGene00012393 | -0.088413554 | 0.6348557 | 0.78338999 | Y6B3B.9 |
| WBGene00012394 | NA | NA | NA | hsd-1 |
| WBGene00012395 | -0.209071257 | 0.08096767 | 0.21275084 | Y6D1A.1 |
| WBGene00012396 | -0.039790355 | 0.7597696 | 0.86405055 | Y6D1A.2 |
| WBGene00012397 | NA | NA | NA | Y6E2A.1 |
| WBGene00012398 | 4.060200724 | 0.00134014 | 0.00893489 | Y6E2A.4 |
| WBGene00012399 | 2.341117405 | 0.00683087 | 0.03341488 | Y6E2A.5 |
| WBGene00012400 | 0.040107227 | 0.58693163 | NA | Y6E2A.7 |
| WBGene00012401 | 0.040107227 | 0.58693163 | NA | irld-57 |
| WBGene00012402 | 0.079249556 | 0.309101 | NA | sfxn-1.3 |
| WBGene00012403 | -0.022930767 | 0.78121253 | NA | NA |
| WBGene00012404 | 2.348523162 | 1.15E-07 | 2.14E-06 | Y6G8.2 |
| WBGene00012405 | 0.489116034 | 0.01155144 | 0.05005181 | ztf-25 |
| WBGene00012406 | 0.020925399 | 0.84643798 | NA | srz-91 |
| WBGene00012407 | 0.221450882 | 0.0412373 | 0.13076787 | Y7A5A.1 |
| WBGene00012408 | 0.03712364 | 0.66489096 | NA | Y7A5A.2 |
| WBGene00012409 | 0.055597606 | 0.44564657 | NA | Y7A5A.3 |
| WBGene00012411 | NA | NA | NA | Y7A5A.5 |
| WBGene00012412 | 0.054509443 | 0.4306858 | NA | Y7A5A.6 |

| | | | | |
|---|---|---|---|---|
| WBGene00012413 | NA | NA | NA | Y7A5A.7 |
| WBGene00012414 | 0.020925399 | 0.84643798 | NA | Y7A5A.8 |
| WBGene00012415 | -0.022930767 | 0.78121253 | NA | Y7A5A.9 |
| WBGene00012416 | 1.007481532 | 0.00147823 | 0.00970961 | Y7A9A.1 |
| WBGene00012418 | 1.143047095 | 0.00172967 | 0.01107791 | NA |
| WBGene00012419 | -0.001488154 | 0.95639912 | NA | NA |
| WBGene00012420 | -0.665021008 | 0.03052888 | 0.1047368 | Y7A9D.1 |
| WBGene00012421 | 1.206596784 | 0.00010663 | 0.00102911 | Y9C2UA.1 |
| WBGene00012423 | 0.010380561 | 0.91756396 | 0.95412315 | Y10G11A.1 |
| WBGene00012424 | 0.399597574 | 0.11170268 | 0.26412127 | dlc-3 |
| WBGene00012425 | 0.015896709 | 0.80422001 | NA | dlc-4 |
| WBGene00012426 | 0.057855329 | 0.58501057 | 0.74805087 | NA |
| WBGene00012428 | -0.205664794 | 0.16215262 | 0.3373226 | Y11D7A.3 |
| WBGene00012429 | 0.092724492 | 0.19640118 | NA | Y11D7A.5 |
| WBGene00012431 | 0.3533952 | 0.0418912 | 0.13221148 | Y11D7A.7 |
| WBGene00012432 | 0.189258199 | 0.1649913 | 0.34123996 | Y11D7A.8 |
| WBGene00012433 | 0.186901046 | 0.23721109 | 0.43352983 | Y11D7A.9 |
| WBGene00012434 | -0.041573793 | 0.73777207 | 0.85261037 | Y11D7A.10 |
| WBGene00012435 | -0.258426154 | 0.00064988 | 0.00485948 | flh-1 |
| WBGene00012436 | -0.384391778 | 0.00122344 | 0.00830428 | flh-3 |
| WBGene00012437 | 1.155646416 | 0.00307181 | 0.0178046 | nmy-3 |
| WBGene00012439 | -0.250902476 | 0.22231053 | 0.41503454 | Y12A6A.1 |
| WBGene00012440 | 0.60818171 | 0.04259205 | 0.13387933 | nlp-53 |
| WBGene00012441 | 0.113207936 | 0.37685042 | 0.5758759 | glb-28 |
| WBGene00012442 | 0.076179833 | 0.65609072 | 0.79670103 | Y15E3A.3 |
| WBGene00012443 | 0.232796332 | 0.1608273 | 0.33544941 | Y15E3A.4 |
| WBGene00012444 | 0.478264519 | 0.05499083 | 0.16141153 | Y15E3A.5 |
| WBGene00012445 | 0.596016384 | 5.82E-08 | 1.14E-06 | Y16B4A.2 |
| WBGene00012446 | -0.008428826 | 0.96489014 | 0.98071026 | nhr-230 |
| WBGene00012448 | 0.263167517 | 0.09544656 | 0.23757788 | cyp-33D3 |
| WBGene00012449 | 0.200733583 | 0.19441544 | 0.3812532 | nhr-231 |
| WBGene00012450 | 0.013204071 | 0.88957758 | NA | Y17D7B.2 |
| WBGene00012451 | 0.040019663 | 0.56356856 | NA | Y17D7B.3 |
| WBGene00012452 | 0.758173443 | 0.00260584 | 0.01555323 | Y17D7B.4 |
| WBGene00012453 | NA | NA | NA | NA |
| WBGene00012454 | -0.023109762 | 0.78121253 | NA | clec-256 |
| WBGene00012455 | 0.020232312 | 0.84643798 | NA | Y17D7C.1 |
| WBGene00012456 | -0.026384613 | 0.67299636 | NA | Y17D7C.2 |
| WBGene00012457 | -0.458026577 | 0.07906552 | 0.20943503 | acbp-6 |
| WBGene00012458 | 0.012961582 | 0.91201396 | 0.95112162 | ash-2 |
| WBGene00012459 | 0.093782373 | 0.6116477 | 0.7663446 | Y17G7B.3 |
| WBGene00012460 | -0.175817149 | 0.22268433 | 0.41543569 | dhps-1 |
| WBGene00012461 | 0.184155824 | 0.20441261 | 0.39347949 | glb-29 |

| | | | | |
|---|---|---|---|---|
| WBGene00012462 | 0.957375089 | 8.61E-05 | 0.00084657 | Y17G7B.8 |
| WBGene00012463 | 0.601317499 | 4.17E-06 | 5.56E-05 | Y17G7B.10 |
| WBGene00012464 | 0.451083727 | 0.0522052 | 0.15551095 | arrd-7 |
| WBGene00012465 | -0.055233028 | 0.6905593 | 0.82232442 | Y17G7B.12 |
| WBGene00012466 | -0.084851918 | 0.45916058 | 0.65047359 | ippk-1 |
| WBGene00012467 | 0.065014541 | 0.37399213 | NA | arrd-8 |
| WBGene00012468 | 0.023422257 | 0.86067478 | 0.92614578 | Y17G7B.17 |
| WBGene00012469 | -0.104256639 | 0.45991946 | 0.65119895 | Y17G7B.18 |
| WBGene00012470 | -0.034792449 | 0.7155449 | NA | Y17G7B.19 |
| WBGene00012471 | -0.166046755 | 0.01645597 | 0.06656443 | Y17G7B.20 |
| WBGene00012472 | -0.031335582 | 0.74557801 | 0.85598121 | Y17G7B.21 |
| WBGene00012473 | -0.049278988 | 0.46988128 | NA | Y17G7B.22 |
| WBGene00012474 | -0.017406491 | 0.86844119 | 0.92992376 | attf-6 |
| WBGene00012475 | NA | NA | NA | Y18D10A.2 |
| WBGene00012476 | -0.156111446 | 0.46338029 | 0.65334658 | Y18D10A.3 |
| WBGene00012477 | NA | NA | NA | Y18D10A.4 |
| WBGene00012478 | 0.712535943 | 0.00119974 | 0.00816996 | mxt-1 |
| WBGene00012479 | -0.063256666 | 0.68092659 | 0.81536323 | Y18D10A.9 |
| WBGene00012480 | -0.08047733 | 0.5076186 | 0.6880885 | clec-104 |
| WBGene00012481 | -0.465495783 | 1.46E-11 | 4.72E-10 | Y18D10A.11 |
| WBGene00012482 | -0.03039785 | 0.70603871 | NA | clec-106 |
| WBGene00012483 | -0.257874014 | 0.09211847 | 0.23149786 | coa-5 |
| WBGene00012484 | -0.101475315 | 0.01932859 | 0.07494437 | car-1 |
| WBGene00012485 | 0.215376828 | 0.32016895 | 0.52007255 | NA |
| WBGene00012486 | 0.057303378 | 0.52744004 | NA | Y18D10A.21 |
| WBGene00012487 | -0.011947027 | 0.93915233 | 0.96586959 | Y18D10A.23 |
| WBGene00012488 | 0.020232312 | 0.84643798 | NA | clec-105 |
| WBGene00012489 | 0.020232312 | 0.84643798 | NA | Y19D2B.1 |
| WBGene00012490 | -0.227974216 | 0.27031377 | 0.46958518 | Y19D2B.2 |
| WBGene00012491 | 1.531622275 | 0.00041226 | 0.00330375 | Y20C6A.1 |
| WBGene00012492 | -0.359090403 | 0.11070055 | 0.26247236 | fbxb-7 |
| WBGene00012493 | NA | NA | NA | NA |
| WBGene00012494 | 0.646084328 | 0.01273478 | 0.0542321 | nhr-232 |
| WBGene00012496 | -0.098911054 | 0.32391458 | 0.52363026 | Y24F12A.1 |
| WBGene00012497 | -0.113026533 | 0.12331463 | 0.28329518 | ragc-1 |
| WBGene00012498 | -0.102967265 | 0.62198047 | 0.7740679 | NA |
| WBGene00012499 | 0.124703695 | 0.55529299 | 0.72706534 | NA |
| WBGene00012500 | 0.032275048 | 0.74536702 | 0.85598121 | hpo-2 |
| WBGene00012501 | 0.078714516 | 0.25916069 | NA | Y26D4A.3 |
| WBGene00012502 | NA | NA | NA | clec-107 |
| WBGene00012503 | NA | NA | NA | Y26D4A.5 |
| WBGene00012504 | NA | NA | NA | clec-108 |
| WBGene00012505 | 0.020925399 | 0.84643798 | NA | pals-13 |

| | | | | |
|---|---|---|---|---|
| WBGene00012509 | 0.020317245 | 0.84643798 | NA | Y26D4A.12 |
| WBGene00012512 | -0.168065589 | 0.16169719 | 0.33656647 | ekl-5 |
| WBGene00012513 | 0.047808857 | 0.81086492 | 0.89531472 | Y26E6A.2 |
| WBGene00012514 | -0.227511907 | 0.27292955 | 0.4730068 | Y26E6A.3 |
| WBGene00012515 | NA | NA | NA | Y26G10.1 |
| WBGene00012516 | NA | NA | NA | sqst-4 |
| WBGene00012517 | NA | NA | NA | NA |
| WBGene00012518 | 0.167544192 | 0.09772238 | 0.24217221 | comt-2 |
| WBGene00012519 | NA | NA | NA | Y32B12A.4 |
| WBGene00012520 | NA | NA | NA | Y32B12A.5 |
| WBGene00012521 | -0.045959069 | 0.50707733 | NA | Y32B12B.1 |
| WBGene00012522 | -0.094204814 | 0.32881105 | 0.52822125 | Y32B12B.2 |
| WBGene00012523 | 0.020232312 | 0.84643798 | NA | srbc-65 |
| WBGene00012524 | -0.04828078 | 0.64699198 | 0.79116425 | Y32B12B.4 |
| WBGene00012525 | 0.353813679 | 0.13591876 | 0.30089333 | Y32B12C.1 |
| WBGene00012526 | -0.087685709 | 0.54616546 | 0.71941196 | Y32B12C.3 |
| WBGene00012527 | -0.298993672 | 0.02144373 | 0.08085439 | set-22 |
| WBGene00012528 | -0.032515812 | 0.68114188 | 0.81548763 | pap-1 |
| WBGene00012529 | 0.189085524 | 0.21974705 | 0.41194834 | Y32F6A.4 |
| WBGene00012530 | 0.211423939 | 0.01617797 | 0.06568542 | Y32F6A.5 |
| WBGene00012531 | 0.220461939 | 0.23230742 | 0.42728806 | Y32F6B.1 |
| WBGene00012532 | 0.610970058 | 0.0338051 | 0.11302591 | crp-1 |
| WBGene00012533 | 0.285518125 | 0.1379423 | 0.3038383 | fbxa-209 |
| WBGene00012535 | -0.759174862 | 0.01345276 | 0.05671232 | Y37A1A.2 |
| WBGene00012536 | 0.201686314 | 0.35699103 | 0.55703993 | Y37A1A.3 |
| WBGene00012537 | NA | NA | NA | Y37A1B.4 |
| WBGene00012538 | 1.784362204 | 1.73E-53 | 4.28E-51 | Y37A1B.5 |
| WBGene00012540 | 0.239907249 | 0.01930642 | 0.07487821 | Y37A1B.7 |
| WBGene00012541 | NA | NA | NA | irld-55 |
| WBGene00012542 | 0.020925399 | 0.84643798 | NA | irld-56 |
| WBGene00012543 | -0.056066704 | 0.52057546 | 0.69924269 | nkcc-1 |
| WBGene00012544 | 0.531254384 | 2.05E-14 | 9.17E-13 | Y37D8A.2 |
| WBGene00012545 | -0.251343612 | 0.20043185 | 0.38900004 | Y37D8A.3 |
| WBGene00012546 | 0.268904984 | 0.02151142 | 0.081047 | Y37D8A.4 |
| WBGene00012547 | 0.31520867 | 0.06166883 | 0.17508681 | Y37D8A.5 |
| WBGene00012548 | 0.658859644 | 0.00806378 | 0.03810905 | Y37D8A.6 |
| WBGene00012549 | 0.120387617 | 0.35614301 | 0.55607228 | Y37D8A.8 |
| WBGene00012550 | -0.158185245 | 0.03555302 | 0.11736907 | spcs-2 |
| WBGene00012551 | 0.140008581 | 0.21171869 | 0.40188946 | cec-7 |
| WBGene00012552 | 0.162916553 | 0.43303129 | 0.62797258 | enu-3.5 |
| WBGene00012553 | -0.096438684 | 0.10947468 | 0.26057856 | cox-5A |
| WBGene00012554 | 1.24664249 | 1.21E-06 | 1.86E-05 | Y37D8A.16 |
| WBGene00012555 | -0.372247067 | 0.00262818 | 0.01564193 | npp-25 |

| | | | | |
|---|---|---|---|---|
| WBGene00012556 | -0.412621042 | 0.02774182 | 0.097919 | mrps-10 |
| WBGene00012557 | 0.438808641 | 4.76E-09 | 1.12E-07 | Y37D8A.19 |
| WBGene00012558 | 0.030337087 | 0.81656586 | 0.89916823 | rbm-7 |
| WBGene00012559 | -0.068834434 | 0.50692823 | 0.6876578 | epg-3 |
| WBGene00012560 | 0.038058219 | 0.85128858 | 0.92034307 | Y37D8A.25 |
| WBGene00012561 | -0.027373974 | 0.68245543 | NA | Y37D8A.26 |
| WBGene00012562 | -0.225508923 | 0.06347969 | 0.17873194 | Y37H2A.1 |
| WBGene00012564 | 1.512896873 | 2.68E-08 | 5.57E-07 | fbxa-107 |
| WBGene00012565 | -0.166601452 | 0.16116472 | 0.33571074 | fbxa-210 |
| WBGene00012566 | -0.287982266 | 0.16124926 | 0.3358251 | fbxa-211 |
| WBGene00012567 | 0.009169979 | 0.96508678 | 0.98071026 | Y37H2A.7 |
| WBGene00012568 | 0.33025668 | 0.15397853 | 0.32715947 | NA |
| WBGene00012569 | 0.352817678 | 0.12464638 | 0.28511378 | NA |
| WBGene00012570 | NA | NA | NA | Y37H2A.10 |
| WBGene00012571 | NA | NA | NA | Y37H2A.11 |
| WBGene00012572 | -0.031417647 | 0.69702864 | NA | NA |
| WBGene00012573 | 2.337795593 | 0.00312436 | 0.01801605 | Y37H2C.1 |
| WBGene00012574 | NA | NA | NA | fbxa-213 |
| WBGene00012575 | NA | NA | NA | Y37H2C.4 |
| WBGene00012576 | 0.135138903 | 0.48514944 | 0.66991162 | Y37H9A.1 |
| WBGene00012577 | 0.010005953 | 0.92153544 | 0.95613845 | Y37H9A.2 |
| WBGene00012578 | 0.22271344 | 0.12102558 | 0.27992663 | Y37H9A.3 |
| WBGene00012579 | NA | NA | NA | Y38E10A.1 |
| WBGene00012580 | NA | NA | NA | Y38E10A.2 |
| WBGene00012581 | -0.107684206 | 0.58962993 | 0.75112054 | Y38E10A.3 |
| WBGene00012582 | 0.488871102 | 0.06442059 | 0.18068618 | clec-8 |
| WBGene00012583 | 0.767782053 | 0.00704132 | 0.03415905 | clec-4 |
| WBGene00012584 | 0.087235659 | 0.39796914 | 0.59623206 | ceh-100 |
| WBGene00012585 | 1.17742476 | 1.25E-17 | 7.76E-16 | lips-15 |
| WBGene00012586 | -0.022930767 | 0.78121253 | NA | Y38E10A.8 |
| WBGene00012587 | 0.014067275 | 0.88561763 | NA | Y38E10A.9 |
| WBGene00012588 | 0.300868522 | 0.17358284 | 0.35400841 | lips-16 |
| WBGene00012589 | 1.09086173 | 0.0123206 | 0.05277527 | nspe-4 |
| WBGene00012590 | NA | NA | NA | nspe-3 |
| WBGene00012591 | -0.073506567 | 0.68411407 | 0.8176383 | nspe-1 |
| WBGene00012592 | -1.074211755 | 7.60E-06 | 9.59E-05 | Y38E10A.14 |
| WBGene00012593 | -2.772593931 | 0.00111388 | 0.00769623 | nspe-7 |
| WBGene00012594 | -0.071242738 | 0.3311583 | NA | nspe-5 |
| WBGene00012595 | 1.630352868 | 0.00253646 | 0.01518245 | Y38E10A.17 |
| WBGene00012596 | -0.002914971 | 0.95718083 | NA | nhr-234 |
| WBGene00012597 | -0.098125101 | 0.31335306 | 0.5138324 | nhr-235 |
| WBGene00012600 | -0.1687441 | 0.07893412 | 0.20924759 | Y38E10A.22 |
| WBGene00012602 | -0.193551928 | 0.10455166 | 0.25284464 | Y38E10A.24 |

| | | | | |
|---|---|---|---|---|
| WBGene00012603 | 0.020232312 | 0.84643798 | NA | nspe-6 |
| WBGene00012604 | 0.232017506 | 0.28715267 | 0.48733074 | nspe-2 |
| WBGene00012605 | 0.158202224 | 0.36917627 | 0.56913865 | Y38F1A.1 |
| WBGene00012606 | 0.311187134 | 0.17873254 | 0.36014632 | Y38F1A.2 |
| WBGene00012607 | 0.167383987 | 0.30012725 | 0.50064522 | Y38F1A.4 |
| WBGene00012608 | -0.016822594 | 0.76202164 | 0.86495294 | hphd-1 |
| WBGene00012609 | -0.023109762 | 0.78121253 | NA | Y38F1A.7 |
| WBGene00012610 | 0.370974111 | 0.12562896 | 0.28666356 | Y38F1A.8 |
| WBGene00012612 | NA | NA | NA | Y38H6A.1 |
| WBGene00012614 | NA | NA | NA | Y38H6A.3 |
| WBGene00012615 | -0.450936942 | 4.78E-16 | 2.55E-14 | dct-16 |
| WBGene00012616 | 0.020232312 | 0.84643798 | NA | dct-14 |
| WBGene00012617 | NA | NA | NA | Y38H6C.4 |
| WBGene00012618 | 0.033463284 | 0.6365037 | NA | dct-10 |
| WBGene00012620 | 0.020317245 | 0.84643798 | NA | NA |
| WBGene00012621 | -0.025226573 | 0.69184128 | NA | Y38H6C.8 |
| WBGene00012622 | -0.102475475 | 0.49356007 | 0.67654836 | Y38H6C.9 |
| WBGene00012623 | 1.234646588 | 0.00102784 | 0.00718998 | fbxa-149 |
| WBGene00012624 | 0.131764043 | 0.53332139 | 0.70899686 | fbxa-150 |
| WBGene00012625 | -0.002950686 | 0.96771748 | NA | Y38H6C.13 |
| WBGene00012626 | 0.190357669 | 0.25036685 | 0.44895131 | famh-161 |
| WBGene00012627 | 0.017141582 | 0.79355725 | NA | Y38H6C.15 |
| WBGene00012628 | 7.73E-05 | 0.99861971 | 0.99923396 | Y38H6C.16 |
| WBGene00012629 | 0.523229723 | 0.03584019 | 0.11799778 | slc-36.3 |
| WBGene00012630 | -0.023109762 | 0.78121253 | NA | Y38H6C.18 |
| WBGene00012631 | 2.125252916 | 3.23E-09 | 7.65E-08 | Y38H6C.19 |
| WBGene00012632 | 0.572569154 | 0.03459033 | 0.11507316 | Y38H6C.20 |
| WBGene00012633 | 1.735060829 | 1.20E-06 | 1.85E-05 | Y38H6C.21 |
| WBGene00012634 | 1.6700878 | 1.88E-07 | 3.37E-06 | Y38H6C.23 |
| WBGene00012635 | -0.051779043 | 0.59330596 | 0.7534436 | Y38H8A.1 |
| WBGene00012636 | 0.018998992 | 0.77286019 | NA | Y38H8A.2 |
| WBGene00012637 | 1.760091679 | 0.01293777 | 0.05495266 | Y38H8A.3 |
| WBGene00012638 | 0.100264625 | 0.38646629 | 0.5851299 | Y38H8A.4 |
| WBGene00012639 | -0.054790336 | 0.58423077 | 0.74778226 | fezf-1 |
| WBGene00012640 | -0.37781052 | 0.03361669 | 0.11255018 | Y38H8A.7 |
| WBGene00012641 | -0.106430147 | 0.24903741 | 0.44749101 | epg-6 |
| WBGene00012642 | 0.036821884 | 0.59344795 | NA | Y39A1A.2 |
| WBGene00012643 | -0.203807036 | 0.14035253 | 0.30785039 | Y39A1A.3 |
| WBGene00012644 | 0.109561741 | 0.33584521 | 0.53516648 | rabx-5 |
| WBGene00012645 | -0.269106897 | 0.10692594 | 0.25676607 | mrpl-22 |
| WBGene00012646 | 0.07290376 | 0.5184404 | 0.69772333 | lron-10 |
| WBGene00012647 | -0.136518851 | 0.51674743 | 0.69614457 | swt-4 |
| WBGene00012648 | 0.20582614 | 0.19176731 | 0.3779482 | Y39A1A.9 |

| | | | | |
|---|---|---|---|---|
| WBGene00012649 | NA | NA | NA | Y39A1A.10 |
| WBGene00012650 | -0.157354475 | 0.04698041 | 0.14380037 | orc-1 |
| WBGene00012651 | -0.019989768 | 0.86459938 | 0.92776221 | orc-4 |
| WBGene00012652 | -0.388798474 | 0.0017577 | 0.01123286 | Y39A1A.14 |
| WBGene00012653 | 0.020232312 | 0.84643798 | NA | NA |
| WBGene00012654 | -0.54085794 | 0.0506926 | 0.15190141 | Y39A1A.16 |
| WBGene00012655 | 0.05112733 | 0.65340503 | 0.79469207 | Y39A1A.17 |
| WBGene00012656 | -0.023056499 | 0.78121253 | NA | Y39A1A.18 |
| WBGene00012657 | -0.169221268 | 0.42879924 | 0.62368862 | Y39A1A.20 |
| WBGene00012658 | -0.459683758 | 0.00703957 | 0.03415905 | Y39A1A.21 |
| WBGene00012659 | -0.134634628 | 0.19485801 | 0.38181426 | Y39A1A.22 |
| WBGene00012660 | NA | NA | NA | Y39A1A.24 |
| WBGene00012661 | -0.022930767 | 0.78121253 | NA | clec-163 |
| WBGene00012662 | -0.036806978 | 0.84092677 | 0.91408863 | Y39A1C.1 |
| WBGene00012663 | -0.017346276 | 0.8550596 | 0.92241889 | hex-3 |
| WBGene00012664 | 2.621529492 | 0 | 0 | Y39B6A.1 |
| WBGene00012665 | -0.237067819 | 0.0106176 | 0.04680887 | pph-5 |
| WBGene00012666 | 0.056863947 | 0.70220797 | 0.83093594 | Y39B6A.3 |
| WBGene00012668 | 0.459186675 | 0.02445703 | 0.08934017 | Y39B6A.5 |
| WBGene00012669 | 0.036935836 | 0.83098026 | 0.90780346 | Y39B6A.7 |
| WBGene00012670 | 0.038562771 | 0.65069375 | NA | Y39B6A.8 |
| WBGene00012671 | -0.129337259 | 0.26928973 | 0.46836195 | Y39B6A.9 |
| WBGene00012672 | -0.19705651 | 0.08809719 | 0.22512414 | Y39B6A.10 |
| WBGene00012673 | -0.09746693 | 0.63733951 | 0.78499939 | dyf-17 |
| WBGene00012674 | -0.122924752 | 0.46107834 | 0.65192158 | bed-1 |
| WBGene00012675 | 0.016720015 | 0.91254899 | 0.95126775 | Y39B6A.13 |
| WBGene00012676 | -0.325957685 | 0.01964888 | 0.07564533 | pro-3 |
| WBGene00012678 | -0.068014835 | 0.71069889 | 0.8370479 | zhp-4 |
| WBGene00012679 | 0.0398788 | 0.58971221 | NA | Y39B6A.18 |
| WBGene00012680 | -0.122004589 | 0.54351416 | 0.71696212 | NA |
| WBGene00012681 | -0.003421551 | 0.95689871 | NA | asp-15 |
| WBGene00012682 | 0.020232312 | 0.84643798 | NA | asp-16 |
| WBGene00012683 | 1.7068543 | 0.0004379 | 0.00346554 | asp-17 |
| WBGene00012684 | -0.007072416 | 0.9244612 | NA | Y39B6A.25 |
| WBGene00012686 | 0.718581909 | 0.01056446 | 0.04663254 | Y39B6A.27 |
| WBGene00012688 | 0.186648832 | 0.19378656 | 0.38068282 | Y39B6A.29 |
| WBGene00012689 | 2.681791036 | 0.00459766 | 0.02454589 | Y39B6A.30 |
| WBGene00012690 | 0.127299467 | 0.16839118 | NA | NA |
| WBGene00012692 | -0.524916563 | 6.29E-05 | 0.00064473 | Y39B6A.33 |
| WBGene00012693 | -0.160645743 | 0.31922539 | 0.51940343 | Y39B6A.34 |
| WBGene00012694 | -0.077279393 | 0.46660045 | 0.65621087 | gtf-2F2 |
| WBGene00012695 | -0.095182118 | 0.59042793 | 0.75162641 | Y39B6A.37 |
| WBGene00012696 | 0.026252451 | 0.89154437 | 0.9434794 | reps-1 |

| | | | | |
|---|---|---|---|---|
| WBGene00012697 | -0.338815118 | 0.00207176 | 0.01285828 | mrps-35 |
| WBGene00012698 | 0.034321451 | 0.8593551 | 0.92479368 | sws-1 |
| WBGene00012699 | 0.212887363 | 0.32356777 | 0.52340689 | Y39B6A.41 |
| WBGene00012700 | -0.062804712 | 0.5801958 | 0.745406 | Y39B6A.42 |
| WBGene00012701 | 0.094834361 | 0.50527041 | 0.68636705 | Y39B6A.43 |
| WBGene00012702 | 0.423987219 | 0.03954183 | 0.12695875 | ztf-20 |
| WBGene00012703 | -0.147204044 | 0.48539573 | 0.66999896 | nhr-145 |
| WBGene00012704 | 0.024098832 | 0.90873956 | 0.95061846 | mlt-3 |
| WBGene00012705 | 0.017279211 | 0.79380376 | NA | sre-18 |
| WBGene00012706 | NA | NA | NA | NA |
| WBGene00012707 | 0.018860755 | 0.77262519 | NA | sre-16 |
| WBGene00012708 | -0.006392486 | 0.92872129 | NA | sre-19 |
| WBGene00012709 | 0.014087259 | 0.85351751 | NA | sre-15 |
| WBGene00012710 | -0.126268522 | 0.51950772 | 0.69857757 | Y39C12A.9 |
| WBGene00012711 | 0.065014541 | 0.37399213 | NA | Y39E4A.1 |
| WBGene00012712 | 0.68210316 | 0.00014106 | 0.00129894 | ttm-1 |
| WBGene00012713 | -0.155343206 | 0.08837498 | 0.22557497 | bckd-1A |
| WBGene00012714 | -0.378492532 | 3.88E-07 | 6.57E-06 | abce-1 |
| WBGene00012715 | -0.336701565 | 0.00018179 | 0.00162381 | snpc-1.2 |
| WBGene00012716 | -0.236081016 | 0.00902623 | 0.04150534 | Y39E4B.5 |
| WBGene00012717 | -0.139164071 | 0.36342112 | 0.56323156 | Y39E4B.6 |
| WBGene00012718 | -0.059419294 | 0.61632781 | 0.76993648 | dhhc-8 |
| WBGene00012719 | -0.20478647 | 0.10028111 | 0.24593229 | Y39E4B.10 |
| WBGene00012720 | 0.179107049 | 0.15052123 | 0.32214315 | Y39E4B.11 |
| WBGene00012721 | 0.094973622 | 0.44189603 | 0.63595638 | Y39E4B.13 |
| WBGene00012722 | -0.133626697 | 0.31354926 | 0.51405222 | Y39G8B.1 |
| WBGene00012723 | -0.071823007 | 0.71898342 | 0.84237564 | Y39G8B.2 |
| WBGene00012724 | NA | NA | NA | sre-48 |
| WBGene00012725 | 0.036687355 | 0.59489568 | NA | sre-47 |
| WBGene00012726 | 0.06111288 | 0.51882185 | NA | Y39G8B.5 |
| WBGene00012727 | -0.027350315 | 0.68023343 | NA | Y39G8B.7 |
| WBGene00012728 | 2.619392793 | 0.00392993 | 0.02158216 | Y39G8B.9 |
| WBGene00012729 | NA | NA | NA | Y39G8B.10 |
| WBGene00012730 | -0.073028493 | 0.3595958 | 0.55958116 | xrn-1 |
| WBGene00012731 | 0.015551528 | 0.84511556 | NA | Y39G8C.2 |
| WBGene00012732 | 0.110867301 | 0.5066609 | 0.68739475 | Y40B1A.1 |
| WBGene00012734 | -0.013093083 | 0.92681879 | 0.9586234 | Y40B1A.3 |
| WBGene00012735 | -0.074053686 | 0.46440146 | 0.6542823 | sptf-3 |
| WBGene00012738 | -0.633948031 | 2.84E-07 | 4.92E-06 | eif-3.J |
| WBGene00012739 | -0.273981313 | 0.01779905 | 0.07060307 | Y40B1B.7 |
| WBGene00012740 | 0.034462747 | 0.75887573 | 0.86363774 | slc-25A46 |
| WBGene00012741 | 0.175525442 | 0.04514366 | 0.13961729 | Y40H4A.2 |
| WBGene00012742 | NA | NA | NA | Y40H7A.2 |

| | | | | |
|---|---|---|---|---|
| WBGene00012742 | NA | NA | NA | Y40H7A.3 |
| WBGene00012743 | 0.020232312 | 0.84643798 | NA | Y40H7A.2 |
| WBGene00012743 | 0.020232312 | 0.84643798 | NA | Y40H7A.3 |
| WBGene00012744 | 0.08687367 | 0.29583168 | NA | Y40H7A.4 |
| WBGene00012745 | 0.084908438 | 0.30232871 | NA | NA |
| WBGene00012746 | NA | NA | NA | Y40H7A.9 |
| WBGene00012747 | 0.519816371 | 3.52E-06 | 4.77E-05 | Y40H7A.10 |
| WBGene00012748 | 0.310827113 | 0.0817468 | 0.21402984 | Y40H7A.11 |
| WBGene00012749 | -0.023109762 | 0.78121253 | NA | NA |
| WBGene00012750 | 0.237114345 | 0.26744288 | 0.46635384 | faah-6 |
| WBGene00012752 | NA | NA | NA | NA |
| WBGene00012753 | 0.498380229 | 0.06585803 | 0.18320871 | Y41C4A.6 |
| WBGene00012754 | 0.042199095 | 0.55922995 | NA | Y41C4A.7 |
| WBGene00012755 | 0.075553414 | 0.71790138 | 0.84178346 | Y41C4A.8 |
| WBGene00012756 | 0.024827989 | 0.85933471 | 0.92479368 | Y41C4A.9 |
| WBGene00012757 | 0.097012966 | 0.43337374 | 0.62840695 | Y41C4A.32 |
| WBGene00012757 | 0.097012966 | 0.43337374 | 0.62840695 | Y41C4A.11 |
| WBGene00012758 | 0.268745011 | 0.23320325 | 0.4286665 | Y41C4A.12 |
| WBGene00012759 | 0.36249857 | 0.06735928 | 0.18611194 | sup-1 |
| WBGene00012760 | 0.034220862 | 0.73084044 | 0.84986592 | Y41C4A.17 |
| WBGene00012761 | 0.042199095 | 0.55922995 | NA | Y41C4A.18 |
| WBGene00012762 | 0.0171241 | 0.82341871 | 0.9034778 | Y41E3.1 |
| WBGene00012763 | 0.486690257 | 0.07135278 | 0.1942138 | atln-2 |
| WBGene00012764 | 0.03555743 | 0.72324271 | 0.84472009 | Y41E3.6 |
| WBGene00012765 | -0.017608354 | 0.86149562 | 0.92662044 | Y41E3.7 |
| WBGene00012766 | 0.167013773 | 0.34329456 | 0.54313812 | Y41E3.8 |
| WBGene00012767 | 0.113465139 | 0.27540697 | 0.47572289 | fcd-2 |
| WBGene00012768 | -0.14266603 | 0.00456465 | 0.02441763 | eef-1B.2 |
| WBGene00012769 | 0.051282414 | 0.42264356 | 0.6186446 | hrpu-1 |
| WBGene00012770 | NA | NA | NA | srt-47 |
| WBGene00012771 | NA | NA | NA | Y41E3.13 |
| WBGene00012772 | 0.0398788 | 0.58971221 | NA | srt-49 |
| WBGene00012773 | NA | NA | NA | srt-48 |
| WBGene00012775 | 0.0398788 | 0.58971221 | NA | srt-50 |
| WBGene00012776 | 1.384099335 | 0.00364639 | 0.02037666 | Y42A5A.1 |
| WBGene00012777 | -0.107703593 | 0.57513174 | 0.742287 | lact-8 |
| WBGene00012778 | 0.102624893 | 0.52468283 | 0.70205458 | Y42A5A.3 |
| WBGene00012779 | 0.152188604 | 0.12115868 | 0.28014596 | cdkl-1 |
| WBGene00012780 | -0.055534994 | 0.78716286 | 0.87949869 | Y42A5A.5 |
| WBGene00012781 | 2.189694268 | 2.57E-06 | 3.62E-05 | nspd-7 |
| WBGene00012782 | 1.925643025 | 6.07E-107 | 3.55E-104 | Y43C5A.2 |
| WBGene00012783 | 1.603922962 | 1.23E-11 | 4.02E-10 | Y43C5A.3 |
| WBGene00012784 | 0.031515541 | 0.7478565 | 0.85763038 | Y43C5A.4 |

| | | | | |
|---|---|---|---|---|
| WBGene00012785 | 0.112161357 | 0.37756611 | 0.57666862 | Y43C5B.2 |
| WBGene00012786 | 0.149443023 | 0.47559756 | 0.66272236 | Y43C5B.3 |
| WBGene00012787 | 0.145661165 | 0.24853685 | 0.44692066 | Y43D4A.1 |
| WBGene00012788 | 0.056472646 | 0.63132828 | 0.78093021 | Y43D4A.2 |
| WBGene00012789 | NA | NA | NA | Y43D4A.3 |
| WBGene00012789 | NA | NA | NA | Y43D4A.4 |
| WBGene00012790 | NA | NA | NA | Y43D4A.3 |
| WBGene00012790 | NA | NA | NA | Y43D4A.4 |
| WBGene00012791 | 0.009107272 | 0.91616128 | NA | Y43D4A.5 |
| WBGene00012792 | -0.148929253 | 0.17799625 | 0.35935509 | Y43D4A.6 |
| WBGene00012793 | NA | NA | NA | NA |
| WBGene00012794 | 1.011658371 | 1.58E-05 | 0.00018497 | Y43E12A.2 |
| WBGene00012795 | 0.009713829 | 0.95337892 | 0.97339057 | Y43E12A.3 |
| WBGene00012796 | -0.028867812 | 0.887219 | 0.94060632 | Y43F4A.1 |
| WBGene00012798 | 0.020317245 | 0.84643798 | NA | Y43F4A.3 |
| WBGene00012799 | NA | NA | NA | Y43F4A.4 |
| WBGene00012802 | -0.138757711 | 0.44589581 | 0.63930108 | set-25 |
| WBGene00012803 | -0.194785022 | 0.01339305 | 0.05652572 | Y43F4B.5 |
| WBGene00012804 | -0.256610646 | 0.03577032 | 0.11792689 | Y43F4B.7 |
| WBGene00012806 | 0.022409978 | 0.88658753 | 0.94033043 | Y43F4B.9 |
| WBGene00012807 | -0.294871946 | 0.02720101 | 0.09645194 | Y43F4B.10 |
| WBGene00012808 | 0.278247912 | 0.06457251 | 0.18104292 | Y43F8A.1 |
| WBGene00012809 | 2.079920439 | 0.00842401 | 0.03935417 | Y43F8A.2 |
| WBGene00012810 | -0.060993816 | 0.75604621 | 0.8618244 | nceh-1 |
| WBGene00012811 | 0.112487263 | 0.58138116 | 0.74581663 | Y43F8A.5 |
| WBGene00012812 | -0.218448937 | 0.08818042 | 0.22527474 | Y43F8B.1 |
| WBGene00012813 | 0.15747488 | 0.26247597 | 0.46050457 | Y43F8B.2 |
| WBGene00012814 | 1.031881419 | 0.01501991 | 0.06191063 | Y43F8B.3 |
| WBGene00012815 | -0.027535543 | 0.68063395 | NA | phy-4 |
| WBGene00012816 | NA | NA | NA | scl-21 |
| WBGene00012817 | NA | NA | NA | NA |
| WBGene00012818 | 0.020232312 | 0.84643798 | NA | Y43F8B.7 |
| WBGene00012819 | 0.445366701 | 0.08626056 | 0.22186041 | Y43F8B.9 |
| WBGene00012820 | NA | NA | NA | Y43F8B.10 |
| WBGene00012821 | 0.08118481 | 0.63952068 | 0.78619899 | Y43F8B.11 |
| WBGene00012822 | 0.028487078 | 0.82208323 | 0.90246087 | Y43F8B.12 |
| WBGene00012823 | 0.008615607 | 0.94220991 | 0.9672483 | Y43F8B.13 |
| WBGene00012824 | 1.399513002 | 0.00560901 | 0.02865368 | Y43F8B.14 |
| WBGene00012825 | 1.508206757 | 6.98E-09 | 1.60E-07 | Y43F8C.3 |
| WBGene00012826 | 0.060160288 | 0.52140784 | NA | dyf-19 |
| WBGene00012827 | 0.100461934 | 0.42314462 | 0.61903081 | Y43F8C.5 |
| WBGene00012828 | -0.215893841 | 0.20175377 | 0.39041461 | Y43F8C.6 |
| WBGene00012829 | 0.00388439 | 0.97520286 | 0.98597553 | Y43F8C.7 |

| | | | | |
|---|---|---|---|---|
| WBGene00012830 | -0.156488521 | 0.42730244 | 0.62252204 | mrps-28 |
| WBGene00012831 | 0.038633797 | 0.57999349 | NA | Y43F8C.9 |
| WBGene00012832 | -0.040794878 | 0.53203165 | NA | dmd-3 |
| WBGene00012833 | 0.127514091 | 0.32280888 | 0.52291098 | Y43F8C.11 |
| WBGene00012834 | -0.030063509 | 0.83896376 | 0.9127791 | Y43F8C.13 |
| WBGene00012835 | 0.017271684 | 0.89160447 | 0.9434794 | ani-3 |
| WBGene00012836 | NA | NA | NA | Y43F8C.15 |
| WBGene00012837 | -0.039636691 | 0.54579661 | NA | Y43F8C.16 |
| WBGene00012838 | 0.020925399 | 0.84643798 | NA | Y43F8C.17 |
| WBGene00012839 | 0.083520471 | 0.30840907 | NA | Y43F8C.18 |
| WBGene00012840 | -0.04812918 | 0.80697133 | 0.89296858 | grsp-1 |
| WBGene00012841 | NA | NA | NA | Y43F11A.1 |
| WBGene00012841 | NA | NA | NA | Y43F11A.4 |
| WBGene00012844 | NA | NA | NA | Y43F11A.1 |
| WBGene00012844 | NA | NA | NA | Y43F11A.4 |
| WBGene00012845 | -0.168351876 | 0.20126799 | 0.38993842 | set-24 |
| WBGene00012846 | 0.018795205 | 0.77532208 | NA | srxa-14 |
| WBGene00012847 | 0.087256674 | 0.23800185 | NA | srxa-15 |
| WBGene00012848 | -0.135962045 | 0.34323066 | 0.54313812 | Y44A6B.3 |
| WBGene00012849 | -0.054932793 | 0.45568274 | NA | ttr-13 |
| WBGene00012850 | 0.539115843 | 0.05436639 | 0.1599806 | Y44A6C.1 |
| WBGene00012851 | -1.011017383 | 1.71E-05 | 0.00019827 | Y44A6C.2 |
| WBGene00012852 | 0.063999456 | 0.32439727 | NA | Y44A6D.1 |
| WBGene00012853 | 0.665399332 | 0.01381487 | 0.05787387 | Y44A6D.2 |
| WBGene00012854 | 0.233359053 | 0.26579336 | 0.46443258 | Y44A6D.3 |
| WBGene00012855 | 0.037002567 | 0.70655889 | 0.83411778 | Y44A6D.5 |
| WBGene00012856 | 0.083922467 | 0.3064294 | NA | Y44A6D.6 |
| WBGene00012857 | 0.205663063 | 0.34658078 | 0.54638625 | pbo-5 |
| WBGene00012858 | -0.395872109 | 0.01001938 | 0.04492917 | Y44F5A.1 |
| WBGene00012859 | 0.049775231 | 0.7984455 | 0.88715493 | Y45F3A.1 |
| WBGene00012860 | -0.028528971 | 0.78571123 | 0.87882597 | acdh-11 |
| WBGene00012861 | 0.377704087 | 0.12290849 | 0.28254094 | Y45F3A.4 |
| WBGene00012862 | NA | NA | NA | Y45F3A.5 |
| WBGene00012863 | 0.0398788 | 0.58971221 | NA | Y45F3A.8 |
| WBGene00012864 | 0.25910331 | 0.17108936 | 0.35007502 | Y45F3A.9 |
| WBGene00012866 | -0.155921665 | 0.46187144 | 0.65228105 | afmd-2 |
| WBGene00012867 | -0.115131236 | 0.54087854 | 0.714905 | seld-1 |
| WBGene00012868 | 0.06054199 | 0.46366406 | 0.6536122 | tbc-9 |
| WBGene00012869 | 0.019263388 | 0.92031799 | 0.95550377 | Y45F10A.7 |
| WBGene00012870 | NA | NA | NA | NA |
| WBGene00012871 | 0.020317245 | 0.84643798 | NA | Y45F10B.3 |
| WBGene00012872 | NA | NA | NA | Y45F10B.8 |
| WBGene00012873 | 0.020317245 | 0.84643798 | NA | Y45F10B.9 |

| | | | | |
|---|---|---|---|---|
| WBGene00012874 | NA | NA | NA | Y45F10B.12 |
| WBGene00012875 | -0.165565916 | 0.36013888 | 0.55995869 | Y45F10B.13 |
| WBGene00012876 | 0.081768833 | 0.41230943 | NA | NA |
| WBGene00012877 | 0.033463284 | 0.6365037 | NA | Y45F10C.1 |
| WBGene00012878 | -0.248122951 | 0.26779697 | 0.4665971 | Y45F10C.2 |
| WBGene00012879 | -0.108700004 | 0.09903519 | 0.24407452 | fbxa-215 |
| WBGene00012880 | -0.673832435 | 0.00018189 | 0.00162381 | Y45F10C.4 |
| WBGene00012883 | 0.124399388 | 0.41169669 | 0.60940766 | Y45F10D.2 |
| WBGene00012885 | 0.017039327 | 0.87023399 | 0.93075432 | Y45F10D.4 |
| WBGene00012886 | 0.101263703 | 0.6059818 | 0.76212732 | Y45F10D.6 |
| WBGene00012887 | -0.44393059 | 0.00045228 | 0.00356587 | Y45F10D.7 |
| WBGene00012888 | 0.049766118 | 0.60409483 | 0.76062542 | sas-6 |
| WBGene00012889 | -0.023653653 | 0.90498182 | 0.94913088 | icap-1 |
| WBGene00012890 | -0.067645116 | 0.69019131 | 0.82195306 | lact-9 |
| WBGene00012891 | 0.971887659 | 0.00255645 | 0.01528336 | sorb-1 |
| WBGene00012892 | -0.048745395 | 0.47417081 | NA | Y45F10D.14 |
| WBGene00012893 | NA | NA | NA | Y45F10D.15 |
| WBGene00012894 | 0.100049001 | 0.28891285 | 0.48912732 | tbc-17 |
| WBGene00012895 | 0.059485535 | 0.76959453 | 0.86887982 | cox-10 |
| WBGene00012896 | 0.01844828 | 0.75685348 | 0.86226195 | snrp-200 |
| WBGene00012897 | -0.055132706 | 0.63685924 | 0.78460587 | pisy-1 |
| WBGene00012898 | 0.077883306 | 0.65098877 | 0.79333499 | NA |
| WBGene00012899 | -1.201938156 | 0.00143736 | 0.00947671 | Y46G5A.7 |
| WBGene00012900 | -0.22090557 | 0.3110535 | 0.51198812 | Y46G5A.8 |
| WBGene00012901 | 0.254903481 | 0.25658271 | 0.45479751 | cnp-2 |
| WBGene00012903 | 0.01788382 | 0.84734946 | 0.91781042 | vps-2 |
| WBGene00012904 | -0.008170231 | 0.94796741 | 0.97044737 | tiar-2 |
| WBGene00012905 | 0.149717874 | 0.0720891 | NA | Y46G5A.14 |
| WBGene00012906 | 0.046352613 | 0.82373121 | 0.9034778 | Y46G5A.15 |
| WBGene00012907 | -0.137038941 | 0.04802969 | 0.14604119 | cpt-1 |
| WBGene00012908 | -0.026551884 | 0.83225152 | 0.9084712 | Y46G5A.18 |
| WBGene00012909 | -0.637087203 | 0.0003442 | 0.0028248 | spds-1 |
| WBGene00012910 | 1.158892459 | 0.0110261 | 0.04828991 | Y46G5A.20 |
| WBGene00012911 | 0.762188392 | 3.18E-08 | 6.52E-07 | acl-7 |
| WBGene00012912 | 0.182023446 | 0.04351057 | NA | Y46G5A.22 |
| WBGene00012913 | NA | NA | NA | Y46G5A.23 |
| WBGene00012914 | 0.097637856 | 0.47774389 | 0.66470097 | bcmo-1 |
| WBGene00012915 | 0.027495845 | 0.86312534 | 0.9270525 | lgc-35 |
| WBGene00012916 | -0.381036569 | 0.08592551 | 0.22135188 | Y46G5A.28 |
| WBGene00012917 | -0.272523303 | 0.18086763 | 0.36309928 | Y46G5A.29 |
| WBGene00012919 | 0.185443564 | 0.38986718 | 0.58786842 | Y46G5A.34 |
| WBGene00012920 | 0.341810035 | 0.07540883 | 0.20220892 | alkb-7 |
| WBGene00012921 | 0.038765322 | 0.66219346 | NA | Y46G5A.36 |

| | | | | |
|---|---|---|---|---|
| WBGene00012922 | -0.87887537 | 0.00920424 | 0.04212545 | Y47D3A.1 |
| WBGene00012923 | 0.478458421 | 0.06973484 | 0.19080318 | fbxa-128 |
| WBGene00012924 | -0.015333811 | 0.89506391 | 0.94517269 | NA |
| WBGene00012925 | 0.043239853 | 0.68908408 | 0.82103516 | wht-8 |
| WBGene00012926 | 0.545633853 | 0.02742023 | 0.0969709 | Y47D3A.13 |
| WBGene00012927 | -0.156014598 | 0.30119255 | 0.50179337 | Y47D3A.14 |
| WBGene00012928 | 0.030552264 | 0.7905445 | 0.88212499 | aakb-2 |
| WBGene00012929 | 0.075396987 | 0.25977821 | 0.45792525 | rsks-1 |
| WBGene00012930 | 0.096508411 | 0.36690272 | 0.5665882 | obr-1 |
| WBGene00012931 | -0.227970294 | 0.24948912 | 0.44781778 | Y47D3A.20 |
| WBGene00012932 | -0.21458051 | 0.06471641 | 0.18127281 | Y47D3A.21 |
| WBGene00012933 | -0.211343066 | 0.19015858 | 0.37607886 | mib-1 |
| WBGene00012934 | 0.044378294 | 0.67395274 | 0.81052588 | gly-9 |
| WBGene00012935 | 0.102944555 | 0.32565276 | 0.52475997 | mcm-10 |
| WBGene00012936 | -0.033854871 | 0.57666788 | 0.7437451 | pola-1 |
| WBGene00012937 | 0.164052509 | 0.16906755 | 0.34728855 | Y47D3A.31 |
| WBGene00012938 | -0.077546968 | 0.7030994 | 0.83144159 | Y47D3A.32 |
| WBGene00012939 | 0.088315715 | 0.49781183 | 0.67976712 | subs-4 |
| WBGene00012940 | 0.013656784 | 0.85751767 | NA | Y47D3B.3 |
| WBGene00012941 | -0.130817644 | 0.49468195 | 0.67742391 | Y47D3B.4 |
| WBGene00012942 | 0.307576943 | 0.05377479 | 0.15882928 | Y47D3B.6 |
| WBGene00012943 | 0.247286237 | 0.07022543 | 0.19171555 | bed-2 |
| WBGene00012945 | 0.020317245 | 0.84643798 | NA | C17D12.3 |
| WBGene00012945 | 0.020317245 | 0.84643798 | NA | Y47H9A.1 |
| WBGene00012946 | 0.03014578 | 0.80353576 | 0.89074999 | Y47H9B.2 |
| WBGene00012947 | 0.220528871 | 0.14311763 | 0.31151213 | Y47H9C.1 |
| WBGene00012948 | 0.38006465 | 0.10891736 | 0.2596319 | dhhc-2 |
| WBGene00012950 | -0.06811578 | 0.56801877 | 0.73707018 | eif-2Bbeta |
| WBGene00012951 | 0.028763232 | 0.8121087 | 0.8958773 | Y47H9C.8 |
| WBGene00012952 | -0.038777867 | 0.73849711 | 0.85304428 | Y47H9C.9 |
| WBGene00012953 | -0.017313551 | 0.93214825 | 0.96185847 | fbxa-216 |
| WBGene00012954 | 0.222686734 | 0.30536559 | 0.50592194 | NA |
| WBGene00012955 | -0.001993815 | 0.95612996 | NA | Y47H9C.12 |
| WBGene00012956 | -0.274017305 | 0.10338691 | 0.25102616 | NA |
| WBGene00012958 | -0.295082063 | 0.20056725 | 0.38919788 | Y47H10A.2 |
| WBGene00012959 | 0.157495019 | 0.11734854 | 0.27380421 | Y47H10A.3 |
| WBGene00012960 | -0.029355792 | 0.87376946 | 0.93276403 | Y47H10A.4 |
| WBGene00012961 | 0.233592217 | 0.29044539 | 0.49092715 | Y47H10A.5 |
| WBGene00012962 | 0.037379395 | 0.80084347 | 0.88894232 | dmsr-5 |
| WBGene00012964 | -0.613614315 | 5.79E-08 | 1.14E-06 | Y48A6B.3 |
| WBGene00012965 | -0.045771477 | 0.50913401 | NA | fipr-17 |
| WBGene00012966 | -0.082950073 | 0.50113675 | 0.68284254 | exos-1 |
| WBGene00012967 | 0.164603708 | 0.34335346 | 0.54313812 | kvs-4 |

| | | | | |
|---|---|---|---|---|
| WBGene00012968 | 1.186983759 | 7.55E-11 | 2.20E-09 | Y48A6B.7 |
| WBGene00012969 | 0.168530234 | 0.27385049 | 0.47404173 | Y48A6B.8 |
| WBGene00012970 | 0.356831825 | 0.10059935 | 0.24650628 | Y48A6B.9 |
| WBGene00012972 | -0.107425426 | 0.1495311 | 0.32088686 | rsa-2 |
| WBGene00012973 | -0.167250815 | 0.02311017 | 0.08534299 | spat-2 |
| WBGene00012974 | -0.195379442 | 0.35843714 | 0.55876045 | NA |
| WBGene00012975 | -0.620764795 | 0.03640128 | 0.11928966 | NA |
| WBGene00012976 | -0.483669439 | 0.01983133 | 0.07608766 | sup-35 |
| WBGene00012977 | -0.188342378 | 0.08568916 | 0.22090578 | Y48A6C.4 |
| WBGene00012978 | -0.181892265 | 0.0578537 | 0.16739844 | Y48B6A.1 |
| WBGene00012979 | 0.409401944 | 0.0045742 | 0.02445975 | Y48B6A.5 |
| WBGene00012980 | 0.403169419 | 0.0290426 | 0.10126524 | efhd-1 |
| WBGene00012981 | -0.417725418 | 0.07648905 | 0.2044689 | Y48B6A.10 |
| WBGene00012982 | -0.070913547 | 0.53106051 | 0.70766758 | jmjd-2 |
| WBGene00012983 | -0.066084017 | 0.37592852 | 0.57500725 | men-1 |
| WBGene00012984 | -0.014499664 | 0.92186663 | 0.95620205 | Y48B6A.13 |
| WBGene00012985 | 0.085309563 | 0.24820057 | NA | Y48C3A.1 |
| WBGene00012987 | -0.085848321 | 0.64625254 | 0.79060565 | Y48C3A.3 |
| WBGene00012988 | 0.279104155 | 0.13524284 | 0.29987738 | ztf-22 |
| WBGene00012989 | 1.449469723 | 3.53E-05 | 0.00038207 | Y48C3A.5 |
| WBGene00012990 | -0.054351937 | 0.62414919 | 0.7755287 | sam-10 |
| WBGene00012992 | -0.229354053 | 0.07962067 | 0.21045789 | mrpl-20 |
| WBGene00012993 | 0.107381446 | 0.58439483 | 0.74778226 | dmsr-3 |
| WBGene00012994 | 0.04694182 | 0.65488283 | 0.79569622 | Y48C3A.12 |
| WBGene00012995 | 0.047767255 | 0.63897663 | 0.78584032 | Y48C3A.14 |
| WBGene00012996 | 0.079346689 | 0.54143676 | 0.7152554 | pinn-4 |
| WBGene00012997 | -0.005033349 | 0.97693452 | 0.98677463 | Y48C3A.18 |
| WBGene00012998 | -0.271658754 | 0.04602186 | 0.14164517 | Y48C3A.20 |
| WBGene00012999 | -0.26579008 | 0.02304068 | 0.0852806 | rpoa-1 |
| WBGene00013000 | -0.220947084 | 0.2257193 | 0.41890687 | ddl-2 |
| WBGene00013001 | -0.043232631 | 0.76615052 | 0.86713124 | noa-1 |
| WBGene00013002 | 0.165537307 | 0.19662155 | 0.3841045 | Y48E1B.3 |
| WBGene00013004 | -0.243795829 | 0.14689688 | 0.31721492 | mrpl-37 |
| WBGene00013006 | -0.225558057 | 0.1889001 | 0.37415135 | lin-38 |
| WBGene00013007 | -3.269860687 | 1.38E-12 | 5.07E-11 | Y48E1B.8 |
| WBGene00013008 | -0.024401669 | 0.89693163 | 0.94582116 | clec-146 |
| WBGene00013009 | 0.352561071 | 0.08156528 | 0.21374571 | fbxc-23 |
| WBGene00013010 | NA | NA | NA | NA |
| WBGene00013011 | 0.164733213 | 0.1300995 | 0.29227968 | snx-14 |
| WBGene00013013 | 0.113975773 | 0.5241373 | 0.70158112 | clec-145 |
| WBGene00013014 | 0.12279158 | 0.37022121 | 0.57006877 | Y48E1C.1 |
| WBGene00013015 | 0.065403378 | 0.63802593 | 0.78552676 | Y48E1C.2 |
| WBGene00013017 | -0.119808674 | 0.44072424 | 0.63491524 | Y48E1C.4 |

| | | | | |
|---|---|---|---|---|
| WBGene00013018 | -0.100003704 | 0.34130642 | 0.54139407 | Y48G10A.1 |
| WBGene00013019 | -0.26468712 | 0.03944901 | 0.12679978 | Y48G10A.2 |
| WBGene00013020 | 0.034449231 | 0.75930708 | 0.86379292 | Y48G10A.3 |
| WBGene00013021 | 0.04076229 | 0.7592577 | 0.86379292 | Y48G10A.4 |
| WBGene00013023 | 0.146352711 | 0.48067607 | 0.66700581 | Y48G10A.6 |
| WBGene00013024 | -0.236740615 | 0.01260134 | 0.05380467 | cept-2 |
| WBGene00013025 | 0.086040495 | 0.10399421 | 0.25210364 | vha-13 |
| WBGene00013026 | 0.123629897 | 0.47203887 | 0.66010393 | Y49A3A.3 |
| WBGene00013027 | 0.010461576 | 0.94167279 | 0.96696811 | Y49A3A.4 |
| WBGene00013028 | 0.111319389 | 0.1344031 | NA | efhc-1 |
| WBGene00013029 | -0.125575875 | 0.37460258 | 0.57340912 | glrx-5 |
| WBGene00013030 | 0.084445234 | 0.4459905 | 0.63930108 | Y49E10.4 |
| WBGene00013031 | 0.031760039 | 0.7704688 | 0.86939759 | NA |
| WBGene00013032 | 0.072952506 | 0.34318498 | NA | NA |
| WBGene00013033 | 0.167641235 | 0.16100856 | 0.33556248 | Y49E10.10 |
| WBGene00013034 | 0.014270203 | 0.85592685 | 0.92266343 | tat-1 |
| WBGene00013035 | 0.43682091 | 0.00309466 | 0.01790864 | Y49E10.16 |
| WBGene00013036 | -0.010877204 | 0.91996234 | 0.95544619 | fbxa-218 |
| WBGene00013037 | 1.498726781 | 1.10E-13 | 4.58E-12 | Y49E10.18 |
| WBGene00013038 | -0.064381973 | 0.2948619 | 0.49581636 | ani-1 |
| WBGene00013039 | 0.044386016 | 0.61408954 | 0.76832036 | scav-3 |
| WBGene00013040 | -0.004343542 | 0.97136731 | 0.98427495 | Y49E10.21 |
| WBGene00013041 | 0.213608322 | 0.32941057 | 0.52894276 | pstk-1 |
| WBGene00013042 | 0.069055409 | 0.7219778 | 0.84401764 | cccp-1 |
| WBGene00013043 | 0.059980168 | 0.38780842 | NA | arrd-16 |
| WBGene00013044 | NA | NA | NA | Y49E10.25 |
| WBGene00013045 | NA | NA | NA | Y49E10.27 |
| WBGene00013046 | 0.260861622 | 0.21528763 | 0.40639906 | Y50E8A.1 |
| WBGene00013047 | NA | NA | NA | Y50E8A.2 |
| WBGene00013048 | 0.047630714 | 0.79571554 | 0.88526377 | Y50E8A.5 |
| WBGene00013049 | 0.033034883 | 0.64219503 | NA | Y50E8A.6 |
| WBGene00013050 | NA | NA | NA | lipl-8 |
| WBGene00013051 | NA | NA | NA | NA |
| WBGene00013052 | 0.063421086 | 0.41095452 | NA | scrm-7 |
| WBGene00013053 | 0.0398788 | 0.58971221 | NA | Y50E8A.10 |
| WBGene00013054 | -0.04942813 | 0.54924154 | NA | Y50E8A.11 |
| WBGene00013055 | -0.087173176 | 0.570686 | 0.7392863 | Y50E8A.12 |
| WBGene00013056 | NA | NA | NA | NA |
| WBGene00013058 | NA | NA | NA | clec-247 |
| WBGene00013059 | NA | NA | NA | NA |
| WBGene00013060 | -0.123519925 | 0.10757993 | NA | Y51A2A.4 |
| WBGene00013061 | 0.020925399 | 0.84643798 | NA | Y51A2A.5 |
| WBGene00013062 | NA | NA | NA | Y51A2A.6 |

| | | | | |
|---|---|---|---|---|
| WBGene00013063 | NA | NA | NA | clec-249 |
| WBGene00013063 | NA | NA | NA | clec-248 |
| WBGene00013064 | NA | NA | NA | clec-249 |
| WBGene00013064 | NA | NA | NA | clec-248 |
| WBGene00013066 | 0.020232312 | 0.84643798 | NA | Y51A2B.2 |
| WBGene00013067 | NA | NA | NA | nhr-288 |
| WBGene00013068 | 0.018657025 | 0.77508599 | NA | Y51A2B.4 |
| WBGene00013069 | 0.040107227 | 0.58693163 | NA | Y51A2B.5 |
| WBGene00013070 | 0.0398788 | 0.58971221 | NA | Y51A2B.6 |
| WBGene00013071 | NA | NA | NA | Y51A2B.8 |
| WBGene00013072 | NA | NA | NA | Y51A2D.1 |
| WBGene00013073 | 0.106083914 | 0.59585646 | 0.75477619 | hmit-1.1 |
| WBGene00013074 | -2.343488002 | 5.56E-09 | 1.29E-07 | hmit-1.2 |
| WBGene00013075 | -0.038018477 | 0.76568131 | 0.86686824 | Y51A2D.7 |
| WBGene00013076 | 0.086470338 | 0.29130368 | NA | Y51A2D.8 |
| WBGene00013077 | 0.029774986 | 0.78618463 | 0.87893473 | ttr-24 |
| WBGene00013078 | -0.391043678 | 0.01486866 | 0.06146019 | ttr-25 |
| WBGene00013079 | 2.466559692 | 1.78E-18 | 1.18E-16 | ttr-26 |
| WBGene00013080 | 0.903379283 | 6.95E-07 | 1.13E-05 | Y51A2D.13 |
| WBGene00013081 | 0.63540062 | 0.00026683 | 0.00226736 | Y51A2D.14 |
| WBGene00013082 | -0.187897263 | 0.3201148 | 0.52007255 | grdn-1 |
| WBGene00013083 | -1.271341665 | 5.09E-09 | 1.19E-07 | Y51A2D.18 |
| WBGene00013085 | 0.008012039 | 0.84942629 | NA | mpz-6 |
| WBGene00013086 | -0.52017803 | 0.0252415 | 0.09139553 | arrd-2 |
| WBGene00013087 | -0.003108065 | 0.95728506 | NA | Y51B9A.5 |
| WBGene00013088 | 0.055461652 | 0.71485914 | 0.8401205 | Y51B9A.6 |
| WBGene00013089 | NA | NA | NA | Y51B9A.7 |
| WBGene00013090 | 0.020925399 | 0.84643798 | NA | Y51B9A.8 |
| WBGene00013091 | 2.928955193 | 1.14E-05 | 0.00013895 | Y51B9A.9 |
| WBGene00013093 | -0.031619997 | 0.76873544 | 0.86844708 | cup-14 |
| WBGene00013094 | -0.048680451 | 0.58270546 | 0.74659933 | Y51H1A.3 |
| WBGene00013095 | 0.079674594 | 0.51137094 | 0.69129317 | ing-3 |
| WBGene00013096 | -0.011461958 | 0.91114061 | 0.95112162 | mcd-1 |
| WBGene00013098 | 0.032136579 | 0.69225183 | NA | Y51H4A.1 |
| WBGene00013100 | -1.599528736 | 0.00261866 | 0.01559799 | zip-7 |
| WBGene00013101 | 2.630870835 | 3.17E-07 | 5.45E-06 | Y51H4A.5 |
| WBGene00013102 | NA | NA | NA | Y51H4A.6 |
| WBGene00013103 | 0.244077183 | 0.11869467 | 0.27614948 | Y51H4A.7 |
| WBGene00013104 | 0.616446697 | 0.03468407 | 0.11525406 | Y51H4A.8 |
| WBGene00013105 | NA | NA | NA | fip-7 |
| WBGene00013106 | -0.030537472 | 0.76057094 | 0.86431035 | set-26 |
| WBGene00013107 | 0.320017313 | 0.05665194 | 0.16499724 | Y51H4A.13 |
| WBGene00013109 | -0.272137557 | 0.05358883 | 0.15850076 | Y51H4A.15 |

| | | | | |
|---|---|---|---|---|
| WBGene00013111 | 0.661513885 | 0.01083368 | 0.04756089 | sta-1 |
| WBGene00013112 | NA | NA | NA | NA |
| WBGene00013113 | NA | NA | NA | NA |
| WBGene00013114 | NA | NA | NA | NA |
| WBGene00013116 | 0.038125098 | 0.61119482 | NA | Y51H4A.22 |
| WBGene00013117 | NA | NA | NA | Y51H4A.23 |
| WBGene00013118 | 0.017604133 | 0.79437959 | NA | Y51H4A.24 |
| WBGene00013119 | 1.361300443 | 0.0001746 | 0.00156926 | Y51H4A.25 |
| WBGene00013120 | 0.038125098 | 0.61119482 | NA | fipr-28 |
| WBGene00013121 | 0.033463284 | 0.6365037 | NA | spe-38 |
| WBGene00013122 | -0.05600714 | 0.69655216 | 0.82690507 | impt-1 |
| WBGene00013123 | 0.052832883 | 0.665714 | 0.80405202 | Y52B11A.3 |
| WBGene00013124 | 0.331046994 | 0.15418818 | 0.32715947 | Y52B11A.4 |
| WBGene00013125 | NA | NA | NA | clec-92 |
| WBGene00013126 | -0.072554423 | 0.63464083 | 0.78332291 | mltn-2 |
| WBGene00013127 | 0.126093133 | 0.18372465 | 0.366923 | Y52B11A.8 |
| WBGene00013128 | -0.084083917 | 0.46541744 | 0.65508332 | dxbp-1 |
| WBGene00013129 | -0.306435802 | 0.02428749 | 0.08880948 | Y52B11A.10 |
| WBGene00013130 | NA | NA | NA | Y52B11B.1 |
| WBGene00013131 | -0.18839207 | 0.26268167 | 0.46064466 | pigl-1 |
| WBGene00013132 | 0.059913012 | 0.64218078 | 0.78811138 | strd-1 |
| WBGene00013134 | 0.332657547 | 0.14854513 | 0.31940802 | NA |
| WBGene00013135 | 0.031992033 | 0.87789403 | 0.93480719 | rga-2 |
| WBGene00013136 | 0.178079333 | 0.17858552 | 0.36004827 | Y53C10A.5 |
| WBGene00013137 | -0.05131226 | 0.66533369 | 0.80372529 | Y53C10A.6 |
| WBGene00013138 | 0.582344324 | 0.01068766 | 0.0470668 | Y53C10A.10 |
| WBGene00013139 | -0.07751712 | 0.52749982 | 0.70472898 | Y53C12A.3 |
| WBGene00013140 | -0.191986384 | 0.05399351 | 0.15921834 | mop-25.2 |
| WBGene00013141 | -0.003085443 | 0.98364809 | 0.99058801 | Y53C12A.6 |
| WBGene00013142 | 0.065014541 | 0.37399213 | NA | Y53C12A.7 |
| WBGene00013143 | -0.285135854 | 0.01856296 | 0.07281926 | Y53C12B.1 |
| WBGene00013144 | -0.162928012 | 0.04520905 | 0.13973098 | pno-1 |
| WBGene00013145 | 0.068906547 | 0.67982161 | 0.81463972 | cutl-2 |
| WBGene00013146 | 0.876518741 | 0.01943385 | 0.07519317 | Y53C12B.7 |
| WBGene00013147 | -0.106690593 | 0.29790363 | 0.49893568 | eyg-1 |
| WBGene00013148 | NA | NA | NA | Y53F4A.2 |
| WBGene00013149 | 0.517349088 | 0.06100186 | 0.17369277 | Y53F4B.1 |
| WBGene00013150 | -0.126067944 | 0.310461 | 0.51126394 | pole-4 |
| WBGene00013151 | -0.112078181 | 0.54359308 | 0.71696212 | nsun-5 |
| WBGene00013152 | 0.020925399 | 0.84643798 | NA | Y53F4B.5 |
| WBGene00013153 | 0.010650806 | 0.9195564 | 0.95531294 | NA |
| WBGene00013154 | -0.045771477 | 0.50913401 | NA | Y53F4B.7 |
| WBGene00013155 | -0.06496591 | 0.59727875 | 0.75574783 | NA |

| | | | | |
|---|---|---|---|---|
| WBGene00013156 | -0.00706155 | 0.9449049 | 0.96892791 | Y53F4B.9 |
| WBGene00013157 | -0.083977 | 0.47488736 | 0.66202117 | Y53F4B.10 |
| WBGene00013158 | 0.25579143 | 0.10016576 | 0.24569055 | Y53F4B.11 |
| WBGene00013159 | 0.114847813 | 0.57405568 | 0.74187918 | Y53F4B.12 |
| WBGene00013160 | 0.047913648 | 0.62302073 | 0.77453057 | Y53F4B.13 |
| WBGene00013161 | 0.350834238 | 0.05393531 | 0.15912311 | Y53F4B.14 |
| WBGene00013162 | 0.053133271 | 0.55059075 | NA | Y53F4B.16 |
| WBGene00013163 | 0.020317245 | 0.84643798 | NA | Y53F4B.17 |
| WBGene00013164 | -0.147743522 | 0.34920887 | 0.54916939 | Y53F4B.18 |
| WBGene00013165 | 0.165163527 | 0.30051825 | 0.50083684 | Y53F4B.19 |
| WBGene00013166 | -0.036742574 | 0.72869215 | 0.84857437 | Y53F4B.20 |
| WBGene00013167 | -0.065290945 | 0.7375893 | 0.85246644 | Y53F4B.21 |
| WBGene00013168 | -0.231994922 | 0.00406126 | 0.02214561 | arp-1 |
| WBGene00013169 | -0.052653409 | 0.78484899 | 0.87855213 | Y53F4B.23 |
| WBGene00013170 | 0.075450426 | 0.33451267 | NA | Y53F4B.24 |
| WBGene00013171 | 0.098837507 | 0.23357279 | NA | Y53F4B.25 |
| WBGene00013172 | NA | NA | NA | Y53F4B.26 |
| WBGene00013173 | 4.060200724 | 0.00134014 | 0.00893489 | Y53F4B.27 |
| WBGene00013175 | 0.093081795 | 0.25403827 | NA | Y53F4B.36 |
| WBGene00013176 | 0.609211717 | 1.59E-05 | 0.00018516 | Y53F4B.39 |
| WBGene00013177 | 0.156168179 | 0.20774792 | 0.3967698 | rsy-1 |
| WBGene00013178 | -0.235356814 | 0.11694616 | 0.27352701 | Y53H1A.2 |
| WBGene00013179 | NA | NA | NA | clec-100 |
| WBGene00013180 | 1.277766994 | 0.00690564 | 0.0336122 | cutl-10 |
| WBGene00013181 | 0.365634742 | 0.00386986 | 0.02132427 | Y53H1B.2 |
| WBGene00013182 | 0.020317245 | 0.84643798 | NA | Y53H1B.4 |
| WBGene00013184 | NA | NA | NA | clpr-3 |
| WBGene00013184 | NA | NA | NA | clpr-4 |
| WBGene00013186 | -0.014272784 | 0.89800511 | 0.94632825 | Y53H1C.3 |
| WBGene00013187 | 0.188290841 | 0.22180762 | 0.41448442 | npr-34 |
| WBGene00013188 | 0.018601266 | 0.92574068 | 0.95816777 | smg-9 |
| WBGene00013189 | -0.007206306 | 0.93204662 | 0.96185847 | Y54E2A.4 |
| WBGene00013190 | 0.107005324 | 0.31506195 | 0.51557662 | Y54E2A.5 |
| WBGene00013192 | 0.126299345 | 0.16983897 | NA | Y54E2A.7 |
| WBGene00013193 | -0.100360465 | 0.48670677 | 0.67130229 | Y54E2A.8 |
| WBGene00013194 | 0.148601991 | 0.12731679 | 0.28871852 | Y54E2A.9 |
| WBGene00013195 | 0.019133773 | 0.92106718 | 0.95586508 | Y54E2A.10 |
| WBGene00013196 | -0.118193118 | 0.21765843 | 0.40939583 | tbc-20 |
| WBGene00013197 | 0.006217309 | 0.97628361 | 0.98666172 | ttm-5 |
| WBGene00013198 | 0.786966111 | 0.01653617 | 0.06677389 | Y54E5A.2 |
| WBGene00013200 | -0.179099396 | 0.10115533 | 0.24733052 | Y54E5A.5 |
| WBGene00013201 | -0.123486573 | 0.43001702 | 0.62496319 | dus-2 |
| WBGene00013202 | 0.080516925 | 0.48399456 | 0.66932694 | Y54E5A.7 |

| | | | | |
|---|---|---|---|---|
| WBGene00013203 | 0.404458068 | 0.10858538 | 0.25913587 | Y54E5A.8 |
| WBGene00013204 | -0.032810315 | 0.81192302 | 0.89587496 | Y54E5B.2 |
| WBGene00013207 | -1.494437065 | 0.00234193 | 0.01419799 | zipt-2.3 |
| WBGene00013208 | 0.260883982 | 0.02348655 | 0.08639863 | Y54G9A.5 |
| WBGene00013209 | -0.251888103 | 0.00362435 | 0.02030772 | bub-3 |
| WBGene00013210 | -0.129867878 | 0.34569541 | 0.54529744 | Y54G9A.7 |
| WBGene00013211 | -0.112875505 | 0.58599206 | 0.74884435 | Y54G9A.9 |
| WBGene00013212 | 0.065756331 | 0.55926674 | 0.73003231 | Y54G11A.1 |
| WBGene00013213 | -0.147603241 | 0.19287319 | 0.37932398 | Y54G11A.2 |
| WBGene00013214 | -0.314772267 | 0.00114339 | 0.007863 | Y54G11A.3 |
| WBGene00013215 | 0.748450038 | 0.02590123 | 0.09315154 | Y54G11A.4 |
| WBGene00013216 | 0.128953994 | 0.45816811 | 0.64973678 | Y54G11A.7 |
| WBGene00013217 | -0.127847552 | 0.27265904 | 0.4726499 | ddl-3 |
| WBGene00013218 | -0.160755203 | 0.3564389 | 0.55641561 | Y54G11A.9 |
| WBGene00013219 | -0.076819873 | 0.46845671 | 0.65738945 | Y54G11A.11 |
| WBGene00013220 | 0.339642624 | 0.15319457 | 0.32633809 | ctl-3 |
| WBGene00013221 | 0.437772843 | 0.07165672 | 0.19475144 | Y54G11A.14 |
| WBGene00013222 | 0.24281132 | 0.19438309 | 0.38124084 | dmsr-6 |
| WBGene00013223 | -0.085723133 | 0.2800547 | 0.47978947 | efa-6 |
| WBGene00013224 | -0.073801251 | 0.49635093 | 0.67886649 | Y55D9A.2 |
| WBGene00013225 | 0.054447204 | 0.50236317 | 0.68381361 | Y56A3A.2 |
| WBGene00013226 | 0.123562325 | 0.30496557 | 0.50562059 | faah-5 |
| WBGene00013227 | 0.296143656 | 0.12968085 | 0.29201626 | Y56A3A.6 |
| WBGene00013228 | 0.181123108 | 0.20911353 | 0.39854611 | Y56A3A.7 |
| WBGene00013230 | -0.874120123 | 0.01669357 | 0.0672381 | fbxb-22 |
| WBGene00013231 | 0.1123147 | 0.4386348 | 0.63346342 | tsen-2 |
| WBGene00013232 | -0.164210849 | 0.09147351 | 0.2302327 | faah-4 |
| WBGene00013233 | -0.8340183 | 0.0075245 | 0.03606095 | sdz-33 |
| WBGene00013234 | -0.868724381 | 0.00872677 | 0.04043314 | fbxb-24 |
| WBGene00013235 | -0.061781165 | 0.49609422 | 0.67868768 | Y56A3A.16 |
| WBGene00013236 | 0.15376839 | 0.34488703 | 0.54459567 | Y56A3A.18 |
| WBGene00013237 | -0.205107191 | 0.01149223 | 0.04983937 | Y56A3A.19 |
| WBGene00013238 | -0.490016574 | 3.15E-17 | 1.88E-15 | trap-4 |
| WBGene00013239 | 0.000254221 | 0.99898588 | 0.99946374 | Y56A3A.22 |
| WBGene00013240 | 0.018522703 | 0.91102265 | 0.95112162 | Y56A3A.28 |
| WBGene00013241 | -0.122865795 | 0.30259219 | 0.50315188 | ung-1 |
| WBGene00013242 | 0.223593753 | 0.28330986 | 0.48327387 | Y56A3A.30 |
| WBGene00013243 | 0.252054304 | 0.08745544 | 0.22393059 | Y56A3A.31 |
| WBGene00013244 | -0.06575753 | 0.74523937 | 0.85590582 | Y56A3A.33 |
| WBGene00013246 | -0.187100567 | 0.19828308 | 0.38640524 | Y57A10A.1 |
| WBGene00013247 | -0.221254657 | 0.19052832 | 0.37655577 | Y57A10A.2 |
| WBGene00013248 | -0.197400377 | 0.30570668 | 0.50637532 | Y57A10A.3 |
| WBGene00013249 | 0.195873703 | 0.35973393 | 0.55964982 | Y57A10A.4 |

| | | | | |
|---|---|---|---|---|
| WBGene00013250 | 0.094483257 | 0.58368226 | 0.7475238 | Y57A10A.5 |
| WBGene00013251 | -0.071796258 | 0.71829188 | 0.8419271 | NA |
| WBGene00013252 | -0.007584536 | 0.96768736 | 0.98210943 | Y57A10A.7 |
| WBGene00013253 | -0.123943468 | 0.44819481 | 0.64088487 | Y57A10A.8 |
| WBGene00013254 | -0.2192104 | 0.30472695 | 0.50545453 | tag-276 |
| WBGene00013255 | 0.045833958 | 0.75690121 | 0.86226195 | tric-1B.1 |
| WBGene00013256 | -0.009306566 | 0.91830014 | 0.95454969 | Y57A10A.13 |
| WBGene00013257 | 0.227439428 | 0.3022496 | 0.50276842 | Y57A10A.14 |
| WBGene00013258 | 0.328840094 | 0.02917092 | 0.10154337 | polg-1 |
| WBGene00013259 | -0.018099977 | 0.91188964 | 0.95112162 | trpp-5 |
| WBGene00013260 | -0.271086869 | 0.00730569 | 0.03515114 | rsr-2 |
| WBGene00013261 | 0.343100178 | 0.036046 | 0.11846229 | sinh-1 |
| WBGene00013263 | -0.740305109 | 0.00036236 | 0.00294907 | txdc-12.1 |
| WBGene00013264 | 0.015234133 | 0.86748955 | NA | Y57A10A.24 |
| WBGene00013265 | -0.156875206 | 0.270559 | 0.46984394 | parn-2 |
| WBGene00013266 | 0.34373049 | 0.02779556 | 0.09799649 | Y57A10A.26 |
| WBGene00013267 | -0.481654769 | 0.00228693 | 0.01390485 | Y57A10A.27 |
| WBGene00013268 | -0.199060815 | 0.11126253 | 0.2633357 | tric-1B.2 |
| WBGene00013269 | -0.015715448 | 0.91021597 | 0.95112162 | Y57A10A.29 |
| WBGene00013270 | 0.109706156 | 0.1305858 | 0.29288894 | Y57A10A.31 |
| WBGene00013272 | 0.015194928 | 0.8086441 | NA | aqp-12 |
| WBGene00013273 | 1.554710805 | 7.79E-05 | 0.00077565 | 5-Mar |
| WBGene00013275 | 0.186837399 | 0.27781195 | 0.47767779 | btb-14 |
| WBGene00013276 | -0.002914971 | 0.95718083 | NA | sre-42 |
| WBGene00013277 | -0.003270206 | 0.95698316 | NA | sre-41 |
| WBGene00013278 | -0.171945439 | 0.42299213 | 0.61886956 | Y57A10B.6 |
| WBGene00013279 | 0.075726029 | 0.27693214 | NA | Y57A10B.7 |
| WBGene00013280 | -0.329984127 | 0.13958868 | 0.3069534 | Y57A10C.1 |
| WBGene00013281 | -0.023056499 | 0.78121253 | NA | sre-27 |
| WBGene00013282 | NA | NA | NA | sre-26 |
| WBGene00013283 | -0.022930767 | 0.78121253 | NA | NA |
| WBGene00013284 | 0.020030979 | 0.85837239 | 0.92421167 | daf-22 |
| WBGene00013285 | NA | NA | NA | dct-12 |
| WBGene00013286 | NA | NA | NA | Y57A10C.8 |
| WBGene00013287 | 0.083520471 | 0.30840907 | NA | Y57A10C.9 |
| WBGene00013288 | -0.008998908 | 0.92213607 | NA | Y57A10C.10 |
| WBGene00013289 | -0.01814637 | 0.90679273 | 0.94953171 | tag-273 |
| WBGene00013290 | 2.335305641 | 0.00018109 | 0.00161868 | Y57G11A.2 |
| WBGene00013292 | 0.517960179 | 0.04004092 | 0.12803875 | Y57G11A.4 |
| WBGene00013293 | 0.020317245 | 0.84643798 | NA | Y57G11A.5 |
| WBGene00013294 | -0.022930767 | 0.78121253 | NA | pals-30 |
| WBGene00013295 | -0.342606132 | 0.15201202 | 0.32451635 | Y57G11B.2 |
| WBGene00013296 | 0.014306166 | 0.87659352 | NA | Y57G11B.3 |

| | | | | |
|---|---|---|---|---|
| WBGene00013297 | -0.81835403 | 2.87E-27 | 3.09E-25 | Y57G11B.5 |
| WBGene00013298 | 0.020317245 | 0.84643798 | NA | Y57G11B.6 |
| WBGene00013299 | 0.042065281 | 0.558937 | NA | irld-18 |
| WBGene00013300 | 0.056411843 | 0.41891328 | NA | lgc-7 |
| WBGene00013301 | -0.141698499 | 0.15930177 | 0.33371545 | Y57G11C.3 |
| WBGene00013302 | -0.105727237 | 0.51726495 | 0.69664944 | vti-1 |
| WBGene00013303 | 0.345783189 | 0.02264258 | 0.08415323 | Y57G11C.5 |
| WBGene00013304 | 0.106997704 | 0.16248985 | NA | ptp-5.2 |
| WBGene00013305 | NA | NA | NA | Y57G11C.7 |
| WBGene00013306 | 0.555900787 | 0.0447036 | 0.13851405 | Y57G11C.8 |
| WBGene00013307 | -0.00278556 | 0.96959346 | 0.98346632 | Y57G11C.9 |
| WBGene00013308 | 0.020317245 | 0.84643798 | NA | Y57G11C.1147 |
| WBGene00013308 | 0.020317245 | 0.84643798 | NA | nuo-3 |
| WBGene00013310 | 0.127996092 | 0.51214744 | 0.69204898 | Y57G11C.14 |
| WBGene00013311 | -0.479106358 | 1.59E-25 | 1.51E-23 | sec-61 |
| WBGene00013312 | -0.008343726 | 0.95315319 | 0.97322797 | hhat-2 |
| WBGene00013313 | 0.108942554 | 0.15594412 | NA | Y57G11C.18 |
| WBGene00013315 | 0.015062379 | 0.80838968 | NA | Y57G11C.20 |
| WBGene00013316 | NA | NA | NA | NA |
| WBGene00013317 | 0.163364275 | 0.44690355 | 0.64010124 | Y57G11C.22 |
| WBGene00013318 | 0.056411843 | 0.41891328 | NA | Y57G11C.23 |
| WBGene00013319 | -0.078356303 | 0.70263666 | 0.83114502 | ccch-5 |
| WBGene00013320 | NA | NA | NA | NA |
| WBGene00013321 | 0.063408649 | 0.74559992 | 0.85598121 | Y57G11C.31 |
| WBGene00013322 | 0.110126488 | 0.13663711 | NA | Y57G11C.32 |
| WBGene00013323 | 0.060474398 | 0.65355659 | 0.79476224 | Y57G11C.33 |
| WBGene00013324 | -0.290265792 | 0.06401177 | 0.17977214 | mrps-7 |
| WBGene00013325 | 0.301821721 | 0.05320967 | 0.15770095 | Y57G11C.36 |
| WBGene00013326 | 0.17869136 | 0.4074066 | 0.60539819 | anoh-2 |
| WBGene00013327 | 0.225104774 | 0.29261243 | 0.49350749 | Y57G11C.38 |
| WBGene00013328 | 0.020232312 | 0.84643798 | NA | Y57G11C.39 |
| WBGene00013329 | 0.367893683 | 0.12860444 | 0.29034659 | Y57G11C.40 |
| WBGene00013330 | -0.029234136 | 0.7166623 | NA | Y57G11C.41 |
| WBGene00013331 | 0.285582086 | 0.13980805 | 0.30722209 | Y57G11C.42 |
| WBGene00013332 | -0.067473239 | 0.61111671 | 0.76604108 | Y57G11C.43 |
| WBGene00013333 | 0.177616853 | 0.0463229 | NA | Y57G11C.44 |
| WBGene00013334 | 0.207718945 | 0.33294063 | 0.53262498 | nlp-56 |
| WBGene00013335 | 0.089911953 | 0.63108163 | 0.78083781 | Y57G11C.46 |
| WBGene00013336 | -0.191423903 | 0.23339932 | 0.42875992 | pnc-2 |
| WBGene00013337 | 0.020925399 | 0.84643798 | NA | Y57G11C.48 |
| WBGene00013338 | NA | NA | NA | lgc-8 |
| WBGene00013339 | -0.077838302 | 0.51243566 | 0.6921829 | phf-14 |
| WBGene00013340 | -0.14314908 | 0.00520363 | 0.0269619 | tcc-1 |

| | | | | |
|---|---|---|---|---|
| WBGene00013341 | NA | NA | NA | NA |
| WBGene00013343 | -0.131479471 | 0.08254442 | 0.21546315 | prp-6 |
| WBGene00013344 | -0.158972363 | 0.38380184 | 0.58327725 | ebp-1 |
| WBGene00013345 | -0.042228557 | 0.82169901 | 0.90237738 | Y59A8B.8 |
| WBGene00013347 | -0.285606121 | 0.00048164 | 0.00377098 | nova-1 |
| WBGene00013348 | 0.113409512 | 0.52032339 | 0.69916842 | fbxa-106 |
| WBGene00013349 | -0.262083415 | 0.02526949 | 0.09143621 | Y59A8B.12 |
| WBGene00013350 | 0.078701316 | 0.6489586 | 0.79237696 | slr-2 |
| WBGene00013351 | NA | NA | NA | Y59A8B.19 |
| WBGene00013352 | 0.460898561 | 0.07892685 | 0.20924759 | lon-8 |
| WBGene00013353 | 0.079983042 | 0.54714476 | 0.71995249 | Y59A8B.21 |
| WBGene00013354 | -0.172254908 | 0.03348219 | 0.11222819 | snx-6 |
| WBGene00013355 | 0.035892531 | 0.71909837 | 0.84237564 | gck-3 |
| WBGene00013356 | NA | NA | NA | Y59A8B.24 |
| WBGene00013357 | 0.020317245 | 0.84643798 | NA | clec-260 |
| WBGene00013358 | -0.142747146 | 0.22228721 | 0.41503454 | abhd-3.1 |
| WBGene00013359 | 0.154856748 | 0.38086484 | 0.58025412 | Y60A3A.8 |
| WBGene00013360 | -0.179556916 | 0.03834769 | 0.12403112 | tmed-4 |
| WBGene00013361 | -0.197314623 | 0.0460141 | 0.14164517 | fars-2 |
| WBGene00013362 | -0.042157314 | 0.77926545 | 0.87514194 | algn-7 |
| WBGene00013363 | 0.017348279 | 0.83364574 | NA | NA |
| WBGene00013364 | 0.18113801 | 0.32250044 | 0.52270521 | Y60A3A.16 |
| WBGene00013365 | 0.020330753 | 0.87677353 | 0.93447196 | Y60A3A.19 |
| WBGene00013366 | 0.313717607 | 0.12727752 | 0.28871852 | Y60A3A.21 |
| WBGene00013367 | 0.020925399 | 0.84643798 | NA | Y60A3A.23 |
| WBGene00013368 | 0.020317245 | 0.84643798 | NA | Y60A9.1 |
| WBGene00013369 | -0.18876098 | 0.27755032 | 0.47751049 | NA |
| WBGene00013370 | -0.202988757 | 0.29366966 | 0.49477762 | Y60A9.3 |
| WBGene00013371 | NA | NA | NA | Y60A9A.1 |
| WBGene00013372 | NA | NA | NA | NA |
| WBGene00013373 | -0.030511598 | 0.74077754 | NA | Y62E10A.2 |
| WBGene00013374 | 0.020232312 | 0.84643798 | NA | Y62E10A.3 |
| WBGene00013375 | 0.060484201 | 0.41382805 | NA | srsx-25 |
| WBGene00013376 | -0.269354774 | 0.07121622 | 0.1939142 | Y62E10A.6 |
| WBGene00013378 | -0.224346456 | 0.0210131 | 0.07956585 | emc-3 |
| WBGene00013379 | -0.038560974 | 0.73892034 | 0.85307041 | Y62E10A.13 |
| WBGene00013380 | -0.309633788 | 0.00064392 | 0.00482728 | memi-2 |
| WBGene00013381 | -0.265431174 | 0.01171419 | 0.05059216 | cyp-31A5 |
| WBGene00013382 | 0.471327875 | 0.04004087 | 0.12803875 | gcl-1 |
| WBGene00013383 | -0.547813759 | 0.00034269 | 0.00281397 | aptf-2 |
| WBGene00013385 | 0.139562744 | 0.41687029 | 0.61390042 | Y62E10A.19 |
| WBGene00013386 | NA | NA | NA | NA |
| WBGene00013387 | 0.208399913 | 0.27680852 | 0.47673845 | Y62F5A.9 |

| | | | | |
|---|---|---|---|---|
| WBGene00013388 | 0.151903689 | 0.41613107 | 0.61330531 | Y62F5A.10 |
| WBGene00013389 | 0.018795205 | 0.77532208 | NA | Y62H9A.1 |
| WBGene00013390 | NA | NA | NA | Y62H9A.2 |
| WBGene00013391 | -1.426882436 | 6.89E-43 | 1.40E-40 | Y62H9A.3 |
| WBGene00013392 | -1.515794881 | 6.04E-46 | 1.30E-43 | Y62H9A.4 |
| WBGene00013393 | 0.264933216 | 2.71E-06 | 3.78E-05 | Y62H9A.5 |
| WBGene00013394 | 0.176774522 | 0.00016173 | 0.00146616 | ule-5 |
| WBGene00013395 | NA | NA | NA | Y62H9A.7 |
| WBGene00013396 | NA | NA | NA | Y62H9A.8 |
| WBGene00013397 | 0.020317245 | 0.84643798 | NA | Y62H9A.9 |
| WBGene00013398 | 0.018860755 | 0.77262519 | NA | Y62H9A.10 |
| WBGene00013399 | -0.095519004 | 0.23723836 | NA | Y62H9A.11 |
| WBGene00013400 | -0.127460451 | 0.27439053 | 0.47475204 | Y62H9A.12 |
| WBGene00013401 | 0.020317245 | 0.84643798 | NA | Y62H9A.13 |
| WBGene00013402 | 0.020232312 | 0.84643798 | NA | Y62H9A.14 |
| WBGene00013403 | -0.223927603 | 0.18799853 | 0.37307424 | fbxb-100 |
| WBGene00013404 | -0.033968392 | 0.74230162 | 0.85420405 | fbxb-101 |
| WBGene00013405 | -0.086564913 | 0.49155374 | 0.67506222 | tdpt-1 |
| WBGene00013406 | -0.257981769 | 0.03214517 | 0.10880816 | Y63D3A.7 |
| WBGene00013407 | -0.238017163 | 0.07702056 | 0.20532738 | fld-1 |
| WBGene00013408 | -1.212036581 | 0.0080218 | 0.03793513 | fbxb-93 |
| WBGene00013409 | -0.037330026 | 0.7388568 | 0.85307041 | fbxb-56 |
| WBGene00013410 | -0.073545765 | 0.64744574 | 0.79141821 | NA |
| WBGene00013411 | NA | NA | NA | Y64G10A.1 |
| WBGene00013412 | NA | NA | NA | Y64G10A.2 |
| WBGene00013415 | 0.841248114 | 0.0021548 | 0.01326676 | ilcr-1 |
| WBGene00013416 | 0.20429507 | 0.35190871 | 0.55181487 | Y64G10A.7 |
| WBGene00013417 | NA | NA | NA | srab-24 |
| WBGene00013418 | -0.221046434 | 0.30154976 | 0.50214187 | Y65A5A.1 |
| WBGene00013419 | 0.039406759 | 0.73697999 | 0.85209856 | Y65A5A.2 |
| WBGene00013421 | 0.015451648 | 0.80337623 | NA | fbxa-220 |
| WBGene00013422 | -0.140581065 | 0.38400222 | 0.58333436 | Y66A7A.2 |
| WBGene00013424 | 0.119325954 | 0.42729532 | 0.62252204 | Y66A7A.4 |
| WBGene00013425 | 0.262485172 | 0.12601274 | 0.28699798 | ceh-91 |
| WBGene00013426 | -0.073606975 | 0.70747288 | 0.83486019 | Y66A7A.7 |
| WBGene00013427 | -0.025049301 | 0.69142686 | NA | Y66D12A.1 |
| WBGene00013429 | 0.254260001 | 0.18232633 | 0.36517643 | Y66D12A.3 |
| WBGene00013431 | 0.051165021 | 0.66520351 | 0.80363434 | ceh-92 |
| WBGene00013432 | 0.189758743 | 0.37076861 | 0.57036433 | Y66D12A.6 |
| WBGene00013433 | 0.070461734 | 0.68375997 | 0.81761922 | Y66D12A.7 |
| WBGene00013434 | 0.134502123 | 0.24271624 | 0.44083966 | Y66D12A.8 |
| WBGene00013435 | -0.253709765 | 0.02976946 | 0.10301459 | Y66D12A.9 |
| WBGene00013436 | -0.069908454 | 0.55254857 | 0.72491534 | Y66D12A.10 |

| | | | | |
|---|---|---|---|---|
| WBGene00013437 | 0.034490003 | 0.73115255 | 0.84986592 | Y66D12A.11 |
| WBGene00013438 | -0.577208419 | 0.00406466 | 0.02215588 | ztf-29 |
| WBGene00013439 | 0.721963698 | 0.00271466 | 0.01608469 | Y66D12A.13 |
| WBGene00013440 | 0.242071282 | 0.22268556 | 0.41543569 | nrps-1 |
| WBGene00013441 | -0.111942288 | 0.23881896 | 0.43592425 | xpb-1 |
| WBGene00013442 | -0.120270438 | 0.51868786 | 0.6979082 | Y66D12A.16 |
| WBGene00013443 | -0.136559581 | 0.16436392 | 0.34037514 | such-1 |
| WBGene00013445 | -0.329047726 | 0.00676769 | 0.03318345 | Y66D12A.19 |
| WBGene00013446 | -0.005041403 | 0.9643435 | 0.98048286 | Y66D12A.21 |
| WBGene00013447 | 0.178100306 | 0.14953972 | 0.32088686 | nars-2 |
| WBGene00013448 | 0.068895128 | 0.73120208 | 0.84986592 | Y66D12A.24 |
| WBGene00013449 | 0.1298478 | 0.10432033 | NA | Y67A6A.1 |
| WBGene00013450 | 0.020232312 | 0.84643798 | NA | oac-56 |
| WBGene00013451 | NA | NA | NA | Y67A10A.2 |
| WBGene00013452 | -0.045771477 | 0.50913401 | NA | Y67A10A.3 |
| WBGene00013453 | -0.37604367 | 0.08938584 | 0.22718866 | fbxa-83 |
| WBGene00013454 | NA | NA | NA | fbxa-126 |
| WBGene00013455 | 0.516431829 | 0.04562604 | 0.1407397 | ssu-2 |
| WBGene00013456 | 0.062364914 | 0.51089023 | NA | Y67A10A.7 |
| WBGene00013457 | -0.173057042 | 0.25697514 | 0.45521755 | paqr-3 |
| WBGene00013458 | 0.010296628 | 0.96162111 | 0.97914283 | Y67A10A.9 |
| WBGene00013459 | -0.124966083 | 0.53287735 | 0.70870744 | Y67A10A.10 |
| WBGene00013460 | -0.096101688 | 0.32020981 | 0.52007255 | cpsf-3 |
| WBGene00013461 | -0.074720407 | 0.4549304 | 0.64691491 | Y67H2A.2 |
| WBGene00013462 | 0.161243922 | 0.10253804 | 0.24950298 | micu-1 |
| WBGene00013463 | -0.228163793 | 3.08E-05 | 0.00033657 | kdp-1 |
| WBGene00013464 | 0.059768375 | 0.62223378 | 0.77414385 | Y67H2A.7 |
| WBGene00013465 | -0.345275931 | 0.00023538 | 0.00202721 | Y67H2A.10 |
| WBGene00013466 | 0.019325396 | 0.77340931 | NA | srz-47 |
| WBGene00013467 | NA | NA | NA | NA |
| WBGene00013468 | NA | NA | NA | srz-96 |
| WBGene00013469 | NA | NA | NA | Y68A4A.10 |
| WBGene00013470 | NA | NA | NA | clec-243 |
| WBGene00013471 | NA | NA | NA | clec-242 |
| WBGene00013472 | NA | NA | NA | Y68A4B.3 |
| WBGene00013473 | 2.121408152 | 0.00057927 | 0.00440573 | Y69E1A.1 |
| WBGene00013474 | 0.619745445 | 0.03880491 | 0.12511401 | Y69E1A.2 |
| WBGene00013475 | 0.018998992 | 0.77286019 | NA | Y69E1A.3 |
| WBGene00013476 | 0.038726127 | 0.57551883 | NA | Y69E1A.4 |
| WBGene00013477 | -1.025970659 | 3.64E-07 | 6.20E-06 | Y69E1A.5 |
| WBGene00013478 | 0.042516415 | 0.55991373 | NA | Y69E1A.8 |
| WBGene00013479 | 0.067086156 | 0.59568752 | 0.75477619 | Y69H2.1 |
| WBGene00013480 | 0.098378334 | 0.61180802 | 0.76638272 | egas-3 |

| | | | | | |
|---|---|---|---|---|---|
| WBGene00013481 | | -0.075279535 | 0.39607814 | 0.59403606 | Y69H2.3 |
| WBGene00013482 | | -0.069438859 | 0.52828608 | 0.70537326 | Y69H2.7 |
| WBGene00013483 | NA | | NA | NA | nhr-241 |
| WBGene00013484 | | 0.056218791 | 0.75218799 | 0.85970214 | Y69H2.9 |
| WBGene00013485 | | 0.079005989 | 0.59660972 | 0.75522171 | Y69H2.10 |
| WBGene00013486 | | 0.193882624 | 0.18857754 | 0.37365866 | egas-1 |
| WBGene00013487 | NA | | NA | NA | egas-2 |
| WBGene00013489 | | -1.947540019 | 5.63E-205 | 6.87E-202 | col-42 |
| WBGene00013490 | | -0.003421551 | 0.95689871 | NA | Y70C5A.2 |
| WBGene00013491 | NA | | NA | NA | Y70C5B.1 |
| WBGene00013491 | NA | | NA | NA | W06G6.21 |
| WBGene00013492 | | 0.070972377 | 0.71258687 | 0.83839476 | Y70C5C.1 |
| WBGene00013493 | | 0.090416984 | 0.26156585 | NA | clec-9 |
| WBGene00013494 | | 0.020232312 | 0.84643798 | NA | irld-58 |
| WBGene00013495 | | -0.00346345 | 0.95708614 | NA | NA |
| WBGene00013496 | | 0.246373656 | 0.10547568 | 0.25420073 | Y70D2A.1 |
| WBGene00013497 | | 0.08277932 | 0.67758289 | 0.81268871 | hex-5 |
| WBGene00013498 | | -0.011426835 | 0.90807442 | 0.950398 | Y70G10A.2 |
| WBGene00013499 | | 1.030038771 | 1.41E-06 | 2.13E-05 | Y70G10A.3 |
| WBGene00013500 | NA | | NA | NA | Y71A12B.2 |
| WBGene00013500 | NA | | NA | NA | Y71A12B.3 |
| WBGene00013501 | NA | | NA | NA | Y71A12B.2 |
| WBGene00013501 | NA | | NA | NA | Y71A12B.3 |
| WBGene00013502 | NA | | NA | NA | clec-111 |
| WBGene00013503 | NA | | NA | NA | clec-112 |
| WBGene00013505 | | -0.027350315 | 0.68023343 | NA | Y71A12B.8 |
| WBGene00013506 | | -0.015018289 | 0.92348042 | 0.95667422 | usp-3 |
| WBGene00013507 | | 0.284992333 | 0.11283482 | 0.26619636 | Y71A12B.10 |
| WBGene00013508 | | -1.380139155 | 4.93E-12 | 1.70E-10 | Y71A12B.11 |
| WBGene00013509 | | -0.023029752 | 0.9090862 | 0.95068624 | Y71A12B.12 |
| WBGene00013510 | | 0.263981135 | 0.2357595 | 0.43157725 | NA |
| WBGene00013511 | | -0.110097519 | 0.33260744 | 0.53237822 | Y71A12B.15 |
| WBGene00013512 | NA | | NA | NA | nhr-276 |
| WBGene00013513 | | 0.074182736 | 0.67283729 | 0.80971649 | Y71A12C.2 |
| WBGene00013514 | | 0.171263083 | 0.37673234 | 0.57581566 | Y73F4A.1 |
| WBGene00013515 | | 1.371793161 | 2.15E-08 | 4.55E-07 | Y73F4A.2 |
| WBGene00013516 | | 2.585170406 | 1.12E-10 | 3.19E-09 | Y73F4A.3 |
| WBGene00013517 | | -0.026384613 | 0.67299636 | NA | lgc-52 |
| WBGene00013518 | | 0.696793598 | 0.0304255 | 0.10444544 | Y73F8A.5 |
| WBGene00013519 | NA | | NA | NA | Y73F8A.10 |
| WBGene00013520 | | -0.317050246 | 0.17655795 | 0.35738765 | best-21 |
| WBGene00013521 | | 0.059980168 | 0.38780842 | NA | Y73F8A.12 |
| WBGene00013522 | | -0.056327531 | 0.78384329 | 0.87839645 | Y73F8A.13 |

| | | | | |
|---|---|---|---|---|
| WBGene00013523 | 0.020925399 | 0.84643798 | NA | Y73F8A.14 |
| WBGene00013524 | 0.0398788 | 0.58971221 | NA | Y73F8A.15 |
| WBGene00013525 | NA | NA | NA | irld-59 |
| WBGene00013526 | 0.131317611 | 0.10121567 | NA | Y73F8A.20 |
| WBGene00013527 | 0.051702513 | 0.65590877 | 0.79661222 | Y73F8A.22 |
| WBGene00013528 | 0.020232312 | 0.84643798 | NA | Y73F8A.23 |
| WBGene00013529 | -0.069670897 | 0.5962174 | 0.75503103 | Y73F8A.24 |
| WBGene00013530 | -0.105672906 | 0.25307784 | 0.45115216 | ntl-11 |
| WBGene00013531 | 0.197950965 | 0.31880279 | 0.51898076 | Y73F8A.26 |
| WBGene00013532 | -0.319407134 | 0.00746747 | 0.03583042 | Y73F8A.27 |
| WBGene00013534 | 0.084818269 | 0.57494977 | 0.74227635 | acr-24 |
| WBGene00013535 | 0.120371441 | 0.53687231 | 0.71185903 | NA |
| WBGene00013536 | 0.010746493 | 0.91649192 | 0.95348243 | Y73F8A.32 |
| WBGene00013537 | 0.042199095 | 0.55922995 | NA | NA |
| WBGene00013538 | -0.072975895 | 0.43810261 | 0.63294458 | ari-1.4 |
| WBGene00013539 | 0.527596706 | 0.0410867 | 0.13048818 | Y73F8A.35 |
| WBGene00013540 | 0.811417691 | 0.0006268 | 0.00472555 | Y75B8A.3 |
| WBGene00013541 | -0.193405412 | 0.26175533 | 0.45968092 | lonp-2 |
| WBGene00013542 | 0.037971159 | 0.84244865 | 0.91507461 | lron-12 |
| WBGene00013543 | 0.197091815 | 0.34034906 | 0.5401272 | Y75B8A.6 |
| WBGene00013544 | -0.266171967 | 0.06654674 | 0.18446648 | Y75B8A.7 |
| WBGene00013545 | 0.024600961 | 0.78121715 | 0.87672852 | Y75B8A.8 |
| WBGene00013547 | 0.017279211 | 0.79380376 | NA | Y75B8A.10 |
| WBGene00013548 | 0.503809859 | 0.06594875 | 0.1833499 | lury-1 |
| WBGene00013549 | -0.026793974 | 0.85146282 | 0.92034307 | Y75B8A.13 |
| WBGene00013550 | -0.141345944 | 0.34059215 | 0.54037817 | Y75B8A.14 |
| WBGene00013551 | -0.095769879 | 0.48126179 | 0.66742323 | Y75B8A.16 |
| WBGene00013552 | -0.129280088 | 0.1306922 | 0.2930716 | gmn-1 |
| WBGene00013553 | -0.171992417 | 0.20179756 | 0.39044774 | Y75B8A.18 |
| WBGene00013554 | -0.150223775 | 0.45175503 | 0.64408856 | Y75B8A.19 |
| WBGene00013555 | 0.106253557 | 0.37088808 | 0.57036433 | fbxa-89 |
| WBGene00013556 | 0.201898175 | 0.21670616 | 0.40823404 | Y75B8A.23 |
| WBGene00013557 | -0.261937959 | 0.00054405 | 0.00416819 | Y75B8A.24 |
| WBGene00013558 | -0.047219207 | 0.67158392 | 0.80894366 | Y75B8A.25 |
| WBGene00013559 | 1.030956716 | 0.00078376 | 0.00570614 | Y75B8A.28 |
| WBGene00013560 | 0.131068514 | 0.46835568 | 0.65738945 | zip-12 |
| WBGene00013561 | -0.106872159 | 0.36247473 | 0.56230055 | Y75B8A.31 |
| WBGene00013562 | 0.211661371 | 0.30926941 | 0.51014122 | Y75B8A.32 |
| WBGene00013563 | -0.031228362 | 0.74290509 | NA | Y75B8A.33 |
| WBGene00013564 | -0.023056499 | 0.78121253 | NA | Y75B8A.34 |
| WBGene00013565 | NA | NA | NA | Y75B8A.37 |
| WBGene00013566 | -0.008989285 | 0.92510764 | 0.95775003 | Y75B12A.2 |
| WBGene00013567 | -0.158158084 | 0.01899517 | 0.07408319 | Y75B12B.1 |

| | | | | | |
|---|---|---|---|---|---|
| WBGene00013568 | | 0.184844347 | 0.30888642 | 0.50968174 | Y75B12B.3 |
| WBGene00013569 | | -0.077602922 | 0.70802074 | 0.83523278 | bath-36 |
| WBGene00013570 | NA | | NA | NA | Y75B12B.8 |
| WBGene00013570 | NA | | NA | NA | C41G6.13 |
| WBGene00013571 | NA | | NA | NA | NA |
| WBGene00013572 | NA | | NA | NA | Y75B12B.10 |
| WBGene00013573 | NA | | NA | NA | Y75B12B.11 |
| WBGene00013574 | | 0.073478574 | 0.67180372 | 0.80894366 | lron-13 |
| WBGene00013575 | | -0.134332946 | 0.1960112 | 0.38361181 | acs-5 |
| WBGene00013576 | | 0.067107961 | 0.73897573 | 0.85307041 | Y76A2B.4 |
| WBGene00013577 | | 0.104126981 | 0.27715047 | 0.4772151 | Y76A2B.5 |
| WBGene00013578 | | -0.051121758 | 0.56999236 | 0.7387023 | scav-2 |
| WBGene00013579 | | -0.117201499 | 0.55788478 | 0.7288294 | Y79H2A.2 |
| WBGene00013580 | | -0.12586561 | 0.20739693 | 0.39646147 | Y79H2A.3 |
| WBGene00013581 | | 0.055325607 | 0.63835786 | 0.78552676 | Y79H2A.4 |
| WBGene00013583 | | -2.035676047 | 0.00026873 | 0.00228083 | ceh-51 |
| WBGene00013584 | | -0.02943488 | 0.88499899 | 0.93940914 | nhr-243 |
| WBGene00013585 | | 0.482526698 | 0.0725216 | 0.19622782 | cyp-42A1 |
| WBGene00013586 | | -0.023109762 | 0.78121253 | NA | Y80D3A.8 |
| WBGene00013587 | | -0.772156377 | 0.00018643 | 0.00166235 | Y80D3A.9 |
| WBGene00013588 | | 1.111596039 | 0.0068294 | 0.03341488 | nlp-42 |
| WBGene00013589 | | -0.162384244 | 0.16380049 | 0.33954605 | NA |
| WBGene00013590 | | 0.023925244 | 0.85303918 | 0.92124708 | Y81G3A.1 |
| WBGene00013591 | | -0.179300792 | 0.12050775 | 0.27934672 | gcn-2 |
| WBGene00013592 | | 0.058702183 | 0.51805387 | NA | Y81G3A.4 |
| WBGene00013593 | | 0.005973769 | 0.9554201 | 0.9749307 | Y87G2A.1 |
| WBGene00013594 | | -0.393785356 | 0.06341462 | 0.17865182 | Y87G2A.2 |
| WBGene00013595 | | -0.033334028 | 0.79857021 | 0.88715885 | atg-4.1 |
| WBGene00013596 | | 0.31859312 | 0.00448734 | 0.02406564 | nyn-2 |
| WBGene00013597 | | -0.055456846 | 0.38701998 | 0.58569567 | gpi-1 |
| WBGene00013598 | | -0.083062933 | 0.4605 | 0.6515443 | vps-28 |
| WBGene00013599 | | -0.057434385 | 0.6988011 | 0.82846224 | eipr-1 |
| WBGene00013601 | | -0.004766758 | 0.96577432 | 0.98082345 | Y87G2A.13 |
| WBGene00013602 | | 0.401231293 | 0.06886162 | 0.18904987 | Y87G2A.16 |
| WBGene00013603 | | 0.144111152 | 0.49948592 | 0.68141757 | soem-1 |
| WBGene00013604 | | -0.087257471 | 0.61677401 | 0.77029671 | Y87G2A.18 |
| WBGene00013605 | | -0.106347677 | 0.48920054 | 0.67323737 | Y95D11A.1 |
| WBGene00013606 | | -0.270305661 | 1.13E-05 | 0.0001376 | cand-1 |
| WBGene00013607 | | 0.21997274 | 0.126272 | 0.28734042 | clec-27 |
| WBGene00013608 | | 2.826664876 | 0.00378801 | 0.0209838 | clec-35 |
| WBGene00013609 | | 0.285170158 | 0.03244867 | 0.10959191 | clec-37 |
| WBGene00013610 | | -0.061424779 | 0.6729225 | 0.8097193 | fbxa-206 |
| WBGene00013611 | | 0.041778173 | 0.81849009 | 0.90020383 | Y102A5C.2 |

| | | | | |
|---|---|---|---|---|
| WBGene00013612 | 0.322212364 | 0.15431574 | 0.32720301 | NA |
| WBGene00013613 | 0.008151406 | 0.84680818 | NA | Y102A5C.4 |
| WBGene00013614 | NA | NA | NA | NA |
| WBGene00013615 | 0.0398788 | 0.58971221 | NA | NA |
| WBGene00013616 | NA | NA | NA | fbxa-148 |
| WBGene00013617 | NA | NA | NA | NA |
| WBGene00013618 | NA | NA | NA | NA |
| WBGene00013619 | NA | NA | NA | fbxa-112 |
| WBGene00013620 | -0.004944805 | 0.94514249 | NA | fbxa-111 |
| WBGene00013621 | NA | NA | NA | clec-239 |
| WBGene00013622 | -0.081849184 | 0.47479968 | 0.66198859 | clec-238 |
| WBGene00013623 | -0.089318869 | 0.64793594 | 0.7915913 | fbxa-90 |
| WBGene00013624 | NA | NA | NA | NA |
| WBGene00013625 | NA | NA | NA | srz-53 |
| WBGene00013626 | NA | NA | NA | srz-54 |
| WBGene00013627 | NA | NA | NA | NA |
| WBGene00013628 | NA | NA | NA | Y102A5C.27 |
| WBGene00013629 | NA | NA | NA | srz-95 |
| WBGene00013631 | NA | NA | NA | Y102F5A.1 |
| WBGene00013632 | -0.410021872 | 0.04236464 | 0.13333206 | Y105C5A.1 |
| WBGene00013633 | -0.306704489 | 0.12564542 | 0.28666356 | Y105C5A.8 |
| WBGene00013634 | NA | NA | NA | Y105C5A.9 |
| WBGene00013635 | -0.022930767 | 0.78121253 | NA | Y105C5A.10 |
| WBGene00013636 | NA | NA | NA | Y105C5A.13 |
| WBGene00013636 | NA | NA | NA | Y105C5A.12 |
| WBGene00013637 | 0.038125098 | 0.61119482 | NA | Y105C5A.13 |
| WBGene00013638 | -0.530767862 | 0.02525066 | 0.09139553 | NA |
| WBGene00013639 | 0.667189958 | 0.00560487 | 0.02864253 | Y105C5A.15 |
| WBGene00013640 | NA | NA | NA | NA |
| WBGene00013642 | 0.125529235 | 0.45835135 | 0.64976513 | daf-38 |
| WBGene00013643 | 1.403451655 | 4.38E-08 | 8.76E-07 | Y105C5A.24 |
| WBGene00013644 | -0.297788091 | 0.14829489 | 0.31901051 | NA |
| WBGene00013645 | NA | NA | NA | Y105C5B.1 |
| WBGene00013646 | 0.221641161 | 0.10040235 | 0.24610595 | Y105C5B.3 |
| WBGene00013647 | 0.098858075 | 0.32817568 | 0.52759636 | Y105C5B.5 |
| WBGene00013648 | 5.039725977 | 0.0001717 | 0.00154986 | NA |
| WBGene00013649 | -0.022930767 | 0.78121253 | NA | Y105C5B.8 |
| WBGene00013650 | -0.476377926 | 0.06327305 | 0.17846903 | Y105C5B.9 |
| WBGene00013651 | -0.032662398 | 0.7706675 | 0.86948778 | Y105C5B.11 |
| WBGene00013652 | -0.485931979 | 0.05748069 | 0.1665825 | NA |
| WBGene00013653 | 0.032270442 | 0.76696213 | 0.86758224 | Y105C5B.14 |
| WBGene00013654 | 0.907617142 | 2.36E-16 | 1.31E-14 | Y105C5B.15 |
| WBGene00013655 | 0.073887753 | 0.28837975 | NA | lgc-19 |

| | | | | |
|---|---|---|---|---|
| WBGene00013656 | 0.020925399 | 0.84643798 | NA | Y105C5B.17 |
| WBGene00013657 | 0.229957869 | 0.02159441 | 0.08119255 | Y105C5B.18 |
| WBGene00013658 | 0.003505105 | 0.98628229 | 0.99217803 | Y105C5B.19 |
| WBGene00013659 | -0.058221182 | 0.74660024 | 0.85672656 | Y105C5B.20 |
| WBGene00013661 | NA | NA | NA | NA |
| WBGene00013662 | 0.074892148 | 0.28555128 | NA | Y105C5B.25 |
| WBGene00013665 | 0.017501802 | 0.83221062 | NA | hlh-32 |
| WBGene00013666 | 0.723473114 | 0.01512975 | 0.06231074 | Y105E8A.1 |
| WBGene00013667 | 0.100058215 | 0.31047386 | 0.51126394 | Y105E8A.2 |
| WBGene00013668 | -0.148789025 | 0.06290292 | 0.17772319 | Y105E8A.3 |
| WBGene00013669 | -0.099236699 | 0.40123954 | 0.59956605 | tcab-1 |
| WBGene00013670 | 0.151028205 | 0.08635452 | 0.22200466 | hpo-13 |
| WBGene00013671 | 0.27977494 | 0.07575261 | 0.20301922 | Y105E8A.11 |
| WBGene00013672 | 0.094538716 | 0.5806653 | 0.74567654 | catp-1 |
| WBGene00013673 | -0.165447005 | 0.28875167 | 0.48896752 | Y105E8A.13 |
| WBGene00013674 | -0.146311816 | 0.1124189 | 0.26542401 | Y105E8A.14 |
| WBGene00013676 | 0.085377123 | 0.4202259 | 0.61691841 | ekl-4 |
| WBGene00013677 | -0.248949176 | 0.00574915 | 0.02924715 | yars-1 |
| WBGene00013678 | -0.01055144 | 0.9309597 | 0.96102242 | Y105E8A.20 |
| WBGene00013679 | 0.033807916 | 0.72666429 | 0.8471289 | Y105E8A.21 |
| WBGene00013680 | -0.079977633 | 0.48959162 | 0.67356802 | rpom-1 |
| WBGene00013682 | 0.26107167 | 0.00160601 | 0.01042266 | Y105E8A.25 |
| WBGene00013683 | 0.215201907 | 0.17395559 | 0.35444388 | zoo-1 |
| WBGene00013684 | 0.309321554 | 0.02842676 | 0.09966385 | Y105E8A.27 |
| WBGene00013685 | 0.374047715 | 0.07120245 | 0.1939142 | Y105E8A.28 |
| WBGene00013687 | -0.082850799 | 0.36372044 | 0.56356544 | exoc-8 |
| WBGene00013688 | -0.062191712 | 0.44764844 | 0.64054152 | riok-2 |
| WBGene00013689 | -0.08813977 | 0.26486335 | 0.46341591 | bath-40 |
| WBGene00013690 | -0.318944451 | 0.00479985 | 0.02531508 | hprt-1 |
| WBGene00013691 | 0.056588823 | 0.7857272 | 0.87882597 | Y105E8B.6 |
| WBGene00013692 | 0.44367446 | 0.07477668 | 0.2010662 | Y105E8B.7 |
| WBGene00013693 | 0.063403701 | 0.75777433 | 0.86298802 | Y105E8B.9 |
| WBGene00013694 | -0.144647887 | 0.2837986 | 0.48371307 | Y106G6A.1 |
| WBGene00013695 | 0.002151486 | 0.98364635 | 0.99058801 | epg-8 |
| WBGene00013696 | 0.102123031 | 0.50888825 | 0.68923523 | Y106G6A.4 |
| WBGene00013697 | -0.155084506 | 0.13609806 | 0.30108972 | dsbn-1 |
| WBGene00013698 | -0.366290511 | 0.09896363 | 0.24405148 | Y106G6D.1 |
| WBGene00013699 | -0.794223876 | 0.00190507 | 0.01201213 | Y106G6D.2 |
| WBGene00013700 | 0.371210144 | 0.01846443 | 0.0725155 | Y106G6D.3 |
| WBGene00013701 | 0.359223346 | 0.0727439 | 0.19672025 | Y106G6D.4 |
| WBGene00013702 | -0.165050326 | 0.39107975 | 0.58907393 | Y106G6D.6 |
| WBGene00013703 | -0.197129344 | 0.04114807 | 0.13057958 | rbm-12 |
| WBGene00013704 | 0.098859193 | 0.60173832 | 0.75908213 | Y106G6D.8 |

| | | | | |
|---|---|---|---|---|
| WBGene00013705 | 0.020232312 | 0.84643798 | NA | Y106G6E.1 |
| WBGene00013706 | NA | NA | NA | NA |
| WBGene00013707 | NA | NA | NA | NA |
| WBGene00013708 | 0.243294756 | 0.2448929 | 0.4429796 | Y106G6E.4 |
| WBGene00013709 | 0.018122852 | 0.8327738 | 0.90874571 | csnk-1 |
| WBGene00013710 | 0.0103327 | 0.86888291 | 0.93000745 | Y106G6G.1 |
| WBGene00013711 | 0.431847025 | 0.09303552 | 0.23312221 | Y106G6G.2 |
| WBGene00013712 | 0.017138317 | 0.82172227 | NA | dlc-6 |
| WBGene00013713 | 1.757124966 | 0.00186839 | 0.01182154 | Y106G6G.4 |
| WBGene00013716 | 0.071272178 | 0.69823792 | 0.82816765 | Y106G6H.1 |
| WBGene00013717 | -0.0391316 | 0.81460962 | 0.897892 | madf-10 |
| WBGene00013718 | 0.004888746 | 0.95737628 | 0.97651847 | pdpr-1 |
| WBGene00013719 | -0.117411034 | 0.21844528 | 0.41031093 | Y106G6H.6 |
| WBGene00013720 | -0.167562036 | 0.19181559 | 0.37797739 | Y106G6H.8 |
| WBGene00013721 | NA | NA | NA | Y106G6H.9 |
| WBGene00013722 | -0.174651877 | 0.21388705 | 0.40474606 | NA |
| WBGene00013723 | 2.756250371 | 0.00099216 | 0.0069987 | Y106G6H.13 |
| WBGene00013724 | -0.088755159 | 0.51514697 | 0.69494764 | Y106G6H.14 |
| WBGene00013725 | -0.183475638 | 0.13672846 | 0.30202798 | ska-1 |
| WBGene00013726 | -0.010723225 | 0.91811339 | 0.95442332 | Y106G6H.16 |
| WBGene00013727 | -0.371332463 | 0.0799996 | 0.21104039 | Y111B2A.1 |
| WBGene00013728 | -0.152096965 | 0.09886579 | 0.24389117 | Y111B2A.2 |
| WBGene00013729 | -0.10366736 | 0.22469688 | 0.41759166 | Y111B2A.3 |
| WBGene00013730 | -0.105995731 | 0.36805026 | 0.56782127 | sql-1 |
| WBGene00013732 | 0.545398509 | 5.86E-07 | 9.69E-06 | aakg-1 |
| WBGene00013734 | 0.038521875 | 0.71750824 | 0.84158547 | Y111B2A.10 |
| WBGene00013735 | -0.152937333 | 0.19535654 | 0.38258628 | Y111B2A.12 |
| WBGene00013736 | 0.335131406 | 0.1302627 | 0.29250015 | gtf-2A2 |
| WBGene00013737 | -0.22034479 | 0.04130055 | 0.13094009 | tpst-1 |
| WBGene00013738 | -0.147494774 | 0.18788535 | 0.37294324 | elpc-2 |
| WBGene00013739 | -0.122865728 | 0.44955075 | 0.64204323 | spin-4 |
| WBGene00013740 | -0.030410358 | 0.78846095 | 0.88033682 | hut-1 |
| WBGene00013741 | 0.129096005 | 0.54262291 | 0.71643448 | Y111B2A.21 |
| WBGene00013742 | -0.057405274 | 0.66437829 | 0.80310126 | sas-1 |
| WBGene00013744 | 0.020317245 | 0.84643798 | NA | Y111B2A.27 |
| WBGene00013746 | 1.105574771 | 0.01427449 | 0.05942359 | lgc-55 |
| WBGene00013747 | 0.106997704 | 0.16248985 | NA | spe-19 |
| WBGene00013748 | -0.024914729 | 0.8631637 | 0.9270525 | ssu-1 |
| WBGene00013749 | 0.020925399 | 0.84643798 | NA | Y113G7A.12 |
| WBGene00013750 | 0.204495506 | 0.27487078 | 0.47516409 | Y113G7A.13 |
| WBGene00013751 | 0.181322802 | 0.1486404 | 0.31951903 | Y113G7A.14 |
| WBGene00013752 | 0.476447151 | 0.06430163 | 0.18038708 | nlp-69 |
| WBGene00013753 | 0.049310307 | 0.64658669 | 0.79080075 | Y113G7A.16 |

| | | | | |
|---|---|---|---|---|
| WBGene00013754 | 0.056411843 | 0.41891328 | NA | fbxa-116 |
| WBGene00013755 | 0.397028191 | 0.11096638 | 0.26296689 | fbxa-115 |
| WBGene00013756 | -0.143164647 | 0.45829466 | 0.64976513 | ftr-1 |
| WBGene00013757 | 0.002302099 | 0.98732238 | 0.99274686 | fbxa-113 |
| WBGene00013758 | -0.150881643 | 0.38870581 | 0.58712904 | fbxa-114 |
| WBGene00013759 | -0.16163644 | 0.38515924 | 0.5843644 | fbxb-59 |
| WBGene00013760 | 0.016059264 | 0.84014476 | NA | srbc-34 |
| WBGene00013761 | 0.03037485 | 0.78455509 | 0.87854069 | Y113G7B.11 |
| WBGene00013762 | 0.115368909 | 0.5661531 | 0.73538965 | Y113G7B.12 |
| WBGene00013763 | 0.393387294 | 0.09910873 | 0.24412027 | NA |
| WBGene00013764 | 0.131894968 | 0.16034585 | NA | Y113G7B.15 |
| WBGene00013765 | -0.206229834 | 0.11018805 | 0.2618821 | cdkr-3 |
| WBGene00013766 | -0.45814154 | 3.45E-24 | 3.10E-22 | prmt-1 |
| WBGene00013768 | -0.006282279 | 0.96515708 | 0.98071026 | sld-5 |
| WBGene00013770 | -0.023109762 | 0.78121253 | NA | NA |
| WBGene00013771 | 2.905156022 | 0.00211725 | 0.01309617 | Y113G7C.1 |
| WBGene00013772 | NA | NA | NA | NA |
| WBGene00013773 | -0.001993815 | 0.95612996 | NA | Y116A8A.2 |
| WBGene00013774 | 0.040107227 | 0.58693163 | NA | clec-193 |
| WBGene00013775 | NA | NA | NA | Y116A8A.4 |
| WBGene00013776 | 0.060177748 | 0.41318197 | NA | Y116A8A.6 |
| WBGene00013777 | NA | NA | NA | Y116A8A.7 |
| WBGene00013778 | 0.110173013 | 0.4178843 | 0.61452833 | clec-194 |
| WBGene00013779 | -0.594095337 | 0.03878861 | 0.12509661 | Y116A8B.1 |
| WBGene00013781 | 0.19260518 | 0.16175989 | 0.33664911 | Y116A8B.4 |
| WBGene00013782 | 0.014067275 | 0.88561763 | NA | npr-32 |
| WBGene00013783 | NA | NA | NA | NA |
| WBGene00013784 | 0.285510179 | 0.21035768 | 0.3999801 | Y116A8C.3 |
| WBGene00013785 | 0.100806964 | 0.4256516 | 0.6215184 | nep-23 |
| WBGene00013786 | 0.080169123 | 0.46912059 | 0.65776619 | nep-24 |
| WBGene00013788 | NA | NA | NA | NA |
| WBGene00013789 | -0.085412565 | 0.56398809 | 0.73404113 | Y116A8C.9 |
| WBGene00013790 | -0.004663515 | 0.9746267 | 0.98580376 | Y116A8C.10 |
| WBGene00013791 | -0.023109762 | 0.78121253 | NA | Y116A8C.11 |
| WBGene00013792 | 0.291519435 | 0.03032318 | 0.104143 | Y116A8C.13 |
| WBGene00013794 | -0.148683362 | 0.47704628 | 0.66410831 | dct-13 |
| WBGene00013795 | 0.020232312 | 0.84643798 | NA | nhr-229 |
| WBGene00013796 | -0.391525536 | 0.09878046 | 0.24380389 | Y116A8C.19 |
| WBGene00013797 | -0.226320757 | 0.28226191 | 0.4820479 | Y116A8C.20 |
| WBGene00013798 | 0.020925399 | 0.84643798 | NA | clec-195 |
| WBGene00013799 | -0.736770091 | 0.0132533 | 0.0560166 | athp-3 |
| WBGene00013800 | 0.096598598 | 0.180451 | NA | Y116A8C.23 |
| WBGene00013801 | 0.020925399 | 0.84643798 | NA | Y116A8C.24 |

| | | | | | |
|---|---|---|---|---|---|
| WBGene00013802 | | 0.177350845 | 0.37137138 | 0.57065999 | Y116A8C.25 |
| WBGene00013803 | | -0.089306078 | 0.3456186 | 0.54528038 | snx-13 |
| WBGene00013804 | | -0.245468718 | 0.11311294 | 0.26680951 | Y116A8C.27 |
| WBGene00013805 | | -0.212647629 | 0.12968527 | 0.29201626 | bca-2 |
| WBGene00013806 | | 0.231885625 | 0.24949454 | 0.44781778 | Y116A8C.29 |
| WBGene00013807 | | 0.105890734 | 0.46257482 | 0.65252509 | Y116A8C.30 |
| WBGene00013808 | | -0.10064589 | 0.22504249 | 0.41796868 | sfa-1 |
| WBGene00013809 | | 0.092748393 | 0.64599832 | 0.79047734 | Y116A8C.33 |
| WBGene00013810 | NA | | NA | NA | Y116A8C.37 |
| WBGene00013811 | | 0.148021412 | 0.07455439 | NA | Y116A8C.38 |
| WBGene00013812 | | 0.056517449 | 0.53618828 | NA | Y116A8C.40 |
| WBGene00013814 | | 0.042065281 | 0.558937 | NA | Y116A8C.43 |
| WBGene00013815 | | 0.017279211 | 0.79380376 | NA | Y116A8C.44 |
| WBGene00013816 | | -2.812460292 | 0.00284413 | 0.01673667 | Y116F11A.1 |
| WBGene00013817 | NA | | NA | NA | Y116F11A.3 |
| WBGene00013819 | | 0.031395149 | 0.87477237 | 0.93322225 | Y116F11A.6 |
| WBGene00013820 | NA | | NA | NA | Y116F11B.2 |
| WBGene00013821 | NA | | NA | NA | NA |
| WBGene00013822 | | 0.155708417 | 0.37716314 | 0.57623349 | Y116F11B.6 |
| WBGene00013823 | | 0.018657025 | 0.77508599 | NA | Y116F11B.7 |
| WBGene00013824 | | 0.020232312 | 0.84643798 | NA | Y116F11B.8 |
| WBGene00013825 | | -0.027350315 | 0.68023343 | NA | Y116F11B.9 |
| WBGene00013826 | | -0.079344577 | 0.67673981 | 0.81237075 | NA |
| WBGene00013827 | | 0.020232312 | 0.84643798 | NA | Y116F11B.11 |
| WBGene00013828 | | -0.055120925 | 0.45606622 | NA | Y116F11B.13 |
| WBGene00013829 | | -0.004864426 | 0.94835335 | NA | Y116F11B.14 |
| WBGene00013830 | | -0.045771477 | 0.50913401 | NA | ZC15.1 |
| WBGene00013831 | NA | | NA | NA | clec-262 |
| WBGene00013832 | | -0.003270206 | 0.95698316 | NA | ZC15.3 |
| WBGene00013833 | | 0.056188364 | 0.42092875 | NA | ZC15.5 |
| WBGene00013834 | NA | | NA | NA | clec-261 |
| WBGene00013835 | | 0.101014341 | 0.61072309 | 0.7658156 | mlt-4 |
| WBGene00013836 | | 0.677110873 | 0.03397083 | 0.11334213 | fbxa-29 |
| WBGene00013837 | | 0.304150879 | 0.00963391 | 0.04368229 | fbxa-30 |
| WBGene00013838 | | 0.135958595 | 0.477722 | 0.66470097 | fbxa-31 |
| WBGene00013839 | | 0.067451448 | 0.71047596 | 0.83691999 | fbxa-32 |
| WBGene00013840 | | -0.022930767 | 0.78121253 | NA | fbxa-151 |
| WBGene00013841 | | 0.135885083 | 0.43827224 | 0.63306471 | NA |
| WBGene00013842 | | 0.18814839 | 0.27962747 | 0.47944787 | NA |
| WBGene00013843 | | 0.020925399 | 0.84643798 | NA | NA |
| WBGene00013844 | | 0.097328453 | 0.33491518 | 0.53414958 | fbxa-50 |
| WBGene00013845 | | 0.316480087 | 0.14819981 | 0.31885282 | fbxa-37 |
| WBGene00013846 | | 0.110677789 | 0.1956955 | NA | ZC84.1 |

| | | | | |
|---|---|---|---|---|
| WBGene00013847 | 0.859802243 | 2.15E-10 | 5.92E-09 | cls-3 |
| WBGene00013848 | 0.205462705 | 0.28091576 | 0.48048327 | npr-29 |
| WBGene00013849 | 0.048633652 | 0.56849329 | NA | ZC84.6 |
| WBGene00013850 | -0.005691347 | 0.93660638 | NA | ZC84.7 |
| WBGene00013852 | 0.188030149 | 0.19394606 | 0.3808686 | hhat-1 |
| WBGene00013853 | 1.378730766 | 9.02E-06 | 0.00011235 | ZC116.1 |
| WBGene00013854 | 1.341741184 | 0.01006191 | 0.04507847 | cyc-2.2 |
| WBGene00013855 | 0.043937347 | 0.82847357 | 0.9062566 | ZC116.3 |
| WBGene00013856 | -0.013563163 | 0.90068549 | 0.94699722 | ZC168.2 |
| WBGene00013857 | -0.09907532 | 0.29034994 | 0.49092715 | orc-5 |
| WBGene00013858 | 0.094392367 | 0.48203425 | 0.66787768 | ssp-34 |
| WBGene00013859 | 0.177257794 | 0.38579477 | 0.58472269 | ZC247.1 |
| WBGene00013860 | 0.361106777 | 0.12207289 | 0.28123828 | ZC247.2 |
| WBGene00013862 | -0.135310979 | 0.12745237 | 0.28874635 | wdr-5.3 |
| WBGene00013863 | -0.177216564 | 0.30617763 | 0.50686869 | ZC302.3 |
| WBGene00013865 | 0.127693253 | 0.10904601 | NA | ZC334.7 |
| WBGene00013866 | -0.518667073 | 0.00154638 | 0.01008041 | cbs-1 |
| WBGene00013867 | 0.436740552 | 8.76E-12 | 2.95E-10 | ZC373.2 |
| WBGene00013868 | -0.029234136 | 0.7166623 | NA | ZC373.3 |
| WBGene00013869 | 0.38389282 | 0.02692131 | 0.09579719 | mlck-1 |
| WBGene00013870 | 0.039228601 | 0.84696617 | 0.91761389 | ZC373.5 |
| WBGene00013871 | -0.062260494 | 0.60871717 | 0.76427649 | gnrr-3 |
| WBGene00013872 | 0.079224272 | 0.70487361 | 0.83266537 | ZC374.2 |
| WBGene00013873 | -0.282511758 | 0.09079328 | 0.22907194 | ZC376.1 |
| WBGene00013874 | -0.523709221 | 0.00396969 | 0.02174347 | ZC376.2 |
| WBGene00013875 | -0.217300151 | 0.30463953 | 0.50545453 | ZC376.3 |
| WBGene00013876 | -0.261330901 | 0.23758718 | 0.43394635 | ceh-74 |
| WBGene00013877 | -0.025427004 | 0.76414104 | 0.86623378 | ZC376.6 |
| WBGene00013878 | -0.123443725 | 0.15387983 | 0.3271315 | atfs-1 |
| WBGene00013879 | 0.075154054 | 0.33389143 | NA | ZC376.8 |
| WBGene00013880 | -0.312721124 | 0.00403539 | 0.02206054 | mppb-1 |
| WBGene00013881 | -0.063554704 | 0.55438349 | 0.7264592 | mans-4 |
| WBGene00013882 | 0.423614092 | 0.01238664 | 0.05301165 | msa-1 |
| WBGene00013883 | 0.01538317 | 0.9179428 | 0.95431375 | npr-13 |
| WBGene00013884 | -0.211900053 | 0.21688267 | 0.40840891 | ZC412.3 |
| WBGene00013886 | 0.077387215 | 0.23878538 | NA | ZC412.5 |
| WBGene00013887 | 0.116824906 | 0.24792566 | 0.44626009 | nspa-5 |
| WBGene00013887 | 0.116824906 | 0.24792566 | 0.44626009 | nspa-4 |
| WBGene00013888 | NA | NA | NA | nspa-1 |
| WBGene00013888 | NA | NA | NA | nspa-3 |
| WBGene00013889 | NA | NA | NA | H12D21.3 |
| WBGene00013889 | NA | NA | NA | ZC412.8 |
| WBGene00013890 | NA | NA | NA | H12D21.2 |

| | | | | |
|---|---|---|---|---|
| WBGene00013890 | NA | NA | NA | ZC412.9 |
| WBGene00013891 | 0.035895112 | 0.72552929 | 0.84627743 | ZC434.3 |
| WBGene00013892 | -0.050373728 | 0.61093462 | 0.76600957 | ZC434.4 |
| WBGene00013893 | -0.007862018 | 0.94122561 | 0.96684409 | coa-6 |
| WBGene00013894 | -0.316638849 | 2.04E-07 | 3.63E-06 | ZC434.8 |
| WBGene00013895 | 0.216882114 | 0.1854973 | 0.36940506 | ZC434.9 |
| WBGene00013896 | 0.58572748 | 0.03010775 | 0.10372113 | ZC443.1 |
| WBGene00013897 | 0.073458642 | 0.29111983 | NA | ZC443.2 |
| WBGene00013898 | 2.123969079 | 6.41E-08 | 1.25E-06 | ZC443.3 |
| WBGene00013899 | 1.817950497 | 1.97E-08 | 4.19E-07 | ZC443.4 |
| WBGene00013900 | 7.686928478 | 8.82E-09 | 1.99E-07 | ugt-18 |
| WBGene00013901 | -0.06855867 | 0.73979883 | 0.8534033 | ugt-16 |
| WBGene00013902 | 1.810184942 | 1.07E-06 | 1.68E-05 | ZC455.1 |
| WBGene00013903 | 0.098237519 | 0.52043047 | 0.69923849 | ugt-3 |
| WBGene00013904 | -0.046017863 | 0.82052714 | 0.90148005 | ugt-6 |
| WBGene00013905 | 0.170414296 | 0.41553512 | 0.61275434 | ugt-4 |
| WBGene00013906 | -0.030695822 | 0.88013101 | 0.93621027 | ugt-5 |
| WBGene00013907 | -0.023056499 | 0.78121253 | NA | sre-20 |
| WBGene00013908 | NA | NA | NA | srbc-43 |
| WBGene00013909 | 0.020317245 | 0.84643798 | NA | srbc-44 |
| WBGene00013910 | NA | NA | NA | srbc-45 |
| WBGene00013911 | 0.020925399 | 0.84643798 | NA | irld-60 |
| WBGene00013912 | NA | NA | NA | irld-19 |
| WBGene00013913 | NA | NA | NA | irld-61 |
| WBGene00013914 | 0.166714343 | 0.09920004 | 0.244263 | lgc-37 |
| WBGene00013915 | NA | NA | NA | irld-20 |
| WBGene00013916 | 0.04909525 | 0.45360729 | NA | ZC504.1 |
| WBGene00013917 | -0.24776506 | 0.07633061 | 0.2040995 | cdk-11.2 |
| WBGene00013918 | NA | NA | NA | NA |
| WBGene00013919 | 0.03558057 | 0.76091332 | 0.86448005 | ZC506.1 |
| WBGene00013920 | 0.450614275 | 0.00131536 | 0.00881383 | pssy-1 |
| WBGene00013921 | 0.332030825 | 0.09175432 | 0.23086011 | best-22 |
| WBGene00013923 | 0.51830498 | 0.06030958 | 0.17202271 | ghi-1 |
| WBGene00013924 | -0.20950938 | 0.00086887 | 0.00625129 | rad-23 |
| WBGene00013925 | 0.006078422 | 0.96341289 | 0.97987691 | coa-7 |
| WBGene00013926 | -0.292487405 | 0.0169933 | 0.06809486 | nep-1 |
| WBGene00013927 | 0.020232312 | 0.84643798 | NA | clec-95 |
| WBGene00013928 | 0.020317245 | 0.84643798 | NA | clec-94 |
| WBGene00013929 | NA | NA | NA | clec-93 |
| WBGene00013930 | NA | NA | NA | clec-96 |
| WBGene00013931 | -1.667536612 | 1.28E-12 | 4.75E-11 | clec-97 |
| WBGene00013932 | 0.020317245 | 0.84643798 | NA | clec-98 |
| WBGene00013933 | 0.074107881 | 0.33444485 | NA | clec-99 |

| | | | | |
|---|---|---|---|---|
| WBGene00013934 | 0.821308234 | 1.36E-05 | 0.00016152 | ZK131.11 |
| WBGene00013935 | 2.37493824 | 4.04E-09 | 9.53E-08 | ZK218.1 |
| WBGene00013937 | 7.788419418 | 5.61E-09 | 1.29E-07 | ZK218.3 |
| WBGene00013938 | 0.019975058 | 0.61254052 | NA | ZK218.4 |
| WBGene00013939 | 7.944997725 | 2.46E-28 | 2.82E-26 | ZK218.5 |
| WBGene00013940 | -0.022930767 | 0.78121253 | NA | nhr-248 |
| WBGene00013941 | 6.558301405 | 1.23E-06 | 1.90E-05 | ZK218.7 |
| WBGene00013942 | 0.020232312 | 0.84643798 | NA | ZK218.8 |
| WBGene00013943 | 7.149694034 | 1.06E-07 | 1.99E-06 | ZK218.11 |
| WBGene00013944 | NA | NA | NA | ZK228.1 |
| WBGene00013946 | -1.06918805 | 1.44E-06 | 2.18E-05 | ZK228.3 |
| WBGene00013947 | -0.585630965 | 0.00019288 | 0.00171249 | ZK228.4 |
| WBGene00013948 | NA | NA | NA | NA |
| WBGene00013949 | 0.549764751 | 0.04719914 | 0.14429788 | ZK262.2 |
| WBGene00013950 | 1.561579046 | 0.00262419 | 0.01562452 | ZK262.3 |
| WBGene00013951 | 0.033034883 | 0.64219503 | NA | ZK262.4 |
| WBGene00013953 | -0.001842784 | 0.95621055 | NA | ZK262.8 |
| WBGene00013954 | 0.035135283 | 0.6810866 | NA | ZK262.9 |
| WBGene00013955 | 0.219535981 | 0.29981765 | 0.50024276 | kri-1 |
| WBGene00013956 | 0.784314505 | 0.02479795 | 0.09029264 | ZK265.3 |
| WBGene00013957 | 0.140566555 | 0.20721908 | 0.39645783 | sre-23 |
| WBGene00013958 | -0.249505192 | 0.03051616 | 0.10473211 | nol-16 |
| WBGene00013959 | 1.706478833 | 0.00323047 | 0.01851168 | ZK265.7 |
| WBGene00013960 | 0.055897183 | 0.44636755 | NA | cutl-8 |
| WBGene00013961 | NA | NA | NA | ZK285.2 |
| WBGene00013962 | 0.513133511 | 0.062528 | 0.17694814 | ZK287.1 |
| WBGene00013963 | 0.304273686 | 0.15374207 | 0.32707608 | sulp-8 |
| WBGene00013964 | 0.387734856 | 0.09102423 | 0.22953245 | nlp-73 |
| WBGene00013965 | 0.044623782 | 0.80751064 | 0.89336279 | ZK287.4 |
| WBGene00013966 | -0.125874268 | 0.45528449 | 0.64722985 | ztf-9 |
| WBGene00013967 | -0.126893052 | 0.38094993 | 0.580263 | ZK287.7 |
| WBGene00013968 | 0.872108506 | 0.0038649 | 0.02130498 | ZK287.9 |
| WBGene00013969 | 0.991928299 | 1.27E-15 | 6.46E-14 | tep-1 |
| WBGene00013970 | 0.186426358 | 0.35926593 | 0.55945595 | klu-1 |
| WBGene00013971 | NA | NA | NA | scl-19 |
| WBGene00013972 | NA | NA | NA | scl-20 |
| WBGene00013973 | NA | NA | NA | asp-18 |
| WBGene00013974 | -0.029029491 | 0.71849588 | NA | npr-9 |
| WBGene00013975 | -1.998888731 | 0.00437546 | 0.02356922 | ZK455.5 |
| WBGene00013976 | 0.101381433 | 0.62956328 | 0.77968456 | hizr-1 |
| WBGene00013978 | -0.049807096 | 0.59052113 | 0.7516797 | ZK507.1 |
| WBGene00013979 | 0.053252768 | 0.6327435 | 0.78177194 | ZK507.3 |
| WBGene00013980 | 0.004860649 | 0.98265372 | 0.98995618 | dos-1 |

| | | | | |
|---|---|---|---|---|
| WBGene00013982 | -0.435116811 | 0.08672771 | 0.22265386 | ZK512.1 |
| WBGene00013983 | -0.477413301 | 2.08E-05 | 0.00023643 | ZK512.2 |
| WBGene00013984 | 0.006482178 | 0.9671831 | 0.98172923 | ZK512.4 |
| WBGene00013985 | 0.087691876 | 0.2977713 | 0.49887512 | sec-16 |
| WBGene00013986 | -1.07233134 | 0.00035716 | 0.00290998 | ZK512.7 |
| WBGene00013987 | 0.135148567 | 0.36533093 | 0.56493559 | ZK512.8 |
| WBGene00013988 | 0.127918313 | 0.10854122 | NA | ZK512.10 |
| WBGene00013989 | -0.068403008 | 0.71452325 | 0.83986311 | ZK512.11 |
| WBGene00013993 | NA | NA | NA | NA |
| WBGene00013994 | -0.159243662 | 0.08092941 | 0.21268849 | ZK524.4 |
| WBGene00013995 | 0.048485426 | 0.45975815 | NA | NA |
| WBGene00013996 | -0.323461034 | 0.109706 | 0.26095947 | ZK550.2 |
| WBGene00013997 | -0.427682048 | 8.33E-08 | 1.59E-06 | ZK550.3 |
| WBGene00013998 | 0.095357104 | 0.37083802 | 0.57036433 | gtf-2E1 |
| WBGene00013999 | 0.379639215 | 0.09824263 | 0.24309114 | ZK550.5 |
| WBGene00014000 | -0.079539485 | 0.54463027 | 0.71804879 | ZK550.6 |
| WBGene00014001 | 0.293721546 | 0.01425305 | 0.05936811 | pyk-2 |
| WBGene00014002 | 0.41863521 | 0.09260406 | 0.23228269 | ZK593.2 |
| WBGene00014003 | 1.785204212 | 1.69E-24 | 1.54E-22 | ZK593.3 |
| WBGene00014004 | -0.18924692 | 0.0621235 | 0.17603061 | fic-1 |
| WBGene00014005 | 0.20611848 | 0.20269619 | 0.39141056 | NA |
| WBGene00014006 | 2.761566238 | 1.71E-53 | 4.28E-51 | ZK596.1 |
| WBGene00014007 | 0.253989901 | 0.07913064 | 0.20946513 | ZK596.2 |
| WBGene00014008 | 0.049257133 | 0.73687681 | 0.85204655 | ZK596.3 |
| WBGene00014009 | 0.015896709 | 0.80422001 | NA | lips-6 |
| WBGene00014011 | -0.054949109 | 0.51476748 | 0.6947365 | ZK632.2 |
| WBGene00014012 | -0.105586186 | 0.32211274 | 0.52226496 | riok-3 |
| WBGene00014013 | 0.369936644 | 1.25E-05 | 0.00014972 | ZK632.4 |
| WBGene00014014 | 0.027720869 | 0.8227951 | 0.90297152 | ZK632.5 |
| WBGene00014015 | -0.144220513 | 0.05295454 | 0.15707123 | panl-3 |
| WBGene00014016 | -0.130837288 | 0.06976782 | 0.19082013 | ZK632.9 |
| WBGene00014017 | -0.008607464 | 0.91578876 | 0.95302176 | ZK632.10 |
| WBGene00014018 | -0.27149784 | 0.01055066 | 0.04661203 | ZK632.11 |
| WBGene00014019 | -0.123984242 | 0.2275404 | 0.42116549 | ZK632.12 |
| WBGene00014020 | -0.183035214 | 0.22414144 | 0.41711331 | ZK632.14 |
| WBGene00014021 | 0.015457248 | 0.94139454 | 0.96688582 | svop-1 |
| WBGene00014022 | -0.09356376 | 0.37967478 | 0.5792246 | ZK637.2 |
| WBGene00014023 | -0.142281421 | 0.12352512 | 0.28360221 | lnkn-1 |
| WBGene00014024 | -0.010404895 | 0.91903511 | 0.95511024 | ZK637.4 |
| WBGene00014025 | -0.156885174 | 0.07471365 | 0.20100745 | asna-1 |
| WBGene00014026 | NA | NA | NA | NA |
| WBGene00014027 | 0.220198644 | 0.30598405 | 0.50666281 | tpk-1 |
| WBGene00014028 | 0.469797739 | 0.03372303 | 0.11280303 | trxr-2 |

| | | | | |
|---|---|---|---|---|
| WBGene00014029 | 0.072255658 | 0.48921333 | 0.67323737 | ZK637.12 |
| WBGene00014030 | -0.242402722 | 0.23608073 | 0.43192937 | glb-1 |
| WBGene00014031 | -0.030717075 | 0.87510087 | 0.93336868 | ZK637.14 |
| WBGene00014032 | -0.101771545 | 0.44599558 | 0.63930108 | ZK637.15 |
| WBGene00014033 | 0.13838871 | 0.41845297 | 0.61517923 | arrd-15 |
| WBGene00014034 | -0.239330251 | 0.09881778 | 0.24383415 | ZK643.2 |
| WBGene00014035 | 1.20282443 | 0.00027361 | 0.00231288 | seb-2 |
| WBGene00014036 | 0.075521853 | 0.38992267 | 0.58786842 | ZK643.5 |
| WBGene00014037 | 0.05787416 | 0.40594086 | NA | ZK643.6 |
| WBGene00014038 | NA | NA | NA | ZK643.7 |
| WBGene00014039 | 0.484775955 | 0.02107992 | 0.07979088 | ZK662.2 |
| WBGene00014040 | -0.053158774 | 0.74905031 | 0.85832711 | ZK662.5 |
| WBGene00014041 | 0.065079126 | 0.68825994 | 0.82045382 | ZK666.1 |
| WBGene00014042 | 0.040107227 | 0.58693163 | NA | ZK666.2 |
| WBGene00014043 | 0.020925399 | 0.84643798 | NA | clec-58 |
| WBGene00014044 | -0.328717488 | 0.04257496 | 0.13387888 | ZK666.4 |
| WBGene00014045 | NA | NA | NA | clec-59 |
| WBGene00014046 | 6.083722645 | 1.38E-09 | 3.41E-08 | clec-60 |
| WBGene00014047 | 7.926184591 | 2.56E-09 | 6.13E-08 | clec-61 |
| WBGene00014048 | 0.058314668 | 0.40208118 | NA | ZK666.8 |
| WBGene00014049 | 0.15038247 | 0.07114191 | NA | ZK666.11 |
| WBGene00014050 | NA | NA | NA | ZK666.12 |
| WBGene00014051 | 0.486493139 | 0.0155757 | 0.06375282 | spv-1 |
| WBGene00014052 | 0.1092394 | 0.60442969 | 0.76077049 | ZK669.2 |
| WBGene00014053 | 0.631589149 | 0.00153779 | 0.01002885 | ZK669.3 |
| WBGene00014054 | -0.279496426 | 0.00036862 | 0.00299665 | dbt-1 |
| WBGene00014055 | 0.333070625 | 0.14238362 | 0.31058875 | ZK669.5 |
| WBGene00014057 | 2.120651773 | 6.42E-24 | 5.66E-22 | ZK673.1 |
| WBGene00014058 | 0.446425989 | 1.05E-07 | 1.98E-06 | ZK673.2 |
| WBGene00014060 | -0.279768177 | 0.21905018 | 0.41095767 | ZK673.4 |
| WBGene00014061 | -0.597869066 | 0.04344587 | 0.13588786 | ZK673.5 |
| WBGene00014062 | 0.143627571 | 0.24513902 | 0.44331526 | ZK673.6 |
| WBGene00014063 | 0.033338513 | 0.73736524 | 0.85234206 | clec-143 |
| WBGene00014064 | -0.288893112 | 0.19788799 | 0.38578935 | sdz-37 |
| WBGene00014065 | 0.122766639 | 0.39384284 | 0.59220016 | ZK673.11 |
| WBGene00014066 | 0.090159675 | 0.50709578 | 0.68770721 | rev-1 |
| WBGene00014067 | 0.145910628 | 0.20595942 | 0.39495113 | ZK675.4 |
| WBGene00014068 | NA | NA | NA | nhr-255 |
| WBGene00014069 | 0.07342307 | 0.54199906 | 0.71586903 | ZK678.3 |
| WBGene00014070 | -0.067905328 | 0.31502275 | NA | srxa-9 |
| WBGene00014073 | 0.063086775 | 0.41202602 | NA | ZK757.1 |
| WBGene00014074 | 0.190152639 | 0.26913528 | 0.46820634 | ZK757.2 |
| WBGene00014075 | -0.072391998 | 0.48458991 | 0.66978766 | dhhc-4 |

| | | | | |
|---|---|---|---|---|
| WBGene00014076 | 0.126967865 | 0.43227394 | 0.62736252 | ZK792.1 |
| WBGene00014077 | 1.653566532 | 0.00041677 | 0.00332892 | ZK792.4 |
| WBGene00014078 | -0.194907991 | 0.0495884 | 0.149325 | ZK792.5 |
| WBGene00014079 | -0.051497731 | 0.80232439 | 0.88993372 | ZK792.7 |
| WBGene00014080 | 0.077804739 | 0.65789547 | 0.79831751 | atg-4.2 |
| WBGene00014081 | 0.044712996 | 0.82814754 | 0.90617864 | ipmk-1 |
| WBGene00014082 | 0.160804925 | 0.24534409 | 0.44352178 | ZK795.2 |
| WBGene00014083 | -0.186646912 | 0.14278077 | 0.31112566 | ZK795.3 |
| WBGene00014084 | -0.003614866 | 0.95700117 | NA | snb-7 |
| WBGene00014085 | 0.050653172 | 0.67767793 | 0.81273613 | ZK809.1 |
| WBGene00014086 | -0.275375327 | 4.35E-05 | 0.00046235 | ZK809.3 |
| WBGene00014087 | -0.298746712 | 0.00034876 | 0.002851 | ZK809.5 |
| WBGene00014088 | 0.27243501 | 0.09345208 | 0.23380732 | ZK809.8 |
| WBGene00014089 | 0.168996703 | 0.37877548 | 0.57815386 | ZK822.1 |
| WBGene00014090 | 0.252899474 | 0.07187316 | 0.19515856 | ZK822.2 |
| WBGene00014091 | 0.905327859 | 1.40E-11 | 4.55E-10 | ZK822.4 |
| WBGene00014092 | 0.174271929 | 0.18505973 | 0.36878448 | ZK822.5 |
| WBGene00014093 | -0.036143964 | 0.7368532 | 0.85204655 | ZK829.1 |
| WBGene00014094 | 0.05787416 | 0.40594086 | NA | ZK829.3 |
| WBGene00014095 | 0.050033421 | 0.33622684 | 0.53559973 | gdh-1 |
| WBGene00014096 | -0.015928828 | 0.8365765 | 0.9111952 | ZK829.7 |
| WBGene00014097 | 0.164119715 | 0.03623899 | 0.11896302 | ZK829.9 |
| WBGene00014098 | -0.029388334 | 0.72264497 | 0.84446045 | ogdh-2 |
| WBGene00014099 | 0.143244284 | 0.25146726 | 0.44991306 | ZK836.3 |
| WBGene00014100 | -0.025226573 | 0.69184128 | NA | ZK849.1 |
| WBGene00014101 | 1.768325106 | 0.00907137 | 0.04161332 | gopc-1 |
| WBGene00014102 | 0.078633594 | 0.26310689 | NA | best-25 |
| WBGene00014103 | 0.325741239 | 0.12465086 | 0.28511378 | best-26 |
| WBGene00014104 | 0.104747489 | 0.16723639 | NA | ZK849.6 |
| WBGene00014105 | 0.071781729 | 0.54750682 | 0.72018982 | ZK856.4 |
| WBGene00014106 | 0.674511553 | 0.00705258 | 0.03420235 | ZK856.5 |
| WBGene00014107 | 0.137938171 | 0.3348676 | 0.53413188 | ZK856.6 |
| WBGene00014108 | 0.131936578 | 0.27158659 | 0.47140478 | ZK856.7 |
| WBGene00014109 | -0.016491466 | 0.8623948 | 0.92672958 | chpf-1 |
| WBGene00014111 | -0.147049268 | 0.29390122 | 0.49505381 | rpc-25 |
| WBGene00014112 | -0.104991732 | 0.43559411 | 0.63075199 | ZK856.11 |
| WBGene00014113 | -0.2009902 | 0.02622137 | 0.09411792 | hpo-40 |
| WBGene00014114 | -0.093744352 | 0.26609173 | 0.46473861 | tftc-3 |
| WBGene00014115 | -0.218718345 | 6.89E-05 | 0.00069749 | gld-4 |
| WBGene00014116 | 0.354554422 | 0.09644801 | 0.23978525 | ZK858.2 |
| WBGene00014117 | -0.067893643 | 0.26356592 | 0.4616976 | clec-91 |
| WBGene00014118 | -0.038343909 | 0.79149658 | 0.88250563 | ZK858.5 |
| WBGene00014119 | -0.092028671 | 0.17617377 | 0.35699717 | ZK858.6 |

| | | | | |
|---|---|---|---|---|
| WBGene00014120 | -0.119236178 | 0.26349907 | 0.46163573 | ZK858.7 |
| WBGene00014121 | 0.335879993 | 0.04797753 | 0.14594619 | ZK858.8 |
| WBGene00014122 | 0.038125098 | 0.61119482 | NA | ZK863.1 |
| WBGene00014123 | 0.05298094 | 0.62758239 | 0.77834351 | elpc-3 |
| WBGene00014124 | -0.077252579 | 0.1899903 | 0.37579682 | usip-1 |
| WBGene00014125 | 1.855921797 | 3.37E-08 | 6.88E-07 | ZK863.8 |
| WBGene00014126 | NA | NA | NA | NA |
| WBGene00014127 | 0.072653761 | 0.34256154 | NA | ZK892.3 |
| WBGene00014128 | -0.022930767 | 0.78121253 | NA | ZK892.4 |
| WBGene00014129 | 0.153594666 | 0.12171927 | 0.28070284 | ZK892.5 |
| WBGene00014130 | 0.106434012 | 0.45244663 | 0.64476067 | ZK892.6 |
| WBGene00014131 | -0.042791409 | 0.80429063 | 0.89116459 | sdz-38 |
| WBGene00014132 | -0.034291257 | 0.77433896 | 0.87215145 | ZK896.1 |
| WBGene00014134 | -0.198472933 | 0.31521678 | 0.51565238 | NA |
| WBGene00014135 | 0.015367421 | 0.94128787 | 0.96684409 | ZK896.4 |
| WBGene00014136 | 1.58491024 | 0.00035643 | 0.00290565 | ZK896.5 |
| WBGene00014137 | 0.957022997 | 9.29E-06 | 0.00011533 | clec-187 |
| WBGene00014138 | -1.177828042 | 1.88E-15 | 9.32E-14 | clec-186 |
| WBGene00014139 | -0.122723106 | 0.22993987 | 0.42426586 | nstp-5 |
| WBGene00014140 | 0.255230207 | 0.25549674 | 0.45380659 | ZK899.1 |
| WBGene00014141 | 0.20952333 | 0.16751831 | 0.3451014 | ZK899.2 |
| WBGene00014142 | -0.084318206 | 0.38493547 | NA | ZK899.3 |
| WBGene00014143 | -0.356320315 | 0.1250795 | 0.28578167 | ZK899.5 |
| WBGene00014144 | -0.217785593 | 0.32022285 | 0.52007255 | ZK899.6 |
| WBGene00014145 | -0.022729726 | 0.70048408 | NA | ZK899.7 |
| WBGene00014147 | NA | NA | NA | NA |
| WBGene00014148 | 0.100002565 | 0.5936577 | 0.75344508 | ZK909.3 |
| WBGene00014149 | -0.02791392 | 0.6763823 | NA | fbxb-70 |
| WBGene00014150 | -0.112954998 | 0.58173793 | 0.74601252 | ZK909.6 |
| WBGene00014151 | -0.121821044 | 0.16330005 | 0.33887458 | vps-15 |
| WBGene00014152 | 1.411275909 | 4.12E-10 | 1.09E-08 | ZK930.2 |
| WBGene00014153 | -0.16820594 | 0.41729823 | 0.61426198 | vab-23 |
| WBGene00014154 | 0.034981685 | 0.68236555 | NA | ZK930.4 |
| WBGene00014155 | -0.157890824 | 0.22008709 | 0.41227192 | ZK930.5 |
| WBGene00014156 | -0.179322428 | 0.27554002 | 0.47572289 | ZK930.6 |
| WBGene00014157 | -0.033151313 | 0.7567589 | 0.86226195 | ZK930.7 |
| WBGene00014158 | 1.893310646 | 0.00055122 | 0.00421655 | ZK938.1 |
| WBGene00014159 | -0.727722331 | 0.01971008 | 0.07578168 | arrd-4 |
| WBGene00014160 | NA | NA | NA | ZK938.3 |
| WBGene00014161 | 0.056411843 | 0.41891328 | NA | arrd-5 |
| WBGene00014162 | -0.0260089 | 0.6772113 | NA | chil-9 |
| WBGene00014163 | -0.021775862 | 0.87833432 | 0.93491899 | rnh-1.2 |
| WBGene00014164 | 0.176048533 | 0.23229223 | 0.42728806 | lact-2 |

| | | | | |
|---|---|---|---|---|
| WBGene00014165 | -0.505307351 | 8.87E-12 | 2.98E-10 | puf-12 |
| WBGene00014166 | 0.254452107 | 0.21176764 | 0.40193025 | ZK945.4 |
| WBGene00014168 | 0.153513723 | 0.06686266 | NA | ZK945.6 |
| WBGene00014169 | 0.335616679 | 0.04807211 | 0.14611246 | ZK945.7 |
| WBGene00014170 | NA | NA | NA | ZK945.8 |
| WBGene00014171 | 1.178620785 | 6.72E-05 | 0.00068366 | nep-26 |
| WBGene00014172 | 0.041654345 | 0.76748559 | 0.86797131 | clpp-1 |
| WBGene00014173 | -1.889398764 | 0.00565299 | 0.02882742 | ZK970.7 |
| WBGene00014174 | 0.034700471 | 0.72961513 | 0.84921656 | ZK970.8 |
| WBGene00014175 | 0.197230943 | 0.26285201 | 0.46079322 | ZK971.1 |
| WBGene00014176 | -0.222458076 | 0.05980305 | 0.17111129 | ZK1010.2 |
| WBGene00014177 | -0.13268461 | 0.24638889 | 0.44469671 | frg-1 |
| WBGene00014178 | 0.077821302 | 0.41957405 | 0.61623018 | ZK1010.4 |
| WBGene00014179 | 0.461386585 | 0.03879097 | 0.12509661 | ZK1010.5 |
| WBGene00014180 | -0.023056499 | 0.78121253 | NA | ZK1010.6 |
| WBGene00014181 | 0.041720845 | 0.80840818 | 0.89382448 | ZK1010.8 |
| WBGene00014182 | NA | NA | NA | ZK1025.2 |
| WBGene00014182 | NA | NA | NA | ZK1025.8 |
| WBGene00014183 | 0.108717649 | 0.15669083 | NA | ZK1025.3 |
| WBGene00014184 | 0.0398788 | 0.58971221 | NA | ZK1025.4 |
| WBGene00014185 | NA | NA | NA | NA |
| WBGene00014186 | NA | NA | NA | nhr-244 |
| WBGene00014188 | NA | NA | NA | ZK1025.2 |
| WBGene00014188 | NA | NA | NA | ZK1025.8 |
| WBGene00014189 | NA | NA | NA | nhr-245 |
| WBGene00014189 | NA | NA | NA | nhr-73 |
| WBGene00014190 | 0.0398788 | 0.58971221 | NA | irld-62 |
| WBGene00014191 | NA | NA | NA | srt-22 |
| WBGene00014192 | -0.023109762 | 0.78121253 | NA | nhr-246 |
| WBGene00014193 | NA | NA | NA | nhr-247 |
| WBGene00014194 | 0.253742757 | 0.25189602 | 0.44993917 | ZK1037.6 |
| WBGene00014195 | NA | NA | NA | srz-10 |
| WBGene00014196 | -0.001842784 | 0.95621055 | NA | ZK1053.1 |
| WBGene00014197 | 0.040218942 | 0.56174727 | NA | ZK1053.2 |
| WBGene00014198 | -0.107957322 | 0.4620466 | 0.65228464 | ZK1053.3 |
| WBGene00014199 | -0.629981712 | 0.00021983 | 0.00191121 | ZK1053.4 |
| WBGene00014200 | 0.316433592 | 0.09918532 | 0.244263 | scrm-2 |
| WBGene00014201 | 0.108942554 | 0.15594412 | NA | ZK1053.6 |
| WBGene00014202 | -0.090595626 | 0.24298633 | 0.44103246 | mmcm-1 |
| WBGene00014203 | 0.021966857 | 0.90136217 | 0.94736852 | ZK1058.3 |
| WBGene00014204 | -0.365089103 | 8.88E-10 | 2.25E-08 | ccdc-47 |
| WBGene00014205 | -0.122484548 | 0.43477934 | 0.63013308 | ZK1058.5 |
| WBGene00014206 | -1.486393856 | 0.00013988 | 0.00129047 | nit-1 |

| | | | | |
|---|---|---|---|---|
| WBGene00014207 | 0.03240537 | 0.79802007 | 0.88691671 | ZK1058.9 |
| WBGene00014208 | -0.052531436 | 0.54715324 | 0.71995249 | znfx-1 |
| WBGene00014209 | -0.135939811 | 0.18872366 | 0.37389758 | ZK1067.3 |
| WBGene00014210 | 0.293153005 | 0.18809642 | 0.37310928 | ZK1067.4 |
| WBGene00014213 | 0.217421804 | 0.15562633 | 0.32912395 | ZK1073.1 |
| WBGene00014214 | 0.370538826 | 0.13008948 | 0.29227968 | ZK1073.2 |
| WBGene00014215 | 0.353513151 | 0.02098336 | 0.07948705 | obr-3 |
| WBGene00014216 | 0.001382454 | 0.99318165 | 0.99586212 | ZK1086.2 |
| WBGene00014217 | NA | NA | NA | ZK1086.3 |
| WBGene00014218 | -0.308095584 | 0.0001328 | 0.00123604 | ZK1098.1 |
| WBGene00014219 | -0.150439136 | 0.11556947 | 0.27123262 | ZK1098.2 |
| WBGene00014220 | -0.125842783 | 0.48328725 | 0.66879098 | ZK1098.3 |
| WBGene00014221 | -0.294991984 | 0.02566479 | 0.09246017 | eif-2Balpha |
| WBGene00014222 | -0.207092297 | 0.10816167 | 0.25869594 | trpp-3 |
| WBGene00014223 | -0.09320539 | 0.59166412 | 0.75228436 | ZK1098.6 |
| WBGene00014224 | -0.502990882 | 0.00054196 | 0.00415439 | mrps-23 |
| WBGene00014225 | 0.040529223 | 0.56245259 | NA | ZK1098.9 |
| WBGene00014226 | 0.029364674 | 0.86141959 | 0.92660673 | ZK1098.11 |
| WBGene00014227 | 0.072706637 | 0.54715603 | 0.71995249 | ZK1128.1 |
| WBGene00014228 | 0.013666139 | 0.92315536 | 0.95651506 | mett-10 |
| WBGene00014229 | 0.052437311 | 0.67746016 | 0.81260807 | ZK1128.3 |
| WBGene00014230 | -0.28681491 | 0.01608831 | 0.06539392 | gtf-2H3 |
| WBGene00014232 | -0.188191683 | 0.03854409 | 0.12446493 | ttll-4 |
| WBGene00014233 | 0.358144671 | 0.08947886 | 0.22726891 | ZK1128.7 |
| WBGene00014234 | -0.176941148 | 0.09793179 | 0.24252696 | vps-29 |
| WBGene00014235 | NA | NA | NA | ZK1225.1 |
| WBGene00014236 | NA | NA | NA | ZK1225.2 |
| WBGene00014238 | 0.294735706 | 0.01008608 | 0.04514532 | ZK1225.4 |
| WBGene00014239 | -0.023056499 | 0.78121253 | NA | ZK1225.5 |
| WBGene00014240 | 0.102731084 | 0.23319132 | NA | htas-1 |
| WBGene00014241 | 0.048485426 | 0.45975815 | NA | ZK1251.3 |
| WBGene00014242 | -0.175358841 | 0.35089175 | 0.5508692 | sdz-36 |
| WBGene00014243 | 0.025498297 | 0.78297086 | 0.87782191 | dcaf-1 |
| WBGene00014244 | -0.432131373 | 0.00669874 | 0.03292253 | ZK1307.1 |
| WBGene00014245 | 0.056411843 | 0.41891328 | NA | ZK1307.2 |
| WBGene00014246 | 0.017348279 | 0.83364574 | NA | ZK1307.3 |
| WBGene00014247 | 0.472686981 | 0.00339953 | 0.01929168 | ZK1307.4 |
| WBGene00014248 | 0.041698592 | 0.73668273 | 0.85204655 | ZK1307.7 |
| WBGene00014249 | -0.036930664 | 0.59878303 | 0.75680097 | ZK1307.8 |
| WBGene00014250 | -0.073507145 | 0.5051338 | 0.68624515 | ZK1307.9 |
| WBGene00014251 | -0.399228972 | 0.01812819 | 0.07155969 | gstk-1 |
| WBGene00014252 | 1.61649091 | 6.72E-55 | 1.73E-52 | ZK1320.2 |
| WBGene00014253 | -0.002559553 | 0.97979005 | 0.98829529 | ZK1320.3 |

| | | | | | |
|---|---|---|---|---|---|
| WBGene00014254 | | 0.232822339 | 0.19425764 | 0.38119905 | cyp-13A10 |
| WBGene00014255 | | 0.036687355 | 0.59489568 | NA | ZK1320.5 |
| WBGene00014256 | | -0.153865376 | 0.28839371 | 0.48863582 | ZK1320.7 |
| WBGene00014258 | | -0.38928937 | 6.84E-07 | 1.11E-05 | ZK1320.9 |
| WBGene00014259 | | 0.335048428 | 0.10242035 | 0.2493824 | ZK1320.11 |
| WBGene00014260 | | -0.03323923 | 0.69122143 | NA | ZK1321.1 |
| WBGene00014261 | | 0.581455553 | 0.020642 | 0.07839678 | shk-1 |
| WBGene00014262 | | 0.149713681 | 0.19748425 | 0.38525874 | ZK1321.4 |
| WBGene00014263 | NA | | NA | NA | NA |
| WBGene00014264 | NA | | NA | NA | NA |
| WBGene00014265 | NA | | NA | NA | NA |
| WBGene00014266 | NA | | NA | NA | NA |
| WBGene00014267 | NA | | NA | NA | NA |
| WBGene00014268 | NA | | NA | NA | NA |
| WBGene00014269 | NA | | NA | NA | NA |
| WBGene00014270 | NA | | NA | NA | NA |
| WBGene00014271 | NA | | NA | NA | NA |
| WBGene00014272 | NA | | NA | NA | NA |
| WBGene00014273 | NA | | NA | NA | NA |
| WBGene00014274 | NA | | NA | NA | NA |
| WBGene00014275 | NA | | NA | NA | NA |
| WBGene00014276 | NA | | NA | NA | NA |
| WBGene00014277 | NA | | NA | NA | NA |
| WBGene00014278 | NA | | NA | NA | NA |
| WBGene00014279 | NA | | NA | NA | NA |
| WBGene00014280 | NA | | NA | NA | NA |
| WBGene00014281 | NA | | NA | NA | NA |
| WBGene00014282 | NA | | NA | NA | NA |
| WBGene00014283 | NA | | NA | NA | NA |
| WBGene00014284 | NA | | NA | NA | NA |
| WBGene00014285 | NA | | NA | NA | NA |
| WBGene00014286 | NA | | NA | NA | NA |
| WBGene00014287 | NA | | NA | NA | NA |
| WBGene00014288 | NA | | NA | NA | NA |
| WBGene00014289 | NA | | NA | NA | NA |
| WBGene00014290 | | 0.038125098 | 0.61119482 | NA | NA |
| WBGene00014291 | NA | | NA | NA | NA |
| WBGene00014292 | NA | | NA | NA | NA |
| WBGene00014293 | NA | | NA | NA | NA |
| WBGene00014294 | | 0.255597446 | 0.15732305 | 0.33127668 | NA |
| WBGene00014295 | NA | | NA | NA | NA |
| WBGene00014296 | NA | | NA | NA | NA |
| WBGene00014297 | NA | | NA | NA | NA |

| | | | | |
|---|---|---|---|---|
| WBGene00014298 | NA | NA | NA | NA |
| WBGene00014299 | NA | NA | NA | NA |
| WBGene00014300 | 1.97088968 | 1.32E-06 | 2.02E-05 | D2023.1 |
| WBGene00014301 | NA | NA | NA | NA |
| WBGene00014302 | NA | NA | NA | NA |
| WBGene00014303 | NA | NA | NA | NA |
| WBGene00014304 | NA | NA | NA | NA |
| WBGene00014305 | NA | NA | NA | NA |
| WBGene00014306 | NA | NA | NA | NA |
| WBGene00014307 | -0.039884454 | 0.6855485 | 0.81848942 | NA |
| WBGene00014308 | 1.019054853 | 6.10E-07 | 1.01E-05 | NA |
| WBGene00014309 | 0.379364125 | 1.98E-13 | 8.06E-12 | NA |
| WBGene00014310 | NA | NA | NA | NA |
| WBGene00014311 | NA | NA | NA | NA |
| WBGene00014312 | NA | NA | NA | NA |
| WBGene00014313 | NA | NA | NA | NA |
| WBGene00014314 | NA | NA | NA | NA |
| WBGene00014315 | NA | NA | NA | NA |
| WBGene00014316 | NA | NA | NA | NA |
| WBGene00014317 | NA | NA | NA | NA |
| WBGene00014318 | NA | NA | NA | NA |
| WBGene00014319 | NA | NA | NA | NA |
| WBGene00014320 | NA | NA | NA | NA |
| WBGene00014321 | NA | NA | NA | NA |
| WBGene00014322 | NA | NA | NA | NA |
| WBGene00014323 | NA | NA | NA | NA |
| WBGene00014324 | NA | NA | NA | NA |
| WBGene00014325 | NA | NA | NA | NA |
| WBGene00014326 | NA | NA | NA | NA |
| WBGene00014327 | NA | NA | NA | NA |
| WBGene00014328 | NA | NA | NA | NA |
| WBGene00014329 | NA | NA | NA | NA |
| WBGene00014330 | NA | NA | NA | NA |
| WBGene00014331 | NA | NA | NA | NA |
| WBGene00014332 | NA | NA | NA | NA |
| WBGene00014333 | NA | NA | NA | NA |
| WBGene00014334 | NA | NA | NA | NA |
| WBGene00014335 | NA | NA | NA | NA |
| WBGene00014336 | NA | NA | NA | NA |
| WBGene00014337 | NA | NA | NA | NA |
| WBGene00014338 | NA | NA | NA | NA |
| WBGene00014339 | NA | NA | NA | NA |
| WBGene00014340 | NA | NA | NA | NA |

| | | | | |
|---|---|---|---|---|
| WBGene00014341 | NA | NA | NA | NA |
| WBGene00014342 | NA | NA | NA | NA |
| WBGene00014343 | NA | NA | NA | NA |
| WBGene00014344 | NA | NA | NA | NA |
| WBGene00014345 | NA | NA | NA | NA |
| WBGene00014346 | NA | NA | NA | NA |
| WBGene00014347 | NA | NA | NA | NA |
| WBGene00014348 | NA | NA | NA | NA |
| WBGene00014349 | NA | NA | NA | NA |
| WBGene00014350 | NA | NA | NA | NA |
| WBGene00014351 | NA | NA | NA | NA |
| WBGene00014352 | NA | NA | NA | NA |
| WBGene00014353 | NA | NA | NA | NA |
| WBGene00014354 | NA | NA | NA | NA |
| WBGene00014355 | NA | NA | NA | NA |
| WBGene00014356 | NA | NA | NA | NA |
| WBGene00014357 | NA | NA | NA | NA |
| WBGene00014358 | NA | NA | NA | NA |
| WBGene00014359 | NA | NA | NA | NA |
| WBGene00014360 | NA | NA | NA | NA |
| WBGene00014361 | NA | NA | NA | NA |
| WBGene00014362 | NA | NA | NA | NA |
| WBGene00014363 | NA | NA | NA | NA |
| WBGene00014364 | NA | NA | NA | NA |
| WBGene00014365 | NA | NA | NA | NA |
| WBGene00014366 | NA | NA | NA | NA |
| WBGene00014367 | NA | NA | NA | NA |
| WBGene00014368 | NA | NA | NA | NA |
| WBGene00014369 | NA | NA | NA | NA |
| WBGene00014370 | NA | NA | NA | NA |
| WBGene00014371 | NA | NA | NA | NA |
| WBGene00014372 | NA | NA | NA | NA |
| WBGene00014373 | NA | NA | NA | NA |
| WBGene00014374 | NA | NA | NA | NA |
| WBGene00014375 | 0.116750398 | 0.18127159 | NA | NA |
| WBGene00014376 | NA | NA | NA | NA |
| WBGene00014377 | NA | NA | NA | NA |
| WBGene00014378 | NA | NA | NA | NA |
| WBGene00014379 | NA | NA | NA | NA |
| WBGene00014380 | NA | NA | NA | NA |
| WBGene00014381 | NA | NA | NA | NA |
| WBGene00014382 | NA | NA | NA | NA |
| WBGene00014384 | NA | NA | NA | NA |

| | | | | |
|---|---|---|---|---|
| WBGene00014385 | NA | NA | NA | NA |
| WBGene00014386 | NA | NA | NA | NA |
| WBGene00014387 | NA | NA | NA | NA |
| WBGene00014388 | NA | NA | NA | NA |
| WBGene00014389 | NA | NA | NA | NA |
| WBGene00014390 | NA | NA | NA | NA |
| WBGene00014391 | NA | NA | NA | NA |
| WBGene00014392 | NA | NA | NA | NA |
| WBGene00014393 | NA | NA | NA | NA |
| WBGene00014394 | NA | NA | NA | NA |
| WBGene00014395 | NA | NA | NA | NA |
| WBGene00014396 | NA | NA | NA | NA |
| WBGene00014397 | NA | NA | NA | NA |
| WBGene00014398 | NA | NA | NA | NA |
| WBGene00014399 | NA | NA | NA | NA |
| WBGene00014400 | NA | NA | NA | NA |
| WBGene00014401 | NA | NA | NA | NA |
| WBGene00014402 | NA | NA | NA | NA |
| WBGene00014403 | NA | NA | NA | NA |
| WBGene00014404 | NA | NA | NA | NA |
| WBGene00014405 | NA | NA | NA | NA |
| WBGene00014406 | NA | NA | NA | NA |
| WBGene00014407 | NA | NA | NA | NA |
| WBGene00014408 | NA | NA | NA | NA |
| WBGene00014409 | NA | NA | NA | NA |
| WBGene00014410 | NA | NA | NA | NA |
| WBGene00014411 | NA | NA | NA | NA |
| WBGene00014413 | NA | NA | NA | NA |
| WBGene00014414 | NA | NA | NA | NA |
| WBGene00014415 | NA | NA | NA | NA |
| WBGene00014416 | NA | NA | NA | NA |
| WBGene00014417 | NA | NA | NA | NA |
| WBGene00014418 | NA | NA | NA | NA |
| WBGene00014419 | NA | NA | NA | NA |
| WBGene00014420 | NA | NA | NA | NA |
| WBGene00014421 | NA | NA | NA | NA |
| WBGene00014422 | NA | NA | NA | NA |
| WBGene00014423 | NA | NA | NA | NA |
| WBGene00014424 | NA | NA | NA | NA |
| WBGene00014425 | NA | NA | NA | NA |
| WBGene00014426 | NA | NA | NA | NA |
| WBGene00014427 | NA | NA | NA | NA |
| WBGene00014428 | NA | NA | NA | NA |

| | | | | |
|---|---|---|---|---|
| WBGene00014429 | NA | NA | NA | NA |
| WBGene00014430 | NA | NA | NA | NA |
| WBGene00014431 | NA | NA | NA | NA |
| WBGene00014432 | NA | NA | NA | NA |
| WBGene00014433 | NA | NA | NA | NA |
| WBGene00014434 | NA | NA | NA | NA |
| WBGene00014435 | NA | NA | NA | NA |
| WBGene00014436 | NA | NA | NA | NA |
| WBGene00014437 | NA | NA | NA | NA |
| WBGene00014438 | NA | NA | NA | NA |
| WBGene00014439 | 0.020317245 | 0.84643798 | NA | NA |
| WBGene00014440 | NA | NA | NA | NA |
| WBGene00014441 | NA | NA | NA | NA |
| WBGene00014442 | NA | NA | NA | NA |
| WBGene00014443 | NA | NA | NA | NA |
| WBGene00014444 | NA | NA | NA | NA |
| WBGene00014445 | NA | NA | NA | NA |
| WBGene00014446 | NA | NA | NA | NA |
| WBGene00014447 | NA | NA | NA | NA |
| WBGene00014448 | NA | NA | NA | NA |
| WBGene00014449 | NA | NA | NA | NA |
| WBGene00014450 | NA | NA | NA | NA |
| WBGene00014451 | NA | NA | NA | NA |
| WBGene00014452 | NA | NA | NA | NA |
| WBGene00014453 | NA | NA | NA | NA |
| WBGene00014454 | 0.685479488 | 0.00229664 | 0.01395229 | NA |
| WBGene00014455 | NA | NA | NA | NA |
| WBGene00014456 | NA | NA | NA | NA |
| WBGene00014457 | NA | NA | NA | NA |
| WBGene00014458 | NA | NA | NA | NA |
| WBGene00014459 | NA | NA | NA | NA |
| WBGene00014460 | NA | NA | NA | NA |
| WBGene00014461 | NA | NA | NA | NA |
| WBGene00014462 | NA | NA | NA | NA |
| WBGene00014463 | NA | NA | NA | NA |
| WBGene00014464 | NA | NA | NA | NA |
| WBGene00014465 | NA | NA | NA | NA |
| WBGene00014466 | NA | NA | NA | NA |
| WBGene00014467 | NA | NA | NA | NA |
| WBGene00014468 | NA | NA | NA | NA |
| WBGene00014469 | NA | NA | NA | NA |
| WBGene00014470 | NA | NA | NA | NA |
| WBGene00014471 | NA | NA | NA | NA |

| | | | | |
|---|---|---|---|---|
| WBGene00014472 | 0.224817616 | 0.25270464 | 0.45081621 | NA |
| WBGene00014473 | NA | NA | NA | NA |
| WBGene00014474 | NA | NA | NA | NA |
| WBGene00014475 | NA | NA | NA | NA |
| WBGene00014476 | NA | NA | NA | NA |
| WBGene00014477 | NA | NA | NA | NA |
| WBGene00014478 | NA | NA | NA | NA |
| WBGene00014479 | NA | NA | NA | NA |
| WBGene00014480 | NA | NA | NA | NA |
| WBGene00014481 | NA | NA | NA | NA |
| WBGene00014482 | NA | NA | NA | NA |
| WBGene00014483 | NA | NA | NA | NA |
| WBGene00014484 | -0.001993815 | 0.95612996 | NA | NA |
| WBGene00014485 | NA | NA | NA | NA |
| WBGene00014486 | NA | NA | NA | NA |
| WBGene00014487 | NA | NA | NA | NA |
| WBGene00014488 | NA | NA | NA | NA |
| WBGene00014489 | NA | NA | NA | NA |
| WBGene00014490 | NA | NA | NA | NA |
| WBGene00014491 | NA | NA | NA | NA |
| WBGene00014492 | NA | NA | NA | NA |
| WBGene00014493 | NA | NA | NA | NA |
| WBGene00014494 | NA | NA | NA | NA |
| WBGene00014495 | NA | NA | NA | NA |
| WBGene00014496 | NA | NA | NA | NA |
| WBGene00014497 | NA | NA | NA | NA |
| WBGene00014498 | NA | NA | NA | NA |
| WBGene00014499 | NA | NA | NA | NA |
| WBGene00014500 | 0.020925399 | 0.84643798 | NA | NA |
| WBGene00014501 | NA | NA | NA | NA |
| WBGene00014502 | NA | NA | NA | NA |
| WBGene00014503 | NA | NA | NA | NA |
| WBGene00014504 | NA | NA | NA | NA |
| WBGene00014505 | 0.019573073 | 0.86550973 | 0.92809516 | NA |
| WBGene00014506 | NA | NA | NA | NA |
| WBGene00014507 | NA | NA | NA | NA |
| WBGene00014508 | NA | NA | NA | NA |
| WBGene00014509 | NA | NA | NA | NA |
| WBGene00014510 | NA | NA | NA | NA |
| WBGene00014511 | NA | NA | NA | NA |
| WBGene00014512 | NA | NA | NA | NA |
| WBGene00014513 | 0.059762547 | 0.38965825 | NA | NA |
| WBGene00014514 | NA | NA | NA | NA |

| | | | | |
|---|---|---|---|---|
| WBGene00014515 | NA | NA | NA | NA |
| WBGene00014516 | NA | NA | NA | NA |
| WBGene00014517 | -0.023109762 | 0.78121253 | NA | NA |
| WBGene00014518 | NA | NA | NA | NA |
| WBGene00014519 | NA | NA | NA | NA |
| WBGene00014520 | NA | NA | NA | NA |
| WBGene00014521 | NA | NA | NA | NA |
| WBGene00014522 | NA | NA | NA | NA |
| WBGene00014523 | NA | NA | NA | NA |
| WBGene00014524 | NA | NA | NA | NA |
| WBGene00014525 | NA | NA | NA | NA |
| WBGene00014526 | NA | NA | NA | NA |
| WBGene00014527 | NA | NA | NA | NA |
| WBGene00014528 | NA | NA | NA | NA |
| WBGene00014529 | NA | NA | NA | NA |
| WBGene00014530 | NA | NA | NA | NA |
| WBGene00014531 | NA | NA | NA | NA |
| WBGene00014532 | NA | NA | NA | NA |
| WBGene00014533 | NA | NA | NA | NA |
| WBGene00014534 | NA | NA | NA | NA |
| WBGene00014535 | NA | NA | NA | NA |
| WBGene00014536 | NA | NA | NA | NA |
| WBGene00014537 | NA | NA | NA | NA |
| WBGene00014538 | NA | NA | NA | NA |
| WBGene00014539 | NA | NA | NA | NA |
| WBGene00014540 | NA | NA | NA | NA |
| WBGene00014541 | NA | NA | NA | NA |
| WBGene00014542 | -0.00346345 | 0.95708614 | NA | NA |
| WBGene00014543 | -0.022930767 | 0.78121253 | NA | NA |
| WBGene00014544 | NA | NA | NA | NA |
| WBGene00014545 | NA | NA | NA | NA |
| WBGene00014546 | NA | NA | NA | NA |
| WBGene00014547 | NA | NA | NA | NA |
| WBGene00014548 | NA | NA | NA | NA |
| WBGene00014549 | NA | NA | NA | NA |
| WBGene00014550 | NA | NA | NA | NA |
| WBGene00014551 | NA | NA | NA | NA |
| WBGene00014552 | NA | NA | NA | NA |
| WBGene00014553 | NA | NA | NA | NA |
| WBGene00014554 | NA | NA | NA | NA |
| WBGene00014555 | NA | NA | NA | NA |
| WBGene00014556 | NA | NA | NA | NA |
| WBGene00014557 | NA | NA | NA | NA |

| | | | | |
|---|---|---|---|---|
| WBGene00014558 | NA | NA | NA | NA |
| WBGene00014559 | NA | NA | NA | NA |
| WBGene00014560 | NA | NA | NA | NA |
| WBGene00014561 | NA | NA | NA | NA |
| WBGene00014562 | 0.295130165 | 0.18219008 | 0.36500341 | NA |
| WBGene00014563 | NA | NA | NA | NA |
| WBGene00014564 | NA | NA | NA | NA |
| WBGene00014565 | NA | NA | NA | NA |
| WBGene00014566 | NA | NA | NA | NA |
| WBGene00014567 | NA | NA | NA | NA |
| WBGene00014568 | NA | NA | NA | NA |
| WBGene00014569 | 0.122842057 | 0.30970911 | 0.5105386 | bkip-1 |
| WBGene00014570 | NA | NA | NA | NA |
| WBGene00014571 | NA | NA | NA | NA |
| WBGene00014572 | NA | NA | NA | NA |
| WBGene00014573 | NA | NA | NA | NA |
| WBGene00014574 | NA | NA | NA | NA |
| WBGene00014575 | NA | NA | NA | NA |
| WBGene00014576 | -0.376908034 | 0.02711077 | 0.0961998 | NA |
| WBGene00014577 | NA | NA | NA | NA |
| WBGene00014578 | NA | NA | NA | NA |
| WBGene00014579 | NA | NA | NA | NA |
| WBGene00014580 | 0.020317245 | 0.84643798 | NA | NA |
| WBGene00014581 | NA | NA | NA | NA |
| WBGene00014582 | NA | NA | NA | NA |
| WBGene00014583 | NA | NA | NA | NA |
| WBGene00014584 | NA | NA | NA | NA |
| WBGene00014585 | NA | NA | NA | NA |
| WBGene00014586 | NA | NA | NA | NA |
| WBGene00014587 | NA | NA | NA | NA |
| WBGene00014588 | NA | NA | NA | NA |
| WBGene00014589 | NA | NA | NA | NA |
| WBGene00014590 | NA | NA | NA | NA |
| WBGene00014591 | NA | NA | NA | NA |
| WBGene00014592 | NA | NA | NA | NA |
| WBGene00014593 | NA | NA | NA | NA |
| WBGene00014594 | NA | NA | NA | NA |
| WBGene00014595 | NA | NA | NA | NA |
| WBGene00014596 | NA | NA | NA | NA |
| WBGene00014597 | NA | NA | NA | NA |
| WBGene00014598 | NA | NA | NA | NA |
| WBGene00014599 | NA | NA | NA | NA |
| WBGene00014600 | NA | NA | NA | NA |

| | | | | |
|---|---|---|---|---|
| WBGene00014601 | NA | NA | NA | NA |
| WBGene00014602 | NA | NA | NA | NA |
| WBGene00014603 | NA | NA | NA | NA |
| WBGene00014604 | NA | NA | NA | NA |
| WBGene00014605 | NA | NA | NA | NA |
| WBGene00014606 | NA | NA | NA | NA |
| WBGene00014607 | NA | NA | NA | NA |
| WBGene00014608 | NA | NA | NA | NA |
| WBGene00014609 | NA | NA | NA | NA |
| WBGene00014610 | NA | NA | NA | NA |
| WBGene00014611 | NA | NA | NA | NA |
| WBGene00014612 | NA | NA | NA | NA |
| WBGene00014613 | -0.029088148 | 0.84627055 | 0.91732043 | NA |
| WBGene00014614 | NA | NA | NA | NA |
| WBGene00014615 | -0.445779487 | 0.02516365 | 0.09123765 | NA |
| WBGene00014616 | NA | NA | NA | NA |
| WBGene00014617 | NA | NA | NA | NA |
| WBGene00014618 | NA | NA | NA | NA |
| WBGene00014619 | NA | NA | NA | NA |
| WBGene00014620 | NA | NA | NA | NA |
| WBGene00014621 | NA | NA | NA | NA |
| WBGene00014622 | NA | NA | NA | NA |
| WBGene00014623 | NA | NA | NA | NA |
| WBGene00014624 | NA | NA | NA | NA |
| WBGene00014625 | NA | NA | NA | NA |
| WBGene00014626 | NA | NA | NA | NA |
| WBGene00014627 | NA | NA | NA | NA |
| WBGene00014628 | NA | NA | NA | NA |
| WBGene00014629 | NA | NA | NA | NA |
| WBGene00014630 | NA | NA | NA | NA |
| WBGene00014631 | NA | NA | NA | NA |
| WBGene00014632 | NA | NA | NA | NA |
| WBGene00014633 | NA | NA | NA | NA |
| WBGene00014634 | NA | NA | NA | NA |
| WBGene00014635 | NA | NA | NA | NA |
| WBGene00014636 | NA | NA | NA | NA |
| WBGene00014637 | NA | NA | NA | NA |
| WBGene00014638 | NA | NA | NA | NA |
| WBGene00014639 | NA | NA | NA | NA |
| WBGene00014640 | NA | NA | NA | NA |
| WBGene00014641 | -0.023109762 | 0.78121253 | NA | NA |
| WBGene00014643 | 0.062064903 | 0.75884324 | 0.86363774 | NA |
| WBGene00014644 | NA | NA | NA | NA |

| | | | | |
|---|---|---|---|---|
| WBGene00014645 | NA | NA | NA | NA |
| WBGene00014646 | NA | NA | NA | NA |
| WBGene00014647 | NA | NA | NA | NA |
| WBGene00014648 | NA | NA | NA | NA |
| WBGene00014649 | NA | NA | NA | NA |
| WBGene00014650 | NA | NA | NA | NA |
| WBGene00014651 | NA | NA | NA | NA |
| WBGene00014652 | 0.020232312 | 0.84643798 | NA | NA |
| WBGene00014653 | NA | NA | NA | NA |
| WBGene00014654 | NA | NA | NA | NA |
| WBGene00014655 | NA | NA | NA | NA |
| WBGene00014656 | NA | NA | NA | NA |
| WBGene00014657 | NA | NA | NA | NA |
| WBGene00014658 | NA | NA | NA | NA |
| WBGene00014659 | NA | NA | NA | NA |
| WBGene00014660 | -0.008998908 | 0.92213607 | NA | NA |
| WBGene00014663 | 0.121499948 | 0.51735841 | 0.69671122 | NA |
| WBGene00014664 | NA | NA | NA | NA |
| WBGene00014665 | NA | NA | NA | NA |
| WBGene00014666 | -0.681443545 | 1.42E-08 | 3.10E-07 | C05D12.3 |
| WBGene00014667 | -0.007512361 | 0.92013177 | 0.95544619 | NA |
| WBGene00014668 | -0.063245565 | 0.60203823 | 0.75927657 | C06C3.9 |
| WBGene00014669 | 0.684433781 | 0.00031478 | 0.00261856 | zipt-2.1 |
| WBGene00014670 | NA | NA | NA | NA |
| WBGene00014671 | NA | NA | NA | NA |
| WBGene00014672 | 0.237995253 | 0.13079134 | 0.29315921 | NA |
| WBGene00014673 | NA | NA | NA | NA |
| WBGene00014674 | NA | NA | NA | NA |
| WBGene00014675 | NA | NA | NA | NA |
| WBGene00014676 | NA | NA | NA | NA |
| WBGene00014677 | NA | NA | NA | NA |
| WBGene00014678 | 0.020925399 | 0.84643798 | NA | NA |
| WBGene00014679 | NA | NA | NA | srz-25 |
| WBGene00014681 | NA | NA | NA | NA |
| WBGene00014682 | NA | NA | NA | NA |
| WBGene00014683 | NA | NA | NA | NA |
| WBGene00014684 | NA | NA | NA | NA |
| WBGene00014685 | NA | NA | NA | NA |
| WBGene00014686 | NA | NA | NA | NA |
| WBGene00014689 | NA | NA | NA | NA |
| WBGene00014691 | NA | NA | NA | NA |
| WBGene00014692 | NA | NA | NA | NA |
| WBGene00014693 | NA | NA | NA | NA |

| | | | | |
|---|---|---|---|---|
| WBGene00014694 | NA | NA | NA | NA |
| WBGene00014695 | NA | NA | NA | NA |
| WBGene00014696 | NA | NA | NA | NA |
| WBGene00014697 | 0.100799411 | 0.61616461 | 0.76979828 | cyp-25A3 |
| WBGene00014698 | -0.137992434 | 0.38131231 | 0.58063375 | C37A5.3 |
| WBGene00014699 | NA | NA | NA | C37A5.5 |
| WBGene00014700 | NA | NA | NA | C37A5.6 |
| WBGene00014701 | -0.048370879 | 0.47720013 | NA | NA |
| WBGene00014702 | NA | NA | NA | NA |
| WBGene00014703 | 0.020232312 | 0.84643798 | NA | NA |
| WBGene00014704 | NA | NA | NA | NA |
| WBGene00014705 | NA | NA | NA | NA |
| WBGene00014706 | NA | NA | NA | NA |
| WBGene00014707 | NA | NA | NA | NA |
| WBGene00014708 | NA | NA | NA | NA |
| WBGene00014710 | NA | NA | NA | NA |
| WBGene00014711 | NA | NA | NA | NA |
| WBGene00014714 | 0.155352187 | 0.0675224 | NA | NA |
| WBGene00014715 | NA | NA | NA | NA |
| WBGene00014716 | NA | NA | NA | NA |
| WBGene00014717 | NA | NA | NA | NA |
| WBGene00014718 | NA | NA | NA | NA |
| WBGene00014719 | NA | NA | NA | NA |
| WBGene00014720 | NA | NA | NA | NA |
| WBGene00014721 | NA | NA | NA | NA |
| WBGene00014722 | NA | NA | NA | NA |
| WBGene00014723 | NA | NA | NA | NA |
| WBGene00014724 | NA | NA | NA | NA |
| WBGene00014727 | NA | NA | NA | NA |
| WBGene00014728 | 0.079249556 | 0.309101 | NA | NA |
| WBGene00014729 | -0.050042403 | 0.47734215 | NA | NA |
| WBGene00014730 | 0.038125098 | 0.61119482 | NA | NA |
| WBGene00014731 | -0.082294594 | 0.47461207 | 0.66192665 | NA |
| WBGene00014732 | 0.141944435 | 0.25537914 | 0.45376287 | NA |
| WBGene00014733 | NA | NA | NA | NA |
| WBGene00014734 | -0.001488154 | 0.95639912 | NA | NA |
| WBGene00014735 | NA | NA | NA | NA |
| WBGene00014738 | NA | NA | NA | NA |
| WBGene00014740 | NA | NA | NA | NA |
| WBGene00014741 | -0.003108065 | 0.95728506 | NA | NA |
| WBGene00014744 | 0.059980168 | 0.38780842 | NA | NA |
| WBGene00014745 | NA | NA | NA | NA |
| WBGene00014746 | -0.429161782 | 0.07484185 | 0.20113061 | NA |

| | | | | |
|---|---|---|---|---|
| WBGene00014748 | NA | NA | NA | NA |
| WBGene00014751 | 0.056844058 | 0.53406644 | NA | NA |
| WBGene00014752 | NA | NA | NA | NA |
| WBGene00014753 | NA | NA | NA | NA |
| WBGene00014754 | 0.038125098 | 0.61119482 | NA | NA |
| WBGene00014755 | 0.040107227 | 0.58693163 | NA | NA |
| WBGene00014757 | NA | NA | NA | NA |
| WBGene00014758 | NA | NA | NA | NA |
| WBGene00014759 | 5.560315607 | 1.03E-19 | 7.58E-18 | NA |
| WBGene00014760 | -0.005017292 | 0.94670066 | NA | NA |
| WBGene00014761 | 2.911929625 | 1.06E-08 | 2.37E-07 | F46A8.11 |
| WBGene00014763 | NA | NA | NA | NA |
| WBGene00014764 | 0.0398788 | 0.58971221 | NA | NA |
| WBGene00014766 | 0.020317245 | 0.84643798 | NA | NA |
| WBGene00014767 | NA | NA | NA | NA |
| WBGene00014768 | NA | NA | NA | NA |
| WBGene00014769 | 0.165260652 | 0.16430727 | 0.34031563 | NA |
| WBGene00014770 | NA | NA | NA | NA |
| WBGene00014771 | 0.0398788 | 0.58971221 | NA | NA |
| WBGene00014772 | 0.249991155 | 0.08866911 | 0.22608926 | NA |
| WBGene00014773 | NA | NA | NA | NA |
| WBGene00014775 | NA | NA | NA | NA |
| WBGene00014776 | NA | NA | NA | NA |
| WBGene00014777 | NA | NA | NA | NA |
| WBGene00014778 | NA | NA | NA | NA |
| WBGene00014779 | NA | NA | NA | NA |
| WBGene00014781 | NA | NA | NA | NA |
| WBGene00014782 | NA | NA | NA | NA |
| WBGene00014783 | NA | NA | NA | NA |
| WBGene00014784 | NA | NA | NA | NA |
| WBGene00014785 | -0.023056499 | 0.78121253 | NA | NA |
| WBGene00014786 | NA | NA | NA | NA |
| WBGene00014787 | NA | NA | NA | NA |
| WBGene00014788 | NA | NA | NA | NA |
| WBGene00014789 | -0.031862882 | 0.8392916 | 0.91293227 | NA |
| WBGene00014790 | 0.064809585 | 0.40075491 | NA | NA |
| WBGene00014793 | NA | NA | NA | H40L08.2 |
| WBGene00014794 | NA | NA | NA | NA |
| WBGene00014796 | NA | NA | NA | NA |
| WBGene00014798 | -0.023056499 | 0.78121253 | NA | NA |
| WBGene00014799 | 0.020925399 | 0.84643798 | NA | NA |
| WBGene00014800 | NA | NA | NA | NA |
| WBGene00014801 | 0.140504992 | 0.50645195 | 0.68733435 | K08E5.1 |

| | | | | |
|---|---|---|---|---|
| WBGene00014803 | NA | NA | NA | NA |
| WBGene00014805 | 0.020232312 | 0.84643798 | NA | NA |
| WBGene00014806 | NA | NA | NA | NA |
| WBGene00014807 | -0.054724347 | 0.76799379 | 0.86834482 | M04D5.1 |
| WBGene00014809 | 0.014378275 | 0.85050389 | NA | NA |
| WBGene00014811 | 0.140586117 | 0.47983375 | 0.66640541 | NA |
| WBGene00014813 | 0.296177177 | 0.05972722 | 0.17105736 | M176.9 |
| WBGene00014814 | -0.055333281 | 0.74512378 | 0.85590582 | NA |
| WBGene00014817 | NA | NA | NA | NA |
| WBGene00014818 | NA | NA | NA | NA |
| WBGene00014819 | NA | NA | NA | NA |
| WBGene00014820 | 0.686694202 | 0.01018746 | 0.04550173 | NA |
| WBGene00014821 | 0.500190862 | 0.057297 | 0.16631353 | NA |
| WBGene00014822 | -0.038932455 | 0.8479186 | 0.9180823 | NA |
| WBGene00014824 | NA | NA | NA | NA |
| WBGene00014825 | 0.017604133 | 0.79437959 | NA | NA |
| WBGene00014826 | -0.114649883 | 0.25183884 | 0.44993917 | R13H4.2 |
| WBGene00014827 | NA | NA | NA | NA |
| WBGene00014828 | NA | NA | NA | NA |
| WBGene00014829 | NA | NA | NA | NA |
| WBGene00014832 | NA | NA | NA | NA |
| WBGene00014833 | NA | NA | NA | NA |
| WBGene00014834 | NA | NA | NA | NA |
| WBGene00014835 | 0.033034883 | 0.64219503 | NA | NA |
| WBGene00014836 | -0.011153545 | 0.9542113 | 0.97396875 | T09F5.12 |
| WBGene00014837 | 0.038125098 | 0.61119482 | NA | NA |
| WBGene00014838 | 3.866079376 | 0.00012886 | 0.00120694 | NA |
| WBGene00014839 | NA | NA | NA | NA |
| WBGene00014841 | NA | NA | NA | NA |
| WBGene00014842 | 0.23608847 | 0.0199448 | 0.07648292 | NA |
| WBGene00014843 | NA | NA | NA | NA |
| WBGene00014844 | 0.020925399 | 0.84643798 | NA | NA |
| WBGene00014845 | NA | NA | NA | NA |
| WBGene00014846 | 0.020317245 | 0.84643798 | NA | NA |
| WBGene00014848 | 0.329856 | 0.16478881 | 0.34096566 | VM106R.1 |
| WBGene00014849 | NA | NA | NA | NA |
| WBGene00014850 | NA | NA | NA | NA |
| WBGene00014851 | NA | NA | NA | NA |
| WBGene00014852 | 0.102066491 | 0.50984924 | 0.689717 | NA |
| WBGene00014853 | NA | NA | NA | NA |
| WBGene00014854 | NA | NA | NA | NA |
| WBGene00014855 | NA | NA | NA | NA |
| WBGene00014856 | NA | NA | NA | NA |

| | | | | |
|---|---|---|---|---|
| WBGene00014857 | NA | NA | NA | NA |
| WBGene00014858 | 0.037992904 | 0.65794732 | NA | NA |
| WBGene00014861 | NA | NA | NA | NA |
| WBGene00014862 | 0.020925399 | 0.84643798 | NA | NA |
| WBGene00014863 | NA | NA | NA | NA |
| WBGene00014864 | 0.112219298 | 0.4596656 | 0.65098856 | Y18D10A.22 |
| WBGene00014865 | NA | NA | NA | NA |
| WBGene00014866 | NA | NA | NA | NA |
| WBGene00014867 | NA | NA | NA | NA |
| WBGene00014868 | NA | NA | NA | NA |
| WBGene00014869 | NA | NA | NA | NA |
| WBGene00014870 | 0.036932429 | 0.82628257 | 0.90477236 | NA |
| WBGene00014871 | 0.223048296 | 0.29855645 | 0.49933345 | Y37H9A.5 |
| WBGene00014872 | NA | NA | NA | NA |
| WBGene00014873 | NA | NA | NA | NA |
| WBGene00014874 | -0.098113324 | 0.55779332 | 0.72877494 | NA |
| WBGene00014875 | NA | NA | NA | NA |
| WBGene00014876 | NA | NA | NA | NA |
| WBGene00014877 | NA | NA | NA | NA |
| WBGene00014878 | NA | NA | NA | NA |
| WBGene00014879 | NA | NA | NA | NA |
| WBGene00014880 | NA | NA | NA | NA |
| WBGene00014881 | NA | NA | NA | NA |
| WBGene00014882 | NA | NA | NA | NA |
| WBGene00014883 | 0.077402455 | 0.41757064 | 0.61440233 | NA |
| WBGene00014884 | 0.235037138 | 0.22766395 | 0.42120558 | NA |
| WBGene00014885 | NA | NA | NA | NA |
| WBGene00014886 | 0.048613912 | 0.47961343 | NA | NA |
| WBGene00014887 | NA | NA | NA | NA |
| WBGene00014888 | -0.053591705 | 0.45278076 | NA | NA |
| WBGene00014889 | NA | NA | NA | NA |
| WBGene00014890 | NA | NA | NA | NA |
| WBGene00014891 | NA | NA | NA | NA |
| WBGene00014892 | NA | NA | NA | NA |
| WBGene00014893 | NA | NA | NA | NA |
| WBGene00014894 | NA | NA | NA | NA |
| WBGene00014895 | NA | NA | NA | NA |
| WBGene00014896 | NA | NA | NA | NA |
| WBGene00014897 | NA | NA | NA | NA |
| WBGene00014898 | NA | NA | NA | NA |
| WBGene00014902 | -0.074967263 | 0.4157793 | NA | NA |
| WBGene00014903 | NA | NA | NA | NA |
| WBGene00014904 | NA | NA | NA | NA |

| | | | | | |
|---|---|---|---|---|---|
| WBGene00014905 | | 0.033463284 | 0.6365037 | NA | NA |
| WBGene00014906 | | -0.023056499 | 0.78121253 | NA | NA |
| WBGene00014907 | | 0.055460599 | 0.59656118 | 0.75522171 | NA |
| WBGene00014908 | NA | | NA | NA | NA |
| WBGene00014909 | | -0.002914971 | 0.95718083 | NA | NA |
| WBGene00014910 | NA | | NA | NA | NA |
| WBGene00014911 | | -0.023109762 | 0.78121253 | NA | NA |
| WBGene00014912 | NA | | NA | NA | NA |
| WBGene00014914 | | 0.001382111 | 0.99503781 | 0.99691196 | Y52B11A.11 |
| WBGene00014915 | | -0.340236429 | 0.0533687 | 0.15810907 | NA |
| WBGene00014916 | | -0.005495226 | 0.97670502 | 0.98670757 | NA |
| WBGene00014917 | | 0.064762459 | 0.75232254 | 0.8597888 | NA |
| WBGene00014918 | | -0.069464566 | 0.71817947 | 0.84186977 | NA |
| WBGene00014920 | | -0.092959704 | 0.5700767 | 0.7387023 | NA |
| WBGene00014921 | | 0.020317245 | 0.84643798 | NA | NA |
| WBGene00014922 | NA | | NA | NA | NA |
| WBGene00014923 | NA | | NA | NA | NA |
| WBGene00014924 | NA | | NA | NA | Y57G11C.28 |
| WBGene00014925 | | 0.020317245 | 0.84643798 | NA | NA |
| WBGene00014926 | | 0.032317057 | 0.61887865 | NA | Y57G11C.30 |
| WBGene00014927 | NA | | NA | NA | NA |
| WBGene00014928 | NA | | NA | NA | NA |
| WBGene00014929 | NA | | NA | NA | NA |
| WBGene00014930 | NA | | NA | NA | NA |
| WBGene00014931 | | 0.033463284 | 0.6365037 | NA | NA |
| WBGene00014932 | | -0.022930767 | 0.78121253 | NA | NA |
| WBGene00014933 | | 0.115146689 | 0.4076507 | 0.60562291 | NA |
| WBGene00014934 | NA | | NA | NA | NA |
| WBGene00014935 | NA | | NA | NA | NA |
| WBGene00014936 | NA | | NA | NA | NA |
| WBGene00014937 | NA | | NA | NA | NA |
| WBGene00014938 | | -0.240123132 | 0.14579502 | 0.31553362 | Y62E10A.23 |
| WBGene00014938 | | -0.240123132 | 0.14579502 | 0.31553362 | mdt-9 |
| WBGene00014939 | NA | | NA | NA | NA |
| WBGene00014941 | | 0.142812335 | 0.40774058 | 0.60569499 | NA |
| WBGene00014943 | NA | | NA | NA | NA |
| WBGene00014944 | | 0.083479586 | 0.30566331 | NA | NA |
| WBGene00014945 | NA | | NA | NA | NA |
| WBGene00014946 | | -0.160203397 | 0.42895522 | 0.62379156 | NA |
| WBGene00014947 | | 0.271222351 | 0.2238375 | 0.4168345 | NA |
| WBGene00014948 | NA | | NA | NA | NA |
| WBGene00014949 | | -0.022930767 | 0.78121253 | NA | NA |
| WBGene00014950 | NA | | NA | NA | NA |

| | | | | |
|---|---|---|---|---|
| WBGene00014951 | NA | NA | NA | NA |
| WBGene00014952 | NA | NA | NA | NA |
| WBGene00014953 | NA | NA | NA | Y81G3A.2 |
| WBGene00014954 | 0.404417828 | 0.0174363 | 0.06946528 | NA |
| WBGene00014955 | 0.289406996 | 0.01587306 | 0.06475271 | NA |
| WBGene00014956 | 0.040107227 | 0.58693163 | NA | NA |
| WBGene00014957 | NA | NA | NA | NA |
| WBGene00014958 | -0.079349983 | 0.44930436 | 0.64190722 | NA |
| WBGene00014961 | NA | NA | NA | NA |
| WBGene00014962 | NA | NA | NA | NA |
| WBGene00014963 | NA | NA | NA | NA |
| WBGene00014964 | NA | NA | NA | NA |
| WBGene00014965 | 0.082816298 | 0.49394277 | 0.67686828 | Y106G6D.5 |
| WBGene00014966 | NA | NA | NA | NA |
| WBGene00014967 | NA | NA | NA | NA |
| WBGene00014969 | NA | NA | NA | NA |
| WBGene00014970 | NA | NA | NA | NA |
| WBGene00014971 | NA | NA | NA | NA |
| WBGene00014972 | NA | NA | NA | NA |
| WBGene00014973 | NA | NA | NA | NA |
| WBGene00014974 | -0.023109762 | 0.78121253 | NA | NA |
| WBGene00014975 | 0.068068345 | 0.70982509 | 0.83662447 | NA |
| WBGene00014976 | -0.008603946 | 0.9233394 | NA | NA |
| WBGene00014977 | 0.095967469 | 0.18297344 | NA | NA |
| WBGene00014978 | 0.123112817 | 0.34971833 | 0.54959362 | NA |
| WBGene00014979 | 0.020232312 | 0.84643798 | NA | NA |
| WBGene00014980 | NA | NA | NA | NA |
| WBGene00014981 | NA | NA | NA | NA |
| WBGene00014983 | 0.020317245 | 0.84643798 | NA | NA |
| WBGene00014984 | NA | NA | NA | NA |
| WBGene00014987 | -0.294272222 | 0.03947979 | 0.12684302 | NA |
| WBGene00014989 | 0.230677762 | 0.25808348 | 0.45629758 | best-23 |
| WBGene00014990 | NA | NA | NA | NA |
| WBGene00014991 | 0.020232312 | 0.84643798 | NA | NA |
| WBGene00014992 | 0.033463284 | 0.6365037 | NA | NA |
| WBGene00014993 | NA | NA | NA | NA |
| WBGene00014994 | 0.10838748 | 0.59571131 | 0.75477619 | NA |
| WBGene00014996 | 0.040107227 | 0.58693163 | NA | NA |
| WBGene00014997 | 0.0398788 | 0.58971221 | NA | AC7.3 |
| WBGene00014998 | -0.032616144 | 0.7322195 | NA | AH9.1 |
| WBGene00014999 | -0.412871787 | 0.02822695 | 0.09909332 | AH9.3 |
| WBGene00015000 | 0.105191732 | 0.16565281 | NA | AH9.4 |
| WBGene00015001 | -0.140740875 | 0.35200922 | 0.55191336 | B0025.4 |

| | | | | |
|---|---|---|---|---|
| WBGene00015002 | 0.127047307 | 0.47977272 | 0.66640541 | B0034.1 |
| WBGene00015003 | NA | NA | NA | B0034.2 |
| WBGene00015004 | NA | NA | NA | B0034.4 |
| WBGene00015005 | 0.04866948 | 0.68243089 | 0.81629663 | B0034.5 |
| WBGene00015006 | 0.033463284 | 0.6365037 | NA | B0041.1 |
| WBGene00015007 | -0.007803671 | 0.905006 | 0.94913088 | ain-2 |
| WBGene00015008 | -0.00226094 | 0.98937808 | 0.99398816 | lmd-2 |
| WBGene00015009 | -0.286007193 | 0.00228493 | 0.01389848 | B0041.5 |
| WBGene00015010 | -0.341034954 | 0.13981427 | 0.30722209 | ptps-1 |
| WBGene00015011 | -0.31926372 | 0.00027113 | 0.00229859 | B0041.8 |
| WBGene00015012 | 0.040664498 | 0.81068817 | 0.89531472 | bath-20 |
| WBGene00015013 | 0.41561011 | 0.10188737 | 0.24858073 | NA |
| WBGene00015014 | 0.848196005 | 0.00314031 | 0.01808705 | bath-24 |
| WBGene00015015 | 0.065014541 | 0.37399213 | NA | math-1 |
| WBGene00015016 | 0.162085497 | 0.44483207 | 0.63821458 | NA |
| WBGene00015017 | -0.185412957 | 0.05928513 | 0.17016145 | B0205.1 |
| WBGene00015018 | 0.05787416 | 0.40594086 | NA | srz-85 |
| WBGene00015019 | 0.018860755 | 0.77262519 | NA | B0205.4 |
| WBGene00015020 | 0.038195847 | 0.57893903 | NA | B0205.5 |
| WBGene00015021 | -0.106927308 | 0.36433633 | 0.56409986 | B0205.6 |
| WBGene00015022 | -0.011053399 | 0.93428909 | 0.96290283 | B0205.8 |
| WBGene00015023 | 0.02986583 | 0.78182139 | 0.87727215 | B0205.9 |
| WBGene00015024 | 2.378764669 | 0.0028021 | 0.01652925 | B0205.10 |
| WBGene00015025 | -0.271496107 | 0.01416664 | 0.05904177 | mrpl-9 |
| WBGene00015026 | 0.065254007 | 0.40163332 | NA | B0207.1 |
| WBGene00015027 | 0.083378938 | 0.54129624 | 0.71522926 | B0207.2 |
| WBGene00015028 | 0.038530293 | 0.79050273 | 0.88212499 | B0207.5 |
| WBGene00015029 | -0.112680862 | 0.37218492 | 0.57146318 | B0207.6 |
| WBGene00015030 | 0.056411843 | 0.41891328 | NA | B0207.7 |
| WBGene00015032 | 0.079587097 | 0.41025132 | 0.60798576 | B0207.9 |
| WBGene00015033 | 0.132442484 | 0.10147609 | NA | B0207.10 |
| WBGene00015034 | 0.020317245 | 0.84643798 | NA | B0207.11 |
| WBGene00015035 | 0.357781079 | 0.10405452 | 0.25210364 | otpl-1 |
| WBGene00015036 | 0.033034883 | 0.64219503 | NA | sre-5 |
| WBGene00015037 | -0.108297507 | 0.5032468 | 0.68450728 | B0212.3 |
| WBGene00015038 | 0.178376061 | 0.409092 | 0.60696351 | nstp-6 |
| WBGene00015040 | 0.293343649 | 0.12158055 | 0.28058968 | cyp-34A5 |
| WBGene00015041 | 0.067040533 | 0.67076634 | 0.80831463 | cyp-34A6 |
| WBGene00015042 | 0.096619834 | 0.33829084 | 0.53789273 | cyp-34A7 |
| WBGene00015043 | -0.872774105 | 1.32E-05 | 0.00015705 | cyp-34A8 |
| WBGene00015044 | 1.07726098 | 0.00110921 | 0.00767407 | cyp-34A9 |
| WBGene00015045 | 2.633722356 | 0.00512305 | 0.02663586 | cyp-34A10 |
| WBGene00015046 | 0.044317259 | 0.23895686 | 0.43601289 | nlp-34 |

| | | | | |
|---|---|---|---|---|
| WBGene00015047 | 0.712740332 | 8.91E-07 | 1.42E-05 | faah-1 |
| WBGene00015048 | 0.170940657 | 0.42692466 | 0.62241028 | faah-2 |
| WBGene00015049 | 0.102280207 | 0.23256935 | NA | B0218.5 |
| WBGene00015050 | -0.239646809 | 0.2076929 | 0.39676823 | clec-51 |
| WBGene00015051 | 0.391469089 | 0.0344932 | 0.11482831 | B0218.7 |
| WBGene00015052 | 0.018529657 | 0.92929477 | 0.96025158 | clec-52 |
| WBGene00015053 | 0.258666451 | 0.14329109 | 0.31177364 | B0222.1 |
| WBGene00015054 | NA | NA | NA | pitr-2 |
| WBGene00015055 | 0.060866418 | 0.77039305 | 0.86937911 | pitr-3 |
| WBGene00015056 | -0.115191825 | 0.50964622 | 0.68965343 | B0222.5 |
| WBGene00015057 | 0.898533494 | 9.21E-08 | 1.75E-06 | gad-3 |
| WBGene00015058 | 0.050163214 | 0.55716828 | NA | NA |
| WBGene00015059 | 0.79396505 | 0.00663812 | 0.03269046 | B0228.1 |
| WBGene00015061 | 0.472957215 | 1.70E-07 | 3.09E-06 | cpna-2 |
| WBGene00015062 | 0.056411843 | 0.41891328 | NA | trx-1 |
| WBGene00015063 | -0.346026587 | 0.1475616 | 0.31829199 | B0228.6 |
| WBGene00015064 | 0.43854232 | 0.00091545 | 0.00653019 | B0228.7 |
| WBGene00015065 | NA | NA | NA | B0228.8 |
| WBGene00015066 | NA | NA | NA | B0228.9 |
| WBGene00015067 | 1.863169583 | 1.67E-07 | 3.04E-06 | B0238.1 |
| WBGene00015068 | NA | NA | NA | srt-68 |
| WBGene00015069 | NA | NA | NA | srt-67 |
| WBGene00015070 | 0.033463284 | 0.6365037 | NA | srt-70 |
| WBGene00015071 | NA | NA | NA | B0238.7 |
| WBGene00015072 | NA | NA | NA | srt-69 |
| WBGene00015073 | -0.05676388 | 0.50484898 | 0.68604918 | B0238.9 |
| WBGene00015074 | 0.023411131 | 0.78463481 | 0.87854069 | natc-2 |
| WBGene00015075 | -0.290746672 | 0.00030569 | 0.00256042 | B0238.11 |
| WBGene00015076 | 2.339696353 | 4.81E-08 | 9.61E-07 | B0238.12 |
| WBGene00015077 | 0.353129196 | 0.00647075 | 0.03201697 | NA |
| WBGene00015078 | NA | NA | NA | NA |
| WBGene00015079 | 0.123251369 | 0.41714005 | 0.61417413 | B0244.4 |
| WBGene00015080 | 1.413806653 | 0.00497048 | 0.02604608 | B0244.5 |
| WBGene00015081 | -0.007625068 | 0.96493104 | 0.98071026 | B0244.6 |
| WBGene00015082 | 0.086908003 | 0.52590744 | 0.70317907 | B0244.7 |
| WBGene00015083 | -0.182205313 | 0.00102196 | 0.00715909 | egg-1 |
| WBGene00015084 | 0.20888338 | 0.02881778 | 0.10067315 | B0244.9 |
| WBGene00015085 | 0.485264366 | 0.07213454 | 0.19557812 | B0244.10 |
| WBGene00015087 | 2.315414297 | 1.58E-12 | 5.78E-11 | B0252.1 |
| WBGene00015088 | -0.353696143 | 0.04133662 | 0.13101029 | B0252.3 |
| WBGene00015089 | 4.677876459 | 0.00042482 | 0.00338403 | B0252.5 |
| WBGene00015091 | -0.058967729 | 0.52876454 | 0.70564594 | B0261.1 |
| WBGene00015092 | -0.242721114 | 0.06041667 | 0.17226103 | mrpl-47 |

| | | | | |
|---|---|---|---|---|
| WBGene00015093 | 0.014452015 | 0.85564601 | NA | B0261.5 |
| WBGene00015094 | 2.186679615 | 0.00127664 | 0.00859766 | B0261.6 |
| WBGene00015095 | -0.14737587 | 0.03529082 | 0.11671402 | B0261.7 |
| WBGene00015096 | 0.236665853 | 0.25914802 | 0.45720647 | B0261.8 |
| WBGene00015097 | 0.044778143 | 0.74936529 | 0.85855366 | B0273.1 |
| WBGene00015098 | -0.658577489 | 0.01113067 | 0.048646 | snpc-3.1 |
| WBGene00015099 | -0.202768195 | 0.07507366 | 0.20153163 | ggtb-1 |
| WBGene00015100 | 0.147551864 | 0.45374454 | 0.64589961 | B0280.2 |
| WBGene00015101 | -0.409702556 | 0.00010361 | 0.00100391 | rpia-1 |
| WBGene00015102 | -0.025437362 | 0.56875343 | 0.73763123 | cpg-2 |
| WBGene00015103 | 0.04909525 | 0.45360729 | NA | B0280.7 |
| WBGene00015104 | -0.269412093 | 0.00970842 | 0.04391134 | B0280.9 |
| WBGene00015105 | -0.07557987 | 0.62618702 | 0.77701535 | pot-1 |
| WBGene00015106 | 0.206575708 | 0.02995553 | 0.10341402 | B0280.11 |
| WBGene00015107 | 0.673160453 | 0.02068699 | 0.07854724 | viro-2 |
| WBGene00015108 | -0.045959069 | 0.50707733 | NA | B0281.1 |
| WBGene00015110 | 0.188503917 | 0.37670114 | 0.57581566 | B0281.3 |
| WBGene00015111 | -0.147688616 | 0.46191462 | 0.65228105 | B0281.4 |
| WBGene00015112 | -0.818268849 | 2.27E-05 | 0.00025556 | B0281.5 |
| WBGene00015113 | 0.278059968 | 0.09290935 | 0.23288577 | B0281.6 |
| WBGene00015114 | 0.015684754 | 0.83758486 | NA | B0281.8 |
| WBGene00015115 | 0.298122043 | 0.19435424 | 0.38124084 | B0286.1 |
| WBGene00015116 | 0.555422532 | 0.00013251 | 0.00123495 | pacs-1 |
| WBGene00015117 | 0.257908402 | 0.15245703 | 0.32514543 | B0294.1 |
| WBGene00015118 | 0.109491735 | 0.21453283 | NA | B0294.3 |
| WBGene00015119 | 0.079578344 | 0.41788044 | NA | B0302.2 |
| WBGene00015121 | 0.037023624 | 0.59153183 | NA | B0302.4 |
| WBGene00015122 | 0.632830293 | 0.00299161 | 0.0174433 | B0302.5 |
| WBGene00015124 | -0.06593898 | 0.74142614 | 0.85407613 | anmt-1 |
| WBGene00015125 | -0.151579139 | 0.0420789 | 0.13262743 | B0303.3 |
| WBGene00015126 | -0.219026084 | 0.00947446 | 0.04314637 | B0303.4 |
| WBGene00015128 | 0.136938808 | 0.40824819 | 0.6061102 | B0303.7 |
| WBGene00015129 | 0.043694866 | 0.78611319 | 0.87892197 | sdz-1 |
| WBGene00015130 | -0.06378479 | 0.57318009 | 0.74127107 | vps-33.1 |
| WBGene00015131 | 0.106932302 | 0.46749132 | 0.65691831 | B0303.11 |
| WBGene00015132 | -0.374923696 | 0.10466001 | 0.25285927 | B0303.14 |
| WBGene00015133 | -0.208766195 | 0.05437694 | 0.1599806 | mrpl-11 |
| WBGene00015134 | -0.071479354 | 0.56185283 | 0.73224918 | B0304.2 |
| WBGene00015135 | 0.144968209 | 0.25015977 | 0.44873673 | cyp-23A1 |
| WBGene00015136 | 0.0333868 | 0.84952157 | 0.91911205 | B0304.4 |
| WBGene00015137 | -0.096728802 | 0.64214573 | 0.78811138 | B0310.1 |
| WBGene00015138 | -0.283971006 | 0.15999483 | 0.33471329 | B0310.2 |
| WBGene00015139 | 0.111471772 | 0.4621517 | 0.65236821 | B0310.3 |

| | | | | |
|---|---|---|---|---|
| WBGene00015141 | -0.464980683 | 0.00089113 | 0.00639248 | ugt-46 |
| WBGene00015142 | 0.065014541 | 0.37399213 | NA | B0310.6 |
| WBGene00015143 | -0.072973707 | 0.28018662 | 0.47984024 | rbm-26 |
| WBGene00015145 | -0.188654104 | 0.08877062 | 0.22611347 | arle-14 |
| WBGene00015146 | -0.08877441 | 0.19389419 | 0.38084316 | abi-1 |
| WBGene00015147 | 0.090618346 | 0.50213596 | 0.68363145 | B0336.7 |
| WBGene00015148 | 0.497357494 | 0.06279574 | 0.17752314 | hpo-28 |
| WBGene00015149 | -0.022930767 | 0.78121253 | NA | B0336.12 |
| WBGene00015150 | -0.086595399 | 0.46372383 | 0.6536122 | B0336.13 |
| WBGene00015151 | 0.078203897 | 0.26561294 | NA | B0348.1 |
| WBGene00015152 | 2.724422382 | 0.00422666 | 0.02291943 | B0348.2 |
| WBGene00015153 | -0.157848695 | 0.1694812 | 0.34787245 | B0348.5 |
| WBGene00015155 | 0.233505869 | 0.16035222 | 0.33484732 | B0353.1 |
| WBGene00015156 | -0.157443057 | 0.12984904 | 0.29221004 | B0361.2 |
| WBGene00015157 | -0.108741154 | 0.26579326 | 0.46443258 | B0361.3 |
| WBGene00015158 | 0.22883574 | 0.20281972 | 0.3914648 | B0361.4 |
| WBGene00015159 | 0.09655046 | 0.37992804 | 0.57949019 | psd-1 |
| WBGene00015160 | -0.116702462 | 0.29442086 | 0.49565557 | B0361.6 |
| WBGene00015161 | 0.143246344 | 0.26855653 | 0.46747547 | pho-5 |
| WBGene00015162 | -0.012074514 | 0.88654716 | 0.94033043 | algn-11 |
| WBGene00015163 | 0.209068606 | 0.20789774 | 0.39679714 | B0361.9 |
| WBGene00015164 | -0.2135239 | 0.03658026 | 0.11978786 | ykt-6 |
| WBGene00015165 | 0.057565879 | 0.43429308 | NA | B0361.11 |
| WBGene00015167 | -0.907595181 | 0.0055359 | 0.02839913 | B0403.3 |
| WBGene00015168 | -0.038653527 | 0.49977482 | 0.68168466 | pdi-6 |
| WBGene00015169 | 0.136310359 | 0.45737448 | 0.64918009 | B0403.5 |
| WBGene00015170 | NA | NA | NA | B0410.1 |
| WBGene00015171 | -0.143501406 | 0.36471275 | 0.56433728 | vang-1 |
| WBGene00015172 | -0.27226899 | 0.14817538 | 0.31884712 | B0410.3 |
| WBGene00015173 | 0.088168945 | 0.34290999 | 0.54313812 | trpp-11 |
| WBGene00015175 | NA | NA | NA | srz-4 |
| WBGene00015176 | -0.311821035 | 0.0015948 | 0.01036373 | vps-51 |
| WBGene00015177 | 0.532664927 | 0.02873026 | 0.10048728 | tmc-2 |
| WBGene00015178 | 2.5954802 | 0.00063742 | 0.00478831 | B0416.2 |
| WBGene00015179 | 0.371587891 | 0.12808825 | 0.28949368 | B0416.3 |
| WBGene00015180 | -0.151432811 | 0.40405871 | 0.60230336 | B0416.4 |
| WBGene00015181 | -0.177992813 | 0.0138309 | 0.05792284 | B0416.5 |
| WBGene00015182 | 0.027204404 | 0.88408806 | 0.93885061 | B0416.7 |
| WBGene00015183 | -0.055763791 | 0.73797982 | 0.85272061 | B0432.1 |
| WBGene00015184 | 0.008966487 | 0.92983537 | 0.9606068 | djr-1.1 |
| WBGene00015185 | -0.192070513 | 0.1721306 | 0.35168352 | mrpl-41 |
| WBGene00015186 | 0.069036857 | 0.45706084 | 0.64899891 | misc-1 |
| WBGene00015188 | -0.018521618 | 0.89170131 | 0.94351368 | B0432.7 |

| | | | | |
|---|---|---|---|---|
| WBGene00015189 | 0.029552714 | 0.85723617 | 0.92325428 | B0432.8 |
| WBGene00015190 | -0.02120243 | 0.91358991 | 0.95181597 | B0432.9 |
| WBGene00015191 | -0.056612321 | 0.7114158 | 0.8376204 | B0432.10 |
| WBGene00015192 | 0.05787416 | 0.40594086 | NA | B0432.11 |
| WBGene00015193 | 0.170886701 | 0.09702423 | 0.24101319 | clec-117 |
| WBGene00015194 | -0.053925767 | 0.66517398 | 0.80363434 | trul-1 |
| WBGene00015196 | -1.198341979 | 0.00033254 | 0.00274137 | B0454.5 |
| WBGene00015197 | 0.796435569 | 0.00075156 | 0.00549082 | B0454.6 |
| WBGene00015198 | 0.149442831 | 0.06876524 | NA | clec-2 |
| WBGene00015199 | 0.020317245 | 0.84643798 | NA | B0454.8 |
| WBGene00015200 | 0.57301028 | 0.00794209 | 0.03761895 | lido-13 |
| WBGene00015202 | 0.051211461 | 0.64976881 | 0.79273174 | B0478.3 |
| WBGene00015203 | -0.043884023 | 0.57794454 | 0.74421161 | cdk-11.1 |
| WBGene00015204 | 0.234155041 | 0.08949958 | 0.22726891 | B0495.5 |
| WBGene00015205 | 0.007531417 | 0.96004829 | 0.97815358 | moa-2 |
| WBGene00015206 | 0.168651009 | 0.01361913 | 0.05724882 | B0495.7 |
| WBGene00015207 | -0.240775179 | 0.01617705 | 0.06568542 | B0495.8 |
| WBGene00015208 | 0.029804055 | 0.81603745 | 0.89891744 | B0495.9 |
| WBGene00015210 | 0.012730745 | 0.86589641 | NA | B0496.1 |
| WBGene00015211 | 0.020232312 | 0.84643798 | NA | B0496.2 |
| WBGene00015214 | -0.045959069 | 0.50707733 | NA | srtx-1 |
| WBGene00015215 | 0.0398788 | 0.58971221 | NA | B0496.6 |
| WBGene00015216 | 0.771319552 | 0.00608606 | 0.03050652 | valv-1 |
| WBGene00015217 | -0.021233117 | 0.89569922 | 0.94523157 | tes-1 |
| WBGene00015218 | 0.099358142 | 0.62012789 | 0.77270574 | B0507.1 |
| WBGene00015219 | 0.177387452 | 0.31867631 | 0.51898076 | B0507.2 |
| WBGene00015220 | 0.200834418 | 0.34339472 | 0.54313812 | B0507.3 |
| WBGene00015221 | 0.0398788 | 0.58971221 | NA | B0507.4 |
| WBGene00015222 | NA | NA | NA | B0507.5 |
| WBGene00015223 | 1.582124936 | 0.00473467 | 0.02506648 | B0507.6 |
| WBGene00015224 | 0.148021412 | 0.07455439 | NA | B0507.7 |
| WBGene00015225 | 4.026141811 | 2.86E-06 | 3.96E-05 | B0507.8 |
| WBGene00015226 | 0.037855293 | 0.65911003 | NA | B0507.9 |
| WBGene00015227 | 3.683480301 | 3.37E-12 | 1.18E-10 | ddn-1 |
| WBGene00015228 | -0.083755272 | 0.68623759 | 0.81911011 | B0511.2 |
| WBGene00015229 | NA | NA | NA | fbxa-125 |
| WBGene00015230 | 0.064809585 | 0.40075491 | NA | tag-344 |
| WBGene00015231 | 0.166965996 | 0.31676094 | 0.51698682 | cutl-14 |
| WBGene00015232 | -0.117044998 | 0.10203749 | 0.2487689 | B0511.6 |
| WBGene00015233 | -0.057699838 | 0.60446985 | 0.76077049 | B0511.7 |
| WBGene00015235 | -0.069443608 | 0.44484528 | 0.63821458 | cdc-26 |
| WBGene00015236 | -0.004755153 | 0.97871751 | 0.98782594 | B0511.11 |
| WBGene00015237 | 0.03520117 | 0.63535933 | 0.78369842 | B0511.12 |

| | | | | |
|---|---|---|---|---|
| WBGene00015238 | -0.279951245 | 0.10168462 | 0.24829294 | mppe-1 |
| WBGene00015241 | 0.036687355 | 0.59489568 | NA | B0524.2 |
| WBGene00015242 | NA | NA | NA | B0524.3 |
| WBGene00015243 | 0.036687355 | 0.59489568 | NA | B0524.4 |
| WBGene00015244 | 0.065014541 | 0.37399213 | NA | B0524.5 |
| WBGene00015245 | -0.230803247 | 0.07602044 | 0.20355389 | B0524.6 |
| WBGene00015246 | 0.056188364 | 0.42092875 | NA | scl-23 |
| WBGene00015247 | 0.040231858 | 0.83895209 | 0.9127791 | B0545.4 |
| WBGene00015248 | -0.109565281 | 0.15417778 | 0.32715947 | mai-2 |
| WBGene00015249 | -0.067045509 | 0.48179025 | 0.66785656 | otub-4 |
| WBGene00015250 | 0.164836901 | 0.27554468 | 0.47572289 | B0546.3 |
| WBGene00015251 | 0.607108157 | 0.00209119 | 0.01295113 | B0546.4 |
| WBGene00015252 | -0.023056499 | 0.78121253 | NA | B0546.5 |
| WBGene00015253 | NA | NA | NA | B0547.2 |
| WBGene00015254 | NA | NA | NA | B0554.1 |
| WBGene00015255 | NA | NA | NA | B0554.2 |
| WBGene00015257 | NA | NA | NA | B0554.4 |
| WBGene00015258 | 0.62728718 | 0.04009403 | 0.12816958 | B0554.5 |
| WBGene00015259 | 0.157098662 | 0.06528165 | NA | dod-20 |
| WBGene00015260 | -0.005581345 | 0.94060131 | NA | B0554.7 |
| WBGene00015261 | 0.039863812 | 0.76440518 | 0.86623548 | B0563.1 |
| WBGene00015262 | 0.900291959 | 0.01997687 | 0.07656581 | B0563.5 |
| WBGene00015263 | 0.208069303 | 0.34142139 | 0.5414344 | B0563.6 |
| WBGene00015264 | 0.172678852 | 0.2632795 | 0.46136148 | B0563.7 |
| WBGene00015265 | -0.135719588 | 0.31879835 | 0.51898076 | B0563.8 |
| WBGene00015266 | -0.132101651 | 0.41027931 | 0.60798576 | BE0003N10.1 |
| WBGene00015267 | -0.114980809 | 0.24878994 | 0.44721099 | chin-1 |
| WBGene00015268 | -0.395873685 | 0.09534861 | 0.23749557 | BE0003N10.3 |
| WBGene00015269 | NA | NA | NA | NA |
| WBGene00015270 | NA | NA | NA | C01B4.6 |
| WBGene00015270 | NA | NA | NA | Y19D10A.16 |
| WBGene00015271 | NA | NA | NA | Y19D10A.4 |
| WBGene00015271 | NA | NA | NA | C01B4.7 |
| WBGene00015272 | NA | NA | NA | C01B4.8 |
| WBGene00015272 | NA | NA | NA | Y19D10A.5 |
| WBGene00015273 | 0.036289481 | 0.71793936 | 0.84178346 | mct-2 |
| WBGene00015273 | 0.036289481 | 0.71793936 | 0.84178346 | mct-1 |
| WBGene00015277 | NA | NA | NA | C01B7.7 |
| WBGene00015277 | NA | NA | NA | F07B7.13 |
| WBGene00015278 | 0.095768074 | 0.53506341 | 0.7102968 | C01B10.3 |
| WBGene00015279 | 0.337892928 | 0.15769691 | 0.33182531 | C01B10.4 |
| WBGene00015280 | 1.226411493 | 3.43E-29 | 4.18E-27 | C01B10.6 |
| WBGene00015281 | -0.039636691 | 0.54579661 | NA | C01B10.7 |

| | | | | | |
|---|---|---|---|---|---|
| WBGene00015282 | | -0.252202383 | 0.00379224 | 0.02099929 | C01B10.8 |
| WBGene00015283 | | -0.186835657 | 0.05394855 | 0.15912311 | C01B10.9 |
| WBGene00015284 | | 0.3455779 | 0.14725388 | 0.31775151 | C01B10.10 |
| WBGene00015285 | | -0.152903348 | 0.37897497 | 0.57833776 | gmeb-1 |
| WBGene00015286 | | 0.173332188 | 0.42242519 | 0.6185751 | best-3 |
| WBGene00015287 | | 0.048324405 | 0.69288477 | 0.82387595 | osta-1 |
| WBGene00015288 | | 0.02696904 | 0.86199934 | 0.92668569 | best-4 |
| WBGene00015291 | | -0.034380001 | 0.8008259 | 0.88894232 | C01B12.8 |
| WBGene00015292 | | 0.059302481 | 0.5133157 | NA | C01C4.2 |
| WBGene00015293 | | 0.537804985 | 0.04747939 | 0.14482202 | C01C4.3 |
| WBGene00015294 | | -0.526804299 | 0.02847027 | 0.09979247 | C01C10.2 |
| WBGene00015295 | | 1.60450624 | 2.15E-12 | 7.67E-11 | acl-12 |
| WBGene00015296 | | -0.068958561 | 0.46072814 | 0.65167817 | gtf-2F1 |
| WBGene00015297 | | -0.16809784 | 0.25323891 | 0.45132939 | sco-1 |
| WBGene00015298 | | -0.309635709 | 0.11357696 | 0.26754164 | C01F1.3 |
| WBGene00015299 | | 0.088670835 | 0.58011611 | 0.745406 | C01F1.4 |
| WBGene00015300 | | 0.036202261 | 0.67244084 | NA | C01F1.5 |
| WBGene00015301 | | 0.046342667 | 0.62964317 | 0.77968456 | C01F1.6 |
| WBGene00015303 | | 0.728585404 | 0.02608254 | 0.09371152 | rga-6 |
| WBGene00015305 | | 0.040218942 | 0.56174727 | NA | C01G5.3 |
| WBGene00015306 | | 0.070721981 | 0.44603293 | NA | C01G5.4 |
| WBGene00015307 | | -0.0914067 | 0.51406808 | 0.69413177 | C01G5.5 |
| WBGene00015308 | | -0.088670401 | 0.26716715 | 0.46599883 | ddi-1 |
| WBGene00015309 | | -0.012235072 | 0.90504857 | 0.94913088 | C01G5.7 |
| WBGene00015310 | | -0.027213567 | 0.78395281 | 0.87845194 | fan-1 |
| WBGene00015311 | | -0.102139554 | 0.59526281 | 0.75477619 | C01G5.9 |
| WBGene00015312 | | 0.018787507 | 0.80608907 | 0.89239698 | C01G8.1 |
| WBGene00015313 | | -0.266972456 | 0.03355889 | 0.11243379 | hpo-32 |
| WBGene00015315 | NA | | NA | NA | srbc-29 |
| WBGene00015316 | NA | | NA | NA | srbc-30 |
| WBGene00015317 | NA | | NA | NA | srbc-32 |
| WBGene00015318 | NA | | NA | NA | srbc-36 |
| WBGene00015320 | | 0.03425478 | 0.84700385 | 0.91761389 | C02B8.1 |
| WBGene00015321 | | -0.003108065 | 0.95728506 | NA | fbxc-38 |
| WBGene00015322 | | 1.034669686 | 0.01237251 | 0.05296672 | C02B8.3 |
| WBGene00015323 | | 0.036100058 | 0.7193253 | 0.84246394 | frpr-1 |
| WBGene00015324 | | 0.035387519 | 0.84811495 | 0.9180823 | C02B8.6 |
| WBGene00015325 | NA | | NA | NA | C02B8.7 |
| WBGene00015326 | | -0.212486221 | 0.00454194 | 0.02431638 | ivd-1 |
| WBGene00015327 | | -0.106904903 | 0.36486181 | 0.56450826 | snpn-1 |
| WBGene00015328 | | 0.015728821 | 0.93299591 | 0.96237798 | C02B10.3 |
| WBGene00015329 | | -0.248137225 | 0.01000049 | 0.04487196 | C02B10.4 |
| WBGene00015330 | | -0.130958185 | 0.2049359 | 0.39402101 | C02B10.5 |

| | | | | |
|---|---|---|---|---|
| WBGene00015331 | 0.020925399 | 0.84643798 | NA | C02B10.6 |
| WBGene00015332 | -0.129773543 | 0.4366351 | 0.63163468 | tyr-1 |
| WBGene00015333 | 0.190060513 | 0.3807747 | 0.58017716 | slc-17.3 |
| WBGene00015334 | 0.211846067 | 0.29954457 | 0.50001506 | sup-18 |
| WBGene00015335 | 0.055827801 | 0.59420606 | 0.7538143 | acdh-6 |
| WBGene00015336 | 0.055429719 | 0.78741909 | 0.87950892 | C02D5.2 |
| WBGene00015337 | -0.014257542 | 0.90128897 | 0.9473596 | gsto-2 |
| WBGene00015338 | NA | NA | NA | catp-2 |
| WBGene00015339 | 3.898972799 | 0.00042304 | 0.00337165 | C02E7.6 |
| WBGene00015340 | 1.466039417 | 0.01537761 | 0.06310077 | C02E7.7 |
| WBGene00015341 | NA | NA | NA | C02E7.8 |
| WBGene00015342 | -0.023056499 | 0.78121253 | NA | C02E7.10 |
| WBGene00015344 | -0.230529297 | 0.00037887 | 0.00306975 | nra-4 |
| WBGene00015345 | 0.102352968 | 0.32744523 | 0.52665337 | C02F5.2 |
| WBGene00015346 | -0.250027477 | 0.0095686 | 0.04350742 | C02F5.3 |
| WBGene00015347 | -0.235123939 | 0.02557116 | 0.09222534 | cids-1 |
| WBGene00015348 | 0.370044478 | 0.01841807 | 0.07241112 | C02F5.5 |
| WBGene00015349 | 0.034468843 | 0.749788 | 0.8587464 | henn-1 |
| WBGene00015350 | 0.694902234 | 3.29E-05 | 0.00035822 | fbxl-1 |
| WBGene00015351 | -0.177981395 | 0.31476625 | 0.5154337 | C02F5.10 |
| WBGene00015352 | 0.083294596 | 0.68237735 | 0.81629663 | C02F5.12 |
| WBGene00015353 | -0.100212339 | 0.48965871 | 0.67359703 | C02F5.13 |
| WBGene00015354 | 1.092161128 | 1.74E-05 | 0.00020062 | snet-1 |
| WBGene00015355 | 0.069103791 | 0.68404189 | 0.8176383 | C02F12.5 |
| WBGene00015356 | 0.183862144 | 0.38423379 | 0.58362558 | tag-278 |
| WBGene00015357 | -0.18598907 | 0.20355032 | 0.39247213 | C02F12.8 |
| WBGene00015359 | -0.099867295 | 0.60173037 | 0.75908213 | C02G6.1 |
| WBGene00015360 | 0.015584271 | 0.80362925 | NA | C02G6.2 |
| WBGene00015361 | 0.058314668 | 0.40208118 | NA | C02H6.1 |
| WBGene00015362 | 0.242811141 | 0.27331229 | 0.47355801 | fbxa-105 |
| WBGene00015364 | 0.036100058 | 0.7193253 | 0.84246394 | npr-19 |
| WBGene00015366 | 0.111598012 | 0.52393687 | 0.70139111 | C03A7.2 |
| WBGene00015367 | 0.020317245 | 0.84643798 | NA | srt-64 |
| WBGene00015368 | 0.05366068 | 0.40968658 | NA | srt-65 |
| WBGene00015369 | -0.037026432 | 0.85506553 | 0.92241889 | ugt-51 |
| WBGene00015371 | 0.22458448 | 0.29792964 | 0.49893568 | C03A7.13 |
| WBGene00015372 | NA | NA | NA | C03B1.1 |
| WBGene00015373 | 0.039292341 | 0.65808492 | NA | C03B1.2 |
| WBGene00015374 | NA | NA | NA | C03B1.3 |
| WBGene00015375 | NA | NA | NA | C03B1.4 |
| WBGene00015376 | -0.028668027 | 0.7217843 | NA | C03B1.5 |
| WBGene00015377 | 0.037838157 | 0.65920403 | NA | C03B1.6 |
| WBGene00015378 | 0.068012149 | 0.69285233 | 0.82387595 | C03B1.7 |

| | | | | |
|---|---|---|---|---|
| WBGene00015379 | NA | NA | NA | C03B1.9 |
| WBGene00015380 | NA | NA | NA | C03B1.10 |
| WBGene00015381 | -0.008471898 | 0.96482725 | 0.98071026 | C03B1.13 |
| WBGene00015382 | 0.093198624 | 0.65626959 | 0.79678605 | C03B1.14 |
| WBGene00015385 | 0.121754221 | 0.55978814 | 0.73020832 | C03B8.2 |
| WBGene00015386 | 0.121340625 | 0.17845998 | NA | C03B8.3 |
| WBGene00015387 | 0.737423026 | 0.02453007 | 0.08956227 | kcnl-3 |
| WBGene00015388 | -0.013442408 | 0.90374339 | 0.94878151 | C03F11.2 |
| WBGene00015389 | 0.127312958 | 0.37791896 | 0.57708713 | scav-1 |
| WBGene00015390 | 0.114771846 | 0.58071197 | 0.74567654 | C03F11.4 |
| WBGene00015391 | 0.061311391 | 0.46061872 | 0.65158635 | sdha-1 |
| WBGene00015392 | 0.116956302 | 0.55326595 | 0.72538433 | nspc-7 |
| WBGene00015393 | 0.262759553 | 0.20224217 | 0.39073999 | C03G6.5 |
| WBGene00015394 | NA | NA | NA | C03G6.6 |
| WBGene00015395 | 0.008770705 | 0.93328612 | 0.96254171 | nhr-147 |
| WBGene00015396 | 0.19469062 | 0.18739332 | 0.37206748 | nhr-148 |
| WBGene00015397 | -0.023056499 | 0.78121253 | NA | nhr-149 |
| WBGene00015398 | 0.020925399 | 0.84643798 | NA | tag-293 |
| WBGene00015399 | 0.075726029 | 0.27693214 | NA | cyp-35A1 |
| WBGene00015400 | 0.718356689 | 1.59E-11 | 5.11E-10 | cyp-35A2 |
| WBGene00015401 | -0.003614866 | 0.95700117 | NA | srsx-24 |
| WBGene00015402 | 1.581244375 | 2.08E-05 | 0.00023624 | C03G6.17 |
| WBGene00015403 | 2.076701296 | 5.57E-41 | 1.03E-38 | clec-10 |
| WBGene00015404 | 0.012770023 | 0.90106584 | 0.94730323 | nstp-4 |
| WBGene00015405 | -0.108750918 | 0.41943964 | 0.61617339 | C03H5.3 |
| WBGene00015406 | 0.020925399 | 0.84643798 | NA | C03H5.4 |
| WBGene00015407 | -0.038007311 | 0.84813889 | 0.9180823 | C03H5.5 |
| WBGene00015408 | 0.402822597 | 0.06574619 | 0.1830715 | C03H5.6 |
| WBGene00015409 | -0.278496572 | 0.13592709 | 0.30089333 | C03H5.7 |
| WBGene00015410 | 0.041916149 | 0.82887531 | 0.90652983 | tbc-6 |
| WBGene00015411 | -0.022930767 | 0.78121253 | NA | srz-60 |
| WBGene00015412 | -0.014817053 | 0.89419142 | 0.94489559 | lgc-9 |
| WBGene00015413 | -0.338603026 | 1.25E-05 | 0.00014998 | pdhb-1 |
| WBGene00015414 | -0.010539732 | 0.91155916 | 0.95112162 | C04C3.4 |
| WBGene00015416 | 0.038325924 | 0.57925535 | NA | C04C3.6 |
| WBGene00015417 | NA | NA | NA | C04C3.7 |
| WBGene00015418 | 0.021584543 | 0.80318186 | 0.89045779 | pac-1 |
| WBGene00015419 | 0.029307384 | 0.78484777 | 0.87855213 | srsx-18 |
| WBGene00015421 | -0.023109762 | 0.78121253 | NA | C04E6.4 |
| WBGene00015422 | -0.006388227 | 0.92874308 | NA | C04E6.5 |
| WBGene00015423 | 0.843137543 | 0.00315029 | 0.01813742 | C04E6.7 |
| WBGene00015424 | 0.099732927 | 0.53353672 | 0.70899686 | C04E6.8 |
| WBGene00015425 | -0.024663339 | 0.82101877 | 0.90190094 | C04E6.11 |

| | | | | |
|---|---|---|---|---|
| WBGene00015426 | 0.241164741 | 0.01705846 | 0.06829447 | C04E6.12 |
| WBGene00015427 | 0.572943653 | 0.04930457 | 0.14871617 | C04E6.13 |
| WBGene00015428 | 0.020925399 | 0.84643798 | NA | C04E7.1 |
| WBGene00015429 | -0.052686892 | 0.75176408 | 0.85959087 | sor-3 |
| WBGene00015430 | 0.633515328 | 0.01331558 | 0.05623114 | C04E7.3 |
| WBGene00015432 | NA | NA | NA | NA |
| WBGene00015433 | 0.246558367 | 0.26915891 | 0.46820634 | C04E12.2 |
| WBGene00015434 | 1.029841069 | 7.34E-05 | 0.00073683 | C04E12.4 |
| WBGene00015435 | -0.277910996 | 0.12603638 | 0.28699798 | C04E12.5 |
| WBGene00015436 | 0.040107227 | 0.58693163 | NA | C04E12.6 |
| WBGene00015437 | -0.045407731 | 0.76488977 | 0.86650813 | scrm-3 |
| WBGene00015438 | NA | NA | NA | srbc-2 |
| WBGene00015439 | 0.19907325 | 0.2277681 | 0.4212729 | C04E12.10 |
| WBGene00015440 | 0.418742958 | 0.06364719 | 0.17910023 | arrd-20 |
| WBGene00015441 | 1.588152997 | 0.00502708 | 0.02623283 | arrd-21 |
| WBGene00015442 | -0.01449588 | 0.89369483 | 0.94468695 | C04F1.1 |
| WBGene00015444 | 0.059980168 | 0.38780842 | NA | C04F5.2 |
| WBGene00015446 | NA | NA | NA | srab-1 |
| WBGene00015447 | NA | NA | NA | srab-2 |
| WBGene00015448 | -0.023109762 | 0.78121253 | NA | srab-3 |
| WBGene00015449 | -0.849236434 | 0.00015444 | 0.00140796 | ugt-63 |
| WBGene00015450 | -0.277357593 | 0.01088687 | 0.04775157 | C04F5.8 |
| WBGene00015451 | 0.039240988 | 0.76394289 | 0.86623378 | C04F5.9 |
| WBGene00015453 | -0.033469054 | 0.7256163 | NA | C04G6.2 |
| WBGene00015454 | -0.010267586 | 0.93588873 | 0.96380369 | C04G6.4 |
| WBGene00015455 | 2.896533439 | 0.00060345 | 0.00457301 | C04G6.5 |
| WBGene00015456 | -0.135258361 | 0.49641574 | 0.67886649 | C04G6.6 |
| WBGene00015457 | 0.05787416 | 0.40594086 | NA | C04G6.7 |
| WBGene00015458 | 0.0398788 | 0.58971221 | NA | C04G6.10 |
| WBGene00015459 | NA | NA | NA | C04G6.11 |
| WBGene00015460 | 0.00523983 | 0.97004782 | 0.98346632 | C05C8.1 |
| WBGene00015461 | -0.211636989 | 0.05479646 | 0.16093781 | krr-1 |
| WBGene00015462 | 0.079603473 | 0.58840556 | 0.75051633 | C05C8.5 |
| WBGene00015463 | -0.06677694 | 0.42874676 | 0.62367425 | hpo-9 |
| WBGene00015464 | -0.685241341 | 0.01465961 | 0.06073325 | C05C8.7 |
| WBGene00015465 | -0.213141608 | 0.14505626 | 0.31445617 | C05C8.8 |
| WBGene00015466 | -0.028056057 | 0.83158072 | 0.90832388 | hyls-1 |
| WBGene00015467 | -0.031060291 | 0.700178 | NA | basl-1 |
| WBGene00015468 | -0.038142382 | 0.72125787 | 0.84362463 | madf-11 |
| WBGene00015469 | 0.299686771 | 0.01552762 | 0.06362715 | C05D2.8 |
| WBGene00015470 | -0.043105104 | 0.60889711 | 0.76432215 | C05D2.10 |
| WBGene00015471 | 0.605923754 | 0.0143944 | 0.05985469 | lmp-2 |
| WBGene00015472 | 0.661226378 | 0.00438342 | 0.02360342 | C05D9.3 |

| | | | | |
|---|---|---|---|---|
| WBGene00015473 | 0.132442484 | 0.10147609 | NA | C05D9.4 |
| WBGene00015474 | -0.240034721 | 0.21212278 | 0.40244779 | C05D9.7 |
| WBGene00015476 | 0.981441939 | 0.00039909 | 0.00320874 | C05D9.9 |
| WBGene00015477 | 0.19154085 | 0.18764353 | 0.37251375 | attf-4 |
| WBGene00015478 | 0.259208826 | 0.21424775 | 0.40511447 | mapk-15 |
| WBGene00015479 | -0.688335963 | 0.02642917 | 0.0946781 | wht-1 |
| WBGene00015480 | 0.09389644 | 0.62291618 | 0.77453057 | C05D10.4 |
| WBGene00015481 | -0.199907218 | 0.00871165 | 0.04040141 | C05D11.1 |
| WBGene00015483 | 1.309124354 | 2.30E-07 | 4.04E-06 | C05D11.5 |
| WBGene00015484 | 0.244933724 | 0.0053278 | 0.02750506 | atgl-1 |
| WBGene00015485 | -0.280390071 | 0.00818241 | 0.03845862 | C05D11.8 |
| WBGene00015486 | -0.24979245 | 0.00962147 | 0.04363938 | C05D11.9 |
| WBGene00015487 | -0.42842533 | 0.00197285 | 0.01233326 | mrps-17 |
| WBGene00015488 | 0.428088464 | 0.07783103 | 0.20682048 | C05D11.13 |
| WBGene00015489 | 0.020925399 | 0.84643798 | NA | C05E4.7 |
| WBGene00015491 | 0.056727431 | 0.44364738 | NA | C05E4.12 |
| WBGene00015492 | -0.141274763 | 0.36412494 | 0.56388417 | lnp-1 |
| WBGene00015493 | NA | NA | NA | C05E11.2 |
| WBGene00015494 | 0.03255121 | 0.87372576 | 0.93276403 | C05E11.3 |
| WBGene00015495 | 0.055634304 | 0.71554816 | 0.84042707 | C05E11.6 |
| WBGene00015496 | -0.023109762 | 0.78121253 | NA | C05E11.7 |
| WBGene00015497 | 0.052673899 | 0.78597246 | 0.87892197 | nhr-76 |
| WBGene00015499 | -0.17866267 | 0.03316274 | 0.11136139 | inst-1 |
| WBGene00015500 | 0.009106066 | 0.93921763 | 0.96586959 | C06A5.2 |
| WBGene00015501 | -0.071801347 | 0.44446082 | 0.63794923 | C06A5.3 |
| WBGene00015504 | -0.092099312 | 0.31882553 | 0.51898076 | C06A5.6 |
| WBGene00015505 | -0.009019852 | 0.94837517 | 0.97044737 | C06A5.8 |
| WBGene00015507 | 0.007637901 | 0.96195481 | 0.97924853 | C06A6.2 |
| WBGene00015508 | -0.150147006 | 0.23341997 | 0.42875992 | mvb-12 |
| WBGene00015509 | -0.077287101 | 0.3327061 | 0.53242432 | C06A6.4 |
| WBGene00015510 | -0.337855805 | 0.02544321 | 0.09184272 | erp-44.2 |
| WBGene00015511 | 0.02727927 | 0.7627639 | NA | C06A6.7 |
| WBGene00015512 | 0.733591537 | 1.26E-33 | 1.77E-31 | mthf-1 |
| WBGene00015513 | -0.039031632 | 0.69869183 | 0.82846224 | snpc-1.1 |
| WBGene00015514 | -0.703977734 | 7.29E-29 | 8.68E-27 | nlp-77 |
| WBGene00015515 | -0.227944835 | 0.09251164 | 0.23212732 | spdl-1 |
| WBGene00015516 | 0.294931643 | 0.06006777 | 0.17173448 | C06A8.6 |
| WBGene00015517 | 0.599552848 | 0.02997482 | 0.10343186 | C06A8.8 |
| WBGene00015518 | -0.455687982 | 0.04449237 | 0.13812825 | C06E1.1 |
| WBGene00015519 | -0.155453426 | 0.46462871 | 0.65449527 | doxa-1 |
| WBGene00015520 | NA | NA | NA | fip-3 |
| WBGene00015521 | 0.017604133 | 0.79437959 | NA | fipr-16 |
| WBGene00015522 | 0.027459475 | 0.82976647 | 0.90708661 | C06E1.7 |

| | | | | | |
|---|---|---|---|---|---|
| WBGene00015523 | | 0.120721076 | 0.31511378 | 0.51559911 | ztf-30 |
| WBGene00015524 | | 0.107808673 | 0.35319829 | 0.55300783 | C06E1.9 |
| WBGene00015525 | | -0.294048292 | 0.06754231 | 0.18653278 | rha-2 |
| WBGene00015526 | | 0.184193711 | 0.36889861 | 0.56883038 | C06E1.11 |
| WBGene00015527 | | 0.112859948 | 0.58114176 | 0.74581663 | C06E2.1 |
| WBGene00015528 | NA | | NA | NA | C06E2.2 |
| WBGene00015529 | | -0.257120697 | 0.19660731 | 0.3841045 | C06E2.5 |
| WBGene00015530 | NA | | NA | NA | C06E4.1 |
| WBGene00015531 | | 0.252361842 | 0.16956106 | 0.34793883 | C06E4.2 |
| WBGene00015532 | | 2.125921977 | 0.0003262 | 0.00270439 | C06E4.3 |
| WBGene00015533 | | 0.020232312 | 0.84643798 | NA | NA |
| WBGene00015534 | NA | | NA | NA | C06E4.5 |
| WBGene00015535 | | 0.074833405 | 0.3346945 | NA | C06E4.6 |
| WBGene00015536 | | 0.124133031 | 0.25369644 | 0.45170492 | glb-2 |
| WBGene00015537 | | 2.922237414 | 0.00014141 | 0.00130126 | C06E4.8 |
| WBGene00015538 | | 0.058785534 | 0.5340317 | 0.70937744 | sams-3 |
| WBGene00015539 | | 0.346912519 | 0.13970866 | 0.3070822 | C06E7.2 |
| WBGene00015540 | | 0.165253761 | 0.10829964 | 0.2588431 | sams-4 |
| WBGene00015541 | | 0.38287062 | 0.11857931 | 0.27592494 | C06E7.4 |
| WBGene00015544 | | 0.418745758 | 0.01499062 | 0.06180728 | C06E8.5 |
| WBGene00015545 | | 0.190638467 | 0.35059889 | 0.55058654 | C06G1.1 |
| WBGene00015546 | | 0.058314668 | 0.40208118 | NA | C06G1.2 |
| WBGene00015547 | | -0.066373434 | 0.47700467 | 0.66410831 | ain-1 |
| WBGene00015549 | | 0.097863341 | 0.38937839 | 0.58773772 | C06G3.3 |
| WBGene00015550 | | 0.038125098 | 0.61119482 | NA | NA |
| WBGene00015551 | | 0.140963311 | 0.31279851 | 0.51333563 | C06G3.5 |
| WBGene00015552 | | -0.203862338 | 0.22418309 | 0.41711331 | C06G3.6 |
| WBGene00015553 | | 0.209837336 | 0.0052637 | 0.02723176 | trxr-1 |
| WBGene00015554 | | -0.11404231 | 0.41940679 | 0.61617339 | C06G3.8 |
| WBGene00015555 | | 0.093124654 | 0.32635712 | 0.52559615 | ufl-1 |
| WBGene00015557 | | -0.00848512 | 0.93423748 | 0.96290283 | C06G4.1 |
| WBGene00015558 | | 0.116830083 | 0.11818778 | NA | C06G4.4 |
| WBGene00015559 | | -0.540545259 | 0.05340325 | 0.15810973 | npr-17 |
| WBGene00015560 | | 0.127121974 | 0.4430062 | 0.63685467 | C07A12.2 |
| WBGene00015561 | | 0.083609116 | 0.61764047 | 0.77097465 | C07A12.7 |
| WBGene00015562 | | 0.065014541 | 0.37399213 | NA | NA |
| WBGene00015564 | NA | | NA | NA | C07D8.5 |
| WBGene00015565 | | -0.182204564 | 0.00200107 | 0.01247242 | C07D8.6 |
| WBGene00015566 | | 0.493666004 | 0.06960864 | 0.19056472 | C07D10.1 |
| WBGene00015567 | | -0.052614785 | 0.52273128 | 0.70044791 | bath-44 |
| WBGene00015568 | | 0.059938442 | 0.52286458 | NA | sre-3 |
| WBGene00015569 | | -0.779016603 | 0.00013396 | 0.00124524 | C07D10.5 |
| WBGene00015570 | | -0.024114341 | 0.86351176 | 0.92715427 | C07F11.2 |

| | | | | |
|---|---|---|---|---|
| WBGene00015572 | 0.086764008 | 0.294364 | NA | C07G1.6 |
| WBGene00015573 | 0.017141582 | 0.79355725 | NA | C07G1.7 |
| WBGene00015574 | 2.200418349 | 0.00413793 | 0.02250501 | irg-1 |
| WBGene00015575 | 0.020317245 | 0.84643798 | NA | NA |
| WBGene00015576 | 0.039888668 | 0.56326497 | NA | C07G3.8 |
| WBGene00015577 | 0.521219454 | 0.00066978 | 0.0049854 | ugt-64 |
| WBGene00015578 | -0.050580107 | 0.47300881 | NA | C07G3.10 |
| WBGene00015579 | NA | NA | NA | C07H6.2 |
| WBGene00015580 | 0.225760394 | 0.14340972 | 0.31198539 | cls-1 |
| WBGene00015581 | -0.029034711 | 0.71151921 | 0.8376204 | C07H6.4 |
| WBGene00015582 | 0.095967469 | 0.18297344 | NA | C07H6.9 |
| WBGene00015583 | -0.023109762 | 0.78121253 | NA | C08A9.3 |
| WBGene00015586 | -0.13988159 | 0.47933323 | 0.66615262 | lido-1 |
| WBGene00015587 | -0.849847328 | 0.00513715 | 0.02669022 | lido-2 |
| WBGene00015588 | 0.065443893 | 0.71714817 | 0.84132743 | lido-3 |
| WBGene00015589 | -0.079107273 | 0.67550176 | 0.81138906 | lido-4 |
| WBGene00015590 | -0.129281952 | 0.34247274 | 0.54280413 | bath-15 |
| WBGene00015591 | 0.024668948 | 0.81185276 | 0.89587496 | cyk-7 |
| WBGene00015592 | 0.195792103 | 0.274261 | 0.47469621 | C08D8.1 |
| WBGene00015593 | NA | NA | NA | C08E3.1 |
| WBGene00015594 | 0.058314668 | 0.40208118 | NA | btb-15 |
| WBGene00015595 | -0.024741105 | 0.85405045 | 0.92193118 | bath-33 |
| WBGene00015596 | NA | NA | NA | NA |
| WBGene00015597 | 0.161417489 | 0.16995875 | 0.34842782 | NA |
| WBGene00015598 | -0.006711909 | 0.92834175 | NA | fbxa-163 |
| WBGene00015599 | 0.085537126 | 0.24698522 | NA | fbxa-164 |
| WBGene00015600 | 0.020317245 | 0.84643798 | NA | fbxa-165 |
| WBGene00015601 | 0.269076775 | 0.03780814 | 0.12271091 | fbxa-166 |
| WBGene00015602 | 4.022678468 | 0.00145912 | 0.00959705 | fbxa-158 |
| WBGene00015603 | NA | NA | NA | fbxa-159 |
| WBGene00015604 | NA | NA | NA | fbxa-160 |
| WBGene00015605 | 0.108538718 | 0.15420469 | NA | C08E3.13 |
| WBGene00015606 | -0.023109762 | 0.78121253 | NA | math-2 |
| WBGene00015607 | -0.205891996 | 0.30729333 | 0.50791168 | fbxb-13 |
| WBGene00015608 | 0.599612488 | 0.00114331 | 0.007863 | math-3 |
| WBGene00015609 | 0.87012109 | 1.35E-05 | 0.00015963 | math-4 |
| WBGene00015610 | -1.269854259 | 2.15E-05 | 0.00024354 | C08F1.6 |
| WBGene00015611 | 0.124326451 | 0.11293212 | NA | C08F1.8 |
| WBGene00015613 | -0.909369206 | 0.00216613 | 0.01330289 | C08F1.10 |
| WBGene00015614 | NA | NA | NA | C08G5.1 |
| WBGene00015615 | NA | NA | NA | fbxc-48 |
| WBGene00015616 | 0.037023624 | 0.59153183 | NA | C08G5.3 |
| WBGene00015618 | 0.133532899 | 0.15682793 | 0.33080502 | C08G5.6 |

| | | | | | |
|---|---|---|---|---|---|
| WBGene00015619 | | 0.921104444 | 0.00033855 | 0.00278782 | C08G9.1 |
| WBGene00015620 | | 0.077791496 | 0.67532191 | 0.81135495 | C08G9.2 |
| WBGene00015621 | | 0.315452814 | 0.10007983 | 0.24556206 | C09B7.2 |
| WBGene00015622 | | -0.057595821 | 0.36232248 | NA | C09B8.3 |
| WBGene00015623 | | -2.866743796 | 7.59E-30 | 9.58E-28 | C09B8.4 |
| WBGene00015624 | | -0.761827273 | 0.00391547 | 0.02152702 | C09B8.5 |
| WBGene00015625 | | 0.177835185 | 0.23600316 | 0.43191528 | C09B8.8 |
| WBGene00015626 | | 0.148467664 | 0.07389511 | NA | C09B9.1 |
| WBGene00015627 | | 0.204923027 | 0.10763935 | 0.25791858 | C09B9.2 |
| WBGene00015628 | | 0.036687355 | 0.59489568 | NA | best-6 |
| WBGene00015629 | | 0.086764008 | 0.294364 | NA | C09B9.4 |
| WBGene00015630 | | 0.056188364 | 0.42092875 | NA | C09B9.7 |
| WBGene00015631 | | 0.255163688 | 0.08892032 | 0.22633561 | clec-89 |
| WBGene00015632 | | 1.846788853 | 3.03E-20 | 2.34E-18 | C09D4.1 |
| WBGene00015633 | | 0.623404857 | 0.00011917 | 0.0011322 | C09D4.2 |
| WBGene00015634 | | 0.242389492 | 0.17437513 | 0.35505649 | C09D4.3 |
| WBGene00015635 | | -0.180363561 | 0.04250343 | 0.13368263 | C09D4.4 |
| WBGene00015636 | NA | | NA | NA | C09D4.6 |
| WBGene00015637 | NA | | NA | NA | srxa-5 |
| WBGene00015638 | | -0.062325426 | 0.71612869 | 0.84075838 | C09E7.4 |
| WBGene00015639 | | 0.07149497 | 0.5434358 | 0.71696212 | C09E7.5 |
| WBGene00015640 | | -0.147235163 | 0.34677168 | 0.54645186 | NA |
| WBGene00015641 | | -0.101760305 | 0.47709112 | 0.66410831 | C09E7.7 |
| WBGene00015642 | | -0.180477176 | 0.16591469 | 0.34245844 | C09E7.8 |
| WBGene00015644 | | 0.095285235 | 0.54359374 | 0.71696212 | C09E8.1 |
| WBGene00015645 | | 0.131317611 | 0.10121567 | NA | lips-7 |
| WBGene00015646 | | 0.051931643 | 0.7977473 | 0.88680697 | mlt-10 |
| WBGene00015647 | | -0.024043683 | 0.89782893 | 0.94623621 | C09F5.1 |
| WBGene00015648 | | 0.124575799 | 0.4194659 | 0.61617339 | orai-1 |
| WBGene00015649 | | 0.059855545 | 0.41422968 | NA | C09F5.3 |
| WBGene00015650 | | -0.191178189 | 0.07532891 | 0.2020316 | pkg-2 |
| WBGene00015651 | | 0.118015953 | 0.17505153 | 0.35564444 | ceh-53 |
| WBGene00015652 | NA | | NA | NA | srz-79 |
| WBGene00015653 | NA | | NA | NA | srz-78 |
| WBGene00015654 | NA | | NA | NA | NA |
| WBGene00015655 | | 0.108942554 | 0.15594412 | NA | C09G12.5 |
| WBGene00015656 | NA | | NA | NA | srz-70 |
| WBGene00015658 | | -0.014574424 | 0.8928179 | 0.9441978 | tsg-101 |
| WBGene00015660 | | -0.709312516 | 0.00579587 | 0.02941328 | catp-3 |
| WBGene00015661 | | 0.092724492 | 0.19640118 | NA | C09H5.7 |
| WBGene00015662 | | -0.25286591 | 0.08946942 | 0.22726891 | C10A4.1 |
| WBGene00015663 | NA | | NA | NA | C10A4.2 |
| WBGene00015664 | | 0.012767728 | 0.88865492 | NA | C10A4.3 |

| | | | | |
|---|---|---|---|---|
| WBGene00015665 | 0.132445395 | 0.5026691 | 0.68397568 | C10A4.4 |
| WBGene00015666 | 0.12103813 | 0.48480955 | 0.66987915 | gad-2 |
| WBGene00015667 | -0.04803032 | 0.76019433 | 0.86416163 | C10A4.6 |
| WBGene00015668 | -0.02419332 | 0.70411432 | NA | C10A4.7 |
| WBGene00015670 | 0.292449115 | 0.13694897 | 0.30241193 | C10B5.1 |
| WBGene00015671 | 0.31176244 | 0.07906437 | 0.20943503 | C10B5.3 |
| WBGene00015672 | 0.117958011 | 0.39536997 | 0.59330482 | C10E2.1 |
| WBGene00015673 | 0.077308817 | 0.68994187 | 0.82175538 | C10E2.2 |
| WBGene00015675 | 0.056913249 | 0.73660655 | 0.85204655 | C10E2.5 |
| WBGene00015676 | -0.042823279 | 0.61784804 | 0.77099816 | mct-6 |
| WBGene00015677 | -0.044391564 | 0.55275247 | 0.72508974 | cpg-4 |
| WBGene00015678 | -0.292182378 | 0.18850052 | 0.37361339 | mcp-1 |
| WBGene00015679 | 4.108437273 | 0.00121786 | 0.00827793 | C10F3.7 |
| WBGene00015680 | 0.152808005 | 0.36829914 | 0.56802567 | egal-1 |
| WBGene00015681 | NA | NA | NA | C10G8.2 |
| WBGene00015682 | 0.042516415 | 0.55991373 | NA | C10G8.3 |
| WBGene00015683 | 0.705811774 | 6.91E-16 | 3.56E-14 | C10G8.4 |
| WBGene00015684 | -0.042023227 | 0.69074144 | 0.82235004 | C10G8.8 |
| WBGene00015685 | -0.027535543 | 0.68063395 | NA | C10G11.1 |
| WBGene00015686 | -0.019131564 | 0.88765343 | 0.94086251 | C10G11.6 |
| WBGene00015687 | -0.452007017 | 1.05E-13 | 4.37E-12 | chdp-1 |
| WBGene00015688 | 2.830566734 | 2.65E-07 | 4.62E-06 | C10G11.8 |
| WBGene00015689 | 0.123674741 | 0.11449035 | NA | spch-2 |
| WBGene00015690 | 0.0398788 | 0.58971221 | NA | C10G11.10 |
| WBGene00015691 | 0.044167339 | 0.64487176 | 0.78968348 | viln-1 |
| WBGene00015692 | 0.27761221 | 0.19591574 | 0.38359238 | ugt-25 |
| WBGene00015693 | -0.007894681 | 0.96203131 | 0.97924853 | ugt-28 |
| WBGene00015694 | 0.035541389 | 0.67519412 | NA | ugt-27 |
| WBGene00015695 | -0.290247728 | 0.0425934 | 0.13387933 | ugt-26 |
| WBGene00015696 | 0.060484201 | 0.41382805 | NA | C10H11.7 |
| WBGene00015697 | 0.044053017 | 0.7023194 | 0.83093594 | mlst-8 |
| WBGene00015698 | -0.401096826 | 2.85E-05 | 0.00031359 | kca-1 |
| WBGene00015700 | 0.032760388 | 0.61273797 | NA | asp-9 |
| WBGene00015701 | 0.09048927 | 0.52321955 | 0.7009294 | C11D2.3 |
| WBGene00015702 | 0.001347166 | 0.99244724 | 0.9955752 | C11D2.4 |
| WBGene00015703 | -0.249455438 | 0.0300267 | 0.10353765 | C11D2.7 |
| WBGene00015704 | 0.565061201 | 0.02807296 | 0.09865967 | tiam-1 |
| WBGene00015705 | 0.018350543 | 0.91643839 | 0.95348243 | nhr-152 |
| WBGene00015706 | 0.300764711 | 0.08877128 | 0.22611347 | C12D5.3 |
| WBGene00015707 | -0.001993815 | 0.95612996 | NA | C12D5.4 |
| WBGene00015708 | 0.034981685 | 0.68236555 | NA | C12D5.5 |
| WBGene00015709 | 0.312569092 | 0.15093844 | 0.32275299 | cyp-33A1 |
| WBGene00015710 | 0.497786023 | 0.05385836 | 0.15899127 | C12D5.9 |

| | | | | |
|---|---|---|---|---|
| WBGene00015711 | 0.155449191 | 0.11965772 | 0.27790428 | C12D5.10 |
| WBGene00015712 | 0.036687355 | 0.59489568 | NA | sre-11 |
| WBGene00015713 | 0.948679205 | 1.83E-14 | 8.28E-13 | C12D12.1 |
| WBGene00015714 | -0.043432003 | 0.49754315 | NA | C12D12.3 |
| WBGene00015716 | 0.777314306 | 0.01581631 | 0.06461122 | sox-4 |
| WBGene00015718 | 0.020317245 | 0.84643798 | NA | C13A2.1 |
| WBGene00015719 | NA | NA | NA | C13A2.2 |
| WBGene00015720 | 0.020232312 | 0.84643798 | NA | C13A2.3 |
| WBGene00015721 | NA | NA | NA | C13A2.4 |
| WBGene00015722 | NA | NA | NA | C13A2.5 |
| WBGene00015723 | NA | NA | NA | C13A2.6 |
| WBGene00015724 | NA | NA | NA | C13A2.7 |
| WBGene00015725 | NA | NA | NA | C13A2.9 |
| WBGene00015726 | NA | NA | NA | C13A2.10 |
| WBGene00015727 | NA | NA | NA | C13A2.11 |
| WBGene00015728 | 0.341629705 | 0.15266649 | 0.32540255 | C13A10.1 |
| WBGene00015729 | -0.008912468 | 0.95310787 | 0.97322797 | C13A10.2 |
| WBGene00015730 | NA | NA | NA | srt-1 |
| WBGene00015730 | NA | NA | NA | srt-2 |
| WBGene00015731 | NA | NA | NA | C13B7.6 |
| WBGene00015731 | NA | NA | NA | Y45G12C.16 |
| WBGene00015733 | 0.742029221 | 0.00016081 | 0.00145968 | C13B9.2 |
| WBGene00015734 | -0.13685378 | 0.0883472 | 0.2255434 | copd-1 |
| WBGene00015735 | 0.23562193 | 0.24670807 | 0.44495002 | pdfr-1 |
| WBGene00015736 | NA | NA | NA | srsx-11 |
| WBGene00015737 | NA | NA | NA | srsx-12 |
| WBGene00015738 | NA | NA | NA | srsx-13 |
| WBGene00015739 | -0.279756062 | 0.22059195 | 0.41295062 | ugt-7 |
| WBGene00015740 | 0.324379471 | 0.16998975 | 0.34842782 | C13E3.1 |
| WBGene00015741 | -0.158582236 | 0.36451231 | 0.56414638 | C13F10.1 |
| WBGene00015742 | -0.172702845 | 0.32307944 | 0.52302147 | kxd-1 |
| WBGene00015743 | 0.099904048 | 0.18697496 | 0.37153914 | soap-1 |
| WBGene00015744 | -0.051870708 | 0.70145639 | 0.83037056 | C13F10.5 |
| WBGene00015745 | -0.179198499 | 0.05925528 | 0.17010911 | C13F10.6 |
| WBGene00015746 | -0.243632152 | 0.03939064 | 0.12666778 | C13F10.7 |
| WBGene00015747 | 0.010016919 | 0.92103222 | 0.95586508 | C13G5.2 |
| WBGene00015748 | 0.081686314 | 0.47979525 | 0.66640541 | C14A11.2 |
| WBGene00015750 | 0.010615449 | 0.90819029 | NA | C14A11.5 |
| WBGene00015751 | -0.306368877 | 0.18119252 | 0.36365178 | C14A11.6 |
| WBGene00015752 | 0.663590384 | 3.11E-07 | 5.36E-06 | C14B9.2 |
| WBGene00015753 | -0.024902545 | 0.89729575 | 0.9459022 | C14B9.3 |
| WBGene00015754 | -0.020319659 | 0.8267388 | 0.90512845 | C14B9.8 |
| WBGene00015755 | -0.091558318 | 0.22017725 | 0.41233245 | C14B9.10 |

| | | | | |
|---|---|---|---|---|
| WBGene00015756 | 2.64166142 | 5.86E-68 | 1.83E-65 | C14C6.2 |
| WBGene00015757 | NA | NA | NA | C14C6.3 |
| WBGene00015758 | NA | NA | NA | nhr-155 |
| WBGene00015759 | 4.240423506 | 6.19E-212 | 9.06E-209 | C14C6.5 |
| WBGene00015760 | NA | NA | NA | C14C6.6 |
| WBGene00015761 | NA | NA | NA | NA |
| WBGene00015762 | NA | NA | NA | C14C6.8 |
| WBGene00015763 | 0.080144773 | 0.41491978 | NA | NA |
| WBGene00015764 | NA | NA | NA | C14C6.13 |
| WBGene00015765 | 0.110922946 | 0.41632966 | 0.61345276 | C14C11.1 |
| WBGene00015766 | -0.414358288 | 0.00169378 | 0.01089569 | C14C11.2 |
| WBGene00015767 | 0.452645441 | 0.00906997 | 0.04161332 | hex-2 |
| WBGene00015768 | 0.101227083 | 0.63048585 | 0.78023357 | C14C11.4 |
| WBGene00015769 | -0.042592326 | 0.82741165 | 0.90562762 | C14C11.7 |
| WBGene00015770 | -0.067905328 | 0.31502275 | NA | C14E2.1 |
| WBGene00015771 | NA | NA | NA | got-1.3 |
| WBGene00015772 | NA | NA | NA | C14E2.3 |
| WBGene00015773 | NA | NA | NA | C14E2.4 |
| WBGene00015774 | NA | NA | NA | C14E2.5 |
| WBGene00015775 | 0.020925399 | 0.84643798 | NA | C14E2.6 |
| WBGene00015776 | 0.762722553 | 3.61E-06 | 4.88E-05 | dct-1 |
| WBGene00015777 | -0.112461658 | 0.56219678 | 0.73230632 | sfxn-2 |
| WBGene00015778 | 0.021463232 | 0.75179709 | 0.85959087 | got-2.2 |
| WBGene00015779 | -0.162645898 | 0.43668078 | 0.63163834 | lrch-1 |
| WBGene00015780 | -0.032739353 | 0.81188078 | 0.89587496 | C14F11.4 |
| WBGene00015781 | 0.186203472 | 0.2937709 | 0.49489123 | C14F11.6 |
| WBGene00015782 | -0.003270206 | 0.95698316 | NA | C14F11.7 |
| WBGene00015783 | 0.853094287 | 0.00347182 | 0.01961824 | C15B12.1 |
| WBGene00015784 | 0.234793841 | 0.05092591 | 0.15237134 | C15B12.2 |
| WBGene00015785 | NA | NA | NA | C15B12.3 |
| WBGene00015786 | 0.286902066 | 0.21274355 | 0.40320754 | C15B12.4 |
| WBGene00015787 | -0.133662577 | 0.52467126 | 0.70205458 | actl-1 |
| WBGene00015788 | 0.09001627 | 0.60569422 | 0.76196945 | C15B12.8 |
| WBGene00015789 | -0.342403886 | 0.1099233 | 0.26139144 | syx-6 |
| WBGene00015790 | 1.566988925 | 0.00022894 | 0.00197637 | C15C7.4 |
| WBGene00015791 | 0.41473623 | 0.00596368 | 0.02998429 | C15C7.5 |
| WBGene00015792 | 0.008078454 | 0.9458634 | 0.96970703 | C15C7.6 |
| WBGene00015793 | 0.386986367 | 0.06606266 | 0.18353384 | C15C7.7 |
| WBGene00015794 | -0.071440472 | 0.70832511 | 0.83527012 | C15F1.1 |
| WBGene00015795 | -0.023841414 | 0.8678971 | 0.92968111 | C15F1.2 |
| WBGene00015796 | -0.058146671 | 0.76897397 | 0.86844708 | C15F1.5 |
| WBGene00015797 | -0.027212241 | 0.88696165 | 0.94040159 | C15F1.8 |
| WBGene00015798 | 0.09167768 | 0.1962711 | NA | C15H9.2 |

| | | | | |
|---|---|---|---|---|
| WBGene00015799 | 0.033034883 | 0.64219503 | NA | C15H9.3 |
| WBGene00015800 | -0.152769792 | 0.25562968 | 0.45398765 | C15H9.4 |
| WBGene00015801 | 0.104127123 | 0.49075163 | 0.6745427 | C15H9.5 |
| WBGene00015802 | -1.361149527 | 6.11E-28 | 6.78E-26 | kynu-1 |
| WBGene00015803 | -0.022733584 | 0.82131533 | 0.90202368 | C15H9.9 |
| WBGene00015805 | 1.253514499 | 0.00011964 | 0.00113596 | C15H9.11 |
| WBGene00015806 | -0.224399565 | 0.06199287 | 0.17572849 | smrc-1 |
| WBGene00015807 | 0.030749725 | 0.84419779 | 0.91642626 | C16A3.2 |
| WBGene00015809 | -0.196357461 | 0.01050258 | 0.04651189 | C16A3.4 |
| WBGene00015810 | -0.178221273 | 0.00756428 | 0.03619236 | C16A3.5 |
| WBGene00015811 | -0.206077914 | 0.04400773 | 0.13709778 | C16A3.6 |
| WBGene00015813 | 0.008475808 | 0.91715498 | 0.95383336 | thoc-2 |
| WBGene00015814 | -0.543639294 | 1.05E-11 | 3.45E-10 | oatr-1 |
| WBGene00015815 | -0.036439303 | 0.73959154 | 0.85329864 | C16A11.2 |
| WBGene00015816 | -0.09231438 | 0.60925325 | 0.76442513 | C16A11.3 |
| WBGene00015817 | -0.25648517 | 0.02220974 | 0.08299459 | C16A11.4 |
| WBGene00015818 | 0.009413858 | 0.94244447 | 0.96728563 | C16A11.5 |
| WBGene00015819 | 0.283386313 | 0.01905857 | 0.07421184 | fbxc-44 |
| WBGene00015820 | 0.077911454 | 0.52353693 | 0.70116211 | C16A11.7 |
| WBGene00015821 | NA | NA | NA | clec-135 |
| WBGene00015822 | 0.209862301 | 0.16259318 | 0.33781058 | C16B8.2 |
| WBGene00015823 | 0.699689553 | 6.53E-06 | 8.32E-05 | C16B8.3 |
| WBGene00015824 | 0.108241952 | 0.60304448 | 0.75988622 | C16B8.4 |
| WBGene00015825 | NA | NA | NA | C16C4.1 |
| WBGene00015826 | NA | NA | NA | NA |
| WBGene00015827 | NA | NA | NA | math-13 |
| WBGene00015828 | 1.722957222 | 3.17E-06 | 4.34E-05 | math-14 |
| WBGene00015829 | 0.042516415 | 0.55991373 | NA | math-15 |
| WBGene00015830 | -1.650842642 | 0.00428848 | 0.02319457 | fbxb-98 |
| WBGene00015832 | 0.020317245 | 0.84643798 | NA | math-16 |
| WBGene00015833 | NA | NA | NA | math-17 |
| WBGene00015834 | NA | NA | NA | math-5 |
| WBGene00015835 | 0.020317245 | 0.84643798 | NA | math-6 |
| WBGene00015836 | 0.040107227 | 0.58693163 | NA | math-7 |
| WBGene00015837 | -0.023109762 | 0.78121253 | NA | math-8 |
| WBGene00015838 | 0.020925399 | 0.84643798 | NA | math-9 |
| WBGene00015839 | 2.127997568 | 0.00512592 | 0.02664133 | math-10 |
| WBGene00015840 | NA | NA | NA | math-11 |
| WBGene00015841 | 0.537807245 | 0.03234089 | 0.10930354 | skpo-2 |
| WBGene00015842 | -0.006920944 | 0.96256337 | 0.97924853 | C16C8.4 |
| WBGene00015843 | -0.110322653 | 0.52905866 | 0.70590976 | C16C8.5 |
| WBGene00015845 | NA | NA | NA | C16C8.7 |
| WBGene00015846 | NA | NA | NA | C16C8.8 |

| | | | | |
|---|---|---|---|---|
| WBGene00015847 | -0.003108065 | 0.95728506 | NA | C16C8.9 |
| WBGene00015848 | NA | NA | NA | C16C8.10 |
| WBGene00015849 | 0.327939177 | 0.09695184 | 0.24087424 | C16C8.11 |
| WBGene00015850 | -0.086204285 | 0.64438314 | 0.78947268 | C16C8.12 |
| WBGene00015851 | 0.13154839 | 0.53576647 | 0.7110367 | C16C8.13 |
| WBGene00015852 | 0.050797608 | 0.79465619 | 0.88455571 | C16C8.14 |
| WBGene00015853 | -0.026713541 | 0.83290618 | 0.90882243 | C16C8.16 |
| WBGene00015854 | NA | NA | NA | C16C8.17 |
| WBGene00015855 | 0.062623394 | 0.50571259 | NA | C16C8.18 |
| WBGene00015856 | NA | NA | NA | C16C8.19 |
| WBGene00015857 | 0.443753021 | 0.05132349 | 0.15332813 | C16D9.1 |
| WBGene00015858 | 0.020232312 | 0.84643798 | NA | C16D9.3 |
| WBGene00015859 | 0.458955837 | 0.08184879 | 0.21422027 | C16D9.4 |
| WBGene00015860 | -0.274479913 | 0.14674022 | 0.31699972 | C16D9.5 |
| WBGene00015861 | -0.163620672 | 0.44199648 | 0.63599709 | C16D9.6 |
| WBGene00015862 | 0.020317245 | 0.84643798 | NA | srt-25 |
| WBGene00015863 | NA | NA | NA | C16D9.8 |
| WBGene00015864 | 0.063290468 | 0.41068021 | NA | C16D9.9 |
| WBGene00015865 | 0.031301586 | 0.8610338 | 0.92639594 | C16E9.1 |
| WBGene00015866 | -0.125224255 | 0.47491348 | 0.66202117 | C16E9.2 |
| WBGene00015867 | 0.062552281 | 0.50975435 | NA | ncs-7 |
| WBGene00015868 | 0.144713255 | 0.44048672 | 0.63469528 | C16H3.3 |
| WBGene00015869 | 0.030337975 | 0.78257109 | 0.87764254 | nhr-257 |
| WBGene00015870 | NA | NA | NA | C17A2.2 |
| WBGene00015871 | NA | NA | NA | C17A2.3 |
| WBGene00015872 | NA | NA | NA | C17A2.4 |
| WBGene00015873 | -0.039636691 | 0.54579661 | NA | oac-4 |
| WBGene00015874 | NA | NA | NA | NA |
| WBGene00015875 | NA | NA | NA | C17A2.7 |
| WBGene00015876 | 0.020317245 | 0.84643798 | NA | C17B7.2 |
| WBGene00015877 | 0.012158447 | 0.89398365 | NA | C17B7.3 |
| WBGene00015878 | 0.656927182 | 0.03151074 | 0.10703384 | C17B7.4 |
| WBGene00015879 | 1.582986314 | 0.00131282 | 0.00880083 | C17B7.5 |
| WBGene00015882 | -0.032681568 | 0.75480939 | 0.86095079 | C17B7.8 |
| WBGene00015883 | NA | NA | NA | C17B7.9 |
| WBGene00015884 | 0.107186719 | 0.31444014 | 0.51495727 | C17B7.10 |
| WBGene00015885 | 0.094020064 | 0.64089609 | 0.7871946 | fbxa-65 |
| WBGene00015886 | NA | NA | NA | C17B7.12 |
| WBGene00015887 | -1.058529155 | 1.42E-14 | 6.48E-13 | C17C3.1 |
| WBGene00015888 | -0.003421551 | 0.95689871 | NA | C17C3.2 |
| WBGene00015888 | -0.003421551 | 0.95689871 | NA | C17C3.13 |
| WBGene00015889 | 0.126476384 | 0.25155872 | 0.44992318 | C17C3.3 |
| WBGene00015890 | -0.023449708 | 0.69192352 | NA | C17C3.5 |

| | | | | |
|---|---|---|---|---|
| WBGene00015891 | 0.018860755 | 0.77262519 | NA | C17C3.6 |
| WBGene00015892 | NA | NA | NA | C17C3.9 |
| WBGene00015893 | 0.038125098 | 0.61119482 | NA | C17C3.11 |
| WBGene00015894 | -1.768219204 | 8.38E-46 | 1.78E-43 | acdh-2 |
| WBGene00015895 | NA | NA | NA | C17C3.2 |
| WBGene00015895 | NA | NA | NA | C17C3.13 |
| WBGene00015896 | -0.349553767 | 0.13793603 | 0.3038383 | C17C3.15 |
| WBGene00015897 | 0.751638959 | 0.00590224 | 0.02978792 | nhr-156 |
| WBGene00015898 | -0.162071554 | 0.19758571 | 0.38540533 | sre-9 |
| WBGene00015899 | -0.029547188 | 0.79662998 | 0.88594452 | C17E7.4 |
| WBGene00015900 | 0.020317245 | 0.84643798 | NA | nhr-157 |
| WBGene00015901 | 0.027295506 | 0.82123517 | 0.90200331 | nhr-158 |
| WBGene00015902 | 0.01912147 | 0.77587372 | NA | nhr-159 |
| WBGene00015903 | -0.183457828 | 0.16800119 | 0.34570701 | C17E7.9 |
| WBGene00015904 | NA | NA | NA | irld-22 |
| WBGene00015905 | -0.29554178 | 0.06008862 | 0.17176054 | C17E7.12 |
| WBGene00015906 | 0.038125098 | 0.61119482 | NA | C17E7.13 |
| WBGene00015907 | 1.705300192 | 0.00562094 | 0.02869462 | C17F3.1 |
| WBGene00015908 | 0.019765023 | 0.61254077 | NA | C17F3.3 |
| WBGene00015909 | NA | NA | NA | clec-124 |
| WBGene00015910 | -0.060814557 | 0.60703579 | 0.76297727 | C17F4.2 |
| WBGene00015911 | 0.020232312 | 0.84643798 | NA | C17F4.3 |
| WBGene00015912 | -0.235569283 | 0.04587521 | 0.14129104 | fbxc-50 |
| WBGene00015913 | 1.152848507 | 4.01E-78 | 1.63E-75 | C17F4.7 |
| WBGene00015914 | -0.29330769 | 0.15026871 | 0.32179087 | C17F4.8 |
| WBGene00015915 | -0.19954832 | 0.02153266 | 0.08110616 | C17G10.1 |
| WBGene00015916 | -0.295894938 | 0.01514243 | 0.06231733 | ttc-4 |
| WBGene00015917 | 0.085537126 | 0.24698522 | NA | C17G10.3 |
| WBGene00015918 | 0.056411843 | 0.41891328 | NA | C17G10.6 |
| WBGene00015919 | 0.293706706 | 0.19665034 | 0.3841045 | C17G10.7 |
| WBGene00015920 | -0.226254744 | 0.0016092 | 0.01043879 | eif-3.L |
| WBGene00015921 | 0.162040002 | 0.32366418 | 0.52344718 | C17H11.1 |
| WBGene00015922 | -0.216737831 | 0.19523099 | 0.38244272 | C17H11.2 |
| WBGene00015924 | NA | NA | NA | C17H11.4 |
| WBGene00015925 | -0.039049916 | 0.73898586 | 0.85307041 | C17H11.5 |
| WBGene00015926 | 0.55099145 | 0.00011091 | 0.00106343 | C17H11.6 |
| WBGene00015927 | -0.203102542 | 0.00846137 | 0.03946545 | dyci-1 |
| WBGene00015928 | -0.136529445 | 0.09111604 | 0.22960929 | C17H12.2 |
| WBGene00015929 | 4.915228234 | 0.00024119 | 0.00207115 | C17H12.3 |
| WBGene00015930 | 0.822029396 | 2.37E-06 | 3.38E-05 | C17H12.4 |
| WBGene00015931 | 0.093758303 | 0.45102099 | 0.64329258 | C17H12.5 |
| WBGene00015932 | 0.321850243 | 0.08624755 | 0.22186041 | C17H12.6 |
| WBGene00015933 | 1.245267778 | 5.02E-13 | 1.95E-11 | C17H12.8 |

| | | | | |
|---|---|---|---|---|
| WBGene00015934 | 0.375350319 | 0.0922736 | 0.23180814 | ceh-48 |
| WBGene00015935 | 0.098532775 | 0.51242477 | 0.6921829 | C17H12.10 |
| WBGene00015936 | 0.293088369 | 0.20000417 | 0.38863533 | C17H12.11 |
| WBGene00015937 | 0.388964998 | 0.05390667 | 0.15909042 | C17H12.12 |
| WBGene00015938 | -0.151796069 | 0.06979527 | 0.19082013 | anat-1 |
| WBGene00015939 | 0.086950279 | 0.59940105 | 0.75732394 | damt-1 |
| WBGene00015940 | -0.072493295 | 0.60836061 | 0.76399837 | zipt-3 |
| WBGene00015941 | -0.538508881 | 5.09E-09 | 1.19E-07 | C18A3.3 |
| WBGene00015942 | 0.114449575 | 0.54121803 | 0.71522926 | osta-2 |
| WBGene00015943 | -0.231667201 | 0.00234138 | 0.01419799 | tiar-1 |
| WBGene00015944 | 0.108717649 | 0.15669083 | NA | C18A3.7 |
| WBGene00015945 | -0.027188909 | 0.68205301 | NA | C18A3.9 |
| WBGene00015946 | -0.146464311 | 0.4683774 | 0.65738945 | C18A3.10 |
| WBGene00015947 | 0.231075222 | 0.29221109 | 0.49305781 | C18A11.1 |
| WBGene00015948 | 0.239911715 | 0.20555874 | 0.39464797 | C18A11.2 |
| WBGene00015949 | 0.877237089 | 0.00233283 | 0.01415459 | C18A11.3 |
| WBGene00015950 | -0.243810447 | 0.1300619 | 0.29227968 | C18A11.4 |
| WBGene00015952 | 0.127771948 | 0.09510488 | NA | C18B2.1 |
| WBGene00015953 | -0.077651285 | 0.40152523 | NA | C18B2.2 |
| WBGene00015954 | 0.102039172 | 0.57770648 | 0.74411462 | C18B2.3 |
| WBGene00015955 | 0.110535435 | 0.37466731 | 0.5734376 | C18B2.4 |
| WBGene00015956 | 0.185300173 | 0.08168042 | 0.21393385 | C18B2.5 |
| WBGene00015957 | 0.520892544 | 0.04413394 | 0.13726797 | del-9 |
| WBGene00015958 | NA | NA | NA | srbc-11 |
| WBGene00015959 | NA | NA | NA | srbc-10 |
| WBGene00015960 | 0.020925399 | 0.84643798 | NA | srbc-9 |
| WBGene00015961 | NA | NA | NA | srbc-7 |
| WBGene00015962 | NA | NA | NA | srbc-8 |
| WBGene00015963 | 0.088099123 | 0.52084837 | 0.69928849 | C18B10.6 |
| WBGene00015964 | 0.086430603 | 0.29509282 | NA | glb-5 |
| WBGene00015965 | 0.290749805 | 0.12959359 | 0.29201626 | ugt-48 |
| WBGene00015968 | 0.288748065 | 0.03771961 | 0.12255944 | C18C4.7 |
| WBGene00015969 | -0.070618238 | 0.65937801 | 0.79896991 | glb-6 |
| WBGene00015970 | 0.129273752 | 0.39803102 | 0.59623206 | C18E3.1 |
| WBGene00015971 | -0.186561943 | 0.06492592 | 0.1816344 | swsn-2.2 |
| WBGene00015972 | 0.002118099 | 0.9910631 | 0.9949664 | C18E3.3 |
| WBGene00015974 | -0.09664085 | 0.49091035 | 0.6746216 | snrp-40.2 |
| WBGene00015975 | -0.133094005 | 0.18838992 | 0.37353966 | cas-2 |
| WBGene00015976 | -0.142772303 | 0.21749914 | 0.40915565 | C18E3.9 |
| WBGene00015977 | 0.186067543 | 0.21477877 | 0.4056994 | C18F10.2 |
| WBGene00015978 | 0.08752666 | 0.28916909 | 0.48944794 | C18F10.7 |
| WBGene00015979 | NA | NA | NA | C18F10.9 |
| WBGene00015980 | 0.366618881 | 0.08273291 | 0.21576282 | C18G1.1 |

| | | | | |
|---|---|---|---|---|
| WBGene00015981 | 0.049430777 | 0.81222334 | 0.89593625 | elt-7 |
| WBGene00015982 | 0.020232312 | 0.84643798 | NA | bgnt-1.4 |
| WBGene00015984 | 0.175498593 | 0.17111077 | 0.35007502 | C18G1.6 |
| WBGene00015985 | 0.014087259 | 0.85351751 | NA | C18G1.7 |
| WBGene00015986 | 0.018795205 | 0.77532208 | NA | C18G1.8 |
| WBGene00015987 | 0.117008449 | 0.27039313 | 0.4696116 | C18G1.9 |
| WBGene00015988 | 0.373635548 | 0.12183152 | 0.28082112 | lido-18 |
| WBGene00015989 | 0.150958691 | 0.33442891 | 0.53378105 | NA |
| WBGene00015990 | 0.020232312 | 0.84643798 | NA | NA |
| WBGene00015991 | 0.110039823 | 0.21624696 | NA | NA |
| WBGene00015992 | 0.117193935 | 0.17873164 | 0.36014632 | C18H2.5 |
| WBGene00015993 | 5.060673616 | 0.00017486 | 0.00157063 | C18H7.1 |
| WBGene00015994 | 0.149813019 | 0.07294962 | NA | C18H7.4 |
| WBGene00015996 | 0.030089642 | 0.79695986 | 0.88624406 | C18H7.6 |
| WBGene00015997 | -0.003421551 | 0.95689871 | NA | C18H7.7 |
| WBGene00015998 | 0.020232312 | 0.84643798 | NA | srt-59 |
| WBGene00015999 | 0.057303378 | 0.52744004 | NA | prmt-4 |
| WBGene00016000 | NA | NA | NA | C18H9.1 |
| WBGene00016002 | -0.04398679 | 0.56786251 | 0.73693272 | C18H9.3 |
| WBGene00016003 | 0.44225889 | 0.04944708 | 0.1490232 | C18H9.5 |
| WBGene00016004 | 1.147240109 | 8.72E-14 | 3.70E-12 | C18H9.6 |
| WBGene00016005 | 0.459012566 | 0.06306585 | 0.17801178 | ift-74 |
| WBGene00016006 | 0.907723569 | 5.80E-19 | 4.04E-17 | fln-2 |
| WBGene00016008 | -0.045959069 | 0.50707733 | NA | C23F12.3 |
| WBGene00016009 | -0.007133501 | 0.97009908 | 0.98346632 | C23F12.4 |
| WBGene00016010 | -0.100526018 | 0.45209745 | 0.64445125 | C23G10.1 |
| WBGene00016011 | -0.19954646 | 0.03841496 | 0.1241575 | C23G10.2 |
| WBGene00016012 | 0.047841874 | 0.79276046 | 0.88365481 | C23G10.5 |
| WBGene00016013 | 0.938396404 | 0.01783682 | 0.07069541 | ugt-66 |
| WBGene00016014 | -0.25214006 | 0.00488236 | 0.02568546 | C23G10.7 |
| WBGene00016015 | -0.245152985 | 0.00086889 | 0.00625129 | C23G10.8 |
| WBGene00016016 | NA | NA | NA | C23G10.10 |
| WBGene00016017 | 5.042505831 | 5.67E-07 | 9.42E-06 | C23G10.11 |
| WBGene00016018 | -0.601290858 | 4.20E-06 | 5.59E-05 | C23H3.2 |
| WBGene00016019 | 0.249951134 | 0.07901617 | 0.20942718 | ascc-1 |
| WBGene00016020 | 0.289034007 | 0.00111643 | 0.00769794 | sptl-1 |
| WBGene00016021 | -0.187926549 | 0.11115181 | 0.2631162 | C23H3.5 |
| WBGene00016022 | 0.437102359 | 0.01411796 | 0.05885565 | C23H3.9 |
| WBGene00016023 | -0.272922581 | 0.22295518 | 0.41556803 | prmt-6 |
| WBGene00016024 | 0.192788807 | 0.16143367 | 0.33606575 | glb-7 |
| WBGene00016025 | -0.360532571 | 0.09930582 | 0.24443111 | xbx-4 |
| WBGene00016027 | -0.057295622 | 0.49370829 | 0.67668818 | C23H5.8 |
| WBGene00016028 | 0.993005742 | 1.99E-05 | 0.00022717 | flp-24 |

| | | | | |
|---|---|---|---|---|
| WBGene00016029 | 0.327087665 | 0.1532711 | 0.32640391 | elf-1 |
| WBGene00016030 | 0.160138116 | 0.41901541 | 0.61579411 | C24A1.3 |
| WBGene00016032 | 0.038125098 | 0.61119482 | NA | C24A3.1 |
| WBGene00016033 | 0.590936179 | 0.00378453 | 0.02097251 | C24A3.2 |
| WBGene00016034 | 0.098246573 | 0.63954853 | 0.78619899 | C24A3.4 |
| WBGene00016037 | 0.14309336 | 0.34826073 | 0.54820829 | dop-6 |
| WBGene00016038 | 0.052406951 | 0.64532359 | 0.78998183 | cst-2 |
| WBGene00016039 | 0.087256674 | 0.23800185 | NA | C24A8.5 |
| WBGene00016040 | 0.282432876 | 0.06333929 | 0.17854265 | C24A11.1 |
| WBGene00016044 | 0.509155353 | 0.05605942 | 0.16382555 | C24B5.1 |
| WBGene00016045 | -0.219996933 | 0.02692476 | 0.09579719 | spas-1 |
| WBGene00016046 | 0.377094919 | 0.10660995 | 0.25615308 | C24B5.4 |
| WBGene00016047 | NA | NA | NA | NA |
| WBGene00016048 | 0.153470436 | 0.13471002 | 0.29901295 | C24B9.3 |
| WBGene00016049 | 0.020925399 | 0.84643798 | NA | srt-28 |
| WBGene00016050 | NA | NA | NA | srt-29 |
| WBGene00016051 | NA | NA | NA | srt-27 |
| WBGene00016052 | 1.234480755 | 0.00130349 | 0.00875031 | dod-3 |
| WBGene00016053 | 0.10817994 | 0.31026735 | 0.51113945 | ptp-5.1 |
| WBGene00016054 | 0.541128652 | 0.01069637 | 0.04708525 | C24D10.2 |
| WBGene00016055 | -0.141839034 | 0.2430703 | 0.44109969 | C24D10.4 |
| WBGene00016056 | -0.585013264 | 0.02822057 | 0.09909332 | C24D10.5 |
| WBGene00016057 | -0.18657581 | 0.08979982 | 0.2277675 | C24D10.6 |
| WBGene00016058 | 0.125264789 | 0.23590542 | 0.43179038 | nspd-3 |
| WBGene00016058 | 0.125264789 | 0.23590542 | 0.43179038 | nspd-6 |
| WBGene00016059 | -0.012137697 | 0.94158034 | 0.96696575 | hir-1 |
| WBGene00016060 | -0.023045382 | 0.80071347 | 0.88893281 | mms-19 |
| WBGene00016061 | -0.321015244 | 0.00198578 | 0.0124035 | hpo-15 |
| WBGene00016062 | 0.0920496 | 0.43973231 | 0.63429761 | C24G6.8 |
| WBGene00016063 | -0.022930767 | 0.78121253 | NA | delm-2 |
| WBGene00016064 | -0.093500375 | 0.56590448 | 0.73520032 | acd-1 |
| WBGene00016066 | -0.419481502 | 0.08114275 | 0.21290521 | acd-2 |
| WBGene00016067 | 0.068602198 | 0.72135351 | 0.84362463 | C24H10.1 |
| WBGene00016068 | 0.102606515 | 0.53347555 | 0.70899686 | C24H10.2 |
| WBGene00016069 | -0.024917643 | 0.69549488 | NA | C24H10.3 |
| WBGene00016070 | 0.05348392 | 0.69310748 | 0.82398087 | C24H10.4 |
| WBGene00016071 | -0.352874664 | 0.11327987 | 0.26694522 | C24H12.1 |
| WBGene00016072 | 0.093211156 | 0.52998923 | 0.70676494 | C24H12.2 |
| WBGene00016073 | -0.28309219 | 0.00581036 | 0.02946638 | C24H12.4 |
| WBGene00016074 | -0.195594104 | 0.07513409 | 0.2016199 | dnsn-1 |
| WBGene00016075 | 0.018860755 | 0.77262519 | NA | C24H12.6 |
| WBGene00016076 | -0.301146446 | 0.02685339 | 0.0956362 | NA |
| WBGene00016077 | -0.09367824 | 0.32438231 | 0.52363682 | C24H12.8 |

| | | | | |
|---|---|---|---|---|
| WBGene00016078 | 0.021883722 | 0.87890034 | 0.93517296 | NA |
| WBGene00016080 | -1.654462498 | 0.00671651 | 0.0329988 | NA |
| WBGene00016081 | 0.195875648 | 0.15759438 | 0.33171373 | C25A6.1 |
| WBGene00016082 | -0.022930767 | 0.78121253 | NA | C25A8.1 |
| WBGene00016083 | 0.350834492 | 0.00645741 | 0.03199229 | C25A8.2 |
| WBGene00016084 | -0.03573523 | 0.54063995 | 0.71476071 | cht-3 |
| WBGene00016085 | 0.110039823 | 0.21624696 | NA | C25A8.5 |
| WBGene00016087 | -0.412862088 | 0.00320766 | 0.01843146 | C25A11.2 |
| WBGene00016088 | -1.880757695 | 3.96E-29 | 4.80E-27 | clec-266 |
| WBGene00016089 | 0.004217117 | 0.96910024 | 0.98327073 | aexr-1 |
| WBGene00016090 | 0.020925399 | 0.84643798 | NA | aexr-2 |
| WBGene00016091 | NA | NA | NA | nhr-30 |
| WBGene00016092 | 0.038125098 | 0.61119482 | NA | cyp-33B1 |
| WBGene00016093 | 0.05866295 | 0.775232 | 0.87262026 | srsx-34 |
| WBGene00016094 | 0.481956283 | 0.07046878 | 0.19218598 | C25E10.4 |
| WBGene00016095 | 0.117772431 | 0.57504108 | 0.742287 | C25E10.5 |
| WBGene00016096 | 0.106997704 | 0.16248985 | NA | C25E10.7 |
| WBGene00016097 | 0.194477532 | 0.34728346 | 0.54690515 | C25E10.8 |
| WBGene00016099 | 0.248869702 | 0.23440331 | 0.43000863 | C25E10.10 |
| WBGene00016100 | 0.040107227 | 0.58693163 | NA | C25E10.11 |
| WBGene00016101 | -0.118015346 | 0.53923458 | 0.71382762 | C25E10.12 |
| WBGene00016102 | 0.095649481 | 0.58609907 | 0.74887137 | C25F6.1 |
| WBGene00016103 | -0.048414948 | 0.6113409 | 0.76616175 | dpyd-1 |
| WBGene00016104 | -0.160683376 | 0.44740895 | 0.64038659 | ddr-1 |
| WBGene00016105 | 0.033034883 | 0.64219503 | NA | C25F6.6 |
| WBGene00016106 | 0.102073462 | 0.46874551 | 0.65749215 | C25F6.7 |
| WBGene00016107 | 0.126299345 | 0.16983897 | NA | C25G6.1 |
| WBGene00016108 | 0.190233663 | 0.22801208 | 0.42150566 | C25G6.3 |
| WBGene00016109 | 0.19680543 | 0.07233714 | 0.19590974 | C25G6.4 |
| WBGene00016110 | -0.293868417 | 0.14981482 | 0.32119473 | npr-11 |
| WBGene00016111 | 0.101508053 | 0.49397839 | 0.67686828 | C25H3.1 |
| WBGene00016112 | 0.035394358 | 0.81635908 | 0.89911384 | C25H3.3 |
| WBGene00016113 | -0.250166469 | 0.0168384 | 0.06765535 | eif-2D |
| WBGene00016114 | 0.935207078 | 0.00242236 | 0.01463107 | flp-27 |
| WBGene00016115 | -0.230029421 | 0.0053425 | 0.02757119 | mdt-26 |
| WBGene00016116 | 0.082329893 | 0.42106917 | 0.61783659 | C25H3.7 |
| WBGene00016118 | -0.229748043 | 0.00059405 | 0.00451113 | C25H3.9 |
| WBGene00016119 | 1.359619422 | 1.38E-05 | 0.00016336 | C25H3.10 |
| WBGene00016120 | 0.037558651 | 0.67232022 | 0.80922725 | C25H3.11 |
| WBGene00016121 | -0.131412085 | 0.47594429 | 0.6631424 | ift-43 |
| WBGene00016123 | -0.197107919 | 0.26393005 | 0.4621143 | C25H3.14 |
| WBGene00016124 | -0.186934919 | 0.17951491 | 0.360928 | dnc-4 |
| WBGene00016125 | 0.086653533 | 0.53323879 | 0.70899686 | C26B2.2 |

| | | | | |
|---|---|---|---|---|
| WBGene00016126 | 0.010489182 | 0.90841558 | NA | nhr-258 |
| WBGene00016128 | -0.151579516 | 0.24638844 | 0.44469671 | elpc-4 |
| WBGene00016129 | -0.147543139 | 0.33252151 | 0.53237822 | C26B2.7 |
| WBGene00016130 | 0.015332469 | 0.91995402 | 0.95544619 | C26B2.8 |
| WBGene00016131 | -0.306093905 | 0.11725567 | 0.27380421 | C26B9.1 |
| WBGene00016132 | NA | NA | NA | C26B9.2 |
| WBGene00016133 | 0.585979646 | 0.04085036 | 0.12993486 | C26B9.3 |
| WBGene00016134 | 0.749649731 | 5.78E-10 | 1.50E-08 | C26B9.5 |
| WBGene00016135 | 0.386141021 | 0.11970927 | 0.27797684 | C26B9.6 |
| WBGene00016136 | -0.048745395 | 0.47417081 | NA | C26B9.7 |
| WBGene00016137 | 0.020232312 | 0.84643798 | NA | C26E6.1 |
| WBGene00016138 | -0.091879889 | 0.50743796 | 0.68801245 | flh-2 |
| WBGene00016139 | -0.328754974 | 0.00126835 | 0.00855422 | ntl-9 |
| WBGene00016140 | -0.299043214 | 8.02E-06 | 0.00010067 | rpb-2 |
| WBGene00016142 | -0.193008982 | 0.04691718 | 0.14367902 | mrps-18C |
| WBGene00016143 | 0.072523519 | 0.50710245 | 0.68770721 | eri-9 |
| WBGene00016144 | -0.038153112 | 0.76992867 | 0.86912303 | mmab-1 |
| WBGene00016145 | -0.256511196 | 0.0411224 | 0.13057321 | C26E6.12 |
| WBGene00016146 | 0.190197613 | 0.35164954 | 0.55169019 | C26F1.1 |
| WBGene00016147 | 0.395668021 | 0.1007355 | 0.24671128 | cyp-32A1 |
| WBGene00016148 | -0.212804981 | 0.06163198 | 0.1750444 | C26F1.3 |
| WBGene00016149 | 0.175920508 | 0.17892444 | 0.36025413 | frpr-3 |
| WBGene00016150 | 0.734816511 | 2.04E-05 | 0.00023212 | hint-3 |
| WBGene00016151 | -0.194730242 | 0.00528064 | 0.02730007 | smc-5 |
| WBGene00016152 | -0.37130892 | 0.08443869 | 0.21880829 | pho-12 |
| WBGene00016153 | 0.020317245 | 0.84643798 | NA | C27A2.5 |
| WBGene00016154 | -0.095193536 | 0.29764224 | 0.49871596 | C27A12.2 |
| WBGene00016155 | 0.031677325 | 0.84529944 | 0.9169413 | C27A12.4 |
| WBGene00016156 | 0.039270568 | 0.6799644 | 0.81473806 | ari-1.3 |
| WBGene00016157 | -0.103996608 | 0.22685928 | 0.42054277 | ari-1.2 |
| WBGene00016158 | -0.051686869 | 0.58281758 | 0.74661232 | ari-1.1 |
| WBGene00016159 | -0.206354046 | 0.09948484 | 0.24461044 | pigo-1 |
| WBGene00016160 | 0.020317245 | 0.84643798 | NA | C27D6.1 |
| WBGene00016161 | 2.358851211 | 0.00673078 | 0.03303563 | C27D6.3 |
| WBGene00016162 | 0.162926886 | 0.07244745 | 0.19617222 | crh-2 |
| WBGene00016163 | -0.145489427 | 0.10013439 | 0.24565477 | C27D9.1 |
| WBGene00016164 | NA | NA | NA | C27D9.2 |
| WBGene00016165 | -0.006025102 | 0.97100413 | 0.98424754 | wdr-60 |
| WBGene00016166 | -0.101093239 | 0.33845684 | 0.53801541 | C27F2.4 |
| WBGene00016167 | -0.151980016 | 0.158971 | 0.33321705 | vps-22 |
| WBGene00016168 | NA | NA | NA | C27F2.6 |
| WBGene00016169 | -0.163529477 | 0.34641493 | 0.54618361 | C27F2.7 |
| WBGene00016170 | 0.100074698 | 0.13183367 | 0.2948185 | tmem-131 |

| | | | | |
|---|---|---|---|---|
| WBGene00016171 | -0.103000361 | 0.28820954 | 0.48849576 | C27F2.10 |
| WBGene00016172 | 0.482681995 | 0.02954401 | 0.1025252 | C27H5.2 |
| WBGene00016173 | -0.301544534 | 0.00012123 | 0.00114955 | fust-1 |
| WBGene00016174 | 0.553051192 | 0.00939779 | 0.04286385 | C27H5.4 |
| WBGene00016175 | 0.020925399 | 0.84643798 | NA | C27H5.6 |
| WBGene00016177 | -0.070499841 | 0.64453478 | 0.78954432 | C28C12.1 |
| WBGene00016178 | 0.002636414 | 0.97168566 | 0.98446127 | mesp-1 |
| WBGene00016179 | 0.095843846 | 0.3367954 | 0.53627204 | C28C12.3 |
| WBGene00016180 | 3.630198402 | 2.12E-19 | 1.53E-17 | C28C12.4 |
| WBGene00016181 | 0.022119505 | 0.87600003 | 0.93385149 | C28C12.11 |
| WBGene00016182 | -0.192630498 | 0.03118178 | 0.10614858 | C28C12.12 |
| WBGene00016183 | NA | NA | NA | C28F5.1 |
| WBGene00016184 | 0.231744861 | 0.18428244 | 0.36768591 | glb-9 |
| WBGene00016185 | 0.020925399 | 0.84643798 | NA | C28F5.4 |
| WBGene00016187 | 0.178656729 | 0.18849109 | 0.37361339 | C28G1.2 |
| WBGene00016188 | -0.041236364 | 0.78602031 | 0.87892197 | sec-15 |
| WBGene00016189 | -0.856849789 | 0.00577906 | 0.02934834 | C28G1.4 |
| WBGene00016190 | 0.807168791 | 0.00090327 | 0.00645737 | C28G1.5 |
| WBGene00016191 | 0.431050155 | 0.09472653 | 0.23614698 | C28G1.6 |
| WBGene00016192 | -0.109939384 | 0.36408531 | 0.56388417 | bcl-7 |
| WBGene00016193 | -0.025253941 | 0.8589993 | 0.92455628 | C28H8.2 |
| WBGene00016194 | -0.28569218 | 2.17E-06 | 3.13E-05 | C28H8.3 |
| WBGene00016195 | -0.257058424 | 0.0060863 | 0.03050652 | erd-2.2 |
| WBGene00016196 | 0.042199095 | 0.55922995 | NA | C28H8.5 |
| WBGene00016197 | 0.23027895 | 0.07912771 | 0.20946513 | pxl-1 |
| WBGene00016198 | 0.015062379 | 0.80838968 | NA | C28H8.7 |
| WBGene00016199 | 0.015584271 | 0.80362925 | NA | C28H8.8 |
| WBGene00016200 | -0.20192605 | 0.03720106 | 0.1213204 | dpff-1 |
| WBGene00016201 | -0.59080857 | 2.34E-05 | 0.00026265 | tdo-2 |
| WBGene00016202 | -0.227745384 | 0.00213039 | 0.0131552 | kle-2 |
| WBGene00016203 | -0.0695237 | 0.65545511 | 0.79619334 | tag-250 |
| WBGene00016204 | 0.654244512 | 0.01949459 | 0.07529356 | gsto-1 |
| WBGene00016206 | NA | NA | NA | C29E4.9 |
| WBGene00016207 | 0.256769372 | 0.15562205 | 0.32912395 | C29E4.10 |
| WBGene00016208 | -0.07332912 | 0.72158614 | 0.84376189 | C29E4.11 |
| WBGene00016209 | -0.16084659 | 0.22082279 | 0.41323231 | C29E4.12 |
| WBGene00016210 | -0.134402071 | 0.40300882 | 0.60141856 | C29F5.1 |
| WBGene00016211 | 0.040107227 | 0.58693163 | NA | NA |
| WBGene00016212 | 0.0398788 | 0.58971221 | NA | C29F5.3 |
| WBGene00016213 | NA | NA | NA | C29F5.5 |
| WBGene00016215 | 0.420588629 | 0.09863109 | 0.24347628 | glb-10 |
| WBGene00016216 | 0.616558606 | 0.0372057 | 0.1213204 | pals-22 |
| WBGene00016217 | 0.216685295 | 0.27585602 | 0.47593777 | C29F9.2 |

| | | | | |
|---|---|---|---|---|
| WBGene00016218 | 1.790333854 | 3.79E-11 | 1.16E-09 | pals-23 |
| WBGene00016219 | 1.572239965 | 1.80E-08 | 3.84E-07 | pals-24 |
| WBGene00016220 | NA | NA | NA | C29F9.5 |
| WBGene00016221 | -0.07794758 | 0.39991875 | NA | C29F9.6 |
| WBGene00016222 | NA | NA | NA | C29F9.8 |
| WBGene00016223 | NA | NA | NA | C29F9.9 |
| WBGene00016224 | -0.001066426 | 0.99264874 | 0.99567339 | fbxa-57 |
| WBGene00016225 | NA | NA | NA | fbxa-58 |
| WBGene00016226 | -0.001993815 | 0.95612996 | NA | C29F9.12 |
| WBGene00016227 | NA | NA | NA | nhr-280 |
| WBGene00016230 | -0.122571112 | 0.50949647 | 0.6895487 | NA |
| WBGene00016231 | -0.02791392 | 0.6763823 | NA | C29G2.3 |
| WBGene00016232 | NA | NA | NA | srt-4 |
| WBGene00016233 | 0.040107227 | 0.58693163 | NA | srt-58 |
| WBGene00016234 | -0.139312611 | 0.44737387 | 0.64038659 | C29G2.6 |
| WBGene00016235 | -0.251362136 | 0.00013455 | 0.00124995 | C29H12.2 |
| WBGene00016236 | 0.068766736 | 0.64016987 | 0.78669859 | heri-1 |
| WBGene00016237 | 1.038968857 | 0.00517588 | 0.0268591 | C29H12.6 |
| WBGene00016238 | -0.095375819 | 0.24447074 | 0.44265348 | mob-4 |
| WBGene00016239 | 0.059980168 | 0.38780842 | NA | C30A5.4 |
| WBGene00016242 | 0.129369107 | 0.09367815 | NA | C30A5.10 |
| WBGene00016244 | -0.228860433 | 0.01795217 | 0.07101802 | C30B5.2 |
| WBGene00016245 | -0.240137905 | 0.02260944 | 0.08410182 | rbmx-2 |
| WBGene00016246 | 0.305287405 | 0.1182346 | 0.27547308 | daf-37 |
| WBGene00016247 | 0.114436995 | 0.43551638 | 0.63071288 | C30B5.6 |
| WBGene00016248 | 0.033142676 | 0.82051904 | 0.90148005 | C30B5.7 |
| WBGene00016249 | -0.013107358 | 0.92198925 | 0.95620205 | mrpl-32 |
| WBGene00016250 | -0.567803441 | 4.20E-26 | 4.18E-24 | hsp-110 |
| WBGene00016253 | 0.008615607 | 0.94220991 | 0.9672483 | C30E1.4 |
| WBGene00016254 | 0.020317245 | 0.84643798 | NA | fbxb-114 |
| WBGene00016255 | 0.020232312 | 0.84643798 | NA | NA |
| WBGene00016257 | NA | NA | NA | C30E1.8 |
| WBGene00016258 | 0.126331955 | 0.03643627 | 0.11936974 | vha-16 |
| WBGene00016259 | 0.017604133 | 0.79437959 | NA | C30F8.3 |
| WBGene00016260 | 0.884312461 | 2.65E-07 | 4.62E-06 | rege-1 |
| WBGene00016261 | -0.093442126 | 0.4611595 | 0.65197338 | C30F12.2 |
| WBGene00016262 | 0.131981137 | 0.46173483 | 0.65228105 | C30F12.3 |
| WBGene00016263 | -0.152924805 | 0.13058815 | 0.29288894 | C30F12.4 |
| WBGene00016264 | 0.155534491 | 0.41736754 | 0.61426198 | C30F12.5 |
| WBGene00016265 | -0.002084059 | 0.99012232 | 0.99446943 | trhr-1 |
| WBGene00016266 | -0.186155157 | 0.22580082 | 0.41900505 | idhg-2 |
| WBGene00016268 | 0.056411843 | 0.41891328 | NA | C30G4.2 |
| WBGene00016269 | 1.054443314 | 3.01E-05 | 0.00032904 | C30G4.4 |

| | | | | |
|---|---|---|---|---|
| WBGene00016271 | 0.03646242 | 0.65175403 | NA | irld-23 |
| WBGene00016273 | 0.05847505 | 0.69806172 | 0.82802573 | C30G12.1 |
| WBGene00016274 | -0.26703359 | 0.06801829 | 0.18759283 | C30G12.2 |
| WBGene00016276 | 0.087098428 | 0.46713809 | 0.65674753 | C30G12.4 |
| WBGene00016277 | -0.184791193 | 0.1023023 | 0.24922229 | C30G12.6 |
| WBGene00016278 | -0.065448854 | 0.46522276 | 0.65493524 | erp-44.1 |
| WBGene00016279 | 0.056411843 | 0.41891328 | NA | C31B8.1 |
| WBGene00016280 | NA | NA | NA | NA |
| WBGene00016281 | 5.683353589 | 6.59E-07 | 1.08E-05 | pals-32 |
| WBGene00016282 | 0.022500406 | 0.90751877 | 0.94995226 | C31B8.7 |
| WBGene00016283 | 0.026944522 | 0.89546913 | 0.94517796 | zmp-3 |
| WBGene00016284 | 0.020232312 | 0.84643798 | NA | C31B8.9 |
| WBGene00016285 | 0.020232312 | 0.84643798 | NA | C31B8.12 |
| WBGene00016286 | 0.148467664 | 0.07389511 | NA | C31H1.1 |
| WBGene00016287 | NA | NA | NA | C31H1.2 |
| WBGene00016288 | 4.562105083 | 0.00050941 | 0.00394824 | C31H1.5 |
| WBGene00016289 | 0.691539486 | 0.00332234 | 0.01894913 | lntl-1 |
| WBGene00016291 | -0.156064002 | 0.04894918 | 0.14797953 | C31H1.8 |
| WBGene00016292 | 0.108775863 | 0.52970175 | 0.70651028 | tbc-7 |
| WBGene00016293 | 0.046719465 | 0.2550594 | 0.45356091 | C31H2.3 |
| WBGene00016294 | -0.209777624 | 0.3337532 | 0.53327122 | C31H2.4 |
| WBGene00016295 | -1.410699142 | 0.00130079 | 0.00873831 | fbxc-40 |
| WBGene00016296 | 0.055078355 | 0.44821561 | NA | fbxc-16 |
| WBGene00016297 | -0.029234136 | 0.7166623 | NA | fbxc-15 |
| WBGene00016298 | -0.448293984 | 0.05885775 | 0.16923337 | fbxc-14 |
| WBGene00016299 | -0.045643056 | 0.74154836 | 0.85407613 | C32B5.6 |
| WBGene00016300 | NA | NA | NA | C32B5.7 |
| WBGene00016301 | 0.059762547 | 0.38965825 | NA | NA |
| WBGene00016302 | 0.020317245 | 0.84643798 | NA | fbxc-7 |
| WBGene00016303 | -0.002318378 | 0.98575493 | 0.99192013 | fbxc-32 |
| WBGene00016304 | 0.181322802 | 0.1486404 | 0.31951903 | NA |
| WBGene00016305 | -0.159644261 | 0.41093193 | 0.60870643 | NA |
| WBGene00016306 | -0.022930767 | 0.78121253 | NA | C32B5.13 |
| WBGene00016307 | -0.720396029 | 0.01381526 | 0.05787387 | fbxc-18 |
| WBGene00016308 | -0.136966401 | 0.29212422 | 0.49296803 | C32B5.15 |
| WBGene00016309 | -0.807650585 | 0.007302 | 0.03515114 | sdz-4 |
| WBGene00016310 | 0.008748879 | 0.96558607 | 0.98082345 | C32D5.1 |
| WBGene00016311 | -0.036117513 | 0.6767745 | 0.81237075 | efr-3 |
| WBGene00016312 | 0.087256674 | 0.23800185 | NA | C32D5.4 |
| WBGene00016313 | -0.077393265 | 0.43606337 | 0.63111939 | set-4 |
| WBGene00016314 | 0.257569778 | 0.15759857 | 0.33171373 | C32D5.6 |
| WBGene00016315 | -0.012320251 | 0.95083272 | 0.97201103 | C32D5.7 |
| WBGene00016316 | 0.178808118 | 0.14804416 | 0.31876649 | C32D5.8 |

| | | | | |
|---|---|---|---|---|
| WBGene00016317 | 0.002058059 | 0.9852088 | 0.99157504 | C32D5.10 |
| WBGene00016318 | -0.05728725 | 0.51083179 | 0.69084502 | C32D5.11 |
| WBGene00016319 | -0.058107175 | 0.73212383 | 0.85011851 | C32D5.12 |
| WBGene00016320 | 0.011913685 | 0.89608828 | NA | C32E8.1 |
| WBGene00016321 | 0.237456111 | 0.03458844 | 0.11507316 | tppp-1 |
| WBGene00016322 | 0.261613773 | 0.0704799 | 0.19218598 | C32E8.4 |
| WBGene00016323 | -0.124804468 | 0.1760489 | 0.35699473 | C32E8.5 |
| WBGene00016324 | 0.225588441 | 0.09968493 | 0.24488039 | C32E8.6 |
| WBGene00016325 | -0.313985776 | 0.07044771 | 0.19218598 | C32E8.9 |
| WBGene00016326 | -0.03960741 | 0.56256783 | 0.73259411 | ubr-1 |
| WBGene00016327 | 0.296494789 | 0.07559869 | 0.20268091 | C32E12.1 |
| WBGene00016328 | -0.026874563 | 0.85287912 | 0.92121012 | pde-5 |
| WBGene00016329 | 0.139794374 | 0.35919499 | 0.55945595 | osr-1 |
| WBGene00016330 | 0.481174314 | 0.01290431 | 0.05482645 | C32E12.4 |
| WBGene00016331 | 0.09181415 | 0.27842739 | 0.47823268 | obr-4 |
| WBGene00016332 | 1.204359914 | 2.82E-10 | 7.69E-09 | C32F10.4 |
| WBGene00016333 | -0.310999048 | 1.30E-07 | 2.40E-06 | C32F10.8 |
| WBGene00016334 | NA | NA | NA | C33C12.1 |
| WBGene00016335 | 0.890226022 | 1.16E-05 | 0.0001399 | gba-1 |
| WBGene00016336 | 0.082422657 | 0.46772265 | 0.65693854 | C33C12.4 |
| WBGene00016337 | -0.745205611 | 0.02694981 | 0.09584857 | fbxb-38 |
| WBGene00016338 | -0.243783417 | 0.07500386 | 0.20145505 | NA |
| WBGene00016339 | 0.026204806 | 0.83982727 | 0.91331137 | C33C12.7 |
| WBGene00016340 | 0.201299616 | 0.27434735 | 0.47473343 | gba-2 |
| WBGene00016341 | 0.271722027 | 0.16588403 | 0.34245805 | mtq-2 |
| WBGene00016342 | 0.203232515 | 0.31291434 | 0.51333563 | C33C12.10 |
| WBGene00016343 | 0.324057804 | 0.08680855 | 0.22278071 | cnnm-3 |
| WBGene00016344 | 0.168508535 | 0.41338522 | 0.6111039 | rsef-1 |
| WBGene00016345 | -0.185175284 | 0.21338493 | 0.40410926 | C33E10.1 |
| WBGene00016346 | 0.116419914 | 0.58160408 | 0.74597156 | fbxa-120 |
| WBGene00016347 | -0.023056499 | 0.78121253 | NA | C33E10.5 |
| WBGene00016349 | 0.006799831 | 0.95114203 | 0.97203268 | C33E10.8 |
| WBGene00016350 | -0.003270206 | 0.95698316 | NA | C33E10.10 |
| WBGene00016351 | 2.217059603 | 6.64E-05 | 0.00067688 | C33F10.1 |
| WBGene00016352 | -0.081749707 | 0.60844348 | 0.76399837 | tbck-1 |
| WBGene00016353 | 0.037999047 | 0.8026876 | 0.89017945 | C33F10.4 |
| WBGene00016354 | 0.673512914 | 2.50E-06 | 3.54E-05 | rig-6 |
| WBGene00016355 | 0.17160912 | 0.42204567 | 0.61846519 | lact-5 |
| WBGene00016356 | 0.154044663 | 0.11873891 | 0.27620852 | C33F10.8 |
| WBGene00016357 | 2.228736123 | 0.00414706 | 0.02254018 | C33F10.11 |
| WBGene00016358 | 0.204237019 | 0.12063035 | 0.27938963 | C33F10.12 |
| WBGene00016359 | -0.272730566 | 0.22214628 | 0.41490543 | fbxb-88 |
| WBGene00016360 | 0.410183359 | 0.06013448 | 0.17176135 | C33G8.2 |

| | | | | |
|---|---|---|---|---|
| WBGene00016361 | -2.299261049 | 0.00192253 | 0.01209618 | drd-10 |
| WBGene00016362 | 0.558759366 | 0.01938727 | 0.07511518 | C33G8.4 |
| WBGene00016363 | 0.137202031 | 0.32987038 | 0.52933278 | srab-4 |
| WBGene00016364 | -0.131738445 | 0.53204094 | 0.70833968 | nhr-161 |
| WBGene00016365 | 0.323977541 | 0.09372864 | 0.23421761 | nhr-139 |
| WBGene00016366 | 0.052315454 | 0.76452428 | 0.86623548 | nhr-140 |
| WBGene00016367 | 0.088592481 | 0.64631917 | 0.79060565 | nhr-162 |
| WBGene00016368 | 0.242948459 | 0.20881482 | 0.39818414 | nhr-163 |
| WBGene00016369 | -0.023109762 | 0.78121253 | NA | C33G8.13 |
| WBGene00016370 | 0.146769602 | 0.20207537 | 0.39052078 | C33H5.1 |
| WBGene00016371 | NA | NA | NA | C33H5.2 |
| WBGene00016373 | 0.136428272 | 0.28903738 | 0.48928157 | swd-2.1 |
| WBGene00016374 | -0.158089186 | 0.10967469 | 0.26095947 | swd-2.2 |
| WBGene00016375 | 0.045103234 | 0.75456504 | 0.86091644 | rpap-3 |
| WBGene00016376 | -0.200513968 | 0.060109 | 0.17176135 | sec-10 |
| WBGene00016378 | 0.219233207 | 0.02677047 | 0.09549848 | imp-3 |
| WBGene00016379 | 0.193404084 | 0.33137796 | 0.53090458 | C33H5.13 |
| WBGene00016380 | 0.037937979 | 0.73905863 | 0.85308715 | ntp-1 |
| WBGene00016381 | -0.250171524 | 0.03820636 | 0.1237291 | sgo-1 |
| WBGene00016382 | 0.040107227 | 0.58693163 | NA | C33H5.16 |
| WBGene00016383 | -0.08269857 | 0.61275648 | 0.7670458 | zgpa-1 |
| WBGene00016384 | 0.055466304 | 0.61266315 | 0.7670458 | cdgs-1 |
| WBGene00016386 | 0.03363967 | 0.7669638 | 0.86758224 | tag-272 |
| WBGene00016387 | -0.245140418 | 0.0206362 | 0.07839508 | kbp-5 |
| WBGene00016388 | 0.239907249 | 0.01930642 | 0.07487821 | C34B2.3 |
| WBGene00016389 | 0.033463284 | 0.6365037 | NA | C34B2.4 |
| WBGene00016390 | 0.041770152 | 0.74868872 | 0.8582013 | ttc-1 |
| WBGene00016391 | -0.224932718 | 0.00544899 | 0.02801219 | lonp-1 |
| WBGene00016392 | -0.226301542 | 0.00220214 | 0.01346766 | sdha-2 |
| WBGene00016393 | -0.218977842 | 0.00143063 | 0.00944717 | C34B2.8 |
| WBGene00016394 | -0.048667429 | 0.79353323 | 0.88399982 | C34B2.9 |
| WBGene00016395 | -0.163757854 | 0.12765636 | 0.28897967 | spcs-1 |
| WBGene00016396 | -0.237334968 | 0.28357345 | 0.48344188 | C34B2.11 |
| WBGene00016397 | 0.03636075 | 0.72437557 | 0.84556796 | maph-9 |
| WBGene00016398 | 0.149717874 | 0.0720891 | NA | C34D4.2 |
| WBGene00016399 | 0.36528309 | 0.05874806 | 0.16896442 | C34D4.3 |
| WBGene00016400 | 0.040271391 | 0.7296888 | 0.8492348 | C34D4.4 |
| WBGene00016402 | -0.004599962 | 0.97909461 | 0.98812489 | C34D4.10 |
| WBGene00016403 | 0.184096451 | 0.31253344 | 0.51332311 | grsp-3 |
| WBGene00016404 | -0.068570902 | 0.60063985 | 0.75836219 | mutd-1 |
| WBGene00016405 | -0.054332829 | 0.32276205 | 0.52291098 | hecd-1 |
| WBGene00016406 | 0.308407464 | 0.0940866 | 0.2349517 | C34D10.1 |
| WBGene00016407 | 1.349223107 | 8.68E-20 | 6.42E-18 | C34D10.2 |

| | | | | |
|---|---|---|---|---|
| WBGene00016408 | -0.102884894 | 0.23476562 | 0.43051146 | prmt-5 |
| WBGene00016409 | -0.101665139 | 0.25208231 | 0.45005241 | sumv-1 |
| WBGene00016410 | 0.077099534 | 0.40991873 | 0.60772325 | C34E10.9 |
| WBGene00016411 | -0.255210665 | 0.1230946 | 0.28287994 | C34E10.10 |
| WBGene00016412 | -0.334782996 | 0.00149787 | 0.00981664 | mrps-26 |
| WBGene00016413 | 0.087256674 | 0.23800185 | NA | C34F11.1 |
| WBGene00016414 | 0.097101621 | 0.42758659 | 0.62266713 | C34F11.2 |
| WBGene00016415 | -0.266780591 | 0.00474045 | 0.02507402 | ampd-1 |
| WBGene00016416 | 0.527257044 | 0.05201721 | 0.15498248 | C34F11.5 |
| WBGene00016417 | 0.494835418 | 0.04143455 | 0.13119448 | C34F11.8 |
| WBGene00016418 | -0.127509527 | 0.10584117 | 0.25483506 | hpo-27 |
| WBGene00016419 | -0.493825964 | 1.06E-05 | 0.00013006 | tyr-4 |
| WBGene00016420 | 0.015551528 | 0.84511556 | NA | C34G6.3 |
| WBGene00016421 | -0.055138908 | 0.58203325 | 0.74619516 | cdc-7 |
| WBGene00016422 | -0.853938486 | 1.45E-05 | 0.00017122 | noah-1 |
| WBGene00016423 | 0.218258226 | 0.31636144 | 0.51694731 | tag-275 |
| WBGene00016424 | 0.088510478 | 0.67148346 | 0.80894366 | C34H4.1 |
| WBGene00016425 | 1.214562463 | 6.35E-07 | 1.04E-05 | C34H4.2 |
| WBGene00016427 | 0.105729685 | 0.59740911 | 0.75583464 | C34H4.5 |
| WBGene00016428 | 0.23168764 | 0.26656983 | 0.46528957 | dmsr-7 |
| WBGene00016429 | -0.003108065 | 0.95728506 | NA | C35A11.2 |
| WBGene00016430 | NA | NA | NA | C35A11.3 |
| WBGene00016431 | -0.407349558 | 0.02542231 | 0.09178991 | C35A11.4 |
| WBGene00016432 | -0.042721979 | 0.71324417 | 0.83883106 | C35B1.2 |
| WBGene00016433 | NA | NA | NA | C35B1.3 |
| WBGene00016434 | 0.208573366 | 0.15825653 | 0.33261394 | C35B1.4 |
| WBGene00016435 | -0.014923349 | 0.883421 | 0.93834641 | C35B1.5 |
| WBGene00016436 | -0.013546528 | 0.90664609 | 0.94944606 | C35B1.7 |
| WBGene00016437 | 0.010354767 | 0.92179475 | 0.95620205 | C35B1.8 |
| WBGene00016438 | 0.189646992 | 0.2873046 | 0.48753206 | C35B8.3 |
| WBGene00016439 | -0.071993048 | 0.46763166 | 0.65691994 | C35D10.1 |
| WBGene00016440 | 2.659719839 | 3.19E-10 | 8.56E-09 | gipc-1 |
| WBGene00016441 | 0.020232312 | 0.84643798 | NA | C35D10.3 |
| WBGene00016442 | -0.329557382 | 0.02061821 | 0.07836741 | C35D10.5 |
| WBGene00016443 | -0.168367054 | 0.116011 | 0.27192405 | C35D10.6 |
| WBGene00016444 | -0.319810704 | 0.0011059 | 0.00765916 | C35D10.7 |
| WBGene00016445 | -0.252319613 | 0.25937195 | 0.45741564 | C35D10.8 |
| WBGene00016446 | 0.090994864 | 0.40244051 | 0.60093131 | C35D10.10 |
| WBGene00016447 | 0.757392295 | 0.01779187 | 0.0705937 | msd-4 |
| WBGene00016448 | -0.012857189 | 0.93429394 | 0.96290283 | C35D10.12 |
| WBGene00016449 | -0.090389839 | 0.27968899 | 0.47944787 | tost-1 |
| WBGene00016450 | 1.410484894 | 1.37E-15 | 6.90E-14 | clec-5 |
| WBGene00016451 | 0.304719846 | 0.02854495 | 0.09998243 | clec-6 |

| | | | | |
|---|---|---|---|---|
| WBGene00016452 | 0.129034485 | 0.43725852 | 0.63224959 | C35D10.17 |
| WBGene00016453 | -0.87763118 | 1.22E-10 | 3.46E-09 | vet-2 |
| WBGene00016454 | 0.209660395 | 0.28607175 | 0.48616975 | C35E7.2 |
| WBGene00016455 | -0.300666537 | 0.18977231 | 0.37546709 | C35E7.3 |
| WBGene00016456 | 1.003136665 | 0.00885661 | 0.040944 | C35E7.4 |
| WBGene00016457 | -1.363460805 | 1.02E-11 | 3.36E-10 | C35E7.5 |
| WBGene00016458 | 0.106481349 | 0.48341985 | 0.66891125 | C35E7.6 |
| WBGene00016459 | -0.003421551 | 0.95689871 | NA | C35E7.7 |
| WBGene00016460 | -0.052479813 | 0.75205277 | 0.85970214 | C35E7.8 |
| WBGene00016461 | 0.264253238 | 0.16234808 | 0.33739454 | C35E7.9 |
| WBGene00016462 | 0.161326597 | 0.34461788 | 0.54434679 | C35E7.10 |
| WBGene00016463 | -0.028017345 | 0.87829476 | 0.93491899 | C35E7.11 |
| WBGene00016464 | 0.017396218 | 0.82745944 | NA | C36B7.1 |
| WBGene00016465 | -0.003108065 | 0.95728506 | NA | C36B7.2 |
| WBGene00016466 | -0.003614866 | 0.95700117 | NA | C36B7.3 |
| WBGene00016467 | 0.020925399 | 0.84643798 | NA | C36B7.4 |
| WBGene00016468 | 0.269489456 | 0.14471506 | 0.31407845 | C36B7.5 |
| WBGene00016469 | -0.077680965 | 0.57790734 | 0.74421161 | C36B7.6 |
| WBGene00016470 | NA | NA | NA | NA |
| WBGene00016471 | NA | NA | NA | srab-8 |
| WBGene00016472 | NA | NA | NA | srbc-6 |
| WBGene00016473 | NA | NA | NA | C36C5.4 |
| WBGene00016474 | 0.559154987 | 0.01098989 | 0.04818898 | C36C5.5 |
| WBGene00016475 | 0.017604133 | 0.79437959 | NA | srab-9 |
| WBGene00016476 | NA | NA | NA | srab-10 |
| WBGene00016477 | -0.023056499 | 0.78121253 | NA | srab-11 |
| WBGene00016478 | -0.080022566 | 0.31274104 | NA | srt-72 |
| WBGene00016479 | NA | NA | NA | srab-6 |
| WBGene00016480 | NA | NA | NA | srab-7 |
| WBGene00016481 | 0.885109814 | 0.02033936 | 0.0775895 | C36C5.12 |
| WBGene00016483 | -0.046394786 | 0.50231372 | NA | C36C5.14 |
| WBGene00016484 | NA | NA | NA | C36C5.15 |
| WBGene00016485 | -0.038864259 | 0.71022306 | 0.83679989 | meg-4 |
| WBGene00016486 | 0.087867436 | 0.55943866 | 0.73003231 | fbxa-170 |
| WBGene00016487 | NA | NA | NA | C36C9.4 |
| WBGene00016488 | 0.015896709 | 0.80422001 | NA | C36C9.5 |
| WBGene00016489 | -0.059955199 | 0.6152966 | 0.76921987 | fubl-3 |
| WBGene00016490 | -0.219706299 | 0.2271934 | 0.42089568 | C36E6.2 |
| WBGene00016491 | 0.032795001 | 0.73365212 | 0.85087141 | acdh-5 |
| WBGene00016492 | 0.315551517 | 0.17915829 | 0.36060716 | C37A2.6 |
| WBGene00016493 | -0.117956641 | 0.01946313 | 0.07526669 | C37A2.7 |
| WBGene00016494 | -0.077655529 | 0.48475399 | 0.66987915 | C37A2.8 |
| WBGene00016495 | -0.007801339 | 0.94369453 | 0.96809358 | C37C3.1 |

| | | | | |
|---|---|---|---|---|
| WBGene00016496 | -0.216577844 | 0.00205794 | 0.01278331 | C37C3.2 |
| WBGene00016497 | NA | NA | NA | vps-32.2 |
| WBGene00016499 | -0.021798135 | 0.91173408 | 0.95112162 | C37C3.7 |
| WBGene00016500 | 0.653417518 | 0.00016753 | 0.00151598 | memo-1 |
| WBGene00016501 | -0.041402714 | 0.58115373 | 0.74581663 | C37C3.9 |
| WBGene00016502 | 0.063217264 | 0.41230173 | NA | C37C3.10 |
| WBGene00016503 | 0.177091107 | 0.19177496 | 0.3779482 | C37C3.11 |
| WBGene00016504 | 0.868185589 | 0.01668561 | 0.06722455 | C37C3.12 |
| WBGene00016505 | 1.488280675 | 1.20E-05 | 0.00014381 | ttr-33 |
| WBGene00016506 | 0.331183764 | 0.161063 | 0.33562814 | abhd-5.1 |
| WBGene00016507 | 0.173901136 | 0.36193645 | 0.56165797 | abhd-5.2 |
| WBGene00016508 | -0.260937556 | 0.01653727 | 0.06677389 | C37H5.5 |
| WBGene00016509 | 0.007373827 | 0.91061432 | 0.95112162 | adss-1 |
| WBGene00016511 | -0.202063446 | 0.18340071 | 0.36639736 | C37H5.13 |
| WBGene00016512 | 0.688071158 | 0.02157843 | 0.08119255 | C38C3.3 |
| WBGene00016513 | 0.497023676 | 0.04457723 | 0.13827441 | C38C3.4 |
| WBGene00016514 | 0.015874474 | 0.83570347 | NA | C38C3.6 |
| WBGene00016516 | 0.018860755 | 0.77262519 | NA | NA |
| WBGene00016517 | 0.098788602 | 0.38991223 | 0.58786842 | nhr-260 |
| WBGene00016518 | -0.080047023 | 0.31391044 | NA | C38C5.1 |
| WBGene00016519 | -0.023056499 | 0.78121253 | NA | srz-56 |
| WBGene00016520 | -0.400122957 | 0.05403777 | 0.15928466 | C39B5.2 |
| WBGene00016521 | -0.017206613 | 0.88401282 | 0.9388388 | fbxa-62 |
| WBGene00016522 | 0.261616216 | 0.20619548 | 0.39530025 | fbxa-38 |
| WBGene00016523 | -0.100990959 | 0.61869623 | 0.77167189 | C39B5.5 |
| WBGene00016524 | 0.185967058 | 0.19202704 | 0.37824134 | C39B5.6 |
| WBGene00016525 | 0.039888668 | 0.56326497 | NA | fbxa-12 |
| WBGene00016526 | NA | NA | NA | fbxa-8 |
| WBGene00016527 | NA | NA | NA | fbxa-47 |
| WBGene00016528 | 0.020232312 | 0.84643798 | NA | C39B5.10 |
| WBGene00016529 | NA | NA | NA | C39D10.1 |
| WBGene00016530 | 0.40019498 | 0.10070571 | 0.24668433 | C39D10.2 |
| WBGene00016531 | 0.436102295 | 0.06117349 | 0.17404608 | pcyt-2.2 |
| WBGene00016532 | 0.099856963 | 0.32407743 | 0.52363682 | C39D10.5 |
| WBGene00016533 | 0.402369138 | 0.09251088 | 0.23212732 | C39D10.6 |
| WBGene00016534 | -0.163915498 | 0.00164071 | 0.01061963 | C39D10.7 |
| WBGene00016535 | -0.060852219 | 0.74970148 | 0.85873723 | C39D10.8 |
| WBGene00016537 | 0.020317245 | 0.84643798 | NA | C39E6.2 |
| WBGene00016538 | 0.029311111 | 0.81054068 | 0.89531472 | C39F7.1 |
| WBGene00016539 | 0.189537883 | 0.32559464 | 0.52475997 | madd-2 |
| WBGene00016540 | 0.03743274 | 0.85729403 | 0.92325428 | C39F7.5 |
| WBGene00016541 | 1.589523431 | 0.00363826 | 0.02034117 | C39H7.1 |
| WBGene00016542 | 0.89081604 | 1.35E-05 | 0.00016023 | C39H7.4 |

| | | | | |
|---|---|---|---|---|
| WBGene00016543 | NA | NA | NA | srsx-14 |
| WBGene00016544 | -0.008036936 | 0.96630733 | 0.9811474 | math-18 |
| WBGene00016545 | -0.231906519 | 0.21403283 | 0.40486495 | C40A11.2 |
| WBGene00016546 | -0.02166915 | 0.89990465 | 0.94665211 | C40A11.3 |
| WBGene00016547 | -0.553409927 | 0.03967034 | 0.12725972 | C40A11.4 |
| WBGene00016548 | 0.020317245 | 0.84643798 | NA | hpo-7 |
| WBGene00016549 | -0.214047754 | 0.22254592 | 0.415281 | C40A11.6 |
| WBGene00016550 | -0.128348974 | 0.28509792 | 0.48519338 | C40A11.7 |
| WBGene00016551 | 0.258891384 | 0.23389173 | 0.42939296 | C40A11.8 |
| WBGene00016553 | -0.214475821 | 0.16380122 | 0.33954605 | NA |
| WBGene00016554 | NA | NA | NA | math-19 |
| WBGene00016555 | 0.203388552 | 0.34978829 | 0.54960832 | math-20 |
| WBGene00016556 | 0.018790812 | 0.76755469 | NA | math-21 |
| WBGene00016557 | 0.020925399 | 0.84643798 | NA | ceh-84 |
| WBGene00016558 | -0.975756567 | 1.72E-05 | 0.00019913 | pks-1 |
| WBGene00016559 | 0.216513804 | 0.29953806 | 0.50001506 | C41A3.2 |
| WBGene00016561 | 0.14143924 | 0.44308954 | 0.63685467 | eri-6 |
| WBGene00016562 | -0.072068175 | 0.42890881 | 0.62378602 | C41D11.3 |
| WBGene00016564 | -0.068357959 | 0.49844508 | 0.68044144 | cmt-1 |
| WBGene00016565 | 0.141564975 | 0.39868738 | 0.59684887 | C41D11.6 |
| WBGene00016566 | -0.399947898 | 0.00028713 | 0.00241876 | eri-7 |
| WBGene00016567 | 0.081080092 | 0.62180452 | 0.77393862 | C41D11.9 |
| WBGene00016568 | 0.774581381 | 4.09E-08 | 8.23E-07 | C41G11.1 |
| WBGene00016570 | 0.788739388 | 0.02185841 | 0.08202975 | gnrr-4 |
| WBGene00016572 | 0.067228392 | 0.7405443 | 0.85345632 | C41H7.2 |
| WBGene00016573 | 0.25459779 | 0.05263639 | 0.15654061 | lido-12 |
| WBGene00016574 | 0.012386206 | 0.94732879 | 0.97044477 | lido-11 |
| WBGene00016575 | 0.196420237 | 0.3639769 | 0.56379451 | lido-10 |
| WBGene00016576 | -0.12528424 | 0.27981761 | 0.47949551 | lido-9 |
| WBGene00016577 | 2.126934229 | 1.68E-05 | 0.00019515 | clec-3 |
| WBGene00016578 | -1.329232248 | 0.00348298 | 0.01966329 | sdz-5 |
| WBGene00016579 | 0.036821884 | 0.59344795 | NA | sre-14 |
| WBGene00016580 | 0.069428001 | 0.61363818 | 0.76788689 | ppm-1.H |
| WBGene00016581 | -0.005914326 | 0.94542075 | NA | C42C1.3 |
| WBGene00016582 | -0.124804734 | 0.2128173 | 0.40329511 | vps-8 |
| WBGene00016583 | -0.104878022 | 0.31151756 | 0.51252147 | tag-335 |
| WBGene00016585 | 0.469982051 | 0.02442812 | 0.08925684 | oac-59 |
| WBGene00016586 | -0.249520251 | 0.00260458 | 0.01555206 | C42C1.8 |
| WBGene00016587 | 0.226303229 | 0.21916924 | 0.41112836 | C42C1.9 |
| WBGene00016588 | -0.05102778 | 0.67376463 | 0.81039694 | C42C1.19 |
| WBGene00016588 | -0.05102778 | 0.67376463 | 0.81039694 | hpo-12 |
| WBGene00016589 | 0.020484615 | 0.82001446 | 0.90113575 | ltah-1.1 |
| WBGene00016590 | 0.046271956 | 0.81832572 | 0.90009067 | C42C1.12 |

| | | | | |
|---|---|---|---|---|
| WBGene00016591 | 0.01414096 | 0.93190837 | 0.96179828 | C42C1.13 |
| WBGene00016592 | 0.058974053 | 0.62903606 | 0.77955959 | erl-1 |
| WBGene00016594 | 0.161306156 | 0.306319 | 0.50701475 | C42D4.1 |
| WBGene00016595 | -1.27232767 | 3.87E-05 | 0.00041588 | C42D4.2 |
| WBGene00016596 | 4.545410174 | 0.00053442 | 0.0041116 | C42D4.3 |
| WBGene00016597 | NA | NA | NA | clec-179 |
| WBGene00016598 | 0.093922628 | 0.25299989 | NA | C42D4.13 |
| WBGene00016599 | -0.018092203 | 0.92739358 | 0.95889615 | C42D8.1 |
| WBGene00016600 | 0.11787239 | 0.46230934 | 0.65252509 | ets-5 |
| WBGene00016601 | -0.098838728 | 0.27654135 | 0.47650251 | acin-1 |
| WBGene00016602 | 0.102731088 | 0.38388637 | 0.58327725 | mus-81 |
| WBGene00016603 | -0.197592097 | 0.01307579 | 0.05545845 | met-1 |
| WBGene00016605 | 0.168014255 | 0.16028436 | 0.33484732 | C43E11.5 |
| WBGene00016606 | -0.139353982 | 0.2456522 | 0.44380477 | exoc-7 |
| WBGene00016607 | -0.660344579 | 1.31E-06 | 2.00E-05 | C43E11.9 |
| WBGene00016608 | -0.011221018 | 0.90023799 | 0.94673068 | cogc-5 |
| WBGene00016609 | 0.008191486 | 0.95050058 | 0.97187506 | C43E11.12 |
| WBGene00016610 | 0.08052964 | 0.33089157 | 0.53051416 | paqr-1 |
| WBGene00016611 | 0.437006743 | 0.00425022 | 0.02301314 | bicd-1 |
| WBGene00016612 | 0.193374628 | 0.07320328 | 0.19770693 | C43G2.3 |
| WBGene00016613 | 0.072049825 | 0.68554988 | 0.81848942 | best-9 |
| WBGene00016614 | -0.140699921 | 0.42428478 | 0.62026494 | trpp-12 |
| WBGene00016616 | -0.044180043 | 0.79826641 | 0.88702326 | C43H6.3 |
| WBGene00016617 | -0.363605937 | 0.06496723 | 0.18169728 | C43H6.4 |
| WBGene00016619 | 0.015507174 | 0.80923808 | NA | C43H6.6 |
| WBGene00016620 | -0.308173403 | 0.12699073 | 0.28830381 | dhhc-9 |
| WBGene00016621 | -0.098704558 | 0.5690714 | 0.73797824 | arch-1 |
| WBGene00016622 | 0.089844393 | 0.43805488 | 0.63293807 | mafr-1 |
| WBGene00016623 | -0.088490168 | 0.4944874 | 0.67741395 | psmd-9 |
| WBGene00016624 | -0.118623892 | 0.35690242 | 0.55699307 | hrpl-1 |
| WBGene00016625 | 0.150167444 | 0.34409659 | 0.54381673 | aff-1 |
| WBGene00016626 | 0.106673312 | 0.56374959 | 0.73387195 | clhm-1 |
| WBGene00016627 | 0.484776576 | 2.83E-17 | 1.72E-15 | C44B7.5 |
| WBGene00016628 | 1.479440788 | 0.00051914 | 0.00400878 | slc-36.5 |
| WBGene00016629 | -0.131026867 | 0.52265109 | 0.70044791 | C44B7.7 |
| WBGene00016630 | 0.421759509 | 3.87E-15 | 1.88E-13 | acer-1 |
| WBGene00016631 | -0.055751434 | 0.72061595 | 0.84343578 | C44B7.11 |
| WBGene00016632 | -0.278893132 | 0.00020523 | 0.00181225 | C44B7.12 |
| WBGene00016633 | NA | NA | NA | C44B11.1 |
| WBGene00016634 | 0.833916979 | 0.01959502 | 0.07554551 | C44B11.4 |
| WBGene00016635 | 0.029240686 | 0.88749384 | 0.94082958 | C44B11.6 |
| WBGene00016636 | 0.081275103 | 0.11333445 | 0.26703085 | perm-2 |
| WBGene00016637 | 0.061828942 | 0.64319315 | 0.78886575 | C44B12.3 |

| | | | | | |
|---|---|---|---|---|---|
| WBGene00016638 | | 0.087903415 | 0.04752232 | 0.1448926 | perm-4 |
| WBGene00016639 | | 0.085597384 | 0.55731545 | 0.72847563 | C44B12.6 |
| WBGene00016641 | | -0.265958166 | 0.15042423 | 0.3220297 | C44C1.1 |
| WBGene00016642 | | 0.167725525 | 0.27793638 | 0.47769842 | chil-10 |
| WBGene00016643 | | -0.078667866 | 0.65599902 | 0.79665575 | vps-45 |
| WBGene00016644 | | -1.437679462 | 3.53E-20 | 2.69E-18 | abhd-3.2 |
| WBGene00016645 | | 0.192304772 | 0.33705944 | 0.53663411 | C44C1.6 |
| WBGene00016646 | NA | | NA | NA | fbxc-5 |
| WBGene00016646 | NA | | NA | NA | fbxc-1 |
| WBGene00016647 | NA | | NA | NA | fbxc-4 |
| WBGene00016647 | NA | | NA | NA | fbxc-2 |
| WBGene00016647 | NA | | NA | NA | fbxc-3 |
| WBGene00016648 | NA | | NA | NA | fbxc-4 |
| WBGene00016648 | NA | | NA | NA | fbxc-2 |
| WBGene00016648 | NA | | NA | NA | fbxc-3 |
| WBGene00016649 | NA | | NA | NA | fbxc-5 |
| WBGene00016649 | NA | | NA | NA | fbxc-1 |
| WBGene00016650 | | 0.149617848 | 0.04810413 | 0.14617943 | ubr-4 |
| WBGene00016652 | | -0.395488879 | 2.25E-05 | 0.00025449 | got-2.1 |
| WBGene00016653 | | -0.494099126 | 1.18E-12 | 4.42E-11 | ssb-1 |
| WBGene00016654 | | 0.112754981 | 0.26751186 | 0.46637784 | C44E4.5 |
| WBGene00016655 | | -0.197988974 | 0.01828792 | 0.07209301 | acbp-1 |
| WBGene00016657 | | 0.426522083 | 0.07650494 | 0.20447405 | C44E12.1 |
| WBGene00016658 | | -0.676574313 | 4.20E-08 | 8.42E-07 | C45B2.1 |
| WBGene00016659 | | -0.068414289 | 0.66983458 | 0.80769627 | C45B2.2 |
| WBGene00016660 | | 0.020925399 | 0.84643798 | NA | C45B2.3 |
| WBGene00016661 | | -0.011679484 | 0.94012218 | 0.96639253 | C45B2.6 |
| WBGene00016662 | | -0.107512443 | 0.45050477 | 0.64280678 | C45B2.8 |
| WBGene00016663 | | 4.042394541 | 9.81E-13 | 3.70E-11 | C45E1.4 |
| WBGene00016664 | | -0.061066657 | 0.74154879 | 0.85407613 | C45E5.1 |
| WBGene00016665 | | 0.05110225 | 0.68533095 | 0.81837876 | chil-11 |
| WBGene00016666 | | 0.20369056 | 0.11913724 | 0.27695909 | NA |
| WBGene00016667 | | 0.195733254 | 0.37033196 | 0.57006877 | C45E5.4 |
| WBGene00016668 | NA | | NA | NA | ilys-1 |
| WBGene00016669 | | 0.606402813 | 0.03503962 | 0.11604052 | ilys-2 |
| WBGene00016670 | | 9.320740292 | 1.74E-12 | 6.29E-11 | ilys-3 |
| WBGene00016671 | | -0.442205798 | 0.08778393 | 0.2244887 | C45G7.4 |
| WBGene00016673 | | -0.023056499 | 0.78121253 | NA | ttbk-6 |
| WBGene00016674 | | -0.120467214 | 0.37256191 | 0.57176928 | C45G9.2 |
| WBGene00016675 | | 0.123674741 | 0.11449035 | NA | C45G9.4 |
| WBGene00016676 | | -0.107529328 | 0.34234683 | 0.54273795 | C45G9.5 |
| WBGene00016677 | | 0.76686468 | 0.00074757 | 0.00546711 | C45G9.6 |
| WBGene00016678 | | -0.306035035 | 0.15297578 | 0.32596686 | C45G9.7 |

| | | | | |
|---|---|---|---|---|
| WBGene00016679 | 0.231555863 | 0.13582655 | 0.30076173 | C45G9.8 |
| WBGene00016680 | 0.196642907 | 0.07227576 | 0.1958522 | C45G9.9 |
| WBGene00016681 | -0.006690551 | 0.92537622 | NA | C45G9.10 |
| WBGene00016682 | 0.117958011 | 0.39536997 | 0.59330482 | C45G9.11 |
| WBGene00016683 | 0.301474232 | 0.10697703 | 0.25682978 | C45G9.12 |
| WBGene00016684 | 1.230655866 | 0.00403662 | 0.02206054 | nlp-43 |
| WBGene00016685 | NA | NA | NA | srbc-16 |
| WBGene00016686 | -0.001993815 | 0.95612996 | NA | cyp-33C1 |
| WBGene00016687 | 0.020317245 | 0.84643798 | NA | srbc-17 |
| WBGene00016688 | 0.020232312 | 0.84643798 | NA | NA |
| WBGene00016689 | NA | NA | NA | srbc-21 |
| WBGene00016690 | NA | NA | NA | srbc-22 |
| WBGene00016691 | NA | NA | NA | srbc-23 |
| WBGene00016692 | 0.020925399 | 0.84643798 | NA | srbc-24 |
| WBGene00016693 | 0.111319389 | 0.1344031 | NA | srbc-20 |
| WBGene00016694 | 0.020232312 | 0.84643798 | NA | C45H4.13 |
| WBGene00016695 | -0.170231488 | 0.42021045 | 0.61691841 | C45H4.14 |
| WBGene00016696 | 0.019325396 | 0.77340931 | NA | srbc-19 |
| WBGene00016697 | 0.025105328 | 0.8457329 | 0.9169413 | cyp-33C2 |
| WBGene00016698 | -0.122912575 | 0.38378658 | 0.58327725 | C46A5.1 |
| WBGene00016699 | 0.026625077 | 0.82402478 | 0.9034778 | del-7 |
| WBGene00016700 | 2.519295689 | 0.00571949 | 0.0291198 | C46A5.4 |
| WBGene00016701 | -0.217250374 | 0.18275642 | 0.36558776 | dsb-3 |
| WBGene00016702 | -0.237844656 | 0.09771292 | 0.24217221 | C46A5.6 |
| WBGene00016703 | 0.232220601 | 0.25454869 | 0.45289189 | rdl-1 |
| WBGene00016704 | 0.508666056 | 0.00083059 | 0.00601118 | hosl-1 |
| WBGene00016705 | -0.293792934 | 0.10173243 | 0.24832686 | C46C11.2 |
| WBGene00016706 | 0.194844606 | 0.37126027 | 0.57063899 | C46C11.3 |
| WBGene00016707 | 0.042809887 | 0.65262876 | 0.79427578 | C46E10.1 |
| WBGene00016708 | NA | NA | NA | C46E10.2 |
| WBGene00016709 | NA | NA | NA | C46E10.3 |
| WBGene00016710 | -0.308703185 | 0.08055411 | 0.21193041 | fbxc-52 |
| WBGene00016711 | 0.059980168 | 0.38780842 | NA | fbxc-53 |
| WBGene00016712 | 0.089163361 | 0.67052973 | 0.80824686 | C46E10.8 |
| WBGene00016713 | -0.012178531 | 0.95126519 | 0.97203268 | C46E10.9 |
| WBGene00016714 | -0.532428222 | 0.04290967 | 0.13464974 | C46F2.1 |
| WBGene00016716 | 1.394598963 | 1.21E-15 | 6.20E-14 | acs-17 |
| WBGene00016717 | 0.271868017 | 0.20935269 | 0.39876248 | math-22 |
| WBGene00016718 | NA | NA | NA | math-23 |
| WBGene00016719 | 0.069028883 | 0.74102752 | 0.85379033 | math-24 |
| WBGene00016720 | -0.003421551 | 0.95689871 | NA | math-25 |
| WBGene00016721 | 0.420434292 | 0.01078821 | 0.04741951 | C46G7.1 |
| WBGene00016722 | -0.270776368 | 0.0214806 | 0.08097257 | C46G7.2 |

| | | | | |
|---|---|---|---|---|
| WBGene00016724 | NA | NA | NA | C46H3.1 |
| WBGene00016725 | 0.081564785 | 0.62786032 | 0.77849788 | C46H3.2 |
| WBGene00016727 | NA | NA | NA | C46H11.1 |
| WBGene00016728 | 0.121704737 | 0.53542705 | 0.71065066 | C46H11.2 |
| WBGene00016729 | 0.033034883 | 0.64219503 | NA | C46H11.3 |
| WBGene00016730 | 0.020232312 | 0.84643798 | NA | C46H11.6 |
| WBGene00016731 | 0.056411843 | 0.41891328 | NA | C46H11.7 |
| WBGene00016732 | 0.040107227 | 0.58693163 | NA | phat-1 |
| WBGene00016733 | 2.329884147 | 0.00078105 | 0.00568925 | phat-2 |
| WBGene00016734 | 0.32212835 | 0.15860691 | 0.33297444 | C46H11.10 |
| WBGene00016735 | 0.633804688 | 0.00050064 | 0.00389084 | fhod-1 |
| WBGene00016737 | NA | NA | NA | miro-2 |
| WBGene00016738 | 0.042684586 | 0.76891656 | 0.86844708 | C47D2.1 |
| WBGene00016739 | -0.198169431 | 0.00413072 | 0.02247412 | pitr-1 |
| WBGene00016740 | -0.337394848 | 0.02191688 | 0.0821939 | C48B6.2 |
| WBGene00016741 | -0.215152278 | 0.01040554 | 0.04630624 | C48B6.3 |
| WBGene00016742 | 0.129876603 | 0.24778679 | 0.44611982 | C48B6.4 |
| WBGene00016743 | -0.054932793 | 0.45568274 | NA | srb-15 |
| WBGene00016744 | 0.016985298 | 0.837043 | NA | bbs-9 |
| WBGene00016745 | -0.347576209 | 0.1117023 | 0.26412127 | C48B6.9 |
| WBGene00016746 | -0.224751384 | 0.04245501 | 0.13355904 | C48B6.10 |
| WBGene00016747 | 0.105279812 | 0.38619217 | 0.5851299 | nmur-1 |
| WBGene00016748 | 0.053667986 | 0.63787411 | 0.78546304 | aexr-3 |
| WBGene00016749 | -0.287841353 | 0.05359861 | 0.15850076 | C48E7.1 |
| WBGene00016750 | -0.120309222 | 0.21603206 | 0.40717372 | let-611 |
| WBGene00016751 | 0.858479908 | 0.00265083 | 0.01575116 | C48E7.6 |
| WBGene00016752 | 0.304452338 | 0.00960023 | 0.04358355 | C48E7.7 |
| WBGene00016753 | 0.127192596 | 0.31937912 | 0.51950113 | oac-9 |
| WBGene00016754 | 0.077853024 | 0.62426125 | 0.77554594 | cebp-2 |
| WBGene00016755 | 0.180439214 | 0.08672871 | 0.22265386 | C49A9.1 |
| WBGene00016756 | 0.029715444 | 0.86488909 | 0.9278846 | C49A9.2 |
| WBGene00016757 | 1.524624614 | 0.00025377 | 0.00216645 | C49A9.3 |
| WBGene00016758 | 0.185230958 | 0.36712491 | 0.56675198 | C49A9.4 |
| WBGene00016759 | -0.148762084 | 0.47406498 | 0.66173431 | C49A9.5 |
| WBGene00016760 | 0.193406902 | 0.29970105 | 0.50010521 | C49A9.6 |
| WBGene00016761 | 0.017279211 | 0.79380376 | NA | tkr-2 |
| WBGene00016762 | 0.437233355 | 0.09180432 | 0.23089202 | ugt-24 |
| WBGene00016763 | 0.117210801 | 0.49720777 | 0.67957608 | C49A9.9 |
| WBGene00016764 | -0.540570026 | 0.00310988 | 0.01794709 | C49A9.10 |
| WBGene00016765 | 0.086430603 | 0.29509282 | NA | C49C8.1 |
| WBGene00016766 | 0.137817966 | 0.42780305 | 0.62281811 | C49C8.2 |
| WBGene00016767 | 0.428051513 | 0.07186125 | 0.19515856 | C49C8.3 |
| WBGene00016768 | 0.920765877 | 0.00119958 | 0.00816996 | cyp-33E1 |

| | | | | |
|---|---|---|---|---|
| WBGene00016769 | 1.008617304 | 2.37E-10 | 6.51E-09 | C49C8.5 |
| WBGene00016770 | 0.020317245 | 0.84643798 | NA | C49C8.6 |
| WBGene00016772 | NA | NA | NA | nhr-166 |
| WBGene00016773 | NA | NA | NA | oac-10 |
| WBGene00016775 | 0.020317245 | 0.84643798 | NA | C49D10.7 |
| WBGene00016776 | -0.023056499 | 0.78121253 | NA | oac-11 |
| WBGene00016777 | -0.022930767 | 0.78121253 | NA | nhr-261 |
| WBGene00016778 | 0.204110432 | 0.06872563 | 0.18885359 | nep-3 |
| WBGene00016779 | -0.1602675 | 0.36761892 | 0.56727537 | sdz-7 |
| WBGene00016781 | 1.412339141 | 1.01E-31 | 1.35E-29 | C49G7.3 |
| WBGene00016782 | 1.685400914 | 8.40E-14 | 3.57E-12 | phat-3 |
| WBGene00016783 | -0.007361385 | 0.96580686 | 0.98082345 | irg-2 |
| WBGene00016785 | -0.102439621 | 0.30201465 | 0.50260002 | C49G7.13 |
| WBGene00016785 | -0.102439621 | 0.30201465 | 0.50260002 | C49G7.7 |
| WBGene00016786 | 0.072198567 | 0.5936189 | 0.75344508 | cyp-35A4 |
| WBGene00016788 | -0.484568054 | 0.04440411 | 0.13791272 | C49G7.10 |
| WBGene00016789 | -0.003421551 | 0.95689871 | NA | djr-1.2 |
| WBGene00016790 | -0.425942848 | 4.16E-06 | 5.56E-05 | C49H3.3 |
| WBGene00016791 | -0.156056452 | 0.37545792 | 0.57446749 | C49H3.4 |
| WBGene00016792 | -0.196017084 | 0.07381041 | 0.19901624 | C49H3.6 |
| WBGene00016793 | -0.081200049 | 0.46482589 | 0.6546283 | arp-11 |
| WBGene00016794 | -0.221218946 | 0.04502466 | 0.13933757 | C49H3.9 |
| WBGene00016795 | 0.046923935 | 0.77954998 | 0.8753272 | C49H3.12 |
| WBGene00016796 | -0.117656419 | 0.27596031 | 0.47597246 | cec-2 |
| WBGene00016797 | -0.166366474 | 0.43493743 | 0.6302998 | C50A2.3 |
| WBGene00016798 | -0.03341184 | 0.74259269 | 0.85444405 | ets-8 |
| WBGene00016799 | -0.293002256 | 0.00552037 | 0.02833933 | C50C3.1 |
| WBGene00016800 | 0.186096614 | 0.19772923 | 0.38553119 | C50C3.2 |
| WBGene00016801 | 0.837542348 | 0.012875 | 0.0547482 | C50C3.5 |
| WBGene00016803 | -0.061292794 | 0.33736681 | 0.53700668 | bath-42 |
| WBGene00016804 | NA | NA | NA | C50C3.12 |
| WBGene00016805 | 0.105082988 | 0.48137828 | 0.66742323 | C50D2.1 |
| WBGene00016806 | 0.396670001 | 0.08484112 | 0.21938866 | C50D2.2 |
| WBGene00016807 | 0.020232312 | 0.84643798 | NA | C50D2.3 |
| WBGene00016808 | -0.048451572 | 0.64617017 | 0.79060565 | C50D2.5 |
| WBGene00016809 | -0.349074467 | 0.11317525 | 0.2668275 | C50D2.6 |
| WBGene00016810 | 0.09469965 | 0.4542599 | 0.64633812 | C50D2.7 |
| WBGene00016811 | 0.105192688 | 0.40826848 | 0.6061102 | C50D2.8 |
| WBGene00016812 | -0.072216308 | 0.54061947 | 0.71476071 | C50D2.9 |
| WBGene00016813 | NA | NA | NA | clec-214 |
| WBGene00016815 | NA | NA | NA | clec-213 |
| WBGene00016816 | -0.093963616 | 0.42809788 | 0.62303203 | C50E3.5 |
| WBGene00016817 | 0.611979707 | 0.04086333 | 0.12994786 | C50E3.6 |

| | | | | |
|---|---|---|---|---|
| WBGene00016818 | 0.040107227 | 0.58693163 | NA | C50E3.7 |
| WBGene00016820 | NA | NA | NA | C50E3.9 |
| WBGene00016821 | NA | NA | NA | scl-26 |
| WBGene00016822 | 0.168768884 | 0.2500367 | 0.44857092 | C50E3.11 |
| WBGene00016823 | -0.186031088 | 0.28777423 | 0.48804087 | C50E3.12 |
| WBGene00016824 | -0.142548198 | 0.2583539 | 0.45636252 | C50E3.13 |
| WBGene00016825 | 0.297945986 | 0.101456 | 0.24788969 | C50E10.1 |
| WBGene00016826 | 0.059980168 | 0.38780842 | NA | C50E10.2 |
| WBGene00016827 | 0.0398788 | 0.58971221 | NA | sre-53 |
| WBGene00016828 | 0.020232312 | 0.84643798 | NA | sre-52 |
| WBGene00016829 | -0.003614866 | 0.95700117 | NA | sre-54 |
| WBGene00016830 | NA | NA | NA | sre-55 |
| WBGene00016831 | 0.020232312 | 0.84643798 | NA | sre-56 |
| WBGene00016832 | NA | NA | NA | sre-49 |
| WBGene00016833 | 0.020232312 | 0.84643798 | NA | sre-51 |
| WBGene00016834 | 0.020925399 | 0.84643798 | NA | sre-50 |
| WBGene00016835 | -0.050363979 | 0.60307413 | 0.75988622 | C50F2.1 |
| WBGene00016836 | 0.059956772 | 0.52206879 | 0.70009244 | C50F2.2 |
| WBGene00016837 | -0.145651767 | 0.09120434 | 0.22975272 | syf-1 |
| WBGene00016838 | -0.105392286 | 0.55305531 | 0.7251731 | C50F2.4 |
| WBGene00016839 | 0.168170932 | 0.05190965 | NA | C50F2.5 |
| WBGene00016840 | 1.414990617 | 0.00107576 | 0.00747161 | C50F2.7 |
| WBGene00016841 | 0.225338047 | 0.19812672 | 0.38615193 | magu-3 |
| WBGene00016842 | 0.074442495 | 0.62823156 | 0.77878363 | npr-35 |
| WBGene00016843 | 0.090723942 | 0.20045316 | NA | C50F7.3 |
| WBGene00016844 | -0.297813118 | 3.95E-05 | 0.00042338 | sucg-1 |
| WBGene00016845 | 5.272308117 | 2.28E-37 | 3.79E-35 | C50F7.5 |
| WBGene00016846 | -0.304530918 | 0.17839285 | 0.35995724 | C50F7.6 |
| WBGene00016847 | -0.023109762 | 0.78121253 | NA | C50F7.9 |
| WBGene00016848 | 0.900529774 | 1.59E-05 | 0.00018533 | klo-1 |
| WBGene00016849 | 0.140203537 | 0.50945607 | 0.6895487 | acs-21 |
| WBGene00016850 | NA | NA | NA | srt-8 |
| WBGene00016851 | NA | NA | NA | NA |
| WBGene00016852 | 0.017501802 | 0.83221062 | NA | srt-7 |
| WBGene00016853 | NA | NA | NA | srt-9 |
| WBGene00016854 | NA | NA | NA | NA |
| WBGene00016855 | 0.079437544 | 0.30813247 | NA | C50H11.8 |
| WBGene00016856 | NA | NA | NA | srt-15 |
| WBGene00016857 | NA | NA | NA | srt-16 |
| WBGene00016858 | 0.038125098 | 0.61119482 | NA | C50H11.13 |
| WBGene00016859 | NA | NA | NA | srt-5 |
| WBGene00016860 | -0.066677103 | 0.73175533 | 0.85001776 | cyp-33C9 |
| WBGene00016861 | NA | NA | NA | C50H11.17 |

| | | | | |
|---|---|---|---|---|
| WBGene00016862 | -0.050136128 | 0.77641881 | 0.87335186 | C52A10.1 |
| WBGene00016863 | -0.008158348 | 0.95995755 | 0.9781292 | C52A10.2 |
| WBGene00016864 | -0.248771006 | 0.26716085 | 0.46599883 | C52A10.3 |
| WBGene00016865 | -0.063633685 | 0.7573155 | 0.86259968 | ets-9 |
| WBGene00016866 | 1.862946431 | 7.52E-11 | 2.19E-09 | coel-1 |
| WBGene00016867 | -0.104260154 | 0.35310205 | 0.55297888 | C52B9.4 |
| WBGene00016868 | 0.320956339 | 0.05046642 | 0.15137859 | pqn-15 |
| WBGene00016868 | 0.320956339 | 0.05046642 | 0.15137859 | C52B9.8 |
| WBGene00016869 | NA | NA | NA | C52B9.10 |
| WBGene00016870 | -0.005581345 | 0.94060131 | NA | ets-10 |
| WBGene00016871 | -0.382654833 | 0.0767188 | 0.20493339 | inso-1 |
| WBGene00016872 | 0.840063528 | 0.01026825 | 0.04580664 | dop-4 |
| WBGene00016873 | 0.087256674 | 0.23800185 | NA | C52B11.4 |
| WBGene00016874 | 0.066012137 | 0.67596989 | 0.81175158 | C52B11.5 |
| WBGene00016877 | -2.730167746 | 5.98E-10 | 1.54E-08 | C52D10.3 |
| WBGene00016879 | -0.285227984 | 0.11732358 | 0.27380421 | cnnm-1 |
| WBGene00016880 | -0.511303591 | 0.06095433 | 0.17359119 | fbxb-95 |
| WBGene00016881 | NA | NA | NA | C52E2.2 |
| WBGene00016882 | -0.002914971 | 0.95718083 | NA | C52E2.3 |
| WBGene00016883 | 2.165978664 | 0.00010101 | 0.00098004 | C52E2.4 |
| WBGene00016884 | 1.482256255 | 0.00336251 | 0.01909638 | C52E2.5 |
| WBGene00016885 | -0.324215861 | 0.16038807 | 0.33484732 | fbxb-97 |
| WBGene00016886 | -0.180722727 | 0.38374267 | 0.58327725 | fbxb-96 |
| WBGene00016888 | -0.15546149 | 0.06733178 | 0.18608862 | C52E12.1 |
| WBGene00016889 | -0.006285946 | 0.94443195 | 0.96851076 | lst-6 |
| WBGene00016890 | 0.374537333 | 0.11691635 | 0.27352701 | lst-5 |
| WBGene00016891 | -0.059276785 | 0.58883414 | 0.75083789 | C53A3.1 |
| WBGene00016892 | 0.115609875 | 0.5812871 | 0.74581663 | pgph-3 |
| WBGene00016893 | 0.551458544 | 0.00038692 | 0.00312461 | C53B7.2 |
| WBGene00016894 | 1.328493379 | 1.91E-33 | 2.66E-31 | C53B7.3 |
| WBGene00016895 | -0.013074031 | 0.91135368 | 0.95112162 | C53B7.6 |
| WBGene00016896 | 0.750267279 | 0.00806004 | 0.03810364 | nep-4 |
| WBGene00016898 | 0.57341744 | 0.00010684 | 0.0010305 | clik-2 |
| WBGene00016899 | -0.149665163 | 0.24860058 | 0.44694251 | C53C11.1 |
| WBGene00016900 | -0.037411149 | 0.84104145 | 0.91408863 | C53C11.2 |
| WBGene00016901 | NA | NA | NA | C53C11.4 |
| WBGene00016902 | -0.642335384 | 0.00013674 | 0.00126552 | C53D5.1 |
| WBGene00016903 | 1.022252547 | 0.01588111 | 0.06476751 | 4-Mar |
| WBGene00016905 | -0.34887245 | 0.05808989 | 0.16778099 | ztf-3 |
| WBGene00016906 | -0.222005803 | 0.20255497 | 0.39129269 | C53D5.5 |
| WBGene00016907 | -0.20007718 | 0.00526014 | 0.02722295 | C53H9.2 |
| WBGene00016909 | -0.043929949 | 0.74934859 | 0.85855366 | frpr-4 |
| WBGene00016911 | 0.16542753 | 0.43744255 | 0.63242192 | drn-1 |

| | | | | |
|---|---|---|---|---|
| WBGene00016912 | 4.090223121 | 5.58E-42 | 1.09E-39 | clec-86 |
| WBGene00016913 | -0.21087773 | 0.04570231 | 0.14087257 | lam-2 |
| WBGene00016914 | 0.0954236 | 0.57899789 | 0.74484738 | C54D1.7 |
| WBGene00016915 | 3.851923037 | 0.0004299 | 0.0034078 | C54D2.1 |
| WBGene00016916 | 0.000958512 | 0.99686414 | 0.9978865 | C54D2.2 |
| WBGene00016917 | -0.029893345 | 0.71077086 | NA | C54E4.1 |
| WBGene00016918 | -0.213827554 | 0.10790224 | 0.25834778 | test-1 |
| WBGene00016919 | 0.435840273 | 0.07847146 | 0.20824734 | C54E4.4 |
| WBGene00016920 | -0.039708612 | 0.83663665 | 0.9111952 | C54E4.5 |
| WBGene00016921 | 0.020925399 | 0.84643798 | NA | C54F6.3 |
| WBGene00016922 | 0.020232312 | 0.84643798 | NA | swt-2 |
| WBGene00016923 | 0.180455236 | 0.29476365 | 0.49577045 | C54F6.5 |
| WBGene00016924 | NA | NA | NA | C54F6.20 |
| WBGene00016924 | NA | NA | NA | C54F6.6 |
| WBGene00016925 | -0.002914971 | 0.95718083 | NA | srb-18 |
| WBGene00016926 | 0.019297131 | 0.88547303 | 0.93970793 | nhr-171 |
| WBGene00016927 | -0.049642808 | 0.55117548 | NA | nhr-172 |
| WBGene00016928 | 0.018795205 | 0.77532208 | NA | srb-19 |
| WBGene00016929 | 1.857771381 | 0.00814663 | 0.03836851 | C54F6.12 |
| WBGene00016930 | 0.132364042 | 0.4240534 | 0.62005052 | madf-6 |
| WBGene00016931 | 0.01104132 | 0.91659459 | 0.95352149 | C54G6.2 |
| WBGene00016932 | 0.009183187 | 0.93015602 | 0.96068422 | C54G6.3 |
| WBGene00016933 | -0.001488154 | 0.95639912 | NA | C54G7.1 |
| WBGene00016934 | -0.339110897 | 0.03342089 | 0.11204838 | mboa-3 |
| WBGene00016935 | 0.099439455 | 0.63448332 | 0.78328899 | ifta-1 |
| WBGene00016937 | -0.428282394 | 0.00354521 | 0.0199713 | tag-294 |
| WBGene00016938 | 0.106997704 | 0.16248985 | NA | C54H2.4 |
| WBGene00016939 | 0.381112722 | 0.00912655 | 0.04180909 | C55B6.1 |
| WBGene00016940 | NA | NA | NA | C55B6.4 |
| WBGene00016941 | 0.020925399 | 0.84643798 | NA | C55B6.5 |
| WBGene00016942 | 0.793359619 | 0.00077252 | 0.00562989 | C55B7.3 |
| WBGene00016943 | 0.26008248 | 1.25E-10 | 3.53E-09 | acdh-1 |
| WBGene00016944 | -0.131138389 | 0.1676441 | 0.34526336 | uri-1 |
| WBGene00016945 | 0.149643218 | 0.22424705 | 0.41712629 | sulp-1 |
| WBGene00016946 | 0.062748649 | 0.50854521 | NA | C55B7.10 |
| WBGene00016947 | -0.30779899 | 0.02536865 | 0.09167501 | C55B7.11 |
| WBGene00016948 | -0.166299148 | 0.22315508 | 0.41583474 | scrt-1 |
| WBGene00016949 | -0.045959069 | 0.50707733 | NA | C55C2.3 |
| WBGene00016950 | 0.033034883 | 0.64219503 | NA | C55C2.4 |
| WBGene00016951 | 0.123297938 | 0.4795094 | 0.6663342 | C55C3.1 |
| WBGene00016953 | 3.026157814 | 2.32E-09 | 5.59E-08 | C55C3.3 |
| WBGene00016954 | 0.167669357 | 0.23691338 | 0.43325616 | C55C3.4 |
| WBGene00016955 | -0.024103293 | 0.74655468 | 0.85672656 | perm-5 |

| | | | | |
|---|---|---|---|---|
| WBGene00016956 | 2.368984981 | 0.00609961 | 0.03056103 | C55C3.6 |
| WBGene00016957 | -0.044162616 | 0.70390736 | 0.83192667 | atic-1 |
| WBGene00016958 | 0.308202427 | 0.18801558 | 0.37307424 | ilys-4 |
| WBGene00016959 | NA | NA | NA | C55H1.1 |
| WBGene00016960 | -0.066022831 | 0.53239427 | 0.70853237 | vps-33.2 |
| WBGene00016961 | 0.014285422 | 0.84922615 | 0.91889144 | vps-32.1 |
| WBGene00016962 | 0.046719465 | 0.2550594 | 0.45356091 | C56C10.4 |
| WBGene00016963 | 0.087256674 | 0.23800185 | NA | C56C10.6 |
| WBGene00016964 | -0.068760877 | 0.51754376 | 0.69689674 | C56C10.7 |
| WBGene00016965 | 0.200538634 | 0.14003387 | 0.30737534 | C56C10.9 |
| WBGene00016966 | -0.103825088 | 0.44432462 | 0.63794923 | aipr-1 |
| WBGene00016968 | 0.06004683 | 0.5428823 | 0.71666297 | epg-5 |
| WBGene00016970 | -0.479043457 | 0.00530698 | 0.02741113 | C56E6.2 |
| WBGene00016971 | -0.061789311 | 0.48374453 | 0.66922913 | toe-2 |
| WBGene00016972 | -0.03391287 | 0.73515281 | 0.85146525 | C56E6.4 |
| WBGene00016973 | 0.054244781 | 0.588583 | 0.75058302 | abch-1 |
| WBGene00016974 | 0.559766043 | 0.05139494 | 0.15344072 | lron-15 |
| WBGene00016975 | 1.173604088 | 8.85E-05 | 0.00086953 | nhr-173 |
| WBGene00016976 | 1.406937792 | 0.00755102 | 0.03616438 | C56E10.3 |
| WBGene00016977 | -0.097077859 | 0.12152862 | 0.28054684 | akap-1 |
| WBGene00016978 | -0.120920616 | 0.50440325 | 0.68557073 | C56G2.3 |
| WBGene00016979 | -0.129047878 | 0.53240478 | 0.70853237 | C56G2.4 |
| WBGene00016980 | -0.385490488 | 0.06654955 | 0.18446648 | C56G2.5 |
| WBGene00016981 | -0.329142549 | 1.56E-06 | 2.33E-05 | rpn-13 |
| WBGene00016982 | 0.115626397 | 0.52222951 | 0.70017723 | C56G2.9 |
| WBGene00016983 | -0.004305471 | 0.98074309 | 0.98891595 | C56G2.15 |
| WBGene00016984 | 0.046896329 | 0.76891552 | 0.86844708 | npr-8 |
| WBGene00016985 | 0.020925399 | 0.84643798 | NA | C56G3.2 |
| WBGene00016987 | 0.227403514 | 0.16893542 | 0.34723902 | CC8.2 |
| WBGene00016988 | 0.216142095 | 0.30421174 | 0.50509913 | CD4.1 |
| WBGene00016989 | 0.123593788 | 0.36769351 | 0.56733066 | mrpl-48 |
| WBGene00016990 | -0.236989687 | 0.05173881 | 0.15427859 | vps-37 |
| WBGene00016992 | 0.370861949 | 0.02196042 | 0.08227289 | zhit-1 |
| WBGene00016993 | -0.086893897 | 0.62598283 | 0.77695953 | CD4.8 |
| WBGene00016995 | -0.22416579 | 0.00199009 | 0.01241451 | acly-1 |
| WBGene00016996 | -0.087814966 | 0.66944659 | 0.80736141 | D1005.2 |
| WBGene00016997 | 0.475756994 | 0.03901147 | 0.12561402 | cebp-1 |
| WBGene00016998 | -0.036572237 | 0.84551879 | 0.9169413 | D1005.4 |
| WBGene00016999 | 0.087256674 | 0.23800185 | NA | D1005.5 |
| WBGene00017000 | 0.038125098 | 0.61119482 | NA | D1005.6 |
| WBGene00017001 | -0.139980998 | 0.38430296 | 0.58367009 | D1007.3 |
| WBGene00017002 | -0.237663608 | 0.17871846 | 0.36014632 | D1007.4 |
| WBGene00017003 | -0.144369119 | 0.17223008 | 0.35183766 | D1007.5 |

| | | | | |
|---|---|---|---|---|
| WBGene00017004 | -0.063335955 | 0.51981377 | 0.69876593 | nrd-1 |
| WBGene00017005 | 0.154864457 | 0.29725449 | 0.49840849 | D1007.8 |
| WBGene00017007 | 0.491544796 | 0.00726286 | 0.03500183 | D1007.10 |
| WBGene00017009 | 0.02821937 | 0.79554702 | 0.88521082 | D1007.13 |
| WBGene00017010 | -0.232509242 | 0.16477277 | 0.34096566 | D1007.15 |
| WBGene00017011 | -0.104015665 | 0.24344646 | 0.441454 | eaf-1 |
| WBGene00017012 | -0.439712505 | 0.00026826 | 0.00227819 | acs-22 |
| WBGene00017013 | 0.314324465 | 0.12101017 | 0.27992663 | ensh-1 |
| WBGene00017014 | -0.003108065 | 0.95728506 | NA | dylt-2 |
| WBGene00017015 | -0.244318706 | 0.27382625 | 0.47404173 | D1014.2 |
| WBGene00017016 | -0.109546527 | 0.18494181 | 0.36859965 | snap-1 |
| WBGene00017017 | -0.026715819 | 0.83593541 | 0.91083801 | D1014.4 |
| WBGene00017018 | 0.107533449 | 0.21907273 | NA | D1014.5 |
| WBGene00017019 | NA | NA | NA | D1014.6 |
| WBGene00017020 | 0.038726127 | 0.57551883 | NA | D1014.7 |
| WBGene00017021 | 0.020317245 | 0.84643798 | NA | D1022.2 |
| WBGene00017022 | 0.048581653 | 0.76863122 | 0.86844708 | D1022.3 |
| WBGene00017023 | -0.235145458 | 0.12521235 | 0.28604057 | D1022.4 |
| WBGene00017024 | -0.0059938 | 0.93322519 | NA | D1022.5 |
| WBGene00017025 | -0.103448403 | 0.38034619 | 0.5797656 | D1037.1 |
| WBGene00017026 | -0.110584818 | 0.44026737 | 0.63450679 | ipla-4 |
| WBGene00017027 | -0.520365635 | 0.05664148 | 0.16499724 | D1044.1 |
| WBGene00017028 | -0.063162621 | 0.75829572 | 0.86329917 | dex-1 |
| WBGene00017029 | NA | NA | NA | D1044.4 |
| WBGene00017030 | NA | NA | NA | D1044.5 |
| WBGene00017031 | -0.163765634 | 0.03627128 | 0.1190227 | D1044.6 |
| WBGene00017032 | 1.784762335 | 0.00179864 | 0.01143458 | D1044.7 |
| WBGene00017033 | 0.223124921 | 0.19109792 | 0.37703623 | nekl-4 |
| WBGene00017034 | 0.078776717 | 0.31677567 | NA | otpl-2 |
| WBGene00017035 | 0.055471206 | 0.44533739 | NA | NA |
| WBGene00017036 | 0.215768954 | 0.10199572 | 0.24872901 | D1069.1 |
| WBGene00017037 | -0.210782015 | 0.074377 | 0.20032261 | D1069.3 |
| WBGene00017038 | -0.078813522 | 0.68996919 | 0.82175538 | dmsr-4 |
| WBGene00017039 | 0.10374391 | 0.57755975 | 0.74411462 | trk-1 |
| WBGene00017040 | 1.439166144 | 0.01137385 | 0.04945781 | D1079.1 |
| WBGene00017041 | 0.085706069 | 0.68198441 | 0.81609593 | D2007.1 |
| WBGene00017042 | 0.127773104 | 0.47980369 | 0.66640541 | D2007.2 |
| WBGene00017044 | -0.151343706 | 0.21475207 | 0.4056994 | mrpl-18 |
| WBGene00017045 | 0.079104636 | 0.46422131 | 0.65409145 | atg-13 |
| WBGene00017046 | -0.399363469 | 0.00581388 | 0.02947402 | utx-1 |
| WBGene00017048 | 0.818205084 | 0.0101224 | 0.04528019 | D2021.4 |
| WBGene00017050 | -0.003614866 | 0.95700117 | NA | D2024.1 |
| WBGene00017051 | -0.058214978 | 0.77466004 | 0.87231177 | afmd-1 |

| | | | | |
|---|---|---|---|---|
| WBGene00017052 | 0.153513723 | 0.06686266 | NA | D2024.4 |
| WBGene00017053 | -0.143367451 | 0.14609389 | 0.31589857 | D2024.5 |
| WBGene00017054 | 0.375861268 | 0.12750121 | 0.28874635 | gstk-2 |
| WBGene00017055 | 0.042199095 | 0.55922995 | NA | D2062.1 |
| WBGene00017056 | 0.056411843 | 0.41891328 | NA | D2062.4 |
| WBGene00017057 | NA | NA | NA | D2062.5 |
| WBGene00017058 | 0.073887753 | 0.28837975 | NA | msrp-6 |
| WBGene00017059 | 0.135783879 | 0.29658086 | 0.49751505 | msrp-5 |
| WBGene00017060 | 0.062318725 | 0.73848669 | 0.85304428 | D2063.1 |
| WBGene00017061 | 0.059762547 | 0.38965825 | NA | oac-12 |
| WBGene00017062 | -0.194116949 | 0.05395035 | 0.15912311 | glrx-3 |
| WBGene00017063 | 0.171651246 | 0.38340271 | 0.58308913 | mctp-1 |
| WBGene00017064 | -0.239866677 | 0.01288078 | 0.0547482 | ppfr-2 |
| WBGene00017065 | -0.133713058 | 0.43657844 | 0.63161511 | D2092.4 |
| WBGene00017066 | -0.133938066 | 0.0608901 | 0.17344203 | maco-1 |
| WBGene00017068 | 0.219871264 | 0.02304865 | 0.0852806 | D2092.8 |
| WBGene00017069 | -0.185368661 | 0.36129523 | 0.5611248 | timm-17B.2 |
| WBGene00017070 | -0.182590341 | 0.0339373 | 0.11331267 | praf-3 |
| WBGene00017071 | 1.543066086 | 1.39E-28 | 1.62E-26 | aagr-1 |
| WBGene00017072 | -0.004944805 | 0.94514249 | NA | D2096.5 |
| WBGene00017073 | 2.826717785 | 0.00390098 | 0.02146348 | D2096.6 |
| WBGene00017074 | -0.188904089 | 0.14196141 | 0.30998613 | D2096.7 |
| WBGene00017075 | -0.5406931 | 8.82E-26 | 8.61E-24 | nap-1 |
| WBGene00017076 | 0.866558591 | 0.02139377 | 0.08072361 | D2096.9 |
| WBGene00017077 | 0.239187197 | 0.28501497 | 0.48510861 | D2096.10 |
| WBGene00017078 | 0.328976459 | 0.16050148 | 0.33498249 | D2096.11 |
| WBGene00017079 | -0.066328675 | 0.51614199 | 0.69567629 | D2096.12 |
| WBGene00017080 | 0.18561138 | 0.09844559 | 0.24330551 | lec-12 |
| WBGene00017082 | -0.493785031 | 0.03533926 | 0.11679506 | DC2.5 |
| WBGene00017083 | -0.243569517 | 0.26449731 | 0.46290011 | kin-33 |
| WBGene00017084 | -0.207204107 | 0.03692097 | 0.12063376 | E01A2.1 |
| WBGene00017085 | -0.194220299 | 0.00107726 | 0.00747847 | E01A2.2 |
| WBGene00017087 | -0.072765965 | 0.64392676 | 0.78931424 | E01A2.5 |
| WBGene00017088 | -0.140292006 | 0.08574752 | 0.2209874 | akir-1 |
| WBGene00017089 | -0.106302976 | 0.60474235 | 0.76104803 | poml-2 |
| WBGene00017090 | 0.042199095 | 0.55922995 | NA | E01A2.8 |
| WBGene00017091 | 0.0493951 | 0.79602544 | 0.88554125 | ttr-40 |
| WBGene00017092 | 1.402649942 | 0.00746906 | 0.03583042 | E02C12.6 |
| WBGene00017093 | 0.426947363 | 0.08477456 | 0.21933343 | E02C12.8 |
| WBGene00017094 | NA | NA | NA | E02C12.9 |
| WBGene00017095 | 0.020232312 | 0.84643798 | NA | E02C12.10 |
| WBGene00017096 | -0.023109762 | 0.78121253 | NA | E02C12.11 |
| WBGene00017097 | -0.022930767 | 0.78121253 | NA | E02C12.12 |

| | | | | |
|---|---|---|---|---|
| WBGene00017098 | -0.318680224 | 1.44E-05 | 0.00017018 | E02D9.1 |
| WBGene00017099 | 0.038125098 | 0.61119482 | NA | E02H9.1 |
| WBGene00017100 | 0.346992633 | 0.13076827 | 0.2931524 | E02H9.2 |
| WBGene00017101 | 0.732780488 | 1.01E-06 | 1.59E-05 | E02H9.3 |
| WBGene00017102 | 0.896096716 | 0.0156523 | 0.06403053 | E02H9.4 |
| WBGene00017103 | 0.805382208 | 0.00041587 | 0.00332538 | klo-2 |
| WBGene00017104 | 0.015551528 | 0.84511556 | NA | E02H9.6 |
| WBGene00017105 | 0.554212477 | 0.00270752 | 0.0160489 | E02H9.7 |
| WBGene00017106 | -0.110715943 | 0.33393921 | 0.53334832 | E02H9.9 |
| WBGene00017107 | -0.003421551 | 0.95689871 | NA | srt-12 |
| WBGene00017108 | -0.248523062 | 0.19255836 | 0.37898198 | cyp-43A1 |
| WBGene00017109 | NA | NA | NA | clec-177 |
| WBGene00017110 | 0.075254164 | 0.33558625 | NA | clec-176 |
| WBGene00017111 | NA | NA | NA | E03H12.4 |
| WBGene00017112 | 3.047271543 | 0.00152657 | 0.00997791 | E03H12.5 |
| WBGene00017113 | 0.0380884 | 0.3427816 | NA | srz-24 |
| WBGene00017114 | 0.386769119 | 0.00501247 | 0.02619953 | E03H12.7 |
| WBGene00017115 | NA | NA | NA | E03H12.9 |
| WBGene00017116 | -0.391200324 | 0.05411713 | 0.15948649 | fbxb-78 |
| WBGene00017117 | -0.046873538 | 0.74035571 | 0.85342581 | NA |
| WBGene00017118 | 0.060728101 | 0.52119783 | NA | NA |
| WBGene00017119 | -0.292226838 | 0.00016828 | 0.00152082 | timm-17B.1 |
| WBGene00017120 | -0.127955638 | 0.32901919 | 0.52843022 | tctn-1 |
| WBGene00017121 | -0.131190512 | 0.00373651 | 0.02074566 | cyc-2.1 |
| WBGene00017122 | 0.057595505 | 0.529074 | NA | pcrg-1 |
| WBGene00017123 | 0.049178285 | 0.6584891 | 0.79870748 | maoc-1 |
| WBGene00017124 | 1.161324975 | 3.57E-05 | 0.00038595 | E04F6.4 |
| WBGene00017125 | -0.251450253 | 0.00011796 | 0.0011222 | acdh-12 |
| WBGene00017126 | 0.005610829 | 0.97851642 | 0.98775916 | E04F6.6 |
| WBGene00017127 | 1.098110585 | 3.26E-13 | 1.29E-11 | E04F6.8 |
| WBGene00017128 | 0.688732009 | 4.16E-05 | 0.00044423 | E04F6.9 |
| WBGene00017129 | 0.080369252 | 0.57480361 | 0.74227635 | E04F6.10 |
| WBGene00017131 | -0.198186642 | 0.35521025 | 0.55506646 | E04F6.15 |
| WBGene00017132 | -0.118596226 | 0.17095927 | 0.34987624 | tofu-6 |
| WBGene00017133 | -0.142320642 | 0.49288862 | 0.67594438 | EEED8.2 |
| WBGene00017134 | -0.156310942 | 0.02658407 | 0.09509341 | EEED8.3 |
| WBGene00017135 | -0.977640248 | 0.0003874 | 0.00312677 | EEED8.4 |
| WBGene00017136 | 0.085309563 | 0.24820057 | NA | ccpp-6 |
| WBGene00017137 | -0.033624923 | 0.74033482 | 0.85342581 | pink-1 |
| WBGene00017138 | -0.093382194 | 0.28566978 | 0.48577294 | EEED8.10 |
| WBGene00017139 | 0.020232312 | 0.84643798 | NA | clec-141 |
| WBGene00017140 | -1.10833767 | 0.00012229 | 0.00115738 | EEED8.12 |
| WBGene00017141 | -0.14777554 | 0.48480449 | 0.66987915 | EEED8.13 |

| | | | | |
|---|---|---|---|---|
| WBGene00017142 | -0.027314313 | 0.84781979 | 0.9180823 | EEED8.14 |
| WBGene00017143 | -1.02277416 | 2.24E-06 | 3.21E-05 | EEED8.15 |
| WBGene00017144 | 0.052829138 | 0.61011431 | 0.76524318 | brap-2 |
| WBGene00017146 | -0.010206773 | 0.91769929 | 0.95418978 | EGAP1.1 |
| WBGene00017147 | -0.126428188 | 0.51619856 | 0.69567629 | EGAP2.1 |
| WBGene00017148 | -0.040818897 | 0.73652314 | 0.85204655 | EGAP2.2 |
| WBGene00017149 | 0.066335133 | 0.55437322 | 0.7264592 | EGAP4.1 |
| WBGene00017150 | 0.030249993 | 0.75326501 | NA | EGAP5.1 |
| WBGene00017151 | -0.047693707 | 0.79111105 | 0.88228647 | EGAP9.3 |
| WBGene00017152 | NA | NA | NA | NA |
| WBGene00017153 | 0.405535198 | 0.05340148 | 0.15810973 | E_BE45912.2 |
| WBGene00017154 | 0.064375598 | 0.68860793 | 0.82066819 | ugt-57 |
| WBGene00017156 | 0.089186376 | 0.55686355 | 0.72827505 | F01E11.3 |
| WBGene00017157 | 0.145997954 | 0.40310444 | 0.60143199 | tyra-2 |
| WBGene00017158 | 0.041888246 | 0.69609322 | 0.82650251 | hpo-10 |
| WBGene00017159 | -0.21223649 | 0.05959512 | 0.17081678 | F01F1.2 |
| WBGene00017160 | 0.176703403 | 0.3730618 | 0.57223655 | F01F1.3 |
| WBGene00017161 | -0.117529665 | 0.30434661 | 0.50526576 | rabn-5 |
| WBGene00017162 | -0.191874004 | 0.0130402 | 0.05533958 | ddx-23 |
| WBGene00017163 | -0.055704613 | 0.54057762 | 0.71476071 | dnpp-1 |
| WBGene00017164 | 0.124636911 | 0.40108321 | 0.59939361 | eng-1 |
| WBGene00017165 | -0.108681671 | 0.26916828 | 0.46820634 | F01F1.11 |
| WBGene00017166 | 0.00625784 | 0.89811017 | 0.94632825 | aldo-2 |
| WBGene00017168 | 0.038125098 | 0.61119482 | NA | F01F1.14 |
| WBGene00017169 | -0.236734018 | 0.15737034 | 0.33132861 | F01F1.15 |
| WBGene00017170 | NA | NA | NA | F01G12.1 |
| WBGene00017172 | 0.020925399 | 0.84643798 | NA | F02C9.1 |
| WBGene00017173 | NA | NA | NA | F02C9.2 |
| WBGene00017174 | 0.235857321 | 0.0507998 | 0.15219151 | tat-6 |
| WBGene00017175 | 0.260786756 | 0.01480087 | 0.06123186 | irld-3 |
| WBGene00017176 | 0.061436742 | 0.66023282 | 0.79960862 | nmur-3 |
| WBGene00017177 | -0.088646927 | 0.37944605 | 0.57899631 | F02E8.4 |
| WBGene00017178 | 0.0510032 | 0.73940756 | 0.85327581 | atg-16.1 |
| WBGene00017179 | 0.151930862 | 0.12247265 | 0.2818045 | wht-4 |
| WBGene00017180 | 1.174442109 | 0.01161642 | 0.05025886 | F02E11.2 |
| WBGene00017181 | 0.093200186 | 0.44344392 | 0.63695304 | F02E11.3 |
| WBGene00017182 | -0.01228186 | 0.9089892 | 0.95068624 | F02E11.4 |
| WBGene00017183 | NA | NA | NA | scl-15 |
| WBGene00017184 | 1.088534859 | 7.37E-09 | 1.68E-07 | ncam-1 |
| WBGene00017185 | NA | NA | NA | F07B7.1 |
| WBGene00017185 | NA | NA | NA | W09B7.1 |
| WBGene00017186 | NA | NA | NA | F07B7.2 |
| WBGene00017186 | NA | NA | NA | W09B7.2 |

| | | | | |
|---|---|---|---|---|
| WBGene00017188 | NA | NA | NA | F07B7.7 |
| WBGene00017188 | NA | NA | NA | EGAP798.1 |
| WBGene00017189 | NA | NA | NA | F07B7.8 |
| WBGene00017189 | NA | NA | NA | K06C4.1 |
| WBGene00017190 | 0.075726029 | 0.27693214 | NA | F07B7.12 |
| WBGene00017191 | NA | NA | NA | C01B7.7 |
| WBGene00017191 | NA | NA | NA | F07B7.13 |
| WBGene00017192 | NA | NA | NA | F07B7.14 |
| WBGene00017193 | 0.652836933 | 0.03659378 | 0.11980534 | F07C3.2 |
| WBGene00017194 | 0.642221779 | 0.0239429 | 0.08779062 | F07C3.3 |
| WBGene00017195 | 0.353288889 | 0.00907245 | 0.04161332 | glo-4 |
| WBGene00017197 | 0.074300834 | 0.66519018 | 0.80363434 | F07C3.9 |
| WBGene00017198 | 1.473958145 | 0.0016776 | 0.01080112 | nhr-36 |
| WBGene00017199 | NA | NA | NA | clec-45 |
| WBGene00017200 | 0.246528622 | 0.1260627 | 0.28699798 | F07C4.6 |
| WBGene00017201 | -1.190894829 | 3.97E-08 | 8.03E-07 | grsp-4 |
| WBGene00017202 | NA | NA | NA | clec-46 |
| WBGene00017203 | NA | NA | NA | F07C4.10 |
| WBGene00017204 | 0.033034883 | 0.64219503 | NA | F07C4.11 |
| WBGene00017205 | 6.604753184 | 1.26E-08 | 2.76E-07 | F07C4.12 |
| WBGene00017207 | -0.11073426 | 0.12651588 | NA | fbxb-6 |
| WBGene00017208 | -0.286761748 | 0.20094909 | 0.38964871 | fbxb-35 |
| WBGene00017209 | NA | NA | NA | F07E5.4 |
| WBGene00017210 | -0.259486233 | 0.05192331 | 0.15476572 | F07E5.5 |
| WBGene00017211 | -0.03786196 | 0.77075201 | 0.86951612 | fbxb-36 |
| WBGene00017212 | 0.038125098 | 0.61119482 | NA | F07E5.7 |
| WBGene00017213 | 0.015194928 | 0.8086441 | NA | F07E5.8 |
| WBGene00017214 | 0.580107879 | 0.0259983 | 0.0934318 | F07E5.9 |
| WBGene00017215 | 0.006773875 | 0.94767696 | 0.97044737 | F07F6.1 |
| WBGene00017216 | -0.117009337 | 0.38631371 | 0.5851299 | F07F6.2 |
| WBGene00017217 | -0.188114809 | 0.0127513 | 0.05427085 | F07F6.4 |
| WBGene00017218 | 0.062748649 | 0.50854521 | NA | dct-5 |
| WBGene00017219 | -0.048947948 | 0.81174432 | 0.89587496 | F07F6.7 |
| WBGene00017220 | 0.062336071 | 0.58428839 | 0.74778226 | F07F6.8 |
| WBGene00017221 | -0.045959069 | 0.50707733 | NA | dgn-3 |
| WBGene00017222 | NA | NA | NA | F07G6.2 |
| WBGene00017223 | NA | NA | NA | F07G6.3 |
| WBGene00017224 | -0.023109762 | 0.78121253 | NA | F07G6.5 |
| WBGene00017225 | -0.023056499 | 0.78121253 | NA | fbxa-52 |
| WBGene00017226 | 0.538076184 | 0.04558009 | 0.14064026 | fbxa-53 |
| WBGene00017227 | 0.020317245 | 0.84643798 | NA | F07G6.8 |
| WBGene00017228 | -0.00346345 | 0.95708614 | NA | F07G11.1 |
| WBGene00017229 | 0.020925399 | 0.84643798 | NA | F07G11.2 |

| | | | | |
|---|---|---|---|---|
| WBGene00017230 | NA | NA | NA | F07G11.3 |
| WBGene00017231 | NA | NA | NA | F07G11.4 |
| WBGene00017232 | NA | NA | NA | NA |
| WBGene00017233 | 0.054259843 | 0.60435562 | 0.76077049 | lmd-4 |
| WBGene00017235 | 0.057567325 | 0.77193185 | 0.87031066 | NA |
| WBGene00017236 | -0.02791392 | 0.6763823 | NA | nlp-60 |
| WBGene00017237 | -0.291959568 | 0.00465254 | 0.02473318 | pole-2 |
| WBGene00017238 | -0.234218246 | 0.02231646 | 0.08326586 | F08B4.7 |
| WBGene00017239 | -0.084872006 | 0.68038174 | 0.81494812 | F08B6.1 |
| WBGene00017240 | -0.290974408 | 0.13771092 | 0.30355699 | calu-2 |
| WBGene00017241 | -0.691199595 | 2.58E-08 | 5.38E-07 | pcyt-1 |
| WBGene00017243 | 0.095583547 | 0.5423048 | 0.71620824 | F08C6.5 |
| WBGene00017244 | 0.352363906 | 0.00068564 | 0.00508017 | apy-1 |
| WBGene00017245 | 0.105363132 | 0.32003686 | 0.52005372 | srpa-72 |
| WBGene00017246 | 0.799924136 | 0.0002347 | 0.0020225 | F08D12.2 |
| WBGene00017247 | 1.336647401 | 7.07E-09 | 1.61E-07 | F08D12.3 |
| WBGene00017248 | 0.016059264 | 0.84014476 | NA | F08D12.4 |
| WBGene00017249 | -0.061546666 | 0.73063075 | 0.84986592 | fbxb-108 |
| WBGene00017250 | 0.759320148 | 0.00629385 | 0.03130005 | F08D12.7 |
| WBGene00017251 | -0.814005769 | 0.01695692 | 0.06800216 | fbxb-105 |
| WBGene00017252 | -0.387875795 | 0.11140856 | 0.2635961 | sdz-10 |
| WBGene00017253 | -0.863782041 | 0.00350515 | 0.01976842 | sdz-9 |
| WBGene00017254 | -1.698980573 | 0.00162477 | 0.01052109 | fbxb-111 |
| WBGene00017255 | 0.034950593 | 0.75619813 | 0.86193047 | F08D12.12 |
| WBGene00017257 | -0.109412219 | 0.57924158 | 0.7449645 | F08F1.3 |
| WBGene00017258 | 0.562615027 | 0.02416821 | 0.08848382 | F08F1.4 |
| WBGene00017259 | -0.040265081 | 0.75682972 | 0.86226195 | F08F1.9 |
| WBGene00017260 | 0.529278074 | 0.02782646 | 0.0980051 | F08F3.1 |
| WBGene00017261 | -0.061221654 | 0.49969089 | 0.68163368 | acl-6 |
| WBGene00017262 | -0.192151984 | 0.34344462 | 0.54313812 | F08F3.4 |
| WBGene00017263 | -0.174799173 | 0.06802338 | 0.18759283 | F08F3.6 |
| WBGene00017264 | -0.010829828 | 0.91280382 | 0.95126775 | F08F3.8 |
| WBGene00017265 | 0.228472238 | 0.27895426 | 0.47880063 | snpc-1.3 |
| WBGene00017266 | 0.142969646 | 0.28497444 | 0.48509601 | F08F3.10 |
| WBGene00017267 | NA | NA | NA | numr-2 |
| WBGene00017268 | -0.140101498 | 0.1675566 | 0.34513171 | hmgr-1 |
| WBGene00017269 | -0.16313799 | 0.09051669 | 0.22884732 | F08F8.4 |
| WBGene00017270 | NA | NA | NA | numr-1 |
| WBGene00017271 | 0.154084801 | 0.2353999 | 0.43118853 | F08F8.6 |
| WBGene00017272 | 0.02578909 | 0.82612118 | 0.90477236 | F08F8.7 |
| WBGene00017273 | -0.038399755 | 0.80298878 | 0.89037039 | gos-28 |
| WBGene00017274 | -0.063068215 | 0.49777271 | 0.67976712 | F08F8.9 |
| WBGene00017275 | -0.262130901 | 0.01081352 | 0.04750098 | F08F8.10 |

| | | | | |
|---|---|---|---|---|
| WBGene00017277 | 4.058412585 | 0.00136667 | 0.00908281 | F09C12.2 |
| WBGene00017278 | 0.020232312 | 0.84643798 | NA | F09C12.6 |
| WBGene00017279 | 0.294412269 | 0.07698791 | 0.20527774 | F09C12.8 |
| WBGene00017280 | -0.110717327 | 0.25334832 | 0.45142246 | usp-39 |
| WBGene00017281 | 0.051012917 | 0.76113004 | 0.86464679 | F09D12.2 |
| WBGene00017282 | -0.123192983 | 0.30461954 | 0.50545453 | algn-2 |
| WBGene00017283 | -0.242814262 | 0.01054138 | 0.04659912 | F09E5.3 |
| WBGene00017284 | 0.042585346 | 0.66469561 | 0.80335219 | sec-6 |
| WBGene00017285 | -0.08264523 | 0.4086963 | 0.60654062 | F09E5.7 |
| WBGene00017286 | -0.296616694 | 0.03569687 | 0.11771124 | F09E5.8 |
| WBGene00017287 | -0.023109762 | 0.78121253 | NA | F09E5.9 |
| WBGene00017288 | 0.294554765 | 0.12138602 | 0.28045725 | F09E5.10 |
| WBGene00017289 | -0.053885322 | 0.67459511 | 0.81103194 | F09E5.11 |
| WBGene00017290 | 0.178711381 | 0.20566443 | 0.39466399 | F09E5.12 |
| WBGene00017291 | 0.434963085 | 0.07667384 | 0.20488806 | F09E5.14 |
| WBGene00017292 | 0.363059212 | 0.13057276 | 0.29288894 | F09E5.16 |
| WBGene00017293 | -0.148801748 | 0.19230305 | 0.37863219 | bmy-1 |
| WBGene00017294 | -0.410421488 | 0.09515276 | 0.23704807 | F09E10.1 |
| WBGene00017295 | -0.023056499 | 0.78121253 | NA | F09E10.5 |
| WBGene00017296 | -0.003421551 | 0.95689871 | NA | F09E10.6 |
| WBGene00017297 | 0.212081917 | 0.32444265 | 0.52363682 | F09E10.7 |
| WBGene00017298 | 0.320049549 | 0.04637026 | 0.14250776 | toca-1 |
| WBGene00017299 | 0.208989511 | 0.23266715 | 0.42778849 | molo-1 |
| WBGene00017300 | -0.419965828 | 2.15E-09 | 5.21E-08 | rpc-2 |
| WBGene00017301 | -0.771219666 | 3.82E-31 | 4.99E-29 | hach-1 |
| WBGene00017302 | 0.893054794 | 9.16E-05 | 0.00089746 | F09F7.5 |
| WBGene00017303 | 0.784773503 | 0.00394297 | 0.02162944 | F09F7.6 |
| WBGene00017304 | 0.380550692 | 0.01401496 | 0.05855964 | nmad-1 |
| WBGene00017305 | 0.153682434 | 0.06760542 | NA | nspb-12 |
| WBGene00017306 | 0.684149124 | 0.02752412 | 0.09726783 | F09F9.1 |
| WBGene00017307 | 0.044589818 | 0.20019866 | 0.3887771 | F09F9.2 |
| WBGene00017308 | 0.038125098 | 0.61119482 | NA | F09F9.3 |
| WBGene00017309 | 0.097988987 | 0.5222276 | 0.70017723 | F09F9.4 |
| WBGene00017310 | 0.4589761 | 0.01324221 | 0.05600207 | F09G2.1 |
| WBGene00017311 | 0.113779829 | 0.2053229 | 0.39439004 | F09G2.2 |
| WBGene00017312 | 5.201287658 | 1.67E-31 | 2.22E-29 | pitr-5 |
| WBGene00017313 | -0.090453828 | 0.32088789 | 0.52085583 | cpsf-2 |
| WBGene00017314 | 0.837273925 | 0.01542747 | 0.06326234 | lgc-39 |
| WBGene00017315 | 0.179480997 | 0.30292197 | 0.50352867 | ugt-36 |
| WBGene00017316 | -0.128636765 | 0.13757883 | 0.30344857 | F09G2.8 |
| WBGene00017317 | -0.144016512 | 0.02047699 | 0.0779926 | attf-2 |
| WBGene00017319 | -0.2623678 | 0.01873515 | 0.07322513 | mrps-9 |
| WBGene00017320 | 2.481032932 | 1.56E-07 | 2.86E-06 | F09G8.5 |

| | | | | |
|---|---|---|---|---|
| WBGene00017321 | 0.031571643 | 0.817554 | 0.89973842 | F09G8.7 |
| WBGene00017322 | 1.371194689 | 4.99E-19 | 3.50E-17 | clec-160 |
| WBGene00017323 | NA | NA | NA | mps-4 |
| WBGene00017324 | NA | NA | NA | F10C1.1 |
| WBGene00017325 | 0.020925399 | 0.84643798 | NA | F10C1.3 |
| WBGene00017326 | -0.047846544 | 0.74830715 | 0.85794758 | dmd-5 |
| WBGene00017327 | 0.355558593 | 0.13006559 | 0.29227968 | F10C1.8 |
| WBGene00017328 | -0.073690525 | 0.27444383 | 0.47478816 | nemp-1 |
| WBGene00017329 | -0.174453071 | 0.40019779 | 0.5984532 | ugt-39 |
| WBGene00017330 | NA | NA | NA | sre-10 |
| WBGene00017331 | 0.369925987 | 0.12911789 | 0.29114663 | ugt-40 |
| WBGene00017332 | -0.022443476 | 0.87454977 | 0.93307197 | ugt-37 |
| WBGene00017333 | NA | NA | NA | ugt-38 |
| WBGene00017334 | 0.012863193 | 0.86580227 | NA | F10D2.8 |
| WBGene00017335 | -0.463173927 | 0.07706714 | 0.20538048 | F10D2.10 |
| WBGene00017336 | -0.544498329 | 0.00620712 | 0.03096369 | ugt-41 |
| WBGene00017338 | 0.303125262 | 0.15499953 | 0.32819427 | F10D7.1 |
| WBGene00017339 | -0.032244915 | 0.85680367 | 0.92320152 | F10D7.2 |
| WBGene00017340 | 0.805596165 | 0.00436666 | 0.02353911 | F10D7.3 |
| WBGene00017341 | 0.042516415 | 0.55991373 | NA | F10D7.10 |
| WBGene00017341 | 0.042516415 | 0.55991373 | NA | F10D7.4 |
| WBGene00017342 | 0.225264804 | 0.08609923 | 0.22169871 | F10D7.5 |
| WBGene00017343 | -0.439322122 | 0.08030024 | 0.21156663 | F10E7.1 |
| WBGene00017344 | -0.065968909 | 0.7042366 | 0.83211428 | F10E7.2 |
| WBGene00017345 | -0.023056499 | 0.78121253 | NA | NA |
| WBGene00017347 | -0.368443601 | 8.41E-06 | 0.0001051 | F10E7.5 |
| WBGene00017348 | -0.174638537 | 0.18443185 | 0.36793388 | F10E7.6 |
| WBGene00017349 | 0.376885223 | 1.09E-09 | 2.74E-08 | farl-11 |
| WBGene00017350 | 0.542359891 | 0.0502616 | 0.1509188 | F10E7.9 |
| WBGene00017351 | 0.075076776 | 0.28090944 | NA | cutl-5 |
| WBGene00017352 | 0.115570766 | 0.56038226 | 0.73080315 | F10E7.11 |
| WBGene00017353 | 0.133532899 | 0.15682793 | 0.33080502 | F10E9.1 |
| WBGene00017354 | -0.049434625 | 0.74674664 | 0.85673854 | F10E9.2 |
| WBGene00017355 | -0.158356785 | 0.10705231 | 0.2569431 | F10E9.3 |
| WBGene00017356 | -0.442541767 | 0.00020116 | 0.00177819 | F10E9.4 |
| WBGene00017357 | -0.169620771 | 0.20068428 | 0.38927869 | F10E9.5 |
| WBGene00017358 | 0.054964518 | 0.6433347 | 0.78886575 | F10E9.7 |
| WBGene00017359 | 0.056411843 | 0.41891328 | NA | F10E9.10 |
| WBGene00017360 | -0.213039702 | 0.09074454 | 0.22898841 | F10E9.11 |
| WBGene00017361 | 1.355084138 | 2.51E-05 | 0.00028036 | F10E9.12 |
| WBGene00017362 | 0.871395335 | 3.24E-11 | 1.00E-09 | F10G2.1 |
| WBGene00017363 | 0.013755544 | 0.88500544 | NA | F10G2.2 |
| WBGene00017364 | 2.928498339 | 1.56E-47 | 3.45E-45 | clec-7 |

| | | | | |
|---|---|---|---|---|
| WBGene00017365 | -0.152632883 | 0.45300974 | 0.64524911 | F10G2.4 |
| WBGene00017366 | -0.007072416 | 0.9244612 | NA | F10G2.7 |
| WBGene00017367 | 0.028241825 | 0.79095698 | 0.88224906 | nhr-263 |
| WBGene00017369 | 0.217473108 | 0.21025048 | 0.3998282 | F10G7.5 |
| WBGene00017370 | 0.055503542 | 0.44925503 | NA | NA |
| WBGene00017371 | 0.059980168 | 0.38780842 | NA | sre-39 |
| WBGene00017372 | 0.059640218 | 0.51913979 | 0.69833937 | F10G7.9 |
| WBGene00017373 | 0.221536055 | 0.02219927 | 0.08297665 | F10G7.10 |
| WBGene00017374 | -0.111338413 | 0.36618066 | 0.56583125 | ttr-41 |
| WBGene00017375 | -0.023987882 | 0.87235651 | 0.93179921 | pbo-6 |
| WBGene00017376 | 0.998482413 | 0.00880093 | 0.04072516 | F11C7.2 |
| WBGene00017378 | 0.298029921 | 0.0302759 | 0.10405388 | F11C7.6 |
| WBGene00017379 | 0.193657017 | 0.03471136 | 0.11525777 | F11C7.7 |
| WBGene00017380 | 0.285881244 | 0.16287745 | 0.33830172 | F11D5.1 |
| WBGene00017381 | -0.001724229 | 0.99212959 | 0.99552939 | ddr-2 |
| WBGene00017382 | 0.200678989 | 0.1848976 | 0.36856171 | F11D5.5 |
| WBGene00017383 | NA | NA | NA | F11D5.6 |
| WBGene00017384 | 0.165655217 | 0.18990326 | 0.37567541 | F11G11.4 |
| WBGene00017385 | 0.117337427 | 0.30506366 | 0.50565347 | F11G11.5 |
| WBGene00017386 | 0.866905836 | 0.01901629 | 0.07412605 | nspd-4 |
| WBGene00017386 | 0.866905836 | 0.01901629 | 0.07412605 | nspd-5 |
| WBGene00017387 | 0.148467664 | 0.07389511 | NA | mpst-4 |
| WBGene00017388 | 0.130463311 | 0.16095178 | 0.33556248 | F11G11.13 |
| WBGene00017389 | 0.280620627 | 0.16911239 | 0.34731021 | lgc-38 |
| WBGene00017390 | -1.672752868 | 0.00053512 | 0.00411266 | F12A10.1 |
| WBGene00017391 | NA | NA | NA | NA |
| WBGene00017392 | 0.020232312 | 0.84643798 | NA | NA |
| WBGene00017393 | NA | NA | NA | nep-5 |
| WBGene00017394 | 0.171584703 | 0.26820926 | 0.46714868 | cal-8 |
| WBGene00017395 | NA | NA | NA | F12A10.6 |
| WBGene00017396 | -0.002914971 | 0.95718083 | NA | F12A10.7 |
| WBGene00017397 | -0.227835223 | 0.11815277 | 0.27532623 | F12A10.8 |
| WBGene00017398 | -0.469135119 | 0.0036911 | 0.02057229 | slc-17.6 |
| WBGene00017399 | 0.020232312 | 0.84643798 | NA | lgc-51 |
| WBGene00017400 | -0.034095757 | 0.76523917 | 0.86670277 | stg-2 |
| WBGene00017401 | 0.020232312 | 0.84643798 | NA | F12D9.2 |
| WBGene00017402 | -0.256584925 | 0.07520728 | 0.20174236 | F12E12.1 |
| WBGene00017403 | -0.04044783 | 0.53612608 | NA | F12E12.2 |
| WBGene00017404 | 0.147507992 | 0.20023247 | 0.3887771 | F12E12.3 |
| WBGene00017405 | -0.603719111 | 0.03175167 | 0.10770997 | bath-31 |
| WBGene00017406 | -0.036980345 | 0.82635175 | 0.90477236 | sdz-12 |
| WBGene00017407 | 0.079434944 | 0.70278955 | 0.83120945 | F12E12.6 |
| WBGene00017408 | -0.999055112 | 0.01583097 | 0.06465306 | sdz-11 |

| | | | | |
|---|---|---|---|---|
| WBGene00017410 | NA | NA | NA | F13A2.1 |
| WBGene00017411 | 0.033463284 | 0.6365037 | NA | F13A2.2 |
| WBGene00017412 | NA | NA | NA | F13A2.3 |
| WBGene00017413 | -0.734811697 | 0.00516452 | 0.02682287 | F13A2.4 |
| WBGene00017414 | NA | NA | NA | F13A2.5 |
| WBGene00017415 | 0.0398788 | 0.58971221 | NA | F13A2.6 |
| WBGene00017416 | -0.022242797 | 0.86840192 | 0.92992376 | F13B6.1 |
| WBGene00017417 | -0.570241932 | 0.03280579 | 0.1105176 | F13B6.2 |
| WBGene00017418 | 0.40778261 | 0.08917826 | 0.22684545 | F13B6.3 |
| WBGene00017419 | -0.088339087 | 0.34414006 | 0.54382672 | F13B9.1 |
| WBGene00017420 | -0.010950308 | 0.91266252 | 0.95126775 | F13B9.2 |
| WBGene00017421 | -0.021936164 | 0.86370877 | 0.9272978 | cutl-29 |
| WBGene00017422 | 0.039876613 | 0.84710535 | 0.91761389 | F13C5.1 |
| WBGene00017423 | -0.788676571 | 1.18E-06 | 1.83E-05 | F13C5.2 |
| WBGene00017424 | NA | NA | NA | F13C5.3 |
| WBGene00017426 | 0.274399993 | 0.20583924 | 0.39487583 | F13C5.5 |
| WBGene00017427 | 0.189140056 | 0.36614116 | 0.56582992 | acp-5 |
| WBGene00017428 | 0.527133571 | 0.0359415 | 0.11822502 | F13D11.3 |
| WBGene00017429 | 0.050273403 | 0.7872789 | 0.87949869 | F13D11.4 |
| WBGene00017430 | 1.168974265 | 3.02E-10 | 8.17E-09 | bcl-11 |
| WBGene00017431 | 0.185201085 | 0.19874986 | 0.38710878 | F13H6.3 |
| WBGene00017432 | 0.407058846 | 0.07924226 | 0.20964688 | NA |
| WBGene00017433 | 0.824703518 | 0.0231434 | 0.08541165 | F13H6.5 |
| WBGene00017434 | 2.439466954 | 8.24E-05 | 0.00081302 | F13H8.1 |
| WBGene00017435 | -0.378320631 | 0.00021345 | 0.00187023 | F13H8.2 |
| WBGene00017436 | -1.453647657 | 2.46E-14 | 1.09E-12 | F13H8.3 |
| WBGene00017437 | 0.043540525 | 0.82372534 | 0.9034778 | nmgp-1 |
| WBGene00017438 | 2.631324824 | 0.00390196 | 0.02146348 | F13H8.5 |
| WBGene00017439 | 0.12301234 | 0.43906134 | 0.63370437 | F13H8.6 |
| WBGene00017440 | -0.04048412 | 0.63499111 | 0.78349105 | upb-1 |
| WBGene00017441 | -0.180429242 | 0.34249195 | 0.54280413 | F13H8.8 |
| WBGene00017442 | 0.113776683 | 0.5896248 | 0.75112054 | F13H8.9 |
| WBGene00017443 | 0.216740706 | 0.3242774 | 0.52363682 | F13H8.11 |
| WBGene00017444 | -0.03039785 | 0.70603871 | NA | F13H8.12 |
| WBGene00017445 | 0.069645922 | 0.73571875 | 0.85171647 | sid-5 |
| WBGene00017446 | -0.39924014 | 0.11028834 | 0.26197918 | F14B8.4 |
| WBGene00017447 | 0.137181806 | 0.49268505 | 0.67574197 | F14B8.5 |
| WBGene00017448 | 0.605808143 | 0.00463566 | 0.02468923 | F14B8.6 |
| WBGene00017449 | 1.615656614 | 2.10E-07 | 3.73E-06 | otpl-3 |
| WBGene00017450 | 0.04278989 | 0.80749557 | 0.89336279 | bath-27 |
| WBGene00017451 | NA | NA | NA | lido-14 |
| WBGene00017452 | -0.043712804 | 0.80214741 | 0.88992108 | bath-29 |
| WBGene00017453 | 0.034634319 | 0.82448673 | 0.90374412 | F14D2.5 |

| | | | | |
|---|---|---|---|---|
| WBGene00017454 | 0.078776717 | 0.31677567 | NA | irld-5 |
| WBGene00017455 | NA | NA | NA | F14D2.7 |
| WBGene00017456 | 0.173060666 | 0.41513309 | 0.6126146 | F14D2.8 |
| WBGene00017459 | 0.077046173 | 0.71277962 | 0.83848678 | F14D2.11 |
| WBGene00017460 | -0.013204245 | 0.91697557 | 0.95371452 | bath-30 |
| WBGene00017461 | 0.325346779 | 0.06882559 | 0.18898639 | bath-28 |
| WBGene00017462 | -0.118235053 | 0.57291855 | 0.74106374 | F14D2.14 |
| WBGene00017463 | 0.411152288 | 0.09543814 | 0.23757788 | F14D12.1 |
| WBGene00017464 | 1.059438406 | 2.81E-06 | 3.90E-05 | sulp-2 |
| WBGene00017465 | NA | NA | NA | F14F9.2 |
| WBGene00017466 | 0.052258977 | 0.67850139 | 0.81332396 | F14F9.3 |
| WBGene00017467 | -0.06398369 | 0.75520467 | 0.86125209 | F14F9.4 |
| WBGene00017468 | 0.013445115 | 0.94803944 | 0.97044737 | F14F9.5 |
| WBGene00017469 | 4.092297353 | 0.00124813 | 0.0084367 | F14F9.6 |
| WBGene00017470 | 0.020232312 | 0.84643798 | NA | F14H12.2 |
| WBGene00017471 | 1.305735758 | 4.15E-05 | 0.00044369 | F14H12.3 |
| WBGene00017472 | 0.076914481 | 0.71000596 | 0.83670293 | cst-1 |
| WBGene00017473 | 0.066203979 | 0.68476767 | 0.81809125 | F14H12.6 |
| WBGene00017474 | -0.006963856 | 0.96230013 | 0.97924853 | F14H12.7 |
| WBGene00017475 | 0.223368022 | 0.161853 | 0.33674717 | F14H12.8 |
| WBGene00017476 | NA | NA | NA | F15A8.1 |
| WBGene00017478 | 0.826423186 | 0.00378278 | 0.02097071 | F15A8.6 |
| WBGene00017480 | -0.012696633 | 0.88559529 | 0.93973915 | nstp-2 |
| WBGene00017481 | -0.54508345 | 0.01941096 | 0.07514433 | F15B10.3 |
| WBGene00017482 | 0.114475595 | 0.22008861 | 0.41227192 | set-9 |
| WBGene00017483 | 0.596305906 | 0.00244392 | 0.01473095 | lgc-22 |
| WBGene00017484 | 0.321653993 | 0.12580543 | 0.28690032 | F15E6.3 |
| WBGene00017485 | -1.59307568 | 2.05E-18 | 1.36E-16 | F15E6.4 |
| WBGene00017486 | NA | NA | NA | F15E6.5 |
| WBGene00017487 | 0.047502162 | 0.79631912 | 0.88573336 | F15E6.6 |
| WBGene00017488 | -0.248871039 | 0.16848161 | 0.34645214 | dct-7 |
| WBGene00017489 | 0.606636906 | 0.04297182 | 0.13469288 | F15E6.9 |
| WBGene00017490 | -0.050007053 | 0.47951277 | NA | pud-2.2 |
| WBGene00017490 | -0.050007053 | 0.47951277 | NA | pud-2.1 |
| WBGene00017491 | NA | NA | NA | irld-28 |
| WBGene00017492 | NA | NA | NA | irld-29 |
| WBGene00017493 | -0.004747605 | 0.9473792 | NA | irld-30 |
| WBGene00017494 | NA | NA | NA | NA |
| WBGene00017496 | NA | NA | NA | NA |
| WBGene00017497 | 0.05787416 | 0.40594086 | NA | irld-32 |
| WBGene00017498 | -0.046394786 | 0.50231372 | NA | pud-4 |
| WBGene00017499 | NA | NA | NA | pud-1.2 |
| WBGene00017499 | NA | NA | NA | pud-1.1 |

| | | | | |
|---|---|---|---|---|
| WBGene00017500 | NA | NA | NA | pud-2.2 |
| WBGene00017500 | NA | NA | NA | pud-2.1 |
| WBGene00017501 | -1.469521209 | 1.60E-29 | 1.98E-27 | pud-3 |
| WBGene00017502 | 0.273612317 | 0.11933341 | 0.27728304 | F16B3.2 |
| WBGene00017503 | 0.26878437 | 0.23604085 | 0.43192937 | nhr-177 |
| WBGene00017504 | 0.050022957 | 0.79304284 | 0.88383499 | F16B4.2 |
| WBGene00017505 | -0.022930767 | 0.78121253 | NA | NA |
| WBGene00017506 | -0.191031239 | 0.08107661 | 0.2128823 | F16B4.4 |
| WBGene00017507 | 0.786164654 | 0.00925426 | 0.04231466 | F16B4.5 |
| WBGene00017508 | 0.108942554 | 0.15594412 | NA | F16B4.6 |
| WBGene00017509 | 0.038633797 | 0.57999349 | NA | F16B4.7 |
| WBGene00017510 | 0.373124659 | 0.03798958 | 0.12319056 | nhr-178 |
| WBGene00017511 | NA | NA | NA | srbc-41 |
| WBGene00017512 | 0.197150025 | 0.34344134 | 0.54313812 | nhr-179 |
| WBGene00017513 | 0.364370957 | 0.1358119 | 0.30076173 | F16F9.1 |
| WBGene00017514 | 0.37584244 | 0.05128195 | 0.15325965 | F16F9.3 |
| WBGene00017515 | 0.242661742 | 0.2657369 | 0.46443258 | F16F9.4 |
| WBGene00017516 | NA | NA | NA | F16G10.1 |
| WBGene00017517 | NA | NA | NA | F16G10.2 |
| WBGene00017518 | NA | NA | NA | F16G10.3 |
| WBGene00017519 | NA | NA | NA | hpo-25 |
| WBGene00017520 | 0.020925399 | 0.84643798 | NA | F16G10.5 |
| WBGene00017521 | -0.023056499 | 0.78121253 | NA | F16G10.6 |
| WBGene00017522 | NA | NA | NA | F16G10.7 |
| WBGene00017523 | NA | NA | NA | F16G10.8 |
| WBGene00017524 | NA | NA | NA | F16G10.9 |
| WBGene00017525 | NA | NA | NA | F16G10.10 |
| WBGene00017526 | -0.001993815 | 0.95612996 | NA | F16G10.11 |
| WBGene00017527 | NA | NA | NA | F16G10.13 |
| WBGene00017528 | -0.001842784 | 0.95621055 | NA | F16G10.14 |
| WBGene00017529 | 2.436044216 | 0.00619911 | 0.03093428 | F16G10.15 |
| WBGene00017530 | -0.05292899 | 0.79618475 | 0.88565119 | F16H11.1 |
| WBGene00017532 | -0.225367522 | 0.15258089 | 0.32531481 | ent-5 |
| WBGene00017533 | -0.037304911 | 0.73101342 | 0.84986592 | NA |
| WBGene00017534 | -0.071455798 | 0.4828831 | 0.66848445 | F17A9.2 |
| WBGene00017535 | 0.055363609 | 0.5974506 | 0.75583464 | atf-8 |
| WBGene00017536 | 0.695593991 | 5.66E-06 | 7.33E-05 | F17A9.4 |
| WBGene00017537 | 0.700245148 | 3.58E-06 | 4.85E-05 | F17A9.5 |
| WBGene00017538 | -0.317029936 | 0.02476726 | 0.09020336 | ceh-49 |
| WBGene00017539 | NA | NA | NA | F17E9.2 |
| WBGene00017540 | 0.020232312 | 0.84643798 | NA | F17E9.3 |
| WBGene00017541 | 0.193212682 | 0.03684703 | 0.12047285 | F17E9.4 |
| WBGene00017542 | 2.018913506 | 0.0009383 | 0.0066785 | F17E9.5 |

| | | | | |
|---|---|---|---|---|
| WBGene00017543 | NA | NA | NA | lgc-5 |
| WBGene00017544 | 0.05787416 | 0.40594086 | NA | lgc-6 |
| WBGene00017545 | 0.146094066 | 0.24671158 | 0.44495002 | F18A1.1 |
| WBGene00017546 | -0.247661893 | 1.66E-06 | 2.46E-05 | rpa-1 |
| WBGene00017547 | -0.053860001 | 0.57468913 | 0.74227635 | alfa-1 |
| WBGene00017548 | -0.072513024 | 0.18269721 | 0.36554136 | F18A1.7 |
| WBGene00017549 | -0.283141748 | 0.05986403 | 0.17125229 | pid-1 |
| WBGene00017550 | 0.063527295 | 0.41294485 | NA | nep-6 |
| WBGene00017551 | NA | NA | NA | F18A12.2 |
| WBGene00017552 | NA | NA | NA | nep-7 |
| WBGene00017553 | 0.166384219 | 0.20404194 | 0.39307402 | nep-8 |
| WBGene00017554 | 0.065014541 | 0.37399213 | NA | nep-9 |
| WBGene00017555 | 0.050680134 | 0.55857002 | NA | nep-10 |
| WBGene00017556 | -0.136622989 | 0.5094931 | 0.6895487 | F18A12.7 |
| WBGene00017557 | 1.752357388 | 2.45E-08 | 5.12E-07 | nep-11 |
| WBGene00017559 | 0.168170932 | 0.05190965 | NA | mpz-3 |
| WBGene00017560 | 0.160503133 | 0.05644121 | NA | F18C5.5 |
| WBGene00017562 | -0.025226573 | 0.69184128 | NA | F18C5.9 |
| WBGene00017563 | 0.324077329 | 0.01930631 | 0.07487821 | F18C5.10 |
| WBGene00017564 | NA | NA | NA | srxa-4 |
| WBGene00017565 | 0.382823707 | 0.00334355 | 0.019033 | ddo-2 |
| WBGene00017566 | NA | NA | NA | F18E3.10 |
| WBGene00017567 | -1.004963841 | 1.17E-05 | 0.00014136 | F18E3.11 |
| WBGene00017568 | 0.12182354 | 0.26074633 | 0.45889975 | F18E9.1 |
| WBGene00017569 | 0.524436878 | 0.01193487 | 0.05137855 | F18E9.3 |
| WBGene00017570 | -0.028337346 | 0.7137484 | NA | F18E9.4 |
| WBGene00017571 | 0.231123395 | 0.22981922 | 0.42425231 | jmjd-3.1 |
| WBGene00017573 | 0.032611377 | 0.61967506 | NA | F18E9.7 |
| WBGene00017574 | 0.260705964 | 0.15723554 | 0.33118768 | F18F11.1 |
| WBGene00017576 | 0.153728711 | 0.45728216 | 0.64918009 | cdh-8 |
| WBGene00017577 | 0.007644371 | 0.94085646 | 0.96674008 | F18F11.4 |
| WBGene00017578 | -0.059452647 | 0.73887479 | 0.85307041 | drl-1 |
| WBGene00017579 | 0.038125098 | 0.61119482 | NA | F18G5.1 |
| WBGene00017580 | 0.523936528 | 0.05975024 | 0.17105736 | lgc-4 |
| WBGene00017581 | NA | NA | NA | F18G5.5 |
| WBGene00017582 | 1.472850604 | 2.86E-13 | 1.14E-11 | F18G5.6 |
| WBGene00017583 | -0.009936598 | 0.95823706 | 0.97705612 | F19B10.1 |
| WBGene00017584 | 0.217206988 | 0.15922098 | 0.33359392 | F19B10.2 |
| WBGene00017585 | NA | NA | NA | F19B10.3 |
| WBGene00017586 | 0.012158447 | 0.89398365 | NA | F19B10.4 |
| WBGene00017587 | -0.026148592 | 0.89605195 | 0.94531608 | F19B10.5 |
| WBGene00017588 | NA | NA | NA | F19B10.6 |
| WBGene00017589 | -0.311286722 | 0.01210971 | 0.05206999 | F19B10.10 |

| | | | | | |
|---|---|---|---|---|---|
| WBGene00017590 | | -0.060782953 | 0.55960335 | 0.73003231 | F19B10.11 |
| WBGene00017591 | | 0.991712433 | 4.89E-23 | 4.28E-21 | F19C7.1 |
| WBGene00017592 | | 3.098296771 | 3.54E-27 | 3.73E-25 | F19C7.2 |
| WBGene00017593 | | 0.056700152 | 0.73363303 | 0.85087141 | F19C7.3 |
| WBGene00017594 | | 1.370556303 | 0.00201896 | 0.01256025 | F19C7.4 |
| WBGene00017597 | | 0.064976307 | 0.74362013 | 0.85511642 | F19C7.8 |
| WBGene00017598 | | -0.198025297 | 0.35970673 | 0.55964982 | F19F10.1 |
| WBGene00017599 | | -0.011147643 | 0.95174154 | 0.97232907 | F19F10.3 |
| WBGene00017600 | | 0.117605973 | 0.57770968 | 0.74411462 | ttr-10 |
| WBGene00017601 | | -0.817965166 | 0.00841124 | 0.03931975 | ets-7 |
| WBGene00017602 | | -0.058752924 | 0.35274012 | NA | clec-211 |
| WBGene00017603 | NA | | NA | NA | clec-212 |
| WBGene00017605 | | -0.082596135 | 0.24525069 | 0.44340768 | F19F10.9 |
| WBGene00017606 | | 0.012667846 | 0.91297552 | 0.95131127 | ets-6 |
| WBGene00017607 | | -0.11451627 | 0.25763894 | 0.45573917 | F19F10.11 |
| WBGene00017608 | | -0.120536019 | 0.2427764 | 0.44089423 | F19F10.12 |
| WBGene00017609 | NA | | NA | NA | F19G12.1 |
| WBGene00017610 | NA | | NA | NA | F19G12.2 |
| WBGene00017611 | | -0.003614866 | 0.95700117 | NA | F19G12.3 |
| WBGene00017612 | NA | | NA | NA | F19G12.4 |
| WBGene00017613 | | 0.747770246 | 4.94E-08 | 9.81E-07 | F20A1.1 |
| WBGene00017614 | | -0.011662311 | 0.90617376 | 0.94916953 | F20A1.2 |
| WBGene00017615 | NA | | NA | NA | srsx-32 |
| WBGene00017616 | | 0.219205335 | 0.15484002 | 0.32795291 | F20A1.4 |
| WBGene00017618 | | 1.63039157 | 2.77E-05 | 0.00030695 | F20A1.6 |
| WBGene00017619 | NA | | NA | NA | F20A1.8 |
| WBGene00017620 | | -0.213829203 | 0.03499341 | 0.11597092 | tofu-2 |
| WBGene00017621 | | 0.298811244 | 0.08779607 | 0.2244887 | F20A1.10 |
| WBGene00017625 | | 0.39895608 | 0.08438134 | 0.21872063 | cgt-2 |
| WBGene00017627 | NA | | NA | NA | F20B6.1 |
| WBGene00017628 | | -0.01154686 | 0.8991333 | 0.94665211 | F20B6.4 |
| WBGene00017629 | | 0.020317245 | 0.84643798 | NA | F20B6.5 |
| WBGene00017630 | NA | | NA | NA | F20B6.6 |
| WBGene00017631 | | -0.067796314 | 0.64330712 | 0.78886575 | F20B6.7 |
| WBGene00017632 | | -0.135323655 | 0.48019275 | 0.66665106 | F20B6.9 |
| WBGene00017633 | NA | | NA | NA | F20D6.1 |
| WBGene00017634 | | 0.015451648 | 0.80337623 | NA | F20D6.2 |
| WBGene00017635 | | -0.214838503 | 0.32346167 | 0.52329307 | F20D6.5 |
| WBGene00017636 | | 0.307064213 | 0.06860304 | 0.18865836 | F20D6.6 |
| WBGene00017637 | | 0.445547901 | 0.05851772 | 0.16849851 | F20D6.8 |
| WBGene00017638 | NA | | NA | NA | F20D6.9 |
| WBGene00017639 | | 0.125642348 | 0.52899617 | 0.70589071 | F20D6.10 |
| WBGene00017640 | | -0.580957547 | 0.00240415 | 0.01452708 | F20D6.11 |

| | | | | |
|---|---|---|---|---|
| WBGene00017641 | -0.33324516 | 2.93E-10 | 7.94E-09 | csr-1 |
| WBGene00017642 | -0.101491456 | 0.20537874 | 0.39440584 | F20D12.2 |
| WBGene00017643 | -0.211916709 | 0.00123653 | 0.00838146 | czw-1 |
| WBGene00017644 | 0.0666735 | 0.74962171 | 0.85873723 | exc-9 |
| WBGene00017645 | 0.020232312 | 0.84643798 | NA | F20D12.7 |
| WBGene00017646 | 0.01344775 | 0.91572428 | 0.95302176 | F20H11.1 |
| WBGene00017647 | 0.081280902 | 0.60159271 | 0.75908213 | F20H11.4 |
| WBGene00017648 | 0.527369781 | 7.52E-11 | 2.19E-09 | ddo-3 |
| WBGene00017650 | 0.035512071 | 0.72830584 | 0.84850209 | F21A9.1 |
| WBGene00017651 | 0.006691586 | 0.97328178 | 0.98532835 | F21A9.2 |
| WBGene00017652 | NA | NA | NA | F21C10.1 |
| WBGene00017653 | 0.368300805 | 0.13112205 | 0.29367568 | F21C10.3 |
| WBGene00017654 | -0.133402025 | 0.46593165 | 0.65553385 | F21C10.4 |
| WBGene00017655 | NA | NA | NA | F21C10.5 |
| WBGene00017656 | NA | NA | NA | F21C10.6 |
| WBGene00017657 | 0.235045299 | 0.09852945 | 0.24347168 | F21C10.7 |
| WBGene00017658 | 1.694804063 | 5.99E-23 | 5.22E-21 | F21C10.9 |
| WBGene00017659 | 0.580526197 | 9.88E-05 | 0.00096161 | F21C10.10 |
| WBGene00017660 | -0.107615698 | 0.29442774 | 0.49565557 | F21C10.11 |
| WBGene00017661 | 0.072999249 | 0.57360821 | 0.74143178 | frpr-6 |
| WBGene00017662 | NA | NA | NA | F21D12.2 |
| WBGene00017663 | 1.32191076 | 1.43E-06 | 2.16E-05 | F21D12.3 |
| WBGene00017664 | 0.027299492 | 0.79328707 | 0.8839726 | npax-1 |
| WBGene00017666 | -0.002914971 | 0.95718083 | NA | F21E9.2 |
| WBGene00017667 | 0.183406933 | 0.39370793 | 0.59205811 | ttr-37 |
| WBGene00017668 | -0.023109762 | 0.78121253 | NA | ins-39 |
| WBGene00017670 | NA | NA | NA | F21E9.6 |
| WBGene00017671 | 1.340550899 | 1.57E-18 | 1.05E-16 | pgal-1 |
| WBGene00017672 | 0.073458642 | 0.29111983 | NA | F21F3.2 |
| WBGene00017673 | 0.073300201 | 0.4402788 | NA | icmt-1 |
| WBGene00017674 | -0.188826534 | 0.29667572 | 0.49760903 | F21F3.4 |
| WBGene00017675 | 0.134230503 | 0.05518804 | 0.16176339 | F21F3.6 |
| WBGene00017676 | 0.109585196 | 0.50571323 | 0.68671373 | F21F3.7 |
| WBGene00017677 | 0.573031913 | 0.02733658 | 0.09679887 | NA |
| WBGene00017678 | 2.931807675 | 3.17E-16 | 1.74E-14 | asp-12 |
| WBGene00017679 | NA | NA | NA | F21F8.5 |
| WBGene00017680 | 0.050433702 | 0.64137111 | 0.78746664 | NA |
| WBGene00017681 | 0.50159435 | 0.04533923 | 0.14004555 | slc-17.5 |
| WBGene00017683 | -0.014920977 | 0.88536592 | 0.93966238 | rbbp-5 |
| WBGene00017684 | NA | NA | NA | F21H12.2 |
| WBGene00017685 | NA | NA | NA | F21H12.3 |
| WBGene00017686 | -0.224909029 | 5.23E-05 | 0.00054569 | tpp-2 |
| WBGene00017687 | 0.674587552 | 1.90E-06 | 2.78E-05 | ets-4 |

| | | | | |
|---|---|---|---|---|
| WBGene00017688 | -0.185421758 | 0.26425532 | 0.46262849 | ttr-35 |
| WBGene00017690 | 0.139476708 | 0.49261455 | 0.67574197 | ceh-60 |
| WBGene00017691 | 0.924873036 | 2.00E-76 | 7.71E-74 | ilys-5 |
| WBGene00017692 | 1.326893767 | 0.00360696 | 0.02023351 | F22B7.1 |
| WBGene00017693 | 0.020232312 | 0.84643798 | NA | flp-23 |
| WBGene00017694 | 0.020925399 | 0.84643798 | NA | F22B7.3 |
| WBGene00017695 | 0.0398788 | 0.58971221 | NA | fip-1 |
| WBGene00017696 | -0.065029576 | 0.47839023 | 0.66528413 | polk-1 |
| WBGene00017698 | 1.218327608 | 7.79E-06 | 9.81E-05 | F22B7.9 |
| WBGene00017699 | -0.247561001 | 0.0024539 | 0.01477281 | flcn-1 |
| WBGene00017700 | 0.137438713 | 0.13428296 | 0.29847226 | F22D3.4 |
| WBGene00017701 | 0.05204179 | 0.67560287 | 0.81139374 | F22D3.5 |
| WBGene00017702 | 0.018053288 | 0.92747811 | 0.95891582 | malt-1 |
| WBGene00017703 | -3.727270606 | 2.74E-13 | 1.10E-11 | F22E5.1 |
| WBGene00017704 | -0.003421551 | 0.95689871 | NA | F22E5.2 |
| WBGene00017705 | 0.042516415 | 0.55991373 | NA | F22E5.6 |
| WBGene00017706 | 0.020925399 | 0.84643798 | NA | F22E5.7 |
| WBGene00017707 | 0.963029128 | 0.01835163 | 0.07220806 | F22E5.8 |
| WBGene00017708 | -0.173141383 | 0.31677959 | 0.51698682 | F22E5.9 |
| WBGene00017709 | NA | NA | NA | F22E5.11 |
| WBGene00017710 | 0.054115925 | 0.54377021 | NA | F22E5.12 |
| WBGene00017711 | 0.270805166 | 0.17368027 | 0.35410846 | F22E5.13 |
| WBGene00017712 | 0.205901914 | 0.13874216 | 0.30527862 | F22E5.17 |
| WBGene00017713 | -0.174390558 | 0.38383162 | 0.58327725 | F22E5.20 |
| WBGene00017714 | -0.323624114 | 0.04273427 | 0.13426449 | F22F1.2 |
| WBGene00017715 | 0.220582197 | 0.12566127 | 0.28666356 | F22F1.3 |
| WBGene00017716 | 0.535925262 | 0.04871827 | 0.14746418 | sups-1 |
| WBGene00017717 | 0.042323707 | 0.78734986 | 0.87949869 | F22F4.4 |
| WBGene00017718 | -0.376539969 | 0.1226189 | 0.28205237 | F22F4.5 |
| WBGene00017719 | -0.157289999 | 0.06551202 | 0.18273223 | ldp-1 |
| WBGene00017720 | 0.062826631 | 0.73362249 | 0.85087141 | F22F7.2 |
| WBGene00017721 | 0.143853672 | 0.18032109 | 0.36220072 | F22F7.3 |
| WBGene00017722 | 0.094392367 | 0.48203425 | 0.66787768 | F22F7.4 |
| WBGene00017723 | 0.020317245 | 0.84643798 | NA | bgnt-1.2 |
| WBGene00017724 | 0.58348306 | 0.01450395 | 0.0602589 | F22F7.7 |
| WBGene00017725 | NA | NA | NA | F22H10.1 |
| WBGene00017726 | -0.077772624 | 0.68783582 | 0.82014857 | F22H10.2 |
| WBGene00017727 | 0.056711677 | 0.49169131 | 0.67507131 | F22H10.3 |
| WBGene00017729 | -0.195200122 | 0.21359541 | 0.40429869 | NA |
| WBGene00017730 | -0.28811786 | 0.20519192 | 0.39430567 | F22H10.6 |
| WBGene00017731 | -0.048370879 | 0.47720013 | NA | NA |
| WBGene00017732 | 0.051836356 | 0.79387898 | 0.88399982 | F23C8.3 |
| WBGene00017733 | -0.115600504 | 0.23914603 | 0.43606953 | ubxn-1 |

| | | | | |
|---|---|---|---|---|
| WBGene00017734 | -0.020640089 | 0.77352677 | 0.87150484 | F23C8.5 |
| WBGene00017735 | 0.071010711 | 0.50862988 | 0.68894903 | did-2 |
| WBGene00017736 | 0.34871602 | 0.14089925 | 0.30860862 | F23C8.7 |
| WBGene00017737 | 0.027525261 | 0.866152 | 0.92862709 | F23C8.8 |
| WBGene00017738 | -0.229866941 | 0.10078241 | 0.24671128 | tipn-1 |
| WBGene00017739 | -0.023756719 | 0.6882956 | NA | NA |
| WBGene00017741 | 0.145197519 | 0.37329172 | 0.57229033 | F23C8.13 |
| WBGene00017742 | -0.276643202 | 0.01705232 | 0.06828858 | nfyc-1 |
| WBGene00017743 | 0.020925399 | 0.84643798 | NA | F23F1.2 |
| WBGene00017744 | 0.002584832 | 0.99116077 | 0.9949664 | fbxc-54 |
| WBGene00017745 | 0.258650687 | 0.23493982 | 0.43077695 | F23F1.4 |
| WBGene00017746 | -0.142511639 | 0.12649195 | 0.28757277 | F23F1.5 |
| WBGene00017747 | 0.386618489 | 0.08687844 | 0.22288194 | F23F1.6 |
| WBGene00017748 | -0.432002124 | 0.09190318 | 0.23107581 | F23F1.7 |
| WBGene00017749 | -0.102010137 | 0.47374062 | 0.66158302 | rpoa-49 |
| WBGene00017750 | 0.05947379 | 0.72510367 | 0.84584837 | F23F1.10 |
| WBGene00017751 | 0.357095376 | 0.14083388 | 0.30856693 | F23F12.3 |
| WBGene00017752 | -0.208470888 | 0.32325414 | 0.52320738 | sdz-15 |
| WBGene00017754 | 0.030337975 | 0.78257109 | 0.87764254 | F23F12.8 |
| WBGene00017755 | -1.36304384 | 2.26E-08 | 4.75E-07 | zip-8 |
| WBGene00017756 | -0.749640792 | 0.00898944 | 0.0414403 | F23F12.12 |
| WBGene00017757 | -0.205413146 | 0.0259851 | 0.0934302 | bra-2 |
| WBGene00017758 | -0.215267636 | 0.08915205 | 0.22684545 | F23H11.2 |
| WBGene00017759 | -0.338919911 | 5.90E-06 | 7.59E-05 | sucl-2 |
| WBGene00017760 | -0.004547385 | 0.96502292 | 0.98071026 | F23H11.4 |
| WBGene00017761 | -0.029136177 | 0.74962119 | NA | F23H11.6 |
| WBGene00017762 | 2.328743873 | 1.01E-05 | 0.0001248 | F23H11.7 |
| WBGene00017763 | 0.294673557 | 0.00593643 | 0.02989207 | crls-1 |
| WBGene00017764 | 7.442183702 | 2.52E-08 | 5.26E-07 | F25A2.1 |
| WBGene00017765 | 0.061625353 | 0.46725454 | 0.65684817 | gcst-1 |
| WBGene00017766 | -0.084382256 | 0.5593593 | 0.73003231 | peli-1 |
| WBGene00017767 | 0.131719055 | 0.52544823 | 0.70269341 | F25B4.4 |
| WBGene00017768 | -0.329825789 | 7.83E-05 | 0.00077921 | F25B4.5 |
| WBGene00017769 | -0.094868626 | 0.5798366 | 0.74533693 | hmgs-1 |
| WBGene00017770 | -0.222860747 | 0.05814614 | 0.16787963 | F25B4.7 |
| WBGene00017771 | -0.367626296 | 0.13108697 | 0.29364202 | F25B4.8 |
| WBGene00017772 | -0.279958953 | 0.00111676 | 0.00769794 | clec-1 |
| WBGene00017773 | -0.117990604 | 0.39456585 | 0.59262392 | dct-6 |
| WBGene00017774 | -0.128937403 | 0.07179749 | 0.19502544 | nop-1 |
| WBGene00017775 | 0.040714684 | 0.52056317 | 0.69924269 | F25B5.3 |
| WBGene00017776 | 0.076398068 | 0.57473114 | 0.74227635 | F25B5.5 |
| WBGene00017777 | 0.070307821 | 0.39061206 | 0.58849055 | F25B5.6 |
| WBGene00017778 | -0.015436309 | 0.86337665 | 0.92712602 | nono-1 |

| | | | | |
|---|---|---|---|---|
| WBGene00017779 | 0.2234044 | 0.20095878 | 0.38964871 | gtr-1 |
| WBGene00017780 | -0.157783506 | 0.1446749 | 0.31403783 | F25E2.2 |
| WBGene00017781 | 1.191099294 | 2.96E-05 | 0.00032419 | F25E2.3 |
| WBGene00017782 | -0.299960652 | 0.0200891 | 0.07689529 | F25E5.1 |
| WBGene00017783 | -0.166658029 | 0.31190593 | 0.51266736 | F25E5.2 |
| WBGene00017784 | 0.04919219 | 0.62443867 | 0.77558589 | F25E5.3 |
| WBGene00017785 | 1.008539197 | 0.00208653 | 0.01293346 | F25E5.4 |
| WBGene00017786 | 0.303912834 | 0.14970532 | 0.321054 | F25E5.5 |
| WBGene00017787 | 0.714642978 | 0.03053184 | 0.1047368 | nhr-141 |
| WBGene00017788 | -0.018764354 | 0.88300075 | 0.93817228 | F25E5.7 |
| WBGene00017789 | 0.371527913 | 0.0156211 | 0.06392075 | F25E5.8 |
| WBGene00017790 | 0.040107227 | 0.58693163 | NA | F25E5.9 |
| WBGene00017791 | -0.050570864 | 0.59666325 | 0.75522171 | try-8 |
| WBGene00017792 | 0.22138764 | 0.14184366 | 0.30983837 | F25E5.16 |
| WBGene00017794 | 0.099211179 | 0.38822106 | 0.58676004 | mltn-13 |
| WBGene00017795 | 0.362622476 | 0.13127194 | 0.29383159 | F25F8.1 |
| WBGene00017796 | 0.287049279 | 0.19985981 | 0.38844384 | F25G6.1 |
| WBGene00017797 | -0.131115962 | 0.21546478 | 0.40652318 | symk-1 |
| WBGene00017798 | 0.251224559 | 0.09952456 | 0.24465061 | slc-17.8 |
| WBGene00017799 | -0.337708829 | 0.00133888 | 0.00893489 | F25G6.8 |
| WBGene00017800 | -0.272289965 | 0.00012527 | 0.00118174 | F25G6.9 |
| WBGene00017801 | -0.063421601 | 0.52227549 | 0.70017723 | F26A1.1 |
| WBGene00017802 | 0.083209383 | 0.30786773 | NA | F26A1.3 |
| WBGene00017803 | 0.020232312 | 0.84643798 | NA | F26A1.4 |
| WBGene00017804 | NA | NA | NA | F26A1.6 |
| WBGene00017805 | NA | NA | NA | F26A1.7 |
| WBGene00017806 | -0.008390518 | 0.91872349 | NA | F26A1.8 |
| WBGene00017807 | 0.876062078 | 0.01951122 | 0.07529356 | F26A1.9 |
| WBGene00017808 | 0.18458729 | 0.13570169 | 0.30057615 | nspd-9 |
| WBGene00017809 | NA | NA | NA | clec-156 |
| WBGene00017810 | -0.050223056 | 0.47776088 | NA | clec-157 |
| WBGene00017811 | 0.054750408 | 0.63113412 | 0.78083781 | F26A1.13 |
| WBGene00017813 | 0.016496914 | 0.93532442 | 0.96342232 | nlp-61 |
| WBGene00017814 | 0.257395924 | 0.2533528 | 0.45142246 | F26A10.2 |
| WBGene00017815 | 0.0398788 | 0.58971221 | NA | F26B1.1 |
| WBGene00017816 | -0.192838213 | 0.000348 | 0.00284637 | hrpk-1 |
| WBGene00017817 | 0.036687355 | 0.59489568 | NA | F26B1.5 |
| WBGene00017818 | -0.077489523 | 0.70830885 | 0.83527012 | F26D11.20 |
| WBGene00017818 | -0.077489523 | 0.70830885 | 0.83527012 | F26D11.1 |
| WBGene00017819 | -0.026384613 | 0.67299636 | NA | F26D11.2 |
| WBGene00017821 | -0.023109762 | 0.78121253 | NA | F26D11.4 |
| WBGene00017822 | 0.020317245 | 0.84643798 | NA | clec-216 |
| WBGene00017823 | 0.063015825 | 0.74419103 | 0.85554724 | pals-34 |

| | | | | |
|---|---|---|---|---|
| WBGene00017824 | NA | NA | NA | clec-217 |
| WBGene00017825 | -0.232534423 | 0.01392813 | 0.05827984 | cee-1 |
| WBGene00017826 | -0.176630333 | 0.06613089 | 0.18362328 | tag-340 |
| WBGene00017827 | 0.144684983 | 0.20195766 | 0.39049949 | F26F4.5 |
| WBGene00017828 | 0.016712302 | 0.88772539 | 0.94087067 | F26F4.6 |
| WBGene00017829 | -0.121081339 | 0.46055446 | 0.6515584 | F26F4.9 |
| WBGene00017830 | -0.372046909 | 5.85E-05 | 0.00060473 | rpb-8 |
| WBGene00017831 | -0.015565515 | 0.90288971 | 0.94829375 | F26F4.12 |
| WBGene00017832 | -0.373910835 | 0.00860755 | 0.03998922 | kbp-2 |
| WBGene00017833 | 0.020232312 | 0.84643798 | NA | F26F12.2 |
| WBGene00017834 | 0.744599304 | 5.88E-05 | 0.00060655 | F26F12.3 |
| WBGene00017835 | 0.219066643 | 0.14285076 | 0.31113924 | F26F12.4 |
| WBGene00017836 | 0.046774719 | 0.71268368 | 0.83844129 | F26F12.5 |
| WBGene00017837 | 0.021197103 | 0.82020665 | 0.90127932 | F26G1.1 |
| WBGene00017838 | 0.850360819 | 0.01375716 | 0.0577296 | F26G1.2 |
| WBGene00017839 | 0.020317245 | 0.84643798 | NA | F26G1.3 |
| WBGene00017840 | 0.593514358 | 0.02362427 | 0.086818 | ttm-2 |
| WBGene00017841 | -0.029454259 | 0.88041709 | 0.93644654 | F26G1.5 |
| WBGene00017842 | 1.047402464 | 0.00575911 | 0.02928763 | nep-12 |
| WBGene00017843 | 0.010428208 | 0.93279671 | 0.9622403 | F26G5.1 |
| WBGene00017845 | 0.020925399 | 0.84643798 | NA | srsx-9 |
| WBGene00017848 | -0.504390913 | 0.06270581 | 0.17733741 | F27B3.6 |
| WBGene00017849 | -0.267445151 | 0.2236413 | 0.41658171 | NA |
| WBGene00017850 | 0.94772544 | 0.01124312 | 0.04904961 | F27B10.1 |
| WBGene00017851 | 1.190490803 | 0.00329844 | 0.01882749 | F27C1.1 |
| WBGene00017852 | 0.004235642 | 0.95791236 | 0.97692915 | F27C1.2 |
| WBGene00017853 | 0.031063203 | 0.70177318 | NA | F27C1.3 |
| WBGene00017854 | -0.13998398 | 0.50654352 | 0.68739475 | F27C1.4 |
| WBGene00017855 | -0.170619806 | 0.07357637 | 0.19853144 | F27C1.6 |
| WBGene00017858 | 0.238876263 | 0.25352521 | 0.45156479 | F27C1.11 |
| WBGene00017860 | NA | NA | NA | F27C1.13 |
| WBGene00017861 | 0.312280305 | 0.14392333 | 0.31287029 | F27D9.2 |
| WBGene00017864 | -0.281505416 | 6.58E-06 | 8.38E-05 | pcca-1 |
| WBGene00017865 | 1.529341913 | 0.00056459 | 0.00429857 | F27D9.7 |
| WBGene00017866 | 0.308905301 | 0.18001108 | 0.36172691 | stn-2 |
| WBGene00017867 | 0.568493721 | 0.04972192 | 0.14960514 | slc-28.1 |
| WBGene00017868 | 0.682568022 | 0.03365526 | 0.11262782 | slc-28.2 |
| WBGene00017869 | 0.104747489 | 0.16723639 | NA | F28A10.1 |
| WBGene00017870 | NA | NA | NA | F28A10.2 |
| WBGene00017871 | 0.020925399 | 0.84643798 | NA | F28A10.3 |
| WBGene00017872 | 0.020317245 | 0.84643798 | NA | F28A10.4 |
| WBGene00017873 | -0.133728773 | 0.14814175 | NA | F28A10.5 |
| WBGene00017874 | -0.792467926 | 3.48E-18 | 2.25E-16 | acdh-9 |

| | | | | |
|---|---|---|---|---|
| WBGene00017875 | 0.036687355 | 0.59489568 | NA | F28A10.7 |
| WBGene00017876 | NA | NA | NA | F28A10.8 |
| WBGene00017877 | 0.020925399 | 0.84643798 | NA | F28A10.9 |
| WBGene00017878 | NA | NA | NA | F28A10.10 |
| WBGene00017879 | 0.20746505 | 0.06510887 | 0.18184975 | acd-4 |
| WBGene00017880 | 0.653371692 | 0.03546198 | 0.11714298 | F28A12.3 |
| WBGene00017881 | 0.293236611 | 4.85E-05 | 0.00051141 | asp-13 |
| WBGene00017885 | 0.142972434 | 0.08695359 | 0.2229252 | dip-2 |
| WBGene00017886 | -0.164558265 | 0.16417553 | 0.34012932 | F28B3.5 |
| WBGene00017887 | -0.107065373 | 0.42252524 | 0.61861921 | F28B3.6 |
| WBGene00017888 | -0.219979034 | 0.17508771 | 0.35564444 | acl-11 |
| WBGene00017889 | -0.02903899 | 0.82517273 | 0.90395465 | F28B3.10 |
| WBGene00017890 | -0.006711909 | 0.92834175 | NA | F28B4.1 |
| WBGene00017891 | 0.065268826 | 0.63260819 | 0.78177194 | rgl-1 |
| WBGene00017892 | 1.149671354 | 1.72E-38 | 3.00E-36 | F28B4.3 |
| WBGene00017893 | 0.243249257 | 0.10611292 | 0.25544107 | F28B4.4 |
| WBGene00017894 | 0.088670189 | 0.60166861 | 0.75908213 | F28B12.1 |
| WBGene00017895 | -0.117316724 | 0.12174462 | 0.28070284 | vrk-1 |
| WBGene00017896 | 0.020317245 | 0.84643798 | NA | F28C10.1 |
| WBGene00017898 | -0.12836299 | 0.44818274 | 0.64088487 | F28C10.3 |
| WBGene00017899 | -0.022930767 | 0.78121253 | NA | F28C10.4 |
| WBGene00017900 | 0.565582642 | 0.01872152 | 0.0731914 | F28E10.1 |
| WBGene00017901 | 0.589442281 | 0.02685132 | 0.0956362 | igeg-1 |
| WBGene00017902 | 0.0398788 | 0.58971221 | NA | F28E10.4 |
| WBGene00017904 | 0.593650965 | 0.00341002 | 0.01933619 | lim-8 |
| WBGene00017905 | -0.115183772 | 0.57101613 | 0.73951536 | F28F5.6 |
| WBGene00017906 | 0.020317245 | 0.84643798 | NA | F28F9.2 |
| WBGene00017907 | 0.036275992 | 0.83743255 | 0.91185854 | F28F9.3 |
| WBGene00017908 | -0.689311656 | 0.00022457 | 0.00194439 | F28H1.1 |
| WBGene00017909 | -0.102372442 | 0.59532935 | 0.75477619 | F28H1.4 |
| WBGene00017910 | 0.020232312 | 0.84643798 | NA | F28H1.5 |
| WBGene00017911 | -1.760335239 | 0.00419434 | 0.02276947 | fbxb-106 |
| WBGene00017912 | -0.775474551 | 0.02427975 | 0.08880948 | fbxb-107 |
| WBGene00017913 | NA | NA | NA | F29A7.3 |
| WBGene00017914 | 0.123000759 | 0.30675717 | 0.50731183 | F29A7.4 |
| WBGene00017916 | -0.03059097 | 0.84422037 | 0.91642626 | F29A7.6 |
| WBGene00017917 | 0.020317245 | 0.84643798 | NA | clec-120 |
| WBGene00017918 | -0.002914971 | 0.95718083 | NA | F29A7.8 |
| WBGene00017919 | -0.050684452 | 0.739496 | 0.85327581 | F29B9.1 |
| WBGene00017920 | -0.146434085 | 0.08365673 | 0.21751343 | jmjd-1.2 |
| WBGene00017921 | -0.019487929 | 0.90925889 | 0.95068624 | F29B9.5 |
| WBGene00017922 | 0.131539016 | 0.10075387 | NA | F29B9.7 |
| WBGene00017923 | -0.236680509 | 0.08257572 | 0.21546804 | F29B9.8 |

| | | | | |
|---|---|---|---|---|
| WBGene00017924 | -0.028198819 | 0.86525538 | 0.92800557 | mrps-21 |
| WBGene00017925 | -0.124845947 | 0.12395125 | 0.28431306 | F29B9.11 |
| WBGene00017926 | -0.145516922 | 0.0632766 | 0.17846903 | cox-6C |
| WBGene00017927 | -0.039636691 | 0.54579661 | NA | F29C4.4 |
| WBGene00017928 | -0.013735366 | 0.91389879 | 0.9519924 | tut-1 |
| WBGene00017929 | -0.199833432 | 0.03587245 | 0.11805091 | grld-1 |
| WBGene00017930 | -0.222208119 | 0.26892468 | 0.46800501 | F29G9.1 |
| WBGene00017931 | -0.047422545 | 0.57100983 | 0.73951536 | picc-1 |
| WBGene00017933 | -0.037086712 | 0.84846756 | 0.91827422 | F29G9.7 |
| WBGene00017934 | 0.313164632 | 0.0093323 | 0.042605 | F30B5.4 |
| WBGene00017935 | 0.020925399 | 0.84643798 | NA | srbc-70 |
| WBGene00017936 | 0.23417208 | 0.29064227 | 0.49114651 | zipt-2.2 |
| WBGene00017937 | 2.9676978 | 0.00051642 | 0.00399339 | F30H5.3 |
| WBGene00017938 | 0.099840818 | 0.50751468 | 0.68801245 | F31A3.3 |
| WBGene00017939 | -0.090353319 | 0.57431792 | 0.74208708 | F31A3.5 |
| WBGene00017941 | -0.030698512 | 0.70341898 | NA | F31A9.2 |
| WBGene00017942 | 0.087447402 | 0.57106708 | 0.73951536 | fbxc-37 |
| WBGene00017944 | 0.07033102 | 0.6968198 | 0.82702162 | F31D5.1 |
| WBGene00017945 | 0.306352661 | 0.16037152 | 0.33484732 | F31D5.2 |
| WBGene00017947 | 0.499809713 | 0.06582353 | 0.18318234 | mth-2 |
| WBGene00017948 | -0.367847634 | 0.12135849 | 0.28045725 | mth-1 |
| WBGene00017949 | -0.090837784 | 0.44901872 | 0.64173691 | F31D5.6 |
| WBGene00017950 | 0.636027276 | 0.03531902 | 0.11675451 | F31E3.2 |
| WBGene00017951 | -0.064765489 | 0.46529185 | 0.65496952 | panl-2 |
| WBGene00017952 | 0.20262512 | 0.2962535 | 0.49737336 | F31E3.6 |
| WBGene00017954 | -0.278751436 | 0.13015265 | 0.29235425 | F31E8.4 |
| WBGene00017955 | 0.206575708 | 0.02995553 | 0.10341402 | F31E8.5 |
| WBGene00017956 | NA | NA | NA | NA |
| WBGene00017957 | 0.252262223 | 0.26102278 | 0.45919656 | F31F4.1 |
| WBGene00017959 | 0.402620694 | 0.06506619 | 0.1818351 | ugt-42 |
| WBGene00017960 | -0.039636691 | 0.54579661 | NA | F31F4.11 |
| WBGene00017961 | 0.603902879 | 0.0431075 | 0.13503143 | nhr-180 |
| WBGene00017962 | -0.371716526 | 0.00719227 | 0.03475312 | fbxa-72 |
| WBGene00017963 | 0.058314668 | 0.40208118 | NA | F31F4.17 |
| WBGene00017964 | 0.789653176 | 7.36E-11 | 2.16E-09 | F31F7.1 |
| WBGene00017965 | 0.23518399 | 0.08691838 | 0.2229252 | F31F7.2 |
| WBGene00017967 | -0.058310724 | 0.54027232 | 0.7146844 | ada-2 |
| WBGene00017968 | 0.171095565 | 0.20477458 | 0.39397451 | skpo-3 |
| WBGene00017969 | 0.432708538 | 0.00110913 | 0.00767407 | ctsa-3.1 |
| WBGene00017970 | 2.177385302 | 1.10E-60 | 2.99E-58 | F32A5.4 |
| WBGene00017971 | -0.271625906 | 0.17212035 | 0.35168352 | F32A5.8 |
| WBGene00017973 | 0.228513489 | 0.08620776 | 0.22186041 | ift-81 |
| WBGene00017974 | -0.032749525 | 0.82633383 | 0.90477236 | vps-41 |

| | | | | |
|---|---|---|---|---|
| WBGene00017975 | 0.260447381 | 0.07142242 | 0.19436131 | F32B5.1 |
| WBGene00017976 | 0.128000389 | 0.16547331 | NA | F32B5.2 |
| WBGene00017977 | NA | NA | NA | F32B5.3 |
| WBGene00017978 | 0.300861436 | 0.05872254 | 0.16894412 | msrp-1 |
| WBGene00017979 | 0.36566275 | 0.00584557 | 0.0295696 | F32B5.6 |
| WBGene00017980 | 0.340978512 | 0.12426282 | 0.28458189 | F32B5.7 |
| WBGene00017981 | -0.201595429 | 0.11376826 | 0.26788052 | figl-1 |
| WBGene00017982 | -0.268793969 | 0.00060639 | 0.00459054 | hpo-18 |
| WBGene00017983 | 0.510850798 | 0.06386977 | 0.17951954 | F32D1.3 |
| WBGene00017984 | -0.304612896 | 4.14E-08 | 8.32E-07 | F32D1.5 |
| WBGene00017985 | -0.01020712 | 0.90764388 | 0.9500153 | neg-1 |
| WBGene00017986 | -0.018426956 | 0.80277183 | 0.89020543 | F32D1.7 |
| WBGene00017987 | 0.0398788 | 0.58971221 | NA | F32D1.8 |
| WBGene00017988 | 0.082354209 | 0.29626344 | 0.49737336 | fipp-1 |
| WBGene00017989 | -0.115514623 | 0.1520661 | 0.32451635 | nol-10 |
| WBGene00017990 | -0.395072661 | 0.00253503 | 0.01518013 | cec-4 |
| WBGene00017991 | 0.030620925 | 0.85055393 | 0.9197276 | clec-180 |
| WBGene00017992 | 0.272952474 | 0.05265947 | 0.15654563 | F32E10.5 |
| WBGene00017993 | -0.285535965 | 2.82E-05 | 0.00031099 | cec-5 |
| WBGene00017995 | -0.109314095 | 0.42266935 | 0.6186446 | F32E10.8 |
| WBGene00017996 | 0.071228933 | 0.41352747 | 0.61119077 | F33D4.4 |
| WBGene00017997 | -0.174151768 | 0.08463994 | 0.21917464 | mrpl-1 |
| WBGene00017998 | 3.243353972 | 0.00095611 | 0.00677561 | F33D4.6 |
| WBGene00017999 | -0.300304953 | 0.02991096 | 0.10340434 | emc-6 |
| WBGene00018000 | NA | NA | NA | F33D11.1 |
| WBGene00018001 | 0.038125098 | 0.61119482 | NA | F33D11.2 |
| WBGene00018002 | 0.042065281 | 0.558937 | NA | twk-47 |
| WBGene00018003 | NA | NA | NA | F33D11.6 |
| WBGene00018004 | 0.153513723 | 0.06686266 | NA | F33D11.7 |
| WBGene00018005 | 0.041009898 | 0.76417308 | 0.86623378 | nlp-62 |
| WBGene00018006 | -0.118556014 | 0.12996578 | 0.29224835 | gpaa-1 |
| WBGene00018007 | -0.211902161 | 0.00335602 | 0.01908564 | F33D11.10 |
| WBGene00018008 | -0.126307999 | 0.06241177 | 0.17667647 | vpr-1 |
| WBGene00018009 | -0.091722184 | 0.38005734 | 0.57962703 | dhhc-3 |
| WBGene00018010 | -0.097873999 | 0.50152937 | 0.68305968 | F33E11.2 |
| WBGene00018011 | 0.046579756 | 0.61504082 | 0.76911622 | F33E11.3 |
| WBGene00018013 | -0.05901204 | 0.65296467 | 0.79443429 | phf-10 |
| WBGene00018014 | -0.124885809 | 0.38525492 | 0.58438845 | F33G12.2 |
| WBGene00018015 | -0.144315445 | 0.16071631 | 0.33528748 | F33G12.3 |
| WBGene00018016 | -0.047064613 | 0.58842817 | 0.75051633 | lrr-1 |
| WBGene00018017 | -0.015920087 | 0.83380516 | 0.90939671 | golg-2 |
| WBGene00018018 | 0.114736086 | 0.51866347 | 0.6979082 | F33G12.6 |
| WBGene00018019 | NA | NA | NA | F33H12.1 |

| | | | | |
|---|---|---|---|---|
| WBGene00018021 | -0.441345551 | 0.08625669 | 0.22186041 | F34D6.1 |
| WBGene00018022 | NA | NA | NA | ceh-87 |
| WBGene00018023 | 0.178877119 | 0.15081028 | 0.32252639 | set-11 |
| WBGene00018024 | 0.500763901 | 0.00214885 | 0.01323571 | F35A5.1 |
| WBGene00018025 | 2.291518569 | 1.89E-08 | 4.04E-07 | F35A5.2 |
| WBGene00018026 | 0.127693253 | 0.10904601 | NA | F35A5.4 |
| WBGene00018027 | 0.040107227 | 0.58693163 | NA | F35A5.5 |
| WBGene00018030 | NA | NA | NA | F35B3.3 |
| WBGene00018031 | 0.118486893 | 0.36313154 | 0.56290195 | F35B3.4 |
| WBGene00018033 | 1.236647486 | 0.00683618 | 0.03341852 | F35B3.7 |
| WBGene00018034 | NA | NA | NA | sek-5 |
| WBGene00018035 | 0.161911593 | 0.45088235 | 0.64322016 | sek-4 |
| WBGene00018036 | -0.070332404 | 0.65896694 | 0.79896991 | F35C8.5 |
| WBGene00018037 | -0.126688306 | 0.20316823 | 0.39190857 | chtl-1 |
| WBGene00018038 | -0.007460734 | 0.96049543 | 0.97847297 | F35C8.8 |
| WBGene00018039 | 0.033463284 | 0.6365037 | NA | F35D2.1 |
| WBGene00018040 | 0.033180193 | 0.61386379 | NA | F35D2.2 |
| WBGene00018041 | -0.205122821 | 0.34514953 | 0.54490539 | F35D2.3 |
| WBGene00018042 | 0.056208621 | 0.59185768 | 0.75246512 | mks-3 |
| WBGene00018043 | -0.033349775 | 0.75000248 | 0.8587464 | F35D11.1 |
| WBGene00018044 | 0.252225684 | 0.14107438 | 0.30869378 | F35D11.3 |
| WBGene00018045 | -0.086913713 | 0.55188535 | 0.72422277 | F35D11.4 |
| WBGene00018046 | -0.330747379 | 0.0571433 | 0.16595987 | coa-3 |
| WBGene00018047 | 0.020317245 | 0.84643798 | NA | clec-136 |
| WBGene00018048 | 0.020232312 | 0.84643798 | NA | clec-137 |
| WBGene00018049 | 0.020232312 | 0.84643798 | NA | clec-138 |
| WBGene00018050 | -0.11419834 | 0.52528182 | 0.70253504 | clec-139 |
| WBGene00018051 | -0.186821815 | 0.27239562 | 0.47247297 | che-10 |
| WBGene00018052 | 1.497525064 | 2.57E-07 | 4.48E-06 | F35F10.1 |
| WBGene00018053 | NA | NA | NA | srbc-3 |
| WBGene00018054 | -0.03991741 | 0.81287416 | 0.89638233 | F35F10.4 |
| WBGene00018055 | -0.303303712 | 0.15038999 | 0.32200348 | F35F10.5 |
| WBGene00018056 | 0.04324155 | 0.81825431 | 0.90007974 | F35F10.6 |
| WBGene00018057 | -0.04044783 | 0.53612608 | NA | F35F10.7 |
| WBGene00018058 | NA | NA | NA | srbc-1 |
| WBGene00018059 | NA | NA | NA | F35F10.10 |
| WBGene00018060 | 0.539485828 | 0.04735185 | 0.14452334 | arrd-18 |
| WBGene00018061 | 0.094268991 | 0.62825035 | 0.77878363 | arrd-19 |
| WBGene00018062 | 0.089404422 | 0.46158874 | 0.65228105 | F35F10.13 |
| WBGene00018063 | -0.024740619 | 0.69507748 | NA | F35F10.14 |
| WBGene00018064 | -0.099691148 | 0.37022011 | 0.57006877 | cdc-73 |
| WBGene00018065 | 0.097875719 | 0.5361549 | 0.71128259 | F35F11.2 |
| WBGene00018066 | -0.003421551 | 0.95689871 | NA | F35F11.3 |

| | | | | |
|---|---|---|---|---|
| WBGene00018067 | 0.193416981 | 0.07497323 | 0.20140975 | npr-7 |
| WBGene00018068 | 0.106233677 | 0.54069224 | 0.71476071 | F35H10.2 |
| WBGene00018069 | 0.070749802 | 0.63656461 | 0.78445295 | F35H10.3 |
| WBGene00018070 | -0.211588212 | 0.25826252 | 0.45633859 | F35H10.5 |
| WBGene00018071 | -0.289636676 | 0.04880927 | 0.14761753 | F35H10.6 |
| WBGene00018072 | 0.03259923 | 0.76544649 | 0.86673775 | nprl-3 |
| WBGene00018073 | 0.0100547 | 0.9443329 | 0.96847702 | F35H10.10 |
| WBGene00018074 | -0.010949441 | 0.94336908 | 0.96789535 | F35H12.1 |
| WBGene00018075 | 0.261832454 | 0.07319925 | 0.19770693 | tbc-11 |
| WBGene00018076 | 0.07625825 | 0.65949543 | 0.79904606 | pifk-1 |
| WBGene00018077 | -0.290623523 | 0.18144203 | 0.36400284 | F35H12.5 |
| WBGene00018078 | 0.231562217 | 0.09622533 | 0.23939422 | F35H12.6 |
| WBGene00018080 | 0.020925399 | 0.84643798 | NA | F36A4.1 |
| WBGene00018081 | 0.127693253 | 0.10904601 | NA | F36A4.2 |
| WBGene00018082 | 0.012386589 | 0.8695009 | NA | F36A4.3 |
| WBGene00018083 | 0.153838603 | 0.19309915 | 0.37948519 | F36A4.4 |
| WBGene00018084 | -0.041653177 | 0.81234345 | 0.89600124 | F36A4.5 |
| WBGene00018085 | 0.186692786 | 0.38907094 | 0.58749873 | ttr-20 |
| WBGene00018087 | 0.106997704 | 0.16248985 | NA | F36D4.1 |
| WBGene00018088 | 0.067837916 | 0.66788529 | 0.8060096 | trpp-4 |
| WBGene00018089 | 0.06057409 | 0.63831116 | 0.78552676 | F36D4.4 |
| WBGene00018090 | -0.25102005 | 0.00105787 | 0.00736132 | F36D4.5 |
| WBGene00018091 | 0.020232312 | 0.84643798 | NA | F36D4.6 |
| WBGene00018092 | -0.010141451 | 0.90973981 | NA | F36F12.1 |
| WBGene00018093 | NA | NA | NA | F36F12.2 |
| WBGene00018094 | -0.042308943 | 0.80873275 | 0.89403928 | F36F12.3 |
| WBGene00018096 | 0.017604133 | 0.79437959 | NA | clec-207 |
| WBGene00018097 | NA | NA | NA | clec-208 |
| WBGene00018098 | 0.074892148 | 0.28555128 | NA | F36F12.7 |
| WBGene00018099 | -0.266406999 | 0.13987418 | 0.30727009 | ztf-28 |
| WBGene00018100 | 0.269833629 | 0.14311255 | 0.31151213 | math-26 |
| WBGene00018101 | 1.05293789 | 1.36E-06 | 2.07E-05 | math-27 |
| WBGene00018102 | -0.035224226 | 0.73465952 | 0.85123061 | math-28 |
| WBGene00018103 | -0.739457767 | 0.02688652 | 0.09573093 | F36H5.4 |
| WBGene00018104 | -0.217721304 | 0.29869827 | 0.49942984 | fbxb-53 |
| WBGene00018105 | NA | NA | NA | clec-19 |
| WBGene00018106 | -0.037596645 | 0.76049334 | 0.86431035 | F36H5.8 |
| WBGene00018107 | -0.108538544 | 0.12871363 | NA | F36H5.9 |
| WBGene00018108 | -0.023056499 | 0.78121253 | NA | F36H5.10 |
| WBGene00018109 | -0.235449552 | 0.17753055 | 0.35876119 | fbxb-12 |
| WBGene00018110 | NA | NA | NA | F36H9.1 |
| WBGene00018111 | 0.074958105 | 0.3349622 | NA | F36H9.2 |
| WBGene00018112 | -0.281211917 | 0.18657271 | 0.37097584 | F36H9.4 |

| | | | | |
|---|---|---|---|---|
| WBGene00018113 | -0.135243383 | 0.38972229 | 0.58786842 | F36H9.5 |
| WBGene00018114 | 0.040107227 | 0.58693163 | NA | srt-41 |
| WBGene00018115 | 0.150770881 | 0.19254261 | 0.37898198 | F36H9.7 |
| WBGene00018117 | 0.904033642 | 0.00047847 | 0.00375216 | nlp-47 |
| WBGene00018118 | 0.031768563 | 0.86278661 | 0.92691949 | F36H12.2 |
| WBGene00018119 | 0.188646571 | 0.13213114 | 0.29530332 | F36H12.3 |
| WBGene00018120 | 0.010018959 | 0.92686789 | 0.9586234 | F36H12.4 |
| WBGene00018121 | 0.019975402 | 0.61487492 | NA | F36H12.5 |
| WBGene00018122 | 0.020317245 | 0.84643798 | NA | ttbk-2 |
| WBGene00018123 | 0.020317245 | 0.84643798 | NA | F36H12.9 |
| WBGene00018124 | 0.038125098 | 0.61119482 | NA | F36H12.10 |
| WBGene00018125 | NA | NA | NA | NA |
| WBGene00018127 | 0.129578469 | 0.09326699 | NA | NA |
| WBGene00018130 | 0.048173932 | 0.76738438 | 0.86792389 | F36H12.17 |
| WBGene00018131 | -0.157988466 | 0.03727014 | 0.12150348 | F37A4.1 |
| WBGene00018132 | -0.057890456 | 0.75784173 | 0.86299765 | F37A4.2 |
| WBGene00018133 | 0.034143018 | 0.68679874 | NA | F37A4.3 |
| WBGene00018134 | 0.020466419 | 0.88484023 | 0.93940914 | F37A4.4 |
| WBGene00018135 | 0.097090099 | 0.3927689 | 0.59095501 | F37A4.5 |
| WBGene00018136 | -0.014502424 | 0.90191734 | 0.94780763 | F37A4.6 |
| WBGene00018137 | -0.099961766 | 0.16013638 | 0.33477934 | bath-41 |
| WBGene00018138 | -1.252161857 | 7.92E-18 | 5.00E-16 | folt-2 |
| WBGene00018139 | 0.020925399 | 0.84643798 | NA | sre-24 |
| WBGene00018140 | -0.017443821 | 0.92211029 | 0.95620205 | F37B4.10 |
| WBGene00018141 | -0.022930767 | 0.78121253 | NA | oac-21 |
| WBGene00018142 | 0.123219986 | 0.25620201 | 0.45460486 | oac-22 |
| WBGene00018143 | NA | NA | NA | oac-23 |
| WBGene00018144 | -0.003785899 | 0.98355316 | 0.99058801 | F37C4.4 |
| WBGene00018145 | -0.132155979 | 0.14073346 | 0.30845488 | F37C4.5 |
| WBGene00018146 | -0.573529166 | 0.0009614 | 0.0068098 | F37C4.6 |
| WBGene00018148 | 0.020925399 | 0.84643798 | NA | F37C4.8 |
| WBGene00018149 | -0.151395655 | 0.13849205 | 0.3048735 | F37C12.1 |
| WBGene00018150 | 0.042984009 | 0.66101122 | 0.80041893 | epg-4 |
| WBGene00018151 | -0.277168254 | 0.00191764 | 0.01208096 | F37C12.3 |
| WBGene00018152 | -0.077740403 | 0.16812074 | 0.34590441 | acs-4 |
| WBGene00018153 | 0.931700731 | 0.01824967 | 0.07198099 | F37C12.10 |
| WBGene00018154 | -0.11791085 | 0.2693994 | 0.4684414 | exos-9 |
| WBGene00018155 | -0.023056499 | 0.78121253 | NA | F37C12.14 |
| WBGene00018156 | -0.247176031 | 0.00259997 | 0.01553683 | ncbp-1 |
| WBGene00018157 | 0.018795205 | 0.77532208 | NA | lron-6 |
| WBGene00018158 | 0.117953933 | 0.36507476 | 0.56462961 | comp-1 |
| WBGene00018159 | -0.180019878 | 0.2301883 | 0.42455339 | F37F2.2 |
| WBGene00018160 | -0.030673769 | 0.6989942 | 0.82858699 | odr-8 |

| | | | | |
|---|---|---|---|---|
| WBGene00018161 | -0.227568449 | 0.01361868 | 0.05724882 | F38A5.2 |
| WBGene00018162 | -0.022930767 | 0.78121253 | NA | nspb-3 |
| WBGene00018163 | -0.034507753 | 0.71774519 | NA | F38A5.6 |
| WBGene00018164 | -0.206193584 | 0.09361957 | 0.23406969 | sup-36 |
| WBGene00018165 | 0.036687355 | 0.59489568 | NA | F38A5.8 |
| WBGene00018166 | -0.023109762 | 0.78121253 | NA | nspb-5 |
| WBGene00018167 | 0.020317245 | 0.84643798 | NA | nspb-4 |
| WBGene00018168 | 0.0398788 | 0.58971221 | NA | irld-7 |
| WBGene00018169 | NA | NA | NA | nspb-2 |
| WBGene00018170 | NA | NA | NA | nspb-1 |
| WBGene00018172 | NA | NA | NA | F38B6.2 |
| WBGene00018173 | 0.13079285 | 0.1612623 | NA | F38B6.3 |
| WBGene00018174 | 0.18529225 | 0.22108459 | 0.41356214 | F38B6.4 |
| WBGene00018175 | 0.154733662 | 0.45296939 | 0.64524911 | F38B6.6 |
| WBGene00018176 | 0.015194928 | 0.8086441 | NA | F38B6.7 |
| WBGene00018178 | 0.059980168 | 0.38780842 | NA | F38E1.3 |
| WBGene00018179 | NA | NA | NA | F38E1.6 |
| WBGene00018180 | -0.045771477 | 0.50913401 | NA | srsx-22 |
| WBGene00018181 | -0.126252475 | 0.22945308 | 0.42368805 | mpdu-1 |
| WBGene00018182 | 0.020232312 | 0.84643798 | NA | F38E1.10 |
| WBGene00018184 | 0.863678279 | 2.02E-06 | 2.93E-05 | F38E9.1 |
| WBGene00018186 | 0.020317245 | 0.84643798 | NA | F38E9.4 |
| WBGene00018187 | -0.023744263 | 0.8619846 | 0.92668569 | twf-2 |
| WBGene00018188 | 0.210022234 | 0.3274238 | 0.52665337 | F38E9.6 |
| WBGene00018189 | -0.109140056 | 0.59266646 | 0.75314295 | nhr-181 |
| WBGene00018190 | NA | NA | NA | F38H12.5 |
| WBGene00018191 | 0.032775352 | 0.85598916 | 0.92266343 | frpr-7 |
| WBGene00018192 | NA | NA | NA | F39B3.3 |
| WBGene00018193 | 0.272426647 | 0.00553177 | 0.02838789 | F39C12.1 |
| WBGene00018194 | NA | NA | NA | NA |
| WBGene00018195 | 0.920357814 | 0.00826117 | 0.03874177 | btb-16 |
| WBGene00018196 | 0.127693253 | 0.10904601 | NA | nep-13 |
| WBGene00018198 | NA | NA | NA | F39E9.6 |
| WBGene00018199 | -0.062432032 | 0.72846203 | 0.84853595 | NA |
| WBGene00018200 | 0.75599166 | 0.01577701 | 0.06446866 | btb-17 |
| WBGene00018201 | 0.038726127 | 0.57551883 | NA | btb-18 |
| WBGene00018202 | -0.16791813 | 0.045681 | NA | F39F10.2 |
| WBGene00018203 | -0.562627733 | 0.03579707 | 0.11794837 | F39F10.3 |
| WBGene00018204 | -0.339865134 | 0.15426991 | 0.32715947 | F39F10.4 |
| WBGene00018205 | 0.020317245 | 0.84643798 | NA | F39F10.5 |
| WBGene00018206 | 0.163010394 | 0.42152667 | 0.61820815 | ugt-61 |
| WBGene00018207 | -0.070748829 | 0.73213198 | 0.85011851 | F39G3.2 |
| WBGene00018208 | -0.014237987 | 0.91845227 | 0.95464005 | F39G3.3 |

| | | | | |
|---|---|---|---|---|
| WBGene00018209 | -0.003108065 | 0.95728506 | NA | F39G3.4 |
| WBGene00018210 | 0.803423579 | 0.01296014 | 0.05503172 | F39G3.5 |
| WBGene00018211 | -0.075452738 | 0.28904131 | NA | oac-24 |
| WBGene00018212 | 0.109034875 | 0.58197401 | 0.74618455 | F39H12.1 |
| WBGene00018213 | 0.117515562 | 0.35988921 | 0.55977244 | F39H12.2 |
| WBGene00018214 | 0.119592331 | 0.48043615 | 0.66679929 | F39H12.3 |
| WBGene00018215 | -0.182054273 | 0.39016206 | 0.58799411 | igcm-1 |
| WBGene00018216 | 0.494509768 | 0.01758979 | 0.06997809 | F40A3.1 |
| WBGene00018217 | -0.23731831 | 0.14953887 | 0.32088686 | F40A3.2 |
| WBGene00018218 | 0.083427241 | 0.32516901 | 0.52443264 | F40A3.3 |
| WBGene00018219 | 0.048027723 | 0.68511652 | 0.81830567 | F40A3.4 |
| WBGene00018221 | 0.435597301 | 0.02493372 | 0.09069681 | F40A3.6 |
| WBGene00018222 | -0.019341412 | 0.92067924 | 0.9557288 | F40A3.7 |
| WBGene00018223 | 0.773274891 | 0.02172054 | 0.08158296 | bath-13 |
| WBGene00018224 | 0.15564525 | 0.19100689 | 0.37703623 | NA |
| WBGene00018225 | 1.17045 | 0.0002243 | 0.00194318 | NA |
| WBGene00018226 | 0.059002333 | 0.71706242 | 0.84132743 | F40B5.2 |
| WBGene00018227 | 0.042065281 | 0.558937 | NA | nep-15 |
| WBGene00018228 | NA | NA | NA | F40C5.1 |
| WBGene00018229 | 0.186507357 | 0.38995821 | 0.58786842 | F40C5.2 |
| WBGene00018229 | 0.186507357 | 0.38995821 | 0.58786842 | T24A6.17 |
| WBGene00018230 | -0.019382952 | 0.86502913 | 0.92790421 | mage-1 |
| WBGene00018231 | -0.022930767 | 0.78121253 | NA | F40E3.3 |
| WBGene00018232 | 2.193115976 | 0.00048348 | 0.00378128 | F40E3.5 |
| WBGene00018233 | 0.175586955 | 0.04633883 | NA | F40E3.6 |
| WBGene00018234 | 0.020232312 | 0.84643798 | NA | F40E12.1 |
| WBGene00018235 | -0.05790816 | 0.62175418 | 0.77393862 | F40E12.2 |
| WBGene00018236 | -0.499297046 | 0.05459007 | 0.16039597 | fbxb-71 |
| WBGene00018237 | 1.18629036 | 1.28E-82 | 5.37E-80 | irg-7 |
| WBGene00018238 | 0.049687035 | 0.70060721 | 0.82976818 | F40F4.7 |
| WBGene00018239 | -0.089833419 | 0.53633143 | 0.71133513 | sec-20 |
| WBGene00018240 | -0.154102615 | 0.46903191 | 0.65776619 | cox-17 |
| WBGene00018241 | -0.091349597 | 0.65824334 | 0.79858641 | F40G9.5 |
| WBGene00018242 | -0.36236061 | 0.04466367 | 0.13848386 | F40G9.6 |
| WBGene00018243 | 0.033034883 | 0.64219503 | NA | F40G9.7 |
| WBGene00018244 | NA | NA | NA | F40G9.8 |
| WBGene00018245 | -0.164011998 | 0.40684132 | 0.60478869 | F40G9.9 |
| WBGene00018246 | -0.001842784 | 0.95621055 | NA | clec-148 |
| WBGene00018247 | -0.196440258 | 0.15558887 | 0.32912395 | F40G9.12 |
| WBGene00018248 | -0.016434121 | 0.88563082 | 0.93973915 | F40G9.14 |
| WBGene00018249 | 0.284552984 | 0.086946 | 0.2229252 | F40H3.1 |
| WBGene00018250 | -1.150945003 | 0.0002217 | 0.00192406 | F40H3.2 |
| WBGene00018251 | 0.009119847 | 0.93123385 | 0.96116988 | F40H3.3 |

| | | | | |
|---|---|---|---|---|
| WBGene00018252 | 0.068914553 | 0.74046448 | 0.85343151 | F40H3.6 |
| WBGene00018253 | 0.120080595 | 0.31712731 | 0.51727506 | F40H6.1 |
| WBGene00018254 | 0.038125098 | 0.61119482 | NA | F40H6.2 |
| WBGene00018255 | NA | NA | NA | F40H6.5 |
| WBGene00018256 | -0.050580107 | 0.47300881 | NA | cutl-28 |
| WBGene00018257 | 0.14575775 | 0.48746642 | 0.67171726 | F41B4.1 |
| WBGene00018258 | 1.025731878 | 0.00072288 | 0.00531311 | F41B4.2 |
| WBGene00018259 | 0.538378571 | 0.05385859 | 0.15899127 | F41B4.3 |
| WBGene00018260 | 0.821434837 | 0.00651322 | 0.03218362 | cyp-33C7 |
| WBGene00018261 | 0.130113876 | 0.4837867 | 0.66922913 | cyp-33C5 |
| WBGene00018262 | 0.780777962 | 0.0099817 | 0.04480138 | cyp-33C3 |
| WBGene00018264 | 0.01912147 | 0.77587372 | NA | cyp-33C6 |
| WBGene00018265 | 0.070279488 | 0.73555818 | 0.85166527 | nhr-182 |
| WBGene00018266 | 0.229946209 | 0.25225568 | 0.45017504 | nhr-183 |
| WBGene00018267 | 2.242068798 | 0.00739423 | 0.03550653 | F41C3.1 |
| WBGene00018268 | 0.246678743 | 0.26845073 | 0.46735451 | F41C3.2 |
| WBGene00018269 | 0.89834307 | 2.76E-16 | 1.53E-14 | acs-11 |
| WBGene00018270 | -0.263218285 | 0.02832588 | 0.09938153 | eas-1 |
| WBGene00018271 | 0.215685495 | 0.0001281 | 0.00120228 | ctsa-1.1 |
| WBGene00018273 | 0.016335094 | 0.91271761 | 0.95126775 | F41C3.7 |
| WBGene00018274 | 0.319138698 | 0.17438776 | 0.35505649 | F41C3.8 |
| WBGene00018275 | 0.194026145 | 0.37392488 | 0.57284024 | F41C3.11 |
| WBGene00018278 | 0.163001738 | 0.39992432 | 0.59821127 | F41C6.4 |
| WBGene00018279 | 0.011573942 | 0.90227566 | NA | F41C6.6 |
| WBGene00018280 | 0.071360176 | 0.72118593 | 0.84362463 | F41C6.7 |
| WBGene00018281 | 0.031988303 | 0.87829163 | 0.93491899 | tbc-18 |
| WBGene00018282 | -0.491082873 | 0.04644301 | 0.14255181 | F41D9.2 |
| WBGene00018283 | 2.336333388 | 0.00143547 | 0.00947123 | sulp-3 |
| WBGene00018284 | 0.056411843 | 0.41891328 | NA | F41E6.1 |
| WBGene00018285 | -0.051918093 | 0.40006665 | 0.59836304 | smk-1 |
| WBGene00018286 | 1.860331819 | 4.39E-13 | 1.72E-11 | F41E6.5 |
| WBGene00018288 | -0.102525887 | 0.35522982 | 0.55506646 | F41E6.7 |
| WBGene00018289 | -0.005017292 | 0.94670066 | NA | F41E6.8 |
| WBGene00018290 | -0.063952076 | 0.58435235 | 0.74778226 | vps-60 |
| WBGene00018291 | 0.020232312 | 0.84643798 | NA | fip-4 |
| WBGene00018292 | 0.205256481 | 0.06161959 | 0.17504316 | F41E6.11 |
| WBGene00018293 | -0.78009514 | 0.00196401 | 0.01231059 | F41E6.12 |
| WBGene00018294 | -0.132671539 | 0.0573389 | 0.16640214 | atg-18 |
| WBGene00018295 | 0.138655625 | 0.29571064 | 0.4967871 | oac-29 |
| WBGene00018296 | 0.020232312 | 0.84643798 | NA | F41F3.1 |
| WBGene00018297 | 5.964761852 | 1.94E-15 | 9.55E-14 | F41F3.3 |
| WBGene00018298 | 0.93189345 | 0.00556449 | 0.02849588 | nlp-50 |
| WBGene00018299 | 0.040107227 | 0.58693163 | NA | F41G3.2 |

| | | | | |
|---|---|---|---|---|
| WBGene00018300 | 0.056188364 | 0.42092875 | NA | F41G3.3 |
| WBGene00018301 | 0.313043815 | 0.02599277 | 0.0934318 | F41G3.5 |
| WBGene00018302 | -0.14445509 | 0.08514649 | 0.21991164 | F41G3.6 |
| WBGene00018303 | 1.824278044 | 5.41E-34 | 7.84E-32 | F41G3.10 |
| WBGene00018304 | 0.721783814 | 0.01771426 | 0.07036203 | agr-1 |
| WBGene00018305 | -0.081326066 | 0.51983871 | 0.69876593 | exos-8 |
| WBGene00018307 | -0.019795209 | 0.90971981 | 0.95103233 | dyla-1 |
| WBGene00018309 | 0.437074932 | 0.06850306 | 0.18847613 | F41G4.5 |
| WBGene00018310 | 0.184952829 | 0.22760142 | 0.42118901 | F41G4.7 |
| WBGene00018311 | 1.041792795 | 0.00291422 | 0.01705318 | F41G4.8 |
| WBGene00018312 | 0.020232312 | 0.84643798 | NA | F41H8.1 |
| WBGene00018313 | 0.020232312 | 0.84643798 | NA | F41H8.2 |
| WBGene00018314 | 3.281630729 | 8.92E-14 | 3.76E-12 | F41H10.1 |
| WBGene00018315 | NA | NA | NA | F41H10.2 |
| WBGene00018316 | -0.241535979 | 0.00370049 | 0.02060816 | F41H10.3 |
| WBGene00018317 | -0.010020512 | 0.93496504 | 0.96327984 | F41H10.4 |
| WBGene00018318 | 0.160484768 | 0.27631716 | 0.47632079 | F41H10.5 |
| WBGene00018319 | 0.046840734 | 0.57497138 | 0.74227635 | hda-6 |
| WBGene00018321 | -0.032471433 | 0.72237634 | 0.84436547 | sand-1 |
| WBGene00018322 | NA | NA | NA | F41H10.12 |
| WBGene00018323 | -0.022930767 | 0.78121253 | NA | F42A6.1 |
| WBGene00018324 | NA | NA | NA | F42A6.2 |
| WBGene00018325 | -0.080472653 | 0.40010183 | NA | NA |
| WBGene00018326 | 0.015896709 | 0.80422001 | NA | NA |
| WBGene00018327 | -0.075452738 | 0.28904131 | NA | F42A6.5 |
| WBGene00018328 | 0.0383613 | 0.81913544 | 0.90050774 | F42A6.6 |
| WBGene00018329 | -0.050921541 | 0.47213035 | NA | NA |
| WBGene00018330 | -0.135598069 | 0.37385596 | 0.57279459 | elks-1 |
| WBGene00018332 | 0.093796813 | 0.25278271 | NA | F42A9.3 |
| WBGene00018333 | 0.000420658 | 0.99875197 | 0.99929799 | cyp-33E3 |
| WBGene00018334 | 0.4482023 | 0.06014756 | 0.17176135 | cyp-33E2 |
| WBGene00018335 | -0.452692196 | 0.00016666 | 0.00150897 | F42A9.6 |
| WBGene00018336 | 0.155143374 | 0.19189259 | 0.37807823 | F42A9.7 |
| WBGene00018337 | -0.100960232 | 0.53599171 | 0.71119246 | F42A9.8 |
| WBGene00018338 | 0.065014541 | 0.37399213 | NA | F42A9.9 |
| WBGene00018339 | -0.147437363 | 0.09964867 | 0.24483238 | abcf-3 |
| WBGene00018340 | -0.659689442 | 3.62E-05 | 0.00039104 | anmt-2 |
| WBGene00018341 | -0.093554928 | 0.20187566 | 0.39045849 | F42A10.5 |
| WBGene00018342 | 0.820395829 | 1.88E-09 | 4.59E-08 | F42A10.6 |
| WBGene00018343 | -1.289211112 | 7.21E-10 | 1.85E-08 | F42A10.7 |
| WBGene00018344 | 0.156841323 | 0.30690944 | 0.50739172 | npr-27 |
| WBGene00018345 | 0.307955169 | 0.06343063 | 0.17866254 | F42C5.3 |
| WBGene00018346 | 1.202496205 | 0.00381694 | 0.02111213 | F42C5.4 |

| | | | | |
|---|---|---|---|---|
| WBGene00018347 | 0.206689679 | 0.11838982 | 0.2757031 | F42C5.5 |
| WBGene00018348 | -0.187272889 | 0.15642652 | 0.33033906 | F42C5.6 |
| WBGene00018349 | 0.212774359 | 0.14398628 | 0.31289979 | F42C5.9 |
| WBGene00018350 | 0.278937906 | 0.05277818 | 0.15677121 | ifo-1 |
| WBGene00018351 | 0.317923994 | 0.16319371 | 0.33876635 | F42G2.2 |
| WBGene00018352 | 0.133275061 | 0.30684081 | 0.50739172 | fbxc-20 |
| WBGene00018353 | 1.946075707 | 1.20E-07 | 2.23E-06 | fbxa-182 |
| WBGene00018354 | 1.600537659 | 0.0004753 | 0.0037293 | F42G2.5 |
| WBGene00018355 | -0.706747938 | 0.00242621 | 0.0146402 | ceh-86 |
| WBGene00018356 | 0.464547542 | 0.05914178 | 0.1698499 | islo-1 |
| WBGene00018357 | 0.002898813 | 0.98401722 | 0.99078441 | moc-3 |
| WBGene00018358 | -0.527258473 | 0.05844245 | 0.16836992 | erp-44.3 |
| WBGene00018359 | 0.145703839 | 0.20637314 | 0.39553725 | F42G8.8 |
| WBGene00018360 | 0.164581923 | 0.16825046 | 0.34609315 | irld-8 |
| WBGene00018361 | -0.066771669 | 0.37379274 | 0.57275766 | F42G8.10 |
| WBGene00018362 | -0.182530106 | 0.00486496 | 0.02561232 | ppm-1.G |
| WBGene00018363 | 0.100982316 | 0.5075155 | 0.68801245 | nlp-52 |
| WBGene00018364 | -0.008726428 | 0.92319081 | 0.95651506 | dagl-2 |
| WBGene00018365 | 0.016985298 | 0.837043 | NA | tpst-2 |
| WBGene00018366 | -0.281247765 | 0.06289264 | 0.17772319 | F42H10.2 |
| WBGene00018367 | -0.301670585 | 0.12694648 | 0.28824805 | F42H10.3 |
| WBGene00018369 | 0.273614925 | 0.21828046 | 0.41024958 | mig-39 |
| WBGene00018370 | -0.115740949 | 0.53488057 | 0.71011847 | F42H10.6 |
| WBGene00018371 | -0.079615971 | 0.42425322 | 0.62026494 | ess-2 |
| WBGene00018373 | 0.594347572 | 0.00261413 | 0.01558364 | F43B10.1 |
| WBGene00018374 | 0.388961185 | 0.07619658 | 0.2038686 | tag-343 |
| WBGene00018375 | 1.157686917 | 0.00435244 | 0.02349709 | F43C9.1 |
| WBGene00018376 | 0.614341322 | 0.0163313 | 0.06617952 | F43C9.2 |
| WBGene00018377 | 0.020925399 | 0.84643798 | NA | hum-10 |
| WBGene00018378 | NA | NA | NA | F43C11.1 |
| WBGene00018379 | NA | NA | NA | F43C11.2 |
| WBGene00018380 | 0.06081927 | 0.72054584 | 0.84342114 | decr-1.1 |
| WBGene00018381 | 0.020925399 | 0.84643798 | NA | F43C11.4 |
| WBGene00018382 | NA | NA | NA | NA |
| WBGene00018383 | -0.141244746 | 0.36684789 | 0.56656328 | F43C11.6 |
| WBGene00018384 | 2.204417051 | 2.13E-18 | 1.40E-16 | F43C11.7 |
| WBGene00018385 | -0.003108065 | 0.95728506 | NA | F43C11.8 |
| WBGene00018386 | -0.118673056 | 0.48263767 | 0.66839748 | F43C11.9 |
| WBGene00018387 | -0.015220816 | 0.89223296 | 0.94374205 | F43C11.10 |
| WBGene00018388 | NA | NA | NA | F43C11.11 |
| WBGene00018389 | NA | NA | NA | F43C11.12 |
| WBGene00018390 | -0.032299407 | 0.82057404 | 0.90148005 | F43E2.1 |
| WBGene00018391 | -0.060107221 | 0.48713551 | 0.67155416 | rpb-4 |

| | | | | |
|---|---|---|---|---|
| WBGene00018392 | 0.719881676 | 0.01658147 | 0.06691546 | insc-1 |
| WBGene00018393 | 1.303310061 | 5.25E-66 | 1.54E-63 | msra-1 |
| WBGene00018394 | 0.265456264 | 0.06566294 | 0.18287448 | F43E2.6 |
| WBGene00018395 | -0.029554546 | 0.60924513 | 0.76442513 | mtch-1 |
| WBGene00018397 | 0.097800217 | 0.43063954 | 0.62563083 | F43E2.11 |
| WBGene00018398 | 0.115364152 | 0.5852114 | 0.74817325 | sptl-2 |
| WBGene00018399 | -0.255897849 | 0.18316138 | 0.36616603 | F43H9.3 |
| WBGene00018400 | -0.07277345 | 0.66619683 | 0.80456881 | F43H9.4 |
| WBGene00018402 | 0.275237303 | 0.10353722 | 0.25126611 | kvs-3 |
| WBGene00018403 | 0.600170197 | 0.01512956 | 0.06231074 | F44A2.3 |
| WBGene00018404 | NA | NA | NA | nhr-39 |
| WBGene00018405 | 0.661450718 | 0.00558291 | 0.02857021 | F44A2.5 |
| WBGene00018406 | -0.006239126 | 0.93041881 | NA | F44A2.7 |
| WBGene00018407 | -0.20348209 | 0.29877312 | 0.49942984 | F44B9.2 |
| WBGene00018408 | 0.000843525 | 0.99454014 | 0.99665038 | F44B9.5 |
| WBGene00018409 | -0.235853899 | 0.00383304 | 0.0211612 | F44B9.8 |
| WBGene00018410 | -0.02791392 | 0.6763823 | NA | F44B9.9 |
| WBGene00018411 | 0.223004298 | 0.14227794 | 0.31044581 | F44B9.10 |
| WBGene00018412 | 0.020925399 | 0.84643798 | NA | nhr-37 |
| WBGene00018413 | 0.314652284 | 0.18066805 | 0.36284785 | cyp-33C4 |
| WBGene00018414 | -0.023056499 | 0.78121253 | NA | nstp-8 |
| WBGene00018415 | 0.034877446 | 0.86196008 | 0.92668569 | nhr-184 |
| WBGene00018417 | -0.090424941 | 0.4405282 | 0.63469528 | F44E2.3 |
| WBGene00018418 | -0.142620268 | 0.45970704 | 0.65098856 | F44E2.4 |
| WBGene00018419 | 0.078191741 | 0.60621487 | 0.76218049 | F44E2.6 |
| WBGene00018420 | -0.254836412 | 0.00250498 | 0.01504324 | F44E2.7 |
| WBGene00018421 | -0.222819718 | 0.01323757 | 0.05599864 | F44E2.8 |
| WBGene00018422 | -0.314375132 | 0.08628326 | 0.22186041 | F44E2.9 |
| WBGene00018423 | -0.107654007 | 0.35177794 | 0.55175334 | F44E2.10 |
| WBGene00018424 | -0.140220476 | 0.3546765 | 0.55451895 | pgph-2 |
| WBGene00018425 | 0.033034883 | 0.64219503 | NA | F44E7.3 |
| WBGene00018426 | -0.251511146 | 7.40E-06 | 9.36E-05 | F44E7.4 |
| WBGene00018427 | -0.136613137 | 0.32476475 | 0.52389607 | F44E7.5 |
| WBGene00018428 | 0.020232312 | 0.84643798 | NA | fbxa-183 |
| WBGene00018429 | 0.053292352 | 0.54948352 | NA | F44E7.7 |
| WBGene00018430 | 0.28580391 | 0.19296751 | 0.37937925 | nhr-142 |
| WBGene00018431 | -0.229882876 | 0.03478806 | 0.11546858 | F44E7.9 |
| WBGene00018432 | NA | NA | NA | irld-33 |
| WBGene00018433 | -0.139607937 | 0.47312638 | 0.66116737 | ceh-82 |
| WBGene00018434 | NA | NA | NA | ceh-81 |
| WBGene00018435 | 0.564715434 | 0.04802521 | 0.14604119 | btb-9 |
| WBGene00018436 | -0.399289368 | 0.09820092 | 0.24302902 | fbxb-11 |
| WBGene00018437 | -0.590087048 | 0.01829607 | 0.07210571 | btb-8 |

| | | | | | |
|---|---|---|---|---|---|
| WBGene00018438 | | -0.536016076 | 0.00145673 | 0.00958563 | btb-6 |
| WBGene00018439 | | 0.06966648 | 0.73563585 | 0.85168784 | fbxa-184 |
| WBGene00018440 | NA | | NA | NA | F45C12.9 |
| WBGene00018441 | | -0.118547189 | 0.37604779 | 0.57512961 | math-29 |
| WBGene00018442 | | -0.160880888 | 0.43237132 | 0.62736252 | sdz-18 |
| WBGene00018443 | | 0.289977068 | 0.1759786 | 0.35695522 | btb-7 |
| WBGene00018444 | | -0.280602326 | 0.19612177 | 0.38373234 | fbxb-48 |
| WBGene00018445 | | -0.17787052 | 0.21116782 | 0.40110379 | fbxb-5 |
| WBGene00018446 | | -0.463049068 | 0.00460739 | 0.0245655 | ceh-83 |
| WBGene00018447 | | 0.033463284 | 0.6365037 | NA | irld-9 |
| WBGene00018448 | | 2.974890507 | 0.0028936 | 0.01697944 | F45D11.1 |
| WBGene00018449 | NA | | NA | NA | NA |
| WBGene00018450 | NA | | NA | NA | NA |
| WBGene00018451 | NA | | NA | NA | NA |
| WBGene00018452 | | -1.481568594 | 0.00787209 | 0.03733571 | F45D11.5 |
| WBGene00018454 | NA | | NA | NA | fbxb-28 |
| WBGene00018455 | | 0.036273277 | 0.85137455 | 0.92034307 | fbxc-42 |
| WBGene00018456 | | -0.871478677 | 0.01730497 | 0.06902885 | fbxc-39 |
| WBGene00018457 | | -0.023056499 | 0.78121253 | NA | fbxb-29 |
| WBGene00018458 | | -0.172943572 | 0.2788166 | 0.47871443 | fbxb-30 |
| WBGene00018459 | NA | | NA | NA | F45D11.16 |
| WBGene00018459 | NA | | NA | NA | F45D11.14 |
| WBGene00018459 | NA | | NA | NA | F45D11.15 |
| WBGene00018460 | NA | | NA | NA | F45D11.16 |
| WBGene00018460 | NA | | NA | NA | F45D11.15 |
| WBGene00018461 | NA | | NA | NA | F45D11.16 |
| WBGene00018461 | NA | | NA | NA | F45D11.15 |
| WBGene00018462 | | -0.084481673 | 0.67743739 | 0.81260807 | F45E1.1 |
| WBGene00018463 | NA | | NA | NA | F45E1.2 |
| WBGene00018465 | | -1.007140192 | 0.00465405 | 0.02473318 | F45E1.4 |
| WBGene00018466 | | -0.111056817 | 0.37073835 | 0.57036433 | F45E1.5 |
| WBGene00018467 | | 0.169191266 | 0.32554562 | 0.52475997 | sdpn-1 |
| WBGene00018468 | | 1.194607389 | 0.00235832 | 0.01427895 | cla-1 |
| WBGene00018470 | | 0.14172135 | 0.45806193 | 0.64973678 | F45E4.5 |
| WBGene00018471 | | -0.014102269 | 0.89579728 | 0.94523157 | F45E4.6 |
| WBGene00018472 | | 0.017141582 | 0.79355725 | NA | nep-16 |
| WBGene00018473 | | -0.42439557 | 0.09068631 | 0.22888093 | cima-1 |
| WBGene00018474 | | -0.156122933 | 0.08933974 | 0.22712679 | cnep-1 |
| WBGene00018475 | | -0.088056845 | 0.49052042 | 0.67440225 | mrpl-14 |
| WBGene00018476 | | 0.100264625 | 0.38646629 | 0.5851299 | F45E12.6 |
| WBGene00018477 | NA | | NA | NA | otpl-4 |
| WBGene00018478 | | 0.01417254 | 0.88154901 | NA | otpl-5 |
| WBGene00018479 | | 0.083000496 | 0.4060317 | 0.60415566 | otpl-6 |

| | | | | |
|---|---|---|---|---|
| WBGene00018480 | 0.017279211 | 0.79380376 | NA | otpl-7 |
| WBGene00018481 | 0.143603334 | 0.32443401 | 0.52363682 | F45F2.9 |
| WBGene00018482 | -0.119633348 | 0.07168948 | 0.19480431 | F45F2.10 |
| WBGene00018483 | -0.333964037 | 0.00011223 | 0.00107537 | F45F2.11 |
| WBGene00018484 | 0.030855962 | 0.84121404 | 0.91420834 | F46C8.1 |
| WBGene00018485 | 0.194428413 | 0.35310427 | 0.55297888 | F46C8.3 |
| WBGene00018486 | 0.021085282 | 0.88229181 | 0.93762317 | glb-16 |
| WBGene00018487 | 2.549572342 | 0.00466192 | 0.02475705 | F46C8.8 |
| WBGene00018488 | -0.61044206 | 0.00032713 | 0.00271051 | acs-1 |
| WBGene00018489 | -0.418213651 | 0.09715815 | 0.24122307 | F46E10.2 |
| WBGene00018490 | 0.020232312 | 0.84643798 | NA | F46E10.3 |
| WBGene00018491 | -0.295721822 | 1.92E-09 | 4.70E-08 | mdh-1 |
| WBGene00018492 | 0.403390551 | 0.10937841 | 0.26039175 | hpo-26 |
| WBGene00018493 | NA | NA | NA | F46F5.1 |
| WBGene00018494 | NA | NA | NA | NA |
| WBGene00018495 | 0.020925399 | 0.84643798 | NA | NA |
| WBGene00018496 | 0.020232312 | 0.84643798 | NA | F46F5.5 |
| WBGene00018497 | 0.14657853 | 0.12995995 | 0.29224835 | msrp-3 |
| WBGene00018498 | NA | NA | NA | F46F5.7 |
| WBGene00018499 | NA | NA | NA | F46F5.8 |
| WBGene00018500 | 0.073458642 | 0.29111983 | NA | msrp-4 |
| WBGene00018501 | NA | NA | NA | F46F5.10 |
| WBGene00018502 | 0.108942554 | 0.15594412 | NA | F46F5.11 |
| WBGene00018503 | NA | NA | NA | NA |
| WBGene00018504 | NA | NA | NA | F46F5.13 |
| WBGene00018505 | 0.020317245 | 0.84643798 | NA | F46F5.14 |
| WBGene00018506 | -0.027728334 | 0.67598605 | NA | F46F5.15 |
| WBGene00018507 | 0.017279211 | 0.79380376 | NA | F46F5.16 |
| WBGene00018508 | 0.162015371 | 0.08152533 | 0.21367925 | F46F11.1 |
| WBGene00018509 | -0.068850643 | 0.41206588 | 0.60964597 | xrep-4 |
| WBGene00018510 | -0.024617809 | 0.90407813 | 0.94889643 | F46F11.7 |
| WBGene00018511 | 0.030384816 | 0.7751599 | 0.87262026 | F46F11.8 |
| WBGene00018512 | -0.175355393 | 0.08119901 | 0.21301463 | trpp-8 |
| WBGene00018513 | 0.016781953 | 0.92261507 | 0.9563868 | F46F11.10 |
| WBGene00018514 | 0.108232392 | 0.47034403 | 0.65840969 | twnk-1 |
| WBGene00018515 | 0.111601271 | 0.49140275 | 0.67504482 | F46G11.2 |
| WBGene00018516 | -0.05923384 | 0.73923142 | 0.85321933 | gakh-1 |
| WBGene00018517 | -0.027315403 | 0.85662138 | 0.92307306 | F46G11.4 |
| WBGene00018518 | 0.092794942 | 0.47327187 | 0.66124376 | F46H5.2 |
| WBGene00018519 | -0.42684554 | 4.18E-14 | 1.82E-12 | F46H5.3 |
| WBGene00018520 | 0.192848076 | 0.14125839 | 0.30896166 | F46H5.4 |
| WBGene00018521 | 0.144683003 | 0.20772647 | 0.3967698 | F46H5.5 |
| WBGene00018522 | 0.271380206 | 0.00439196 | 0.02363198 | F46H5.7 |

| | | | | | |
|---|---|---|---|---|---|
| WBGene00018523 | | -0.289281508 | 0.20566232 | 0.39466399 | lact-1 |
| WBGene00018525 | NA | | NA | NA | F47B3.6 |
| WBGene00018525 | NA | | NA | NA | F47B3.1 |
| WBGene00018526 | NA | | NA | NA | F47B3.2 |
| WBGene00018527 | NA | | NA | NA | F47B3.3 |
| WBGene00018528 | | 0.055503542 | 0.44925503 | NA | F47B3.4 |
| WBGene00018529 | NA | | NA | NA | F47B3.5 |
| WBGene00018530 | NA | | NA | NA | F47B3.6 |
| WBGene00018530 | NA | | NA | NA | F47B3.1 |
| WBGene00018531 | | 0.100692133 | 0.38370663 | 0.58327725 | F47B3.7 |
| WBGene00018532 | | -0.311440401 | 0.02053915 | 0.07815019 | F47B7.1 |
| WBGene00018533 | | -0.314361712 | 0.0049548 | 0.02597321 | F47B7.2 |
| WBGene00018534 | NA | | NA | NA | F47B7.3 |
| WBGene00018535 | | -0.022930767 | 0.78121253 | NA | F47B7.4 |
| WBGene00018536 | NA | | NA | NA | F47B7.5 |
| WBGene00018537 | | -0.005221332 | 0.94448901 | NA | F47B7.6 |
| WBGene00018538 | | 0.862197766 | 0.00150192 | 0.00982992 | marg-1 |
| WBGene00018539 | | 0.020925399 | 0.84643798 | NA | nhr-185 |
| WBGene00018540 | | 1.4811537 | 0.00102003 | 0.00715578 | NA |
| WBGene00018541 | | 0.092724492 | 0.19640118 | NA | nhr-186 |
| WBGene00018542 | | 0.01417254 | 0.88154901 | NA | nhr-187 |
| WBGene00018543 | | 0.220773028 | 0.30289711 | 0.50352867 | ugt-32 |
| WBGene00018544 | | 0.020232312 | 0.84643798 | NA | nhr-188 |
| WBGene00018545 | NA | | NA | NA | nhr-189 |
| WBGene00018547 | | 0.205003558 | 0.34529552 | 0.54496048 | clec-78 |
| WBGene00018548 | | 2.665852878 | 0.00473188 | 0.02506493 | clec-79 |
| WBGene00018549 | NA | | NA | NA | F47C12.6 |
| WBGene00018550 | NA | | NA | NA | F47C12.7 |
| WBGene00018551 | | 0.020317245 | 0.84643798 | NA | F47C12.8 |
| WBGene00018552 | | 0.020925399 | 0.84643798 | NA | srt-19 |
| WBGene00018553 | NA | | NA | NA | NA |
| WBGene00018554 | NA | | NA | NA | srt-3 |
| WBGene00018555 | NA | | NA | NA | srt-35 |
| WBGene00018556 | NA | | NA | NA | srt-36 |
| WBGene00018557 | NA | | NA | NA | srt-38 |
| WBGene00018558 | NA | | NA | NA | srt-20 |
| WBGene00018559 | NA | | NA | NA | srt-37 |
| WBGene00018560 | | 0.058772436 | 0.73880008 | 0.85307041 | F47D12.3 |
| WBGene00018561 | | -0.575551516 | 0.03917008 | 0.12603156 | gadr-3 |
| WBGene00018562 | | 0.229736655 | 0.2940626 | 0.4952117 | F47D12.6 |
| WBGene00018563 | | 0.108564604 | 0.38554444 | 0.5846459 | F47D12.7 |
| WBGene00018564 | | 0.015510664 | 0.84215011 | 0.91488609 | F47D12.9 |
| WBGene00018565 | | 0.020925399 | 0.84643798 | NA | F47E1.1 |

| | | | | |
|---|---|---|---|---|
| WBGene00018566 | 0.109440096 | 0.59907885 | 0.75710949 | F47E1.2 |
| WBGene00018567 | NA | NA | NA | F47E1.3 |
| WBGene00018568 | 0.101109079 | 0.62138937 | 0.7736192 | F47E1.4 |
| WBGene00018569 | 0.572085638 | 0.01938958 | 0.07511518 | F47F2.1 |
| WBGene00018571 | 0.020925399 | 0.84643798 | NA | F47F2.3 |
| WBGene00018572 | 0.559741565 | 0.04286794 | 0.13456893 | lin-42 |
| WBGene00018573 | NA | NA | NA | oac-30 |
| WBGene00018575 | 0.071413559 | 0.63657802 | 0.78445295 | clec-119 |
| WBGene00018576 | -0.139216675 | 0.51029615 | 0.69018438 | F47G3.1 |
| WBGene00018577 | 0.072344087 | 0.72891497 | 0.84861661 | asna-2 |
| WBGene00018578 | -0.225875161 | 0.17741744 | 0.3586316 | F47G3.3 |
| WBGene00018579 | 0.227161397 | 0.30450445 | 0.50541317 | acs-16 |
| WBGene00018580 | 0.092938702 | 0.19548342 | NA | F47G6.3 |
| WBGene00018582 | -0.059402382 | 0.7623558 | 0.86502767 | F48A9.2 |
| WBGene00018585 | -0.033405687 | 0.75063716 | 0.85919975 | F48A11.4 |
| WBGene00018586 | -0.158887673 | 0.01055902 | 0.04663254 | ubxn-3 |
| WBGene00018587 | 0.378775402 | 0.10185016 | 0.24853626 | F48B9.1 |
| WBGene00018589 | -0.047129293 | 0.81031113 | 0.89517582 | F48B9.3 |
| WBGene00018590 | 1.44097124 | 4.46E-05 | 0.0004731 | pdf-2 |
| WBGene00018591 | 0.087256674 | 0.23800185 | NA | npax-2 |
| WBGene00018593 | 0.222414255 | 0.26033882 | 0.45840351 | F48B9.8 |
| WBGene00018594 | 1.3397815 | 6.12E-05 | 0.00062938 | aman-3 |
| WBGene00018595 | 0.113657359 | 0.589458 | 0.75111006 | fbxa-145 |
| WBGene00018596 | -0.023056499 | 0.78121253 | NA | F48C1.3 |
| WBGene00018598 | -0.258673322 | 0.08345049 | 0.21713111 | F48C1.5 |
| WBGene00018599 | -0.047002638 | 0.70798122 | 0.83523278 | F48C1.6 |
| WBGene00018600 | 0.037023624 | 0.59153183 | NA | F48C1.8 |
| WBGene00018601 | 0.05787416 | 0.40594086 | NA | F48C1.9 |
| WBGene00018602 | -1.105118086 | 2.05E-21 | 1.68E-19 | F48D6.4 |
| WBGene00018603 | 0.2558867 | 0.24370669 | 0.44163915 | F48E3.2 |
| WBGene00018604 | -0.203465024 | 0.00646823 | 0.03201533 | uggt-1 |
| WBGene00018605 | 0.283870752 | 0.00010839 | 0.00104334 | F48E3.4 |
| WBGene00018606 | -1.708344736 | 1.73E-05 | 0.00020004 | F48E3.6 |
| WBGene00018607 | 0.91532194 | 0.00168049 | 0.01081494 | F48E3.8 |
| WBGene00018608 | 0.083652064 | 0.57978382 | 0.74533452 | F48E3.9 |
| WBGene00018609 | -0.123166326 | 0.1963196 | 0.38390747 | F48E8.2 |
| WBGene00018610 | -0.126912892 | 0.13840731 | 0.30477085 | F48E8.3 |
| WBGene00018611 | 0.035513447 | 0.71658805 | 0.84107469 | F48E8.4 |
| WBGene00018612 | -0.28988935 | 0.0269523 | 0.09584857 | disl-2 |
| WBGene00018613 | -0.145076044 | 0.07192 | 0.19521332 | skpt-1 |
| WBGene00018614 | 0.243369537 | 0.12803107 | 0.28940913 | pals-31 |
| WBGene00018615 | -0.098728361 | 0.63576031 | 0.7840437 | F48G7.4 |
| WBGene00018616 | 0.394058249 | 0.00488566 | 0.02569357 | F48G7.5 |

| | | | | |
|---|---|---|---|---|
| WBGene00018617 | NA | NA | NA | srt-17 |
| WBGene00018618 | 0.020925399 | 0.84643798 | NA | F48G7.7 |
| WBGene00018619 | 0.048485426 | 0.45975815 | NA | F48G7.8 |
| WBGene00018620 | -0.003614866 | 0.95700117 | NA | F48G7.9 |
| WBGene00018621 | 0.10313377 | 0.32392045 | 0.52363026 | F48G7.10 |
| WBGene00018622 | 0.455022315 | 0.08192085 | 0.21437054 | nhr-190 |
| WBGene00018623 | 0.020317245 | 0.84643798 | NA | F48G7.12 |
| WBGene00018624 | 0.108614162 | 0.30880061 | 0.50965502 | F48G7.13 |
| WBGene00018625 | -0.260319509 | 0.00141097 | 0.0093475 | prp-17 |
| WBGene00018626 | 0.034981685 | 0.68236555 | NA | F49D11.2 |
| WBGene00018627 | -0.009288933 | 0.92571861 | 0.95816777 | F49D11.3 |
| WBGene00018628 | NA | NA | NA | F49D11.4 |
| WBGene00018630 | 0.05787416 | 0.40594086 | NA | F49D11.6 |
| WBGene00018631 | NA | NA | NA | F49D11.7 |
| WBGene00018632 | -0.157372449 | 0.13059074 | 0.29288894 | tag-296 |
| WBGene00018634 | -0.116855799 | 0.56743335 | 0.73659312 | pigx-1 |
| WBGene00018635 | -0.010976791 | 0.91993668 | 0.95544619 | nprl-2 |
| WBGene00018636 | -0.264664054 | 0.00466785 | 0.02477956 | oef-1 |
| WBGene00018637 | -0.299899548 | 0.18250128 | 0.36537689 | F49E8.6 |
| WBGene00018638 | -0.159377097 | 0.16981579 | 0.34826628 | F49E8.7 |
| WBGene00018639 | 0.350312439 | 0.10843793 | 0.25903731 | tbc-16 |
| WBGene00018640 | 0.297999526 | 0.01001424 | 0.04491988 | F49E10.2 |
| WBGene00018641 | 0.778065319 | 0.02211269 | 0.08271637 | F49E10.4 |
| WBGene00018643 | -0.311094877 | 0.15857688 | 0.33295915 | drd-50 |
| WBGene00018644 | NA | NA | NA | NA |
| WBGene00018645 | 8.347750352 | 3.03E-10 | 8.18E-09 | F49F1.5 |
| WBGene00018646 | 0.294099092 | 0.16083976 | 0.33544941 | mul-1 |
| WBGene00018647 | 0.181502243 | 0.35469246 | 0.55451895 | F49F1.7 |
| WBGene00018648 | 0.134614265 | 0.27004599 | 0.46923955 | F49F1.8 |
| WBGene00018649 | NA | NA | NA | F49F1.9 |
| WBGene00018650 | NA | NA | NA | F49F1.10 |
| WBGene00018651 | NA | NA | NA | F49F1.11 |
| WBGene00018652 | 0.020317245 | 0.84643798 | NA | F49F1.12 |
| WBGene00018653 | 0.063217264 | 0.41230173 | NA | F49H12.2 |
| WBGene00018654 | 0.313150022 | 0.09663767 | 0.24021597 | F49H12.3 |
| WBGene00018655 | 0.0627133 | 0.50876942 | NA | F49H12.4 |
| WBGene00018656 | -0.845835323 | 4.35E-06 | 5.76E-05 | txdc-12.2 |
| WBGene00018657 | 0.242428649 | 0.00451504 | 0.02420531 | acl-4 |
| WBGene00018658 | 0.3965005 | 0.10425254 | 0.25244404 | NA |
| WBGene00018659 | -0.140081034 | 0.35445778 | 0.5544462 | F52C6.2 |
| WBGene00018660 | 0.194072979 | 0.37322223 | 0.57229033 | F52C6.3 |
| WBGene00018661 | 0.092172396 | 0.65296291 | 0.79443429 | F52C6.4 |
| WBGene00018662 | 0.020317245 | 0.84643798 | NA | math-30 |

| | | | | |
|---|---|---|---|---|
| WBGene00018663 | -0.023109762 | 0.78121253 | NA | math-31 |
| WBGene00018664 | 0.674546305 | 0.02703781 | 0.09603604 | NA |
| WBGene00018665 | -0.093522779 | 0.63130868 | 0.78093021 | bath-4 |
| WBGene00018666 | 0.060861088 | 0.76980602 | 0.86905158 | NA |
| WBGene00018667 | 0.064558723 | 0.75013593 | 0.85883173 | bath-7 |
| WBGene00018668 | -0.093750394 | 0.56838929 | 0.73735492 | bath-2 |
| WBGene00018669 | 0.954503602 | 0.01130801 | 0.04924465 | F52C6.12 |
| WBGene00018670 | 0.166089675 | 0.27890129 | 0.47876583 | F52C6.13 |
| WBGene00018671 | 0.030765213 | 0.70446454 | NA | F52C6.14 |
| WBGene00018672 | 0.170497822 | 0.1696411 | 0.34800552 | sorf-2 |
| WBGene00018674 | -0.125173851 | 0.39303283 | 0.59122507 | F52C9.3 |
| WBGene00018675 | -0.082336446 | 0.39430161 | NA | F52C9.5 |
| WBGene00018678 | 0.015560945 | 0.90435405 | 0.94901431 | F52C12.1 |
| WBGene00018679 | -0.220380751 | 0.12822701 | 0.28971392 | F52C12.2 |
| WBGene00018680 | -0.044640635 | 0.80971616 | 0.89476527 | tsen-54 |
| WBGene00018681 | -0.109107012 | 0.33763511 | 0.53731692 | denn-4 |
| WBGene00018682 | -0.339347363 | 0.03925987 | 0.126275 | aagr-4 |
| WBGene00018683 | 0.364232966 | 0.06773459 | 0.18693725 | F52D1.2 |
| WBGene00018684 | NA | NA | NA | fbxa-41 |
| WBGene00018687 | -0.03147554 | 0.70637395 | NA | F52D2.5 |
| WBGene00018688 | -0.048825734 | 0.75513221 | 0.8612519 | F52D2.6 |
| WBGene00018689 | -0.310968996 | 0.08318909 | 0.2166438 | F52D2.7 |
| WBGene00018690 | -0.129763116 | 0.3126595 | 0.51333563 | fbxa-46 |
| WBGene00018691 | NA | NA | NA | fbxa-17 |
| WBGene00018692 | 0.364393396 | 0.1300808 | 0.29227968 | F52E1.2 |
| WBGene00018693 | -0.002914971 | 0.95718083 | NA | F52E1.3 |
| WBGene00018694 | 0.247187107 | 0.22789394 | 0.42139368 | F52E1.5 |
| WBGene00018696 | 0.2905668 | 0.15575684 | 0.32930481 | pho-6 |
| WBGene00018697 | 0.236469572 | 0.28752014 | 0.48778475 | F52E1.9 |
| WBGene00018698 | -0.139046218 | 0.14808202 | 0.31878678 | vha-18 |
| WBGene00018699 | 0.05787416 | 0.40594086 | NA | NA |
| WBGene00018700 | 0.13397748 | 0.03824429 | 0.12378446 | lmd-3 |
| WBGene00018701 | -0.027666327 | 0.63362755 | 0.78260005 | pccb-1 |
| WBGene00018702 | 0.160857408 | 0.45093779 | 0.64323657 | F52E4.5 |
| WBGene00018703 | -0.02857628 | 0.84178899 | 0.9146295 | sec-3 |
| WBGene00018704 | -0.288310482 | 0.18174306 | 0.36443528 | ztf-13 |
| WBGene00018706 | 0.207676267 | 0.06493234 | 0.1816344 | F52F10.2 |
| WBGene00018707 | -0.215748808 | 0.22458802 | 0.41754834 | oac-31 |
| WBGene00018708 | 0.265056598 | 0.2155555 | 0.40653717 | oac-32 |
| WBGene00018709 | -0.022930767 | 0.78121253 | NA | srsx-8 |
| WBGene00018710 | 0.107025413 | 0.35838916 | 0.55876045 | F52G3.1 |
| WBGene00018711 | -0.133386211 | 0.26309007 | 0.46113362 | F52G3.3 |
| WBGene00018712 | -0.004944805 | 0.94514249 | NA | F52G3.4 |

| | | | | |
|---|---|---|---|---|
| WBGene00018713 | -0.123830601 | 0.43796005 | 0.63286351 | F52G3.5 |
| WBGene00018714 | -0.063981495 | 0.75245784 | 0.85987631 | F52H2.1 |
| WBGene00018715 | 0.167270207 | 0.40872412 | 0.60654062 | F52H2.3 |
| WBGene00018716 | 0.162301498 | 0.43748168 | 0.63242192 | F52H2.4 |
| WBGene00018717 | 0.029934296 | 0.87905637 | 0.93527101 | F52H2.5 |
| WBGene00018718 | 0.199060785 | 0.19198401 | 0.37820747 | F52H2.6 |
| WBGene00018719 | 0.185449876 | 0.3198361 | 0.51991615 | F52H2.7 |
| WBGene00018720 | 0.159887966 | 0.34719177 | 0.54681957 | F53A3.1 |
| WBGene00018721 | 0.094692813 | 0.30656783 | 0.50711327 | polh-1 |
| WBGene00018723 | 0.335664605 | 0.1185546 | 0.27591129 | F53A3.7 |
| WBGene00018724 | 0.287105622 | 0.21269842 | 0.40320754 | F53A9.1 |
| WBGene00018725 | -0.280284213 | 0.19764706 | 0.38547364 | kreg-1 |
| WBGene00018726 | 0.075726029 | 0.27693214 | NA | F53A9.3 |
| WBGene00018727 | -0.001488154 | 0.95639912 | NA | jbts-14 |
| WBGene00018728 | 0.126230452 | 0.45583044 | 0.64769152 | frpr-8 |
| WBGene00018729 | 1.757520919 | 3.65E-10 | 9.70E-09 | F53A9.6 |
| WBGene00018730 | 2.581465363 | 1.66E-11 | 5.32E-10 | F53A9.7 |
| WBGene00018731 | 4.026524366 | 2.70E-38 | 4.65E-36 | F53A9.8 |
| WBGene00018732 | 2.559343855 | 1.82E-07 | 3.27E-06 | F53A9.9 |
| WBGene00018734 | 0.305358771 | 0.06994528 | 0.19109846 | F53A10.2 |
| WBGene00018735 | -0.291446529 | 0.03384929 | 0.1131433 | F53B1.2 |
| WBGene00018736 | 0.153838603 | 0.19309915 | 0.37948519 | F53B1.3 |
| WBGene00018737 | 2.3948127 | 0.00132578 | 0.00886335 | F53B1.4 |
| WBGene00018738 | 0.211652051 | 0.23408784 | 0.42959139 | swip-10 |
| WBGene00018739 | -0.165470437 | 0.32380257 | 0.52355531 | F53B1.8 |
| WBGene00018740 | -0.216141733 | 0.09997746 | 0.24539492 | tra-4 |
| WBGene00018741 | 0.058314668 | 0.40208118 | NA | comt-1 |
| WBGene00018742 | 0.344784925 | 0.11192838 | 0.26456948 | F53B3.3 |
| WBGene00018743 | -0.694190165 | 0.00020634 | 0.00181659 | F53B3.5 |
| WBGene00018744 | -0.57471044 | 0.04312662 | 0.13506244 | F53B3.6 |
| WBGene00018745 | 0.129818876 | 0.14983923 | 0.32120003 | F53C3.1 |
| WBGene00018746 | -0.501133665 | 0.05819528 | 0.16792217 | fbxb-103 |
| WBGene00018747 | -0.031763797 | 0.75218228 | 0.85970214 | F53C3.3 |
| WBGene00018748 | 0.085486216 | 0.53592025 | 0.71117633 | F53C3.4 |
| WBGene00018749 | NA | NA | NA | F53C3.5 |
| WBGene00018750 | 0.073695829 | 0.43812771 | NA | F53C3.6 |
| WBGene00018751 | -0.045771477 | 0.50913401 | NA | F53C3.7 |
| WBGene00018752 | -0.02989064 | 0.74575433 | NA | F53C3.8 |
| WBGene00018754 | -0.537292927 | 0.04927891 | 0.14866937 | F53C3.11 |
| WBGene00018755 | 0.791547006 | 0.00283936 | 0.01671536 | bcmo-2 |
| WBGene00018756 | 0.105449885 | 0.21739024 | 0.40904904 | plpp-1.1 |
| WBGene00018757 | -0.007430444 | 0.96107789 | 0.97890942 | tag-304 |
| WBGene00018758 | 0.340728928 | 0.08527366 | 0.22015369 | F53E10.1 |

| | | | | |
|---|---|---|---|---|
| WBGene00018759 | NA | NA | NA | NA |
| WBGene00018760 | 0.454130429 | 0.0582772 | 0.16812541 | irg-3 |
| WBGene00018761 | -0.014893655 | 0.89933725 | 0.94665211 | F53E10.5 |
| WBGene00018762 | -0.360373031 | 0.00835981 | 0.03911664 | F53E10.6 |
| WBGene00018763 | 0.677972319 | 0.03242276 | 0.10952967 | F53F10.1 |
| WBGene00018764 | 0.200662143 | 0.0035721 | 0.02007644 | F53F10.2 |
| WBGene00018765 | 0.210100436 | 0.08543803 | 0.22030613 | mpc-2 |
| WBGene00018766 | -0.297245696 | 0.14169791 | 0.30966242 | F53G2.1 |
| WBGene00018767 | -0.031720952 | 0.73904865 | NA | F53G2.2 |
| WBGene00018768 | -0.022930767 | 0.78121253 | NA | F53G2.3 |
| WBGene00018769 | -0.537897529 | 3.18E-07 | 5.45E-06 | mnat-1 |
| WBGene00018771 | 0.037127706 | 0.59421403 | NA | duox-2 |
| WBGene00018772 | 0.524073075 | 0.04606111 | 0.14173616 | F53G12.4 |
| WBGene00018773 | 0.166646557 | 0.31624929 | 0.51687942 | F53G12.8 |
| WBGene00018774 | 0.032962945 | 0.83678515 | 0.91128915 | F53G12.9 |
| WBGene00018776 | 0.002520325 | 0.97431152 | 0.98568926 | F53H1.1 |
| WBGene00018777 | -0.103743751 | 0.49129147 | 0.67495527 | F53H1.3 |
| WBGene00018778 | -0.097543647 | 0.1726349 | 0.35261152 | F53H1.4 |
| WBGene00018779 | -0.844879733 | 0.00142308 | 0.00940917 | F53H8.3 |
| WBGene00018781 | -0.086022667 | 0.56643889 | 0.73547639 | F54A3.2 |
| WBGene00018782 | -0.454574745 | 4.82E-15 | 2.32E-13 | cct-3 |
| WBGene00018783 | 0.319652108 | 0.07250629 | 0.19622782 | cbs-2 |
| WBGene00018784 | -0.279464885 | 0.00953403 | 0.04337716 | F54A3.5 |
| WBGene00018785 | -0.141739088 | 0.29837853 | 0.49925925 | F54A3.6 |
| WBGene00018786 | -0.359853116 | 0.05802704 | 0.16770111 | hmbx-1 |
| WBGene00018787 | 0.402346323 | 0.06950565 | 0.19038957 | cutl-20 |
| WBGene00018788 | 0.20085754 | 0.32156788 | 0.52161514 | shc-1 |
| WBGene00018789 | -0.055564708 | 0.73484235 | 0.85124035 | F54C1.1 |
| WBGene00018791 | -0.027728334 | 0.67598605 | NA | F54C1.6 |
| WBGene00018792 | 0.1161173 | 0.12124304 | NA | F54C1.8 |
| WBGene00018793 | -0.09816825 | 0.43552402 | 0.63071288 | mrpl-40 |
| WBGene00018794 | 0.210697331 | 0.07488082 | 0.2011984 | attf-3 |
| WBGene00018795 | 0.009302745 | 0.96421939 | 0.98045715 | F54C4.4 |
| WBGene00018797 | -0.137711177 | 0.12580802 | 0.28690032 | ept-1 |
| WBGene00018798 | 0.204385101 | 0.06841036 | 0.18834313 | gnrr-1 |
| WBGene00018802 | -0.161929353 | 0.06133852 | 0.17444778 | F54D8.6 |
| WBGene00018803 | 3.917744712 | 1.73E-07 | 3.14E-06 | fbxa-24 |
| WBGene00018804 | 0.270276953 | 0.22468646 | 0.41759166 | lido-15 |
| WBGene00018805 | 0.318738412 | 0.17667391 | 0.35749508 | F54D10.4 |
| WBGene00018806 | -0.401602131 | 0.00167601 | 0.01079563 | F54D10.5 |
| WBGene00018807 | 0.54338802 | 0.05132581 | 0.15332813 | NA |
| WBGene00018808 | -0.020798608 | 0.8882677 | 0.94130917 | F54D10.7 |
| WBGene00018809 | 0.020232312 | 0.84643798 | NA | F54D10.8 |

| | | | | | |
|---|---|---|---|---|---|
| WBGene00018810 | | -0.022930767 | 0.78121253 | NA | F54D10.9 |
| WBGene00018811 | | 0.011816894 | 0.82695986 | 0.90530277 | pmt-2 |
| WBGene00018812 | | -0.02052286 | 0.83021256 | 0.90730329 | sumv-2 |
| WBGene00018813 | | -0.301879966 | 0.12955192 | 0.29199041 | F54D11.3 |
| WBGene00018814 | | 0.038451716 | 0.84292826 | 0.91545974 | F54D11.4 |
| WBGene00018815 | NA | | NA | NA | F54D12.1 |
| WBGene00018816 | NA | | NA | NA | F54D12.2 |
| WBGene00018819 | | 0.107297512 | 0.26266261 | 0.46064466 | rog-1 |
| WBGene00018820 | NA | | NA | NA | F54D12.7 |
| WBGene00018821 | | 0.036687355 | 0.59489568 | NA | F54D12.8 |
| WBGene00018822 | | 0.020232312 | 0.84643798 | NA | F54D12.9 |
| WBGene00018823 | | -0.836271167 | 3.10E-07 | 5.35E-06 | F54E2.1 |
| WBGene00018824 | NA | | NA | NA | F54E2.2 |
| WBGene00018825 | NA | | NA | NA | F54E2.5 |
| WBGene00018826 | NA | | NA | NA | srt-34 |
| WBGene00018827 | | 0.030334142 | 0.83211504 | 0.9084712 | pst-2 |
| WBGene00018828 | | -1.289540428 | 0.0006367 | 0.00478786 | sdz-21 |
| WBGene00018829 | NA | | NA | NA | F54E7.6 |
| WBGene00018830 | | -0.183784152 | 0.08924937 | 0.22697585 | ska-3 |
| WBGene00018831 | | -0.103051657 | 0.56576579 | 0.73512398 | F54E7.9 |
| WBGene00018833 | | -0.378786418 | 0.03736765 | 0.12174004 | ztf-1 |
| WBGene00018834 | NA | | NA | NA | F54F2.6 |
| WBGene00018835 | | -0.173018771 | 0.38450074 | 0.58378879 | F54F2.7 |
| WBGene00018836 | | -0.093688336 | 0.46605979 | 0.65560147 | F54F2.9 |
| WBGene00018837 | | 0.523975162 | 0.01157533 | 0.05011068 | F54G2.1 |
| WBGene00018838 | | 0.206003443 | 0.20015622 | 0.3887619 | F54G2.2 |
| WBGene00018839 | | 0.085537126 | 0.24698522 | NA | F54H5.2 |
| WBGene00018840 | | 0.116737817 | 0.11962812 | NA | F54H5.3 |
| WBGene00018841 | | 0.109036101 | 0.21396767 | NA | F54H5.5 |
| WBGene00018842 | | 0.012876793 | 0.86436819 | NA | F54H12.2 |
| WBGene00018844 | | 0.018860755 | 0.77262519 | NA | F54H12.4 |
| WBGene00018845 | | -0.509094012 | 0.01125406 | 0.0490681 | F54H12.5 |
| WBGene00018846 | | 0.019081625 | 0.7159199 | 0.84069484 | eef-1B.1 |
| WBGene00018847 | | -0.009852926 | 0.8934574 | 0.94462086 | 6-Mar |
| WBGene00018848 | | -0.067816854 | 0.60824316 | 0.76399837 | F55A3.2 |
| WBGene00018849 | | -0.180982512 | 0.00055236 | 0.00422305 | spt-16 |
| WBGene00018850 | | 0.221576165 | 0.18588011 | 0.37006672 | gpx-8 |
| WBGene00018851 | | -0.017080194 | 0.88641162 | 0.94033043 | F55A3.6 |
| WBGene00018852 | | 0.775660256 | 0.00161902 | 0.01049318 | F55A3.7 |
| WBGene00018853 | | -0.399519236 | 0.02558442 | 0.09223873 | sec-22 |
| WBGene00018854 | | 0.144338594 | 0.49658352 | 0.67886649 | nlf-1 |
| WBGene00018855 | | -0.023056499 | 0.78121253 | NA | F55A4.3 |
| WBGene00018856 | | 0.012101543 | 0.92831319 | 0.95971144 | NA |

| | | | | |
|---|---|---|---|---|
| WBGene00018857 | -0.201969336 | 0.19472972 | 0.38171613 | stau-1 |
| WBGene00018859 | 0.077911454 | 0.52353693 | 0.70116211 | F55A4.7 |
| WBGene00018860 | -0.093544222 | 0.61238241 | 0.7668267 | vpat-1 |
| WBGene00018861 | 0.118954819 | 0.42848268 | 0.62341398 | cutl-21 |
| WBGene00018862 | 0.050032281 | 0.46003352 | 0.65119895 | wago-2 |
| WBGene00018863 | -0.164405782 | 0.33852444 | 0.53803042 | F55A12.2 |
| WBGene00018864 | -0.030855043 | 0.70118875 | 0.83025511 | F55A12.5 |
| WBGene00018865 | 0.013869033 | 0.8872176 | NA | F55A12.6 |
| WBGene00018866 | -0.217819153 | 0.06374843 | 0.1792959 | nath-10 |
| WBGene00018867 | -0.058481123 | 0.71400018 | 0.83958531 | zhp-1 |
| WBGene00018868 | NA | NA | NA | F55C7.2 |
| WBGene00018869 | 0.343467451 | 0.00977268 | 0.04410661 | rfip-1 |
| WBGene00018870 | NA | NA | NA | F55C12.2 |
| WBGene00018871 | 0.189078077 | 0.35226952 | 0.552085 | F55C12.4 |
| WBGene00018872 | 0.078266152 | 0.31674333 | 0.51698682 | F55C12.5 |
| WBGene00018873 | 0.119971181 | 0.4228963 | 0.6187912 | F55C12.6 |
| WBGene00018874 | 0.334752644 | 0.13703492 | 0.30247674 | tag-234 |
| WBGene00018875 | NA | NA | NA | F55D1.1 |
| WBGene00018877 | -0.181412949 | 0.20763828 | 0.39671566 | aman-1 |
| WBGene00018878 | -0.434019891 | 0.0146641 | 0.06073466 | glit-1 |
| WBGene00018879 | 0.218385837 | 0.24477368 | 0.44292812 | F55D10.4 |
| WBGene00018880 | 0.038569885 | 0.77101525 | 0.86974607 | acc-3 |
| WBGene00018881 | NA | NA | NA | F55E10.1 |
| WBGene00018882 | NA | NA | NA | F55E10.2 |
| WBGene00018883 | NA | NA | NA | F55E10.4 |
| WBGene00018884 | NA | NA | NA | F55E10.5 |
| WBGene00018885 | -0.285742418 | 0.13310461 | 0.29668448 | drd-5 |
| WBGene00018886 | 0.417946616 | 0.08932409 | 0.22712642 | npr-28 |
| WBGene00018888 | NA | NA | NA | F55F1.2 |
| WBGene00018890 | -0.355420459 | 0.00030966 | 0.00258483 | F55F8.2 |
| WBGene00018891 | -0.352881841 | 0.0015363 | 0.01002357 | F55F8.3 |
| WBGene00018892 | -0.161634602 | 0.05102165 | 0.15257515 | cir-1 |
| WBGene00018893 | -0.199907795 | 0.05341239 | 0.15810973 | wdr-12 |
| WBGene00018894 | NA | NA | NA | mff-2 |
| WBGene00018895 | 0.149813019 | 0.07294962 | NA | F55F8.7 |
| WBGene00018896 | -0.008657085 | 0.92287164 | NA | F55F8.8 |
| WBGene00018897 | -0.229670671 | 0.01007909 | 0.0451295 | zipt-2.4 |
| WBGene00018898 | -0.166718117 | 0.01410908 | 0.05883538 | F55F10.1 |
| WBGene00018899 | 0.115495999 | 0.4007278 | 0.59903477 | F55G1.1 |
| WBGene00018900 | -0.135453493 | 0.01403287 | 0.05860104 | rod-1 |
| WBGene00018901 | -0.03389328 | 0.84016716 | 0.91361314 | slc-25A18.2 |
| WBGene00018902 | -0.239583252 | 0.03981651 | 0.12750514 | F55G1.6 |
| WBGene00018903 | 1.033017826 | 0.00054469 | 0.00417094 | F55G1.7 |

| | | | | | |
|---|---|---|---|---|---|
| WBGene00018904 | | -0.08467363 | 0.54471268 | 0.71809278 | F55G1.9 |
| WBGene00018906 | | 0.187630164 | 0.25113956 | 0.44967521 | egas-4 |
| WBGene00018907 | | 1.403287618 | 1.24E-05 | 0.00014915 | F55G1.15 |
| WBGene00018908 | | -0.078296355 | 0.5347503 | 0.71000989 | fncm-1 |
| WBGene00018909 | | -0.039546067 | 0.70386886 | 0.83192667 | slx-1 |
| WBGene00018910 | NA | | NA | NA | clec-209 |
| WBGene00018910 | NA | | NA | NA | F56A4.2 |
| WBGene00018911 | NA | | NA | NA | F56A4.3 |
| WBGene00018912 | NA | | NA | NA | F56A4.4 |
| WBGene00018913 | NA | | NA | NA | F56A4.5 |
| WBGene00018914 | NA | | NA | NA | NA |
| WBGene00018915 | NA | | NA | NA | str-267 |
| WBGene00018915 | NA | | NA | NA | str-154 |
| WBGene00018917 | NA | | NA | NA | irld-35 |
| WBGene00018918 | NA | | NA | NA | F56A4.10 |
| WBGene00018919 | NA | | NA | NA | NA |
| WBGene00018920 | NA | | NA | NA | F56A4.12 |
| WBGene00018921 | | 0.231202686 | 0.2776537 | 0.47761423 | sago-2 |
| WBGene00018923 | | 0.061384931 | 0.59572663 | 0.75477619 | eme-1 |
| WBGene00018924 | | 0.111272776 | 0.28369516 | 0.48359306 | F56A11.4 |
| WBGene00018925 | | -0.08102854 | 0.46046786 | 0.6515443 | F56A11.5 |
| WBGene00018926 | | 0.104881425 | 0.42162007 | 0.61824826 | F56A11.6 |
| WBGene00018928 | | 0.001458058 | 0.99492469 | 0.99689928 | F56B3.2 |
| WBGene00018929 | | 0.003456717 | 0.97812168 | 0.98749686 | F56B3.3 |
| WBGene00018930 | | 1.0882402 | 0.01589809 | 0.06481871 | F56B3.6 |
| WBGene00018931 | | 0.165815219 | 0.38279534 | 0.5823469 | ugt-52 |
| WBGene00018932 | | -0.292947166 | 0.04047921 | 0.12911897 | mrpl-2 |
| WBGene00018933 | NA | | NA | NA | F56B3.9 |
| WBGene00018934 | | -0.295812289 | 0.009409 | 0.04290165 | F56B3.11 |
| WBGene00018935 | | -0.274050554 | 0.01646721 | 0.06656443 | skr-18 |
| WBGene00018937 | | 0.052425971 | 0.65297236 | 0.79443429 | F56B6.6 |
| WBGene00018938 | NA | | NA | NA | F56C3.1 |
| WBGene00018939 | | -0.088405548 | 0.64496639 | 0.78968348 | fbxa-190 |
| WBGene00018940 | NA | | NA | NA | F56C3.3 |
| WBGene00018941 | | 0.0328853 | 0.6130758 | NA | F56C3.4 |
| WBGene00018942 | NA | | NA | NA | F56C3.5 |
| WBGene00018943 | | -0.005221332 | 0.94448901 | NA | dgn-2 |
| WBGene00018944 | NA | | NA | NA | F56C3.7 |
| WBGene00018945 | | 0.015896709 | 0.80422001 | NA | F56C3.8 |
| WBGene00018946 | | 0.025105328 | 0.8457329 | 0.9169413 | F56C3.9 |
| WBGene00018948 | | -0.161461618 | 0.03709105 | 0.12105439 | F56C9.3 |
| WBGene00018949 | | 0.034400479 | 0.75985204 | 0.8640619 | acbp-4 |
| WBGene00018950 | | -0.21552752 | 0.00305178 | 0.01771796 | F56C9.6 |

| | | | | |
|---|---|---|---|---|
| WBGene00018951 | 0.838607234 | 4.65E-17 | 2.74E-15 | F56C9.7 |
| WBGene00018952 | 0.699120015 | 0.0066034 | 0.03255231 | F56C9.8 |
| WBGene00018953 | -0.080061371 | 0.15421147 | 0.32715947 | F56C9.10 |
| WBGene00018954 | -0.273676941 | 0.09003218 | 0.22813449 | F56C9.11 |
| WBGene00018955 | 0.017567764 | 0.92890291 | 0.95998218 | F56C11.3 |
| WBGene00018957 | -0.069199135 | 0.63862198 | 0.78565367 | F56C11.5 |
| WBGene00018958 | 0.345083885 | 0.13158373 | 0.29439872 | F56C11.6 |
| WBGene00018959 | -0.116658429 | 0.20534357 | 0.39439004 | F56D1.1 |
| WBGene00018960 | -0.240554599 | 0.00957593 | 0.04351372 | ilcr-2 |
| WBGene00018961 | -0.287925563 | 0.03365429 | 0.11262782 | mrps-16 |
| WBGene00018963 | -0.104432864 | 0.09947926 | 0.24461044 | ucr-1 |
| WBGene00018964 | -0.088824232 | 0.43619634 | 0.63124945 | F56D2.2 |
| WBGene00018965 | 0.080388946 | 0.40624029 | 0.60428981 | F56D2.3 |
| WBGene00018966 | 0.794188145 | 0.01867731 | 0.07305758 | F56D2.5 |
| WBGene00018967 | -0.361390303 | 5.25E-06 | 6.84E-05 | ddx-15 |
| WBGene00018968 | NA | NA | NA | F56D2.8 |
| WBGene00018969 | 2.071548249 | 0.00334317 | 0.019033 | F56D3.1 |
| WBGene00018970 | -0.027350315 | 0.68023343 | NA | clec-68 |
| WBGene00018971 | 1.797079741 | 2.78E-05 | 0.00030761 | clec-67 |
| WBGene00018972 | -0.003421551 | 0.95689871 | NA | srz-19 |
| WBGene00018973 | NA | NA | NA | srz-20 |
| WBGene00018974 | -0.058311164 | 0.48259026 | 0.66839748 | fcho-1 |
| WBGene00018975 | -0.060840329 | 0.48261263 | 0.66839748 | F56E10.1 |
| WBGene00018976 | -0.010524949 | 0.95181351 | 0.97233475 | daam-1 |
| WBGene00018978 | 0.013755544 | 0.88500544 | NA | ttr-58 |
| WBGene00018979 | -0.001488154 | 0.95639912 | NA | ttr-55 |
| WBGene00018980 | -0.003448773 | 0.96209218 | NA | F56F4.3 |
| WBGene00018981 | 0.020317245 | 0.84643798 | NA | F56F4.4 |
| WBGene00018982 | NA | NA | NA | abt-6 |
| WBGene00018984 | 0.429370616 | 0.00020797 | 0.0018266 | F56F10.1 |
| WBGene00018985 | 0.172943961 | 0.4017238 | 0.60010592 | F56F10.2 |
| WBGene00018986 | -0.130050609 | 0.43879833 | 0.6335804 | cdo-1 |
| WBGene00018987 | -0.057320952 | 0.73405017 | 0.85093534 | lgl-1 |
| WBGene00018988 | 0.177616853 | 0.0463229 | NA | F56F11.1 |
| WBGene00018989 | -0.136852922 | 0.50840505 | 0.68889943 | F56F11.2 |
| WBGene00018990 | 1.068594891 | 2.36E-06 | 3.36E-05 | klf-1 |
| WBGene00018991 | -0.171473069 | 0.04401779 | 0.13709778 | F56F11.4 |
| WBGene00018992 | 0.207440471 | 0.26374733 | 0.46190487 | NA |
| WBGene00018993 | 0.020925399 | 0.84643798 | NA | nhr-266 |
| WBGene00018994 | NA | NA | NA | F56H1.3 |
| WBGene00018995 | -0.141027649 | 0.12063804 | 0.27938963 | ccpp-1 |
| WBGene00018996 | -0.020370431 | 0.89576927 | 0.94523157 | F57B9.1 |
| WBGene00018997 | 0.01912147 | 0.77587372 | NA | F57B9.3 |

| | | | | |
|---|---|---|---|---|
| WBGene00018998 | -0.195121566 | 0.07690199 | 0.20523552 | flap-1 |
| WBGene00018999 | 0.094173262 | 0.55800934 | 0.72887515 | F57B9.8 |
| WBGene00019001 | -0.043720748 | 0.50281943 | 0.68411665 | ipgm-1 |
| WBGene00019002 | -0.151195759 | 0.10246635 | 0.2494529 | F57B10.4 |
| WBGene00019003 | -0.130865335 | 0.19972139 | 0.38822634 | tmed-3 |
| WBGene00019004 | 0.093185969 | 0.38154465 | 0.58074595 | xpg-1 |
| WBGene00019005 | -0.387648437 | 0.01021637 | 0.0456169 | F57B10.8 |
| WBGene00019006 | 0.22547065 | 0.15654001 | 0.33048339 | spg-20 |
| WBGene00019007 | -0.200373636 | 0.01554407 | 0.06365895 | ucr-11 |
| WBGene00019008 | -0.075200874 | 0.54437268 | 0.71783846 | pdxk-1 |
| WBGene00019009 | 0.163048515 | 0.41176785 | 0.60945138 | clec-90 |
| WBGene00019010 | -0.062508836 | 0.73401499 | 0.85093534 | glo-2 |
| WBGene00019011 | -0.109665104 | 0.27595644 | 0.47597246 | F57C9.4 |
| WBGene00019012 | -0.131352738 | 0.34964705 | 0.54956334 | F57C9.6 |
| WBGene00019013 | -0.129941207 | 0.44025055 | 0.63450679 | tofu-4 |
| WBGene00019014 | 0.444760158 | 0.08649477 | 0.22217016 | F57C12.2 |
| WBGene00019015 | 0.169787267 | 0.27502886 | 0.4753509 | F57F4.1 |
| WBGene00019016 | -0.045771477 | 0.50913401 | NA | F57F4.2 |
| WBGene00019017 | 0.465162168 | 0.00271628 | 0.01608782 | F57F4.4 |
| WBGene00019018 | 0.485337468 | 0.01924383 | 0.0747543 | abts-3 |
| WBGene00019019 | -0.006084888 | 0.94102417 | NA | frpr-10 |
| WBGene00019020 | 0.178768928 | 0.09068156 | 0.22888093 | F57H12.5 |
| WBGene00019021 | -0.00946502 | 0.91279816 | NA | F57H12.6 |
| WBGene00019022 | 0.422045966 | 0.04064391 | 0.12954359 | F58A6.1 |
| WBGene00019023 | 0.106065124 | 0.22635466 | NA | F58A6.2 |
| WBGene00019024 | 0.197193241 | 0.07248009 | 0.1962243 | F58A6.5 |
| WBGene00019025 | 0.059892517 | 0.69610016 | 0.82650251 | srb-16 |
| WBGene00019026 | 2.292093509 | 0.00710298 | 0.0343898 | F58A6.9 |
| WBGene00019027 | 0.020317245 | 0.84643798 | NA | srb-12 |
| WBGene00019028 | NA | NA | NA | srb-13 |
| WBGene00019029 | -0.071490547 | 0.65037334 | 0.79304706 | F58B6.1 |
| WBGene00019030 | 0.394474343 | 0.10078156 | 0.24671128 | exc-6 |
| WBGene00019032 | -0.023056499 | 0.78121253 | NA | srsx-17 |
| WBGene00019034 | -0.220126429 | 0.12605256 | 0.28699798 | fbxb-25 |
| WBGene00019035 | -0.469878003 | 0.07685254 | 0.20521578 | fbxb-26 |
| WBGene00019036 | -0.022930767 | 0.78121253 | NA | irld-36 |
| WBGene00019037 | -0.982841358 | 0.00216045 | 0.01329049 | fbxb-17 |
| WBGene00019038 | 0.020925399 | 0.84643798 | NA | irld-37 |
| WBGene00019039 | -0.323241267 | 0.12766344 | 0.28897967 | fbxb-18 |
| WBGene00019040 | -0.071737948 | 0.65401053 | 0.7949984 | fbxb-19 |
| WBGene00019041 | -0.141317545 | 0.42707153 | 0.62241233 | fbxc-22 |
| WBGene00019042 | -1.011555474 | 0.01258567 | 0.05375343 | fbxc-21 |
| WBGene00019043 | -0.26895114 | 0.21626519 | 0.40756065 | fbxc-26 |

| | | | | | |
|---|---|---|---|---|---|
| WBGene00019044 | | 0.016338669 | 0.93367219 | 0.96273643 | F58E1.13 |
| WBGene00019045 | | 0.007407455 | 0.96942906 | 0.98346632 | fbxb-47 |
| WBGene00019046 | | 0.020232312 | 0.84643798 | NA | F58E2.2 |
| WBGene00019047 | | 0.249901714 | 0.22845172 | 0.42211145 | F58E2.3 |
| WBGene00019048 | | 0.766349698 | 0.02639616 | 0.09461484 | F58E2.4 |
| WBGene00019049 | NA | | NA | NA | F58E2.5 |
| WBGene00019050 | NA | | NA | NA | NA |
| WBGene00019051 | NA | | NA | NA | srz-23 |
| WBGene00019053 | NA | | NA | NA | F58F6.5 |
| WBGene00019054 | | 0.020925399 | 0.84643798 | NA | F58F6.6 |
| WBGene00019055 | | 0.204737186 | 0.03089539 | 0.105465 | F58F9.1 |
| WBGene00019057 | | 0.309530409 | 0.16519125 | 0.34141236 | F58F9.3 |
| WBGene00019058 | | 0.26769639 | 0.23658145 | 0.43270319 | F58F9.4 |
| WBGene00019059 | NA | | NA | NA | F58F9.6 |
| WBGene00019060 | | 0.027501208 | 0.84995557 | 0.91926207 | acox-3 |
| WBGene00019061 | | 0.033976521 | 0.61858811 | 0.77164084 | F58F12.1 |
| WBGene00019062 | NA | | NA | NA | F58F12.2 |
| WBGene00019063 | | 0.078203897 | 0.26561294 | NA | F58F12.3 |
| WBGene00019064 | | 0.202412836 | 0.18475449 | 0.36847712 | myo-5 |
| WBGene00019065 | NA | | NA | NA | F58G4.3 |
| WBGene00019066 | | 0.029316356 | 0.78959001 | 0.8813958 | sdz-23 |
| WBGene00019067 | | 0.110641221 | 0.44172106 | 0.63591328 | F58H7.1 |
| WBGene00019068 | | -0.278797204 | 0.11986776 | 0.2781274 | faah-3 |
| WBGene00019069 | | 0.199703294 | 0.21451468 | 0.40546209 | lgc-30 |
| WBGene00019070 | | -0.040167252 | 0.7594393 | 0.86387621 | F58H7.5 |
| WBGene00019072 | NA | | NA | NA | fbxc-8 |
| WBGene00019073 | NA | | NA | NA | fbxc-4 |
| WBGene00019073 | NA | | NA | NA | fbxc-2 |
| WBGene00019073 | NA | | NA | NA | fbxc-3 |
| WBGene00019074 | | -0.074638063 | 0.59328957 | 0.7534436 | cpd-1 |
| WBGene00019075 | | -0.234032443 | 0.12138908 | 0.28045725 | F59A3.2 |
| WBGene00019076 | | -0.437533933 | 0.00117234 | 0.0080244 | mrpl-24 |
| WBGene00019077 | | -0.222203376 | 0.01717014 | 0.06864773 | zipt-11 |
| WBGene00019079 | | -0.25724325 | 0.20147939 | 0.39024763 | rpa-4 |
| WBGene00019080 | | 0.03732969 | 0.59229443 | NA | F59A3.7 |
| WBGene00019081 | | -0.010443884 | 0.92014147 | 0.95544619 | F59A3.8 |
| WBGene00019082 | | -0.326100583 | 0.11732733 | 0.27380421 | pup-3 |
| WBGene00019083 | NA | | NA | NA | F59A3.10 |
| WBGene00019084 | | 0.105750925 | 0.22592505 | NA | F59A6.2 |
| WBGene00019085 | NA | | NA | NA | F59A6.3 |
| WBGene00019086 | | 0.22585412 | 0.02141775 | 0.08077723 | F59A6.4 |
| WBGene00019087 | | -0.24788091 | 0.00111443 | 0.00769638 | F59A6.5 |
| WBGene00019089 | | 0.020317245 | 0.84643798 | NA | clec-206 |

| | | | | |
|---|---|---|---|---|
| WBGene00019090 | -0.461493969 | 0.0791568 | 0.20949652 | F59A7.2 |
| WBGene00019091 | NA | NA | NA | srab-13 |
| WBGene00019092 | 1.735916967 | 2.83E-05 | 0.00031099 | F59A7.5 |
| WBGene00019094 | -0.095328367 | 0.55568042 | 0.72744251 | F59A7.7 |
| WBGene00019095 | 0.007169742 | 0.962694 | 0.97924853 | NA |
| WBGene00019096 | 0.063731418 | 0.41159439 | NA | cysl-4 |
| WBGene00019097 | 0.428563452 | 5.09E-05 | 0.00053315 | F59B1.2 |
| WBGene00019098 | 0.056753339 | 0.53102759 | NA | F59B1.4 |
| WBGene00019099 | 0.020925399 | 0.84643798 | NA | F59B1.6 |
| WBGene00019100 | 0.106228733 | 0.58014871 | 0.745406 | F59B1.8 |
| WBGene00019101 | 0.295423432 | 0.17850175 | 0.36000072 | F59B1.10 |
| WBGene00019102 | 0.027691389 | 0.87371036 | 0.93276403 | F59C12.3 |
| WBGene00019103 | 0.008290134 | 0.94759413 | 0.97044737 | F59D6.1 |
| WBGene00019104 | 0.020232312 | 0.84643798 | NA | asp-7 |
| WBGene00019105 | 1.544735431 | 3.19E-06 | 4.36E-05 | asp-8 |
| WBGene00019106 | -0.003421551 | 0.95689871 | NA | irld-38 |
| WBGene00019107 | NA | NA | NA | NA |
| WBGene00019108 | 0.648684484 | 0.03634187 | 0.11918312 | chpf-2 |
| WBGene00019109 | 0.218233794 | 0.2084291 | 0.39762802 | F59E11.2 |
| WBGene00019111 | 0.755998049 | 0.00553847 | 0.02840236 | F59E11.5 |
| WBGene00019112 | 0.020317245 | 0.84643798 | NA | F59E11.6 |
| WBGene00019113 | 0.420140415 | 0.05097562 | 0.15246866 | F59E11.7 |
| WBGene00019114 | -0.096923728 | 0.5764155 | 0.7435506 | nhr-194 |
| WBGene00019115 | 0.280212879 | 0.19949373 | 0.38803905 | nhr-195 |
| WBGene00019116 | 0.593967618 | 0.02375885 | 0.08722501 | nhr-143 |
| WBGene00019117 | 0.715133671 | 0.02021552 | 0.07727819 | NA |
| WBGene00019118 | -0.199237264 | 0.0106023 | 0.04675549 | F59E12.1 |
| WBGene00019119 | 0.020232312 | 0.84643798 | NA | F59E12.3 |
| WBGene00019120 | 0.096665086 | 0.57762838 | 0.74411462 | npl-4.1 |
| WBGene00019121 | -0.178917214 | 0.25167041 | 0.44993917 | npl-4.2 |
| WBGene00019122 | -0.00587957 | 0.97506921 | 0.98591235 | F59E12.6 |
| WBGene00019123 | NA | NA | NA | F59E12.8 |
| WBGene00019124 | 0.088744194 | 0.23108353 | 0.42600025 | F59E12.9 |
| WBGene00019125 | -0.250569973 | 0.13106814 | 0.29364202 | ddl-1 |
| WBGene00019126 | NA | NA | NA | sam-4 |
| WBGene00019127 | -0.076691572 | 0.30917263 | 0.51003904 | cgt-3 |
| WBGene00019128 | 0.261497906 | 0.19154843 | 0.37775601 | F59G1.4 |
| WBGene00019129 | -0.180044446 | 0.21948042 | 0.41160662 | F59G1.8 |
| WBGene00019130 | 0.369831775 | 0.00277288 | 0.01638989 | gbas-1 |
| WBGene00019131 | 1.367177496 | 6.30E-06 | 8.05E-05 | NA |
| WBGene00019132 | 0.508313028 | 0.01845945 | 0.07251537 | bath-19 |
| WBGene00019133 | -0.024891995 | 0.88675232 | 0.94033043 | F59H6.2 |
| WBGene00019134 | 0.494615422 | 0.06801589 | 0.18759283 | F59H6.3 |

| | | | | |
|---|---|---|---|---|
| WBGene00019135 | NA | NA | NA | math-32 |
| WBGene00019137 | NA | NA | NA | ceh-85 |
| WBGene00019138 | 1.046801027 | 0.00501239 | 0.02619953 | bath-21 |
| WBGene00019139 | 1.181854274 | 1.03E-05 | 0.0001268 | bath-1 |
| WBGene00019140 | 0.087861741 | 0.67147002 | 0.80894366 | bath-3 |
| WBGene00019141 | 0.084956964 | 0.60076312 | 0.75845242 | bath-5 |
| WBGene00019142 | 0.154471412 | 0.44686492 | 0.64010124 | btb-4 |
| WBGene00019143 | -0.205038937 | 0.33975587 | 0.53946055 | H01M10.1 |
| WBGene00019144 | 0.033034883 | 0.64219503 | NA | H01M10.2 |
| WBGene00019145 | 0.063999456 | 0.32439727 | NA | H02F09.2 |
| WBGene00019146 | 3.6018405 | 1.67E-12 | 6.08E-11 | H02F09.3 |
| WBGene00019148 | -0.182102376 | 0.39271942 | 0.59095501 | H03E18.1 |
| WBGene00019149 | 0.194017805 | 0.18712382 | 0.37163319 | H03E18.2 |
| WBGene00019150 | 0.049950653 | 0.56395944 | NA | H04J21.1 |
| WBGene00019151 | 1.628071784 | 0.01395102 | 0.05834243 | pck-3 |
| WBGene00019152 | 0.102413642 | 0.23275609 | NA | nspb-6 |
| WBGene00019153 | 0.127961806 | 0.3536781 | 0.55358147 | H04M03.3 |
| WBGene00019154 | 4.173512293 | 0.00017347 | 0.00156195 | glf-1 |
| WBGene00019155 | NA | NA | NA | H04M03.11 |
| WBGene00019156 | 0.51326612 | 0.05513323 | 0.16163509 | H04M03.12 |
| WBGene00019157 | -0.107759893 | 0.22145355 | 0.41403415 | H05C05.1 |
| WBGene00019158 | 0.189726608 | 0.1250083 | 0.28566356 | H05C05.2 |
| WBGene00019159 | 0.079128521 | 0.30759726 | NA | H05C05.3 |
| WBGene00019160 | 1.343475594 | 7.28E-13 | 2.77E-11 | nlp-49 |
| WBGene00019162 | -0.158868573 | 0.00539067 | 0.02773477 | eif-1.A |
| WBGene00019163 | 0.093795983 | 0.31792002 | 0.51816397 | ubxn-6 |
| WBGene00019164 | 0.423112302 | 0.00436605 | 0.02353911 | H06H21.8 |
| WBGene00019165 | 0.02821937 | 0.79554702 | 0.88521082 | mpz-4 |
| WBGene00019166 | 1.300261618 | 6.24E-07 | 1.03E-05 | tat-2 |
| WBGene00019167 | 0.28804109 | 0.03260758 | 0.10999882 | H06I04.1 |
| WBGene00019168 | -0.450857953 | 4.85E-08 | 9.68E-07 | H06I04.3 |
| WBGene00019169 | 0.086632749 | 0.29414331 | NA | H06I04.5 |
| WBGene00019170 | 0.485573625 | 0.03541321 | 0.11701304 | H06I04.6 |
| WBGene00019171 | 0.053037263 | 0.64766425 | 0.79141821 | fipr-29 |
| WBGene00019174 | -0.104347022 | 0.29885287 | 0.49942984 | H09G03.1 |
| WBGene00019176 | 0.347048881 | 0.1480228 | 0.31876649 | NA |
| WBGene00019177 | 0.139055123 | 0.13773188 | 0.3035575 | H10D18.1 |
| WBGene00019178 | NA | NA | NA | scl-12 |
| WBGene00019179 | NA | NA | NA | scl-13 |
| WBGene00019180 | 0.24808328 | 0.25257637 | 0.45064229 | H10D18.5 |
| WBGene00019181 | 0.065014541 | 0.37399213 | NA | H10D18.6 |
| WBGene00019182 | 0.822157789 | 0.00846401 | 0.03946545 | H10E21.1 |
| WBGene00019183 | 0.205478269 | 0.29000881 | 0.49054729 | npr-30 |

| | | | | |
|---|---|---|---|---|
| WBGene00019184 | 1.593661338 | 0.01459507 | 0.06051727 | H10E21.4 |
| WBGene00019185 | 1.172006097 | 0.00173955 | 0.01113144 | H10E21.5 |
| WBGene00019186 | NA | NA | NA | fbxb-58 |
| WBGene00019187 | 0.250831721 | 0.25100483 | 0.44956215 | H11E01.2 |
| WBGene00019188 | 0.909563403 | 7.85E-05 | 0.0007805 | H11E01.3 |
| WBGene00019189 | 0.630176159 | 0.01529001 | 0.06281175 | H11L12.1 |
| WBGene00019190 | -0.02419332 | 0.70411432 | NA | H12I13.1 |
| WBGene00019192 | NA | NA | NA | H12I13.3 |
| WBGene00019193 | NA | NA | NA | NA |
| WBGene00019194 | NA | NA | NA | H12I13.6 |
| WBGene00019196 | -0.070588531 | 0.56530652 | 0.73491979 | H14A12.3 |
| WBGene00019197 | 0.016458072 | 0.93790474 | 0.96517113 | H14A12.5 |
| WBGene00019198 | 1.750941036 | 0.0004093 | 0.00328178 | strm-1 |
| WBGene00019199 | 0.056048518 | 0.59213766 | 0.75269035 | H14E04.2 |
| WBGene00019200 | 0.082009015 | 0.63267399 | 0.78177194 | H14E04.3 |
| WBGene00019202 | NA | NA | NA | H14N18.2 |
| WBGene00019203 | -0.072799822 | 0.4791797 | 0.66615262 | ttr-47 |
| WBGene00019204 | 1.033536597 | 1.39E-06 | 2.11E-05 | H14N18.4 |
| WBGene00019205 | 0.350056443 | 0.00349098 | 0.0196961 | kcc-2 |
| WBGene00019207 | -0.198605887 | 0.03628156 | 0.1190227 | fgt-1 |
| WBGene00019208 | -5.983781857 | 1.92E-07 | 3.44E-06 | lips-14 |
| WBGene00019209 | -0.153663006 | 0.02416418 | 0.08848382 | emc-1 |
| WBGene00019211 | 0.021843841 | 0.87751194 | 0.93473466 | H18N23.2 |
| WBGene00019212 | -0.157525288 | 0.35814329 | 0.55854025 | zmp-2 |
| WBGene00019213 | 0.054898452 | 0.7718126 | 0.87031066 | H20E11.1 |
| WBGene00019214 | 0.04909525 | 0.45360729 | NA | H20E11.2 |
| WBGene00019215 | -0.178690511 | 0.40666818 | 0.60471122 | H20E11.3 |
| WBGene00019216 | 0.033034883 | 0.64219503 | NA | H20J04.1 |
| WBGene00019217 | -0.014167487 | 0.86228434 | 0.92672958 | athp-2 |
| WBGene00019218 | -0.10640698 | 0.32241807 | 0.52270214 | madf-3 |
| WBGene00019219 | -0.333810963 | 0.06234313 | 0.17655045 | H20J04.4 |
| WBGene00019220 | -0.358592777 | 0.00398498 | 0.02181905 | pfd-2 |
| WBGene00019221 | 0.028993486 | 0.85720946 | 0.92325428 | H20J04.6 |
| WBGene00019222 | NA | NA | NA | H20J04.7 |
| WBGene00019224 | 0.23904313 | 0.21977867 | 0.41195487 | gnrr-5 |
| WBGene00019225 | 0.272826326 | 0.14591333 | 0.31569634 | bgal-2 |
| WBGene00019226 | 0.247583149 | 0.10309561 | 0.25058216 | H22K11.3 |
| WBGene00019227 | 0.557311757 | 0.02945346 | 0.10233226 | sgca-1 |
| WBGene00019228 | -0.006766984 | 0.93963666 | NA | H23L24.1 |
| WBGene00019229 | 0.019325396 | 0.77340931 | NA | ipla-5 |
| WBGene00019230 | 0.173857204 | 0.15827648 | 0.33261394 | ttll-11 |
| WBGene00019231 | 0.108942554 | 0.15594412 | NA | H23L24.4 |
| WBGene00019232 | 0.226451031 | 0.2146314 | 0.40560951 | ugt-13 |

| | | | | |
|---|---|---|---|---|
| WBGene00019233 | 0.448289654 | 0.02966751 | 0.10282354 | ugt-14 |
| WBGene00019234 | 2.041496968 | 1.06E-09 | 2.68E-08 | ugt-8 |
| WBGene00019235 | -0.001842784 | 0.95621055 | NA | H23N18.4 |
| WBGene00019236 | 0.003036824 | 0.98176769 | 0.98947207 | H23N18.5 |
| WBGene00019237 | 0.417204374 | 0.00500957 | 0.02619953 | H24G06.1 |
| WBGene00019239 | -0.991720914 | 0.01526559 | 0.06272904 | H24K24.2 |
| WBGene00019240 | -0.015470651 | 0.84160933 | 0.91450217 | H24K24.3 |
| WBGene00019241 | -0.055003877 | 0.64402349 | 0.78931424 | trm-2A |
| WBGene00019242 | 0.038125098 | 0.61119482 | NA | srbc-40 |
| WBGene00019243 | -0.094304235 | 0.54638554 | 0.71958539 | fbxb-72 |
| WBGene00019244 | 0.262665871 | 0.12528123 | 0.28610864 | H25P19.1 |
| WBGene00019245 | -0.13071711 | 0.16431194 | 0.34031563 | sacy-1 |
| WBGene00019246 | -0.260502917 | 0.00089585 | 0.00641691 | rpb-5 |
| WBGene00019247 | -0.093450274 | 0.25669304 | 0.45488293 | syp-4 |
| WBGene00019248 | 0.072135779 | 0.42199099 | 0.61846519 | H27M09.5 |
| WBGene00019249 | 0.069500743 | 0.49122743 | 0.67495527 | H28G03.1 |
| WBGene00019250 | 0.121139343 | 0.36593493 | 0.56563063 | H28G03.2 |
| WBGene00019251 | NA | NA | NA | H28G03.3 |
| WBGene00019253 | NA | NA | NA | H31G24.1 |
| WBGene00019254 | -0.128864841 | 0.44216541 | 0.63611514 | H31G24.3 |
| WBGene00019255 | 0.07984558 | 0.3112552 | NA | H32C10.1 |
| WBGene00019257 | NA | NA | NA | dhhc-13 |
| WBGene00019259 | -0.416409095 | 1.44E-14 | 6.55E-13 | H34C03.2 |
| WBGene00019260 | 0.123679789 | 0.26051292 | 0.45854408 | H34I24.1 |
| WBGene00019261 | 1.567460308 | 3.49E-42 | 6.94E-40 | H34I24.2 |
| WBGene00019262 | 3.805748335 | 1.12E-05 | 0.00013692 | dmsr-12 |
| WBGene00019263 | 0.449333379 | 0.02679061 | 0.09550554 | H35B03.1 |
| WBGene00019264 | -0.200307724 | 0.09429658 | 0.23535565 | H35B03.2 |
| WBGene00019265 | 0.019325396 | 0.77340931 | NA | H35N09.1 |
| WBGene00019266 | -0.11108296 | 0.59137172 | 0.75215398 | H35N09.2 |
| WBGene00019267 | -0.081737357 | 0.47313108 | 0.66116737 | H39E20.1 |
| WBGene00019268 | 0.494826938 | 0.00823185 | 0.03864062 | H41C03.1 |
| WBGene00019269 | -0.004864426 | 0.94835335 | NA | H41C03.2 |
| WBGene00019270 | 0.227645604 | 0.2628608 | 0.46079322 | H41C03.3 |
| WBGene00019272 | 0.425672234 | 0.05047917 | 0.15138581 | H42K12.3 |
| WBGene00019273 | 1.829795219 | 1.67E-16 | 9.44E-15 | H43E16.1 |
| WBGene00019274 | -0.009927241 | 0.94282132 | 0.96758451 | H43I07.1 |
| WBGene00019275 | -0.058920144 | 0.63757048 | 0.78521781 | rpac-40 |
| WBGene00019276 | -0.136214264 | 0.31305603 | 0.51343714 | algn-5 |
| WBGene00019277 | 0.076207694 | 0.71521901 | 0.84034359 | sgcb-1 |
| WBGene00019278 | -0.026531149 | 0.8836563 | 0.93852825 | NA |
| WBGene00019279 | 0.106158671 | 0.57122824 | 0.73953069 | K01A2.4 |
| WBGene00019280 | 0.218361663 | 0.25379058 | 0.45176267 | K01A2.5 |

| | | | | |
|---|---|---|---|---|
| WBGene00019281 | NA | NA | NA | K01A2.6 |
| WBGene00019282 | NA | NA | NA | mps-2 |
| WBGene00019283 | NA | NA | NA | K01A2.9 |
| WBGene00019283 | NA | NA | NA | K01A2.12 |
| WBGene00019284 | 1.96925666 | 1.77E-06 | 2.61E-05 | K01A2.10 |
| WBGene00019285 | 0.795696435 | 0.02429876 | 0.08882851 | cbn-1 |
| WBGene00019286 | 0.087256674 | 0.23800185 | NA | K01A12.2 |
| WBGene00019287 | 0.028414915 | 0.78722706 | 0.87949869 | K01A12.3 |
| WBGene00019288 | -0.06730507 | 0.31931151 | NA | K02A2.1 |
| WBGene00019289 | -0.103164013 | 0.61730558 | 0.7706319 | kcc-3 |
| WBGene00019290 | -0.011865107 | 0.90063048 | 0.94699722 | K02A2.5 |
| WBGene00019292 | NA | NA | NA | K02A6.1 |
| WBGene00019293 | 0.073315368 | 0.43833996 | NA | K02A6.2 |
| WBGene00019294 | 1.058467767 | 2.34E-05 | 0.00026297 | K02A6.3 |
| WBGene00019295 | -0.173430804 | 0.07723661 | 0.20566135 | pfkb-1.2 |
| WBGene00019296 | -0.136251814 | 0.14071296 | 0.30845488 | mcu-1 |
| WBGene00019297 | 0.020232312 | 0.84643798 | NA | K02B2.6 |
| WBGene00019298 | 1.135802031 | 3.62E-22 | 3.01E-20 | K02D7.1 |
| WBGene00019299 | -0.006239126 | 0.93041881 | NA | snai-1 |
| WBGene00019300 | 1.763669824 | 5.14E-05 | 0.00053703 | swt-1 |
| WBGene00019301 | 0.002610778 | 0.97574662 | 0.98632327 | K02D10.1 |
| WBGene00019302 | 0.05891516 | 0.51827007 | NA | K02D10.2 |
| WBGene00019303 | 0.013755544 | 0.88500544 | NA | K02D10.3 |
| WBGene00019304 | 0.69949623 | 0.00850577 | 0.03962231 | K02D10.4 |
| WBGene00019305 | -0.066585328 | 0.55879454 | 0.72969237 | snap-29 |
| WBGene00019306 | -0.045959069 | 0.50707733 | NA | K02E7.1 |
| WBGene00019309 | 0.020317245 | 0.84643798 | NA | K02E7.4 |
| WBGene00019310 | -0.188866329 | 0.12349483 | 0.28357714 | NA |
| WBGene00019311 | -0.522305434 | 0.04795929 | 0.14592104 | K02E7.6 |
| WBGene00019312 | 0.020317245 | 0.84643798 | NA | K02E7.7 |
| WBGene00019313 | -1.270975384 | 7.78E-05 | 0.00077527 | btb-10 |
| WBGene00019314 | 1.776659209 | 0.00373398 | 0.02073946 | K02E7.10 |
| WBGene00019315 | -0.033175302 | 0.78006802 | 0.87577455 | K02E7.11 |
| WBGene00019316 | -0.029194423 | 0.73736399 | NA | K02E7.12 |
| WBGene00019317 | 0.034998565 | 0.86721463 | 0.92929001 | K02E10.1 |
| WBGene00019318 | 0.154359758 | 0.3392587 | 0.53902188 | K02E10.4 |
| WBGene00019319 | -0.011535809 | 0.91042331 | 0.95112162 | K02E10.5 |
| WBGene00019320 | -0.130266292 | 0.48092869 | 0.66716666 | K02E10.6 |
| WBGene00019321 | 0.248806235 | 0.25184797 | 0.44993917 | K02E10.7 |
| WBGene00019322 | 0.191039058 | 2.58E-06 | 3.63E-05 | ahcy-1 |
| WBGene00019323 | -0.291648926 | 9.34E-05 | 0.00091398 | teg-4 |
| WBGene00019324 | 1.285403238 | 8.06E-07 | 1.30E-05 | K02F2.5 |
| WBGene00019326 | 0.242297392 | 0.02293172 | 0.08506292 | K02F3.2 |

| | | | | |
|---|---|---|---|---|
| WBGene00019327 | 0.703945021 | 2.81E-14 | 1.24E-12 | zip-2 |
| WBGene00019328 | 0.003012278 | 0.99063419 | 0.9947106 | clec-149 |
| WBGene00019329 | -0.13382308 | 0.48500245 | 0.6698982 | ceeh-1 |
| WBGene00019331 | 0.058693962 | 0.518099 | NA | dos-3 |
| WBGene00019332 | -0.102223817 | 0.50697209 | 0.6876578 | K02F3.9 |
| WBGene00019333 | -0.159856203 | 0.06294678 | 0.17781282 | moma-1 |
| WBGene00019334 | -0.030176175 | 0.76612622 | 0.86713124 | K02F3.12 |
| WBGene00019335 | -0.003270206 | 0.95698316 | NA | K02F6.1 |
| WBGene00019337 | 0.450514127 | 0.08101866 | 0.21280844 | K02F6.3 |
| WBGene00019338 | -0.022930767 | 0.78121253 | NA | K02F6.4 |
| WBGene00019339 | NA | NA | NA | K02F6.5 |
| WBGene00019340 | -0.027188909 | 0.68205301 | NA | K02F6.6 |
| WBGene00019341 | -0.147782324 | 0.25043594 | 0.44901207 | K02F6.7 |
| WBGene00019342 | NA | NA | NA | K02F6.8 |
| WBGene00019343 | 0.12383152 | 0.53780063 | 0.71258851 | nep-18 |
| WBGene00019344 | 0.192911844 | 0.30689521 | 0.50739172 | dhhc-10 |
| WBGene00019345 | 0.371346262 | 0.0574645 | 0.16656854 | tmem-135 |
| WBGene00019346 | 0.008725183 | 0.96588433 | 0.98082345 | K02G10.5 |
| WBGene00019347 | 0.040735483 | 0.77909993 | 0.87502317 | mbl-1 |
| WBGene00019348 | 0.036687355 | 0.59489568 | NA | K02H11.4 |
| WBGene00019349 | 0.020232312 | 0.84643798 | NA | trx-5 |
| WBGene00019350 | NA | NA | NA | K02H11.9 |
| WBGene00019351 | 0.253629831 | 0.00288861 | 0.01697118 | lron-7 |
| WBGene00019352 | 0.531728614 | 0.02684972 | 0.0956362 | K03A1.4 |
| WBGene00019353 | 0.086663543 | 0.45673111 | 0.64869855 | K03B4.1 |
| WBGene00019354 | NA | NA | NA | K03B4.2 |
| WBGene00019355 | 0.116885593 | 0.56530747 | 0.73491979 | K03B4.4 |
| WBGene00019356 | NA | NA | NA | K03B4.6 |
| WBGene00019357 | 0.039576682 | 0.62972269 | 0.77968456 | cpg-8 |
| WBGene00019359 | NA | NA | NA | K03C7.3 |
| WBGene00019360 | 0.036946674 | 0.80304221 | 0.89037039 | neto-1 |
| WBGene00019361 | 0.25298423 | 0.09309911 | 0.23320702 | clik-3 |
| WBGene00019362 | -0.024931353 | 0.84100775 | 0.91408863 | cdk-2 |
| WBGene00019364 | 0.028432716 | 0.81082345 | 0.89531472 | mksr-1 |
| WBGene00019365 | 0.375745242 | 0.02765367 | 0.09767529 | rei-2 |
| WBGene00019367 | 0.133275529 | 0.30952779 | 0.51040962 | K03H6.1 |
| WBGene00019368 | -0.110829446 | 0.56150697 | 0.73186357 | K03H6.2 |
| WBGene00019369 | 0.020232312 | 0.84643798 | NA | K03H6.4 |
| WBGene00019370 | 1.137621707 | 0.011459 | 0.04975423 | K03H6.5 |
| WBGene00019371 | NA | NA | NA | K03H9.1 |
| WBGene00019372 | -0.024016858 | 0.7036901 | NA | K03H9.3 |
| WBGene00019373 | 0.429373018 | 0.03548511 | 0.11717129 | K04A8.1 |
| WBGene00019374 | 0.036687355 | 0.59489568 | NA | K04A8.2 |

| | | | | |
|---|---|---|---|---|
| WBGene00019376 | 7.010582796 | 3.30E-22 | 2.78E-20 | lipl-4 |
| WBGene00019379 | 0.139470123 | 0.21332115 | 0.40404075 | K04A8.10 |
| WBGene00019380 | -0.503880935 | 2.93E-08 | 6.06E-07 | K04C2.2 |
| WBGene00019381 | -0.184398662 | 0.11975146 | 0.27797684 | K04C2.3 |
| WBGene00019382 | 0.013724068 | 0.88156189 | NA | K04C2.5 |
| WBGene00019383 | 0.033463284 | 0.6365037 | NA | K04E7.1 |
| WBGene00019384 | NA | NA | NA | NA |
| WBGene00019385 | NA | NA | NA | NA |
| WBGene00019386 | 0.020317245 | 0.84643798 | NA | irld-42 |
| WBGene00019387 | NA | NA | NA | K04F1.8 |
| WBGene00019388 | -0.149432937 | 0.46464202 | 0.65449527 | K04F1.9 |
| WBGene00019389 | NA | NA | NA | irld-40 |
| WBGene00019390 | NA | NA | NA | NA |
| WBGene00019391 | 0.056411843 | 0.41891328 | NA | irld-11 |
| WBGene00019392 | NA | NA | NA | irld-44 |
| WBGene00019393 | 0.05366068 | 0.40968658 | NA | irld-45 |
| WBGene00019394 | 0.126517799 | 0.55031903 | 0.72288004 | K04F10.1 |
| WBGene00019395 | -0.184279407 | 0.30539841 | 0.50592194 | K04F10.2 |
| WBGene00019396 | -0.154854986 | 0.11275854 | 0.26605926 | K04F10.3 |
| WBGene00019399 | -0.249027238 | 0.01845402 | 0.0725135 | K04F10.7 |
| WBGene00019400 | -0.270358989 | 0.0050779 | 0.02646633 | K04G7.1 |
| WBGene00019401 | -0.102685567 | 0.11358779 | 0.26754164 | nuo-4 |
| WBGene00019402 | -0.245872169 | 0.00745511 | 0.03578695 | syf-2 |
| WBGene00019403 | 0.089174571 | 0.63012782 | 0.77998829 | K05B2.2 |
| WBGene00019404 | 0.003370003 | 0.98081646 | 0.98892182 | K05B2.4 |
| WBGene00019405 | 0.105191732 | 0.16565281 | NA | K05F1.1 |
| WBGene00019406 | 0.137077131 | 0.40577789 | 0.60394369 | acdh-8 |
| WBGene00019407 | -0.177815072 | 0.39409632 | 0.59233798 | lect-2 |
| WBGene00019408 | 0.757152493 | 0.00153213 | 0.01000532 | K05F1.6 |
| WBGene00019409 | NA | NA | NA | K05F1.8 |
| WBGene00019410 | 2.14635982 | 0.00478484 | 0.0252541 | K05F1.9 |
| WBGene00019411 | 1.427107417 | 1.09E-09 | 2.74E-08 | K05F1.10 |
| WBGene00019412 | 0.032432121 | 0.8735949 | 0.93276403 | fbxb-49 |
| WBGene00019413 | -0.54850407 | 0.05274151 | 0.1567259 | fbxb-50 |
| WBGene00019414 | -0.843026153 | 0.02129028 | 0.08040005 | fbxb-51 |
| WBGene00019415 | -0.907996622 | 0.00179523 | 0.01142283 | K05F6.4 |
| WBGene00019416 | -0.259931341 | 0.18635893 | 0.37072776 | fbxb-44 |
| WBGene00019417 | -0.047440791 | 0.7931249 | 0.88385916 | fbxb-52 |
| WBGene00019418 | -0.300454012 | 0.19415452 | 0.38110708 | fbxb-54 |
| WBGene00019419 | -0.046394786 | 0.50231372 | NA | K05F6.8 |
| WBGene00019420 | -0.648215266 | 0.03637424 | 0.11924659 | fbxb-46 |
| WBGene00019421 | 0.331890264 | 0.00729139 | 0.03511618 | K05F6.10 |
| WBGene00019422 | 0.033034883 | 0.64219503 | NA | K05G3.1 |

| | | | | |
|---|---|---|---|---|
| WBGene00019423 | -0.005242408 | 0.94179184 | NA | K05G3.2 |
| WBGene00019424 | -0.02780059 | 0.7786177 | 0.87468286 | aptf-1 |
| WBGene00019425 | 0.104747489 | 0.16723639 | NA | K06A1.2 |
| WBGene00019426 | -0.056421686 | 0.76627922 | 0.86720987 | cutl-16 |
| WBGene00019427 | 0.042545583 | 0.75422678 | 0.86082276 | atg-16.2 |
| WBGene00019428 | 0.951167427 | 6.88E-05 | 0.00069738 | dgk-5 |
| WBGene00019429 | -0.207548748 | 0.05739964 | 0.16643091 | K06A5.1 |
| WBGene00019430 | 2.02681625 | 5.37E-05 | 0.00055787 | K06A5.2 |
| WBGene00019431 | 0.023310283 | 0.85561574 | 0.92262213 | K06A5.3 |
| WBGene00019432 | -0.042937595 | 0.57723348 | 0.74411462 | knl-2 |
| WBGene00019433 | -0.217579566 | 0.00160262 | 0.01040992 | acdh-3 |
| WBGene00019434 | -0.022640462 | 0.90601824 | 0.94916953 | nmtn-1 |
| WBGene00019435 | 3.404161284 | 7.91E-37 | 1.26E-34 | K06A9.1 |
| WBGene00019436 | 0.859407087 | 0.01767615 | 0.07024877 | K06A9.2 |
| WBGene00019437 | NA | NA | NA | K06A9.3 |
| WBGene00019438 | NA | NA | NA | cyp-25A6 |
| WBGene00019439 | 0.451464192 | 0.08541072 | 0.22030613 | NA |
| WBGene00019441 | 0.146653878 | 0.38547534 | 0.58460167 | NA |
| WBGene00019442 | NA | NA | NA | F07B7.8 |
| WBGene00019442 | NA | NA | NA | K06C4.1 |
| WBGene00019444 | 0.263168864 | 0.23508211 | 0.43087597 | frpr-11 |
| WBGene00019445 | -0.112664652 | 0.47264477 | 0.66074593 | frpr-12 |
| WBGene00019448 | 1.222140673 | 0.00943188 | 0.04299255 | frpr-13 |
| WBGene00019449 | NA | NA | NA | K06H6.1 |
| WBGene00019450 | NA | NA | NA | K06H6.2 |
| WBGene00019451 | NA | NA | NA | nstp-7 |
| WBGene00019452 | NA | NA | NA | K06H6.4 |
| WBGene00019453 | NA | NA | NA | K06H6.5 |
| WBGene00019454 | NA | NA | NA | NA |
| WBGene00019455 | 0.033145653 | 0.76091485 | 0.86448005 | K06H7.1 |
| WBGene00019456 | -0.100527727 | 0.6241944 | 0.7755287 | K06H7.2 |
| WBGene00019457 | -0.08896386 | 0.3192481 | 0.51940343 | vms-1 |
| WBGene00019458 | -0.292811871 | 0.06130868 | 0.17439681 | K06H7.7 |
| WBGene00019459 | 0.032491831 | 0.73597967 | 0.85181646 | K06H7.8 |
| WBGene00019460 | -0.029926892 | 0.80689665 | 0.89296858 | idi-1 |
| WBGene00019461 | 0.221293577 | 0.09961163 | 0.24478246 | K07A3.3 |
| WBGene00019462 | 0.178180073 | 0.40297083 | 0.60141856 | ccm-2 |
| WBGene00019464 | -0.226211657 | 0.12034381 | 0.27905506 | K07B1.4 |
| WBGene00019465 | -0.02128682 | 0.83170757 | 0.90832691 | acl-14 |
| WBGene00019466 | 1.102540808 | 1.41E-45 | 2.95E-43 | tos-1 |
| WBGene00019467 | -0.150168744 | 0.30843475 | 0.50933828 | K07B1.7 |
| WBGene00019469 | 0.020317245 | 0.84643798 | NA | NA |
| WBGene00019470 | 0.020232312 | 0.84643798 | NA | cyp-35B3 |

| | | | | |
|---|---|---|---|---|
| WBGene00019471 | 0.020925399 | 0.84643798 | NA | cyp-35B2 |
| WBGene00019472 | -0.008998908 | 0.92213607 | NA | cyp-35B1 |
| WBGene00019473 | -0.934232201 | 0.01074996 | 0.04730694 | cyp-35A5 |
| WBGene00019476 | -0.069250997 | 0.70567263 | 0.83334029 | timp-1 |
| WBGene00019477 | 0.010415971 | 0.95975148 | 0.97798729 | K07C11.4 |
| WBGene00019478 | 0.02709111 | 0.87973709 | 0.93585924 | cri-2 |
| WBGene00019479 | 0.270478938 | 0.03559457 | 0.11747974 | K07C11.7 |
| WBGene00019480 | 0.402179347 | 0.11033774 | 0.26203648 | K07C11.8 |
| WBGene00019481 | -0.044379621 | 0.67014424 | 0.80796767 | cogc-6 |
| WBGene00019482 | 0.036687355 | 0.59489568 | NA | K07C11.10 |
| WBGene00019483 | -0.022930767 | 0.78121253 | NA | K07D4.2 |
| WBGene00019484 | -0.012675 | 0.90920875 | 0.95068624 | K07D4.4 |
| WBGene00019485 | 0.059281883 | 0.42158309 | NA | K07D4.5 |
| WBGene00019486 | 0.036687355 | 0.59489568 | NA | K07D4.6 |
| WBGene00019487 | 0.336982053 | 0.02923081 | 0.10165518 | ephx-1 |
| WBGene00019488 | 1.443791058 | 0.0002794 | 0.00235641 | K07D4.9 |
| WBGene00019489 | -0.354086871 | 0.03399899 | 0.11336414 | K07E1.1 |
| WBGene00019490 | -0.448562446 | 0.00280762 | 0.01655513 | K07E3.1 |
| WBGene00019491 | 0.036027941 | 0.84425636 | 0.91642626 | K07E3.2 |
| WBGene00019492 | 0.350145606 | 0.00012751 | 0.00119977 | K07E3.4 |
| WBGene00019493 | 0.165911671 | 0.36020019 | 0.55995869 | catp-5 |
| WBGene00019494 | NA | NA | NA | NA |
| WBGene00019495 | 0.187865031 | 0.37966804 | 0.5792246 | sdz-24 |
| WBGene00019496 | 0.203993328 | 0.06827547 | 0.18811088 | frpr-14 |
| WBGene00019497 | 0.311499734 | 0.07307434 | 0.19746796 | K07E8.6 |
| WBGene00019498 | -0.218028084 | 0.14569827 | 0.31541748 | K07E8.7 |
| WBGene00019501 | 0.05787416 | 0.40594086 | NA | K07E12.2 |
| WBGene00019502 | NA | NA | NA | K07G6.1 |
| WBGene00019503 | -0.068112638 | 0.55843745 | 0.72929111 | tbce-1 |
| WBGene00019504 | -0.1739867 | 0.0105188 | 0.04652743 | K07H8.2 |
| WBGene00019506 | 0.127403933 | 0.16699325 | NA | K07H8.5 |
| WBGene00019507 | 0.062394992 | 0.6753379 | 0.81135495 | K07H8.7 |
| WBGene00019508 | -0.023109762 | 0.78121253 | NA | K07H8.8 |
| WBGene00019509 | 0.296265557 | 0.05281318 | 0.15682423 | K07H8.9 |
| WBGene00019510 | -0.461482017 | 6.47E-22 | 5.36E-20 | K07H8.10 |
| WBGene00019511 | -0.100237363 | 0.47387108 | 0.66163899 | lido-8 |
| WBGene00019512 | 0.201839397 | 0.15427286 | 0.32715947 | K08A2.2 |
| WBGene00019513 | 0.06250988 | 0.70533467 | 0.83307559 | lido-7 |
| WBGene00019514 | -0.11079177 | 0.12635372 | NA | K08B4.2 |
| WBGene00019515 | 0.625490536 | 0.01729267 | 0.06900571 | ugt-19 |
| WBGene00019516 | -0.338659524 | 0.14511865 | 0.31448819 | ugt-20 |
| WBGene00019517 | -0.064322954 | 0.64717529 | 0.79127191 | NA |
| WBGene00019518 | 0.048219258 | 0.73663732 | 0.85204655 | K08B5.1 |

| | | | | | |
|---|---|---|---|---|---|
| WBGene00019519 | | -0.139229829 | 0.50093208 | 0.68269142 | K08B5.2 |
| WBGene00019520 | | 1.744902581 | 0.00943954 | 0.0430141 | K08B12.1 |
| WBGene00019521 | | -0.260865986 | 0.04795137 | 0.14592104 | dmd-7 |
| WBGene00019522 | | -0.129172479 | 0.53983695 | 0.71445958 | K08B12.3 |
| WBGene00019524 | | -0.003108065 | 0.95728506 | NA | K08D9.2 |
| WBGene00019525 | | 1.411508362 | 0.00471502 | 0.02499369 | K08D9.4 |
| WBGene00019526 | NA | | NA | NA | K08D9.5 |
| WBGene00019527 | | -0.012687209 | 0.90571599 | 0.94916953 | K08D9.6 |
| WBGene00019529 | | -0.023056499 | 0.78121253 | NA | K08D10.5 |
| WBGene00019530 | | 0.090103632 | 0.20323472 | NA | scrm-8 |
| WBGene00019531 | | 0.131042911 | 0.31624036 | 0.51687942 | scrm-5 |
| WBGene00019532 | | 1.557123739 | 0.00076183 | 0.00555475 | NA |
| WBGene00019533 | | -0.00346345 | 0.95708614 | NA | K08D10.10 |
| WBGene00019534 | | 0.020317245 | 0.84643798 | NA | K08D10.11 |
| WBGene00019535 | | 0.235804662 | 0.25226978 | 0.45017504 | tsen-34 |
| WBGene00019536 | | 0.298723055 | 0.13762776 | 0.30351077 | rpi-2 |
| WBGene00019537 | | -0.124231104 | 0.02024861 | 0.0773441 | K08D12.3 |
| WBGene00019538 | | -0.008563965 | 0.95510463 | 0.97467672 | K08D12.4 |
| WBGene00019540 | | -0.072902588 | 0.55547011 | 0.72723222 | K08D12.6 |
| WBGene00019541 | | -0.086014004 | 0.49451543 | 0.67741395 | K08F11.1 |
| WBGene00019543 | | -0.329734641 | 1.76E-06 | 2.60E-05 | cif-1 |
| WBGene00019544 | | -0.076233133 | 0.43574819 | 0.63091271 | miro-1 |
| WBGene00019545 | | 0.036687355 | 0.59489568 | NA | K09B3.1 |
| WBGene00019546 | | 1.248997013 | 8.30E-06 | 0.00010391 | glct-6 |
| WBGene00019547 | | 0.268763647 | 0.23719591 | 0.43352983 | K09C4.1 |
| WBGene00019548 | NA | | NA | NA | K09C4.2 |
| WBGene00019549 | | 0.056727431 | 0.44364738 | NA | K09C4.4 |
| WBGene00019550 | | -0.254144031 | 0.22011688 | 0.41227213 | K09C4.5 |
| WBGene00019552 | | 0.056844058 | 0.53406644 | NA | K09C4.7 |
| WBGene00019553 | NA | | NA | NA | K09C4.9 |
| WBGene00019554 | | -0.197091386 | 0.25984474 | 0.4579683 | K09C4.10 |
| WBGene00019556 | | 0.056430134 | 0.44291702 | NA | K09C6.2 |
| WBGene00019557 | NA | | NA | NA | K09C6.3 |
| WBGene00019558 | | 0.020232312 | 0.84643798 | NA | srbc-12 |
| WBGene00019559 | NA | | NA | NA | srbc-13 |
| WBGene00019560 | | -0.023056499 | 0.78121253 | NA | K09C6.6 |
| WBGene00019561 | | 0.354351423 | 0.04158679 | 0.13150589 | K09C6.7 |
| WBGene00019562 | | 0.018998992 | 0.77286019 | NA | K09C6.8 |
| WBGene00019563 | | 0.247496628 | 0.25515829 | 0.45364572 | K09C6.9 |
| WBGene00019564 | | -0.135568764 | 0.51139924 | 0.69129317 | K09D9.1 |
| WBGene00019565 | | 0.103622993 | 0.62201412 | 0.7740679 | cyp-35A3 |
| WBGene00019566 | | -0.023056499 | 0.78121253 | NA | K09D9.3 |
| WBGene00019567 | | 0.038125098 | 0.61119482 | NA | K09D9.9 |

| | | | | |
|---|---|---|---|---|
| WBGene00019568 | 0.153682434 | 0.06760542 | NA | K09D9.11 |
| WBGene00019569 | 0.08115332 | 0.67503748 | 0.81123072 | NA |
| WBGene00019570 | -0.050953199 | 0.55282647 | NA | K09E2.1 |
| WBGene00019571 | 0.175488315 | 0.09096736 | 0.22943207 | K09E2.2 |
| WBGene00019572 | 0.113353412 | 0.39675449 | 0.59468258 | K09E2.3 |
| WBGene00019574 | NA | NA | NA | K09E3.2 |
| WBGene00019575 | -0.023056499 | 0.78121253 | NA | K09E3.4 |
| WBGene00019576 | -0.071916023 | 0.71451128 | 0.83986311 | lido-5 |
| WBGene00019577 | 0.172673684 | 0.40794813 | 0.60581891 | lido-6 |
| WBGene00019578 | -0.261851799 | 0.15707403 | 0.33108565 | K09E3.7 |
| WBGene00019579 | 0.616739648 | 0.04141413 | 0.13118657 | oac-57 |
| WBGene00019580 | 1.070784709 | 0.00654958 | 0.03231965 | oac-58 |
| WBGene00019582 | NA | NA | NA | K09F5.1 |
| WBGene00019583 | NA | NA | NA | K09F5.4 |
| WBGene00019584 | -0.360801763 | 0.13257338 | 0.29580404 | set-12 |
| WBGene00019586 | 0.166262928 | 0.36116651 | 0.56098439 | K09F6.3 |
| WBGene00019587 | -1.119027047 | 0.00725362 | 0.03498034 | K09F6.4 |
| WBGene00019588 | -0.006116961 | 0.96502422 | 0.98071026 | K09F6.5 |
| WBGene00019589 | 1.064711731 | 0.00033971 | 0.00279579 | K09F6.6 |
| WBGene00019590 | 0.753955312 | 0.00698558 | 0.03393078 | K09F6.7 |
| WBGene00019591 | -0.375732055 | 0.10439726 | 0.25259961 | fbxc-25 |
| WBGene00019592 | 1.338088883 | 1.44E-08 | 3.13E-07 | K09F6.9 |
| WBGene00019593 | 1.512540254 | 4.29E-06 | 5.70E-05 | K09F6.10 |
| WBGene00019594 | -0.00346345 | 0.95708614 | NA | K09H9.1 |
| WBGene00019595 | 0.166226702 | 0.12564494 | 0.28666356 | dscc-1 |
| WBGene00019597 | 0.032412747 | 0.87537262 | 0.93344293 | K09H9.5 |
| WBGene00019598 | 0.610560315 | 0.00085492 | 0.0061629 | K09H9.7 |
| WBGene00019599 | 0.197910052 | 0.15226902 | 0.32480061 | acds-10 |
| WBGene00019600 | -0.073627727 | 0.33351631 | 0.53304196 | rga-3 |
| WBGene00019601 | NA | NA | NA | K09H11.4 |
| WBGene00019603 | 0.54537232 | 0.01032523 | 0.04604681 | K09H11.6 |
| WBGene00019604 | NA | NA | NA | pgph-1 |
| WBGene00019605 | 0.395566961 | 0.04183111 | 0.13207877 | ctsa-4.1 |
| WBGene00019606 | -0.085301401 | 0.08849865 | 0.22573322 | clec-88 |
| WBGene00019607 | -0.141900641 | 0.28034042 | 0.47994201 | K10B2.4 |
| WBGene00019608 | -0.227440588 | 0.00068104 | 0.00505376 | ani-2 |
| WBGene00019609 | 0.105392323 | 0.6155169 | 0.7693173 | K10B3.1 |
| WBGene00019611 | -0.487991905 | 0.01863526 | 0.07297107 | K10B3.5 |
| WBGene00019612 | 0.395142006 | 0.07765466 | 0.20649143 | gpcp-1 |
| WBGene00019613 | 0.038125098 | 0.61119482 | NA | K10B4.1 |
| WBGene00019614 | 0.038595855 | 0.57520533 | NA | srt-54 |
| WBGene00019615 | 0.473112055 | 0.01631925 | 0.0661509 | K10B4.3 |
| WBGene00019616 | -0.00504514 | 0.94402823 | NA | nmur-2 |

| | | | | |
|---|---|---|---|---|
| WBGene00019617 | 0.699986799 | 1.20E-11 | 3.93E-10 | ctsa-2 |
| WBGene00019618 | 0.063217264 | 0.41230173 | NA | K10C2.2 |
| WBGene00019619 | 0.179620393 | 0.11211521 | 0.26492556 | asp-14 |
| WBGene00019620 | -0.698233386 | 2.45E-11 | 7.69E-10 | fah-1 |
| WBGene00019621 | NA | NA | NA | K10C2.6 |
| WBGene00019622 | -1.034633897 | 0.0147871 | 0.06119218 | K10C2.7 |
| WBGene00019623 | 0.265383618 | 0.08787533 | 0.22465212 | K10C9.1 |
| WBGene00019624 | -0.144621366 | 0.32332844 | 0.52325099 | rnst-2 |
| WBGene00019625 | 0.020925399 | 0.84643798 | NA | K10C9.4 |
| WBGene00019626 | 0.00542986 | 0.96540615 | 0.980746 | K10C9.7 |
| WBGene00019627 | -0.202538151 | 0.01980283 | 0.07601814 | hira-1 |
| WBGene00019628 | -0.271775773 | 1.97E-07 | 3.53E-06 | pup-2 |
| WBGene00019629 | -0.086711284 | 0.22651117 | 0.42005699 | cid-1 |
| WBGene00019630 | -0.189470882 | 0.14869047 | 0.31957974 | emb-1 |
| WBGene00019631 | -0.339161055 | 0.0144269 | 0.05995578 | K10D2.5 |
| WBGene00019633 | 0.115486913 | 0.3603355 | 0.56010957 | K10D2.7 |
| WBGene00019636 | 1.162483398 | 8.03E-05 | 0.00079414 | gsto-3 |
| WBGene00019638 | -0.010113687 | 0.92007878 | 0.95544619 | K10F12.6 |
| WBGene00019639 | 0.020232312 | 0.84643798 | NA | K10F12.7 |
| WBGene00019640 | 0.020317245 | 0.84643798 | NA | dos-2 |
| WBGene00019641 | -0.012173249 | 0.9106505 | 0.95112162 | K10G6.4 |
| WBGene00019642 | 0.379352847 | 0.01861379 | 0.07290651 | K11C4.1 |
| WBGene00019643 | 0.243548939 | 0.24994496 | 0.44846129 | K11C4.2 |
| WBGene00019644 | 1.269666041 | 0.00712749 | 0.03447426 | cpt-4 |
| WBGene00019645 | 0.346904857 | 0.01124978 | 0.04906404 | swt-7 |
| WBGene00019646 | 0.201651104 | 0.15458457 | 0.32763602 | K11D12.6 |
| WBGene00019647 | 0.085433766 | 0.57289886 | 0.74106374 | K11D12.7 |
| WBGene00019648 | 0.17013281 | 0.29074922 | 0.49121385 | K11D12.8 |
| WBGene00019649 | 0.148947424 | 0.45040928 | 0.64279069 | K11D12.9 |
| WBGene00019650 | 0.358041904 | 0.13956887 | 0.3069534 | K11D12.11 |
| WBGene00019651 | 0.136400924 | 0.46752878 | 0.65691831 | K11D12.12 |
| WBGene00019652 | 0.190385575 | 0.28623081 | 0.4862171 | K11G9.1 |
| WBGene00019653 | 0.03234015 | 0.87706969 | 0.9345835 | K11G9.2 |
| WBGene00019654 | -0.498717558 | 0.06515976 | 0.18191261 | K11G9.3 |
| WBGene00019655 | -0.143855945 | 0.46204795 | 0.65228464 | K11G9.5 |
| WBGene00019656 | -0.223757741 | 0.07999766 | 0.21104039 | slc-25A10 |
| WBGene00019657 | -0.046421884 | 0.73974766 | 0.8534033 | K11G12.6 |
| WBGene00019658 | -0.137446089 | 0.40452854 | 0.60263556 | K11H12.1 |
| WBGene00019659 | -0.154242656 | 0.24193307 | 0.43996299 | K11H12.3 |
| WBGene00019660 | -2.063143699 | 3.30E-06 | 4.49E-05 | K11H12.4 |
| WBGene00019661 | 0.169324683 | 0.35999239 | 0.55987345 | K11H12.5 |
| WBGene00019662 | -0.104347022 | 0.29885287 | 0.49942984 | K11H12.6 |
| WBGene00019663 | -0.197556177 | 0.26085996 | 0.45904454 | K11H12.7 |

| | | | | |
|---|---|---|---|---|
| WBGene00019664 | -0.109265445 | 0.24289049 | 0.44097731 | K11H12.8 |
| WBGene00019665 | 0.481098838 | 0.06340789 | 0.17865182 | K11H12.9 |
| WBGene00019666 | 0.260272351 | 0.0216338 | 0.0812989 | sago-1 |
| WBGene00019667 | 0.0398788 | 0.58971221 | NA | spe-49 |
| WBGene00019668 | 3.691833351 | 0.00228974 | 0.01391621 | fil-2 |
| WBGene00019669 | -0.123020751 | 0.31494955 | 0.51557662 | K12B6.4 |
| WBGene00019672 | -0.0231085 | 0.90501469 | 0.94913088 | K12B6.8 |
| WBGene00019673 | 0.13400701 | 0.18175715 | 0.36443528 | K12C11.1 |
| WBGene00019674 | -0.108407985 | 0.50811933 | 0.68863974 | K12C11.3 |
| WBGene00019675 | 0.033034883 | 0.64219503 | NA | K12D9.1 |
| WBGene00019676 | NA | NA | NA | NA |
| WBGene00019677 | 0.026955476 | 0.86759827 | 0.92947386 | K12H4.2 |
| WBGene00019678 | -0.388466639 | 0.00024413 | 0.0020915 | K12H4.3 |
| WBGene00019679 | -0.122165705 | 0.23727082 | 0.43353077 | spcs-3 |
| WBGene00019680 | -0.245543562 | 0.00796867 | 0.03773264 | K12H4.5 |
| WBGene00019681 | NA | NA | NA | K12H4.6 |
| WBGene00019682 | 0.243709074 | 0.00048402 | 0.00378348 | K12H4.7 |
| WBGene00019683 | -0.048168134 | 0.81076492 | 0.89531472 | K12H6.2 |
| WBGene00019684 | 0.020232312 | 0.84643798 | NA | K12H6.4 |
| WBGene00019685 | NA | NA | NA | K12H6.5 |
| WBGene00019686 | -0.188287779 | 0.17489188 | 0.35564444 | K12H6.6 |
| WBGene00019687 | -0.908894803 | 0.01003996 | 0.04500449 | K12H6.7 |
| WBGene00019688 | 0.020925399 | 0.84643798 | NA | K12H6.8 |
| WBGene00019689 | 0.104747489 | 0.16723639 | NA | K12H6.9 |
| WBGene00019690 | -0.221747417 | 0.27660996 | 0.47656466 | set-13 |
| WBGene00019691 | 0.195948603 | 0.36637525 | 0.56601245 | K12H6.12 |
| WBGene00019692 | -0.124415789 | 0.30022044 | 0.50066452 | fubl-4 |
| WBGene00019693 | -0.158319974 | 0.02711797 | 0.0961998 | ostd-1 |
| WBGene00019697 | 0.04439038 | 0.58545656 | 0.74840426 | M01B12.4 |
| WBGene00019698 | -0.013079016 | 0.92273818 | 0.9564467 | riok-1 |
| WBGene00019699 | 3.966189701 | 1.73E-08 | 3.72E-07 | math-34 |
| WBGene00019700 | -1.039104431 | 0.00051195 | 0.00396375 | btb-11 |
| WBGene00019702 | 0.033034883 | 0.64219503 | NA | fbxb-39 |
| WBGene00019703 | -0.940565553 | 0.00520418 | 0.0269619 | fbxb-41 |
| WBGene00019704 | -0.57494724 | 0.04685266 | 0.14356841 | fbxb-40 |
| WBGene00019705 | -1.10196266 | 0.00910889 | 0.04175429 | fbxb-45 |
| WBGene00019706 | 0.059779397 | 0.52382254 | NA | NA |
| WBGene00019709 | NA | NA | NA | M01E10.3 |
| WBGene00019710 | -0.036138042 | 0.79532223 | 0.8851625 | M01E11.1 |
| WBGene00019711 | -0.222964483 | 0.00939211 | 0.04285129 | M01E11.2 |
| WBGene00019712 | 0.065414424 | 0.58549432 | 0.74840426 | rmh-1 |
| WBGene00019713 | 0.093310266 | 0.15266478 | 0.32540255 | pqn-52 |
| WBGene00019714 | NA | NA | NA | M01G4.1 |

| | | | | |
|---|---|---|---|---|
| WBGene00019715 | -0.16787315 | 0.0859343 | 0.22135188 | M01G5.1 |
| WBGene00019716 | 0.055893576 | 0.66257682 | 0.80169186 | M01G5.3 |
| WBGene00019717 | 1.575667374 | 0.00208496 | 0.01292923 | trx-3 |
| WBGene00019718 | 1.653740524 | 0.01048076 | 0.04648337 | M01H9.2 |
| WBGene00019719 | 0.290910722 | 0.00011626 | 0.00110809 | M01H9.3 |
| WBGene00019720 | 0.254308152 | 0.25978602 | 0.45792525 | M01H9.4 |
| WBGene00019721 | 0.018342935 | 0.76675379 | NA | M01H9.5 |
| WBGene00019722 | -0.029327418 | 0.71586712 | NA | M02A10.1 |
| WBGene00019723 | NA | NA | NA | fbxc-47 |
| WBGene00019724 | -0.026152034 | 0.87784003 | 0.93480719 | M02B7.2 |
| WBGene00019725 | 0.227853258 | 0.16963549 | 0.34800552 | algn-14 |
| WBGene00019726 | 0.288154494 | 0.01215798 | 0.05226219 | bris-1 |
| WBGene00019727 | -0.354215455 | 0.00451848 | 0.02421452 | zig-12 |
| WBGene00019728 | 0.135401195 | 0.50917867 | 0.68943724 | M02D8.2 |
| WBGene00019729 | -0.408630963 | 0.01785657 | 0.07073539 | M02D8.3 |
| WBGene00019730 | 0.472715035 | 0.00014395 | 0.0013214 | asns-2 |
| WBGene00019731 | -0.542168494 | 0.03808214 | 0.12340873 | M02D8.5 |
| WBGene00019732 | 0.642875345 | 0.02700323 | 0.095975 | M02D8.6 |
| WBGene00019733 | 0.253988561 | 0.11717744 | 0.27380421 | M02E1.1 |
| WBGene00019734 | 0.133673795 | 0.46750873 | 0.65691831 | nlp-64 |
| WBGene00019735 | -0.023587884 | 0.86682275 | 0.92907407 | M02F4.1 |
| WBGene00019736 | -0.003614866 | 0.95700117 | NA | M02F4.2 |
| WBGene00019738 | -1.511950263 | 1.39E-07 | 2.55E-06 | clec-265 |
| WBGene00019739 | NA | NA | NA | NA |
| WBGene00019740 | NA | NA | NA | srt-31 |
| WBGene00019741 | 0.222092167 | 0.28251475 | 0.48236717 | nhr-201 |
| WBGene00019742 | 0.351878298 | 0.14567028 | 0.31540351 | nhr-202 |
| WBGene00019743 | 0.410128509 | 0.09774622 | 0.24219028 | nhr-203 |
| WBGene00019744 | 0.645539029 | 0.00900762 | 0.04147188 | M02H5.8 |
| WBGene00019745 | NA | NA | NA | srbc-57 |
| WBGene00019746 | -0.010398296 | 0.95310662 | 0.97322797 | M03A1.3 |
| WBGene00019747 | 0.0914353 | 0.26746624 | 0.46635384 | ipla-1 |
| WBGene00019748 | 0.167781679 | 0.22288851 | 0.41556803 | atg-2 |
| WBGene00019749 | 0.110064003 | 0.60113609 | 0.75866162 | M03A8.3 |
| WBGene00019750 | 0.433583251 | 0.05267881 | 0.15657134 | M03D4.3 |
| WBGene00019751 | 0.231280466 | 0.27637082 | 0.47632079 | M03D4.4 |
| WBGene00019753 | 0.320933863 | 0.10460215 | 0.25284464 | M03E7.1 |
| WBGene00019754 | 0.036085749 | 0.67342551 | NA | M03E7.2 |
| WBGene00019755 | NA | NA | NA | M03E7.3 |
| WBGene00019756 | -0.027728334 | 0.67598605 | NA | M03E7.4 |
| WBGene00019757 | -0.185569407 | 0.14802328 | 0.31876649 | memb-2 |
| WBGene00019758 | NA | NA | NA | M03F4.4 |
| WBGene00019759 | -0.050863974 | 0.73490199 | 0.85124209 | M03F4.6 |

| | | | | |
|---|---|---|---|---|
| WBGene00019760 | -0.745150538 | 8.99E-21 | 7.07E-19 | calu-1 |
| WBGene00019761 | 0.151587434 | 0.12223267 | 0.28134075 | M03F8.1 |
| WBGene00019762 | -0.191694176 | 0.01237253 | 0.05296672 | M03F8.3 |
| WBGene00019763 | 1.251804068 | 0.00579045 | 0.02939597 | galt-1 |
| WBGene00019764 | 0.178387348 | 0.28612214 | 0.48616975 | M03F8.5 |
| WBGene00019765 | -0.06961745 | 0.65871065 | 0.79882238 | M03F8.6 |
| WBGene00019766 | -0.269331586 | 0.00212706 | 0.01314573 | M4.1 |
| WBGene00019767 | -0.142166112 | 0.01894569 | 0.07392961 | rpa-2 |
| WBGene00019768 | -0.038024303 | 0.83767857 | 0.91205859 | M04F3.2 |
| WBGene00019769 | -0.468703819 | 0.07593903 | 0.20340713 | M04F3.3 |
| WBGene00019770 | 0.049085195 | 0.77007858 | 0.86915823 | M04F3.4 |
| WBGene00019771 | 0.023322146 | 0.8249726 | 0.90387067 | M04F3.5 |
| WBGene00019772 | 2.167750524 | 2.63E-18 | 1.73E-16 | M04G7.1 |
| WBGene00019773 | 0.05787416 | 0.40594086 | NA | M04G7.2 |
| WBGene00019774 | 0.089971942 | 0.66214783 | 0.8014937 | M04G7.3 |
| WBGene00019776 | 0.00776142 | 0.9393658 | 0.96589373 | M6.4 |
| WBGene00019777 | 0.467648573 | 0.02085252 | 0.07913473 | M57.1 |
| WBGene00019778 | -0.178528653 | 0.0650202 | 0.18177597 | M57.2 |
| WBGene00019779 | 0.084463956 | 0.41201192 | 0.60964575 | endu-2 |
| WBGene00019780 | 0.62357563 | 5.76E-07 | 9.57E-06 | M60.4 |
| WBGene00019781 | 0.321284979 | 0.15176209 | 0.32424969 | M60.6 |
| WBGene00019782 | 1.186743917 | 0.00598272 | 0.03004906 | M60.7 |
| WBGene00019783 | 0.148649225 | 0.20159765 | 0.39027971 | M70.1 |
| WBGene00019784 | NA | NA | NA | M70.2 |
| WBGene00019785 | 0.953553441 | 0.01822136 | 0.07188869 | M70.3 |
| WBGene00019786 | 0.184103833 | 0.09332315 | 0.23360431 | M70.4 |
| WBGene00019787 | 0.071610072 | 0.66237513 | 0.80166643 | M70.5 |
| WBGene00019788 | NA | NA | NA | sdz-25 |
| WBGene00019788 | NA | NA | NA | M116.1 |
| WBGene00019789 | 1.253052246 | 1.33E-05 | 0.00015827 | NA |
| WBGene00019791 | NA | NA | NA | sdz-25 |
| WBGene00019791 | NA | NA | NA | M116.1 |
| WBGene00019792 | -0.098138435 | 0.53645698 | 0.71143719 | M116.5 |
| WBGene00019793 | -0.216557118 | 0.15897282 | 0.33321705 | M151.1 |
| WBGene00019794 | -0.079092944 | 0.69198691 | 0.82320708 | M151.2 |
| WBGene00019795 | 0.136701384 | 0.51964569 | 0.69869893 | M151.3 |
| WBGene00019796 | 0.173256543 | 0.42227157 | 0.61854675 | M151.4 |
| WBGene00019797 | -0.465088122 | 0.0757372 | 0.20301507 | fbxb-31 |
| WBGene00019798 | -0.137672455 | 0.38021782 | 0.5797656 | fbxb-32 |
| WBGene00019799 | -0.07223649 | 0.67228243 | 0.80922725 | M151.7 |
| WBGene00019800 | -0.326716233 | 0.00035132 | 0.00286877 | mtss-1 |
| WBGene00019801 | -0.074886538 | 0.53882062 | 0.71366679 | aak-1 |
| WBGene00019803 | 0.484899918 | 0.00965135 | 0.04373425 | PDB1.1 |

| | | | | |
|---|---|---|---|---|
| WBGene00019804 | -0.008303499 | 0.92600956 | NA | R01B10.2 |
| WBGene00019805 | NA | NA | NA | R01B10.3 |
| WBGene00019806 | 0.072582149 | 0.72025481 | 0.84328945 | pgap-3 |
| WBGene00019807 | -0.141396447 | 0.20948985 | 0.39895172 | jamp-1 |
| WBGene00019808 | 0.230180148 | 0.06674892 | 0.18487909 | R01B10.6 |
| WBGene00019809 | NA | NA | NA | R01H2.1 |
| WBGene00019810 | 0.085429064 | 0.39813282 | 0.59632354 | R01H2.2 |
| WBGene00019811 | -0.031369802 | 0.6590341 | 0.79896991 | egg-2 |
| WBGene00019812 | -0.00504514 | 0.94402823 | NA | R01H2.4 |
| WBGene00019813 | 0.099541136 | 0.56243021 | 0.73248005 | ger-1 |
| WBGene00019814 | -0.600589811 | 0.03405294 | 0.11346588 | R02C2.1 |
| WBGene00019815 | -0.257532422 | 0.25114365 | 0.44967521 | kin-34 |
| WBGene00019816 | 0.055389999 | 0.78343565 | 0.87807405 | nhr-204 |
| WBGene00019817 | -0.678796473 | 0.03294063 | 0.11079341 | R02C2.6 |
| WBGene00019818 | -0.033100131 | 0.83517839 | 0.91041969 | R02C2.7 |
| WBGene00019819 | -0.272394165 | 0.00309822 | 0.01791512 | aass-1 |
| WBGene00019820 | -0.045212129 | 0.6500983 | 0.79283982 | cogc-8 |
| WBGene00019821 | -0.197381154 | 0.08832319 | 0.22552142 | gtf-2H1 |
| WBGene00019822 | 0.039049124 | 0.66444643 | 0.80311733 | R02D3.4 |
| WBGene00019823 | -0.091537316 | 0.36198379 | 0.56165797 | fnta-1 |
| WBGene00019824 | -0.073892553 | 0.41579748 | 0.61306051 | R02D3.7 |
| WBGene00019825 | 0.052058788 | 0.70134638 | 0.83030746 | R02D3.8 |
| WBGene00019826 | 0.010168758 | 0.92577292 | 0.95816777 | R02E4.1 |
| WBGene00019827 | 0.112984106 | 0.2604145 | 0.45848108 | mop-25.1 |
| WBGene00019828 | 0.146091726 | 0.24340539 | 0.44143421 | R02E12.4 |
| WBGene00019829 | 0.412711382 | 0.08151666 | 0.21367925 | R02E12.5 |
| WBGene00019830 | -0.669377899 | 0.01217537 | 0.05232155 | hrg-1 |
| WBGene00019831 | 0.263756851 | 0.00023958 | 0.00205846 | R02F2.1 |
| WBGene00019832 | 0.301515164 | 0.19599923 | 0.38361181 | osg-1 |
| WBGene00019833 | -0.166772498 | 0.15515349 | 0.32836112 | R02F2.4 |
| WBGene00019834 | 0.058314668 | 0.40208118 | NA | R02F2.5 |
| WBGene00019835 | NA | NA | NA | R02F2.6 |
| WBGene00019836 | -0.221358603 | 0.14083368 | 0.30856693 | R02F2.7 |
| WBGene00019837 | 0.086786644 | 0.65933601 | 0.79896991 | R02F2.8 |
| WBGene00019838 | -0.255928648 | 0.09005548 | 0.22815407 | R02F2.9 |
| WBGene00019839 | 4.015081074 | 0.000278 | 0.0023473 | R02F11.1 |
| WBGene00019840 | 0.459258192 | 0.04538591 | 0.14015926 | R02F11.2 |
| WBGene00019841 | 0.023616554 | 0.90243196 | 0.94802873 | R02F11.3 |
| WBGene00019842 | -0.07816385 | 0.43274638 | 0.62769807 | R02F11.4 |
| WBGene00019843 | -0.416835202 | 0.09984546 | 0.24519245 | R03E9.2 |
| WBGene00019844 | 0.687044124 | 1.16E-05 | 0.00014014 | abts-4 |
| WBGene00019845 | 0.040107227 | 0.58693163 | NA | irld-13 |
| WBGene00019846 | 1.297152967 | 3.88E-06 | 5.20E-05 | gpx-7 |

| | | | | |
|---|---|---|---|---|
| WBGene00019847 | -0.14180954 | 0.49655079 | 0.67886649 | R03G5.6 |
| WBGene00019849 | -0.00346345 | 0.95708614 | NA | oac-40 |
| WBGene00019850 | NA | NA | NA | srt-18 |
| WBGene00019851 | NA | NA | NA | oac-41 |
| WBGene00019853 | 0.054509443 | 0.4306858 | NA | NA |
| WBGene00019854 | -0.069564184 | 0.73168143 | 0.85001776 | R03H10.1 |
| WBGene00019855 | -0.327649016 | 0.13675548 | 0.30204209 | R03H10.2 |
| WBGene00019856 | NA | NA | NA | R03H10.4 |
| WBGene00019857 | NA | NA | NA | R03H10.5 |
| WBGene00019858 | 0.023852276 | 0.87056491 | 0.93091263 | R03H10.6 |
| WBGene00019859 | 0.122340125 | 0.26157228 | 0.45963512 | R03H10.7 |
| WBGene00019861 | 0.020317245 | 0.84643798 | NA | R04A9.1 |
| WBGene00019862 | -0.22543152 | 0.02723461 | 0.09652431 | nrde-3 |
| WBGene00019863 | 0.325322453 | 0.15484751 | 0.32795291 | R04A9.3 |
| WBGene00019864 | -0.142724352 | 0.41431416 | 0.61181736 | ceh-93 |
| WBGene00019865 | -0.249748341 | 0.24368122 | 0.44163915 | R04A9.6 |
| WBGene00019866 | 0.106997704 | 0.16248985 | NA | R04B3.1 |
| WBGene00019867 | 0.29331832 | 0.17993399 | 0.36162163 | R04B3.2 |
| WBGene00019868 | 0.15888422 | 0.28762454 | 0.48790533 | R04B3.3 |
| WBGene00019869 | -0.029288295 | 0.87062115 | 0.93091263 | cnnm-2 |
| WBGene00019870 | 0.136504554 | 0.29911873 | 0.49958329 | R04E5.7 |
| WBGene00019871 | 0.422419315 | 0.00082413 | 0.00596737 | R04E5.8 |
| WBGene00019872 | 0.095178505 | 0.64995231 | 0.79273174 | R04E5.9 |
| WBGene00019873 | 0.18472607 | 0.3252682 | 0.52447706 | arrd-26 |
| WBGene00019874 | -0.006388227 | 0.92874308 | NA | R05C11.2 |
| WBGene00019875 | 0.573732004 | 2.32E-06 | 3.32E-05 | mca-2 |
| WBGene00019877 | 0.018383274 | 0.83258267 | 0.90860486 | R05D3.2 |
| WBGene00019878 | 0.019718458 | 0.92319611 | 0.95651506 | R05D3.3 |
| WBGene00019879 | NA | NA | NA | R05D3.5 |
| WBGene00019880 | NA | NA | NA | R05D3.6 |
| WBGene00019880 | NA | NA | NA | ZC262.5 |
| WBGene00019881 | -0.15983515 | 0.26841473 | 0.46735451 | R05D3.8 |
| WBGene00019882 | -0.260449763 | 0.11041338 | 0.26217357 | R05D3.9 |
| WBGene00019883 | -0.147162813 | 0.0415506 | 0.13147662 | met-2 |
| WBGene00019884 | -0.026002146 | 0.80689723 | 0.89296858 | R05D3.12 |
| WBGene00019885 | 1.919999181 | 0.00207793 | 0.01289111 | R05D8.7 |
| WBGene00019886 | -0.04044783 | 0.53612608 | NA | R05D8.9 |
| WBGene00019887 | 0.020925399 | 0.84643798 | NA | R05D8.11 |
| WBGene00019888 | -0.08183197 | 0.34340906 | 0.54313812 | btbd-10 |
| WBGene00019890 | -0.13572561 | 0.06504032 | 0.18179751 | R05F9.6 |
| WBGene00019891 | NA | NA | NA | R05F9.7 |
| WBGene00019892 | -0.104834882 | 0.41104122 | 0.60880673 | R05F9.9 |
| WBGene00019893 | -0.39660903 | 4.95E-09 | 1.16E-07 | sgt-1 |

| | | | | | |
|---|---|---|---|---|---|
| WBGene00019894 | | -0.016915164 | 0.93039556 | 0.96077877 | R05F9.11 |
| WBGene00019895 | | 0.23288469 | 0.00055365 | 0.0042307 | aagr-2 |
| WBGene00019896 | | 0.178630669 | 0.19962303 | 0.38808669 | R05G6.1 |
| WBGene00019898 | | -0.224921799 | 0.12149134 | 0.28051644 | R05G6.4 |
| WBGene00019899 | | 0.135237889 | 0.40955264 | 0.60752382 | R05G6.5 |
| WBGene00019900 | | -0.144248068 | 0.00687067 | 0.03350883 | vdac-1 |
| WBGene00019901 | NA | | NA | NA | R05G6.9 |
| WBGene00019902 | | 1.673375553 | 2.23E-10 | 6.13E-09 | R05G6.10 |
| WBGene00019903 | NA | | NA | NA | R05G9.1 |
| WBGene00019904 | | 0.999503598 | 0.0038309 | 0.0211574 | twk-49 |
| WBGene00019905 | | -0.294594183 | 0.04793808 | 0.14591713 | R05G9.3 |
| WBGene00019906 | | 0.078387035 | 0.47443094 | 0.6618896 | R05G9R.1 |
| WBGene00019907 | | -0.120972496 | 0.50720416 | 0.68778143 | R05H11.1 |
| WBGene00019908 | | 0.079578344 | 0.41788044 | NA | R05H11.2 |
| WBGene00019909 | | 0.042242919 | 0.75306696 | 0.86007127 | R06A10.1 |
| WBGene00019911 | | 0.547835415 | 0.04574052 | 0.14095706 | R06A10.4 |
| WBGene00019912 | | 0.357613672 | 0.05946866 | 0.17062133 | NA |
| WBGene00019913 | | 0.119290655 | 0.55256166 | 0.72491534 | R06B10.2 |
| WBGene00019914 | | -0.339557789 | 0.00124629 | 0.00842816 | clec-150 |
| WBGene00019915 | | 0.073547312 | 0.68478877 | 0.81809125 | tbc-10 |
| WBGene00019916 | NA | | NA | NA | R06B10.7 |
| WBGene00019917 | | 1.712189433 | 8.44E-06 | 0.00010538 | clec-43 |
| WBGene00019918 | | -0.809506413 | 0.00312552 | 0.01801605 | fbxc-36 |
| WBGene00019919 | NA | | NA | NA | R07C3.3 |
| WBGene00019920 | | 0.180353161 | 0.08874952 | 0.22611347 | acs-15 |
| WBGene00019921 | | -0.207003335 | 0.18514074 | 0.36889569 | fbxc-34 |
| WBGene00019922 | | -0.82219872 | 0.00095242 | 0.00675601 | fbxc-28 |
| WBGene00019923 | | -0.325834718 | 0.16467633 | 0.34084094 | fbxc-29 |
| WBGene00019924 | | -0.004914234 | 0.97958526 | 0.98820677 | fbxc-33 |
| WBGene00019925 | | -0.256764473 | 0.18973417 | 0.37544235 | fbxc-31 |
| WBGene00019926 | | -0.025848508 | 0.67901638 | NA | fbxc-27 |
| WBGene00019927 | | -0.54938449 | 0.05248191 | 0.15611289 | fbxc-24 |
| WBGene00019928 | | 0.244377767 | 0.10629635 | 0.25568726 | clec-44 |
| WBGene00019929 | | 0.042132183 | 0.71222909 | 0.83815617 | R07C3.13 |
| WBGene00019930 | | -0.027535543 | 0.68063395 | NA | fbxc-35 |
| WBGene00019931 | | 8.889129511 | 1.71E-11 | 5.46E-10 | R07C12.1 |
| WBGene00019932 | | 2.207387993 | 1.19E-30 | 1.53E-28 | NA |
| WBGene00019933 | | 3.881638042 | 0.00041628 | 0.00332684 | R07C12.3 |
| WBGene00019934 | | 0.142381686 | 0.2554642 | 0.45380659 | NA |
| WBGene00019935 | | 0.480672897 | 0.02023548 | 0.07731411 | R07E4.1 |
| WBGene00019937 | | -0.07019187 | 0.66714485 | 0.80531517 | R07E4.3 |
| WBGene00019938 | | 0.014601088 | 0.91088072 | 0.95112162 | R07E4.5 |
| WBGene00019939 | | -0.121040235 | 0.56400517 | 0.73404113 | lips-17 |

| | | | | |
|---|---|---|---|---|
| WBGene00019940 | 0.125353415 | 0.02962872 | 0.10272178 | npp-21 |
| WBGene00019941 | -0.325877824 | 5.34E-05 | 0.00055445 | pgam-5 |
| WBGene00019943 | -0.402472227 | 1.45E-05 | 0.00017122 | R07G3.7 |
| WBGene00019944 | 0.456899032 | 0.06907327 | 0.1894888 | R07G3.8 |
| WBGene00019945 | 0.05787416 | 0.40594086 | NA | R08C7.1 |
| WBGene00019946 | 0.159841335 | 0.08646039 | 0.22215979 | chat-1 |
| WBGene00019947 | -0.104671719 | 0.06887783 | 0.18905892 | htz-1 |
| WBGene00019948 | NA | NA | NA | R08C7.4 |
| WBGene00019949 | 0.04032971 | 0.564277 | NA | R08C7.5 |
| WBGene00019950 | 0.083610696 | 0.30589327 | NA | clec-175 |
| WBGene00019951 | 0.106065124 | 0.22635466 | NA | R08C7.8 |
| WBGene00019952 | -0.801194164 | 0.02301244 | 0.08521119 | fbxb-74 |
| WBGene00019953 | -0.102252859 | 0.16468198 | 0.34084094 | wapl-1 |
| WBGene00019954 | NA | NA | NA | R08C7.11 |
| WBGene00019955 | 0.728814498 | 0.0015642 | 0.01017879 | R08C7.12 |
| WBGene00019956 | -1.26656935 | 0.00025941 | 0.00221197 | fbxb-75 |
| WBGene00019957 | 1.232293421 | 1.83E-50 | 4.38E-48 | R08E3.1 |
| WBGene00019959 | -0.026638715 | 0.8456952 | 0.9169413 | R08E3.3 |
| WBGene00019960 | -0.09346205 | 0.64338815 | 0.78886575 | ztf-16 |
| WBGene00019961 | 0.125980438 | 0.53169157 | 0.70813211 | R08E5.1 |
| WBGene00019962 | 0.183896953 | 0.27677294 | 0.47673325 | cysl-3 |
| WBGene00019963 | 0.150806003 | 0.35071399 | 0.55069792 | R08E5.3 |
| WBGene00019964 | NA | NA | NA | R08E5.4 |
| WBGene00019965 | -0.037400748 | 0.80823122 | 0.89375478 | R08F11.1 |
| WBGene00019966 | 0.033463284 | 0.6365037 | NA | srt-30 |
| WBGene00019967 | 1.073198772 | 0.00034599 | 0.00283474 | cyp-33C8 |
| WBGene00019968 | 1.385077017 | 0.00814751 | 0.03836851 | R08F11.4 |
| WBGene00019969 | 0.168951373 | 0.19642853 | 0.38396663 | fpn-1.3 |
| WBGene00019970 | 3.264407994 | 5.64E-06 | 7.31E-05 | R08F11.7 |
| WBGene00019971 | -0.138981911 | 0.02057168 | 0.07821085 | ergo-1 |
| WBGene00019972 | 0.177521067 | 0.40601947 | 0.60415566 | R09A1.2 |
| WBGene00019973 | 0.667280587 | 0.03108855 | 0.10595147 | R09A1.3 |
| WBGene00019975 | 2.024603109 | 0.00025442 | 0.00217075 | flp-34 |
| WBGene00019976 | NA | NA | NA | NA |
| WBGene00019977 | 1.300639328 | 4.69E-05 | 0.00049661 | fpn-1.2 |
| WBGene00019978 | 0.717409733 | 9.56E-11 | 2.75E-09 | hacd-1 |
| WBGene00019979 | -0.208978924 | 0.31882378 | 0.51898076 | R09B5.11 |
| WBGene00019980 | 0.047995428 | 0.23236376 | 0.42733801 | chil-14 |
| WBGene00019981 | 0.020317245 | 0.84643798 | NA | srbc-59 |
| WBGene00019982 | NA | NA | NA | srbc-58 |
| WBGene00019983 | -0.224400478 | 0.00255167 | 0.01526102 | sti-1 |
| WBGene00019986 | 0.838706144 | 1.19E-06 | 1.85E-05 | R09F10.1 |
| WBGene00019987 | -0.28762183 | 0.09481581 | 0.23632929 | R09F10.3 |

| | | | | |
|---|---|---|---|---|
| WBGene00019988 | 0.161839292 | 0.44432498 | 0.63794923 | R09F10.5 |
| WBGene00019989 | -0.297889594 | 0.00049174 | 0.00383159 | R09F10.8 |
| WBGene00019991 | 0.015395676 | 0.86609647 | NA | R09H3.3 |
| WBGene00019992 | NA | NA | NA | R10A10.1 |
| WBGene00019993 | 0.051933388 | 0.74376915 | 0.85522062 | rbx-2 |
| WBGene00019994 | 0.216594157 | 0.29968138 | 0.50010521 | cdh-1 |
| WBGene00019995 | 0.167697175 | 0.09780429 | 0.24229315 | R10F2.4 |
| WBGene00019996 | -0.004864426 | 0.94835335 | NA | R10F2.5 |
| WBGene00019997 | 0.078283472 | 0.7083996 | 0.83528054 | R10F2.6 |
| WBGene00019998 | 0.598722187 | 0.02032626 | 0.07757997 | R10H1.1 |
| WBGene00019999 | 0.017708609 | 0.82438962 | NA | srab-14 |
| WBGene00020003 | 0.40050424 | 0.07294273 | 0.19714869 | R11E3.1 |
| WBGene00020004 | 1.556219005 | 1.97E-06 | 2.88E-05 | R11E3.2 |
| WBGene00020006 | -0.102105328 | 0.60178211 | 0.75908213 | set-15 |
| WBGene00020007 | 0.744453199 | 8.79E-08 | 1.67E-06 | R11F4.1 |
| WBGene00020008 | 0.041065091 | 0.74438723 | 0.85556581 | R11F4.2 |
| WBGene00020010 | -0.044468772 | 0.75280805 | 0.86007127 | R11G1.1 |
| WBGene00020011 | 0.016311432 | 0.92544324 | 0.95802973 | R11G1.2 |
| WBGene00020012 | 0.20522493 | 0.28684681 | 0.48698101 | R11G1.6 |
| WBGene00020013 | NA | NA | NA | R11G11.3 |
| WBGene00020014 | 0.020232312 | 0.84643798 | NA | R11G11.6 |
| WBGene00020015 | 0.569258301 | 0.03493126 | 0.11583886 | nhr-210 |
| WBGene00020016 | 5.999276816 | 4.71E-16 | 2.52E-14 | lipl-3 |
| WBGene00020017 | 0.121012255 | 0.17811699 | NA | R12A1.3 |
| WBGene00020018 | -0.095716246 | 0.6230107 | 0.77453057 | R12B2.2 |
| WBGene00020019 | 0.085186218 | 0.39527467 | 0.59330482 | R12B2.3 |
| WBGene00020020 | 0.04032971 | 0.564277 | NA | R12B2.6 |
| WBGene00020021 | 0.015507174 | 0.80923808 | NA | R12B2.7 |
| WBGene00020022 | -0.388722334 | 0.00012443 | 0.00117459 | gldc-1 |
| WBGene00020023 | 0.076416372 | 0.53138767 | 0.70779174 | frpr-16 |
| WBGene00020025 | 0.062126925 | 0.5923772 | 0.75292947 | R12C12.5 |
| WBGene00020026 | -0.131771018 | 0.30725574 | 0.50790688 | R12C12.6 |
| WBGene00020027 | 0.047485087 | 0.69192536 | 0.82320708 | R12C12.7 |
| WBGene00020028 | -0.185918307 | 0.14930279 | 0.32056638 | R12C12.8 |
| WBGene00020029 | 0.17794369 | 0.3266453 | 0.52574947 | R12C12.9 |
| WBGene00020030 | 0.073474819 | 0.52807739 | 0.70521324 | kel-20 |
| WBGene00020031 | 0.028411467 | 0.66119718 | 0.80057789 | suco-1 |
| WBGene00020032 | NA | NA | NA | R12E2.6 |
| WBGene00020033 | 0.261189988 | 0.07602163 | 0.20355389 | R12E2.7 |
| WBGene00020034 | 0.06560776 | 0.524929 | 0.70219146 | R12E2.8 |
| WBGene00020035 | -0.281764886 | 0.000347 | 0.00284054 | egg-5 |
| WBGene00020036 | NA | NA | NA | R12E2.11 |
| WBGene00020037 | -0.366848645 | 0.00501596 | 0.02620003 | mrps-6 |

| | | | | |
|---|---|---|---|---|
| WBGene00020038 | 0.000155267 | 0.99953953 | 0.99981268 | R12E2.13 |
| WBGene00020039 | 3.190427978 | 0.00198957 | 0.01241451 | R12E2.14 |
| WBGene00020040 | 2.399352301 | 0.00582915 | 0.02953099 | R12E2.15 |
| WBGene00020041 | -0.001993815 | 0.95612996 | NA | R13A1.1 |
| WBGene00020042 | 0.020232312 | 0.84643798 | NA | R13A1.3 |
| WBGene00020043 | 0.460508308 | 0.04958335 | 0.149325 | R13A1.5 |
| WBGene00020045 | -0.013074031 | 0.91135368 | 0.95112162 | glb-23 |
| WBGene00020046 | 0.18768487 | 0.29156135 | 0.49225357 | best-16 |
| WBGene00020047 | -0.156302527 | 0.45999771 | 0.65119895 | ttr-32 |
| WBGene00020048 | -0.003448773 | 0.96209218 | NA | lgc-12 |
| WBGene00020049 | -0.401721219 | 0.097417 | 0.2417018 | ttr-8 |
| WBGene00020050 | 0.119968624 | 0.4956512 | 0.67825155 | R13A5.7 |
| WBGene00020051 | 0.475299723 | 0.06018446 | 0.1718332 | mfsd-6 |
| WBGene00020052 | -0.160514709 | 0.45032335 | 0.64273583 | R13A5.10 |
| WBGene00020053 | 0.123301879 | 0.34912712 | 0.5491588 | R13A5.11 |
| WBGene00020054 | 0.093250216 | 0.65430003 | 0.79525209 | R13D7.2 |
| WBGene00020055 | NA | NA | NA | srt-6 |
| WBGene00020056 | NA | NA | NA | R13D11.1 |
| WBGene00020057 | NA | NA | NA | R13D11.3 |
| WBGene00020058 | 0.190239985 | 0.37423351 | 0.57307319 | R13D11.4 |
| WBGene00020059 | NA | NA | NA | NA |
| WBGene00020060 | 0.015062379 | 0.80838968 | NA | srab-16 |
| WBGene00020062 | 0.131282588 | 0.15838945 | 0.33270946 | nhr-270 |
| WBGene00020063 | NA | NA | NA | R13D11.10 |
| WBGene00020064 | -0.119603117 | 0.5247689 | 0.70210559 | kbp-1 |
| WBGene00020065 | 0.020232312 | 0.84643798 | NA | clec-159 |
| WBGene00020066 | -0.003421551 | 0.95689871 | NA | dhhc-5 |
| WBGene00020067 | 0.040107227 | 0.58693163 | NA | clec-158 |
| WBGene00020068 | -0.454142969 | 4.11E-11 | 1.25E-09 | cra-1 |
| WBGene00020069 | 1.396115229 | 0.00889624 | 0.04110126 | R13H7.2 |
| WBGene00020070 | 0.124326451 | 0.11293212 | NA | rmd-6 |
| WBGene00020071 | 0.040107227 | 0.58693163 | NA | R13H9.5 |
| WBGene00020072 | NA | NA | NA | R13H9.6 |
| WBGene00020073 | -1.109141642 | 7.20E-07 | 1.17E-05 | sdz-28 |
| WBGene00020075 | 0.536525813 | 0.03835045 | 0.12403112 | math-35 |
| WBGene00020076 | NA | NA | NA | R52.4 |
| WBGene00020077 | -0.003270206 | 0.95698316 | NA | R52.5 |
| WBGene00020078 | -0.131194822 | 0.15149574 | NA | R52.6 |
| WBGene00020080 | 0.055670063 | 0.72169114 | 0.84380644 | math-37 |
| WBGene00020081 | -0.520203973 | 0.05663059 | 0.16499724 | NA |
| WBGene00020082 | -0.157250779 | 0.17932342 | 0.36082247 | gcp-2.1 |
| WBGene00020083 | -0.034547387 | 0.83133788 | 0.90812638 | R57.2 |
| WBGene00020084 | 1.553119839 | 0.00971273 | 0.04391726 | R105.1 |

| | | | | |
|---|---|---|---|---|
| WBGene00020086 | 0.269625789 | 0.2188923 | 0.41076674 | npr-24 |
| WBGene00020087 | 0.063115168 | 0.70067151 | 0.82977688 | NA |
| WBGene00020088 | -0.20905346 | 0.16000672 | 0.33471329 | R119.2 |
| WBGene00020089 | -0.172993628 | 0.31292402 | 0.51333563 | R119.3 |
| WBGene00020090 | 0.096067513 | 0.6145891 | 0.76874831 | R119.5 |
| WBGene00020091 | -0.013047251 | 0.8693958 | 0.93006166 | rnp-8 |
| WBGene00020092 | -0.013085104 | 0.87834238 | 0.93491899 | pcf-11 |
| WBGene00020093 | -0.050263476 | 0.7324102 | 0.85023927 | R144.3 |
| WBGene00020094 | -0.046863448 | 0.65753716 | 0.79799416 | wip-1 |
| WBGene00020095 | -0.069914616 | 0.57852427 | 0.74449976 | smcr-8 |
| WBGene00020096 | -0.042289579 | 0.75294185 | 0.86007127 | R144.6 |
| WBGene00020097 | -0.186939285 | 0.00075616 | 0.00551893 | larp-1 |
| WBGene00020098 | -0.312134419 | 0.05815933 | 0.1678846 | R144.10 |
| WBGene00020099 | -0.081473248 | 0.61965783 | 0.77258029 | R144.11 |
| WBGene00020100 | 0.143682932 | 0.32969934 | 0.52923232 | mks-1 |
| WBGene00020101 | 0.071450508 | 0.56032512 | 0.73080315 | lmtr-5 |
| WBGene00020102 | -0.0390962 | 0.65249255 | 0.79424205 | R148.3 |
| WBGene00020103 | 0.022711003 | 0.85162225 | 0.92039576 | R148.4 |
| WBGene00020104 | -0.022224631 | 0.89925404 | 0.94665211 | R148.5 |
| WBGene00020105 | 0.51463428 | 0.02457587 | 0.08967287 | R148.7 |
| WBGene00020106 | 0.302558373 | 0.19479559 | 0.38179413 | R151.1 |
| WBGene00020107 | -0.088870695 | 0.11059102 | 0.26229761 | R151.2 |
| WBGene00020109 | -0.219550041 | 0.06591295 | 0.18332232 | R151.6 |
| WBGene00020110 | -0.125613107 | 0.09245263 | 0.2320976 | hsp-75 |
| WBGene00020111 | 0.000960103 | 0.99084247 | 0.9948515 | R151.8 |
| WBGene00020112 | -0.285878624 | 0.00301909 | 0.01756155 | pfd-5 |
| WBGene00020113 | -0.163544317 | 0.21305195 | 0.40365788 | R151.10 |
| WBGene00020114 | 0.775656464 | 3.41E-07 | 5.84E-06 | pde-4 |
| WBGene00020115 | -0.294926983 | 2.69E-05 | 0.00029804 | mboa-6 |
| WBGene00020116 | 0.372348972 | 0.04919441 | 0.14853688 | moa-1 |
| WBGene00020117 | 0.125899713 | 0.17088208 | NA | R155.3 |
| WBGene00020118 | -0.002914971 | 0.95718083 | NA | R155.4 |
| WBGene00020119 | NA | NA | NA | R160.2 |
| WBGene00020120 | 0.004898077 | 0.97863106 | 0.98780678 | R160.3 |
| WBGene00020121 | 0.224105962 | 0.1003294 | 0.2459683 | R160.4 |
| WBGene00020122 | 0.038125098 | 0.61119482 | NA | R160.5 |
| WBGene00020123 | -0.023109762 | 0.78121253 | NA | R160.6 |
| WBGene00020125 | -0.018906196 | 0.88683031 | 0.94033043 | R173.3 |
| WBGene00020126 | 0.848594883 | 0.01063937 | 0.04689075 | flp-26 |
| WBGene00020127 | -0.023109762 | 0.78121253 | NA | R193.1 |
| WBGene00020128 | -0.190194979 | 0.17320293 | 0.35347679 | R193.2 |
| WBGene00020129 | 0.020925399 | 0.84643798 | NA | R193.3 |
| WBGene00020130 | -0.040047473 | 0.81208829 | 0.8958773 | igcm-2 |

| | | | | |
|---|---|---|---|---|
| WBGene00020131 | 0.129027369 | 0.12333334 | 0.28329518 | gcy-28 |
| WBGene00020134 | 0.036805092 | 0.83050845 | 0.90749117 | T01B6.1 |
| WBGene00020135 | 0.860299098 | 0.00371394 | 0.0206595 | aakg-4 |
| WBGene00020136 | 0.004209733 | 0.97124536 | 0.98427495 | nlp-45 |
| WBGene00020137 | 0.027633105 | 0.8175753 | 0.89973842 | T01B10.5 |
| WBGene00020138 | 0.059530286 | 0.58049605 | 0.74553006 | T01B11.1 |
| WBGene00020139 | 0.029442912 | 0.82819318 | 0.90617864 | eppl-1 |
| WBGene00020140 | 0.472967109 | 0.01244179 | 0.0532321 | ant-1.4 |
| WBGene00020141 | 4.808338701 | 1.95E-06 | 2.84E-05 | lmd-5 |
| WBGene00020142 | 0.16865532 | 0.18328419 | 0.36634318 | aak-2 |
| WBGene00020143 | -0.090697185 | 0.63156788 | 0.78099644 | T01C8.2 |
| WBGene00020144 | -0.174866321 | 0.36921837 | 0.56914362 | T01C8.3 |
| WBGene00020145 | 0.033463284 | 0.6365037 | NA | got-1.1 |
| WBGene00020146 | 0.352625824 | 0.01169529 | 0.05054036 | got-1.2 |
| WBGene00020148 | -1.170285201 | 0.00045955 | 0.00361544 | T01D1.3 |
| WBGene00020149 | -0.223551258 | 0.20735808 | 0.39645783 | T01D1.4 |
| WBGene00020150 | NA | NA | NA | T01D1.5 |
| WBGene00020151 | 0.055204521 | 0.44852756 | NA | T01G6.1 |
| WBGene00020152 | 0.457217737 | 0.0810904 | 0.2128823 | nhr-211 |
| WBGene00020153 | 0.903394687 | 0.0038732 | 0.02133464 | nhr-212 |
| WBGene00020154 | 0.020317245 | 0.84643798 | NA | T01G6.10 |
| WBGene00020155 | 1.088961767 | 8.79E-27 | 8.87E-25 | T02B11.3 |
| WBGene00020156 | 0.132122826 | 0.36648507 | 0.56612238 | T02B11.4 |
| WBGene00020157 | 0.205323007 | 0.1972154 | 0.38483681 | T02B11.6 |
| WBGene00020158 | 0.358153659 | 0.00040847 | 0.00327874 | T02C5.1 |
| WBGene00020160 | -0.6000226 | 2.42E-05 | 0.00027041 | igcm-3 |
| WBGene00020163 | -1.304177271 | 0.0001518 | 0.00138472 | T02G5.3 |
| WBGene00020164 | 0.596847289 | 0.00046463 | 0.00365261 | NA |
| WBGene00020166 | -0.305396964 | 5.71E-06 | 7.37E-05 | T02G5.7 |
| WBGene00020167 | 0.672996607 | 0.03025201 | 0.10404504 | NA |
| WBGene00020168 | -0.36247158 | 0.00031578 | 0.00262542 | mct-5 |
| WBGene00020169 | 0.525352716 | 7.48E-10 | 1.92E-08 | mmaa-1 |
| WBGene00020170 | -0.192722189 | 0.33062291 | 0.53019167 | T02G5.14 |
| WBGene00020171 | -0.276256144 | 0.02640488 | 0.09461484 | T02H6.1 |
| WBGene00020172 | -0.146173446 | 0.23021169 | 0.42455339 | thoc-1 |
| WBGene00020173 | 0.033034883 | 0.64219503 | NA | T02H6.3 |
| WBGene00020174 | NA | NA | NA | T02H6.4 |
| WBGene00020175 | -0.205312346 | 0.2175028 | 0.40915565 | T02H6.5 |
| WBGene00020176 | NA | NA | NA | T02H6.6 |
| WBGene00020177 | -0.003108065 | 0.95728506 | NA | T02H6.7 |
| WBGene00020178 | 0.020925399 | 0.84643798 | NA | T02H6.8 |
| WBGene00020179 | 0.025227316 | 0.77846045 | NA | T02H6.9 |
| WBGene00020180 | NA | NA | NA | T02H6.10 |

| | | | | |
|---|---|---|---|---|
| WBGene00020181 | -0.096234447 | 0.12154283 | 0.28054684 | T02H6.11 |
| WBGene00020182 | -3.081491569 | 4.89E-20 | 3.67E-18 | ugt-53 |
| WBGene00020183 | 0.020529042 | 0.91253035 | 0.95126775 | T03D3.5 |
| WBGene00020184 | -0.272066546 | 0.00756261 | 0.03619236 | uba-5 |
| WBGene00020185 | -0.249350212 | 0.00101799 | 0.00714839 | pgk-1 |
| WBGene00020187 | 1.347339692 | 0.00665036 | 0.03272871 | gsp-4 |
| WBGene00020188 | 0.311492305 | 0.17613488 | 0.35699717 | T03F1.6 |
| WBGene00020189 | -0.035166213 | 0.83605703 | 0.91090274 | tfbm-1 |
| WBGene00020190 | -0.074003116 | 0.53906741 | 0.71373539 | guk-1 |
| WBGene00020191 | -0.223212126 | 0.29913317 | 0.49958329 | clec-53 |
| WBGene00020192 | 0.041945952 | 0.84060457 | 0.91391904 | T03F1.11 |
| WBGene00020193 | -0.227414211 | 0.1519531 | 0.32444879 | T03F1.12 |
| WBGene00020194 | 2.135466593 | 0.00811454 | 0.03827205 | T03G6.1 |
| WBGene00020195 | 0.217194313 | 0.14211675 | 0.31014031 | T03G6.3 |
| WBGene00020196 | 0.107005324 | 0.31506195 | 0.51557662 | T03G11.3 |
| WBGene00020197 | 0.005471932 | 0.97231782 | 0.98476102 | mans-3 |
| WBGene00020199 | -0.263955897 | 0.04533952 | 0.14004555 | T03G11.6 |
| WBGene00020200 | 1.165899023 | 4.86E-07 | 8.14E-06 | T04A6.1 |
| WBGene00020201 | 0.104095495 | 0.43114873 | 0.62617285 | T04A6.2 |
| WBGene00020202 | 0.11411957 | 0.12533078 | NA | T04A6.3 |
| WBGene00020203 | NA | NA | NA | T04B8.1 |
| WBGene00020204 | -1.261841354 | 0.01268805 | 0.05409667 | NA |
| WBGene00020205 | -1.194723267 | 1.23E-06 | 1.89E-05 | arrd-1 |
| WBGene00020207 | -0.097333473 | 0.35959671 | 0.55958116 | T04B8.5 |
| WBGene00020208 | 0.16599564 | 0.07474821 | 0.2010635 | T04C4.1 |
| WBGene00020209 | 0.140618303 | 0.1240794 | 0.28442955 | T04C9.1 |
| WBGene00020210 | NA | NA | NA | T04C9.2 |
| WBGene00020211 | NA | NA | NA | T04C9.3 |
| WBGene00020212 | -0.113518066 | 0.49188241 | 0.67512065 | T04D1.2 |
| WBGene00020214 | -0.420549685 | 0.01970144 | 0.07578168 | attf-5 |
| WBGene00020215 | -0.366576053 | 0.08767866 | 0.22426713 | T04G9.4 |
| WBGene00020216 | -0.280376084 | 0.00012667 | 0.00119344 | trap-2 |
| WBGene00020217 | NA | NA | NA | T04G9.6 |
| WBGene00020218 | 0.380388164 | 1.73E-06 | 2.57E-05 | T04G9.7 |
| WBGene00020219 | 0.102824752 | 0.48135423 | 0.66742323 | T05A7.1 |
| WBGene00020220 | 0.103693905 | 0.32163492 | 0.52161514 | clec-140 |
| WBGene00020221 | NA | NA | NA | T05A7.3 |
| WBGene00020222 | 0.241592003 | 0.27767587 | 0.47761423 | fut-6 |
| WBGene00020223 | 2.776984398 | 0.00098817 | 0.00697578 | T05A7.6 |
| WBGene00020224 | 0.053344873 | 0.73448265 | 0.85122777 | T05A7.7 |
| WBGene00020225 | 0.020317245 | 0.84643798 | NA | sre-40 |
| WBGene00020227 | 0.0398788 | 0.58971221 | NA | T05A8.1 |
| WBGene00020228 | 0.020317245 | 0.84643798 | NA | T05A8.2 |

| | | | | | |
|---|---|---|---|---|---|
| WBGene00020229 | | 0.706091868 | 0.00459729 | 0.02454589 | T05A8.3 |
| WBGene00020230 | | 0.877437775 | 4.40E-06 | 5.81E-05 | nep-2 |
| WBGene00020231 | | -0.035362779 | 0.74196272 | 0.85412964 | T05A8.5 |
| WBGene00020232 | NA | | NA | NA | T05A8.6 |
| WBGene00020233 | | -0.022930767 | 0.78121253 | NA | T05A8.7 |
| WBGene00020234 | | -0.158849943 | 0.08348451 | 0.21714277 | T05A12.3 |
| WBGene00020235 | | -0.190949199 | 0.18521434 | 0.36899213 | T05A12.4 |
| WBGene00020236 | | 0.169373515 | 0.04734603 | 0.14452334 | lgc-1 |
| WBGene00020237 | | -0.15074667 | 0.47814712 | 0.66507239 | phat-4 |
| WBGene00020238 | NA | | NA | NA | T05B4.4 |
| WBGene00020239 | NA | | NA | NA | T05B4.8 |
| WBGene00020240 | | 0.038125098 | 0.61119482 | NA | T05B4.9 |
| WBGene00020241 | NA | | NA | NA | T05B4.10 |
| WBGene00020242 | | 4.900412263 | 0.00030139 | 0.00252583 | phat-5 |
| WBGene00020243 | NA | | NA | NA | T05B4.12 |
| WBGene00020244 | NA | | NA | NA | T05B4.13 |
| WBGene00020245 | | -0.099356841 | 0.6295171 | 0.77968456 | T05B11.1 |
| WBGene00020246 | | -0.134235742 | 0.0810112 | 0.21280844 | clic-1 |
| WBGene00020247 | | -0.813473042 | 0.0006847 | 0.00507837 | T05B11.4 |
| WBGene00020248 | | -0.080021475 | 0.5984837 | 0.75668392 | T05C1.1 |
| WBGene00020249 | | 0.154629375 | 0.05747417 | NA | T05C1.2 |
| WBGene00020250 | | 0.065014541 | 0.37399213 | NA | T05C1.3 |
| WBGene00020251 | | 0.578748612 | 0.00020105 | 0.00177819 | camt-1 |
| WBGene00020252 | | 0.020232312 | 0.84643798 | NA | T05C1.5 |
| WBGene00020254 | | 0.012622424 | 0.93680209 | 0.96440159 | T05C3.2 |
| WBGene00020256 | | 0.100918934 | 0.57995445 | 0.745406 | T05C3.6 |
| WBGene00020258 | | -0.165166015 | 0.41304481 | 0.61066233 | T05E7.1 |
| WBGene00020259 | | 0.086979454 | 0.30640438 | 0.50701475 | T05E7.3 |
| WBGene00020260 | | 0.42005619 | 0.0373604 | 0.12174004 | T05E7.4 |
| WBGene00020261 | | -0.836204754 | 1.99E-06 | 2.89E-05 | vet-1 |
| WBGene00020262 | | 0.117176072 | 0.51768311 | 0.69702027 | ferl-1 |
| WBGene00020263 | | -0.194144565 | 0.02881007 | 0.10067023 | let-355 |
| WBGene00020264 | | -0.091400391 | 0.35044136 | 0.55039819 | acl-8 |
| WBGene00020265 | | 0.020232312 | 0.84643798 | NA | fbxa-196 |
| WBGene00020266 | | 0.09179788 | 0.35502997 | 0.55492622 | T05H4.3 |
| WBGene00020267 | | 0.090921299 | 0.49770536 | 0.67976712 | T05H4.4 |
| WBGene00020268 | | -0.108523728 | 0.13505954 | 0.29956169 | hpo-19 |
| WBGene00020269 | | -0.423591947 | 9.79E-14 | 4.09E-12 | erfa-1 |
| WBGene00020270 | | 0.115914333 | 0.27363721 | 0.47384061 | T05H4.7 |
| WBGene00020271 | | 0.018476367 | 0.76699386 | NA | NA |
| WBGene00020273 | | -0.105909881 | 0.41390442 | 0.61154897 | T05H4.10 |
| WBGene00020274 | | -0.287215809 | 0.00136155 | 0.00905698 | T05H4.11 |
| WBGene00020275 | | -0.174470611 | 0.01186672 | 0.05114532 | atp-4 |

| | | | | |
|---|---|---|---|---|
| WBGene00020276 | 0.210045425 | 0.24935646 | 0.447792 | T05H4.15 |
| WBGene00020277 | 0.249308295 | 0.16829349 | 0.34611391 | T06A1.1 |
| WBGene00020279 | 0.330590118 | 0.11624349 | 0.2721767 | glb-25 |
| WBGene00020280 | 0.155369088 | 0.40425886 | 0.60246181 | T06A1.5 |
| WBGene00020281 | 0.176238726 | 0.14992525 | 0.32125686 | T06A4.1 |
| WBGene00020282 | 0.042516415 | 0.55991373 | NA | mps-3 |
| WBGene00020283 | 0.189117931 | 0.35731965 | 0.55743388 | T06A4.3 |
| WBGene00020284 | -0.116347437 | 0.27543434 | 0.47572289 | mel-46 |
| WBGene00020285 | -0.010851459 | 0.91937786 | 0.95526302 | T06A10.2 |
| WBGene00020286 | -0.016892602 | 0.89432366 | 0.94489559 | gtsf-1 |
| WBGene00020287 | -0.07075857 | 0.50971517 | 0.68965343 | lsy-13 |
| WBGene00020288 | NA | NA | NA | srt-73 |
| WBGene00020289 | 0.258278784 | 0.09204602 | 0.23139523 | T06C10.3 |
| WBGene00020290 | -0.127668598 | 0.19955174 | 0.38803905 | T06D4.1 |
| WBGene00020291 | -0.022930767 | 0.78121253 | NA | T06D4.2 |
| WBGene00020292 | 0.015896709 | 0.80422001 | NA | nep-19 |
| WBGene00020293 | 0.061973585 | 0.53921241 | 0.71382762 | nep-20 |
| WBGene00020294 | 0.563511295 | 0.04461118 | 0.1383504 | T06F4.1 |
| WBGene00020295 | -0.34545184 | 0.05635027 | 0.16447948 | pqbp-1.2 |
| WBGene00020296 | -0.047420833 | 0.69424841 | 0.82485238 | rrp-8 |
| WBGene00020297 | -0.077187749 | 0.29933947 | 0.4998228 | nog-1 |
| WBGene00020298 | -0.033890276 | 0.73951347 | 0.85327581 | T07A9.10 |
| WBGene00020299 | -0.097485747 | 0.63450634 | 0.78328899 | T07A9.12 |
| WBGene00020300 | -0.334669211 | 0.00472166 | 0.02501984 | tag-261 |
| WBGene00020301 | 0.420771395 | 0.02586282 | 0.09305912 | T07D1.2 |
| WBGene00020302 | 0.027734128 | 0.79804936 | 0.88691671 | T07D1.3 |
| WBGene00020304 | 0.063738357 | 0.75924606 | 0.86379292 | NA |
| WBGene00020305 | -0.009020277 | 0.91686706 | NA | T07D3.2 |
| WBGene00020306 | -0.050545243 | 0.47515706 | NA | T07D3.3 |
| WBGene00020307 | 0.059483468 | 0.77579539 | 0.87314926 | T07D3.4 |
| WBGene00020308 | NA | NA | NA | T07D3.5 |
| WBGene00020309 | 0.065014541 | 0.37399213 | NA | T07D3.6 |
| WBGene00020311 | -0.350431266 | 0.00694646 | 0.03377717 | T07D3.9 |
| WBGene00020313 | -0.014638923 | 0.89450925 | 0.94489559 | T07E3.2 |
| WBGene00020314 | -0.242684871 | 0.01125949 | 0.04907717 | T07E3.3 |
| WBGene00020315 | -0.503474757 | 0.01386447 | 0.05804682 | T07E3.4 |
| WBGene00020316 | 0.039514574 | 0.76531035 | 0.86671634 | brc-2 |
| WBGene00020317 | 1.530994853 | 3.08E-20 | 2.36E-18 | pdf-1 |
| WBGene00020318 | 0.02766226 | 0.81885014 | 0.90042384 | T07F8.1 |
| WBGene00020319 | -0.071611806 | 0.64521235 | 0.7899117 | npr-31 |
| WBGene00020320 | 0.139382837 | 0.42163847 | 0.61824826 | T07F8.4 |
| WBGene00020321 | 0.211967466 | 0.33516361 | 0.53432502 | T07F12.1 |
| WBGene00020322 | 0.079781354 | 0.59794941 | 0.75608389 | T07F12.2 |

| | | | | |
|---|---|---|---|---|
| WBGene00020323 | -0.001705491 | 0.98911322 | 0.99398816 | arrd-23 |
| WBGene00020324 | 1.537223117 | 0.00027903 | 0.00235466 | T07F12.4 |
| WBGene00020325 | 0.085309563 | 0.24820057 | NA | bath-46 |
| WBGene00020326 | 6.876871763 | 6.35E-12 | 2.18E-10 | math-38 |
| WBGene00020327 | NA | NA | NA | clec-21 |
| WBGene00020328 | 0.020925399 | 0.84643798 | NA | clec-20 |
| WBGene00020329 | 4.233826956 | 0.00014239 | 0.00130866 | bath-26 |
| WBGene00020330 | 0.062023591 | 0.73121788 | 0.84986592 | T07H6.1 |
| WBGene00020333 | -0.023056499 | 0.78121253 | NA | sre-12 |
| WBGene00020334 | -0.199023522 | 0.06949628 | 0.19038957 | epg-7 |
| WBGene00020335 | -0.174742682 | 0.41024475 | 0.60798576 | ttr-30 |
| WBGene00020338 | 0.157937701 | 0.46183634 | 0.65228105 | NA |
| WBGene00020339 | -0.353326896 | 0.06578914 | 0.18315627 | ttr-59 |
| WBGene00020340 | 0.708697934 | 2.05E-07 | 3.65E-06 | T08B1.1 |
| WBGene00020341 | 0.400360135 | 0.1093057 | 0.26039175 | T08B1.4 |
| WBGene00020342 | 0.0398788 | 0.58971221 | NA | fbxa-201 |
| WBGene00020343 | 0.096432954 | 0.61822598 | 0.77125483 | acs-3 |
| WBGene00020345 | -0.141885284 | 0.50202667 | 0.68359084 | T08B2.4 |
| WBGene00020346 | -0.166159853 | 0.06613834 | 0.18362328 | rbm-5 |
| WBGene00020347 | -0.114385423 | 0.02626341 | 0.09424574 | ech-1.2 |
| WBGene00020348 | -0.31020027 | 0.01976845 | 0.07594591 | mrpl-23 |
| WBGene00020349 | -0.19549499 | 0.34908991 | 0.5491588 | T08B2.11 |
| WBGene00020350 | 0.123301879 | 0.34912712 | 0.5491588 | T08B2.12 |
| WBGene00020351 | 0.071678115 | 0.54236457 | 0.71622257 | T08B6.1 |
| WBGene00020352 | 0.159699531 | 0.27529111 | 0.47569186 | NA |
| WBGene00020353 | 2.084009954 | 0.00318832 | 0.01833471 | T08B6.4 |
| WBGene00020354 | -0.338937531 | 0.15559222 | 0.32912395 | sgnh-1 |
| WBGene00020356 | 0.497918821 | 0.00315884 | 0.01817947 | T08B6.9 |
| WBGene00020357 | 3.271098562 | 0.00172751 | 0.01107373 | T08E11.1 |
| WBGene00020358 | 0.356850723 | 0.14168665 | 0.30966242 | math-39 |
| WBGene00020359 | 0.269060995 | 0.2374311 | 0.4337695 | math-40 |
| WBGene00020360 | 0.453835974 | 0.06619031 | 0.18367937 | math-41 |
| WBGene00020361 | -0.191468339 | 0.33089647 | 0.53051416 | fbxc-19 |
| WBGene00020362 | -0.284527513 | 0.18478124 | 0.36848027 | fbxb-10 |
| WBGene00020363 | 0.456698717 | 0.05031931 | 0.15100924 | fbxa-3 |
| WBGene00020364 | 5.927221469 | 1.15E-05 | 0.00013981 | T08E11.8 |
| WBGene00020365 | -0.035576297 | 0.82486212 | 0.90387067 | T08G2.2 |
| WBGene00020366 | -0.306614469 | 0.00045742 | 0.00360248 | acdh-10 |
| WBGene00020368 | 0.128610589 | 0.51417679 | 0.69421453 | ast-1 |
| WBGene00020369 | 0.407676081 | 3.65E-05 | 0.00039416 | T08H10.1 |
| WBGene00020370 | 0.037127706 | 0.59421403 | NA | T08H10.3 |
| WBGene00020371 | -0.003614866 | 0.95700117 | NA | oac-60 |
| WBGene00020372 | 0.209341215 | 0.06615756 | 0.18362328 | T09A12.1 |

| | | | | |
|---|---|---|---|---|
| WBGene00020373 | 0.041863657 | 0.80470169 | 0.89140028 | gpx-6 |
| WBGene00020374 | 0.267305311 | 0.05810051 | 0.16778099 | T09A12.5 |
| WBGene00020375 | -0.091341949 | 0.12238146 | 0.28163894 | pigv-1 |
| WBGene00020376 | -0.09099252 | 0.43053472 | 0.62559139 | T09B4.2 |
| WBGene00020377 | -0.062497681 | 0.63938188 | 0.78619217 | T09B4.3 |
| WBGene00020378 | -0.048983422 | 0.80568409 | 0.89208355 | T09B4.4 |
| WBGene00020379 | -1.030123039 | 7.38E-15 | 3.47E-13 | T09B4.5 |
| WBGene00020380 | 3.701637942 | 0.00227318 | 0.01383852 | T09B4.6 |
| WBGene00020381 | 0.265926596 | 0.074455 | 0.20049579 | T09B4.7 |
| WBGene00020382 | -0.527473756 | 0.00026011 | 0.00221665 | T09B4.8 |
| WBGene00020383 | -0.307587534 | 0.0033606 | 0.01909296 | tin-44 |
| WBGene00020384 | 0.036687355 | 0.59489568 | NA | T09D3.3 |
| WBGene00020385 | -0.01515683 | 0.89178711 | 0.94353629 | nhr-216 |
| WBGene00020386 | 1.701524586 | 7.66E-07 | 1.24E-05 | cyp-34A4 |
| WBGene00020388 | 0.228300653 | 0.12409917 | 0.28442955 | NA |
| WBGene00020389 | -0.392665166 | 0.00045686 | 0.00360005 | T10B5.3 |
| WBGene00020390 | 0.310991116 | 0.18408174 | 0.36743568 | T10B5.4 |
| WBGene00020391 | -0.274280235 | 5.38E-06 | 6.98E-05 | cct-7 |
| WBGene00020392 | -0.1713839 | 0.07363505 | 0.19865317 | knl-3 |
| WBGene00020393 | 0.078878961 | 0.58266613 | 0.74659933 | T10B5.7 |
| WBGene00020394 | 0.998542494 | 0.00226706 | 0.01381847 | T10B5.8 |
| WBGene00020396 | 0.018184259 | 0.89842296 | 0.94652159 | T10B5.10 |
| WBGene00020397 | -0.465291926 | 0.00829118 | 0.0388576 | pcbd-1 |
| WBGene00020398 | 0.118889379 | 0.24769539 | 0.44607264 | cerk-1 |
| WBGene00020399 | -0.349203739 | 0.00021678 | 0.00188924 | ztf-4 |
| WBGene00020400 | NA | NA | NA | T10B11.4 |
| WBGene00020401 | 0.009939852 | 0.96272819 | 0.97924853 | T10B11.5 |
| WBGene00020402 | -0.08202732 | 0.52718454 | 0.70443633 | T10B11.6 |
| WBGene00020403 | -0.252425864 | 0.01967008 | 0.07570707 | hrde-4 |
| WBGene00020404 | -0.112225542 | 0.41734641 | 0.61426198 | T10B11.8 |
| WBGene00020406 | NA | NA | NA | T10D4.1 |
| WBGene00020407 | 0.085832948 | 0.68215022 | 0.8161756 | chil-24 |
| WBGene00020408 | -0.293576484 | 0.04816695 | 0.1463202 | T10D4.6 |
| WBGene00020409 | NA | NA | NA | T10D4.7 |
| WBGene00020410 | NA | NA | NA | T10D4.11 |
| WBGene00020411 | 0.036840012 | 0.76789095 | 0.86834482 | T10E9.1 |
| WBGene00020412 | -0.24670958 | 0.0087174 | 0.0404025 | T10E9.2 |
| WBGene00020413 | -0.309511423 | 0.17941313 | 0.36082247 | T10E9.3 |
| WBGene00020414 | 2.692191956 | 1.25E-05 | 0.00014968 | T10E9.4 |
| WBGene00020415 | 0.017141582 | 0.79355725 | NA | T10E9.5 |
| WBGene00020416 | 0.018860755 | 0.77262519 | NA | T10E9.6 |
| WBGene00020417 | -0.162998315 | 0.05740492 | 0.16643091 | nuo-2 |
| WBGene00020418 | -0.055586347 | 0.74524195 | 0.85590582 | T10E9.8 |

| | | | | |
|---|---|---|---|---|
| WBGene00020419 | 0.668573884 | 0.03501729 | 0.11599279 | acdh-4 |
| WBGene00020420 | 0.014602567 | 0.85416114 | NA | T10E10.3 |
| WBGene00020421 | 0.051828392 | 0.7389691 | 0.85307041 | T10E10.4 |
| WBGene00020422 | -0.263532032 | 0.00965442 | 0.04373463 | T10F2.2 |
| WBGene00020423 | -0.358259525 | 1.80E-07 | 3.24E-06 | prp-19 |
| WBGene00020424 | -0.005394017 | 0.97450861 | 0.98575242 | T10H9.1 |
| WBGene00020425 | 0.111124896 | 0.32101477 | 0.52094627 | syx-18 |
| WBGene00020426 | 0.329788164 | 0.11508916 | 0.27038196 | T10H10.2 |
| WBGene00020427 | 0.119954914 | 0.50912854 | 0.68943312 | sav-1 |
| WBGene00020428 | -0.023109762 | 0.78121253 | NA | irld-48 |
| WBGene00020430 | NA | NA | NA | irld-49 |
| WBGene00020431 | NA | NA | NA | irld-50 |
| WBGene00020432 | NA | NA | NA | hpa-2 |
| WBGene00020433 | 0.043817472 | 0.77188916 | 0.87031066 | T11F8.1 |
| WBGene00020434 | -0.143723185 | 0.47033482 | 0.65840969 | NA |
| WBGene00020435 | 0.063086775 | 0.41202602 | NA | T11F8.4 |
| WBGene00020436 | -0.203450149 | 0.32084504 | 0.52084403 | T12A2.1 |
| WBGene00020437 | -0.251699923 | 5.09E-06 | 6.64E-05 | stt-3 |
| WBGene00020438 | -0.406181414 | 0.10418029 | 0.25236659 | T12A2.3 |
| WBGene00020439 | 0.054259843 | 0.60435562 | 0.76077049 | T12A2.5 |
| WBGene00020440 | -0.001842784 | 0.95621055 | NA | T12A2.6 |
| WBGene00020441 | -0.163689509 | 0.09896412 | 0.24405148 | bcas-2 |
| WBGene00020442 | -0.102947065 | 0.28590259 | 0.48599789 | gen-1 |
| WBGene00020443 | -0.100893266 | 0.47649029 | 0.66371366 | esyt-2 |
| WBGene00020444 | 0.456631102 | 0.01402779 | 0.05859654 | T12B3.1 |
| WBGene00020445 | -0.389951495 | 0.05083637 | 0.15226991 | T12B3.2 |
| WBGene00020446 | 0.170766361 | 0.34461043 | 0.54434679 | T12B3.3 |
| WBGene00020447 | 0.069401995 | 0.66411012 | 0.80296004 | mob-3 |
| WBGene00020448 | 0.537586249 | 0.05292338 | 0.1570598 | fbxa-51 |
| WBGene00020449 | 0.161207082 | 0.44147354 | 0.63568195 | fbxa-54 |
| WBGene00020450 | 0.16317096 | 0.36895425 | 0.56885627 | fbxa-10 |
| WBGene00020451 | 0.270516646 | 0.22178603 | 0.41448442 | fbxa-11 |
| WBGene00020452 | NA | NA | NA | fbxa-22 |
| WBGene00020453 | 0.735326139 | 0.00956315 | 0.04349615 | fbxa-55 |
| WBGene00020454 | 0.018795205 | 0.77532208 | NA | fbxa-56 |
| WBGene00020455 | 1.559383032 | 9.23E-08 | 1.75E-06 | fbxa-59 |
| WBGene00020456 | 1.450368163 | 3.57E-12 | 1.25E-10 | fbxa-60 |
| WBGene00020457 | 0.73771812 | 0.00861636 | 0.03999752 | fbxa-67 |
| WBGene00020458 | 0.17504235 | 0.28473495 | 0.4847447 | NA |
| WBGene00020459 | 0.08357617 | 0.40331318 | 0.60162081 | fbxa-23 |
| WBGene00020460 | -0.031682333 | 0.8434806 | 0.91599165 | nhr-273 |
| WBGene00020461 | -0.07507442 | 0.61611526 | 0.76979828 | T12C9.7 |
| WBGene00020462 | -0.030120241 | 0.76560344 | 0.86684711 | ari-2 |

| | | | | |
|---|---|---|---|---|
| WBGene00020463 | -0.062946533 | 0.49564907 | 0.67825155 | cec-6 |
| WBGene00020464 | -0.171228185 | 0.19168648 | 0.37787555 | T12E12.3 |
| WBGene00020465 | 0.036687355 | 0.59489568 | NA | T12E12.6 |
| WBGene00020466 | -0.128692252 | 0.27220487 | 0.47230997 | sld-2 |
| WBGene00020467 | -0.467649499 | 0.00059906 | 0.0045421 | T12F5.2 |
| WBGene00020468 | 0.034909354 | 0.71754056 | 0.84158547 | T13A10.1 |
| WBGene00020469 | 0.085347488 | 0.3030966 | NA | T13A10.2 |
| WBGene00020471 | 0.61494337 | 9.69E-07 | 1.54E-05 | pho-14 |
| WBGene00020472 | 0.040107227 | 0.58693163 | NA | lips-11 |
| WBGene00020473 | -0.035761389 | 0.73076692 | 0.84986592 | lips-12 |
| WBGene00020474 | 0.017604133 | 0.79437959 | NA | lips-13 |
| WBGene00020475 | 0.083236723 | 0.57735823 | 0.74411462 | sut-1 |
| WBGene00020476 | 0.126118189 | 0.53897109 | 0.71373539 | T13B5.9 |
| WBGene00020478 | 0.074910648 | 0.55630987 | 0.72794301 | T13C2.2 |
| WBGene00020479 | 0.269224091 | 0.22209557 | 0.41486365 | T13C2.3 |
| WBGene00020480 | -0.296756519 | 0.04557839 | 0.14064026 | ssup-72 |
| WBGene00020481 | -0.035906287 | 0.48138715 | 0.66742323 | T13C2.6 |
| WBGene00020482 | 0.041995817 | 0.78681617 | 0.87943931 | T13C2.7 |
| WBGene00020483 | NA | NA | NA | T13C5.2 |
| WBGene00020484 | 0.016453512 | 0.83636998 | NA | T13C5.3 |
| WBGene00020485 | 0.085309563 | 0.24820057 | NA | ceh-54 |
| WBGene00020486 | 0.406607352 | 0.07100865 | 0.19349295 | T13C5.6 |
| WBGene00020487 | -0.035554943 | 0.7427605 | 0.85444405 | T13C5.7 |
| WBGene00020488 | -0.023056499 | 0.78121253 | NA | T13G4.1 |
| WBGene00020489 | -0.059828008 | 0.75973729 | 0.86405055 | hot-8 |
| WBGene00020490 | 0.477326982 | 0.06402103 | 0.17977214 | tmc-1 |
| WBGene00020491 | -0.074582689 | 0.7161774 | 0.84075838 | T13G4.4 |
| WBGene00020492 | 0.020317245 | 0.84643798 | NA | T13G4.5 |
| WBGene00020494 | 0.020232312 | 0.84643798 | NA | T13G4.7 |
| WBGene00020496 | 0.012882159 | 0.87753076 | 0.93473466 | spat-3 |
| WBGene00020497 | 0.504139627 | 0.05963613 | 0.17090088 | T14A8.2 |
| WBGene00020498 | -0.053610607 | 0.62507449 | 0.77616111 | T14B4.1 |
| WBGene00020499 | -0.285205991 | 0.06554186 | 0.18274583 | mrps-18.C |
| WBGene00020500 | -0.069168883 | 0.51362431 | 0.69359652 | T14B4.3 |
| WBGene00020501 | -0.269553164 | 0.19677072 | 0.38427639 | T14B4.5 |
| WBGene00020502 | -0.1096078 | 0.59363063 | 0.75344508 | T14B4.8 |
| WBGene00020503 | -0.036383874 | 0.78272454 | 0.87768017 | T14B4.9 |
| WBGene00020504 | 0.273410425 | 0.03354931 | 0.11242742 | svh-2 |
| WBGene00020505 | 0.05787416 | 0.40594086 | NA | T14E8.2 |
| WBGene00020506 | 0.882523617 | 0.01971017 | 0.07578168 | dop-3 |
| WBGene00020507 | 0.114914524 | 0.0830605 | 0.21637781 | vha-15 |
| WBGene00020508 | 0.052553175 | 0.76211191 | 0.86496748 | T14F9.2 |
| WBGene00020509 | 0.283934784 | 0.0170932 | 0.06841485 | hex-1 |

| | | | | | |
|---|---|---|---|---|---|
| WBGene00020510 | | 0.2941387 | 0.17782034 | 0.35909899 | T14G11.1 |
| WBGene00020511 | | -0.003194923 | 0.97450377 | 0.98575242 | immt-1 |
| WBGene00020515 | NA | | NA | NA | T14G12.6 |
| WBGene00020516 | | -0.217233878 | 0.02961338 | 0.10271723 | T15B7.1 |
| WBGene00020517 | | 0.191793112 | 0.00871704 | 0.0404025 | hpo-8 |
| WBGene00020520 | NA | | NA | NA | T15B7.8 |
| WBGene00020522 | | -0.004598644 | 0.97369751 | 0.98547665 | T15B7.10 |
| WBGene00020523 | NA | | NA | NA | dmsr-14 |
| WBGene00020524 | | 0.020925399 | 0.84643798 | NA | dmsr-13 |
| WBGene00020525 | NA | | NA | NA | dmsr-15 |
| WBGene00020526 | | -0.494546439 | 0.04295341 | 0.13469288 | T15B7.14 |
| WBGene00020527 | | -0.011042939 | 0.91191786 | 0.95112162 | T15B7.15 |
| WBGene00020528 | | -0.287957859 | 0.20747007 | 0.39654951 | lgc-54 |
| WBGene00020530 | | 0.525841216 | 0.00235799 | 0.01427895 | T15B12.1 |
| WBGene00020531 | | 0.050144643 | 0.55745549 | NA | T15B12.2 |
| WBGene00020532 | | 0.179129814 | 0.09050034 | 0.2288455 | math-42 |
| WBGene00020533 | | 0.020232312 | 0.84643798 | NA | T16A1.2 |
| WBGene00020534 | | 0.020925399 | 0.84643798 | NA | fbxc-49 |
| WBGene00020535 | NA | | NA | NA | T16A1.4 |
| WBGene00020536 | NA | | NA | NA | T16A1.5 |
| WBGene00020537 | | -0.679037118 | 0.01542995 | 0.06326234 | fbxb-37 |
| WBGene00020539 | NA | | NA | NA | T16D1.1 |
| WBGene00020540 | | 0.290077816 | 0.20917982 | 0.39862055 | pals-25 |
| WBGene00020541 | | -0.97299024 | 0.01275021 | 0.05427085 | fbxb-81 |
| WBGene00020542 | | -0.571688896 | 0.04832627 | 0.14658068 | fbxb-82 |
| WBGene00020543 | | -0.318731088 | 0.17469482 | 0.35538514 | fbxb-83 |
| WBGene00020544 | | -0.995739862 | 0.0028263 | 0.01665188 | fbxb-84 |
| WBGene00020545 | | -0.005691347 | 0.93660638 | NA | adbp-2 |
| WBGene00020546 | | 0.035177603 | 0.67823902 | NA | zig-13 |
| WBGene00020547 | | -0.001993815 | 0.95612996 | NA | NA |
| WBGene00020548 | | -0.105834181 | 0.21658577 | NA | T17A3.12 |
| WBGene00020549 | | -0.644409415 | 2.30E-17 | 1.41E-15 | nmt-1 |
| WBGene00020550 | | -0.182605719 | 0.39653151 | 0.59453332 | T17H7.1 |
| WBGene00020551 | | 0.019429395 | 0.91607613 | 0.95318532 | T17H7.7 |
| WBGene00020553 | | -0.021918395 | 0.76194329 | 0.86495294 | ttc-17 |
| WBGene00020554 | | 0.992437953 | 0.01695758 | 0.06800216 | T19A5.3 |
| WBGene00020555 | | 0.019325396 | 0.77340931 | NA | nhr-219 |
| WBGene00020556 | | 0.114616758 | 0.44113327 | 0.63531696 | pamn-1 |
| WBGene00020557 | | -0.088451021 | 0.42009069 | 0.61684362 | T19B4.3 |
| WBGene00020558 | | -0.068414745 | 0.45316755 | 0.64528841 | T19B4.5 |
| WBGene00020559 | | 0.029820668 | 0.8860653 | 0.94006391 | best-17 |
| WBGene00020560 | | 0.306703639 | 0.02889657 | 0.10087619 | T19C3.2 |
| WBGene00020561 | | 0.229584169 | 0.23916827 | 0.43606953 | T19C3.3 |

| | | | | |
|---|---|---|---|---|
| WBGene00020562 | 0.601724038 | 0.0005166 | 0.00399339 | T19C3.4 |
| WBGene00020563 | 0.048909231 | 0.7989684 | 0.88739921 | T19C3.5 |
| WBGene00020564 | 0.04913709 | 0.56495507 | NA | T19C3.6 |
| WBGene00020565 | 0.063086775 | 0.41202602 | NA | T19C3.7 |
| WBGene00020566 | 0.026415684 | 0.89048389 | 0.94297517 | ttr-7 |
| WBGene00020567 | 0.094999169 | 0.63825408 | 0.78552676 | adt-3 |
| WBGene00020568 | -0.034867888 | 0.82993196 | 0.9071919 | T19D2.3 |
| WBGene00020569 | -0.085322767 | 0.67159417 | 0.80894366 | lgc-32 |
| WBGene00020571 | 0.015896709 | 0.80422001 | NA | lpr-7 |
| WBGene00020572 | 0.136815004 | 0.51620977 | 0.69567629 | rund-1 |
| WBGene00020573 | 0.015516711 | 0.87392635 | NA | T19D7.5 |
| WBGene00020574 | 0.380653795 | 0.1063019 | 0.25568726 | T19D7.6 |
| WBGene00020575 | 0.216642551 | 0.16531711 | 0.34162424 | T19D7.7 |
| WBGene00020576 | 1.248609869 | 1.65E-12 | 6.00E-11 | T19D12.1 |
| WBGene00020577 | 0.569477264 | 0.00369231 | 0.02057229 | T19D12.2 |
| WBGene00020578 | 0.059762547 | 0.38965825 | NA | T19D12.3 |
| WBGene00020579 | 0.204969385 | 0.21021854 | 0.39981938 | T19D12.4 |
| WBGene00020580 | 0.038221642 | 0.65613466 | NA | T19D12.5 |
| WBGene00020581 | 0.478385221 | 3.91E-05 | 0.00041984 | T19D12.6 |
| WBGene00020582 | 0.108116047 | 0.44969821 | 0.64204323 | oig-8 |
| WBGene00020583 | 0.745330575 | 0.02228211 | 0.08315891 | T19D12.9 |
| WBGene00020584 | 0.038765322 | 0.66219346 | NA | T19D12.10 |
| WBGene00020585 | 0.046423366 | 0.8236864 | 0.9034778 | clec-178 |
| WBGene00020586 | 0.373560654 | 0.04010568 | 0.12817885 | frpr-18 |
| WBGene00020587 | 0.016007174 | 0.93916666 | 0.96586959 | ugt-9 |
| WBGene00020588 | 0.051250263 | 0.31673338 | 0.51698682 | T19H12.2 |
| WBGene00020589 | 0.647921782 | 0.00660978 | 0.03257066 | T19H12.3 |
| WBGene00020590 | NA | NA | NA | T19H12.6 |
| WBGene00020591 | -0.002295056 | 0.98566488 | 0.99189769 | NA |
| WBGene00020592 | -0.676368228 | 2.86E-06 | 3.96E-05 | ugt-12 |
| WBGene00020593 | 0.047608179 | 0.81922897 | 0.90054293 | ugt-11 |
| WBGene00020594 | NA | NA | NA | ugt-10 |
| WBGene00020595 | 0.934035011 | 0.01616266 | 0.06565969 | T20B5.2 |
| WBGene00020596 | 0.11290185 | 0.4751123 | 0.66217223 | oga-1 |
| WBGene00020597 | NA | NA | NA | T20B6.1 |
| WBGene00020598 | 0.038125098 | 0.61119482 | NA | T20B6.2 |
| WBGene00020599 | -0.023411554 | 0.8744067 | 0.93303611 | T20B6.3 |
| WBGene00020600 | -0.293898132 | 0.01248886 | 0.0533867 | trd-1 |
| WBGene00020601 | -0.087248958 | 0.46165769 | 0.65228105 | T20B12.3 |
| WBGene00020602 | 0.020925399 | 0.84643798 | NA | T20B12.4 |
| WBGene00020603 | NA | NA | NA | T20B12.5 |
| WBGene00020604 | 0.018525853 | 0.77805821 | 0.87435671 | T20B12.7 |
| WBGene00020605 | 0.610068554 | 0.04004436 | 0.12803875 | lgc-50 |

| | | | | |
|---|---|---|---|---|
| WBGene00020607 | NA | NA | NA | srab-20 |
| WBGene00020608 | NA | NA | NA | srab-22 |
| WBGene00020609 | 0.016864519 | 0.82560414 | NA | T20D4.3 |
| WBGene00020611 | 0.099305307 | 0.57128478 | 0.7395385 | T20D4.5 |
| WBGene00020612 | 0.183869046 | 0.39362595 | 0.59205646 | arrd-22 |
| WBGene00020613 | 3.186218402 | 0.00100921 | 0.00710038 | T20D4.7 |
| WBGene00020614 | -0.023756719 | 0.6882956 | NA | T20D4.8 |
| WBGene00020615 | 0.098674686 | 0.39052766 | 0.58842394 | T20D4.9 |
| WBGene00020616 | 0.13684499 | 0.38688979 | 0.58559427 | T20D4.10 |
| WBGene00020617 | 0.224982256 | 0.30470463 | 0.50545453 | T20D4.11 |
| WBGene00020618 | -0.022930767 | 0.78121253 | NA | T20D4.12 |
| WBGene00020619 | 0.236586887 | 0.01979246 | 0.07599826 | T20D4.13 |
| WBGene00020620 | 0.156042338 | 0.21860529 | 0.41042776 | T20D4.15 |
| WBGene00020622 | NA | NA | NA | T20D4.17 |
| WBGene00020623 | NA | NA | NA | srab-21 |
| WBGene00020624 | 0.007004473 | 0.95172663 | 0.97232907 | T20D4.19 |
| WBGene00020625 | 0.160422812 | 0.36176873 | 0.56162188 | mrrf-1 |
| WBGene00020626 | 1.546760495 | 0.00977863 | 0.04411987 | T20F5.4 |
| WBGene00020627 | 0.438258028 | 0.07367644 | 0.1986916 | T20F5.5 |
| WBGene00020628 | -0.063048233 | 0.42585865 | 0.62160822 | T20F5.6 |
| WBGene00020629 | -0.186556158 | 0.0633039 | 0.17847856 | T20F5.7 |
| WBGene00020630 | -0.056635481 | 0.66655189 | 0.80470744 | T20F7.1 |
| WBGene00020631 | 0.117113962 | 0.26279598 | 0.46078993 | T20F7.3 |
| WBGene00020632 | 0.266136757 | 0.24175219 | 0.4397433 | NA |
| WBGene00020633 | -0.016099065 | 0.91152829 | 0.95112162 | aakg-2 |
| WBGene00020634 | 0.10343811 | 0.38053245 | 0.57992875 | acs-9 |
| WBGene00020635 | 0.100264625 | 0.38646629 | 0.5851299 | T20H4.2 |
| WBGene00020636 | -0.32832985 | 6.97E-05 | 0.00070428 | T20H4.5 |
| WBGene00020637 | 0.132442484 | 0.10147609 | NA | fbxa-7 |
| WBGene00020638 | -0.005017292 | 0.94670066 | NA | fbxa-43 |
| WBGene00020639 | 0.350381882 | 0.13993437 | 0.30730149 | fbxa-71 |
| WBGene00020640 | 0.108717649 | 0.15669083 | NA | fbxa-73 |
| WBGene00020641 | 0.065014541 | 0.37399213 | NA | NA |
| WBGene00020642 | -0.003614866 | 0.95700117 | NA | T20H9.6 |
| WBGene00020645 | NA | NA | NA | T21D9.2 |
| WBGene00020646 | -0.001488154 | 0.95639912 | NA | zmp-6 |
| WBGene00020647 | -0.116901078 | 0.24474161 | 0.44292812 | pqbp-1.1 |
| WBGene00020648 | 0.11501967 | 0.26873634 | 0.46773285 | T21D12.7 |
| WBGene00020649 | 1.0441216 | 1.33E-09 | 3.31E-08 | sma-10 |
| WBGene00020650 | 0.017604133 | 0.79437959 | NA | T21D12.11 |
| WBGene00020651 | 0.249723681 | 0.21154645 | 0.40161457 | T21D12.12 |
| WBGene00020652 | -0.129852377 | 0.11245305 | 0.26542401 | egg-4 |
| WBGene00020653 | -0.022930767 | 0.78121253 | NA | T21E3.2 |

| | | | | |
|---|---|---|---|---|
| WBGene00020655 | -0.023109762 | 0.78121253 | NA | T21E12.3 |
| WBGene00020656 | 0.020232312 | 0.84643798 | NA | T21E12.5 |
| WBGene00020657 | 1.747284549 | 0.01309692 | 0.05548382 | lgc-53 |
| WBGene00020658 | -0.061149225 | 0.583807 | 0.74761814 | T21F4.1 |
| WBGene00020659 | 0.101779524 | 0.32286833 | 0.52291098 | T21G5.1 |
| WBGene00020660 | 0.060105893 | 0.41982798 | NA | T21G5.2 |
| WBGene00020661 | 0.434085154 | 0.0751094 | 0.20159059 | smz-2 |
| WBGene00020662 | 0.646842203 | 5.80E-18 | 3.71E-16 | T21H3.1 |
| WBGene00020663 | -0.004864426 | 0.94835335 | NA | cutl-22 |
| WBGene00020664 | 0.018998992 | 0.77286019 | NA | T21H3.5 |
| WBGene00020665 | -0.524076228 | 0.00823432 | 0.03864062 | T22B2.1 |
| WBGene00020666 | -0.136826863 | 0.29257571 | 0.49350242 | T22B2.2 |
| WBGene00020667 | -0.00346345 | 0.95708614 | NA | T22B2.3 |
| WBGene00020669 | 0.020232312 | 0.84643798 | NA | T22B2.5 |
| WBGene00020670 | 0.020232312 | 0.84643798 | NA | T22B2.6 |
| WBGene00020672 | -0.183512333 | 0.35978409 | 0.55966839 | T22B7.3 |
| WBGene00020673 | -1.147623142 | 3.48E-05 | 0.00037747 | T22B7.4 |
| WBGene00020674 | -1.801970074 | 1.03E-06 | 1.61E-05 | T22B7.7 |
| WBGene00020675 | 0.050601671 | 0.68421308 | 0.8176383 | T22B11.1 |
| WBGene00020678 | 1.391568863 | 8.18E-17 | 4.77E-15 | T22B11.4 |
| WBGene00020679 | 0.048922319 | 0.39657617 | 0.5945394 | ogdh-1 |
| WBGene00020680 | 0.020232312 | 0.84643798 | NA | T22D1.1 |
| WBGene00020682 | -0.067991047 | 0.35384421 | 0.5536639 | T22D1.3 |
| WBGene00020683 | -0.258070266 | 9.74E-07 | 1.54E-05 | ribo-1 |
| WBGene00020684 | -0.133215402 | 0.04979675 | 0.14973797 | T22D1.5 |
| WBGene00020686 | -0.003421551 | 0.95689871 | NA | T22D1.8 |
| WBGene00020687 | -0.391606655 | 7.09E-07 | 1.15E-05 | ruvb-2 |
| WBGene00020688 | -0.332686708 | 0.16117142 | 0.33571074 | T22D1.11 |
| WBGene00020689 | 0.647647866 | 0.03691543 | 0.12063376 | npr-12 |
| WBGene00020690 | -0.201237857 | 0.31930199 | 0.51943339 | T22E5.1 |
| WBGene00020691 | -0.037152691 | 0.7475597 | 0.85735716 | igdb-3 |
| WBGene00020692 | 0.282089734 | 0.20485813 | 0.39397451 | T22E5.6 |
| WBGene00020693 | -0.068129594 | 0.73164287 | 0.85001776 | lron-8 |
| WBGene00020694 | -0.02791392 | 0.6763823 | NA | dhhc-11 |
| WBGene00020695 | -0.189747002 | 0.06199148 | 0.17572849 | T22F3.2 |
| WBGene00020696 | 0.012263599 | 0.76869394 | 0.86844708 | pygl-1 |
| WBGene00020697 | 0.092322524 | 0.57010084 | 0.7387023 | T22F3.7 |
| WBGene00020698 | 0.204978633 | 0.03076956 | 0.10520716 | T22F3.8 |
| WBGene00020699 | 0.463127696 | 0.03321175 | 0.1115004 | T22F3.10 |
| WBGene00020700 | 7.127475263 | 1.14E-17 | 7.08E-16 | T22F3.11 |
| WBGene00020701 | 1.715775828 | 7.14E-15 | 3.37E-13 | T22F7.1 |
| WBGene00020702 | 0.366943365 | 0.09836689 | 0.2432018 | T22F7.3 |
| WBGene00020703 | 0.347813374 | 0.11098097 | 0.26296689 | T22F7.4 |

| | | | | |
|---|---|---|---|---|
| WBGene00020704 | 0.195703989 | 0.22424591 | 0.41712629 | T22F7.5 |
| WBGene00020705 | -0.219290304 | 0.0777876 | 0.20676983 | T22H9.1 |
| WBGene00020706 | 0.26099443 | 0.01250346 | 0.05341791 | atg-9 |
| WBGene00020707 | 0.085309563 | 0.24820057 | NA | wago-10 |
| WBGene00020708 | 0.010693481 | 0.91044081 | 0.95112162 | dmd-8 |
| WBGene00020709 | -0.116032084 | 0.20154515 | 0.39027971 | T23B3.1 |
| WBGene00020710 | 0.318803962 | 0.10678492 | 0.25651158 | T23B3.2 |
| WBGene00020711 | -0.059168106 | 0.58774465 | 0.75003656 | NA |
| WBGene00020712 | 0.158421825 | 0.2514916 | 0.44991306 | ckr-1 |
| WBGene00020713 | 0.546047142 | 0.03037092 | 0.1042825 | T23B3.5 |
| WBGene00020714 | 0.144749483 | 0.37320851 | 0.57229033 | T23B3.6 |
| WBGene00020715 | 0.700911833 | 0.02016503 | 0.07716576 | nspd-4 |
| WBGene00020715 | 0.700911833 | 0.02016503 | 0.07716576 | nspd-5 |
| WBGene00020716 | -0.157556413 | 0.21716133 | 0.4087973 | phf-30 |
| WBGene00020717 | -0.074812003 | 0.59334777 | 0.7534436 | mrpl-4 |
| WBGene00020718 | -0.183788032 | 0.11733385 | 0.27380421 | mrps-2 |
| WBGene00020719 | -0.162496628 | 0.02750561 | 0.09722587 | natc-1 |
| WBGene00020720 | 0.017279211 | 0.79380376 | NA | T23B12.5 |
| WBGene00020721 | 0.157304252 | 0.13650108 | 0.30161672 | T23B12.6 |
| WBGene00020722 | 0.098970981 | 0.63026476 | 0.78009183 | T23B12.8 |
| WBGene00020725 | -0.210139536 | 0.18830706 | 0.37342593 | cnp-3 |
| WBGene00020726 | -0.287136188 | 0.15227317 | 0.32480061 | T23C6.4 |
| WBGene00020727 | 0.18747054 | 0.31767368 | 0.51787779 | npr-21 |
| WBGene00020728 | 0.065250956 | 0.56039389 | 0.73080315 | T23E1.1 |
| WBGene00020729 | -0.192666755 | 0.37350372 | 0.572527 | NA |
| WBGene00020730 | 0.075450426 | 0.33451267 | NA | T23E1.3 |
| WBGene00020732 | 0.031268349 | 0.76419839 | 0.86623378 | T23E7.2 |
| WBGene00020735 | 0.472990588 | 0.06023319 | 0.17193881 | T23F2.2 |
| WBGene00020736 | 1.215716718 | 2.58E-06 | 3.63E-05 | T23F2.3 |
| WBGene00020737 | 0.033463284 | 0.6365037 | NA | T23F2.4 |
| WBGene00020738 | -0.110348414 | 0.24556533 | 0.44370259 | T23F2.5 |
| WBGene00020739 | NA | NA | NA | T23F4.1 |
| WBGene00020740 | 0.678945894 | 0.02050738 | 0.07806773 | T23F4.2 |
| WBGene00020741 | 0.193525222 | 0.15975817 | 0.33441858 | dlhd-1 |
| WBGene00020742 | -0.135855174 | 0.10735973 | 0.25742774 | T23H2.3 |
| WBGene00020743 | 0.204978633 | 0.03076956 | 0.10520716 | mam-8 |
| WBGene00020744 | 0.164860012 | 0.42922382 | 0.62405819 | T24A6.1 |
| WBGene00020746 | NA | NA | NA | srbc-66 |
| WBGene00020747 | 0.020925399 | 0.84643798 | NA | T24A6.7 |
| WBGene00020748 | -0.023056499 | 0.78121253 | NA | nhr-221 |
| WBGene00020749 | 0.020232312 | 0.84643798 | NA | srbc-67 |
| WBGene00020750 | -0.003421551 | 0.95689871 | NA | nhr-222 |
| WBGene00020751 | NA | NA | NA | NA |

| | | | | |
|---|---|---|---|---|
| WBGene00020752 | NA | NA | NA | srbc-68 |
| WBGene00020753 | NA | NA | NA | T24A6.16 |
| WBGene00020754 | 0.013553178 | 0.74301546 | NA | F40C5.2 |
| WBGene00020754 | 0.013553178 | 0.74301546 | NA | T24A6.17 |
| WBGene00020755 | NA | NA | NA | grl-31 |
| WBGene00020756 | -0.256881302 | 0.15777137 | 0.33193429 | T24A6.20 |
| WBGene00020757 | -0.232462947 | 0.00244679 | 0.01474215 | ucr-2.3 |
| WBGene00020758 | -0.950267544 | 2.13E-06 | 3.09E-05 | T24C4.2 |
| WBGene00020759 | 0.032136579 | 0.69225183 | NA | T24C4.3 |
| WBGene00020760 | -1.041957865 | 0.01140837 | 0.04954906 | T24C4.4 |
| WBGene00020761 | -0.308205388 | 0.00911953 | 0.04179 | T24C4.5 |
| WBGene00020762 | -0.062161318 | 0.39440537 | 0.59249945 | zer-1 |
| WBGene00020763 | -0.151765055 | 0.14098487 | 0.30863628 | ztf-18 |
| WBGene00020764 | 0.186439191 | 0.27264466 | 0.4726499 | T24C12.1 |
| WBGene00020765 | -0.44688778 | 0.00986166 | 0.04439868 | T24C12.3 |
| WBGene00020766 | 0.037023624 | 0.59153183 | NA | T24C12.4 |
| WBGene00020767 | -0.042343348 | 0.83011103 | 0.90726004 | lgc-40 |
| WBGene00020769 | NA | NA | NA | T24D8.6 |
| WBGene00020770 | 0.033256964 | 0.80895655 | 0.89421916 | eat-17 |
| WBGene00020771 | -0.046864186 | 0.79375226 | 0.88399982 | T24E12.1 |
| WBGene00020772 | -0.003108065 | 0.95728506 | NA | T24E12.2 |
| WBGene00020773 | NA | NA | NA | T24E12.3 |
| WBGene00020774 | 1.777445596 | 0.00148106 | 0.00971951 | T24E12.5 |
| WBGene00020775 | -0.022930767 | 0.78121253 | NA | T24E12.6 |
| WBGene00020777 | 0.02460575 | 0.90369401 | 0.94878151 | T24E12.10 |
| WBGene00020778 | 0.149976618 | 0.46374725 | 0.6536122 | T24E12.11 |
| WBGene00020779 | -0.107998266 | 0.18174419 | 0.36443528 | T24G10.2 |
| WBGene00020781 | -0.208068257 | 0.00085302 | 0.00615226 | T24H7.2 |
| WBGene00020782 | -0.146074664 | 0.17564157 | 0.35637032 | T24H7.3 |
| WBGene00020783 | -0.046884725 | 0.76356048 | 0.86600735 | blos-4 |
| WBGene00020784 | 0.363359621 | 0.08024191 | 0.21152716 | tat-4 |
| WBGene00020785 | 0.038125098 | 0.61119482 | NA | T25B2.1 |
| WBGene00020786 | 0.709519231 | 0.00123037 | 0.00834746 | T25B2.2 |
| WBGene00020787 | NA | NA | NA | T25B6.1 |
| WBGene00020788 | 0.104844688 | 0.41397741 | 0.61154897 | nep-22 |
| WBGene00020789 | 0.020317245 | 0.84643798 | NA | T25B6.3 |
| WBGene00020790 | 0.069987627 | 0.55165619 | 0.72405185 | T25B6.4 |
| WBGene00020791 | NA | NA | NA | T25B6.5 |
| WBGene00020792 | NA | NA | NA | T25B6.6 |
| WBGene00020793 | -0.056659509 | 0.52915738 | NA | T25D1.1 |
| WBGene00020794 | 0.001333159 | 0.99231666 | 0.9955752 | T25D1.2 |
| WBGene00020795 | -0.008657085 | 0.92287164 | NA | T25D1.3 |
| WBGene00020796 | -0.307591953 | 0.01327314 | 0.05608429 | mrpl-28 |

| | | | | |
|---|---|---|---|---|
| WBGene00020797 | 0.062077014 | 0.76460531 | 0.86625291 | T25D3.3 |
| WBGene00020798 | -0.074006163 | 0.38975571 | 0.58786842 | T25D3.4 |
| WBGene00020799 | 0.748623165 | 0.00759782 | 0.03628169 | T25D10.1 |
| WBGene00020800 | 0.15218237 | 0.34342626 | 0.54313812 | zig-10 |
| WBGene00020801 | 0.092461635 | 0.64269419 | 0.78841103 | T25D10.4 |
| WBGene00020802 | -0.006464466 | 0.93734527 | NA | btb-2 |
| WBGene00020803 | 0.200945507 | 0.03188128 | 0.10804951 | T25E4.1 |
| WBGene00020804 | 0.020317245 | 0.84643798 | NA | T25E4.2 |
| WBGene00020805 | NA | NA | NA | T25F10.1 |
| WBGene00020806 | 0.516945586 | 0.03083429 | 0.10537928 | T25F10.3 |
| WBGene00020807 | 0.086561712 | 0.29531459 | NA | T25F10.4 |
| WBGene00020808 | -0.378904434 | 6.42E-10 | 1.65E-08 | clik-1 |
| WBGene00020809 | -0.224974136 | 0.13563027 | 0.30048096 | T25G12.1 |
| WBGene00020811 | 0.110937945 | 0.59795747 | 0.75608389 | T25G12.3 |
| WBGene00020812 | -0.187328633 | 0.05834647 | 0.16822582 | acdh-7 |
| WBGene00020813 | 1.054210929 | 9.97E-05 | 0.00096938 | T25G12.6 |
| WBGene00020817 | -0.061406079 | 0.72916584 | 0.84882858 | T25G12.11 |
| WBGene00020819 | -0.035447805 | 0.7530984 | 0.86007127 | T26A5.2 |
| WBGene00020820 | -0.041099115 | 0.68700498 | 0.81969194 | algn-1 |
| WBGene00020821 | -0.117529335 | 0.0850465 | 0.21977511 | jhdm-1 |
| WBGene00020822 | -0.118420834 | 0.31113558 | 0.51206565 | T26A5.6 |
| WBGene00020823 | -0.109034714 | 0.40978721 | 0.60768708 | T26A5.8 |
| WBGene00020824 | 0.04734063 | 0.58722545 | 0.74981371 | T26A8.1 |
| WBGene00020825 | 0.105191732 | 0.16565281 | NA | tmem-231 |
| WBGene00020826 | 0.0398788 | 0.58971221 | NA | T26A8.3 |
| WBGene00020827 | -0.315474601 | 0.00059799 | 0.00453874 | T26A8.4 |
| WBGene00020828 | NA | NA | NA | T26C11.2 |
| WBGene00020829 | 0.020232312 | 0.84643798 | NA | T26C11.3 |
| WBGene00020830 | 0.155905369 | 0.38942993 | 0.58773772 | T26C11.4 |
| WBGene00020831 | -0.092850127 | 0.23802433 | 0.43469056 | T26C12.1 |
| WBGene00020832 | 0.053246302 | 0.54058394 | NA | T26C12.2 |
| WBGene00020833 | -0.271276872 | 0.21101003 | 0.40096257 | T26C12.3 |
| WBGene00020834 | 0.231471588 | 0.29794377 | 0.49893568 | T27A1.2 |
| WBGene00020835 | -0.230085645 | 0.21845323 | 0.41031093 | T27A1.3 |
| WBGene00020836 | 0.218787001 | 0.26512396 | 0.46365025 | lgc-34 |
| WBGene00020837 | -0.076405218 | 0.70833371 | 0.83527012 | slc-36.2 |
| WBGene00020838 | 0.166270599 | 0.09788772 | 0.24245883 | trak-1 |
| WBGene00020839 | -0.132273446 | 0.08482586 | 0.21938866 | usp-5 |
| WBGene00020840 | 2.43807243 | 0.00110159 | 0.00763295 | spch-3 |
| WBGene00020841 | 0.251563532 | 0.04219116 | 0.13284316 | T27A3.5 |
| WBGene00020842 | 0.169358749 | 0.20529087 | 0.39439004 | nmrk-1 |
| WBGene00020843 | -0.146905105 | 0.2163831 | 0.40773037 | T27A3.7 |
| WBGene00020844 | -0.004658348 | 0.97221349 | 0.98476102 | T27A10.2 |

| | | | | |
|---|---|---|---|---|
| WBGene00020845 | NA | NA | NA | T27A10.5 |
| WBGene00020846 | 0.204391769 | 0.35074504 | 0.55069792 | T27A10.6 |
| WBGene00020847 | 0.18719994 | 0.28106009 | 0.48055596 | cgr-1 |
| WBGene00020849 | 0.178840338 | 0.25842658 | 0.45636252 | nhr-225 |
| WBGene00020850 | 0.147017577 | 0.46968009 | 0.6581095 | nhr-226 |
| WBGene00020851 | 0.071716943 | 0.58035149 | 0.74546945 | nhr-227 |
| WBGene00020852 | -0.002935336 | 0.98675991 | 0.99231759 | nhr-228 |
| WBGene00020853 | 0.0398788 | 0.58971221 | NA | fbxa-137 |
| WBGene00020854 | 1.768290015 | 5.51E-11 | 1.65E-09 | T27C4.1 |
| WBGene00020855 | -0.469479829 | 0.06973146 | 0.19080318 | hpo-36 |
| WBGene00020856 | -0.001842784 | 0.95621055 | NA | srt-61 |
| WBGene00020857 | NA | NA | NA | srt-62 |
| WBGene00020858 | 0.159882618 | 0.23223135 | 0.42728806 | mop-25.3 |
| WBGene00020859 | 0.157008394 | 0.11701758 | 0.27352701 | bath-25 |
| WBGene00020860 | 0.486713781 | 0.04653177 | 0.14276439 | nspd-10 |
| WBGene00020861 | 0.281999517 | 0.19029337 | 0.37614219 | T27E4.1 |
| WBGene00020862 | 1.597833842 | 9.76E-27 | 9.79E-25 | fip-2 |
| WBGene00020863 | 0.059762547 | 0.38965825 | NA | T27E4.5 |
| WBGene00020864 | 0.05787416 | 0.40594086 | NA | oac-50 |
| WBGene00020865 | 1.054651989 | 0.00010319 | 0.00100054 | T27E4.7 |
| WBGene00020866 | -0.193975978 | 0.04081891 | 0.12989126 | vps-24 |
| WBGene00020867 | 0.128171136 | 0.51557221 | 0.69531934 | shc-2 |
| WBGene00020868 | -0.321984886 | 1.02E-05 | 0.00012538 | eif-1 |
| WBGene00020869 | 0.971381063 | 1.13E-07 | 2.10E-06 | T28A11.2 |
| WBGene00020870 | 0.020232312 | 0.84643798 | NA | T28A11.3 |
| WBGene00020871 | 0.020925399 | 0.84643798 | NA | T28A11.4 |
| WBGene00020872 | 0.078203897 | 0.26561294 | NA | T28A11.5 |
| WBGene00020873 | 0.038125098 | 0.61119482 | NA | T28A11.6 |
| WBGene00020874 | NA | NA | NA | srbc-5 |
| WBGene00020875 | NA | NA | NA | T28A11.13 |
| WBGene00020877 | -0.09036188 | 0.19485829 | NA | srt-63 |
| WBGene00020878 | 0.015005315 | 0.93069083 | 0.96081261 | T28A11.16 |
| WBGene00020879 | 0.045392335 | 0.73169103 | 0.85001776 | T28A11.17 |
| WBGene00020880 | 0.09777701 | 0.17585057 | NA | T28A11.18 |
| WBGene00020881 | 4.569719654 | 4.96E-05 | 0.00052104 | T28A11.19 |
| WBGene00020882 | 0.090723942 | 0.20045316 | NA | T28A11.20 |
| WBGene00020883 | -0.028179567 | 0.88674952 | 0.94033043 | fbxa-64 |
| WBGene00020884 | 0.283337417 | 0.14131775 | 0.3089954 | T28B4.1 |
| WBGene00020885 | 0.063527295 | 0.41294485 | NA | NA |
| WBGene00020886 | 0.086576897 | 0.45539071 | 0.64725516 | ttr-6 |
| WBGene00020887 | 1.162263124 | 0.01015712 | 0.04539389 | hic-1 |
| WBGene00020888 | NA | NA | NA | T28C12.1 |
| WBGene00020889 | 1.687426619 | 0.00072045 | 0.00530301 | T28C12.2 |

| | | | | |
|---|---|---|---|---|
| WBGene00020890 | 0.07797108 | 0.4146317 | 0.61208134 | fbxa-202 |
| WBGene00020891 | -0.095585907 | 0.54752833 | 0.72018982 | T28C12.4 |
| WBGene00020892 | -0.001993815 | 0.95612996 | NA | T28C12.5 |
| WBGene00020893 | 0.006171253 | 0.96633737 | 0.9811474 | T28C12.6 |
| WBGene00020894 | 0.066544425 | 0.74888569 | 0.85820565 | T28D9.1 |
| WBGene00020895 | 1.800482583 | 5.03E-15 | 2.42E-13 | plpp-1.2 |
| WBGene00020896 | -0.187437406 | 0.14460475 | 0.31397867 | T28D9.16 |
| WBGene00020896 | -0.187437406 | 0.14460475 | 0.31397867 | T28D9.4 |
| WBGene00020897 | 0.330859901 | 0.13272427 | 0.29608656 | del-10 |
| WBGene00020898 | 0.207056196 | 0.02971382 | 0.10287067 | lido-16 |
| WBGene00020899 | NA | NA | NA | T28D9.11 |
| WBGene00020900 | -0.040794878 | 0.53203165 | NA | T28F2.1 |
| WBGene00020901 | 0.018132227 | 0.92899433 | 0.96000889 | T28F2.2 |
| WBGene00020902 | -0.308559693 | 0.12187194 | 0.2808637 | jmjc-1 |
| WBGene00020903 | -0.122054254 | 0.54924998 | 0.72192916 | acd-5 |
| WBGene00020904 | 0.053914589 | 0.60183221 | 0.75908213 | T28F12.1 |
| WBGene00020905 | 0.347142525 | 0.05418019 | 0.15960807 | T28H11.7 |
| WBGene00020906 | 1.009907989 | 0.01323642 | 0.05599864 | T28H11.8 |
| WBGene00020908 | 0.554401804 | 0.00518843 | 0.02690888 | VC5.2 |
| WBGene00020909 | -0.094239777 | 0.29913757 | 0.49958329 | W01A11.1 |
| WBGene00020910 | -0.069350425 | 0.55397208 | 0.72611506 | dgtr-1 |
| WBGene00020911 | -0.720620776 | 0.00014193 | 0.00130528 | cpt-6 |
| WBGene00020912 | 0.326879133 | 0.07163099 | 0.19471765 | W01A11.7 |
| WBGene00020913 | 0.017604133 | 0.79437959 | NA | W01B11.1 |
| WBGene00020914 | 0.126263134 | 0.3576116 | 0.55777048 | sulp-6 |
| WBGene00020915 | -0.648093356 | 5.61E-30 | 7.14E-28 | nol-58 |
| WBGene00020917 | -0.036333501 | 0.83090405 | 0.90780346 | W01B11.6 |
| WBGene00020918 | NA | NA | NA | W01C8.1 |
| WBGene00020919 | 0.165026462 | 0.34610117 | 0.54574768 | set-19 |
| WBGene00020920 | 0.082315756 | 0.67488938 | 0.8111554 | set-20 |
| WBGene00020921 | 0.051008413 | 0.73235958 | 0.85023927 | W01C8.5 |
| WBGene00020922 | 0.0398788 | 0.58971221 | NA | W01H2.2 |
| WBGene00020923 | NA | NA | NA | W02B3.3 |
| WBGene00020924 | -0.089401349 | 0.65502094 | 0.795798 | W02B3.4 |
| WBGene00020927 | -0.023109762 | 0.78121253 | NA | W02B3.7 |
| WBGene00020929 | 0.802121533 | 0.02306399 | 0.08529423 | W02C12.2 |
| WBGene00020930 | 0.828636851 | 2.37E-12 | 8.43E-11 | hlh-30 |
| WBGene00020931 | 0.017748311 | 0.8460236 | 0.91712065 | cytb-5.2 |
| WBGene00020932 | -0.227746103 | 0.01533214 | 0.06294947 | dhod-1 |
| WBGene00020933 | -0.215900108 | 0.0302664 | 0.10405388 | W02D3.4 |
| WBGene00020935 | -0.159118739 | 0.10465331 | 0.25285927 | fnci-1 |
| WBGene00020936 | -0.138227424 | 0.13970543 | 0.3070822 | hrpf-1 |
| WBGene00020937 | 0.022882923 | 0.89507679 | 0.94517269 | W02D3.12 |

| | | | | |
|---|---|---|---|---|
| WBGene00020938 | -0.243374931 | 0.16511066 | 0.34135557 | clec-218 |
| WBGene00020939 | 0.067587245 | 0.67194665 | 0.80897713 | W02D7.3 |
| WBGene00020940 | 0.106949217 | 0.60685886 | 0.76292466 | W02D7.4 |
| WBGene00020941 | 0.039362348 | 0.84386715 | 0.91621455 | W02D7.5 |
| WBGene00020942 | 0.204025841 | 0.11186051 | 0.26445175 | W02D7.6 |
| WBGene00020943 | 1.975860822 | 0.01041644 | 0.04632376 | W02D7.8 |
| WBGene00020944 | 0.039034462 | 0.57625038 | NA | W02D7.9 |
| WBGene00020945 | 0.074892148 | 0.28555128 | NA | clec-219 |
| WBGene00020946 | -0.075452738 | 0.28904131 | NA | W02D7.11 |
| WBGene00020947 | 0.132552616 | 0.52262352 | 0.70044791 | W02F12.2 |
| WBGene00020948 | -0.078905988 | 0.25609656 | 0.45448602 | era-1 |
| WBGene00020949 | -0.056172192 | 0.61160444 | 0.7663446 | W02F12.4 |
| WBGene00020950 | -0.224691311 | 0.00190163 | 0.01200074 | dlst-1 |
| WBGene00020951 | -0.220661134 | 0.03156009 | 0.1071594 | sna-1 |
| WBGene00020952 | 0.556438147 | 0.00038983 | 0.00314464 | kel-8 |
| WBGene00020953 | 0.028617328 | 0.86831562 | 0.92992376 | W02G9.3 |
| WBGene00020954 | 2.207748177 | 2.24E-34 | 3.28E-32 | W02G9.4 |
| WBGene00020955 | 0.045819525 | 0.74678623 | 0.85673854 | angl-1 |
| WBGene00020957 | -0.125974239 | 0.3437078 | 0.54349561 | W02H5.2 |
| WBGene00020961 | -0.001842784 | 0.95621055 | NA | sknr-1 |
| WBGene00020962 | 0.317709831 | 0.02766136 | 0.09767529 | W02H5.8 |
| WBGene00020964 | 0.103918 | 0.29478393 | 0.49577045 | polq-1 |
| WBGene00020966 | 0.111535092 | 0.41417029 | 0.61170859 | W03A5.1 |
| WBGene00020967 | -0.243091945 | 0.25843192 | 0.45636252 | W03A5.2 |
| WBGene00020970 | 0.088140323 | 0.47028247 | 0.65840969 | W03A5.6 |
| WBGene00020971 | NA | NA | NA | NA |
| WBGene00020972 | 0.211532234 | 0.32831399 | 0.52776077 | trpl-2 |
| WBGene00020973 | 0.072780805 | 0.44132469 | NA | W03B1.3 |
| WBGene00020974 | NA | NA | NA | W03B1.5 |
| WBGene00020976 | 0.110517154 | 0.48617477 | 0.67069489 | oac-51 |
| WBGene00020977 | 0.028835113 | 0.81116185 | 0.89550752 | oac-52 |
| WBGene00020978 | 0.020925399 | 0.84643798 | NA | W03B1.9 |
| WBGene00020979 | 0.035893496 | 0.67498843 | NA | fbxb-55 |
| WBGene00020981 | 2.232723258 | 6.53E-27 | 6.73E-25 | W03D2.6 |
| WBGene00020982 | 0.014452015 | 0.85564601 | NA | ilys-6 |
| WBGene00020983 | 0.122115126 | 0.17630691 | NA | W03D2.9 |
| WBGene00020984 | 0.059980168 | 0.38780842 | NA | W03D8.1 |
| WBGene00020985 | 0.124500775 | 0.41394845 | 0.61154897 | W03D8.2 |
| WBGene00020986 | 0.01912147 | 0.77587372 | NA | W03D8.3 |
| WBGene00020987 | 0.053957331 | 0.54487037 | NA | W03D8.5 |
| WBGene00020988 | -0.291202115 | 0.12983941 | 0.29221004 | W03D8.7 |
| WBGene00020989 | 2.492937197 | 5.02E-07 | 8.38E-06 | W03D8.8 |
| WBGene00020990 | 1.285775724 | 0.0067462 | 0.03310022 | W03D8.9 |

| | | | | |
|---|---|---|---|---|
| WBGene00020991 | 0.104747489 | 0.16723639 | NA | NA |
| WBGene00020992 | -0.320250504 | 0.15722957 | 0.33118768 | NA |
| WBGene00020993 | 0.073092156 | 0.57340206 | 0.74142714 | W03F8.3 |
| WBGene00020994 | -0.188620191 | 0.0385801 | 0.12455375 | W03F8.4 |
| WBGene00020995 | 1.582195688 | 0.00017443 | 0.00156866 | W03F8.6 |
| WBGene00020997 | 0.094296888 | 0.43676652 | 0.63169995 | gpa-18 |
| WBGene00020999 | -0.167874656 | 0.06103836 | 0.1737629 | W03F9.1 |
| WBGene00021000 | 0.118725776 | 0.57348232 | 0.74143178 | W03F9.2 |
| WBGene00021001 | NA | NA | NA | NA |
| WBGene00021002 | -0.70709032 | 0.02849921 | 0.09987003 | W03F9.4 |
| WBGene00021003 | -0.123519925 | 0.10757993 | NA | W03F9.9 |
| WBGene00021004 | -0.168641741 | 0.02123302 | 0.08026661 | sftb-2 |
| WBGene00021005 | -0.186171522 | 0.01787616 | 0.07077935 | ule-1 |
| WBGene00021006 | 0.075726029 | 0.27693214 | NA | dct-9 |
| WBGene00021007 | 2.361094891 | 0.00614402 | 0.03071169 | W03F11.4 |
| WBGene00021008 | 0.087212725 | 0.59341312 | 0.75344508 | W03F11.5 |
| WBGene00021009 | 0.040617991 | 0.67866582 | 0.81345447 | afd-1 |
| WBGene00021010 | 0.085309563 | 0.24820057 | NA | W03G1.2 |
| WBGene00021011 | 0.292659763 | 0.11162589 | 0.26406765 | W03G1.5 |
| WBGene00021012 | -0.106541516 | 0.21987477 | 0.41199107 | pig-1 |
| WBGene00021013 | 0.0398788 | 0.58971221 | NA | NA |
| WBGene00021014 | 0.095280942 | 0.42476347 | 0.62055982 | pid-4 |
| WBGene00021015 | 0.041852112 | 0.82410764 | 0.9034778 | enu-3.3 |
| WBGene00021016 | 0.020232312 | 0.84643798 | NA | W03G9.5 |
| WBGene00021017 | 0.161418676 | 0.05530083 | NA | cil-7 |
| WBGene00021018 | -0.14635151 | 0.46406576 | 0.65393521 | NA |
| WBGene00021019 | 0.765191137 | 0.01886917 | 0.0737096 | W04B5.2 |
| WBGene00021020 | 0.606157464 | 7.07E-05 | 0.00071347 | W04B5.3 |
| WBGene00021021 | -0.015723373 | 0.92239812 | 0.95629733 | mrpl-30 |
| WBGene00021022 | -0.054255229 | 0.7199459 | 0.8430559 | W04B5.5 |
| WBGene00021023 | NA | NA | NA | W04B5.6 |
| WBGene00021024 | -0.313781181 | 0.00446373 | 0.02395657 | W04C9.2 |
| WBGene00021025 | 0.084908438 | 0.30232871 | NA | cutl-13 |
| WBGene00021026 | -0.488959566 | 6.04E-05 | 0.00062327 | W04C9.4 |
| WBGene00021027 | -0.158890521 | 0.23161624 | 0.42654268 | W04C9.5 |
| WBGene00021028 | 0.246015716 | 0.1807174 | 0.36289719 | W04C9.6 |
| WBGene00021029 | NA | NA | NA | W04D12.1 |
| WBGene00021030 | -0.023756719 | 0.6882956 | NA | W04H10.1 |
| WBGene00021031 | -0.026384613 | 0.67299636 | NA | W04H10.2 |
| WBGene00021032 | -0.007072416 | 0.9244612 | NA | clec-118 |
| WBGene00021033 | NA | NA | NA | srz-32 |
| WBGene00021034 | 1.401268958 | 0.00097446 | 0.00689231 | enu-3.4 |
| WBGene00021035 | -0.054201219 | 0.31289816 | 0.51333563 | W05F2.3 |

| | | | | |
|---|---|---|---|---|
| WBGene00021036 | 0.786747594 | 2.17E-06 | 3.13E-05 | W05F2.4 |
| WBGene00021037 | 0.031410033 | 0.87128043 | 0.93126126 | NA |
| WBGene00021038 | -0.031673288 | 0.77006594 | 0.86915823 | W05F2.6 |
| WBGene00021039 | 1.042679406 | 0.00366088 | 0.02043424 | W05F2.7 |
| WBGene00021040 | 0.118564696 | 0.57008613 | 0.7387023 | W05G11.2 |
| WBGene00021041 | NA | NA | NA | clp-10 |
| WBGene00021042 | -0.008099777 | 0.92782023 | NA | btb-5 |
| WBGene00021043 | 0.139926145 | 0.13476541 | 0.29909055 | pck-1 |
| WBGene00021044 | -0.005597223 | 0.97724625 | 0.98702141 | W05H7.1 |
| WBGene00021045 | 0.234398651 | 0.14591204 | 0.31569634 | W05H7.2 |
| WBGene00021046 | -0.080679793 | 0.69177369 | 0.82310107 | sedl-1 |
| WBGene00021047 | 0.010866368 | 0.93578795 | 0.96380369 | zfp-3 |
| WBGene00021048 | 1.030302178 | 3.04E-20 | 2.34E-18 | W05H9.1 |
| WBGene00021049 | -0.073948877 | 0.58987665 | 0.75112054 | W05H9.2 |
| WBGene00021050 | -0.171424481 | 0.17602919 | 0.35699473 | NA |
| WBGene00021051 | -0.271889519 | 0.04712953 | 0.14411519 | W05H9.4 |
| WBGene00021052 | -0.00738099 | 0.96958667 | 0.98346632 | W06A11.1 |
| WBGene00021053 | -0.269354006 | 0.20200463 | 0.39052078 | gadr-4 |
| WBGene00021054 | -0.212548318 | 0.20936324 | 0.39876248 | fbxb-118 |
| WBGene00021055 | 0.53775001 | 0.04209546 | 0.13262743 | W06A11.4 |
| WBGene00021056 | -0.168224781 | 0.0089166 | 0.04114337 | W06B4.1 |
| WBGene00021057 | 0.486416517 | 0.00122313 | 0.00830428 | W06B4.2 |
| WBGene00021058 | -0.127696287 | 0.31371124 | 0.5141086 | vps-18 |
| WBGene00021059 | 0.053591844 | 0.78407914 | 0.87847449 | atat-2 |
| WBGene00021060 | -0.030538533 | 0.84993018 | 0.91926207 | dct-11 |
| WBGene00021061 | -0.011291203 | 0.91318181 | 0.9514585 | W06E11.1 |
| WBGene00021063 | -0.155838801 | 0.11745448 | 0.27391546 | sbds-1 |
| WBGene00021066 | 0.057929007 | 0.76917533 | 0.86854049 | W06H8.2 |
| WBGene00021067 | NA | NA | NA | W06H8.4 |
| WBGene00021068 | 0.432171502 | 0.00290421 | 0.01701059 | W06H8.6 |
| WBGene00021070 | -0.008484127 | 0.96463095 | 0.98070702 | cpr-8 |
| WBGene00021072 | 5.798064543 | 1.82E-05 | 0.00020968 | W07B8.4 |
| WBGene00021073 | -0.540576915 | 9.31E-09 | 2.09E-07 | nol-2 |
| WBGene00021074 | -0.248619799 | 0.08451692 | 0.21892209 | W07E6.2 |
| WBGene00021075 | 0.274956278 | 0.1284228 | 0.29002595 | sms-5 |
| WBGene00021076 | -0.227887885 | 0.28063447 | 0.48033309 | W07E6.5 |
| WBGene00021078 | 0.161119638 | 0.2229464 | 0.41556803 | unc-132 |
| WBGene00021079 | 0.420789337 | 0.08055312 | 0.21193041 | W08A12.2 |
| WBGene00021080 | -0.050364839 | 0.78753546 | 0.8795718 | W08A12.3 |
| WBGene00021081 | 0.266950212 | 0.03876463 | 0.12506676 | pals-33 |
| WBGene00021082 | 0.011718217 | 0.95551921 | 0.97496388 | zip-11 |
| WBGene00021083 | 0.107031567 | 0.14107589 | NA | W08E12.2 |
| WBGene00021084 | 0.040107227 | 0.58693163 | NA | W08E12.3 |

| | | | | |
|---|---|---|---|---|
| WBGene00021085 | 0.020232312 | 0.84643798 | NA | W08E12.4 |
| WBGene00021085 | 0.020232312 | 0.84643798 | NA | W08E12.5 |
| WBGene00021086 | NA | NA | NA | W08E12.4 |
| WBGene00021086 | NA | NA | NA | W08E12.5 |
| WBGene00021087 | 0.128000389 | 0.16547331 | NA | W08E12.6 |
| WBGene00021088 | -0.341943752 | 3.31E-13 | 1.31E-11 | W08E12.7 |
| WBGene00021089 | -0.215033116 | 0.17755868 | 0.35876852 | W08E12.8 |
| WBGene00021090 | NA | NA | NA | W08E12.9 |
| WBGene00021091 | 0.03732969 | 0.59229443 | NA | W08F4.1 |
| WBGene00021092 | -0.099416824 | 0.57806626 | 0.74423743 | fbxb-34 |
| WBGene00021093 | -0.200498344 | 0.00344103 | 0.01948897 | W08F4.3 |
| WBGene00021094 | 0.038282788 | 0.76011509 | 0.86416163 | W08F4.5 |
| WBGene00021095 | -0.208467911 | 0.33343288 | 0.53304196 | mlt-8 |
| WBGene00021096 | 0.249632919 | 0.13230996 | 0.29556762 | W08F4.7 |
| WBGene00021097 | -0.095497193 | 0.25831892 | 0.45636252 | cdc-37 |
| WBGene00021098 | NA | NA | NA | fbxb-14 |
| WBGene00021099 | -0.029327418 | 0.71586712 | NA | W08F4.10 |
| WBGene00021100 | NA | NA | NA | NA |
| WBGene00021101 | -0.202553664 | 0.24769965 | 0.44607264 | W08F4.12 |
| WBGene00021103 | -0.195890573 | 0.08430949 | 0.21862829 | eri-3 |
| WBGene00021104 | -0.153614668 | 0.36276368 | 0.56256996 | W09B6.4 |
| WBGene00021106 | NA | NA | NA | F07B7.1 |
| WBGene00021106 | NA | NA | NA | W09B7.1 |
| WBGene00021107 | NA | NA | NA | F07B7.2 |
| WBGene00021107 | NA | NA | NA | W09B7.2 |
| WBGene00021108 | NA | NA | NA | W09B7.3 |
| WBGene00021109 | 0.020925399 | 0.84643798 | NA | W09C3.1 |
| WBGene00021110 | -0.007713692 | 0.92886983 | NA | W09C3.2 |
| WBGene00021111 | 0.080169123 | 0.46912059 | 0.65776619 | W09C3.3 |
| WBGene00021112 | -0.248430635 | 0.08939511 | 0.22718866 | W09C3.4 |
| WBGene00021113 | 1.577975473 | 0.00584649 | 0.0295696 | gsp-3 |
| WBGene00021114 | -0.016185326 | 0.92136547 | 0.95610687 | W09C3.7 |
| WBGene00021115 | 0.020317245 | 0.84643798 | NA | W09C3.8 |
| WBGene00021116 | 0.052276084 | 0.74020096 | 0.85342581 | ncs-6 |
| WBGene00021117 | 0.160695095 | 0.27585154 | 0.47593777 | clec-126 |
| WBGene00021118 | 0.240122814 | 0.10384866 | 0.25189665 | clec-125 |
| WBGene00021120 | NA | NA | NA | W09G12.6 |
| WBGene00021121 | 0.862064942 | 0.01056486 | 0.04663254 | W09G12.7 |
| WBGene00021122 | NA | NA | NA | W09G12.8 |
| WBGene00021123 | 0.135577623 | 0.52190931 | 0.70005196 | W09G12.9 |
| WBGene00021124 | 0.057634798 | 0.69534183 | 0.82587001 | W10C4.1 |
| WBGene00021125 | 0.033843731 | 0.73364668 | 0.85087141 | ccb-2 |
| WBGene00021127 | 0.688075335 | 0.02862796 | 0.10024922 | W10C8.4 |

| | | | | |
|---|---|---|---|---|
| WBGene00021128 | 0.262656297 | 0.04403342 | 0.13709778 | W10C8.5 |
| WBGene00021129 | -0.02791392 | 0.6763823 | NA | W10D9.1 |
| WBGene00021130 | -0.008786608 | 0.93354413 | 0.96267218 | NA |
| WBGene00021131 | 0.013457052 | 0.94697461 | 0.97031324 | W10D9.3 |
| WBGene00021132 | -0.215276462 | 0.10321872 | 0.25078415 | nfyb-1 |
| WBGene00021133 | -0.452297986 | 0.00031068 | 0.00259062 | tomm-22 |
| WBGene00021134 | 0.008770835 | 0.94116496 | 0.96684409 | W10G11.1 |
| WBGene00021135 | 0.018861788 | 0.92698551 | 0.95867732 | W10G11.2 |
| WBGene00021136 | -0.266049046 | 0.22478682 | 0.41761662 | W10G11.3 |
| WBGene00021137 | -0.063068723 | 0.58146895 | 0.74586357 | W10G11.4 |
| WBGene00021138 | NA | NA | NA | clec-127 |
| WBGene00021139 | NA | NA | NA | clec-128 |
| WBGene00021140 | NA | NA | NA | clec-131 |
| WBGene00021141 | NA | NA | NA | clec-134 |
| WBGene00021142 | 0.020232312 | 0.84643798 | NA | clec-133 |
| WBGene00021143 | NA | NA | NA | clec-132 |
| WBGene00021144 | NA | NA | NA | clec-130 |
| WBGene00021145 | NA | NA | NA | clec-129 |
| WBGene00021146 | 0.076672338 | 0.6971398 | 0.82733434 | lgc-45 |
| WBGene00021148 | 0.797325619 | 0.00958418 | 0.0435377 | W10G11.19 |
| WBGene00021149 | -0.23527948 | 0.08346519 | 0.21713111 | dnc-3 |
| WBGene00021150 | NA | NA | NA | Y1B5A.1 |
| WBGene00021151 | 0.017279211 | 0.79380376 | NA | NA |
| WBGene00021152 | 0.269510511 | 0.20890336 | 0.39824923 | mgl-3 |
| WBGene00021153 | -0.090332411 | 0.39799806 | 0.59623206 | Y4C6A.3 |
| WBGene00021154 | 0.127266368 | 0.1668541 | NA | Y4C6A.4 |
| WBGene00021155 | -0.159260925 | 0.11265202 | 0.26585077 | Y4C6B.1 |
| WBGene00021156 | 0.328254124 | 0.15783095 | 0.33196421 | Y4C6B.2 |
| WBGene00021157 | 0.971703643 | 0.00651233 | 0.03218362 | Y4C6B.3 |
| WBGene00021158 | 1.288486384 | 0.00196806 | 0.01231909 | Y4C6B.4 |
| WBGene00021159 | -0.307644457 | 0.1216601 | 0.28068477 | Y4C6B.5 |
| WBGene00021160 | 0.239447822 | 0.09794941 | 0.24252957 | gba-4 |
| WBGene00021161 | 0.031165256 | 0.8667256 | 0.92904485 | Y4C6B.7 |
| WBGene00021162 | NA | NA | NA | Y5H2A.1 |
| WBGene00021163 | 0.262264889 | 0.11942836 | 0.27743563 | nhr-275 |
| WBGene00021164 | 0.085537126 | 0.24698522 | NA | Y5H2B.1 |
| WBGene00021165 | -0.361982564 | 0.13547534 | 0.30025649 | Y5H2B.3 |
| WBGene00021166 | NA | NA | NA | srbc-42 |
| WBGene00021167 | 3.252648065 | 2.62E-05 | 0.00029149 | cyp-32B1 |
| WBGene00021168 | 0.283388711 | 0.01115116 | 0.04870648 | cyp-33C12 |
| WBGene00021169 | 0.0398788 | 0.58971221 | NA | srt-40 |
| WBGene00021170 | 0.026407777 | 0.78268632 | 0.87768017 | arx-4 |
| WBGene00021171 | 2.795633741 | 3.20E-09 | 7.60E-08 | Y8A9A.2 |

| | | | | |
|---|---|---|---|---|
| WBGene00021172 | NA | NA | NA | Y8A9A.3 |
| WBGene00021173 | NA | NA | NA | Y8A9A.4 |
| WBGene00021174 | -0.108518519 | 0.44502689 | 0.63835003 | fbxb-99 |
| WBGene00021175 | 0.038195847 | 0.57893903 | NA | math-46 |
| WBGene00021176 | 0.404754335 | 0.00452007 | 0.02421452 | Y9C9A.1 |
| WBGene00021177 | NA | NA | NA | Y9C9A.5 |
| WBGene00021178 | 0.191342153 | 0.20482584 | 0.39397451 | Y9C9A.8 |
| WBGene00021179 | 0.020232312 | 0.84643798 | NA | fbxa-86 |
| WBGene00021180 | -0.900899001 | 0.01918544 | 0.07458685 | Y9C9A.13 |
| WBGene00021181 | -0.005068434 | 0.94614153 | NA | NA |
| WBGene00021182 | 0.020925399 | 0.84643798 | NA | srz-28 |
| WBGene00021183 | 0.013399338 | 0.88795937 | NA | Y9C9A.16 |
| WBGene00021184 | NA | NA | NA | Y9C9A.17 |
| WBGene00021185 | 0.101225005 | 0.62101414 | 0.77341511 | Y9C12A.1 |
| WBGene00021186 | 0.071816598 | 0.69726927 | 0.82742093 | Y9D1A.1 |
| WBGene00021187 | 0.277905924 | 0.22399594 | 0.41697706 | Y9D1A.2 |
| WBGene00021188 | -0.094106979 | 0.62969964 | 0.77968456 | Y13C8A.1 |
| WBGene00021190 | 0.032649045 | 0.87507928 | 0.93336868 | Y14H12A.1 |
| WBGene00021191 | -0.329949934 | 0.00028335 | 0.00238836 | Y14H12B.1 |
| WBGene00021192 | -0.325353541 | 0.00141257 | 0.00935391 | Y14H12B.2 |
| WBGene00021193 | 0.02028958 | 0.87791809 | 0.93480719 | math-44 |
| WBGene00021195 | -0.001212288 | 0.99125552 | 0.99499329 | Y17G9A.2 |
| WBGene00021196 | 0.128826435 | 0.24887962 | 0.44726236 | Y17G9A.3 |
| WBGene00021197 | 1.77212102 | 0.00019844 | 0.00175763 | Y17G9A.4 |
| WBGene00021198 | -0.164687516 | 0.33154705 | 0.53109194 | npp-26 |
| WBGene00021199 | -0.206250987 | 0.0576603 | 0.16706996 | Y17G9B.2 |
| WBGene00021200 | -0.170185322 | 0.09835797 | 0.2432018 | cyp-31A3 |
| WBGene00021201 | 0.264240655 | 0.17012961 | 0.34864471 | Y17G9B.4 |
| WBGene00021202 | 0.054556904 | 0.62949402 | 0.77968456 | Y17G9B.5 |
| WBGene00021204 | -0.167260285 | 0.28418546 | 0.48420335 | fbxb-77 |
| WBGene00021205 | 0.14101667 | 0.36610877 | 0.56582992 | Y17G9B.8 |
| WBGene00021207 | 0.239091413 | 0.10855775 | 0.25911216 | ptp-5.3 |
| WBGene00021208 | 0.110553036 | 0.4143693 | 0.61181736 | Y18H1A.2 |
| WBGene00021209 | 0.004351428 | 0.96424614 | 0.98045715 | hgap-1 |
| WBGene00021210 | 0.213486839 | 0.1695077 | 0.34787807 | Y18H1A.4 |
| WBGene00021212 | NA | NA | NA | Y18H1A.8 |
| WBGene00021213 | 0.209959862 | 0.25279502 | 0.4509225 | Y18H1A.9 |
| WBGene00021214 | 0.155125059 | 0.25077165 | 0.44933884 | hasp-2 |
| WBGene00021215 | 0.256261337 | 0.04134953 | 0.13101029 | Y18H1A.11 |
| WBGene00021216 | NA | NA | NA | Y19D10A.1 |
| WBGene00021217 | NA | NA | NA | str-258 |
| WBGene00021217 | NA | NA | NA | Y19D10A.2 |
| WBGene00021218 | NA | NA | NA | NA |

| | | | | |
|---|---|---|---|---|
| WBGene00021219 | NA | NA | NA | Y19D10A.4 |
| WBGene00021219 | NA | NA | NA | C01B4.7 |
| WBGene00021220 | NA | NA | NA | C01B4.8 |
| WBGene00021220 | NA | NA | NA | Y19D10A.5 |
| WBGene00021221 | NA | NA | NA | nas-2 |
| WBGene00021221 | NA | NA | NA | Y19D10A.6 |
| WBGene00021222 | NA | NA | NA | irld-35 |
| WBGene00021222 | NA | NA | NA | irld-53 |
| WBGene00021223 | NA | NA | NA | Y19D10A.8 |
| WBGene00021224 | NA | NA | NA | clec-209 |
| WBGene00021224 | NA | NA | NA | F56A4.2 |
| WBGene00021225 | NA | NA | NA | NA |
| WBGene00021226 | NA | NA | NA | F56A4.12 |
| WBGene00021226 | NA | NA | NA | Y19D10A.11 |
| WBGene00021227 | NA | NA | NA | mct-2 |
| WBGene00021227 | NA | NA | NA | mct-1 |
| WBGene00021228 | NA | NA | NA | str-265 |
| WBGene00021230 | NA | NA | NA | NA |
| WBGene00021232 | 0.011705027 | 0.89401441 | NA | irld-15 |
| WBGene00021233 | -0.059246987 | 0.34871219 | NA | NA |
| WBGene00021234 | 0.04032971 | 0.564277 | NA | NA |
| WBGene00021235 | 0.850079192 | 0.0110235 | 0.04828991 | Y19D10B.6 |
| WBGene00021236 | NA | NA | NA | pud-1.2 |
| WBGene00021236 | NA | NA | NA | pud-1.1 |
| WBGene00021237 | 1.008776268 | 1.07E-05 | 0.00013104 | crtc-1 |
| WBGene00021238 | 0.243448639 | 0.06110293 | 0.17387911 | gap-3 |
| WBGene00021239 | 1.279713747 | 1.34E-10 | 3.77E-09 | Y20F4.4 |
| WBGene00021241 | NA | NA | NA | Y22D7AL.1 |
| WBGene00021243 | NA | NA | NA | Y22D7AL.3 |
| WBGene00021244 | -0.291714955 | 0.0543524 | 0.1599806 | Y22D7AL.4 |
| WBGene00021245 | 0.048048547 | 0.71882416 | 0.84228128 | Y22D7AL.6 |
| WBGene00021246 | 0.183107086 | 0.32093044 | 0.52086715 | Y22D7AL.7 |
| WBGene00021247 | 0.548836867 | 0.01322036 | 0.05595818 | Y22D7AL.9 |
| WBGene00021248 | -0.371088409 | 5.02E-07 | 8.38E-06 | Y22D7AL.10 |
| WBGene00021249 | 0.056742221 | 0.78308056 | 0.87787767 | Y22D7AL.11 |
| WBGene00021250 | NA | NA | NA | Y22D7AL.12 |
| WBGene00021251 | NA | NA | NA | Y22D7AL.13 |
| WBGene00021252 | 0.028858446 | 0.79092535 | 0.88224906 | cht-4 |
| WBGene00021253 | 1.055782819 | 2.31E-06 | 3.31E-05 | Y22D7AL.15 |
| WBGene00021254 | 0.077941203 | 0.47428272 | 0.66187389 | Y22D7AL.16 |
| WBGene00021256 | -0.112395546 | 0.57849615 | 0.74449976 | Y22D7AR.2 |
| WBGene00021257 | NA | NA | NA | Y22D7AR.3 |
| WBGene00021259 | 0.157701754 | 0.11700178 | 0.27352701 | glb-30 |

| | | | | |
|---|---|---|---|---|
| WBGene00021260 | 0.393976673 | 0.10612991 | 0.25544107 | Y22D7AR.6 |
| WBGene00021261 | -0.086623065 | 0.54826084 | 0.72092678 | Y22D7AR.7 |
| WBGene00021262 | 0.278522319 | 0.21832955 | 0.41028917 | fbxa-74 |
| WBGene00021263 | 0.801187619 | 2.85E-09 | 6.79E-08 | Y22D7AR.10 |
| WBGene00021264 | -0.233270777 | 0.26826807 | 0.46719552 | fbxa-13 |
| WBGene00021265 | 0.107582918 | 0.22104628 | NA | NA |
| WBGene00021266 | -0.009168143 | 0.91551225 | NA | Y22D7AR.14 |
| WBGene00021267 | NA | NA | NA | Y23B4A.1 |
| WBGene00021268 | 0.340567345 | 0.12631872 | 0.28740208 | capa-1 |
| WBGene00021269 | 0.05362107 | 0.7907154 | 0.88218122 | Y23H5A.2 |
| WBGene00021270 | -0.304671749 | 9.57E-05 | 0.00093259 | pid-3 |
| WBGene00021271 | 0.012581561 | 0.90489012 | 0.94913088 | spe-47 |
| WBGene00021272 | -0.216130994 | 0.30488715 | 0.50558985 | Y23H5B.1 |
| WBGene00021273 | NA | NA | NA | NA |
| WBGene00021274 | 0.21794342 | 0.20439665 | 0.39347949 | Y23H5B.3 |
| WBGene00021275 | 1.492741095 | 0.00033203 | 0.00274137 | dmsr-2 |
| WBGene00021276 | -0.508120207 | 1.78E-07 | 3.21E-06 | Y23H5B.5 |
| WBGene00021277 | -0.070298134 | 0.45845734 | 0.64984741 | Y23H5B.6 |
| WBGene00021278 | -0.23413361 | 0.01715451 | 0.06861036 | NA |
| WBGene00021279 | 0.143715068 | 0.42176564 | 0.61831088 | Y23H5B.8 |
| WBGene00021280 | NA | NA | NA | nspd-8 |
| WBGene00021281 | 0.273920376 | 0.01147998 | 0.04983052 | ell-1 |
| WBGene00021282 | -0.259050288 | 0.11855397 | 0.27591129 | set-21 |
| WBGene00021284 | NA | NA | NA | Y24D9A.6 |
| WBGene00021285 | 0.0398788 | 0.58971221 | NA | Y24D9A.7 |
| WBGene00021286 | 0.006942898 | 0.93498871 | 0.96327984 | tald-1 |
| WBGene00021287 | 1.420162675 | 0.00236159 | 0.01428759 | Y24D9B.1 |
| WBGene00021288 | 0.255826417 | 0.15410016 | 0.32715947 | clec-123 |
| WBGene00021289 | -0.006558791 | 0.92999162 | NA | Y25C1A.2 |
| WBGene00021290 | -0.022930767 | 0.78121253 | NA | clec-122 |
| WBGene00021291 | NA | NA | NA | clec-121 |
| WBGene00021292 | 0.014887007 | 0.82556451 | 0.90431617 | copb-1 |
| WBGene00021293 | -0.81818786 | 0.00283582 | 0.01670124 | Y25C1A.6 |
| WBGene00021294 | -0.073386253 | 0.49745519 | 0.67972388 | Y25C1A.7 |
| WBGene00021295 | -0.078795209 | 0.5646733 | 0.73455191 | Y25C1A.8 |
| WBGene00021296 | 0.025235922 | 0.86091363 | 0.92633472 | Y25C1A.13 |
| WBGene00021297 | 0.015451648 | 0.80337623 | NA | fbxc-55 |
| WBGene00021298 | 0.036250435 | 0.83528431 | 0.91046736 | Y27F2A.6 |
| WBGene00021299 | -0.220285486 | 0.2954446 | 0.49651107 | Y27F2A.8 |
| WBGene00021300 | 0.020317245 | 0.84643798 | NA | Y27F2A.9 |
| WBGene00021301 | -0.027728334 | 0.67598605 | NA | Y32G9A.2 |
| WBGene00021302 | 0.017279211 | 0.79380376 | NA | Y32G9A.3 |
| WBGene00021303 | 0.020925399 | 0.84643798 | NA | Y32G9A.5 |

| | | | | |
|---|---|---|---|---|
| WBGene00021304 | 0.205231528 | 0.29743932 | 0.49843299 | nphp-2 |
| WBGene00021305 | 0.238286175 | 0.03415028 | 0.11376434 | zig-11 |
| WBGene00021309 | 0.092117212 | 0.60324177 | 0.75988622 | Y32G9B.1 |
| WBGene00021310 | -0.029467247 | 0.7339295 | NA | Y32H12A.1 |
| WBGene00021311 | -0.240676092 | 0.01166463 | 0.05042274 | thoc-5 |
| WBGene00021312 | -0.067897906 | 0.49320646 | 0.67612921 | szy-2 |
| WBGene00021313 | -0.208770426 | 0.0577245 | 0.16712627 | paqr-2 |
| WBGene00021314 | 0.05041318 | 0.79104503 | 0.88228006 | Y32H12A.6 |
| WBGene00021315 | -0.150649693 | 0.15218037 | 0.3246973 | Y32H12A.7 |
| WBGene00021316 | -0.017752577 | 0.8223309 | 0.90266507 | Y32H12A.8 |
| WBGene00021317 | NA | NA | NA | Y32H12A.9 |
| WBGene00021319 | -0.081724596 | 0.64429793 | 0.78947268 | Y34B4A.2 |
| WBGene00021320 | 0.411248636 | 0.08088997 | 0.212623 | Y34B4A.4 |
| WBGene00021321 | 1.514312141 | 4.86E-08 | 9.68E-07 | Y34B4A.5 |
| WBGene00021322 | 0.818756262 | 1.22E-18 | 8.18E-17 | Y34B4A.6 |
| WBGene00021323 | -0.023626663 | 0.88299983 | 0.93817228 | Y34B4A.7 |
| WBGene00021324 | -0.295598461 | 0.09130364 | 0.22991608 | rga-8 |
| WBGene00021325 | 0.613417701 | 2.02E-10 | 5.59E-09 | Y34B4A.9 |
| WBGene00021326 | 0.348853682 | 0.09754078 | 0.24192874 | Y34B4A.10 |
| WBGene00021327 | -0.318946657 | 0.0046354 | 0.02468923 | mrpl-38 |
| WBGene00021328 | 0.383313571 | 0.10508555 | 0.25347201 | npr-23 |
| WBGene00021329 | -0.07312731 | 0.55513094 | 0.72703012 | Y34D9A.3 |
| WBGene00021331 | 0.003129956 | 0.98170471 | 0.98947207 | glrx-10 |
| WBGene00021332 | -0.123808398 | 0.33422407 | 0.53367105 | Y34D9A.7 |
| WBGene00021333 | 0.217717791 | 0.3211877 | 0.52116913 | Y34D9A.8 |
| WBGene00021334 | -0.014189202 | 0.86871649 | 0.93000745 | vps-4 |
| WBGene00021335 | 1.461971559 | 4.69E-10 | 1.23E-08 | spp-23 |
| WBGene00021336 | 2.426896904 | 7.78E-18 | 4.93E-16 | Y34F4.1 |
| WBGene00021337 | 2.672217286 | 3.95E-17 | 2.34E-15 | Y34F4.2 |
| WBGene00021338 | -0.032936777 | 0.86927963 | 0.93000745 | Y34F4.3 |
| WBGene00021339 | 0.153738106 | 0.38129479 | 0.58063375 | Y34F4.4 |
| WBGene00021340 | 0.774439606 | 0.00048737 | 0.00380562 | Y34F4.5 |
| WBGene00021342 | -0.081102108 | 0.66217251 | 0.8014937 | NA |
| WBGene00021343 | 0.316356342 | 0.13649581 | 0.30161672 | cutl-23 |
| WBGene00021344 | 0.247134547 | 0.12379016 | 0.28407706 | rev-3 |
| WBGene00021345 | -0.039976722 | 0.81930524 | 0.90055916 | Y37B11A.3 |
| WBGene00021346 | 0.112501827 | 0.44034923 | 0.6345623 | Y37E3.1 |
| WBGene00021347 | -0.187743628 | 0.12693819 | 0.28824805 | rpb-10 |
| WBGene00021348 | -0.055638708 | 0.52115236 | 0.69956833 | moag-4 |
| WBGene00021349 | 0.089349279 | 0.55587942 | 0.72763797 | arl-13 |
| WBGene00021350 | -0.279056377 | 4.53E-10 | 1.19E-08 | Y37E3.8 |
| WBGene00021351 | -0.319104672 | 2.96E-07 | 5.12E-06 | eif-2alpha |
| WBGene00021352 | 0.002706658 | 0.97746803 | 0.98710921 | pcyt-2.1 |

| | | | | |
|---|---|---|---|---|
| WBGene00021353 | 0.666163668 | 0.02193386 | 0.08220176 | iglr-3 |
| WBGene00021354 | -0.634186385 | 0.00219025 | 0.01341174 | fpn-1.1 |
| WBGene00021355 | -0.270770286 | 6.67E-05 | 0.00067887 | Y37E3.17 |
| WBGene00021357 | 0.167948754 | 0.20206198 | 0.39052078 | Y37E11AL.1 |
| WBGene00021358 | 0.553596715 | 0.03917558 | 0.12603156 | Y37E11AL.2 |
| WBGene00021359 | -0.156872649 | 0.12280274 | 0.28238651 | Y37E11AL.3 |
| WBGene00021360 | NA | NA | NA | NA |
| WBGene00021362 | 0.604351947 | 0.04390892 | 0.13692661 | fmil-1 |
| WBGene00021363 | -0.19593541 | 0.09959134 | 0.24477367 | taf-6.2 |
| WBGene00021364 | 0.029550419 | 0.78609414 | 0.87892197 | Y37E11AL.9 |
| WBGene00021365 | 0.122674653 | 0.23873403 | 0.43587791 | smgl-2 |
| WBGene00021366 | 0.00961793 | 0.94649812 | 0.97008603 | Y37E11AM.2 |
| WBGene00021367 | -0.497377055 | 0.0051688 | 0.02683559 | Y37E11AM.4 |
| WBGene00021367 | -0.497377055 | 0.0051688 | 0.02683559 | Y37E11AM.3 |
| WBGene00021368 | 0.948208629 | 4.97E-05 | 0.00052166 | best-20 |
| WBGene00021369 | 0.364483493 | 0.13411637 | 0.29820707 | siah-1 |
| WBGene00021370 | 0.550631254 | 0.00758467 | 0.03626621 | nape-2 |
| WBGene00021371 | 0.633555229 | 0.02050423 | 0.07806773 | nape-1 |
| WBGene00021372 | -0.003769594 | 0.97942736 | 0.98820177 | ugt-45 |
| WBGene00021373 | 0.108971426 | 0.21480919 | NA | Y37E11AR.7 |
| WBGene00021374 | 0.063554114 | 0.74668702 | 0.85673854 | Y37E11B.1 |
| WBGene00021375 | 0.02415023 | 0.88996366 | 0.942697 | Y37E11B.2 |
| WBGene00021376 | 0.118329422 | 0.49227905 | 0.67536147 | blos-7 |
| WBGene00021377 | -0.090997487 | 0.55611109 | 0.72787614 | Y37E11B.5 |
| WBGene00021378 | -0.137696544 | 0.43525288 | 0.63045722 | Y37E11B.6 |
| WBGene00021379 | -0.280162283 | 0.15350922 | 0.32677064 | Y37E11B.7 |
| WBGene00021380 | 0.020317245 | 0.84643798 | NA | NA |
| WBGene00021381 | 0.261341384 | 0.12764418 | 0.28897967 | Y37E11B.10 |
| WBGene00021382 | 0.147548069 | 0.32469786 | 0.52384588 | Y37F4.1 |
| WBGene00021383 | NA | NA | NA | Y37F4.2 |
| WBGene00021384 | NA | NA | NA | Y37F4.3 |
| WBGene00021385 | NA | NA | NA | Y37F4.4 |
| WBGene00021386 | 0.256724347 | 0.04113935 | 0.13057958 | Y37F4.5 |
| WBGene00021387 | -0.150051352 | 0.46150141 | 0.6522678 | Y37F4.6 |
| WBGene00021388 | 0.76143247 | 0.02336498 | 0.08608119 | Y37F4.7 |
| WBGene00021389 | 0.129591746 | 0.31645742 | 0.51698682 | Y38A10A.2 |
| WBGene00021390 | 0.040107227 | 0.58693163 | NA | fbxa-214 |
| WBGene00021391 | 0.079374187 | 0.41591412 | 0.613109 | smut-1 |
| WBGene00021392 | -0.16362767 | 0.13045883 | 0.29281738 | Y38A10A.7 |
| WBGene00021394 | -0.05220419 | 0.68663092 | 0.81942482 | Y38C1AA.1 |
| WBGene00021395 | 0.170810731 | 0.24372955 | 0.44163915 | pnc-1 |
| WBGene00021396 | 0.301491409 | 0.15355331 | 0.32681698 | cutl-24 |
| WBGene00021397 | 0.331067853 | 0.13460512 | 0.29882541 | Y38C1AA.6 |

| | | | | | |
|---|---|---|---|---|---|
| WBGene00021398 | | 0.982805962 | 0.01102154 | 0.04828991 | Y38C1AA.7 |
| WBGene00021400 | | 1.271050781 | 0.00712252 | 0.03446158 | Y38C1AA.9 |
| WBGene00021401 | | 0.000726343 | 0.99807185 | 0.99882228 | prdx-6 |
| WBGene00021402 | | -0.222542977 | 0.14346535 | 0.31206006 | Y38C1AA.12 |
| WBGene00021403 | NA | | NA | NA | Y38C1AB.5 |
| WBGene00021403 | NA | | NA | NA | Y38C1AB.1 |
| WBGene00021404 | | 0.020232312 | 0.84643798 | NA | trpl-3 |
| WBGene00021404 | | 0.020232312 | 0.84643798 | NA | trpl-4 |
| WBGene00021405 | NA | | NA | NA | Y38C1AB.3 |
| WBGene00021405 | NA | | NA | NA | Y38C1AB.7 |
| WBGene00021406 | | -0.004242534 | 0.97382054 | 0.98551763 | frm-5.2 |
| WBGene00021407 | NA | | NA | NA | Y38C1AB.5 |
| WBGene00021407 | NA | | NA | NA | Y38C1AB.1 |
| WBGene00021408 | | 0.020232312 | 0.84643798 | NA | trpl-3 |
| WBGene00021408 | | 0.020232312 | 0.84643798 | NA | trpl-4 |
| WBGene00021409 | NA | | NA | NA | Y38C1AB.3 |
| WBGene00021409 | NA | | NA | NA | Y38C1AB.7 |
| WBGene00021410 | | -0.122996209 | 0.53448992 | 0.7099217 | Y38C1BA.1 |
| WBGene00021412 | | -0.030364153 | 0.8805152 | 0.93648286 | cyp-29A3 |
| WBGene00021413 | | -0.045959069 | 0.50707733 | NA | srt-52 |
| WBGene00021414 | NA | | NA | NA | Y38C9B.3 |
| WBGene00021415 | | 0.368980536 | 0.13003432 | 0.29227968 | nsy-4 |
| WBGene00021416 | | 0.010778738 | 0.92477379 | 0.95760755 | mksr-2 |
| WBGene00021417 | | 0.182977979 | 0.2037876 | 0.39279281 | nhr-236 |
| WBGene00021418 | | 0.161414913 | 0.44798595 | 0.64077396 | NA |
| WBGene00021419 | | 0.143944107 | 0.37068596 | 0.57036433 | eri-5 |
| WBGene00021420 | | -0.343709035 | 6.67E-05 | 0.00067887 | trap-3 |
| WBGene00021421 | | -0.09309157 | 0.50644977 | 0.68733435 | Y38F2AR.3 |
| WBGene00021423 | | -0.119529446 | 0.36335745 | 0.56319251 | tftc-1 |
| WBGene00021425 | | 0.023289466 | 0.83871289 | 0.9127791 | ppgn-1 |
| WBGene00021427 | | -0.483873885 | 6.38E-06 | 8.14E-05 | Y38F2AR.9 |
| WBGene00021428 | | 0.170581473 | 0.15838979 | 0.33270946 | Y38F2AR.10 |
| WBGene00021430 | | 0.14763563 | 0.34920733 | 0.54916939 | Y38F2AR.12 |
| WBGene00021433 | NA | | NA | NA | Y39A3A.2 |
| WBGene00021434 | | 0.078351211 | 0.48317535 | 0.66876255 | Y39A3A.3 |
| WBGene00021435 | | 0.105751795 | 0.42780503 | 0.62281811 | Y39A3A.4 |
| WBGene00021436 | | 0.040107227 | 0.58693163 | NA | Y39A3B.1 |
| WBGene00021437 | | 0.018599829 | 0.91488207 | 0.95275542 | lgc-42 |
| WBGene00021438 | | -0.072145491 | 0.47763203 | 0.66470097 | Y39A3B.3 |
| WBGene00021439 | | 1.11761121 | 0.01133109 | 0.04931585 | ckr-2 |
| WBGene00021440 | | 0.140198009 | 0.14737295 | 0.3179616 | Y39A3CL.1 |
| WBGene00021441 | | -0.023771824 | 0.8612341 | 0.92654132 | Y39A3CL.3 |
| WBGene00021442 | | 0.119590863 | 0.31275853 | 0.51333563 | Y39A3CL.4 |

| | | | | | |
|---|---|---|---|---|---|
| WBGene00021443 | | 0.047268195 | 0.6515691 | 0.79364585 | Y39A3CL.7 |
| WBGene00021444 | | -0.05796181 | 0.61989004 | 0.77264865 | ttc-7 |
| WBGene00021446 | | 0.023658411 | 0.9105004 | 0.95112162 | hlh-33 |
| WBGene00021447 | | 0.104747489 | 0.16723639 | NA | Y39A3CR.8 |
| WBGene00021448 | | 0.079102063 | 0.69293139 | 0.82387595 | Y39D8A.1 |
| WBGene00021449 | | -0.032864604 | 0.76821099 | 0.86844708 | mltn-7 |
| WBGene00021450 | | 0.040107227 | 0.58693163 | NA | mltn-6 |
| WBGene00021451 | | 0.063503222 | 0.60329372 | 0.75988622 | mltn-8 |
| WBGene00021452 | | -0.131728267 | 0.44998893 | 0.64232114 | Y39F10A.1 |
| WBGene00021453 | NA | | NA | NA | Y39F10A.2 |
| WBGene00021454 | NA | | NA | NA | Y39F10A.3 |
| WBGene00021456 | | 0.058862203 | 0.52860185 | 0.70562177 | rfip-2 |
| WBGene00021457 | | 0.035234593 | 0.66618922 | NA | Y39F10C.1 |
| WBGene00021458 | | -1.218259274 | 2.96E-06 | 4.08E-05 | Y39G10AL.1 |
| WBGene00021460 | | -0.177898193 | 0.02823019 | 0.09909332 | zwl-1 |
| WBGene00021461 | | 0.211272535 | 0.23192349 | 0.42690368 | nekl-1 |
| WBGene00021463 | | -0.646112215 | 0.0253717 | 0.09167501 | zeel-1 |
| WBGene00021464 | | 0.987633217 | 0.00098142 | 0.00693152 | ugt-31 |
| WBGene00021465 | | -0.129649804 | 0.3228514 | 0.52291098 | ekl-7 |
| WBGene00021466 | | -0.488161558 | 1.84E-11 | 5.85E-10 | eif-2gamma |
| WBGene00021467 | | -0.107269222 | 0.21436356 | 0.40522878 | Y39G10AR.9 |
| WBGene00021468 | | -1.269076114 | 9.16E-34 | 1.32E-31 | epg-2 |
| WBGene00021469 | | 0.639513382 | 1.48E-06 | 2.22E-05 | Y39G10AR.11 |
| WBGene00021470 | | -0.23840273 | 0.00057957 | 0.00440576 | tpxl-1 |
| WBGene00021471 | | 0.026758925 | 0.76705959 | NA | Y39G10AR.15 |
| WBGene00021472 | | 0.060553361 | 0.64442238 | 0.78947268 | Y39G10AR.16 |
| WBGene00021473 | | 0.19944119 | 0.35903434 | 0.55927458 | sydn-1 |
| WBGene00021474 | | 0.00379525 | 0.96829961 | 0.98266269 | dot-1.1 |
| WBGene00021475 | | -0.194244555 | 0.19079296 | 0.37682446 | tbca-1 |
| WBGene00021476 | | -0.029839699 | 0.83548735 | 0.91048532 | nsun-4 |
| WBGene00021477 | NA | | NA | NA | srbc-64 |
| WBGene00021478 | NA | | NA | NA | Y39H10A.1 |
| WBGene00021479 | | -0.043953102 | 0.75647898 | 0.86211635 | oac-55 |
| WBGene00021482 | | 0.17856612 | 0.28439822 | 0.48434039 | Y39H10A.6 |
| WBGene00021483 | NA | | NA | NA | NA |
| WBGene00021484 | | 0.133035765 | 0.27244002 | 0.47249247 | Y40A1A.2 |
| WBGene00021485 | | 0.182590025 | 0.33965552 | 0.5393597 | Y40A1A.3 |
| WBGene00021486 | | -0.170375189 | 0.00081921 | 0.00593472 | lbp-9 |
| WBGene00021487 | | -0.167576153 | 0.42616335 | 0.62179031 | comt-3 |
| WBGene00021488 | | 0.020232312 | 0.84643798 | NA | srab-23 |
| WBGene00021489 | | 0.020925399 | 0.84643798 | NA | NA |
| WBGene00021490 | | -0.00346345 | 0.95708614 | NA | Y40B10A.5 |
| WBGene00021491 | | -0.087157727 | 0.2060916 | NA | comt-4 |

| | | | | |
|---|---|---|---|---|
| WBGene00021492 | 0.060534915 | 0.72853478 | 0.84853595 | comt-5 |
| WBGene00021493 | 0.246168681 | 0.22739846 | 0.42103879 | Y40B10A.9 |
| WBGene00021494 | NA | NA | NA | Y40B10B.1 |
| WBGene00021495 | 0.518253307 | 0.03788476 | 0.12293237 | sqst-2 |
| WBGene00021496 | -0.005581345 | 0.94060131 | NA | Y40C5A.3 |
| WBGene00021497 | 0.378464052 | 0.0842238 | 0.21851129 | Y40C5A.4 |
| WBGene00021500 | 0.019325396 | 0.77340931 | NA | Y40C7B.4 |
| WBGene00021502 | -0.101201689 | 0.44868326 | 0.6413954 | Y40D12A.1 |
| WBGene00021503 | 0.543499483 | 0.00039894 | 0.00320874 | ctsa-4.2 |
| WBGene00021504 | 0.056411843 | 0.41891328 | NA | Y41D4A.1 |
| WBGene00021505 | -0.666630461 | 0.03369224 | 0.11272578 | Y41D4A.3 |
| WBGene00021506 | -0.085289308 | 0.24910242 | 0.44751957 | Y41D4A.4 |
| WBGene00021507 | -0.11429524 | 0.1584211 | 0.3327275 | Y41D4A.5 |
| WBGene00021508 | 0.092854015 | 0.51130253 | 0.69129309 | Y41D4A.6 |
| WBGene00021509 | 0.537511659 | 0.05642585 | 0.16456789 | Y41D4A.7 |
| WBGene00021510 | 0.115538027 | 0.56627163 | 0.73538965 | frpr-19 |
| WBGene00021511 | 0.130877758 | 0.34943567 | 0.54940803 | Y41D4B.1 |
| WBGene00021512 | 0.032855688 | 0.76449512 | 0.86623548 | Y41D4B.4 |
| WBGene00021513 | 0.065254007 | 0.40163332 | NA | Y41D4B.6 |
| WBGene00021514 | -0.242358929 | 0.03074179 | 0.10518593 | Y41D4B.11 |
| WBGene00021515 | 0.129928931 | 0.4028221 | 0.60131712 | set-23 |
| WBGene00021516 | -0.23780635 | 0.25770539 | 0.45573917 | Y41D4B.14 |
| WBGene00021517 | 0.204737186 | 0.03089539 | 0.105465 | Y41D4B.15 |
| WBGene00021518 | 1.24507217 | 1.90E-08 | 4.04E-07 | hpo-6 |
| WBGene00021519 | 4.054497961 | 2.55E-06 | 3.59E-05 | Y41D4B.17 |
| WBGene00021520 | 0.127777177 | 0.49205152 | 0.67515197 | NA |
| WBGene00021521 | -0.007949484 | 0.92915667 | NA | nhr-287 |
| WBGene00021522 | 0.176277565 | 0.15191609 | 0.32441708 | nhr-274 |
| WBGene00021524 | 0.03732969 | 0.59229443 | NA | Y41D4B.24 |
| WBGene00021525 | -0.318929437 | 0.09814076 | 0.24292119 | Y41D4B.26 |
| WBGene00021526 | 0.05787416 | 0.40594086 | NA | Y41G9A.2 |
| WBGene00021527 | 0.652562495 | 0.03401394 | 0.11338755 | aakg-3 |
| WBGene00021528 | 0.859382282 | 0.00590839 | 0.02980864 | gbb-1 |
| WBGene00021529 | 0.02007495 | 0.88901265 | 0.94194334 | Y41G9A.5 |
| WBGene00021530 | -0.0260089 | 0.6772113 | NA | Y41G9A.6 |
| WBGene00021531 | 0.243473094 | 0.27787705 | 0.47767968 | Y42G9A.1 |
| WBGene00021532 | 0.761862465 | 0.0068474 | 0.03346222 | Y42G9A.2 |
| WBGene00021533 | 0.071424916 | 0.70961946 | 0.83644944 | Y42G9A.3 |
| WBGene00021534 | 0.118143958 | 0.14375959 | 0.31256075 | mvk-1 |
| WBGene00021535 | -0.054274206 | 0.65207758 | 0.7939756 | wht-7 |
| WBGene00021536 | -0.281400776 | 0.00981966 | 0.04426407 | Y42H9AR.1 |
| WBGene00021537 | -0.059340233 | 0.59553729 | 0.75477619 | Y42H9AR.2 |
| WBGene00021538 | -0.141924471 | 0.3517941 | 0.55175334 | rabs-5 |

| | | | | |
|---|---|---|---|---|
| WBGene00021539 | -0.067575327 | 0.45342321 | 0.64558682 | Y42H9AR.4 |
| WBGene00021540 | -0.14161434 | 0.4200777 | 0.61684362 | Y42H9AR.5 |
| WBGene00021541 | -0.237488931 | 0.00691688 | 0.03365571 | Y42H9B.3 |
| WBGene00021542 | -1.232978963 | 0.00118945 | 0.00811117 | Y43B11AL.1 |
| WBGene00021543 | 0.349472483 | 0.04784387 | 0.14569094 | Y43B11AR.1 |
| WBGene00021544 | 0.117295054 | 0.54347435 | 0.71696212 | Y43B11AR.3 |
| WBGene00021545 | -0.068637798 | 0.42272976 | 0.61864479 | Y43H11AL.1 |
| WBGene00021546 | -0.01239108 | 0.90915892 | 0.95068624 | laat-1 |
| WBGene00021547 | 0.03375759 | 0.86464827 | 0.92776221 | Y44E3A.1 |
| WBGene00021548 | 0.002368253 | 0.98613961 | 0.99217084 | trx-4 |
| WBGene00021549 | 0.001030126 | 0.99268114 | 0.99567339 | cdap-2 |
| WBGene00021551 | -0.155820357 | 0.07207733 | 0.1954954 | edc-4 |
| WBGene00021552 | -0.024937974 | 0.82455409 | 0.90375029 | zip-4 |
| WBGene00021553 | -0.065297811 | 0.68164277 | 0.81588716 | tyr-5 |
| WBGene00021554 | 0.028767395 | 0.78201412 | 0.87735392 | Y45G5AL.1 |
| WBGene00021555 | -0.320923497 | 0.00303566 | 0.01764397 | pid-5 |
| WBGene00021556 | 0.006936155 | 0.97214025 | 0.98476102 | Y45G5AM.3 |
| WBGene00021557 | -1.198893469 | 0.00111046 | 0.00767622 | Y45G5AM.5 |
| WBGene00021558 | 0.244361542 | 0.25099279 | 0.44956215 | Y45G5AM.6 |
| WBGene00021559 | -0.034413237 | 0.7116883 | 0.83774147 | Y45G5AM.7 |
| WBGene00021560 | -0.160330409 | 0.29936112 | 0.4998228 | coh-4 |
| WBGene00021561 | -0.00158694 | 0.98446735 | 0.99110132 | Y45G5AM.9 |
| WBGene00021562 | -0.06122429 | 0.31850456 | 0.51882792 | nuo-5 |
| WBGene00021563 | -0.279481973 | 0.11324837 | 0.26692598 | Y45G12B.2 |
| WBGene00021564 | 0.216768683 | 0.14759131 | 0.31829199 | Y45G12B.3 |
| WBGene00021565 | -1.33615435 | 0.00616642 | 0.03081317 | Y45G12C.1 |
| WBGene00021566 | NA | NA | NA | Y45G12C.3 |
| WBGene00021567 | NA | NA | NA | F56A4.5 |
| WBGene00021567 | NA | NA | NA | Y45G12C.4 |
| WBGene00021568 | NA | NA | NA | srt-1 |
| WBGene00021568 | NA | NA | NA | srt-2 |
| WBGene00021569 | NA | NA | NA | Y45G12C.6 |
| WBGene00021570 | NA | NA | NA | srd-73 |
| WBGene00021570 | NA | NA | NA | srd-60 |
| WBGene00021571 | NA | NA | NA | srbc-81 |
| WBGene00021571 | NA | NA | NA | Y19D10A.17 |
| WBGene00021572 | NA | NA | NA | srd-71 |
| WBGene00021572 | NA | NA | NA | srd-72 |
| WBGene00021573 | NA | NA | NA | Y45G12C.10 |
| WBGene00021573 | NA | NA | NA | str-119 |
| WBGene00021574 | NA | NA | NA | NA |
| WBGene00021576 | -1.646747962 | 3.71E-11 | 1.14E-09 | fbxc-51 |
| WBGene00021578 | NA | NA | NA | Y46B2A.3 |

| | | | | |
|---|---|---|---|---|
| WBGene00021579 | 2.223965973 | 0.00217022 | 0.01331485 | clec-73 |
| WBGene00021580 | 0.108717649 | 0.15669083 | NA | clec-174 |
| WBGene00021581 | 2.626270291 | 0.00061611 | 0.0046593 | clec-70 |
| WBGene00021582 | 6.384827118 | 2.22E-06 | 3.20E-05 | clec-71 |
| WBGene00021583 | 1.538299582 | 0.00161505 | 0.01047208 | clec-72 |
| WBGene00021585 | -0.116352194 | 0.46740063 | 0.65691831 | clec-74 |
| WBGene00021586 | 0.042065281 | 0.558937 | NA | clec-75 |
| WBGene00021587 | 4.776194425 | 2.33E-06 | 3.33E-05 | clec-76 |
| WBGene00021589 | -0.139193654 | 0.28809305 | 0.48841713 | Y46D2A.1 |
| WBGene00021590 | -2.804671548 | 2.69E-08 | 5.60E-07 | Y46D2A.2 |
| WBGene00021591 | NA | NA | NA | Y46D2A.3 |
| WBGene00021592 | -0.023109762 | 0.78121253 | NA | Y46E12A.2 |
| WBGene00021593 | NA | NA | NA | Y46E12A.3 |
| WBGene00021594 | 0.038640051 | 0.8344408 | 0.90975112 | tig-3 |
| WBGene00021595 | -0.065928555 | 0.60386042 | 0.76039566 | Y46E12BL.2 |
| WBGene00021596 | -0.066496573 | 0.43927794 | 0.63389201 | spsb-2 |
| WBGene00021597 | 0.152804458 | 0.1747447 | 0.35543723 | spsb-1 |
| WBGene00021599 | 0.01319506 | 0.88869344 | NA | srt-42 |
| WBGene00021600 | 1.675039661 | 0.00147695 | 0.0097056 | Y46H3A.4 |
| WBGene00021601 | 0.035695256 | 0.67391046 | NA | Y46H3A.5 |
| WBGene00021602 | 0.687500408 | 0.02092693 | 0.07937594 | clec-205 |
| WBGene00021603 | -0.110383369 | 0.59378965 | 0.7534819 | clec-204 |
| WBGene00021604 | NA | NA | NA | NA |
| WBGene00021605 | -0.47487918 | 0.00708582 | 0.03431807 | Y46H3C.5 |
| WBGene00021606 | 0.277702973 | 0.19665647 | 0.3841045 | NA |
| WBGene00021607 | -0.396360093 | 0.0384979 | 0.12437064 | Y46H3C.7 |
| WBGene00021608 | 0.018998992 | 0.77286019 | NA | Y46H3D.1 |
| WBGene00021609 | 0.020317245 | 0.84643798 | NA | irld-17 |
| WBGene00021610 | -0.003717304 | 0.98133251 | 0.98926637 | nhr-237 |
| WBGene00021611 | 0.094280228 | 0.63409211 | 0.78290965 | nhr-238 |
| WBGene00021612 | NA | NA | NA | Y46H3D.8 |
| WBGene00021613 | -0.233301967 | 0.08047096 | 0.21182584 | ttc-37 |
| WBGene00021614 | 0.250242994 | 0.20593461 | 0.39495113 | Y47A7.2 |
| WBGene00021615 | NA | NA | NA | Y47C4A.1 |
| WBGene00021615 | NA | NA | NA | Y108F1.1 |
| WBGene00021616 | NA | NA | NA | Y47D7A.2 |
| WBGene00021617 | NA | NA | NA | Y47D7A.3 |
| WBGene00021619 | 0.134501521 | 0.43014923 | 0.62509328 | Y47D7A.6 |
| WBGene00021620 | NA | NA | NA | Y47D7A.7 |
| WBGene00021621 | 0.020317245 | 0.84643798 | NA | Y47D7A.9 |
| WBGene00021622 | NA | NA | NA | hpo-37 |
| WBGene00021623 | 0.020317245 | 0.84643798 | NA | Y47D7A.11 |
| WBGene00021624 | 0.127693253 | 0.10904601 | NA | Y47D7A.12 |

| | | | | |
|---|---|---|---|---|
| WBGene00021625 | 8.220391215 | 1.19E-09 | 2.99E-08 | Y47D7A.13 |
| WBGene00021626 | -0.036727213 | 0.82000854 | 0.90113575 | rft-2 |
| WBGene00021627 | 0.384651938 | 0.04080752 | 0.12988324 | Y47D7A.15 |
| WBGene00021628 | -0.068345027 | 0.42781789 | 0.62281811 | Y47D9A.1 |
| WBGene00021629 | 0.212753408 | 0.23164487 | 0.42654268 | scpl-3 |
| WBGene00021630 | 0.038125098 | 0.61119482 | NA | Y47D9A.3 |
| WBGene00021631 | -0.022930767 | 0.78121253 | NA | Y47D9A.4 |
| WBGene00021632 | -0.005068434 | 0.94614153 | NA | Y47D9A.5 |
| WBGene00021633 | 0.039071363 | 0.81177867 | 0.89587496 | Y47G6A.3 |
| WBGene00021634 | -0.152150164 | 0.28353007 | 0.48342422 | rde-10 |
| WBGene00021635 | 0.777767027 | 0.00026648 | 0.00226568 | Y47G6A.5 |
| WBGene00021636 | -0.120971462 | 0.24370353 | 0.44163915 | pcaf-1 |
| WBGene00021637 | 0.167747069 | 0.30357054 | 0.50432047 | Y47G6A.7 |
| WBGene00021638 | -0.089944766 | 0.52097302 | 0.6993917 | Y47G6A.9 |
| WBGene00021639 | 0.168599116 | 0.29542224 | 0.49651107 | Y47G6A.13 |
| WBGene00021640 | -0.279167185 | 0.05127094 | 0.15325803 | Y47G6A.14 |
| WBGene00021641 | 0.270145001 | 0.21494972 | 0.40587776 | Y47G6A.15 |
| WBGene00021643 | 0.095082031 | 0.53114945 | 0.70766758 | ccep-290 |
| WBGene00021644 | -0.209942217 | 0.12408734 | 0.28442955 | Y47G6A.18 |
| WBGene00021645 | -0.908675415 | 4.87E-07 | 8.15E-06 | Y47G6A.19 |
| WBGene00021646 | 0.124589977 | 0.45835492 | 0.64976513 | Y47G6A.21 |
| WBGene00021647 | -0.207581351 | 0.0913163 | 0.22991608 | Y47G6A.22 |
| WBGene00021648 | -0.169775891 | 0.20734593 | 0.39645783 | mis-12 |
| WBGene00021649 | -0.199474411 | 0.14427378 | 0.31335298 | Y47G6A.25 |
| WBGene00021650 | 0.155352187 | 0.0675224 | NA | Y47G6A.26 |
| WBGene00021651 | 0.02274406 | 0.9118234 | 0.95112162 | NA |
| WBGene00021653 | 0.017348279 | 0.83364574 | NA | Y47G6A.30 |
| WBGene00021654 | 0.076286008 | 0.7137579 | 0.83936783 | Y47G7B.2 |
| WBGene00021655 | -0.034621659 | 0.79881336 | 0.88729432 | Y48A5A.1 |
| WBGene00021656 | -0.126391773 | 0.27510907 | 0.47543342 | Y48D7A.1 |
| WBGene00021657 | 0.000198995 | 0.99976838 | 0.99983667 | Y48G1A.1 |
| WBGene00021658 | 0.132507018 | 0.26595032 | 0.46459594 | Y48G1A.2 |
| WBGene00021660 | -0.298900053 | 0.01959034 | 0.07554551 | nol-14 |
| WBGene00021661 | 0.106917325 | 0.45044959 | 0.64279069 | mbtr-1 |
| WBGene00021662 | 0.113771128 | 0.58771347 | 0.75003656 | snpc-3.3 |
| WBGene00021664 | 1.055394093 | 0.00012784 | 0.00120134 | NA |
| WBGene00021666 | -0.020431082 | 0.87765373 | 0.93479763 | Y48G1BL.7 |
| WBGene00021667 | -0.227394069 | 0.29733657 | 0.49843299 | snpc-3.2 |
| WBGene00021668 | -0.005343958 | 0.9373808 | NA | NA |
| WBGene00021671 | 0.560707796 | 0.04471087 | 0.13851405 | Y48G1BM.6 |
| WBGene00021672 | NA | NA | NA | Y48G1BM.7 |
| WBGene00021673 | -0.076631788 | 0.69882581 | 0.82846224 | NA |
| WBGene00021675 | -0.015487121 | 0.93374324 | 0.96274188 | Y48G1BR.1 |

| | | | | | |
|---|---|---|---|---|---|
| WBGene00021676 | | -0.184443324 | 0.17940498 | 0.36082247 | pid-2 |
| WBGene00021677 | | 0.094043326 | 0.46948742 | 0.65798188 | pgs-1 |
| WBGene00021678 | | 0.086812998 | 0.52968645 | 0.70651028 | Y48G1C.5 |
| WBGene00021679 | | -0.018364707 | 0.92736548 | 0.95889615 | Y48G1C.6 |
| WBGene00021681 | | -0.041823801 | 0.56946476 | 0.7382268 | Y48G1C.8 |
| WBGene00021682 | | 0.026785325 | 0.85795994 | 0.92383552 | Y48G1C.9 |
| WBGene00021683 | | -0.065596938 | 0.65459735 | 0.79548139 | Y48G1C.10 |
| WBGene00021685 | | -0.111976283 | 0.19161024 | 0.3778203 | herc-1 |
| WBGene00021686 | | -0.242926635 | 0.00245739 | 0.01478779 | Y48G8AL.5 |
| WBGene00021687 | | 0.47827209 | 0.0348654 | 0.11569906 | Y48G8AL.7 |
| WBGene00021689 | | 0.015604342 | 0.89897205 | 0.94663864 | Y48G8AL.10 |
| WBGene00021690 | | 0.095967469 | 0.18297344 | NA | Y48G8AL.12 |
| WBGene00021691 | | 0.421467847 | 2.66E-05 | 0.0002952 | Y48G8AL.13 |
| WBGene00021692 | | 0.189998377 | 0.35131559 | 0.55135723 | Y48G8AL.15 |
| WBGene00021693 | | 0.019074982 | 0.88679847 | 0.94033043 | Y48G8AR.2 |
| WBGene00021694 | | -0.023109762 | 0.78121253 | NA | Y48G8AR.3 |
| WBGene00021697 | | -0.089923466 | 0.27475458 | 0.47510119 | gcn-1 |
| WBGene00021698 | | 0.175109696 | 0.26199982 | 0.45988963 | frl-1 |
| WBGene00021700 | | -0.027350315 | 0.68023343 | NA | Y48G9A.6 |
| WBGene00021701 | | 0.068706873 | 0.55842168 | 0.72929111 | Y48G9A.7 |
| WBGene00021702 | | 0.670511109 | 0.00095504 | 0.00677132 | Y48G9A.9 |
| WBGene00021703 | | -0.085062742 | 0.43071529 | 0.62566748 | cpt-3 |
| WBGene00021704 | | 0.225910943 | 0.22234883 | 0.41503454 | Y48G9A.11 |
| WBGene00021705 | NA | | NA | NA | Y48G9A.12 |
| WBGene00021706 | NA | | NA | NA | srbc-39 |
| WBGene00021707 | NA | | NA | NA | srbc-38 |
| WBGene00021708 | | 0.015194928 | 0.8086441 | NA | srbc-37 |
| WBGene00021709 | | 0.054590615 | 0.74818338 | 0.85793804 | ugt-29 |
| WBGene00021710 | | 0.015896709 | 0.80422001 | NA | cyp-33C11 |
| WBGene00021711 | | 0.040037202 | 0.84387359 | 0.91621455 | wago-11 |
| WBGene00021713 | | -0.958462305 | 0.01419773 | 0.05915451 | Y49F6A.5 |
| WBGene00021714 | | -0.254925706 | 0.04726711 | 0.14440365 | cyh-1 |
| WBGene00021715 | | -0.189560988 | 0.18618604 | 0.37052463 | Y49F6B.2 |
| WBGene00021716 | | 0.004038446 | 0.97420074 | 0.98568926 | fbxb-112 |
| WBGene00021717 | | -0.190982736 | 0.07352057 | 0.19845402 | sdz-32 |
| WBGene00021718 | NA | | NA | NA | Y49F6B.6 |
| WBGene00021719 | | -1.305542141 | 0.00106666 | 0.00741193 | fbxb-104 |
| WBGene00021720 | | 0.24126326 | 0.04516034 | 0.13963939 | Y49F6B.8 |
| WBGene00021721 | | -0.033915454 | 0.8076896 | 0.89346646 | Y49F6B.9 |
| WBGene00021722 | NA | | NA | NA | srz-5 |
| WBGene00021723 | | -0.358706511 | 0.08040748 | 0.21171511 | NA |
| WBGene00021724 | NA | | NA | NA | Y49F6C.2 |
| WBGene00021725 | | -1.009947333 | 0.00117689 | 0.00804803 | bath-9 |

| | | | | |
|---|---|---|---|---|
| WBGene00021726 | -1.164465651 | 0.00601624 | 0.03019669 | bath-10 |
| WBGene00021727 | -0.089324451 | 0.54956842 | 0.72221805 | bath-23 |
| WBGene00021728 | NA | NA | NA | Y49F6C.6 |
| WBGene00021729 | 0.017604133 | 0.79437959 | NA | Y49F6C.7 |
| WBGene00021730 | 0.134939349 | 0.35099388 | 0.55097046 | Y49F6C.8 |
| WBGene00021731 | -0.202652332 | 0.25840925 | 0.45636252 | Y49G5A.1 |
| WBGene00021732 | 0.331468433 | 0.005177 | 0.0268591 | Y49G5B.1 |
| WBGene00021733 | 0.0083797 | 0.94606742 | 0.96978037 | otub-2 |
| WBGene00021734 | -0.027648852 | 0.73121158 | NA | Y50C1A.2 |
| WBGene00021735 | 0.048933848 | 0.68341751 | 0.81741 | Y50D4A.1 |
| WBGene00021736 | -0.069394287 | 0.62769119 | 0.77835408 | wrb-1 |
| WBGene00021737 | NA | NA | NA | NA |
| WBGene00021738 | -0.156180336 | 0.25713754 | 0.45539154 | Y50D4A.4 |
| WBGene00021739 | 0.135846218 | 0.51844355 | 0.69772333 | Y50D4A.5 |
| WBGene00021740 | 0.513454595 | 0.04765717 | 0.14524326 | Y50D4B.1 |
| WBGene00021741 | 0.162205727 | 0.43064732 | 0.62563083 | Y50D4B.2 |
| WBGene00021743 | 0.121035315 | 0.32631236 | 0.52559615 | Y50D4B.4 |
| WBGene00021744 | NA | NA | NA | clec-203 |
| WBGene00021745 | -0.188368889 | 0.32035382 | 0.52021964 | Y50D4B.6 |
| WBGene00021746 | 0.015896709 | 0.80422001 | NA | Y50D4B.7 |
| WBGene00021747 | -0.143156368 | 0.46734017 | 0.65690548 | NA |
| WBGene00021748 | 0.275940871 | 0.03770179 | 0.12255593 | Y50D4C.3 |
| WBGene00021749 | 0.112098895 | 0.39162 | 0.58970569 | Y50D4C.5 |
| WBGene00021750 | 0.206428824 | 0.28290242 | 0.48274758 | Y50D4C.6 |
| WBGene00021751 | 0.044490289 | 0.80421207 | 0.89116459 | Y50D7A.1 |
| WBGene00021752 | -0.160146927 | 0.14757785 | 0.31829199 | xpd-1 |
| WBGene00021753 | -0.090524537 | 0.3670243 | 0.56666208 | Y50D7A.3 |
| WBGene00021754 | -0.474053918 | 1.81E-07 | 3.25E-06 | hpo-29 |
| WBGene00021755 | -0.053370558 | 0.75205058 | 0.85970214 | hpo-38 |
| WBGene00021756 | 0.007307828 | 0.94832993 | 0.97044737 | Y50D7A.8 |
| WBGene00021757 | -0.196674569 | 0.12735088 | 0.28871852 | taco-1 |
| WBGene00021758 | 0.108992413 | 0.50000977 | 0.68187807 | Y50D7A.10 |
| WBGene00021763 | 0.03923301 | 0.58912771 | 0.751016 | Y51F10.2 |
| WBGene00021764 | -0.20535402 | 0.03025074 | 0.10404504 | Y51F10.3 |
| WBGene00021765 | 0.023236063 | 0.84530622 | 0.9169413 | Y51F10.4 |
| WBGene00021766 | 0.119794408 | 0.31967787 | 0.51987478 | hex-4 |
| WBGene00021768 | 1.08437866 | 3.51E-49 | 8.03E-47 | Y51F10.7 |
| WBGene00021771 | -0.023149854 | 0.87737214 | 0.9347017 | spe-48 |
| WBGene00021772 | 2.966887188 | 1.74E-09 | 4.27E-08 | math-45 |
| WBGene00021773 | -0.321930678 | 0.16181795 | 0.33672208 | fbxb-42 |
| WBGene00021774 | -0.899305122 | 0.01664769 | 0.06710872 | fbxb-43 |
| WBGene00021776 | 3.873054621 | 0.00178052 | 0.01134678 | Y51H7BR.4 |
| WBGene00021778 | 0.036687355 | 0.59489568 | NA | Y51H7BR.7 |

| | | | | |
|---|---|---|---|---|
| WBGene00021779 | -0.998429147 | 0.00014547 | 0.00133277 | Y51H7C.1 |
| WBGene00021780 | 0.020925399 | 0.84643798 | NA | scl-17 |
| WBGene00021781 | -0.006549512 | 0.94966534 | 0.97122452 | Y51H7C.3 |
| WBGene00021783 | -0.094093254 | 0.46194394 | 0.65228105 | alkb-1 |
| WBGene00021784 | 0.085104625 | 0.39173624 | 0.58975942 | cogc-4 |
| WBGene00021785 | -0.172083732 | 0.17861509 | 0.36005832 | Y51H7C.7 |
| WBGene00021786 | 0.020232312 | 0.84643798 | NA | Y51H7C.8 |
| WBGene00021787 | 0.153576436 | 0.21929823 | 0.41131762 | Y51H7C.9 |
| WBGene00021788 | -0.007913497 | 0.92703506 | NA | Y51H7C.10 |
| WBGene00021789 | -0.374620604 | 0.0177377 | 0.07043602 | nol-6 |
| WBGene00021790 | -0.029986232 | 0.77833643 | 0.87450109 | Y51H7C.12 |
| WBGene00021791 | 0.11782495 | 0.55485833 | 0.72686037 | Y51H7C.13 |
| WBGene00021792 | 0.020317245 | 0.84643798 | NA | NA |
| WBGene00021793 | NA | NA | NA | Y52D5A.2 |
| WBGene00021794 | 0.017501802 | 0.83221062 | NA | Y52E8A.1 |
| WBGene00021795 | -0.131263148 | 0.38304728 | 0.58260911 | Y52E8A.2 |
| WBGene00021796 | 0.388897881 | 0.09024894 | 0.22852106 | Y52E8A.3 |
| WBGene00021797 | 0.033896333 | 0.86872336 | 0.93000745 | plep-1 |
| WBGene00021798 | 0.239907249 | 0.01930642 | 0.07487821 | Y52E8A.6 |
| WBGene00021799 | 0.052621217 | 0.76079648 | 0.86447973 | Y53G8AL.1 |
| WBGene00021800 | -0.269644871 | 0.00024434 | 0.00209204 | Y53G8AL.2 |
| WBGene00021801 | 0.059762547 | 0.38965825 | NA | Y53G8AL.3 |
| WBGene00021803 | 0.020317245 | 0.84643798 | NA | Y53G8AM.2 |
| WBGene00021804 | -0.023056499 | 0.78121253 | NA | Y53G8AM.4 |
| WBGene00021805 | NA | NA | NA | Y53G8AM.5 |
| WBGene00021806 | NA | NA | NA | Y53G8AM.6 |
| WBGene00021807 | 0.106997704 | 0.16248985 | NA | Y53G8AM.7 |
| WBGene00021808 | 0.206094031 | 0.2579592 | 0.45613294 | Y53G8AM.8 |
| WBGene00021809 | -0.005377232 | 0.94281242 | NA | Y53G8AR.1 |
| WBGene00021810 | -0.022446615 | 0.88324572 | 0.9382283 | phf-15 |
| WBGene00021811 | -0.275743047 | 0.02378471 | 0.08729804 | ral-1 |
| WBGene00021812 | -0.058540327 | 0.72388083 | 0.84536486 | Y53G8AR.5 |
| WBGene00021813 | -0.332804405 | 0.00253321 | 0.01517542 | Y53G8AR.6 |
| WBGene00021814 | 0.34323269 | 0.10847192 | 0.25907624 | Y53G8AR.7 |
| WBGene00021815 | -0.105307701 | 0.48374528 | 0.66922913 | Y53G8AR.8 |
| WBGene00021816 | -0.309342973 | 0.00242687 | 0.0146402 | Y53G8AR.9 |
| WBGene00021817 | 0.094835281 | 0.65145709 | 0.79364585 | Y53G8B.1 |
| WBGene00021818 | -0.284100005 | 0.11000006 | 0.26148903 | Y53G8B.2 |
| WBGene00021819 | 0.118550079 | 0.47431056 | 0.66187389 | NA |
| WBGene00021820 | 0.616450369 | 6.87E-05 | 0.00069738 | nipa-1 |
| WBGene00021821 | NA | NA | NA | Y54B9A.1 |
| WBGene00021822 | NA | NA | NA | Y54C5A.1 |
| WBGene00021826 | -0.277788032 | 0.00039379 | 0.00317307 | txl-1 |

| | | | | |
|---|---|---|---|---|
| WBGene00021827 | -0.026070939 | 0.833277 | 0.90902382 | dnc-6 |
| WBGene00021828 | -0.089834843 | 0.46488874 | 0.65465384 | Y54E10A.6 |
| WBGene00021829 | -0.202979899 | 0.16993494 | 0.34842782 | mrpl-17 |
| WBGene00021830 | -0.444104455 | 0.00037484 | 0.00304513 | Y54E10A.10 |
| WBGene00021831 | -0.110151072 | 0.22517775 | 0.41816684 | Y54E10A.11 |
| WBGene00021832 | -0.178858618 | 0.09947937 | 0.24461044 | Y54E10A.12 |
| WBGene00021833 | 0.020925399 | 0.84643798 | NA | Y54E10A.13 |
| WBGene00021834 | 0.843491571 | 1.13E-07 | 2.11E-06 | mab-31 |
| WBGene00021835 | -0.519001461 | 0.0036315 | 0.0203322 | Y54E10A.17 |
| WBGene00021837 | 0.0398788 | 0.58971221 | NA | Y54E10BL.1 |
| WBGene00021838 | 0.034398412 | 0.72141905 | 0.84363389 | Y54E10BL.3 |
| WBGene00021839 | -0.100500498 | 0.28780445 | 0.48804087 | nduf-5 |
| WBGene00021840 | -0.341053236 | 0.00555644 | 0.02847768 | pign-1 |
| WBGene00021841 | -0.137342136 | 0.25172653 | 0.44993917 | Y54E10BR.2 |
| WBGene00021842 | 0.18738378 | 0.08612966 | 0.22169907 | Y54E10BR.3 |
| WBGene00021843 | -0.164353797 | 0.11051848 | 0.26225301 | Y54E10BR.4 |
| WBGene00021844 | -0.156916246 | 0.03940928 | 0.12669987 | sec-11 |
| WBGene00021845 | -0.144267484 | 0.13210578 | 0.29529699 | rpb-7 |
| WBGene00021846 | 0.146454836 | 0.26030611 | 0.45840351 | ztf-23 |
| WBGene00021847 | -0.111798672 | 0.29914943 | 0.49958329 | Y54F10AL.1 |
| WBGene00021848 | 0.05447302 | 0.54638635 | NA | nhr-239 |
| WBGene00021849 | -0.231882108 | 0.01033711 | 0.0460577 | Y54F10AM.5 |
| WBGene00021850 | 0.280844156 | 0.14327842 | 0.31177364 | Y54F10AM.6 |
| WBGene00021852 | -0.10296942 | 0.37281433 | 0.57195734 | Y54F10AM.8 |
| WBGene00021853 | -0.084878454 | 0.54196834 | 0.71586903 | Y54F10AM.11 |
| WBGene00021854 | -0.174916879 | 0.09942018 | 0.24461044 | Y54F10AR.1 |
| WBGene00021855 | 0.312845818 | 0.03397709 | 0.11334213 | Y54F10AR.2 |
| WBGene00021856 | -0.111483232 | 0.29459059 | 0.49569181 | ppm-1.D |
| WBGene00021857 | -0.336075282 | 6.05E-06 | 7.76E-05 | iffb-1 |
| WBGene00021858 | 0.203606308 | 0.17072654 | 0.34949765 | Y54F10BM.3 |
| WBGene00021859 | 0.245065684 | 0.10401449 | 0.25210364 | fbxa-39 |
| WBGene00021860 | 0.108717649 | 0.15669083 | NA | fbxa-9 |
| WBGene00021861 | -0.451041791 | 0.0660597 | 0.18353384 | Y54F10BM.6 |
| WBGene00021862 | 0.090782289 | 0.39629677 | 0.59430308 | fbxa-48 |
| WBGene00021863 | -0.044983226 | 0.77801224 | 0.87435671 | Y54F10BM.9 |
| WBGene00021864 | 0.037023624 | 0.59153183 | NA | fbxa-1 |
| WBGene00021865 | 0.063217264 | 0.41230173 | NA | fbxa-66 |
| WBGene00021866 | 0.086128422 | 0.63966683 | 0.78627841 | Y54F10BM.12 |
| WBGene00021867 | -0.295123363 | 0.19948305 | 0.38803905 | Y54F10BM.13 |
| WBGene00021868 | -0.039796036 | 0.60913567 | 0.76442513 | atln-1 |
| WBGene00021869 | -0.030725551 | 0.73068227 | 0.84986592 | vrp-1 |
| WBGene00021870 | -0.059596853 | 0.76058839 | 0.86431035 | samt-1 |
| WBGene00021871 | 0.136055298 | 0.34961372 | 0.54956334 | dml-1 |

| | | | | |
|---|---|---|---|---|
| WBGene00021872 | 1.931496388 | 2.61E-32 | 3.58E-30 | clec-85 |
| WBGene00021873 | 6.505443287 | 1.16E-08 | 2.56E-07 | clec-82 |
| WBGene00021875 | 0.288199872 | 0.21205431 | 0.40241359 | Y54G2A.10 |
| WBGene00021876 | 1.744700375 | 2.18E-24 | 1.97E-22 | Y54G2A.11 |
| WBGene00021877 | -0.041434975 | 0.76908408 | 0.86850443 | Y54G2A.12 |
| WBGene00021878 | 0.345506161 | 0.0067906 | 0.03328463 | Y54G2A.13 |
| WBGene00021879 | 1.193483388 | 2.26E-19 | 1.62E-17 | clec-83 |
| WBGene00021880 | 0.193036759 | 0.0748227 | 0.20111604 | Y54G2A.15 |
| WBGene00021881 | 0.173800876 | 0.39849201 | 0.5966784 | Y54G2A.16 |
| WBGene00021882 | 0.226487296 | 0.02991741 | 0.10340434 | nprt-1 |
| WBGene00021883 | -0.150236007 | 0.05676712 | 0.16520123 | Y54G2A.18 |
| WBGene00021884 | -0.013247841 | 0.94660088 | 0.97012344 | Y54G2A.19 |
| WBGene00021885 | -0.027373974 | 0.68245543 | NA | NA |
| WBGene00021886 | 0.117809727 | 0.57246749 | 0.74067661 | NA |
| WBGene00021887 | 0.142019813 | 0.36195221 | 0.56165797 | fbxc-45 |
| WBGene00021888 | -0.344601776 | 7.98E-06 | 0.00010026 | manf-1 |
| WBGene00021889 | 0.020232312 | 0.84643798 | NA | NA |
| WBGene00021890 | 0.366521308 | 0.00043738 | 0.00346329 | Y54G2A.26 |
| WBGene00021892 | 0.118929849 | 0.36167745 | 0.56153972 | Y54G2A.28 |
| WBGene00021893 | 0.602148975 | 0.02730535 | 0.09674195 | ndnf-1 |
| WBGene00021894 | 0.371974845 | 0.12678662 | 0.28806376 | Y54G2A.32 |
| WBGene00021895 | 1.657299099 | 1.52E-21 | 1.25E-19 | clec-84 |
| WBGene00021897 | 0.248341314 | 0.24846237 | 0.4468965 | ser-6 |
| WBGene00021898 | 0.421084496 | 0.02199995 | 0.08239993 | Y54G2A.36 |
| WBGene00021899 | -0.324532586 | 6.80E-05 | 0.00069003 | Y54H5A.1 |
| WBGene00021900 | -0.135406259 | 0.24207299 | 0.44005348 | Y54H5A.2 |
| WBGene00021901 | -0.255890735 | 0.0022305 | 0.01361832 | rbm-42 |
| WBGene00021902 | -0.027493333 | 0.85319222 | 0.92127645 | oxy-4 |
| WBGene00021904 | 0.003112091 | 0.98495871 | 0.99139148 | gtf-2H5 |
| WBGene00021905 | -0.14535449 | 0.36929895 | 0.56920791 | helq-1 |
| WBGene00021906 | 0.020199342 | 0.87192921 | 0.9314787 | Y55B1AR.2 |
| WBGene00021907 | -0.04044783 | 0.53612608 | NA | Y55B1AR.3 |
| WBGene00021908 | 0.131496369 | 0.24300313 | 0.44103246 | Y55B1AR.4 |
| WBGene00021909 | 1.215275935 | 2.15E-06 | 3.10E-05 | Y55B1BL.1 |
| WBGene00021910 | -0.126037033 | 0.21968012 | 0.41187561 | stim-1 |
| WBGene00021911 | NA | NA | NA | Y55B1BR.1 |
| WBGene00021912 | 0.081319772 | 0.66132551 | 0.80066707 | Y55B1BR.2 |
| WBGene00021913 | -1.013942804 | 2.13E-06 | 3.08E-05 | cec-8 |
| WBGene00021914 | -0.197574768 | 0.2610718 | 0.45919656 | magu-1 |
| WBGene00021915 | 0.124012378 | 0.54459915 | 0.71804879 | Y55D5A.1 |
| WBGene00021916 | -0.001488154 | 0.95639912 | NA | Y55D5A.2 |
| WBGene00021917 | 0.671901221 | 0.03299266 | 0.11091745 | Y55D5A.3 |
| WBGene00021918 | 0.608877457 | 0.0414828 | 0.13130013 | Y55D5A.4 |

| | | | | |
|---|---|---|---|---|
| WBGene00021919 | -0.296212453 | 0.08570076 | 0.22090578 | cutl-25 |
| WBGene00021920 | -0.235303981 | 0.0534295 | 0.15812844 | mrps-25 |
| WBGene00021921 | -0.308014769 | 0.00101362 | 0.00712451 | rbm-39 |
| WBGene00021922 | 0.032895483 | 0.79154517 | 0.88250563 | atg-3 |
| WBGene00021923 | -0.092529152 | 0.6025746 | 0.75975669 | Y55F3AM.5 |
| WBGene00021925 | 0.284398529 | 0.16008281 | 0.33477681 | immp-2 |
| WBGene00021926 | 0.052377149 | 0.64718811 | 0.79127191 | Y55F3AM.9 |
| WBGene00021927 | 0.28013391 | 0.20761726 | 0.39671566 | abhd-14 |
| WBGene00021928 | 1.045762638 | 0.00026115 | 0.00222426 | Y55F3AM.11 |
| WBGene00021929 | -0.056372776 | 0.60978465 | 0.76489522 | dcap-1 |
| WBGene00021930 | -0.100105538 | 0.30187658 | 0.50253269 | Y55F3AM.13 |
| WBGene00021931 | -0.012374601 | 0.9071207 | 0.94967137 | Y55F3AM.14 |
| WBGene00021932 | -0.26482348 | 0.15067179 | 0.32241824 | cox-18 |
| WBGene00021933 | -0.220989788 | 0.11371564 | 0.26779968 | Y55F3AR.2 |
| WBGene00021934 | -0.395390601 | 2.66E-11 | 8.29E-10 | cct-8 |
| WBGene00021935 | -0.190832571 | 0.15888846 | 0.33321705 | mrpl-46 |
| WBGene00021936 | -0.121103532 | 0.23240693 | 0.42736371 | zipt-15 |
| WBGene00021938 | 0.127586374 | 0.22111635 | 0.41356214 | Y55F3BR.1 |
| WBGene00021939 | 0.206546826 | 0.24471222 | 0.44292812 | Y55F3BR.2 |
| WBGene00021941 | 0.02616051 | 0.87569999 | 0.93359962 | lgc-33 |
| WBGene00021942 | 0.079324102 | 0.70050479 | 0.82976818 | madf-1 |
| WBGene00021943 | 0.064714772 | 0.67546451 | 0.81138906 | Y55F3BR.6 |
| WBGene00021945 | -0.006757146 | 0.95844787 | 0.97720301 | lem-4 |
| WBGene00021947 | 0.018790812 | 0.76755469 | NA | srt-23 |
| WBGene00021948 | 0.081477177 | 0.67169209 | 0.80894366 | kvs-5 |
| WBGene00021949 | NA | NA | NA | clec-164 |
| WBGene00021950 | 0.355514989 | 0.13170553 | 0.29457694 | cutl-26 |
| WBGene00021951 | NA | NA | NA | srt-24 |
| WBGene00021952 | 0.047391751 | 0.51874555 | 0.6979082 | vha-19 |
| WBGene00021953 | -0.05239914 | 0.54259756 | NA | Y55H10A.2 |
| WBGene00021956 | -0.06197892 | 0.37456456 | 0.57340912 | Y57E12AL.1 |
| WBGene00021957 | NA | NA | NA | Y57E12AL.2 |
| WBGene00021958 | 0.020317245 | 0.84643798 | NA | Y57E12AL.4 |
| WBGene00021959 | -0.243314982 | 0.00217079 | 0.01331485 | Y57E12AL.6 |
| WBGene00021960 | -0.050045707 | 0.59447361 | 0.75407219 | Y57E12AM.1 |
| WBGene00021961 | NA | NA | NA | Y57E12B.1 |
| WBGene00021963 | 0.056550688 | 0.43750329 | NA | lipl-6 |
| WBGene00021964 | 0.020317245 | 0.84643798 | NA | Y57E12B.4 |
| WBGene00021965 | 0.043065537 | 0.83205279 | 0.9084712 | Y57G7A.1 |
| WBGene00021966 | -0.356398523 | 0.13641896 | 0.30153552 | Y57G7A.2 |
| WBGene00021967 | 0.020232312 | 0.84643798 | NA | Y57G7A.3 |
| WBGene00021968 | 0.08963136 | 0.47024696 | 0.65840969 | Y57G7A.5 |
| WBGene00021969 | 0.494948423 | 0.03913427 | 0.12595403 | Y57G7A.6 |

| | | | | |
|---|---|---|---|---|
| WBGene00021970 | -0.027373974 | 0.68245543 | NA | srz-64 |
| WBGene00021971 | 0.163523892 | 0.39674975 | 0.59468258 | NA |
| WBGene00021972 | 0.04909525 | 0.45360729 | NA | glb-31 |
| WBGene00021973 | -0.231372539 | 0.05032045 | 0.15100924 | emc-2 |
| WBGene00021974 | 0.014067275 | 0.88561763 | NA | Y57G7A.11 |
| WBGene00021975 | -0.878443255 | 0.00044577 | 0.00352402 | Y58A7A.1 |
| WBGene00021976 | -0.168051168 | 0.26545033 | 0.46405472 | Y58A7A.2 |
| WBGene00021977 | 0.316740702 | 0.13607574 | 0.30108582 | Y58A7A.3 |
| WBGene00021978 | 0.175586955 | 0.04633883 | NA | Y58A7A.4 |
| WBGene00021979 | -0.147519964 | 0.45470437 | 0.64678155 | Y58A7A.5 |
| WBGene00021980 | 0.129292902 | 0.42668086 | 0.62233856 | glb-32 |
| WBGene00021981 | 0.712120848 | 0.01803461 | 0.07120946 | lgc-26 |
| WBGene00021983 | 0.613808837 | 0.02428384 | 0.08880948 | npr-5 |
| WBGene00021984 | 1.219877629 | 3.68E-06 | 4.97E-05 | Y59C2A.1 |
| WBGene00021985 | -0.110511577 | 0.49756683 | 0.67974955 | Y59C2A.2 |
| WBGene00021986 | 0.13803036 | 0.25345118 | 0.45154279 | Y59C2A.3 |
| WBGene00021987 | NA | NA | NA | fbxa-40 |
| WBGene00021988 | NA | NA | NA | rgs-8.1 |
| WBGene00021988 | NA | NA | NA | rgs-8.2 |
| WBGene00021990 | -0.030550281 | 0.70468744 | NA | Y59E1B.1 |
| WBGene00021992 | 0.020317245 | 0.84643798 | NA | thn-6 |
| WBGene00021993 | 0.210221437 | 0.10759814 | 0.25791493 | Y59E9AL.3 |
| WBGene00021994 | -0.015502984 | 0.93872458 | 0.96565974 | Y59E9AL.4 |
| WBGene00021995 | NA | NA | NA | NA |
| WBGene00021996 | 0.696185066 | 0.02962045 | 0.10271744 | Y59E9AL.6 |
| WBGene00021997 | NA | NA | NA | Y59E9AR.7 |
| WBGene00021997 | NA | NA | NA | Y59E9AR.1 |
| WBGene00021998 | NA | NA | NA | Y59E9AR.2 |
| WBGene00021999 | NA | NA | NA | thn-5 |
| WBGene00022000 | NA | NA | NA | tbx-30 |
| WBGene00022000 | NA | NA | NA | tbx-42 |
| WBGene00022001 | NA | NA | NA | NA |
| WBGene00022002 | NA | NA | NA | Y59E9AR.7 |
| WBGene00022002 | NA | NA | NA | Y59E9AR.1 |
| WBGene00022003 | NA | NA | NA | Y59E9AR.2 |
| WBGene00022003 | NA | NA | NA | Y59E9AR.8 |
| WBGene00022004 | 1.998252552 | 0.00112943 | 0.00778166 | npr-22 |
| WBGene00022005 | 0.020317245 | 0.84643798 | NA | Y59H11AM.1 |
| WBGene00022006 | 0.020232312 | 0.84643798 | NA | mpst-6 |
| WBGene00022006 | 0.020232312 | 0.84643798 | NA | mpst-5 |
| WBGene00022007 | NA | NA | NA | mpst-6 |
| WBGene00022007 | NA | NA | NA | mpst-5 |
| WBGene00022008 | 0.033034883 | 0.64219503 | NA | Y59H11AM.4 |

| | | | | |
|---|---|---|---|---|
| WBGene00022010 | 0.990112463 | 9.56E-17 | 5.53E-15 | catp-7 |
| WBGene00022011 | 0.020925399 | 0.84643798 | NA | Y59H11AR.3 |
| WBGene00022012 | 0.412404473 | 0.05371674 | 0.15868983 | Y59H11AR.4 |
| WBGene00022013 | 0.020232312 | 0.84643798 | NA | Y60C6A.1 |
| WBGene00022014 | 0.052638593 | 0.68835364 | 0.82049871 | Y61A9LA.1 |
| WBGene00022015 | -0.059491009 | 0.60889382 | 0.76432215 | Y61A9LA.3 |
| WBGene00022016 | -0.007072416 | 0.9244612 | NA | Y61A9LA.4 |
| WBGene00022017 | 0.054307791 | 0.78469207 | 0.87854069 | NA |
| WBGene00022018 | 0.017396218 | 0.82745944 | NA | Y61A9LA.7 |
| WBGene00022019 | -0.070559512 | 0.40966834 | 0.60757234 | sut-2 |
| WBGene00022020 | -0.072889732 | 0.33672315 | NA | hpo-39 |
| WBGene00022021 | -0.331370141 | 0.00020755 | 0.00182401 | Y61A9LA.10 |
| WBGene00022022 | 0.18589038 | 0.29806565 | 0.49896857 | Y61A9LA.11 |
| WBGene00022023 | NA | NA | NA | NA |
| WBGene00022024 | 0.194885556 | 0.03436775 | 0.1144367 | Y64H9A.2 |
| WBGene00022025 | 0.11777392 | 0.27247141 | 0.47249247 | cus-2 |
| WBGene00022026 | 0.540366637 | 0.04983038 | 0.14980835 | Y65B4A.2 |
| WBGene00022027 | -0.133926551 | 0.23571921 | 0.43155745 | vps-20 |
| WBGene00022028 | 0.011730136 | 0.94293394 | 0.96758451 | Y65B4A.4 |
| WBGene00022029 | -0.038166674 | 0.68024854 | 0.81488454 | Y65B4A.6 |
| WBGene00022030 | 0.020317245 | 0.84643798 | NA | Y65B4A.7 |
| WBGene00022031 | 0.254072903 | 0.08244923 | 0.21537824 | Y65B4A.8 |
| WBGene00022032 | 0.09167768 | 0.1962711 | NA | Y65B4A.9 |
| WBGene00022033 | 0.013204071 | 0.88957758 | NA | Y65B4BL.1 |
| WBGene00022034 | -0.309413964 | 3.43E-05 | 0.00037255 | deps-1 |
| WBGene00022035 | 0.117308937 | 0.48532076 | 0.66999896 | Y65B4BL.3 |
| WBGene00022036 | -0.023171074 | 0.8719095 | 0.9314787 | Y65B4BL.4 |
| WBGene00022037 | -0.11063767 | 0.0247581 | 0.09019242 | acs-13 |
| WBGene00022038 | 0.039135871 | 0.65930174 | NA | Y65B4BL.6 |
| WBGene00022039 | 0.114836815 | 0.47471005 | 0.66192665 | Y65B4BL.7 |
| WBGene00022040 | 0.685999473 | 0.03286477 | 0.1106146 | Y65B4BR.1 |
| WBGene00022042 | -0.246529637 | 3.69E-07 | 6.28E-06 | icd-2 |
| WBGene00022043 | -0.030119851 | 0.78439294 | 0.87847449 | psf-3 |
| WBGene00022044 | -0.173923536 | 0.14161603 | 0.30960136 | dpm-1 |
| WBGene00022045 | -0.196121276 | 0.25228374 | 0.45017504 | mrpl-55 |
| WBGene00022046 | -0.567754131 | 1.35E-10 | 3.79E-09 | Y66H1A.4 |
| WBGene00022047 | 0.435897336 | 0.02734513 | 0.09679887 | Y66H1A.5 |
| WBGene00022048 | 0.418499093 | 1.88E-14 | 8.47E-13 | fln-1 |
| WBGene00022051 | 0.247145411 | 0.11087262 | 0.26283776 | rep-1 |
| WBGene00022052 | -0.103326139 | 0.36352891 | 0.56333899 | Y67D2.2 |
| WBGene00022053 | -0.276555833 | 0.00219236 | 0.01341902 | cisd-3.2 |
| WBGene00022054 | -0.02156459 | 0.89877275 | 0.94659373 | Y67D2.4 |
| WBGene00022055 | -0.09241258 | 0.6255005 | 0.77662422 | Y67D2.5 |

| | | | | | |
|---|---|---|---|---|---|
| WBGene00022056 | | 0.026856163 | 0.83904011 | 0.91279435 | ddx-35 |
| WBGene00022057 | | -0.020710663 | 0.85557336 | 0.92262213 | Y67D2.7 |
| WBGene00022059 | | 0.091609967 | 0.34753862 | 0.54724811 | Y67D8A.2 |
| WBGene00022060 | | 1.397692016 | 8.29E-10 | 2.12E-08 | dmd-9 |
| WBGene00022061 | | 0.017348279 | 0.83364574 | NA | Y67D8B.1 |
| WBGene00022062 | | 0.048422943 | 0.69491855 | 0.82543425 | Y67D8B.2 |
| WBGene00022063 | | 0.054620117 | 0.71913483 | 0.84237564 | nlp-81 |
| WBGene00022064 | NA | | NA | NA | Y67D8B.5 |
| WBGene00022066 | NA | | NA | NA | Y67D8C.2 |
| WBGene00022067 | | -0.029893044 | 0.84817753 | 0.9180823 | Y67D8C.3 |
| WBGene00022069 | | 0.011902649 | 0.83493479 | 0.91022191 | eel-1 |
| WBGene00022070 | | -0.058648864 | 0.51609488 | NA | Y67D8C.6 |
| WBGene00022071 | | -0.023056499 | 0.78121253 | NA | Y67D8C.7 |
| WBGene00022072 | | -0.506394064 | 0.0007157 | 0.00527088 | cpg-9 |
| WBGene00022073 | | 0.151809777 | 0.47689732 | 0.66409108 | Y67D8C.9 |
| WBGene00022074 | | -0.134142133 | 0.14838875 | 0.31916552 | Y69A2AR.1 |
| WBGene00022075 | | -0.260401902 | 0.01962375 | 0.07562583 | Y69A2AR.3 |
| WBGene00022076 | | -0.073776905 | 0.71712824 | 0.84132743 | daao-1 |
| WBGene00022077 | | -0.182328645 | 0.28584061 | 0.48594895 | vamp-7 |
| WBGene00022078 | | 0.349372952 | 0.13361521 | 0.2973944 | epg-9 |
| WBGene00022079 | | 0.193198565 | 0.03499485 | 0.11597092 | NA |
| WBGene00022082 | | 0.453919003 | 0.04979297 | 0.14973797 | NA |
| WBGene00022083 | | 1.022000795 | 0.00779001 | 0.03701836 | Y69A2AR.12 |
| WBGene00022085 | | 0.05787416 | 0.40594086 | NA | NA |
| WBGene00022086 | | 0.012386589 | 0.8695009 | NA | sprr-3 |
| WBGene00022087 | | -0.256407649 | 0.01101387 | 0.04827968 | Y69A2AR.16 |
| WBGene00022089 | | -0.084827998 | 0.09316584 | 0.23328904 | Y69A2AR.18 |
| WBGene00022090 | | 1.239757119 | 0.01324922 | 0.05601554 | Y69A2AR.19 |
| WBGene00022092 | | -0.039292859 | 0.69258238 | 0.82366166 | Y69A2AR.21 |
| WBGene00022093 | | 0.575959459 | 0.00510144 | 0.0265518 | Y69A2AR.22 |
| WBGene00022094 | | 0.148467664 | 0.07389511 | NA | Y69A2AR.23 |
| WBGene00022095 | NA | | NA | NA | Y69A2AR.24 |
| WBGene00022096 | | 0.218558388 | 0.09862053 | 0.24347628 | Y69A2AR.25 |
| WBGene00022097 | NA | | NA | NA | nhr-242 |
| WBGene00022099 | | -0.129989534 | 0.37224536 | 0.57146318 | Y69A2AR.28 |
| WBGene00022100 | | 0.265125268 | 0.2438973 | 0.44183375 | Y69A2AR.31 |
| WBGene00022101 | | -0.080707559 | 0.64154551 | 0.78759583 | Y69A2AR.32 |
| WBGene00022102 | | 0.098261066 | 0.23575476 | NA | Y69F12A.1 |
| WBGene00022103 | | -0.018864281 | 0.92405038 | 0.95699382 | cdh-12 |
| WBGene00022104 | | -0.757220677 | 0.01148861 | 0.04983846 | acsd-1 |
| WBGene00022106 | | 0.500843508 | 0.02569049 | 0.09253001 | lgc-46 |
| WBGene00022107 | | -0.340396045 | 0.03640368 | 0.11928966 | Y71F9AL.1 |
| WBGene00022108 | | 0.4970562 | 0.04414964 | 0.13726797 | Y71F9AL.2 |

| | | | | |
|---|---|---|---|---|
| WBGene00022110 | -0.12067992 | 0.54980643 | 0.72233632 | Y71F9AL.4 |
| WBGene00022111 | 0.083761825 | 0.56455965 | 0.73446933 | Y71F9AL.6 |
| WBGene00022112 | -0.695570737 | 0.03096518 | 0.10565399 | Y71F9AL.7 |
| WBGene00022113 | 0.166536609 | 0.31184447 | 0.51266736 | Y71F9AL.8 |
| WBGene00022114 | -0.187706488 | 0.00355629 | 0.02001063 | Y71F9AL.9 |
| WBGene00022115 | 0.1124627 | 0.39958404 | 0.59782443 | Y71F9AL.10 |
| WBGene00022116 | -0.023056499 | 0.78121253 | NA | Y71F9AL.11 |
| WBGene00022117 | -0.311535854 | 0.00310968 | 0.01794709 | Y71F9AL.12 |
| WBGene00022118 | -0.03349962 | 0.82606832 | 0.90477236 | dnc-5 |
| WBGene00022119 | -0.107306184 | 0.05093268 | 0.15237134 | copa-1 |
| WBGene00022121 | -0.202459769 | 0.06826994 | 0.18811088 | cogc-3 |
| WBGene00022122 | -0.363617251 | 2.90E-10 | 7.85E-09 | trap-1 |
| WBGene00022123 | -0.25619393 | 0.16015667 | 0.33477934 | Y71F9AR.2 |
| WBGene00022124 | 0.010107413 | 0.94056559 | 0.96666778 | supr-1 |
| WBGene00022125 | -0.070335385 | 0.72460537 | 0.84567136 | Y71F9B.1 |
| WBGene00022126 | 0.051056763 | 0.69846041 | 0.82829733 | Y71F9B.2 |
| WBGene00022127 | -0.166498468 | 0.00682044 | 0.03339732 | yop-1 |
| WBGene00022128 | 0.039818838 | 0.75643948 | 0.86211635 | Y71F9B.6 |
| WBGene00022129 | -0.047878695 | 0.80497957 | 0.89157318 | lron-11 |
| WBGene00022130 | -0.01842476 | 0.92666292 | 0.9586234 | Y71F9B.9 |
| WBGene00022131 | 0.55287525 | 0.00878986 | 0.04069967 | Y71F9B.13 |
| WBGene00022133 | 0.18285475 | 0.08713723 | 0.22327199 | Y71F9B.15 |
| WBGene00022134 | -0.166353221 | 0.02836631 | 0.09949957 | hrpu-2 |
| WBGene00022138 | 0.173539298 | 0.24180508 | 0.43978486 | trpp-10 |
| WBGene00022139 | 0.616846487 | 2.41E-07 | 4.22E-06 | tub-2 |
| WBGene00022140 | -0.074719773 | 0.68001496 | 0.81473806 | Y71G12A.4 |
| WBGene00022141 | -0.206116587 | 0.02053971 | 0.07815019 | chaf-2 |
| WBGene00022142 | 0.214956244 | 0.10395465 | 0.25207022 | Y71G12B.2 |
| WBGene00022143 | 0.038125098 | 0.61119482 | NA | Y71G12B.3 |
| WBGene00022144 | 1.174890354 | 6.59E-09 | 1.52E-07 | pghm-1 |
| WBGene00022145 | -0.10962359 | 0.40041289 | 0.59869742 | Y71G12B.5 |
| WBGene00022146 | -0.672439545 | 0.00016012 | 0.0014543 | Y71G12B.6 |
| WBGene00022148 | -0.272575295 | 0.01115967 | 0.0487291 | ddx-27 |
| WBGene00022149 | -0.216488497 | 0.07143352 | 0.19436131 | lin-65 |
| WBGene00022150 | -0.143661325 | 0.26115313 | 0.45922927 | Y71G12B.10 |
| WBGene00022152 | -0.07713773 | 0.55700815 | 0.72833405 | atg-5 |
| WBGene00022153 | -0.085259443 | 0.48399005 | 0.66932694 | Y71G12B.13 |
| WBGene00022154 | 0.101276993 | 0.60051302 | 0.75826745 | drag-1 |
| WBGene00022155 | -0.000238269 | 0.99710505 | 0.99805942 | Y71G12B.17 |
| WBGene00022156 | NA | NA | NA | Y71G12B.18 |
| WBGene00022157 | 0.010572984 | 0.90770248 | NA | Y71G12B.22 |
| WBGene00022158 | 0.10656138 | 0.60917908 | 0.76442513 | Y71G12B.23 |
| WBGene00022159 | -0.351525315 | 0.00981682 | 0.04426407 | mppa-1 |

| | | | | |
|---|---|---|---|---|
| WBGene00022160 | -0.015628375 | 0.93801106 | 0.96517113 | Y71G12B.25 |
| WBGene00022161 | 0.233523966 | 0.22717137 | 0.42089568 | Y71G12B.26 |
| WBGene00022162 | 0.175586955 | 0.04633883 | NA | Y71G12B.27 |
| WBGene00022163 | 0.010623157 | 0.91671291 | 0.95357682 | NA |
| WBGene00022165 | -0.097690919 | 0.4221666 | 0.61846519 | Y71H2AL.2 |
| WBGene00022166 | -0.081766901 | 0.57577882 | 0.74292568 | Y71H2AM.1 |
| WBGene00022167 | -0.203859049 | 0.03471667 | 0.11525777 | Y71H2AM.2 |
| WBGene00022168 | 0.027301256 | 0.81392919 | 0.8972771 | Y71H2AM.3 |
| WBGene00022169 | 0.163650311 | 0.14084781 | 0.30856693 | Y71H2AM.4 |
| WBGene00022170 | -0.073205968 | 0.19452472 | 0.38141641 | cox-6B |
| WBGene00022171 | 0.213779226 | 0.31763793 | 0.51787716 | Y71H2AM.6 |
| WBGene00022172 | -0.116835943 | 0.41554797 | 0.61275434 | cosa-1 |
| WBGene00022173 | -0.007235886 | 0.96670975 | 0.98138937 | set-27 |
| WBGene00022174 | -0.034948898 | 0.75461052 | 0.86091644 | Y71H2AM.9 |
| WBGene00022175 | 0.046197674 | 0.8227133 | 0.90294943 | Y71H2AM.10 |
| WBGene00022176 | -0.214672854 | 0.04413515 | 0.13726797 | Y71H2AM.11 |
| WBGene00022177 | 0.81134034 | 0.01361638 | 0.05724882 | Y71H2AM.12 |
| WBGene00022178 | 0.070073019 | 0.52159067 | 0.69999149 | Y71H2AM.13 |
| WBGene00022179 | 0.073887753 | 0.28837975 | NA | Y71H2AM.14 |
| WBGene00022180 | 0.685809006 | 0.0328648 | 0.1106146 | Y71H2AM.15 |
| WBGene00022181 | 1.792064131 | 0.00034077 | 0.00280295 | pho-9 |
| WBGene00022182 | -0.055129194 | 0.62127414 | 0.7735436 | swsn-3 |
| WBGene00022185 | -0.039230231 | 0.74422851 | 0.85554724 | Y71H2AM.20 |
| WBGene00022188 | 0.012128498 | 0.91519957 | 0.95281497 | Y71H2AR.1 |
| WBGene00022189 | 0.042516415 | 0.55991373 | NA | Y71H2AR.2 |
| WBGene00022190 | -0.049378039 | 0.80975616 | 0.89476527 | josd-1 |
| WBGene00022191 | -0.66031782 | 0.02426891 | 0.08880806 | Y71H2B.1 |
| WBGene00022192 | -0.158188983 | 0.08979167 | 0.2277675 | Y71H2B.2 |
| WBGene00022193 | 0.081748318 | 0.47215314 | 0.66018471 | ppfr-4 |
| WBGene00022194 | 0.11906692 | 0.4067066 | 0.60471122 | Y71H2B.4 |
| WBGene00022195 | 0.051141474 | 0.69458029 | 0.82516634 | Y71H2B.5 |
| WBGene00022196 | 0.349976814 | 0.1322112 | 0.29543714 | gpa-17 |
| WBGene00022197 | -0.164502526 | 0.43988208 | 0.63432616 | Y71H2B.8 |
| WBGene00022198 | 0.086662932 | 0.64768616 | 0.79141821 | Y71H2B.11 |
| WBGene00022199 | -0.179566405 | 0.05717363 | 0.16598832 | pfk-1.1 |
| WBGene00022200 | 0.013415987 | 0.91099194 | 0.95112162 | fard-1 |
| WBGene00022201 | -0.254292252 | 0.00584687 | 0.0295696 | Y71H10B.1 |
| WBGene00022202 | 0.537884999 | 0.05688667 | 0.16542055 | Y72A10A.1 |
| WBGene00022203 | 0.240818402 | 0.03064995 | 0.10501895 | Y73B3A.1 |
| WBGene00022204 | -0.003270206 | 0.95698316 | NA | NA |
| WBGene00022205 | 0.388616959 | 0.04377018 | 0.13663949 | NA |
| WBGene00022206 | -0.088561214 | 0.56592497 | 0.73520032 | Y73B3A.4 |
| WBGene00022207 | 0.020317245 | 0.84643798 | NA | elk-2 |

| | | | | |
|---|---|---|---|---|
| WBGene00022209 | NA | NA | NA | Y73B3A.7 |
| WBGene00022210 | NA | NA | NA | Y73B3A.8 |
| WBGene00022211 | 0.071495183 | 0.57771249 | 0.74411462 | NA |
| WBGene00022213 | 0.05366068 | 0.40968658 | NA | Y73B3A.11 |
| WBGene00022214 | 0.01912147 | 0.77587372 | NA | cal-6 |
| WBGene00022215 | 0.214693348 | 0.12968391 | 0.29201626 | Y73B3A.13 |
| WBGene00022216 | 0.020232312 | 0.84643798 | NA | fbxa-16 |
| WBGene00022217 | NA | NA | NA | fbxa-221 |
| WBGene00022218 | -0.027535543 | 0.68063395 | NA | Y73B3A.16 |
| WBGene00022219 | 0.45187014 | 0.04548494 | 0.14040585 | Y73B3A.18 |
| WBGene00022221 | -0.058752347 | 0.75177583 | 0.85959087 | Y73B3A.20 |
| WBGene00022222 | -0.057638897 | 0.77985598 | 0.87560364 | NA |
| WBGene00022223 | 0.868485053 | 0.02132558 | 0.08049185 | Y73B3B.1 |
| WBGene00022224 | 0.003144905 | 0.98219148 | 0.98969481 | set-28 |
| WBGene00022225 | 0.039888668 | 0.56326497 | NA | Y73B3B.3 |
| WBGene00022227 | 0.106039833 | 0.54609416 | 0.71941196 | NA |
| WBGene00022228 | 0.106267305 | 0.22564025 | NA | Y73B6A.1 |
| WBGene00022229 | 0.063086775 | 0.41202602 | NA | NA |
| WBGene00022230 | 0.049192156 | 0.75417626 | 0.86082276 | Y73B6A.3 |
| WBGene00022231 | -0.065338335 | 0.72105696 | 0.84361475 | tyr-6 |
| WBGene00022232 | -0.232596568 | 0.1010707 | 0.24716488 | exos-2 |
| WBGene00022233 | 0.303208368 | 0.00983072 | 0.04430026 | ipla-6 |
| WBGene00022235 | -0.25637655 | 8.34E-08 | 1.59E-06 | sqd-1 |
| WBGene00022236 | 0.639510528 | 0.03763811 | 0.12240486 | Y73B6BL.12 |
| WBGene00022237 | -0.131174895 | 0.46385758 | 0.65370477 | Y73B6BL.14 |
| WBGene00022240 | 0.168407257 | 0.38650306 | 0.5851299 | shl-1 |
| WBGene00022242 | 0.30355105 | 0.19068057 | 0.37675496 | sfrp-1 |
| WBGene00022243 | NA | NA | NA | Y73B6BL.22 |
| WBGene00022244 | -0.334320621 | 0.00020564 | 0.00181384 | Y73B6BL.23 |
| WBGene00022245 | -0.565033921 | 4.80E-14 | 2.08E-12 | acp-6 |
| WBGene00022246 | 0.015644614 | 0.93494131 | 0.96327984 | acp-7 |
| WBGene00022247 | -0.082704109 | 0.64028557 | 0.78675996 | lgc-10 |
| WBGene00022248 | -0.021501642 | 0.90007936 | 0.94668216 | Y73B6BL.27 |
| WBGene00022249 | -0.007072416 | 0.9244612 | NA | NA |
| WBGene00022250 | -0.197919606 | 0.26311797 | 0.46113362 | Y73B6BL.29 |
| WBGene00022251 | -0.194272642 | 0.10875455 | 0.25941805 | blos-2 |
| WBGene00022252 | -0.312736786 | 0.08110937 | 0.21289391 | Y73B6BL.31 |
| WBGene00022253 | -0.281976463 | 0.00032463 | 0.00269283 | hrpf-2 |
| WBGene00022254 | 2.144165372 | 6.74E-07 | 1.10E-05 | nlp-71 |
| WBGene00022255 | 0.231657286 | 0.19948108 | 0.38803905 | tmed-13 |
| WBGene00022256 | 0.01409643 | 0.85915526 | NA | Y73B6BL.37 |
| WBGene00022257 | -0.162703047 | 0.00099861 | 0.00702912 | puf-11 |
| WBGene00022258 | -0.067645366 | 0.71566626 | 0.8404644 | Y73C8B.1 |

| | | | | |
|---|---|---|---|---|
| WBGene00022259 | -0.223070029 | 0.293402 | 0.49444046 | Y73C8B.2 |
| WBGene00022260 | 0.161405194 | 0.37869096 | 0.57808511 | Y73C8B.3 |
| WBGene00022261 | 2.157862258 | 0.00051607 | 0.00399339 | clec-210 |
| WBGene00022262 | -0.312213827 | 0.17557194 | 0.35627842 | Y73C8C.3 |
| WBGene00022263 | 0.015507174 | 0.80923808 | NA | Y73C8C.4 |
| WBGene00022264 | NA | NA | NA | sdz-34 |
| WBGene00022265 | -0.670831009 | 0.00330581 | 0.01886218 | Y73C8C.8 |
| WBGene00022266 | NA | NA | NA | srt-56 |
| WBGene00022267 | 0.095967469 | 0.18297344 | NA | Y73C8C.10 |
| WBGene00022268 | -0.038736524 | 0.72112164 | 0.843623 | Y73E7A.1 |
| WBGene00022269 | -0.209553215 | 0.23404304 | 0.42956302 | Y73E7A.2 |
| WBGene00022270 | -0.158195667 | 0.44515574 | 0.63844021 | Y73E7A.3 |
| WBGene00022271 | 0.701779215 | 0.01178887 | 0.0508547 | cpx-1 |
| WBGene00022272 | 0.015452195 | 0.83957939 | NA | NA |
| WBGene00022273 | -0.227330948 | 0.12612918 | 0.28708764 | Y73E7A.6 |
| WBGene00022274 | 0.063533602 | 0.65336588 | 0.79469207 | Y73E7A.8 |
| WBGene00022275 | 0.113630647 | 0.33263929 | 0.53237822 | Y74C9A.1 |
| WBGene00022276 | 0.239346101 | 0.01077517 | 0.04738939 | nlp-40 |
| WBGene00022277 | -0.102381717 | 0.47118408 | 0.65914448 | homt-1 |
| WBGene00022278 | -0.088534269 | 0.3118462 | 0.51266736 | rcor-1 |
| WBGene00022279 | -0.283350446 | 0.07251902 | 0.19622782 | sesn-1 |
| WBGene00022280 | -0.096535808 | 0.47402837 | 0.66173431 | Y74C10AL.2 |
| WBGene00022281 | -0.106847453 | 0.540098 | 0.71451837 | abtm-1 |
| WBGene00022282 | -0.006239126 | 0.93041881 | NA | Y74C10AR.2 |
| WBGene00022283 | 0.396949169 | 0.1021783 | 0.24900008 | lgc-27 |
| WBGene00022284 | 0.110179411 | 0.60095451 | 0.75861226 | glb-33 |
| WBGene00022285 | 0.037502531 | 0.8234947 | 0.9034778 | Y75B7AL.2 |
| WBGene00022286 | -0.06367715 | 0.38830771 | 0.58683043 | rga-4 |
| WBGene00022287 | 0.237687726 | 0.17889267 | 0.36025413 | Y75B7AR.1 |
| WBGene00022288 | 0.056430134 | 0.44291702 | NA | Y75B7B.1 |
| WBGene00022289 | NA | NA | NA | Y75B7B.2 |
| WBGene00022290 | 0.020232312 | 0.84643798 | NA | Y75D11A.1 |
| WBGene00022291 | 0.020232312 | 0.84643798 | NA | Y75D11A.2 |
| WBGene00022292 | -0.120548814 | 0.4808427 | 0.66716666 | Y75D11A.3 |
| WBGene00022295 | 0.133829313 | 0.25201253 | 0.45002018 | cng-2 |
| WBGene00022296 | 0.044100214 | 0.59157906 | 0.75224898 | xpc-1 |
| WBGene00022297 | -0.246813917 | 0.04292127 | 0.13464974 | Y76B12C.3 |
| WBGene00022298 | -0.082604507 | 0.64089596 | 0.7871946 | Y76B12C.4 |
| WBGene00022300 | -0.047150721 | 0.67379017 | 0.81039694 | Y76B12C.6 |
| WBGene00022301 | -0.093823332 | 0.35692295 | 0.55699307 | cpsf-1 |
| WBGene00022302 | 0.125795252 | 0.48992097 | 0.67384467 | Y76B12C.8 |
| WBGene00022303 | 0.059281883 | 0.42158309 | NA | Y76B12C.9 |
| WBGene00022306 | 0.098471607 | 0.3128288 | 0.51333563 | hxk-3 |

| | | | | |
|---|---|---|---|---|
| WBGene00022307 | 0.587530182 | 0.00056633 | 0.0043096 | Y77E11A.2 |
| WBGene00022308 | NA | NA | NA | Y77E11A.3 |
| WBGene00022309 | -0.407653758 | 0.0618964 | 0.17559101 | rpc-11 |
| WBGene00022310 | -0.216295463 | 0.01522775 | 0.06259108 | Y77E11A.7 |
| WBGene00022311 | 0.020317245 | 0.84643798 | NA | Y77E11A.8 |
| WBGene00022312 | 0.018998992 | 0.77286019 | NA | clec-171 |
| WBGene00022313 | 0.081914551 | 0.67333682 | 0.80998475 | Y77E11A.12 |
| WBGene00022314 | 0.166606288 | 0.20364034 | 0.39256065 | Y77E11A.14 |
| WBGene00022315 | -0.023056499 | 0.78121253 | NA | Y81B9A.1 |
| WBGene00022316 | NA | NA | NA | Y81B9A.2 |
| WBGene00022317 | NA | NA | NA | Y81B9A.3 |
| WBGene00022318 | NA | NA | NA | Y82E9BL.1 |
| WBGene00022319 | -0.05998906 | 0.61167291 | 0.7663446 | Y82E9BL.2 |
| WBGene00022320 | 0.052960832 | 0.64985033 | 0.79273174 | Y82E9BL.3 |
| WBGene00022321 | 0.020317245 | 0.84643798 | NA | fbxa-25 |
| WBGene00022322 | -0.126664939 | 0.3339206 | 0.53334832 | Y82E9BL.5 |
| WBGene00022323 | -0.105006899 | 0.50668426 | 0.68739475 | Y82E9BL.6 |
| WBGene00022324 | 0.096669932 | 0.60368886 | 0.76024502 | fbxa-26 |
| WBGene00022325 | -0.037453375 | 0.76413055 | 0.86623378 | fbxa-5 |
| WBGene00022326 | 0.031908494 | 0.87877231 | 0.93510469 | fbxa-14 |
| WBGene00022327 | -0.016801818 | 0.89128093 | 0.94340978 | fbxa-15 |
| WBGene00022328 | 0.0398788 | 0.58971221 | NA | Y82E9BL.12 |
| WBGene00022329 | 0.276112532 | 0.22558537 | 0.41876447 | fbxa-79 |
| WBGene00022330 | -0.140180139 | 0.46891682 | 0.65766943 | fbxa-80 |
| WBGene00022331 | -0.024099226 | 0.89877904 | 0.94659373 | fbxa-19 |
| WBGene00022332 | 0.075407067 | 0.52000167 | 0.69892081 | fbxa-20 |
| WBGene00022333 | -0.019311943 | 0.88319803 | 0.9382283 | fbxa-27 |
| WBGene00022334 | -0.609750814 | 0.02677558 | 0.09549848 | Y82E9BR.1 |
| WBGene00022335 | -0.204762308 | 0.12658526 | 0.287682 | Y82E9BR.2 |
| WBGene00022336 | -0.021479181 | 0.63787689 | 0.78546304 | Y82E9BR.3 |
| WBGene00022337 | 0.217462994 | 0.30220163 | 0.50276842 | pals-16 |
| WBGene00022338 | 0.519175772 | 0.04888006 | 0.1478011 | Y82E9BR.5 |
| WBGene00022339 | 0.106997704 | 0.16248985 | NA | Y82E9BR.6 |
| WBGene00022340 | -0.022930767 | 0.78121253 | NA | Y82E9BR.7 |
| WBGene00022341 | NA | NA | NA | Y82E9BR.8 |
| WBGene00022342 | NA | NA | NA | Y82E9BR.9 |
| WBGene00022343 | NA | NA | NA | Y82E9BR.10 |
| WBGene00022344 | -0.067666074 | 0.31672907 | NA | pals-40 |
| WBGene00022345 | 0.336374653 | 0.05655287 | 0.16483986 | fbxa-138 |
| WBGene00022346 | 0.592267749 | 0.02655099 | 0.09500297 | pals-17 |
| WBGene00022347 | -0.170735145 | 0.13439481 | 0.29867553 | Y82E9BR.14 |
| WBGene00022348 | -0.016878909 | 0.87309895 | 0.93245617 | Y82E9BR.16 |
| WBGene00022349 | -1.227592922 | 1.40E-06 | 2.12E-05 | Y82E9BR.17 |

| | | | | |
|---|---|---|---|---|
| WBGene00022350 | -0.109504998 | 0.25059656 | 0.44909133 | Y82E9BR.18 |
| WBGene00022351 | 0.109815432 | 0.37988224 | 0.57948072 | Y82E9BR.19 |
| WBGene00022352 | 0.020232312 | 0.84643798 | NA | Y82E9BR.20 |
| WBGene00022353 | 0.865155953 | 0.02148819 | 0.08098033 | pals-18 |
| WBGene00022354 | 0.361312039 | 0.05641617 | 0.16456789 | Y82E9BR.22 |
| WBGene00022356 | 0.020232312 | 0.84643798 | NA | eak-3 |
| WBGene00022357 | 0.144867011 | 0.262531 | 0.46054593 | kbp-4 |
| WBGene00022358 | 0.118431157 | 0.43381782 | 0.62886402 | Y92H12A.2 |
| WBGene00022360 | -0.294158746 | 0.01726573 | 0.06895461 | Y92H12A.4 |
| WBGene00022361 | -0.179754978 | 0.06516847 | 0.18191261 | Y92H12A.5 |
| WBGene00022363 | -0.167980985 | 0.22731261 | 0.42095673 | Y92H12BL.1 |
| WBGene00022365 | 0.300083782 | 0.19698975 | 0.38449899 | Y92H12BL.4 |
| WBGene00022366 | 0.154999802 | 0.43233222 | 0.62736252 | Y92H12BL.5 |
| WBGene00022367 | -0.121036166 | 0.5637444 | 0.73387195 | Y92H12BM.1 |
| WBGene00022368 | -0.315138696 | 0.00862359 | 0.0400063 | Y92H12BR.2 |
| WBGene00022369 | -0.158852243 | 0.01222031 | 0.0524685 | Y92H12BR.3 |
| WBGene00022370 | -0.093578744 | 0.55679154 | 0.72824592 | NA |
| WBGene00022371 | 0.035981444 | 0.8165062 | 0.89916823 | set-29 |
| WBGene00022372 | 0.020850884 | 0.88764304 | 0.94086251 | Y92H12BR.7 |
| WBGene00022373 | -0.258850916 | 0.04422825 | 0.13745399 | mrpl-15 |
| WBGene00022374 | 0.050954652 | 0.67929817 | 0.81414575 | nhr-277 |
| WBGene00022375 | 0.198655298 | 0.22679177 | 0.42047086 | Y94H6A.2 |
| WBGene00022376 | -0.134276656 | 0.34901461 | 0.5491588 | Y94H6A.3 |
| WBGene00022377 | 0.385208964 | 0.11774346 | 0.27441611 | gpx-4 |
| WBGene00022378 | -0.8633009 | 1.06E-08 | 2.36E-07 | Y94H6A.5 |
| WBGene00022379 | 0.111418843 | 0.59553021 | 0.75477619 | Y94H6A.7 |
| WBGene00022380 | -0.158445027 | 0.08261689 | 0.21553705 | Y94H6A.8 |
| WBGene00022381 | -0.247082349 | 0.00414902 | 0.02254018 | ubxn-2 |
| WBGene00022382 | -0.00214591 | 0.98644096 | 0.99221025 | Y94H6A.10 |
| WBGene00022385 | 0.067139527 | 0.64746283 | 0.79141821 | Y95B8A.4 |
| WBGene00022386 | 1.526329688 | 1.08E-05 | 0.00013173 | Y95B8A.6 |
| WBGene00022388 | 0.002122699 | 0.9795452 | 0.98820677 | Y95B8A.8 |
| WBGene00022389 | 0.755728087 | 5.34E-06 | 6.94E-05 | pde-6 |
| WBGene00022390 | -0.06242786 | 0.74201502 | 0.85412964 | him-19 |
| WBGene00022391 | 0.560386805 | 0.05164376 | 0.15402653 | prhg-1 |
| WBGene00022392 | 0.035402431 | 0.72436751 | 0.84556796 | Y97E10AL.1 |
| WBGene00022393 | -0.099339568 | 0.29652554 | 0.49751505 | abhd-12 |
| WBGene00022394 | -0.411150968 | 0.00052688 | 0.00405792 | natb-1 |
| WBGene00022395 | -0.050042403 | 0.47734215 | NA | ceh-76 |
| WBGene00022396 | 1.912351898 | 3.09E-05 | 0.00033696 | Y97E10AR.1 |
| WBGene00022397 | 0.153923898 | 0.43262025 | 0.62762516 | Y97E10AR.2 |
| WBGene00022398 | -0.094183121 | 0.44025464 | 0.63450679 | Y97E10AR.3 |
| WBGene00022399 | -0.11361737 | 0.33264093 | 0.53237822 | Y97E10AR.4 |

| | | | | |
|---|---|---|---|---|
| WBGene00022400 | -0.26405179 | 0.02766611 | 0.09767529 | rpb-9 |
| WBGene00022401 | -0.028859527 | 0.74216684 | 0.85412964 | Y97E10AR.6 |
| WBGene00022402 | -0.231021184 | 0.06273607 | 0.17738872 | lmtr-2 |
| WBGene00022403 | -0.022930767 | 0.78121253 | NA | Y97E10B.1 |
| WBGene00022404 | -0.023056499 | 0.78121253 | NA | srsx-5 |
| WBGene00022405 | NA | NA | NA | srsx-6 |
| WBGene00022406 | -0.04044783 | 0.53612608 | NA | srsx-7 |
| WBGene00022407 | NA | NA | NA | NA |
| WBGene00022408 | NA | NA | NA | srsx-3 |
| WBGene00022409 | 0.106997704 | 0.16248985 | NA | srt-53 |
| WBGene00022410 | 1.683726198 | 2.13E-14 | 9.53E-13 | Y97E10C.1 |
| WBGene00022411 | -0.008760898 | 0.95393023 | 0.97374974 | Y102A11A.1 |
| WBGene00022412 | 0.158241241 | 0.43940553 | 0.63401363 | Y102A11A.2 |
| WBGene00022413 | 0.921364791 | 6.36E-11 | 1.89E-09 | Y102A11A.3 |
| WBGene00022415 | 1.708588435 | 0.00563244 | 0.02874329 | Y102A11A.5 |
| WBGene00022416 | -0.110972833 | 0.58281576 | 0.74661232 | Y102A11A.6 |
| WBGene00022417 | 0.106493795 | 0.36759306 | 0.56727537 | Y102A11A.7 |
| WBGene00022418 | -0.443473776 | 0.05712335 | 0.16595987 | igcm-4 |
| WBGene00022419 | -0.160162758 | 0.37671041 | 0.57581566 | Y102A11A.9 |
| WBGene00022420 | -0.207293162 | 0.00307945 | 0.0178418 | wdr-4 |
| WBGene00022421 | NA | NA | NA | Y102E9.3 |
| WBGene00022423 | 0.744547215 | 0.01171255 | 0.05059216 | nhr-41 |
| WBGene00022425 | -0.102734472 | 0.17191262 | 0.35133623 | plst-1 |
| WBGene00022426 | -0.038152924 | 0.76635788 | 0.86723186 | Y104H12D.2 |
| WBGene00022427 | 0.047142791 | 0.66247948 | 0.80166643 | spl-3 |
| WBGene00022428 | -0.103358954 | 0.58165602 | 0.74597282 | Y104H12D.4 |
| WBGene00022429 | NA | NA | NA | Y47C4A.1 |
| WBGene00022429 | NA | NA | NA | Y108F1.1 |
| WBGene00022431 | 0.074892148 | 0.28555128 | NA | set-33 |
| WBGene00022432 | 0.040107227 | 0.58693163 | NA | NA |
| WBGene00022433 | NA | NA | NA | Y108F1.5 |
| WBGene00022434 | -0.01524123 | 0.89068127 | 0.9430654 | Y108G3AL.2 |
| WBGene00022435 | 0.296438284 | 0.19640897 | 0.38396663 | cfap-36 |
| WBGene00022438 | -0.082765362 | 0.44296292 | 0.63685467 | Y110A2AL.1 |
| WBGene00022439 | 0.079578344 | 0.41788044 | NA | Y110A2AL.2 |
| WBGene00022440 | 2.795582546 | 3.26E-16 | 1.79E-14 | Y110A2AL.3 |
| WBGene00022441 | 0.641870816 | 0.03719079 | 0.1213204 | Y110A2AL.4 |
| WBGene00022442 | 0.020232312 | 0.84643798 | NA | Y110A2AL.5 |
| WBGene00022443 | NA | NA | NA | Y110A2AL.6 |
| WBGene00022444 | 0.033034883 | 0.64219503 | NA | Y110A2AL.7 |
| WBGene00022445 | 2.449577011 | 1.58E-12 | 5.78E-11 | Y110A2AL.9 |
| WBGene00022446 | 0.020317245 | 0.84643798 | NA | Y110A2AL.10 |
| WBGene00022447 | -0.14486503 | 0.45520113 | 0.64717418 | pigw-1 |

| | | | | | |
|---|---|---|---|---|---|
| WBGene00022448 | | -0.210998325 | 0.09119355 | 0.22975272 | pinn-1 |
| WBGene00022450 | | 0.076352835 | 0.60574649 | 0.76196945 | ubc-26 |
| WBGene00022452 | | 0.011553092 | 0.89861475 | 0.94658742 | Y110A2AR.1 |
| WBGene00022453 | | -0.493134873 | 0.00178095 | 0.01134678 | ncap-1 |
| WBGene00022454 | | 0.109861508 | 0.27238593 | 0.47247297 | trf-2 |
| WBGene00022455 | | 0.324480552 | 0.00094725 | 0.00672582 | tyms-1 |
| WBGene00022456 | | -0.037027364 | 0.57116588 | 0.73951536 | pfkb-1.1 |
| WBGene00022457 | | 0.116618711 | 0.53800675 | 0.71275968 | Y110A7A.7 |
| WBGene00022458 | | -0.216415931 | 0.00240058 | 0.01451151 | prp-31 |
| WBGene00022459 | | -0.098491566 | 0.23393353 | 0.42941585 | Y110A7A.9 |
| WBGene00022460 | | -0.035681377 | 0.79391528 | 0.88399982 | use-1 |
| WBGene00022462 | | -0.331505757 | 0.00523465 | 0.02710062 | Y110A7A.15 |
| WBGene00022463 | | -0.180036485 | 0.04990224 | 0.14996978 | elpc-1 |
| WBGene00022465 | | -0.005221332 | 0.94448901 | NA | ift-20 |
| WBGene00022466 | | -0.019052288 | 0.85773706 | 0.92366345 | Y119C1A.1 |
| WBGene00022467 | | 0.038633797 | 0.57999349 | NA | Y119C1B.1 |
| WBGene00022469 | | 0.31793095 | 0.08952739 | 0.22728812 | Y119C1B.3 |
| WBGene00022470 | | 0.04332264 | 0.76140058 | 0.86476346 | mrpl-19 |
| WBGene00022471 | | 0.380899361 | 0.00320192 | 0.01840569 | rnf-145 |
| WBGene00022472 | | 0.122725279 | 0.5007015 | 0.68269142 | Y119C1B.6 |
| WBGene00022473 | | -0.125759037 | 0.10496455 | 0.25334286 | bet-1 |
| WBGene00022474 | | 0.071527255 | 0.70352363 | 0.83180889 | zig-9 |
| WBGene00022475 | | -0.217483307 | 0.2345492 | 0.43020249 | Y119C1B.10 |
| WBGene00022476 | | 0.020925399 | 0.84643798 | NA | fbxa-75 |
| WBGene00022477 | NA | | NA | NA | fbxa-28 |
| WBGene00022478 | NA | | NA | NA | fbxa-35 |
| WBGene00022479 | | 0.0398788 | 0.58971221 | NA | fbxa-36 |
| WBGene00022481 | NA | | NA | NA | srz-9 |
| WBGene00022481 | NA | | NA | NA | srz-8 |
| WBGene00022482 | NA | | NA | NA | fbxa-208 |
| WBGene00022483 | NA | | NA | NA | srz-9 |
| WBGene00022483 | NA | | NA | NA | srz-8 |
| WBGene00022484 | | 0.01912147 | 0.77587372 | NA | fbxa-77 |
| WBGene00022485 | | 0.247930087 | 0.21985848 | 0.41199107 | fbxa-63 |
| WBGene00022486 | | 1.149151503 | 0.00172114 | 0.01103774 | fbxa-61 |
| WBGene00022487 | | 3.061763351 | 2.35E-09 | 5.65E-08 | fbxa-21 |
| WBGene00022488 | | 0.070250284 | 0.49397441 | 0.67686828 | orc-3 |
| WBGene00022489 | | -0.013887331 | 0.9044381 | 0.94903449 | Y119D3B.12 |
| WBGene00022490 | | 1.471532623 | 2.94E-17 | 1.78E-15 | Y119D3B.13 |
| WBGene00022491 | | 0.143223729 | 0.25770054 | 0.45573917 | Y119D3B.14 |
| WBGene00022492 | | -0.172409118 | 0.03759599 | 0.12232064 | dss-1 |
| WBGene00022493 | | -0.288692978 | 0.02959172 | 0.10266645 | mrpl-45 |
| WBGene00022494 | NA | | NA | NA | fbxa-91 |

| | | | | |
|---|---|---|---|---|
| WBGene00022495 | NA | NA | NA | fbxa-78 |
| WBGene00022496 | NA | NA | NA | fbxa-92 |
| WBGene00022497 | -0.946821888 | 3.23E-29 | 3.97E-27 | Y119D3B.21 |
| WBGene00022498 | 0.098736425 | 0.4730723 | 0.66116737 | hsd-2 |
| WBGene00022499 | 0.704157135 | 0.00783434 | 0.03718074 | set-30 |
| WBGene00022500 | -0.50546313 | 2.27E-11 | 7.15E-10 | lfi-1 |
| WBGene00022501 | -0.058438715 | 0.72571904 | 0.84643133 | ZC8.6 |
| WBGene00022502 | -0.120227129 | 0.47641856 | 0.6636769 | ZC13.1 |
| WBGene00022503 | -0.005581345 | 0.94060131 | NA | ZC13.2 |
| WBGene00022504 | -0.013417424 | 0.90475368 | 0.94913088 | mam-1 |
| WBGene00022505 | 0.120282145 | 0.50360703 | 0.68467923 | ZC21.3 |
| WBGene00022506 | 0.059762547 | 0.38965825 | NA | ZC21.6 |
| WBGene00022507 | 0.015584271 | 0.80362925 | NA | ZC21.8 |
| WBGene00022508 | -0.400939479 | 0.03012241 | 0.10372113 | ZC53.1 |
| WBGene00022509 | NA | NA | NA | ZC53.2 |
| WBGene00022511 | NA | NA | NA | ZC53.4 |
| WBGene00022512 | -0.418941899 | 0.09754152 | 0.24192874 | dot-1.5 |
| WBGene00022515 | -0.021909038 | 0.86198872 | 0.92668569 | ttr-48 |
| WBGene00022516 | -0.057813906 | 0.58943945 | 0.75111006 | mtx-2 |
| WBGene00022517 | 0.633658395 | 0.03306207 | 0.11109979 | ZC123.1 |
| WBGene00022518 | 0.321616788 | 0.04693549 | 0.14370192 | zfh-2 |
| WBGene00022519 | 0.357000619 | 0.11724749 | 0.27380421 | ZC123.4 |
| WBGene00022521 | NA | NA | NA | ZC132.2 |
| WBGene00022522 | 0.078347594 | 0.31595553 | NA | ZC132.11 |
| WBGene00022522 | 0.078347594 | 0.31595553 | NA | ZC132.3 |
| WBGene00022526 | NA | NA | NA | ZC132.8 |
| WBGene00022527 | 0.03502957 | 0.67055915 | NA | ZC132.9 |
| WBGene00022528 | -0.022930767 | 0.78121253 | NA | srt-13 |
| WBGene00022529 | -0.001488154 | 0.95639912 | NA | srt-14 |
| WBGene00022530 | 0.127731691 | 0.16731752 | NA | ZC155.2 |
| WBGene00022531 | 0.022094124 | 0.79756344 | 0.88678055 | morc-1 |
| WBGene00022532 | -0.1827109 | 0.03700226 | 0.12082912 | ZC155.4 |
| WBGene00022533 | 0.153934188 | 0.12196186 | 0.2810267 | cutl-19 |
| WBGene00022534 | -0.147639566 | 0.2166908 | 0.40823404 | syx-16 |
| WBGene00022536 | -0.022930767 | 0.78121253 | NA | ZC178.2 |
| WBGene00022537 | 0.020925399 | 0.84643798 | NA | ZC190.2 |
| WBGene00022538 | 0.271389853 | 0.21495637 | 0.40587776 | ZC190.4 |
| WBGene00022539 | 0.020317245 | 0.84643798 | NA | ZC190.5 |
| WBGene00022540 | NA | NA | NA | ZC190.6 |
| WBGene00022541 | -0.002914971 | 0.95718083 | NA | ZC190.7 |
| WBGene00022542 | 0.108717649 | 0.15669083 | NA | ZC190.8 |
| WBGene00022544 | 0.149563834 | 0.32252707 | 0.52270521 | ZC190.10 |
| WBGene00022545 | 0.522667958 | 0.02470294 | 0.09005871 | NA |

| | | | | |
|---|---|---|---|---|
| WBGene00022546 | -0.141496145 | 0.38093446 | 0.580263 | ZC196.2 |
| WBGene00022547 | 0.078617024 | 0.41549642 | 0.61275434 | ZC196.3 |
| WBGene00022548 | 2.262851829 | 0.00027175 | 0.00230177 | ZC196.4 |
| WBGene00022549 | 0.845151066 | 0.00348382 | 0.01966329 | ZC196.5 |
| WBGene00022550 | 0.395470409 | 0.01759848 | 0.06997809 | droe-8 |
| WBGene00022551 | NA | NA | NA | ZC196.8 |
| WBGene00022552 | NA | NA | NA | ZC196.9 |
| WBGene00022553 | -0.058752924 | 0.35274012 | NA | ZC204.1 |
| WBGene00022554 | 0.06840532 | 0.68387582 | 0.8176383 | duxl-1 |
| WBGene00022555 | -0.001885728 | 0.99043033 | 0.99471055 | btb-12 |
| WBGene00022556 | NA | NA | NA | ZC204.6 |
| WBGene00022557 | -0.37431767 | 0.12594953 | 0.28697495 | fbxb-15 |
| WBGene00022558 | -0.076683892 | 0.54242118 | 0.71623273 | fbxb-21 |
| WBGene00022559 | -1.101688719 | 0.00494602 | 0.02594577 | fbxb-20 |
| WBGene00022560 | -0.135033468 | 0.33844742 | 0.53801541 | fbxb-16 |
| WBGene00022561 | NA | NA | NA | btb-13 |
| WBGene00022562 | 0.368669252 | 0.12939536 | 0.29168245 | ZC204.12 |
| WBGene00022563 | NA | NA | NA | ZC204.13 |
| WBGene00022564 | 0.582596617 | 0.03262939 | 0.10999998 | ZC204.14 |
| WBGene00022565 | 0.020232312 | 0.84643798 | NA | ZC239.2 |
| WBGene00022566 | -0.006558791 | 0.92999162 | NA | ZC239.3 |
| WBGene00022567 | 0.036692668 | 0.59311964 | NA | ZC239.4 |
| WBGene00022568 | -0.037309845 | 0.74106738 | 0.85379033 | ZC239.5 |
| WBGene00022569 | 0.236828759 | 0.28186829 | 0.48154418 | ZC239.6 |
| WBGene00022570 | NA | NA | NA | sdz-35 |
| WBGene00022571 | 0.222997264 | 0.12500083 | 0.28566356 | ZC239.13 |
| WBGene00022572 | 0.35036515 | 0.0641266 | 0.17996504 | ZC239.14 |
| WBGene00022573 | 0.181864499 | 0.29698656 | 0.49801628 | ZC239.15 |
| WBGene00022574 | 0.302748702 | 0.18643326 | 0.37076465 | ZC239.16 |
| WBGene00022575 | 0.108538718 | 0.15420469 | NA | ZC239.17 |
| WBGene00022576 | 0.008351447 | 0.96917853 | 0.98328202 | ZC250.2 |
| WBGene00022577 | 0.038125098 | 0.61119482 | NA | nstp-3 |
| WBGene00022578 | NA | NA | NA | ZC262.1 |
| WBGene00022579 | 0.103724844 | 0.32616342 | 0.52545759 | ZC262.2 |
| WBGene00022580 | -0.148280716 | 0.04879218 | 0.14761753 | iglr-2 |
| WBGene00022581 | 0.020232312 | 0.84643798 | NA | ZC262.4 |
| WBGene00022582 | NA | NA | NA | R05D3.6 |
| WBGene00022582 | NA | NA | NA | ZC262.5 |
| WBGene00022583 | -0.207560477 | 0.08542657 | 0.22030613 | mrps-18A |
| WBGene00022584 | -0.284090124 | 0.18135732 | 0.36388277 | ZC266.1 |
| WBGene00022586 | -0.188719155 | 0.01515381 | 0.06232215 | ZC308.4 |
| WBGene00022587 | -0.005689931 | 0.94487179 | NA | ZC317.1 |
| WBGene00022588 | -0.02791392 | 0.6763823 | NA | ZC317.2 |

| | | | | |
|---|---|---|---|---|
| WBGene00022589 | 0.096253655 | 0.43861735 | 0.63346342 | ZC317.6 |
| WBGene00022590 | 0.064756509 | 0.71779893 | 0.84175363 | ZC317.7 |
| WBGene00022591 | -0.14071593 | 0.3650945 | 0.56462961 | cuti-1 |
| WBGene00022592 | 1.300042444 | 0.00011322 | 0.00108416 | klu-2 |
| WBGene00022593 | 1.211387444 | 1.52E-05 | 0.00017848 | ZC328.3 |
| WBGene00022594 | 0.109724155 | 0.21583934 | NA | ZC328.5 |
| WBGene00022596 | -0.006711909 | 0.92834175 | NA | ZC395.4 |
| WBGene00022597 | 1.140518583 | 0.00047098 | 0.00369739 | ZC395.5 |
| WBGene00022598 | 0.010130051 | 0.90541279 | 0.94916953 | ztf-8 |
| WBGene00022599 | -0.224337348 | 1.57E-05 | 0.00018368 | daf-41 |
| WBGene00022600 | NA | NA | NA | ZC395.11 |
| WBGene00022601 | 0.118871522 | 0.43239637 | 0.62736252 | ZC404.1 |
| WBGene00022603 | -0.238478657 | 0.00672719 | 0.03302911 | gck-2 |
| WBGene00022604 | 0.020317245 | 0.84643798 | NA | dmsr-10 |
| WBGene00022605 | 2.95945005 | 0.00188222 | 0.01189879 | dmsr-11 |
| WBGene00022606 | 0.020317245 | 0.84643798 | NA | dmsr-9 |
| WBGene00022608 | 0.579823031 | 0.0239372 | 0.08779062 | madf-9 |
| WBGene00022609 | 0.051516243 | 0.77222037 | 0.87056891 | ZC416.2 |
| WBGene00022610 | 0.075482451 | 0.46830301 | 0.65738945 | ltah-1.2 |
| WBGene00022611 | 0.148021412 | 0.07455439 | NA | ZC449.1 |
| WBGene00022612 | 0.06604503 | 0.62657453 | 0.77729856 | ZC449.2 |
| WBGene00022613 | 0.061101924 | 0.67017003 | 0.80796767 | sek-3 |
| WBGene00022614 | 0.416750008 | 0.10084967 | 0.24678924 | ZC449.4 |
| WBGene00022615 | 0.287090803 | 0.13118969 | 0.2937373 | ZC449.5 |
| WBGene00022616 | NA | NA | NA | hsd-3 |
| WBGene00022617 | 0.011886376 | 0.89631667 | NA | ZC477.2 |
| WBGene00022618 | -0.090441088 | 0.34578727 | 0.54537018 | ZC477.3 |
| WBGene00022619 | NA | NA | NA | ZC477.4 |
| WBGene00022620 | -0.064058407 | 0.53103205 | 0.70766758 | rde-8 |
| WBGene00022621 | 0.716545182 | 0.02351585 | 0.0864847 | ZC477.7 |
| WBGene00022622 | 0.056188364 | 0.42092875 | NA | NA |
| WBGene00022623 | 0.17722574 | 0.34540485 | 0.54500188 | ZC487.1 |
| WBGene00022624 | NA | NA | NA | ZC487.2 |
| WBGene00022627 | 0.178629006 | 0.18703843 | 0.3715644 | ZC513.2 |
| WBGene00022628 | 0.056188364 | 0.42092875 | NA | ZC513.3 |
| WBGene00022629 | -0.063085576 | 0.49730827 | 0.67958655 | algn-12 |
| WBGene00022630 | NA | NA | NA | fbxa-223 |
| WBGene00022631 | -0.171515305 | 0.38834991 | 0.58683363 | nekl-2 |
| WBGene00022632 | 0.035512071 | 0.72830584 | 0.84850209 | ZC581.2 |
| WBGene00022633 | 0.572298845 | 0.00249642 | 0.01500415 | ZC581.3 |
| WBGene00022634 | 0.229899396 | 0.05281745 | 0.15682423 | ZC581.7 |
| WBGene00022635 | 0.845537097 | 0.01148764 | 0.04983846 | ZC581.9 |
| WBGene00022636 | 0.020232312 | 0.84643798 | NA | NA |

| | | | | |
|---|---|---|---|---|
| WBGene00022637 | 0.020925399 | 0.84643798 | NA | nhr-252 |
| WBGene00022638 | 0.00492204 | 0.97001696 | 0.98346632 | scl-27 |
| WBGene00022639 | -0.055120925 | 0.45606622 | NA | nhr-253 |
| WBGene00022640 | NA | NA | NA | nhr-254 |
| WBGene00022641 | 0.085420076 | 0.67180835 | 0.80894366 | ZK6.6 |
| WBGene00022642 | 0.946022229 | 5.72E-59 | 1.50E-56 | lipl-5 |
| WBGene00022643 | 0.842367739 | 0.00530774 | 0.02741113 | ZK6.8 |
| WBGene00022644 | 1.21838294 | 5.79E-37 | 9.32E-35 | dod-19 |
| WBGene00022645 | -0.308204087 | 0.00027182 | 0.00230177 | ZK6.11 |
| WBGene00022646 | 0.082547838 | 0.35570412 | 0.55568332 | acl-9 |
| WBGene00022647 | 0.302306532 | 0.17507896 | 0.35564444 | slc-17.1 |
| WBGene00022648 | -0.013434131 | 0.89447308 | 0.94489559 | ZK54.3 |
| WBGene00022649 | 0.573581735 | 0.04701985 | 0.14383988 | ZK84.1 |
| WBGene00022650 | 0.390047521 | 0.03703267 | 0.12089082 | ZK84.2 |
| WBGene00022651 | -0.027535543 | 0.68063395 | NA | ZK84.4 |
| WBGene00022652 | -0.005221332 | 0.94448901 | NA | ZK84.5 |
| WBGene00022653 | 0.77309437 | 3.11E-12 | 1.09E-10 | ZK105.1 |
| WBGene00022654 | 0.040107227 | 0.58693163 | NA | ZK105.3 |
| WBGene00022655 | 0.169792106 | 0.0983704 | 0.2432018 | srsx-40 |
| WBGene00022656 | -0.046394786 | 0.50231372 | NA | ZK105.5 |
| WBGene00022657 | -0.066966092 | 0.64574676 | 0.79023557 | ZK105.6 |
| WBGene00022658 | 0.020925399 | 0.84643798 | NA | NA |
| WBGene00022660 | NA | NA | NA | ZK112.4 |
| WBGene00022661 | 0.073226744 | 0.33932533 | NA | ZK112.5 |
| WBGene00022662 | -0.001488154 | 0.95639912 | NA | ZK112.6 |
| WBGene00022663 | -0.271596217 | 0.10795549 | 0.25837987 | glrx-21 |
| WBGene00022664 | 0.558162347 | 0.00879698 | 0.04071973 | ZK121.2 |
| WBGene00022665 | 0.529504055 | 0.02272324 | 0.08441791 | ZK154.1 |
| WBGene00022666 | 0.227612425 | 0.22496755 | 0.4178825 | ZK154.4 |
| WBGene00022667 | -0.157822895 | 0.21720062 | 0.4087973 | ZK154.5 |
| WBGene00022668 | 0.380341241 | 0.08252058 | 0.21546315 | ZK154.6 |
| WBGene00022669 | -0.010502654 | 0.93049495 | 0.96081261 | ZK177.1 |
| WBGene00022670 | -0.023056499 | 0.78121253 | NA | ZK177.2 |
| WBGene00022671 | 0.038740512 | 0.64922655 | NA | ZK177.3 |
| WBGene00022672 | -0.092156505 | 0.38148152 | 0.58074595 | npp-24 |
| WBGene00022673 | -0.044683205 | 0.64161775 | 0.78761847 | ZK177.8 |
| WBGene00022674 | 0.020232312 | 0.84643798 | NA | ZK177.9 |
| WBGene00022675 | 1.298589747 | 0.00013022 | 0.00121665 | gbb-2 |
| WBGene00022677 | -0.082067675 | 0.55154976 | 0.72397705 | rfth-1 |
| WBGene00022678 | -0.079943155 | 0.22608628 | 0.41948159 | sar-1 |
| WBGene00022679 | 4.854123889 | 1.41E-10 | 3.94E-09 | ZK180.5 |
| WBGene00022680 | 0.630048201 | 0.02294417 | 0.08507051 | ZK180.6 |
| WBGene00022681 | -0.220851284 | 0.21143765 | 0.40156423 | ZK185.1 |

| | | | | | |
|---|---|---|---|---|---|
| WBGene00022682 | | -0.062240949 | 0.73113185 | 0.84986592 | ZK185.2 |
| WBGene00022683 | | -0.961625249 | 0.00060797 | 0.00460015 | ZK185.3 |
| WBGene00022685 | | 0.063290468 | 0.41068021 | NA | ZK250.2 |
| WBGene00022688 | | 0.056304773 | 0.44260385 | NA | NA |
| WBGene00022689 | | 0.042516415 | 0.55991373 | NA | math-48 |
| WBGene00022690 | | -0.003614866 | 0.95700117 | NA | math-49 |
| WBGene00022691 | NA | | NA | NA | math-50 |
| WBGene00022693 | NA | | NA | NA | NA |
| WBGene00022694 | | -0.108698191 | 0.23963071 | 0.43670695 | gcna-1 |
| WBGene00022695 | | 0.083827478 | 0.68482524 | 0.81809125 | ZK328.6 |
| WBGene00022696 | | 0.167242032 | 0.23230648 | 0.42728806 | ift-139 |
| WBGene00022697 | | -0.018130561 | 0.85513117 | 0.92241889 | cyy-1 |
| WBGene00022698 | | 0.20360585 | 0.1687739 | 0.34695573 | ZK353.2 |
| WBGene00022699 | | 0.0398788 | 0.58971221 | NA | ZK353.3 |
| WBGene00022700 | | -0.00346345 | 0.95708614 | NA | ZK353.4 |
| WBGene00022702 | | -0.115028613 | 0.24600899 | 0.4442849 | cutc-1 |
| WBGene00022703 | | -0.251861683 | 0.00101559 | 0.00713495 | ubxn-4 |
| WBGene00022704 | | 0.000717374 | 0.99655161 | 0.99784654 | ZK353.9 |
| WBGene00022705 | | 0.24890981 | 0.10047081 | 0.24623254 | ZK354.2 |
| WBGene00022706 | | 0.081973927 | 0.58955362 | 0.75112054 | ZK354.3 |
| WBGene00022707 | | 0.169280425 | 0.15423391 | 0.32715947 | ZK354.6 |
| WBGene00022708 | | 0.19934579 | 0.06804369 | 0.18761349 | ZK354.7 |
| WBGene00022709 | | 0.262652301 | 0.03980871 | 0.12750514 | ZK354.8 |
| WBGene00022710 | | -0.003421551 | 0.95689871 | NA | ZK354.9 |
| WBGene00022711 | NA | | NA | NA | irld-63 |
| WBGene00022712 | | -0.316938869 | 0.05511134 | 0.16160326 | ZK355.2 |
| WBGene00022713 | | 0.020232312 | 0.84643798 | NA | ZK355.3 |
| WBGene00022714 | NA | | NA | NA | irld-64 |
| WBGene00022715 | | -0.023109762 | 0.78121253 | NA | irld-65 |
| WBGene00022716 | NA | | NA | NA | irld-66 |
| WBGene00022717 | | -0.033483072 | 0.69354288 | 0.82426831 | hipr-1 |
| WBGene00022718 | | -0.0557734 | 0.38938691 | 0.58773772 | ZK370.4 |
| WBGene00022719 | | 0.00714057 | 0.93756611 | 0.96498457 | pdhk-2 |
| WBGene00022720 | | -0.15442744 | 0.37840671 | 0.57771144 | ZK370.6 |
| WBGene00022721 | | 0.185070935 | 0.17622811 | 0.35701616 | ugtp-1 |
| WBGene00022722 | | -0.257417321 | 0.00376108 | 0.0208662 | tomm-70 |
| WBGene00022723 | | 0.020925399 | 0.84643798 | NA | ZK380.2 |
| WBGene00022724 | | -0.028819752 | 0.7204202 | NA | ZK380.4 |
| WBGene00022725 | | 0.038125098 | 0.61119482 | NA | ZK381.2 |
| WBGene00022726 | | -0.027535543 | 0.68063395 | NA | nhr-249 |
| WBGene00022727 | | 0.127596356 | 0.53193182 | 0.70832322 | prkl-1 |
| WBGene00022728 | NA | | NA | NA | ZK402.1 |
| WBGene00022729 | NA | | NA | NA | ZK402.2 |

| | | | | |
|---|---|---|---|---|
| WBGene00022730 | 0.028882636 | 0.80486432 | 0.89151297 | ZK402.3 |
| WBGene00022731 | 0.083479586 | 0.30566331 | NA | ZK402.5 |
| WBGene00022732 | 0.106997704 | 0.16248985 | NA | ZK418.2 |
| WBGene00022733 | NA | NA | NA | tmem-17 |
| WBGene00022734 | -0.303933971 | 0.00070842 | 0.00522783 | ZK418.5 |
| WBGene00022735 | 0.12779741 | 0.5187716 | 0.6979082 | ZK418.6 |
| WBGene00022736 | 0.198828134 | 0.36197499 | 0.56165797 | ZK418.7 |
| WBGene00022737 | -0.293915402 | 0.02779724 | 0.09799649 | tofu-7 |
| WBGene00022738 | -0.185212218 | 0.01802232 | 0.07120946 | fubl-2 |
| WBGene00022739 | -0.466635196 | 4.76E-07 | 8.00E-06 | toe-1 |
| WBGene00022741 | -0.628716112 | 0.01406586 | 0.0587053 | ZK430.5 |
| WBGene00022742 | -0.035623218 | 0.7483089 | 0.85794758 | ZK430.7 |
| WBGene00022743 | -0.171924501 | 0.24450604 | 0.44266266 | mlt-7 |
| WBGene00022744 | 0.055597606 | 0.44564657 | NA | ZK470.1 |
| WBGene00022745 | 0.387367857 | 0.00237815 | 0.01438184 | ZK470.2 |
| WBGene00022746 | NA | NA | NA | ZK470.4 |
| WBGene00022747 | -0.001488154 | 0.95639912 | NA | ZK470.6 |
| WBGene00022748 | 0.237516861 | 0.00013383 | 0.00124482 | oaz-1 |
| WBGene00022749 | -0.132881359 | 0.19228374 | 0.37863219 | ZK484.3 |
| WBGene00022750 | -0.072610547 | 0.29210312 | 0.49296803 | ZK484.4 |
| WBGene00022751 | 2.359940239 | 2.27E-07 | 4.00E-06 | msrp-2 |
| WBGene00022752 | -0.020834105 | 0.87559585 | 0.93355657 | ZK484.6 |
| WBGene00022753 | 0.180007984 | 0.0451335 | NA | ZK484.7 |
| WBGene00022754 | 1.64785948 | 0.00042259 | 0.00336986 | nspd-1 |
| WBGene00022755 | 0.584878809 | 0.0105353 | 0.04658632 | NA |
| WBGene00022756 | NA | NA | NA | nhr-251 |
| WBGene00022757 | NA | NA | NA | ZK488.5 |
| WBGene00022758 | NA | NA | NA | ZK488.6 |
| WBGene00022759 | -0.106389613 | 0.19617432 | 0.38376959 | ZK546.2 |
| WBGene00022760 | 0.170624885 | 0.2428728 | 0.44097731 | NA |
| WBGene00022761 | 0.033492732 | 0.73622334 | 0.85189646 | ZK546.4 |
| WBGene00022762 | 0.003490283 | 0.97800472 | 0.98744687 | ZK546.5 |
| WBGene00022763 | 0.020925399 | 0.84643798 | NA | NA |
| WBGene00022765 | -0.333645396 | 7.29E-05 | 0.00073257 | ZK546.14 |
| WBGene00022766 | 0.371086131 | 0.00142413 | 0.00940917 | cblc-1 |
| WBGene00022767 | 0.290028151 | 0.137666 | 0.30354939 | ZK563.2 |
| WBGene00022769 | -0.197671663 | 0.25349145 | 0.4515596 | sorf-1 |
| WBGene00022770 | 0.024089169 | 0.90843965 | 0.95050846 | acp-1 |
| WBGene00022771 | -0.312215598 | 0.133218 | 0.29682617 | ZK616.1 |
| WBGene00022772 | -0.021266517 | 0.91597648 | 0.95314937 | ZK616.2 |
| WBGene00022773 | 0.07941194 | 0.64929917 | 0.79266148 | ZK616.3 |
| WBGene00022774 | -0.303827727 | 0.0004129 | 0.00330702 | swsn-6 |
| WBGene00022775 | 0.043724012 | 0.64901199 | 0.79237696 | ZK616.5 |

| | | | | |
|---|---|---|---|---|
| WBGene00022776 | -0.23575933 | 0.00862372 | 0.0400063 | perm-3 |
| WBGene00022778 | 0.078203897 | 0.26561294 | NA | ZK616.8 |
| WBGene00022780 | 0.205728946 | 0.13960814 | 0.3069534 | ZK622.1 |
| WBGene00022781 | 0.893945613 | 6.86E-50 | 1.59E-47 | pmt-1 |
| WBGene00022782 | 0.114253242 | 0.58828416 | 0.75051633 | ZK622.4 |
| WBGene00022783 | -0.341122375 | 0.09043732 | 0.22876518 | tomm-7 |
| WBGene00022785 | -0.09198727 | 0.36019117 | 0.55995869 | ZK652.6 |
| WBGene00022786 | 0.203424848 | 0.35205939 | 0.55193292 | ZK652.8 |
| WBGene00022787 | 0.079292702 | 0.46236451 | 0.65252509 | tag-307 |
| WBGene00022788 | -0.048655502 | 0.75806979 | 0.86312309 | slc-17.7 |
| WBGene00022789 | 1.841688754 | 0.00584411 | 0.0295696 | lron-2 |
| WBGene00022790 | 0.035889078 | 0.72085409 | 0.84351221 | ZK682.7 |
| WBGene00022791 | -0.254496776 | 0.03247758 | 0.10966427 | ZK686.1 |
| WBGene00022792 | -0.356712139 | 0.00026466 | 0.00225152 | ZK686.2 |
| WBGene00022793 | -0.228476486 | 0.00224294 | 0.01368285 | ZK686.3 |
| WBGene00022794 | 0.043038804 | 0.72452559 | 0.84564565 | snu-23 |
| WBGene00022795 | -0.006738828 | 0.92480778 | NA | ZK686.5 |
| WBGene00022796 | 0.052960832 | 0.64985033 | 0.79273174 | ZK688.1 |
| WBGene00022797 | 0.818729264 | 8.18E-05 | 0.00080774 | best-24 |
| WBGene00022798 | 0.355745373 | 0.01165212 | 0.05038357 | fahd-1 |
| WBGene00022799 | -0.023109762 | 0.78121253 | NA | ZK688.4 |
| WBGene00022800 | 0.026847413 | 0.70157131 | 0.83043945 | ZK688.5 |
| WBGene00022801 | 0.122008166 | 0.31699154 | 0.51716884 | pcp-5 |
| WBGene00022802 | -0.216942183 | 0.28565755 | 0.48577294 | ZK688.7 |
| WBGene00022803 | 0.072997001 | 0.56565728 | 0.73508275 | ZK688.9 |
| WBGene00022804 | 0.020317245 | 0.84643798 | NA | ZK697.1 |
| WBGene00022805 | -0.0341551 | 0.85187857 | 0.92048984 | nhr-256 |
| WBGene00022806 | NA | NA | NA | NA |
| WBGene00022807 | -0.022930767 | 0.78121253 | NA | srab-25 |
| WBGene00022808 | 0.020232312 | 0.84643798 | NA | ZK697.8 |
| WBGene00022809 | 0.015452195 | 0.83957939 | NA | ZK697.14 |
| WBGene00022810 | 0.163393119 | 0.37060064 | 0.57036433 | ZK721.3 |
| WBGene00022811 | 0.047846954 | 0.76120168 | 0.86464679 | ZK721.4 |
| WBGene00022812 | -0.213657166 | 0.07923383 | 0.20964688 | uvs-1 |
| WBGene00022813 | -0.331461923 | 0.14929262 | 0.32056638 | ZK742.3 |
| WBGene00022814 | -0.023109762 | 0.78121253 | NA | ZK742.4 |
| WBGene00022815 | 0.041781973 | 0.74456948 | 0.85560331 | asic-1 |
| WBGene00022816 | -1.120497386 | 1.74E-05 | 0.00020129 | fbn-1 |
| WBGene00022817 | 0.049919234 | 0.57576642 | 0.74292568 | upp-1 |
| WBGene00022818 | NA | NA | NA | ZK783.3 |
| WBGene00022820 | 0.143278645 | 0.0278164 | 0.0980051 | ZK813.1 |
| WBGene00022821 | -0.259462664 | 0.01651827 | 0.06673398 | ZK813.2 |
| WBGene00022822 | 0.055423694 | 0.46954407 | 0.65798188 | ZK813.3 |

| | | | | |
|---|---|---|---|---|
| WBGene00022823 | 0.28048758 | 0.22218851 | 0.41493138 | ZK813.4 |
| WBGene00022824 | 0.13169099 | 0.16083826 | NA | ZK813.5 |
| WBGene00022825 | -0.233624077 | 0.09860808 | 0.24347628 | ZK816.1 |
| WBGene00022826 | NA | NA | NA | ZK816.3 |
| WBGene00022827 | 0.087256674 | 0.23800185 | NA | ZK816.4 |
| WBGene00022829 | 0.0398788 | 0.58971221 | NA | ZK867.2 |
| WBGene00022830 | -0.108947824 | 0.08643893 | 0.22214886 | ZK973.1 |
| WBGene00022831 | -0.158355423 | 0.04975925 | 0.14968669 | cec-10 |
| WBGene00022832 | -0.177499259 | 0.03329389 | 0.11172491 | pdp-1 |
| WBGene00022833 | -0.025848508 | 0.67901638 | NA | ZK973.4 |
| WBGene00022834 | -0.176370602 | 0.38798398 | 0.58658338 | ZK973.8 |
| WBGene00022835 | -0.194889682 | 0.11114184 | 0.2631162 | ZK973.9 |
| WBGene00022836 | -0.026981404 | 0.83647967 | 0.9111598 | ZK973.11 |
| WBGene00022837 | -0.166124132 | 0.28688747 | 0.48699356 | ceh-45 |
| WBGene00022838 | 0.059762547 | 0.38965825 | NA | ZK993.2 |
| WBGene00022840 | NA | NA | NA | ZK994.1 |
| WBGene00022842 | 0.193657017 | 0.03471136 | 0.11525777 | ZK994.6 |
| WBGene00022843 | -0.029216896 | 0.74896678 | NA | ZK1055.2 |
| WBGene00022845 | 0.042065281 | 0.558937 | NA | ZK1055.4 |
| WBGene00022846 | 0.01476227 | 0.84668595 | NA | ZK1055.5 |
| WBGene00022847 | -0.061583366 | 0.76365573 | 0.86603661 | ZK1055.6 |
| WBGene00022848 | 0.152625496 | 0.42660652 | 0.62229213 | ZK1055.7 |
| WBGene00022849 | -0.092835266 | 0.59787667 | 0.75608389 | acs-6 |
| WBGene00022850 | -0.261296125 | 0.01900082 | 0.07408547 | ZK1127.3 |
| WBGene00022851 | -0.342417649 | 0.00391913 | 0.02153903 | ZK1127.4 |
| WBGene00022852 | -0.549095451 | 0.00010412 | 0.00100821 | ZK1127.5 |
| WBGene00022853 | 0.07300667 | 0.63845032 | 0.78556509 | ZK1127.6 |
| WBGene00022854 | -0.071679992 | 0.5243493 | 0.70180073 | cin-4 |
| WBGene00022855 | 0.017968393 | 0.79767581 | 0.88680697 | tcer-1 |
| WBGene00022856 | -0.942153686 | 1.15E-08 | 2.54E-07 | cth-2 |
| WBGene00022857 | -0.148068667 | 0.41958874 | 0.61623018 | ZK1127.12 |
| WBGene00022858 | 0.898297798 | 5.76E-13 | 2.22E-11 | ZK1193.2 |
| WBGene00022861 | 0.521357264 | 0.00617864 | 0.03086371 | dve-1 |
| WBGene00022862 | -0.024836555 | 0.85519527 | 0.92241889 | ZK1236.1 |
| WBGene00022864 | -0.060913368 | 0.66505423 | 0.80363434 | ZK1236.5 |
| WBGene00022865 | 0.081350753 | 0.46596139 | 0.65553385 | ufbp-1 |
| WBGene00022866 | -0.03147554 | 0.70637395 | NA | ZK1240.1 |
| WBGene00022867 | 0.530984044 | 0.03467184 | 0.11523958 | ZK1240.2 |
| WBGene00022868 | 0.597002898 | 0.00596161 | 0.02998416 | ZK1240.3 |
| WBGene00022869 | 0.018476367 | 0.76699386 | NA | ZK1240.4 |
| WBGene00022870 | 0.332291283 | 0.10817134 | 0.25869594 | ZK1240.5 |
| WBGene00022871 | 0.020317245 | 0.84643798 | NA | ZK1240.6 |
| WBGene00022872 | 0.020925399 | 0.84643798 | NA | ZK1240.8 |

| | | | | | |
|---|---|---|---|---|---|
| WBGene00022873 | | 0.512637194 | 0.05788633 | 0.16740658 | ZK1240.9 |
| WBGene00022874 | | 0.020317245 | 0.84643798 | NA | nep-25 |
| WBGene00022875 | NA | | NA | NA | F58A6.9 |
| WBGene00022875 | NA | | NA | NA | ZK1248.4 |
| WBGene00022876 | | 0.131539016 | 0.10075387 | NA | ZK1248.5 |
| WBGene00022877 | | -0.002914971 | 0.95718083 | NA | wago-5 |
| WBGene00022880 | | -0.209909435 | 0.00841396 | 0.03931975 | tbc-2 |
| WBGene00022881 | | -0.213515801 | 0.0854051 | 0.22030613 | ZK1248.11 |
| WBGene00022882 | | 0.054996767 | 0.59371268 | 0.75344954 | ZK1248.13 |
| WBGene00022883 | | -0.21820225 | 0.04372164 | 0.13655812 | ZK1248.15 |
| WBGene00022884 | | 0.020925399 | 0.84643798 | NA | ZK1248.17 |
| WBGene00022885 | | -0.065384577 | 0.72426276 | 0.84556796 | ZK1248.19 |
| WBGene00022886 | NA | | NA | NA | ZK1290.1 |
| WBGene00022887 | | -0.593411507 | 0.03191343 | 0.10810841 | ZK1290.5 |
| WBGene00022888 | | 0.148539892 | 0.12780549 | 0.2891224 | rnh-1.1 |
| WBGene00022889 | | 0.020317245 | 0.84643798 | NA | ZK1290.7 |
| WBGene00022890 | | 0.0398788 | 0.58971221 | NA | fbxa-224 |
| WBGene00022891 | | 0.177882915 | 0.26135732 | 0.4594228 | ZK1290.10 |
| WBGene00022892 | | 0.063731418 | 0.41159439 | NA | ZK1290.11 |
| WBGene00022893 | | 0.677740472 | 0.00793297 | 0.03759793 | ZK1290.13 |
| WBGene00022894 | NA | | NA | NA | NA |
| WBGene00022895 | NA | | NA | NA | NA |
| WBGene00022896 | NA | | NA | NA | NA |
| WBGene00022897 | NA | | NA | NA | NA |
| WBGene00022898 | NA | | NA | NA | NA |
| WBGene00022899 | NA | | NA | NA | NA |
| WBGene00022900 | NA | | NA | NA | NA |
| WBGene00022901 | NA | | NA | NA | NA |
| WBGene00022902 | NA | | NA | NA | NA |
| WBGene00022903 | NA | | NA | NA | NA |
| WBGene00022904 | NA | | NA | NA | NA |
| WBGene00022905 | NA | | NA | NA | NA |
| WBGene00022906 | NA | | NA | NA | NA |
| WBGene00022907 | NA | | NA | NA | NA |
| WBGene00022908 | NA | | NA | NA | NA |
| WBGene00022909 | NA | | NA | NA | NA |
| WBGene00022910 | | 0.477539831 | 0.02636618 | 0.09454499 | NA |
| WBGene00022911 | NA | | NA | NA | NA |
| WBGene00022912 | NA | | NA | NA | NA |
| WBGene00022913 | NA | | NA | NA | NA |
| WBGene00022914 | NA | | NA | NA | NA |
| WBGene00022915 | NA | | NA | NA | NA |
| WBGene00022916 | NA | | NA | NA | NA |

| | | | | |
|---|---|---|---|---|
| WBGene00022917 | NA | NA | NA | NA |
| WBGene00022918 | NA | NA | NA | NA |
| WBGene00022919 | NA | NA | NA | NA |
| WBGene00022920 | NA | NA | NA | NA |
| WBGene00022921 | NA | NA | NA | NA |
| WBGene00022922 | NA | NA | NA | NA |
| WBGene00022923 | NA | NA | NA | NA |
| WBGene00022924 | NA | NA | NA | NA |
| WBGene00022925 | NA | NA | NA | NA |
| WBGene00022926 | NA | NA | NA | NA |
| WBGene00022927 | 0.020925399 | 0.84643798 | NA | NA |
| WBGene00022928 | NA | NA | NA | NA |
| WBGene00022929 | NA | NA | NA | NA |
| WBGene00022930 | NA | NA | NA | NA |
| WBGene00022931 | NA | NA | NA | NA |
| WBGene00022932 | 0.020925399 | 0.84643798 | NA | NA |
| WBGene00022933 | NA | NA | NA | NA |
| WBGene00022934 | NA | NA | NA | NA |
| WBGene00022935 | NA | NA | NA | NA |
| WBGene00022936 | NA | NA | NA | NA |
| WBGene00022937 | NA | NA | NA | NA |
| WBGene00022938 | NA | NA | NA | NA |
| WBGene00022939 | NA | NA | NA | NA |
| WBGene00022940 | NA | NA | NA | NA |
| WBGene00022941 | NA | NA | NA | NA |
| WBGene00022942 | NA | NA | NA | NA |
| WBGene00022943 | NA | NA | NA | NA |
| WBGene00022944 | NA | NA | NA | NA |
| WBGene00022945 | NA | NA | NA | NA |
| WBGene00022946 | NA | NA | NA | NA |
| WBGene00022947 | NA | NA | NA | NA |
| WBGene00022948 | NA | NA | NA | NA |
| WBGene00022949 | NA | NA | NA | NA |
| WBGene00022950 | NA | NA | NA | NA |
| WBGene00022951 | 0.809162151 | 3.43E-14 | 1.50E-12 | NA |
| WBGene00022952 | NA | NA | NA | NA |
| WBGene00022953 | NA | NA | NA | NA |
| WBGene00022954 | NA | NA | NA | NA |
| WBGene00022955 | NA | NA | NA | NA |
| WBGene00022956 | NA | NA | NA | NA |
| WBGene00022957 | NA | NA | NA | NA |
| WBGene00022958 | NA | NA | NA | NA |
| WBGene00022959 | NA | NA | NA | NA |

| | | | | |
|---|---|---|---|---|
| WBGene00022960 | NA | NA | NA | NA |
| WBGene00022961 | NA | NA | NA | NA |
| WBGene00022962 | NA | NA | NA | NA |
| WBGene00022963 | NA | NA | NA | NA |
| WBGene00022964 | NA | NA | NA | NA |
| WBGene00022965 | NA | NA | NA | NA |
| WBGene00022966 | NA | NA | NA | NA |
| WBGene00022967 | NA | NA | NA | NA |
| WBGene00022968 | NA | NA | NA | NA |
| WBGene00022969 | NA | NA | NA | NA |
| WBGene00022970 | -0.293016193 | 0.12641844 | 0.28753959 | NA |
| WBGene00022971 | NA | NA | NA | NA |
| WBGene00022972 | NA | NA | NA | NA |
| WBGene00022973 | NA | NA | NA | NA |
| WBGene00022974 | NA | NA | NA | NA |
| WBGene00022975 | NA | NA | NA | NA |
| WBGene00022976 | NA | NA | NA | NA |
| WBGene00022977 | NA | NA | NA | NA |
| WBGene00022978 | NA | NA | NA | NA |
| WBGene00022979 | NA | NA | NA | NA |
| WBGene00022980 | NA | NA | NA | NA |
| WBGene00022981 | NA | NA | NA | NA |
| WBGene00022982 | -0.022930767 | 0.78121253 | NA | NA |
| WBGene00022983 | NA | NA | NA | NA |
| WBGene00022984 | NA | NA | NA | NA |
| WBGene00022985 | NA | NA | NA | NA |
| WBGene00022986 | NA | NA | NA | NA |
| WBGene00022987 | NA | NA | NA | NA |
| WBGene00022988 | NA | NA | NA | NA |
| WBGene00022989 | NA | NA | NA | NA |
| WBGene00022990 | NA | NA | NA | NA |
| WBGene00022991 | NA | NA | NA | NA |
| WBGene00022992 | NA | NA | NA | NA |
| WBGene00022993 | NA | NA | NA | NA |
| WBGene00022994 | NA | NA | NA | NA |
| WBGene00022995 | NA | NA | NA | NA |
| WBGene00022996 | NA | NA | NA | NA |
| WBGene00022997 | NA | NA | NA | NA |
| WBGene00022998 | NA | NA | NA | NA |
| WBGene00022999 | NA | NA | NA | NA |
| WBGene00023000 | NA | NA | NA | NA |
| WBGene00023001 | NA | NA | NA | NA |
| WBGene00023002 | NA | NA | NA | NA |

| | | | | |
|---|---|---|---|---|
| WBGene00023003 | NA | NA | NA | NA |
| WBGene00023004 | NA | NA | NA | NA |
| WBGene00023005 | NA | NA | NA | NA |
| WBGene00023006 | NA | NA | NA | NA |
| WBGene00023007 | NA | NA | NA | NA |
| WBGene00023008 | NA | NA | NA | NA |
| WBGene00023009 | NA | NA | NA | NA |
| WBGene00023010 | NA | NA | NA | NA |
| WBGene00023011 | NA | NA | NA | NA |
| WBGene00023012 | NA | NA | NA | NA |
| WBGene00023013 | NA | NA | NA | NA |
| WBGene00023014 | NA | NA | NA | NA |
| WBGene00023015 | NA | NA | NA | NA |
| WBGene00023016 | NA | NA | NA | NA |
| WBGene00023017 | NA | NA | NA | NA |
| WBGene00023018 | NA | NA | NA | NA |
| WBGene00023019 | NA | NA | NA | NA |
| WBGene00023020 | -0.025086968 | 0.74513755 | NA | NA |
| WBGene00023021 | -0.150477144 | 0.4709828 | 0.65898884 | NA |
| WBGene00023022 | NA | NA | NA | NA |
| WBGene00023023 | NA | NA | NA | NA |
| WBGene00023024 | NA | NA | NA | NA |
| WBGene00023025 | NA | NA | NA | NA |
| WBGene00023026 | NA | NA | NA | NA |
| WBGene00023027 | NA | NA | NA | NA |
| WBGene00023028 | NA | NA | NA | NA |
| WBGene00023029 | NA | NA | NA | NA |
| WBGene00023030 | NA | NA | NA | NA |
| WBGene00023031 | NA | NA | NA | NA |
| WBGene00023032 | NA | NA | NA | NA |
| WBGene00023033 | NA | NA | NA | NA |
| WBGene00023034 | NA | NA | NA | NA |
| WBGene00023035 | NA | NA | NA | NA |
| WBGene00023036 | 0.063096179 | 0.76192582 | 0.86495294 | NA |
| WBGene00023037 | NA | NA | NA | NA |
| WBGene00023038 | NA | NA | NA | NA |
| WBGene00023039 | -0.098129057 | 0.56484805 | 0.73464869 | NA |
| WBGene00023040 | NA | NA | NA | NA |
| WBGene00023041 | NA | NA | NA | NA |
| WBGene00023042 | NA | NA | NA | NA |
| WBGene00023043 | NA | NA | NA | NA |
| WBGene00023044 | NA | NA | NA | NA |
| WBGene00023045 | NA | NA | NA | NA |

| | | | | |
|---|---|---|---|---|
| WBGene00023046 | NA | NA | NA | NA |
| WBGene00023047 | NA | NA | NA | NA |
| WBGene00023048 | NA | NA | NA | NA |
| WBGene00023049 | NA | NA | NA | NA |
| WBGene00023050 | NA | NA | NA | NA |
| WBGene00023051 | NA | NA | NA | NA |
| WBGene00023052 | NA | NA | NA | NA |
| WBGene00023053 | NA | NA | NA | NA |
| WBGene00023054 | NA | NA | NA | NA |
| WBGene00023055 | NA | NA | NA | NA |
| WBGene00023056 | NA | NA | NA | NA |
| WBGene00023057 | NA | NA | NA | NA |
| WBGene00023058 | NA | NA | NA | NA |
| WBGene00023059 | NA | NA | NA | NA |
| WBGene00023060 | NA | NA | NA | NA |
| WBGene00023061 | NA | NA | NA | NA |
| WBGene00023062 | NA | NA | NA | NA |
| WBGene00023063 | NA | NA | NA | NA |
| WBGene00023064 | NA | NA | NA | NA |
| WBGene00023065 | NA | NA | NA | NA |
| WBGene00023066 | NA | NA | NA | NA |
| WBGene00023067 | -0.606175045 | 1.48E-05 | 0.00017459 | rpl-41.1 |
| WBGene00023068 | -0.032974911 | 0.72944715 | NA | rpl-41.2 |
| WBGene00023069 | 0.48908824 | 0.01557133 | 0.06375275 | NA |
| WBGene00023070 | NA | NA | NA | NA |
| WBGene00023071 | -0.050223056 | 0.47776088 | NA | NA |
| WBGene00023072 | NA | NA | NA | NA |
| WBGene00023073 | NA | NA | NA | NA |
| WBGene00023074 | NA | NA | NA | NA |
| WBGene00023075 | NA | NA | NA | NA |
| WBGene00023076 | NA | NA | NA | NA |
| WBGene00023077 | NA | NA | NA | NA |
| WBGene00023078 | NA | NA | NA | NA |
| WBGene00023079 | NA | NA | NA | NA |
| WBGene00023080 | NA | NA | NA | NA |
| WBGene00023081 | NA | NA | NA | NA |
| WBGene00023082 | NA | NA | NA | NA |
| WBGene00023083 | NA | NA | NA | NA |
| WBGene00023084 | NA | NA | NA | NA |
| WBGene00023085 | NA | NA | NA | NA |
| WBGene00023086 | NA | NA | NA | NA |
| WBGene00023087 | NA | NA | NA | NA |
| WBGene00023088 | NA | NA | NA | NA |

| | | | | |
|---|---|---|---|---|
| WBGene00023089 | NA | NA | NA | NA |
| WBGene00023090 | NA | NA | NA | NA |
| WBGene00023091 | NA | NA | NA | NA |
| WBGene00023092 | NA | NA | NA | NA |
| WBGene00023093 | NA | NA | NA | NA |
| WBGene00023094 | NA | NA | NA | NA |
| WBGene00023095 | NA | NA | NA | NA |
| WBGene00023096 | 0.020925399 | 0.84643798 | NA | NA |
| WBGene00023097 | NA | NA | NA | NA |
| WBGene00023098 | NA | NA | NA | NA |
| WBGene00023099 | NA | NA | NA | NA |
| WBGene00023100 | NA | NA | NA | NA |
| WBGene00023101 | NA | NA | NA | NA |
| WBGene00023102 | NA | NA | NA | NA |
| WBGene00023103 | NA | NA | NA | NA |
| WBGene00023104 | 0.033034883 | 0.64219503 | NA | NA |
| WBGene00023105 | NA | NA | NA | NA |
| WBGene00023106 | NA | NA | NA | NA |
| WBGene00023107 | NA | NA | NA | NA |
| WBGene00023108 | NA | NA | NA | NA |
| WBGene00023109 | NA | NA | NA | NA |
| WBGene00023110 | NA | NA | NA | NA |
| WBGene00023111 | NA | NA | NA | NA |
| WBGene00023112 | NA | NA | NA | NA |
| WBGene00023113 | NA | NA | NA | NA |
| WBGene00023115 | NA | NA | NA | NA |
| WBGene00023116 | 0.020232312 | 0.84643798 | NA | NA |
| WBGene00023117 | 0.020317245 | 0.84643798 | NA | NA |
| WBGene00023118 | NA | NA | NA | NA |
| WBGene00023119 | NA | NA | NA | NA |
| WBGene00023120 | NA | NA | NA | NA |
| WBGene00023121 | NA | NA | NA | NA |
| WBGene00023122 | NA | NA | NA | NA |
| WBGene00023123 | NA | NA | NA | NA |
| WBGene00023124 | NA | NA | NA | NA |
| WBGene00023125 | NA | NA | NA | NA |
| WBGene00023126 | NA | NA | NA | NA |
| WBGene00023127 | NA | NA | NA | NA |
| WBGene00023128 | NA | NA | NA | NA |
| WBGene00023129 | NA | NA | NA | NA |
| WBGene00023130 | NA | NA | NA | NA |
| WBGene00023131 | NA | NA | NA | NA |
| WBGene00023132 | NA | NA | NA | NA |

| | | | | |
|---|---|---|---|---|
| WBGene00023133 | NA | NA | NA | NA |
| WBGene00023134 | NA | NA | NA | NA |
| WBGene00023135 | NA | NA | NA | NA |
| WBGene00023136 | NA | NA | NA | NA |
| WBGene00023137 | 0.020317245 | 0.84643798 | NA | NA |
| WBGene00023138 | NA | NA | NA | NA |
| WBGene00023139 | NA | NA | NA | NA |
| WBGene00023140 | NA | NA | NA | NA |
| WBGene00023141 | NA | NA | NA | NA |
| WBGene00023142 | NA | NA | NA | NA |
| WBGene00023143 | NA | NA | NA | NA |
| WBGene00023144 | NA | NA | NA | NA |
| WBGene00023145 | NA | NA | NA | NA |
| WBGene00023146 | NA | NA | NA | NA |
| WBGene00023147 | NA | NA | NA | NA |
| WBGene00023148 | NA | NA | NA | NA |
| WBGene00023149 | NA | NA | NA | NA |
| WBGene00023150 | NA | NA | NA | NA |
| WBGene00023151 | NA | NA | NA | NA |
| WBGene00023152 | NA | NA | NA | NA |
| WBGene00023153 | NA | NA | NA | NA |
| WBGene00023154 | NA | NA | NA | NA |
| WBGene00023155 | NA | NA | NA | NA |
| WBGene00023156 | NA | NA | NA | NA |
| WBGene00023157 | NA | NA | NA | NA |
| WBGene00023158 | NA | NA | NA | NA |
| WBGene00023159 | NA | NA | NA | NA |
| WBGene00023160 | NA | NA | NA | NA |
| WBGene00023161 | NA | NA | NA | NA |
| WBGene00023162 | NA | NA | NA | NA |
| WBGene00023163 | -0.196148184 | 0.19415871 | 0.38110708 | NA |
| WBGene00023164 | NA | NA | NA | NA |
| WBGene00023165 | NA | NA | NA | NA |
| WBGene00023166 | NA | NA | NA | NA |
| WBGene00023167 | NA | NA | NA | NA |
| WBGene00023168 | NA | NA | NA | NA |
| WBGene00023169 | NA | NA | NA | NA |
| WBGene00023170 | NA | NA | NA | NA |
| WBGene00023171 | NA | NA | NA | NA |
| WBGene00023172 | 0.242743183 | 0.17396931 | 0.35444388 | Y23H5A.8 |
| WBGene00023173 | NA | NA | NA | NA |
| WBGene00023174 | NA | NA | NA | NA |
| WBGene00023175 | NA | NA | NA | NA |

| | | | | |
|---|---|---|---|---|
| WBGene00023176 | NA | NA | NA | NA |
| WBGene00023177 | NA | NA | NA | NA |
| WBGene00023178 | NA | NA | NA | NA |
| WBGene00023179 | NA | NA | NA | NA |
| WBGene00023180 | NA | NA | NA | NA |
| WBGene00023181 | NA | NA | NA | NA |
| WBGene00023182 | NA | NA | NA | NA |
| WBGene00023183 | NA | NA | NA | NA |
| WBGene00023184 | NA | NA | NA | NA |
| WBGene00023185 | NA | NA | NA | NA |
| WBGene00023186 | NA | NA | NA | Y67D8B.3 |
| WBGene00023187 | 0.020232312 | 0.84643798 | NA | NA |
| WBGene00023188 | NA | NA | NA | NA |
| WBGene00023189 | NA | NA | NA | NA |
| WBGene00023190 | NA | NA | NA | NA |
| WBGene00023191 | -0.004747605 | 0.9473792 | NA | NA |
| WBGene00023192 | NA | NA | NA | NA |
| WBGene00023193 | -0.003270206 | 0.95698316 | NA | NA |
| WBGene00023194 | NA | NA | NA | NA |
| WBGene00023195 | NA | NA | NA | NA |
| WBGene00023196 | NA | NA | NA | NA |
| WBGene00023197 | NA | NA | NA | NA |
| WBGene00023198 | NA | NA | NA | NA |
| WBGene00023199 | NA | NA | NA | NA |
| WBGene00023200 | 0.020925399 | 0.84643798 | NA | NA |
| WBGene00023201 | 0.020925399 | 0.84643798 | NA | NA |
| WBGene00023202 | NA | NA | NA | NA |
| WBGene00023203 | NA | NA | NA | NA |
| WBGene00023204 | NA | NA | NA | NA |
| WBGene00023205 | NA | NA | NA | NA |
| WBGene00023206 | NA | NA | NA | NA |
| WBGene00023207 | NA | NA | NA | NA |
| WBGene00023208 | NA | NA | NA | NA |
| WBGene00023209 | 0.135221628 | 0.0937172 | 0.23421761 | NA |
| WBGene00023210 | NA | NA | NA | NA |
| WBGene00023211 | NA | NA | NA | NA |
| WBGene00023212 | NA | NA | NA | NA |
| WBGene00023213 | NA | NA | NA | NA |
| WBGene00023214 | NA | NA | NA | NA |
| WBGene00023215 | NA | NA | NA | NA |
| WBGene00023216 | NA | NA | NA | NA |
| WBGene00023217 | NA | NA | NA | NA |
| WBGene00023218 | NA | NA | NA | NA |

| | | | | |
|---|---|---|---|---|
| WBGene00023219 | NA | NA | NA | NA |
| WBGene00023220 | NA | NA | NA | NA |
| WBGene00023221 | NA | NA | NA | NA |
| WBGene00023222 | NA | NA | NA | NA |
| WBGene00023223 | 0.467292038 | 0.0307048 | 0.10513307 | NA |
| WBGene00023224 | NA | NA | NA | NA |
| WBGene00023225 | 0.0398788 | 0.58971221 | NA | NA |
| WBGene00023226 | NA | NA | NA | NA |
| WBGene00023227 | NA | NA | NA | NA |
| WBGene00023228 | NA | NA | NA | NA |
| WBGene00023229 | NA | NA | NA | NA |
| WBGene00023230 | NA | NA | NA | NA |
| WBGene00023231 | NA | NA | NA | NA |
| WBGene00023232 | NA | NA | NA | NA |
| WBGene00023233 | NA | NA | NA | NA |
| WBGene00023234 | NA | NA | NA | NA |
| WBGene00023235 | 0.020232312 | 0.84643798 | NA | NA |
| WBGene00023236 | NA | NA | NA | NA |
| WBGene00023237 | NA | NA | NA | NA |
| WBGene00023238 | NA | NA | NA | NA |
| WBGene00023239 | NA | NA | NA | NA |
| WBGene00023240 | NA | NA | NA | NA |
| WBGene00023241 | NA | NA | NA | NA |
| WBGene00023242 | NA | NA | NA | NA |
| WBGene00023243 | NA | NA | NA | NA |
| WBGene00023244 | -0.215761687 | 0.12049293 | 0.27934672 | NA |
| WBGene00023245 | -0.110326467 | 0.49191907 | 0.67512065 | NA |
| WBGene00023246 | -0.027350315 | 0.68023343 | NA | NA |
| WBGene00023247 | -0.002914971 | 0.95718083 | NA | NA |
| WBGene00023248 | NA | NA | NA | NA |
| WBGene00023249 | NA | NA | NA | NA |
| WBGene00023250 | NA | NA | NA | NA |
| WBGene00023251 | NA | NA | NA | NA |
| WBGene00023252 | NA | NA | NA | NA |
| WBGene00023253 | 0.020317245 | 0.84643798 | NA | NA |
| WBGene00023254 | NA | NA | NA | NA |
| WBGene00023255 | NA | NA | NA | NA |
| WBGene00023256 | NA | NA | NA | NA |
| WBGene00023257 | NA | NA | NA | NA |
| WBGene00023259 | -0.136778718 | 0.28736355 | 0.48757558 | NA |
| WBGene00023260 | NA | NA | NA | NA |
| WBGene00023262 | NA | NA | NA | NA |
| WBGene00023263 | 0.195051885 | 0.37157809 | 0.57085779 | NA |

| | | | | |
|---|---|---|---|---|
| WBGene00023264 | NA | NA | NA | NA |
| WBGene00023265 | NA | NA | NA | NA |
| WBGene00023266 | NA | NA | NA | NA |
| WBGene00023267 | -0.023109762 | 0.78121253 | NA | NA |
| WBGene00023268 | NA | NA | NA | NA |
| WBGene00023269 | NA | NA | NA | NA |
| WBGene00023270 | NA | NA | NA | NA |
| WBGene00023271 | -0.266245142 | 0.10580179 | 0.25483506 | NA |
| WBGene00023272 | NA | NA | NA | NA |
| WBGene00023273 | NA | NA | NA | NA |
| WBGene00023274 | NA | NA | NA | NA |
| WBGene00023275 | NA | NA | NA | NA |
| WBGene00023276 | -0.022930767 | 0.78121253 | NA | NA |
| WBGene00023277 | -0.023109762 | 0.78121253 | NA | NA |
| WBGene00023278 | NA | NA | NA | NA |
| WBGene00023279 | NA | NA | NA | NA |
| WBGene00023281 | NA | NA | NA | NA |
| WBGene00023282 | 0.038125098 | 0.61119482 | NA | NA |
| WBGene00023283 | NA | NA | NA | NA |
| WBGene00023284 | 0.364280698 | 0.13459122 | 0.29882541 | F41B5.1 |
| WBGene00023285 | NA | NA | NA | NA |
| WBGene00023287 | NA | NA | NA | NA |
| WBGene00023288 | NA | NA | NA | NA |
| WBGene00023289 | NA | NA | NA | NA |
| WBGene00023290 | NA | NA | NA | NA |
| WBGene00023291 | -0.061343715 | 0.58835742 | 0.75051633 | NA |
| WBGene00023293 | NA | NA | NA | NA |
| WBGene00023294 | NA | NA | NA | NA |
| WBGene00023295 | NA | NA | NA | NA |
| WBGene00023296 | NA | NA | NA | NA |
| WBGene00023297 | NA | NA | NA | NA |
| WBGene00023298 | NA | NA | NA | NA |
| WBGene00023299 | NA | NA | NA | NA |
| WBGene00023300 | -0.023109762 | 0.78121253 | NA | NA |
| WBGene00023301 | NA | NA | NA | NA |
| WBGene00023302 | NA | NA | NA | NA |
| WBGene00023303 | NA | NA | NA | NA |
| WBGene00023304 | NA | NA | NA | NA |
| WBGene00023305 | NA | NA | NA | NA |
| WBGene00023306 | NA | NA | NA | NA |
| WBGene00023307 | NA | NA | NA | NA |
| WBGene00023308 | -0.023109762 | 0.78121253 | NA | NA |
| WBGene00023309 | NA | NA | NA | NA |

| | | | | |
|---|---|---|---|---|
| WBGene00023310 | NA | NA | NA | NA |
| WBGene00023311 | NA | NA | NA | NA |
| WBGene00023312 | NA | NA | NA | NA |
| WBGene00023313 | NA | NA | NA | NA |
| WBGene00023314 | -0.00346345 | 0.95708614 | NA | NA |
| WBGene00023315 | NA | NA | NA | NA |
| WBGene00023316 | NA | NA | NA | NA |
| WBGene00023317 | NA | NA | NA | NA |
| WBGene00023318 | -0.297319311 | 0.1911062 | 0.37703623 | NA |
| WBGene00023319 | NA | NA | NA | NA |
| WBGene00023320 | 0.020317245 | 0.84643798 | NA | NA |
| WBGene00023321 | 0.020925399 | 0.84643798 | NA | NA |
| WBGene00023322 | 0.005645504 | 0.96072387 | 0.9786376 | NA |
| WBGene00023323 | 0.012907434 | 0.94951224 | 0.97122452 | NA |
| WBGene00023324 | NA | NA | NA | NA |
| WBGene00023325 | 0.020232312 | 0.84643798 | NA | NA |
| WBGene00023326 | NA | NA | NA | NA |
| WBGene00023327 | NA | NA | NA | NA |
| WBGene00023328 | NA | NA | NA | NA |
| WBGene00023329 | NA | NA | NA | NA |
| WBGene00023330 | NA | NA | NA | NA |
| WBGene00023331 | NA | NA | NA | NA |
| WBGene00023332 | NA | NA | NA | NA |
| WBGene00023333 | NA | NA | NA | NA |
| WBGene00023334 | NA | NA | NA | NA |
| WBGene00023335 | NA | NA | NA | NA |
| WBGene00023336 | NA | NA | NA | NA |
| WBGene00023337 | NA | NA | NA | NA |
| WBGene00023338 | NA | NA | NA | NA |
| WBGene00023339 | NA | NA | NA | NA |
| WBGene00023340 | NA | NA | NA | NA |
| WBGene00023341 | NA | NA | NA | NA |
| WBGene00023342 | NA | NA | NA | NA |
| WBGene00023343 | NA | NA | NA | NA |
| WBGene00023344 | NA | NA | NA | NA |
| WBGene00023345 | 0.094874781 | 0.40302901 | 0.60141856 | NA |
| WBGene00023346 | 0.059980168 | 0.38780842 | NA | NA |
| WBGene00023348 | NA | NA | NA | NA |
| WBGene00023349 | NA | NA | NA | NA |
| WBGene00023351 | NA | NA | NA | NA |
| WBGene00023352 | NA | NA | NA | NA |
| WBGene00023353 | NA | NA | NA | NA |
| WBGene00023354 | NA | NA | NA | NA |

| | | | | |
|---|---|---|---|---|
| WBGene00023355 | -0.075020616 | 0.71488891 | 0.8401205 | NA |
| WBGene00023356 | NA | NA | NA | NA |
| WBGene00023357 | NA | NA | NA | NA |
| WBGene00023358 | NA | NA | NA | NA |
| WBGene00023359 | NA | NA | NA | NA |
| WBGene00023360 | NA | NA | NA | NA |
| WBGene00023361 | NA | NA | NA | NA |
| WBGene00023362 | NA | NA | NA | NA |
| WBGene00023363 | NA | NA | NA | NA |
| WBGene00023364 | NA | NA | NA | NA |
| WBGene00023365 | NA | NA | NA | NA |
| WBGene00023366 | NA | NA | NA | NA |
| WBGene00023367 | NA | NA | NA | NA |
| WBGene00023368 | NA | NA | NA | NA |
| WBGene00023369 | NA | NA | NA | NA |
| WBGene00023370 | NA | NA | NA | NA |
| WBGene00023371 | NA | NA | NA | NA |
| WBGene00023372 | NA | NA | NA | NA |
| WBGene00023373 | NA | NA | NA | NA |
| WBGene00023375 | NA | NA | NA | NA |
| WBGene00023376 | NA | NA | NA | NA |
| WBGene00023377 | NA | NA | NA | NA |
| WBGene00023378 | NA | NA | NA | NA |
| WBGene00023379 | NA | NA | NA | NA |
| WBGene00023380 | NA | NA | NA | NA |
| WBGene00023381 | NA | NA | NA | NA |
| WBGene00023382 | NA | NA | NA | NA |
| WBGene00023383 | NA | NA | NA | NA |
| WBGene00023384 | NA | NA | NA | NA |
| WBGene00023385 | NA | NA | NA | NA |
| WBGene00023386 | -0.023109762 | 0.78121253 | NA | NA |
| WBGene00023387 | NA | NA | NA | NA |
| WBGene00023388 | NA | NA | NA | NA |
| WBGene00023389 | NA | NA | NA | NA |
| WBGene00023390 | NA | NA | NA | NA |
| WBGene00023391 | NA | NA | NA | NA |
| WBGene00023392 | NA | NA | NA | NA |
| WBGene00023393 | NA | NA | NA | NA |
| WBGene00023394 | NA | NA | NA | NA |
| WBGene00023395 | 0.038125098 | 0.61119482 | NA | NA |
| WBGene00023396 | NA | NA | NA | NA |
| WBGene00023397 | NA | NA | NA | NA |
| WBGene00023398 | NA | NA | NA | NA |

| | | | | |
|---|---|---|---|---|
| WBGene00023399 | NA | NA | NA | NA |
| WBGene00023400 | -0.045771477 | 0.50913401 | NA | NA |
| WBGene00023401 | -0.741850009 | 0.00020601 | 0.0018159 | NA |
| WBGene00023402 | NA | NA | NA | NA |
| WBGene00023404 | -0.386672636 | 0.06564535 | 0.18286028 | plr-1 |
| WBGene00023405 | -0.060195382 | 0.64038101 | 0.78675996 | sor-1 |
| WBGene00023407 | -0.019806875 | 0.92183447 | 0.95620205 | cex-1 |
| WBGene00023408 | 0.597897528 | 0.02316035 | 0.08543478 | cex-2 |
| WBGene00023409 | -0.133964628 | 0.28823953 | 0.48849576 | mbd-2 |
| WBGene00023410 | NA | NA | NA | clpr-2 |
| WBGene00023411 | NA | NA | NA | srz-3 |
| WBGene00023413 | NA | NA | NA | H31B20.2 |
| WBGene00023414 | -0.110295553 | 0.59210973 | 0.75269035 | R01H2.7 |
| WBGene00023415 | 0.212944783 | 0.32540744 | 0.5246438 | F16C3.3 |
| WBGene00023416 | 0.107377299 | 0.31369484 | 0.5141086 | spi-1 |
| WBGene00023417 | -0.055443334 | 0.76229675 | 0.86502767 | swm-1 |
| WBGene00023418 | -0.075183631 | 0.4812579 | 0.66742323 | R06F6.12 |
| WBGene00023419 | -0.28411153 | 0.14522037 | 0.31460967 | F53F4.16 |
| WBGene00023420 | 0.58163194 | 0.01051771 | 0.04652743 | C26D10.7 |
| WBGene00023421 | 0.064938948 | 0.59587843 | 0.75477619 | rde-11 |
| WBGene00023422 | -0.236815473 | 0.08591283 | 0.22135188 | mrpl-21 |
| WBGene00023423 | 0.020317245 | 0.84643798 | NA | F43C1.7 |
| WBGene00023424 | 0.378131582 | 0.0454674 | 0.14038132 | C53D6.10 |
| WBGene00023425 | 0.064941241 | 0.4010185 | NA | try-9 |
| WBGene00023427 | 0.381418346 | 0.04825111 | 0.14641337 | F14D2.15 |
| WBGene00023428 | -0.023109762 | 0.78121253 | NA | NA |
| WBGene00023429 | -0.022930767 | 0.78121253 | NA | NA |
| WBGene00023430 | -0.055809923 | 0.57791914 | 0.74421161 | NA |
| WBGene00023431 | 0.034505823 | 0.68373529 | NA | C25E10.13 |
| WBGene00023432 | 0.957587209 | 0.00249777 | 0.01500608 | K12B6.9 |
| WBGene00023450 | 0.029477762 | 0.85583728 | 0.92263556 | T12D8.10 |
| WBGene00023451 | 0.308033497 | 0.07406858 | 0.19956705 | mlc-7 |
| WBGene00023452 | -0.023056499 | 0.78121253 | NA | NA |
| WBGene00023453 | NA | NA | NA | NA |
| WBGene00023454 | NA | NA | NA | NA |
| WBGene00023455 | 0.020317245 | 0.84643798 | NA | NA |
| WBGene00023456 | NA | NA | NA | NA |
| WBGene00023457 | NA | NA | NA | NA |
| WBGene00023458 | 0.029576845 | 0.85187466 | 0.92048984 | M02E1.3 |
| WBGene00023459 | NA | NA | NA | Y18D10A.26 |
| WBGene00023460 | -0.025577507 | 0.74049476 | NA | F46H6.5 |
| WBGene00023461 | NA | NA | NA | NA |
| WBGene00023462 | 0.020317245 | 0.84643798 | NA | srz-83 |

| | | | | | |
|---|---|---|---|---|---|
| WBGene00023463 | | -0.023109762 | 0.78121253 | NA | srz-82 |
| WBGene00023464 | | 0.036687355 | 0.59489568 | NA | NA |
| WBGene00023465 | | -0.023056499 | 0.78121253 | NA | NA |
| WBGene00023466 | NA | | NA | NA | NA |
| WBGene00023467 | | 0.020232312 | 0.84643798 | NA | NA |
| WBGene00023468 | NA | | NA | NA | NA |
| WBGene00023469 | | 0.015551528 | 0.84511556 | NA | NA |
| WBGene00023470 | | -0.048531013 | 0.47590296 | NA | NA |
| WBGene00023471 | | 0.018860755 | 0.77262519 | NA | NA |
| WBGene00023472 | | -0.00901083 | 0.91276867 | NA | NA |
| WBGene00023474 | NA | | NA | NA | srz-37 |
| WBGene00023475 | | 0.020925399 | 0.84643798 | NA | srz-69 |
| WBGene00023476 | NA | | NA | NA | NA |
| WBGene00023477 | | 0.083053927 | 0.69074932 | 0.82235004 | nspc-1 |
| WBGene00023478 | NA | | NA | NA | srz-62 |
| WBGene00023479 | NA | | NA | NA | srz-67 |
| WBGene00023480 | NA | | NA | NA | NA |
| WBGene00023481 | NA | | NA | NA | NA |
| WBGene00023482 | NA | | NA | NA | NA |
| WBGene00023483 | | 0.15038247 | 0.07114191 | NA | Y75B8A.39 |
| WBGene00023484 | | 0.521279479 | 0.00279475 | 0.01649249 | Y54G2A.37 |
| WBGene00023485 | | -0.10870849 | 0.59289875 | 0.75314295 | ttr-49 |
| WBGene00023486 | | -0.022719221 | 0.89730042 | 0.9459022 | K08B4.7 |
| WBGene00023487 | | -0.166521021 | 0.300344 | 0.50066452 | mrps-24 |
| WBGene00023488 | NA | | NA | NA | NA |
| WBGene00023489 | | -0.091006486 | 0.61697529 | 0.77048236 | F42A10.9 |
| WBGene00023490 | | 0.091046425 | 0.35523343 | 0.55506646 | F36A2.14 |
| WBGene00023491 | | -0.246307644 | 0.20986151 | 0.39929595 | tsp-21 |
| WBGene00023492 | | -0.362312144 | 0.13237839 | 0.2955991 | T24D3.2 |
| WBGene00023493 | | 0.137372735 | 0.12646195 | 0.28757277 | nspc-12 |
| WBGene00023497 | | -0.029853837 | 0.76423344 | 0.86623378 | lin-15B |
| WBGene00023498 | | -0.077363124 | 0.58422115 | 0.74778226 | lin-15A |
| WBGene00023499 | | -0.023056499 | 0.78121253 | NA | srz-63 |
| WBGene00023500 | | -0.887508972 | 0.00348125 | 0.01966329 | C11H1.9 |
| WBGene00023501 | | 2.352843574 | 3.06E-05 | 0.000335 | F11E6.11 |
| WBGene00023502 | | 0.11573919 | 0.53035599 | 0.70706083 | clec-184 |
| WBGene00023503 | | -0.00783928 | 0.95763941 | 0.97671882 | F17C11.13 |
| WBGene00023504 | | 0.142678735 | 0.40985751 | 0.60772325 | F26F2.8 |
| WBGene00023505 | NA | | NA | NA | srz-102 |
| WBGene00023506 | NA | | NA | NA | srz-11 |
| WBGene00023507 | NA | | NA | NA | NA |
| WBGene00023508 | NA | | NA | NA | srz-38 |
| WBGene00023509 | NA | | NA | NA | NA |

| | | | | |
|---|---|---|---|---|
| WBGene00023510 | 0.020317245 | 0.84643798 | NA | NA |
| WBGene00023511 | NA | NA | NA | NA |
| WBGene00023512 | NA | NA | NA | NA |
| WBGene00023513 | NA | NA | NA | srz-71 |
| WBGene00023514 | NA | NA | NA | srz-72 |
| WBGene00023515 | 0.017631418 | 0.81668032 | NA | srz-97 |
| WBGene00023516 | NA | NA | NA | NA |
| WBGene00023517 | NA | NA | NA | srz-75 |
| WBGene00023518 | NA | NA | NA | srz-76 |
| WBGene00023519 | 0.14692852 | 0.37313086 | 0.57228251 | Y60A3A.24 |
| WBGene00023520 | 0.269646712 | 0.19344887 | 0.38007045 | oig-5 |
| WBGene00043049 | 0.035695256 | 0.67391046 | NA | F32H2.11 |
| WBGene00043050 | 0.15930175 | 0.24437246 | 0.44253027 | oig-4 |
| WBGene00043051 | -0.249120666 | 0.06204125 | 0.17583159 | C29E4.13 |
| WBGene00043052 | 0.020232312 | 0.84643798 | NA | srz-104 |
| WBGene00043053 | 0.372629881 | 0.04644156 | 0.14255181 | NA |
| WBGene00043054 | 0.093607072 | 0.48225698 | 0.66812305 | T21B4.15 |
| WBGene00043055 | NA | NA | NA | NA |
| WBGene00043056 | 0.000340583 | 0.99935204 | 0.99976175 | nfya-2 |
| WBGene00043057 | -0.171887677 | 0.20432747 | 0.39341905 | M142.8 |
| WBGene00043058 | NA | NA | NA | sre-34 |
| WBGene00043059 | 0.018998992 | 0.77286019 | NA | srab-26 |
| WBGene00043060 | -0.111517582 | 0.46806943 | 0.65729949 | R144.12 |
| WBGene00043061 | 0.040107227 | 0.58693163 | NA | Y54G2A.38 |
| WBGene00043062 | 3.530563554 | 5.41E-64 | 1.55E-61 | clec-80 |
| WBGene00043064 | -0.233014599 | 0.06396051 | 0.17974009 | nbet-1 |
| WBGene00043065 | 0.382035559 | 0.0049366 | 0.02591493 | C55C3.7 |
| WBGene00043066 | -0.010875694 | 0.94213577 | 0.9672483 | acr-25 |
| WBGene00043067 | 0.319205642 | 0.09361138 | 0.23406969 | dlc-5 |
| WBGene00043068 | -0.569663864 | 0.01159358 | 0.05017486 | F32E10.9 |
| WBGene00043069 | NA | NA | NA | F20D6.12 |
| WBGene00043097 | 0.295907392 | 0.01705079 | 0.06828858 | C02D5.4 |
| WBGene00043120 | -0.023056499 | 0.78121253 | NA | C09H10.4 |
| WBGene00043147 | 0.157590572 | 0.27378165 | 0.47403467 | nspd-3 |
| WBGene00043147 | 0.157590572 | 0.27378165 | 0.47403467 | nspd-6 |
| WBGene00043156 | -0.334200536 | 0.00721521 | 0.03484098 | C27F2.9 |
| WBGene00043187 | NA | NA | NA | nspa-9 |
| WBGene00043239 | -0.003108065 | 0.95728506 | NA | NA |
| WBGene00043279 | 0.077033784 | 0.7107854 | 0.83708245 | F22D6.8 |
| WBGene00043283 | NA | NA | NA | NA |
| WBGene00043297 | 0.052453824 | 0.61124345 | 0.76613435 | F28F5.4 |
| WBGene00043302 | NA | NA | NA | F32D8.11 |
| WBGene00043308 | NA | NA | NA | tmem-107 |

| | | | | |
|---|---|---|---|---|
| WBGene00043327 | -0.023056499 | 0.78121253 | NA | F45D11.7 |
| WBGene00043328 | NA | NA | NA | F45D11.11 |
| WBGene00043466 | NA | NA | NA | R05G6.3 |
| WBGene00043469 | -0.022930767 | 0.78121253 | NA | R09H3.2 |
| WBGene00043534 | -0.100425495 | 0.5675002 | 0.73659312 | W02B12.13 |
| WBGene00043666 | NA | NA | NA | Y45G5AM.4 |
| WBGene00043702 | 2.434738106 | 1.00E-11 | 3.32E-10 | Y54G2A.7 |
| WBGene00043705 | -0.155873068 | 0.45730753 | 0.64918009 | egrh-2 |
| WBGene00043743 | 0.09777701 | 0.17585057 | NA | Y59E9AL.2 |
| WBGene00043948 | NA | NA | NA | ZC262.7 |
| WBGene00043952 | -0.08242813 | 0.39390996 | NA | ZC404.2 |
| WBGene00043980 | 0.141071593 | 0.49069187 | 0.6745427 | F11D5.7 |
| WBGene00043981 | 0.024279323 | 0.81248486 | 0.89608971 | T14E8.4 |
| WBGene00043982 | 0.536596403 | 0.05571405 | 0.16301149 | B0403.6 |
| WBGene00043983 | 1.01105163 | 0.00101867 | 0.00714973 | C39D10.11 |
| WBGene00043984 | NA | NA | NA | C56E6.7 |
| WBGene00043985 | -0.124816705 | 0.55130291 | 0.72373276 | C44C10.11 |
| WBGene00043986 | -0.023056499 | 0.78121253 | NA | fut-5 |
| WBGene00043987 | NA | NA | NA | NA |
| WBGene00043988 | NA | NA | NA | AC8.3 |
| WBGene00043988 | NA | NA | NA | AC8.10 |
| WBGene00043989 | NA | NA | NA | AC8.4 |
| WBGene00043989 | NA | NA | NA | AC8.11 |
| WBGene00043990 | NA | NA | NA | AC8.7 |
| WBGene00043990 | NA | NA | NA | AC8.12 |
| WBGene00043991 | NA | NA | NA | NA |
| WBGene00043992 | -0.003116542 | 0.98093731 | 0.98897556 | bbs-4 |
| WBGene00043993 | NA | NA | NA | NA |
| WBGene00043994 | 0.242484499 | 0.24745434 | 0.44591953 | C07A9.12 |
| WBGene00043995 | 0.957791156 | 0.01762517 | 0.0700652 | F40F12.9 |
| WBGene00043996 | -0.12206767 | 0.28021627 | 0.47984024 | fbxb-87 |
| WBGene00043997 | 0.103693905 | 0.32163492 | 0.52161514 | Y79H2A.12 |
| WBGene00043998 | 0.133167519 | 0.48834132 | 0.67241655 | T21C12.8 |
| WBGene00044000 | NA | NA | NA | NA |
| WBGene00044001 | -0.132676953 | 0.34889458 | 0.54902897 | C09G4.4 |
| WBGene00044002 | NA | NA | NA | srt-74 |
| WBGene00044003 | -0.025049301 | 0.69142686 | NA | Y41E3.18 |
| WBGene00044004 | 0.014087259 | 0.85351751 | NA | C24A8.6 |
| WBGene00044005 | 0.02981377 | 0.8058743 | 0.89222668 | F37C12.18 |
| WBGene00044006 | 0.019248146 | 0.89538084 | 0.94517796 | nlp-82 |
| WBGene00044007 | 0.541781583 | 0.03158426 | 0.1071938 | ZK1320.13 |
| WBGene00044008 | 0.011259363 | 0.90828167 | 0.95041108 | gip-2 |
| WBGene00044009 | 0.243774512 | 0.17265706 | 0.35261152 | ZK822.6 |

| | | | | |
|---|---|---|---|---|
| WBGene00044011 | 0.087256674 | 0.23800185 | NA | T06E4.12 |
| WBGene00044012 | NA | NA | NA | srxa-19 |
| WBGene00044013 | NA | NA | NA | F15H10.9 |
| WBGene00044014 | NA | NA | NA | F15H10.10 |
| WBGene00044015 | 0.065014541 | 0.37399213 | NA | NA |
| WBGene00044016 | 0.037023624 | 0.59153183 | NA | C35A5.10 |
| WBGene00044018 | 0.020925399 | 0.84643798 | NA | T04B2.8 |
| WBGene00044019 | 0.083589196 | 0.6320162 | 0.78140088 | C55A6.12 |
| WBGene00044020 | -0.019880188 | 0.87392987 | 0.93286725 | ZK856.14 |
| WBGene00044021 | 0.041666597 | 0.76192104 | 0.86495294 | R02E4.2 |
| WBGene00044022 | 0.036687355 | 0.59489568 | NA | NA |
| WBGene00044024 | 0.356202027 | 0.00628979 | 0.0312908 | C35A5.11 |
| WBGene00044025 | 0.135826349 | 0.31388227 | 0.51433132 | T01G1.4 |
| WBGene00044026 | -0.045490149 | 0.6543688 | 0.79526966 | cisd-1 |
| WBGene00044027 | 0.020317245 | 0.84643798 | NA | F27E5.8 |
| WBGene00044028 | 0.121530836 | 0.43521555 | 0.63045722 | nlp-63 |
| WBGene00044029 | 0.316204249 | 0.17632249 | 0.35715793 | C05D12.7 |
| WBGene00044030 | -0.00346345 | 0.95708614 | NA | Y17G7B.23 |
| WBGene00044031 | NA | NA | NA | F49C5.9 |
| WBGene00044032 | 0.167550193 | 0.39037422 | 0.58825329 | ceh-99 |
| WBGene00044034 | 0.007687992 | 0.96255151 | 0.97924853 | Y38E10A.28 |
| WBGene00044035 | -0.087854351 | 0.59282738 | 0.75314295 | F27E5.7 |
| WBGene00044036 | -0.022930767 | 0.78121253 | NA | NA |
| WBGene00044037 | NA | NA | NA | NA |
| WBGene00044038 | -0.028246693 | 0.71442752 | NA | NA |
| WBGene00044040 | NA | NA | NA | NA |
| WBGene00044041 | 0.137421262 | 0.30773026 | 0.50846165 | NA |
| WBGene00044042 | 0.020317245 | 0.84643798 | NA | NA |
| WBGene00044043 | 0.090723942 | 0.20045316 | NA | Y46G5A.38 |
| WBGene00044044 | 0.020232312 | 0.84643798 | NA | C08B6.14 |
| WBGene00044045 | -0.034819511 | 0.77344997 | 0.87148538 | R11G10.3 |
| WBGene00044046 | NA | NA | NA | NA |
| WBGene00044047 | -0.16372506 | 0.35465032 | 0.55451895 | Y57A10C.11 |
| WBGene00044048 | 0.020317245 | 0.84643798 | NA | F57G9.7 |
| WBGene00044049 | 0.630697566 | 0.00329563 | 0.01881878 | sfxn-5 |
| WBGene00044051 | NA | NA | NA | NA |
| WBGene00044052 | NA | NA | NA | NA |
| WBGene00044053 | -0.331355495 | 0.160599 | 0.33512594 | NA |
| WBGene00044054 | 0.020232312 | 0.84643798 | NA | NA |
| WBGene00044056 | NA | NA | NA | NA |
| WBGene00044057 | 0.020232312 | 0.84643798 | NA | NA |
| WBGene00044058 | 0.05787416 | 0.40594086 | NA | F17B5.6 |
| WBGene00044059 | NA | NA | NA | T27C5.12 |

| | | | | |
|---|---|---|---|---|
| WBGene00044061 | -0.209986269 | 0.1305401 | 0.29288894 | tbc-12 |
| WBGene00044062 | 0.037472146 | 0.66218673 | NA | snb-6 |
| WBGene00044063 | -0.104066679 | 0.37590636 | 0.57500725 | bpnt-1 |
| WBGene00044065 | -0.035014186 | 0.82359213 | 0.9034778 | tag-229 |
| WBGene00044066 | -0.799990759 | 0.01728018 | 0.06899349 | fbxb-66 |
| WBGene00044067 | -0.412685801 | 2.94E-05 | 0.00032253 | zipt-7.1 |
| WBGene00044068 | -0.025359402 | 0.89718689 | 0.9459022 | syd-9 |
| WBGene00044069 | -0.042799924 | 0.55634523 | 0.72794301 | hat-1 |
| WBGene00044070 | 1.083186147 | 0.00111616 | 0.00769794 | set-18 |
| WBGene00044071 | -0.014371354 | 0.91872446 | 0.95485517 | dhhc-14 |
| WBGene00044072 | -0.115771314 | 0.34202057 | 0.54229187 | ham-3 |
| WBGene00044073 | 1.444840334 | 3.16E-05 | 0.00034383 | tag-244 |
| WBGene00044074 | -0.00346345 | 0.95708614 | NA | mltn-5 |
| WBGene00044077 | -0.134720823 | 0.10332065 | 0.2509069 | tag-232 |
| WBGene00044078 | 0.313321632 | 0.18248724 | 0.36537689 | tag-243 |
| WBGene00044079 | 0.929889508 | 9.36E-05 | 0.000915 | tag-241 |
| WBGene00044080 | -0.90793001 | 2.97E-31 | 3.91E-29 | D1086.10 |
| WBGene00044081 | 0.040530812 | 0.58729646 | 0.74981371 | D1086.11 |
| WBGene00044082 | 0.968375344 | 9.04E-06 | 0.00011257 | D1086.12 |
| WBGene00044083 | NA | NA | NA | tin-9.2 |
| WBGene00044084 | -0.027373974 | 0.68245543 | NA | sre-46 |
| WBGene00044085 | NA | NA | NA | NA |
| WBGene00044086 | -0.001993815 | 0.95612996 | NA | clec-228 |
| WBGene00044087 | NA | NA | NA | C45H4.18 |
| WBGene00044088 | 0.017604133 | 0.79437959 | NA | oac-2 |
| WBGene00044089 | NA | NA | NA | srbc-31 |
| WBGene00044090 | 0.040107227 | 0.58693163 | NA | srbc-33 |
| WBGene00044091 | -0.023056499 | 0.78121253 | NA | srbc-60 |
| WBGene00044092 | NA | NA | NA | srbc-71 |
| WBGene00044093 | 0.020317245 | 0.84643798 | NA | NA |
| WBGene00044094 | -0.096655194 | 0.38841785 | 0.58687573 | fitm-2 |
| WBGene00044095 | 0.020317245 | 0.84643798 | NA | NA |
| WBGene00044096 | NA | NA | NA | NA |
| WBGene00044097 | NA | NA | NA | NA |
| WBGene00044099 | NA | NA | NA | srd-61 |
| WBGene00044099 | NA | NA | NA | srd-75 |
| WBGene00044100 | NA | NA | NA | srbc-27 |
| WBGene00044101 | NA | NA | NA | NA |
| WBGene00044102 | NA | NA | NA | NA |
| WBGene00044103 | 0.020232312 | 0.84643798 | NA | NA |
| WBGene00044104 | NA | NA | NA | NA |
| WBGene00044105 | NA | NA | NA | srbc-55 |
| WBGene00044106 | 0.020232312 | 0.84643798 | NA | srbc-84 |

| | | | | |
|---|---|---|---|---|
| WBGene00044107 | 0.086553653 | 0.61662723 | 0.77017909 | F58G6.9 |
| WBGene00044109 | 0.27487826 | 0.01138934 | 0.04951048 | K02E11.10 |
| WBGene00044112 | -0.009725633 | 0.92224477 | 0.95626565 | srm-5 |
| WBGene00044113 | NA | NA | NA | C54F6.15 |
| WBGene00044114 | NA | NA | NA | srxa-11 |
| WBGene00044115 | 0.020317245 | 0.84643798 | NA | srxa-12 |
| WBGene00044116 | 0.033034883 | 0.64219503 | NA | srxa-13 |
| WBGene00044117 | NA | NA | NA | srsx-10 |
| WBGene00044119 | -0.023056499 | 0.78121253 | NA | NA |
| WBGene00044120 | 0.181723676 | 0.22797698 | 0.421494 | C36B1.13 |
| WBGene00044121 | 0.079160218 | 0.57509343 | 0.742287 | gcp-2.3 |
| WBGene00044122 | 0.106997704 | 0.16248985 | NA | T28B8.6 |
| WBGene00044124 | NA | NA | NA | srsx-15 |
| WBGene00044125 | NA | NA | NA | srsx-16 |
| WBGene00044126 | NA | NA | NA | NA |
| WBGene00044127 | NA | NA | NA | BE10.5 |
| WBGene00044128 | 0.020925399 | 0.84643798 | NA | NA |
| WBGene00044129 | -0.001488154 | 0.95639912 | NA | NA |
| WBGene00044130 | NA | NA | NA | NA |
| WBGene00044131 | NA | NA | NA | Y45G12C.16 |
| WBGene00044132 | 0.110775655 | 0.48754635 | 0.67176418 | C23H4.8 |
| WBGene00044133 | NA | NA | NA | C44C10.12 |
| WBGene00044134 | -0.0096926 | 0.95917177 | 0.97766875 | D1025.10 |
| WBGene00044135 | -0.003270206 | 0.95698316 | NA | F08G12.11 |
| WBGene00044136 | -0.022930767 | 0.78121253 | NA | F23D12.8 |
| WBGene00044137 | 0.020925399 | 0.84643798 | NA | F28H6.8 |
| WBGene00044138 | 0.020232312 | 0.84643798 | NA | sssh-1 |
| WBGene00044139 | 0.038726127 | 0.57551883 | NA | F42D1.4 |
| WBGene00044140 | NA | NA | NA | F42F12.11 |
| WBGene00044141 | NA | NA | NA | H03G16.6 |
| WBGene00044142 | 0.033463284 | 0.6365037 | NA | H13N06.7 |
| WBGene00044143 | 0.118013047 | 0.54838024 | 0.72098017 | K09E9.4 |
| WBGene00044144 | 0.032208757 | 0.85564918 | 0.92262213 | M153.4 |
| WBGene00044145 | NA | NA | NA | M163.9 |
| WBGene00044146 | NA | NA | NA | M163.10 |
| WBGene00044147 | NA | NA | NA | M163.11 |
| WBGene00044148 | -0.07867744 | 0.31038252 | NA | T08D10.4 |
| WBGene00044149 | -0.128104259 | 0.53896214 | 0.71373539 | T24D5.6 |
| WBGene00044150 | -0.373501248 | 0.1256125 | 0.28666356 | VB0395L.1 |
| WBGene00044151 | 0.020232312 | 0.84643798 | NA | W04G3.12 |
| WBGene00044152 | NA | NA | NA | W04G3.10 |
| WBGene00044153 | 0.01091784 | 0.90563006 | NA | W04G3.11 |
| WBGene00044154 | 0.105191732 | 0.16565281 | NA | Y7A5A.10 |

| | | | | |
|---|---|---|---|---|
| WBGene00044155 | 0.197712019 | 0.07031294 | 0.1918829 | NA |
| WBGene00044156 | -0.022088885 | 0.89918339 | 0.94665211 | Y66C5A.1 |
| WBGene00044157 | NA | NA | NA | Y66C5A.2 |
| WBGene00044158 | -0.003108065 | 0.95728506 | NA | Y53C12A.8 |
| WBGene00044159 | NA | NA | NA | F26C11.4 |
| WBGene00044160 | NA | NA | NA | W10C6.2 |
| WBGene00044161 | -0.014064954 | 0.9025887 | 0.94811315 | C47G2.8 |
| WBGene00044162 | NA | NA | NA | F49C5.10 |
| WBGene00044163 | 0.15038247 | 0.07114191 | NA | F49C5.11 |
| WBGene00044164 | -0.00346345 | 0.95708614 | NA | Y46G5A.39 |
| WBGene00044166 | 0.919575158 | 0.01374244 | 0.05768436 | C54C6.7 |
| WBGene00044167 | -0.076823182 | 0.65797335 | 0.79832501 | C36A4.11 |
| WBGene00044168 | 0.020925399 | 0.84643798 | NA | NA |
| WBGene00044169 | -0.049065066 | 0.47159718 | NA | C06C3.10 |
| WBGene00044170 | NA | NA | NA | NA |
| WBGene00044171 | NA | NA | NA | NA |
| WBGene00044174 | -0.247081524 | 0.2654853 | 0.46406044 | fipr-5 |
| WBGene00044175 | -0.000991934 | 0.99413819 | 0.99638399 | fipr-9 |
| WBGene00044176 | 0.72199683 | 0.01847749 | 0.0725473 | C30G7.2 |
| WBGene00044177 | 1.724547408 | 0.00545838 | 0.02805058 | C30G7.3 |
| WBGene00044178 | 0.126525176 | 0.54970665 | 0.72232362 | C30G7.4 |
| WBGene00044179 | NA | NA | NA | C30G7.5 |
| WBGene00044180 | 0.038595855 | 0.57520533 | NA | C47E8.11 |
| WBGene00044181 | 0.020317245 | 0.84643798 | NA | F09C6.11 |
| WBGene00044182 | 0.456054642 | 0.00360108 | 0.02021606 | F09C6.12 |
| WBGene00044183 | 0.06594186 | 0.5947082 | 0.75425526 | F09C6.13 |
| WBGene00044184 | NA | NA | NA | F11A5.16 |
| WBGene00044185 | NA | NA | NA | F14F8.14 |
| WBGene00044187 | 0.01912147 | 0.77587372 | NA | ttll-9 |
| WBGene00044188 | -0.084723833 | 0.53741687 | 0.71232305 | F28F8.9 |
| WBGene00044189 | NA | NA | NA | F36D3.14 |
| WBGene00044191 | -0.181784406 | 0.27964895 | 0.47944787 | F43D2.6 |
| WBGene00044192 | 0.033463284 | 0.6365037 | NA | F46B6.13 |
| WBGene00044193 | NA | NA | NA | F47B8.13 |
| WBGene00044194 | NA | NA | NA | F47B8.14 |
| WBGene00044195 | 0.020317245 | 0.84643798 | NA | F49A5.10 |
| WBGene00044196 | -0.003270206 | 0.95698316 | NA | NA |
| WBGene00044197 | -0.34791693 | 0.14303363 | 0.31144488 | fbxb-63 |
| WBGene00044198 | 0.016985298 | 0.837043 | NA | F56H9.6 |
| WBGene00044199 | NA | NA | NA | F57B1.8 |
| WBGene00044200 | -0.213652977 | 0.23066085 | 0.4252746 | H37A05.4 |
| WBGene00044201 | NA | NA | NA | H39E23.3 |
| WBGene00044202 | 0.048802683 | 0.71232684 | 0.83815617 | K10G4.10 |

| | | | | |
|---|---|---|---|---|
| WBGene00044203 | NA | NA | NA | T02E9.6 |
| WBGene00044204 | -0.147204197 | 0.25293644 | 0.4510337 | suds-3 |
| WBGene00044205 | -0.013730065 | 0.90947463 | 0.9508439 | T13F3.8 |
| WBGene00044206 | 4.815723991 | 1.35E-16 | 7.68E-15 | T26H5.9 |
| WBGene00044207 | 0.149442831 | 0.06876524 | NA | Y6G8.5 |
| WBGene00044208 | 0.020925399 | 0.84643798 | NA | Y6G8.6 |
| WBGene00044209 | -0.930333471 | 0.00381977 | 0.02111981 | NA |
| WBGene00044210 | -0.13078516 | 0.53282989 | 0.70870744 | Y51A2D.21 |
| WBGene00044211 | 0.088289671 | 0.63136882 | 0.78093021 | Y60A3A.25 |
| WBGene00044212 | 6.694246595 | 6.53E-07 | 1.07E-05 | Y68A4A.13 |
| WBGene00044213 | 0.318539771 | 0.17014319 | 0.34864471 | Y102A5C.36 |
| WBGene00044215 | 0.020232312 | 0.84643798 | NA | ZK228.10 |
| WBGene00044216 | NA | NA | NA | NA |
| WBGene00044217 | 0.097567973 | 0.43970529 | 0.63429761 | NA |
| WBGene00044218 | 0.046446288 | 0.82340593 | 0.9034778 | F23H12.10 |
| WBGene00044219 | -0.197650096 | 0.36084949 | 0.56083865 | NA |
| WBGene00044220 | 0.194083313 | 0.18711526 | 0.37163319 | NA |
| WBGene00044221 | 0.409952077 | 0.05975603 | 0.17105736 | F41G3.18 |
| WBGene00044222 | -0.022930767 | 0.78121253 | NA | T16G12.10 |
| WBGene00044224 | 0.04969721 | 0.67790397 | 0.81292048 | NA |
| WBGene00044225 | -0.050837788 | 0.78366327 | 0.87826193 | C38D4.10 |
| WBGene00044226 | NA | NA | NA | NA |
| WBGene00044227 | 0.065254007 | 0.40163332 | NA | C07A9.13 |
| WBGene00044228 | NA | NA | NA | T03F6.7 |
| WBGene00044229 | 0.036821884 | 0.59344795 | NA | NA |
| WBGene00044230 | NA | NA | NA | C36E8.6 |
| WBGene00044232 | 0.046154497 | 0.6664957 | 0.80470744 | C18D11.7 |
| WBGene00044233 | 0.125999388 | 0.50406316 | 0.68523572 | dolk-1 |
| WBGene00044235 | -0.023056499 | 0.78121253 | NA | C06A1.7 |
| WBGene00044236 | 0.186920603 | 0.34998467 | 0.54973989 | C17E4.11 |
| WBGene00044237 | 0.065014541 | 0.37399213 | NA | pals-9 |
| WBGene00044238 | 0.065014541 | 0.37399213 | NA | C30H6.12 |
| WBGene00044239 | -0.002914971 | 0.95718083 | NA | C33D9.10 |
| WBGene00044240 | 0.020925399 | 0.84643798 | NA | C47E12.14 |
| WBGene00044241 | 0.148722629 | 0.44652482 | 0.63980915 | C53D6.11 |
| WBGene00044242 | NA | NA | NA | C54C8.12 |
| WBGene00044243 | 0.085537126 | 0.24698522 | NA | F16C3.4 |
| WBGene00044244 | NA | NA | NA | F26D10.13 |
| WBGene00044245 | NA | NA | NA | F32A7.8 |
| WBGene00044246 | NA | NA | NA | F36D1.10 |
| WBGene00044247 | 0.020232312 | 0.84643798 | NA | F36H2.4 |
| WBGene00044249 | 0.040087849 | 0.56144502 | NA | F55B11.6 |
| WBGene00044250 | 0.056411843 | 0.41891328 | NA | F56G4.7 |

| | | | | | |
|---|---|---|---|---|---|
| WBGene00044251 | | 0.287357137 | 0.20785341 | 0.39677283 | F59C6.12 |
| WBGene00044253 | | 0.150607842 | 0.4778461 | 0.66478001 | M117.6 |
| WBGene00044256 | | 0.06375261 | 0.63596542 | 0.78416454 | Y45F10C.6 |
| WBGene00044257 | | 0.040218942 | 0.56174727 | NA | Y45F10D.16 |
| WBGene00044258 | | 0.305398921 | 0.11022268 | 0.26189065 | Y57G11C.51 |
| WBGene00044259 | | -0.108739123 | 0.36964675 | 0.56950417 | Y73F8A.36 |
| WBGene00044260 | | 0.242259451 | 0.2512779 | 0.44986057 | Y87G2A.19 |
| WBGene00044261 | | 0.087256674 | 0.23800185 | NA | Y87G2A.20 |
| WBGene00044262 | | 0.133726375 | 0.51480065 | 0.6947365 | NA |
| WBGene00044263 | NA | | NA | NA | Y105C5A.26 |
| WBGene00044264 | | 0.020232312 | 0.84643798 | NA | Y105E8A.32 |
| WBGene00044265 | NA | | NA | NA | Y105E8B.11 |
| WBGene00044266 | NA | | NA | NA | ZK39.9 |
| WBGene00044267 | NA | | NA | NA | NA |
| WBGene00044268 | | -0.067940784 | 0.63163574 | 0.78099644 | NA |
| WBGene00044270 | | 0.085537126 | 0.24698522 | NA | NA |
| WBGene00044271 | NA | | NA | NA | nspa-2 |
| WBGene00044272 | NA | | NA | NA | nspa-5 |
| WBGene00044272 | NA | | NA | NA | nspa-4 |
| WBGene00044273 | NA | | NA | NA | nspa-7 |
| WBGene00044273 | NA | | NA | NA | nspa-6 |
| WBGene00044274 | NA | | NA | NA | nspa-7 |
| WBGene00044274 | NA | | NA | NA | nspa-6 |
| WBGene00044275 | NA | | NA | NA | nspa-10 |
| WBGene00044276 | NA | | NA | NA | NA |
| WBGene00044277 | NA | | NA | NA | NA |
| WBGene00044278 | | 0.075726029 | 0.27693214 | NA | F09C6.14 |
| WBGene00044279 | | 0.015584271 | 0.80362925 | NA | NA |
| WBGene00044280 | | 2.039326358 | 9.68E-08 | 1.83E-06 | W02A2.9 |
| WBGene00044281 | | -0.623239729 | 0.03148538 | 0.10698016 | R04A9.7 |
| WBGene00044282 | NA | | NA | NA | T19H12.12 |
| WBGene00044283 | | 0.052819602 | 0.80001766 | 0.88826172 | spp-22 |
| WBGene00044284 | NA | | NA | NA | spp-21 |
| WBGene00044285 | | 0.033034883 | 0.64219503 | NA | NA |
| WBGene00044286 | NA | | NA | NA | ugt-35 |
| WBGene00044287 | | 0.072424635 | 0.53990717 | 0.71445958 | F21H12.7 |
| WBGene00044288 | | -0.116558739 | 0.17232192 | NA | F30H5.5 |
| WBGene00044289 | | 1.769578303 | 0.00464672 | 0.02471217 | Y37F4.8 |
| WBGene00044290 | | -0.104387478 | 0.52632366 | 0.70360708 | B0035.18 |
| WBGene00044291 | | 0.106516128 | 0.594081 | 0.75372096 | C25F9.10 |
| WBGene00044292 | | 0.038125098 | 0.61119482 | NA | F56D6.8 |
| WBGene00044293 | | 0.033034883 | 0.64219503 | NA | K08D12.7 |
| WBGene00044294 | | -0.246480367 | 0.10849069 | 0.25907881 | C01B10.11 |

| | | | | |
|---|---|---|---|---|
| WBGene00044295 | 0.018790812 | 0.76755469 | NA | C09E7.10 |
| WBGene00044296 | 0.961551805 | 8.51E-07 | 1.37E-05 | F10C1.9 |
| WBGene00044297 | 0.054390058 | 0.532651 | NA | C25B8.8 |
| WBGene00044298 | 0.038125098 | 0.61119482 | NA | T23B7.2 |
| WBGene00044299 | 0.058314668 | 0.40208118 | NA | C42D8.9 |
| WBGene00044300 | 0.157815695 | 0.10232055 | 0.24922229 | D1022.9 |
| WBGene00044301 | 0.670423042 | 0.03179246 | 0.10782335 | lgc-28 |
| WBGene00044302 | 2.97753672 | 0.00171452 | 0.01100012 | F28E10.5 |
| WBGene00044303 | -0.031854812 | 0.84063573 | 0.91391904 | hst-3.2 |
| WBGene00044304 | -0.067499821 | 0.71406939 | 0.83959925 | F52B10.3 |
| WBGene00044305 | -0.020007185 | 0.86252588 | 0.92672958 | rad-8 |
| WBGene00044306 | -0.022930767 | 0.78121253 | NA | F58F9.8 |
| WBGene00044307 | NA | NA | NA | F58F9.9 |
| WBGene00044308 | 0.020317245 | 0.84643798 | NA | NA |
| WBGene00044309 | NA | NA | NA | NA |
| WBGene00044310 | -0.044978524 | 0.8062723 | 0.89246487 | H20J04.9 |
| WBGene00044311 | 0.04818669 | 0.62629569 | 0.77708433 | K04C2.7 |
| WBGene00044312 | -0.13984316 | 0.25182083 | 0.44993917 | K04C2.8 |
| WBGene00044313 | 0.060484201 | 0.41382805 | NA | M02D8.7 |
| WBGene00044314 | -0.581319261 | 0.03064136 | 0.10501409 | fbxb-33 |
| WBGene00044315 | 0.411556161 | 0.00442461 | 0.0237902 | F41G3.20 |
| WBGene00044316 | 1.570303125 | 6.55E-05 | 0.0006688 | F41G3.21 |
| WBGene00044317 | -0.004747605 | 0.9473792 | NA | F41G3.19 |
| WBGene00044318 | -0.568821822 | 0.00165946 | 0.01071256 | tag-267 |
| WBGene00044319 | -0.085791648 | 0.55727902 | 0.72847563 | tag-266 |
| WBGene00044321 | -0.208179068 | 0.07216496 | 0.19562436 | mrps-30 |
| WBGene00044322 | -0.119816237 | 0.53019294 | 0.70690782 | tag-280 |
| WBGene00044323 | -0.150198961 | 0.47176202 | 0.65988991 | tag-281 |
| WBGene00044324 | -0.0041533 | 0.97348809 | 0.98538191 | ufm-1 |
| WBGene00044325 | 0.122507161 | 0.28445311 | 0.48437753 | tag-321 |
| WBGene00044326 | -0.130158314 | 0.22021356 | 0.41234771 | tag-322 |
| WBGene00044329 | -0.214025794 | 0.02257686 | 0.08400197 | cpsf-4 |
| WBGene00044330 | -0.07655404 | 0.71118174 | 0.8374145 | alr-1 |
| WBGene00044331 | 1.884597359 | 0.00755705 | 0.03618143 | clec-25 |
| WBGene00044332 | 3.488932363 | 3.93E-10 | 1.04E-08 | clec-36 |
| WBGene00044333 | 0.048485426 | 0.45975815 | NA | C01G12.13 |
| WBGene00044337 | NA | NA | NA | nspc-2 |
| WBGene00044337 | NA | NA | NA | nspc-6 |
| WBGene00044338 | 1.374299475 | 0.00521514 | 0.02700915 | nspc-3 |
| WBGene00044339 | -0.876984625 | 1.16E-05 | 0.00014014 | nspc-4 |
| WBGene00044340 | 0.015896709 | 0.80422001 | NA | nspc-5 |
| WBGene00044341 | -0.072491415 | 0.72132278 | 0.84362463 | ave-1 |
| WBGene00044342 | 0.136637339 | 0.33638191 | 0.53578846 | T05B11.7 |

| | | | | |
|---|---|---|---|---|
| WBGene00044343 | NA | NA | NA | NA |
| WBGene00044344 | -0.05535379 | 0.67075782 | 0.80831463 | mrpl-39 |
| WBGene00044345 | -0.139693584 | 0.31130472 | 0.5122686 | Y48G1C.12 |
| WBGene00044346 | 0.040107227 | 0.58693163 | NA | Y59E9AL.8 |
| WBGene00044347 | 0.068765814 | 0.58514502 | 0.74815372 | Y71G12B.30 |
| WBGene00044348 | 0.121788739 | 0.42121463 | 0.61793621 | Y71G12B.31 |
| WBGene00044349 | 0.111696043 | 0.33558103 | 0.53480372 | Y71G12B.32 |
| WBGene00044350 | 0.027524151 | 0.8951458 | 0.94517739 | Y73B6BL.44 |
| WBGene00044352 | NA | NA | NA | ZK381.8 |
| WBGene00044353 | -0.022930767 | 0.78121253 | NA | ZK418.10 |
| WBGene00044354 | 0.056770337 | 0.59158492 | 0.75224898 | ZK418.11 |
| WBGene00044355 | 0.093181679 | 0.65573762 | 0.79647042 | ZK563.7 |
| WBGene00044356 | -0.023056499 | 0.78121253 | NA | W02D7.12 |
| WBGene00044357 | NA | NA | NA | nspc-2 |
| WBGene00044357 | NA | NA | NA | nspc-6 |
| WBGene00044358 | 0.020925399 | 0.84643798 | NA | Y71G12B.33 |
| WBGene00044359 | -0.658967577 | 0.01800333 | 0.07118842 | C16C4.17 |
| WBGene00044360 | 0.00981207 | 0.94859697 | 0.97060649 | C33C12.11 |
| WBGene00044361 | NA | NA | NA | K09F6.11 |
| WBGene00044362 | 0.020317245 | 0.84643798 | NA | F42G2.7 |
| WBGene00044363 | 0.077128902 | 0.52194298 | 0.70005196 | F22E5.21 |
| WBGene00044364 | NA | NA | NA | nspf-1 |
| WBGene00044365 | NA | NA | NA | nspf-2 |
| WBGene00044366 | NA | NA | NA | nspf-3 |
| WBGene00044367 | -0.0550775 | 0.58232579 | 0.74624349 | T05A8.8 |
| WBGene00044368 | NA | NA | NA | K12H6.10 |
| WBGene00044369 | NA | NA | NA | irld-67 |
| WBGene00044370 | NA | NA | NA | C41H7.9 |
| WBGene00044371 | 0.001029307 | 0.99761053 | 0.99849711 | ZC239.20 |
| WBGene00044372 | -0.21090971 | 0.28842376 | 0.48863582 | ZC239.21 |
| WBGene00044373 | NA | NA | NA | C17F4.11 |
| WBGene00044374 | NA | NA | NA | C17F4.12 |
| WBGene00044375 | 0.020232312 | 0.84643798 | NA | F39E9.14 |
| WBGene00044376 | NA | NA | NA | Y49F6B.12 |
| WBGene00044377 | 0.020317245 | 0.84643798 | NA | Y49F6B.13 |
| WBGene00044378 | NA | NA | NA | Y49F6B.14 |
| WBGene00044379 | -0.023109762 | 0.78121253 | NA | F40H7.12 |
| WBGene00044380 | NA | NA | NA | T24E12.12 |
| WBGene00044381 | 0.032192384 | 0.61853721 | NA | K10G6.5 |
| WBGene00044382 | 0.033463284 | 0.6365037 | NA | Y14H12A.2 |
| WBGene00044383 | 0.085309563 | 0.24820057 | NA | T05A7.11 |
| WBGene00044384 | 0.038125098 | 0.61119482 | NA | F11G11.14 |
| WBGene00044385 | 0.106997704 | 0.16248985 | NA | F59A6.10 |

| | | | | |
|---|---|---|---|---|
| WBGene00044386 | 0.033463284 | 0.6365037 | NA | C27A2.7 |
| WBGene00044387 | 0.083479586 | 0.30566331 | NA | C27A2.8 |
| WBGene00044388 | 0.059776078 | 0.52388083 | NA | C27D6.11 |
| WBGene00044389 | 1.208327545 | 0.00322065 | 0.01848433 | C27D6.12 |
| WBGene00044390 | NA | NA | NA | ZK177.11 |
| WBGene00044391 | -0.003108065 | 0.95728506 | NA | C25H3.15 |
| WBGene00044392 | 0.065014541 | 0.37399213 | NA | C25H3.16 |
| WBGene00044393 | 0.038125098 | 0.61119482 | NA | ZK1248.20 |
| WBGene00044394 | 0.013399338 | 0.88795937 | NA | B0034.7 |
| WBGene00044395 | 0.013156723 | 0.8627609 | NA | R12C12.10 |
| WBGene00044396 | 0.425487802 | 0.09000387 | 0.22810224 | ZK1290.14 |
| WBGene00044397 | NA | NA | NA | NA |
| WBGene00044398 | NA | NA | NA | NA |
| WBGene00044399 | -0.034785937 | 0.72283628 | NA | F11F1.8 |
| WBGene00044401 | 0.183888072 | 0.27903198 | 0.47882175 | Y55B1BR.6 |
| WBGene00044402 | -0.276352709 | 0.10317103 | 0.25070988 | Y82E9BL.18 |
| WBGene00044403 | 0.31303662 | 0.1601828 | 0.33477934 | fbxa-219 |
| WBGene00044405 | NA | NA | NA | NA |
| WBGene00044406 | 0.086647583 | 0.55123073 | 0.72373276 | F02A9.7 |
| WBGene00044409 | NA | NA | NA | C45G9.14 |
| WBGene00044410 | -0.067263114 | 0.71543324 | 0.84039298 | F48E8.8 |
| WBGene00044411 | 0.079596829 | 0.48303725 | 0.66863462 | R12B2.8 |
| WBGene00044412 | -0.027493884 | 0.89025095 | 0.94282725 | F40H6.6 |
| WBGene00044413 | -0.038323147 | 0.85057276 | 0.9197276 | B0205.12 |
| WBGene00044414 | 0.304078173 | 0.19284222 | 0.37932398 | C06A5.12 |
| WBGene00044415 | NA | NA | NA | C08E3.14 |
| WBGene00044416 | NA | NA | NA | C08E3.15 |
| WBGene00044417 | -0.036106717 | 0.76600275 | 0.86713124 | C08F1.11 |
| WBGene00044418 | 1.263414489 | 0.00461316 | 0.0245873 | C08G5.7 |
| WBGene00044419 | 0.017604133 | 0.79437959 | NA | fbxc-17 |
| WBGene00044420 | 0.092784318 | 0.61439167 | 0.7686327 | C50D2.10 |
| WBGene00044421 | 1.676194741 | 0.00344361 | 0.01948897 | D1007.18 |
| WBGene00044423 | 0.199512413 | 0.33402768 | 0.53343143 | F53F10.8 |
| WBGene00044424 | NA | NA | NA | F54D12.10 |
| WBGene00044425 | NA | NA | NA | F54D12.11 |
| WBGene00044427 | 0.048485426 | 0.45975815 | NA | F56A6.5 |
| WBGene00044428 | -0.046394786 | 0.50231372 | NA | T01D1.7 |
| WBGene00044429 | NA | NA | NA | T07H3.7 |
| WBGene00044430 | NA | NA | NA | NA |
| WBGene00044431 | -0.296230376 | 0.03108356 | 0.10595147 | mzt-1 |
| WBGene00044432 | -0.022930767 | 0.78121253 | NA | W03G9.9 |
| WBGene00044433 | 0.224179964 | 0.30054999 | 0.50083684 | W10C8.6 |
| WBGene00044434 | 0.020925399 | 0.84643798 | NA | Y18H1A.15 |

| | | | | |
|---|---|---|---|---|
| WBGene00044435 | 0.196550163 | 0.20159707 | 0.39027971 | Y20F4.8 |
| WBGene00044436 | -0.274756052 | 0.21474493 | 0.4056994 | Y47G6A.31 |
| WBGene00044437 | -0.006239126 | 0.93041881 | NA | Y47G6A.32 |
| WBGene00044438 | 0.00170114 | 0.99338358 | 0.99586212 | Y51F10.11 |
| WBGene00044439 | -0.221566686 | 0.2056357 | 0.39466399 | Y51H7C.15 |
| WBGene00044440 | 0.228642282 | 0.10310135 | 0.25058216 | adpr-1 |
| WBGene00044441 | 0.020232312 | 0.84643798 | NA | Y119C1B.11 |
| WBGene00044442 | 0.083520471 | 0.30840907 | NA | T08A9.13 |
| WBGene00044443 | -0.239345871 | 0.14528285 | 0.31461118 | ZC21.10 |
| WBGene00044444 | -0.002914971 | 0.95718083 | NA | F09G8.10 |
| WBGene00044445 | 0.065707583 | 0.7499283 | 0.8587464 | C02F5.14 |
| WBGene00044446 | 0.040107227 | 0.58693163 | NA | C06G4.6 |
| WBGene00044447 | 0.389956889 | 0.0322606 | 0.10910588 | ZK688.10 |
| WBGene00044448 | 0.033034883 | 0.64219503 | NA | ZK783.6 |
| WBGene00044449 | NA | NA | NA | fipr-20 |
| WBGene00044450 | 0.0398788 | 0.58971221 | NA | fipr-18 |
| WBGene00044451 | 0.020232312 | 0.84643798 | NA | fipr-19 |
| WBGene00044452 | -0.027310994 | 0.89292211 | 0.9441978 | Y102E9.5 |
| WBGene00044453 | 0.020232312 | 0.84643798 | NA | Y102E9.6 |
| WBGene00044454 | -0.001993815 | 0.95612996 | NA | F27B3.8 |
| WBGene00044455 | 0.475852977 | 0.07231446 | 0.19588535 | F23F12.13 |
| WBGene00044457 | -0.577950217 | 0.0409959 | 0.13025628 | C18H7.11 |
| WBGene00044458 | NA | NA | NA | K11H12.10 |
| WBGene00044459 | -0.552760083 | 0.04564115 | 0.1407397 | Y55F3C.9 |
| WBGene00044460 | -0.086533417 | 0.42974627 | 0.6246317 | Y55F3C.10 |
| WBGene00044461 | 0.063217264 | 0.41230173 | NA | Y18H1A.14 |
| WBGene00044462 | 0.059409224 | 0.42187437 | NA | C44B12.9 |
| WBGene00044463 | -0.027058162 | 0.72388304 | NA | W09G12.10 |
| WBGene00044464 | 0.056727431 | 0.44364738 | NA | Y77E11A.16 |
| WBGene00044466 | 0.145166861 | 0.45257818 | 0.6448226 | Y48A5A.3 |
| WBGene00044467 | 0.077697904 | 0.68050942 | 0.81496025 | R05C11.4 |
| WBGene00044468 | 0.237102807 | 0.21889164 | 0.41076674 | Y69A2AL.2 |
| WBGene00044469 | 1.252935055 | 1.65E-08 | 3.56E-07 | clec-173 |
| WBGene00044470 | NA | NA | NA | C09G12.17 |
| WBGene00044471 | NA | NA | NA | F56D6.9 |
| WBGene00044472 | 0.034123182 | 0.68690178 | NA | dct-8 |
| WBGene00044473 | NA | NA | NA | F56D6.11 |
| WBGene00044474 | 0.059762547 | 0.38965825 | NA | F56D6.12 |
| WBGene00044475 | 0.090103632 | 0.20323472 | NA | F56D6.13 |
| WBGene00044476 | 0.09777701 | 0.17585057 | NA | F56D6.14 |
| WBGene00044477 | 0.020925399 | 0.84643798 | NA | F49F1.14 |
| WBGene00044478 | 0.017604133 | 0.79437959 | NA | K06B9.6 |
| WBGene00044479 | 0.036894305 | 0.59120501 | NA | F15E6.10 |

| | | | | |
|---|---|---|---|---|
| WBGene00044480 | 0.020317245 | 0.84643798 | NA | ZK185.4 |
| WBGene00044481 | 0.631852191 | 0.00335642 | 0.01908564 | ZK185.5 |
| WBGene00044482 | 0.146122617 | 0.32837116 | 0.52779472 | F29B9.12 |
| WBGene00044483 | -0.093633366 | 0.49881108 | 0.68081411 | H06H21.11 |
| WBGene00044484 | 0.209898521 | 0.1977044 | 0.38553119 | hot-9 |
| WBGene00044485 | 0.237531969 | 0.28070736 | 0.48036693 | Y54G2A.41 |
| WBGene00044486 | 0.015896709 | 0.80422001 | NA | Y54G2A.42 |
| WBGene00044487 | 0.017141582 | 0.79355725 | NA | Y54G2A.44 |
| WBGene00044488 | 0.126043471 | 0.47758872 | 0.66470097 | Y54G2A.45 |
| WBGene00044489 | 0.084975776 | 0.67799822 | 0.81292048 | pudl-1 |
| WBGene00044490 | 0.132098125 | 0.43203428 | 0.62721011 | pudl-2 |
| WBGene00044491 | 0.078475818 | 0.52084483 | 0.69928849 | Y54G2A.48 |
| WBGene00044492 | 0.91156773 | 0.01993797 | 0.07647677 | Y54G2A.49 |
| WBGene00044493 | 0.192336727 | 0.22990355 | 0.42425231 | Y54G2A.50 |
| WBGene00044495 | 0.456148977 | 0.06345801 | 0.17870526 | Y54G2A.40 |
| WBGene00044498 | NA | NA | NA | Y59E9AR.10 |
| WBGene00044499 | -0.048732643 | 0.59290524 | 0.75314295 | T11F8.5 |
| WBGene00044500 | 0.68277203 | 0.01746795 | 0.06953461 | T19E7.6 |
| WBGene00044501 | 0.040107227 | 0.58693163 | NA | C55C3.8 |
| WBGene00044502 | 0.329372606 | 0.14490504 | 0.3142999 | C46G7.5 |
| WBGene00044503 | -0.015341772 | 0.8879423 | 0.94103245 | NA |
| WBGene00044504 | 0.125592892 | 0.42948585 | 0.62431517 | R13A1.10 |
| WBGene00044505 | -0.022930767 | 0.78121253 | NA | D2024.10 |
| WBGene00044508 | 0.115307647 | 0.52740392 | 0.70466516 | nsy-7 |
| WBGene00044509 | 0.342977242 | 0.11164995 | 0.26408189 | K07H8.12 |
| WBGene00044510 | 0.236429133 | 0.09297852 | 0.23301926 | K07H8.11 |
| WBGene00044511 | NA | NA | NA | clec-181 |
| WBGene00044512 | 0.205126475 | 0.23456776 | 0.43020249 | W03F9.11 |
| WBGene00044513 | 0.480794376 | 0.04093263 | 0.13008348 | nlp-70 |
| WBGene00044514 | 1.254627814 | 0.00393393 | 0.02159606 | R09E12.9 |
| WBGene00044515 | 0.005979718 | 0.96159783 | 0.97914283 | NA |
| WBGene00044516 | NA | NA | NA | Y32G9A.12 |
| WBGene00044517 | 0.132459662 | 0.49320811 | 0.67612921 | NA |
| WBGene00044518 | NA | NA | NA | fipr-25 |
| WBGene00044519 | NA | NA | NA | F59A7.11 |
| WBGene00044520 | -0.028297421 | 0.84955558 | 0.91911205 | F59A7.12 |
| WBGene00044521 | -0.022930767 | 0.78121253 | NA | Y5H2A.4 |
| WBGene00044523 | 0.01912147 | 0.77587372 | NA | NA |
| WBGene00044524 | NA | NA | NA | Y73C8C.12 |
| WBGene00044525 | -0.023056499 | 0.78121253 | NA | Y73C8B.5 |
| WBGene00044526 | -0.355283825 | 0.09312525 | 0.2332273 | T28A11.22 |
| WBGene00044527 | 0.208726275 | 0.28342311 | 0.48335442 | Y39H10B.2 |
| WBGene00044528 | 0.057855329 | 0.58501057 | 0.74805087 | lgc-29 |

| | | | | |
|---|---|---|---|---|
| WBGene00044529 | 0.102555946 | 0.30149619 | 0.50212782 | irld-1 |
| WBGene00044530 | 0.611141475 | 0.04051667 | 0.12921032 | F32D1.11 |
| WBGene00044531 | NA | NA | NA | F13A2.9 |
| WBGene00044533 | 0.155930036 | 0.12000031 | 0.27834673 | F41F3.8 |
| WBGene00044534 | 0.060160288 | 0.52140784 | NA | H23N18.6 |
| WBGene00044535 | 0.501559706 | 1.08E-07 | 2.03E-06 | K11D12.13 |
| WBGene00044536 | 0.063527295 | 0.41294485 | NA | F14F9.8 |
| WBGene00044537 | 0.09266566 | 0.60618724 | 0.76218049 | B0238.15 |
| WBGene00044538 | 0.040107227 | 0.58693163 | NA | T09D3.8 |
| WBGene00044539 | -0.055731908 | 0.5783904 | 0.7444901 | K09H11.9 |
| WBGene00044541 | 0.014602567 | 0.85416114 | NA | K09H11.11 |
| WBGene00044542 | 0.020925399 | 0.84643798 | NA | CD4.11 |
| WBGene00044543 | NA | NA | NA | C53A3.3 |
| WBGene00044544 | 0.177428927 | 0.34406781 | 0.54381673 | F26F12.8 |
| WBGene00044545 | NA | NA | NA | EGAP9.4 |
| WBGene00044546 | NA | NA | NA | T01C4.8 |
| WBGene00044547 | -0.002914971 | 0.95718083 | NA | C13A2.12 |
| WBGene00044548 | NA | NA | NA | cnc-9 |
| WBGene00044549 | NA | NA | NA | clec-215 |
| WBGene00044550 | NA | NA | NA | C50E3.15 |
| WBGene00044551 | NA | NA | NA | ZK105.8 |
| WBGene00044552 | 0.020232312 | 0.84643798 | NA | Y97E10AR.8 |
| WBGene00044553 | -0.10074509 | 0.61915214 | 0.77208129 | T23B12.11 |
| WBGene00044554 | -0.325339041 | 0.13780279 | 0.30365277 | F41E6.15 |
| WBGene00044556 | 0.231543092 | 0.27431134 | 0.47472723 | F38G1.3 |
| WBGene00044557 | 0.065014541 | 0.37399213 | NA | ZC13.10 |
| WBGene00044558 | 0.020317245 | 0.84643798 | NA | F56C3.10 |
| WBGene00044559 | -0.004944805 | 0.94514249 | NA | ZK380.5 |
| WBGene00044560 | NA | NA | NA | C36C9.6 |
| WBGene00044561 | NA | NA | NA | Y75D11A.6 |
| WBGene00044562 | NA | NA | NA | C14E2.7 |
| WBGene00044563 | -0.012142019 | 0.90244358 | 0.94802873 | T26C11.8 |
| WBGene00044564 | 0.038125098 | 0.61119482 | NA | ZK813.6 |
| WBGene00044565 | NA | NA | NA | F53B1.9 |
| WBGene00044567 | -0.269674308 | 0.14980992 | 0.32119473 | C46C11.4 |
| WBGene00044568 | 0.349909333 | 0.10185217 | 0.24853626 | ntc-1 |
| WBGene00044569 | 0.020232312 | 0.84643798 | NA | C25F6.8 |
| WBGene00044570 | 0.274849728 | 0.20511185 | 0.39425529 | T22B7.8 |
| WBGene00044571 | -0.022930767 | 0.78121253 | NA | C46F4.3 |
| WBGene00044573 | -0.001993815 | 0.95612996 | NA | C15B12.9 |
| WBGene00044574 | 0.049227885 | 0.75989762 | 0.8640619 | F23G4.1 |
| WBGene00044575 | NA | NA | NA | F38B6.8 |
| WBGene00044577 | NA | NA | NA | F43E12.1 |

| | | | | |
|---|---|---|---|---|
| WBGene00044578 | NA | NA | NA | M02D8.6 |
| WBGene00044578 | NA | NA | NA | K01A12.4 |
| WBGene00044579 | -0.046394786 | 0.50231372 | NA | C06E2.9 |
| WBGene00044580 | NA | NA | NA | C24A3.9 |
| WBGene00044581 | NA | NA | NA | Y81B9A.4 |
| WBGene00044582 | 0.224834386 | 0.20643188 | 0.39559805 | B0222.11 |
| WBGene00044583 | NA | NA | NA | F07G6.9 |
| WBGene00044584 | 0.183370727 | 0.08738565 | 0.22379103 | T03G11.9 |
| WBGene00044585 | 0.009909198 | 0.95217727 | 0.97257063 | NA |
| WBGene00044586 | 0.257112251 | 0.16901128 | 0.34728789 | Y61A9LA.12 |
| WBGene00044587 | NA | NA | NA | srz-105 |
| WBGene00044588 | 0.033167789 | 0.82892956 | 0.90652983 | ZK1236.9 |
| WBGene00044589 | 0.020925399 | 0.84643798 | NA | C29F5.8 |
| WBGene00044602 | 0.020232312 | 0.84643798 | NA | C31B8.16 |
| WBGene00044603 | -0.00346345 | 0.95708614 | NA | acbp-7 |
| WBGene00044604 | -0.213521413 | 0.22727789 | 0.42094567 | T10F2.5 |
| WBGene00044605 | -0.257376153 | 0.11492741 | 0.27017536 | ttr-36 |
| WBGene00044606 | NA | NA | NA | NA |
| WBGene00044607 | 0.950957355 | 0.00281676 | 0.01660235 | K02A11.4 |
| WBGene00044608 | -0.150430065 | 0.29555452 | 0.49663878 | glrx-22 |
| WBGene00044609 | 0.197707572 | 0.24213438 | 0.44011042 | tbcc-1 |
| WBGene00044610 | NA | NA | NA | D2092.10 |
| WBGene00044611 | 0.203653921 | 0.14312834 | 0.31151213 | T27A3.8 |
| WBGene00044612 | 0.737118287 | 0.02496121 | 0.09072917 | Y57E12AR.1 |
| WBGene00044614 | -0.119734266 | 0.42801244 | 0.62297744 | C36B1.14 |
| WBGene00044615 | 0.82504174 | 0.02342878 | 0.08622945 | B0252.8 |
| WBGene00044616 | -0.064153509 | 0.64758829 | 0.79141821 | M01A12.4 |
| WBGene00044617 | 0.078203897 | 0.26561294 | NA | bus-1 |
| WBGene00044618 | 0.078387035 | 0.47443094 | 0.6618896 | bus-2 |
| WBGene00044620 | 0.03732969 | 0.59229443 | NA | bus-4 |
| WBGene00044623 | 0.062302706 | 0.63461401 | 0.78332291 | bus-8 |
| WBGene00044630 | 0.00057415 | 0.99806931 | 0.99882228 | bus-17 |
| WBGene00044631 | 0.081768833 | 0.41230943 | NA | bus-18 |
| WBGene00044632 | 0.084706123 | 0.6276295 | 0.77834351 | NA |
| WBGene00044633 | 0.106764072 | 0.31580329 | 0.51638105 | F54H12.7 |
| WBGene00044634 | -0.002914971 | 0.95718083 | NA | C29E4.14 |
| WBGene00044636 | 0.235789599 | 0.23696955 | 0.43330475 | B0393.9 |
| WBGene00044637 | 0.003184887 | 0.98877847 | 0.99380144 | rft-1 |
| WBGene00044638 | -1.353366619 | 6.33E-35 | 9.46E-33 | F23A7.8 |
| WBGene00044639 | -0.099452571 | 0.17595618 | NA | NA |
| WBGene00044640 | 0.241909766 | 0.26050357 | 0.45854408 | B0432.14 |
| WBGene00044641 | 0.073766927 | 0.57454872 | 0.74225428 | T01D1.8 |
| WBGene00044642 | 0.337727758 | 0.02339408 | 0.08616672 | NA |

| | | | | |
|---|---|---|---|---|
| WBGene00044643 | 0.231410419 | 0.08886609 | 0.22627625 | H17B01.5 |
| WBGene00044644 | 1.175734727 | 0.00097769 | 0.00691183 | B0205.13 |
| WBGene00044645 | -0.022930767 | 0.78121253 | NA | Y51H7BR.8 |
| WBGene00044646 | 0.086764008 | 0.294364 | NA | B0205.14 |
| WBGene00044648 | 0.057298498 | 0.75741822 | 0.86264957 | R144.13 |
| WBGene00044649 | -0.07942555 | 0.55521643 | 0.72703012 | C23H5.11 |
| WBGene00044650 | 0.169623337 | 0.11592948 | 0.27191983 | Y71G10AR.4 |
| WBGene00044651 | -0.018222174 | 0.89103873 | 0.94328981 | egrh-3 |
| WBGene00044653 | NA | NA | NA | T13F3.9 |
| WBGene00044654 | 0.09059338 | 0.49208023 | 0.67515197 | fbxa-85 |
| WBGene00044655 | NA | NA | NA | fbxa-194 |
| WBGene00044657 | NA | NA | NA | NA |
| WBGene00044658 | 6.287299971 | 3.13E-06 | 4.29E-05 | C01G10.17 |
| WBGene00044659 | 0.020317245 | 0.84643798 | NA | fbxa-142 |
| WBGene00044660 | 0.296939398 | 0.06851103 | 0.18847613 | NA |
| WBGene00044661 | 0.208291566 | 0.26521431 | 0.4637486 | fbxa-143 |
| WBGene00044662 | NA | NA | NA | F26G1.9 |
| WBGene00044663 | NA | NA | NA | F26G1.10 |
| WBGene00044664 | NA | NA | NA | F59A6.11 |
| WBGene00044665 | 0.27499971 | 0.13949929 | 0.30689844 | F59A6.12 |
| WBGene00044666 | 0.070823658 | 0.70624553 | 0.83388232 | F33G12.7 |
| WBGene00044667 | 0.093296287 | 0.6525987 | 0.79427578 | ZK622.5 |
| WBGene00044668 | NA | NA | NA | C18A3.11 |
| WBGene00044670 | NA | NA | NA | NA |
| WBGene00044672 | 0.172491974 | 0.4122296 | 0.60976496 | fbxa-4 |
| WBGene00044673 | NA | NA | NA | C07D10.6 |
| WBGene00044674 | NA | NA | NA | B0280.17 |
| WBGene00044675 | -0.084602292 | 0.65713319 | 0.79763613 | fbxa-76 |
| WBGene00044676 | -0.117340069 | 0.48723253 | 0.67155416 | fbxa-42 |
| WBGene00044677 | 0.061389344 | 0.64188352 | 0.78787866 | ZK353.10 |
| WBGene00044678 | -0.02791392 | 0.6763823 | NA | B0303.16 |
| WBGene00044679 | 0.055508127 | 0.63725864 | 0.78496583 | K02B12.9 |
| WBGene00044680 | -0.045269523 | 0.82391583 | 0.9034778 | F30A10.13 |
| WBGene00044681 | 0.067686538 | 0.55755777 | 0.7287273 | T24C4.8 |
| WBGene00044683 | -0.001488154 | 0.95639912 | NA | C36E6.8 |
| WBGene00044684 | 0.070327578 | 0.44827578 | NA | T08G11.2 |
| WBGene00044686 | 0.861496569 | 1.46E-06 | 2.20E-05 | flp-28 |
| WBGene00044687 | NA | NA | NA | B0563.9 |
| WBGene00044689 | -0.044433245 | 0.49495692 | 0.67757497 | arid-1 |
| WBGene00044690 | 0.173538763 | 0.41817484 | 0.61483208 | M02B7.7 |
| WBGene00044691 | NA | NA | NA | NA |
| WBGene00044692 | -1.392508158 | 4.28E-05 | 0.00045585 | fbxb-91 |
| WBGene00044693 | -0.326954529 | 0.14606433 | 0.31589857 | fbxb-92 |

| | | | | | |
|---|---|---|---|---|---|
| WBGene00044694 | | -0.440201055 | 0.08875196 | 0.22611347 | fbxb-90 |
| WBGene00044695 | | -0.135302469 | 0.31990848 | 0.51995782 | NA |
| WBGene00044696 | | 1.038987008 | 3.89E-35 | 5.93E-33 | F52E1.14 |
| WBGene00044697 | | 3.377720531 | 1.27E-05 | 0.00015153 | poml-4 |
| WBGene00044698 | | 0.013981771 | 0.94709564 | 0.97031324 | K09F5.6 |
| WBGene00044699 | | 0.915768498 | 0.01047276 | 0.04648337 | nhr-286 |
| WBGene00044700 | | -0.241298913 | 0.27677236 | 0.47673325 | NA |
| WBGene00044701 | | 0.250509902 | 0.1983413 | 0.38646726 | AH9.6 |
| WBGene00044702 | | -0.091421884 | 0.64766885 | 0.79141821 | NA |
| WBGene00044703 | | 0.00027486 | 0.99963959 | 0.99983667 | NA |
| WBGene00044704 | NA | | NA | NA | NA |
| WBGene00044705 | | 0.123867178 | 0.25990428 | 0.4579683 | T22F3.12 |
| WBGene00044706 | NA | | NA | NA | NA |
| WBGene00044707 | | 0.017141582 | 0.79355725 | NA | pals-15 |
| WBGene00044708 | | 0.085309563 | 0.24820057 | NA | pals-11 |
| WBGene00044709 | NA | | NA | NA | pals-12 |
| WBGene00044710 | NA | | NA | NA | NA |
| WBGene00044711 | NA | | NA | NA | pals-10 |
| WBGene00044712 | | 0.506305468 | 0.05911887 | 0.16981742 | F22G12.8 |
| WBGene00044713 | NA | | NA | NA | NA |
| WBGene00044714 | NA | | NA | NA | NA |
| WBGene00044716 | NA | | NA | NA | NA |
| WBGene00044718 | NA | | NA | NA | T20D4.20 |
| WBGene00044719 | | 0.358294277 | 0.05966571 | 0.17095218 | clec-172 |
| WBGene00044720 | | -0.282037106 | 0.1941863 | 0.38111014 | metl-2 |
| WBGene00044722 | NA | | NA | NA | NA |
| WBGene00044723 | | -0.054802699 | 0.60102331 | 0.75861226 | K11H12.11 |
| WBGene00044724 | | 0.464315228 | 0.01278981 | 0.05441896 | R02E4.3 |
| WBGene00044725 | | -0.226624954 | 0.26933375 | 0.46838288 | R11G1.7 |
| WBGene00044726 | | 0.017348279 | 0.83364574 | NA | Y53F4B.43 |
| WBGene00044727 | NA | | NA | NA | NA |
| WBGene00044728 | | 3.375135092 | 3.42E-21 | 2.77E-19 | NA |
| WBGene00044729 | | -0.001993815 | 0.95612996 | NA | NA |
| WBGene00044732 | | 0.027068195 | 0.83441834 | 0.90975112 | T03G11.10 |
| WBGene00044733 | | -0.456433948 | 0.04570349 | 0.14087257 | W08F4.13 |
| WBGene00044734 | NA | | NA | NA | C01B4.6 |
| WBGene00044734 | NA | | NA | NA | Y19D10A.16 |
| WBGene00044735 | NA | | NA | NA | srbc-81 |
| WBGene00044735 | NA | | NA | NA | Y19D10A.17 |
| WBGene00044736 | | -0.016116384 | 0.93606062 | 0.96386107 | gfrp-1 |
| WBGene00044737 | | -0.465684311 | 0.00024473 | 0.00209296 | ttr-45 |
| WBGene00044738 | | -0.096280646 | 0.23426011 | NA | folt-3 |
| WBGene00044739 | NA | | NA | NA | NA |

| | | | | | |
|---|---|---|---|---|---|
| WBGene00044741 | | 0.0398788 | 0.58971221 | NA | Y57G11C.52 |
| WBGene00044742 | | -0.145620914 | 0.48887535 | 0.67296202 | Y55F3BR.10 |
| WBGene00044743 | | 0.472430669 | 0.0689245 | 0.18911611 | crb-3 |
| WBGene00044744 | | 0.0398788 | 0.58971221 | NA | F10G7.12 |
| WBGene00044745 | | -0.102107353 | 0.58984444 | 0.75112054 | Y62E10A.20 |
| WBGene00044746 | | -0.206817671 | 0.23146028 | 0.42642631 | C53H9.3 |
| WBGene00044747 | NA | | NA | NA | NA |
| WBGene00044748 | | -0.046394786 | 0.50231372 | NA | NA |
| WBGene00044749 | NA | | NA | NA | NA |
| WBGene00044750 | NA | | NA | NA | NA |
| WBGene00044751 | NA | | NA | NA | NA |
| WBGene00044752 | NA | | NA | NA | NA |
| WBGene00044753 | | 0.139887606 | 0.29519663 | 0.49626522 | F49H12.7 |
| WBGene00044754 | | 0.061275255 | 0.51782541 | NA | Y119C1B.12 |
| WBGene00044755 | NA | | NA | NA | C32D5.14 |
| WBGene00044756 | | 0.119197413 | 0.46589463 | 0.65553385 | F58F12.4 |
| WBGene00044757 | | 0.622050843 | 0.04072053 | 0.12966273 | Y55F3BR.11 |
| WBGene00044758 | | -0.104134361 | 0.55493786 | 0.72686037 | Y37H2A.12 |
| WBGene00044759 | | -0.005068434 | 0.94614153 | NA | T06E6.15 |
| WBGene00044760 | | 0.146150641 | 0.25476187 | 0.45316105 | Y71H2AM.25 |
| WBGene00044761 | | 0.389595226 | 0.06339713 | 0.17865182 | pals-19 |
| WBGene00044762 | | 0.035959371 | 0.80859432 | 0.89395375 | ZK688.11 |
| WBGene00044763 | | 0.055259634 | 0.63569622 | 0.78403069 | ZK688.12 |
| WBGene00044764 | | 0.098430752 | 0.39151046 | 0.58960138 | NA |
| WBGene00044767 | | 0.1166417 | 0.53186624 | 0.70830031 | T23B7.3 |
| WBGene00044768 | | 0.039749717 | 0.80544448 | 0.89195315 | NA |
| WBGene00044769 | | -0.023056499 | 0.78121253 | NA | NA |
| WBGene00044770 | | 0.235104661 | 0.29109498 | 0.49174127 | C53C11.5 |
| WBGene00044771 | | 0.03732969 | 0.59229443 | NA | tmem-218 |
| WBGene00044772 | | -0.011313224 | 0.93857441 | 0.96561506 | T23E7.6 |
| WBGene00044773 | | 0.033279134 | 0.81608033 | 0.89891744 | C08A9.10 |
| WBGene00044774 | | 0.68292931 | 0.02852439 | 0.09993435 | F16B3.3 |
| WBGene00044775 | NA | | NA | NA | ZK1290.15 |
| WBGene00044776 | | 1.63770722 | 0.00302381 | 0.01758206 | T02B11.8 |
| WBGene00044777 | | 0.038325924 | 0.57925535 | NA | T02B11.9 |
| WBGene00044778 | NA | | NA | NA | K09C6.10 |
| WBGene00044779 | | 0.379171765 | 0.12430588 | 0.2845917 | T10H9.8 |
| WBGene00044780 | | 0.060105893 | 0.41982798 | NA | F18E9.8 |
| WBGene00044781 | | 0.020232312 | 0.84643798 | NA | NA |
| WBGene00044782 | NA | | NA | NA | NA |
| WBGene00044783 | | 3.687098275 | 0.00230266 | 0.01397727 | T26H5.10 |
| WBGene00044784 | | -0.071475789 | 0.53349285 | 0.70899686 | F13H10.8 |
| WBGene00044786 | | 0.020317245 | 0.84643798 | NA | ZK1053.7 |

| | | | | |
|---|---|---|---|---|
| WBGene00044787 | 0.073887753 | 0.28837975 | NA | pals-36 |
| WBGene00044788 | -0.103712433 | 0.58785036 | 0.75010608 | ttr-38 |
| WBGene00044789 | -0.366945171 | 0.00993782 | 0.04467289 | T07A9.14 |
| WBGene00044790 | 0.321305441 | 0.06868529 | 0.18884617 | clec-224 |
| WBGene00044791 | 0.037127706 | 0.59421403 | NA | bnc-1 |
| WBGene00044792 | 0.120174015 | 0.5661784 | 0.73538965 | F59C6.14 |
| WBGene00044794 | NA | NA | NA | Y59A8B.25 |
| WBGene00044795 | -0.00888715 | 0.96221207 | 0.97924853 | C06H5.8 |
| WBGene00044796 | -0.164387262 | 0.27229276 | 0.47240648 | W03D8.11 |
| WBGene00044797 | -0.276023391 | 0.22293759 | 0.41556803 | F13E9.13 |
| WBGene00044798 | -0.48117872 | 0.05688768 | 0.16542055 | tbx-43 |
| WBGene00044799 | 0.091751247 | 0.48722461 | 0.67155416 | ZK686.6 |
| WBGene00044800 | NA | NA | NA | NA |
| WBGene00044801 | NA | NA | NA | ZC262.9 |
| WBGene00044801 | NA | NA | NA | ZC262.10 |
| WBGene00044802 | NA | NA | NA | ZC262.9 |
| WBGene00044802 | NA | NA | NA | ZC262.10 |
| WBGene00044803 | NA | NA | NA | C02G6.3 |
| WBGene00044804 | NA | NA | NA | NA |
| WBGene00044805 | 0.020232312 | 0.84643798 | NA | Y53C10A.15 |
| WBGene00044806 | 0.020925399 | 0.84643798 | NA | NA |
| WBGene00044807 | 1.383186731 | 0.00811396 | 0.03827205 | T19H5.6 |
| WBGene00044808 | NA | NA | NA | NA |
| WBGene00044809 | NA | NA | NA | NA |
| WBGene00044810 | 0.040218942 | 0.56174727 | NA | Y37E3.19 |
| WBGene00044811 | 2.738950381 | 2.22E-19 | 1.59E-17 | F12E12.11 |
| WBGene00044889 | NA | NA | NA | T10D4.14 |
| WBGene00044891 | 0.173614898 | 0.32207235 | 0.52225731 | T07C4.12 |
| WBGene00044892 | NA | NA | NA | clec-257 |
| WBGene00044894 | -0.097132393 | 0.62217571 | 0.77413738 | Y71F9AR.4 |
| WBGene00044895 | NA | NA | NA | R06A10.5 |
| WBGene00044896 | 0.020317245 | 0.84643798 | NA | K12C11.5 |
| WBGene00044897 | -0.053591705 | 0.45278076 | NA | M01D7.9 |
| WBGene00044898 | -0.61862441 | 0.03673695 | 0.1202203 | let-383 |
| WBGene00044900 | 0.033463284 | 0.6365037 | NA | cnc-11 |
| WBGene00044901 | -0.083598104 | 0.68666915 | 0.81942482 | Y41G9A.10 |
| WBGene00044902 | NA | NA | NA | NA |
| WBGene00044903 | -0.216054471 | 0.3119914 | 0.51266736 | K03H1.13 |
| WBGene00044905 | -0.41945482 | 0.0603688 | 0.17215809 | NA |
| WBGene00044906 | NA | NA | NA | Y67A10A.11 |
| WBGene00044907 | -0.099326088 | 0.44280478 | 0.63678468 | NA |
| WBGene00044908 | NA | NA | NA | F38A1.15 |
| WBGene00044909 | NA | NA | NA | NA |

| | | | | | |
|---|---|---|---|---|---|
| WBGene00044911 | | 0.039034462 | 0.57625038 | NA | H38K22.6 |
| WBGene00044912 | NA | | NA | NA | H38K22.7 |
| WBGene00044914 | NA | | NA | NA | NA |
| WBGene00044916 | | -0.165570607 | 0.04837287 | 0.14666126 | F40F8.11 |
| WBGene00044917 | | 0.020925399 | 0.84643798 | NA | F40F8.12 |
| WBGene00044920 | | -0.117731018 | 0.51003818 | 0.6898992 | NA |
| WBGene00044921 | | 0.285358996 | 0.21574281 | 0.40673326 | F53C11.9 |
| WBGene00044922 | | -2.712876836 | 6.53E-08 | 1.27E-06 | Y43C5A.7 |
| WBGene00044923 | NA | | NA | NA | NA |
| WBGene00044924 | | -0.015399996 | 0.91440441 | 0.95232572 | M106.7 |
| WBGene00044925 | | -0.023056499 | 0.78121253 | NA | NA |
| WBGene00044926 | NA | | NA | NA | NA |
| WBGene00044927 | | 0.219821287 | 0.09468512 | 0.23608393 | NA |
| WBGene00044928 | NA | | NA | NA | NA |
| WBGene00044929 | | -0.023109762 | 0.78121253 | NA | NA |
| WBGene00044930 | NA | | NA | NA | NA |
| WBGene00044931 | NA | | NA | NA | NA |
| WBGene00044932 | | -0.023056499 | 0.78121253 | NA | NA |
| WBGene00044933 | NA | | NA | NA | NA |
| WBGene00044934 | NA | | NA | NA | NA |
| WBGene00044935 | NA | | NA | NA | NA |
| WBGene00044936 | | 0.020925399 | 0.84643798 | NA | NA |
| WBGene00044937 | NA | | NA | NA | NA |
| WBGene00044938 | NA | | NA | NA | NA |
| WBGene00044939 | | 0.018790812 | 0.76755469 | NA | NA |
| WBGene00044940 | | -0.073738544 | 0.42233186 | NA | NA |
| WBGene00044941 | NA | | NA | NA | NA |
| WBGene00044942 | | -0.023756719 | 0.6882956 | NA | NA |
| WBGene00044943 | | -0.003108065 | 0.95728506 | NA | NA |
| WBGene00044944 | | -0.022930767 | 0.78121253 | NA | NA |
| WBGene00044945 | NA | | NA | NA | NA |
| WBGene00044946 | NA | | NA | NA | NA |
| WBGene00044947 | NA | | NA | NA | NA |
| WBGene00044948 | | -0.005017292 | 0.94670066 | NA | NA |
| WBGene00044949 | | -0.108135811 | 0.59760582 | 0.75590037 | Y49A10A.2 |
| WBGene00044951 | | -0.022930767 | 0.78121253 | NA | NA |
| WBGene00044952 | | 0.020925399 | 0.84643798 | NA | NA |
| WBGene00044953 | | -0.044105179 | 0.80262551 | 0.89017803 | D2096.13 |
| WBGene00044954 | NA | | NA | NA | NA |
| WBGene00044955 | NA | | NA | NA | NA |
| WBGene00044956 | | -0.003614866 | 0.95700117 | NA | NA |
| WBGene00044957 | | 0.020232312 | 0.84643798 | NA | NA |
| WBGene00044958 | NA | | NA | NA | NA |

| | | | | | |
|---|---|---|---|---|---|
| WBGene00044960 | | 0.020232312 | 0.84643798 | NA | NA |
| WBGene00044961 | | 0.020232312 | 0.84643798 | NA | NA |
| WBGene00044962 | | -0.023056499 | 0.78121253 | NA | NA |
| WBGene00044963 | NA | | NA | NA | NA |
| WBGene00044964 | NA | | NA | NA | NA |
| WBGene00044965 | | -0.022930767 | 0.78121253 | NA | NA |
| WBGene00044966 | NA | | NA | NA | NA |
| WBGene00044967 | NA | | NA | NA | NA |
| WBGene00044968 | NA | | NA | NA | NA |
| WBGene00044969 | NA | | NA | NA | NA |
| WBGene00044970 | NA | | NA | NA | NA |
| WBGene00044971 | NA | | NA | NA | NA |
| WBGene00044972 | | -0.023109762 | 0.78121253 | NA | NA |
| WBGene00044973 | | 0.020232312 | 0.84643798 | NA | NA |
| WBGene00044974 | NA | | NA | NA | NA |
| WBGene00044975 | NA | | NA | NA | NA |
| WBGene00044976 | | -0.540151029 | 0.04622251 | 0.14214188 | NA |
| WBGene00044977 | | 0.232811011 | 0.20171699 | 0.39039503 | F56C4.4 |
| WBGene00044978 | | -0.227520763 | 0.11240567 | 0.26542401 | F58D2.3 |
| WBGene00044979 | | -0.003614866 | 0.95700117 | NA | Y41E3.19 |
| WBGene00044980 | NA | | NA | NA | NA |
| WBGene00044981 | | -0.023109762 | 0.78121253 | NA | NA |
| WBGene00044982 | NA | | NA | NA | NA |
| WBGene00044983 | NA | | NA | NA | NA |
| WBGene00044984 | NA | | NA | NA | NA |
| WBGene00044985 | | 0.020925399 | 0.84643798 | NA | NA |
| WBGene00044986 | NA | | NA | NA | NA |
| WBGene00044987 | | -0.148423409 | 0.18209116 | 0.36488466 | K07A1.17 |
| WBGene00044988 | | -0.511148549 | 0.05500586 | 0.16141321 | W01A8.8 |
| WBGene00044989 | | 0.024298218 | 0.83953481 | 0.91312898 | Y37A1B.17 |
| WBGene00044990 | | 0.010181931 | 0.947008 | 0.97031324 | T25B9.11 |
| WBGene00044991 | | 0.032016602 | 0.77614564 | 0.87325354 | T25B9.12 |
| WBGene00044995 | | -0.022930767 | 0.78121253 | NA | NA |
| WBGene00044997 | NA | | NA | NA | NA |
| WBGene00044998 | | -0.141080065 | 0.39277257 | 0.59095501 | M199.9 |
| WBGene00045000 | NA | | NA | NA | NA |
| WBGene00045002 | NA | | NA | NA | NA |
| WBGene00045003 | NA | | NA | NA | NA |
| WBGene00045006 | NA | | NA | NA | NA |
| WBGene00045007 | NA | | NA | NA | NA |
| WBGene00045009 | NA | | NA | NA | NA |
| WBGene00045011 | NA | | NA | NA | NA |
| WBGene00045012 | NA | | NA | NA | NA |

| | | | | |
|---|---|---|---|---|
| WBGene00045013 | 0.416044759 | 0.10161405 | 0.24816202 | F08A7.1 |
| WBGene00045016 | NA | NA | NA | NA |
| WBGene00045017 | NA | NA | NA | NA |
| WBGene00045018 | NA | NA | NA | NA |
| WBGene00045019 | NA | NA | NA | NA |
| WBGene00045022 | NA | NA | NA | NA |
| WBGene00045024 | 0.020317245 | 0.84643798 | NA | C43F9.11 |
| WBGene00045025 | NA | NA | NA | NA |
| WBGene00045026 | NA | NA | NA | NA |
| WBGene00045027 | NA | NA | NA | NA |
| WBGene00045028 | NA | NA | NA | NA |
| WBGene00045029 | NA | NA | NA | NA |
| WBGene00045036 | 0.033061864 | 0.76892385 | 0.86844708 | K08E4.7 |
| WBGene00045037 | NA | NA | NA | NA |
| WBGene00045038 | 0.440769816 | 1.10E-07 | 2.06E-06 | ule-2 |
| WBGene00045040 | NA | NA | NA | NA |
| WBGene00045043 | 0.165585509 | 0.3481695 | 0.54812362 | NA |
| WBGene00045045 | NA | NA | NA | NA |
| WBGene00045046 | NA | NA | NA | NA |
| WBGene00045048 | -0.320417299 | 0.07728811 | 0.20570801 | D1007.19 |
| WBGene00045050 | 0.077039485 | 0.57968921 | 0.74527834 | ZC15.10 |
| WBGene00045051 | NA | NA | NA | NA |
| WBGene00045052 | 1.61695132 | 0.00121656 | 0.00827292 | K12C11.6 |
| WBGene00045053 | 0.357535191 | 0.1349316 | 0.29932326 | K12C11.7 |
| WBGene00045055 | NA | NA | NA | NA |
| WBGene00045056 | 0.111583235 | 0.4466794 | 0.63996801 | F13H10.9 |
| WBGene00045057 | NA | NA | NA | NA |
| WBGene00045058 | -0.79879624 | 0.00659286 | 0.03251129 | gadr-5 |
| WBGene00045059 | NA | NA | NA | NA |
| WBGene00045060 | 0.047133413 | 0.70006286 | 0.82952576 | F55B11.7 |
| WBGene00045061 | 0.102989984 | 0.32666778 | 0.52574947 | Y57G11B.8 |
| WBGene00045062 | -0.001480842 | 0.98929087 | 0.99398816 | NA |
| WBGene00045063 | 0.093996533 | 0.61550527 | 0.7693173 | Y37A1A.4 |
| WBGene00045065 | NA | NA | NA | NA |
| WBGene00045066 | NA | NA | NA | NA |
| WBGene00045067 | -0.158154199 | 0.26122486 | 0.45924511 | F07C6.6 |
| WBGene00045068 | NA | NA | NA | NA |
| WBGene00045069 | NA | NA | NA | C08F11.17 |
| WBGene00045071 | NA | NA | NA | NA |
| WBGene00045072 | NA | NA | NA | NA |
| WBGene00045074 | 0.0398788 | 0.58971221 | NA | NA |
| WBGene00045075 | NA | NA | NA | NA |
| WBGene00045076 | 0.020317245 | 0.84643798 | NA | NA |

| | | | | |
|---|---|---|---|---|
| WBGene00045077 | -0.051353384 | 0.78626117 | 0.87895318 | NA |
| WBGene00045078 | -0.321780002 | 0.1080477 | 0.25852695 | C46C2.7 |
| WBGene00045082 | -0.133034304 | 0.22186138 | 0.41453195 | NA |
| WBGene00045083 | NA | NA | NA | NA |
| WBGene00045084 | NA | NA | NA | NA |
| WBGene00045085 | NA | NA | NA | NA |
| WBGene00045086 | NA | NA | NA | NA |
| WBGene00045087 | NA | NA | NA | NA |
| WBGene00045088 | 0.0398788 | 0.58971221 | NA | NA |
| WBGene00045089 | NA | NA | NA | NA |
| WBGene00045090 | NA | NA | NA | NA |
| WBGene00045091 | NA | NA | NA | NA |
| WBGene00045092 | NA | NA | NA | NA |
| WBGene00045093 | NA | NA | NA | NA |
| WBGene00045094 | NA | NA | NA | NA |
| WBGene00045095 | -0.004244673 | 0.95303937 | NA | NA |
| WBGene00045096 | 0.038125098 | 0.61119482 | NA | K01A6.7 |
| WBGene00045097 | -0.023109762 | 0.78121253 | NA | NA |
| WBGene00045098 | -0.022930767 | 0.78121253 | NA | NA |
| WBGene00045099 | -0.023056499 | 0.78121253 | NA | NA |
| WBGene00045100 | NA | NA | NA | NA |
| WBGene00045101 | 0.020317245 | 0.84643798 | NA | C35D6.13 |
| WBGene00045102 | NA | NA | NA | NA |
| WBGene00045103 | NA | NA | NA | T09E11.12 |
| WBGene00045104 | NA | NA | NA | NA |
| WBGene00045105 | NA | NA | NA | NA |
| WBGene00045106 | NA | NA | NA | NA |
| WBGene00045107 | NA | NA | NA | NA |
| WBGene00045108 | 0.020232312 | 0.84643798 | NA | H01G02.4 |
| WBGene00045109 | 0.065014541 | 0.37399213 | NA | NA |
| WBGene00045110 | NA | NA | NA | NA |
| WBGene00045111 | NA | NA | NA | NA |
| WBGene00045112 | NA | NA | NA | NA |
| WBGene00045113 | 0.020925399 | 0.84643798 | NA | NA |
| WBGene00045114 | NA | NA | NA | NA |
| WBGene00045115 | NA | NA | NA | NA |
| WBGene00045116 | NA | NA | NA | NA |
| WBGene00045117 | NA | NA | NA | NA |
| WBGene00045118 | NA | NA | NA | NA |
| WBGene00045119 | NA | NA | NA | NA |
| WBGene00045120 | -0.008657085 | 0.92287164 | NA | NA |
| WBGene00045121 | 0.020925399 | 0.84643798 | NA | NA |
| WBGene00045122 | NA | NA | NA | F22B3.10 |

| | | | | | |
|---|---|---|---|---|---|
| WBGene00045123 | | -0.022930767 | 0.78121253 | NA | NA |
| WBGene00045124 | | 0.020925399 | 0.84643798 | NA | NA |
| WBGene00045125 | | 0.020925399 | 0.84643798 | NA | NA |
| WBGene00045126 | NA | | NA | NA | NA |
| WBGene00045127 | NA | | NA | NA | NA |
| WBGene00045128 | NA | | NA | NA | Y116A8C.49 |
| WBGene00045129 | NA | | NA | NA | NA |
| WBGene00045130 | | 0.020317245 | 0.84643798 | NA | NA |
| WBGene00045131 | NA | | NA | NA | NA |
| WBGene00045132 | | 0.020317245 | 0.84643798 | NA | NA |
| WBGene00045133 | NA | | NA | NA | NA |
| WBGene00045134 | NA | | NA | NA | NA |
| WBGene00045136 | NA | | NA | NA | NA |
| WBGene00045137 | | -0.023109762 | 0.78121253 | NA | NA |
| WBGene00045138 | | 0.01052236 | 0.92318938 | 0.95651506 | NA |
| WBGene00045139 | NA | | NA | NA | NA |
| WBGene00045140 | NA | | NA | NA | NA |
| WBGene00045141 | NA | | NA | NA | NA |
| WBGene00045142 | NA | | NA | NA | NA |
| WBGene00045143 | NA | | NA | NA | NA |
| WBGene00045144 | NA | | NA | NA | Y64G10A.10 |
| WBGene00045145 | NA | | NA | NA | NA |
| WBGene00045146 | | 0.194715235 | 0.15908078 | 0.3333956 | Y41E3.22 |
| WBGene00045147 | NA | | NA | NA | NA |
| WBGene00045148 | | 0.036687355 | 0.59489568 | NA | NA |
| WBGene00045149 | NA | | NA | NA | NA |
| WBGene00045150 | | 0.020317245 | 0.84643798 | NA | NA |
| WBGene00045151 | NA | | NA | NA | NA |
| WBGene00045152 | NA | | NA | NA | NA |
| WBGene00045153 | NA | | NA | NA | NA |
| WBGene00045154 | NA | | NA | NA | NA |
| WBGene00045155 | NA | | NA | NA | NA |
| WBGene00045156 | NA | | NA | NA | NA |
| WBGene00045157 | NA | | NA | NA | NA |
| WBGene00045158 | NA | | NA | NA | NA |
| WBGene00045159 | NA | | NA | NA | NA |
| WBGene00045160 | NA | | NA | NA | NA |
| WBGene00045161 | | -0.022930767 | 0.78121253 | NA | NA |
| WBGene00045162 | | 0.059762547 | 0.38965825 | NA | NA |
| WBGene00045163 | | -0.007072416 | 0.9244612 | NA | NA |
| WBGene00045164 | NA | | NA | NA | NA |
| WBGene00045165 | NA | | NA | NA | NA |
| WBGene00045167 | NA | | NA | NA | NA |

| | | | | |
|---|---|---|---|---|
| WBGene00045168 | NA | NA | NA | NA |
| WBGene00045169 | NA | NA | NA | NA |
| WBGene00045170 | NA | NA | NA | NA |
| WBGene00045171 | -0.046394786 | 0.50231372 | NA | NA |
| WBGene00045172 | NA | NA | NA | NA |
| WBGene00045174 | NA | NA | NA | NA |
| WBGene00045175 | -0.397493817 | 0.03255155 | 0.10983805 | NA |
| WBGene00045176 | NA | NA | NA | NA |
| WBGene00045177 | 0.244759531 | 0.08243182 | 0.21537824 | T21D12.14 |
| WBGene00045178 | NA | NA | NA | fipr-27 |
| WBGene00045179 | NA | NA | NA | NA |
| WBGene00045180 | 0.038125098 | 0.61119482 | NA | K07F5.17 |
| WBGene00045182 | NA | NA | NA | NA |
| WBGene00045183 | -0.035762513 | 0.77595522 | 0.87322248 | T28C6.10 |
| WBGene00045184 | NA | NA | NA | NA |
| WBGene00045185 | NA | NA | NA | NA |
| WBGene00045188 | -0.003421551 | 0.95689871 | NA | NA |
| WBGene00045189 | NA | NA | NA | NA |
| WBGene00045190 | -0.00346345 | 0.95708614 | NA | R01H2.8 |
| WBGene00045192 | -0.251427076 | 0.00095154 | 0.00675303 | aho-3 |
| WBGene00045193 | NA | NA | NA | NA |
| WBGene00045194 | 0.055519824 | 0.53935207 | NA | NA |
| WBGene00045195 | -0.015946848 | 0.89462076 | 0.94489559 | F53F4.17 |
| WBGene00045197 | NA | NA | NA | NA |
| WBGene00045198 | -0.001488154 | 0.95639912 | NA | K10D11.7 |
| WBGene00045199 | NA | NA | NA | NA |
| WBGene00045200 | -0.023109762 | 0.78121253 | NA | NA |
| WBGene00045201 | NA | NA | NA | NA |
| WBGene00045202 | 0.020317245 | 0.84643798 | NA | NA |
| WBGene00045203 | NA | NA | NA | NA |
| WBGene00045204 | NA | NA | NA | NA |
| WBGene00045205 | NA | NA | NA | NA |
| WBGene00045207 | 0.083078564 | 0.30763554 | NA | F13E9.14 |
| WBGene00045208 | 1.381939786 | 7.83E-05 | 0.00077916 | F13E9.15 |
| WBGene00045209 | NA | NA | NA | F13E9.16 |
| WBGene00045210 | 0.122833479 | 0.25726816 | 0.45540601 | C54D2.6 |
| WBGene00045211 | NA | NA | NA | C18D11.9 |
| WBGene00045212 | 0.183002387 | 0.34983119 | 0.54961674 | Y110A7A.21 |
| WBGene00045215 | 0.146558499 | 0.28429772 | 0.4842819 | ceh-63 |
| WBGene00045218 | 0.23830156 | 0.26429627 | 0.46264487 | NA |
| WBGene00045234 | NA | NA | NA | Y26D4A.17 |
| WBGene00045235 | NA | NA | NA | NA |
| WBGene00045236 | NA | NA | NA | NA |

| | | | | |
|---|---|---|---|---|
| WBGene00045237 | 0.172806247 | 0.189681 | 0.37543862 | mrt-1 |
| WBGene00045238 | NA | NA | NA | NA |
| WBGene00045239 | -0.284605643 | 0.18912664 | 0.37454392 | Y49E10.29 |
| WBGene00045242 | 0.033463284 | 0.6365037 | NA | C05B5.11 |
| WBGene00045245 | -0.04864578 | 0.79360873 | 0.88399982 | ttr-34 |
| WBGene00045246 | 0.087909695 | 0.67536253 | 0.81135495 | C29E4.15 |
| WBGene00045247 | 0.12863238 | 0.24937397 | 0.447792 | F54H12.8 |
| WBGene00045248 | 0.030572857 | 0.77880322 | 0.87482414 | ZK180.7 |
| WBGene00045249 | -0.044083573 | 0.81987693 | 0.9011199 | T07A9.15 |
| WBGene00045250 | 0.055519824 | 0.53935207 | NA | Y73B6BL.47 |
| WBGene00045251 | -0.011662311 | 0.90617376 | 0.94916953 | F54F7.9 |
| WBGene00045252 | -0.303255269 | 0.13732385 | 0.30297747 | Y62H9A.15 |
| WBGene00045253 | 0.244642537 | 0.23142639 | 0.42641754 | F46C3.6 |
| WBGene00045254 | -0.002914971 | 0.95718083 | NA | NA |
| WBGene00045255 | 0.159627108 | 0.45425282 | 0.64633812 | nhr-146 |
| WBGene00045261 | -0.431011932 | 0.00196365 | 0.01231059 | H29C22.1 |
| WBGene00045265 | -2.197673614 | 2.69E-15 | 1.32E-13 | K10C2.8 |
| WBGene00045268 | -0.434960183 | 0.0005581 | 0.00425803 | ttr-42 |
| WBGene00045269 | 0.015062379 | 0.80838968 | NA | F46G11.6 |
| WBGene00045270 | 0.156387729 | 0.3099499 | 0.51074581 | C04E7.5 |
| WBGene00045271 | 0.084263254 | 0.68685273 | 0.81957708 | F07G6.10 |
| WBGene00045272 | -0.331998983 | 0.03700025 | 0.12082912 | F59C12.4 |
| WBGene00045273 | 0.09117635 | 0.59948228 | 0.75735785 | NA |
| WBGene00045275 | -2.950257162 | 9.93E-12 | 3.30E-10 | F18E3.12 |
| WBGene00045276 | -1.337441675 | 1.67E-06 | 2.48E-05 | F18E3.13 |
| WBGene00045277 | 0.03187317 | 0.79154866 | 0.88250563 | ZK742.6 |
| WBGene00045278 | NA | NA | NA | NA |
| WBGene00045279 | NA | NA | NA | NA |
| WBGene00045280 | NA | NA | NA | NA |
| WBGene00045281 | NA | NA | NA | NA |
| WBGene00045282 | 0.020317245 | 0.84643798 | NA | NA |
| WBGene00045283 | NA | NA | NA | K08E4.8 |
| WBGene00045285 | NA | NA | NA | NA |
| WBGene00045286 | NA | NA | NA | NA |
| WBGene00045287 | NA | NA | NA | NA |
| WBGene00045289 | 0.016616329 | 0.9055977 | 0.94916953 | fbxa-169 |
| WBGene00045290 | NA | NA | NA | NA |
| WBGene00045291 | -0.11305787 | 0.58722657 | 0.74981371 | F23D12.10 |
| WBGene00045292 | -0.093696363 | 0.55284606 | 0.72508974 | fbxa-179 |
| WBGene00045293 | 0.257916091 | 0.17324814 | 0.35347679 | NA |
| WBGene00045294 | NA | NA | NA | NA |
| WBGene00045295 | 0.056411843 | 0.41891328 | NA | fbxa-222 |
| WBGene00045297 | 1.516869258 | 0.00030589 | 0.00256062 | fbxc-43 |

| | | | | | |
|---|---|---|---|---|---|
| WBGene00045298 | | 0.020317245 | 0.84643798 | NA | fbxc-12 |
| WBGene00045299 | NA | | NA | NA | fbxc-11 |
| WBGene00045299 | NA | | NA | NA | fbxc-10 |
| WBGene00045299 | NA | | NA | NA | fbxc-9 |
| WBGene00045300 | NA | | NA | NA | fbxc-11 |
| WBGene00045300 | NA | | NA | NA | fbxc-10 |
| WBGene00045300 | NA | | NA | NA | fbxc-9 |
| WBGene00045301 | NA | | NA | NA | fbxc-11 |
| WBGene00045301 | NA | | NA | NA | fbxc-10 |
| WBGene00045301 | NA | | NA | NA | fbxc-9 |
| WBGene00045302 | | -0.104347022 | 0.29885287 | 0.49942984 | fbxc-30 |
| WBGene00045303 | NA | | NA | NA | Y27F2A.10 |
| WBGene00045304 | | -0.001842784 | 0.95621055 | NA | Y27F2A.11 |
| WBGene00045305 | | -1.429819356 | 9.91E-09 | 2.22E-07 | ZC250.4 |
| WBGene00045306 | | -0.101609559 | 0.22055508 | NA | ZC250.5 |
| WBGene00045308 | | -0.050728434 | 0.78515522 | 0.87870478 | C37H5.14 |
| WBGene00045309 | NA | | NA | NA | NA |
| WBGene00045311 | | -0.050223056 | 0.47776088 | NA | NA |
| WBGene00045313 | NA | | NA | NA | NA |
| WBGene00045314 | | -0.023056499 | 0.78121253 | NA | NA |
| WBGene00045315 | NA | | NA | NA | NA |
| WBGene00045316 | | -0.021600209 | 0.91516461 | 0.95281497 | NA |
| WBGene00045319 | NA | | NA | NA | NA |
| WBGene00045321 | | 0.020232312 | 0.84643798 | NA | NA |
| WBGene00045322 | | 0.033034883 | 0.64219503 | NA | NA |
| WBGene00045323 | NA | | NA | NA | NA |
| WBGene00045324 | NA | | NA | NA | NA |
| WBGene00045325 | NA | | NA | NA | NA |
| WBGene00045326 | NA | | NA | NA | NA |
| WBGene00045327 | NA | | NA | NA | NA |
| WBGene00045328 | NA | | NA | NA | NA |
| WBGene00045329 | NA | | NA | NA | NA |
| WBGene00045330 | NA | | NA | NA | NA |
| WBGene00045331 | | -0.031119177 | 0.73352595 | NA | F14D7.12 |
| WBGene00045332 | NA | | NA | NA | NA |
| WBGene00045336 | | 0.085488282 | 0.61466334 | 0.7687755 | R11D1.12 |
| WBGene00045337 | NA | | NA | NA | M01B2.12 |
| WBGene00045338 | | 1.718424669 | 0.00973231 | 0.04399218 | M01B2.13 |
| WBGene00045339 | | 0.020925399 | 0.84643798 | NA | T10H4.13 |
| WBGene00045340 | | 0.122326112 | 0.30956314 | 0.51040962 | T10G3.8 |
| WBGene00045341 | NA | | NA | NA | NA |
| WBGene00045342 | NA | | NA | NA | NA |
| WBGene00045343 | NA | | NA | NA | NA |

| | | | | |
|---|---|---|---|---|
| WBGene00045344 | NA | NA | NA | Y43F8C.23 |
| WBGene00045345 | NA | NA | NA | NA |
| WBGene00045346 | NA | NA | NA | NA |
| WBGene00045347 | NA | NA | NA | NA |
| WBGene00045348 | NA | NA | NA | NA |
| WBGene00045350 | NA | NA | NA | NA |
| WBGene00045351 | NA | NA | NA | NA |
| WBGene00045352 | NA | NA | NA | NA |
| WBGene00045353 | NA | NA | NA | NA |
| WBGene00045354 | NA | NA | NA | NA |
| WBGene00045355 | 0.310899088 | 0.11769847 | 0.27435492 | D1086.17 |
| WBGene00045356 | -0.003108065 | 0.95728506 | NA | T03E6.9 |
| WBGene00045357 | -0.071049844 | 0.53976591 | 0.71445958 | F13A7.14 |
| WBGene00045358 | NA | NA | NA | NA |
| WBGene00045359 | NA | NA | NA | NA |
| WBGene00045360 | NA | NA | NA | NA |
| WBGene00045361 | 0.033034883 | 0.64219503 | NA | clec-253 |
| WBGene00045362 | 0.020925399 | 0.84643798 | NA | F11D11.12 |
| WBGene00045363 | NA | NA | NA | NA |
| WBGene00045364 | NA | NA | NA | clec-250 |
| WBGene00045365 | -0.347099556 | 0.1423903 | 0.31058875 | ZC449.7 |
| WBGene00045366 | 0.088716647 | 0.63709579 | 0.78483125 | NA |
| WBGene00045367 | -0.001993815 | 0.95612996 | NA | NA |
| WBGene00045368 | NA | NA | NA | NA |
| WBGene00045369 | NA | NA | NA | NA |
| WBGene00045370 | NA | NA | NA | NA |
| WBGene00045371 | NA | NA | NA | NA |
| WBGene00045372 | NA | NA | NA | NA |
| WBGene00045373 | NA | NA | NA | NA |
| WBGene00045374 | NA | NA | NA | NA |
| WBGene00045375 | -0.003108065 | 0.95728506 | NA | NA |
| WBGene00045376 | NA | NA | NA | NA |
| WBGene00045377 | NA | NA | NA | NA |
| WBGene00045378 | NA | NA | NA | NA |
| WBGene00045379 | 0.298661451 | 0.05846993 | 0.16841593 | C45B11.6 |
| WBGene00045380 | NA | NA | NA | W06G6.15 |
| WBGene00045381 | NA | NA | NA | F28B1.9 |
| WBGene00045382 | 0.172240472 | 0.32416859 | 0.52363682 | T05E12.8 |
| WBGene00045383 | -0.306052314 | 0.00010466 | 0.00101145 | sup-46 |
| WBGene00045385 | 0.105191732 | 0.16565281 | NA | lgc-24 |
| WBGene00045386 | 0.144165731 | 0.28426153 | 0.48427659 | nlp-76 |
| WBGene00045387 | 0.746761718 | 0.01368031 | 0.05746165 | K09H9.8 |
| WBGene00045388 | 0.036687355 | 0.59489568 | NA | NA |

| | | | | |
|---|---|---|---|---|
| WBGene00045389 | NA | NA | NA | NA |
| WBGene00045390 | -0.071081044 | 0.65515705 | 0.79589731 | C38C3.10 |
| WBGene00045391 | NA | NA | NA | NA |
| WBGene00045392 | 0.207946895 | 0.2870265 | 0.48717308 | F26D11.12 |
| WBGene00045393 | 0.056411843 | 0.41891328 | NA | F26D11.13 |
| WBGene00045394 | 0.483363098 | 5.08E-15 | 2.43E-13 | ZK813.7 |
| WBGene00045395 | -0.001993815 | 0.95612996 | NA | F09F9.5 |
| WBGene00045396 | 0.038125098 | 0.61119482 | NA | ZC334.12 |
| WBGene00045397 | 0.505118162 | 0.05639518 | 0.164544 | Y54G2A.52 |
| WBGene00045398 | NA | NA | NA | Y71F9AM.8 |
| WBGene00045399 | -0.799764893 | 4.76E-09 | 1.12E-07 | Y47G6A.33 |
| WBGene00045400 | 0.192190606 | 0.30656765 | 0.50711327 | C54D10.13 |
| WBGene00045401 | 1.143593301 | 0.0164092 | 0.06644002 | eol-1 |
| WBGene00045402 | NA | NA | NA | Y75B12B.13 |
| WBGene00045403 | 0.52783243 | 0.0338677 | 0.11315768 | K10H10.12 |
| WBGene00045404 | -0.007241538 | 0.96112444 | 0.97890942 | NA |
| WBGene00045405 | 0.950083873 | 0.01790694 | 0.07087742 | D1086.18 |
| WBGene00045406 | -0.109699439 | 0.50816657 | 0.68864002 | D1086.19 |
| WBGene00045407 | -0.262182998 | 0.06146869 | 0.17468228 | T07D4.5 |
| WBGene00045408 | 0.038125098 | 0.61119482 | NA | NA |
| WBGene00045410 | 0.839994316 | 0.00023406 | 0.0020182 | NA |
| WBGene00045411 | 3.978314734 | 0.00030942 | 0.00258423 | C25F9.11 |
| WBGene00045412 | 0.336219997 | 0.00672504 | 0.03302911 | C25F9.12 |
| WBGene00045413 | 0.020317245 | 0.84643798 | NA | C25F9.13 |
| WBGene00045414 | 0.011191389 | 0.90553779 | NA | T19H5.7 |
| WBGene00045415 | 0.127758703 | 0.16886741 | NA | Y43F8B.15 |
| WBGene00045416 | 3.164166749 | 1.31E-10 | 3.68E-09 | Y37H2A.14 |
| WBGene00045417 | 4.800353348 | 2.18E-05 | 0.00024672 | Y37H2A.13 |
| WBGene00045418 | -0.005447915 | 0.96223306 | 0.97924853 | F15B9.10 |
| WBGene00045419 | 0.024968169 | 0.80388827 | 0.89090365 | lsy-12 |
| WBGene00045433 | -0.499492706 | 0.00530258 | 0.02740385 | F49D11.10 |
| WBGene00045434 | -0.21385749 | 0.08171765 | 0.2139918 | Y95D11A.3 |
| WBGene00045455 | 0.038911608 | 0.73676145 | 0.85204655 | F26G1.11 |
| WBGene00045456 | 0.226194845 | 0.05419888 | 0.15963102 | C10A4.9 |
| WBGene00045457 | 0.379629012 | 0.03829531 | 0.12390755 | F33H12.7 |
| WBGene00045458 | -0.016082418 | 0.89371347 | 0.94468695 | F17B5.8 |
| WBGene00045459 | NA | NA | NA | Y59A8B.26 |
| WBGene00045460 | 0.040107227 | 0.58693163 | NA | C15C8.8 |
| WBGene00045461 | 1.487924982 | 0.00078415 | 0.00570614 | C25F9.14 |
| WBGene00045468 | 0.152593967 | 0.27564702 | 0.47574538 | F08A8.8 |
| WBGene00045473 | -0.166748323 | 0.40528089 | 0.60332664 | F30A10.14 |
| WBGene00045474 | 0.093904075 | 0.4036632 | 0.60202027 | C49G9.2 |
| WBGene00045475 | 0.392895022 | 0.09726235 | 0.24139991 | T07A5.7 |

| | | | | | |
|---|---|---|---|---|---|
| WBGene00045480 | | -0.015100905 | 0.892162 | 0.94374205 | F14F3.5 |
| WBGene00045481 | | 0.038125098 | 0.61119482 | NA | M153.5 |
| WBGene00045482 | | 0.133358339 | 0.34036019 | 0.5401272 | T03F6.9 |
| WBGene00045483 | | -0.223308382 | 0.02331257 | 0.08590971 | oxy-5 |
| WBGene00045484 | | 0.127602834 | 0.50303917 | 0.68435203 | F34D10.9 |
| WBGene00045486 | | -0.084578794 | 0.55650697 | 0.72797554 | K05F6.12 |
| WBGene00045487 | NA | | NA | NA | NA |
| WBGene00045488 | | -0.4008533 | 0.10969278 | 0.26095947 | F57B1.9 |
| WBGene00045490 | NA | | NA | NA | F14D7.14 |
| WBGene00045491 | NA | | NA | NA | NA |
| WBGene00045492 | NA | | NA | NA | NA |
| WBGene00045493 | | 0.084908438 | 0.30232871 | NA | F55G7.4 |
| WBGene00045494 | | -0.004968869 | 0.94188684 | NA | ZK662.6 |
| WBGene00045495 | | -0.096825462 | 0.56568386 | 0.73508275 | F11C1.7 |
| WBGene00045496 | NA | | NA | NA | NA |
| WBGene00045497 | | -0.045771477 | 0.50913401 | NA | K03A11.6 |
| WBGene00045498 | | -0.508975999 | 0.0420953 | 0.13262743 | F56F4.8 |
| WBGene00045500 | | -0.027373974 | 0.68245543 | NA | T05E12.9 |
| WBGene00045501 | NA | | NA | NA | Y6G8.8 |
| WBGene00045502 | NA | | NA | NA | NA |
| WBGene00045506 | NA | | NA | NA | F57G4.10 |
| WBGene00045507 | | 0.137423788 | 0.4418591 | 0.63595638 | F10A3.17 |
| WBGene00045508 | NA | | NA | NA | D1081.10 |
| WBGene00045509 | | -0.085871677 | 0.6684478 | 0.80655548 | NA |
| WBGene00045510 | | 0.033034883 | 0.64219503 | NA | F11A5.17 |
| WBGene00045511 | | -0.06247386 | 0.60508778 | 0.7614173 | F11A5.18 |
| WBGene00045513 | | 0.020232312 | 0.84643798 | NA | F14H3.14 |
| WBGene00045514 | | 0.020232312 | 0.84643798 | NA | NA |
| WBGene00045515 | NA | | NA | NA | ZK1037.13 |
| WBGene00045516 | | 0.201254814 | 0.30616095 | 0.50686869 | F43D2.7 |
| WBGene00045517 | | 0.065254007 | 0.40163332 | NA | R11D1.13 |
| WBGene00045518 | NA | | NA | NA | NA |
| WBGene00045519 | NA | | NA | NA | NA |
| WBGene00045520 | NA | | NA | NA | NA |
| WBGene00045521 | NA | | NA | NA | NA |
| WBGene00045522 | NA | | NA | NA | NA |
| WBGene00045523 | NA | | NA | NA | NA |
| WBGene00045524 | NA | | NA | NA | NA |
| WBGene00045525 | NA | | NA | NA | NA |
| WBGene00045526 | NA | | NA | NA | NA |
| WBGene00045527 | NA | | NA | NA | NA |
| WBGene00045528 | NA | | NA | NA | NA |
| WBGene00045529 | NA | | NA | NA | NA |

| | | | | |
|---|---|---|---|---|
| WBGene00045530 | NA | NA | NA | NA |
| WBGene00045531 | NA | NA | NA | NA |
| WBGene00045532 | NA | NA | NA | NA |
| WBGene00045533 | NA | NA | NA | NA |
| WBGene00045534 | NA | NA | NA | NA |
| WBGene00045535 | NA | NA | NA | NA |
| WBGene00045536 | NA | NA | NA | NA |
| WBGene00045537 | NA | NA | NA | NA |
| WBGene00045538 | NA | NA | NA | NA |
| WBGene00045539 | NA | NA | NA | NA |
| WBGene00045540 | NA | NA | NA | NA |
| WBGene00045541 | NA | NA | NA | NA |
| WBGene00045542 | NA | NA | NA | NA |
| WBGene00045543 | NA | NA | NA | NA |
| WBGene00045544 | NA | NA | NA | NA |
| WBGene00045545 | NA | NA | NA | NA |
| WBGene00045546 | NA | NA | NA | NA |
| WBGene00045547 | NA | NA | NA | NA |
| WBGene00045548 | NA | NA | NA | NA |
| WBGene00045549 | NA | NA | NA | NA |
| WBGene00045550 | NA | NA | NA | NA |
| WBGene00045551 | NA | NA | NA | NA |
| WBGene00045552 | NA | NA | NA | NA |
| WBGene00045553 | NA | NA | NA | NA |
| WBGene00045554 | NA | NA | NA | NA |
| WBGene00045555 | NA | NA | NA | NA |
| WBGene00045556 | NA | NA | NA | NA |
| WBGene00045557 | NA | NA | NA | NA |
| WBGene00045558 | NA | NA | NA | NA |
| WBGene00045559 | NA | NA | NA | NA |
| WBGene00045560 | NA | NA | NA | NA |
| WBGene00045561 | NA | NA | NA | NA |
| WBGene00045562 | NA | NA | NA | NA |
| WBGene00045563 | NA | NA | NA | NA |
| WBGene00045564 | NA | NA | NA | NA |
| WBGene00045565 | NA | NA | NA | NA |
| WBGene00045566 | NA | NA | NA | NA |
| WBGene00045567 | NA | NA | NA | NA |
| WBGene00045568 | NA | NA | NA | NA |
| WBGene00045569 | NA | NA | NA | NA |
| WBGene00045570 | NA | NA | NA | NA |
| WBGene00045571 | NA | NA | NA | NA |
| WBGene00045572 | NA | NA | NA | NA |

| | | | | |
|---|---|---|---|---|
| WBGene00045573 | NA | NA | NA | NA |
| WBGene00045574 | NA | NA | NA | NA |
| WBGene00045575 | NA | NA | NA | NA |
| WBGene00045576 | NA | NA | NA | NA |
| WBGene00045577 | NA | NA | NA | NA |
| WBGene00045578 | NA | NA | NA | NA |
| WBGene00045579 | NA | NA | NA | NA |
| WBGene00045580 | NA | NA | NA | NA |
| WBGene00045581 | NA | NA | NA | NA |
| WBGene00045582 | NA | NA | NA | NA |
| WBGene00045583 | NA | NA | NA | NA |
| WBGene00045584 | NA | NA | NA | NA |
| WBGene00045585 | NA | NA | NA | NA |
| WBGene00045586 | NA | NA | NA | NA |
| WBGene00045587 | NA | NA | NA | NA |
| WBGene00045588 | NA | NA | NA | NA |
| WBGene00045589 | NA | NA | NA | NA |
| WBGene00045590 | NA | NA | NA | NA |
| WBGene00045591 | NA | NA | NA | NA |
| WBGene00045592 | 0.020232312 | 0.84643798 | NA | NA |
| WBGene00045593 | NA | NA | NA | NA |
| WBGene00045594 | NA | NA | NA | NA |
| WBGene00045595 | NA | NA | NA | NA |
| WBGene00045596 | NA | NA | NA | NA |
| WBGene00045597 | NA | NA | NA | NA |
| WBGene00045598 | NA | NA | NA | NA |
| WBGene00045599 | NA | NA | NA | NA |
| WBGene00045600 | NA | NA | NA | NA |
| WBGene00045601 | NA | NA | NA | NA |
| WBGene00045602 | NA | NA | NA | NA |
| WBGene00045603 | NA | NA | NA | NA |
| WBGene00045604 | NA | NA | NA | NA |
| WBGene00045605 | NA | NA | NA | NA |
| WBGene00045606 | NA | NA | NA | NA |
| WBGene00045607 | -0.022930767 | 0.78121253 | NA | NA |
| WBGene00045608 | NA | NA | NA | NA |
| WBGene00045609 | NA | NA | NA | NA |
| WBGene00045610 | NA | NA | NA | NA |
| WBGene00045611 | NA | NA | NA | NA |
| WBGene00045612 | NA | NA | NA | NA |
| WBGene00045613 | NA | NA | NA | NA |
| WBGene00045614 | NA | NA | NA | NA |
| WBGene00045615 | NA | NA | NA | NA |

| | | | | |
|---|---|---|---|---|
| WBGene00045616 | NA | NA | NA | NA |
| WBGene00045617 | NA | NA | NA | NA |
| WBGene00045618 | NA | NA | NA | NA |
| WBGene00045619 | NA | NA | NA | NA |
| WBGene00045620 | NA | NA | NA | NA |
| WBGene00045621 | NA | NA | NA | NA |
| WBGene00045622 | NA | NA | NA | NA |
| WBGene00045623 | NA | NA | NA | NA |
| WBGene00045624 | NA | NA | NA | NA |
| WBGene00045625 | NA | NA | NA | NA |
| WBGene00045626 | NA | NA | NA | NA |
| WBGene00045627 | NA | NA | NA | NA |
| WBGene00045628 | NA | NA | NA | NA |
| WBGene00045629 | NA | NA | NA | NA |
| WBGene00045630 | NA | NA | NA | NA |
| WBGene00045631 | NA | NA | NA | NA |
| WBGene00045632 | NA | NA | NA | NA |
| WBGene00045633 | NA | NA | NA | NA |
| WBGene00045634 | NA | NA | NA | NA |
| WBGene00045635 | NA | NA | NA | NA |
| WBGene00045636 | NA | NA | NA | NA |
| WBGene00045637 | NA | NA | NA | NA |
| WBGene00045638 | NA | NA | NA | NA |
| WBGene00045639 | NA | NA | NA | NA |
| WBGene00045640 | NA | NA | NA | NA |
| WBGene00045641 | NA | NA | NA | NA |
| WBGene00045642 | NA | NA | NA | NA |
| WBGene00045643 | NA | NA | NA | NA |
| WBGene00045644 | NA | NA | NA | NA |
| WBGene00045645 | NA | NA | NA | NA |
| WBGene00045646 | NA | NA | NA | NA |
| WBGene00045647 | NA | NA | NA | NA |
| WBGene00045648 | NA | NA | NA | NA |
| WBGene00045649 | NA | NA | NA | NA |
| WBGene00045650 | NA | NA | NA | NA |
| WBGene00045651 | NA | NA | NA | NA |
| WBGene00045652 | NA | NA | NA | NA |
| WBGene00045653 | NA | NA | NA | NA |
| WBGene00045654 | NA | NA | NA | NA |
| WBGene00045655 | NA | NA | NA | NA |
| WBGene00045656 | NA | NA | NA | NA |
| WBGene00045657 | NA | NA | NA | NA |
| WBGene00045658 | NA | NA | NA | NA |

| | | | | |
|---|---|---|---|---|
| WBGene00045659 | NA | NA | NA | NA |
| WBGene00045660 | NA | NA | NA | NA |
| WBGene00045661 | NA | NA | NA | NA |
| WBGene00045662 | NA | NA | NA | NA |
| WBGene00045663 | NA | NA | NA | NA |
| WBGene00045664 | NA | NA | NA | NA |
| WBGene00045665 | NA | NA | NA | NA |
| WBGene00045666 | NA | NA | NA | NA |
| WBGene00045667 | NA | NA | NA | NA |
| WBGene00045668 | NA | NA | NA | NA |
| WBGene00045669 | NA | NA | NA | NA |
| WBGene00045670 | NA | NA | NA | NA |
| WBGene00045671 | NA | NA | NA | NA |
| WBGene00045672 | NA | NA | NA | NA |
| WBGene00045673 | NA | NA | NA | NA |
| WBGene00045674 | NA | NA | NA | NA |
| WBGene00045675 | NA | NA | NA | NA |
| WBGene00045676 | NA | NA | NA | NA |
| WBGene00045677 | NA | NA | NA | NA |
| WBGene00045678 | NA | NA | NA | NA |
| WBGene00045679 | NA | NA | NA | NA |
| WBGene00045680 | NA | NA | NA | NA |
| WBGene00045681 | NA | NA | NA | NA |
| WBGene00045682 | NA | NA | NA | NA |
| WBGene00045683 | NA | NA | NA | NA |
| WBGene00045684 | NA | NA | NA | NA |
| WBGene00045685 | NA | NA | NA | NA |
| WBGene00045686 | NA | NA | NA | NA |
| WBGene00045687 | NA | NA | NA | NA |
| WBGene00045688 | NA | NA | NA | NA |
| WBGene00045689 | NA | NA | NA | NA |
| WBGene00045690 | NA | NA | NA | NA |
| WBGene00045691 | NA | NA | NA | NA |
| WBGene00045692 | NA | NA | NA | NA |
| WBGene00045693 | NA | NA | NA | NA |
| WBGene00045694 | NA | NA | NA | NA |
| WBGene00045695 | NA | NA | NA | NA |
| WBGene00045696 | NA | NA | NA | NA |
| WBGene00045697 | NA | NA | NA | NA |
| WBGene00045698 | NA | NA | NA | NA |
| WBGene00045699 | NA | NA | NA | NA |
| WBGene00045700 | NA | NA | NA | NA |
| WBGene00045701 | NA | NA | NA | NA |

| | | | | |
|---|---|---|---|---|
| WBGene00045702 | NA | NA | NA | NA |
| WBGene00045703 | NA | NA | NA | NA |
| WBGene00045704 | NA | NA | NA | NA |
| WBGene00045705 | NA | NA | NA | NA |
| WBGene00045706 | NA | NA | NA | NA |
| WBGene00045707 | NA | NA | NA | NA |
| WBGene00045708 | NA | NA | NA | NA |
| WBGene00045709 | NA | NA | NA | NA |
| WBGene00045710 | NA | NA | NA | NA |
| WBGene00045711 | NA | NA | NA | NA |
| WBGene00045712 | NA | NA | NA | NA |
| WBGene00045713 | NA | NA | NA | NA |
| WBGene00045714 | NA | NA | NA | NA |
| WBGene00045715 | NA | NA | NA | NA |
| WBGene00045716 | NA | NA | NA | NA |
| WBGene00045717 | NA | NA | NA | NA |
| WBGene00045718 | NA | NA | NA | NA |
| WBGene00045719 | NA | NA | NA | NA |
| WBGene00045720 | NA | NA | NA | NA |
| WBGene00045721 | NA | NA | NA | NA |
| WBGene00045722 | NA | NA | NA | NA |
| WBGene00045723 | NA | NA | NA | NA |
| WBGene00045724 | NA | NA | NA | NA |
| WBGene00045725 | NA | NA | NA | NA |
| WBGene00045726 | NA | NA | NA | NA |
| WBGene00045727 | -0.002914971 | 0.95718083 | NA | NA |
| WBGene00045728 | NA | NA | NA | NA |
| WBGene00045729 | NA | NA | NA | NA |
| WBGene00045730 | NA | NA | NA | NA |
| WBGene00045731 | NA | NA | NA | NA |
| WBGene00045732 | NA | NA | NA | NA |
| WBGene00045733 | NA | NA | NA | NA |
| WBGene00045734 | NA | NA | NA | NA |
| WBGene00045735 | NA | NA | NA | NA |
| WBGene00045736 | NA | NA | NA | NA |
| WBGene00045737 | NA | NA | NA | NA |
| WBGene00045738 | NA | NA | NA | NA |
| WBGene00045739 | NA | NA | NA | NA |
| WBGene00045740 | NA | NA | NA | NA |
| WBGene00045741 | NA | NA | NA | NA |
| WBGene00045743 | NA | NA | NA | NA |
| WBGene00045744 | NA | NA | NA | NA |
| WBGene00045745 | NA | NA | NA | NA |

| | | | | |
|---|---|---|---|---|
| WBGene00045746 | NA | NA | NA | NA |
| WBGene00045747 | NA | NA | NA | NA |
| WBGene00045748 | NA | NA | NA | NA |
| WBGene00045749 | NA | NA | NA | NA |
| WBGene00045750 | NA | NA | NA | NA |
| WBGene00045751 | NA | NA | NA | NA |
| WBGene00045752 | NA | NA | NA | NA |
| WBGene00045753 | NA | NA | NA | NA |
| WBGene00045754 | NA | NA | NA | NA |
| WBGene00045755 | NA | NA | NA | NA |
| WBGene00045756 | NA | NA | NA | NA |
| WBGene00045757 | NA | NA | NA | NA |
| WBGene00045758 | NA | NA | NA | NA |
| WBGene00045759 | NA | NA | NA | NA |
| WBGene00045760 | NA | NA | NA | NA |
| WBGene00045761 | NA | NA | NA | NA |
| WBGene00045762 | NA | NA | NA | NA |
| WBGene00045763 | NA | NA | NA | NA |
| WBGene00045764 | NA | NA | NA | NA |
| WBGene00045765 | NA | NA | NA | NA |
| WBGene00045766 | NA | NA | NA | NA |
| WBGene00045767 | NA | NA | NA | NA |
| WBGene00045768 | NA | NA | NA | NA |
| WBGene00045769 | NA | NA | NA | NA |
| WBGene00045770 | NA | NA | NA | NA |
| WBGene00045771 | NA | NA | NA | NA |
| WBGene00045772 | NA | NA | NA | NA |
| WBGene00045773 | NA | NA | NA | NA |
| WBGene00045774 | NA | NA | NA | NA |
| WBGene00045775 | NA | NA | NA | NA |
| WBGene00045776 | NA | NA | NA | NA |
| WBGene00045777 | NA | NA | NA | NA |
| WBGene00045778 | NA | NA | NA | NA |
| WBGene00045779 | NA | NA | NA | NA |
| WBGene00045780 | NA | NA | NA | NA |
| WBGene00045781 | NA | NA | NA | NA |
| WBGene00045782 | NA | NA | NA | NA |
| WBGene00045783 | NA | NA | NA | NA |
| WBGene00045784 | NA | NA | NA | NA |
| WBGene00045785 | NA | NA | NA | NA |
| WBGene00045786 | NA | NA | NA | NA |
| WBGene00045787 | NA | NA | NA | NA |
| WBGene00045788 | NA | NA | NA | NA |

| | | | | |
|---|---|---|---|---|
| WBGene00045789 | NA | NA | NA | NA |
| WBGene00045790 | NA | NA | NA | NA |
| WBGene00045791 | NA | NA | NA | NA |
| WBGene00045792 | NA | NA | NA | NA |
| WBGene00045793 | NA | NA | NA | NA |
| WBGene00045794 | NA | NA | NA | NA |
| WBGene00045795 | NA | NA | NA | NA |
| WBGene00045796 | NA | NA | NA | NA |
| WBGene00045797 | NA | NA | NA | NA |
| WBGene00045798 | NA | NA | NA | NA |
| WBGene00045799 | NA | NA | NA | NA |
| WBGene00045800 | NA | NA | NA | NA |
| WBGene00045801 | NA | NA | NA | NA |
| WBGene00045802 | NA | NA | NA | NA |
| WBGene00045803 | NA | NA | NA | NA |
| WBGene00045804 | NA | NA | NA | NA |
| WBGene00045805 | NA | NA | NA | NA |
| WBGene00045806 | NA | NA | NA | NA |
| WBGene00045807 | NA | NA | NA | NA |
| WBGene00045808 | 0.033463284 | 0.6365037 | NA | NA |
| WBGene00045809 | NA | NA | NA | NA |
| WBGene00045810 | NA | NA | NA | NA |
| WBGene00045811 | NA | NA | NA | NA |
| WBGene00045812 | NA | NA | NA | NA |
| WBGene00045813 | NA | NA | NA | NA |
| WBGene00045814 | NA | NA | NA | NA |
| WBGene00045815 | NA | NA | NA | NA |
| WBGene00045816 | NA | NA | NA | NA |
| WBGene00045817 | NA | NA | NA | NA |
| WBGene00045818 | NA | NA | NA | NA |
| WBGene00045819 | NA | NA | NA | NA |
| WBGene00045820 | NA | NA | NA | NA |
| WBGene00045821 | NA | NA | NA | NA |
| WBGene00045822 | NA | NA | NA | NA |
| WBGene00045823 | NA | NA | NA | NA |
| WBGene00045824 | NA | NA | NA | NA |
| WBGene00045825 | NA | NA | NA | NA |
| WBGene00045826 | NA | NA | NA | NA |
| WBGene00045827 | NA | NA | NA | NA |
| WBGene00045828 | NA | NA | NA | NA |
| WBGene00045829 | NA | NA | NA | NA |
| WBGene00045830 | NA | NA | NA | NA |
| WBGene00045831 | NA | NA | NA | NA |

| | | | | |
|---|---|---|---|---|
| WBGene00045832 | NA | NA | NA | NA |
| WBGene00045833 | NA | NA | NA | NA |
| WBGene00045834 | NA | NA | NA | NA |
| WBGene00045835 | NA | NA | NA | NA |
| WBGene00045836 | NA | NA | NA | NA |
| WBGene00045837 | NA | NA | NA | NA |
| WBGene00045838 | NA | NA | NA | NA |
| WBGene00045839 | NA | NA | NA | NA |
| WBGene00045840 | NA | NA | NA | NA |
| WBGene00045841 | NA | NA | NA | NA |
| WBGene00045842 | NA | NA | NA | NA |
| WBGene00045843 | NA | NA | NA | NA |
| WBGene00045844 | NA | NA | NA | NA |
| WBGene00045845 | NA | NA | NA | NA |
| WBGene00045846 | NA | NA | NA | NA |
| WBGene00045847 | NA | NA | NA | NA |
| WBGene00045848 | NA | NA | NA | NA |
| WBGene00045849 | NA | NA | NA | NA |
| WBGene00045850 | NA | NA | NA | NA |
| WBGene00045851 | NA | NA | NA | NA |
| WBGene00045852 | NA | NA | NA | NA |
| WBGene00045853 | NA | NA | NA | NA |
| WBGene00045854 | NA | NA | NA | NA |
| WBGene00045855 | NA | NA | NA | NA |
| WBGene00045856 | NA | NA | NA | NA |
| WBGene00045857 | NA | NA | NA | NA |
| WBGene00045858 | NA | NA | NA | NA |
| WBGene00045859 | NA | NA | NA | NA |
| WBGene00045860 | NA | NA | NA | NA |
| WBGene00045861 | NA | NA | NA | NA |
| WBGene00045862 | NA | NA | NA | NA |
| WBGene00045863 | NA | NA | NA | NA |
| WBGene00045864 | NA | NA | NA | NA |
| WBGene00045865 | NA | NA | NA | NA |
| WBGene00045866 | NA | NA | NA | NA |
| WBGene00045867 | NA | NA | NA | NA |
| WBGene00045868 | NA | NA | NA | NA |
| WBGene00045869 | NA | NA | NA | NA |
| WBGene00045870 | NA | NA | NA | NA |
| WBGene00045871 | NA | NA | NA | NA |
| WBGene00045872 | NA | NA | NA | NA |
| WBGene00045873 | NA | NA | NA | NA |
| WBGene00045874 | NA | NA | NA | NA |

| | | | | |
|---|---|---|---|---|
| WBGene00045875 | NA | NA | NA | NA |
| WBGene00045876 | NA | NA | NA | NA |
| WBGene00045877 | NA | NA | NA | NA |
| WBGene00045878 | NA | NA | NA | NA |
| WBGene00045879 | NA | NA | NA | NA |
| WBGene00045880 | NA | NA | NA | NA |
| WBGene00045881 | NA | NA | NA | NA |
| WBGene00045882 | NA | NA | NA | NA |
| WBGene00045883 | NA | NA | NA | NA |
| WBGene00045884 | NA | NA | NA | NA |
| WBGene00045885 | NA | NA | NA | NA |
| WBGene00045886 | NA | NA | NA | NA |
| WBGene00045887 | NA | NA | NA | NA |
| WBGene00045888 | NA | NA | NA | NA |
| WBGene00045889 | NA | NA | NA | NA |
| WBGene00045890 | NA | NA | NA | NA |
| WBGene00045891 | NA | NA | NA | NA |
| WBGene00045892 | NA | NA | NA | NA |
| WBGene00045893 | NA | NA | NA | NA |
| WBGene00045894 | NA | NA | NA | NA |
| WBGene00045895 | NA | NA | NA | NA |
| WBGene00045896 | NA | NA | NA | NA |
| WBGene00045897 | NA | NA | NA | NA |
| WBGene00045898 | NA | NA | NA | NA |
| WBGene00045899 | NA | NA | NA | NA |
| WBGene00045900 | NA | NA | NA | NA |
| WBGene00045901 | NA | NA | NA | NA |
| WBGene00045902 | NA | NA | NA | NA |
| WBGene00045903 | NA | NA | NA | NA |
| WBGene00045904 | NA | NA | NA | NA |
| WBGene00045905 | NA | NA | NA | NA |
| WBGene00045906 | NA | NA | NA | NA |
| WBGene00045907 | NA | NA | NA | NA |
| WBGene00045908 | NA | NA | NA | NA |
| WBGene00045909 | NA | NA | NA | NA |
| WBGene00045910 | NA | NA | NA | NA |
| WBGene00045911 | NA | NA | NA | NA |
| WBGene00045912 | NA | NA | NA | NA |
| WBGene00045913 | NA | NA | NA | NA |
| WBGene00045914 | NA | NA | NA | NA |
| WBGene00045915 | NA | NA | NA | NA |
| WBGene00045916 | NA | NA | NA | NA |
| WBGene00045917 | NA | NA | NA | NA |

| | | | | |
|---|---|---|---|---|
| WBGene00045918 | NA | NA | NA | NA |
| WBGene00045919 | NA | NA | NA | NA |
| WBGene00045920 | NA | NA | NA | NA |
| WBGene00045921 | NA | NA | NA | NA |
| WBGene00045922 | NA | NA | NA | NA |
| WBGene00045923 | NA | NA | NA | NA |
| WBGene00045924 | NA | NA | NA | NA |
| WBGene00045925 | NA | NA | NA | NA |
| WBGene00045926 | NA | NA | NA | NA |
| WBGene00045927 | NA | NA | NA | NA |
| WBGene00045928 | NA | NA | NA | NA |
| WBGene00045929 | NA | NA | NA | NA |
| WBGene00045930 | NA | NA | NA | NA |
| WBGene00045931 | NA | NA | NA | NA |
| WBGene00045932 | NA | NA | NA | NA |
| WBGene00045933 | NA | NA | NA | NA |
| WBGene00045934 | NA | NA | NA | NA |
| WBGene00045935 | NA | NA | NA | NA |
| WBGene00045936 | NA | NA | NA | NA |
| WBGene00045937 | NA | NA | NA | NA |
| WBGene00045938 | NA | NA | NA | NA |
| WBGene00045939 | NA | NA | NA | NA |
| WBGene00045940 | NA | NA | NA | NA |
| WBGene00045941 | NA | NA | NA | NA |
| WBGene00045942 | NA | NA | NA | NA |
| WBGene00045943 | NA | NA | NA | NA |
| WBGene00045944 | NA | NA | NA | NA |
| WBGene00045945 | NA | NA | NA | NA |
| WBGene00045946 | NA | NA | NA | NA |
| WBGene00045947 | NA | NA | NA | NA |
| WBGene00045948 | NA | NA | NA | NA |
| WBGene00045949 | NA | NA | NA | NA |
| WBGene00045950 | NA | NA | NA | NA |
| WBGene00045951 | NA | NA | NA | NA |
| WBGene00045952 | NA | NA | NA | NA |
| WBGene00045953 | NA | NA | NA | NA |
| WBGene00045954 | NA | NA | NA | NA |
| WBGene00045955 | NA | NA | NA | NA |
| WBGene00045956 | NA | NA | NA | NA |
| WBGene00045957 | NA | NA | NA | NA |
| WBGene00045958 | -0.023056499 | 0.78121253 | NA | NA |
| WBGene00045959 | NA | NA | NA | NA |
| WBGene00045960 | NA | NA | NA | NA |

| | | | | | |
|---|---|---|---|---|---|
| WBGene00045961 | NA | | NA | NA | NA |
| WBGene00045962 | NA | | NA | NA | NA |
| WBGene00045963 | NA | | NA | NA | NA |
| WBGene00045964 | NA | | NA | NA | NA |
| WBGene00045965 | NA | | NA | NA | NA |
| WBGene00045966 | NA | | NA | NA | NA |
| WBGene00045967 | NA | | NA | NA | NA |
| WBGene00045968 | NA | | NA | NA | NA |
| WBGene00045969 | NA | | NA | NA | NA |
| WBGene00045970 | NA | | NA | NA | NA |
| WBGene00045971 | NA | | NA | NA | NA |
| WBGene00045972 | NA | | NA | NA | NA |
| WBGene00045973 | NA | | NA | NA | NA |
| WBGene00045974 | NA | | NA | NA | NA |
| WBGene00045975 | NA | | NA | NA | NA |
| WBGene00045976 | NA | | NA | NA | NA |
| WBGene00045977 | NA | | NA | NA | NA |
| WBGene00045978 | NA | | NA | NA | NA |
| WBGene00045979 | NA | | NA | NA | NA |
| WBGene00045980 | NA | | NA | NA | NA |
| WBGene00045981 | NA | | NA | NA | NA |
| WBGene00045982 | NA | | NA | NA | NA |
| WBGene00045983 | NA | | NA | NA | NA |
| WBGene00045984 | NA | | NA | NA | NA |
| WBGene00045985 | NA | | NA | NA | NA |
| WBGene00045986 | | 0.020925399 | 0.84643798 | NA | NA |
| WBGene00045987 | | 0.020232312 | 0.84643798 | NA | NA |
| WBGene00045988 | NA | | NA | NA | NA |
| WBGene00045989 | NA | | NA | NA | NA |
| WBGene00045990 | NA | | NA | NA | NA |
| WBGene00045991 | NA | | NA | NA | NA |
| WBGene00045992 | NA | | NA | NA | NA |
| WBGene00045993 | NA | | NA | NA | NA |
| WBGene00045994 | NA | | NA | NA | NA |
| WBGene00045995 | NA | | NA | NA | NA |
| WBGene00045996 | NA | | NA | NA | NA |
| WBGene00045997 | NA | | NA | NA | NA |
| WBGene00045998 | NA | | NA | NA | NA |
| WBGene00045999 | NA | | NA | NA | NA |
| WBGene00046000 | NA | | NA | NA | NA |
| WBGene00046001 | NA | | NA | NA | NA |
| WBGene00046002 | NA | | NA | NA | NA |
| WBGene00046003 | NA | | NA | NA | NA |

| | | | | |
|---|---|---|---|---|
| WBGene00046004 | NA | NA | NA | NA |
| WBGene00046005 | NA | NA | NA | NA |
| WBGene00046006 | NA | NA | NA | NA |
| WBGene00046007 | NA | NA | NA | NA |
| WBGene00046008 | NA | NA | NA | NA |
| WBGene00046009 | NA | NA | NA | NA |
| WBGene00046010 | NA | NA | NA | NA |
| WBGene00046011 | NA | NA | NA | NA |
| WBGene00046012 | NA | NA | NA | NA |
| WBGene00046013 | NA | NA | NA | NA |
| WBGene00046014 | NA | NA | NA | NA |
| WBGene00046015 | NA | NA | NA | NA |
| WBGene00046016 | NA | NA | NA | NA |
| WBGene00046017 | NA | NA | NA | NA |
| WBGene00046018 | NA | NA | NA | NA |
| WBGene00046019 | NA | NA | NA | NA |
| WBGene00046020 | NA | NA | NA | NA |
| WBGene00046021 | NA | NA | NA | NA |
| WBGene00046022 | NA | NA | NA | NA |
| WBGene00046023 | NA | NA | NA | NA |
| WBGene00046024 | NA | NA | NA | NA |
| WBGene00046025 | NA | NA | NA | NA |
| WBGene00046026 | NA | NA | NA | NA |
| WBGene00046027 | NA | NA | NA | NA |
| WBGene00046028 | NA | NA | NA | NA |
| WBGene00046029 | NA | NA | NA | NA |
| WBGene00046030 | NA | NA | NA | NA |
| WBGene00046031 | NA | NA | NA | NA |
| WBGene00046032 | NA | NA | NA | NA |
| WBGene00046033 | NA | NA | NA | NA |
| WBGene00046034 | NA | NA | NA | NA |
| WBGene00046035 | NA | NA | NA | NA |
| WBGene00046036 | NA | NA | NA | NA |
| WBGene00046037 | NA | NA | NA | NA |
| WBGene00046038 | NA | NA | NA | NA |
| WBGene00046039 | NA | NA | NA | NA |
| WBGene00046040 | NA | NA | NA | NA |
| WBGene00046041 | NA | NA | NA | NA |
| WBGene00046042 | NA | NA | NA | NA |
| WBGene00046043 | NA | NA | NA | NA |
| WBGene00046044 | NA | NA | NA | NA |
| WBGene00046045 | NA | NA | NA | NA |
| WBGene00046046 | NA | NA | NA | NA |

| | | | | |
|---|---|---|---|---|
| WBGene00046047 | NA | NA | NA | NA |
| WBGene00046048 | NA | NA | NA | NA |
| WBGene00046049 | NA | NA | NA | NA |
| WBGene00046050 | NA | NA | NA | NA |
| WBGene00046051 | NA | NA | NA | NA |
| WBGene00046052 | NA | NA | NA | NA |
| WBGene00046053 | NA | NA | NA | NA |
| WBGene00046054 | NA | NA | NA | NA |
| WBGene00046055 | NA | NA | NA | NA |
| WBGene00046056 | NA | NA | NA | NA |
| WBGene00046057 | NA | NA | NA | NA |
| WBGene00046058 | NA | NA | NA | NA |
| WBGene00046059 | NA | NA | NA | NA |
| WBGene00046060 | NA | NA | NA | NA |
| WBGene00046061 | NA | NA | NA | NA |
| WBGene00046062 | NA | NA | NA | NA |
| WBGene00046063 | NA | NA | NA | NA |
| WBGene00046064 | NA | NA | NA | NA |
| WBGene00046065 | NA | NA | NA | NA |
| WBGene00046066 | NA | NA | NA | NA |
| WBGene00046067 | NA | NA | NA | NA |
| WBGene00046068 | NA | NA | NA | NA |
| WBGene00046069 | NA | NA | NA | NA |
| WBGene00046070 | NA | NA | NA | NA |
| WBGene00046071 | NA | NA | NA | NA |
| WBGene00046072 | NA | NA | NA | NA |
| WBGene00046073 | NA | NA | NA | NA |
| WBGene00046074 | NA | NA | NA | NA |
| WBGene00046075 | NA | NA | NA | NA |
| WBGene00046076 | NA | NA | NA | NA |
| WBGene00046077 | NA | NA | NA | NA |
| WBGene00046078 | NA | NA | NA | NA |
| WBGene00046079 | NA | NA | NA | NA |
| WBGene00046080 | NA | NA | NA | NA |
| WBGene00046081 | NA | NA | NA | NA |
| WBGene00046082 | NA | NA | NA | NA |
| WBGene00046083 | NA | NA | NA | NA |
| WBGene00046084 | NA | NA | NA | NA |
| WBGene00046085 | NA | NA | NA | NA |
| WBGene00046086 | NA | NA | NA | NA |
| WBGene00046087 | NA | NA | NA | NA |
| WBGene00046088 | NA | NA | NA | NA |
| WBGene00046089 | NA | NA | NA | NA |

| | | | | |
|---|---|---|---|---|
| WBGene00046090 | NA | NA | NA | NA |
| WBGene00046091 | NA | NA | NA | NA |
| WBGene00046092 | NA | NA | NA | NA |
| WBGene00046093 | NA | NA | NA | NA |
| WBGene00046094 | NA | NA | NA | NA |
| WBGene00046095 | NA | NA | NA | NA |
| WBGene00046096 | NA | NA | NA | NA |
| WBGene00046097 | NA | NA | NA | NA |
| WBGene00046098 | NA | NA | NA | NA |
| WBGene00046099 | NA | NA | NA | NA |
| WBGene00046100 | NA | NA | NA | NA |
| WBGene00046101 | NA | NA | NA | NA |
| WBGene00046102 | NA | NA | NA | NA |
| WBGene00046103 | NA | NA | NA | NA |
| WBGene00046104 | NA | NA | NA | NA |
| WBGene00046105 | NA | NA | NA | NA |
| WBGene00046106 | NA | NA | NA | NA |
| WBGene00046107 | NA | NA | NA | NA |
| WBGene00046108 | NA | NA | NA | NA |
| WBGene00046109 | NA | NA | NA | NA |
| WBGene00046110 | NA | NA | NA | NA |
| WBGene00046111 | NA | NA | NA | NA |
| WBGene00046112 | NA | NA | NA | NA |
| WBGene00046113 | NA | NA | NA | NA |
| WBGene00046114 | NA | NA | NA | NA |
| WBGene00046115 | NA | NA | NA | NA |
| WBGene00046116 | NA | NA | NA | NA |
| WBGene00046117 | NA | NA | NA | NA |
| WBGene00046118 | NA | NA | NA | NA |
| WBGene00046119 | NA | NA | NA | NA |
| WBGene00046120 | NA | NA | NA | NA |
| WBGene00046121 | NA | NA | NA | NA |
| WBGene00046122 | NA | NA | NA | NA |
| WBGene00046123 | NA | NA | NA | NA |
| WBGene00046124 | NA | NA | NA | NA |
| WBGene00046125 | NA | NA | NA | NA |
| WBGene00046126 | NA | NA | NA | NA |
| WBGene00046127 | NA | NA | NA | NA |
| WBGene00046128 | NA | NA | NA | NA |
| WBGene00046129 | NA | NA | NA | NA |
| WBGene00046130 | NA | NA | NA | NA |
| WBGene00046131 | NA | NA | NA | NA |
| WBGene00046132 | NA | NA | NA | NA |

| | | | | |
|---|---|---|---|---|
| WBGene00046133 | NA | NA | NA | NA |
| WBGene00046134 | NA | NA | NA | NA |
| WBGene00046135 | NA | NA | NA | NA |
| WBGene00046136 | NA | NA | NA | NA |
| WBGene00046137 | NA | NA | NA | NA |
| WBGene00046138 | NA | NA | NA | NA |
| WBGene00046139 | NA | NA | NA | NA |
| WBGene00046140 | NA | NA | NA | NA |
| WBGene00046141 | NA | NA | NA | NA |
| WBGene00046142 | NA | NA | NA | NA |
| WBGene00046143 | NA | NA | NA | NA |
| WBGene00046144 | NA | NA | NA | NA |
| WBGene00046145 | NA | NA | NA | NA |
| WBGene00046146 | NA | NA | NA | NA |
| WBGene00046147 | -0.022930767 | 0.78121253 | NA | NA |
| WBGene00046148 | NA | NA | NA | NA |
| WBGene00046149 | NA | NA | NA | NA |
| WBGene00046150 | NA | NA | NA | NA |
| WBGene00046151 | NA | NA | NA | NA |
| WBGene00046152 | NA | NA | NA | NA |
| WBGene00046153 | NA | NA | NA | NA |
| WBGene00046154 | NA | NA | NA | NA |
| WBGene00046155 | NA | NA | NA | NA |
| WBGene00046156 | NA | NA | NA | NA |
| WBGene00046157 | NA | NA | NA | NA |
| WBGene00046158 | NA | NA | NA | NA |
| WBGene00046159 | NA | NA | NA | NA |
| WBGene00046160 | NA | NA | NA | NA |
| WBGene00046161 | NA | NA | NA | NA |
| WBGene00046162 | NA | NA | NA | NA |
| WBGene00046163 | NA | NA | NA | NA |
| WBGene00046164 | NA | NA | NA | NA |
| WBGene00046165 | NA | NA | NA | NA |
| WBGene00046166 | NA | NA | NA | NA |
| WBGene00046167 | NA | NA | NA | NA |
| WBGene00046168 | NA | NA | NA | NA |
| WBGene00046169 | NA | NA | NA | NA |
| WBGene00046170 | NA | NA | NA | NA |
| WBGene00046171 | NA | NA | NA | NA |
| WBGene00046172 | NA | NA | NA | NA |
| WBGene00046173 | NA | NA | NA | NA |
| WBGene00046174 | NA | NA | NA | NA |
| WBGene00046175 | NA | NA | NA | NA |

| | | | | |
|---|---|---|---|---|
| WBGene00046176 | NA | NA | NA | NA |
| WBGene00046177 | NA | NA | NA | NA |
| WBGene00046178 | NA | NA | NA | NA |
| WBGene00046179 | NA | NA | NA | NA |
| WBGene00046180 | NA | NA | NA | NA |
| WBGene00046181 | NA | NA | NA | NA |
| WBGene00046182 | NA | NA | NA | NA |
| WBGene00046183 | NA | NA | NA | NA |
| WBGene00046184 | NA | NA | NA | NA |
| WBGene00046185 | NA | NA | NA | NA |
| WBGene00046186 | NA | NA | NA | NA |
| WBGene00046187 | NA | NA | NA | NA |
| WBGene00046188 | NA | NA | NA | NA |
| WBGene00046189 | NA | NA | NA | NA |
| WBGene00046190 | NA | NA | NA | NA |
| WBGene00046191 | NA | NA | NA | NA |
| WBGene00046192 | NA | NA | NA | NA |
| WBGene00046193 | NA | NA | NA | NA |
| WBGene00046194 | NA | NA | NA | NA |
| WBGene00046195 | NA | NA | NA | NA |
| WBGene00046196 | NA | NA | NA | NA |
| WBGene00046197 | NA | NA | NA | NA |
| WBGene00046198 | NA | NA | NA | NA |
| WBGene00046199 | NA | NA | NA | NA |
| WBGene00046200 | NA | NA | NA | NA |
| WBGene00046201 | NA | NA | NA | NA |
| WBGene00046202 | NA | NA | NA | NA |
| WBGene00046203 | NA | NA | NA | NA |
| WBGene00046204 | NA | NA | NA | NA |
| WBGene00046205 | NA | NA | NA | NA |
| WBGene00046206 | NA | NA | NA | NA |
| WBGene00046207 | NA | NA | NA | NA |
| WBGene00046208 | NA | NA | NA | NA |
| WBGene00046209 | NA | NA | NA | NA |
| WBGene00046210 | NA | NA | NA | NA |
| WBGene00046211 | NA | NA | NA | NA |
| WBGene00046212 | NA | NA | NA | NA |
| WBGene00046213 | NA | NA | NA | NA |
| WBGene00046214 | NA | NA | NA | NA |
| WBGene00046215 | NA | NA | NA | NA |
| WBGene00046216 | NA | NA | NA | NA |
| WBGene00046217 | NA | NA | NA | NA |
| WBGene00046218 | NA | NA | NA | NA |

| | | | | |
|---|---|---|---|---|
| WBGene00046219 | NA | NA | NA | NA |
| WBGene00046220 | NA | NA | NA | NA |
| WBGene00046221 | NA | NA | NA | NA |
| WBGene00046222 | NA | NA | NA | NA |
| WBGene00046223 | NA | NA | NA | NA |
| WBGene00046224 | NA | NA | NA | NA |
| WBGene00046225 | NA | NA | NA | NA |
| WBGene00046226 | NA | NA | NA | NA |
| WBGene00046227 | NA | NA | NA | NA |
| WBGene00046228 | NA | NA | NA | NA |
| WBGene00046229 | NA | NA | NA | NA |
| WBGene00046230 | NA | NA | NA | NA |
| WBGene00046231 | NA | NA | NA | NA |
| WBGene00046232 | NA | NA | NA | NA |
| WBGene00046233 | NA | NA | NA | NA |
| WBGene00046234 | NA | NA | NA | NA |
| WBGene00046235 | NA | NA | NA | NA |
| WBGene00046236 | NA | NA | NA | NA |
| WBGene00046237 | NA | NA | NA | NA |
| WBGene00046238 | NA | NA | NA | NA |
| WBGene00046239 | NA | NA | NA | NA |
| WBGene00046240 | NA | NA | NA | NA |
| WBGene00046241 | NA | NA | NA | NA |
| WBGene00046242 | NA | NA | NA | NA |
| WBGene00046243 | NA | NA | NA | NA |
| WBGene00046244 | NA | NA | NA | NA |
| WBGene00046245 | NA | NA | NA | NA |
| WBGene00046246 | NA | NA | NA | NA |
| WBGene00046247 | NA | NA | NA | NA |
| WBGene00046248 | NA | NA | NA | NA |
| WBGene00046249 | NA | NA | NA | NA |
| WBGene00046250 | NA | NA | NA | NA |
| WBGene00046251 | NA | NA | NA | NA |
| WBGene00046252 | NA | NA | NA | NA |
| WBGene00046253 | NA | NA | NA | NA |
| WBGene00046254 | NA | NA | NA | NA |
| WBGene00046255 | NA | NA | NA | NA |
| WBGene00046256 | NA | NA | NA | NA |
| WBGene00046257 | NA | NA | NA | NA |
| WBGene00046258 | NA | NA | NA | NA |
| WBGene00046259 | NA | NA | NA | NA |
| WBGene00046260 | NA | NA | NA | NA |
| WBGene00046261 | NA | NA | NA | NA |

| | | | | |
|---|---|---|---|---|
| WBGene00046262 | NA | NA | NA | NA |
| WBGene00046263 | NA | NA | NA | NA |
| WBGene00046264 | NA | NA | NA | NA |
| WBGene00046265 | NA | NA | NA | NA |
| WBGene00046266 | NA | NA | NA | NA |
| WBGene00046267 | NA | NA | NA | NA |
| WBGene00046268 | NA | NA | NA | NA |
| WBGene00046269 | NA | NA | NA | NA |
| WBGene00046270 | NA | NA | NA | NA |
| WBGene00046271 | NA | NA | NA | NA |
| WBGene00046272 | NA | NA | NA | NA |
| WBGene00046273 | NA | NA | NA | NA |
| WBGene00046274 | NA | NA | NA | NA |
| WBGene00046275 | NA | NA | NA | NA |
| WBGene00046276 | NA | NA | NA | NA |
| WBGene00046277 | NA | NA | NA | NA |
| WBGene00046278 | NA | NA | NA | NA |
| WBGene00046279 | NA | NA | NA | NA |
| WBGene00046280 | NA | NA | NA | NA |
| WBGene00046281 | NA | NA | NA | NA |
| WBGene00046282 | NA | NA | NA | NA |
| WBGene00046283 | NA | NA | NA | NA |
| WBGene00046284 | NA | NA | NA | NA |
| WBGene00046285 | NA | NA | NA | NA |
| WBGene00046286 | NA | NA | NA | NA |
| WBGene00046287 | NA | NA | NA | NA |
| WBGene00046288 | NA | NA | NA | NA |
| WBGene00046289 | NA | NA | NA | NA |
| WBGene00046290 | NA | NA | NA | NA |
| WBGene00046291 | NA | NA | NA | NA |
| WBGene00046292 | NA | NA | NA | NA |
| WBGene00046293 | NA | NA | NA | NA |
| WBGene00046294 | NA | NA | NA | NA |
| WBGene00046295 | NA | NA | NA | NA |
| WBGene00046296 | NA | NA | NA | NA |
| WBGene00046297 | NA | NA | NA | NA |
| WBGene00046298 | NA | NA | NA | NA |
| WBGene00046299 | NA | NA | NA | NA |
| WBGene00046300 | NA | NA | NA | NA |
| WBGene00046301 | NA | NA | NA | NA |
| WBGene00046302 | NA | NA | NA | NA |
| WBGene00046303 | NA | NA | NA | NA |
| WBGene00046304 | NA | NA | NA | NA |

| | | | | |
|---|---|---|---|---|
| WBGene00046305 | NA | NA | NA | NA |
| WBGene00046306 | NA | NA | NA | NA |
| WBGene00046307 | NA | NA | NA | NA |
| WBGene00046308 | NA | NA | NA | NA |
| WBGene00046309 | NA | NA | NA | NA |
| WBGene00046310 | NA | NA | NA | NA |
| WBGene00046311 | NA | NA | NA | NA |
| WBGene00046312 | NA | NA | NA | NA |
| WBGene00046313 | -0.023056499 | 0.78121253 | NA | NA |
| WBGene00046314 | NA | NA | NA | NA |
| WBGene00046315 | NA | NA | NA | NA |
| WBGene00046316 | NA | NA | NA | NA |
| WBGene00046317 | NA | NA | NA | NA |
| WBGene00046318 | NA | NA | NA | NA |
| WBGene00046319 | NA | NA | NA | NA |
| WBGene00046320 | NA | NA | NA | NA |
| WBGene00046321 | NA | NA | NA | NA |
| WBGene00046322 | NA | NA | NA | NA |
| WBGene00046323 | NA | NA | NA | NA |
| WBGene00046324 | NA | NA | NA | NA |
| WBGene00046325 | NA | NA | NA | NA |
| WBGene00046326 | NA | NA | NA | NA |
| WBGene00046327 | NA | NA | NA | NA |
| WBGene00046328 | NA | NA | NA | NA |
| WBGene00046329 | NA | NA | NA | NA |
| WBGene00046330 | NA | NA | NA | NA |
| WBGene00046331 | NA | NA | NA | NA |
| WBGene00046332 | NA | NA | NA | NA |
| WBGene00046333 | NA | NA | NA | NA |
| WBGene00046334 | NA | NA | NA | NA |
| WBGene00046335 | 0.020232312 | 0.84643798 | NA | NA |
| WBGene00046336 | NA | NA | NA | NA |
| WBGene00046337 | NA | NA | NA | NA |
| WBGene00046338 | NA | NA | NA | NA |
| WBGene00046339 | NA | NA | NA | NA |
| WBGene00046340 | NA | NA | NA | NA |
| WBGene00046341 | NA | NA | NA | NA |
| WBGene00046342 | NA | NA | NA | NA |
| WBGene00046343 | NA | NA | NA | NA |
| WBGene00046344 | NA | NA | NA | NA |
| WBGene00046345 | NA | NA | NA | NA |
| WBGene00046346 | NA | NA | NA | NA |
| WBGene00046347 | NA | NA | NA | NA |

| | | | | | |
|---|---|---|---|---|---|
| WBGene00046348 | NA | | NA | NA | NA |
| WBGene00046349 | NA | | NA | NA | NA |
| WBGene00046350 | NA | | NA | NA | NA |
| WBGene00046351 | NA | | NA | NA | NA |
| WBGene00046352 | NA | | NA | NA | NA |
| WBGene00046353 | NA | | NA | NA | NA |
| WBGene00046354 | NA | | NA | NA | NA |
| WBGene00046355 | NA | | NA | NA | NA |
| WBGene00046356 | NA | | NA | NA | NA |
| WBGene00046357 | NA | | NA | NA | NA |
| WBGene00046358 | NA | | NA | NA | NA |
| WBGene00046359 | NA | | NA | NA | NA |
| WBGene00046360 | NA | | NA | NA | NA |
| WBGene00046361 | NA | | NA | NA | NA |
| WBGene00046362 | NA | | NA | NA | NA |
| WBGene00046363 | NA | | NA | NA | NA |
| WBGene00046364 | | 0.020925399 | 0.84643798 | NA | NA |
| WBGene00046365 | NA | | NA | NA | NA |
| WBGene00046366 | NA | | NA | NA | NA |
| WBGene00046367 | NA | | NA | NA | NA |
| WBGene00046368 | NA | | NA | NA | NA |
| WBGene00046369 | NA | | NA | NA | NA |
| WBGene00046370 | NA | | NA | NA | NA |
| WBGene00046371 | NA | | NA | NA | NA |
| WBGene00046372 | | 0.020232312 | 0.84643798 | NA | NA |
| WBGene00046373 | NA | | NA | NA | NA |
| WBGene00046374 | NA | | NA | NA | NA |
| WBGene00046375 | NA | | NA | NA | NA |
| WBGene00046376 | NA | | NA | NA | NA |
| WBGene00046377 | NA | | NA | NA | NA |
| WBGene00046378 | NA | | NA | NA | NA |
| WBGene00046379 | NA | | NA | NA | NA |
| WBGene00046380 | NA | | NA | NA | NA |
| WBGene00046381 | NA | | NA | NA | NA |
| WBGene00046382 | NA | | NA | NA | NA |
| WBGene00046383 | NA | | NA | NA | NA |
| WBGene00046384 | NA | | NA | NA | NA |
| WBGene00046385 | NA | | NA | NA | NA |
| WBGene00046386 | NA | | NA | NA | NA |
| WBGene00046387 | NA | | NA | NA | NA |
| WBGene00046388 | NA | | NA | NA | NA |
| WBGene00046389 | NA | | NA | NA | NA |
| WBGene00046390 | NA | | NA | NA | NA |

| | | | | | |
|---|---|---|---|---|---|
| WBGene00046391 | NA | | NA | NA | NA |
| WBGene00046392 | NA | | NA | NA | NA |
| WBGene00046393 | NA | | NA | NA | NA |
| WBGene00046394 | NA | | NA | NA | NA |
| WBGene00046395 | NA | | NA | NA | NA |
| WBGene00046396 | NA | | NA | NA | NA |
| WBGene00046397 | NA | | NA | NA | NA |
| WBGene00046398 | NA | | NA | NA | NA |
| WBGene00046399 | NA | | NA | NA | NA |
| WBGene00046400 | NA | | NA | NA | NA |
| WBGene00046401 | NA | | NA | NA | NA |
| WBGene00046402 | NA | | NA | NA | NA |
| WBGene00046403 | NA | | NA | NA | NA |
| WBGene00046404 | NA | | NA | NA | NA |
| WBGene00046405 | NA | | NA | NA | NA |
| WBGene00046406 | NA | | NA | NA | NA |
| WBGene00046407 | NA | | NA | NA | NA |
| WBGene00046408 | NA | | NA | NA | NA |
| WBGene00046409 | NA | | NA | NA | NA |
| WBGene00046410 | NA | | NA | NA | NA |
| WBGene00046411 | NA | | NA | NA | NA |
| WBGene00046412 | NA | | NA | NA | NA |
| WBGene00046413 | NA | | NA | NA | NA |
| WBGene00046414 | NA | | NA | NA | NA |
| WBGene00046415 | NA | | NA | NA | NA |
| WBGene00046416 | NA | | NA | NA | NA |
| WBGene00046417 | NA | | NA | NA | NA |
| WBGene00046418 | NA | | NA | NA | NA |
| WBGene00046419 | NA | | NA | NA | NA |
| WBGene00046420 | NA | | NA | NA | NA |
| WBGene00046421 | NA | | NA | NA | NA |
| WBGene00046422 | NA | | NA | NA | NA |
| WBGene00046423 | NA | | NA | NA | NA |
| WBGene00046424 | NA | | NA | NA | NA |
| WBGene00046425 | NA | | NA | NA | NA |
| WBGene00046426 | NA | | NA | NA | NA |
| WBGene00046427 | NA | | NA | NA | NA |
| WBGene00046428 | NA | | NA | NA | NA |
| WBGene00046429 | NA | | NA | NA | NA |
| WBGene00046430 | NA | | NA | NA | NA |
| WBGene00046431 | NA | | NA | NA | NA |
| WBGene00046432 | NA | | NA | NA | NA |
| WBGene00046433 | | 0.020925399 | 0.84643798 | NA | NA |

| | | | | |
|---|---|---|---|---|
| WBGene00046434 | NA | NA | NA | NA |
| WBGene00046435 | NA | NA | NA | NA |
| WBGene00046436 | NA | NA | NA | NA |
| WBGene00046437 | NA | NA | NA | NA |
| WBGene00046438 | NA | NA | NA | NA |
| WBGene00046439 | NA | NA | NA | NA |
| WBGene00046440 | NA | NA | NA | NA |
| WBGene00046441 | NA | NA | NA | NA |
| WBGene00046442 | NA | NA | NA | NA |
| WBGene00046443 | NA | NA | NA | NA |
| WBGene00046444 | NA | NA | NA | NA |
| WBGene00046445 | NA | NA | NA | NA |
| WBGene00046446 | NA | NA | NA | NA |
| WBGene00046447 | NA | NA | NA | NA |
| WBGene00046448 | NA | NA | NA | NA |
| WBGene00046449 | NA | NA | NA | NA |
| WBGene00046450 | NA | NA | NA | NA |
| WBGene00046451 | NA | NA | NA | NA |
| WBGene00046452 | NA | NA | NA | NA |
| WBGene00046453 | NA | NA | NA | NA |
| WBGene00046454 | NA | NA | NA | NA |
| WBGene00046455 | NA | NA | NA | NA |
| WBGene00046456 | NA | NA | NA | NA |
| WBGene00046457 | NA | NA | NA | NA |
| WBGene00046458 | NA | NA | NA | NA |
| WBGene00046459 | NA | NA | NA | NA |
| WBGene00046460 | NA | NA | NA | NA |
| WBGene00046461 | NA | NA | NA | NA |
| WBGene00046462 | NA | NA | NA | NA |
| WBGene00046463 | NA | NA | NA | NA |
| WBGene00046464 | NA | NA | NA | NA |
| WBGene00046465 | NA | NA | NA | NA |
| WBGene00046466 | NA | NA | NA | NA |
| WBGene00046467 | NA | NA | NA | NA |
| WBGene00046468 | NA | NA | NA | NA |
| WBGene00046469 | NA | NA | NA | NA |
| WBGene00046470 | NA | NA | NA | NA |
| WBGene00046471 | NA | NA | NA | NA |
| WBGene00046472 | NA | NA | NA | NA |
| WBGene00046473 | NA | NA | NA | NA |
| WBGene00046474 | NA | NA | NA | NA |
| WBGene00046475 | NA | NA | NA | NA |
| WBGene00046476 | NA | NA | NA | NA |

| | | | | |
|---|---|---|---|---|
| WBGene00046477 | NA | NA | NA | NA |
| WBGene00046478 | NA | NA | NA | NA |
| WBGene00046479 | NA | NA | NA | NA |
| WBGene00046480 | NA | NA | NA | NA |
| WBGene00046481 | NA | NA | NA | NA |
| WBGene00046482 | NA | NA | NA | NA |
| WBGene00046483 | NA | NA | NA | NA |
| WBGene00046484 | NA | NA | NA | NA |
| WBGene00046485 | NA | NA | NA | NA |
| WBGene00046486 | NA | NA | NA | NA |
| WBGene00046487 | NA | NA | NA | NA |
| WBGene00046488 | NA | NA | NA | NA |
| WBGene00046489 | NA | NA | NA | NA |
| WBGene00046490 | NA | NA | NA | NA |
| WBGene00046491 | NA | NA | NA | NA |
| WBGene00046492 | NA | NA | NA | NA |
| WBGene00046493 | NA | NA | NA | NA |
| WBGene00046494 | NA | NA | NA | NA |
| WBGene00046495 | NA | NA | NA | NA |
| WBGene00046496 | NA | NA | NA | NA |
| WBGene00046497 | NA | NA | NA | NA |
| WBGene00046498 | NA | NA | NA | NA |
| WBGene00046499 | NA | NA | NA | NA |
| WBGene00046500 | NA | NA | NA | NA |
| WBGene00046501 | NA | NA | NA | NA |
| WBGene00046502 | NA | NA | NA | NA |
| WBGene00046503 | NA | NA | NA | NA |
| WBGene00046504 | NA | NA | NA | NA |
| WBGene00046505 | NA | NA | NA | NA |
| WBGene00046506 | NA | NA | NA | NA |
| WBGene00046507 | NA | NA | NA | NA |
| WBGene00046508 | NA | NA | NA | NA |
| WBGene00046509 | NA | NA | NA | NA |
| WBGene00046510 | NA | NA | NA | NA |
| WBGene00046511 | NA | NA | NA | NA |
| WBGene00046512 | NA | NA | NA | NA |
| WBGene00046513 | NA | NA | NA | NA |
| WBGene00046514 | NA | NA | NA | NA |
| WBGene00046515 | NA | NA | NA | NA |
| WBGene00046516 | NA | NA | NA | NA |
| WBGene00046517 | NA | NA | NA | NA |
| WBGene00046518 | NA | NA | NA | NA |
| WBGene00046519 | NA | NA | NA | NA |

| | | | | |
|---|---|---|---|---|
| WBGene00046520 | NA | NA | NA | NA |
| WBGene00046521 | NA | NA | NA | NA |
| WBGene00046522 | NA | NA | NA | NA |
| WBGene00046523 | NA | NA | NA | NA |
| WBGene00046524 | NA | NA | NA | NA |
| WBGene00046525 | NA | NA | NA | NA |
| WBGene00046526 | NA | NA | NA | NA |
| WBGene00046527 | 0.020317245 | 0.84643798 | NA | NA |
| WBGene00046528 | NA | NA | NA | NA |
| WBGene00046529 | NA | NA | NA | NA |
| WBGene00046530 | NA | NA | NA | NA |
| WBGene00046531 | NA | NA | NA | NA |
| WBGene00046532 | NA | NA | NA | NA |
| WBGene00046533 | NA | NA | NA | NA |
| WBGene00046534 | NA | NA | NA | NA |
| WBGene00046535 | NA | NA | NA | NA |
| WBGene00046536 | NA | NA | NA | NA |
| WBGene00046537 | NA | NA | NA | NA |
| WBGene00046538 | NA | NA | NA | NA |
| WBGene00046539 | NA | NA | NA | NA |
| WBGene00046540 | NA | NA | NA | NA |
| WBGene00046541 | NA | NA | NA | NA |
| WBGene00046542 | NA | NA | NA | NA |
| WBGene00046543 | NA | NA | NA | NA |
| WBGene00046544 | NA | NA | NA | NA |
| WBGene00046545 | NA | NA | NA | NA |
| WBGene00046546 | NA | NA | NA | NA |
| WBGene00046547 | 0.020317245 | 0.84643798 | NA | NA |
| WBGene00046548 | NA | NA | NA | NA |
| WBGene00046549 | NA | NA | NA | NA |
| WBGene00046550 | NA | NA | NA | NA |
| WBGene00046551 | NA | NA | NA | NA |
| WBGene00046552 | NA | NA | NA | NA |
| WBGene00046553 | NA | NA | NA | NA |
| WBGene00046554 | NA | NA | NA | NA |
| WBGene00046555 | NA | NA | NA | NA |
| WBGene00046556 | NA | NA | NA | NA |
| WBGene00046557 | NA | NA | NA | NA |
| WBGene00046558 | NA | NA | NA | NA |
| WBGene00046559 | NA | NA | NA | NA |
| WBGene00046560 | NA | NA | NA | NA |
| WBGene00046561 | NA | NA | NA | NA |
| WBGene00046562 | NA | NA | NA | NA |

| | | | | |
|---|---|---|---|---|
| WBGene00046563 | NA | NA | NA | NA |
| WBGene00046564 | NA | NA | NA | NA |
| WBGene00046565 | NA | NA | NA | NA |
| WBGene00046566 | NA | NA | NA | NA |
| WBGene00046567 | NA | NA | NA | NA |
| WBGene00046568 | NA | NA | NA | NA |
| WBGene00046569 | NA | NA | NA | NA |
| WBGene00046570 | NA | NA | NA | NA |
| WBGene00046571 | NA | NA | NA | NA |
| WBGene00046572 | NA | NA | NA | NA |
| WBGene00046573 | NA | NA | NA | NA |
| WBGene00046574 | NA | NA | NA | NA |
| WBGene00046575 | NA | NA | NA | NA |
| WBGene00046576 | NA | NA | NA | NA |
| WBGene00046577 | NA | NA | NA | NA |
| WBGene00046578 | NA | NA | NA | NA |
| WBGene00046579 | NA | NA | NA | NA |
| WBGene00046580 | NA | NA | NA | NA |
| WBGene00046581 | NA | NA | NA | NA |
| WBGene00046582 | NA | NA | NA | NA |
| WBGene00046583 | NA | NA | NA | NA |
| WBGene00046584 | NA | NA | NA | NA |
| WBGene00046585 | NA | NA | NA | NA |
| WBGene00046586 | NA | NA | NA | NA |
| WBGene00046587 | NA | NA | NA | NA |
| WBGene00046588 | NA | NA | NA | NA |
| WBGene00046589 | NA | NA | NA | NA |
| WBGene00046590 | NA | NA | NA | NA |
| WBGene00046591 | NA | NA | NA | NA |
| WBGene00046592 | NA | NA | NA | NA |
| WBGene00046593 | NA | NA | NA | NA |
| WBGene00046594 | NA | NA | NA | NA |
| WBGene00046595 | NA | NA | NA | NA |
| WBGene00046596 | NA | NA | NA | NA |
| WBGene00046597 | NA | NA | NA | NA |
| WBGene00046598 | NA | NA | NA | NA |
| WBGene00046599 | NA | NA | NA | NA |
| WBGene00046600 | NA | NA | NA | NA |
| WBGene00046601 | NA | NA | NA | NA |
| WBGene00046602 | NA | NA | NA | NA |
| WBGene00046603 | NA | NA | NA | NA |
| WBGene00046604 | NA | NA | NA | NA |
| WBGene00046605 | NA | NA | NA | NA |

| | | | | |
|---|---|---|---|---|
| WBGene00046606 | NA | NA | NA | NA |
| WBGene00046607 | NA | NA | NA | NA |
| WBGene00046608 | NA | NA | NA | NA |
| WBGene00046609 | NA | NA | NA | NA |
| WBGene00046610 | NA | NA | NA | NA |
| WBGene00046611 | NA | NA | NA | NA |
| WBGene00046612 | NA | NA | NA | NA |
| WBGene00046613 | NA | NA | NA | NA |
| WBGene00046614 | NA | NA | NA | NA |
| WBGene00046615 | NA | NA | NA | NA |
| WBGene00046616 | NA | NA | NA | NA |
| WBGene00046617 | -0.023109762 | 0.78121253 | NA | NA |
| WBGene00046618 | NA | NA | NA | NA |
| WBGene00046619 | NA | NA | NA | NA |
| WBGene00046620 | NA | NA | NA | NA |
| WBGene00046621 | NA | NA | NA | NA |
| WBGene00046622 | NA | NA | NA | NA |
| WBGene00046623 | NA | NA | NA | NA |
| WBGene00046624 | NA | NA | NA | NA |
| WBGene00046625 | NA | NA | NA | NA |
| WBGene00046626 | NA | NA | NA | NA |
| WBGene00046627 | NA | NA | NA | NA |
| WBGene00046628 | NA | NA | NA | NA |
| WBGene00046629 | NA | NA | NA | NA |
| WBGene00046630 | NA | NA | NA | NA |
| WBGene00046631 | NA | NA | NA | NA |
| WBGene00046632 | NA | NA | NA | NA |
| WBGene00046633 | NA | NA | NA | NA |
| WBGene00046634 | NA | NA | NA | NA |
| WBGene00046635 | NA | NA | NA | NA |
| WBGene00046636 | NA | NA | NA | NA |
| WBGene00046637 | NA | NA | NA | NA |
| WBGene00046638 | NA | NA | NA | NA |
| WBGene00046639 | NA | NA | NA | NA |
| WBGene00046640 | NA | NA | NA | NA |
| WBGene00046641 | NA | NA | NA | NA |
| WBGene00046642 | NA | NA | NA | NA |
| WBGene00046643 | NA | NA | NA | NA |
| WBGene00046644 | NA | NA | NA | NA |
| WBGene00046645 | NA | NA | NA | NA |
| WBGene00046646 | NA | NA | NA | NA |
| WBGene00046647 | NA | NA | NA | NA |
| WBGene00046648 | NA | NA | NA | NA |

| | | | | |
|---|---|---|---|---|
| WBGene00046649 | NA | NA | NA | NA |
| WBGene00046650 | NA | NA | NA | NA |
| WBGene00046651 | NA | NA | NA | NA |
| WBGene00046652 | NA | NA | NA | NA |
| WBGene00046653 | NA | NA | NA | NA |
| WBGene00046654 | NA | NA | NA | NA |
| WBGene00046655 | NA | NA | NA | NA |
| WBGene00046656 | NA | NA | NA | NA |
| WBGene00046657 | NA | NA | NA | NA |
| WBGene00046658 | NA | NA | NA | NA |
| WBGene00046659 | NA | NA | NA | NA |
| WBGene00046660 | NA | NA | NA | NA |
| WBGene00046661 | NA | NA | NA | NA |
| WBGene00046662 | NA | NA | NA | NA |
| WBGene00046663 | NA | NA | NA | NA |
| WBGene00046664 | NA | NA | NA | NA |
| WBGene00046665 | NA | NA | NA | NA |
| WBGene00046666 | NA | NA | NA | NA |
| WBGene00046667 | NA | NA | NA | NA |
| WBGene00046668 | NA | NA | NA | NA |
| WBGene00046669 | NA | NA | NA | NA |
| WBGene00046670 | NA | NA | NA | NA |
| WBGene00046671 | NA | NA | NA | NA |
| WBGene00046672 | NA | NA | NA | NA |
| WBGene00046673 | NA | NA | NA | NA |
| WBGene00046674 | NA | NA | NA | NA |
| WBGene00046675 | NA | NA | NA | NA |
| WBGene00046676 | NA | NA | NA | NA |
| WBGene00046677 | NA | NA | NA | NA |
| WBGene00046678 | NA | NA | NA | NA |
| WBGene00046679 | NA | NA | NA | NA |
| WBGene00046680 | NA | NA | NA | NA |
| WBGene00046681 | NA | NA | NA | NA |
| WBGene00046682 | NA | NA | NA | NA |
| WBGene00046683 | NA | NA | NA | NA |
| WBGene00046684 | NA | NA | NA | NA |
| WBGene00046685 | NA | NA | NA | NA |
| WBGene00046686 | NA | NA | NA | NA |
| WBGene00046687 | NA | NA | NA | NA |
| WBGene00046688 | NA | NA | NA | NA |
| WBGene00046689 | NA | NA | NA | NA |
| WBGene00046690 | NA | NA | NA | NA |
| WBGene00046691 | NA | NA | NA | NA |

| | | | | |
|---|---|---|---|---|
| WBGene00046692 | NA | NA | NA | NA |
| WBGene00046693 | NA | NA | NA | NA |
| WBGene00046694 | 0.038125098 | 0.61119482 | NA | NA |
| WBGene00046695 | NA | NA | NA | NA |
| WBGene00046696 | NA | NA | NA | NA |
| WBGene00046697 | NA | NA | NA | NA |
| WBGene00046698 | NA | NA | NA | NA |
| WBGene00046699 | NA | NA | NA | NA |
| WBGene00046700 | NA | NA | NA | NA |
| WBGene00046701 | NA | NA | NA | NA |
| WBGene00046702 | NA | NA | NA | NA |
| WBGene00046703 | NA | NA | NA | NA |
| WBGene00046704 | NA | NA | NA | NA |
| WBGene00046705 | NA | NA | NA | NA |
| WBGene00046706 | NA | NA | NA | NA |
| WBGene00046707 | NA | NA | NA | NA |
| WBGene00046708 | NA | NA | NA | NA |
| WBGene00046709 | NA | NA | NA | NA |
| WBGene00046710 | NA | NA | NA | NA |
| WBGene00046711 | NA | NA | NA | NA |
| WBGene00046712 | NA | NA | NA | NA |
| WBGene00046713 | NA | NA | NA | NA |
| WBGene00046714 | NA | NA | NA | NA |
| WBGene00046715 | NA | NA | NA | NA |
| WBGene00046716 | NA | NA | NA | NA |
| WBGene00046717 | NA | NA | NA | NA |
| WBGene00046718 | NA | NA | NA | NA |
| WBGene00046719 | NA | NA | NA | NA |
| WBGene00046720 | NA | NA | NA | NA |
| WBGene00046721 | NA | NA | NA | NA |
| WBGene00046722 | NA | NA | NA | NA |
| WBGene00046723 | NA | NA | NA | NA |
| WBGene00046724 | NA | NA | NA | NA |
| WBGene00046725 | NA | NA | NA | NA |
| WBGene00046726 | NA | NA | NA | NA |
| WBGene00046727 | NA | NA | NA | NA |
| WBGene00046728 | NA | NA | NA | NA |
| WBGene00046729 | NA | NA | NA | NA |
| WBGene00046730 | NA | NA | NA | NA |
| WBGene00046731 | NA | NA | NA | NA |
| WBGene00046732 | NA | NA | NA | NA |
| WBGene00046733 | NA | NA | NA | NA |
| WBGene00046734 | NA | NA | NA | NA |

| | | | | |
|---|---|---|---|---|
| WBGene00046735 | NA | NA | NA | NA |
| WBGene00046736 | NA | NA | NA | NA |
| WBGene00046737 | NA | NA | NA | NA |
| WBGene00046738 | NA | NA | NA | NA |
| WBGene00046739 | NA | NA | NA | NA |
| WBGene00046740 | NA | NA | NA | NA |
| WBGene00046741 | NA | NA | NA | NA |
| WBGene00046742 | NA | NA | NA | NA |
| WBGene00046743 | NA | NA | NA | NA |
| WBGene00046744 | NA | NA | NA | NA |
| WBGene00046745 | NA | NA | NA | NA |
| WBGene00046746 | NA | NA | NA | NA |
| WBGene00046747 | NA | NA | NA | NA |
| WBGene00046748 | NA | NA | NA | NA |
| WBGene00046749 | NA | NA | NA | NA |
| WBGene00046750 | NA | NA | NA | NA |
| WBGene00046751 | NA | NA | NA | NA |
| WBGene00046752 | NA | NA | NA | NA |
| WBGene00046753 | NA | NA | NA | NA |
| WBGene00046754 | NA | NA | NA | NA |
| WBGene00046755 | NA | NA | NA | NA |
| WBGene00046756 | NA | NA | NA | NA |
| WBGene00046757 | NA | NA | NA | NA |
| WBGene00046758 | NA | NA | NA | NA |
| WBGene00046759 | NA | NA | NA | NA |
| WBGene00046760 | NA | NA | NA | NA |
| WBGene00046761 | NA | NA | NA | NA |
| WBGene00046762 | NA | NA | NA | NA |
| WBGene00046763 | NA | NA | NA | NA |
| WBGene00046764 | NA | NA | NA | NA |
| WBGene00046765 | NA | NA | NA | NA |
| WBGene00046766 | NA | NA | NA | NA |
| WBGene00046767 | NA | NA | NA | NA |
| WBGene00046768 | NA | NA | NA | NA |
| WBGene00046769 | NA | NA | NA | NA |
| WBGene00046770 | NA | NA | NA | NA |
| WBGene00046771 | NA | NA | NA | NA |
| WBGene00046772 | NA | NA | NA | NA |
| WBGene00046773 | NA | NA | NA | NA |
| WBGene00046774 | NA | NA | NA | NA |
| WBGene00046775 | NA | NA | NA | NA |
| WBGene00046776 | NA | NA | NA | NA |
| WBGene00046777 | NA | NA | NA | NA |

| | | | | |
|---|---|---|---|---|
| WBGene00046778 | NA | NA | NA | NA |
| WBGene00046779 | NA | NA | NA | NA |
| WBGene00046780 | NA | NA | NA | NA |
| WBGene00046781 | NA | NA | NA | NA |
| WBGene00046782 | NA | NA | NA | NA |
| WBGene00046783 | NA | NA | NA | NA |
| WBGene00046784 | NA | NA | NA | NA |
| WBGene00046785 | NA | NA | NA | NA |
| WBGene00046786 | NA | NA | NA | NA |
| WBGene00046787 | NA | NA | NA | NA |
| WBGene00046788 | NA | NA | NA | NA |
| WBGene00046789 | NA | NA | NA | NA |
| WBGene00046790 | NA | NA | NA | NA |
| WBGene00046791 | NA | NA | NA | NA |
| WBGene00046792 | NA | NA | NA | NA |
| WBGene00046793 | NA | NA | NA | NA |
| WBGene00046794 | NA | NA | NA | NA |
| WBGene00046795 | NA | NA | NA | NA |
| WBGene00046796 | NA | NA | NA | NA |
| WBGene00046797 | NA | NA | NA | NA |
| WBGene00046798 | NA | NA | NA | NA |
| WBGene00046799 | NA | NA | NA | NA |
| WBGene00046800 | NA | NA | NA | NA |
| WBGene00046801 | NA | NA | NA | NA |
| WBGene00046802 | NA | NA | NA | NA |
| WBGene00046803 | NA | NA | NA | NA |
| WBGene00046804 | NA | NA | NA | NA |
| WBGene00046805 | NA | NA | NA | NA |
| WBGene00046806 | NA | NA | NA | NA |
| WBGene00046807 | NA | NA | NA | NA |
| WBGene00046808 | NA | NA | NA | NA |
| WBGene00046809 | NA | NA | NA | NA |
| WBGene00046810 | NA | NA | NA | NA |
| WBGene00046811 | NA | NA | NA | NA |
| WBGene00046812 | NA | NA | NA | NA |
| WBGene00046813 | NA | NA | NA | NA |
| WBGene00046814 | NA | NA | NA | NA |
| WBGene00046815 | NA | NA | NA | NA |
| WBGene00046816 | NA | NA | NA | NA |
| WBGene00046817 | NA | NA | NA | NA |
| WBGene00046818 | NA | NA | NA | NA |
| WBGene00046819 | NA | NA | NA | NA |
| WBGene00046820 | NA | NA | NA | NA |

| | | | | |
|---|---|---|---|---|
| WBGene00046821 | NA | NA | NA | NA |
| WBGene00046822 | NA | NA | NA | NA |
| WBGene00046823 | NA | NA | NA | NA |
| WBGene00046824 | NA | NA | NA | NA |
| WBGene00046825 | NA | NA | NA | NA |
| WBGene00046826 | NA | NA | NA | NA |
| WBGene00046827 | NA | NA | NA | NA |
| WBGene00046828 | NA | NA | NA | NA |
| WBGene00046829 | NA | NA | NA | NA |
| WBGene00046830 | NA | NA | NA | NA |
| WBGene00046831 | NA | NA | NA | NA |
| WBGene00046832 | NA | NA | NA | NA |
| WBGene00046833 | NA | NA | NA | NA |
| WBGene00046834 | NA | NA | NA | NA |
| WBGene00046835 | NA | NA | NA | NA |
| WBGene00046836 | NA | NA | NA | NA |
| WBGene00046837 | NA | NA | NA | NA |
| WBGene00046838 | NA | NA | NA | NA |
| WBGene00046839 | NA | NA | NA | NA |
| WBGene00046840 | NA | NA | NA | NA |
| WBGene00046841 | NA | NA | NA | NA |
| WBGene00046842 | NA | NA | NA | NA |
| WBGene00046843 | NA | NA | NA | NA |
| WBGene00046844 | NA | NA | NA | NA |
| WBGene00046845 | NA | NA | NA | NA |
| WBGene00046846 | 0.020232312 | 0.84643798 | NA | NA |
| WBGene00046847 | NA | NA | NA | NA |
| WBGene00046848 | NA | NA | NA | NA |
| WBGene00046849 | NA | NA | NA | NA |
| WBGene00046850 | NA | NA | NA | NA |
| WBGene00046851 | NA | NA | NA | NA |
| WBGene00046852 | NA | NA | NA | NA |
| WBGene00046853 | NA | NA | NA | NA |
| WBGene00046854 | NA | NA | NA | NA |
| WBGene00046855 | NA | NA | NA | NA |
| WBGene00046856 | NA | NA | NA | NA |
| WBGene00046857 | NA | NA | NA | NA |
| WBGene00046858 | NA | NA | NA | NA |
| WBGene00046859 | 0.020232312 | 0.84643798 | NA | NA |
| WBGene00046860 | NA | NA | NA | NA |
| WBGene00046861 | NA | NA | NA | NA |
| WBGene00046862 | NA | NA | NA | NA |
| WBGene00046863 | NA | NA | NA | NA |

| | | | | |
|---|---|---|---|---|
| WBGene00046864 | NA | NA | NA | NA |
| WBGene00046865 | NA | NA | NA | NA |
| WBGene00046866 | NA | NA | NA | NA |
| WBGene00046867 | NA | NA | NA | NA |
| WBGene00046868 | NA | NA | NA | NA |
| WBGene00046869 | NA | NA | NA | NA |
| WBGene00046870 | NA | NA | NA | NA |
| WBGene00046871 | NA | NA | NA | NA |
| WBGene00046872 | NA | NA | NA | NA |
| WBGene00046873 | NA | NA | NA | NA |
| WBGene00046874 | NA | NA | NA | NA |
| WBGene00046875 | NA | NA | NA | NA |
| WBGene00046876 | NA | NA | NA | NA |
| WBGene00046877 | NA | NA | NA | NA |
| WBGene00046878 | NA | NA | NA | NA |
| WBGene00046879 | NA | NA | NA | NA |
| WBGene00046880 | NA | NA | NA | NA |
| WBGene00046881 | NA | NA | NA | NA |
| WBGene00046882 | NA | NA | NA | NA |
| WBGene00046883 | NA | NA | NA | NA |
| WBGene00046884 | NA | NA | NA | NA |
| WBGene00046885 | NA | NA | NA | NA |
| WBGene00046886 | NA | NA | NA | NA |
| WBGene00046887 | NA | NA | NA | NA |
| WBGene00046888 | NA | NA | NA | NA |
| WBGene00046889 | NA | NA | NA | NA |
| WBGene00046890 | NA | NA | NA | NA |
| WBGene00046891 | NA | NA | NA | NA |
| WBGene00046892 | NA | NA | NA | NA |
| WBGene00046893 | NA | NA | NA | NA |
| WBGene00046894 | NA | NA | NA | NA |
| WBGene00046895 | NA | NA | NA | NA |
| WBGene00046896 | NA | NA | NA | NA |
| WBGene00046897 | NA | NA | NA | NA |
| WBGene00046898 | NA | NA | NA | NA |
| WBGene00046899 | NA | NA | NA | NA |
| WBGene00046900 | NA | NA | NA | NA |
| WBGene00046901 | NA | NA | NA | NA |
| WBGene00046902 | NA | NA | NA | NA |
| WBGene00046903 | NA | NA | NA | NA |
| WBGene00046904 | NA | NA | NA | NA |
| WBGene00046905 | NA | NA | NA | NA |
| WBGene00046906 | NA | NA | NA | NA |

| | | | | |
|---|---|---|---|---|
| WBGene00046907 | NA | NA | NA | NA |
| WBGene00046908 | NA | NA | NA | NA |
| WBGene00046909 | NA | NA | NA | NA |
| WBGene00046910 | NA | NA | NA | NA |
| WBGene00046911 | NA | NA | NA | NA |
| WBGene00046912 | NA | NA | NA | NA |
| WBGene00046913 | NA | NA | NA | NA |
| WBGene00046914 | NA | NA | NA | NA |
| WBGene00046915 | NA | NA | NA | NA |
| WBGene00046916 | NA | NA | NA | NA |
| WBGene00046917 | NA | NA | NA | NA |
| WBGene00046918 | NA | NA | NA | NA |
| WBGene00046919 | NA | NA | NA | NA |
| WBGene00046920 | NA | NA | NA | NA |
| WBGene00046921 | NA | NA | NA | NA |
| WBGene00046922 | NA | NA | NA | NA |
| WBGene00046923 | NA | NA | NA | NA |
| WBGene00046924 | NA | NA | NA | NA |
| WBGene00046925 | NA | NA | NA | NA |
| WBGene00046926 | NA | NA | NA | NA |
| WBGene00046927 | NA | NA | NA | NA |
| WBGene00046928 | NA | NA | NA | NA |
| WBGene00046929 | NA | NA | NA | NA |
| WBGene00046930 | NA | NA | NA | NA |
| WBGene00046931 | NA | NA | NA | NA |
| WBGene00046932 | NA | NA | NA | NA |
| WBGene00046933 | NA | NA | NA | NA |
| WBGene00046934 | NA | NA | NA | NA |
| WBGene00046935 | NA | NA | NA | NA |
| WBGene00046936 | NA | NA | NA | NA |
| WBGene00046937 | NA | NA | NA | NA |
| WBGene00046938 | NA | NA | NA | NA |
| WBGene00046939 | NA | NA | NA | NA |
| WBGene00046940 | NA | NA | NA | NA |
| WBGene00046941 | NA | NA | NA | NA |
| WBGene00046942 | NA | NA | NA | NA |
| WBGene00046943 | NA | NA | NA | NA |
| WBGene00046944 | NA | NA | NA | NA |
| WBGene00046945 | NA | NA | NA | NA |
| WBGene00046946 | NA | NA | NA | NA |
| WBGene00046947 | NA | NA | NA | NA |
| WBGene00046948 | NA | NA | NA | NA |
| WBGene00046949 | NA | NA | NA | NA |

| | | | | |
|---|---|---|---|---|
| WBGene00046950 | NA | NA | NA | NA |
| WBGene00046951 | NA | NA | NA | NA |
| WBGene00046952 | NA | NA | NA | NA |
| WBGene00046953 | NA | NA | NA | NA |
| WBGene00046954 | NA | NA | NA | NA |
| WBGene00046955 | NA | NA | NA | NA |
| WBGene00046956 | NA | NA | NA | NA |
| WBGene00046957 | NA | NA | NA | NA |
| WBGene00046958 | NA | NA | NA | NA |
| WBGene00046959 | NA | NA | NA | NA |
| WBGene00046960 | NA | NA | NA | NA |
| WBGene00046961 | NA | NA | NA | NA |
| WBGene00046962 | NA | NA | NA | NA |
| WBGene00046963 | NA | NA | NA | NA |
| WBGene00046964 | NA | NA | NA | NA |
| WBGene00046965 | NA | NA | NA | NA |
| WBGene00046966 | NA | NA | NA | NA |
| WBGene00046967 | NA | NA | NA | NA |
| WBGene00046968 | NA | NA | NA | NA |
| WBGene00046969 | NA | NA | NA | NA |
| WBGene00046970 | NA | NA | NA | NA |
| WBGene00046971 | NA | NA | NA | NA |
| WBGene00046972 | NA | NA | NA | NA |
| WBGene00046973 | NA | NA | NA | NA |
| WBGene00046974 | NA | NA | NA | NA |
| WBGene00046975 | NA | NA | NA | NA |
| WBGene00046976 | NA | NA | NA | NA |
| WBGene00046977 | NA | NA | NA | NA |
| WBGene00046978 | NA | NA | NA | NA |
| WBGene00046979 | NA | NA | NA | NA |
| WBGene00046980 | NA | NA | NA | NA |
| WBGene00046981 | NA | NA | NA | NA |
| WBGene00046982 | NA | NA | NA | NA |
| WBGene00046983 | NA | NA | NA | NA |
| WBGene00046984 | NA | NA | NA | NA |
| WBGene00046985 | NA | NA | NA | NA |
| WBGene00046986 | NA | NA | NA | NA |
| WBGene00046987 | NA | NA | NA | NA |
| WBGene00046988 | NA | NA | NA | NA |
| WBGene00046989 | NA | NA | NA | NA |
| WBGene00046990 | NA | NA | NA | NA |
| WBGene00046991 | NA | NA | NA | NA |
| WBGene00046992 | NA | NA | NA | NA |

| | | | | |
|---|---|---|---|---|
| WBGene00046993 | NA | NA | NA | NA |
| WBGene00046994 | NA | NA | NA | NA |
| WBGene00046995 | NA | NA | NA | NA |
| WBGene00046996 | NA | NA | NA | NA |
| WBGene00046997 | NA | NA | NA | NA |
| WBGene00046998 | NA | NA | NA | NA |
| WBGene00046999 | NA | NA | NA | NA |
| WBGene00047000 | NA | NA | NA | NA |
| WBGene00047001 | NA | NA | NA | NA |
| WBGene00047002 | NA | NA | NA | NA |
| WBGene00047003 | NA | NA | NA | NA |
| WBGene00047004 | NA | NA | NA | NA |
| WBGene00047005 | NA | NA | NA | NA |
| WBGene00047006 | NA | NA | NA | NA |
| WBGene00047007 | NA | NA | NA | NA |
| WBGene00047008 | NA | NA | NA | NA |
| WBGene00047009 | NA | NA | NA | NA |
| WBGene00047010 | NA | NA | NA | NA |
| WBGene00047011 | NA | NA | NA | NA |
| WBGene00047012 | NA | NA | NA | NA |
| WBGene00047013 | NA | NA | NA | NA |
| WBGene00047014 | NA | NA | NA | NA |
| WBGene00047015 | NA | NA | NA | NA |
| WBGene00047016 | NA | NA | NA | NA |
| WBGene00047017 | NA | NA | NA | NA |
| WBGene00047018 | NA | NA | NA | NA |
| WBGene00047019 | NA | NA | NA | NA |
| WBGene00047020 | NA | NA | NA | NA |
| WBGene00047021 | NA | NA | NA | NA |
| WBGene00047022 | NA | NA | NA | NA |
| WBGene00047023 | NA | NA | NA | NA |
| WBGene00047024 | NA | NA | NA | NA |
| WBGene00047025 | NA | NA | NA | NA |
| WBGene00047026 | NA | NA | NA | NA |
| WBGene00047027 | NA | NA | NA | NA |
| WBGene00047028 | NA | NA | NA | NA |
| WBGene00047029 | NA | NA | NA | NA |
| WBGene00047030 | NA | NA | NA | NA |
| WBGene00047031 | NA | NA | NA | NA |
| WBGene00047032 | NA | NA | NA | NA |
| WBGene00047033 | NA | NA | NA | NA |
| WBGene00047034 | NA | NA | NA | NA |
| WBGene00047035 | NA | NA | NA | NA |

| | | | | |
|---|---|---|---|---|
| WBGene00047036 | NA | NA | NA | NA |
| WBGene00047037 | NA | NA | NA | NA |
| WBGene00047038 | NA | NA | NA | NA |
| WBGene00047039 | NA | NA | NA | NA |
| WBGene00047040 | NA | NA | NA | NA |
| WBGene00047041 | NA | NA | NA | NA |
| WBGene00047042 | NA | NA | NA | NA |
| WBGene00047043 | NA | NA | NA | NA |
| WBGene00047044 | NA | NA | NA | NA |
| WBGene00047045 | NA | NA | NA | NA |
| WBGene00047046 | NA | NA | NA | NA |
| WBGene00047047 | NA | NA | NA | NA |
| WBGene00047048 | NA | NA | NA | NA |
| WBGene00047049 | NA | NA | NA | NA |
| WBGene00047050 | NA | NA | NA | NA |
| WBGene00047051 | NA | NA | NA | NA |
| WBGene00047052 | NA | NA | NA | NA |
| WBGene00047053 | NA | NA | NA | NA |
| WBGene00047054 | NA | NA | NA | NA |
| WBGene00047055 | NA | NA | NA | NA |
| WBGene00047056 | NA | NA | NA | NA |
| WBGene00047057 | NA | NA | NA | NA |
| WBGene00047058 | NA | NA | NA | NA |
| WBGene00047059 | NA | NA | NA | NA |
| WBGene00047060 | NA | NA | NA | NA |
| WBGene00047061 | NA | NA | NA | NA |
| WBGene00047062 | NA | NA | NA | NA |
| WBGene00047063 | NA | NA | NA | NA |
| WBGene00047064 | NA | NA | NA | NA |
| WBGene00047065 | NA | NA | NA | NA |
| WBGene00047066 | NA | NA | NA | NA |
| WBGene00047067 | NA | NA | NA | NA |
| WBGene00047068 | NA | NA | NA | NA |
| WBGene00047069 | NA | NA | NA | NA |
| WBGene00047070 | NA | NA | NA | NA |
| WBGene00047071 | NA | NA | NA | NA |
| WBGene00047072 | NA | NA | NA | NA |
| WBGene00047073 | NA | NA | NA | NA |
| WBGene00047074 | NA | NA | NA | NA |
| WBGene00047075 | NA | NA | NA | NA |
| WBGene00047076 | NA | NA | NA | NA |
| WBGene00047077 | NA | NA | NA | NA |
| WBGene00047078 | NA | NA | NA | NA |

| | | | | |
|---|---|---|---|---|
| WBGene00047079 | NA | NA | NA | NA |
| WBGene00047080 | NA | NA | NA | NA |
| WBGene00047081 | NA | NA | NA | NA |
| WBGene00047082 | NA | NA | NA | NA |
| WBGene00047083 | NA | NA | NA | NA |
| WBGene00047084 | NA | NA | NA | NA |
| WBGene00047085 | NA | NA | NA | NA |
| WBGene00047086 | NA | NA | NA | NA |
| WBGene00047087 | NA | NA | NA | NA |
| WBGene00047088 | NA | NA | NA | NA |
| WBGene00047089 | NA | NA | NA | NA |
| WBGene00047090 | NA | NA | NA | NA |
| WBGene00047091 | NA | NA | NA | NA |
| WBGene00047092 | NA | NA | NA | NA |
| WBGene00047093 | NA | NA | NA | NA |
| WBGene00047094 | NA | NA | NA | NA |
| WBGene00047095 | NA | NA | NA | NA |
| WBGene00047096 | NA | NA | NA | NA |
| WBGene00047097 | NA | NA | NA | NA |
| WBGene00047098 | NA | NA | NA | NA |
| WBGene00047099 | NA | NA | NA | NA |
| WBGene00047100 | NA | NA | NA | NA |
| WBGene00047101 | NA | NA | NA | NA |
| WBGene00047102 | NA | NA | NA | NA |
| WBGene00047103 | NA | NA | NA | NA |
| WBGene00047104 | NA | NA | NA | NA |
| WBGene00047105 | NA | NA | NA | NA |
| WBGene00047106 | NA | NA | NA | NA |
| WBGene00047107 | NA | NA | NA | NA |
| WBGene00047108 | NA | NA | NA | NA |
| WBGene00047109 | NA | NA | NA | NA |
| WBGene00047110 | NA | NA | NA | NA |
| WBGene00047111 | NA | NA | NA | NA |
| WBGene00047112 | NA | NA | NA | NA |
| WBGene00047113 | NA | NA | NA | NA |
| WBGene00047114 | NA | NA | NA | NA |
| WBGene00047115 | NA | NA | NA | NA |
| WBGene00047116 | NA | NA | NA | NA |
| WBGene00047117 | NA | NA | NA | NA |
| WBGene00047118 | NA | NA | NA | NA |
| WBGene00047119 | NA | NA | NA | NA |
| WBGene00047120 | NA | NA | NA | NA |
| WBGene00047121 | NA | NA | NA | NA |

| | | | | |
|---|---|---|---|---|
| WBGene00047122 | NA | NA | NA | NA |
| WBGene00047123 | NA | NA | NA | NA |
| WBGene00047124 | NA | NA | NA | NA |
| WBGene00047125 | NA | NA | NA | NA |
| WBGene00047126 | NA | NA | NA | NA |
| WBGene00047127 | NA | NA | NA | NA |
| WBGene00047128 | NA | NA | NA | NA |
| WBGene00047129 | NA | NA | NA | NA |
| WBGene00047130 | -0.022930767 | 0.78121253 | NA | NA |
| WBGene00047131 | NA | NA | NA | NA |
| WBGene00047132 | NA | NA | NA | NA |
| WBGene00047133 | NA | NA | NA | NA |
| WBGene00047134 | NA | NA | NA | NA |
| WBGene00047135 | NA | NA | NA | NA |
| WBGene00047136 | NA | NA | NA | NA |
| WBGene00047137 | NA | NA | NA | NA |
| WBGene00047138 | NA | NA | NA | NA |
| WBGene00047139 | NA | NA | NA | NA |
| WBGene00047140 | NA | NA | NA | NA |
| WBGene00047141 | NA | NA | NA | NA |
| WBGene00047142 | NA | NA | NA | NA |
| WBGene00047143 | NA | NA | NA | NA |
| WBGene00047144 | NA | NA | NA | NA |
| WBGene00047145 | NA | NA | NA | NA |
| WBGene00047146 | NA | NA | NA | NA |
| WBGene00047147 | NA | NA | NA | NA |
| WBGene00047148 | NA | NA | NA | NA |
| WBGene00047149 | NA | NA | NA | NA |
| WBGene00047150 | NA | NA | NA | NA |
| WBGene00047151 | NA | NA | NA | NA |
| WBGene00047152 | NA | NA | NA | NA |
| WBGene00047153 | NA | NA | NA | NA |
| WBGene00047154 | NA | NA | NA | NA |
| WBGene00047155 | NA | NA | NA | NA |
| WBGene00047156 | NA | NA | NA | NA |
| WBGene00047157 | NA | NA | NA | NA |
| WBGene00047158 | NA | NA | NA | NA |
| WBGene00047159 | NA | NA | NA | NA |
| WBGene00047160 | NA | NA | NA | NA |
| WBGene00047161 | NA | NA | NA | NA |
| WBGene00047162 | NA | NA | NA | NA |
| WBGene00047163 | NA | NA | NA | NA |
| WBGene00047164 | NA | NA | NA | NA |

| | | | | |
|---|---|---|---|---|
| WBGene00047165 | NA | NA | NA | NA |
| WBGene00047166 | NA | NA | NA | NA |
| WBGene00047167 | NA | NA | NA | NA |
| WBGene00047168 | NA | NA | NA | NA |
| WBGene00047169 | NA | NA | NA | NA |
| WBGene00047170 | NA | NA | NA | NA |
| WBGene00047171 | NA | NA | NA | NA |
| WBGene00047172 | NA | NA | NA | NA |
| WBGene00047173 | NA | NA | NA | NA |
| WBGene00047174 | NA | NA | NA | NA |
| WBGene00047175 | NA | NA | NA | NA |
| WBGene00047176 | NA | NA | NA | NA |
| WBGene00047177 | NA | NA | NA | NA |
| WBGene00047178 | NA | NA | NA | NA |
| WBGene00047179 | NA | NA | NA | NA |
| WBGene00047180 | NA | NA | NA | NA |
| WBGene00047181 | NA | NA | NA | NA |
| WBGene00047182 | NA | NA | NA | NA |
| WBGene00047183 | NA | NA | NA | NA |
| WBGene00047184 | NA | NA | NA | NA |
| WBGene00047185 | NA | NA | NA | NA |
| WBGene00047186 | NA | NA | NA | NA |
| WBGene00047187 | NA | NA | NA | NA |
| WBGene00047188 | NA | NA | NA | NA |
| WBGene00047189 | NA | NA | NA | NA |
| WBGene00047190 | NA | NA | NA | NA |
| WBGene00047191 | NA | NA | NA | NA |
| WBGene00047192 | NA | NA | NA | NA |
| WBGene00047193 | NA | NA | NA | NA |
| WBGene00047194 | NA | NA | NA | NA |
| WBGene00047195 | NA | NA | NA | NA |
| WBGene00047196 | NA | NA | NA | NA |
| WBGene00047197 | NA | NA | NA | NA |
| WBGene00047198 | NA | NA | NA | NA |
| WBGene00047199 | NA | NA | NA | NA |
| WBGene00047200 | NA | NA | NA | NA |
| WBGene00047201 | NA | NA | NA | NA |
| WBGene00047202 | NA | NA | NA | NA |
| WBGene00047203 | NA | NA | NA | NA |
| WBGene00047204 | NA | NA | NA | NA |
| WBGene00047205 | NA | NA | NA | NA |
| WBGene00047206 | NA | NA | NA | NA |
| WBGene00047207 | NA | NA | NA | NA |

| | | | | |
|---|---|---|---|---|
| WBGene00047208 | NA | NA | NA | NA |
| WBGene00047209 | NA | NA | NA | NA |
| WBGene00047210 | NA | NA | NA | NA |
| WBGene00047211 | NA | NA | NA | NA |
| WBGene00047212 | NA | NA | NA | NA |
| WBGene00047213 | NA | NA | NA | NA |
| WBGene00047214 | NA | NA | NA | NA |
| WBGene00047215 | NA | NA | NA | NA |
| WBGene00047216 | NA | NA | NA | NA |
| WBGene00047217 | NA | NA | NA | NA |
| WBGene00047218 | NA | NA | NA | NA |
| WBGene00047219 | NA | NA | NA | NA |
| WBGene00047220 | NA | NA | NA | NA |
| WBGene00047221 | NA | NA | NA | NA |
| WBGene00047222 | NA | NA | NA | NA |
| WBGene00047223 | NA | NA | NA | NA |
| WBGene00047224 | NA | NA | NA | NA |
| WBGene00047225 | NA | NA | NA | NA |
| WBGene00047226 | NA | NA | NA | NA |
| WBGene00047227 | NA | NA | NA | NA |
| WBGene00047228 | NA | NA | NA | NA |
| WBGene00047229 | NA | NA | NA | NA |
| WBGene00047230 | NA | NA | NA | NA |
| WBGene00047231 | NA | NA | NA | NA |
| WBGene00047232 | -0.023056499 | 0.78121253 | NA | NA |
| WBGene00047233 | NA | NA | NA | NA |
| WBGene00047234 | NA | NA | NA | NA |
| WBGene00047235 | NA | NA | NA | NA |
| WBGene00047236 | NA | NA | NA | NA |
| WBGene00047237 | NA | NA | NA | NA |
| WBGene00047238 | NA | NA | NA | NA |
| WBGene00047239 | NA | NA | NA | NA |
| WBGene00047240 | NA | NA | NA | NA |
| WBGene00047241 | NA | NA | NA | NA |
| WBGene00047242 | NA | NA | NA | NA |
| WBGene00047243 | NA | NA | NA | NA |
| WBGene00047244 | NA | NA | NA | NA |
| WBGene00047245 | NA | NA | NA | NA |
| WBGene00047246 | NA | NA | NA | NA |
| WBGene00047247 | NA | NA | NA | NA |
| WBGene00047248 | NA | NA | NA | NA |
| WBGene00047249 | NA | NA | NA | NA |
| WBGene00047250 | NA | NA | NA | NA |

| | | | | |
|---|---|---|---|---|
| WBGene00047251 | NA | NA | NA | NA |
| WBGene00047252 | NA | NA | NA | NA |
| WBGene00047253 | NA | NA | NA | NA |
| WBGene00047254 | NA | NA | NA | NA |
| WBGene00047255 | NA | NA | NA | NA |
| WBGene00047256 | NA | NA | NA | NA |
| WBGene00047257 | NA | NA | NA | NA |
| WBGene00047258 | NA | NA | NA | NA |
| WBGene00047259 | NA | NA | NA | NA |
| WBGene00047260 | NA | NA | NA | NA |
| WBGene00047261 | NA | NA | NA | NA |
| WBGene00047262 | NA | NA | NA | NA |
| WBGene00047263 | NA | NA | NA | NA |
| WBGene00047264 | NA | NA | NA | NA |
| WBGene00047265 | NA | NA | NA | NA |
| WBGene00047266 | NA | NA | NA | NA |
| WBGene00047267 | NA | NA | NA | NA |
| WBGene00047268 | NA | NA | NA | NA |
| WBGene00047269 | NA | NA | NA | NA |
| WBGene00047270 | NA | NA | NA | NA |
| WBGene00047271 | NA | NA | NA | NA |
| WBGene00047272 | NA | NA | NA | NA |
| WBGene00047273 | NA | NA | NA | NA |
| WBGene00047274 | NA | NA | NA | NA |
| WBGene00047275 | NA | NA | NA | NA |
| WBGene00047276 | NA | NA | NA | NA |
| WBGene00047277 | NA | NA | NA | NA |
| WBGene00047278 | NA | NA | NA | NA |
| WBGene00047279 | NA | NA | NA | NA |
| WBGene00047280 | NA | NA | NA | NA |
| WBGene00047281 | NA | NA | NA | NA |
| WBGene00047282 | NA | NA | NA | NA |
| WBGene00047283 | NA | NA | NA | NA |
| WBGene00047284 | NA | NA | NA | NA |
| WBGene00047285 | NA | NA | NA | NA |
| WBGene00047286 | NA | NA | NA | NA |
| WBGene00047287 | NA | NA | NA | NA |
| WBGene00047288 | NA | NA | NA | NA |
| WBGene00047289 | NA | NA | NA | NA |
| WBGene00047290 | NA | NA | NA | NA |
| WBGene00047291 | NA | NA | NA | NA |
| WBGene00047292 | NA | NA | NA | NA |
| WBGene00047293 | NA | NA | NA | NA |

| | | | | |
|---|---|---|---|---|
| WBGene00047294 | NA | NA | NA | NA |
| WBGene00047295 | NA | NA | NA | NA |
| WBGene00047296 | NA | NA | NA | NA |
| WBGene00047297 | NA | NA | NA | NA |
| WBGene00047298 | NA | NA | NA | NA |
| WBGene00047299 | NA | NA | NA | NA |
| WBGene00047300 | NA | NA | NA | NA |
| WBGene00047301 | NA | NA | NA | NA |
| WBGene00047302 | NA | NA | NA | NA |
| WBGene00047303 | NA | NA | NA | NA |
| WBGene00047304 | NA | NA | NA | NA |
| WBGene00047305 | NA | NA | NA | NA |
| WBGene00047306 | NA | NA | NA | NA |
| WBGene00047307 | NA | NA | NA | NA |
| WBGene00047308 | NA | NA | NA | NA |
| WBGene00047309 | NA | NA | NA | NA |
| WBGene00047310 | NA | NA | NA | NA |
| WBGene00047311 | NA | NA | NA | NA |
| WBGene00047312 | NA | NA | NA | NA |
| WBGene00047313 | NA | NA | NA | NA |
| WBGene00047314 | NA | NA | NA | NA |
| WBGene00047315 | NA | NA | NA | NA |
| WBGene00047316 | NA | NA | NA | NA |
| WBGene00047317 | NA | NA | NA | NA |
| WBGene00047318 | NA | NA | NA | NA |
| WBGene00047319 | NA | NA | NA | NA |
| WBGene00047320 | NA | NA | NA | NA |
| WBGene00047321 | NA | NA | NA | NA |
| WBGene00047322 | NA | NA | NA | NA |
| WBGene00047323 | NA | NA | NA | NA |
| WBGene00047324 | NA | NA | NA | NA |
| WBGene00047325 | NA | NA | NA | NA |
| WBGene00047326 | NA | NA | NA | NA |
| WBGene00047327 | NA | NA | NA | NA |
| WBGene00047328 | NA | NA | NA | NA |
| WBGene00047329 | NA | NA | NA | NA |
| WBGene00047330 | NA | NA | NA | NA |
| WBGene00047331 | NA | NA | NA | NA |
| WBGene00047332 | NA | NA | NA | NA |
| WBGene00047333 | NA | NA | NA | NA |
| WBGene00047334 | NA | NA | NA | NA |
| WBGene00047335 | NA | NA | NA | NA |
| WBGene00047336 | NA | NA | NA | NA |

| | | | | |
|---|---|---|---|---|
| WBGene00047337 | NA | NA | NA | NA |
| WBGene00047338 | NA | NA | NA | NA |
| WBGene00047339 | NA | NA | NA | NA |
| WBGene00047340 | NA | NA | NA | NA |
| WBGene00047341 | NA | NA | NA | NA |
| WBGene00047342 | NA | NA | NA | NA |
| WBGene00047343 | NA | NA | NA | NA |
| WBGene00047344 | NA | NA | NA | NA |
| WBGene00047345 | NA | NA | NA | NA |
| WBGene00047346 | NA | NA | NA | NA |
| WBGene00047347 | NA | NA | NA | NA |
| WBGene00047348 | NA | NA | NA | NA |
| WBGene00047349 | NA | NA | NA | NA |
| WBGene00047350 | NA | NA | NA | NA |
| WBGene00047351 | NA | NA | NA | NA |
| WBGene00047352 | NA | NA | NA | NA |
| WBGene00047353 | NA | NA | NA | NA |
| WBGene00047354 | NA | NA | NA | NA |
| WBGene00047355 | NA | NA | NA | NA |
| WBGene00047356 | NA | NA | NA | NA |
| WBGene00047357 | NA | NA | NA | NA |
| WBGene00047358 | NA | NA | NA | NA |
| WBGene00047359 | NA | NA | NA | NA |
| WBGene00047360 | NA | NA | NA | NA |
| WBGene00047361 | NA | NA | NA | NA |
| WBGene00047362 | NA | NA | NA | NA |
| WBGene00047363 | NA | NA | NA | NA |
| WBGene00047364 | NA | NA | NA | NA |
| WBGene00047365 | NA | NA | NA | NA |
| WBGene00047366 | NA | NA | NA | NA |
| WBGene00047367 | NA | NA | NA | NA |
| WBGene00047368 | NA | NA | NA | NA |
| WBGene00047369 | NA | NA | NA | NA |
| WBGene00047370 | NA | NA | NA | NA |
| WBGene00047371 | NA | NA | NA | NA |
| WBGene00047372 | NA | NA | NA | NA |
| WBGene00047373 | NA | NA | NA | NA |
| WBGene00047374 | NA | NA | NA | NA |
| WBGene00047375 | NA | NA | NA | NA |
| WBGene00047376 | NA | NA | NA | NA |
| WBGene00047377 | NA | NA | NA | NA |
| WBGene00047378 | NA | NA | NA | NA |
| WBGene00047379 | NA | NA | NA | NA |

| | | | | |
|---|---|---|---|---|
| WBGene00047380 | NA | NA | NA | NA |
| WBGene00047381 | NA | NA | NA | NA |
| WBGene00047382 | NA | NA | NA | NA |
| WBGene00047383 | NA | NA | NA | NA |
| WBGene00047384 | NA | NA | NA | NA |
| WBGene00047385 | NA | NA | NA | NA |
| WBGene00047386 | NA | NA | NA | NA |
| WBGene00047387 | NA | NA | NA | NA |
| WBGene00047388 | NA | NA | NA | NA |
| WBGene00047389 | NA | NA | NA | NA |
| WBGene00047390 | NA | NA | NA | NA |
| WBGene00047391 | NA | NA | NA | NA |
| WBGene00047392 | NA | NA | NA | NA |
| WBGene00047393 | NA | NA | NA | NA |
| WBGene00047394 | NA | NA | NA | NA |
| WBGene00047395 | NA | NA | NA | NA |
| WBGene00047396 | NA | NA | NA | NA |
| WBGene00047397 | NA | NA | NA | NA |
| WBGene00047398 | NA | NA | NA | NA |
| WBGene00047399 | NA | NA | NA | NA |
| WBGene00047400 | NA | NA | NA | NA |
| WBGene00047401 | NA | NA | NA | NA |
| WBGene00047402 | NA | NA | NA | NA |
| WBGene00047403 | NA | NA | NA | NA |
| WBGene00047404 | NA | NA | NA | NA |
| WBGene00047405 | NA | NA | NA | NA |
| WBGene00047406 | NA | NA | NA | NA |
| WBGene00047407 | NA | NA | NA | NA |
| WBGene00047408 | NA | NA | NA | NA |
| WBGene00047409 | NA | NA | NA | NA |
| WBGene00047410 | NA | NA | NA | NA |
| WBGene00047411 | NA | NA | NA | NA |
| WBGene00047412 | NA | NA | NA | NA |
| WBGene00047413 | NA | NA | NA | NA |
| WBGene00047414 | NA | NA | NA | NA |
| WBGene00047415 | NA | NA | NA | NA |
| WBGene00047416 | NA | NA | NA | NA |
| WBGene00047417 | 0.020232312 | 0.84643798 | NA | NA |
| WBGene00047418 | NA | NA | NA | NA |
| WBGene00047419 | NA | NA | NA | NA |
| WBGene00047420 | NA | NA | NA | NA |
| WBGene00047421 | NA | NA | NA | NA |
| WBGene00047422 | NA | NA | NA | NA |

| | | | | |
|---|---|---|---|---|
| WBGene00047423 | NA | NA | NA | NA |
| WBGene00047424 | NA | NA | NA | NA |
| WBGene00047425 | NA | NA | NA | NA |
| WBGene00047426 | NA | NA | NA | NA |
| WBGene00047427 | NA | NA | NA | NA |
| WBGene00047428 | NA | NA | NA | NA |
| WBGene00047429 | NA | NA | NA | NA |
| WBGene00047430 | NA | NA | NA | NA |
| WBGene00047431 | NA | NA | NA | NA |
| WBGene00047432 | NA | NA | NA | NA |
| WBGene00047433 | NA | NA | NA | NA |
| WBGene00047434 | NA | NA | NA | NA |
| WBGene00047435 | NA | NA | NA | NA |
| WBGene00047436 | NA | NA | NA | NA |
| WBGene00047437 | NA | NA | NA | NA |
| WBGene00047438 | NA | NA | NA | NA |
| WBGene00047439 | NA | NA | NA | NA |
| WBGene00047440 | NA | NA | NA | NA |
| WBGene00047441 | NA | NA | NA | NA |
| WBGene00047442 | NA | NA | NA | NA |
| WBGene00047443 | NA | NA | NA | NA |
| WBGene00047444 | NA | NA | NA | NA |
| WBGene00047445 | NA | NA | NA | NA |
| WBGene00047446 | NA | NA | NA | NA |
| WBGene00047447 | NA | NA | NA | NA |
| WBGene00047448 | NA | NA | NA | NA |
| WBGene00047449 | NA | NA | NA | NA |
| WBGene00047450 | NA | NA | NA | NA |
| WBGene00047451 | NA | NA | NA | NA |
| WBGene00047452 | NA | NA | NA | NA |
| WBGene00047453 | NA | NA | NA | NA |
| WBGene00047454 | NA | NA | NA | NA |
| WBGene00047455 | NA | NA | NA | NA |
| WBGene00047456 | NA | NA | NA | NA |
| WBGene00047457 | NA | NA | NA | NA |
| WBGene00047458 | NA | NA | NA | NA |
| WBGene00047459 | NA | NA | NA | NA |
| WBGene00047460 | NA | NA | NA | NA |
| WBGene00047461 | NA | NA | NA | NA |
| WBGene00047462 | NA | NA | NA | NA |
| WBGene00047463 | NA | NA | NA | NA |
| WBGene00047464 | NA | NA | NA | NA |
| WBGene00047465 | NA | NA | NA | NA |

| | | | | |
|---|---|---|---|---|
| WBGene00047466 | NA | NA | NA | NA |
| WBGene00047467 | NA | NA | NA | NA |
| WBGene00047468 | NA | NA | NA | NA |
| WBGene00047469 | NA | NA | NA | NA |
| WBGene00047470 | NA | NA | NA | NA |
| WBGene00047471 | NA | NA | NA | NA |
| WBGene00047472 | NA | NA | NA | NA |
| WBGene00047473 | NA | NA | NA | NA |
| WBGene00047474 | NA | NA | NA | NA |
| WBGene00047475 | NA | NA | NA | NA |
| WBGene00047476 | NA | NA | NA | NA |
| WBGene00047477 | NA | NA | NA | NA |
| WBGene00047478 | NA | NA | NA | NA |
| WBGene00047479 | NA | NA | NA | NA |
| WBGene00047480 | NA | NA | NA | NA |
| WBGene00047481 | NA | NA | NA | NA |
| WBGene00047482 | NA | NA | NA | NA |
| WBGene00047483 | NA | NA | NA | NA |
| WBGene00047484 | NA | NA | NA | NA |
| WBGene00047485 | NA | NA | NA | NA |
| WBGene00047486 | NA | NA | NA | NA |
| WBGene00047487 | NA | NA | NA | NA |
| WBGene00047488 | NA | NA | NA | NA |
| WBGene00047489 | NA | NA | NA | NA |
| WBGene00047490 | NA | NA | NA | NA |
| WBGene00047491 | NA | NA | NA | NA |
| WBGene00047492 | NA | NA | NA | NA |
| WBGene00047493 | NA | NA | NA | NA |
| WBGene00047494 | NA | NA | NA | NA |
| WBGene00047495 | NA | NA | NA | NA |
| WBGene00047496 | NA | NA | NA | NA |
| WBGene00047497 | NA | NA | NA | NA |
| WBGene00047498 | NA | NA | NA | NA |
| WBGene00047499 | NA | NA | NA | NA |
| WBGene00047500 | NA | NA | NA | NA |
| WBGene00047501 | NA | NA | NA | NA |
| WBGene00047502 | NA | NA | NA | NA |
| WBGene00047503 | NA | NA | NA | NA |
| WBGene00047504 | NA | NA | NA | NA |
| WBGene00047505 | NA | NA | NA | NA |
| WBGene00047506 | NA | NA | NA | NA |
| WBGene00047507 | NA | NA | NA | NA |
| WBGene00047508 | NA | NA | NA | NA |

| | | | | |
|---|---|---|---|---|
| WBGene00047509 | NA | NA | NA | NA |
| WBGene00047510 | NA | NA | NA | NA |
| WBGene00047511 | NA | NA | NA | NA |
| WBGene00047512 | NA | NA | NA | NA |
| WBGene00047513 | NA | NA | NA | NA |
| WBGene00047514 | NA | NA | NA | NA |
| WBGene00047515 | NA | NA | NA | NA |
| WBGene00047516 | NA | NA | NA | NA |
| WBGene00047517 | NA | NA | NA | NA |
| WBGene00047518 | NA | NA | NA | NA |
| WBGene00047519 | NA | NA | NA | NA |
| WBGene00047520 | NA | NA | NA | NA |
| WBGene00047521 | NA | NA | NA | NA |
| WBGene00047522 | NA | NA | NA | NA |
| WBGene00047523 | NA | NA | NA | NA |
| WBGene00047524 | NA | NA | NA | NA |
| WBGene00047525 | NA | NA | NA | NA |
| WBGene00047526 | NA | NA | NA | NA |
| WBGene00047527 | NA | NA | NA | NA |
| WBGene00047528 | NA | NA | NA | NA |
| WBGene00047529 | NA | NA | NA | NA |
| WBGene00047530 | NA | NA | NA | NA |
| WBGene00047531 | NA | NA | NA | NA |
| WBGene00047532 | NA | NA | NA | NA |
| WBGene00047533 | NA | NA | NA | NA |
| WBGene00047534 | NA | NA | NA | NA |
| WBGene00047535 | NA | NA | NA | NA |
| WBGene00047536 | NA | NA | NA | NA |
| WBGene00047537 | NA | NA | NA | NA |
| WBGene00047538 | NA | NA | NA | NA |
| WBGene00047539 | NA | NA | NA | NA |
| WBGene00047540 | NA | NA | NA | NA |
| WBGene00047541 | NA | NA | NA | NA |
| WBGene00047542 | NA | NA | NA | NA |
| WBGene00047543 | NA | NA | NA | NA |
| WBGene00047544 | NA | NA | NA | NA |
| WBGene00047545 | NA | NA | NA | NA |
| WBGene00047546 | NA | NA | NA | NA |
| WBGene00047547 | NA | NA | NA | NA |
| WBGene00047548 | NA | NA | NA | NA |
| WBGene00047549 | NA | NA | NA | NA |
| WBGene00047550 | NA | NA | NA | NA |
| WBGene00047551 | NA | NA | NA | NA |

| | | | | |
|---|---|---|---|---|
| WBGene00047552 | NA | NA | NA | NA |
| WBGene00047553 | NA | NA | NA | NA |
| WBGene00047554 | NA | NA | NA | NA |
| WBGene00047555 | NA | NA | NA | NA |
| WBGene00047556 | NA | NA | NA | NA |
| WBGene00047557 | NA | NA | NA | NA |
| WBGene00047558 | NA | NA | NA | NA |
| WBGene00047559 | NA | NA | NA | NA |
| WBGene00047560 | NA | NA | NA | NA |
| WBGene00047561 | NA | NA | NA | NA |
| WBGene00047562 | NA | NA | NA | NA |
| WBGene00047563 | NA | NA | NA | NA |
| WBGene00047564 | NA | NA | NA | NA |
| WBGene00047565 | NA | NA | NA | NA |
| WBGene00047566 | NA | NA | NA | NA |
| WBGene00047567 | NA | NA | NA | NA |
| WBGene00047568 | NA | NA | NA | NA |
| WBGene00047569 | NA | NA | NA | NA |
| WBGene00047570 | NA | NA | NA | NA |
| WBGene00047571 | NA | NA | NA | NA |
| WBGene00047572 | NA | NA | NA | NA |
| WBGene00047573 | NA | NA | NA | NA |
| WBGene00047574 | NA | NA | NA | NA |
| WBGene00047575 | NA | NA | NA | NA |
| WBGene00047576 | NA | NA | NA | NA |
| WBGene00047577 | NA | NA | NA | NA |
| WBGene00047578 | NA | NA | NA | NA |
| WBGene00047579 | NA | NA | NA | NA |
| WBGene00047580 | NA | NA | NA | NA |
| WBGene00047581 | NA | NA | NA | NA |
| WBGene00047582 | NA | NA | NA | NA |
| WBGene00047583 | NA | NA | NA | NA |
| WBGene00047584 | NA | NA | NA | NA |
| WBGene00047585 | NA | NA | NA | NA |
| WBGene00047586 | NA | NA | NA | NA |
| WBGene00047587 | NA | NA | NA | NA |
| WBGene00047588 | NA | NA | NA | NA |
| WBGene00047589 | NA | NA | NA | NA |
| WBGene00047590 | NA | NA | NA | NA |
| WBGene00047591 | NA | NA | NA | NA |
| WBGene00047592 | NA | NA | NA | NA |
| WBGene00047593 | NA | NA | NA | NA |
| WBGene00047594 | NA | NA | NA | NA |

| | | | | |
|---|---|---|---|---|
| WBGene00047595 | NA | NA | NA | NA |
| WBGene00047596 | NA | NA | NA | NA |
| WBGene00047597 | NA | NA | NA | NA |
| WBGene00047598 | NA | NA | NA | NA |
| WBGene00047599 | NA | NA | NA | NA |
| WBGene00047600 | NA | NA | NA | NA |
| WBGene00047601 | NA | NA | NA | NA |
| WBGene00047602 | NA | NA | NA | NA |
| WBGene00047603 | NA | NA | NA | NA |
| WBGene00047604 | NA | NA | NA | NA |
| WBGene00047605 | NA | NA | NA | NA |
| WBGene00047606 | NA | NA | NA | NA |
| WBGene00047607 | NA | NA | NA | NA |
| WBGene00047608 | NA | NA | NA | NA |
| WBGene00047609 | NA | NA | NA | NA |
| WBGene00047610 | NA | NA | NA | NA |
| WBGene00047611 | NA | NA | NA | NA |
| WBGene00047612 | NA | NA | NA | NA |
| WBGene00047613 | NA | NA | NA | NA |
| WBGene00047614 | NA | NA | NA | NA |
| WBGene00047615 | -0.023109762 | 0.78121253 | NA | NA |
| WBGene00047616 | NA | NA | NA | NA |
| WBGene00047617 | NA | NA | NA | NA |
| WBGene00047618 | NA | NA | NA | NA |
| WBGene00047619 | NA | NA | NA | NA |
| WBGene00047620 | NA | NA | NA | NA |
| WBGene00047621 | NA | NA | NA | NA |
| WBGene00047622 | NA | NA | NA | NA |
| WBGene00047623 | NA | NA | NA | NA |
| WBGene00047624 | NA | NA | NA | NA |
| WBGene00047625 | NA | NA | NA | NA |
| WBGene00047626 | NA | NA | NA | NA |
| WBGene00047627 | NA | NA | NA | NA |
| WBGene00047628 | NA | NA | NA | NA |
| WBGene00047629 | NA | NA | NA | NA |
| WBGene00047630 | NA | NA | NA | NA |
| WBGene00047631 | NA | NA | NA | NA |
| WBGene00047632 | NA | NA | NA | NA |
| WBGene00047633 | NA | NA | NA | NA |
| WBGene00047634 | NA | NA | NA | NA |
| WBGene00047635 | NA | NA | NA | NA |
| WBGene00047636 | NA | NA | NA | NA |
| WBGene00047637 | NA | NA | NA | NA |

| | | | | |
|---|---|---|---|---|
| WBGene00047638 | NA | NA | NA | NA |
| WBGene00047639 | NA | NA | NA | NA |
| WBGene00047640 | NA | NA | NA | NA |
| WBGene00047641 | NA | NA | NA | NA |
| WBGene00047642 | NA | NA | NA | NA |
| WBGene00047643 | NA | NA | NA | NA |
| WBGene00047644 | NA | NA | NA | NA |
| WBGene00047645 | NA | NA | NA | NA |
| WBGene00047646 | NA | NA | NA | NA |
| WBGene00047647 | NA | NA | NA | NA |
| WBGene00047648 | NA | NA | NA | NA |
| WBGene00047649 | NA | NA | NA | NA |
| WBGene00047650 | NA | NA | NA | NA |
| WBGene00047651 | NA | NA | NA | NA |
| WBGene00047652 | NA | NA | NA | NA |
| WBGene00047653 | NA | NA | NA | NA |
| WBGene00047654 | NA | NA | NA | NA |
| WBGene00047655 | NA | NA | NA | NA |
| WBGene00047656 | NA | NA | NA | NA |
| WBGene00047657 | NA | NA | NA | NA |
| WBGene00047658 | NA | NA | NA | NA |
| WBGene00047659 | NA | NA | NA | NA |
| WBGene00047660 | NA | NA | NA | NA |
| WBGene00047661 | NA | NA | NA | NA |
| WBGene00047662 | NA | NA | NA | NA |
| WBGene00047663 | NA | NA | NA | NA |
| WBGene00047664 | NA | NA | NA | NA |
| WBGene00047665 | NA | NA | NA | NA |
| WBGene00047666 | NA | NA | NA | NA |
| WBGene00047667 | NA | NA | NA | NA |
| WBGene00047668 | NA | NA | NA | NA |
| WBGene00047669 | NA | NA | NA | NA |
| WBGene00047670 | NA | NA | NA | NA |
| WBGene00047671 | NA | NA | NA | NA |
| WBGene00047672 | NA | NA | NA | NA |
| WBGene00047673 | NA | NA | NA | NA |
| WBGene00047674 | NA | NA | NA | NA |
| WBGene00047675 | NA | NA | NA | NA |
| WBGene00047676 | NA | NA | NA | NA |
| WBGene00047677 | NA | NA | NA | NA |
| WBGene00047678 | NA | NA | NA | NA |
| WBGene00047679 | NA | NA | NA | NA |
| WBGene00047680 | NA | NA | NA | NA |

| | | | | |
|---|---|---|---|---|
| WBGene00047681 | NA | NA | NA | NA |
| WBGene00047682 | NA | NA | NA | NA |
| WBGene00047683 | NA | NA | NA | NA |
| WBGene00047684 | NA | NA | NA | NA |
| WBGene00047685 | NA | NA | NA | NA |
| WBGene00047686 | NA | NA | NA | NA |
| WBGene00047687 | NA | NA | NA | NA |
| WBGene00047688 | NA | NA | NA | NA |
| WBGene00047689 | NA | NA | NA | NA |
| WBGene00047690 | NA | NA | NA | NA |
| WBGene00047691 | NA | NA | NA | NA |
| WBGene00047692 | NA | NA | NA | NA |
| WBGene00047693 | NA | NA | NA | NA |
| WBGene00047694 | NA | NA | NA | NA |
| WBGene00047695 | NA | NA | NA | NA |
| WBGene00047696 | NA | NA | NA | NA |
| WBGene00047697 | NA | NA | NA | NA |
| WBGene00047698 | NA | NA | NA | NA |
| WBGene00047699 | NA | NA | NA | NA |
| WBGene00047700 | NA | NA | NA | NA |
| WBGene00047701 | NA | NA | NA | NA |
| WBGene00047702 | NA | NA | NA | NA |
| WBGene00047703 | NA | NA | NA | NA |
| WBGene00047704 | NA | NA | NA | NA |
| WBGene00047705 | NA | NA | NA | NA |
| WBGene00047706 | NA | NA | NA | NA |
| WBGene00047707 | NA | NA | NA | NA |
| WBGene00047708 | NA | NA | NA | NA |
| WBGene00047709 | NA | NA | NA | NA |
| WBGene00047710 | NA | NA | NA | NA |
| WBGene00047711 | NA | NA | NA | NA |
| WBGene00047712 | NA | NA | NA | NA |
| WBGene00047713 | NA | NA | NA | NA |
| WBGene00047714 | NA | NA | NA | NA |
| WBGene00047715 | NA | NA | NA | NA |
| WBGene00047716 | NA | NA | NA | NA |
| WBGene00047717 | NA | NA | NA | NA |
| WBGene00047718 | NA | NA | NA | NA |
| WBGene00047719 | NA | NA | NA | NA |
| WBGene00047720 | NA | NA | NA | NA |
| WBGene00047721 | NA | NA | NA | NA |
| WBGene00047722 | NA | NA | NA | NA |
| WBGene00047723 | NA | NA | NA | NA |

| | | | | | |
|---|---|---|---|---|---|
| WBGene00047724 | NA | | NA | NA | NA |
| WBGene00047725 | NA | | NA | NA | NA |
| WBGene00047726 | | -0.023056499 | 0.78121253 | NA | NA |
| WBGene00047727 | NA | | NA | NA | NA |
| WBGene00047728 | NA | | NA | NA | NA |
| WBGene00047729 | NA | | NA | NA | NA |
| WBGene00047730 | NA | | NA | NA | NA |
| WBGene00047731 | NA | | NA | NA | NA |
| WBGene00047732 | NA | | NA | NA | NA |
| WBGene00047733 | NA | | NA | NA | NA |
| WBGene00047734 | NA | | NA | NA | NA |
| WBGene00047735 | NA | | NA | NA | NA |
| WBGene00047736 | NA | | NA | NA | NA |
| WBGene00047737 | NA | | NA | NA | NA |
| WBGene00047738 | NA | | NA | NA | NA |
| WBGene00047739 | NA | | NA | NA | NA |
| WBGene00047740 | NA | | NA | NA | NA |
| WBGene00047741 | NA | | NA | NA | NA |
| WBGene00047742 | NA | | NA | NA | NA |
| WBGene00047743 | NA | | NA | NA | NA |
| WBGene00047744 | NA | | NA | NA | NA |
| WBGene00047745 | NA | | NA | NA | NA |
| WBGene00047746 | NA | | NA | NA | NA |
| WBGene00047747 | NA | | NA | NA | NA |
| WBGene00047748 | NA | | NA | NA | NA |
| WBGene00047749 | NA | | NA | NA | NA |
| WBGene00047750 | NA | | NA | NA | NA |
| WBGene00047751 | NA | | NA | NA | NA |
| WBGene00047752 | NA | | NA | NA | NA |
| WBGene00047753 | NA | | NA | NA | NA |
| WBGene00047754 | NA | | NA | NA | NA |
| WBGene00047755 | NA | | NA | NA | NA |
| WBGene00047756 | NA | | NA | NA | NA |
| WBGene00047757 | NA | | NA | NA | NA |
| WBGene00047758 | NA | | NA | NA | NA |
| WBGene00047759 | NA | | NA | NA | NA |
| WBGene00047760 | NA | | NA | NA | NA |
| WBGene00047761 | NA | | NA | NA | NA |
| WBGene00047762 | NA | | NA | NA | NA |
| WBGene00047763 | NA | | NA | NA | NA |
| WBGene00047764 | NA | | NA | NA | NA |
| WBGene00047765 | NA | | NA | NA | NA |
| WBGene00047766 | NA | | NA | NA | NA |

| | | | | |
|---|---|---|---|---|
| WBGene00047767 | NA | NA | NA | NA |
| WBGene00047768 | NA | NA | NA | NA |
| WBGene00047769 | NA | NA | NA | NA |
| WBGene00047770 | NA | NA | NA | NA |
| WBGene00047771 | NA | NA | NA | NA |
| WBGene00047772 | NA | NA | NA | NA |
| WBGene00047773 | NA | NA | NA | NA |
| WBGene00047774 | NA | NA | NA | NA |
| WBGene00047775 | NA | NA | NA | NA |
| WBGene00047776 | 0.020317245 | 0.84643798 | NA | NA |
| WBGene00047777 | NA | NA | NA | NA |
| WBGene00047778 | NA | NA | NA | NA |
| WBGene00047779 | NA | NA | NA | NA |
| WBGene00047780 | NA | NA | NA | NA |
| WBGene00047781 | NA | NA | NA | NA |
| WBGene00047782 | NA | NA | NA | NA |
| WBGene00047783 | NA | NA | NA | NA |
| WBGene00047784 | NA | NA | NA | NA |
| WBGene00047785 | NA | NA | NA | NA |
| WBGene00047786 | NA | NA | NA | NA |
| WBGene00047787 | NA | NA | NA | NA |
| WBGene00047788 | NA | NA | NA | NA |
| WBGene00047789 | NA | NA | NA | NA |
| WBGene00047790 | NA | NA | NA | NA |
| WBGene00047791 | NA | NA | NA | NA |
| WBGene00047792 | NA | NA | NA | NA |
| WBGene00047793 | NA | NA | NA | NA |
| WBGene00047794 | NA | NA | NA | NA |
| WBGene00047795 | NA | NA | NA | NA |
| WBGene00047796 | NA | NA | NA | NA |
| WBGene00047797 | NA | NA | NA | NA |
| WBGene00047798 | NA | NA | NA | NA |
| WBGene00047799 | NA | NA | NA | NA |
| WBGene00047800 | NA | NA | NA | NA |
| WBGene00047801 | NA | NA | NA | NA |
| WBGene00047802 | NA | NA | NA | NA |
| WBGene00047803 | NA | NA | NA | NA |
| WBGene00047804 | NA | NA | NA | NA |
| WBGene00047805 | NA | NA | NA | NA |
| WBGene00047806 | NA | NA | NA | NA |
| WBGene00047807 | NA | NA | NA | NA |
| WBGene00047808 | NA | NA | NA | NA |
| WBGene00047809 | NA | NA | NA | NA |

| | | | | |
|---|---|---|---|---|
| WBGene00047810 | NA | NA | NA | NA |
| WBGene00047811 | NA | NA | NA | NA |
| WBGene00047812 | NA | NA | NA | NA |
| WBGene00047813 | NA | NA | NA | NA |
| WBGene00047814 | NA | NA | NA | NA |
| WBGene00047815 | NA | NA | NA | NA |
| WBGene00047816 | NA | NA | NA | NA |
| WBGene00047817 | NA | NA | NA | NA |
| WBGene00047818 | NA | NA | NA | NA |
| WBGene00047819 | NA | NA | NA | NA |
| WBGene00047820 | NA | NA | NA | NA |
| WBGene00047821 | NA | NA | NA | NA |
| WBGene00047822 | NA | NA | NA | NA |
| WBGene00047823 | NA | NA | NA | NA |
| WBGene00047824 | NA | NA | NA | NA |
| WBGene00047825 | NA | NA | NA | NA |
| WBGene00047826 | NA | NA | NA | NA |
| WBGene00047827 | NA | NA | NA | NA |
| WBGene00047828 | NA | NA | NA | NA |
| WBGene00047829 | NA | NA | NA | NA |
| WBGene00047830 | NA | NA | NA | NA |
| WBGene00047831 | NA | NA | NA | NA |
| WBGene00047832 | NA | NA | NA | NA |
| WBGene00047833 | NA | NA | NA | NA |
| WBGene00047834 | NA | NA | NA | NA |
| WBGene00047835 | 0.0398788 | 0.58971221 | NA | NA |
| WBGene00047836 | NA | NA | NA | NA |
| WBGene00047837 | NA | NA | NA | NA |
| WBGene00047838 | NA | NA | NA | NA |
| WBGene00047839 | NA | NA | NA | NA |
| WBGene00047840 | NA | NA | NA | NA |
| WBGene00047841 | NA | NA | NA | NA |
| WBGene00047842 | NA | NA | NA | NA |
| WBGene00047843 | NA | NA | NA | NA |
| WBGene00047844 | NA | NA | NA | NA |
| WBGene00047845 | NA | NA | NA | NA |
| WBGene00047846 | NA | NA | NA | NA |
| WBGene00047847 | NA | NA | NA | NA |
| WBGene00047848 | NA | NA | NA | NA |
| WBGene00047849 | NA | NA | NA | NA |
| WBGene00047850 | NA | NA | NA | NA |
| WBGene00047851 | NA | NA | NA | NA |
| WBGene00047852 | NA | NA | NA | NA |

| | | | | |
|---|---|---|---|---|
| WBGene00047853 | NA | NA | NA | NA |
| WBGene00047854 | NA | NA | NA | NA |
| WBGene00047855 | NA | NA | NA | NA |
| WBGene00047856 | NA | NA | NA | NA |
| WBGene00047857 | NA | NA | NA | NA |
| WBGene00047858 | NA | NA | NA | NA |
| WBGene00047859 | NA | NA | NA | NA |
| WBGene00047860 | NA | NA | NA | NA |
| WBGene00047861 | NA | NA | NA | NA |
| WBGene00047862 | NA | NA | NA | NA |
| WBGene00047863 | NA | NA | NA | NA |
| WBGene00047864 | NA | NA | NA | NA |
| WBGene00047865 | NA | NA | NA | NA |
| WBGene00047866 | NA | NA | NA | NA |
| WBGene00047867 | NA | NA | NA | NA |
| WBGene00047868 | NA | NA | NA | NA |
| WBGene00047869 | NA | NA | NA | NA |
| WBGene00047870 | NA | NA | NA | NA |
| WBGene00047871 | NA | NA | NA | NA |
| WBGene00047872 | NA | NA | NA | NA |
| WBGene00047873 | NA | NA | NA | NA |
| WBGene00047874 | NA | NA | NA | NA |
| WBGene00047875 | NA | NA | NA | NA |
| WBGene00047876 | NA | NA | NA | NA |
| WBGene00047877 | NA | NA | NA | NA |
| WBGene00047878 | NA | NA | NA | NA |
| WBGene00047879 | NA | NA | NA | NA |
| WBGene00047880 | NA | NA | NA | NA |
| WBGene00047881 | NA | NA | NA | NA |
| WBGene00047882 | NA | NA | NA | NA |
| WBGene00047883 | NA | NA | NA | NA |
| WBGene00047884 | NA | NA | NA | NA |
| WBGene00047885 | NA | NA | NA | NA |
| WBGene00047886 | NA | NA | NA | NA |
| WBGene00047887 | NA | NA | NA | NA |
| WBGene00047888 | NA | NA | NA | NA |
| WBGene00047889 | NA | NA | NA | NA |
| WBGene00047890 | NA | NA | NA | NA |
| WBGene00047891 | NA | NA | NA | NA |
| WBGene00047892 | NA | NA | NA | NA |
| WBGene00047893 | NA | NA | NA | NA |
| WBGene00047894 | NA | NA | NA | NA |
| WBGene00047895 | NA | NA | NA | NA |

| | | | | |
|---|---|---|---|---|
| WBGene00047896 | NA | NA | NA | NA |
| WBGene00047897 | NA | NA | NA | NA |
| WBGene00047898 | NA | NA | NA | NA |
| WBGene00047899 | NA | NA | NA | NA |
| WBGene00047900 | NA | NA | NA | NA |
| WBGene00047901 | NA | NA | NA | NA |
| WBGene00047902 | NA | NA | NA | NA |
| WBGene00047903 | NA | NA | NA | NA |
| WBGene00047905 | NA | NA | NA | NA |
| WBGene00047906 | NA | NA | NA | NA |
| WBGene00047907 | 0.020317245 | 0.84643798 | NA | NA |
| WBGene00047908 | NA | NA | NA | NA |
| WBGene00047909 | NA | NA | NA | NA |
| WBGene00047910 | NA | NA | NA | NA |
| WBGene00047911 | NA | NA | NA | NA |
| WBGene00047912 | NA | NA | NA | NA |
| WBGene00047913 | NA | NA | NA | NA |
| WBGene00047914 | NA | NA | NA | NA |
| WBGene00047915 | NA | NA | NA | NA |
| WBGene00047916 | NA | NA | NA | NA |
| WBGene00047917 | NA | NA | NA | NA |
| WBGene00047918 | NA | NA | NA | NA |
| WBGene00047919 | NA | NA | NA | NA |
| WBGene00047920 | NA | NA | NA | NA |
| WBGene00047921 | NA | NA | NA | NA |
| WBGene00047922 | NA | NA | NA | NA |
| WBGene00047923 | NA | NA | NA | NA |
| WBGene00047924 | NA | NA | NA | NA |
| WBGene00047925 | NA | NA | NA | NA |
| WBGene00047926 | NA | NA | NA | NA |
| WBGene00047927 | NA | NA | NA | NA |
| WBGene00047928 | NA | NA | NA | NA |
| WBGene00047929 | NA | NA | NA | NA |
| WBGene00047930 | NA | NA | NA | NA |
| WBGene00047931 | NA | NA | NA | NA |
| WBGene00047932 | NA | NA | NA | NA |
| WBGene00047933 | NA | NA | NA | NA |
| WBGene00047934 | NA | NA | NA | NA |
| WBGene00047935 | NA | NA | NA | NA |
| WBGene00047936 | NA | NA | NA | NA |
| WBGene00047937 | NA | NA | NA | NA |
| WBGene00047938 | NA | NA | NA | NA |
| WBGene00047939 | NA | NA | NA | NA |

| | | | | |
|---|---|---|---|---|
| WBGene00047940 | NA | NA | NA | NA |
| WBGene00047941 | NA | NA | NA | NA |
| WBGene00047942 | NA | NA | NA | NA |
| WBGene00047943 | NA | NA | NA | NA |
| WBGene00047944 | NA | NA | NA | NA |
| WBGene00047945 | NA | NA | NA | NA |
| WBGene00047946 | NA | NA | NA | NA |
| WBGene00047947 | NA | NA | NA | NA |
| WBGene00047948 | NA | NA | NA | NA |
| WBGene00047949 | NA | NA | NA | NA |
| WBGene00047950 | NA | NA | NA | NA |
| WBGene00047951 | NA | NA | NA | NA |
| WBGene00047952 | NA | NA | NA | NA |
| WBGene00047953 | NA | NA | NA | NA |
| WBGene00047954 | NA | NA | NA | NA |
| WBGene00047955 | NA | NA | NA | NA |
| WBGene00047956 | NA | NA | NA | NA |
| WBGene00047957 | NA | NA | NA | NA |
| WBGene00047958 | NA | NA | NA | NA |
| WBGene00047959 | NA | NA | NA | NA |
| WBGene00047960 | NA | NA | NA | NA |
| WBGene00047961 | NA | NA | NA | NA |
| WBGene00047962 | NA | NA | NA | NA |
| WBGene00047963 | NA | NA | NA | NA |
| WBGene00047964 | NA | NA | NA | NA |
| WBGene00047965 | NA | NA | NA | NA |
| WBGene00047966 | NA | NA | NA | NA |
| WBGene00047967 | NA | NA | NA | NA |
| WBGene00047968 | NA | NA | NA | NA |
| WBGene00047969 | NA | NA | NA | NA |
| WBGene00047970 | NA | NA | NA | NA |
| WBGene00047971 | NA | NA | NA | NA |
| WBGene00047972 | NA | NA | NA | NA |
| WBGene00047973 | NA | NA | NA | NA |
| WBGene00047974 | NA | NA | NA | NA |
| WBGene00047975 | NA | NA | NA | NA |
| WBGene00047976 | NA | NA | NA | NA |
| WBGene00047977 | NA | NA | NA | NA |
| WBGene00047978 | NA | NA | NA | NA |
| WBGene00047979 | NA | NA | NA | NA |
| WBGene00047980 | NA | NA | NA | NA |
| WBGene00047981 | NA | NA | NA | NA |
| WBGene00047982 | NA | NA | NA | NA |

| | | | | |
|---|---|---|---|---|
| WBGene00047983 | NA | NA | NA | NA |
| WBGene00047984 | NA | NA | NA | NA |
| WBGene00047985 | NA | NA | NA | NA |
| WBGene00047986 | NA | NA | NA | NA |
| WBGene00047987 | NA | NA | NA | NA |
| WBGene00047988 | NA | NA | NA | NA |
| WBGene00047989 | NA | NA | NA | NA |
| WBGene00047990 | NA | NA | NA | NA |
| WBGene00047991 | NA | NA | NA | NA |
| WBGene00047992 | NA | NA | NA | NA |
| WBGene00047993 | NA | NA | NA | NA |
| WBGene00047994 | NA | NA | NA | NA |
| WBGene00047995 | NA | NA | NA | NA |
| WBGene00047996 | NA | NA | NA | NA |
| WBGene00047997 | NA | NA | NA | NA |
| WBGene00047998 | NA | NA | NA | NA |
| WBGene00047999 | NA | NA | NA | NA |
| WBGene00048000 | NA | NA | NA | NA |
| WBGene00048001 | NA | NA | NA | NA |
| WBGene00048002 | NA | NA | NA | NA |
| WBGene00048003 | NA | NA | NA | NA |
| WBGene00048004 | NA | NA | NA | NA |
| WBGene00048005 | NA | NA | NA | NA |
| WBGene00048006 | NA | NA | NA | NA |
| WBGene00048007 | NA | NA | NA | NA |
| WBGene00048008 | NA | NA | NA | NA |
| WBGene00048009 | NA | NA | NA | NA |
| WBGene00048010 | NA | NA | NA | NA |
| WBGene00048011 | NA | NA | NA | NA |
| WBGene00048012 | NA | NA | NA | NA |
| WBGene00048013 | NA | NA | NA | NA |
| WBGene00048014 | NA | NA | NA | NA |
| WBGene00048015 | NA | NA | NA | NA |
| WBGene00048016 | NA | NA | NA | NA |
| WBGene00048017 | NA | NA | NA | NA |
| WBGene00048018 | NA | NA | NA | NA |
| WBGene00048019 | NA | NA | NA | NA |
| WBGene00048020 | NA | NA | NA | NA |
| WBGene00048021 | NA | NA | NA | NA |
| WBGene00048022 | NA | NA | NA | NA |
| WBGene00048023 | NA | NA | NA | NA |
| WBGene00048024 | NA | NA | NA | NA |
| WBGene00048025 | NA | NA | NA | NA |

| | | | | |
|---|---|---|---|---|
| WBGene00048026 | NA | NA | NA | NA |
| WBGene00048027 | NA | NA | NA | NA |
| WBGene00048028 | NA | NA | NA | NA |
| WBGene00048029 | NA | NA | NA | NA |
| WBGene00048030 | NA | NA | NA | NA |
| WBGene00048031 | NA | NA | NA | NA |
| WBGene00048032 | NA | NA | NA | NA |
| WBGene00048033 | NA | NA | NA | NA |
| WBGene00048034 | NA | NA | NA | NA |
| WBGene00048035 | NA | NA | NA | NA |
| WBGene00048036 | NA | NA | NA | NA |
| WBGene00048037 | NA | NA | NA | NA |
| WBGene00048038 | NA | NA | NA | NA |
| WBGene00048039 | NA | NA | NA | NA |
| WBGene00048040 | NA | NA | NA | NA |
| WBGene00048041 | NA | NA | NA | NA |
| WBGene00048042 | NA | NA | NA | NA |
| WBGene00048043 | NA | NA | NA | NA |
| WBGene00048044 | NA | NA | NA | NA |
| WBGene00048045 | NA | NA | NA | NA |
| WBGene00048046 | NA | NA | NA | NA |
| WBGene00048047 | NA | NA | NA | NA |
| WBGene00048048 | NA | NA | NA | NA |
| WBGene00048049 | NA | NA | NA | NA |
| WBGene00048050 | NA | NA | NA | NA |
| WBGene00048051 | NA | NA | NA | NA |
| WBGene00048052 | NA | NA | NA | NA |
| WBGene00048053 | NA | NA | NA | NA |
| WBGene00048054 | NA | NA | NA | NA |
| WBGene00048055 | NA | NA | NA | NA |
| WBGene00048056 | NA | NA | NA | NA |
| WBGene00048057 | NA | NA | NA | NA |
| WBGene00048058 | NA | NA | NA | NA |
| WBGene00048059 | NA | NA | NA | NA |
| WBGene00048060 | NA | NA | NA | NA |
| WBGene00048061 | NA | NA | NA | NA |
| WBGene00048062 | NA | NA | NA | NA |
| WBGene00048063 | NA | NA | NA | NA |
| WBGene00048064 | NA | NA | NA | NA |
| WBGene00048065 | NA | NA | NA | NA |
| WBGene00048066 | NA | NA | NA | NA |
| WBGene00048067 | NA | NA | NA | NA |
| WBGene00048068 | NA | NA | NA | NA |

| | | | | |
|---|---|---|---|---|
| WBGene00048069 | NA | NA | NA | NA |
| WBGene00048070 | NA | NA | NA | NA |
| WBGene00048071 | NA | NA | NA | NA |
| WBGene00048072 | NA | NA | NA | NA |
| WBGene00048073 | NA | NA | NA | NA |
| WBGene00048074 | NA | NA | NA | NA |
| WBGene00048075 | NA | NA | NA | NA |
| WBGene00048076 | NA | NA | NA | NA |
| WBGene00048077 | NA | NA | NA | NA |
| WBGene00048078 | NA | NA | NA | NA |
| WBGene00048079 | NA | NA | NA | NA |
| WBGene00048080 | NA | NA | NA | NA |
| WBGene00048081 | NA | NA | NA | NA |
| WBGene00048082 | NA | NA | NA | NA |
| WBGene00048083 | NA | NA | NA | NA |
| WBGene00048084 | NA | NA | NA | NA |
| WBGene00048085 | NA | NA | NA | NA |
| WBGene00048086 | NA | NA | NA | NA |
| WBGene00048087 | NA | NA | NA | NA |
| WBGene00048088 | NA | NA | NA | NA |
| WBGene00048089 | NA | NA | NA | NA |
| WBGene00048090 | NA | NA | NA | NA |
| WBGene00048091 | NA | NA | NA | NA |
| WBGene00048092 | NA | NA | NA | NA |
| WBGene00048093 | NA | NA | NA | NA |
| WBGene00048094 | -0.023109762 | 0.78121253 | NA | NA |
| WBGene00048095 | NA | NA | NA | NA |
| WBGene00048096 | NA | NA | NA | NA |
| WBGene00048097 | NA | NA | NA | NA |
| WBGene00048098 | NA | NA | NA | NA |
| WBGene00048099 | NA | NA | NA | NA |
| WBGene00048100 | NA | NA | NA | NA |
| WBGene00048101 | NA | NA | NA | NA |
| WBGene00048102 | NA | NA | NA | NA |
| WBGene00048103 | NA | NA | NA | NA |
| WBGene00048104 | NA | NA | NA | NA |
| WBGene00048105 | NA | NA | NA | NA |
| WBGene00048106 | NA | NA | NA | NA |
| WBGene00048107 | NA | NA | NA | NA |
| WBGene00048108 | NA | NA | NA | NA |
| WBGene00048109 | NA | NA | NA | NA |
| WBGene00048110 | NA | NA | NA | NA |
| WBGene00048111 | NA | NA | NA | NA |

| | | | | |
|---|---|---|---|---|
| WBGene00048112 | NA | NA | NA | NA |
| WBGene00048113 | NA | NA | NA | NA |
| WBGene00048114 | NA | NA | NA | NA |
| WBGene00048115 | NA | NA | NA | NA |
| WBGene00048116 | NA | NA | NA | NA |
| WBGene00048117 | NA | NA | NA | NA |
| WBGene00048118 | NA | NA | NA | NA |
| WBGene00048119 | NA | NA | NA | NA |
| WBGene00048120 | NA | NA | NA | NA |
| WBGene00048121 | NA | NA | NA | NA |
| WBGene00048122 | NA | NA | NA | NA |
| WBGene00048123 | NA | NA | NA | NA |
| WBGene00048124 | NA | NA | NA | NA |
| WBGene00048125 | NA | NA | NA | NA |
| WBGene00048126 | NA | NA | NA | NA |
| WBGene00048127 | NA | NA | NA | NA |
| WBGene00048128 | NA | NA | NA | NA |
| WBGene00048129 | NA | NA | NA | NA |
| WBGene00048130 | NA | NA | NA | NA |
| WBGene00048131 | NA | NA | NA | NA |
| WBGene00048132 | NA | NA | NA | NA |
| WBGene00048133 | NA | NA | NA | NA |
| WBGene00048134 | NA | NA | NA | NA |
| WBGene00048135 | NA | NA | NA | NA |
| WBGene00048136 | NA | NA | NA | NA |
| WBGene00048137 | NA | NA | NA | NA |
| WBGene00048138 | NA | NA | NA | NA |
| WBGene00048139 | NA | NA | NA | NA |
| WBGene00048140 | NA | NA | NA | NA |
| WBGene00048141 | NA | NA | NA | NA |
| WBGene00048142 | NA | NA | NA | NA |
| WBGene00048143 | NA | NA | NA | NA |
| WBGene00048144 | NA | NA | NA | NA |
| WBGene00048145 | NA | NA | NA | NA |
| WBGene00048146 | NA | NA | NA | NA |
| WBGene00048147 | NA | NA | NA | NA |
| WBGene00048148 | NA | NA | NA | NA |
| WBGene00048149 | NA | NA | NA | NA |
| WBGene00048150 | NA | NA | NA | NA |
| WBGene00048151 | NA | NA | NA | NA |
| WBGene00048152 | NA | NA | NA | NA |
| WBGene00048153 | NA | NA | NA | NA |
| WBGene00048154 | NA | NA | NA | NA |

| | | | | |
|---|---|---|---|---|
| WBGene00048155 | NA | NA | NA | NA |
| WBGene00048156 | NA | NA | NA | NA |
| WBGene00048157 | NA | NA | NA | NA |
| WBGene00048158 | NA | NA | NA | NA |
| WBGene00048159 | NA | NA | NA | NA |
| WBGene00048160 | NA | NA | NA | NA |
| WBGene00048161 | NA | NA | NA | NA |
| WBGene00048162 | NA | NA | NA | NA |
| WBGene00048163 | NA | NA | NA | NA |
| WBGene00048164 | NA | NA | NA | NA |
| WBGene00048165 | NA | NA | NA | NA |
| WBGene00048166 | NA | NA | NA | NA |
| WBGene00048167 | NA | NA | NA | NA |
| WBGene00048168 | NA | NA | NA | NA |
| WBGene00048169 | NA | NA | NA | NA |
| WBGene00048170 | NA | NA | NA | NA |
| WBGene00048171 | NA | NA | NA | NA |
| WBGene00048172 | NA | NA | NA | NA |
| WBGene00048173 | NA | NA | NA | NA |
| WBGene00048174 | NA | NA | NA | NA |
| WBGene00048175 | NA | NA | NA | NA |
| WBGene00048176 | NA | NA | NA | NA |
| WBGene00048177 | NA | NA | NA | NA |
| WBGene00048178 | NA | NA | NA | NA |
| WBGene00048179 | NA | NA | NA | NA |
| WBGene00048180 | NA | NA | NA | NA |
| WBGene00048181 | NA | NA | NA | NA |
| WBGene00048182 | NA | NA | NA | NA |
| WBGene00048183 | NA | NA | NA | NA |
| WBGene00048184 | NA | NA | NA | NA |
| WBGene00048185 | NA | NA | NA | NA |
| WBGene00048186 | NA | NA | NA | NA |
| WBGene00048187 | NA | NA | NA | NA |
| WBGene00048188 | NA | NA | NA | NA |
| WBGene00048189 | NA | NA | NA | NA |
| WBGene00048190 | NA | NA | NA | NA |
| WBGene00048191 | NA | NA | NA | NA |
| WBGene00048192 | NA | NA | NA | NA |
| WBGene00048193 | NA | NA | NA | NA |
| WBGene00048194 | NA | NA | NA | NA |
| WBGene00048195 | NA | NA | NA | NA |
| WBGene00048196 | NA | NA | NA | NA |
| WBGene00048197 | NA | NA | NA | NA |

| | | | | |
|---|---|---|---|---|
| WBGene00048198 | NA | NA | NA | NA |
| WBGene00048199 | NA | NA | NA | NA |
| WBGene00048200 | NA | NA | NA | NA |
| WBGene00048201 | NA | NA | NA | NA |
| WBGene00048202 | NA | NA | NA | NA |
| WBGene00048203 | NA | NA | NA | NA |
| WBGene00048204 | NA | NA | NA | NA |
| WBGene00048205 | NA | NA | NA | NA |
| WBGene00048206 | NA | NA | NA | NA |
| WBGene00048207 | NA | NA | NA | NA |
| WBGene00048208 | NA | NA | NA | NA |
| WBGene00048209 | NA | NA | NA | NA |
| WBGene00048210 | NA | NA | NA | NA |
| WBGene00048211 | NA | NA | NA | NA |
| WBGene00048212 | NA | NA | NA | NA |
| WBGene00048213 | NA | NA | NA | NA |
| WBGene00048214 | NA | NA | NA | NA |
| WBGene00048215 | NA | NA | NA | NA |
| WBGene00048216 | NA | NA | NA | NA |
| WBGene00048217 | NA | NA | NA | NA |
| WBGene00048218 | NA | NA | NA | NA |
| WBGene00048219 | NA | NA | NA | NA |
| WBGene00048220 | NA | NA | NA | NA |
| WBGene00048221 | NA | NA | NA | NA |
| WBGene00048222 | 0.020317245 | 0.84643798 | NA | NA |
| WBGene00048223 | NA | NA | NA | NA |
| WBGene00048224 | NA | NA | NA | NA |
| WBGene00048225 | NA | NA | NA | NA |
| WBGene00048226 | NA | NA | NA | NA |
| WBGene00048227 | NA | NA | NA | NA |
| WBGene00048228 | NA | NA | NA | NA |
| WBGene00048229 | NA | NA | NA | NA |
| WBGene00048230 | NA | NA | NA | NA |
| WBGene00048231 | NA | NA | NA | NA |
| WBGene00048232 | NA | NA | NA | NA |
| WBGene00048233 | NA | NA | NA | NA |
| WBGene00048234 | NA | NA | NA | NA |
| WBGene00048235 | NA | NA | NA | NA |
| WBGene00048236 | NA | NA | NA | NA |
| WBGene00048237 | NA | NA | NA | NA |
| WBGene00048238 | NA | NA | NA | NA |
| WBGene00048239 | NA | NA | NA | NA |
| WBGene00048240 | NA | NA | NA | NA |

| | | | | |
|---|---|---|---|---|
| WBGene00048241 | NA | NA | NA | NA |
| WBGene00048242 | NA | NA | NA | NA |
| WBGene00048243 | NA | NA | NA | NA |
| WBGene00048244 | NA | NA | NA | NA |
| WBGene00048245 | NA | NA | NA | NA |
| WBGene00048246 | NA | NA | NA | NA |
| WBGene00048247 | NA | NA | NA | NA |
| WBGene00048248 | NA | NA | NA | NA |
| WBGene00048249 | NA | NA | NA | NA |
| WBGene00048250 | NA | NA | NA | NA |
| WBGene00048251 | NA | NA | NA | NA |
| WBGene00048252 | NA | NA | NA | NA |
| WBGene00048253 | NA | NA | NA | NA |
| WBGene00048254 | NA | NA | NA | NA |
| WBGene00048255 | NA | NA | NA | NA |
| WBGene00048256 | NA | NA | NA | NA |
| WBGene00048257 | NA | NA | NA | NA |
| WBGene00048258 | NA | NA | NA | NA |
| WBGene00048259 | NA | NA | NA | NA |
| WBGene00048260 | NA | NA | NA | NA |
| WBGene00048261 | NA | NA | NA | NA |
| WBGene00048262 | NA | NA | NA | NA |
| WBGene00048263 | NA | NA | NA | NA |
| WBGene00048264 | NA | NA | NA | NA |
| WBGene00048265 | NA | NA | NA | NA |
| WBGene00048266 | NA | NA | NA | NA |
| WBGene00048267 | NA | NA | NA | NA |
| WBGene00048268 | NA | NA | NA | NA |
| WBGene00048269 | NA | NA | NA | NA |
| WBGene00048270 | NA | NA | NA | NA |
| WBGene00048271 | NA | NA | NA | NA |
| WBGene00048272 | NA | NA | NA | NA |
| WBGene00048273 | NA | NA | NA | NA |
| WBGene00048274 | NA | NA | NA | NA |
| WBGene00048275 | NA | NA | NA | NA |
| WBGene00048276 | NA | NA | NA | NA |
| WBGene00048277 | NA | NA | NA | NA |
| WBGene00048278 | NA | NA | NA | NA |
| WBGene00048279 | NA | NA | NA | NA |
| WBGene00048280 | NA | NA | NA | NA |
| WBGene00048281 | NA | NA | NA | NA |
| WBGene00048282 | NA | NA | NA | NA |
| WBGene00048283 | NA | NA | NA | NA |

| | | | | |
|---|---|---|---|---|
| WBGene00048284 | NA | NA | NA | NA |
| WBGene00048285 | NA | NA | NA | NA |
| WBGene00048286 | NA | NA | NA | NA |
| WBGene00048287 | NA | NA | NA | NA |
| WBGene00048288 | NA | NA | NA | NA |
| WBGene00048289 | NA | NA | NA | NA |
| WBGene00048290 | NA | NA | NA | NA |
| WBGene00048291 | NA | NA | NA | NA |
| WBGene00048292 | NA | NA | NA | NA |
| WBGene00048293 | NA | NA | NA | NA |
| WBGene00048294 | NA | NA | NA | NA |
| WBGene00048295 | NA | NA | NA | NA |
| WBGene00048296 | NA | NA | NA | NA |
| WBGene00048297 | NA | NA | NA | NA |
| WBGene00048298 | NA | NA | NA | NA |
| WBGene00048299 | NA | NA | NA | NA |
| WBGene00048300 | NA | NA | NA | NA |
| WBGene00048301 | NA | NA | NA | NA |
| WBGene00048302 | NA | NA | NA | NA |
| WBGene00048303 | NA | NA | NA | NA |
| WBGene00048304 | NA | NA | NA | NA |
| WBGene00048305 | NA | NA | NA | NA |
| WBGene00048306 | NA | NA | NA | NA |
| WBGene00048307 | NA | NA | NA | NA |
| WBGene00048308 | NA | NA | NA | NA |
| WBGene00048309 | NA | NA | NA | NA |
| WBGene00048310 | NA | NA | NA | NA |
| WBGene00048311 | NA | NA | NA | NA |
| WBGene00048312 | NA | NA | NA | NA |
| WBGene00048313 | NA | NA | NA | NA |
| WBGene00048314 | NA | NA | NA | NA |
| WBGene00048315 | NA | NA | NA | NA |
| WBGene00048316 | NA | NA | NA | NA |
| WBGene00048317 | NA | NA | NA | NA |
| WBGene00048318 | NA | NA | NA | NA |
| WBGene00048319 | NA | NA | NA | NA |
| WBGene00048320 | NA | NA | NA | NA |
| WBGene00048321 | NA | NA | NA | NA |
| WBGene00048322 | NA | NA | NA | NA |
| WBGene00048323 | NA | NA | NA | NA |
| WBGene00048324 | NA | NA | NA | NA |
| WBGene00048325 | NA | NA | NA | NA |
| WBGene00048326 | NA | NA | NA | NA |

| | | | | |
|---|---|---|---|---|
| WBGene00048327 | NA | NA | NA | NA |
| WBGene00048328 | NA | NA | NA | NA |
| WBGene00048329 | NA | NA | NA | NA |
| WBGene00048330 | NA | NA | NA | NA |
| WBGene00048331 | NA | NA | NA | NA |
| WBGene00048332 | NA | NA | NA | NA |
| WBGene00048333 | NA | NA | NA | NA |
| WBGene00048334 | NA | NA | NA | NA |
| WBGene00048335 | NA | NA | NA | NA |
| WBGene00048336 | NA | NA | NA | NA |
| WBGene00048337 | NA | NA | NA | NA |
| WBGene00048338 | NA | NA | NA | NA |
| WBGene00048339 | NA | NA | NA | NA |
| WBGene00048340 | NA | NA | NA | NA |
| WBGene00048341 | NA | NA | NA | NA |
| WBGene00048342 | NA | NA | NA | NA |
| WBGene00048343 | NA | NA | NA | NA |
| WBGene00048344 | NA | NA | NA | NA |
| WBGene00048345 | NA | NA | NA | NA |
| WBGene00048346 | NA | NA | NA | NA |
| WBGene00048347 | NA | NA | NA | NA |
| WBGene00048348 | NA | NA | NA | NA |
| WBGene00048349 | NA | NA | NA | NA |
| WBGene00048350 | NA | NA | NA | NA |
| WBGene00048351 | NA | NA | NA | NA |
| WBGene00048352 | NA | NA | NA | NA |
| WBGene00048353 | NA | NA | NA | NA |
| WBGene00048354 | NA | NA | NA | NA |
| WBGene00048355 | NA | NA | NA | NA |
| WBGene00048356 | NA | NA | NA | NA |
| WBGene00048357 | NA | NA | NA | NA |
| WBGene00048358 | NA | NA | NA | NA |
| WBGene00048359 | NA | NA | NA | NA |
| WBGene00048360 | NA | NA | NA | NA |
| WBGene00048361 | NA | NA | NA | NA |
| WBGene00048362 | NA | NA | NA | NA |
| WBGene00048363 | NA | NA | NA | NA |
| WBGene00048364 | NA | NA | NA | NA |
| WBGene00048365 | NA | NA | NA | NA |
| WBGene00048366 | NA | NA | NA | NA |
| WBGene00048367 | NA | NA | NA | NA |
| WBGene00048368 | NA | NA | NA | NA |
| WBGene00048369 | NA | NA | NA | NA |

| | | | | |
|---|---|---|---|---|
| WBGene00048370 | NA | NA | NA | NA |
| WBGene00048371 | NA | NA | NA | NA |
| WBGene00048372 | NA | NA | NA | NA |
| WBGene00048373 | NA | NA | NA | NA |
| WBGene00048374 | NA | NA | NA | NA |
| WBGene00048375 | NA | NA | NA | NA |
| WBGene00048376 | NA | NA | NA | NA |
| WBGene00048377 | NA | NA | NA | NA |
| WBGene00048378 | NA | NA | NA | NA |
| WBGene00048379 | NA | NA | NA | NA |
| WBGene00048380 | NA | NA | NA | NA |
| WBGene00048381 | NA | NA | NA | NA |
| WBGene00048382 | NA | NA | NA | NA |
| WBGene00048383 | NA | NA | NA | NA |
| WBGene00048384 | NA | NA | NA | NA |
| WBGene00048385 | NA | NA | NA | NA |
| WBGene00048386 | NA | NA | NA | NA |
| WBGene00048387 | NA | NA | NA | NA |
| WBGene00048388 | NA | NA | NA | NA |
| WBGene00048389 | NA | NA | NA | NA |
| WBGene00048390 | NA | NA | NA | NA |
| WBGene00048391 | NA | NA | NA | NA |
| WBGene00048392 | NA | NA | NA | NA |
| WBGene00048393 | NA | NA | NA | NA |
| WBGene00048394 | NA | NA | NA | NA |
| WBGene00048395 | NA | NA | NA | NA |
| WBGene00048396 | NA | NA | NA | NA |
| WBGene00048397 | NA | NA | NA | NA |
| WBGene00048398 | NA | NA | NA | NA |
| WBGene00048399 | NA | NA | NA | NA |
| WBGene00048400 | NA | NA | NA | NA |
| WBGene00048401 | NA | NA | NA | NA |
| WBGene00048402 | NA | NA | NA | NA |
| WBGene00048403 | NA | NA | NA | NA |
| WBGene00048404 | NA | NA | NA | NA |
| WBGene00048405 | NA | NA | NA | NA |
| WBGene00048406 | NA | NA | NA | NA |
| WBGene00048407 | NA | NA | NA | NA |
| WBGene00048408 | NA | NA | NA | NA |
| WBGene00048409 | NA | NA | NA | NA |
| WBGene00048410 | NA | NA | NA | NA |
| WBGene00048411 | NA | NA | NA | NA |
| WBGene00048412 | NA | NA | NA | NA |

| | | | | |
|---|---|---|---|---|
| WBGene00048413 | NA | NA | NA | NA |
| WBGene00048414 | NA | NA | NA | NA |
| WBGene00048415 | NA | NA | NA | NA |
| WBGene00048416 | NA | NA | NA | NA |
| WBGene00048417 | NA | NA | NA | NA |
| WBGene00048418 | NA | NA | NA | NA |
| WBGene00048419 | NA | NA | NA | NA |
| WBGene00048420 | NA | NA | NA | NA |
| WBGene00048421 | NA | NA | NA | NA |
| WBGene00048422 | NA | NA | NA | NA |
| WBGene00048423 | NA | NA | NA | NA |
| WBGene00048424 | NA | NA | NA | NA |
| WBGene00048425 | NA | NA | NA | NA |
| WBGene00048426 | NA | NA | NA | NA |
| WBGene00048427 | NA | NA | NA | NA |
| WBGene00048428 | NA | NA | NA | NA |
| WBGene00048429 | NA | NA | NA | NA |
| WBGene00048430 | NA | NA | NA | NA |
| WBGene00048431 | NA | NA | NA | NA |
| WBGene00048432 | NA | NA | NA | NA |
| WBGene00048433 | NA | NA | NA | NA |
| WBGene00048434 | NA | NA | NA | NA |
| WBGene00048435 | NA | NA | NA | NA |
| WBGene00048436 | NA | NA | NA | NA |
| WBGene00048437 | NA | NA | NA | NA |
| WBGene00048438 | NA | NA | NA | NA |
| WBGene00048439 | NA | NA | NA | NA |
| WBGene00048440 | NA | NA | NA | NA |
| WBGene00048441 | NA | NA | NA | NA |
| WBGene00048442 | NA | NA | NA | NA |
| WBGene00048443 | NA | NA | NA | NA |
| WBGene00048444 | NA | NA | NA | NA |
| WBGene00048445 | NA | NA | NA | NA |
| WBGene00048446 | NA | NA | NA | NA |
| WBGene00048447 | NA | NA | NA | NA |
| WBGene00048448 | NA | NA | NA | NA |
| WBGene00048449 | NA | NA | NA | NA |
| WBGene00048450 | NA | NA | NA | NA |
| WBGene00048451 | NA | NA | NA | NA |
| WBGene00048452 | NA | NA | NA | NA |
| WBGene00048453 | NA | NA | NA | NA |
| WBGene00048454 | NA | NA | NA | NA |
| WBGene00048455 | NA | NA | NA | NA |

| | | | | |
|---|---|---|---|---|
| WBGene00048456 | NA | NA | NA | NA |
| WBGene00048457 | NA | NA | NA | NA |
| WBGene00048458 | NA | NA | NA | NA |
| WBGene00048459 | NA | NA | NA | NA |
| WBGene00048460 | NA | NA | NA | NA |
| WBGene00048461 | NA | NA | NA | NA |
| WBGene00048462 | NA | NA | NA | NA |
| WBGene00048463 | NA | NA | NA | NA |
| WBGene00048464 | NA | NA | NA | NA |
| WBGene00048465 | NA | NA | NA | NA |
| WBGene00048466 | NA | NA | NA | NA |
| WBGene00048467 | NA | NA | NA | NA |
| WBGene00048468 | NA | NA | NA | NA |
| WBGene00048469 | NA | NA | NA | NA |
| WBGene00048470 | NA | NA | NA | NA |
| WBGene00048471 | NA | NA | NA | NA |
| WBGene00048472 | NA | NA | NA | NA |
| WBGene00048473 | NA | NA | NA | NA |
| WBGene00048474 | NA | NA | NA | NA |
| WBGene00048475 | NA | NA | NA | NA |
| WBGene00048476 | NA | NA | NA | NA |
| WBGene00048477 | NA | NA | NA | NA |
| WBGene00048478 | NA | NA | NA | NA |
| WBGene00048479 | NA | NA | NA | NA |
| WBGene00048480 | NA | NA | NA | NA |
| WBGene00048481 | NA | NA | NA | NA |
| WBGene00048482 | NA | NA | NA | NA |
| WBGene00048483 | NA | NA | NA | NA |
| WBGene00048484 | NA | NA | NA | NA |
| WBGene00048485 | NA | NA | NA | NA |
| WBGene00048486 | NA | NA | NA | NA |
| WBGene00048487 | NA | NA | NA | NA |
| WBGene00048488 | NA | NA | NA | NA |
| WBGene00048489 | NA | NA | NA | NA |
| WBGene00048490 | NA | NA | NA | NA |
| WBGene00048491 | NA | NA | NA | NA |
| WBGene00048492 | NA | NA | NA | NA |
| WBGene00048493 | NA | NA | NA | NA |
| WBGene00048494 | NA | NA | NA | NA |
| WBGene00048495 | NA | NA | NA | NA |
| WBGene00048496 | NA | NA | NA | NA |
| WBGene00048497 | NA | NA | NA | NA |
| WBGene00048498 | NA | NA | NA | NA |

| | | | | |
|---|---|---|---|---|
| WBGene00048499 | NA | NA | NA | NA |
| WBGene00048500 | NA | NA | NA | NA |
| WBGene00048501 | NA | NA | NA | NA |
| WBGene00048502 | NA | NA | NA | NA |
| WBGene00048503 | NA | NA | NA | NA |
| WBGene00048504 | NA | NA | NA | NA |
| WBGene00048505 | NA | NA | NA | NA |
| WBGene00048506 | NA | NA | NA | NA |
| WBGene00048507 | NA | NA | NA | NA |
| WBGene00048508 | NA | NA | NA | NA |
| WBGene00048509 | NA | NA | NA | NA |
| WBGene00048510 | NA | NA | NA | NA |
| WBGene00048511 | NA | NA | NA | NA |
| WBGene00048512 | NA | NA | NA | NA |
| WBGene00048513 | NA | NA | NA | NA |
| WBGene00048514 | NA | NA | NA | NA |
| WBGene00048515 | NA | NA | NA | NA |
| WBGene00048516 | NA | NA | NA | NA |
| WBGene00048517 | NA | NA | NA | NA |
| WBGene00048518 | NA | NA | NA | NA |
| WBGene00048519 | NA | NA | NA | NA |
| WBGene00048520 | NA | NA | NA | NA |
| WBGene00048521 | NA | NA | NA | NA |
| WBGene00048522 | NA | NA | NA | NA |
| WBGene00048523 | NA | NA | NA | NA |
| WBGene00048524 | NA | NA | NA | NA |
| WBGene00048525 | NA | NA | NA | NA |
| WBGene00048526 | NA | NA | NA | NA |
| WBGene00048527 | NA | NA | NA | NA |
| WBGene00048528 | NA | NA | NA | NA |
| WBGene00048529 | NA | NA | NA | NA |
| WBGene00048530 | NA | NA | NA | NA |
| WBGene00048531 | NA | NA | NA | NA |
| WBGene00048532 | NA | NA | NA | NA |
| WBGene00048533 | NA | NA | NA | NA |
| WBGene00048534 | NA | NA | NA | NA |
| WBGene00048535 | NA | NA | NA | NA |
| WBGene00048536 | NA | NA | NA | NA |
| WBGene00048537 | NA | NA | NA | NA |
| WBGene00048538 | NA | NA | NA | NA |
| WBGene00048539 | NA | NA | NA | NA |
| WBGene00048540 | NA | NA | NA | NA |
| WBGene00048541 | NA | NA | NA | NA |

| | | | | |
|---|---|---|---|---|
| WBGene00048542 | NA | NA | NA | NA |
| WBGene00048543 | NA | NA | NA | NA |
| WBGene00048544 | NA | NA | NA | NA |
| WBGene00048545 | NA | NA | NA | NA |
| WBGene00048546 | NA | NA | NA | NA |
| WBGene00048547 | NA | NA | NA | NA |
| WBGene00048548 | NA | NA | NA | NA |
| WBGene00048549 | NA | NA | NA | NA |
| WBGene00048550 | 0.020232312 | 0.84643798 | NA | NA |
| WBGene00048551 | NA | NA | NA | NA |
| WBGene00048552 | NA | NA | NA | NA |
| WBGene00048553 | NA | NA | NA | NA |
| WBGene00048554 | NA | NA | NA | NA |
| WBGene00048555 | NA | NA | NA | NA |
| WBGene00048556 | NA | NA | NA | NA |
| WBGene00048557 | NA | NA | NA | NA |
| WBGene00048558 | NA | NA | NA | NA |
| WBGene00048559 | NA | NA | NA | NA |
| WBGene00048560 | NA | NA | NA | NA |
| WBGene00048561 | NA | NA | NA | NA |
| WBGene00048562 | NA | NA | NA | NA |
| WBGene00048563 | NA | NA | NA | NA |
| WBGene00048564 | NA | NA | NA | NA |
| WBGene00048565 | NA | NA | NA | NA |
| WBGene00048566 | NA | NA | NA | NA |
| WBGene00048567 | NA | NA | NA | NA |
| WBGene00048568 | NA | NA | NA | NA |
| WBGene00048569 | NA | NA | NA | NA |
| WBGene00048570 | NA | NA | NA | NA |
| WBGene00048571 | NA | NA | NA | NA |
| WBGene00048572 | NA | NA | NA | NA |
| WBGene00048573 | NA | NA | NA | NA |
| WBGene00048574 | NA | NA | NA | NA |
| WBGene00048575 | NA | NA | NA | NA |
| WBGene00048576 | NA | NA | NA | NA |
| WBGene00048577 | NA | NA | NA | NA |
| WBGene00048578 | NA | NA | NA | NA |
| WBGene00048579 | NA | NA | NA | NA |
| WBGene00048580 | NA | NA | NA | NA |
| WBGene00048581 | NA | NA | NA | NA |
| WBGene00048582 | NA | NA | NA | NA |
| WBGene00048583 | NA | NA | NA | NA |
| WBGene00048584 | NA | NA | NA | NA |

| | | | | |
|---|---|---|---|---|
| WBGene00048585 | NA | NA | NA | NA |
| WBGene00048586 | NA | NA | NA | NA |
| WBGene00048587 | NA | NA | NA | NA |
| WBGene00048588 | NA | NA | NA | NA |
| WBGene00048589 | NA | NA | NA | NA |
| WBGene00048590 | 0.020317245 | 0.84643798 | NA | NA |
| WBGene00048591 | NA | NA | NA | NA |
| WBGene00048592 | NA | NA | NA | NA |
| WBGene00048593 | NA | NA | NA | NA |
| WBGene00048594 | NA | NA | NA | NA |
| WBGene00048595 | NA | NA | NA | NA |
| WBGene00048596 | NA | NA | NA | NA |
| WBGene00048597 | NA | NA | NA | NA |
| WBGene00048598 | NA | NA | NA | NA |
| WBGene00048599 | NA | NA | NA | NA |
| WBGene00048600 | NA | NA | NA | NA |
| WBGene00048601 | NA | NA | NA | NA |
| WBGene00048602 | NA | NA | NA | NA |
| WBGene00048603 | NA | NA | NA | NA |
| WBGene00048604 | NA | NA | NA | NA |
| WBGene00048605 | NA | NA | NA | NA |
| WBGene00048606 | NA | NA | NA | NA |
| WBGene00048607 | NA | NA | NA | NA |
| WBGene00048608 | NA | NA | NA | NA |
| WBGene00048609 | NA | NA | NA | NA |
| WBGene00048610 | NA | NA | NA | NA |
| WBGene00048611 | NA | NA | NA | NA |
| WBGene00048612 | NA | NA | NA | NA |
| WBGene00048613 | NA | NA | NA | NA |
| WBGene00048614 | NA | NA | NA | NA |
| WBGene00048615 | NA | NA | NA | NA |
| WBGene00048616 | NA | NA | NA | NA |
| WBGene00048617 | NA | NA | NA | NA |
| WBGene00048618 | NA | NA | NA | NA |
| WBGene00048619 | NA | NA | NA | NA |
| WBGene00048620 | NA | NA | NA | NA |
| WBGene00048621 | NA | NA | NA | NA |
| WBGene00048622 | NA | NA | NA | NA |
| WBGene00048623 | NA | NA | NA | NA |
| WBGene00048624 | NA | NA | NA | NA |
| WBGene00048625 | NA | NA | NA | NA |
| WBGene00048626 | NA | NA | NA | NA |
| WBGene00048627 | NA | NA | NA | NA |

| | | | | |
|---|---|---|---|---|
| WBGene00048628 | NA | NA | NA | NA |
| WBGene00048629 | NA | NA | NA | NA |
| WBGene00048630 | NA | NA | NA | NA |
| WBGene00048631 | NA | NA | NA | NA |
| WBGene00048632 | NA | NA | NA | NA |
| WBGene00048633 | NA | NA | NA | NA |
| WBGene00048634 | NA | NA | NA | NA |
| WBGene00048635 | NA | NA | NA | NA |
| WBGene00048636 | NA | NA | NA | NA |
| WBGene00048637 | NA | NA | NA | NA |
| WBGene00048638 | NA | NA | NA | NA |
| WBGene00048639 | NA | NA | NA | NA |
| WBGene00048640 | NA | NA | NA | NA |
| WBGene00048641 | NA | NA | NA | NA |
| WBGene00048642 | NA | NA | NA | NA |
| WBGene00048643 | NA | NA | NA | NA |
| WBGene00048644 | NA | NA | NA | NA |
| WBGene00048645 | NA | NA | NA | NA |
| WBGene00048646 | NA | NA | NA | NA |
| WBGene00048647 | NA | NA | NA | NA |
| WBGene00048648 | NA | NA | NA | NA |
| WBGene00048649 | NA | NA | NA | NA |
| WBGene00048650 | NA | NA | NA | NA |
| WBGene00048651 | NA | NA | NA | NA |
| WBGene00048652 | NA | NA | NA | NA |
| WBGene00048653 | NA | NA | NA | NA |
| WBGene00048654 | NA | NA | NA | NA |
| WBGene00048655 | NA | NA | NA | NA |
| WBGene00048656 | NA | NA | NA | NA |
| WBGene00048657 | NA | NA | NA | NA |
| WBGene00048658 | NA | NA | NA | NA |
| WBGene00048659 | NA | NA | NA | NA |
| WBGene00048660 | NA | NA | NA | NA |
| WBGene00048661 | NA | NA | NA | NA |
| WBGene00048662 | NA | NA | NA | NA |
| WBGene00048663 | NA | NA | NA | NA |
| WBGene00048664 | NA | NA | NA | NA |
| WBGene00048665 | NA | NA | NA | NA |
| WBGene00048666 | NA | NA | NA | NA |
| WBGene00048667 | NA | NA | NA | NA |
| WBGene00048668 | NA | NA | NA | NA |
| WBGene00048669 | NA | NA | NA | NA |
| WBGene00048670 | NA | NA | NA | NA |

| | | | | |
|---|---|---|---|---|
| WBGene00048671 | NA | NA | NA | NA |
| WBGene00048672 | NA | NA | NA | NA |
| WBGene00048673 | NA | NA | NA | NA |
| WBGene00048674 | NA | NA | NA | NA |
| WBGene00048675 | NA | NA | NA | NA |
| WBGene00048676 | NA | NA | NA | NA |
| WBGene00048677 | NA | NA | NA | NA |
| WBGene00048678 | NA | NA | NA | NA |
| WBGene00048679 | NA | NA | NA | NA |
| WBGene00048680 | NA | NA | NA | NA |
| WBGene00048681 | NA | NA | NA | NA |
| WBGene00048682 | NA | NA | NA | NA |
| WBGene00048683 | NA | NA | NA | NA |
| WBGene00048684 | NA | NA | NA | NA |
| WBGene00048685 | NA | NA | NA | NA |
| WBGene00048686 | NA | NA | NA | NA |
| WBGene00048687 | NA | NA | NA | NA |
| WBGene00048688 | NA | NA | NA | NA |
| WBGene00048689 | NA | NA | NA | NA |
| WBGene00048690 | NA | NA | NA | NA |
| WBGene00048691 | NA | NA | NA | NA |
| WBGene00048692 | NA | NA | NA | NA |
| WBGene00048693 | NA | NA | NA | NA |
| WBGene00048694 | NA | NA | NA | NA |
| WBGene00048695 | NA | NA | NA | NA |
| WBGene00048696 | NA | NA | NA | NA |
| WBGene00048697 | NA | NA | NA | NA |
| WBGene00048698 | NA | NA | NA | NA |
| WBGene00048699 | NA | NA | NA | NA |
| WBGene00048700 | NA | NA | NA | NA |
| WBGene00048701 | NA | NA | NA | NA |
| WBGene00048702 | NA | NA | NA | NA |
| WBGene00048703 | NA | NA | NA | NA |
| WBGene00048704 | NA | NA | NA | NA |
| WBGene00048705 | NA | NA | NA | NA |
| WBGene00048706 | NA | NA | NA | NA |
| WBGene00048707 | NA | NA | NA | NA |
| WBGene00048708 | NA | NA | NA | NA |
| WBGene00048709 | NA | NA | NA | NA |
| WBGene00048710 | NA | NA | NA | NA |
| WBGene00048711 | NA | NA | NA | NA |
| WBGene00048712 | NA | NA | NA | NA |
| WBGene00048713 | NA | NA | NA | NA |

| | | | | |
|---|---|---|---|---|
| WBGene00048714 | NA | NA | NA | NA |
| WBGene00048715 | NA | NA | NA | NA |
| WBGene00048716 | NA | NA | NA | NA |
| WBGene00048717 | NA | NA | NA | NA |
| WBGene00048718 | NA | NA | NA | NA |
| WBGene00048719 | NA | NA | NA | NA |
| WBGene00048720 | NA | NA | NA | NA |
| WBGene00048721 | 0.020317245 | 0.84643798 | NA | NA |
| WBGene00048722 | NA | NA | NA | NA |
| WBGene00048723 | NA | NA | NA | NA |
| WBGene00048724 | NA | NA | NA | NA |
| WBGene00048725 | NA | NA | NA | NA |
| WBGene00048726 | NA | NA | NA | NA |
| WBGene00048727 | NA | NA | NA | NA |
| WBGene00048728 | NA | NA | NA | NA |
| WBGene00048729 | NA | NA | NA | NA |
| WBGene00048730 | NA | NA | NA | NA |
| WBGene00048731 | NA | NA | NA | NA |
| WBGene00048732 | NA | NA | NA | NA |
| WBGene00048733 | NA | NA | NA | NA |
| WBGene00048734 | NA | NA | NA | NA |
| WBGene00048735 | NA | NA | NA | NA |
| WBGene00048736 | NA | NA | NA | NA |
| WBGene00048737 | NA | NA | NA | NA |
| WBGene00048738 | NA | NA | NA | NA |
| WBGene00048739 | NA | NA | NA | NA |
| WBGene00048740 | NA | NA | NA | NA |
| WBGene00048741 | NA | NA | NA | NA |
| WBGene00048742 | NA | NA | NA | NA |
| WBGene00048743 | NA | NA | NA | NA |
| WBGene00048744 | NA | NA | NA | NA |
| WBGene00048745 | NA | NA | NA | NA |
| WBGene00048746 | NA | NA | NA | NA |
| WBGene00048747 | NA | NA | NA | NA |
| WBGene00048748 | NA | NA | NA | NA |
| WBGene00048749 | NA | NA | NA | NA |
| WBGene00048750 | NA | NA | NA | NA |
| WBGene00048751 | NA | NA | NA | NA |
| WBGene00048752 | NA | NA | NA | NA |
| WBGene00048753 | NA | NA | NA | NA |
| WBGene00048754 | NA | NA | NA | NA |
| WBGene00048755 | NA | NA | NA | NA |
| WBGene00048756 | NA | NA | NA | NA |

| | | | | |
|---|---|---|---|---|
| WBGene00048757 | NA | NA | NA | NA |
| WBGene00048758 | NA | NA | NA | NA |
| WBGene00048759 | NA | NA | NA | NA |
| WBGene00048760 | NA | NA | NA | NA |
| WBGene00048761 | NA | NA | NA | NA |
| WBGene00048762 | NA | NA | NA | NA |
| WBGene00048763 | 0.020232312 | 0.84643798 | NA | NA |
| WBGene00048764 | NA | NA | NA | NA |
| WBGene00048765 | NA | NA | NA | NA |
| WBGene00048766 | NA | NA | NA | NA |
| WBGene00048767 | NA | NA | NA | NA |
| WBGene00048768 | NA | NA | NA | NA |
| WBGene00048769 | NA | NA | NA | NA |
| WBGene00048770 | NA | NA | NA | NA |
| WBGene00048771 | NA | NA | NA | NA |
| WBGene00048772 | NA | NA | NA | NA |
| WBGene00048773 | NA | NA | NA | NA |
| WBGene00048774 | NA | NA | NA | NA |
| WBGene00048775 | NA | NA | NA | NA |
| WBGene00048776 | NA | NA | NA | NA |
| WBGene00048777 | NA | NA | NA | NA |
| WBGene00048778 | NA | NA | NA | NA |
| WBGene00048779 | NA | NA | NA | NA |
| WBGene00048780 | NA | NA | NA | NA |
| WBGene00048781 | NA | NA | NA | NA |
| WBGene00048782 | NA | NA | NA | NA |
| WBGene00048783 | NA | NA | NA | NA |
| WBGene00048784 | NA | NA | NA | NA |
| WBGene00048785 | NA | NA | NA | NA |
| WBGene00048786 | NA | NA | NA | NA |
| WBGene00048787 | NA | NA | NA | NA |
| WBGene00048788 | NA | NA | NA | NA |
| WBGene00048789 | NA | NA | NA | NA |
| WBGene00048790 | NA | NA | NA | NA |
| WBGene00048791 | NA | NA | NA | NA |
| WBGene00048792 | NA | NA | NA | NA |
| WBGene00048793 | NA | NA | NA | NA |
| WBGene00048794 | NA | NA | NA | NA |
| WBGene00048795 | NA | NA | NA | NA |
| WBGene00048796 | NA | NA | NA | NA |
| WBGene00048797 | NA | NA | NA | NA |
| WBGene00048798 | NA | NA | NA | NA |
| WBGene00048799 | NA | NA | NA | NA |

| | | | | |
|---|---|---|---|---|
| WBGene00048800 | NA | NA | NA | NA |
| WBGene00048801 | NA | NA | NA | NA |
| WBGene00048802 | NA | NA | NA | NA |
| WBGene00048803 | NA | NA | NA | NA |
| WBGene00048804 | NA | NA | NA | NA |
| WBGene00048805 | NA | NA | NA | NA |
| WBGene00048806 | NA | NA | NA | NA |
| WBGene00048807 | NA | NA | NA | NA |
| WBGene00048808 | NA | NA | NA | NA |
| WBGene00048809 | NA | NA | NA | NA |
| WBGene00048810 | NA | NA | NA | NA |
| WBGene00048811 | NA | NA | NA | NA |
| WBGene00048812 | NA | NA | NA | NA |
| WBGene00048813 | NA | NA | NA | NA |
| WBGene00048814 | NA | NA | NA | NA |
| WBGene00048815 | NA | NA | NA | NA |
| WBGene00048816 | NA | NA | NA | NA |
| WBGene00048817 | NA | NA | NA | NA |
| WBGene00048819 | NA | NA | NA | NA |
| WBGene00048820 | NA | NA | NA | NA |
| WBGene00048821 | NA | NA | NA | NA |
| WBGene00048822 | NA | NA | NA | NA |
| WBGene00048823 | NA | NA | NA | NA |
| WBGene00048824 | NA | NA | NA | NA |
| WBGene00048825 | NA | NA | NA | NA |
| WBGene00048826 | NA | NA | NA | NA |
| WBGene00048827 | NA | NA | NA | NA |
| WBGene00048828 | NA | NA | NA | NA |
| WBGene00048829 | NA | NA | NA | NA |
| WBGene00048830 | NA | NA | NA | NA |
| WBGene00048831 | NA | NA | NA | NA |
| WBGene00048832 | NA | NA | NA | NA |
| WBGene00048833 | NA | NA | NA | NA |
| WBGene00048834 | NA | NA | NA | NA |
| WBGene00048835 | NA | NA | NA | NA |
| WBGene00048836 | NA | NA | NA | NA |
| WBGene00048837 | NA | NA | NA | NA |
| WBGene00048838 | NA | NA | NA | NA |
| WBGene00048839 | NA | NA | NA | NA |
| WBGene00048840 | NA | NA | NA | NA |
| WBGene00048841 | NA | NA | NA | NA |
| WBGene00048842 | NA | NA | NA | NA |
| WBGene00048843 | NA | NA | NA | NA |

| | | | | |
|---|---|---|---|---|
| WBGene00048844 | NA | NA | NA | NA |
| WBGene00048845 | NA | NA | NA | NA |
| WBGene00048846 | NA | NA | NA | NA |
| WBGene00048847 | NA | NA | NA | NA |
| WBGene00048848 | NA | NA | NA | NA |
| WBGene00048849 | NA | NA | NA | NA |
| WBGene00048850 | NA | NA | NA | NA |
| WBGene00048851 | NA | NA | NA | NA |
| WBGene00048852 | NA | NA | NA | NA |
| WBGene00048853 | NA | NA | NA | NA |
| WBGene00048854 | NA | NA | NA | NA |
| WBGene00048855 | NA | NA | NA | NA |
| WBGene00048856 | NA | NA | NA | NA |
| WBGene00048857 | NA | NA | NA | NA |
| WBGene00048858 | NA | NA | NA | NA |
| WBGene00048859 | NA | NA | NA | NA |
| WBGene00048860 | NA | NA | NA | NA |
| WBGene00048861 | NA | NA | NA | NA |
| WBGene00048862 | NA | NA | NA | NA |
| WBGene00048863 | NA | NA | NA | NA |
| WBGene00048864 | NA | NA | NA | NA |
| WBGene00048865 | NA | NA | NA | NA |
| WBGene00048866 | NA | NA | NA | NA |
| WBGene00048867 | NA | NA | NA | NA |
| WBGene00048868 | NA | NA | NA | NA |
| WBGene00048869 | NA | NA | NA | NA |
| WBGene00048870 | NA | NA | NA | NA |
| WBGene00048871 | NA | NA | NA | NA |
| WBGene00048872 | NA | NA | NA | NA |
| WBGene00048873 | NA | NA | NA | NA |
| WBGene00048874 | NA | NA | NA | NA |
| WBGene00048875 | NA | NA | NA | NA |
| WBGene00048876 | NA | NA | NA | NA |
| WBGene00048877 | NA | NA | NA | NA |
| WBGene00048878 | NA | NA | NA | NA |
| WBGene00048879 | NA | NA | NA | NA |
| WBGene00048880 | NA | NA | NA | NA |
| WBGene00048881 | NA | NA | NA | NA |
| WBGene00048882 | NA | NA | NA | NA |
| WBGene00048883 | NA | NA | NA | NA |
| WBGene00048884 | NA | NA | NA | NA |
| WBGene00048885 | NA | NA | NA | NA |
| WBGene00048886 | NA | NA | NA | NA |

| | | | | |
|---|---|---|---|---|
| WBGene00048887 | NA | NA | NA | NA |
| WBGene00048888 | NA | NA | NA | NA |
| WBGene00048889 | NA | NA | NA | NA |
| WBGene00048890 | NA | NA | NA | NA |
| WBGene00048891 | NA | NA | NA | NA |
| WBGene00048892 | NA | NA | NA | NA |
| WBGene00048893 | NA | NA | NA | NA |
| WBGene00048894 | NA | NA | NA | NA |
| WBGene00048895 | NA | NA | NA | NA |
| WBGene00048896 | NA | NA | NA | NA |
| WBGene00048897 | NA | NA | NA | NA |
| WBGene00048898 | NA | NA | NA | NA |
| WBGene00048899 | NA | NA | NA | NA |
| WBGene00048900 | NA | NA | NA | NA |
| WBGene00048901 | NA | NA | NA | NA |
| WBGene00048902 | NA | NA | NA | NA |
| WBGene00048903 | NA | NA | NA | NA |
| WBGene00048904 | NA | NA | NA | NA |
| WBGene00048905 | NA | NA | NA | NA |
| WBGene00048906 | NA | NA | NA | NA |
| WBGene00048907 | NA | NA | NA | NA |
| WBGene00048908 | NA | NA | NA | NA |
| WBGene00048909 | NA | NA | NA | NA |
| WBGene00048910 | NA | NA | NA | NA |
| WBGene00048911 | NA | NA | NA | NA |
| WBGene00048912 | NA | NA | NA | NA |
| WBGene00048913 | NA | NA | NA | NA |
| WBGene00048914 | NA | NA | NA | NA |
| WBGene00048915 | NA | NA | NA | NA |
| WBGene00048916 | NA | NA | NA | NA |
| WBGene00048917 | NA | NA | NA | NA |
| WBGene00048918 | NA | NA | NA | NA |
| WBGene00048919 | NA | NA | NA | NA |
| WBGene00048920 | NA | NA | NA | NA |
| WBGene00048921 | NA | NA | NA | NA |
| WBGene00048922 | NA | NA | NA | NA |
| WBGene00048923 | NA | NA | NA | NA |
| WBGene00048924 | NA | NA | NA | NA |
| WBGene00048925 | NA | NA | NA | NA |
| WBGene00048926 | NA | NA | NA | NA |
| WBGene00048927 | NA | NA | NA | NA |
| WBGene00048928 | NA | NA | NA | NA |
| WBGene00048929 | NA | NA | NA | NA |

| | | | | |
|---|---|---|---|---|
| WBGene00048930 | NA | NA | NA | NA |
| WBGene00048931 | NA | NA | NA | NA |
| WBGene00048932 | NA | NA | NA | NA |
| WBGene00048933 | NA | NA | NA | NA |
| WBGene00048934 | NA | NA | NA | NA |
| WBGene00048935 | NA | NA | NA | NA |
| WBGene00048936 | NA | NA | NA | NA |
| WBGene00048937 | NA | NA | NA | NA |
| WBGene00048938 | 0.020317245 | 0.84643798 | NA | NA |
| WBGene00048939 | NA | NA | NA | NA |
| WBGene00048940 | NA | NA | NA | NA |
| WBGene00048941 | NA | NA | NA | NA |
| WBGene00048942 | NA | NA | NA | NA |
| WBGene00048943 | NA | NA | NA | NA |
| WBGene00048944 | NA | NA | NA | NA |
| WBGene00048945 | NA | NA | NA | NA |
| WBGene00048946 | NA | NA | NA | NA |
| WBGene00048947 | NA | NA | NA | NA |
| WBGene00048948 | NA | NA | NA | NA |
| WBGene00048949 | NA | NA | NA | NA |
| WBGene00048950 | NA | NA | NA | NA |
| WBGene00048951 | NA | NA | NA | NA |
| WBGene00048952 | NA | NA | NA | NA |
| WBGene00048953 | NA | NA | NA | NA |
| WBGene00048954 | NA | NA | NA | NA |
| WBGene00048955 | NA | NA | NA | NA |
| WBGene00048956 | NA | NA | NA | NA |
| WBGene00048957 | NA | NA | NA | NA |
| WBGene00048958 | NA | NA | NA | NA |
| WBGene00048959 | NA | NA | NA | NA |
| WBGene00048960 | NA | NA | NA | NA |
| WBGene00048961 | NA | NA | NA | NA |
| WBGene00048962 | NA | NA | NA | NA |
| WBGene00048963 | NA | NA | NA | NA |
| WBGene00048964 | NA | NA | NA | NA |
| WBGene00048965 | NA | NA | NA | NA |
| WBGene00048966 | NA | NA | NA | NA |
| WBGene00048967 | NA | NA | NA | NA |
| WBGene00048968 | NA | NA | NA | NA |
| WBGene00048969 | NA | NA | NA | NA |
| WBGene00048970 | NA | NA | NA | NA |
| WBGene00048971 | NA | NA | NA | NA |
| WBGene00048972 | NA | NA | NA | NA |

| | | | | |
|---|---|---|---|---|
| WBGene00048973 | NA | NA | NA | NA |
| WBGene00048974 | NA | NA | NA | NA |
| WBGene00048975 | NA | NA | NA | NA |
| WBGene00048976 | NA | NA | NA | NA |
| WBGene00048977 | NA | NA | NA | NA |
| WBGene00048978 | NA | NA | NA | NA |
| WBGene00048979 | NA | NA | NA | NA |
| WBGene00048980 | NA | NA | NA | NA |
| WBGene00048981 | NA | NA | NA | NA |
| WBGene00048982 | NA | NA | NA | NA |
| WBGene00048983 | NA | NA | NA | NA |
| WBGene00048984 | NA | NA | NA | NA |
| WBGene00048985 | NA | NA | NA | NA |
| WBGene00048986 | NA | NA | NA | NA |
| WBGene00048987 | NA | NA | NA | NA |
| WBGene00048988 | NA | NA | NA | NA |
| WBGene00048989 | NA | NA | NA | NA |
| WBGene00048990 | NA | NA | NA | NA |
| WBGene00048991 | NA | NA | NA | NA |
| WBGene00048992 | NA | NA | NA | NA |
| WBGene00048993 | NA | NA | NA | NA |
| WBGene00048994 | NA | NA | NA | NA |
| WBGene00048995 | NA | NA | NA | NA |
| WBGene00048996 | NA | NA | NA | NA |
| WBGene00048997 | NA | NA | NA | NA |
| WBGene00048998 | NA | NA | NA | NA |
| WBGene00048999 | NA | NA | NA | NA |
| WBGene00049000 | NA | NA | NA | NA |
| WBGene00049001 | NA | NA | NA | NA |
| WBGene00049002 | NA | NA | NA | NA |
| WBGene00049003 | NA | NA | NA | NA |
| WBGene00049004 | NA | NA | NA | NA |
| WBGene00049005 | NA | NA | NA | NA |
| WBGene00049006 | NA | NA | NA | NA |
| WBGene00049007 | NA | NA | NA | NA |
| WBGene00049008 | NA | NA | NA | NA |
| WBGene00049009 | NA | NA | NA | NA |
| WBGene00049010 | NA | NA | NA | NA |
| WBGene00049011 | NA | NA | NA | NA |
| WBGene00049012 | NA | NA | NA | NA |
| WBGene00049013 | NA | NA | NA | NA |
| WBGene00049014 | NA | NA | NA | NA |
| WBGene00049015 | NA | NA | NA | NA |

| | | | | |
|---|---|---|---|---|
| WBGene00049016 | NA | NA | NA | NA |
| WBGene00049017 | NA | NA | NA | NA |
| WBGene00049018 | NA | NA | NA | NA |
| WBGene00049019 | NA | NA | NA | NA |
| WBGene00049020 | NA | NA | NA | NA |
| WBGene00049021 | NA | NA | NA | NA |
| WBGene00049022 | 0.020232312 | 0.84643798 | NA | NA |
| WBGene00049023 | NA | NA | NA | NA |
| WBGene00049024 | NA | NA | NA | NA |
| WBGene00049025 | NA | NA | NA | NA |
| WBGene00049026 | NA | NA | NA | NA |
| WBGene00049027 | NA | NA | NA | NA |
| WBGene00049028 | NA | NA | NA | NA |
| WBGene00049029 | NA | NA | NA | NA |
| WBGene00049030 | NA | NA | NA | NA |
| WBGene00049031 | NA | NA | NA | NA |
| WBGene00049032 | NA | NA | NA | NA |
| WBGene00049033 | NA | NA | NA | NA |
| WBGene00049034 | NA | NA | NA | NA |
| WBGene00049035 | NA | NA | NA | NA |
| WBGene00049036 | NA | NA | NA | NA |
| WBGene00049037 | NA | NA | NA | NA |
| WBGene00049038 | NA | NA | NA | NA |
| WBGene00049039 | NA | NA | NA | NA |
| WBGene00049040 | NA | NA | NA | NA |
| WBGene00049041 | NA | NA | NA | NA |
| WBGene00049042 | NA | NA | NA | NA |
| WBGene00049043 | NA | NA | NA | NA |
| WBGene00049044 | NA | NA | NA | NA |
| WBGene00049045 | NA | NA | NA | NA |
| WBGene00049046 | NA | NA | NA | NA |
| WBGene00049047 | NA | NA | NA | NA |
| WBGene00049048 | NA | NA | NA | NA |
| WBGene00049049 | -0.023056499 | 0.78121253 | NA | NA |
| WBGene00049050 | NA | NA | NA | NA |
| WBGene00049051 | NA | NA | NA | NA |
| WBGene00049052 | NA | NA | NA | NA |
| WBGene00049053 | NA | NA | NA | NA |
| WBGene00049054 | NA | NA | NA | NA |
| WBGene00049055 | NA | NA | NA | NA |
| WBGene00049056 | NA | NA | NA | NA |
| WBGene00049057 | NA | NA | NA | NA |
| WBGene00049058 | NA | NA | NA | NA |

| | | | | |
|---|---|---|---|---|
| WBGene00049059 | NA | NA | NA | NA |
| WBGene00049060 | NA | NA | NA | NA |
| WBGene00049061 | NA | NA | NA | NA |
| WBGene00049062 | NA | NA | NA | NA |
| WBGene00049063 | NA | NA | NA | NA |
| WBGene00049064 | NA | NA | NA | NA |
| WBGene00049065 | NA | NA | NA | NA |
| WBGene00049066 | NA | NA | NA | NA |
| WBGene00049067 | NA | NA | NA | NA |
| WBGene00049068 | NA | NA | NA | NA |
| WBGene00049069 | NA | NA | NA | NA |
| WBGene00049070 | NA | NA | NA | NA |
| WBGene00049071 | NA | NA | NA | NA |
| WBGene00049072 | NA | NA | NA | NA |
| WBGene00049073 | NA | NA | NA | NA |
| WBGene00049074 | NA | NA | NA | NA |
| WBGene00049075 | NA | NA | NA | NA |
| WBGene00049076 | NA | NA | NA | NA |
| WBGene00049077 | NA | NA | NA | NA |
| WBGene00049078 | NA | NA | NA | NA |
| WBGene00049079 | NA | NA | NA | NA |
| WBGene00049080 | NA | NA | NA | NA |
| WBGene00049081 | NA | NA | NA | NA |
| WBGene00049082 | NA | NA | NA | NA |
| WBGene00049083 | NA | NA | NA | NA |
| WBGene00049084 | NA | NA | NA | NA |
| WBGene00049085 | NA | NA | NA | NA |
| WBGene00049086 | NA | NA | NA | NA |
| WBGene00049087 | NA | NA | NA | NA |
| WBGene00049088 | NA | NA | NA | NA |
| WBGene00049089 | NA | NA | NA | NA |
| WBGene00049090 | NA | NA | NA | NA |
| WBGene00049091 | NA | NA | NA | NA |
| WBGene00049092 | NA | NA | NA | NA |
| WBGene00049093 | NA | NA | NA | NA |
| WBGene00049094 | NA | NA | NA | NA |
| WBGene00049095 | NA | NA | NA | NA |
| WBGene00049096 | NA | NA | NA | NA |
| WBGene00049097 | NA | NA | NA | NA |
| WBGene00049098 | NA | NA | NA | NA |
| WBGene00049099 | NA | NA | NA | NA |
| WBGene00049100 | NA | NA | NA | NA |
| WBGene00049101 | NA | NA | NA | NA |

| | | | | |
|---|---|---|---|---|
| WBGene00049102 | NA | NA | NA | NA |
| WBGene00049103 | NA | NA | NA | NA |
| WBGene00049104 | NA | NA | NA | NA |
| WBGene00049105 | NA | NA | NA | NA |
| WBGene00049106 | NA | NA | NA | NA |
| WBGene00049107 | NA | NA | NA | NA |
| WBGene00049108 | NA | NA | NA | NA |
| WBGene00049109 | NA | NA | NA | NA |
| WBGene00049110 | NA | NA | NA | NA |
| WBGene00049111 | NA | NA | NA | NA |
| WBGene00049112 | NA | NA | NA | NA |
| WBGene00049113 | NA | NA | NA | NA |
| WBGene00049114 | NA | NA | NA | NA |
| WBGene00049115 | NA | NA | NA | NA |
| WBGene00049116 | NA | NA | NA | NA |
| WBGene00049117 | NA | NA | NA | NA |
| WBGene00049118 | NA | NA | NA | NA |
| WBGene00049119 | NA | NA | NA | NA |
| WBGene00049120 | NA | NA | NA | NA |
| WBGene00049121 | NA | NA | NA | NA |
| WBGene00049122 | NA | NA | NA | NA |
| WBGene00049123 | NA | NA | NA | NA |
| WBGene00049124 | NA | NA | NA | NA |
| WBGene00049125 | NA | NA | NA | NA |
| WBGene00049126 | NA | NA | NA | NA |
| WBGene00049127 | NA | NA | NA | NA |
| WBGene00049128 | NA | NA | NA | NA |
| WBGene00049129 | NA | NA | NA | NA |
| WBGene00049130 | NA | NA | NA | NA |
| WBGene00049131 | NA | NA | NA | NA |
| WBGene00049132 | NA | NA | NA | NA |
| WBGene00049133 | NA | NA | NA | NA |
| WBGene00049134 | NA | NA | NA | NA |
| WBGene00049135 | NA | NA | NA | NA |
| WBGene00049136 | NA | NA | NA | NA |
| WBGene00049137 | NA | NA | NA | NA |
| WBGene00049138 | NA | NA | NA | NA |
| WBGene00049139 | NA | NA | NA | NA |
| WBGene00049140 | NA | NA | NA | NA |
| WBGene00049141 | NA | NA | NA | NA |
| WBGene00049142 | NA | NA | NA | NA |
| WBGene00049143 | NA | NA | NA | NA |
| WBGene00049144 | NA | NA | NA | NA |

| | | | | |
|---|---|---|---|---|
| WBGene00049145 | NA | NA | NA | NA |
| WBGene00049146 | NA | NA | NA | NA |
| WBGene00049147 | NA | NA | NA | NA |
| WBGene00049148 | NA | NA | NA | NA |
| WBGene00049149 | NA | NA | NA | NA |
| WBGene00049150 | NA | NA | NA | NA |
| WBGene00049151 | NA | NA | NA | NA |
| WBGene00049152 | NA | NA | NA | NA |
| WBGene00049153 | NA | NA | NA | NA |
| WBGene00049154 | NA | NA | NA | NA |
| WBGene00049155 | NA | NA | NA | NA |
| WBGene00049156 | NA | NA | NA | NA |
| WBGene00049157 | NA | NA | NA | NA |
| WBGene00049158 | NA | NA | NA | NA |
| WBGene00049159 | NA | NA | NA | NA |
| WBGene00049160 | NA | NA | NA | NA |
| WBGene00049161 | NA | NA | NA | NA |
| WBGene00049162 | NA | NA | NA | NA |
| WBGene00049163 | NA | NA | NA | NA |
| WBGene00049164 | NA | NA | NA | NA |
| WBGene00049165 | NA | NA | NA | NA |
| WBGene00049166 | NA | NA | NA | NA |
| WBGene00049167 | NA | NA | NA | NA |
| WBGene00049168 | NA | NA | NA | NA |
| WBGene00049169 | NA | NA | NA | NA |
| WBGene00049170 | NA | NA | NA | NA |
| WBGene00049171 | NA | NA | NA | NA |
| WBGene00049172 | NA | NA | NA | NA |
| WBGene00049173 | NA | NA | NA | NA |
| WBGene00049174 | NA | NA | NA | NA |
| WBGene00049175 | NA | NA | NA | NA |
| WBGene00049176 | NA | NA | NA | NA |
| WBGene00049177 | NA | NA | NA | NA |
| WBGene00049178 | NA | NA | NA | NA |
| WBGene00049179 | NA | NA | NA | NA |
| WBGene00049180 | NA | NA | NA | NA |
| WBGene00049181 | NA | NA | NA | NA |
| WBGene00049182 | NA | NA | NA | NA |
| WBGene00049183 | NA | NA | NA | NA |
| WBGene00049184 | NA | NA | NA | NA |
| WBGene00049185 | NA | NA | NA | NA |
| WBGene00049186 | NA | NA | NA | NA |
| WBGene00049187 | NA | NA | NA | NA |

| | | | | |
|---|---|---|---|---|
| WBGene00049188 | NA | NA | NA | NA |
| WBGene00049189 | NA | NA | NA | NA |
| WBGene00049190 | NA | NA | NA | NA |
| WBGene00049191 | NA | NA | NA | NA |
| WBGene00049192 | NA | NA | NA | NA |
| WBGene00049193 | NA | NA | NA | NA |
| WBGene00049194 | NA | NA | NA | NA |
| WBGene00049195 | NA | NA | NA | NA |
| WBGene00049196 | NA | NA | NA | NA |
| WBGene00049197 | NA | NA | NA | NA |
| WBGene00049198 | NA | NA | NA | NA |
| WBGene00049199 | NA | NA | NA | NA |
| WBGene00049200 | NA | NA | NA | NA |
| WBGene00049201 | NA | NA | NA | NA |
| WBGene00049202 | NA | NA | NA | NA |
| WBGene00049203 | NA | NA | NA | NA |
| WBGene00049204 | NA | NA | NA | NA |
| WBGene00049205 | NA | NA | NA | NA |
| WBGene00049206 | NA | NA | NA | NA |
| WBGene00049207 | NA | NA | NA | NA |
| WBGene00049208 | NA | NA | NA | NA |
| WBGene00049209 | NA | NA | NA | NA |
| WBGene00049210 | NA | NA | NA | NA |
| WBGene00049211 | NA | NA | NA | NA |
| WBGene00049212 | NA | NA | NA | NA |
| WBGene00049213 | NA | NA | NA | NA |
| WBGene00049214 | NA | NA | NA | NA |
| WBGene00049215 | NA | NA | NA | NA |
| WBGene00049216 | NA | NA | NA | NA |
| WBGene00049217 | NA | NA | NA | NA |
| WBGene00049218 | NA | NA | NA | NA |
| WBGene00049219 | NA | NA | NA | NA |
| WBGene00049220 | NA | NA | NA | NA |
| WBGene00049221 | NA | NA | NA | NA |
| WBGene00049222 | NA | NA | NA | NA |
| WBGene00049223 | NA | NA | NA | NA |
| WBGene00049224 | NA | NA | NA | NA |
| WBGene00049225 | NA | NA | NA | NA |
| WBGene00049226 | NA | NA | NA | NA |
| WBGene00049227 | NA | NA | NA | NA |
| WBGene00049228 | NA | NA | NA | NA |
| WBGene00049229 | NA | NA | NA | NA |
| WBGene00049230 | NA | NA | NA | NA |

| | | | | |
|---|---|---|---|---|
| WBGene00049231 | NA | NA | NA | NA |
| WBGene00049232 | NA | NA | NA | NA |
| WBGene00049233 | NA | NA | NA | NA |
| WBGene00049234 | NA | NA | NA | NA |
| WBGene00049235 | NA | NA | NA | NA |
| WBGene00049236 | NA | NA | NA | NA |
| WBGene00049237 | NA | NA | NA | NA |
| WBGene00049238 | NA | NA | NA | NA |
| WBGene00049239 | NA | NA | NA | NA |
| WBGene00049240 | NA | NA | NA | NA |
| WBGene00049241 | NA | NA | NA | NA |
| WBGene00049242 | NA | NA | NA | NA |
| WBGene00049243 | NA | NA | NA | NA |
| WBGene00049244 | NA | NA | NA | NA |
| WBGene00049245 | NA | NA | NA | NA |
| WBGene00049246 | NA | NA | NA | NA |
| WBGene00049247 | NA | NA | NA | NA |
| WBGene00049248 | NA | NA | NA | NA |
| WBGene00049249 | NA | NA | NA | NA |
| WBGene00049250 | NA | NA | NA | NA |
| WBGene00049251 | NA | NA | NA | NA |
| WBGene00049252 | NA | NA | NA | NA |
| WBGene00049253 | NA | NA | NA | NA |
| WBGene00049254 | NA | NA | NA | NA |
| WBGene00049255 | NA | NA | NA | NA |
| WBGene00049256 | NA | NA | NA | NA |
| WBGene00049257 | NA | NA | NA | NA |
| WBGene00049258 | NA | NA | NA | NA |
| WBGene00049259 | NA | NA | NA | NA |
| WBGene00049260 | NA | NA | NA | NA |
| WBGene00049261 | NA | NA | NA | NA |
| WBGene00049262 | NA | NA | NA | NA |
| WBGene00049263 | NA | NA | NA | NA |
| WBGene00049264 | NA | NA | NA | NA |
| WBGene00049265 | NA | NA | NA | NA |
| WBGene00049266 | NA | NA | NA | NA |
| WBGene00049267 | NA | NA | NA | NA |
| WBGene00049268 | NA | NA | NA | NA |
| WBGene00049269 | NA | NA | NA | NA |
| WBGene00049270 | NA | NA | NA | NA |
| WBGene00049271 | NA | NA | NA | NA |
| WBGene00049272 | NA | NA | NA | NA |
| WBGene00049273 | NA | NA | NA | NA |

| | | | | |
|---|---|---|---|---|
| WBGene00049274 | NA | NA | NA | NA |
| WBGene00049275 | NA | NA | NA | NA |
| WBGene00049276 | NA | NA | NA | NA |
| WBGene00049277 | NA | NA | NA | NA |
| WBGene00049278 | NA | NA | NA | NA |
| WBGene00049279 | NA | NA | NA | NA |
| WBGene00049280 | NA | NA | NA | NA |
| WBGene00049281 | NA | NA | NA | NA |
| WBGene00049282 | 0.020317245 | 0.84643798 | NA | NA |
| WBGene00049283 | NA | NA | NA | NA |
| WBGene00049284 | NA | NA | NA | NA |
| WBGene00049285 | NA | NA | NA | NA |
| WBGene00049286 | NA | NA | NA | NA |
| WBGene00049287 | NA | NA | NA | NA |
| WBGene00049288 | NA | NA | NA | NA |
| WBGene00049289 | NA | NA | NA | NA |
| WBGene00049290 | NA | NA | NA | NA |
| WBGene00049291 | NA | NA | NA | NA |
| WBGene00049292 | NA | NA | NA | NA |
| WBGene00049293 | NA | NA | NA | NA |
| WBGene00049294 | NA | NA | NA | NA |
| WBGene00049295 | NA | NA | NA | NA |
| WBGene00049296 | NA | NA | NA | NA |
| WBGene00049297 | NA | NA | NA | NA |
| WBGene00049298 | NA | NA | NA | NA |
| WBGene00049299 | NA | NA | NA | NA |
| WBGene00049300 | NA | NA | NA | NA |
| WBGene00049301 | NA | NA | NA | NA |
| WBGene00049302 | NA | NA | NA | NA |
| WBGene00049303 | NA | NA | NA | NA |
| WBGene00049304 | NA | NA | NA | NA |
| WBGene00049305 | NA | NA | NA | NA |
| WBGene00049306 | NA | NA | NA | NA |
| WBGene00049307 | NA | NA | NA | NA |
| WBGene00049308 | NA | NA | NA | NA |
| WBGene00049309 | NA | NA | NA | NA |
| WBGene00049310 | NA | NA | NA | NA |
| WBGene00049311 | NA | NA | NA | NA |
| WBGene00049312 | NA | NA | NA | NA |
| WBGene00049313 | NA | NA | NA | NA |
| WBGene00049314 | NA | NA | NA | NA |
| WBGene00049315 | NA | NA | NA | NA |
| WBGene00049316 | NA | NA | NA | NA |

| | | | | |
|---|---|---|---|---|
| WBGene00049317 | NA | NA | NA | NA |
| WBGene00049318 | NA | NA | NA | NA |
| WBGene00049319 | NA | NA | NA | NA |
| WBGene00049320 | NA | NA | NA | NA |
| WBGene00049321 | NA | NA | NA | NA |
| WBGene00049322 | NA | NA | NA | NA |
| WBGene00049323 | -0.046394786 | 0.50231372 | NA | NA |
| WBGene00049324 | NA | NA | NA | NA |
| WBGene00049325 | NA | NA | NA | NA |
| WBGene00049326 | NA | NA | NA | NA |
| WBGene00049327 | NA | NA | NA | NA |
| WBGene00049328 | NA | NA | NA | NA |
| WBGene00049329 | NA | NA | NA | NA |
| WBGene00049330 | NA | NA | NA | NA |
| WBGene00049331 | NA | NA | NA | NA |
| WBGene00049332 | NA | NA | NA | NA |
| WBGene00049333 | NA | NA | NA | NA |
| WBGene00049334 | NA | NA | NA | NA |
| WBGene00049335 | NA | NA | NA | NA |
| WBGene00049336 | NA | NA | NA | NA |
| WBGene00049337 | NA | NA | NA | NA |
| WBGene00049338 | NA | NA | NA | NA |
| WBGene00049339 | NA | NA | NA | NA |
| WBGene00049340 | NA | NA | NA | NA |
| WBGene00049341 | NA | NA | NA | NA |
| WBGene00049342 | NA | NA | NA | NA |
| WBGene00049343 | NA | NA | NA | NA |
| WBGene00049344 | NA | NA | NA | NA |
| WBGene00049345 | NA | NA | NA | NA |
| WBGene00049346 | NA | NA | NA | NA |
| WBGene00049347 | NA | NA | NA | NA |
| WBGene00049348 | NA | NA | NA | NA |
| WBGene00049349 | NA | NA | NA | NA |
| WBGene00049350 | NA | NA | NA | NA |
| WBGene00049351 | NA | NA | NA | NA |
| WBGene00049352 | NA | NA | NA | NA |
| WBGene00049353 | NA | NA | NA | NA |
| WBGene00049354 | NA | NA | NA | NA |
| WBGene00049355 | NA | NA | NA | NA |
| WBGene00049356 | NA | NA | NA | NA |
| WBGene00049357 | NA | NA | NA | NA |
| WBGene00049358 | NA | NA | NA | NA |
| WBGene00049359 | NA | NA | NA | NA |

| | | | | |
|---|---|---|---|---|
| WBGene00049360 | NA | NA | NA | NA |
| WBGene00049361 | NA | NA | NA | NA |
| WBGene00049362 | NA | NA | NA | NA |
| WBGene00049363 | NA | NA | NA | NA |
| WBGene00049364 | NA | NA | NA | NA |
| WBGene00049365 | NA | NA | NA | NA |
| WBGene00049366 | NA | NA | NA | NA |
| WBGene00049367 | NA | NA | NA | NA |
| WBGene00049368 | NA | NA | NA | NA |
| WBGene00049369 | NA | NA | NA | NA |
| WBGene00049370 | NA | NA | NA | NA |
| WBGene00049371 | NA | NA | NA | NA |
| WBGene00049372 | NA | NA | NA | NA |
| WBGene00049373 | NA | NA | NA | NA |
| WBGene00049374 | NA | NA | NA | NA |
| WBGene00049375 | NA | NA | NA | NA |
| WBGene00049376 | NA | NA | NA | NA |
| WBGene00049377 | NA | NA | NA | NA |
| WBGene00049378 | NA | NA | NA | NA |
| WBGene00049379 | NA | NA | NA | NA |
| WBGene00049380 | NA | NA | NA | NA |
| WBGene00049381 | NA | NA | NA | NA |
| WBGene00049382 | NA | NA | NA | NA |
| WBGene00049383 | NA | NA | NA | NA |
| WBGene00049384 | NA | NA | NA | NA |
| WBGene00049385 | NA | NA | NA | NA |
| WBGene00049386 | NA | NA | NA | NA |
| WBGene00049387 | NA | NA | NA | NA |
| WBGene00049388 | NA | NA | NA | NA |
| WBGene00049389 | NA | NA | NA | NA |
| WBGene00049390 | NA | NA | NA | NA |
| WBGene00049391 | NA | NA | NA | NA |
| WBGene00049392 | NA | NA | NA | NA |
| WBGene00049393 | NA | NA | NA | NA |
| WBGene00049394 | NA | NA | NA | NA |
| WBGene00049395 | NA | NA | NA | NA |
| WBGene00049396 | NA | NA | NA | NA |
| WBGene00049397 | NA | NA | NA | NA |
| WBGene00049398 | NA | NA | NA | NA |
| WBGene00049399 | NA | NA | NA | NA |
| WBGene00049400 | NA | NA | NA | NA |
| WBGene00049401 | NA | NA | NA | NA |
| WBGene00049402 | NA | NA | NA | NA |

| | | | | |
|---|---|---|---|---|
| WBGene00049403 | NA | NA | NA | NA |
| WBGene00049404 | NA | NA | NA | NA |
| WBGene00049405 | NA | NA | NA | NA |
| WBGene00049406 | NA | NA | NA | NA |
| WBGene00049407 | NA | NA | NA | NA |
| WBGene00049408 | NA | NA | NA | NA |
| WBGene00049409 | NA | NA | NA | NA |
| WBGene00049410 | NA | NA | NA | NA |
| WBGene00049411 | NA | NA | NA | NA |
| WBGene00049412 | NA | NA | NA | NA |
| WBGene00049413 | NA | NA | NA | NA |
| WBGene00049414 | 0.020317245 | 0.84643798 | NA | NA |
| WBGene00049415 | NA | NA | NA | NA |
| WBGene00049416 | NA | NA | NA | NA |
| WBGene00049417 | NA | NA | NA | NA |
| WBGene00049418 | NA | NA | NA | NA |
| WBGene00049419 | NA | NA | NA | NA |
| WBGene00049420 | NA | NA | NA | NA |
| WBGene00049421 | NA | NA | NA | NA |
| WBGene00049422 | NA | NA | NA | NA |
| WBGene00049423 | NA | NA | NA | NA |
| WBGene00049424 | NA | NA | NA | NA |
| WBGene00049425 | NA | NA | NA | NA |
| WBGene00049426 | NA | NA | NA | NA |
| WBGene00049427 | NA | NA | NA | NA |
| WBGene00049428 | NA | NA | NA | NA |
| WBGene00049429 | NA | NA | NA | NA |
| WBGene00049430 | NA | NA | NA | NA |
| WBGene00049431 | NA | NA | NA | NA |
| WBGene00049432 | NA | NA | NA | NA |
| WBGene00049433 | NA | NA | NA | NA |
| WBGene00049434 | NA | NA | NA | NA |
| WBGene00049435 | NA | NA | NA | NA |
| WBGene00049436 | NA | NA | NA | NA |
| WBGene00049437 | NA | NA | NA | NA |
| WBGene00049438 | NA | NA | NA | NA |
| WBGene00049439 | NA | NA | NA | NA |
| WBGene00049440 | NA | NA | NA | NA |
| WBGene00049441 | NA | NA | NA | NA |
| WBGene00049443 | NA | NA | NA | NA |
| WBGene00049444 | NA | NA | NA | NA |
| WBGene00049445 | NA | NA | NA | NA |
| WBGene00049446 | NA | NA | NA | NA |

| | | | | |
|---|---|---|---|---|
| WBGene00049447 | NA | NA | NA | NA |
| WBGene00049448 | NA | NA | NA | NA |
| WBGene00049449 | NA | NA | NA | NA |
| WBGene00049450 | NA | NA | NA | NA |
| WBGene00049451 | NA | NA | NA | NA |
| WBGene00049452 | NA | NA | NA | NA |
| WBGene00049453 | NA | NA | NA | NA |
| WBGene00049454 | NA | NA | NA | NA |
| WBGene00049455 | NA | NA | NA | NA |
| WBGene00049456 | NA | NA | NA | NA |
| WBGene00049457 | NA | NA | NA | NA |
| WBGene00049458 | NA | NA | NA | NA |
| WBGene00049459 | NA | NA | NA | NA |
| WBGene00049460 | NA | NA | NA | NA |
| WBGene00049461 | NA | NA | NA | NA |
| WBGene00049462 | NA | NA | NA | NA |
| WBGene00049463 | NA | NA | NA | NA |
| WBGene00049464 | NA | NA | NA | NA |
| WBGene00049465 | NA | NA | NA | NA |
| WBGene00049466 | NA | NA | NA | NA |
| WBGene00049467 | NA | NA | NA | NA |
| WBGene00049468 | NA | NA | NA | NA |
| WBGene00049469 | NA | NA | NA | NA |
| WBGene00049470 | NA | NA | NA | NA |
| WBGene00049471 | NA | NA | NA | NA |
| WBGene00049472 | NA | NA | NA | NA |
| WBGene00049473 | NA | NA | NA | NA |
| WBGene00049474 | NA | NA | NA | NA |
| WBGene00049475 | NA | NA | NA | NA |
| WBGene00049476 | NA | NA | NA | NA |
| WBGene00049477 | NA | NA | NA | NA |
| WBGene00049478 | NA | NA | NA | NA |
| WBGene00049479 | NA | NA | NA | NA |
| WBGene00049480 | NA | NA | NA | NA |
| WBGene00049481 | NA | NA | NA | NA |
| WBGene00049482 | NA | NA | NA | NA |
| WBGene00049483 | NA | NA | NA | NA |
| WBGene00049484 | NA | NA | NA | NA |
| WBGene00049485 | NA | NA | NA | NA |
| WBGene00049486 | NA | NA | NA | NA |
| WBGene00049487 | NA | NA | NA | NA |
| WBGene00049488 | NA | NA | NA | NA |
| WBGene00049489 | NA | NA | NA | NA |

| | | | | |
|---|---|---|---|---|
| WBGene00049490 | NA | NA | NA | NA |
| WBGene00049491 | NA | NA | NA | NA |
| WBGene00049492 | NA | NA | NA | NA |
| WBGene00049493 | NA | NA | NA | NA |
| WBGene00049494 | NA | NA | NA | NA |
| WBGene00049495 | NA | NA | NA | NA |
| WBGene00049496 | NA | NA | NA | NA |
| WBGene00049497 | NA | NA | NA | NA |
| WBGene00049498 | NA | NA | NA | NA |
| WBGene00049499 | NA | NA | NA | NA |
| WBGene00049500 | NA | NA | NA | NA |
| WBGene00049501 | NA | NA | NA | NA |
| WBGene00049502 | NA | NA | NA | NA |
| WBGene00049503 | NA | NA | NA | NA |
| WBGene00049504 | NA | NA | NA | NA |
| WBGene00049505 | NA | NA | NA | NA |
| WBGene00049506 | NA | NA | NA | NA |
| WBGene00049507 | NA | NA | NA | NA |
| WBGene00049508 | NA | NA | NA | NA |
| WBGene00049509 | NA | NA | NA | NA |
| WBGene00049510 | NA | NA | NA | NA |
| WBGene00049511 | NA | NA | NA | NA |
| WBGene00049512 | NA | NA | NA | NA |
| WBGene00049513 | NA | NA | NA | NA |
| WBGene00049514 | NA | NA | NA | NA |
| WBGene00049515 | NA | NA | NA | NA |
| WBGene00049516 | NA | NA | NA | NA |
| WBGene00049517 | NA | NA | NA | NA |
| WBGene00049518 | NA | NA | NA | NA |
| WBGene00049519 | NA | NA | NA | NA |
| WBGene00049520 | NA | NA | NA | NA |
| WBGene00049521 | NA | NA | NA | NA |
| WBGene00049522 | NA | NA | NA | NA |
| WBGene00049523 | NA | NA | NA | NA |
| WBGene00049524 | NA | NA | NA | NA |
| WBGene00049525 | NA | NA | NA | NA |
| WBGene00049526 | NA | NA | NA | NA |
| WBGene00049527 | NA | NA | NA | NA |
| WBGene00049528 | NA | NA | NA | NA |
| WBGene00049529 | NA | NA | NA | NA |
| WBGene00049530 | NA | NA | NA | NA |
| WBGene00049531 | NA | NA | NA | NA |
| WBGene00049532 | NA | NA | NA | NA |

| | | | | |
|---|---|---|---|---|
| WBGene00049533 | NA | NA | NA | NA |
| WBGene00049534 | NA | NA | NA | NA |
| WBGene00049535 | NA | NA | NA | NA |
| WBGene00049536 | NA | NA | NA | NA |
| WBGene00049537 | NA | NA | NA | NA |
| WBGene00049538 | NA | NA | NA | NA |
| WBGene00049539 | NA | NA | NA | NA |
| WBGene00049540 | NA | NA | NA | NA |
| WBGene00049541 | NA | NA | NA | NA |
| WBGene00049542 | NA | NA | NA | NA |
| WBGene00049543 | NA | NA | NA | NA |
| WBGene00049544 | NA | NA | NA | NA |
| WBGene00049545 | NA | NA | NA | NA |
| WBGene00049546 | NA | NA | NA | NA |
| WBGene00049547 | NA | NA | NA | NA |
| WBGene00049548 | NA | NA | NA | NA |
| WBGene00049549 | NA | NA | NA | NA |
| WBGene00049550 | NA | NA | NA | NA |
| WBGene00049551 | NA | NA | NA | NA |
| WBGene00049552 | NA | NA | NA | NA |
| WBGene00049553 | NA | NA | NA | NA |
| WBGene00049554 | NA | NA | NA | NA |
| WBGene00049555 | NA | NA | NA | NA |
| WBGene00049556 | NA | NA | NA | NA |
| WBGene00049557 | NA | NA | NA | NA |
| WBGene00049558 | NA | NA | NA | NA |
| WBGene00049559 | NA | NA | NA | NA |
| WBGene00049560 | NA | NA | NA | NA |
| WBGene00049561 | NA | NA | NA | NA |
| WBGene00049562 | NA | NA | NA | NA |
| WBGene00049563 | NA | NA | NA | NA |
| WBGene00049564 | NA | NA | NA | NA |
| WBGene00049565 | NA | NA | NA | NA |
| WBGene00049566 | NA | NA | NA | NA |
| WBGene00049567 | NA | NA | NA | NA |
| WBGene00049568 | NA | NA | NA | NA |
| WBGene00049569 | NA | NA | NA | NA |
| WBGene00049570 | NA | NA | NA | NA |
| WBGene00049571 | 0.020925399 | 0.84643798 | NA | NA |
| WBGene00049572 | NA | NA | NA | NA |
| WBGene00049573 | NA | NA | NA | NA |
| WBGene00049574 | NA | NA | NA | NA |
| WBGene00049575 | NA | NA | NA | NA |

| | | | | |
|---|---|---|---|---|
| WBGene00049576 | NA | NA | NA | NA |
| WBGene00049577 | NA | NA | NA | NA |
| WBGene00049578 | NA | NA | NA | NA |
| WBGene00049579 | NA | NA | NA | NA |
| WBGene00049580 | NA | NA | NA | NA |
| WBGene00049581 | NA | NA | NA | NA |
| WBGene00049582 | NA | NA | NA | NA |
| WBGene00049583 | NA | NA | NA | NA |
| WBGene00049584 | NA | NA | NA | NA |
| WBGene00049585 | NA | NA | NA | NA |
| WBGene00049586 | NA | NA | NA | NA |
| WBGene00049587 | NA | NA | NA | NA |
| WBGene00049588 | NA | NA | NA | NA |
| WBGene00049589 | NA | NA | NA | NA |
| WBGene00049590 | NA | NA | NA | NA |
| WBGene00049591 | NA | NA | NA | NA |
| WBGene00049592 | NA | NA | NA | NA |
| WBGene00049593 | NA | NA | NA | NA |
| WBGene00049594 | NA | NA | NA | NA |
| WBGene00049595 | NA | NA | NA | NA |
| WBGene00049596 | NA | NA | NA | NA |
| WBGene00049597 | NA | NA | NA | NA |
| WBGene00049598 | NA | NA | NA | NA |
| WBGene00049599 | NA | NA | NA | NA |
| WBGene00049600 | NA | NA | NA | NA |
| WBGene00049601 | NA | NA | NA | NA |
| WBGene00049602 | NA | NA | NA | NA |
| WBGene00049603 | NA | NA | NA | NA |
| WBGene00049604 | NA | NA | NA | NA |
| WBGene00049605 | NA | NA | NA | NA |
| WBGene00049606 | NA | NA | NA | NA |
| WBGene00049607 | NA | NA | NA | NA |
| WBGene00049608 | NA | NA | NA | NA |
| WBGene00049609 | NA | NA | NA | NA |
| WBGene00049610 | NA | NA | NA | NA |
| WBGene00049611 | NA | NA | NA | NA |
| WBGene00049612 | NA | NA | NA | NA |
| WBGene00049613 | NA | NA | NA | NA |
| WBGene00049614 | NA | NA | NA | NA |
| WBGene00049615 | NA | NA | NA | NA |
| WBGene00049616 | NA | NA | NA | NA |
| WBGene00049617 | NA | NA | NA | NA |
| WBGene00049618 | NA | NA | NA | NA |

| | | | | |
|---|---|---|---|---|
| WBGene00049619 | NA | NA | NA | NA |
| WBGene00049620 | NA | NA | NA | NA |
| WBGene00049621 | NA | NA | NA | NA |
| WBGene00049622 | NA | NA | NA | NA |
| WBGene00049623 | NA | NA | NA | NA |
| WBGene00049624 | NA | NA | NA | NA |
| WBGene00049625 | NA | NA | NA | NA |
| WBGene00049626 | NA | NA | NA | NA |
| WBGene00049627 | NA | NA | NA | NA |
| WBGene00049628 | NA | NA | NA | NA |
| WBGene00049629 | NA | NA | NA | NA |
| WBGene00049630 | NA | NA | NA | NA |
| WBGene00049631 | NA | NA | NA | NA |
| WBGene00049632 | NA | NA | NA | NA |
| WBGene00049633 | NA | NA | NA | NA |
| WBGene00049634 | NA | NA | NA | NA |
| WBGene00049635 | NA | NA | NA | NA |
| WBGene00049636 | NA | NA | NA | NA |
| WBGene00049637 | NA | NA | NA | NA |
| WBGene00049638 | NA | NA | NA | NA |
| WBGene00049639 | NA | NA | NA | NA |
| WBGene00049640 | NA | NA | NA | NA |
| WBGene00049641 | NA | NA | NA | NA |
| WBGene00049642 | NA | NA | NA | NA |
| WBGene00049643 | NA | NA | NA | NA |
| WBGene00049644 | NA | NA | NA | NA |
| WBGene00049645 | NA | NA | NA | NA |
| WBGene00049646 | NA | NA | NA | NA |
| WBGene00049647 | NA | NA | NA | NA |
| WBGene00049648 | NA | NA | NA | NA |
| WBGene00049649 | NA | NA | NA | NA |
| WBGene00049650 | NA | NA | NA | NA |
| WBGene00049651 | NA | NA | NA | NA |
| WBGene00049652 | NA | NA | NA | NA |
| WBGene00049653 | NA | NA | NA | NA |
| WBGene00049654 | NA | NA | NA | NA |
| WBGene00049655 | NA | NA | NA | NA |
| WBGene00049656 | NA | NA | NA | NA |
| WBGene00049657 | NA | NA | NA | NA |
| WBGene00049658 | NA | NA | NA | NA |
| WBGene00049659 | NA | NA | NA | NA |
| WBGene00049660 | NA | NA | NA | NA |
| WBGene00049661 | NA | NA | NA | NA |

| | | | | |
|---|---|---|---|---|
| WBGene00049662 | NA | NA | NA | NA |
| WBGene00049663 | NA | NA | NA | NA |
| WBGene00049664 | NA | NA | NA | NA |
| WBGene00049665 | NA | NA | NA | NA |
| WBGene00049666 | NA | NA | NA | NA |
| WBGene00049667 | NA | NA | NA | NA |
| WBGene00049668 | NA | NA | NA | NA |
| WBGene00049669 | NA | NA | NA | NA |
| WBGene00049670 | NA | NA | NA | NA |
| WBGene00049671 | NA | NA | NA | NA |
| WBGene00049672 | NA | NA | NA | NA |
| WBGene00049673 | NA | NA | NA | NA |
| WBGene00049674 | 0.020232312 | 0.84643798 | NA | NA |
| WBGene00049675 | NA | NA | NA | NA |
| WBGene00049676 | NA | NA | NA | NA |
| WBGene00049677 | NA | NA | NA | NA |
| WBGene00049678 | NA | NA | NA | NA |
| WBGene00049679 | NA | NA | NA | NA |
| WBGene00049680 | NA | NA | NA | NA |
| WBGene00049681 | NA | NA | NA | NA |
| WBGene00049682 | NA | NA | NA | NA |
| WBGene00049683 | NA | NA | NA | NA |
| WBGene00049684 | NA | NA | NA | NA |
| WBGene00049685 | NA | NA | NA | NA |
| WBGene00049686 | NA | NA | NA | NA |
| WBGene00049687 | NA | NA | NA | NA |
| WBGene00049688 | NA | NA | NA | NA |
| WBGene00049689 | NA | NA | NA | NA |
| WBGene00049690 | NA | NA | NA | NA |
| WBGene00049691 | NA | NA | NA | NA |
| WBGene00049692 | NA | NA | NA | NA |
| WBGene00049693 | NA | NA | NA | NA |
| WBGene00049694 | NA | NA | NA | NA |
| WBGene00049695 | NA | NA | NA | NA |
| WBGene00049696 | NA | NA | NA | NA |
| WBGene00049697 | NA | NA | NA | NA |
| WBGene00049698 | NA | NA | NA | NA |
| WBGene00049699 | NA | NA | NA | NA |
| WBGene00049700 | NA | NA | NA | NA |
| WBGene00049701 | NA | NA | NA | NA |
| WBGene00049702 | NA | NA | NA | NA |
| WBGene00049703 | NA | NA | NA | NA |
| WBGene00049704 | NA | NA | NA | NA |

| | | | | |
|---|---|---|---|---|
| WBGene00049705 | NA | NA | NA | NA |
| WBGene00049706 | NA | NA | NA | NA |
| WBGene00049707 | NA | NA | NA | NA |
| WBGene00049708 | NA | NA | NA | NA |
| WBGene00049709 | NA | NA | NA | NA |
| WBGene00049710 | NA | NA | NA | NA |
| WBGene00049711 | NA | NA | NA | NA |
| WBGene00049712 | NA | NA | NA | NA |
| WBGene00049713 | NA | NA | NA | NA |
| WBGene00049714 | NA | NA | NA | NA |
| WBGene00049715 | NA | NA | NA | NA |
| WBGene00049716 | NA | NA | NA | NA |
| WBGene00049717 | NA | NA | NA | NA |
| WBGene00049718 | NA | NA | NA | NA |
| WBGene00049719 | NA | NA | NA | NA |
| WBGene00049720 | NA | NA | NA | NA |
| WBGene00049721 | NA | NA | NA | NA |
| WBGene00049722 | NA | NA | NA | NA |
| WBGene00049723 | NA | NA | NA | NA |
| WBGene00049724 | NA | NA | NA | NA |
| WBGene00049725 | NA | NA | NA | NA |
| WBGene00049726 | NA | NA | NA | NA |
| WBGene00049727 | NA | NA | NA | NA |
| WBGene00049728 | NA | NA | NA | NA |
| WBGene00049729 | NA | NA | NA | NA |
| WBGene00049730 | NA | NA | NA | NA |
| WBGene00049731 | NA | NA | NA | NA |
| WBGene00049732 | NA | NA | NA | NA |
| WBGene00049733 | NA | NA | NA | NA |
| WBGene00049734 | NA | NA | NA | NA |
| WBGene00049735 | NA | NA | NA | NA |
| WBGene00049736 | NA | NA | NA | NA |
| WBGene00049737 | NA | NA | NA | NA |
| WBGene00049738 | NA | NA | NA | NA |
| WBGene00049739 | NA | NA | NA | NA |
| WBGene00049740 | NA | NA | NA | NA |
| WBGene00049741 | NA | NA | NA | NA |
| WBGene00049742 | NA | NA | NA | NA |
| WBGene00049743 | NA | NA | NA | NA |
| WBGene00049744 | NA | NA | NA | NA |
| WBGene00049745 | NA | NA | NA | NA |
| WBGene00049746 | NA | NA | NA | NA |
| WBGene00049747 | NA | NA | NA | NA |

| | | | | |
|---|---|---|---|---|
| WBGene00049748 | NA | NA | NA | NA |
| WBGene00049749 | NA | NA | NA | NA |
| WBGene00049750 | NA | NA | NA | NA |
| WBGene00049751 | NA | NA | NA | NA |
| WBGene00049752 | NA | NA | NA | NA |
| WBGene00049753 | NA | NA | NA | NA |
| WBGene00049754 | NA | NA | NA | NA |
| WBGene00049755 | NA | NA | NA | NA |
| WBGene00049756 | NA | NA | NA | NA |
| WBGene00049757 | NA | NA | NA | NA |
| WBGene00049758 | NA | NA | NA | NA |
| WBGene00049759 | NA | NA | NA | NA |
| WBGene00049760 | NA | NA | NA | NA |
| WBGene00049761 | NA | NA | NA | NA |
| WBGene00049762 | 0.020232312 | 0.84643798 | NA | NA |
| WBGene00049763 | NA | NA | NA | NA |
| WBGene00049764 | NA | NA | NA | NA |
| WBGene00049765 | NA | NA | NA | NA |
| WBGene00049766 | NA | NA | NA | NA |
| WBGene00049767 | NA | NA | NA | NA |
| WBGene00049768 | NA | NA | NA | NA |
| WBGene00049769 | NA | NA | NA | NA |
| WBGene00049770 | NA | NA | NA | NA |
| WBGene00049771 | NA | NA | NA | NA |
| WBGene00049772 | NA | NA | NA | NA |
| WBGene00049773 | NA | NA | NA | NA |
| WBGene00049774 | NA | NA | NA | NA |
| WBGene00049775 | NA | NA | NA | NA |
| WBGene00049776 | NA | NA | NA | NA |
| WBGene00049777 | NA | NA | NA | NA |
| WBGene00049778 | NA | NA | NA | NA |
| WBGene00049779 | NA | NA | NA | NA |
| WBGene00049780 | NA | NA | NA | NA |
| WBGene00049781 | NA | NA | NA | NA |
| WBGene00049782 | NA | NA | NA | NA |
| WBGene00049783 | NA | NA | NA | NA |
| WBGene00049784 | NA | NA | NA | NA |
| WBGene00049785 | NA | NA | NA | NA |
| WBGene00049786 | NA | NA | NA | NA |
| WBGene00049787 | NA | NA | NA | NA |
| WBGene00049788 | NA | NA | NA | NA |
| WBGene00049789 | NA | NA | NA | NA |
| WBGene00049790 | NA | NA | NA | NA |

| | | | | |
|---|---|---|---|---|
| WBGene00049791 | NA | NA | NA | NA |
| WBGene00049792 | NA | NA | NA | NA |
| WBGene00049793 | NA | NA | NA | NA |
| WBGene00049794 | NA | NA | NA | NA |
| WBGene00049795 | NA | NA | NA | NA |
| WBGene00049796 | NA | NA | NA | NA |
| WBGene00049797 | NA | NA | NA | NA |
| WBGene00049798 | NA | NA | NA | NA |
| WBGene00049799 | NA | NA | NA | NA |
| WBGene00049800 | NA | NA | NA | NA |
| WBGene00049801 | NA | NA | NA | NA |
| WBGene00049802 | NA | NA | NA | NA |
| WBGene00049803 | NA | NA | NA | NA |
| WBGene00049804 | NA | NA | NA | NA |
| WBGene00049805 | NA | NA | NA | NA |
| WBGene00049806 | NA | NA | NA | NA |
| WBGene00049807 | NA | NA | NA | NA |
| WBGene00049808 | NA | NA | NA | NA |
| WBGene00049809 | NA | NA | NA | NA |
| WBGene00049810 | NA | NA | NA | NA |
| WBGene00049811 | NA | NA | NA | NA |
| WBGene00049812 | NA | NA | NA | NA |
| WBGene00049813 | NA | NA | NA | NA |
| WBGene00049814 | NA | NA | NA | NA |
| WBGene00049815 | NA | NA | NA | NA |
| WBGene00049816 | NA | NA | NA | NA |
| WBGene00049817 | NA | NA | NA | NA |
| WBGene00049818 | NA | NA | NA | NA |
| WBGene00049819 | NA | NA | NA | NA |
| WBGene00049820 | NA | NA | NA | NA |
| WBGene00049821 | NA | NA | NA | NA |
| WBGene00049822 | NA | NA | NA | NA |
| WBGene00049823 | NA | NA | NA | NA |
| WBGene00049824 | NA | NA | NA | NA |
| WBGene00049825 | NA | NA | NA | NA |
| WBGene00049826 | NA | NA | NA | NA |
| WBGene00049827 | NA | NA | NA | NA |
| WBGene00049828 | NA | NA | NA | NA |
| WBGene00049829 | NA | NA | NA | NA |
| WBGene00049830 | NA | NA | NA | NA |
| WBGene00049831 | NA | NA | NA | NA |
| WBGene00049832 | NA | NA | NA | NA |
| WBGene00049833 | NA | NA | NA | NA |

| | | | | |
|---|---|---|---|---|
| WBGene00049834 | NA | NA | NA | NA |
| WBGene00049835 | NA | NA | NA | NA |
| WBGene00049836 | NA | NA | NA | NA |
| WBGene00049837 | NA | NA | NA | NA |
| WBGene00049838 | NA | NA | NA | NA |
| WBGene00049839 | NA | NA | NA | NA |
| WBGene00049840 | NA | NA | NA | NA |
| WBGene00049841 | NA | NA | NA | NA |
| WBGene00049842 | NA | NA | NA | NA |
| WBGene00049843 | NA | NA | NA | NA |
| WBGene00049844 | NA | NA | NA | NA |
| WBGene00049845 | NA | NA | NA | NA |
| WBGene00049846 | NA | NA | NA | NA |
| WBGene00049847 | NA | NA | NA | NA |
| WBGene00049848 | NA | NA | NA | NA |
| WBGene00049849 | NA | NA | NA | NA |
| WBGene00049850 | NA | NA | NA | NA |
| WBGene00049851 | NA | NA | NA | NA |
| WBGene00049852 | NA | NA | NA | NA |
| WBGene00049853 | NA | NA | NA | NA |
| WBGene00049854 | NA | NA | NA | NA |
| WBGene00049855 | NA | NA | NA | NA |
| WBGene00049856 | NA | NA | NA | NA |
| WBGene00049857 | NA | NA | NA | NA |
| WBGene00049858 | NA | NA | NA | NA |
| WBGene00049859 | NA | NA | NA | NA |
| WBGene00049860 | NA | NA | NA | NA |
| WBGene00049861 | NA | NA | NA | NA |
| WBGene00049862 | NA | NA | NA | NA |
| WBGene00049863 | NA | NA | NA | NA |
| WBGene00049864 | NA | NA | NA | NA |
| WBGene00049865 | NA | NA | NA | NA |
| WBGene00049866 | NA | NA | NA | NA |
| WBGene00049867 | NA | NA | NA | NA |
| WBGene00049868 | NA | NA | NA | NA |
| WBGene00049869 | NA | NA | NA | NA |
| WBGene00049870 | NA | NA | NA | NA |
| WBGene00049871 | NA | NA | NA | NA |
| WBGene00049872 | NA | NA | NA | NA |
| WBGene00049873 | NA | NA | NA | NA |
| WBGene00049874 | NA | NA | NA | NA |
| WBGene00049875 | NA | NA | NA | NA |
| WBGene00049876 | NA | NA | NA | NA |

| | | | | |
|---|---|---|---|---|
| WBGene00049877 | NA | NA | NA | NA |
| WBGene00049878 | NA | NA | NA | NA |
| WBGene00049879 | NA | NA | NA | NA |
| WBGene00049880 | NA | NA | NA | NA |
| WBGene00049881 | NA | NA | NA | NA |
| WBGene00049882 | NA | NA | NA | NA |
| WBGene00049883 | NA | NA | NA | NA |
| WBGene00049884 | NA | NA | NA | NA |
| WBGene00049885 | NA | NA | NA | NA |
| WBGene00049886 | 0.020317245 | 0.84643798 | NA | NA |
| WBGene00049887 | NA | NA | NA | NA |
| WBGene00049888 | 0.020317245 | 0.84643798 | NA | NA |
| WBGene00049889 | NA | NA | NA | NA |
| WBGene00049890 | NA | NA | NA | NA |
| WBGene00049891 | NA | NA | NA | NA |
| WBGene00049892 | NA | NA | NA | NA |
| WBGene00049893 | NA | NA | NA | NA |
| WBGene00049894 | NA | NA | NA | NA |
| WBGene00049895 | NA | NA | NA | NA |
| WBGene00049896 | NA | NA | NA | NA |
| WBGene00049897 | NA | NA | NA | NA |
| WBGene00049898 | NA | NA | NA | NA |
| WBGene00049899 | NA | NA | NA | NA |
| WBGene00049900 | NA | NA | NA | NA |
| WBGene00049901 | NA | NA | NA | NA |
| WBGene00049902 | NA | NA | NA | NA |
| WBGene00049903 | NA | NA | NA | NA |
| WBGene00049904 | NA | NA | NA | NA |
| WBGene00049905 | NA | NA | NA | NA |
| WBGene00049906 | NA | NA | NA | NA |
| WBGene00049907 | NA | NA | NA | NA |
| WBGene00049908 | NA | NA | NA | NA |
| WBGene00049909 | NA | NA | NA | NA |
| WBGene00049910 | NA | NA | NA | NA |
| WBGene00049911 | NA | NA | NA | NA |
| WBGene00049912 | NA | NA | NA | NA |
| WBGene00049913 | NA | NA | NA | NA |
| WBGene00049914 | NA | NA | NA | NA |
| WBGene00049915 | NA | NA | NA | NA |
| WBGene00049916 | NA | NA | NA | NA |
| WBGene00049917 | NA | NA | NA | NA |
| WBGene00049918 | NA | NA | NA | NA |
| WBGene00049919 | NA | NA | NA | NA |

| | | | | |
|---|---|---|---|---|
| WBGene00049920 | NA | NA | NA | NA |
| WBGene00049921 | NA | NA | NA | NA |
| WBGene00049922 | NA | NA | NA | NA |
| WBGene00049923 | NA | NA | NA | NA |
| WBGene00049924 | NA | NA | NA | NA |
| WBGene00049925 | NA | NA | NA | NA |
| WBGene00049926 | NA | NA | NA | NA |
| WBGene00049927 | NA | NA | NA | NA |
| WBGene00049928 | NA | NA | NA | NA |
| WBGene00049929 | NA | NA | NA | NA |
| WBGene00049930 | NA | NA | NA | NA |
| WBGene00049931 | NA | NA | NA | NA |
| WBGene00049932 | NA | NA | NA | NA |
| WBGene00049933 | NA | NA | NA | NA |
| WBGene00049934 | NA | NA | NA | NA |
| WBGene00049935 | NA | NA | NA | NA |
| WBGene00049936 | NA | NA | NA | NA |
| WBGene00049937 | NA | NA | NA | NA |
| WBGene00049938 | NA | NA | NA | NA |
| WBGene00049939 | NA | NA | NA | NA |
| WBGene00049940 | NA | NA | NA | NA |
| WBGene00049941 | NA | NA | NA | NA |
| WBGene00049942 | NA | NA | NA | NA |
| WBGene00049943 | NA | NA | NA | NA |
| WBGene00049944 | NA | NA | NA | NA |
| WBGene00049945 | -0.003270206 | 0.95698316 | NA | NA |
| WBGene00049946 | NA | NA | NA | NA |
| WBGene00049947 | NA | NA | NA | NA |
| WBGene00049948 | NA | NA | NA | NA |
| WBGene00049949 | NA | NA | NA | NA |
| WBGene00049950 | NA | NA | NA | NA |
| WBGene00049951 | NA | NA | NA | NA |
| WBGene00049952 | NA | NA | NA | NA |
| WBGene00049953 | NA | NA | NA | NA |
| WBGene00049954 | NA | NA | NA | NA |
| WBGene00049955 | NA | NA | NA | NA |
| WBGene00049956 | NA | NA | NA | NA |
| WBGene00049957 | NA | NA | NA | NA |
| WBGene00049958 | NA | NA | NA | NA |
| WBGene00049959 | NA | NA | NA | NA |
| WBGene00049960 | NA | NA | NA | NA |
| WBGene00049961 | NA | NA | NA | NA |
| WBGene00049962 | NA | NA | NA | NA |

| | | | | | |
|---|---|---|---|---|---|
| WBGene00049963 | NA | | NA | NA | NA |
| WBGene00049964 | NA | | NA | NA | NA |
| WBGene00049965 | NA | | NA | NA | NA |
| WBGene00049966 | NA | | NA | NA | NA |
| WBGene00049967 | NA | | NA | NA | NA |
| WBGene00049968 | NA | | NA | NA | NA |
| WBGene00049969 | NA | | NA | NA | NA |
| WBGene00049970 | NA | | NA | NA | NA |
| WBGene00049971 | NA | | NA | NA | NA |
| WBGene00049972 | NA | | NA | NA | NA |
| WBGene00049973 | NA | | NA | NA | NA |
| WBGene00049974 | NA | | NA | NA | NA |
| WBGene00049975 | NA | | NA | NA | NA |
| WBGene00049976 | NA | | NA | NA | NA |
| WBGene00049977 | NA | | NA | NA | NA |
| WBGene00049978 | NA | | NA | NA | NA |
| WBGene00049979 | NA | | NA | NA | NA |
| WBGene00049980 | NA | | NA | NA | NA |
| WBGene00049981 | NA | | NA | NA | NA |
| WBGene00049982 | NA | | NA | NA | NA |
| WBGene00049983 | NA | | NA | NA | NA |
| WBGene00049984 | NA | | NA | NA | NA |
| WBGene00049985 | NA | | NA | NA | NA |
| WBGene00049986 | NA | | NA | NA | NA |
| WBGene00049987 | NA | | NA | NA | NA |
| WBGene00049988 | | 0.020232312 | 0.84643798 | NA | NA |
| WBGene00049989 | NA | | NA | NA | NA |
| WBGene00049990 | NA | | NA | NA | NA |
| WBGene00049991 | NA | | NA | NA | NA |
| WBGene00049992 | NA | | NA | NA | NA |
| WBGene00049993 | NA | | NA | NA | NA |
| WBGene00049994 | NA | | NA | NA | NA |
| WBGene00049995 | NA | | NA | NA | NA |
| WBGene00049996 | NA | | NA | NA | NA |
| WBGene00049997 | NA | | NA | NA | NA |
| WBGene00049998 | NA | | NA | NA | NA |
| WBGene00049999 | NA | | NA | NA | NA |
| WBGene00050000 | NA | | NA | NA | NA |
| WBGene00050001 | NA | | NA | NA | NA |
| WBGene00050002 | NA | | NA | NA | NA |
| WBGene00050003 | NA | | NA | NA | NA |
| WBGene00050004 | NA | | NA | NA | NA |
| WBGene00050005 | NA | | NA | NA | NA |

| | | | | |
|---|---|---|---|---|
| WBGene00050006 | NA | NA | NA | NA |
| WBGene00050007 | NA | NA | NA | NA |
| WBGene00050008 | NA | NA | NA | NA |
| WBGene00050009 | NA | NA | NA | NA |
| WBGene00050010 | NA | NA | NA | NA |
| WBGene00050011 | NA | NA | NA | NA |
| WBGene00050012 | NA | NA | NA | NA |
| WBGene00050013 | NA | NA | NA | NA |
| WBGene00050014 | NA | NA | NA | NA |
| WBGene00050015 | NA | NA | NA | NA |
| WBGene00050016 | NA | NA | NA | NA |
| WBGene00050017 | NA | NA | NA | NA |
| WBGene00050018 | -0.023056499 | 0.78121253 | NA | NA |
| WBGene00050019 | NA | NA | NA | NA |
| WBGene00050020 | NA | NA | NA | NA |
| WBGene00050021 | NA | NA | NA | NA |
| WBGene00050022 | NA | NA | NA | NA |
| WBGene00050023 | NA | NA | NA | NA |
| WBGene00050024 | NA | NA | NA | NA |
| WBGene00050025 | NA | NA | NA | NA |
| WBGene00050026 | NA | NA | NA | NA |
| WBGene00050027 | NA | NA | NA | NA |
| WBGene00050028 | NA | NA | NA | NA |
| WBGene00050029 | NA | NA | NA | NA |
| WBGene00050030 | NA | NA | NA | NA |
| WBGene00050031 | NA | NA | NA | NA |
| WBGene00050032 | NA | NA | NA | NA |
| WBGene00050033 | NA | NA | NA | NA |
| WBGene00050034 | NA | NA | NA | NA |
| WBGene00050035 | NA | NA | NA | NA |
| WBGene00050036 | NA | NA | NA | NA |
| WBGene00050037 | NA | NA | NA | NA |
| WBGene00050038 | NA | NA | NA | NA |
| WBGene00050039 | NA | NA | NA | NA |
| WBGene00050040 | NA | NA | NA | NA |
| WBGene00050041 | NA | NA | NA | NA |
| WBGene00050042 | NA | NA | NA | NA |
| WBGene00050043 | NA | NA | NA | NA |
| WBGene00050044 | NA | NA | NA | NA |
| WBGene00050045 | NA | NA | NA | NA |
| WBGene00050046 | NA | NA | NA | NA |
| WBGene00050047 | NA | NA | NA | NA |
| WBGene00050048 | NA | NA | NA | NA |

| | | | | |
|---|---|---|---|---|
| WBGene00050049 | NA | NA | NA | NA |
| WBGene00050050 | NA | NA | NA | NA |
| WBGene00050051 | NA | NA | NA | NA |
| WBGene00050052 | NA | NA | NA | NA |
| WBGene00050053 | NA | NA | NA | NA |
| WBGene00050054 | NA | NA | NA | NA |
| WBGene00050055 | NA | NA | NA | NA |
| WBGene00050056 | NA | NA | NA | NA |
| WBGene00050057 | NA | NA | NA | NA |
| WBGene00050058 | NA | NA | NA | NA |
| WBGene00050059 | NA | NA | NA | NA |
| WBGene00050060 | NA | NA | NA | NA |
| WBGene00050061 | NA | NA | NA | NA |
| WBGene00050062 | NA | NA | NA | NA |
| WBGene00050063 | NA | NA | NA | NA |
| WBGene00050064 | NA | NA | NA | NA |
| WBGene00050065 | NA | NA | NA | NA |
| WBGene00050066 | NA | NA | NA | NA |
| WBGene00050067 | NA | NA | NA | NA |
| WBGene00050068 | NA | NA | NA | NA |
| WBGene00050069 | NA | NA | NA | NA |
| WBGene00050070 | NA | NA | NA | NA |
| WBGene00050071 | NA | NA | NA | NA |
| WBGene00050072 | NA | NA | NA | NA |
| WBGene00050073 | NA | NA | NA | NA |
| WBGene00050074 | NA | NA | NA | NA |
| WBGene00050075 | NA | NA | NA | NA |
| WBGene00050076 | NA | NA | NA | NA |
| WBGene00050077 | NA | NA | NA | NA |
| WBGene00050078 | NA | NA | NA | NA |
| WBGene00050079 | NA | NA | NA | NA |
| WBGene00050080 | NA | NA | NA | NA |
| WBGene00050081 | NA | NA | NA | NA |
| WBGene00050082 | NA | NA | NA | NA |
| WBGene00050083 | NA | NA | NA | NA |
| WBGene00050084 | NA | NA | NA | NA |
| WBGene00050085 | NA | NA | NA | NA |
| WBGene00050086 | NA | NA | NA | NA |
| WBGene00050087 | NA | NA | NA | NA |
| WBGene00050088 | NA | NA | NA | NA |
| WBGene00050089 | NA | NA | NA | NA |
| WBGene00050090 | NA | NA | NA | NA |
| WBGene00050091 | NA | NA | NA | NA |

| | | | | |
|---|---|---|---|---|
| WBGene00050092 | NA | NA | NA | NA |
| WBGene00050093 | NA | NA | NA | NA |
| WBGene00050094 | NA | NA | NA | NA |
| WBGene00050095 | NA | NA | NA | NA |
| WBGene00050096 | NA | NA | NA | NA |
| WBGene00050097 | NA | NA | NA | NA |
| WBGene00050098 | NA | NA | NA | NA |
| WBGene00050099 | NA | NA | NA | NA |
| WBGene00050100 | NA | NA | NA | NA |
| WBGene00050101 | NA | NA | NA | NA |
| WBGene00050102 | NA | NA | NA | NA |
| WBGene00050103 | NA | NA | NA | NA |
| WBGene00050104 | NA | NA | NA | NA |
| WBGene00050105 | NA | NA | NA | NA |
| WBGene00050106 | NA | NA | NA | NA |
| WBGene00050107 | NA | NA | NA | NA |
| WBGene00050108 | NA | NA | NA | NA |
| WBGene00050109 | NA | NA | NA | NA |
| WBGene00050110 | NA | NA | NA | NA |
| WBGene00050111 | NA | NA | NA | NA |
| WBGene00050112 | NA | NA | NA | NA |
| WBGene00050113 | NA | NA | NA | NA |
| WBGene00050114 | NA | NA | NA | NA |
| WBGene00050115 | NA | NA | NA | NA |
| WBGene00050116 | NA | NA | NA | NA |
| WBGene00050117 | NA | NA | NA | NA |
| WBGene00050118 | NA | NA | NA | NA |
| WBGene00050119 | NA | NA | NA | NA |
| WBGene00050120 | NA | NA | NA | NA |
| WBGene00050121 | NA | NA | NA | NA |
| WBGene00050122 | NA | NA | NA | NA |
| WBGene00050123 | NA | NA | NA | NA |
| WBGene00050124 | NA | NA | NA | NA |
| WBGene00050125 | NA | NA | NA | NA |
| WBGene00050126 | NA | NA | NA | NA |
| WBGene00050127 | NA | NA | NA | NA |
| WBGene00050128 | NA | NA | NA | NA |
| WBGene00050129 | NA | NA | NA | NA |
| WBGene00050130 | NA | NA | NA | NA |
| WBGene00050131 | NA | NA | NA | NA |
| WBGene00050132 | NA | NA | NA | NA |
| WBGene00050133 | NA | NA | NA | NA |
| WBGene00050134 | NA | NA | NA | NA |

| | | | | |
|---|---|---|---|---|
| WBGene00050135 | NA | NA | NA | NA |
| WBGene00050136 | NA | NA | NA | NA |
| WBGene00050137 | NA | NA | NA | NA |
| WBGene00050138 | NA | NA | NA | NA |
| WBGene00050139 | NA | NA | NA | NA |
| WBGene00050140 | NA | NA | NA | NA |
| WBGene00050141 | NA | NA | NA | NA |
| WBGene00050142 | NA | NA | NA | NA |
| WBGene00050143 | NA | NA | NA | NA |
| WBGene00050144 | NA | NA | NA | NA |
| WBGene00050145 | NA | NA | NA | NA |
| WBGene00050146 | NA | NA | NA | NA |
| WBGene00050147 | NA | NA | NA | NA |
| WBGene00050148 | NA | NA | NA | NA |
| WBGene00050149 | NA | NA | NA | NA |
| WBGene00050150 | NA | NA | NA | NA |
| WBGene00050151 | NA | NA | NA | NA |
| WBGene00050152 | NA | NA | NA | NA |
| WBGene00050153 | NA | NA | NA | NA |
| WBGene00050154 | NA | NA | NA | NA |
| WBGene00050155 | NA | NA | NA | NA |
| WBGene00050156 | NA | NA | NA | NA |
| WBGene00050157 | NA | NA | NA | NA |
| WBGene00050158 | NA | NA | NA | NA |
| WBGene00050159 | NA | NA | NA | NA |
| WBGene00050160 | NA | NA | NA | NA |
| WBGene00050161 | NA | NA | NA | NA |
| WBGene00050162 | NA | NA | NA | NA |
| WBGene00050163 | NA | NA | NA | NA |
| WBGene00050164 | NA | NA | NA | NA |
| WBGene00050165 | NA | NA | NA | NA |
| WBGene00050166 | NA | NA | NA | NA |
| WBGene00050167 | NA | NA | NA | NA |
| WBGene00050168 | NA | NA | NA | NA |
| WBGene00050169 | NA | NA | NA | NA |
| WBGene00050170 | NA | NA | NA | NA |
| WBGene00050171 | NA | NA | NA | NA |
| WBGene00050172 | NA | NA | NA | NA |
| WBGene00050173 | NA | NA | NA | NA |
| WBGene00050174 | NA | NA | NA | NA |
| WBGene00050175 | NA | NA | NA | NA |
| WBGene00050176 | NA | NA | NA | NA |
| WBGene00050177 | NA | NA | NA | NA |

| | | | | |
|---|---|---|---|---|
| WBGene00050178 | NA | NA | NA | NA |
| WBGene00050179 | NA | NA | NA | NA |
| WBGene00050180 | NA | NA | NA | NA |
| WBGene00050181 | NA | NA | NA | NA |
| WBGene00050182 | NA | NA | NA | NA |
| WBGene00050183 | NA | NA | NA | NA |
| WBGene00050184 | NA | NA | NA | NA |
| WBGene00050185 | NA | NA | NA | NA |
| WBGene00050186 | NA | NA | NA | NA |
| WBGene00050187 | NA | NA | NA | NA |
| WBGene00050188 | NA | NA | NA | NA |
| WBGene00050189 | NA | NA | NA | NA |
| WBGene00050190 | NA | NA | NA | NA |
| WBGene00050191 | NA | NA | NA | NA |
| WBGene00050192 | NA | NA | NA | NA |
| WBGene00050193 | NA | NA | NA | NA |
| WBGene00050194 | NA | NA | NA | NA |
| WBGene00050195 | NA | NA | NA | NA |
| WBGene00050196 | NA | NA | NA | NA |
| WBGene00050197 | NA | NA | NA | NA |
| WBGene00050198 | NA | NA | NA | NA |
| WBGene00050199 | NA | NA | NA | NA |
| WBGene00050200 | NA | NA | NA | NA |
| WBGene00050201 | NA | NA | NA | NA |
| WBGene00050202 | NA | NA | NA | NA |
| WBGene00050203 | NA | NA | NA | NA |
| WBGene00050204 | NA | NA | NA | NA |
| WBGene00050205 | NA | NA | NA | NA |
| WBGene00050206 | NA | NA | NA | NA |
| WBGene00050207 | NA | NA | NA | NA |
| WBGene00050208 | NA | NA | NA | NA |
| WBGene00050209 | NA | NA | NA | NA |
| WBGene00050210 | NA | NA | NA | NA |
| WBGene00050211 | NA | NA | NA | NA |
| WBGene00050212 | NA | NA | NA | NA |
| WBGene00050213 | NA | NA | NA | NA |
| WBGene00050214 | NA | NA | NA | NA |
| WBGene00050215 | NA | NA | NA | NA |
| WBGene00050216 | NA | NA | NA | NA |
| WBGene00050217 | NA | NA | NA | NA |
| WBGene00050218 | NA | NA | NA | NA |
| WBGene00050219 | NA | NA | NA | NA |
| WBGene00050220 | NA | NA | NA | NA |

| | | | | |
|---|---|---|---|---|
| WBGene00050221 | NA | NA | NA | NA |
| WBGene00050222 | NA | NA | NA | NA |
| WBGene00050223 | NA | NA | NA | NA |
| WBGene00050224 | NA | NA | NA | NA |
| WBGene00050225 | NA | NA | NA | NA |
| WBGene00050226 | NA | NA | NA | NA |
| WBGene00050227 | NA | NA | NA | NA |
| WBGene00050228 | NA | NA | NA | NA |
| WBGene00050229 | NA | NA | NA | NA |
| WBGene00050230 | NA | NA | NA | NA |
| WBGene00050231 | NA | NA | NA | NA |
| WBGene00050232 | NA | NA | NA | NA |
| WBGene00050233 | NA | NA | NA | NA |
| WBGene00050234 | NA | NA | NA | NA |
| WBGene00050235 | NA | NA | NA | NA |
| WBGene00050236 | NA | NA | NA | NA |
| WBGene00050237 | NA | NA | NA | NA |
| WBGene00050238 | NA | NA | NA | NA |
| WBGene00050239 | NA | NA | NA | NA |
| WBGene00050240 | NA | NA | NA | NA |
| WBGene00050241 | NA | NA | NA | NA |
| WBGene00050242 | NA | NA | NA | NA |
| WBGene00050243 | NA | NA | NA | NA |
| WBGene00050244 | NA | NA | NA | NA |
| WBGene00050245 | NA | NA | NA | NA |
| WBGene00050246 | NA | NA | NA | NA |
| WBGene00050247 | NA | NA | NA | NA |
| WBGene00050248 | NA | NA | NA | NA |
| WBGene00050249 | NA | NA | NA | NA |
| WBGene00050250 | NA | NA | NA | NA |
| WBGene00050251 | NA | NA | NA | NA |
| WBGene00050252 | NA | NA | NA | NA |
| WBGene00050253 | NA | NA | NA | NA |
| WBGene00050254 | NA | NA | NA | NA |
| WBGene00050255 | NA | NA | NA | NA |
| WBGene00050256 | NA | NA | NA | NA |
| WBGene00050257 | NA | NA | NA | NA |
| WBGene00050258 | NA | NA | NA | NA |
| WBGene00050259 | NA | NA | NA | NA |
| WBGene00050260 | 0.020317245 | 0.84643798 | NA | NA |
| WBGene00050261 | NA | NA | NA | NA |
| WBGene00050262 | NA | NA | NA | NA |
| WBGene00050263 | NA | NA | NA | NA |

| | | | | |
|---|---|---|---|---|
| WBGene00050264 | NA | NA | NA | NA |
| WBGene00050265 | NA | NA | NA | NA |
| WBGene00050266 | NA | NA | NA | NA |
| WBGene00050267 | NA | NA | NA | NA |
| WBGene00050268 | NA | NA | NA | NA |
| WBGene00050269 | 0.020317245 | 0.84643798 | NA | NA |
| WBGene00050270 | NA | NA | NA | NA |
| WBGene00050271 | NA | NA | NA | NA |
| WBGene00050272 | NA | NA | NA | NA |
| WBGene00050273 | NA | NA | NA | NA |
| WBGene00050274 | NA | NA | NA | NA |
| WBGene00050275 | NA | NA | NA | NA |
| WBGene00050276 | NA | NA | NA | NA |
| WBGene00050277 | NA | NA | NA | NA |
| WBGene00050278 | NA | NA | NA | NA |
| WBGene00050279 | NA | NA | NA | NA |
| WBGene00050280 | NA | NA | NA | NA |
| WBGene00050281 | NA | NA | NA | NA |
| WBGene00050282 | NA | NA | NA | NA |
| WBGene00050283 | NA | NA | NA | NA |
| WBGene00050284 | NA | NA | NA | NA |
| WBGene00050285 | NA | NA | NA | NA |
| WBGene00050286 | NA | NA | NA | NA |
| WBGene00050287 | NA | NA | NA | NA |
| WBGene00050288 | 0.020232312 | 0.84643798 | NA | NA |
| WBGene00050289 | NA | NA | NA | NA |
| WBGene00050290 | NA | NA | NA | NA |
| WBGene00050291 | NA | NA | NA | NA |
| WBGene00050292 | NA | NA | NA | NA |
| WBGene00050293 | NA | NA | NA | NA |
| WBGene00050294 | NA | NA | NA | NA |
| WBGene00050295 | NA | NA | NA | NA |
| WBGene00050296 | NA | NA | NA | NA |
| WBGene00050297 | NA | NA | NA | NA |
| WBGene00050298 | NA | NA | NA | NA |
| WBGene00050299 | NA | NA | NA | NA |
| WBGene00050300 | NA | NA | NA | NA |
| WBGene00050301 | NA | NA | NA | NA |
| WBGene00050302 | NA | NA | NA | NA |
| WBGene00050303 | NA | NA | NA | NA |
| WBGene00050304 | NA | NA | NA | NA |
| WBGene00050305 | NA | NA | NA | NA |
| WBGene00050306 | NA | NA | NA | NA |

| | | | | |
|---|---|---|---|---|
| WBGene00050307 | NA | NA | NA | NA |
| WBGene00050308 | NA | NA | NA | NA |
| WBGene00050309 | NA | NA | NA | NA |
| WBGene00050310 | NA | NA | NA | NA |
| WBGene00050311 | NA | NA | NA | NA |
| WBGene00050312 | NA | NA | NA | NA |
| WBGene00050313 | NA | NA | NA | NA |
| WBGene00050314 | NA | NA | NA | NA |
| WBGene00050315 | NA | NA | NA | NA |
| WBGene00050316 | NA | NA | NA | NA |
| WBGene00050317 | NA | NA | NA | NA |
| WBGene00050318 | NA | NA | NA | NA |
| WBGene00050319 | NA | NA | NA | NA |
| WBGene00050320 | NA | NA | NA | NA |
| WBGene00050321 | NA | NA | NA | NA |
| WBGene00050322 | NA | NA | NA | NA |
| WBGene00050323 | NA | NA | NA | NA |
| WBGene00050324 | NA | NA | NA | NA |
| WBGene00050325 | NA | NA | NA | NA |
| WBGene00050326 | NA | NA | NA | NA |
| WBGene00050327 | NA | NA | NA | NA |
| WBGene00050328 | NA | NA | NA | NA |
| WBGene00050329 | NA | NA | NA | NA |
| WBGene00050330 | NA | NA | NA | NA |
| WBGene00050331 | NA | NA | NA | NA |
| WBGene00050332 | NA | NA | NA | NA |
| WBGene00050333 | NA | NA | NA | NA |
| WBGene00050334 | NA | NA | NA | NA |
| WBGene00050335 | NA | NA | NA | NA |
| WBGene00050336 | NA | NA | NA | NA |
| WBGene00050337 | NA | NA | NA | NA |
| WBGene00050338 | NA | NA | NA | NA |
| WBGene00050339 | NA | NA | NA | NA |
| WBGene00050340 | NA | NA | NA | NA |
| WBGene00050341 | NA | NA | NA | NA |
| WBGene00050342 | NA | NA | NA | NA |
| WBGene00050343 | NA | NA | NA | NA |
| WBGene00050344 | NA | NA | NA | NA |
| WBGene00050345 | NA | NA | NA | NA |
| WBGene00050346 | NA | NA | NA | NA |
| WBGene00050347 | NA | NA | NA | NA |
| WBGene00050348 | NA | NA | NA | NA |
| WBGene00050349 | 0.020317245 | 0.84643798 | NA | NA |

| | | | | |
|---|---|---|---|---|
| WBGene00050350 | NA | NA | NA | NA |
| WBGene00050351 | NA | NA | NA | NA |
| WBGene00050352 | NA | NA | NA | NA |
| WBGene00050353 | NA | NA | NA | NA |
| WBGene00050354 | NA | NA | NA | NA |
| WBGene00050355 | NA | NA | NA | NA |
| WBGene00050356 | NA | NA | NA | NA |
| WBGene00050357 | NA | NA | NA | NA |
| WBGene00050358 | NA | NA | NA | NA |
| WBGene00050359 | NA | NA | NA | NA |
| WBGene00050360 | NA | NA | NA | NA |
| WBGene00050361 | NA | NA | NA | NA |
| WBGene00050362 | NA | NA | NA | NA |
| WBGene00050363 | NA | NA | NA | NA |
| WBGene00050364 | NA | NA | NA | NA |
| WBGene00050365 | NA | NA | NA | NA |
| WBGene00050366 | NA | NA | NA | NA |
| WBGene00050367 | NA | NA | NA | NA |
| WBGene00050368 | NA | NA | NA | NA |
| WBGene00050369 | NA | NA | NA | NA |
| WBGene00050370 | NA | NA | NA | NA |
| WBGene00050371 | NA | NA | NA | NA |
| WBGene00050372 | NA | NA | NA | NA |
| WBGene00050373 | NA | NA | NA | NA |
| WBGene00050374 | NA | NA | NA | NA |
| WBGene00050375 | NA | NA | NA | NA |
| WBGene00050376 | NA | NA | NA | NA |
| WBGene00050377 | NA | NA | NA | NA |
| WBGene00050378 | NA | NA | NA | NA |
| WBGene00050379 | NA | NA | NA | NA |
| WBGene00050380 | NA | NA | NA | NA |
| WBGene00050381 | NA | NA | NA | NA |
| WBGene00050382 | NA | NA | NA | NA |
| WBGene00050383 | NA | NA | NA | NA |
| WBGene00050384 | NA | NA | NA | NA |
| WBGene00050385 | NA | NA | NA | NA |
| WBGene00050386 | NA | NA | NA | NA |
| WBGene00050387 | NA | NA | NA | NA |
| WBGene00050388 | NA | NA | NA | NA |
| WBGene00050389 | NA | NA | NA | NA |
| WBGene00050390 | NA | NA | NA | NA |
| WBGene00050391 | NA | NA | NA | NA |
| WBGene00050392 | NA | NA | NA | NA |

| | | | | |
|---|---|---|---|---|
| WBGene00050393 | NA | NA | NA | NA |
| WBGene00050394 | NA | NA | NA | NA |
| WBGene00050395 | NA | NA | NA | NA |
| WBGene00050396 | NA | NA | NA | NA |
| WBGene00050397 | 0.040107227 | 0.58693163 | NA | NA |
| WBGene00050398 | NA | NA | NA | NA |
| WBGene00050399 | NA | NA | NA | NA |
| WBGene00050400 | NA | NA | NA | NA |
| WBGene00050401 | NA | NA | NA | NA |
| WBGene00050402 | NA | NA | NA | NA |
| WBGene00050403 | NA | NA | NA | NA |
| WBGene00050404 | NA | NA | NA | NA |
| WBGene00050405 | NA | NA | NA | NA |
| WBGene00050406 | NA | NA | NA | NA |
| WBGene00050407 | NA | NA | NA | NA |
| WBGene00050408 | NA | NA | NA | NA |
| WBGene00050409 | NA | NA | NA | NA |
| WBGene00050410 | NA | NA | NA | NA |
| WBGene00050411 | NA | NA | NA | NA |
| WBGene00050412 | NA | NA | NA | NA |
| WBGene00050413 | NA | NA | NA | NA |
| WBGene00050414 | NA | NA | NA | NA |
| WBGene00050415 | NA | NA | NA | NA |
| WBGene00050416 | NA | NA | NA | NA |
| WBGene00050417 | -0.023109762 | 0.78121253 | NA | NA |
| WBGene00050418 | NA | NA | NA | NA |
| WBGene00050419 | NA | NA | NA | NA |
| WBGene00050420 | NA | NA | NA | NA |
| WBGene00050421 | NA | NA | NA | NA |
| WBGene00050422 | NA | NA | NA | NA |
| WBGene00050423 | NA | NA | NA | NA |
| WBGene00050424 | NA | NA | NA | NA |
| WBGene00050425 | NA | NA | NA | NA |
| WBGene00050426 | NA | NA | NA | NA |
| WBGene00050427 | NA | NA | NA | NA |
| WBGene00050428 | NA | NA | NA | NA |
| WBGene00050429 | NA | NA | NA | NA |
| WBGene00050430 | NA | NA | NA | NA |
| WBGene00050431 | NA | NA | NA | NA |
| WBGene00050432 | NA | NA | NA | NA |
| WBGene00050433 | NA | NA | NA | NA |
| WBGene00050434 | NA | NA | NA | NA |
| WBGene00050435 | NA | NA | NA | NA |

| | | | | |
|---|---|---|---|---|
| WBGene00050436 | NA | NA | NA | NA |
| WBGene00050437 | NA | NA | NA | NA |
| WBGene00050438 | NA | NA | NA | NA |
| WBGene00050439 | NA | NA | NA | NA |
| WBGene00050440 | NA | NA | NA | NA |
| WBGene00050441 | NA | NA | NA | NA |
| WBGene00050442 | NA | NA | NA | NA |
| WBGene00050443 | NA | NA | NA | NA |
| WBGene00050444 | NA | NA | NA | NA |
| WBGene00050445 | NA | NA | NA | NA |
| WBGene00050446 | NA | NA | NA | NA |
| WBGene00050447 | NA | NA | NA | NA |
| WBGene00050448 | NA | NA | NA | NA |
| WBGene00050449 | NA | NA | NA | NA |
| WBGene00050450 | NA | NA | NA | NA |
| WBGene00050451 | NA | NA | NA | NA |
| WBGene00050452 | NA | NA | NA | NA |
| WBGene00050453 | NA | NA | NA | NA |
| WBGene00050454 | NA | NA | NA | NA |
| WBGene00050455 | NA | NA | NA | NA |
| WBGene00050456 | NA | NA | NA | NA |
| WBGene00050457 | NA | NA | NA | NA |
| WBGene00050458 | NA | NA | NA | NA |
| WBGene00050459 | NA | NA | NA | NA |
| WBGene00050460 | NA | NA | NA | NA |
| WBGene00050461 | NA | NA | NA | NA |
| WBGene00050462 | NA | NA | NA | NA |
| WBGene00050463 | NA | NA | NA | NA |
| WBGene00050464 | NA | NA | NA | NA |
| WBGene00050465 | NA | NA | NA | NA |
| WBGene00050466 | NA | NA | NA | NA |
| WBGene00050467 | NA | NA | NA | NA |
| WBGene00050468 | NA | NA | NA | NA |
| WBGene00050469 | NA | NA | NA | NA |
| WBGene00050470 | NA | NA | NA | NA |
| WBGene00050471 | NA | NA | NA | NA |
| WBGene00050472 | NA | NA | NA | NA |
| WBGene00050473 | NA | NA | NA | NA |
| WBGene00050474 | NA | NA | NA | NA |
| WBGene00050475 | NA | NA | NA | NA |
| WBGene00050476 | NA | NA | NA | NA |
| WBGene00050477 | NA | NA | NA | NA |
| WBGene00050478 | NA | NA | NA | NA |

| | | | | |
|---|---|---|---|---|
| WBGene00050479 | NA | NA | NA | NA |
| WBGene00050480 | NA | NA | NA | NA |
| WBGene00050481 | NA | NA | NA | NA |
| WBGene00050482 | NA | NA | NA | NA |
| WBGene00050483 | NA | NA | NA | NA |
| WBGene00050484 | NA | NA | NA | NA |
| WBGene00050485 | NA | NA | NA | NA |
| WBGene00050486 | NA | NA | NA | NA |
| WBGene00050487 | NA | NA | NA | NA |
| WBGene00050488 | NA | NA | NA | NA |
| WBGene00050489 | NA | NA | NA | NA |
| WBGene00050490 | NA | NA | NA | NA |
| WBGene00050491 | NA | NA | NA | NA |
| WBGene00050492 | NA | NA | NA | NA |
| WBGene00050493 | NA | NA | NA | NA |
| WBGene00050494 | NA | NA | NA | NA |
| WBGene00050495 | NA | NA | NA | NA |
| WBGene00050496 | NA | NA | NA | NA |
| WBGene00050497 | NA | NA | NA | NA |
| WBGene00050498 | NA | NA | NA | NA |
| WBGene00050499 | NA | NA | NA | NA |
| WBGene00050500 | NA | NA | NA | NA |
| WBGene00050501 | NA | NA | NA | NA |
| WBGene00050502 | NA | NA | NA | NA |
| WBGene00050503 | NA | NA | NA | NA |
| WBGene00050504 | NA | NA | NA | NA |
| WBGene00050505 | NA | NA | NA | NA |
| WBGene00050506 | NA | NA | NA | NA |
| WBGene00050507 | NA | NA | NA | NA |
| WBGene00050508 | NA | NA | NA | NA |
| WBGene00050509 | NA | NA | NA | NA |
| WBGene00050510 | NA | NA | NA | NA |
| WBGene00050511 | NA | NA | NA | NA |
| WBGene00050512 | NA | NA | NA | NA |
| WBGene00050513 | NA | NA | NA | NA |
| WBGene00050514 | NA | NA | NA | NA |
| WBGene00050515 | NA | NA | NA | NA |
| WBGene00050516 | NA | NA | NA | NA |
| WBGene00050517 | NA | NA | NA | NA |
| WBGene00050518 | NA | NA | NA | NA |
| WBGene00050519 | NA | NA | NA | NA |
| WBGene00050520 | NA | NA | NA | NA |
| WBGene00050521 | NA | NA | NA | NA |

| | | | | |
|---|---|---|---|---|
| WBGene00050522 | NA | NA | NA | NA |
| WBGene00050523 | NA | NA | NA | NA |
| WBGene00050524 | NA | NA | NA | NA |
| WBGene00050525 | NA | NA | NA | NA |
| WBGene00050526 | NA | NA | NA | NA |
| WBGene00050527 | NA | NA | NA | NA |
| WBGene00050528 | NA | NA | NA | NA |
| WBGene00050529 | NA | NA | NA | NA |
| WBGene00050530 | NA | NA | NA | NA |
| WBGene00050531 | NA | NA | NA | NA |
| WBGene00050532 | NA | NA | NA | NA |
| WBGene00050533 | NA | NA | NA | NA |
| WBGene00050534 | NA | NA | NA | NA |
| WBGene00050535 | NA | NA | NA | NA |
| WBGene00050536 | NA | NA | NA | NA |
| WBGene00050537 | NA | NA | NA | NA |
| WBGene00050538 | NA | NA | NA | NA |
| WBGene00050539 | NA | NA | NA | NA |
| WBGene00050540 | NA | NA | NA | NA |
| WBGene00050541 | NA | NA | NA | NA |
| WBGene00050542 | NA | NA | NA | NA |
| WBGene00050543 | NA | NA | NA | NA |
| WBGene00050544 | NA | NA | NA | NA |
| WBGene00050545 | NA | NA | NA | NA |
| WBGene00050546 | NA | NA | NA | NA |
| WBGene00050547 | NA | NA | NA | NA |
| WBGene00050548 | NA | NA | NA | NA |
| WBGene00050549 | NA | NA | NA | NA |
| WBGene00050550 | NA | NA | NA | NA |
| WBGene00050551 | NA | NA | NA | NA |
| WBGene00050552 | NA | NA | NA | NA |
| WBGene00050553 | NA | NA | NA | NA |
| WBGene00050554 | NA | NA | NA | NA |
| WBGene00050555 | NA | NA | NA | NA |
| WBGene00050556 | NA | NA | NA | NA |
| WBGene00050557 | NA | NA | NA | NA |
| WBGene00050558 | NA | NA | NA | NA |
| WBGene00050559 | NA | NA | NA | NA |
| WBGene00050560 | NA | NA | NA | NA |
| WBGene00050561 | NA | NA | NA | NA |
| WBGene00050562 | NA | NA | NA | NA |
| WBGene00050563 | NA | NA | NA | NA |
| WBGene00050564 | NA | NA | NA | NA |

| | | | | | |
|---|---|---|---|---|---|
| WBGene00050565 | NA | | NA | NA | NA |
| WBGene00050566 | NA | | NA | NA | NA |
| WBGene00050567 | NA | | NA | NA | NA |
| WBGene00050568 | NA | | NA | NA | NA |
| WBGene00050569 | NA | | NA | NA | NA |
| WBGene00050570 | NA | | NA | NA | NA |
| WBGene00050571 | NA | | NA | NA | NA |
| WBGene00050572 | NA | | NA | NA | NA |
| WBGene00050573 | NA | | NA | NA | NA |
| WBGene00050574 | NA | | NA | NA | NA |
| WBGene00050575 | NA | | NA | NA | NA |
| WBGene00050576 | NA | | NA | NA | NA |
| WBGene00050577 | NA | | NA | NA | NA |
| WBGene00050578 | NA | | NA | NA | NA |
| WBGene00050579 | NA | | NA | NA | NA |
| WBGene00050580 | NA | | NA | NA | NA |
| WBGene00050581 | NA | | NA | NA | NA |
| WBGene00050582 | NA | | NA | NA | NA |
| WBGene00050583 | NA | | NA | NA | NA |
| WBGene00050584 | NA | | NA | NA | NA |
| WBGene00050585 | NA | | NA | NA | NA |
| WBGene00050586 | NA | | NA | NA | NA |
| WBGene00050587 | NA | | NA | NA | NA |
| WBGene00050588 | NA | | NA | NA | NA |
| WBGene00050589 | NA | | NA | NA | NA |
| WBGene00050590 | | -0.023056499 | 0.78121253 | NA | NA |
| WBGene00050591 | NA | | NA | NA | NA |
| WBGene00050592 | NA | | NA | NA | NA |
| WBGene00050593 | NA | | NA | NA | NA |
| WBGene00050594 | NA | | NA | NA | NA |
| WBGene00050595 | NA | | NA | NA | NA |
| WBGene00050596 | NA | | NA | NA | NA |
| WBGene00050597 | NA | | NA | NA | NA |
| WBGene00050598 | NA | | NA | NA | NA |
| WBGene00050599 | NA | | NA | NA | NA |
| WBGene00050600 | NA | | NA | NA | NA |
| WBGene00050601 | NA | | NA | NA | NA |
| WBGene00050602 | NA | | NA | NA | NA |
| WBGene00050603 | NA | | NA | NA | NA |
| WBGene00050604 | NA | | NA | NA | NA |
| WBGene00050605 | NA | | NA | NA | NA |
| WBGene00050606 | NA | | NA | NA | NA |
| WBGene00050607 | | -0.023056499 | 0.78121253 | NA | NA |

| | | | | |
|---|---|---|---|---|
| WBGene00050608 | NA | NA | NA | NA |
| WBGene00050609 | NA | NA | NA | NA |
| WBGene00050610 | NA | NA | NA | NA |
| WBGene00050611 | NA | NA | NA | NA |
| WBGene00050612 | NA | NA | NA | NA |
| WBGene00050613 | NA | NA | NA | NA |
| WBGene00050614 | NA | NA | NA | NA |
| WBGene00050615 | NA | NA | NA | NA |
| WBGene00050616 | NA | NA | NA | NA |
| WBGene00050617 | NA | NA | NA | NA |
| WBGene00050618 | NA | NA | NA | NA |
| WBGene00050619 | NA | NA | NA | NA |
| WBGene00050620 | NA | NA | NA | NA |
| WBGene00050621 | NA | NA | NA | NA |
| WBGene00050622 | NA | NA | NA | NA |
| WBGene00050623 | NA | NA | NA | NA |
| WBGene00050624 | NA | NA | NA | NA |
| WBGene00050625 | NA | NA | NA | NA |
| WBGene00050626 | NA | NA | NA | NA |
| WBGene00050627 | NA | NA | NA | NA |
| WBGene00050628 | NA | NA | NA | NA |
| WBGene00050629 | NA | NA | NA | NA |
| WBGene00050630 | NA | NA | NA | NA |
| WBGene00050631 | NA | NA | NA | NA |
| WBGene00050632 | NA | NA | NA | NA |
| WBGene00050633 | NA | NA | NA | NA |
| WBGene00050634 | NA | NA | NA | NA |
| WBGene00050635 | NA | NA | NA | NA |
| WBGene00050636 | NA | NA | NA | NA |
| WBGene00050637 | NA | NA | NA | NA |
| WBGene00050638 | NA | NA | NA | NA |
| WBGene00050639 | NA | NA | NA | NA |
| WBGene00050640 | NA | NA | NA | NA |
| WBGene00050641 | NA | NA | NA | NA |
| WBGene00050642 | NA | NA | NA | NA |
| WBGene00050643 | NA | NA | NA | NA |
| WBGene00050644 | NA | NA | NA | NA |
| WBGene00050645 | NA | NA | NA | NA |
| WBGene00050646 | NA | NA | NA | NA |
| WBGene00050647 | NA | NA | NA | NA |
| WBGene00050648 | NA | NA | NA | NA |
| WBGene00050649 | NA | NA | NA | NA |
| WBGene00050650 | NA | NA | NA | NA |

| | | | | |
|---|---|---|---|---|
| WBGene00050651 | NA | NA | NA | NA |
| WBGene00050652 | NA | NA | NA | NA |
| WBGene00050653 | NA | NA | NA | NA |
| WBGene00050654 | NA | NA | NA | NA |
| WBGene00050655 | NA | NA | NA | NA |
| WBGene00050656 | NA | NA | NA | NA |
| WBGene00050657 | NA | NA | NA | NA |
| WBGene00050658 | NA | NA | NA | NA |
| WBGene00050659 | NA | NA | NA | NA |
| WBGene00050660 | NA | NA | NA | NA |
| WBGene00050661 | NA | NA | NA | NA |
| WBGene00050662 | NA | NA | NA | NA |
| WBGene00050663 | NA | NA | NA | NA |
| WBGene00050664 | NA | NA | NA | NA |
| WBGene00050665 | NA | NA | NA | NA |
| WBGene00050666 | NA | NA | NA | NA |
| WBGene00050667 | NA | NA | NA | NA |
| WBGene00050668 | NA | NA | NA | NA |
| WBGene00050669 | NA | NA | NA | NA |
| WBGene00050670 | NA | NA | NA | NA |
| WBGene00050671 | NA | NA | NA | NA |
| WBGene00050672 | 0.020232312 | 0.84643798 | NA | NA |
| WBGene00050673 | NA | NA | NA | NA |
| WBGene00050674 | NA | NA | NA | NA |
| WBGene00050675 | NA | NA | NA | NA |
| WBGene00050676 | NA | NA | NA | NA |
| WBGene00050677 | NA | NA | NA | NA |
| WBGene00050678 | NA | NA | NA | NA |
| WBGene00050679 | NA | NA | NA | NA |
| WBGene00050680 | NA | NA | NA | NA |
| WBGene00050681 | NA | NA | NA | NA |
| WBGene00050682 | NA | NA | NA | NA |
| WBGene00050683 | NA | NA | NA | NA |
| WBGene00050684 | NA | NA | NA | NA |
| WBGene00050685 | NA | NA | NA | NA |
| WBGene00050686 | NA | NA | NA | NA |
| WBGene00050687 | NA | NA | NA | NA |
| WBGene00050688 | NA | NA | NA | NA |
| WBGene00050689 | NA | NA | NA | NA |
| WBGene00050690 | NA | NA | NA | NA |
| WBGene00050691 | NA | NA | NA | NA |
| WBGene00050692 | NA | NA | NA | NA |
| WBGene00050693 | NA | NA | NA | NA |

| | | | | |
|---|---|---|---|---|
| WBGene00050694 | NA | NA | NA | NA |
| WBGene00050695 | NA | NA | NA | NA |
| WBGene00050696 | NA | NA | NA | NA |
| WBGene00050697 | NA | NA | NA | NA |
| WBGene00050698 | NA | NA | NA | NA |
| WBGene00050699 | NA | NA | NA | NA |
| WBGene00050700 | NA | NA | NA | NA |
| WBGene00050701 | NA | NA | NA | NA |
| WBGene00050702 | NA | NA | NA | NA |
| WBGene00050703 | NA | NA | NA | NA |
| WBGene00050704 | NA | NA | NA | NA |
| WBGene00050705 | NA | NA | NA | NA |
| WBGene00050706 | NA | NA | NA | NA |
| WBGene00050707 | NA | NA | NA | NA |
| WBGene00050708 | NA | NA | NA | NA |
| WBGene00050709 | NA | NA | NA | NA |
| WBGene00050710 | NA | NA | NA | NA |
| WBGene00050711 | NA | NA | NA | NA |
| WBGene00050712 | -0.023109762 | 0.78121253 | NA | NA |
| WBGene00050713 | NA | NA | NA | NA |
| WBGene00050714 | NA | NA | NA | NA |
| WBGene00050715 | NA | NA | NA | NA |
| WBGene00050716 | NA | NA | NA | NA |
| WBGene00050717 | NA | NA | NA | NA |
| WBGene00050718 | NA | NA | NA | NA |
| WBGene00050719 | NA | NA | NA | NA |
| WBGene00050720 | NA | NA | NA | NA |
| WBGene00050721 | NA | NA | NA | NA |
| WBGene00050722 | NA | NA | NA | NA |
| WBGene00050723 | NA | NA | NA | NA |
| WBGene00050724 | NA | NA | NA | NA |
| WBGene00050725 | NA | NA | NA | NA |
| WBGene00050726 | NA | NA | NA | NA |
| WBGene00050727 | NA | NA | NA | NA |
| WBGene00050728 | NA | NA | NA | NA |
| WBGene00050729 | NA | NA | NA | NA |
| WBGene00050730 | NA | NA | NA | NA |
| WBGene00050731 | NA | NA | NA | NA |
| WBGene00050732 | NA | NA | NA | NA |
| WBGene00050733 | NA | NA | NA | NA |
| WBGene00050734 | NA | NA | NA | NA |
| WBGene00050735 | NA | NA | NA | NA |
| WBGene00050736 | NA | NA | NA | NA |

| | | | | |
|---|---|---|---|---|
| WBGene00050737 | NA | NA | NA | NA |
| WBGene00050738 | NA | NA | NA | NA |
| WBGene00050739 | NA | NA | NA | NA |
| WBGene00050740 | NA | NA | NA | NA |
| WBGene00050741 | NA | NA | NA | NA |
| WBGene00050742 | NA | NA | NA | NA |
| WBGene00050743 | NA | NA | NA | NA |
| WBGene00050744 | NA | NA | NA | NA |
| WBGene00050745 | NA | NA | NA | NA |
| WBGene00050746 | NA | NA | NA | NA |
| WBGene00050747 | NA | NA | NA | NA |
| WBGene00050748 | NA | NA | NA | NA |
| WBGene00050749 | NA | NA | NA | NA |
| WBGene00050750 | NA | NA | NA | NA |
| WBGene00050751 | NA | NA | NA | NA |
| WBGene00050752 | NA | NA | NA | NA |
| WBGene00050753 | NA | NA | NA | NA |
| WBGene00050754 | NA | NA | NA | NA |
| WBGene00050755 | NA | NA | NA | NA |
| WBGene00050756 | NA | NA | NA | NA |
| WBGene00050757 | NA | NA | NA | NA |
| WBGene00050758 | NA | NA | NA | NA |
| WBGene00050759 | NA | NA | NA | NA |
| WBGene00050760 | NA | NA | NA | NA |
| WBGene00050761 | NA | NA | NA | NA |
| WBGene00050762 | NA | NA | NA | NA |
| WBGene00050763 | NA | NA | NA | NA |
| WBGene00050764 | NA | NA | NA | NA |
| WBGene00050765 | NA | NA | NA | NA |
| WBGene00050766 | NA | NA | NA | NA |
| WBGene00050767 | NA | NA | NA | NA |
| WBGene00050768 | NA | NA | NA | NA |
| WBGene00050769 | NA | NA | NA | NA |
| WBGene00050770 | NA | NA | NA | NA |
| WBGene00050771 | NA | NA | NA | NA |
| WBGene00050772 | NA | NA | NA | NA |
| WBGene00050773 | NA | NA | NA | NA |
| WBGene00050774 | NA | NA | NA | NA |
| WBGene00050775 | NA | NA | NA | NA |
| WBGene00050776 | NA | NA | NA | NA |
| WBGene00050777 | NA | NA | NA | NA |
| WBGene00050778 | NA | NA | NA | NA |
| WBGene00050779 | NA | NA | NA | NA |

| | | | | |
|---|---|---|---|---|
| WBGene00050780 | NA | NA | NA | NA |
| WBGene00050781 | NA | NA | NA | NA |
| WBGene00050782 | NA | NA | NA | NA |
| WBGene00050783 | NA | NA | NA | NA |
| WBGene00050784 | NA | NA | NA | NA |
| WBGene00050785 | NA | NA | NA | NA |
| WBGene00050786 | NA | NA | NA | NA |
| WBGene00050787 | NA | NA | NA | NA |
| WBGene00050788 | NA | NA | NA | NA |
| WBGene00050789 | NA | NA | NA | NA |
| WBGene00050790 | NA | NA | NA | NA |
| WBGene00050791 | NA | NA | NA | NA |
| WBGene00050792 | NA | NA | NA | NA |
| WBGene00050793 | NA | NA | NA | NA |
| WBGene00050794 | NA | NA | NA | NA |
| WBGene00050795 | NA | NA | NA | NA |
| WBGene00050796 | NA | NA | NA | NA |
| WBGene00050797 | NA | NA | NA | NA |
| WBGene00050798 | NA | NA | NA | NA |
| WBGene00050799 | NA | NA | NA | NA |
| WBGene00050800 | NA | NA | NA | NA |
| WBGene00050801 | NA | NA | NA | NA |
| WBGene00050802 | NA | NA | NA | NA |
| WBGene00050803 | NA | NA | NA | NA |
| WBGene00050804 | NA | NA | NA | NA |
| WBGene00050805 | NA | NA | NA | NA |
| WBGene00050806 | NA | NA | NA | NA |
| WBGene00050807 | NA | NA | NA | NA |
| WBGene00050808 | NA | NA | NA | NA |
| WBGene00050809 | NA | NA | NA | NA |
| WBGene00050810 | NA | NA | NA | NA |
| WBGene00050811 | NA | NA | NA | NA |
| WBGene00050812 | NA | NA | NA | NA |
| WBGene00050813 | NA | NA | NA | NA |
| WBGene00050814 | NA | NA | NA | NA |
| WBGene00050815 | NA | NA | NA | NA |
| WBGene00050816 | NA | NA | NA | NA |
| WBGene00050817 | NA | NA | NA | NA |
| WBGene00050818 | -0.023109762 | 0.78121253 | NA | NA |
| WBGene00050819 | NA | NA | NA | NA |
| WBGene00050820 | NA | NA | NA | NA |
| WBGene00050821 | NA | NA | NA | NA |
| WBGene00050822 | NA | NA | NA | NA |

| | | | | |
|---|---|---|---|---|
| WBGene00050823 | NA | NA | NA | NA |
| WBGene00050824 | NA | NA | NA | NA |
| WBGene00050825 | NA | NA | NA | NA |
| WBGene00050826 | NA | NA | NA | NA |
| WBGene00050827 | NA | NA | NA | NA |
| WBGene00050828 | NA | NA | NA | NA |
| WBGene00050829 | NA | NA | NA | NA |
| WBGene00050830 | NA | NA | NA | NA |
| WBGene00050831 | NA | NA | NA | NA |
| WBGene00050832 | NA | NA | NA | NA |
| WBGene00050833 | NA | NA | NA | NA |
| WBGene00050834 | NA | NA | NA | NA |
| WBGene00050835 | NA | NA | NA | NA |
| WBGene00050836 | NA | NA | NA | NA |
| WBGene00050837 | NA | NA | NA | NA |
| WBGene00050838 | NA | NA | NA | NA |
| WBGene00050839 | NA | NA | NA | NA |
| WBGene00050840 | NA | NA | NA | NA |
| WBGene00050841 | NA | NA | NA | NA |
| WBGene00050842 | NA | NA | NA | NA |
| WBGene00050843 | NA | NA | NA | NA |
| WBGene00050844 | NA | NA | NA | NA |
| WBGene00050845 | NA | NA | NA | NA |
| WBGene00050846 | NA | NA | NA | NA |
| WBGene00050847 | NA | NA | NA | NA |
| WBGene00050848 | NA | NA | NA | NA |
| WBGene00050849 | NA | NA | NA | NA |
| WBGene00050850 | NA | NA | NA | NA |
| WBGene00050851 | NA | NA | NA | NA |
| WBGene00050852 | NA | NA | NA | NA |
| WBGene00050853 | NA | NA | NA | NA |
| WBGene00050854 | NA | NA | NA | NA |
| WBGene00050855 | NA | NA | NA | NA |
| WBGene00050856 | NA | NA | NA | NA |
| WBGene00050857 | NA | NA | NA | NA |
| WBGene00050858 | NA | NA | NA | NA |
| WBGene00050859 | 0.020317245 | 0.84643798 | NA | NA |
| WBGene00050860 | NA | NA | NA | NA |
| WBGene00050861 | NA | NA | NA | NA |
| WBGene00050862 | NA | NA | NA | NA |
| WBGene00050863 | NA | NA | NA | NA |
| WBGene00050864 | NA | NA | NA | NA |
| WBGene00050865 | NA | NA | NA | NA |

| | | | | |
|---|---|---|---|---|
| WBGene00050866 | NA | NA | NA | NA |
| WBGene00050867 | NA | NA | NA | NA |
| WBGene00050868 | NA | NA | NA | NA |
| WBGene00050869 | NA | NA | NA | NA |
| WBGene00050870 | NA | NA | NA | NA |
| WBGene00050871 | NA | NA | NA | NA |
| WBGene00050873 | NA | NA | NA | NA |
| WBGene00050874 | NA | NA | NA | NA |
| WBGene00050875 | 0.020925399 | 0.84643798 | NA | bah-1 |
| WBGene00050878 | -0.022930767 | 0.78121253 | NA | NA |
| WBGene00050879 | 0.058314668 | 0.40208118 | NA | D1081.11 |
| WBGene00050880 | -0.003614866 | 0.95700117 | NA | F32H2.12 |
| WBGene00050881 | -0.023056499 | 0.78121253 | NA | ZK39.10 |
| WBGene00050883 | NA | NA | NA | NA |
| WBGene00050884 | 0.46427828 | 0.07433442 | 0.20024476 | Y69H2.15 |
| WBGene00050885 | 0.169732125 | 0.41810164 | 0.6147862 | Y94H6A.12 |
| WBGene00050889 | 0.059762547 | 0.38965825 | NA | T04C12.9 |
| WBGene00050890 | 0.063731418 | 0.41159439 | NA | NA |
| WBGene00050891 | 0.020232312 | 0.84643798 | NA | C08E8.9 |
| WBGene00050892 | NA | NA | NA | ttr-57 |
| WBGene00050893 | NA | NA | NA | Y94A7B.11 |
| WBGene00050894 | NA | NA | NA | F49H6.15 |
| WBGene00050896 | 0.020925399 | 0.84643798 | NA | ttr-56 |
| WBGene00050897 | NA | NA | NA | NA |
| WBGene00050898 | -0.022930767 | 0.78121253 | NA | T19B10.12 |
| WBGene00050899 | 0.372822128 | 0.06561139 | 0.18286028 | T10C6.15 |
| WBGene00050900 | NA | NA | NA | NA |
| WBGene00050902 | NA | NA | NA | NA |
| WBGene00050903 | -0.062834629 | 0.6043851 | 0.76077049 | NA |
| WBGene00050904 | 0.315587732 | 0.14400101 | 0.31289979 | Y70C5A.3 |
| WBGene00050906 | 0.020925399 | 0.84643798 | NA | F20E11.17 |
| WBGene00050907 | NA | NA | NA | Y51A2B.9 |
| WBGene00050910 | NA | NA | NA | Y43F8B.17 |
| WBGene00050911 | -0.045959069 | 0.50707733 | NA | Y43F8B.18 |
| WBGene00050912 | NA | NA | NA | NA |
| WBGene00050913 | -0.131976046 | 0.47501532 | 0.66210009 | T12B5.14 |
| WBGene00050914 | -0.225301961 | 0.18270826 | 0.36554136 | T12B5.15 |
| WBGene00050915 | 0.056411843 | 0.41891328 | NA | Y6E2A.10 |
| WBGene00050916 | NA | NA | NA | F55F10.3 |
| WBGene00050917 | 0.002527202 | 0.98663161 | 0.99225672 | Y53G8AL.4 |
| WBGene00050919 | -0.006766984 | 0.93963666 | NA | C45G9.15 |
| WBGene00050920 | 0.17233687 | 0.09395797 | 0.23471056 | NA |
| WBGene00050921 | NA | NA | NA | NA |

| | | | | |
|---|---|---|---|---|
| WBGene00050922 | NA | NA | NA | NA |
| WBGene00050923 | NA | NA | NA | NA |
| WBGene00050924 | NA | NA | NA | NA |
| WBGene00050926 | -0.0059938 | 0.93322519 | NA | NA |
| WBGene00050927 | 0.04319202 | 0.81605735 | 0.89891744 | NA |
| WBGene00050928 | NA | NA | NA | NA |
| WBGene00050929 | NA | NA | NA | T08G5.7 |
| WBGene00050929 | NA | NA | NA | F58D12.5 |
| WBGene00050930 | NA | NA | NA | NA |
| WBGene00050931 | NA | NA | NA | F59B2.14 |
| WBGene00050932 | -0.001842784 | 0.95621055 | NA | NA |
| WBGene00050933 | NA | NA | NA | NA |
| WBGene00050934 | 0.038125098 | 0.61119482 | NA | T19C9.10 |
| WBGene00050935 | 0.212711649 | 0.13344536 | 0.29715186 | NA |
| WBGene00050936 | NA | NA | NA | ZK856.16 |
| WBGene00050937 | 0.04670115 | 0.80375049 | 0.89088582 | R02D5.8 |
| WBGene00050938 | NA | NA | NA | NA |
| WBGene00050939 | 0.020317245 | 0.84643798 | NA | C05G5.7 |
| WBGene00050940 | -0.0550775 | 0.58232579 | 0.74624349 | C25D7.16 |
| WBGene00050941 | 0.090827668 | 0.46993704 | 0.65834352 | C33F10.14 |
| WBGene00050943 | 0.512626975 | 0.05487528 | 0.16113697 | ZC412.10 |
| WBGene00050944 | 0.090831671 | 0.60961512 | 0.76480853 | C14A11.9 |
| WBGene00050945 | 0.184894095 | 0.27775531 | 0.47767779 | NA |
| WBGene00050946 | -0.339186675 | 0.15482084 | 0.32795291 | NA |
| WBGene00050947 | -0.009669557 | 0.95056768 | 0.97187579 | K07C5.13 |
| WBGene00050948 | -0.142292786 | 0.47016548 | 0.65840969 | NA |
| WBGene00050949 | NA | NA | NA | NA |
| WBGene00050950 | NA | NA | NA | NA |
| WBGene00050951 | 0.040107227 | 0.58693163 | NA | NA |
| WBGene00050954 | NA | NA | NA | NA |
| WBGene00050955 | NA | NA | NA | C14A6.12 |
| WBGene00050956 | 0.092790431 | 0.40230482 | 0.60078997 | C14A6.13 |
| WBGene00050957 | 0.24160414 | 0.27743277 | 0.47747657 | C10A4.10 |
| WBGene00050958 | 0.129401398 | 0.10528579 | NA | NA |
| WBGene00050959 | NA | NA | NA | F26F2.2 |
| WBGene00050959 | NA | NA | NA | F26F2.9 |
| WBGene00050960 | NA | NA | NA | NA |
| WBGene00050961 | NA | NA | NA | NA |
| WBGene00050962 | NA | NA | NA | NA |
| WBGene00050963 | NA | NA | NA | NA |
| WBGene00050964 | NA | NA | NA | NA |
| WBGene00050965 | NA | NA | NA | F49H6.17 |
| WBGene00050967 | 0.126299345 | 0.16983897 | NA | C25F9.15 |

| | | | | |
|---|---|---|---|---|
| WBGene00050968 | NA | NA | NA | R11G10.4 |
| WBGene00050970 | -0.201733842 | 0.19083624 | 0.37685911 | NA |
| WBGene00050971 | 0.03741491 | 0.57187985 | NA | NA |
| WBGene00050972 | NA | NA | NA | NA |
| WBGene00050973 | NA | NA | NA | NA |
| WBGene00050974 | 0.016453512 | 0.83636998 | NA | Y6G8.9 |
| WBGene00050976 | NA | NA | NA | NA |
| WBGene00050977 | -0.023056499 | 0.78121253 | NA | NA |
| WBGene00077434 | NA | NA | NA | NA |
| WBGene00077436 | 0.020317245 | 0.84643798 | NA | NA |
| WBGene00077437 | 0.392490288 | 0.08849142 | 0.22573322 | NA |
| WBGene00077438 | 0.015062379 | 0.80838968 | NA | NA |
| WBGene00077439 | 0.036687355 | 0.59489568 | NA | F55B12.10 |
| WBGene00077440 | NA | NA | NA | NA |
| WBGene00077441 | 0.038955509 | 0.78295772 | 0.87782191 | NA |
| WBGene00077442 | NA | NA | NA | NA |
| WBGene00077443 | 1.618721567 | 0.0010617 | 0.00738096 | NA |
| WBGene00077444 | 0.0398788 | 0.58971221 | NA | NA |
| WBGene00077445 | NA | NA | NA | Y51A2A.12 |
| WBGene00077446 | NA | NA | NA | NA |
| WBGene00077447 | NA | NA | NA | NA |
| WBGene00077448 | 1.119091799 | 0.01603839 | 0.06520911 | NA |
| WBGene00077449 | -0.079472169 | 0.3156132 | NA | NA |
| WBGene00077450 | 0.015507174 | 0.80923808 | NA | K06B4.15 |
| WBGene00077451 | NA | NA | NA | NA |
| WBGene00077452 | NA | NA | NA | NA |
| WBGene00077453 | -0.260536439 | 0.05680297 | 0.1652727 | Y62F5A.12 |
| WBGene00077454 | NA | NA | NA | NA |
| WBGene00077455 | NA | NA | NA | Y69H2.17 |
| WBGene00077456 | NA | NA | NA | NA |
| WBGene00077457 | -0.066169458 | 0.73477136 | 0.85123781 | NA |
| WBGene00077459 | NA | NA | NA | NA |
| WBGene00077460 | NA | NA | NA | NA |
| WBGene00077461 | NA | NA | NA | NA |
| WBGene00077462 | 0.020925399 | 0.84643798 | NA | NA |
| WBGene00077463 | NA | NA | NA | NA |
| WBGene00077464 | NA | NA | NA | Y70C5B.2 |
| WBGene00077465 | NA | NA | NA | NA |
| WBGene00077466 | NA | NA | NA | NA |
| WBGene00077467 | NA | NA | NA | NA |
| WBGene00077468 | NA | NA | NA | NA |
| WBGene00077469 | NA | NA | NA | NA |
| WBGene00077470 | NA | NA | NA | NA |

| | | | | |
|---|---|---|---|---|
| WBGene00077471 | NA | NA | NA | NA |
| WBGene00077472 | NA | NA | NA | NA |
| WBGene00077473 | NA | NA | NA | NA |
| WBGene00077474 | NA | NA | NA | NA |
| WBGene00077475 | NA | NA | NA | NA |
| WBGene00077476 | NA | NA | NA | NA |
| WBGene00077477 | NA | NA | NA | NA |
| WBGene00077478 | NA | NA | NA | NA |
| WBGene00077479 | NA | NA | NA | NA |
| WBGene00077480 | NA | NA | NA | NA |
| WBGene00077481 | NA | NA | NA | NA |
| WBGene00077482 | NA | NA | NA | NA |
| WBGene00077483 | NA | NA | NA | NA |
| WBGene00077484 | NA | NA | NA | NA |
| WBGene00077485 | NA | NA | NA | NA |
| WBGene00077486 | NA | NA | NA | NA |
| WBGene00077487 | NA | NA | NA | NA |
| WBGene00077488 | 0.014323743 | 0.88029836 | NA | NA |
| WBGene00077489 | NA | NA | NA | C04G6.13 |
| WBGene00077490 | 0.058148341 | 0.65231153 | 0.79408772 | M03A1.8 |
| WBGene00077491 | 0.012581561 | 0.90489012 | 0.94913088 | K08H10.10 |
| WBGene00077492 | NA | NA | NA | F09C6.15 |
| WBGene00077493 | NA | NA | NA | F09C6.16 |
| WBGene00077494 | NA | NA | NA | NA |
| WBGene00077495 | 0.020317245 | 0.84643798 | NA | Y20C6A.4 |
| WBGene00077496 | NA | NA | NA | NA |
| WBGene00077497 | 0.015584271 | 0.80362925 | NA | K06A4.10 |
| WBGene00077498 | NA | NA | NA | NA |
| WBGene00077499 | 0.020317245 | 0.84643798 | NA | NA |
| WBGene00077500 | 0.10913774 | 0.41517284 | 0.6126146 | C27H6.9 |
| WBGene00077501 | NA | NA | NA | NA |
| WBGene00077502 | NA | NA | NA | NA |
| WBGene00077503 | 0.020317245 | 0.84643798 | NA | AC3.12 |
| WBGene00077504 | NA | NA | NA | NA |
| WBGene00077506 | NA | NA | NA | NA |
| WBGene00077508 | -0.062008946 | 0.58438726 | 0.74778226 | T27F6.10 |
| WBGene00077510 | NA | NA | NA | NA |
| WBGene00077514 | 0.020317245 | 0.84643798 | NA | F07B10.7 |
| WBGene00077515 | NA | NA | NA | NA |
| WBGene00077516 | NA | NA | NA | NA |
| WBGene00077517 | NA | NA | NA | NA |
| WBGene00077519 | 0.109724155 | 0.21583934 | NA | T27C10.8 |
| WBGene00077520 | 0.142712883 | 0.2530174 | 0.45109934 | C37A5.11 |

| | | | | | |
|---|---|---|---|---|---|
| WBGene00077521 | | 0.108062642 | 0.56493214 | 0.73469279 | maf-1 |
| WBGene00077522 | | 0.160104831 | 0.23165166 | 0.42654268 | T10D4.15 |
| WBGene00077523 | | -0.023056499 | 0.78121253 | NA | C23H5.12 |
| WBGene00077524 | NA | | NA | NA | NA |
| WBGene00077525 | | 0.382857145 | 0.11596872 | 0.27192405 | C41G7.8 |
| WBGene00077526 | | -0.503697084 | 6.26E-05 | 0.00064226 | C25A1.16 |
| WBGene00077527 | NA | | NA | NA | NA |
| WBGene00077531 | | -0.305111756 | 0.12142875 | 0.28048552 | F40G9.15 |
| WBGene00077534 | NA | | NA | NA | NA |
| WBGene00077535 | | -0.023056499 | 0.78121253 | NA | NA |
| WBGene00077536 | | -0.03304608 | 0.72886299 | NA | F38B7.10 |
| WBGene00077537 | | -0.132114892 | 0.52968636 | 0.70651028 | NA |
| WBGene00077538 | | -0.001842784 | 0.95621055 | NA | T04F3.5 |
| WBGene00077539 | | 1.538393511 | 0.00984414 | 0.04433344 | Y52B11A.12 |
| WBGene00077543 | | 0.270822119 | 0.15704755 | 0.33108565 | F27D4.8 |
| WBGene00077546 | NA | | NA | NA | C01A2.9 |
| WBGene00077548 | | 0.040107227 | 0.58693163 | NA | F21C3.7 |
| WBGene00077549 | | 0.175008625 | 0.15653285 | 0.33048339 | NA |
| WBGene00077551 | NA | | NA | NA | NA |
| WBGene00077552 | NA | | NA | NA | NA |
| WBGene00077553 | | 0.052960832 | 0.64985033 | 0.79273174 | C26C6.10 |
| WBGene00077555 | | 0.020317245 | 0.84643798 | NA | NA |
| WBGene00077556 | | 0.020925399 | 0.84643798 | NA | NA |
| WBGene00077557 | | -0.00346345 | 0.95708614 | NA | F15D3.9 |
| WBGene00077558 | | -0.156985938 | 0.16770155 | 0.34527078 | T06D10.3 |
| WBGene00077559 | | -0.156713779 | 0.4428862 | 0.63683922 | R05D7.7 |
| WBGene00077561 | | 0.033463284 | 0.6365037 | NA | NA |
| WBGene00077562 | NA | | NA | NA | ZC443.7 |
| WBGene00077563 | | 0.019325396 | 0.77340931 | NA | peel-1 |
| WBGene00077564 | | 0.039292341 | 0.65808492 | NA | NA |
| WBGene00077565 | NA | | NA | NA | W07G4.7 |
| WBGene00077566 | NA | | NA | NA | NA |
| WBGene00077567 | NA | | NA | NA | NA |
| WBGene00077568 | | -0.056836877 | 0.64669574 | 0.79086805 | NA |
| WBGene00077569 | | 0.020925399 | 0.84643798 | NA | F53F4.18 |
| WBGene00077570 | | -0.127689698 | 0.3047365 | 0.50545453 | NA |
| WBGene00077571 | NA | | NA | NA | NA |
| WBGene00077572 | NA | | NA | NA | NA |
| WBGene00077573 | | 0.036692668 | 0.59311964 | NA | T08G5.15 |
| WBGene00077574 | | 0.020317245 | 0.84643798 | NA | NA |
| WBGene00077575 | NA | | NA | NA | NA |
| WBGene00077576 | NA | | NA | NA | C15H11.13 |
| WBGene00077577 | NA | | NA | NA | R02D5.9 |

| | | | | |
|---|---|---|---|---|
| WBGene00077579 | 0.040107227 | 0.58693163 | NA | NA |
| WBGene00077580 | NA | NA | NA | NA |
| WBGene00077581 | 0.059762547 | 0.38965825 | NA | NA |
| WBGene00077582 | -0.023109762 | 0.78121253 | NA | F14H8.8 |
| WBGene00077583 | NA | NA | NA | C01G10.18 |
| WBGene00077584 | NA | NA | NA | NA |
| WBGene00077585 | NA | NA | NA | T01G5.8 |
| WBGene00077586 | -0.023056499 | 0.78121253 | NA | NA |
| WBGene00077588 | NA | NA | NA | T26H8.5 |
| WBGene00077589 | NA | NA | NA | NA |
| WBGene00077590 | NA | NA | NA | NA |
| WBGene00077592 | -0.023056499 | 0.78121253 | NA | T25B2.3 |
| WBGene00077593 | 0.234030257 | 0.15447712 | 0.32749776 | C49G7.12 |
| WBGene00077594 | 0.245314565 | 0.21584949 | 0.40688198 | NA |
| WBGene00077595 | NA | NA | NA | NA |
| WBGene00077596 | -0.046394786 | 0.50231372 | NA | NA |
| WBGene00077597 | NA | NA | NA | NA |
| WBGene00077598 | NA | NA | NA | NA |
| WBGene00077599 | 0.013848069 | 0.85560057 | NA | T20F5.8 |
| WBGene00077600 | NA | NA | NA | NA |
| WBGene00077601 | NA | NA | NA | NA |
| WBGene00077602 | NA | NA | NA | NA |
| WBGene00077603 | NA | NA | NA | NA |
| WBGene00077604 | NA | NA | NA | NA |
| WBGene00077605 | NA | NA | NA | NA |
| WBGene00077606 | -0.003421551 | 0.95689871 | NA | NA |
| WBGene00077607 | 0.055348204 | 0.72475673 | 0.84572464 | T10C6.16 |
| WBGene00077608 | NA | NA | NA | NA |
| WBGene00077609 | NA | NA | NA | NA |
| WBGene00077610 | NA | NA | NA | NA |
| WBGene00077611 | NA | NA | NA | NA |
| WBGene00077612 | NA | NA | NA | NA |
| WBGene00077613 | 0.020232312 | 0.84643798 | NA | NA |
| WBGene00077614 | NA | NA | NA | NA |
| WBGene00077615 | 0.020232312 | 0.84643798 | NA | NA |
| WBGene00077616 | -0.001993815 | 0.95612996 | NA | F57E7.4 |
| WBGene00077617 | NA | NA | NA | NA |
| WBGene00077618 | NA | NA | NA | NA |
| WBGene00077619 | NA | NA | NA | NA |
| WBGene00077620 | NA | NA | NA | NA |
| WBGene00077621 | NA | NA | NA | NA |
| WBGene00077622 | 0.040107227 | 0.58693163 | NA | T25E12.14 |
| WBGene00077624 | NA | NA | NA | Y102A5C.40 |

| | | | | |
|---|---|---|---|---|
| WBGene00077625 | NA | NA | NA | NA |
| WBGene00077626 | NA | NA | NA | NA |
| WBGene00077627 | NA | NA | NA | NA |
| WBGene00077629 | 0.153452331 | 0.12161515 | 0.28062528 | NA |
| WBGene00077630 | -0.023056499 | 0.78121253 | NA | NA |
| WBGene00077631 | NA | NA | NA | NA |
| WBGene00077632 | NA | NA | NA | NA |
| WBGene00077633 | NA | NA | NA | NA |
| WBGene00077634 | -0.023109762 | 0.78121253 | NA | NA |
| WBGene00077635 | 0.020232312 | 0.84643798 | NA | NA |
| WBGene00077636 | 0.249120396 | 0.25824181 | 0.45633859 | NA |
| WBGene00077637 | NA | NA | NA | NA |
| WBGene00077638 | NA | NA | NA | NA |
| WBGene00077640 | NA | NA | NA | Y26G10.4 |
| WBGene00077641 | NA | NA | NA | NA |
| WBGene00077642 | NA | NA | NA | NA |
| WBGene00077643 | -0.133601674 | 0.49483065 | 0.67746545 | trpp-1 |
| WBGene00077644 | NA | NA | NA | NA |
| WBGene00077646 | NA | NA | NA | NA |
| WBGene00077647 | NA | NA | NA | NA |
| WBGene00077648 | NA | NA | NA | NA |
| WBGene00077649 | NA | NA | NA | NA |
| WBGene00077650 | NA | NA | NA | NA |
| WBGene00077651 | NA | NA | NA | NA |
| WBGene00077652 | -0.029508487 | 0.75403176 | 0.86073437 | NA |
| WBGene00077653 | 0.111571299 | 0.40320121 | 0.60151508 | NA |
| WBGene00077654 | NA | NA | NA | NA |
| WBGene00077655 | NA | NA | NA | asp-19 |
| WBGene00077656 | NA | NA | NA | NA |
| WBGene00077657 | NA | NA | NA | NA |
| WBGene00077658 | -0.073369501 | 0.59863272 | 0.75674164 | M106.8 |
| WBGene00077659 | NA | NA | NA | NA |
| WBGene00077660 | NA | NA | NA | NA |
| WBGene00077662 | -0.035285237 | 0.71906987 | NA | NA |
| WBGene00077663 | -0.020670548 | 0.90165616 | 0.9476095 | F55C9.14 |
| WBGene00077664 | -0.152604505 | 0.31217701 | 0.51291478 | NA |
| WBGene00077666 | NA | NA | NA | NA |
| WBGene00077667 | NA | NA | NA | NA |
| WBGene00077669 | 0.093671727 | 0.44322154 | 0.6368667 | Y38H6A.5 |
| WBGene00077670 | -0.027373974 | 0.68245543 | NA | NA |
| WBGene00077671 | NA | NA | NA | NA |
| WBGene00077673 | -0.022930767 | 0.78121253 | NA | B0563.10 |
| WBGene00077674 | 0.020232312 | 0.84643798 | NA | K02A6.4 |

| | | | | |
|---|---|---|---|---|
| WBGene00077679 | -0.023109762 | 0.78121253 | NA | K11E4.6 |
| WBGene00077680 | NA | NA | NA | F54F7.10 |
| WBGene00077681 | -0.001488154 | 0.95639912 | NA | Y44A6D.7 |
| WBGene00077682 | 0.020925399 | 0.84643798 | NA | F58B4.7 |
| WBGene00077684 | -0.034056522 | 0.77393929 | 0.87183544 | Y26D4A.21 |
| WBGene00077685 | 0.036692668 | 0.59311964 | NA | C04F12.12 |
| WBGene00077686 | NA | NA | NA | NA |
| WBGene00077687 | 0.01161837 | 0.90275923 | NA | NA |
| WBGene00077688 | NA | NA | NA | Y43F8B.19 |
| WBGene00077689 | 0.033463284 | 0.6365037 | NA | NA |
| WBGene00077690 | -0.024500043 | 0.90386874 | 0.94884507 | mtp-18 |
| WBGene00077691 | 0.237708622 | 0.01995706 | 0.0765099 | T06E4.14 |
| WBGene00077692 | -0.221911104 | 0.24198912 | 0.44001027 | C08E8.10 |
| WBGene00077693 | 0.007595709 | 0.97006532 | 0.98346632 | T04C12.11 |
| WBGene00077694 | NA | NA | NA | NA |
| WBGene00077695 | NA | NA | NA | NA |
| WBGene00077696 | 0.020317245 | 0.84643798 | NA | 1-Mar |
| WBGene00077697 | 2.048481377 | 0.00214376 | 0.0132155 | F58E6.13 |
| WBGene00077698 | 0.783388048 | 0.02312162 | 0.08535643 | lurp-4 |
| WBGene00077699 | 0.015062379 | 0.80838968 | NA | C45B11.8 |
| WBGene00077701 | 0.31067508 | 0.09015573 | 0.22832902 | poml-3 |
| WBGene00077703 | -0.023056499 | 0.78121253 | NA | K08E5.5 |
| WBGene00077704 | NA | NA | NA | F12A10.9 |
| WBGene00077705 | 0.017604133 | 0.79437959 | NA | NA |
| WBGene00077706 | -0.003108065 | 0.95728506 | NA | NA |
| WBGene00077707 | NA | NA | NA | NA |
| WBGene00077708 | NA | NA | NA | NA |
| WBGene00077709 | NA | NA | NA | NA |
| WBGene00077712 | 0.117748482 | 0.56625539 | 0.73538965 | nipi-4 |
| WBGene00077714 | -0.128727831 | 0.27103892 | 0.47056806 | R102.11 |
| WBGene00077715 | -0.023056499 | 0.78121253 | NA | NA |
| WBGene00077716 | NA | NA | NA | NA |
| WBGene00077717 | NA | NA | NA | NA |
| WBGene00077718 | 0.020317245 | 0.84643798 | NA | NA |
| WBGene00077719 | NA | NA | NA | NA |
| WBGene00077720 | NA | NA | NA | NA |
| WBGene00077721 | NA | NA | NA | NA |
| WBGene00077722 | NA | NA | NA | NA |
| WBGene00077723 | NA | NA | NA | NA |
| WBGene00077724 | NA | NA | NA | NA |
| WBGene00077725 | NA | NA | NA | NA |
| WBGene00077726 | NA | NA | NA | NA |
| WBGene00077727 | NA | NA | NA | NA |

| | | | | |
|---|---|---|---|---|
| WBGene00077728 | NA | NA | NA | T23G11.11 |
| WBGene00077732 | -0.097656189 | 0.16259485 | 0.33781058 | szy-4 |
| WBGene00077750 | NA | NA | NA | NA |
| WBGene00077751 | -0.009169485 | 0.91550466 | NA | irld-27 |
| WBGene00077752 | 0.033463284 | 0.6365037 | NA | Y105C5A.508 |
| WBGene00077753 | NA | NA | NA | NA |
| WBGene00077754 | NA | NA | NA | NA |
| WBGene00077755 | NA | NA | NA | NA |
| WBGene00077756 | -0.030092673 | 0.77124503 | 0.86993825 | Y11D7A.19 |
| WBGene00077757 | 0.402968953 | 0.07728979 | 0.20570801 | E02H4.7 |
| WBGene00077758 | 0.020925399 | 0.84643798 | NA | irld-12 |
| WBGene00077759 | 0.020232312 | 0.84643798 | NA | irld-51 |
| WBGene00077761 | 0.957788288 | 1.49E-11 | 4.82E-10 | zip-9 |
| WBGene00077762 | 0.018998992 | 0.77286019 | NA | Y57G11C.499 |
| WBGene00077763 | 0.059675786 | 0.4188661 | NA | glb-34 |
| WBGene00077764 | 0.806182967 | 0.00014095 | 0.00129875 | F59C6.16 |
| WBGene00077765 | -0.156376353 | 0.10719009 | 0.25718947 | NA |
| WBGene00077766 | NA | NA | NA | rgba-1 |
| WBGene00077767 | NA | NA | NA | NA |
| WBGene00077768 | 0.018860755 | 0.77262519 | NA | K04G2.12 |
| WBGene00077769 | 0.1298478 | 0.10432033 | NA | F33E2.10 |
| WBGene00077770 | 1.25082706 | 7.19E-05 | 0.00072398 | M04D5.3 |
| WBGene00077771 | -0.025792645 | 0.76453076 | 0.86623548 | C41G7.9 |
| WBGene00077772 | -0.461891105 | 0.07980821 | 0.21076336 | NA |
| WBGene00077773 | -0.079472169 | 0.3156132 | NA | F25H5.10 |
| WBGene00077775 | 2.117779954 | 4.88E-05 | 0.00051448 | nlp-58 |
| WBGene00077781 | -0.118865677 | 0.4871132 | 0.67155416 | NA |
| WBGene00077782 | 0.020232312 | 0.84643798 | NA | T26H10.2 |
| WBGene00077783 | 0.085537126 | 0.24698522 | NA | D1054.18 |
| WBGene00077784 | NA | NA | NA | NA |
| WBGene00077785 | -0.039636691 | 0.54579661 | NA | NA |
| WBGene00077786 | 0.018256332 | 0.90119032 | 0.94732391 | F53F8.7 |
| WBGene00077787 | 0.013292061 | 0.94931114 | 0.97120148 | NA |
| WBGene00077788 | 0.020925399 | 0.84643798 | NA | W07G4.8 |
| WBGene00077789 | NA | NA | NA | NA |
| WBGene00086546 | 0.057502245 | 0.62040318 | 0.7729172 | cil-1 |
| WBGene00086551 | -0.298323982 | 0.01494031 | 0.06165193 | D2005.7 |
| WBGene00086552 | NA | NA | NA | C17D12.9 |
| WBGene00086553 | -0.045959069 | 0.50707733 | NA | ZC334.13 |
| WBGene00086554 | 0.056430134 | 0.44291702 | NA | D1081.12 |
| WBGene00086555 | -0.134964184 | 0.34666586 | 0.54639908 | F49B2.7 |
| WBGene00086556 | 0.151890614 | 0.19731044 | 0.38497098 | F36D1.12 |
| WBGene00086557 | 0.370043486 | 0.11910824 | 0.27693563 | K07A12.8 |

| | | | | |
|---|---|---|---|---|
| WBGene00086558 | 0.300785675 | 0.02950964 | 0.10243197 | F12E12.12 |
| WBGene00086559 | 0.104747489 | 0.16723639 | NA | Y43F8B.20 |
| WBGene00086560 | 0.020925399 | 0.84643798 | NA | ZK355.8 |
| WBGene00086561 | NA | NA | NA | K01A2.9 |
| WBGene00086561 | NA | NA | NA | K01A2.12 |
| WBGene00086562 | -0.201746294 | 0.1520734 | 0.32451635 | T02G6.11 |
| WBGene00086563 | -0.023109762 | 0.78121253 | NA | K08C9.9 |
| WBGene00086564 | NA | NA | NA | K08C9.10 |
| WBGene00086565 | 0.246061673 | 0.11610549 | 0.2720276 | W06G6.17 |
| WBGene00086566 | 0.072474879 | 0.54136812 | 0.71522926 | C08H9.16 |
| WBGene00086567 | 0.020925399 | 0.84643798 | NA | C05C10.8 |
| WBGene00086568 | 0.457833528 | 0.04492037 | 0.13907362 | Y2H9A.6 |
| WBGene00086569 | 0.652448376 | 0.01567863 | 0.06410244 | F38A6.5 |
| WBGene00119201 | NA | NA | NA | C08E8.11 |
| WBGene00119203 | 0.135412618 | 0.48120778 | 0.66742323 | T04A8.18 |
| WBGene00119204 | 0.105820647 | 0.22502924 | NA | ZC116.5 |
| WBGene00138711 | 0.039135871 | 0.65930174 | NA | R07B1.13 |
| WBGene00138714 | NA | NA | NA | T02D1.9 |
| WBGene00138715 | 0.020232312 | 0.84643798 | NA | Y6G8.14 |
| WBGene00138716 | NA | NA | NA | F17B5.10 |
| WBGene00138717 | -0.128629727 | 0.39460813 | 0.59262392 | F52C12.6 |
| WBGene00138719 | NA | NA | NA | F26F2.10 |
| WBGene00138720 | -0.166140451 | 0.31280933 | 0.51333563 | F58H1.8 |
| WBGene00138721 | 5.452653057 | 5.26E-05 | 0.00054787 | pals-37 |
| WBGene00138724 | -0.023449708 | 0.69192352 | NA | F15B9.11 |
| WBGene00164968 | -0.041018366 | 0.83096958 | 0.90780346 | K08H2.10 |
| WBGene00164969 | 0.078633594 | 0.26310689 | NA | K09C8.9 |
| WBGene00164971 | 0.020232312 | 0.84643798 | NA | T21E8.6 |
| WBGene00164972 | NA | NA | NA | T21E8.7 |
| WBGene00164973 | 0.249687257 | 0.07974061 | 0.21066082 | C39B10.7 |
| WBGene00164974 | -0.004864426 | 0.94835335 | NA | F46C3.7 |
| WBGene00164983 | -0.000370622 | 0.99632306 | 0.99769349 | NA |
| WBGene00164985 | -0.37484398 | 0.12173034 | 0.28070284 | ZK1010.10 |
| WBGene00164986 | 0.017708609 | 0.82438962 | NA | Y48A6C.6 |
| WBGene00164991 | NA | NA | NA | NA |
| WBGene00164992 | NA | NA | NA | NA |
| WBGene00164993 | NA | NA | NA | NA |
| WBGene00164994 | NA | NA | NA | NA |
| WBGene00164995 | NA | NA | NA | NA |
| WBGene00164996 | NA | NA | NA | NA |
| WBGene00164997 | NA | NA | NA | NA |
| WBGene00164998 | NA | NA | NA | NA |
| WBGene00164999 | NA | NA | NA | NA |

| | | | | |
|---|---|---|---|---|
| WBGene00165000 | NA | NA | NA | NA |
| WBGene00165001 | NA | NA | NA | NA |
| WBGene00165002 | NA | NA | NA | NA |
| WBGene00165003 | NA | NA | NA | NA |
| WBGene00165004 | NA | NA | NA | NA |
| WBGene00165005 | NA | NA | NA | NA |
| WBGene00165006 | NA | NA | NA | NA |
| WBGene00165007 | -0.023056499 | 0.78121253 | NA | NA |
| WBGene00165009 | NA | NA | NA | NA |
| WBGene00165010 | NA | NA | NA | NA |
| WBGene00165011 | NA | NA | NA | NA |
| WBGene00165012 | NA | NA | NA | NA |
| WBGene00165013 | NA | NA | NA | NA |
| WBGene00165014 | NA | NA | NA | NA |
| WBGene00165015 | NA | NA | NA | NA |
| WBGene00165016 | NA | NA | NA | NA |
| WBGene00165017 | NA | NA | NA | NA |
| WBGene00165018 | NA | NA | NA | NA |
| WBGene00165019 | NA | NA | NA | NA |
| WBGene00165020 | NA | NA | NA | NA |
| WBGene00165021 | NA | NA | NA | NA |
| WBGene00165022 | NA | NA | NA | NA |
| WBGene00165023 | NA | NA | NA | NA |
| WBGene00165024 | NA | NA | NA | NA |
| WBGene00165025 | NA | NA | NA | NA |
| WBGene00165026 | NA | NA | NA | NA |
| WBGene00165027 | NA | NA | NA | NA |
| WBGene00165028 | NA | NA | NA | NA |
| WBGene00165029 | NA | NA | NA | NA |
| WBGene00165030 | NA | NA | NA | NA |
| WBGene00165031 | NA | NA | NA | NA |
| WBGene00165032 | NA | NA | NA | NA |
| WBGene00165033 | NA | NA | NA | NA |
| WBGene00165034 | NA | NA | NA | NA |
| WBGene00165035 | NA | NA | NA | NA |
| WBGene00165036 | NA | NA | NA | NA |
| WBGene00165037 | NA | NA | NA | NA |
| WBGene00165038 | NA | NA | NA | NA |
| WBGene00165039 | NA | NA | NA | NA |
| WBGene00165040 | NA | NA | NA | NA |
| WBGene00165041 | NA | NA | NA | NA |
| WBGene00165042 | NA | NA | NA | NA |
| WBGene00165043 | NA | NA | NA | NA |

| | | | | |
|---|---|---|---|---|
| WBGene00165044 | NA | NA | NA | NA |
| WBGene00165045 | NA | NA | NA | NA |
| WBGene00165046 | NA | NA | NA | NA |
| WBGene00165047 | NA | NA | NA | NA |
| WBGene00165048 | NA | NA | NA | NA |
| WBGene00165049 | NA | NA | NA | NA |
| WBGene00165050 | NA | NA | NA | NA |
| WBGene00165051 | NA | NA | NA | NA |
| WBGene00165052 | NA | NA | NA | NA |
| WBGene00165053 | NA | NA | NA | NA |
| WBGene00165054 | NA | NA | NA | NA |
| WBGene00165055 | NA | NA | NA | NA |
| WBGene00165056 | NA | NA | NA | NA |
| WBGene00165057 | NA | NA | NA | NA |
| WBGene00165058 | NA | NA | NA | NA |
| WBGene00165059 | NA | NA | NA | NA |
| WBGene00165060 | NA | NA | NA | NA |
| WBGene00165061 | NA | NA | NA | NA |
| WBGene00165062 | NA | NA | NA | NA |
| WBGene00165063 | NA | NA | NA | NA |
| WBGene00165064 | NA | NA | NA | NA |
| WBGene00165065 | NA | NA | NA | NA |
| WBGene00165066 | NA | NA | NA | NA |
| WBGene00165067 | NA | NA | NA | NA |
| WBGene00165068 | NA | NA | NA | NA |
| WBGene00165069 | NA | NA | NA | NA |
| WBGene00165070 | NA | NA | NA | NA |
| WBGene00165071 | NA | NA | NA | NA |
| WBGene00165072 | NA | NA | NA | NA |
| WBGene00165073 | NA | NA | NA | NA |
| WBGene00165074 | NA | NA | NA | NA |
| WBGene00165075 | NA | NA | NA | NA |
| WBGene00165076 | NA | NA | NA | NA |
| WBGene00165077 | NA | NA | NA | NA |
| WBGene00165078 | NA | NA | NA | NA |
| WBGene00165079 | NA | NA | NA | NA |
| WBGene00165080 | NA | NA | NA | NA |
| WBGene00165081 | NA | NA | NA | NA |
| WBGene00165082 | NA | NA | NA | NA |
| WBGene00165083 | NA | NA | NA | NA |
| WBGene00165084 | NA | NA | NA | NA |
| WBGene00165085 | NA | NA | NA | NA |
| WBGene00165086 | NA | NA | NA | NA |

| | | | | |
|---|---|---|---|---|
| WBGene00165087 | NA | NA | NA | NA |
| WBGene00165088 | NA | NA | NA | NA |
| WBGene00165089 | NA | NA | NA | NA |
| WBGene00165090 | NA | NA | NA | NA |
| WBGene00165091 | NA | NA | NA | NA |
| WBGene00165092 | NA | NA | NA | NA |
| WBGene00165093 | NA | NA | NA | NA |
| WBGene00165094 | NA | NA | NA | NA |
| WBGene00165095 | NA | NA | NA | NA |
| WBGene00165096 | NA | NA | NA | NA |
| WBGene00165097 | NA | NA | NA | NA |
| WBGene00165098 | NA | NA | NA | NA |
| WBGene00165099 | NA | NA | NA | NA |
| WBGene00165100 | NA | NA | NA | NA |
| WBGene00165101 | NA | NA | NA | NA |
| WBGene00165102 | NA | NA | NA | NA |
| WBGene00165103 | NA | NA | NA | NA |
| WBGene00165104 | NA | NA | NA | NA |
| WBGene00165105 | NA | NA | NA | NA |
| WBGene00165106 | NA | NA | NA | NA |
| WBGene00165107 | NA | NA | NA | NA |
| WBGene00165108 | NA | NA | NA | NA |
| WBGene00165109 | NA | NA | NA | NA |
| WBGene00165110 | NA | NA | NA | NA |
| WBGene00165111 | NA | NA | NA | NA |
| WBGene00165112 | NA | NA | NA | NA |
| WBGene00165113 | NA | NA | NA | NA |
| WBGene00165114 | NA | NA | NA | NA |
| WBGene00165115 | NA | NA | NA | NA |
| WBGene00165116 | NA | NA | NA | NA |
| WBGene00165117 | NA | NA | NA | NA |
| WBGene00165118 | NA | NA | NA | NA |
| WBGene00165119 | NA | NA | NA | NA |
| WBGene00165120 | NA | NA | NA | NA |
| WBGene00165121 | NA | NA | NA | NA |
| WBGene00165122 | NA | NA | NA | NA |
| WBGene00165123 | NA | NA | NA | NA |
| WBGene00165124 | NA | NA | NA | NA |
| WBGene00165125 | NA | NA | NA | NA |
| WBGene00165126 | -0.023056499 | 0.78121253 | NA | NA |
| WBGene00165127 | NA | NA | NA | NA |
| WBGene00165128 | NA | NA | NA | NA |
| WBGene00165129 | NA | NA | NA | NA |

| | | | | |
|---|---|---|---|---|
| WBGene00165130 | NA | NA | NA | NA |
| WBGene00165131 | NA | NA | NA | NA |
| WBGene00165132 | NA | NA | NA | NA |
| WBGene00165133 | NA | NA | NA | NA |
| WBGene00165134 | NA | NA | NA | NA |
| WBGene00165135 | NA | NA | NA | NA |
| WBGene00165136 | NA | NA | NA | NA |
| WBGene00165137 | NA | NA | NA | NA |
| WBGene00165138 | NA | NA | NA | NA |
| WBGene00165139 | NA | NA | NA | NA |
| WBGene00165140 | NA | NA | NA | NA |
| WBGene00165141 | NA | NA | NA | NA |
| WBGene00165142 | NA | NA | NA | NA |
| WBGene00165143 | NA | NA | NA | NA |
| WBGene00165144 | NA | NA | NA | NA |
| WBGene00165146 | NA | NA | NA | NA |
| WBGene00165147 | NA | NA | NA | NA |
| WBGene00165148 | NA | NA | NA | NA |
| WBGene00165149 | NA | NA | NA | NA |
| WBGene00165150 | NA | NA | NA | NA |
| WBGene00165151 | NA | NA | NA | NA |
| WBGene00165152 | NA | NA | NA | NA |
| WBGene00165153 | NA | NA | NA | NA |
| WBGene00165154 | NA | NA | NA | NA |
| WBGene00165155 | NA | NA | NA | NA |
| WBGene00165156 | NA | NA | NA | NA |
| WBGene00165157 | NA | NA | NA | NA |
| WBGene00165158 | NA | NA | NA | NA |
| WBGene00165159 | NA | NA | NA | NA |
| WBGene00165160 | NA | NA | NA | NA |
| WBGene00165161 | NA | NA | NA | NA |
| WBGene00165162 | NA | NA | NA | NA |
| WBGene00165163 | NA | NA | NA | NA |
| WBGene00165164 | NA | NA | NA | NA |
| WBGene00165165 | NA | NA | NA | NA |
| WBGene00165166 | NA | NA | NA | NA |
| WBGene00165167 | NA | NA | NA | NA |
| WBGene00165168 | NA | NA | NA | NA |
| WBGene00165169 | NA | NA | NA | NA |
| WBGene00165170 | NA | NA | NA | NA |
| WBGene00165171 | NA | NA | NA | NA |
| WBGene00165172 | NA | NA | NA | NA |
| WBGene00165173 | NA | NA | NA | NA |

| | | | | |
|---|---|---|---|---|
| WBGene00165174 | NA | NA | NA | NA |
| WBGene00165175 | NA | NA | NA | NA |
| WBGene00165176 | NA | NA | NA | NA |
| WBGene00165177 | NA | NA | NA | NA |
| WBGene00165178 | NA | NA | NA | NA |
| WBGene00165179 | NA | NA | NA | NA |
| WBGene00165180 | NA | NA | NA | NA |
| WBGene00165181 | NA | NA | NA | NA |
| WBGene00165182 | NA | NA | NA | NA |
| WBGene00165183 | NA | NA | NA | NA |
| WBGene00165184 | NA | NA | NA | NA |
| WBGene00165185 | NA | NA | NA | NA |
| WBGene00165186 | NA | NA | NA | NA |
| WBGene00165187 | NA | NA | NA | NA |
| WBGene00165188 | NA | NA | NA | NA |
| WBGene00165189 | NA | NA | NA | NA |
| WBGene00165190 | NA | NA | NA | NA |
| WBGene00165191 | NA | NA | NA | NA |
| WBGene00165192 | NA | NA | NA | NA |
| WBGene00165193 | NA | NA | NA | NA |
| WBGene00165194 | NA | NA | NA | NA |
| WBGene00165195 | NA | NA | NA | NA |
| WBGene00165196 | NA | NA | NA | NA |
| WBGene00165197 | NA | NA | NA | NA |
| WBGene00165198 | NA | NA | NA | NA |
| WBGene00165199 | NA | NA | NA | NA |
| WBGene00165200 | NA | NA | NA | NA |
| WBGene00165201 | NA | NA | NA | NA |
| WBGene00165202 | NA | NA | NA | NA |
| WBGene00165203 | NA | NA | NA | NA |
| WBGene00165204 | NA | NA | NA | NA |
| WBGene00165205 | NA | NA | NA | NA |
| WBGene00165206 | NA | NA | NA | NA |
| WBGene00165207 | NA | NA | NA | NA |
| WBGene00165208 | NA | NA | NA | NA |
| WBGene00165209 | NA | NA | NA | NA |
| WBGene00165210 | NA | NA | NA | NA |
| WBGene00165211 | NA | NA | NA | NA |
| WBGene00165212 | NA | NA | NA | NA |
| WBGene00165213 | NA | NA | NA | NA |
| WBGene00165214 | NA | NA | NA | NA |
| WBGene00165215 | NA | NA | NA | NA |
| WBGene00165217 | NA | NA | NA | NA |

| | | | | |
|---|---|---|---|---|
| WBGene00165218 | NA | NA | NA | NA |
| WBGene00165219 | NA | NA | NA | NA |
| WBGene00165220 | NA | NA | NA | NA |
| WBGene00165221 | NA | NA | NA | NA |
| WBGene00165222 | NA | NA | NA | NA |
| WBGene00165223 | NA | NA | NA | NA |
| WBGene00165224 | NA | NA | NA | NA |
| WBGene00165225 | NA | NA | NA | NA |
| WBGene00165226 | NA | NA | NA | NA |
| WBGene00165227 | NA | NA | NA | NA |
| WBGene00165228 | NA | NA | NA | NA |
| WBGene00165229 | NA | NA | NA | NA |
| WBGene00165230 | NA | NA | NA | NA |
| WBGene00165231 | NA | NA | NA | NA |
| WBGene00165232 | NA | NA | NA | NA |
| WBGene00165233 | NA | NA | NA | NA |
| WBGene00165234 | NA | NA | NA | NA |
| WBGene00165235 | NA | NA | NA | NA |
| WBGene00165236 | NA | NA | NA | NA |
| WBGene00165237 | NA | NA | NA | NA |
| WBGene00165238 | NA | NA | NA | NA |
| WBGene00165239 | NA | NA | NA | NA |
| WBGene00165240 | NA | NA | NA | NA |
| WBGene00165241 | NA | NA | NA | NA |
| WBGene00165242 | NA | NA | NA | NA |
| WBGene00165243 | NA | NA | NA | NA |
| WBGene00165244 | NA | NA | NA | NA |
| WBGene00165245 | NA | NA | NA | NA |
| WBGene00165246 | NA | NA | NA | NA |
| WBGene00165247 | NA | NA | NA | NA |
| WBGene00165248 | NA | NA | NA | NA |
| WBGene00165249 | NA | NA | NA | NA |
| WBGene00165250 | NA | NA | NA | NA |
| WBGene00165251 | NA | NA | NA | NA |
| WBGene00165252 | NA | NA | NA | NA |
| WBGene00165253 | NA | NA | NA | NA |
| WBGene00165254 | NA | NA | NA | NA |
| WBGene00165255 | NA | NA | NA | NA |
| WBGene00165256 | NA | NA | NA | NA |
| WBGene00165257 | NA | NA | NA | NA |
| WBGene00165258 | NA | NA | NA | NA |
| WBGene00165259 | NA | NA | NA | NA |
| WBGene00165260 | NA | NA | NA | NA |

| | | | | |
|---|---|---|---|---|
| WBGene00165261 | NA | NA | NA | NA |
| WBGene00165262 | NA | NA | NA | NA |
| WBGene00165263 | NA | NA | NA | NA |
| WBGene00165264 | NA | NA | NA | NA |
| WBGene00165265 | NA | NA | NA | NA |
| WBGene00165266 | NA | NA | NA | NA |
| WBGene00165267 | NA | NA | NA | NA |
| WBGene00165268 | NA | NA | NA | NA |
| WBGene00165269 | NA | NA | NA | NA |
| WBGene00165270 | NA | NA | NA | NA |
| WBGene00165271 | NA | NA | NA | NA |
| WBGene00165272 | NA | NA | NA | NA |
| WBGene00165273 | NA | NA | NA | NA |
| WBGene00165274 | NA | NA | NA | NA |
| WBGene00165275 | NA | NA | NA | NA |
| WBGene00165276 | 0.020317245 | 0.84643798 | NA | NA |
| WBGene00165277 | NA | NA | NA | NA |
| WBGene00165278 | NA | NA | NA | NA |
| WBGene00165279 | NA | NA | NA | NA |
| WBGene00165280 | NA | NA | NA | NA |
| WBGene00165281 | NA | NA | NA | NA |
| WBGene00165282 | NA | NA | NA | NA |
| WBGene00165283 | NA | NA | NA | NA |
| WBGene00165284 | NA | NA | NA | NA |
| WBGene00165285 | NA | NA | NA | NA |
| WBGene00165286 | NA | NA | NA | NA |
| WBGene00165287 | NA | NA | NA | NA |
| WBGene00165288 | NA | NA | NA | NA |
| WBGene00165289 | NA | NA | NA | NA |
| WBGene00165290 | NA | NA | NA | NA |
| WBGene00165291 | NA | NA | NA | NA |
| WBGene00165292 | NA | NA | NA | NA |
| WBGene00165293 | NA | NA | NA | NA |
| WBGene00165294 | NA | NA | NA | NA |
| WBGene00165295 | NA | NA | NA | NA |
| WBGene00165296 | NA | NA | NA | NA |
| WBGene00165297 | NA | NA | NA | NA |
| WBGene00165298 | NA | NA | NA | NA |
| WBGene00165299 | NA | NA | NA | NA |
| WBGene00165300 | NA | NA | NA | NA |
| WBGene00165301 | NA | NA | NA | NA |
| WBGene00165302 | NA | NA | NA | NA |
| WBGene00165303 | NA | NA | NA | NA |

| | | | | |
|---|---|---|---|---|
| WBGene00165304 | NA | NA | NA | NA |
| WBGene00165305 | NA | NA | NA | NA |
| WBGene00165306 | NA | NA | NA | NA |
| WBGene00165307 | NA | NA | NA | NA |
| WBGene00165308 | NA | NA | NA | NA |
| WBGene00165309 | NA | NA | NA | NA |
| WBGene00165310 | NA | NA | NA | NA |
| WBGene00165311 | NA | NA | NA | NA |
| WBGene00165312 | NA | NA | NA | NA |
| WBGene00165313 | NA | NA | NA | NA |
| WBGene00165314 | NA | NA | NA | NA |
| WBGene00165315 | NA | NA | NA | NA |
| WBGene00165316 | NA | NA | NA | NA |
| WBGene00165317 | NA | NA | NA | NA |
| WBGene00165318 | NA | NA | NA | NA |
| WBGene00165319 | NA | NA | NA | NA |
| WBGene00165320 | NA | NA | NA | NA |
| WBGene00165321 | NA | NA | NA | NA |
| WBGene00165322 | NA | NA | NA | NA |
| WBGene00165323 | NA | NA | NA | NA |
| WBGene00165324 | NA | NA | NA | NA |
| WBGene00165325 | NA | NA | NA | NA |
| WBGene00165326 | NA | NA | NA | NA |
| WBGene00165327 | NA | NA | NA | NA |
| WBGene00165328 | NA | NA | NA | NA |
| WBGene00165329 | NA | NA | NA | NA |
| WBGene00165330 | NA | NA | NA | NA |
| WBGene00165331 | NA | NA | NA | NA |
| WBGene00165332 | NA | NA | NA | NA |
| WBGene00165333 | NA | NA | NA | NA |
| WBGene00165334 | NA | NA | NA | NA |
| WBGene00165335 | NA | NA | NA | NA |
| WBGene00165336 | NA | NA | NA | NA |
| WBGene00165337 | NA | NA | NA | NA |
| WBGene00165338 | NA | NA | NA | NA |
| WBGene00165339 | NA | NA | NA | NA |
| WBGene00165340 | NA | NA | NA | NA |
| WBGene00165341 | NA | NA | NA | NA |
| WBGene00165342 | NA | NA | NA | NA |
| WBGene00165343 | NA | NA | NA | NA |
| WBGene00165344 | NA | NA | NA | NA |
| WBGene00165345 | NA | NA | NA | NA |
| WBGene00165346 | NA | NA | NA | NA |

| | | | | |
|---|---|---|---|---|
| WBGene00165347 | NA | NA | NA | NA |
| WBGene00165348 | NA | NA | NA | NA |
| WBGene00165349 | NA | NA | NA | NA |
| WBGene00165350 | NA | NA | NA | NA |
| WBGene00165351 | NA | NA | NA | NA |
| WBGene00165352 | NA | NA | NA | NA |
| WBGene00165353 | NA | NA | NA | NA |
| WBGene00165354 | NA | NA | NA | NA |
| WBGene00165355 | NA | NA | NA | NA |
| WBGene00165356 | NA | NA | NA | NA |
| WBGene00165357 | NA | NA | NA | NA |
| WBGene00165358 | NA | NA | NA | NA |
| WBGene00165359 | NA | NA | NA | NA |
| WBGene00165360 | NA | NA | NA | NA |
| WBGene00165361 | NA | NA | NA | NA |
| WBGene00165362 | NA | NA | NA | NA |
| WBGene00165363 | NA | NA | NA | NA |
| WBGene00165364 | NA | NA | NA | NA |
| WBGene00165365 | NA | NA | NA | NA |
| WBGene00165366 | NA | NA | NA | NA |
| WBGene00165367 | NA | NA | NA | NA |
| WBGene00165368 | NA | NA | NA | NA |
| WBGene00165369 | NA | NA | NA | NA |
| WBGene00165370 | NA | NA | NA | NA |
| WBGene00165371 | NA | NA | NA | NA |
| WBGene00165372 | NA | NA | NA | NA |
| WBGene00165373 | NA | NA | NA | NA |
| WBGene00165374 | NA | NA | NA | NA |
| WBGene00165375 | NA | NA | NA | NA |
| WBGene00165376 | NA | NA | NA | NA |
| WBGene00165377 | NA | NA | NA | NA |
| WBGene00165378 | NA | NA | NA | NA |
| WBGene00165379 | NA | NA | NA | NA |
| WBGene00165380 | NA | NA | NA | NA |
| WBGene00165381 | NA | NA | NA | NA |
| WBGene00165382 | NA | NA | NA | NA |
| WBGene00165383 | NA | NA | NA | NA |
| WBGene00165384 | NA | NA | NA | NA |
| WBGene00165385 | NA | NA | NA | NA |
| WBGene00165386 | NA | NA | NA | NA |
| WBGene00165387 | NA | NA | NA | NA |
| WBGene00165388 | NA | NA | NA | NA |
| WBGene00165389 | NA | NA | NA | NA |

| | | | | |
|---|---|---|---|---|
| WBGene00165390 | NA | NA | NA | NA |
| WBGene00165391 | NA | NA | NA | NA |
| WBGene00165392 | NA | NA | NA | NA |
| WBGene00165393 | NA | NA | NA | NA |
| WBGene00165394 | NA | NA | NA | NA |
| WBGene00165395 | NA | NA | NA | NA |
| WBGene00165396 | NA | NA | NA | NA |
| WBGene00165397 | NA | NA | NA | NA |
| WBGene00165398 | NA | NA | NA | NA |
| WBGene00165399 | NA | NA | NA | NA |
| WBGene00165400 | NA | NA | NA | NA |
| WBGene00165401 | NA | NA | NA | NA |
| WBGene00165402 | NA | NA | NA | NA |
| WBGene00165403 | NA | NA | NA | NA |
| WBGene00165404 | NA | NA | NA | NA |
| WBGene00165405 | NA | NA | NA | NA |
| WBGene00165406 | NA | NA | NA | NA |
| WBGene00165407 | NA | NA | NA | NA |
| WBGene00165408 | NA | NA | NA | NA |
| WBGene00165409 | NA | NA | NA | NA |
| WBGene00165410 | NA | NA | NA | NA |
| WBGene00165411 | NA | NA | NA | NA |
| WBGene00165412 | NA | NA | NA | NA |
| WBGene00165413 | NA | NA | NA | NA |
| WBGene00165414 | NA | NA | NA | NA |
| WBGene00165415 | NA | NA | NA | NA |
| WBGene00165416 | NA | NA | NA | NA |
| WBGene00165417 | NA | NA | NA | NA |
| WBGene00165418 | NA | NA | NA | NA |
| WBGene00165419 | NA | NA | NA | NA |
| WBGene00165420 | NA | NA | NA | NA |
| WBGene00165421 | NA | NA | NA | NA |
| WBGene00165422 | NA | NA | NA | NA |
| WBGene00165423 | NA | NA | NA | NA |
| WBGene00165424 | NA | NA | NA | NA |
| WBGene00165425 | NA | NA | NA | NA |
| WBGene00165426 | NA | NA | NA | NA |
| WBGene00165427 | NA | NA | NA | NA |
| WBGene00165428 | NA | NA | NA | NA |
| WBGene00165429 | NA | NA | NA | NA |
| WBGene00165430 | NA | NA | NA | NA |
| WBGene00165431 | NA | NA | NA | NA |
| WBGene00165432 | NA | NA | NA | NA |

| | | | | |
|---|---|---|---|---|
| WBGene00165433 | NA | NA | NA | NA |
| WBGene00165434 | NA | NA | NA | NA |
| WBGene00165435 | NA | NA | NA | NA |
| WBGene00165436 | NA | NA | NA | NA |
| WBGene00165438 | NA | NA | NA | NA |
| WBGene00165439 | NA | NA | NA | NA |
| WBGene00165440 | NA | NA | NA | NA |
| WBGene00165441 | NA | NA | NA | NA |
| WBGene00165442 | NA | NA | NA | NA |
| WBGene00165443 | NA | NA | NA | NA |
| WBGene00165444 | NA | NA | NA | NA |
| WBGene00165445 | NA | NA | NA | NA |
| WBGene00165446 | NA | NA | NA | NA |
| WBGene00165447 | NA | NA | NA | NA |
| WBGene00165448 | NA | NA | NA | NA |
| WBGene00165449 | NA | NA | NA | NA |
| WBGene00165450 | NA | NA | NA | NA |
| WBGene00165451 | NA | NA | NA | NA |
| WBGene00165452 | NA | NA | NA | NA |
| WBGene00165453 | NA | NA | NA | NA |
| WBGene00165454 | NA | NA | NA | NA |
| WBGene00165455 | NA | NA | NA | NA |
| WBGene00165456 | NA | NA | NA | NA |
| WBGene00165457 | NA | NA | NA | NA |
| WBGene00165458 | NA | NA | NA | NA |
| WBGene00165459 | NA | NA | NA | NA |
| WBGene00165460 | NA | NA | NA | NA |
| WBGene00165461 | NA | NA | NA | NA |
| WBGene00165462 | NA | NA | NA | NA |
| WBGene00165463 | NA | NA | NA | NA |
| WBGene00165464 | NA | NA | NA | NA |
| WBGene00165465 | NA | NA | NA | NA |
| WBGene00165466 | NA | NA | NA | NA |
| WBGene00165467 | NA | NA | NA | NA |
| WBGene00165468 | NA | NA | NA | NA |
| WBGene00165469 | NA | NA | NA | NA |
| WBGene00165470 | NA | NA | NA | NA |
| WBGene00165471 | NA | NA | NA | NA |
| WBGene00165472 | NA | NA | NA | NA |
| WBGene00165473 | NA | NA | NA | NA |
| WBGene00165474 | NA | NA | NA | NA |
| WBGene00165475 | NA | NA | NA | NA |
| WBGene00165476 | NA | NA | NA | NA |

| | | | | |
|---|---|---|---|---|
| WBGene00165477 | NA | NA | NA | NA |
| WBGene00165478 | NA | NA | NA | NA |
| WBGene00165479 | NA | NA | NA | NA |
| WBGene00165480 | NA | NA | NA | NA |
| WBGene00165481 | NA | NA | NA | NA |
| WBGene00165482 | NA | NA | NA | NA |
| WBGene00165483 | NA | NA | NA | NA |
| WBGene00165484 | NA | NA | NA | NA |
| WBGene00165485 | NA | NA | NA | NA |
| WBGene00165486 | NA | NA | NA | NA |
| WBGene00165487 | NA | NA | NA | NA |
| WBGene00165488 | NA | NA | NA | NA |
| WBGene00165489 | NA | NA | NA | NA |
| WBGene00165490 | NA | NA | NA | NA |
| WBGene00165491 | NA | NA | NA | NA |
| WBGene00165492 | NA | NA | NA | NA |
| WBGene00165493 | NA | NA | NA | NA |
| WBGene00165494 | NA | NA | NA | NA |
| WBGene00165495 | NA | NA | NA | NA |
| WBGene00165496 | NA | NA | NA | NA |
| WBGene00165497 | NA | NA | NA | NA |
| WBGene00165498 | NA | NA | NA | NA |
| WBGene00165499 | 0.020232312 | 0.84643798 | NA | NA |
| WBGene00165500 | NA | NA | NA | NA |
| WBGene00165501 | NA | NA | NA | NA |
| WBGene00165502 | NA | NA | NA | NA |
| WBGene00165503 | NA | NA | NA | NA |
| WBGene00165504 | NA | NA | NA | NA |
| WBGene00165505 | NA | NA | NA | NA |
| WBGene00165506 | NA | NA | NA | NA |
| WBGene00165507 | NA | NA | NA | NA |
| WBGene00165508 | NA | NA | NA | NA |
| WBGene00165509 | NA | NA | NA | NA |
| WBGene00165510 | NA | NA | NA | NA |
| WBGene00165511 | NA | NA | NA | NA |
| WBGene00165512 | NA | NA | NA | NA |
| WBGene00165513 | NA | NA | NA | NA |
| WBGene00165514 | NA | NA | NA | NA |
| WBGene00165515 | NA | NA | NA | NA |
| WBGene00165516 | NA | NA | NA | NA |
| WBGene00165517 | NA | NA | NA | NA |
| WBGene00165518 | NA | NA | NA | NA |
| WBGene00165519 | NA | NA | NA | NA |

| | | | | |
|---|---|---|---|---|
| WBGene00165520 | NA | NA | NA | NA |
| WBGene00165521 | NA | NA | NA | NA |
| WBGene00165522 | NA | NA | NA | NA |
| WBGene00165523 | NA | NA | NA | NA |
| WBGene00165524 | NA | NA | NA | NA |
| WBGene00165525 | NA | NA | NA | NA |
| WBGene00165526 | NA | NA | NA | NA |
| WBGene00165527 | NA | NA | NA | NA |
| WBGene00165528 | NA | NA | NA | NA |
| WBGene00165529 | NA | NA | NA | NA |
| WBGene00165530 | NA | NA | NA | NA |
| WBGene00165531 | NA | NA | NA | NA |
| WBGene00165532 | NA | NA | NA | NA |
| WBGene00165533 | NA | NA | NA | NA |
| WBGene00165534 | NA | NA | NA | NA |
| WBGene00165535 | NA | NA | NA | NA |
| WBGene00165536 | NA | NA | NA | NA |
| WBGene00165537 | NA | NA | NA | NA |
| WBGene00165538 | NA | NA | NA | NA |
| WBGene00165539 | NA | NA | NA | NA |
| WBGene00165540 | NA | NA | NA | NA |
| WBGene00165541 | NA | NA | NA | NA |
| WBGene00165542 | NA | NA | NA | NA |
| WBGene00165543 | NA | NA | NA | NA |
| WBGene00165544 | NA | NA | NA | NA |
| WBGene00165545 | NA | NA | NA | NA |
| WBGene00165546 | NA | NA | NA | NA |
| WBGene00165547 | NA | NA | NA | NA |
| WBGene00165548 | NA | NA | NA | NA |
| WBGene00165549 | NA | NA | NA | NA |
| WBGene00165550 | NA | NA | NA | NA |
| WBGene00165551 | NA | NA | NA | NA |
| WBGene00165552 | NA | NA | NA | NA |
| WBGene00165553 | NA | NA | NA | NA |
| WBGene00165554 | NA | NA | NA | NA |
| WBGene00165555 | NA | NA | NA | NA |
| WBGene00165556 | NA | NA | NA | NA |
| WBGene00165557 | NA | NA | NA | NA |
| WBGene00165558 | NA | NA | NA | NA |
| WBGene00165559 | NA | NA | NA | NA |
| WBGene00165560 | NA | NA | NA | NA |
| WBGene00165561 | NA | NA | NA | NA |
| WBGene00165562 | NA | NA | NA | NA |

| | | | | |
|---|---|---|---|---|
| WBGene00165563 | NA | NA | NA | NA |
| WBGene00165564 | NA | NA | NA | NA |
| WBGene00165565 | NA | NA | NA | NA |
| WBGene00165566 | NA | NA | NA | NA |
| WBGene00165567 | NA | NA | NA | NA |
| WBGene00165568 | NA | NA | NA | NA |
| WBGene00165569 | NA | NA | NA | NA |
| WBGene00165570 | NA | NA | NA | NA |
| WBGene00165571 | NA | NA | NA | NA |
| WBGene00165572 | NA | NA | NA | NA |
| WBGene00165573 | NA | NA | NA | NA |
| WBGene00165574 | NA | NA | NA | NA |
| WBGene00165575 | 0.040107227 | 0.58693163 | NA | NA |
| WBGene00165576 | NA | NA | NA | NA |
| WBGene00165577 | NA | NA | NA | NA |
| WBGene00165578 | NA | NA | NA | NA |
| WBGene00165579 | NA | NA | NA | NA |
| WBGene00165580 | NA | NA | NA | NA |
| WBGene00165581 | NA | NA | NA | NA |
| WBGene00165582 | NA | NA | NA | NA |
| WBGene00165583 | NA | NA | NA | NA |
| WBGene00165584 | NA | NA | NA | NA |
| WBGene00165585 | NA | NA | NA | NA |
| WBGene00165586 | NA | NA | NA | NA |
| WBGene00165587 | NA | NA | NA | NA |
| WBGene00165588 | NA | NA | NA | NA |
| WBGene00165589 | NA | NA | NA | NA |
| WBGene00165590 | NA | NA | NA | NA |
| WBGene00165591 | NA | NA | NA | NA |
| WBGene00165592 | NA | NA | NA | NA |
| WBGene00165593 | NA | NA | NA | NA |
| WBGene00165594 | NA | NA | NA | NA |
| WBGene00165595 | NA | NA | NA | NA |
| WBGene00165596 | NA | NA | NA | NA |
| WBGene00165597 | NA | NA | NA | NA |
| WBGene00165598 | NA | NA | NA | NA |
| WBGene00165599 | NA | NA | NA | NA |
| WBGene00165600 | NA | NA | NA | NA |
| WBGene00165601 | NA | NA | NA | NA |
| WBGene00165602 | NA | NA | NA | NA |
| WBGene00165603 | NA | NA | NA | NA |
| WBGene00165604 | NA | NA | NA | NA |
| WBGene00165605 | NA | NA | NA | NA |

| | | | | |
|---|---|---|---|---|
| WBGene00165606 | NA | NA | NA | NA |
| WBGene00165607 | NA | NA | NA | NA |
| WBGene00165608 | NA | NA | NA | NA |
| WBGene00165609 | NA | NA | NA | NA |
| WBGene00165610 | NA | NA | NA | NA |
| WBGene00165611 | NA | NA | NA | NA |
| WBGene00165612 | NA | NA | NA | NA |
| WBGene00165613 | NA | NA | NA | NA |
| WBGene00165614 | NA | NA | NA | NA |
| WBGene00165615 | NA | NA | NA | NA |
| WBGene00165616 | NA | NA | NA | NA |
| WBGene00165617 | NA | NA | NA | NA |
| WBGene00165618 | NA | NA | NA | NA |
| WBGene00165619 | NA | NA | NA | NA |
| WBGene00165620 | NA | NA | NA | NA |
| WBGene00165621 | NA | NA | NA | NA |
| WBGene00165622 | NA | NA | NA | NA |
| WBGene00165623 | NA | NA | NA | NA |
| WBGene00165624 | NA | NA | NA | NA |
| WBGene00165625 | NA | NA | NA | NA |
| WBGene00165626 | NA | NA | NA | NA |
| WBGene00165627 | NA | NA | NA | NA |
| WBGene00165628 | NA | NA | NA | NA |
| WBGene00165629 | NA | NA | NA | NA |
| WBGene00165630 | NA | NA | NA | NA |
| WBGene00165631 | NA | NA | NA | NA |
| WBGene00165632 | NA | NA | NA | NA |
| WBGene00165633 | NA | NA | NA | NA |
| WBGene00165634 | NA | NA | NA | NA |
| WBGene00165635 | NA | NA | NA | NA |
| WBGene00165636 | NA | NA | NA | NA |
| WBGene00165637 | NA | NA | NA | NA |
| WBGene00165638 | NA | NA | NA | NA |
| WBGene00165639 | NA | NA | NA | NA |
| WBGene00165640 | NA | NA | NA | NA |
| WBGene00165641 | NA | NA | NA | NA |
| WBGene00165642 | NA | NA | NA | NA |
| WBGene00165643 | NA | NA | NA | NA |
| WBGene00165644 | NA | NA | NA | NA |
| WBGene00165645 | NA | NA | NA | NA |
| WBGene00165646 | NA | NA | NA | NA |
| WBGene00165647 | NA | NA | NA | NA |
| WBGene00165648 | NA | NA | NA | NA |

| | | | | |
|---|---|---|---|---|
| WBGene00165649 | NA | NA | NA | NA |
| WBGene00165650 | NA | NA | NA | NA |
| WBGene00165651 | NA | NA | NA | NA |
| WBGene00165652 | NA | NA | NA | NA |
| WBGene00165653 | NA | NA | NA | NA |
| WBGene00165654 | NA | NA | NA | NA |
| WBGene00165655 | NA | NA | NA | NA |
| WBGene00165656 | NA | NA | NA | NA |
| WBGene00165657 | NA | NA | NA | NA |
| WBGene00165658 | NA | NA | NA | NA |
| WBGene00165659 | NA | NA | NA | NA |
| WBGene00165660 | NA | NA | NA | NA |
| WBGene00165661 | NA | NA | NA | NA |
| WBGene00165662 | NA | NA | NA | NA |
| WBGene00165663 | NA | NA | NA | NA |
| WBGene00165664 | NA | NA | NA | NA |
| WBGene00165665 | NA | NA | NA | NA |
| WBGene00165666 | NA | NA | NA | NA |
| WBGene00165667 | NA | NA | NA | NA |
| WBGene00165668 | NA | NA | NA | NA |
| WBGene00165669 | NA | NA | NA | NA |
| WBGene00165670 | NA | NA | NA | NA |
| WBGene00165671 | NA | NA | NA | NA |
| WBGene00165672 | NA | NA | NA | NA |
| WBGene00165673 | NA | NA | NA | NA |
| WBGene00165674 | NA | NA | NA | NA |
| WBGene00165675 | NA | NA | NA | NA |
| WBGene00165676 | NA | NA | NA | NA |
| WBGene00165677 | NA | NA | NA | NA |
| WBGene00165678 | NA | NA | NA | NA |
| WBGene00165679 | NA | NA | NA | NA |
| WBGene00165680 | NA | NA | NA | NA |
| WBGene00165681 | NA | NA | NA | NA |
| WBGene00165682 | NA | NA | NA | NA |
| WBGene00165683 | NA | NA | NA | NA |
| WBGene00165684 | NA | NA | NA | NA |
| WBGene00165685 | NA | NA | NA | NA |
| WBGene00165686 | NA | NA | NA | NA |
| WBGene00165687 | NA | NA | NA | NA |
| WBGene00165688 | NA | NA | NA | NA |
| WBGene00165689 | NA | NA | NA | NA |
| WBGene00165690 | NA | NA | NA | NA |
| WBGene00165691 | NA | NA | NA | NA |

| | | | | |
|---|---|---|---|---|
| WBGene00165692 | NA | NA | NA | NA |
| WBGene00165693 | NA | NA | NA | NA |
| WBGene00165694 | NA | NA | NA | NA |
| WBGene00165695 | NA | NA | NA | NA |
| WBGene00165696 | NA | NA | NA | NA |
| WBGene00165697 | NA | NA | NA | NA |
| WBGene00165698 | NA | NA | NA | NA |
| WBGene00165699 | NA | NA | NA | NA |
| WBGene00165700 | NA | NA | NA | NA |
| WBGene00165701 | NA | NA | NA | NA |
| WBGene00165702 | NA | NA | NA | NA |
| WBGene00165703 | NA | NA | NA | NA |
| WBGene00165704 | NA | NA | NA | NA |
| WBGene00165705 | NA | NA | NA | NA |
| WBGene00165706 | NA | NA | NA | NA |
| WBGene00165707 | NA | NA | NA | NA |
| WBGene00165708 | NA | NA | NA | NA |
| WBGene00165709 | NA | NA | NA | NA |
| WBGene00165710 | NA | NA | NA | NA |
| WBGene00165711 | NA | NA | NA | NA |
| WBGene00165712 | NA | NA | NA | NA |
| WBGene00165713 | NA | NA | NA | NA |
| WBGene00165714 | NA | NA | NA | NA |
| WBGene00165715 | NA | NA | NA | NA |
| WBGene00165716 | NA | NA | NA | NA |
| WBGene00165717 | NA | NA | NA | NA |
| WBGene00165718 | NA | NA | NA | NA |
| WBGene00165719 | NA | NA | NA | NA |
| WBGene00165720 | NA | NA | NA | NA |
| WBGene00165721 | NA | NA | NA | NA |
| WBGene00165722 | NA | NA | NA | NA |
| WBGene00165723 | NA | NA | NA | NA |
| WBGene00165724 | NA | NA | NA | NA |
| WBGene00165725 | NA | NA | NA | NA |
| WBGene00165726 | NA | NA | NA | NA |
| WBGene00165727 | NA | NA | NA | NA |
| WBGene00165728 | NA | NA | NA | NA |
| WBGene00165729 | NA | NA | NA | NA |
| WBGene00165730 | NA | NA | NA | NA |
| WBGene00165731 | NA | NA | NA | NA |
| WBGene00165732 | NA | NA | NA | NA |
| WBGene00165733 | NA | NA | NA | NA |
| WBGene00165734 | NA | NA | NA | NA |

| | | | | |
|---|---|---|---|---|
| WBGene00165735 | NA | NA | NA | NA |
| WBGene00165736 | NA | NA | NA | NA |
| WBGene00165737 | NA | NA | NA | NA |
| WBGene00165738 | 0.020232312 | 0.84643798 | NA | NA |
| WBGene00165739 | NA | NA | NA | NA |
| WBGene00165740 | NA | NA | NA | NA |
| WBGene00165741 | NA | NA | NA | NA |
| WBGene00165742 | NA | NA | NA | NA |
| WBGene00165743 | NA | NA | NA | NA |
| WBGene00165744 | NA | NA | NA | NA |
| WBGene00165745 | NA | NA | NA | NA |
| WBGene00165746 | NA | NA | NA | NA |
| WBGene00165747 | NA | NA | NA | NA |
| WBGene00165748 | NA | NA | NA | NA |
| WBGene00165749 | NA | NA | NA | NA |
| WBGene00165750 | NA | NA | NA | NA |
| WBGene00165751 | NA | NA | NA | NA |
| WBGene00165752 | NA | NA | NA | NA |
| WBGene00165753 | NA | NA | NA | NA |
| WBGene00165754 | NA | NA | NA | NA |
| WBGene00165755 | NA | NA | NA | NA |
| WBGene00165756 | NA | NA | NA | NA |
| WBGene00165757 | NA | NA | NA | NA |
| WBGene00165758 | NA | NA | NA | NA |
| WBGene00165759 | NA | NA | NA | NA |
| WBGene00165760 | NA | NA | NA | NA |
| WBGene00165761 | NA | NA | NA | NA |
| WBGene00165762 | NA | NA | NA | NA |
| WBGene00165763 | NA | NA | NA | NA |
| WBGene00165764 | NA | NA | NA | NA |
| WBGene00165765 | NA | NA | NA | NA |
| WBGene00165766 | NA | NA | NA | NA |
| WBGene00165767 | NA | NA | NA | NA |
| WBGene00165768 | NA | NA | NA | NA |
| WBGene00165769 | NA | NA | NA | NA |
| WBGene00165770 | NA | NA | NA | NA |
| WBGene00165771 | NA | NA | NA | NA |
| WBGene00165772 | NA | NA | NA | NA |
| WBGene00165773 | NA | NA | NA | NA |
| WBGene00165774 | NA | NA | NA | NA |
| WBGene00165775 | NA | NA | NA | NA |
| WBGene00165776 | NA | NA | NA | NA |
| WBGene00165777 | NA | NA | NA | NA |

| | | | | |
|---|---|---|---|---|
| WBGene00165778 | NA | NA | NA | NA |
| WBGene00165779 | NA | NA | NA | NA |
| WBGene00165780 | NA | NA | NA | NA |
| WBGene00165781 | NA | NA | NA | NA |
| WBGene00165782 | NA | NA | NA | NA |
| WBGene00165783 | NA | NA | NA | NA |
| WBGene00165784 | NA | NA | NA | NA |
| WBGene00165785 | NA | NA | NA | NA |
| WBGene00165786 | NA | NA | NA | NA |
| WBGene00165787 | NA | NA | NA | NA |
| WBGene00165788 | NA | NA | NA | NA |
| WBGene00165789 | NA | NA | NA | NA |
| WBGene00165790 | NA | NA | NA | NA |
| WBGene00165791 | NA | NA | NA | NA |
| WBGene00165792 | NA | NA | NA | NA |
| WBGene00165793 | NA | NA | NA | NA |
| WBGene00165794 | NA | NA | NA | NA |
| WBGene00165795 | NA | NA | NA | NA |
| WBGene00165796 | NA | NA | NA | NA |
| WBGene00165797 | NA | NA | NA | NA |
| WBGene00165798 | NA | NA | NA | NA |
| WBGene00165799 | NA | NA | NA | NA |
| WBGene00165800 | NA | NA | NA | NA |
| WBGene00165801 | NA | NA | NA | NA |
| WBGene00165802 | NA | NA | NA | NA |
| WBGene00165803 | NA | NA | NA | NA |
| WBGene00165804 | NA | NA | NA | NA |
| WBGene00165805 | NA | NA | NA | NA |
| WBGene00165806 | NA | NA | NA | NA |
| WBGene00165807 | NA | NA | NA | NA |
| WBGene00165808 | NA | NA | NA | NA |
| WBGene00165809 | NA | NA | NA | NA |
| WBGene00165810 | NA | NA | NA | NA |
| WBGene00165811 | 0.020317245 | 0.84643798 | NA | NA |
| WBGene00165812 | NA | NA | NA | NA |
| WBGene00165813 | NA | NA | NA | NA |
| WBGene00165814 | NA | NA | NA | NA |
| WBGene00165815 | NA | NA | NA | NA |
| WBGene00165816 | NA | NA | NA | NA |
| WBGene00165817 | NA | NA | NA | NA |
| WBGene00165818 | NA | NA | NA | NA |
| WBGene00165819 | NA | NA | NA | NA |
| WBGene00165820 | NA | NA | NA | NA |

| | | | | |
|---|---|---|---|---|
| WBGene00165821 | NA | NA | NA | NA |
| WBGene00165822 | NA | NA | NA | NA |
| WBGene00165823 | NA | NA | NA | NA |
| WBGene00165824 | NA | NA | NA | NA |
| WBGene00165825 | NA | NA | NA | NA |
| WBGene00165826 | NA | NA | NA | NA |
| WBGene00165827 | NA | NA | NA | NA |
| WBGene00165828 | NA | NA | NA | NA |
| WBGene00165829 | NA | NA | NA | NA |
| WBGene00165830 | NA | NA | NA | NA |
| WBGene00165831 | NA | NA | NA | NA |
| WBGene00165832 | NA | NA | NA | NA |
| WBGene00165833 | NA | NA | NA | NA |
| WBGene00165834 | NA | NA | NA | NA |
| WBGene00165835 | NA | NA | NA | NA |
| WBGene00165836 | NA | NA | NA | NA |
| WBGene00165837 | NA | NA | NA | NA |
| WBGene00165838 | NA | NA | NA | NA |
| WBGene00165839 | NA | NA | NA | NA |
| WBGene00165840 | NA | NA | NA | NA |
| WBGene00165841 | NA | NA | NA | NA |
| WBGene00165842 | NA | NA | NA | NA |
| WBGene00165843 | NA | NA | NA | NA |
| WBGene00165844 | NA | NA | NA | NA |
| WBGene00165845 | NA | NA | NA | NA |
| WBGene00165846 | NA | NA | NA | NA |
| WBGene00165847 | NA | NA | NA | NA |
| WBGene00165848 | NA | NA | NA | NA |
| WBGene00165849 | NA | NA | NA | NA |
| WBGene00165850 | NA | NA | NA | NA |
| WBGene00165851 | NA | NA | NA | NA |
| WBGene00165852 | NA | NA | NA | NA |
| WBGene00165853 | NA | NA | NA | NA |
| WBGene00165854 | NA | NA | NA | NA |
| WBGene00165855 | NA | NA | NA | NA |
| WBGene00165856 | NA | NA | NA | NA |
| WBGene00165857 | NA | NA | NA | NA |
| WBGene00165858 | NA | NA | NA | NA |
| WBGene00165859 | NA | NA | NA | NA |
| WBGene00165860 | NA | NA | NA | NA |
| WBGene00165861 | NA | NA | NA | NA |
| WBGene00165862 | NA | NA | NA | NA |
| WBGene00165863 | NA | NA | NA | NA |

| | | | | |
|---|---|---|---|---|
| WBGene00165864 | NA | NA | NA | NA |
| WBGene00165865 | NA | NA | NA | NA |
| WBGene00165866 | NA | NA | NA | NA |
| WBGene00165867 | NA | NA | NA | NA |
| WBGene00165868 | NA | NA | NA | NA |
| WBGene00165869 | NA | NA | NA | NA |
| WBGene00165870 | NA | NA | NA | NA |
| WBGene00165871 | NA | NA | NA | NA |
| WBGene00165872 | NA | NA | NA | NA |
| WBGene00165873 | NA | NA | NA | NA |
| WBGene00165874 | NA | NA | NA | NA |
| WBGene00165875 | NA | NA | NA | NA |
| WBGene00165876 | NA | NA | NA | NA |
| WBGene00165877 | NA | NA | NA | NA |
| WBGene00165878 | NA | NA | NA | NA |
| WBGene00165879 | NA | NA | NA | NA |
| WBGene00165880 | NA | NA | NA | NA |
| WBGene00165881 | NA | NA | NA | NA |
| WBGene00165882 | NA | NA | NA | NA |
| WBGene00165883 | NA | NA | NA | NA |
| WBGene00165884 | NA | NA | NA | NA |
| WBGene00165885 | NA | NA | NA | NA |
| WBGene00165886 | NA | NA | NA | NA |
| WBGene00165887 | NA | NA | NA | NA |
| WBGene00165888 | NA | NA | NA | NA |
| WBGene00165889 | NA | NA | NA | NA |
| WBGene00165890 | NA | NA | NA | NA |
| WBGene00165891 | NA | NA | NA | NA |
| WBGene00165892 | NA | NA | NA | NA |
| WBGene00165893 | NA | NA | NA | NA |
| WBGene00165894 | NA | NA | NA | NA |
| WBGene00165895 | NA | NA | NA | NA |
| WBGene00165896 | NA | NA | NA | NA |
| WBGene00165897 | NA | NA | NA | NA |
| WBGene00165898 | NA | NA | NA | NA |
| WBGene00165899 | NA | NA | NA | NA |
| WBGene00165900 | NA | NA | NA | NA |
| WBGene00165901 | NA | NA | NA | NA |
| WBGene00165902 | NA | NA | NA | NA |
| WBGene00165903 | NA | NA | NA | NA |
| WBGene00165904 | NA | NA | NA | NA |
| WBGene00165905 | NA | NA | NA | NA |
| WBGene00165906 | NA | NA | NA | NA |

| | | | | |
|---|---|---|---|---|
| WBGene00165907 | NA | NA | NA | NA |
| WBGene00165908 | NA | NA | NA | NA |
| WBGene00165909 | NA | NA | NA | NA |
| WBGene00165910 | NA | NA | NA | NA |
| WBGene00165911 | NA | NA | NA | NA |
| WBGene00165912 | NA | NA | NA | NA |
| WBGene00165913 | NA | NA | NA | NA |
| WBGene00165914 | NA | NA | NA | NA |
| WBGene00165915 | NA | NA | NA | NA |
| WBGene00165916 | NA | NA | NA | NA |
| WBGene00165917 | NA | NA | NA | NA |
| WBGene00165918 | NA | NA | NA | NA |
| WBGene00165919 | NA | NA | NA | NA |
| WBGene00165920 | NA | NA | NA | NA |
| WBGene00165921 | NA | NA | NA | NA |
| WBGene00165922 | NA | NA | NA | NA |
| WBGene00165923 | NA | NA | NA | NA |
| WBGene00165924 | NA | NA | NA | NA |
| WBGene00165925 | NA | NA | NA | NA |
| WBGene00165926 | NA | NA | NA | NA |
| WBGene00165927 | NA | NA | NA | NA |
| WBGene00165928 | NA | NA | NA | NA |
| WBGene00165929 | NA | NA | NA | NA |
| WBGene00165930 | NA | NA | NA | NA |
| WBGene00165931 | NA | NA | NA | NA |
| WBGene00165932 | NA | NA | NA | NA |
| WBGene00165933 | NA | NA | NA | NA |
| WBGene00165934 | NA | NA | NA | NA |
| WBGene00165935 | NA | NA | NA | NA |
| WBGene00165936 | NA | NA | NA | NA |
| WBGene00165937 | NA | NA | NA | NA |
| WBGene00165938 | NA | NA | NA | NA |
| WBGene00165939 | NA | NA | NA | NA |
| WBGene00165940 | NA | NA | NA | NA |
| WBGene00165941 | NA | NA | NA | NA |
| WBGene00165942 | NA | NA | NA | NA |
| WBGene00165943 | NA | NA | NA | NA |
| WBGene00165944 | NA | NA | NA | NA |
| WBGene00165945 | NA | NA | NA | NA |
| WBGene00165946 | NA | NA | NA | NA |
| WBGene00165947 | NA | NA | NA | NA |
| WBGene00165948 | NA | NA | NA | NA |
| WBGene00165949 | NA | NA | NA | NA |

| | | | | |
|---|---|---|---|---|
| WBGene00165950 | NA | NA | NA | NA |
| WBGene00165951 | NA | NA | NA | NA |
| WBGene00165952 | NA | NA | NA | NA |
| WBGene00165953 | NA | NA | NA | NA |
| WBGene00165954 | NA | NA | NA | NA |
| WBGene00165955 | NA | NA | NA | NA |
| WBGene00165956 | NA | NA | NA | NA |
| WBGene00165957 | NA | NA | NA | NA |
| WBGene00165958 | NA | NA | NA | NA |
| WBGene00165959 | NA | NA | NA | NA |
| WBGene00165960 | NA | NA | NA | NA |
| WBGene00165961 | NA | NA | NA | NA |
| WBGene00165962 | NA | NA | NA | NA |
| WBGene00165963 | NA | NA | NA | NA |
| WBGene00165964 | NA | NA | NA | NA |
| WBGene00165965 | NA | NA | NA | NA |
| WBGene00165966 | NA | NA | NA | NA |
| WBGene00165967 | NA | NA | NA | NA |
| WBGene00165968 | NA | NA | NA | NA |
| WBGene00165969 | NA | NA | NA | NA |
| WBGene00165970 | NA | NA | NA | NA |
| WBGene00165971 | NA | NA | NA | NA |
| WBGene00165972 | NA | NA | NA | NA |
| WBGene00165973 | NA | NA | NA | NA |
| WBGene00165974 | NA | NA | NA | NA |
| WBGene00165975 | NA | NA | NA | NA |
| WBGene00165976 | NA | NA | NA | NA |
| WBGene00165977 | NA | NA | NA | NA |
| WBGene00165978 | NA | NA | NA | NA |
| WBGene00165979 | NA | NA | NA | NA |
| WBGene00165980 | NA | NA | NA | NA |
| WBGene00165981 | NA | NA | NA | NA |
| WBGene00165982 | NA | NA | NA | NA |
| WBGene00165983 | NA | NA | NA | NA |
| WBGene00165984 | NA | NA | NA | NA |
| WBGene00165985 | NA | NA | NA | NA |
| WBGene00165986 | NA | NA | NA | NA |
| WBGene00165987 | NA | NA | NA | NA |
| WBGene00165988 | NA | NA | NA | NA |
| WBGene00165989 | NA | NA | NA | NA |
| WBGene00165990 | NA | NA | NA | NA |
| WBGene00165991 | NA | NA | NA | NA |
| WBGene00165992 | NA | NA | NA | NA |

| | | | | |
|---|---|---|---|---|
| WBGene00165993 | NA | NA | NA | NA |
| WBGene00165994 | NA | NA | NA | NA |
| WBGene00165995 | NA | NA | NA | NA |
| WBGene00165996 | NA | NA | NA | NA |
| WBGene00165997 | NA | NA | NA | NA |
| WBGene00165998 | NA | NA | NA | NA |
| WBGene00165999 | NA | NA | NA | NA |
| WBGene00166000 | NA | NA | NA | NA |
| WBGene00166001 | NA | NA | NA | NA |
| WBGene00166002 | NA | NA | NA | NA |
| WBGene00166003 | NA | NA | NA | NA |
| WBGene00166004 | NA | NA | NA | NA |
| WBGene00166005 | NA | NA | NA | NA |
| WBGene00166006 | NA | NA | NA | NA |
| WBGene00166007 | NA | NA | NA | NA |
| WBGene00166008 | NA | NA | NA | NA |
| WBGene00166009 | NA | NA | NA | NA |
| WBGene00166010 | NA | NA | NA | NA |
| WBGene00166011 | NA | NA | NA | NA |
| WBGene00166012 | NA | NA | NA | NA |
| WBGene00166013 | NA | NA | NA | NA |
| WBGene00166014 | NA | NA | NA | NA |
| WBGene00166015 | NA | NA | NA | NA |
| WBGene00166016 | NA | NA | NA | NA |
| WBGene00166017 | NA | NA | NA | NA |
| WBGene00166018 | NA | NA | NA | NA |
| WBGene00166019 | NA | NA | NA | NA |
| WBGene00166020 | NA | NA | NA | NA |
| WBGene00166021 | NA | NA | NA | NA |
| WBGene00166022 | NA | NA | NA | NA |
| WBGene00166023 | NA | NA | NA | NA |
| WBGene00166024 | NA | NA | NA | NA |
| WBGene00166025 | NA | NA | NA | NA |
| WBGene00166026 | NA | NA | NA | NA |
| WBGene00166027 | NA | NA | NA | NA |
| WBGene00166028 | NA | NA | NA | NA |
| WBGene00166029 | NA | NA | NA | NA |
| WBGene00166030 | NA | NA | NA | NA |
| WBGene00166031 | NA | NA | NA | NA |
| WBGene00166032 | NA | NA | NA | NA |
| WBGene00166033 | NA | NA | NA | NA |
| WBGene00166034 | NA | NA | NA | NA |
| WBGene00166035 | NA | NA | NA | NA |

| | | | | |
|---|---|---|---|---|
| WBGene00166036 | NA | NA | NA | NA |
| WBGene00166037 | NA | NA | NA | NA |
| WBGene00166038 | NA | NA | NA | NA |
| WBGene00166039 | NA | NA | NA | NA |
| WBGene00166040 | NA | NA | NA | NA |
| WBGene00166041 | NA | NA | NA | NA |
| WBGene00166042 | NA | NA | NA | NA |
| WBGene00166043 | NA | NA | NA | NA |
| WBGene00166044 | NA | NA | NA | NA |
| WBGene00166045 | NA | NA | NA | NA |
| WBGene00166046 | NA | NA | NA | NA |
| WBGene00166047 | NA | NA | NA | NA |
| WBGene00166048 | NA | NA | NA | NA |
| WBGene00166049 | NA | NA | NA | NA |
| WBGene00166050 | NA | NA | NA | NA |
| WBGene00166051 | NA | NA | NA | NA |
| WBGene00166052 | NA | NA | NA | NA |
| WBGene00166053 | NA | NA | NA | NA |
| WBGene00166054 | NA | NA | NA | NA |
| WBGene00166055 | NA | NA | NA | NA |
| WBGene00166057 | NA | NA | NA | NA |
| WBGene00166058 | NA | NA | NA | NA |
| WBGene00166059 | NA | NA | NA | NA |
| WBGene00166060 | NA | NA | NA | NA |
| WBGene00166062 | NA | NA | NA | NA |
| WBGene00166063 | NA | NA | NA | NA |
| WBGene00166064 | NA | NA | NA | NA |
| WBGene00166065 | NA | NA | NA | NA |
| WBGene00166066 | NA | NA | NA | NA |
| WBGene00166067 | NA | NA | NA | NA |
| WBGene00166068 | NA | NA | NA | NA |
| WBGene00166069 | NA | NA | NA | NA |
| WBGene00166070 | NA | NA | NA | NA |
| WBGene00166071 | NA | NA | NA | NA |
| WBGene00166072 | NA | NA | NA | NA |
| WBGene00166073 | NA | NA | NA | NA |
| WBGene00166074 | NA | NA | NA | NA |
| WBGene00166075 | NA | NA | NA | NA |
| WBGene00166076 | NA | NA | NA | NA |
| WBGene00166077 | NA | NA | NA | NA |
| WBGene00166078 | NA | NA | NA | NA |
| WBGene00166079 | NA | NA | NA | NA |
| WBGene00166080 | NA | NA | NA | NA |

| | | | | |
|---|---|---|---|---|
| WBGene00166081 | NA | NA | NA | NA |
| WBGene00166082 | NA | NA | NA | NA |
| WBGene00166083 | NA | NA | NA | NA |
| WBGene00166084 | NA | NA | NA | NA |
| WBGene00166085 | NA | NA | NA | NA |
| WBGene00166086 | NA | NA | NA | NA |
| WBGene00166087 | NA | NA | NA | NA |
| WBGene00166088 | NA | NA | NA | NA |
| WBGene00166089 | NA | NA | NA | NA |
| WBGene00166090 | NA | NA | NA | NA |
| WBGene00166091 | NA | NA | NA | NA |
| WBGene00166092 | NA | NA | NA | NA |
| WBGene00166093 | NA | NA | NA | NA |
| WBGene00166094 | NA | NA | NA | NA |
| WBGene00166095 | NA | NA | NA | NA |
| WBGene00166096 | NA | NA | NA | NA |
| WBGene00166097 | NA | NA | NA | NA |
| WBGene00166098 | NA | NA | NA | NA |
| WBGene00166099 | NA | NA | NA | NA |
| WBGene00166100 | NA | NA | NA | NA |
| WBGene00166101 | NA | NA | NA | NA |
| WBGene00166102 | NA | NA | NA | NA |
| WBGene00166103 | NA | NA | NA | NA |
| WBGene00166104 | NA | NA | NA | NA |
| WBGene00166105 | NA | NA | NA | NA |
| WBGene00166106 | NA | NA | NA | NA |
| WBGene00166107 | NA | NA | NA | NA |
| WBGene00166108 | NA | NA | NA | NA |
| WBGene00166109 | NA | NA | NA | NA |
| WBGene00166110 | NA | NA | NA | NA |
| WBGene00166111 | NA | NA | NA | NA |
| WBGene00166112 | NA | NA | NA | NA |
| WBGene00166113 | NA | NA | NA | NA |
| WBGene00166114 | NA | NA | NA | NA |
| WBGene00166115 | NA | NA | NA | NA |
| WBGene00166116 | NA | NA | NA | NA |
| WBGene00166117 | NA | NA | NA | NA |
| WBGene00166118 | NA | NA | NA | NA |
| WBGene00166119 | NA | NA | NA | NA |
| WBGene00166120 | NA | NA | NA | NA |
| WBGene00166121 | NA | NA | NA | NA |
| WBGene00166122 | NA | NA | NA | NA |
| WBGene00166123 | NA | NA | NA | NA |

| | | | | |
|---|---|---|---|---|
| WBGene00166124 | NA | NA | NA | NA |
| WBGene00166125 | NA | NA | NA | NA |
| WBGene00166126 | NA | NA | NA | NA |
| WBGene00166127 | NA | NA | NA | NA |
| WBGene00166128 | NA | NA | NA | NA |
| WBGene00166129 | NA | NA | NA | NA |
| WBGene00166130 | NA | NA | NA | NA |
| WBGene00166131 | NA | NA | NA | NA |
| WBGene00166132 | NA | NA | NA | NA |
| WBGene00166133 | NA | NA | NA | NA |
| WBGene00166134 | NA | NA | NA | NA |
| WBGene00166135 | NA | NA | NA | NA |
| WBGene00166136 | NA | NA | NA | NA |
| WBGene00166137 | NA | NA | NA | NA |
| WBGene00166138 | NA | NA | NA | NA |
| WBGene00166139 | NA | NA | NA | NA |
| WBGene00166140 | NA | NA | NA | NA |
| WBGene00166141 | NA | NA | NA | NA |
| WBGene00166142 | NA | NA | NA | NA |
| WBGene00166143 | NA | NA | NA | NA |
| WBGene00166144 | NA | NA | NA | NA |
| WBGene00166145 | NA | NA | NA | NA |
| WBGene00166146 | NA | NA | NA | NA |
| WBGene00166147 | NA | NA | NA | NA |
| WBGene00166148 | NA | NA | NA | NA |
| WBGene00166149 | NA | NA | NA | NA |
| WBGene00166150 | NA | NA | NA | NA |
| WBGene00166151 | NA | NA | NA | NA |
| WBGene00166152 | NA | NA | NA | NA |
| WBGene00166153 | NA | NA | NA | NA |
| WBGene00166154 | NA | NA | NA | NA |
| WBGene00166155 | NA | NA | NA | NA |
| WBGene00166156 | NA | NA | NA | NA |
| WBGene00166157 | NA | NA | NA | NA |
| WBGene00166158 | -0.023109762 | 0.78121253 | NA | NA |
| WBGene00166159 | NA | NA | NA | NA |
| WBGene00166160 | NA | NA | NA | NA |
| WBGene00166161 | NA | NA | NA | NA |
| WBGene00166162 | NA | NA | NA | NA |
| WBGene00166163 | NA | NA | NA | NA |
| WBGene00166164 | NA | NA | NA | NA |
| WBGene00166165 | NA | NA | NA | NA |
| WBGene00166166 | NA | NA | NA | NA |

| | | | | |
|---|---|---|---|---|
| WBGene00166167 | NA | NA | NA | NA |
| WBGene00166168 | -0.022930767 | 0.78121253 | NA | NA |
| WBGene00166169 | NA | NA | NA | NA |
| WBGene00166170 | NA | NA | NA | NA |
| WBGene00166171 | NA | NA | NA | NA |
| WBGene00166172 | NA | NA | NA | NA |
| WBGene00166173 | NA | NA | NA | NA |
| WBGene00166174 | NA | NA | NA | NA |
| WBGene00166175 | NA | NA | NA | NA |
| WBGene00166176 | NA | NA | NA | NA |
| WBGene00166177 | NA | NA | NA | NA |
| WBGene00166178 | NA | NA | NA | NA |
| WBGene00166179 | NA | NA | NA | NA |
| WBGene00166180 | NA | NA | NA | NA |
| WBGene00166181 | NA | NA | NA | NA |
| WBGene00166182 | NA | NA | NA | NA |
| WBGene00166183 | NA | NA | NA | NA |
| WBGene00166184 | NA | NA | NA | NA |
| WBGene00166185 | NA | NA | NA | NA |
| WBGene00166186 | NA | NA | NA | NA |
| WBGene00166187 | NA | NA | NA | NA |
| WBGene00166188 | NA | NA | NA | NA |
| WBGene00166189 | NA | NA | NA | NA |
| WBGene00166190 | NA | NA | NA | NA |
| WBGene00166191 | NA | NA | NA | NA |
| WBGene00166192 | NA | NA | NA | NA |
| WBGene00166193 | NA | NA | NA | NA |
| WBGene00166194 | NA | NA | NA | NA |
| WBGene00166195 | NA | NA | NA | NA |
| WBGene00166196 | NA | NA | NA | NA |
| WBGene00166197 | NA | NA | NA | NA |
| WBGene00166198 | NA | NA | NA | NA |
| WBGene00166199 | NA | NA | NA | NA |
| WBGene00166200 | NA | NA | NA | NA |
| WBGene00166201 | NA | NA | NA | NA |
| WBGene00166202 | NA | NA | NA | NA |
| WBGene00166203 | NA | NA | NA | NA |
| WBGene00166204 | NA | NA | NA | NA |
| WBGene00166205 | NA | NA | NA | NA |
| WBGene00166206 | NA | NA | NA | NA |
| WBGene00166207 | NA | NA | NA | NA |
| WBGene00166208 | NA | NA | NA | NA |
| WBGene00166209 | NA | NA | NA | NA |

| | | | | |
|---|---|---|---|---|
| WBGene00166210 | NA | NA | NA | NA |
| WBGene00166211 | NA | NA | NA | NA |
| WBGene00166212 | NA | NA | NA | NA |
| WBGene00166213 | NA | NA | NA | NA |
| WBGene00166214 | NA | NA | NA | NA |
| WBGene00166215 | NA | NA | NA | NA |
| WBGene00166216 | NA | NA | NA | NA |
| WBGene00166217 | NA | NA | NA | NA |
| WBGene00166218 | NA | NA | NA | NA |
| WBGene00166219 | NA | NA | NA | NA |
| WBGene00166220 | NA | NA | NA | NA |
| WBGene00166221 | NA | NA | NA | NA |
| WBGene00166222 | NA | NA | NA | NA |
| WBGene00166223 | NA | NA | NA | NA |
| WBGene00166224 | NA | NA | NA | NA |
| WBGene00166225 | NA | NA | NA | NA |
| WBGene00166226 | NA | NA | NA | NA |
| WBGene00166227 | NA | NA | NA | NA |
| WBGene00166228 | NA | NA | NA | NA |
| WBGene00166229 | NA | NA | NA | NA |
| WBGene00166230 | NA | NA | NA | NA |
| WBGene00166231 | NA | NA | NA | NA |
| WBGene00166232 | NA | NA | NA | NA |
| WBGene00166233 | NA | NA | NA | NA |
| WBGene00166234 | NA | NA | NA | NA |
| WBGene00166235 | -0.023109762 | 0.78121253 | NA | NA |
| WBGene00166236 | NA | NA | NA | NA |
| WBGene00166237 | NA | NA | NA | NA |
| WBGene00166238 | NA | NA | NA | NA |
| WBGene00166239 | NA | NA | NA | NA |
| WBGene00166240 | NA | NA | NA | NA |
| WBGene00166241 | NA | NA | NA | NA |
| WBGene00166242 | NA | NA | NA | NA |
| WBGene00166243 | NA | NA | NA | NA |
| WBGene00166244 | NA | NA | NA | NA |
| WBGene00166245 | NA | NA | NA | NA |
| WBGene00166246 | NA | NA | NA | NA |
| WBGene00166247 | NA | NA | NA | NA |
| WBGene00166248 | NA | NA | NA | NA |
| WBGene00166249 | NA | NA | NA | NA |
| WBGene00166250 | NA | NA | NA | NA |
| WBGene00166251 | NA | NA | NA | NA |
| WBGene00166252 | NA | NA | NA | NA |

| | | | | |
|---|---|---|---|---|
| WBGene00166253 | NA | NA | NA | NA |
| WBGene00166254 | NA | NA | NA | NA |
| WBGene00166255 | NA | NA | NA | NA |
| WBGene00166256 | NA | NA | NA | NA |
| WBGene00166257 | NA | NA | NA | NA |
| WBGene00166258 | NA | NA | NA | NA |
| WBGene00166259 | NA | NA | NA | NA |
| WBGene00166260 | NA | NA | NA | NA |
| WBGene00166261 | NA | NA | NA | NA |
| WBGene00166262 | NA | NA | NA | NA |
| WBGene00166263 | NA | NA | NA | NA |
| WBGene00166264 | NA | NA | NA | NA |
| WBGene00166265 | NA | NA | NA | NA |
| WBGene00166266 | NA | NA | NA | NA |
| WBGene00166267 | NA | NA | NA | NA |
| WBGene00166268 | NA | NA | NA | NA |
| WBGene00166269 | NA | NA | NA | NA |
| WBGene00166270 | NA | NA | NA | NA |
| WBGene00166271 | NA | NA | NA | NA |
| WBGene00166272 | NA | NA | NA | NA |
| WBGene00166273 | NA | NA | NA | NA |
| WBGene00166274 | NA | NA | NA | NA |
| WBGene00166275 | NA | NA | NA | NA |
| WBGene00166276 | NA | NA | NA | NA |
| WBGene00166277 | NA | NA | NA | NA |
| WBGene00166278 | NA | NA | NA | NA |
| WBGene00166279 | NA | NA | NA | NA |
| WBGene00166280 | NA | NA | NA | NA |
| WBGene00166281 | NA | NA | NA | NA |
| WBGene00166282 | NA | NA | NA | NA |
| WBGene00166283 | NA | NA | NA | NA |
| WBGene00166284 | NA | NA | NA | NA |
| WBGene00166285 | NA | NA | NA | NA |
| WBGene00166286 | NA | NA | NA | NA |
| WBGene00166287 | NA | NA | NA | NA |
| WBGene00166288 | NA | NA | NA | NA |
| WBGene00166289 | NA | NA | NA | NA |
| WBGene00166290 | NA | NA | NA | NA |
| WBGene00166291 | NA | NA | NA | NA |
| WBGene00166292 | NA | NA | NA | NA |
| WBGene00166293 | NA | NA | NA | NA |
| WBGene00166294 | NA | NA | NA | NA |
| WBGene00166295 | NA | NA | NA | NA |

| | | | | |
|---|---|---|---|---|
| WBGene00166296 | NA | NA | NA | NA |
| WBGene00166297 | NA | NA | NA | NA |
| WBGene00166298 | NA | NA | NA | NA |
| WBGene00166299 | NA | NA | NA | NA |
| WBGene00166300 | NA | NA | NA | NA |
| WBGene00166301 | NA | NA | NA | NA |
| WBGene00166302 | NA | NA | NA | NA |
| WBGene00166303 | NA | NA | NA | NA |
| WBGene00166304 | NA | NA | NA | NA |
| WBGene00166305 | NA | NA | NA | NA |
| WBGene00166306 | NA | NA | NA | NA |
| WBGene00166307 | NA | NA | NA | NA |
| WBGene00166308 | NA | NA | NA | NA |
| WBGene00166309 | NA | NA | NA | NA |
| WBGene00166310 | NA | NA | NA | NA |
| WBGene00166311 | NA | NA | NA | NA |
| WBGene00166312 | NA | NA | NA | NA |
| WBGene00166313 | NA | NA | NA | NA |
| WBGene00166314 | NA | NA | NA | NA |
| WBGene00166315 | NA | NA | NA | NA |
| WBGene00166316 | NA | NA | NA | NA |
| WBGene00166317 | NA | NA | NA | NA |
| WBGene00166318 | NA | NA | NA | NA |
| WBGene00166319 | NA | NA | NA | NA |
| WBGene00166320 | NA | NA | NA | NA |
| WBGene00166321 | NA | NA | NA | NA |
| WBGene00166322 | NA | NA | NA | NA |
| WBGene00166323 | NA | NA | NA | NA |
| WBGene00166324 | NA | NA | NA | NA |
| WBGene00166325 | NA | NA | NA | NA |
| WBGene00166326 | NA | NA | NA | NA |
| WBGene00166327 | NA | NA | NA | NA |
| WBGene00166328 | NA | NA | NA | NA |
| WBGene00166329 | NA | NA | NA | NA |
| WBGene00166330 | NA | NA | NA | NA |
| WBGene00166331 | NA | NA | NA | NA |
| WBGene00166332 | NA | NA | NA | NA |
| WBGene00166333 | NA | NA | NA | NA |
| WBGene00166334 | NA | NA | NA | NA |
| WBGene00166335 | NA | NA | NA | NA |
| WBGene00166336 | NA | NA | NA | NA |
| WBGene00166337 | NA | NA | NA | NA |
| WBGene00166338 | NA | NA | NA | NA |

| | | | | |
|---|---|---|---|---|
| WBGene00166339 | NA | NA | NA | NA |
| WBGene00166340 | NA | NA | NA | NA |
| WBGene00166341 | NA | NA | NA | NA |
| WBGene00166342 | NA | NA | NA | NA |
| WBGene00166343 | NA | NA | NA | NA |
| WBGene00166344 | NA | NA | NA | NA |
| WBGene00166345 | NA | NA | NA | NA |
| WBGene00166346 | NA | NA | NA | NA |
| WBGene00166347 | NA | NA | NA | NA |
| WBGene00166348 | NA | NA | NA | NA |
| WBGene00166349 | NA | NA | NA | NA |
| WBGene00166350 | NA | NA | NA | NA |
| WBGene00166351 | NA | NA | NA | NA |
| WBGene00166352 | NA | NA | NA | NA |
| WBGene00166353 | NA | NA | NA | NA |
| WBGene00166354 | NA | NA | NA | NA |
| WBGene00166355 | NA | NA | NA | NA |
| WBGene00166356 | NA | NA | NA | NA |
| WBGene00166357 | NA | NA | NA | NA |
| WBGene00166358 | NA | NA | NA | NA |
| WBGene00166359 | NA | NA | NA | NA |
| WBGene00166360 | NA | NA | NA | NA |
| WBGene00166361 | NA | NA | NA | NA |
| WBGene00166362 | NA | NA | NA | NA |
| WBGene00166363 | NA | NA | NA | NA |
| WBGene00166364 | NA | NA | NA | NA |
| WBGene00166365 | NA | NA | NA | NA |
| WBGene00166366 | NA | NA | NA | NA |
| WBGene00166367 | NA | NA | NA | NA |
| WBGene00166368 | NA | NA | NA | NA |
| WBGene00166369 | NA | NA | NA | NA |
| WBGene00166370 | NA | NA | NA | NA |
| WBGene00166371 | NA | NA | NA | NA |
| WBGene00166372 | NA | NA | NA | NA |
| WBGene00166373 | NA | NA | NA | NA |
| WBGene00166374 | NA | NA | NA | NA |
| WBGene00166375 | NA | NA | NA | NA |
| WBGene00166376 | NA | NA | NA | NA |
| WBGene00166377 | NA | NA | NA | NA |
| WBGene00166378 | NA | NA | NA | NA |
| WBGene00166379 | NA | NA | NA | NA |
| WBGene00166380 | NA | NA | NA | NA |
| WBGene00166381 | NA | NA | NA | NA |

| | | | | |
|---|---|---|---|---|
| WBGene00166382 | NA | NA | NA | NA |
| WBGene00166383 | NA | NA | NA | NA |
| WBGene00166384 | NA | NA | NA | NA |
| WBGene00166385 | NA | NA | NA | NA |
| WBGene00166386 | NA | NA | NA | NA |
| WBGene00166387 | NA | NA | NA | NA |
| WBGene00166388 | NA | NA | NA | NA |
| WBGene00166389 | NA | NA | NA | NA |
| WBGene00166390 | NA | NA | NA | NA |
| WBGene00166391 | NA | NA | NA | NA |
| WBGene00166392 | NA | NA | NA | NA |
| WBGene00166393 | NA | NA | NA | NA |
| WBGene00166394 | NA | NA | NA | NA |
| WBGene00166395 | NA | NA | NA | NA |
| WBGene00166396 | NA | NA | NA | NA |
| WBGene00166397 | NA | NA | NA | NA |
| WBGene00166398 | NA | NA | NA | NA |
| WBGene00166399 | NA | NA | NA | NA |
| WBGene00166400 | NA | NA | NA | NA |
| WBGene00166401 | NA | NA | NA | NA |
| WBGene00166402 | NA | NA | NA | NA |
| WBGene00166403 | NA | NA | NA | NA |
| WBGene00166404 | NA | NA | NA | NA |
| WBGene00166405 | NA | NA | NA | NA |
| WBGene00166406 | NA | NA | NA | NA |
| WBGene00166407 | NA | NA | NA | NA |
| WBGene00166408 | NA | NA | NA | NA |
| WBGene00166409 | NA | NA | NA | NA |
| WBGene00166410 | NA | NA | NA | NA |
| WBGene00166411 | NA | NA | NA | NA |
| WBGene00166412 | NA | NA | NA | NA |
| WBGene00166413 | NA | NA | NA | NA |
| WBGene00166414 | NA | NA | NA | NA |
| WBGene00166415 | NA | NA | NA | NA |
| WBGene00166416 | NA | NA | NA | NA |
| WBGene00166417 | NA | NA | NA | NA |
| WBGene00166418 | -0.022930767 | 0.78121253 | NA | NA |
| WBGene00166419 | NA | NA | NA | NA |
| WBGene00166420 | NA | NA | NA | NA |
| WBGene00166421 | NA | NA | NA | NA |
| WBGene00166422 | NA | NA | NA | NA |
| WBGene00166423 | NA | NA | NA | NA |
| WBGene00166424 | NA | NA | NA | NA |

| | | | | |
|---|---|---|---|---|
| WBGene00166425 | NA | NA | NA | NA |
| WBGene00166426 | NA | NA | NA | NA |
| WBGene00166427 | NA | NA | NA | NA |
| WBGene00166428 | NA | NA | NA | NA |
| WBGene00166429 | NA | NA | NA | NA |
| WBGene00166430 | NA | NA | NA | NA |
| WBGene00166431 | NA | NA | NA | NA |
| WBGene00166432 | NA | NA | NA | NA |
| WBGene00166433 | NA | NA | NA | NA |
| WBGene00166434 | NA | NA | NA | NA |
| WBGene00166435 | NA | NA | NA | NA |
| WBGene00166436 | NA | NA | NA | NA |
| WBGene00166437 | NA | NA | NA | NA |
| WBGene00166438 | NA | NA | NA | NA |
| WBGene00166439 | NA | NA | NA | NA |
| WBGene00166440 | NA | NA | NA | NA |
| WBGene00166441 | NA | NA | NA | NA |
| WBGene00166442 | NA | NA | NA | NA |
| WBGene00166443 | NA | NA | NA | NA |
| WBGene00166444 | NA | NA | NA | NA |
| WBGene00166445 | NA | NA | NA | NA |
| WBGene00166446 | NA | NA | NA | NA |
| WBGene00166447 | NA | NA | NA | NA |
| WBGene00166448 | NA | NA | NA | NA |
| WBGene00166449 | NA | NA | NA | NA |
| WBGene00166450 | NA | NA | NA | NA |
| WBGene00166451 | NA | NA | NA | NA |
| WBGene00166452 | NA | NA | NA | NA |
| WBGene00166453 | NA | NA | NA | NA |
| WBGene00166454 | NA | NA | NA | NA |
| WBGene00166455 | NA | NA | NA | NA |
| WBGene00166456 | NA | NA | NA | NA |
| WBGene00166457 | NA | NA | NA | NA |
| WBGene00166458 | NA | NA | NA | NA |
| WBGene00166459 | NA | NA | NA | NA |
| WBGene00166460 | NA | NA | NA | NA |
| WBGene00166461 | NA | NA | NA | NA |
| WBGene00166462 | NA | NA | NA | NA |
| WBGene00166463 | NA | NA | NA | NA |
| WBGene00166464 | NA | NA | NA | NA |
| WBGene00166465 | NA | NA | NA | NA |
| WBGene00166466 | NA | NA | NA | NA |
| WBGene00166467 | NA | NA | NA | NA |

| | | | | |
|---|---|---|---|---|
| WBGene00166468 | NA | NA | NA | NA |
| WBGene00166469 | NA | NA | NA | NA |
| WBGene00166470 | NA | NA | NA | NA |
| WBGene00166471 | NA | NA | NA | NA |
| WBGene00166472 | NA | NA | NA | NA |
| WBGene00166473 | NA | NA | NA | NA |
| WBGene00166474 | NA | NA | NA | NA |
| WBGene00166475 | NA | NA | NA | NA |
| WBGene00166476 | NA | NA | NA | NA |
| WBGene00166477 | NA | NA | NA | NA |
| WBGene00166478 | NA | NA | NA | NA |
| WBGene00166479 | NA | NA | NA | NA |
| WBGene00166480 | NA | NA | NA | NA |
| WBGene00166481 | NA | NA | NA | NA |
| WBGene00166482 | NA | NA | NA | NA |
| WBGene00166483 | NA | NA | NA | NA |
| WBGene00166484 | NA | NA | NA | NA |
| WBGene00166485 | NA | NA | NA | NA |
| WBGene00166486 | NA | NA | NA | NA |
| WBGene00166487 | NA | NA | NA | NA |
| WBGene00166488 | NA | NA | NA | NA |
| WBGene00166489 | NA | NA | NA | NA |
| WBGene00166490 | NA | NA | NA | NA |
| WBGene00166491 | NA | NA | NA | NA |
| WBGene00166492 | NA | NA | NA | NA |
| WBGene00166493 | NA | NA | NA | NA |
| WBGene00166494 | NA | NA | NA | NA |
| WBGene00166495 | NA | NA | NA | NA |
| WBGene00166496 | NA | NA | NA | NA |
| WBGene00166497 | NA | NA | NA | NA |
| WBGene00166498 | NA | NA | NA | NA |
| WBGene00166499 | NA | NA | NA | NA |
| WBGene00166500 | NA | NA | NA | NA |
| WBGene00166501 | NA | NA | NA | NA |
| WBGene00166502 | NA | NA | NA | NA |
| WBGene00166503 | NA | NA | NA | NA |
| WBGene00166504 | NA | NA | NA | NA |
| WBGene00166505 | NA | NA | NA | NA |
| WBGene00166506 | NA | NA | NA | NA |
| WBGene00166507 | NA | NA | NA | NA |
| WBGene00166508 | NA | NA | NA | NA |
| WBGene00166509 | NA | NA | NA | NA |
| WBGene00166510 | NA | NA | NA | NA |

| | | | | |
|---|---|---|---|---|
| WBGene00166511 | NA | NA | NA | NA |
| WBGene00166512 | NA | NA | NA | NA |
| WBGene00166513 | NA | NA | NA | NA |
| WBGene00166514 | NA | NA | NA | NA |
| WBGene00166515 | NA | NA | NA | NA |
| WBGene00166516 | NA | NA | NA | NA |
| WBGene00166517 | NA | NA | NA | NA |
| WBGene00166518 | NA | NA | NA | NA |
| WBGene00166519 | NA | NA | NA | NA |
| WBGene00166520 | NA | NA | NA | NA |
| WBGene00166521 | NA | NA | NA | NA |
| WBGene00166522 | NA | NA | NA | NA |
| WBGene00166523 | NA | NA | NA | NA |
| WBGene00166524 | NA | NA | NA | NA |
| WBGene00166525 | NA | NA | NA | NA |
| WBGene00166526 | NA | NA | NA | NA |
| WBGene00166527 | NA | NA | NA | NA |
| WBGene00166528 | NA | NA | NA | NA |
| WBGene00166529 | NA | NA | NA | NA |
| WBGene00166530 | NA | NA | NA | NA |
| WBGene00166531 | NA | NA | NA | NA |
| WBGene00166532 | NA | NA | NA | NA |
| WBGene00166533 | NA | NA | NA | NA |
| WBGene00166534 | NA | NA | NA | NA |
| WBGene00166535 | NA | NA | NA | NA |
| WBGene00166536 | NA | NA | NA | NA |
| WBGene00166537 | NA | NA | NA | NA |
| WBGene00166538 | NA | NA | NA | NA |
| WBGene00166539 | NA | NA | NA | NA |
| WBGene00166540 | NA | NA | NA | NA |
| WBGene00166541 | NA | NA | NA | NA |
| WBGene00166542 | NA | NA | NA | NA |
| WBGene00166543 | NA | NA | NA | NA |
| WBGene00166544 | NA | NA | NA | NA |
| WBGene00166545 | NA | NA | NA | NA |
| WBGene00166546 | NA | NA | NA | NA |
| WBGene00166547 | NA | NA | NA | NA |
| WBGene00166548 | NA | NA | NA | NA |
| WBGene00166549 | NA | NA | NA | NA |
| WBGene00166550 | NA | NA | NA | NA |
| WBGene00166551 | NA | NA | NA | NA |
| WBGene00166552 | NA | NA | NA | NA |
| WBGene00166553 | NA | NA | NA | NA |

| | | | | |
|---|---|---|---|---|
| WBGene00166554 | NA | NA | NA | NA |
| WBGene00166555 | NA | NA | NA | NA |
| WBGene00166556 | NA | NA | NA | NA |
| WBGene00166557 | NA | NA | NA | NA |
| WBGene00166558 | NA | NA | NA | NA |
| WBGene00166559 | NA | NA | NA | NA |
| WBGene00166560 | NA | NA | NA | NA |
| WBGene00166561 | NA | NA | NA | NA |
| WBGene00166562 | NA | NA | NA | NA |
| WBGene00166563 | NA | NA | NA | NA |
| WBGene00166564 | NA | NA | NA | NA |
| WBGene00166565 | NA | NA | NA | NA |
| WBGene00166566 | NA | NA | NA | NA |
| WBGene00166567 | 0.038125098 | 0.61119482 | NA | NA |
| WBGene00166568 | NA | NA | NA | NA |
| WBGene00166569 | NA | NA | NA | NA |
| WBGene00166570 | NA | NA | NA | NA |
| WBGene00166571 | NA | NA | NA | NA |
| WBGene00166572 | NA | NA | NA | NA |
| WBGene00166573 | NA | NA | NA | NA |
| WBGene00166574 | NA | NA | NA | NA |
| WBGene00166575 | NA | NA | NA | NA |
| WBGene00166576 | NA | NA | NA | NA |
| WBGene00166577 | NA | NA | NA | NA |
| WBGene00166578 | NA | NA | NA | NA |
| WBGene00166579 | NA | NA | NA | NA |
| WBGene00166580 | NA | NA | NA | NA |
| WBGene00166581 | NA | NA | NA | NA |
| WBGene00166582 | NA | NA | NA | NA |
| WBGene00166583 | NA | NA | NA | NA |
| WBGene00166584 | NA | NA | NA | NA |
| WBGene00166585 | NA | NA | NA | NA |
| WBGene00166586 | NA | NA | NA | NA |
| WBGene00166587 | NA | NA | NA | NA |
| WBGene00166588 | NA | NA | NA | NA |
| WBGene00166589 | NA | NA | NA | NA |
| WBGene00166590 | NA | NA | NA | NA |
| WBGene00166591 | NA | NA | NA | NA |
| WBGene00166592 | NA | NA | NA | NA |
| WBGene00166593 | NA | NA | NA | NA |
| WBGene00166594 | NA | NA | NA | NA |
| WBGene00166595 | NA | NA | NA | NA |
| WBGene00166596 | NA | NA | NA | NA |

| | | | | |
|---|---|---|---|---|
| WBGene00166597 | NA | NA | NA | NA |
| WBGene00166598 | NA | NA | NA | NA |
| WBGene00166599 | NA | NA | NA | NA |
| WBGene00166600 | NA | NA | NA | NA |
| WBGene00166601 | NA | NA | NA | NA |
| WBGene00166602 | NA | NA | NA | NA |
| WBGene00166603 | NA | NA | NA | NA |
| WBGene00166604 | NA | NA | NA | NA |
| WBGene00166605 | NA | NA | NA | NA |
| WBGene00166606 | NA | NA | NA | NA |
| WBGene00166607 | NA | NA | NA | NA |
| WBGene00166608 | NA | NA | NA | NA |
| WBGene00166609 | NA | NA | NA | NA |
| WBGene00166610 | NA | NA | NA | NA |
| WBGene00166611 | NA | NA | NA | NA |
| WBGene00166612 | NA | NA | NA | NA |
| WBGene00166613 | NA | NA | NA | NA |
| WBGene00166614 | NA | NA | NA | NA |
| WBGene00166615 | NA | NA | NA | NA |
| WBGene00166616 | NA | NA | NA | NA |
| WBGene00166617 | NA | NA | NA | NA |
| WBGene00166618 | NA | NA | NA | NA |
| WBGene00166619 | NA | NA | NA | NA |
| WBGene00166620 | NA | NA | NA | NA |
| WBGene00166621 | NA | NA | NA | NA |
| WBGene00166622 | NA | NA | NA | NA |
| WBGene00166623 | NA | NA | NA | NA |
| WBGene00166624 | NA | NA | NA | NA |
| WBGene00166625 | NA | NA | NA | NA |
| WBGene00166626 | NA | NA | NA | NA |
| WBGene00166627 | NA | NA | NA | NA |
| WBGene00166628 | NA | NA | NA | NA |
| WBGene00166629 | NA | NA | NA | NA |
| WBGene00166630 | NA | NA | NA | NA |
| WBGene00166631 | NA | NA | NA | NA |
| WBGene00166632 | NA | NA | NA | NA |
| WBGene00166633 | NA | NA | NA | NA |
| WBGene00166634 | NA | NA | NA | NA |
| WBGene00166635 | NA | NA | NA | NA |
| WBGene00166636 | NA | NA | NA | NA |
| WBGene00166637 | NA | NA | NA | NA |
| WBGene00166638 | NA | NA | NA | NA |
| WBGene00166639 | NA | NA | NA | NA |

| | | | | |
|---|---|---|---|---|
| WBGene00166640 | NA | NA | NA | NA |
| WBGene00166641 | NA | NA | NA | NA |
| WBGene00166642 | NA | NA | NA | NA |
| WBGene00166643 | NA | NA | NA | NA |
| WBGene00166644 | NA | NA | NA | NA |
| WBGene00166645 | NA | NA | NA | NA |
| WBGene00166646 | NA | NA | NA | NA |
| WBGene00166647 | NA | NA | NA | NA |
| WBGene00166648 | NA | NA | NA | NA |
| WBGene00166649 | NA | NA | NA | NA |
| WBGene00166650 | NA | NA | NA | NA |
| WBGene00166651 | NA | NA | NA | NA |
| WBGene00166652 | NA | NA | NA | NA |
| WBGene00166653 | NA | NA | NA | NA |
| WBGene00166654 | NA | NA | NA | NA |
| WBGene00166655 | NA | NA | NA | NA |
| WBGene00166656 | NA | NA | NA | NA |
| WBGene00166657 | NA | NA | NA | NA |
| WBGene00166658 | NA | NA | NA | NA |
| WBGene00166659 | NA | NA | NA | NA |
| WBGene00166660 | NA | NA | NA | NA |
| WBGene00166661 | NA | NA | NA | NA |
| WBGene00166662 | NA | NA | NA | NA |
| WBGene00166663 | NA | NA | NA | NA |
| WBGene00166664 | NA | NA | NA | NA |
| WBGene00166665 | NA | NA | NA | NA |
| WBGene00166666 | NA | NA | NA | NA |
| WBGene00166667 | NA | NA | NA | NA |
| WBGene00166668 | NA | NA | NA | NA |
| WBGene00166669 | NA | NA | NA | NA |
| WBGene00166670 | NA | NA | NA | NA |
| WBGene00166671 | NA | NA | NA | NA |
| WBGene00166672 | NA | NA | NA | NA |
| WBGene00166673 | NA | NA | NA | NA |
| WBGene00166674 | NA | NA | NA | NA |
| WBGene00166675 | NA | NA | NA | NA |
| WBGene00166676 | NA | NA | NA | NA |
| WBGene00166677 | NA | NA | NA | NA |
| WBGene00166678 | NA | NA | NA | NA |
| WBGene00166679 | NA | NA | NA | NA |
| WBGene00166680 | NA | NA | NA | NA |
| WBGene00166681 | NA | NA | NA | NA |
| WBGene00166682 | NA | NA | NA | NA |

| | | | | |
|---|---|---|---|---|
| WBGene00166683 | NA | NA | NA | NA |
| WBGene00166684 | NA | NA | NA | NA |
| WBGene00166685 | NA | NA | NA | NA |
| WBGene00166686 | NA | NA | NA | NA |
| WBGene00166687 | NA | NA | NA | NA |
| WBGene00166688 | NA | NA | NA | NA |
| WBGene00166689 | NA | NA | NA | NA |
| WBGene00166690 | NA | NA | NA | NA |
| WBGene00166691 | NA | NA | NA | NA |
| WBGene00166692 | NA | NA | NA | NA |
| WBGene00166693 | NA | NA | NA | NA |
| WBGene00166694 | NA | NA | NA | NA |
| WBGene00166695 | NA | NA | NA | NA |
| WBGene00166696 | NA | NA | NA | NA |
| WBGene00166697 | NA | NA | NA | NA |
| WBGene00166698 | NA | NA | NA | NA |
| WBGene00166699 | NA | NA | NA | NA |
| WBGene00166700 | NA | NA | NA | NA |
| WBGene00166701 | NA | NA | NA | NA |
| WBGene00166702 | NA | NA | NA | NA |
| WBGene00166703 | NA | NA | NA | NA |
| WBGene00166704 | NA | NA | NA | NA |
| WBGene00166705 | NA | NA | NA | NA |
| WBGene00166706 | NA | NA | NA | NA |
| WBGene00166707 | NA | NA | NA | NA |
| WBGene00166708 | NA | NA | NA | NA |
| WBGene00166709 | NA | NA | NA | NA |
| WBGene00166710 | NA | NA | NA | NA |
| WBGene00166711 | NA | NA | NA | NA |
| WBGene00166712 | NA | NA | NA | NA |
| WBGene00166713 | NA | NA | NA | NA |
| WBGene00166714 | NA | NA | NA | NA |
| WBGene00166715 | NA | NA | NA | NA |
| WBGene00166716 | NA | NA | NA | NA |
| WBGene00166717 | NA | NA | NA | NA |
| WBGene00166718 | NA | NA | NA | NA |
| WBGene00166719 | NA | NA | NA | NA |
| WBGene00166720 | NA | NA | NA | NA |
| WBGene00166721 | NA | NA | NA | NA |
| WBGene00166722 | NA | NA | NA | NA |
| WBGene00166723 | NA | NA | NA | NA |
| WBGene00166724 | NA | NA | NA | NA |
| WBGene00166725 | NA | NA | NA | NA |

| | | | | |
|---|---|---|---|---|
| WBGene00166726 | NA | NA | NA | NA |
| WBGene00166727 | NA | NA | NA | NA |
| WBGene00166728 | NA | NA | NA | NA |
| WBGene00166729 | NA | NA | NA | NA |
| WBGene00166730 | NA | NA | NA | NA |
| WBGene00166731 | NA | NA | NA | NA |
| WBGene00166732 | NA | NA | NA | NA |
| WBGene00166733 | NA | NA | NA | NA |
| WBGene00166734 | NA | NA | NA | NA |
| WBGene00166735 | NA | NA | NA | NA |
| WBGene00166736 | NA | NA | NA | NA |
| WBGene00166737 | NA | NA | NA | NA |
| WBGene00166738 | NA | NA | NA | NA |
| WBGene00166739 | NA | NA | NA | NA |
| WBGene00166740 | NA | NA | NA | NA |
| WBGene00166741 | NA | NA | NA | NA |
| WBGene00166742 | NA | NA | NA | NA |
| WBGene00166743 | NA | NA | NA | NA |
| WBGene00166744 | NA | NA | NA | NA |
| WBGene00166745 | NA | NA | NA | NA |
| WBGene00166746 | NA | NA | NA | NA |
| WBGene00166747 | NA | NA | NA | NA |
| WBGene00166748 | NA | NA | NA | NA |
| WBGene00166749 | NA | NA | NA | NA |
| WBGene00166750 | NA | NA | NA | NA |
| WBGene00166751 | NA | NA | NA | NA |
| WBGene00166752 | NA | NA | NA | NA |
| WBGene00166753 | NA | NA | NA | NA |
| WBGene00166754 | NA | NA | NA | NA |
| WBGene00166755 | NA | NA | NA | NA |
| WBGene00166756 | NA | NA | NA | NA |
| WBGene00166757 | NA | NA | NA | NA |
| WBGene00166758 | NA | NA | NA | NA |
| WBGene00166759 | NA | NA | NA | NA |
| WBGene00166760 | NA | NA | NA | NA |
| WBGene00166761 | NA | NA | NA | NA |
| WBGene00166762 | NA | NA | NA | NA |
| WBGene00166763 | NA | NA | NA | NA |
| WBGene00166764 | NA | NA | NA | NA |
| WBGene00166765 | NA | NA | NA | NA |
| WBGene00166766 | NA | NA | NA | NA |
| WBGene00166767 | NA | NA | NA | NA |
| WBGene00166768 | NA | NA | NA | NA |

| | | | | |
|---|---|---|---|---|
| WBGene00166769 | NA | NA | NA | NA |
| WBGene00166770 | NA | NA | NA | NA |
| WBGene00166771 | NA | NA | NA | NA |
| WBGene00166772 | NA | NA | NA | NA |
| WBGene00166773 | NA | NA | NA | NA |
| WBGene00166774 | NA | NA | NA | NA |
| WBGene00166775 | NA | NA | NA | NA |
| WBGene00166776 | NA | NA | NA | NA |
| WBGene00166777 | NA | NA | NA | NA |
| WBGene00166778 | NA | NA | NA | NA |
| WBGene00166779 | NA | NA | NA | NA |
| WBGene00166780 | NA | NA | NA | NA |
| WBGene00166781 | NA | NA | NA | NA |
| WBGene00166782 | NA | NA | NA | NA |
| WBGene00166783 | NA | NA | NA | NA |
| WBGene00166784 | NA | NA | NA | NA |
| WBGene00166785 | NA | NA | NA | NA |
| WBGene00166786 | NA | NA | NA | NA |
| WBGene00166787 | NA | NA | NA | NA |
| WBGene00166788 | NA | NA | NA | NA |
| WBGene00166789 | NA | NA | NA | NA |
| WBGene00166790 | NA | NA | NA | NA |
| WBGene00166791 | NA | NA | NA | NA |
| WBGene00166792 | NA | NA | NA | NA |
| WBGene00166793 | NA | NA | NA | NA |
| WBGene00166794 | NA | NA | NA | NA |
| WBGene00166795 | NA | NA | NA | NA |
| WBGene00166796 | NA | NA | NA | NA |
| WBGene00166797 | NA | NA | NA | NA |
| WBGene00166798 | NA | NA | NA | NA |
| WBGene00166799 | NA | NA | NA | NA |
| WBGene00166800 | NA | NA | NA | NA |
| WBGene00166801 | NA | NA | NA | NA |
| WBGene00166802 | NA | NA | NA | NA |
| WBGene00166803 | NA | NA | NA | NA |
| WBGene00166804 | NA | NA | NA | NA |
| WBGene00166805 | NA | NA | NA | NA |
| WBGene00166806 | NA | NA | NA | NA |
| WBGene00166807 | NA | NA | NA | NA |
| WBGene00166808 | NA | NA | NA | NA |
| WBGene00166809 | NA | NA | NA | NA |
| WBGene00166810 | NA | NA | NA | NA |
| WBGene00166811 | NA | NA | NA | NA |

| | | | | |
|---|---|---|---|---|
| WBGene00166812 | NA | NA | NA | NA |
| WBGene00166813 | NA | NA | NA | NA |
| WBGene00166814 | NA | NA | NA | NA |
| WBGene00166815 | NA | NA | NA | NA |
| WBGene00166816 | NA | NA | NA | NA |
| WBGene00166817 | NA | NA | NA | NA |
| WBGene00166819 | NA | NA | NA | NA |
| WBGene00166820 | NA | NA | NA | NA |
| WBGene00166821 | NA | NA | NA | NA |
| WBGene00166822 | NA | NA | NA | NA |
| WBGene00166823 | NA | NA | NA | NA |
| WBGene00166824 | NA | NA | NA | NA |
| WBGene00166825 | NA | NA | NA | NA |
| WBGene00166826 | NA | NA | NA | NA |
| WBGene00166827 | NA | NA | NA | NA |
| WBGene00166828 | NA | NA | NA | NA |
| WBGene00166829 | NA | NA | NA | NA |
| WBGene00166830 | NA | NA | NA | NA |
| WBGene00166831 | NA | NA | NA | NA |
| WBGene00166832 | NA | NA | NA | NA |
| WBGene00166833 | NA | NA | NA | NA |
| WBGene00166834 | NA | NA | NA | NA |
| WBGene00166835 | NA | NA | NA | NA |
| WBGene00166836 | NA | NA | NA | NA |
| WBGene00166837 | NA | NA | NA | NA |
| WBGene00166838 | NA | NA | NA | NA |
| WBGene00166839 | NA | NA | NA | NA |
| WBGene00166840 | NA | NA | NA | NA |
| WBGene00166841 | NA | NA | NA | NA |
| WBGene00166842 | NA | NA | NA | NA |
| WBGene00166843 | NA | NA | NA | NA |
| WBGene00166844 | NA | NA | NA | NA |
| WBGene00166845 | NA | NA | NA | NA |
| WBGene00166846 | NA | NA | NA | NA |
| WBGene00166847 | NA | NA | NA | NA |
| WBGene00166848 | NA | NA | NA | NA |
| WBGene00166849 | NA | NA | NA | NA |
| WBGene00166850 | NA | NA | NA | NA |
| WBGene00166851 | NA | NA | NA | NA |
| WBGene00166852 | NA | NA | NA | NA |
| WBGene00166853 | NA | NA | NA | NA |
| WBGene00166854 | NA | NA | NA | NA |
| WBGene00166855 | NA | NA | NA | NA |

| | | | | |
|---|---|---|---|---|
| WBGene00166856 | NA | NA | NA | NA |
| WBGene00166857 | NA | NA | NA | NA |
| WBGene00166858 | NA | NA | NA | NA |
| WBGene00166859 | NA | NA | NA | NA |
| WBGene00166860 | NA | NA | NA | NA |
| WBGene00166861 | NA | NA | NA | NA |
| WBGene00166862 | NA | NA | NA | NA |
| WBGene00166863 | NA | NA | NA | NA |
| WBGene00166864 | NA | NA | NA | NA |
| WBGene00166865 | NA | NA | NA | NA |
| WBGene00166866 | NA | NA | NA | NA |
| WBGene00166867 | NA | NA | NA | NA |
| WBGene00166868 | NA | NA | NA | NA |
| WBGene00166869 | NA | NA | NA | NA |
| WBGene00166870 | NA | NA | NA | NA |
| WBGene00166871 | NA | NA | NA | NA |
| WBGene00166872 | NA | NA | NA | NA |
| WBGene00166873 | NA | NA | NA | NA |
| WBGene00166874 | NA | NA | NA | NA |
| WBGene00166875 | NA | NA | NA | NA |
| WBGene00166876 | NA | NA | NA | NA |
| WBGene00166877 | NA | NA | NA | NA |
| WBGene00166878 | NA | NA | NA | NA |
| WBGene00166879 | NA | NA | NA | NA |
| WBGene00166880 | NA | NA | NA | NA |
| WBGene00166881 | NA | NA | NA | NA |
| WBGene00166882 | NA | NA | NA | NA |
| WBGene00166883 | NA | NA | NA | NA |
| WBGene00166884 | NA | NA | NA | NA |
| WBGene00166885 | NA | NA | NA | NA |
| WBGene00166886 | NA | NA | NA | NA |
| WBGene00166887 | NA | NA | NA | NA |
| WBGene00166888 | NA | NA | NA | NA |
| WBGene00166889 | NA | NA | NA | NA |
| WBGene00166890 | NA | NA | NA | NA |
| WBGene00166891 | NA | NA | NA | NA |
| WBGene00166892 | NA | NA | NA | NA |
| WBGene00166893 | NA | NA | NA | NA |
| WBGene00166894 | NA | NA | NA | NA |
| WBGene00166895 | NA | NA | NA | NA |
| WBGene00166896 | NA | NA | NA | NA |
| WBGene00166897 | NA | NA | NA | NA |
| WBGene00166898 | NA | NA | NA | NA |

| | | | | |
|---|---|---|---|---|
| WBGene00166899 | NA | NA | NA | NA |
| WBGene00166900 | NA | NA | NA | NA |
| WBGene00166901 | NA | NA | NA | NA |
| WBGene00166902 | NA | NA | NA | NA |
| WBGene00166903 | NA | NA | NA | NA |
| WBGene00166904 | NA | NA | NA | NA |
| WBGene00166905 | NA | NA | NA | NA |
| WBGene00166906 | NA | NA | NA | NA |
| WBGene00166907 | NA | NA | NA | NA |
| WBGene00166908 | NA | NA | NA | NA |
| WBGene00166909 | NA | NA | NA | NA |
| WBGene00166910 | NA | NA | NA | NA |
| WBGene00166911 | NA | NA | NA | NA |
| WBGene00166912 | NA | NA | NA | NA |
| WBGene00166913 | NA | NA | NA | NA |
| WBGene00166914 | NA | NA | NA | NA |
| WBGene00166915 | NA | NA | NA | NA |
| WBGene00166916 | NA | NA | NA | NA |
| WBGene00166917 | NA | NA | NA | NA |
| WBGene00166918 | NA | NA | NA | NA |
| WBGene00166919 | NA | NA | NA | NA |
| WBGene00166920 | NA | NA | NA | NA |
| WBGene00166921 | NA | NA | NA | NA |
| WBGene00166922 | NA | NA | NA | NA |
| WBGene00166923 | NA | NA | NA | NA |
| WBGene00166924 | NA | NA | NA | NA |
| WBGene00166925 | NA | NA | NA | NA |
| WBGene00166926 | NA | NA | NA | NA |
| WBGene00166927 | NA | NA | NA | NA |
| WBGene00166928 | NA | NA | NA | NA |
| WBGene00166929 | NA | NA | NA | NA |
| WBGene00166930 | NA | NA | NA | NA |
| WBGene00166931 | NA | NA | NA | NA |
| WBGene00166932 | NA | NA | NA | NA |
| WBGene00166933 | NA | NA | NA | NA |
| WBGene00166934 | NA | NA | NA | NA |
| WBGene00166935 | NA | NA | NA | NA |
| WBGene00166936 | NA | NA | NA | NA |
| WBGene00166937 | NA | NA | NA | NA |
| WBGene00166938 | NA | NA | NA | NA |
| WBGene00166939 | NA | NA | NA | NA |
| WBGene00166940 | NA | NA | NA | NA |
| WBGene00166941 | NA | NA | NA | NA |

| | | | | |
|---|---|---|---|---|
| WBGene00166942 | NA | NA | NA | NA |
| WBGene00166943 | NA | NA | NA | NA |
| WBGene00166944 | NA | NA | NA | NA |
| WBGene00166945 | NA | NA | NA | NA |
| WBGene00166946 | NA | NA | NA | NA |
| WBGene00166947 | NA | NA | NA | NA |
| WBGene00166948 | NA | NA | NA | NA |
| WBGene00166949 | NA | NA | NA | NA |
| WBGene00166950 | NA | NA | NA | NA |
| WBGene00166951 | NA | NA | NA | NA |
| WBGene00166952 | NA | NA | NA | NA |
| WBGene00166953 | NA | NA | NA | NA |
| WBGene00166954 | NA | NA | NA | NA |
| WBGene00166955 | NA | NA | NA | NA |
| WBGene00166956 | NA | NA | NA | NA |
| WBGene00166957 | NA | NA | NA | NA |
| WBGene00166958 | NA | NA | NA | NA |
| WBGene00166959 | NA | NA | NA | NA |
| WBGene00166960 | NA | NA | NA | NA |
| WBGene00166961 | NA | NA | NA | NA |
| WBGene00166962 | NA | NA | NA | NA |
| WBGene00166963 | NA | NA | NA | NA |
| WBGene00166964 | NA | NA | NA | NA |
| WBGene00166965 | NA | NA | NA | NA |
| WBGene00166966 | NA | NA | NA | NA |
| WBGene00166967 | NA | NA | NA | NA |
| WBGene00166968 | NA | NA | NA | NA |
| WBGene00166969 | NA | NA | NA | NA |
| WBGene00166970 | NA | NA | NA | NA |
| WBGene00166971 | NA | NA | NA | NA |
| WBGene00166972 | NA | NA | NA | NA |
| WBGene00166973 | NA | NA | NA | NA |
| WBGene00166974 | NA | NA | NA | NA |
| WBGene00166975 | NA | NA | NA | NA |
| WBGene00166976 | NA | NA | NA | NA |
| WBGene00166977 | NA | NA | NA | NA |
| WBGene00166978 | NA | NA | NA | NA |
| WBGene00166979 | NA | NA | NA | NA |
| WBGene00166980 | NA | NA | NA | NA |
| WBGene00166981 | NA | NA | NA | NA |
| WBGene00166982 | NA | NA | NA | NA |
| WBGene00166983 | NA | NA | NA | NA |
| WBGene00166984 | NA | NA | NA | NA |

| | | | | |
|---|---|---|---|---|
| WBGene00166985 | NA | NA | NA | NA |
| WBGene00166986 | NA | NA | NA | NA |
| WBGene00166987 | NA | NA | NA | NA |
| WBGene00166988 | NA | NA | NA | NA |
| WBGene00166989 | NA | NA | NA | NA |
| WBGene00166990 | NA | NA | NA | NA |
| WBGene00166991 | -0.003270206 | 0.95698316 | NA | NA |
| WBGene00166992 | NA | NA | NA | NA |
| WBGene00166993 | NA | NA | NA | NA |
| WBGene00166994 | NA | NA | NA | NA |
| WBGene00166995 | NA | NA | NA | NA |
| WBGene00166996 | NA | NA | NA | NA |
| WBGene00166997 | NA | NA | NA | NA |
| WBGene00166998 | NA | NA | NA | NA |
| WBGene00166999 | NA | NA | NA | NA |
| WBGene00167000 | NA | NA | NA | NA |
| WBGene00167001 | NA | NA | NA | NA |
| WBGene00167002 | NA | NA | NA | NA |
| WBGene00167003 | NA | NA | NA | NA |
| WBGene00167004 | NA | NA | NA | NA |
| WBGene00167005 | NA | NA | NA | NA |
| WBGene00167006 | NA | NA | NA | NA |
| WBGene00167007 | NA | NA | NA | NA |
| WBGene00167008 | NA | NA | NA | NA |
| WBGene00167009 | 0.020925399 | 0.84643798 | NA | NA |
| WBGene00167010 | NA | NA | NA | NA |
| WBGene00167011 | NA | NA | NA | NA |
| WBGene00167012 | NA | NA | NA | NA |
| WBGene00167013 | NA | NA | NA | NA |
| WBGene00167014 | NA | NA | NA | NA |
| WBGene00167015 | NA | NA | NA | NA |
| WBGene00167016 | NA | NA | NA | NA |
| WBGene00167017 | NA | NA | NA | NA |
| WBGene00167018 | NA | NA | NA | NA |
| WBGene00167019 | NA | NA | NA | NA |
| WBGene00167020 | NA | NA | NA | NA |
| WBGene00167021 | NA | NA | NA | NA |
| WBGene00167022 | NA | NA | NA | NA |
| WBGene00167023 | NA | NA | NA | NA |
| WBGene00167024 | NA | NA | NA | NA |
| WBGene00167025 | NA | NA | NA | NA |
| WBGene00167026 | NA | NA | NA | NA |
| WBGene00167027 | NA | NA | NA | NA |

| | | | | |
|---|---|---|---|---|
| WBGene00167028 | NA | NA | NA | NA |
| WBGene00167029 | NA | NA | NA | NA |
| WBGene00167030 | NA | NA | NA | NA |
| WBGene00167031 | NA | NA | NA | NA |
| WBGene00167032 | NA | NA | NA | NA |
| WBGene00167033 | NA | NA | NA | NA |
| WBGene00167034 | NA | NA | NA | NA |
| WBGene00167035 | NA | NA | NA | NA |
| WBGene00167036 | NA | NA | NA | NA |
| WBGene00167037 | NA | NA | NA | NA |
| WBGene00167038 | NA | NA | NA | NA |
| WBGene00167039 | NA | NA | NA | NA |
| WBGene00167040 | NA | NA | NA | NA |
| WBGene00167041 | NA | NA | NA | NA |
| WBGene00167042 | NA | NA | NA | NA |
| WBGene00167043 | NA | NA | NA | NA |
| WBGene00167044 | NA | NA | NA | NA |
| WBGene00167045 | NA | NA | NA | NA |
| WBGene00167046 | NA | NA | NA | NA |
| WBGene00167047 | NA | NA | NA | NA |
| WBGene00167048 | NA | NA | NA | NA |
| WBGene00167049 | NA | NA | NA | NA |
| WBGene00167050 | NA | NA | NA | NA |
| WBGene00167051 | NA | NA | NA | NA |
| WBGene00167052 | NA | NA | NA | NA |
| WBGene00167053 | NA | NA | NA | NA |
| WBGene00167054 | NA | NA | NA | NA |
| WBGene00167055 | NA | NA | NA | NA |
| WBGene00167056 | NA | NA | NA | NA |
| WBGene00167057 | NA | NA | NA | NA |
| WBGene00167058 | NA | NA | NA | NA |
| WBGene00167059 | NA | NA | NA | NA |
| WBGene00167060 | NA | NA | NA | NA |
| WBGene00167061 | NA | NA | NA | NA |
| WBGene00167062 | NA | NA | NA | NA |
| WBGene00167063 | NA | NA | NA | NA |
| WBGene00167064 | NA | NA | NA | NA |
| WBGene00167065 | NA | NA | NA | NA |
| WBGene00167066 | NA | NA | NA | NA |
| WBGene00167067 | NA | NA | NA | NA |
| WBGene00167068 | NA | NA | NA | NA |
| WBGene00167069 | NA | NA | NA | NA |
| WBGene00167070 | NA | NA | NA | NA |

| | | | | |
|---|---|---|---|---|
| WBGene00167071 | NA | NA | NA | NA |
| WBGene00167072 | NA | NA | NA | NA |
| WBGene00167073 | NA | NA | NA | NA |
| WBGene00167074 | NA | NA | NA | NA |
| WBGene00167075 | NA | NA | NA | NA |
| WBGene00167076 | NA | NA | NA | NA |
| WBGene00167077 | NA | NA | NA | NA |
| WBGene00167078 | NA | NA | NA | NA |
| WBGene00167079 | NA | NA | NA | NA |
| WBGene00167080 | NA | NA | NA | NA |
| WBGene00167081 | NA | NA | NA | NA |
| WBGene00167082 | NA | NA | NA | NA |
| WBGene00167083 | NA | NA | NA | NA |
| WBGene00167084 | NA | NA | NA | NA |
| WBGene00167085 | NA | NA | NA | NA |
| WBGene00167086 | NA | NA | NA | NA |
| WBGene00167087 | NA | NA | NA | NA |
| WBGene00167088 | NA | NA | NA | NA |
| WBGene00167089 | NA | NA | NA | NA |
| WBGene00167090 | NA | NA | NA | NA |
| WBGene00167091 | NA | NA | NA | NA |
| WBGene00167092 | NA | NA | NA | NA |
| WBGene00167093 | NA | NA | NA | NA |
| WBGene00167094 | NA | NA | NA | NA |
| WBGene00167095 | NA | NA | NA | NA |
| WBGene00167096 | NA | NA | NA | NA |
| WBGene00167097 | NA | NA | NA | NA |
| WBGene00167098 | NA | NA | NA | NA |
| WBGene00167099 | NA | NA | NA | NA |
| WBGene00167100 | NA | NA | NA | NA |
| WBGene00167101 | NA | NA | NA | NA |
| WBGene00167102 | NA | NA | NA | NA |
| WBGene00167103 | NA | NA | NA | NA |
| WBGene00167104 | NA | NA | NA | NA |
| WBGene00167105 | NA | NA | NA | NA |
| WBGene00167106 | NA | NA | NA | NA |
| WBGene00167107 | NA | NA | NA | NA |
| WBGene00167108 | NA | NA | NA | NA |
| WBGene00167109 | NA | NA | NA | NA |
| WBGene00167110 | NA | NA | NA | NA |
| WBGene00167111 | NA | NA | NA | NA |
| WBGene00167112 | NA | NA | NA | NA |
| WBGene00167113 | NA | NA | NA | NA |

| | | | | |
|---|---|---|---|---|
| WBGene00167114 | NA | NA | NA | NA |
| WBGene00167115 | NA | NA | NA | NA |
| WBGene00167116 | NA | NA | NA | NA |
| WBGene00167117 | NA | NA | NA | NA |
| WBGene00167118 | NA | NA | NA | NA |
| WBGene00167119 | NA | NA | NA | NA |
| WBGene00167120 | NA | NA | NA | NA |
| WBGene00167121 | NA | NA | NA | NA |
| WBGene00167122 | NA | NA | NA | NA |
| WBGene00167123 | NA | NA | NA | NA |
| WBGene00167124 | NA | NA | NA | NA |
| WBGene00167125 | NA | NA | NA | NA |
| WBGene00167126 | NA | NA | NA | NA |
| WBGene00167127 | NA | NA | NA | NA |
| WBGene00167128 | NA | NA | NA | NA |
| WBGene00167129 | NA | NA | NA | NA |
| WBGene00167130 | NA | NA | NA | NA |
| WBGene00167131 | NA | NA | NA | NA |
| WBGene00167132 | NA | NA | NA | NA |
| WBGene00167133 | NA | NA | NA | NA |
| WBGene00167134 | NA | NA | NA | NA |
| WBGene00167135 | NA | NA | NA | NA |
| WBGene00167136 | NA | NA | NA | NA |
| WBGene00167137 | NA | NA | NA | NA |
| WBGene00167138 | NA | NA | NA | NA |
| WBGene00167139 | NA | NA | NA | NA |
| WBGene00167140 | NA | NA | NA | NA |
| WBGene00167141 | NA | NA | NA | NA |
| WBGene00167142 | NA | NA | NA | NA |
| WBGene00167143 | NA | NA | NA | NA |
| WBGene00167144 | NA | NA | NA | NA |
| WBGene00167145 | NA | NA | NA | NA |
| WBGene00167146 | NA | NA | NA | NA |
| WBGene00167147 | -0.001842784 | 0.95621055 | NA | NA |
| WBGene00167148 | NA | NA | NA | NA |
| WBGene00167149 | NA | NA | NA | NA |
| WBGene00167150 | NA | NA | NA | NA |
| WBGene00167151 | NA | NA | NA | NA |
| WBGene00167152 | NA | NA | NA | NA |
| WBGene00167153 | NA | NA | NA | NA |
| WBGene00167154 | NA | NA | NA | NA |
| WBGene00167155 | NA | NA | NA | NA |
| WBGene00167156 | NA | NA | NA | NA |

| | | | | |
|---|---|---|---|---|
| WBGene00167157 | NA | NA | NA | NA |
| WBGene00167158 | NA | NA | NA | NA |
| WBGene00167159 | 0.036894305 | 0.59120501 | NA | NA |
| WBGene00167160 | NA | NA | NA | NA |
| WBGene00167161 | NA | NA | NA | NA |
| WBGene00167162 | NA | NA | NA | NA |
| WBGene00167163 | NA | NA | NA | NA |
| WBGene00167164 | NA | NA | NA | NA |
| WBGene00167165 | NA | NA | NA | NA |
| WBGene00167166 | NA | NA | NA | NA |
| WBGene00167167 | NA | NA | NA | NA |
| WBGene00167168 | NA | NA | NA | NA |
| WBGene00167169 | NA | NA | NA | NA |
| WBGene00167170 | NA | NA | NA | NA |
| WBGene00167171 | NA | NA | NA | NA |
| WBGene00167172 | NA | NA | NA | NA |
| WBGene00167173 | NA | NA | NA | NA |
| WBGene00167174 | NA | NA | NA | NA |
| WBGene00167175 | NA | NA | NA | NA |
| WBGene00167176 | NA | NA | NA | NA |
| WBGene00167177 | NA | NA | NA | NA |
| WBGene00167178 | NA | NA | NA | NA |
| WBGene00167179 | NA | NA | NA | NA |
| WBGene00167180 | NA | NA | NA | NA |
| WBGene00167181 | NA | NA | NA | NA |
| WBGene00167182 | NA | NA | NA | NA |
| WBGene00167183 | NA | NA | NA | NA |
| WBGene00167184 | NA | NA | NA | NA |
| WBGene00167185 | NA | NA | NA | NA |
| WBGene00167186 | NA | NA | NA | NA |
| WBGene00167187 | NA | NA | NA | NA |
| WBGene00167188 | NA | NA | NA | NA |
| WBGene00167189 | NA | NA | NA | NA |
| WBGene00167190 | NA | NA | NA | NA |
| WBGene00167191 | NA | NA | NA | NA |
| WBGene00167192 | NA | NA | NA | NA |
| WBGene00167193 | NA | NA | NA | NA |
| WBGene00167194 | NA | NA | NA | NA |
| WBGene00167195 | NA | NA | NA | NA |
| WBGene00167196 | NA | NA | NA | NA |
| WBGene00167197 | NA | NA | NA | NA |
| WBGene00167198 | NA | NA | NA | NA |
| WBGene00167199 | NA | NA | NA | NA |

| | | | | |
|---|---|---|---|---|
| WBGene00167200 | NA | NA | NA | NA |
| WBGene00167201 | NA | NA | NA | NA |
| WBGene00167202 | NA | NA | NA | NA |
| WBGene00167203 | NA | NA | NA | NA |
| WBGene00167204 | NA | NA | NA | NA |
| WBGene00167205 | NA | NA | NA | NA |
| WBGene00167206 | NA | NA | NA | NA |
| WBGene00167207 | NA | NA | NA | NA |
| WBGene00167208 | NA | NA | NA | NA |
| WBGene00167209 | NA | NA | NA | NA |
| WBGene00167210 | NA | NA | NA | NA |
| WBGene00167211 | NA | NA | NA | NA |
| WBGene00167212 | NA | NA | NA | NA |
| WBGene00167213 | NA | NA | NA | NA |
| WBGene00167214 | NA | NA | NA | NA |
| WBGene00167215 | NA | NA | NA | NA |
| WBGene00167216 | NA | NA | NA | NA |
| WBGene00167217 | NA | NA | NA | NA |
| WBGene00167218 | NA | NA | NA | NA |
| WBGene00167219 | NA | NA | NA | NA |
| WBGene00167220 | NA | NA | NA | NA |
| WBGene00167221 | NA | NA | NA | NA |
| WBGene00167222 | NA | NA | NA | NA |
| WBGene00167223 | NA | NA | NA | NA |
| WBGene00167224 | NA | NA | NA | NA |
| WBGene00167225 | NA | NA | NA | NA |
| WBGene00167226 | NA | NA | NA | NA |
| WBGene00167227 | NA | NA | NA | NA |
| WBGene00167228 | NA | NA | NA | NA |
| WBGene00167229 | NA | NA | NA | NA |
| WBGene00167230 | NA | NA | NA | NA |
| WBGene00167231 | NA | NA | NA | NA |
| WBGene00167232 | NA | NA | NA | NA |
| WBGene00167233 | NA | NA | NA | NA |
| WBGene00167234 | NA | NA | NA | NA |
| WBGene00167235 | NA | NA | NA | NA |
| WBGene00167236 | NA | NA | NA | NA |
| WBGene00167237 | NA | NA | NA | NA |
| WBGene00167238 | NA | NA | NA | NA |
| WBGene00167239 | NA | NA | NA | NA |
| WBGene00167240 | NA | NA | NA | NA |
| WBGene00167241 | NA | NA | NA | NA |
| WBGene00167242 | NA | NA | NA | NA |

| | | | | |
|---|---|---|---|---|
| WBGene00167243 | NA | NA | NA | NA |
| WBGene00167244 | NA | NA | NA | NA |
| WBGene00167245 | NA | NA | NA | NA |
| WBGene00167246 | NA | NA | NA | NA |
| WBGene00167247 | NA | NA | NA | NA |
| WBGene00167248 | NA | NA | NA | NA |
| WBGene00167249 | NA | NA | NA | NA |
| WBGene00167250 | NA | NA | NA | NA |
| WBGene00167251 | NA | NA | NA | NA |
| WBGene00167252 | NA | NA | NA | NA |
| WBGene00167253 | NA | NA | NA | NA |
| WBGene00167254 | NA | NA | NA | NA |
| WBGene00167255 | NA | NA | NA | NA |
| WBGene00167256 | NA | NA | NA | NA |
| WBGene00167257 | NA | NA | NA | NA |
| WBGene00167258 | NA | NA | NA | NA |
| WBGene00167259 | NA | NA | NA | NA |
| WBGene00167260 | NA | NA | NA | NA |
| WBGene00167261 | NA | NA | NA | NA |
| WBGene00167262 | NA | NA | NA | NA |
| WBGene00167263 | NA | NA | NA | NA |
| WBGene00167264 | NA | NA | NA | NA |
| WBGene00167265 | NA | NA | NA | NA |
| WBGene00167266 | NA | NA | NA | NA |
| WBGene00167267 | NA | NA | NA | NA |
| WBGene00167268 | NA | NA | NA | NA |
| WBGene00167269 | NA | NA | NA | NA |
| WBGene00167270 | NA | NA | NA | NA |
| WBGene00167271 | NA | NA | NA | NA |
| WBGene00167272 | NA | NA | NA | NA |
| WBGene00167273 | NA | NA | NA | NA |
| WBGene00167274 | NA | NA | NA | NA |
| WBGene00167275 | NA | NA | NA | NA |
| WBGene00167276 | NA | NA | NA | NA |
| WBGene00167277 | NA | NA | NA | NA |
| WBGene00167278 | NA | NA | NA | NA |
| WBGene00167279 | NA | NA | NA | NA |
| WBGene00167280 | NA | NA | NA | NA |
| WBGene00167281 | NA | NA | NA | NA |
| WBGene00167282 | NA | NA | NA | NA |
| WBGene00167283 | NA | NA | NA | NA |
| WBGene00167284 | NA | NA | NA | NA |
| WBGene00167285 | NA | NA | NA | NA |

| | | | | |
|---|---|---|---|---|
| WBGene00167286 | NA | NA | NA | NA |
| WBGene00167287 | NA | NA | NA | NA |
| WBGene00167288 | NA | NA | NA | NA |
| WBGene00167289 | NA | NA | NA | NA |
| WBGene00167290 | NA | NA | NA | NA |
| WBGene00167291 | NA | NA | NA | NA |
| WBGene00167292 | NA | NA | NA | NA |
| WBGene00167293 | NA | NA | NA | NA |
| WBGene00167294 | NA | NA | NA | NA |
| WBGene00167295 | NA | NA | NA | NA |
| WBGene00167296 | NA | NA | NA | NA |
| WBGene00167297 | NA | NA | NA | NA |
| WBGene00167298 | NA | NA | NA | NA |
| WBGene00167299 | NA | NA | NA | NA |
| WBGene00167300 | NA | NA | NA | NA |
| WBGene00167301 | NA | NA | NA | NA |
| WBGene00167302 | NA | NA | NA | NA |
| WBGene00167303 | NA | NA | NA | NA |
| WBGene00167304 | NA | NA | NA | NA |
| WBGene00167305 | NA | NA | NA | NA |
| WBGene00167306 | NA | NA | NA | NA |
| WBGene00167307 | NA | NA | NA | NA |
| WBGene00167308 | NA | NA | NA | NA |
| WBGene00167309 | NA | NA | NA | NA |
| WBGene00167310 | NA | NA | NA | NA |
| WBGene00167311 | NA | NA | NA | NA |
| WBGene00167312 | NA | NA | NA | NA |
| WBGene00167313 | NA | NA | NA | NA |
| WBGene00167314 | NA | NA | NA | NA |
| WBGene00167315 | NA | NA | NA | NA |
| WBGene00167316 | NA | NA | NA | NA |
| WBGene00167317 | NA | NA | NA | NA |
| WBGene00167318 | NA | NA | NA | NA |
| WBGene00167319 | NA | NA | NA | NA |
| WBGene00167320 | NA | NA | NA | NA |
| WBGene00167321 | NA | NA | NA | NA |
| WBGene00167322 | NA | NA | NA | NA |
| WBGene00167323 | NA | NA | NA | NA |
| WBGene00167324 | NA | NA | NA | NA |
| WBGene00167325 | NA | NA | NA | NA |
| WBGene00167326 | NA | NA | NA | NA |
| WBGene00167327 | NA | NA | NA | NA |
| WBGene00167328 | NA | NA | NA | NA |

| | | | | |
|---|---|---|---|---|
| WBGene00167329 | NA | NA | NA | NA |
| WBGene00167330 | NA | NA | NA | NA |
| WBGene00167331 | NA | NA | NA | NA |
| WBGene00167332 | NA | NA | NA | NA |
| WBGene00167333 | NA | NA | NA | NA |
| WBGene00167334 | NA | NA | NA | NA |
| WBGene00167335 | NA | NA | NA | NA |
| WBGene00167336 | NA | NA | NA | NA |
| WBGene00167337 | NA | NA | NA | NA |
| WBGene00167338 | NA | NA | NA | NA |
| WBGene00167339 | NA | NA | NA | NA |
| WBGene00167340 | NA | NA | NA | NA |
| WBGene00167341 | NA | NA | NA | NA |
| WBGene00167342 | NA | NA | NA | NA |
| WBGene00167343 | NA | NA | NA | NA |
| WBGene00167344 | NA | NA | NA | NA |
| WBGene00167345 | NA | NA | NA | NA |
| WBGene00167346 | NA | NA | NA | NA |
| WBGene00167347 | NA | NA | NA | NA |
| WBGene00167348 | NA | NA | NA | NA |
| WBGene00167349 | NA | NA | NA | NA |
| WBGene00167350 | NA | NA | NA | NA |
| WBGene00167351 | NA | NA | NA | NA |
| WBGene00167352 | NA | NA | NA | NA |
| WBGene00167353 | NA | NA | NA | NA |
| WBGene00167354 | NA | NA | NA | NA |
| WBGene00167355 | NA | NA | NA | NA |
| WBGene00167356 | NA | NA | NA | NA |
| WBGene00167357 | NA | NA | NA | NA |
| WBGene00167358 | NA | NA | NA | NA |
| WBGene00167359 | NA | NA | NA | NA |
| WBGene00167360 | NA | NA | NA | NA |
| WBGene00167361 | NA | NA | NA | NA |
| WBGene00167362 | NA | NA | NA | NA |
| WBGene00167363 | NA | NA | NA | NA |
| WBGene00167364 | NA | NA | NA | NA |
| WBGene00167365 | NA | NA | NA | NA |
| WBGene00167366 | NA | NA | NA | NA |
| WBGene00167367 | NA | NA | NA | NA |
| WBGene00167368 | NA | NA | NA | NA |
| WBGene00167369 | NA | NA | NA | NA |
| WBGene00167370 | NA | NA | NA | NA |
| WBGene00167371 | NA | NA | NA | NA |

| | | | | |
|---|---|---|---|---|
| WBGene00167372 | NA | NA | NA | NA |
| WBGene00167373 | NA | NA | NA | NA |
| WBGene00167374 | NA | NA | NA | NA |
| WBGene00167375 | NA | NA | NA | NA |
| WBGene00167376 | NA | NA | NA | NA |
| WBGene00167377 | NA | NA | NA | NA |
| WBGene00167378 | NA | NA | NA | NA |
| WBGene00167379 | NA | NA | NA | NA |
| WBGene00167380 | NA | NA | NA | NA |
| WBGene00167381 | NA | NA | NA | NA |
| WBGene00167382 | NA | NA | NA | NA |
| WBGene00167383 | NA | NA | NA | NA |
| WBGene00167384 | NA | NA | NA | NA |
| WBGene00167385 | NA | NA | NA | NA |
| WBGene00167386 | NA | NA | NA | NA |
| WBGene00167387 | NA | NA | NA | NA |
| WBGene00167388 | NA | NA | NA | NA |
| WBGene00167389 | NA | NA | NA | NA |
| WBGene00167390 | NA | NA | NA | NA |
| WBGene00167391 | NA | NA | NA | NA |
| WBGene00167392 | NA | NA | NA | NA |
| WBGene00167393 | NA | NA | NA | NA |
| WBGene00167394 | NA | NA | NA | NA |
| WBGene00167395 | NA | NA | NA | NA |
| WBGene00167396 | NA | NA | NA | NA |
| WBGene00167397 | NA | NA | NA | NA |
| WBGene00167398 | NA | NA | NA | NA |
| WBGene00167399 | NA | NA | NA | NA |
| WBGene00167400 | NA | NA | NA | NA |
| WBGene00167401 | NA | NA | NA | NA |
| WBGene00167402 | NA | NA | NA | NA |
| WBGene00167403 | NA | NA | NA | NA |
| WBGene00167404 | NA | NA | NA | NA |
| WBGene00167405 | NA | NA | NA | NA |
| WBGene00167406 | NA | NA | NA | NA |
| WBGene00167407 | NA | NA | NA | NA |
| WBGene00167408 | NA | NA | NA | NA |
| WBGene00167409 | NA | NA | NA | NA |
| WBGene00167410 | NA | NA | NA | NA |
| WBGene00167411 | NA | NA | NA | NA |
| WBGene00167412 | NA | NA | NA | NA |
| WBGene00167413 | NA | NA | NA | NA |
| WBGene00167414 | NA | NA | NA | NA |

| | | | | |
|---|---|---|---|---|
| WBGene00167415 | NA | NA | NA | NA |
| WBGene00167416 | NA | NA | NA | NA |
| WBGene00167417 | NA | NA | NA | NA |
| WBGene00167418 | NA | NA | NA | NA |
| WBGene00167419 | NA | NA | NA | NA |
| WBGene00167420 | NA | NA | NA | NA |
| WBGene00167421 | NA | NA | NA | NA |
| WBGene00167422 | NA | NA | NA | NA |
| WBGene00167423 | NA | NA | NA | NA |
| WBGene00167424 | NA | NA | NA | NA |
| WBGene00167425 | NA | NA | NA | NA |
| WBGene00167426 | NA | NA | NA | NA |
| WBGene00167427 | NA | NA | NA | NA |
| WBGene00167428 | NA | NA | NA | NA |
| WBGene00167429 | NA | NA | NA | NA |
| WBGene00167430 | NA | NA | NA | NA |
| WBGene00167431 | NA | NA | NA | NA |
| WBGene00167432 | NA | NA | NA | NA |
| WBGene00167433 | NA | NA | NA | NA |
| WBGene00167434 | NA | NA | NA | NA |
| WBGene00167435 | NA | NA | NA | NA |
| WBGene00167436 | NA | NA | NA | NA |
| WBGene00167437 | NA | NA | NA | NA |
| WBGene00167438 | NA | NA | NA | NA |
| WBGene00167439 | NA | NA | NA | NA |
| WBGene00167440 | NA | NA | NA | NA |
| WBGene00167441 | NA | NA | NA | NA |
| WBGene00167442 | NA | NA | NA | NA |
| WBGene00167443 | NA | NA | NA | NA |
| WBGene00167444 | NA | NA | NA | NA |
| WBGene00167445 | NA | NA | NA | NA |
| WBGene00167446 | NA | NA | NA | NA |
| WBGene00167447 | NA | NA | NA | NA |
| WBGene00167448 | NA | NA | NA | NA |
| WBGene00167449 | NA | NA | NA | NA |
| WBGene00167450 | NA | NA | NA | NA |
| WBGene00167451 | NA | NA | NA | NA |
| WBGene00167452 | NA | NA | NA | NA |
| WBGene00167453 | NA | NA | NA | NA |
| WBGene00167454 | NA | NA | NA | NA |
| WBGene00167455 | NA | NA | NA | NA |
| WBGene00167456 | NA | NA | NA | NA |
| WBGene00167457 | NA | NA | NA | NA |

| | | | | |
|---|---|---|---|---|
| WBGene00167458 | NA | NA | NA | NA |
| WBGene00167459 | NA | NA | NA | NA |
| WBGene00167460 | NA | NA | NA | NA |
| WBGene00167461 | NA | NA | NA | NA |
| WBGene00167462 | NA | NA | NA | NA |
| WBGene00167463 | NA | NA | NA | NA |
| WBGene00167464 | NA | NA | NA | NA |
| WBGene00167465 | NA | NA | NA | NA |
| WBGene00167466 | NA | NA | NA | NA |
| WBGene00167467 | NA | NA | NA | NA |
| WBGene00167468 | NA | NA | NA | NA |
| WBGene00167469 | NA | NA | NA | NA |
| WBGene00167470 | NA | NA | NA | NA |
| WBGene00167471 | NA | NA | NA | NA |
| WBGene00167472 | NA | NA | NA | NA |
| WBGene00167473 | NA | NA | NA | NA |
| WBGene00167474 | NA | NA | NA | NA |
| WBGene00167475 | NA | NA | NA | NA |
| WBGene00167476 | NA | NA | NA | NA |
| WBGene00167477 | NA | NA | NA | NA |
| WBGene00167478 | NA | NA | NA | NA |
| WBGene00167479 | NA | NA | NA | NA |
| WBGene00167480 | NA | NA | NA | NA |
| WBGene00167481 | NA | NA | NA | NA |
| WBGene00167482 | NA | NA | NA | NA |
| WBGene00167483 | NA | NA | NA | NA |
| WBGene00167484 | NA | NA | NA | NA |
| WBGene00167485 | NA | NA | NA | NA |
| WBGene00167487 | NA | NA | NA | NA |
| WBGene00167488 | NA | NA | NA | NA |
| WBGene00167489 | NA | NA | NA | NA |
| WBGene00167490 | NA | NA | NA | NA |
| WBGene00167491 | NA | NA | NA | NA |
| WBGene00167492 | NA | NA | NA | NA |
| WBGene00167493 | NA | NA | NA | NA |
| WBGene00167494 | NA | NA | NA | NA |
| WBGene00167495 | NA | NA | NA | NA |
| WBGene00167496 | NA | NA | NA | NA |
| WBGene00167497 | NA | NA | NA | NA |
| WBGene00167498 | NA | NA | NA | NA |
| WBGene00167499 | NA | NA | NA | NA |
| WBGene00167500 | NA | NA | NA | NA |
| WBGene00167501 | NA | NA | NA | NA |

| | | | | |
|---|---|---|---|---|
| WBGene00167502 | NA | NA | NA | NA |
| WBGene00167503 | NA | NA | NA | NA |
| WBGene00167504 | NA | NA | NA | NA |
| WBGene00167505 | NA | NA | NA | NA |
| WBGene00167506 | NA | NA | NA | NA |
| WBGene00167507 | NA | NA | NA | NA |
| WBGene00167508 | NA | NA | NA | NA |
| WBGene00167509 | NA | NA | NA | NA |
| WBGene00167510 | NA | NA | NA | NA |
| WBGene00167511 | NA | NA | NA | NA |
| WBGene00167512 | NA | NA | NA | NA |
| WBGene00167513 | NA | NA | NA | NA |
| WBGene00167514 | NA | NA | NA | NA |
| WBGene00167515 | NA | NA | NA | NA |
| WBGene00167516 | NA | NA | NA | NA |
| WBGene00167517 | NA | NA | NA | NA |
| WBGene00167518 | NA | NA | NA | NA |
| WBGene00167519 | NA | NA | NA | NA |
| WBGene00167520 | NA | NA | NA | NA |
| WBGene00167521 | NA | NA | NA | NA |
| WBGene00167522 | NA | NA | NA | NA |
| WBGene00167523 | NA | NA | NA | NA |
| WBGene00167524 | NA | NA | NA | NA |
| WBGene00167525 | NA | NA | NA | NA |
| WBGene00167526 | NA | NA | NA | NA |
| WBGene00167527 | NA | NA | NA | NA |
| WBGene00167528 | NA | NA | NA | NA |
| WBGene00167529 | NA | NA | NA | NA |
| WBGene00167530 | NA | NA | NA | NA |
| WBGene00167531 | NA | NA | NA | NA |
| WBGene00167532 | NA | NA | NA | NA |
| WBGene00167533 | NA | NA | NA | NA |
| WBGene00167534 | NA | NA | NA | NA |
| WBGene00167535 | NA | NA | NA | NA |
| WBGene00167536 | NA | NA | NA | NA |
| WBGene00167537 | NA | NA | NA | NA |
| WBGene00167538 | NA | NA | NA | NA |
| WBGene00167539 | NA | NA | NA | NA |
| WBGene00167540 | NA | NA | NA | NA |
| WBGene00167541 | NA | NA | NA | NA |
| WBGene00167542 | NA | NA | NA | NA |
| WBGene00167543 | NA | NA | NA | NA |
| WBGene00167544 | NA | NA | NA | NA |

| | | | | |
|---|---|---|---|---|
| WBGene00167545 | NA | NA | NA | NA |
| WBGene00167546 | NA | NA | NA | NA |
| WBGene00167547 | NA | NA | NA | NA |
| WBGene00167548 | NA | NA | NA | NA |
| WBGene00167549 | NA | NA | NA | NA |
| WBGene00167550 | NA | NA | NA | NA |
| WBGene00167551 | NA | NA | NA | NA |
| WBGene00167552 | NA | NA | NA | NA |
| WBGene00167553 | NA | NA | NA | NA |
| WBGene00167554 | NA | NA | NA | NA |
| WBGene00167555 | NA | NA | NA | NA |
| WBGene00167556 | NA | NA | NA | NA |
| WBGene00167557 | NA | NA | NA | NA |
| WBGene00167558 | NA | NA | NA | NA |
| WBGene00167559 | NA | NA | NA | NA |
| WBGene00167560 | NA | NA | NA | NA |
| WBGene00167561 | NA | NA | NA | NA |
| WBGene00167562 | NA | NA | NA | NA |
| WBGene00167563 | NA | NA | NA | NA |
| WBGene00167564 | NA | NA | NA | NA |
| WBGene00167565 | NA | NA | NA | NA |
| WBGene00167566 | NA | NA | NA | NA |
| WBGene00167567 | NA | NA | NA | NA |
| WBGene00167568 | NA | NA | NA | NA |
| WBGene00167569 | NA | NA | NA | NA |
| WBGene00167570 | NA | NA | NA | NA |
| WBGene00167571 | NA | NA | NA | NA |
| WBGene00167572 | NA | NA | NA | NA |
| WBGene00167573 | NA | NA | NA | NA |
| WBGene00167574 | NA | NA | NA | NA |
| WBGene00167575 | NA | NA | NA | NA |
| WBGene00167576 | NA | NA | NA | NA |
| WBGene00167577 | NA | NA | NA | NA |
| WBGene00167578 | NA | NA | NA | NA |
| WBGene00167579 | NA | NA | NA | NA |
| WBGene00167580 | NA | NA | NA | NA |
| WBGene00167581 | NA | NA | NA | NA |
| WBGene00167582 | NA | NA | NA | NA |
| WBGene00167583 | NA | NA | NA | NA |
| WBGene00167584 | NA | NA | NA | NA |
| WBGene00167585 | NA | NA | NA | NA |
| WBGene00167586 | NA | NA | NA | NA |
| WBGene00167587 | NA | NA | NA | NA |

| | | | | |
|---|---|---|---|---|
| WBGene00167588 | NA | NA | NA | NA |
| WBGene00167589 | NA | NA | NA | NA |
| WBGene00167590 | NA | NA | NA | NA |
| WBGene00167591 | NA | NA | NA | NA |
| WBGene00167592 | NA | NA | NA | NA |
| WBGene00167593 | NA | NA | NA | NA |
| WBGene00167594 | NA | NA | NA | NA |
| WBGene00167595 | NA | NA | NA | NA |
| WBGene00167596 | NA | NA | NA | NA |
| WBGene00167597 | NA | NA | NA | NA |
| WBGene00167598 | NA | NA | NA | NA |
| WBGene00167599 | NA | NA | NA | NA |
| WBGene00167600 | NA | NA | NA | NA |
| WBGene00167601 | NA | NA | NA | NA |
| WBGene00167602 | NA | NA | NA | NA |
| WBGene00167603 | NA | NA | NA | NA |
| WBGene00167604 | NA | NA | NA | NA |
| WBGene00167605 | NA | NA | NA | NA |
| WBGene00167606 | NA | NA | NA | NA |
| WBGene00167607 | NA | NA | NA | NA |
| WBGene00167608 | NA | NA | NA | NA |
| WBGene00167609 | NA | NA | NA | NA |
| WBGene00167610 | NA | NA | NA | NA |
| WBGene00167611 | NA | NA | NA | NA |
| WBGene00167612 | NA | NA | NA | NA |
| WBGene00167613 | NA | NA | NA | NA |
| WBGene00167614 | NA | NA | NA | NA |
| WBGene00167615 | NA | NA | NA | NA |
| WBGene00167616 | NA | NA | NA | NA |
| WBGene00167617 | NA | NA | NA | NA |
| WBGene00167618 | NA | NA | NA | NA |
| WBGene00167619 | NA | NA | NA | NA |
| WBGene00167620 | NA | NA | NA | NA |
| WBGene00167621 | NA | NA | NA | NA |
| WBGene00167622 | NA | NA | NA | NA |
| WBGene00167623 | NA | NA | NA | NA |
| WBGene00167624 | NA | NA | NA | NA |
| WBGene00167625 | NA | NA | NA | NA |
| WBGene00167626 | NA | NA | NA | NA |
| WBGene00167627 | NA | NA | NA | NA |
| WBGene00167628 | NA | NA | NA | NA |
| WBGene00167629 | NA | NA | NA | NA |
| WBGene00167630 | NA | NA | NA | NA |

| | | | | |
|---|---|---|---|---|
| WBGene00167631 | NA | NA | NA | NA |
| WBGene00167632 | NA | NA | NA | NA |
| WBGene00167633 | NA | NA | NA | NA |
| WBGene00167634 | NA | NA | NA | NA |
| WBGene00167635 | NA | NA | NA | NA |
| WBGene00167636 | NA | NA | NA | NA |
| WBGene00167637 | NA | NA | NA | NA |
| WBGene00167638 | NA | NA | NA | NA |
| WBGene00167639 | NA | NA | NA | NA |
| WBGene00167640 | NA | NA | NA | NA |
| WBGene00167641 | NA | NA | NA | NA |
| WBGene00167642 | NA | NA | NA | NA |
| WBGene00167643 | NA | NA | NA | NA |
| WBGene00167644 | NA | NA | NA | NA |
| WBGene00167645 | NA | NA | NA | NA |
| WBGene00167646 | NA | NA | NA | NA |
| WBGene00167647 | NA | NA | NA | NA |
| WBGene00167648 | NA | NA | NA | NA |
| WBGene00167649 | NA | NA | NA | NA |
| WBGene00167650 | NA | NA | NA | NA |
| WBGene00167651 | NA | NA | NA | NA |
| WBGene00167652 | NA | NA | NA | NA |
| WBGene00167653 | NA | NA | NA | NA |
| WBGene00167654 | NA | NA | NA | NA |
| WBGene00167655 | NA | NA | NA | NA |
| WBGene00167656 | NA | NA | NA | NA |
| WBGene00167657 | NA | NA | NA | NA |
| WBGene00167658 | NA | NA | NA | NA |
| WBGene00167659 | NA | NA | NA | NA |
| WBGene00167660 | NA | NA | NA | NA |
| WBGene00167661 | NA | NA | NA | NA |
| WBGene00167662 | NA | NA | NA | NA |
| WBGene00167663 | NA | NA | NA | NA |
| WBGene00167664 | NA | NA | NA | NA |
| WBGene00167665 | NA | NA | NA | NA |
| WBGene00167666 | NA | NA | NA | NA |
| WBGene00167667 | NA | NA | NA | NA |
| WBGene00167668 | NA | NA | NA | NA |
| WBGene00167669 | NA | NA | NA | NA |
| WBGene00167670 | NA | NA | NA | NA |
| WBGene00167671 | NA | NA | NA | NA |
| WBGene00167672 | NA | NA | NA | NA |
| WBGene00167673 | NA | NA | NA | NA |

| | | | | |
|---|---|---|---|---|
| WBGene00167674 | NA | NA | NA | NA |
| WBGene00167675 | NA | NA | NA | NA |
| WBGene00167676 | -0.00346345 | 0.95708614 | NA | NA |
| WBGene00167677 | NA | NA | NA | NA |
| WBGene00167678 | NA | NA | NA | NA |
| WBGene00167679 | NA | NA | NA | NA |
| WBGene00167680 | NA | NA | NA | NA |
| WBGene00167681 | NA | NA | NA | NA |
| WBGene00167682 | NA | NA | NA | NA |
| WBGene00167683 | NA | NA | NA | NA |
| WBGene00167684 | NA | NA | NA | NA |
| WBGene00167685 | NA | NA | NA | NA |
| WBGene00167686 | NA | NA | NA | NA |
| WBGene00167687 | NA | NA | NA | NA |
| WBGene00167688 | NA | NA | NA | NA |
| WBGene00167689 | NA | NA | NA | NA |
| WBGene00167690 | NA | NA | NA | NA |
| WBGene00167691 | NA | NA | NA | NA |
| WBGene00167692 | NA | NA | NA | NA |
| WBGene00167693 | NA | NA | NA | NA |
| WBGene00167694 | NA | NA | NA | NA |
| WBGene00167695 | NA | NA | NA | NA |
| WBGene00167696 | NA | NA | NA | NA |
| WBGene00167697 | NA | NA | NA | NA |
| WBGene00167698 | NA | NA | NA | NA |
| WBGene00167699 | NA | NA | NA | NA |
| WBGene00167700 | NA | NA | NA | NA |
| WBGene00167701 | NA | NA | NA | NA |
| WBGene00167702 | NA | NA | NA | NA |
| WBGene00167703 | NA | NA | NA | NA |
| WBGene00167704 | NA | NA | NA | NA |
| WBGene00167705 | NA | NA | NA | NA |
| WBGene00167706 | NA | NA | NA | NA |
| WBGene00167707 | NA | NA | NA | NA |
| WBGene00167708 | NA | NA | NA | NA |
| WBGene00167709 | NA | NA | NA | NA |
| WBGene00167710 | NA | NA | NA | NA |
| WBGene00167711 | NA | NA | NA | NA |
| WBGene00167712 | NA | NA | NA | NA |
| WBGene00167713 | NA | NA | NA | NA |
| WBGene00167714 | NA | NA | NA | NA |
| WBGene00167715 | NA | NA | NA | NA |
| WBGene00167716 | NA | NA | NA | NA |

| | | | | |
|---|---|---|---|---|
| WBGene00167717 | NA | NA | NA | NA |
| WBGene00167718 | NA | NA | NA | NA |
| WBGene00167719 | NA | NA | NA | NA |
| WBGene00167720 | NA | NA | NA | NA |
| WBGene00167721 | NA | NA | NA | NA |
| WBGene00167722 | NA | NA | NA | NA |
| WBGene00167723 | NA | NA | NA | NA |
| WBGene00167724 | NA | NA | NA | NA |
| WBGene00167725 | NA | NA | NA | NA |
| WBGene00167726 | NA | NA | NA | NA |
| WBGene00167727 | NA | NA | NA | NA |
| WBGene00167728 | NA | NA | NA | NA |
| WBGene00167729 | NA | NA | NA | NA |
| WBGene00167730 | NA | NA | NA | NA |
| WBGene00167731 | NA | NA | NA | NA |
| WBGene00167732 | NA | NA | NA | NA |
| WBGene00167733 | NA | NA | NA | NA |
| WBGene00167734 | NA | NA | NA | NA |
| WBGene00167735 | NA | NA | NA | NA |
| WBGene00167736 | NA | NA | NA | NA |
| WBGene00167737 | NA | NA | NA | NA |
| WBGene00167738 | NA | NA | NA | NA |
| WBGene00167739 | NA | NA | NA | NA |
| WBGene00167740 | NA | NA | NA | NA |
| WBGene00167741 | NA | NA | NA | NA |
| WBGene00167742 | NA | NA | NA | NA |
| WBGene00167743 | NA | NA | NA | NA |
| WBGene00167744 | NA | NA | NA | NA |
| WBGene00167745 | NA | NA | NA | NA |
| WBGene00167746 | NA | NA | NA | NA |
| WBGene00167747 | NA | NA | NA | NA |
| WBGene00167748 | NA | NA | NA | NA |
| WBGene00167749 | NA | NA | NA | NA |
| WBGene00167750 | NA | NA | NA | NA |
| WBGene00167751 | NA | NA | NA | NA |
| WBGene00167752 | NA | NA | NA | NA |
| WBGene00167753 | NA | NA | NA | NA |
| WBGene00167754 | NA | NA | NA | NA |
| WBGene00167755 | NA | NA | NA | NA |
| WBGene00167756 | NA | NA | NA | NA |
| WBGene00167757 | NA | NA | NA | NA |
| WBGene00167758 | NA | NA | NA | NA |
| WBGene00167759 | NA | NA | NA | NA |

| | | | | |
|---|---|---|---|---|
| WBGene00167760 | NA | NA | NA | NA |
| WBGene00167761 | NA | NA | NA | NA |
| WBGene00167762 | NA | NA | NA | NA |
| WBGene00167763 | NA | NA | NA | NA |
| WBGene00167764 | NA | NA | NA | NA |
| WBGene00167765 | NA | NA | NA | NA |
| WBGene00167766 | NA | NA | NA | NA |
| WBGene00167767 | NA | NA | NA | NA |
| WBGene00167768 | NA | NA | NA | NA |
| WBGene00167769 | NA | NA | NA | NA |
| WBGene00167770 | NA | NA | NA | NA |
| WBGene00167771 | NA | NA | NA | NA |
| WBGene00167772 | NA | NA | NA | NA |
| WBGene00167773 | NA | NA | NA | NA |
| WBGene00167774 | NA | NA | NA | NA |
| WBGene00167775 | NA | NA | NA | NA |
| WBGene00167776 | NA | NA | NA | NA |
| WBGene00167777 | NA | NA | NA | NA |
| WBGene00167778 | NA | NA | NA | NA |
| WBGene00167779 | NA | NA | NA | NA |
| WBGene00167780 | NA | NA | NA | NA |
| WBGene00167781 | NA | NA | NA | NA |
| WBGene00167782 | NA | NA | NA | NA |
| WBGene00167783 | NA | NA | NA | NA |
| WBGene00167784 | NA | NA | NA | NA |
| WBGene00167785 | NA | NA | NA | NA |
| WBGene00167786 | NA | NA | NA | NA |
| WBGene00167787 | NA | NA | NA | NA |
| WBGene00167788 | NA | NA | NA | NA |
| WBGene00167789 | NA | NA | NA | NA |
| WBGene00167790 | NA | NA | NA | NA |
| WBGene00167791 | NA | NA | NA | NA |
| WBGene00167792 | NA | NA | NA | NA |
| WBGene00167793 | NA | NA | NA | NA |
| WBGene00167794 | NA | NA | NA | NA |
| WBGene00167795 | NA | NA | NA | NA |
| WBGene00167796 | NA | NA | NA | NA |
| WBGene00167797 | NA | NA | NA | NA |
| WBGene00167799 | NA | NA | NA | NA |
| WBGene00167800 | NA | NA | NA | NA |
| WBGene00167801 | NA | NA | NA | NA |
| WBGene00167802 | NA | NA | NA | NA |
| WBGene00167803 | NA | NA | NA | NA |

| | | | | |
|---|---|---|---|---|
| WBGene00167804 | NA | NA | NA | NA |
| WBGene00167805 | NA | NA | NA | NA |
| WBGene00167806 | NA | NA | NA | NA |
| WBGene00167807 | NA | NA | NA | NA |
| WBGene00167808 | NA | NA | NA | NA |
| WBGene00167809 | NA | NA | NA | NA |
| WBGene00167810 | NA | NA | NA | NA |
| WBGene00167811 | NA | NA | NA | NA |
| WBGene00167812 | NA | NA | NA | NA |
| WBGene00167813 | NA | NA | NA | NA |
| WBGene00167814 | NA | NA | NA | NA |
| WBGene00167815 | NA | NA | NA | NA |
| WBGene00167816 | NA | NA | NA | NA |
| WBGene00167817 | NA | NA | NA | NA |
| WBGene00167818 | NA | NA | NA | NA |
| WBGene00167819 | NA | NA | NA | NA |
| WBGene00167820 | NA | NA | NA | NA |
| WBGene00167821 | NA | NA | NA | NA |
| WBGene00167822 | NA | NA | NA | NA |
| WBGene00167823 | NA | NA | NA | NA |
| WBGene00167824 | NA | NA | NA | NA |
| WBGene00167825 | NA | NA | NA | NA |
| WBGene00167826 | NA | NA | NA | NA |
| WBGene00167827 | NA | NA | NA | NA |
| WBGene00167828 | NA | NA | NA | NA |
| WBGene00167829 | NA | NA | NA | NA |
| WBGene00167830 | NA | NA | NA | NA |
| WBGene00167831 | NA | NA | NA | NA |
| WBGene00167832 | NA | NA | NA | NA |
| WBGene00167833 | NA | NA | NA | NA |
| WBGene00167834 | NA | NA | NA | NA |
| WBGene00167835 | NA | NA | NA | NA |
| WBGene00167836 | NA | NA | NA | NA |
| WBGene00167837 | NA | NA | NA | NA |
| WBGene00167838 | NA | NA | NA | NA |
| WBGene00167839 | NA | NA | NA | NA |
| WBGene00167840 | NA | NA | NA | NA |
| WBGene00167841 | NA | NA | NA | NA |
| WBGene00167842 | NA | NA | NA | NA |
| WBGene00167843 | NA | NA | NA | NA |
| WBGene00167845 | NA | NA | NA | NA |
| WBGene00167846 | NA | NA | NA | NA |
| WBGene00167847 | NA | NA | NA | NA |

| | | | | |
|---|---|---|---|---|
| WBGene00167848 | NA | NA | NA | NA |
| WBGene00167849 | NA | NA | NA | NA |
| WBGene00167850 | NA | NA | NA | NA |
| WBGene00167851 | NA | NA | NA | NA |
| WBGene00167852 | NA | NA | NA | NA |
| WBGene00167853 | NA | NA | NA | NA |
| WBGene00167854 | NA | NA | NA | NA |
| WBGene00167855 | NA | NA | NA | NA |
| WBGene00167856 | NA | NA | NA | NA |
| WBGene00167857 | NA | NA | NA | NA |
| WBGene00167858 | NA | NA | NA | NA |
| WBGene00167859 | NA | NA | NA | NA |
| WBGene00167860 | NA | NA | NA | NA |
| WBGene00167861 | NA | NA | NA | NA |
| WBGene00167862 | NA | NA | NA | NA |
| WBGene00167863 | NA | NA | NA | NA |
| WBGene00167864 | NA | NA | NA | NA |
| WBGene00167865 | NA | NA | NA | NA |
| WBGene00167866 | NA | NA | NA | NA |
| WBGene00167867 | NA | NA | NA | NA |
| WBGene00167868 | NA | NA | NA | NA |
| WBGene00167869 | NA | NA | NA | NA |
| WBGene00167870 | NA | NA | NA | NA |
| WBGene00167871 | NA | NA | NA | NA |
| WBGene00167872 | NA | NA | NA | NA |
| WBGene00167873 | NA | NA | NA | NA |
| WBGene00167874 | NA | NA | NA | NA |
| WBGene00167875 | NA | NA | NA | NA |
| WBGene00167876 | NA | NA | NA | NA |
| WBGene00167877 | NA | NA | NA | NA |
| WBGene00167878 | NA | NA | NA | NA |
| WBGene00167879 | NA | NA | NA | NA |
| WBGene00167880 | NA | NA | NA | NA |
| WBGene00167881 | NA | NA | NA | NA |
| WBGene00167882 | NA | NA | NA | NA |
| WBGene00167883 | NA | NA | NA | NA |
| WBGene00167884 | NA | NA | NA | NA |
| WBGene00167885 | NA | NA | NA | NA |
| WBGene00167886 | NA | NA | NA | NA |
| WBGene00167887 | NA | NA | NA | NA |
| WBGene00167888 | NA | NA | NA | NA |
| WBGene00167889 | NA | NA | NA | NA |
| WBGene00167890 | NA | NA | NA | NA |

| | | | | |
|---|---|---|---|---|
| WBGene00167891 | NA | NA | NA | NA |
| WBGene00167892 | NA | NA | NA | NA |
| WBGene00167893 | NA | NA | NA | NA |
| WBGene00167894 | NA | NA | NA | NA |
| WBGene00167895 | NA | NA | NA | NA |
| WBGene00167896 | NA | NA | NA | NA |
| WBGene00167897 | NA | NA | NA | NA |
| WBGene00167898 | NA | NA | NA | NA |
| WBGene00167899 | NA | NA | NA | NA |
| WBGene00167900 | NA | NA | NA | NA |
| WBGene00167901 | NA | NA | NA | NA |
| WBGene00167902 | NA | NA | NA | NA |
| WBGene00167903 | NA | NA | NA | NA |
| WBGene00167904 | NA | NA | NA | NA |
| WBGene00167905 | NA | NA | NA | NA |
| WBGene00167906 | NA | NA | NA | NA |
| WBGene00167907 | NA | NA | NA | NA |
| WBGene00167908 | NA | NA | NA | NA |
| WBGene00167909 | 0.020232312 | 0.84643798 | NA | NA |
| WBGene00167910 | NA | NA | NA | NA |
| WBGene00167911 | NA | NA | NA | NA |
| WBGene00167912 | NA | NA | NA | NA |
| WBGene00167913 | NA | NA | NA | NA |
| WBGene00167914 | NA | NA | NA | NA |
| WBGene00167915 | NA | NA | NA | NA |
| WBGene00167916 | NA | NA | NA | NA |
| WBGene00167917 | NA | NA | NA | NA |
| WBGene00167918 | NA | NA | NA | NA |
| WBGene00167919 | NA | NA | NA | NA |
| WBGene00167920 | NA | NA | NA | NA |
| WBGene00167921 | NA | NA | NA | NA |
| WBGene00167922 | NA | NA | NA | NA |
| WBGene00167923 | NA | NA | NA | NA |
| WBGene00167924 | NA | NA | NA | NA |
| WBGene00167925 | NA | NA | NA | NA |
| WBGene00167926 | NA | NA | NA | NA |
| WBGene00167927 | NA | NA | NA | NA |
| WBGene00167928 | NA | NA | NA | NA |
| WBGene00167929 | NA | NA | NA | NA |
| WBGene00167930 | NA | NA | NA | NA |
| WBGene00167931 | NA | NA | NA | NA |
| WBGene00167932 | NA | NA | NA | NA |
| WBGene00167933 | NA | NA | NA | NA |

| | | | | |
|---|---|---|---|---|
| WBGene00167934 | NA | NA | NA | NA |
| WBGene00167935 | NA | NA | NA | NA |
| WBGene00167936 | NA | NA | NA | NA |
| WBGene00167937 | NA | NA | NA | NA |
| WBGene00167938 | NA | NA | NA | NA |
| WBGene00167939 | NA | NA | NA | NA |
| WBGene00167940 | NA | NA | NA | NA |
| WBGene00167941 | NA | NA | NA | NA |
| WBGene00167942 | NA | NA | NA | NA |
| WBGene00167943 | NA | NA | NA | NA |
| WBGene00167944 | NA | NA | NA | NA |
| WBGene00167945 | NA | NA | NA | NA |
| WBGene00167946 | NA | NA | NA | NA |
| WBGene00167947 | NA | NA | NA | NA |
| WBGene00167948 | NA | NA | NA | NA |
| WBGene00167949 | NA | NA | NA | NA |
| WBGene00167950 | NA | NA | NA | NA |
| WBGene00167951 | NA | NA | NA | NA |
| WBGene00167952 | NA | NA | NA | NA |
| WBGene00167953 | NA | NA | NA | NA |
| WBGene00167954 | NA | NA | NA | NA |
| WBGene00167955 | NA | NA | NA | NA |
| WBGene00167956 | NA | NA | NA | NA |
| WBGene00167957 | 0.020317245 | 0.84643798 | NA | NA |
| WBGene00167958 | NA | NA | NA | NA |
| WBGene00167959 | NA | NA | NA | NA |
| WBGene00167960 | NA | NA | NA | NA |
| WBGene00167961 | NA | NA | NA | NA |
| WBGene00167962 | NA | NA | NA | NA |
| WBGene00167963 | NA | NA | NA | NA |
| WBGene00167964 | NA | NA | NA | NA |
| WBGene00167965 | NA | NA | NA | NA |
| WBGene00167966 | NA | NA | NA | NA |
| WBGene00167967 | NA | NA | NA | NA |
| WBGene00167968 | NA | NA | NA | NA |
| WBGene00167969 | NA | NA | NA | NA |
| WBGene00167970 | NA | NA | NA | NA |
| WBGene00167971 | NA | NA | NA | NA |
| WBGene00167972 | NA | NA | NA | NA |
| WBGene00167973 | NA | NA | NA | NA |
| WBGene00167974 | NA | NA | NA | NA |
| WBGene00167975 | NA | NA | NA | NA |
| WBGene00167976 | NA | NA | NA | NA |

| | | | | |
|---|---|---|---|---|
| WBGene00167977 | NA | NA | NA | NA |
| WBGene00167978 | NA | NA | NA | NA |
| WBGene00167979 | NA | NA | NA | NA |
| WBGene00167980 | NA | NA | NA | NA |
| WBGene00167981 | NA | NA | NA | NA |
| WBGene00167982 | NA | NA | NA | NA |
| WBGene00167983 | NA | NA | NA | NA |
| WBGene00167984 | NA | NA | NA | NA |
| WBGene00167985 | NA | NA | NA | NA |
| WBGene00167986 | NA | NA | NA | NA |
| WBGene00167987 | NA | NA | NA | NA |
| WBGene00167988 | NA | NA | NA | NA |
| WBGene00167989 | NA | NA | NA | NA |
| WBGene00167990 | NA | NA | NA | NA |
| WBGene00167991 | NA | NA | NA | NA |
| WBGene00167992 | NA | NA | NA | NA |
| WBGene00167993 | NA | NA | NA | NA |
| WBGene00167994 | NA | NA | NA | NA |
| WBGene00167995 | NA | NA | NA | NA |
| WBGene00167996 | NA | NA | NA | NA |
| WBGene00167997 | NA | NA | NA | NA |
| WBGene00167998 | NA | NA | NA | NA |
| WBGene00167999 | NA | NA | NA | NA |
| WBGene00168000 | NA | NA | NA | NA |
| WBGene00168001 | NA | NA | NA | NA |
| WBGene00168002 | NA | NA | NA | NA |
| WBGene00168003 | NA | NA | NA | NA |
| WBGene00168004 | NA | NA | NA | NA |
| WBGene00168005 | NA | NA | NA | NA |
| WBGene00168006 | NA | NA | NA | NA |
| WBGene00168007 | NA | NA | NA | NA |
| WBGene00168008 | NA | NA | NA | NA |
| WBGene00168009 | NA | NA | NA | NA |
| WBGene00168010 | NA | NA | NA | NA |
| WBGene00168011 | NA | NA | NA | NA |
| WBGene00168012 | NA | NA | NA | NA |
| WBGene00168013 | 0.020317245 | 0.84643798 | NA | NA |
| WBGene00168014 | NA | NA | NA | NA |
| WBGene00168015 | NA | NA | NA | NA |
| WBGene00168016 | NA | NA | NA | NA |
| WBGene00168017 | NA | NA | NA | NA |
| WBGene00168018 | NA | NA | NA | NA |
| WBGene00168019 | NA | NA | NA | NA |

| | | | | |
|---|---|---|---|---|
| WBGene00168020 | NA | NA | NA | NA |
| WBGene00168021 | NA | NA | NA | NA |
| WBGene00168022 | NA | NA | NA | NA |
| WBGene00168023 | NA | NA | NA | NA |
| WBGene00168024 | NA | NA | NA | NA |
| WBGene00168025 | NA | NA | NA | NA |
| WBGene00168026 | NA | NA | NA | NA |
| WBGene00168027 | NA | NA | NA | NA |
| WBGene00168028 | NA | NA | NA | NA |
| WBGene00168029 | NA | NA | NA | NA |
| WBGene00168030 | NA | NA | NA | NA |
| WBGene00168031 | NA | NA | NA | NA |
| WBGene00168032 | NA | NA | NA | NA |
| WBGene00168033 | NA | NA | NA | NA |
| WBGene00168034 | NA | NA | NA | NA |
| WBGene00168035 | NA | NA | NA | NA |
| WBGene00168036 | NA | NA | NA | NA |
| WBGene00168037 | NA | NA | NA | NA |
| WBGene00168038 | NA | NA | NA | NA |
| WBGene00168039 | NA | NA | NA | NA |
| WBGene00168040 | NA | NA | NA | NA |
| WBGene00168041 | NA | NA | NA | NA |
| WBGene00168042 | NA | NA | NA | NA |
| WBGene00168043 | NA | NA | NA | NA |
| WBGene00168044 | NA | NA | NA | NA |
| WBGene00168045 | NA | NA | NA | NA |
| WBGene00168046 | NA | NA | NA | NA |
| WBGene00168047 | NA | NA | NA | NA |
| WBGene00168048 | NA | NA | NA | NA |
| WBGene00168049 | NA | NA | NA | NA |
| WBGene00168050 | NA | NA | NA | NA |
| WBGene00168051 | NA | NA | NA | NA |
| WBGene00168052 | NA | NA | NA | NA |
| WBGene00168053 | NA | NA | NA | NA |
| WBGene00168054 | NA | NA | NA | NA |
| WBGene00168055 | NA | NA | NA | NA |
| WBGene00168056 | NA | NA | NA | NA |
| WBGene00168057 | NA | NA | NA | NA |
| WBGene00168058 | NA | NA | NA | NA |
| WBGene00168059 | NA | NA | NA | NA |
| WBGene00168060 | NA | NA | NA | NA |
| WBGene00168061 | NA | NA | NA | NA |
| WBGene00168062 | NA | NA | NA | NA |

| | | | | |
|---|---|---|---|---|
| WBGene00168063 | NA | NA | NA | NA |
| WBGene00168064 | NA | NA | NA | NA |
| WBGene00168065 | NA | NA | NA | NA |
| WBGene00168066 | NA | NA | NA | NA |
| WBGene00168067 | NA | NA | NA | NA |
| WBGene00168068 | NA | NA | NA | NA |
| WBGene00168069 | NA | NA | NA | NA |
| WBGene00168070 | NA | NA | NA | NA |
| WBGene00168071 | NA | NA | NA | NA |
| WBGene00168072 | NA | NA | NA | NA |
| WBGene00168073 | NA | NA | NA | NA |
| WBGene00168074 | NA | NA | NA | NA |
| WBGene00168075 | NA | NA | NA | NA |
| WBGene00168076 | NA | NA | NA | NA |
| WBGene00168077 | NA | NA | NA | NA |
| WBGene00168078 | NA | NA | NA | NA |
| WBGene00168079 | NA | NA | NA | NA |
| WBGene00168080 | NA | NA | NA | NA |
| WBGene00168081 | NA | NA | NA | NA |
| WBGene00168082 | NA | NA | NA | NA |
| WBGene00168083 | NA | NA | NA | NA |
| WBGene00168084 | NA | NA | NA | NA |
| WBGene00168085 | NA | NA | NA | NA |
| WBGene00168086 | NA | NA | NA | NA |
| WBGene00168087 | NA | NA | NA | NA |
| WBGene00168088 | NA | NA | NA | NA |
| WBGene00168089 | NA | NA | NA | NA |
| WBGene00168090 | NA | NA | NA | NA |
| WBGene00168091 | NA | NA | NA | NA |
| WBGene00168092 | NA | NA | NA | NA |
| WBGene00168093 | NA | NA | NA | NA |
| WBGene00168094 | NA | NA | NA | NA |
| WBGene00168095 | NA | NA | NA | NA |
| WBGene00168096 | NA | NA | NA | NA |
| WBGene00168097 | NA | NA | NA | NA |
| WBGene00168098 | NA | NA | NA | NA |
| WBGene00168099 | NA | NA | NA | NA |
| WBGene00168100 | NA | NA | NA | NA |
| WBGene00168101 | NA | NA | NA | NA |
| WBGene00168102 | NA | NA | NA | NA |
| WBGene00168103 | NA | NA | NA | NA |
| WBGene00168104 | NA | NA | NA | NA |
| WBGene00168105 | NA | NA | NA | NA |

| | | | | |
|---|---|---|---|---|
| WBGene00168106 | NA | NA | NA | NA |
| WBGene00168107 | NA | NA | NA | NA |
| WBGene00168108 | NA | NA | NA | NA |
| WBGene00168109 | NA | NA | NA | NA |
| WBGene00168110 | NA | NA | NA | NA |
| WBGene00168111 | NA | NA | NA | NA |
| WBGene00168112 | NA | NA | NA | NA |
| WBGene00168113 | NA | NA | NA | NA |
| WBGene00168114 | NA | NA | NA | NA |
| WBGene00168115 | NA | NA | NA | NA |
| WBGene00168116 | NA | NA | NA | NA |
| WBGene00168117 | NA | NA | NA | NA |
| WBGene00168118 | NA | NA | NA | NA |
| WBGene00168119 | NA | NA | NA | NA |
| WBGene00168120 | NA | NA | NA | NA |
| WBGene00168121 | NA | NA | NA | NA |
| WBGene00168122 | NA | NA | NA | NA |
| WBGene00168123 | NA | NA | NA | NA |
| WBGene00168124 | NA | NA | NA | NA |
| WBGene00168125 | NA | NA | NA | NA |
| WBGene00168126 | NA | NA | NA | NA |
| WBGene00168127 | NA | NA | NA | NA |
| WBGene00168128 | NA | NA | NA | NA |
| WBGene00168129 | NA | NA | NA | NA |
| WBGene00168130 | NA | NA | NA | NA |
| WBGene00168131 | NA | NA | NA | NA |
| WBGene00168132 | NA | NA | NA | NA |
| WBGene00168133 | NA | NA | NA | NA |
| WBGene00168134 | NA | NA | NA | NA |
| WBGene00168135 | NA | NA | NA | NA |
| WBGene00168136 | NA | NA | NA | NA |
| WBGene00168137 | NA | NA | NA | NA |
| WBGene00168138 | NA | NA | NA | NA |
| WBGene00168139 | NA | NA | NA | NA |
| WBGene00168140 | NA | NA | NA | NA |
| WBGene00168141 | NA | NA | NA | NA |
| WBGene00168142 | NA | NA | NA | NA |
| WBGene00168143 | NA | NA | NA | NA |
| WBGene00168144 | NA | NA | NA | NA |
| WBGene00168145 | NA | NA | NA | NA |
| WBGene00168146 | NA | NA | NA | NA |
| WBGene00168147 | NA | NA | NA | NA |
| WBGene00168148 | NA | NA | NA | NA |

| | | | | |
|---|---|---|---|---|
| WBGene00168149 | NA | NA | NA | NA |
| WBGene00168150 | NA | NA | NA | NA |
| WBGene00168151 | NA | NA | NA | NA |
| WBGene00168152 | NA | NA | NA | NA |
| WBGene00168153 | NA | NA | NA | NA |
| WBGene00168154 | NA | NA | NA | NA |
| WBGene00168155 | NA | NA | NA | NA |
| WBGene00168156 | NA | NA | NA | NA |
| WBGene00168157 | NA | NA | NA | NA |
| WBGene00168158 | NA | NA | NA | NA |
| WBGene00168159 | NA | NA | NA | NA |
| WBGene00168160 | NA | NA | NA | NA |
| WBGene00168161 | NA | NA | NA | NA |
| WBGene00168162 | NA | NA | NA | NA |
| WBGene00168163 | NA | NA | NA | NA |
| WBGene00168164 | NA | NA | NA | NA |
| WBGene00168165 | NA | NA | NA | NA |
| WBGene00168166 | NA | NA | NA | NA |
| WBGene00168167 | NA | NA | NA | NA |
| WBGene00168168 | NA | NA | NA | NA |
| WBGene00168169 | NA | NA | NA | NA |
| WBGene00168170 | NA | NA | NA | NA |
| WBGene00168171 | NA | NA | NA | NA |
| WBGene00168172 | NA | NA | NA | NA |
| WBGene00168173 | NA | NA | NA | NA |
| WBGene00168174 | NA | NA | NA | NA |
| WBGene00168175 | NA | NA | NA | NA |
| WBGene00168176 | NA | NA | NA | NA |
| WBGene00168177 | NA | NA | NA | NA |
| WBGene00168178 | NA | NA | NA | NA |
| WBGene00168179 | NA | NA | NA | NA |
| WBGene00168180 | NA | NA | NA | NA |
| WBGene00168181 | NA | NA | NA | NA |
| WBGene00168182 | NA | NA | NA | NA |
| WBGene00168183 | NA | NA | NA | NA |
| WBGene00168184 | NA | NA | NA | NA |
| WBGene00168185 | NA | NA | NA | NA |
| WBGene00168186 | NA | NA | NA | NA |
| WBGene00168187 | NA | NA | NA | NA |
| WBGene00168188 | NA | NA | NA | NA |
| WBGene00168189 | NA | NA | NA | NA |
| WBGene00168190 | NA | NA | NA | NA |
| WBGene00168191 | NA | NA | NA | NA |

| | | | | |
|---|---|---|---|---|
| WBGene00168192 | NA | NA | NA | NA |
| WBGene00168193 | NA | NA | NA | NA |
| WBGene00168194 | NA | NA | NA | NA |
| WBGene00168195 | NA | NA | NA | NA |
| WBGene00168196 | NA | NA | NA | NA |
| WBGene00168197 | NA | NA | NA | NA |
| WBGene00168198 | NA | NA | NA | NA |
| WBGene00168199 | NA | NA | NA | NA |
| WBGene00168200 | NA | NA | NA | NA |
| WBGene00168201 | NA | NA | NA | NA |
| WBGene00168202 | NA | NA | NA | NA |
| WBGene00168203 | NA | NA | NA | NA |
| WBGene00168204 | NA | NA | NA | NA |
| WBGene00168205 | NA | NA | NA | NA |
| WBGene00168206 | NA | NA | NA | NA |
| WBGene00168207 | NA | NA | NA | NA |
| WBGene00168208 | NA | NA | NA | NA |
| WBGene00168209 | NA | NA | NA | NA |
| WBGene00168210 | NA | NA | NA | NA |
| WBGene00168211 | NA | NA | NA | NA |
| WBGene00168212 | NA | NA | NA | NA |
| WBGene00168213 | NA | NA | NA | NA |
| WBGene00168214 | NA | NA | NA | NA |
| WBGene00168215 | NA | NA | NA | NA |
| WBGene00168216 | NA | NA | NA | NA |
| WBGene00168217 | NA | NA | NA | NA |
| WBGene00168218 | NA | NA | NA | NA |
| WBGene00168219 | NA | NA | NA | NA |
| WBGene00168220 | NA | NA | NA | NA |
| WBGene00168221 | NA | NA | NA | NA |
| WBGene00168222 | NA | NA | NA | NA |
| WBGene00168223 | NA | NA | NA | NA |
| WBGene00168224 | NA | NA | NA | NA |
| WBGene00168225 | NA | NA | NA | NA |
| WBGene00168226 | NA | NA | NA | NA |
| WBGene00168227 | NA | NA | NA | NA |
| WBGene00168228 | NA | NA | NA | NA |
| WBGene00168229 | NA | NA | NA | NA |
| WBGene00168230 | NA | NA | NA | NA |
| WBGene00168231 | NA | NA | NA | NA |
| WBGene00168232 | NA | NA | NA | NA |
| WBGene00168233 | NA | NA | NA | NA |
| WBGene00168234 | NA | NA | NA | NA |

| | | | | |
|---|---|---|---|---|
| WBGene00168235 | NA | NA | NA | NA |
| WBGene00168236 | 0.020317245 | 0.84643798 | NA | NA |
| WBGene00168237 | NA | NA | NA | NA |
| WBGene00168238 | NA | NA | NA | NA |
| WBGene00168239 | NA | NA | NA | NA |
| WBGene00168240 | NA | NA | NA | NA |
| WBGene00168241 | NA | NA | NA | NA |
| WBGene00168242 | NA | NA | NA | NA |
| WBGene00168243 | NA | NA | NA | NA |
| WBGene00168244 | NA | NA | NA | NA |
| WBGene00168245 | NA | NA | NA | NA |
| WBGene00168246 | NA | NA | NA | NA |
| WBGene00168247 | NA | NA | NA | NA |
| WBGene00168248 | NA | NA | NA | NA |
| WBGene00168249 | NA | NA | NA | NA |
| WBGene00168250 | NA | NA | NA | NA |
| WBGene00168251 | NA | NA | NA | NA |
| WBGene00168252 | NA | NA | NA | NA |
| WBGene00168253 | NA | NA | NA | NA |
| WBGene00168254 | NA | NA | NA | NA |
| WBGene00168255 | NA | NA | NA | NA |
| WBGene00168256 | NA | NA | NA | NA |
| WBGene00168257 | NA | NA | NA | NA |
| WBGene00168258 | NA | NA | NA | NA |
| WBGene00168259 | NA | NA | NA | NA |
| WBGene00168260 | NA | NA | NA | NA |
| WBGene00168261 | NA | NA | NA | NA |
| WBGene00168262 | NA | NA | NA | NA |
| WBGene00168263 | NA | NA | NA | NA |
| WBGene00168264 | NA | NA | NA | NA |
| WBGene00168265 | NA | NA | NA | NA |
| WBGene00168266 | NA | NA | NA | NA |
| WBGene00168267 | NA | NA | NA | NA |
| WBGene00168268 | NA | NA | NA | NA |
| WBGene00168269 | NA | NA | NA | NA |
| WBGene00168270 | NA | NA | NA | NA |
| WBGene00168271 | NA | NA | NA | NA |
| WBGene00168272 | NA | NA | NA | NA |
| WBGene00168273 | NA | NA | NA | NA |
| WBGene00168274 | NA | NA | NA | NA |
| WBGene00168275 | NA | NA | NA | NA |
| WBGene00168276 | NA | NA | NA | NA |
| WBGene00168277 | NA | NA | NA | NA |

| | | | | |
|---|---|---|---|---|
| WBGene00168278 | NA | NA | NA | NA |
| WBGene00168279 | NA | NA | NA | NA |
| WBGene00168280 | NA | NA | NA | NA |
| WBGene00168281 | NA | NA | NA | NA |
| WBGene00168282 | NA | NA | NA | NA |
| WBGene00168283 | NA | NA | NA | NA |
| WBGene00168284 | NA | NA | NA | NA |
| WBGene00168285 | NA | NA | NA | NA |
| WBGene00168286 | NA | NA | NA | NA |
| WBGene00168287 | NA | NA | NA | NA |
| WBGene00168288 | NA | NA | NA | NA |
| WBGene00168289 | NA | NA | NA | NA |
| WBGene00168290 | NA | NA | NA | NA |
| WBGene00168291 | NA | NA | NA | NA |
| WBGene00168292 | NA | NA | NA | NA |
| WBGene00168293 | NA | NA | NA | NA |
| WBGene00168294 | NA | NA | NA | NA |
| WBGene00168295 | NA | NA | NA | NA |
| WBGene00168296 | NA | NA | NA | NA |
| WBGene00168297 | NA | NA | NA | NA |
| WBGene00168298 | NA | NA | NA | NA |
| WBGene00168299 | NA | NA | NA | NA |
| WBGene00168300 | NA | NA | NA | NA |
| WBGene00168301 | NA | NA | NA | NA |
| WBGene00168302 | NA | NA | NA | NA |
| WBGene00168303 | NA | NA | NA | NA |
| WBGene00168304 | NA | NA | NA | NA |
| WBGene00168305 | NA | NA | NA | NA |
| WBGene00168306 | NA | NA | NA | NA |
| WBGene00168307 | NA | NA | NA | NA |
| WBGene00168308 | NA | NA | NA | NA |
| WBGene00168309 | NA | NA | NA | NA |
| WBGene00168310 | NA | NA | NA | NA |
| WBGene00168311 | NA | NA | NA | NA |
| WBGene00168312 | NA | NA | NA | NA |
| WBGene00168313 | NA | NA | NA | NA |
| WBGene00168314 | NA | NA | NA | NA |
| WBGene00168315 | NA | NA | NA | NA |
| WBGene00168316 | NA | NA | NA | NA |
| WBGene00168317 | NA | NA | NA | NA |
| WBGene00168318 | NA | NA | NA | NA |
| WBGene00168319 | NA | NA | NA | NA |
| WBGene00168320 | NA | NA | NA | NA |

| | | | | |
|---|---|---|---|---|
| WBGene00168321 | NA | NA | NA | NA |
| WBGene00168322 | NA | NA | NA | NA |
| WBGene00168323 | NA | NA | NA | NA |
| WBGene00168324 | NA | NA | NA | NA |
| WBGene00168325 | NA | NA | NA | NA |
| WBGene00168326 | NA | NA | NA | NA |
| WBGene00168327 | NA | NA | NA | NA |
| WBGene00168328 | NA | NA | NA | NA |
| WBGene00168329 | NA | NA | NA | NA |
| WBGene00168330 | NA | NA | NA | NA |
| WBGene00168331 | NA | NA | NA | NA |
| WBGene00168332 | NA | NA | NA | NA |
| WBGene00168333 | NA | NA | NA | NA |
| WBGene00168334 | NA | NA | NA | NA |
| WBGene00168335 | NA | NA | NA | NA |
| WBGene00168336 | NA | NA | NA | NA |
| WBGene00168337 | NA | NA | NA | NA |
| WBGene00168338 | NA | NA | NA | NA |
| WBGene00168339 | NA | NA | NA | NA |
| WBGene00168340 | NA | NA | NA | NA |
| WBGene00168341 | NA | NA | NA | NA |
| WBGene00168342 | NA | NA | NA | NA |
| WBGene00168343 | NA | NA | NA | NA |
| WBGene00168344 | NA | NA | NA | NA |
| WBGene00168345 | NA | NA | NA | NA |
| WBGene00168346 | NA | NA | NA | NA |
| WBGene00168347 | NA | NA | NA | NA |
| WBGene00168348 | NA | NA | NA | NA |
| WBGene00168349 | NA | NA | NA | NA |
| WBGene00168350 | NA | NA | NA | NA |
| WBGene00168351 | NA | NA | NA | NA |
| WBGene00168352 | NA | NA | NA | NA |
| WBGene00168353 | NA | NA | NA | NA |
| WBGene00168354 | NA | NA | NA | NA |
| WBGene00168355 | NA | NA | NA | NA |
| WBGene00168356 | NA | NA | NA | NA |
| WBGene00168357 | NA | NA | NA | NA |
| WBGene00168358 | NA | NA | NA | NA |
| WBGene00168359 | NA | NA | NA | NA |
| WBGene00168360 | NA | NA | NA | NA |
| WBGene00168361 | NA | NA | NA | NA |
| WBGene00168362 | NA | NA | NA | NA |
| WBGene00168363 | NA | NA | NA | NA |

| | | | | |
|---|---|---|---|---|
| WBGene00168364 | NA | NA | NA | NA |
| WBGene00168365 | NA | NA | NA | NA |
| WBGene00168366 | NA | NA | NA | NA |
| WBGene00168367 | NA | NA | NA | NA |
| WBGene00168368 | NA | NA | NA | NA |
| WBGene00168369 | NA | NA | NA | NA |
| WBGene00168370 | NA | NA | NA | NA |
| WBGene00168371 | NA | NA | NA | NA |
| WBGene00168372 | NA | NA | NA | NA |
| WBGene00168373 | NA | NA | NA | NA |
| WBGene00168374 | NA | NA | NA | NA |
| WBGene00168375 | NA | NA | NA | NA |
| WBGene00168376 | NA | NA | NA | NA |
| WBGene00168377 | NA | NA | NA | NA |
| WBGene00168378 | NA | NA | NA | NA |
| WBGene00168379 | NA | NA | NA | NA |
| WBGene00168380 | NA | NA | NA | NA |
| WBGene00168381 | NA | NA | NA | NA |
| WBGene00168382 | NA | NA | NA | NA |
| WBGene00168383 | NA | NA | NA | NA |
| WBGene00168384 | NA | NA | NA | NA |
| WBGene00168385 | NA | NA | NA | NA |
| WBGene00168386 | NA | NA | NA | NA |
| WBGene00168387 | NA | NA | NA | NA |
| WBGene00168388 | NA | NA | NA | NA |
| WBGene00168389 | NA | NA | NA | NA |
| WBGene00168390 | NA | NA | NA | NA |
| WBGene00168391 | NA | NA | NA | NA |
| WBGene00168392 | NA | NA | NA | NA |
| WBGene00168393 | NA | NA | NA | NA |
| WBGene00168394 | NA | NA | NA | NA |
| WBGene00168395 | NA | NA | NA | NA |
| WBGene00168396 | NA | NA | NA | NA |
| WBGene00168397 | NA | NA | NA | NA |
| WBGene00168398 | NA | NA | NA | NA |
| WBGene00168399 | NA | NA | NA | NA |
| WBGene00168400 | NA | NA | NA | NA |
| WBGene00168401 | NA | NA | NA | NA |
| WBGene00168402 | NA | NA | NA | NA |
| WBGene00168403 | NA | NA | NA | NA |
| WBGene00168404 | NA | NA | NA | NA |
| WBGene00168405 | NA | NA | NA | NA |
| WBGene00168406 | NA | NA | NA | NA |

| | | | | |
|---|---|---|---|---|
| WBGene00168407 | NA | NA | NA | NA |
| WBGene00168408 | NA | NA | NA | NA |
| WBGene00168409 | NA | NA | NA | NA |
| WBGene00168410 | NA | NA | NA | NA |
| WBGene00168411 | NA | NA | NA | NA |
| WBGene00168412 | NA | NA | NA | NA |
| WBGene00168413 | NA | NA | NA | NA |
| WBGene00168414 | NA | NA | NA | NA |
| WBGene00168415 | NA | NA | NA | NA |
| WBGene00168416 | NA | NA | NA | NA |
| WBGene00168417 | NA | NA | NA | NA |
| WBGene00168418 | NA | NA | NA | NA |
| WBGene00168419 | NA | NA | NA | NA |
| WBGene00168420 | NA | NA | NA | NA |
| WBGene00168421 | NA | NA | NA | NA |
| WBGene00168422 | NA | NA | NA | NA |
| WBGene00168423 | NA | NA | NA | NA |
| WBGene00168424 | NA | NA | NA | NA |
| WBGene00168425 | NA | NA | NA | NA |
| WBGene00168426 | NA | NA | NA | NA |
| WBGene00168427 | NA | NA | NA | NA |
| WBGene00168428 | NA | NA | NA | NA |
| WBGene00168429 | NA | NA | NA | NA |
| WBGene00168430 | NA | NA | NA | NA |
| WBGene00168431 | NA | NA | NA | NA |
| WBGene00168432 | NA | NA | NA | NA |
| WBGene00168433 | NA | NA | NA | NA |
| WBGene00168434 | NA | NA | NA | NA |
| WBGene00168435 | NA | NA | NA | NA |
| WBGene00168436 | NA | NA | NA | NA |
| WBGene00168437 | NA | NA | NA | NA |
| WBGene00168438 | NA | NA | NA | NA |
| WBGene00168439 | NA | NA | NA | NA |
| WBGene00168440 | NA | NA | NA | NA |
| WBGene00168441 | NA | NA | NA | NA |
| WBGene00168442 | NA | NA | NA | NA |
| WBGene00168443 | NA | NA | NA | NA |
| WBGene00168444 | NA | NA | NA | NA |
| WBGene00168445 | NA | NA | NA | NA |
| WBGene00168446 | NA | NA | NA | NA |
| WBGene00168447 | NA | NA | NA | NA |
| WBGene00168448 | NA | NA | NA | NA |
| WBGene00168449 | NA | NA | NA | NA |

| | | | | |
|---|---|---|---|---|
| WBGene00168450 | NA | NA | NA | NA |
| WBGene00168451 | NA | NA | NA | NA |
| WBGene00168452 | NA | NA | NA | NA |
| WBGene00168453 | NA | NA | NA | NA |
| WBGene00168454 | NA | NA | NA | NA |
| WBGene00168455 | NA | NA | NA | NA |
| WBGene00168456 | NA | NA | NA | NA |
| WBGene00168457 | NA | NA | NA | NA |
| WBGene00168458 | NA | NA | NA | NA |
| WBGene00168459 | NA | NA | NA | NA |
| WBGene00168460 | NA | NA | NA | NA |
| WBGene00168461 | NA | NA | NA | NA |
| WBGene00168462 | NA | NA | NA | NA |
| WBGene00168463 | NA | NA | NA | NA |
| WBGene00168464 | NA | NA | NA | NA |
| WBGene00168465 | NA | NA | NA | NA |
| WBGene00168466 | NA | NA | NA | NA |
| WBGene00168467 | NA | NA | NA | NA |
| WBGene00168468 | NA | NA | NA | NA |
| WBGene00168469 | NA | NA | NA | NA |
| WBGene00168470 | NA | NA | NA | NA |
| WBGene00168472 | NA | NA | NA | NA |
| WBGene00168473 | NA | NA | NA | NA |
| WBGene00168474 | NA | NA | NA | NA |
| WBGene00168475 | NA | NA | NA | NA |
| WBGene00168476 | NA | NA | NA | NA |
| WBGene00168477 | NA | NA | NA | NA |
| WBGene00168478 | NA | NA | NA | NA |
| WBGene00168479 | NA | NA | NA | NA |
| WBGene00168480 | NA | NA | NA | NA |
| WBGene00168481 | NA | NA | NA | NA |
| WBGene00168482 | NA | NA | NA | NA |
| WBGene00168483 | NA | NA | NA | NA |
| WBGene00168484 | NA | NA | NA | NA |
| WBGene00168485 | 0.020232312 | 0.84643798 | NA | NA |
| WBGene00168486 | NA | NA | NA | NA |
| WBGene00168487 | NA | NA | NA | NA |
| WBGene00168488 | NA | NA | NA | NA |
| WBGene00168489 | NA | NA | NA | NA |
| WBGene00168490 | NA | NA | NA | NA |
| WBGene00168491 | NA | NA | NA | NA |
| WBGene00168492 | NA | NA | NA | NA |
| WBGene00168493 | NA | NA | NA | NA |

| | | | | |
|---|---|---|---|---|
| WBGene00168494 | NA | NA | NA | NA |
| WBGene00168495 | NA | NA | NA | NA |
| WBGene00168496 | NA | NA | NA | NA |
| WBGene00168497 | NA | NA | NA | NA |
| WBGene00168498 | NA | NA | NA | NA |
| WBGene00168499 | NA | NA | NA | NA |
| WBGene00168500 | NA | NA | NA | NA |
| WBGene00168501 | NA | NA | NA | NA |
| WBGene00168502 | NA | NA | NA | NA |
| WBGene00168503 | NA | NA | NA | NA |
| WBGene00168504 | NA | NA | NA | NA |
| WBGene00168505 | NA | NA | NA | NA |
| WBGene00168506 | NA | NA | NA | NA |
| WBGene00168507 | NA | NA | NA | NA |
| WBGene00168508 | NA | NA | NA | NA |
| WBGene00168509 | NA | NA | NA | NA |
| WBGene00168510 | NA | NA | NA | NA |
| WBGene00168511 | NA | NA | NA | NA |
| WBGene00168512 | NA | NA | NA | NA |
| WBGene00168513 | NA | NA | NA | NA |
| WBGene00168514 | NA | NA | NA | NA |
| WBGene00168515 | NA | NA | NA | NA |
| WBGene00168516 | NA | NA | NA | NA |
| WBGene00168517 | NA | NA | NA | NA |
| WBGene00168518 | NA | NA | NA | NA |
| WBGene00168519 | NA | NA | NA | NA |
| WBGene00168520 | NA | NA | NA | NA |
| WBGene00168521 | NA | NA | NA | NA |
| WBGene00168522 | NA | NA | NA | NA |
| WBGene00168523 | NA | NA | NA | NA |
| WBGene00168524 | NA | NA | NA | NA |
| WBGene00168525 | NA | NA | NA | NA |
| WBGene00168526 | NA | NA | NA | NA |
| WBGene00168527 | NA | NA | NA | NA |
| WBGene00168528 | NA | NA | NA | NA |
| WBGene00168529 | NA | NA | NA | NA |
| WBGene00168530 | NA | NA | NA | NA |
| WBGene00168531 | NA | NA | NA | NA |
| WBGene00168532 | NA | NA | NA | NA |
| WBGene00168533 | NA | NA | NA | NA |
| WBGene00168534 | NA | NA | NA | NA |
| WBGene00168535 | NA | NA | NA | NA |
| WBGene00168536 | NA | NA | NA | NA |

| | | | | |
|---|---|---|---|---|
| WBGene00168537 | NA | NA | NA | NA |
| WBGene00168538 | NA | NA | NA | NA |
| WBGene00168539 | NA | NA | NA | NA |
| WBGene00168540 | NA | NA | NA | NA |
| WBGene00168541 | NA | NA | NA | NA |
| WBGene00168542 | NA | NA | NA | NA |
| WBGene00168543 | NA | NA | NA | NA |
| WBGene00168544 | 0.020317245 | 0.84643798 | NA | NA |
| WBGene00168545 | NA | NA | NA | NA |
| WBGene00168546 | NA | NA | NA | NA |
| WBGene00168547 | NA | NA | NA | NA |
| WBGene00168548 | NA | NA | NA | NA |
| WBGene00168549 | NA | NA | NA | NA |
| WBGene00168550 | NA | NA | NA | NA |
| WBGene00168551 | NA | NA | NA | NA |
| WBGene00168552 | NA | NA | NA | NA |
| WBGene00168553 | NA | NA | NA | NA |
| WBGene00168554 | NA | NA | NA | NA |
| WBGene00168555 | NA | NA | NA | NA |
| WBGene00168556 | NA | NA | NA | NA |
| WBGene00168557 | NA | NA | NA | NA |
| WBGene00168558 | NA | NA | NA | NA |
| WBGene00168559 | NA | NA | NA | NA |
| WBGene00168560 | NA | NA | NA | NA |
| WBGene00168561 | NA | NA | NA | NA |
| WBGene00168562 | NA | NA | NA | NA |
| WBGene00168563 | NA | NA | NA | NA |
| WBGene00168564 | NA | NA | NA | NA |
| WBGene00168565 | NA | NA | NA | NA |
| WBGene00168566 | NA | NA | NA | NA |
| WBGene00168567 | NA | NA | NA | NA |
| WBGene00168568 | NA | NA | NA | NA |
| WBGene00168569 | NA | NA | NA | NA |
| WBGene00168570 | NA | NA | NA | NA |
| WBGene00168571 | NA | NA | NA | NA |
| WBGene00168572 | NA | NA | NA | NA |
| WBGene00168573 | NA | NA | NA | NA |
| WBGene00168574 | NA | NA | NA | NA |
| WBGene00168575 | NA | NA | NA | NA |
| WBGene00168576 | NA | NA | NA | NA |
| WBGene00168577 | NA | NA | NA | NA |
| WBGene00168578 | NA | NA | NA | NA |
| WBGene00168579 | NA | NA | NA | NA |

| | | | | |
|---|---|---|---|---|
| WBGene00168580 | NA | NA | NA | NA |
| WBGene00168581 | NA | NA | NA | NA |
| WBGene00168582 | NA | NA | NA | NA |
| WBGene00168583 | NA | NA | NA | NA |
| WBGene00168584 | NA | NA | NA | NA |
| WBGene00168585 | NA | NA | NA | NA |
| WBGene00168586 | NA | NA | NA | NA |
| WBGene00168587 | NA | NA | NA | NA |
| WBGene00168588 | NA | NA | NA | NA |
| WBGene00168589 | 0.020317245 | 0.84643798 | NA | NA |
| WBGene00168590 | NA | NA | NA | NA |
| WBGene00168591 | NA | NA | NA | NA |
| WBGene00168592 | NA | NA | NA | NA |
| WBGene00168593 | NA | NA | NA | NA |
| WBGene00168594 | NA | NA | NA | NA |
| WBGene00168595 | NA | NA | NA | NA |
| WBGene00168596 | NA | NA | NA | NA |
| WBGene00168597 | 0.056188364 | 0.42092875 | NA | NA |
| WBGene00168598 | NA | NA | NA | NA |
| WBGene00168599 | NA | NA | NA | NA |
| WBGene00168600 | NA | NA | NA | NA |
| WBGene00168601 | NA | NA | NA | NA |
| WBGene00168602 | NA | NA | NA | NA |
| WBGene00168603 | NA | NA | NA | NA |
| WBGene00168604 | NA | NA | NA | NA |
| WBGene00168605 | -0.023056499 | 0.78121253 | NA | NA |
| WBGene00168606 | NA | NA | NA | NA |
| WBGene00168607 | NA | NA | NA | NA |
| WBGene00168608 | NA | NA | NA | NA |
| WBGene00168609 | NA | NA | NA | NA |
| WBGene00168610 | NA | NA | NA | NA |
| WBGene00168611 | NA | NA | NA | NA |
| WBGene00168612 | NA | NA | NA | NA |
| WBGene00168613 | NA | NA | NA | NA |
| WBGene00168614 | NA | NA | NA | NA |
| WBGene00168615 | NA | NA | NA | NA |
| WBGene00168616 | NA | NA | NA | NA |
| WBGene00168617 | NA | NA | NA | NA |
| WBGene00168618 | NA | NA | NA | NA |
| WBGene00168619 | NA | NA | NA | NA |
| WBGene00168620 | NA | NA | NA | NA |
| WBGene00168621 | NA | NA | NA | NA |
| WBGene00168622 | NA | NA | NA | NA |

| | | | | |
|---|---|---|---|---|
| WBGene00168623 | NA | NA | NA | NA |
| WBGene00168624 | NA | NA | NA | NA |
| WBGene00168625 | NA | NA | NA | NA |
| WBGene00168626 | NA | NA | NA | NA |
| WBGene00168627 | NA | NA | NA | NA |
| WBGene00168628 | NA | NA | NA | NA |
| WBGene00168629 | NA | NA | NA | NA |
| WBGene00168630 | NA | NA | NA | NA |
| WBGene00168631 | NA | NA | NA | NA |
| WBGene00168632 | NA | NA | NA | NA |
| WBGene00168633 | NA | NA | NA | NA |
| WBGene00168634 | NA | NA | NA | NA |
| WBGene00168635 | NA | NA | NA | NA |
| WBGene00168636 | NA | NA | NA | NA |
| WBGene00168637 | NA | NA | NA | NA |
| WBGene00168638 | NA | NA | NA | NA |
| WBGene00168639 | NA | NA | NA | NA |
| WBGene00168640 | NA | NA | NA | NA |
| WBGene00168641 | NA | NA | NA | NA |
| WBGene00168642 | -0.001842784 | 0.95621055 | NA | NA |
| WBGene00168643 | NA | NA | NA | NA |
| WBGene00168644 | NA | NA | NA | NA |
| WBGene00168645 | NA | NA | NA | NA |
| WBGene00168646 | NA | NA | NA | NA |
| WBGene00168647 | NA | NA | NA | NA |
| WBGene00168648 | NA | NA | NA | NA |
| WBGene00168649 | NA | NA | NA | NA |
| WBGene00168650 | NA | NA | NA | NA |
| WBGene00168651 | NA | NA | NA | NA |
| WBGene00168652 | NA | NA | NA | NA |
| WBGene00168653 | NA | NA | NA | NA |
| WBGene00168654 | NA | NA | NA | NA |
| WBGene00168655 | NA | NA | NA | NA |
| WBGene00168656 | NA | NA | NA | NA |
| WBGene00168657 | NA | NA | NA | NA |
| WBGene00168658 | NA | NA | NA | NA |
| WBGene00168659 | NA | NA | NA | NA |
| WBGene00168660 | NA | NA | NA | NA |
| WBGene00168661 | NA | NA | NA | NA |
| WBGene00168662 | NA | NA | NA | NA |
| WBGene00168663 | NA | NA | NA | NA |
| WBGene00168664 | NA | NA | NA | NA |
| WBGene00168665 | NA | NA | NA | NA |

| | | | | |
|---|---|---|---|---|
| WBGene00168666 | NA | NA | NA | NA |
| WBGene00168667 | NA | NA | NA | NA |
| WBGene00168668 | NA | NA | NA | NA |
| WBGene00168669 | NA | NA | NA | NA |
| WBGene00168670 | NA | NA | NA | NA |
| WBGene00168671 | NA | NA | NA | NA |
| WBGene00168672 | NA | NA | NA | NA |
| WBGene00168673 | NA | NA | NA | NA |
| WBGene00168674 | NA | NA | NA | NA |
| WBGene00168675 | NA | NA | NA | NA |
| WBGene00168676 | NA | NA | NA | NA |
| WBGene00168677 | NA | NA | NA | NA |
| WBGene00168678 | NA | NA | NA | NA |
| WBGene00168679 | NA | NA | NA | NA |
| WBGene00168680 | NA | NA | NA | NA |
| WBGene00168681 | NA | NA | NA | NA |
| WBGene00168682 | NA | NA | NA | NA |
| WBGene00168683 | NA | NA | NA | NA |
| WBGene00168684 | NA | NA | NA | NA |
| WBGene00168685 | NA | NA | NA | NA |
| WBGene00168686 | NA | NA | NA | NA |
| WBGene00168687 | NA | NA | NA | NA |
| WBGene00168688 | NA | NA | NA | NA |
| WBGene00168689 | NA | NA | NA | NA |
| WBGene00168690 | NA | NA | NA | NA |
| WBGene00168691 | NA | NA | NA | NA |
| WBGene00168692 | NA | NA | NA | NA |
| WBGene00168693 | NA | NA | NA | NA |
| WBGene00168694 | NA | NA | NA | NA |
| WBGene00168695 | NA | NA | NA | NA |
| WBGene00168696 | NA | NA | NA | NA |
| WBGene00168697 | NA | NA | NA | NA |
| WBGene00168698 | NA | NA | NA | NA |
| WBGene00168699 | NA | NA | NA | NA |
| WBGene00168700 | NA | NA | NA | NA |
| WBGene00168701 | NA | NA | NA | NA |
| WBGene00168702 | NA | NA | NA | NA |
| WBGene00168703 | NA | NA | NA | NA |
| WBGene00168704 | NA | NA | NA | NA |
| WBGene00168705 | NA | NA | NA | NA |
| WBGene00168706 | NA | NA | NA | NA |
| WBGene00168707 | NA | NA | NA | NA |
| WBGene00168708 | NA | NA | NA | NA |

| | | | | |
|---|---|---|---|---|
| WBGene00168709 | NA | NA | NA | NA |
| WBGene00168710 | NA | NA | NA | NA |
| WBGene00168711 | NA | NA | NA | NA |
| WBGene00168712 | NA | NA | NA | NA |
| WBGene00168713 | NA | NA | NA | NA |
| WBGene00168714 | NA | NA | NA | NA |
| WBGene00168715 | NA | NA | NA | NA |
| WBGene00168716 | NA | NA | NA | NA |
| WBGene00168717 | NA | NA | NA | NA |
| WBGene00168718 | NA | NA | NA | NA |
| WBGene00168719 | NA | NA | NA | NA |
| WBGene00168720 | NA | NA | NA | NA |
| WBGene00168721 | NA | NA | NA | NA |
| WBGene00168722 | NA | NA | NA | NA |
| WBGene00168723 | NA | NA | NA | NA |
| WBGene00168724 | NA | NA | NA | NA |
| WBGene00168725 | NA | NA | NA | NA |
| WBGene00168726 | NA | NA | NA | NA |
| WBGene00168727 | NA | NA | NA | NA |
| WBGene00168728 | NA | NA | NA | NA |
| WBGene00168729 | NA | NA | NA | NA |
| WBGene00168730 | NA | NA | NA | NA |
| WBGene00168731 | NA | NA | NA | NA |
| WBGene00168732 | NA | NA | NA | NA |
| WBGene00168733 | NA | NA | NA | NA |
| WBGene00168734 | NA | NA | NA | NA |
| WBGene00168735 | NA | NA | NA | NA |
| WBGene00168736 | NA | NA | NA | NA |
| WBGene00168737 | NA | NA | NA | NA |
| WBGene00168738 | NA | NA | NA | NA |
| WBGene00168739 | NA | NA | NA | NA |
| WBGene00168740 | NA | NA | NA | NA |
| WBGene00168741 | NA | NA | NA | NA |
| WBGene00168742 | NA | NA | NA | NA |
| WBGene00168743 | NA | NA | NA | NA |
| WBGene00168744 | NA | NA | NA | NA |
| WBGene00168745 | NA | NA | NA | NA |
| WBGene00168746 | NA | NA | NA | NA |
| WBGene00168747 | NA | NA | NA | NA |
| WBGene00168748 | NA | NA | NA | NA |
| WBGene00168749 | NA | NA | NA | NA |
| WBGene00168750 | NA | NA | NA | NA |
| WBGene00168751 | NA | NA | NA | NA |

| | | | | |
|---|---|---|---|---|
| WBGene00168752 | NA | NA | NA | NA |
| WBGene00168753 | NA | NA | NA | NA |
| WBGene00168754 | NA | NA | NA | NA |
| WBGene00168755 | NA | NA | NA | NA |
| WBGene00168756 | NA | NA | NA | NA |
| WBGene00168757 | NA | NA | NA | NA |
| WBGene00168758 | NA | NA | NA | NA |
| WBGene00168759 | NA | NA | NA | NA |
| WBGene00168760 | NA | NA | NA | NA |
| WBGene00168761 | NA | NA | NA | NA |
| WBGene00168762 | NA | NA | NA | NA |
| WBGene00168763 | NA | NA | NA | NA |
| WBGene00168764 | NA | NA | NA | NA |
| WBGene00168765 | NA | NA | NA | NA |
| WBGene00168766 | NA | NA | NA | NA |
| WBGene00168767 | NA | NA | NA | NA |
| WBGene00168768 | NA | NA | NA | NA |
| WBGene00168769 | NA | NA | NA | NA |
| WBGene00168770 | NA | NA | NA | NA |
| WBGene00168771 | NA | NA | NA | NA |
| WBGene00168772 | NA | NA | NA | NA |
| WBGene00168773 | NA | NA | NA | NA |
| WBGene00168774 | NA | NA | NA | NA |
| WBGene00168775 | NA | NA | NA | NA |
| WBGene00168776 | NA | NA | NA | NA |
| WBGene00168777 | NA | NA | NA | NA |
| WBGene00168778 | NA | NA | NA | NA |
| WBGene00168779 | NA | NA | NA | NA |
| WBGene00168780 | NA | NA | NA | NA |
| WBGene00168781 | NA | NA | NA | NA |
| WBGene00168782 | NA | NA | NA | NA |
| WBGene00168783 | NA | NA | NA | NA |
| WBGene00168784 | NA | NA | NA | NA |
| WBGene00168785 | NA | NA | NA | NA |
| WBGene00168786 | NA | NA | NA | NA |
| WBGene00168787 | NA | NA | NA | NA |
| WBGene00168788 | NA | NA | NA | NA |
| WBGene00168789 | NA | NA | NA | NA |
| WBGene00168790 | NA | NA | NA | NA |
| WBGene00168791 | NA | NA | NA | NA |
| WBGene00168792 | NA | NA | NA | NA |
| WBGene00168793 | NA | NA | NA | NA |
| WBGene00168794 | NA | NA | NA | NA |

| | | | | |
|---|---|---|---|---|
| WBGene00168795 | NA | NA | NA | NA |
| WBGene00168796 | NA | NA | NA | NA |
| WBGene00168797 | NA | NA | NA | NA |
| WBGene00168798 | NA | NA | NA | NA |
| WBGene00168799 | NA | NA | NA | NA |
| WBGene00168800 | NA | NA | NA | NA |
| WBGene00168801 | NA | NA | NA | NA |
| WBGene00168802 | NA | NA | NA | NA |
| WBGene00168803 | NA | NA | NA | NA |
| WBGene00168804 | NA | NA | NA | NA |
| WBGene00168805 | NA | NA | NA | NA |
| WBGene00168806 | NA | NA | NA | NA |
| WBGene00168807 | NA | NA | NA | NA |
| WBGene00168808 | NA | NA | NA | NA |
| WBGene00168809 | NA | NA | NA | NA |
| WBGene00168810 | NA | NA | NA | NA |
| WBGene00168811 | NA | NA | NA | NA |
| WBGene00168812 | NA | NA | NA | NA |
| WBGene00168813 | NA | NA | NA | NA |
| WBGene00168814 | NA | NA | NA | NA |
| WBGene00168815 | NA | NA | NA | NA |
| WBGene00168816 | NA | NA | NA | NA |
| WBGene00168817 | NA | NA | NA | NA |
| WBGene00168818 | NA | NA | NA | NA |
| WBGene00168819 | NA | NA | NA | NA |
| WBGene00168820 | NA | NA | NA | NA |
| WBGene00168821 | NA | NA | NA | NA |
| WBGene00168822 | NA | NA | NA | NA |
| WBGene00168823 | NA | NA | NA | NA |
| WBGene00168824 | NA | NA | NA | NA |
| WBGene00168825 | NA | NA | NA | NA |
| WBGene00168826 | NA | NA | NA | NA |
| WBGene00168827 | NA | NA | NA | NA |
| WBGene00168828 | NA | NA | NA | NA |
| WBGene00168829 | NA | NA | NA | NA |
| WBGene00168830 | NA | NA | NA | NA |
| WBGene00168831 | NA | NA | NA | NA |
| WBGene00168832 | NA | NA | NA | NA |
| WBGene00168833 | NA | NA | NA | NA |
| WBGene00168834 | NA | NA | NA | NA |
| WBGene00168835 | NA | NA | NA | NA |
| WBGene00168836 | NA | NA | NA | NA |
| WBGene00168837 | NA | NA | NA | NA |

| | | | | |
|---|---|---|---|---|
| WBGene00168838 | NA | NA | NA | NA |
| WBGene00168839 | NA | NA | NA | NA |
| WBGene00168840 | NA | NA | NA | NA |
| WBGene00168841 | NA | NA | NA | NA |
| WBGene00168842 | NA | NA | NA | NA |
| WBGene00168843 | NA | NA | NA | NA |
| WBGene00168844 | NA | NA | NA | NA |
| WBGene00168845 | NA | NA | NA | NA |
| WBGene00168846 | NA | NA | NA | NA |
| WBGene00168847 | NA | NA | NA | NA |
| WBGene00168848 | NA | NA | NA | NA |
| WBGene00168849 | NA | NA | NA | NA |
| WBGene00168850 | NA | NA | NA | NA |
| WBGene00168851 | NA | NA | NA | NA |
| WBGene00168852 | NA | NA | NA | NA |
| WBGene00168853 | NA | NA | NA | NA |
| WBGene00168854 | NA | NA | NA | NA |
| WBGene00168855 | NA | NA | NA | NA |
| WBGene00168856 | NA | NA | NA | NA |
| WBGene00168857 | NA | NA | NA | NA |
| WBGene00168858 | NA | NA | NA | NA |
| WBGene00168859 | NA | NA | NA | NA |
| WBGene00168860 | NA | NA | NA | NA |
| WBGene00168861 | NA | NA | NA | NA |
| WBGene00168862 | NA | NA | NA | NA |
| WBGene00168863 | NA | NA | NA | NA |
| WBGene00168864 | NA | NA | NA | NA |
| WBGene00168865 | NA | NA | NA | NA |
| WBGene00168866 | NA | NA | NA | NA |
| WBGene00168867 | NA | NA | NA | NA |
| WBGene00168868 | NA | NA | NA | NA |
| WBGene00168869 | NA | NA | NA | NA |
| WBGene00168870 | NA | NA | NA | NA |
| WBGene00168871 | NA | NA | NA | NA |
| WBGene00168872 | NA | NA | NA | NA |
| WBGene00168873 | NA | NA | NA | NA |
| WBGene00168874 | NA | NA | NA | NA |
| WBGene00168875 | NA | NA | NA | NA |
| WBGene00168876 | NA | NA | NA | NA |
| WBGene00168877 | NA | NA | NA | NA |
| WBGene00168878 | NA | NA | NA | NA |
| WBGene00168879 | NA | NA | NA | NA |
| WBGene00168880 | NA | NA | NA | NA |

| | | | | |
|---|---|---|---|---|
| WBGene00168881 | NA | NA | NA | NA |
| WBGene00168882 | NA | NA | NA | NA |
| WBGene00168883 | NA | NA | NA | NA |
| WBGene00168884 | NA | NA | NA | NA |
| WBGene00168885 | NA | NA | NA | NA |
| WBGene00168886 | NA | NA | NA | NA |
| WBGene00168887 | NA | NA | NA | NA |
| WBGene00168888 | NA | NA | NA | NA |
| WBGene00168889 | NA | NA | NA | NA |
| WBGene00168890 | NA | NA | NA | NA |
| WBGene00168891 | NA | NA | NA | NA |
| WBGene00168892 | 0.020925399 | 0.84643798 | NA | NA |
| WBGene00168893 | NA | NA | NA | NA |
| WBGene00168894 | NA | NA | NA | NA |
| WBGene00168895 | NA | NA | NA | NA |
| WBGene00168896 | NA | NA | NA | NA |
| WBGene00168897 | NA | NA | NA | NA |
| WBGene00168898 | NA | NA | NA | NA |
| WBGene00168899 | NA | NA | NA | NA |
| WBGene00168900 | NA | NA | NA | NA |
| WBGene00168901 | NA | NA | NA | NA |
| WBGene00168902 | 0.038125098 | 0.61119482 | NA | NA |
| WBGene00168903 | NA | NA | NA | NA |
| WBGene00168904 | NA | NA | NA | NA |
| WBGene00168905 | NA | NA | NA | NA |
| WBGene00168906 | NA | NA | NA | NA |
| WBGene00168907 | NA | NA | NA | NA |
| WBGene00168908 | NA | NA | NA | NA |
| WBGene00168909 | NA | NA | NA | NA |
| WBGene00168910 | NA | NA | NA | NA |
| WBGene00168911 | NA | NA | NA | NA |
| WBGene00168912 | NA | NA | NA | NA |
| WBGene00168913 | NA | NA | NA | NA |
| WBGene00168914 | NA | NA | NA | NA |
| WBGene00168915 | NA | NA | NA | NA |
| WBGene00168916 | NA | NA | NA | NA |
| WBGene00168917 | NA | NA | NA | NA |
| WBGene00168918 | NA | NA | NA | NA |
| WBGene00168919 | NA | NA | NA | NA |
| WBGene00168920 | NA | NA | NA | NA |
| WBGene00168921 | NA | NA | NA | NA |
| WBGene00168922 | NA | NA | NA | NA |
| WBGene00168923 | NA | NA | NA | NA |

| | | | | |
|---|---|---|---|---|
| WBGene00168924 | NA | NA | NA | NA |
| WBGene00168925 | NA | NA | NA | NA |
| WBGene00168926 | NA | NA | NA | NA |
| WBGene00168927 | NA | NA | NA | NA |
| WBGene00168928 | NA | NA | NA | NA |
| WBGene00168929 | NA | NA | NA | NA |
| WBGene00168930 | NA | NA | NA | NA |
| WBGene00168931 | NA | NA | NA | NA |
| WBGene00168932 | NA | NA | NA | NA |
| WBGene00168933 | NA | NA | NA | NA |
| WBGene00168934 | NA | NA | NA | NA |
| WBGene00168935 | NA | NA | NA | NA |
| WBGene00168936 | NA | NA | NA | NA |
| WBGene00168937 | NA | NA | NA | NA |
| WBGene00168938 | NA | NA | NA | NA |
| WBGene00168939 | NA | NA | NA | NA |
| WBGene00168940 | NA | NA | NA | NA |
| WBGene00168941 | NA | NA | NA | NA |
| WBGene00168942 | NA | NA | NA | NA |
| WBGene00168943 | NA | NA | NA | NA |
| WBGene00168944 | NA | NA | NA | NA |
| WBGene00168945 | NA | NA | NA | NA |
| WBGene00168946 | NA | NA | NA | NA |
| WBGene00168947 | NA | NA | NA | NA |
| WBGene00168948 | NA | NA | NA | NA |
| WBGene00168949 | NA | NA | NA | NA |
| WBGene00168950 | NA | NA | NA | NA |
| WBGene00168951 | NA | NA | NA | NA |
| WBGene00168952 | NA | NA | NA | NA |
| WBGene00168953 | NA | NA | NA | NA |
| WBGene00168954 | NA | NA | NA | NA |
| WBGene00168955 | NA | NA | NA | NA |
| WBGene00168956 | NA | NA | NA | NA |
| WBGene00168957 | NA | NA | NA | NA |
| WBGene00168958 | NA | NA | NA | NA |
| WBGene00168959 | NA | NA | NA | NA |
| WBGene00168960 | NA | NA | NA | NA |
| WBGene00168961 | NA | NA | NA | NA |
| WBGene00168962 | NA | NA | NA | NA |
| WBGene00168963 | NA | NA | NA | NA |
| WBGene00168964 | NA | NA | NA | NA |
| WBGene00168965 | NA | NA | NA | NA |
| WBGene00168966 | NA | NA | NA | NA |

| | | | | |
|---|---|---|---|---|
| WBGene00168967 | NA | NA | NA | NA |
| WBGene00168968 | NA | NA | NA | NA |
| WBGene00168969 | NA | NA | NA | NA |
| WBGene00168970 | NA | NA | NA | NA |
| WBGene00168971 | NA | NA | NA | NA |
| WBGene00168972 | NA | NA | NA | NA |
| WBGene00168973 | NA | NA | NA | NA |
| WBGene00168974 | NA | NA | NA | NA |
| WBGene00168975 | NA | NA | NA | NA |
| WBGene00168976 | NA | NA | NA | NA |
| WBGene00168977 | NA | NA | NA | NA |
| WBGene00168978 | NA | NA | NA | NA |
| WBGene00168979 | NA | NA | NA | NA |
| WBGene00168980 | NA | NA | NA | NA |
| WBGene00168981 | NA | NA | NA | NA |
| WBGene00168982 | NA | NA | NA | NA |
| WBGene00168983 | NA | NA | NA | NA |
| WBGene00168984 | NA | NA | NA | NA |
| WBGene00168985 | NA | NA | NA | NA |
| WBGene00168986 | NA | NA | NA | NA |
| WBGene00168987 | NA | NA | NA | NA |
| WBGene00168988 | NA | NA | NA | NA |
| WBGene00168989 | NA | NA | NA | NA |
| WBGene00168990 | NA | NA | NA | NA |
| WBGene00168991 | NA | NA | NA | NA |
| WBGene00168992 | NA | NA | NA | NA |
| WBGene00168993 | NA | NA | NA | NA |
| WBGene00168994 | NA | NA | NA | NA |
| WBGene00168995 | NA | NA | NA | NA |
| WBGene00168996 | NA | NA | NA | NA |
| WBGene00168997 | NA | NA | NA | NA |
| WBGene00168998 | NA | NA | NA | NA |
| WBGene00168999 | NA | NA | NA | NA |
| WBGene00169000 | NA | NA | NA | NA |
| WBGene00169001 | NA | NA | NA | NA |
| WBGene00169002 | NA | NA | NA | NA |
| WBGene00169003 | NA | NA | NA | NA |
| WBGene00169004 | NA | NA | NA | NA |
| WBGene00169005 | NA | NA | NA | NA |
| WBGene00169006 | 0.020925399 | 0.84643798 | NA | NA |
| WBGene00169007 | NA | NA | NA | NA |
| WBGene00169008 | NA | NA | NA | NA |
| WBGene00169009 | NA | NA | NA | NA |

| | | | | |
|---|---|---|---|---|
| WBGene00169010 | NA | NA | NA | NA |
| WBGene00169011 | NA | NA | NA | NA |
| WBGene00169012 | NA | NA | NA | NA |
| WBGene00169013 | NA | NA | NA | NA |
| WBGene00169014 | NA | NA | NA | NA |
| WBGene00169015 | NA | NA | NA | NA |
| WBGene00169016 | NA | NA | NA | NA |
| WBGene00169017 | NA | NA | NA | NA |
| WBGene00169018 | NA | NA | NA | NA |
| WBGene00169019 | NA | NA | NA | NA |
| WBGene00169020 | NA | NA | NA | NA |
| WBGene00169021 | NA | NA | NA | NA |
| WBGene00169022 | NA | NA | NA | NA |
| WBGene00169023 | NA | NA | NA | NA |
| WBGene00169024 | NA | NA | NA | NA |
| WBGene00169025 | NA | NA | NA | NA |
| WBGene00169026 | NA | NA | NA | NA |
| WBGene00169027 | NA | NA | NA | NA |
| WBGene00169028 | NA | NA | NA | NA |
| WBGene00169029 | -0.244713 | 0.14396507 | 0.31289979 | NA |
| WBGene00169030 | NA | NA | NA | NA |
| WBGene00169031 | NA | NA | NA | NA |
| WBGene00169032 | NA | NA | NA | NA |
| WBGene00169033 | NA | NA | NA | NA |
| WBGene00169034 | NA | NA | NA | NA |
| WBGene00169035 | NA | NA | NA | NA |
| WBGene00169036 | NA | NA | NA | NA |
| WBGene00169037 | NA | NA | NA | NA |
| WBGene00169038 | NA | NA | NA | NA |
| WBGene00169039 | NA | NA | NA | NA |
| WBGene00169040 | NA | NA | NA | NA |
| WBGene00169041 | NA | NA | NA | NA |
| WBGene00169042 | NA | NA | NA | NA |
| WBGene00169043 | NA | NA | NA | NA |
| WBGene00169044 | NA | NA | NA | NA |
| WBGene00169045 | NA | NA | NA | NA |
| WBGene00169046 | NA | NA | NA | NA |
| WBGene00169047 | NA | NA | NA | NA |
| WBGene00169048 | NA | NA | NA | NA |
| WBGene00169049 | NA | NA | NA | NA |
| WBGene00169050 | NA | NA | NA | NA |
| WBGene00169051 | NA | NA | NA | NA |
| WBGene00169052 | NA | NA | NA | NA |

| | | | | |
|---|---|---|---|---|
| WBGene00169053 | NA | NA | NA | NA |
| WBGene00169054 | NA | NA | NA | NA |
| WBGene00169055 | NA | NA | NA | NA |
| WBGene00169056 | NA | NA | NA | NA |
| WBGene00169057 | NA | NA | NA | NA |
| WBGene00169058 | NA | NA | NA | NA |
| WBGene00169059 | NA | NA | NA | NA |
| WBGene00169060 | NA | NA | NA | NA |
| WBGene00169061 | NA | NA | NA | NA |
| WBGene00169062 | NA | NA | NA | NA |
| WBGene00169063 | NA | NA | NA | NA |
| WBGene00169064 | NA | NA | NA | NA |
| WBGene00169065 | NA | NA | NA | NA |
| WBGene00169066 | NA | NA | NA | NA |
| WBGene00169067 | NA | NA | NA | NA |
| WBGene00169068 | NA | NA | NA | NA |
| WBGene00169069 | NA | NA | NA | NA |
| WBGene00169070 | NA | NA | NA | NA |
| WBGene00169071 | NA | NA | NA | NA |
| WBGene00169072 | NA | NA | NA | NA |
| WBGene00169073 | NA | NA | NA | NA |
| WBGene00169074 | NA | NA | NA | NA |
| WBGene00169075 | NA | NA | NA | NA |
| WBGene00169076 | NA | NA | NA | NA |
| WBGene00169077 | NA | NA | NA | NA |
| WBGene00169078 | NA | NA | NA | NA |
| WBGene00169079 | NA | NA | NA | NA |
| WBGene00169080 | NA | NA | NA | NA |
| WBGene00169081 | NA | NA | NA | NA |
| WBGene00169082 | NA | NA | NA | NA |
| WBGene00169083 | NA | NA | NA | NA |
| WBGene00169084 | NA | NA | NA | NA |
| WBGene00169085 | NA | NA | NA | NA |
| WBGene00169086 | NA | NA | NA | NA |
| WBGene00169087 | NA | NA | NA | NA |
| WBGene00169088 | NA | NA | NA | NA |
| WBGene00169089 | 0.036687355 | 0.59489568 | NA | NA |
| WBGene00169090 | NA | NA | NA | NA |
| WBGene00169091 | NA | NA | NA | NA |
| WBGene00169092 | NA | NA | NA | NA |
| WBGene00169093 | NA | NA | NA | NA |
| WBGene00169094 | NA | NA | NA | NA |
| WBGene00169095 | NA | NA | NA | NA |

| | | | | |
|---|---|---|---|---|
| WBGene00169096 | NA | NA | NA | NA |
| WBGene00169097 | NA | NA | NA | NA |
| WBGene00169098 | NA | NA | NA | NA |
| WBGene00169099 | NA | NA | NA | NA |
| WBGene00169100 | NA | NA | NA | NA |
| WBGene00169101 | NA | NA | NA | NA |
| WBGene00169102 | NA | NA | NA | NA |
| WBGene00169103 | NA | NA | NA | NA |
| WBGene00169104 | NA | NA | NA | NA |
| WBGene00169105 | NA | NA | NA | NA |
| WBGene00169106 | NA | NA | NA | NA |
| WBGene00169107 | NA | NA | NA | NA |
| WBGene00169108 | NA | NA | NA | NA |
| WBGene00169109 | NA | NA | NA | NA |
| WBGene00169110 | NA | NA | NA | NA |
| WBGene00169111 | NA | NA | NA | NA |
| WBGene00169112 | NA | NA | NA | NA |
| WBGene00169113 | NA | NA | NA | NA |
| WBGene00169114 | NA | NA | NA | NA |
| WBGene00169115 | NA | NA | NA | NA |
| WBGene00169116 | NA | NA | NA | NA |
| WBGene00169117 | NA | NA | NA | NA |
| WBGene00169118 | NA | NA | NA | NA |
| WBGene00169119 | NA | NA | NA | NA |
| WBGene00169120 | NA | NA | NA | NA |
| WBGene00169121 | NA | NA | NA | NA |
| WBGene00169122 | NA | NA | NA | NA |
| WBGene00169123 | NA | NA | NA | NA |
| WBGene00169124 | NA | NA | NA | NA |
| WBGene00169125 | NA | NA | NA | NA |
| WBGene00169126 | NA | NA | NA | NA |
| WBGene00169127 | NA | NA | NA | NA |
| WBGene00169128 | NA | NA | NA | NA |
| WBGene00169129 | NA | NA | NA | NA |
| WBGene00169130 | NA | NA | NA | NA |
| WBGene00169131 | NA | NA | NA | NA |
| WBGene00169132 | NA | NA | NA | NA |
| WBGene00169133 | NA | NA | NA | NA |
| WBGene00169134 | NA | NA | NA | NA |
| WBGene00169135 | NA | NA | NA | NA |
| WBGene00169136 | NA | NA | NA | NA |
| WBGene00169137 | NA | NA | NA | NA |
| WBGene00169138 | NA | NA | NA | NA |

| | | | | |
|---|---|---|---|---|
| WBGene00169139 | NA | NA | NA | NA |
| WBGene00169140 | NA | NA | NA | NA |
| WBGene00169141 | NA | NA | NA | NA |
| WBGene00169142 | NA | NA | NA | NA |
| WBGene00169143 | NA | NA | NA | NA |
| WBGene00169144 | NA | NA | NA | NA |
| WBGene00169145 | NA | NA | NA | NA |
| WBGene00169146 | NA | NA | NA | NA |
| WBGene00169147 | NA | NA | NA | NA |
| WBGene00169148 | NA | NA | NA | NA |
| WBGene00169149 | NA | NA | NA | NA |
| WBGene00169150 | NA | NA | NA | NA |
| WBGene00169151 | NA | NA | NA | NA |
| WBGene00169152 | NA | NA | NA | NA |
| WBGene00169153 | NA | NA | NA | NA |
| WBGene00169154 | NA | NA | NA | NA |
| WBGene00169155 | NA | NA | NA | NA |
| WBGene00169156 | NA | NA | NA | NA |
| WBGene00169157 | NA | NA | NA | NA |
| WBGene00169158 | NA | NA | NA | NA |
| WBGene00169159 | NA | NA | NA | NA |
| WBGene00169160 | NA | NA | NA | NA |
| WBGene00169161 | NA | NA | NA | NA |
| WBGene00169162 | NA | NA | NA | NA |
| WBGene00169163 | NA | NA | NA | NA |
| WBGene00169164 | NA | NA | NA | NA |
| WBGene00169165 | NA | NA | NA | NA |
| WBGene00169166 | NA | NA | NA | NA |
| WBGene00169167 | NA | NA | NA | NA |
| WBGene00169168 | NA | NA | NA | NA |
| WBGene00169169 | NA | NA | NA | NA |
| WBGene00169170 | NA | NA | NA | NA |
| WBGene00169171 | NA | NA | NA | NA |
| WBGene00169172 | NA | NA | NA | NA |
| WBGene00169173 | NA | NA | NA | NA |
| WBGene00169174 | NA | NA | NA | NA |
| WBGene00169175 | NA | NA | NA | NA |
| WBGene00169176 | NA | NA | NA | NA |
| WBGene00169177 | NA | NA | NA | NA |
| WBGene00169178 | NA | NA | NA | NA |
| WBGene00169179 | NA | NA | NA | NA |
| WBGene00169180 | NA | NA | NA | NA |
| WBGene00169181 | NA | NA | NA | NA |

| | | | | |
|---|---|---|---|---|
| WBGene00169182 | NA | NA | NA | NA |
| WBGene00169183 | NA | NA | NA | NA |
| WBGene00169184 | NA | NA | NA | NA |
| WBGene00169185 | NA | NA | NA | NA |
| WBGene00169186 | NA | NA | NA | NA |
| WBGene00169187 | NA | NA | NA | NA |
| WBGene00169188 | NA | NA | NA | NA |
| WBGene00169189 | NA | NA | NA | NA |
| WBGene00169190 | NA | NA | NA | NA |
| WBGene00169191 | NA | NA | NA | NA |
| WBGene00169192 | NA | NA | NA | NA |
| WBGene00169193 | NA | NA | NA | NA |
| WBGene00169194 | NA | NA | NA | NA |
| WBGene00169195 | NA | NA | NA | NA |
| WBGene00169196 | NA | NA | NA | NA |
| WBGene00169197 | NA | NA | NA | NA |
| WBGene00169198 | NA | NA | NA | NA |
| WBGene00169199 | NA | NA | NA | NA |
| WBGene00169200 | NA | NA | NA | NA |
| WBGene00169201 | NA | NA | NA | NA |
| WBGene00169202 | NA | NA | NA | NA |
| WBGene00169203 | NA | NA | NA | NA |
| WBGene00169204 | NA | NA | NA | NA |
| WBGene00169205 | NA | NA | NA | NA |
| WBGene00169206 | NA | NA | NA | NA |
| WBGene00169207 | NA | NA | NA | NA |
| WBGene00169208 | NA | NA | NA | NA |
| WBGene00169209 | NA | NA | NA | NA |
| WBGene00169210 | NA | NA | NA | NA |
| WBGene00169211 | NA | NA | NA | NA |
| WBGene00169212 | NA | NA | NA | NA |
| WBGene00169213 | NA | NA | NA | NA |
| WBGene00169214 | NA | NA | NA | NA |
| WBGene00169215 | NA | NA | NA | NA |
| WBGene00169216 | NA | NA | NA | NA |
| WBGene00169217 | NA | NA | NA | NA |
| WBGene00169218 | NA | NA | NA | NA |
| WBGene00169219 | NA | NA | NA | NA |
| WBGene00169220 | NA | NA | NA | NA |
| WBGene00169221 | NA | NA | NA | NA |
| WBGene00169222 | NA | NA | NA | NA |
| WBGene00169223 | NA | NA | NA | NA |
| WBGene00169224 | NA | NA | NA | NA |

| | | | | |
|---|---|---|---|---|
| WBGene00169225 | NA | NA | NA | NA |
| WBGene00169226 | NA | NA | NA | NA |
| WBGene00169227 | NA | NA | NA | NA |
| WBGene00169228 | NA | NA | NA | NA |
| WBGene00169229 | NA | NA | NA | NA |
| WBGene00169230 | NA | NA | NA | NA |
| WBGene00169231 | NA | NA | NA | NA |
| WBGene00169232 | NA | NA | NA | NA |
| WBGene00169233 | NA | NA | NA | NA |
| WBGene00169234 | NA | NA | NA | NA |
| WBGene00169235 | NA | NA | NA | NA |
| WBGene00169236 | NA | NA | NA | NA |
| WBGene00169237 | NA | NA | NA | NA |
| WBGene00169238 | NA | NA | NA | NA |
| WBGene00169239 | NA | NA | NA | NA |
| WBGene00169240 | NA | NA | NA | NA |
| WBGene00169242 | NA | NA | NA | NA |
| WBGene00169243 | NA | NA | NA | NA |
| WBGene00169244 | NA | NA | NA | NA |
| WBGene00169245 | NA | NA | NA | NA |
| WBGene00169246 | NA | NA | NA | NA |
| WBGene00169247 | NA | NA | NA | NA |
| WBGene00169248 | NA | NA | NA | NA |
| WBGene00169249 | NA | NA | NA | NA |
| WBGene00169250 | NA | NA | NA | NA |
| WBGene00169251 | NA | NA | NA | NA |
| WBGene00169252 | NA | NA | NA | NA |
| WBGene00169253 | NA | NA | NA | NA |
| WBGene00169254 | NA | NA | NA | NA |
| WBGene00169255 | NA | NA | NA | NA |
| WBGene00169256 | NA | NA | NA | NA |
| WBGene00169257 | NA | NA | NA | NA |
| WBGene00169258 | NA | NA | NA | NA |
| WBGene00169259 | NA | NA | NA | NA |
| WBGene00169260 | NA | NA | NA | NA |
| WBGene00169261 | NA | NA | NA | NA |
| WBGene00169262 | NA | NA | NA | NA |
| WBGene00169263 | NA | NA | NA | NA |
| WBGene00169264 | NA | NA | NA | NA |
| WBGene00169265 | NA | NA | NA | NA |
| WBGene00169266 | NA | NA | NA | NA |
| WBGene00169267 | NA | NA | NA | NA |
| WBGene00169268 | NA | NA | NA | NA |

| | | | | |
|---|---|---|---|---|
| WBGene00169269 | NA | NA | NA | NA |
| WBGene00169270 | NA | NA | NA | NA |
| WBGene00169271 | NA | NA | NA | NA |
| WBGene00169272 | NA | NA | NA | NA |
| WBGene00169273 | NA | NA | NA | NA |
| WBGene00169274 | NA | NA | NA | NA |
| WBGene00169275 | NA | NA | NA | NA |
| WBGene00169276 | NA | NA | NA | NA |
| WBGene00169277 | NA | NA | NA | NA |
| WBGene00169278 | NA | NA | NA | NA |
| WBGene00169279 | NA | NA | NA | NA |
| WBGene00169280 | NA | NA | NA | NA |
| WBGene00169281 | NA | NA | NA | NA |
| WBGene00169282 | NA | NA | NA | NA |
| WBGene00169283 | NA | NA | NA | NA |
| WBGene00169284 | NA | NA | NA | NA |
| WBGene00169285 | NA | NA | NA | NA |
| WBGene00169286 | NA | NA | NA | NA |
| WBGene00169287 | NA | NA | NA | NA |
| WBGene00169288 | NA | NA | NA | NA |
| WBGene00169289 | NA | NA | NA | NA |
| WBGene00169290 | NA | NA | NA | NA |
| WBGene00169291 | NA | NA | NA | NA |
| WBGene00169292 | NA | NA | NA | NA |
| WBGene00169293 | NA | NA | NA | NA |
| WBGene00169294 | NA | NA | NA | NA |
| WBGene00169295 | NA | NA | NA | NA |
| WBGene00169296 | NA | NA | NA | NA |
| WBGene00169297 | NA | NA | NA | NA |
| WBGene00169298 | NA | NA | NA | NA |
| WBGene00169299 | NA | NA | NA | NA |
| WBGene00169300 | NA | NA | NA | NA |
| WBGene00169301 | NA | NA | NA | NA |
| WBGene00169302 | NA | NA | NA | NA |
| WBGene00169303 | NA | NA | NA | NA |
| WBGene00169304 | NA | NA | NA | NA |
| WBGene00169305 | NA | NA | NA | NA |
| WBGene00169306 | NA | NA | NA | NA |
| WBGene00169307 | NA | NA | NA | NA |
| WBGene00169308 | NA | NA | NA | NA |
| WBGene00169309 | NA | NA | NA | NA |
| WBGene00169310 | NA | NA | NA | NA |
| WBGene00169311 | NA | NA | NA | NA |

| | | | | |
|---|---|---|---|---|
| WBGene00169312 | NA | NA | NA | NA |
| WBGene00169313 | NA | NA | NA | NA |
| WBGene00169314 | NA | NA | NA | NA |
| WBGene00169315 | NA | NA | NA | NA |
| WBGene00169316 | NA | NA | NA | NA |
| WBGene00169317 | NA | NA | NA | NA |
| WBGene00169318 | NA | NA | NA | NA |
| WBGene00169319 | NA | NA | NA | NA |
| WBGene00169320 | NA | NA | NA | NA |
| WBGene00169321 | NA | NA | NA | NA |
| WBGene00169322 | NA | NA | NA | NA |
| WBGene00169323 | NA | NA | NA | NA |
| WBGene00169324 | NA | NA | NA | NA |
| WBGene00169325 | NA | NA | NA | NA |
| WBGene00169326 | NA | NA | NA | NA |
| WBGene00169327 | NA | NA | NA | NA |
| WBGene00169328 | NA | NA | NA | NA |
| WBGene00169329 | NA | NA | NA | NA |
| WBGene00169330 | NA | NA | NA | NA |
| WBGene00169331 | NA | NA | NA | NA |
| WBGene00169332 | NA | NA | NA | NA |
| WBGene00169333 | NA | NA | NA | NA |
| WBGene00169334 | NA | NA | NA | NA |
| WBGene00169335 | NA | NA | NA | NA |
| WBGene00169336 | NA | NA | NA | NA |
| WBGene00169337 | NA | NA | NA | NA |
| WBGene00169338 | NA | NA | NA | NA |
| WBGene00169339 | NA | NA | NA | NA |
| WBGene00169340 | NA | NA | NA | NA |
| WBGene00169341 | NA | NA | NA | NA |
| WBGene00169342 | NA | NA | NA | NA |
| WBGene00169343 | NA | NA | NA | NA |
| WBGene00169344 | NA | NA | NA | NA |
| WBGene00169345 | NA | NA | NA | NA |
| WBGene00169346 | NA | NA | NA | NA |
| WBGene00169347 | NA | NA | NA | NA |
| WBGene00169348 | NA | NA | NA | NA |
| WBGene00169349 | NA | NA | NA | NA |
| WBGene00169350 | NA | NA | NA | NA |
| WBGene00169351 | NA | NA | NA | NA |
| WBGene00169352 | NA | NA | NA | NA |
| WBGene00169353 | NA | NA | NA | NA |
| WBGene00169354 | NA | NA | NA | NA |

| | | | | |
|---|---|---|---|---|
| WBGene00169355 | NA | NA | NA | NA |
| WBGene00169356 | NA | NA | NA | NA |
| WBGene00169357 | NA | NA | NA | NA |
| WBGene00169358 | NA | NA | NA | NA |
| WBGene00169359 | NA | NA | NA | NA |
| WBGene00169360 | NA | NA | NA | NA |
| WBGene00169361 | NA | NA | NA | NA |
| WBGene00169362 | NA | NA | NA | NA |
| WBGene00169363 | NA | NA | NA | NA |
| WBGene00169364 | NA | NA | NA | NA |
| WBGene00169365 | NA | NA | NA | NA |
| WBGene00169366 | NA | NA | NA | NA |
| WBGene00169367 | NA | NA | NA | NA |
| WBGene00169368 | NA | NA | NA | NA |
| WBGene00169369 | NA | NA | NA | NA |
| WBGene00169370 | NA | NA | NA | NA |
| WBGene00169371 | NA | NA | NA | NA |
| WBGene00169372 | NA | NA | NA | NA |
| WBGene00169373 | NA | NA | NA | NA |
| WBGene00169374 | NA | NA | NA | NA |
| WBGene00169375 | NA | NA | NA | NA |
| WBGene00169376 | NA | NA | NA | NA |
| WBGene00169377 | NA | NA | NA | NA |
| WBGene00169378 | NA | NA | NA | NA |
| WBGene00169379 | NA | NA | NA | NA |
| WBGene00169380 | NA | NA | NA | NA |
| WBGene00169381 | NA | NA | NA | NA |
| WBGene00169382 | NA | NA | NA | NA |
| WBGene00169383 | NA | NA | NA | NA |
| WBGene00169384 | NA | NA | NA | NA |
| WBGene00169385 | NA | NA | NA | NA |
| WBGene00169386 | NA | NA | NA | NA |
| WBGene00169387 | NA | NA | NA | NA |
| WBGene00169388 | NA | NA | NA | NA |
| WBGene00169389 | NA | NA | NA | NA |
| WBGene00169390 | NA | NA | NA | NA |
| WBGene00169391 | NA | NA | NA | NA |
| WBGene00169392 | NA | NA | NA | NA |
| WBGene00169393 | NA | NA | NA | NA |
| WBGene00169394 | NA | NA | NA | NA |
| WBGene00169395 | NA | NA | NA | NA |
| WBGene00169396 | NA | NA | NA | NA |
| WBGene00169397 | NA | NA | NA | NA |

| | | | | |
|---|---|---|---|---|
| WBGene00169398 | NA | NA | NA | NA |
| WBGene00169399 | NA | NA | NA | NA |
| WBGene00169401 | NA | NA | NA | NA |
| WBGene00169402 | NA | NA | NA | NA |
| WBGene00169403 | NA | NA | NA | NA |
| WBGene00169404 | NA | NA | NA | NA |
| WBGene00169405 | NA | NA | NA | NA |
| WBGene00169406 | NA | NA | NA | NA |
| WBGene00169407 | NA | NA | NA | NA |
| WBGene00169408 | NA | NA | NA | NA |
| WBGene00169409 | NA | NA | NA | NA |
| WBGene00169410 | NA | NA | NA | NA |
| WBGene00169411 | NA | NA | NA | NA |
| WBGene00169412 | NA | NA | NA | NA |
| WBGene00169413 | NA | NA | NA | NA |
| WBGene00169414 | NA | NA | NA | NA |
| WBGene00169415 | NA | NA | NA | NA |
| WBGene00169416 | NA | NA | NA | NA |
| WBGene00169417 | NA | NA | NA | NA |
| WBGene00169418 | NA | NA | NA | NA |
| WBGene00169419 | NA | NA | NA | NA |
| WBGene00169420 | NA | NA | NA | NA |
| WBGene00169421 | NA | NA | NA | NA |
| WBGene00169422 | NA | NA | NA | NA |
| WBGene00169423 | NA | NA | NA | NA |
| WBGene00169424 | NA | NA | NA | NA |
| WBGene00169425 | NA | NA | NA | NA |
| WBGene00169426 | NA | NA | NA | NA |
| WBGene00169427 | NA | NA | NA | NA |
| WBGene00169428 | NA | NA | NA | NA |
| WBGene00169429 | NA | NA | NA | NA |
| WBGene00169430 | NA | NA | NA | NA |
| WBGene00169431 | NA | NA | NA | NA |
| WBGene00169432 | NA | NA | NA | NA |
| WBGene00169433 | NA | NA | NA | NA |
| WBGene00169434 | NA | NA | NA | NA |
| WBGene00169435 | NA | NA | NA | NA |
| WBGene00169436 | NA | NA | NA | NA |
| WBGene00169437 | NA | NA | NA | NA |
| WBGene00169438 | NA | NA | NA | NA |
| WBGene00169439 | NA | NA | NA | NA |
| WBGene00169440 | NA | NA | NA | NA |
| WBGene00169441 | NA | NA | NA | NA |

| | | | | |
|---|---|---|---|---|
| WBGene00169442 | NA | NA | NA | NA |
| WBGene00169443 | NA | NA | NA | NA |
| WBGene00169444 | NA | NA | NA | NA |
| WBGene00169445 | NA | NA | NA | NA |
| WBGene00169446 | NA | NA | NA | NA |
| WBGene00169447 | NA | NA | NA | NA |
| WBGene00169448 | NA | NA | NA | NA |
| WBGene00169449 | NA | NA | NA | NA |
| WBGene00169450 | NA | NA | NA | NA |
| WBGene00169451 | NA | NA | NA | NA |
| WBGene00169452 | NA | NA | NA | NA |
| WBGene00169453 | NA | NA | NA | NA |
| WBGene00169454 | NA | NA | NA | NA |
| WBGene00169455 | NA | NA | NA | NA |
| WBGene00169456 | NA | NA | NA | NA |
| WBGene00169457 | NA | NA | NA | NA |
| WBGene00169458 | NA | NA | NA | NA |
| WBGene00169459 | NA | NA | NA | NA |
| WBGene00169460 | NA | NA | NA | NA |
| WBGene00169461 | NA | NA | NA | NA |
| WBGene00169462 | NA | NA | NA | NA |
| WBGene00169463 | NA | NA | NA | NA |
| WBGene00169464 | NA | NA | NA | NA |
| WBGene00169465 | NA | NA | NA | NA |
| WBGene00169466 | NA | NA | NA | NA |
| WBGene00169467 | NA | NA | NA | NA |
| WBGene00169468 | NA | NA | NA | NA |
| WBGene00169469 | NA | NA | NA | NA |
| WBGene00169470 | NA | NA | NA | NA |
| WBGene00169471 | NA | NA | NA | NA |
| WBGene00169472 | NA | NA | NA | NA |
| WBGene00169473 | NA | NA | NA | NA |
| WBGene00169474 | NA | NA | NA | NA |
| WBGene00169475 | NA | NA | NA | NA |
| WBGene00169476 | NA | NA | NA | NA |
| WBGene00169477 | NA | NA | NA | NA |
| WBGene00169478 | NA | NA | NA | NA |
| WBGene00169479 | NA | NA | NA | NA |
| WBGene00169480 | NA | NA | NA | NA |
| WBGene00169481 | NA | NA | NA | NA |
| WBGene00169482 | NA | NA | NA | NA |
| WBGene00169483 | NA | NA | NA | NA |
| WBGene00169484 | NA | NA | NA | NA |

| | | | | |
|---|---|---|---|---|
| WBGene00169485 | NA | NA | NA | NA |
| WBGene00169486 | -0.003108065 | 0.95728506 | NA | NA |
| WBGene00169487 | NA | NA | NA | NA |
| WBGene00169488 | NA | NA | NA | NA |
| WBGene00169489 | NA | NA | NA | NA |
| WBGene00169490 | NA | NA | NA | NA |
| WBGene00169491 | NA | NA | NA | NA |
| WBGene00169492 | NA | NA | NA | NA |
| WBGene00169493 | NA | NA | NA | NA |
| WBGene00169494 | NA | NA | NA | NA |
| WBGene00169495 | NA | NA | NA | NA |
| WBGene00169496 | NA | NA | NA | NA |
| WBGene00169497 | NA | NA | NA | NA |
| WBGene00169498 | NA | NA | NA | NA |
| WBGene00169499 | NA | NA | NA | NA |
| WBGene00169500 | NA | NA | NA | NA |
| WBGene00169501 | NA | NA | NA | NA |
| WBGene00169502 | NA | NA | NA | NA |
| WBGene00169503 | NA | NA | NA | NA |
| WBGene00169504 | NA | NA | NA | NA |
| WBGene00169505 | NA | NA | NA | NA |
| WBGene00169506 | NA | NA | NA | NA |
| WBGene00169507 | NA | NA | NA | NA |
| WBGene00169508 | NA | NA | NA | NA |
| WBGene00169509 | NA | NA | NA | NA |
| WBGene00169510 | NA | NA | NA | NA |
| WBGene00169511 | NA | NA | NA | NA |
| WBGene00169512 | NA | NA | NA | NA |
| WBGene00169513 | NA | NA | NA | NA |
| WBGene00169514 | NA | NA | NA | NA |
| WBGene00169515 | NA | NA | NA | NA |
| WBGene00169516 | NA | NA | NA | NA |
| WBGene00169517 | NA | NA | NA | NA |
| WBGene00169518 | NA | NA | NA | NA |
| WBGene00169519 | NA | NA | NA | NA |
| WBGene00169520 | NA | NA | NA | NA |
| WBGene00169521 | NA | NA | NA | NA |
| WBGene00169522 | NA | NA | NA | NA |
| WBGene00169523 | NA | NA | NA | NA |
| WBGene00169524 | NA | NA | NA | NA |
| WBGene00169525 | NA | NA | NA | NA |
| WBGene00169526 | NA | NA | NA | NA |
| WBGene00169527 | NA | NA | NA | NA |

| | | | | | |
|---|---|---|---|---|---|
| WBGene00169528 | | 0.020232312 | 0.84643798 | NA | NA |
| WBGene00169529 | NA | | NA | NA | NA |
| WBGene00169530 | NA | | NA | NA | NA |
| WBGene00169531 | NA | | NA | NA | NA |
| WBGene00169532 | NA | | NA | NA | NA |
| WBGene00169533 | NA | | NA | NA | NA |
| WBGene00169534 | NA | | NA | NA | NA |
| WBGene00169535 | NA | | NA | NA | NA |
| WBGene00169536 | NA | | NA | NA | NA |
| WBGene00169537 | NA | | NA | NA | NA |
| WBGene00169538 | | 0.0398788 | 0.58971221 | NA | NA |
| WBGene00169539 | NA | | NA | NA | NA |
| WBGene00169540 | NA | | NA | NA | NA |
| WBGene00169541 | NA | | NA | NA | NA |
| WBGene00169542 | NA | | NA | NA | NA |
| WBGene00169543 | NA | | NA | NA | NA |
| WBGene00169544 | NA | | NA | NA | NA |
| WBGene00169545 | NA | | NA | NA | NA |
| WBGene00169546 | NA | | NA | NA | NA |
| WBGene00169547 | NA | | NA | NA | NA |
| WBGene00169548 | NA | | NA | NA | NA |
| WBGene00169549 | NA | | NA | NA | NA |
| WBGene00169550 | NA | | NA | NA | NA |
| WBGene00169551 | NA | | NA | NA | NA |
| WBGene00169552 | NA | | NA | NA | NA |
| WBGene00169553 | NA | | NA | NA | NA |
| WBGene00169554 | NA | | NA | NA | NA |
| WBGene00169555 | NA | | NA | NA | NA |
| WBGene00169556 | NA | | NA | NA | NA |
| WBGene00169557 | NA | | NA | NA | NA |
| WBGene00169558 | NA | | NA | NA | NA |
| WBGene00169559 | NA | | NA | NA | NA |
| WBGene00169560 | NA | | NA | NA | NA |
| WBGene00169561 | NA | | NA | NA | NA |
| WBGene00169562 | NA | | NA | NA | NA |
| WBGene00169563 | NA | | NA | NA | NA |
| WBGene00169564 | NA | | NA | NA | NA |
| WBGene00169565 | NA | | NA | NA | NA |
| WBGene00169566 | NA | | NA | NA | NA |
| WBGene00169567 | NA | | NA | NA | NA |
| WBGene00169568 | NA | | NA | NA | NA |
| WBGene00169569 | NA | | NA | NA | NA |
| WBGene00169570 | NA | | NA | NA | NA |

| | | | | |
|---|---|---|---|---|
| WBGene00169571 | NA | NA | NA | NA |
| WBGene00169572 | NA | NA | NA | NA |
| WBGene00169573 | NA | NA | NA | NA |
| WBGene00169574 | NA | NA | NA | NA |
| WBGene00169575 | NA | NA | NA | NA |
| WBGene00169576 | NA | NA | NA | NA |
| WBGene00169577 | NA | NA | NA | NA |
| WBGene00169578 | NA | NA | NA | NA |
| WBGene00169579 | NA | NA | NA | NA |
| WBGene00169580 | NA | NA | NA | NA |
| WBGene00169581 | NA | NA | NA | NA |
| WBGene00169582 | NA | NA | NA | NA |
| WBGene00169583 | NA | NA | NA | NA |
| WBGene00169584 | NA | NA | NA | NA |
| WBGene00169585 | NA | NA | NA | NA |
| WBGene00169586 | NA | NA | NA | NA |
| WBGene00169587 | NA | NA | NA | NA |
| WBGene00169588 | NA | NA | NA | NA |
| WBGene00169589 | NA | NA | NA | NA |
| WBGene00169590 | NA | NA | NA | NA |
| WBGene00169591 | NA | NA | NA | NA |
| WBGene00169592 | NA | NA | NA | NA |
| WBGene00169593 | NA | NA | NA | NA |
| WBGene00169594 | NA | NA | NA | NA |
| WBGene00169595 | NA | NA | NA | NA |
| WBGene00169596 | NA | NA | NA | NA |
| WBGene00169597 | NA | NA | NA | NA |
| WBGene00169598 | NA | NA | NA | NA |
| WBGene00169599 | NA | NA | NA | NA |
| WBGene00169600 | NA | NA | NA | NA |
| WBGene00169601 | NA | NA | NA | NA |
| WBGene00169602 | NA | NA | NA | NA |
| WBGene00169603 | NA | NA | NA | NA |
| WBGene00169604 | NA | NA | NA | NA |
| WBGene00169605 | NA | NA | NA | NA |
| WBGene00169606 | NA | NA | NA | NA |
| WBGene00169607 | NA | NA | NA | NA |
| WBGene00169608 | NA | NA | NA | NA |
| WBGene00169609 | NA | NA | NA | NA |
| WBGene00169610 | NA | NA | NA | NA |
| WBGene00169611 | NA | NA | NA | NA |
| WBGene00169612 | NA | NA | NA | NA |
| WBGene00169613 | NA | NA | NA | NA |

| | | | | |
|---|---|---|---|---|
| WBGene00169614 | NA | NA | NA | NA |
| WBGene00169615 | NA | NA | NA | NA |
| WBGene00169617 | NA | NA | NA | NA |
| WBGene00169618 | NA | NA | NA | NA |
| WBGene00169619 | NA | NA | NA | NA |
| WBGene00169620 | NA | NA | NA | NA |
| WBGene00169621 | NA | NA | NA | NA |
| WBGene00169622 | 0.020925399 | 0.84643798 | NA | NA |
| WBGene00169623 | NA | NA | NA | NA |
| WBGene00169624 | NA | NA | NA | NA |
| WBGene00169625 | NA | NA | NA | NA |
| WBGene00169626 | NA | NA | NA | NA |
| WBGene00169627 | NA | NA | NA | NA |
| WBGene00169628 | NA | NA | NA | NA |
| WBGene00169629 | NA | NA | NA | NA |
| WBGene00169630 | NA | NA | NA | NA |
| WBGene00169631 | NA | NA | NA | NA |
| WBGene00169632 | NA | NA | NA | NA |
| WBGene00169633 | NA | NA | NA | NA |
| WBGene00169634 | NA | NA | NA | NA |
| WBGene00169635 | NA | NA | NA | NA |
| WBGene00169636 | NA | NA | NA | NA |
| WBGene00169637 | NA | NA | NA | NA |
| WBGene00169638 | NA | NA | NA | NA |
| WBGene00169639 | NA | NA | NA | NA |
| WBGene00169640 | NA | NA | NA | NA |
| WBGene00169641 | NA | NA | NA | NA |
| WBGene00169642 | NA | NA | NA | NA |
| WBGene00169643 | NA | NA | NA | NA |
| WBGene00169644 | NA | NA | NA | NA |
| WBGene00169645 | NA | NA | NA | NA |
| WBGene00169646 | NA | NA | NA | NA |
| WBGene00169647 | NA | NA | NA | NA |
| WBGene00169648 | NA | NA | NA | NA |
| WBGene00169649 | NA | NA | NA | NA |
| WBGene00169650 | NA | NA | NA | NA |
| WBGene00169651 | NA | NA | NA | NA |
| WBGene00169652 | NA | NA | NA | NA |
| WBGene00169653 | NA | NA | NA | NA |
| WBGene00169654 | NA | NA | NA | NA |
| WBGene00169655 | NA | NA | NA | NA |
| WBGene00169656 | NA | NA | NA | NA |
| WBGene00169657 | NA | NA | NA | NA |

| | | | | | |
|---|---|---|---|---|---|
| WBGene00169658 | NA | | NA | NA | NA |
| WBGene00169659 | NA | | NA | NA | NA |
| WBGene00169660 | NA | | NA | NA | NA |
| WBGene00169661 | NA | | NA | NA | NA |
| WBGene00169662 | NA | | NA | NA | NA |
| WBGene00169663 | NA | | NA | NA | NA |
| WBGene00169664 | | 0.020232312 | 0.84643798 | NA | NA |
| WBGene00169665 | NA | | NA | NA | NA |
| WBGene00169666 | NA | | NA | NA | NA |
| WBGene00169667 | NA | | NA | NA | NA |
| WBGene00169668 | NA | | NA | NA | NA |
| WBGene00169669 | NA | | NA | NA | NA |
| WBGene00169670 | NA | | NA | NA | NA |
| WBGene00169671 | NA | | NA | NA | NA |
| WBGene00169672 | NA | | NA | NA | NA |
| WBGene00169673 | NA | | NA | NA | NA |
| WBGene00169674 | NA | | NA | NA | NA |
| WBGene00169675 | NA | | NA | NA | NA |
| WBGene00169676 | NA | | NA | NA | NA |
| WBGene00169677 | NA | | NA | NA | NA |
| WBGene00169678 | NA | | NA | NA | NA |
| WBGene00169679 | NA | | NA | NA | NA |
| WBGene00169680 | NA | | NA | NA | NA |
| WBGene00169681 | NA | | NA | NA | NA |
| WBGene00169682 | NA | | NA | NA | NA |
| WBGene00169683 | NA | | NA | NA | NA |
| WBGene00169684 | NA | | NA | NA | NA |
| WBGene00169685 | NA | | NA | NA | NA |
| WBGene00169686 | NA | | NA | NA | NA |
| WBGene00169687 | NA | | NA | NA | NA |
| WBGene00169688 | | 0.020317245 | 0.84643798 | NA | NA |
| WBGene00169689 | NA | | NA | NA | NA |
| WBGene00169690 | NA | | NA | NA | NA |
| WBGene00169691 | NA | | NA | NA | NA |
| WBGene00169692 | NA | | NA | NA | NA |
| WBGene00169693 | NA | | NA | NA | NA |
| WBGene00169694 | NA | | NA | NA | NA |
| WBGene00169695 | NA | | NA | NA | NA |
| WBGene00169696 | NA | | NA | NA | NA |
| WBGene00169697 | NA | | NA | NA | NA |
| WBGene00169698 | NA | | NA | NA | NA |
| WBGene00169699 | NA | | NA | NA | NA |
| WBGene00169700 | NA | | NA | NA | NA |

| | | | | |
|---|---|---|---|---|
| WBGene00169701 | NA | NA | NA | NA |
| WBGene00169702 | NA | NA | NA | NA |
| WBGene00169703 | NA | NA | NA | NA |
| WBGene00169704 | NA | NA | NA | NA |
| WBGene00169705 | NA | NA | NA | NA |
| WBGene00169706 | NA | NA | NA | NA |
| WBGene00169707 | NA | NA | NA | NA |
| WBGene00169708 | NA | NA | NA | NA |
| WBGene00169709 | NA | NA | NA | NA |
| WBGene00169710 | NA | NA | NA | NA |
| WBGene00169711 | NA | NA | NA | NA |
| WBGene00169712 | NA | NA | NA | NA |
| WBGene00169713 | NA | NA | NA | NA |
| WBGene00169714 | NA | NA | NA | NA |
| WBGene00169715 | NA | NA | NA | NA |
| WBGene00169716 | NA | NA | NA | NA |
| WBGene00169717 | NA | NA | NA | NA |
| WBGene00169718 | NA | NA | NA | NA |
| WBGene00169719 | NA | NA | NA | NA |
| WBGene00169720 | NA | NA | NA | NA |
| WBGene00169721 | NA | NA | NA | NA |
| WBGene00169722 | NA | NA | NA | NA |
| WBGene00169723 | NA | NA | NA | NA |
| WBGene00169724 | NA | NA | NA | NA |
| WBGene00169725 | NA | NA | NA | NA |
| WBGene00169726 | NA | NA | NA | NA |
| WBGene00169727 | NA | NA | NA | NA |
| WBGene00169728 | NA | NA | NA | NA |
| WBGene00169729 | NA | NA | NA | NA |
| WBGene00169730 | NA | NA | NA | NA |
| WBGene00169731 | NA | NA | NA | NA |
| WBGene00169732 | NA | NA | NA | NA |
| WBGene00169733 | NA | NA | NA | NA |
| WBGene00169734 | -0.046394786 | 0.50231372 | NA | NA |
| WBGene00169735 | NA | NA | NA | NA |
| WBGene00169736 | NA | NA | NA | NA |
| WBGene00169737 | NA | NA | NA | NA |
| WBGene00169738 | NA | NA | NA | NA |
| WBGene00169739 | NA | NA | NA | NA |
| WBGene00169740 | NA | NA | NA | NA |
| WBGene00169741 | NA | NA | NA | NA |
| WBGene00169742 | NA | NA | NA | NA |
| WBGene00169743 | NA | NA | NA | NA |

| | | | | |
|---|---|---|---|---|
| WBGene00169744 | NA | NA | NA | NA |
| WBGene00169745 | NA | NA | NA | NA |
| WBGene00169746 | NA | NA | NA | NA |
| WBGene00169747 | NA | NA | NA | NA |
| WBGene00169748 | NA | NA | NA | NA |
| WBGene00169749 | NA | NA | NA | NA |
| WBGene00169750 | NA | NA | NA | NA |
| WBGene00169751 | NA | NA | NA | NA |
| WBGene00169752 | NA | NA | NA | NA |
| WBGene00169753 | NA | NA | NA | NA |
| WBGene00169754 | NA | NA | NA | NA |
| WBGene00169755 | NA | NA | NA | NA |
| WBGene00169756 | NA | NA | NA | NA |
| WBGene00169757 | NA | NA | NA | NA |
| WBGene00169758 | NA | NA | NA | NA |
| WBGene00169759 | NA | NA | NA | NA |
| WBGene00169760 | NA | NA | NA | NA |
| WBGene00169761 | NA | NA | NA | NA |
| WBGene00169762 | NA | NA | NA | NA |
| WBGene00169763 | NA | NA | NA | NA |
| WBGene00169764 | NA | NA | NA | NA |
| WBGene00169765 | NA | NA | NA | NA |
| WBGene00169766 | NA | NA | NA | NA |
| WBGene00169767 | NA | NA | NA | NA |
| WBGene00169768 | NA | NA | NA | NA |
| WBGene00169769 | NA | NA | NA | NA |
| WBGene00169770 | NA | NA | NA | NA |
| WBGene00169771 | NA | NA | NA | NA |
| WBGene00169772 | NA | NA | NA | NA |
| WBGene00169773 | NA | NA | NA | NA |
| WBGene00169774 | NA | NA | NA | NA |
| WBGene00169775 | NA | NA | NA | NA |
| WBGene00169776 | NA | NA | NA | NA |
| WBGene00169777 | NA | NA | NA | NA |
| WBGene00169778 | NA | NA | NA | NA |
| WBGene00169779 | NA | NA | NA | NA |
| WBGene00169780 | NA | NA | NA | NA |
| WBGene00169781 | NA | NA | NA | NA |
| WBGene00169782 | NA | NA | NA | NA |
| WBGene00169783 | NA | NA | NA | NA |
| WBGene00169784 | NA | NA | NA | NA |
| WBGene00169785 | NA | NA | NA | NA |
| WBGene00169786 | NA | NA | NA | NA |

| | | | | |
|---|---|---|---|---|
| WBGene00169787 | NA | NA | NA | NA |
| WBGene00169788 | NA | NA | NA | NA |
| WBGene00169789 | NA | NA | NA | NA |
| WBGene00169790 | NA | NA | NA | NA |
| WBGene00169791 | NA | NA | NA | NA |
| WBGene00169792 | NA | NA | NA | NA |
| WBGene00169793 | NA | NA | NA | NA |
| WBGene00169794 | NA | NA | NA | NA |
| WBGene00169795 | NA | NA | NA | NA |
| WBGene00169796 | NA | NA | NA | NA |
| WBGene00169797 | NA | NA | NA | NA |
| WBGene00169798 | NA | NA | NA | NA |
| WBGene00169799 | NA | NA | NA | NA |
| WBGene00169800 | NA | NA | NA | NA |
| WBGene00169801 | NA | NA | NA | NA |
| WBGene00169802 | NA | NA | NA | NA |
| WBGene00169803 | NA | NA | NA | NA |
| WBGene00169804 | NA | NA | NA | NA |
| WBGene00169805 | NA | NA | NA | NA |
| WBGene00169806 | NA | NA | NA | NA |
| WBGene00169807 | NA | NA | NA | NA |
| WBGene00169808 | NA | NA | NA | NA |
| WBGene00169809 | NA | NA | NA | NA |
| WBGene00169810 | NA | NA | NA | NA |
| WBGene00169811 | NA | NA | NA | NA |
| WBGene00169812 | NA | NA | NA | NA |
| WBGene00169813 | NA | NA | NA | NA |
| WBGene00169814 | NA | NA | NA | NA |
| WBGene00169815 | NA | NA | NA | NA |
| WBGene00169816 | NA | NA | NA | NA |
| WBGene00169817 | NA | NA | NA | NA |
| WBGene00169818 | NA | NA | NA | NA |
| WBGene00169819 | NA | NA | NA | NA |
| WBGene00169820 | NA | NA | NA | NA |
| WBGene00169821 | NA | NA | NA | NA |
| WBGene00169822 | NA | NA | NA | NA |
| WBGene00169823 | NA | NA | NA | NA |
| WBGene00169824 | NA | NA | NA | NA |
| WBGene00169825 | NA | NA | NA | NA |
| WBGene00169826 | NA | NA | NA | NA |
| WBGene00169827 | NA | NA | NA | NA |
| WBGene00169828 | NA | NA | NA | NA |
| WBGene00169829 | NA | NA | NA | NA |

| | | | | |
|---|---|---|---|---|
| WBGene00169830 | NA | NA | NA | NA |
| WBGene00169831 | NA | NA | NA | NA |
| WBGene00169832 | NA | NA | NA | NA |
| WBGene00169833 | NA | NA | NA | NA |
| WBGene00169834 | NA | NA | NA | NA |
| WBGene00169835 | -0.022930767 | 0.78121253 | NA | NA |
| WBGene00169836 | NA | NA | NA | NA |
| WBGene00169837 | NA | NA | NA | NA |
| WBGene00169838 | NA | NA | NA | NA |
| WBGene00169839 | NA | NA | NA | NA |
| WBGene00169840 | NA | NA | NA | NA |
| WBGene00169841 | NA | NA | NA | NA |
| WBGene00169842 | NA | NA | NA | NA |
| WBGene00169843 | NA | NA | NA | NA |
| WBGene00169844 | NA | NA | NA | NA |
| WBGene00169845 | NA | NA | NA | NA |
| WBGene00169846 | 0.020232312 | 0.84643798 | NA | NA |
| WBGene00169847 | NA | NA | NA | NA |
| WBGene00169848 | NA | NA | NA | NA |
| WBGene00169849 | NA | NA | NA | NA |
| WBGene00169850 | NA | NA | NA | NA |
| WBGene00169851 | NA | NA | NA | NA |
| WBGene00169852 | NA | NA | NA | NA |
| WBGene00169853 | NA | NA | NA | NA |
| WBGene00169854 | NA | NA | NA | NA |
| WBGene00169855 | NA | NA | NA | NA |
| WBGene00169856 | NA | NA | NA | NA |
| WBGene00169857 | NA | NA | NA | NA |
| WBGene00169858 | NA | NA | NA | NA |
| WBGene00169859 | NA | NA | NA | NA |
| WBGene00169860 | NA | NA | NA | NA |
| WBGene00169861 | NA | NA | NA | NA |
| WBGene00169862 | NA | NA | NA | NA |
| WBGene00169863 | NA | NA | NA | NA |
| WBGene00169864 | NA | NA | NA | NA |
| WBGene00169865 | NA | NA | NA | NA |
| WBGene00169866 | NA | NA | NA | NA |
| WBGene00169867 | NA | NA | NA | NA |
| WBGene00169868 | NA | NA | NA | NA |
| WBGene00169869 | NA | NA | NA | NA |
| WBGene00169870 | NA | NA | NA | NA |
| WBGene00169871 | NA | NA | NA | NA |
| WBGene00169872 | NA | NA | NA | NA |

| | | | | |
|---|---|---|---|---|
| WBGene00169873 | NA | NA | NA | NA |
| WBGene00169874 | NA | NA | NA | NA |
| WBGene00169875 | NA | NA | NA | NA |
| WBGene00169876 | NA | NA | NA | NA |
| WBGene00169877 | NA | NA | NA | NA |
| WBGene00169878 | NA | NA | NA | NA |
| WBGene00169879 | NA | NA | NA | NA |
| WBGene00169880 | NA | NA | NA | NA |
| WBGene00169881 | NA | NA | NA | NA |
| WBGene00169882 | NA | NA | NA | NA |
| WBGene00169883 | NA | NA | NA | NA |
| WBGene00169884 | NA | NA | NA | NA |
| WBGene00169885 | NA | NA | NA | NA |
| WBGene00169886 | NA | NA | NA | NA |
| WBGene00169887 | NA | NA | NA | NA |
| WBGene00169888 | NA | NA | NA | NA |
| WBGene00169889 | NA | NA | NA | NA |
| WBGene00169890 | NA | NA | NA | NA |
| WBGene00169891 | NA | NA | NA | NA |
| WBGene00169892 | NA | NA | NA | NA |
| WBGene00169893 | NA | NA | NA | NA |
| WBGene00169894 | NA | NA | NA | NA |
| WBGene00169895 | NA | NA | NA | NA |
| WBGene00169896 | NA | NA | NA | NA |
| WBGene00169897 | NA | NA | NA | NA |
| WBGene00169898 | NA | NA | NA | NA |
| WBGene00169899 | NA | NA | NA | NA |
| WBGene00169900 | NA | NA | NA | NA |
| WBGene00169901 | NA | NA | NA | NA |
| WBGene00169902 | NA | NA | NA | NA |
| WBGene00169903 | NA | NA | NA | NA |
| WBGene00169904 | NA | NA | NA | NA |
| WBGene00169905 | NA | NA | NA | NA |
| WBGene00169906 | NA | NA | NA | NA |
| WBGene00169907 | NA | NA | NA | NA |
| WBGene00169908 | NA | NA | NA | NA |
| WBGene00169909 | NA | NA | NA | NA |
| WBGene00169910 | NA | NA | NA | NA |
| WBGene00169911 | NA | NA | NA | NA |
| WBGene00169912 | NA | NA | NA | NA |
| WBGene00169913 | NA | NA | NA | NA |
| WBGene00169914 | NA | NA | NA | NA |
| WBGene00169915 | NA | NA | NA | NA |

| | | | | |
|---|---|---|---|---|
| WBGene00169916 | NA | NA | NA | NA |
| WBGene00169917 | NA | NA | NA | NA |
| WBGene00169918 | NA | NA | NA | NA |
| WBGene00169919 | NA | NA | NA | NA |
| WBGene00169920 | NA | NA | NA | NA |
| WBGene00169921 | NA | NA | NA | NA |
| WBGene00169922 | NA | NA | NA | NA |
| WBGene00169923 | NA | NA | NA | NA |
| WBGene00169924 | NA | NA | NA | NA |
| WBGene00169925 | NA | NA | NA | NA |
| WBGene00169926 | NA | NA | NA | NA |
| WBGene00169927 | NA | NA | NA | NA |
| WBGene00169928 | NA | NA | NA | NA |
| WBGene00169929 | NA | NA | NA | NA |
| WBGene00169930 | NA | NA | NA | NA |
| WBGene00169931 | NA | NA | NA | NA |
| WBGene00169932 | NA | NA | NA | NA |
| WBGene00169933 | NA | NA | NA | NA |
| WBGene00169934 | NA | NA | NA | NA |
| WBGene00169935 | NA | NA | NA | NA |
| WBGene00169936 | NA | NA | NA | NA |
| WBGene00169937 | NA | NA | NA | NA |
| WBGene00169938 | NA | NA | NA | NA |
| WBGene00169939 | NA | NA | NA | NA |
| WBGene00169940 | -0.022930767 | 0.78121253 | NA | NA |
| WBGene00169941 | NA | NA | NA | NA |
| WBGene00169942 | NA | NA | NA | NA |
| WBGene00169943 | NA | NA | NA | NA |
| WBGene00169944 | NA | NA | NA | NA |
| WBGene00169946 | NA | NA | NA | NA |
| WBGene00169947 | NA | NA | NA | NA |
| WBGene00169948 | NA | NA | NA | NA |
| WBGene00169949 | NA | NA | NA | NA |
| WBGene00169950 | NA | NA | NA | NA |
| WBGene00169951 | NA | NA | NA | NA |
| WBGene00169952 | NA | NA | NA | NA |
| WBGene00169953 | NA | NA | NA | NA |
| WBGene00169954 | NA | NA | NA | NA |
| WBGene00169955 | NA | NA | NA | NA |
| WBGene00169956 | 0.020317245 | 0.84643798 | NA | NA |
| WBGene00169957 | NA | NA | NA | NA |
| WBGene00169958 | NA | NA | NA | NA |
| WBGene00169959 | NA | NA | NA | NA |

| | | | | |
|---|---|---|---|---|
| WBGene00169960 | NA | NA | NA | NA |
| WBGene00169961 | NA | NA | NA | NA |
| WBGene00169962 | NA | NA | NA | NA |
| WBGene00169963 | NA | NA | NA | NA |
| WBGene00169964 | NA | NA | NA | NA |
| WBGene00169965 | NA | NA | NA | NA |
| WBGene00169966 | NA | NA | NA | NA |
| WBGene00169967 | NA | NA | NA | NA |
| WBGene00169968 | NA | NA | NA | NA |
| WBGene00169969 | NA | NA | NA | NA |
| WBGene00169970 | NA | NA | NA | NA |
| WBGene00169971 | NA | NA | NA | NA |
| WBGene00169972 | NA | NA | NA | NA |
| WBGene00169973 | -0.022930767 | 0.78121253 | NA | NA |
| WBGene00169974 | NA | NA | NA | NA |
| WBGene00169975 | NA | NA | NA | NA |
| WBGene00169976 | NA | NA | NA | NA |
| WBGene00169977 | NA | NA | NA | NA |
| WBGene00169978 | NA | NA | NA | NA |
| WBGene00169979 | NA | NA | NA | NA |
| WBGene00169980 | NA | NA | NA | NA |
| WBGene00169981 | NA | NA | NA | NA |
| WBGene00169982 | NA | NA | NA | NA |
| WBGene00169983 | NA | NA | NA | NA |
| WBGene00169984 | NA | NA | NA | NA |
| WBGene00169985 | NA | NA | NA | NA |
| WBGene00169986 | NA | NA | NA | NA |
| WBGene00169987 | NA | NA | NA | NA |
| WBGene00169988 | NA | NA | NA | NA |
| WBGene00169989 | NA | NA | NA | NA |
| WBGene00169990 | NA | NA | NA | NA |
| WBGene00169991 | NA | NA | NA | NA |
| WBGene00169992 | NA | NA | NA | NA |
| WBGene00169993 | NA | NA | NA | NA |
| WBGene00169994 | NA | NA | NA | NA |
| WBGene00169995 | NA | NA | NA | NA |
| WBGene00169996 | NA | NA | NA | NA |
| WBGene00169997 | NA | NA | NA | NA |
| WBGene00169998 | NA | NA | NA | NA |
| WBGene00169999 | NA | NA | NA | NA |
| WBGene00170000 | NA | NA | NA | NA |
| WBGene00170001 | NA | NA | NA | NA |
| WBGene00170002 | NA | NA | NA | NA |

| | | | | |
|---|---|---|---|---|
| WBGene00170003 | NA | NA | NA | NA |
| WBGene00170004 | NA | NA | NA | NA |
| WBGene00170005 | NA | NA | NA | NA |
| WBGene00170006 | NA | NA | NA | NA |
| WBGene00170007 | NA | NA | NA | NA |
| WBGene00170008 | NA | NA | NA | NA |
| WBGene00170009 | NA | NA | NA | NA |
| WBGene00170010 | NA | NA | NA | NA |
| WBGene00170011 | NA | NA | NA | NA |
| WBGene00170012 | NA | NA | NA | NA |
| WBGene00170013 | NA | NA | NA | NA |
| WBGene00170014 | NA | NA | NA | NA |
| WBGene00170015 | NA | NA | NA | NA |
| WBGene00170016 | NA | NA | NA | NA |
| WBGene00170017 | NA | NA | NA | NA |
| WBGene00170018 | NA | NA | NA | NA |
| WBGene00170019 | NA | NA | NA | NA |
| WBGene00170020 | NA | NA | NA | NA |
| WBGene00170021 | NA | NA | NA | NA |
| WBGene00170022 | NA | NA | NA | NA |
| WBGene00170023 | NA | NA | NA | NA |
| WBGene00170024 | NA | NA | NA | NA |
| WBGene00170025 | NA | NA | NA | NA |
| WBGene00170026 | NA | NA | NA | NA |
| WBGene00170027 | NA | NA | NA | NA |
| WBGene00170028 | NA | NA | NA | NA |
| WBGene00170029 | NA | NA | NA | NA |
| WBGene00170030 | NA | NA | NA | NA |
| WBGene00170031 | NA | NA | NA | NA |
| WBGene00170032 | NA | NA | NA | NA |
| WBGene00170033 | NA | NA | NA | NA |
| WBGene00170034 | NA | NA | NA | NA |
| WBGene00170035 | NA | NA | NA | NA |
| WBGene00170036 | NA | NA | NA | NA |
| WBGene00170037 | NA | NA | NA | NA |
| WBGene00170038 | NA | NA | NA | NA |
| WBGene00170039 | NA | NA | NA | NA |
| WBGene00170040 | NA | NA | NA | NA |
| WBGene00170041 | NA | NA | NA | NA |
| WBGene00170042 | NA | NA | NA | NA |
| WBGene00170043 | NA | NA | NA | NA |
| WBGene00170044 | NA | NA | NA | NA |
| WBGene00170045 | NA | NA | NA | NA |

| | | | | |
|---|---|---|---|---|
| WBGene00170046 | NA | NA | NA | NA |
| WBGene00170047 | NA | NA | NA | NA |
| WBGene00170048 | NA | NA | NA | NA |
| WBGene00170049 | NA | NA | NA | NA |
| WBGene00170050 | NA | NA | NA | NA |
| WBGene00170051 | NA | NA | NA | NA |
| WBGene00170052 | NA | NA | NA | NA |
| WBGene00170053 | NA | NA | NA | NA |
| WBGene00170054 | NA | NA | NA | NA |
| WBGene00170055 | NA | NA | NA | NA |
| WBGene00170056 | NA | NA | NA | NA |
| WBGene00170057 | NA | NA | NA | NA |
| WBGene00170058 | NA | NA | NA | NA |
| WBGene00170059 | NA | NA | NA | NA |
| WBGene00170060 | NA | NA | NA | NA |
| WBGene00170061 | NA | NA | NA | NA |
| WBGene00170062 | NA | NA | NA | NA |
| WBGene00170063 | NA | NA | NA | NA |
| WBGene00170064 | NA | NA | NA | NA |
| WBGene00170065 | NA | NA | NA | NA |
| WBGene00170066 | NA | NA | NA | NA |
| WBGene00170067 | NA | NA | NA | NA |
| WBGene00170068 | NA | NA | NA | NA |
| WBGene00170069 | NA | NA | NA | NA |
| WBGene00170070 | NA | NA | NA | NA |
| WBGene00170071 | NA | NA | NA | NA |
| WBGene00170072 | NA | NA | NA | NA |
| WBGene00170073 | NA | NA | NA | NA |
| WBGene00170074 | NA | NA | NA | NA |
| WBGene00170075 | NA | NA | NA | NA |
| WBGene00170076 | NA | NA | NA | NA |
| WBGene00170077 | NA | NA | NA | NA |
| WBGene00170078 | NA | NA | NA | NA |
| WBGene00170079 | NA | NA | NA | NA |
| WBGene00170080 | NA | NA | NA | NA |
| WBGene00170081 | NA | NA | NA | NA |
| WBGene00170082 | NA | NA | NA | NA |
| WBGene00170083 | NA | NA | NA | NA |
| WBGene00170084 | NA | NA | NA | NA |
| WBGene00170085 | NA | NA | NA | NA |
| WBGene00170086 | NA | NA | NA | NA |
| WBGene00170087 | NA | NA | NA | NA |
| WBGene00170088 | NA | NA | NA | NA |

| | | | | |
|---|---|---|---|---|
| WBGene00170089 | NA | NA | NA | NA |
| WBGene00170090 | NA | NA | NA | NA |
| WBGene00170091 | NA | NA | NA | NA |
| WBGene00170092 | NA | NA | NA | NA |
| WBGene00170093 | NA | NA | NA | NA |
| WBGene00170094 | NA | NA | NA | NA |
| WBGene00170095 | NA | NA | NA | NA |
| WBGene00170096 | NA | NA | NA | NA |
| WBGene00170097 | NA | NA | NA | NA |
| WBGene00170098 | NA | NA | NA | NA |
| WBGene00170099 | NA | NA | NA | NA |
| WBGene00170100 | NA | NA | NA | NA |
| WBGene00170101 | NA | NA | NA | NA |
| WBGene00170102 | NA | NA | NA | NA |
| WBGene00170103 | NA | NA | NA | NA |
| WBGene00170104 | NA | NA | NA | NA |
| WBGene00170105 | NA | NA | NA | NA |
| WBGene00170106 | NA | NA | NA | NA |
| WBGene00170107 | NA | NA | NA | NA |
| WBGene00170108 | NA | NA | NA | NA |
| WBGene00170109 | NA | NA | NA | NA |
| WBGene00170110 | NA | NA | NA | NA |
| WBGene00170111 | NA | NA | NA | NA |
| WBGene00170112 | NA | NA | NA | NA |
| WBGene00170113 | NA | NA | NA | NA |
| WBGene00170114 | NA | NA | NA | NA |
| WBGene00170115 | NA | NA | NA | NA |
| WBGene00170116 | NA | NA | NA | NA |
| WBGene00170117 | NA | NA | NA | NA |
| WBGene00170118 | NA | NA | NA | NA |
| WBGene00170119 | NA | NA | NA | NA |
| WBGene00170120 | NA | NA | NA | NA |
| WBGene00170121 | NA | NA | NA | NA |
| WBGene00170122 | NA | NA | NA | NA |
| WBGene00170123 | NA | NA | NA | NA |
| WBGene00170124 | NA | NA | NA | NA |
| WBGene00170125 | NA | NA | NA | NA |
| WBGene00170126 | NA | NA | NA | NA |
| WBGene00170127 | NA | NA | NA | NA |
| WBGene00170128 | NA | NA | NA | NA |
| WBGene00170129 | NA | NA | NA | NA |
| WBGene00170130 | NA | NA | NA | NA |
| WBGene00170131 | NA | NA | NA | NA |

| | | | | |
|---|---|---|---|---|
| WBGene00170132 | NA | NA | NA | NA |
| WBGene00170133 | NA | NA | NA | NA |
| WBGene00170134 | NA | NA | NA | NA |
| WBGene00170135 | NA | NA | NA | NA |
| WBGene00170136 | NA | NA | NA | NA |
| WBGene00170137 | NA | NA | NA | NA |
| WBGene00170138 | NA | NA | NA | NA |
| WBGene00170139 | NA | NA | NA | NA |
| WBGene00170140 | NA | NA | NA | NA |
| WBGene00170141 | NA | NA | NA | NA |
| WBGene00170142 | NA | NA | NA | NA |
| WBGene00170143 | -0.027188909 | 0.68205301 | NA | NA |
| WBGene00170144 | NA | NA | NA | NA |
| WBGene00170145 | NA | NA | NA | NA |
| WBGene00170146 | NA | NA | NA | NA |
| WBGene00170147 | NA | NA | NA | NA |
| WBGene00170148 | NA | NA | NA | NA |
| WBGene00170149 | NA | NA | NA | NA |
| WBGene00170150 | NA | NA | NA | NA |
| WBGene00170151 | NA | NA | NA | NA |
| WBGene00170152 | NA | NA | NA | NA |
| WBGene00170153 | NA | NA | NA | NA |
| WBGene00170154 | NA | NA | NA | NA |
| WBGene00170155 | NA | NA | NA | NA |
| WBGene00170156 | NA | NA | NA | NA |
| WBGene00170157 | NA | NA | NA | NA |
| WBGene00170158 | NA | NA | NA | NA |
| WBGene00170159 | NA | NA | NA | NA |
| WBGene00170160 | NA | NA | NA | NA |
| WBGene00170161 | NA | NA | NA | NA |
| WBGene00170162 | NA | NA | NA | NA |
| WBGene00170163 | NA | NA | NA | NA |
| WBGene00170164 | NA | NA | NA | NA |
| WBGene00170165 | NA | NA | NA | NA |
| WBGene00170166 | NA | NA | NA | NA |
| WBGene00170167 | NA | NA | NA | NA |
| WBGene00170168 | NA | NA | NA | NA |
| WBGene00170169 | NA | NA | NA | NA |
| WBGene00170170 | NA | NA | NA | NA |
| WBGene00170171 | NA | NA | NA | NA |
| WBGene00170172 | NA | NA | NA | NA |
| WBGene00170173 | NA | NA | NA | NA |
| WBGene00170174 | NA | NA | NA | NA |

| | | | | |
|---|---|---|---|---|
| WBGene00170175 | NA | NA | NA | NA |
| WBGene00170176 | NA | NA | NA | NA |
| WBGene00170177 | NA | NA | NA | NA |
| WBGene00170178 | NA | NA | NA | NA |
| WBGene00170179 | NA | NA | NA | NA |
| WBGene00170180 | NA | NA | NA | NA |
| WBGene00170181 | NA | NA | NA | NA |
| WBGene00170182 | NA | NA | NA | NA |
| WBGene00170183 | NA | NA | NA | NA |
| WBGene00170184 | NA | NA | NA | NA |
| WBGene00170185 | NA | NA | NA | NA |
| WBGene00170186 | NA | NA | NA | NA |
| WBGene00170187 | NA | NA | NA | NA |
| WBGene00170188 | NA | NA | NA | NA |
| WBGene00170189 | NA | NA | NA | NA |
| WBGene00170190 | NA | NA | NA | NA |
| WBGene00170191 | NA | NA | NA | NA |
| WBGene00170192 | NA | NA | NA | NA |
| WBGene00170193 | NA | NA | NA | NA |
| WBGene00170194 | NA | NA | NA | NA |
| WBGene00170195 | NA | NA | NA | NA |
| WBGene00170196 | NA | NA | NA | NA |
| WBGene00170197 | NA | NA | NA | NA |
| WBGene00170198 | NA | NA | NA | NA |
| WBGene00170199 | NA | NA | NA | NA |
| WBGene00170200 | NA | NA | NA | NA |
| WBGene00170201 | NA | NA | NA | NA |
| WBGene00170202 | NA | NA | NA | NA |
| WBGene00170203 | NA | NA | NA | NA |
| WBGene00170204 | NA | NA | NA | NA |
| WBGene00170205 | NA | NA | NA | NA |
| WBGene00170206 | NA | NA | NA | NA |
| WBGene00170207 | NA | NA | NA | NA |
| WBGene00170208 | NA | NA | NA | NA |
| WBGene00170209 | NA | NA | NA | NA |
| WBGene00170210 | NA | NA | NA | NA |
| WBGene00170211 | NA | NA | NA | NA |
| WBGene00170212 | NA | NA | NA | NA |
| WBGene00170213 | NA | NA | NA | NA |
| WBGene00170214 | NA | NA | NA | NA |
| WBGene00170215 | NA | NA | NA | NA |
| WBGene00170216 | NA | NA | NA | NA |
| WBGene00170217 | NA | NA | NA | NA |

| | | | | |
|---|---|---|---|---|
| WBGene00170218 | NA | NA | NA | NA |
| WBGene00170219 | NA | NA | NA | NA |
| WBGene00170220 | NA | NA | NA | NA |
| WBGene00170221 | NA | NA | NA | NA |
| WBGene00170222 | NA | NA | NA | NA |
| WBGene00170223 | NA | NA | NA | NA |
| WBGene00170224 | NA | NA | NA | NA |
| WBGene00170225 | NA | NA | NA | NA |
| WBGene00170226 | NA | NA | NA | NA |
| WBGene00170227 | NA | NA | NA | NA |
| WBGene00170228 | NA | NA | NA | NA |
| WBGene00170229 | NA | NA | NA | NA |
| WBGene00170230 | NA | NA | NA | NA |
| WBGene00170231 | NA | NA | NA | NA |
| WBGene00170232 | NA | NA | NA | NA |
| WBGene00170233 | NA | NA | NA | NA |
| WBGene00170234 | NA | NA | NA | NA |
| WBGene00170235 | NA | NA | NA | NA |
| WBGene00170236 | NA | NA | NA | NA |
| WBGene00170237 | NA | NA | NA | NA |
| WBGene00170238 | NA | NA | NA | NA |
| WBGene00170239 | NA | NA | NA | NA |
| WBGene00170240 | NA | NA | NA | NA |
| WBGene00170241 | NA | NA | NA | NA |
| WBGene00170242 | NA | NA | NA | NA |
| WBGene00170243 | NA | NA | NA | NA |
| WBGene00170244 | NA | NA | NA | NA |
| WBGene00170245 | NA | NA | NA | NA |
| WBGene00170246 | NA | NA | NA | NA |
| WBGene00170247 | NA | NA | NA | NA |
| WBGene00170248 | NA | NA | NA | NA |
| WBGene00170249 | NA | NA | NA | NA |
| WBGene00170250 | NA | NA | NA | NA |
| WBGene00170251 | NA | NA | NA | NA |
| WBGene00170252 | NA | NA | NA | NA |
| WBGene00170253 | NA | NA | NA | NA |
| WBGene00170254 | NA | NA | NA | NA |
| WBGene00170255 | NA | NA | NA | NA |
| WBGene00170256 | NA | NA | NA | NA |
| WBGene00170257 | NA | NA | NA | NA |
| WBGene00170258 | NA | NA | NA | NA |
| WBGene00170259 | NA | NA | NA | NA |
| WBGene00170260 | NA | NA | NA | NA |

| | | | | |
|---|---|---|---|---|
| WBGene00170261 | NA | NA | NA | NA |
| WBGene00170262 | NA | NA | NA | NA |
| WBGene00170263 | NA | NA | NA | NA |
| WBGene00170264 | NA | NA | NA | NA |
| WBGene00170265 | NA | NA | NA | NA |
| WBGene00170266 | NA | NA | NA | NA |
| WBGene00170267 | NA | NA | NA | NA |
| WBGene00170268 | NA | NA | NA | NA |
| WBGene00170269 | NA | NA | NA | NA |
| WBGene00170270 | NA | NA | NA | NA |
| WBGene00170271 | NA | NA | NA | NA |
| WBGene00170272 | NA | NA | NA | NA |
| WBGene00170273 | NA | NA | NA | NA |
| WBGene00170274 | NA | NA | NA | NA |
| WBGene00170275 | NA | NA | NA | NA |
| WBGene00170276 | NA | NA | NA | NA |
| WBGene00170277 | NA | NA | NA | NA |
| WBGene00170278 | NA | NA | NA | NA |
| WBGene00170279 | NA | NA | NA | NA |
| WBGene00170280 | NA | NA | NA | NA |
| WBGene00170281 | NA | NA | NA | NA |
| WBGene00170282 | NA | NA | NA | NA |
| WBGene00170283 | NA | NA | NA | NA |
| WBGene00170284 | NA | NA | NA | NA |
| WBGene00170285 | NA | NA | NA | NA |
| WBGene00170286 | NA | NA | NA | NA |
| WBGene00170287 | NA | NA | NA | NA |
| WBGene00170288 | NA | NA | NA | NA |
| WBGene00170289 | NA | NA | NA | NA |
| WBGene00170290 | NA | NA | NA | NA |
| WBGene00170291 | NA | NA | NA | NA |
| WBGene00170292 | NA | NA | NA | NA |
| WBGene00170293 | NA | NA | NA | NA |
| WBGene00170294 | NA | NA | NA | NA |
| WBGene00170295 | NA | NA | NA | NA |
| WBGene00170296 | NA | NA | NA | NA |
| WBGene00170297 | NA | NA | NA | NA |
| WBGene00170298 | NA | NA | NA | NA |
| WBGene00170299 | NA | NA | NA | NA |
| WBGene00170300 | NA | NA | NA | NA |
| WBGene00170301 | NA | NA | NA | NA |
| WBGene00170302 | NA | NA | NA | NA |
| WBGene00170303 | NA | NA | NA | NA |

| | | | | |
|---|---|---|---|---|
| WBGene00170304 | NA | NA | NA | NA |
| WBGene00170305 | NA | NA | NA | NA |
| WBGene00170306 | NA | NA | NA | NA |
| WBGene00170307 | NA | NA | NA | NA |
| WBGene00170308 | NA | NA | NA | NA |
| WBGene00170309 | NA | NA | NA | NA |
| WBGene00170310 | NA | NA | NA | NA |
| WBGene00170311 | NA | NA | NA | NA |
| WBGene00170312 | NA | NA | NA | NA |
| WBGene00170313 | NA | NA | NA | NA |
| WBGene00170314 | NA | NA | NA | NA |
| WBGene00170315 | NA | NA | NA | NA |
| WBGene00170316 | NA | NA | NA | NA |
| WBGene00170317 | NA | NA | NA | NA |
| WBGene00170318 | NA | NA | NA | NA |
| WBGene00170319 | NA | NA | NA | NA |
| WBGene00170320 | NA | NA | NA | NA |
| WBGene00170321 | NA | NA | NA | NA |
| WBGene00170322 | NA | NA | NA | NA |
| WBGene00170323 | NA | NA | NA | NA |
| WBGene00170324 | NA | NA | NA | NA |
| WBGene00170325 | NA | NA | NA | NA |
| WBGene00170326 | NA | NA | NA | NA |
| WBGene00170327 | NA | NA | NA | NA |
| WBGene00170328 | NA | NA | NA | NA |
| WBGene00170329 | NA | NA | NA | NA |
| WBGene00170330 | NA | NA | NA | NA |
| WBGene00170331 | NA | NA | NA | NA |
| WBGene00170332 | NA | NA | NA | NA |
| WBGene00170333 | NA | NA | NA | NA |
| WBGene00170334 | NA | NA | NA | NA |
| WBGene00170335 | NA | NA | NA | NA |
| WBGene00170336 | NA | NA | NA | NA |
| WBGene00170337 | NA | NA | NA | NA |
| WBGene00170338 | NA | NA | NA | NA |
| WBGene00170339 | NA | NA | NA | NA |
| WBGene00170340 | NA | NA | NA | NA |
| WBGene00170341 | NA | NA | NA | NA |
| WBGene00170342 | NA | NA | NA | NA |
| WBGene00170343 | NA | NA | NA | NA |
| WBGene00170344 | NA | NA | NA | NA |
| WBGene00170345 | NA | NA | NA | NA |
| WBGene00170346 | NA | NA | NA | NA |

| | | | | |
|---|---|---|---|---|
| WBGene00170347 | NA | NA | NA | NA |
| WBGene00170348 | NA | NA | NA | NA |
| WBGene00170349 | NA | NA | NA | NA |
| WBGene00170350 | NA | NA | NA | NA |
| WBGene00170351 | NA | NA | NA | NA |
| WBGene00170352 | NA | NA | NA | NA |
| WBGene00170353 | NA | NA | NA | NA |
| WBGene00170354 | 0.020232312 | 0.84643798 | NA | NA |
| WBGene00170355 | NA | NA | NA | NA |
| WBGene00170356 | NA | NA | NA | NA |
| WBGene00170357 | NA | NA | NA | NA |
| WBGene00170358 | NA | NA | NA | NA |
| WBGene00170359 | NA | NA | NA | NA |
| WBGene00170360 | NA | NA | NA | NA |
| WBGene00170361 | NA | NA | NA | NA |
| WBGene00170362 | NA | NA | NA | NA |
| WBGene00170363 | NA | NA | NA | NA |
| WBGene00170364 | NA | NA | NA | NA |
| WBGene00170365 | NA | NA | NA | NA |
| WBGene00170366 | NA | NA | NA | NA |
| WBGene00170367 | NA | NA | NA | NA |
| WBGene00170368 | NA | NA | NA | NA |
| WBGene00170369 | NA | NA | NA | NA |
| WBGene00170370 | NA | NA | NA | NA |
| WBGene00170371 | NA | NA | NA | NA |
| WBGene00170372 | NA | NA | NA | NA |
| WBGene00170373 | NA | NA | NA | NA |
| WBGene00170374 | NA | NA | NA | NA |
| WBGene00170375 | NA | NA | NA | NA |
| WBGene00170376 | NA | NA | NA | NA |
| WBGene00170377 | NA | NA | NA | NA |
| WBGene00170378 | NA | NA | NA | NA |
| WBGene00170379 | NA | NA | NA | NA |
| WBGene00170380 | NA | NA | NA | NA |
| WBGene00170381 | NA | NA | NA | NA |
| WBGene00170382 | NA | NA | NA | NA |
| WBGene00170383 | NA | NA | NA | NA |
| WBGene00170384 | NA | NA | NA | NA |
| WBGene00170385 | NA | NA | NA | NA |
| WBGene00170386 | NA | NA | NA | NA |
| WBGene00170387 | NA | NA | NA | NA |
| WBGene00170388 | NA | NA | NA | NA |
| WBGene00170389 | NA | NA | NA | NA |

| | | | | |
|---|---|---|---|---|
| WBGene00170390 | NA | NA | NA | NA |
| WBGene00170391 | NA | NA | NA | NA |
| WBGene00170392 | NA | NA | NA | NA |
| WBGene00170393 | NA | NA | NA | NA |
| WBGene00170394 | NA | NA | NA | NA |
| WBGene00170395 | NA | NA | NA | NA |
| WBGene00170396 | NA | NA | NA | NA |
| WBGene00170397 | NA | NA | NA | NA |
| WBGene00170398 | NA | NA | NA | NA |
| WBGene00170399 | NA | NA | NA | NA |
| WBGene00170400 | NA | NA | NA | NA |
| WBGene00170401 | NA | NA | NA | NA |
| WBGene00170402 | NA | NA | NA | NA |
| WBGene00170403 | NA | NA | NA | NA |
| WBGene00170404 | NA | NA | NA | NA |
| WBGene00170405 | NA | NA | NA | NA |
| WBGene00170406 | NA | NA | NA | NA |
| WBGene00170407 | NA | NA | NA | NA |
| WBGene00170408 | NA | NA | NA | NA |
| WBGene00170409 | NA | NA | NA | NA |
| WBGene00170410 | NA | NA | NA | NA |
| WBGene00170411 | NA | NA | NA | NA |
| WBGene00170412 | NA | NA | NA | NA |
| WBGene00170413 | NA | NA | NA | NA |
| WBGene00170414 | NA | NA | NA | NA |
| WBGene00170415 | NA | NA | NA | NA |
| WBGene00170416 | NA | NA | NA | NA |
| WBGene00170417 | NA | NA | NA | NA |
| WBGene00170418 | NA | NA | NA | NA |
| WBGene00170419 | NA | NA | NA | NA |
| WBGene00170420 | NA | NA | NA | NA |
| WBGene00170421 | NA | NA | NA | NA |
| WBGene00170422 | NA | NA | NA | NA |
| WBGene00170423 | NA | NA | NA | NA |
| WBGene00170424 | NA | NA | NA | NA |
| WBGene00170425 | NA | NA | NA | NA |
| WBGene00170426 | NA | NA | NA | NA |
| WBGene00170427 | NA | NA | NA | NA |
| WBGene00170428 | NA | NA | NA | NA |
| WBGene00170429 | NA | NA | NA | NA |
| WBGene00170430 | NA | NA | NA | NA |
| WBGene00170431 | NA | NA | NA | NA |
| WBGene00170432 | NA | NA | NA | NA |

| | | | | |
|---|---|---|---|---|
| WBGene00170433 | NA | NA | NA | NA |
| WBGene00170434 | NA | NA | NA | NA |
| WBGene00170435 | NA | NA | NA | NA |
| WBGene00170436 | NA | NA | NA | NA |
| WBGene00170437 | NA | NA | NA | NA |
| WBGene00170438 | NA | NA | NA | NA |
| WBGene00170439 | NA | NA | NA | NA |
| WBGene00170440 | NA | NA | NA | NA |
| WBGene00170441 | NA | NA | NA | NA |
| WBGene00170442 | NA | NA | NA | NA |
| WBGene00170443 | NA | NA | NA | NA |
| WBGene00170444 | NA | NA | NA | NA |
| WBGene00170445 | NA | NA | NA | NA |
| WBGene00170446 | NA | NA | NA | NA |
| WBGene00170447 | NA | NA | NA | NA |
| WBGene00170448 | NA | NA | NA | NA |
| WBGene00170449 | NA | NA | NA | NA |
| WBGene00170450 | NA | NA | NA | NA |
| WBGene00170451 | NA | NA | NA | NA |
| WBGene00170452 | 0.020317245 | 0.84643798 | NA | NA |
| WBGene00170453 | NA | NA | NA | NA |
| WBGene00170454 | NA | NA | NA | NA |
| WBGene00170455 | NA | NA | NA | NA |
| WBGene00170456 | NA | NA | NA | NA |
| WBGene00170457 | NA | NA | NA | NA |
| WBGene00170458 | NA | NA | NA | NA |
| WBGene00170459 | NA | NA | NA | NA |
| WBGene00170460 | NA | NA | NA | NA |
| WBGene00170461 | NA | NA | NA | NA |
| WBGene00170462 | NA | NA | NA | NA |
| WBGene00170463 | NA | NA | NA | NA |
| WBGene00170464 | NA | NA | NA | NA |
| WBGene00170465 | NA | NA | NA | NA |
| WBGene00170466 | NA | NA | NA | NA |
| WBGene00170467 | NA | NA | NA | NA |
| WBGene00170468 | NA | NA | NA | NA |
| WBGene00170469 | NA | NA | NA | NA |
| WBGene00170470 | NA | NA | NA | NA |
| WBGene00170471 | NA | NA | NA | NA |
| WBGene00170472 | NA | NA | NA | NA |
| WBGene00170473 | NA | NA | NA | NA |
| WBGene00170474 | NA | NA | NA | NA |
| WBGene00170475 | NA | NA | NA | NA |

| | | | | |
|---|---|---|---|---|
| WBGene00170476 | NA | NA | NA | NA |
| WBGene00170477 | NA | NA | NA | NA |
| WBGene00170478 | NA | NA | NA | NA |
| WBGene00170479 | NA | NA | NA | NA |
| WBGene00170480 | NA | NA | NA | NA |
| WBGene00170481 | NA | NA | NA | NA |
| WBGene00170482 | NA | NA | NA | NA |
| WBGene00170483 | NA | NA | NA | NA |
| WBGene00170484 | NA | NA | NA | NA |
| WBGene00170485 | NA | NA | NA | NA |
| WBGene00170486 | NA | NA | NA | NA |
| WBGene00170487 | NA | NA | NA | NA |
| WBGene00170488 | NA | NA | NA | NA |
| WBGene00170489 | NA | NA | NA | NA |
| WBGene00170490 | NA | NA | NA | NA |
| WBGene00170491 | NA | NA | NA | NA |
| WBGene00170492 | NA | NA | NA | NA |
| WBGene00170493 | NA | NA | NA | NA |
| WBGene00170494 | NA | NA | NA | NA |
| WBGene00170495 | NA | NA | NA | NA |
| WBGene00170496 | NA | NA | NA | NA |
| WBGene00170497 | NA | NA | NA | NA |
| WBGene00170498 | NA | NA | NA | NA |
| WBGene00170499 | NA | NA | NA | NA |
| WBGene00170500 | NA | NA | NA | NA |
| WBGene00170501 | NA | NA | NA | NA |
| WBGene00170502 | NA | NA | NA | NA |
| WBGene00170503 | NA | NA | NA | NA |
| WBGene00170504 | NA | NA | NA | NA |
| WBGene00170505 | NA | NA | NA | NA |
| WBGene00170506 | NA | NA | NA | NA |
| WBGene00170507 | NA | NA | NA | NA |
| WBGene00170508 | NA | NA | NA | NA |
| WBGene00170509 | NA | NA | NA | NA |
| WBGene00170510 | NA | NA | NA | NA |
| WBGene00170511 | NA | NA | NA | NA |
| WBGene00170512 | NA | NA | NA | NA |
| WBGene00170513 | NA | NA | NA | NA |
| WBGene00170514 | NA | NA | NA | NA |
| WBGene00170515 | NA | NA | NA | NA |
| WBGene00170516 | NA | NA | NA | NA |
| WBGene00170517 | NA | NA | NA | NA |
| WBGene00170518 | NA | NA | NA | NA |

| | | | | |
|---|---|---|---|---|
| WBGene00170519 | NA | NA | NA | NA |
| WBGene00170520 | NA | NA | NA | NA |
| WBGene00170521 | NA | NA | NA | NA |
| WBGene00170522 | NA | NA | NA | NA |
| WBGene00170523 | NA | NA | NA | NA |
| WBGene00170524 | NA | NA | NA | NA |
| WBGene00170525 | NA | NA | NA | NA |
| WBGene00170526 | NA | NA | NA | NA |
| WBGene00170527 | NA | NA | NA | NA |
| WBGene00170528 | NA | NA | NA | NA |
| WBGene00170529 | NA | NA | NA | NA |
| WBGene00170530 | NA | NA | NA | NA |
| WBGene00170531 | NA | NA | NA | NA |
| WBGene00170532 | NA | NA | NA | NA |
| WBGene00170533 | NA | NA | NA | NA |
| WBGene00170534 | NA | NA | NA | NA |
| WBGene00170535 | NA | NA | NA | NA |
| WBGene00170536 | NA | NA | NA | NA |
| WBGene00170537 | NA | NA | NA | NA |
| WBGene00170538 | NA | NA | NA | NA |
| WBGene00170539 | 0.01409643 | 0.85915526 | NA | NA |
| WBGene00170540 | NA | NA | NA | NA |
| WBGene00170541 | NA | NA | NA | NA |
| WBGene00170542 | NA | NA | NA | NA |
| WBGene00170543 | NA | NA | NA | NA |
| WBGene00170544 | NA | NA | NA | NA |
| WBGene00170545 | NA | NA | NA | NA |
| WBGene00170546 | NA | NA | NA | NA |
| WBGene00170547 | NA | NA | NA | NA |
| WBGene00170548 | NA | NA | NA | NA |
| WBGene00170549 | NA | NA | NA | NA |
| WBGene00170550 | NA | NA | NA | NA |
| WBGene00170551 | NA | NA | NA | NA |
| WBGene00170552 | NA | NA | NA | NA |
| WBGene00170553 | NA | NA | NA | NA |
| WBGene00170554 | NA | NA | NA | NA |
| WBGene00170555 | NA | NA | NA | NA |
| WBGene00170556 | NA | NA | NA | NA |
| WBGene00170557 | NA | NA | NA | NA |
| WBGene00170558 | NA | NA | NA | NA |
| WBGene00170559 | NA | NA | NA | NA |
| WBGene00170560 | NA | NA | NA | NA |
| WBGene00170561 | NA | NA | NA | NA |

| | | | | |
|---|---|---|---|---|
| WBGene00170562 | NA | NA | NA | NA |
| WBGene00170563 | NA | NA | NA | NA |
| WBGene00170564 | NA | NA | NA | NA |
| WBGene00170565 | NA | NA | NA | NA |
| WBGene00170566 | NA | NA | NA | NA |
| WBGene00170567 | NA | NA | NA | NA |
| WBGene00170568 | NA | NA | NA | NA |
| WBGene00170569 | NA | NA | NA | NA |
| WBGene00170570 | NA | NA | NA | NA |
| WBGene00170571 | NA | NA | NA | NA |
| WBGene00170572 | NA | NA | NA | NA |
| WBGene00170573 | NA | NA | NA | NA |
| WBGene00170574 | NA | NA | NA | NA |
| WBGene00170575 | NA | NA | NA | NA |
| WBGene00170576 | NA | NA | NA | NA |
| WBGene00170577 | NA | NA | NA | NA |
| WBGene00170578 | NA | NA | NA | NA |
| WBGene00170579 | NA | NA | NA | NA |
| WBGene00170580 | 0.020232312 | 0.84643798 | NA | NA |
| WBGene00170581 | NA | NA | NA | NA |
| WBGene00170582 | NA | NA | NA | NA |
| WBGene00170583 | NA | NA | NA | NA |
| WBGene00170584 | NA | NA | NA | NA |
| WBGene00170585 | NA | NA | NA | NA |
| WBGene00170586 | NA | NA | NA | NA |
| WBGene00170587 | NA | NA | NA | NA |
| WBGene00170588 | NA | NA | NA | NA |
| WBGene00170589 | NA | NA | NA | NA |
| WBGene00170590 | NA | NA | NA | NA |
| WBGene00170591 | NA | NA | NA | NA |
| WBGene00170592 | NA | NA | NA | NA |
| WBGene00170593 | NA | NA | NA | NA |
| WBGene00170594 | NA | NA | NA | NA |
| WBGene00170595 | NA | NA | NA | NA |
| WBGene00170596 | NA | NA | NA | NA |
| WBGene00170597 | NA | NA | NA | NA |
| WBGene00170598 | NA | NA | NA | NA |
| WBGene00170599 | NA | NA | NA | NA |
| WBGene00170600 | NA | NA | NA | NA |
| WBGene00170601 | NA | NA | NA | NA |
| WBGene00170602 | NA | NA | NA | NA |
| WBGene00170603 | NA | NA | NA | NA |
| WBGene00170604 | NA | NA | NA | NA |

| | | | | |
|---|---|---|---|---|
| WBGene00170605 | NA | NA | NA | NA |
| WBGene00170606 | NA | NA | NA | NA |
| WBGene00170607 | NA | NA | NA | NA |
| WBGene00170608 | NA | NA | NA | NA |
| WBGene00170609 | NA | NA | NA | NA |
| WBGene00170610 | NA | NA | NA | NA |
| WBGene00170611 | NA | NA | NA | NA |
| WBGene00170612 | NA | NA | NA | NA |
| WBGene00170613 | NA | NA | NA | NA |
| WBGene00170614 | NA | NA | NA | NA |
| WBGene00170615 | NA | NA | NA | NA |
| WBGene00170616 | NA | NA | NA | NA |
| WBGene00170617 | NA | NA | NA | NA |
| WBGene00170618 | NA | NA | NA | NA |
| WBGene00170619 | NA | NA | NA | NA |
| WBGene00170620 | NA | NA | NA | NA |
| WBGene00170621 | NA | NA | NA | NA |
| WBGene00170622 | NA | NA | NA | NA |
| WBGene00170623 | NA | NA | NA | NA |
| WBGene00170624 | NA | NA | NA | NA |
| WBGene00170625 | NA | NA | NA | NA |
| WBGene00170626 | NA | NA | NA | NA |
| WBGene00170627 | NA | NA | NA | NA |
| WBGene00170628 | NA | NA | NA | NA |
| WBGene00170629 | NA | NA | NA | NA |
| WBGene00170630 | NA | NA | NA | NA |
| WBGene00170631 | NA | NA | NA | NA |
| WBGene00170632 | NA | NA | NA | NA |
| WBGene00170633 | NA | NA | NA | NA |
| WBGene00170634 | NA | NA | NA | NA |
| WBGene00170635 | NA | NA | NA | NA |
| WBGene00170636 | NA | NA | NA | NA |
| WBGene00170637 | NA | NA | NA | NA |
| WBGene00170638 | NA | NA | NA | NA |
| WBGene00170639 | NA | NA | NA | NA |
| WBGene00170640 | NA | NA | NA | NA |
| WBGene00170641 | NA | NA | NA | NA |
| WBGene00170642 | NA | NA | NA | NA |
| WBGene00170643 | NA | NA | NA | NA |
| WBGene00170644 | NA | NA | NA | NA |
| WBGene00170645 | NA | NA | NA | NA |
| WBGene00170646 | NA | NA | NA | NA |
| WBGene00170647 | NA | NA | NA | NA |

| | | | | |
|---|---|---|---|---|
| WBGene00170648 | NA | NA | NA | NA |
| WBGene00170649 | NA | NA | NA | NA |
| WBGene00170650 | NA | NA | NA | NA |
| WBGene00170651 | NA | NA | NA | NA |
| WBGene00170652 | NA | NA | NA | NA |
| WBGene00170653 | NA | NA | NA | NA |
| WBGene00170654 | NA | NA | NA | NA |
| WBGene00170655 | NA | NA | NA | NA |
| WBGene00170656 | NA | NA | NA | NA |
| WBGene00170657 | NA | NA | NA | NA |
| WBGene00170658 | NA | NA | NA | NA |
| WBGene00170659 | NA | NA | NA | NA |
| WBGene00170660 | NA | NA | NA | NA |
| WBGene00170661 | NA | NA | NA | NA |
| WBGene00170662 | NA | NA | NA | NA |
| WBGene00170663 | NA | NA | NA | NA |
| WBGene00170664 | NA | NA | NA | NA |
| WBGene00170665 | NA | NA | NA | NA |
| WBGene00170666 | NA | NA | NA | NA |
| WBGene00170667 | NA | NA | NA | NA |
| WBGene00170668 | NA | NA | NA | NA |
| WBGene00170669 | NA | NA | NA | NA |
| WBGene00170670 | NA | NA | NA | NA |
| WBGene00170671 | NA | NA | NA | NA |
| WBGene00170672 | NA | NA | NA | NA |
| WBGene00170673 | NA | NA | NA | NA |
| WBGene00170674 | 0.020232312 | 0.84643798 | NA | NA |
| WBGene00170675 | NA | NA | NA | NA |
| WBGene00170676 | NA | NA | NA | NA |
| WBGene00170677 | NA | NA | NA | NA |
| WBGene00170678 | NA | NA | NA | NA |
| WBGene00170679 | NA | NA | NA | NA |
| WBGene00170680 | NA | NA | NA | NA |
| WBGene00170681 | NA | NA | NA | NA |
| WBGene00170682 | NA | NA | NA | NA |
| WBGene00170683 | NA | NA | NA | NA |
| WBGene00170684 | NA | NA | NA | NA |
| WBGene00170685 | NA | NA | NA | NA |
| WBGene00170686 | NA | NA | NA | NA |
| WBGene00170687 | NA | NA | NA | NA |
| WBGene00170688 | NA | NA | NA | NA |
| WBGene00170689 | NA | NA | NA | NA |
| WBGene00170690 | NA | NA | NA | NA |

| | | | | |
|---|---|---|---|---|
| WBGene00170691 | NA | NA | NA | NA |
| WBGene00170692 | NA | NA | NA | NA |
| WBGene00170693 | NA | NA | NA | NA |
| WBGene00170694 | NA | NA | NA | NA |
| WBGene00170695 | NA | NA | NA | NA |
| WBGene00170697 | NA | NA | NA | NA |
| WBGene00170698 | NA | NA | NA | NA |
| WBGene00170699 | NA | NA | NA | NA |
| WBGene00170700 | NA | NA | NA | NA |
| WBGene00170701 | NA | NA | NA | NA |
| WBGene00170702 | NA | NA | NA | NA |
| WBGene00170703 | NA | NA | NA | NA |
| WBGene00170704 | NA | NA | NA | NA |
| WBGene00170705 | NA | NA | NA | NA |
| WBGene00170706 | NA | NA | NA | NA |
| WBGene00170707 | NA | NA | NA | NA |
| WBGene00170708 | NA | NA | NA | NA |
| WBGene00170709 | NA | NA | NA | NA |
| WBGene00170710 | NA | NA | NA | NA |
| WBGene00170711 | NA | NA | NA | NA |
| WBGene00170712 | NA | NA | NA | NA |
| WBGene00170713 | NA | NA | NA | NA |
| WBGene00170714 | NA | NA | NA | NA |
| WBGene00170715 | NA | NA | NA | NA |
| WBGene00170716 | NA | NA | NA | NA |
| WBGene00170717 | NA | NA | NA | NA |
| WBGene00170718 | NA | NA | NA | NA |
| WBGene00170719 | NA | NA | NA | NA |
| WBGene00170720 | NA | NA | NA | NA |
| WBGene00170721 | NA | NA | NA | NA |
| WBGene00170722 | NA | NA | NA | NA |
| WBGene00170723 | NA | NA | NA | NA |
| WBGene00170724 | NA | NA | NA | NA |
| WBGene00170725 | NA | NA | NA | NA |
| WBGene00170726 | NA | NA | NA | NA |
| WBGene00170727 | NA | NA | NA | NA |
| WBGene00170728 | NA | NA | NA | NA |
| WBGene00170729 | NA | NA | NA | NA |
| WBGene00170730 | NA | NA | NA | NA |
| WBGene00170731 | NA | NA | NA | NA |
| WBGene00170732 | NA | NA | NA | NA |
| WBGene00170733 | NA | NA | NA | NA |
| WBGene00170734 | NA | NA | NA | NA |

| | | | | |
|---|---|---|---|---|
| WBGene00170735 | NA | NA | NA | NA |
| WBGene00170736 | NA | NA | NA | NA |
| WBGene00170737 | NA | NA | NA | NA |
| WBGene00170738 | NA | NA | NA | NA |
| WBGene00170739 | NA | NA | NA | NA |
| WBGene00170740 | NA | NA | NA | NA |
| WBGene00170741 | NA | NA | NA | NA |
| WBGene00170742 | NA | NA | NA | NA |
| WBGene00170743 | NA | NA | NA | NA |
| WBGene00170744 | -0.023109762 | 0.78121253 | NA | NA |
| WBGene00170745 | NA | NA | NA | NA |
| WBGene00170746 | NA | NA | NA | NA |
| WBGene00170747 | NA | NA | NA | NA |
| WBGene00170748 | NA | NA | NA | NA |
| WBGene00170749 | NA | NA | NA | NA |
| WBGene00170750 | NA | NA | NA | NA |
| WBGene00170751 | NA | NA | NA | NA |
| WBGene00170752 | NA | NA | NA | NA |
| WBGene00170753 | NA | NA | NA | NA |
| WBGene00170754 | NA | NA | NA | NA |
| WBGene00170755 | NA | NA | NA | NA |
| WBGene00170756 | NA | NA | NA | NA |
| WBGene00170757 | NA | NA | NA | NA |
| WBGene00170758 | NA | NA | NA | NA |
| WBGene00170759 | NA | NA | NA | NA |
| WBGene00170760 | NA | NA | NA | NA |
| WBGene00170761 | NA | NA | NA | NA |
| WBGene00170762 | NA | NA | NA | NA |
| WBGene00170763 | NA | NA | NA | NA |
| WBGene00170764 | NA | NA | NA | NA |
| WBGene00170765 | NA | NA | NA | NA |
| WBGene00170766 | NA | NA | NA | NA |
| WBGene00170767 | NA | NA | NA | NA |
| WBGene00170768 | NA | NA | NA | NA |
| WBGene00170769 | NA | NA | NA | NA |
| WBGene00170770 | NA | NA | NA | NA |
| WBGene00170771 | NA | NA | NA | NA |
| WBGene00170772 | NA | NA | NA | NA |
| WBGene00170773 | NA | NA | NA | NA |
| WBGene00170774 | NA | NA | NA | NA |
| WBGene00170775 | NA | NA | NA | NA |
| WBGene00170776 | NA | NA | NA | NA |
| WBGene00170777 | NA | NA | NA | NA |

| | | | | |
|---|---|---|---|---|
| WBGene00170778 | NA | NA | NA | NA |
| WBGene00170779 | NA | NA | NA | NA |
| WBGene00170780 | NA | NA | NA | NA |
| WBGene00170781 | NA | NA | NA | NA |
| WBGene00170782 | NA | NA | NA | NA |
| WBGene00170783 | NA | NA | NA | NA |
| WBGene00170784 | NA | NA | NA | NA |
| WBGene00170785 | NA | NA | NA | NA |
| WBGene00170786 | NA | NA | NA | NA |
| WBGene00170787 | NA | NA | NA | NA |
| WBGene00170788 | NA | NA | NA | NA |
| WBGene00170789 | NA | NA | NA | NA |
| WBGene00170790 | NA | NA | NA | NA |
| WBGene00170791 | NA | NA | NA | NA |
| WBGene00170792 | NA | NA | NA | NA |
| WBGene00170793 | NA | NA | NA | NA |
| WBGene00170794 | NA | NA | NA | NA |
| WBGene00170795 | NA | NA | NA | NA |
| WBGene00170796 | NA | NA | NA | NA |
| WBGene00170797 | NA | NA | NA | NA |
| WBGene00170798 | NA | NA | NA | NA |
| WBGene00170799 | NA | NA | NA | NA |
| WBGene00170800 | NA | NA | NA | NA |
| WBGene00170801 | NA | NA | NA | NA |
| WBGene00170802 | NA | NA | NA | NA |
| WBGene00170803 | NA | NA | NA | NA |
| WBGene00170804 | NA | NA | NA | NA |
| WBGene00170805 | NA | NA | NA | NA |
| WBGene00170806 | NA | NA | NA | NA |
| WBGene00170807 | NA | NA | NA | NA |
| WBGene00170808 | NA | NA | NA | NA |
| WBGene00170809 | NA | NA | NA | NA |
| WBGene00170810 | NA | NA | NA | NA |
| WBGene00170811 | NA | NA | NA | NA |
| WBGene00170812 | NA | NA | NA | NA |
| WBGene00170813 | NA | NA | NA | NA |
| WBGene00170814 | NA | NA | NA | NA |
| WBGene00170815 | NA | NA | NA | NA |
| WBGene00170816 | NA | NA | NA | NA |
| WBGene00170817 | NA | NA | NA | NA |
| WBGene00170818 | NA | NA | NA | NA |
| WBGene00170819 | -0.022930767 | 0.78121253 | NA | NA |
| WBGene00170820 | NA | NA | NA | NA |

| | | | | |
|---|---|---|---|---|
| WBGene00170821 | NA | NA | NA | NA |
| WBGene00170822 | NA | NA | NA | NA |
| WBGene00170823 | NA | NA | NA | NA |
| WBGene00170824 | NA | NA | NA | NA |
| WBGene00170825 | NA | NA | NA | NA |
| WBGene00170826 | NA | NA | NA | NA |
| WBGene00170827 | NA | NA | NA | NA |
| WBGene00170828 | NA | NA | NA | NA |
| WBGene00170829 | NA | NA | NA | NA |
| WBGene00170830 | NA | NA | NA | NA |
| WBGene00170831 | NA | NA | NA | NA |
| WBGene00170832 | NA | NA | NA | NA |
| WBGene00170833 | NA | NA | NA | NA |
| WBGene00170834 | NA | NA | NA | NA |
| WBGene00170835 | 0.036687355 | 0.59489568 | NA | NA |
| WBGene00170836 | NA | NA | NA | NA |
| WBGene00170837 | NA | NA | NA | NA |
| WBGene00170838 | NA | NA | NA | NA |
| WBGene00170839 | NA | NA | NA | NA |
| WBGene00170840 | NA | NA | NA | NA |
| WBGene00170841 | NA | NA | NA | NA |
| WBGene00170842 | NA | NA | NA | NA |
| WBGene00170843 | NA | NA | NA | NA |
| WBGene00170844 | NA | NA | NA | NA |
| WBGene00170845 | NA | NA | NA | NA |
| WBGene00170846 | 0.020317245 | 0.84643798 | NA | NA |
| WBGene00170847 | NA | NA | NA | NA |
| WBGene00170848 | NA | NA | NA | NA |
| WBGene00170849 | NA | NA | NA | NA |
| WBGene00170850 | NA | NA | NA | NA |
| WBGene00170851 | NA | NA | NA | NA |
| WBGene00170852 | NA | NA | NA | NA |
| WBGene00170853 | NA | NA | NA | NA |
| WBGene00170854 | NA | NA | NA | NA |
| WBGene00170855 | NA | NA | NA | NA |
| WBGene00170856 | NA | NA | NA | NA |
| WBGene00170857 | NA | NA | NA | NA |
| WBGene00170858 | NA | NA | NA | NA |
| WBGene00170859 | NA | NA | NA | NA |
| WBGene00170860 | NA | NA | NA | NA |
| WBGene00170861 | NA | NA | NA | NA |
| WBGene00170862 | NA | NA | NA | NA |
| WBGene00170863 | NA | NA | NA | NA |

| | | | | |
|---|---|---|---|---|
| WBGene00170864 | NA | NA | NA | NA |
| WBGene00170865 | NA | NA | NA | NA |
| WBGene00170866 | NA | NA | NA | NA |
| WBGene00170867 | NA | NA | NA | NA |
| WBGene00170868 | NA | NA | NA | NA |
| WBGene00170869 | NA | NA | NA | NA |
| WBGene00170870 | NA | NA | NA | NA |
| WBGene00170871 | NA | NA | NA | NA |
| WBGene00170872 | NA | NA | NA | NA |
| WBGene00170873 | NA | NA | NA | NA |
| WBGene00170874 | NA | NA | NA | NA |
| WBGene00170875 | NA | NA | NA | NA |
| WBGene00170876 | NA | NA | NA | NA |
| WBGene00170877 | NA | NA | NA | NA |
| WBGene00170878 | NA | NA | NA | NA |
| WBGene00170879 | NA | NA | NA | NA |
| WBGene00170880 | NA | NA | NA | NA |
| WBGene00170881 | NA | NA | NA | NA |
| WBGene00170882 | NA | NA | NA | NA |
| WBGene00170883 | NA | NA | NA | NA |
| WBGene00170884 | NA | NA | NA | NA |
| WBGene00170885 | NA | NA | NA | NA |
| WBGene00170886 | NA | NA | NA | NA |
| WBGene00170887 | NA | NA | NA | NA |
| WBGene00170888 | NA | NA | NA | NA |
| WBGene00170889 | NA | NA | NA | NA |
| WBGene00170890 | NA | NA | NA | NA |
| WBGene00170891 | NA | NA | NA | NA |
| WBGene00170892 | NA | NA | NA | NA |
| WBGene00170893 | NA | NA | NA | NA |
| WBGene00170894 | NA | NA | NA | NA |
| WBGene00170895 | NA | NA | NA | NA |
| WBGene00170896 | NA | NA | NA | NA |
| WBGene00170897 | NA | NA | NA | NA |
| WBGene00170898 | NA | NA | NA | NA |
| WBGene00170899 | NA | NA | NA | NA |
| WBGene00170900 | NA | NA | NA | NA |
| WBGene00170901 | NA | NA | NA | NA |
| WBGene00170902 | NA | NA | NA | NA |
| WBGene00170903 | 0.020925399 | 0.84643798 | NA | NA |
| WBGene00170904 | NA | NA | NA | NA |
| WBGene00170905 | NA | NA | NA | NA |
| WBGene00170906 | NA | NA | NA | NA |

| | | | | |
|---|---|---|---|---|
| WBGene00170907 | NA | NA | NA | NA |
| WBGene00170908 | NA | NA | NA | NA |
| WBGene00170909 | NA | NA | NA | NA |
| WBGene00170910 | NA | NA | NA | NA |
| WBGene00170911 | NA | NA | NA | NA |
| WBGene00170912 | NA | NA | NA | NA |
| WBGene00170913 | 0.020925399 | 0.84643798 | NA | NA |
| WBGene00170914 | NA | NA | NA | NA |
| WBGene00170915 | NA | NA | NA | NA |
| WBGene00170916 | NA | NA | NA | NA |
| WBGene00170917 | NA | NA | NA | NA |
| WBGene00170918 | NA | NA | NA | NA |
| WBGene00170919 | NA | NA | NA | NA |
| WBGene00170920 | NA | NA | NA | NA |
| WBGene00170921 | NA | NA | NA | NA |
| WBGene00170922 | NA | NA | NA | NA |
| WBGene00170923 | NA | NA | NA | NA |
| WBGene00170924 | NA | NA | NA | NA |
| WBGene00170925 | NA | NA | NA | NA |
| WBGene00170926 | NA | NA | NA | NA |
| WBGene00170927 | NA | NA | NA | NA |
| WBGene00170928 | NA | NA | NA | NA |
| WBGene00170929 | NA | NA | NA | NA |
| WBGene00170930 | NA | NA | NA | NA |
| WBGene00170931 | NA | NA | NA | NA |
| WBGene00170932 | NA | NA | NA | NA |
| WBGene00170933 | NA | NA | NA | NA |
| WBGene00170934 | NA | NA | NA | NA |
| WBGene00170935 | NA | NA | NA | NA |
| WBGene00170936 | NA | NA | NA | NA |
| WBGene00170937 | NA | NA | NA | NA |
| WBGene00170938 | NA | NA | NA | NA |
| WBGene00170939 | NA | NA | NA | NA |
| WBGene00170940 | NA | NA | NA | NA |
| WBGene00170941 | NA | NA | NA | NA |
| WBGene00170942 | NA | NA | NA | NA |
| WBGene00170943 | NA | NA | NA | NA |
| WBGene00170944 | NA | NA | NA | NA |
| WBGene00170945 | NA | NA | NA | NA |
| WBGene00170946 | NA | NA | NA | NA |
| WBGene00170947 | NA | NA | NA | NA |
| WBGene00170948 | NA | NA | NA | NA |
| WBGene00170949 | NA | NA | NA | NA |

| | | | | |
|---|---|---|---|---|
| WBGene00170950 | 0.020232312 | 0.84643798 | NA | NA |
| WBGene00170951 | NA | NA | NA | NA |
| WBGene00170952 | NA | NA | NA | NA |
| WBGene00170953 | NA | NA | NA | NA |
| WBGene00170954 | NA | NA | NA | NA |
| WBGene00170955 | NA | NA | NA | NA |
| WBGene00170956 | NA | NA | NA | NA |
| WBGene00170957 | NA | NA | NA | NA |
| WBGene00170958 | NA | NA | NA | NA |
| WBGene00170959 | NA | NA | NA | NA |
| WBGene00170960 | NA | NA | NA | NA |
| WBGene00170961 | NA | NA | NA | NA |
| WBGene00170962 | NA | NA | NA | NA |
| WBGene00170963 | NA | NA | NA | NA |
| WBGene00170964 | NA | NA | NA | NA |
| WBGene00170965 | NA | NA | NA | NA |
| WBGene00170966 | NA | NA | NA | NA |
| WBGene00170967 | NA | NA | NA | NA |
| WBGene00170968 | NA | NA | NA | NA |
| WBGene00170969 | NA | NA | NA | NA |
| WBGene00170970 | NA | NA | NA | NA |
| WBGene00170971 | NA | NA | NA | NA |
| WBGene00170972 | NA | NA | NA | NA |
| WBGene00170973 | NA | NA | NA | NA |
| WBGene00170974 | NA | NA | NA | NA |
| WBGene00170975 | NA | NA | NA | NA |
| WBGene00170976 | NA | NA | NA | NA |
| WBGene00170977 | NA | NA | NA | NA |
| WBGene00170978 | NA | NA | NA | NA |
| WBGene00170979 | NA | NA | NA | NA |
| WBGene00170980 | NA | NA | NA | NA |
| WBGene00170981 | NA | NA | NA | NA |
| WBGene00170982 | NA | NA | NA | NA |
| WBGene00170983 | NA | NA | NA | NA |
| WBGene00170984 | NA | NA | NA | NA |
| WBGene00170985 | NA | NA | NA | NA |
| WBGene00170986 | NA | NA | NA | NA |
| WBGene00170987 | NA | NA | NA | NA |
| WBGene00170988 | NA | NA | NA | NA |
| WBGene00170989 | NA | NA | NA | NA |
| WBGene00170990 | NA | NA | NA | NA |
| WBGene00170991 | NA | NA | NA | NA |
| WBGene00170992 | NA | NA | NA | NA |

| | | | | |
|---|---|---|---|---|
| WBGene00170993 | NA | NA | NA | NA |
| WBGene00170994 | NA | NA | NA | NA |
| WBGene00170995 | NA | NA | NA | NA |
| WBGene00170996 | NA | NA | NA | NA |
| WBGene00170997 | NA | NA | NA | NA |
| WBGene00170998 | NA | NA | NA | NA |
| WBGene00170999 | NA | NA | NA | NA |
| WBGene00171000 | NA | NA | NA | NA |
| WBGene00171001 | NA | NA | NA | NA |
| WBGene00171002 | NA | NA | NA | NA |
| WBGene00171003 | NA | NA | NA | NA |
| WBGene00171004 | NA | NA | NA | NA |
| WBGene00171005 | NA | NA | NA | NA |
| WBGene00171006 | NA | NA | NA | NA |
| WBGene00171007 | NA | NA | NA | NA |
| WBGene00171008 | NA | NA | NA | NA |
| WBGene00171009 | NA | NA | NA | NA |
| WBGene00171010 | NA | NA | NA | NA |
| WBGene00171011 | NA | NA | NA | NA |
| WBGene00171012 | NA | NA | NA | NA |
| WBGene00171013 | NA | NA | NA | NA |
| WBGene00171014 | NA | NA | NA | NA |
| WBGene00171015 | NA | NA | NA | NA |
| WBGene00171016 | NA | NA | NA | NA |
| WBGene00171017 | NA | NA | NA | NA |
| WBGene00171018 | NA | NA | NA | NA |
| WBGene00171019 | NA | NA | NA | NA |
| WBGene00171020 | NA | NA | NA | NA |
| WBGene00171021 | NA | NA | NA | NA |
| WBGene00171022 | NA | NA | NA | NA |
| WBGene00171023 | NA | NA | NA | NA |
| WBGene00171024 | NA | NA | NA | NA |
| WBGene00171025 | NA | NA | NA | NA |
| WBGene00171026 | NA | NA | NA | NA |
| WBGene00171027 | 0.056411843 | 0.41891328 | NA | NA |
| WBGene00171028 | NA | NA | NA | NA |
| WBGene00171029 | NA | NA | NA | NA |
| WBGene00171030 | NA | NA | NA | NA |
| WBGene00171031 | NA | NA | NA | NA |
| WBGene00171032 | NA | NA | NA | NA |
| WBGene00171033 | NA | NA | NA | NA |
| WBGene00171034 | NA | NA | NA | NA |
| WBGene00171035 | NA | NA | NA | NA |

| | | | | |
|---|---|---|---|---|
| WBGene00171036 | NA | NA | NA | NA |
| WBGene00171037 | NA | NA | NA | NA |
| WBGene00171038 | NA | NA | NA | NA |
| WBGene00171039 | NA | NA | NA | NA |
| WBGene00171040 | NA | NA | NA | NA |
| WBGene00171041 | NA | NA | NA | NA |
| WBGene00171042 | NA | NA | NA | NA |
| WBGene00171043 | NA | NA | NA | NA |
| WBGene00171044 | NA | NA | NA | NA |
| WBGene00171045 | NA | NA | NA | NA |
| WBGene00171046 | NA | NA | NA | NA |
| WBGene00171047 | NA | NA | NA | NA |
| WBGene00171048 | NA | NA | NA | NA |
| WBGene00171049 | NA | NA | NA | NA |
| WBGene00171050 | NA | NA | NA | NA |
| WBGene00171051 | NA | NA | NA | NA |
| WBGene00171052 | NA | NA | NA | NA |
| WBGene00171053 | NA | NA | NA | NA |
| WBGene00171054 | NA | NA | NA | NA |
| WBGene00171055 | NA | NA | NA | NA |
| WBGene00171056 | NA | NA | NA | NA |
| WBGene00171057 | NA | NA | NA | NA |
| WBGene00171058 | NA | NA | NA | NA |
| WBGene00171059 | NA | NA | NA | NA |
| WBGene00171060 | NA | NA | NA | NA |
| WBGene00171061 | NA | NA | NA | NA |
| WBGene00171062 | NA | NA | NA | NA |
| WBGene00171063 | NA | NA | NA | NA |
| WBGene00171064 | 0.0398788 | 0.58971221 | NA | NA |
| WBGene00171065 | NA | NA | NA | NA |
| WBGene00171066 | NA | NA | NA | NA |
| WBGene00171067 | NA | NA | NA | NA |
| WBGene00171068 | NA | NA | NA | NA |
| WBGene00171069 | NA | NA | NA | NA |
| WBGene00171070 | NA | NA | NA | NA |
| WBGene00171071 | NA | NA | NA | NA |
| WBGene00171072 | NA | NA | NA | NA |
| WBGene00171073 | NA | NA | NA | NA |
| WBGene00171074 | NA | NA | NA | NA |
| WBGene00171075 | NA | NA | NA | NA |
| WBGene00171076 | NA | NA | NA | NA |
| WBGene00171077 | NA | NA | NA | NA |
| WBGene00171078 | NA | NA | NA | NA |

| | | | | |
|---|---|---|---|---|
| WBGene00171079 | NA | NA | NA | NA |
| WBGene00171080 | NA | NA | NA | NA |
| WBGene00171081 | NA | NA | NA | NA |
| WBGene00171082 | NA | NA | NA | NA |
| WBGene00171083 | NA | NA | NA | NA |
| WBGene00171084 | NA | NA | NA | NA |
| WBGene00171085 | NA | NA | NA | NA |
| WBGene00171086 | NA | NA | NA | NA |
| WBGene00171087 | NA | NA | NA | NA |
| WBGene00171088 | NA | NA | NA | NA |
| WBGene00171089 | NA | NA | NA | NA |
| WBGene00171090 | NA | NA | NA | NA |
| WBGene00171091 | NA | NA | NA | NA |
| WBGene00171092 | NA | NA | NA | NA |
| WBGene00171093 | NA | NA | NA | NA |
| WBGene00171094 | NA | NA | NA | NA |
| WBGene00171095 | NA | NA | NA | NA |
| WBGene00171096 | NA | NA | NA | NA |
| WBGene00171097 | NA | NA | NA | NA |
| WBGene00171098 | NA | NA | NA | NA |
| WBGene00171099 | NA | NA | NA | NA |
| WBGene00171100 | NA | NA | NA | NA |
| WBGene00171101 | NA | NA | NA | NA |
| WBGene00171102 | NA | NA | NA | NA |
| WBGene00171103 | NA | NA | NA | NA |
| WBGene00171104 | NA | NA | NA | NA |
| WBGene00171105 | NA | NA | NA | NA |
| WBGene00171106 | NA | NA | NA | NA |
| WBGene00171107 | NA | NA | NA | NA |
| WBGene00171108 | NA | NA | NA | NA |
| WBGene00171109 | NA | NA | NA | NA |
| WBGene00171110 | NA | NA | NA | NA |
| WBGene00171111 | NA | NA | NA | NA |
| WBGene00171112 | NA | NA | NA | NA |
| WBGene00171113 | NA | NA | NA | NA |
| WBGene00171114 | NA | NA | NA | NA |
| WBGene00171115 | NA | NA | NA | NA |
| WBGene00171116 | NA | NA | NA | NA |
| WBGene00171117 | NA | NA | NA | NA |
| WBGene00171118 | NA | NA | NA | NA |
| WBGene00171119 | NA | NA | NA | NA |
| WBGene00171120 | NA | NA | NA | NA |
| WBGene00171121 | NA | NA | NA | NA |

| | | | | |
|---|---|---|---|---|
| WBGene00171122 | NA | NA | NA | NA |
| WBGene00171123 | NA | NA | NA | NA |
| WBGene00171124 | NA | NA | NA | NA |
| WBGene00171125 | NA | NA | NA | NA |
| WBGene00171126 | NA | NA | NA | NA |
| WBGene00171127 | NA | NA | NA | NA |
| WBGene00171128 | NA | NA | NA | NA |
| WBGene00171129 | NA | NA | NA | NA |
| WBGene00171130 | NA | NA | NA | NA |
| WBGene00171131 | NA | NA | NA | NA |
| WBGene00171132 | NA | NA | NA | NA |
| WBGene00171133 | NA | NA | NA | NA |
| WBGene00171134 | NA | NA | NA | NA |
| WBGene00171135 | NA | NA | NA | NA |
| WBGene00171136 | NA | NA | NA | NA |
| WBGene00171137 | NA | NA | NA | NA |
| WBGene00171138 | NA | NA | NA | NA |
| WBGene00171139 | NA | NA | NA | NA |
| WBGene00171140 | NA | NA | NA | NA |
| WBGene00171141 | NA | NA | NA | NA |
| WBGene00171142 | NA | NA | NA | NA |
| WBGene00171143 | NA | NA | NA | NA |
| WBGene00171144 | NA | NA | NA | NA |
| WBGene00171145 | NA | NA | NA | NA |
| WBGene00171146 | NA | NA | NA | NA |
| WBGene00171147 | NA | NA | NA | NA |
| WBGene00171148 | NA | NA | NA | NA |
| WBGene00171149 | NA | NA | NA | NA |
| WBGene00171150 | NA | NA | NA | NA |
| WBGene00171151 | NA | NA | NA | NA |
| WBGene00171152 | NA | NA | NA | NA |
| WBGene00171153 | NA | NA | NA | NA |
| WBGene00171154 | NA | NA | NA | NA |
| WBGene00171155 | NA | NA | NA | NA |
| WBGene00171156 | NA | NA | NA | NA |
| WBGene00171157 | NA | NA | NA | NA |
| WBGene00171158 | NA | NA | NA | NA |
| WBGene00171159 | NA | NA | NA | NA |
| WBGene00171160 | NA | NA | NA | NA |
| WBGene00171161 | NA | NA | NA | NA |
| WBGene00171162 | NA | NA | NA | NA |
| WBGene00171163 | NA | NA | NA | NA |
| WBGene00171164 | NA | NA | NA | NA |

| | | | | |
|---|---|---|---|---|
| WBGene00171165 | NA | NA | NA | NA |
| WBGene00171166 | NA | NA | NA | NA |
| WBGene00171167 | NA | NA | NA | NA |
| WBGene00171168 | NA | NA | NA | NA |
| WBGene00171169 | NA | NA | NA | NA |
| WBGene00171170 | NA | NA | NA | NA |
| WBGene00171171 | NA | NA | NA | NA |
| WBGene00171172 | NA | NA | NA | NA |
| WBGene00171173 | NA | NA | NA | NA |
| WBGene00171174 | NA | NA | NA | NA |
| WBGene00171175 | NA | NA | NA | NA |
| WBGene00171176 | NA | NA | NA | NA |
| WBGene00171177 | NA | NA | NA | NA |
| WBGene00171178 | NA | NA | NA | NA |
| WBGene00171179 | NA | NA | NA | NA |
| WBGene00171180 | NA | NA | NA | NA |
| WBGene00171181 | NA | NA | NA | NA |
| WBGene00171182 | NA | NA | NA | NA |
| WBGene00171183 | NA | NA | NA | NA |
| WBGene00171184 | NA | NA | NA | NA |
| WBGene00171185 | NA | NA | NA | NA |
| WBGene00171186 | NA | NA | NA | NA |
| WBGene00171187 | NA | NA | NA | NA |
| WBGene00171188 | NA | NA | NA | NA |
| WBGene00171189 | NA | NA | NA | NA |
| WBGene00171190 | NA | NA | NA | NA |
| WBGene00171191 | NA | NA | NA | NA |
| WBGene00171192 | NA | NA | NA | NA |
| WBGene00171193 | NA | NA | NA | NA |
| WBGene00171194 | NA | NA | NA | NA |
| WBGene00171195 | NA | NA | NA | NA |
| WBGene00171196 | NA | NA | NA | NA |
| WBGene00171197 | NA | NA | NA | NA |
| WBGene00171198 | NA | NA | NA | NA |
| WBGene00171199 | NA | NA | NA | NA |
| WBGene00171200 | NA | NA | NA | NA |
| WBGene00171201 | NA | NA | NA | NA |
| WBGene00171202 | NA | NA | NA | NA |
| WBGene00171203 | NA | NA | NA | NA |
| WBGene00171204 | NA | NA | NA | NA |
| WBGene00171205 | NA | NA | NA | NA |
| WBGene00171206 | NA | NA | NA | NA |
| WBGene00171207 | NA | NA | NA | NA |

| | | | | |
|---|---|---|---|---|
| WBGene00171208 | NA | NA | NA | NA |
| WBGene00171209 | NA | NA | NA | NA |
| WBGene00171210 | NA | NA | NA | NA |
| WBGene00171211 | NA | NA | NA | NA |
| WBGene00171212 | NA | NA | NA | NA |
| WBGene00171213 | NA | NA | NA | NA |
| WBGene00171214 | NA | NA | NA | NA |
| WBGene00171215 | NA | NA | NA | NA |
| WBGene00171216 | NA | NA | NA | NA |
| WBGene00171217 | NA | NA | NA | NA |
| WBGene00171218 | NA | NA | NA | NA |
| WBGene00171219 | NA | NA | NA | NA |
| WBGene00171220 | NA | NA | NA | NA |
| WBGene00171221 | NA | NA | NA | NA |
| WBGene00171222 | NA | NA | NA | NA |
| WBGene00171223 | NA | NA | NA | NA |
| WBGene00171224 | NA | NA | NA | NA |
| WBGene00171225 | NA | NA | NA | NA |
| WBGene00171226 | NA | NA | NA | NA |
| WBGene00171227 | NA | NA | NA | NA |
| WBGene00171228 | NA | NA | NA | NA |
| WBGene00171229 | NA | NA | NA | NA |
| WBGene00171230 | NA | NA | NA | NA |
| WBGene00171231 | NA | NA | NA | NA |
| WBGene00171232 | NA | NA | NA | NA |
| WBGene00171233 | 0.020925399 | 0.84643798 | NA | NA |
| WBGene00171234 | NA | NA | NA | NA |
| WBGene00171235 | NA | NA | NA | NA |
| WBGene00171236 | NA | NA | NA | NA |
| WBGene00171237 | NA | NA | NA | NA |
| WBGene00171238 | NA | NA | NA | NA |
| WBGene00171239 | NA | NA | NA | NA |
| WBGene00171240 | NA | NA | NA | NA |
| WBGene00171241 | NA | NA | NA | NA |
| WBGene00171242 | NA | NA | NA | NA |
| WBGene00171243 | NA | NA | NA | NA |
| WBGene00171244 | NA | NA | NA | NA |
| WBGene00171245 | NA | NA | NA | NA |
| WBGene00171246 | NA | NA | NA | NA |
| WBGene00171247 | NA | NA | NA | NA |
| WBGene00171248 | NA | NA | NA | NA |
| WBGene00171249 | NA | NA | NA | NA |
| WBGene00171250 | NA | NA | NA | NA |

| | | | | |
|---|---|---|---|---|
| WBGene00171251 | NA | NA | NA | NA |
| WBGene00171252 | NA | NA | NA | NA |
| WBGene00171253 | NA | NA | NA | NA |
| WBGene00171254 | NA | NA | NA | NA |
| WBGene00171255 | NA | NA | NA | NA |
| WBGene00171256 | NA | NA | NA | NA |
| WBGene00171257 | NA | NA | NA | NA |
| WBGene00171258 | NA | NA | NA | NA |
| WBGene00171259 | NA | NA | NA | NA |
| WBGene00171260 | NA | NA | NA | NA |
| WBGene00171261 | NA | NA | NA | NA |
| WBGene00171262 | NA | NA | NA | NA |
| WBGene00171263 | NA | NA | NA | NA |
| WBGene00171264 | NA | NA | NA | NA |
| WBGene00171265 | NA | NA | NA | NA |
| WBGene00171266 | NA | NA | NA | NA |
| WBGene00171267 | NA | NA | NA | NA |
| WBGene00171268 | NA | NA | NA | NA |
| WBGene00171269 | NA | NA | NA | NA |
| WBGene00171270 | NA | NA | NA | NA |
| WBGene00171271 | NA | NA | NA | NA |
| WBGene00171272 | NA | NA | NA | NA |
| WBGene00171273 | NA | NA | NA | NA |
| WBGene00171274 | NA | NA | NA | NA |
| WBGene00171275 | NA | NA | NA | NA |
| WBGene00171276 | NA | NA | NA | NA |
| WBGene00171277 | NA | NA | NA | NA |
| WBGene00171278 | NA | NA | NA | NA |
| WBGene00171279 | NA | NA | NA | NA |
| WBGene00171280 | NA | NA | NA | NA |
| WBGene00171281 | NA | NA | NA | NA |
| WBGene00171282 | NA | NA | NA | NA |
| WBGene00171283 | NA | NA | NA | NA |
| WBGene00171284 | NA | NA | NA | NA |
| WBGene00171285 | NA | NA | NA | NA |
| WBGene00171286 | NA | NA | NA | NA |
| WBGene00171287 | NA | NA | NA | NA |
| WBGene00171288 | NA | NA | NA | NA |
| WBGene00171289 | NA | NA | NA | NA |
| WBGene00171290 | NA | NA | NA | NA |
| WBGene00171291 | NA | NA | NA | NA |
| WBGene00171292 | NA | NA | NA | NA |
| WBGene00171293 | NA | NA | NA | NA |

| | | | | |
|---|---|---|---|---|
| WBGene00171294 | NA | NA | NA | NA |
| WBGene00171295 | NA | NA | NA | NA |
| WBGene00171296 | NA | NA | NA | NA |
| WBGene00171297 | NA | NA | NA | NA |
| WBGene00171298 | NA | NA | NA | NA |
| WBGene00171299 | NA | NA | NA | NA |
| WBGene00171300 | NA | NA | NA | NA |
| WBGene00171301 | NA | NA | NA | NA |
| WBGene00171302 | NA | NA | NA | NA |
| WBGene00171303 | NA | NA | NA | NA |
| WBGene00171304 | NA | NA | NA | NA |
| WBGene00171305 | NA | NA | NA | NA |
| WBGene00171306 | NA | NA | NA | NA |
| WBGene00171307 | NA | NA | NA | NA |
| WBGene00171308 | NA | NA | NA | NA |
| WBGene00171309 | NA | NA | NA | NA |
| WBGene00171310 | NA | NA | NA | NA |
| WBGene00171311 | NA | NA | NA | NA |
| WBGene00171312 | NA | NA | NA | NA |
| WBGene00171313 | NA | NA | NA | NA |
| WBGene00171314 | NA | NA | NA | NA |
| WBGene00171315 | NA | NA | NA | NA |
| WBGene00171316 | NA | NA | NA | NA |
| WBGene00171317 | NA | NA | NA | NA |
| WBGene00171318 | NA | NA | NA | NA |
| WBGene00171319 | NA | NA | NA | NA |
| WBGene00171320 | NA | NA | NA | NA |
| WBGene00171321 | NA | NA | NA | NA |
| WBGene00171322 | NA | NA | NA | NA |
| WBGene00171323 | NA | NA | NA | NA |
| WBGene00171324 | NA | NA | NA | NA |
| WBGene00171325 | NA | NA | NA | NA |
| WBGene00171326 | NA | NA | NA | NA |
| WBGene00171327 | NA | NA | NA | NA |
| WBGene00171328 | NA | NA | NA | NA |
| WBGene00171329 | NA | NA | NA | NA |
| WBGene00171330 | NA | NA | NA | NA |
| WBGene00171331 | NA | NA | NA | NA |
| WBGene00171332 | NA | NA | NA | NA |
| WBGene00171333 | NA | NA | NA | NA |
| WBGene00171334 | NA | NA | NA | NA |
| WBGene00171335 | NA | NA | NA | NA |
| WBGene00171336 | NA | NA | NA | NA |

| | | | | |
|---|---|---|---|---|
| WBGene00171337 | NA | NA | NA | NA |
| WBGene00171338 | NA | NA | NA | NA |
| WBGene00171339 | NA | NA | NA | NA |
| WBGene00171340 | NA | NA | NA | NA |
| WBGene00171341 | NA | NA | NA | NA |
| WBGene00171342 | NA | NA | NA | NA |
| WBGene00171343 | NA | NA | NA | NA |
| WBGene00171344 | NA | NA | NA | NA |
| WBGene00171345 | NA | NA | NA | NA |
| WBGene00171346 | NA | NA | NA | NA |
| WBGene00171347 | NA | NA | NA | NA |
| WBGene00171348 | NA | NA | NA | NA |
| WBGene00171349 | NA | NA | NA | NA |
| WBGene00171350 | NA | NA | NA | NA |
| WBGene00171351 | NA | NA | NA | NA |
| WBGene00171352 | NA | NA | NA | NA |
| WBGene00171353 | NA | NA | NA | NA |
| WBGene00171354 | NA | NA | NA | NA |
| WBGene00171355 | NA | NA | NA | NA |
| WBGene00171356 | NA | NA | NA | NA |
| WBGene00171357 | NA | NA | NA | NA |
| WBGene00171358 | NA | NA | NA | NA |
| WBGene00171359 | NA | NA | NA | NA |
| WBGene00171360 | NA | NA | NA | NA |
| WBGene00171361 | NA | NA | NA | NA |
| WBGene00171362 | NA | NA | NA | NA |
| WBGene00171363 | NA | NA | NA | NA |
| WBGene00171364 | NA | NA | NA | NA |
| WBGene00171365 | NA | NA | NA | NA |
| WBGene00171366 | NA | NA | NA | NA |
| WBGene00171367 | NA | NA | NA | NA |
| WBGene00171368 | NA | NA | NA | NA |
| WBGene00171369 | NA | NA | NA | NA |
| WBGene00171370 | NA | NA | NA | NA |
| WBGene00171371 | NA | NA | NA | NA |
| WBGene00171372 | NA | NA | NA | NA |
| WBGene00171373 | NA | NA | NA | NA |
| WBGene00171374 | NA | NA | NA | NA |
| WBGene00171375 | NA | NA | NA | NA |
| WBGene00171376 | NA | NA | NA | NA |
| WBGene00171377 | NA | NA | NA | NA |
| WBGene00171378 | NA | NA | NA | NA |
| WBGene00171379 | NA | NA | NA | NA |

| | | | | |
|---|---|---|---|---|
| WBGene00171380 | NA | NA | NA | NA |
| WBGene00171381 | NA | NA | NA | NA |
| WBGene00171382 | NA | NA | NA | NA |
| WBGene00171383 | NA | NA | NA | NA |
| WBGene00171384 | NA | NA | NA | NA |
| WBGene00171385 | NA | NA | NA | NA |
| WBGene00171386 | NA | NA | NA | NA |
| WBGene00171387 | NA | NA | NA | NA |
| WBGene00171388 | NA | NA | NA | NA |
| WBGene00171389 | NA | NA | NA | NA |
| WBGene00171390 | NA | NA | NA | NA |
| WBGene00171391 | NA | NA | NA | NA |
| WBGene00171392 | NA | NA | NA | NA |
| WBGene00171393 | NA | NA | NA | NA |
| WBGene00171394 | NA | NA | NA | NA |
| WBGene00171395 | NA | NA | NA | NA |
| WBGene00171396 | NA | NA | NA | NA |
| WBGene00171397 | NA | NA | NA | NA |
| WBGene00171398 | NA | NA | NA | NA |
| WBGene00171399 | NA | NA | NA | NA |
| WBGene00171400 | NA | NA | NA | NA |
| WBGene00171401 | NA | NA | NA | NA |
| WBGene00171402 | NA | NA | NA | NA |
| WBGene00171403 | NA | NA | NA | NA |
| WBGene00171404 | NA | NA | NA | NA |
| WBGene00171405 | NA | NA | NA | NA |
| WBGene00171406 | NA | NA | NA | NA |
| WBGene00171407 | NA | NA | NA | NA |
| WBGene00171408 | NA | NA | NA | NA |
| WBGene00171409 | NA | NA | NA | NA |
| WBGene00171410 | NA | NA | NA | NA |
| WBGene00171411 | NA | NA | NA | NA |
| WBGene00171412 | NA | NA | NA | NA |
| WBGene00171413 | NA | NA | NA | NA |
| WBGene00171414 | NA | NA | NA | NA |
| WBGene00171415 | NA | NA | NA | NA |
| WBGene00171416 | NA | NA | NA | NA |
| WBGene00171417 | NA | NA | NA | NA |
| WBGene00171418 | NA | NA | NA | NA |
| WBGene00171419 | 0.038125098 | 0.61119482 | NA | NA |
| WBGene00171420 | NA | NA | NA | NA |
| WBGene00171421 | NA | NA | NA | NA |
| WBGene00171422 | NA | NA | NA | NA |

| | | | | |
|---|---|---|---|---|
| WBGene00171423 | NA | NA | NA | NA |
| WBGene00171424 | NA | NA | NA | NA |
| WBGene00171425 | NA | NA | NA | NA |
| WBGene00171426 | NA | NA | NA | NA |
| WBGene00171427 | NA | NA | NA | NA |
| WBGene00171428 | NA | NA | NA | NA |
| WBGene00171429 | NA | NA | NA | NA |
| WBGene00171430 | NA | NA | NA | NA |
| WBGene00171431 | NA | NA | NA | NA |
| WBGene00171432 | NA | NA | NA | NA |
| WBGene00171433 | NA | NA | NA | NA |
| WBGene00171434 | NA | NA | NA | NA |
| WBGene00171435 | NA | NA | NA | NA |
| WBGene00171436 | NA | NA | NA | NA |
| WBGene00171437 | NA | NA | NA | NA |
| WBGene00171438 | NA | NA | NA | NA |
| WBGene00171439 | NA | NA | NA | NA |
| WBGene00171440 | NA | NA | NA | NA |
| WBGene00171441 | NA | NA | NA | NA |
| WBGene00171442 | NA | NA | NA | NA |
| WBGene00171443 | NA | NA | NA | NA |
| WBGene00171444 | NA | NA | NA | NA |
| WBGene00171445 | NA | NA | NA | NA |
| WBGene00171446 | NA | NA | NA | NA |
| WBGene00171447 | NA | NA | NA | NA |
| WBGene00171448 | NA | NA | NA | NA |
| WBGene00171449 | NA | NA | NA | NA |
| WBGene00171450 | NA | NA | NA | NA |
| WBGene00171451 | NA | NA | NA | NA |
| WBGene00171452 | NA | NA | NA | NA |
| WBGene00171453 | NA | NA | NA | NA |
| WBGene00171454 | 0.020232312 | 0.84643798 | NA | NA |
| WBGene00171455 | NA | NA | NA | NA |
| WBGene00171456 | NA | NA | NA | NA |
| WBGene00171457 | NA | NA | NA | NA |
| WBGene00171458 | NA | NA | NA | NA |
| WBGene00171459 | NA | NA | NA | NA |
| WBGene00171460 | NA | NA | NA | NA |
| WBGene00171461 | NA | NA | NA | NA |
| WBGene00171462 | NA | NA | NA | NA |
| WBGene00171463 | NA | NA | NA | NA |
| WBGene00171464 | NA | NA | NA | NA |
| WBGene00171465 | NA | NA | NA | NA |

| | | | | |
|---|---|---|---|---|
| WBGene00171466 | NA | NA | NA | NA |
| WBGene00171467 | NA | NA | NA | NA |
| WBGene00171468 | NA | NA | NA | NA |
| WBGene00171469 | NA | NA | NA | NA |
| WBGene00171470 | NA | NA | NA | NA |
| WBGene00171471 | NA | NA | NA | NA |
| WBGene00171472 | NA | NA | NA | NA |
| WBGene00171473 | NA | NA | NA | NA |
| WBGene00171474 | NA | NA | NA | NA |
| WBGene00171475 | NA | NA | NA | NA |
| WBGene00171476 | NA | NA | NA | NA |
| WBGene00171477 | NA | NA | NA | NA |
| WBGene00171478 | NA | NA | NA | NA |
| WBGene00171479 | -0.003270206 | 0.95698316 | NA | NA |
| WBGene00171480 | NA | NA | NA | NA |
| WBGene00171481 | NA | NA | NA | NA |
| WBGene00171482 | NA | NA | NA | NA |
| WBGene00171483 | NA | NA | NA | NA |
| WBGene00171484 | 0.020925399 | 0.84643798 | NA | NA |
| WBGene00171485 | NA | NA | NA | NA |
| WBGene00171486 | NA | NA | NA | NA |
| WBGene00171487 | NA | NA | NA | NA |
| WBGene00171488 | NA | NA | NA | NA |
| WBGene00171489 | NA | NA | NA | NA |
| WBGene00171490 | NA | NA | NA | NA |
| WBGene00171491 | NA | NA | NA | NA |
| WBGene00171492 | NA | NA | NA | NA |
| WBGene00171493 | NA | NA | NA | NA |
| WBGene00171494 | NA | NA | NA | NA |
| WBGene00171495 | NA | NA | NA | NA |
| WBGene00171496 | NA | NA | NA | NA |
| WBGene00171497 | NA | NA | NA | NA |
| WBGene00171498 | NA | NA | NA | NA |
| WBGene00171499 | NA | NA | NA | NA |
| WBGene00171500 | NA | NA | NA | NA |
| WBGene00171501 | NA | NA | NA | NA |
| WBGene00171502 | NA | NA | NA | NA |
| WBGene00171503 | NA | NA | NA | NA |
| WBGene00171504 | NA | NA | NA | NA |
| WBGene00171505 | NA | NA | NA | NA |
| WBGene00171506 | NA | NA | NA | NA |
| WBGene00171507 | NA | NA | NA | NA |
| WBGene00171508 | NA | NA | NA | NA |

| | | | | |
|---|---|---|---|---|
| WBGene00171509 | NA | NA | NA | NA |
| WBGene00171510 | NA | NA | NA | NA |
| WBGene00171511 | NA | NA | NA | NA |
| WBGene00171512 | NA | NA | NA | NA |
| WBGene00171513 | NA | NA | NA | NA |
| WBGene00171514 | NA | NA | NA | NA |
| WBGene00171515 | NA | NA | NA | NA |
| WBGene00171516 | NA | NA | NA | NA |
| WBGene00171517 | NA | NA | NA | NA |
| WBGene00171518 | -0.023109762 | 0.78121253 | NA | NA |
| WBGene00171519 | NA | NA | NA | NA |
| WBGene00171520 | NA | NA | NA | NA |
| WBGene00171521 | NA | NA | NA | NA |
| WBGene00171522 | NA | NA | NA | NA |
| WBGene00171523 | NA | NA | NA | NA |
| WBGene00171524 | NA | NA | NA | NA |
| WBGene00171525 | NA | NA | NA | NA |
| WBGene00171526 | NA | NA | NA | NA |
| WBGene00171527 | NA | NA | NA | NA |
| WBGene00171528 | NA | NA | NA | NA |
| WBGene00171529 | NA | NA | NA | NA |
| WBGene00171530 | NA | NA | NA | NA |
| WBGene00171531 | NA | NA | NA | NA |
| WBGene00171532 | NA | NA | NA | NA |
| WBGene00171533 | NA | NA | NA | NA |
| WBGene00171534 | NA | NA | NA | NA |
| WBGene00171535 | NA | NA | NA | NA |
| WBGene00171536 | NA | NA | NA | NA |
| WBGene00171537 | NA | NA | NA | NA |
| WBGene00171538 | NA | NA | NA | NA |
| WBGene00171539 | NA | NA | NA | NA |
| WBGene00171540 | NA | NA | NA | NA |
| WBGene00171541 | NA | NA | NA | NA |
| WBGene00171542 | NA | NA | NA | NA |
| WBGene00171543 | NA | NA | NA | NA |
| WBGene00171544 | NA | NA | NA | NA |
| WBGene00171545 | NA | NA | NA | NA |
| WBGene00171546 | NA | NA | NA | NA |
| WBGene00171547 | NA | NA | NA | NA |
| WBGene00171548 | NA | NA | NA | NA |
| WBGene00171549 | NA | NA | NA | NA |
| WBGene00171550 | NA | NA | NA | NA |
| WBGene00171551 | NA | NA | NA | NA |

| | | | | |
|---|---|---|---|---|
| WBGene00171552 | NA | NA | NA | NA |
| WBGene00171553 | NA | NA | NA | NA |
| WBGene00171554 | NA | NA | NA | NA |
| WBGene00171555 | NA | NA | NA | NA |
| WBGene00171556 | NA | NA | NA | NA |
| WBGene00171557 | NA | NA | NA | NA |
| WBGene00171558 | NA | NA | NA | NA |
| WBGene00171559 | NA | NA | NA | NA |
| WBGene00171560 | NA | NA | NA | NA |
| WBGene00171561 | NA | NA | NA | NA |
| WBGene00171562 | NA | NA | NA | NA |
| WBGene00171563 | NA | NA | NA | NA |
| WBGene00171564 | NA | NA | NA | NA |
| WBGene00171565 | NA | NA | NA | NA |
| WBGene00171566 | NA | NA | NA | NA |
| WBGene00171567 | NA | NA | NA | NA |
| WBGene00171568 | -0.003614866 | 0.95700117 | NA | NA |
| WBGene00171569 | NA | NA | NA | NA |
| WBGene00171570 | NA | NA | NA | NA |
| WBGene00171571 | NA | NA | NA | NA |
| WBGene00171572 | NA | NA | NA | NA |
| WBGene00171573 | NA | NA | NA | NA |
| WBGene00171574 | NA | NA | NA | NA |
| WBGene00171575 | NA | NA | NA | NA |
| WBGene00171576 | -0.003614866 | 0.95700117 | NA | NA |
| WBGene00171577 | NA | NA | NA | NA |
| WBGene00171578 | NA | NA | NA | NA |
| WBGene00171579 | NA | NA | NA | NA |
| WBGene00171580 | NA | NA | NA | NA |
| WBGene00171581 | NA | NA | NA | NA |
| WBGene00171582 | NA | NA | NA | NA |
| WBGene00171583 | NA | NA | NA | NA |
| WBGene00171584 | NA | NA | NA | NA |
| WBGene00171585 | NA | NA | NA | NA |
| WBGene00171586 | NA | NA | NA | NA |
| WBGene00171587 | NA | NA | NA | NA |
| WBGene00171588 | NA | NA | NA | NA |
| WBGene00171589 | NA | NA | NA | NA |
| WBGene00171590 | -0.022930767 | 0.78121253 | NA | NA |
| WBGene00171591 | NA | NA | NA | NA |
| WBGene00171592 | NA | NA | NA | NA |
| WBGene00171593 | NA | NA | NA | NA |
| WBGene00171594 | NA | NA | NA | NA |

| | | | | |
|---|---|---|---|---|
| WBGene00171595 | NA | NA | NA | NA |
| WBGene00171596 | NA | NA | NA | NA |
| WBGene00171597 | NA | NA | NA | NA |
| WBGene00171598 | NA | NA | NA | NA |
| WBGene00171599 | NA | NA | NA | NA |
| WBGene00171600 | NA | NA | NA | NA |
| WBGene00171601 | NA | NA | NA | NA |
| WBGene00171602 | NA | NA | NA | NA |
| WBGene00171603 | NA | NA | NA | NA |
| WBGene00171604 | NA | NA | NA | NA |
| WBGene00171605 | NA | NA | NA | NA |
| WBGene00171606 | NA | NA | NA | NA |
| WBGene00171607 | NA | NA | NA | NA |
| WBGene00171608 | NA | NA | NA | NA |
| WBGene00171609 | NA | NA | NA | NA |
| WBGene00171610 | NA | NA | NA | NA |
| WBGene00171611 | NA | NA | NA | NA |
| WBGene00171612 | NA | NA | NA | NA |
| WBGene00171613 | NA | NA | NA | NA |
| WBGene00171614 | NA | NA | NA | NA |
| WBGene00171615 | NA | NA | NA | NA |
| WBGene00171616 | NA | NA | NA | NA |
| WBGene00171617 | NA | NA | NA | NA |
| WBGene00171618 | NA | NA | NA | NA |
| WBGene00171619 | NA | NA | NA | NA |
| WBGene00171620 | NA | NA | NA | NA |
| WBGene00171621 | NA | NA | NA | NA |
| WBGene00171622 | NA | NA | NA | NA |
| WBGene00171623 | NA | NA | NA | NA |
| WBGene00171624 | NA | NA | NA | NA |
| WBGene00171625 | NA | NA | NA | NA |
| WBGene00171626 | NA | NA | NA | NA |
| WBGene00171627 | NA | NA | NA | NA |
| WBGene00171628 | NA | NA | NA | NA |
| WBGene00171629 | NA | NA | NA | NA |
| WBGene00171630 | NA | NA | NA | NA |
| WBGene00171631 | NA | NA | NA | NA |
| WBGene00171632 | NA | NA | NA | NA |
| WBGene00171633 | NA | NA | NA | NA |
| WBGene00171634 | NA | NA | NA | NA |
| WBGene00171635 | NA | NA | NA | NA |
| WBGene00171636 | NA | NA | NA | NA |
| WBGene00171637 | NA | NA | NA | NA |

| | | | | |
|---|---|---|---|---|
| WBGene00171638 | NA | NA | NA | NA |
| WBGene00171639 | NA | NA | NA | NA |
| WBGene00171640 | NA | NA | NA | NA |
| WBGene00171641 | NA | NA | NA | NA |
| WBGene00171642 | NA | NA | NA | NA |
| WBGene00171643 | NA | NA | NA | NA |
| WBGene00171644 | NA | NA | NA | NA |
| WBGene00171645 | NA | NA | NA | NA |
| WBGene00171646 | NA | NA | NA | NA |
| WBGene00171647 | NA | NA | NA | NA |
| WBGene00171648 | NA | NA | NA | NA |
| WBGene00171649 | NA | NA | NA | NA |
| WBGene00171650 | NA | NA | NA | NA |
| WBGene00171651 | NA | NA | NA | NA |
| WBGene00171652 | NA | NA | NA | NA |
| WBGene00171653 | 0.020925399 | 0.84643798 | NA | NA |
| WBGene00171654 | NA | NA | NA | NA |
| WBGene00171655 | NA | NA | NA | NA |
| WBGene00171656 | NA | NA | NA | NA |
| WBGene00171657 | NA | NA | NA | NA |
| WBGene00171658 | NA | NA | NA | NA |
| WBGene00171659 | NA | NA | NA | NA |
| WBGene00171660 | NA | NA | NA | NA |
| WBGene00171661 | NA | NA | NA | NA |
| WBGene00171662 | NA | NA | NA | NA |
| WBGene00171663 | NA | NA | NA | NA |
| WBGene00171664 | NA | NA | NA | NA |
| WBGene00171665 | NA | NA | NA | NA |
| WBGene00171666 | NA | NA | NA | NA |
| WBGene00171667 | NA | NA | NA | NA |
| WBGene00171668 | NA | NA | NA | NA |
| WBGene00171669 | NA | NA | NA | NA |
| WBGene00171670 | NA | NA | NA | NA |
| WBGene00171671 | NA | NA | NA | NA |
| WBGene00171672 | NA | NA | NA | NA |
| WBGene00171673 | NA | NA | NA | NA |
| WBGene00171674 | NA | NA | NA | NA |
| WBGene00171675 | NA | NA | NA | NA |
| WBGene00171676 | NA | NA | NA | NA |
| WBGene00171677 | NA | NA | NA | NA |
| WBGene00171678 | NA | NA | NA | NA |
| WBGene00171679 | NA | NA | NA | NA |
| WBGene00171680 | NA | NA | NA | NA |

| | | | | |
|---|---|---|---|---|
| WBGene00171681 | NA | NA | NA | NA |
| WBGene00171682 | NA | NA | NA | NA |
| WBGene00171683 | NA | NA | NA | NA |
| WBGene00171684 | NA | NA | NA | NA |
| WBGene00171685 | NA | NA | NA | NA |
| WBGene00171686 | NA | NA | NA | NA |
| WBGene00171687 | NA | NA | NA | NA |
| WBGene00171688 | NA | NA | NA | NA |
| WBGene00171689 | NA | NA | NA | NA |
| WBGene00171690 | NA | NA | NA | NA |
| WBGene00171691 | NA | NA | NA | NA |
| WBGene00171692 | NA | NA | NA | NA |
| WBGene00171693 | NA | NA | NA | NA |
| WBGene00171694 | NA | NA | NA | NA |
| WBGene00171695 | NA | NA | NA | NA |
| WBGene00171696 | NA | NA | NA | NA |
| WBGene00171697 | NA | NA | NA | NA |
| WBGene00171698 | NA | NA | NA | NA |
| WBGene00171699 | NA | NA | NA | NA |
| WBGene00171700 | NA | NA | NA | NA |
| WBGene00171701 | NA | NA | NA | NA |
| WBGene00171702 | NA | NA | NA | NA |
| WBGene00171703 | NA | NA | NA | NA |
| WBGene00171704 | NA | NA | NA | NA |
| WBGene00171705 | NA | NA | NA | NA |
| WBGene00171706 | NA | NA | NA | NA |
| WBGene00171707 | NA | NA | NA | NA |
| WBGene00171708 | NA | NA | NA | NA |
| WBGene00171709 | NA | NA | NA | NA |
| WBGene00171710 | NA | NA | NA | NA |
| WBGene00171712 | NA | NA | NA | NA |
| WBGene00171713 | NA | NA | NA | NA |
| WBGene00171714 | NA | NA | NA | NA |
| WBGene00171715 | NA | NA | NA | NA |
| WBGene00171716 | NA | NA | NA | NA |
| WBGene00171717 | NA | NA | NA | NA |
| WBGene00171718 | NA | NA | NA | NA |
| WBGene00171719 | NA | NA | NA | NA |
| WBGene00171720 | NA | NA | NA | NA |
| WBGene00171721 | NA | NA | NA | NA |
| WBGene00171722 | NA | NA | NA | NA |
| WBGene00171723 | NA | NA | NA | NA |
| WBGene00171724 | NA | NA | NA | NA |

| | | | | |
|---|---|---|---|---|
| WBGene00171725 | NA | NA | NA | NA |
| WBGene00171726 | NA | NA | NA | NA |
| WBGene00171727 | NA | NA | NA | NA |
| WBGene00171728 | NA | NA | NA | NA |
| WBGene00171729 | NA | NA | NA | NA |
| WBGene00171730 | NA | NA | NA | NA |
| WBGene00171731 | NA | NA | NA | NA |
| WBGene00171732 | NA | NA | NA | NA |
| WBGene00171733 | NA | NA | NA | NA |
| WBGene00171734 | NA | NA | NA | NA |
| WBGene00171735 | NA | NA | NA | NA |
| WBGene00171736 | NA | NA | NA | NA |
| WBGene00171737 | NA | NA | NA | NA |
| WBGene00171738 | NA | NA | NA | NA |
| WBGene00171739 | NA | NA | NA | NA |
| WBGene00171740 | NA | NA | NA | NA |
| WBGene00171741 | NA | NA | NA | NA |
| WBGene00171742 | NA | NA | NA | NA |
| WBGene00171743 | NA | NA | NA | NA |
| WBGene00171744 | NA | NA | NA | NA |
| WBGene00171745 | NA | NA | NA | NA |
| WBGene00171746 | NA | NA | NA | NA |
| WBGene00171747 | NA | NA | NA | NA |
| WBGene00171748 | NA | NA | NA | NA |
| WBGene00171749 | NA | NA | NA | NA |
| WBGene00171750 | NA | NA | NA | NA |
| WBGene00171751 | NA | NA | NA | NA |
| WBGene00171752 | NA | NA | NA | NA |
| WBGene00171753 | NA | NA | NA | NA |
| WBGene00171754 | NA | NA | NA | NA |
| WBGene00171755 | NA | NA | NA | NA |
| WBGene00171756 | NA | NA | NA | NA |
| WBGene00171757 | NA | NA | NA | NA |
| WBGene00171758 | NA | NA | NA | NA |
| WBGene00171759 | NA | NA | NA | NA |
| WBGene00171760 | NA | NA | NA | NA |
| WBGene00171761 | NA | NA | NA | NA |
| WBGene00171762 | NA | NA | NA | NA |
| WBGene00171763 | NA | NA | NA | NA |
| WBGene00171764 | NA | NA | NA | NA |
| WBGene00171765 | NA | NA | NA | NA |
| WBGene00171766 | NA | NA | NA | NA |
| WBGene00171767 | NA | NA | NA | NA |

| | | | | |
|---|---|---|---|---|
| WBGene00171768 | NA | NA | NA | NA |
| WBGene00171769 | NA | NA | NA | NA |
| WBGene00171770 | NA | NA | NA | NA |
| WBGene00171771 | NA | NA | NA | NA |
| WBGene00171772 | NA | NA | NA | NA |
| WBGene00171773 | NA | NA | NA | NA |
| WBGene00171774 | NA | NA | NA | NA |
| WBGene00171775 | NA | NA | NA | NA |
| WBGene00171776 | NA | NA | NA | NA |
| WBGene00171777 | NA | NA | NA | NA |
| WBGene00171778 | NA | NA | NA | NA |
| WBGene00171779 | NA | NA | NA | NA |
| WBGene00171780 | NA | NA | NA | NA |
| WBGene00171781 | NA | NA | NA | NA |
| WBGene00171782 | NA | NA | NA | NA |
| WBGene00171783 | NA | NA | NA | NA |
| WBGene00171784 | NA | NA | NA | NA |
| WBGene00171785 | NA | NA | NA | NA |
| WBGene00171786 | NA | NA | NA | NA |
| WBGene00171787 | NA | NA | NA | NA |
| WBGene00171788 | NA | NA | NA | NA |
| WBGene00171789 | NA | NA | NA | NA |
| WBGene00171790 | NA | NA | NA | NA |
| WBGene00171791 | NA | NA | NA | NA |
| WBGene00171792 | NA | NA | NA | NA |
| WBGene00171793 | NA | NA | NA | NA |
| WBGene00171794 | NA | NA | NA | NA |
| WBGene00171795 | NA | NA | NA | NA |
| WBGene00171796 | NA | NA | NA | NA |
| WBGene00171797 | NA | NA | NA | NA |
| WBGene00171798 | NA | NA | NA | NA |
| WBGene00171799 | NA | NA | NA | NA |
| WBGene00171800 | NA | NA | NA | NA |
| WBGene00171801 | NA | NA | NA | NA |
| WBGene00171802 | NA | NA | NA | NA |
| WBGene00171803 | NA | NA | NA | NA |
| WBGene00171804 | NA | NA | NA | NA |
| WBGene00171805 | NA | NA | NA | NA |
| WBGene00171806 | NA | NA | NA | NA |
| WBGene00171807 | NA | NA | NA | NA |
| WBGene00171809 | NA | NA | NA | NA |
| WBGene00171810 | NA | NA | NA | NA |
| WBGene00171811 | -0.022930767 | 0.78121253 | NA | NA |

| | | | | |
|---|---|---|---|---|
| WBGene00171812 | NA | NA | NA | NA |
| WBGene00171813 | NA | NA | NA | NA |
| WBGene00171814 | NA | NA | NA | NA |
| WBGene00171815 | NA | NA | NA | NA |
| WBGene00171816 | NA | NA | NA | NA |
| WBGene00171817 | NA | NA | NA | NA |
| WBGene00171818 | NA | NA | NA | NA |
| WBGene00171819 | NA | NA | NA | NA |
| WBGene00171820 | NA | NA | NA | NA |
| WBGene00171821 | NA | NA | NA | NA |
| WBGene00171822 | NA | NA | NA | NA |
| WBGene00171823 | NA | NA | NA | NA |
| WBGene00171824 | NA | NA | NA | NA |
| WBGene00171825 | NA | NA | NA | NA |
| WBGene00171826 | NA | NA | NA | NA |
| WBGene00171827 | NA | NA | NA | NA |
| WBGene00171828 | NA | NA | NA | NA |
| WBGene00171829 | NA | NA | NA | NA |
| WBGene00171830 | NA | NA | NA | NA |
| WBGene00171831 | NA | NA | NA | NA |
| WBGene00171832 | NA | NA | NA | NA |
| WBGene00171833 | NA | NA | NA | NA |
| WBGene00171834 | NA | NA | NA | NA |
| WBGene00171835 | NA | NA | NA | NA |
| WBGene00171836 | NA | NA | NA | NA |
| WBGene00171837 | NA | NA | NA | NA |
| WBGene00171838 | NA | NA | NA | NA |
| WBGene00171839 | NA | NA | NA | NA |
| WBGene00171840 | NA | NA | NA | NA |
| WBGene00171841 | NA | NA | NA | NA |
| WBGene00171842 | NA | NA | NA | NA |
| WBGene00171843 | NA | NA | NA | NA |
| WBGene00171844 | NA | NA | NA | NA |
| WBGene00171845 | NA | NA | NA | NA |
| WBGene00171846 | NA | NA | NA | NA |
| WBGene00171847 | NA | NA | NA | NA |
| WBGene00171848 | NA | NA | NA | NA |
| WBGene00171849 | NA | NA | NA | NA |
| WBGene00171850 | NA | NA | NA | NA |
| WBGene00171851 | NA | NA | NA | NA |
| WBGene00171852 | NA | NA | NA | NA |
| WBGene00171853 | NA | NA | NA | NA |
| WBGene00171854 | NA | NA | NA | NA |

| | | | | |
|---|---|---|---|---|
| WBGene00171855 | NA | NA | NA | NA |
| WBGene00171856 | NA | NA | NA | NA |
| WBGene00171857 | NA | NA | NA | NA |
| WBGene00171858 | NA | NA | NA | NA |
| WBGene00171859 | NA | NA | NA | NA |
| WBGene00171860 | NA | NA | NA | NA |
| WBGene00171861 | NA | NA | NA | NA |
| WBGene00171862 | NA | NA | NA | NA |
| WBGene00171863 | NA | NA | NA | NA |
| WBGene00171864 | NA | NA | NA | NA |
| WBGene00171865 | -0.022930767 | 0.78121253 | NA | NA |
| WBGene00171866 | NA | NA | NA | NA |
| WBGene00171867 | NA | NA | NA | NA |
| WBGene00171868 | NA | NA | NA | NA |
| WBGene00171869 | NA | NA | NA | NA |
| WBGene00171870 | NA | NA | NA | NA |
| WBGene00171871 | NA | NA | NA | NA |
| WBGene00171872 | NA | NA | NA | NA |
| WBGene00171873 | NA | NA | NA | NA |
| WBGene00171875 | NA | NA | NA | NA |
| WBGene00171876 | NA | NA | NA | NA |
| WBGene00171877 | NA | NA | NA | NA |
| WBGene00171878 | NA | NA | NA | NA |
| WBGene00171879 | NA | NA | NA | NA |
| WBGene00171880 | NA | NA | NA | NA |
| WBGene00171881 | NA | NA | NA | NA |
| WBGene00171882 | NA | NA | NA | NA |
| WBGene00171883 | NA | NA | NA | NA |
| WBGene00171884 | NA | NA | NA | NA |
| WBGene00171885 | NA | NA | NA | NA |
| WBGene00171886 | NA | NA | NA | NA |
| WBGene00171887 | NA | NA | NA | NA |
| WBGene00171888 | NA | NA | NA | NA |
| WBGene00171889 | NA | NA | NA | NA |
| WBGene00171890 | NA | NA | NA | NA |
| WBGene00171891 | NA | NA | NA | NA |
| WBGene00171892 | NA | NA | NA | NA |
| WBGene00171893 | NA | NA | NA | NA |
| WBGene00171894 | NA | NA | NA | NA |
| WBGene00171895 | NA | NA | NA | NA |
| WBGene00171896 | NA | NA | NA | NA |
| WBGene00171897 | NA | NA | NA | NA |
| WBGene00171898 | NA | NA | NA | NA |

| | | | | |
|---|---|---|---|---|
| WBGene00171899 | NA | NA | NA | NA |
| WBGene00171900 | NA | NA | NA | NA |
| WBGene00171901 | NA | NA | NA | NA |
| WBGene00171902 | NA | NA | NA | NA |
| WBGene00171903 | NA | NA | NA | NA |
| WBGene00171904 | NA | NA | NA | NA |
| WBGene00171905 | NA | NA | NA | NA |
| WBGene00171906 | NA | NA | NA | NA |
| WBGene00171907 | NA | NA | NA | NA |
| WBGene00171908 | NA | NA | NA | NA |
| WBGene00171909 | NA | NA | NA | NA |
| WBGene00171910 | NA | NA | NA | NA |
| WBGene00171911 | NA | NA | NA | NA |
| WBGene00171912 | NA | NA | NA | NA |
| WBGene00171913 | NA | NA | NA | NA |
| WBGene00171914 | NA | NA | NA | NA |
| WBGene00171915 | NA | NA | NA | NA |
| WBGene00171916 | NA | NA | NA | NA |
| WBGene00171917 | NA | NA | NA | NA |
| WBGene00171918 | NA | NA | NA | NA |
| WBGene00171919 | NA | NA | NA | NA |
| WBGene00171920 | NA | NA | NA | NA |
| WBGene00171921 | NA | NA | NA | NA |
| WBGene00171922 | NA | NA | NA | NA |
| WBGene00171923 | NA | NA | NA | NA |
| WBGene00171924 | NA | NA | NA | NA |
| WBGene00171925 | NA | NA | NA | NA |
| WBGene00171926 | NA | NA | NA | NA |
| WBGene00171927 | NA | NA | NA | NA |
| WBGene00171928 | NA | NA | NA | NA |
| WBGene00171929 | NA | NA | NA | NA |
| WBGene00171930 | NA | NA | NA | NA |
| WBGene00171931 | NA | NA | NA | NA |
| WBGene00171932 | NA | NA | NA | NA |
| WBGene00171933 | NA | NA | NA | NA |
| WBGene00171934 | NA | NA | NA | NA |
| WBGene00171935 | NA | NA | NA | NA |
| WBGene00171936 | NA | NA | NA | NA |
| WBGene00171937 | NA | NA | NA | NA |
| WBGene00171938 | NA | NA | NA | NA |
| WBGene00171939 | NA | NA | NA | NA |
| WBGene00171940 | NA | NA | NA | NA |
| WBGene00171941 | NA | NA | NA | NA |

| | | | | |
|---|---|---|---|---|
| WBGene00171942 | NA | NA | NA | NA |
| WBGene00171943 | NA | NA | NA | NA |
| WBGene00171944 | NA | NA | NA | NA |
| WBGene00171945 | NA | NA | NA | NA |
| WBGene00171946 | NA | NA | NA | NA |
| WBGene00171947 | NA | NA | NA | NA |
| WBGene00171948 | NA | NA | NA | NA |
| WBGene00171949 | NA | NA | NA | NA |
| WBGene00171950 | NA | NA | NA | NA |
| WBGene00171951 | NA | NA | NA | NA |
| WBGene00171952 | NA | NA | NA | NA |
| WBGene00171953 | NA | NA | NA | NA |
| WBGene00171954 | NA | NA | NA | NA |
| WBGene00171955 | NA | NA | NA | NA |
| WBGene00171956 | NA | NA | NA | NA |
| WBGene00171957 | NA | NA | NA | NA |
| WBGene00171958 | NA | NA | NA | NA |
| WBGene00171959 | NA | NA | NA | NA |
| WBGene00171960 | NA | NA | NA | NA |
| WBGene00171961 | NA | NA | NA | NA |
| WBGene00171962 | NA | NA | NA | NA |
| WBGene00171963 | NA | NA | NA | NA |
| WBGene00171964 | NA | NA | NA | NA |
| WBGene00171965 | NA | NA | NA | NA |
| WBGene00171966 | NA | NA | NA | NA |
| WBGene00171967 | NA | NA | NA | NA |
| WBGene00171968 | NA | NA | NA | NA |
| WBGene00171969 | NA | NA | NA | NA |
| WBGene00171970 | NA | NA | NA | NA |
| WBGene00171971 | NA | NA | NA | NA |
| WBGene00171972 | NA | NA | NA | NA |
| WBGene00171973 | NA | NA | NA | NA |
| WBGene00171974 | NA | NA | NA | NA |
| WBGene00171975 | NA | NA | NA | NA |
| WBGene00171976 | NA | NA | NA | NA |
| WBGene00171977 | NA | NA | NA | NA |
| WBGene00171978 | NA | NA | NA | NA |
| WBGene00171979 | NA | NA | NA | NA |
| WBGene00171980 | NA | NA | NA | NA |
| WBGene00171981 | NA | NA | NA | NA |
| WBGene00171982 | NA | NA | NA | NA |
| WBGene00171983 | NA | NA | NA | NA |
| WBGene00171984 | NA | NA | NA | NA |

| | | | | |
|---|---|---|---|---|
| WBGene00171985 | NA | NA | NA | NA |
| WBGene00171986 | NA | NA | NA | NA |
| WBGene00171987 | NA | NA | NA | NA |
| WBGene00171988 | NA | NA | NA | NA |
| WBGene00171989 | NA | NA | NA | NA |
| WBGene00171990 | NA | NA | NA | NA |
| WBGene00171991 | NA | NA | NA | NA |
| WBGene00171992 | NA | NA | NA | NA |
| WBGene00171993 | NA | NA | NA | NA |
| WBGene00171994 | NA | NA | NA | NA |
| WBGene00171995 | NA | NA | NA | NA |
| WBGene00171996 | NA | NA | NA | NA |
| WBGene00171997 | NA | NA | NA | NA |
| WBGene00171998 | NA | NA | NA | NA |
| WBGene00171999 | NA | NA | NA | NA |
| WBGene00172000 | NA | NA | NA | NA |
| WBGene00172001 | NA | NA | NA | NA |
| WBGene00172002 | NA | NA | NA | NA |
| WBGene00172003 | NA | NA | NA | NA |
| WBGene00172004 | NA | NA | NA | NA |
| WBGene00172006 | NA | NA | NA | NA |
| WBGene00172007 | NA | NA | NA | NA |
| WBGene00172008 | NA | NA | NA | NA |
| WBGene00172009 | NA | NA | NA | NA |
| WBGene00172010 | NA | NA | NA | NA |
| WBGene00172011 | NA | NA | NA | NA |
| WBGene00172012 | NA | NA | NA | NA |
| WBGene00172013 | NA | NA | NA | NA |
| WBGene00172014 | NA | NA | NA | NA |
| WBGene00172015 | NA | NA | NA | NA |
| WBGene00172016 | NA | NA | NA | NA |
| WBGene00172017 | NA | NA | NA | NA |
| WBGene00172018 | NA | NA | NA | NA |
| WBGene00172019 | NA | NA | NA | NA |
| WBGene00172020 | NA | NA | NA | NA |
| WBGene00172021 | NA | NA | NA | NA |
| WBGene00172022 | NA | NA | NA | NA |
| WBGene00172023 | NA | NA | NA | NA |
| WBGene00172024 | NA | NA | NA | NA |
| WBGene00172025 | NA | NA | NA | NA |
| WBGene00172026 | NA | NA | NA | NA |
| WBGene00172027 | NA | NA | NA | NA |
| WBGene00172028 | NA | NA | NA | NA |

| | | | | |
|---|---|---|---|---|
| WBGene00172029 | NA | NA | NA | NA |
| WBGene00172030 | NA | NA | NA | NA |
| WBGene00172031 | NA | NA | NA | NA |
| WBGene00172032 | NA | NA | NA | NA |
| WBGene00172033 | NA | NA | NA | NA |
| WBGene00172034 | NA | NA | NA | NA |
| WBGene00172035 | NA | NA | NA | NA |
| WBGene00172036 | NA | NA | NA | NA |
| WBGene00172037 | NA | NA | NA | NA |
| WBGene00172038 | NA | NA | NA | NA |
| WBGene00172039 | NA | NA | NA | NA |
| WBGene00172040 | NA | NA | NA | NA |
| WBGene00172041 | NA | NA | NA | NA |
| WBGene00172042 | NA | NA | NA | NA |
| WBGene00172043 | NA | NA | NA | NA |
| WBGene00172044 | NA | NA | NA | NA |
| WBGene00172045 | NA | NA | NA | NA |
| WBGene00172046 | NA | NA | NA | NA |
| WBGene00172047 | NA | NA | NA | NA |
| WBGene00172048 | NA | NA | NA | NA |
| WBGene00172049 | NA | NA | NA | NA |
| WBGene00172050 | NA | NA | NA | NA |
| WBGene00172051 | NA | NA | NA | NA |
| WBGene00172052 | NA | NA | NA | NA |
| WBGene00172053 | NA | NA | NA | NA |
| WBGene00172054 | NA | NA | NA | NA |
| WBGene00172055 | NA | NA | NA | NA |
| WBGene00172056 | NA | NA | NA | NA |
| WBGene00172057 | NA | NA | NA | NA |
| WBGene00172058 | NA | NA | NA | NA |
| WBGene00172059 | NA | NA | NA | NA |
| WBGene00172060 | NA | NA | NA | NA |
| WBGene00172061 | NA | NA | NA | NA |
| WBGene00172062 | NA | NA | NA | NA |
| WBGene00172063 | NA | NA | NA | NA |
| WBGene00172064 | NA | NA | NA | NA |
| WBGene00172065 | NA | NA | NA | NA |
| WBGene00172066 | NA | NA | NA | NA |
| WBGene00172067 | NA | NA | NA | NA |
| WBGene00172068 | NA | NA | NA | NA |
| WBGene00172069 | NA | NA | NA | NA |
| WBGene00172070 | NA | NA | NA | NA |
| WBGene00172071 | NA | NA | NA | NA |

| | | | | |
|---|---|---|---|---|
| WBGene00172072 | NA | NA | NA | NA |
| WBGene00172073 | NA | NA | NA | NA |
| WBGene00172074 | NA | NA | NA | NA |
| WBGene00172075 | NA | NA | NA | NA |
| WBGene00172076 | NA | NA | NA | NA |
| WBGene00172077 | NA | NA | NA | NA |
| WBGene00172078 | NA | NA | NA | NA |
| WBGene00172079 | NA | NA | NA | NA |
| WBGene00172080 | NA | NA | NA | NA |
| WBGene00172081 | NA | NA | NA | NA |
| WBGene00172082 | NA | NA | NA | NA |
| WBGene00172083 | NA | NA | NA | NA |
| WBGene00172084 | NA | NA | NA | NA |
| WBGene00172085 | NA | NA | NA | NA |
| WBGene00172086 | NA | NA | NA | NA |
| WBGene00172087 | NA | NA | NA | NA |
| WBGene00172088 | NA | NA | NA | NA |
| WBGene00172089 | NA | NA | NA | NA |
| WBGene00172090 | NA | NA | NA | NA |
| WBGene00172091 | NA | NA | NA | NA |
| WBGene00172092 | NA | NA | NA | NA |
| WBGene00172093 | NA | NA | NA | NA |
| WBGene00172094 | NA | NA | NA | NA |
| WBGene00172095 | NA | NA | NA | NA |
| WBGene00172096 | NA | NA | NA | NA |
| WBGene00172097 | NA | NA | NA | NA |
| WBGene00172098 | NA | NA | NA | NA |
| WBGene00172099 | NA | NA | NA | NA |
| WBGene00172100 | NA | NA | NA | NA |
| WBGene00172101 | NA | NA | NA | NA |
| WBGene00172102 | NA | NA | NA | NA |
| WBGene00172103 | NA | NA | NA | NA |
| WBGene00172104 | NA | NA | NA | NA |
| WBGene00172105 | NA | NA | NA | NA |
| WBGene00172106 | NA | NA | NA | NA |
| WBGene00172107 | NA | NA | NA | NA |
| WBGene00172108 | NA | NA | NA | NA |
| WBGene00172109 | NA | NA | NA | NA |
| WBGene00172110 | NA | NA | NA | NA |
| WBGene00172111 | NA | NA | NA | NA |
| WBGene00172112 | NA | NA | NA | NA |
| WBGene00172113 | NA | NA | NA | NA |
| WBGene00172114 | NA | NA | NA | NA |

| | | | | |
|---|---|---|---|---|
| WBGene00172115 | NA | NA | NA | NA |
| WBGene00172116 | NA | NA | NA | NA |
| WBGene00172117 | NA | NA | NA | NA |
| WBGene00172118 | NA | NA | NA | NA |
| WBGene00172119 | NA | NA | NA | NA |
| WBGene00172120 | NA | NA | NA | NA |
| WBGene00172121 | NA | NA | NA | NA |
| WBGene00172122 | NA | NA | NA | NA |
| WBGene00172123 | NA | NA | NA | NA |
| WBGene00172124 | NA | NA | NA | NA |
| WBGene00172125 | NA | NA | NA | NA |
| WBGene00172126 | NA | NA | NA | NA |
| WBGene00172127 | NA | NA | NA | NA |
| WBGene00172128 | NA | NA | NA | NA |
| WBGene00172129 | NA | NA | NA | NA |
| WBGene00172130 | NA | NA | NA | NA |
| WBGene00172131 | NA | NA | NA | NA |
| WBGene00172132 | NA | NA | NA | NA |
| WBGene00172133 | NA | NA | NA | NA |
| WBGene00172134 | NA | NA | NA | NA |
| WBGene00172135 | NA | NA | NA | NA |
| WBGene00172136 | NA | NA | NA | NA |
| WBGene00172137 | NA | NA | NA | NA |
| WBGene00172138 | NA | NA | NA | NA |
| WBGene00172139 | NA | NA | NA | NA |
| WBGene00172140 | NA | NA | NA | NA |
| WBGene00172141 | NA | NA | NA | NA |
| WBGene00172142 | NA | NA | NA | NA |
| WBGene00172143 | NA | NA | NA | NA |
| WBGene00172144 | 0.020232312 | 0.84643798 | NA | NA |
| WBGene00172145 | NA | NA | NA | NA |
| WBGene00172146 | NA | NA | NA | NA |
| WBGene00172147 | NA | NA | NA | NA |
| WBGene00172148 | NA | NA | NA | NA |
| WBGene00172149 | NA | NA | NA | NA |
| WBGene00172150 | NA | NA | NA | NA |
| WBGene00172151 | NA | NA | NA | NA |
| WBGene00172152 | NA | NA | NA | NA |
| WBGene00172153 | NA | NA | NA | NA |
| WBGene00172154 | NA | NA | NA | NA |
| WBGene00172155 | NA | NA | NA | NA |
| WBGene00172156 | NA | NA | NA | NA |
| WBGene00172157 | NA | NA | NA | NA |

| | | | | |
|---|---|---|---|---|
| WBGene00172158 | NA | NA | NA | NA |
| WBGene00172159 | NA | NA | NA | NA |
| WBGene00172160 | NA | NA | NA | NA |
| WBGene00172161 | NA | NA | NA | NA |
| WBGene00172162 | NA | NA | NA | NA |
| WBGene00172163 | NA | NA | NA | NA |
| WBGene00172164 | NA | NA | NA | NA |
| WBGene00172165 | NA | NA | NA | NA |
| WBGene00172166 | NA | NA | NA | NA |
| WBGene00172167 | NA | NA | NA | NA |
| WBGene00172168 | NA | NA | NA | NA |
| WBGene00172169 | 0.020232312 | 0.84643798 | NA | NA |
| WBGene00172170 | NA | NA | NA | NA |
| WBGene00172171 | NA | NA | NA | NA |
| WBGene00172172 | NA | NA | NA | NA |
| WBGene00172173 | NA | NA | NA | NA |
| WBGene00172174 | NA | NA | NA | NA |
| WBGene00172175 | NA | NA | NA | NA |
| WBGene00172176 | NA | NA | NA | NA |
| WBGene00172177 | NA | NA | NA | NA |
| WBGene00172178 | NA | NA | NA | NA |
| WBGene00172179 | NA | NA | NA | NA |
| WBGene00172180 | NA | NA | NA | NA |
| WBGene00172181 | NA | NA | NA | NA |
| WBGene00172182 | NA | NA | NA | NA |
| WBGene00172183 | NA | NA | NA | NA |
| WBGene00172184 | NA | NA | NA | NA |
| WBGene00172185 | NA | NA | NA | NA |
| WBGene00172186 | NA | NA | NA | NA |
| WBGene00172187 | NA | NA | NA | NA |
| WBGene00172188 | NA | NA | NA | NA |
| WBGene00172189 | NA | NA | NA | NA |
| WBGene00172190 | NA | NA | NA | NA |
| WBGene00172191 | NA | NA | NA | NA |
| WBGene00172192 | NA | NA | NA | NA |
| WBGene00172193 | NA | NA | NA | NA |
| WBGene00172194 | NA | NA | NA | NA |
| WBGene00172195 | NA | NA | NA | NA |
| WBGene00172196 | NA | NA | NA | NA |
| WBGene00172197 | NA | NA | NA | NA |
| WBGene00172198 | NA | NA | NA | NA |
| WBGene00172199 | NA | NA | NA | NA |
| WBGene00172200 | NA | NA | NA | NA |

| | | | | |
|---|---|---|---|---|
| WBGene00172201 | NA | NA | NA | NA |
| WBGene00172202 | NA | NA | NA | NA |
| WBGene00172203 | NA | NA | NA | NA |
| WBGene00172204 | NA | NA | NA | NA |
| WBGene00172205 | NA | NA | NA | NA |
| WBGene00172206 | NA | NA | NA | NA |
| WBGene00172207 | 0.020232312 | 0.84643798 | NA | NA |
| WBGene00172208 | NA | NA | NA | NA |
| WBGene00172209 | NA | NA | NA | NA |
| WBGene00172210 | NA | NA | NA | NA |
| WBGene00172211 | NA | NA | NA | NA |
| WBGene00172212 | NA | NA | NA | NA |
| WBGene00172213 | NA | NA | NA | NA |
| WBGene00172214 | NA | NA | NA | NA |
| WBGene00172215 | NA | NA | NA | NA |
| WBGene00172216 | NA | NA | NA | NA |
| WBGene00172217 | NA | NA | NA | NA |
| WBGene00172218 | NA | NA | NA | NA |
| WBGene00172219 | NA | NA | NA | NA |
| WBGene00172220 | NA | NA | NA | NA |
| WBGene00172221 | NA | NA | NA | NA |
| WBGene00172222 | NA | NA | NA | NA |
| WBGene00172223 | NA | NA | NA | NA |
| WBGene00172224 | NA | NA | NA | NA |
| WBGene00172225 | NA | NA | NA | NA |
| WBGene00172226 | NA | NA | NA | NA |
| WBGene00172227 | NA | NA | NA | NA |
| WBGene00172228 | NA | NA | NA | NA |
| WBGene00172229 | NA | NA | NA | NA |
| WBGene00172230 | NA | NA | NA | NA |
| WBGene00172231 | NA | NA | NA | NA |
| WBGene00172232 | NA | NA | NA | NA |
| WBGene00172233 | NA | NA | NA | NA |
| WBGene00172234 | NA | NA | NA | NA |
| WBGene00172235 | NA | NA | NA | NA |
| WBGene00172236 | NA | NA | NA | NA |
| WBGene00172237 | NA | NA | NA | NA |
| WBGene00172238 | NA | NA | NA | NA |
| WBGene00172239 | NA | NA | NA | NA |
| WBGene00172240 | NA | NA | NA | NA |
| WBGene00172241 | NA | NA | NA | NA |
| WBGene00172242 | NA | NA | NA | NA |
| WBGene00172243 | NA | NA | NA | NA |

| | | | | |
|---|---|---|---|---|
| WBGene00172244 | NA | NA | NA | NA |
| WBGene00172245 | NA | NA | NA | NA |
| WBGene00172246 | NA | NA | NA | NA |
| WBGene00172247 | NA | NA | NA | NA |
| WBGene00172248 | NA | NA | NA | NA |
| WBGene00172249 | NA | NA | NA | NA |
| WBGene00172250 | NA | NA | NA | NA |
| WBGene00172251 | NA | NA | NA | NA |
| WBGene00172252 | NA | NA | NA | NA |
| WBGene00172253 | NA | NA | NA | NA |
| WBGene00172254 | NA | NA | NA | NA |
| WBGene00172255 | NA | NA | NA | NA |
| WBGene00172256 | NA | NA | NA | NA |
| WBGene00172257 | NA | NA | NA | NA |
| WBGene00172258 | NA | NA | NA | NA |
| WBGene00172259 | NA | NA | NA | NA |
| WBGene00172260 | NA | NA | NA | NA |
| WBGene00172261 | NA | NA | NA | NA |
| WBGene00172262 | NA | NA | NA | NA |
| WBGene00172263 | NA | NA | NA | NA |
| WBGene00172264 | NA | NA | NA | NA |
| WBGene00172265 | NA | NA | NA | NA |
| WBGene00172266 | NA | NA | NA | NA |
| WBGene00172267 | NA | NA | NA | NA |
| WBGene00172268 | NA | NA | NA | NA |
| WBGene00172269 | NA | NA | NA | NA |
| WBGene00172270 | NA | NA | NA | NA |
| WBGene00172271 | NA | NA | NA | NA |
| WBGene00172272 | NA | NA | NA | NA |
| WBGene00172273 | NA | NA | NA | NA |
| WBGene00172274 | NA | NA | NA | NA |
| WBGene00172275 | NA | NA | NA | NA |
| WBGene00172276 | NA | NA | NA | NA |
| WBGene00172277 | NA | NA | NA | NA |
| WBGene00172278 | NA | NA | NA | NA |
| WBGene00172279 | NA | NA | NA | NA |
| WBGene00172280 | NA | NA | NA | NA |
| WBGene00172281 | NA | NA | NA | NA |
| WBGene00172282 | NA | NA | NA | NA |
| WBGene00172283 | NA | NA | NA | NA |
| WBGene00172284 | NA | NA | NA | NA |
| WBGene00172285 | NA | NA | NA | NA |
| WBGene00172286 | NA | NA | NA | NA |

| | | | | |
|---|---|---|---|---|
| WBGene00172287 | NA | NA | NA | NA |
| WBGene00172288 | NA | NA | NA | NA |
| WBGene00172289 | NA | NA | NA | NA |
| WBGene00172290 | NA | NA | NA | NA |
| WBGene00172291 | NA | NA | NA | NA |
| WBGene00172292 | NA | NA | NA | NA |
| WBGene00172293 | NA | NA | NA | NA |
| WBGene00172294 | NA | NA | NA | NA |
| WBGene00172295 | NA | NA | NA | NA |
| WBGene00172296 | NA | NA | NA | NA |
| WBGene00172297 | NA | NA | NA | NA |
| WBGene00172298 | NA | NA | NA | NA |
| WBGene00172299 | NA | NA | NA | NA |
| WBGene00172300 | NA | NA | NA | NA |
| WBGene00172301 | NA | NA | NA | NA |
| WBGene00172302 | NA | NA | NA | NA |
| WBGene00172303 | NA | NA | NA | NA |
| WBGene00172304 | NA | NA | NA | NA |
| WBGene00172305 | NA | NA | NA | NA |
| WBGene00172306 | NA | NA | NA | NA |
| WBGene00172307 | NA | NA | NA | NA |
| WBGene00172308 | NA | NA | NA | NA |
| WBGene00172309 | NA | NA | NA | NA |
| WBGene00172310 | NA | NA | NA | NA |
| WBGene00172311 | 0.020317245 | 0.84643798 | NA | NA |
| WBGene00172312 | NA | NA | NA | NA |
| WBGene00172313 | NA | NA | NA | NA |
| WBGene00172314 | NA | NA | NA | NA |
| WBGene00172315 | NA | NA | NA | NA |
| WBGene00172316 | NA | NA | NA | NA |
| WBGene00172317 | NA | NA | NA | NA |
| WBGene00172318 | NA | NA | NA | NA |
| WBGene00172319 | NA | NA | NA | NA |
| WBGene00172320 | NA | NA | NA | NA |
| WBGene00172321 | NA | NA | NA | NA |
| WBGene00172322 | NA | NA | NA | NA |
| WBGene00172323 | NA | NA | NA | NA |
| WBGene00172324 | NA | NA | NA | NA |
| WBGene00172325 | NA | NA | NA | NA |
| WBGene00172326 | NA | NA | NA | NA |
| WBGene00172327 | NA | NA | NA | NA |
| WBGene00172328 | NA | NA | NA | NA |
| WBGene00172329 | NA | NA | NA | NA |

| | | | | |
|---|---|---|---|---|
| WBGene00172330 | NA | NA | NA | NA |
| WBGene00172331 | NA | NA | NA | NA |
| WBGene00172332 | NA | NA | NA | NA |
| WBGene00172333 | NA | NA | NA | NA |
| WBGene00172334 | NA | NA | NA | NA |
| WBGene00172335 | NA | NA | NA | NA |
| WBGene00172336 | NA | NA | NA | NA |
| WBGene00172337 | NA | NA | NA | NA |
| WBGene00172338 | NA | NA | NA | NA |
| WBGene00172339 | NA | NA | NA | NA |
| WBGene00172340 | NA | NA | NA | NA |
| WBGene00172341 | NA | NA | NA | NA |
| WBGene00172342 | NA | NA | NA | NA |
| WBGene00172343 | NA | NA | NA | NA |
| WBGene00172344 | NA | NA | NA | NA |
| WBGene00172345 | NA | NA | NA | NA |
| WBGene00172346 | NA | NA | NA | NA |
| WBGene00172347 | NA | NA | NA | NA |
| WBGene00172348 | NA | NA | NA | NA |
| WBGene00172349 | NA | NA | NA | NA |
| WBGene00172350 | NA | NA | NA | NA |
| WBGene00172351 | NA | NA | NA | NA |
| WBGene00172352 | NA | NA | NA | NA |
| WBGene00172353 | NA | NA | NA | NA |
| WBGene00172354 | NA | NA | NA | NA |
| WBGene00172355 | NA | NA | NA | NA |
| WBGene00172356 | NA | NA | NA | NA |
| WBGene00172357 | NA | NA | NA | NA |
| WBGene00172358 | NA | NA | NA | NA |
| WBGene00172359 | NA | NA | NA | NA |
| WBGene00172360 | NA | NA | NA | NA |
| WBGene00172361 | NA | NA | NA | NA |
| WBGene00172362 | NA | NA | NA | NA |
| WBGene00172363 | NA | NA | NA | NA |
| WBGene00172364 | NA | NA | NA | NA |
| WBGene00172365 | NA | NA | NA | NA |
| WBGene00172366 | NA | NA | NA | NA |
| WBGene00172367 | NA | NA | NA | NA |
| WBGene00172368 | NA | NA | NA | NA |
| WBGene00172369 | NA | NA | NA | NA |
| WBGene00172370 | NA | NA | NA | NA |
| WBGene00172371 | -0.023109762 | 0.78121253 | NA | NA |
| WBGene00172372 | NA | NA | NA | NA |

| | | | | |
|---|---|---|---|---|
| WBGene00172373 | NA | NA | NA | NA |
| WBGene00172374 | NA | NA | NA | NA |
| WBGene00172375 | NA | NA | NA | NA |
| WBGene00172376 | NA | NA | NA | NA |
| WBGene00172377 | NA | NA | NA | NA |
| WBGene00172378 | NA | NA | NA | NA |
| WBGene00172379 | NA | NA | NA | NA |
| WBGene00172380 | NA | NA | NA | NA |
| WBGene00172381 | NA | NA | NA | NA |
| WBGene00172382 | NA | NA | NA | NA |
| WBGene00172383 | NA | NA | NA | NA |
| WBGene00172384 | NA | NA | NA | NA |
| WBGene00172385 | NA | NA | NA | NA |
| WBGene00172386 | NA | NA | NA | NA |
| WBGene00172387 | NA | NA | NA | NA |
| WBGene00172388 | NA | NA | NA | NA |
| WBGene00172389 | NA | NA | NA | NA |
| WBGene00172390 | NA | NA | NA | NA |
| WBGene00172391 | NA | NA | NA | NA |
| WBGene00172392 | NA | NA | NA | NA |
| WBGene00172393 | NA | NA | NA | NA |
| WBGene00172394 | NA | NA | NA | NA |
| WBGene00172395 | NA | NA | NA | NA |
| WBGene00172396 | NA | NA | NA | NA |
| WBGene00172397 | NA | NA | NA | NA |
| WBGene00172398 | NA | NA | NA | NA |
| WBGene00172399 | NA | NA | NA | NA |
| WBGene00172400 | NA | NA | NA | NA |
| WBGene00172401 | NA | NA | NA | NA |
| WBGene00172402 | NA | NA | NA | NA |
| WBGene00172403 | NA | NA | NA | NA |
| WBGene00172404 | NA | NA | NA | NA |
| WBGene00172405 | NA | NA | NA | NA |
| WBGene00172406 | NA | NA | NA | NA |
| WBGene00172407 | NA | NA | NA | NA |
| WBGene00172408 | NA | NA | NA | NA |
| WBGene00172409 | NA | NA | NA | NA |
| WBGene00172410 | NA | NA | NA | NA |
| WBGene00172411 | NA | NA | NA | NA |
| WBGene00172412 | -0.023056499 | 0.78121253 | NA | NA |
| WBGene00172413 | NA | NA | NA | NA |
| WBGene00172414 | NA | NA | NA | NA |
| WBGene00172415 | NA | NA | NA | NA |

| | | | | |
|---|---|---|---|---|
| WBGene00172416 | NA | NA | NA | NA |
| WBGene00172417 | NA | NA | NA | NA |
| WBGene00172418 | NA | NA | NA | NA |
| WBGene00172419 | NA | NA | NA | NA |
| WBGene00172420 | NA | NA | NA | NA |
| WBGene00172421 | NA | NA | NA | NA |
| WBGene00172422 | NA | NA | NA | NA |
| WBGene00172423 | NA | NA | NA | NA |
| WBGene00172424 | NA | NA | NA | NA |
| WBGene00172425 | NA | NA | NA | NA |
| WBGene00172426 | NA | NA | NA | NA |
| WBGene00172427 | NA | NA | NA | NA |
| WBGene00172428 | NA | NA | NA | NA |
| WBGene00172429 | NA | NA | NA | NA |
| WBGene00172430 | NA | NA | NA | NA |
| WBGene00172431 | NA | NA | NA | NA |
| WBGene00172432 | NA | NA | NA | NA |
| WBGene00172433 | NA | NA | NA | NA |
| WBGene00172434 | NA | NA | NA | NA |
| WBGene00172435 | NA | NA | NA | NA |
| WBGene00172436 | NA | NA | NA | NA |
| WBGene00172437 | NA | NA | NA | NA |
| WBGene00172438 | NA | NA | NA | NA |
| WBGene00172439 | NA | NA | NA | NA |
| WBGene00172440 | NA | NA | NA | NA |
| WBGene00172441 | NA | NA | NA | NA |
| WBGene00172442 | NA | NA | NA | NA |
| WBGene00172443 | NA | NA | NA | NA |
| WBGene00172444 | NA | NA | NA | NA |
| WBGene00172445 | NA | NA | NA | NA |
| WBGene00172446 | NA | NA | NA | NA |
| WBGene00172447 | NA | NA | NA | NA |
| WBGene00172448 | NA | NA | NA | NA |
| WBGene00172449 | NA | NA | NA | NA |
| WBGene00172450 | NA | NA | NA | NA |
| WBGene00172451 | NA | NA | NA | NA |
| WBGene00172452 | NA | NA | NA | NA |
| WBGene00172453 | NA | NA | NA | NA |
| WBGene00172454 | NA | NA | NA | NA |
| WBGene00172455 | NA | NA | NA | NA |
| WBGene00172456 | NA | NA | NA | NA |
| WBGene00172457 | 0.020232312 | 0.84643798 | NA | NA |
| WBGene00172458 | NA | NA | NA | NA |

| | | | | |
|---|---|---|---|---|
| WBGene00172459 | NA | NA | NA | NA |
| WBGene00172460 | NA | NA | NA | NA |
| WBGene00172462 | NA | NA | NA | NA |
| WBGene00172463 | NA | NA | NA | NA |
| WBGene00172464 | NA | NA | NA | NA |
| WBGene00172465 | NA | NA | NA | NA |
| WBGene00172466 | NA | NA | NA | NA |
| WBGene00172467 | NA | NA | NA | NA |
| WBGene00172468 | NA | NA | NA | NA |
| WBGene00172469 | 0.020317245 | 0.84643798 | NA | NA |
| WBGene00172470 | NA | NA | NA | NA |
| WBGene00172471 | NA | NA | NA | NA |
| WBGene00172472 | NA | NA | NA | NA |
| WBGene00172473 | NA | NA | NA | NA |
| WBGene00172474 | NA | NA | NA | NA |
| WBGene00172475 | NA | NA | NA | NA |
| WBGene00172476 | NA | NA | NA | NA |
| WBGene00172477 | NA | NA | NA | NA |
| WBGene00172478 | NA | NA | NA | NA |
| WBGene00172479 | NA | NA | NA | NA |
| WBGene00172480 | NA | NA | NA | NA |
| WBGene00172481 | NA | NA | NA | NA |
| WBGene00172482 | NA | NA | NA | NA |
| WBGene00172483 | NA | NA | NA | NA |
| WBGene00172484 | NA | NA | NA | NA |
| WBGene00172485 | NA | NA | NA | NA |
| WBGene00172486 | NA | NA | NA | NA |
| WBGene00172487 | NA | NA | NA | NA |
| WBGene00172488 | NA | NA | NA | NA |
| WBGene00172489 | NA | NA | NA | NA |
| WBGene00172490 | NA | NA | NA | NA |
| WBGene00172491 | NA | NA | NA | NA |
| WBGene00172492 | NA | NA | NA | NA |
| WBGene00172493 | NA | NA | NA | NA |
| WBGene00172494 | NA | NA | NA | NA |
| WBGene00172495 | NA | NA | NA | NA |
| WBGene00172496 | NA | NA | NA | NA |
| WBGene00172497 | NA | NA | NA | NA |
| WBGene00172498 | NA | NA | NA | NA |
| WBGene00172499 | NA | NA | NA | NA |
| WBGene00172500 | NA | NA | NA | NA |
| WBGene00172501 | NA | NA | NA | NA |
| WBGene00172502 | NA | NA | NA | NA |

| | | | | |
|---|---|---|---|---|
| WBGene00172503 | NA | NA | NA | NA |
| WBGene00172504 | NA | NA | NA | NA |
| WBGene00172505 | NA | NA | NA | NA |
| WBGene00172506 | NA | NA | NA | NA |
| WBGene00172507 | NA | NA | NA | NA |
| WBGene00172508 | NA | NA | NA | NA |
| WBGene00172509 | NA | NA | NA | NA |
| WBGene00172510 | NA | NA | NA | NA |
| WBGene00172511 | NA | NA | NA | NA |
| WBGene00172512 | NA | NA | NA | NA |
| WBGene00172513 | NA | NA | NA | NA |
| WBGene00172514 | NA | NA | NA | NA |
| WBGene00172515 | NA | NA | NA | NA |
| WBGene00172516 | NA | NA | NA | NA |
| WBGene00172517 | NA | NA | NA | NA |
| WBGene00172518 | NA | NA | NA | NA |
| WBGene00172519 | NA | NA | NA | NA |
| WBGene00172520 | NA | NA | NA | NA |
| WBGene00172521 | NA | NA | NA | NA |
| WBGene00172522 | NA | NA | NA | NA |
| WBGene00172523 | NA | NA | NA | NA |
| WBGene00172524 | NA | NA | NA | NA |
| WBGene00172525 | NA | NA | NA | NA |
| WBGene00172526 | NA | NA | NA | NA |
| WBGene00172527 | NA | NA | NA | NA |
| WBGene00172528 | NA | NA | NA | NA |
| WBGene00172529 | NA | NA | NA | NA |
| WBGene00172530 | NA | NA | NA | NA |
| WBGene00172531 | NA | NA | NA | NA |
| WBGene00172532 | NA | NA | NA | NA |
| WBGene00172533 | NA | NA | NA | NA |
| WBGene00172534 | NA | NA | NA | NA |
| WBGene00172535 | NA | NA | NA | NA |
| WBGene00172536 | NA | NA | NA | NA |
| WBGene00172537 | NA | NA | NA | NA |
| WBGene00172538 | NA | NA | NA | NA |
| WBGene00172539 | NA | NA | NA | NA |
| WBGene00172540 | NA | NA | NA | NA |
| WBGene00172541 | NA | NA | NA | NA |
| WBGene00172542 | NA | NA | NA | NA |
| WBGene00172543 | NA | NA | NA | NA |
| WBGene00172544 | NA | NA | NA | NA |
| WBGene00172545 | NA | NA | NA | NA |

| | | | | |
|---|---|---|---|---|
| WBGene00172546 | NA | NA | NA | NA |
| WBGene00172547 | NA | NA | NA | NA |
| WBGene00172548 | NA | NA | NA | NA |
| WBGene00172549 | NA | NA | NA | NA |
| WBGene00172550 | NA | NA | NA | NA |
| WBGene00172551 | NA | NA | NA | NA |
| WBGene00172552 | NA | NA | NA | NA |
| WBGene00172553 | NA | NA | NA | NA |
| WBGene00172554 | NA | NA | NA | NA |
| WBGene00172555 | NA | NA | NA | NA |
| WBGene00172556 | NA | NA | NA | NA |
| WBGene00172557 | NA | NA | NA | NA |
| WBGene00172558 | NA | NA | NA | NA |
| WBGene00172559 | NA | NA | NA | NA |
| WBGene00172560 | NA | NA | NA | NA |
| WBGene00172561 | NA | NA | NA | NA |
| WBGene00172562 | NA | NA | NA | NA |
| WBGene00172563 | NA | NA | NA | NA |
| WBGene00172564 | NA | NA | NA | NA |
| WBGene00172565 | NA | NA | NA | NA |
| WBGene00172566 | NA | NA | NA | NA |
| WBGene00172567 | NA | NA | NA | NA |
| WBGene00172568 | NA | NA | NA | NA |
| WBGene00172569 | NA | NA | NA | NA |
| WBGene00172570 | NA | NA | NA | NA |
| WBGene00172571 | NA | NA | NA | NA |
| WBGene00172572 | NA | NA | NA | NA |
| WBGene00172573 | NA | NA | NA | NA |
| WBGene00172574 | NA | NA | NA | NA |
| WBGene00172575 | NA | NA | NA | NA |
| WBGene00172576 | NA | NA | NA | NA |
| WBGene00172577 | NA | NA | NA | NA |
| WBGene00172578 | NA | NA | NA | NA |
| WBGene00172579 | NA | NA | NA | NA |
| WBGene00172580 | NA | NA | NA | NA |
| WBGene00172581 | NA | NA | NA | NA |
| WBGene00172582 | NA | NA | NA | NA |
| WBGene00172583 | NA | NA | NA | NA |
| WBGene00172584 | NA | NA | NA | NA |
| WBGene00172585 | 0.020925399 | 0.84643798 | NA | NA |
| WBGene00172586 | NA | NA | NA | NA |
| WBGene00172587 | NA | NA | NA | NA |
| WBGene00172588 | NA | NA | NA | NA |

| | | | | |
|---|---|---|---|---|
| WBGene00172589 | NA | NA | NA | NA |
| WBGene00172590 | NA | NA | NA | NA |
| WBGene00172591 | NA | NA | NA | NA |
| WBGene00172592 | NA | NA | NA | NA |
| WBGene00172593 | NA | NA | NA | NA |
| WBGene00172594 | NA | NA | NA | NA |
| WBGene00172595 | NA | NA | NA | NA |
| WBGene00172596 | NA | NA | NA | NA |
| WBGene00172597 | NA | NA | NA | NA |
| WBGene00172598 | 0.020925399 | 0.84643798 | NA | NA |
| WBGene00172599 | NA | NA | NA | NA |
| WBGene00172600 | NA | NA | NA | NA |
| WBGene00172601 | NA | NA | NA | NA |
| WBGene00172602 | NA | NA | NA | NA |
| WBGene00172603 | NA | NA | NA | NA |
| WBGene00172604 | NA | NA | NA | NA |
| WBGene00172605 | NA | NA | NA | NA |
| WBGene00172606 | NA | NA | NA | NA |
| WBGene00172607 | NA | NA | NA | NA |
| WBGene00172608 | NA | NA | NA | NA |
| WBGene00172609 | NA | NA | NA | NA |
| WBGene00172610 | NA | NA | NA | NA |
| WBGene00172611 | NA | NA | NA | NA |
| WBGene00172612 | NA | NA | NA | NA |
| WBGene00172613 | NA | NA | NA | NA |
| WBGene00172614 | NA | NA | NA | NA |
| WBGene00172615 | NA | NA | NA | NA |
| WBGene00172616 | NA | NA | NA | NA |
| WBGene00172617 | NA | NA | NA | NA |
| WBGene00172618 | NA | NA | NA | NA |
| WBGene00172619 | NA | NA | NA | NA |
| WBGene00172620 | NA | NA | NA | NA |
| WBGene00172621 | NA | NA | NA | NA |
| WBGene00172622 | NA | NA | NA | NA |
| WBGene00172623 | NA | NA | NA | NA |
| WBGene00172624 | NA | NA | NA | NA |
| WBGene00172625 | NA | NA | NA | NA |
| WBGene00172626 | NA | NA | NA | NA |
| WBGene00172627 | NA | NA | NA | NA |
| WBGene00172628 | NA | NA | NA | NA |
| WBGene00172629 | NA | NA | NA | NA |
| WBGene00172630 | NA | NA | NA | NA |
| WBGene00172631 | NA | NA | NA | NA |

| | | | | |
|---|---|---|---|---|
| WBGene00172632 | NA | NA | NA | NA |
| WBGene00172633 | NA | NA | NA | NA |
| WBGene00172634 | NA | NA | NA | NA |
| WBGene00172635 | NA | NA | NA | NA |
| WBGene00172636 | NA | NA | NA | NA |
| WBGene00172637 | NA | NA | NA | NA |
| WBGene00172638 | NA | NA | NA | NA |
| WBGene00172639 | NA | NA | NA | NA |
| WBGene00172640 | NA | NA | NA | NA |
| WBGene00172641 | NA | NA | NA | NA |
| WBGene00172642 | NA | NA | NA | NA |
| WBGene00172643 | NA | NA | NA | NA |
| WBGene00172644 | NA | NA | NA | NA |
| WBGene00172645 | NA | NA | NA | NA |
| WBGene00172646 | NA | NA | NA | NA |
| WBGene00172647 | NA | NA | NA | NA |
| WBGene00172648 | NA | NA | NA | NA |
| WBGene00172649 | NA | NA | NA | NA |
| WBGene00172650 | NA | NA | NA | NA |
| WBGene00172651 | NA | NA | NA | NA |
| WBGene00172652 | NA | NA | NA | NA |
| WBGene00172653 | NA | NA | NA | NA |
| WBGene00172654 | NA | NA | NA | NA |
| WBGene00172655 | NA | NA | NA | NA |
| WBGene00172656 | NA | NA | NA | NA |
| WBGene00172657 | NA | NA | NA | NA |
| WBGene00172658 | NA | NA | NA | NA |
| WBGene00172659 | NA | NA | NA | NA |
| WBGene00172660 | NA | NA | NA | NA |
| WBGene00172661 | NA | NA | NA | NA |
| WBGene00172662 | NA | NA | NA | NA |
| WBGene00172663 | NA | NA | NA | NA |
| WBGene00172664 | NA | NA | NA | NA |
| WBGene00172665 | NA | NA | NA | NA |
| WBGene00172666 | NA | NA | NA | NA |
| WBGene00172667 | NA | NA | NA | NA |
| WBGene00172668 | NA | NA | NA | NA |
| WBGene00172669 | NA | NA | NA | NA |
| WBGene00172670 | NA | NA | NA | NA |
| WBGene00172671 | NA | NA | NA | NA |
| WBGene00172672 | NA | NA | NA | NA |
| WBGene00172673 | NA | NA | NA | NA |
| WBGene00172674 | NA | NA | NA | NA |

| | | | | |
|---|---|---|---|---|
| WBGene00172675 | NA | NA | NA | NA |
| WBGene00172676 | NA | NA | NA | NA |
| WBGene00172677 | NA | NA | NA | NA |
| WBGene00172678 | NA | NA | NA | NA |
| WBGene00172679 | NA | NA | NA | NA |
| WBGene00172680 | NA | NA | NA | NA |
| WBGene00172681 | NA | NA | NA | NA |
| WBGene00172682 | NA | NA | NA | NA |
| WBGene00172683 | NA | NA | NA | NA |
| WBGene00172684 | NA | NA | NA | NA |
| WBGene00172685 | NA | NA | NA | NA |
| WBGene00172686 | NA | NA | NA | NA |
| WBGene00172687 | NA | NA | NA | NA |
| WBGene00172688 | NA | NA | NA | NA |
| WBGene00172689 | NA | NA | NA | NA |
| WBGene00172690 | NA | NA | NA | NA |
| WBGene00172691 | NA | NA | NA | NA |
| WBGene00172692 | NA | NA | NA | NA |
| WBGene00172693 | NA | NA | NA | NA |
| WBGene00172694 | NA | NA | NA | NA |
| WBGene00172695 | NA | NA | NA | NA |
| WBGene00172696 | NA | NA | NA | NA |
| WBGene00172697 | NA | NA | NA | NA |
| WBGene00172698 | NA | NA | NA | NA |
| WBGene00172699 | NA | NA | NA | NA |
| WBGene00172700 | NA | NA | NA | NA |
| WBGene00172701 | NA | NA | NA | NA |
| WBGene00172702 | NA | NA | NA | NA |
| WBGene00172703 | NA | NA | NA | NA |
| WBGene00172704 | NA | NA | NA | NA |
| WBGene00172705 | NA | NA | NA | NA |
| WBGene00172706 | NA | NA | NA | NA |
| WBGene00172707 | NA | NA | NA | NA |
| WBGene00172708 | NA | NA | NA | NA |
| WBGene00172709 | NA | NA | NA | NA |
| WBGene00172710 | NA | NA | NA | NA |
| WBGene00172711 | NA | NA | NA | NA |
| WBGene00172712 | NA | NA | NA | NA |
| WBGene00172713 | NA | NA | NA | NA |
| WBGene00172714 | NA | NA | NA | NA |
| WBGene00172715 | NA | NA | NA | NA |
| WBGene00172716 | NA | NA | NA | NA |
| WBGene00172717 | NA | NA | NA | NA |

| | | | | |
|---|---|---|---|---|
| WBGene00172718 | NA | NA | NA | NA |
| WBGene00172719 | NA | NA | NA | NA |
| WBGene00172720 | NA | NA | NA | NA |
| WBGene00172721 | NA | NA | NA | NA |
| WBGene00172722 | NA | NA | NA | NA |
| WBGene00172723 | NA | NA | NA | NA |
| WBGene00172724 | NA | NA | NA | NA |
| WBGene00172725 | NA | NA | NA | NA |
| WBGene00172726 | NA | NA | NA | NA |
| WBGene00172727 | NA | NA | NA | NA |
| WBGene00172728 | NA | NA | NA | NA |
| WBGene00172729 | NA | NA | NA | NA |
| WBGene00172730 | NA | NA | NA | NA |
| WBGene00172731 | NA | NA | NA | NA |
| WBGene00172732 | NA | NA | NA | NA |
| WBGene00172733 | NA | NA | NA | NA |
| WBGene00172734 | NA | NA | NA | NA |
| WBGene00172735 | NA | NA | NA | NA |
| WBGene00172736 | NA | NA | NA | NA |
| WBGene00172737 | NA | NA | NA | NA |
| WBGene00172738 | NA | NA | NA | NA |
| WBGene00172739 | NA | NA | NA | NA |
| WBGene00172740 | NA | NA | NA | NA |
| WBGene00172741 | NA | NA | NA | NA |
| WBGene00172742 | NA | NA | NA | NA |
| WBGene00172743 | NA | NA | NA | NA |
| WBGene00172744 | NA | NA | NA | NA |
| WBGene00172745 | NA | NA | NA | NA |
| WBGene00172746 | NA | NA | NA | NA |
| WBGene00172747 | NA | NA | NA | NA |
| WBGene00172748 | NA | NA | NA | NA |
| WBGene00172749 | NA | NA | NA | NA |
| WBGene00172750 | NA | NA | NA | NA |
| WBGene00172751 | NA | NA | NA | NA |
| WBGene00172752 | NA | NA | NA | NA |
| WBGene00172753 | NA | NA | NA | NA |
| WBGene00172754 | NA | NA | NA | NA |
| WBGene00172755 | NA | NA | NA | NA |
| WBGene00172756 | NA | NA | NA | NA |
| WBGene00172757 | NA | NA | NA | NA |
| WBGene00172758 | NA | NA | NA | NA |
| WBGene00172759 | NA | NA | NA | NA |
| WBGene00172760 | NA | NA | NA | NA |

| | | | | |
|---|---|---|---|---|
| WBGene00172761 | NA | NA | NA | NA |
| WBGene00172762 | NA | NA | NA | NA |
| WBGene00172763 | NA | NA | NA | NA |
| WBGene00172764 | NA | NA | NA | NA |
| WBGene00172765 | NA | NA | NA | NA |
| WBGene00172766 | NA | NA | NA | NA |
| WBGene00172767 | NA | NA | NA | NA |
| WBGene00172768 | NA | NA | NA | NA |
| WBGene00172769 | NA | NA | NA | NA |
| WBGene00172770 | NA | NA | NA | NA |
| WBGene00172771 | NA | NA | NA | NA |
| WBGene00172772 | NA | NA | NA | NA |
| WBGene00172773 | NA | NA | NA | NA |
| WBGene00172774 | NA | NA | NA | NA |
| WBGene00172775 | NA | NA | NA | NA |
| WBGene00172776 | NA | NA | NA | NA |
| WBGene00172777 | NA | NA | NA | NA |
| WBGene00172778 | NA | NA | NA | NA |
| WBGene00172779 | NA | NA | NA | NA |
| WBGene00172780 | NA | NA | NA | NA |
| WBGene00172781 | NA | NA | NA | NA |
| WBGene00172782 | NA | NA | NA | NA |
| WBGene00172783 | NA | NA | NA | NA |
| WBGene00172784 | NA | NA | NA | NA |
| WBGene00172785 | NA | NA | NA | NA |
| WBGene00172786 | NA | NA | NA | NA |
| WBGene00172787 | NA | NA | NA | NA |
| WBGene00172788 | NA | NA | NA | NA |
| WBGene00172789 | NA | NA | NA | NA |
| WBGene00172790 | NA | NA | NA | NA |
| WBGene00172791 | NA | NA | NA | NA |
| WBGene00172792 | NA | NA | NA | NA |
| WBGene00172793 | NA | NA | NA | NA |
| WBGene00172794 | NA | NA | NA | NA |
| WBGene00172795 | NA | NA | NA | NA |
| WBGene00172796 | NA | NA | NA | NA |
| WBGene00172797 | NA | NA | NA | NA |
| WBGene00172798 | NA | NA | NA | NA |
| WBGene00172799 | NA | NA | NA | NA |
| WBGene00172800 | NA | NA | NA | NA |
| WBGene00172801 | NA | NA | NA | NA |
| WBGene00172802 | NA | NA | NA | NA |
| WBGene00172803 | NA | NA | NA | NA |

| | | | | |
|---|---|---|---|---|
| WBGene00172804 | NA | NA | NA | NA |
| WBGene00172805 | NA | NA | NA | NA |
| WBGene00172806 | NA | NA | NA | NA |
| WBGene00172807 | NA | NA | NA | NA |
| WBGene00172808 | NA | NA | NA | NA |
| WBGene00172809 | NA | NA | NA | NA |
| WBGene00172810 | NA | NA | NA | NA |
| WBGene00172811 | NA | NA | NA | NA |
| WBGene00172812 | NA | NA | NA | NA |
| WBGene00172813 | NA | NA | NA | NA |
| WBGene00172814 | NA | NA | NA | NA |
| WBGene00172815 | NA | NA | NA | NA |
| WBGene00172816 | NA | NA | NA | NA |
| WBGene00172817 | NA | NA | NA | NA |
| WBGene00172818 | NA | NA | NA | NA |
| WBGene00172819 | NA | NA | NA | NA |
| WBGene00172820 | NA | NA | NA | NA |
| WBGene00172821 | NA | NA | NA | NA |
| WBGene00172822 | NA | NA | NA | NA |
| WBGene00172823 | NA | NA | NA | NA |
| WBGene00172824 | NA | NA | NA | NA |
| WBGene00172825 | NA | NA | NA | NA |
| WBGene00172826 | NA | NA | NA | NA |
| WBGene00172827 | NA | NA | NA | NA |
| WBGene00172828 | NA | NA | NA | NA |
| WBGene00172829 | NA | NA | NA | NA |
| WBGene00172830 | NA | NA | NA | NA |
| WBGene00172831 | NA | NA | NA | NA |
| WBGene00172832 | NA | NA | NA | NA |
| WBGene00172833 | NA | NA | NA | NA |
| WBGene00172834 | NA | NA | NA | NA |
| WBGene00172835 | NA | NA | NA | NA |
| WBGene00172836 | NA | NA | NA | NA |
| WBGene00172837 | NA | NA | NA | NA |
| WBGene00172838 | NA | NA | NA | NA |
| WBGene00172839 | NA | NA | NA | NA |
| WBGene00172840 | NA | NA | NA | NA |
| WBGene00172841 | NA | NA | NA | NA |
| WBGene00172842 | NA | NA | NA | NA |
| WBGene00172843 | NA | NA | NA | NA |
| WBGene00172844 | NA | NA | NA | NA |
| WBGene00172845 | NA | NA | NA | NA |
| WBGene00172846 | NA | NA | NA | NA |

| | | | | |
|---|---|---|---|---|
| WBGene00172847 | NA | NA | NA | NA |
| WBGene00172848 | NA | NA | NA | NA |
| WBGene00172849 | NA | NA | NA | NA |
| WBGene00172850 | NA | NA | NA | NA |
| WBGene00172851 | NA | NA | NA | NA |
| WBGene00172852 | NA | NA | NA | NA |
| WBGene00172853 | NA | NA | NA | NA |
| WBGene00172854 | NA | NA | NA | NA |
| WBGene00172855 | NA | NA | NA | NA |
| WBGene00172856 | NA | NA | NA | NA |
| WBGene00172857 | NA | NA | NA | NA |
| WBGene00172858 | NA | NA | NA | NA |
| WBGene00172859 | NA | NA | NA | NA |
| WBGene00172860 | NA | NA | NA | NA |
| WBGene00172861 | NA | NA | NA | NA |
| WBGene00172862 | NA | NA | NA | NA |
| WBGene00172863 | NA | NA | NA | NA |
| WBGene00172864 | NA | NA | NA | NA |
| WBGene00172865 | NA | NA | NA | NA |
| WBGene00172866 | NA | NA | NA | NA |
| WBGene00172867 | NA | NA | NA | NA |
| WBGene00172868 | NA | NA | NA | NA |
| WBGene00172869 | NA | NA | NA | NA |
| WBGene00172870 | NA | NA | NA | NA |
| WBGene00172871 | NA | NA | NA | NA |
| WBGene00172872 | NA | NA | NA | NA |
| WBGene00172873 | NA | NA | NA | NA |
| WBGene00172874 | NA | NA | NA | NA |
| WBGene00172875 | NA | NA | NA | NA |
| WBGene00172876 | NA | NA | NA | NA |
| WBGene00172877 | NA | NA | NA | NA |
| WBGene00172878 | NA | NA | NA | NA |
| WBGene00172879 | NA | NA | NA | NA |
| WBGene00172880 | NA | NA | NA | NA |
| WBGene00172881 | NA | NA | NA | NA |
| WBGene00172882 | NA | NA | NA | NA |
| WBGene00172883 | NA | NA | NA | NA |
| WBGene00172884 | NA | NA | NA | NA |
| WBGene00172885 | NA | NA | NA | NA |
| WBGene00172886 | NA | NA | NA | NA |
| WBGene00172887 | NA | NA | NA | NA |
| WBGene00172888 | NA | NA | NA | NA |
| WBGene00172889 | NA | NA | NA | NA |

| | | | | |
|---|---|---|---|---|
| WBGene00172890 | NA | NA | NA | NA |
| WBGene00172891 | NA | NA | NA | NA |
| WBGene00172892 | NA | NA | NA | NA |
| WBGene00172893 | NA | NA | NA | NA |
| WBGene00172894 | NA | NA | NA | NA |
| WBGene00172895 | NA | NA | NA | NA |
| WBGene00172896 | NA | NA | NA | NA |
| WBGene00172897 | NA | NA | NA | NA |
| WBGene00172898 | NA | NA | NA | NA |
| WBGene00172899 | NA | NA | NA | NA |
| WBGene00172900 | NA | NA | NA | NA |
| WBGene00172901 | NA | NA | NA | NA |
| WBGene00172902 | NA | NA | NA | NA |
| WBGene00172903 | NA | NA | NA | NA |
| WBGene00172904 | NA | NA | NA | NA |
| WBGene00172905 | NA | NA | NA | NA |
| WBGene00172906 | NA | NA | NA | NA |
| WBGene00172907 | NA | NA | NA | NA |
| WBGene00172908 | NA | NA | NA | NA |
| WBGene00172909 | NA | NA | NA | NA |
| WBGene00172910 | NA | NA | NA | NA |
| WBGene00172911 | NA | NA | NA | NA |
| WBGene00172912 | NA | NA | NA | NA |
| WBGene00172913 | NA | NA | NA | NA |
| WBGene00172914 | NA | NA | NA | NA |
| WBGene00172915 | NA | NA | NA | NA |
| WBGene00172916 | NA | NA | NA | NA |
| WBGene00172917 | NA | NA | NA | NA |
| WBGene00172918 | NA | NA | NA | NA |
| WBGene00172919 | NA | NA | NA | NA |
| WBGene00172920 | NA | NA | NA | NA |
| WBGene00172921 | NA | NA | NA | NA |
| WBGene00172922 | NA | NA | NA | NA |
| WBGene00172923 | NA | NA | NA | NA |
| WBGene00172924 | NA | NA | NA | NA |
| WBGene00172925 | NA | NA | NA | NA |
| WBGene00172926 | NA | NA | NA | NA |
| WBGene00172927 | NA | NA | NA | NA |
| WBGene00172928 | NA | NA | NA | NA |
| WBGene00172929 | NA | NA | NA | NA |
| WBGene00172930 | NA | NA | NA | NA |
| WBGene00172931 | NA | NA | NA | NA |
| WBGene00172932 | NA | NA | NA | NA |

| | | | | |
|---|---|---|---|---|
| WBGene00172933 | NA | NA | NA | NA |
| WBGene00172934 | NA | NA | NA | NA |
| WBGene00172935 | NA | NA | NA | NA |
| WBGene00172936 | NA | NA | NA | NA |
| WBGene00172937 | NA | NA | NA | NA |
| WBGene00172938 | NA | NA | NA | NA |
| WBGene00172939 | NA | NA | NA | NA |
| WBGene00172940 | NA | NA | NA | NA |
| WBGene00172941 | NA | NA | NA | NA |
| WBGene00172942 | NA | NA | NA | NA |
| WBGene00172943 | NA | NA | NA | NA |
| WBGene00172944 | NA | NA | NA | NA |
| WBGene00172945 | NA | NA | NA | NA |
| WBGene00172946 | NA | NA | NA | NA |
| WBGene00172947 | NA | NA | NA | NA |
| WBGene00172948 | NA | NA | NA | NA |
| WBGene00172949 | NA | NA | NA | NA |
| WBGene00172950 | NA | NA | NA | NA |
| WBGene00172951 | NA | NA | NA | NA |
| WBGene00172952 | NA | NA | NA | NA |
| WBGene00172953 | NA | NA | NA | NA |
| WBGene00172954 | NA | NA | NA | NA |
| WBGene00172955 | NA | NA | NA | NA |
| WBGene00172956 | NA | NA | NA | NA |
| WBGene00172957 | NA | NA | NA | NA |
| WBGene00172958 | NA | NA | NA | NA |
| WBGene00172959 | NA | NA | NA | NA |
| WBGene00172960 | NA | NA | NA | NA |
| WBGene00172961 | NA | NA | NA | NA |
| WBGene00172962 | NA | NA | NA | NA |
| WBGene00172963 | NA | NA | NA | NA |
| WBGene00172964 | NA | NA | NA | NA |
| WBGene00172965 | NA | NA | NA | NA |
| WBGene00172966 | NA | NA | NA | NA |
| WBGene00172967 | NA | NA | NA | NA |
| WBGene00172968 | NA | NA | NA | NA |
| WBGene00172969 | NA | NA | NA | NA |
| WBGene00172970 | NA | NA | NA | NA |
| WBGene00172971 | NA | NA | NA | NA |
| WBGene00172972 | NA | NA | NA | NA |
| WBGene00172973 | NA | NA | NA | NA |
| WBGene00172974 | NA | NA | NA | NA |
| WBGene00172975 | NA | NA | NA | NA |

| | | | | |
|---|---|---|---|---|
| WBGene00172976 | NA | NA | NA | NA |
| WBGene00172977 | NA | NA | NA | NA |
| WBGene00172978 | NA | NA | NA | NA |
| WBGene00172979 | NA | NA | NA | NA |
| WBGene00172980 | NA | NA | NA | NA |
| WBGene00172981 | NA | NA | NA | NA |
| WBGene00172982 | NA | NA | NA | NA |
| WBGene00172983 | NA | NA | NA | NA |
| WBGene00172984 | NA | NA | NA | NA |
| WBGene00172985 | NA | NA | NA | NA |
| WBGene00172986 | NA | NA | NA | NA |
| WBGene00172987 | NA | NA | NA | NA |
| WBGene00172988 | NA | NA | NA | NA |
| WBGene00172989 | NA | NA | NA | NA |
| WBGene00172990 | NA | NA | NA | NA |
| WBGene00172991 | NA | NA | NA | NA |
| WBGene00172992 | NA | NA | NA | NA |
| WBGene00172993 | NA | NA | NA | NA |
| WBGene00172994 | NA | NA | NA | NA |
| WBGene00172995 | NA | NA | NA | NA |
| WBGene00172996 | NA | NA | NA | NA |
| WBGene00172997 | NA | NA | NA | NA |
| WBGene00172998 | NA | NA | NA | NA |
| WBGene00172999 | NA | NA | NA | NA |
| WBGene00173000 | NA | NA | NA | NA |
| WBGene00173001 | NA | NA | NA | NA |
| WBGene00173002 | NA | NA | NA | NA |
| WBGene00173003 | NA | NA | NA | NA |
| WBGene00173004 | NA | NA | NA | NA |
| WBGene00173005 | NA | NA | NA | NA |
| WBGene00173006 | NA | NA | NA | NA |
| WBGene00173007 | NA | NA | NA | NA |
| WBGene00173008 | NA | NA | NA | NA |
| WBGene00173009 | NA | NA | NA | NA |
| WBGene00173010 | NA | NA | NA | NA |
| WBGene00173011 | NA | NA | NA | NA |
| WBGene00173012 | NA | NA | NA | NA |
| WBGene00173013 | NA | NA | NA | NA |
| WBGene00173014 | NA | NA | NA | NA |
| WBGene00173015 | NA | NA | NA | NA |
| WBGene00173016 | NA | NA | NA | NA |
| WBGene00173017 | NA | NA | NA | NA |
| WBGene00173018 | NA | NA | NA | NA |

| | | | | |
|---|---|---|---|---|
| WBGene00173019 | NA | NA | NA | NA |
| WBGene00173020 | NA | NA | NA | NA |
| WBGene00173021 | NA | NA | NA | NA |
| WBGene00173022 | NA | NA | NA | NA |
| WBGene00173023 | NA | NA | NA | NA |
| WBGene00173024 | NA | NA | NA | NA |
| WBGene00173025 | NA | NA | NA | NA |
| WBGene00173026 | NA | NA | NA | NA |
| WBGene00173027 | NA | NA | NA | NA |
| WBGene00173028 | NA | NA | NA | NA |
| WBGene00173029 | NA | NA | NA | NA |
| WBGene00173030 | NA | NA | NA | NA |
| WBGene00173031 | NA | NA | NA | NA |
| WBGene00173032 | NA | NA | NA | NA |
| WBGene00173033 | NA | NA | NA | NA |
| WBGene00173034 | NA | NA | NA | NA |
| WBGene00173035 | NA | NA | NA | NA |
| WBGene00173036 | NA | NA | NA | NA |
| WBGene00173037 | NA | NA | NA | NA |
| WBGene00173038 | NA | NA | NA | NA |
| WBGene00173039 | NA | NA | NA | NA |
| WBGene00173040 | NA | NA | NA | NA |
| WBGene00173041 | NA | NA | NA | NA |
| WBGene00173042 | NA | NA | NA | NA |
| WBGene00173043 | NA | NA | NA | NA |
| WBGene00173044 | NA | NA | NA | NA |
| WBGene00173045 | NA | NA | NA | NA |
| WBGene00173046 | NA | NA | NA | NA |
| WBGene00173047 | NA | NA | NA | NA |
| WBGene00173048 | NA | NA | NA | NA |
| WBGene00173049 | NA | NA | NA | NA |
| WBGene00173050 | NA | NA | NA | NA |
| WBGene00173051 | NA | NA | NA | NA |
| WBGene00173052 | NA | NA | NA | NA |
| WBGene00173053 | NA | NA | NA | NA |
| WBGene00173054 | NA | NA | NA | NA |
| WBGene00173055 | NA | NA | NA | NA |
| WBGene00173056 | NA | NA | NA | NA |
| WBGene00173057 | NA | NA | NA | NA |
| WBGene00173058 | NA | NA | NA | NA |
| WBGene00173059 | NA | NA | NA | NA |
| WBGene00173060 | NA | NA | NA | NA |
| WBGene00173061 | NA | NA | NA | NA |

| | | | | |
|---|---|---|---|---|
| WBGene00173062 | NA | NA | NA | NA |
| WBGene00173063 | NA | NA | NA | NA |
| WBGene00173064 | NA | NA | NA | NA |
| WBGene00173065 | NA | NA | NA | NA |
| WBGene00173066 | NA | NA | NA | NA |
| WBGene00173067 | NA | NA | NA | NA |
| WBGene00173068 | NA | NA | NA | NA |
| WBGene00173069 | NA | NA | NA | NA |
| WBGene00173070 | NA | NA | NA | NA |
| WBGene00173071 | NA | NA | NA | NA |
| WBGene00173072 | NA | NA | NA | NA |
| WBGene00173073 | NA | NA | NA | NA |
| WBGene00173074 | NA | NA | NA | NA |
| WBGene00173075 | NA | NA | NA | NA |
| WBGene00173076 | NA | NA | NA | NA |
| WBGene00173077 | NA | NA | NA | NA |
| WBGene00173078 | NA | NA | NA | NA |
| WBGene00173079 | NA | NA | NA | NA |
| WBGene00173080 | NA | NA | NA | NA |
| WBGene00173081 | NA | NA | NA | NA |
| WBGene00173082 | NA | NA | NA | NA |
| WBGene00173083 | NA | NA | NA | NA |
| WBGene00173084 | NA | NA | NA | NA |
| WBGene00173085 | NA | NA | NA | NA |
| WBGene00173086 | NA | NA | NA | NA |
| WBGene00173087 | NA | NA | NA | NA |
| WBGene00173088 | NA | NA | NA | NA |
| WBGene00173089 | NA | NA | NA | NA |
| WBGene00173090 | NA | NA | NA | NA |
| WBGene00173091 | NA | NA | NA | NA |
| WBGene00173092 | NA | NA | NA | NA |
| WBGene00173093 | NA | NA | NA | NA |
| WBGene00173094 | NA | NA | NA | NA |
| WBGene00173095 | NA | NA | NA | NA |
| WBGene00173096 | NA | NA | NA | NA |
| WBGene00173097 | NA | NA | NA | NA |
| WBGene00173098 | NA | NA | NA | NA |
| WBGene00173099 | NA | NA | NA | NA |
| WBGene00173100 | NA | NA | NA | NA |
| WBGene00173101 | NA | NA | NA | NA |
| WBGene00173102 | NA | NA | NA | NA |
| WBGene00173103 | NA | NA | NA | NA |
| WBGene00173104 | NA | NA | NA | NA |

| | | | | |
|---|---|---|---|---|
| WBGene00173105 | NA | NA | NA | NA |
| WBGene00173106 | NA | NA | NA | NA |
| WBGene00173107 | NA | NA | NA | NA |
| WBGene00173108 | NA | NA | NA | NA |
| WBGene00173109 | NA | NA | NA | NA |
| WBGene00173110 | NA | NA | NA | NA |
| WBGene00173111 | NA | NA | NA | NA |
| WBGene00173112 | NA | NA | NA | NA |
| WBGene00173113 | NA | NA | NA | NA |
| WBGene00173114 | NA | NA | NA | NA |
| WBGene00173115 | NA | NA | NA | NA |
| WBGene00173116 | NA | NA | NA | NA |
| WBGene00173117 | NA | NA | NA | NA |
| WBGene00173118 | NA | NA | NA | NA |
| WBGene00173119 | NA | NA | NA | NA |
| WBGene00173120 | NA | NA | NA | NA |
| WBGene00173121 | NA | NA | NA | NA |
| WBGene00173122 | NA | NA | NA | NA |
| WBGene00173123 | NA | NA | NA | NA |
| WBGene00173124 | NA | NA | NA | NA |
| WBGene00173125 | NA | NA | NA | NA |
| WBGene00173126 | NA | NA | NA | NA |
| WBGene00173127 | NA | NA | NA | NA |
| WBGene00173128 | NA | NA | NA | NA |
| WBGene00173129 | NA | NA | NA | NA |
| WBGene00173130 | NA | NA | NA | NA |
| WBGene00173131 | NA | NA | NA | NA |
| WBGene00173132 | NA | NA | NA | NA |
| WBGene00173133 | NA | NA | NA | NA |
| WBGene00173134 | NA | NA | NA | NA |
| WBGene00173135 | -0.022930767 | 0.78121253 | NA | NA |
| WBGene00173136 | NA | NA | NA | NA |
| WBGene00173137 | NA | NA | NA | NA |
| WBGene00173138 | NA | NA | NA | NA |
| WBGene00173139 | NA | NA | NA | NA |
| WBGene00173140 | NA | NA | NA | NA |
| WBGene00173141 | NA | NA | NA | NA |
| WBGene00173142 | NA | NA | NA | NA |
| WBGene00173143 | NA | NA | NA | NA |
| WBGene00173144 | NA | NA | NA | NA |
| WBGene00173145 | NA | NA | NA | NA |
| WBGene00173146 | NA | NA | NA | NA |
| WBGene00173147 | NA | NA | NA | NA |

| | | | | |
|---|---|---|---|---|
| WBGene00173148 | NA | NA | NA | NA |
| WBGene00173149 | NA | NA | NA | NA |
| WBGene00173150 | NA | NA | NA | NA |
| WBGene00173151 | NA | NA | NA | NA |
| WBGene00173152 | NA | NA | NA | NA |
| WBGene00173153 | NA | NA | NA | NA |
| WBGene00173154 | NA | NA | NA | NA |
| WBGene00173155 | NA | NA | NA | NA |
| WBGene00173156 | NA | NA | NA | NA |
| WBGene00173157 | NA | NA | NA | NA |
| WBGene00173158 | NA | NA | NA | NA |
| WBGene00173159 | 0.020317245 | 0.84643798 | NA | NA |
| WBGene00173160 | NA | NA | NA | NA |
| WBGene00173161 | NA | NA | NA | NA |
| WBGene00173162 | NA | NA | NA | NA |
| WBGene00173163 | NA | NA | NA | NA |
| WBGene00173164 | NA | NA | NA | NA |
| WBGene00173165 | NA | NA | NA | NA |
| WBGene00173166 | NA | NA | NA | NA |
| WBGene00173167 | NA | NA | NA | NA |
| WBGene00173168 | NA | NA | NA | NA |
| WBGene00173169 | NA | NA | NA | NA |
| WBGene00173170 | NA | NA | NA | NA |
| WBGene00173171 | NA | NA | NA | NA |
| WBGene00173172 | NA | NA | NA | NA |
| WBGene00173173 | NA | NA | NA | NA |
| WBGene00173174 | NA | NA | NA | NA |
| WBGene00173175 | NA | NA | NA | NA |
| WBGene00173176 | NA | NA | NA | NA |
| WBGene00173177 | NA | NA | NA | NA |
| WBGene00173178 | NA | NA | NA | NA |
| WBGene00173179 | NA | NA | NA | NA |
| WBGene00173180 | NA | NA | NA | NA |
| WBGene00173181 | NA | NA | NA | NA |
| WBGene00173182 | NA | NA | NA | NA |
| WBGene00173183 | NA | NA | NA | NA |
| WBGene00173184 | NA | NA | NA | NA |
| WBGene00173185 | NA | NA | NA | NA |
| WBGene00173186 | NA | NA | NA | NA |
| WBGene00173187 | NA | NA | NA | NA |
| WBGene00173188 | NA | NA | NA | NA |
| WBGene00173189 | NA | NA | NA | NA |
| WBGene00173190 | NA | NA | NA | NA |

| | | | | |
|---|---|---|---|---|
| WBGene00173191 | NA | NA | NA | NA |
| WBGene00173192 | NA | NA | NA | NA |
| WBGene00173193 | NA | NA | NA | NA |
| WBGene00173194 | NA | NA | NA | NA |
| WBGene00173195 | NA | NA | NA | NA |
| WBGene00173196 | NA | NA | NA | NA |
| WBGene00173197 | NA | NA | NA | NA |
| WBGene00173198 | NA | NA | NA | NA |
| WBGene00173199 | NA | NA | NA | NA |
| WBGene00173200 | NA | NA | NA | NA |
| WBGene00173201 | NA | NA | NA | NA |
| WBGene00173202 | NA | NA | NA | NA |
| WBGene00173203 | NA | NA | NA | NA |
| WBGene00173204 | NA | NA | NA | NA |
| WBGene00173205 | NA | NA | NA | NA |
| WBGene00173206 | NA | NA | NA | NA |
| WBGene00173207 | NA | NA | NA | NA |
| WBGene00173208 | NA | NA | NA | NA |
| WBGene00173209 | NA | NA | NA | NA |
| WBGene00173210 | NA | NA | NA | NA |
| WBGene00173211 | NA | NA | NA | NA |
| WBGene00173212 | NA | NA | NA | NA |
| WBGene00173213 | NA | NA | NA | NA |
| WBGene00173214 | NA | NA | NA | NA |
| WBGene00173215 | NA | NA | NA | NA |
| WBGene00173216 | NA | NA | NA | NA |
| WBGene00173217 | NA | NA | NA | NA |
| WBGene00173218 | NA | NA | NA | NA |
| WBGene00173219 | NA | NA | NA | NA |
| WBGene00173220 | NA | NA | NA | NA |
| WBGene00173221 | NA | NA | NA | NA |
| WBGene00173222 | NA | NA | NA | NA |
| WBGene00173223 | NA | NA | NA | NA |
| WBGene00173224 | NA | NA | NA | NA |
| WBGene00173225 | NA | NA | NA | NA |
| WBGene00173226 | NA | NA | NA | NA |
| WBGene00173227 | NA | NA | NA | NA |
| WBGene00173228 | NA | NA | NA | NA |
| WBGene00173229 | NA | NA | NA | NA |
| WBGene00173230 | NA | NA | NA | NA |
| WBGene00173231 | NA | NA | NA | NA |
| WBGene00173232 | NA | NA | NA | NA |
| WBGene00173233 | NA | NA | NA | NA |

| | | | | |
|---|---|---|---|---|
| WBGene00173234 | NA | NA | NA | NA |
| WBGene00173235 | NA | NA | NA | NA |
| WBGene00173236 | NA | NA | NA | NA |
| WBGene00173237 | NA | NA | NA | NA |
| WBGene00173238 | NA | NA | NA | NA |
| WBGene00173239 | NA | NA | NA | NA |
| WBGene00173240 | NA | NA | NA | NA |
| WBGene00173241 | NA | NA | NA | NA |
| WBGene00173242 | NA | NA | NA | NA |
| WBGene00173243 | NA | NA | NA | NA |
| WBGene00173244 | NA | NA | NA | NA |
| WBGene00173245 | NA | NA | NA | NA |
| WBGene00173246 | -0.023056499 | 0.78121253 | NA | NA |
| WBGene00173247 | NA | NA | NA | NA |
| WBGene00173248 | NA | NA | NA | NA |
| WBGene00173249 | NA | NA | NA | NA |
| WBGene00173250 | NA | NA | NA | NA |
| WBGene00173251 | NA | NA | NA | NA |
| WBGene00173252 | NA | NA | NA | NA |
| WBGene00173253 | NA | NA | NA | NA |
| WBGene00173254 | NA | NA | NA | NA |
| WBGene00173255 | NA | NA | NA | NA |
| WBGene00173256 | NA | NA | NA | NA |
| WBGene00173257 | NA | NA | NA | NA |
| WBGene00173258 | NA | NA | NA | NA |
| WBGene00173259 | NA | NA | NA | NA |
| WBGene00173260 | NA | NA | NA | NA |
| WBGene00173261 | NA | NA | NA | NA |
| WBGene00173262 | NA | NA | NA | NA |
| WBGene00173263 | NA | NA | NA | NA |
| WBGene00173264 | NA | NA | NA | NA |
| WBGene00173265 | NA | NA | NA | NA |
| WBGene00173266 | NA | NA | NA | NA |
| WBGene00173267 | NA | NA | NA | NA |
| WBGene00173268 | NA | NA | NA | NA |
| WBGene00173269 | NA | NA | NA | NA |
| WBGene00173270 | NA | NA | NA | NA |
| WBGene00173271 | NA | NA | NA | NA |
| WBGene00173272 | NA | NA | NA | NA |
| WBGene00173273 | NA | NA | NA | NA |
| WBGene00173274 | NA | NA | NA | NA |
| WBGene00173275 | NA | NA | NA | NA |
| WBGene00173276 | NA | NA | NA | NA |

| | | | | |
|---|---|---|---|---|
| WBGene00173277 | NA | NA | NA | NA |
| WBGene00173278 | NA | NA | NA | NA |
| WBGene00173279 | NA | NA | NA | NA |
| WBGene00173280 | NA | NA | NA | NA |
| WBGene00173281 | NA | NA | NA | NA |
| WBGene00173282 | NA | NA | NA | NA |
| WBGene00173283 | NA | NA | NA | NA |
| WBGene00173284 | NA | NA | NA | NA |
| WBGene00173285 | NA | NA | NA | NA |
| WBGene00173286 | NA | NA | NA | NA |
| WBGene00173287 | NA | NA | NA | NA |
| WBGene00173288 | NA | NA | NA | NA |
| WBGene00173289 | NA | NA | NA | NA |
| WBGene00173290 | NA | NA | NA | NA |
| WBGene00173291 | NA | NA | NA | NA |
| WBGene00173292 | NA | NA | NA | NA |
| WBGene00173293 | NA | NA | NA | NA |
| WBGene00173294 | NA | NA | NA | NA |
| WBGene00173295 | NA | NA | NA | NA |
| WBGene00173296 | NA | NA | NA | NA |
| WBGene00173297 | NA | NA | NA | NA |
| WBGene00173298 | NA | NA | NA | NA |
| WBGene00173299 | NA | NA | NA | NA |
| WBGene00173300 | NA | NA | NA | NA |
| WBGene00173301 | NA | NA | NA | NA |
| WBGene00173302 | NA | NA | NA | NA |
| WBGene00173303 | NA | NA | NA | NA |
| WBGene00173304 | NA | NA | NA | NA |
| WBGene00173305 | NA | NA | NA | NA |
| WBGene00173306 | NA | NA | NA | NA |
| WBGene00173307 | NA | NA | NA | NA |
| WBGene00173308 | NA | NA | NA | NA |
| WBGene00173309 | NA | NA | NA | NA |
| WBGene00173310 | NA | NA | NA | NA |
| WBGene00173311 | NA | NA | NA | NA |
| WBGene00173312 | NA | NA | NA | NA |
| WBGene00173313 | NA | NA | NA | NA |
| WBGene00173314 | NA | NA | NA | NA |
| WBGene00173315 | NA | NA | NA | NA |
| WBGene00173316 | NA | NA | NA | NA |
| WBGene00173317 | NA | NA | NA | NA |
| WBGene00173318 | NA | NA | NA | NA |
| WBGene00173319 | NA | NA | NA | NA |

| | | | | |
|---|---|---|---|---|
| WBGene00173320 | NA | NA | NA | NA |
| WBGene00173321 | NA | NA | NA | NA |
| WBGene00173322 | NA | NA | NA | NA |
| WBGene00173323 | 0.020232312 | 0.84643798 | NA | NA |
| WBGene00173324 | NA | NA | NA | NA |
| WBGene00173325 | NA | NA | NA | NA |
| WBGene00173326 | NA | NA | NA | NA |
| WBGene00173327 | NA | NA | NA | NA |
| WBGene00173328 | NA | NA | NA | NA |
| WBGene00173329 | NA | NA | NA | NA |
| WBGene00173330 | NA | NA | NA | NA |
| WBGene00173331 | NA | NA | NA | NA |
| WBGene00173332 | NA | NA | NA | NA |
| WBGene00173333 | NA | NA | NA | NA |
| WBGene00173334 | NA | NA | NA | NA |
| WBGene00173335 | NA | NA | NA | NA |
| WBGene00173336 | NA | NA | NA | NA |
| WBGene00173337 | NA | NA | NA | NA |
| WBGene00173338 | NA | NA | NA | NA |
| WBGene00173339 | NA | NA | NA | NA |
| WBGene00173340 | NA | NA | NA | NA |
| WBGene00173341 | NA | NA | NA | NA |
| WBGene00173342 | NA | NA | NA | NA |
| WBGene00173343 | NA | NA | NA | NA |
| WBGene00173344 | NA | NA | NA | NA |
| WBGene00173345 | NA | NA | NA | NA |
| WBGene00173346 | NA | NA | NA | NA |
| WBGene00173347 | NA | NA | NA | NA |
| WBGene00173348 | NA | NA | NA | NA |
| WBGene00173349 | NA | NA | NA | NA |
| WBGene00173350 | NA | NA | NA | NA |
| WBGene00173351 | NA | NA | NA | NA |
| WBGene00173352 | NA | NA | NA | NA |
| WBGene00173353 | NA | NA | NA | NA |
| WBGene00173354 | NA | NA | NA | NA |
| WBGene00173355 | NA | NA | NA | NA |
| WBGene00173356 | NA | NA | NA | NA |
| WBGene00173357 | NA | NA | NA | NA |
| WBGene00173358 | NA | NA | NA | NA |
| WBGene00173359 | NA | NA | NA | NA |
| WBGene00173360 | NA | NA | NA | NA |
| WBGene00173361 | NA | NA | NA | NA |
| WBGene00173362 | NA | NA | NA | NA |

| | | | | |
|---|---|---|---|---|
| WBGene00173363 | NA | NA | NA | NA |
| WBGene00173364 | NA | NA | NA | NA |
| WBGene00173365 | NA | NA | NA | NA |
| WBGene00173366 | NA | NA | NA | NA |
| WBGene00173367 | NA | NA | NA | NA |
| WBGene00173368 | NA | NA | NA | NA |
| WBGene00173369 | NA | NA | NA | NA |
| WBGene00173370 | NA | NA | NA | NA |
| WBGene00173371 | NA | NA | NA | NA |
| WBGene00173372 | NA | NA | NA | NA |
| WBGene00173373 | NA | NA | NA | NA |
| WBGene00173374 | NA | NA | NA | NA |
| WBGene00173375 | NA | NA | NA | NA |
| WBGene00173376 | NA | NA | NA | NA |
| WBGene00173377 | NA | NA | NA | NA |
| WBGene00173378 | NA | NA | NA | NA |
| WBGene00173379 | NA | NA | NA | NA |
| WBGene00173380 | NA | NA | NA | NA |
| WBGene00173381 | NA | NA | NA | NA |
| WBGene00173382 | NA | NA | NA | NA |
| WBGene00173383 | NA | NA | NA | NA |
| WBGene00173384 | NA | NA | NA | NA |
| WBGene00173385 | NA | NA | NA | NA |
| WBGene00173386 | NA | NA | NA | NA |
| WBGene00173387 | NA | NA | NA | NA |
| WBGene00173388 | NA | NA | NA | NA |
| WBGene00173389 | NA | NA | NA | NA |
| WBGene00173390 | NA | NA | NA | NA |
| WBGene00173391 | NA | NA | NA | NA |
| WBGene00173392 | NA | NA | NA | NA |
| WBGene00173393 | NA | NA | NA | NA |
| WBGene00173394 | NA | NA | NA | NA |
| WBGene00173395 | NA | NA | NA | NA |
| WBGene00173396 | NA | NA | NA | NA |
| WBGene00173397 | NA | NA | NA | NA |
| WBGene00173398 | NA | NA | NA | NA |
| WBGene00173399 | NA | NA | NA | NA |
| WBGene00173400 | NA | NA | NA | NA |
| WBGene00173401 | NA | NA | NA | NA |
| WBGene00173402 | NA | NA | NA | NA |
| WBGene00173403 | NA | NA | NA | NA |
| WBGene00173404 | NA | NA | NA | NA |
| WBGene00173405 | NA | NA | NA | NA |

| | | | | |
|---|---|---|---|---|
| WBGene00173406 | NA | NA | NA | NA |
| WBGene00173407 | NA | NA | NA | NA |
| WBGene00173408 | NA | NA | NA | NA |
| WBGene00173409 | NA | NA | NA | NA |
| WBGene00173410 | NA | NA | NA | NA |
| WBGene00173411 | NA | NA | NA | NA |
| WBGene00173412 | NA | NA | NA | NA |
| WBGene00173413 | NA | NA | NA | NA |
| WBGene00173414 | NA | NA | NA | NA |
| WBGene00173415 | NA | NA | NA | NA |
| WBGene00173416 | NA | NA | NA | NA |
| WBGene00173417 | NA | NA | NA | NA |
| WBGene00173418 | NA | NA | NA | NA |
| WBGene00173419 | NA | NA | NA | NA |
| WBGene00173420 | NA | NA | NA | NA |
| WBGene00173421 | NA | NA | NA | NA |
| WBGene00173422 | NA | NA | NA | NA |
| WBGene00173423 | NA | NA | NA | NA |
| WBGene00173424 | NA | NA | NA | NA |
| WBGene00173425 | NA | NA | NA | NA |
| WBGene00173426 | NA | NA | NA | NA |
| WBGene00173427 | NA | NA | NA | NA |
| WBGene00173428 | NA | NA | NA | NA |
| WBGene00173429 | NA | NA | NA | NA |
| WBGene00173430 | NA | NA | NA | NA |
| WBGene00173431 | NA | NA | NA | NA |
| WBGene00173432 | NA | NA | NA | NA |
| WBGene00173433 | NA | NA | NA | NA |
| WBGene00173434 | NA | NA | NA | NA |
| WBGene00173435 | NA | NA | NA | NA |
| WBGene00173436 | NA | NA | NA | NA |
| WBGene00173437 | NA | NA | NA | NA |
| WBGene00173438 | NA | NA | NA | NA |
| WBGene00173439 | NA | NA | NA | NA |
| WBGene00173440 | NA | NA | NA | NA |
| WBGene00173441 | NA | NA | NA | NA |
| WBGene00173442 | NA | NA | NA | NA |
| WBGene00173443 | NA | NA | NA | NA |
| WBGene00173444 | NA | NA | NA | NA |
| WBGene00173445 | NA | NA | NA | NA |
| WBGene00173446 | NA | NA | NA | NA |
| WBGene00173447 | NA | NA | NA | NA |
| WBGene00173448 | NA | NA | NA | NA |

| | | | | |
|---|---|---|---|---|
| WBGene00173449 | NA | NA | NA | NA |
| WBGene00173450 | NA | NA | NA | NA |
| WBGene00173451 | NA | NA | NA | NA |
| WBGene00173452 | NA | NA | NA | NA |
| WBGene00173453 | NA | NA | NA | NA |
| WBGene00173454 | NA | NA | NA | NA |
| WBGene00173455 | NA | NA | NA | NA |
| WBGene00173456 | NA | NA | NA | NA |
| WBGene00173457 | NA | NA | NA | NA |
| WBGene00173458 | NA | NA | NA | NA |
| WBGene00173459 | NA | NA | NA | NA |
| WBGene00173460 | NA | NA | NA | NA |
| WBGene00173461 | NA | NA | NA | NA |
| WBGene00173462 | NA | NA | NA | NA |
| WBGene00173463 | NA | NA | NA | NA |
| WBGene00173464 | NA | NA | NA | NA |
| WBGene00173465 | NA | NA | NA | NA |
| WBGene00173466 | NA | NA | NA | NA |
| WBGene00173467 | NA | NA | NA | NA |
| WBGene00173468 | NA | NA | NA | NA |
| WBGene00173469 | NA | NA | NA | NA |
| WBGene00173470 | NA | NA | NA | NA |
| WBGene00173471 | NA | NA | NA | NA |
| WBGene00173472 | NA | NA | NA | NA |
| WBGene00173473 | NA | NA | NA | NA |
| WBGene00173474 | NA | NA | NA | NA |
| WBGene00173475 | NA | NA | NA | NA |
| WBGene00173476 | NA | NA | NA | NA |
| WBGene00173477 | NA | NA | NA | NA |
| WBGene00173478 | NA | NA | NA | NA |
| WBGene00173480 | NA | NA | NA | NA |
| WBGene00173481 | NA | NA | NA | NA |
| WBGene00173482 | NA | NA | NA | NA |
| WBGene00173483 | NA | NA | NA | NA |
| WBGene00173484 | NA | NA | NA | NA |
| WBGene00173485 | NA | NA | NA | NA |
| WBGene00173486 | NA | NA | NA | NA |
| WBGene00173487 | NA | NA | NA | NA |
| WBGene00173488 | NA | NA | NA | NA |
| WBGene00173489 | NA | NA | NA | NA |
| WBGene00173490 | NA | NA | NA | NA |
| WBGene00173491 | NA | NA | NA | NA |
| WBGene00173492 | NA | NA | NA | NA |

| | | | | |
|---|---|---|---|---|
| WBGene00173493 | NA | NA | NA | NA |
| WBGene00173494 | NA | NA | NA | NA |
| WBGene00173495 | NA | NA | NA | NA |
| WBGene00173496 | NA | NA | NA | NA |
| WBGene00173497 | NA | NA | NA | NA |
| WBGene00173498 | -0.022930767 | 0.78121253 | NA | NA |
| WBGene00173499 | NA | NA | NA | NA |
| WBGene00173500 | NA | NA | NA | NA |
| WBGene00173501 | NA | NA | NA | NA |
| WBGene00173502 | NA | NA | NA | NA |
| WBGene00173503 | NA | NA | NA | NA |
| WBGene00173504 | NA | NA | NA | NA |
| WBGene00173505 | 0.020232312 | 0.84643798 | NA | NA |
| WBGene00173506 | NA | NA | NA | NA |
| WBGene00173507 | NA | NA | NA | NA |
| WBGene00173508 | NA | NA | NA | NA |
| WBGene00173509 | -0.023056499 | 0.78121253 | NA | NA |
| WBGene00173510 | NA | NA | NA | NA |
| WBGene00173511 | NA | NA | NA | NA |
| WBGene00173512 | NA | NA | NA | NA |
| WBGene00173513 | NA | NA | NA | NA |
| WBGene00173514 | NA | NA | NA | NA |
| WBGene00173515 | NA | NA | NA | NA |
| WBGene00173516 | NA | NA | NA | NA |
| WBGene00173517 | NA | NA | NA | NA |
| WBGene00173518 | NA | NA | NA | NA |
| WBGene00173519 | NA | NA | NA | NA |
| WBGene00173520 | NA | NA | NA | NA |
| WBGene00173521 | NA | NA | NA | NA |
| WBGene00173522 | NA | NA | NA | NA |
| WBGene00173523 | NA | NA | NA | NA |
| WBGene00173524 | NA | NA | NA | NA |
| WBGene00173525 | NA | NA | NA | NA |
| WBGene00173526 | NA | NA | NA | NA |
| WBGene00173527 | NA | NA | NA | NA |
| WBGene00173528 | NA | NA | NA | NA |
| WBGene00173529 | NA | NA | NA | NA |
| WBGene00173530 | NA | NA | NA | NA |
| WBGene00173531 | NA | NA | NA | NA |
| WBGene00173532 | NA | NA | NA | NA |
| WBGene00173533 | NA | NA | NA | NA |
| WBGene00173534 | NA | NA | NA | NA |
| WBGene00173535 | NA | NA | NA | NA |

| | | | | |
|---|---|---|---|---|
| WBGene00173536 | NA | NA | NA | NA |
| WBGene00173537 | NA | NA | NA | NA |
| WBGene00173538 | NA | NA | NA | NA |
| WBGene00173539 | NA | NA | NA | NA |
| WBGene00173540 | NA | NA | NA | NA |
| WBGene00173541 | NA | NA | NA | NA |
| WBGene00173542 | NA | NA | NA | NA |
| WBGene00173543 | NA | NA | NA | NA |
| WBGene00173544 | NA | NA | NA | NA |
| WBGene00173545 | NA | NA | NA | NA |
| WBGene00173546 | NA | NA | NA | NA |
| WBGene00173547 | NA | NA | NA | NA |
| WBGene00173548 | NA | NA | NA | NA |
| WBGene00173550 | NA | NA | NA | NA |
| WBGene00173551 | NA | NA | NA | NA |
| WBGene00173552 | NA | NA | NA | NA |
| WBGene00173553 | NA | NA | NA | NA |
| WBGene00173554 | NA | NA | NA | NA |
| WBGene00173555 | NA | NA | NA | NA |
| WBGene00173556 | NA | NA | NA | NA |
| WBGene00173557 | -0.003614866 | 0.95700117 | NA | NA |
| WBGene00173558 | NA | NA | NA | NA |
| WBGene00173559 | NA | NA | NA | NA |
| WBGene00173560 | -0.023109762 | 0.78121253 | NA | NA |
| WBGene00173561 | NA | NA | NA | NA |
| WBGene00173562 | NA | NA | NA | NA |
| WBGene00173563 | NA | NA | NA | NA |
| WBGene00173564 | NA | NA | NA | NA |
| WBGene00173565 | NA | NA | NA | NA |
| WBGene00173566 | NA | NA | NA | NA |
| WBGene00173567 | NA | NA | NA | NA |
| WBGene00173568 | NA | NA | NA | NA |
| WBGene00173569 | NA | NA | NA | NA |
| WBGene00173570 | NA | NA | NA | NA |
| WBGene00173571 | NA | NA | NA | NA |
| WBGene00173572 | NA | NA | NA | NA |
| WBGene00173573 | NA | NA | NA | NA |
| WBGene00173574 | NA | NA | NA | NA |
| WBGene00173575 | NA | NA | NA | NA |
| WBGene00173576 | NA | NA | NA | NA |
| WBGene00173577 | NA | NA | NA | NA |
| WBGene00173578 | NA | NA | NA | NA |
| WBGene00173579 | NA | NA | NA | NA |

| | | | | |
|---|---|---|---|---|
| WBGene00173580 | NA | NA | NA | NA |
| WBGene00173581 | NA | NA | NA | NA |
| WBGene00173582 | NA | NA | NA | NA |
| WBGene00173583 | NA | NA | NA | NA |
| WBGene00173584 | NA | NA | NA | NA |
| WBGene00173585 | NA | NA | NA | NA |
| WBGene00173586 | NA | NA | NA | NA |
| WBGene00173587 | NA | NA | NA | NA |
| WBGene00173588 | NA | NA | NA | NA |
| WBGene00173589 | NA | NA | NA | NA |
| WBGene00173590 | NA | NA | NA | NA |
| WBGene00173591 | NA | NA | NA | NA |
| WBGene00173592 | NA | NA | NA | NA |
| WBGene00173593 | NA | NA | NA | NA |
| WBGene00173594 | NA | NA | NA | NA |
| WBGene00173595 | NA | NA | NA | NA |
| WBGene00173596 | NA | NA | NA | NA |
| WBGene00173597 | NA | NA | NA | NA |
| WBGene00173598 | NA | NA | NA | NA |
| WBGene00173599 | NA | NA | NA | NA |
| WBGene00173600 | NA | NA | NA | NA |
| WBGene00173601 | NA | NA | NA | NA |
| WBGene00173602 | NA | NA | NA | NA |
| WBGene00173603 | NA | NA | NA | NA |
| WBGene00173604 | NA | NA | NA | NA |
| WBGene00173605 | NA | NA | NA | NA |
| WBGene00173606 | NA | NA | NA | NA |
| WBGene00173607 | NA | NA | NA | NA |
| WBGene00173608 | NA | NA | NA | NA |
| WBGene00173609 | NA | NA | NA | NA |
| WBGene00173610 | NA | NA | NA | NA |
| WBGene00173611 | NA | NA | NA | NA |
| WBGene00173612 | NA | NA | NA | NA |
| WBGene00173613 | 0.020232312 | 0.84643798 | NA | NA |
| WBGene00173614 | NA | NA | NA | NA |
| WBGene00173615 | NA | NA | NA | NA |
| WBGene00173616 | NA | NA | NA | NA |
| WBGene00173617 | NA | NA | NA | NA |
| WBGene00173618 | NA | NA | NA | NA |
| WBGene00173619 | NA | NA | NA | NA |
| WBGene00173620 | NA | NA | NA | NA |
| WBGene00173621 | NA | NA | NA | NA |
| WBGene00173622 | NA | NA | NA | NA |

| | | | | |
|---|---|---|---|---|
| WBGene00173623 | NA | NA | NA | NA |
| WBGene00173624 | NA | NA | NA | NA |
| WBGene00173625 | NA | NA | NA | NA |
| WBGene00173626 | NA | NA | NA | NA |
| WBGene00173627 | NA | NA | NA | NA |
| WBGene00173628 | NA | NA | NA | NA |
| WBGene00173629 | NA | NA | NA | NA |
| WBGene00173630 | NA | NA | NA | NA |
| WBGene00173631 | NA | NA | NA | NA |
| WBGene00173632 | NA | NA | NA | NA |
| WBGene00173633 | NA | NA | NA | NA |
| WBGene00173634 | NA | NA | NA | NA |
| WBGene00173635 | NA | NA | NA | NA |
| WBGene00173636 | NA | NA | NA | NA |
| WBGene00173637 | NA | NA | NA | NA |
| WBGene00173638 | NA | NA | NA | NA |
| WBGene00173639 | NA | NA | NA | NA |
| WBGene00173640 | NA | NA | NA | NA |
| WBGene00173641 | NA | NA | NA | NA |
| WBGene00173642 | NA | NA | NA | NA |
| WBGene00173643 | NA | NA | NA | NA |
| WBGene00173644 | NA | NA | NA | NA |
| WBGene00173645 | NA | NA | NA | NA |
| WBGene00173646 | NA | NA | NA | NA |
| WBGene00173647 | NA | NA | NA | NA |
| WBGene00173648 | NA | NA | NA | NA |
| WBGene00173649 | NA | NA | NA | NA |
| WBGene00173650 | NA | NA | NA | NA |
| WBGene00173651 | NA | NA | NA | NA |
| WBGene00173652 | NA | NA | NA | NA |
| WBGene00173653 | NA | NA | NA | NA |
| WBGene00173654 | NA | NA | NA | NA |
| WBGene00173655 | NA | NA | NA | NA |
| WBGene00173656 | NA | NA | NA | NA |
| WBGene00173657 | NA | NA | NA | NA |
| WBGene00173658 | NA | NA | NA | NA |
| WBGene00173659 | NA | NA | NA | NA |
| WBGene00173660 | NA | NA | NA | NA |
| WBGene00173661 | NA | NA | NA | NA |
| WBGene00173662 | NA | NA | NA | NA |
| WBGene00173663 | NA | NA | NA | NA |
| WBGene00173664 | NA | NA | NA | NA |
| WBGene00173665 | NA | NA | NA | NA |

| | | | | |
|---|---|---|---|---|
| WBGene00173666 | NA | NA | NA | NA |
| WBGene00173667 | NA | NA | NA | NA |
| WBGene00173668 | NA | NA | NA | NA |
| WBGene00173669 | NA | NA | NA | NA |
| WBGene00173670 | NA | NA | NA | NA |
| WBGene00173671 | NA | NA | NA | NA |
| WBGene00173672 | NA | NA | NA | NA |
| WBGene00173673 | NA | NA | NA | NA |
| WBGene00173674 | NA | NA | NA | NA |
| WBGene00173675 | NA | NA | NA | NA |
| WBGene00173676 | NA | NA | NA | NA |
| WBGene00173677 | NA | NA | NA | NA |
| WBGene00173678 | NA | NA | NA | NA |
| WBGene00173679 | NA | NA | NA | NA |
| WBGene00173680 | NA | NA | NA | NA |
| WBGene00173681 | NA | NA | NA | NA |
| WBGene00173682 | NA | NA | NA | NA |
| WBGene00173683 | NA | NA | NA | NA |
| WBGene00173684 | NA | NA | NA | NA |
| WBGene00173685 | NA | NA | NA | NA |
| WBGene00173686 | NA | NA | NA | NA |
| WBGene00173687 | NA | NA | NA | NA |
| WBGene00173688 | NA | NA | NA | NA |
| WBGene00173689 | NA | NA | NA | NA |
| WBGene00173690 | NA | NA | NA | NA |
| WBGene00173691 | NA | NA | NA | NA |
| WBGene00173692 | NA | NA | NA | NA |
| WBGene00173693 | NA | NA | NA | NA |
| WBGene00173694 | NA | NA | NA | NA |
| WBGene00173695 | NA | NA | NA | NA |
| WBGene00173696 | NA | NA | NA | NA |
| WBGene00173697 | NA | NA | NA | NA |
| WBGene00173698 | NA | NA | NA | NA |
| WBGene00173699 | NA | NA | NA | NA |
| WBGene00173700 | NA | NA | NA | NA |
| WBGene00173701 | NA | NA | NA | NA |
| WBGene00173702 | NA | NA | NA | NA |
| WBGene00173703 | NA | NA | NA | NA |
| WBGene00173704 | NA | NA | NA | NA |
| WBGene00173705 | NA | NA | NA | NA |
| WBGene00173706 | NA | NA | NA | NA |
| WBGene00173707 | NA | NA | NA | NA |
| WBGene00173708 | NA | NA | NA | NA |

| | | | | |
|---|---|---|---|---|
| WBGene00173709 | NA | NA | NA | NA |
| WBGene00173710 | NA | NA | NA | NA |
| WBGene00173711 | NA | NA | NA | NA |
| WBGene00173712 | NA | NA | NA | NA |
| WBGene00173713 | NA | NA | NA | NA |
| WBGene00173714 | NA | NA | NA | NA |
| WBGene00173715 | NA | NA | NA | NA |
| WBGene00173716 | NA | NA | NA | NA |
| WBGene00173717 | NA | NA | NA | NA |
| WBGene00173718 | NA | NA | NA | NA |
| WBGene00173719 | NA | NA | NA | NA |
| WBGene00173720 | NA | NA | NA | NA |
| WBGene00173721 | NA | NA | NA | NA |
| WBGene00173722 | NA | NA | NA | NA |
| WBGene00173723 | NA | NA | NA | NA |
| WBGene00173724 | NA | NA | NA | NA |
| WBGene00173725 | NA | NA | NA | NA |
| WBGene00173726 | NA | NA | NA | NA |
| WBGene00173727 | NA | NA | NA | NA |
| WBGene00173728 | NA | NA | NA | NA |
| WBGene00173729 | NA | NA | NA | NA |
| WBGene00173730 | NA | NA | NA | NA |
| WBGene00173731 | NA | NA | NA | NA |
| WBGene00173732 | NA | NA | NA | NA |
| WBGene00173733 | NA | NA | NA | NA |
| WBGene00173734 | NA | NA | NA | NA |
| WBGene00173735 | NA | NA | NA | NA |
| WBGene00173736 | NA | NA | NA | NA |
| WBGene00173737 | NA | NA | NA | NA |
| WBGene00173738 | NA | NA | NA | NA |
| WBGene00173739 | NA | NA | NA | NA |
| WBGene00173740 | NA | NA | NA | NA |
| WBGene00173741 | NA | NA | NA | NA |
| WBGene00173742 | NA | NA | NA | NA |
| WBGene00173743 | NA | NA | NA | NA |
| WBGene00173744 | NA | NA | NA | NA |
| WBGene00173745 | NA | NA | NA | NA |
| WBGene00173746 | NA | NA | NA | NA |
| WBGene00173747 | NA | NA | NA | NA |
| WBGene00173748 | NA | NA | NA | NA |
| WBGene00173749 | NA | NA | NA | NA |
| WBGene00173750 | NA | NA | NA | NA |
| WBGene00173751 | NA | NA | NA | NA |

| | | | | |
|---|---|---|---|---|
| WBGene00173752 | NA | NA | NA | NA |
| WBGene00173753 | NA | NA | NA | NA |
| WBGene00173754 | NA | NA | NA | NA |
| WBGene00173755 | NA | NA | NA | NA |
| WBGene00173756 | NA | NA | NA | NA |
| WBGene00173757 | NA | NA | NA | NA |
| WBGene00173758 | NA | NA | NA | NA |
| WBGene00173759 | NA | NA | NA | NA |
| WBGene00173760 | NA | NA | NA | NA |
| WBGene00173761 | NA | NA | NA | NA |
| WBGene00173762 | NA | NA | NA | NA |
| WBGene00173763 | NA | NA | NA | NA |
| WBGene00173764 | NA | NA | NA | NA |
| WBGene00173765 | NA | NA | NA | NA |
| WBGene00173766 | NA | NA | NA | NA |
| WBGene00173767 | NA | NA | NA | NA |
| WBGene00173768 | NA | NA | NA | NA |
| WBGene00173769 | NA | NA | NA | NA |
| WBGene00173770 | NA | NA | NA | NA |
| WBGene00173771 | NA | NA | NA | NA |
| WBGene00173772 | NA | NA | NA | NA |
| WBGene00173773 | NA | NA | NA | NA |
| WBGene00173774 | NA | NA | NA | NA |
| WBGene00173775 | NA | NA | NA | NA |
| WBGene00173776 | 0.020232312 | 0.84643798 | NA | NA |
| WBGene00173777 | NA | NA | NA | NA |
| WBGene00173778 | NA | NA | NA | NA |
| WBGene00173779 | NA | NA | NA | NA |
| WBGene00173780 | NA | NA | NA | NA |
| WBGene00173781 | NA | NA | NA | NA |
| WBGene00173782 | NA | NA | NA | NA |
| WBGene00173783 | NA | NA | NA | NA |
| WBGene00173784 | NA | NA | NA | NA |
| WBGene00173785 | NA | NA | NA | NA |
| WBGene00173786 | NA | NA | NA | NA |
| WBGene00173787 | NA | NA | NA | NA |
| WBGene00173788 | NA | NA | NA | NA |
| WBGene00173789 | NA | NA | NA | NA |
| WBGene00173790 | NA | NA | NA | NA |
| WBGene00173791 | NA | NA | NA | NA |
| WBGene00173792 | NA | NA | NA | NA |
| WBGene00173793 | NA | NA | NA | NA |
| WBGene00173794 | NA | NA | NA | NA |

| | | | | |
|---|---|---|---|---|
| WBGene00173795 | NA | NA | NA | NA |
| WBGene00173796 | NA | NA | NA | NA |
| WBGene00173797 | NA | NA | NA | NA |
| WBGene00173798 | NA | NA | NA | NA |
| WBGene00173799 | NA | NA | NA | NA |
| WBGene00173800 | NA | NA | NA | NA |
| WBGene00173801 | NA | NA | NA | NA |
| WBGene00173802 | NA | NA | NA | NA |
| WBGene00173803 | NA | NA | NA | NA |
| WBGene00173804 | NA | NA | NA | NA |
| WBGene00173805 | NA | NA | NA | NA |
| WBGene00173806 | NA | NA | NA | NA |
| WBGene00173807 | NA | NA | NA | NA |
| WBGene00173808 | NA | NA | NA | NA |
| WBGene00173809 | NA | NA | NA | NA |
| WBGene00173810 | NA | NA | NA | NA |
| WBGene00173811 | NA | NA | NA | NA |
| WBGene00173812 | NA | NA | NA | NA |
| WBGene00173813 | NA | NA | NA | NA |
| WBGene00173814 | NA | NA | NA | NA |
| WBGene00173815 | NA | NA | NA | NA |
| WBGene00173816 | NA | NA | NA | NA |
| WBGene00173817 | NA | NA | NA | NA |
| WBGene00173818 | NA | NA | NA | NA |
| WBGene00173819 | NA | NA | NA | NA |
| WBGene00173820 | NA | NA | NA | NA |
| WBGene00173821 | NA | NA | NA | NA |
| WBGene00173822 | NA | NA | NA | NA |
| WBGene00173823 | NA | NA | NA | NA |
| WBGene00173824 | NA | NA | NA | NA |
| WBGene00173825 | NA | NA | NA | NA |
| WBGene00173826 | NA | NA | NA | NA |
| WBGene00173827 | NA | NA | NA | NA |
| WBGene00173828 | NA | NA | NA | NA |
| WBGene00173829 | NA | NA | NA | NA |
| WBGene00173830 | NA | NA | NA | NA |
| WBGene00173831 | NA | NA | NA | NA |
| WBGene00173832 | NA | NA | NA | NA |
| WBGene00173833 | NA | NA | NA | NA |
| WBGene00173834 | NA | NA | NA | NA |
| WBGene00173835 | NA | NA | NA | NA |
| WBGene00173836 | NA | NA | NA | NA |
| WBGene00173837 | NA | NA | NA | NA |

| | | | | |
|---|---|---|---|---|
| WBGene00173838 | NA | NA | NA | NA |
| WBGene00173839 | NA | NA | NA | NA |
| WBGene00173840 | NA | NA | NA | NA |
| WBGene00173841 | NA | NA | NA | NA |
| WBGene00173842 | NA | NA | NA | NA |
| WBGene00173843 | NA | NA | NA | NA |
| WBGene00173844 | NA | NA | NA | NA |
| WBGene00173845 | NA | NA | NA | NA |
| WBGene00173846 | NA | NA | NA | NA |
| WBGene00173847 | NA | NA | NA | NA |
| WBGene00173848 | NA | NA | NA | NA |
| WBGene00173849 | NA | NA | NA | NA |
| WBGene00173850 | NA | NA | NA | NA |
| WBGene00173851 | NA | NA | NA | NA |
| WBGene00173852 | NA | NA | NA | NA |
| WBGene00173853 | NA | NA | NA | NA |
| WBGene00173854 | NA | NA | NA | NA |
| WBGene00173855 | NA | NA | NA | NA |
| WBGene00173856 | NA | NA | NA | NA |
| WBGene00173857 | NA | NA | NA | NA |
| WBGene00173858 | NA | NA | NA | NA |
| WBGene00173859 | NA | NA | NA | NA |
| WBGene00173860 | NA | NA | NA | NA |
| WBGene00173861 | NA | NA | NA | NA |
| WBGene00173862 | NA | NA | NA | NA |
| WBGene00173863 | NA | NA | NA | NA |
| WBGene00173864 | NA | NA | NA | NA |
| WBGene00173865 | NA | NA | NA | NA |
| WBGene00173866 | NA | NA | NA | NA |
| WBGene00173867 | NA | NA | NA | NA |
| WBGene00173868 | NA | NA | NA | NA |
| WBGene00173869 | NA | NA | NA | NA |
| WBGene00173870 | NA | NA | NA | NA |
| WBGene00173871 | NA | NA | NA | NA |
| WBGene00173872 | NA | NA | NA | NA |
| WBGene00173873 | NA | NA | NA | NA |
| WBGene00173874 | NA | NA | NA | NA |
| WBGene00173875 | NA | NA | NA | NA |
| WBGene00173876 | NA | NA | NA | NA |
| WBGene00173877 | NA | NA | NA | NA |
| WBGene00173878 | NA | NA | NA | NA |
| WBGene00173879 | NA | NA | NA | NA |
| WBGene00173880 | NA | NA | NA | NA |

| | | | | |
|---|---|---|---|---|
| WBGene00173881 | NA | NA | NA | NA |
| WBGene00173882 | NA | NA | NA | NA |
| WBGene00173883 | NA | NA | NA | NA |
| WBGene00173884 | NA | NA | NA | NA |
| WBGene00173885 | NA | NA | NA | NA |
| WBGene00173886 | NA | NA | NA | NA |
| WBGene00173887 | NA | NA | NA | NA |
| WBGene00173888 | NA | NA | NA | NA |
| WBGene00173889 | NA | NA | NA | NA |
| WBGene00173890 | NA | NA | NA | NA |
| WBGene00173891 | NA | NA | NA | NA |
| WBGene00173892 | NA | NA | NA | NA |
| WBGene00173893 | NA | NA | NA | NA |
| WBGene00173894 | NA | NA | NA | NA |
| WBGene00173895 | NA | NA | NA | NA |
| WBGene00173896 | NA | NA | NA | NA |
| WBGene00173897 | NA | NA | NA | NA |
| WBGene00173898 | NA | NA | NA | NA |
| WBGene00173899 | NA | NA | NA | NA |
| WBGene00173900 | NA | NA | NA | NA |
| WBGene00173901 | NA | NA | NA | NA |
| WBGene00173902 | -0.022930767 | 0.78121253 | NA | NA |
| WBGene00173903 | NA | NA | NA | NA |
| WBGene00173904 | NA | NA | NA | NA |
| WBGene00173905 | NA | NA | NA | NA |
| WBGene00173906 | NA | NA | NA | NA |
| WBGene00173907 | NA | NA | NA | NA |
| WBGene00173908 | NA | NA | NA | NA |
| WBGene00173909 | NA | NA | NA | NA |
| WBGene00173910 | NA | NA | NA | NA |
| WBGene00173911 | NA | NA | NA | NA |
| WBGene00173912 | NA | NA | NA | NA |
| WBGene00173913 | NA | NA | NA | NA |
| WBGene00173914 | NA | NA | NA | NA |
| WBGene00173915 | NA | NA | NA | NA |
| WBGene00173916 | NA | NA | NA | NA |
| WBGene00173917 | NA | NA | NA | NA |
| WBGene00173918 | NA | NA | NA | NA |
| WBGene00173919 | NA | NA | NA | NA |
| WBGene00173920 | NA | NA | NA | NA |
| WBGene00173921 | NA | NA | NA | NA |
| WBGene00173922 | NA | NA | NA | NA |
| WBGene00173923 | NA | NA | NA | NA |

| | | | | |
|---|---|---|---|---|
| WBGene00173924 | NA | NA | NA | NA |
| WBGene00173925 | NA | NA | NA | NA |
| WBGene00173926 | NA | NA | NA | NA |
| WBGene00173927 | NA | NA | NA | NA |
| WBGene00173928 | NA | NA | NA | NA |
| WBGene00173929 | NA | NA | NA | NA |
| WBGene00173930 | NA | NA | NA | NA |
| WBGene00173931 | NA | NA | NA | NA |
| WBGene00173932 | NA | NA | NA | NA |
| WBGene00173933 | NA | NA | NA | NA |
| WBGene00173934 | NA | NA | NA | NA |
| WBGene00173935 | NA | NA | NA | NA |
| WBGene00173936 | NA | NA | NA | NA |
| WBGene00173937 | NA | NA | NA | NA |
| WBGene00173938 | NA | NA | NA | NA |
| WBGene00173939 | NA | NA | NA | NA |
| WBGene00173940 | NA | NA | NA | NA |
| WBGene00173941 | NA | NA | NA | NA |
| WBGene00173942 | NA | NA | NA | NA |
| WBGene00173943 | NA | NA | NA | NA |
| WBGene00173944 | NA | NA | NA | NA |
| WBGene00173945 | NA | NA | NA | NA |
| WBGene00173946 | NA | NA | NA | NA |
| WBGene00173947 | NA | NA | NA | NA |
| WBGene00173948 | NA | NA | NA | NA |
| WBGene00173949 | NA | NA | NA | NA |
| WBGene00173950 | NA | NA | NA | NA |
| WBGene00173951 | NA | NA | NA | NA |
| WBGene00173952 | NA | NA | NA | NA |
| WBGene00173953 | NA | NA | NA | NA |
| WBGene00173954 | NA | NA | NA | NA |
| WBGene00173955 | NA | NA | NA | NA |
| WBGene00173956 | NA | NA | NA | NA |
| WBGene00173957 | NA | NA | NA | NA |
| WBGene00173958 | NA | NA | NA | NA |
| WBGene00173959 | NA | NA | NA | NA |
| WBGene00173960 | NA | NA | NA | NA |
| WBGene00173961 | NA | NA | NA | NA |
| WBGene00173962 | NA | NA | NA | NA |
| WBGene00173963 | NA | NA | NA | NA |
| WBGene00173964 | NA | NA | NA | NA |
| WBGene00173965 | NA | NA | NA | NA |
| WBGene00173966 | NA | NA | NA | NA |

| | | | | |
|---|---|---|---|---|
| WBGene00173967 | NA | NA | NA | NA |
| WBGene00173968 | NA | NA | NA | NA |
| WBGene00173969 | NA | NA | NA | NA |
| WBGene00173970 | NA | NA | NA | NA |
| WBGene00173971 | NA | NA | NA | NA |
| WBGene00173972 | NA | NA | NA | NA |
| WBGene00173973 | NA | NA | NA | NA |
| WBGene00173974 | NA | NA | NA | NA |
| WBGene00173975 | NA | NA | NA | NA |
| WBGene00173976 | NA | NA | NA | NA |
| WBGene00173977 | NA | NA | NA | NA |
| WBGene00173978 | NA | NA | NA | NA |
| WBGene00173979 | NA | NA | NA | NA |
| WBGene00173980 | NA | NA | NA | NA |
| WBGene00173981 | NA | NA | NA | NA |
| WBGene00173982 | NA | NA | NA | NA |
| WBGene00173983 | NA | NA | NA | NA |
| WBGene00173984 | NA | NA | NA | NA |
| WBGene00173985 | NA | NA | NA | NA |
| WBGene00173986 | NA | NA | NA | NA |
| WBGene00173987 | NA | NA | NA | NA |
| WBGene00173988 | NA | NA | NA | NA |
| WBGene00173989 | NA | NA | NA | NA |
| WBGene00173990 | NA | NA | NA | NA |
| WBGene00173991 | NA | NA | NA | NA |
| WBGene00173992 | NA | NA | NA | NA |
| WBGene00173993 | NA | NA | NA | NA |
| WBGene00173994 | NA | NA | NA | NA |
| WBGene00173995 | NA | NA | NA | NA |
| WBGene00173996 | NA | NA | NA | NA |
| WBGene00173997 | NA | NA | NA | NA |
| WBGene00173998 | NA | NA | NA | NA |
| WBGene00173999 | NA | NA | NA | NA |
| WBGene00174000 | NA | NA | NA | NA |
| WBGene00174001 | NA | NA | NA | NA |
| WBGene00174002 | NA | NA | NA | NA |
| WBGene00174003 | NA | NA | NA | NA |
| WBGene00174004 | NA | NA | NA | NA |
| WBGene00174005 | NA | NA | NA | NA |
| WBGene00174006 | NA | NA | NA | NA |
| WBGene00174007 | NA | NA | NA | NA |
| WBGene00174008 | NA | NA | NA | NA |
| WBGene00174009 | NA | NA | NA | NA |

| | | | | |
|---|---|---|---|---|
| WBGene00174010 | NA | NA | NA | NA |
| WBGene00174011 | NA | NA | NA | NA |
| WBGene00174012 | NA | NA | NA | NA |
| WBGene00174013 | NA | NA | NA | NA |
| WBGene00174014 | NA | NA | NA | NA |
| WBGene00174015 | NA | NA | NA | NA |
| WBGene00174016 | NA | NA | NA | NA |
| WBGene00174017 | NA | NA | NA | NA |
| WBGene00174018 | NA | NA | NA | NA |
| WBGene00174019 | NA | NA | NA | NA |
| WBGene00174020 | NA | NA | NA | NA |
| WBGene00174021 | NA | NA | NA | NA |
| WBGene00174022 | NA | NA | NA | NA |
| WBGene00174023 | NA | NA | NA | NA |
| WBGene00174024 | NA | NA | NA | NA |
| WBGene00174025 | NA | NA | NA | NA |
| WBGene00174026 | NA | NA | NA | NA |
| WBGene00174027 | NA | NA | NA | NA |
| WBGene00174028 | NA | NA | NA | NA |
| WBGene00174029 | NA | NA | NA | NA |
| WBGene00174030 | NA | NA | NA | NA |
| WBGene00174031 | NA | NA | NA | NA |
| WBGene00174032 | NA | NA | NA | NA |
| WBGene00174033 | NA | NA | NA | NA |
| WBGene00174034 | NA | NA | NA | NA |
| WBGene00174035 | NA | NA | NA | NA |
| WBGene00174036 | NA | NA | NA | NA |
| WBGene00174037 | NA | NA | NA | NA |
| WBGene00174038 | -0.023056499 | 0.78121253 | NA | NA |
| WBGene00174039 | NA | NA | NA | NA |
| WBGene00174040 | NA | NA | NA | NA |
| WBGene00174041 | NA | NA | NA | NA |
| WBGene00174042 | NA | NA | NA | NA |
| WBGene00174043 | NA | NA | NA | NA |
| WBGene00174044 | NA | NA | NA | NA |
| WBGene00174045 | NA | NA | NA | NA |
| WBGene00174046 | NA | NA | NA | NA |
| WBGene00174047 | NA | NA | NA | NA |
| WBGene00174048 | NA | NA | NA | NA |
| WBGene00174049 | NA | NA | NA | NA |
| WBGene00174050 | NA | NA | NA | NA |
| WBGene00174051 | NA | NA | NA | NA |
| WBGene00174052 | NA | NA | NA | NA |

| | | | | |
|---|---|---|---|---|
| WBGene00174053 | NA | NA | NA | NA |
| WBGene00174054 | NA | NA | NA | NA |
| WBGene00174055 | NA | NA | NA | NA |
| WBGene00174056 | NA | NA | NA | NA |
| WBGene00174057 | NA | NA | NA | NA |
| WBGene00174058 | NA | NA | NA | NA |
| WBGene00174059 | NA | NA | NA | NA |
| WBGene00174060 | NA | NA | NA | NA |
| WBGene00174061 | NA | NA | NA | NA |
| WBGene00174062 | NA | NA | NA | NA |
| WBGene00174063 | NA | NA | NA | NA |
| WBGene00174064 | NA | NA | NA | NA |
| WBGene00174065 | NA | NA | NA | NA |
| WBGene00174066 | NA | NA | NA | NA |
| WBGene00174067 | NA | NA | NA | NA |
| WBGene00174068 | NA | NA | NA | NA |
| WBGene00174069 | NA | NA | NA | NA |
| WBGene00174070 | NA | NA | NA | NA |
| WBGene00174071 | NA | NA | NA | NA |
| WBGene00174072 | NA | NA | NA | NA |
| WBGene00174073 | NA | NA | NA | NA |
| WBGene00174074 | NA | NA | NA | NA |
| WBGene00174075 | NA | NA | NA | NA |
| WBGene00174076 | NA | NA | NA | NA |
| WBGene00174077 | NA | NA | NA | NA |
| WBGene00174078 | NA | NA | NA | NA |
| WBGene00174079 | NA | NA | NA | NA |
| WBGene00174080 | NA | NA | NA | NA |
| WBGene00174081 | NA | NA | NA | NA |
| WBGene00174082 | NA | NA | NA | NA |
| WBGene00174083 | NA | NA | NA | NA |
| WBGene00174084 | NA | NA | NA | NA |
| WBGene00174085 | NA | NA | NA | NA |
| WBGene00174086 | NA | NA | NA | NA |
| WBGene00174087 | NA | NA | NA | NA |
| WBGene00174088 | NA | NA | NA | NA |
| WBGene00174089 | NA | NA | NA | NA |
| WBGene00174090 | NA | NA | NA | NA |
| WBGene00174091 | NA | NA | NA | NA |
| WBGene00174092 | NA | NA | NA | NA |
| WBGene00174093 | NA | NA | NA | NA |
| WBGene00174094 | NA | NA | NA | NA |
| WBGene00174095 | NA | NA | NA | NA |

| | | | | |
|---|---|---|---|---|
| WBGene00174096 | NA | NA | NA | NA |
| WBGene00174097 | NA | NA | NA | NA |
| WBGene00174098 | NA | NA | NA | NA |
| WBGene00174099 | NA | NA | NA | NA |
| WBGene00174100 | NA | NA | NA | NA |
| WBGene00174101 | NA | NA | NA | NA |
| WBGene00174102 | NA | NA | NA | NA |
| WBGene00174103 | NA | NA | NA | NA |
| WBGene00174104 | NA | NA | NA | NA |
| WBGene00174105 | NA | NA | NA | NA |
| WBGene00174106 | NA | NA | NA | NA |
| WBGene00174107 | NA | NA | NA | NA |
| WBGene00174108 | NA | NA | NA | NA |
| WBGene00174109 | NA | NA | NA | NA |
| WBGene00174110 | NA | NA | NA | NA |
| WBGene00174111 | NA | NA | NA | NA |
| WBGene00174112 | NA | NA | NA | NA |
| WBGene00174113 | NA | NA | NA | NA |
| WBGene00174114 | NA | NA | NA | NA |
| WBGene00174115 | NA | NA | NA | NA |
| WBGene00174116 | NA | NA | NA | NA |
| WBGene00174117 | NA | NA | NA | NA |
| WBGene00174118 | NA | NA | NA | NA |
| WBGene00174119 | NA | NA | NA | NA |
| WBGene00174120 | NA | NA | NA | NA |
| WBGene00174121 | NA | NA | NA | NA |
| WBGene00174122 | NA | NA | NA | NA |
| WBGene00174123 | NA | NA | NA | NA |
| WBGene00174124 | NA | NA | NA | NA |
| WBGene00174125 | NA | NA | NA | NA |
| WBGene00174126 | NA | NA | NA | NA |
| WBGene00174127 | NA | NA | NA | NA |
| WBGene00174128 | NA | NA | NA | NA |
| WBGene00174129 | NA | NA | NA | NA |
| WBGene00174130 | NA | NA | NA | NA |
| WBGene00174131 | NA | NA | NA | NA |
| WBGene00174132 | NA | NA | NA | NA |
| WBGene00174133 | NA | NA | NA | NA |
| WBGene00174134 | NA | NA | NA | NA |
| WBGene00174135 | NA | NA | NA | NA |
| WBGene00174136 | NA | NA | NA | NA |
| WBGene00174137 | NA | NA | NA | NA |
| WBGene00174138 | NA | NA | NA | NA |

| | | | | |
|---|---|---|---|---|
| WBGene00174139 | NA | NA | NA | NA |
| WBGene00174140 | NA | NA | NA | NA |
| WBGene00174141 | NA | NA | NA | NA |
| WBGene00174142 | NA | NA | NA | NA |
| WBGene00174143 | NA | NA | NA | NA |
| WBGene00174144 | NA | NA | NA | NA |
| WBGene00174145 | NA | NA | NA | NA |
| WBGene00174146 | NA | NA | NA | NA |
| WBGene00174147 | NA | NA | NA | NA |
| WBGene00174148 | NA | NA | NA | NA |
| WBGene00174149 | NA | NA | NA | NA |
| WBGene00174150 | NA | NA | NA | NA |
| WBGene00174151 | NA | NA | NA | NA |
| WBGene00174152 | NA | NA | NA | NA |
| WBGene00174153 | NA | NA | NA | NA |
| WBGene00174154 | NA | NA | NA | NA |
| WBGene00174155 | NA | NA | NA | NA |
| WBGene00174156 | NA | NA | NA | NA |
| WBGene00174157 | NA | NA | NA | NA |
| WBGene00174158 | NA | NA | NA | NA |
| WBGene00174159 | NA | NA | NA | NA |
| WBGene00174160 | NA | NA | NA | NA |
| WBGene00174161 | NA | NA | NA | NA |
| WBGene00174162 | NA | NA | NA | NA |
| WBGene00174163 | NA | NA | NA | NA |
| WBGene00174164 | NA | NA | NA | NA |
| WBGene00174165 | NA | NA | NA | NA |
| WBGene00174166 | NA | NA | NA | NA |
| WBGene00174167 | NA | NA | NA | NA |
| WBGene00174168 | NA | NA | NA | NA |
| WBGene00174169 | NA | NA | NA | NA |
| WBGene00174170 | NA | NA | NA | NA |
| WBGene00174171 | NA | NA | NA | NA |
| WBGene00174172 | NA | NA | NA | NA |
| WBGene00174173 | NA | NA | NA | NA |
| WBGene00174174 | NA | NA | NA | NA |
| WBGene00174175 | NA | NA | NA | NA |
| WBGene00174176 | NA | NA | NA | NA |
| WBGene00174177 | NA | NA | NA | NA |
| WBGene00174178 | NA | NA | NA | NA |
| WBGene00174179 | NA | NA | NA | NA |
| WBGene00174180 | NA | NA | NA | NA |
| WBGene00174181 | NA | NA | NA | NA |

| | | | | |
|---|---|---|---|---|
| WBGene00174182 | NA | NA | NA | NA |
| WBGene00174183 | NA | NA | NA | NA |
| WBGene00174184 | 0.020925399 | 0.84643798 | NA | NA |
| WBGene00174185 | NA | NA | NA | NA |
| WBGene00174186 | NA | NA | NA | NA |
| WBGene00174187 | NA | NA | NA | NA |
| WBGene00174188 | NA | NA | NA | NA |
| WBGene00174189 | NA | NA | NA | NA |
| WBGene00174190 | NA | NA | NA | NA |
| WBGene00174191 | NA | NA | NA | NA |
| WBGene00174192 | NA | NA | NA | NA |
| WBGene00174193 | NA | NA | NA | NA |
| WBGene00174194 | NA | NA | NA | NA |
| WBGene00174195 | NA | NA | NA | NA |
| WBGene00174196 | NA | NA | NA | NA |
| WBGene00174197 | NA | NA | NA | NA |
| WBGene00174198 | NA | NA | NA | NA |
| WBGene00174199 | NA | NA | NA | NA |
| WBGene00174200 | NA | NA | NA | NA |
| WBGene00174201 | NA | NA | NA | NA |
| WBGene00174202 | NA | NA | NA | NA |
| WBGene00174203 | NA | NA | NA | NA |
| WBGene00174204 | NA | NA | NA | NA |
| WBGene00174205 | NA | NA | NA | NA |
| WBGene00174206 | NA | NA | NA | NA |
| WBGene00174207 | NA | NA | NA | NA |
| WBGene00174208 | NA | NA | NA | NA |
| WBGene00174209 | NA | NA | NA | NA |
| WBGene00174210 | NA | NA | NA | NA |
| WBGene00174211 | NA | NA | NA | NA |
| WBGene00174212 | NA | NA | NA | NA |
| WBGene00174213 | NA | NA | NA | NA |
| WBGene00174214 | NA | NA | NA | NA |
| WBGene00174215 | NA | NA | NA | NA |
| WBGene00174216 | NA | NA | NA | NA |
| WBGene00174217 | NA | NA | NA | NA |
| WBGene00174218 | NA | NA | NA | NA |
| WBGene00174219 | NA | NA | NA | NA |
| WBGene00174220 | NA | NA | NA | NA |
| WBGene00174221 | NA | NA | NA | NA |
| WBGene00174222 | NA | NA | NA | NA |
| WBGene00174223 | NA | NA | NA | NA |
| WBGene00174224 | NA | NA | NA | NA |

| | | | | |
|---|---|---|---|---|
| WBGene00174225 | NA | NA | NA | NA |
| WBGene00174226 | NA | NA | NA | NA |
| WBGene00174227 | NA | NA | NA | NA |
| WBGene00174228 | NA | NA | NA | NA |
| WBGene00174229 | NA | NA | NA | NA |
| WBGene00174230 | NA | NA | NA | NA |
| WBGene00174231 | NA | NA | NA | NA |
| WBGene00174232 | NA | NA | NA | NA |
| WBGene00174233 | NA | NA | NA | NA |
| WBGene00174234 | NA | NA | NA | NA |
| WBGene00174235 | NA | NA | NA | NA |
| WBGene00174236 | NA | NA | NA | NA |
| WBGene00174237 | NA | NA | NA | NA |
| WBGene00174238 | NA | NA | NA | NA |
| WBGene00174239 | NA | NA | NA | NA |
| WBGene00174240 | NA | NA | NA | NA |
| WBGene00174241 | NA | NA | NA | NA |
| WBGene00174242 | NA | NA | NA | NA |
| WBGene00174243 | NA | NA | NA | NA |
| WBGene00174244 | NA | NA | NA | NA |
| WBGene00174245 | NA | NA | NA | NA |
| WBGene00174246 | NA | NA | NA | NA |
| WBGene00174247 | NA | NA | NA | NA |
| WBGene00174248 | NA | NA | NA | NA |
| WBGene00174249 | NA | NA | NA | NA |
| WBGene00174250 | NA | NA | NA | NA |
| WBGene00174251 | NA | NA | NA | NA |
| WBGene00174252 | NA | NA | NA | NA |
| WBGene00174253 | NA | NA | NA | NA |
| WBGene00174254 | NA | NA | NA | NA |
| WBGene00174255 | NA | NA | NA | NA |
| WBGene00174256 | NA | NA | NA | NA |
| WBGene00174257 | NA | NA | NA | NA |
| WBGene00174258 | NA | NA | NA | NA |
| WBGene00174259 | NA | NA | NA | NA |
| WBGene00174260 | NA | NA | NA | NA |
| WBGene00174261 | NA | NA | NA | NA |
| WBGene00174262 | NA | NA | NA | NA |
| WBGene00174263 | NA | NA | NA | NA |
| WBGene00174264 | NA | NA | NA | NA |
| WBGene00174265 | NA | NA | NA | NA |
| WBGene00174266 | NA | NA | NA | NA |
| WBGene00174267 | NA | NA | NA | NA |

| | | | | |
|---|---|---|---|---|
| WBGene00174268 | NA | NA | NA | NA |
| WBGene00174269 | NA | NA | NA | NA |
| WBGene00174270 | 0.171941111 | 0.04983501 | NA | NA |
| WBGene00174271 | NA | NA | NA | NA |
| WBGene00174272 | NA | NA | NA | NA |
| WBGene00174273 | NA | NA | NA | NA |
| WBGene00174274 | NA | NA | NA | NA |
| WBGene00174275 | NA | NA | NA | NA |
| WBGene00174276 | NA | NA | NA | NA |
| WBGene00174277 | NA | NA | NA | NA |
| WBGene00174278 | NA | NA | NA | NA |
| WBGene00174279 | NA | NA | NA | NA |
| WBGene00174280 | NA | NA | NA | NA |
| WBGene00174281 | NA | NA | NA | NA |
| WBGene00174282 | NA | NA | NA | NA |
| WBGene00174283 | NA | NA | NA | NA |
| WBGene00174284 | NA | NA | NA | NA |
| WBGene00174285 | NA | NA | NA | NA |
| WBGene00174286 | NA | NA | NA | NA |
| WBGene00174287 | NA | NA | NA | NA |
| WBGene00174288 | NA | NA | NA | NA |
| WBGene00174289 | NA | NA | NA | NA |
| WBGene00174290 | NA | NA | NA | NA |
| WBGene00174291 | NA | NA | NA | NA |
| WBGene00174292 | NA | NA | NA | NA |
| WBGene00174293 | NA | NA | NA | NA |
| WBGene00174294 | NA | NA | NA | NA |
| WBGene00174295 | NA | NA | NA | NA |
| WBGene00174296 | NA | NA | NA | NA |
| WBGene00174297 | NA | NA | NA | NA |
| WBGene00174298 | NA | NA | NA | NA |
| WBGene00174299 | NA | NA | NA | NA |
| WBGene00174300 | NA | NA | NA | NA |
| WBGene00174301 | NA | NA | NA | NA |
| WBGene00174302 | NA | NA | NA | NA |
| WBGene00174303 | NA | NA | NA | NA |
| WBGene00174304 | NA | NA | NA | NA |
| WBGene00174305 | NA | NA | NA | NA |
| WBGene00174306 | NA | NA | NA | NA |
| WBGene00174307 | NA | NA | NA | NA |
| WBGene00174308 | NA | NA | NA | NA |
| WBGene00174309 | NA | NA | NA | NA |
| WBGene00174310 | NA | NA | NA | NA |

| | | | | |
|---|---|---|---|---|
| WBGene00174311 | -0.023056499 | 0.78121253 | NA | NA |
| WBGene00174312 | NA | NA | NA | NA |
| WBGene00174313 | NA | NA | NA | NA |
| WBGene00174314 | NA | NA | NA | NA |
| WBGene00174315 | -0.023056499 | 0.78121253 | NA | NA |
| WBGene00174316 | NA | NA | NA | NA |
| WBGene00174317 | NA | NA | NA | NA |
| WBGene00174318 | NA | NA | NA | NA |
| WBGene00174319 | NA | NA | NA | NA |
| WBGene00174320 | NA | NA | NA | NA |
| WBGene00174321 | NA | NA | NA | NA |
| WBGene00174322 | -0.04044783 | 0.53612608 | NA | NA |
| WBGene00174323 | NA | NA | NA | NA |
| WBGene00174324 | NA | NA | NA | NA |
| WBGene00174325 | NA | NA | NA | NA |
| WBGene00174326 | NA | NA | NA | NA |
| WBGene00174327 | NA | NA | NA | NA |
| WBGene00174328 | NA | NA | NA | NA |
| WBGene00174329 | NA | NA | NA | NA |
| WBGene00174330 | NA | NA | NA | NA |
| WBGene00174331 | NA | NA | NA | NA |
| WBGene00174332 | NA | NA | NA | NA |
| WBGene00174333 | NA | NA | NA | NA |
| WBGene00174334 | NA | NA | NA | NA |
| WBGene00174335 | NA | NA | NA | NA |
| WBGene00174336 | NA | NA | NA | NA |
| WBGene00174337 | NA | NA | NA | NA |
| WBGene00174338 | NA | NA | NA | NA |
| WBGene00174339 | NA | NA | NA | NA |
| WBGene00174340 | NA | NA | NA | NA |
| WBGene00174341 | NA | NA | NA | NA |
| WBGene00174342 | NA | NA | NA | NA |
| WBGene00174343 | NA | NA | NA | NA |
| WBGene00174344 | NA | NA | NA | NA |
| WBGene00174345 | NA | NA | NA | NA |
| WBGene00174346 | NA | NA | NA | NA |
| WBGene00174347 | NA | NA | NA | NA |
| WBGene00174348 | NA | NA | NA | NA |
| WBGene00174349 | NA | NA | NA | NA |
| WBGene00174350 | NA | NA | NA | NA |
| WBGene00174351 | NA | NA | NA | NA |
| WBGene00174352 | NA | NA | NA | NA |
| WBGene00174353 | NA | NA | NA | NA |

| | | | | |
|---|---|---|---|---|
| WBGene00174354 | NA | NA | NA | NA |
| WBGene00174355 | NA | NA | NA | NA |
| WBGene00174356 | NA | NA | NA | NA |
| WBGene00174357 | NA | NA | NA | NA |
| WBGene00174358 | NA | NA | NA | NA |
| WBGene00174359 | NA | NA | NA | NA |
| WBGene00174360 | NA | NA | NA | NA |
| WBGene00174361 | NA | NA | NA | NA |
| WBGene00174362 | NA | NA | NA | NA |
| WBGene00174363 | NA | NA | NA | NA |
| WBGene00174364 | NA | NA | NA | NA |
| WBGene00174365 | NA | NA | NA | NA |
| WBGene00174366 | NA | NA | NA | NA |
| WBGene00174367 | NA | NA | NA | NA |
| WBGene00174368 | NA | NA | NA | NA |
| WBGene00174369 | NA | NA | NA | NA |
| WBGene00174370 | NA | NA | NA | NA |
| WBGene00174371 | NA | NA | NA | NA |
| WBGene00174372 | NA | NA | NA | NA |
| WBGene00174373 | NA | NA | NA | NA |
| WBGene00174374 | NA | NA | NA | NA |
| WBGene00174375 | NA | NA | NA | NA |
| WBGene00174376 | NA | NA | NA | NA |
| WBGene00174377 | NA | NA | NA | NA |
| WBGene00174378 | NA | NA | NA | NA |
| WBGene00174379 | NA | NA | NA | NA |
| WBGene00174380 | NA | NA | NA | NA |
| WBGene00174381 | NA | NA | NA | NA |
| WBGene00174382 | NA | NA | NA | NA |
| WBGene00174383 | NA | NA | NA | NA |
| WBGene00174384 | NA | NA | NA | NA |
| WBGene00174385 | NA | NA | NA | NA |
| WBGene00174386 | NA | NA | NA | NA |
| WBGene00174387 | NA | NA | NA | NA |
| WBGene00174388 | NA | NA | NA | NA |
| WBGene00174389 | NA | NA | NA | NA |
| WBGene00174390 | NA | NA | NA | NA |
| WBGene00174391 | NA | NA | NA | NA |
| WBGene00174392 | NA | NA | NA | NA |
| WBGene00174393 | NA | NA | NA | NA |
| WBGene00174394 | NA | NA | NA | NA |
| WBGene00174395 | NA | NA | NA | NA |
| WBGene00174396 | NA | NA | NA | NA |

| | | | | |
|---|---|---|---|---|
| WBGene00174397 | NA | NA | NA | NA |
| WBGene00174398 | NA | NA | NA | NA |
| WBGene00174399 | NA | NA | NA | NA |
| WBGene00174400 | NA | NA | NA | NA |
| WBGene00174401 | NA | NA | NA | NA |
| WBGene00174402 | NA | NA | NA | NA |
| WBGene00174403 | NA | NA | NA | NA |
| WBGene00174404 | NA | NA | NA | NA |
| WBGene00174405 | NA | NA | NA | NA |
| WBGene00174406 | NA | NA | NA | NA |
| WBGene00174407 | NA | NA | NA | NA |
| WBGene00174408 | NA | NA | NA | NA |
| WBGene00174409 | NA | NA | NA | NA |
| WBGene00174410 | NA | NA | NA | NA |
| WBGene00174411 | NA | NA | NA | NA |
| WBGene00174412 | NA | NA | NA | NA |
| WBGene00174413 | NA | NA | NA | NA |
| WBGene00174414 | NA | NA | NA | NA |
| WBGene00174415 | NA | NA | NA | NA |
| WBGene00174416 | NA | NA | NA | NA |
| WBGene00174417 | NA | NA | NA | NA |
| WBGene00174418 | NA | NA | NA | NA |
| WBGene00174419 | NA | NA | NA | NA |
| WBGene00174420 | NA | NA | NA | NA |
| WBGene00174421 | NA | NA | NA | NA |
| WBGene00174422 | NA | NA | NA | NA |
| WBGene00174423 | NA | NA | NA | NA |
| WBGene00174424 | NA | NA | NA | NA |
| WBGene00174425 | NA | NA | NA | NA |
| WBGene00174426 | NA | NA | NA | NA |
| WBGene00174427 | NA | NA | NA | NA |
| WBGene00174428 | NA | NA | NA | NA |
| WBGene00174429 | NA | NA | NA | NA |
| WBGene00174430 | NA | NA | NA | NA |
| WBGene00174431 | NA | NA | NA | NA |
| WBGene00174432 | NA | NA | NA | NA |
| WBGene00174433 | NA | NA | NA | NA |
| WBGene00174434 | NA | NA | NA | NA |
| WBGene00174435 | NA | NA | NA | NA |
| WBGene00174436 | NA | NA | NA | NA |
| WBGene00174437 | NA | NA | NA | NA |
| WBGene00174438 | NA | NA | NA | NA |
| WBGene00174439 | NA | NA | NA | NA |

| | | | | |
|---|---|---|---|---|
| WBGene00174440 | NA | NA | NA | NA |
| WBGene00174441 | NA | NA | NA | NA |
| WBGene00174442 | NA | NA | NA | NA |
| WBGene00174443 | NA | NA | NA | NA |
| WBGene00174444 | NA | NA | NA | NA |
| WBGene00174445 | NA | NA | NA | NA |
| WBGene00174446 | NA | NA | NA | NA |
| WBGene00174447 | NA | NA | NA | NA |
| WBGene00174448 | NA | NA | NA | NA |
| WBGene00174449 | NA | NA | NA | NA |
| WBGene00174450 | NA | NA | NA | NA |
| WBGene00174451 | NA | NA | NA | NA |
| WBGene00174452 | NA | NA | NA | NA |
| WBGene00174453 | NA | NA | NA | NA |
| WBGene00174454 | NA | NA | NA | NA |
| WBGene00174455 | NA | NA | NA | NA |
| WBGene00174456 | NA | NA | NA | NA |
| WBGene00174457 | NA | NA | NA | NA |
| WBGene00174458 | NA | NA | NA | NA |
| WBGene00174459 | NA | NA | NA | NA |
| WBGene00174460 | NA | NA | NA | NA |
| WBGene00174461 | NA | NA | NA | NA |
| WBGene00174462 | NA | NA | NA | NA |
| WBGene00174463 | NA | NA | NA | NA |
| WBGene00174464 | NA | NA | NA | NA |
| WBGene00174465 | NA | NA | NA | NA |
| WBGene00174466 | NA | NA | NA | NA |
| WBGene00174467 | NA | NA | NA | NA |
| WBGene00174468 | NA | NA | NA | NA |
| WBGene00174469 | NA | NA | NA | NA |
| WBGene00174470 | NA | NA | NA | NA |
| WBGene00174471 | NA | NA | NA | NA |
| WBGene00174472 | NA | NA | NA | NA |
| WBGene00174473 | NA | NA | NA | NA |
| WBGene00174474 | NA | NA | NA | NA |
| WBGene00174475 | NA | NA | NA | NA |
| WBGene00174476 | NA | NA | NA | NA |
| WBGene00174477 | NA | NA | NA | NA |
| WBGene00174478 | NA | NA | NA | NA |
| WBGene00174479 | NA | NA | NA | NA |
| WBGene00174480 | NA | NA | NA | NA |
| WBGene00174481 | NA | NA | NA | NA |
| WBGene00174482 | NA | NA | NA | NA |

| | | | | |
|---|---|---|---|---|
| WBGene00174483 | NA | NA | NA | NA |
| WBGene00174484 | NA | NA | NA | NA |
| WBGene00174485 | NA | NA | NA | NA |
| WBGene00174486 | NA | NA | NA | NA |
| WBGene00174487 | NA | NA | NA | NA |
| WBGene00174488 | NA | NA | NA | NA |
| WBGene00174489 | NA | NA | NA | NA |
| WBGene00174490 | NA | NA | NA | NA |
| WBGene00174491 | NA | NA | NA | NA |
| WBGene00174492 | NA | NA | NA | NA |
| WBGene00174493 | NA | NA | NA | NA |
| WBGene00174494 | NA | NA | NA | NA |
| WBGene00174495 | NA | NA | NA | NA |
| WBGene00174496 | NA | NA | NA | NA |
| WBGene00174497 | NA | NA | NA | NA |
| WBGene00174498 | NA | NA | NA | NA |
| WBGene00174499 | NA | NA | NA | NA |
| WBGene00174500 | NA | NA | NA | NA |
| WBGene00174501 | NA | NA | NA | NA |
| WBGene00174502 | NA | NA | NA | NA |
| WBGene00174503 | NA | NA | NA | NA |
| WBGene00174504 | NA | NA | NA | NA |
| WBGene00174505 | NA | NA | NA | NA |
| WBGene00174506 | NA | NA | NA | NA |
| WBGene00174507 | NA | NA | NA | NA |
| WBGene00174508 | NA | NA | NA | NA |
| WBGene00174509 | NA | NA | NA | NA |
| WBGene00174510 | NA | NA | NA | NA |
| WBGene00174511 | NA | NA | NA | NA |
| WBGene00174512 | NA | NA | NA | NA |
| WBGene00174513 | NA | NA | NA | NA |
| WBGene00174514 | NA | NA | NA | NA |
| WBGene00174515 | NA | NA | NA | NA |
| WBGene00174516 | NA | NA | NA | NA |
| WBGene00174517 | NA | NA | NA | NA |
| WBGene00174518 | NA | NA | NA | NA |
| WBGene00174519 | NA | NA | NA | NA |
| WBGene00174520 | NA | NA | NA | NA |
| WBGene00174521 | NA | NA | NA | NA |
| WBGene00174522 | NA | NA | NA | NA |
| WBGene00174523 | NA | NA | NA | NA |
| WBGene00174524 | NA | NA | NA | NA |
| WBGene00174525 | NA | NA | NA | NA |

| | | | | |
|---|---|---|---|---|
| WBGene00174526 | NA | NA | NA | NA |
| WBGene00174527 | NA | NA | NA | NA |
| WBGene00174528 | NA | NA | NA | NA |
| WBGene00174529 | NA | NA | NA | NA |
| WBGene00174530 | NA | NA | NA | NA |
| WBGene00174531 | NA | NA | NA | NA |
| WBGene00174532 | NA | NA | NA | NA |
| WBGene00174533 | NA | NA | NA | NA |
| WBGene00174534 | NA | NA | NA | NA |
| WBGene00174535 | NA | NA | NA | NA |
| WBGene00174536 | NA | NA | NA | NA |
| WBGene00174537 | NA | NA | NA | NA |
| WBGene00174538 | NA | NA | NA | NA |
| WBGene00174539 | NA | NA | NA | NA |
| WBGene00174540 | NA | NA | NA | NA |
| WBGene00174541 | NA | NA | NA | NA |
| WBGene00174542 | NA | NA | NA | NA |
| WBGene00174543 | NA | NA | NA | NA |
| WBGene00174544 | NA | NA | NA | NA |
| WBGene00174545 | NA | NA | NA | NA |
| WBGene00174546 | NA | NA | NA | NA |
| WBGene00174547 | NA | NA | NA | NA |
| WBGene00174548 | NA | NA | NA | NA |
| WBGene00174549 | NA | NA | NA | NA |
| WBGene00174550 | NA | NA | NA | NA |
| WBGene00174551 | NA | NA | NA | NA |
| WBGene00174552 | NA | NA | NA | NA |
| WBGene00174553 | NA | NA | NA | NA |
| WBGene00174554 | NA | NA | NA | NA |
| WBGene00174555 | NA | NA | NA | NA |
| WBGene00174556 | NA | NA | NA | NA |
| WBGene00174557 | NA | NA | NA | NA |
| WBGene00174558 | NA | NA | NA | NA |
| WBGene00174559 | NA | NA | NA | NA |
| WBGene00174560 | NA | NA | NA | NA |
| WBGene00174561 | NA | NA | NA | NA |
| WBGene00174562 | NA | NA | NA | NA |
| WBGene00174563 | NA | NA | NA | NA |
| WBGene00174564 | NA | NA | NA | NA |
| WBGene00174565 | NA | NA | NA | NA |
| WBGene00174566 | NA | NA | NA | NA |
| WBGene00174567 | NA | NA | NA | NA |
| WBGene00174568 | NA | NA | NA | NA |

| | | | | |
|---|---|---|---|---|
| WBGene00174569 | NA | NA | NA | NA |
| WBGene00174570 | 0.04909525 | 0.45360729 | NA | NA |
| WBGene00174571 | -0.023109762 | 0.78121253 | NA | NA |
| WBGene00174572 | NA | NA | NA | NA |
| WBGene00174573 | NA | NA | NA | NA |
| WBGene00174574 | NA | NA | NA | NA |
| WBGene00174575 | NA | NA | NA | NA |
| WBGene00174576 | NA | NA | NA | NA |
| WBGene00174577 | NA | NA | NA | NA |
| WBGene00174578 | NA | NA | NA | NA |
| WBGene00174579 | NA | NA | NA | NA |
| WBGene00174580 | NA | NA | NA | NA |
| WBGene00174581 | NA | NA | NA | NA |
| WBGene00174582 | NA | NA | NA | NA |
| WBGene00174583 | NA | NA | NA | NA |
| WBGene00174584 | NA | NA | NA | NA |
| WBGene00174585 | NA | NA | NA | NA |
| WBGene00174586 | NA | NA | NA | NA |
| WBGene00174587 | NA | NA | NA | NA |
| WBGene00174588 | NA | NA | NA | NA |
| WBGene00174589 | NA | NA | NA | NA |
| WBGene00174590 | NA | NA | NA | NA |
| WBGene00174591 | NA | NA | NA | NA |
| WBGene00174592 | NA | NA | NA | NA |
| WBGene00174593 | NA | NA | NA | NA |
| WBGene00174594 | NA | NA | NA | NA |
| WBGene00174595 | NA | NA | NA | NA |
| WBGene00174596 | NA | NA | NA | NA |
| WBGene00174597 | NA | NA | NA | NA |
| WBGene00174598 | NA | NA | NA | NA |
| WBGene00174599 | NA | NA | NA | NA |
| WBGene00174600 | NA | NA | NA | NA |
| WBGene00174601 | NA | NA | NA | NA |
| WBGene00174602 | NA | NA | NA | NA |
| WBGene00174603 | NA | NA | NA | NA |
| WBGene00174604 | NA | NA | NA | NA |
| WBGene00174605 | NA | NA | NA | NA |
| WBGene00174606 | NA | NA | NA | NA |
| WBGene00174607 | NA | NA | NA | NA |
| WBGene00174608 | NA | NA | NA | NA |
| WBGene00174609 | NA | NA | NA | NA |
| WBGene00174610 | NA | NA | NA | NA |
| WBGene00174611 | NA | NA | NA | NA |

| | | | | |
|---|---|---|---|---|
| WBGene00174612 | NA | NA | NA | NA |
| WBGene00174613 | NA | NA | NA | NA |
| WBGene00174614 | NA | NA | NA | NA |
| WBGene00174615 | NA | NA | NA | NA |
| WBGene00174616 | NA | NA | NA | NA |
| WBGene00174617 | NA | NA | NA | NA |
| WBGene00174618 | NA | NA | NA | NA |
| WBGene00174619 | NA | NA | NA | NA |
| WBGene00174620 | NA | NA | NA | NA |
| WBGene00174621 | NA | NA | NA | NA |
| WBGene00174622 | NA | NA | NA | NA |
| WBGene00174623 | NA | NA | NA | NA |
| WBGene00174624 | NA | NA | NA | NA |
| WBGene00174625 | NA | NA | NA | NA |
| WBGene00174626 | NA | NA | NA | NA |
| WBGene00174627 | NA | NA | NA | NA |
| WBGene00174628 | NA | NA | NA | NA |
| WBGene00174629 | NA | NA | NA | NA |
| WBGene00174630 | NA | NA | NA | NA |
| WBGene00174631 | NA | NA | NA | NA |
| WBGene00174632 | NA | NA | NA | NA |
| WBGene00174633 | NA | NA | NA | NA |
| WBGene00174634 | NA | NA | NA | NA |
| WBGene00174635 | NA | NA | NA | NA |
| WBGene00174636 | NA | NA | NA | NA |
| WBGene00174637 | NA | NA | NA | NA |
| WBGene00174638 | NA | NA | NA | NA |
| WBGene00174639 | NA | NA | NA | NA |
| WBGene00174640 | NA | NA | NA | NA |
| WBGene00174641 | NA | NA | NA | NA |
| WBGene00174642 | NA | NA | NA | NA |
| WBGene00174643 | NA | NA | NA | NA |
| WBGene00174644 | NA | NA | NA | NA |
| WBGene00174645 | NA | NA | NA | NA |
| WBGene00174646 | 0.020317245 | 0.84643798 | NA | NA |
| WBGene00174647 | NA | NA | NA | NA |
| WBGene00174648 | NA | NA | NA | NA |
| WBGene00174649 | NA | NA | NA | NA |
| WBGene00174650 | NA | NA | NA | NA |
| WBGene00174651 | NA | NA | NA | NA |
| WBGene00174652 | NA | NA | NA | NA |
| WBGene00174653 | NA | NA | NA | NA |
| WBGene00174654 | NA | NA | NA | NA |

| | | | | |
|---|---|---|---|---|
| WBGene00174655 | NA | NA | NA | NA |
| WBGene00174656 | NA | NA | NA | NA |
| WBGene00174657 | NA | NA | NA | NA |
| WBGene00174658 | NA | NA | NA | NA |
| WBGene00174659 | NA | NA | NA | NA |
| WBGene00174660 | NA | NA | NA | NA |
| WBGene00174661 | NA | NA | NA | NA |
| WBGene00174662 | NA | NA | NA | NA |
| WBGene00174663 | NA | NA | NA | NA |
| WBGene00174664 | NA | NA | NA | NA |
| WBGene00174665 | NA | NA | NA | NA |
| WBGene00174666 | NA | NA | NA | NA |
| WBGene00174667 | NA | NA | NA | NA |
| WBGene00174668 | NA | NA | NA | NA |
| WBGene00174669 | NA | NA | NA | NA |
| WBGene00174670 | NA | NA | NA | NA |
| WBGene00174671 | NA | NA | NA | NA |
| WBGene00174672 | NA | NA | NA | NA |
| WBGene00174673 | NA | NA | NA | NA |
| WBGene00174674 | NA | NA | NA | NA |
| WBGene00174675 | NA | NA | NA | NA |
| WBGene00174676 | NA | NA | NA | NA |
| WBGene00174677 | NA | NA | NA | NA |
| WBGene00174678 | NA | NA | NA | NA |
| WBGene00174679 | NA | NA | NA | NA |
| WBGene00174680 | NA | NA | NA | NA |
| WBGene00174681 | NA | NA | NA | NA |
| WBGene00174682 | NA | NA | NA | NA |
| WBGene00174683 | NA | NA | NA | NA |
| WBGene00174684 | NA | NA | NA | NA |
| WBGene00174685 | NA | NA | NA | NA |
| WBGene00174686 | NA | NA | NA | NA |
| WBGene00174687 | NA | NA | NA | NA |
| WBGene00174688 | NA | NA | NA | NA |
| WBGene00174689 | NA | NA | NA | NA |
| WBGene00174690 | NA | NA | NA | NA |
| WBGene00174691 | NA | NA | NA | NA |
| WBGene00174692 | NA | NA | NA | NA |
| WBGene00174693 | NA | NA | NA | NA |
| WBGene00174694 | NA | NA | NA | NA |
| WBGene00174695 | NA | NA | NA | NA |
| WBGene00174696 | NA | NA | NA | NA |
| WBGene00174697 | NA | NA | NA | NA |

| | | | | |
|---|---|---|---|---|
| WBGene00174698 | NA | NA | NA | NA |
| WBGene00174699 | NA | NA | NA | NA |
| WBGene00174700 | NA | NA | NA | NA |
| WBGene00174701 | NA | NA | NA | NA |
| WBGene00174702 | NA | NA | NA | NA |
| WBGene00174703 | NA | NA | NA | NA |
| WBGene00174704 | NA | NA | NA | NA |
| WBGene00174705 | NA | NA | NA | NA |
| WBGene00174706 | NA | NA | NA | NA |
| WBGene00174707 | NA | NA | NA | NA |
| WBGene00174708 | NA | NA | NA | NA |
| WBGene00174709 | NA | NA | NA | NA |
| WBGene00174710 | NA | NA | NA | NA |
| WBGene00174711 | NA | NA | NA | NA |
| WBGene00174712 | NA | NA | NA | NA |
| WBGene00174713 | NA | NA | NA | NA |
| WBGene00174714 | NA | NA | NA | NA |
| WBGene00174715 | NA | NA | NA | NA |
| WBGene00174716 | NA | NA | NA | NA |
| WBGene00174717 | NA | NA | NA | NA |
| WBGene00174718 | NA | NA | NA | NA |
| WBGene00174719 | NA | NA | NA | NA |
| WBGene00174720 | NA | NA | NA | NA |
| WBGene00174721 | NA | NA | NA | NA |
| WBGene00174722 | NA | NA | NA | NA |
| WBGene00174723 | NA | NA | NA | NA |
| WBGene00174724 | NA | NA | NA | NA |
| WBGene00174725 | NA | NA | NA | NA |
| WBGene00174726 | NA | NA | NA | NA |
| WBGene00174727 | NA | NA | NA | NA |
| WBGene00174728 | NA | NA | NA | NA |
| WBGene00174729 | NA | NA | NA | NA |
| WBGene00174730 | NA | NA | NA | NA |
| WBGene00174731 | NA | NA | NA | NA |
| WBGene00174732 | NA | NA | NA | NA |
| WBGene00174733 | NA | NA | NA | NA |
| WBGene00174734 | NA | NA | NA | NA |
| WBGene00174735 | NA | NA | NA | NA |
| WBGene00174736 | NA | NA | NA | NA |
| WBGene00174737 | NA | NA | NA | NA |
| WBGene00174738 | NA | NA | NA | NA |
| WBGene00174739 | NA | NA | NA | NA |
| WBGene00174740 | NA | NA | NA | NA |

| | | | | |
|---|---|---|---|---|
| WBGene00174741 | NA | NA | NA | NA |
| WBGene00174742 | NA | NA | NA | NA |
| WBGene00174743 | NA | NA | NA | NA |
| WBGene00174744 | NA | NA | NA | NA |
| WBGene00174745 | NA | NA | NA | NA |
| WBGene00174746 | NA | NA | NA | NA |
| WBGene00174747 | NA | NA | NA | NA |
| WBGene00174748 | NA | NA | NA | NA |
| WBGene00174749 | NA | NA | NA | NA |
| WBGene00174750 | NA | NA | NA | NA |
| WBGene00174751 | NA | NA | NA | NA |
| WBGene00174752 | NA | NA | NA | NA |
| WBGene00174753 | -0.023056499 | 0.78121253 | NA | NA |
| WBGene00174754 | NA | NA | NA | NA |
| WBGene00174755 | NA | NA | NA | NA |
| WBGene00174756 | NA | NA | NA | NA |
| WBGene00174757 | NA | NA | NA | NA |
| WBGene00174758 | NA | NA | NA | NA |
| WBGene00174759 | NA | NA | NA | NA |
| WBGene00174760 | NA | NA | NA | NA |
| WBGene00174761 | NA | NA | NA | NA |
| WBGene00174762 | NA | NA | NA | NA |
| WBGene00174763 | NA | NA | NA | NA |
| WBGene00174764 | NA | NA | NA | NA |
| WBGene00174765 | NA | NA | NA | NA |
| WBGene00174766 | NA | NA | NA | NA |
| WBGene00174767 | NA | NA | NA | NA |
| WBGene00174768 | NA | NA | NA | NA |
| WBGene00174769 | NA | NA | NA | NA |
| WBGene00174770 | NA | NA | NA | NA |
| WBGene00174771 | -0.023056499 | 0.78121253 | NA | NA |
| WBGene00174772 | NA | NA | NA | NA |
| WBGene00174773 | NA | NA | NA | NA |
| WBGene00174774 | NA | NA | NA | NA |
| WBGene00174775 | NA | NA | NA | NA |
| WBGene00174776 | NA | NA | NA | NA |
| WBGene00174777 | NA | NA | NA | NA |
| WBGene00174778 | NA | NA | NA | NA |
| WBGene00174779 | NA | NA | NA | NA |
| WBGene00174780 | NA | NA | NA | NA |
| WBGene00174781 | NA | NA | NA | NA |
| WBGene00174782 | NA | NA | NA | NA |
| WBGene00174783 | NA | NA | NA | NA |

| | | | | |
|---|---|---|---|---|
| WBGene00174784 | NA | NA | NA | NA |
| WBGene00174785 | NA | NA | NA | NA |
| WBGene00174786 | NA | NA | NA | NA |
| WBGene00174787 | NA | NA | NA | NA |
| WBGene00174788 | NA | NA | NA | NA |
| WBGene00174789 | NA | NA | NA | NA |
| WBGene00174790 | NA | NA | NA | NA |
| WBGene00174791 | NA | NA | NA | NA |
| WBGene00174792 | NA | NA | NA | NA |
| WBGene00174793 | NA | NA | NA | NA |
| WBGene00174794 | NA | NA | NA | NA |
| WBGene00174795 | NA | NA | NA | NA |
| WBGene00174796 | NA | NA | NA | NA |
| WBGene00174797 | NA | NA | NA | NA |
| WBGene00174798 | NA | NA | NA | NA |
| WBGene00174799 | NA | NA | NA | NA |
| WBGene00174800 | NA | NA | NA | NA |
| WBGene00174801 | NA | NA | NA | NA |
| WBGene00174802 | NA | NA | NA | NA |
| WBGene00174803 | 0.020317245 | 0.84643798 | NA | NA |
| WBGene00174804 | NA | NA | NA | NA |
| WBGene00174805 | NA | NA | NA | NA |
| WBGene00174806 | NA | NA | NA | NA |
| WBGene00174807 | NA | NA | NA | NA |
| WBGene00174808 | NA | NA | NA | NA |
| WBGene00174809 | NA | NA | NA | NA |
| WBGene00174810 | NA | NA | NA | NA |
| WBGene00174811 | NA | NA | NA | NA |
| WBGene00174812 | NA | NA | NA | NA |
| WBGene00174813 | NA | NA | NA | NA |
| WBGene00174814 | NA | NA | NA | NA |
| WBGene00174815 | NA | NA | NA | NA |
| WBGene00174816 | NA | NA | NA | NA |
| WBGene00174817 | NA | NA | NA | NA |
| WBGene00174818 | NA | NA | NA | NA |
| WBGene00174819 | NA | NA | NA | NA |
| WBGene00174820 | NA | NA | NA | NA |
| WBGene00174821 | NA | NA | NA | NA |
| WBGene00174822 | NA | NA | NA | NA |
| WBGene00174823 | NA | NA | NA | NA |
| WBGene00174824 | NA | NA | NA | NA |
| WBGene00174825 | NA | NA | NA | NA |
| WBGene00174826 | NA | NA | NA | NA |

| | | | | |
|---|---|---|---|---|
| WBGene00174827 | NA | NA | NA | NA |
| WBGene00174828 | NA | NA | NA | NA |
| WBGene00174829 | NA | NA | NA | NA |
| WBGene00174830 | NA | NA | NA | NA |
| WBGene00174831 | NA | NA | NA | NA |
| WBGene00174832 | NA | NA | NA | NA |
| WBGene00174833 | NA | NA | NA | NA |
| WBGene00174834 | NA | NA | NA | NA |
| WBGene00174835 | NA | NA | NA | NA |
| WBGene00174836 | NA | NA | NA | NA |
| WBGene00174837 | 0.020232312 | 0.84643798 | NA | NA |
| WBGene00174838 | NA | NA | NA | NA |
| WBGene00174839 | NA | NA | NA | NA |
| WBGene00174840 | NA | NA | NA | NA |
| WBGene00174841 | NA | NA | NA | NA |
| WBGene00174842 | NA | NA | NA | NA |
| WBGene00174843 | NA | NA | NA | NA |
| WBGene00174844 | NA | NA | NA | NA |
| WBGene00174845 | NA | NA | NA | NA |
| WBGene00174846 | NA | NA | NA | NA |
| WBGene00174847 | NA | NA | NA | NA |
| WBGene00174848 | NA | NA | NA | NA |
| WBGene00174849 | NA | NA | NA | NA |
| WBGene00174850 | NA | NA | NA | NA |
| WBGene00174851 | NA | NA | NA | NA |
| WBGene00174852 | NA | NA | NA | NA |
| WBGene00174853 | NA | NA | NA | NA |
| WBGene00174854 | NA | NA | NA | NA |
| WBGene00174855 | NA | NA | NA | NA |
| WBGene00174856 | NA | NA | NA | NA |
| WBGene00174857 | NA | NA | NA | NA |
| WBGene00174858 | NA | NA | NA | NA |
| WBGene00174859 | NA | NA | NA | NA |
| WBGene00174860 | NA | NA | NA | NA |
| WBGene00174861 | NA | NA | NA | NA |
| WBGene00174862 | NA | NA | NA | NA |
| WBGene00174863 | NA | NA | NA | NA |
| WBGene00174864 | NA | NA | NA | NA |
| WBGene00174865 | NA | NA | NA | NA |
| WBGene00174866 | NA | NA | NA | NA |
| WBGene00174867 | NA | NA | NA | NA |
| WBGene00174868 | NA | NA | NA | NA |
| WBGene00174869 | NA | NA | NA | NA |

| | | | | |
|---|---|---|---|---|
| WBGene00174870 | NA | NA | NA | NA |
| WBGene00174871 | NA | NA | NA | NA |
| WBGene00174872 | NA | NA | NA | NA |
| WBGene00174873 | NA | NA | NA | NA |
| WBGene00174874 | NA | NA | NA | NA |
| WBGene00174875 | NA | NA | NA | NA |
| WBGene00174876 | NA | NA | NA | NA |
| WBGene00174877 | NA | NA | NA | NA |
| WBGene00174878 | NA | NA | NA | NA |
| WBGene00174879 | NA | NA | NA | NA |
| WBGene00174880 | NA | NA | NA | NA |
| WBGene00174881 | NA | NA | NA | NA |
| WBGene00174882 | NA | NA | NA | NA |
| WBGene00174883 | NA | NA | NA | NA |
| WBGene00174884 | NA | NA | NA | NA |
| WBGene00174885 | NA | NA | NA | NA |
| WBGene00174886 | NA | NA | NA | NA |
| WBGene00174887 | NA | NA | NA | NA |
| WBGene00174888 | NA | NA | NA | NA |
| WBGene00174889 | NA | NA | NA | NA |
| WBGene00174890 | NA | NA | NA | NA |
| WBGene00174891 | NA | NA | NA | NA |
| WBGene00174892 | NA | NA | NA | NA |
| WBGene00174893 | NA | NA | NA | NA |
| WBGene00174894 | NA | NA | NA | NA |
| WBGene00174895 | NA | NA | NA | NA |
| WBGene00174896 | NA | NA | NA | NA |
| WBGene00174897 | NA | NA | NA | NA |
| WBGene00174898 | NA | NA | NA | NA |
| WBGene00174899 | NA | NA | NA | NA |
| WBGene00174900 | NA | NA | NA | NA |
| WBGene00174901 | NA | NA | NA | NA |
| WBGene00174902 | NA | NA | NA | NA |
| WBGene00174903 | NA | NA | NA | NA |
| WBGene00174904 | NA | NA | NA | NA |
| WBGene00174905 | NA | NA | NA | NA |
| WBGene00174906 | NA | NA | NA | NA |
| WBGene00174907 | NA | NA | NA | NA |
| WBGene00174908 | NA | NA | NA | NA |
| WBGene00174909 | NA | NA | NA | NA |
| WBGene00174910 | NA | NA | NA | NA |
| WBGene00174911 | NA | NA | NA | NA |
| WBGene00174912 | NA | NA | NA | NA |

| | | | | |
|---|---|---|---|---|
| WBGene00174913 | NA | NA | NA | NA |
| WBGene00174914 | NA | NA | NA | NA |
| WBGene00174915 | NA | NA | NA | NA |
| WBGene00174916 | NA | NA | NA | NA |
| WBGene00174917 | NA | NA | NA | NA |
| WBGene00174918 | NA | NA | NA | NA |
| WBGene00174919 | NA | NA | NA | NA |
| WBGene00174920 | NA | NA | NA | NA |
| WBGene00174921 | NA | NA | NA | NA |
| WBGene00174922 | NA | NA | NA | NA |
| WBGene00174923 | NA | NA | NA | NA |
| WBGene00174924 | NA | NA | NA | NA |
| WBGene00174925 | NA | NA | NA | NA |
| WBGene00174926 | NA | NA | NA | NA |
| WBGene00174927 | NA | NA | NA | NA |
| WBGene00174928 | NA | NA | NA | NA |
| WBGene00174929 | NA | NA | NA | NA |
| WBGene00174930 | NA | NA | NA | NA |
| WBGene00174931 | NA | NA | NA | NA |
| WBGene00174932 | NA | NA | NA | NA |
| WBGene00174933 | NA | NA | NA | NA |
| WBGene00174934 | NA | NA | NA | NA |
| WBGene00174935 | NA | NA | NA | NA |
| WBGene00174936 | 0.020317245 | 0.84643798 | NA | NA |
| WBGene00174937 | NA | NA | NA | NA |
| WBGene00174938 | NA | NA | NA | NA |
| WBGene00174939 | NA | NA | NA | NA |
| WBGene00174940 | NA | NA | NA | NA |
| WBGene00174941 | NA | NA | NA | NA |
| WBGene00174942 | NA | NA | NA | NA |
| WBGene00174943 | NA | NA | NA | NA |
| WBGene00174944 | NA | NA | NA | NA |
| WBGene00174945 | NA | NA | NA | NA |
| WBGene00174946 | NA | NA | NA | NA |
| WBGene00174947 | NA | NA | NA | NA |
| WBGene00174948 | NA | NA | NA | NA |
| WBGene00174949 | NA | NA | NA | NA |
| WBGene00174950 | NA | NA | NA | NA |
| WBGene00174951 | NA | NA | NA | NA |
| WBGene00174952 | NA | NA | NA | NA |
| WBGene00174953 | NA | NA | NA | NA |
| WBGene00174954 | NA | NA | NA | NA |
| WBGene00174955 | NA | NA | NA | NA |

| | | | | |
|---|---|---|---|---|
| WBGene00174956 | NA | NA | NA | NA |
| WBGene00174957 | NA | NA | NA | NA |
| WBGene00174958 | NA | NA | NA | NA |
| WBGene00174959 | NA | NA | NA | NA |
| WBGene00174960 | NA | NA | NA | NA |
| WBGene00174961 | NA | NA | NA | NA |
| WBGene00174962 | NA | NA | NA | NA |
| WBGene00174963 | NA | NA | NA | NA |
| WBGene00174964 | NA | NA | NA | NA |
| WBGene00174965 | NA | NA | NA | NA |
| WBGene00174966 | NA | NA | NA | NA |
| WBGene00174967 | NA | NA | NA | NA |
| WBGene00174968 | NA | NA | NA | NA |
| WBGene00174969 | NA | NA | NA | NA |
| WBGene00174970 | NA | NA | NA | NA |
| WBGene00174971 | NA | NA | NA | NA |
| WBGene00174972 | NA | NA | NA | NA |
| WBGene00174973 | NA | NA | NA | NA |
| WBGene00174974 | NA | NA | NA | NA |
| WBGene00174975 | NA | NA | NA | NA |
| WBGene00174976 | NA | NA | NA | NA |
| WBGene00174977 | NA | NA | NA | NA |
| WBGene00174978 | NA | NA | NA | NA |
| WBGene00174979 | NA | NA | NA | NA |
| WBGene00174980 | NA | NA | NA | NA |
| WBGene00174981 | NA | NA | NA | NA |
| WBGene00174982 | NA | NA | NA | NA |
| WBGene00174983 | NA | NA | NA | NA |
| WBGene00174984 | NA | NA | NA | NA |
| WBGene00174985 | NA | NA | NA | NA |
| WBGene00174986 | NA | NA | NA | NA |
| WBGene00174987 | NA | NA | NA | NA |
| WBGene00174988 | NA | NA | NA | NA |
| WBGene00174989 | NA | NA | NA | NA |
| WBGene00174990 | NA | NA | NA | NA |
| WBGene00174991 | NA | NA | NA | NA |
| WBGene00174992 | NA | NA | NA | NA |
| WBGene00174993 | NA | NA | NA | NA |
| WBGene00174994 | NA | NA | NA | NA |
| WBGene00174995 | NA | NA | NA | NA |
| WBGene00174996 | NA | NA | NA | NA |
| WBGene00174997 | NA | NA | NA | NA |
| WBGene00174998 | NA | NA | NA | NA |

| | | | | |
|---|---|---|---|---|
| WBGene00174999 | NA | NA | NA | NA |
| WBGene00175000 | NA | NA | NA | NA |
| WBGene00175001 | NA | NA | NA | NA |
| WBGene00175002 | NA | NA | NA | NA |
| WBGene00175003 | NA | NA | NA | NA |
| WBGene00175004 | NA | NA | NA | NA |
| WBGene00175005 | NA | NA | NA | NA |
| WBGene00175006 | NA | NA | NA | NA |
| WBGene00175007 | NA | NA | NA | NA |
| WBGene00175008 | NA | NA | NA | NA |
| WBGene00175009 | NA | NA | NA | NA |
| WBGene00175010 | NA | NA | NA | NA |
| WBGene00175011 | NA | NA | NA | NA |
| WBGene00175012 | NA | NA | NA | NA |
| WBGene00175013 | NA | NA | NA | NA |
| WBGene00175014 | NA | NA | NA | NA |
| WBGene00175015 | NA | NA | NA | NA |
| WBGene00175016 | NA | NA | NA | NA |
| WBGene00175017 | NA | NA | NA | NA |
| WBGene00175018 | NA | NA | NA | NA |
| WBGene00175019 | NA | NA | NA | NA |
| WBGene00175020 | NA | NA | NA | NA |
| WBGene00175021 | NA | NA | NA | NA |
| WBGene00175022 | NA | NA | NA | NA |
| WBGene00175023 | NA | NA | NA | NA |
| WBGene00175024 | NA | NA | NA | NA |
| WBGene00175025 | NA | NA | NA | NA |
| WBGene00175027 | 0.048485426 | 0.45975815 | NA | W02F12.8 |
| WBGene00175028 | -0.022930767 | 0.78121253 | NA | nspe-8 |
| WBGene00175029 | NA | NA | NA | NA |
| WBGene00175030 | -0.078571677 | 0.59987914 | 0.75772845 | Y53C12A.10 |
| WBGene00175031 | 0.004783976 | 0.97838285 | 0.98769243 | C06C3.12 |
| WBGene00175034 | -0.003108065 | 0.95728506 | NA | F54B3.4 |
| WBGene00175035 | 0.003450034 | 0.97917496 | 0.98812489 | F54C9.14 |
| WBGene00175036 | -0.027535543 | 0.68063395 | NA | NA |
| WBGene00184956 | NA | NA | NA | F49C5.12 |
| WBGene00184990 | 0.037426138 | 0.79823094 | 0.88702326 | C14B1.12 |
| WBGene00185001 | -0.392782465 | 0.09060939 | 0.22888093 | F56A8.9 |
| WBGene00185002 | -0.293459654 | 0.0226388 | 0.08415323 | K11H3.8 |
| WBGene00185005 | -0.658616008 | 0.03359003 | 0.11248666 | T03F6.10 |
| WBGene00185006 | 0.020317245 | 0.84643798 | NA | F59B2.15 |
| WBGene00185014 | -0.2531091 | 0.17892164 | 0.36025413 | F17C8.9 |
| WBGene00185048 | -0.087665835 | 0.65368542 | 0.794835 | urm-1 |

| | | | | |
|---|---|---|---|---|
| WBGene00185055 | 0.005252628 | 0.98030388 | 0.98860925 | Y47D3B.12 |
| WBGene00185067 | -0.269025035 | 0.07724416 | 0.20566135 | Y54G11A.17 |
| WBGene00185076 | 0.121439489 | 0.26182825 | 0.45969869 | NA |
| WBGene00185077 | 0.119185029 | 0.36207125 | 0.56173415 | F58D2.4 |
| WBGene00185078 | -0.035644833 | 0.81600385 | 0.89891744 | F40G9.17 |
| WBGene00185079 | 0.093688625 | 0.58971985 | 0.75112054 | W10D9.6 |
| WBGene00185086 | -0.286598647 | 0.00997489 | 0.04478454 | F59A3.12 |
| WBGene00185088 | -0.307211872 | 0.16232658 | 0.33739454 | Y55F3BL.4 |
| WBGene00185089 | 0.155188708 | 0.44752126 | 0.64048473 | C16A11.10 |
| WBGene00185092 | 0.310875315 | 0.14805083 | 0.31876649 | K08D8.7 |
| WBGene00185093 | NA | NA | NA | NA |
| WBGene00185095 | 0.033463284 | 0.6365037 | NA | W09C2.7 |
| WBGene00185096 | NA | NA | NA | W09C2.9 |
| WBGene00185097 | -0.009020277 | 0.91686706 | NA | C49C3.20 |
| WBGene00185098 | -0.222598497 | 0.31196066 | 0.51266736 | Y71G12B.35 |
| WBGene00185100 | NA | NA | NA | T12A7.9 |
| WBGene00185101 | NA | NA | NA | Y65A5A.24 |
| WBGene00185102 | -0.050223056 | 0.47776088 | NA | Y105C5A.1268 |
| WBGene00185116 | NA | NA | NA | W09C2.8 |
| WBGene00185117 | -0.023056499 | 0.78121253 | NA | C45B11.9 |
| WBGene00185118 | 0.201839397 | 0.15427286 | 0.32715947 | K02E2.11 |
| WBGene00185119 | -0.005691347 | 0.93660638 | NA | NA |
| WBGene00189930 | NA | NA | NA | Y26G10.5 |
| WBGene00189931 | NA | NA | NA | Y26G10.6 |
| WBGene00189932 | NA | NA | NA | Y26G10.7 |
| WBGene00189933 | 0.015062379 | 0.80838968 | NA | F38B7.11 |
| WBGene00189937 | 0.036687355 | 0.59489568 | NA | Y6G8.15 |
| WBGene00189943 | NA | NA | NA | Y116F11B.17 |
| WBGene00189944 | 0.085537126 | 0.24698522 | NA | NA |
| WBGene00189946 | NA | NA | NA | NA |
| WBGene00189947 | 0.018790812 | 0.76755469 | NA | C06B3.16 |
| WBGene00189948 | -0.023056499 | 0.78121253 | NA | F56H9.8 |
| WBGene00189949 | 0.535136728 | 0.0127181 | 0.05420835 | Y48G1C.13 |
| WBGene00189950 | 0.040019663 | 0.56356856 | NA | C39B5.14 |
| WBGene00189951 | 0.239408657 | 0.27022568 | 0.46948786 | NA |
| WBGene00189952 | 0.223258487 | 0.12476038 | 0.28523059 | K10D2.8 |
| WBGene00189953 | NA | NA | NA | R155.5 |
| WBGene00189954 | -0.164316497 | 0.34334879 | 0.54313812 | F40G9.18 |
| WBGene00189955 | -0.022930767 | 0.78121253 | NA | F40G9.19 |
| WBGene00189956 | 2.54970279 | 0.00535223 | 0.02761171 | Y34F4.6 |
| WBGene00189957 | -0.022930767 | 0.78121253 | NA | M03F8.7 |
| WBGene00189958 | -0.20903565 | 0.29855201 | 0.49933345 | F45H11.8 |
| WBGene00189961 | NA | NA | NA | Y54G2A.57 |

| | | | | |
|---|---|---|---|---|
| WBGene00189964 | NA | NA | NA | NA |
| WBGene00189966 | NA | NA | NA | NA |
| WBGene00189967 | NA | NA | NA | NA |
| WBGene00189968 | NA | NA | NA | NA |
| WBGene00189969 | NA | NA | NA | NA |
| WBGene00189970 | NA | NA | NA | NA |
| WBGene00189971 | NA | NA | NA | NA |
| WBGene00189972 | NA | NA | NA | NA |
| WBGene00189973 | NA | NA | NA | NA |
| WBGene00189974 | NA | NA | NA | NA |
| WBGene00189975 | NA | NA | NA | NA |
| WBGene00189976 | NA | NA | NA | NA |
| WBGene00189977 | NA | NA | NA | NA |
| WBGene00189978 | NA | NA | NA | NA |
| WBGene00189979 | -0.050007053 | 0.47951277 | NA | NA |
| WBGene00189980 | NA | NA | NA | NA |
| WBGene00189981 | NA | NA | NA | NA |
| WBGene00189982 | NA | NA | NA | NA |
| WBGene00189983 | NA | NA | NA | NA |
| WBGene00189984 | NA | NA | NA | NA |
| WBGene00189985 | NA | NA | NA | NA |
| WBGene00189986 | NA | NA | NA | NA |
| WBGene00189987 | NA | NA | NA | C18H7.12 |
| WBGene00189993 | -0.023056499 | 0.78121253 | NA | C49C3.21 |
| WBGene00189995 | 0.036692668 | 0.59311964 | NA | C06E7.88 |
| WBGene00194641 | 0.39756 | 0.09567834 | 0.23807385 | F47G3.4 |
| WBGene00194642 | 0.219794743 | 0.27558553 | 0.47572289 | C25H3.17 |
| WBGene00194643 | -0.027373974 | 0.68245543 | NA | C48D1.9 |
| WBGene00194644 | 0.020232312 | 0.84643798 | NA | NA |
| WBGene00194645 | NA | NA | NA | Y105C5A.1269 |
| WBGene00194646 | 0.020317245 | 0.84643798 | NA | Y105C5A.1270 |
| WBGene00194647 | 0.056188364 | 0.42092875 | NA | Y51H4A.935 |
| WBGene00194649 | 0.119946747 | 0.48003614 | 0.66658152 | Y57G11C.1130 |
| WBGene00194650 | -0.023056499 | 0.78121253 | NA | NA |
| WBGene00194651 | -0.100061045 | 0.63102327 | 0.78083263 | C33D9.13 |
| WBGene00194652 | -0.077397239 | 0.49126487 | 0.67495527 | NA |
| WBGene00194653 | NA | NA | NA | F36H2.6 |
| WBGene00194654 | 0.056430134 | 0.44291702 | NA | M05B5.7 |
| WBGene00194656 | NA | NA | NA | T05F1.15 |
| WBGene00194657 | 0.0398788 | 0.58971221 | NA | NA |
| WBGene00194658 | 0.017141582 | 0.79355725 | NA | R09E10.13 |
| WBGene00194659 | 0.066255937 | 0.61997653 | 0.77264865 | NA |
| WBGene00194660 | 0.076301704 | 0.42782167 | NA | W09C5.12 |

| | | | | |
|---|---|---|---|---|
| WBGene00194661 | NA | NA | NA | Y57G11C.1131 |
| WBGene00194663 | 0.440797278 | 0.08956757 | 0.22735069 | F38B2.6 |
| WBGene00194664 | 0.106997704 | 0.16248985 | NA | K09E9.7 |
| WBGene00194665 | -0.064037742 | 0.36180097 | NA | NA |
| WBGene00194666 | NA | NA | NA | NA |
| WBGene00194667 | -0.118808254 | 0.16692368 | NA | NA |
| WBGene00194668 | NA | NA | NA | NA |
| WBGene00194669 | 0.020232312 | 0.84643798 | NA | NA |
| WBGene00194670 | 0.052087141 | 0.64257817 | 0.78833475 | K07E3.9 |
| WBGene00194672 | NA | NA | NA | Y116A8C.463 |
| WBGene00194673 | NA | NA | NA | NA |
| WBGene00194674 | -0.653706406 | 2.54E-08 | 5.30E-07 | F23D12.11 |
| WBGene00194675 | -0.103565832 | 0.41252049 | 0.6101336 | NA |
| WBGene00194676 | NA | NA | NA | NA |
| WBGene00194677 | -0.014735285 | 0.89564173 | 0.94523157 | D1054.19 |
| WBGene00194678 | -0.079857863 | 0.31359805 | NA | NA |
| WBGene00194679 | 0.395427712 | 0.05740565 | 0.16643091 | T26C11.9 |
| WBGene00194680 | 0.017604133 | 0.79437959 | NA | Y17D7C.6 |
| WBGene00194681 | 1.620868419 | 0.00686228 | 0.03348286 | NA |
| WBGene00194682 | -0.083404677 | 0.68758295 | 0.81998681 | F40D4.17 |
| WBGene00194683 | 0.013204071 | 0.88957758 | NA | NA |
| WBGene00194684 | NA | NA | NA | Y48A6B.14 |
| WBGene00194686 | NA | NA | NA | NA |
| WBGene00194687 | 0.257774783 | 0.18257089 | 0.36541631 | F57C12.6 |
| WBGene00194689 | NA | NA | NA | Y38H6C.24 |
| WBGene00194690 | NA | NA | NA | NA |
| WBGene00194691 | -0.00346345 | 0.95708614 | NA | ZK1098.12 |
| WBGene00194692 | 0.976033562 | 0.01778979 | 0.0705937 | T02C12.5 |
| WBGene00194695 | 3.365143575 | 0.00064227 | 0.00481736 | B0462.5 |
| WBGene00194696 | NA | NA | NA | EEED8.18 |
| WBGene00194697 | 0.019325396 | 0.77340931 | NA | F59A3.13 |
| WBGene00194698 | NA | NA | NA | F49D11.11 |
| WBGene00194699 | 0.019325396 | 0.77340931 | NA | Y43F8B.22 |
| WBGene00194700 | 0.195017896 | 0.12460389 | 0.28511378 | B0025.5 |
| WBGene00194701 | 0.020925399 | 0.84643798 | NA | Y23H5B.12 |
| WBGene00194702 | NA | NA | NA | C32E8.12 |
| WBGene00194703 | 1.955339255 | 0.00015644 | 0.00142532 | F26B1.8 |
| WBGene00194704 | NA | NA | NA | T07H3.8 |
| WBGene00194706 | 0.04814143 | 0.73187104 | 0.85001776 | C53A5.16 |
| WBGene00194707 | -0.068102233 | 0.65050857 | 0.79307985 | C53A5.17 |
| WBGene00194708 | NA | NA | NA | Y36E3A.2 |
| WBGene00194710 | 0.015584271 | 0.80362925 | NA | mks-2 |
| WBGene00194711 | 0.020232312 | 0.84643798 | NA | D2062.13 |

| | | | | |
|---|---|---|---|---|
| WBGene00194712 | NA | NA | NA | T11F9.21 |
| WBGene00194713 | 0.049863756 | 0.69102395 | 0.8224095 | F19B10.13 |
| WBGene00194714 | NA | NA | NA | ZC204.17 |
| WBGene00194715 | NA | NA | NA | T25E12.16 |
| WBGene00194716 | -0.060665711 | 0.6685444 | 0.80660557 | NA |
| WBGene00194717 | 4.580866897 | 1.90E-16 | 1.07E-14 | ZC239.22 |
| WBGene00194725 | -0.008533167 | 0.91735298 | NA | T16H12.13 |
| WBGene00194726 | 0.677918337 | 0.02139802 | 0.08072361 | R13A5.15 |
| WBGene00194727 | 0.108614162 | 0.30880061 | 0.50965502 | C18D11.10 |
| WBGene00194729 | 0.020925399 | 0.84643798 | NA | ZK783.7 |
| WBGene00194730 | -0.003270206 | 0.95698316 | NA | C29E4.17 |
| WBGene00194731 | 0.020232312 | 0.84643798 | NA | Y39A1A.27 |
| WBGene00194733 | 0.020317245 | 0.84643798 | NA | Y41C4A.21 |
| WBGene00194734 | NA | NA | NA | T04A6.4 |
| WBGene00194735 | NA | NA | NA | Y82E9BL.19 |
| WBGene00194737 | 0.020232312 | 0.84643798 | NA | Y50D7A.13 |
| WBGene00194738 | -0.023109762 | 0.78121253 | NA | Y46E12A.5 |
| WBGene00194742 | 0.082289338 | 0.54816819 | 0.72092678 | F37C12.21 |
| WBGene00194744 | NA | NA | NA | Y38H6C.25 |
| WBGene00194746 | -0.003421551 | 0.95689871 | NA | pals-20 |
| WBGene00194747 | 0.030154492 | 0.86190318 | 0.92668569 | H05C05.4 |
| WBGene00194748 | 0.020317245 | 0.84643798 | NA | NA |
| WBGene00194759 | NA | NA | NA | NA |
| WBGene00194768 | NA | NA | NA | NA |
| WBGene00194769 | 2.21659445 | 1.27E-08 | 2.79E-07 | NA |
| WBGene00194771 | NA | NA | NA | NA |
| WBGene00194780 | NA | NA | NA | C12D8.20 |
| WBGene00194783 | -0.094856784 | 0.23887585 | NA | K03B8.14 |
| WBGene00194784 | 0.060177748 | 0.41318197 | NA | T11F9.22 |
| WBGene00194785 | -0.003270206 | 0.95698316 | NA | C52E4.12 |
| WBGene00194786 | NA | NA | NA | NA |
| WBGene00194787 | 0.0398788 | 0.58971221 | NA | T16G1.13 |
| WBGene00194788 | NA | NA | NA | R02D5.10 |
| WBGene00194792 | 0.036687355 | 0.59489568 | NA | NA |
| WBGene00194793 | 0.01736423 | 0.90604415 | 0.94916953 | W04G3.13 |
| WBGene00194794 | 0.081897742 | 0.57947321 | 0.74518754 | NA |
| WBGene00194795 | 0.020925399 | 0.84643798 | NA | F42F12.14 |
| WBGene00194799 | NA | NA | NA | Y70C5A.4 |
| WBGene00194800 | 0.019325396 | 0.77340931 | NA | NA |
| WBGene00194801 | 0.020232312 | 0.84643798 | NA | Y6G8.16 |
| WBGene00194802 | 0.020232312 | 0.84643798 | NA | NA |
| WBGene00194803 | 0.180007984 | 0.0451335 | NA | C25F9.16 |
| WBGene00194804 | NA | NA | NA | NA |

| | | | | | |
|---|---|---|---|---|---|
| WBGene00194805 | NA | | NA | NA | NA |
| WBGene00194806 | NA | | NA | NA | NA |
| WBGene00194807 | NA | | NA | NA | NA |
| WBGene00194808 | NA | | NA | NA | NA |
| WBGene00194809 | NA | | NA | NA | NA |
| WBGene00194810 | NA | | NA | NA | NA |
| WBGene00194811 | | 0.017279211 | 0.79380376 | NA | NA |
| WBGene00194812 | | 0.079182682 | 0.70181564 | 0.8306615 | NA |
| WBGene00194814 | | -0.030716767 | 0.73679148 | NA | NA |
| WBGene00194815 | | 5.189440523 | 0.00011754 | 0.00111888 | Y43F8B.23 |
| WBGene00194816 | | 0.178535255 | 0.04064772 | 0.12954359 | ZK616.61 |
| WBGene00194821 | | -0.033040371 | 0.77019441 | 0.86922195 | dmsr-16 |
| WBGene00194822 | NA | | NA | NA | NA |
| WBGene00194823 | NA | | NA | NA | NA |
| WBGene00194824 | NA | | NA | NA | NA |
| WBGene00194825 | NA | | NA | NA | NA |
| WBGene00194826 | NA | | NA | NA | NA |
| WBGene00194827 | NA | | NA | NA | NA |
| WBGene00194828 | NA | | NA | NA | NA |
| WBGene00194829 | NA | | NA | NA | NA |
| WBGene00194830 | NA | | NA | NA | NA |
| WBGene00194831 | NA | | NA | NA | NA |
| WBGene00194832 | NA | | NA | NA | Y73B6BL.270 |
| WBGene00194833 | | 0.020232312 | 0.84643798 | NA | F17E9.15 |
| WBGene00194834 | NA | | NA | NA | NA |
| WBGene00194835 | | 0.016985298 | 0.837043 | NA | Y17G9B.11 |
| WBGene00194836 | | -0.023109762 | 0.78121253 | NA | NA |
| WBGene00194839 | | 0.020317245 | 0.84643798 | NA | C01B10.44 |
| WBGene00194840 | NA | | NA | NA | NA |
| WBGene00194841 | NA | | NA | NA | NA |
| WBGene00194842 | | 0.020925399 | 0.84643798 | NA | NA |
| WBGene00194843 | NA | | NA | NA | NA |
| WBGene00194844 | | 0.085537126 | 0.24698522 | NA | NA |
| WBGene00194847 | NA | | NA | NA | NA |
| WBGene00194848 | | 0.056595066 | 0.62610877 | 0.77700176 | T27D12.6 |
| WBGene00194849 | | 0.490264264 | 0.06818193 | 0.18792388 | K09A9.8 |
| WBGene00194850 | | 0.020232312 | 0.84643798 | NA | F44F4.15 |
| WBGene00194851 | | -0.062042028 | 0.58426481 | 0.74778226 | NA |
| WBGene00194852 | | -0.251232981 | 0.25209206 | 0.45005241 | T28A8.8 |
| WBGene00194853 | NA | | NA | NA | NA |
| WBGene00194854 | NA | | NA | NA | NA |
| WBGene00194855 | | 1.547517658 | 0.00571563 | 0.02911711 | Y75B8A.44 |
| WBGene00194856 | NA | | NA | NA | NA |

| | | | | |
|---|---|---|---|---|
| WBGene00194861 | 0.25115933 | 0.25291591 | 0.4510337 | H04D03.6 |
| WBGene00194864 | 0.085309563 | 0.24820057 | NA | Y66A7A.9 |
| WBGene00194865 | NA | NA | NA | NA |
| WBGene00194866 | 0.018657025 | 0.77508599 | NA | Y41C4A.22 |
| WBGene00194867 | NA | NA | NA | W09C2.10 |
| WBGene00194868 | 0.020317245 | 0.84643798 | NA | M03C11.9 |
| WBGene00194869 | -0.0260089 | 0.6772113 | NA | K01A6.8 |
| WBGene00194870 | 0.017348279 | 0.83364574 | NA | F53A2.11 |
| WBGene00194871 | NA | NA | NA | R107.9 |
| WBGene00194883 | 0.080478428 | 0.4100328 | 0.60780503 | B0496.11 |
| WBGene00194884 | NA | NA | NA | NA |
| WBGene00194885 | NA | NA | NA | NA |
| WBGene00194886 | NA | NA | NA | C05B5.12 |
| WBGene00194888 | NA | NA | NA | K10H10.13 |
| WBGene00194890 | NA | NA | NA | C01B9.4 |
| WBGene00194891 | 0.020317245 | 0.84643798 | NA | NA |
| WBGene00194892 | 0.096060922 | 0.46777497 | 0.65694901 | dsb-2 |
| WBGene00194894 | 0.04909525 | 0.45360729 | NA | T05B4.14 |
| WBGene00194895 | 0.020925399 | 0.84643798 | NA | ZK105.11 |
| WBGene00194896 | -0.00346345 | 0.95708614 | NA | W02H5.11 |
| WBGene00194898 | NA | NA | NA | C17B7.14 |
| WBGene00194902 | 0.154005181 | 0.12099973 | 0.27992663 | Y39H10B.3 |
| WBGene00194905 | 0.052437311 | 0.67746016 | 0.81260807 | Y53F4B.51 |
| WBGene00194906 | -0.046813543 | 0.80436719 | 0.89116459 | Y53C12A.11 |
| WBGene00194907 | NA | NA | NA | E02H4.8 |
| WBGene00194908 | -1.795099112 | 2.55E-09 | 6.10E-08 | F22F7.8 |
| WBGene00194909 | 0.049166068 | 0.71037278 | 0.83686577 | F41E6.17 |
| WBGene00194910 | 0.059675786 | 0.4188661 | NA | ZC404.15 |
| WBGene00194912 | 0.196684894 | 0.03290959 | 0.11071444 | ZK856.18 |
| WBGene00194913 | NA | NA | NA | Y105C5A.1272 |
| WBGene00194914 | NA | NA | NA | F44D12.16 |
| WBGene00194915 | 0.012672266 | 0.86772563 | NA | JC8.16 |
| WBGene00194916 | 0.020317245 | 0.84643798 | NA | F35D2.6 |
| WBGene00194917 | -0.023109762 | 0.78121253 | NA | Y45F10B.59 |
| WBGene00194918 | 0.020232312 | 0.84643798 | NA | K08D8.9 |
| WBGene00194919 | NA | NA | NA | K08D8.10 |
| WBGene00194920 | NA | NA | NA | H12I19.115 |
| WBGene00194921 | 0.436550064 | 0.0547458 | 0.16082127 | ZK742.7 |
| WBGene00194922 | NA | NA | NA | F13E9.19 |
| WBGene00194923 | -0.06730507 | 0.31931151 | NA | NA |
| WBGene00194924 | NA | NA | NA | Y51H4A.936 |
| WBGene00194925 | NA | NA | NA | K08D8.11 |
| WBGene00194926 | 0.062819936 | 0.57356367 | 0.74143178 | K08D8.12 |

| | | | | |
|---|---|---|---|---|
| WBGene00194927 | NA | NA | NA | Y51H4A.937 |
| WBGene00194928 | -0.067666074 | 0.31672907 | NA | ZC513.14 |
| WBGene00194929 | 0.017138317 | 0.82172227 | NA | Y105C5B.1413 |
| WBGene00194930 | NA | NA | NA | Y73F8A.1159 |
| WBGene00194931 | NA | NA | NA | Y105C5A.1273 |
| WBGene00194932 | 0.033034883 | 0.64219503 | NA | NA |
| WBGene00194933 | NA | NA | NA | T12A7.10 |
| WBGene00194934 | 0.077127579 | 0.6765328 | 0.81229431 | ZK809.9 |
| WBGene00194935 | 0.020317245 | 0.84643798 | NA | M02G9.4 |
| WBGene00194939 | -0.105834181 | 0.21658577 | NA | NA |
| WBGene00194940 | NA | NA | NA | C34C6.9 |
| WBGene00194945 | NA | NA | NA | NA |
| WBGene00194951 | NA | NA | NA | Y105C5A.1274 |
| WBGene00194952 | -0.003421551 | 0.95689871 | NA | ZK666.14 |
| WBGene00194955 | -0.003421551 | 0.95689871 | NA | F37H8.6 |
| WBGene00194957 | NA | NA | NA | Y116A8C.465 |
| WBGene00194981 | -0.567041381 | 0.04832359 | 0.14658068 | Y49E10.30 |
| WBGene00194982 | -0.045959069 | 0.50707733 | NA | Y17D7B.10 |
| WBGene00194983 | 0.037818748 | 0.65938735 | NA | T25B2.4 |
| WBGene00194984 | NA | NA | NA | NA |
| WBGene00194985 | 0.005800696 | 0.97742521 | 0.98710921 | Y25C1A.14 |
| WBGene00194986 | 0.093035954 | 0.52202924 | 0.70009244 | pigb-1 |
| WBGene00194987 | NA | NA | NA | NA |
| WBGene00194988 | NA | NA | NA | NA |
| WBGene00194989 | NA | NA | NA | NA |
| WBGene00194990 | NA | NA | NA | NA |
| WBGene00194991 | NA | NA | NA | NA |
| WBGene00194992 | NA | NA | NA | NA |
| WBGene00194993 | NA | NA | NA | NA |
| WBGene00194994 | NA | NA | NA | NA |
| WBGene00194995 | NA | NA | NA | NA |
| WBGene00194996 | NA | NA | NA | NA |
| WBGene00194997 | -0.003270206 | 0.95698316 | NA | NA |
| WBGene00194998 | -0.005523802 | 0.97668004 | 0.98670757 | C16C4.18 |
| WBGene00194999 | NA | NA | NA | NA |
| WBGene00195000 | NA | NA | NA | NA |
| WBGene00195001 | NA | NA | NA | NA |
| WBGene00195002 | NA | NA | NA | NA |
| WBGene00195003 | NA | NA | NA | Y46C8AL.11 |
| WBGene00195004 | 0.059208151 | 0.51856536 | NA | K08D12.8 |
| WBGene00195005 | 0.112443425 | 0.19418407 | NA | NA |
| WBGene00195006 | -0.023056499 | 0.78121253 | NA | C35B1.9 |
| WBGene00195007 | NA | NA | NA | NA |

| | | | | |
|---|---|---|---|---|
| WBGene00195008 | NA | NA | NA | NA |
| WBGene00195009 | NA | NA | NA | F17E9.16 |
| WBGene00195010 | NA | NA | NA | Y37E11AL.12 |
| WBGene00195011 | -0.0260089 | 0.6772113 | NA | ZK180.8 |
| WBGene00195012 | NA | NA | NA | Y38C1BA.4 |
| WBGene00195016 | -0.00346345 | 0.95708614 | NA | NA |
| WBGene00195017 | 0.020925399 | 0.84643798 | NA | NA |
| WBGene00195018 | NA | NA | NA | NA |
| WBGene00195019 | NA | NA | NA | NA |
| WBGene00195020 | NA | NA | NA | NA |
| WBGene00195021 | NA | NA | NA | NA |
| WBGene00195022 | NA | NA | NA | NA |
| WBGene00195023 | NA | NA | NA | NA |
| WBGene00195024 | -0.027188909 | 0.68205301 | NA | NA |
| WBGene00195025 | NA | NA | NA | NA |
| WBGene00195026 | NA | NA | NA | NA |
| WBGene00195027 | NA | NA | NA | NA |
| WBGene00195028 | NA | NA | NA | NA |
| WBGene00195029 | NA | NA | NA | NA |
| WBGene00195030 | NA | NA | NA | NA |
| WBGene00195031 | NA | NA | NA | NA |
| WBGene00195032 | NA | NA | NA | NA |
| WBGene00195033 | NA | NA | NA | NA |
| WBGene00195034 | -0.019569613 | 0.8786347 | 0.93503852 | NA |
| WBGene00195035 | NA | NA | NA | NA |
| WBGene00195036 | NA | NA | NA | NA |
| WBGene00195037 | NA | NA | NA | NA |
| WBGene00195038 | NA | NA | NA | NA |
| WBGene00195039 | NA | NA | NA | NA |
| WBGene00195040 | NA | NA | NA | NA |
| WBGene00195043 | NA | NA | NA | NA |
| WBGene00195045 | 0.04913709 | 0.56495507 | NA | W05F2.8 |
| WBGene00195046 | 0.033463284 | 0.6365037 | NA | F42A9.11 |
| WBGene00195048 | NA | NA | NA | K08D10.13 |
| WBGene00195049 | -0.001993815 | 0.95612996 | NA | CC8.3 |
| WBGene00195050 | NA | NA | NA | T22D1.17 |
| WBGene00195051 | NA | NA | NA | T22D1.18 |
| WBGene00195056 | 0.054099237 | 0.60535803 | 0.76162644 | C45E1.5 |
| WBGene00195057 | -0.001842784 | 0.95621055 | NA | B0261.9 |
| WBGene00195058 | NA | NA | NA | C54E10.7 |
| WBGene00195060 | 0.012506684 | 0.93027172 | 0.96071865 | NA |
| WBGene00195061 | -0.045959069 | 0.50707733 | NA | ZK228.13 |
| WBGene00195063 | -0.207371243 | 0.26494801 | 0.46350864 | coa-1 |

| | | | | |
|---|---|---|---|---|
| WBGene00195064 | NA | NA | NA | NA |
| WBGene00195065 | NA | NA | NA | NA |
| WBGene00195067 | NA | NA | NA | C04C3.9 |
| WBGene00195068 | NA | NA | NA | Y48G8AL.16 |
| WBGene00195069 | -0.003270206 | 0.95698316 | NA | W03G9.10 |
| WBGene00195070 | 0.187540592 | 0.34681272 | 0.54645772 | C16C8.20 |
| WBGene00195071 | -0.192596064 | 0.26160988 | 0.45964601 | C16C8.21 |
| WBGene00195072 | 0.127758703 | 0.16886741 | NA | C16C8.22 |
| WBGene00195073 | 0.23365545 | 0.15392995 | 0.32715947 | F02E11.7 |
| WBGene00195077 | -0.484329231 | 0.04414902 | 0.13726797 | T24E12.13 |
| WBGene00195081 | 0.059762547 | 0.38965825 | NA | C04F6.7 |
| WBGene00195084 | 0.11294248 | 0.42363646 | 0.61965737 | C43C3.4 |
| WBGene00195086 | 0.098926403 | 0.23155549 | NA | F21C10.13 |
| WBGene00195093 | -0.096839557 | 0.35563809 | 0.55563945 | Y59E9AL.36 |
| WBGene00195117 | NA | NA | NA | NA |
| WBGene00195142 | 0.282867873 | 0.18391647 | 0.36725601 | M110.10 |
| WBGene00195143 | -0.002914971 | 0.95718083 | NA | nlp-78 |
| WBGene00195144 | NA | NA | NA | C16D2.3 |
| WBGene00195145 | 0.074892148 | 0.28555128 | NA | ZK666.15 |
| WBGene00195146 | -0.007708341 | 0.95671055 | 0.97597542 | F54D5.17 |
| WBGene00195147 | 0.01409643 | 0.85915526 | NA | F14E5.8 |
| WBGene00195148 | NA | NA | NA | W02B12.16 |
| WBGene00195149 | -0.046394786 | 0.50231372 | NA | F26H11.8 |
| WBGene00195150 | 0.038125098 | 0.61119482 | NA | Y81G3A.6 |
| WBGene00195151 | NA | NA | NA | NA |
| WBGene00195154 | NA | NA | NA | F13A2.10 |
| WBGene00195155 | -0.02609845 | 0.85357667 | 0.92153958 | W04A8.9 |
| WBGene00195157 | -0.023109762 | 0.78121253 | NA | C12D8.21 |
| WBGene00195158 | NA | NA | NA | F26F2.11 |
| WBGene00195159 | 0.042199095 | 0.55922995 | NA | NA |
| WBGene00195160 | NA | NA | NA | Y47D7A.17 |
| WBGene00195164 | -0.148185327 | 0.11602837 | 0.27192405 | K12B6.11 |
| WBGene00195165 | 4.37471588 | 9.13E-05 | 0.00089547 | F57G4.11 |
| WBGene00195166 | 0.085537126 | 0.24698522 | NA | W05B10.6 |
| WBGene00195168 | 0.020317245 | 0.84643798 | NA | T13C5.9 |
| WBGene00195169 | 0.022700991 | 0.87184544 | 0.9314787 | Y50D4A.6 |
| WBGene00195170 | 0.149354056 | 0.20024371 | 0.3887771 | NA |
| WBGene00195171 | NA | NA | NA | T09D3.9 |
| WBGene00195172 | NA | NA | NA | NA |
| WBGene00195174 | 0.020232312 | 0.84643798 | NA | K08E5.6 |
| WBGene00195175 | NA | NA | NA | NA |
| WBGene00195176 | 2.83528913 | 7.90E-06 | 9.94E-05 | Y43F8B.24 |
| WBGene00195177 | 0.020925399 | 0.84643798 | NA | Y43F8B.25 |

| | | | | |
|---|---|---|---|---|
| WBGene00195178 | NA | NA | NA | ZC581.10 |
| WBGene00195179 | -0.039636691 | 0.54579661 | NA | F48C1.11 |
| WBGene00195180 | NA | NA | NA | C41D11.10 |
| WBGene00195181 | 0.038125098 | 0.61119482 | NA | F15H10.12 |
| WBGene00195182 | -0.023109762 | 0.78121253 | NA | C17G10.10 |
| WBGene00195183 | -0.386491421 | 0.11596973 | 0.27192405 | R04A9.9 |
| WBGene00195184 | -0.026014132 | 0.85891664 | 0.92455628 | F19G12.8 |
| WBGene00195185 | 0.231966365 | 0.25719778 | 0.45539154 | F19G12.9 |
| WBGene00195186 | 0.47102138 | 0.07189693 | 0.19518689 | F19G12.10 |
| WBGene00195188 | NA | NA | NA | F19G12.11 |
| WBGene00195189 | -0.626178063 | 0.02730923 | 0.09674195 | ZC449.8 |
| WBGene00195192 | NA | NA | NA | NA |
| WBGene00195208 | 0.020317245 | 0.84643798 | NA | W02H3.3 |
| WBGene00195209 | 0.059711191 | 0.42255368 | NA | C07B5.8 |
| WBGene00195210 | NA | NA | NA | Y7A5A.13 |
| WBGene00195211 | 0.065014541 | 0.37399213 | NA | C34F6.12 |
| WBGene00195212 | -0.032974911 | 0.72944715 | NA | F09B12.7 |
| WBGene00195213 | NA | NA | NA | F11C1.9 |
| WBGene00195214 | 0.040107227 | 0.58693163 | NA | F11C1.10 |
| WBGene00195215 | 0.020317245 | 0.84643798 | NA | F55G7.5 |
| WBGene00195226 | -0.046467764 | 0.75840133 | 0.86329917 | ZC404.16 |
| WBGene00195227 | 0.038125098 | 0.61119482 | NA | Y71A12B.23 |
| WBGene00195228 | NA | NA | NA | Y71A12B.24 |
| WBGene00195229 | NA | NA | NA | Y71A12B.25 |
| WBGene00195230 | NA | NA | NA | F36D1.13 |
| WBGene00195230 | NA | NA | NA | F36D1.14 |
| WBGene00195231 | NA | NA | NA | F36D1.13 |
| WBGene00195231 | NA | NA | NA | F36D1.14 |
| WBGene00195232 | NA | NA | NA | F36D1.15 |
| WBGene00195233 | 0.40079545 | 0.10792927 | 0.25837025 | NA |
| WBGene00195234 | NA | NA | NA | NA |
| WBGene00195235 | NA | NA | NA | NA |
| WBGene00195236 | NA | NA | NA | NA |
| WBGene00195237 | NA | NA | NA | NA |
| WBGene00195238 | -0.022729726 | 0.70048408 | NA | NA |
| WBGene00195239 | NA | NA | NA | ZK1127.13 |
| WBGene00195240 | 0.020925399 | 0.84643798 | NA | Y39B6A.69 |
| WBGene00195241 | NA | NA | NA | Y113G7B.27 |
| WBGene00195242 | NA | NA | NA | Y113G7B.28 |
| WBGene00195243 | NA | NA | NA | F46B3.23 |
| WBGene00195244 | -0.116325205 | 0.53064384 | 0.70725139 | F30A10.15 |
| WBGene00195245 | NA | NA | NA | NA |
| WBGene00195246 | 0.191737311 | 0.12829736 | 0.28978736 | ZK546.19 |

| | | | | | |
|---|---|---|---|---|---|
| WBGene00195247 | | -0.003614866 | 0.95700117 | NA | NA |
| WBGene00195248 | | -0.227339988 | 0.20406773 | 0.39307402 | emc-5 |
| WBGene00195250 | NA | | NA | NA | C13C4.8 |
| WBGene00195252 | NA | | NA | NA | NA |
| WBGene00195253 | | 0.123609679 | 0.51348311 | 0.69346981 | NA |
| WBGene00195254 | | -0.163548736 | 0.17745738 | 0.35866283 | NA |
| WBGene00195255 | | 0.032086008 | 0.74678542 | NA | NA |
| WBGene00195256 | | 0.020232312 | 0.84643798 | NA | F25D1.8 |
| WBGene00195257 | | -0.022930767 | 0.78121253 | NA | NA |
| WBGene00195258 | NA | | NA | NA | NA |
| WBGene00195259 | NA | | NA | NA | NA |
| WBGene00195260 | NA | | NA | NA | NA |
| WBGene00195261 | NA | | NA | NA | NA |
| WBGene00195262 | NA | | NA | NA | NA |
| WBGene00195263 | NA | | NA | NA | NA |
| WBGene00195264 | NA | | NA | NA | NA |
| WBGene00195265 | NA | | NA | NA | NA |
| WBGene00195266 | NA | | NA | NA | NA |
| WBGene00195267 | | 0.040019663 | 0.56356856 | NA | NA |
| WBGene00195268 | NA | | NA | NA | NA |
| WBGene00195269 | NA | | NA | NA | NA |
| WBGene00195270 | NA | | NA | NA | NA |
| WBGene00195271 | NA | | NA | NA | NA |
| WBGene00195272 | NA | | NA | NA | NA |
| WBGene00195273 | NA | | NA | NA | NA |
| WBGene00195274 | NA | | NA | NA | NA |
| WBGene00195275 | NA | | NA | NA | NA |
| WBGene00195276 | NA | | NA | NA | NA |
| WBGene00195277 | NA | | NA | NA | NA |
| WBGene00195278 | NA | | NA | NA | NA |
| WBGene00195279 | NA | | NA | NA | NA |
| WBGene00195280 | NA | | NA | NA | NA |
| WBGene00195281 | NA | | NA | NA | NA |
| WBGene00195282 | NA | | NA | NA | NA |
| WBGene00195283 | NA | | NA | NA | NA |
| WBGene00195284 | NA | | NA | NA | NA |
| WBGene00195285 | NA | | NA | NA | NA |
| WBGene00195286 | NA | | NA | NA | NA |
| WBGene00195287 | NA | | NA | NA | NA |
| WBGene00195288 | NA | | NA | NA | NA |
| WBGene00195289 | NA | | NA | NA | NA |
| WBGene00195290 | NA | | NA | NA | NA |
| WBGene00195291 | NA | | NA | NA | NA |

| | | | | |
|---|---|---|---|---|
| WBGene00195292 | NA | NA | NA | NA |
| WBGene00195293 | NA | NA | NA | NA |
| WBGene00195294 | NA | NA | NA | NA |
| WBGene00195295 | 0.020232312 | 0.84643798 | NA | NA |
| WBGene00195296 | NA | NA | NA | NA |
| WBGene00195297 | NA | NA | NA | NA |
| WBGene00195298 | NA | NA | NA | NA |
| WBGene00195299 | NA | NA | NA | NA |
| WBGene00195300 | NA | NA | NA | NA |
| WBGene00195301 | -0.023056499 | 0.78121253 | NA | NA |
| WBGene00195302 | NA | NA | NA | NA |
| WBGene00195303 | NA | NA | NA | NA |
| WBGene00195304 | NA | NA | NA | NA |
| WBGene00195305 | NA | NA | NA | NA |
| WBGene00195306 | NA | NA | NA | NA |
| WBGene00195307 | NA | NA | NA | NA |
| WBGene00195308 | NA | NA | NA | NA |
| WBGene00195309 | NA | NA | NA | NA |
| WBGene00195310 | NA | NA | NA | NA |
| WBGene00195311 | -0.002914971 | 0.95718083 | NA | NA |
| WBGene00195312 | NA | NA | NA | NA |
| WBGene00195313 | NA | NA | NA | NA |
| WBGene00195314 | NA | NA | NA | NA |
| WBGene00195315 | NA | NA | NA | NA |
| WBGene00195316 | NA | NA | NA | NA |
| WBGene00195317 | NA | NA | NA | NA |
| WBGene00195318 | NA | NA | NA | NA |
| WBGene00195319 | NA | NA | NA | NA |
| WBGene00195320 | NA | NA | NA | NA |
| WBGene00195321 | NA | NA | NA | NA |
| WBGene00195322 | NA | NA | NA | NA |
| WBGene00195323 | NA | NA | NA | NA |
| WBGene00195324 | NA | NA | NA | NA |
| WBGene00195325 | NA | NA | NA | NA |
| WBGene00195326 | NA | NA | NA | NA |
| WBGene00195327 | NA | NA | NA | NA |
| WBGene00195328 | NA | NA | NA | NA |
| WBGene00195329 | NA | NA | NA | NA |
| WBGene00195330 | NA | NA | NA | NA |
| WBGene00195331 | NA | NA | NA | NA |
| WBGene00195332 | NA | NA | NA | NA |
| WBGene00195333 | NA | NA | NA | NA |
| WBGene00195334 | NA | NA | NA | NA |

| | | | | |
|---|---|---|---|---|
| WBGene00195335 | NA | NA | NA | NA |
| WBGene00195336 | NA | NA | NA | NA |
| WBGene00195337 | NA | NA | NA | NA |
| WBGene00195338 | -0.003108065 | 0.95728506 | NA | NA |
| WBGene00195339 | NA | NA | NA | NA |
| WBGene00195340 | NA | NA | NA | NA |
| WBGene00195341 | NA | NA | NA | NA |
| WBGene00195342 | NA | NA | NA | NA |
| WBGene00195343 | NA | NA | NA | NA |
| WBGene00195344 | NA | NA | NA | NA |
| WBGene00195345 | NA | NA | NA | NA |
| WBGene00195346 | NA | NA | NA | NA |
| WBGene00195347 | NA | NA | NA | NA |
| WBGene00195348 | NA | NA | NA | NA |
| WBGene00195349 | NA | NA | NA | NA |
| WBGene00195350 | NA | NA | NA | NA |
| WBGene00195351 | NA | NA | NA | NA |
| WBGene00195352 | NA | NA | NA | NA |
| WBGene00195353 | NA | NA | NA | NA |
| WBGene00195354 | NA | NA | NA | NA |
| WBGene00195355 | NA | NA | NA | NA |
| WBGene00195356 | NA | NA | NA | NA |
| WBGene00195357 | NA | NA | NA | NA |
| WBGene00195358 | NA | NA | NA | NA |
| WBGene00195359 | NA | NA | NA | NA |
| WBGene00195360 | NA | NA | NA | NA |
| WBGene00195361 | NA | NA | NA | NA |
| WBGene00195362 | NA | NA | NA | NA |
| WBGene00195363 | NA | NA | NA | NA |
| WBGene00195364 | NA | NA | NA | NA |
| WBGene00195365 | NA | NA | NA | NA |
| WBGene00195366 | NA | NA | NA | NA |
| WBGene00195367 | NA | NA | NA | NA |
| WBGene00195368 | NA | NA | NA | NA |
| WBGene00195369 | NA | NA | NA | NA |
| WBGene00195370 | -0.003421551 | 0.95689871 | NA | NA |
| WBGene00195371 | NA | NA | NA | NA |
| WBGene00195372 | NA | NA | NA | NA |
| WBGene00195373 | NA | NA | NA | NA |
| WBGene00195374 | 0.018860755 | 0.77262519 | NA | NA |
| WBGene00195375 | NA | NA | NA | NA |
| WBGene00195376 | NA | NA | NA | NA |
| WBGene00195377 | NA | NA | NA | NA |

| | | | | |
|---|---|---|---|---|
| WBGene00195378 | NA | NA | NA | NA |
| WBGene00195379 | NA | NA | NA | NA |
| WBGene00195380 | NA | NA | NA | NA |
| WBGene00195381 | NA | NA | NA | NA |
| WBGene00195382 | NA | NA | NA | NA |
| WBGene00195383 | NA | NA | NA | NA |
| WBGene00195384 | NA | NA | NA | NA |
| WBGene00195385 | -0.023056499 | 0.78121253 | NA | NA |
| WBGene00195386 | 0.020925399 | 0.84643798 | NA | NA |
| WBGene00195387 | NA | NA | NA | NA |
| WBGene00195388 | NA | NA | NA | NA |
| WBGene00195389 | NA | NA | NA | NA |
| WBGene00195390 | 0.020232312 | 0.84643798 | NA | NA |
| WBGene00195391 | NA | NA | NA | NA |
| WBGene00195392 | NA | NA | NA | NA |
| WBGene00195393 | NA | NA | NA | NA |
| WBGene00195394 | NA | NA | NA | NA |
| WBGene00195395 | NA | NA | NA | NA |
| WBGene00195396 | NA | NA | NA | NA |
| WBGene00195397 | NA | NA | NA | NA |
| WBGene00195398 | NA | NA | NA | NA |
| WBGene00195399 | NA | NA | NA | NA |
| WBGene00195400 | NA | NA | NA | NA |
| WBGene00195401 | 0.020925399 | 0.84643798 | NA | NA |
| WBGene00195402 | NA | NA | NA | NA |
| WBGene00195403 | NA | NA | NA | NA |
| WBGene00195404 | NA | NA | NA | NA |
| WBGene00195405 | NA | NA | NA | NA |
| WBGene00195406 | NA | NA | NA | NA |
| WBGene00195407 | NA | NA | NA | NA |
| WBGene00195408 | NA | NA | NA | NA |
| WBGene00195409 | NA | NA | NA | NA |
| WBGene00195410 | 0.020317245 | 0.84643798 | NA | NA |
| WBGene00195411 | NA | NA | NA | NA |
| WBGene00195412 | NA | NA | NA | NA |
| WBGene00195413 | NA | NA | NA | NA |
| WBGene00195414 | 0.020232312 | 0.84643798 | NA | NA |
| WBGene00195415 | NA | NA | NA | NA |
| WBGene00195416 | NA | NA | NA | NA |
| WBGene00195417 | NA | NA | NA | NA |
| WBGene00195418 | NA | NA | NA | NA |
| WBGene00195419 | NA | NA | NA | NA |
| WBGene00195420 | NA | NA | NA | NA |

| | | | | |
|---|---|---|---|---|
| WBGene00195421 | NA | NA | NA | NA |
| WBGene00195422 | NA | NA | NA | NA |
| WBGene00195423 | NA | NA | NA | NA |
| WBGene00195424 | NA | NA | NA | NA |
| WBGene00195425 | -0.023109762 | 0.78121253 | NA | NA |
| WBGene00195426 | NA | NA | NA | NA |
| WBGene00195427 | NA | NA | NA | NA |
| WBGene00195428 | NA | NA | NA | NA |
| WBGene00195429 | NA | NA | NA | NA |
| WBGene00195430 | NA | NA | NA | NA |
| WBGene00195431 | NA | NA | NA | NA |
| WBGene00195432 | NA | NA | NA | NA |
| WBGene00195433 | NA | NA | NA | NA |
| WBGene00195434 | NA | NA | NA | NA |
| WBGene00195435 | NA | NA | NA | NA |
| WBGene00195436 | NA | NA | NA | NA |
| WBGene00195437 | NA | NA | NA | NA |
| WBGene00195438 | NA | NA | NA | NA |
| WBGene00195439 | NA | NA | NA | NA |
| WBGene00195440 | NA | NA | NA | NA |
| WBGene00195441 | NA | NA | NA | NA |
| WBGene00195442 | NA | NA | NA | NA |
| WBGene00195443 | NA | NA | NA | NA |
| WBGene00195444 | NA | NA | NA | NA |
| WBGene00195445 | NA | NA | NA | NA |
| WBGene00195446 | NA | NA | NA | NA |
| WBGene00195447 | NA | NA | NA | NA |
| WBGene00195448 | NA | NA | NA | NA |
| WBGene00195449 | NA | NA | NA | NA |
| WBGene00195450 | NA | NA | NA | NA |
| WBGene00195451 | NA | NA | NA | NA |
| WBGene00195452 | NA | NA | NA | NA |
| WBGene00195453 | 0.020317245 | 0.84643798 | NA | NA |
| WBGene00195454 | NA | NA | NA | NA |
| WBGene00195455 | -0.023056499 | 0.78121253 | NA | NA |
| WBGene00195456 | NA | NA | NA | NA |
| WBGene00195457 | NA | NA | NA | NA |
| WBGene00195458 | 0.020317245 | 0.84643798 | NA | NA |
| WBGene00195459 | NA | NA | NA | NA |
| WBGene00195460 | NA | NA | NA | NA |
| WBGene00195461 | NA | NA | NA | NA |
| WBGene00195462 | NA | NA | NA | NA |
| WBGene00195463 | NA | NA | NA | NA |

| | | | | |
|---|---|---|---|---|
| WBGene00195464 | NA | NA | NA | NA |
| WBGene00195465 | NA | NA | NA | NA |
| WBGene00195466 | NA | NA | NA | NA |
| WBGene00195467 | NA | NA | NA | NA |
| WBGene00195468 | NA | NA | NA | NA |
| WBGene00195469 | NA | NA | NA | NA |
| WBGene00195470 | NA | NA | NA | NA |
| WBGene00195471 | NA | NA | NA | NA |
| WBGene00195472 | NA | NA | NA | NA |
| WBGene00195473 | NA | NA | NA | NA |
| WBGene00195474 | NA | NA | NA | NA |
| WBGene00195475 | NA | NA | NA | NA |
| WBGene00195476 | NA | NA | NA | NA |
| WBGene00195477 | NA | NA | NA | NA |
| WBGene00195478 | NA | NA | NA | NA |
| WBGene00195479 | NA | NA | NA | NA |
| WBGene00195480 | NA | NA | NA | NA |
| WBGene00195481 | NA | NA | NA | NA |
| WBGene00195482 | NA | NA | NA | NA |
| WBGene00195483 | NA | NA | NA | NA |
| WBGene00195484 | NA | NA | NA | NA |
| WBGene00195485 | 0.017604133 | 0.79437959 | NA | NA |
| WBGene00195486 | NA | NA | NA | NA |
| WBGene00195487 | NA | NA | NA | NA |
| WBGene00195488 | NA | NA | NA | NA |
| WBGene00195489 | NA | NA | NA | NA |
| WBGene00195490 | NA | NA | NA | NA |
| WBGene00195491 | NA | NA | NA | NA |
| WBGene00195492 | NA | NA | NA | NA |
| WBGene00195493 | NA | NA | NA | NA |
| WBGene00195494 | 0.020232312 | 0.84643798 | NA | NA |
| WBGene00195495 | NA | NA | NA | NA |
| WBGene00195496 | NA | NA | NA | NA |
| WBGene00195497 | NA | NA | NA | NA |
| WBGene00195498 | NA | NA | NA | NA |
| WBGene00195499 | 0.020925399 | 0.84643798 | NA | NA |
| WBGene00195500 | NA | NA | NA | NA |
| WBGene00195501 | NA | NA | NA | NA |
| WBGene00195502 | NA | NA | NA | NA |
| WBGene00195503 | NA | NA | NA | NA |
| WBGene00195504 | NA | NA | NA | NA |
| WBGene00195505 | NA | NA | NA | NA |
| WBGene00195506 | NA | NA | NA | NA |

| | | | | |
|---|---|---|---|---|
| WBGene00195507 | NA | NA | NA | NA |
| WBGene00195508 | NA | NA | NA | NA |
| WBGene00195509 | NA | NA | NA | NA |
| WBGene00195510 | NA | NA | NA | NA |
| WBGene00195511 | NA | NA | NA | NA |
| WBGene00195512 | -0.022930767 | 0.78121253 | NA | NA |
| WBGene00195513 | NA | NA | NA | NA |
| WBGene00195514 | NA | NA | NA | NA |
| WBGene00195515 | NA | NA | NA | NA |
| WBGene00195516 | NA | NA | NA | NA |
| WBGene00195517 | NA | NA | NA | NA |
| WBGene00195518 | NA | NA | NA | NA |
| WBGene00195519 | NA | NA | NA | NA |
| WBGene00195520 | NA | NA | NA | NA |
| WBGene00195521 | NA | NA | NA | NA |
| WBGene00195522 | NA | NA | NA | NA |
| WBGene00195523 | NA | NA | NA | NA |
| WBGene00195524 | NA | NA | NA | NA |
| WBGene00195525 | NA | NA | NA | NA |
| WBGene00195526 | NA | NA | NA | NA |
| WBGene00195527 | NA | NA | NA | NA |
| WBGene00195528 | 0.038125098 | 0.61119482 | NA | NA |
| WBGene00195529 | NA | NA | NA | NA |
| WBGene00195530 | -0.025049301 | 0.69142686 | NA | NA |
| WBGene00195531 | NA | NA | NA | NA |
| WBGene00195532 | NA | NA | NA | NA |
| WBGene00195533 | NA | NA | NA | NA |
| WBGene00195534 | NA | NA | NA | NA |
| WBGene00195535 | NA | NA | NA | NA |
| WBGene00195536 | NA | NA | NA | NA |
| WBGene00195537 | NA | NA | NA | NA |
| WBGene00195538 | NA | NA | NA | NA |
| WBGene00195539 | NA | NA | NA | NA |
| WBGene00195540 | NA | NA | NA | NA |
| WBGene00195541 | NA | NA | NA | NA |
| WBGene00195542 | NA | NA | NA | NA |
| WBGene00195543 | NA | NA | NA | NA |
| WBGene00195544 | NA | NA | NA | NA |
| WBGene00195545 | NA | NA | NA | NA |
| WBGene00195546 | NA | NA | NA | NA |
| WBGene00195547 | NA | NA | NA | NA |
| WBGene00195548 | 0.020232312 | 0.84643798 | NA | NA |
| WBGene00195549 | NA | NA | NA | NA |

| | | | | |
|---|---|---|---|---|
| WBGene00195550 | NA | NA | NA | NA |
| WBGene00195551 | NA | NA | NA | NA |
| WBGene00195552 | NA | NA | NA | NA |
| WBGene00195553 | NA | NA | NA | NA |
| WBGene00195554 | NA | NA | NA | NA |
| WBGene00195555 | NA | NA | NA | NA |
| WBGene00195556 | NA | NA | NA | NA |
| WBGene00195557 | NA | NA | NA | NA |
| WBGene00195558 | NA | NA | NA | NA |
| WBGene00195559 | NA | NA | NA | NA |
| WBGene00195560 | NA | NA | NA | NA |
| WBGene00195561 | NA | NA | NA | NA |
| WBGene00195562 | NA | NA | NA | NA |
| WBGene00195563 | NA | NA | NA | NA |
| WBGene00195564 | NA | NA | NA | NA |
| WBGene00195565 | NA | NA | NA | NA |
| WBGene00195566 | NA | NA | NA | NA |
| WBGene00195567 | NA | NA | NA | NA |
| WBGene00195568 | NA | NA | NA | NA |
| WBGene00195569 | NA | NA | NA | NA |
| WBGene00195570 | NA | NA | NA | NA |
| WBGene00195571 | NA | NA | NA | NA |
| WBGene00195572 | NA | NA | NA | NA |
| WBGene00195573 | NA | NA | NA | NA |
| WBGene00195574 | NA | NA | NA | NA |
| WBGene00195575 | NA | NA | NA | NA |
| WBGene00195576 | NA | NA | NA | NA |
| WBGene00195577 | NA | NA | NA | NA |
| WBGene00195578 | NA | NA | NA | NA |
| WBGene00195579 | NA | NA | NA | NA |
| WBGene00195580 | -0.023056499 | 0.78121253 | NA | NA |
| WBGene00195581 | NA | NA | NA | NA |
| WBGene00195582 | NA | NA | NA | NA |
| WBGene00195583 | NA | NA | NA | NA |
| WBGene00195584 | NA | NA | NA | NA |
| WBGene00195585 | NA | NA | NA | NA |
| WBGene00195586 | NA | NA | NA | NA |
| WBGene00195587 | NA | NA | NA | NA |
| WBGene00195588 | NA | NA | NA | NA |
| WBGene00195589 | NA | NA | NA | NA |
| WBGene00195590 | NA | NA | NA | NA |
| WBGene00195591 | NA | NA | NA | NA |
| WBGene00195592 | NA | NA | NA | NA |

| | | | | |
|---|---|---|---|---|
| WBGene00195593 | NA | NA | NA | NA |
| WBGene00195594 | NA | NA | NA | NA |
| WBGene00195595 | NA | NA | NA | NA |
| WBGene00195596 | NA | NA | NA | NA |
| WBGene00195597 | NA | NA | NA | NA |
| WBGene00195598 | NA | NA | NA | NA |
| WBGene00195599 | NA | NA | NA | NA |
| WBGene00195600 | NA | NA | NA | NA |
| WBGene00195601 | NA | NA | NA | NA |
| WBGene00195602 | NA | NA | NA | NA |
| WBGene00195603 | NA | NA | NA | NA |
| WBGene00195604 | NA | NA | NA | NA |
| WBGene00195605 | NA | NA | NA | NA |
| WBGene00195606 | NA | NA | NA | NA |
| WBGene00195607 | NA | NA | NA | NA |
| WBGene00195608 | NA | NA | NA | NA |
| WBGene00195609 | NA | NA | NA | NA |
| WBGene00195610 | NA | NA | NA | NA |
| WBGene00195611 | NA | NA | NA | NA |
| WBGene00195612 | NA | NA | NA | NA |
| WBGene00195613 | NA | NA | NA | NA |
| WBGene00195614 | NA | NA | NA | NA |
| WBGene00195615 | NA | NA | NA | NA |
| WBGene00195616 | NA | NA | NA | NA |
| WBGene00195617 | NA | NA | NA | NA |
| WBGene00195618 | NA | NA | NA | NA |
| WBGene00195619 | NA | NA | NA | NA |
| WBGene00195620 | NA | NA | NA | NA |
| WBGene00195621 | NA | NA | NA | NA |
| WBGene00195622 | NA | NA | NA | NA |
| WBGene00195623 | -0.02791392 | 0.6763823 | NA | NA |
| WBGene00195624 | NA | NA | NA | NA |
| WBGene00195625 | NA | NA | NA | NA |
| WBGene00195626 | NA | NA | NA | NA |
| WBGene00195627 | NA | NA | NA | NA |
| WBGene00195628 | NA | NA | NA | NA |
| WBGene00195629 | NA | NA | NA | NA |
| WBGene00195630 | NA | NA | NA | NA |
| WBGene00195631 | NA | NA | NA | NA |
| WBGene00195632 | 0.020232312 | 0.84643798 | NA | NA |
| WBGene00195633 | NA | NA | NA | NA |
| WBGene00195634 | NA | NA | NA | NA |
| WBGene00195635 | NA | NA | NA | NA |

| | | | | |
|---|---|---|---|---|
| WBGene00195636 | NA | NA | NA | NA |
| WBGene00195637 | NA | NA | NA | NA |
| WBGene00195638 | NA | NA | NA | NA |
| WBGene00195639 | NA | NA | NA | NA |
| WBGene00195640 | NA | NA | NA | NA |
| WBGene00195641 | -0.023056499 | 0.78121253 | NA | NA |
| WBGene00195642 | NA | NA | NA | NA |
| WBGene00195643 | NA | NA | NA | NA |
| WBGene00195644 | NA | NA | NA | NA |
| WBGene00195645 | NA | NA | NA | NA |
| WBGene00195646 | NA | NA | NA | NA |
| WBGene00195647 | NA | NA | NA | NA |
| WBGene00195648 | NA | NA | NA | NA |
| WBGene00195649 | NA | NA | NA | NA |
| WBGene00195650 | NA | NA | NA | NA |
| WBGene00195651 | NA | NA | NA | NA |
| WBGene00195652 | NA | NA | NA | NA |
| WBGene00195653 | NA | NA | NA | NA |
| WBGene00195654 | NA | NA | NA | NA |
| WBGene00195655 | NA | NA | NA | NA |
| WBGene00195656 | NA | NA | NA | NA |
| WBGene00195657 | NA | NA | NA | NA |
| WBGene00195658 | NA | NA | NA | NA |
| WBGene00195659 | NA | NA | NA | NA |
| WBGene00195660 | NA | NA | NA | NA |
| WBGene00195661 | NA | NA | NA | NA |
| WBGene00195662 | NA | NA | NA | NA |
| WBGene00195663 | NA | NA | NA | NA |
| WBGene00195664 | NA | NA | NA | NA |
| WBGene00195665 | NA | NA | NA | NA |
| WBGene00195666 | NA | NA | NA | NA |
| WBGene00195667 | NA | NA | NA | NA |
| WBGene00195668 | NA | NA | NA | NA |
| WBGene00195669 | NA | NA | NA | NA |
| WBGene00195670 | NA | NA | NA | NA |
| WBGene00195671 | NA | NA | NA | NA |
| WBGene00195672 | NA | NA | NA | NA |
| WBGene00195673 | NA | NA | NA | NA |
| WBGene00195674 | NA | NA | NA | NA |
| WBGene00195675 | NA | NA | NA | NA |
| WBGene00195676 | NA | NA | NA | NA |
| WBGene00195677 | NA | NA | NA | NA |
| WBGene00195678 | NA | NA | NA | NA |

| | | | | |
|---|---|---|---|---|
| WBGene00195679 | NA | NA | NA | NA |
| WBGene00195680 | NA | NA | NA | NA |
| WBGene00195681 | NA | NA | NA | NA |
| WBGene00195682 | NA | NA | NA | NA |
| WBGene00195683 | 0.020925399 | 0.84643798 | NA | NA |
| WBGene00195684 | NA | NA | NA | NA |
| WBGene00195685 | NA | NA | NA | NA |
| WBGene00195686 | NA | NA | NA | NA |
| WBGene00195687 | NA | NA | NA | NA |
| WBGene00195688 | NA | NA | NA | NA |
| WBGene00195689 | NA | NA | NA | NA |
| WBGene00195690 | NA | NA | NA | NA |
| WBGene00195691 | NA | NA | NA | NA |
| WBGene00195692 | NA | NA | NA | NA |
| WBGene00195693 | NA | NA | NA | NA |
| WBGene00195694 | NA | NA | NA | NA |
| WBGene00195695 | NA | NA | NA | NA |
| WBGene00195696 | NA | NA | NA | NA |
| WBGene00195697 | NA | NA | NA | NA |
| WBGene00195698 | NA | NA | NA | NA |
| WBGene00195699 | NA | NA | NA | NA |
| WBGene00195700 | 0.020317245 | 0.84643798 | NA | NA |
| WBGene00195701 | NA | NA | NA | NA |
| WBGene00195702 | NA | NA | NA | NA |
| WBGene00195703 | NA | NA | NA | NA |
| WBGene00195704 | NA | NA | NA | NA |
| WBGene00195705 | NA | NA | NA | NA |
| WBGene00195706 | NA | NA | NA | NA |
| WBGene00195707 | NA | NA | NA | NA |
| WBGene00195708 | NA | NA | NA | NA |
| WBGene00195709 | NA | NA | NA | NA |
| WBGene00195710 | NA | NA | NA | NA |
| WBGene00195711 | NA | NA | NA | NA |
| WBGene00195712 | NA | NA | NA | NA |
| WBGene00195713 | NA | NA | NA | NA |
| WBGene00195714 | NA | NA | NA | NA |
| WBGene00195715 | NA | NA | NA | NA |
| WBGene00195716 | NA | NA | NA | NA |
| WBGene00195717 | NA | NA | NA | NA |
| WBGene00195718 | NA | NA | NA | NA |
| WBGene00195719 | NA | NA | NA | NA |
| WBGene00195720 | NA | NA | NA | NA |
| WBGene00195721 | NA | NA | NA | NA |

| | | | | |
|---|---|---|---|---|
| WBGene00195722 | NA | NA | NA | NA |
| WBGene00195723 | NA | NA | NA | NA |
| WBGene00195724 | NA | NA | NA | NA |
| WBGene00195725 | NA | NA | NA | NA |
| WBGene00195726 | NA | NA | NA | NA |
| WBGene00195727 | NA | NA | NA | NA |
| WBGene00195728 | NA | NA | NA | NA |
| WBGene00195729 | NA | NA | NA | NA |
| WBGene00195730 | NA | NA | NA | NA |
| WBGene00195731 | NA | NA | NA | NA |
| WBGene00195732 | NA | NA | NA | NA |
| WBGene00195733 | NA | NA | NA | NA |
| WBGene00195734 | NA | NA | NA | NA |
| WBGene00195735 | NA | NA | NA | NA |
| WBGene00195736 | NA | NA | NA | NA |
| WBGene00195737 | NA | NA | NA | NA |
| WBGene00195738 | NA | NA | NA | NA |
| WBGene00195739 | NA | NA | NA | NA |
| WBGene00195740 | NA | NA | NA | NA |
| WBGene00195741 | NA | NA | NA | NA |
| WBGene00195742 | NA | NA | NA | NA |
| WBGene00195743 | NA | NA | NA | NA |
| WBGene00195744 | NA | NA | NA | NA |
| WBGene00195745 | NA | NA | NA | NA |
| WBGene00195746 | NA | NA | NA | NA |
| WBGene00195747 | NA | NA | NA | NA |
| WBGene00195748 | NA | NA | NA | NA |
| WBGene00195749 | NA | NA | NA | NA |
| WBGene00195750 | NA | NA | NA | NA |
| WBGene00195751 | NA | NA | NA | NA |
| WBGene00195752 | NA | NA | NA | NA |
| WBGene00195753 | NA | NA | NA | NA |
| WBGene00195754 | NA | NA | NA | NA |
| WBGene00195755 | NA | NA | NA | NA |
| WBGene00195756 | NA | NA | NA | NA |
| WBGene00195757 | NA | NA | NA | NA |
| WBGene00195758 | NA | NA | NA | NA |
| WBGene00195759 | NA | NA | NA | NA |
| WBGene00195760 | NA | NA | NA | NA |
| WBGene00195761 | NA | NA | NA | NA |
| WBGene00195762 | NA | NA | NA | NA |
| WBGene00195763 | NA | NA | NA | NA |
| WBGene00195764 | NA | NA | NA | NA |

| | | | | |
|---|---|---|---|---|
| WBGene00195765 | NA | NA | NA | NA |
| WBGene00195766 | NA | NA | NA | NA |
| WBGene00195767 | NA | NA | NA | NA |
| WBGene00195768 | NA | NA | NA | NA |
| WBGene00195769 | NA | NA | NA | NA |
| WBGene00195770 | NA | NA | NA | NA |
| WBGene00195771 | NA | NA | NA | NA |
| WBGene00195772 | NA | NA | NA | NA |
| WBGene00195773 | NA | NA | NA | NA |
| WBGene00195774 | NA | NA | NA | NA |
| WBGene00195775 | NA | NA | NA | NA |
| WBGene00195776 | NA | NA | NA | NA |
| WBGene00195777 | NA | NA | NA | NA |
| WBGene00195778 | NA | NA | NA | NA |
| WBGene00195779 | NA | NA | NA | NA |
| WBGene00195780 | NA | NA | NA | NA |
| WBGene00195781 | NA | NA | NA | NA |
| WBGene00195782 | NA | NA | NA | NA |
| WBGene00195783 | NA | NA | NA | NA |
| WBGene00195784 | NA | NA | NA | NA |
| WBGene00195785 | NA | NA | NA | NA |
| WBGene00195786 | NA | NA | NA | NA |
| WBGene00195787 | NA | NA | NA | NA |
| WBGene00195788 | NA | NA | NA | NA |
| WBGene00195789 | NA | NA | NA | NA |
| WBGene00195790 | NA | NA | NA | NA |
| WBGene00195791 | NA | NA | NA | NA |
| WBGene00195792 | NA | NA | NA | NA |
| WBGene00195793 | NA | NA | NA | NA |
| WBGene00195794 | NA | NA | NA | NA |
| WBGene00195795 | 0.020925399 | 0.84643798 | NA | NA |
| WBGene00195796 | NA | NA | NA | NA |
| WBGene00195797 | NA | NA | NA | NA |
| WBGene00195798 | NA | NA | NA | NA |
| WBGene00195799 | NA | NA | NA | NA |
| WBGene00195800 | NA | NA | NA | NA |
| WBGene00195801 | NA | NA | NA | NA |
| WBGene00195802 | NA | NA | NA | NA |
| WBGene00195803 | -0.023056499 | 0.78121253 | NA | NA |
| WBGene00195804 | NA | NA | NA | NA |
| WBGene00195805 | NA | NA | NA | NA |
| WBGene00195806 | NA | NA | NA | NA |
| WBGene00195807 | NA | NA | NA | NA |

| | | | | |
|---|---|---|---|---|
| WBGene00195808 | NA | NA | NA | NA |
| WBGene00195809 | NA | NA | NA | NA |
| WBGene00195810 | NA | NA | NA | NA |
| WBGene00195811 | NA | NA | NA | NA |
| WBGene00195812 | NA | NA | NA | NA |
| WBGene00195813 | NA | NA | NA | NA |
| WBGene00195814 | NA | NA | NA | NA |
| WBGene00195815 | NA | NA | NA | NA |
| WBGene00195816 | NA | NA | NA | NA |
| WBGene00195817 | NA | NA | NA | NA |
| WBGene00195818 | NA | NA | NA | NA |
| WBGene00195819 | -0.023109762 | 0.78121253 | NA | NA |
| WBGene00195820 | -0.027535543 | 0.68063395 | NA | NA |
| WBGene00195821 | NA | NA | NA | NA |
| WBGene00195822 | NA | NA | NA | NA |
| WBGene00195823 | NA | NA | NA | NA |
| WBGene00195824 | NA | NA | NA | NA |
| WBGene00195825 | NA | NA | NA | NA |
| WBGene00195826 | NA | NA | NA | NA |
| WBGene00195827 | NA | NA | NA | NA |
| WBGene00195828 | NA | NA | NA | NA |
| WBGene00195829 | NA | NA | NA | NA |
| WBGene00195830 | NA | NA | NA | NA |
| WBGene00195831 | NA | NA | NA | NA |
| WBGene00195832 | NA | NA | NA | NA |
| WBGene00195833 | NA | NA | NA | NA |
| WBGene00195834 | NA | NA | NA | NA |
| WBGene00195835 | NA | NA | NA | NA |
| WBGene00195836 | NA | NA | NA | NA |
| WBGene00195837 | NA | NA | NA | NA |
| WBGene00195838 | NA | NA | NA | NA |
| WBGene00195839 | NA | NA | NA | NA |
| WBGene00195840 | NA | NA | NA | NA |
| WBGene00195841 | NA | NA | NA | NA |
| WBGene00195842 | NA | NA | NA | NA |
| WBGene00195843 | NA | NA | NA | NA |
| WBGene00195844 | NA | NA | NA | NA |
| WBGene00195845 | NA | NA | NA | NA |
| WBGene00195846 | NA | NA | NA | NA |
| WBGene00195847 | NA | NA | NA | NA |
| WBGene00195848 | NA | NA | NA | NA |
| WBGene00195849 | NA | NA | NA | NA |
| WBGene00195850 | NA | NA | NA | NA |

| | | | | |
|---|---|---|---|---|
| WBGene00195851 | NA | NA | NA | NA |
| WBGene00195852 | NA | NA | NA | NA |
| WBGene00195853 | NA | NA | NA | NA |
| WBGene00195854 | NA | NA | NA | NA |
| WBGene00195855 | NA | NA | NA | NA |
| WBGene00195856 | NA | NA | NA | NA |
| WBGene00195857 | NA | NA | NA | NA |
| WBGene00195858 | NA | NA | NA | NA |
| WBGene00195859 | NA | NA | NA | NA |
| WBGene00195860 | NA | NA | NA | NA |
| WBGene00195861 | NA | NA | NA | NA |
| WBGene00195862 | NA | NA | NA | NA |
| WBGene00195863 | NA | NA | NA | NA |
| WBGene00195864 | NA | NA | NA | NA |
| WBGene00195865 | NA | NA | NA | NA |
| WBGene00195866 | NA | NA | NA | NA |
| WBGene00195867 | NA | NA | NA | NA |
| WBGene00195868 | NA | NA | NA | NA |
| WBGene00195869 | NA | NA | NA | NA |
| WBGene00195870 | NA | NA | NA | NA |
| WBGene00195871 | NA | NA | NA | NA |
| WBGene00195872 | NA | NA | NA | NA |
| WBGene00195873 | NA | NA | NA | NA |
| WBGene00195874 | NA | NA | NA | NA |
| WBGene00195875 | NA | NA | NA | NA |
| WBGene00195876 | NA | NA | NA | NA |
| WBGene00195877 | 0.033034883 | 0.64219503 | NA | NA |
| WBGene00195878 | NA | NA | NA | NA |
| WBGene00195879 | NA | NA | NA | NA |
| WBGene00195880 | NA | NA | NA | NA |
| WBGene00195881 | NA | NA | NA | NA |
| WBGene00195882 | NA | NA | NA | NA |
| WBGene00195883 | NA | NA | NA | NA |
| WBGene00195884 | NA | NA | NA | NA |
| WBGene00195885 | NA | NA | NA | NA |
| WBGene00195886 | NA | NA | NA | NA |
| WBGene00195887 | NA | NA | NA | NA |
| WBGene00195888 | 0.020925399 | 0.84643798 | NA | NA |
| WBGene00195889 | NA | NA | NA | NA |
| WBGene00195890 | NA | NA | NA | NA |
| WBGene00195891 | NA | NA | NA | NA |
| WBGene00195892 | NA | NA | NA | NA |
| WBGene00195893 | NA | NA | NA | NA |

| | | | | |
|---|---|---|---|---|
| WBGene00195894 | NA | NA | NA | NA |
| WBGene00195895 | NA | NA | NA | NA |
| WBGene00195896 | NA | NA | NA | NA |
| WBGene00195897 | NA | NA | NA | NA |
| WBGene00195898 | NA | NA | NA | NA |
| WBGene00195899 | NA | NA | NA | NA |
| WBGene00195900 | NA | NA | NA | NA |
| WBGene00195901 | NA | NA | NA | NA |
| WBGene00195902 | NA | NA | NA | NA |
| WBGene00195903 | NA | NA | NA | NA |
| WBGene00195904 | NA | NA | NA | NA |
| WBGene00195905 | NA | NA | NA | NA |
| WBGene00195906 | NA | NA | NA | NA |
| WBGene00195907 | NA | NA | NA | NA |
| WBGene00195908 | NA | NA | NA | NA |
| WBGene00195909 | NA | NA | NA | NA |
| WBGene00195910 | NA | NA | NA | NA |
| WBGene00195911 | NA | NA | NA | NA |
| WBGene00195912 | NA | NA | NA | NA |
| WBGene00195913 | NA | NA | NA | NA |
| WBGene00195914 | NA | NA | NA | NA |
| WBGene00195915 | NA | NA | NA | NA |
| WBGene00195916 | NA | NA | NA | NA |
| WBGene00195917 | NA | NA | NA | NA |
| WBGene00195918 | NA | NA | NA | NA |
| WBGene00195919 | NA | NA | NA | NA |
| WBGene00195920 | NA | NA | NA | NA |
| WBGene00195921 | NA | NA | NA | NA |
| WBGene00195922 | NA | NA | NA | NA |
| WBGene00195923 | NA | NA | NA | NA |
| WBGene00195924 | NA | NA | NA | NA |
| WBGene00195925 | NA | NA | NA | NA |
| WBGene00195926 | NA | NA | NA | NA |
| WBGene00195927 | NA | NA | NA | NA |
| WBGene00195928 | NA | NA | NA | NA |
| WBGene00195929 | NA | NA | NA | NA |
| WBGene00195930 | NA | NA | NA | NA |
| WBGene00195931 | NA | NA | NA | NA |
| WBGene00195932 | NA | NA | NA | NA |
| WBGene00195933 | NA | NA | NA | NA |
| WBGene00195934 | NA | NA | NA | NA |
| WBGene00195935 | NA | NA | NA | NA |
| WBGene00195936 | NA | NA | NA | NA |

| | | | | |
|---|---|---|---|---|
| WBGene00195937 | NA | NA | NA | NA |
| WBGene00195938 | NA | NA | NA | NA |
| WBGene00195939 | NA | NA | NA | NA |
| WBGene00195940 | NA | NA | NA | NA |
| WBGene00195941 | NA | NA | NA | NA |
| WBGene00195942 | NA | NA | NA | NA |
| WBGene00195943 | 0.017141582 | 0.79355725 | NA | NA |
| WBGene00195944 | NA | NA | NA | NA |
| WBGene00195945 | NA | NA | NA | NA |
| WBGene00195946 | NA | NA | NA | NA |
| WBGene00195947 | NA | NA | NA | NA |
| WBGene00195948 | NA | NA | NA | NA |
| WBGene00195949 | NA | NA | NA | NA |
| WBGene00195950 | NA | NA | NA | NA |
| WBGene00195951 | NA | NA | NA | NA |
| WBGene00195952 | NA | NA | NA | NA |
| WBGene00195953 | NA | NA | NA | NA |
| WBGene00195954 | NA | NA | NA | NA |
| WBGene00195955 | NA | NA | NA | NA |
| WBGene00195956 | NA | NA | NA | NA |
| WBGene00195957 | NA | NA | NA | NA |
| WBGene00195958 | NA | NA | NA | NA |
| WBGene00195959 | NA | NA | NA | NA |
| WBGene00195960 | NA | NA | NA | NA |
| WBGene00195961 | NA | NA | NA | NA |
| WBGene00195962 | NA | NA | NA | NA |
| WBGene00195963 | NA | NA | NA | NA |
| WBGene00195964 | NA | NA | NA | NA |
| WBGene00195965 | NA | NA | NA | NA |
| WBGene00195966 | NA | NA | NA | NA |
| WBGene00195967 | NA | NA | NA | NA |
| WBGene00195968 | NA | NA | NA | NA |
| WBGene00195969 | NA | NA | NA | NA |
| WBGene00195970 | NA | NA | NA | NA |
| WBGene00195971 | NA | NA | NA | NA |
| WBGene00195972 | NA | NA | NA | NA |
| WBGene00195973 | NA | NA | NA | NA |
| WBGene00195974 | NA | NA | NA | NA |
| WBGene00195975 | NA | NA | NA | NA |
| WBGene00195976 | NA | NA | NA | NA |
| WBGene00195977 | NA | NA | NA | NA |
| WBGene00195978 | NA | NA | NA | NA |
| WBGene00195979 | NA | NA | NA | NA |

| | | | | |
|---|---|---|---|---|
| WBGene00195980 | NA | NA | NA | NA |
| WBGene00195981 | 0.020232312 | 0.84643798 | NA | NA |
| WBGene00195982 | NA | NA | NA | NA |
| WBGene00195983 | NA | NA | NA | NA |
| WBGene00195984 | NA | NA | NA | NA |
| WBGene00195985 | NA | NA | NA | NA |
| WBGene00195986 | NA | NA | NA | NA |
| WBGene00195987 | NA | NA | NA | NA |
| WBGene00195988 | NA | NA | NA | NA |
| WBGene00195989 | NA | NA | NA | NA |
| WBGene00195990 | NA | NA | NA | NA |
| WBGene00195991 | NA | NA | NA | NA |
| WBGene00195992 | NA | NA | NA | NA |
| WBGene00195993 | NA | NA | NA | NA |
| WBGene00195994 | 0.020232312 | 0.84643798 | NA | NA |
| WBGene00195995 | NA | NA | NA | NA |
| WBGene00195996 | NA | NA | NA | NA |
| WBGene00195997 | NA | NA | NA | NA |
| WBGene00195998 | NA | NA | NA | NA |
| WBGene00195999 | NA | NA | NA | NA |
| WBGene00196000 | 0.020232312 | 0.84643798 | NA | NA |
| WBGene00196001 | NA | NA | NA | NA |
| WBGene00196002 | NA | NA | NA | NA |
| WBGene00196003 | NA | NA | NA | NA |
| WBGene00196004 | NA | NA | NA | NA |
| WBGene00196005 | NA | NA | NA | NA |
| WBGene00196006 | NA | NA | NA | NA |
| WBGene00196007 | NA | NA | NA | NA |
| WBGene00196008 | NA | NA | NA | NA |
| WBGene00196009 | NA | NA | NA | NA |
| WBGene00196010 | NA | NA | NA | NA |
| WBGene00196011 | NA | NA | NA | NA |
| WBGene00196012 | NA | NA | NA | NA |
| WBGene00196013 | NA | NA | NA | NA |
| WBGene00196014 | NA | NA | NA | NA |
| WBGene00196015 | NA | NA | NA | NA |
| WBGene00196016 | -0.022930767 | 0.78121253 | NA | NA |
| WBGene00196017 | NA | NA | NA | NA |
| WBGene00196018 | NA | NA | NA | NA |
| WBGene00196019 | NA | NA | NA | NA |
| WBGene00196020 | -0.023109762 | 0.78121253 | NA | NA |
| WBGene00196021 | NA | NA | NA | NA |
| WBGene00196022 | NA | NA | NA | NA |

| | | | | |
|---|---|---|---|---|
| WBGene00196023 | NA | NA | NA | NA |
| WBGene00196024 | NA | NA | NA | NA |
| WBGene00196025 | NA | NA | NA | NA |
| WBGene00196026 | NA | NA | NA | NA |
| WBGene00196027 | 0.038633797 | 0.57999349 | NA | NA |
| WBGene00196028 | NA | NA | NA | NA |
| WBGene00196029 | NA | NA | NA | NA |
| WBGene00196030 | NA | NA | NA | NA |
| WBGene00196031 | NA | NA | NA | NA |
| WBGene00196032 | NA | NA | NA | NA |
| WBGene00196033 | NA | NA | NA | NA |
| WBGene00196034 | NA | NA | NA | NA |
| WBGene00196035 | NA | NA | NA | NA |
| WBGene00196036 | NA | NA | NA | NA |
| WBGene00196037 | NA | NA | NA | NA |
| WBGene00196038 | NA | NA | NA | NA |
| WBGene00196039 | NA | NA | NA | NA |
| WBGene00196040 | NA | NA | NA | NA |
| WBGene00196041 | NA | NA | NA | NA |
| WBGene00196042 | NA | NA | NA | NA |
| WBGene00196043 | NA | NA | NA | NA |
| WBGene00196044 | NA | NA | NA | NA |
| WBGene00196045 | NA | NA | NA | NA |
| WBGene00196046 | 0.020925399 | 0.84643798 | NA | NA |
| WBGene00196047 | NA | NA | NA | NA |
| WBGene00196048 | NA | NA | NA | NA |
| WBGene00196049 | NA | NA | NA | NA |
| WBGene00196050 | NA | NA | NA | NA |
| WBGene00196051 | NA | NA | NA | NA |
| WBGene00196052 | NA | NA | NA | NA |
| WBGene00196053 | NA | NA | NA | NA |
| WBGene00196054 | NA | NA | NA | NA |
| WBGene00196055 | -0.023056499 | 0.78121253 | NA | NA |
| WBGene00196056 | NA | NA | NA | NA |
| WBGene00196057 | NA | NA | NA | NA |
| WBGene00196058 | NA | NA | NA | NA |
| WBGene00196059 | NA | NA | NA | NA |
| WBGene00196060 | NA | NA | NA | NA |
| WBGene00196061 | NA | NA | NA | NA |
| WBGene00196062 | NA | NA | NA | NA |
| WBGene00196063 | NA | NA | NA | NA |
| WBGene00196064 | NA | NA | NA | NA |
| WBGene00196065 | NA | NA | NA | NA |

| | | | | |
|---|---|---|---|---|
| WBGene00196066 | NA | NA | NA | NA |
| WBGene00196067 | NA | NA | NA | NA |
| WBGene00196068 | NA | NA | NA | NA |
| WBGene00196069 | NA | NA | NA | NA |
| WBGene00196070 | NA | NA | NA | NA |
| WBGene00196071 | NA | NA | NA | NA |
| WBGene00196072 | NA | NA | NA | NA |
| WBGene00196073 | NA | NA | NA | NA |
| WBGene00196074 | NA | NA | NA | NA |
| WBGene00196075 | NA | NA | NA | NA |
| WBGene00196076 | NA | NA | NA | NA |
| WBGene00196077 | NA | NA | NA | NA |
| WBGene00196078 | NA | NA | NA | NA |
| WBGene00196079 | -0.023056499 | 0.78121253 | NA | NA |
| WBGene00196080 | NA | NA | NA | NA |
| WBGene00196081 | 0.020925399 | 0.84643798 | NA | NA |
| WBGene00196082 | NA | NA | NA | NA |
| WBGene00196083 | NA | NA | NA | NA |
| WBGene00196084 | NA | NA | NA | NA |
| WBGene00196085 | NA | NA | NA | NA |
| WBGene00196086 | 0.020232312 | 0.84643798 | NA | NA |
| WBGene00196087 | NA | NA | NA | NA |
| WBGene00196088 | NA | NA | NA | NA |
| WBGene00196089 | NA | NA | NA | NA |
| WBGene00196090 | NA | NA | NA | NA |
| WBGene00196091 | NA | NA | NA | NA |
| WBGene00196092 | NA | NA | NA | NA |
| WBGene00196093 | NA | NA | NA | NA |
| WBGene00196094 | NA | NA | NA | NA |
| WBGene00196095 | NA | NA | NA | NA |
| WBGene00196096 | NA | NA | NA | NA |
| WBGene00196097 | NA | NA | NA | NA |
| WBGene00196098 | NA | NA | NA | NA |
| WBGene00196099 | NA | NA | NA | NA |
| WBGene00196100 | NA | NA | NA | NA |
| WBGene00196101 | NA | NA | NA | NA |
| WBGene00196102 | NA | NA | NA | NA |
| WBGene00196103 | NA | NA | NA | NA |
| WBGene00196104 | NA | NA | NA | NA |
| WBGene00196105 | NA | NA | NA | NA |
| WBGene00196106 | NA | NA | NA | NA |
| WBGene00196107 | NA | NA | NA | NA |
| WBGene00196108 | NA | NA | NA | NA |

| | | | | |
|---|---|---|---|---|
| WBGene00196109 | NA | NA | NA | NA |
| WBGene00196110 | NA | NA | NA | NA |
| WBGene00196111 | NA | NA | NA | NA |
| WBGene00196112 | NA | NA | NA | NA |
| WBGene00196113 | NA | NA | NA | NA |
| WBGene00196114 | NA | NA | NA | NA |
| WBGene00196115 | NA | NA | NA | NA |
| WBGene00196116 | NA | NA | NA | NA |
| WBGene00196117 | NA | NA | NA | NA |
| WBGene00196118 | NA | NA | NA | NA |
| WBGene00196119 | NA | NA | NA | NA |
| WBGene00196120 | NA | NA | NA | NA |
| WBGene00196121 | NA | NA | NA | NA |
| WBGene00196122 | NA | NA | NA | NA |
| WBGene00196123 | NA | NA | NA | NA |
| WBGene00196124 | NA | NA | NA | NA |
| WBGene00196125 | NA | NA | NA | NA |
| WBGene00196126 | NA | NA | NA | NA |
| WBGene00196127 | NA | NA | NA | NA |
| WBGene00196128 | NA | NA | NA | NA |
| WBGene00196129 | NA | NA | NA | NA |
| WBGene00196130 | NA | NA | NA | NA |
| WBGene00196131 | NA | NA | NA | NA |
| WBGene00196132 | NA | NA | NA | NA |
| WBGene00196133 | NA | NA | NA | NA |
| WBGene00196134 | NA | NA | NA | NA |
| WBGene00196135 | NA | NA | NA | NA |
| WBGene00196136 | NA | NA | NA | NA |
| WBGene00196137 | NA | NA | NA | NA |
| WBGene00196138 | 0.020925399 | 0.84643798 | NA | NA |
| WBGene00196139 | NA | NA | NA | NA |
| WBGene00196140 | NA | NA | NA | NA |
| WBGene00196141 | NA | NA | NA | NA |
| WBGene00196142 | NA | NA | NA | NA |
| WBGene00196143 | NA | NA | NA | NA |
| WBGene00196144 | NA | NA | NA | NA |
| WBGene00196145 | NA | NA | NA | NA |
| WBGene00196146 | NA | NA | NA | NA |
| WBGene00196147 | NA | NA | NA | NA |
| WBGene00196148 | NA | NA | NA | NA |
| WBGene00196149 | NA | NA | NA | NA |
| WBGene00196150 | NA | NA | NA | NA |
| WBGene00196151 | NA | NA | NA | NA |

| | | | | |
|---|---|---|---|---|
| WBGene00196152 | NA | NA | NA | NA |
| WBGene00196153 | NA | NA | NA | NA |
| WBGene00196154 | NA | NA | NA | NA |
| WBGene00196155 | NA | NA | NA | NA |
| WBGene00196156 | NA | NA | NA | NA |
| WBGene00196157 | NA | NA | NA | NA |
| WBGene00196158 | NA | NA | NA | NA |
| WBGene00196159 | NA | NA | NA | NA |
| WBGene00196160 | NA | NA | NA | NA |
| WBGene00196161 | NA | NA | NA | NA |
| WBGene00196162 | -0.045959069 | 0.50707733 | NA | NA |
| WBGene00196163 | NA | NA | NA | NA |
| WBGene00196164 | NA | NA | NA | NA |
| WBGene00196165 | NA | NA | NA | NA |
| WBGene00196166 | NA | NA | NA | NA |
| WBGene00196167 | NA | NA | NA | NA |
| WBGene00196168 | NA | NA | NA | NA |
| WBGene00196169 | NA | NA | NA | NA |
| WBGene00196170 | NA | NA | NA | NA |
| WBGene00196171 | NA | NA | NA | NA |
| WBGene00196172 | NA | NA | NA | NA |
| WBGene00196173 | NA | NA | NA | NA |
| WBGene00196174 | NA | NA | NA | NA |
| WBGene00196175 | NA | NA | NA | NA |
| WBGene00196176 | NA | NA | NA | NA |
| WBGene00196177 | NA | NA | NA | NA |
| WBGene00196178 | NA | NA | NA | NA |
| WBGene00196179 | NA | NA | NA | NA |
| WBGene00196180 | 0.020317245 | 0.84643798 | NA | NA |
| WBGene00196181 | NA | NA | NA | NA |
| WBGene00196182 | 0.020317245 | 0.84643798 | NA | NA |
| WBGene00196183 | NA | NA | NA | NA |
| WBGene00196184 | -0.075452738 | 0.28904131 | NA | NA |
| WBGene00196185 | NA | NA | NA | NA |
| WBGene00196186 | NA | NA | NA | NA |
| WBGene00196187 | NA | NA | NA | NA |
| WBGene00196188 | NA | NA | NA | NA |
| WBGene00196189 | NA | NA | NA | NA |
| WBGene00196190 | NA | NA | NA | NA |
| WBGene00196191 | NA | NA | NA | NA |
| WBGene00196192 | NA | NA | NA | NA |
| WBGene00196193 | NA | NA | NA | NA |
| WBGene00196194 | NA | NA | NA | NA |

| | | | | |
|---|---|---|---|---|
| WBGene00196195 | NA | NA | NA | NA |
| WBGene00196196 | NA | NA | NA | NA |
| WBGene00196197 | NA | NA | NA | NA |
| WBGene00196198 | NA | NA | NA | NA |
| WBGene00196199 | NA | NA | NA | NA |
| WBGene00196200 | NA | NA | NA | NA |
| WBGene00196201 | NA | NA | NA | NA |
| WBGene00196202 | NA | NA | NA | NA |
| WBGene00196203 | NA | NA | NA | NA |
| WBGene00196204 | NA | NA | NA | NA |
| WBGene00196205 | NA | NA | NA | NA |
| WBGene00196206 | NA | NA | NA | NA |
| WBGene00196207 | -0.027373974 | 0.68245543 | NA | NA |
| WBGene00196208 | NA | NA | NA | NA |
| WBGene00196209 | NA | NA | NA | NA |
| WBGene00196210 | NA | NA | NA | NA |
| WBGene00196211 | NA | NA | NA | NA |
| WBGene00196212 | NA | NA | NA | NA |
| WBGene00196213 | NA | NA | NA | NA |
| WBGene00196214 | NA | NA | NA | NA |
| WBGene00196215 | NA | NA | NA | NA |
| WBGene00196216 | NA | NA | NA | NA |
| WBGene00196217 | NA | NA | NA | NA |
| WBGene00196218 | NA | NA | NA | NA |
| WBGene00196219 | NA | NA | NA | NA |
| WBGene00196220 | -0.046394786 | 0.50231372 | NA | NA |
| WBGene00196221 | NA | NA | NA | NA |
| WBGene00196222 | NA | NA | NA | NA |
| WBGene00196223 | NA | NA | NA | NA |
| WBGene00196224 | NA | NA | NA | NA |
| WBGene00196225 | NA | NA | NA | NA |
| WBGene00196226 | NA | NA | NA | NA |
| WBGene00196227 | 0.040107227 | 0.58693163 | NA | NA |
| WBGene00196228 | NA | NA | NA | NA |
| WBGene00196229 | NA | NA | NA | NA |
| WBGene00196230 | NA | NA | NA | NA |
| WBGene00196231 | NA | NA | NA | NA |
| WBGene00196232 | NA | NA | NA | NA |
| WBGene00196233 | NA | NA | NA | NA |
| WBGene00196234 | NA | NA | NA | NA |
| WBGene00196235 | NA | NA | NA | NA |
| WBGene00196236 | NA | NA | NA | NA |
| WBGene00196237 | 0.020925399 | 0.84643798 | NA | NA |

| | | | | |
|---|---|---|---|---|
| WBGene00196238 | NA | NA | NA | NA |
| WBGene00196239 | NA | NA | NA | NA |
| WBGene00196240 | NA | NA | NA | NA |
| WBGene00196241 | NA | NA | NA | NA |
| WBGene00196242 | NA | NA | NA | NA |
| WBGene00196243 | NA | NA | NA | NA |
| WBGene00196244 | NA | NA | NA | NA |
| WBGene00196245 | NA | NA | NA | NA |
| WBGene00196246 | NA | NA | NA | NA |
| WBGene00196247 | NA | NA | NA | NA |
| WBGene00196248 | NA | NA | NA | NA |
| WBGene00196249 | NA | NA | NA | NA |
| WBGene00196250 | NA | NA | NA | NA |
| WBGene00196251 | 0.020317245 | 0.84643798 | NA | NA |
| WBGene00196252 | NA | NA | NA | NA |
| WBGene00196253 | NA | NA | NA | NA |
| WBGene00196254 | NA | NA | NA | NA |
| WBGene00196255 | NA | NA | NA | NA |
| WBGene00196256 | NA | NA | NA | NA |
| WBGene00196257 | NA | NA | NA | NA |
| WBGene00196258 | NA | NA | NA | NA |
| WBGene00196259 | NA | NA | NA | NA |
| WBGene00196260 | NA | NA | NA | NA |
| WBGene00196261 | NA | NA | NA | NA |
| WBGene00196262 | NA | NA | NA | NA |
| WBGene00196263 | NA | NA | NA | NA |
| WBGene00196264 | NA | NA | NA | NA |
| WBGene00196265 | NA | NA | NA | NA |
| WBGene00196266 | NA | NA | NA | NA |
| WBGene00196267 | NA | NA | NA | NA |
| WBGene00196268 | NA | NA | NA | NA |
| WBGene00196269 | NA | NA | NA | NA |
| WBGene00196270 | NA | NA | NA | NA |
| WBGene00196271 | NA | NA | NA | NA |
| WBGene00196272 | NA | NA | NA | NA |
| WBGene00196273 | NA | NA | NA | NA |
| WBGene00196274 | NA | NA | NA | NA |
| WBGene00196275 | NA | NA | NA | NA |
| WBGene00196276 | NA | NA | NA | NA |
| WBGene00196277 | NA | NA | NA | NA |
| WBGene00196278 | NA | NA | NA | NA |
| WBGene00196279 | NA | NA | NA | NA |
| WBGene00196280 | -0.001993815 | 0.95612996 | NA | NA |

| | | | | |
|---|---|---|---|---|
| WBGene00196281 | NA | NA | NA | NA |
| WBGene00196282 | NA | NA | NA | NA |
| WBGene00196283 | NA | NA | NA | NA |
| WBGene00196284 | NA | NA | NA | NA |
| WBGene00196285 | NA | NA | NA | NA |
| WBGene00196286 | NA | NA | NA | NA |
| WBGene00196287 | NA | NA | NA | NA |
| WBGene00196288 | NA | NA | NA | NA |
| WBGene00196289 | NA | NA | NA | NA |
| WBGene00196290 | NA | NA | NA | NA |
| WBGene00196291 | NA | NA | NA | NA |
| WBGene00196292 | NA | NA | NA | NA |
| WBGene00196293 | NA | NA | NA | NA |
| WBGene00196294 | NA | NA | NA | NA |
| WBGene00196295 | 0.020925399 | 0.84643798 | NA | NA |
| WBGene00196296 | NA | NA | NA | NA |
| WBGene00196297 | NA | NA | NA | NA |
| WBGene00196298 | NA | NA | NA | NA |
| WBGene00196299 | NA | NA | NA | NA |
| WBGene00196300 | NA | NA | NA | NA |
| WBGene00196301 | NA | NA | NA | NA |
| WBGene00196302 | NA | NA | NA | NA |
| WBGene00196303 | NA | NA | NA | NA |
| WBGene00196304 | NA | NA | NA | NA |
| WBGene00196305 | NA | NA | NA | NA |
| WBGene00196306 | NA | NA | NA | NA |
| WBGene00196307 | NA | NA | NA | NA |
| WBGene00196308 | NA | NA | NA | NA |
| WBGene00196309 | NA | NA | NA | NA |
| WBGene00196310 | NA | NA | NA | NA |
| WBGene00196311 | NA | NA | NA | NA |
| WBGene00196312 | NA | NA | NA | NA |
| WBGene00196313 | NA | NA | NA | NA |
| WBGene00196314 | NA | NA | NA | NA |
| WBGene00196315 | NA | NA | NA | NA |
| WBGene00196316 | NA | NA | NA | NA |
| WBGene00196317 | NA | NA | NA | NA |
| WBGene00196318 | NA | NA | NA | NA |
| WBGene00196319 | 0.020232312 | 0.84643798 | NA | NA |
| WBGene00196320 | NA | NA | NA | NA |
| WBGene00196321 | NA | NA | NA | NA |
| WBGene00196322 | NA | NA | NA | NA |
| WBGene00196323 | NA | NA | NA | NA |

| | | | | |
|---|---|---|---|---|
| WBGene00196324 | NA | NA | NA | NA |
| WBGene00196325 | NA | NA | NA | NA |
| WBGene00196326 | NA | NA | NA | NA |
| WBGene00196327 | NA | NA | NA | NA |
| WBGene00196328 | 0.020317245 | 0.84643798 | NA | NA |
| WBGene00196329 | NA | NA | NA | NA |
| WBGene00196330 | NA | NA | NA | NA |
| WBGene00196331 | 0.020925399 | 0.84643798 | NA | NA |
| WBGene00196332 | NA | NA | NA | NA |
| WBGene00196333 | NA | NA | NA | NA |
| WBGene00196334 | 0.020232312 | 0.84643798 | NA | NA |
| WBGene00196335 | NA | NA | NA | NA |
| WBGene00196336 | 0.020925399 | 0.84643798 | NA | NA |
| WBGene00196337 | NA | NA | NA | NA |
| WBGene00196338 | 0.020317245 | 0.84643798 | NA | NA |
| WBGene00196339 | NA | NA | NA | NA |
| WBGene00196340 | NA | NA | NA | NA |
| WBGene00196341 | NA | NA | NA | NA |
| WBGene00196342 | NA | NA | NA | NA |
| WBGene00196343 | NA | NA | NA | NA |
| WBGene00196344 | NA | NA | NA | NA |
| WBGene00196345 | NA | NA | NA | NA |
| WBGene00196346 | NA | NA | NA | NA |
| WBGene00196347 | NA | NA | NA | NA |
| WBGene00196348 | NA | NA | NA | NA |
| WBGene00196349 | NA | NA | NA | NA |
| WBGene00196350 | NA | NA | NA | NA |
| WBGene00196351 | NA | NA | NA | NA |
| WBGene00196352 | NA | NA | NA | NA |
| WBGene00196353 | NA | NA | NA | NA |
| WBGene00196354 | NA | NA | NA | NA |
| WBGene00196355 | NA | NA | NA | NA |
| WBGene00196356 | NA | NA | NA | NA |
| WBGene00196357 | NA | NA | NA | NA |
| WBGene00196358 | NA | NA | NA | NA |
| WBGene00196359 | NA | NA | NA | NA |
| WBGene00196360 | NA | NA | NA | NA |
| WBGene00196361 | NA | NA | NA | NA |
| WBGene00196362 | NA | NA | NA | NA |
| WBGene00196363 | NA | NA | NA | NA |
| WBGene00196364 | NA | NA | NA | NA |
| WBGene00196365 | NA | NA | NA | NA |
| WBGene00196366 | NA | NA | NA | NA |

| | | | | |
|---|---|---|---|---|
| WBGene00196367 | NA | NA | NA | NA |
| WBGene00196368 | NA | NA | NA | NA |
| WBGene00196369 | NA | NA | NA | NA |
| WBGene00196370 | NA | NA | NA | NA |
| WBGene00196371 | NA | NA | NA | NA |
| WBGene00196372 | NA | NA | NA | NA |
| WBGene00196373 | NA | NA | NA | NA |
| WBGene00196374 | 0.020232312 | 0.84643798 | NA | NA |
| WBGene00196375 | NA | NA | NA | NA |
| WBGene00196376 | NA | NA | NA | NA |
| WBGene00196377 | NA | NA | NA | NA |
| WBGene00196378 | NA | NA | NA | NA |
| WBGene00196379 | NA | NA | NA | NA |
| WBGene00196380 | NA | NA | NA | NA |
| WBGene00196381 | NA | NA | NA | NA |
| WBGene00196382 | NA | NA | NA | NA |
| WBGene00196383 | NA | NA | NA | NA |
| WBGene00196384 | NA | NA | NA | NA |
| WBGene00196385 | NA | NA | NA | NA |
| WBGene00196386 | NA | NA | NA | NA |
| WBGene00196387 | NA | NA | NA | NA |
| WBGene00196388 | NA | NA | NA | NA |
| WBGene00196389 | NA | NA | NA | NA |
| WBGene00196390 | NA | NA | NA | NA |
| WBGene00196391 | 0.020232312 | 0.84643798 | NA | NA |
| WBGene00196392 | NA | NA | NA | NA |
| WBGene00196393 | NA | NA | NA | NA |
| WBGene00196394 | NA | NA | NA | NA |
| WBGene00196395 | NA | NA | NA | NA |
| WBGene00196396 | NA | NA | NA | NA |
| WBGene00196397 | NA | NA | NA | NA |
| WBGene00196398 | NA | NA | NA | NA |
| WBGene00196399 | NA | NA | NA | NA |
| WBGene00196400 | NA | NA | NA | NA |
| WBGene00196401 | NA | NA | NA | NA |
| WBGene00196402 | NA | NA | NA | NA |
| WBGene00196403 | NA | NA | NA | NA |
| WBGene00196404 | NA | NA | NA | NA |
| WBGene00196405 | NA | NA | NA | NA |
| WBGene00196406 | NA | NA | NA | NA |
| WBGene00196407 | NA | NA | NA | NA |
| WBGene00196408 | NA | NA | NA | NA |
| WBGene00196409 | 0.056979086 | 0.43850646 | NA | NA |

| | | | | |
|---|---|---|---|---|
| WBGene00196410 | NA | NA | NA | NA |
| WBGene00196411 | NA | NA | NA | NA |
| WBGene00196412 | NA | NA | NA | NA |
| WBGene00196413 | NA | NA | NA | NA |
| WBGene00196414 | NA | NA | NA | NA |
| WBGene00196415 | NA | NA | NA | NA |
| WBGene00196416 | NA | NA | NA | NA |
| WBGene00196417 | NA | NA | NA | NA |
| WBGene00196418 | NA | NA | NA | NA |
| WBGene00196419 | NA | NA | NA | NA |
| WBGene00196420 | 0.020232312 | 0.84643798 | NA | NA |
| WBGene00196421 | NA | NA | NA | NA |
| WBGene00196422 | NA | NA | NA | NA |
| WBGene00196423 | NA | NA | NA | NA |
| WBGene00196424 | NA | NA | NA | NA |
| WBGene00196425 | NA | NA | NA | NA |
| WBGene00196426 | NA | NA | NA | NA |
| WBGene00196427 | NA | NA | NA | NA |
| WBGene00196428 | NA | NA | NA | NA |
| WBGene00196429 | NA | NA | NA | NA |
| WBGene00196430 | NA | NA | NA | NA |
| WBGene00196431 | NA | NA | NA | NA |
| WBGene00196432 | NA | NA | NA | NA |
| WBGene00196433 | NA | NA | NA | NA |
| WBGene00196434 | NA | NA | NA | NA |
| WBGene00196435 | NA | NA | NA | NA |
| WBGene00196436 | NA | NA | NA | NA |
| WBGene00196437 | NA | NA | NA | NA |
| WBGene00196438 | NA | NA | NA | NA |
| WBGene00196439 | NA | NA | NA | NA |
| WBGene00196440 | NA | NA | NA | NA |
| WBGene00196441 | NA | NA | NA | NA |
| WBGene00196442 | 0.020925399 | 0.84643798 | NA | NA |
| WBGene00196443 | NA | NA | NA | NA |
| WBGene00196444 | NA | NA | NA | NA |
| WBGene00196445 | NA | NA | NA | NA |
| WBGene00196446 | NA | NA | NA | NA |
| WBGene00196447 | NA | NA | NA | NA |
| WBGene00196448 | NA | NA | NA | NA |
| WBGene00196449 | NA | NA | NA | NA |
| WBGene00196450 | NA | NA | NA | NA |
| WBGene00196451 | NA | NA | NA | NA |
| WBGene00196452 | NA | NA | NA | NA |

| | | | | |
|---|---|---|---|---|
| WBGene00196453 | NA | NA | NA | NA |
| WBGene00196454 | NA | NA | NA | NA |
| WBGene00196455 | NA | NA | NA | NA |
| WBGene00196456 | NA | NA | NA | NA |
| WBGene00196457 | NA | NA | NA | NA |
| WBGene00196458 | NA | NA | NA | NA |
| WBGene00196459 | NA | NA | NA | NA |
| WBGene00196460 | NA | NA | NA | NA |
| WBGene00196461 | NA | NA | NA | NA |
| WBGene00196462 | NA | NA | NA | NA |
| WBGene00196463 | NA | NA | NA | NA |
| WBGene00196464 | NA | NA | NA | NA |
| WBGene00196465 | NA | NA | NA | NA |
| WBGene00196466 | NA | NA | NA | NA |
| WBGene00196467 | NA | NA | NA | NA |
| WBGene00196468 | NA | NA | NA | NA |
| WBGene00196469 | NA | NA | NA | NA |
| WBGene00196470 | 0.038125098 | 0.61119482 | NA | NA |
| WBGene00196471 | -0.023056499 | 0.78121253 | NA | NA |
| WBGene00196472 | NA | NA | NA | NA |
| WBGene00196473 | NA | NA | NA | NA |
| WBGene00196474 | NA | NA | NA | NA |
| WBGene00196475 | NA | NA | NA | NA |
| WBGene00196476 | NA | NA | NA | NA |
| WBGene00196477 | NA | NA | NA | NA |
| WBGene00196478 | NA | NA | NA | NA |
| WBGene00196479 | NA | NA | NA | NA |
| WBGene00196480 | NA | NA | NA | NA |
| WBGene00196481 | NA | NA | NA | NA |
| WBGene00196482 | NA | NA | NA | NA |
| WBGene00196483 | NA | NA | NA | NA |
| WBGene00196484 | NA | NA | NA | NA |
| WBGene00196485 | NA | NA | NA | NA |
| WBGene00196486 | NA | NA | NA | NA |
| WBGene00196487 | NA | NA | NA | NA |
| WBGene00196488 | NA | NA | NA | NA |
| WBGene00196489 | NA | NA | NA | NA |
| WBGene00196490 | NA | NA | NA | NA |
| WBGene00196491 | NA | NA | NA | NA |
| WBGene00196492 | NA | NA | NA | NA |
| WBGene00196493 | NA | NA | NA | NA |
| WBGene00196494 | NA | NA | NA | NA |
| WBGene00196495 | NA | NA | NA | NA |

| | | | | |
|---|---|---|---|---|
| WBGene00196496 | NA | NA | NA | NA |
| WBGene00196497 | NA | NA | NA | NA |
| WBGene00196498 | NA | NA | NA | NA |
| WBGene00196499 | NA | NA | NA | NA |
| WBGene00196500 | NA | NA | NA | NA |
| WBGene00196501 | NA | NA | NA | NA |
| WBGene00196502 | NA | NA | NA | NA |
| WBGene00196503 | NA | NA | NA | NA |
| WBGene00196504 | NA | NA | NA | NA |
| WBGene00196505 | NA | NA | NA | NA |
| WBGene00196506 | NA | NA | NA | NA |
| WBGene00196507 | NA | NA | NA | NA |
| WBGene00196508 | NA | NA | NA | NA |
| WBGene00196509 | NA | NA | NA | NA |
| WBGene00196510 | NA | NA | NA | NA |
| WBGene00196511 | NA | NA | NA | NA |
| WBGene00196512 | NA | NA | NA | NA |
| WBGene00196513 | NA | NA | NA | NA |
| WBGene00196514 | NA | NA | NA | NA |
| WBGene00196515 | NA | NA | NA | NA |
| WBGene00196516 | NA | NA | NA | NA |
| WBGene00196517 | NA | NA | NA | NA |
| WBGene00196518 | NA | NA | NA | NA |
| WBGene00196519 | NA | NA | NA | NA |
| WBGene00196520 | NA | NA | NA | NA |
| WBGene00196521 | NA | NA | NA | NA |
| WBGene00196522 | NA | NA | NA | NA |
| WBGene00196523 | NA | NA | NA | NA |
| WBGene00196524 | NA | NA | NA | NA |
| WBGene00196525 | NA | NA | NA | NA |
| WBGene00196526 | NA | NA | NA | NA |
| WBGene00196527 | NA | NA | NA | NA |
| WBGene00196528 | NA | NA | NA | NA |
| WBGene00196529 | NA | NA | NA | NA |
| WBGene00196530 | NA | NA | NA | NA |
| WBGene00196531 | NA | NA | NA | NA |
| WBGene00196532 | NA | NA | NA | NA |
| WBGene00196533 | NA | NA | NA | NA |
| WBGene00196534 | NA | NA | NA | NA |
| WBGene00196535 | NA | NA | NA | NA |
| WBGene00196536 | NA | NA | NA | NA |
| WBGene00196537 | NA | NA | NA | NA |
| WBGene00196538 | NA | NA | NA | NA |

| | | | | |
|---|---|---|---|---|
| WBGene00196539 | NA | NA | NA | NA |
| WBGene00196540 | NA | NA | NA | NA |
| WBGene00196541 | NA | NA | NA | NA |
| WBGene00196542 | NA | NA | NA | NA |
| WBGene00196543 | NA | NA | NA | NA |
| WBGene00196544 | NA | NA | NA | NA |
| WBGene00196545 | NA | NA | NA | NA |
| WBGene00196546 | 0.020925399 | 0.84643798 | NA | NA |
| WBGene00196547 | NA | NA | NA | NA |
| WBGene00196548 | NA | NA | NA | NA |
| WBGene00196549 | NA | NA | NA | NA |
| WBGene00196550 | NA | NA | NA | NA |
| WBGene00196551 | NA | NA | NA | NA |
| WBGene00196552 | NA | NA | NA | NA |
| WBGene00196553 | NA | NA | NA | NA |
| WBGene00196554 | NA | NA | NA | NA |
| WBGene00196555 | NA | NA | NA | NA |
| WBGene00196556 | NA | NA | NA | NA |
| WBGene00196557 | NA | NA | NA | NA |
| WBGene00196558 | NA | NA | NA | NA |
| WBGene00196559 | 0.020317245 | 0.84643798 | NA | NA |
| WBGene00196560 | 0.020232312 | 0.84643798 | NA | NA |
| WBGene00196561 | NA | NA | NA | NA |
| WBGene00196562 | NA | NA | NA | NA |
| WBGene00196563 | NA | NA | NA | NA |
| WBGene00196564 | NA | NA | NA | NA |
| WBGene00196565 | NA | NA | NA | NA |
| WBGene00196566 | NA | NA | NA | NA |
| WBGene00196567 | NA | NA | NA | NA |
| WBGene00196568 | NA | NA | NA | NA |
| WBGene00196569 | NA | NA | NA | NA |
| WBGene00196570 | NA | NA | NA | NA |
| WBGene00196571 | NA | NA | NA | NA |
| WBGene00196572 | 0.020925399 | 0.84643798 | NA | NA |
| WBGene00196573 | NA | NA | NA | NA |
| WBGene00196574 | NA | NA | NA | NA |
| WBGene00196575 | NA | NA | NA | NA |
| WBGene00196576 | NA | NA | NA | NA |
| WBGene00196577 | NA | NA | NA | NA |
| WBGene00196578 | NA | NA | NA | NA |
| WBGene00196579 | NA | NA | NA | NA |
| WBGene00196580 | NA | NA | NA | NA |
| WBGene00196581 | NA | NA | NA | NA |

| | | | | |
|---|---|---|---|---|
| WBGene00196582 | NA | NA | NA | NA |
| WBGene00196583 | NA | NA | NA | NA |
| WBGene00196584 | NA | NA | NA | NA |
| WBGene00196585 | NA | NA | NA | NA |
| WBGene00196586 | NA | NA | NA | NA |
| WBGene00196587 | NA | NA | NA | NA |
| WBGene00196588 | NA | NA | NA | NA |
| WBGene00196589 | NA | NA | NA | NA |
| WBGene00196590 | NA | NA | NA | NA |
| WBGene00196591 | -0.026384613 | 0.67299636 | NA | NA |
| WBGene00196592 | NA | NA | NA | NA |
| WBGene00196593 | NA | NA | NA | NA |
| WBGene00196594 | NA | NA | NA | NA |
| WBGene00196595 | NA | NA | NA | NA |
| WBGene00196596 | NA | NA | NA | NA |
| WBGene00196597 | NA | NA | NA | NA |
| WBGene00196598 | NA | NA | NA | NA |
| WBGene00196599 | NA | NA | NA | NA |
| WBGene00196600 | NA | NA | NA | NA |
| WBGene00196601 | -0.022930767 | 0.78121253 | NA | NA |
| WBGene00196602 | NA | NA | NA | NA |
| WBGene00196603 | NA | NA | NA | NA |
| WBGene00196604 | NA | NA | NA | NA |
| WBGene00196605 | NA | NA | NA | NA |
| WBGene00196606 | 0.038125098 | 0.61119482 | NA | NA |
| WBGene00196607 | NA | NA | NA | NA |
| WBGene00196608 | NA | NA | NA | NA |
| WBGene00196609 | NA | NA | NA | NA |
| WBGene00196610 | NA | NA | NA | NA |
| WBGene00196611 | NA | NA | NA | NA |
| WBGene00196612 | NA | NA | NA | NA |
| WBGene00196613 | NA | NA | NA | NA |
| WBGene00196614 | NA | NA | NA | NA |
| WBGene00196615 | NA | NA | NA | NA |
| WBGene00196616 | NA | NA | NA | NA |
| WBGene00196617 | NA | NA | NA | NA |
| WBGene00196618 | NA | NA | NA | NA |
| WBGene00196619 | NA | NA | NA | NA |
| WBGene00196620 | NA | NA | NA | NA |
| WBGene00196621 | NA | NA | NA | NA |
| WBGene00196622 | NA | NA | NA | NA |
| WBGene00196623 | NA | NA | NA | NA |
| WBGene00196624 | NA | NA | NA | NA |

| | | | | |
|---|---|---|---|---|
| WBGene00196625 | NA | NA | NA | NA |
| WBGene00196626 | NA | NA | NA | NA |
| WBGene00196627 | NA | NA | NA | NA |
| WBGene00196628 | NA | NA | NA | NA |
| WBGene00196629 | NA | NA | NA | NA |
| WBGene00196630 | NA | NA | NA | NA |
| WBGene00196631 | -0.022930767 | 0.78121253 | NA | NA |
| WBGene00196632 | NA | NA | NA | NA |
| WBGene00196633 | NA | NA | NA | NA |
| WBGene00196634 | NA | NA | NA | NA |
| WBGene00196635 | NA | NA | NA | NA |
| WBGene00196636 | NA | NA | NA | NA |
| WBGene00196637 | NA | NA | NA | NA |
| WBGene00196638 | NA | NA | NA | NA |
| WBGene00196639 | NA | NA | NA | NA |
| WBGene00196640 | NA | NA | NA | NA |
| WBGene00196641 | NA | NA | NA | NA |
| WBGene00196642 | NA | NA | NA | NA |
| WBGene00196643 | NA | NA | NA | NA |
| WBGene00196644 | NA | NA | NA | NA |
| WBGene00196645 | -0.023109762 | 0.78121253 | NA | NA |
| WBGene00196646 | NA | NA | NA | NA |
| WBGene00196647 | NA | NA | NA | NA |
| WBGene00196648 | NA | NA | NA | NA |
| WBGene00196649 | NA | NA | NA | NA |
| WBGene00196650 | NA | NA | NA | NA |
| WBGene00196651 | NA | NA | NA | NA |
| WBGene00196652 | NA | NA | NA | NA |
| WBGene00196653 | NA | NA | NA | NA |
| WBGene00196654 | NA | NA | NA | NA |
| WBGene00196655 | NA | NA | NA | NA |
| WBGene00196656 | NA | NA | NA | NA |
| WBGene00196657 | NA | NA | NA | NA |
| WBGene00196658 | NA | NA | NA | NA |
| WBGene00196659 | 0.020925399 | 0.84643798 | NA | NA |
| WBGene00196660 | NA | NA | NA | NA |
| WBGene00196661 | NA | NA | NA | NA |
| WBGene00196662 | NA | NA | NA | NA |
| WBGene00196663 | NA | NA | NA | NA |
| WBGene00196664 | NA | NA | NA | NA |
| WBGene00196665 | NA | NA | NA | NA |
| WBGene00196666 | NA | NA | NA | NA |
| WBGene00196667 | NA | NA | NA | NA |

| | | | | |
|---|---|---|---|---|
| WBGene00196668 | NA | NA | NA | NA |
| WBGene00196669 | NA | NA | NA | NA |
| WBGene00196670 | NA | NA | NA | NA |
| WBGene00196671 | NA | NA | NA | NA |
| WBGene00196672 | NA | NA | NA | NA |
| WBGene00196673 | NA | NA | NA | NA |
| WBGene00196674 | NA | NA | NA | NA |
| WBGene00196675 | NA | NA | NA | NA |
| WBGene00196676 | NA | NA | NA | NA |
| WBGene00196677 | NA | NA | NA | NA |
| WBGene00196678 | NA | NA | NA | NA |
| WBGene00196679 | -0.023109762 | 0.78121253 | NA | NA |
| WBGene00196680 | NA | NA | NA | NA |
| WBGene00196681 | -0.023109762 | 0.78121253 | NA | NA |
| WBGene00196682 | NA | NA | NA | NA |
| WBGene00196683 | NA | NA | NA | NA |
| WBGene00196684 | NA | NA | NA | NA |
| WBGene00196685 | NA | NA | NA | NA |
| WBGene00196686 | NA | NA | NA | NA |
| WBGene00196687 | -0.023109762 | 0.78121253 | NA | NA |
| WBGene00196688 | NA | NA | NA | NA |
| WBGene00196689 | NA | NA | NA | NA |
| WBGene00196690 | NA | NA | NA | NA |
| WBGene00196691 | NA | NA | NA | NA |
| WBGene00196692 | NA | NA | NA | NA |
| WBGene00196693 | NA | NA | NA | NA |
| WBGene00196694 | NA | NA | NA | NA |
| WBGene00196695 | -0.022930767 | 0.78121253 | NA | NA |
| WBGene00196696 | NA | NA | NA | NA |
| WBGene00196697 | NA | NA | NA | NA |
| WBGene00196698 | NA | NA | NA | NA |
| WBGene00196699 | NA | NA | NA | NA |
| WBGene00196700 | NA | NA | NA | NA |
| WBGene00196701 | NA | NA | NA | NA |
| WBGene00196702 | NA | NA | NA | NA |
| WBGene00196703 | NA | NA | NA | NA |
| WBGene00196704 | NA | NA | NA | NA |
| WBGene00196705 | NA | NA | NA | NA |
| WBGene00196706 | -0.023109762 | 0.78121253 | NA | NA |
| WBGene00196707 | NA | NA | NA | NA |
| WBGene00196708 | NA | NA | NA | NA |
| WBGene00196709 | NA | NA | NA | NA |
| WBGene00196710 | NA | NA | NA | NA |

| | | | | |
|---|---|---|---|---|
| WBGene00196711 | NA | NA | NA | NA |
| WBGene00196712 | NA | NA | NA | NA |
| WBGene00196713 | NA | NA | NA | NA |
| WBGene00196714 | NA | NA | NA | NA |
| WBGene00196715 | NA | NA | NA | NA |
| WBGene00196716 | NA | NA | NA | NA |
| WBGene00196717 | NA | NA | NA | NA |
| WBGene00196718 | NA | NA | NA | NA |
| WBGene00196719 | NA | NA | NA | NA |
| WBGene00196720 | 0.020925399 | 0.84643798 | NA | NA |
| WBGene00196721 | NA | NA | NA | NA |
| WBGene00196722 | NA | NA | NA | NA |
| WBGene00196723 | NA | NA | NA | NA |
| WBGene00196724 | NA | NA | NA | NA |
| WBGene00196725 | NA | NA | NA | NA |
| WBGene00196726 | NA | NA | NA | NA |
| WBGene00196727 | NA | NA | NA | NA |
| WBGene00196728 | NA | NA | NA | NA |
| WBGene00196729 | NA | NA | NA | NA |
| WBGene00196730 | NA | NA | NA | NA |
| WBGene00196731 | NA | NA | NA | NA |
| WBGene00196732 | NA | NA | NA | NA |
| WBGene00196733 | NA | NA | NA | NA |
| WBGene00196734 | NA | NA | NA | NA |
| WBGene00196735 | NA | NA | NA | NA |
| WBGene00196736 | NA | NA | NA | NA |
| WBGene00196737 | NA | NA | NA | NA |
| WBGene00196738 | NA | NA | NA | NA |
| WBGene00196739 | NA | NA | NA | NA |
| WBGene00196740 | NA | NA | NA | NA |
| WBGene00196741 | NA | NA | NA | NA |
| WBGene00196742 | NA | NA | NA | NA |
| WBGene00196743 | NA | NA | NA | NA |
| WBGene00196744 | NA | NA | NA | NA |
| WBGene00196745 | NA | NA | NA | NA |
| WBGene00196746 | NA | NA | NA | NA |
| WBGene00196747 | NA | NA | NA | NA |
| WBGene00196748 | NA | NA | NA | NA |
| WBGene00196749 | NA | NA | NA | NA |
| WBGene00196750 | NA | NA | NA | NA |
| WBGene00196751 | NA | NA | NA | NA |
| WBGene00196752 | NA | NA | NA | NA |
| WBGene00196753 | NA | NA | NA | NA |

| | | | | |
|---|---|---|---|---|
| WBGene00196754 | NA | NA | NA | NA |
| WBGene00196755 | NA | NA | NA | NA |
| WBGene00196756 | NA | NA | NA | NA |
| WBGene00196757 | NA | NA | NA | NA |
| WBGene00196758 | 0.020232312 | 0.84643798 | NA | NA |
| WBGene00196759 | NA | NA | NA | NA |
| WBGene00196760 | NA | NA | NA | NA |
| WBGene00196761 | NA | NA | NA | NA |
| WBGene00196762 | NA | NA | NA | NA |
| WBGene00196763 | NA | NA | NA | NA |
| WBGene00196764 | NA | NA | NA | NA |
| WBGene00196765 | NA | NA | NA | NA |
| WBGene00196766 | NA | NA | NA | NA |
| WBGene00196767 | NA | NA | NA | NA |
| WBGene00196768 | NA | NA | NA | NA |
| WBGene00196769 | NA | NA | NA | NA |
| WBGene00196770 | NA | NA | NA | NA |
| WBGene00196771 | NA | NA | NA | NA |
| WBGene00196772 | NA | NA | NA | NA |
| WBGene00196773 | NA | NA | NA | NA |
| WBGene00196774 | NA | NA | NA | NA |
| WBGene00196775 | NA | NA | NA | NA |
| WBGene00196776 | NA | NA | NA | NA |
| WBGene00196777 | NA | NA | NA | NA |
| WBGene00196778 | NA | NA | NA | NA |
| WBGene00196779 | NA | NA | NA | NA |
| WBGene00196780 | 0.020317245 | 0.84643798 | NA | NA |
| WBGene00196781 | NA | NA | NA | NA |
| WBGene00196782 | NA | NA | NA | NA |
| WBGene00196783 | NA | NA | NA | NA |
| WBGene00196784 | NA | NA | NA | NA |
| WBGene00196785 | NA | NA | NA | NA |
| WBGene00196786 | NA | NA | NA | NA |
| WBGene00196787 | NA | NA | NA | NA |
| WBGene00196788 | NA | NA | NA | NA |
| WBGene00196789 | NA | NA | NA | NA |
| WBGene00196790 | NA | NA | NA | NA |
| WBGene00196791 | NA | NA | NA | NA |
| WBGene00196792 | NA | NA | NA | NA |
| WBGene00196793 | NA | NA | NA | NA |
| WBGene00196794 | NA | NA | NA | NA |
| WBGene00196795 | NA | NA | NA | NA |
| WBGene00196796 | NA | NA | NA | NA |

| | | | | |
|---|---|---|---|---|
| WBGene00196797 | NA | NA | NA | NA |
| WBGene00196798 | NA | NA | NA | NA |
| WBGene00196799 | NA | NA | NA | NA |
| WBGene00196800 | NA | NA | NA | NA |
| WBGene00196801 | NA | NA | NA | NA |
| WBGene00196802 | -0.003270206 | 0.95698316 | NA | NA |
| WBGene00196803 | NA | NA | NA | NA |
| WBGene00196804 | 0.020925399 | 0.84643798 | NA | NA |
| WBGene00196805 | NA | NA | NA | NA |
| WBGene00196806 | NA | NA | NA | NA |
| WBGene00196807 | NA | NA | NA | NA |
| WBGene00196808 | NA | NA | NA | NA |
| WBGene00196809 | NA | NA | NA | NA |
| WBGene00196810 | NA | NA | NA | NA |
| WBGene00196811 | NA | NA | NA | NA |
| WBGene00196812 | NA | NA | NA | NA |
| WBGene00196813 | NA | NA | NA | NA |
| WBGene00196814 | -0.023109762 | 0.78121253 | NA | NA |
| WBGene00196815 | NA | NA | NA | NA |
| WBGene00196816 | NA | NA | NA | NA |
| WBGene00196817 | NA | NA | NA | NA |
| WBGene00196818 | NA | NA | NA | NA |
| WBGene00196819 | NA | NA | NA | NA |
| WBGene00196820 | NA | NA | NA | NA |
| WBGene00196821 | NA | NA | NA | NA |
| WBGene00196822 | NA | NA | NA | NA |
| WBGene00196823 | NA | NA | NA | NA |
| WBGene00196824 | NA | NA | NA | NA |
| WBGene00196825 | NA | NA | NA | NA |
| WBGene00196826 | NA | NA | NA | NA |
| WBGene00196827 | NA | NA | NA | NA |
| WBGene00196828 | NA | NA | NA | NA |
| WBGene00196829 | NA | NA | NA | NA |
| WBGene00196830 | 0.056188364 | 0.42092875 | NA | NA |
| WBGene00196831 | NA | NA | NA | NA |
| WBGene00196832 | NA | NA | NA | NA |
| WBGene00196833 | NA | NA | NA | NA |
| WBGene00196834 | NA | NA | NA | NA |
| WBGene00196835 | NA | NA | NA | NA |
| WBGene00196836 | NA | NA | NA | NA |
| WBGene00196837 | NA | NA | NA | NA |
| WBGene00196838 | NA | NA | NA | NA |
| WBGene00196839 | NA | NA | NA | NA |

| | | | | |
|---|---|---|---|---|
| WBGene00196840 | NA | NA | NA | NA |
| WBGene00196841 | NA | NA | NA | NA |
| WBGene00196842 | NA | NA | NA | NA |
| WBGene00196843 | NA | NA | NA | NA |
| WBGene00196844 | NA | NA | NA | NA |
| WBGene00196845 | NA | NA | NA | NA |
| WBGene00196846 | NA | NA | NA | NA |
| WBGene00196847 | NA | NA | NA | NA |
| WBGene00196848 | 0.040107227 | 0.58693163 | NA | NA |
| WBGene00196849 | NA | NA | NA | NA |
| WBGene00196850 | NA | NA | NA | NA |
| WBGene00196851 | NA | NA | NA | NA |
| WBGene00196852 | 0.020925399 | 0.84643798 | NA | NA |
| WBGene00196853 | NA | NA | NA | NA |
| WBGene00196854 | NA | NA | NA | NA |
| WBGene00196855 | NA | NA | NA | NA |
| WBGene00196856 | NA | NA | NA | NA |
| WBGene00196857 | NA | NA | NA | NA |
| WBGene00196858 | NA | NA | NA | NA |
| WBGene00196859 | NA | NA | NA | NA |
| WBGene00196860 | NA | NA | NA | NA |
| WBGene00196861 | NA | NA | NA | NA |
| WBGene00196862 | NA | NA | NA | NA |
| WBGene00196863 | NA | NA | NA | NA |
| WBGene00196864 | NA | NA | NA | NA |
| WBGene00196865 | NA | NA | NA | NA |
| WBGene00196866 | NA | NA | NA | NA |
| WBGene00196867 | NA | NA | NA | NA |
| WBGene00196868 | NA | NA | NA | NA |
| WBGene00196869 | NA | NA | NA | NA |
| WBGene00196870 | NA | NA | NA | NA |
| WBGene00196871 | NA | NA | NA | NA |
| WBGene00196872 | NA | NA | NA | NA |
| WBGene00196873 | NA | NA | NA | NA |
| WBGene00196874 | NA | NA | NA | NA |
| WBGene00196875 | NA | NA | NA | NA |
| WBGene00196876 | NA | NA | NA | NA |
| WBGene00196877 | NA | NA | NA | NA |
| WBGene00196878 | NA | NA | NA | NA |
| WBGene00196879 | -0.023056499 | 0.78121253 | NA | NA |
| WBGene00196880 | NA | NA | NA | NA |
| WBGene00196881 | NA | NA | NA | NA |
| WBGene00196882 | NA | NA | NA | NA |

| | | | | |
|---|---|---|---|---|
| WBGene00196883 | NA | NA | NA | NA |
| WBGene00196884 | NA | NA | NA | NA |
| WBGene00196885 | -0.003270206 | 0.95698316 | NA | NA |
| WBGene00196886 | NA | NA | NA | NA |
| WBGene00196887 | NA | NA | NA | NA |
| WBGene00196888 | -0.023109762 | 0.78121253 | NA | NA |
| WBGene00196889 | NA | NA | NA | NA |
| WBGene00196890 | NA | NA | NA | NA |
| WBGene00196891 | NA | NA | NA | NA |
| WBGene00196892 | NA | NA | NA | NA |
| WBGene00196893 | NA | NA | NA | NA |
| WBGene00196894 | NA | NA | NA | NA |
| WBGene00196895 | NA | NA | NA | NA |
| WBGene00196896 | NA | NA | NA | NA |
| WBGene00196897 | NA | NA | NA | NA |
| WBGene00196898 | NA | NA | NA | NA |
| WBGene00196899 | NA | NA | NA | NA |
| WBGene00196900 | NA | NA | NA | NA |
| WBGene00196901 | NA | NA | NA | NA |
| WBGene00196902 | NA | NA | NA | NA |
| WBGene00196903 | NA | NA | NA | NA |
| WBGene00196904 | NA | NA | NA | NA |
| WBGene00196905 | NA | NA | NA | NA |
| WBGene00196906 | NA | NA | NA | NA |
| WBGene00196907 | NA | NA | NA | NA |
| WBGene00196908 | NA | NA | NA | NA |
| WBGene00196909 | 0.020232312 | 0.84643798 | NA | NA |
| WBGene00196910 | NA | NA | NA | NA |
| WBGene00196911 | NA | NA | NA | NA |
| WBGene00196912 | NA | NA | NA | NA |
| WBGene00196913 | NA | NA | NA | NA |
| WBGene00196914 | 0.056188364 | 0.42092875 | NA | NA |
| WBGene00196915 | NA | NA | NA | NA |
| WBGene00196916 | NA | NA | NA | NA |
| WBGene00196917 | NA | NA | NA | NA |
| WBGene00196918 | NA | NA | NA | NA |
| WBGene00196919 | NA | NA | NA | NA |
| WBGene00196920 | NA | NA | NA | NA |
| WBGene00196921 | NA | NA | NA | NA |
| WBGene00196922 | NA | NA | NA | NA |
| WBGene00196923 | NA | NA | NA | NA |
| WBGene00196924 | NA | NA | NA | NA |
| WBGene00196925 | NA | NA | NA | NA |

| | | | | |
|---|---|---|---|---|
| WBGene00196926 | NA | NA | NA | NA |
| WBGene00196927 | NA | NA | NA | NA |
| WBGene00196928 | NA | NA | NA | NA |
| WBGene00196929 | NA | NA | NA | NA |
| WBGene00196930 | NA | NA | NA | NA |
| WBGene00196931 | NA | NA | NA | NA |
| WBGene00196932 | NA | NA | NA | NA |
| WBGene00196933 | NA | NA | NA | NA |
| WBGene00196934 | NA | NA | NA | NA |
| WBGene00196935 | NA | NA | NA | NA |
| WBGene00196936 | NA | NA | NA | NA |
| WBGene00196937 | NA | NA | NA | NA |
| WBGene00196938 | NA | NA | NA | NA |
| WBGene00196939 | NA | NA | NA | NA |
| WBGene00196940 | NA | NA | NA | NA |
| WBGene00196941 | NA | NA | NA | NA |
| WBGene00196942 | NA | NA | NA | NA |
| WBGene00196943 | NA | NA | NA | NA |
| WBGene00196944 | 0.063217264 | 0.41230173 | NA | NA |
| WBGene00196945 | NA | NA | NA | NA |
| WBGene00196946 | -0.003421551 | 0.95689871 | NA | NA |
| WBGene00196947 | NA | NA | NA | NA |
| WBGene00196948 | NA | NA | NA | NA |
| WBGene00196949 | 0.020232312 | 0.84643798 | NA | NA |
| WBGene00196950 | 0.020317245 | 0.84643798 | NA | NA |
| WBGene00196951 | NA | NA | NA | NA |
| WBGene00196952 | NA | NA | NA | NA |
| WBGene00196953 | NA | NA | NA | NA |
| WBGene00196954 | NA | NA | NA | NA |
| WBGene00196955 | 0.020232312 | 0.84643798 | NA | NA |
| WBGene00196956 | NA | NA | NA | NA |
| WBGene00196957 | NA | NA | NA | NA |
| WBGene00196958 | NA | NA | NA | NA |
| WBGene00196959 | NA | NA | NA | NA |
| WBGene00196960 | NA | NA | NA | NA |
| WBGene00196961 | NA | NA | NA | NA |
| WBGene00196962 | NA | NA | NA | NA |
| WBGene00196963 | NA | NA | NA | NA |
| WBGene00196964 | NA | NA | NA | NA |
| WBGene00196965 | NA | NA | NA | NA |
| WBGene00196966 | NA | NA | NA | NA |
| WBGene00196967 | NA | NA | NA | NA |
| WBGene00196968 | NA | NA | NA | NA |

| | | | | |
|---|---|---|---|---|
| WBGene00196969 | NA | NA | NA | NA |
| WBGene00196970 | NA | NA | NA | NA |
| WBGene00196971 | NA | NA | NA | NA |
| WBGene00196972 | NA | NA | NA | NA |
| WBGene00196973 | NA | NA | NA | NA |
| WBGene00196974 | NA | NA | NA | NA |
| WBGene00196975 | NA | NA | NA | NA |
| WBGene00196976 | NA | NA | NA | NA |
| WBGene00196977 | NA | NA | NA | NA |
| WBGene00196978 | NA | NA | NA | NA |
| WBGene00196979 | NA | NA | NA | NA |
| WBGene00196980 | NA | NA | NA | NA |
| WBGene00196981 | NA | NA | NA | NA |
| WBGene00196982 | NA | NA | NA | NA |
| WBGene00196983 | NA | NA | NA | NA |
| WBGene00196984 | NA | NA | NA | NA |
| WBGene00196985 | NA | NA | NA | NA |
| WBGene00196986 | NA | NA | NA | NA |
| WBGene00196987 | NA | NA | NA | NA |
| WBGene00196988 | NA | NA | NA | NA |
| WBGene00196989 | NA | NA | NA | NA |
| WBGene00196990 | NA | NA | NA | NA |
| WBGene00196991 | NA | NA | NA | NA |
| WBGene00196992 | NA | NA | NA | NA |
| WBGene00196993 | NA | NA | NA | NA |
| WBGene00196994 | 0.020232312 | 0.84643798 | NA | NA |
| WBGene00196995 | 0.020232312 | 0.84643798 | NA | NA |
| WBGene00196996 | NA | NA | NA | NA |
| WBGene00196997 | NA | NA | NA | NA |
| WBGene00196998 | NA | NA | NA | NA |
| WBGene00196999 | NA | NA | NA | NA |
| WBGene00197000 | NA | NA | NA | NA |
| WBGene00197001 | NA | NA | NA | NA |
| WBGene00197002 | NA | NA | NA | NA |
| WBGene00197003 | NA | NA | NA | NA |
| WBGene00197004 | NA | NA | NA | NA |
| WBGene00197005 | NA | NA | NA | NA |
| WBGene00197006 | NA | NA | NA | NA |
| WBGene00197007 | NA | NA | NA | NA |
| WBGene00197008 | NA | NA | NA | NA |
| WBGene00197009 | NA | NA | NA | NA |
| WBGene00197010 | NA | NA | NA | NA |
| WBGene00197011 | NA | NA | NA | NA |

| | | | | |
|---|---|---|---|---|
| WBGene00197012 | NA | NA | NA | NA |
| WBGene00197013 | NA | NA | NA | NA |
| WBGene00197014 | NA | NA | NA | NA |
| WBGene00197015 | NA | NA | NA | NA |
| WBGene00197016 | NA | NA | NA | NA |
| WBGene00197017 | NA | NA | NA | NA |
| WBGene00197018 | NA | NA | NA | NA |
| WBGene00197019 | NA | NA | NA | NA |
| WBGene00197020 | NA | NA | NA | NA |
| WBGene00197021 | NA | NA | NA | NA |
| WBGene00197022 | NA | NA | NA | NA |
| WBGene00197023 | NA | NA | NA | NA |
| WBGene00197024 | NA | NA | NA | NA |
| WBGene00197025 | -0.045771477 | 0.50913401 | NA | NA |
| WBGene00197026 | NA | NA | NA | NA |
| WBGene00197027 | NA | NA | NA | NA |
| WBGene00197028 | NA | NA | NA | NA |
| WBGene00197029 | NA | NA | NA | NA |
| WBGene00197030 | NA | NA | NA | NA |
| WBGene00197031 | NA | NA | NA | NA |
| WBGene00197032 | NA | NA | NA | NA |
| WBGene00197033 | 0.018342935 | 0.76675379 | NA | NA |
| WBGene00197034 | NA | NA | NA | NA |
| WBGene00197035 | NA | NA | NA | NA |
| WBGene00197036 | NA | NA | NA | NA |
| WBGene00197037 | NA | NA | NA | NA |
| WBGene00197038 | NA | NA | NA | NA |
| WBGene00197039 | NA | NA | NA | NA |
| WBGene00197040 | NA | NA | NA | NA |
| WBGene00197041 | -0.003421551 | 0.95689871 | NA | NA |
| WBGene00197042 | NA | NA | NA | NA |
| WBGene00197043 | NA | NA | NA | NA |
| WBGene00197044 | NA | NA | NA | NA |
| WBGene00197045 | 0.01912147 | 0.77587372 | NA | NA |
| WBGene00197046 | NA | NA | NA | NA |
| WBGene00197047 | NA | NA | NA | NA |
| WBGene00197048 | NA | NA | NA | NA |
| WBGene00197049 | NA | NA | NA | NA |
| WBGene00197050 | 0.020232312 | 0.84643798 | NA | NA |
| WBGene00197051 | NA | NA | NA | NA |
| WBGene00197052 | NA | NA | NA | NA |
| WBGene00197053 | NA | NA | NA | NA |
| WBGene00197054 | NA | NA | NA | NA |

| | | | | |
|---|---|---|---|---|
| WBGene00197055 | NA | NA | NA | NA |
| WBGene00197056 | NA | NA | NA | NA |
| WBGene00197057 | NA | NA | NA | NA |
| WBGene00197058 | NA | NA | NA | NA |
| WBGene00197059 | NA | NA | NA | NA |
| WBGene00197060 | -0.022930767 | 0.78121253 | NA | NA |
| WBGene00197061 | NA | NA | NA | NA |
| WBGene00197062 | NA | NA | NA | NA |
| WBGene00197063 | NA | NA | NA | NA |
| WBGene00197064 | NA | NA | NA | NA |
| WBGene00197065 | NA | NA | NA | NA |
| WBGene00197066 | NA | NA | NA | NA |
| WBGene00197067 | NA | NA | NA | NA |
| WBGene00197068 | NA | NA | NA | NA |
| WBGene00197069 | NA | NA | NA | NA |
| WBGene00197070 | NA | NA | NA | NA |
| WBGene00197071 | NA | NA | NA | NA |
| WBGene00197072 | NA | NA | NA | NA |
| WBGene00197073 | NA | NA | NA | NA |
| WBGene00197074 | NA | NA | NA | NA |
| WBGene00197075 | NA | NA | NA | NA |
| WBGene00197076 | NA | NA | NA | NA |
| WBGene00197077 | NA | NA | NA | NA |
| WBGene00197078 | NA | NA | NA | NA |
| WBGene00197079 | NA | NA | NA | NA |
| WBGene00197080 | 0.059980168 | 0.38780842 | NA | NA |
| WBGene00197081 | NA | NA | NA | NA |
| WBGene00197082 | NA | NA | NA | NA |
| WBGene00197083 | NA | NA | NA | NA |
| WBGene00197084 | NA | NA | NA | NA |
| WBGene00197085 | NA | NA | NA | NA |
| WBGene00197086 | NA | NA | NA | NA |
| WBGene00197087 | NA | NA | NA | NA |
| WBGene00197088 | NA | NA | NA | NA |
| WBGene00197089 | NA | NA | NA | NA |
| WBGene00197090 | NA | NA | NA | NA |
| WBGene00197091 | NA | NA | NA | NA |
| WBGene00197092 | NA | NA | NA | NA |
| WBGene00197093 | NA | NA | NA | NA |
| WBGene00197094 | NA | NA | NA | NA |
| WBGene00197095 | NA | NA | NA | NA |
| WBGene00197096 | NA | NA | NA | NA |
| WBGene00197097 | NA | NA | NA | NA |

| | | | | |
|---|---|---|---|---|
| WBGene00197098 | NA | NA | NA | NA |
| WBGene00197099 | -0.022930767 | 0.78121253 | NA | NA |
| WBGene00197100 | NA | NA | NA | NA |
| WBGene00197101 | NA | NA | NA | NA |
| WBGene00197102 | NA | NA | NA | NA |
| WBGene00197103 | NA | NA | NA | NA |
| WBGene00197104 | NA | NA | NA | NA |
| WBGene00197105 | NA | NA | NA | NA |
| WBGene00197106 | NA | NA | NA | NA |
| WBGene00197107 | NA | NA | NA | NA |
| WBGene00197108 | NA | NA | NA | NA |
| WBGene00197109 | NA | NA | NA | NA |
| WBGene00197110 | NA | NA | NA | NA |
| WBGene00197111 | NA | NA | NA | NA |
| WBGene00197112 | NA | NA | NA | NA |
| WBGene00197113 | NA | NA | NA | NA |
| WBGene00197114 | NA | NA | NA | NA |
| WBGene00197115 | NA | NA | NA | NA |
| WBGene00197116 | NA | NA | NA | NA |
| WBGene00197117 | NA | NA | NA | NA |
| WBGene00197118 | NA | NA | NA | NA |
| WBGene00197119 | NA | NA | NA | NA |
| WBGene00197120 | NA | NA | NA | NA |
| WBGene00197121 | NA | NA | NA | NA |
| WBGene00197122 | NA | NA | NA | NA |
| WBGene00197123 | NA | NA | NA | NA |
| WBGene00197124 | NA | NA | NA | NA |
| WBGene00197125 | NA | NA | NA | NA |
| WBGene00197126 | 0.020317245 | 0.84643798 | NA | NA |
| WBGene00197127 | NA | NA | NA | NA |
| WBGene00197128 | NA | NA | NA | NA |
| WBGene00197129 | NA | NA | NA | NA |
| WBGene00197130 | NA | NA | NA | NA |
| WBGene00197131 | NA | NA | NA | NA |
| WBGene00197132 | NA | NA | NA | NA |
| WBGene00197133 | NA | NA | NA | NA |
| WBGene00197134 | NA | NA | NA | NA |
| WBGene00197135 | NA | NA | NA | NA |
| WBGene00197136 | NA | NA | NA | NA |
| WBGene00197137 | NA | NA | NA | NA |
| WBGene00197138 | NA | NA | NA | NA |
| WBGene00197139 | NA | NA | NA | NA |
| WBGene00197140 | 0.020232312 | 0.84643798 | NA | NA |

| | | | | |
|---|---|---|---|---|
| WBGene00197141 | NA | NA | NA | NA |
| WBGene00197142 | NA | NA | NA | NA |
| WBGene00197143 | NA | NA | NA | NA |
| WBGene00197144 | NA | NA | NA | NA |
| WBGene00197145 | -0.001842784 | 0.95621055 | NA | NA |
| WBGene00197146 | NA | NA | NA | NA |
| WBGene00197147 | NA | NA | NA | NA |
| WBGene00197148 | NA | NA | NA | NA |
| WBGene00197149 | NA | NA | NA | NA |
| WBGene00197150 | NA | NA | NA | NA |
| WBGene00197151 | NA | NA | NA | NA |
| WBGene00197152 | NA | NA | NA | NA |
| WBGene00197153 | 0.020232312 | 0.84643798 | NA | NA |
| WBGene00197154 | NA | NA | NA | NA |
| WBGene00197155 | NA | NA | NA | NA |
| WBGene00197156 | NA | NA | NA | NA |
| WBGene00197157 | NA | NA | NA | NA |
| WBGene00197158 | NA | NA | NA | NA |
| WBGene00197159 | NA | NA | NA | NA |
| WBGene00197160 | NA | NA | NA | NA |
| WBGene00197161 | NA | NA | NA | NA |
| WBGene00197162 | NA | NA | NA | NA |
| WBGene00197163 | NA | NA | NA | NA |
| WBGene00197164 | NA | NA | NA | NA |
| WBGene00197165 | NA | NA | NA | NA |
| WBGene00197166 | NA | NA | NA | NA |
| WBGene00197167 | NA | NA | NA | NA |
| WBGene00197168 | NA | NA | NA | NA |
| WBGene00197169 | 0.020317245 | 0.84643798 | NA | NA |
| WBGene00197170 | NA | NA | NA | NA |
| WBGene00197171 | NA | NA | NA | NA |
| WBGene00197172 | NA | NA | NA | NA |
| WBGene00197173 | NA | NA | NA | NA |
| WBGene00197174 | NA | NA | NA | NA |
| WBGene00197175 | NA | NA | NA | NA |
| WBGene00197176 | NA | NA | NA | NA |
| WBGene00197177 | NA | NA | NA | NA |
| WBGene00197178 | NA | NA | NA | NA |
| WBGene00197179 | NA | NA | NA | NA |
| WBGene00197180 | NA | NA | NA | NA |
| WBGene00197181 | NA | NA | NA | NA |
| WBGene00197182 | NA | NA | NA | NA |
| WBGene00197183 | 0.020232312 | 0.84643798 | NA | NA |

| | | | | |
|---|---|---|---|---|
| WBGene00197184 | NA | NA | NA | NA |
| WBGene00197185 | NA | NA | NA | NA |
| WBGene00197186 | NA | NA | NA | NA |
| WBGene00197187 | NA | NA | NA | NA |
| WBGene00197188 | NA | NA | NA | NA |
| WBGene00197189 | NA | NA | NA | NA |
| WBGene00197190 | NA | NA | NA | NA |
| WBGene00197191 | NA | NA | NA | NA |
| WBGene00197192 | NA | NA | NA | NA |
| WBGene00197193 | NA | NA | NA | NA |
| WBGene00197194 | NA | NA | NA | NA |
| WBGene00197195 | NA | NA | NA | NA |
| WBGene00197196 | NA | NA | NA | NA |
| WBGene00197197 | NA | NA | NA | NA |
| WBGene00197198 | NA | NA | NA | NA |
| WBGene00197199 | NA | NA | NA | NA |
| WBGene00197200 | NA | NA | NA | NA |
| WBGene00197201 | NA | NA | NA | NA |
| WBGene00197202 | NA | NA | NA | NA |
| WBGene00197203 | NA | NA | NA | NA |
| WBGene00197204 | NA | NA | NA | NA |
| WBGene00197205 | NA | NA | NA | NA |
| WBGene00197206 | NA | NA | NA | NA |
| WBGene00197207 | NA | NA | NA | NA |
| WBGene00197208 | 0.0398788 | 0.58971221 | NA | NA |
| WBGene00197209 | NA | NA | NA | NA |
| WBGene00197210 | NA | NA | NA | NA |
| WBGene00197211 | NA | NA | NA | NA |
| WBGene00197212 | NA | NA | NA | NA |
| WBGene00197213 | NA | NA | NA | NA |
| WBGene00197214 | NA | NA | NA | NA |
| WBGene00197215 | NA | NA | NA | NA |
| WBGene00197216 | NA | NA | NA | NA |
| WBGene00197217 | NA | NA | NA | NA |
| WBGene00197218 | NA | NA | NA | NA |
| WBGene00197219 | NA | NA | NA | NA |
| WBGene00197220 | NA | NA | NA | NA |
| WBGene00197221 | NA | NA | NA | NA |
| WBGene00197222 | NA | NA | NA | NA |
| WBGene00197223 | NA | NA | NA | NA |
| WBGene00197224 | NA | NA | NA | NA |
| WBGene00197225 | NA | NA | NA | NA |
| WBGene00197226 | NA | NA | NA | NA |

| | | | | |
|---|---|---|---|---|
| WBGene00197227 | NA | NA | NA | NA |
| WBGene00197228 | 0.020232312 | 0.84643798 | NA | NA |
| WBGene00197229 | NA | NA | NA | NA |
| WBGene00197230 | NA | NA | NA | NA |
| WBGene00197231 | NA | NA | NA | NA |
| WBGene00197232 | NA | NA | NA | NA |
| WBGene00197233 | NA | NA | NA | NA |
| WBGene00197234 | NA | NA | NA | NA |
| WBGene00197235 | NA | NA | NA | NA |
| WBGene00197236 | NA | NA | NA | NA |
| WBGene00197237 | NA | NA | NA | NA |
| WBGene00197238 | NA | NA | NA | NA |
| WBGene00197239 | NA | NA | NA | NA |
| WBGene00197240 | NA | NA | NA | NA |
| WBGene00197241 | NA | NA | NA | NA |
| WBGene00197242 | NA | NA | NA | NA |
| WBGene00197243 | NA | NA | NA | NA |
| WBGene00197244 | NA | NA | NA | NA |
| WBGene00197245 | NA | NA | NA | NA |
| WBGene00197246 | NA | NA | NA | NA |
| WBGene00197247 | NA | NA | NA | NA |
| WBGene00197248 | NA | NA | NA | NA |
| WBGene00197249 | NA | NA | NA | NA |
| WBGene00197250 | NA | NA | NA | NA |
| WBGene00197251 | NA | NA | NA | NA |
| WBGene00197252 | NA | NA | NA | NA |
| WBGene00197253 | NA | NA | NA | NA |
| WBGene00197254 | NA | NA | NA | NA |
| WBGene00197255 | NA | NA | NA | NA |
| WBGene00197256 | NA | NA | NA | NA |
| WBGene00197257 | NA | NA | NA | NA |
| WBGene00197258 | NA | NA | NA | NA |
| WBGene00197259 | NA | NA | NA | NA |
| WBGene00197260 | NA | NA | NA | NA |
| WBGene00197261 | NA | NA | NA | NA |
| WBGene00197262 | NA | NA | NA | NA |
| WBGene00197263 | NA | NA | NA | NA |
| WBGene00197264 | NA | NA | NA | NA |
| WBGene00197265 | NA | NA | NA | NA |
| WBGene00197266 | NA | NA | NA | NA |
| WBGene00197267 | NA | NA | NA | NA |
| WBGene00197268 | NA | NA | NA | NA |
| WBGene00197269 | NA | NA | NA | NA |

| | | | | |
|---|---|---|---|---|
| WBGene00197270 | NA | NA | NA | NA |
| WBGene00197271 | NA | NA | NA | NA |
| WBGene00197272 | NA | NA | NA | NA |
| WBGene00197273 | NA | NA | NA | NA |
| WBGene00197274 | NA | NA | NA | NA |
| WBGene00197275 | NA | NA | NA | NA |
| WBGene00197276 | NA | NA | NA | NA |
| WBGene00197277 | NA | NA | NA | NA |
| WBGene00197278 | NA | NA | NA | NA |
| WBGene00197279 | NA | NA | NA | NA |
| WBGene00197280 | NA | NA | NA | NA |
| WBGene00197281 | NA | NA | NA | NA |
| WBGene00197282 | NA | NA | NA | NA |
| WBGene00197283 | NA | NA | NA | NA |
| WBGene00197284 | NA | NA | NA | NA |
| WBGene00197285 | NA | NA | NA | NA |
| WBGene00197286 | 0.020317245 | 0.84643798 | NA | NA |
| WBGene00197287 | NA | NA | NA | NA |
| WBGene00197288 | NA | NA | NA | NA |
| WBGene00197289 | NA | NA | NA | NA |
| WBGene00197290 | NA | NA | NA | NA |
| WBGene00197291 | NA | NA | NA | NA |
| WBGene00197292 | NA | NA | NA | NA |
| WBGene00197293 | NA | NA | NA | NA |
| WBGene00197294 | NA | NA | NA | NA |
| WBGene00197295 | NA | NA | NA | NA |
| WBGene00197296 | -0.023056499 | 0.78121253 | NA | NA |
| WBGene00197297 | NA | NA | NA | NA |
| WBGene00197298 | NA | NA | NA | NA |
| WBGene00197299 | NA | NA | NA | NA |
| WBGene00197300 | 0.017604133 | 0.79437959 | NA | NA |
| WBGene00197301 | NA | NA | NA | NA |
| WBGene00197302 | NA | NA | NA | NA |
| WBGene00197303 | NA | NA | NA | NA |
| WBGene00197304 | NA | NA | NA | NA |
| WBGene00197305 | NA | NA | NA | NA |
| WBGene00197306 | NA | NA | NA | NA |
| WBGene00197307 | NA | NA | NA | NA |
| WBGene00197308 | NA | NA | NA | NA |
| WBGene00197309 | NA | NA | NA | NA |
| WBGene00197310 | NA | NA | NA | NA |
| WBGene00197311 | NA | NA | NA | NA |
| WBGene00197312 | NA | NA | NA | NA |

| | | | | |
|---|---|---|---|---|
| WBGene00197313 | NA | NA | NA | NA |
| WBGene00197314 | NA | NA | NA | NA |
| WBGene00197315 | NA | NA | NA | NA |
| WBGene00197316 | NA | NA | NA | NA |
| WBGene00197317 | NA | NA | NA | NA |
| WBGene00197318 | NA | NA | NA | NA |
| WBGene00197319 | NA | NA | NA | NA |
| WBGene00197320 | NA | NA | NA | NA |
| WBGene00197321 | NA | NA | NA | NA |
| WBGene00197322 | NA | NA | NA | NA |
| WBGene00197323 | NA | NA | NA | NA |
| WBGene00197324 | NA | NA | NA | NA |
| WBGene00197325 | NA | NA | NA | NA |
| WBGene00197326 | NA | NA | NA | NA |
| WBGene00197327 | NA | NA | NA | NA |
| WBGene00197328 | -0.023056499 | 0.78121253 | NA | NA |
| WBGene00197329 | NA | NA | NA | NA |
| WBGene00197330 | NA | NA | NA | NA |
| WBGene00197331 | NA | NA | NA | NA |
| WBGene00197332 | NA | NA | NA | NA |
| WBGene00197333 | NA | NA | NA | NA |
| WBGene00197334 | NA | NA | NA | NA |
| WBGene00197335 | NA | NA | NA | NA |
| WBGene00197336 | NA | NA | NA | NA |
| WBGene00197337 | NA | NA | NA | NA |
| WBGene00197338 | NA | NA | NA | NA |
| WBGene00197339 | NA | NA | NA | NA |
| WBGene00197340 | NA | NA | NA | NA |
| WBGene00197341 | NA | NA | NA | NA |
| WBGene00197342 | NA | NA | NA | NA |
| WBGene00197343 | NA | NA | NA | NA |
| WBGene00197344 | NA | NA | NA | NA |
| WBGene00197345 | NA | NA | NA | NA |
| WBGene00197346 | NA | NA | NA | NA |
| WBGene00197347 | 0.020232312 | 0.84643798 | NA | NA |
| WBGene00197348 | NA | NA | NA | NA |
| WBGene00197349 | NA | NA | NA | NA |
| WBGene00197350 | NA | NA | NA | NA |
| WBGene00197351 | NA | NA | NA | NA |
| WBGene00197352 | NA | NA | NA | NA |
| WBGene00197353 | NA | NA | NA | NA |
| WBGene00197354 | NA | NA | NA | NA |
| WBGene00197355 | NA | NA | NA | NA |

| | | | | |
|---|---|---|---|---|
| WBGene00197356 | NA | NA | NA | NA |
| WBGene00197357 | NA | NA | NA | NA |
| WBGene00197358 | NA | NA | NA | NA |
| WBGene00197359 | NA | NA | NA | NA |
| WBGene00197360 | NA | NA | NA | NA |
| WBGene00197361 | NA | NA | NA | NA |
| WBGene00197362 | NA | NA | NA | NA |
| WBGene00197363 | NA | NA | NA | NA |
| WBGene00197364 | NA | NA | NA | NA |
| WBGene00197365 | NA | NA | NA | NA |
| WBGene00197366 | NA | NA | NA | NA |
| WBGene00197367 | NA | NA | NA | NA |
| WBGene00197368 | NA | NA | NA | NA |
| WBGene00197369 | NA | NA | NA | NA |
| WBGene00197370 | NA | NA | NA | NA |
| WBGene00197371 | NA | NA | NA | NA |
| WBGene00197372 | NA | NA | NA | NA |
| WBGene00197373 | NA | NA | NA | NA |
| WBGene00197374 | NA | NA | NA | NA |
| WBGene00197375 | NA | NA | NA | NA |
| WBGene00197376 | NA | NA | NA | NA |
| WBGene00197377 | NA | NA | NA | NA |
| WBGene00197378 | NA | NA | NA | NA |
| WBGene00197379 | NA | NA | NA | NA |
| WBGene00197380 | NA | NA | NA | NA |
| WBGene00197381 | NA | NA | NA | NA |
| WBGene00197382 | NA | NA | NA | NA |
| WBGene00197383 | NA | NA | NA | NA |
| WBGene00197384 | NA | NA | NA | NA |
| WBGene00197385 | NA | NA | NA | NA |
| WBGene00197386 | NA | NA | NA | NA |
| WBGene00197387 | NA | NA | NA | NA |
| WBGene00197388 | NA | NA | NA | NA |
| WBGene00197389 | NA | NA | NA | NA |
| WBGene00197390 | NA | NA | NA | NA |
| WBGene00197391 | NA | NA | NA | NA |
| WBGene00197392 | NA | NA | NA | NA |
| WBGene00197393 | NA | NA | NA | NA |
| WBGene00197394 | NA | NA | NA | NA |
| WBGene00197395 | NA | NA | NA | NA |
| WBGene00197396 | NA | NA | NA | NA |
| WBGene00197397 | NA | NA | NA | NA |
| WBGene00197398 | NA | NA | NA | NA |

| | | | | |
|---|---|---|---|---|
| WBGene00197399 | NA | NA | NA | NA |
| WBGene00197400 | NA | NA | NA | NA |
| WBGene00197401 | NA | NA | NA | NA |
| WBGene00197402 | NA | NA | NA | NA |
| WBGene00197403 | NA | NA | NA | NA |
| WBGene00197404 | NA | NA | NA | NA |
| WBGene00197405 | 0.056188364 | 0.42092875 | NA | NA |
| WBGene00197406 | NA | NA | NA | NA |
| WBGene00197407 | NA | NA | NA | NA |
| WBGene00197408 | NA | NA | NA | NA |
| WBGene00197409 | NA | NA | NA | NA |
| WBGene00197410 | NA | NA | NA | NA |
| WBGene00197411 | NA | NA | NA | NA |
| WBGene00197412 | NA | NA | NA | NA |
| WBGene00197413 | NA | NA | NA | NA |
| WBGene00197414 | NA | NA | NA | NA |
| WBGene00197415 | NA | NA | NA | NA |
| WBGene00197416 | 0.020317245 | 0.84643798 | NA | NA |
| WBGene00197417 | NA | NA | NA | NA |
| WBGene00197418 | NA | NA | NA | NA |
| WBGene00197419 | NA | NA | NA | NA |
| WBGene00197420 | NA | NA | NA | NA |
| WBGene00197421 | NA | NA | NA | NA |
| WBGene00197422 | NA | NA | NA | NA |
| WBGene00197423 | NA | NA | NA | NA |
| WBGene00197424 | NA | NA | NA | NA |
| WBGene00197425 | NA | NA | NA | NA |
| WBGene00197426 | NA | NA | NA | NA |
| WBGene00197427 | NA | NA | NA | NA |
| WBGene00197428 | NA | NA | NA | NA |
| WBGene00197429 | NA | NA | NA | NA |
| WBGene00197430 | NA | NA | NA | NA |
| WBGene00197431 | NA | NA | NA | NA |
| WBGene00197432 | NA | NA | NA | NA |
| WBGene00197433 | NA | NA | NA | NA |
| WBGene00197434 | NA | NA | NA | NA |
| WBGene00197435 | NA | NA | NA | NA |
| WBGene00197436 | NA | NA | NA | NA |
| WBGene00197437 | NA | NA | NA | NA |
| WBGene00197438 | NA | NA | NA | NA |
| WBGene00197439 | NA | NA | NA | NA |
| WBGene00197440 | NA | NA | NA | NA |
| WBGene00197441 | NA | NA | NA | NA |

| | | | | |
|---|---|---|---|---|
| WBGene00197442 | NA | NA | NA | NA |
| WBGene00197443 | NA | NA | NA | NA |
| WBGene00197444 | 0.038125098 | 0.61119482 | NA | NA |
| WBGene00197445 | NA | NA | NA | NA |
| WBGene00197446 | NA | NA | NA | NA |
| WBGene00197447 | NA | NA | NA | NA |
| WBGene00197448 | NA | NA | NA | NA |
| WBGene00197449 | NA | NA | NA | NA |
| WBGene00197450 | NA | NA | NA | NA |
| WBGene00197451 | NA | NA | NA | NA |
| WBGene00197452 | 0.020925399 | 0.84643798 | NA | NA |
| WBGene00197453 | NA | NA | NA | NA |
| WBGene00197454 | NA | NA | NA | NA |
| WBGene00197455 | NA | NA | NA | NA |
| WBGene00197456 | NA | NA | NA | NA |
| WBGene00197457 | 0.020925399 | 0.84643798 | NA | NA |
| WBGene00197458 | NA | NA | NA | NA |
| WBGene00197459 | NA | NA | NA | NA |
| WBGene00197460 | NA | NA | NA | NA |
| WBGene00197461 | NA | NA | NA | NA |
| WBGene00197462 | NA | NA | NA | NA |
| WBGene00197463 | NA | NA | NA | NA |
| WBGene00197464 | NA | NA | NA | NA |
| WBGene00197465 | NA | NA | NA | NA |
| WBGene00197466 | NA | NA | NA | NA |
| WBGene00197467 | NA | NA | NA | NA |
| WBGene00197468 | NA | NA | NA | NA |
| WBGene00197469 | NA | NA | NA | NA |
| WBGene00197470 | NA | NA | NA | NA |
| WBGene00197471 | NA | NA | NA | NA |
| WBGene00197472 | NA | NA | NA | NA |
| WBGene00197473 | NA | NA | NA | NA |
| WBGene00197474 | NA | NA | NA | NA |
| WBGene00197475 | NA | NA | NA | NA |
| WBGene00197476 | NA | NA | NA | NA |
| WBGene00197477 | 0.020925399 | 0.84643798 | NA | NA |
| WBGene00197478 | NA | NA | NA | NA |
| WBGene00197479 | 0.020925399 | 0.84643798 | NA | NA |
| WBGene00197480 | NA | NA | NA | NA |
| WBGene00197481 | NA | NA | NA | NA |
| WBGene00197482 | NA | NA | NA | NA |
| WBGene00197483 | NA | NA | NA | NA |
| WBGene00197484 | NA | NA | NA | NA |

| | | | | |
|---|---|---|---|---|
| WBGene00197485 | NA | NA | NA | NA |
| WBGene00197486 | NA | NA | NA | NA |
| WBGene00197487 | NA | NA | NA | NA |
| WBGene00197488 | NA | NA | NA | NA |
| WBGene00197489 | NA | NA | NA | NA |
| WBGene00197490 | NA | NA | NA | NA |
| WBGene00197491 | NA | NA | NA | NA |
| WBGene00197492 | NA | NA | NA | NA |
| WBGene00197493 | NA | NA | NA | NA |
| WBGene00197494 | NA | NA | NA | NA |
| WBGene00197495 | NA | NA | NA | NA |
| WBGene00197496 | NA | NA | NA | NA |
| WBGene00197497 | NA | NA | NA | NA |
| WBGene00197498 | NA | NA | NA | NA |
| WBGene00197499 | NA | NA | NA | NA |
| WBGene00197500 | NA | NA | NA | NA |
| WBGene00197501 | NA | NA | NA | NA |
| WBGene00197502 | NA | NA | NA | NA |
| WBGene00197503 | 0.020232312 | 0.84643798 | NA | NA |
| WBGene00197504 | NA | NA | NA | NA |
| WBGene00197505 | NA | NA | NA | NA |
| WBGene00197506 | NA | NA | NA | NA |
| WBGene00197507 | NA | NA | NA | NA |
| WBGene00197508 | NA | NA | NA | NA |
| WBGene00197509 | NA | NA | NA | NA |
| WBGene00197510 | NA | NA | NA | NA |
| WBGene00197511 | NA | NA | NA | NA |
| WBGene00197512 | NA | NA | NA | NA |
| WBGene00197513 | NA | NA | NA | NA |
| WBGene00197514 | NA | NA | NA | NA |
| WBGene00197515 | NA | NA | NA | NA |
| WBGene00197516 | NA | NA | NA | NA |
| WBGene00197517 | NA | NA | NA | NA |
| WBGene00197518 | NA | NA | NA | NA |
| WBGene00197519 | NA | NA | NA | NA |
| WBGene00197520 | NA | NA | NA | NA |
| WBGene00197521 | NA | NA | NA | NA |
| WBGene00197522 | NA | NA | NA | NA |
| WBGene00197523 | -0.003108065 | 0.95728506 | NA | NA |
| WBGene00197524 | NA | NA | NA | NA |
| WBGene00197525 | NA | NA | NA | NA |
| WBGene00197526 | NA | NA | NA | NA |
| WBGene00197527 | NA | NA | NA | NA |

| | | | | |
|---|---|---|---|---|
| WBGene00197528 | NA | NA | NA | NA |
| WBGene00197529 | NA | NA | NA | NA |
| WBGene00197530 | NA | NA | NA | NA |
| WBGene00197531 | NA | NA | NA | NA |
| WBGene00197532 | NA | NA | NA | NA |
| WBGene00197533 | NA | NA | NA | NA |
| WBGene00197534 | NA | NA | NA | NA |
| WBGene00197535 | NA | NA | NA | NA |
| WBGene00197536 | NA | NA | NA | NA |
| WBGene00197537 | NA | NA | NA | NA |
| WBGene00197538 | NA | NA | NA | NA |
| WBGene00197539 | -0.023056499 | 0.78121253 | NA | NA |
| WBGene00197540 | NA | NA | NA | NA |
| WBGene00197541 | NA | NA | NA | NA |
| WBGene00197542 | NA | NA | NA | NA |
| WBGene00197543 | -0.023056499 | 0.78121253 | NA | NA |
| WBGene00197544 | NA | NA | NA | NA |
| WBGene00197545 | NA | NA | NA | NA |
| WBGene00197546 | NA | NA | NA | NA |
| WBGene00197547 | NA | NA | NA | NA |
| WBGene00197548 | NA | NA | NA | NA |
| WBGene00197549 | NA | NA | NA | NA |
| WBGene00197550 | NA | NA | NA | NA |
| WBGene00197551 | NA | NA | NA | NA |
| WBGene00197552 | NA | NA | NA | NA |
| WBGene00197553 | NA | NA | NA | NA |
| WBGene00197554 | NA | NA | NA | NA |
| WBGene00197555 | NA | NA | NA | NA |
| WBGene00197556 | NA | NA | NA | NA |
| WBGene00197557 | NA | NA | NA | NA |
| WBGene00197558 | NA | NA | NA | NA |
| WBGene00197559 | NA | NA | NA | NA |
| WBGene00197560 | NA | NA | NA | NA |
| WBGene00197561 | NA | NA | NA | NA |
| WBGene00197562 | NA | NA | NA | NA |
| WBGene00197563 | 0.020317245 | 0.84643798 | NA | NA |
| WBGene00197564 | NA | NA | NA | NA |
| WBGene00197565 | -0.023109762 | 0.78121253 | NA | NA |
| WBGene00197566 | NA | NA | NA | NA |
| WBGene00197567 | NA | NA | NA | NA |
| WBGene00197568 | NA | NA | NA | NA |
| WBGene00197569 | NA | NA | NA | NA |
| WBGene00197570 | NA | NA | NA | NA |

| | | | | |
|---|---|---|---|---|
| WBGene00197571 | NA | NA | NA | NA |
| WBGene00197572 | NA | NA | NA | NA |
| WBGene00197573 | -0.023109762 | 0.78121253 | NA | NA |
| WBGene00197574 | NA | NA | NA | NA |
| WBGene00197575 | NA | NA | NA | NA |
| WBGene00197576 | NA | NA | NA | NA |
| WBGene00197577 | NA | NA | NA | NA |
| WBGene00197578 | NA | NA | NA | NA |
| WBGene00197579 | 0.020925399 | 0.84643798 | NA | NA |
| WBGene00197580 | NA | NA | NA | NA |
| WBGene00197581 | NA | NA | NA | NA |
| WBGene00197582 | NA | NA | NA | NA |
| WBGene00197583 | NA | NA | NA | NA |
| WBGene00197584 | NA | NA | NA | NA |
| WBGene00197585 | NA | NA | NA | NA |
| WBGene00197586 | NA | NA | NA | NA |
| WBGene00197587 | NA | NA | NA | NA |
| WBGene00197588 | NA | NA | NA | NA |
| WBGene00197589 | NA | NA | NA | NA |
| WBGene00197590 | NA | NA | NA | NA |
| WBGene00197591 | NA | NA | NA | NA |
| WBGene00197592 | NA | NA | NA | NA |
| WBGene00197593 | NA | NA | NA | NA |
| WBGene00197594 | NA | NA | NA | NA |
| WBGene00197595 | NA | NA | NA | NA |
| WBGene00197596 | NA | NA | NA | NA |
| WBGene00197597 | -0.023109762 | 0.78121253 | NA | NA |
| WBGene00197598 | -0.003270206 | 0.95698316 | NA | NA |
| WBGene00197599 | NA | NA | NA | NA |
| WBGene00197600 | NA | NA | NA | NA |
| WBGene00197601 | NA | NA | NA | NA |
| WBGene00197602 | NA | NA | NA | NA |
| WBGene00197603 | NA | NA | NA | NA |
| WBGene00197604 | -0.023109762 | 0.78121253 | NA | NA |
| WBGene00197605 | NA | NA | NA | NA |
| WBGene00197606 | NA | NA | NA | NA |
| WBGene00197607 | NA | NA | NA | NA |
| WBGene00197608 | NA | NA | NA | NA |
| WBGene00197609 | NA | NA | NA | NA |
| WBGene00197610 | NA | NA | NA | NA |
| WBGene00197611 | NA | NA | NA | NA |
| WBGene00197612 | NA | NA | NA | NA |
| WBGene00197613 | NA | NA | NA | NA |

| | | | | |
|---|---|---|---|---|
| WBGene00197614 | NA | NA | NA | NA |
| WBGene00197615 | NA | NA | NA | NA |
| WBGene00197616 | NA | NA | NA | NA |
| WBGene00197617 | NA | NA | NA | NA |
| WBGene00197618 | NA | NA | NA | NA |
| WBGene00197619 | NA | NA | NA | NA |
| WBGene00197620 | NA | NA | NA | NA |
| WBGene00197621 | NA | NA | NA | NA |
| WBGene00197622 | NA | NA | NA | NA |
| WBGene00197623 | NA | NA | NA | NA |
| WBGene00197624 | NA | NA | NA | NA |
| WBGene00197625 | NA | NA | NA | NA |
| WBGene00197626 | NA | NA | NA | NA |
| WBGene00197627 | NA | NA | NA | NA |
| WBGene00197628 | NA | NA | NA | NA |
| WBGene00197629 | NA | NA | NA | NA |
| WBGene00197630 | 0.020317245 | 0.84643798 | NA | NA |
| WBGene00197631 | NA | NA | NA | NA |
| WBGene00197632 | NA | NA | NA | NA |
| WBGene00197633 | 0.020232312 | 0.84643798 | NA | NA |
| WBGene00197634 | NA | NA | NA | NA |
| WBGene00197635 | NA | NA | NA | NA |
| WBGene00197636 | 0.020925399 | 0.84643798 | NA | NA |
| WBGene00197637 | NA | NA | NA | NA |
| WBGene00197638 | NA | NA | NA | NA |
| WBGene00197639 | NA | NA | NA | NA |
| WBGene00197640 | NA | NA | NA | NA |
| WBGene00197641 | NA | NA | NA | NA |
| WBGene00197642 | NA | NA | NA | NA |
| WBGene00197643 | NA | NA | NA | NA |
| WBGene00197644 | NA | NA | NA | NA |
| WBGene00197645 | NA | NA | NA | NA |
| WBGene00197646 | NA | NA | NA | NA |
| WBGene00197647 | NA | NA | NA | NA |
| WBGene00197648 | NA | NA | NA | NA |
| WBGene00197649 | NA | NA | NA | NA |
| WBGene00197650 | NA | NA | NA | NA |
| WBGene00197651 | -0.023056499 | 0.78121253 | NA | NA |
| WBGene00197652 | NA | NA | NA | NA |
| WBGene00197653 | NA | NA | NA | NA |
| WBGene00197654 | NA | NA | NA | NA |
| WBGene00197655 | NA | NA | NA | NA |
| WBGene00197656 | NA | NA | NA | NA |

| | | | | |
|---|---|---|---|---|
| WBGene00197657 | NA | NA | NA | NA |
| WBGene00197658 | NA | NA | NA | NA |
| WBGene00197659 | NA | NA | NA | NA |
| WBGene00197660 | NA | NA | NA | NA |
| WBGene00197661 | NA | NA | NA | NA |
| WBGene00197662 | NA | NA | NA | NA |
| WBGene00197663 | 0.020925399 | 0.84643798 | NA | NA |
| WBGene00197664 | NA | NA | NA | NA |
| WBGene00197665 | NA | NA | NA | NA |
| WBGene00197666 | NA | NA | NA | NA |
| WBGene00197667 | NA | NA | NA | NA |
| WBGene00197668 | NA | NA | NA | NA |
| WBGene00197669 | NA | NA | NA | NA |
| WBGene00197670 | 0.033463284 | 0.6365037 | NA | NA |
| WBGene00197671 | NA | NA | NA | NA |
| WBGene00197672 | NA | NA | NA | NA |
| WBGene00197673 | NA | NA | NA | NA |
| WBGene00197674 | NA | NA | NA | NA |
| WBGene00197675 | NA | NA | NA | NA |
| WBGene00197676 | NA | NA | NA | NA |
| WBGene00197677 | NA | NA | NA | NA |
| WBGene00197678 | NA | NA | NA | NA |
| WBGene00197679 | NA | NA | NA | NA |
| WBGene00197680 | NA | NA | NA | NA |
| WBGene00197681 | NA | NA | NA | NA |
| WBGene00197682 | NA | NA | NA | NA |
| WBGene00197683 | NA | NA | NA | NA |
| WBGene00197684 | NA | NA | NA | NA |
| WBGene00197685 | NA | NA | NA | NA |
| WBGene00197686 | NA | NA | NA | NA |
| WBGene00197687 | NA | NA | NA | NA |
| WBGene00197688 | NA | NA | NA | NA |
| WBGene00197689 | NA | NA | NA | NA |
| WBGene00197690 | NA | NA | NA | NA |
| WBGene00197691 | NA | NA | NA | NA |
| WBGene00197692 | NA | NA | NA | NA |
| WBGene00197693 | NA | NA | NA | NA |
| WBGene00197694 | NA | NA | NA | NA |
| WBGene00197695 | NA | NA | NA | NA |
| WBGene00197696 | NA | NA | NA | NA |
| WBGene00197697 | NA | NA | NA | NA |
| WBGene00197698 | NA | NA | NA | NA |
| WBGene00197699 | NA | NA | NA | NA |

| | | | | |
|---|---|---|---|---|
| WBGene00197700 | NA | NA | NA | NA |
| WBGene00197701 | NA | NA | NA | NA |
| WBGene00197702 | NA | NA | NA | NA |
| WBGene00197703 | NA | NA | NA | NA |
| WBGene00197704 | NA | NA | NA | NA |
| WBGene00197705 | NA | NA | NA | NA |
| WBGene00197706 | NA | NA | NA | NA |
| WBGene00197707 | NA | NA | NA | NA |
| WBGene00197708 | NA | NA | NA | NA |
| WBGene00197709 | NA | NA | NA | NA |
| WBGene00197710 | NA | NA | NA | NA |
| WBGene00197711 | NA | NA | NA | NA |
| WBGene00197712 | NA | NA | NA | NA |
| WBGene00197713 | NA | NA | NA | NA |
| WBGene00197714 | 0.020317245 | 0.84643798 | NA | NA |
| WBGene00197715 | NA | NA | NA | NA |
| WBGene00197716 | NA | NA | NA | NA |
| WBGene00197717 | NA | NA | NA | NA |
| WBGene00197718 | NA | NA | NA | NA |
| WBGene00197719 | NA | NA | NA | NA |
| WBGene00197720 | NA | NA | NA | NA |
| WBGene00197721 | NA | NA | NA | NA |
| WBGene00197722 | NA | NA | NA | NA |
| WBGene00197723 | NA | NA | NA | NA |
| WBGene00197724 | NA | NA | NA | NA |
| WBGene00197725 | NA | NA | NA | NA |
| WBGene00197726 | NA | NA | NA | NA |
| WBGene00197727 | 0.020232312 | 0.84643798 | NA | NA |
| WBGene00197728 | NA | NA | NA | NA |
| WBGene00197729 | NA | NA | NA | NA |
| WBGene00197730 | NA | NA | NA | NA |
| WBGene00197731 | NA | NA | NA | NA |
| WBGene00197732 | NA | NA | NA | NA |
| WBGene00197733 | -0.003614866 | 0.95700117 | NA | NA |
| WBGene00197734 | NA | NA | NA | NA |
| WBGene00197735 | NA | NA | NA | NA |
| WBGene00197736 | NA | NA | NA | NA |
| WBGene00197737 | NA | NA | NA | NA |
| WBGene00197738 | NA | NA | NA | NA |
| WBGene00197739 | NA | NA | NA | NA |
| WBGene00197740 | NA | NA | NA | NA |
| WBGene00197741 | NA | NA | NA | NA |
| WBGene00197742 | 0.015070939 | 0.84337138 | NA | NA |

| | | | | |
|---|---|---|---|---|
| WBGene00197743 | NA | NA | NA | NA |
| WBGene00197744 | NA | NA | NA | NA |
| WBGene00197745 | NA | NA | NA | NA |
| WBGene00197746 | NA | NA | NA | NA |
| WBGene00197747 | NA | NA | NA | NA |
| WBGene00197748 | NA | NA | NA | NA |
| WBGene00197749 | NA | NA | NA | NA |
| WBGene00197750 | NA | NA | NA | NA |
| WBGene00197751 | NA | NA | NA | NA |
| WBGene00197752 | NA | NA | NA | NA |
| WBGene00197753 | NA | NA | NA | NA |
| WBGene00197754 | NA | NA | NA | NA |
| WBGene00197755 | NA | NA | NA | NA |
| WBGene00197756 | NA | NA | NA | NA |
| WBGene00197757 | NA | NA | NA | NA |
| WBGene00197758 | NA | NA | NA | NA |
| WBGene00197759 | NA | NA | NA | NA |
| WBGene00197760 | NA | NA | NA | NA |
| WBGene00197761 | NA | NA | NA | NA |
| WBGene00197762 | NA | NA | NA | NA |
| WBGene00197763 | NA | NA | NA | NA |
| WBGene00197764 | NA | NA | NA | NA |
| WBGene00197765 | NA | NA | NA | NA |
| WBGene00197766 | NA | NA | NA | NA |
| WBGene00197767 | NA | NA | NA | NA |
| WBGene00197768 | NA | NA | NA | NA |
| WBGene00197769 | NA | NA | NA | NA |
| WBGene00197770 | NA | NA | NA | NA |
| WBGene00197771 | NA | NA | NA | NA |
| WBGene00197772 | NA | NA | NA | NA |
| WBGene00197773 | NA | NA | NA | NA |
| WBGene00197774 | NA | NA | NA | NA |
| WBGene00197775 | NA | NA | NA | NA |
| WBGene00197776 | NA | NA | NA | NA |
| WBGene00197777 | NA | NA | NA | NA |
| WBGene00197778 | NA | NA | NA | NA |
| WBGene00197779 | NA | NA | NA | NA |
| WBGene00197780 | NA | NA | NA | NA |
| WBGene00197781 | NA | NA | NA | NA |
| WBGene00197782 | NA | NA | NA | NA |
| WBGene00197783 | NA | NA | NA | NA |
| WBGene00197784 | NA | NA | NA | NA |
| WBGene00197785 | NA | NA | NA | NA |

| | | | | |
|---|---|---|---|---|
| WBGene00197786 | NA | NA | NA | NA |
| WBGene00197787 | NA | NA | NA | NA |
| WBGene00197788 | NA | NA | NA | NA |
| WBGene00197789 | NA | NA | NA | NA |
| WBGene00197790 | NA | NA | NA | NA |
| WBGene00197791 | NA | NA | NA | NA |
| WBGene00197792 | NA | NA | NA | NA |
| WBGene00197793 | NA | NA | NA | NA |
| WBGene00197794 | NA | NA | NA | NA |
| WBGene00197795 | NA | NA | NA | NA |
| WBGene00197796 | 0.020232312 | 0.84643798 | NA | NA |
| WBGene00197797 | NA | NA | NA | NA |
| WBGene00197798 | NA | NA | NA | NA |
| WBGene00197799 | NA | NA | NA | NA |
| WBGene00197800 | NA | NA | NA | NA |
| WBGene00197801 | NA | NA | NA | NA |
| WBGene00197802 | NA | NA | NA | NA |
| WBGene00197803 | NA | NA | NA | NA |
| WBGene00197804 | NA | NA | NA | NA |
| WBGene00197805 | NA | NA | NA | NA |
| WBGene00197806 | NA | NA | NA | NA |
| WBGene00197807 | NA | NA | NA | NA |
| WBGene00197808 | NA | NA | NA | NA |
| WBGene00197809 | NA | NA | NA | NA |
| WBGene00197810 | NA | NA | NA | NA |
| WBGene00197811 | NA | NA | NA | NA |
| WBGene00197812 | NA | NA | NA | NA |
| WBGene00197813 | NA | NA | NA | NA |
| WBGene00197814 | 0.020232312 | 0.84643798 | NA | NA |
| WBGene00197815 | NA | NA | NA | NA |
| WBGene00197816 | NA | NA | NA | NA |
| WBGene00197817 | -0.022930767 | 0.78121253 | NA | NA |
| WBGene00197818 | NA | NA | NA | NA |
| WBGene00197819 | NA | NA | NA | NA |
| WBGene00197820 | NA | NA | NA | NA |
| WBGene00197821 | NA | NA | NA | NA |
| WBGene00197822 | NA | NA | NA | NA |
| WBGene00197823 | NA | NA | NA | NA |
| WBGene00197824 | NA | NA | NA | NA |
| WBGene00197825 | NA | NA | NA | NA |
| WBGene00197826 | NA | NA | NA | NA |
| WBGene00197827 | 0.020925399 | 0.84643798 | NA | NA |
| WBGene00197828 | NA | NA | NA | NA |

| | | | | |
|---|---|---|---|---|
| WBGene00197829 | NA | NA | NA | NA |
| WBGene00197830 | NA | NA | NA | NA |
| WBGene00197831 | NA | NA | NA | NA |
| WBGene00197832 | -0.023056499 | 0.78121253 | NA | NA |
| WBGene00197833 | NA | NA | NA | NA |
| WBGene00197834 | 0.020232312 | 0.84643798 | NA | NA |
| WBGene00197835 | NA | NA | NA | NA |
| WBGene00197836 | NA | NA | NA | NA |
| WBGene00197837 | NA | NA | NA | NA |
| WBGene00197838 | NA | NA | NA | NA |
| WBGene00197839 | NA | NA | NA | NA |
| WBGene00197840 | -0.022930767 | 0.78121253 | NA | NA |
| WBGene00197841 | NA | NA | NA | NA |
| WBGene00197842 | NA | NA | NA | NA |
| WBGene00197843 | NA | NA | NA | NA |
| WBGene00197844 | NA | NA | NA | NA |
| WBGene00197845 | NA | NA | NA | NA |
| WBGene00197846 | NA | NA | NA | NA |
| WBGene00197847 | NA | NA | NA | NA |
| WBGene00197848 | NA | NA | NA | NA |
| WBGene00197849 | NA | NA | NA | NA |
| WBGene00197850 | NA | NA | NA | NA |
| WBGene00197851 | NA | NA | NA | NA |
| WBGene00197852 | NA | NA | NA | NA |
| WBGene00197853 | NA | NA | NA | NA |
| WBGene00197854 | NA | NA | NA | NA |
| WBGene00197855 | NA | NA | NA | NA |
| WBGene00197856 | NA | NA | NA | NA |
| WBGene00197857 | NA | NA | NA | NA |
| WBGene00197858 | NA | NA | NA | NA |
| WBGene00197859 | NA | NA | NA | NA |
| WBGene00197860 | NA | NA | NA | NA |
| WBGene00197861 | NA | NA | NA | NA |
| WBGene00197862 | NA | NA | NA | NA |
| WBGene00197863 | NA | NA | NA | NA |
| WBGene00197864 | NA | NA | NA | NA |
| WBGene00197865 | NA | NA | NA | NA |
| WBGene00197866 | NA | NA | NA | NA |
| WBGene00197867 | NA | NA | NA | NA |
| WBGene00197868 | NA | NA | NA | NA |
| WBGene00197869 | NA | NA | NA | NA |
| WBGene00197870 | NA | NA | NA | NA |
| WBGene00197871 | NA | NA | NA | NA |

| | | | | |
|---|---|---|---|---|
| WBGene00197872 | NA | NA | NA | NA |
| WBGene00197873 | NA | NA | NA | NA |
| WBGene00197874 | NA | NA | NA | NA |
| WBGene00197875 | NA | NA | NA | NA |
| WBGene00197876 | NA | NA | NA | NA |
| WBGene00197877 | NA | NA | NA | NA |
| WBGene00197878 | NA | NA | NA | NA |
| WBGene00197879 | -0.022930767 | 0.78121253 | NA | NA |
| WBGene00197880 | NA | NA | NA | NA |
| WBGene00197881 | NA | NA | NA | NA |
| WBGene00197882 | NA | NA | NA | NA |
| WBGene00197883 | NA | NA | NA | NA |
| WBGene00197884 | NA | NA | NA | NA |
| WBGene00197885 | NA | NA | NA | NA |
| WBGene00197886 | NA | NA | NA | NA |
| WBGene00197887 | -0.022729726 | 0.70048408 | NA | NA |
| WBGene00197888 | 0.020232312 | 0.84643798 | NA | NA |
| WBGene00197889 | NA | NA | NA | NA |
| WBGene00197890 | NA | NA | NA | NA |
| WBGene00197891 | NA | NA | NA | NA |
| WBGene00197892 | NA | NA | NA | NA |
| WBGene00197893 | NA | NA | NA | NA |
| WBGene00197894 | NA | NA | NA | NA |
| WBGene00197895 | NA | NA | NA | NA |
| WBGene00197896 | NA | NA | NA | NA |
| WBGene00197897 | NA | NA | NA | NA |
| WBGene00197898 | NA | NA | NA | NA |
| WBGene00197899 | NA | NA | NA | NA |
| WBGene00197900 | NA | NA | NA | NA |
| WBGene00197901 | NA | NA | NA | NA |
| WBGene00197902 | NA | NA | NA | NA |
| WBGene00197903 | NA | NA | NA | NA |
| WBGene00197904 | NA | NA | NA | NA |
| WBGene00197905 | NA | NA | NA | NA |
| WBGene00197906 | NA | NA | NA | NA |
| WBGene00197907 | NA | NA | NA | NA |
| WBGene00197908 | NA | NA | NA | NA |
| WBGene00197909 | NA | NA | NA | NA |
| WBGene00197910 | NA | NA | NA | NA |
| WBGene00197911 | NA | NA | NA | NA |
| WBGene00197912 | NA | NA | NA | NA |
| WBGene00197913 | NA | NA | NA | NA |
| WBGene00197914 | NA | NA | NA | NA |

| | | | | |
|---|---|---|---|---|
| WBGene00197915 | NA | NA | NA | NA |
| WBGene00197916 | NA | NA | NA | NA |
| WBGene00197917 | NA | NA | NA | NA |
| WBGene00197918 | NA | NA | NA | NA |
| WBGene00197919 | NA | NA | NA | NA |
| WBGene00197920 | NA | NA | NA | NA |
| WBGene00197921 | NA | NA | NA | NA |
| WBGene00197922 | NA | NA | NA | NA |
| WBGene00197923 | NA | NA | NA | NA |
| WBGene00197924 | NA | NA | NA | NA |
| WBGene00197925 | NA | NA | NA | NA |
| WBGene00197926 | NA | NA | NA | NA |
| WBGene00197927 | NA | NA | NA | NA |
| WBGene00197928 | 0.020925399 | 0.84643798 | NA | NA |
| WBGene00197929 | NA | NA | NA | NA |
| WBGene00197930 | -0.022930767 | 0.78121253 | NA | NA |
| WBGene00197931 | NA | NA | NA | NA |
| WBGene00197932 | NA | NA | NA | NA |
| WBGene00197933 | NA | NA | NA | NA |
| WBGene00197934 | NA | NA | NA | NA |
| WBGene00197935 | NA | NA | NA | NA |
| WBGene00197936 | NA | NA | NA | NA |
| WBGene00197937 | NA | NA | NA | NA |
| WBGene00197938 | NA | NA | NA | NA |
| WBGene00197939 | NA | NA | NA | NA |
| WBGene00197940 | 0.108942554 | 0.15594412 | NA | NA |
| WBGene00197941 | NA | NA | NA | NA |
| WBGene00197942 | NA | NA | NA | NA |
| WBGene00197943 | NA | NA | NA | NA |
| WBGene00197944 | -0.003614866 | 0.95700117 | NA | NA |
| WBGene00197945 | NA | NA | NA | NA |
| WBGene00197946 | NA | NA | NA | NA |
| WBGene00197947 | NA | NA | NA | NA |
| WBGene00197948 | NA | NA | NA | NA |
| WBGene00197949 | NA | NA | NA | NA |
| WBGene00197950 | -0.023109762 | 0.78121253 | NA | NA |
| WBGene00197951 | NA | NA | NA | NA |
| WBGene00197952 | NA | NA | NA | NA |
| WBGene00197953 | NA | NA | NA | NA |
| WBGene00197954 | NA | NA | NA | NA |
| WBGene00197955 | NA | NA | NA | NA |
| WBGene00197956 | NA | NA | NA | NA |
| WBGene00197957 | NA | NA | NA | NA |

| | | | | |
|---|---|---|---|---|
| WBGene00197958 | NA | NA | NA | NA |
| WBGene00197959 | NA | NA | NA | NA |
| WBGene00197960 | NA | NA | NA | NA |
| WBGene00197961 | NA | NA | NA | NA |
| WBGene00197962 | NA | NA | NA | NA |
| WBGene00197963 | NA | NA | NA | NA |
| WBGene00197964 | NA | NA | NA | NA |
| WBGene00197965 | NA | NA | NA | NA |
| WBGene00197966 | NA | NA | NA | NA |
| WBGene00197967 | NA | NA | NA | NA |
| WBGene00197968 | NA | NA | NA | NA |
| WBGene00197969 | NA | NA | NA | NA |
| WBGene00197970 | NA | NA | NA | NA |
| WBGene00197971 | NA | NA | NA | NA |
| WBGene00197972 | NA | NA | NA | NA |
| WBGene00197973 | NA | NA | NA | NA |
| WBGene00197974 | NA | NA | NA | NA |
| WBGene00197975 | NA | NA | NA | NA |
| WBGene00197976 | NA | NA | NA | NA |
| WBGene00197977 | NA | NA | NA | NA |
| WBGene00197978 | NA | NA | NA | NA |
| WBGene00197979 | NA | NA | NA | NA |
| WBGene00197980 | NA | NA | NA | NA |
| WBGene00197981 | 0.020317245 | 0.84643798 | NA | NA |
| WBGene00197982 | NA | NA | NA | NA |
| WBGene00197983 | NA | NA | NA | NA |
| WBGene00197984 | NA | NA | NA | NA |
| WBGene00197985 | NA | NA | NA | NA |
| WBGene00197986 | NA | NA | NA | NA |
| WBGene00197987 | NA | NA | NA | NA |
| WBGene00197988 | NA | NA | NA | NA |
| WBGene00197989 | NA | NA | NA | NA |
| WBGene00197990 | NA | NA | NA | NA |
| WBGene00197991 | NA | NA | NA | NA |
| WBGene00197992 | NA | NA | NA | NA |
| WBGene00197993 | NA | NA | NA | NA |
| WBGene00197994 | NA | NA | NA | NA |
| WBGene00197995 | NA | NA | NA | NA |
| WBGene00197996 | 0.017141582 | 0.79355725 | NA | NA |
| WBGene00197997 | 0.020317245 | 0.84643798 | NA | NA |
| WBGene00197998 | -0.023056499 | 0.78121253 | NA | NA |
| WBGene00197999 | NA | NA | NA | NA |
| WBGene00198000 | NA | NA | NA | NA |

| | | | | |
|---|---|---|---|---|
| WBGene00198001 | NA | NA | NA | NA |
| WBGene00198002 | NA | NA | NA | NA |
| WBGene00198003 | NA | NA | NA | NA |
| WBGene00198004 | NA | NA | NA | NA |
| WBGene00198005 | NA | NA | NA | NA |
| WBGene00198006 | NA | NA | NA | NA |
| WBGene00198007 | NA | NA | NA | NA |
| WBGene00198008 | NA | NA | NA | NA |
| WBGene00198009 | NA | NA | NA | NA |
| WBGene00198010 | NA | NA | NA | NA |
| WBGene00198011 | NA | NA | NA | NA |
| WBGene00198012 | NA | NA | NA | NA |
| WBGene00198013 | NA | NA | NA | NA |
| WBGene00198014 | NA | NA | NA | NA |
| WBGene00198015 | NA | NA | NA | NA |
| WBGene00198016 | NA | NA | NA | NA |
| WBGene00198017 | NA | NA | NA | NA |
| WBGene00198018 | NA | NA | NA | NA |
| WBGene00198019 | NA | NA | NA | NA |
| WBGene00198020 | NA | NA | NA | NA |
| WBGene00198021 | NA | NA | NA | NA |
| WBGene00198022 | NA | NA | NA | NA |
| WBGene00198023 | NA | NA | NA | NA |
| WBGene00198024 | NA | NA | NA | NA |
| WBGene00198025 | NA | NA | NA | NA |
| WBGene00198026 | NA | NA | NA | NA |
| WBGene00198027 | NA | NA | NA | NA |
| WBGene00198028 | NA | NA | NA | NA |
| WBGene00198029 | 0.020925399 | 0.84643798 | NA | NA |
| WBGene00198030 | NA | NA | NA | NA |
| WBGene00198031 | NA | NA | NA | NA |
| WBGene00198032 | NA | NA | NA | NA |
| WBGene00198033 | NA | NA | NA | NA |
| WBGene00198034 | NA | NA | NA | NA |
| WBGene00198035 | NA | NA | NA | NA |
| WBGene00198036 | NA | NA | NA | NA |
| WBGene00198037 | NA | NA | NA | NA |
| WBGene00198038 | NA | NA | NA | NA |
| WBGene00198039 | NA | NA | NA | NA |
| WBGene00198040 | NA | NA | NA | NA |
| WBGene00198041 | NA | NA | NA | NA |
| WBGene00198042 | -0.022930767 | 0.78121253 | NA | NA |
| WBGene00198043 | NA | NA | NA | NA |

| | | | | |
|---|---|---|---|---|
| WBGene00198044 | NA | NA | NA | NA |
| WBGene00198045 | NA | NA | NA | NA |
| WBGene00198046 | NA | NA | NA | NA |
| WBGene00198047 | NA | NA | NA | NA |
| WBGene00198048 | NA | NA | NA | NA |
| WBGene00198049 | NA | NA | NA | NA |
| WBGene00198050 | NA | NA | NA | NA |
| WBGene00198051 | NA | NA | NA | NA |
| WBGene00198052 | NA | NA | NA | NA |
| WBGene00198053 | NA | NA | NA | NA |
| WBGene00198054 | NA | NA | NA | NA |
| WBGene00198055 | NA | NA | NA | NA |
| WBGene00198056 | NA | NA | NA | NA |
| WBGene00198057 | 0.020925399 | 0.84643798 | NA | NA |
| WBGene00198058 | NA | NA | NA | NA |
| WBGene00198059 | NA | NA | NA | NA |
| WBGene00198060 | NA | NA | NA | NA |
| WBGene00198061 | NA | NA | NA | NA |
| WBGene00198062 | NA | NA | NA | NA |
| WBGene00198063 | 0.020232312 | 0.84643798 | NA | NA |
| WBGene00198064 | NA | NA | NA | NA |
| WBGene00198065 | NA | NA | NA | NA |
| WBGene00198066 | NA | NA | NA | NA |
| WBGene00198067 | NA | NA | NA | NA |
| WBGene00198068 | NA | NA | NA | NA |
| WBGene00198069 | NA | NA | NA | NA |
| WBGene00198070 | NA | NA | NA | NA |
| WBGene00198071 | NA | NA | NA | NA |
| WBGene00198072 | NA | NA | NA | NA |
| WBGene00198073 | 0.020232312 | 0.84643798 | NA | NA |
| WBGene00198074 | NA | NA | NA | NA |
| WBGene00198075 | NA | NA | NA | NA |
| WBGene00198076 | NA | NA | NA | NA |
| WBGene00198077 | NA | NA | NA | NA |
| WBGene00198078 | NA | NA | NA | NA |
| WBGene00198079 | NA | NA | NA | NA |
| WBGene00198080 | NA | NA | NA | NA |
| WBGene00198081 | -0.023109762 | 0.78121253 | NA | NA |
| WBGene00198082 | 0.020925399 | 0.84643798 | NA | NA |
| WBGene00198083 | NA | NA | NA | NA |
| WBGene00198084 | NA | NA | NA | NA |
| WBGene00198085 | NA | NA | NA | NA |
| WBGene00198086 | NA | NA | NA | NA |

| | | | | |
|---|---|---|---|---|
| WBGene00198087 | NA | NA | NA | NA |
| WBGene00198088 | NA | NA | NA | NA |
| WBGene00198089 | NA | NA | NA | NA |
| WBGene00198090 | NA | NA | NA | NA |
| WBGene00198091 | NA | NA | NA | NA |
| WBGene00198092 | NA | NA | NA | NA |
| WBGene00198093 | NA | NA | NA | NA |
| WBGene00198094 | NA | NA | NA | NA |
| WBGene00198095 | NA | NA | NA | NA |
| WBGene00198096 | NA | NA | NA | NA |
| WBGene00198097 | NA | NA | NA | NA |
| WBGene00198098 | NA | NA | NA | NA |
| WBGene00198099 | NA | NA | NA | NA |
| WBGene00198100 | NA | NA | NA | NA |
| WBGene00198101 | NA | NA | NA | NA |
| WBGene00198102 | NA | NA | NA | NA |
| WBGene00198103 | 0.020317245 | 0.84643798 | NA | NA |
| WBGene00198104 | NA | NA | NA | NA |
| WBGene00198105 | NA | NA | NA | NA |
| WBGene00198106 | NA | NA | NA | NA |
| WBGene00198107 | NA | NA | NA | NA |
| WBGene00198108 | NA | NA | NA | NA |
| WBGene00198109 | NA | NA | NA | NA |
| WBGene00198110 | NA | NA | NA | NA |
| WBGene00198111 | NA | NA | NA | NA |
| WBGene00198112 | NA | NA | NA | NA |
| WBGene00198113 | NA | NA | NA | NA |
| WBGene00198114 | NA | NA | NA | NA |
| WBGene00198115 | NA | NA | NA | NA |
| WBGene00198116 | NA | NA | NA | NA |
| WBGene00198117 | NA | NA | NA | NA |
| WBGene00198118 | NA | NA | NA | NA |
| WBGene00198119 | NA | NA | NA | NA |
| WBGene00198120 | NA | NA | NA | NA |
| WBGene00198121 | NA | NA | NA | NA |
| WBGene00198122 | NA | NA | NA | NA |
| WBGene00198123 | NA | NA | NA | NA |
| WBGene00198124 | NA | NA | NA | NA |
| WBGene00198125 | NA | NA | NA | NA |
| WBGene00198126 | 0.020232312 | 0.84643798 | NA | NA |
| WBGene00198127 | NA | NA | NA | NA |
| WBGene00198128 | NA | NA | NA | NA |
| WBGene00198129 | NA | NA | NA | NA |

| | | | | |
|---|---|---|---|---|
| WBGene00198130 | NA | NA | NA | NA |
| WBGene00198131 | NA | NA | NA | NA |
| WBGene00198132 | NA | NA | NA | NA |
| WBGene00198133 | NA | NA | NA | NA |
| WBGene00198134 | NA | NA | NA | NA |
| WBGene00198135 | NA | NA | NA | NA |
| WBGene00198136 | NA | NA | NA | NA |
| WBGene00198137 | NA | NA | NA | NA |
| WBGene00198138 | NA | NA | NA | NA |
| WBGene00198139 | NA | NA | NA | NA |
| WBGene00198140 | NA | NA | NA | NA |
| WBGene00198141 | NA | NA | NA | NA |
| WBGene00198142 | NA | NA | NA | NA |
| WBGene00198143 | NA | NA | NA | NA |
| WBGene00198144 | NA | NA | NA | NA |
| WBGene00198145 | NA | NA | NA | NA |
| WBGene00198146 | NA | NA | NA | NA |
| WBGene00198147 | NA | NA | NA | NA |
| WBGene00198148 | NA | NA | NA | NA |
| WBGene00198149 | NA | NA | NA | NA |
| WBGene00198150 | NA | NA | NA | NA |
| WBGene00198151 | NA | NA | NA | NA |
| WBGene00198152 | NA | NA | NA | NA |
| WBGene00198153 | NA | NA | NA | NA |
| WBGene00198154 | NA | NA | NA | NA |
| WBGene00198155 | NA | NA | NA | NA |
| WBGene00198156 | 0.036687355 | 0.59489568 | NA | NA |
| WBGene00198157 | NA | NA | NA | NA |
| WBGene00198158 | NA | NA | NA | NA |
| WBGene00198159 | NA | NA | NA | NA |
| WBGene00198160 | NA | NA | NA | NA |
| WBGene00198161 | NA | NA | NA | NA |
| WBGene00198162 | NA | NA | NA | NA |
| WBGene00198163 | NA | NA | NA | NA |
| WBGene00198164 | NA | NA | NA | NA |
| WBGene00198165 | NA | NA | NA | NA |
| WBGene00198166 | NA | NA | NA | NA |
| WBGene00198167 | NA | NA | NA | NA |
| WBGene00198168 | 0.020925399 | 0.84643798 | NA | NA |
| WBGene00198169 | NA | NA | NA | NA |
| WBGene00198170 | NA | NA | NA | NA |
| WBGene00198171 | NA | NA | NA | NA |
| WBGene00198172 | NA | NA | NA | NA |

| | | | | |
|---|---|---|---|---|
| WBGene00198173 | NA | NA | NA | NA |
| WBGene00198174 | NA | NA | NA | NA |
| WBGene00198175 | NA | NA | NA | NA |
| WBGene00198176 | NA | NA | NA | NA |
| WBGene00198177 | NA | NA | NA | NA |
| WBGene00198178 | NA | NA | NA | NA |
| WBGene00198179 | NA | NA | NA | NA |
| WBGene00198180 | NA | NA | NA | NA |
| WBGene00198181 | NA | NA | NA | NA |
| WBGene00198182 | NA | NA | NA | NA |
| WBGene00198183 | NA | NA | NA | NA |
| WBGene00198184 | NA | NA | NA | NA |
| WBGene00198185 | NA | NA | NA | NA |
| WBGene00198186 | NA | NA | NA | NA |
| WBGene00198187 | NA | NA | NA | NA |
| WBGene00198188 | NA | NA | NA | NA |
| WBGene00198189 | NA | NA | NA | NA |
| WBGene00198190 | NA | NA | NA | NA |
| WBGene00198191 | NA | NA | NA | NA |
| WBGene00198192 | NA | NA | NA | NA |
| WBGene00198193 | NA | NA | NA | NA |
| WBGene00198194 | NA | NA | NA | NA |
| WBGene00198195 | NA | NA | NA | NA |
| WBGene00198196 | NA | NA | NA | NA |
| WBGene00198197 | NA | NA | NA | NA |
| WBGene00198198 | NA | NA | NA | NA |
| WBGene00198199 | NA | NA | NA | NA |
| WBGene00198200 | NA | NA | NA | NA |
| WBGene00198201 | NA | NA | NA | NA |
| WBGene00198202 | NA | NA | NA | NA |
| WBGene00198203 | NA | NA | NA | NA |
| WBGene00198204 | NA | NA | NA | NA |
| WBGene00198205 | NA | NA | NA | NA |
| WBGene00198206 | NA | NA | NA | NA |
| WBGene00198207 | NA | NA | NA | NA |
| WBGene00198208 | NA | NA | NA | NA |
| WBGene00198209 | NA | NA | NA | NA |
| WBGene00198210 | NA | NA | NA | NA |
| WBGene00198211 | NA | NA | NA | NA |
| WBGene00198212 | NA | NA | NA | NA |
| WBGene00198213 | NA | NA | NA | NA |
| WBGene00198214 | NA | NA | NA | NA |
| WBGene00198215 | NA | NA | NA | NA |

| | | | | |
|---|---|---|---|---|
| WBGene00198216 | NA | NA | NA | NA |
| WBGene00198217 | NA | NA | NA | NA |
| WBGene00198218 | NA | NA | NA | NA |
| WBGene00198219 | NA | NA | NA | NA |
| WBGene00198220 | 0.036687355 | 0.59489568 | NA | NA |
| WBGene00198221 | NA | NA | NA | NA |
| WBGene00198222 | NA | NA | NA | NA |
| WBGene00198223 | NA | NA | NA | NA |
| WBGene00198224 | NA | NA | NA | NA |
| WBGene00198225 | NA | NA | NA | NA |
| WBGene00198226 | NA | NA | NA | NA |
| WBGene00198227 | NA | NA | NA | NA |
| WBGene00198228 | NA | NA | NA | NA |
| WBGene00198229 | NA | NA | NA | NA |
| WBGene00198230 | NA | NA | NA | NA |
| WBGene00198231 | NA | NA | NA | NA |
| WBGene00198232 | NA | NA | NA | NA |
| WBGene00198233 | NA | NA | NA | NA |
| WBGene00198234 | NA | NA | NA | NA |
| WBGene00198235 | 0.033463284 | 0.6365037 | NA | NA |
| WBGene00198236 | NA | NA | NA | NA |
| WBGene00198237 | NA | NA | NA | NA |
| WBGene00198238 | 0.020232312 | 0.84643798 | NA | NA |
| WBGene00198239 | NA | NA | NA | NA |
| WBGene00198240 | NA | NA | NA | NA |
| WBGene00198241 | NA | NA | NA | NA |
| WBGene00198242 | NA | NA | NA | NA |
| WBGene00198243 | NA | NA | NA | NA |
| WBGene00198244 | NA | NA | NA | NA |
| WBGene00198245 | NA | NA | NA | NA |
| WBGene00198246 | NA | NA | NA | NA |
| WBGene00198247 | NA | NA | NA | NA |
| WBGene00198248 | NA | NA | NA | NA |
| WBGene00198249 | NA | NA | NA | NA |
| WBGene00198250 | NA | NA | NA | NA |
| WBGene00198251 | NA | NA | NA | NA |
| WBGene00198252 | NA | NA | NA | NA |
| WBGene00198253 | NA | NA | NA | NA |
| WBGene00198254 | NA | NA | NA | NA |
| WBGene00198255 | NA | NA | NA | NA |
| WBGene00198256 | NA | NA | NA | NA |
| WBGene00198257 | 0.020317245 | 0.84643798 | NA | NA |
| WBGene00198258 | NA | NA | NA | NA |

| | | | | |
|---|---|---|---|---|
| WBGene00198259 | NA | NA | NA | NA |
| WBGene00198260 | NA | NA | NA | NA |
| WBGene00198261 | NA | NA | NA | NA |
| WBGene00198262 | NA | NA | NA | NA |
| WBGene00198263 | NA | NA | NA | NA |
| WBGene00198264 | NA | NA | NA | NA |
| WBGene00198265 | -0.023109762 | 0.78121253 | NA | NA |
| WBGene00198266 | NA | NA | NA | NA |
| WBGene00198267 | NA | NA | NA | NA |
| WBGene00198268 | NA | NA | NA | NA |
| WBGene00198269 | NA | NA | NA | NA |
| WBGene00198270 | NA | NA | NA | NA |
| WBGene00198271 | NA | NA | NA | NA |
| WBGene00198272 | NA | NA | NA | NA |
| WBGene00198273 | NA | NA | NA | NA |
| WBGene00198274 | NA | NA | NA | NA |
| WBGene00198275 | NA | NA | NA | NA |
| WBGene00198276 | 0.020232312 | 0.84643798 | NA | NA |
| WBGene00198277 | NA | NA | NA | NA |
| WBGene00198278 | NA | NA | NA | NA |
| WBGene00198279 | NA | NA | NA | NA |
| WBGene00198280 | NA | NA | NA | NA |
| WBGene00198281 | NA | NA | NA | NA |
| WBGene00198282 | NA | NA | NA | NA |
| WBGene00198283 | NA | NA | NA | NA |
| WBGene00198284 | NA | NA | NA | NA |
| WBGene00198285 | NA | NA | NA | NA |
| WBGene00198286 | NA | NA | NA | NA |
| WBGene00198287 | NA | NA | NA | NA |
| WBGene00198288 | NA | NA | NA | NA |
| WBGene00198289 | NA | NA | NA | NA |
| WBGene00198290 | NA | NA | NA | NA |
| WBGene00198291 | NA | NA | NA | NA |
| WBGene00198292 | 0.020317245 | 0.84643798 | NA | NA |
| WBGene00198293 | NA | NA | NA | NA |
| WBGene00198294 | NA | NA | NA | NA |
| WBGene00198295 | NA | NA | NA | NA |
| WBGene00198296 | 0.020925399 | 0.84643798 | NA | NA |
| WBGene00198297 | NA | NA | NA | NA |
| WBGene00198298 | NA | NA | NA | NA |
| WBGene00198299 | NA | NA | NA | NA |
| WBGene00198300 | NA | NA | NA | NA |
| WBGene00198301 | NA | NA | NA | NA |

| | | | | |
|---|---|---|---|---|
| WBGene00198302 | NA | NA | NA | NA |
| WBGene00198303 | NA | NA | NA | NA |
| WBGene00198304 | NA | NA | NA | NA |
| WBGene00198305 | NA | NA | NA | NA |
| WBGene00198306 | NA | NA | NA | NA |
| WBGene00198307 | -0.022930767 | 0.78121253 | NA | NA |
| WBGene00198308 | NA | NA | NA | NA |
| WBGene00198309 | -0.022930767 | 0.78121253 | NA | NA |
| WBGene00198310 | NA | NA | NA | NA |
| WBGene00198311 | NA | NA | NA | NA |
| WBGene00198312 | NA | NA | NA | NA |
| WBGene00198313 | 0.020317245 | 0.84643798 | NA | NA |
| WBGene00198314 | NA | NA | NA | NA |
| WBGene00198315 | 0.020925399 | 0.84643798 | NA | NA |
| WBGene00198316 | NA | NA | NA | NA |
| WBGene00198317 | NA | NA | NA | NA |
| WBGene00198318 | NA | NA | NA | NA |
| WBGene00198319 | NA | NA | NA | NA |
| WBGene00198320 | NA | NA | NA | NA |
| WBGene00198321 | NA | NA | NA | NA |
| WBGene00198322 | NA | NA | NA | NA |
| WBGene00198323 | NA | NA | NA | NA |
| WBGene00198324 | NA | NA | NA | NA |
| WBGene00198325 | 0.020925399 | 0.84643798 | NA | NA |
| WBGene00198326 | NA | NA | NA | NA |
| WBGene00198327 | NA | NA | NA | NA |
| WBGene00198328 | NA | NA | NA | NA |
| WBGene00198329 | NA | NA | NA | NA |
| WBGene00198330 | NA | NA | NA | NA |
| WBGene00198331 | NA | NA | NA | NA |
| WBGene00198332 | NA | NA | NA | NA |
| WBGene00198333 | NA | NA | NA | NA |
| WBGene00198334 | NA | NA | NA | NA |
| WBGene00198335 | NA | NA | NA | NA |
| WBGene00198336 | NA | NA | NA | NA |
| WBGene00198337 | NA | NA | NA | NA |
| WBGene00198338 | NA | NA | NA | NA |
| WBGene00198339 | NA | NA | NA | NA |
| WBGene00198340 | NA | NA | NA | NA |
| WBGene00198341 | NA | NA | NA | NA |
| WBGene00198342 | NA | NA | NA | NA |
| WBGene00198343 | NA | NA | NA | NA |
| WBGene00198344 | NA | NA | NA | NA |

| | | | | |
|---|---|---|---|---|
| WBGene00198345 | NA | NA | NA | NA |
| WBGene00198346 | NA | NA | NA | NA |
| WBGene00198347 | NA | NA | NA | NA |
| WBGene00198348 | NA | NA | NA | NA |
| WBGene00198349 | NA | NA | NA | NA |
| WBGene00198350 | 0.033034883 | 0.64219503 | NA | NA |
| WBGene00198351 | NA | NA | NA | NA |
| WBGene00198352 | NA | NA | NA | NA |
| WBGene00198353 | NA | NA | NA | NA |
| WBGene00198354 | NA | NA | NA | NA |
| WBGene00198355 | NA | NA | NA | NA |
| WBGene00198356 | NA | NA | NA | NA |
| WBGene00198357 | NA | NA | NA | NA |
| WBGene00198358 | NA | NA | NA | NA |
| WBGene00198359 | NA | NA | NA | NA |
| WBGene00198360 | NA | NA | NA | NA |
| WBGene00198361 | NA | NA | NA | NA |
| WBGene00198362 | NA | NA | NA | NA |
| WBGene00198363 | NA | NA | NA | NA |
| WBGene00198364 | NA | NA | NA | NA |
| WBGene00198365 | -0.023056499 | 0.78121253 | NA | NA |
| WBGene00198366 | NA | NA | NA | NA |
| WBGene00198367 | NA | NA | NA | NA |
| WBGene00198368 | NA | NA | NA | NA |
| WBGene00198369 | NA | NA | NA | NA |
| WBGene00198370 | NA | NA | NA | NA |
| WBGene00198371 | NA | NA | NA | NA |
| WBGene00198372 | NA | NA | NA | NA |
| WBGene00198373 | NA | NA | NA | NA |
| WBGene00198374 | NA | NA | NA | NA |
| WBGene00198375 | NA | NA | NA | NA |
| WBGene00198376 | NA | NA | NA | NA |
| WBGene00198377 | NA | NA | NA | NA |
| WBGene00198378 | NA | NA | NA | NA |
| WBGene00198379 | NA | NA | NA | NA |
| WBGene00198380 | NA | NA | NA | NA |
| WBGene00198381 | NA | NA | NA | NA |
| WBGene00198382 | NA | NA | NA | NA |
| WBGene00198383 | -0.022930767 | 0.78121253 | NA | NA |
| WBGene00198384 | 0.020317245 | 0.84643798 | NA | NA |
| WBGene00198385 | NA | NA | NA | NA |
| WBGene00198386 | NA | NA | NA | NA |
| WBGene00198387 | NA | NA | NA | NA |

| | | | | |
|---|---|---|---|---|
| WBGene00198388 | NA | NA | NA | NA |
| WBGene00198389 | NA | NA | NA | NA |
| WBGene00198390 | NA | NA | NA | NA |
| WBGene00198391 | NA | NA | NA | NA |
| WBGene00198392 | NA | NA | NA | NA |
| WBGene00198393 | NA | NA | NA | NA |
| WBGene00198394 | NA | NA | NA | NA |
| WBGene00198395 | NA | NA | NA | NA |
| WBGene00198396 | NA | NA | NA | NA |
| WBGene00198397 | NA | NA | NA | NA |
| WBGene00198398 | NA | NA | NA | NA |
| WBGene00198399 | NA | NA | NA | NA |
| WBGene00198400 | NA | NA | NA | NA |
| WBGene00198401 | NA | NA | NA | NA |
| WBGene00198402 | NA | NA | NA | NA |
| WBGene00198403 | NA | NA | NA | NA |
| WBGene00198404 | NA | NA | NA | NA |
| WBGene00198405 | NA | NA | NA | NA |
| WBGene00198406 | NA | NA | NA | NA |
| WBGene00198407 | NA | NA | NA | NA |
| WBGene00198408 | NA | NA | NA | NA |
| WBGene00198409 | NA | NA | NA | NA |
| WBGene00198410 | NA | NA | NA | NA |
| WBGene00198411 | NA | NA | NA | NA |
| WBGene00198412 | NA | NA | NA | NA |
| WBGene00198413 | NA | NA | NA | NA |
| WBGene00198414 | 0.020317245 | 0.84643798 | NA | NA |
| WBGene00198415 | NA | NA | NA | NA |
| WBGene00198416 | NA | NA | NA | NA |
| WBGene00198417 | NA | NA | NA | NA |
| WBGene00198418 | NA | NA | NA | NA |
| WBGene00198419 | NA | NA | NA | NA |
| WBGene00198420 | NA | NA | NA | NA |
| WBGene00198421 | NA | NA | NA | NA |
| WBGene00198422 | NA | NA | NA | NA |
| WBGene00198423 | NA | NA | NA | NA |
| WBGene00198424 | NA | NA | NA | NA |
| WBGene00198425 | NA | NA | NA | NA |
| WBGene00198426 | NA | NA | NA | NA |
| WBGene00198427 | NA | NA | NA | NA |
| WBGene00198428 | NA | NA | NA | NA |
| WBGene00198429 | -0.023109762 | 0.78121253 | NA | NA |
| WBGene00198430 | NA | NA | NA | NA |

| | | | | |
|---|---|---|---|---|
| WBGene00198431 | NA | NA | NA | NA |
| WBGene00198432 | NA | NA | NA | NA |
| WBGene00198433 | -0.023056499 | 0.78121253 | NA | NA |
| WBGene00198434 | NA | NA | NA | NA |
| WBGene00198435 | NA | NA | NA | NA |
| WBGene00198436 | NA | NA | NA | NA |
| WBGene00198437 | NA | NA | NA | NA |
| WBGene00198438 | NA | NA | NA | NA |
| WBGene00198439 | NA | NA | NA | NA |
| WBGene00198440 | NA | NA | NA | NA |
| WBGene00198441 | NA | NA | NA | NA |
| WBGene00198442 | NA | NA | NA | NA |
| WBGene00198443 | NA | NA | NA | NA |
| WBGene00198444 | NA | NA | NA | NA |
| WBGene00198445 | 0.020925399 | 0.84643798 | NA | NA |
| WBGene00198446 | 0.056411843 | 0.41891328 | NA | NA |
| WBGene00198447 | NA | NA | NA | NA |
| WBGene00198448 | NA | NA | NA | NA |
| WBGene00198449 | NA | NA | NA | NA |
| WBGene00198450 | NA | NA | NA | NA |
| WBGene00198451 | NA | NA | NA | NA |
| WBGene00198452 | NA | NA | NA | NA |
| WBGene00198453 | NA | NA | NA | NA |
| WBGene00198454 | 0.020317245 | 0.84643798 | NA | NA |
| WBGene00198455 | NA | NA | NA | NA |
| WBGene00198456 | NA | NA | NA | NA |
| WBGene00198457 | 0.038125098 | 0.61119482 | NA | NA |
| WBGene00198458 | NA | NA | NA | NA |
| WBGene00198459 | NA | NA | NA | NA |
| WBGene00198460 | NA | NA | NA | NA |
| WBGene00198461 | NA | NA | NA | NA |
| WBGene00198462 | NA | NA | NA | NA |
| WBGene00198463 | NA | NA | NA | NA |
| WBGene00198464 | NA | NA | NA | NA |
| WBGene00198465 | NA | NA | NA | NA |
| WBGene00198466 | NA | NA | NA | NA |
| WBGene00198467 | NA | NA | NA | NA |
| WBGene00198468 | NA | NA | NA | NA |
| WBGene00198469 | NA | NA | NA | NA |
| WBGene00198470 | NA | NA | NA | NA |
| WBGene00198471 | NA | NA | NA | NA |
| WBGene00198472 | NA | NA | NA | NA |
| WBGene00198473 | NA | NA | NA | NA |

| | | | | | |
|---|---|---|---|---|---|
| WBGene00198474 | NA | | NA | NA | NA |
| WBGene00198475 | NA | | NA | NA | NA |
| WBGene00198476 | NA | | NA | NA | NA |
| WBGene00198477 | | 0.020317245 | 0.84643798 | NA | NA |
| WBGene00198478 | NA | | NA | NA | NA |
| WBGene00198479 | NA | | NA | NA | NA |
| WBGene00198480 | | 0.020232312 | 0.84643798 | NA | NA |
| WBGene00198481 | NA | | NA | NA | NA |
| WBGene00198482 | NA | | NA | NA | NA |
| WBGene00198483 | NA | | NA | NA | NA |
| WBGene00198484 | NA | | NA | NA | NA |
| WBGene00198485 | NA | | NA | NA | NA |
| WBGene00198486 | NA | | NA | NA | NA |
| WBGene00198487 | NA | | NA | NA | NA |
| WBGene00198488 | NA | | NA | NA | NA |
| WBGene00198489 | NA | | NA | NA | NA |
| WBGene00198490 | NA | | NA | NA | NA |
| WBGene00198491 | NA | | NA | NA | NA |
| WBGene00198492 | NA | | NA | NA | NA |
| WBGene00198493 | NA | | NA | NA | NA |
| WBGene00198494 | NA | | NA | NA | NA |
| WBGene00198495 | NA | | NA | NA | NA |
| WBGene00198496 | NA | | NA | NA | NA |
| WBGene00198497 | | 0.056188364 | 0.42092875 | NA | NA |
| WBGene00198498 | NA | | NA | NA | NA |
| WBGene00198499 | NA | | NA | NA | NA |
| WBGene00198500 | NA | | NA | NA | NA |
| WBGene00198501 | NA | | NA | NA | NA |
| WBGene00198502 | NA | | NA | NA | NA |
| WBGene00198503 | NA | | NA | NA | NA |
| WBGene00198504 | NA | | NA | NA | NA |
| WBGene00198505 | NA | | NA | NA | NA |
| WBGene00198506 | NA | | NA | NA | NA |
| WBGene00198507 | NA | | NA | NA | NA |
| WBGene00198508 | NA | | NA | NA | NA |
| WBGene00198509 | NA | | NA | NA | NA |
| WBGene00198510 | NA | | NA | NA | NA |
| WBGene00198511 | NA | | NA | NA | NA |
| WBGene00198512 | NA | | NA | NA | NA |
| WBGene00198513 | NA | | NA | NA | NA |
| WBGene00198514 | NA | | NA | NA | NA |
| WBGene00198515 | NA | | NA | NA | NA |
| WBGene00198516 | NA | | NA | NA | NA |

| | | | | |
|---|---|---|---|---|
| WBGene00198517 | NA | NA | NA | NA |
| WBGene00198518 | NA | NA | NA | NA |
| WBGene00198519 | NA | NA | NA | NA |
| WBGene00198520 | NA | NA | NA | NA |
| WBGene00198521 | 0.040107227 | 0.58693163 | NA | NA |
| WBGene00198522 | NA | NA | NA | NA |
| WBGene00198523 | NA | NA | NA | NA |
| WBGene00198524 | NA | NA | NA | NA |
| WBGene00198525 | NA | NA | NA | NA |
| WBGene00198526 | NA | NA | NA | NA |
| WBGene00198527 | NA | NA | NA | NA |
| WBGene00198528 | 0.040107227 | 0.58693163 | NA | NA |
| WBGene00198529 | NA | NA | NA | NA |
| WBGene00198530 | -0.022930767 | 0.78121253 | NA | NA |
| WBGene00198531 | NA | NA | NA | NA |
| WBGene00198532 | NA | NA | NA | NA |
| WBGene00198533 | NA | NA | NA | NA |
| WBGene00198534 | NA | NA | NA | NA |
| WBGene00198535 | NA | NA | NA | NA |
| WBGene00198536 | NA | NA | NA | NA |
| WBGene00198537 | NA | NA | NA | NA |
| WBGene00198538 | -0.022930767 | 0.78121253 | NA | NA |
| WBGene00198539 | NA | NA | NA | NA |
| WBGene00198540 | NA | NA | NA | NA |
| WBGene00198541 | NA | NA | NA | NA |
| WBGene00198542 | NA | NA | NA | NA |
| WBGene00198543 | NA | NA | NA | NA |
| WBGene00198544 | NA | NA | NA | NA |
| WBGene00198545 | NA | NA | NA | NA |
| WBGene00198546 | NA | NA | NA | NA |
| WBGene00198547 | NA | NA | NA | NA |
| WBGene00198548 | NA | NA | NA | NA |
| WBGene00198549 | NA | NA | NA | NA |
| WBGene00198550 | NA | NA | NA | NA |
| WBGene00198551 | NA | NA | NA | NA |
| WBGene00198552 | NA | NA | NA | NA |
| WBGene00198553 | NA | NA | NA | NA |
| WBGene00198554 | NA | NA | NA | NA |
| WBGene00198555 | NA | NA | NA | NA |
| WBGene00198556 | NA | NA | NA | NA |
| WBGene00198557 | NA | NA | NA | NA |
| WBGene00198558 | NA | NA | NA | NA |
| WBGene00198559 | NA | NA | NA | NA |

| | | | | |
|---|---|---|---|---|
| WBGene00198560 | NA | NA | NA | NA |
| WBGene00198561 | NA | NA | NA | NA |
| WBGene00198562 | NA | NA | NA | NA |
| WBGene00198563 | NA | NA | NA | NA |
| WBGene00198564 | NA | NA | NA | NA |
| WBGene00198565 | NA | NA | NA | NA |
| WBGene00198566 | NA | NA | NA | NA |
| WBGene00198567 | NA | NA | NA | NA |
| WBGene00198568 | NA | NA | NA | NA |
| WBGene00198569 | NA | NA | NA | NA |
| WBGene00198570 | NA | NA | NA | NA |
| WBGene00198571 | NA | NA | NA | NA |
| WBGene00198572 | NA | NA | NA | NA |
| WBGene00198573 | NA | NA | NA | NA |
| WBGene00198574 | NA | NA | NA | NA |
| WBGene00198575 | NA | NA | NA | NA |
| WBGene00198576 | NA | NA | NA | NA |
| WBGene00198577 | NA | NA | NA | NA |
| WBGene00198578 | NA | NA | NA | NA |
| WBGene00198579 | NA | NA | NA | NA |
| WBGene00198580 | NA | NA | NA | NA |
| WBGene00198581 | NA | NA | NA | NA |
| WBGene00198582 | NA | NA | NA | NA |
| WBGene00198583 | NA | NA | NA | NA |
| WBGene00198584 | NA | NA | NA | NA |
| WBGene00198585 | NA | NA | NA | NA |
| WBGene00198586 | NA | NA | NA | NA |
| WBGene00198587 | NA | NA | NA | NA |
| WBGene00198588 | NA | NA | NA | NA |
| WBGene00198589 | NA | NA | NA | NA |
| WBGene00198590 | NA | NA | NA | NA |
| WBGene00198591 | NA | NA | NA | NA |
| WBGene00198592 | NA | NA | NA | NA |
| WBGene00198593 | NA | NA | NA | NA |
| WBGene00198594 | NA | NA | NA | NA |
| WBGene00198595 | NA | NA | NA | NA |
| WBGene00198596 | NA | NA | NA | NA |
| WBGene00198597 | NA | NA | NA | NA |
| WBGene00198598 | NA | NA | NA | NA |
| WBGene00198599 | NA | NA | NA | NA |
| WBGene00198600 | NA | NA | NA | NA |
| WBGene00198601 | NA | NA | NA | NA |
| WBGene00198602 | NA | NA | NA | NA |

| | | | | |
|---|---|---|---|---|
| WBGene00198603 | NA | NA | NA | NA |
| WBGene00198604 | NA | NA | NA | NA |
| WBGene00198605 | NA | NA | NA | NA |
| WBGene00198606 | NA | NA | NA | NA |
| WBGene00198607 | NA | NA | NA | NA |
| WBGene00198608 | NA | NA | NA | NA |
| WBGene00198609 | NA | NA | NA | NA |
| WBGene00198610 | NA | NA | NA | NA |
| WBGene00198611 | NA | NA | NA | NA |
| WBGene00198612 | NA | NA | NA | NA |
| WBGene00198613 | NA | NA | NA | NA |
| WBGene00198614 | NA | NA | NA | NA |
| WBGene00198615 | NA | NA | NA | NA |
| WBGene00198616 | NA | NA | NA | NA |
| WBGene00198617 | NA | NA | NA | NA |
| WBGene00198618 | NA | NA | NA | NA |
| WBGene00198619 | NA | NA | NA | NA |
| WBGene00198620 | 0.020925399 | 0.84643798 | NA | NA |
| WBGene00198621 | NA | NA | NA | NA |
| WBGene00198622 | NA | NA | NA | NA |
| WBGene00198623 | NA | NA | NA | NA |
| WBGene00198624 | -0.022930767 | 0.78121253 | NA | NA |
| WBGene00198625 | NA | NA | NA | NA |
| WBGene00198626 | NA | NA | NA | NA |
| WBGene00198627 | NA | NA | NA | NA |
| WBGene00198628 | NA | NA | NA | NA |
| WBGene00198629 | NA | NA | NA | NA |
| WBGene00198630 | NA | NA | NA | NA |
| WBGene00198631 | NA | NA | NA | NA |
| WBGene00198632 | NA | NA | NA | NA |
| WBGene00198633 | NA | NA | NA | NA |
| WBGene00198634 | NA | NA | NA | NA |
| WBGene00198635 | NA | NA | NA | NA |
| WBGene00198636 | -0.023056499 | 0.78121253 | NA | NA |
| WBGene00198637 | NA | NA | NA | NA |
| WBGene00198638 | NA | NA | NA | NA |
| WBGene00198639 | NA | NA | NA | NA |
| WBGene00198640 | NA | NA | NA | NA |
| WBGene00198641 | NA | NA | NA | NA |
| WBGene00198642 | NA | NA | NA | NA |
| WBGene00198643 | NA | NA | NA | NA |
| WBGene00198644 | NA | NA | NA | NA |
| WBGene00198645 | NA | NA | NA | NA |

| | | | | |
|---|---|---|---|---|
| WBGene00198646 | NA | NA | NA | NA |
| WBGene00198647 | NA | NA | NA | NA |
| WBGene00198648 | NA | NA | NA | NA |
| WBGene00198649 | NA | NA | NA | NA |
| WBGene00198650 | NA | NA | NA | NA |
| WBGene00198651 | NA | NA | NA | NA |
| WBGene00198652 | 0.020317245 | 0.84643798 | NA | NA |
| WBGene00198653 | 0.020317245 | 0.84643798 | NA | NA |
| WBGene00198654 | NA | NA | NA | NA |
| WBGene00198655 | NA | NA | NA | NA |
| WBGene00198656 | -0.001842784 | 0.95621055 | NA | NA |
| WBGene00198657 | NA | NA | NA | NA |
| WBGene00198658 | NA | NA | NA | NA |
| WBGene00198659 | NA | NA | NA | NA |
| WBGene00198660 | NA | NA | NA | NA |
| WBGene00198661 | NA | NA | NA | NA |
| WBGene00198662 | NA | NA | NA | NA |
| WBGene00198663 | NA | NA | NA | NA |
| WBGene00198664 | NA | NA | NA | NA |
| WBGene00198665 | NA | NA | NA | NA |
| WBGene00198666 | NA | NA | NA | NA |
| WBGene00198667 | NA | NA | NA | NA |
| WBGene00198668 | -0.022930767 | 0.78121253 | NA | NA |
| WBGene00198669 | NA | NA | NA | NA |
| WBGene00198670 | NA | NA | NA | NA |
| WBGene00198671 | NA | NA | NA | NA |
| WBGene00198672 | NA | NA | NA | NA |
| WBGene00198673 | NA | NA | NA | NA |
| WBGene00198674 | NA | NA | NA | NA |
| WBGene00198675 | NA | NA | NA | NA |
| WBGene00198676 | NA | NA | NA | NA |
| WBGene00198677 | 0.020232312 | 0.84643798 | NA | NA |
| WBGene00198678 | NA | NA | NA | NA |
| WBGene00198679 | NA | NA | NA | NA |
| WBGene00198680 | NA | NA | NA | NA |
| WBGene00198681 | NA | NA | NA | NA |
| WBGene00198682 | NA | NA | NA | NA |
| WBGene00198683 | NA | NA | NA | NA |
| WBGene00198684 | NA | NA | NA | NA |
| WBGene00198685 | NA | NA | NA | NA |
| WBGene00198686 | NA | NA | NA | NA |
| WBGene00198687 | NA | NA | NA | NA |
| WBGene00198688 | NA | NA | NA | NA |

| | | | | |
|---|---|---|---|---|
| WBGene00198689 | NA | NA | NA | NA |
| WBGene00198690 | NA | NA | NA | NA |
| WBGene00198691 | NA | NA | NA | NA |
| WBGene00198692 | 0.015507174 | 0.80923808 | NA | NA |
| WBGene00198693 | NA | NA | NA | NA |
| WBGene00198694 | NA | NA | NA | NA |
| WBGene00198695 | NA | NA | NA | NA |
| WBGene00198696 | NA | NA | NA | NA |
| WBGene00198697 | NA | NA | NA | NA |
| WBGene00198698 | NA | NA | NA | NA |
| WBGene00198699 | NA | NA | NA | NA |
| WBGene00198700 | NA | NA | NA | NA |
| WBGene00198701 | NA | NA | NA | NA |
| WBGene00198702 | NA | NA | NA | NA |
| WBGene00198703 | NA | NA | NA | NA |
| WBGene00198704 | NA | NA | NA | NA |
| WBGene00198705 | NA | NA | NA | NA |
| WBGene00198706 | NA | NA | NA | NA |
| WBGene00198707 | NA | NA | NA | NA |
| WBGene00198708 | NA | NA | NA | NA |
| WBGene00198709 | -0.023056499 | 0.78121253 | NA | NA |
| WBGene00198710 | NA | NA | NA | NA |
| WBGene00198711 | NA | NA | NA | NA |
| WBGene00198712 | NA | NA | NA | NA |
| WBGene00198713 | NA | NA | NA | NA |
| WBGene00198714 | NA | NA | NA | NA |
| WBGene00198715 | NA | NA | NA | NA |
| WBGene00198716 | NA | NA | NA | NA |
| WBGene00198717 | NA | NA | NA | NA |
| WBGene00198718 | NA | NA | NA | NA |
| WBGene00198719 | NA | NA | NA | NA |
| WBGene00198720 | NA | NA | NA | NA |
| WBGene00198721 | NA | NA | NA | NA |
| WBGene00198722 | NA | NA | NA | NA |
| WBGene00198723 | NA | NA | NA | NA |
| WBGene00198724 | NA | NA | NA | NA |
| WBGene00198725 | NA | NA | NA | NA |
| WBGene00198726 | NA | NA | NA | NA |
| WBGene00198727 | NA | NA | NA | NA |
| WBGene00198728 | NA | NA | NA | NA |
| WBGene00198729 | NA | NA | NA | NA |
| WBGene00198730 | NA | NA | NA | NA |
| WBGene00198731 | NA | NA | NA | NA |

| | | | | |
|---|---|---|---|---|
| WBGene00198732 | NA | NA | NA | NA |
| WBGene00198733 | NA | NA | NA | NA |
| WBGene00198734 | NA | NA | NA | NA |
| WBGene00198735 | NA | NA | NA | NA |
| WBGene00198736 | NA | NA | NA | NA |
| WBGene00198737 | 0.020317245 | 0.84643798 | NA | NA |
| WBGene00198738 | NA | NA | NA | NA |
| WBGene00198739 | NA | NA | NA | NA |
| WBGene00198740 | NA | NA | NA | NA |
| WBGene00198741 | NA | NA | NA | NA |
| WBGene00198742 | NA | NA | NA | NA |
| WBGene00198743 | NA | NA | NA | NA |
| WBGene00198744 | NA | NA | NA | NA |
| WBGene00198745 | NA | NA | NA | NA |
| WBGene00198746 | 0.020317245 | 0.84643798 | NA | NA |
| WBGene00198747 | NA | NA | NA | NA |
| WBGene00198748 | NA | NA | NA | NA |
| WBGene00198749 | NA | NA | NA | NA |
| WBGene00198750 | 0.020317245 | 0.84643798 | NA | NA |
| WBGene00198751 | NA | NA | NA | NA |
| WBGene00198752 | NA | NA | NA | NA |
| WBGene00198753 | NA | NA | NA | NA |
| WBGene00198754 | NA | NA | NA | NA |
| WBGene00198755 | 0.020232312 | 0.84643798 | NA | NA |
| WBGene00198756 | NA | NA | NA | NA |
| WBGene00198757 | NA | NA | NA | NA |
| WBGene00198758 | NA | NA | NA | NA |
| WBGene00198759 | NA | NA | NA | NA |
| WBGene00198760 | NA | NA | NA | NA |
| WBGene00198761 | NA | NA | NA | NA |
| WBGene00198762 | NA | NA | NA | NA |
| WBGene00198763 | NA | NA | NA | NA |
| WBGene00198764 | NA | NA | NA | NA |
| WBGene00198765 | NA | NA | NA | NA |
| WBGene00198766 | NA | NA | NA | NA |
| WBGene00198767 | NA | NA | NA | NA |
| WBGene00198768 | NA | NA | NA | NA |
| WBGene00198769 | NA | NA | NA | NA |
| WBGene00198770 | NA | NA | NA | NA |
| WBGene00198771 | NA | NA | NA | NA |
| WBGene00198772 | NA | NA | NA | NA |
| WBGene00198773 | NA | NA | NA | NA |
| WBGene00198774 | NA | NA | NA | NA |

| | | | | |
|---|---|---|---|---|
| WBGene00198775 | NA | NA | NA | NA |
| WBGene00198776 | NA | NA | NA | NA |
| WBGene00198777 | NA | NA | NA | NA |
| WBGene00198778 | NA | NA | NA | NA |
| WBGene00198779 | NA | NA | NA | NA |
| WBGene00198780 | NA | NA | NA | NA |
| WBGene00198781 | NA | NA | NA | NA |
| WBGene00198782 | NA | NA | NA | NA |
| WBGene00198783 | NA | NA | NA | NA |
| WBGene00198784 | NA | NA | NA | NA |
| WBGene00198785 | NA | NA | NA | NA |
| WBGene00198786 | NA | NA | NA | NA |
| WBGene00198787 | NA | NA | NA | NA |
| WBGene00198788 | NA | NA | NA | NA |
| WBGene00198789 | NA | NA | NA | NA |
| WBGene00198790 | NA | NA | NA | NA |
| WBGene00198791 | NA | NA | NA | NA |
| WBGene00198792 | NA | NA | NA | NA |
| WBGene00198793 | NA | NA | NA | NA |
| WBGene00198794 | NA | NA | NA | NA |
| WBGene00198795 | NA | NA | NA | NA |
| WBGene00198796 | NA | NA | NA | NA |
| WBGene00198797 | NA | NA | NA | NA |
| WBGene00198798 | NA | NA | NA | NA |
| WBGene00198799 | NA | NA | NA | NA |
| WBGene00198800 | NA | NA | NA | NA |
| WBGene00198801 | NA | NA | NA | NA |
| WBGene00198802 | NA | NA | NA | NA |
| WBGene00198803 | NA | NA | NA | NA |
| WBGene00198804 | NA | NA | NA | NA |
| WBGene00198805 | NA | NA | NA | NA |
| WBGene00198806 | NA | NA | NA | NA |
| WBGene00198807 | NA | NA | NA | NA |
| WBGene00198808 | NA | NA | NA | NA |
| WBGene00198809 | NA | NA | NA | NA |
| WBGene00198810 | NA | NA | NA | NA |
| WBGene00198811 | NA | NA | NA | NA |
| WBGene00198812 | NA | NA | NA | NA |
| WBGene00198813 | NA | NA | NA | NA |
| WBGene00198814 | NA | NA | NA | NA |
| WBGene00198815 | NA | NA | NA | NA |
| WBGene00198816 | NA | NA | NA | NA |
| WBGene00198817 | NA | NA | NA | NA |

| | | | | |
|---|---|---|---|---|
| WBGene00198818 | NA | NA | NA | NA |
| WBGene00198819 | NA | NA | NA | NA |
| WBGene00198820 | NA | NA | NA | NA |
| WBGene00198821 | NA | NA | NA | NA |
| WBGene00198822 | NA | NA | NA | NA |
| WBGene00198823 | NA | NA | NA | NA |
| WBGene00198824 | NA | NA | NA | NA |
| WBGene00198825 | NA | NA | NA | NA |
| WBGene00198826 | -0.00346345 | 0.95708614 | NA | NA |
| WBGene00198827 | NA | NA | NA | NA |
| WBGene00198828 | NA | NA | NA | NA |
| WBGene00198829 | NA | NA | NA | NA |
| WBGene00198830 | NA | NA | NA | NA |
| WBGene00198831 | NA | NA | NA | NA |
| WBGene00198832 | NA | NA | NA | NA |
| WBGene00198833 | NA | NA | NA | NA |
| WBGene00198834 | NA | NA | NA | NA |
| WBGene00198835 | NA | NA | NA | NA |
| WBGene00198836 | NA | NA | NA | NA |
| WBGene00198837 | NA | NA | NA | NA |
| WBGene00198838 | NA | NA | NA | NA |
| WBGene00198839 | NA | NA | NA | NA |
| WBGene00198840 | NA | NA | NA | NA |
| WBGene00198841 | NA | NA | NA | NA |
| WBGene00198842 | -0.023056499 | 0.78121253 | NA | NA |
| WBGene00198843 | NA | NA | NA | NA |
| WBGene00198844 | NA | NA | NA | NA |
| WBGene00198845 | NA | NA | NA | NA |
| WBGene00198846 | NA | NA | NA | NA |
| WBGene00198847 | NA | NA | NA | NA |
| WBGene00198848 | NA | NA | NA | NA |
| WBGene00198849 | NA | NA | NA | NA |
| WBGene00198850 | NA | NA | NA | NA |
| WBGene00198851 | -0.023056499 | 0.78121253 | NA | NA |
| WBGene00198852 | NA | NA | NA | NA |
| WBGene00198853 | NA | NA | NA | NA |
| WBGene00198854 | NA | NA | NA | NA |
| WBGene00198855 | 0.020232312 | 0.84643798 | NA | NA |
| WBGene00198856 | 0.058314668 | 0.40208118 | NA | NA |
| WBGene00198857 | NA | NA | NA | NA |
| WBGene00198858 | NA | NA | NA | NA |
| WBGene00198859 | NA | NA | NA | NA |
| WBGene00198860 | NA | NA | NA | NA |

| | | | | |
|---|---|---|---|---|
| WBGene00198861 | NA | NA | NA | NA |
| WBGene00198862 | NA | NA | NA | NA |
| WBGene00198863 | NA | NA | NA | NA |
| WBGene00198864 | NA | NA | NA | NA |
| WBGene00198865 | NA | NA | NA | NA |
| WBGene00198866 | NA | NA | NA | NA |
| WBGene00198867 | NA | NA | NA | NA |
| WBGene00198868 | NA | NA | NA | NA |
| WBGene00198869 | NA | NA | NA | NA |
| WBGene00198870 | NA | NA | NA | NA |
| WBGene00198871 | NA | NA | NA | NA |
| WBGene00198872 | NA | NA | NA | NA |
| WBGene00198873 | NA | NA | NA | NA |
| WBGene00198874 | NA | NA | NA | NA |
| WBGene00198875 | NA | NA | NA | NA |
| WBGene00198876 | NA | NA | NA | NA |
| WBGene00198877 | NA | NA | NA | NA |
| WBGene00198878 | NA | NA | NA | NA |
| WBGene00198879 | NA | NA | NA | NA |
| WBGene00198880 | NA | NA | NA | NA |
| WBGene00198881 | NA | NA | NA | NA |
| WBGene00198882 | NA | NA | NA | NA |
| WBGene00198883 | NA | NA | NA | NA |
| WBGene00198884 | NA | NA | NA | NA |
| WBGene00198885 | NA | NA | NA | NA |
| WBGene00198886 | NA | NA | NA | NA |
| WBGene00198887 | NA | NA | NA | NA |
| WBGene00198888 | NA | NA | NA | NA |
| WBGene00198889 | NA | NA | NA | NA |
| WBGene00198890 | 0.020925399 | 0.84643798 | NA | NA |
| WBGene00198891 | NA | NA | NA | NA |
| WBGene00198892 | NA | NA | NA | NA |
| WBGene00198893 | NA | NA | NA | NA |
| WBGene00198894 | NA | NA | NA | NA |
| WBGene00198895 | NA | NA | NA | NA |
| WBGene00198896 | NA | NA | NA | NA |
| WBGene00198897 | NA | NA | NA | NA |
| WBGene00198898 | NA | NA | NA | NA |
| WBGene00198899 | NA | NA | NA | NA |
| WBGene00198900 | NA | NA | NA | NA |
| WBGene00198901 | NA | NA | NA | NA |
| WBGene00198902 | NA | NA | NA | NA |
| WBGene00198903 | NA | NA | NA | NA |

| | | | | |
|---|---|---|---|---|
| WBGene00198904 | NA | NA | NA | NA |
| WBGene00198905 | NA | NA | NA | NA |
| WBGene00198906 | NA | NA | NA | NA |
| WBGene00198907 | NA | NA | NA | NA |
| WBGene00198908 | NA | NA | NA | NA |
| WBGene00198909 | NA | NA | NA | NA |
| WBGene00198910 | NA | NA | NA | NA |
| WBGene00198911 | NA | NA | NA | NA |
| WBGene00198912 | NA | NA | NA | NA |
| WBGene00198913 | NA | NA | NA | NA |
| WBGene00198914 | NA | NA | NA | NA |
| WBGene00198915 | NA | NA | NA | NA |
| WBGene00198916 | NA | NA | NA | NA |
| WBGene00198917 | NA | NA | NA | NA |
| WBGene00198918 | NA | NA | NA | NA |
| WBGene00198919 | NA | NA | NA | NA |
| WBGene00198920 | NA | NA | NA | NA |
| WBGene00198921 | NA | NA | NA | NA |
| WBGene00198922 | NA | NA | NA | NA |
| WBGene00198923 | NA | NA | NA | NA |
| WBGene00198924 | NA | NA | NA | NA |
| WBGene00198925 | -0.001488154 | 0.95639912 | NA | NA |
| WBGene00198926 | NA | NA | NA | NA |
| WBGene00198927 | NA | NA | NA | NA |
| WBGene00198928 | NA | NA | NA | NA |
| WBGene00198929 | NA | NA | NA | NA |
| WBGene00198930 | NA | NA | NA | NA |
| WBGene00198931 | NA | NA | NA | NA |
| WBGene00198932 | 0.020925399 | 0.84643798 | NA | NA |
| WBGene00198933 | NA | NA | NA | NA |
| WBGene00198934 | NA | NA | NA | NA |
| WBGene00198935 | NA | NA | NA | NA |
| WBGene00198936 | -0.022930767 | 0.78121253 | NA | NA |
| WBGene00198937 | NA | NA | NA | NA |
| WBGene00198938 | NA | NA | NA | NA |
| WBGene00198939 | NA | NA | NA | NA |
| WBGene00198940 | NA | NA | NA | NA |
| WBGene00198941 | NA | NA | NA | NA |
| WBGene00198942 | NA | NA | NA | NA |
| WBGene00198943 | NA | NA | NA | NA |
| WBGene00198944 | NA | NA | NA | NA |
| WBGene00198945 | NA | NA | NA | NA |
| WBGene00198946 | NA | NA | NA | NA |

| | | | | |
|---|---|---|---|---|
| WBGene00198947 | NA | NA | NA | NA |
| WBGene00198948 | NA | NA | NA | NA |
| WBGene00198949 | NA | NA | NA | NA |
| WBGene00198950 | NA | NA | NA | NA |
| WBGene00198951 | NA | NA | NA | NA |
| WBGene00198952 | NA | NA | NA | NA |
| WBGene00198953 | -0.023056499 | 0.78121253 | NA | NA |
| WBGene00198954 | NA | NA | NA | NA |
| WBGene00198955 | 0.020317245 | 0.84643798 | NA | NA |
| WBGene00198956 | NA | NA | NA | NA |
| WBGene00198957 | NA | NA | NA | NA |
| WBGene00198958 | NA | NA | NA | NA |
| WBGene00198959 | NA | NA | NA | NA |
| WBGene00198960 | NA | NA | NA | NA |
| WBGene00198961 | NA | NA | NA | NA |
| WBGene00198962 | NA | NA | NA | NA |
| WBGene00198963 | NA | NA | NA | NA |
| WBGene00198964 | NA | NA | NA | NA |
| WBGene00198965 | NA | NA | NA | NA |
| WBGene00198966 | NA | NA | NA | NA |
| WBGene00198967 | NA | NA | NA | NA |
| WBGene00198968 | NA | NA | NA | NA |
| WBGene00198969 | NA | NA | NA | NA |
| WBGene00198970 | NA | NA | NA | NA |
| WBGene00198971 | NA | NA | NA | NA |
| WBGene00198972 | NA | NA | NA | NA |
| WBGene00198973 | NA | NA | NA | NA |
| WBGene00198974 | NA | NA | NA | NA |
| WBGene00198975 | -0.022930767 | 0.78121253 | NA | NA |
| WBGene00198976 | NA | NA | NA | NA |
| WBGene00198977 | NA | NA | NA | NA |
| WBGene00198978 | NA | NA | NA | NA |
| WBGene00198979 | NA | NA | NA | NA |
| WBGene00198980 | NA | NA | NA | NA |
| WBGene00198981 | NA | NA | NA | NA |
| WBGene00198982 | NA | NA | NA | NA |
| WBGene00198983 | NA | NA | NA | NA |
| WBGene00198984 | NA | NA | NA | NA |
| WBGene00198985 | NA | NA | NA | NA |
| WBGene00198986 | NA | NA | NA | NA |
| WBGene00198987 | NA | NA | NA | NA |
| WBGene00198988 | NA | NA | NA | NA |
| WBGene00198989 | NA | NA | NA | NA |

| | | | | |
|---|---|---|---|---|
| WBGene00198990 | NA | NA | NA | NA |
| WBGene00198991 | NA | NA | NA | NA |
| WBGene00198992 | NA | NA | NA | NA |
| WBGene00198993 | NA | NA | NA | NA |
| WBGene00198994 | NA | NA | NA | NA |
| WBGene00198995 | NA | NA | NA | NA |
| WBGene00198996 | NA | NA | NA | NA |
| WBGene00198997 | NA | NA | NA | NA |
| WBGene00198998 | NA | NA | NA | NA |
| WBGene00198999 | NA | NA | NA | NA |
| WBGene00199000 | NA | NA | NA | NA |
| WBGene00199001 | NA | NA | NA | NA |
| WBGene00199002 | NA | NA | NA | NA |
| WBGene00199003 | -0.023109762 | 0.78121253 | NA | NA |
| WBGene00199004 | NA | NA | NA | NA |
| WBGene00199005 | NA | NA | NA | NA |
| WBGene00199006 | NA | NA | NA | NA |
| WBGene00199007 | NA | NA | NA | NA |
| WBGene00199008 | NA | NA | NA | NA |
| WBGene00199009 | NA | NA | NA | NA |
| WBGene00199010 | 0.020925399 | 0.84643798 | NA | NA |
| WBGene00199011 | NA | NA | NA | NA |
| WBGene00199012 | NA | NA | NA | NA |
| WBGene00199013 | -0.023109762 | 0.78121253 | NA | NA |
| WBGene00199014 | NA | NA | NA | NA |
| WBGene00199015 | NA | NA | NA | NA |
| WBGene00199016 | NA | NA | NA | NA |
| WBGene00199017 | NA | NA | NA | NA |
| WBGene00199018 | -0.008998908 | 0.92213607 | NA | NA |
| WBGene00199019 | NA | NA | NA | NA |
| WBGene00199020 | NA | NA | NA | NA |
| WBGene00199021 | NA | NA | NA | NA |
| WBGene00199022 | NA | NA | NA | NA |
| WBGene00199023 | NA | NA | NA | NA |
| WBGene00199024 | NA | NA | NA | NA |
| WBGene00199025 | NA | NA | NA | NA |
| WBGene00199026 | NA | NA | NA | NA |
| WBGene00199027 | NA | NA | NA | NA |
| WBGene00199028 | NA | NA | NA | NA |
| WBGene00199029 | NA | NA | NA | NA |
| WBGene00199030 | NA | NA | NA | NA |
| WBGene00199031 | NA | NA | NA | NA |
| WBGene00199032 | NA | NA | NA | NA |

| | | | | |
|---|---|---|---|---|
| WBGene00199033 | NA | NA | NA | NA |
| WBGene00199034 | NA | NA | NA | NA |
| WBGene00199035 | NA | NA | NA | NA |
| WBGene00199036 | NA | NA | NA | NA |
| WBGene00199037 | NA | NA | NA | NA |
| WBGene00199038 | NA | NA | NA | NA |
| WBGene00199039 | NA | NA | NA | NA |
| WBGene00199040 | NA | NA | NA | NA |
| WBGene00199041 | NA | NA | NA | NA |
| WBGene00199042 | 0.020232312 | 0.84643798 | NA | NA |
| WBGene00199043 | NA | NA | NA | NA |
| WBGene00199044 | NA | NA | NA | NA |
| WBGene00199045 | NA | NA | NA | NA |
| WBGene00199046 | NA | NA | NA | NA |
| WBGene00199047 | NA | NA | NA | NA |
| WBGene00199048 | NA | NA | NA | NA |
| WBGene00199049 | NA | NA | NA | NA |
| WBGene00199050 | NA | NA | NA | NA |
| WBGene00199051 | NA | NA | NA | NA |
| WBGene00199052 | NA | NA | NA | NA |
| WBGene00199053 | NA | NA | NA | NA |
| WBGene00199054 | NA | NA | NA | NA |
| WBGene00199055 | NA | NA | NA | NA |
| WBGene00199056 | NA | NA | NA | NA |
| WBGene00199057 | NA | NA | NA | NA |
| WBGene00199058 | NA | NA | NA | NA |
| WBGene00199059 | NA | NA | NA | NA |
| WBGene00199060 | NA | NA | NA | NA |
| WBGene00199061 | NA | NA | NA | NA |
| WBGene00199062 | NA | NA | NA | NA |
| WBGene00199063 | NA | NA | NA | NA |
| WBGene00199064 | NA | NA | NA | NA |
| WBGene00199065 | NA | NA | NA | NA |
| WBGene00199066 | NA | NA | NA | NA |
| WBGene00199067 | NA | NA | NA | NA |
| WBGene00199068 | NA | NA | NA | NA |
| WBGene00199069 | NA | NA | NA | NA |
| WBGene00199070 | NA | NA | NA | NA |
| WBGene00199071 | NA | NA | NA | NA |
| WBGene00199072 | NA | NA | NA | NA |
| WBGene00199073 | NA | NA | NA | NA |
| WBGene00199074 | NA | NA | NA | NA |
| WBGene00199075 | NA | NA | NA | NA |

| | | | | |
|---|---|---|---|---|
| WBGene00199076 | NA | NA | NA | NA |
| WBGene00199077 | NA | NA | NA | NA |
| WBGene00199078 | 0.020232312 | 0.84643798 | NA | NA |
| WBGene00199079 | NA | NA | NA | NA |
| WBGene00199080 | NA | NA | NA | NA |
| WBGene00199081 | NA | NA | NA | NA |
| WBGene00199082 | -0.025848508 | 0.67901638 | NA | NA |
| WBGene00199083 | NA | NA | NA | NA |
| WBGene00199084 | NA | NA | NA | NA |
| WBGene00199085 | NA | NA | NA | NA |
| WBGene00199086 | NA | NA | NA | NA |
| WBGene00199087 | NA | NA | NA | NA |
| WBGene00199088 | NA | NA | NA | NA |
| WBGene00199089 | NA | NA | NA | NA |
| WBGene00199090 | NA | NA | NA | NA |
| WBGene00199091 | -0.023056499 | 0.78121253 | NA | NA |
| WBGene00199092 | NA | NA | NA | NA |
| WBGene00199093 | NA | NA | NA | NA |
| WBGene00199094 | NA | NA | NA | NA |
| WBGene00199095 | NA | NA | NA | NA |
| WBGene00199096 | NA | NA | NA | NA |
| WBGene00199097 | NA | NA | NA | NA |
| WBGene00199098 | NA | NA | NA | NA |
| WBGene00199099 | NA | NA | NA | NA |
| WBGene00199100 | NA | NA | NA | NA |
| WBGene00199101 | -0.023109762 | 0.78121253 | NA | NA |
| WBGene00199102 | NA | NA | NA | NA |
| WBGene00199103 | NA | NA | NA | NA |
| WBGene00199104 | 0.020232312 | 0.84643798 | NA | NA |
| WBGene00199105 | NA | NA | NA | NA |
| WBGene00199106 | NA | NA | NA | NA |
| WBGene00199107 | NA | NA | NA | NA |
| WBGene00199108 | NA | NA | NA | NA |
| WBGene00199109 | NA | NA | NA | NA |
| WBGene00199110 | NA | NA | NA | NA |
| WBGene00199111 | NA | NA | NA | NA |
| WBGene00199112 | NA | NA | NA | NA |
| WBGene00199113 | NA | NA | NA | NA |
| WBGene00199114 | NA | NA | NA | NA |
| WBGene00199115 | NA | NA | NA | NA |
| WBGene00199116 | -0.023056499 | 0.78121253 | NA | NA |
| WBGene00199117 | NA | NA | NA | NA |
| WBGene00199118 | NA | NA | NA | NA |

| | | | | |
|---|---|---|---|---|
| WBGene00199119 | NA | NA | NA | NA |
| WBGene00199120 | NA | NA | NA | NA |
| WBGene00199121 | NA | NA | NA | NA |
| WBGene00199122 | NA | NA | NA | NA |
| WBGene00199123 | NA | NA | NA | NA |
| WBGene00199124 | NA | NA | NA | NA |
| WBGene00199125 | NA | NA | NA | NA |
| WBGene00199126 | NA | NA | NA | NA |
| WBGene00199127 | NA | NA | NA | NA |
| WBGene00199128 | NA | NA | NA | NA |
| WBGene00199129 | NA | NA | NA | NA |
| WBGene00199130 | NA | NA | NA | NA |
| WBGene00199131 | NA | NA | NA | NA |
| WBGene00199132 | NA | NA | NA | NA |
| WBGene00199133 | NA | NA | NA | NA |
| WBGene00199134 | NA | NA | NA | NA |
| WBGene00199135 | NA | NA | NA | NA |
| WBGene00199136 | NA | NA | NA | NA |
| WBGene00199137 | NA | NA | NA | NA |
| WBGene00199138 | NA | NA | NA | NA |
| WBGene00199139 | NA | NA | NA | NA |
| WBGene00199140 | NA | NA | NA | NA |
| WBGene00199141 | NA | NA | NA | NA |
| WBGene00199142 | NA | NA | NA | NA |
| WBGene00199143 | NA | NA | NA | NA |
| WBGene00199144 | NA | NA | NA | NA |
| WBGene00199145 | NA | NA | NA | NA |
| WBGene00199146 | NA | NA | NA | NA |
| WBGene00199147 | NA | NA | NA | NA |
| WBGene00199148 | NA | NA | NA | NA |
| WBGene00199149 | NA | NA | NA | NA |
| WBGene00199150 | NA | NA | NA | NA |
| WBGene00199151 | NA | NA | NA | NA |
| WBGene00199152 | NA | NA | NA | NA |
| WBGene00199153 | NA | NA | NA | NA |
| WBGene00199154 | NA | NA | NA | NA |
| WBGene00199155 | NA | NA | NA | NA |
| WBGene00199156 | NA | NA | NA | NA |
| WBGene00199157 | NA | NA | NA | NA |
| WBGene00199158 | NA | NA | NA | NA |
| WBGene00199159 | NA | NA | NA | NA |
| WBGene00199160 | NA | NA | NA | NA |
| WBGene00199161 | NA | NA | NA | NA |

| | | | | |
|---|---|---|---|---|
| WBGene00199162 | NA | NA | NA | NA |
| WBGene00199163 | NA | NA | NA | NA |
| WBGene00199164 | NA | NA | NA | NA |
| WBGene00199165 | NA | NA | NA | NA |
| WBGene00199166 | NA | NA | NA | NA |
| WBGene00199167 | NA | NA | NA | NA |
| WBGene00199168 | NA | NA | NA | NA |
| WBGene00199169 | NA | NA | NA | NA |
| WBGene00199170 | NA | NA | NA | NA |
| WBGene00199171 | NA | NA | NA | NA |
| WBGene00199172 | NA | NA | NA | NA |
| WBGene00199173 | NA | NA | NA | NA |
| WBGene00199174 | NA | NA | NA | NA |
| WBGene00199175 | 0.020925399 | 0.84643798 | NA | NA |
| WBGene00199176 | NA | NA | NA | NA |
| WBGene00199177 | NA | NA | NA | NA |
| WBGene00199178 | NA | NA | NA | NA |
| WBGene00199179 | -0.023056499 | 0.78121253 | NA | NA |
| WBGene00199180 | NA | NA | NA | NA |
| WBGene00199181 | NA | NA | NA | NA |
| WBGene00199182 | NA | NA | NA | NA |
| WBGene00199183 | NA | NA | NA | NA |
| WBGene00199184 | NA | NA | NA | NA |
| WBGene00199185 | NA | NA | NA | NA |
| WBGene00199186 | NA | NA | NA | NA |
| WBGene00199187 | NA | NA | NA | NA |
| WBGene00199188 | NA | NA | NA | NA |
| WBGene00199189 | NA | NA | NA | NA |
| WBGene00199190 | NA | NA | NA | NA |
| WBGene00199191 | NA | NA | NA | NA |
| WBGene00199192 | NA | NA | NA | NA |
| WBGene00199193 | NA | NA | NA | NA |
| WBGene00199194 | NA | NA | NA | NA |
| WBGene00199195 | NA | NA | NA | NA |
| WBGene00199196 | NA | NA | NA | NA |
| WBGene00199197 | NA | NA | NA | NA |
| WBGene00199198 | NA | NA | NA | NA |
| WBGene00199199 | NA | NA | NA | NA |
| WBGene00199200 | NA | NA | NA | NA |
| WBGene00199201 | NA | NA | NA | NA |
| WBGene00199202 | NA | NA | NA | NA |
| WBGene00199203 | NA | NA | NA | NA |
| WBGene00199204 | NA | NA | NA | NA |

| | | | | |
|---|---|---|---|---|
| WBGene00199205 | NA | NA | NA | NA |
| WBGene00199206 | NA | NA | NA | NA |
| WBGene00199207 | NA | NA | NA | NA |
| WBGene00199208 | NA | NA | NA | NA |
| WBGene00199209 | NA | NA | NA | NA |
| WBGene00199210 | NA | NA | NA | NA |
| WBGene00199211 | NA | NA | NA | NA |
| WBGene00199212 | NA | NA | NA | NA |
| WBGene00199213 | NA | NA | NA | NA |
| WBGene00199214 | NA | NA | NA | NA |
| WBGene00199215 | NA | NA | NA | NA |
| WBGene00199216 | NA | NA | NA | NA |
| WBGene00199217 | NA | NA | NA | NA |
| WBGene00199218 | NA | NA | NA | NA |
| WBGene00199219 | NA | NA | NA | NA |
| WBGene00199220 | NA | NA | NA | NA |
| WBGene00199221 | NA | NA | NA | NA |
| WBGene00199222 | NA | NA | NA | NA |
| WBGene00199223 | NA | NA | NA | NA |
| WBGene00199224 | NA | NA | NA | NA |
| WBGene00199225 | NA | NA | NA | NA |
| WBGene00199226 | NA | NA | NA | NA |
| WBGene00199227 | NA | NA | NA | NA |
| WBGene00199228 | NA | NA | NA | NA |
| WBGene00199229 | NA | NA | NA | NA |
| WBGene00199230 | NA | NA | NA | NA |
| WBGene00199231 | -0.022930767 | 0.78121253 | NA | NA |
| WBGene00199232 | NA | NA | NA | NA |
| WBGene00199233 | NA | NA | NA | NA |
| WBGene00199234 | NA | NA | NA | NA |
| WBGene00199235 | 0.020317245 | 0.84643798 | NA | NA |
| WBGene00199236 | NA | NA | NA | NA |
| WBGene00199237 | NA | NA | NA | NA |
| WBGene00199238 | NA | NA | NA | NA |
| WBGene00199239 | NA | NA | NA | NA |
| WBGene00199240 | NA | NA | NA | NA |
| WBGene00199241 | NA | NA | NA | NA |
| WBGene00199242 | NA | NA | NA | NA |
| WBGene00199243 | NA | NA | NA | NA |
| WBGene00199244 | NA | NA | NA | NA |
| WBGene00199245 | NA | NA | NA | NA |
| WBGene00199246 | NA | NA | NA | NA |
| WBGene00199247 | NA | NA | NA | NA |

| | | | | |
|---|---|---|---|---|
| WBGene00199248 | NA | NA | NA | NA |
| WBGene00199249 | NA | NA | NA | NA |
| WBGene00199250 | NA | NA | NA | NA |
| WBGene00199251 | 0.020317245 | 0.84643798 | NA | NA |
| WBGene00199252 | NA | NA | NA | NA |
| WBGene00199253 | NA | NA | NA | NA |
| WBGene00199254 | NA | NA | NA | NA |
| WBGene00199255 | NA | NA | NA | NA |
| WBGene00199256 | NA | NA | NA | NA |
| WBGene00199257 | NA | NA | NA | NA |
| WBGene00199258 | NA | NA | NA | NA |
| WBGene00199259 | NA | NA | NA | NA |
| WBGene00199260 | NA | NA | NA | NA |
| WBGene00199261 | NA | NA | NA | NA |
| WBGene00199262 | NA | NA | NA | NA |
| WBGene00199263 | NA | NA | NA | NA |
| WBGene00199264 | NA | NA | NA | NA |
| WBGene00199265 | NA | NA | NA | NA |
| WBGene00199266 | NA | NA | NA | NA |
| WBGene00199267 | NA | NA | NA | NA |
| WBGene00199268 | NA | NA | NA | NA |
| WBGene00199269 | NA | NA | NA | NA |
| WBGene00199270 | NA | NA | NA | NA |
| WBGene00199271 | NA | NA | NA | NA |
| WBGene00199272 | NA | NA | NA | NA |
| WBGene00199273 | NA | NA | NA | NA |
| WBGene00199274 | NA | NA | NA | NA |
| WBGene00199275 | NA | NA | NA | NA |
| WBGene00199276 | NA | NA | NA | NA |
| WBGene00199277 | NA | NA | NA | NA |
| WBGene00199278 | NA | NA | NA | NA |
| WBGene00199279 | NA | NA | NA | NA |
| WBGene00199280 | NA | NA | NA | NA |
| WBGene00199281 | NA | NA | NA | NA |
| WBGene00199282 | NA | NA | NA | NA |
| WBGene00199283 | NA | NA | NA | NA |
| WBGene00199284 | NA | NA | NA | NA |
| WBGene00199285 | NA | NA | NA | NA |
| WBGene00199286 | NA | NA | NA | NA |
| WBGene00199287 | NA | NA | NA | NA |
| WBGene00199288 | NA | NA | NA | NA |
| WBGene00199289 | NA | NA | NA | NA |
| WBGene00199290 | NA | NA | NA | NA |

| | | | | |
|---|---|---|---|---|
| WBGene00199291 | NA | NA | NA | NA |
| WBGene00199292 | NA | NA | NA | NA |
| WBGene00199293 | NA | NA | NA | NA |
| WBGene00199294 | NA | NA | NA | NA |
| WBGene00199295 | NA | NA | NA | NA |
| WBGene00199296 | 0.033463284 | 0.6365037 | NA | NA |
| WBGene00199297 | NA | NA | NA | NA |
| WBGene00199298 | NA | NA | NA | NA |
| WBGene00199299 | NA | NA | NA | NA |
| WBGene00199300 | 0.020317245 | 0.84643798 | NA | NA |
| WBGene00199301 | -0.005221332 | 0.94448901 | NA | NA |
| WBGene00199302 | NA | NA | NA | NA |
| WBGene00199303 | NA | NA | NA | NA |
| WBGene00199304 | NA | NA | NA | NA |
| WBGene00199305 | NA | NA | NA | NA |
| WBGene00199306 | NA | NA | NA | NA |
| WBGene00199307 | NA | NA | NA | NA |
| WBGene00199308 | NA | NA | NA | NA |
| WBGene00199309 | NA | NA | NA | NA |
| WBGene00199310 | NA | NA | NA | NA |
| WBGene00199311 | NA | NA | NA | NA |
| WBGene00199312 | NA | NA | NA | NA |
| WBGene00199313 | NA | NA | NA | NA |
| WBGene00199314 | NA | NA | NA | NA |
| WBGene00199315 | NA | NA | NA | NA |
| WBGene00199316 | NA | NA | NA | NA |
| WBGene00199317 | NA | NA | NA | NA |
| WBGene00199318 | NA | NA | NA | NA |
| WBGene00199319 | NA | NA | NA | NA |
| WBGene00199320 | NA | NA | NA | NA |
| WBGene00199321 | NA | NA | NA | NA |
| WBGene00199322 | NA | NA | NA | NA |
| WBGene00199323 | NA | NA | NA | NA |
| WBGene00199324 | NA | NA | NA | NA |
| WBGene00199325 | NA | NA | NA | NA |
| WBGene00199326 | NA | NA | NA | NA |
| WBGene00199327 | NA | NA | NA | NA |
| WBGene00199328 | NA | NA | NA | NA |
| WBGene00199329 | 0.020232312 | 0.84643798 | NA | NA |
| WBGene00199330 | NA | NA | NA | NA |
| WBGene00199331 | NA | NA | NA | NA |
| WBGene00199332 | NA | NA | NA | NA |
| WBGene00199333 | NA | NA | NA | NA |

| | | | | |
|---|---|---|---|---|
| WBGene00199334 | NA | NA | NA | NA |
| WBGene00199335 | NA | NA | NA | NA |
| WBGene00199336 | -0.022930767 | 0.78121253 | NA | NA |
| WBGene00199337 | NA | NA | NA | NA |
| WBGene00199338 | NA | NA | NA | NA |
| WBGene00199339 | NA | NA | NA | NA |
| WBGene00199340 | NA | NA | NA | NA |
| WBGene00199341 | NA | NA | NA | NA |
| WBGene00199342 | NA | NA | NA | NA |
| WBGene00199343 | NA | NA | NA | NA |
| WBGene00199344 | NA | NA | NA | NA |
| WBGene00199345 | NA | NA | NA | NA |
| WBGene00199346 | NA | NA | NA | NA |
| WBGene00199347 | NA | NA | NA | NA |
| WBGene00199348 | NA | NA | NA | NA |
| WBGene00199349 | NA | NA | NA | NA |
| WBGene00199350 | NA | NA | NA | NA |
| WBGene00199351 | NA | NA | NA | NA |
| WBGene00199352 | NA | NA | NA | NA |
| WBGene00199353 | NA | NA | NA | NA |
| WBGene00199354 | NA | NA | NA | NA |
| WBGene00199355 | NA | NA | NA | NA |
| WBGene00199356 | NA | NA | NA | NA |
| WBGene00199357 | NA | NA | NA | NA |
| WBGene00199358 | NA | NA | NA | NA |
| WBGene00199359 | NA | NA | NA | NA |
| WBGene00199360 | NA | NA | NA | NA |
| WBGene00199361 | NA | NA | NA | NA |
| WBGene00199362 | -0.023056499 | 0.78121253 | NA | NA |
| WBGene00199363 | NA | NA | NA | NA |
| WBGene00199364 | NA | NA | NA | NA |
| WBGene00199365 | NA | NA | NA | NA |
| WBGene00199366 | NA | NA | NA | NA |
| WBGene00199367 | NA | NA | NA | NA |
| WBGene00199368 | 0.020232312 | 0.84643798 | NA | NA |
| WBGene00199369 | NA | NA | NA | NA |
| WBGene00199370 | NA | NA | NA | NA |
| WBGene00199371 | 0.020925399 | 0.84643798 | NA | NA |
| WBGene00199372 | NA | NA | NA | NA |
| WBGene00199373 | NA | NA | NA | NA |
| WBGene00199374 | NA | NA | NA | NA |
| WBGene00199375 | NA | NA | NA | NA |
| WBGene00199376 | NA | NA | NA | NA |

| | | | | |
|---|---|---|---|---|
| WBGene00199377 | NA | NA | NA | NA |
| WBGene00199378 | NA | NA | NA | NA |
| WBGene00199379 | NA | NA | NA | NA |
| WBGene00199380 | NA | NA | NA | NA |
| WBGene00199381 | NA | NA | NA | NA |
| WBGene00199382 | NA | NA | NA | NA |
| WBGene00199383 | NA | NA | NA | NA |
| WBGene00199384 | NA | NA | NA | NA |
| WBGene00199385 | NA | NA | NA | NA |
| WBGene00199386 | NA | NA | NA | NA |
| WBGene00199387 | NA | NA | NA | NA |
| WBGene00199388 | NA | NA | NA | NA |
| WBGene00199389 | NA | NA | NA | NA |
| WBGene00199390 | NA | NA | NA | NA |
| WBGene00199391 | NA | NA | NA | NA |
| WBGene00199392 | NA | NA | NA | NA |
| WBGene00199393 | NA | NA | NA | NA |
| WBGene00199394 | NA | NA | NA | NA |
| WBGene00199395 | NA | NA | NA | NA |
| WBGene00199396 | NA | NA | NA | NA |
| WBGene00199397 | NA | NA | NA | NA |
| WBGene00199398 | 0.020232312 | 0.84643798 | NA | NA |
| WBGene00199399 | NA | NA | NA | NA |
| WBGene00199400 | NA | NA | NA | NA |
| WBGene00199401 | NA | NA | NA | NA |
| WBGene00199402 | -0.023109762 | 0.78121253 | NA | NA |
| WBGene00199403 | NA | NA | NA | NA |
| WBGene00199404 | NA | NA | NA | NA |
| WBGene00199405 | NA | NA | NA | NA |
| WBGene00199406 | 0.016985298 | 0.837043 | NA | NA |
| WBGene00199407 | NA | NA | NA | NA |
| WBGene00199408 | NA | NA | NA | NA |
| WBGene00199409 | NA | NA | NA | NA |
| WBGene00199410 | NA | NA | NA | NA |
| WBGene00199411 | NA | NA | NA | NA |
| WBGene00199412 | NA | NA | NA | NA |
| WBGene00199413 | NA | NA | NA | NA |
| WBGene00199414 | NA | NA | NA | NA |
| WBGene00199415 | NA | NA | NA | NA |
| WBGene00199416 | 0.020925399 | 0.84643798 | NA | NA |
| WBGene00199417 | NA | NA | NA | NA |
| WBGene00199418 | NA | NA | NA | NA |
| WBGene00199419 | NA | NA | NA | NA |

| | | | | |
|---|---|---|---|---|
| WBGene00199420 | NA | NA | NA | NA |
| WBGene00199421 | NA | NA | NA | NA |
| WBGene00199422 | NA | NA | NA | NA |
| WBGene00199423 | NA | NA | NA | NA |
| WBGene00199424 | NA | NA | NA | NA |
| WBGene00199425 | NA | NA | NA | NA |
| WBGene00199426 | NA | NA | NA | NA |
| WBGene00199427 | NA | NA | NA | NA |
| WBGene00199428 | NA | NA | NA | NA |
| WBGene00199429 | NA | NA | NA | NA |
| WBGene00199430 | NA | NA | NA | NA |
| WBGene00199431 | NA | NA | NA | NA |
| WBGene00199432 | NA | NA | NA | NA |
| WBGene00199433 | NA | NA | NA | NA |
| WBGene00199434 | NA | NA | NA | NA |
| WBGene00199435 | NA | NA | NA | NA |
| WBGene00199436 | NA | NA | NA | NA |
| WBGene00199437 | 0.011886678 | 0.86820775 | NA | NA |
| WBGene00199438 | -0.022930767 | 0.78121253 | NA | NA |
| WBGene00199439 | NA | NA | NA | NA |
| WBGene00199440 | NA | NA | NA | NA |
| WBGene00199441 | NA | NA | NA | NA |
| WBGene00199442 | NA | NA | NA | NA |
| WBGene00199443 | NA | NA | NA | NA |
| WBGene00199444 | NA | NA | NA | NA |
| WBGene00199445 | NA | NA | NA | NA |
| WBGene00199446 | NA | NA | NA | NA |
| WBGene00199447 | NA | NA | NA | NA |
| WBGene00199448 | 0.020925399 | 0.84643798 | NA | NA |
| WBGene00199449 | NA | NA | NA | NA |
| WBGene00199450 | NA | NA | NA | NA |
| WBGene00199451 | NA | NA | NA | NA |
| WBGene00199452 | NA | NA | NA | NA |
| WBGene00199453 | NA | NA | NA | NA |
| WBGene00199454 | NA | NA | NA | NA |
| WBGene00199455 | NA | NA | NA | NA |
| WBGene00199456 | NA | NA | NA | NA |
| WBGene00199457 | NA | NA | NA | NA |
| WBGene00199458 | NA | NA | NA | NA |
| WBGene00199459 | NA | NA | NA | NA |
| WBGene00199460 | NA | NA | NA | NA |
| WBGene00199461 | NA | NA | NA | NA |
| WBGene00199462 | NA | NA | NA | NA |

| | | | | |
|---|---|---|---|---|
| WBGene00199463 | NA | NA | NA | NA |
| WBGene00199464 | NA | NA | NA | NA |
| WBGene00199465 | NA | NA | NA | NA |
| WBGene00199466 | NA | NA | NA | NA |
| WBGene00199467 | NA | NA | NA | NA |
| WBGene00199468 | NA | NA | NA | NA |
| WBGene00199469 | NA | NA | NA | NA |
| WBGene00199470 | NA | NA | NA | NA |
| WBGene00199471 | NA | NA | NA | NA |
| WBGene00199472 | NA | NA | NA | NA |
| WBGene00199473 | NA | NA | NA | NA |
| WBGene00199474 | NA | NA | NA | NA |
| WBGene00199475 | NA | NA | NA | NA |
| WBGene00199476 | NA | NA | NA | NA |
| WBGene00199477 | NA | NA | NA | NA |
| WBGene00199478 | 0.020925399 | 0.84643798 | NA | NA |
| WBGene00199479 | NA | NA | NA | NA |
| WBGene00199480 | NA | NA | NA | NA |
| WBGene00199481 | NA | NA | NA | NA |
| WBGene00199482 | NA | NA | NA | NA |
| WBGene00199483 | NA | NA | NA | NA |
| WBGene00199484 | NA | NA | NA | NA |
| WBGene00199485 | NA | NA | NA | NA |
| WBGene00199486 | NA | NA | NA | NA |
| WBGene00199487 | NA | NA | NA | NA |
| WBGene00199488 | NA | NA | NA | NA |
| WBGene00199489 | NA | NA | NA | NA |
| WBGene00199490 | NA | NA | NA | NA |
| WBGene00199491 | NA | NA | NA | NA |
| WBGene00199492 | NA | NA | NA | NA |
| WBGene00199493 | NA | NA | NA | NA |
| WBGene00199494 | NA | NA | NA | NA |
| WBGene00199495 | NA | NA | NA | NA |
| WBGene00199496 | NA | NA | NA | NA |
| WBGene00199497 | NA | NA | NA | NA |
| WBGene00199498 | NA | NA | NA | NA |
| WBGene00199499 | NA | NA | NA | NA |
| WBGene00199500 | NA | NA | NA | NA |
| WBGene00199501 | NA | NA | NA | NA |
| WBGene00199502 | NA | NA | NA | NA |
| WBGene00199503 | NA | NA | NA | NA |
| WBGene00199504 | 0.020925399 | 0.84643798 | NA | NA |
| WBGene00199505 | NA | NA | NA | NA |

| | | | | |
|---|---|---|---|---|
| WBGene00199506 | NA | NA | NA | NA |
| WBGene00199507 | -0.023056499 | 0.78121253 | NA | NA |
| WBGene00199508 | NA | NA | NA | NA |
| WBGene00199509 | NA | NA | NA | NA |
| WBGene00199510 | NA | NA | NA | NA |
| WBGene00199511 | NA | NA | NA | NA |
| WBGene00199512 | NA | NA | NA | NA |
| WBGene00199513 | NA | NA | NA | NA |
| WBGene00199514 | NA | NA | NA | NA |
| WBGene00199515 | NA | NA | NA | NA |
| WBGene00199516 | NA | NA | NA | NA |
| WBGene00199517 | NA | NA | NA | NA |
| WBGene00199518 | NA | NA | NA | NA |
| WBGene00199519 | NA | NA | NA | NA |
| WBGene00199520 | NA | NA | NA | NA |
| WBGene00199521 | NA | NA | NA | NA |
| WBGene00199522 | NA | NA | NA | NA |
| WBGene00199523 | NA | NA | NA | NA |
| WBGene00199524 | NA | NA | NA | NA |
| WBGene00199525 | 0.020925399 | 0.84643798 | NA | NA |
| WBGene00199526 | NA | NA | NA | NA |
| WBGene00199527 | NA | NA | NA | NA |
| WBGene00199528 | NA | NA | NA | NA |
| WBGene00199529 | NA | NA | NA | NA |
| WBGene00199530 | NA | NA | NA | NA |
| WBGene00199531 | NA | NA | NA | NA |
| WBGene00199532 | NA | NA | NA | NA |
| WBGene00199533 | NA | NA | NA | NA |
| WBGene00199534 | NA | NA | NA | NA |
| WBGene00199535 | NA | NA | NA | NA |
| WBGene00199536 | NA | NA | NA | NA |
| WBGene00199537 | NA | NA | NA | NA |
| WBGene00199538 | NA | NA | NA | NA |
| WBGene00199539 | NA | NA | NA | NA |
| WBGene00199540 | NA | NA | NA | NA |
| WBGene00199541 | NA | NA | NA | NA |
| WBGene00199542 | NA | NA | NA | NA |
| WBGene00199543 | NA | NA | NA | NA |
| WBGene00199544 | NA | NA | NA | NA |
| WBGene00199545 | NA | NA | NA | NA |
| WBGene00199546 | NA | NA | NA | NA |
| WBGene00199547 | NA | NA | NA | NA |
| WBGene00199548 | NA | NA | NA | NA |

| | | | | |
|---|---|---|---|---|
| WBGene00199549 | NA | NA | NA | NA |
| WBGene00199550 | NA | NA | NA | NA |
| WBGene00199551 | NA | NA | NA | NA |
| WBGene00199552 | NA | NA | NA | NA |
| WBGene00199553 | NA | NA | NA | NA |
| WBGene00199554 | NA | NA | NA | NA |
| WBGene00199555 | NA | NA | NA | NA |
| WBGene00199556 | NA | NA | NA | NA |
| WBGene00199557 | NA | NA | NA | NA |
| WBGene00199558 | NA | NA | NA | NA |
| WBGene00199559 | NA | NA | NA | NA |
| WBGene00199560 | NA | NA | NA | NA |
| WBGene00199561 | NA | NA | NA | NA |
| WBGene00199562 | NA | NA | NA | NA |
| WBGene00199563 | NA | NA | NA | NA |
| WBGene00199564 | NA | NA | NA | NA |
| WBGene00199565 | NA | NA | NA | NA |
| WBGene00199566 | NA | NA | NA | NA |
| WBGene00199567 | NA | NA | NA | NA |
| WBGene00199568 | NA | NA | NA | NA |
| WBGene00199569 | NA | NA | NA | NA |
| WBGene00199570 | NA | NA | NA | NA |
| WBGene00199571 | NA | NA | NA | NA |
| WBGene00199572 | NA | NA | NA | NA |
| WBGene00199573 | NA | NA | NA | NA |
| WBGene00199574 | NA | NA | NA | NA |
| WBGene00199575 | NA | NA | NA | NA |
| WBGene00199576 | NA | NA | NA | NA |
| WBGene00199577 | NA | NA | NA | NA |
| WBGene00199578 | NA | NA | NA | NA |
| WBGene00199579 | NA | NA | NA | NA |
| WBGene00199580 | NA | NA | NA | NA |
| WBGene00199581 | NA | NA | NA | NA |
| WBGene00199582 | NA | NA | NA | NA |
| WBGene00199583 | NA | NA | NA | NA |
| WBGene00199584 | NA | NA | NA | NA |
| WBGene00199585 | NA | NA | NA | NA |
| WBGene00199586 | NA | NA | NA | NA |
| WBGene00199587 | NA | NA | NA | NA |
| WBGene00199588 | NA | NA | NA | NA |
| WBGene00199589 | NA | NA | NA | NA |
| WBGene00199590 | NA | NA | NA | NA |
| WBGene00199591 | NA | NA | NA | NA |

| | | | | |
|---|---|---|---|---|
| WBGene00199592 | NA | NA | NA | NA |
| WBGene00199593 | NA | NA | NA | NA |
| WBGene00199594 | NA | NA | NA | NA |
| WBGene00199595 | NA | NA | NA | NA |
| WBGene00199596 | NA | NA | NA | NA |
| WBGene00199597 | NA | NA | NA | NA |
| WBGene00199598 | NA | NA | NA | NA |
| WBGene00199599 | NA | NA | NA | NA |
| WBGene00199600 | NA | NA | NA | NA |
| WBGene00199601 | NA | NA | NA | NA |
| WBGene00199602 | NA | NA | NA | NA |
| WBGene00199603 | NA | NA | NA | NA |
| WBGene00199604 | NA | NA | NA | NA |
| WBGene00199605 | NA | NA | NA | NA |
| WBGene00199606 | -0.023109762 | 0.78121253 | NA | NA |
| WBGene00199607 | NA | NA | NA | NA |
| WBGene00199608 | NA | NA | NA | NA |
| WBGene00199609 | NA | NA | NA | NA |
| WBGene00199610 | NA | NA | NA | NA |
| WBGene00199611 | NA | NA | NA | NA |
| WBGene00199612 | NA | NA | NA | NA |
| WBGene00199613 | NA | NA | NA | NA |
| WBGene00199614 | NA | NA | NA | NA |
| WBGene00199615 | NA | NA | NA | NA |
| WBGene00199616 | NA | NA | NA | NA |
| WBGene00199617 | NA | NA | NA | NA |
| WBGene00199618 | NA | NA | NA | NA |
| WBGene00199619 | NA | NA | NA | NA |
| WBGene00199620 | NA | NA | NA | NA |
| WBGene00199621 | NA | NA | NA | NA |
| WBGene00199622 | NA | NA | NA | NA |
| WBGene00199623 | NA | NA | NA | NA |
| WBGene00199624 | NA | NA | NA | NA |
| WBGene00199625 | NA | NA | NA | NA |
| WBGene00199626 | NA | NA | NA | NA |
| WBGene00199627 | NA | NA | NA | NA |
| WBGene00199628 | NA | NA | NA | NA |
| WBGene00199629 | NA | NA | NA | NA |
| WBGene00199630 | NA | NA | NA | NA |
| WBGene00199631 | NA | NA | NA | NA |
| WBGene00199632 | NA | NA | NA | NA |
| WBGene00199633 | NA | NA | NA | NA |
| WBGene00199634 | NA | NA | NA | NA |

| | | | | |
|---|---|---|---|---|
| WBGene00199635 | NA | NA | NA | NA |
| WBGene00199636 | -0.001488154 | 0.95639912 | NA | NA |
| WBGene00199637 | NA | NA | NA | NA |
| WBGene00199638 | NA | NA | NA | NA |
| WBGene00199639 | NA | NA | NA | NA |
| WBGene00199640 | NA | NA | NA | NA |
| WBGene00199641 | NA | NA | NA | NA |
| WBGene00199642 | NA | NA | NA | NA |
| WBGene00199643 | NA | NA | NA | NA |
| WBGene00199644 | NA | NA | NA | NA |
| WBGene00199645 | NA | NA | NA | NA |
| WBGene00199646 | NA | NA | NA | NA |
| WBGene00199647 | NA | NA | NA | NA |
| WBGene00199648 | NA | NA | NA | NA |
| WBGene00199649 | NA | NA | NA | NA |
| WBGene00199650 | NA | NA | NA | NA |
| WBGene00199651 | NA | NA | NA | NA |
| WBGene00199652 | NA | NA | NA | NA |
| WBGene00199653 | NA | NA | NA | NA |
| WBGene00199654 | NA | NA | NA | NA |
| WBGene00199655 | NA | NA | NA | NA |
| WBGene00199656 | NA | NA | NA | NA |
| WBGene00199657 | NA | NA | NA | NA |
| WBGene00199658 | 0.020317245 | 0.84643798 | NA | NA |
| WBGene00199659 | NA | NA | NA | NA |
| WBGene00199660 | NA | NA | NA | NA |
| WBGene00199661 | -0.003108065 | 0.95728506 | NA | NA |
| WBGene00199662 | NA | NA | NA | NA |
| WBGene00199663 | NA | NA | NA | NA |
| WBGene00199664 | NA | NA | NA | NA |
| WBGene00199665 | NA | NA | NA | NA |
| WBGene00199666 | NA | NA | NA | NA |
| WBGene00199667 | NA | NA | NA | NA |
| WBGene00199668 | NA | NA | NA | NA |
| WBGene00199669 | -0.022930767 | 0.78121253 | NA | NA |
| WBGene00199670 | -0.00346345 | 0.95708614 | NA | NA |
| WBGene00199671 | NA | NA | NA | NA |
| WBGene00199672 | NA | NA | NA | NA |
| WBGene00199673 | NA | NA | NA | NA |
| WBGene00199674 | NA | NA | NA | NA |
| WBGene00199675 | NA | NA | NA | NA |
| WBGene00199676 | NA | NA | NA | NA |
| WBGene00199677 | NA | NA | NA | NA |

| | | | | |
|---|---|---|---|---|
| WBGene00199678 | NA | NA | NA | NA |
| WBGene00199679 | NA | NA | NA | NA |
| WBGene00199680 | NA | NA | NA | NA |
| WBGene00199681 | NA | NA | NA | NA |
| WBGene00199682 | NA | NA | NA | NA |
| WBGene00199683 | NA | NA | NA | NA |
| WBGene00199684 | NA | NA | NA | NA |
| WBGene00199685 | NA | NA | NA | NA |
| WBGene00199686 | NA | NA | NA | NA |
| WBGene00199687 | -0.002914971 | 0.95718083 | NA | NA |
| WBGene00199688 | NA | NA | NA | NA |
| WBGene00199689 | NA | NA | NA | NA |
| WBGene00199690 | NA | NA | NA | NA |
| WBGene00199691 | NA | NA | NA | NA |
| WBGene00199692 | NA | NA | NA | NA |
| WBGene00199693 | NA | NA | NA | NA |
| WBGene00199694 | NA | NA | NA | NA |
| WBGene00199695 | NA | NA | NA | NA |
| WBGene00199696 | NA | NA | NA | NA |
| WBGene00199697 | NA | NA | NA | NA |
| WBGene00199698 | NA | NA | NA | NA |
| WBGene00199699 | NA | NA | NA | NA |
| WBGene00199700 | NA | NA | NA | NA |
| WBGene00199701 | NA | NA | NA | NA |
| WBGene00199702 | NA | NA | NA | NA |
| WBGene00199703 | NA | NA | NA | NA |
| WBGene00199704 | NA | NA | NA | NA |
| WBGene00199705 | NA | NA | NA | NA |
| WBGene00199706 | NA | NA | NA | NA |
| WBGene00199707 | NA | NA | NA | NA |
| WBGene00199708 | NA | NA | NA | NA |
| WBGene00199709 | NA | NA | NA | NA |
| WBGene00199710 | NA | NA | NA | NA |
| WBGene00199711 | NA | NA | NA | NA |
| WBGene00199712 | NA | NA | NA | NA |
| WBGene00199713 | NA | NA | NA | NA |
| WBGene00199714 | NA | NA | NA | NA |
| WBGene00199715 | NA | NA | NA | NA |
| WBGene00199716 | 0.020317245 | 0.84643798 | NA | NA |
| WBGene00199717 | NA | NA | NA | NA |
| WBGene00199718 | NA | NA | NA | NA |
| WBGene00199719 | NA | NA | NA | NA |
| WBGene00199720 | 0.017604133 | 0.79437959 | NA | NA |

| | | | | |
|---|---|---|---|---|
| WBGene00199721 | NA | NA | NA | NA |
| WBGene00199722 | NA | NA | NA | NA |
| WBGene00199723 | NA | NA | NA | NA |
| WBGene00199724 | NA | NA | NA | NA |
| WBGene00199725 | NA | NA | NA | NA |
| WBGene00199726 | NA | NA | NA | NA |
| WBGene00199727 | NA | NA | NA | NA |
| WBGene00199728 | NA | NA | NA | NA |
| WBGene00199729 | NA | NA | NA | NA |
| WBGene00199730 | NA | NA | NA | NA |
| WBGene00199731 | NA | NA | NA | NA |
| WBGene00199732 | NA | NA | NA | NA |
| WBGene00199733 | NA | NA | NA | NA |
| WBGene00199734 | NA | NA | NA | NA |
| WBGene00199735 | NA | NA | NA | NA |
| WBGene00199736 | NA | NA | NA | NA |
| WBGene00199737 | NA | NA | NA | NA |
| WBGene00199738 | NA | NA | NA | NA |
| WBGene00199739 | NA | NA | NA | NA |
| WBGene00199740 | -0.002914971 | 0.95718083 | NA | NA |
| WBGene00199741 | NA | NA | NA | NA |
| WBGene00199742 | NA | NA | NA | NA |
| WBGene00199743 | NA | NA | NA | NA |
| WBGene00199744 | NA | NA | NA | NA |
| WBGene00199745 | NA | NA | NA | NA |
| WBGene00199746 | NA | NA | NA | NA |
| WBGene00199747 | NA | NA | NA | NA |
| WBGene00199748 | NA | NA | NA | NA |
| WBGene00199749 | NA | NA | NA | NA |
| WBGene00199750 | NA | NA | NA | NA |
| WBGene00199751 | NA | NA | NA | NA |
| WBGene00199752 | NA | NA | NA | NA |
| WBGene00199753 | NA | NA | NA | NA |
| WBGene00199754 | NA | NA | NA | NA |
| WBGene00199755 | NA | NA | NA | NA |
| WBGene00199756 | NA | NA | NA | NA |
| WBGene00199757 | NA | NA | NA | NA |
| WBGene00199758 | NA | NA | NA | NA |
| WBGene00199759 | NA | NA | NA | NA |
| WBGene00199760 | NA | NA | NA | NA |
| WBGene00199761 | 0.0398788 | 0.58971221 | NA | NA |
| WBGene00199762 | NA | NA | NA | NA |
| WBGene00199763 | NA | NA | NA | NA |

| | | | | |
|---|---|---|---|---|
| WBGene00199764 | NA | NA | NA | NA |
| WBGene00199765 | NA | NA | NA | NA |
| WBGene00199766 | NA | NA | NA | NA |
| WBGene00199767 | NA | NA | NA | NA |
| WBGene00199768 | NA | NA | NA | NA |
| WBGene00199769 | NA | NA | NA | NA |
| WBGene00199770 | NA | NA | NA | NA |
| WBGene00199771 | NA | NA | NA | NA |
| WBGene00199772 | NA | NA | NA | NA |
| WBGene00199773 | NA | NA | NA | NA |
| WBGene00199774 | NA | NA | NA | NA |
| WBGene00199775 | NA | NA | NA | NA |
| WBGene00199776 | NA | NA | NA | NA |
| WBGene00199777 | NA | NA | NA | NA |
| WBGene00199778 | NA | NA | NA | NA |
| WBGene00199779 | NA | NA | NA | NA |
| WBGene00199780 | NA | NA | NA | NA |
| WBGene00199781 | NA | NA | NA | NA |
| WBGene00199782 | NA | NA | NA | NA |
| WBGene00199783 | NA | NA | NA | NA |
| WBGene00199784 | NA | NA | NA | NA |
| WBGene00199785 | NA | NA | NA | NA |
| WBGene00199786 | NA | NA | NA | NA |
| WBGene00199787 | NA | NA | NA | NA |
| WBGene00199788 | NA | NA | NA | NA |
| WBGene00199789 | NA | NA | NA | NA |
| WBGene00199790 | NA | NA | NA | NA |
| WBGene00199791 | NA | NA | NA | NA |
| WBGene00199792 | NA | NA | NA | NA |
| WBGene00199793 | NA | NA | NA | NA |
| WBGene00199794 | NA | NA | NA | NA |
| WBGene00199795 | NA | NA | NA | NA |
| WBGene00199796 | NA | NA | NA | NA |
| WBGene00199797 | NA | NA | NA | NA |
| WBGene00199798 | NA | NA | NA | NA |
| WBGene00199799 | NA | NA | NA | NA |
| WBGene00199800 | NA | NA | NA | NA |
| WBGene00199801 | NA | NA | NA | NA |
| WBGene00199802 | NA | NA | NA | NA |
| WBGene00199803 | NA | NA | NA | NA |
| WBGene00199804 | NA | NA | NA | NA |
| WBGene00199805 | NA | NA | NA | NA |
| WBGene00199806 | NA | NA | NA | NA |

| | | | | | |
|---|---|---|---|---|---|
| WBGene00199807 | NA | | NA | NA | NA |
| WBGene00199808 | NA | | NA | NA | NA |
| WBGene00199809 | NA | | NA | NA | NA |
| WBGene00199810 | NA | | NA | NA | NA |
| WBGene00199811 | NA | | NA | NA | NA |
| WBGene00199812 | NA | | NA | NA | NA |
| WBGene00199813 | NA | | NA | NA | NA |
| WBGene00199814 | NA | | NA | NA | NA |
| WBGene00199815 | NA | | NA | NA | NA |
| WBGene00199816 | NA | | NA | NA | NA |
| WBGene00199817 | NA | | NA | NA | NA |
| WBGene00199818 | NA | | NA | NA | NA |
| WBGene00199819 | NA | | NA | NA | NA |
| WBGene00199820 | NA | | NA | NA | NA |
| WBGene00199821 | NA | | NA | NA | NA |
| WBGene00199822 | NA | | NA | NA | NA |
| WBGene00199823 | NA | | NA | NA | NA |
| WBGene00199824 | NA | | NA | NA | NA |
| WBGene00199825 | NA | | NA | NA | NA |
| WBGene00199826 | NA | | NA | NA | NA |
| WBGene00199827 | NA | | NA | NA | NA |
| WBGene00199828 | NA | | NA | NA | NA |
| WBGene00199829 | NA | | NA | NA | NA |
| WBGene00199830 | NA | | NA | NA | NA |
| WBGene00199831 | | -0.023109762 | 0.78121253 | NA | NA |
| WBGene00199832 | NA | | NA | NA | NA |
| WBGene00199833 | | 0.033034883 | 0.64219503 | NA | NA |
| WBGene00199834 | NA | | NA | NA | NA |
| WBGene00199835 | NA | | NA | NA | NA |
| WBGene00199836 | NA | | NA | NA | NA |
| WBGene00199837 | NA | | NA | NA | NA |
| WBGene00199838 | NA | | NA | NA | NA |
| WBGene00199839 | NA | | NA | NA | NA |
| WBGene00199840 | NA | | NA | NA | NA |
| WBGene00199841 | NA | | NA | NA | NA |
| WBGene00199842 | NA | | NA | NA | NA |
| WBGene00199843 | NA | | NA | NA | NA |
| WBGene00199844 | NA | | NA | NA | NA |
| WBGene00199845 | NA | | NA | NA | NA |
| WBGene00199846 | NA | | NA | NA | NA |
| WBGene00199847 | NA | | NA | NA | NA |
| WBGene00199848 | NA | | NA | NA | NA |
| WBGene00199849 | NA | | NA | NA | NA |

| | | | | |
|---|---|---|---|---|
| WBGene00199850 | NA | NA | NA | NA |
| WBGene00199851 | NA | NA | NA | NA |
| WBGene00199852 | NA | NA | NA | NA |
| WBGene00199853 | 0.036687355 | 0.59489568 | NA | NA |
| WBGene00199854 | NA | NA | NA | NA |
| WBGene00199855 | NA | NA | NA | NA |
| WBGene00199856 | NA | NA | NA | NA |
| WBGene00199857 | NA | NA | NA | NA |
| WBGene00199858 | NA | NA | NA | NA |
| WBGene00199859 | NA | NA | NA | NA |
| WBGene00199860 | NA | NA | NA | NA |
| WBGene00199861 | NA | NA | NA | NA |
| WBGene00199862 | NA | NA | NA | NA |
| WBGene00199863 | NA | NA | NA | NA |
| WBGene00199864 | NA | NA | NA | NA |
| WBGene00199865 | NA | NA | NA | NA |
| WBGene00199866 | 0.020925399 | 0.84643798 | NA | NA |
| WBGene00199867 | NA | NA | NA | NA |
| WBGene00199868 | NA | NA | NA | NA |
| WBGene00199869 | NA | NA | NA | NA |
| WBGene00199870 | NA | NA | NA | NA |
| WBGene00199871 | NA | NA | NA | NA |
| WBGene00199872 | NA | NA | NA | NA |
| WBGene00199873 | NA | NA | NA | NA |
| WBGene00199874 | NA | NA | NA | NA |
| WBGene00199875 | NA | NA | NA | NA |
| WBGene00199876 | NA | NA | NA | NA |
| WBGene00199877 | NA | NA | NA | NA |
| WBGene00199878 | NA | NA | NA | NA |
| WBGene00199879 | NA | NA | NA | NA |
| WBGene00199880 | NA | NA | NA | NA |
| WBGene00199881 | NA | NA | NA | NA |
| WBGene00199882 | -0.023109762 | 0.78121253 | NA | NA |
| WBGene00199883 | NA | NA | NA | NA |
| WBGene00199884 | NA | NA | NA | NA |
| WBGene00199885 | NA | NA | NA | NA |
| WBGene00199886 | NA | NA | NA | NA |
| WBGene00199887 | NA | NA | NA | NA |
| WBGene00199888 | NA | NA | NA | NA |
| WBGene00199889 | NA | NA | NA | NA |
| WBGene00199890 | NA | NA | NA | NA |
| WBGene00199891 | NA | NA | NA | NA |
| WBGene00199892 | NA | NA | NA | NA |

| | | | | |
|---|---|---|---|---|
| WBGene00199893 | NA | NA | NA | NA |
| WBGene00199894 | NA | NA | NA | NA |
| WBGene00199895 | NA | NA | NA | NA |
| WBGene00199896 | NA | NA | NA | NA |
| WBGene00199897 | NA | NA | NA | NA |
| WBGene00199898 | NA | NA | NA | NA |
| WBGene00199899 | -0.023056499 | 0.78121253 | NA | NA |
| WBGene00199900 | NA | NA | NA | NA |
| WBGene00199901 | NA | NA | NA | NA |
| WBGene00199902 | NA | NA | NA | NA |
| WBGene00199903 | NA | NA | NA | NA |
| WBGene00199904 | NA | NA | NA | NA |
| WBGene00199905 | NA | NA | NA | NA |
| WBGene00199906 | NA | NA | NA | NA |
| WBGene00199907 | NA | NA | NA | NA |
| WBGene00199908 | NA | NA | NA | NA |
| WBGene00199909 | NA | NA | NA | NA |
| WBGene00199910 | NA | NA | NA | NA |
| WBGene00199911 | NA | NA | NA | NA |
| WBGene00199912 | NA | NA | NA | NA |
| WBGene00199913 | NA | NA | NA | NA |
| WBGene00199914 | NA | NA | NA | NA |
| WBGene00199915 | NA | NA | NA | NA |
| WBGene00199916 | NA | NA | NA | NA |
| WBGene00199917 | NA | NA | NA | NA |
| WBGene00199918 | NA | NA | NA | NA |
| WBGene00199919 | NA | NA | NA | NA |
| WBGene00199920 | NA | NA | NA | NA |
| WBGene00199921 | NA | NA | NA | NA |
| WBGene00199922 | NA | NA | NA | NA |
| WBGene00199923 | NA | NA | NA | NA |
| WBGene00199924 | NA | NA | NA | NA |
| WBGene00199925 | NA | NA | NA | NA |
| WBGene00199926 | NA | NA | NA | NA |
| WBGene00199927 | NA | NA | NA | NA |
| WBGene00199928 | NA | NA | NA | NA |
| WBGene00199929 | NA | NA | NA | NA |
| WBGene00199930 | NA | NA | NA | NA |
| WBGene00199931 | NA | NA | NA | NA |
| WBGene00199932 | NA | NA | NA | NA |
| WBGene00199933 | NA | NA | NA | NA |
| WBGene00199934 | NA | NA | NA | NA |
| WBGene00199935 | NA | NA | NA | NA |

| | | | | |
|---|---|---|---|---|
| WBGene00199936 | NA | NA | NA | NA |
| WBGene00199937 | -0.003421551 | 0.95689871 | NA | NA |
| WBGene00199938 | NA | NA | NA | NA |
| WBGene00199939 | NA | NA | NA | NA |
| WBGene00199940 | NA | NA | NA | NA |
| WBGene00199941 | NA | NA | NA | NA |
| WBGene00199942 | NA | NA | NA | NA |
| WBGene00199943 | NA | NA | NA | NA |
| WBGene00199944 | NA | NA | NA | NA |
| WBGene00199945 | NA | NA | NA | NA |
| WBGene00199946 | NA | NA | NA | NA |
| WBGene00199947 | NA | NA | NA | NA |
| WBGene00199948 | NA | NA | NA | NA |
| WBGene00199949 | NA | NA | NA | NA |
| WBGene00199950 | NA | NA | NA | NA |
| WBGene00199951 | NA | NA | NA | NA |
| WBGene00199952 | NA | NA | NA | NA |
| WBGene00199953 | NA | NA | NA | NA |
| WBGene00199954 | NA | NA | NA | NA |
| WBGene00199955 | NA | NA | NA | NA |
| WBGene00199956 | NA | NA | NA | NA |
| WBGene00199957 | NA | NA | NA | NA |
| WBGene00199958 | NA | NA | NA | NA |
| WBGene00199959 | NA | NA | NA | NA |
| WBGene00199960 | NA | NA | NA | NA |
| WBGene00199961 | NA | NA | NA | NA |
| WBGene00199962 | NA | NA | NA | NA |
| WBGene00199963 | NA | NA | NA | NA |
| WBGene00199964 | NA | NA | NA | NA |
| WBGene00199965 | NA | NA | NA | NA |
| WBGene00199966 | NA | NA | NA | NA |
| WBGene00199967 | NA | NA | NA | NA |
| WBGene00199968 | NA | NA | NA | NA |
| WBGene00199969 | NA | NA | NA | NA |
| WBGene00199970 | NA | NA | NA | NA |
| WBGene00199971 | NA | NA | NA | NA |
| WBGene00199972 | NA | NA | NA | NA |
| WBGene00199973 | NA | NA | NA | NA |
| WBGene00199974 | NA | NA | NA | NA |
| WBGene00199975 | NA | NA | NA | NA |
| WBGene00199976 | NA | NA | NA | NA |
| WBGene00199977 | NA | NA | NA | NA |
| WBGene00199978 | 0.040107227 | 0.58693163 | NA | NA |

| | | | | |
|---|---|---|---|---|
| WBGene00199979 | NA | NA | NA | NA |
| WBGene00199980 | NA | NA | NA | NA |
| WBGene00199981 | NA | NA | NA | NA |
| WBGene00199982 | NA | NA | NA | NA |
| WBGene00199983 | NA | NA | NA | NA |
| WBGene00199984 | NA | NA | NA | NA |
| WBGene00199985 | NA | NA | NA | NA |
| WBGene00199986 | NA | NA | NA | NA |
| WBGene00199987 | NA | NA | NA | NA |
| WBGene00199988 | NA | NA | NA | NA |
| WBGene00199989 | NA | NA | NA | NA |
| WBGene00199990 | NA | NA | NA | NA |
| WBGene00199991 | NA | NA | NA | NA |
| WBGene00199992 | NA | NA | NA | NA |
| WBGene00199993 | NA | NA | NA | NA |
| WBGene00199994 | NA | NA | NA | NA |
| WBGene00199995 | NA | NA | NA | NA |
| WBGene00199996 | NA | NA | NA | NA |
| WBGene00199997 | -0.023056499 | 0.78121253 | NA | NA |
| WBGene00199998 | NA | NA | NA | NA |
| WBGene00199999 | NA | NA | NA | NA |
| WBGene00200000 | NA | NA | NA | NA |
| WBGene00200001 | NA | NA | NA | NA |
| WBGene00200002 | NA | NA | NA | NA |
| WBGene00200003 | NA | NA | NA | NA |
| WBGene00200004 | NA | NA | NA | NA |
| WBGene00200005 | 0.020317245 | 0.84643798 | NA | NA |
| WBGene00200006 | NA | NA | NA | NA |
| WBGene00200007 | NA | NA | NA | NA |
| WBGene00200008 | 0.020232312 | 0.84643798 | NA | NA |
| WBGene00200009 | -0.003614866 | 0.95700117 | NA | NA |
| WBGene00200010 | NA | NA | NA | NA |
| WBGene00200011 | NA | NA | NA | NA |
| WBGene00200012 | NA | NA | NA | NA |
| WBGene00200013 | NA | NA | NA | NA |
| WBGene00200014 | NA | NA | NA | NA |
| WBGene00200015 | 0.077894142 | 0.31923424 | NA | NA |
| WBGene00200016 | NA | NA | NA | NA |
| WBGene00200017 | NA | NA | NA | NA |
| WBGene00200018 | 0.020317245 | 0.84643798 | NA | NA |
| WBGene00200019 | NA | NA | NA | NA |
| WBGene00200020 | NA | NA | NA | NA |
| WBGene00200021 | NA | NA | NA | NA |

| | | | | |
|---|---|---|---|---|
| WBGene00200022 | NA | NA | NA | NA |
| WBGene00200023 | NA | NA | NA | NA |
| WBGene00200024 | NA | NA | NA | NA |
| WBGene00200025 | NA | NA | NA | NA |
| WBGene00200026 | NA | NA | NA | NA |
| WBGene00200027 | NA | NA | NA | NA |
| WBGene00200028 | NA | NA | NA | NA |
| WBGene00200029 | NA | NA | NA | NA |
| WBGene00200030 | NA | NA | NA | NA |
| WBGene00200031 | NA | NA | NA | NA |
| WBGene00200032 | NA | NA | NA | NA |
| WBGene00200033 | NA | NA | NA | NA |
| WBGene00200034 | NA | NA | NA | NA |
| WBGene00200035 | NA | NA | NA | NA |
| WBGene00200036 | NA | NA | NA | NA |
| WBGene00200037 | NA | NA | NA | NA |
| WBGene00200038 | NA | NA | NA | NA |
| WBGene00200039 | NA | NA | NA | NA |
| WBGene00200040 | 0.020232312 | 0.84643798 | NA | NA |
| WBGene00200041 | 0.018860755 | 0.77262519 | NA | NA |
| WBGene00200042 | NA | NA | NA | NA |
| WBGene00200043 | NA | NA | NA | NA |
| WBGene00200044 | NA | NA | NA | NA |
| WBGene00200045 | NA | NA | NA | NA |
| WBGene00200046 | NA | NA | NA | NA |
| WBGene00200047 | NA | NA | NA | NA |
| WBGene00200048 | NA | NA | NA | NA |
| WBGene00200049 | NA | NA | NA | NA |
| WBGene00200050 | NA | NA | NA | NA |
| WBGene00200051 | NA | NA | NA | NA |
| WBGene00200052 | NA | NA | NA | NA |
| WBGene00200053 | NA | NA | NA | NA |
| WBGene00200054 | NA | NA | NA | NA |
| WBGene00200055 | NA | NA | NA | NA |
| WBGene00200056 | 0.020232312 | 0.84643798 | NA | NA |
| WBGene00200057 | NA | NA | NA | NA |
| WBGene00200058 | NA | NA | NA | NA |
| WBGene00200059 | NA | NA | NA | NA |
| WBGene00200060 | NA | NA | NA | NA |
| WBGene00200061 | NA | NA | NA | NA |
| WBGene00200062 | NA | NA | NA | NA |
| WBGene00200063 | NA | NA | NA | NA |
| WBGene00200064 | NA | NA | NA | NA |

| | | | | |
|---|---|---|---|---|
| WBGene00200065 | NA | NA | NA | NA |
| WBGene00200066 | NA | NA | NA | NA |
| WBGene00200067 | NA | NA | NA | NA |
| WBGene00200068 | NA | NA | NA | NA |
| WBGene00200069 | NA | NA | NA | NA |
| WBGene00200070 | NA | NA | NA | NA |
| WBGene00200071 | NA | NA | NA | NA |
| WBGene00200072 | NA | NA | NA | NA |
| WBGene00200073 | NA | NA | NA | NA |
| WBGene00200074 | NA | NA | NA | NA |
| WBGene00200075 | NA | NA | NA | NA |
| WBGene00200076 | NA | NA | NA | NA |
| WBGene00200077 | NA | NA | NA | NA |
| WBGene00200078 | NA | NA | NA | NA |
| WBGene00200079 | NA | NA | NA | NA |
| WBGene00200080 | NA | NA | NA | NA |
| WBGene00200081 | NA | NA | NA | NA |
| WBGene00200082 | NA | NA | NA | NA |
| WBGene00200083 | NA | NA | NA | NA |
| WBGene00200084 | NA | NA | NA | NA |
| WBGene00200085 | NA | NA | NA | NA |
| WBGene00200086 | NA | NA | NA | NA |
| WBGene00200087 | NA | NA | NA | NA |
| WBGene00200088 | NA | NA | NA | NA |
| WBGene00200089 | NA | NA | NA | NA |
| WBGene00200090 | NA | NA | NA | NA |
| WBGene00200091 | NA | NA | NA | NA |
| WBGene00200092 | NA | NA | NA | NA |
| WBGene00200093 | NA | NA | NA | NA |
| WBGene00200094 | NA | NA | NA | NA |
| WBGene00200095 | NA | NA | NA | NA |
| WBGene00200096 | NA | NA | NA | NA |
| WBGene00200097 | NA | NA | NA | NA |
| WBGene00200098 | -0.002914971 | 0.95718083 | NA | NA |
| WBGene00200099 | -0.040794878 | 0.53203165 | NA | NA |
| WBGene00200100 | NA | NA | NA | NA |
| WBGene00200101 | 0.05998475 | 0.41450796 | NA | NA |
| WBGene00200102 | NA | NA | NA | NA |
| WBGene00200103 | NA | NA | NA | NA |
| WBGene00200104 | NA | NA | NA | NA |
| WBGene00200105 | NA | NA | NA | NA |
| WBGene00200106 | NA | NA | NA | NA |
| WBGene00200107 | NA | NA | NA | NA |

| | | | | |
|---|---|---|---|---|
| WBGene00200108 | NA | NA | NA | NA |
| WBGene00200109 | NA | NA | NA | NA |
| WBGene00200110 | NA | NA | NA | NA |
| WBGene00200111 | NA | NA | NA | NA |
| WBGene00200112 | NA | NA | NA | NA |
| WBGene00200113 | NA | NA | NA | NA |
| WBGene00200114 | NA | NA | NA | NA |
| WBGene00200115 | NA | NA | NA | NA |
| WBGene00200116 | NA | NA | NA | NA |
| WBGene00200117 | NA | NA | NA | NA |
| WBGene00200118 | NA | NA | NA | NA |
| WBGene00200119 | NA | NA | NA | NA |
| WBGene00200120 | NA | NA | NA | NA |
| WBGene00200121 | 0.020232312 | 0.84643798 | NA | NA |
| WBGene00200122 | NA | NA | NA | NA |
| WBGene00200123 | NA | NA | NA | NA |
| WBGene00200124 | NA | NA | NA | NA |
| WBGene00200125 | NA | NA | NA | NA |
| WBGene00200126 | NA | NA | NA | NA |
| WBGene00200127 | NA | NA | NA | NA |
| WBGene00200128 | NA | NA | NA | NA |
| WBGene00200129 | NA | NA | NA | NA |
| WBGene00200130 | NA | NA | NA | NA |
| WBGene00200131 | NA | NA | NA | NA |
| WBGene00200132 | NA | NA | NA | NA |
| WBGene00200133 | NA | NA | NA | NA |
| WBGene00200134 | NA | NA | NA | NA |
| WBGene00200135 | NA | NA | NA | NA |
| WBGene00200136 | NA | NA | NA | NA |
| WBGene00200137 | NA | NA | NA | NA |
| WBGene00200138 | NA | NA | NA | NA |
| WBGene00200139 | NA | NA | NA | NA |
| WBGene00200140 | NA | NA | NA | NA |
| WBGene00200141 | NA | NA | NA | NA |
| WBGene00200142 | NA | NA | NA | NA |
| WBGene00200143 | NA | NA | NA | NA |
| WBGene00200144 | -0.023056499 | 0.78121253 | NA | NA |
| WBGene00200145 | NA | NA | NA | NA |
| WBGene00200146 | NA | NA | NA | NA |
| WBGene00200147 | NA | NA | NA | NA |
| WBGene00200148 | NA | NA | NA | NA |
| WBGene00200149 | NA | NA | NA | NA |
| WBGene00200150 | -0.022930767 | 0.78121253 | NA | NA |

| | | | | |
|---|---|---|---|---|
| WBGene00200151 | NA | NA | NA | NA |
| WBGene00200152 | NA | NA | NA | NA |
| WBGene00200153 | NA | NA | NA | NA |
| WBGene00200154 | NA | NA | NA | NA |
| WBGene00200155 | NA | NA | NA | NA |
| WBGene00200156 | NA | NA | NA | NA |
| WBGene00200157 | NA | NA | NA | NA |
| WBGene00200158 | NA | NA | NA | NA |
| WBGene00200159 | NA | NA | NA | NA |
| WBGene00200160 | NA | NA | NA | NA |
| WBGene00200161 | NA | NA | NA | NA |
| WBGene00200162 | NA | NA | NA | NA |
| WBGene00200163 | NA | NA | NA | NA |
| WBGene00200164 | -0.026384613 | 0.67299636 | NA | NA |
| WBGene00200165 | NA | NA | NA | NA |
| WBGene00200166 | NA | NA | NA | NA |
| WBGene00200167 | NA | NA | NA | NA |
| WBGene00200168 | NA | NA | NA | NA |
| WBGene00200169 | 0.040107227 | 0.58693163 | NA | NA |
| WBGene00200170 | NA | NA | NA | NA |
| WBGene00200171 | NA | NA | NA | NA |
| WBGene00200172 | NA | NA | NA | NA |
| WBGene00200173 | NA | NA | NA | NA |
| WBGene00200174 | NA | NA | NA | NA |
| WBGene00200175 | NA | NA | NA | NA |
| WBGene00200176 | NA | NA | NA | NA |
| WBGene00200177 | NA | NA | NA | NA |
| WBGene00200178 | NA | NA | NA | NA |
| WBGene00200179 | NA | NA | NA | NA |
| WBGene00200180 | NA | NA | NA | NA |
| WBGene00200181 | NA | NA | NA | NA |
| WBGene00200182 | NA | NA | NA | NA |
| WBGene00200183 | NA | NA | NA | NA |
| WBGene00200184 | NA | NA | NA | NA |
| WBGene00200185 | NA | NA | NA | NA |
| WBGene00200186 | NA | NA | NA | NA |
| WBGene00200187 | NA | NA | NA | NA |
| WBGene00200188 | NA | NA | NA | NA |
| WBGene00200189 | NA | NA | NA | NA |
| WBGene00200190 | NA | NA | NA | NA |
| WBGene00200191 | NA | NA | NA | NA |
| WBGene00200192 | NA | NA | NA | NA |
| WBGene00200193 | NA | NA | NA | NA |

| | | | | |
|---|---|---|---|---|
| WBGene00200194 | NA | NA | NA | NA |
| WBGene00200195 | NA | NA | NA | NA |
| WBGene00200196 | NA | NA | NA | NA |
| WBGene00200197 | NA | NA | NA | NA |
| WBGene00200198 | NA | NA | NA | NA |
| WBGene00200199 | NA | NA | NA | NA |
| WBGene00200200 | NA | NA | NA | NA |
| WBGene00200201 | NA | NA | NA | NA |
| WBGene00200202 | NA | NA | NA | NA |
| WBGene00200203 | NA | NA | NA | NA |
| WBGene00200204 | NA | NA | NA | NA |
| WBGene00200205 | NA | NA | NA | NA |
| WBGene00200206 | NA | NA | NA | NA |
| WBGene00200207 | NA | NA | NA | NA |
| WBGene00200208 | NA | NA | NA | NA |
| WBGene00200209 | NA | NA | NA | NA |
| WBGene00200210 | NA | NA | NA | NA |
| WBGene00200211 | NA | NA | NA | NA |
| WBGene00200212 | 0.148021412 | 0.07455439 | NA | NA |
| WBGene00200213 | NA | NA | NA | NA |
| WBGene00200214 | NA | NA | NA | NA |
| WBGene00200215 | NA | NA | NA | NA |
| WBGene00200216 | NA | NA | NA | NA |
| WBGene00200217 | NA | NA | NA | NA |
| WBGene00200218 | NA | NA | NA | NA |
| WBGene00200219 | NA | NA | NA | NA |
| WBGene00200220 | NA | NA | NA | NA |
| WBGene00200221 | NA | NA | NA | NA |
| WBGene00200222 | NA | NA | NA | NA |
| WBGene00200223 | NA | NA | NA | NA |
| WBGene00200224 | NA | NA | NA | NA |
| WBGene00200225 | NA | NA | NA | NA |
| WBGene00200226 | NA | NA | NA | NA |
| WBGene00200227 | NA | NA | NA | NA |
| WBGene00200228 | NA | NA | NA | NA |
| WBGene00200229 | NA | NA | NA | NA |
| WBGene00200230 | NA | NA | NA | NA |
| WBGene00200231 | NA | NA | NA | NA |
| WBGene00200232 | NA | NA | NA | NA |
| WBGene00200233 | NA | NA | NA | NA |
| WBGene00200234 | NA | NA | NA | NA |
| WBGene00200235 | NA | NA | NA | NA |
| WBGene00200236 | NA | NA | NA | NA |

| | | | | |
|---|---|---|---|---|
| WBGene00200237 | NA | NA | NA | NA |
| WBGene00200238 | NA | NA | NA | NA |
| WBGene00200239 | NA | NA | NA | NA |
| WBGene00200240 | NA | NA | NA | NA |
| WBGene00200241 | NA | NA | NA | NA |
| WBGene00200242 | NA | NA | NA | NA |
| WBGene00200243 | NA | NA | NA | NA |
| WBGene00200244 | NA | NA | NA | NA |
| WBGene00200245 | NA | NA | NA | NA |
| WBGene00200246 | NA | NA | NA | NA |
| WBGene00200247 | NA | NA | NA | NA |
| WBGene00200248 | NA | NA | NA | NA |
| WBGene00200249 | NA | NA | NA | NA |
| WBGene00200250 | NA | NA | NA | NA |
| WBGene00200251 | NA | NA | NA | NA |
| WBGene00200252 | NA | NA | NA | NA |
| WBGene00200253 | NA | NA | NA | NA |
| WBGene00200254 | NA | NA | NA | NA |
| WBGene00200255 | NA | NA | NA | NA |
| WBGene00200256 | NA | NA | NA | NA |
| WBGene00200257 | NA | NA | NA | NA |
| WBGene00200258 | NA | NA | NA | NA |
| WBGene00200259 | NA | NA | NA | NA |
| WBGene00200260 | NA | NA | NA | NA |
| WBGene00200261 | NA | NA | NA | NA |
| WBGene00200262 | 0.020317245 | 0.84643798 | NA | NA |
| WBGene00200263 | NA | NA | NA | NA |
| WBGene00200264 | -0.022930767 | 0.78121253 | NA | NA |
| WBGene00200265 | NA | NA | NA | NA |
| WBGene00200266 | NA | NA | NA | NA |
| WBGene00200267 | NA | NA | NA | NA |
| WBGene00200268 | NA | NA | NA | NA |
| WBGene00200269 | NA | NA | NA | NA |
| WBGene00200270 | NA | NA | NA | NA |
| WBGene00200271 | NA | NA | NA | NA |
| WBGene00200272 | NA | NA | NA | NA |
| WBGene00200273 | NA | NA | NA | NA |
| WBGene00200274 | NA | NA | NA | NA |
| WBGene00200275 | NA | NA | NA | NA |
| WBGene00200276 | NA | NA | NA | NA |
| WBGene00200277 | NA | NA | NA | NA |
| WBGene00200278 | NA | NA | NA | NA |
| WBGene00200279 | NA | NA | NA | NA |

| | | | | |
|---|---|---|---|---|
| WBGene00200280 | NA | NA | NA | NA |
| WBGene00200281 | NA | NA | NA | NA |
| WBGene00200282 | NA | NA | NA | NA |
| WBGene00200283 | NA | NA | NA | NA |
| WBGene00200284 | 0.033034883 | 0.64219503 | NA | NA |
| WBGene00200285 | NA | NA | NA | NA |
| WBGene00200286 | NA | NA | NA | NA |
| WBGene00200287 | NA | NA | NA | NA |
| WBGene00200288 | NA | NA | NA | NA |
| WBGene00200289 | -0.022930767 | 0.78121253 | NA | NA |
| WBGene00200290 | -0.023056499 | 0.78121253 | NA | NA |
| WBGene00200291 | NA | NA | NA | NA |
| WBGene00200292 | NA | NA | NA | NA |
| WBGene00200293 | NA | NA | NA | NA |
| WBGene00200294 | NA | NA | NA | NA |
| WBGene00200295 | NA | NA | NA | NA |
| WBGene00200296 | NA | NA | NA | NA |
| WBGene00200297 | NA | NA | NA | NA |
| WBGene00200298 | 0.020317245 | 0.84643798 | NA | NA |
| WBGene00200299 | NA | NA | NA | NA |
| WBGene00200300 | -0.023109762 | 0.78121253 | NA | NA |
| WBGene00200301 | NA | NA | NA | NA |
| WBGene00200302 | NA | NA | NA | NA |
| WBGene00200303 | NA | NA | NA | NA |
| WBGene00200304 | NA | NA | NA | NA |
| WBGene00200305 | NA | NA | NA | NA |
| WBGene00200306 | 0.020925399 | 0.84643798 | NA | NA |
| WBGene00200307 | NA | NA | NA | NA |
| WBGene00200308 | NA | NA | NA | NA |
| WBGene00200309 | NA | NA | NA | NA |
| WBGene00200310 | NA | NA | NA | NA |
| WBGene00200311 | NA | NA | NA | NA |
| WBGene00200312 | NA | NA | NA | NA |
| WBGene00200313 | NA | NA | NA | NA |
| WBGene00200314 | NA | NA | NA | NA |
| WBGene00200315 | NA | NA | NA | NA |
| WBGene00200316 | NA | NA | NA | NA |
| WBGene00200317 | NA | NA | NA | NA |
| WBGene00200318 | NA | NA | NA | NA |
| WBGene00200319 | NA | NA | NA | NA |
| WBGene00200320 | NA | NA | NA | NA |
| WBGene00200321 | NA | NA | NA | NA |
| WBGene00200322 | NA | NA | NA | NA |

| | | | | |
|---|---|---|---|---|
| WBGene00200323 | NA | NA | NA | NA |
| WBGene00200324 | NA | NA | NA | NA |
| WBGene00200325 | NA | NA | NA | NA |
| WBGene00200326 | NA | NA | NA | NA |
| WBGene00200327 | NA | NA | NA | NA |
| WBGene00200328 | NA | NA | NA | NA |
| WBGene00200329 | NA | NA | NA | NA |
| WBGene00200330 | -0.04044783 | 0.53612608 | NA | NA |
| WBGene00200331 | NA | NA | NA | NA |
| WBGene00200332 | NA | NA | NA | NA |
| WBGene00200333 | NA | NA | NA | NA |
| WBGene00200334 | NA | NA | NA | NA |
| WBGene00200335 | NA | NA | NA | NA |
| WBGene00200336 | NA | NA | NA | NA |
| WBGene00200337 | NA | NA | NA | NA |
| WBGene00200338 | NA | NA | NA | NA |
| WBGene00200339 | NA | NA | NA | NA |
| WBGene00200340 | NA | NA | NA | NA |
| WBGene00200341 | NA | NA | NA | NA |
| WBGene00200342 | NA | NA | NA | NA |
| WBGene00200343 | NA | NA | NA | NA |
| WBGene00200344 | NA | NA | NA | NA |
| WBGene00200345 | NA | NA | NA | NA |
| WBGene00200346 | NA | NA | NA | NA |
| WBGene00200347 | NA | NA | NA | NA |
| WBGene00200348 | NA | NA | NA | NA |
| WBGene00200349 | NA | NA | NA | NA |
| WBGene00200350 | NA | NA | NA | NA |
| WBGene00200351 | NA | NA | NA | NA |
| WBGene00200352 | NA | NA | NA | NA |
| WBGene00200353 | NA | NA | NA | NA |
| WBGene00200354 | NA | NA | NA | NA |
| WBGene00200355 | 0.018657025 | 0.77508599 | NA | NA |
| WBGene00200356 | NA | NA | NA | NA |
| WBGene00200357 | NA | NA | NA | NA |
| WBGene00200358 | NA | NA | NA | NA |
| WBGene00200359 | NA | NA | NA | NA |
| WBGene00200360 | NA | NA | NA | NA |
| WBGene00200361 | NA | NA | NA | NA |
| WBGene00200362 | NA | NA | NA | NA |
| WBGene00200363 | NA | NA | NA | NA |
| WBGene00200364 | NA | NA | NA | NA |
| WBGene00200365 | NA | NA | NA | NA |

| | | | | |
|---|---|---|---|---|
| WBGene00200366 | NA | NA | NA | NA |
| WBGene00200367 | NA | NA | NA | NA |
| WBGene00200368 | NA | NA | NA | NA |
| WBGene00200369 | NA | NA | NA | NA |
| WBGene00200370 | NA | NA | NA | NA |
| WBGene00200371 | NA | NA | NA | NA |
| WBGene00200372 | NA | NA | NA | NA |
| WBGene00200373 | NA | NA | NA | NA |
| WBGene00200374 | NA | NA | NA | NA |
| WBGene00200375 | NA | NA | NA | NA |
| WBGene00200376 | NA | NA | NA | NA |
| WBGene00200377 | NA | NA | NA | NA |
| WBGene00200378 | NA | NA | NA | NA |
| WBGene00200379 | NA | NA | NA | NA |
| WBGene00200380 | NA | NA | NA | NA |
| WBGene00200381 | NA | NA | NA | NA |
| WBGene00200382 | NA | NA | NA | NA |
| WBGene00200383 | NA | NA | NA | NA |
| WBGene00200384 | NA | NA | NA | NA |
| WBGene00200385 | NA | NA | NA | NA |
| WBGene00200386 | NA | NA | NA | NA |
| WBGene00200387 | NA | NA | NA | NA |
| WBGene00200388 | NA | NA | NA | NA |
| WBGene00200389 | NA | NA | NA | NA |
| WBGene00200390 | NA | NA | NA | NA |
| WBGene00200391 | NA | NA | NA | NA |
| WBGene00200392 | NA | NA | NA | NA |
| WBGene00200393 | NA | NA | NA | NA |
| WBGene00200394 | NA | NA | NA | NA |
| WBGene00200395 | NA | NA | NA | NA |
| WBGene00200396 | NA | NA | NA | NA |
| WBGene00200397 | 0.020232312 | 0.84643798 | NA | NA |
| WBGene00200398 | NA | NA | NA | NA |
| WBGene00200399 | NA | NA | NA | NA |
| WBGene00200400 | NA | NA | NA | NA |
| WBGene00200401 | NA | NA | NA | NA |
| WBGene00200402 | NA | NA | NA | NA |
| WBGene00200403 | NA | NA | NA | NA |
| WBGene00200404 | NA | NA | NA | NA |
| WBGene00200405 | NA | NA | NA | NA |
| WBGene00200406 | NA | NA | NA | NA |
| WBGene00200407 | NA | NA | NA | NA |
| WBGene00200408 | NA | NA | NA | NA |

| | | | | |
|---|---|---|---|---|
| WBGene00200409 | NA | NA | NA | NA |
| WBGene00200410 | NA | NA | NA | NA |
| WBGene00200411 | NA | NA | NA | NA |
| WBGene00200412 | NA | NA | NA | NA |
| WBGene00200413 | NA | NA | NA | NA |
| WBGene00200414 | NA | NA | NA | NA |
| WBGene00200415 | NA | NA | NA | NA |
| WBGene00200416 | NA | NA | NA | NA |
| WBGene00200417 | NA | NA | NA | NA |
| WBGene00200418 | NA | NA | NA | NA |
| WBGene00200419 | NA | NA | NA | NA |
| WBGene00200420 | NA | NA | NA | NA |
| WBGene00200421 | NA | NA | NA | NA |
| WBGene00200422 | NA | NA | NA | NA |
| WBGene00200423 | NA | NA | NA | NA |
| WBGene00200424 | NA | NA | NA | NA |
| WBGene00200425 | NA | NA | NA | NA |
| WBGene00200426 | NA | NA | NA | NA |
| WBGene00200427 | NA | NA | NA | NA |
| WBGene00200428 | -0.023056499 | 0.78121253 | NA | NA |
| WBGene00200429 | NA | NA | NA | NA |
| WBGene00200430 | NA | NA | NA | NA |
| WBGene00200431 | NA | NA | NA | NA |
| WBGene00200432 | NA | NA | NA | NA |
| WBGene00200433 | -0.022930767 | 0.78121253 | NA | NA |
| WBGene00200434 | NA | NA | NA | NA |
| WBGene00200435 | NA | NA | NA | NA |
| WBGene00200436 | NA | NA | NA | NA |
| WBGene00200437 | NA | NA | NA | NA |
| WBGene00200438 | NA | NA | NA | NA |
| WBGene00200439 | NA | NA | NA | NA |
| WBGene00200440 | NA | NA | NA | NA |
| WBGene00200441 | -0.023056499 | 0.78121253 | NA | NA |
| WBGene00200442 | NA | NA | NA | NA |
| WBGene00200443 | NA | NA | NA | NA |
| WBGene00200444 | NA | NA | NA | NA |
| WBGene00200445 | NA | NA | NA | NA |
| WBGene00200446 | -0.022930767 | 0.78121253 | NA | NA |
| WBGene00200447 | NA | NA | NA | NA |
| WBGene00200448 | NA | NA | NA | NA |
| WBGene00200449 | NA | NA | NA | NA |
| WBGene00200450 | NA | NA | NA | NA |
| WBGene00200451 | NA | NA | NA | NA |

| | | | | |
|---|---|---|---|---|
| WBGene00200452 | NA | NA | NA | NA |
| WBGene00200453 | NA | NA | NA | NA |
| WBGene00200454 | NA | NA | NA | NA |
| WBGene00200455 | NA | NA | NA | NA |
| WBGene00200456 | NA | NA | NA | NA |
| WBGene00200457 | NA | NA | NA | NA |
| WBGene00200458 | 0.020232312 | 0.84643798 | NA | NA |
| WBGene00200459 | 0.018657025 | 0.77508599 | NA | NA |
| WBGene00200460 | NA | NA | NA | NA |
| WBGene00200461 | NA | NA | NA | NA |
| WBGene00200462 | NA | NA | NA | NA |
| WBGene00200463 | NA | NA | NA | NA |
| WBGene00200464 | NA | NA | NA | NA |
| WBGene00200465 | NA | NA | NA | NA |
| WBGene00200466 | NA | NA | NA | NA |
| WBGene00200467 | NA | NA | NA | NA |
| WBGene00200468 | NA | NA | NA | NA |
| WBGene00200469 | NA | NA | NA | NA |
| WBGene00200470 | NA | NA | NA | NA |
| WBGene00200471 | NA | NA | NA | NA |
| WBGene00200472 | NA | NA | NA | NA |
| WBGene00200473 | NA | NA | NA | NA |
| WBGene00200474 | NA | NA | NA | NA |
| WBGene00200475 | NA | NA | NA | NA |
| WBGene00200476 | NA | NA | NA | NA |
| WBGene00200477 | NA | NA | NA | NA |
| WBGene00200478 | NA | NA | NA | NA |
| WBGene00200479 | NA | NA | NA | NA |
| WBGene00200480 | NA | NA | NA | NA |
| WBGene00200481 | NA | NA | NA | NA |
| WBGene00200482 | NA | NA | NA | NA |
| WBGene00200483 | NA | NA | NA | NA |
| WBGene00200484 | NA | NA | NA | NA |
| WBGene00200485 | NA | NA | NA | NA |
| WBGene00200486 | NA | NA | NA | NA |
| WBGene00200487 | NA | NA | NA | NA |
| WBGene00200488 | NA | NA | NA | NA |
| WBGene00200489 | NA | NA | NA | NA |
| WBGene00200490 | NA | NA | NA | NA |
| WBGene00200491 | NA | NA | NA | NA |
| WBGene00200492 | NA | NA | NA | NA |
| WBGene00200493 | NA | NA | NA | NA |
| WBGene00200494 | NA | NA | NA | NA |

| | | | | |
|---|---|---|---|---|
| WBGene00200495 | NA | NA | NA | NA |
| WBGene00200496 | NA | NA | NA | NA |
| WBGene00200497 | NA | NA | NA | NA |
| WBGene00200498 | NA | NA | NA | NA |
| WBGene00200499 | -0.022930767 | 0.78121253 | NA | NA |
| WBGene00200500 | NA | NA | NA | NA |
| WBGene00200501 | NA | NA | NA | NA |
| WBGene00200502 | NA | NA | NA | NA |
| WBGene00200503 | NA | NA | NA | NA |
| WBGene00200504 | NA | NA | NA | NA |
| WBGene00200505 | NA | NA | NA | NA |
| WBGene00200506 | NA | NA | NA | NA |
| WBGene00200507 | NA | NA | NA | NA |
| WBGene00200508 | NA | NA | NA | NA |
| WBGene00200509 | NA | NA | NA | NA |
| WBGene00200510 | NA | NA | NA | NA |
| WBGene00200511 | NA | NA | NA | NA |
| WBGene00200512 | NA | NA | NA | NA |
| WBGene00200513 | NA | NA | NA | NA |
| WBGene00200514 | NA | NA | NA | NA |
| WBGene00200515 | NA | NA | NA | NA |
| WBGene00200516 | NA | NA | NA | NA |
| WBGene00200517 | NA | NA | NA | NA |
| WBGene00200518 | NA | NA | NA | NA |
| WBGene00200519 | NA | NA | NA | NA |
| WBGene00200520 | NA | NA | NA | NA |
| WBGene00200521 | NA | NA | NA | NA |
| WBGene00200522 | NA | NA | NA | NA |
| WBGene00200523 | NA | NA | NA | NA |
| WBGene00200524 | NA | NA | NA | NA |
| WBGene00200525 | NA | NA | NA | NA |
| WBGene00200526 | NA | NA | NA | NA |
| WBGene00200527 | NA | NA | NA | NA |
| WBGene00200528 | 0.020925399 | 0.84643798 | NA | NA |
| WBGene00200529 | NA | NA | NA | NA |
| WBGene00200530 | NA | NA | NA | NA |
| WBGene00200531 | NA | NA | NA | NA |
| WBGene00200532 | NA | NA | NA | NA |
| WBGene00200533 | NA | NA | NA | NA |
| WBGene00200534 | NA | NA | NA | NA |
| WBGene00200535 | NA | NA | NA | NA |
| WBGene00200536 | NA | NA | NA | NA |
| WBGene00200537 | NA | NA | NA | NA |

| | | | | | |
|---|---|---|---|---|---|
| WBGene00200538 | NA | | NA | NA | NA |
| WBGene00200539 | NA | | NA | NA | NA |
| WBGene00200540 | NA | | NA | NA | NA |
| WBGene00200541 | NA | | NA | NA | NA |
| WBGene00200542 | NA | | NA | NA | NA |
| WBGene00200543 | NA | | NA | NA | NA |
| WBGene00200544 | NA | | NA | NA | NA |
| WBGene00200545 | NA | | NA | NA | NA |
| WBGene00200546 | NA | | NA | NA | NA |
| WBGene00200547 | NA | | NA | NA | NA |
| WBGene00200548 | NA | | NA | NA | NA |
| WBGene00200549 | NA | | NA | NA | NA |
| WBGene00200550 | NA | | NA | NA | NA |
| WBGene00200551 | NA | | NA | NA | NA |
| WBGene00200552 | NA | | NA | NA | NA |
| WBGene00200553 | NA | | NA | NA | NA |
| WBGene00200554 | NA | | NA | NA | NA |
| WBGene00200555 | NA | | NA | NA | NA |
| WBGene00200556 | NA | | NA | NA | NA |
| WBGene00200557 | NA | | NA | NA | NA |
| WBGene00200558 | NA | | NA | NA | NA |
| WBGene00200559 | NA | | NA | NA | NA |
| WBGene00200560 | NA | | NA | NA | NA |
| WBGene00200561 | NA | | NA | NA | NA |
| WBGene00200562 | | 0.020317245 | 0.84643798 | NA | NA |
| WBGene00200563 | NA | | NA | NA | NA |
| WBGene00200564 | NA | | NA | NA | NA |
| WBGene00200565 | NA | | NA | NA | NA |
| WBGene00200566 | NA | | NA | NA | NA |
| WBGene00200567 | | 0.020925399 | 0.84643798 | NA | NA |
| WBGene00200568 | NA | | NA | NA | NA |
| WBGene00200569 | NA | | NA | NA | NA |
| WBGene00200570 | NA | | NA | NA | NA |
| WBGene00200571 | NA | | NA | NA | NA |
| WBGene00200572 | NA | | NA | NA | NA |
| WBGene00200573 | NA | | NA | NA | NA |
| WBGene00200574 | NA | | NA | NA | NA |
| WBGene00200575 | NA | | NA | NA | NA |
| WBGene00200576 | NA | | NA | NA | NA |
| WBGene00200577 | NA | | NA | NA | NA |
| WBGene00200578 | NA | | NA | NA | NA |
| WBGene00200579 | NA | | NA | NA | NA |
| WBGene00200580 | NA | | NA | NA | NA |

| | | | | |
|---|---|---|---|---|
| WBGene00200581 | NA | NA | NA | NA |
| WBGene00200582 | NA | NA | NA | NA |
| WBGene00200583 | NA | NA | NA | NA |
| WBGene00200584 | NA | NA | NA | NA |
| WBGene00200585 | NA | NA | NA | NA |
| WBGene00200586 | NA | NA | NA | NA |
| WBGene00200587 | NA | NA | NA | NA |
| WBGene00200588 | NA | NA | NA | NA |
| WBGene00200589 | NA | NA | NA | NA |
| WBGene00200590 | -0.022930767 | 0.78121253 | NA | NA |
| WBGene00200591 | NA | NA | NA | NA |
| WBGene00200592 | NA | NA | NA | NA |
| WBGene00200593 | NA | NA | NA | NA |
| WBGene00200594 | NA | NA | NA | NA |
| WBGene00200595 | NA | NA | NA | NA |
| WBGene00200596 | NA | NA | NA | NA |
| WBGene00200597 | NA | NA | NA | NA |
| WBGene00200598 | 0.056188364 | 0.42092875 | NA | NA |
| WBGene00200599 | NA | NA | NA | NA |
| WBGene00200600 | 0.01912147 | 0.77587372 | NA | NA |
| WBGene00200601 | NA | NA | NA | NA |
| WBGene00200602 | NA | NA | NA | NA |
| WBGene00200603 | NA | NA | NA | NA |
| WBGene00200604 | NA | NA | NA | NA |
| WBGene00200605 | NA | NA | NA | NA |
| WBGene00200606 | -0.001993815 | 0.95612996 | NA | NA |
| WBGene00200607 | NA | NA | NA | NA |
| WBGene00200608 | NA | NA | NA | NA |
| WBGene00200609 | NA | NA | NA | NA |
| WBGene00200610 | NA | NA | NA | NA |
| WBGene00200611 | NA | NA | NA | NA |
| WBGene00200612 | NA | NA | NA | NA |
| WBGene00200613 | NA | NA | NA | NA |
| WBGene00200614 | NA | NA | NA | NA |
| WBGene00200615 | NA | NA | NA | NA |
| WBGene00200616 | NA | NA | NA | NA |
| WBGene00200617 | NA | NA | NA | NA |
| WBGene00200618 | NA | NA | NA | NA |
| WBGene00200619 | NA | NA | NA | NA |
| WBGene00200620 | -0.023056499 | 0.78121253 | NA | NA |
| WBGene00200621 | NA | NA | NA | NA |
| WBGene00200622 | NA | NA | NA | NA |
| WBGene00200623 | NA | NA | NA | NA |

| | | | | |
|---|---|---|---|---|
| WBGene00200624 | NA | NA | NA | NA |
| WBGene00200625 | NA | NA | NA | NA |
| WBGene00200626 | -0.022930767 | 0.78121253 | NA | NA |
| WBGene00200627 | NA | NA | NA | NA |
| WBGene00200628 | NA | NA | NA | NA |
| WBGene00200629 | NA | NA | NA | NA |
| WBGene00200630 | NA | NA | NA | NA |
| WBGene00200631 | NA | NA | NA | NA |
| WBGene00200632 | NA | NA | NA | NA |
| WBGene00200633 | NA | NA | NA | NA |
| WBGene00200634 | NA | NA | NA | NA |
| WBGene00200635 | NA | NA | NA | NA |
| WBGene00200636 | NA | NA | NA | NA |
| WBGene00200637 | NA | NA | NA | NA |
| WBGene00200638 | NA | NA | NA | NA |
| WBGene00200639 | NA | NA | NA | NA |
| WBGene00200640 | NA | NA | NA | NA |
| WBGene00200641 | NA | NA | NA | NA |
| WBGene00200642 | NA | NA | NA | NA |
| WBGene00200643 | 0.020925399 | 0.84643798 | NA | NA |
| WBGene00200644 | NA | NA | NA | NA |
| WBGene00200645 | NA | NA | NA | NA |
| WBGene00200646 | NA | NA | NA | NA |
| WBGene00200647 | NA | NA | NA | NA |
| WBGene00200648 | NA | NA | NA | NA |
| WBGene00200649 | 0.020232312 | 0.84643798 | NA | NA |
| WBGene00200650 | NA | NA | NA | NA |
| WBGene00200651 | 0.020232312 | 0.84643798 | NA | NA |
| WBGene00200652 | NA | NA | NA | NA |
| WBGene00200653 | NA | NA | NA | NA |
| WBGene00200654 | NA | NA | NA | NA |
| WBGene00200655 | NA | NA | NA | NA |
| WBGene00200656 | NA | NA | NA | NA |
| WBGene00200657 | NA | NA | NA | NA |
| WBGene00200658 | NA | NA | NA | NA |
| WBGene00200659 | NA | NA | NA | NA |
| WBGene00200660 | NA | NA | NA | NA |
| WBGene00200661 | NA | NA | NA | NA |
| WBGene00200662 | NA | NA | NA | NA |
| WBGene00200663 | NA | NA | NA | NA |
| WBGene00200664 | NA | NA | NA | NA |
| WBGene00200665 | NA | NA | NA | NA |
| WBGene00200666 | NA | NA | NA | NA |

| | | | | |
|---|---|---|---|---|
| WBGene00200667 | NA | NA | NA | NA |
| WBGene00200668 | NA | NA | NA | NA |
| WBGene00200669 | NA | NA | NA | NA |
| WBGene00200670 | NA | NA | NA | NA |
| WBGene00200671 | NA | NA | NA | NA |
| WBGene00200672 | NA | NA | NA | NA |
| WBGene00200673 | NA | NA | NA | NA |
| WBGene00200674 | NA | NA | NA | NA |
| WBGene00200675 | NA | NA | NA | NA |
| WBGene00200676 | NA | NA | NA | NA |
| WBGene00200677 | NA | NA | NA | NA |
| WBGene00200678 | NA | NA | NA | NA |
| WBGene00200679 | NA | NA | NA | NA |
| WBGene00200680 | 0.020317245 | 0.84643798 | NA | NA |
| WBGene00200681 | NA | NA | NA | NA |
| WBGene00200682 | NA | NA | NA | NA |
| WBGene00200683 | NA | NA | NA | NA |
| WBGene00200684 | NA | NA | NA | NA |
| WBGene00200685 | 0.020925399 | 0.84643798 | NA | NA |
| WBGene00200686 | NA | NA | NA | NA |
| WBGene00200687 | NA | NA | NA | NA |
| WBGene00200688 | NA | NA | NA | NA |
| WBGene00200689 | NA | NA | NA | NA |
| WBGene00200690 | NA | NA | NA | NA |
| WBGene00200691 | NA | NA | NA | NA |
| WBGene00200692 | NA | NA | NA | NA |
| WBGene00200693 | NA | NA | NA | NA |
| WBGene00200694 | NA | NA | NA | NA |
| WBGene00200695 | NA | NA | NA | NA |
| WBGene00200696 | NA | NA | NA | NA |
| WBGene00200697 | NA | NA | NA | NA |
| WBGene00200698 | NA | NA | NA | NA |
| WBGene00200699 | NA | NA | NA | NA |
| WBGene00200700 | NA | NA | NA | NA |
| WBGene00200701 | NA | NA | NA | NA |
| WBGene00200702 | NA | NA | NA | NA |
| WBGene00200703 | NA | NA | NA | NA |
| WBGene00200704 | NA | NA | NA | NA |
| WBGene00200705 | NA | NA | NA | NA |
| WBGene00200706 | NA | NA | NA | NA |
| WBGene00200707 | NA | NA | NA | NA |
| WBGene00200708 | NA | NA | NA | NA |
| WBGene00200709 | NA | NA | NA | NA |

| | | | | |
|---|---|---|---|---|
| WBGene00200710 | NA | NA | NA | NA |
| WBGene00200711 | NA | NA | NA | NA |
| WBGene00200712 | NA | NA | NA | NA |
| WBGene00200713 | 0.020317245 | 0.84643798 | NA | NA |
| WBGene00200714 | NA | NA | NA | NA |
| WBGene00200715 | NA | NA | NA | NA |
| WBGene00200716 | NA | NA | NA | NA |
| WBGene00200717 | NA | NA | NA | NA |
| WBGene00200718 | NA | NA | NA | NA |
| WBGene00200719 | NA | NA | NA | NA |
| WBGene00200720 | NA | NA | NA | NA |
| WBGene00200721 | 0.020232312 | 0.84643798 | NA | NA |
| WBGene00200722 | 0.020232312 | 0.84643798 | NA | NA |
| WBGene00200723 | -0.001993815 | 0.95612996 | NA | NA |
| WBGene00200724 | NA | NA | NA | NA |
| WBGene00200725 | NA | NA | NA | NA |
| WBGene00200726 | NA | NA | NA | NA |
| WBGene00200727 | NA | NA | NA | NA |
| WBGene00200728 | NA | NA | NA | NA |
| WBGene00200729 | NA | NA | NA | NA |
| WBGene00200730 | 0.020317245 | 0.84643798 | NA | NA |
| WBGene00200731 | NA | NA | NA | NA |
| WBGene00200732 | NA | NA | NA | NA |
| WBGene00200733 | NA | NA | NA | NA |
| WBGene00200734 | -0.023056499 | 0.78121253 | NA | NA |
| WBGene00200735 | NA | NA | NA | NA |
| WBGene00200736 | NA | NA | NA | NA |
| WBGene00200737 | NA | NA | NA | NA |
| WBGene00200738 | NA | NA | NA | NA |
| WBGene00200739 | NA | NA | NA | NA |
| WBGene00200740 | NA | NA | NA | NA |
| WBGene00200741 | NA | NA | NA | NA |
| WBGene00200742 | NA | NA | NA | NA |
| WBGene00200743 | NA | NA | NA | NA |
| WBGene00200744 | NA | NA | NA | NA |
| WBGene00200745 | NA | NA | NA | NA |
| WBGene00200746 | NA | NA | NA | NA |
| WBGene00200747 | NA | NA | NA | NA |
| WBGene00200748 | NA | NA | NA | NA |
| WBGene00200749 | NA | NA | NA | NA |
| WBGene00200750 | NA | NA | NA | NA |
| WBGene00200751 | NA | NA | NA | NA |
| WBGene00200752 | NA | NA | NA | NA |

| | | | | |
|---|---|---|---|---|
| WBGene00200753 | NA | NA | NA | NA |
| WBGene00200754 | NA | NA | NA | NA |
| WBGene00200755 | NA | NA | NA | NA |
| WBGene00200756 | NA | NA | NA | NA |
| WBGene00200757 | NA | NA | NA | NA |
| WBGene00200758 | NA | NA | NA | NA |
| WBGene00200759 | NA | NA | NA | NA |
| WBGene00200760 | NA | NA | NA | NA |
| WBGene00200761 | NA | NA | NA | NA |
| WBGene00200762 | NA | NA | NA | NA |
| WBGene00200763 | NA | NA | NA | NA |
| WBGene00200764 | NA | NA | NA | NA |
| WBGene00200765 | NA | NA | NA | NA |
| WBGene00200766 | NA | NA | NA | NA |
| WBGene00200767 | NA | NA | NA | NA |
| WBGene00200768 | NA | NA | NA | NA |
| WBGene00200769 | NA | NA | NA | NA |
| WBGene00200770 | NA | NA | NA | NA |
| WBGene00200771 | NA | NA | NA | NA |
| WBGene00200772 | NA | NA | NA | NA |
| WBGene00200773 | NA | NA | NA | NA |
| WBGene00200774 | NA | NA | NA | NA |
| WBGene00200775 | NA | NA | NA | NA |
| WBGene00200776 | NA | NA | NA | NA |
| WBGene00200777 | NA | NA | NA | NA |
| WBGene00200778 | NA | NA | NA | NA |
| WBGene00200779 | NA | NA | NA | NA |
| WBGene00200780 | NA | NA | NA | NA |
| WBGene00200781 | NA | NA | NA | NA |
| WBGene00200782 | -0.023056499 | 0.78121253 | NA | NA |
| WBGene00200783 | NA | NA | NA | NA |
| WBGene00200784 | NA | NA | NA | NA |
| WBGene00200785 | NA | NA | NA | NA |
| WBGene00200786 | NA | NA | NA | NA |
| WBGene00200787 | NA | NA | NA | NA |
| WBGene00200788 | NA | NA | NA | NA |
| WBGene00200789 | NA | NA | NA | NA |
| WBGene00200790 | NA | NA | NA | NA |
| WBGene00200791 | NA | NA | NA | NA |
| WBGene00200792 | NA | NA | NA | NA |
| WBGene00200793 | NA | NA | NA | NA |
| WBGene00200794 | NA | NA | NA | NA |
| WBGene00200795 | 0.020232312 | 0.84643798 | NA | NA |

| | | | | |
|---|---|---|---|---|
| WBGene00200796 | NA | NA | NA | NA |
| WBGene00200797 | NA | NA | NA | NA |
| WBGene00200798 | NA | NA | NA | NA |
| WBGene00200799 | NA | NA | NA | NA |
| WBGene00200800 | NA | NA | NA | NA |
| WBGene00200801 | NA | NA | NA | NA |
| WBGene00200802 | NA | NA | NA | NA |
| WBGene00200803 | NA | NA | NA | NA |
| WBGene00200804 | NA | NA | NA | NA |
| WBGene00200805 | NA | NA | NA | NA |
| WBGene00200806 | NA | NA | NA | NA |
| WBGene00200807 | NA | NA | NA | NA |
| WBGene00200808 | NA | NA | NA | NA |
| WBGene00200809 | -0.023109762 | 0.78121253 | NA | NA |
| WBGene00200810 | NA | NA | NA | NA |
| WBGene00200811 | NA | NA | NA | NA |
| WBGene00200812 | NA | NA | NA | NA |
| WBGene00200813 | NA | NA | NA | NA |
| WBGene00200814 | NA | NA | NA | NA |
| WBGene00200815 | NA | NA | NA | NA |
| WBGene00200816 | NA | NA | NA | NA |
| WBGene00200817 | NA | NA | NA | NA |
| WBGene00200818 | NA | NA | NA | NA |
| WBGene00200819 | NA | NA | NA | NA |
| WBGene00200820 | NA | NA | NA | NA |
| WBGene00200821 | NA | NA | NA | NA |
| WBGene00200822 | NA | NA | NA | NA |
| WBGene00200823 | NA | NA | NA | NA |
| WBGene00200824 | NA | NA | NA | NA |
| WBGene00200825 | NA | NA | NA | NA |
| WBGene00200826 | NA | NA | NA | NA |
| WBGene00200827 | NA | NA | NA | NA |
| WBGene00200828 | NA | NA | NA | NA |
| WBGene00200829 | NA | NA | NA | NA |
| WBGene00200830 | NA | NA | NA | NA |
| WBGene00200831 | NA | NA | NA | NA |
| WBGene00200832 | NA | NA | NA | NA |
| WBGene00200833 | NA | NA | NA | NA |
| WBGene00200834 | NA | NA | NA | NA |
| WBGene00200835 | NA | NA | NA | NA |
| WBGene00200836 | NA | NA | NA | NA |
| WBGene00200837 | NA | NA | NA | NA |
| WBGene00200838 | NA | NA | NA | NA |

| | | | | |
|---|---|---|---|---|
| WBGene00200839 | NA | NA | NA | NA |
| WBGene00200840 | NA | NA | NA | NA |
| WBGene00200841 | NA | NA | NA | NA |
| WBGene00200842 | NA | NA | NA | NA |
| WBGene00200843 | NA | NA | NA | NA |
| WBGene00200844 | NA | NA | NA | NA |
| WBGene00200845 | NA | NA | NA | NA |
| WBGene00200846 | NA | NA | NA | NA |
| WBGene00200847 | NA | NA | NA | NA |
| WBGene00200848 | NA | NA | NA | NA |
| WBGene00200849 | NA | NA | NA | NA |
| WBGene00200850 | NA | NA | NA | NA |
| WBGene00200851 | NA | NA | NA | NA |
| WBGene00200852 | NA | NA | NA | NA |
| WBGene00200853 | NA | NA | NA | NA |
| WBGene00200854 | NA | NA | NA | NA |
| WBGene00200855 | NA | NA | NA | NA |
| WBGene00200856 | NA | NA | NA | NA |
| WBGene00200857 | NA | NA | NA | NA |
| WBGene00200858 | NA | NA | NA | NA |
| WBGene00200859 | NA | NA | NA | NA |
| WBGene00200860 | NA | NA | NA | NA |
| WBGene00200861 | NA | NA | NA | NA |
| WBGene00200862 | NA | NA | NA | NA |
| WBGene00200863 | 0.020925399 | 0.84643798 | NA | NA |
| WBGene00200864 | NA | NA | NA | NA |
| WBGene00200865 | NA | NA | NA | NA |
| WBGene00200866 | NA | NA | NA | NA |
| WBGene00200867 | NA | NA | NA | NA |
| WBGene00200868 | NA | NA | NA | NA |
| WBGene00200869 | NA | NA | NA | NA |
| WBGene00200870 | -0.023056499 | 0.78121253 | NA | NA |
| WBGene00200871 | NA | NA | NA | NA |
| WBGene00200872 | NA | NA | NA | NA |
| WBGene00200873 | NA | NA | NA | NA |
| WBGene00200874 | NA | NA | NA | NA |
| WBGene00200875 | NA | NA | NA | NA |
| WBGene00200876 | NA | NA | NA | NA |
| WBGene00200877 | NA | NA | NA | NA |
| WBGene00200878 | NA | NA | NA | NA |
| WBGene00200879 | NA | NA | NA | NA |
| WBGene00200880 | NA | NA | NA | NA |
| WBGene00200881 | NA | NA | NA | NA |

| | | | | |
|---|---|---|---|---|
| WBGene00200882 | NA | NA | NA | NA |
| WBGene00200883 | NA | NA | NA | NA |
| WBGene00200884 | -0.003421551 | 0.95689871 | NA | NA |
| WBGene00200885 | NA | NA | NA | NA |
| WBGene00200886 | NA | NA | NA | NA |
| WBGene00200887 | NA | NA | NA | NA |
| WBGene00200888 | NA | NA | NA | NA |
| WBGene00200889 | 0.033034883 | 0.64219503 | NA | NA |
| WBGene00200890 | NA | NA | NA | NA |
| WBGene00200891 | NA | NA | NA | NA |
| WBGene00200892 | NA | NA | NA | NA |
| WBGene00200893 | NA | NA | NA | NA |
| WBGene00200894 | NA | NA | NA | NA |
| WBGene00200895 | NA | NA | NA | NA |
| WBGene00200896 | NA | NA | NA | NA |
| WBGene00200897 | NA | NA | NA | NA |
| WBGene00200898 | NA | NA | NA | NA |
| WBGene00200899 | NA | NA | NA | NA |
| WBGene00200900 | NA | NA | NA | NA |
| WBGene00200901 | NA | NA | NA | NA |
| WBGene00200902 | NA | NA | NA | NA |
| WBGene00200903 | NA | NA | NA | NA |
| WBGene00200904 | NA | NA | NA | NA |
| WBGene00200905 | NA | NA | NA | NA |
| WBGene00200906 | 0.020925399 | 0.84643798 | NA | NA |
| WBGene00200907 | NA | NA | NA | NA |
| WBGene00200908 | NA | NA | NA | NA |
| WBGene00200909 | NA | NA | NA | NA |
| WBGene00200910 | 0.085038347 | 0.30255919 | NA | NA |
| WBGene00200911 | NA | NA | NA | NA |
| WBGene00200912 | NA | NA | NA | NA |
| WBGene00200913 | NA | NA | NA | NA |
| WBGene00200914 | NA | NA | NA | NA |
| WBGene00200915 | NA | NA | NA | NA |
| WBGene00200916 | NA | NA | NA | NA |
| WBGene00200917 | NA | NA | NA | NA |
| WBGene00200918 | NA | NA | NA | NA |
| WBGene00200919 | NA | NA | NA | NA |
| WBGene00200920 | NA | NA | NA | NA |
| WBGene00200921 | NA | NA | NA | NA |
| WBGene00200922 | NA | NA | NA | NA |
| WBGene00200923 | NA | NA | NA | NA |
| WBGene00200924 | NA | NA | NA | NA |

| | | | | |
|---|---|---|---|---|
| WBGene00200925 | NA | NA | NA | NA |
| WBGene00200926 | NA | NA | NA | NA |
| WBGene00200927 | NA | NA | NA | NA |
| WBGene00200928 | NA | NA | NA | NA |
| WBGene00200929 | NA | NA | NA | NA |
| WBGene00200930 | NA | NA | NA | NA |
| WBGene00200931 | 0.020317245 | 0.84643798 | NA | NA |
| WBGene00200932 | NA | NA | NA | NA |
| WBGene00200933 | NA | NA | NA | NA |
| WBGene00200934 | NA | NA | NA | NA |
| WBGene00200935 | NA | NA | NA | NA |
| WBGene00200936 | NA | NA | NA | NA |
| WBGene00200937 | NA | NA | NA | NA |
| WBGene00200938 | NA | NA | NA | NA |
| WBGene00200939 | 0.020925399 | 0.84643798 | NA | NA |
| WBGene00200940 | NA | NA | NA | NA |
| WBGene00200941 | NA | NA | NA | NA |
| WBGene00200942 | NA | NA | NA | NA |
| WBGene00200943 | NA | NA | NA | NA |
| WBGene00200944 | NA | NA | NA | NA |
| WBGene00200945 | NA | NA | NA | NA |
| WBGene00200946 | NA | NA | NA | NA |
| WBGene00200947 | NA | NA | NA | NA |
| WBGene00200948 | NA | NA | NA | NA |
| WBGene00200949 | NA | NA | NA | NA |
| WBGene00200950 | NA | NA | NA | NA |
| WBGene00200951 | NA | NA | NA | NA |
| WBGene00200952 | NA | NA | NA | NA |
| WBGene00200953 | NA | NA | NA | NA |
| WBGene00200954 | 0.020317245 | 0.84643798 | NA | NA |
| WBGene00200955 | NA | NA | NA | NA |
| WBGene00200956 | NA | NA | NA | NA |
| WBGene00200957 | NA | NA | NA | NA |
| WBGene00200958 | NA | NA | NA | NA |
| WBGene00200959 | NA | NA | NA | NA |
| WBGene00200960 | NA | NA | NA | NA |
| WBGene00200961 | NA | NA | NA | NA |
| WBGene00200962 | NA | NA | NA | NA |
| WBGene00200963 | NA | NA | NA | NA |
| WBGene00200964 | NA | NA | NA | NA |
| WBGene00200965 | NA | NA | NA | NA |
| WBGene00200966 | NA | NA | NA | NA |
| WBGene00200967 | NA | NA | NA | NA |

| | | | | |
|---|---|---|---|---|
| WBGene00200968 | 0.020925399 | 0.84643798 | NA | NA |
| WBGene00200969 | NA | NA | NA | NA |
| WBGene00200970 | NA | NA | NA | NA |
| WBGene00200971 | NA | NA | NA | NA |
| WBGene00200972 | NA | NA | NA | NA |
| WBGene00200973 | NA | NA | NA | NA |
| WBGene00200974 | NA | NA | NA | NA |
| WBGene00200975 | NA | NA | NA | NA |
| WBGene00200976 | NA | NA | NA | NA |
| WBGene00200977 | 0.020925399 | 0.84643798 | NA | NA |
| WBGene00200978 | NA | NA | NA | NA |
| WBGene00200979 | NA | NA | NA | NA |
| WBGene00200980 | NA | NA | NA | NA |
| WBGene00200981 | NA | NA | NA | NA |
| WBGene00200982 | NA | NA | NA | NA |
| WBGene00200983 | NA | NA | NA | NA |
| WBGene00200984 | NA | NA | NA | NA |
| WBGene00200985 | NA | NA | NA | NA |
| WBGene00200986 | -0.022930767 | 0.78121253 | NA | NA |
| WBGene00200987 | NA | NA | NA | NA |
| WBGene00200988 | NA | NA | NA | NA |
| WBGene00200989 | NA | NA | NA | NA |
| WBGene00200990 | NA | NA | NA | NA |
| WBGene00200991 | NA | NA | NA | NA |
| WBGene00200992 | NA | NA | NA | NA |
| WBGene00200993 | NA | NA | NA | NA |
| WBGene00200994 | NA | NA | NA | NA |
| WBGene00200995 | NA | NA | NA | NA |
| WBGene00200996 | NA | NA | NA | NA |
| WBGene00200997 | NA | NA | NA | NA |
| WBGene00200998 | -0.014709656 | 0.88496452 | 0.93940914 | NA |
| WBGene00200999 | NA | NA | NA | NA |
| WBGene00201000 | NA | NA | NA | NA |
| WBGene00201001 | NA | NA | NA | NA |
| WBGene00201002 | NA | NA | NA | NA |
| WBGene00201003 | NA | NA | NA | NA |
| WBGene00201004 | NA | NA | NA | NA |
| WBGene00201005 | NA | NA | NA | NA |
| WBGene00201006 | NA | NA | NA | NA |
| WBGene00201007 | NA | NA | NA | NA |
| WBGene00201008 | NA | NA | NA | NA |
| WBGene00201009 | NA | NA | NA | NA |
| WBGene00201010 | NA | NA | NA | NA |

| | | | | |
|---|---|---|---|---|
| WBGene00201011 | NA | NA | NA | NA |
| WBGene00201012 | NA | NA | NA | NA |
| WBGene00201013 | NA | NA | NA | NA |
| WBGene00201014 | 0.020232312 | 0.84643798 | NA | NA |
| WBGene00201015 | NA | NA | NA | NA |
| WBGene00201016 | NA | NA | NA | NA |
| WBGene00201017 | NA | NA | NA | NA |
| WBGene00201018 | NA | NA | NA | NA |
| WBGene00201019 | NA | NA | NA | NA |
| WBGene00201020 | NA | NA | NA | NA |
| WBGene00201021 | NA | NA | NA | NA |
| WBGene00201022 | NA | NA | NA | NA |
| WBGene00201023 | NA | NA | NA | NA |
| WBGene00201024 | NA | NA | NA | NA |
| WBGene00201025 | NA | NA | NA | NA |
| WBGene00201026 | NA | NA | NA | NA |
| WBGene00201027 | NA | NA | NA | NA |
| WBGene00201028 | NA | NA | NA | NA |
| WBGene00201029 | NA | NA | NA | NA |
| WBGene00201030 | NA | NA | NA | NA |
| WBGene00201031 | 0.020925399 | 0.84643798 | NA | NA |
| WBGene00201032 | NA | NA | NA | NA |
| WBGene00201033 | NA | NA | NA | NA |
| WBGene00201034 | NA | NA | NA | NA |
| WBGene00201035 | NA | NA | NA | NA |
| WBGene00201036 | NA | NA | NA | NA |
| WBGene00201037 | NA | NA | NA | NA |
| WBGene00201038 | NA | NA | NA | NA |
| WBGene00201039 | NA | NA | NA | NA |
| WBGene00201040 | NA | NA | NA | NA |
| WBGene00201041 | NA | NA | NA | NA |
| WBGene00201042 | NA | NA | NA | NA |
| WBGene00201043 | NA | NA | NA | NA |
| WBGene00201044 | NA | NA | NA | NA |
| WBGene00201045 | NA | NA | NA | NA |
| WBGene00201046 | NA | NA | NA | NA |
| WBGene00201047 | 0.038125098 | 0.61119482 | NA | NA |
| WBGene00201048 | 0.020317245 | 0.84643798 | NA | NA |
| WBGene00201049 | NA | NA | NA | NA |
| WBGene00201050 | NA | NA | NA | NA |
| WBGene00201051 | NA | NA | NA | NA |
| WBGene00201052 | NA | NA | NA | NA |
| WBGene00201053 | NA | NA | NA | NA |

| | | | | |
|---|---|---|---|---|
| WBGene00201054 | NA | NA | NA | NA |
| WBGene00201055 | NA | NA | NA | NA |
| WBGene00201056 | NA | NA | NA | NA |
| WBGene00201057 | NA | NA | NA | NA |
| WBGene00201058 | NA | NA | NA | NA |
| WBGene00201059 | NA | NA | NA | NA |
| WBGene00201060 | NA | NA | NA | NA |
| WBGene00201061 | NA | NA | NA | NA |
| WBGene00201062 | NA | NA | NA | NA |
| WBGene00201063 | NA | NA | NA | NA |
| WBGene00201064 | 0.038125098 | 0.61119482 | NA | NA |
| WBGene00201065 | NA | NA | NA | NA |
| WBGene00201066 | NA | NA | NA | NA |
| WBGene00201067 | NA | NA | NA | NA |
| WBGene00201068 | NA | NA | NA | NA |
| WBGene00201069 | NA | NA | NA | NA |
| WBGene00201070 | NA | NA | NA | NA |
| WBGene00201071 | NA | NA | NA | NA |
| WBGene00201072 | NA | NA | NA | NA |
| WBGene00201073 | NA | NA | NA | NA |
| WBGene00201074 | NA | NA | NA | NA |
| WBGene00201075 | NA | NA | NA | NA |
| WBGene00201076 | NA | NA | NA | NA |
| WBGene00201077 | NA | NA | NA | NA |
| WBGene00201078 | NA | NA | NA | NA |
| WBGene00201079 | NA | NA | NA | NA |
| WBGene00201080 | NA | NA | NA | NA |
| WBGene00201081 | NA | NA | NA | NA |
| WBGene00201082 | NA | NA | NA | NA |
| WBGene00201083 | NA | NA | NA | NA |
| WBGene00201084 | NA | NA | NA | NA |
| WBGene00201085 | NA | NA | NA | NA |
| WBGene00201086 | NA | NA | NA | NA |
| WBGene00201087 | NA | NA | NA | NA |
| WBGene00201088 | NA | NA | NA | NA |
| WBGene00201089 | NA | NA | NA | NA |
| WBGene00201090 | NA | NA | NA | NA |
| WBGene00201091 | NA | NA | NA | NA |
| WBGene00201092 | NA | NA | NA | NA |
| WBGene00201093 | NA | NA | NA | NA |
| WBGene00201094 | NA | NA | NA | NA |
| WBGene00201095 | NA | NA | NA | NA |
| WBGene00201096 | NA | NA | NA | NA |

| | | | | |
|---|---|---|---|---|
| WBGene00201097 | NA | NA | NA | NA |
| WBGene00201098 | NA | NA | NA | NA |
| WBGene00201099 | NA | NA | NA | NA |
| WBGene00201100 | NA | NA | NA | NA |
| WBGene00201101 | NA | NA | NA | NA |
| WBGene00201102 | NA | NA | NA | NA |
| WBGene00201103 | NA | NA | NA | NA |
| WBGene00201104 | NA | NA | NA | NA |
| WBGene00201105 | NA | NA | NA | NA |
| WBGene00201106 | NA | NA | NA | NA |
| WBGene00201107 | NA | NA | NA | NA |
| WBGene00201108 | NA | NA | NA | NA |
| WBGene00201109 | NA | NA | NA | NA |
| WBGene00201110 | NA | NA | NA | NA |
| WBGene00201111 | NA | NA | NA | NA |
| WBGene00201112 | 0.020232312 | 0.84643798 | NA | NA |
| WBGene00201113 | NA | NA | NA | NA |
| WBGene00201114 | NA | NA | NA | NA |
| WBGene00201115 | NA | NA | NA | NA |
| WBGene00201116 | NA | NA | NA | NA |
| WBGene00201117 | NA | NA | NA | NA |
| WBGene00201118 | NA | NA | NA | NA |
| WBGene00201119 | NA | NA | NA | NA |
| WBGene00201120 | NA | NA | NA | NA |
| WBGene00201121 | NA | NA | NA | NA |
| WBGene00201122 | NA | NA | NA | NA |
| WBGene00201123 | NA | NA | NA | NA |
| WBGene00201124 | NA | NA | NA | NA |
| WBGene00201125 | NA | NA | NA | NA |
| WBGene00201126 | NA | NA | NA | NA |
| WBGene00201127 | NA | NA | NA | NA |
| WBGene00201128 | NA | NA | NA | NA |
| WBGene00201129 | NA | NA | NA | NA |
| WBGene00201130 | NA | NA | NA | NA |
| WBGene00201131 | NA | NA | NA | NA |
| WBGene00201132 | NA | NA | NA | NA |
| WBGene00201133 | NA | NA | NA | NA |
| WBGene00201134 | -0.045959069 | 0.50707733 | NA | NA |
| WBGene00201135 | NA | NA | NA | NA |
| WBGene00201136 | NA | NA | NA | NA |
| WBGene00201137 | NA | NA | NA | NA |
| WBGene00201138 | NA | NA | NA | NA |
| WBGene00201139 | NA | NA | NA | NA |

| | | | | |
|---|---|---|---|---|
| WBGene00201140 | NA | NA | NA | NA |
| WBGene00201141 | -0.022930767 | 0.78121253 | NA | NA |
| WBGene00201142 | NA | NA | NA | NA |
| WBGene00201143 | 0.020317245 | 0.84643798 | NA | NA |
| WBGene00201145 | NA | NA | NA | NA |
| WBGene00201146 | NA | NA | NA | NA |
| WBGene00201147 | NA | NA | NA | NA |
| WBGene00201148 | NA | NA | NA | NA |
| WBGene00201149 | NA | NA | NA | NA |
| WBGene00201150 | NA | NA | NA | NA |
| WBGene00201151 | NA | NA | NA | NA |
| WBGene00201152 | NA | NA | NA | NA |
| WBGene00201153 | 0.0398788 | 0.58971221 | NA | NA |
| WBGene00201154 | NA | NA | NA | NA |
| WBGene00201155 | NA | NA | NA | NA |
| WBGene00201156 | NA | NA | NA | NA |
| WBGene00201157 | NA | NA | NA | NA |
| WBGene00201158 | NA | NA | NA | NA |
| WBGene00201159 | NA | NA | NA | NA |
| WBGene00201160 | NA | NA | NA | NA |
| WBGene00201161 | NA | NA | NA | NA |
| WBGene00201162 | NA | NA | NA | NA |
| WBGene00201163 | NA | NA | NA | NA |
| WBGene00201164 | NA | NA | NA | NA |
| WBGene00201165 | NA | NA | NA | NA |
| WBGene00201166 | NA | NA | NA | NA |
| WBGene00201167 | NA | NA | NA | NA |
| WBGene00201168 | 0.020232312 | 0.84643798 | NA | NA |
| WBGene00201169 | NA | NA | NA | NA |
| WBGene00201170 | NA | NA | NA | NA |
| WBGene00201171 | NA | NA | NA | NA |
| WBGene00201172 | NA | NA | NA | NA |
| WBGene00201173 | NA | NA | NA | NA |
| WBGene00201174 | NA | NA | NA | NA |
| WBGene00201175 | NA | NA | NA | NA |
| WBGene00201176 | 0.020232312 | 0.84643798 | NA | NA |
| WBGene00201177 | NA | NA | NA | NA |
| WBGene00201178 | NA | NA | NA | NA |
| WBGene00201179 | NA | NA | NA | NA |
| WBGene00201180 | NA | NA | NA | NA |
| WBGene00201181 | NA | NA | NA | NA |
| WBGene00201182 | NA | NA | NA | NA |
| WBGene00201183 | NA | NA | NA | NA |

| | | | | |
|---|---|---|---|---|
| WBGene00201184 | NA | NA | NA | NA |
| WBGene00201185 | NA | NA | NA | NA |
| WBGene00201186 | NA | NA | NA | NA |
| WBGene00201187 | NA | NA | NA | NA |
| WBGene00201188 | NA | NA | NA | NA |
| WBGene00201189 | NA | NA | NA | NA |
| WBGene00201190 | NA | NA | NA | NA |
| WBGene00201191 | NA | NA | NA | NA |
| WBGene00201192 | NA | NA | NA | NA |
| WBGene00201193 | NA | NA | NA | NA |
| WBGene00201194 | NA | NA | NA | NA |
| WBGene00201195 | NA | NA | NA | NA |
| WBGene00201196 | NA | NA | NA | NA |
| WBGene00201197 | NA | NA | NA | NA |
| WBGene00201198 | NA | NA | NA | NA |
| WBGene00201199 | NA | NA | NA | NA |
| WBGene00201200 | NA | NA | NA | NA |
| WBGene00201201 | NA | NA | NA | NA |
| WBGene00201202 | -0.003270206 | 0.95698316 | NA | NA |
| WBGene00201203 | 0.020925399 | 0.84643798 | NA | NA |
| WBGene00201204 | -0.022930767 | 0.78121253 | NA | NA |
| WBGene00201205 | NA | NA | NA | NA |
| WBGene00201206 | NA | NA | NA | NA |
| WBGene00201207 | 0.020317245 | 0.84643798 | NA | NA |
| WBGene00201208 | NA | NA | NA | NA |
| WBGene00201209 | NA | NA | NA | NA |
| WBGene00201210 | NA | NA | NA | NA |
| WBGene00201211 | NA | NA | NA | NA |
| WBGene00201212 | NA | NA | NA | NA |
| WBGene00201213 | NA | NA | NA | NA |
| WBGene00201214 | NA | NA | NA | NA |
| WBGene00201215 | NA | NA | NA | NA |
| WBGene00201216 | NA | NA | NA | NA |
| WBGene00201217 | NA | NA | NA | NA |
| WBGene00201218 | NA | NA | NA | NA |
| WBGene00201219 | 0.020317245 | 0.84643798 | NA | NA |
| WBGene00201220 | NA | NA | NA | NA |
| WBGene00201221 | NA | NA | NA | NA |
| WBGene00201222 | NA | NA | NA | NA |
| WBGene00201223 | NA | NA | NA | NA |
| WBGene00201224 | NA | NA | NA | NA |
| WBGene00201225 | NA | NA | NA | NA |
| WBGene00201226 | NA | NA | NA | NA |

| | | | | |
|---|---|---|---|---|
| WBGene00201227 | NA | NA | NA | NA |
| WBGene00201228 | NA | NA | NA | NA |
| WBGene00201229 | NA | NA | NA | NA |
| WBGene00201230 | NA | NA | NA | NA |
| WBGene00201231 | NA | NA | NA | NA |
| WBGene00201232 | NA | NA | NA | NA |
| WBGene00201233 | NA | NA | NA | NA |
| WBGene00201234 | NA | NA | NA | NA |
| WBGene00201235 | NA | NA | NA | NA |
| WBGene00201236 | NA | NA | NA | NA |
| WBGene00201237 | NA | NA | NA | NA |
| WBGene00201238 | NA | NA | NA | NA |
| WBGene00201239 | NA | NA | NA | NA |
| WBGene00201240 | NA | NA | NA | NA |
| WBGene00201241 | 0.020317245 | 0.84643798 | NA | NA |
| WBGene00201242 | NA | NA | NA | NA |
| WBGene00201243 | NA | NA | NA | NA |
| WBGene00201244 | NA | NA | NA | NA |
| WBGene00201245 | NA | NA | NA | NA |
| WBGene00201246 | -0.023109762 | 0.78121253 | NA | NA |
| WBGene00201247 | NA | NA | NA | NA |
| WBGene00201248 | NA | NA | NA | NA |
| WBGene00201249 | NA | NA | NA | NA |
| WBGene00201250 | NA | NA | NA | NA |
| WBGene00201251 | NA | NA | NA | NA |
| WBGene00201252 | NA | NA | NA | NA |
| WBGene00201253 | 0.020232312 | 0.84643798 | NA | NA |
| WBGene00201254 | NA | NA | NA | NA |
| WBGene00201255 | NA | NA | NA | NA |
| WBGene00201256 | NA | NA | NA | NA |
| WBGene00201257 | NA | NA | NA | NA |
| WBGene00201258 | NA | NA | NA | NA |
| WBGene00201259 | NA | NA | NA | NA |
| WBGene00201260 | NA | NA | NA | NA |
| WBGene00201261 | NA | NA | NA | NA |
| WBGene00201262 | NA | NA | NA | NA |
| WBGene00201263 | NA | NA | NA | NA |
| WBGene00201264 | NA | NA | NA | NA |
| WBGene00201265 | NA | NA | NA | NA |
| WBGene00201266 | NA | NA | NA | NA |
| WBGene00201267 | NA | NA | NA | NA |
| WBGene00201268 | NA | NA | NA | NA |
| WBGene00201269 | NA | NA | NA | NA |

| | | | | |
|---|---|---|---|---|
| WBGene00201270 | NA | NA | NA | NA |
| WBGene00201271 | NA | NA | NA | NA |
| WBGene00201272 | NA | NA | NA | NA |
| WBGene00201273 | NA | NA | NA | NA |
| WBGene00201274 | NA | NA | NA | NA |
| WBGene00201275 | NA | NA | NA | NA |
| WBGene00201276 | NA | NA | NA | NA |
| WBGene00201277 | NA | NA | NA | NA |
| WBGene00201278 | NA | NA | NA | NA |
| WBGene00201279 | NA | NA | NA | NA |
| WBGene00201280 | NA | NA | NA | NA |
| WBGene00201281 | NA | NA | NA | NA |
| WBGene00201282 | NA | NA | NA | NA |
| WBGene00201283 | NA | NA | NA | NA |
| WBGene00201284 | NA | NA | NA | NA |
| WBGene00201285 | NA | NA | NA | NA |
| WBGene00201286 | NA | NA | NA | NA |
| WBGene00201287 | NA | NA | NA | NA |
| WBGene00201288 | NA | NA | NA | NA |
| WBGene00201289 | NA | NA | NA | NA |
| WBGene00201290 | 0.020232312 | 0.84643798 | NA | NA |
| WBGene00201291 | -0.023109762 | 0.78121253 | NA | NA |
| WBGene00201292 | NA | NA | NA | NA |
| WBGene00201293 | NA | NA | NA | NA |
| WBGene00201294 | 0.038125098 | 0.61119482 | NA | NA |
| WBGene00201295 | NA | NA | NA | NA |
| WBGene00201296 | NA | NA | NA | NA |
| WBGene00201297 | NA | NA | NA | NA |
| WBGene00201298 | NA | NA | NA | NA |
| WBGene00201299 | NA | NA | NA | NA |
| WBGene00201300 | NA | NA | NA | NA |
| WBGene00201301 | NA | NA | NA | NA |
| WBGene00201302 | NA | NA | NA | NA |
| WBGene00201303 | NA | NA | NA | NA |
| WBGene00201304 | NA | NA | NA | NA |
| WBGene00201305 | NA | NA | NA | NA |
| WBGene00201306 | NA | NA | NA | NA |
| WBGene00201307 | NA | NA | NA | NA |
| WBGene00201308 | 0.020317245 | 0.84643798 | NA | NA |
| WBGene00201309 | NA | NA | NA | NA |
| WBGene00201310 | NA | NA | NA | NA |
| WBGene00201311 | NA | NA | NA | NA |
| WBGene00201312 | NA | NA | NA | NA |

| | | | | |
|---|---|---|---|---|
| WBGene00201313 | NA | NA | NA | NA |
| WBGene00201314 | NA | NA | NA | NA |
| WBGene00201315 | NA | NA | NA | NA |
| WBGene00201316 | NA | NA | NA | NA |
| WBGene00201317 | NA | NA | NA | NA |
| WBGene00201318 | NA | NA | NA | NA |
| WBGene00201319 | NA | NA | NA | NA |
| WBGene00201320 | NA | NA | NA | NA |
| WBGene00201321 | NA | NA | NA | NA |
| WBGene00201322 | NA | NA | NA | NA |
| WBGene00201323 | NA | NA | NA | NA |
| WBGene00201324 | NA | NA | NA | NA |
| WBGene00201325 | NA | NA | NA | NA |
| WBGene00201326 | NA | NA | NA | NA |
| WBGene00201327 | NA | NA | NA | NA |
| WBGene00201328 | NA | NA | NA | NA |
| WBGene00201329 | NA | NA | NA | NA |
| WBGene00201330 | NA | NA | NA | NA |
| WBGene00201331 | NA | NA | NA | NA |
| WBGene00201332 | NA | NA | NA | NA |
| WBGene00201333 | NA | NA | NA | NA |
| WBGene00201334 | NA | NA | NA | NA |
| WBGene00201335 | NA | NA | NA | NA |
| WBGene00201336 | NA | NA | NA | NA |
| WBGene00201337 | NA | NA | NA | NA |
| WBGene00201338 | NA | NA | NA | NA |
| WBGene00201339 | NA | NA | NA | NA |
| WBGene00201340 | NA | NA | NA | NA |
| WBGene00201341 | NA | NA | NA | NA |
| WBGene00201342 | NA | NA | NA | NA |
| WBGene00201343 | NA | NA | NA | NA |
| WBGene00201344 | NA | NA | NA | NA |
| WBGene00201345 | NA | NA | NA | NA |
| WBGene00201346 | NA | NA | NA | NA |
| WBGene00201347 | NA | NA | NA | NA |
| WBGene00201348 | NA | NA | NA | NA |
| WBGene00201349 | NA | NA | NA | NA |
| WBGene00201350 | NA | NA | NA | NA |
| WBGene00201351 | NA | NA | NA | NA |
| WBGene00201352 | NA | NA | NA | NA |
| WBGene00201353 | NA | NA | NA | NA |
| WBGene00201354 | NA | NA | NA | NA |
| WBGene00201355 | NA | NA | NA | NA |

| | | | | |
|---|---|---|---|---|
| WBGene00201356 | NA | NA | NA | NA |
| WBGene00201357 | NA | NA | NA | NA |
| WBGene00201358 | NA | NA | NA | NA |
| WBGene00201359 | NA | NA | NA | NA |
| WBGene00201360 | NA | NA | NA | NA |
| WBGene00201361 | NA | NA | NA | NA |
| WBGene00201362 | NA | NA | NA | NA |
| WBGene00201363 | NA | NA | NA | NA |
| WBGene00201364 | NA | NA | NA | NA |
| WBGene00201365 | NA | NA | NA | NA |
| WBGene00201366 | NA | NA | NA | NA |
| WBGene00201367 | NA | NA | NA | NA |
| WBGene00201368 | NA | NA | NA | NA |
| WBGene00201369 | NA | NA | NA | NA |
| WBGene00201370 | NA | NA | NA | NA |
| WBGene00201371 | NA | NA | NA | NA |
| WBGene00201372 | -0.023056499 | 0.78121253 | NA | NA |
| WBGene00201373 | NA | NA | NA | NA |
| WBGene00201374 | NA | NA | NA | NA |
| WBGene00201375 | NA | NA | NA | NA |
| WBGene00201376 | NA | NA | NA | NA |
| WBGene00201377 | NA | NA | NA | NA |
| WBGene00201378 | NA | NA | NA | NA |
| WBGene00201379 | NA | NA | NA | NA |
| WBGene00201380 | NA | NA | NA | NA |
| WBGene00201381 | NA | NA | NA | NA |
| WBGene00201382 | NA | NA | NA | NA |
| WBGene00201383 | NA | NA | NA | NA |
| WBGene00201384 | NA | NA | NA | NA |
| WBGene00201385 | NA | NA | NA | NA |
| WBGene00201386 | NA | NA | NA | NA |
| WBGene00201387 | NA | NA | NA | NA |
| WBGene00201388 | NA | NA | NA | NA |
| WBGene00201389 | NA | NA | NA | NA |
| WBGene00201390 | NA | NA | NA | NA |
| WBGene00201391 | NA | NA | NA | NA |
| WBGene00201392 | NA | NA | NA | NA |
| WBGene00201393 | NA | NA | NA | NA |
| WBGene00201394 | NA | NA | NA | NA |
| WBGene00201395 | NA | NA | NA | NA |
| WBGene00201396 | -0.046394786 | 0.50231372 | NA | NA |
| WBGene00201397 | NA | NA | NA | NA |
| WBGene00201398 | NA | NA | NA | NA |

| | | | | |
|---|---|---|---|---|
| WBGene00201399 | NA | NA | NA | NA |
| WBGene00201400 | NA | NA | NA | NA |
| WBGene00201401 | NA | NA | NA | NA |
| WBGene00201402 | NA | NA | NA | NA |
| WBGene00201403 | NA | NA | NA | NA |
| WBGene00201404 | NA | NA | NA | NA |
| WBGene00201405 | NA | NA | NA | NA |
| WBGene00201406 | NA | NA | NA | NA |
| WBGene00201407 | NA | NA | NA | NA |
| WBGene00201408 | NA | NA | NA | NA |
| WBGene00201409 | NA | NA | NA | NA |
| WBGene00201410 | NA | NA | NA | NA |
| WBGene00201411 | NA | NA | NA | NA |
| WBGene00201412 | NA | NA | NA | NA |
| WBGene00201413 | NA | NA | NA | NA |
| WBGene00201414 | NA | NA | NA | NA |
| WBGene00201415 | NA | NA | NA | NA |
| WBGene00201416 | NA | NA | NA | NA |
| WBGene00201417 | NA | NA | NA | NA |
| WBGene00201418 | NA | NA | NA | NA |
| WBGene00201419 | NA | NA | NA | NA |
| WBGene00201420 | NA | NA | NA | NA |
| WBGene00201421 | NA | NA | NA | NA |
| WBGene00201422 | NA | NA | NA | NA |
| WBGene00201423 | -0.023056499 | 0.78121253 | NA | NA |
| WBGene00201424 | NA | NA | NA | NA |
| WBGene00201425 | NA | NA | NA | NA |
| WBGene00201426 | NA | NA | NA | NA |
| WBGene00201427 | NA | NA | NA | NA |
| WBGene00201428 | NA | NA | NA | NA |
| WBGene00201429 | NA | NA | NA | NA |
| WBGene00201430 | NA | NA | NA | NA |
| WBGene00201431 | NA | NA | NA | NA |
| WBGene00201432 | NA | NA | NA | NA |
| WBGene00201433 | NA | NA | NA | NA |
| WBGene00201434 | NA | NA | NA | NA |
| WBGene00201435 | NA | NA | NA | NA |
| WBGene00201436 | NA | NA | NA | NA |
| WBGene00201437 | NA | NA | NA | NA |
| WBGene00201438 | NA | NA | NA | NA |
| WBGene00201439 | NA | NA | NA | NA |
| WBGene00201440 | NA | NA | NA | NA |
| WBGene00201441 | NA | NA | NA | NA |

| | | | | |
|---|---|---|---|---|
| WBGene00201442 | NA | NA | NA | NA |
| WBGene00201443 | NA | NA | NA | NA |
| WBGene00201444 | NA | NA | NA | NA |
| WBGene00201445 | NA | NA | NA | NA |
| WBGene00201446 | NA | NA | NA | NA |
| WBGene00201447 | NA | NA | NA | NA |
| WBGene00201448 | NA | NA | NA | NA |
| WBGene00201449 | NA | NA | NA | NA |
| WBGene00201450 | NA | NA | NA | NA |
| WBGene00201451 | NA | NA | NA | NA |
| WBGene00201452 | NA | NA | NA | NA |
| WBGene00201453 | NA | NA | NA | NA |
| WBGene00201454 | NA | NA | NA | NA |
| WBGene00201455 | NA | NA | NA | NA |
| WBGene00201456 | NA | NA | NA | NA |
| WBGene00201457 | NA | NA | NA | NA |
| WBGene00201458 | -0.022930767 | 0.78121253 | NA | NA |
| WBGene00201459 | NA | NA | NA | NA |
| WBGene00201460 | NA | NA | NA | NA |
| WBGene00201461 | NA | NA | NA | NA |
| WBGene00201462 | NA | NA | NA | NA |
| WBGene00201463 | NA | NA | NA | NA |
| WBGene00201464 | NA | NA | NA | NA |
| WBGene00201465 | NA | NA | NA | NA |
| WBGene00201466 | 0.015899775 | 0.84174438 | NA | NA |
| WBGene00201467 | NA | NA | NA | NA |
| WBGene00201468 | NA | NA | NA | NA |
| WBGene00201469 | NA | NA | NA | NA |
| WBGene00201470 | NA | NA | NA | NA |
| WBGene00201471 | NA | NA | NA | NA |
| WBGene00201472 | NA | NA | NA | NA |
| WBGene00201473 | NA | NA | NA | NA |
| WBGene00201474 | NA | NA | NA | NA |
| WBGene00201475 | NA | NA | NA | NA |
| WBGene00201476 | NA | NA | NA | NA |
| WBGene00201477 | NA | NA | NA | NA |
| WBGene00201478 | NA | NA | NA | NA |
| WBGene00201479 | NA | NA | NA | NA |
| WBGene00201480 | NA | NA | NA | NA |
| WBGene00201481 | NA | NA | NA | NA |
| WBGene00201482 | NA | NA | NA | NA |
| WBGene00201483 | NA | NA | NA | NA |
| WBGene00201484 | NA | NA | NA | NA |

| | | | | |
|---|---|---|---|---|
| WBGene00201485 | NA | NA | NA | NA |
| WBGene00201486 | NA | NA | NA | NA |
| WBGene00201487 | NA | NA | NA | NA |
| WBGene00201488 | NA | NA | NA | NA |
| WBGene00201489 | NA | NA | NA | NA |
| WBGene00201490 | NA | NA | NA | NA |
| WBGene00201491 | NA | NA | NA | NA |
| WBGene00201492 | NA | NA | NA | NA |
| WBGene00201493 | NA | NA | NA | NA |
| WBGene00201494 | NA | NA | NA | NA |
| WBGene00201495 | NA | NA | NA | NA |
| WBGene00201496 | NA | NA | NA | NA |
| WBGene00201497 | NA | NA | NA | NA |
| WBGene00201498 | NA | NA | NA | NA |
| WBGene00201499 | NA | NA | NA | NA |
| WBGene00201500 | NA | NA | NA | NA |
| WBGene00201501 | NA | NA | NA | NA |
| WBGene00201502 | NA | NA | NA | NA |
| WBGene00201503 | -0.001842784 | 0.95621055 | NA | NA |
| WBGene00201504 | NA | NA | NA | NA |
| WBGene00201505 | NA | NA | NA | NA |
| WBGene00201506 | NA | NA | NA | NA |
| WBGene00201507 | NA | NA | NA | NA |
| WBGene00201508 | NA | NA | NA | NA |
| WBGene00201509 | NA | NA | NA | NA |
| WBGene00201510 | NA | NA | NA | NA |
| WBGene00201511 | NA | NA | NA | NA |
| WBGene00201512 | NA | NA | NA | NA |
| WBGene00201513 | NA | NA | NA | NA |
| WBGene00201514 | NA | NA | NA | NA |
| WBGene00201515 | NA | NA | NA | NA |
| WBGene00201516 | NA | NA | NA | NA |
| WBGene00201517 | NA | NA | NA | NA |
| WBGene00201518 | NA | NA | NA | NA |
| WBGene00201519 | NA | NA | NA | NA |
| WBGene00201520 | NA | NA | NA | NA |
| WBGene00201521 | NA | NA | NA | NA |
| WBGene00201522 | NA | NA | NA | NA |
| WBGene00201523 | 0.0398788 | 0.58971221 | NA | NA |
| WBGene00201524 | NA | NA | NA | NA |
| WBGene00201525 | NA | NA | NA | NA |
| WBGene00201526 | NA | NA | NA | NA |
| WBGene00201527 | NA | NA | NA | NA |

| | | | | |
|---|---|---|---|---|
| WBGene00201528 | NA | NA | NA | NA |
| WBGene00201529 | NA | NA | NA | NA |
| WBGene00201530 | NA | NA | NA | NA |
| WBGene00201531 | NA | NA | NA | NA |
| WBGene00201532 | NA | NA | NA | NA |
| WBGene00201533 | NA | NA | NA | NA |
| WBGene00201534 | NA | NA | NA | NA |
| WBGene00201535 | NA | NA | NA | NA |
| WBGene00201536 | NA | NA | NA | NA |
| WBGene00201537 | NA | NA | NA | NA |
| WBGene00201538 | -0.00346345 | 0.95708614 | NA | NA |
| WBGene00201539 | 0.020925399 | 0.84643798 | NA | NA |
| WBGene00201540 | NA | NA | NA | NA |
| WBGene00201541 | NA | NA | NA | NA |
| WBGene00201542 | NA | NA | NA | NA |
| WBGene00201543 | NA | NA | NA | NA |
| WBGene00201544 | NA | NA | NA | NA |
| WBGene00201545 | NA | NA | NA | NA |
| WBGene00201546 | NA | NA | NA | NA |
| WBGene00201547 | NA | NA | NA | NA |
| WBGene00201548 | NA | NA | NA | NA |
| WBGene00201549 | NA | NA | NA | NA |
| WBGene00201550 | NA | NA | NA | NA |
| WBGene00201551 | NA | NA | NA | NA |
| WBGene00201552 | NA | NA | NA | NA |
| WBGene00201553 | NA | NA | NA | NA |
| WBGene00201554 | NA | NA | NA | NA |
| WBGene00201555 | NA | NA | NA | NA |
| WBGene00201556 | NA | NA | NA | NA |
| WBGene00201557 | NA | NA | NA | NA |
| WBGene00201558 | NA | NA | NA | NA |
| WBGene00201559 | NA | NA | NA | NA |
| WBGene00201560 | NA | NA | NA | NA |
| WBGene00201561 | NA | NA | NA | NA |
| WBGene00201562 | NA | NA | NA | NA |
| WBGene00201563 | NA | NA | NA | NA |
| WBGene00201564 | NA | NA | NA | NA |
| WBGene00201565 | NA | NA | NA | NA |
| WBGene00201566 | NA | NA | NA | NA |
| WBGene00201567 | NA | NA | NA | NA |
| WBGene00201568 | NA | NA | NA | NA |
| WBGene00201569 | NA | NA | NA | NA |
| WBGene00201570 | NA | NA | NA | NA |

| | | | | |
|---|---|---|---|---|
| WBGene00201571 | NA | NA | NA | NA |
| WBGene00201572 | NA | NA | NA | NA |
| WBGene00201573 | NA | NA | NA | NA |
| WBGene00201574 | NA | NA | NA | NA |
| WBGene00201575 | NA | NA | NA | NA |
| WBGene00201576 | NA | NA | NA | NA |
| WBGene00201577 | NA | NA | NA | NA |
| WBGene00201578 | NA | NA | NA | NA |
| WBGene00201579 | NA | NA | NA | NA |
| WBGene00201580 | NA | NA | NA | NA |
| WBGene00201581 | NA | NA | NA | NA |
| WBGene00201582 | -0.022930767 | 0.78121253 | NA | NA |
| WBGene00201583 | NA | NA | NA | NA |
| WBGene00201584 | NA | NA | NA | NA |
| WBGene00201585 | NA | NA | NA | NA |
| WBGene00201586 | NA | NA | NA | NA |
| WBGene00201587 | NA | NA | NA | NA |
| WBGene00201588 | NA | NA | NA | NA |
| WBGene00201589 | -0.04044783 | 0.53612608 | NA | NA |
| WBGene00201590 | NA | NA | NA | NA |
| WBGene00201591 | -0.023056499 | 0.78121253 | NA | NA |
| WBGene00201592 | NA | NA | NA | NA |
| WBGene00201593 | 0.020925399 | 0.84643798 | NA | NA |
| WBGene00201594 | -0.023056499 | 0.78121253 | NA | NA |
| WBGene00201595 | NA | NA | NA | NA |
| WBGene00201596 | NA | NA | NA | NA |
| WBGene00201597 | NA | NA | NA | NA |
| WBGene00201598 | NA | NA | NA | NA |
| WBGene00201599 | NA | NA | NA | NA |
| WBGene00201600 | NA | NA | NA | NA |
| WBGene00201601 | NA | NA | NA | NA |
| WBGene00201602 | NA | NA | NA | NA |
| WBGene00201603 | NA | NA | NA | NA |
| WBGene00201604 | NA | NA | NA | NA |
| WBGene00201605 | NA | NA | NA | NA |
| WBGene00201606 | NA | NA | NA | NA |
| WBGene00201607 | NA | NA | NA | NA |
| WBGene00201608 | NA | NA | NA | NA |
| WBGene00201609 | NA | NA | NA | NA |
| WBGene00201610 | NA | NA | NA | NA |
| WBGene00201611 | NA | NA | NA | NA |
| WBGene00201612 | NA | NA | NA | NA |
| WBGene00201613 | NA | NA | NA | NA |

| | | | | |
|---|---|---|---|---|
| WBGene00201614 | NA | NA | NA | NA |
| WBGene00201615 | NA | NA | NA | NA |
| WBGene00201616 | NA | NA | NA | NA |
| WBGene00201617 | NA | NA | NA | NA |
| WBGene00201618 | NA | NA | NA | NA |
| WBGene00201619 | NA | NA | NA | NA |
| WBGene00201620 | -0.022930767 | 0.78121253 | NA | NA |
| WBGene00201621 | NA | NA | NA | NA |
| WBGene00201622 | NA | NA | NA | NA |
| WBGene00201623 | NA | NA | NA | NA |
| WBGene00201624 | NA | NA | NA | NA |
| WBGene00201625 | 0.020925399 | 0.84643798 | NA | NA |
| WBGene00201626 | NA | NA | NA | NA |
| WBGene00201627 | NA | NA | NA | NA |
| WBGene00201628 | NA | NA | NA | NA |
| WBGene00201629 | NA | NA | NA | NA |
| WBGene00201630 | NA | NA | NA | NA |
| WBGene00201631 | NA | NA | NA | NA |
| WBGene00201632 | NA | NA | NA | NA |
| WBGene00201633 | NA | NA | NA | NA |
| WBGene00201634 | -0.023056499 | 0.78121253 | NA | NA |
| WBGene00201636 | NA | NA | NA | NA |
| WBGene00201637 | NA | NA | NA | NA |
| WBGene00201638 | 0.020925399 | 0.84643798 | NA | NA |
| WBGene00201639 | 0.020925399 | 0.84643798 | NA | NA |
| WBGene00201640 | -0.075452738 | 0.28904131 | NA | NA |
| WBGene00201641 | NA | NA | NA | NA |
| WBGene00201642 | NA | NA | NA | NA |
| WBGene00201643 | NA | NA | NA | NA |
| WBGene00201644 | NA | NA | NA | NA |
| WBGene00201645 | NA | NA | NA | NA |
| WBGene00201646 | NA | NA | NA | NA |
| WBGene00201647 | NA | NA | NA | NA |
| WBGene00201648 | NA | NA | NA | NA |
| WBGene00201649 | NA | NA | NA | NA |
| WBGene00201650 | NA | NA | NA | NA |
| WBGene00201651 | NA | NA | NA | NA |
| WBGene00201652 | NA | NA | NA | NA |
| WBGene00201653 | NA | NA | NA | NA |
| WBGene00201654 | NA | NA | NA | NA |
| WBGene00201655 | NA | NA | NA | NA |
| WBGene00201656 | NA | NA | NA | NA |
| WBGene00201657 | NA | NA | NA | NA |

| | | | | |
|---|---|---|---|---|
| WBGene00201658 | NA | NA | NA | NA |
| WBGene00201659 | NA | NA | NA | NA |
| WBGene00201660 | NA | NA | NA | NA |
| WBGene00201661 | NA | NA | NA | NA |
| WBGene00201662 | NA | NA | NA | NA |
| WBGene00201663 | NA | NA | NA | NA |
| WBGene00201664 | NA | NA | NA | NA |
| WBGene00201665 | NA | NA | NA | NA |
| WBGene00201666 | NA | NA | NA | NA |
| WBGene00201667 | NA | NA | NA | NA |
| WBGene00201668 | NA | NA | NA | NA |
| WBGene00201669 | NA | NA | NA | NA |
| WBGene00201670 | 0.017631418 | 0.81668032 | NA | NA |
| WBGene00201671 | NA | NA | NA | NA |
| WBGene00201672 | NA | NA | NA | NA |
| WBGene00201673 | -0.023109762 | 0.78121253 | NA | NA |
| WBGene00201674 | NA | NA | NA | NA |
| WBGene00201675 | NA | NA | NA | NA |
| WBGene00201676 | NA | NA | NA | NA |
| WBGene00201677 | NA | NA | NA | NA |
| WBGene00201678 | 0.020317245 | 0.84643798 | NA | NA |
| WBGene00201679 | NA | NA | NA | NA |
| WBGene00201680 | NA | NA | NA | NA |
| WBGene00201681 | NA | NA | NA | NA |
| WBGene00201682 | NA | NA | NA | NA |
| WBGene00201683 | 0.020925399 | 0.84643798 | NA | NA |
| WBGene00201684 | NA | NA | NA | NA |
| WBGene00201685 | NA | NA | NA | NA |
| WBGene00201686 | NA | NA | NA | NA |
| WBGene00201687 | NA | NA | NA | NA |
| WBGene00201688 | NA | NA | NA | NA |
| WBGene00201689 | NA | NA | NA | NA |
| WBGene00201690 | NA | NA | NA | NA |
| WBGene00201691 | NA | NA | NA | NA |
| WBGene00201692 | NA | NA | NA | NA |
| WBGene00201693 | NA | NA | NA | NA |
| WBGene00201694 | NA | NA | NA | NA |
| WBGene00201695 | NA | NA | NA | NA |
| WBGene00201696 | -0.023109762 | 0.78121253 | NA | NA |
| WBGene00201697 | NA | NA | NA | NA |
| WBGene00201698 | NA | NA | NA | NA |
| WBGene00201699 | NA | NA | NA | NA |
| WBGene00201700 | NA | NA | NA | NA |

| | | | | |
|---|---|---|---|---|
| WBGene00201701 | NA | NA | NA | NA |
| WBGene00201702 | NA | NA | NA | NA |
| WBGene00201703 | NA | NA | NA | NA |
| WBGene00201704 | NA | NA | NA | NA |
| WBGene00201705 | NA | NA | NA | NA |
| WBGene00201706 | NA | NA | NA | NA |
| WBGene00201707 | NA | NA | NA | NA |
| WBGene00201708 | NA | NA | NA | NA |
| WBGene00201709 | NA | NA | NA | NA |
| WBGene00201710 | NA | NA | NA | NA |
| WBGene00201711 | NA | NA | NA | NA |
| WBGene00201712 | NA | NA | NA | NA |
| WBGene00201713 | NA | NA | NA | NA |
| WBGene00201714 | NA | NA | NA | NA |
| WBGene00201715 | NA | NA | NA | NA |
| WBGene00201716 | NA | NA | NA | NA |
| WBGene00201717 | 0.020925399 | 0.84643798 | NA | NA |
| WBGene00201718 | NA | NA | NA | NA |
| WBGene00201719 | NA | NA | NA | NA |
| WBGene00201720 | NA | NA | NA | NA |
| WBGene00201721 | NA | NA | NA | NA |
| WBGene00201722 | NA | NA | NA | NA |
| WBGene00201723 | NA | NA | NA | NA |
| WBGene00201724 | NA | NA | NA | NA |
| WBGene00201725 | NA | NA | NA | NA |
| WBGene00201726 | NA | NA | NA | NA |
| WBGene00201727 | NA | NA | NA | NA |
| WBGene00201728 | NA | NA | NA | NA |
| WBGene00201729 | 0.056411843 | 0.41891328 | NA | NA |
| WBGene00201730 | NA | NA | NA | NA |
| WBGene00201731 | NA | NA | NA | NA |
| WBGene00201732 | NA | NA | NA | NA |
| WBGene00201733 | NA | NA | NA | NA |
| WBGene00201734 | NA | NA | NA | NA |
| WBGene00201735 | NA | NA | NA | NA |
| WBGene00201736 | NA | NA | NA | NA |
| WBGene00201737 | NA | NA | NA | NA |
| WBGene00201738 | NA | NA | NA | NA |
| WBGene00201739 | 0.020925399 | 0.84643798 | NA | NA |
| WBGene00201740 | NA | NA | NA | NA |
| WBGene00201741 | NA | NA | NA | NA |
| WBGene00201742 | NA | NA | NA | NA |
| WBGene00201743 | NA | NA | NA | NA |

| | | | | |
|---|---|---|---|---|
| WBGene00201744 | NA | NA | NA | NA |
| WBGene00201745 | NA | NA | NA | NA |
| WBGene00201746 | NA | NA | NA | NA |
| WBGene00201747 | NA | NA | NA | NA |
| WBGene00201748 | NA | NA | NA | NA |
| WBGene00201749 | NA | NA | NA | NA |
| WBGene00201750 | NA | NA | NA | NA |
| WBGene00201751 | NA | NA | NA | NA |
| WBGene00201752 | NA | NA | NA | NA |
| WBGene00201753 | NA | NA | NA | NA |
| WBGene00201754 | NA | NA | NA | NA |
| WBGene00201755 | NA | NA | NA | NA |
| WBGene00201756 | NA | NA | NA | NA |
| WBGene00201757 | NA | NA | NA | NA |
| WBGene00201758 | NA | NA | NA | NA |
| WBGene00201759 | NA | NA | NA | NA |
| WBGene00201760 | NA | NA | NA | NA |
| WBGene00201761 | NA | NA | NA | NA |
| WBGene00201762 | NA | NA | NA | NA |
| WBGene00201763 | NA | NA | NA | NA |
| WBGene00201764 | NA | NA | NA | NA |
| WBGene00201765 | NA | NA | NA | NA |
| WBGene00201766 | NA | NA | NA | NA |
| WBGene00201767 | NA | NA | NA | NA |
| WBGene00201768 | NA | NA | NA | NA |
| WBGene00201769 | NA | NA | NA | NA |
| WBGene00201770 | NA | NA | NA | NA |
| WBGene00201771 | NA | NA | NA | NA |
| WBGene00201772 | NA | NA | NA | NA |
| WBGene00201773 | NA | NA | NA | NA |
| WBGene00201774 | NA | NA | NA | NA |
| WBGene00201775 | NA | NA | NA | NA |
| WBGene00201776 | NA | NA | NA | NA |
| WBGene00201777 | NA | NA | NA | NA |
| WBGene00201778 | 0.033463284 | 0.6365037 | NA | NA |
| WBGene00201779 | NA | NA | NA | NA |
| WBGene00201780 | NA | NA | NA | NA |
| WBGene00201781 | NA | NA | NA | NA |
| WBGene00201782 | NA | NA | NA | NA |
| WBGene00201783 | NA | NA | NA | NA |
| WBGene00201784 | NA | NA | NA | NA |
| WBGene00201785 | NA | NA | NA | NA |
| WBGene00201786 | NA | NA | NA | NA |

| | | | | |
|---|---|---|---|---|
| WBGene00201787 | NA | NA | NA | NA |
| WBGene00201788 | NA | NA | NA | NA |
| WBGene00201789 | NA | NA | NA | NA |
| WBGene00201790 | NA | NA | NA | NA |
| WBGene00201791 | NA | NA | NA | NA |
| WBGene00201792 | NA | NA | NA | NA |
| WBGene00201793 | NA | NA | NA | NA |
| WBGene00201794 | NA | NA | NA | NA |
| WBGene00201795 | NA | NA | NA | NA |
| WBGene00201796 | NA | NA | NA | NA |
| WBGene00201797 | NA | NA | NA | NA |
| WBGene00201798 | NA | NA | NA | NA |
| WBGene00201799 | NA | NA | NA | NA |
| WBGene00201800 | NA | NA | NA | NA |
| WBGene00201801 | NA | NA | NA | NA |
| WBGene00201802 | NA | NA | NA | NA |
| WBGene00201803 | NA | NA | NA | NA |
| WBGene00201804 | NA | NA | NA | NA |
| WBGene00201805 | NA | NA | NA | NA |
| WBGene00201806 | NA | NA | NA | NA |
| WBGene00201807 | NA | NA | NA | NA |
| WBGene00201808 | NA | NA | NA | NA |
| WBGene00201809 | NA | NA | NA | NA |
| WBGene00201810 | NA | NA | NA | NA |
| WBGene00201811 | NA | NA | NA | NA |
| WBGene00201812 | NA | NA | NA | NA |
| WBGene00201813 | NA | NA | NA | NA |
| WBGene00201814 | NA | NA | NA | NA |
| WBGene00201815 | NA | NA | NA | NA |
| WBGene00201816 | NA | NA | NA | NA |
| WBGene00201817 | NA | NA | NA | NA |
| WBGene00201818 | NA | NA | NA | NA |
| WBGene00201819 | NA | NA | NA | NA |
| WBGene00201820 | 0.020317245 | 0.84643798 | NA | NA |
| WBGene00201821 | NA | NA | NA | NA |
| WBGene00201822 | NA | NA | NA | NA |
| WBGene00201823 | NA | NA | NA | NA |
| WBGene00201824 | NA | NA | NA | NA |
| WBGene00201825 | NA | NA | NA | NA |
| WBGene00201826 | NA | NA | NA | NA |
| WBGene00201827 | NA | NA | NA | NA |
| WBGene00201828 | NA | NA | NA | NA |
| WBGene00201829 | NA | NA | NA | NA |

| | | | | |
|---|---|---|---|---|
| WBGene00201830 | NA | NA | NA | NA |
| WBGene00201831 | NA | NA | NA | NA |
| WBGene00201832 | NA | NA | NA | NA |
| WBGene00201833 | NA | NA | NA | NA |
| WBGene00201834 | NA | NA | NA | NA |
| WBGene00201835 | NA | NA | NA | NA |
| WBGene00201836 | NA | NA | NA | NA |
| WBGene00201837 | NA | NA | NA | NA |
| WBGene00201838 | NA | NA | NA | NA |
| WBGene00201839 | NA | NA | NA | NA |
| WBGene00201840 | NA | NA | NA | NA |
| WBGene00201841 | NA | NA | NA | NA |
| WBGene00201842 | NA | NA | NA | NA |
| WBGene00201843 | NA | NA | NA | NA |
| WBGene00201844 | NA | NA | NA | NA |
| WBGene00201845 | NA | NA | NA | NA |
| WBGene00201846 | NA | NA | NA | NA |
| WBGene00201847 | NA | NA | NA | NA |
| WBGene00201848 | NA | NA | NA | NA |
| WBGene00201849 | NA | NA | NA | NA |
| WBGene00201850 | NA | NA | NA | NA |
| WBGene00201851 | NA | NA | NA | NA |
| WBGene00201852 | NA | NA | NA | NA |
| WBGene00201853 | NA | NA | NA | NA |
| WBGene00201854 | NA | NA | NA | NA |
| WBGene00201855 | NA | NA | NA | NA |
| WBGene00201856 | NA | NA | NA | NA |
| WBGene00201857 | NA | NA | NA | NA |
| WBGene00201858 | NA | NA | NA | NA |
| WBGene00201859 | NA | NA | NA | NA |
| WBGene00201860 | NA | NA | NA | NA |
| WBGene00201861 | NA | NA | NA | NA |
| WBGene00201862 | NA | NA | NA | NA |
| WBGene00201863 | NA | NA | NA | NA |
| WBGene00201864 | NA | NA | NA | NA |
| WBGene00201865 | NA | NA | NA | NA |
| WBGene00201866 | NA | NA | NA | NA |
| WBGene00201867 | NA | NA | NA | NA |
| WBGene00201868 | NA | NA | NA | NA |
| WBGene00201869 | NA | NA | NA | NA |
| WBGene00201870 | NA | NA | NA | NA |
| WBGene00201871 | -0.003421551 | 0.95689871 | NA | NA |
| WBGene00201872 | NA | NA | NA | NA |

| | | | | |
|---|---|---|---|---|
| WBGene00201873 | NA | NA | NA | NA |
| WBGene00201874 | NA | NA | NA | NA |
| WBGene00201875 | NA | NA | NA | NA |
| WBGene00201876 | NA | NA | NA | NA |
| WBGene00201877 | NA | NA | NA | NA |
| WBGene00201878 | NA | NA | NA | NA |
| WBGene00201879 | NA | NA | NA | NA |
| WBGene00201880 | NA | NA | NA | NA |
| WBGene00201881 | NA | NA | NA | NA |
| WBGene00201882 | NA | NA | NA | NA |
| WBGene00201883 | NA | NA | NA | NA |
| WBGene00201884 | NA | NA | NA | NA |
| WBGene00201885 | NA | NA | NA | NA |
| WBGene00201886 | NA | NA | NA | NA |
| WBGene00201887 | NA | NA | NA | NA |
| WBGene00201888 | NA | NA | NA | NA |
| WBGene00201889 | NA | NA | NA | NA |
| WBGene00201890 | NA | NA | NA | NA |
| WBGene00201891 | NA | NA | NA | NA |
| WBGene00201892 | NA | NA | NA | NA |
| WBGene00201893 | NA | NA | NA | NA |
| WBGene00201894 | NA | NA | NA | NA |
| WBGene00201895 | NA | NA | NA | NA |
| WBGene00201896 | NA | NA | NA | NA |
| WBGene00201897 | NA | NA | NA | NA |
| WBGene00201898 | NA | NA | NA | NA |
| WBGene00201899 | 0.020317245 | 0.84643798 | NA | NA |
| WBGene00201900 | 0.017501802 | 0.83221062 | NA | NA |
| WBGene00201901 | NA | NA | NA | NA |
| WBGene00201902 | NA | NA | NA | NA |
| WBGene00201903 | NA | NA | NA | NA |
| WBGene00201904 | 0.020232312 | 0.84643798 | NA | NA |
| WBGene00201905 | NA | NA | NA | NA |
| WBGene00201906 | NA | NA | NA | NA |
| WBGene00201907 | NA | NA | NA | NA |
| WBGene00201908 | NA | NA | NA | NA |
| WBGene00201909 | NA | NA | NA | NA |
| WBGene00201910 | NA | NA | NA | NA |
| WBGene00201911 | NA | NA | NA | NA |
| WBGene00201912 | NA | NA | NA | NA |
| WBGene00201913 | NA | NA | NA | NA |
| WBGene00201914 | NA | NA | NA | NA |
| WBGene00201915 | 0.020925399 | 0.84643798 | NA | NA |

| | | | | |
|---|---|---|---|---|
| WBGene00201916 | NA | NA | NA | NA |
| WBGene00201917 | NA | NA | NA | NA |
| WBGene00201918 | NA | NA | NA | NA |
| WBGene00201919 | NA | NA | NA | NA |
| WBGene00201920 | 0.038125098 | 0.61119482 | NA | NA |
| WBGene00201921 | NA | NA | NA | NA |
| WBGene00201922 | NA | NA | NA | NA |
| WBGene00201923 | NA | NA | NA | NA |
| WBGene00201924 | NA | NA | NA | NA |
| WBGene00201925 | NA | NA | NA | NA |
| WBGene00201926 | NA | NA | NA | NA |
| WBGene00201927 | NA | NA | NA | NA |
| WBGene00201928 | NA | NA | NA | NA |
| WBGene00201929 | NA | NA | NA | NA |
| WBGene00201930 | NA | NA | NA | NA |
| WBGene00201931 | NA | NA | NA | NA |
| WBGene00201932 | NA | NA | NA | NA |
| WBGene00201933 | NA | NA | NA | NA |
| WBGene00201934 | NA | NA | NA | NA |
| WBGene00201935 | NA | NA | NA | NA |
| WBGene00201936 | 0.020317245 | 0.84643798 | NA | NA |
| WBGene00201937 | NA | NA | NA | NA |
| WBGene00201938 | NA | NA | NA | NA |
| WBGene00201939 | NA | NA | NA | NA |
| WBGene00201940 | NA | NA | NA | NA |
| WBGene00201941 | 0.020232312 | 0.84643798 | NA | NA |
| WBGene00201942 | NA | NA | NA | NA |
| WBGene00201943 | NA | NA | NA | NA |
| WBGene00201944 | NA | NA | NA | NA |
| WBGene00201945 | NA | NA | NA | NA |
| WBGene00201946 | NA | NA | NA | NA |
| WBGene00201947 | NA | NA | NA | NA |
| WBGene00201948 | NA | NA | NA | NA |
| WBGene00201949 | NA | NA | NA | NA |
| WBGene00201950 | NA | NA | NA | NA |
| WBGene00201951 | NA | NA | NA | NA |
| WBGene00201952 | NA | NA | NA | NA |
| WBGene00201953 | -0.023109762 | 0.78121253 | NA | NA |
| WBGene00201954 | NA | NA | NA | NA |
| WBGene00201955 | NA | NA | NA | NA |
| WBGene00201956 | NA | NA | NA | NA |
| WBGene00201957 | NA | NA | NA | NA |
| WBGene00201958 | NA | NA | NA | NA |

| | | | | |
|---|---|---|---|---|
| WBGene00201959 | NA | NA | NA | NA |
| WBGene00201960 | 0.020317245 | 0.84643798 | NA | NA |
| WBGene00201961 | NA | NA | NA | NA |
| WBGene00201962 | NA | NA | NA | NA |
| WBGene00201963 | NA | NA | NA | NA |
| WBGene00201964 | 0.015908114 | 0.8416225 | NA | NA |
| WBGene00201965 | NA | NA | NA | NA |
| WBGene00201966 | NA | NA | NA | NA |
| WBGene00201967 | -0.023109762 | 0.78121253 | NA | NA |
| WBGene00201968 | NA | NA | NA | NA |
| WBGene00201969 | NA | NA | NA | NA |
| WBGene00201970 | NA | NA | NA | NA |
| WBGene00201971 | NA | NA | NA | NA |
| WBGene00201972 | NA | NA | NA | NA |
| WBGene00201973 | NA | NA | NA | NA |
| WBGene00201974 | NA | NA | NA | NA |
| WBGene00201975 | NA | NA | NA | NA |
| WBGene00201976 | NA | NA | NA | NA |
| WBGene00201977 | NA | NA | NA | NA |
| WBGene00201978 | NA | NA | NA | NA |
| WBGene00201979 | NA | NA | NA | NA |
| WBGene00201980 | NA | NA | NA | NA |
| WBGene00201981 | NA | NA | NA | NA |
| WBGene00201982 | NA | NA | NA | NA |
| WBGene00201983 | NA | NA | NA | NA |
| WBGene00201984 | NA | NA | NA | NA |
| WBGene00201985 | NA | NA | NA | NA |
| WBGene00201986 | NA | NA | NA | NA |
| WBGene00201987 | NA | NA | NA | NA |
| WBGene00201988 | NA | NA | NA | NA |
| WBGene00201989 | NA | NA | NA | NA |
| WBGene00201990 | NA | NA | NA | NA |
| WBGene00201991 | 0.020317245 | 0.84643798 | NA | NA |
| WBGene00201992 | NA | NA | NA | NA |
| WBGene00201993 | NA | NA | NA | NA |
| WBGene00201994 | -0.045959069 | 0.50707733 | NA | NA |
| WBGene00201995 | NA | NA | NA | NA |
| WBGene00201996 | NA | NA | NA | NA |
| WBGene00201997 | 0.020925399 | 0.84643798 | NA | NA |
| WBGene00201998 | NA | NA | NA | NA |
| WBGene00201999 | NA | NA | NA | NA |
| WBGene00202000 | NA | NA | NA | NA |
| WBGene00202001 | NA | NA | NA | NA |

| | | | | |
|---|---|---|---|---|
| WBGene00202002 | NA | NA | NA | NA |
| WBGene00202003 | NA | NA | NA | NA |
| WBGene00202004 | NA | NA | NA | NA |
| WBGene00202005 | NA | NA | NA | NA |
| WBGene00202006 | NA | NA | NA | NA |
| WBGene00202007 | NA | NA | NA | NA |
| WBGene00202008 | NA | NA | NA | NA |
| WBGene00202009 | NA | NA | NA | NA |
| WBGene00202010 | NA | NA | NA | NA |
| WBGene00202011 | NA | NA | NA | NA |
| WBGene00202012 | NA | NA | NA | NA |
| WBGene00202013 | NA | NA | NA | NA |
| WBGene00202014 | NA | NA | NA | NA |
| WBGene00202015 | NA | NA | NA | NA |
| WBGene00202016 | NA | NA | NA | NA |
| WBGene00202017 | NA | NA | NA | NA |
| WBGene00202018 | NA | NA | NA | NA |
| WBGene00202019 | NA | NA | NA | NA |
| WBGene00202020 | NA | NA | NA | NA |
| WBGene00202021 | NA | NA | NA | NA |
| WBGene00202022 | NA | NA | NA | NA |
| WBGene00202023 | NA | NA | NA | NA |
| WBGene00202024 | NA | NA | NA | NA |
| WBGene00202025 | NA | NA | NA | NA |
| WBGene00202026 | NA | NA | NA | NA |
| WBGene00202027 | NA | NA | NA | NA |
| WBGene00202028 | NA | NA | NA | NA |
| WBGene00202029 | NA | NA | NA | NA |
| WBGene00202030 | 0.020925399 | 0.84643798 | NA | NA |
| WBGene00202031 | NA | NA | NA | NA |
| WBGene00202032 | NA | NA | NA | NA |
| WBGene00202033 | 0.015062379 | 0.80838968 | NA | NA |
| WBGene00202034 | NA | NA | NA | NA |
| WBGene00202035 | NA | NA | NA | NA |
| WBGene00202036 | NA | NA | NA | NA |
| WBGene00202037 | NA | NA | NA | NA |
| WBGene00202038 | NA | NA | NA | NA |
| WBGene00202039 | NA | NA | NA | NA |
| WBGene00202040 | NA | NA | NA | NA |
| WBGene00202041 | NA | NA | NA | NA |
| WBGene00202042 | NA | NA | NA | NA |
| WBGene00202043 | NA | NA | NA | NA |
| WBGene00202044 | NA | NA | NA | NA |

| | | | | |
|---|---|---|---|---|
| WBGene00202045 | NA | NA | NA | NA |
| WBGene00202046 | NA | NA | NA | NA |
| WBGene00202047 | NA | NA | NA | NA |
| WBGene00202048 | NA | NA | NA | NA |
| WBGene00202049 | NA | NA | NA | NA |
| WBGene00202050 | NA | NA | NA | NA |
| WBGene00202051 | NA | NA | NA | NA |
| WBGene00202052 | NA | NA | NA | NA |
| WBGene00202053 | NA | NA | NA | NA |
| WBGene00202054 | NA | NA | NA | NA |
| WBGene00202055 | NA | NA | NA | NA |
| WBGene00202056 | -0.023056499 | 0.78121253 | NA | NA |
| WBGene00202057 | NA | NA | NA | NA |
| WBGene00202058 | NA | NA | NA | NA |
| WBGene00202059 | NA | NA | NA | NA |
| WBGene00202060 | NA | NA | NA | NA |
| WBGene00202061 | NA | NA | NA | NA |
| WBGene00202062 | NA | NA | NA | NA |
| WBGene00202063 | NA | NA | NA | NA |
| WBGene00202064 | NA | NA | NA | NA |
| WBGene00202065 | NA | NA | NA | NA |
| WBGene00202066 | -0.023056499 | 0.78121253 | NA | NA |
| WBGene00202067 | NA | NA | NA | NA |
| WBGene00202068 | NA | NA | NA | NA |
| WBGene00202069 | NA | NA | NA | NA |
| WBGene00202070 | NA | NA | NA | NA |
| WBGene00202071 | NA | NA | NA | NA |
| WBGene00202072 | NA | NA | NA | NA |
| WBGene00202073 | NA | NA | NA | NA |
| WBGene00202074 | NA | NA | NA | NA |
| WBGene00202075 | NA | NA | NA | NA |
| WBGene00202076 | NA | NA | NA | NA |
| WBGene00202077 | NA | NA | NA | NA |
| WBGene00202078 | NA | NA | NA | NA |
| WBGene00202079 | NA | NA | NA | NA |
| WBGene00202080 | NA | NA | NA | NA |
| WBGene00202081 | 0.020317245 | 0.84643798 | NA | NA |
| WBGene00202082 | NA | NA | NA | NA |
| WBGene00202083 | NA | NA | NA | NA |
| WBGene00202084 | NA | NA | NA | NA |
| WBGene00202085 | NA | NA | NA | NA |
| WBGene00202086 | NA | NA | NA | NA |
| WBGene00202087 | NA | NA | NA | NA |

| | | | | |
|---|---|---|---|---|
| WBGene00202088 | NA | NA | NA | NA |
| WBGene00202089 | 0.020925399 | 0.84643798 | NA | NA |
| WBGene00202090 | NA | NA | NA | NA |
| WBGene00202091 | NA | NA | NA | NA |
| WBGene00202092 | NA | NA | NA | NA |
| WBGene00202093 | NA | NA | NA | NA |
| WBGene00202094 | NA | NA | NA | NA |
| WBGene00202095 | NA | NA | NA | NA |
| WBGene00202096 | NA | NA | NA | NA |
| WBGene00202097 | NA | NA | NA | NA |
| WBGene00202098 | NA | NA | NA | NA |
| WBGene00202099 | NA | NA | NA | NA |
| WBGene00202100 | NA | NA | NA | NA |
| WBGene00202101 | NA | NA | NA | NA |
| WBGene00202102 | NA | NA | NA | NA |
| WBGene00202103 | NA | NA | NA | NA |
| WBGene00202104 | NA | NA | NA | NA |
| WBGene00202105 | NA | NA | NA | NA |
| WBGene00202106 | NA | NA | NA | NA |
| WBGene00202107 | NA | NA | NA | NA |
| WBGene00202108 | -0.022930767 | 0.78121253 | NA | NA |
| WBGene00202109 | NA | NA | NA | NA |
| WBGene00202110 | NA | NA | NA | NA |
| WBGene00202111 | NA | NA | NA | NA |
| WBGene00202112 | NA | NA | NA | NA |
| WBGene00202113 | NA | NA | NA | NA |
| WBGene00202114 | NA | NA | NA | NA |
| WBGene00202115 | NA | NA | NA | NA |
| WBGene00202116 | NA | NA | NA | NA |
| WBGene00202117 | NA | NA | NA | NA |
| WBGene00202118 | NA | NA | NA | NA |
| WBGene00202119 | NA | NA | NA | NA |
| WBGene00202120 | NA | NA | NA | NA |
| WBGene00202121 | NA | NA | NA | NA |
| WBGene00202122 | -0.075452738 | 0.28904131 | NA | NA |
| WBGene00202123 | NA | NA | NA | NA |
| WBGene00202124 | NA | NA | NA | NA |
| WBGene00202125 | NA | NA | NA | NA |
| WBGene00202126 | NA | NA | NA | NA |
| WBGene00202127 | NA | NA | NA | NA |
| WBGene00202128 | 0.020317245 | 0.84643798 | NA | NA |
| WBGene00202129 | NA | NA | NA | NA |
| WBGene00202130 | NA | NA | NA | NA |

| | | | | |
|---|---|---|---|---|
| WBGene00202131 | NA | NA | NA | NA |
| WBGene00202132 | NA | NA | NA | NA |
| WBGene00202133 | NA | NA | NA | NA |
| WBGene00202134 | NA | NA | NA | NA |
| WBGene00202135 | NA | NA | NA | NA |
| WBGene00202136 | NA | NA | NA | NA |
| WBGene00202137 | NA | NA | NA | NA |
| WBGene00202138 | NA | NA | NA | NA |
| WBGene00202139 | NA | NA | NA | NA |
| WBGene00202140 | NA | NA | NA | NA |
| WBGene00202141 | NA | NA | NA | NA |
| WBGene00202142 | 0.020232312 | 0.84643798 | NA | NA |
| WBGene00202143 | NA | NA | NA | NA |
| WBGene00202144 | NA | NA | NA | NA |
| WBGene00202145 | NA | NA | NA | NA |
| WBGene00202146 | NA | NA | NA | NA |
| WBGene00202147 | NA | NA | NA | NA |
| WBGene00202148 | NA | NA | NA | NA |
| WBGene00202149 | NA | NA | NA | NA |
| WBGene00202150 | NA | NA | NA | NA |
| WBGene00202151 | NA | NA | NA | NA |
| WBGene00202152 | NA | NA | NA | NA |
| WBGene00202153 | NA | NA | NA | NA |
| WBGene00202154 | NA | NA | NA | NA |
| WBGene00202155 | NA | NA | NA | NA |
| WBGene00202156 | NA | NA | NA | NA |
| WBGene00202157 | NA | NA | NA | NA |
| WBGene00202158 | NA | NA | NA | NA |
| WBGene00202159 | NA | NA | NA | NA |
| WBGene00202160 | NA | NA | NA | NA |
| WBGene00202161 | NA | NA | NA | NA |
| WBGene00202162 | NA | NA | NA | NA |
| WBGene00202163 | NA | NA | NA | NA |
| WBGene00202164 | NA | NA | NA | NA |
| WBGene00202165 | NA | NA | NA | NA |
| WBGene00202166 | NA | NA | NA | NA |
| WBGene00202167 | NA | NA | NA | NA |
| WBGene00202168 | 0.020232312 | 0.84643798 | NA | NA |
| WBGene00202169 | NA | NA | NA | NA |
| WBGene00202170 | NA | NA | NA | NA |
| WBGene00202171 | NA | NA | NA | NA |
| WBGene00202172 | NA | NA | NA | NA |
| WBGene00202173 | NA | NA | NA | NA |

| | | | | |
|---|---|---|---|---|
| WBGene00202174 | NA | NA | NA | NA |
| WBGene00202175 | NA | NA | NA | NA |
| WBGene00202176 | NA | NA | NA | NA |
| WBGene00202177 | NA | NA | NA | NA |
| WBGene00202178 | -0.023056499 | 0.78121253 | NA | NA |
| WBGene00202179 | NA | NA | NA | NA |
| WBGene00202180 | NA | NA | NA | NA |
| WBGene00202181 | NA | NA | NA | NA |
| WBGene00202182 | NA | NA | NA | NA |
| WBGene00202183 | NA | NA | NA | NA |
| WBGene00202184 | NA | NA | NA | NA |
| WBGene00202185 | NA | NA | NA | NA |
| WBGene00202186 | NA | NA | NA | NA |
| WBGene00202187 | NA | NA | NA | NA |
| WBGene00202188 | NA | NA | NA | NA |
| WBGene00202189 | NA | NA | NA | NA |
| WBGene00202190 | NA | NA | NA | NA |
| WBGene00202191 | NA | NA | NA | NA |
| WBGene00202192 | NA | NA | NA | NA |
| WBGene00202193 | NA | NA | NA | NA |
| WBGene00202194 | NA | NA | NA | NA |
| WBGene00202195 | NA | NA | NA | NA |
| WBGene00202196 | NA | NA | NA | NA |
| WBGene00202197 | NA | NA | NA | NA |
| WBGene00202198 | NA | NA | NA | NA |
| WBGene00202199 | NA | NA | NA | NA |
| WBGene00202200 | NA | NA | NA | NA |
| WBGene00202201 | NA | NA | NA | NA |
| WBGene00202202 | NA | NA | NA | NA |
| WBGene00202203 | NA | NA | NA | NA |
| WBGene00202204 | NA | NA | NA | NA |
| WBGene00202205 | NA | NA | NA | NA |
| WBGene00202206 | NA | NA | NA | NA |
| WBGene00202207 | NA | NA | NA | NA |
| WBGene00202208 | NA | NA | NA | NA |
| WBGene00202209 | NA | NA | NA | NA |
| WBGene00202210 | NA | NA | NA | NA |
| WBGene00202211 | NA | NA | NA | NA |
| WBGene00202212 | NA | NA | NA | NA |
| WBGene00202213 | NA | NA | NA | NA |
| WBGene00202214 | NA | NA | NA | NA |
| WBGene00202215 | NA | NA | NA | NA |
| WBGene00202216 | NA | NA | NA | NA |

| | | | | |
|---|---|---|---|---|
| WBGene00202217 | NA | NA | NA | NA |
| WBGene00202218 | -0.00346345 | 0.95708614 | NA | NA |
| WBGene00202219 | NA | NA | NA | NA |
| WBGene00202220 | NA | NA | NA | NA |
| WBGene00202221 | NA | NA | NA | NA |
| WBGene00202222 | NA | NA | NA | NA |
| WBGene00202223 | NA | NA | NA | NA |
| WBGene00202224 | NA | NA | NA | NA |
| WBGene00202225 | NA | NA | NA | NA |
| WBGene00202226 | NA | NA | NA | NA |
| WBGene00202227 | NA | NA | NA | NA |
| WBGene00202228 | NA | NA | NA | NA |
| WBGene00202229 | NA | NA | NA | NA |
| WBGene00202230 | NA | NA | NA | NA |
| WBGene00202231 | 0.020232312 | 0.84643798 | NA | NA |
| WBGene00202232 | NA | NA | NA | NA |
| WBGene00202233 | NA | NA | NA | NA |
| WBGene00202234 | NA | NA | NA | NA |
| WBGene00202235 | NA | NA | NA | NA |
| WBGene00202236 | NA | NA | NA | NA |
| WBGene00202237 | NA | NA | NA | NA |
| WBGene00202238 | NA | NA | NA | NA |
| WBGene00202239 | NA | NA | NA | NA |
| WBGene00202240 | NA | NA | NA | NA |
| WBGene00202241 | NA | NA | NA | NA |
| WBGene00202242 | NA | NA | NA | NA |
| WBGene00202243 | NA | NA | NA | NA |
| WBGene00202244 | NA | NA | NA | NA |
| WBGene00202245 | NA | NA | NA | NA |
| WBGene00202246 | NA | NA | NA | NA |
| WBGene00202247 | -0.023109762 | 0.78121253 | NA | NA |
| WBGene00202248 | NA | NA | NA | NA |
| WBGene00202249 | NA | NA | NA | NA |
| WBGene00202250 | NA | NA | NA | NA |
| WBGene00202251 | NA | NA | NA | NA |
| WBGene00202252 | NA | NA | NA | NA |
| WBGene00202253 | NA | NA | NA | NA |
| WBGene00202254 | NA | NA | NA | NA |
| WBGene00202255 | NA | NA | NA | NA |
| WBGene00202256 | NA | NA | NA | NA |
| WBGene00202257 | NA | NA | NA | NA |
| WBGene00202258 | NA | NA | NA | NA |
| WBGene00202259 | NA | NA | NA | NA |

| | | | | |
|---|---|---|---|---|
| WBGene00202260 | NA | NA | NA | NA |
| WBGene00202261 | NA | NA | NA | NA |
| WBGene00202262 | NA | NA | NA | NA |
| WBGene00202263 | NA | NA | NA | NA |
| WBGene00202264 | NA | NA | NA | NA |
| WBGene00202265 | NA | NA | NA | NA |
| WBGene00202266 | NA | NA | NA | NA |
| WBGene00202267 | NA | NA | NA | NA |
| WBGene00202268 | NA | NA | NA | NA |
| WBGene00202269 | NA | NA | NA | NA |
| WBGene00202270 | NA | NA | NA | NA |
| WBGene00202271 | NA | NA | NA | NA |
| WBGene00202272 | NA | NA | NA | NA |
| WBGene00202273 | NA | NA | NA | NA |
| WBGene00202274 | NA | NA | NA | NA |
| WBGene00202275 | NA | NA | NA | NA |
| WBGene00202276 | NA | NA | NA | NA |
| WBGene00202277 | NA | NA | NA | NA |
| WBGene00202278 | NA | NA | NA | NA |
| WBGene00202279 | NA | NA | NA | NA |
| WBGene00202280 | NA | NA | NA | NA |
| WBGene00202281 | NA | NA | NA | NA |
| WBGene00202282 | NA | NA | NA | NA |
| WBGene00202283 | NA | NA | NA | NA |
| WBGene00202284 | NA | NA | NA | NA |
| WBGene00202285 | NA | NA | NA | NA |
| WBGene00202286 | NA | NA | NA | NA |
| WBGene00202287 | NA | NA | NA | NA |
| WBGene00202288 | NA | NA | NA | NA |
| WBGene00202289 | NA | NA | NA | NA |
| WBGene00202290 | NA | NA | NA | NA |
| WBGene00202291 | NA | NA | NA | NA |
| WBGene00202292 | NA | NA | NA | NA |
| WBGene00202293 | NA | NA | NA | NA |
| WBGene00202294 | NA | NA | NA | NA |
| WBGene00202295 | NA | NA | NA | NA |
| WBGene00202296 | NA | NA | NA | NA |
| WBGene00202297 | 0.020232312 | 0.84643798 | NA | NA |
| WBGene00202298 | NA | NA | NA | NA |
| WBGene00202299 | NA | NA | NA | NA |
| WBGene00202300 | -0.023109762 | 0.78121253 | NA | NA |
| WBGene00202301 | NA | NA | NA | NA |
| WBGene00202302 | NA | NA | NA | NA |

| | | | | |
|---|---|---|---|---|
| WBGene00202303 | NA | NA | NA | NA |
| WBGene00202304 | NA | NA | NA | NA |
| WBGene00202305 | NA | NA | NA | NA |
| WBGene00202306 | NA | NA | NA | NA |
| WBGene00202307 | NA | NA | NA | NA |
| WBGene00202308 | NA | NA | NA | NA |
| WBGene00202309 | NA | NA | NA | NA |
| WBGene00202310 | NA | NA | NA | NA |
| WBGene00202311 | NA | NA | NA | NA |
| WBGene00202312 | NA | NA | NA | NA |
| WBGene00202313 | NA | NA | NA | NA |
| WBGene00202314 | NA | NA | NA | NA |
| WBGene00202315 | NA | NA | NA | NA |
| WBGene00202316 | NA | NA | NA | NA |
| WBGene00202317 | NA | NA | NA | NA |
| WBGene00202318 | NA | NA | NA | NA |
| WBGene00202319 | NA | NA | NA | NA |
| WBGene00202320 | NA | NA | NA | NA |
| WBGene00202321 | NA | NA | NA | NA |
| WBGene00202322 | NA | NA | NA | NA |
| WBGene00202323 | NA | NA | NA | NA |
| WBGene00202324 | NA | NA | NA | NA |
| WBGene00202325 | NA | NA | NA | NA |
| WBGene00202326 | NA | NA | NA | NA |
| WBGene00202327 | 0.040107227 | 0.58693163 | NA | NA |
| WBGene00202328 | NA | NA | NA | NA |
| WBGene00202329 | NA | NA | NA | NA |
| WBGene00202330 | NA | NA | NA | NA |
| WBGene00202331 | NA | NA | NA | NA |
| WBGene00202332 | NA | NA | NA | NA |
| WBGene00202333 | NA | NA | NA | NA |
| WBGene00202334 | -0.022930767 | 0.78121253 | NA | NA |
| WBGene00202335 | NA | NA | NA | NA |
| WBGene00202336 | NA | NA | NA | NA |
| WBGene00202337 | NA | NA | NA | NA |
| WBGene00202338 | NA | NA | NA | NA |
| WBGene00202339 | -0.023109762 | 0.78121253 | NA | NA |
| WBGene00202340 | NA | NA | NA | NA |
| WBGene00202341 | NA | NA | NA | NA |
| WBGene00202342 | NA | NA | NA | NA |
| WBGene00202343 | NA | NA | NA | NA |
| WBGene00202344 | NA | NA | NA | NA |
| WBGene00202345 | NA | NA | NA | NA |

| | | | | |
|---|---|---|---|---|
| WBGene00202346 | NA | NA | NA | NA |
| WBGene00202347 | NA | NA | NA | NA |
| WBGene00202348 | NA | NA | NA | NA |
| WBGene00202349 | -0.002914971 | 0.95718083 | NA | NA |
| WBGene00202350 | NA | NA | NA | NA |
| WBGene00202351 | NA | NA | NA | NA |
| WBGene00202352 | NA | NA | NA | NA |
| WBGene00202353 | NA | NA | NA | NA |
| WBGene00202354 | NA | NA | NA | NA |
| WBGene00202355 | NA | NA | NA | NA |
| WBGene00202356 | NA | NA | NA | NA |
| WBGene00202357 | NA | NA | NA | NA |
| WBGene00202358 | NA | NA | NA | NA |
| WBGene00202359 | NA | NA | NA | NA |
| WBGene00202360 | NA | NA | NA | NA |
| WBGene00202361 | NA | NA | NA | NA |
| WBGene00202362 | NA | NA | NA | NA |
| WBGene00202363 | NA | NA | NA | NA |
| WBGene00202364 | NA | NA | NA | NA |
| WBGene00202365 | NA | NA | NA | NA |
| WBGene00202366 | NA | NA | NA | NA |
| WBGene00202367 | NA | NA | NA | NA |
| WBGene00202368 | NA | NA | NA | NA |
| WBGene00202369 | NA | NA | NA | NA |
| WBGene00202370 | NA | NA | NA | NA |
| WBGene00202371 | NA | NA | NA | NA |
| WBGene00202372 | 0.036687355 | 0.59489568 | NA | NA |
| WBGene00202373 | NA | NA | NA | NA |
| WBGene00202374 | NA | NA | NA | NA |
| WBGene00202375 | NA | NA | NA | NA |
| WBGene00202376 | NA | NA | NA | NA |
| WBGene00202377 | NA | NA | NA | NA |
| WBGene00202378 | NA | NA | NA | NA |
| WBGene00202379 | NA | NA | NA | NA |
| WBGene00202380 | NA | NA | NA | NA |
| WBGene00202381 | NA | NA | NA | NA |
| WBGene00202382 | NA | NA | NA | NA |
| WBGene00202383 | NA | NA | NA | NA |
| WBGene00202384 | 0.036113109 | 0.67051389 | NA | NA |
| WBGene00202385 | NA | NA | NA | NA |
| WBGene00202386 | NA | NA | NA | NA |
| WBGene00202387 | NA | NA | NA | NA |
| WBGene00202388 | NA | NA | NA | NA |

| | | | | |
|---|---|---|---|---|
| WBGene00202389 | NA | NA | NA | NA |
| WBGene00202390 | NA | NA | NA | NA |
| WBGene00202391 | NA | NA | NA | NA |
| WBGene00202392 | NA | NA | NA | NA |
| WBGene00202393 | NA | NA | NA | NA |
| WBGene00202394 | NA | NA | NA | NA |
| WBGene00202395 | -0.023056499 | 0.78121253 | NA | NA |
| WBGene00202396 | NA | NA | NA | NA |
| WBGene00202397 | NA | NA | NA | NA |
| WBGene00202398 | NA | NA | NA | NA |
| WBGene00202399 | NA | NA | NA | NA |
| WBGene00202400 | 0.074409796 | 0.33504472 | NA | NA |
| WBGene00202401 | NA | NA | NA | NA |
| WBGene00202402 | NA | NA | NA | NA |
| WBGene00202403 | NA | NA | NA | NA |
| WBGene00202404 | NA | NA | NA | NA |
| WBGene00202405 | NA | NA | NA | NA |
| WBGene00202406 | NA | NA | NA | NA |
| WBGene00202407 | NA | NA | NA | NA |
| WBGene00202408 | NA | NA | NA | NA |
| WBGene00202409 | NA | NA | NA | NA |
| WBGene00202410 | NA | NA | NA | NA |
| WBGene00202411 | NA | NA | NA | NA |
| WBGene00202412 | NA | NA | NA | NA |
| WBGene00202413 | NA | NA | NA | NA |
| WBGene00202414 | NA | NA | NA | NA |
| WBGene00202415 | NA | NA | NA | NA |
| WBGene00202416 | NA | NA | NA | NA |
| WBGene00202417 | NA | NA | NA | NA |
| WBGene00202418 | NA | NA | NA | NA |
| WBGene00202419 | NA | NA | NA | NA |
| WBGene00202420 | NA | NA | NA | NA |
| WBGene00202421 | NA | NA | NA | NA |
| WBGene00202422 | 0.0398788 | 0.58971221 | NA | NA |
| WBGene00202423 | 0.055204521 | 0.44852756 | NA | NA |
| WBGene00202424 | NA | NA | NA | NA |
| WBGene00202425 | NA | NA | NA | NA |
| WBGene00202426 | NA | NA | NA | NA |
| WBGene00202427 | NA | NA | NA | NA |
| WBGene00202428 | NA | NA | NA | NA |
| WBGene00202429 | NA | NA | NA | NA |
| WBGene00202430 | NA | NA | NA | NA |
| WBGene00202431 | NA | NA | NA | NA |

| | | | | |
|---|---|---|---|---|
| WBGene00202432 | NA | NA | NA | NA |
| WBGene00202433 | NA | NA | NA | NA |
| WBGene00202434 | NA | NA | NA | NA |
| WBGene00202435 | NA | NA | NA | NA |
| WBGene00202436 | NA | NA | NA | NA |
| WBGene00202437 | NA | NA | NA | NA |
| WBGene00202438 | NA | NA | NA | NA |
| WBGene00202439 | NA | NA | NA | NA |
| WBGene00202440 | NA | NA | NA | NA |
| WBGene00202441 | 0.020317245 | 0.84643798 | NA | NA |
| WBGene00202442 | NA | NA | NA | NA |
| WBGene00202443 | NA | NA | NA | NA |
| WBGene00202444 | NA | NA | NA | NA |
| WBGene00202445 | NA | NA | NA | NA |
| WBGene00202446 | NA | NA | NA | NA |
| WBGene00202447 | NA | NA | NA | NA |
| WBGene00202448 | NA | NA | NA | NA |
| WBGene00202449 | NA | NA | NA | NA |
| WBGene00202450 | NA | NA | NA | NA |
| WBGene00202451 | NA | NA | NA | NA |
| WBGene00202452 | NA | NA | NA | NA |
| WBGene00202453 | NA | NA | NA | NA |
| WBGene00202454 | NA | NA | NA | NA |
| WBGene00202455 | NA | NA | NA | NA |
| WBGene00202456 | NA | NA | NA | NA |
| WBGene00202457 | NA | NA | NA | NA |
| WBGene00202458 | NA | NA | NA | NA |
| WBGene00202459 | NA | NA | NA | NA |
| WBGene00202460 | NA | NA | NA | NA |
| WBGene00202461 | NA | NA | NA | NA |
| WBGene00202462 | NA | NA | NA | NA |
| WBGene00202463 | NA | NA | NA | NA |
| WBGene00202464 | NA | NA | NA | NA |
| WBGene00202465 | NA | NA | NA | NA |
| WBGene00202466 | NA | NA | NA | NA |
| WBGene00202467 | NA | NA | NA | NA |
| WBGene00202468 | NA | NA | NA | NA |
| WBGene00202469 | NA | NA | NA | NA |
| WBGene00202470 | NA | NA | NA | NA |
| WBGene00202471 | NA | NA | NA | NA |
| WBGene00202472 | NA | NA | NA | NA |
| WBGene00202473 | NA | NA | NA | NA |
| WBGene00202474 | NA | NA | NA | NA |

| | | | | |
|---|---|---|---|---|
| WBGene00202475 | NA | NA | NA | NA |
| WBGene00202476 | NA | NA | NA | NA |
| WBGene00202477 | NA | NA | NA | NA |
| WBGene00202478 | NA | NA | NA | NA |
| WBGene00202479 | NA | NA | NA | NA |
| WBGene00202480 | NA | NA | NA | NA |
| WBGene00202481 | NA | NA | NA | NA |
| WBGene00202482 | NA | NA | NA | NA |
| WBGene00202483 | 0.020317245 | 0.84643798 | NA | NA |
| WBGene00202484 | NA | NA | NA | NA |
| WBGene00202485 | NA | NA | NA | NA |
| WBGene00202486 | NA | NA | NA | NA |
| WBGene00202487 | NA | NA | NA | NA |
| WBGene00202488 | NA | NA | NA | NA |
| WBGene00202489 | NA | NA | NA | NA |
| WBGene00202490 | NA | NA | NA | NA |
| WBGene00202491 | NA | NA | NA | NA |
| WBGene00202492 | NA | NA | NA | NA |
| WBGene00202493 | NA | NA | NA | NA |
| WBGene00202494 | NA | NA | NA | NA |
| WBGene00202495 | NA | NA | NA | T26H5.11 |
| WBGene00202496 | 0.020317245 | 0.84643798 | NA | R02F11.9 |
| WBGene00202497 | NA | NA | NA | R02F11.12 |
| WBGene00202497 | NA | NA | NA | R02F11.10 |
| WBGene00202498 | NA | NA | NA | Y60C6A.2 |
| WBGene00202499 | NA | NA | NA | B0507.15 |
| WBGene00202500 | NA | NA | NA | K09H11.15 |
| WBGene00202501 | 0.039292341 | 0.65808492 | NA | Y57E12B.10 |
| WBGene00202502 | NA | NA | NA | F41B5.12 |
| WBGene00202504 | -0.022930767 | 0.78121253 | NA | Y32B12B.8 |
| WBGene00202511 | NA | NA | NA | F07C3.16 |
| WBGene00202512 | -0.048370879 | 0.47720013 | NA | C25E10.16 |
| WBGene00202513 | -0.023109762 | 0.78121253 | NA | ZK938.10 |
| WBGene00202514 | 0.020317245 | 0.84643798 | NA | T09F5.20 |
| WBGene00206354 | 0.007418874 | 0.9638064 | 0.98020905 | F27E5.9 |
| WBGene00206355 | 0.436742974 | 0.01465496 | 0.06073117 | C34C6.10 |
| WBGene00206356 | 0.016059264 | 0.84014476 | NA | T08B2.15 |
| WBGene00206357 | -0.046394786 | 0.50231372 | NA | C09D4.9 |
| WBGene00206358 | -0.0260089 | 0.6772113 | NA | F46E10.18 |
| WBGene00206359 | -0.085381841 | 0.49464546 | 0.67742391 | C05E4.15 |
| WBGene00206360 | -0.00346345 | 0.95708614 | NA | T13C5.10 |
| WBGene00206361 | 0.032961943 | 0.7401812 | 0.85342581 | K02G10.15 |
| WBGene00206362 | -0.077339924 | 0.67561649 | 0.81139374 | B0041.11 |

| | | | | |
|---|---|---|---|---|
| WBGene00206363 | NA | NA | NA | F56H1.10 |
| WBGene00206364 | 0.020925399 | 0.84643798 | NA | C07A12.18 |
| WBGene00206365 | 0.020232312 | 0.84643798 | NA | B0563.18 |
| WBGene00206366 | 0.055201863 | 0.70223679 | 0.83093594 | C03H5.9 |
| WBGene00206368 | 0.033034883 | 0.64219503 | NA | Y39F10C.3 |
| WBGene00206369 | NA | NA | NA | NA |
| WBGene00206370 | NA | NA | NA | F02D8.9 |
| WBGene00206371 | 0.020232312 | 0.84643798 | NA | F28B12.6 |
| WBGene00206372 | 0.0398788 | 0.58971221 | NA | B0454.26 |
| WBGene00206373 | 0.38122275 | 0.10120659 | 0.24737325 | C27A2.12 |
| WBGene00206374 | -0.011975515 | 0.91432153 | 0.95230714 | K10G6.9 |
| WBGene00206375 | -0.022930767 | 0.78121253 | NA | NA |
| WBGene00206377 | -0.114688067 | 0.44906046 | 0.64173691 | F44E5.15 |
| WBGene00206378 | -0.052776296 | 0.53974799 | NA | C14E2.11 |
| WBGene00206379 | NA | NA | NA | C55B6.7 |
| WBGene00206380 | -0.091800229 | 0.6348064 | 0.78338999 | C42D4.18 |
| WBGene00206381 | 0.145427569 | 0.20160409 | 0.39027971 | F52H2.15 |
| WBGene00206382 | 0.075029222 | 0.33362495 | NA | F01G12.18 |
| WBGene00206383 | -0.023056499 | 0.78121253 | NA | C32C4.16 |
| WBGene00206384 | -0.103089982 | 0.30595116 | 0.50666281 | D1005.10 |
| WBGene00206385 | -0.06730507 | 0.31931151 | NA | T06D4.6 |
| WBGene00206386 | -0.163315737 | 0.39440609 | 0.59249945 | K10C2.12 |
| WBGene00206388 | -0.871416017 | 0.02062374 | 0.07836806 | K10C2.14 |
| WBGene00206389 | -0.342199432 | 0.12812077 | 0.28952248 | C29F7.10 |
| WBGene00206390 | NA | NA | NA | F14F7.7 |
| WBGene00206391 | NA | NA | NA | NA |
| WBGene00206392 | -0.003614866 | 0.95700117 | NA | F47D2.11 |
| WBGene00206393 | -0.023109762 | 0.78121253 | NA | W04G5.16 |
| WBGene00206394 | NA | NA | NA | NA |
| WBGene00206395 | 0.040107227 | 0.58693163 | NA | C08H9.18 |
| WBGene00206411 | NA | NA | NA | NA |
| WBGene00206412 | -0.058819927 | 0.64250184 | 0.78833475 | T24H7.8 |
| WBGene00206413 | -0.039799734 | 0.74173477 | 0.85408877 | ZK1240.11 |
| WBGene00206414 | 0.015052905 | 0.91935465 | 0.95526302 | F31D5.7 |
| WBGene00206415 | 0.174913048 | 0.04692711 | NA | F59H6.14 |
| WBGene00206416 | 0.042199095 | 0.55922995 | NA | W04H10.5 |
| WBGene00206417 | -0.067666074 | 0.31672907 | NA | W04H10.6 |
| WBGene00206418 | 0.509324255 | 0.05241564 | 0.15597915 | M04C3.5 |
| WBGene00206420 | -0.235110995 | 0.28860262 | 0.48877166 | NA |
| WBGene00206421 | NA | NA | NA | T26G10.9 |
| WBGene00206421 | NA | NA | NA | T26G10.8 |
| WBGene00206422 | NA | NA | NA | T26G10.9 |
| WBGene00206422 | NA | NA | NA | T26G10.8 |

| | | | | |
|---|---|---|---|---|
| WBGene00206423 | NA | NA | NA | NA |
| WBGene00206459 | -0.322077585 | 0.07686734 | 0.20521785 | NA |
| WBGene00206460 | -0.022930767 | 0.78121253 | NA | F56H9.9 |
| WBGene00206461 | 0.108717649 | 0.15669083 | NA | Y47D3A.34 |
| WBGene00206462 | -0.02791392 | 0.6763823 | NA | B0391.14 |
| WBGene00206464 | 0.030221926 | 0.78559125 | 0.87882516 | F59C6.18 |
| WBGene00206466 | 0.038125098 | 0.61119482 | NA | T08D10.5 |
| WBGene00206468 | NA | NA | NA | ZC262.11 |
| WBGene00206468 | NA | NA | NA | ZC262.12 |
| WBGene00206469 | 0.017501802 | 0.83221062 | NA | F58D2.5 |
| WBGene00206470 | -0.068085205 | 0.31374805 | NA | NA |
| WBGene00206471 | NA | NA | NA | Y22F5A.11 |
| WBGene00206473 | -0.003614866 | 0.95700117 | NA | B0273.115 |
| WBGene00206474 | -0.00963504 | 0.92298606 | 0.95651506 | T22A3.12 |
| WBGene00206475 | -0.045959069 | 0.50707733 | NA | F27D9.12 |
| WBGene00206477 | NA | NA | NA | ZC262.11 |
| WBGene00206477 | NA | NA | NA | ZC262.12 |
| WBGene00206478 | NA | NA | NA | M6.11 |
| WBGene00206479 | 0.020317245 | 0.84643798 | NA | C44C11.6 |
| WBGene00206481 | 0.020232312 | 0.84643798 | NA | Y51B9A.14 |
| WBGene00206482 | -0.01651397 | 0.93259344 | 0.9620984 | B0416.10 |
| WBGene00206483 | 0.017950189 | 0.90418573 | 0.94890569 | B0416.11 |
| WBGene00206484 | 0.004660483 | 0.98192785 | 0.98949726 | C31H2.14 |
| WBGene00206485 | 0.248307939 | 0.14875142 | 0.31960586 | C10A4.13 |
| WBGene00206486 | NA | NA | NA | Y54F10BM.20 |
| WBGene00206487 | -0.029492044 | 0.87611156 | 0.9339024 | best-19 |
| WBGene00206488 | NA | NA | NA | ZK757.10 |
| WBGene00206489 | NA | NA | NA | NA |
| WBGene00206490 | NA | NA | NA | Y57G11C.1142 |
| WBGene00206491 | -0.004851975 | 0.97922593 | 0.98812489 | NA |
| WBGene00206492 | -0.046394786 | 0.50231372 | NA | F11D11.21 |
| WBGene00206493 | 0.036821884 | 0.59344795 | NA | Y38H8A.12 |
| WBGene00206494 | NA | NA | NA | NA |
| WBGene00206495 | NA | NA | NA | F20E11.18 |
| WBGene00206496 | NA | NA | NA | Y51A2B.16 |
| WBGene00206497 | 0.032439478 | 0.68988446 | NA | Y52B11A.18 |
| WBGene00206498 | NA | NA | NA | C29A12.22 |
| WBGene00206499 | NA | NA | NA | C06H5.11 |
| WBGene00206500 | -0.02791392 | 0.6763823 | NA | F55C9.15 |
| WBGene00206501 | 0.020232312 | 0.84643798 | NA | F14D7.18 |
| WBGene00206502 | 0.033698991 | 0.67889023 | NA | T01B7.13 |
| WBGene00206503 | NA | NA | NA | T26E4.18 |
| WBGene00206504 | NA | NA | NA | K10G4.16 |

| | | | | |
|---|---|---|---|---|
| WBGene00206504 | NA | NA | NA | K10G4.15 |
| WBGene00206505 | -0.00346345 | 0.95708614 | NA | B0250.18 |
| WBGene00206506 | NA | NA | NA | F59A1.18 |
| WBGene00206507 | -0.159798892 | 0.09065013 | 0.22888093 | Y51A2A.16 |
| WBGene00206508 | NA | NA | NA | ZK262.19 |
| WBGene00206509 | NA | NA | NA | NA |
| WBGene00206510 | NA | NA | NA | Y113G7A.22 |
| WBGene00206511 | NA | NA | NA | W02H5.16 |
| WBGene00206512 | -0.046394786 | 0.50231372 | NA | Y39A3B.7 |
| WBGene00206513 | NA | NA | NA | B0244.15 |
| WBGene00206514 | -0.027373974 | 0.68245543 | NA | R05G9.5 |
| WBGene00206515 | NA | NA | NA | T28C6.11 |
| WBGene00206516 | 0.033034883 | 0.64219503 | NA | F53E4.2 |
| WBGene00206517 | -0.001842784 | 0.95621055 | NA | T27F7.12 |
| WBGene00206518 | NA | NA | NA | F36H5.12 |
| WBGene00206519 | -0.342178647 | 0.08295698 | 0.21618298 | F36H5.13 |
| WBGene00206520 | 0.020317245 | 0.84643798 | NA | ZK897.10 |
| WBGene00206521 | -0.050520088 | 0.73719273 | 0.85227723 | T02G5.15 |
| WBGene00206522 | 0.19684369 | 0.2763571 | 0.47632079 | F14D2.19 |
| WBGene00206523 | NA | NA | NA | C54E10.10 |
| WBGene00206524 | 0.020232312 | 0.84643798 | NA | C54E10.11 |
| WBGene00206525 | 1.879406923 | 0.00017526 | 0.00157329 | R74.10 |
| WBGene00206526 | NA | NA | NA | NA |
| WBGene00206527 | 0.040107227 | 0.58693163 | NA | T27A8.7 |
| WBGene00206528 | NA | NA | NA | Y53H1A.7 |
| WBGene00206529 | 0.039888668 | 0.56326497 | NA | C04A2.15 |
| WBGene00206530 | -0.028536222 | 0.71191204 | NA | Y52B11A.19 |
| WBGene00206531 | -0.003421551 | 0.95689871 | NA | NA |
| WBGene00206532 | 0.166645955 | 0.20118447 | 0.38993141 | F11A6.15 |
| WBGene00206533 | 0.0398788 | 0.58971221 | NA | C30A5.16 |
| WBGene00206534 | -0.023056499 | 0.78121253 | NA | C31A11.13 |
| WBGene00206535 | -0.002914971 | 0.95718083 | NA | M162.15 |
| WBGene00206536 | 0.110545227 | 0.1354043 | NA | NA |
| WBGene00206537 | 0.020317245 | 0.84643798 | NA | Y71H9A.10 |
| WBGene00206538 | 0.0398788 | 0.58971221 | NA | C09G5.13 |
| WBGene00215616 | 0.020317245 | 0.84643798 | NA | NA |
| WBGene00219205 | 0.020925399 | 0.84643798 | NA | R03D7.15 |
| WBGene00219206 | -0.009139816 | 0.95860788 | 0.97725557 | W05H12.4 |
| WBGene00219207 | 0.139177552 | 0.44452961 | 0.63794923 | C01F6.14 |
| WBGene00219208 | -0.046447909 | 0.81060632 | 0.89531472 | B0365.9 |
| WBGene00219209 | NA | NA | NA | F29F11.20 |
| WBGene00219210 | -0.163306443 | 0.43275626 | 0.62769807 | Y37H2A.18 |
| WBGene00219212 | NA | NA | NA | K10G4.16 |

| | | | | |
|---|---|---|---|---|
| WBGene00219212 | NA | NA | NA | K10G4.15 |
| WBGene00219214 | 0.059980168 | 0.38780842 | NA | T10B9.14 |
| WBGene00219215 | NA | NA | NA | NA |
| WBGene00219216 | 4.246986134 | 7.12E-27 | 7.29E-25 | K08D10.14 |
| WBGene00219217 | -0.974733602 | 0.01381243 | 0.05787387 | C08F8.15 |
| WBGene00219218 | -0.022350167 | 0.91184453 | 0.95112162 | Y73B6BL.288 |
| WBGene00219219 | NA | NA | NA | NA |
| WBGene00219220 | NA | NA | NA | NA |
| WBGene00219221 | NA | NA | NA | NA |
| WBGene00219222 | NA | NA | NA | NA |
| WBGene00219223 | 0.056411843 | 0.41891328 | NA | NA |
| WBGene00219224 | NA | NA | NA | NA |
| WBGene00219225 | NA | NA | NA | NA |
| WBGene00219226 | -0.003421551 | 0.95689871 | NA | NA |
| WBGene00219227 | NA | NA | NA | NA |
| WBGene00219228 | NA | NA | NA | NA |
| WBGene00219229 | NA | NA | NA | NA |
| WBGene00219230 | NA | NA | NA | NA |
| WBGene00219231 | NA | NA | NA | NA |
| WBGene00219232 | NA | NA | NA | NA |
| WBGene00219233 | 0.020317245 | 0.84643798 | NA | NA |
| WBGene00219234 | NA | NA | NA | B0511.17 |
| WBGene00219235 | 0.055597606 | 0.44564657 | NA | K03B4.8 |
| WBGene00219236 | -0.011440004 | 0.94206245 | 0.9672483 | K03B4.9 |
| WBGene00219238 | -0.180635116 | 0.24911444 | 0.44751957 | M151.9 |
| WBGene00219239 | -0.030550281 | 0.70468744 | NA | K05F6.14 |
| WBGene00219241 | -0.006690551 | 0.92537622 | NA | Y116A8A.150 |
| WBGene00219242 | 0.020232312 | 0.84643798 | NA | NA |
| WBGene00219243 | NA | NA | NA | T23G4.127 |
| WBGene00219244 | NA | NA | NA | Y51H4A.938 |
| WBGene00219245 | NA | NA | NA | NA |
| WBGene00219246 | NA | NA | NA | NA |
| WBGene00219247 | NA | NA | NA | NA |
| WBGene00219248 | NA | NA | NA | NA |
| WBGene00219249 | NA | NA | NA | NA |
| WBGene00219250 | NA | NA | NA | NA |
| WBGene00219251 | NA | NA | NA | NA |
| WBGene00219252 | -0.003421551 | 0.95689871 | NA | NA |
| WBGene00219253 | NA | NA | NA | NA |
| WBGene00219254 | NA | NA | NA | NA |
| WBGene00219255 | NA | NA | NA | NA |
| WBGene00219256 | NA | NA | NA | NA |
| WBGene00219257 | NA | NA | NA | NA |

| | | | | |
|---|---|---|---|---|
| WBGene00219258 | NA | NA | NA | NA |
| WBGene00219259 | NA | NA | NA | NA |
| WBGene00219260 | NA | NA | NA | NA |
| WBGene00219261 | NA | NA | NA | NA |
| WBGene00219262 | NA | NA | NA | NA |
| WBGene00219263 | NA | NA | NA | NA |
| WBGene00219264 | NA | NA | NA | NA |
| WBGene00219265 | NA | NA | NA | NA |
| WBGene00219266 | NA | NA | NA | NA |
| WBGene00219267 | NA | NA | NA | NA |
| WBGene00219269 | NA | NA | NA | NA |
| WBGene00219270 | NA | NA | NA | NA |
| WBGene00219271 | 0.063112098 | 0.73091819 | 0.84986592 | F36H5.14 |
| WBGene00219272 | NA | NA | NA | C17C3.24 |
| WBGene00219274 | -0.13732995 | 0.38769987 | 0.58621438 | T25G12.13 |
| WBGene00219275 | -0.523558304 | 0.00130044 | 0.00873831 | Y39G10AR.32 |
| WBGene00219278 | -0.027959859 | 0.85298421 | 0.92124708 | R09A8.9 |
| WBGene00219279 | 0.120872687 | 0.52301527 | 0.70071985 | NA |
| WBGene00219280 | 0.020232312 | 0.84643798 | NA | C05B5.16 |
| WBGene00219281 | NA | NA | NA | K10D3.8 |
| WBGene00219282 | NA | NA | NA | F43G9.21 |
| WBGene00219283 | 0.153645126 | 0.11943698 | 0.27743563 | F55H12.7 |
| WBGene00219284 | NA | NA | NA | C41G7.15 |
| WBGene00219285 | 0.039888668 | 0.56326497 | NA | K07A1.20 |
| WBGene00219286 | -0.03748849 | 0.76224224 | 0.86502767 | C04F12.16 |
| WBGene00219287 | 0.030854145 | 0.73726291 | 0.85229106 | F36D1.23 |
| WBGene00219288 | -0.070528244 | 0.65955484 | 0.79905192 | Y87G2A.25 |
| WBGene00219289 | -0.128640004 | 0.35376985 | 0.55364577 | T05C12.15 |
| WBGene00219290 | NA | NA | NA | AH6.17 |
| WBGene00219291 | NA | NA | NA | F35H8.12 |
| WBGene00219292 | NA | NA | NA | NA |
| WBGene00219293 | NA | NA | NA | B0334.20 |
| WBGene00219294 | NA | NA | NA | F43G6.16 |
| WBGene00219295 | 0.048220471 | 0.66873757 | 0.80677215 | C34C12.12 |
| WBGene00219296 | 0.020317245 | 0.84643798 | NA | C54C6.11 |
| WBGene00219297 | 0.361927665 | 0.04495961 | 0.13916568 | T23F11.11 |
| WBGene00219298 | 0.020317245 | 0.84643798 | NA | F55H2.8 |
| WBGene00219299 | 0.015684754 | 0.83758486 | NA | C01G12.16 |
| WBGene00219300 | -0.006738828 | 0.92480778 | NA | W10G6.6 |
| WBGene00219301 | NA | NA | NA | C44C10.15 |
| WBGene00219302 | NA | NA | NA | Y43F8B.28 |
| WBGene00219303 | -0.148946342 | 0.48028984 | 0.66672262 | NA |
| WBGene00219304 | 0.294921369 | 0.17328073 | 0.35349075 | NA |

| | | | | | |
|---|---|---|---|---|---|
| WBGene00219306 | | -0.133496872 | 0.52527532 | 0.70253504 | NA |
| WBGene00219307 | | 0.960387224 | 0.00891647 | 0.04114337 | NA |
| WBGene00219308 | | 0.517815638 | 0.03717665 | 0.12130675 | W06G6.20 |
| WBGene00219309 | | 0.109291922 | 0.21520653 | NA | F26F2.13 |
| WBGene00219310 | | 0.020232312 | 0.84643798 | NA | C17E4.19 |
| WBGene00219311 | NA | | NA | NA | NA |
| WBGene00219312 | NA | | NA | NA | NA |
| WBGene00219313 | | -0.490915882 | 0.00071348 | 0.00525723 | C17E4.20 |
| WBGene00219314 | NA | | NA | NA | C05B5.17 |
| WBGene00219315 | | -0.068557401 | 0.66014035 | 0.79959405 | F07E5.12 |
| WBGene00219316 | | 1.557851146 | 4.06E-12 | 1.41E-10 | F21A3.11 |
| WBGene00219317 | | 0.109591702 | 0.21566448 | NA | F31E3.12 |
| WBGene00219318 | | 0.020925399 | 0.84643798 | NA | R08A2.9 |
| WBGene00219319 | | 0.108545822 | 0.40506514 | 0.60312812 | H25P06.5 |
| WBGene00219321 | NA | | NA | NA | T07H3.10 |
| WBGene00219322 | | 0.1161173 | 0.12124304 | NA | C33C12.12 |
| WBGene00219323 | NA | | NA | NA | NA |
| WBGene00219324 | NA | | NA | NA | F47B8.18 |
| WBGene00219325 | | 0.065014541 | 0.37399213 | NA | F17C11.22 |
| WBGene00219326 | | 0.132442484 | 0.10147609 | NA | F36D3.16 |
| WBGene00219327 | NA | | NA | NA | ZK822.9 |
| WBGene00219329 | | 0.16127196 | 0.34427146 | 0.54397566 | K02D10.8 |
| WBGene00219330 | | -0.023109762 | 0.78121253 | NA | F18E2.15 |
| WBGene00219331 | NA | | NA | NA | NA |
| WBGene00219332 | NA | | NA | NA | Y116A8C.467 |
| WBGene00219337 | | -0.006690551 | 0.92537622 | NA | T16G1.18 |
| WBGene00219342 | | -0.050921541 | 0.47213035 | NA | Y57G11C.1143 |
| WBGene00219343 | | 0.020232312 | 0.84643798 | NA | NA |
| WBGene00219351 | NA | | NA | NA | Y7A5A.20 |
| WBGene00219352 | | 0.020317245 | 0.84643798 | NA | NA |
| WBGene00219362 | | 0.015451648 | 0.80337623 | NA | ZK470.14 |
| WBGene00219368 | | -0.003270206 | 0.95698316 | NA | F14H12.17 |
| WBGene00219369 | | 0.178768928 | 0.09068156 | 0.22888093 | R160.10 |
| WBGene00219370 | NA | | NA | NA | ZK262.20 |
| WBGene00219371 | | -0.022930767 | 0.78121253 | NA | C03G5.14 |
| WBGene00219373 | | 0.200211695 | 0.15329115 | 0.32640391 | F01E11.18 |
| WBGene00219374 | | -0.192379976 | 0.27823997 | 0.47802292 | C25A11.10 |
| WBGene00219375 | | 0.114418898 | 0.52066316 | 0.6992609 | C02B8.12 |
| WBGene00219376 | | 1.089850822 | 0.00583431 | 0.02954693 | C28G1.10 |
| WBGene00219377 | | 0.020925399 | 0.84643798 | NA | NA |
| WBGene00219378 | NA | | NA | NA | F43G6.17 |
| WBGene00219381 | NA | | NA | NA | C47A10.16 |
| WBGene00219383 | NA | | NA | NA | Y61B8A.6 |

| | | | | |
|---|---|---|---|---|
| WBGene00219394 | NA | NA | NA | Y48A6C.8 |
| WBGene00219395 | NA | NA | NA | M199.134 |
| WBGene00219399 | -0.18920017 | 0.37172111 | 0.57098881 | Y55F3AM.21 |
| WBGene00219410 | 0.163299072 | 0.16825975 | 0.34609315 | C04C3.14 |
| WBGene00219411 | -0.023109762 | 0.78121253 | NA | T14G12.11 |
| WBGene00219412 | NA | NA | NA | ZK381.62 |
| WBGene00219413 | 0.046892931 | 0.77338651 | 0.87148095 | Y38F2AL.12 |
| WBGene00219414 | 0.020317245 | 0.84643798 | NA | F14H3.15 |
| WBGene00219415 | 0.017279211 | 0.79380376 | NA | NA |
| WBGene00219416 | NA | NA | NA | NA |
| WBGene00219417 | 0.020232312 | 0.84643798 | NA | NA |
| WBGene00219418 | 0.063421086 | 0.41095452 | NA | F44D12.19 |
| WBGene00219420 | 0.127792695 | 0.25173677 | 0.44993917 | T19C4.24 |
| WBGene00219421 | -0.00346345 | 0.95708614 | NA | Y66D12A.30 |
| WBGene00219422 | -0.161976723 | 0.32419871 | 0.52363682 | Y52B11A.20 |
| WBGene00219423 | 0.484958701 | 0.06423711 | 0.18024062 | C23H5.15 |
| WBGene00219424 | 0.074892148 | 0.28555128 | NA | Y54G2A.72 |
| WBGene00219425 | 0.019171305 | 0.91395432 | 0.9519924 | Y54G2A.73 |
| WBGene00219434 | -0.049665371 | 0.48040244 | NA | F54D5.23 |
| WBGene00219436 | 0.165221751 | 0.10135336 | 0.24769062 | C49H3.16 |
| WBGene00219437 | NA | NA | NA | C50A2.7 |
| WBGene00219442 | NA | NA | NA | NA |
| WBGene00219443 | -0.226108783 | 0.09149039 | 0.2302356 | Y57G11C.1144 |
| WBGene00219444 | 0.020925399 | 0.84643798 | NA | NA |
| WBGene00219445 | NA | NA | NA | C06H5.12 |
| WBGene00219446 | 0.020925399 | 0.84643798 | NA | C06H5.13 |
| WBGene00219447 | NA | NA | NA | C06H5.14 |
| WBGene00219448 | NA | NA | NA | NA |
| WBGene00219449 | -0.030550281 | 0.70468744 | NA | Y38A10A.11 |
| WBGene00219450 | -0.032465584 | 0.75303242 | 0.86007127 | F53B6.13 |
| WBGene00219451 | -0.340838805 | 0.03979758 | 0.12750514 | F52D2.12 |
| WBGene00219452 | -0.253868589 | 0.15000116 | 0.32135893 | F21D9.11 |
| WBGene00219453 | 0.018795205 | 0.77532208 | NA | NA |
| WBGene00219454 | NA | NA | NA | C26H9A.3 |
| WBGene00219454 | NA | NA | NA | T23B5.6 |
| WBGene00219455 | NA | NA | NA | Y43F8B.29 |
| WBGene00219456 | 0.10726575 | 0.22063221 | NA | ZK822.10 |
| WBGene00219458 | NA | NA | NA | H21P03.10 |
| WBGene00219460 | 0.020925399 | 0.84643798 | NA | K09H9.9 |
| WBGene00219476 | NA | NA | NA | D1046.16 |
| WBGene00219477 | NA | NA | NA | NA |
| WBGene00219478 | 0.018790812 | 0.76755469 | NA | Y105C5B.1418 |
| WBGene00219483 | NA | NA | NA | NA |

| | | | | | |
|---|---|---|---|---|---|
| WBGene00219489 | | -0.023109762 | 0.78121253 | NA | C10C6.13 |
| WBGene00219490 | | -0.023109762 | 0.78121253 | NA | NA |
| WBGene00219491 | | -0.104863495 | 0.52179407 | 0.70004463 | Y53F4B.62 |
| WBGene00219492 | | -0.023109762 | 0.78121253 | NA | NA |
| WBGene00219493 | | 0.001350812 | 0.99224991 | 0.9955752 | C18D4.12 |
| WBGene00219497 | NA | | NA | NA | NA |
| WBGene00219498 | NA | | NA | NA | Y106G6E.8 |
| WBGene00219544 | NA | | NA | NA | C17G1.14 |
| WBGene00219545 | NA | | NA | NA | NA |
| WBGene00219546 | NA | | NA | NA | NA |
| WBGene00219547 | | -0.003108065 | 0.95728506 | NA | T06A10.106 |
| WBGene00219548 | | 0.009560321 | 0.92725366 | 0.95888691 | NA |
| WBGene00219549 | | 0.020317245 | 0.84643798 | NA | NA |
| WBGene00219550 | NA | | NA | NA | NA |
| WBGene00219551 | NA | | NA | NA | NA |
| WBGene00219552 | NA | | NA | NA | NA |
| WBGene00219553 | NA | | NA | NA | NA |
| WBGene00219554 | | -0.030698512 | 0.70341898 | NA | NA |
| WBGene00219555 | NA | | NA | NA | NA |
| WBGene00219556 | NA | | NA | NA | NA |
| WBGene00219557 | NA | | NA | NA | NA |
| WBGene00219558 | NA | | NA | NA | NA |
| WBGene00219559 | NA | | NA | NA | NA |
| WBGene00219560 | NA | | NA | NA | NA |
| WBGene00219561 | | 0.073887753 | 0.28837975 | NA | NA |
| WBGene00219562 | NA | | NA | NA | NA |
| WBGene00219563 | | 0.05998475 | 0.41450796 | NA | NA |
| WBGene00219564 | NA | | NA | NA | NA |
| WBGene00219565 | NA | | NA | NA | NA |
| WBGene00219566 | NA | | NA | NA | NA |
| WBGene00219567 | NA | | NA | NA | NA |
| WBGene00219568 | NA | | NA | NA | NA |
| WBGene00219569 | | 0.020317245 | 0.84643798 | NA | NA |
| WBGene00219570 | | 0.056411843 | 0.41891328 | NA | NA |
| WBGene00219571 | | -0.015717668 | 0.92505406 | 0.95775003 | NA |
| WBGene00219572 | NA | | NA | NA | NA |
| WBGene00219573 | NA | | NA | NA | NA |
| WBGene00219574 | | 0.020317245 | 0.84643798 | NA | NA |
| WBGene00219575 | | 0.192871103 | 0.37012201 | 0.57005643 | NA |
| WBGene00219576 | NA | | NA | NA | NA |
| WBGene00219577 | NA | | NA | NA | NA |
| WBGene00219578 | NA | | NA | NA | NA |
| WBGene00219579 | NA | | NA | NA | NA |

| | | | | |
|---|---|---|---|---|
| WBGene00219580 | NA | NA | NA | NA |
| WBGene00219581 | 0.034618355 | 0.68540017 | NA | NA |
| WBGene00219582 | NA | NA | NA | NA |
| WBGene00219583 | 0.020232312 | 0.84643798 | NA | NA |
| WBGene00219584 | NA | NA | NA | NA |
| WBGene00219585 | NA | NA | NA | NA |
| WBGene00219586 | NA | NA | NA | NA |
| WBGene00219587 | NA | NA | NA | NA |
| WBGene00219588 | -0.022930767 | 0.78121253 | NA | NA |
| WBGene00219589 | -0.005873038 | 0.97122029 | 0.98427495 | NA |
| WBGene00219590 | -0.050740777 | 0.47171914 | NA | NA |
| WBGene00219591 | 0.020232312 | 0.84643798 | NA | NA |
| WBGene00219592 | 0.011886678 | 0.86820775 | NA | NA |
| WBGene00219593 | NA | NA | NA | NA |
| WBGene00219594 | NA | NA | NA | NA |
| WBGene00219596 | -0.003421551 | 0.95689871 | NA | NA |
| WBGene00219597 | NA | NA | NA | NA |
| WBGene00219598 | -0.003421551 | 0.95689871 | NA | NA |
| WBGene00219599 | 0.065014541 | 0.37399213 | NA | NA |
| WBGene00219600 | NA | NA | NA | NA |
| WBGene00219601 | NA | NA | NA | NA |
| WBGene00219602 | 0.020232312 | 0.84643798 | NA | NA |
| WBGene00219603 | NA | NA | NA | NA |
| WBGene00219604 | NA | NA | NA | NA |
| WBGene00219605 | -0.023109762 | 0.78121253 | NA | NA |
| WBGene00219606 | NA | NA | NA | NA |
| WBGene00219607 | NA | NA | NA | NA |
| WBGene00219608 | NA | NA | NA | NA |
| WBGene00219609 | -0.808717383 | 6.93E-08 | 1.34E-06 | NA |
| WBGene00219610 | NA | NA | NA | NA |
| WBGene00219611 | NA | NA | NA | NA |
| WBGene00219612 | -0.023056499 | 0.78121253 | NA | NA |
| WBGene00219613 | NA | NA | NA | NA |
| WBGene00219614 | 0.020232312 | 0.84643798 | NA | NA |
| WBGene00219615 | -0.022930767 | 0.78121253 | NA | NA |
| WBGene00219616 | NA | NA | NA | NA |
| WBGene00219617 | NA | NA | NA | NA |
| WBGene00219618 | -0.027535543 | 0.68063395 | NA | NA |
| WBGene00219619 | 0.016241841 | 0.91147605 | 0.95112162 | NA |
| WBGene00219620 | 0.059779397 | 0.52382254 | NA | NA |
| WBGene00219621 | NA | NA | NA | NA |
| WBGene00219622 | NA | NA | NA | NA |
| WBGene00219623 | NA | NA | NA | NA |

| | | | | |
|---|---|---|---|---|
| WBGene00219624 | NA | NA | NA | NA |
| WBGene00219625 | NA | NA | NA | NA |
| WBGene00219626 | NA | NA | NA | NA |
| WBGene00219627 | NA | NA | NA | NA |
| WBGene00219628 | NA | NA | NA | NA |
| WBGene00219629 | -0.00346345 | 0.95708614 | NA | NA |
| WBGene00219630 | NA | NA | NA | NA |
| WBGene00219631 | NA | NA | NA | NA |
| WBGene00219632 | NA | NA | NA | NA |
| WBGene00219633 | NA | NA | NA | NA |
| WBGene00219634 | NA | NA | NA | NA |
| WBGene00219635 | NA | NA | NA | NA |
| WBGene00219636 | -0.058556085 | 0.62645154 | 0.77721185 | NA |
| WBGene00219637 | NA | NA | NA | NA |
| WBGene00219638 | NA | NA | NA | NA |
| WBGene00219639 | NA | NA | NA | NA |
| WBGene00219640 | NA | NA | NA | NA |
| WBGene00219641 | 0.020925399 | 0.84643798 | NA | NA |
| WBGene00219642 | NA | NA | NA | NA |
| WBGene00219643 | NA | NA | NA | NA |
| WBGene00219644 | 0.077417255 | 0.47929224 | 0.66615262 | NA |
| WBGene00219645 | NA | NA | NA | NA |
| WBGene00219646 | -0.023756719 | 0.6882956 | NA | NA |
| WBGene00219647 | -0.005691347 | 0.93660638 | NA | NA |
| WBGene00219648 | 0.022527562 | 0.85645312 | 0.92295967 | NA |
| WBGene00219649 | 0.063290468 | 0.41068021 | NA | NA |
| WBGene00219650 | 0.065014541 | 0.37399213 | NA | NA |
| WBGene00219651 | 0.013656784 | 0.85751767 | NA | NA |
| WBGene00219652 | -0.023056499 | 0.78121253 | NA | NA |
| WBGene00219653 | NA | NA | NA | NA |
| WBGene00219654 | -0.001993815 | 0.95612996 | NA | NA |
| WBGene00219655 | 0.030454968 | 0.75166778 | NA | NA |
| WBGene00219656 | 0.017279211 | 0.79380376 | NA | NA |
| WBGene00219657 | -0.0260089 | 0.6772113 | NA | NA |
| WBGene00219658 | -0.003421551 | 0.95689871 | NA | NA |
| WBGene00219659 | NA | NA | NA | NA |
| WBGene00219660 | -0.00504514 | 0.94402823 | NA | NA |
| WBGene00219661 | 0.059980168 | 0.38780842 | NA | NA |
| WBGene00219662 | -0.022930767 | 0.78121253 | NA | NA |
| WBGene00219663 | NA | NA | NA | NA |
| WBGene00219664 | 0.033034883 | 0.64219503 | NA | NA |
| WBGene00219665 | 0.01912147 | 0.77587372 | NA | NA |
| WBGene00219666 | NA | NA | NA | NA |

| | | | | |
|---|---|---|---|---|
| WBGene00219667 | NA | NA | NA | NA |
| WBGene00219668 | NA | NA | NA | NA |
| WBGene00219669 | NA | NA | NA | NA |
| WBGene00219670 | NA | NA | NA | NA |
| WBGene00219671 | NA | NA | NA | NA |
| WBGene00219672 | NA | NA | NA | NA |
| WBGene00219673 | 0.01912147 | 0.77587372 | NA | NA |
| WBGene00219674 | NA | NA | NA | NA |
| WBGene00219675 | -0.032501778 | 0.75294777 | 0.86007127 | NA |
| WBGene00219676 | 0.0398788 | 0.58971221 | NA | NA |
| WBGene00219677 | 0.011113023 | 0.90389313 | NA | NA |
| WBGene00219678 | -0.032039727 | 0.87429848 | 0.93298863 | NA |
| WBGene00219679 | NA | NA | NA | NA |
| WBGene00219680 | 0.064207016 | 0.57325152 | 0.74129796 | NA |
| WBGene00219681 | NA | NA | NA | NA |
| WBGene00219682 | 0.020232312 | 0.84643798 | NA | NA |
| WBGene00219683 | 0.054115925 | 0.54377021 | NA | NA |
| WBGene00219684 | NA | NA | NA | NA |
| WBGene00219685 | 0.040107227 | 0.58693163 | NA | NA |
| WBGene00219686 | -0.028717298 | 0.75525005 | NA | NA |
| WBGene00219687 | -0.242337869 | 0.14463961 | 0.31400779 | NA |
| WBGene00219688 | 0.008044468 | 0.9441175 | 0.96832395 | NA |
| WBGene00219689 | -0.00346345 | 0.95708614 | NA | NA |
| WBGene00219690 | NA | NA | NA | NA |
| WBGene00219691 | NA | NA | NA | NA |
| WBGene00219692 | NA | NA | NA | NA |
| WBGene00219693 | NA | NA | NA | NA |
| WBGene00219694 | NA | NA | NA | NA |
| WBGene00219695 | 0.05787416 | 0.40594086 | NA | NA |
| WBGene00219696 | 0.014602567 | 0.85416114 | NA | NA |
| WBGene00219697 | NA | NA | NA | NA |
| WBGene00219698 | NA | NA | NA | NA |
| WBGene00219699 | 0.130819452 | 0.53619466 | 0.71128259 | NA |
| WBGene00219700 | -0.094484395 | 0.24037165 | NA | NA |
| WBGene00219701 | -0.05087301 | 0.55686612 | NA | NA |
| WBGene00219702 | -0.001488154 | 0.95639912 | NA | NA |
| WBGene00219703 | 0.040107227 | 0.58693163 | NA | NA |
| WBGene00219704 | NA | NA | NA | NA |
| WBGene00219705 | 0.033463284 | 0.6365037 | NA | NA |
| WBGene00219706 | NA | NA | NA | NA |
| WBGene00219707 | 0.837128088 | 0.01035375 | 0.04610376 | NA |
| WBGene00219708 | 0.020232312 | 0.84643798 | NA | NA |
| WBGene00219709 | 0.020317245 | 0.84643798 | NA | NA |

| | | | | | |
|---|---|---|---|---|---|
| WBGene00219710 | | -0.002914971 | 0.95718083 | NA | NA |
| WBGene00219711 | | 0.020232312 | 0.84643798 | NA | NA |
| WBGene00219712 | NA | | NA | NA | NA |
| WBGene00219713 | | 0.072527804 | 0.34229409 | NA | NA |
| WBGene00219714 | NA | | NA | NA | NA |
| WBGene00219715 | | 0.038125098 | 0.61119482 | NA | NA |
| WBGene00219716 | NA | | NA | NA | NA |
| WBGene00219717 | | 0.020317245 | 0.84643798 | NA | NA |
| WBGene00219718 | | -0.001488154 | 0.95639912 | NA | NA |
| WBGene00219719 | NA | | NA | NA | NA |
| WBGene00219720 | | -0.029683071 | 0.71268666 | NA | NA |
| WBGene00219721 | | 0.056411843 | 0.41891328 | NA | NA |
| WBGene00219722 | NA | | NA | NA | NA |
| WBGene00219723 | | 0.020232312 | 0.84643798 | NA | NA |
| WBGene00219724 | NA | | NA | NA | NA |
| WBGene00219725 | | -0.20707524 | 0.24816018 | 0.44657242 | NA |
| WBGene00219726 | | -0.005691347 | 0.93660638 | NA | NA |
| WBGene00219727 | | -0.022930767 | 0.78121253 | NA | NA |
| WBGene00219728 | | -0.278077887 | 0.15622975 | 0.32997112 | NA |
| WBGene00219729 | | 0.020317245 | 0.84643798 | NA | NA |
| WBGene00219730 | NA | | NA | NA | NA |
| WBGene00219731 | | 0.040107227 | 0.58693163 | NA | NA |
| WBGene00219732 | | -0.001993815 | 0.95612996 | NA | NA |
| WBGene00219733 | | -0.023109762 | 0.78121253 | NA | NA |
| WBGene00219734 | NA | | NA | NA | NA |
| WBGene00219735 | | 0.020925399 | 0.84643798 | NA | NA |
| WBGene00219736 | | 0.167740353 | 0.38144007 | 0.58074595 | NA |
| WBGene00219737 | NA | | NA | NA | NA |
| WBGene00219738 | NA | | NA | NA | NA |
| WBGene00219739 | NA | | NA | NA | NA |
| WBGene00219740 | | 0.033034883 | 0.64219503 | NA | NA |
| WBGene00219741 | | 0.040107227 | 0.58693163 | NA | NA |
| WBGene00219742 | NA | | NA | NA | NA |
| WBGene00219743 | | 0.056411843 | 0.41891328 | NA | NA |
| WBGene00219744 | NA | | NA | NA | NA |
| WBGene00219745 | NA | | NA | NA | NA |
| WBGene00219746 | | 0.090555256 | 0.45782412 | 0.64957873 | NA |
| WBGene00219747 | NA | | NA | NA | NA |
| WBGene00219748 | | 0.167112319 | 0.16550044 | 0.34195163 | NA |
| WBGene00219749 | | -0.004747605 | 0.9473792 | NA | NA |
| WBGene00219750 | NA | | NA | NA | Y38C1AA.19 |
| WBGene00219751 | | -0.040794878 | 0.53203165 | NA | NA |
| WBGene00219752 | NA | | NA | NA | NA |

| | | | | |
|---|---|---|---|---|
| WBGene00219753 | NA | NA | NA | NA |
| WBGene00219754 | 0.169792106 | 0.0983704 | 0.2432018 | NA |
| WBGene00219755 | -0.037563819 | 0.74062602 | 0.85348332 | NA |
| WBGene00219756 | NA | NA | NA | NA |
| WBGene00219757 | NA | NA | NA | NA |
| WBGene00219758 | -0.054089928 | 0.73593831 | 0.85181646 | NA |
| WBGene00219759 | NA | NA | NA | NA |
| WBGene00219760 | NA | NA | NA | NA |
| WBGene00219761 | NA | NA | NA | NA |
| WBGene00219762 | NA | NA | NA | NA |
| WBGene00219763 | -0.022930767 | 0.78121253 | NA | NA |
| WBGene00219764 | NA | NA | NA | NA |
| WBGene00219765 | NA | NA | NA | NA |
| WBGene00219766 | NA | NA | NA | NA |
| WBGene00219767 | NA | NA | NA | NA |
| WBGene00219768 | 0.020317245 | 0.84643798 | NA | NA |
| WBGene00219769 | 0.155352187 | 0.0675224 | NA | NA |
| WBGene00219770 | -0.096677039 | 0.36415032 | 0.56388417 | NA |
| WBGene00219771 | -0.007316781 | 0.93252041 | NA | NA |
| WBGene00219772 | 0.020925399 | 0.84643798 | NA | NA |
| WBGene00219773 | 0.020317245 | 0.84643798 | NA | NA |
| WBGene00219774 | NA | NA | NA | NA |
| WBGene00219775 | NA | NA | NA | NA |
| WBGene00219776 | -0.003421551 | 0.95689871 | NA | NA |
| WBGene00219777 | NA | NA | NA | NA |
| WBGene00219778 | NA | NA | NA | NA |
| WBGene00219779 | NA | NA | NA | NA |
| WBGene00219780 | 0.076900573 | 0.64490541 | 0.78968348 | NA |
| WBGene00219781 | NA | NA | NA | NA |
| WBGene00219782 | 0.020925399 | 0.84643798 | NA | NA |
| WBGene00219783 | NA | NA | NA | NA |
| WBGene00219784 | 0.0398788 | 0.58971221 | NA | NA |
| WBGene00219785 | NA | NA | NA | NA |
| WBGene00219786 | NA | NA | NA | NA |
| WBGene00219787 | NA | NA | NA | NA |
| WBGene00219788 | NA | NA | NA | NA |
| WBGene00219789 | NA | NA | NA | NA |
| WBGene00219790 | NA | NA | NA | NA |
| WBGene00219791 | NA | NA | NA | NA |
| WBGene00219792 | NA | NA | NA | NA |
| WBGene00219793 | NA | NA | NA | NA |
| WBGene00219794 | NA | NA | NA | NA |
| WBGene00219795 | NA | NA | NA | NA |

| | | | | | |
|---|---|---|---|---|---|
| WBGene00219796 | | -0.71959169 | 0.02291771 | 0.0850325 | C03D6.9 |
| WBGene00219797 | | 0.040107227 | 0.58693163 | NA | NA |
| WBGene00219798 | NA | | NA | NA | NA |
| WBGene00219799 | NA | | NA | NA | NA |
| WBGene00219800 | | -0.13586275 | 0.47426688 | 0.66187389 | NA |
| WBGene00219801 | NA | | NA | NA | NA |
| WBGene00219803 | | -0.078358442 | 0.67727164 | 0.81260807 | NA |
| WBGene00219804 | NA | | NA | NA | NA |
| WBGene00219805 | | 0.039489191 | 0.65655312 | NA | NA |
| WBGene00219806 | NA | | NA | NA | NA |
| WBGene00219807 | NA | | NA | NA | NA |
| WBGene00219808 | NA | | NA | NA | NA |
| WBGene00219809 | NA | | NA | NA | NA |
| WBGene00219810 | NA | | NA | NA | NA |
| WBGene00219811 | NA | | NA | NA | NA |
| WBGene00219812 | NA | | NA | NA | NA |
| WBGene00219813 | NA | | NA | NA | NA |
| WBGene00219814 | NA | | NA | NA | NA |
| WBGene00219815 | NA | | NA | NA | NA |
| WBGene00219816 | | -1.241797541 | 0.00401244 | 0.02194477 | NA |
| WBGene00219817 | NA | | NA | NA | NA |
| WBGene00219818 | NA | | NA | NA | NA |
| WBGene00219819 | NA | | NA | NA | NA |
| WBGene00219820 | NA | | NA | NA | NA |
| WBGene00219821 | NA | | NA | NA | NA |
| WBGene00219822 | NA | | NA | NA | NA |
| WBGene00219823 | NA | | NA | NA | NA |
| WBGene00219824 | | -0.001993815 | 0.95612996 | NA | NA |
| WBGene00219825 | | 0.020317245 | 0.84643798 | NA | NA |
| WBGene00219826 | NA | | NA | NA | NA |
| WBGene00219827 | | 0.040107227 | 0.58693163 | NA | NA |
| WBGene00219828 | NA | | NA | NA | NA |
| WBGene00219829 | NA | | NA | NA | NA |
| WBGene00219830 | NA | | NA | NA | NA |
| WBGene00219831 | NA | | NA | NA | NA |
| WBGene00219832 | NA | | NA | NA | NA |
| WBGene00219833 | | -0.045959069 | 0.50707733 | NA | NA |
| WBGene00219834 | | -0.023109762 | 0.78121253 | NA | NA |
| WBGene00219835 | | 0.020925399 | 0.84643798 | NA | NA |
| WBGene00219836 | NA | | NA | NA | NA |
| WBGene00219837 | | 0.020232312 | 0.84643798 | NA | NA |
| WBGene00219838 | | 0.481119677 | 0.02918142 | 0.10155578 | NA |
| WBGene00219839 | NA | | NA | NA | NA |

| | | | | |
|---|---|---|---|---|
| WBGene00219840 | NA | NA | NA | NA |
| WBGene00219841 | NA | NA | NA | NA |
| WBGene00219842 | NA | NA | NA | NA |
| WBGene00219843 | NA | NA | NA | NA |
| WBGene00219844 | NA | NA | NA | NA |
| WBGene00219845 | 0.020232312 | 0.84643798 | NA | NA |
| WBGene00219846 | NA | NA | NA | NA |
| WBGene00219847 | NA | NA | NA | NA |
| WBGene00219848 | NA | NA | NA | NA |
| WBGene00219849 | -0.925943164 | 0.00508044 | 0.02646633 | NA |
| WBGene00219850 | NA | NA | NA | NA |
| WBGene00219851 | NA | NA | NA | NA |
| WBGene00219852 | NA | NA | NA | NA |
| WBGene00219853 | NA | NA | NA | NA |
| WBGene00219854 | NA | NA | NA | NA |
| WBGene00219855 | NA | NA | NA | NA |
| WBGene00219856 | NA | NA | NA | NA |
| WBGene00219857 | 0.020317245 | 0.84643798 | NA | NA |
| WBGene00219858 | NA | NA | NA | NA |
| WBGene00219859 | 0.020252628 | 0.8890894 | 0.94194334 | NA |
| WBGene00219860 | NA | NA | NA | NA |
| WBGene00219861 | 0.020317245 | 0.84643798 | NA | NA |
| WBGene00219862 | NA | NA | NA | NA |
| WBGene00219863 | 0.020925399 | 0.84643798 | NA | NA |
| WBGene00219864 | NA | NA | NA | NA |
| WBGene00219865 | 0.020232312 | 0.84643798 | NA | NA |
| WBGene00219866 | NA | NA | NA | NA |
| WBGene00219867 | NA | NA | NA | NA |
| WBGene00219868 | NA | NA | NA | NA |
| WBGene00219869 | NA | NA | NA | NA |
| WBGene00219870 | NA | NA | NA | NA |
| WBGene00219871 | NA | NA | NA | NA |
| WBGene00219872 | NA | NA | NA | NA |
| WBGene00219873 | 0.020232312 | 0.84643798 | NA | NA |
| WBGene00219874 | NA | NA | NA | NA |
| WBGene00219875 | NA | NA | NA | NA |
| WBGene00219876 | NA | NA | NA | NA |
| WBGene00219877 | NA | NA | NA | NA |
| WBGene00219878 | NA | NA | NA | NA |
| WBGene00219879 | NA | NA | NA | NA |
| WBGene00219880 | NA | NA | NA | NA |
| WBGene00219881 | NA | NA | NA | NA |
| WBGene00219882 | -0.022930767 | 0.78121253 | NA | NA |

| | | | | |
|---|---|---|---|---|
| WBGene00219883 | NA | NA | NA | NA |
| WBGene00219884 | NA | NA | NA | NA |
| WBGene00219885 | NA | NA | NA | NA |
| WBGene00219886 | -0.144131037 | 0.48052933 | 0.6668654 | NA |
| WBGene00219887 | NA | NA | NA | NA |
| WBGene00219888 | -0.022930767 | 0.78121253 | NA | NA |
| WBGene00219889 | NA | NA | NA | NA |
| WBGene00219890 | NA | NA | NA | NA |
| WBGene00219891 | 0.020317245 | 0.84643798 | NA | NA |
| WBGene00219892 | NA | NA | NA | NA |
| WBGene00219893 | -0.022930767 | 0.78121253 | NA | NA |
| WBGene00219894 | NA | NA | NA | NA |
| WBGene00219895 | NA | NA | NA | NA |
| WBGene00219896 | -0.023109762 | 0.78121253 | NA | NA |
| WBGene00219897 | NA | NA | NA | NA |
| WBGene00219898 | NA | NA | NA | NA |
| WBGene00219899 | NA | NA | NA | NA |
| WBGene00219900 | NA | NA | NA | NA |
| WBGene00219901 | NA | NA | NA | NA |
| WBGene00219902 | NA | NA | NA | NA |
| WBGene00219903 | NA | NA | NA | NA |
| WBGene00219904 | 0.032917009 | 0.86254655 | 0.92672958 | NA |
| WBGene00219905 | 0.05787416 | 0.40594086 | NA | NA |
| WBGene00219906 | NA | NA | NA | NA |
| WBGene00219907 | 0.020317245 | 0.84643798 | NA | NA |
| WBGene00219908 | NA | NA | NA | NA |
| WBGene00219909 | NA | NA | NA | NA |
| WBGene00219910 | NA | NA | NA | NA |
| WBGene00219911 | -0.023109762 | 0.78121253 | NA | NA |
| WBGene00219912 | -0.021550488 | 0.8699792 | 0.93061778 | NA |
| WBGene00219913 | NA | NA | NA | NA |
| WBGene00219914 | NA | NA | NA | NA |
| WBGene00219915 | NA | NA | NA | NA |
| WBGene00219916 | NA | NA | NA | NA |
| WBGene00219917 | 0.040107227 | 0.58693163 | NA | NA |
| WBGene00219918 | NA | NA | NA | NA |
| WBGene00219919 | NA | NA | NA | NA |
| WBGene00219920 | NA | NA | NA | NA |
| WBGene00219921 | NA | NA | NA | NA |
| WBGene00219922 | NA | NA | NA | NA |
| WBGene00219923 | NA | NA | NA | NA |
| WBGene00219924 | NA | NA | NA | NA |
| WBGene00219925 | NA | NA | NA | NA |

| | | | | | |
|---|---|---|---|---|---|
| WBGene00219926 | NA | | NA | NA | NA |
| WBGene00219927 | NA | | NA | NA | NA |
| WBGene00219928 | NA | | NA | NA | NA |
| WBGene00219929 | NA | | NA | NA | NA |
| WBGene00219930 | | 0.020232312 | 0.84643798 | NA | NA |
| WBGene00219931 | NA | | NA | NA | NA |
| WBGene00219932 | | -0.10317501 | 0.22262224 | NA | NA |
| WBGene00219933 | NA | | NA | NA | NA |
| WBGene00219934 | NA | | NA | NA | NA |
| WBGene00219935 | | 0.020232312 | 0.84643798 | NA | NA |
| WBGene00219936 | NA | | NA | NA | NA |
| WBGene00219937 | NA | | NA | NA | NA |
| WBGene00219938 | NA | | NA | NA | NA |
| WBGene00219939 | NA | | NA | NA | NA |
| WBGene00219940 | NA | | NA | NA | NA |
| WBGene00219941 | | 0.020232312 | 0.84643798 | NA | NA |
| WBGene00219942 | NA | | NA | NA | NA |
| WBGene00219943 | NA | | NA | NA | NA |
| WBGene00219944 | NA | | NA | NA | NA |
| WBGene00219945 | NA | | NA | NA | NA |
| WBGene00219946 | NA | | NA | NA | NA |
| WBGene00219947 | NA | | NA | NA | NA |
| WBGene00219948 | NA | | NA | NA | NA |
| WBGene00219949 | | -0.056791266 | 0.52823312 | NA | NA |
| WBGene00219950 | | 2.834074357 | 0.00248857 | 0.01496312 | NA |
| WBGene00219951 | | 0.03732969 | 0.59229443 | NA | NA |
| WBGene00219952 | NA | | NA | NA | NA |
| WBGene00219953 | NA | | NA | NA | NA |
| WBGene00219954 | NA | | NA | NA | NA |
| WBGene00219955 | | -0.006558791 | 0.92999162 | NA | NA |
| WBGene00219956 | NA | | NA | NA | NA |
| WBGene00219957 | NA | | NA | NA | NA |
| WBGene00219958 | NA | | NA | NA | NA |
| WBGene00219959 | NA | | NA | NA | NA |
| WBGene00219960 | NA | | NA | NA | NA |
| WBGene00219961 | | 0.056411843 | 0.41891328 | NA | NA |
| WBGene00219962 | NA | | NA | NA | NA |
| WBGene00219963 | NA | | NA | NA | NA |
| WBGene00219964 | NA | | NA | NA | NA |
| WBGene00219965 | | -0.022930767 | 0.78121253 | NA | NA |
| WBGene00219966 | NA | | NA | NA | NA |
| WBGene00219967 | NA | | NA | NA | NA |
| WBGene00219968 | NA | | NA | NA | NA |

| | | | | |
|---|---|---|---|---|
| WBGene00219969 | NA | NA | NA | NA |
| WBGene00219970 | NA | NA | NA | NA |
| WBGene00219971 | -0.054932793 | 0.45568274 | NA | NA |
| WBGene00219972 | 0.015896709 | 0.80422001 | NA | NA |
| WBGene00219973 | NA | NA | NA | NA |
| WBGene00219974 | NA | NA | NA | NA |
| WBGene00219975 | 0.065014541 | 0.37399213 | NA | NA |
| WBGene00219976 | 0.078195096 | 0.4175888 | 0.61440233 | NA |
| WBGene00219977 | NA | NA | NA | NA |
| WBGene00219978 | 0.020232312 | 0.84643798 | NA | NA |
| WBGene00219979 | -0.003108065 | 0.95728506 | NA | NA |
| WBGene00219980 | NA | NA | NA | NA |
| WBGene00219981 | NA | NA | NA | NA |
| WBGene00219982 | NA | NA | NA | NA |
| WBGene00219983 | NA | NA | NA | NA |
| WBGene00219984 | NA | NA | NA | NA |
| WBGene00219985 | NA | NA | NA | NA |
| WBGene00219986 | NA | NA | NA | NA |
| WBGene00219987 | 0.020925399 | 0.84643798 | NA | NA |
| WBGene00219988 | NA | NA | NA | NA |
| WBGene00219989 | NA | NA | NA | NA |
| WBGene00219990 | NA | NA | NA | NA |
| WBGene00219991 | NA | NA | NA | NA |
| WBGene00219992 | NA | NA | NA | NA |
| WBGene00219994 | NA | NA | NA | NA |
| WBGene00219995 | NA | NA | NA | NA |
| WBGene00219996 | NA | NA | NA | NA |
| WBGene00219997 | NA | NA | NA | NA |
| WBGene00219998 | NA | NA | NA | NA |
| WBGene00219999 | NA | NA | NA | NA |
| WBGene00220000 | 0.123938409 | 0.30637787 | 0.50701475 | NA |
| WBGene00220001 | 0.020925399 | 0.84643798 | NA | NA |
| WBGene00220002 | -0.022930767 | 0.78121253 | NA | NA |
| WBGene00220003 | NA | NA | NA | NA |
| WBGene00220004 | NA | NA | NA | NA |
| WBGene00220005 | NA | NA | NA | NA |
| WBGene00220006 | -0.065572498 | 0.66507655 | 0.80363434 | NA |
| WBGene00220007 | 0.016277737 | 0.9126034 | 0.95126775 | NA |
| WBGene00220008 | 0.040107227 | 0.58693163 | NA | NA |
| WBGene00220009 | -0.045959069 | 0.50707733 | NA | NA |
| WBGene00220010 | NA | NA | NA | NA |
| WBGene00220011 | -0.022930767 | 0.78121253 | NA | NA |
| WBGene00220012 | NA | NA | NA | NA |

| | | | | |
|---|---|---|---|---|
| WBGene00220013 | NA | NA | NA | NA |
| WBGene00220014 | NA | NA | NA | NA |
| WBGene00220015 | NA | NA | NA | NA |
| WBGene00220016 | NA | NA | NA | NA |
| WBGene00220017 | 0.040937706 | 0.64530356 | NA | NA |
| WBGene00220018 | NA | NA | NA | NA |
| WBGene00220019 | NA | NA | NA | NA |
| WBGene00220020 | NA | NA | NA | NA |
| WBGene00220021 | NA | NA | NA | NA |
| WBGene00220022 | NA | NA | NA | NA |
| WBGene00220023 | NA | NA | NA | NA |
| WBGene00220024 | NA | NA | NA | NA |
| WBGene00220025 | -0.005017292 | 0.94670066 | NA | NA |
| WBGene00220026 | -0.049826614 | 0.47909173 | NA | NA |
| WBGene00220027 | NA | NA | NA | NA |
| WBGene00220028 | NA | NA | NA | NA |
| WBGene00220029 | 0.020232312 | 0.84643798 | NA | NA |
| WBGene00220030 | NA | NA | NA | NA |
| WBGene00220031 | NA | NA | NA | NA |
| WBGene00220032 | -0.001993815 | 0.95612996 | NA | NA |
| WBGene00220033 | 0.020317245 | 0.84643798 | NA | NA |
| WBGene00220034 | NA | NA | NA | NA |
| WBGene00220035 | 0.033463284 | 0.6365037 | NA | NA |
| WBGene00220036 | NA | NA | NA | NA |
| WBGene00220037 | NA | NA | NA | NA |
| WBGene00220038 | NA | NA | NA | NA |
| WBGene00220039 | NA | NA | NA | NA |
| WBGene00220040 | NA | NA | NA | NA |
| WBGene00220041 | NA | NA | NA | NA |
| WBGene00220042 | 0.129596382 | 0.27625721 | 0.47632079 | NA |
| WBGene00220043 | NA | NA | NA | NA |
| WBGene00220044 | NA | NA | NA | NA |
| WBGene00220045 | NA | NA | NA | NA |
| WBGene00220046 | -0.001488154 | 0.95639912 | NA | NA |
| WBGene00220047 | -0.001842784 | 0.95621055 | NA | NA |
| WBGene00220048 | NA | NA | NA | NA |
| WBGene00220049 | NA | NA | NA | NA |
| WBGene00220050 | NA | NA | NA | NA |
| WBGene00220051 | NA | NA | NA | NA |
| WBGene00220052 | NA | NA | NA | NA |
| WBGene00220053 | 0.020232312 | 0.84643798 | NA | NA |
| WBGene00220054 | 0.038125098 | 0.61119482 | NA | NA |
| WBGene00220055 | NA | NA | NA | NA |

| | | | | |
|---|---|---|---|---|
| WBGene00220056 | NA | NA | NA | NA |
| WBGene00220057 | NA | NA | NA | NA |
| WBGene00220058 | -0.023056499 | 0.78121253 | NA | NA |
| WBGene00220059 | NA | NA | NA | NA |
| WBGene00220060 | NA | NA | NA | NA |
| WBGene00220061 | NA | NA | NA | NA |
| WBGene00220062 | NA | NA | NA | NA |
| WBGene00220063 | NA | NA | NA | NA |
| WBGene00220064 | NA | NA | NA | NA |
| WBGene00220065 | NA | NA | NA | NA |
| WBGene00220066 | NA | NA | NA | NA |
| WBGene00220067 | NA | NA | NA | NA |
| WBGene00220068 | NA | NA | NA | NA |
| WBGene00220069 | NA | NA | NA | NA |
| WBGene00220070 | NA | NA | NA | NA |
| WBGene00220071 | 0.020925399 | 0.84643798 | NA | NA |
| WBGene00220072 | NA | NA | NA | NA |
| WBGene00220073 | 0.020232312 | 0.84643798 | NA | NA |
| WBGene00220074 | NA | NA | NA | NA |
| WBGene00220075 | 0.020317245 | 0.84643798 | NA | NA |
| WBGene00220076 | 0.970047494 | 0.001444 | 0.00951039 | NA |
| WBGene00220077 | NA | NA | NA | NA |
| WBGene00220078 | NA | NA | NA | NA |
| WBGene00220079 | NA | NA | NA | NA |
| WBGene00220080 | NA | NA | NA | NA |
| WBGene00220081 | 0.059980168 | 0.38780842 | NA | NA |
| WBGene00220082 | NA | NA | NA | NA |
| WBGene00220083 | NA | NA | NA | NA |
| WBGene00220084 | 0.050219891 | 0.68891964 | 0.82090603 | NA |
| WBGene00220085 | NA | NA | NA | NA |
| WBGene00220086 | NA | NA | NA | NA |
| WBGene00220087 | NA | NA | NA | NA |
| WBGene00220088 | NA | NA | NA | NA |
| WBGene00220089 | NA | NA | NA | NA |
| WBGene00220090 | -0.010141451 | 0.90973981 | NA | NA |
| WBGene00220091 | NA | NA | NA | NA |
| WBGene00220092 | 0.033034883 | 0.64219503 | NA | NA |
| WBGene00220093 | NA | NA | NA | NA |
| WBGene00220094 | NA | NA | NA | NA |
| WBGene00220095 | NA | NA | NA | NA |
| WBGene00220096 | NA | NA | NA | NA |
| WBGene00220097 | NA | NA | NA | NA |
| WBGene00220098 | NA | NA | NA | NA |

| | | | | |
|---|---|---|---|---|
| WBGene00220099 | NA | NA | NA | NA |
| WBGene00220100 | NA | NA | NA | NA |
| WBGene00220101 | NA | NA | NA | NA |
| WBGene00220102 | NA | NA | NA | NA |
| WBGene00220104 | NA | NA | NA | NA |
| WBGene00220105 | NA | NA | NA | NA |
| WBGene00220106 | NA | NA | NA | NA |
| WBGene00220107 | NA | NA | NA | NA |
| WBGene00220108 | 0.020232312 | 0.84643798 | NA | NA |
| WBGene00220109 | NA | NA | NA | NA |
| WBGene00220110 | NA | NA | NA | NA |
| WBGene00220111 | NA | NA | NA | NA |
| WBGene00220112 | NA | NA | NA | NA |
| WBGene00220113 | -0.003421551 | 0.95689871 | NA | NA |
| WBGene00220114 | NA | NA | NA | NA |
| WBGene00220115 | NA | NA | NA | NA |
| WBGene00220116 | NA | NA | NA | NA |
| WBGene00220117 | NA | NA | NA | NA |
| WBGene00220118 | NA | NA | NA | NA |
| WBGene00220119 | NA | NA | NA | NA |
| WBGene00220120 | NA | NA | NA | NA |
| WBGene00220121 | 0.087256674 | 0.23800185 | NA | NA |
| WBGene00220122 | NA | NA | NA | NA |
| WBGene00220123 | -0.026384613 | 0.67299636 | NA | NA |
| WBGene00220124 | NA | NA | NA | NA |
| WBGene00220125 | NA | NA | NA | NA |
| WBGene00220126 | -0.045771477 | 0.50913401 | NA | NA |
| WBGene00220127 | -0.03323923 | 0.69122143 | NA | NA |
| WBGene00220128 | NA | NA | NA | NA |
| WBGene00220129 | 0.020232312 | 0.84643798 | NA | NA |
| WBGene00220130 | NA | NA | NA | NA |
| WBGene00220131 | NA | NA | NA | NA |
| WBGene00220132 | NA | NA | NA | NA |
| WBGene00220133 | 0.038125098 | 0.61119482 | NA | NA |
| WBGene00220134 | -0.003421551 | 0.95689871 | NA | NA |
| WBGene00220135 | 0.036687355 | 0.59489568 | NA | NA |
| WBGene00220136 | NA | NA | NA | NA |
| WBGene00220137 | NA | NA | NA | NA |
| WBGene00220138 | -0.027535543 | 0.68063395 | NA | NA |
| WBGene00220139 | NA | NA | NA | NA |
| WBGene00220140 | 0.092724492 | 0.19640118 | NA | NA |
| WBGene00220141 | NA | NA | NA | NA |
| WBGene00220142 | NA | NA | NA | NA |

| | | | | |
|---|---|---|---|---|
| WBGene00220143 | NA | NA | NA | NA |
| WBGene00220144 | NA | NA | NA | NA |
| WBGene00220145 | NA | NA | NA | NA |
| WBGene00220146 | NA | NA | NA | NA |
| WBGene00220147 | -0.022930767 | 0.78121253 | NA | NA |
| WBGene00220148 | -0.023056499 | 0.78121253 | NA | NA |
| WBGene00220149 | NA | NA | NA | NA |
| WBGene00220150 | NA | NA | NA | NA |
| WBGene00220151 | NA | NA | NA | NA |
| WBGene00220152 | NA | NA | NA | NA |
| WBGene00220153 | NA | NA | NA | NA |
| WBGene00220154 | NA | NA | NA | NA |
| WBGene00220155 | NA | NA | NA | NA |
| WBGene00220156 | NA | NA | NA | NA |
| WBGene00220157 | -0.023109762 | 0.78121253 | NA | NA |
| WBGene00220158 | NA | NA | NA | NA |
| WBGene00220159 | NA | NA | NA | NA |
| WBGene00220160 | NA | NA | NA | NA |
| WBGene00220161 | NA | NA | NA | NA |
| WBGene00220162 | NA | NA | NA | NA |
| WBGene00220163 | NA | NA | NA | NA |
| WBGene00220164 | NA | NA | NA | NA |
| WBGene00220165 | 0.051784019 | 0.80385651 | 0.89090365 | NA |
| WBGene00220166 | NA | NA | NA | NA |
| WBGene00220167 | -0.022930767 | 0.78121253 | NA | NA |
| WBGene00220168 | 0.032317057 | 0.61887865 | NA | NA |
| WBGene00220169 | NA | NA | NA | NA |
| WBGene00220170 | NA | NA | NA | NA |
| WBGene00220171 | NA | NA | NA | NA |
| WBGene00220172 | -0.023109762 | 0.78121253 | NA | NA |
| WBGene00220173 | NA | NA | NA | NA |
| WBGene00220174 | NA | NA | NA | NA |
| WBGene00220175 | -0.022930767 | 0.78121253 | NA | NA |
| WBGene00220176 | NA | NA | NA | NA |
| WBGene00220177 | 0.020232312 | 0.84643798 | NA | NA |
| WBGene00220178 | NA | NA | NA | NA |
| WBGene00220179 | -0.023109762 | 0.78121253 | NA | NA |
| WBGene00220180 | NA | NA | NA | NA |
| WBGene00220181 | NA | NA | NA | NA |
| WBGene00220182 | NA | NA | NA | NA |
| WBGene00220183 | NA | NA | NA | NA |
| WBGene00220184 | NA | NA | NA | NA |
| WBGene00220185 | NA | NA | NA | NA |

| | | | | |
|---|---|---|---|---|
| WBGene00220186 | NA | NA | NA | NA |
| WBGene00220187 | NA | NA | NA | NA |
| WBGene00220188 | NA | NA | NA | NA |
| WBGene00220189 | NA | NA | NA | NA |
| WBGene00220190 | NA | NA | NA | NA |
| WBGene00220191 | -0.023109762 | 0.78121253 | NA | NA |
| WBGene00220192 | NA | NA | NA | NA |
| WBGene00220193 | NA | NA | NA | NA |
| WBGene00220194 | NA | NA | NA | NA |
| WBGene00220195 | NA | NA | NA | NA |
| WBGene00220196 | NA | NA | NA | NA |
| WBGene00220197 | NA | NA | NA | NA |
| WBGene00220198 | -0.046394786 | 0.50231372 | NA | NA |
| WBGene00220199 | NA | NA | NA | NA |
| WBGene00220200 | NA | NA | NA | NA |
| WBGene00220201 | -0.022930767 | 0.78121253 | NA | NA |
| WBGene00220202 | NA | NA | NA | NA |
| WBGene00220203 | NA | NA | NA | NA |
| WBGene00220204 | NA | NA | NA | NA |
| WBGene00220205 | NA | NA | NA | NA |
| WBGene00220206 | NA | NA | NA | NA |
| WBGene00220207 | NA | NA | NA | NA |
| WBGene00220208 | NA | NA | NA | NA |
| WBGene00220209 | NA | NA | NA | NA |
| WBGene00220210 | NA | NA | NA | NA |
| WBGene00220211 | NA | NA | NA | NA |
| WBGene00220212 | NA | NA | NA | NA |
| WBGene00220213 | NA | NA | NA | NA |
| WBGene00220214 | NA | NA | NA | NA |
| WBGene00220215 | NA | NA | NA | NA |
| WBGene00220216 | NA | NA | NA | NA |
| WBGene00220217 | NA | NA | NA | NA |
| WBGene00220218 | NA | NA | NA | NA |
| WBGene00220219 | NA | NA | NA | NA |
| WBGene00220220 | NA | NA | NA | NA |
| WBGene00220221 | NA | NA | NA | NA |
| WBGene00220222 | NA | NA | NA | NA |
| WBGene00220223 | NA | NA | NA | NA |
| WBGene00220224 | NA | NA | NA | NA |
| WBGene00220225 | NA | NA | NA | NA |
| WBGene00220226 | NA | NA | NA | NA |
| WBGene00220227 | NA | NA | NA | NA |
| WBGene00220228 | NA | NA | NA | NA |

| | | | | |
|---|---|---|---|---|
| WBGene00220229 | NA | NA | NA | NA |
| WBGene00220230 | NA | NA | NA | NA |
| WBGene00220231 | NA | NA | NA | NA |
| WBGene00220232 | NA | NA | NA | NA |
| WBGene00220233 | NA | NA | NA | NA |
| WBGene00220234 | NA | NA | NA | NA |
| WBGene00220235 | NA | NA | NA | NA |
| WBGene00220236 | 0.020317245 | 0.84643798 | NA | NA |
| WBGene00220237 | NA | NA | NA | NA |
| WBGene00220238 | 0.016613161 | 0.83477339 | NA | NA |
| WBGene00220239 | NA | NA | NA | NA |
| WBGene00220240 | 0.020232312 | 0.84643798 | NA | NA |
| WBGene00220241 | NA | NA | NA | NA |
| WBGene00220242 | NA | NA | NA | NA |
| WBGene00220243 | 0.01912147 | 0.77587372 | NA | NA |
| WBGene00220244 | NA | NA | NA | NA |
| WBGene00220245 | NA | NA | NA | NA |
| WBGene00220246 | NA | NA | NA | NA |
| WBGene00220247 | NA | NA | NA | NA |
| WBGene00220248 | 0.020925399 | 0.84643798 | NA | NA |
| WBGene00220249 | -0.002914971 | 0.95718083 | NA | R144.19 |
| WBGene00220250 | NA | NA | NA | C09B9.85 |
| WBGene00220251 | 0.020232312 | 0.84643798 | NA | NA |
| WBGene00220252 | 0.020232312 | 0.84643798 | NA | NA |
| WBGene00220253 | NA | NA | NA | C05A2.4 |
| WBGene00220254 | 0.020232312 | 0.84643798 | NA | Y105C5A.1285 |
| WBGene00220255 | NA | NA | NA | NA |
| WBGene00220256 | NA | NA | NA | NA |
| WBGene00220257 | NA | NA | NA | C02H6.3 |
| WBGene00220258 | -0.003108065 | 0.95728506 | NA | C50C3.19 |
| WBGene00220259 | -0.007128374 | 0.96159491 | 0.97914283 | ZK418.13 |
| WBGene00220260 | 0.020925399 | 0.84643798 | NA | NA |
| WBGene00220261 | 0.131317611 | 0.10121567 | NA | ZC434.10 |
| WBGene00235003 | NA | NA | NA | NA |
| WBGene00235080 | NA | NA | NA | NA |
| WBGene00235081 | NA | NA | NA | Y70C5B.1 |
| WBGene00235081 | NA | NA | NA | W06G6.21 |
| WBGene00235084 | NA | NA | NA | NA |
| WBGene00235085 | NA | NA | NA | NA |
| WBGene00235086 | 0.036687355 | 0.59489568 | NA | NA |
| WBGene00235087 | NA | NA | NA | NA |
| WBGene00235088 | NA | NA | NA | NA |
| WBGene00235089 | NA | NA | NA | NA |

| | | | | |
|---|---|---|---|---|
| WBGene00235090 | 0.055503542 | 0.44925503 | NA | C38D9.13 |
| WBGene00235091 | NA | NA | NA | NA |
| WBGene00235092 | 0.020232312 | 0.84643798 | NA | C54F6.17 |
| WBGene00235093 | -0.007072416 | 0.9244612 | NA | C54F6.18 |
| WBGene00235094 | -0.250556247 | 0.25507961 | 0.45356091 | K08D9.9 |
| WBGene00235095 | 0.05787416 | 0.40594086 | NA | B0035.21 |
| WBGene00235102 | 0.016311866 | 0.86300826 | 0.9270525 | Y54G2A.75 |
| WBGene00235106 | 0.155122056 | 0.06782466 | NA | NA |
| WBGene00235107 | NA | NA | NA | NA |
| WBGene00235108 | NA | NA | NA | NA |
| WBGene00235109 | NA | NA | NA | NA |
| WBGene00235110 | NA | NA | NA | T14G12.12 |
| WBGene00235111 | NA | NA | NA | NA |
| WBGene00235112 | NA | NA | NA | C26H9A.3 |
| WBGene00235112 | NA | NA | NA | T23B5.6 |
| WBGene00235113 | 0.117508794 | 0.17909081 | 0.3605209 | ZK105.12 |
| WBGene00235114 | 0.102413642 | 0.23275609 | NA | ZK105.13 |
| WBGene00235115 | NA | NA | NA | F47C12.13 |
| WBGene00235116 | NA | NA | NA | C03G6.21 |
| WBGene00235117 | NA | NA | NA | Y19D10B.8 |
| WBGene00235118 | NA | NA | NA | K04A8.18 |
| WBGene00235119 | NA | NA | NA | NA |
| WBGene00235120 | -0.023109762 | 0.78121253 | NA | R105.5 |
| WBGene00235121 | NA | NA | NA | NA |
| WBGene00235128 | -0.022930767 | 0.78121253 | NA | NA |
| WBGene00235129 | NA | NA | NA | ZC518.8 |
| WBGene00235132 | 0.018476367 | 0.76699386 | NA | ZK792.12 |
| WBGene00235133 | 0.182879948 | 0.22641527 | 0.42003877 | T26H5.14 |
| WBGene00235134 | 0.020925399 | 0.84643798 | NA | R07H5.16 |
| WBGene00235152 | NA | NA | NA | NA |
| WBGene00235153 | NA | NA | NA | NA |
| WBGene00235157 | NA | NA | NA | NA |
| WBGene00235158 | -0.274347073 | 0.19903143 | 0.38750254 | Y37E3.30 |
| WBGene00235159 | NA | NA | NA | NA |
| WBGene00235160 | NA | NA | NA | NA |
| WBGene00235161 | NA | NA | NA | NA |
| WBGene00235162 | -0.022930767 | 0.78121253 | NA | NA |
| WBGene00235163 | NA | NA | NA | NA |
| WBGene00235164 | 0.327828137 | 0.1105092 | 0.26225301 | K09F6.13 |
| WBGene00235165 | NA | NA | NA | Y47D7A.18 |
| WBGene00235166 | NA | NA | NA | NA |
| WBGene00235167 | NA | NA | NA | NA |
| WBGene00235175 | NA | NA | NA | NA |

| | | | | |
|---|---|---|---|---|
| WBGene00235179 | NA | NA | NA | NA |
| WBGene00235180 | NA | NA | NA | NA |
| WBGene00235181 | NA | NA | NA | NA |
| WBGene00235183 | NA | NA | NA | NA |
| WBGene00235188 | NA | NA | NA | NA |
| WBGene00235190 | NA | NA | NA | NA |
| WBGene00235197 | NA | NA | NA | NA |
| WBGene00235198 | NA | NA | NA | NA |
| WBGene00235200 | NA | NA | NA | NA |
| WBGene00235201 | NA | NA | NA | NA |
| WBGene00235206 | NA | NA | NA | NA |
| WBGene00235209 | NA | NA | NA | NA |
| WBGene00235235 | NA | NA | NA | NA |
| WBGene00235237 | NA | NA | NA | NA |
| WBGene00235247 | NA | NA | NA | NA |
| WBGene00235250 | NA | NA | NA | NA |
| WBGene00235251 | NA | NA | NA | NA |
| WBGene00235252 | NA | NA | NA | NA |
| WBGene00235254 | -0.022930767 | 0.78121253 | NA | F49D11.14 |
| WBGene00235255 | NA | NA | NA | NA |
| WBGene00235256 | -0.270471155 | 0.22434768 | 0.41720746 | Y54G11A.19 |
| WBGene00235257 | -0.027535543 | 0.68063395 | NA | T05C7.4 |
| WBGene00235258 | NA | NA | NA | T05C7.5 |
| WBGene00235259 | 0.054438138 | 0.58727981 | 0.74981371 | NA |
| WBGene00235260 | 0.020232312 | 0.84643798 | NA | T05C7.7 |
| WBGene00235261 | NA | NA | NA | T05C7.8 |
| WBGene00235262 | NA | NA | NA | T05C7.9 |
| WBGene00235263 | 0.106905496 | 0.30908553 | 0.50995281 | Y66H1A.8 |
| WBGene00235264 | NA | NA | NA | M04G7.5 |
| WBGene00235265 | NA | NA | NA | NA |
| WBGene00235266 | -0.045959069 | 0.50707733 | NA | NA |
| WBGene00235267 | -0.807731718 | 0.02245609 | 0.08361638 | Y77E11A.23 |
| WBGene00235268 | 0.204737186 | 0.03089539 | 0.105465 | Y54G2A.76 |
| WBGene00235269 | 0.043643477 | 0.71534859 | 0.84039298 | F59E12.15 |
| WBGene00235270 | NA | NA | NA | NA |
| WBGene00235271 | 0.106997704 | 0.16248985 | NA | C49C8.8 |
| WBGene00235272 | -0.184668252 | 0.35720257 | 0.55731062 | NA |
| WBGene00235273 | 0.131539016 | 0.10075387 | NA | T11F8.12 |
| WBGene00235274 | 0.061794036 | 0.58114697 | 0.74581663 | C54E4.12 |
| WBGene00235275 | 0.065014541 | 0.37399213 | NA | T09A12.8 |
| WBGene00235276 | NA | NA | NA | NA |
| WBGene00235277 | 0.020925399 | 0.84643798 | NA | T04C4.3 |
| WBGene00235278 | NA | NA | NA | NA |

| | | | | | |
|---|---|---|---|---|---|
| WBGene00235279 | | -0.046394786 | 0.50231372 | NA | Y67D8C.22 |
| WBGene00235280 | | -0.003421551 | 0.95689871 | NA | F42A9.17 |
| WBGene00235281 | | 0.020317245 | 0.84643798 | NA | NA |
| WBGene00235282 | NA | | NA | NA | F17E9.18 |
| WBGene00235283 | NA | | NA | NA | Y55F3BL.6 |
| WBGene00235284 | | -0.003614866 | 0.95700117 | NA | T12G3.11 |
| WBGene00235285 | | 0.020925399 | 0.84643798 | NA | F56D6.18 |
| WBGene00235286 | | -0.040794878 | 0.53203165 | NA | NA |
| WBGene00235287 | NA | | NA | NA | Y55F3C.17 |
| WBGene00235288 | | 0.0398788 | 0.58971221 | NA | Y69A2AR.46 |
| WBGene00235289 | NA | | NA | NA | Y73B6BL.289 |
| WBGene00235290 | NA | | NA | NA | Y59H11AR.10 |
| WBGene00235291 | | 0.549956903 | 0.0484081 | 0.14670169 | C01G5.25 |
| WBGene00235292 | | -0.016029059 | 0.89319957 | 0.94441647 | NA |
| WBGene00235293 | | 0.020925399 | 0.84643798 | NA | F56D6.19 |
| WBGene00235294 | | 0.020232312 | 0.84643798 | NA | Y69A2AR.47 |
| WBGene00235295 | | 0.076064012 | 0.48487322 | 0.66987915 | C42D4.19 |
| WBGene00235296 | | -0.050042403 | 0.47734215 | NA | C07G1.15 |
| WBGene00235297 | | 0.056411843 | 0.41891328 | NA | F56D6.20 |
| WBGene00235298 | NA | | NA | NA | F19B6.10 |
| WBGene00235299 | | 0.020232312 | 0.84643798 | NA | NA |
| WBGene00235300 | | 0.040107227 | 0.58693163 | NA | T06A10.107 |
| WBGene00235301 | NA | | NA | NA | B0513.90 |
| WBGene00235302 | NA | | NA | NA | C07C7.3 |
| WBGene00235303 | | 0.020925399 | 0.84643798 | NA | T13A10.64 |
| WBGene00235304 | NA | | NA | NA | Y116A8C.468 |
| WBGene00235305 | | 0.020925399 | 0.84643798 | NA | Y116A8C.469 |
| WBGene00235307 | | 0.038125098 | 0.61119482 | NA | NA |
| WBGene00235308 | | 0.0398788 | 0.58971221 | NA | C37C3.17 |
| WBGene00235309 | | 0.005436156 | 0.89690084 | NA | F21C10.17 |
| WBGene00235310 | | -0.262231713 | 0.21112701 | 0.40107832 | R02D1.2 |
| WBGene00235311 | | 3.896562938 | 0.00170886 | 0.01097677 | Y57E12B.11 |
| WBGene00235312 | NA | | NA | NA | F28A12.10 |
| WBGene00235313 | | 0.040107227 | 0.58693163 | NA | C25E10.17 |
| WBGene00235314 | | 0.065014541 | 0.37399213 | NA | NA |
| WBGene00235315 | | 0.106014872 | 0.56816832 | 0.73716355 | Y50D4A.10 |
| WBGene00235316 | | 0.020317245 | 0.84643798 | NA | B0222.15 |
| WBGene00235317 | NA | | NA | NA | K04A8.19 |
| WBGene00235318 | | -0.005221332 | 0.94448901 | NA | NA |
| WBGene00235319 | | -0.186227704 | 0.33912708 | 0.53887123 | NA |
| WBGene00235320 | | 0.020925399 | 0.84643798 | NA | R02F11.11 |
| WBGene00235321 | NA | | NA | NA | Y73C8B.9 |
| WBGene00235322 | NA | | NA | NA | F32F2.4 |

| | | | | |
|---|---|---|---|---|
| WBGene00235323 | NA | NA | NA | NA |
| WBGene00235324 | NA | NA | NA | F10G2.10 |
| WBGene00235325 | 0.012634358 | 0.93607954 | 0.96386107 | C08D8.7 |
| WBGene00235326 | NA | NA | NA | F22F7.11 |
| WBGene00235327 | NA | NA | NA | T02B11.10 |
| WBGene00235328 | -0.0260089 | 0.6772113 | NA | T06E4.21 |
| WBGene00235329 | NA | NA | NA | F37B4.15 |
| WBGene00235330 | -0.034050838 | 0.7451853 | 0.85590582 | F21C10.18 |
| WBGene00235331 | NA | NA | NA | NA |
| WBGene00235332 | 5.555353159 | 1.04E-06 | 1.63E-05 | K04A8.20 |
| WBGene00235333 | 0.015507174 | 0.80923808 | NA | C24H12.18 |
| WBGene00235334 | NA | NA | NA | NA |
| WBGene00235335 | -0.045771477 | 0.50913401 | NA | R02D5.24 |
| WBGene00235336 | NA | NA | NA | C54F6.19 |
| WBGene00235338 | -0.032838509 | 0.74430088 | 0.85556326 | NA |
| WBGene00235339 | NA | NA | NA | W06G6.12 |
| WBGene00235339 | NA | NA | NA | F14F8.15 |
| WBGene00235340 | 0.189218374 | 0.36153483 | 0.56137782 | C45G7.13 |
| WBGene00235341 | -0.023056499 | 0.78121253 | NA | NA |
| WBGene00235342 | NA | NA | NA | Y69A2AR.50 |
| WBGene00235343 | 0.017279211 | 0.79380376 | NA | Y69A2AR.51 |
| WBGene00235346 | NA | NA | NA | NA |
| WBGene00235348 | NA | NA | NA | NA |
| WBGene00235350 | NA | NA | NA | Y54G2A.77 |
| WBGene00235351 | 0.038125098 | 0.61119482 | NA | C32B5.18 |
| WBGene00235352 | 1.442035167 | 0.0001587 | 0.00144227 | ZK250.13 |
| WBGene00235353 | 0.19278513 | 0.31827268 | 0.51856558 | NA |
| WBGene00235354 | -0.008998908 | 0.92213607 | NA | ZK250.14 |
| WBGene00235355 | -0.235232833 | 0.17673563 | 0.35754958 | NA |
| WBGene00235356 | NA | NA | NA | NA |
| WBGene00235357 | NA | NA | NA | NA |
| WBGene00235358 | NA | NA | NA | F56D6.21 |
| WBGene00235359 | 0.065014541 | 0.37399213 | NA | F56D6.22 |
| WBGene00235361 | NA | NA | NA | F56D6.23 |
| WBGene00235362 | NA | NA | NA | NA |
| WBGene00235363 | NA | NA | NA | F49F1.2 |
| WBGene00235364 | NA | NA | NA | F49F1.4 |
| WBGene00235365 | NA | NA | NA | F49F1.15 |
| WBGene00235366 | NA | NA | NA | NA |
| WBGene00235367 | NA | NA | NA | F49F1.17 |
| WBGene00235368 | 0.038125098 | 0.61119482 | NA | F49F1.18 |
| WBGene00235369 | 0.020925399 | 0.84643798 | NA | K08D10.18 |
| WBGene00235370 | -0.022930767 | 0.78121253 | NA | NA |

| | | | | | |
|---|---|---|---|---|---|
| WBGene00235371 | | -0.023109762 | 0.78121253 | NA | F30B5.11 |
| WBGene00235372 | | -0.003614866 | 0.95700117 | NA | NA |
| WBGene00235373 | NA | | NA | NA | NA |
| WBGene00235374 | | 5.96878484 | 1.05E-05 | 0.00012844 | R11E3.19 |
| WBGene00235375 | | 0.029901071 | 0.7449032 | 0.85585245 | NA |
| WBGene00235376 | NA | | NA | NA | NA |
| WBGene00235377 | NA | | NA | NA | NA |
| WBGene00235378 | NA | | NA | NA | NA |
| WBGene00235379 | NA | | NA | NA | NA |
| WBGene00235380 | | -0.040794878 | 0.53203165 | NA | C24D10.49 |
| WBGene00235381 | | 0.023021378 | 0.90817744 | 0.95041108 | NA |
| WBGene00235382 | NA | | NA | NA | NA |
| WBGene00235383 | | 0.124326451 | 0.11293212 | NA | ZK616.65 |
| WBGene00235385 | | 0.04785768 | 0.81537062 | 0.89862521 | NA |
| WBGene00235386 | NA | | NA | NA | NA |
| WBGene00235387 | | 0.050891708 | 0.65487245 | 0.79569622 | NA |
| WBGene00235388 | NA | | NA | NA | W04C9.9 |
| WBGene00235389 | NA | | NA | NA | NA |
| WBGene00235390 | NA | | NA | NA | F56A6.8 |
| WBGene00235391 | | 0.539757072 | 0.05264905 | 0.15654563 | F38A5.22 |
| WBGene00236785 | | 0.058314668 | 0.40208118 | NA | NA |
| WBGene00236786 | NA | | NA | NA | C17H12.37 |
| WBGene00236787 | NA | | NA | NA | NA |
| WBGene00236788 | NA | | NA | NA | NA |
| WBGene00236789 | | 0.013306624 | 0.88406782 | NA | Y18H1A.18 |
| WBGene00236797 | NA | | NA | NA | NA |
| WBGene00236798 | | 0.020317245 | 0.84643798 | NA | NA |
| WBGene00236799 | NA | | NA | NA | NA |
| WBGene00236800 | NA | | NA | NA | NA |
| WBGene00236806 | NA | | NA | NA | D2024.18 |
| WBGene00236807 | NA | | NA | NA | NA |
| WBGene00236808 | | -0.099452571 | 0.17595618 | NA | C54G6.8 |
| WBGene00236809 | | 0.056411843 | 0.41891328 | NA | ZK993.5 |
| WBGene00249808 | NA | | NA | NA | NA |
| WBGene00249812 | NA | | NA | NA | NA |
| WBGene00249813 | NA | | NA | NA | NA |
| WBGene00249814 | NA | | NA | NA | NA |
| WBGene00249815 | | -0.188229011 | 0.2914 | 0.49208621 | F20D12.12 |
| WBGene00249816 | NA | | NA | NA | NA |
| WBGene00249817 | | 1.785437504 | 0.00444974 | 0.02390777 | F42G8.19 |
| WBGene00249825 | NA | | NA | NA | NA |
| WBGene00249826 | | 0.072952506 | 0.34318498 | NA | NA |
| WBGene00255375 | | 1.941140893 | 0.00013251 | 0.00123495 | F08B4.8 |

| | | | | |
|---|---|---|---|---|
| WBGene00255420 | NA | NA | NA | Y51F10.15 |
| WBGene00255426 | -0.089291045 | 0.6507991 | 0.79325861 | NA |
| WBGene00255435 | NA | NA | NA | NA |
| WBGene00255441 | NA | NA | NA | NA |
| WBGene00255457 | 0.033829031 | 0.83199189 | 0.9084712 | F23C8.14 |
| WBGene00255469 | -0.001842784 | 0.95621055 | NA | C25G4.17 |
| WBGene00255522 | NA | NA | NA | Y57G11B.223 |
| WBGene00255543 | NA | NA | NA | NA |
| WBGene00255545 | NA | NA | NA | M199.135 |
| WBGene00255546 | NA | NA | NA | M199.136 |
| WBGene00255549 | 0.020317245 | 0.84643798 | NA | NA |
| WBGene00255550 | NA | NA | NA | W03D8.14 |
| WBGene00255553 | NA | NA | NA | Y51H4A.941 |
| WBGene00255560 | NA | NA | NA | NA |
| WBGene00255561 | NA | NA | NA | NA |
| WBGene00255562 | NA | NA | NA | NA |
| WBGene00255563 | NA | NA | NA | NA |
| WBGene00255564 | NA | NA | NA | NA |
| WBGene00255565 | NA | NA | NA | NA |
| WBGene00255566 | NA | NA | NA | NA |
| WBGene00255567 | 0.020232312 | 0.84643798 | NA | NA |
| WBGene00255568 | NA | NA | NA | NA |
| WBGene00255569 | NA | NA | NA | NA |
| WBGene00255570 | 0.020232312 | 0.84643798 | NA | NA |
| WBGene00255571 | NA | NA | NA | NA |
| WBGene00255572 | NA | NA | NA | NA |
| WBGene00255573 | NA | NA | NA | NA |
| WBGene00255574 | NA | NA | NA | NA |
| WBGene00255575 | NA | NA | NA | NA |
| WBGene00255576 | NA | NA | NA | NA |
| WBGene00255577 | NA | NA | NA | NA |
| WBGene00255578 | NA | NA | NA | NA |
| WBGene00255579 | NA | NA | NA | NA |
| WBGene00255580 | NA | NA | NA | NA |
| WBGene00255581 | NA | NA | NA | NA |
| WBGene00255582 | NA | NA | NA | NA |
| WBGene00255583 | NA | NA | NA | NA |
| WBGene00255584 | -0.054932793 | 0.45568274 | NA | NA |
| WBGene00255586 | -0.023056499 | 0.78121253 | NA | NA |
| WBGene00255587 | NA | NA | NA | NA |
| WBGene00255588 | NA | NA | NA | NA |
| WBGene00255594 | 0.207013528 | 0.23712389 | 0.43347869 | Y48G1BL.8 |
| WBGene00255595 | NA | NA | NA | D1046.18 |

| | | | | |
|---|---|---|---|---|
| WBGene00255596 | -0.26853996 | 0.19076287 | 0.37682446 | NA |
| WBGene00255597 | NA | NA | NA | pals-21 |
| WBGene00255598 | 0.118977087 | 0.49162731 | 0.67507131 | BE0003N10.6 |
| WBGene00255600 | 0.052272507 | 0.78715687 | 0.87949869 | F54H5.14 |
| WBGene00255605 | NA | NA | NA | T23B3.8 |
| WBGene00255606 | -0.027188909 | 0.68205301 | NA | NA |
| WBGene00255607 | 0.020232312 | 0.84643798 | NA | F22H10.11 |
| WBGene00255611 | NA | NA | NA | Y116A8C.470 |
| WBGene00255702 | 0.086632749 | 0.29414331 | NA | C06A12.19 |
| WBGene00255703 | NA | NA | NA | NA |
| WBGene00255704 | NA | NA | NA | B0348.10 |
| WBGene00255709 | 0.020925399 | 0.84643798 | NA | NA |
| WBGene00255714 | -0.254174938 | 0.03215179 | 0.10880816 | B0511.19 |
| WBGene00255728 | 0.040107227 | 0.58693163 | NA | NA |
| WBGene00255729 | NA | NA | NA | R11G11.16 |
| WBGene00255730 | -0.023756719 | 0.6882956 | NA | NA |
| WBGene00255731 | NA | NA | NA | T02B11.11 |
| WBGene00255732 | NA | NA | NA | T02B11.12 |
| WBGene00255733 | NA | NA | NA | T02B11.13 |
| WBGene00255736 | 0.108614162 | 0.30880061 | 0.50965502 | NA |
| WBGene00268189 | 0.092938702 | 0.19548342 | NA | NA |
| WBGene00268190 | 0.020317245 | 0.84643798 | NA | NA |
| WBGene00268191 | NA | NA | NA | NA |
| WBGene00268192 | -0.001842784 | 0.95621055 | NA | NA |
| WBGene00268193 | NA | NA | NA | NA |
| WBGene00268194 | NA | NA | NA | F36H9.10 |
| WBGene00268196 | 0.102989984 | 0.32666778 | 0.52574947 | F59A7.14 |
| WBGene00268197 | NA | NA | NA | Y40B10A.11 |
| WBGene00268198 | NA | NA | NA | F22F7.12 |
| WBGene00268199 | NA | NA | NA | C45H4.21 |
| WBGene00268202 | NA | NA | NA | NA |
| WBGene00268203 | NA | NA | NA | NA |
| WBGene00268204 | NA | NA | NA | NA |
| WBGene00268205 | NA | NA | NA | NA |
| WBGene00268206 | NA | NA | NA | F54E2.9 |
| WBGene00268207 | -0.104657486 | 0.21322776 | NA | NA |
| WBGene00268208 | NA | NA | NA | NA |
| WBGene00268209 | -0.002914971 | 0.95718083 | NA | NA |
| WBGene00268210 | 0.020317245 | 0.84643798 | NA | NA |
| WBGene00268211 | NA | NA | NA | NA |
| WBGene00268212 | -0.556832618 | 0.04389493 | 0.13691215 | NA |
| WBGene00268213 | 0.018998992 | 0.77286019 | NA | C17B7.15 |
| WBGene00268214 | NA | NA | NA | NA |

| | | | | |
|---|---|---|---|---|
| WBGene00268215 | -0.023109762 | 0.78121253 | NA | C49F5.13 |
| WBGene00268216 | 0.040107227 | 0.58693163 | NA | F49B2.9 |
| WBGene00269331 | NA | NA | NA | NA |
| WBGene00269332 | NA | NA | NA | Y39F10A.6 |
| WBGene00269333 | 0.020232312 | 0.84643798 | NA | NA |
| WBGene00269336 | NA | NA | NA | NA |
| WBGene00269337 | 0.059762547 | 0.38965825 | NA | NA |
| WBGene00269338 | 0.020232312 | 0.84643798 | NA | NA |
| WBGene00269346 | 0.0398788 | 0.58971221 | NA | NA |
| WBGene00269352 | NA | NA | NA | K08D9.10 |
| WBGene00269355 | -0.044780899 | 0.80507741 | 0.89161409 | T28A11.25 |
| WBGene00269373 | -0.03039785 | 0.70603871 | NA | M01D1.12 |
| WBGene00269376 | NA | NA | NA | NA |
| WBGene00269378 | 0.036894305 | 0.59120501 | NA | C07G3.15 |
| WBGene00269379 | 0.075519207 | 0.42603099 | 0.62170069 | C30B5.17 |
| WBGene00269381 | NA | NA | NA | F08F8.11 |
| WBGene00269382 | 0.196370465 | 0.14874752 | 0.31960586 | F33D11.16 |
| WBGene00269383 | 0.242349331 | 0.25645077 | 0.45467374 | NA |
| WBGene00269384 | -0.097822242 | 0.45487637 | 0.64690092 | NA |
| WBGene00269385 | NA | NA | NA | F41F3.9 |
| WBGene00269386 | -0.031690166 | 0.70451177 | NA | NA |
| WBGene00269387 | -0.025848508 | 0.67901638 | NA | btb-22 |
| WBGene00269388 | -0.942769609 | 0.01009256 | 0.04516048 | F45C12.17 |
| WBGene00269389 | 0.046567441 | 0.70369999 | 0.8318388 | C08F1.12 |
| WBGene00269390 | -0.075958958 | 0.41028869 | NA | NA |
| WBGene00269391 | NA | NA | NA | F54F2.14 |
| WBGene00269392 | 0.020317245 | 0.84643798 | NA | F54F7.14 |
| WBGene00269393 | -0.026384613 | 0.67299636 | NA | NA |
| WBGene00269394 | 0.056677809 | 0.43780437 | NA | F59H6.15 |
| WBGene00269419 | -0.14515815 | 0.46028777 | 0.65136993 | NA |
| WBGene00269420 | NA | NA | NA | NA |
| WBGene00269421 | 2.453671294 | 3.93E-17 | 2.34E-15 | R06F6.14 |
| WBGene00269422 | 0.020925399 | 0.84643798 | NA | T01H10.11 |
| WBGene00269423 | NA | NA | NA | NA |
| WBGene00269424 | NA | NA | NA | K09D9.14 |
| WBGene00269425 | 0.098194087 | 0.17424046 | NA | C49G7.13 |
| WBGene00269426 | 0.020317245 | 0.84643798 | NA | NA |
| WBGene00269427 | NA | NA | NA | NA |
| WBGene00269428 | NA | NA | NA | NA |
| WBGene00269429 | 0.020925399 | 0.84643798 | NA | H43I07.4 |
| WBGene00269430 | NA | NA | NA | NA |
| WBGene00269431 | -0.150977363 | 0.06525693 | NA | T16A1.10 |
| WBGene00269432 | 0.016036363 | 0.86497314 | NA | ZC132.11 |

| | | | | |
|---|---|---|---|---|
| WBGene00269433 | 0.084701974 | 0.52975989 | 0.70652346 | T16G12.12 |
| WBGene00269434 | 0.745356907 | 0.02521591 | 0.0913601 | sqst-5 |
| WBGene00269435 | NA | NA | NA | T27C10.9 |
| WBGene00269436 | 0.063086775 | 0.41202602 | NA | NA |
| WBGene00269437 | -0.001993815 | 0.95612996 | NA | Y47D7A.19 |
| WBGene00269438 | 0.042199095 | 0.55922995 | NA | Y56A3A.41 |
| WBGene00270290 | -0.068622032 | 0.67829208 | 0.81313963 | ZK938.11 |
| WBGene00270291 | -0.050580107 | 0.47300881 | NA | NA |
| WBGene00270292 | -0.023109762 | 0.78121253 | NA | NA |
| WBGene00270293 | -0.240985533 | 0.17388578 | 0.3543794 | F36H5.17 |
| WBGene00270303 | -0.001993815 | 0.95612996 | NA | T28D9.16 |
| WBGene00270309 | 0.048485426 | 0.45975815 | NA | K05F1.13 |
| WBGene00270318 | 0.127403933 | 0.16699325 | NA | F26G1.15 |
| WBGene00270319 | 0.380672199 | 0.07870309 | 0.20874853 | F26A1.19 |
| WBGene00270320 | -0.037432977 | 0.73008835 | 0.84956474 | F27D9.13 |
| WBGene00270321 | 2.318243745 | 1.38E-06 | 2.10E-05 | Y41C4A.32 |
| WBGene00270322 | -0.148114095 | 0.31901332 | 0.51919453 | C17G10.13 |
| WBGene00271414 | 0.020232312 | 0.84643798 | NA | T05A7.12 |
| WBGene00271415 | -0.029531685 | 0.71404005 | NA | T05A7.13 |
| WBGene00271437 | 0.015945659 | 0.84133371 | NA | F10D2.16 |
| WBGene00271444 | 0.036894305 | 0.59120501 | NA | T04C12.33 |
| WBGene00271567 | 0.020317245 | 0.84643798 | NA | T28F4.8 |
| WBGene00271577 | 0.181789487 | 0.04369581 | NA | B0244.16 |
| WBGene00271598 | -0.01462739 | 0.89230388 | 0.94374205 | NA |
| WBGene00271605 | -0.001993815 | 0.95612996 | NA | H05L14.3 |
| WBGene00271606 | 0.241627593 | 0.14781877 | 0.31864173 | F22F4.9 |
| WBGene00271632 | 0.018795205 | 0.77532208 | NA | NA |
| WBGene00271633 | NA | NA | NA | NA |
| WBGene00271634 | 0.020925399 | 0.84643798 | NA | NA |
| WBGene00271635 | NA | NA | NA | NA |
| WBGene00271636 | 3.54274657 | 4.73E-06 | 6.23E-05 | NA |
| WBGene00271637 | NA | NA | NA | NA |
| WBGene00271638 | NA | NA | NA | NA |
| WBGene00271639 | NA | NA | NA | NA |
| WBGene00271640 | NA | NA | NA | NA |
| WBGene00271641 | NA | NA | NA | NA |
| WBGene00271642 | 0.510234033 | 0.00081208 | 0.00589447 | NA |
| WBGene00271643 | -0.022930767 | 0.78121253 | NA | NA |
| WBGene00271644 | NA | NA | NA | NA |
| WBGene00271645 | NA | NA | NA | NA |
| WBGene00271646 | NA | NA | NA | NA |
| WBGene00271647 | 0.122015251 | 0.44002191 | 0.63446531 | NA |
| WBGene00271648 | NA | NA | NA | NA |

| | | | | |
|---|---|---|---|---|
| WBGene00271649 | NA | NA | NA | NA |
| WBGene00271650 | NA | NA | NA | NA |
| WBGene00271651 | -0.045959069 | 0.50707733 | NA | NA |
| WBGene00271652 | -0.022930767 | 0.78121253 | NA | NA |
| WBGene00271653 | NA | NA | NA | NA |
| WBGene00271654 | 0.020925399 | 0.84643798 | NA | NA |
| WBGene00271655 | NA | NA | NA | NA |
| WBGene00271656 | NA | NA | NA | NA |
| WBGene00271657 | NA | NA | NA | Y38E10A.33 |
| WBGene00271694 | -0.483958732 | 0.02674532 | 0.09546083 | T10E9.14 |
| WBGene00271701 | 0.261167849 | 0.16552225 | 0.34195163 | F10D7.10 |
| WBGene00271703 | NA | NA | NA | ZK1010.12 |
| WBGene00271706 | -0.060738075 | 0.76512821 | 0.86670277 | D2089.8 |
| WBGene00271707 | NA | NA | NA | ZK105.14 |
| WBGene00271715 | 0.527373325 | 0.04690051 | 0.14367902 | B0244.17 |
| WBGene00271716 | 0.010180195 | 0.95906264 | 0.97762557 | NA |
| WBGene00271744 | 0.138296718 | 0.51561322 | 0.69531934 | K09F6.15 |
| WBGene00271774 | NA | NA | NA | R02F2.10 |
| WBGene00271775 | -0.164196603 | 0.16218585 | 0.3373438 | EGAP1.5 |
| WBGene00271776 | NA | NA | NA | F53G12.18 |
| WBGene00271777 | NA | NA | NA | Y48G8AR.8 |
| WBGene00271778 | -0.024740619 | 0.69507748 | NA | T01A4.9 |
| WBGene00271779 | NA | NA | NA | C35E7.12 |
| WBGene00271780 | NA | NA | NA | F11A6.16 |
| WBGene00271781 | -0.102949619 | 0.35340968 | 0.55327963 | NA |
| WBGene00271786 | -0.05813386 | 0.56632684 | 0.73539612 | NA |
| WBGene00271788 | 0.287633732 | 0.03144426 | 0.10690908 | M01H9.10 |
| WBGene00271789 | 0.020317245 | 0.84643798 | NA | K01G12.22 |
| WBGene00271790 | NA | NA | NA | NA |
| WBGene00271791 | -0.282741653 | 0.05611391 | 0.16395205 | NA |
| WBGene00271792 | NA | NA | NA | NA |
| WBGene00271795 | 1.220949489 | 0.01135518 | 0.04939131 | Y46C8AM.1 |
| WBGene00271796 | 0.078776717 | 0.31677567 | NA | C54F6.20 |
| WBGene00271797 | NA | NA | NA | E02C12.19 |
| WBGene00271798 | -0.022930767 | 0.78121253 | NA | F54D5.24 |
| WBGene00271799 | NA | NA | NA | NA |
| WBGene00271800 | NA | NA | NA | NA |
| WBGene00271801 | NA | NA | NA | NA |
| WBGene00271802 | 0.015062379 | 0.80838968 | NA | NA |
| WBGene00271803 | -0.155683077 | 0.3317596 | 0.53137429 | C09G5.14 |
| WBGene00271804 | NA | NA | NA | F11D11.22 |
| WBGene00271805 | 0.038125098 | 0.61119482 | NA | NA |
| WBGene00271806 | 0.020232312 | 0.84643798 | NA | ZK938.12 |

| | | | | | |
|---|---|---|---|---|---|
| WBGene00271807 | | -0.003614866 | 0.95700117 | NA | F19F10.14 |
| WBGene00271808 | | 0.038125098 | 0.61119482 | NA | F21C10.19 |
| WBGene00271809 | | 0.057565879 | 0.43429308 | NA | NA |
| WBGene00271810 | | 0.006971683 | 0.95127038 | 0.97203268 | C27H5.12 |
| WBGene00271811 | | 0.020925399 | 0.84643798 | NA | NA |
| WBGene00271812 | | 2.077807212 | 0.00328671 | 0.01877513 | F58F12.12 |
| WBGene00271813 | NA | | NA | NA | NA |
| WBGene00271815 | | 0.020317245 | 0.84643798 | NA | K04A8.21 |
| WBGene00271816 | NA | | NA | NA | R02F11.12 |
| WBGene00271817 | | -0.13047768 | 0.15321597 | NA | F55C12.19 |
| WBGene00271818 | | 0.071340251 | 0.58210087 | 0.74621652 | K05F1.14 |
| WBGene00271819 | | 0.17338789 | 0.09011165 | 0.22825685 | F10C1.23 |
| WBGene00271820 | | 0.075076776 | 0.28090944 | NA | NA |
| WBGene00271821 | | 0.017279211 | 0.79380376 | NA | T28C12.15 |
| WBGene00271822 | | -0.09479125 | 0.54553335 | 0.71904517 | F11G11.15 |
| WBGene00271823 | NA | | NA | NA | NA |
| WBGene00271824 | | -0.244787593 | 0.23342977 | 0.42875992 | R03H10.11 |
| WBGene00271827 | | 2.297115913 | 2.18E-05 | 0.00024672 | T19H12.15 |
| WBGene00271830 | | -0.026384613 | 0.67299636 | NA | W09G10.9 |
| WBGene00271831 | NA | | NA | NA | NA |
| WBGene00271832 | NA | | NA | NA | Y60C6A.3 |
| WBGene00271837 | NA | | NA | NA | NA |
| WBGene00284844 | | 0.038125098 | 0.61119482 | NA | H38K22.9 |
| WBGene00284845 | NA | | NA | NA | Y71F9AR.10 |
| WBGene00284846 | | 0.220894503 | 0.10004178 | 0.24550984 | ZC262.13 |
| WBGene00284847 | | -6.730693184 | 8.66E-07 | 1.39E-05 | F54D8.10 |
| WBGene00284848 | | -0.023109762 | 0.78121253 | NA | M02E1.4 |
| WBGene00284849 | NA | | NA | NA | C03A3.9 |
| WBGene00284850 | | 0.110126488 | 0.13663711 | NA | Y46B2A.4 |
| WBGene00284851 | | 0.116830083 | 0.11818778 | NA | F28A10.13 |
| WBGene00284852 | | 0.031202807 | 0.8692816 | 0.93000745 | NA |
| WBGene00284853 | NA | | NA | NA | C32E8.13 |
| WBGene00284854 | NA | | NA | NA | Y95D11A.5 |
| WBGene00284855 | | 0.068135573 | 0.6096633 | 0.76480853 | C04C11.25 |
| WBGene00284856 | | -0.022930767 | 0.78121253 | NA | NA |
| WBGene00284857 | NA | | NA | NA | NA |
| WBGene00284858 | NA | | NA | NA | NA |
| WBGene00284859 | | -0.377940131 | 0.04047533 | 0.12911897 | NA |
| WBGene00284861 | | -0.067064794 | 0.32104386 | NA | NA |
| WBGene00284867 | | 0.017604133 | 0.79437959 | NA | T22B7.22 |
| WBGene00302964 | | 0.128556704 | 0.48789289 | 0.67198869 | Y62E10A.23 |
| WBGene00302968 | NA | | NA | NA | Y62E10A.24 |
| WBGene00302971 | NA | | NA | NA | F59E12.17 |

| | | | | | |
|---|---|---|---|---|---|
| WBGene00302974 | | 0.119570007 | 0.49013581 | 0.67393674 | Y65B4BM.3 |
| WBGene00302976 | NA | | NA | NA | F49E8.8 |
| WBGene00302978 | | 0.062194465 | 0.67695793 | 0.8124716 | C42C1.19 |
| WBGene00302980 | | -0.240457591 | 2.36E-06 | 3.36E-05 | unc-60 |
| WBGene00302982 | | 0.057170847 | 0.77842141 | 0.87452946 | W02H5.19 |
| WBGene00302984 | NA | | NA | NA | Y51H7C.41 |
| WBGene00302986 | | -0.180648497 | 0.39401073 | 0.59231709 | Y37E11AM.4 |
| WBGene00302988 | NA | | NA | NA | NA |
| WBGene00302991 | | 0.262150586 | 0.22416246 | 0.41711331 | cox-16 |
| WBGene00302993 | | 0.161652037 | 0.31177329 | 0.51266736 | T14B4.19 |
| WBGene00302995 | | -0.166067568 | 0.34869109 | 0.54882674 | H06H21.37 |
| WBGene00302998 | | -0.123080227 | 0.52810313 | 0.70521324 | F53G2.12 |
| WBGene00303000 | NA | | NA | NA | C37C3.18 |
| WBGene00303002 | NA | | NA | NA | Y48E1B.17 |
| WBGene00303005 | NA | | NA | NA | C34E10.12 |
| WBGene00303007 | NA | | NA | NA | Y57G11C.1147 |
| WBGene00303009 | | -0.092463938 | 0.52161103 | 0.69999149 | mrpl-42 |
| WBGene00303011 | | 0.634089495 | 0.00056399 | 0.0042962 | F17C11.26 |
| WBGene00303013 | | 0.037472146 | 0.66218673 | NA | T04H1.12 |
| WBGene00303015 | | 0.439160479 | 0.08799443 | 0.22491734 | F20C5.11 |
| WBGene00303019 | NA | | NA | NA | Y70G10A.4 |
| WBGene00303021 | | -0.507313605 | 9.63E-12 | 3.22E-10 | F02A9.10 |
| WBGene00303023 | | 0.047463612 | 0.78437976 | 0.87847449 | K07F5.22 |
| WBGene00303024 | NA | | NA | NA | H40L08.7 |
| WBGene00303026 | | -0.204110235 | 0.33351704 | 0.53304196 | T20D3.14 |
| WBGene00303028 | NA | | NA | NA | F45H11.11 |
| WBGene00303030 | | -0.069356039 | 0.72290396 | 0.84469568 | C25A1.20 |
| WBGene00303031 | NA | | NA | NA | B0035.22 |
| WBGene00303032 | NA | | NA | NA | B0041.12 |
| WBGene00303033 | NA | | NA | NA | B0361.14 |
| WBGene00303034 | NA | | NA | NA | C02C6.14 |
| WBGene00303035 | NA | | NA | NA | C07G2.5 |
| WBGene00303036 | NA | | NA | NA | C08B6.18 |
| WBGene00303037 | NA | | NA | NA | C09E8.6 |
| WBGene00303038 | NA | | NA | NA | C09E10.3 |
| WBGene00303039 | NA | | NA | NA | C13F10.8 |
| WBGene00303040 | NA | | NA | NA | C13G3.12 |
| WBGene00303041 | NA | | NA | NA | C14C10.10 |
| WBGene00303042 | NA | | NA | NA | C26H9A.4 |
| WBGene00303043 | NA | | NA | NA | C28D4.18 |
| WBGene00303044 | NA | | NA | NA | C34D10.5 |
| WBGene00303045 | NA | | NA | NA | C43H6.12 |
| WBGene00303046 | NA | | NA | NA | C48A7.16 |

| | | | | |
|---|---|---|---|---|
| WBGene00303047 | NA | NA | NA | C49F8.14 |
| WBGene00303048 | NA | NA | NA | F55C5.17 |
| WBGene00303049 | NA | NA | NA | C55C3.73 |
| WBGene00303050 | NA | NA | NA | C56G2.22 |
| WBGene00303051 | NA | NA | NA | F01E11.20 |
| WBGene00303052 | NA | NA | NA | F08B12.11 |
| WBGene00303053 | NA | NA | NA | F08C6.18 |
| WBGene00303054 | NA | NA | NA | F44E5.17 |
| WBGene00303055 | NA | NA | NA | F13G3.15 |
| WBGene00303056 | NA | NA | NA | F27D4.12 |
| WBGene00303057 | NA | NA | NA | F33D11.17 |
| WBGene00303058 | NA | NA | NA | F47G4.14 |
| WBGene00303059 | NA | NA | NA | F48F5.8 |
| WBGene00303060 | NA | NA | NA | F52G3.7 |
| WBGene00303061 | NA | NA | NA | F52H3.11 |
| WBGene00303062 | NA | NA | NA | F55A4.13 |
| WBGene00303063 | NA | NA | NA | T06E8.8 |
| WBGene00303064 | NA | NA | NA | F59A6.13 |
| WBGene00303065 | NA | NA | NA | H03A11.13 |
| WBGene00303066 | NA | NA | NA | H40L08.8 |
| WBGene00303067 | NA | NA | NA | K01D12.20 |
| WBGene00303068 | NA | NA | NA | K08F8.15 |
| WBGene00303069 | NA | NA | NA | K11D12.14 |
| WBGene00303070 | NA | NA | NA | C53H9.4 |
| WBGene00303071 | NA | NA | NA | M176.16 |
| WBGene00303072 | NA | NA | NA | R10E9.4 |
| WBGene00303073 | NA | NA | NA | R12B2.13 |
| WBGene00303074 | NA | NA | NA | T03G6.7 |
| WBGene00303075 | NA | NA | NA | T04C10.11 |
| WBGene00303076 | NA | NA | NA | T04H1.13 |
| WBGene00303077 | NA | NA | NA | T05A8.11 |
| WBGene00303078 | NA | NA | NA | T06D10.8 |
| WBGene00303079 | NA | NA | NA | T09F3.7 |
| WBGene00303080 | NA | NA | NA | T14B4.20 |
| WBGene00303081 | NA | NA | NA | T23F2.13 |
| WBGene00303082 | NA | NA | NA | T24D1.7 |
| WBGene00303083 | NA | NA | NA | T28F3.11 |
| WBGene00303085 | NA | NA | NA | Y32F6B.9 |
| WBGene00303086 | NA | NA | NA | Y37D8A.29 |
| WBGene00303087 | NA | NA | NA | Y39G8B.13 |
| WBGene00303088 | NA | NA | NA | Y44E3A.7 |
| WBGene00303089 | NA | NA | NA | Y46G5A.47 |
| WBGene00303090 | NA | NA | NA | Y47D9A.7 |

| | | | | |
|---|---|---|---|---|
| WBGene00303091 | -0.092289776 | 0.64133352 | 0.78746664 | Y50D4C.12 |
| WBGene00303092 | NA | NA | NA | Y54F10AM.15 |
| WBGene00303093 | NA | NA | NA | Y55B1BL.2 |
| WBGene00303094 | NA | NA | NA | Y71G12A.5 |
| WBGene00303095 | NA | NA | NA | Y71H2AL.4 |
| WBGene00303096 | NA | NA | NA | Y75B8A.63 |
| WBGene00303097 | NA | NA | NA | Y92H12BL.6 |
| WBGene00303098 | NA | NA | NA | ZK377.15 |
| WBGene00303099 | NA | NA | NA | ZK617.26 |
| WBGene00303100 | NA | NA | NA | ZK856.19 |
| WBGene00303101 | NA | NA | NA | F20B6.10 |
| WBGene00303102 | NA | NA | NA | T19D2.13 |
| WBGene00303103 | NA | NA | NA | W02D3.13 |
| WBGene00303104 | NA | NA | NA | ZC328.8 |
| WBGene00303105 | -0.003108065 | 0.95728506 | NA | R07E5.19 |
| WBGene00303113 | NA | NA | NA | NA |
| WBGene00303114 | NA | NA | NA | NA |
| WBGene00303115 | NA | NA | NA | NA |
| WBGene00303208 | NA | NA | NA | Y71H2AM.29 |
| WBGene00303231 | 0.497560198 | 0.06006235 | 0.17173448 | C53A3.4 |
| WBGene00303233 | 0.042065281 | 0.558937 | NA | D1014.13 |
| WBGene00303359 | NA | NA | NA | ZK177.12 |
| WBGene00303360 | NA | NA | NA | NA |
| WBGene00303362 | NA | NA | NA | C27D6.15 |
| WBGene00303363 | 0.233276312 | 0.13336063 | 0.29700836 | NA |
| WBGene00303364 | 0.292986828 | 0.0779546 | 0.20706337 | Y14H12A.5 |
| WBGene00303365 | -0.109190711 | 0.28932382 | 0.4895758 | M151.10 |
| WBGene00303366 | 0.016471116 | 0.91160016 | 0.95112162 | Y49F6A.11 |
| WBGene00303367 | NA | NA | NA | Y49F6A.12 |
| WBGene00303368 | NA | NA | NA | Y49F6B.19 |
| WBGene00303369 | NA | NA | NA | Y49F6B.20 |
| WBGene00303370 | 0.015584271 | 0.80362925 | NA | Y59A8A.8 |
| WBGene00303371 | NA | NA | NA | NA |
| WBGene00303378 | -0.003421551 | 0.95689871 | NA | Y61B8B.3 |
| WBGene00303398 | 0.033034883 | 0.64219503 | NA | F46H5.11 |
| WBGene00303421 | NA | NA | NA | F20D6.15 |
| WBGene00303423 | 0.124316213 | 0.46925718 | 0.65783171 | F10D7.11 |
| WBGene00303425 | 0.093320698 | 0.44732956 | 0.64038659 | R02E12.12 |
| WBGene00304173 | -1.235164168 | 0.0111066 | 0.04858432 | F07E5.13 |
| WBGene00304175 | NA | NA | NA | C37E2.10 |
| WBGene00304181 | NA | NA | NA | F38B7.17 |
| WBGene00304182 | NA | NA | NA | NA |
| WBGene00304183 | NA | NA | NA | NA |

| | | | | | |
|---|---|---|---|---|---|
| WBGene00304189 | | 0.124815915 | 0.55297497 | 0.72513269 | NA |
| WBGene00304194 | NA | | NA | NA | M163.21 |
| WBGene00304195 | NA | | NA | NA | M163.22 |
| WBGene00304201 | | 7.569052085 | 3.72E-11 | 1.14E-09 | NA |
| WBGene00304202 | | 0.040529223 | 0.56245259 | NA | NA |
| WBGene00304203 | | -0.004944805 | 0.94514249 | NA | NA |
| WBGene00304215 | | -0.071124496 | 0.3419818 | NA | NA |
| WBGene00304219 | | 0.478477097 | 0.05976084 | 0.17105736 | NA |
| WBGene00304220 | | -0.050740777 | 0.47171914 | NA | NA |
| WBGene00304237 | NA | | NA | NA | NA |
| WBGene00304238 | NA | | NA | NA | NA |
| WBGene00304775 | | -0.03117915 | 0.73302896 | NA | NA |
| WBGene00304776 | | 0.018860755 | 0.77262519 | NA | NA |
| WBGene00304782 | | 0.052594298 | 0.67659718 | 0.81230499 | NA |
| WBGene00304783 | | 0.062968892 | 0.63385307 | 0.78268051 | NA |
| WBGene00304785 | NA | | NA | NA | NA |
| WBGene00304786 | NA | | NA | NA | NA |
| WBGene00304787 | | 0.109724155 | 0.21583934 | NA | NA |
| WBGene00304788 | | -0.032749713 | 0.76895765 | 0.86844708 | NA |
| WBGene00304789 | | 0.226517075 | 0.25638087 | 0.45460486 | NA |
| WBGene00304790 | | -0.022930767 | 0.78121253 | NA | NA |
| WBGene00304791 | | 0.176461773 | 0.09037499 | 0.22872606 | NA |
| WBGene00304792 | | -0.898235924 | 0.01781622 | 0.07065201 | NA |
| WBGene00304793 | | -0.008998908 | 0.92213607 | NA | NA |
| WBGene00304794 | | 0.020317245 | 0.84643798 | NA | NA |
| WBGene00304797 | | -0.023109762 | 0.78121253 | NA | NA |
| WBGene00304798 | NA | | NA | NA | NA |
| WBGene00304806 | | 0.175541666 | 0.09345266 | 0.23380732 | NA |
| WBGene00304808 | | 0.038726127 | 0.57551883 | NA | NA |
| WBGene00304811 | | 0.165853693 | 0.42039113 | 0.61703724 | NA |
| WBGene00304812 | | -0.772886744 | 4.68E-11 | 1.42E-09 | NA |
| WBGene00304813 | | -0.562709291 | 0.04971841 | 0.14960514 | NA |
| WBGene00304814 | | 0.014087259 | 0.85351751 | NA | NA |
| WBGene00304815 | | -0.67545706 | 1.14E-05 | 0.00013823 | NA |
| WBGene00304816 | | -0.023109762 | 0.78121253 | NA | NA |
| WBGene00304817 | NA | | NA | NA | NA |
| WBGene00304818 | NA | | NA | NA | NA |
| WBGene00304819 | NA | | NA | NA | NA |
| WBGene00304820 | | -0.00346345 | 0.95708614 | NA | NA |
| WBGene00304822 | | -0.053591705 | 0.45278076 | NA | NA |
| WBGene00304823 | | 0.020317245 | 0.84643798 | NA | NA |
| WBGene00304824 | | -0.099557361 | 0.39916515 | 0.59738086 | NA |
| WBGene00304825 | | -0.101609559 | 0.22055508 | NA | NA |

| | | | | | |
|---|---|---|---|---|---|
| WBGene00304826 | | 0.020925399 | 0.84643798 | NA | NA |
| WBGene00304827 | NA | | NA | NA | NA |
| WBGene00304828 | NA | | NA | NA | NA |
| WBGene00304895 | | -0.418859061 | 0.0803884 | 0.21171511 | NA |
| WBGene00304901 | | -0.028446343 | 0.71258617 | NA | NA |
| WBGene00304906 | | 0.018795205 | 0.77532208 | NA | NA |
| WBGene00304907 | NA | | NA | NA | NA |
| WBGene00304908 | | 0.051546931 | 0.80002599 | 0.88826172 | NA |
| WBGene00304930 | NA | | NA | NA | NA |
| WBGene00304981 | NA | | NA | NA | NA |
| WBGene00304985 | | -0.053570221 | 0.53767408 | NA | NA |
| WBGene00304992 | | -0.732632284 | 0.02284524 | 0.08480109 | NA |
| WBGene00304996 | | -0.025853814 | 0.76517134 | NA | NA |
| WBGene00305018 | | -0.056791266 | 0.52823312 | NA | NA |
| WBGene00305047 | NA | | NA | NA | NA |
| WBGene00305048 | NA | | NA | NA | NA |
| WBGene00305049 | | 0.038765322 | 0.66219346 | NA | NA |
| WBGene00305050 | | -0.240484081 | 0.2802482 | 0.47984024 | NA |
| WBGene00305065 | | -0.006711909 | 0.92834175 | NA | NA |
| WBGene00305066 | | 0.020232312 | 0.84643798 | NA | NA |
| WBGene00305067 | | 0.329719306 | 0.08162191 | 0.21385583 | NA |
| WBGene00305069 | NA | | NA | NA | NA |
| WBGene00305086 | | 0.038125098 | 0.61119482 | NA | NA |
| WBGene00305090 | NA | | NA | NA | NA |
| WBGene00305091 | NA | | NA | NA | NA |
| WBGene00305092 | | 0.06678304 | 0.68739362 | 0.81995518 | NA |
| WBGene00305093 | | -0.023109762 | 0.78121253 | NA | NA |
| WBGene00305095 | | -0.090268704 | 0.56219598 | 0.73230632 | NA |
| WBGene00305102 | NA | | NA | NA | NA |
| WBGene00305103 | | 0.126763858 | 0.38578002 | 0.58472269 | NA |
| WBGene00305104 | | 0.040107227 | 0.58693163 | NA | NA |
| WBGene00305109 | | -0.023109762 | 0.78121253 | NA | NA |
| WBGene00305110 | | -0.079824684 | 0.4472736 | 0.64038659 | NA |
| WBGene00305111 | | -0.003270206 | 0.95698316 | NA | NA |
| WBGene00305112 | | 0.044157924 | 0.71225742 | 0.83815617 | NA |
| WBGene00305113 | | -0.022930767 | 0.78121253 | NA | NA |
| WBGene00305114 | NA | | NA | NA | NA |
| WBGene00305130 | NA | | NA | NA | NA |
| WBGene00305131 | | 0.336948984 | 0.08686201 | 0.22287885 | NA |
| WBGene00305132 | NA | | NA | NA | NA |
| WBGene00305157 | NA | | NA | NA | NA |
| WBGene00305158 | | 0.754301303 | 0.01835726 | 0.07221082 | NA |
| WBGene00305159 | NA | | NA | NA | NA |

| | | | | |
|---|---|---|---|---|
| WBGene00305173 | 0.034658478 | 0.67366584 | NA | NA |
