## Supplementary Table 2 - Transcriptome analysis of WT and lbp-3 Tg animals for "Lysosome Lipid Signaling from the Periphery to Neurons Regulates Longevity"

| ID | log2FoldChange | pvalue | padj | SYMBOL |
|---|---|---|---|---|
| WBGene00000001 | -0.300450611 | 0.10633685 | 0.26575456 | aap-1 |
| WBGene00000002 | 0.504813784 | 0.21631946 | 0.41787345 | aat-1 |
| WBGene00000003 | 0.806532085 | 0.04066217 | 0.13906461 | aat-2 |
| WBGene00000004 | 0.150762319 | 0.60198658 | 0.76683654 | aat-3 |
| WBGene00000005 | 0.26223662 | 0.50173713 | 0.68948393 | aat-4 |
| WBGene00000006 | 1.130303134 | 0.00159945 | 0.01315642 | aat-5 |
| WBGene00000007 | 1.124486016 | 0.00600855 | 0.0356374 | aat-6 |
| WBGene00000008 | 0.698847757 | 0.61017687 | 1 | aat-7 |
| WBGene00000009 | 2.178953833 | 0.04775041 | 0.15609525 | aat-8 |
| WBGene00000010 | -0.242829476 | 0.29684366 | 0.50717406 | aat-9 |
| WBGene00000012 | 1.712479368 | 0.58529868 | 1 | abf-1 |
| WBGene00000013 | 1.026909457 | 0.25386023 | 0.46180957 | abf-2 |
| WBGene00000015 | 2.093642713 | 0.24945587 | 1 | abf-4 |
| WBGene00000016 | 0.63503133 | 0.14972666 | 0.3312375 | abf-5 |
| WBGene00000017 | 0.306348595 | 0.40085147 | 0.60833979 | abf-6 |
| WBGene00000018 | 0.12284015 | 0.60611655 | 0.76927253 | abl-1 |
| WBGene00000019 | 2.102898496 | 0.14717924 | 1 | abt-1 |
| WBGene00000020 | 1.286866132 | 7.5817E-05 | 0.0012252 | abt-2 |
| WBGene00000022 | 1.13913733 | 3.7105E-05 | 0.00067238 | abt-4 |
| WBGene00000023 | 0.593855373 | 0.71573207 | 1 | abt-5 |
| WBGene00000024 | -0.861206134 | 0.70037023 | 1 | abu-1 |
| WBGene00000027 | -2.705999569 | 0.42423051 | 1 | abu-4 |
| WBGene00000029 | -0.156269293 | 0.96924949 | 1 | abu-6 |
| WBGene00000031 | -0.465832989 | 0.35245225 | 0.56141732 | abu-8 |
| WBGene00000033 | -0.183677144 | 0.84745953 | 0.91633779 | abu-10 |
| WBGene00000034 | -0.183594323 | 0.83792469 | 0.91143626 | abu-11 |
| WBGene00000035 | 0.258580679 | 0.50846477 | 0.69510987 | ace-1 |
| WBGene00000036 | 0.835678445 | 0.00720115 | 0.040616 | ace-2 |
| WBGene00000037 | 0.280414365 | 0.44364463 | 0.64321584 | ace-3 |
| WBGene00000038 | 0.929916491 | 0.2789422 | 0.48770698 | ace-4 |
| WBGene00000039 | 0.277420899 | 0.41901513 | 0.62348775 | acn-1 |
| WBGene00000040 | 0.519206897 | 0.00136849 | 0.01169165 | aco-1 |
| WBGene00000041 | 0.018325779 | 0.91139359 | 0.95398134 | aco-2 |
| WBGene00000042 | 0.763546346 | 0.30047188 | 0.51007117 | acr-2 |
| WBGene00000043 | 0.878184137 | 0.523167 | 1 | acr-3 |
| WBGene00000044 | 0.15546908 | 0.84934678 | 0.91751216 | acr-5 |
| WBGene00000045 | -0.988317891 | 0.59464747 | 1 | acr-6 |
| WBGene00000046 | -0.188352933 | 0.88395112 | 1 | acr-7 |
| WBGene00000047 | 0.697492737 | 0.18444739 | 0.37693584 | acr-8 |
| WBGene00000048 | -0.372640048 | 0.66625019 | 0.80861938 | acr-9 |
| WBGene00000049 | 1.251616604 | 0.41368807 | 1 | acr-10 |
| WBGene00000050 | 0.520074244 | 0.57453862 | 0.74795819 | acr-11 |

| | | | | |
|---|---|---|---|---|
| WBGene00000051 | 0.659810976 | 0.26650329 | 0.47484833 | acr-12 |
| WBGene00000053 | 1.25857425 | 0.46917901 | 1 | acr-14 |
| WBGene00000054 | 0.380722639 | 0.53185965 | 0.71541193 | acr-15 |
| WBGene00000055 | -0.359454572 | 0.568782 | 0.74341002 | acr-16 |
| WBGene00000056 | 0.480880519 | 0.69175662 | 1 | acr-17 |
| WBGene00000057 | 0.113286124 | 0.88627959 | 0.93995577 | acr-18 |
| WBGene00000058 | 2.661602148 | 0.00873979 | 0.04674442 | acr-19 |
| WBGene00000059 | 0.962764182 | 0.31222778 | 0.5204225 | acr-20 |
| WBGene00000060 | 2.019308854 | 0.10056278 | 1 | acr-21 |
| WBGene00000061 | 0.697764096 | 0.35682764 | 0.56521999 | lgc-11 |
| WBGene00000062 | 0.557322029 | 0.64044244 | 1 | acr-23 |
| WBGene00000063 | 0.200509607 | 0.14266424 | 0.32052915 | act-1 |
| WBGene00000064 | -0.789411977 | 5.79E-10 | 4.89E-08 | act-2 |
| WBGene00000065 | 0.200509607 | 0.14266424 | 0.32052915 | act-3 |
| WBGene00000066 | 0.210592588 | 0.17972675 | 0.37204817 | act-4 |
| WBGene00000067 | 0.422646134 | 0.00628496 | 0.03692426 | act-5 |
| WBGene00000068 | 1.499812174 | 1.1539E-05 | 0.00025611 | acy-1 |
| WBGene00000069 | 0.957954869 | 0.15516383 | 0.33888182 | acy-2 |
| WBGene00000070 | 0.937872205 | 0.00442801 | 0.02848992 | acy-3 |
| WBGene00000071 | 1.410352305 | 0.00059476 | 0.00609501 | acy-4 |
| WBGene00000072 | 0.827382845 | 0.04971197 | 0.1598816 | add-1 |
| WBGene00000073 | 0.187885122 | 0.49332313 | 0.68209043 | add-2 |
| WBGene00000074 | 0.092791087 | 0.75668199 | 0.86407788 | adm-2 |
| WBGene00000075 | -0.355605336 | 0.22079891 | 0.42382918 | adm-4 |
| WBGene00000079 | 0.033458117 | 0.88934658 | 0.94186643 | adr-1 |
| WBGene00000080 | -0.256883241 | 0.15689497 | 0.34118964 | adr-2 |
| WBGene00000081 | 0.456107356 | 0.00298069 | 0.02112586 | ads-1 |
| WBGene00000082 | 0.152207756 | 0.83221094 | 0.90915448 | adt-1 |
| WBGene00000083 | 0.497091628 | 0.05939601 | 0.18127877 | adt-2 |
| WBGene00000084 | 0.974007168 | 0.02225003 | 0.09270337 | aex-1 |
| WBGene00000085 | 1.39428765 | 0.22060383 | 1 | aex-2 |
| WBGene00000086 | 1.216501002 | 0.00217671 | 0.0166106 | aex-3 |
| WBGene00000088 | 2.143019531 | 3.22E-15 | 6.40E-13 | aex-5 |
| WBGene00000089 | 0.391424466 | 0.36402393 | 0.57174854 | aex-6 |
| WBGene00000090 | 0.028186994 | 0.87983613 | 0.93574569 | age-1 |
| WBGene00000092 | -0.134937611 | 0.56982668 | 0.74425222 | ags-3 |
| WBGene00000093 | -0.411106945 | 0.1499274 | 0.33131935 | agt-1 |
| WBGene00000094 | -0.316522807 | 0.26542291 | 0.47425189 | agt-2 |
| WBGene00000095 | 0.043145232 | 0.87228061 | 0.93134806 | aha-1 |
| WBGene00000096 | -0.343783753 | 0.6885529 | 0.82264522 | ahr-1 |
| WBGene00000097 | -0.327189109 | 0.19172921 | 0.3866281 | aip-1 |
| WBGene00000098 | -0.279275886 | 0.05832673 | 0.17896002 | air-1 |
| WBGene00000099 | -0.585757557 | 0.00157197 | 0.01297433 | air-2 |

| | | | | |
|---|---|---|---|---|
| WBGene00000100 | 0.775783981 | 0.00027636 | 0.00338931 | ajm-1 |
| WBGene00000101 | 0.437151222 | 0.00559705 | 0.03389214 | aka-1 |
| WBGene00000102 | -0.064506091 | 0.65472654 | 0.80163724 | akt-1 |
| WBGene00000103 | 0.365478071 | 0.28532968 | 0.49509613 | akt-2 |
| WBGene00000105 | 0.633555074 | 0.01514791 | 0.07019452 | alg-1 |
| WBGene00000106 | -0.358466444 | 0.07294136 | 0.20807825 | alg-2 |
| WBGene00000107 | 0.246667181 | 0.09938914 | 0.25393171 | alh-1 |
| WBGene00000109 | 0.284901248 | 0.0825705 | 0.22548097 | alh-3 |
| WBGene00000110 | 0.195799991 | 0.27113818 | 0.47974773 | alh-4 |
| WBGene00000111 | 1.914442742 | 4.5798E-06 | 0.00011633 | alh-5 |
| WBGene00000112 | -0.203490694 | 0.3139171 | 0.52177554 | alh-6 |
| WBGene00000113 | -0.060168383 | 0.81046384 | 0.8965687 | alh-7 |
| WBGene00000114 | 0.308622887 | 0.04926564 | 0.15933378 | alh-8 |
| WBGene00000115 | -0.744472092 | 2.7467E-05 | 0.00052187 | alh-9 |
| WBGene00000116 | -0.311082982 | 0.23954515 | 0.44640333 | alh-10 |
| WBGene00000117 | 0.038338124 | 0.88784496 | 0.94084004 | alh-11 |
| WBGene00000118 | 0.62917819 | 0.00058483 | 0.00601315 | alh-12 |
| WBGene00000120 | -1.079450938 | 3.7157E-06 | 9.8713E-05 | aly-1 |
| WBGene00000121 | -0.622298119 | 1.2387E-05 | 0.00027147 | aly-2 |
| WBGene00000122 | -0.825607824 | 8.34E-08 | 3.9842E-06 | aly-3 |
| WBGene00000123 | -0.143043548 | 0.50750479 | 0.69418834 | ama-1 |
| WBGene00000133 | -2.603968543 | 0.24486328 | 1 | amt-1 |
| WBGene00000134 | 1.451058156 | 0.28227533 | 1 | amt-2 |
| WBGene00000135 | 1.216099164 | 0.08663449 | 0.23265413 | amt-3 |
| WBGene00000136 | 0.234514147 | 0.32780489 | 0.53644415 | amt-4 |
| WBGene00000137 | -0.026882052 | 0.92354021 | 0.95953374 | amx-1 |
| WBGene00000138 | 1.237205925 | 0.0012948 | 0.0111728 | amx-2 |
| WBGene00000139 | 0.549655142 | 0.46134225 | 0.65666477 | amx-3 |
| WBGene00000140 | 1.745891746 | 4.81E-12 | 5.96E-10 | anc-1 |
| WBGene00000142 | -0.527670081 | 0.00633357 | 0.03712981 | aos-1 |
| WBGene00000143 | -0.123802557 | 0.51489982 | 0.70153728 | apc-2 |
| WBGene00000144 | -0.972380675 | 0.00057003 | 0.00591513 | apc-10 |
| WBGene00000145 | -1.280194246 | 2.90E-07 | 1.1721E-05 | apc-11 |
| WBGene00000146 | -0.13341101 | 0.37316932 | 0.58115042 | ape-1 |
| WBGene00000147 | 0.02511552 | 0.92519152 | 0.96044148 | aph-1 |
| WBGene00000148 | -0.281519956 | 0.14451599 | 0.32307998 | aph-2 |
| WBGene00000149 | 0.971832788 | 3.11E-09 | 2.14E-07 | apl-1 |
| WBGene00000150 | 0.490758272 | 0.05887445 | 0.18023064 | apm-1 |
| WBGene00000151 | -0.078406682 | 0.6769349 | 0.81495175 | apn-1 |
| WBGene00000155 | -0.730868122 | 1.92E-07 | 8.1039E-06 | app-1 |
| WBGene00000157 | -0.637777676 | 0.01979362 | 0.08494676 | aps-2 |
| WBGene00000158 | 0.098073292 | 0.64175891 | 0.79287626 | apg-1 |
| WBGene00000159 | -0.268712923 | 0.27262641 | 0.48137831 | aps-1 |

| | | | | |
|---|---|---|---|---|
| WBGene00000160 | 0.213898798 | 0.27890635 | 0.48770698 | apb-1 |
| WBGene00000161 | 0.260785505 | 0.23579505 | 0.44209298 | apa-2 |
| WBGene00000162 | -0.106551599 | 0.5965664 | 0.7626563 | apd-3 |
| WBGene00000163 | -0.127960836 | 0.40647062 | 0.61334325 | apb-3 |
| WBGene00000164 | -0.683426479 | 0.01185208 | 0.0587811 | apm-3 |
| WBGene00000165 | -0.750280342 | 0.00218126 | 0.0166244 | aps-3 |
| WBGene00000166 | 0.984690205 | 0.07611352 | 0.21413202 | apt-9 |
| WBGene00000168 | -0.061838678 | 0.81637391 | 0.90039932 | apx-1 |
| WBGene00000169 | 0.705166086 | 0.0027899 | 0.02017462 | aqp-1 |
| WBGene00000170 | 0.371819167 | 0.02302241 | 0.09475086 | aqp-2 |
| WBGene00000171 | 0.92795125 | 0.4466972 | 1 | aqp-3 |
| WBGene00000172 | 0.583778601 | 0.06203269 | 0.18642052 | aqp-4 |
| WBGene00000173 | 1.299498743 | 0.33175818 | 1 | aqp-5 |
| WBGene00000174 | 1.824823365 | 0.13169181 | 1 | aqp-6 |
| WBGene00000175 | 0.864418322 | 8.80E-07 | 3.0084E-05 | aqp-7 |
| WBGene00000176 | 0.126394322 | 0.6012953 | 0.76643845 | aqp-8 |
| WBGene00000177 | 1.472386254 | 0.22425479 | 1 | aqp-9 |
| WBGene00000178 | 0.061137041 | 0.81219931 | 0.89750722 | aqp-10 |
| WBGene00000179 | 0.141741618 | 0.50909811 | 0.69566237 | aqp-11 |
| WBGene00000180 | 0.093773504 | 0.75033362 | 0.86087701 | arc-1 |
| WBGene00000181 | -0.48848112 | 0.00216146 | 0.016515 | ard-1 |
| WBGene00000182 | -0.324101574 | 0.03631855 | 0.12893084 | arf-1.2 |
| WBGene00000183 | -0.243990079 | 0.09205859 | 0.24150461 | arf-3 |
| WBGene00000184 | -0.217463673 | 0.39113448 | 0.59953881 | arf-6 |
| WBGene00000185 | 1.529802032 | 0.64971375 | 1 | arg-1 |
| WBGene00000186 | 0.213426984 | 0.44168494 | 0.64137027 | ark-1 |
| WBGene00000187 | -0.570581749 | 0.02100749 | 0.08871372 | arl-1 |
| WBGene00000188 | 0.19312178 | 0.75431028 | 0.86280319 | arl-3 |
| WBGene00000189 | -0.719983685 | 0.03943443 | 0.13609252 | arl-5 |
| WBGene00000190 | 0.557163635 | 0.41574466 | 0.62082449 | arf-1.1 |
| WBGene00000192 | -0.321831347 | 0.05384586 | 0.16905678 | arl-8 |
| WBGene00000193 | -0.399747356 | 0.66717535 | 0.80920839 | arl-6 |
| WBGene00000195 | -0.269269345 | 0.44034997 | 0.64035082 | arr-1 |
| WBGene00000196 | 0.26084744 | 0.28437712 | 0.49417901 | aars-1 |
| WBGene00000197 | -0.312950782 | 0.09290583 | 0.24304454 | aars-2 |
| WBGene00000198 | 0.088034992 | 0.58813527 | 0.75642763 | art-1 |
| WBGene00000199 | -0.113846932 | 0.53237156 | 0.71586256 | arx-1 |
| WBGene00000200 | 0.042239693 | 0.80327806 | 0.89239902 | arx-2 |
| WBGene00000201 | -0.264187612 | 0.18117631 | 0.37323727 | arx-3 |
| WBGene00000202 | -0.731362217 | 0.00073249 | 0.00721852 | exos-4.2 |
| WBGene00000203 | -0.590672446 | 0.00709475 | 0.04013854 | arx-5 |
| WBGene00000204 | -0.414641942 | 0.04197847 | 0.14229939 | arx-6 |
| WBGene00000205 | -0.492295882 | 0.07232747 | 0.20700797 | arx-7 |

| | | | | |
|---|---|---|---|---|
| WBGene00000206 | -0.720504479 | 8.55E-07 | 2.9321E-05 | asb-1 |
| WBGene00000207 | 0.338651188 | 0.04342374 | 0.14535638 | asb-2 |
| WBGene00000208 | -0.212992556 | 0.36452471 | 0.57203053 | asc-1 |
| WBGene00000209 | -0.741594656 | 2.1511E-06 | 6.2329E-05 | asg-1 |
| WBGene00000210 | -0.07593162 | 0.66177612 | 0.80528531 | asg-2 |
| WBGene00000211 | -0.406669346 | 0.24912858 | 0.45684766 | asm-1 |
| WBGene00000212 | 2.458429036 | 6.41E-09 | 4.11E-07 | asm-2 |
| WBGene00000213 | 2.878221121 | 0.02625227 | 1 | asm-3 |
| WBGene00000214 | 1.354308268 | 1.33E-20 | 4.62E-18 | asp-1 |
| WBGene00000215 | 0.766441526 | 1.74E-07 | 7.5103E-06 | asp-2 |
| WBGene00000216 | 1.042142681 | 1.20E-12 | 1.66E-10 | asp-3 |
| WBGene00000217 | -0.609849984 | 0.00014933 | 0.00209238 | asp-4 |
| WBGene00000218 | 1.10190234 | 1.63E-12 | 2.18E-10 | asp-5 |
| WBGene00000219 | 1.352085686 | 2.44E-26 | 1.56E-23 | asp-6 |
| WBGene00000220 | -0.117825156 | 0.79269546 | 0.88562765 | atf-2 |
| WBGene00000221 | 0.895164743 | 1.1151E-05 | 0.00024933 | atf-5 |
| WBGene00000222 | -0.002226703 | 0.99457407 | 0.99728475 | atf-6 |
| WBGene00000223 | 0.270362172 | 0.09855541 | 0.25265229 | atf-7 |
| WBGene00000224 | 0.698409604 | 0.00189216 | 0.01492707 | atgp-1 |
| WBGene00000225 | 0.795337251 | 0.00476141 | 0.0301295 | atgp-2 |
| WBGene00000226 | 0.814761763 | 1.6282E-05 | 0.00034201 | atl-1 |
| WBGene00000227 | 0.249410042 | 0.32485645 | 0.53327805 | atm-1 |
| WBGene00000228 | 0.401289846 | 0.05686649 | 0.17612653 | atn-1 |
| WBGene00000229 | -0.247814911 | 0.10588066 | 0.26498244 | atp-2 |
| WBGene00000230 | -0.018665255 | 0.89464354 | 0.94453253 | atp-3 |
| WBGene00000231 | 0.372220668 | 0.10614344 | 0.26549649 | atx-2 |
| WBGene00000232 | 0.834138089 | 0.03071405 | 0.11552024 | avr-14 |
| WBGene00000233 | 0.110430524 | 0.7322611 | 0.85026607 | avr-15 |
| WBGene00000235 | -0.727557493 | 3.3552E-05 | 0.00061165 | baf-1 |
| WBGene00000236 | -0.843657269 | 4.9969E-05 | 0.00086544 | bag-1 |
| WBGene00000237 | 0.9042274 | 0.00301492 | 0.02128743 | bam-2 |
| WBGene00000238 | 0.956480973 | 0.00858738 | 0.04617335 | bar-1 |
| WBGene00000239 | -0.133092625 | 0.64407354 | 0.79428104 | bas-1 |
| WBGene00000240 | -0.082691017 | 0.67869579 | 0.81641653 | pah-1 |
| WBGene00000241 | 1.355260724 | 0.1124478 | 0.27474919 | bbs-1 |
| WBGene00000242 | 0.157197587 | 0.82413846 | 0.90501674 | bbs-2 |
| WBGene00000244 | 0.764608182 | 0.31009385 | 0.518332 | bbs-8 |
| WBGene00000245 | -0.172094185 | 0.5443009 | 0.7236484 | bca-1 |
| WBGene00000246 | -0.413307714 | 0.41565124 | 0.62082449 | bcc-1 |
| WBGene00000247 | -0.206638071 | 0.37155565 | 0.57908308 | bec-1 |
| WBGene00000248 | 0.195130373 | 0.42577098 | 0.62970697 | ben-1 |
| WBGene00000249 | -0.769283351 | 0.00130763 | 0.01125956 | bir-1 |
| WBGene00000250 | 0.101914458 | 0.6906606 | 0.82370435 | bir-2 |

| | | | | |
|---|---|---|---|---|
| WBGene00000251 | 0.09697944 | 0.97612634 | 1 | bli-1 |
| WBGene00000252 | -1.094201388 | 0.7816772 | 1 | bli-2 |
| WBGene00000253 | 0.677254934 | 0.02606072 | 0.10302938 | bli-3 |
| WBGene00000254 | -0.069515051 | 0.64835598 | 0.79737183 | bli-4 |
| WBGene00000255 | 0.28530093 | 0.92084641 | 1 | bli-5 |
| WBGene00000256 | 0.970263619 | 0.55793369 | 1 | bli-6 |
| WBGene00000257 | -0.383803893 | 0.06932433 | 0.20101902 | bmk-1 |
| WBGene00000259 | -0.104622562 | 0.63250272 | 0.78720815 | bpl-1 |
| WBGene00000262 | -0.930116849 | 0.0123733 | 0.06059872 | bra-1 |
| WBGene00000263 | -0.35147055 | 0.03860265 | 0.13432927 | F23H11.5 |
| WBGene00000264 | 0.101411313 | 0.66824371 | 0.81001866 | brc-1 |
| WBGene00000265 | -0.133993867 | 0.54757149 | 0.72633896 | brd-1 |
| WBGene00000266 | -0.10579149 | 0.66568139 | 0.8082846 | bre-1 |
| WBGene00000267 | 0.621533787 | 0.31824547 | 0.52611905 | bre-2 |
| WBGene00000268 | 0.343722019 | 0.06882307 | 0.19999543 | bre-3 |
| WBGene00000269 | -0.102872704 | 0.75632541 | 0.86391446 | bre-4 |
| WBGene00000270 | -0.649347919 | 0.03550355 | 0.1271155 | bre-5 |
| WBGene00000271 | -0.018179046 | 0.93068971 | 0.96384889 | brf-1 |
| WBGene00000272 | 0.713482268 | 0.53197236 | 1 | bro-1 |
| WBGene00000273 | 0.007910526 | 0.96061289 | 0.9796557 | brp-1 |
| WBGene00000274 | -0.073656075 | 0.73526834 | 0.852128 | btf-1 |
| WBGene00000275 | -0.408766974 | 0.025187 | 0.10072312 | bub-1 |
| WBGene00000276 | -0.033735044 | 0.83484147 | 0.91032502 | byn-1 |
| WBGene00000277 | 1.00499668 | 4.28E-07 | 1.6421E-05 | cab-1 |
| WBGene00000279 | 0.616623379 | 0.29456033 | 0.50490709 | cah-1 |
| WBGene00000280 | 0.879308496 | 0.15574204 | 0.33959979 | cah-2 |
| WBGene00000281 | 0.946717698 | 1.709E-05 | 0.00035635 | cah-3 |
| WBGene00000282 | 0.253628654 | 0.1295712 | 0.30084603 | cah-4 |
| WBGene00000283 | 0.468644337 | 0.08832607 | 0.23562642 | cah-5 |
| WBGene00000284 | 0.633397785 | 0.12547665 | 0.29536855 | cah-6 |
| WBGene00000285 | 0.586884148 | 0.15259477 | 0.33490055 | cal-1 |
| WBGene00000286 | 0.942601836 | 0.00804487 | 0.04407614 | cal-2 |
| WBGene00000287 | 0.526894812 | 0.47909191 | 0.67113235 | cal-3 |
| WBGene00000288 | 1.226102426 | 0.00122707 | 0.01073304 | cal-4 |
| WBGene00000289 | -0.075280659 | 0.69002011 | 0.82353771 | cam-1 |
| WBGene00000292 | -0.177762537 | 0.26361056 | 0.4721438 | cap-1 |
| WBGene00000293 | -0.258898671 | 0.10759881 | 0.26759651 | cap-2 |
| WBGene00000294 | 0.280985971 | 0.16303099 | 0.34903179 | cas-1 |
| WBGene00000295 | 0.674409926 | 0.36564739 | 0.5734821 | cat-1 |
| WBGene00000296 | 0.684767922 | 0.5048665 | 1 | cat-2 |
| WBGene00000298 | 0.314602668 | 0.09062679 | 0.23935172 | cat-4 |
| WBGene00000301 | -1.431460036 | 8.16E-15 | 1.50E-12 | cav-1 |
| WBGene00000302 | 0.744521622 | 0.26157436 | 0.47006725 | cav-2 |

| | | | | |
|---|---|---|---|---|
| WBGene00000366 | 0.517741973 | 0.02207766 | 0.09223842 | cbp-1 |
| WBGene00000367 | 1.628227545 | 4.4039E-05 | 0.00077941 | cca-1 |
| WBGene00000368 | 0.676842912 | 0.03331534 | 0.12190523 | ccb-1 |
| WBGene00000369 | -0.5422898 | 0.00253626 | 0.0187875 | ccf-1 |
| WBGene00000370 | 0.249578767 | 0.16273211 | 0.34858311 | ccg-1 |
| WBGene00000371 | -0.375091219 | 0.01292579 | 0.06263017 | cco-1 |
| WBGene00000372 | 1.410232066 | 0.10731709 | 0.26726908 | cyp-13A7 |
| WBGene00000373 | 0.530695168 | 0.1663014 | 0.35329226 | cyp-14A5 |
| WBGene00000375 | -0.390664831 | 0.06647746 | 0.19523533 | cyp-44A1 |
| WBGene00000376 | -0.292404701 | 0.0826321 | 0.22549704 | ccr-4 |
| WBGene00000377 | -0.406401982 | 0.00659474 | 0.03812704 | cct-1 |
| WBGene00000378 | -0.432534508 | 0.00733538 | 0.04113904 | cct-2 |
| WBGene00000379 | -0.22540893 | 0.13623858 | 0.31161881 | cct-4 |
| WBGene00000380 | -0.33629935 | 0.01882944 | 0.08212916 | cct-5 |
| WBGene00000381 | -0.234472735 | 0.14123101 | 0.31918944 | cct-6 |
| WBGene00000382 | -0.310058978 | 0.07974375 | 0.22043917 | cdc-6 |
| WBGene00000383 | 0.201306813 | 0.23107029 | 0.43640702 | cdc-14 |
| WBGene00000386 | -0.425650458 | 0.01724533 | 0.07691976 | cdc-25.1 |
| WBGene00000387 | -0.72564099 | 0.00188786 | 0.01490281 | cdc-25.2 |
| WBGene00000388 | -0.670781934 | 0.0029512 | 0.02095355 | cdc-25.3 |
| WBGene00000389 | -1.969805064 | 0.62862569 | 1 | cdc-25.4 |
| WBGene00000390 | -0.123811006 | 0.49346961 | 0.68212223 | cdc-42 |
| WBGene00000391 | -0.072625003 | 0.86095719 | 0.9251156 | cdd-1 |
| WBGene00000392 | -0.538206895 | 0.04756381 | 0.15561095 | cdd-2 |
| WBGene00000393 | 0.336120961 | 0.20539302 | 0.40488336 | cdf-1 |
| WBGene00000395 | 0.487274457 | 0.18566665 | 0.3782814 | cdh-3 |
| WBGene00000396 | 0.272882444 | 0.23058414 | 0.4360994 | cdh-4 |
| WBGene00000397 | -0.625472677 | 0.2671495 | 0.47544153 | cdh-5 |
| WBGene00000399 | 1.401955864 | 0.14073647 | 0.31848676 | cdh-7 |
| WBGene00000401 | -0.691090412 | 0.26931376 | 0.47783631 | cdh-9 |
| WBGene00000402 | 0.725321499 | 0.40761535 | 0.6141434 | cdh-10 |
| WBGene00000403 | 0.785471542 | 2.5057E-05 | 0.00048923 | casy-1 |
| WBGene00000405 | -0.582055893 | 7.0368E-05 | 0.00115921 | cdk-1 |
| WBGene00000406 | -0.81534081 | 0.00802233 | 0.04399239 | cdk-4 |
| WBGene00000407 | -0.054421298 | 0.80167431 | 0.89148362 | cdk-5 |
| WBGene00000408 | -0.442186173 | 0.03736299 | 0.13121898 | cdk-7 |
| WBGene00000409 | 0.058646177 | 0.7561512 | 0.86387802 | cdk-8 |
| WBGene00000410 | 0.045323573 | 0.85543824 | 0.92113008 | cdk-9 |
| WBGene00000411 | -0.600122005 | 0.00082759 | 0.0079179 | cdl-1 |
| WBGene00000412 | 0.444307195 | 0.66004764 | 1 | cdr-1 |
| WBGene00000413 | -0.310838023 | 0.16340194 | 0.34957937 | cdt-1 |
| WBGene00000414 | -0.355815753 | 0.10289895 | 0.25967495 | cec-1 |
| WBGene00000415 | -0.004134358 | 0.98350818 | 0.99200572 | ced-1 |

| | | | | |
|---|---|---|---|---|
| WBGene00000416 | -0.521983802 | 0.01019998 | 0.0525628 | ced-2 |
| WBGene00000417 | 0.054566154 | 0.78914195 | 0.88377202 | ced-3 |
| WBGene00000418 | -0.204831178 | 0.41381244 | 0.61934132 | ced-4 |
| WBGene00000419 | 0.513752832 | 0.00781041 | 0.04304295 | ced-5 |
| WBGene00000420 | -0.342456891 | 0.11729815 | 0.28233879 | ced-6 |
| WBGene00000421 | 0.856836534 | 2.6597E-06 | 7.4575E-05 | ced-7 |
| WBGene00000422 | 0.056835123 | 0.92206497 | 0.95919722 | ced-8 |
| WBGene00000423 | -0.793285953 | 0.00017359 | 0.00235221 | ced-9 |
| WBGene00000424 | -0.317040687 | 0.15153938 | 0.33336244 | ced-10 |
| WBGene00000425 | 0.46177396 | 0.32533842 | 0.5338183 | ced-11 |
| WBGene00000426 | -0.135522738 | 0.46542289 | 0.65974005 | ced-12 |
| WBGene00000428 | 0.224523095 | 0.92090023 | 1 | ceh-1 |
| WBGene00000429 | -0.044534513 | 0.97429035 | 1 | ceh-2 |
| WBGene00000430 | 0.635387686 | 0.82210541 | 1 | ceh-5 |
| WBGene00000431 | 0.648061841 | 0.2999526 | 0.50968853 | ceh-6 |
| WBGene00000433 | 0.278363362 | 0.78990626 | 1 | ceh-8 |
| WBGene00000434 | -0.947931206 | 0.45677197 | 1 | ceh-9 |
| WBGene00000435 | 0.628157342 | 0.48898522 | 0.67872065 | ceh-10 |
| WBGene00000436 | 0.574508983 | 0.8551971 | 1 | ceh-12 |
| WBGene00000437 | 0.084110705 | 0.82082387 | 0.90284677 | ceh-13 |
| WBGene00000438 | 0.022233908 | 0.97339692 | 0.9861492 | ceh-14 |
| WBGene00000439 | -0.944767306 | 0.27321009 | 0.48186715 | ceh-16 |
| WBGene00000440 | 0.138724036 | 0.88990066 | 0.94196731 | ceh-17 |
| WBGene00000441 | 1.096222314 | 0.00109715 | 0.00983787 | ceh-18 |
| WBGene00000442 | 0.085173975 | 0.93718725 | 1 | ceh-19 |
| WBGene00000443 | 0.242797095 | 0.32039154 | 0.52858726 | ceh-20 |
| WBGene00000444 | -0.409451291 | 0.28063034 | 0.48974169 | ceh-21 |
| WBGene00000445 | 0.037709469 | 0.95619035 | 0.97777369 | ceh-22 |
| WBGene00000446 | 0.569829665 | 0.5411282 | 0.72203951 | ceh-23 |
| WBGene00000447 | 0.208243434 | 0.82401417 | 0.90496496 | ceh-24 |
| WBGene00000448 | 0.625363197 | 0.13219054 | 0.30481857 | pros-1 |
| WBGene00000449 | -0.681282352 | 0.17664032 | 0.36810678 | ceh-27 |
| WBGene00000451 | 1.661953952 | 0.18503666 | 1 | ceh-30 |
| WBGene00000452 | 1.433216648 | 0.16088159 | 1 | ceh-31 |
| WBGene00000453 | -0.212029995 | 0.45904578 | 0.65487177 | ceh-32 |
| WBGene00000454 | -0.006475189 | 0.99557144 | 1 | ceh-33 |
| WBGene00000455 | 0.823339874 | 0.10786488 | 0.26759651 | ceh-34 |
| WBGene00000457 | -0.847822256 | 0.24851321 | 0.45656791 | ceh-36 |
| WBGene00000458 | 0.469865023 | 0.29267177 | 0.5028057 | ceh-37 |
| WBGene00000459 | -0.059810944 | 0.76144034 | 0.86657736 | ceh-38 |
| WBGene00000460 | -0.546076575 | 0.06174559 | 0.18592589 | ceh-39 |
| WBGene00000461 | -1.031400495 | 0.02635639 | 0.10373661 | ceh-40 |
| WBGene00000462 | -0.405144075 | 0.30936943 | 0.51807644 | ceh-41 |

| | | | | |
|---|---|---|---|---|
| WBGene00000463 | -0.796317316 | 0.01946607 | 0.0839565 | ceh-43 |
| WBGene00000464 | -0.170941526 | 0.31691109 | 0.52491372 | ceh-44 |
| WBGene00000465 | -0.373075583 | 0.02538198 | 0.10120283 | cpg-1 |
| WBGene00000466 | -0.189636338 | 0.29923535 | 0.50906705 | cel-1 |
| WBGene00000467 | -0.476458327 | 0.026149 | 0.1032287 | cep-1 |
| WBGene00000468 | 1.046325029 | 0.47829595 | 1 | ces-1 |
| WBGene00000469 | -0.191468335 | 0.75295442 | 0.86217294 | ces-2 |
| WBGene00000472 | -0.273594386 | 0.06736397 | 0.1971139 | cey-1 |
| WBGene00000473 | -0.696340259 | 5.58E-07 | 2.0791E-05 | cey-2 |
| WBGene00000474 | -0.882981668 | 2.20E-09 | 1.57E-07 | cey-3 |
| WBGene00000475 | -0.350773966 | 0.01322241 | 0.06355238 | cey-4 |
| WBGene00000476 | 1.072849474 | 0.01178027 | 0.05854454 | cfi-1 |
| WBGene00000477 | 0.992292283 | 0.0702889 | 0.20260624 | cft-1 |
| WBGene00000478 | 0.088134656 | 0.82605729 | 0.90625042 | cfz-2 |
| WBGene00000479 | -0.106444318 | 0.60168062 | 0.76670734 | cgh-1 |
| WBGene00000480 | -0.192246656 | 0.26810246 | 0.47664841 | cgp-1 |
| WBGene00000481 | 1.443191319 | 0.00532771 | 0.03265207 | cha-1 |
| WBGene00000482 | 0.157377761 | 0.43984595 | 0.64007787 | chd-3 |
| WBGene00000483 | -1.14715222 | 0.68932198 | 1 | che-1 |
| WBGene00000484 | 0.128594534 | 0.87986763 | 0.93574569 | che-2 |
| WBGene00000485 | 1.842984844 | 0.00109003 | 0.00981195 | che-3 |
| WBGene00000487 | -0.019206579 | 0.99153961 | 1 | che-6 |
| WBGene00000488 | 0.464297498 | 0.45062855 | 0.64900133 | che-7 |
| WBGene00000490 | 0.741736497 | 0.21902627 | 0.42176018 | che-11 |
| WBGene00000491 | 1.208972059 | 0.02251806 | 0.09327592 | che-12 |
| WBGene00000492 | 0.53914761 | 0.5096255 | 0.69630466 | che-13 |
| WBGene00000493 | -1.090035098 | 0.16369366 | 0.3498336 | che-14 |
| WBGene00000496 | 0.016469419 | 0.91617407 | 0.95593584 | chs-1 |
| WBGene00000497 | 0.425807684 | 0.3172004 | 0.52493219 | chs-2 |
| WBGene00000498 | -0.548383573 | 0.00612093 | 0.03618024 | chk-1 |
| WBGene00000499 | 0.097919665 | 0.86027166 | 0.92454265 | chk-2 |
| WBGene00000500 | -0.504464007 | 0.01126339 | 0.05674224 | chn-1 |
| WBGene00000501 | 1.090577193 | 0.03053514 | 0.11502549 | cho-1 |
| WBGene00000502 | -0.544366124 | 0.00563596 | 0.0340768 | chp-1 |
| WBGene00000503 | -1.843991391 | 1.07E-14 | 1.93E-12 | cht-1 |
| WBGene00000506 | 0.105453169 | 0.70534825 | 0.83345073 | cic-1 |
| WBGene00000507 | 0.312256267 | 0.23196619 | 0.43758569 | cit-1.1 |
| WBGene00000508 | 0.188744879 | 0.40665028 | 0.61346186 | cit-1.2 |
| WBGene00000509 | 0.322060552 | 0.1063207 | 0.26575456 | cka-1 |
| WBGene00000510 | 0.415209984 | 0.28919747 | 0.49938081 | cka-2 |
| WBGene00000511 | 1.259617189 | 0.05690827 | 0.17616605 | ckb-1 |
| WBGene00000512 | -0.938517875 | 0.00126128 | 0.01096932 | ckb-2 |
| WBGene00000513 | 1.158132015 | 0.62662247 | 1 | ckb-3 |

| | | | | |
|---|---|---|---|---|
| WBGene00000514 | 0.079619475 | 0.84231662 | 0.91337104 | ckb-4 |
| WBGene00000515 | -0.235516446 | 0.45016165 | 0.6488677 | ckc-1 |
| WBGene00000516 | -0.085037567 | 0.84584972 | 0.91552657 | cki-1 |
| WBGene00000517 | 0.077131051 | 0.75100916 | 0.86140786 | cki-2 |
| WBGene00000518 | 0.887647744 | 0.0677701 | 0.19774952 | ckk-1 |
| WBGene00000519 | 0.035918577 | 0.8643588 | 0.92704772 | cku-70 |
| WBGene00000520 | -0.211959971 | 0.26735132 | 0.47566125 | cku-80 |
| WBGene00000521 | 0.518740847 | 0.68093333 | 1 | cky-1 |
| WBGene00000522 | 0.421849455 | 0.11575524 | 0.28023616 | clc-1 |
| WBGene00000523 | -0.040273731 | 0.95331824 | 0.97623454 | clc-2 |
| WBGene00000524 | 0.145373994 | 0.7921723 | 0.88527422 | clc-3 |
| WBGene00000525 | -0.558804446 | 0.87818675 | 1 | clc-4 |
| WBGene00000526 | -0.015463173 | 0.97368986 | 0.98628138 | clc-5 |
| WBGene00000527 | 0.694761036 | 0.00271321 | 0.01974886 | cle-1 |
| WBGene00000528 | 1.051621643 | 6.018E-06 | 0.00014673 | clh-1 |
| WBGene00000529 | 0.832782579 | 0.13101827 | 0.30345151 | clh-2 |
| WBGene00000530 | 0.245725555 | 0.39738478 | 0.60512336 | clh-3 |
| WBGene00000531 | 0.799997511 | 0.32147721 | 0.52965027 | clh-4 |
| WBGene00000532 | -0.089425422 | 0.62503608 | 0.78190379 | clh-5 |
| WBGene00000533 | -0.186927909 | 0.42166868 | 0.62631238 | clh-6 |
| WBGene00000534 | -0.255654002 | 0.09463993 | 0.24636008 | cpi-2 |
| WBGene00000535 | -2.312842038 | 1.01E-19 | 3.22E-17 | cpi-1 |
| WBGene00000536 | -0.139209957 | 0.61603166 | 0.77593531 | clk-1 |
| WBGene00000537 | 0.07788777 | 0.71739498 | 0.84015553 | clk-2 |
| WBGene00000539 | 1.644408806 | 3.8709E-06 | 0.0001015 | cln-3.1 |
| WBGene00000540 | 0.176552237 | 0.58223933 | 0.75273854 | cln-3.2 |
| WBGene00000541 | 0.352650251 | 0.54409038 | 0.72360624 | cln-3.3 |
| WBGene00000542 | 0.551115686 | 0.0007055 | 0.00702395 | clp-1 |
| WBGene00000543 | 0.505501379 | 0.02766028 | 0.10729185 | clp-2 |
| WBGene00000544 | -0.755237416 | 0.60992977 | 1 | clp-3 |
| WBGene00000545 | 0.450563828 | 0.13754578 | 0.31389911 | clp-4 |
| WBGene00000546 | 2.223658251 | 0.25271431 | 1 | clp-6 |
| WBGene00000547 | 0.154129099 | 0.38835667 | 0.59713208 | clp-7 |
| WBGene00000548 | 0.339404315 | 0.06358221 | 0.189529 | clr-1 |
| WBGene00000549 | 0.154682239 | 0.5627908 | 0.7382069 | cls-2 |
| WBGene00000550 | 0.110715357 | 0.59656922 | 0.7626563 | clu-1 |
| WBGene00000552 | 0.089393474 | 0.56838208 | 0.74321237 | cmd-1 |
| WBGene00000553 | 1.467362443 | 0.00212251 | 0.01629942 | cmk-1 |
| WBGene00000554 | 0.259091021 | 0.26402959 | 0.47252111 | cnb-1 |
| WBGene00000556 | -3.053966537 | 0.01304671 | 1 | cnc-2 |
| WBGene00000557 | -0.394502164 | 0.57501392 | 0.74815787 | cnc-3 |
| WBGene00000558 | -0.612416984 | 0.49865756 | 0.68695347 | cnc-4 |
| WBGene00000560 | -2.051235694 | 0.34377405 | 1 | cnc-6 |

| | | | | |
|---|---|---|---|---|
| WBGene00000561 | -0.631193206 | 0.54543579 | 1 | cnd-1 |
| WBGene00000562 | 0.118260432 | 0.89095372 | 0.9424176 | cng-1 |
| WBGene00000563 | 1.611773326 | 0.30382351 | 1 | cng-3 |
| WBGene00000564 | 0.120212629 | 0.62078518 | 0.77893271 | cnk-1 |
| WBGene00000565 | -0.086496895 | 0.70602804 | 0.83366276 | cnt-1 |
| WBGene00000566 | -0.146515246 | 0.4468439 | 0.64584638 | cnt-2 |
| WBGene00000567 | -0.162121431 | 0.25438732 | 0.46207607 | cnx-1 |
| WBGene00000583 | 0.251058074 | 0.69621347 | 0.82821161 | cof-2 |
| WBGene00000584 | 0.545661025 | 0.59193243 | 1 | cog-1 |
| WBGene00000585 | 0.113598073 | 0.54048131 | 0.72174866 | cogc-2 |
| WBGene00000591 | -0.098659844 | 0.73750114 | 0.85334533 | coh-1 |
| WBGene00000592 | -0.190966663 | 0.47863527 | 0.67083387 | coh-3 |
| WBGene00000593 | -1.094201388 | 0.7816772 | 1 | col-2 |
| WBGene00000594 | 0.501728336 | 0.0762945 | 0.21436971 | col-3 |
| WBGene00000596 | -0.604396148 | 0.82723443 | 1 | col-7 |
| WBGene00000597 | -1.824661231 | 2.40E-26 | 1.56E-23 | col-8 |
| WBGene00000598 | -0.156269293 | 0.96924949 | 1 | col-9 |
| WBGene00000599 | 0.315856664 | 0.13768831 | 0.31416534 | col-10 |
| WBGene00000601 | 0.126643766 | 0.8828213 | 0.93749089 | col-12 |
| WBGene00000602 | -0.284202651 | 0.88416022 | 1 | col-13 |
| WBGene00000603 | -1.074526991 | 0.00566176 | 0.03416471 | col-14 |
| WBGene00000606 | -0.182667178 | 0.83470648 | 0.91032502 | col-17 |
| WBGene00000608 | -0.311846595 | 0.09602587 | 0.24863513 | col-19 |
| WBGene00000609 | -0.938862224 | 5.5563E-06 | 0.00013683 | col-20 |
| WBGene00000610 | 0.688793783 | 0.86378755 | 1 | col-33 |
| WBGene00000611 | -0.777993272 | 0.17780569 | 0.36971037 | col-34 |
| WBGene00000612 | -1.094201388 | 0.7816772 | 1 | col-35 |
| WBGene00000614 | -0.156269293 | 0.96924949 | 1 | col-37 |
| WBGene00000615 | 0.991814351 | 0.29547591 | 0.50569115 | col-38 |
| WBGene00000616 | 0.853690933 | 0.00052627 | 0.00558035 | col-39 |
| WBGene00000617 | 0.281454817 | 0.94435481 | 1 | col-40 |
| WBGene00000618 | -0.977566462 | 0.35381171 | 1 | col-41 |
| WBGene00000620 | -3.625582342 | 0.00907565 | 1 | col-43 |
| WBGene00000622 | 1.216957619 | 0.74819621 | 1 | col-45 |
| WBGene00000623 | -0.228167292 | 0.87186929 | 1 | col-46 |
| WBGene00000624 | 0.778594282 | 0.70344619 | 1 | col-47 |
| WBGene00000625 | -1.455714105 | 0.00904139 | 0.04793514 | col-48 |
| WBGene00000626 | 1.584577408 | 0.63258867 | 1 | col-49 |
| WBGene00000629 | 0.281454817 | 0.94435481 | 1 | col-52 |
| WBGene00000630 | -0.192831384 | 0.77927277 | 0.87749497 | col-53 |
| WBGene00000631 | -0.818621145 | 0.42215259 | 1 | col-54 |
| WBGene00000633 | -0.753364595 | 0.71590156 | 1 | col-56 |
| WBGene00000634 | -0.573209976 | 0.40071326 | 0.60828453 | col-58 |

| | | | | |
|---|---|---|---|---|
| WBGene00000636 | 2.521428441 | 0.15463352 | 1 | col-60 |
| WBGene00000637 | -1.259309624 | 0.25874952 | 1 | col-61 |
| WBGene00000638 | 0.421206838 | 0.79609137 | 1 | col-62 |
| WBGene00000639 | 1.747998126 | 0.61126322 | 1 | col-63 |
| WBGene00000640 | 1.448645387 | 0.51537893 | 1 | col-64 |
| WBGene00000641 | -1.155231515 | 0.00580753 | 0.03480218 | col-65 |
| WBGene00000644 | -1.396388689 | 0.1757668 | 1 | col-68 |
| WBGene00000645 | 0.842324357 | 0.60400222 | 1 | col-69 |
| WBGene00000646 | -1.14715222 | 0.68932198 | 1 | H17B01.2 |
| WBGene00000647 | 0.202462917 | 0.86786543 | 1 | col-71 |
| WBGene00000648 | -0.156269293 | 0.96924949 | 1 | col-72 |
| WBGene00000649 | -0.054284561 | 0.96500281 | 1 | col-73 |
| WBGene00000650 | -1.06107064 | 0.00013196 | 0.00190953 | col-74 |
| WBGene00000651 | -3.81679611 | 0.07116215 | 1 | col-75 |
| WBGene00000652 | 0.438167738 | 0.18724044 | 0.38072118 | col-76 |
| WBGene00000653 | -0.861830037 | 0.02906114 | 0.11077905 | col-77 |
| WBGene00000655 | 0.962938242 | 0.55520017 | 1 | col-79 |
| WBGene00000656 | -1.108210519 | 1.0243E-06 | 3.4258E-05 | col-80 |
| WBGene00000657 | -0.194986594 | 0.83886517 | 0.91202483 | col-81 |
| WBGene00000658 | -0.420517409 | 0.78174857 | 1 | col-83 |
| WBGene00000659 | -1.366565036 | 0.6506483 | 1 | col-84 |
| WBGene00000660 | 0.377828795 | 0.92526585 | 1 | col-85 |
| WBGene00000661 | -0.298265327 | 0.89527345 | 1 | col-86 |
| WBGene00000662 | 1.529802032 | 0.64971375 | 1 | col-87 |
| WBGene00000663 | 1.584577408 | 0.63258867 | 1 | col-88 |
| WBGene00000664 | 0.204681671 | 0.82487803 | 0.90566506 | col-89 |
| WBGene00000665 | -0.605024566 | 0.47344454 | 0.66645288 | col-90 |
| WBGene00000666 | 0.39107888 | 0.63796904 | 0.79024747 | col-91 |
| WBGene00000667 | 1.947221036 | 0.07169825 | 0.2057406 | col-92 |
| WBGene00000668 | -1.264302164 | 0.01641872 | 0.07418999 | col-93 |
| WBGene00000669 | 0.774277223 | 0.02170308 | 0.09098658 | col-94 |
| WBGene00000670 | -1.051644588 | 1.46E-08 | 8.60E-07 | col-95 |
| WBGene00000671 | -1.038388431 | 0.34347786 | 1 | col-96 |
| WBGene00000672 | -1.31451832 | 0.00374265 | 0.02498046 | col-97 |
| WBGene00000673 | -0.547274581 | 0.01151711 | 0.05763776 | col-98 |
| WBGene00000674 | -0.268885255 | 0.58945954 | 0.75742266 | col-99 |
| WBGene00000675 | -1.815166669 | 5.85E-22 | 2.29E-19 | col-101 |
| WBGene00000677 | -0.406953076 | 0.04868649 | 0.15821805 | col-103 |
| WBGene00000678 | -0.899156291 | 0.04411711 | 0.14694809 | col-104 |
| WBGene00000679 | -1.082680044 | 0.67762556 | 1 | col-105 |
| WBGene00000680 | -1.387036273 | 3.11E-17 | 7.70E-15 | col-106 |
| WBGene00000681 | 0.023314718 | 0.92351412 | 0.95953374 | col-107 |
| WBGene00000683 | -0.942645934 | 0.06462294 | 0.19164316 | col-109 |

| | | | | |
|---|---|---|---|---|
| WBGene00000685 | -0.650106892 | 0.54725837 | 1 | col-111 |
| WBGene00000687 | -1.2100486 | 0.0617998 | 0.18595076 | col-113 |
| WBGene00000689 | -0.450439867 | 0.49107492 | 0.6802214 | col-115 |
| WBGene00000690 | -1.992503642 | 0.08296638 | 1 | col-116 |
| WBGene00000691 | 0.501728336 | 0.0762945 | 0.21436971 | col-117 |
| WBGene00000692 | 0.059429741 | 0.86608095 | 0.92799221 | col-118 |
| WBGene00000693 | -0.536123935 | 0.0028221 | 0.020326 | col-119 |
| WBGene00000694 | 0.056106392 | 0.97371196 | 1 | col-120 |
| WBGene00000695 | -0.887349564 | 0.04798544 | 0.15661054 | col-121 |
| WBGene00000696 | -1.512664243 | 1.63E-18 | 4.70E-16 | col-122 |
| WBGene00000697 | 0.270496667 | 0.92869767 | 1 | col-123 |
| WBGene00000698 | 0.233271951 | 0.56890063 | 0.74341002 | col-124 |
| WBGene00000699 | -0.61969226 | 0.04917759 | 0.15921737 | col-125 |
| WBGene00000700 | -0.263229581 | 0.93409872 | 1 | col-126 |
| WBGene00000701 | -0.263229581 | 0.93409872 | 1 | col-127 |
| WBGene00000702 | 0.23765919 | 0.83508882 | 1 | col-128 |
| WBGene00000703 | 0.202808749 | 0.7292954 | 0.84847759 | col-129 |
| WBGene00000704 | -0.582507698 | 0.2648672 | 0.47360128 | col-130 |
| WBGene00000705 | -0.882730389 | 0.36132254 | 0.56912519 | col-131 |
| WBGene00000706 | -0.62146648 | 0.85925752 | 1 | col-132 |
| WBGene00000707 | 3.225472789 | 0.13593768 | 1 | col-133 |
| WBGene00000708 | 0.890714601 | 0.00078949 | 0.00766199 | col-135 |
| WBGene00000710 | 0.747845955 | 0.85215965 | 1 | col-137 |
| WBGene00000711 | 2.132995841 | 0.26474404 | 1 | col-138 |
| WBGene00000712 | -0.036084025 | 0.96053966 | 0.9796557 | col-139 |
| WBGene00000713 | -0.611240081 | 0.00058598 | 0.00601899 | col-140 |
| WBGene00000714 | 1.166283287 | 0.42407091 | 1 | col-141 |
| WBGene00000715 | -0.262852539 | 0.49598211 | 0.68438489 | col-142 |
| WBGene00000716 | -1.16797646 | 2.69E-09 | 1.90E-07 | col-143 |
| WBGene00000717 | 0.328694508 | 0.14069262 | 0.31844689 | col-144 |
| WBGene00000718 | 0.218252028 | 0.85202216 | 1 | col-145 |
| WBGene00000719 | 1.70532274 | 0.20868167 | 1 | col-146 |
| WBGene00000720 | 0.640197382 | 0.00074312 | 0.00729365 | col-147 |
| WBGene00000722 | 0.104897692 | 0.82187119 | 0.90342582 | col-149 |
| WBGene00000723 | -1.754636245 | 0.04312333 | 0.14481318 | col-150 |
| WBGene00000724 | 0.564510346 | 0.78300443 | 1 | col-151 |
| WBGene00000725 | 0.281454817 | 0.94435481 | 1 | col-152 |
| WBGene00000726 | 0.760835545 | 0.27887463 | 0.48770698 | col-153 |
| WBGene00000727 | 1.080083342 | 0.07326386 | 0.20855723 | col-154 |
| WBGene00000728 | 0.401945397 | 0.31269246 | 0.52091097 | col-155 |
| WBGene00000729 | 0.32595502 | 0.91626577 | 1 | col-156 |
| WBGene00000730 | 1.962133156 | 0.28921013 | 1 | col-157 |
| WBGene00000731 | -2.793287338 | 3.8513E-06 | 0.00010121 | col-158 |

| | | | | |
|---|---|---|---|---|
| WBGene00000732 | -0.746422506 | 0.23629385 | 0.44244693 | col-159 |
| WBGene00000733 | 0.149049274 | 0.53331005 | 0.71646737 | col-160 |
| WBGene00000734 | 1.953641176 | 0.30191136 | 1 | col-161 |
| WBGene00000735 | -1.115761498 | 0.44847094 | 1 | col-162 |
| WBGene00000736 | -0.48070356 | 0.82169194 | 1 | col-163 |
| WBGene00000738 | -1.208817026 | 2.8909E-06 | 8.05E-05 | col-165 |
| WBGene00000739 | 0.630621229 | 0.00193778 | 0.01519809 | col-166 |
| WBGene00000740 | 0.570032147 | 0.00787774 | 0.04335616 | col-167 |
| WBGene00000741 | 0.565228071 | 0.10776562 | 0.26759651 | col-168 |
| WBGene00000742 | 1.270226175 | 0.01053634 | 0.05395266 | col-169 |
| WBGene00000743 | 0.099400265 | 0.90055108 | 0.9476979 | col-170 |
| WBGene00000745 | -0.094483547 | 0.94321804 | 1 | col-172 |
| WBGene00000746 | -1.330302244 | 0.49220638 | 1 | col-173 |
| WBGene00000747 | -0.033793778 | 0.98074606 | 1 | col-174 |
| WBGene00000748 | 0.403207732 | 0.86402224 | 1 | col-175 |
| WBGene00000749 | 1.073653728 | 0.17443347 | 0.36457165 | col-176 |
| WBGene00000750 | 0.27355185 | 0.70364601 | 0.83263838 | col-177 |
| WBGene00000751 | -0.519496448 | 0.00447089 | 0.0286898 | col-178 |
| WBGene00000752 | 0.188528831 | 0.31891349 | 0.52680654 | col-179 |
| WBGene00000753 | 0.08697668 | 0.93685828 | 1 | col-180 |
| WBGene00000754 | -1.659025542 | 1.03E-24 | 5.43E-22 | col-181 |
| WBGene00000755 | -0.096378115 | 0.95471465 | 1 | col-182 |
| WBGene00000757 | -1.285362183 | 1.48E-15 | 3.05E-13 | col-184 |
| WBGene00000759 | 0.26417221 | 0.77752316 | 0.87617493 | col-186 |
| WBGene00000760 | -0.70576507 | 0.76841669 | 1 | col-187 |
| WBGene00000761 | -0.009632103 | 0.9665494 | 0.9830824 | coq-1 |
| WBGene00000762 | 0.017680554 | 0.94628946 | 0.972918 | coq-2 |
| WBGene00000763 | -0.279654203 | 0.4006517 | 0.60826713 | coq-3 |
| WBGene00000764 | -0.259260533 | 0.42780893 | 0.63072596 | coq-4 |
| WBGene00000765 | -0.610144338 | 0.00266 | 0.01945487 | coq-5 |
| WBGene00000766 | 0.026672192 | 0.9248851 | 0.96034913 | coq-6 |
| WBGene00000767 | 0.063401681 | 0.75454249 | 0.86283239 | coq-8 |
| WBGene00000768 | 0.657496157 | 0.04407578 | 0.14685071 | cor-1 |
| WBGene00000770 | 0.076617369 | 0.79298278 | 0.88575825 | cpb-1 |
| WBGene00000771 | -0.440144501 | 0.78503374 | 1 | cpb-2 |
| WBGene00000772 | -0.118884142 | 0.55046368 | 0.72904762 | cpb-3 |
| WBGene00000773 | -0.651712467 | 0.00034956 | 0.0040966 | cpf-1 |
| WBGene00000774 | -0.623813241 | 0.00266679 | 0.01949279 | cpf-2 |
| WBGene00000775 | 0.287885625 | 0.5863012 | 0.75557614 | cpin-1 |
| WBGene00000776 | 0.634912501 | 1.5215E-05 | 0.00032238 | cpl-1 |
| WBGene00000777 | -0.513586646 | 0.00101594 | 0.00935145 | cpn-1 |
| WBGene00000778 | -0.684009739 | 0.22095616 | 0.42397327 | cpn-2 |
| WBGene00000779 | -0.086040355 | 0.57267693 | 0.7465769 | cpn-3 |

| | | | | |
|---|---|---|---|---|
| WBGene00000780 | 0.432474585 | 0.06921837 | 0.20080771 | cpn-4 |
| WBGene00000781 | 2.83242639 | 1.10E-63 | 1.34E-59 | cpr-1 |
| WBGene00000782 | 1.451745273 | 0.51709377 | 1 | cpr-2 |
| WBGene00000783 | 0.67345448 | 0.00684325 | 0.03911669 | cpr-3 |
| WBGene00000784 | 1.724980355 | 1.12E-30 | 8.52E-28 | cpr-4 |
| WBGene00000785 | -0.590715824 | 0.00043988 | 0.00482436 | cpr-5 |
| WBGene00000786 | -0.379935264 | 0.00655876 | 0.03798877 | cpr-6 |
| WBGene00000787 | 0.143737381 | 0.56136683 | 0.73709014 | cps-6 |
| WBGene00000788 | -0.356561609 | 0.04041114 | 0.13851827 | cpz-1 |
| WBGene00000789 | 0.229822735 | 0.21388696 | 0.41449348 | cpz-2 |
| WBGene00000792 | 0.409320733 | 0.31385728 | 0.52177554 | crb-1 |
| WBGene00000793 | 0.156707214 | 0.55354559 | 0.73137414 | crh-1 |
| WBGene00000794 | -0.698453298 | 8.2996E-05 | 0.001321 | crn-1 |
| WBGene00000795 | -0.250404503 | 0.48061073 | 0.67247838 | crn-2 |
| WBGene00000796 | 0.062580399 | 0.7015833 | 0.83157723 | crn-3 |
| WBGene00000797 | -0.644746643 | 0.12015946 | 0.28644335 | crn-4 |
| WBGene00000798 | -0.802261016 | 0.00951789 | 0.04995964 | crn-5 |
| WBGene00000799 | 1.131625899 | 0.00052589 | 0.00558035 | crn-6 |
| WBGene00000800 | -0.161238697 | 0.36821306 | 0.57588021 | cars-1 |
| WBGene00000802 | -0.143618177 | 0.27270809 | 0.48138251 | crt-1 |
| WBGene00000803 | -0.348973729 | 0.28552583 | 0.49536569 | csb-1 |
| WBGene00000804 | -0.979954081 | 1.4798E-06 | 4.5485E-05 | csc-1 |
| WBGene00000812 | 0.006459309 | 0.97422128 | 0.98649644 | csk-1 |
| WBGene00000813 | -0.378431234 | 0.03490975 | 0.12590615 | csn-1 |
| WBGene00000814 | -0.131845036 | 0.58149642 | 0.75233887 | csn-2 |
| WBGene00000815 | -0.406460481 | 0.01648698 | 0.07432888 | csn-3 |
| WBGene00000816 | -0.358809121 | 0.0432747 | 0.1451376 | csn-4 |
| WBGene00000817 | -0.462707155 | 0.00931949 | 0.04915201 | csn-5 |
| WBGene00000818 | -0.287910165 | 0.16686124 | 0.35418459 | csn-6 |
| WBGene00000819 | 0.056677218 | 0.94660341 | 0.97304242 | csp-1 |
| WBGene00000820 | 2.209112991 | 0.10360733 | 1 | csp-2 |
| WBGene00000822 | -0.296419377 | 0.07459855 | 0.21102571 | csq-1 |
| WBGene00000830 | 0.506532115 | 0.40298277 | 0.61081321 | ctl-1 |
| WBGene00000831 | 0.576172214 | 0.00201627 | 0.01564191 | ctl-2 |
| WBGene00000832 | 0.19249187 | 0.48331357 | 0.67431741 | ctn-1 |
| WBGene00000833 | 0.100400583 | 0.47848852 | 0.67082322 | cts-1 |
| WBGene00000834 | 0.052134778 | 0.76278557 | 0.86721412 | cua-1 |
| WBGene00000835 | -0.858029323 | 0.00073112 | 0.00721082 | cuc-1 |
| WBGene00000836 | -0.587336299 | 0.00057243 | 0.00592991 | cul-1 |
| WBGene00000837 | -0.007296954 | 0.96846496 | 0.98386325 | cul-2 |
| WBGene00000838 | -0.203623867 | 0.23054179 | 0.4360994 | cul-3 |
| WBGene00000839 | 0.09074436 | 0.66669001 | 0.80890254 | cul-4 |
| WBGene00000840 | 0.259446047 | 0.17318334 | 0.36296651 | cul-5 |

| | | | | |
|---|---|---|---|---|
| WBGene00000841 | 0.764254968 | 0.1519516 | 0.33384807 | cul-6 |
| WBGene00000843 | -0.599451116 | 0.00075443 | 0.00738672 | cup-2 |
| WBGene00000845 | 0.593768185 | 0.2524754 | 0.46053243 | cup-4 |
| WBGene00000846 | 0.109144199 | 0.57506047 | 0.74815787 | cup-5 |
| WBGene00000853 | -1.100488969 | 0.08934421 | 0.2371834 | cut-6 |
| WBGene00000854 | -0.15494293 | 0.47967146 | 0.67161331 | cux-7 |
| WBGene00000857 | -0.451426857 | 0.18747834 | 0.38098404 | cwn-1 |
| WBGene00000858 | 0.26864006 | 0.4642445 | 0.65886864 | cwn-2 |
| WBGene00000862 | 0.962883245 | 0.17914341 | 0.37122037 | cwp-4 |
| WBGene00000863 | -0.34235068 | 0.07444239 | 0.2108738 | cya-1 |
| WBGene00000864 | 2.312670972 | 0.31200728 | 1 | cya-2 |
| WBGene00000865 | -0.622959095 | 0.00038699 | 0.00440757 | cyb-1 |
| WBGene00000866 | -0.54459962 | 0.0005653 | 0.00587615 | cyb-2.1 |
| WBGene00000867 | -0.561650806 | 6.1397E-05 | 0.00103057 | cyb-2.2 |
| WBGene00000868 | -0.662445619 | 4.3958E-06 | 0.00011294 | cyb-3 |
| WBGene00000869 | -0.265802167 | 0.06155757 | 0.1858206 | cyc-1 |
| WBGene00000870 | -0.494328777 | 0.17116473 | 0.36015788 | cyd-1 |
| WBGene00000871 | -0.007576846 | 0.96899697 | 0.98399257 | cye-1 |
| WBGene00000872 | 0.119669445 | 0.584495 | 0.75436671 | cyk-1 |
| WBGene00000874 | -0.044426282 | 0.80905929 | 0.89607634 | cyk-3 |
| WBGene00000875 | -0.509997422 | 0.00558842 | 0.03385679 | cyk-4 |
| WBGene00000876 | -0.141903034 | 0.45902223 | 0.65487177 | cyl-1 |
| WBGene00000877 | -0.766081198 | 1.1139E-05 | 0.00024933 | cyn-1 |
| WBGene00000878 | 0.549192711 | 0.64227402 | 1 | cyn-2 |
| WBGene00000879 | -0.503693152 | 0.00034957 | 0.0040966 | cyn-3 |
| WBGene00000880 | -0.031782598 | 0.87104646 | 0.93043947 | cyn-4 |
| WBGene00000881 | -0.591406374 | 7.2747E-05 | 0.00118872 | cyn-5 |
| WBGene00000882 | -0.72949096 | 0.0105491 | 0.05397883 | cyn-6 |
| WBGene00000883 | -0.666494257 | 8.06E-07 | 2.779E-05 | cyn-7 |
| WBGene00000884 | -0.695098092 | 0.00554795 | 0.033662 | cyn-8 |
| WBGene00000885 | -0.265252985 | 0.24686351 | 0.45494383 | cyn-9 |
| WBGene00000886 | -0.323724114 | 0.18062142 | 0.37259848 | cyn-10 |
| WBGene00000887 | -0.66626628 | 0.0058555 | 0.03496885 | cyn-11 |
| WBGene00000888 | -0.859701522 | 3.8352E-05 | 0.0006929 | cyn-12 |
| WBGene00000889 | -0.644471766 | 0.00087814 | 0.00832283 | cyn-13 |
| WBGene00000890 | -0.224774432 | 0.23057184 | 0.4360994 | sig-7 |
| WBGene00000891 | -0.379735236 | 0.04302526 | 0.14462062 | cyn-15 |
| WBGene00000892 | -0.149197223 | 0.49213775 | 0.68122728 | cyn-16 |
| WBGene00000893 | -0.384452159 | 0.52902753 | 0.7133411 | cyn-17 |
| WBGene00000894 | -0.243713344 | 0.14271027 | 0.32052915 | dab-1 |
| WBGene00000895 | 0.500485596 | 0.59967405 | 0.76517526 | dac-1 |
| WBGene00000896 | -0.308732982 | 0.23570322 | 0.44202545 | dad-1 |
| WBGene00000897 | -0.452390712 | 0.06723552 | 0.19693762 | daf-1 |

| | | | | |
|---|---|---|---|---|
| WBGene00000898 | 0.890848876 | 0.00035338 | 0.0041282 | daf-2 |
| WBGene00000899 | 0.587033411 | 0.02331752 | 0.09557663 | daf-3 |
| WBGene00000900 | 0.321881528 | 0.18172282 | 0.37382024 | daf-4 |
| WBGene00000901 | 0.665546813 | 0.00662235 | 0.03822359 | daf-5 |
| WBGene00000902 | -1.150817143 | 0.15521244 | 0.33892703 | daf-6 |
| WBGene00000903 | 0.867501746 | 0.0049459 | 0.0309367 | daf-7 |
| WBGene00000904 | 0.264373059 | 0.47702391 | 0.66938827 | daf-8 |
| WBGene00000905 | -0.811295972 | 0.53278583 | 1 | daf-9 |
| WBGene00000906 | 1.272803963 | 0.1159429 | 0.28046677 | daf-10 |
| WBGene00000907 | 0.057219869 | 0.94453991 | 0.97134161 | daf-11 |
| WBGene00000908 | 1.062804602 | 0.00428498 | 0.02770178 | daf-12 |
| WBGene00000910 | 0.621879364 | 0.03391486 | 0.12365175 | daf-14 |
| WBGene00000911 | 0.34476734 | 0.07992159 | 0.2208805 | daf-15 |
| WBGene00000912 | 0.792390054 | 0.0003946 | 0.00446076 | daf-16 |
| WBGene00000913 | -0.158061339 | 0.31073262 | 0.51873445 | daf-18 |
| WBGene00000914 | 0.191133301 | 0.47683832 | 0.66928255 | daf-19 |
| WBGene00000915 | -0.409250358 | 0.01440834 | 0.06772423 | daf-21 |
| WBGene00000917 | -0.686971667 | 0.00451217 | 0.02892412 | daf-25 |
| WBGene00000920 | -0.327593762 | 0.59618396 | 0.76232432 | daf-28 |
| WBGene00000923 | -0.234181841 | 0.24131993 | 0.4489527 | daf-31 |
| WBGene00000928 | -0.948161301 | 0.00823833 | 0.04477642 | dao-2 |
| WBGene00000929 | -0.416764798 | 0.13927733 | 0.31642329 | dao-3 |
| WBGene00000930 | 0.747845955 | 0.85215965 | 1 | dao-4 |
| WBGene00000931 | -0.13532266 | 0.48893042 | 0.67872065 | dao-5 |
| WBGene00000932 | -0.861206134 | 0.70037023 | 1 | dao-6 |
| WBGene00000933 | -0.143429224 | 0.46653376 | 0.66083793 | dap-3 |
| WBGene00000934 | 1.139188783 | 0.04954251 | 0.1594963 | dat-1 |
| WBGene00000935 | -0.119822273 | 0.47788732 | 0.67028996 | daz-1 |
| WBGene00000936 | 0.9485205 | 0.06281246 | 0.18801925 | dbl-1 |
| WBGene00000937 | 0.009898701 | 0.96586632 | 0.98286673 | dbr-1 |
| WBGene00000938 | 0.392303994 | 0.03200348 | 0.1186885 | dcp-66 |
| WBGene00000939 | 0.435606349 | 0.01694909 | 0.07590517 | dcr-1 |
| WBGene00000940 | -0.454174365 | 0.04027419 | 0.13817557 | dcs-1 |
| WBGene00000941 | -0.654879249 | 0.00320765 | 0.02226645 | ddp-1 |
| WBGene00000942 | 1.124599284 | 1.32E-09 | 1.01E-07 | deb-1 |
| WBGene00000950 | 0.377828795 | 0.92526585 | 1 | C47C12.2 |
| WBGene00000950 | 0.728487164 | 0.28950774 | 0.49967941 | deg-1 |
| WBGene00000951 | 1.403730765 | 0.02868472 | 0.10965404 | deg-3 |
| WBGene00000952 | 0.953100521 | 0.38531959 | 1 | del-1 |
| WBGene00000953 | 0.48346977 | 0.58814569 | 0.75642763 | del-2 |
| WBGene00000955 | 1.585708242 | 0.00630005 | 0.03696903 | des-2 |
| WBGene00000958 | 1.116267506 | 0.00104457 | 0.0095358 | dgk-1 |
| WBGene00000959 | 0.105672838 | 0.65791768 | 0.80303357 | dgk-2 |

| | | | | |
|---|---|---|---|---|
| WBGene00000960 | 1.382346278 | 0.10589694 | 0.26498244 | dgk-3 |
| WBGene00000961 | 0.320498315 | 0.31414309 | 0.52190056 | dgn-1 |
| WBGene00000962 | 0.783582638 | 0.0004305 | 0.00475153 | dhc-1 |
| WBGene00000963 | 1.260484839 | 3.4078E-06 | 9.1943E-05 | dhp-1 |
| WBGene00000964 | 0.790533718 | 0.00683243 | 0.03909169 | dhp-2 |
| WBGene00000965 | -0.554258223 | 0.00927546 | 0.04898362 | dhs-1 |
| WBGene00000966 | -0.587302793 | 0.30165835 | 0.51115619 | dhs-2 |
| WBGene00000967 | -0.911256165 | 1.7659E-05 | 0.00036571 | dhs-3 |
| WBGene00000968 | 1.712765709 | 0.01078317 | 0.05491545 | dhs-4 |
| WBGene00000969 | 0.132999747 | 0.71857919 | 0.84097455 | dhs-5 |
| WBGene00000970 | -0.222834439 | 0.1836905 | 0.37589521 | dhs-6 |
| WBGene00000971 | -0.843015906 | 0.29400094 | 0.50437549 | dhs-7 |
| WBGene00000972 | 0.050306222 | 0.96843654 | 1 | dhs-8 |
| WBGene00000973 | 0.35496512 | 0.21150207 | 0.41127027 | dhs-9 |
| WBGene00000974 | -0.882263636 | 0.0047725 | 0.03017863 | dhs-11 |
| WBGene00000975 | -0.567985751 | 0.0102474 | 0.05278476 | dhs-12 |
| WBGene00000976 | -0.464478822 | 0.0249041 | 0.09974122 | dhs-13 |
| WBGene00000977 | -0.737945519 | 0.01639667 | 0.07414228 | dhs-14 |
| WBGene00000978 | 0.80677977 | 0.26013498 | 0.46838185 | dhs-15 |
| WBGene00000979 | 1.399559096 | 0.32283475 | 1 | dhs-16 |
| WBGene00000980 | -0.089818121 | 0.78361202 | 0.88074562 | dhs-17 |
| WBGene00000981 | 0.429865956 | 0.13136616 | 0.30385151 | dhs-18 |
| WBGene00000982 | -0.509388805 | 0.05187115 | 0.1645161 | dhs-19 |
| WBGene00000983 | -0.554106223 | 0.10952459 | 0.27005137 | dhs-20 |
| WBGene00000984 | 0.793088286 | 0.00956456 | 0.05013963 | dhs-21 |
| WBGene00000985 | -0.331877645 | 0.05257818 | 0.16602124 | dhs-22 |
| WBGene00000986 | 2.132974887 | 0.15028077 | 1 | dhs-23 |
| WBGene00000987 | -0.383838021 | 0.08454283 | 0.22901261 | dhs-24 |
| WBGene00000988 | -1.99979362 | 7.17E-17 | 1.64E-14 | dhs-25 |
| WBGene00000989 | -1.643308501 | 0.17476521 | 1 | dhs-26 |
| WBGene00000990 | -0.25259263 | 0.46939962 | 0.66298055 | dhs-27 |
| WBGene00000991 | 0.041435751 | 0.79192122 | 0.88508842 | dhs-28 |
| WBGene00000992 | 0.172070377 | 0.70188047 | 0.83169343 | dhs-29 |
| WBGene00000993 | -0.427646356 | 0.13448698 | 0.30877579 | dhs-30 |
| WBGene00000994 | 0.028398398 | 0.89020146 | 0.94196731 | dic-1 |
| WBGene00000995 | -0.528747948 | 0.01826491 | 0.08025853 | die-1 |
| WBGene00000996 | -0.560078584 | 0.00104348 | 0.0095358 | dif-1 |
| WBGene00000998 | 2.300406281 | 0.00060226 | 0.00616009 | dig-1 |
| WBGene00001000 | 0.54886008 | 0.00195108 | 0.01526291 | dim-1 |
| WBGene00001001 | -0.14145965 | 0.39953183 | 0.60727919 | dis-3 |
| WBGene00001002 | -0.689176135 | 0.00016191 | 0.00222879 | div-1 |
| WBGene00001005 | -0.622686052 | 2.961E-05 | 0.00055391 | dlc-1 |
| WBGene00001006 | 0.386238895 | 0.08585688 | 0.23133342 | dlg-1 |

| | | | | |
|---|---|---|---|---|
| WBGene00001007 | -0.241306506 | 0.22848773 | 0.43373241 | dli-1 |
| WBGene00001008 | 0.916983009 | 0.00010201 | 0.00155001 | dlk-1 |
| WBGene00001016 | 0.043592448 | 0.83508763 | 0.91032502 | dna-2 |
| WBGene00001017 | 0.251384403 | 0.14009135 | 0.3175003 | dnc-1 |
| WBGene00001018 | -0.748586156 | 0.00035776 | 0.00416052 | dnc-2 |
| WBGene00001019 | 0.135123654 | 0.48472567 | 0.67527875 | dnj-1 |
| WBGene00001020 | -0.803618144 | 0.00054869 | 0.00573418 | dnj-2 |
| WBGene00001021 | -0.013252112 | 0.97549465 | 0.98720352 | dnj-3 |
| WBGene00001022 | 0.13239976 | 0.65323555 | 0.80061911 | dnj-4 |
| WBGene00001023 | -0.055975925 | 0.71392831 | 0.83829608 | dnj-5 |
| WBGene00001025 | 0.11489182 | 0.57817363 | 0.74963756 | dnj-7 |
| WBGene00001026 | 0.309760575 | 0.08462537 | 0.22913394 | dnj-8 |
| WBGene00001027 | -0.136040594 | 0.52052529 | 0.70555963 | dnj-9 |
| WBGene00001028 | -0.144720717 | 0.48138251 | 0.67289112 | dnj-10 |
| WBGene00001029 | -0.204104546 | 0.15988566 | 0.34521994 | dnj-11 |
| WBGene00001030 | -0.475197425 | 0.00300893 | 0.02126396 | dnj-12 |
| WBGene00001031 | -0.36264336 | 0.02459418 | 0.09910319 | dnj-13 |
| WBGene00001032 | -0.427177456 | 0.13993411 | 0.31729081 | dnj-14 |
| WBGene00001033 | -0.206764908 | 0.44146023 | 0.64137027 | dnj-15 |
| WBGene00001034 | -0.468051908 | 0.01329295 | 0.0637904 | dnj-16 |
| WBGene00001035 | -0.192140589 | 0.36082828 | 0.56849418 | dnj-17 |
| WBGene00001036 | 0.05140894 | 0.85296078 | 0.91927977 | dnj-18 |
| WBGene00001037 | -0.343406261 | 0.01319724 | 0.06348166 | dnj-19 |
| WBGene00001038 | 0.208202826 | 0.26254103 | 0.4712464 | dnj-20 |
| WBGene00001039 | -0.838433198 | 0.00585984 | 0.03497752 | dnj-21 |
| WBGene00001040 | -0.284491255 | 0.13349371 | 0.30701782 | dnj-22 |
| WBGene00001041 | -0.565006462 | 0.00118399 | 0.01043165 | dnj-23 |
| WBGene00001042 | 0.102051093 | 0.8038812 | 0.892856 | dnj-24 |
| WBGene00001043 | 0.090588615 | 0.65953486 | 0.80368078 | dnj-25 |
| WBGene00001044 | -0.115627056 | 0.77887365 | 0.87725216 | dnj-26 |
| WBGene00001045 | -0.584218189 | 0.01416279 | 0.06680455 | dnj-27 |
| WBGene00001046 | -0.424564612 | 0.08939938 | 0.2371834 | dnj-28 |
| WBGene00001047 | 0.121765037 | 0.48076959 | 0.67254563 | dnj-29 |
| WBGene00001048 | -0.668647841 | 0.00425144 | 0.02760253 | dnj-30 |
| WBGene00001049 | -0.077895559 | 0.72201321 | 0.84328642 | dog-1 |
| WBGene00001050 | -0.044357684 | 0.86458588 | 0.92720937 | dom-3 |
| WBGene00001051 | -0.403135777 | 0.01340234 | 0.06393839 | cks-1 |
| WBGene00001052 | 1.855112419 | 0.00156431 | 0.01294256 | dop-1 |
| WBGene00001053 | 1.016618439 | 0.04146009 | 0.14107818 | dop-2 |
| WBGene00001054 | 0.614021225 | 0.17065076 | 0.35951255 | dpf-1 |
| WBGene00001055 | 0.449822199 | 0.08825841 | 0.23550447 | dpf-2 |
| WBGene00001056 | 0.06201371 | 0.70774147 | 0.83480343 | dpf-3 |
| WBGene00001057 | -0.130043007 | 0.47028727 | 0.66356557 | dpf-4 |

| | | | | |
|---|---|---|---|---|
| WBGene00001058 | -0.203949504 | 0.19418281 | 0.39032673 | dpf-5 |
| WBGene00001059 | -0.120691298 | 0.83067369 | 0.90824619 | dpf-6 |
| WBGene00001060 | -0.070885772 | 0.72576971 | 0.84579072 | dpf-7 |
| WBGene00001061 | -0.309098538 | 0.12002693 | 0.28624902 | dpl-1 |
| WBGene00001063 | -0.616422377 | 0.34185198 | 0.55023529 | dpy-1 |
| WBGene00001064 | -1.03567058 | 0.01959465 | 0.08421189 | dpy-2 |
| WBGene00001065 | -0.752442126 | 0.15810302 | 0.34299953 | dpy-3 |
| WBGene00001066 | -1.567261688 | 0.12893758 | 0.30000597 | dpy-4 |
| WBGene00001067 | 0.640964248 | 0.46441186 | 0.65892471 | dpy-5 |
| WBGene00001068 | 1.241726742 | 0.00020626 | 0.0026898 | dpy-6 |
| WBGene00001069 | -0.669958606 | 0.09188519 | 0.24130927 | dpy-7 |
| WBGene00001070 | -1.469015504 | 0.04279972 | 0.14406193 | dpy-8 |
| WBGene00001071 | -1.07423425 | 0.08729395 | 0.23376757 | dpy-9 |
| WBGene00001072 | -0.485212518 | 0.13095949 | 0.30337616 | dpy-10 |
| WBGene00001073 | 0.075242487 | 0.69015704 | 0.82353771 | dpy-11 |
| WBGene00001074 | 0.486485256 | 0.64157518 | 0.79272992 | dpy-13 |
| WBGene00001075 | -0.649064174 | 0.00690321 | 0.03938528 | dpy-14 |
| WBGene00001076 | -0.79757511 | 0.0004243 | 0.00470019 | dpy-17 |
| WBGene00001077 | -0.100649012 | 0.68997103 | 0.82353771 | dpy-18 |
| WBGene00001078 | 0.654325406 | 0.11435901 | 0.27821121 | dpy-19 |
| WBGene00001079 | 1.732908132 | 0.28460632 | 1 | dpy-20 |
| WBGene00001080 | 0.228944283 | 0.34892141 | 0.55740195 | dpy-21 |
| WBGene00001081 | 0.749033939 | 0.00258131 | 0.01900857 | dpy-22 |
| WBGene00001082 | 0.023315058 | 0.91460214 | 0.95571968 | dpy-23 |
| WBGene00001085 | 0.123444784 | 0.43530823 | 0.63637293 | dpy-26 |
| WBGene00001086 | 0.236047766 | 0.29213487 | 0.50223867 | dpy-27 |
| WBGene00001087 | 0.113763263 | 0.56686627 | 0.74186951 | dpy-28 |
| WBGene00001088 | -0.618716784 | 0.00022747 | 0.00290705 | dpy-30 |
| WBGene00001089 | 0.395920126 | 0.05371764 | 0.16869784 | dre-1 |
| WBGene00001090 | 0.141202167 | 0.47044022 | 0.66356557 | drh-1 |
| WBGene00001092 | -0.576630067 | 0.00509134 | 0.03156996 | dro-1 |
| WBGene00001093 | -0.224968963 | 0.16509643 | 0.35168723 | drp-1 |
| WBGene00001094 | -0.198258456 | 0.15712211 | 0.34156125 | dars-1 |
| WBGene00001095 | 0.002576406 | 0.99230713 | 0.99624583 | dars-2 |
| WBGene00001096 | 1.487047283 | 0.50914254 | 1 | dsc-1 |
| WBGene00001099 | 0.862370114 | 9.1016E-05 | 0.001424 | dsc-4 |
| WBGene00001101 | 0.300649293 | 0.20813251 | 0.40776775 | dsh-1 |
| WBGene00001102 | -0.702925862 | 3.0982E-05 | 0.0005734 | dsh-2 |
| WBGene00001103 | -2.608330477 | 0.0001097 | 0.00164226 | dsl-1 |
| WBGene00001104 | -2.080131177 | 3.84E-11 | 3.96E-09 | dsl-2 |
| WBGene00001105 | -0.037009784 | 0.96113482 | 0.97977647 | dsl-3 |
| WBGene00001106 | -2.934840138 | 0.2654436 | 1 | dsl-4 |
| WBGene00001107 | 1.395728942 | 0.51364788 | 1 | dsl-5 |

| | | | | |
|---|---|---|---|---|
| WBGene00001108 | -0.542325745 | 0.46045118 | 0.6560508 | dsl-6 |
| WBGene00001109 | 0.761396302 | 0.58843477 | 1 | dsl-7 |
| WBGene00001110 | -0.107329713 | 0.64853728 | 0.79743682 | duo-1 |
| WBGene00001112 | 0.105433909 | 0.56093279 | 0.73692733 | duo-3 |
| WBGene00001113 | 0.653371507 | 0.01274167 | 0.06192817 | dur-1 |
| WBGene00001115 | 0.432933257 | 0.28510515 | 0.49491874 | dyb-1 |
| WBGene00001116 | 1.218997862 | 0.00235592 | 0.01767814 | dyc-1 |
| WBGene00001117 | 0.456752984 | 0.46083586 | 0.65630483 | dyf-1 |
| WBGene00001118 | 0.902630191 | 0.13491857 | 0.3093066 | dyf-2 |
| WBGene00001119 | -0.013886031 | 0.98232691 | 0.99089656 | dyf-3 |
| WBGene00001121 | -0.893826314 | 0.27953988 | 0.4885409 | dyf-5 |
| WBGene00001122 | 0.872546367 | 0.26808299 | 0.47664841 | dyf-6 |
| WBGene00001123 | 0.579312553 | 0.53159016 | 0.71520791 | dyf-7 |
| WBGene00001127 | 0.220444988 | 0.81177223 | 0.89724443 | dyf-11 |
| WBGene00001129 | 0.540153299 | 0.592985 | 1 | dyf-13 |
| WBGene00001130 | 0.416180761 | 0.06611464 | 0.1944049 | dyn-1 |
| WBGene00001131 | 0.934500291 | 3.3158E-05 | 0.00060628 | dys-1 |
| WBGene00001132 | 0.498648208 | 0.01909142 | 0.08295699 | alp-1 |
| WBGene00001133 | 1.58450535 | 0.19380325 | 1 | eat-2 |
| WBGene00001134 | 0.036145767 | 0.82381347 | 0.90490539 | eat-3 |
| WBGene00001135 | 0.433704295 | 0.23964431 | 0.44645107 | eat-4 |
| WBGene00001136 | 1.204798034 | 0.3026498 | 1 | eat-5 |
| WBGene00001137 | 0.539485917 | 0.00127109 | 0.01103881 | eat-6 |
| WBGene00001145 | 0.490949537 | 0.07013303 | 0.20243772 | eat-16 |
| WBGene00001147 | 0.905690628 | 0.3062901 | 0.51512233 | eat-18 |
| WBGene00001148 | -0.350233269 | 0.25145235 | 0.4595639 | eat-20 |
| WBGene00001149 | 0.698375099 | 0.00012176 | 0.00178326 | bcat-1 |
| WBGene00001150 | -0.322645079 | 0.58657613 | 0.75578084 | ech-1.1 |
| WBGene00001151 | -0.258275589 | 0.30190557 | 0.51136099 | ech-2 |
| WBGene00001152 | -0.109585263 | 0.7331215 | 0.85085825 | ech-3 |
| WBGene00001153 | -0.226710853 | 0.20982212 | 0.41007351 | ech-4 |
| WBGene00001154 | -0.44944822 | 0.03187552 | 0.11847911 | ech-5 |
| WBGene00001155 | -0.33224533 | 0.03484545 | 0.12579797 | ech-6 |
| WBGene00001156 | -0.255467204 | 0.15831521 | 0.34305991 | ech-7 |
| WBGene00001157 | 0.575992696 | 0.05138069 | 0.16368748 | ech-8 |
| WBGene00001158 | 2.167130122 | 1.75E-10 | 1.61E-08 | ech-9 |
| WBGene00001159 | 0.62290106 | 0.22297766 | 0.42693137 | eff-1 |
| WBGene00001160 | -0.092764736 | 0.59688395 | 0.76297832 | efk-1 |
| WBGene00001161 | -0.508220764 | 0.00752639 | 0.04191648 | efl-1 |
| WBGene00001162 | -0.101312741 | 0.70264644 | 0.83218963 | efl-2 |
| WBGene00001163 | 0.105831333 | 0.79943161 | 0.88979641 | efn-2 |
| WBGene00001164 | -1.221030377 | 0.06784139 | 0.1979006 | efn-3 |
| WBGene00001165 | -0.427060561 | 0.2715364 | 0.48029187 | efn-4 |

| | | | | |
|---|---|---|---|---|
| WBGene00001166 | -0.244407455 | 0.09975554 | 0.25443821 | eftu-2 |
| WBGene00001167 | 0.099950186 | 0.55320723 | 0.73100664 | eef-2 |
| WBGene00001168 | -0.25504144 | 0.09639728 | 0.24913639 | eef-1A.1 |
| WBGene00001169 | 0.538725964 | 0.00013851 | 0.00198313 | eef-1A.2 |
| WBGene00001170 | 0.528147961 | 0.81036933 | 1 | egl-1 |
| WBGene00001171 | 1.107179731 | 0.03132074 | 0.11711274 | egl-2 |
| WBGene00001172 | 1.242791371 | 6.43E-09 | 4.11E-07 | egl-3 |
| WBGene00001173 | 0.093662742 | 0.60400384 | 0.76858292 | egl-4 |
| WBGene00001174 | 0.110571394 | 0.90964867 | 1 | egl-5 |
| WBGene00001175 | -0.391379296 | 0.45148181 | 0.64930592 | egl-6 |
| WBGene00001177 | 0.619429902 | 0.00497133 | 0.0310478 | egl-8 |
| WBGene00001178 | -0.124433481 | 0.67611026 | 0.81466397 | egl-9 |
| WBGene00001179 | 0.624015305 | 0.02468898 | 0.09930834 | egl-10 |
| WBGene00001182 | 0.887962843 | 0.00939437 | 0.04943953 | egl-13 |
| WBGene00001184 | 1.441706699 | 3.2342E-06 | 8.8042E-05 | egl-15 |
| WBGene00001185 | 0.912641807 | 0.51376582 | 1 | egl-17 |
| WBGene00001186 | -0.40815396 | 0.11598381 | 0.28048782 | egl-18 |
| WBGene00001187 | 1.222619149 | 9.73E-07 | 3.3106E-05 | egl-19 |
| WBGene00001188 | 0.826496615 | 0.0609918 | 0.18466371 | egl-20 |
| WBGene00001189 | 0.932302696 | 1.62E-07 | 7.0843E-06 | egl-21 |
| WBGene00001190 | 0.72883681 | 0.1954769 | 0.3919546 | egl-23 |
| WBGene00001193 | -0.28731895 | 0.39228766 | 0.60022237 | egl-26 |
| WBGene00001194 | 0.175004803 | 0.29261201 | 0.50277419 | egl-27 |
| WBGene00001196 | 0.557701147 | 0.03497847 | 0.12601589 | egl-30 |
| WBGene00001202 | 1.219074115 | 0.09050702 | 0.23924447 | egl-36 |
| WBGene00001204 | 0.115195624 | 0.87411538 | 0.93213743 | egl-38 |
| WBGene00001207 | 0.50738893 | 0.24964904 | 0.45743873 | egl-43 |
| WBGene00001208 | 1.369165101 | 3.80E-08 | 1.964E-06 | egl-44 |
| WBGene00001209 | -0.045501264 | 0.81067075 | 0.89671589 | egl-45 |
| WBGene00001210 | 0.246678519 | 0.6576237 | 0.80291727 | egl-46 |
| WBGene00001211 | 1.020947257 | 0.29773931 | 0.5078408 | egl-47 |
| WBGene00001214 | 0.063508488 | 0.70755287 | 0.83477788 | ego-1 |
| WBGene00001215 | 0.194748658 | 0.40424821 | 0.61135744 | ego-2 |
| WBGene00001223 | 1.38322681 | 0.69550305 | 1 | ehn-3 |
| WBGene00001224 | 0.266047532 | 0.27550601 | 0.48411714 | ehs-1 |
| WBGene00001225 | -0.253543831 | 0.13841997 | 0.3153802 | eif-3.B |
| WBGene00001226 | -0.120538203 | 0.37521016 | 0.58305729 | eif-3.C |
| WBGene00001227 | -0.313413254 | 0.05132463 | 0.16359475 | eif-3.D |
| WBGene00001228 | 0.068953351 | 0.65917033 | 0.80368078 | eif-3.E |
| WBGene00001229 | -0.582131884 | 0.00103295 | 0.00947315 | eif-3.F |
| WBGene00001230 | -0.391908214 | 0.02488827 | 0.09974122 | eif-3.G |
| WBGene00001231 | -0.126001006 | 0.44216041 | 0.64190715 | eif-3.H |
| WBGene00001232 | -0.30148859 | 0.08447211 | 0.22890007 | eif-3.I |

| | | | | |
|---|---|---|---|---|
| WBGene00001233 | -0.448426156 | 0.00520763 | 0.03215838 | eif-3.K |
| WBGene00001234 | -0.595237519 | 0.00114259 | 0.01016281 | eif-6 |
| WBGene00001235 | -0.644460668 | 0.01269167 | 0.06174859 | elb-1 |
| WBGene00001236 | 0.067523457 | 0.81747514 | 0.90086942 | elc-1 |
| WBGene00001237 | 1.77510668 | 0.36223503 | 1 | elc-2 |
| WBGene00001239 | 0.43070281 | 0.00260741 | 0.01912786 | elo-1 |
| WBGene00001240 | 0.699824363 | 6.73E-07 | 2.4538E-05 | elo-2 |
| WBGene00001241 | 0.297155393 | 0.07528133 | 0.2123091 | elo-3 |
| WBGene00001242 | -0.589598164 | 0.45076197 | 0.64903951 | elo-4 |
| WBGene00001243 | -0.094767448 | 0.67464257 | 0.81399855 | elo-5 |
| WBGene00001244 | 0.101985939 | 0.59144261 | 0.75865871 | elo-6 |
| WBGene00001245 | -0.263229581 | 0.93409872 | 1 | elo-7 |
| WBGene00001246 | -0.394320382 | 0.8739308 | 1 | elo-8 |
| WBGene00001247 | 0.18118929 | 0.63421167 | 0.78783996 | elo-9 |
| WBGene00001248 | 0.340822661 | 0.43336222 | 0.63529827 | elp-1 |
| WBGene00001249 | -0.675678415 | 0.02671907 | 0.10473002 | elt-1 |
| WBGene00001250 | 1.304734608 | 0.00017489 | 0.00236184 | elt-2 |
| WBGene00001251 | 0.446300335 | 0.13655691 | 0.31211172 | elt-3 |
| WBGene00001252 | 1.219060141 | 0.46174049 | 1 | elt-4 |
| WBGene00001253 | 0.645039605 | 0.21967551 | 0.42247433 | elt-6 |
| WBGene00001258 | 0.103145025 | 0.61599322 | 0.77593531 | emb-4 |
| WBGene00001259 | -0.157032722 | 0.31317566 | 0.52122318 | emb-5 |
| WBGene00001262 | 0.173567029 | 0.32992773 | 0.53846654 | emb-8 |
| WBGene00001263 | 0.938935312 | 1.2307E-05 | 0.0002702 | emb-9 |
| WBGene00001281 | -0.194967774 | 0.27191947 | 0.48056732 | emb-27 |
| WBGene00001284 | 0.289930322 | 0.11911509 | 0.28484891 | emb-30 |
| WBGene00001303 | -0.562609977 | 0.00073001 | 0.00721082 | emo-1 |
| WBGene00001309 | -0.425138599 | 0.02565333 | 0.1020543 | emr-1 |
| WBGene00001310 | -1.676353087 | 0.16399405 | 1 | end-1 |
| WBGene00001311 | -1.859732522 | 0.27647017 | 1 | end-3 |
| WBGene00001320 | 0.050360454 | 0.77648547 | 0.87541184 | ent-1 |
| WBGene00001324 | 0.305224905 | 0.13476373 | 0.30911892 | eor-1 |
| WBGene00001325 | 0.024511414 | 0.93350288 | 0.96489515 | eor-2 |
| WBGene00001328 | 0.801377901 | 0.07240682 | 0.20703984 | epi-1 |
| WBGene00001329 | 0.364378143 | 0.06446383 | 0.19135827 | epn-1 |
| WBGene00001330 | 0.640584801 | 0.00071012 | 0.00704379 | eps-8 |
| WBGene00001331 | -0.137547171 | 0.49351468 | 0.68212223 | erd-2 |
| WBGene00001332 | -0.458964152 | 0.02023269 | 0.08622153 | eri-1 |
| WBGene00001333 | 0.252677842 | 0.12792233 | 0.29864109 | erm-1 |
| WBGene00001334 | -0.211794226 | 0.23243274 | 0.43798943 | ero-1 |
| WBGene00001335 | -0.387980103 | 0.16938387 | 0.35748616 | F35A5.8 |
| WBGene00001336 | -0.404602583 | 0.01719772 | 0.07676381 | qars-1 |
| WBGene00001337 | 0.024460864 | 0.89558126 | 0.94507369 | ears-1 |

| | | | | |
|---|---|---|---|---|
| WBGene00001338 | 0.237882676 | 0.40619144 | 0.61299816 | ears-2 |
| WBGene00001340 | 0.185912911 | 0.32958081 | 0.5381944 | etr-1 |
| WBGene00001345 | 0.832921731 | 0.00375599 | 0.02502825 | fos-1 |
| WBGene00001352 | 0.471706791 | 0.00592224 | 0.03528061 | evl-14 |
| WBGene00001358 | -0.250345373 | 0.43423464 | 0.63564538 | evl-20 |
| WBGene00001362 | 1.440130638 | 0.09853 | 0.2526405 | C46E1.3 |
| WBGene00001365 | -0.155130987 | 0.48969771 | 0.67924367 | exc-4 |
| WBGene00001366 | 1.133848476 | 0.00010136 | 0.00154605 | exc-5 |
| WBGene00001368 | 0.714474804 | 0.04064615 | 0.13904901 | exc-7 |
| WBGene00001371 | -0.381904732 | 0.18387844 | 0.37602631 | exl-1 |
| WBGene00001372 | -0.651902472 | 0.00084774 | 0.00809155 | exo-3 |
| WBGene00001373 | 0.940623821 | 0.4052443 | 1 | exp-1 |
| WBGene00001374 | 0.635421831 | 0.08622025 | 0.23191957 | exp-2 |
| WBGene00001377 | -0.583493717 | 0.16932007 | 0.35745349 | eya-1 |
| WBGene00001385 | -0.045060036 | 0.75814392 | 0.8646088 | far-1 |
| WBGene00001386 | -0.411400162 | 0.00250532 | 0.0186151 | far-2 |
| WBGene00001387 | -3.46631196 | 6.12E-08 | 3.0336E-06 | far-3 |
| WBGene00001388 | 1.111404431 | 0.53076615 | 1 | far-4 |
| WBGene00001389 | -0.050092452 | 0.93964973 | 0.96893897 | far-5 |
| WBGene00001390 | 0.198282949 | 0.33163205 | 0.53979686 | far-6 |
| WBGene00001391 | 0.956929102 | 0.04495793 | 0.14905359 | far-7 |
| WBGene00001392 | 0.532042137 | 0.00151596 | 0.01266707 | far-8 |
| WBGene00001393 | 0.191697818 | 0.12452898 | 0.29397363 | fat-1 |
| WBGene00001394 | 0.89673263 | 3.00E-12 | 3.79E-10 | fat-2 |
| WBGene00001395 | 1.2700722 | 1.50E-12 | 2.02E-10 | fat-3 |
| WBGene00001396 | 0.643331471 | 2.8142E-05 | 0.00053221 | fat-4 |
| WBGene00001397 | -0.674158233 | 0.00037687 | 0.00431654 | fat-5 |
| WBGene00001398 | 0.098293076 | 0.50090044 | 0.68880193 | fat-6 |
| WBGene00001399 | -1.313667105 | 5.75E-14 | 9.56E-12 | fat-7 |
| WBGene00001400 | 1.072617024 | 0.23692879 | 0.44302363 | fax-1 |
| WBGene00001401 | 0.049174397 | 0.89978647 | 0.94730381 | fbf-1 |
| WBGene00001402 | -0.26967093 | 0.48657364 | 0.67622374 | fbf-2 |
| WBGene00001403 | 0.550673507 | 0.04377308 | 0.14608272 | fbl-1 |
| WBGene00001404 | -0.625099127 | 9.2771E-05 | 0.00144401 | fbp-1 |
| WBGene00001405 | 0.03530317 | 0.8500179 | 0.91807378 | fce-1 |
| WBGene00001406 | 0.006425068 | 0.98222088 | 0.99087193 | fce-2 |
| WBGene00001410 | 0.86154388 | 0.01123211 | 0.05660816 | feh-1 |
| WBGene00001411 | -0.110045166 | 0.56069557 | 0.73673213 | fem-1 |
| WBGene00001412 | -0.584928621 | 0.00180469 | 0.01437712 | fem-2 |
| WBGene00001413 | -0.090623734 | 0.71935668 | 0.84172219 | fem-3 |
| WBGene00001414 | 0.864899468 | 0.6533431 | 1 | fer-1 |
| WBGene00001423 | -0.425665648 | 0.00205623 | 0.01590109 | fib-1 |
| WBGene00001424 | 1.477253415 | 0.58357327 | 1 | fis-1 |

| | | | | |
|---|---|---|---|---|
| WBGene00001425 | -0.31997023 | 0.2799738 | 0.4890343 | fis-2 |
| WBGene00001426 | -0.250023649 | 0.1365521 | 0.31211172 | fkb-1 |
| WBGene00001427 | -0.790025335 | 3.85E-08 | 1.9724E-06 | fkb-2 |
| WBGene00001428 | -0.378456375 | 0.17318714 | 0.36296651 | fkb-3 |
| WBGene00001429 | -0.100317667 | 0.84711765 | 0.91624547 | fkb-4 |
| WBGene00001430 | -0.766131367 | 0.00317834 | 0.02210432 | fkb-5 |
| WBGene00001431 | -0.36134058 | 0.01879562 | 0.08205594 | fkb-6 |
| WBGene00001432 | -0.861485674 | 0.09754674 | 0.25102055 | fkb-7 |
| WBGene00001433 | 0.090243594 | 0.87047837 | 0.93024187 | fkb-8 |
| WBGene00001434 | -0.722039885 | 0.36921363 | 0.57676566 | fkh-2 |
| WBGene00001435 | -1.32953625 | 0.01074901 | 0.05476447 | fkh-3 |
| WBGene00001436 | -1.323697109 | 0.00952627 | 0.04998204 | fkh-4 |
| WBGene00001437 | -2.989623253 | 0.00627245 | 0.0368962 | fkh-5 |
| WBGene00001438 | 1.851857589 | 0.06973441 | 0.20177441 | fkh-6 |
| WBGene00001439 | 0.653380181 | 0.02378895 | 0.09691687 | fkh-7 |
| WBGene00001440 | 0.135967355 | 0.87116831 | 0.93048777 | fkh-8 |
| WBGene00001441 | 1.185269097 | 1.6654E-05 | 0.00034922 | fkh-9 |
| WBGene00001442 | 1.734818937 | 0.23499288 | 1 | fkh-10 |
| WBGene00001443 | 0.318856859 | 0.08197921 | 0.22442154 | fli-1 |
| WBGene00001444 | 0.755712997 | 0.00219354 | 0.01668656 | flp-1 |
| WBGene00001445 | 1.365690073 | 0.02297417 | 0.09461854 | flp-2 |
| WBGene00001446 | 0.87725685 | 0.02167327 | 0.09089296 | flp-3 |
| WBGene00001447 | 1.598855223 | 0.00479442 | 0.03028568 | flp-4 |
| WBGene00001448 | 0.701026184 | 0.06177301 | 0.18594442 | flp-5 |
| WBGene00001449 | 0.20442326 | 0.45886672 | 0.65487177 | flp-6 |
| WBGene00001450 | 0.717227829 | 0.14059644 | 0.31834517 | flp-7 |
| WBGene00001451 | 0.293368226 | 0.35010832 | 0.5586365 | flp-8 |
| WBGene00001452 | 1.045650575 | 0.00021603 | 0.00279021 | flp-9 |
| WBGene00001453 | 1.057143155 | 0.12771271 | 0.29839611 | flp-10 |
| WBGene00001454 | 1.405732463 | 0.0004943 | 0.0053219 | flp-11 |
| WBGene00001455 | 0.600246473 | 0.0673271 | 0.19711075 | flp-12 |
| WBGene00001456 | 1.151512563 | 0.00473518 | 0.03000511 | flp-13 |
| WBGene00001457 | 0.547195172 | 0.03625713 | 0.12882581 | flp-14 |
| WBGene00001458 | 0.725021555 | 0.10660726 | 0.2661907 | flp-15 |
| WBGene00001459 | 1.349125507 | 0.00132877 | 0.01140922 | flp-16 |
| WBGene00001460 | 0.919414149 | 0.01562465 | 0.07169268 | flp-17 |
| WBGene00001461 | 0.760299689 | 0.01417171 | 0.06681893 | flp-18 |
| WBGene00001462 | 0.409341517 | 0.29893697 | 0.50881729 | flp-19 |
| WBGene00001463 | 0.348925418 | 0.58402735 | 0.7540432 | flp-20 |
| WBGene00001464 | 1.585160149 | 0.052169 | 0.1651157 | flp-21 |
| WBGene00001465 | 0.846976755 | 0.07485222 | 0.21159041 | flr-1 |
| WBGene00001466 | 0.200012973 | 0.82240466 | 0.90384856 | flr-2 |
| WBGene00001468 | 0.830306314 | 0.12885483 | 0.2998709 | flr-4 |

| | | | | |
|---|---|---|---|---|
| WBGene00001470 | 0.040565047 | 0.83664959 | 0.9111967 | baz-2 |
| WBGene00001475 | 0.430671049 | 0.12772956 | 0.29839611 | fmi-1 |
| WBGene00001476 | 0.06429945 | 0.85619902 | 0.92147082 | fmo-1 |
| WBGene00001477 | 2.350498832 | 0.09137509 | 1 | fmo-2 |
| WBGene00001478 | -0.306758857 | 0.42587299 | 0.62970697 | fmo-3 |
| WBGene00001479 | 0.526822186 | 0.06269699 | 0.18776618 | fmo-4 |
| WBGene00001480 | 0.275173743 | 0.48631943 | 0.67622374 | fmo-5 |
| WBGene00001481 | -0.311176849 | 0.33393056 | 0.54215712 | fog-1 |
| WBGene00001482 | 0.194727848 | 0.503082 | 0.69016029 | fog-2 |
| WBGene00001483 | 2.415035119 | 0.09407257 | 1 | fog-3 |
| WBGene00001484 | -0.187482595 | 0.54649984 | 0.72543326 | fox-1 |
| WBGene00001485 | -0.252460159 | 0.24617779 | 0.45416267 | fre-1 |
| WBGene00001486 | -0.216889862 | 0.57721581 | 0.74903703 | frh-1 |
| WBGene00001487 | 0.937196423 | 0.55866782 | 1 | frk-1 |
| WBGene00001488 | 0.409145117 | 0.04382086 | 0.14620199 | frm-1 |
| WBGene00001489 | 0.280871687 | 0.27496433 | 0.48355295 | frm-2 |
| WBGene00001490 | 0.94319434 | 0.00317463 | 0.02210224 | frm-3 |
| WBGene00001491 | 0.214191377 | 0.25474862 | 0.46230167 | frm-4 |
| WBGene00001492 | -0.293025303 | 0.51720039 | 0.703015 | frm-5.1 |
| WBGene00001493 | 1.032643462 | 1.4954E-06 | 4.5616E-05 | frm-7 |
| WBGene00001494 | 0.634403426 | 0.07674096 | 0.21490008 | frm-8 |
| WBGene00001496 | 0.684460048 | 0.00014885 | 0.0020892 | frm-10 |
| WBGene00001497 | -0.496766053 | 0.00298859 | 0.02115713 | fars-1 |
| WBGene00001498 | -0.136235708 | 0.41944134 | 0.62392028 | fars-3 |
| WBGene00001499 | -0.324135867 | 0.08368272 | 0.22732355 | fsn-1 |
| WBGene00001500 | 1.656887122 | 0.29148686 | 1 | ftn-1 |
| WBGene00001501 | -0.070244582 | 0.68914685 | 0.82299402 | ftn-2 |
| WBGene00001502 | -0.056734548 | 0.70271743 | 0.83218963 | ftt-2 |
| WBGene00001503 | 0.054858837 | 0.73648781 | 0.85280864 | fum-1 |
| WBGene00001505 | 0.113522346 | 0.75302863 | 0.8621766 | fut-1 |
| WBGene00001506 | 0.889786899 | 0.38620912 | 1 | fut-2 |
| WBGene00001507 | 0.256327779 | 0.85584911 | 1 | fut-4 |
| WBGene00001508 | 0.406686032 | 0.21151236 | 0.41127027 | fut-8 |
| WBGene00001509 | -0.085388399 | 0.62035678 | 0.77878286 | fzo-1 |
| WBGene00001510 | 0.050359657 | 0.82895126 | 0.90762341 | fzr-1 |
| WBGene00001511 | -0.519652993 | 0.00129089 | 0.01115495 | fzy-1 |
| WBGene00001512 | 1.62140433 | 0.03883739 | 0.13475287 | gab-1 |
| WBGene00001513 | -0.322251764 | 0.04338248 | 0.14529838 | gad-1 |
| WBGene00001514 | -0.021141803 | 0.91199359 | 0.9544448 | xnd-1 |
| WBGene00001515 | 1.299605261 | 0.00383625 | 0.02536812 | gap-1 |
| WBGene00001516 | 0.112065551 | 0.60809769 | 0.77041783 | gap-2 |
| WBGene00001517 | 1.698784084 | 0.00508589 | 0.03155229 | gar-1 |
| WBGene00001518 | 0.965375629 | 0.01056258 | 0.05401866 | gar-2 |

| | | | | |
|---|---|---|---|---|
| WBGene00001519 | 0.693781007 | 0.10015578 | 0.25508523 | gar-3 |
| WBGene00001520 | 0.221861113 | 0.20406859 | 0.40345168 | gas-1 |
| WBGene00001522 | -0.703167451 | 0.01479348 | 0.06901623 | gbh-1 |
| WBGene00001523 | 0.839792399 | 0.05074667 | 0.16230646 | gbh-2 |
| WBGene00001526 | 0.468699022 | 0.09205353 | 0.24150461 | gck-1 |
| WBGene00001527 | 0.421676729 | 0.02249479 | 0.09326547 | gcs-1 |
| WBGene00001528 | 0.281454817 | 0.94435481 | 1 | gcy-1 |
| WBGene00001530 | 0.993321677 | 0.45380417 | 1 | gcy-3 |
| WBGene00001531 | 1.389979168 | 0.39397789 | 1 | gcy-4 |
| WBGene00001532 | 1.924548786 | 0.34794862 | 1 | gcy-5 |
| WBGene00001533 | 0.370728191 | 0.83463145 | 1 | gcy-6 |
| WBGene00001534 | 1.708484285 | 0.16539049 | 1 | gcy-7 |
| WBGene00001535 | 0.421478679 | 0.63838428 | 0.79036023 | gcy-8 |
| WBGene00001536 | 0.293405867 | 0.72184174 | 0.84322534 | gcy-9 |
| WBGene00001537 | 0.675018486 | 0.20137389 | 0.40040623 | gcy-11 |
| WBGene00001538 | 0.464853908 | 0.07691776 | 0.21525651 | gcy-12 |
| WBGene00001539 | 0.742628056 | 0.76901841 | 1 | gcy-13 |
| WBGene00001540 | 0.096169744 | 0.95008442 | 1 | gcy-14 |
| WBGene00001541 | 1.743666163 | 0.01034079 | 0.0531756 | gcy-15 |
| WBGene00001542 | 1.187732978 | 0.58705413 | 1 | gcy-17 |
| WBGene00001543 | 1.220367297 | 0.02856758 | 0.10951656 | gcy-18 |
| WBGene00001544 | 1.46443947 | 0.03321002 | 0.12177674 | gcy-19 |
| WBGene00001545 | 0.410637484 | 0.77490091 | 1 | gcy-20 |
| WBGene00001546 | 0.969807753 | 0.14845005 | 0.32925321 | gcy-21 |
| WBGene00001547 | 0.932651616 | 0.19391679 | 0.38998571 | gcy-22 |
| WBGene00001548 | 0.523890266 | 0.46914319 | 0.6628497 | gcy-23 |
| WBGene00001549 | 2.250043379 | 0.19197259 | 1 | gcy-25 |
| WBGene00001550 | 0.273191754 | 0.85346958 | 1 | gcy-27 |
| WBGene00001551 | -0.337634122 | 0.70780549 | 0.83480343 | gcy-31 |
| WBGene00001552 | 0.770138371 | 0.29110161 | 0.50117196 | gcy-32 |
| WBGene00001553 | 0.042145706 | 0.95950965 | 0.97910629 | gcy-33 |
| WBGene00001554 | 1.273053973 | 0.2676078 | 1 | gcy-34 |
| WBGene00001555 | 0.27934748 | 0.69687977 | 0.82851717 | gcy-35 |
| WBGene00001556 | 0.96241819 | 0.2181632 | 0.42071361 | gcy-36 |
| WBGene00001557 | 0.707100279 | 0.43949219 | 0.63986984 | gcy-37 |
| WBGene00001558 | -0.139667742 | 0.39268826 | 0.60045695 | gdi-1 |
| WBGene00001559 | 0.513960118 | 0.04748249 | 0.15547149 | gei-1 |
| WBGene00001561 | -0.264148324 | 0.12544848 | 0.29536855 | gei-4 |
| WBGene00001562 | 0.53276221 | 0.03402219 | 0.12389423 | lin-66 |
| WBGene00001563 | -0.117986978 | 0.58554354 | 0.75492026 | gei-6 |
| WBGene00001564 | 0.167730849 | 0.32094999 | 0.52915674 | icl-1 |
| WBGene00001565 | 0.261764021 | 0.31288157 | 0.52103307 | gei-8 |
| WBGene00001566 | -0.180775109 | 0.40380755 | 0.6111802 | acdh-13 |

| | | | | |
|---|---|---|---|---|
| WBGene00001568 | -0.238408544 | 0.1787929 | 0.3708553 | snpc-4 |
| WBGene00001569 | -0.227728309 | 0.23482121 | 0.44071175 | meg-3 |
| WBGene00001570 | 0.314139063 | 0.32109715 | 0.52931982 | gei-13 |
| WBGene00001571 | -0.073115856 | 0.70809657 | 0.8349845 | gei-14 |
| WBGene00001572 | 0.419511778 | 0.1387847 | 0.3156585 | gei-15 |
| WBGene00001574 | 0.314408193 | 0.08091354 | 0.2224296 | gei-17 |
| WBGene00001575 | 0.797267857 | 0.55104665 | 1 | gei-18 |
| WBGene00001577 | 0.280872445 | 0.51739126 | 0.7030383 | gem-4 |
| WBGene00001578 | 0.809093506 | 0.02566842 | 0.10207673 | ges-1 |
| WBGene00001579 | -0.294825655 | 0.17684849 | 0.36841412 | gex-2 |
| WBGene00001580 | 0.039449189 | 0.82678671 | 0.90653097 | gex-3 |
| WBGene00001581 | 1.401872071 | 0.03174758 | 0.11812001 | gfi-1 |
| WBGene00001582 | 0.056167868 | 0.75805941 | 0.86459364 | gfi-2 |
| WBGene00001583 | -0.151451318 | 0.61037782 | 0.77201762 | gfi-3 |
| WBGene00001584 | 0.115937749 | 0.53899541 | 0.72067849 | nrde-4 |
| WBGene00001585 | -0.40105024 | 0.09640415 | 0.24913639 | gfl-1 |
| WBGene00001586 | 1.096560246 | 0.19810918 | 0.39572944 | ggr-1 |
| WBGene00001587 | 1.612444614 | 0.07710196 | 0.21544186 | ggr-2 |
| WBGene00001588 | 0.426533844 | 0.51625801 | 0.70236313 | ggr-3 |
| WBGene00001589 | 0.032919245 | 0.86150221 | 0.92537365 | gip-1 |
| WBGene00001590 | -0.202347611 | 0.55769604 | 0.73485866 | C45G3.3 |
| WBGene00001591 | 0.966471377 | 0.00305681 | 0.02151462 | glc-1 |
| WBGene00001592 | 0.480753737 | 0.25748876 | 0.46548109 | glc-2 |
| WBGene00001593 | 1.004953827 | 0.18801859 | 0.38164405 | glc-3 |
| WBGene00001594 | 0.598842977 | 0.10903811 | 0.26929042 | glc-4 |
| WBGene00001595 | -0.009297684 | 0.96052317 | 0.9796557 | gld-1 |
| WBGene00001596 | 0.385866051 | 0.06774785 | 0.19774952 | gld-2 |
| WBGene00001597 | -0.031975587 | 0.84569679 | 0.91544265 | gld-3 |
| WBGene00001598 | -0.402612878 | 0.013603 | 0.06474087 | glh-1 |
| WBGene00001599 | -0.038710707 | 0.85034449 | 0.91824825 | glh-2 |
| WBGene00001600 | -0.28000824 | 0.18998045 | 0.38435739 | glh-3 |
| WBGene00001601 | -0.278798291 | 0.12954686 | 0.30084603 | glh-4 |
| WBGene00001602 | -0.280998115 | 0.32124672 | 0.52942261 | gln-1 |
| WBGene00001603 | 2.876611589 | 0.15205148 | 1 | gln-2 |
| WBGene00001604 | 0.292924307 | 0.10280389 | 0.25948899 | gln-3 |
| WBGene00001606 | -0.551690792 | 0.00014478 | 0.00205591 | gln-5 |
| WBGene00001607 | -0.519450885 | 0.00015064 | 0.0021071 | gln-6 |
| WBGene00001608 | 0.786366838 | 0.40480547 | 0.61174299 | R07B1.8 |
| WBGene00001609 | -0.095230169 | 0.61172242 | 0.77323497 | glp-1 |
| WBGene00001612 | 1.051398612 | 0.08397164 | 0.22787208 | glr-1 |
| WBGene00001613 | 1.184822838 | 0.10920035 | 0.26952662 | glr-2 |
| WBGene00001614 | -0.050697871 | 0.97187944 | 1 | glr-3 |
| WBGene00001615 | 0.935913969 | 0.01897187 | 0.08261744 | glr-4 |

| | | | | |
|---|---|---|---|---|
| WBGene00001616 | 0.147167345 | 0.72672304 | 0.84642598 | glr-5 |
| WBGene00001617 | 0.778835126 | 0.06301258 | 0.18843244 | glr-6 |
| WBGene00001618 | 1.037045989 | 0.43789003 | 1 | glr-7 |
| WBGene00001619 | 1.957271014 | 0.49743227 | 1 | glr-8 |
| WBGene00001620 | 0.640240028 | 0.00184877 | 0.01468011 | glt-1 |
| WBGene00001621 | 1.135018504 | 0.00365528 | 0.02455145 | glt-3 |
| WBGene00001622 | 0.989374989 | 0.00769033 | 0.04249806 | glt-4 |
| WBGene00001623 | 1.193457339 | 0.0262849 | 0.1035946 | glt-5 |
| WBGene00001624 | 0.584133519 | 0.30342342 | 0.51235935 | glt-6 |
| WBGene00001625 | 0.31329442 | 0.70746424 | 0.83477788 | glt-7 |
| WBGene00001626 | 1.270678414 | 0.5135606 | 1 | gly-1 |
| WBGene00001627 | 0.66187664 | 0.15485074 | 0.33852997 | gly-2 |
| WBGene00001628 | -0.213391305 | 0.23691985 | 0.44302363 | gly-3 |
| WBGene00001629 | -0.171712639 | 0.47747064 | 0.66993771 | gly-4 |
| WBGene00001630 | 0.09342063 | 0.60670598 | 0.76973843 | gly-5 |
| WBGene00001631 | 0.186868991 | 0.43884775 | 0.63931493 | gly-6 |
| WBGene00001632 | -0.345421074 | 0.08925662 | 0.23707387 | gly-7 |
| WBGene00001633 | -0.552318928 | 0.10664312 | 0.26619123 | gly-8 |
| WBGene00001635 | -0.162019036 | 0.40064479 | 0.60826713 | gly-10 |
| WBGene00001636 | -0.149827751 | 0.72837181 | 0.84781793 | gly-11 |
| WBGene00001637 | 0.034427441 | 0.93805696 | 0.96795424 | gly-12 |
| WBGene00001638 | -0.076621059 | 0.7880022 | 0.88339194 | gly-13 |
| WBGene00001639 | 0.619965011 | 0.44669632 | 0.6457926 | gly-14 |
| WBGene00001642 | -0.156269293 | 0.96924949 | 1 | gly-17 |
| WBGene00001643 | 2.367081327 | 0.03847577 | 0.1339509 | gly-18 |
| WBGene00001644 | 0.547520753 | 0.35860112 | 0.56682414 | gly-19 |
| WBGene00001645 | -0.12365432 | 0.48141616 | 0.67289112 | gly-20 |
| WBGene00001646 | 0.356819207 | 0.5536789 | 0.73145475 | gna-1 |
| WBGene00001647 | -0.32418388 | 0.0452622 | 0.14985775 | gna-2 |
| WBGene00001648 | 0.202302936 | 0.26712118 | 0.47544153 | goa-1 |
| WBGene00001649 | -0.080862511 | 0.72935052 | 0.84847759 | gob-1 |
| WBGene00001650 | 1.09572823 | 8.3673E-05 | 0.00132447 | gon-1 |
| WBGene00001651 | 0.763576573 | 0.00053488 | 0.00565183 | gon-2 |
| WBGene00001653 | 0.241918036 | 0.15259615 | 0.33490055 | gon-4 |
| WBGene00001660 | 0.320317627 | 0.21593702 | 0.41759977 | gop-1 |
| WBGene00001661 | -0.489754545 | 0.02310673 | 0.09490485 | gop-2 |
| WBGene00001662 | 0.024185448 | 0.89672138 | 0.94596353 | gop-3 |
| WBGene00001663 | 1.302576327 | 0.14935922 | 0.33078626 | gpa-1 |
| WBGene00001664 | 0.342747198 | 0.61307559 | 0.77405075 | gpa-2 |
| WBGene00001665 | 0.619210825 | 0.32655293 | 0.53515658 | gpa-3 |
| WBGene00001666 | -0.303396526 | 0.63046795 | 0.78556152 | gpa-4 |
| WBGene00001667 | 1.557897343 | 0.63813005 | 1 | gpa-5 |
| WBGene00001668 | 0.75945276 | 0.50255169 | 1 | gpa-6 |

| | | | | |
|---|---|---|---|---|
| WBGene00001669 | 0.147259445 | 0.69368702 | 0.82625862 | gpa-7 |
| WBGene00001670 | 1.655400498 | 0.39459006 | 1 | gpa-8 |
| WBGene00001671 | 1.643120425 | 0.15300206 | 1 | gpa-9 |
| WBGene00001672 | 0.316126866 | 0.78528406 | 1 | gpa-10 |
| WBGene00001673 | 1.014268055 | 0.52746164 | 1 | gpa-11 |
| WBGene00001674 | -0.111862382 | 0.80166872 | 0.89148362 | gpa-12 |
| WBGene00001675 | 0.832900307 | 0.46265096 | 1 | gpa-13 |
| WBGene00001676 | 2.544277474 | 0.02926953 | 1 | gpa-14 |
| WBGene00001677 | 1.363343267 | 0.34848545 | 1 | gpa-15 |
| WBGene00001678 | -0.167148091 | 0.45236791 | 0.65010241 | gpa-16 |
| WBGene00001679 | 0.300997006 | 0.12155214 | 0.28868634 | gpb-1 |
| WBGene00001680 | 0.74624723 | 0.0098086 | 0.05113433 | gpb-2 |
| WBGene00001681 | -0.333918784 | 0.68835456 | 0.82257015 | gpc-1 |
| WBGene00001682 | -0.373343413 | 0.06728893 | 0.19704651 | gpc-2 |
| WBGene00001683 | -0.752570889 | 1.47E-07 | 6.5253E-06 | gpd-1 |
| WBGene00001684 | 0.094798826 | 0.55625169 | 0.73343308 | gpd-2 |
| WBGene00001685 | -0.071800985 | 0.63480294 | 0.78805138 | gpd-3 |
| WBGene00001686 | -1.025798711 | 3.09E-09 | 2.14E-07 | gpd-4 |
| WBGene00001687 | -0.283512492 | 0.65128674 | 0.79895649 | gpn-1 |
| WBGene00001688 | -0.792978748 | 0.00579135 | 0.03472234 | gpr-1 |
| WBGene00001689 | -0.603620918 | 0.01939622 | 0.08371473 | gpr-2 |
| WBGene00001690 | 0.364319412 | 0.63171539 | 0.78652714 | grd-1 |
| WBGene00001691 | 1.487288669 | 0.49618504 | 1 | grd-2 |
| WBGene00001692 | -1.198125674 | 3.2367E-05 | 0.00059361 | grd-3 |
| WBGene00001693 | -0.774842772 | 0.78069886 | 1 | grd-4 |
| WBGene00001694 | -2.403569131 | 6.61E-19 | 2.01E-16 | grd-5 |
| WBGene00001695 | 0.405264827 | 0.76796163 | 1 | grd-6 |
| WBGene00001696 | 1.876192691 | 0.43542138 | 1 | grd-7 |
| WBGene00001698 | 1.851444818 | 0.53669729 | 1 | grd-9 |
| WBGene00001699 | 0.763659326 | 0.0098675 | 0.05133254 | grd-10 |
| WBGene00001700 | 1.569713482 | 0.2905918 | 1 | grd-11 |
| WBGene00001701 | -0.237997406 | 0.83325628 | 1 | grd-12 |
| WBGene00001702 | 0.625027386 | 0.0576578 | 0.17785146 | grd-13 |
| WBGene00001703 | -2.055921171 | 0.00040029 | 0.00451601 | grd-14 |
| WBGene00001704 | -0.838076201 | 0.60277622 | 1 | grd-15 |
| WBGene00001705 | -1.335154934 | 0.17250416 | 0.36203509 | grd-16 |
| WBGene00001707 | 0.043976703 | 0.90706526 | 0.95141938 | grh-1 |
| WBGene00001708 | 0.112317825 | 0.64637368 | 0.79566286 | grk-1 |
| WBGene00001709 | 0.64087702 | 0.03879906 | 0.13466535 | grk-2 |
| WBGene00001710 | -1.081596404 | 0.20734401 | 0.40685432 | grl-1 |
| WBGene00001711 | -0.180165717 | 0.90534843 | 1 | grl-2 |
| WBGene00001712 | -0.156269293 | 0.96924949 | 1 | grl-3 |
| WBGene00001713 | -0.655896705 | 0.03976937 | 0.13696105 | grl-4 |

| | | | | |
|---|---|---|---|---|
| WBGene00001714 | -0.810792369 | 0.20991738 | 0.41007351 | grl-5 |
| WBGene00001715 | -1.187477906 | 0.47721276 | 1 | grl-6 |
| WBGene00001716 | -1.145884371 | 0.00898141 | 0.04764648 | grl-7 |
| WBGene00001717 | -0.477834054 | 0.68409934 | 1 | grl-8 |
| WBGene00001718 | 1.270678414 | 0.5135606 | 1 | grl-9 |
| WBGene00001719 | -0.667092806 | 0.31336981 | 0.52132404 | grl-10 |
| WBGene00001720 | -1.159714244 | 0.75790634 | 1 | grl-11 |
| WBGene00001721 | -2.051235694 | 0.34377405 | 1 | grl-12 |
| WBGene00001722 | 0.688793783 | 0.86378755 | 1 | grl-13 |
| WBGene00001723 | 1.170775457 | 0.37229477 | 1 | grl-14 |
| WBGene00001724 | -1.607478039 | 0.00318738 | 0.02215108 | grl-15 |
| WBGene00001725 | -0.953284076 | 0.01609786 | 0.07317263 | grl-16 |
| WBGene00001726 | -1.027355405 | 0.68015432 | 1 | grl-17 |
| WBGene00001727 | -0.023374929 | 0.9903626 | 1 | grl-18 |
| WBGene00001728 | 0.281454817 | 0.94435481 | 1 | grl-19 |
| WBGene00001729 | 2.365623541 | 0.29275878 | 1 | grl-20 |
| WBGene00001730 | 0.056944093 | 0.93453726 | 0.96559519 | grl-21 |
| WBGene00001731 | -1.041300001 | 0.35003325 | 1 | grl-22 |
| WBGene00001734 | -1.094201388 | 0.7816772 | 1 | grl-25 |
| WBGene00001735 | 0.03940405 | 0.99001443 | 1 | grl-26 |
| WBGene00001736 | -0.559340474 | 0.78183513 | 1 | grl-27 |
| WBGene00001737 | 0.57991002 | 0.82402153 | 1 | grl-28 |
| WBGene00001739 | -0.156269293 | 0.96924949 | 1 | grl-30 |
| WBGene00001740 | 0.347522235 | 0.26853696 | 0.47707159 | gro-1 |
| WBGene00001743 | -0.308902736 | 0.13328801 | 0.30677721 | grp-1 |
| WBGene00001744 | -0.431224511 | 0.00961849 | 0.0503354 | gars-1 |
| WBGene00001745 | 0.090580803 | 0.78292766 | 0.8803042 | gsa-1 |
| WBGene00001746 | 0.208752818 | 0.29109305 | 0.50117196 | gsk-3 |
| WBGene00001747 | 0.036028688 | 0.8218169 | 0.90342582 | gsp-1 |
| WBGene00001748 | -0.310064164 | 0.04951875 | 0.15947139 | gsp-2 |
| WBGene00001749 | -0.307185638 | 0.05057564 | 0.16205827 | gst-1 |
| WBGene00001750 | 1.847274755 | 0.08470456 | 1 | gst-2 |
| WBGene00001751 | 1.099135066 | 0.15141284 | 0.33332315 | gst-3 |
| WBGene00001752 | 0.117777019 | 0.74863517 | 0.85982212 | gst-4 |
| WBGene00001753 | 0.023976891 | 0.93128829 | 0.96397892 | gst-5 |
| WBGene00001754 | 0.807107184 | 0.02895795 | 0.11055928 | gst-6 |
| WBGene00001755 | -0.632661577 | 6.3607E-05 | 0.00106246 | gst-7 |
| WBGene00001756 | 0.720066222 | 0.75158338 | 1 | gst-8 |
| WBGene00001757 | -0.751325212 | 0.47366714 | 1 | gst-9 |
| WBGene00001758 | -0.895771742 | 0.03260054 | 0.12034149 | gst-10 |
| WBGene00001759 | 0.576059267 | 0.43380759 | 0.63555665 | gst-11 |
| WBGene00001760 | -1.031906454 | 0.26582879 | 0.47448212 | gst-12 |
| WBGene00001761 | 0.256669283 | 0.22476968 | 0.42887455 | gst-13 |

| | | | | |
|---|---|---|---|---|
| WBGene00001762 | 2.748584877 | 0.098936 | 1 | gst-14 |
| WBGene00001763 | 0.855037229 | 0.22116565 | 0.42406383 | gst-15 |
| WBGene00001764 | -0.076886952 | 0.91272206 | 0.95467585 | gst-16 |
| WBGene00001766 | 0.281454817 | 0.94435481 | 1 | gst-18 |
| WBGene00001767 | -0.345553352 | 0.70003451 | 0.83055986 | gst-19 |
| WBGene00001768 | 0.071816319 | 0.82102322 | 0.90291441 | gst-20 |
| WBGene00001769 | 0.123786728 | 0.90984828 | 1 | gst-21 |
| WBGene00001770 | 0.280732049 | 0.47943643 | 0.67137689 | gst-22 |
| WBGene00001771 | 0.891993454 | 0.49642359 | 1 | gst-23 |
| WBGene00001772 | 0.972643135 | 0.01548933 | 0.07126889 | gst-24 |
| WBGene00001773 | 1.025181888 | 0.79790347 | 1 | gst-25 |
| WBGene00001774 | 0.137176687 | 0.58009243 | 0.75092251 | gst-26 |
| WBGene00001775 | 0.018244317 | 0.927847 | 0.96207963 | gst-27 |
| WBGene00001776 | 0.113398689 | 0.76501528 | 0.86852913 | gst-28 |
| WBGene00001777 | 2.133182101 | 0.3921092 | 1 | gst-29 |
| WBGene00001778 | -0.912437437 | 0.51463941 | 1 | gst-30 |
| WBGene00001779 | 0.553362071 | 0.70600558 | 1 | gst-31 |
| WBGene00001780 | 1.036568105 | 0.67105582 | 1 | gst-32 |
| WBGene00001781 | 2.223671006 | 0.04025904 | 1 | gst-33 |
| WBGene00001782 | 0.042994488 | 0.97752318 | 1 | gst-34 |
| WBGene00001783 | 1.208523505 | 0.21242207 | 0.4123787 | gst-35 |
| WBGene00001784 | 0.428007368 | 0.03999358 | 0.13751404 | gst-36 |
| WBGene00001786 | 1.66604429 | 0.00475967 | 0.0301295 | gst-38 |
| WBGene00001787 | 0.162319093 | 0.7620367 | 0.86701223 | gst-39 |
| WBGene00001788 | 0.046746622 | 0.9490444 | 0.97411952 | gst-40 |
| WBGene00001789 | 0.525798739 | 0.33667584 | 0.54486555 | gst-41 |
| WBGene00001790 | -0.33800946 | 0.33511758 | 0.54330693 | gst-42 |
| WBGene00001791 | 0.446025225 | 0.29655758 | 0.50697066 | gst-43 |
| WBGene00001792 | -1.254449518 | 0.22604759 | 1 | gst-44 |
| WBGene00001793 | -0.280256976 | 0.11443759 | 0.27821121 | gsy-1 |
| WBGene00001794 | -0.354079759 | 0.02774965 | 0.10743254 | gta-1 |
| WBGene00001795 | 1.045397467 | 1.2574E-06 | 4.0332E-05 | gtl-1 |
| WBGene00001796 | 0.843366875 | 0.03890275 | 0.13484361 | gtl-2 |
| WBGene00001803 | 2.673005303 | 0.00590624 | 0.03521989 | lite-1 |
| WBGene00001804 | -0.492788182 | 0.75077029 | 1 | gur-3 |
| WBGene00001805 | 0.264604285 | 0.73904287 | 0.85446788 | gur-4 |
| WBGene00001806 | 0.749727563 | 0.69881416 | 1 | gur-5 |
| WBGene00001808 | -0.96560263 | 2.9493E-05 | 0.00055344 | gut-2 |
| WBGene00001811 | -0.649632721 | 0.11974073 | 0.28595048 | haf-1 |
| WBGene00001812 | 0.400839616 | 0.06534319 | 0.19297778 | haf-2 |
| WBGene00001813 | -0.222230726 | 0.24093495 | 0.44858017 | haf-3 |
| WBGene00001814 | 0.651097827 | 0.00422087 | 0.02743343 | haf-4 |
| WBGene00001815 | 0.192282861 | 0.36995963 | 0.5774852 | hmt-1 |

| | | | | |
|---|---|---|---|---|
| WBGene00001816 | -0.264315751 | 0.30880211 | 0.51762617 | haf-6 |
| WBGene00001817 | 0.828269156 | 0.01483729 | 0.0690984 | haf-7 |
| WBGene00001819 | 0.493021765 | 0.01589738 | 0.07250566 | haf-9 |
| WBGene00001820 | -0.579206303 | 0.21611559 | 0.41761252 | ham-1 |
| WBGene00001821 | 1.097420571 | 0.08836233 | 0.23562642 | ham-2 |
| WBGene00001823 | -0.695750386 | 0.01238659 | 0.06061845 | hap-1 |
| WBGene00001824 | 0.209983769 | 0.44440377 | 0.64377833 | hbl-1 |
| WBGene00001827 | -0.341220649 | 0.03171339 | 0.11812001 | hcf-1 |
| WBGene00001828 | -0.337829826 | 0.45791787 | 0.65476162 | hch-1 |
| WBGene00001829 | -0.293322408 | 0.12641156 | 0.29662983 | hcp-1 |
| WBGene00001830 | -0.204691229 | 0.22549428 | 0.42964942 | hcp-2 |
| WBGene00001831 | -0.669939612 | 0.00093331 | 0.00874327 | hcp-3 |
| WBGene00001832 | -0.281007739 | 0.0970903 | 0.25037668 | hcp-4 |
| WBGene00001833 | -0.184953094 | 0.28167734 | 0.49104955 | hcp-6 |
| WBGene00001834 | -0.225580843 | 0.21263704 | 0.41273002 | hda-1 |
| WBGene00001835 | -0.170248696 | 0.42958412 | 0.63219161 | hda-2 |
| WBGene00001836 | -0.301326157 | 0.11176048 | 0.27384136 | hda-3 |
| WBGene00001837 | 0.71931551 | 0.00476225 | 0.0301295 | hda-4 |
| WBGene00001838 | -0.265194587 | 0.33183441 | 0.53982957 | hda-10 |
| WBGene00001839 | -0.668355569 | 0.74739695 | 1 | hdl-1 |
| WBGene00001840 | -0.613863906 | 4.1753E-06 | 0.00010809 | hel-1 |
| WBGene00001841 | -0.33900259 | 0.79641528 | 1 | hen-1 |
| WBGene00001842 | -0.634206263 | 0.6152287 | 1 | her-1 |
| WBGene00001843 | -0.072353907 | 0.71754312 | 0.84024798 | hgo-1 |
| WBGene00001844 | 0.098876192 | 0.7158089 | 0.83991842 | hid-1 |
| WBGene00001851 | 0.633365422 | 0.00090066 | 0.00848328 | hif-1 |
| WBGene00001852 | 1.062369884 | 0.0479825 | 0.15661054 | hil-1 |
| WBGene00001853 | -0.8955599 | 2.6416E-05 | 0.00050915 | hil-2 |
| WBGene00001854 | -0.578878022 | 0.02738647 | 0.10652467 | hil-3 |
| WBGene00001855 | -0.971398656 | 1.64E-08 | 9.45E-07 | hil-4 |
| WBGene00001856 | -0.67902389 | 1.1626E-05 | 0.00025757 | hil-5 |
| WBGene00001857 | -1.344429361 | 0.30469171 | 1 | hil-6 |
| WBGene00001858 | -0.472148055 | 0.15856579 | 0.34334711 | hil-7 |
| WBGene00001859 | -2.563099533 | 0.4751044 | 1 | hil-8 |
| WBGene00001860 | 0.152460962 | 0.36015122 | 0.5682386 | him-1 |
| WBGene00001862 | -0.28732869 | 0.14173523 | 0.31991215 | him-3 |
| WBGene00001863 | 1.205670416 | 7.29E-07 | 2.58E-05 | him-4 |
| WBGene00001864 | -0.148915035 | 0.65856234 | 0.8035709 | him-5 |
| WBGene00001865 | 0.019214448 | 0.93215506 | 0.96406579 | him-6 |
| WBGene00001867 | -0.392072086 | 0.10790578 | 0.26759651 | him-8 |
| WBGene00001869 | -0.362493868 | 0.0334199 | 0.12217734 | him-10 |
| WBGene00001872 | -0.169015323 | 0.50273694 | 0.68999878 | him-14 |
| WBGene00001874 | -0.229824492 | 0.18115479 | 0.37323727 | him-17 |

| | | | | |
|---|---|---|---|---|
| WBGene00001875 | 0.471935218 | 0.81492399 | 1 | his-1 |
| WBGene00001876 | 0.351523238 | 0.82866109 | 1 | his-2 |
| WBGene00001877 | 0.15375832 | 0.92579988 | 1 | his-3 |
| WBGene00001878 | 1.239458001 | 0.42143596 | 1 | his-4 |
| WBGene00001879 | -1.559877735 | 0.2439388 | 1 | his-5 |
| WBGene00001880 | -0.775430817 | 0.56537786 | 1 | his-6 |
| WBGene00001881 | -2.247685785 | 0.20276942 | 1 | his-7 |
| WBGene00001882 | -1.890926751 | 0.30009887 | 1 | his-8 |
| WBGene00001883 | -0.156269293 | 0.96924949 | 1 | his-9 |
| WBGene00001886 | -0.156269293 | 0.96924949 | 1 | his-12 |
| WBGene00001887 | -0.156269293 | 0.96924949 | 1 | his-13 |
| WBGene00001890 | -0.156269293 | 0.96924949 | 1 | his-16 |
| WBGene00001891 | -1.094201388 | 0.7816772 | 1 | his-17 |
| WBGene00001892 | -1.22259901 | 0.68385967 | 1 | his-18 |
| WBGene00001898 | -1.002158962 | 1.4869E-06 | 4.5587E-05 | his-24 |
| WBGene00001899 | -0.156269293 | 0.96924949 | 1 | his-25 |
| WBGene00001905 | -0.075918738 | 0.96511751 | 1 | his-31 |
| WBGene00001906 | -1.57933985 | 0.04374758 | 0.14608272 | his-32 |
| WBGene00001909 | -0.001524873 | 0.99506546 | 0.99756601 | his-35 |
| WBGene00001910 | -1.019966439 | 7.0591E-05 | 0.00116131 | C50F4.6 |
| WBGene00001911 | -0.109846055 | 0.62697292 | 0.78313562 | his-37 |
| WBGene00001912 | -0.993541924 | 0.32841422 | 1 | his-38 |
| WBGene00001914 | -1.740416632 | 0.06893397 | 0.20017397 | his-40 |
| WBGene00001915 | 0.06103756 | 0.78905659 | 0.8837593 | his-41 |
| WBGene00001916 | -0.156269293 | 0.96924949 | 1 | his-42 |
| WBGene00001917 | -0.156269293 | 0.96924949 | 1 | his-43 |
| WBGene00001919 | -0.462562375 | 0.80026554 | 1 | his-45 |
| WBGene00001920 | 0.688793783 | 0.86378755 | 1 | his-46 |
| WBGene00001921 | -0.937629543 | 0.45419155 | 1 | his-47 |
| WBGene00001922 | -1.332051688 | 0.08411365 | 0.22815545 | his-48 |
| WBGene00001929 | -0.462562375 | 0.80026554 | 1 | his-55 |
| WBGene00001930 | 0.688793783 | 0.86378755 | 1 | his-56 |
| WBGene00001931 | 0.342629161 | 0.5368574 | 0.71886905 | his-57 |
| WBGene00001932 | -1.479200062 | 0.22816067 | 1 | his-58 |
| WBGene00001933 | -1.290650632 | 0.07693254 | 0.21525651 | his-59 |
| WBGene00001934 | -1.059201162 | 0.02901088 | 0.11065695 | his-60 |
| WBGene00001935 | -0.60641256 | 0.3853499 | 0.5938732 | his-61 |
| WBGene00001936 | -0.792298544 | 0.22628 | 0.43060846 | his-62 |
| WBGene00001937 | -0.043881252 | 0.95755392 | 0.97850563 | his-63 |
| WBGene00001938 | -0.112601668 | 0.92523287 | 1 | his-64 |
| WBGene00001939 | -0.937629543 | 0.45419155 | 1 | his-65 |
| WBGene00001940 | -1.479200062 | 0.22816067 | 1 | his-66 |
| WBGene00001941 | -0.774082565 | 0.21118289 | 0.4111419 | his-67 |

| | | | | |
|---|---|---|---|---|
| WBGene00001942 | -1.149702914 | 0.00200461 | 0.01557134 | his-68 |
| WBGene00001944 | -0.333693239 | 0.89573179 | 1 | his-70 |
| WBGene00001945 | 0.452352113 | 0.05588061 | 0.17389475 | his-71 |
| WBGene00001946 | -0.578622328 | 0.00014686 | 0.00207432 | his-72 |
| WBGene00001947 | -0.682164043 | 0.63656507 | 1 | his-73 |
| WBGene00001948 | -0.289629167 | 0.52432304 | 0.70891225 | hlh-1 |
| WBGene00001949 | -0.69993315 | 0.00282145 | 0.020326 | hlh-2 |
| WBGene00001950 | -1.157786745 | 0.03218295 | 0.1191463 | hlh-3 |
| WBGene00001951 | 1.851444818 | 0.53669729 | 1 | hlh-4 |
| WBGene00001952 | 0.661934724 | 0.50472135 | 1 | hlh-6 |
| WBGene00001953 | -0.73089135 | 0.72186459 | 1 | hlh-8 |
| WBGene00001954 | 2.332054728 | 0.29723791 | 1 | hlh-10 |
| WBGene00001955 | 1.032384922 | 2.6395E-06 | 7.4182E-05 | hlh-11 |
| WBGene00001956 | 0.510884577 | 0.74133419 | 1 | hlh-12 |
| WBGene00001957 | 1.470575353 | 0.51153906 | 1 | hlh-13 |
| WBGene00001958 | -0.001064539 | 0.99909889 | 0.99975765 | hlh-14 |
| WBGene00001959 | -1.14715222 | 0.68932198 | 1 | hlh-15 |
| WBGene00001960 | 0.018830311 | 0.98644685 | 1 | hlh-16 |
| WBGene00001961 | 1.897929842 | 0.32022294 | 1 | hlh-17 |
| WBGene00001962 | -0.01455644 | 0.9964743 | 1 | hlh-19 |
| WBGene00001964 | -1.279666666 | 0.55824702 | 1 | hlh-25 |
| WBGene00001965 | -1.159714244 | 0.75790634 | 1 | hlh-26 |
| WBGene00001966 | 1.017496938 | 0.63763677 | 1 | hlh-27 |
| WBGene00001967 | -0.156269293 | 0.96924949 | 1 | hlh-28 |
| WBGene00001968 | -0.323868354 | 0.91634405 | 1 | hlh-29 |
| WBGene00001971 | -0.578789166 | 0.00019812 | 0.00262017 | hmg-1.1 |
| WBGene00001972 | -0.17973475 | 0.33646951 | 0.54486555 | hmg-1.2 |
| WBGene00001973 | -0.369066421 | 0.011613 | 0.05790286 | hmg-3 |
| WBGene00001974 | -0.456085917 | 0.00247998 | 0.01848341 | hmg-4 |
| WBGene00001975 | -0.295006157 | 0.12183455 | 0.28924389 | hmg-5 |
| WBGene00001976 | -0.80528728 | 0.00037288 | 0.00428295 | hmg-11 |
| WBGene00001977 | -0.868214264 | 1.56E-07 | 6.8548E-06 | hmg-12 |
| WBGene00001978 | 0.132038604 | 0.46620256 | 0.66046269 | hmp-1 |
| WBGene00001979 | -0.381244163 | 0.03205122 | 0.11874696 | hmp-2 |
| WBGene00001980 | -0.028624293 | 0.88795221 | 0.94084004 | hmr-1 |
| WBGene00001981 | -0.821336945 | 0.19004179 | 0.38435739 | hnd-1 |
| WBGene00001983 | -0.028470697 | 0.87076607 | 0.93030157 | hoe-1 |
| WBGene00001984 | -0.635986856 | 0.48832101 | 0.67820992 | hog-1 |
| WBGene00001985 | -0.230783358 | 0.355553 | 0.5638413 | hop-1 |
| WBGene00001986 | 0.352592004 | 0.53069828 | 0.71443643 | hot-1 |
| WBGene00001987 | 0.237228786 | 0.74608802 | 0.85839618 | hot-2 |
| WBGene00001988 | 0.525885669 | 0.48249606 | 0.67353221 | hot-3 |
| WBGene00001989 | 0.008915196 | 0.98942319 | 0.99484196 | hot-4 |

| | | | | |
|---|---|---|---|---|
| WBGene00001990 | -0.288028661 | 0.7046916 | 0.83302778 | hot-5 |
| WBGene00001991 | 0.66862108 | 0.30145382 | 0.51102357 | hot-6 |
| WBGene00001992 | -0.060388924 | 0.94847919 | 0.97374309 | hot-7 |
| WBGene00001993 | -0.280792364 | 0.13798734 | 0.31463007 | hpd-1 |
| WBGene00001994 | 0.506071353 | 0.05049067 | 0.16187147 | hpk-1 |
| WBGene00001995 | 0.064473426 | 0.90622435 | 0.95109561 | hpl-1 |
| WBGene00001996 | -0.256491896 | 0.22482147 | 0.42890595 | hpl-2 |
| WBGene00001997 | -0.28434107 | 0.32679865 | 0.53544701 | hpr-9 |
| WBGene00001998 | -0.232383671 | 0.32150934 | 0.52965027 | hpr-17 |
| WBGene00001999 | -0.537512668 | 0.00038314 | 0.00437599 | hrp-1 |
| WBGene00002000 | -0.302508623 | 0.07571738 | 0.21329111 | hrp-2 |
| WBGene00002001 | -0.431995058 | 0.01452727 | 0.06809869 | hars-1 |
| WBGene00002002 | -0.527407955 | 0.01173312 | 0.058358 | hsb-1 |
| WBGene00002003 | -0.294073808 | 0.12119449 | 0.28817515 | hse-5 |
| WBGene00002004 | -0.089296569 | 0.61431348 | 0.77489662 | hsf-1 |
| WBGene00002005 | -0.242362153 | 0.12332086 | 0.29203015 | hsp-1 |
| WBGene00002007 | -0.168665737 | 0.2758527 | 0.48446621 | hsp-3 |
| WBGene00002008 | -0.53151522 | 0.00086925 | 0.0082514 | hsp-4 |
| WBGene00002010 | -0.254121587 | 0.25384364 | 0.46180957 | hsp-6 |
| WBGene00002011 | 0.271383201 | 0.21220283 | 0.41207318 | hsp-12.2 |
| WBGene00002012 | 1.772131847 | 0.01361831 | 0.06478836 | hsp-12.3 |
| WBGene00002013 | 1.72767182 | 0.30669144 | 1 | hsp-12.6 |
| WBGene00002015 | 0.688793783 | 0.86378755 | 1 | hsp-16.1 |
| WBGene00002016 | 1.89686344 | 0.00230842 | 0.01739695 | hsp-16.2 |
| WBGene00002017 | 0.688793783 | 0.86378755 | 1 | hsp-16.11 |
| WBGene00002018 | 2.437100798 | 0.0017058 | 0.01375174 | hsp-16.41 |
| WBGene00002019 | 0.747845955 | 0.85215965 | 1 | hsp-16.48 |
| WBGene00002020 | 0.747845955 | 0.85215965 | 1 | hsp-16.49 |
| WBGene00002021 | 0.554648404 | 0.01499219 | 0.06968613 | hsp-17 |
| WBGene00002023 | 0.010207681 | 0.95948668 | 0.97910629 | hsp-25 |
| WBGene00002024 | -0.171295732 | 0.39824369 | 0.60593292 | hsp-43 |
| WBGene00002025 | -0.559843617 | 0.00109483 | 0.00982431 | hsp-60 |
| WBGene00002026 | 3.013210945 | 0.05718607 | 1 | hsp-70 |
| WBGene00002027 | -0.175264603 | 0.33009238 | 0.53851809 | hsr-9 |
| WBGene00002028 | -0.109162665 | 0.61194904 | 0.77336039 | hst-1 |
| WBGene00002029 | -0.317376695 | 0.4209343 | 0.62552794 | hst-2 |
| WBGene00002030 | 0.871223796 | 0.31000181 | 0.51830255 | hst-3.1 |
| WBGene00002031 | -0.674919915 | 0.05618014 | 0.17462444 | hst-6 |
| WBGene00002032 | -0.558588005 | 0.00692773 | 0.03945103 | htp-1 |
| WBGene00002033 | -0.538721717 | 0.01301872 | 0.06294757 | htp-2 |
| WBGene00002034 | -0.457404965 | 0.00896838 | 0.04761045 | htp-3 |
| WBGene00002035 | 0.309245818 | 0.07930508 | 0.21977698 | hum-1 |
| WBGene00002036 | 0.151533826 | 0.40330664 | 0.61092276 | hum-2 |

| | | | | |
|---|---|---|---|---|
| WBGene00002037 | 0.973441296 | 4.6383E-05 | 0.00081026 | hum-4 |
| WBGene00002038 | 0.715424088 | 0.00049781 | 0.0053446 | hum-5 |
| WBGene00002039 | 0.62913068 | 0.03310735 | 0.12151039 | hum-6 |
| WBGene00002040 | 0.991763347 | 0.00270146 | 0.01967513 | hum-7 |
| WBGene00002041 | 1.313660795 | 1.61E-08 | 9.34E-07 | hum-8 |
| WBGene00002042 | 0.13787713 | 0.70740665 | 0.83477788 | hus-1 |
| WBGene00002043 | 0.159013398 | 0.43624727 | 0.63705534 | hyl-1 |
| WBGene00002044 | 0.242673418 | 0.28263783 | 0.49211328 | hyl-2 |
| WBGene00002045 | -0.416292179 | 0.00392135 | 0.02586047 | icd-1 |
| WBGene00002046 | -0.415078687 | 0.03604359 | 0.12832998 | icln-1 |
| WBGene00002047 | -0.680726279 | 0.00027911 | 0.0034056 | icp-1 |
| WBGene00002048 | 1.054500092 | 5.07E-07 | 1.9109E-05 | ida-1 |
| WBGene00002050 | 1.041657094 | 1.7261E-06 | 5.1115E-05 | ifa-1 |
| WBGene00002051 | -0.675003548 | 0.15847852 | 0.34328059 | ifa-3 |
| WBGene00002052 | -0.451783504 | 0.04414344 | 0.14696081 | ifa-4 |
| WBGene00002053 | 0.40522365 | 0.02230711 | 0.09282554 | ifb-1 |
| WBGene00002054 | 0.786415204 | 1.3934E-06 | 4.3826E-05 | ifb-2 |
| WBGene00002055 | 1.027389156 | 5.3271E-06 | 0.00013308 | ifc-1 |
| WBGene00002056 | 0.931534061 | 8.34E-09 | 5.24E-07 | ifc-2 |
| WBGene00002057 | 0.94249042 | 0.00085372 | 0.0081358 | ifd-1 |
| WBGene00002058 | 1.001185109 | 0.00023775 | 0.00299432 | ifd-2 |
| WBGene00002059 | -0.734510784 | 4.5201E-06 | 0.00011505 | ife-1 |
| WBGene00002060 | -0.478507771 | 0.04506249 | 0.14927796 | ife-2 |
| WBGene00002061 | -0.578321985 | 0.00248878 | 0.01851489 | ife-3 |
| WBGene00002062 | -0.511821902 | 0.2157824 | 0.41750027 | ife-4 |
| WBGene00002063 | -0.671532004 | 0.13933973 | 0.31650583 | ife-5 |
| WBGene00002064 | -0.834677487 | 8.49E-10 | 6.87E-08 | iff-1 |
| WBGene00002065 | 0.240891969 | 0.11738418 | 0.28243387 | iff-2 |
| WBGene00002066 | 0.294367394 | 0.12689608 | 0.29736447 | ifg-1 |
| WBGene00002067 | 0.839793258 | 0.00017173 | 0.00233738 | ifp-1 |
| WBGene00002068 | -0.798513363 | 2.43E-07 | 1.004E-05 | ify-1 |
| WBGene00002069 | 1.229440368 | 0.00981328 | 0.05113433 | ikb-1 |
| WBGene00002070 | -0.143784293 | 0.36403238 | 0.57174854 | ile-1 |
| WBGene00002071 | -0.368615537 | 0.10200249 | 0.25778773 | ile-2 |
| WBGene00002072 | -0.264152174 | 0.20174552 | 0.40055476 | ima-1 |
| WBGene00002073 | -0.418401616 | 0.00786241 | 0.04329141 | ima-2 |
| WBGene00002074 | 0.108901067 | 0.59499846 | 0.76134495 | ima-3 |
| WBGene00002075 | -0.165385538 | 0.41649351 | 0.62148715 | imb-1 |
| WBGene00002076 | -0.148633543 | 0.45907072 | 0.65487177 | imb-2 |
| WBGene00002077 | -0.016641085 | 0.9286514 | 0.96274927 | imb-3 |
| WBGene00002078 | 0.03854621 | 0.85291065 | 0.91927977 | xpo-1 |
| WBGene00002079 | -0.484797181 | 0.00466041 | 0.0296707 | xpo-2 |
| WBGene00002080 | 0.319428799 | 0.10039763 | 0.25537978 | xpo-3 |

| | | | | |
|---|---|---|---|---|
| WBGene00002081 | 0.023464525 | 0.90819246 | 0.95194377 | ina-1 |
| WBGene00002083 | -0.286773157 | 0.05729558 | 0.17718433 | inf-1 |
| WBGene00002084 | 1.676799393 | 0.01492261 | 0.06938927 | ins-1 |
| WBGene00002085 | -0.73553186 | 0.59937787 | 1 | ins-2 |
| WBGene00002086 | 0.933365627 | 0.29543014 | 0.50569115 | ins-3 |
| WBGene00002087 | 2.0691839 | 0.09893214 | 1 | ins-4 |
| WBGene00002088 | -0.733928226 | 0.39012451 | 0.59849655 | ins-5 |
| WBGene00002089 | 1.097571182 | 0.12625305 | 0.29631885 | ins-6 |
| WBGene00002090 | 2.177932153 | 0.00676312 | 0.03884153 | ins-7 |
| WBGene00002091 | 1.413734769 | 0.68804531 | 1 | ins-8 |
| WBGene00002092 | 1.060949945 | 0.61866902 | 1 | ins-9 |
| WBGene00002093 | 0.371543842 | 0.76771749 | 1 | ins-10 |
| WBGene00002094 | -0.107268465 | 0.96117914 | 1 | ins-11 |
| WBGene00002095 | 3.568124035 | 0.04525241 | 1 | ins-12 |
| WBGene00002096 | -1.969805064 | 0.62862569 | 1 | ins-13 |
| WBGene00002097 | 0.438680766 | 0.71416247 | 1 | ins-14 |
| WBGene00002099 | 0.281454817 | 0.94435481 | 1 | ins-16 |
| WBGene00002100 | 0.59967397 | 0.15315617 | 0.33572094 | ins-17 |
| WBGene00002101 | 1.625729004 | 0.07949376 | 0.22014597 | ins-18 |
| WBGene00002102 | -3.733374421 | 0.11147094 | 1 | ins-19 |
| WBGene00002103 | 0.688793783 | 0.86378755 | 1 | ins-20 |
| WBGene00002104 | -1.094201388 | 0.7816772 | 1 | ins-21 |
| WBGene00002105 | 0.554062667 | 0.58132699 | 1 | ins-22 |
| WBGene00002106 | 2.254752291 | 0.12157854 | 1 | ins-23 |
| WBGene00002107 | 0.593355387 | 0.63867162 | 1 | ins-24 |
| WBGene00002108 | 0.983488853 | 0.80601565 | 1 | ins-25 |
| WBGene00002109 | -0.121822376 | 0.83123172 | 0.90861273 | ins-26 |
| WBGene00002110 | -0.000409304 | 0.99953361 | 0.99989047 | ins-27 |
| WBGene00002111 | 0.953609603 | 0.27092012 | 0.47962106 | ins-28 |
| WBGene00002112 | 0.663847008 | 0.589747 | 1 | ins-29 |
| WBGene00002113 | 1.606143423 | 0.09377429 | 0.24457865 | ins-30 |
| WBGene00002114 | 0.39851023 | 0.84358283 | 1 | ins-31 |
| WBGene00002115 | -0.629292503 | 0.72518963 | 1 | ins-32 |
| WBGene00002116 | 2.793858195 | 0.21955872 | 1 | ins-33 |
| WBGene00002117 | -1.137655086 | 0.05290225 | 0.16682759 | ins-34 |
| WBGene00002118 | 1.03015083 | 0.15027635 | 0.33184889 | ins-35 |
| WBGene00002119 | 0.747845955 | 0.85215965 | 1 | ins-36 |
| WBGene00002120 | -0.934013985 | 0.67351079 | 1 | ins-37 |
| WBGene00002123 | 0.818453507 | 0.01777322 | 0.07863871 | inx-1 |
| WBGene00002124 | -0.898318469 | 0.00010938 | 0.00164131 | inx-2 |
| WBGene00002125 | -0.722048372 | 9.6697E-05 | 0.00149174 | inx-3 |
| WBGene00002127 | 0.462531711 | 0.06217754 | 0.18680279 | inx-5 |
| WBGene00002128 | 0.962441304 | 0.00640824 | 0.03735114 | inx-6 |

| | | | | |
|---|---|---|---|---|
| WBGene00002129 | 0.212364847 | 0.54298538 | 0.7232458 | inx-7 |
| WBGene00002130 | 0.409373982 | 0.02873433 | 0.10980909 | inx-8 |
| WBGene00002131 | 0.098775151 | 0.76745432 | 0.86967173 | inx-9 |
| WBGene00002132 | 0.44697481 | 0.26116815 | 0.46968486 | inx-10 |
| WBGene00002133 | -0.393994617 | 0.38544017 | 0.59393693 | inx-11 |
| WBGene00002134 | 0.42549835 | 0.0257954 | 0.10231361 | inx-12 |
| WBGene00002135 | 0.404876183 | 0.05148679 | 0.16381056 | inx-13 |
| WBGene00002136 | -0.123356016 | 0.49398705 | 0.68261972 | inx-14 |
| WBGene00002137 | 0.093964256 | 0.70289796 | 0.83224928 | inx-15 |
| WBGene00002138 | 0.058121754 | 0.80394546 | 0.892856 | inx-16 |
| WBGene00002139 | 0.535478765 | 0.11281959 | 0.27526982 | inx-17 |
| WBGene00002140 | 0.31463398 | 0.6121397 | 0.77344032 | inx-18 |
| WBGene00002141 | 1.043553179 | 0.30605295 | 1 | inx-19 |
| WBGene00002142 | 0.537575548 | 0.49636864 | 0.68474169 | inx-20 |
| WBGene00002143 | 0.030065931 | 0.90702486 | 0.95141938 | inx-21 |
| WBGene00002144 | -0.320334114 | 0.07750336 | 0.21596701 | inx-22 |
| WBGene00002146 | 0.30957329 | 0.35534853 | 0.5636643 | ipp-5 |
| WBGene00002147 | 0.166015123 | 0.34998446 | 0.55851227 | ire-1 |
| WBGene00002148 | 0.033030732 | 0.85566746 | 0.92113008 | gon-14 |
| WBGene00002149 | 0.76968033 | 0.35769969 | 0.56591502 | irk-1 |
| WBGene00002150 | 0.665295152 | 0.43893581 | 0.63936649 | irk-2 |
| WBGene00002151 | 0.126502535 | 0.79881232 | 0.88951485 | irk-3 |
| WBGene00002152 | 0.226121339 | 0.16857308 | 0.35643429 | iars-1 |
| WBGene00002153 | 0.08763318 | 0.62223192 | 0.7801836 | iars-2 |
| WBGene00002162 | -0.122731878 | 0.4269032 | 0.63044468 | isp-1 |
| WBGene00002163 | 1.35469208 | 0.00317489 | 0.02210224 | ist-1 |
| WBGene00002169 | -0.122617634 | 0.47911936 | 0.67113235 | isw-1 |
| WBGene00002173 | 1.259478999 | 1.25E-09 | 9.66E-08 | itr-1 |
| WBGene00002174 | 1.201270791 | 2.9297E-06 | 8.1396E-05 | itx-1 |
| WBGene00002175 | 0.343805532 | 0.1957898 | 0.39242892 | jac-1 |
| WBGene00002176 | 1.560597489 | 0.00043549 | 0.00479785 | jip-1 |
| WBGene00002177 | 1.691171725 | 0.0820798 | 0.22459565 | jkk-1 |
| WBGene00002178 | 0.037498613 | 0.9208314 | 0.95857104 | jnk-1 |
| WBGene00002179 | 1.146164577 | 1.4788E-06 | 4.5485E-05 | jph-1 |
| WBGene00002180 | -0.832604595 | 0.00284635 | 0.02043612 | jtr-1 |
| WBGene00002181 | 0.095413999 | 0.80994423 | 0.89629222 | kal-1 |
| WBGene00002182 | -0.364977954 | 0.51501727 | 0.70161856 | kap-1 |
| WBGene00002183 | -0.25765494 | 0.13346321 | 0.30700585 | kat-1 |
| WBGene00002184 | 0.548264068 | 0.36872158 | 0.57621943 | kel-1 |
| WBGene00002185 | 0.050915902 | 0.81824385 | 0.9014626 | kel-3 |
| WBGene00002186 | 2.191582552 | 0.22563181 | 1 | kel-10 |
| WBGene00002187 | 0.667568876 | 0.00248856 | 0.01851489 | kgb-1 |
| WBGene00002188 | 1.697403741 | 0.02677712 | 0.10483748 | kgb-2 |

| | | | | |
|---|---|---|---|---|
| WBGene00002189 | 1.07359497 | 1.43E-07 | 6.4127E-06 | kin-1 |
| WBGene00002190 | 0.871974518 | 0.00029711 | 0.0035786 | kin-2 |
| WBGene00002191 | -0.342215233 | 0.03787123 | 0.13258205 | kin-3 |
| WBGene00002192 | 0.088705011 | 0.63971325 | 0.79154394 | kin-4 |
| WBGene00002193 | 0.298149064 | 0.8636957 | 1 | kin-5 |
| WBGene00002194 | -0.156269293 | 0.96924949 | 1 | kin-6 |
| WBGene00002195 | 0.975366077 | 0.11139839 | 0.27322986 | kin-9 |
| WBGene00002196 | -0.490155521 | 0.00443801 | 0.02852401 | kin-10 |
| WBGene00002198 | 0.184118991 | 0.78631855 | 0.88223764 | kin-14 |
| WBGene00002199 | 0.193714194 | 0.6589548 | 0.80357275 | kin-15 |
| WBGene00002200 | 0.514501176 | 0.24065813 | 0.44815063 | kin-16 |
| WBGene00002201 | -0.049152862 | 0.75387293 | 0.86257386 | kin-18 |
| WBGene00002202 | -0.220260776 | 0.23450963 | 0.44049946 | kin-19 |
| WBGene00002203 | 0.043988422 | 0.85194018 | 0.91924443 | kin-20 |
| WBGene00002205 | 0.335597573 | 0.8313888 | 1 | kin-23 |
| WBGene00002206 | 0.747845955 | 0.85215965 | 1 | kin-24 |
| WBGene00002207 | 0.072285888 | 0.77215321 | 0.87263447 | sid-3 |
| WBGene00002208 | 0.377828795 | 0.92526585 | 1 | kin-26 |
| WBGene00002210 | 0.737771299 | 0.00789734 | 0.0434443 | kin-29 |
| WBGene00002211 | 2.020606521 | 0.11672878 | 1 | kin-30 |
| WBGene00002213 | 0.212295176 | 0.5234637 | 0.7083563 | kin-32 |
| WBGene00002214 | -0.4860861 | 0.00109264 | 0.00981195 | klc-1 |
| WBGene00002215 | 0.150813928 | 0.44956736 | 0.64840361 | klc-2 |
| WBGene00002216 | 0.873707216 | 0.00748064 | 0.04169991 | klp-3 |
| WBGene00002217 | 0.957610178 | 0.00012345 | 0.00180183 | klp-4 |
| WBGene00002218 | 1.607437665 | 0.25301208 | 1 | klp-6 |
| WBGene00002219 | -0.319026247 | 0.1514399 | 0.33332315 | klp-7 |
| WBGene00002220 | -0.028186247 | 0.94281326 | 0.97038791 | gcc-1 |
| WBGene00002222 | 0.336040185 | 0.24307631 | 0.4507009 | klp-11 |
| WBGene00002223 | 0.653815229 | 8.0318E-05 | 0.00128308 | klp-12 |
| WBGene00002224 | 0.413805024 | 0.57285146 | 0.74664408 | klp-13 |
| WBGene00002225 | -0.91589041 | 6.11E-07 | 2.269E-05 | klp-15 |
| WBGene00002226 | -0.621603465 | 0.00027138 | 0.00334503 | klp-16 |
| WBGene00002227 | 2.566123675 | 0.23268183 | 1 | klp-17 |
| WBGene00002228 | -0.363771371 | 0.03533258 | 0.12675342 | klp-18 |
| WBGene00002229 | -0.393597347 | 0.02298105 | 0.09461854 | klp-19 |
| WBGene00002230 | 0.243875091 | 0.51477814 | 0.70152895 | klp-20 |
| WBGene00002231 | 0.017104002 | 0.9153436 | 0.95580617 | knl-1 |
| WBGene00002232 | 0.278211346 | 0.27796047 | 0.4868318 | kpc-1 |
| WBGene00002233 | 0.550473646 | 0.40041735 | 0.60821556 | kqt-1 |
| WBGene00002234 | 0.668616537 | 0.13543807 | 0.31002141 | kqt-2 |
| WBGene00002235 | 0.634694441 | 0.23108976 | 0.43640702 | kqt-3 |
| WBGene00002238 | -0.226510444 | 0.17152472 | 0.36066532 | kars-1 |

| | | | | |
|---|---|---|---|---|
| WBGene00002239 | -0.045636778 | 0.87511272 | 0.93273141 | ksr-1 |
| WBGene00002240 | 0.218034413 | 0.38121461 | 0.58944556 | ksr-2 |
| WBGene00002241 | -0.291677334 | 0.11511942 | 0.27930954 | kup-1 |
| WBGene00002242 | 1.438409382 | 0.09258043 | 0.24245449 | kvs-1 |
| WBGene00002243 | 0.562888813 | 0.1899765 | 0.38435739 | lad-2 |
| WBGene00002244 | -0.025224354 | 0.89762718 | 0.94634211 | laf-1 |
| WBGene00002245 | -0.086168708 | 0.66623941 | 0.80861938 | lag-1 |
| WBGene00002246 | -0.387368567 | 0.73027673 | 1 | lag-2 |
| WBGene00002247 | 0.102065476 | 0.62437805 | 0.78151093 | lam-1 |
| WBGene00002248 | 0.372577983 | 0.24196023 | 0.44959276 | lam-3 |
| WBGene00002249 | -0.316670598 | 0.03749344 | 0.13156297 | lap-1 |
| WBGene00002250 | 0.05307195 | 0.76619651 | 0.86897515 | lap-2 |
| WBGene00002251 | -0.001616572 | 0.99240143 | 0.99625813 | lat-1 |
| WBGene00002252 | 0.281727145 | 0.70164543 | 0.83157723 | lat-2 |
| WBGene00002253 | -0.352302458 | 0.05824427 | 0.17884261 | lbp-1 |
| WBGene00002254 | -0.715373545 | 2.1001E-06 | 6.129E-05 | lbp-2 |
| WBGene00002255 | 2.269110776 | 3.30E-36 | 3.64E-33 | lbp-3 |
| WBGene00002256 | -0.95942446 | 1.5967E-05 | 0.00033654 | lbp-4 |
| WBGene00002257 | 0.247704601 | 0.37450417 | 0.58248208 | lbp-5 |
| WBGene00002258 | -0.179006237 | 0.17172457 | 0.36096054 | lbp-6 |
| WBGene00002259 | -0.049233389 | 0.87202907 | 0.93116139 | lbp-7 |
| WBGene00002260 | -1.488391716 | 0.50775624 | 1 | lbp-8 |
| WBGene00002261 | 0.099049572 | 0.67775737 | 0.81560633 | ldb-1 |
| WBGene00002262 | 0.009397175 | 0.95836164 | 0.97852807 | ldh-1 |
| WBGene00002263 | 0.868581417 | 1.1741E-05 | 0.00025965 | lea-1 |
| WBGene00002264 | 0.075935366 | 0.64936816 | 0.79813513 | lec-1 |
| WBGene00002265 | 0.882048765 | 8.00E-08 | 3.8553E-06 | lec-2 |
| WBGene00002266 | 0.446500343 | 0.03934698 | 0.13590659 | lec-3 |
| WBGene00002267 | -0.093368684 | 0.66797788 | 0.80985814 | lec-4 |
| WBGene00002268 | -0.51306978 | 0.0053397 | 0.03270902 | lec-5 |
| WBGene00002269 | -0.32083894 | 0.02257486 | 0.09341561 | lec-6 |
| WBGene00002270 | 1.180904559 | 0.14052012 | 0.31829379 | lec-7 |
| WBGene00002271 | -0.006934643 | 0.96580183 | 0.98286673 | lec-8 |
| WBGene00002272 | 0.062629483 | 0.68795463 | 0.82242734 | lec-9 |
| WBGene00002273 | 0.299885443 | 0.05191719 | 0.16458845 | lec-10 |
| WBGene00002274 | 0.791747385 | 0.00841404 | 0.04544254 | lec-11 |
| WBGene00002275 | -0.720007514 | 0.00038401 | 0.0043818 | lem-2 |
| WBGene00002276 | -0.037147035 | 0.85041172 | 0.91824825 | lem-3 |
| WBGene00002278 | -0.209553873 | 0.45113064 | 0.64911342 | lep-2 |
| WBGene00002280 | 1.124405197 | 0.01222942 | 0.060088 | let-2 |
| WBGene00002282 | 0.529975103 | 0.36450307 | 0.57203053 | let-4 |
| WBGene00002285 | 1.116613983 | 0.69067248 | 1 | let-7 |
| WBGene00002295 | 0.624079234 | 0.00478196 | 0.03022267 | let-19 |

| | | | | |
|---|---|---|---|---|
| WBGene00002297 | -0.331374782 | 0.08146951 | 0.22335581 | ect-2 |
| WBGene00002299 | 1.241430005 | 3.1243E-05 | 0.00057735 | let-23 |
| WBGene00002324 | -0.635593045 | 0.01611093 | 0.07320462 | let-49 |
| WBGene00002335 | -0.153566417 | 0.4319205 | 0.6341955 | let-60 |
| WBGene00002344 | -0.286599554 | 0.04209277 | 0.14263141 | let-70 |
| WBGene00002348 | 1.259089288 | 2.95E-07 | 1.1874E-05 | myo-1 |
| WBGene00002363 | -0.188216752 | 0.29720808 | 0.50751102 | let-92 |
| WBGene00002368 | -0.3919945 | 0.028564 | 0.10951656 | let-99 |
| WBGene00002393 | -0.107661183 | 0.8931797 | 0.94370331 | lpr-1 |
| WBGene00002497 | 0.517816737 | 0.13850389 | 0.31549263 | let-268 |
| WBGene00002583 | 0.561340338 | 0.00310369 | 0.0217187 | let-363 |
| WBGene00002601 | -0.730691012 | 0.31394204 | 0.52177554 | let-381 |
| WBGene00002632 | 0.510722073 | 0.02673242 | 0.10473002 | let-413 |
| WBGene00002637 | 0.157367827 | 0.34645046 | 0.55524371 | let-418 |
| WBGene00002694 | -0.048099322 | 0.82103411 | 0.90291441 | let-502 |
| WBGene00002696 | -0.222432664 | 0.19678667 | 0.39393024 | let-504 |
| WBGene00002717 | 0.422835652 | 0.05805153 | 0.17854141 | let-526 |
| WBGene00002783 | 0.396494331 | 0.03445364 | 0.12486742 | let-607 |
| WBGene00002804 | -0.187367041 | 0.33590952 | 0.54427832 | let-630 |
| WBGene00002827 | 0.609952413 | 0.14990084 | 0.33131935 | let-653 |
| WBGene00002845 | 0.256464868 | 0.23927527 | 0.44625615 | let-711 |
| WBGene00002850 | 0.300726517 | 0.07436481 | 0.21075238 | let-716 |
| WBGene00002855 | -0.304675247 | 0.03454907 | 0.12506268 | let-721 |
| WBGene00002879 | -0.308696593 | 0.07861575 | 0.21836509 | let-754 |
| WBGene00002881 | 1.204998118 | 0.00177088 | 0.01416357 | let-756 |
| WBGene00002889 | -0.051419916 | 0.78339305 | 0.88066436 | let-765 |
| WBGene00002891 | 0.086754032 | 0.57313141 | 0.74692877 | let-767 |
| WBGene00002915 | 1.369561357 | 9.52E-09 | 5.87E-07 | let-805 |
| WBGene00002957 | -0.211517855 | 0.22347845 | 0.42746592 | let-858 |
| WBGene00002974 | 0.606383283 | 0.23946142 | 0.44640333 | lev-1 |
| WBGene00002975 | -0.206832623 | 0.80929695 | 0.89609686 | lev-8 |
| WBGene00002976 | 0.728294903 | 0.15379999 | 0.336812 | lev-9 |
| WBGene00002977 | 0.787522536 | 0.00348763 | 0.02372175 | lev-10 |
| WBGene00002978 | 0.176616874 | 0.24652431 | 0.45445668 | lev-11 |
| WBGene00002979 | 0.754495748 | 0.00218007 | 0.0166244 | lfe-2 |
| WBGene00002980 | -0.556822972 | 0.0001791 | 0.0024107 | lgg-1 |
| WBGene00002981 | 0.17668568 | 0.31775905 | 0.52560117 | lgg-2 |
| WBGene00002982 | -0.511285847 | 0.09583653 | 0.24825077 | lgg-3 |
| WBGene00002983 | 0.070930117 | 0.89869518 | 0.94694691 | lgx-1 |
| WBGene00002985 | -0.36323116 | 0.01715909 | 0.07664771 | lig-1 |
| WBGene00002986 | 0.201349576 | 0.39970343 | 0.60743514 | lig-4 |
| WBGene00002987 | 1.456564815 | 0.07889593 | 0.21883972 | lim-4 |
| WBGene00002988 | 0.93505505 | 0.25089379 | 0.45885383 | lim-6 |

| | | | | |
|---|---|---|---|---|
| WBGene00002989 | 0.627246911 | 0.15948278 | 0.34449938 | lim-7 |
| WBGene00002990 | 1.060019356 | 0.01433417 | 0.06742782 | lin-1 |
| WBGene00002991 | 0.707837194 | 0.03566579 | 0.12738276 | lin-2 |
| WBGene00002992 | -0.118854997 | 0.61190308 | 0.77336039 | lin-3 |
| WBGene00002993 | -1.831477136 | 0.52194793 | 1 | lin-4 |
| WBGene00002994 | -0.442749983 | 0.00766168 | 0.04244005 | lin-5 |
| WBGene00002996 | -0.253517131 | 0.41840462 | 0.62314162 | lin-7 |
| WBGene00002997 | -0.502317821 | 0.1913939 | 0.38619135 | lin-8 |
| WBGene00002998 | 0.099080005 | 0.61629059 | 0.77610041 | lin-9 |
| WBGene00002999 | 0.114072715 | 0.59925988 | 0.76480755 | lin-10 |
| WBGene00003000 | 2.006664523 | 0.00639265 | 0.03727819 | lin-11 |
| WBGene00003001 | -0.545328715 | 0.16252932 | 0.34847392 | lin-12 |
| WBGene00003002 | 0.442895538 | 0.03669291 | 0.12978406 | lin-13 |
| WBGene00003003 | 1.185390173 | 2.5496E-05 | 0.00049528 | lin-14 |
| WBGene00003006 | 0.225300028 | 0.58688337 | 0.75584502 | lin-17 |
| WBGene00003007 | 0.183257646 | 0.62775047 | 0.7837037 | lin-18 |
| WBGene00003008 | 0.91442609 | 0.41564173 | 1 | lin-22 |
| WBGene00003009 | -0.075209904 | 0.68163215 | 0.81759493 | lin-23 |
| WBGene00003010 | -0.18964547 | 0.37751027 | 0.58512604 | lin-24 |
| WBGene00003011 | 0.173748893 | 0.5105929 | 0.69715569 | lin-25 |
| WBGene00003012 | -0.053091467 | 0.76783432 | 0.86985878 | lin-26 |
| WBGene00003014 | -0.454480074 | 0.01907603 | 0.08292788 | lin-28 |
| WBGene00003015 | 0.729443302 | 0.24369854 | 0.45114272 | lin-29 |
| WBGene00003017 | 1.710383207 | 0.19834414 | 1 | lin-31 |
| WBGene00003018 | 0.828439456 | 0.52426463 | 1 | lin-32 |
| WBGene00003019 | 0.2500954 | 0.41723919 | 0.62193997 | lin-33 |
| WBGene00003020 | -0.000980908 | 0.99534755 | 0.99756601 | lin-35 |
| WBGene00003021 | 0.064628134 | 0.75632232 | 0.86391446 | lin-36 |
| WBGene00003022 | -0.554647207 | 0.00304782 | 0.02146383 | lin-37 |
| WBGene00003024 | 0.863789612 | 0.04951212 | 0.15947139 | lin-39 |
| WBGene00003025 | -0.266486181 | 0.20698103 | 0.40665706 | lin-40 |
| WBGene00003026 | 0.246669985 | 0.27249874 | 0.48129287 | lin-41 |
| WBGene00003029 | 0.05265095 | 0.95453636 | 0.97697363 | lin-44 |
| WBGene00003030 | 0.330573098 | 0.0990916 | 0.25339764 | lin-45 |
| WBGene00003031 | 3.223997694 | 0.05140171 | 1 | lin-46 |
| WBGene00003033 | 0.461516258 | 0.67163007 | 1 | lin-48 |
| WBGene00003034 | 0.168219628 | 0.38528985 | 0.59385604 | lin-49 |
| WBGene00003035 | -0.48654281 | 0.04797664 | 0.15661054 | lin-52 |
| WBGene00003036 | -0.547270752 | 0.0003521 | 0.0041184 | lin-53 |
| WBGene00003037 | -0.580381805 | 0.00113337 | 0.01008813 | lin-54 |
| WBGene00003038 | -0.111459961 | 0.73324275 | 0.85091763 | lin-56 |
| WBGene00003040 | 0.472630824 | 0.01141005 | 0.05724354 | lin-59 |
| WBGene00003041 | -0.249999832 | 0.18456913 | 0.37699425 | lin-61 |

| | | | | |
|---|---|---|---|---|
| WBGene00003043 | 0.392589994 | 0.07614673 | 0.21413202 | lip-1 |
| WBGene00003044 | -0.336281712 | 0.07997388 | 0.22093964 | lir-1 |
| WBGene00003045 | -0.265766385 | 0.30211472 | 0.51162795 | lir-2 |
| WBGene00003046 | -0.703106452 | 0.30782247 | 0.51641185 | lir-3 |
| WBGene00003047 | -0.134237176 | 0.51605051 | 0.70217251 | lis-1 |
| WBGene00003048 | 0.295358289 | 0.26830517 | 0.47693895 | lit-1 |
| WBGene00003052 | -0.178423048 | 0.25159269 | 0.45969985 | lmn-1 |
| WBGene00003053 | -0.256121021 | 0.11480425 | 0.27887924 | lmp-1 |
| WBGene00003055 | -0.738516349 | 0.00420799 | 0.0273643 | lon-1 |
| WBGene00003056 | -0.094888295 | 0.75528744 | 0.86329738 | lon-2 |
| WBGene00003057 | -1.784783115 | 0.06770056 | 0.19774952 | lon-3 |
| WBGene00003058 | -0.245531902 | 0.94070824 | 1 | lov-1 |
| WBGene00003059 | -0.526442721 | 0.01753289 | 0.07803036 | lpd-2 |
| WBGene00003060 | 0.631550344 | 0.00382877 | 0.0253473 | lpd-3 |
| WBGene00003061 | -0.469546921 | 0.00625472 | 0.03682764 | lpd-5 |
| WBGene00003062 | -0.146444552 | 0.39166425 | 0.60002469 | lpd-6 |
| WBGene00003063 | -0.091891975 | 0.56618597 | 0.74153871 | lpd-7 |
| WBGene00003064 | -0.721424758 | 0.0003238 | 0.00384285 | lpd-8 |
| WBGene00003065 | -0.344081431 | 0.06229314 | 0.18697183 | lpd-9 |
| WBGene00003066 | -0.176959743 | 0.37264952 | 0.58048985 | lpl-1 |
| WBGene00003068 | 0.280211327 | 0.20960995 | 0.40996353 | lrk-1 |
| WBGene00003071 | 1.320786268 | 1.28E-11 | 1.42E-09 | lrp-1 |
| WBGene00003072 | 0.644558208 | 0.00825904 | 0.04484483 | lrp-2 |
| WBGene00003073 | 0.161810645 | 0.28186498 | 0.49119028 | lars-1 |
| WBGene00003074 | -0.165789753 | 0.46393908 | 0.65886864 | lars-2 |
| WBGene00003075 | -0.461948368 | 0.62881548 | 1 | lrx-1 |
| WBGene00003076 | -1.089059772 | 7.39E-07 | 2.6097E-05 | lsm-1 |
| WBGene00003077 | -0.871070427 | 0.00038844 | 0.00441167 | lsm-3 |
| WBGene00003078 | -0.574748564 | 0.01208364 | 0.05958873 | lsm-4 |
| WBGene00003079 | -0.550417939 | 0.04115006 | 0.14038458 | lsm-5 |
| WBGene00003080 | -1.118866788 | 0.00013801 | 0.00197826 | lsm-6 |
| WBGene00003081 | -1.021550762 | 1.9366E-05 | 0.00039122 | lsm-7 |
| WBGene00003082 | -0.82527921 | 0.00022529 | 0.00289132 | lsm-8 |
| WBGene00003083 | -0.267198296 | 0.16403352 | 0.35019009 | lst-1 |
| WBGene00003084 | -0.088958946 | 0.7305008 | 0.84919669 | lst-2 |
| WBGene00003085 | -0.032821912 | 0.84325099 | 0.91377278 | ccar-1 |
| WBGene00003086 | 0.024709091 | 0.9175858 | 0.95646284 | lst-4 |
| WBGene00003087 | -0.654508308 | 0.02666112 | 0.10458568 | lsy-2 |
| WBGene00003089 | 0.50373956 | 0.14943245 | 0.33082775 | ltd-1 |
| WBGene00003090 | -0.018760959 | 0.90476283 | 0.95064695 | lys-1 |
| WBGene00003091 | -0.648417776 | 0.00041347 | 0.00462241 | lys-2 |
| WBGene00003092 | -0.159847599 | 0.86576328 | 0.92781574 | lys-3 |
| WBGene00003093 | 1.358152639 | 4.18E-16 | 9.24E-14 | lys-4 |

| | | | | |
|---|---|---|---|---|
| WBGene00003094 | 1.722078092 | 0.03128109 | 0.11703659 | lys-5 |
| WBGene00003095 | 2.423045592 | 4.89E-08 | 2.4762E-06 | lys-6 |
| WBGene00003096 | 2.128884603 | 5.20E-58 | 3.16E-54 | lys-7 |
| WBGene00003097 | 0.978663619 | 1.99E-08 | 1.1158E-06 | lys-8 |
| WBGene00003098 | 0.377828795 | 0.92526585 | 1 | lys-9 |
| WBGene00003099 | 1.055788746 | 1.4495E-05 | 0.00030929 | lys-10 |
| WBGene00003100 | -2.95057565 | 0.13733704 | 1 | mab-3 |
| WBGene00003102 | 0.741908977 | 0.33680572 | 0.5450031 | mab-5 |
| WBGene00003104 | -0.145049012 | 0.84666507 | 0.91591915 | mab-7 |
| WBGene00003106 | 1.276569476 | 0.22826538 | 1 | mab-9 |
| WBGene00003107 | 0.122562367 | 0.82542555 | 0.9058566 | mab-10 |
| WBGene00003111 | -0.55394962 | 0.02335566 | 0.09563613 | mab-20 |
| WBGene00003112 | 0.844782267 | 0.20493402 | 0.40437249 | mab-21 |
| WBGene00003114 | 1.38322681 | 0.69550305 | 1 | mab-23 |
| WBGene00003119 | -0.397282212 | 0.10764944 | 0.26759651 | mac-1 |
| WBGene00003123 | -0.790350517 | 1.3397E-06 | 4.247E-05 | mag-1 |
| WBGene00003124 | -0.133230985 | 0.80208715 | 0.89160777 | mai-1 |
| WBGene00003129 | -0.524989974 | 0.04992228 | 0.16047296 | map-1 |
| WBGene00003130 | -0.566963401 | 0.00122895 | 0.01073432 | map-2 |
| WBGene00003131 | -0.156269293 | 0.96924949 | 1 | mar-1 |
| WBGene00003132 | -0.170286299 | 0.32122872 | 0.52942261 | mat-1 |
| WBGene00003133 | 0.052680244 | 0.76896247 | 0.87048702 | mat-2 |
| WBGene00003134 | -0.193698774 | 0.32702115 | 0.5355225 | mat-3 |
| WBGene00003136 | 0.195790525 | 0.36029461 | 0.56837101 | mau-2 |
| WBGene00003142 | -0.079203978 | 0.76848168 | 0.87028947 | mau-8 |
| WBGene00003143 | 1.075538322 | 0.00129403 | 0.0111728 | max-1 |
| WBGene00003144 | 0.087104385 | 0.70915108 | 0.83533552 | max-2 |
| WBGene00003148 | -0.55126601 | 0.00173711 | 0.01394593 | mbf-1 |
| WBGene00003149 | 0.225956692 | 0.30124987 | 0.51098347 | mbk-1 |
| WBGene00003150 | -0.01595671 | 0.93042386 | 0.96368163 | mbk-2 |
| WBGene00003151 | 0.481781884 | 0.0402586 | 0.13817557 | mca-1 |
| WBGene00003153 | 0.523920304 | 0.01547435 | 0.07126889 | mca-3 |
| WBGene00003154 | -0.49576248 | 0.00456088 | 0.02918804 | mcm-2 |
| WBGene00003155 | -0.307547971 | 0.03174965 | 0.11812001 | mcm-3 |
| WBGene00003156 | -0.574939168 | 0.00103307 | 0.00947315 | mcm-4 |
| WBGene00003157 | -0.475606246 | 0.00043329 | 0.00477795 | mcm-5 |
| WBGene00003158 | -0.397097118 | 0.00661837 | 0.03822359 | mcm-6 |
| WBGene00003159 | -0.589387953 | 0.00028239 | 0.0034354 | mcm-7 |
| WBGene00003160 | -0.335843075 | 0.03723352 | 0.13106761 | mdf-1 |
| WBGene00003161 | -0.764037576 | 1.3161E-05 | 0.00028483 | mdf-2 |
| WBGene00003162 | -0.266950785 | 0.10529367 | 0.26412613 | mdh-2 |
| WBGene00003163 | 1.180793071 | 4.79E-09 | 3.14E-07 | mdl-1 |
| WBGene00003164 | -0.403534826 | 0.0332265 | 0.12180039 | mdt-6 |

| | | | | |
|---|---|---|---|---|
| WBGene00003165 | 1.474580068 | 0.00026989 | 0.00333263 | mec-1 |
| WBGene00003166 | 1.140608178 | 0.01510734 | 0.0701022 | mec-2 |
| WBGene00003167 | -0.253605996 | 0.76553971 | 0.86871835 | mec-3 |
| WBGene00003168 | 2.180378641 | 0.02487691 | 0.09974122 | mec-4 |
| WBGene00003169 | 0.257242192 | 0.39216698 | 0.60022237 | mec-5 |
| WBGene00003170 | 0.954203474 | 0.31896507 | 0.52680654 | mec-6 |
| WBGene00003171 | 0.840636602 | 0.00285944 | 0.02051799 | mec-7 |
| WBGene00003172 | 0.167371653 | 0.53441396 | 0.71755507 | mec-8 |
| WBGene00003173 | 1.075638313 | 0.05046466 | 0.16187147 | mec-9 |
| WBGene00003174 | 1.027803128 | 0.23092107 | 0.43640702 | mec-10 |
| WBGene00003175 | 1.000097648 | 0.00020497 | 0.00267591 | mec-12 |
| WBGene00003176 | 0.263652632 | 0.86760974 | 1 | mec-14 |
| WBGene00003177 | -0.422494945 | 0.02255806 | 0.09337791 | mec-15 |
| WBGene00003178 | 0.584925248 | 0.01507832 | 0.07000609 | mec-17 |
| WBGene00003179 | 1.155059074 | 0.0293643 | 0.11148501 | mec-18 |
| WBGene00003181 | 0.952350203 | 0.67617448 | 1 | med-2 |
| WBGene00003182 | 1.208777443 | 0.00047824 | 0.00519815 | mef-2 |
| WBGene00003183 | -0.589812929 | 0.00067156 | 0.00674397 | mei-1 |
| WBGene00003184 | -0.377372512 | 0.04283759 | 0.14410951 | mei-2 |
| WBGene00003185 | -0.322689322 | 0.10182962 | 0.25747525 | mek-1 |
| WBGene00003186 | -0.115251742 | 0.58412564 | 0.7540432 | mek-2 |
| WBGene00003196 | 0.166752965 | 0.36189146 | 0.56967189 | mel-11 |
| WBGene00003209 | -0.073238992 | 0.64580263 | 0.79536368 | mel-26 |
| WBGene00003210 | 0.042383238 | 0.80165972 | 0.89148362 | mel-28 |
| WBGene00003214 | 0.078236801 | 0.56227169 | 0.73781645 | mel-32 |
| WBGene00003216 | -0.395005153 | 0.05403975 | 0.169534 | memi-1 |
| WBGene00003218 | -0.102206934 | 0.55147053 | 0.72990338 | mep-1 |
| WBGene00003219 | 0.040759281 | 0.88981258 | 0.94196731 | mes-1 |
| WBGene00003220 | 0.105889631 | 0.61579074 | 0.77579282 | mes-2 |
| WBGene00003221 | 0.128972898 | 0.52004123 | 0.70534444 | mes-3 |
| WBGene00003222 | -0.204522093 | 0.36677051 | 0.57427918 | mes-4 |
| WBGene00003224 | -0.148454675 | 0.41894479 | 0.62348722 | mes-6 |
| WBGene00003225 | -0.343483534 | 0.05993309 | 0.18213959 | mev-1 |
| WBGene00003228 | -0.382952474 | 0.03035067 | 0.11443713 | mex-1 |
| WBGene00003229 | -0.241098829 | 0.24591075 | 0.45380794 | mex-3 |
| WBGene00003230 | -0.230691073 | 0.18136561 | 0.37346673 | mex-5 |
| WBGene00003231 | -0.296960484 | 0.0807271 | 0.22222266 | mex-6 |
| WBGene00003232 | 0.749703249 | 0.23882794 | 0.44554549 | mgl-1 |
| WBGene00003233 | 0.789157954 | 0.03825359 | 0.13342054 | mgl-2 |
| WBGene00003234 | 0.038066915 | 0.89722086 | 0.94634211 | mif-1 |
| WBGene00003235 | -0.215448403 | 0.40165098 | 0.60925094 | mif-2 |
| WBGene00003236 | -0.324329072 | 0.31279853 | 0.52101618 | mif-3 |
| WBGene00003237 | -0.378571493 | 0.72970295 | 0.84851293 | mif-4 |

| | | | | |
|---|---|---|---|---|
| WBGene00003238 | 0.8124496 | 0.03275658 | 0.12073394 | mig-1 |
| WBGene00003239 | -0.70912235 | 0.01928475 | 0.0834592 | mig-2 |
| WBGene00003241 | -0.376431224 | 0.02365653 | 0.09657496 | mig-5 |
| WBGene00003242 | 1.065463621 | 2.81E-08 | 1.5104E-06 | mig-6 |
| WBGene00003243 | 0.347847154 | 0.17114156 | 0.36015788 | mig-10 |
| WBGene00003245 | 0.911360757 | 0.07727617 | 0.21568044 | mig-13 |
| WBGene00003246 | -0.224256201 | 0.31636532 | 0.52436742 | mig-14 |
| WBGene00003247 | -0.029219098 | 0.89998721 | 0.94735085 | mig-15 |
| WBGene00003248 | -0.098699227 | 0.86645254 | 0.92799221 | mig-17 |
| WBGene00003249 | 0.566761598 | 0.43494998 | 0.63619337 | mig-18 |
| WBGene00003252 | 0.704551429 | 0.51564407 | 1 | mig-21 |
| WBGene00003253 | 0.006710766 | 0.97056005 | 0.98468356 | mig-22 |
| WBGene00003254 | -0.221860714 | 0.41891372 | 0.62348722 | mig-23 |
| WBGene00003258 | -0.061610836 | 0.76406656 | 0.86806496 | cogc-1 |
| WBGene00003270 | -0.156269293 | 0.96924949 | 1 | mir-42 |
| WBGene00003271 | -0.156269293 | 0.96924949 | 1 | mir-43 |
| WBGene00003290 | 0.846183542 | 0.65777268 | 1 | mir-62 |
| WBGene00003298 | 0.983488853 | 0.80601565 | 1 | mir-70 |
| WBGene00003305 | -0.156269293 | 0.96924949 | 1 | mir-77 |
| WBGene00003308 | -1.760055813 | 0.45626364 | 1 | mir-80 |
| WBGene00003314 | 1.181686151 | 0.7579682 | 1 | mir-86 |
| WBGene00003316 | 0.688793783 | 0.86378755 | 1 | C32E12.6 |
| WBGene00003323 | -0.156269293 | 0.96924949 | 1 | mir-230 |
| WBGene00003325 | -0.156269293 | 0.96924949 | 1 | mir-232 |
| WBGene00003331 | 1.025181888 | 0.79790347 | 1 | mir-238 |
| WBGene00003337 | 1.025181888 | 0.79790347 | 1 | mir-243 |
| WBGene00003349 | 0.742628056 | 0.76901841 | 1 | mir-255 |
| WBGene00003350 | -0.156269293 | 0.96924949 | 1 | mir-256 |
| WBGene00003354 | 1.377972441 | 0.36464582 | 1 | mir-260 |
| WBGene00003355 | -0.558804446 | 0.87818675 | 1 | mir-261 |
| WBGene00003356 | 3.055829411 | 0.05484431 | 1 | mir-262 |
| WBGene00003363 | -1.094201388 | 0.7816772 | 1 | mir-270 |
| WBGene00003366 | -0.156269293 | 0.96924949 | 1 | mir-273 |
| WBGene00003367 | 0.235829511 | 0.1449468 | 0.32367097 | mix-1 |
| WBGene00003368 | 0.268897653 | 0.43393384 | 0.63557216 | mkk-4 |
| WBGene00003369 | 0.347194629 | 0.04520833 | 0.14972022 | mlc-1 |
| WBGene00003370 | -0.079294493 | 0.59297 | 0.75981517 | mlc-2 |
| WBGene00003371 | 0.206296338 | 0.19787779 | 0.39539734 | mlc-3 |
| WBGene00003372 | -0.292660447 | 0.09232593 | 0.24194455 | mlc-4 |
| WBGene00003373 | -0.239764183 | 0.2414635 | 0.44908217 | mlh-1 |
| WBGene00003374 | 1.403487479 | 1.3086E-05 | 0.00028422 | mlk-1 |
| WBGene00003375 | -0.433751411 | 0.09382555 | 0.24460725 | mlp-1 |
| WBGene00003376 | -0.27166705 | 0.83141994 | 1 | mls-1 |

| | | | | |
|---|---|---|---|---|
| WBGene00003377 | -0.544474177 | 0.48417133 | 0.67489369 | mls-2 |
| WBGene00003378 | 0.413969953 | 0.10094797 | 0.25601242 | mml-1 |
| WBGene00003380 | 0.058712139 | 0.95441013 | 1 | mnm-2 |
| WBGene00003384 | -0.206723659 | 0.57400801 | 0.74766991 | moc-1 |
| WBGene00003385 | 0.356050503 | 0.42836671 | 0.63100978 | moc-2 |
| WBGene00003386 | 1.800036314 | 0.01091287 | 0.05545967 | mod-1 |
| WBGene00003387 | 1.255638708 | 0.04789358 | 0.15647899 | mod-5 |
| WBGene00003388 | -0.012657068 | 0.97422719 | 0.98649644 | moe-3 |
| WBGene00003389 | -0.03789647 | 0.80286969 | 0.89223258 | mog-1 |
| WBGene00003390 | -0.618878457 | 0.0008581 | 0.00816568 | mog-2 |
| WBGene00003391 | -0.393802715 | 0.06403602 | 0.1906477 | mog-3 |
| WBGene00003392 | -0.129463623 | 0.52359635 | 0.70845684 | mog-4 |
| WBGene00003393 | 0.065177283 | 0.68286554 | 0.81858911 | mog-5 |
| WBGene00003394 | -0.105210313 | 0.83001492 | 0.90810229 | mom-1 |
| WBGene00003395 | -0.415521669 | 0.03417571 | 0.12430365 | mom-2 |
| WBGene00003396 | 0.127560664 | 0.53247536 | 0.71592285 | mom-4 |
| WBGene00003397 | 0.168831581 | 0.3480349 | 0.55657162 | mom-5 |
| WBGene00003400 | 0.487805499 | 0.05775559 | 0.17810786 | dapk-1 |
| WBGene00003401 | -0.029953341 | 0.85695446 | 0.92179358 | mpk-1 |
| WBGene00003402 | 1.247939242 | 0.00092027 | 0.00864782 | mpk-2 |
| WBGene00003403 | 0.437370235 | 0.76963385 | 1 | mps-1 |
| WBGene00003404 | 1.318699142 | 1.3398E-06 | 4.247E-05 | mpz-1 |
| WBGene00003405 | 0.028608611 | 0.86857338 | 0.9289571 | mre-11 |
| WBGene00003406 | -0.621728872 | 0.00011287 | 0.00168245 | mrg-1 |
| WBGene00003407 | 0.836494898 | 3.2536E-05 | 0.0005958 | mrp-1 |
| WBGene00003408 | 1.042699941 | 0.00049976 | 0.00536005 | mrp-2 |
| WBGene00003409 | 1.300961693 | 0.00029084 | 0.00352054 | mrp-3 |
| WBGene00003410 | -0.058125966 | 0.75273561 | 0.86217294 | mrp-4 |
| WBGene00003411 | 0.499432871 | 0.01404677 | 0.06643956 | mrp-5 |
| WBGene00003412 | 0.816927073 | 0.06980545 | 0.20187165 | mrp-6 |
| WBGene00003413 | 0.400390292 | 0.03644456 | 0.12918931 | mrp-7 |
| WBGene00003414 | 1.244074973 | 0.003514 | 0.0238524 | mrp-8 |
| WBGene00003415 | -0.034015977 | 0.8353082 | 0.91036596 | mars-1 |
| WBGene00003417 | -0.271860235 | 0.39438553 | 0.6021422 | mrt-2 |
| WBGene00003418 | -0.466041647 | 0.00403667 | 0.02646288 | msh-2 |
| WBGene00003421 | -0.15262097 | 0.5724579 | 0.7463715 | msh-5 |
| WBGene00003422 | -0.07254053 | 0.67208186 | 0.81248091 | msh-6 |
| WBGene00003423 | 0.700728571 | 0.00757691 | 0.04208198 | msi-1 |
| WBGene00003424 | 1.977885804 | 0.49204652 | 1 | msp-3 |
| WBGene00003425 | -0.18557176 | 0.95664217 | 1 | msp-10 |
| WBGene00003426 | 0.972483005 | 0.33065939 | 0.53886384 | msp-19 |
| WBGene00003429 | 0.8114576 | 0.45018242 | 1 | msp-31 |
| WBGene00003431 | 2.845798441 | 0.08303931 | 1 | msp-33 |

| | | | | |
|---|---|---|---|---|
| WBGene00003432 | 1.424985016 | 0.33176275 | 1 | msp-36 |
| WBGene00003434 | 0.76774009 | 0.54413037 | 1 | msp-38 |
| WBGene00003435 | 2.595051674 | 0.1573287 | 1 | msp-40 |
| WBGene00003438 | 2.415207537 | 0.08219082 | 1 | msp-45 |
| WBGene00003442 | 1.127937484 | 0.55876044 | 1 | msp-49 |
| WBGene00003443 | 1.218433787 | 0.43632226 | 1 | msp-50 |
| WBGene00003444 | 0.922655255 | 0.59312081 | 1 | msp-51 |
| WBGene00003446 | 3.794818473 | 0.02487495 | 1 | msp-53 |
| WBGene00003448 | 0.743013877 | 0.5733175 | 1 | msp-55 |
| WBGene00003449 | 1.996315266 | 0.49147447 | 1 | msp-56 |
| WBGene00003450 | 1.433099883 | 0.39185372 | 1 | msp-57 |
| WBGene00003451 | -1.969805064 | 0.62862569 | 1 | msp-58 |
| WBGene00003452 | 1.581673404 | 0.13579525 | 0.31078041 | msp-59 |
| WBGene00003456 | -0.156269293 | 0.96924949 | 1 | msp-63 |
| WBGene00003457 | 1.384352628 | 0.36184168 | 1 | msp-64 |
| WBGene00003458 | 1.584577408 | 0.63258867 | 1 | msp-65 |
| WBGene00003463 | 1.459759188 | 0.21410202 | 1 | msp-76 |
| WBGene00003464 | 1.731602358 | 0.2487004 | 1 | msp-77 |
| WBGene00003465 | 0.16936395 | 0.92512149 | 1 | msp-78 |
| WBGene00003466 | 2.215950884 | 0.41557421 | 1 | msp-79 |
| WBGene00003467 | 0.673816101 | 0.56207811 | 1 | msp-81 |
| WBGene00003468 | 1.413734769 | 0.68804531 | 1 | msp-113 |
| WBGene00003469 | 0.533180584 | 0.75735363 | 1 | msp-142 |
| WBGene00003470 | 0.393897498 | 0.73658207 | 1 | msp-152 |
| WBGene00003471 | 0.368089963 | 0.6231829 | 0.78057453 | mtd-1 |
| WBGene00003472 | 0.526783508 | 0.0030678 | 0.02154205 | mtk-1 |
| WBGene00003473 | -0.4046795 | 0.77815015 | 1 | mtl-1 |
| WBGene00003474 | 0.263651978 | 0.59504178 | 0.76134495 | mtl-2 |
| WBGene00003475 | 0.124866097 | 0.64567683 | 0.79536368 | mtm-1 |
| WBGene00003476 | 0.1140697 | 0.45297409 | 0.65060434 | mtm-3 |
| WBGene00003477 | 0.181181864 | 0.49315232 | 0.68200961 | mtm-5 |
| WBGene00003478 | -0.157039047 | 0.40710375 | 0.61394145 | mtm-6 |
| WBGene00003479 | -0.340256393 | 0.16582481 | 0.35258827 | mtm-9 |
| WBGene00003480 | 0.450082482 | 0.23459925 | 0.44049946 | klf-3 |
| WBGene00003482 | 0.524928144 | 0.0377853 | 0.13235755 | mua-3 |
| WBGene00003485 | 0.28327101 | 0.1417804 | 0.31995461 | mua-6 |
| WBGene00003495 | 0.382681238 | 0.03610124 | 0.12845988 | mup-2 |
| WBGene00003497 | 0.940912072 | 9.114E-06 | 0.00021077 | mup-4 |
| WBGene00003499 | 0.012932458 | 0.95047348 | 0.97468105 | mut-2 |
| WBGene00003504 | -0.438514616 | 0.01439285 | 0.06767763 | mut-7 |
| WBGene00003507 | -0.651372558 | 0.00396344 | 0.0260814 | mut-14 |
| WBGene00003508 | -0.049149832 | 0.79124104 | 0.88470671 | mut-16 |
| WBGene00003509 | -0.720545512 | 0.00320203 | 0.02224021 | mxl-1 |

| | | | | |
|---|---|---|---|---|
| WBGene00003510 | -0.479835964 | 0.06937319 | 0.20106467 | mxl-2 |
| WBGene00003511 | 0.445586564 | 0.05654868 | 0.17559016 | mxl-3 |
| WBGene00003514 | 1.16220041 | 6.79E-07 | 2.4622E-05 | myo-2 |
| WBGene00003515 | 1.442087101 | 7.60E-14 | 1.25E-11 | myo-3 |
| WBGene00003516 | 0.109821195 | 0.67486598 | 0.81406091 | nab-1 |
| WBGene00003517 | 0.652677685 | 0.05079115 | 0.16240593 | nac-1 |
| WBGene00003518 | 0.203039273 | 0.62438481 | 0.78151093 | nac-2 |
| WBGene00003519 | 1.458360909 | 1.3591E-06 | 4.2894E-05 | nac-3 |
| WBGene00003520 | -1.352073912 | 0.29655201 | 1 | nas-1 |
| WBGene00003522 | -0.468500985 | 0.56881132 | 0.74341002 | nas-3 |
| WBGene00003523 | -0.55592286 | 0.47517844 | 0.66814989 | nas-4 |
| WBGene00003524 | -0.005744587 | 0.9942523 | 0.99712681 | nas-5 |
| WBGene00003525 | 1.167297104 | 0.39064847 | 1 | nas-6 |
| WBGene00003526 | -0.254585669 | 0.62686925 | 0.78309735 | nas-7 |
| WBGene00003527 | -0.01991848 | 0.98346386 | 1 | nas-8 |
| WBGene00003528 | 0.233296096 | 0.63915795 | 0.79095064 | nas-9 |
| WBGene00003529 | -1.094201388 | 0.7816772 | 1 | nas-10 |
| WBGene00003530 | -0.336827109 | 0.16234798 | 0.34830656 | nas-11 |
| WBGene00003531 | 1.244164731 | 0.17944235 | 0.37164953 | nas-12 |
| WBGene00003532 | -0.540005146 | 0.71409829 | 1 | nas-13 |
| WBGene00003533 | -0.327794031 | 0.79421475 | 1 | nas-14 |
| WBGene00003534 | 1.846934612 | 0.15711059 | 1 | nas-15 |
| WBGene00003536 | -2.256128098 | 0.25895524 | 1 | nas-17 |
| WBGene00003538 | -1.159714244 | 0.75790634 | 1 | nas-19 |
| WBGene00003539 | -0.440473649 | 0.38998739 | 0.59836179 | nas-20 |
| WBGene00003540 | 0.839158393 | 0.55523389 | 1 | nas-21 |
| WBGene00003541 | -0.986506794 | 0.6369201 | 1 | nas-22 |
| WBGene00003542 | -2.026187751 | 0.18975352 | 1 | nas-23 |
| WBGene00003543 | 0.465758074 | 0.87314503 | 1 | nas-24 |
| WBGene00003544 | 1.90207306 | 0.03445399 | 0.12486742 | nas-25 |
| WBGene00003545 | 0.281454817 | 0.94435481 | 1 | nas-27 |
| WBGene00003546 | 0.534682108 | 0.63998659 | 1 | nas-28 |
| WBGene00003547 | 2.633697959 | 0.04395463 | 1 | nas-29 |
| WBGene00003548 | -0.995178474 | 0.11674257 | 0.28150377 | Y95B8A.2 |
| WBGene00003548 | 0.017483337 | 0.98842241 | 1 | nas-30 |
| WBGene00003549 | 1.278341087 | 0.01528029 | 0.07063015 | nas-31 |
| WBGene00003550 | 1.127347542 | 0.13153934 | 0.30402039 | nas-32 |
| WBGene00003551 | 0.472363238 | 0.69260485 | 1 | nas-33 |
| WBGene00003552 | 1.8412028 | 0.0864217 | 1 | nas-36 |
| WBGene00003553 | 1.111709496 | 0.00032977 | 0.00390985 | nas-37 |
| WBGene00003554 | -0.235215153 | 0.75950584 | 0.86543035 | nas-38 |
| WBGene00003555 | 1.426582257 | 0.00635602 | 0.0371681 | nas-39 |
| WBGene00003558 | 1.140397171 | 0.02713904 | 0.10574296 | nca-2 |

| | | | | |
|---|---|---|---|---|
| WBGene00003559 | 0.331859238 | 0.58658454 | 0.75578084 | ncl-1 |
| WBGene00003561 | 0.922652649 | 0.00116233 | 0.01029311 | ncr-1 |
| WBGene00003562 | 3.368690017 | 0.00587612 | 0.03505749 | ncr-2 |
| WBGene00003563 | 0.328861473 | 0.70703198 | 0.83453969 | ncs-1 |
| WBGene00003564 | 0.767515287 | 1.7922E-05 | 0.00036836 | ncs-2 |
| WBGene00003565 | 0.689981805 | 0.21379311 | 0.41437773 | ncs-3 |
| WBGene00003566 | 0.69702448 | 0.00379159 | 0.02521516 | ncx-1 |
| WBGene00003567 | 0.961307302 | 8.7123E-06 | 0.00020264 | ncx-2 |
| WBGene00003568 | 0.476258224 | 0.07734384 | 0.21573609 | ncx-3 |
| WBGene00003569 | 1.344535093 | 0.00137884 | 0.01174769 | ncx-4 |
| WBGene00003570 | 1.208752296 | 0.4056585 | 1 | ncx-5 |
| WBGene00003571 | -0.084116928 | 0.90875685 | 0.95237091 | ncx-6 |
| WBGene00003572 | 0.115066182 | 0.72078562 | 0.8428256 | ncx-7 |
| WBGene00003573 | 0.959172488 | 0.13468591 | 0.30899879 | ncx-8 |
| WBGene00003574 | -0.064522962 | 0.96594488 | 1 | ncx-9 |
| WBGene00003575 | 2.892012672 | 0.01050617 | 0.05382084 | ncx-10 |
| WBGene00003576 | -0.273224272 | 0.08480786 | 0.22947452 | ndc-80 |
| WBGene00003577 | 0.162626984 | 0.50255413 | 0.68999878 | ndg-4 |
| WBGene00003578 | 0.078606652 | 0.87333309 | 0.93182731 | ndx-1 |
| WBGene00003579 | -0.574101543 | 0.45907369 | 0.65487177 | ndx-2 |
| WBGene00003580 | -0.288900494 | 0.42463759 | 0.62892251 | ndx-3 |
| WBGene00003581 | -0.780151077 | 0.01321721 | 0.06355238 | ndx-4 |
| WBGene00003582 | 0.220035197 | 0.3160786 | 0.52410684 | dcap-2 |
| WBGene00003583 | -0.396202436 | 0.16122706 | 0.34688245 | ndx-6 |
| WBGene00003584 | -0.120248196 | 0.68230213 | 0.81815607 | ndx-7 |
| WBGene00003585 | 0.403114611 | 0.5217505 | 0.70690467 | ndx-8 |
| WBGene00003586 | -0.301386499 | 0.12092821 | 0.28776048 | ndx-9 |
| WBGene00003587 | -0.520093292 | 0.00847834 | 0.04570849 | ned-8 |
| WBGene00003588 | -0.323818892 | 0.08432398 | 0.22867488 | nex-1 |
| WBGene00003589 | -0.069862847 | 0.80208362 | 0.89160777 | nex-2 |
| WBGene00003590 | -0.738447315 | 0.02008727 | 0.08569202 | nex-3 |
| WBGene00003591 | 0.751978014 | 0.20702802 | 0.40665706 | nex-4 |
| WBGene00003592 | -0.200839015 | 0.27851201 | 0.48751649 | nfi-1 |
| WBGene00003593 | 0.50492173 | 0.11222964 | 0.27460243 | nfm-1 |
| WBGene00003594 | 0.058201526 | 0.81382842 | 0.8986238 | nft-1 |
| WBGene00003595 | -2.540549033 | 0.0219395 | 0.09178756 | ngn-1 |
| WBGene00003596 | -0.104759496 | 0.54530125 | 0.72434382 | ngp-1 |
| WBGene00003597 | 1.099620998 | 0.00095174 | 0.00887485 | nhl-1 |
| WBGene00003598 | 0.6277213 | 0.23372885 | 0.43981741 | nhl-2 |
| WBGene00003599 | -0.024328406 | 0.93611315 | 0.96668791 | nhl-3 |
| WBGene00003600 | 0.91539982 | 0.00053206 | 0.00562694 | nhr-1 |
| WBGene00003600, W | 1.060070137 | 1.8344E-05 | 0.00037495 | crf-1 |
| WBGene00003601 | -1.256423969 | 2.006E-05 | 0.0004019 | nhr-2 |

| | | | | |
|---|---|---|---|---|
| WBGene00003602 | 0.416002566 | 0.10550004 | 0.26442527 | nhr-3 |
| WBGene00003603 | 0.132627618 | 0.68043037 | 0.8167559 | nhr-4 |
| WBGene00003604 | 0.759107543 | 0.102572 | 0.25901136 | nhr-5 |
| WBGene00003605 | 1.148993491 | 0.04152915 | 0.14123401 | nhr-6 |
| WBGene00003606 | 1.273276208 | 0.00063461 | 0.00643679 | nhr-7 |
| WBGene00003607 | 0.773019154 | 0.00426401 | 0.02762505 | nhr-8 |
| WBGene00003609 | 0.061415307 | 0.83294024 | 0.90960882 | nhr-10 |
| WBGene00003610 | -0.268805319 | 0.61319662 | 0.77405075 | nhr-11 |
| WBGene00003611 | -0.301966405 | 0.38150302 | 0.58977507 | nhr-12 |
| WBGene00003612 | -0.260962059 | 0.40550036 | 0.61226986 | nhr-13 |
| WBGene00003613 | 1.154800038 | 0.00069645 | 0.00695929 | nhr-14 |
| WBGene00003614 | 1.155552588 | 0.19024849 | 0.38471135 | nhr-15 |
| WBGene00003615 | 0.91657459 | 0.3594425 | 0.56763676 | nhr-16 |
| WBGene00003616 | -0.153802489 | 0.63424894 | 0.78783996 | nhr-17 |
| WBGene00003617 | 0.820602066 | 0.12616801 | 0.29630283 | nhr-18 |
| WBGene00003618 | 1.148052382 | 0.00068519 | 0.0068694 | nhr-19 |
| WBGene00003619 | 0.135719368 | 0.63093772 | 0.78581631 | nhr-20 |
| WBGene00003620 | 1.310157606 | 0.00031687 | 0.00378283 | nhr-21 |
| WBGene00003621 | 0.272043451 | 0.30484852 | 0.51397943 | nhr-22 |
| WBGene00003622 | -0.112720848 | 0.66995771 | 0.8111245 | nhr-23 |
| WBGene00003623 | 0.749279927 | 0.04452896 | 0.14783895 | nhr-25 |
| WBGene00003624 | 0.355649467 | 0.28242627 | 0.49195657 | nhr-28 |
| WBGene00003625 | 0.774060747 | 0.00010292 | 0.0015567 | nhr-31 |
| WBGene00003626 | 0.835025524 | 0.04248699 | 0.14360649 | nhr-32 |
| WBGene00003627 | 0.844570571 | 0.00058492 | 0.00601315 | nhr-34 |
| WBGene00003628 | 1.087950256 | 0.00136767 | 0.01169165 | nhr-35 |
| WBGene00003629 | 0.680743233 | 0.76740144 | 1 | nhr-38 |
| WBGene00003630 | 0.759352011 | 0.00729858 | 0.04098614 | nhr-40 |
| WBGene00003632 | 0.609435984 | 0.12911525 | 0.30030427 | nhr-42 |
| WBGene00003633 | -0.313877379 | 0.62189023 | 0.77999682 | nhr-43 |
| WBGene00003634 | 0.723054137 | 0.1638252 | 0.34999151 | nhr-44 |
| WBGene00003635 | 0.785819002 | 0.01259497 | 0.06143653 | nhr-45 |
| WBGene00003636 | 0.224943062 | 0.2922986 | 0.50237787 | nhr-46 |
| WBGene00003637 | -0.068888782 | 0.82525289 | 0.90583088 | nhr-47 |
| WBGene00003638 | 0.05341517 | 0.857178 | 0.92195234 | nhr-48 |
| WBGene00003639 | 0.485037959 | 0.01758228 | 0.07810589 | nhr-49 |
| WBGene00003640 | 0.953210817 | 0.05922442 | 0.1808915 | nhr-50 |
| WBGene00003641 | -1.070706988 | 0.48553658 | 1 | nhr-51 |
| WBGene00003642 | -0.741863642 | 0.68909365 | 1 | nhr-52 |
| WBGene00003643 | 0.079704485 | 0.91179022 | 0.95431423 | nhr-53 |
| WBGene00003644 | 0.390803764 | 0.66297945 | 0.80637483 | nhr-54 |
| WBGene00003645 | 1.231260357 | 0.0072059 | 0.040616 | nhr-55 |
| WBGene00003646 | 0.263404986 | 0.65804966 | 0.80303357 | nhr-56 |

| | | | | |
|---|---|---|---|---|
| WBGene00003647 | 0.381349427 | 0.49244236 | 0.68149352 | nhr-57 |
| WBGene00003648 | 2.037398456 | 0.0238748 | 0.09707432 | nhr-58 |
| WBGene00003649 | 0.598335997 | 0.42493375 | 0.62893095 | nhr-59 |
| WBGene00003650 | 0.743468639 | 0.52792973 | 1 | nhr-60 |
| WBGene00003651 | -0.249216838 | 0.49726491 | 0.68558861 | nhr-61 |
| WBGene00003652 | 0.210467162 | 0.62365414 | 0.78099896 | nhr-62 |
| WBGene00003653 | 0.086403265 | 0.90512821 | 0.95070175 | nhr-63 |
| WBGene00003654 | 0.100610553 | 0.82157714 | 0.90326615 | nhr-64 |
| WBGene00003655 | 0.266554493 | 0.60701466 | 0.76984905 | nhr-65 |
| WBGene00003656 | 0.122140589 | 0.50814699 | 0.69491018 | nhr-66 |
| WBGene00003657 | 0.719356577 | 0.35980306 | 0.56805837 | nhr-67 |
| WBGene00003658 | 0.690606704 | 0.0012013 | 0.01055349 | nhr-68 |
| WBGene00003659 | 0.682054533 | 0.03464261 | 0.12525192 | nhr-69 |
| WBGene00003660 | 0.470914509 | 0.23668177 | 0.44283456 | nhr-70 |
| WBGene00003661 | -0.111828084 | 0.66913516 | 0.81045191 | nhr-71 |
| WBGene00003662 | 0.09697944 | 0.97612634 | 1 | nhr-72 |
| WBGene00003663 | -1.082680044 | 0.67762556 | 1 | nhr-73 |
| WBGene00003664 | -1.225663127 | 0.22483593 | 1 | nhr-74 |
| WBGene00003668 | -0.528209627 | 0.35728229 | 0.56561018 | nhr-78 |
| WBGene00003669 | 0.732684946 | 0.09671249 | 0.24972061 | nhr-79 |
| WBGene00003670 | 0.977737078 | 0.02298246 | 0.09461854 | nhr-80 |
| WBGene00003671 | -1.901856996 | 0.64072034 | 1 | nhr-81 |
| WBGene00003674 | 0.12092777 | 0.76410594 | 0.86806496 | nhr-84 |
| WBGene00003675 | 0.472848044 | 0.3533116 | 0.56208019 | nhr-85 |
| WBGene00003676 | -0.022017842 | 0.93247734 | 0.96416346 | nhr-86 |
| WBGene00003677 | 0.831069723 | 0.18573775 | 0.37836275 | nhr-87 |
| WBGene00003678 | 1.13004394 | 0.00063882 | 0.00647404 | nhr-88 |
| WBGene00003680 | 1.449186169 | 0.00381225 | 0.02528609 | nhr-90 |
| WBGene00003681 | -0.14391845 | 0.74800212 | 0.85972621 | nhr-91 |
| WBGene00003682 | 0.102981687 | 0.88189594 | 0.93708197 | nhr-92 |
| WBGene00003683 | 1.501307361 | 0.39583573 | 1 | nhr-93 |
| WBGene00003684 | 1.185494438 | 0.22474156 | 0.42887455 | nhr-94 |
| WBGene00003685 | -0.038714673 | 0.91867772 | 0.95698552 | nhr-95 |
| WBGene00003686 | 0.637442796 | 0.29314077 | 0.50325538 | nhr-96 |
| WBGene00003687 | 0.517956967 | 0.00259874 | 0.01907574 | nhr-97 |
| WBGene00003688 | 1.572038656 | 0.04270387 | 0.14385895 | nhr-98 |
| WBGene00003689 | 1.664730189 | 0.01920601 | 0.08332046 | nhr-99 |
| WBGene00003690 | 0.736354167 | 0.00598006 | 0.03552053 | nhr-100 |
| WBGene00003691 | 0.937336957 | 0.01047781 | 0.05370108 | nhr-101 |
| WBGene00003692 | -0.327503771 | 0.54363668 | 0.72337864 | nhr-102 |
| WBGene00003693 | 0.853413376 | 0.23773671 | 0.44395533 | nhr-103 |
| WBGene00003694 | 0.835008504 | 0.06858229 | 0.199539 | nhr-104 |
| WBGene00003695 | 0.39676488 | 0.25449729 | 0.46216317 | nhr-105 |

| | | | | |
|---|---|---|---|---|
| WBGene00003696 | 0.400090227 | 0.53539546 | 0.7178681 | nhr-106 |
| WBGene00003697 | 0.87656263 | 0.08143637 | 0.22333837 | nhr-107 |
| WBGene00003698 | 0.52853078 | 0.58595362 | 1 | nhr-108 |
| WBGene00003699 | -0.100736636 | 0.85295286 | 0.91927977 | nhr-109 |
| WBGene00003700 | 1.802265266 | 0.01002009 | 0.05194443 | nhr-110 |
| WBGene00003702 | 0.488681396 | 0.45815428 | 0.65477918 | nhr-112 |
| WBGene00003703 | 0.001742171 | 0.99899207 | 1 | nhr-113 |
| WBGene00003704 | 0.926459007 | 2.529E-06 | 7.1572E-05 | nhr-114 |
| WBGene00003705 | 1.729317662 | 0.01129511 | 0.05683132 | nhr-115 |
| WBGene00003706 | 0.073290113 | 0.91840716 | 0.95686787 | nhr-116 |
| WBGene00003707 | 0.750443753 | 0.20783351 | 0.40737918 | nhr-117 |
| WBGene00003709 | 0.299563615 | 0.41721349 | 0.62193997 | nhr-119 |
| WBGene00003710 | 0.587666048 | 0.15081835 | 0.33250147 | nhr-120 |
| WBGene00003711 | 1.826749395 | 7.3124E-05 | 0.00119327 | nhr-121 |
| WBGene00003712 | 0.442720938 | 0.31444704 | 0.52218596 | nhr-122 |
| WBGene00003713 | -0.11967713 | 0.89769345 | 1 | nhr-123 |
| WBGene00003714 | 0.480011494 | 0.77590664 | 1 | nhr-124 |
| WBGene00003715 | 1.157328866 | 0.15555284 | 0.33932513 | nhr-125 |
| WBGene00003716 | 0.323096265 | 0.60306782 | 0.76781108 | nhr-126 |
| WBGene00003717 | -0.265633899 | 0.86436221 | 1 | nhr-127 |
| WBGene00003718 | 0.89701186 | 0.23493002 | 0.44084782 | nhr-128 |
| WBGene00003719 | 1.65644977 | 0.04622209 | 0.15228831 | nhr-129 |
| WBGene00003720 | 0.995490573 | 0.02799973 | 0.10819373 | nhr-130 |
| WBGene00003721 | 0.767769199 | 0.17848712 | 0.37055611 | nhr-131 |
| WBGene00003722 | 0.183051707 | 0.75728239 | 0.86429398 | nhr-132 |
| WBGene00003723 | 0.161833132 | 0.73713893 | 0.85315574 | nhr-133 |
| WBGene00003724 | 0.244313383 | 0.6175613 | 0.77712375 | nhr-134 |
| WBGene00003725 | 0.646541632 | 0.29057091 | 0.50059826 | nhr-135 |
| WBGene00003726 | 0.628081278 | 0.23407262 | 0.4401914 | nhr-136 |
| WBGene00003727 | 0.344401527 | 0.41482806 | 0.62017539 | nhr-137 |
| WBGene00003728 | 0.291338434 | 0.31043881 | 0.5186511 | nhr-138 |
| WBGene00003729 | 0.70997188 | 0.28524801 | 0.49502517 | nhx-1 |
| WBGene00003730 | 1.526030592 | 4.00E-14 | 6.74E-12 | nhx-2 |
| WBGene00003731 | 1.237287119 | 0.02739231 | 0.10652467 | nhx-3 |
| WBGene00003732 | 1.343453783 | 0.00881543 | 0.04700445 | nhx-4 |
| WBGene00003733 | 0.544558537 | 0.08790851 | 0.23501017 | nhx-5 |
| WBGene00003734 | 1.046239859 | 0.51262061 | 1 | nhx-6 |
| WBGene00003735 | 0.584423816 | 0.06569638 | 0.19353487 | nhx-8 |
| WBGene00003736 | 0.038302025 | 0.91097791 | 0.9539162 | nhx-9 |
| WBGene00003738 | 0.541931233 | 0.00988091 | 0.05133254 | nid-1 |
| WBGene00003739 | 0.225594762 | 0.60500613 | 0.76897128 | nlp-1 |
| WBGene00003740 | 2.785013157 | 0.03821618 | 1 | nlp-2 |
| WBGene00003741 | 0.740529367 | 0.05588723 | 0.17389475 | nlp-3 |

| | | | | |
|---|---|---|---|---|
| WBGene00003742 | 0.820015846 | 0.7518774 | 1 | nlp-4 |
| WBGene00003743 | 0.905189753 | 0.20749569 | 0.40703925 | nlp-5 |
| WBGene00003744 | 0.548862028 | 0.15640169 | 0.34042182 | nlp-6 |
| WBGene00003745 | 0.772033993 | 0.177701 | 0.36961342 | nlp-7 |
| WBGene00003746 | 1.081520541 | 0.00052828 | 0.00559317 | nlp-8 |
| WBGene00003747 | 1.005836467 | 0.02216054 | 0.09248923 | nlp-9 |
| WBGene00003748 | 0.933317633 | 0.0316955 | 0.11812001 | nlp-10 |
| WBGene00003749 | 0.591640374 | 0.12706949 | 0.29759852 | nlp-11 |
| WBGene00003750 | -0.140528348 | 0.81124846 | 0.89692775 | nlp-12 |
| WBGene00003751 | 0.756677413 | 0.03317374 | 0.12168047 | nlp-13 |
| WBGene00003752 | 0.866053114 | 0.0281472 | 0.10851365 | nlp-14 |
| WBGene00003753 | 0.686218995 | 0.08680597 | 0.23301156 | nlp-15 |
| WBGene00003754 | 0.656366372 | 0.05978489 | 0.18182573 | nlp-16 |
| WBGene00003755 | 0.802587467 | 0.12509894 | 0.29486046 | nlp-17 |
| WBGene00003756 | 1.015188921 | 0.00496133 | 0.03100132 | nlp-18 |
| WBGene00003757 | 0.98234419 | 0.28751388 | 0.4974059 | nlp-19 |
| WBGene00003758 | 0.42209235 | 0.36576492 | 0.57350663 | nlp-20 |
| WBGene00003759 | 0.745142565 | 0.02594673 | 0.10284664 | nlp-21 |
| WBGene00003761 | -1.901856996 | 0.64072034 | 1 | nlp-23 |
| WBGene00003762 | -0.410760304 | 0.12450343 | 0.29397363 | nlp-24 |
| WBGene00003763 | -0.429656971 | 0.77604656 | 1 | nlp-25 |
| WBGene00003764 | -0.934929804 | 0.13113277 | 0.30360086 | nlp-26 |
| WBGene00003765 | -1.039793605 | 0.0001374 | 0.00197186 | nlp-27 |
| WBGene00003766 | -1.571134344 | 3.8282E-06 | 0.00010103 | nlp-28 |
| WBGene00003767 | -0.958808221 | 0.01652124 | 0.07432888 | nlp-29 |
| WBGene00003768 | -1.537589574 | 0.05528583 | 0.17237425 | nlp-30 |
| WBGene00003769 | -0.446601391 | 0.21605087 | 0.41761252 | nlp-31 |
| WBGene00003770 | -0.156269293 | 0.96924949 | 1 | nlp-32 |
| WBGene00003771 | -0.12385684 | 0.65623751 | 0.80231895 | nlp-33 |
| WBGene00003772 | 0.589694674 | 0.06736028 | 0.1971139 | nlr-1 |
| WBGene00003773 | 0.272711929 | 0.16637936 | 0.35339605 | nlt-1 |
| WBGene00003774 | 1.870866998 | 0.12343464 | 1 | nmr-1 |
| WBGene00003775 | 1.226794432 | 0.24328965 | 1 | nmr-2 |
| WBGene00003776 | 0.532106603 | 0.00483393 | 0.03044023 | nmy-1 |
| WBGene00003777 | 0.149273525 | 0.39826577 | 0.60593292 | nmy-2 |
| WBGene00003778 | 1.058450237 | 0.07436256 | 0.21075238 | nnt-1 |
| WBGene00003779 | 1.006022701 | 0.08589171 | 0.23137592 | nob-1 |
| WBGene00003783 | -0.653733494 | 0.11574973 | 0.28023616 | nos-1 |
| WBGene00003784 | -0.516176817 | 0.00349457 | 0.02375567 | nos-2 |
| WBGene00003785 | -0.050557099 | 0.80944685 | 0.89609686 | nos-3 |
| WBGene00003786 | 1.379771129 | 3.25E-13 | 4.75E-11 | npa-1 |
| WBGene00003787 | -0.315873304 | 0.06005918 | 0.18234021 | npp-1 |
| WBGene00003788 | -0.595323112 | 9.6128E-05 | 0.00148485 | npp-2 |

| | | | | |
|---|---|---|---|---|
| WBGene00003789 | -0.039910353 | 0.82335561 | 0.90456615 | npp-3 |
| WBGene00003790 | -0.131914244 | 0.49677992 | 0.68507553 | npp-4 |
| WBGene00003791 | -0.530137784 | 0.00088307 | 0.00835001 | npp-5 |
| WBGene00003792 | 0.061886854 | 0.72779408 | 0.84742955 | npp-6 |
| WBGene00003793 | 0.153490517 | 0.43301806 | 0.63493626 | npp-7 |
| WBGene00003794 | -0.261053166 | 0.18277175 | 0.37489979 | npp-8 |
| WBGene00003795 | -0.094432967 | 0.60405389 | 0.76858292 | npp-9 |
| WBGene00003796 | -0.190045157 | 0.27389832 | 0.48243138 | npp-10 |
| WBGene00003797 | -0.32334182 | 0.11815773 | 0.28345248 | npp-11 |
| WBGene00003798 | 0.354946336 | 0.09556119 | 0.24783919 | npp-12 |
| WBGene00003799 | -0.214575795 | 0.26113683 | 0.46968486 | npp-13 |
| WBGene00003800 | 0.134662187 | 0.40463486 | 0.61163743 | npp-14 |
| WBGene00003801 | -0.211514172 | 0.2051354 | 0.40457259 | npp-15 |
| WBGene00003802 | -0.719030026 | 6.362E-05 | 0.00106246 | npp-16 |
| WBGene00003803 | -0.22554649 | 0.19336445 | 0.38900377 | rae-1 |
| WBGene00003804 | -0.67333562 | 0.000152 | 0.00211546 | npp-18 |
| WBGene00003805 | -0.413237721 | 0.02490145 | 0.09974122 | npp-19 |
| WBGene00003806 | -0.450378842 | 0.02064651 | 0.087418 | npp-20 |
| WBGene00003807 | 0.375501373 | 0.78102239 | 1 | npr-1 |
| WBGene00003808 | 1.082732261 | 0.19164712 | 0.38657379 | npr-2 |
| WBGene00003812 | 0.350758067 | 0.15228749 | 0.33446498 | nrf-5 |
| WBGene00003813 | 0.804789717 | 0.0006993 | 0.00697632 | nrf-6 |
| WBGene00003815 | -0.426750351 | 0.00754459 | 0.04196006 | nars-1 |
| WBGene00003816 | -0.261432045 | 0.66579229 | 0.80833842 | asns-1 |
| WBGene00003818 | 0.168638902 | 0.41803566 | 0.62282133 | nsf-1 |
| WBGene00003821 | -0.152941005 | 0.3017132 | 0.51117778 | nst-1 |
| WBGene00003822 | 0.797532507 | 0.00161689 | 0.01327293 | nsy-1 |
| WBGene00003825 | -0.068428859 | 0.70544534 | 0.83347722 | ntl-2 |
| WBGene00003826 | -0.041477107 | 0.81091936 | 0.89678607 | ntl-3 |
| WBGene00003827 | 0.271725623 | 0.14527171 | 0.32397939 | ntl-4 |
| WBGene00003828 | 1.066184393 | 3.96E-08 | 2.0205E-06 | nuc-1 |
| WBGene00003829 | -0.596489768 | 0.00092274 | 0.00866436 | nud-1 |
| WBGene00003830 | -0.199920782 | 0.309684 | 0.51818605 | num-1 |
| WBGene00003831 | -0.412401113 | 0.00776383 | 0.04284573 | nuo-1 |
| WBGene00003834 | 0.26743018 | 0.14929669 | 0.33076844 | nxf-1 |
| WBGene00003835 | -0.52709769 | 0.02738023 | 0.10652467 | nxf-2 |
| WBGene00003836 | -0.496043441 | 0.01763376 | 0.07822123 | nxt-1 |
| WBGene00003837 | 0.680767907 | 0.08593996 | 0.23145453 | oat-1 |
| WBGene00003838 | 0.780075108 | 0.49091814 | 1 | ocr-1 |
| WBGene00003839 | 2.602991927 | 0.02621501 | 0.10340367 | ocr-2 |
| WBGene00003840 | 2.007104841 | 0.31492229 | 1 | ocr-3 |
| WBGene00003841 | 0.07688797 | 0.94522541 | 1 | ocr-4 |
| WBGene00003842 | 0.831423551 | 0.18451231 | 0.37694161 | oct-1 |

| | | | | |
|---|---|---|---|---|
| WBGene00003843 | 1.283185584 | 0.02220123 | 0.09256358 | oct-2 |
| WBGene00003844 | -0.974745837 | 0.00034918 | 0.0040966 | odc-1 |
| WBGene00003845 | -0.1543483 | 0.88670749 | 1 | odd-1 |
| WBGene00003846 | 1.041941342 | 0.09936681 | 0.25393171 | odd-2 |
| WBGene00003847 | 0.211916244 | 0.48435503 | 0.67499477 | blmp-1 |
| WBGene00003848 | 1.344563648 | 0.34068438 | 1 | odr-1 |
| WBGene00003849 | 0.391949327 | 0.51432067 | 0.70122035 | odr-2 |
| WBGene00003850 | 0.130085476 | 0.83961965 | 0.91211511 | odr-3 |
| WBGene00003851 | -0.302875017 | 0.10862871 | 0.26875696 | odr-4 |
| WBGene00003854 | 1.270678414 | 0.5135606 | 1 | odr-7 |
| WBGene00003856 | 1.001765124 | 0.17809706 | 0.37012606 | odr-10 |
| WBGene00003858 | 0.368202638 | 0.05328608 | 0.16764609 | ogt-1 |
| WBGene00003859 | 0.427997781 | 0.5798592 | 0.75070264 | oig-1 |
| WBGene00003860 | -1.010163454 | 0.00428116 | 0.02769183 | oig-2 |
| WBGene00003861 | 0.543681994 | 0.14825906 | 0.32891394 | oig-3 |
| WBGene00003862 | 2.012315008 | 0.01312136 | 0.06326707 | old-1 |
| WBGene00003864 | -0.181654762 | 0.26790742 | 0.47644118 | oma-1 |
| WBGene00003865 | -0.354015759 | 0.02634185 | 0.10373545 | oma-2 |
| WBGene00003868 | -0.094787701 | 0.62248369 | 0.7803381 | ooc-3 |
| WBGene00003870 | -0.607170854 | 0.00047936 | 0.00520567 | ooc-5 |
| WBGene00003876 | 0.912497532 | 0.00385616 | 0.02548593 | pept-2 |
| WBGene00003877 | 1.24189593 | 3.92E-15 | 7.67E-13 | pept-1 |
| WBGene00003878 | 1.036833627 | 0.01826418 | 0.08025853 | pept-3 |
| WBGene00003879 | -0.828765424 | 0.69034289 | 1 | ora-1 |
| WBGene00003882 | 0.0869265 | 0.68498368 | 0.81999476 | orc-2 |
| WBGene00003883 | 0.326616897 | 0.38551333 | 0.59397428 | osm-1 |
| WBGene00003884 | 0.736746158 | 0.27850093 | 0.48751649 | osm-3 |
| WBGene00003885 | 1.228611349 | 0.10058468 | 0.25569485 | osm-5 |
| WBGene00003886 | 0.795058093 | 0.38240895 | 0.59076562 | osm-6 |
| WBGene00003887 | 0.196998201 | 0.53127312 | 0.71493169 | osm-7 |
| WBGene00003888 | -1.108887062 | 0.31696487 | 1 | osm-8 |
| WBGene00003889 | -0.667801608 | 0.42376848 | 0.62806093 | osm-9 |
| WBGene00003890 | 0.233738896 | 0.68374274 | 0.81883218 | osm-10 |
| WBGene00003891 | 0.248580915 | 0.24281218 | 0.45048634 | osm-11 |
| WBGene00003892 | 1.163996705 | 0.21051985 | 0.41041347 | osm-12 |
| WBGene00003893 | 0.222367045 | 0.1267658 | 0.29717389 | ost-1 |
| WBGene00003898 | 0.24841244 | 0.2844348 | 0.49417901 | oxi-1 |
| WBGene00003901 | -0.230250414 | 0.15490574 | 0.33852997 | paa-1 |
| WBGene00003902 | -0.011711548 | 0.93220923 | 0.96406579 | pab-1 |
| WBGene00003903 | 0.187136816 | 0.46579067 | 0.65995619 | pab-2 |
| WBGene00003904 | -0.588022972 | 0.0005623 | 0.00585995 | pabp-2 |
| WBGene00003905 | 0.653353426 | 0.00153705 | 0.0127993 | pad-1 |
| WBGene00003906 | -0.22461868 | 0.93331196 | 1 | paf-1 |

| | | | | |
|---|---|---|---|---|
| WBGene00003907 | 0.043362339 | 0.91817655 | 0.95686787 | paf-2 |
| WBGene00003909 | 2.623840182 | 0.01886102 | 0.08216422 | pag-3 |
| WBGene00003911 | -0.281012837 | 0.2576702 | 0.46553184 | pak-1 |
| WBGene00003912 | -0.661465787 | 8.3645E-05 | 0.00132447 | pal-1 |
| WBGene00003914 | -0.375194387 | 0.02582576 | 0.10240059 | pam-1 |
| WBGene00003915 | 0.770849644 | 0.00045084 | 0.00493566 | pan-1 |
| WBGene00003916 | 0.430522532 | 0.62678656 | 0.78309735 | AH10.3 |
| WBGene00003916 | 0.465380149 | 0.01421237 | 0.06695739 | par-1 |
| WBGene00003917 | -0.286473678 | 0.24437575 | 0.45193694 | par-2 |
| WBGene00003918 | 0.010043739 | 0.95467232 | 0.97697363 | par-3 |
| WBGene00003919 | -0.084372063 | 0.63540568 | 0.788316 | par-4 |
| WBGene00003920 | -0.534602436 | 9.2224E-05 | 0.00143734 | par-5 |
| WBGene00003921 | -0.365388424 | 0.10774445 | 0.26759651 | par-6 |
| WBGene00003922 | -0.323797373 | 0.07130741 | 0.20486117 | pas-1 |
| WBGene00003923 | -0.913446538 | 1.35E-09 | 1.02E-07 | pas-2 |
| WBGene00003924 | -0.656105134 | 3.1576E-06 | 8.6538E-05 | pas-3 |
| WBGene00003925 | -0.573038545 | 0.00040098 | 0.00451601 | pas-4 |
| WBGene00003926 | -0.946042956 | 4.18E-10 | 3.62E-08 | pas-5 |
| WBGene00003927 | -0.474601719 | 0.00357399 | 0.02418716 | pas-6 |
| WBGene00003928 | -0.701257704 | 9.1645E-06 | 0.00021153 | pas-7 |
| WBGene00003929 | 0.985234879 | 1.0814E-06 | 3.5798E-05 | pat-2 |
| WBGene00003930 | 0.194212761 | 0.24365664 | 0.45114272 | pat-3 |
| WBGene00003931 | -0.056066262 | 0.83652317 | 0.9111967 | pat-4 |
| WBGene00003932 | 0.069315747 | 0.76102935 | 0.86650332 | pat-6 |
| WBGene00003933 | 0.726174024 | 0.09700752 | 0.25021635 | pat-9 |
| WBGene00003934 | -0.517099864 | 0.00025567 | 0.00318372 | pat-10 |
| WBGene00003936 | 0.614958135 | 0.00853276 | 0.04596105 | pat-12 |
| WBGene00003937 | -1.895791037 | 0.52793577 | 1 | pax-1 |
| WBGene00003938 | 2.07531578 | 0.45514895 | 1 | pax-2 |
| WBGene00003939 | 0.326610492 | 0.65511299 | 0.80180033 | pax-3 |
| WBGene00003941 | 0.411633195 | 0.089381 | 0.2371834 | pbo-1 |
| WBGene00003943 | 0.391950908 | 0.19863078 | 0.39657321 | pbo-4 |
| WBGene00003947 | -0.908419329 | 8.09E-08 | 3.8843E-06 | pbs-1 |
| WBGene00003948 | -0.64375766 | 6.7356E-05 | 0.00111717 | pbs-2 |
| WBGene00003949 | -0.538564021 | 0.00027418 | 0.0033692 | pbs-3 |
| WBGene00003950 | -0.860262611 | 2.05E-08 | 1.1456E-06 | pbs-4 |
| WBGene00003951 | -0.802574016 | 1.2466E-06 | 4.0145E-05 | pbs-5 |
| WBGene00003952 | -0.504859273 | 0.00154027 | 0.01281727 | pbs-6 |
| WBGene00003953 | -0.665804584 | 1.2754E-06 | 4.075E-05 | pbs-7 |
| WBGene00003954 | -0.359035965 | 0.32352793 | 0.53202663 | pcm-1 |
| WBGene00003955 | -0.752844564 | 3.71E-07 | 1.4546E-05 | pcn-1 |
| WBGene00003956 | 0.279208663 | 0.18522258 | 0.377757 | pcp-1 |
| WBGene00003957 | 1.284533456 | 1.47E-09 | 1.10E-07 | pcp-2 |

| | | | | |
|---|---|---|---|---|
| WBGene00003958 | 1.374962501 | 8.67E-10 | 6.97E-08 | pcp-3 |
| WBGene00003959 | 0.910900024 | 0.00825327 | 0.04483353 | pcp-4 |
| WBGene00003960 | -0.236100152 | 0.47273853 | 0.66583741 | pcs-1 |
| WBGene00003961 | -0.116483894 | 0.50037143 | 0.68823038 | pct-1 |
| WBGene00003962 | -0.371447466 | 0.00931658 | 0.04915201 | pdi-1 |
| WBGene00003963 | -0.099248293 | 0.53177489 | 0.71537717 | pdi-2 |
| WBGene00003964 | -0.062321305 | 0.66536582 | 0.80808083 | pdi-3 |
| WBGene00003965 | -0.607259079 | 0.07957623 | 0.22027702 | pdk-1 |
| WBGene00003966 | 0.272606957 | 0.59987837 | 0.76535554 | pdl-1 |
| WBGene00003967 | -0.233331018 | 0.24249766 | 0.45004036 | pdr-1 |
| WBGene00003968 | 0.724867663 | 0.23598088 | 0.44215444 | peb-1 |
| WBGene00003969 | 0.995145988 | 0.03158355 | 0.11791386 | pef-1 |
| WBGene00003970 | 0.005424645 | 0.978217 | 0.98873753 | pek-1 |
| WBGene00003975 | -0.802680005 | 0.01331217 | 0.06379465 | pen-2 |
| WBGene00003976 | -0.770024802 | 0.27714244 | 0.4859599 | pes-1 |
| WBGene00003977 | -2.50408321 | 0.09643964 | 1 | pes-2.1 |
| WBGene00003978 | 0.731079408 | 0.02007369 | 0.08569202 | pes-4 |
| WBGene00003979 | -0.908631441 | 0.00879959 | 0.04700214 | pes-5 |
| WBGene00003980 | 1.142870269 | 7.5086E-06 | 0.0001777 | pes-7 |
| WBGene00003981 | -1.088572391 | 0.18209698 | 0.37384831 | pes-8 |
| WBGene00003982 | -0.475292592 | 0.00142481 | 0.01205586 | pes-9 |
| WBGene00003983 | -2.086867335 | 2.51E-07 | 1.0315E-05 | pes-10 |
| WBGene00003986 | 0.579438659 | 0.00819972 | 0.04460253 | mct-4 |
| WBGene00003987 | -0.093513447 | 0.8610403 | 0.92512303 | pes-23 |
| WBGene00003989 | -0.483409197 | 0.00142451 | 0.01205586 | pfn-1 |
| WBGene00003990 | 0.264772816 | 0.35320436 | 0.56202545 | pfn-2 |
| WBGene00003991 | -0.361677903 | 0.29201548 | 0.50210451 | pfn-3 |
| WBGene00003992 | -0.292588568 | 0.05656555 | 0.17559765 | pgl-1 |
| WBGene00003993 | -0.000323768 | 0.99894074 | 0.99968476 | pgl-2 |
| WBGene00003994 | -0.414355302 | 0.03175552 | 0.11812001 | pgl-3 |
| WBGene00003995 | 0.774459915 | 0.05786744 | 0.17822644 | pgp-1 |
| WBGene00003996 | 0.138112874 | 0.5884866 | 0.75678592 | pgp-2 |
| WBGene00003997 | 0.928043446 | 0.00524757 | 0.03232409 | pgp-3 |
| WBGene00003998 | -0.313847346 | 0.47598795 | 0.66870743 | pgp-4 |
| WBGene00003999 | 1.822789544 | 1.0074E-05 | 0.00022904 | pgp-5 |
| WBGene00004000 | 1.373883095 | 0.01336856 | 0.06389871 | pgp-6 |
| WBGene00004001 | 0.28530093 | 0.92084641 | 1 | pgp-7 |
| WBGene00004002 | 1.665650054 | 0.1016464 | 0.25717336 | pgp-8 |
| WBGene00004003 | 1.484038903 | 7.4669E-06 | 0.00017706 | pgp-9 |
| WBGene00004004 | 0.813422747 | 0.04949641 | 0.15947139 | pgp-10 |
| WBGene00004005 | 1.669471637 | 9.6268E-06 | 0.0002205 | pgp-11 |
| WBGene00004006 | 0.228344069 | 0.69387486 | 0.82632023 | pgp-12 |
| WBGene00004007 | 0.069153844 | 0.93982617 | 0.9689509 | pgp-13 |

| | | | | |
|---|---|---|---|---|
| WBGene00004008 | 1.075601753 | 0.00332501 | 0.02283312 | pgp-14 |
| WBGene00004010 | 1.032739122 | 0.01203331 | 0.0594371 | pha-1 |
| WBGene00004011 | 1.072877033 | 0.41052173 | 1 | pha-2 |
| WBGene00004013 | 0.585676533 | 0.07028616 | 0.20260624 | pha-4 |
| WBGene00004014 | -0.423954431 | 0.00541284 | 0.03304034 | phb-1 |
| WBGene00004015 | -0.41881828 | 0.00256528 | 0.01892167 | phb-2 |
| WBGene00004016 | -0.666930423 | 0.01548776 | 0.07126889 | phf-5 |
| WBGene00004017 | -0.453702271 | 0.47199037 | 0.66486078 | phg-1 |
| WBGene00004020 | 0.86405885 | 6.78E-07 | 2.4622E-05 | pho-1 |
| WBGene00004023 | 1.281635751 | 2.8563E-05 | 0.00053933 | pho-4 |
| WBGene00004024 | 0.788095195 | 0.0782134 | 0.21746833 | php-3 |
| WBGene00004025 | -0.545577747 | 0.07301199 | 0.20813303 | phy-2 |
| WBGene00004026 | -1.10932516 | 0.49911886 | 1 | phy-3 |
| WBGene00004027 | -0.583965191 | 0.00482719 | 0.03041352 | pie-1 |
| WBGene00004028 | -0.542735072 | 0.01852088 | 0.08108981 | pif-1 |
| WBGene00004029 | -0.274926364 | 0.3363012 | 0.54476756 | pik-1 |
| WBGene00004030 | 0.725084524 | 0.26668982 | 0.47511094 | pin-2 |
| WBGene00004031 | 0.396068721 | 0.08238344 | 0.22517275 | pis-1 |
| WBGene00004032 | 0.592916824 | 0.00056664 | 0.00588501 | pkc-1 |
| WBGene00004033 | 0.565994573 | 0.0100164 | 0.05194443 | pkc-2 |
| WBGene00004034 | 0.04330678 | 0.81325495 | 0.89826495 | pkc-3 |
| WBGene00004035 | 2.697061674 | 0.24723036 | 1 | pkd-2 |
| WBGene00004036 | 0.456515182 | 0.34926451 | 0.55765656 | frm-9 |
| WBGene00004036 | 1.027848371 | 0.00314465 | 0.02196735 | plc-1 |
| WBGene00004037 | 0.993568273 | 0.00646765 | 0.03754592 | plc-2 |
| WBGene00004038 | 1.167599074 | 0.00011378 | 0.00169089 | plc-3 |
| WBGene00004039 | 0.606936118 | 0.00295724 | 0.02098417 | plc-4 |
| WBGene00004040 | 0.443812658 | 0.0336806 | 0.1229823 | pld-1 |
| WBGene00004042 | -0.537715324 | 0.00022899 | 0.00292033 | plk-1 |
| WBGene00004043 | -0.526145073 | 0.00861754 | 0.04627403 | plk-2 |
| WBGene00004044 | -0.039000871 | 0.78318776 | 0.8805151 | plk-3 |
| WBGene00004045 | 0.882269668 | 0.01133341 | 0.05690611 | pll-1 |
| WBGene00004046 | -0.535206773 | 0.00019901 | 0.00262339 | plp-1 |
| WBGene00004047 | 0.445325351 | 0.06771661 | 0.19774952 | plx-1 |
| WBGene00004048 | 0.850123065 | 0.01368291 | 0.06499384 | plx-2 |
| WBGene00004049 | 0.088730414 | 0.6528449 | 0.80022109 | parp-1 |
| WBGene00004050 | -0.489959442 | 0.00209805 | 0.01614222 | parp-2 |
| WBGene00004051 | 0.1374642 | 0.44530935 | 0.64463117 | parg-1 |
| WBGene00004052 | 2.200255556 | 0.06633037 | 0.19494482 | parg-2 |
| WBGene00004053 | 0.826140396 | 0.00038829 | 0.00441167 | tank-1 |
| WBGene00004055 | 0.172880104 | 0.52527734 | 0.70985766 | pmk-1 |
| WBGene00004056 | 0.349094951 | 0.37349339 | 0.5812823 | pmk-2 |
| WBGene00004057 | 0.563090526 | 0.07628655 | 0.21436971 | pmk-3 |

| | | | | |
|---|---|---|---|---|
| WBGene00004058 | 1.400800198 | 1.21E-11 | 1.36E-09 | pmp-1 |
| WBGene00004059 | 0.580879432 | 0.00872888 | 0.0467404 | pmp-2 |
| WBGene00004060 | 0.373021695 | 0.03727284 | 0.13111312 | pmp-3 |
| WBGene00004061 | 0.704692099 | 0.05091349 | 0.16271141 | pmp-4 |
| WBGene00004062 | 1.200098356 | 1.61E-15 | 3.26E-13 | pmp-5 |
| WBGene00004063 | 0.097722226 | 0.62603318 | 0.78260593 | pmr-1 |
| WBGene00004064 | 0.343604522 | 0.17082731 | 0.35975964 | pms-2 |
| WBGene00004068 | 0.210119085 | 0.23102 | 0.43640702 | pnk-1 |
| WBGene00004075 | -0.156550308 | 0.33653648 | 0.54486555 | pod-1 |
| WBGene00004076 | 0.786144019 | 0.0001474 | 0.00207843 | pod-2 |
| WBGene00004077 | -0.683333846 | 0.00375345 | 0.02502506 | pop-1 |
| WBGene00004078 | -0.342415086 | 0.02250506 | 0.09326547 | pos-1 |
| WBGene00004083 | 0.700272272 | 0.03635058 | 0.12900683 | pph-1 |
| WBGene00004084 | -0.156269293 | 0.96924949 | 1 | pph-2 |
| WBGene00004085 | -0.286099655 | 0.2678687 | 0.47644118 | pph-4.1 |
| WBGene00004086 | -0.671977571 | 0.00217412 | 0.01660129 | pph-4.2 |
| WBGene00004087 | 0.306637126 | 0.11187414 | 0.27394993 | ppk-1 |
| WBGene00004088 | -0.13961077 | 0.55595596 | 0.73328206 | ppk-2 |
| WBGene00004089 | 0.162444402 | 0.50641992 | 0.69322684 | ppk-3 |
| WBGene00004090 | -0.167958187 | 0.49870751 | 0.68695347 | eif-2Bgamma |
| WBGene00004091 | -0.210596868 | 0.24171658 | 0.44927756 | pps-1 |
| WBGene00004092 | -1.040663402 | 0.05147511 | 0.16381056 | ppt-1 |
| WBGene00004093 | -0.023863085 | 0.90559382 | 0.95088303 | ppw-1 |
| WBGene00004094 | 0.547092888 | 0.05168482 | 0.16413951 | ppw-2 |
| WBGene00004095 | 0.199070745 | 0.27668769 | 0.48544296 | pqe-1 |
| WBGene00004096 | 1.473432776 | 4.19E-12 | 5.25E-10 | pqm-1 |
| WBGene00004098 | -1.901856996 | 0.64072034 | 1 | pqn-2 |
| WBGene00004099 | 0.041803775 | 0.98890197 | 1 | abu-15 |
| WBGene00004100 | 0.940300818 | 0.21200737 | 0.412025 | pqn-8 |
| WBGene00004101 | 0.128189639 | 0.45560401 | 0.65284446 | hgrs-1 |
| WBGene00004102 | -1.604557294 | 0.15553655 | 1 | pqn-10 |
| WBGene00004103 | 0.143798628 | 0.94792951 | 1 | pqn-11 |
| WBGene00004104 | -0.429913232 | 0.40660482 | 0.61346186 | pqn-13 |
| WBGene00004105 | -0.074787394 | 0.70884778 | 0.83525966 | szy-20 |
| WBGene00004106 | 0.284397702 | 0.45826602 | 0.65477918 | pqn-15 |
| WBGene00004107 | 0.041803775 | 0.98890197 | 1 | pqn-16 |
| WBGene00004108 | 1.125197788 | 0.00039044 | 0.00442696 | pqn-18 |
| WBGene00004109 | -0.094752199 | 0.56042817 | 0.73673213 | stam-1 |
| WBGene00004110 | 0.001670252 | 0.99374821 | 0.99678594 | pqn-20 |
| WBGene00004111 | 0.208822519 | 0.37369048 | 0.58147277 | pqn-21 |
| WBGene00004112 | 0.581625842 | 0.00696943 | 0.0396327 | pqn-22 |
| WBGene00004113 | 0.232571534 | 0.45881316 | 0.65487177 | pqn-24 |
| WBGene00004114 | 1.375694503 | 0.00361941 | 0.02442494 | pqn-25 |

| | | | | |
|---|---|---|---|---|
| WBGene00004115 | -0.112382762 | 0.87084074 | 0.93030157 | pqn-26 |
| WBGene00004116 | -0.282968183 | 0.21817457 | 0.42071361 | pqn-27 |
| WBGene00004117 | 0.446329064 | 0.00412295 | 0.02691224 | sin-3 |
| WBGene00004118 | -0.511884107 | 0.61374786 | 1 | pqn-29 |
| WBGene00004119 | 1.826295239 | 9.25E-09 | 5.73E-07 | pqn-31 |
| WBGene00004120 | -0.40910806 | 0.40885567 | 0.61457432 | pqn-32 |
| WBGene00004121 | 0.583665612 | 0.03747613 | 0.13154026 | ptrn-1 |
| WBGene00004122 | 0.545521715 | 0.10208762 | 0.2578955 | pqn-35 |
| WBGene00004123 | 1.000938873 | 0.10673547 | 0.26636697 | pqn-36 |
| WBGene00004124 | 0.882139991 | 0.07615699 | 0.21413202 | pqn-37 |
| WBGene00004125 | -0.102737615 | 0.62904215 | 0.78453054 | mdt-30 |
| WBGene00004126 | 0.149249931 | 0.57986075 | 0.75070264 | pqn-39 |
| WBGene00004127 | -0.233797509 | 0.77008757 | 0.87135445 | abu-12 |
| WBGene00004128 | -0.009109855 | 0.97230464 | 0.98553604 | pqn-41 |
| WBGene00004129 | 1.180870117 | 0.10356033 | 0.26074781 | pqn-42 |
| WBGene00004130 | 1.656792018 | 0.00926157 | 0.04895288 | ketn-1 |
| WBGene00004131 | 1.20040238 | 4.25E-11 | 4.33E-09 | pqn-44 |
| WBGene00004132 | 0.018628664 | 0.9253832 | 0.96044148 | ifet-1 |
| WBGene00004133 | -0.129880392 | 0.79108792 | 0.88470671 | abu-13 |
| WBGene00004134 | 0.948736956 | 0.00227334 | 0.0172074 | myrf-1 |
| WBGene00004135 | 0.816353833 | 0.02262158 | 0.09351331 | pqn-48 |
| WBGene00004136 | -0.391261421 | 0.02305936 | 0.09480655 | pqn-51 |
| WBGene00004138 | 0.248334731 | 0.11609021 | 0.28048782 | pqn-53 |
| WBGene00004139 | 1.555655533 | 0.445513 | 1 | pqn-54 |
| WBGene00004140 | 0.239495512 | 0.41882647 | 0.62346398 | ebax-1 |
| WBGene00004143 | -0.227809044 | 0.1468282 | 0.32655086 | pqn-59 |
| WBGene00004144 | -0.112328776 | 0.7901178 | 0.88437542 | pqn-60 |
| WBGene00004145 | 1.511350264 | 0.00021562 | 0.00278792 | pqn-62 |
| WBGene00004146 | 0.687259521 | 0.81162667 | 1 | pqn-63 |
| WBGene00004147 | 0.009626936 | 0.96054662 | 0.9796557 | larp-5 |
| WBGene00004148 | 0.498816742 | 0.02647763 | 0.10396665 | pqn-65 |
| WBGene00004149 | 0.972820844 | 0.64960696 | 1 | trpl-5 |
| WBGene00004150 | 0.626538234 | 0.51902855 | 1 | pqn-67 |
| WBGene00004151 | -0.454831436 | 0.29735851 | 0.50755372 | pqn-68 |
| WBGene00004152 | -0.671930279 | 0.00021202 | 0.00275443 | pqn-70 |
| WBGene00004153 | -1.184628337 | 0.07292545 | 0.20807825 | pqn-71 |
| WBGene00004154 | 0.006790156 | 0.99096648 | 0.99555847 | pqn-72 |
| WBGene00004155 | -0.08323573 | 0.83709135 | 0.91143626 | pqn-73 |
| WBGene00004156 | -0.20993014 | 0.70523901 | 0.83345073 | pqn-74 |
| WBGene00004157 | 0.954293104 | 0.07597855 | 0.21397718 | pqn-75 |
| WBGene00004161 | 0.185438072 | 0.45854288 | 0.65483658 | pqn-80 |
| WBGene00004163 | -0.290380492 | 0.46782879 | 0.66183982 | pqn-82 |
| WBGene00004164 | -0.293436875 | 0.09165925 | 0.24108209 | pqn-83 |

| | | | | |
|---|---|---|---|---|
| WBGene00004166 | 0.366923025 | 0.08061775 | 0.22209668 | pqn-85 |
| WBGene00004167 | 0.193716764 | 0.28789154 | 0.4978338 | pqn-87 |
| WBGene00004168 | 1.579485857 | 0.0068271 | 0.03909169 | pqn-88 |
| WBGene00004169 | 0.94094351 | 0.00072029 | 0.00713295 | pqn-89 |
| WBGene00004172 | 0.514574896 | 0.05823502 | 0.17884261 | pqn-92 |
| WBGene00004173 | 1.024519647 | 0.00016365 | 0.00224761 | pqn-94 |
| WBGene00004174 | -0.608207948 | 0.05431699 | 0.1701839 | abu-14 |
| WBGene00004175 | -0.477379635 | 0.05455976 | 0.17072422 | pqn-96 |
| WBGene00004176 | 0.642250106 | 0.63591954 | 1 | pqn-97 |
| WBGene00004177 | 0.528572344 | 0.61903194 | 1 | pqn-98 |
| WBGene00004178 | -0.170061382 | 0.34031672 | 0.54870987 | prg-1 |
| WBGene00004179 | 0.201396234 | 0.24378952 | 0.45119643 | prg-2 |
| WBGene00004180 | -0.468988091 | 0.01948249 | 0.08396019 | pri-1 |
| WBGene00004181 | -0.510979668 | 0.00860006 | 0.04622106 | pri-2 |
| WBGene00004182 | 0.702807992 | 0.01018357 | 0.05252284 | prk-1 |
| WBGene00004183 | -0.067865561 | 0.90628656 | 0.95109561 | prk-2 |
| WBGene00004184 | 0.929551444 | 0.00965225 | 0.05046857 | T19D2.2 |
| WBGene00004185 | 0.104444315 | 0.7056255 | 0.83352339 | pro-1 |
| WBGene00004186 | -0.091063312 | 0.56008375 | 0.73664574 | prpf-4 |
| WBGene00004187 | 0.314637459 | 0.13982741 | 0.31713891 | prp-8 |
| WBGene00004188 | -0.624602561 | 0.0011192 | 0.00999868 | prp-21 |
| WBGene00004189 | -0.460892237 | 0.00107543 | 0.00973657 | pars-1 |
| WBGene00004190 | -0.60678453 | 0.02097814 | 0.08862061 | pars-2 |
| WBGene00004191 | 0.962415395 | 0.02733297 | 0.10639615 | prx-1 |
| WBGene00004192 | -0.049900151 | 0.81830322 | 0.9014626 | prx-2 |
| WBGene00004193 | 0.172150345 | 0.62142965 | 0.77957942 | prx-3 |
| WBGene00004194 | 0.325555665 | 0.28809367 | 0.4981124 | prx-5 |
| WBGene00004195 | 0.800102122 | 0.04870977 | 0.15822286 | prx-6 |
| WBGene00004196 | 0.172154488 | 0.58144159 | 0.75233887 | prx-11 |
| WBGene00004197 | 0.204301131 | 0.65664147 | 0.80236354 | prx-12 |
| WBGene00004198 | 0.526648849 | 0.15038283 | 0.33196327 | prx-13 |
| WBGene00004199 | -0.423670459 | 0.0273286 | 0.10639615 | prx-14 |
| WBGene00004201 | -0.182022097 | 0.33887005 | 0.54703115 | prx-19 |
| WBGene00004202 | 0.562018923 | 0.11254571 | 0.27491036 | pry-1 |
| WBGene00004203 | -0.484046391 | 0.02212375 | 0.09239918 | swsn-1 |
| WBGene00004204 | 0.042700062 | 0.79636425 | 0.88760289 | swsn-4 |
| WBGene00004205 | -0.630214825 | 0.00739027 | 0.04130997 | psr-1 |
| WBGene00004206 | 0.179560642 | 0.46446567 | 0.65892471 | pst-1 |
| WBGene00004207 | 0.600284297 | 0.07075484 | 0.20375581 | ptb-1 |
| WBGene00004208 | 0.20575344 | 0.34988986 | 0.55850812 | ptc-1 |
| WBGene00004209 | 0.932931886 | 0.70926986 | 1 | ptc-2 |
| WBGene00004210 | 0.154964874 | 0.53132597 | 0.71493169 | ptc-3 |
| WBGene00004211 | 1.012954158 | 0.52787261 | 1 | ptd-2 |

| | | | | |
|---|---|---|---|---|
| WBGene00004212 | 0.737590418 | 0.04473514 | 0.14843654 | ptl-1 |
| WBGene00004213 | 0.719442927 | 0.01650228 | 0.07432888 | ptp-1 |
| WBGene00004214 | 0.124274331 | 0.48917261 | 0.67890314 | ptp-2 |
| WBGene00004215 | 0.330183184 | 0.07613621 | 0.21413202 | ptp-3 |
| WBGene00004216 | 0.115648431 | 0.83032928 | 0.90819324 | ptr-1 |
| WBGene00004217 | -0.160921188 | 0.37951372 | 0.58756389 | ptr-2 |
| WBGene00004218 | 1.40634393 | 0.06971049 | 0.20175329 | ptr-3 |
| WBGene00004219 | -0.127920502 | 0.73084645 | 0.84935453 | ptr-4 |
| WBGene00004220 | 0.768124649 | 0.0579097 | 0.17827225 | ptr-5 |
| WBGene00004221 | 0.203690887 | 0.67960932 | 0.81647553 | ptr-6 |
| WBGene00004222 | 0.833471181 | 0.01609161 | 0.07317161 | ptr-8 |
| WBGene00004223 | 1.025181888 | 0.79790347 | 1 | ptr-9 |
| WBGene00004224 | 3.03258634 | 0.01964201 | 1 | ptr-10 |
| WBGene00004225 | 0.480046003 | 0.25657831 | 0.46441757 | ptr-11 |
| WBGene00004226 | 1.02344896 | 0.00541052 | 0.03304034 | ptr-12 |
| WBGene00004227 | 0.313832917 | 0.41740731 | 0.62203783 | ptr-13 |
| WBGene00004228 | 0.538252263 | 0.10312518 | 0.26008368 | ptr-14 |
| WBGene00004229 | -0.317578261 | 0.65958134 | 0.80368078 | ptr-15 |
| WBGene00004230 | -0.42221894 | 0.54491417 | 0.7240673 | ptr-16 |
| WBGene00004231 | 0.367375289 | 0.85290706 | 1 | ptr-17 |
| WBGene00004232 | -0.404838174 | 0.32657731 | 0.53515658 | ptr-18 |
| WBGene00004233 | -0.129779465 | 0.88505092 | 0.9392835 | ptr-19 |
| WBGene00004234 | 0.58337399 | 0.14216924 | 0.3202963 | ptr-20 |
| WBGene00004235 | 0.58652567 | 0.01213506 | 0.05974526 | ptr-21 |
| WBGene00004236 | 2.012505705 | 0.03714853 | 0.13095828 | ptr-22 |
| WBGene00004237 | 1.083834871 | 0.00013375 | 0.00193084 | ptr-23 |
| WBGene00004238 | 0.786427524 | 0.00127682 | 0.01107273 | ptr-24 |
| WBGene00004239 | -0.292745092 | 0.0577704 | 0.17810829 | puf-3 |
| WBGene00004241 | -0.353452818 | 0.03275599 | 0.12073394 | puf-5 |
| WBGene00004242 | -0.293741985 | 0.18182553 | 0.37382024 | puf-6 |
| WBGene00004243 | -0.30859307 | 0.15334477 | 0.33605756 | puf-7 |
| WBGene00004244 | -0.155628886 | 0.4005402 | 0.60826713 | puf-8 |
| WBGene00004245 | 0.276517744 | 0.286654 | 0.49685225 | puf-9 |
| WBGene00004248 | 0.054590871 | 0.81095579 | 0.89678607 | pus-1 |
| WBGene00004249 | 0.258890522 | 0.4269431 | 0.63044468 | pvf-1 |
| WBGene00004254 | 0.303280062 | 0.13197165 | 0.30444001 | pxf-1 |
| WBGene00004255 | 0.483677207 | 0.53577724 | 0.7178681 | epac-1 |
| WBGene00004256 | 0.719282678 | 0.01019983 | 0.0525628 | pxn-1 |
| WBGene00004257 | 0.772633261 | 0.08657531 | 0.23254665 | pxn-2 |
| WBGene00004258 | 0.226515841 | 0.2189587 | 0.42169695 | pyc-1 |
| WBGene00004259 | -0.113079126 | 0.64396107 | 0.79428104 | pyr-1 |
| WBGene00004264 | 1.208318798 | 0.0017988 | 0.014349 | qua-1 |
| WBGene00004265 | 0.935828021 | 0.03041085 | 0.11462843 | qui-1 |

| | | | | |
|---|---|---|---|---|
| WBGene00004266 | -0.237797656 | 0.11023912 | 0.27120833 | rab-1 |
| WBGene00004267 | 0.483213306 | 0.06693277 | 0.19624021 | rab-3 |
| WBGene00004268 | -0.023182003 | 0.89770382 | 0.94634211 | rab-5 |
| WBGene00004269 | -0.379528289 | 0.04578662 | 0.15114066 | rab-6.1 |
| WBGene00004270 | -0.445010062 | 0.11825717 | 0.28357895 | rab-6.2 |
| WBGene00004271 | -0.147328451 | 0.47436223 | 0.66742749 | rab-7 |
| WBGene00004272 | -0.457568302 | 0.02962029 | 0.11210187 | rab-8 |
| WBGene00004273 | -0.295135356 | 0.07300638 | 0.20813303 | rab-10 |
| WBGene00004274 | -0.336547582 | 0.02115222 | 0.08907704 | rab-11.1 |
| WBGene00004275 | 1.368756254 | 0.39524415 | 1 | rab-11.2 |
| WBGene00004276 | -0.289046519 | 0.24530076 | 0.45309546 | rab-14 |
| WBGene00004277 | -0.194453484 | 0.42692699 | 0.63044468 | rab-18 |
| WBGene00004278 | -0.469125794 | 0.0612895 | 0.1852878 | rab-19 |
| WBGene00004279 | -0.534043356 | 0.02537012 | 0.10119011 | rab-21 |
| WBGene00004281 | -0.388228939 | 0.57760948 | 0.74906609 | rab-28 |
| WBGene00004282 | -0.747674594 | 0.01487077 | 0.06918667 | rab-30 |
| WBGene00004283 | -0.123520393 | 0.63707451 | 0.78967829 | rab-33 |
| WBGene00004284 | -0.26064388 | 0.24913832 | 0.45684766 | rab-35 |
| WBGene00004285 | 0.479304831 | 0.45681626 | 0.65384393 | rab-37 |
| WBGene00004286 | -0.22005798 | 0.429574 | 0.63219161 | rab-39 |
| WBGene00004287 | -0.156269293 | 0.96924949 | 1 | rac-2 |
| WBGene00004296 | 0.810431143 | 0.00010184 | 0.00154942 | rad-50 |
| WBGene00004297 | -0.400469691 | 0.02578803 | 0.10231361 | rad-51 |
| WBGene00004298 | -0.146540959 | 0.5457431 | 0.72477215 | rad-54 |
| WBGene00004300 | 0.113999328 | 0.63269654 | 0.78722871 | ram-2 |
| WBGene00004301 | 1.242193114 | 0.38564611 | 1 | ram-5 |
| WBGene00004302 | -0.636327005 | 7.62E-07 | 2.6665E-05 | ran-1 |
| WBGene00004303 | -0.118234084 | 0.449029 | 0.6479551 | ran-2 |
| WBGene00004304 | -0.136914482 | 0.41852349 | 0.62320835 | ran-3 |
| WBGene00004305 | -0.351344008 | 0.06497027 | 0.19229743 | ran-4 |
| WBGene00004306 | -0.489334459 | 0.00605892 | 0.03586386 | ran-5 |
| WBGene00004307 | -0.434413405 | 0.0133182 | 0.06379465 | rap-1 |
| WBGene00004308 | -0.182302269 | 0.42564989 | 0.62968384 | rap-2 |
| WBGene00004309 | -0.253586303 | 0.78089761 | 0.8788112 | rap-3 |
| WBGene00004310 | 0.962612215 | 0.00918818 | 0.04860727 | ras-1 |
| WBGene00004311 | 0.311054272 | 0.25828465 | 0.46636436 | ras-2 |
| WBGene00004312 | -0.352417345 | 0.03720681 | 0.13103218 | rba-1 |
| WBGene00004313 | 0.901744973 | 9.4333E-05 | 0.0014627 | rbc-1 |
| WBGene00004314 | 0.067083261 | 0.77129792 | 0.87215499 | rbc-2 |
| WBGene00004315 | -0.131030001 | 0.50834964 | 0.69503074 | rbd-1 |
| WBGene00004316 | 1.137873712 | 0.00106609 | 0.00967868 | rbf-1 |
| WBGene00004317 | 0.102846428 | 0.67961812 | 0.81647553 | rbg-1 |
| WBGene00004318 | 0.25095701 | 0.23806397 | 0.44432509 | rbg-2 |

| | | | | |
|---|---|---|---|---|
| WBGene00004319 | 0.325044545 | 0.09082696 | 0.23962958 | rbr-2 |
| WBGene00004320 | -0.839061472 | 1.4156E-06 | 4.4181E-05 | rbx-1 |
| WBGene00004321 | 1.560784496 | 6.0133E-06 | 0.00014673 | rcan-1 |
| WBGene00004322 | 0.045069772 | 0.8342103 | 0.91023162 | rcq-5 |
| WBGene00004323 | 0.415309284 | 0.03201588 | 0.1186885 | rde-1 |
| WBGene00004324 | 0.08345706 | 0.67492522 | 0.81406091 | rde-2 |
| WBGene00004326 | -0.259262047 | 0.134879 | 0.3093066 | rde-4 |
| WBGene00004328 | 0.658508476 | 0.61162289 | 1 | rdy-2 |
| WBGene00004332 | -0.130318179 | 0.70759694 | 0.83477788 | rec-1 |
| WBGene00004333 | 0.031219136 | 0.8987907 | 0.94694691 | rec-8 |
| WBGene00004334 | -1.661583473 | 0.00040644 | 0.00456479 | ref-1 |
| WBGene00004335 | -0.319045135 | 0.56458195 | 0.73975712 | ref-2 |
| WBGene00004336 | -0.08194798 | 0.58881204 | 0.75696389 | ret-1 |
| WBGene00004337 | -0.265508564 | 0.09347342 | 0.24402558 | rfc-1 |
| WBGene00004338 | -0.720262532 | 0.00025334 | 0.00316442 | rfc-2 |
| WBGene00004339 | -0.538115788 | 0.00394084 | 0.02594674 | rfc-3 |
| WBGene00004340 | -0.618152328 | 0.00022737 | 0.00290705 | rfc-4 |
| WBGene00004341 | -0.389250384 | 0.03919061 | 0.13548213 | rfl-1 |
| WBGene00004342 | 0.417597269 | 0.3172249 | 0.52493219 | rfs-1 |
| WBGene00004343 | 0.076800818 | 0.70230115 | 0.83202949 | rgr-1 |
| WBGene00004344 | -0.010932558 | 0.98426945 | 0.99236135 | rgs-1 |
| WBGene00004345 | 1.174664193 | 0.01788834 | 0.07903286 | rgs-2 |
| WBGene00004346 | 0.979380106 | 0.0695362 | 0.20148901 | rgs-3 |
| WBGene00004347 | 1.639141671 | 0.06896848 | 0.20022628 | Y38E10A.20 |
| WBGene00004347 | 0.780288667 | 0.09106025 | 0.2400266 | rgs-4 |
| WBGene00004348 | 0.118979335 | 0.72870262 | 0.84808076 | rgs-5 |
| WBGene00004349 | 0.739556643 | 0.03457595 | 0.1251227 | rgs-6 |
| WBGene00004350 | -0.043670372 | 0.85951809 | 0.92397814 | rgs-7 |
| WBGene00004352 | -0.712869062 | 0.0128104 | 0.06221244 | rgs-9 |
| WBGene00004353 | -0.394130576 | 0.65723553 | 0.80268551 | rgs-10 |
| WBGene00004354 | -0.477871299 | 0.1074019 | 0.26742544 | rgs-11 |
| WBGene00004355 | 0.092374076 | 0.61020035 | 0.77201762 | rha-1 |
| WBGene00004356 | -0.281667592 | 0.23383971 | 0.43988967 | rhi-1 |
| WBGene00004357 | 0.107001667 | 0.56142398 | 0.73709014 | rho-1 |
| WBGene00004358 | 0.532827701 | 0.00098344 | 0.00913536 | rhr-1 |
| WBGene00004359 | 0.075034619 | 0.91467921 | 0.95571968 | rhr-2 |
| WBGene00004360 | -0.235578605 | 0.37937415 | 0.58747827 | rib-1 |
| WBGene00004361 | -0.298950669 | 0.2362716 | 0.44244693 | rib-2 |
| WBGene00004363 | 0.203706915 | 0.58051883 | 0.75131426 | ric-3 |
| WBGene00004364 | 0.50793246 | 0.05166829 | 0.16412997 | ric-4 |
| WBGene00004367 | 0.003777927 | 0.98820131 | 0.994756 | ric-8 |
| WBGene00004368 | 1.120827571 | 0.0546559 | 0.17089294 | ric-19 |
| WBGene00004369 | 1.052612062 | 0.00267447 | 0.01953718 | rig-1 |

| | | | | |
|---|---|---|---|---|
| WBGene00004370 | 0.890139785 | 0.00472231 | 0.02998555 | rig-3 |
| WBGene00004371 | 0.096438048 | 0.56835595 | 0.74321237 | rig-4 |
| WBGene00004372 | 0.303970234 | 0.63460623 | 0.78788774 | rig-5 |
| WBGene00004373 | -0.072338001 | 0.66672593 | 0.80890254 | rme-1 |
| WBGene00004374 | -0.47018654 | 0.00547483 | 0.03335168 | rme-2 |
| WBGene00004375 | 0.142953036 | 0.61355443 | 0.77426092 | rme-4 |
| WBGene00004377 | 0.171655883 | 0.52817568 | 0.71250899 | rme-6 |
| WBGene00004378 | 0.386635031 | 0.04991779 | 0.16047296 | rme-8 |
| WBGene00004380 | -0.512089442 | 0.02407737 | 0.09763639 | rnf-1 |
| WBGene00004381 | -0.630424011 | 0.0029965 | 0.02118839 | rnf-5 |
| WBGene00004382 | -0.046601496 | 0.84652548 | 0.91591915 | rnh-1.0 |
| WBGene00004383 | -0.409919453 | 0.03920393 | 0.1354896 | rnh-2 |
| WBGene00004384 | -0.239759103 | 0.1806717 | 0.37259848 | rnp-1 |
| WBGene00004385 | -0.785397622 | 0.0010916 | 0.00981195 | rnp-2 |
| WBGene00004386 | -0.381508415 | 0.08793767 | 0.23501017 | rnp-3 |
| WBGene00004387 | -0.79168719 | 1.3522E-05 | 0.00029161 | rnp-4 |
| WBGene00004388 | -0.580832852 | 0.00436 | 0.02812683 | rnp-5 |
| WBGene00004389 | -0.414364655 | 0.01220082 | 0.059996 | rnp-6 |
| WBGene00004390 | -0.387464801 | 0.03884647 | 0.13475287 | rnp-7 |
| WBGene00004391 | -0.404395957 | 0.01224414 | 0.06011166 | rnr-1 |
| WBGene00004392 | -0.858563106 | 5.86E-12 | 7.18E-10 | rnr-2 |
| WBGene00004393 | 1.300278319 | 0.2213834 | 0.42434731 | rnt-1 |
| WBGene00004394 | 1.880100277 | 0.56749367 | 1 | rol-1 |
| WBGene00004395 | 2.603586522 | 1.885E-05 | 0.0003827 | rol-3 |
| WBGene00004397 | -0.445252015 | 0.61489634 | 0.77514863 | rol-6 |
| WBGene00004398 | -1.552550942 | 0.09545977 | 0.24772303 | rol-8 |
| WBGene00004400 | 0.011470786 | 0.97699667 | 0.98807881 | rom-1 |
| WBGene00004401 | 1.023080089 | 0.241571 | 0.44921328 | rom-2 |
| WBGene00004402 | 2.918540233 | 0.18670406 | 1 | rom-3 |
| WBGene00004403 | 0.24628263 | 0.32435597 | 0.53273888 | rom-4 |
| WBGene00004404 | -0.088930849 | 0.74038036 | 0.85519532 | rom-5 |
| WBGene00004405 | 0.06396058 | 0.72626131 | 0.84605053 | rop-1 |
| WBGene00004408 | -0.310396898 | 0.0131671 | 0.06338691 | rla-0 |
| WBGene00004409 | -0.520980704 | 2.0518E-05 | 0.0004104 | rla-1 |
| WBGene00004410 | -0.687989363 | 1.1683E-06 | 3.8277E-05 | rla-2 |
| WBGene00004411 | 0.060862063 | 0.74163814 | 0.85569007 | rpc-1 |
| WBGene00004412 | -0.398113691 | 0.00136245 | 0.01165715 | rpl-1 |
| WBGene00004413 | -0.318396186 | 0.01218797 | 0.05998139 | rpl-2 |
| WBGene00004414 | -0.059848876 | 0.70399938 | 0.83266549 | rpl-3 |
| WBGene00004415 | -0.233294754 | 0.09786274 | 0.25151828 | rpl-4 |
| WBGene00004416 | 0.03532572 | 0.79038456 | 0.8845213 | rpl-5 |
| WBGene00004417 | -0.280052571 | 0.04256413 | 0.14374718 | rpl-6 |
| WBGene00004418 | -0.001959818 | 0.98927393 | 0.99484196 | rpl-7 |

| | | | | |
|---|---|---|---|---|
| WBGene00004419 | -0.00491759 | 0.96965824 | 0.98437435 | rpl-7A |
| WBGene00004420 | -0.19575758 | 0.11771365 | 0.28277829 | rpl-9 |
| WBGene00004421 | -0.391455958 | 0.00143814 | 0.01213378 | rpl-10 |
| WBGene00004422 | -0.712968722 | 2.62E-08 | 1.4127E-06 | rpl-11.1 |
| WBGene00004423 | 0.483065477 | 0.00142494 | 0.01205586 | rpl-11.2 |
| WBGene00004424 | -0.303446177 | 0.02957693 | 0.11207609 | rpl-12 |
| WBGene00004425 | -0.379898741 | 0.00531183 | 0.03258762 | rpl-13 |
| WBGene00004426 | -0.202517243 | 0.12939992 | 0.30073592 | rpl-14 |
| WBGene00004427 | -0.46433053 | 0.00060358 | 0.0061684 | rpl-15 |
| WBGene00004428 | -0.403655397 | 0.00080147 | 0.00774734 | rpl-16 |
| WBGene00004429 | -0.380182495 | 0.00227565 | 0.01721411 | rpl-17 |
| WBGene00004430 | -0.292044287 | 0.03365125 | 0.1229121 | rpl-18 |
| WBGene00004431 | -0.247990384 | 0.04045315 | 0.13858398 | rpl-19 |
| WBGene00004432 | -0.377789426 | 0.00417358 | 0.02718215 | rpl-20 |
| WBGene00004433 | -0.325547815 | 0.00763051 | 0.04232164 | rpl-21 |
| WBGene00004434 | -0.530457811 | 4.764E-05 | 0.00082838 | rpl-22 |
| WBGene00004435 | -0.555455956 | 1.7729E-05 | 0.00036606 | rpl-23 |
| WBGene00004436 | -0.319606659 | 0.01952917 | 0.08404952 | rpl-24.1 |
| WBGene00004437 | 0.021864317 | 0.91101708 | 0.9539162 | rpl-24.2 |
| WBGene00004438 | 0.25518084 | 0.12696847 | 0.2974767 | rpl-25.1 |
| WBGene00004439 | -0.602041982 | 7.29E-07 | 2.58E-05 | rpl-25.2 |
| WBGene00004440 | -0.440791969 | 0.00115064 | 0.01022685 | rpl-26 |
| WBGene00004441 | -0.373161714 | 0.0054019 | 0.0330234 | rpl-27 |
| WBGene00004442 | -0.54297088 | 0.00011772 | 0.00173663 | rpl-28 |
| WBGene00004443 | -0.508952893 | 0.00095341 | 0.00888359 | rpl-29 |
| WBGene00004444 | -0.263415036 | 0.03669786 | 0.12978406 | rpl-30 |
| WBGene00004445 | -0.343347013 | 0.01157002 | 0.05775973 | rpl-31 |
| WBGene00004446 | -0.50567442 | 2.6283E-05 | 0.00050812 | rpl-32 |
| WBGene00004447 | -0.197810533 | 0.11344119 | 0.27656416 | rpl-33 |
| WBGene00004448 | -0.402047057 | 0.00213434 | 0.01637995 | rpl-34 |
| WBGene00004449 | -0.316420316 | 0.01030756 | 0.05302716 | rpl-35 |
| WBGene00004450 | -0.415886595 | 0.00182968 | 0.01454759 | rpl-36 |
| WBGene00004451 | -0.64144198 | 0.01169238 | 0.05817917 | rpl-37 |
| WBGene00004452 | -0.171803435 | 0.17432302 | 0.3644752 | rpl-38 |
| WBGene00004453 | -0.531430084 | 0.00123141 | 0.01074804 | rpl-39 |
| WBGene00004454 | -0.481680873 | 0.00014438 | 0.0020526 | rpl-36.A |
| WBGene00004455 | 0.747845955 | 0.85215965 | 1 | rpl-42 |
| WBGene00004456 | -0.497545294 | 0.00013928 | 0.00199182 | rpl-43 |
| WBGene00004457 | 1.325801526 | 0.00107382 | 0.00972929 | rpm-1 |
| WBGene00004458 | -0.072719201 | 0.65578848 | 0.8022265 | rpn-1 |
| WBGene00004459 | -0.198820253 | 0.26331483 | 0.47200729 | rpn-2 |
| WBGene00004460 | -0.132642444 | 0.33403224 | 0.54224969 | rpn-3 |
| WBGene00004461 | -0.395458385 | 0.00380135 | 0.02523355 | rpn-5 |

| | | | | |
|---|---|---|---|---|
| WBGene00004462 | -0.209935602 | 0.17536551 | 0.36582693 | rpn-6.1 |
| WBGene00004463 | -0.44011112 | 0.00079399 | 0.0076995 | rpn-7 |
| WBGene00004464 | -0.356427458 | 0.01390219 | 0.06582937 | rpn-8 |
| WBGene00004465 | -0.678535524 | 1.2296E-06 | 3.9703E-05 | rpn-9 |
| WBGene00004466 | -0.517465718 | 0.00065414 | 0.00660173 | rpn-10 |
| WBGene00004467 | -0.619486444 | 0.00020176 | 0.00264604 | rpn-11 |
| WBGene00004468 | -0.520818487 | 0.00133375 | 0.0114439 | rpn-12 |
| WBGene00004469 | -0.279645606 | 0.02250785 | 0.09326547 | rps-0 |
| WBGene00004470 | -0.131005395 | 0.30183797 | 0.51131782 | rps-1 |
| WBGene00004471 | -0.505977021 | 6.8498E-05 | 0.00113302 | rps-2 |
| WBGene00004472 | -0.21118991 | 0.11087401 | 0.27238394 | rps-3 |
| WBGene00004473 | -0.273450038 | 0.04251934 | 0.14367585 | rps-4 |
| WBGene00004474 | -0.390105653 | 0.00454255 | 0.02908812 | rps-5 |
| WBGene00004475 | -0.392520987 | 0.00257081 | 0.01895092 | rps-6 |
| WBGene00004476 | -0.284812756 | 0.04030785 | 0.13824225 | rps-7 |
| WBGene00004477 | -0.138952579 | 0.25032613 | 0.45833352 | rps-8 |
| WBGene00004478 | -0.373416837 | 0.00262561 | 0.01924973 | rps-9 |
| WBGene00004479 | -0.445901044 | 0.00064782 | 0.00654891 | rps-10 |
| WBGene00004480 | -0.351808779 | 0.00460992 | 0.02944188 | rps-11 |
| WBGene00004481 | -0.530484522 | 0.0001136 | 0.00169022 | rps-12 |
| WBGene00004482 | -0.297768549 | 0.04027702 | 0.13817557 | rps-13 |
| WBGene00004483 | -0.262898091 | 0.05345955 | 0.1680612 | rps-14 |
| WBGene00004484 | -0.483771245 | 0.00027637 | 0.00338931 | rps-15 |
| WBGene00004485 | -0.420683482 | 0.00080263 | 0.00775234 | rps-16 |
| WBGene00004486 | -0.503587304 | 0.0001977 | 0.00261751 | rps-17 |
| WBGene00004487 | -0.248400752 | 0.05207125 | 0.16493531 | rps-18 |
| WBGene00004488 | -0.400192228 | 0.00323392 | 0.02240559 | rps-19 |
| WBGene00004489 | -0.16650066 | 0.17822258 | 0.37026016 | rps-20 |
| WBGene00004490 | -0.402143408 | 0.00128023 | 0.0110944 | rps-21 |
| WBGene00004491 | -0.314210279 | 0.01770576 | 0.07837169 | rps-22 |
| WBGene00004492 | -0.400465795 | 0.00170403 | 0.01374658 | rps-23 |
| WBGene00004493 | -0.451161804 | 0.00089078 | 0.00840325 | rps-24 |
| WBGene00004494 | -0.382130391 | 0.00166446 | 0.01354434 | rps-25 |
| WBGene00004495 | -0.33852814 | 0.00565162 | 0.03415444 | rps-26 |
| WBGene00004496 | -0.453583856 | 0.00183916 | 0.01461337 | rps-27 |
| WBGene00004497 | -0.422384988 | 0.00156401 | 0.01294256 | rps-28 |
| WBGene00004498 | -0.591632968 | 6.2898E-05 | 0.00105331 | rps-29 |
| WBGene00004499 | -0.484605085 | 0.00040186 | 0.00452176 | rps-30 |
| WBGene00004501 | -0.398767356 | 0.00340326 | 0.02321083 | rpt-1 |
| WBGene00004502 | -0.341683807 | 0.02705506 | 0.10556882 | rpt-2 |
| WBGene00004503 | -0.515254327 | 0.00025805 | 0.00320341 | rpt-3 |
| WBGene00004504 | -0.417101768 | 0.00247986 | 0.01848341 | rpt-4 |
| WBGene00004505 | -0.500994967 | 0.00078234 | 0.00761385 | rpt-5 |

| | | | | |
|---|---|---|---|---|
| WBGene00004506 | -0.397037468 | 0.01088326 | 0.05535553 | rpt-6 |
| WBGene00004507 | 0.807822989 | 0.01070796 | 0.05460115 | rpy-1 |
| WBGene00004508 | 0.144789936 | 0.42245745 | 0.62710061 | rrf-1 |
| WBGene00004509 | 0.53701726 | 0.19460297 | 0.39059777 | rrf-2 |
| WBGene00004510 | 0.115406855 | 0.53335245 | 0.71646737 | rrf-3 |
| WBGene00004512 | -0.058401084 | 0.97750667 | 0.98825319 | rrn-1.1 |
| WBGene00004513 | -0.058401084 | 0.97750667 | 0.98825319 | rrn-1.2 |
| WBGene00004567 | -0.93027164 | NaN | 1 | rrn-2.1 |
| WBGene00004622 | 0.510070289 | NaN | 1 | rrn-3.1 |
| WBGene00004679 | -0.076827625 | 0.65042815 | 0.79854871 | rars-1 |
| WBGene00004680 | -0.365459499 | 0.06141878 | 0.18554004 | rars-2 |
| WBGene00004681 | 0.198438303 | 0.3134181 | 0.52133294 | rsd-2 |
| WBGene00004682 | 0.258137566 | 0.32268519 | 0.53107238 | rsd-3 |
| WBGene00004684 | -0.039818525 | 0.82647135 | 0.90626704 | rsd-6 |
| WBGene00004698 | -0.664216799 | 2.2667E-05 | 0.00044895 | rsp-1 |
| WBGene00004699 | -0.518696203 | 0.0018654 | 0.01478316 | rsp-2 |
| WBGene00004700 | -1.075125558 | 2.00E-11 | 2.15E-09 | rsp-3 |
| WBGene00004701 | -0.781871255 | 5.4702E-06 | 0.00013582 | rsp-4 |
| WBGene00004702 | -0.65670419 | 0.00038257 | 0.00437366 | rsp-5 |
| WBGene00004703 | -0.89232551 | 1.00E-08 | 6.11E-07 | rsp-6 |
| WBGene00004704 | -0.54688776 | 0.0012135 | 0.01063766 | rsp-7 |
| WBGene00004705 | -0.482931102 | 0.00629958 | 0.03696903 | rsp-8 |
| WBGene00004706 | -0.599647598 | 0.01884795 | 0.08215101 | rsr-1 |
| WBGene00004719 | 0.765881845 | 0.04235638 | 0.14336459 | sad-1 |
| WBGene00004721 | -0.517871428 | 0.01266626 | 0.06166041 | san-1 |
| WBGene00004723 | -0.101728814 | 0.63338807 | 0.78758343 | sap-49 |
| WBGene00004726 | 0.018191035 | 0.93219557 | 0.96406579 | sas-4 |
| WBGene00004727 | 0.025193221 | 0.92763342 | 0.96200873 | sax-1 |
| WBGene00004728 | 0.269398548 | 0.1293296 | 0.30068761 | sax-2 |
| WBGene00004729 | 0.09395348 | 0.66848429 | 0.8101485 | sax-3 |
| WBGene00004732 | 0.337235402 | 0.11231988 | 0.27460243 | sax-7 |
| WBGene00004735 | 0.347411292 | 0.20032692 | 0.39884703 | sbp-1 |
| WBGene00004736 | 0.564480422 | 0.00215608 | 0.01648427 | sca-1 |
| WBGene00004737 | -0.281696111 | 0.26444765 | 0.47312981 | scc-1 |
| WBGene00004738 | 0.219680657 | 0.24716782 | 0.45509016 | scc-3 |
| WBGene00004739 | 1.221939521 | 0.39841193 | 1 | C16D6.1 |
| WBGene00004739 | 1.11830908 | 3.3302E-06 | 9.0161E-05 | scd-1 |
| WBGene00004740 | 1.28999501 | 0.00160737 | 0.01321267 | scd-2 |
| WBGene00004742 | -1.901856996 | 0.64072034 | 1 | scl-1 |
| WBGene00004743 | -0.04560104 | 0.84078585 | 0.9124319 | scm-1 |
| WBGene00004744 | 0.457186176 | 0.03916015 | 0.13545069 | scp-1 |
| WBGene00004745 | -0.414081778 | 0.09832799 | 0.25233568 | sdc-1 |
| WBGene00004746 | 0.460215853 | 0.11749305 | 0.28245932 | sdc-2 |

| | | | | |
|---|---|---|---|---|
| WBGene00004747 | 0.435458505 | 0.10408093 | 0.26178715 | sdc-3 |
| WBGene00004748 | 1.004470897 | 0.58499715 | 1 | sdf-9 |
| WBGene00004749 | -0.095018293 | 0.64712992 | 0.79635155 | sdn-1 |
| WBGene00004750 | -1.276556426 | 2.9728E-06 | 8.218E-05 | sea-1 |
| WBGene00004751 | 0.72605698 | 0.00402337 | 0.02640207 | sea-2 |
| WBGene00004752 | 0.142470761 | 0.44893473 | 0.6479551 | sec-5 |
| WBGene00004753 | 0.50039716 | 0.01986274 | 0.08507334 | sec-8 |
| WBGene00004754 | 0.016506022 | 0.9392765 | 0.96880629 | sec-23 |
| WBGene00004755 | 0.213275944 | 0.23285491 | 0.43851271 | sec-24.1 |
| WBGene00004756 | 0.431000692 | 0.03047475 | 0.11483361 | sec-24.2 |
| WBGene00004758 | 0.368097827 | 0.1680962 | 0.35573574 | sek-1 |
| WBGene00004759 | -0.111485503 | 0.48299063 | 0.67409924 | sel-1 |
| WBGene00004760 | 0.583087754 | 0.01758846 | 0.07810589 | sel-2 |
| WBGene00004762 | 0.101582407 | 0.57664032 | 0.74903703 | sel-5 |
| WBGene00004764 | -0.470268562 | 0.51862211 | 0.7042379 | sel-7 |
| WBGene00004765 | 0.144343938 | 0.60648266 | 0.76957629 | sel-8 |
| WBGene00004766 | -0.617069657 | 0.00038677 | 0.00440757 | sel-9 |
| WBGene00004767 | -0.334686346 | 0.06444601 | 0.19135217 | sel-10 |
| WBGene00004768 | -0.090787594 | 0.60721523 | 0.77002299 | sel-11 |
| WBGene00004769 | -0.553041677 | 0.05784227 | 0.17819413 | sel-12 |
| WBGene00004771 | -0.141046516 | 0.67571921 | 0.81451193 | sem-2 |
| WBGene00004773 | 1.125302786 | 4.5944E-05 | 0.00080492 | sem-4 |
| WBGene00004774 | -0.242752058 | 0.3365627 | 0.54486555 | sem-5 |
| WBGene00004776 | 1.211135809 | 0.08602719 | 0.23158671 | ser-1 |
| WBGene00004777 | 0.596678212 | 0.37666238 | 0.58441636 | ser-2 |
| WBGene00004778 | 1.108345567 | 0.02303321 | 0.0947632 | ser-3 |
| WBGene00004779 | 0.426921872 | 0.46003583 | 0.65578196 | ser-4 |
| WBGene00004780 | 0.474715976 | 0.45983108 | 0.65564231 | ser-7 |
| WBGene00004781 | -0.765903883 | 1.1815E-05 | 0.00026082 | set-1 |
| WBGene00004782 | 0.243879268 | 0.30243447 | 0.51175705 | set-2 |
| WBGene00004783 | -0.074461052 | 0.72165015 | 0.84318685 | seu-1 |
| WBGene00004786 | -0.723446895 | 0.00409309 | 0.02674606 | sex-1 |
| WBGene00004787 | 0.090507132 | 0.82905392 | 0.90762341 | sft-1 |
| WBGene00004788 | -0.294747513 | 0.09909661 | 0.25339764 | sft-4 |
| WBGene00004789 | 1.001374338 | 0.00019298 | 0.00257038 | sgk-1 |
| WBGene00004790 | 0.216845799 | 0.63738391 | 0.78988242 | sgn-1 |
| WBGene00004793 | 0.694777541 | 0.24127717 | 0.44894193 | shw-3 |
| WBGene00004795 | 0.58259935 | 0.01410353 | 0.06665279 | sid-1 |
| WBGene00004796 | 0.127970232 | 0.71141354 | 0.83670171 | sid-2 |
| WBGene00004798 | -0.434933136 | 0.00162518 | 0.01331395 | sip-1 |
| WBGene00004800 | -0.185008879 | 0.38186731 | 0.59007904 | sir-2.1 |
| WBGene00004801 | 0.43400549 | 0.16266188 | 0.34857801 | sir-2.2 |
| WBGene00004802 | 0.035600533 | 0.94703679 | 0.97304242 | sir-2.3 |

| | | | | |
|---|---|---|---|---|
| WBGene00004803 | -0.345035471 | 0.25954485 | 0.46780493 | sir-2.4 |
| WBGene00004804 | 0.343264738 | 0.08519177 | 0.23019943 | skn-1 |
| WBGene00004806 | -0.390374175 | 0.01121358 | 0.05653826 | skp-1 |
| WBGene00004807 | -0.080421996 | 0.60782902 | 0.77036604 | skr-1 |
| WBGene00004808 | -0.673194664 | 6.4962E-05 | 0.00108201 | skr-2 |
| WBGene00004809 | 0.207702083 | 0.39574664 | 0.60353725 | skr-3 |
| WBGene00004810 | -0.068216989 | 0.8335735 | 0.90998018 | skr-4 |
| WBGene00004811 | 2.858884662 | 0.00767832 | 0.04245474 | skr-5 |
| WBGene00004813 | -1.677541958 | 1.11E-13 | 1.77E-11 | skr-7 |
| WBGene00004814 | -2.332479414 | 3.25E-13 | 4.75E-11 | skr-8 |
| WBGene00004815 | -2.174527291 | 2.55E-08 | 1.39E-06 | skr-9 |
| WBGene00004816 | -2.23940903 | 6.82E-13 | 9.85E-11 | skr-10 |
| WBGene00004818 | -2.403613759 | 7.19E-10 | 6.02E-08 | skr-12 |
| WBGene00004819 | -1.562989366 | 7.6996E-05 | 0.00123653 | skr-13 |
| WBGene00004820 | -1.740604866 | 7.24E-10 | 6.02E-08 | skr-14 |
| WBGene00004821 | -1.372973452 | 0.00121013 | 0.01061573 | skr-15 |
| WBGene00004822 | -0.382103532 | 0.31586004 | 0.52395912 | skr-16 |
| WBGene00004823 | -0.708548581 | 0.00347465 | 0.02364673 | skr-17 |
| WBGene00004824 | -0.169700037 | 0.45520889 | 0.65257895 | F56B3.4 |
| WBGene00004825 | -0.151302837 | 0.63394328 | 0.78783996 | skr-19 |
| WBGene00004826 | -0.27971624 | 0.51682031 | 0.70281311 | skr-20 |
| WBGene00004827 | -1.326216174 | 0.00179422 | 0.01432188 | skr-21 |
| WBGene00004829 | -0.158645273 | 0.53580156 | 0.7178681 | sli-1 |
| WBGene00004830 | 0.568553443 | 0.13121621 | 0.30367823 | slo-1 |
| WBGene00004831 | 0.997232692 | 0.00027709 | 0.00339446 | slo-2 |
| WBGene00004833 | 0.377828795 | 0.92526585 | 1 | sls-2.1 |
| WBGene00004834 | 0.4828626 | 0.81641781 | 1 | sls-2.2 |
| WBGene00004835 | -0.18557176 | 0.95664217 | 1 | sls-2.3 |
| WBGene00004836 | 0.12332541 | 0.94831365 | 1 | sls-2.4 |
| WBGene00004837 | -0.340763753 | 0.79892838 | 1 | sls-2.5 |
| WBGene00004838 | -0.156269293 | 0.96924949 | 1 | sls-2.6 |
| WBGene00004839 | -0.156269293 | 0.96924949 | 1 | sls-2.7 |
| WBGene00004840 | -1.443360033 | 0.37063936 | 1 | sls-2.8 |
| WBGene00004841 | -1.159714244 | 0.75790634 | 1 | sls-2.9 |
| WBGene00004842 | 0.041803775 | 0.98890197 | 1 | sls-2.10 |
| WBGene00004843 | -1.690101699 | 0.3296611 | 1 | sls-2.11 |
| WBGene00004844 | -0.156269293 | 0.96924949 | 1 | sls-2.12 |
| WBGene00004846 | 0.377828795 | 0.92526585 | 1 | sls-2.14 |
| WBGene00004847 | 0.233739675 | 0.93575889 | 1 | sls-2.15 |
| WBGene00004848 | -0.337470324 | 0.76472132 | 1 | sls-2.16 |
| WBGene00004849 | -0.156269293 | 0.96924949 | 1 | sls-2.17 |
| WBGene00004850 | 0.688793783 | 0.86378755 | 1 | sls-2.18 |
| WBGene00004854 | 1.275309759 | 0.0001739 | 0.00235374 | slt-1 |

| | | | | |
|---|---|---|---|---|
| WBGene00004855 | 0.62451163 | 0.0051867 | 0.0320467 | sma-1 |
| WBGene00004856 | 0.061490292 | 0.88832113 | 0.94102668 | sma-2 |
| WBGene00004857 | 0.527104015 | 0.19098955 | 0.38569596 | sma-3 |
| WBGene00004858 | 1.289665219 | 5.9083E-06 | 0.00014491 | sma-4 |
| WBGene00004859 | 1.00168795 | 0.00030195 | 0.00361531 | sma-5 |
| WBGene00004860 | 0.397424649 | 0.21670278 | 0.41848075 | sma-6 |
| WBGene00004862 | 1.061414267 | 1.4435E-06 | 4.4823E-05 | sma-9 |
| WBGene00004873 | -0.05046932 | 0.78628136 | 0.88223764 | smc-3 |
| WBGene00004874 | 0.204638583 | 0.23239007 | 0.437977 | smc-4 |
| WBGene00004875 | 0.284473242 | 0.27195978 | 0.48056732 | smd-1 |
| WBGene00004876 | -0.439895255 | 0.2067576 | 0.40655714 | smf-1 |
| WBGene00004877 | -1.964653846 | 0.53599202 | 1 | smf-2 |
| WBGene00004878 | 1.262272695 | 0.05461052 | 0.17079681 | smf-3 |
| WBGene00004879 | 0.852392742 | 5.5213E-06 | 0.00013637 | smg-1 |
| WBGene00004880 | 0.438753966 | 0.07855173 | 0.2182372 | smg-2 |
| WBGene00004881 | 0.174602143 | 0.31975477 | 0.52791721 | smg-3 |
| WBGene00004882 | -0.299076656 | 0.12424187 | 0.29363841 | smg-4 |
| WBGene00004883 | -0.093289627 | 0.71406155 | 0.83835424 | smg-5 |
| WBGene00004884 | -0.090965352 | 0.64711107 | 0.79635155 | smg-6 |
| WBGene00004885 | -0.042430001 | 0.86960305 | 0.9297975 | smg-7 |
| WBGene00004886 | -0.661179327 | 0.06641953 | 0.19515961 | smi-1 |
| WBGene00004887 | -0.787519106 | 0.00138194 | 0.01175936 | smn-1 |
| WBGene00004888 | -0.668106517 | 5.5264E-06 | 0.00013637 | smo-1 |
| WBGene00004889 | 0.373566342 | 0.10990946 | 0.27056179 | smp-1 |
| WBGene00004890 | 0.808656653 | 0.006345 | 0.0371585 | smp-2 |
| WBGene00004891 | -0.827035347 | 0.00023306 | 0.00295673 | smr-1 |
| WBGene00004892 | -0.210276001 | 0.29119889 | 0.50121253 | sms-1 |
| WBGene00004893 | -0.437122263 | 0.21003925 | 0.41014101 | sms-2 |
| WBGene00004894 | -0.17411402 | 0.65059297 | 0.79859638 | sms-3 |
| WBGene00004895 | -0.096547947 | 0.6303143 | 0.78556152 | smu-1 |
| WBGene00004896 | -0.343635823 | 0.05557582 | 0.17318943 | smu-2 |
| WBGene00004897 | -0.190346207 | 0.27505992 | 0.4835712 | snb-1 |
| WBGene00004898 | 0.606680726 | 0.10627798 | 0.26571684 | snb-2 |
| WBGene00004899 | 0.281454817 | 0.94435481 | 1 | snb-5 |
| WBGene00004900 | 0.327626249 | 0.15300667 | 0.33549829 | snf-1 |
| WBGene00004901 | 0.747845955 | 0.85215965 | 1 | snf-2 |
| WBGene00004902 | 0.561971616 | 0.04024669 | 0.13817557 | snf-3 |
| WBGene00004904 | 1.051217365 | 0.47132162 | 1 | snf-5 |
| WBGene00004905 | 0.031517257 | 0.89463259 | 0.94453253 | snf-6 |
| WBGene00004906 | 0.03940405 | 0.99001443 | 1 | snf-7 |
| WBGene00004908 | 0.289691386 | 0.55491192 | 0.73238519 | snf-9 |
| WBGene00004909 | -0.356102232 | 0.88046231 | 1 | snf-10 |
| WBGene00004910 | 0.722000009 | 0.01459313 | 0.06832302 | snf-11 |

| | | | | |
|---|---|---|---|---|
| WBGene00004911 | 0.305834269 | 0.63897012 | 0.7907988 | snf-12 |
| WBGene00004912 | 0.406895288 | 0.40305004 | 0.61082973 | sng-1 |
| WBGene00004913 | 0.808774586 | 0.12733701 | 0.29788027 | snn-1 |
| WBGene00004914 | -0.756261832 | 4.747E-06 | 0.00012007 | snr-1 |
| WBGene00004915 | -0.768472112 | 7.67E-07 | 2.6736E-05 | snr-2 |
| WBGene00004916 | -0.953191335 | 1.06E-07 | 4.9607E-06 | snr-3 |
| WBGene00004917 | -0.454114006 | 0.00258176 | 0.01900857 | snr-4 |
| WBGene00004918 | -0.819229794 | 5.2654E-06 | 0.00013208 | snr-5 |
| WBGene00004919 | -0.645757461 | 0.00010158 | 0.00154743 | snr-6 |
| WBGene00004920 | -0.999005866 | 1.30E-07 | 5.867E-06 | snr-7 |
| WBGene00004921 | 0.282651455 | 0.32347536 | 0.53201223 | snt-1 |
| WBGene00004922 | -0.402165233 | 0.2050696 | 0.40450853 | snt-2 |
| WBGene00004923 | 0.850399632 | 0.07136026 | 0.20496451 | snt-3 |
| WBGene00004924 | 1.030368952 | 2.1449E-06 | 6.2329E-05 | snt-4 |
| WBGene00004925 | 0.519915735 | 0.19967769 | 0.39794596 | snt-5 |
| WBGene00004926 | 1.121526124 | 0.03325874 | 0.12180825 | snt-6 |
| WBGene00004927 | -0.153224616 | 0.49607591 | 0.68441563 | snx-1 |
| WBGene00004928 | -0.121109848 | 0.7651616 | 0.86853281 | soc-1 |
| WBGene00004929 | 0.816088952 | 0.00025711 | 0.00319837 | soc-2 |
| WBGene00004930 | -0.343360243 | 0.0246284 | 0.09920818 | sod-1 |
| WBGene00004931 | -0.722367938 | 1.3721E-05 | 0.00029537 | sod-2 |
| WBGene00004932 | 0.1107192 | 0.89402175 | 0.9442643 | sod-3 |
| WBGene00004933 | 0.557401833 | 0.08147952 | 0.22335581 | sod-4 |
| WBGene00004944 | 1.18421521 | 0.03203567 | 0.11872561 | sol-1 |
| WBGene00004945 | 0.013326732 | 0.96784788 | 0.98372885 | sop-2 |
| WBGene00004946 | -0.256855229 | 0.18119269 | 0.37323727 | sop-3 |
| WBGene00004947 | 0.564381581 | 0.01405797 | 0.06646331 | sos-1 |
| WBGene00004949 | 0.451378427 | 0.18749524 | 0.38098404 | sox-2 |
| WBGene00004950 | -0.086660043 | 0.88181674 | 0.93708197 | sox-3 |
| WBGene00004951 | 0.872289306 | 7.4065E-05 | 0.00120217 | spc-1 |
| WBGene00004952 | -0.890638386 | 8.1155E-06 | 0.00019021 | spd-1 |
| WBGene00004953 | -0.09832373 | 0.59222483 | 0.75950161 | spd-2 |
| WBGene00004954 | -0.664780586 | 0.04347671 | 0.1454936 | spd-3 |
| WBGene00004955 | -0.199870744 | 0.19444711 | 0.3905455 | spd-5 |
| WBGene00004958 | 0.919196192 | 0.62466396 | 1 | spe-4 |
| WBGene00004959 | 0.026126153 | 0.94631321 | 0.972918 | spe-5 |
| WBGene00004960 | -0.550214255 | 0.43979443 | 0.64007787 | spe-6 |
| WBGene00004962 | -0.62146648 | 0.85925752 | 1 | spe-8 |
| WBGene00004963 | 1.687239501 | 0.59403242 | 1 | spe-9 |
| WBGene00004964 | -0.245531902 | 0.94070824 | 1 | spe-10 |
| WBGene00004965 | 1.217579795 | 0.4558521 | 1 | spe-11 |
| WBGene00004966 | 0.747845955 | 0.85215965 | 1 | spe-12 |
| WBGene00004969 | 0.288120921 | 0.54015545 | 0.72145516 | spe-15 |

| | | | | |
|---|---|---|---|---|
| WBGene00004972 | 0.434004084 | 0.71647135 | 1 | spe-26 |
| WBGene00004973 | -2.097890267 | 0.34816065 | 1 | spe-27 |
| WBGene00004975 | -0.09395824 | 0.65782551 | 0.80303357 | spe-39 |
| WBGene00004976 | 1.02028718 | 0.17335025 | 0.36304655 | spe-41 |
| WBGene00004978 | -0.163411999 | 0.45788392 | 0.65476162 | spg-7 |
| WBGene00004979 | -0.104239523 | 0.7645742 | 0.86822555 | sph-1 |
| WBGene00004980 | 0.096107552 | 0.63511209 | 0.78811283 | spk-1 |
| WBGene00004981 | 0.112075982 | 0.52036364 | 0.70549804 | spl-1 |
| WBGene00004984 | -0.15636576 | 0.29582121 | 0.50613942 | spn-4 |
| WBGene00004985 | -0.287507059 | 0.35461985 | 0.56294974 | spo-11 |
| WBGene00004986 | 1.371474033 | 1.52E-08 | 8.86E-07 | spp-1 |
| WBGene00004987 | 2.432042688 | 2.63E-31 | 2.13E-28 | spp-2 |
| WBGene00004988 | 0.409645162 | 0.03076106 | 0.1155896 | spp-3 |
| WBGene00004989 | 0.690006475 | 0.00987781 | 0.05133254 | spp-4 |
| WBGene00004990 | 0.130850404 | 0.4348366 | 0.63614305 | spp-5 |
| WBGene00004991 | 0.685293525 | 0.73391023 | 1 | spp-6 |
| WBGene00004992 | -0.693814354 | 0.75039651 | 1 | spp-7 |
| WBGene00004993 | 2.366532077 | 1.07E-33 | 1.08E-30 | spp-8 |
| WBGene00004994 | -0.774842772 | 0.78069886 | 1 | spp-9 |
| WBGene00004995 | 0.384173111 | 0.01311821 | 0.06326707 | spp-10 |
| WBGene00004996 | -0.156269293 | 0.96924949 | 1 | spp-11 |
| WBGene00004997 | 2.524956457 | 0.06642507 | 1 | spp-12 |
| WBGene00004998 | -1.48205703 | 0.08892952 | 0.23639701 | spp-13 |
| WBGene00004999 | 0.151174392 | 0.31121524 | 0.51944793 | spp-14 |
| WBGene00005000 | 0.372684449 | 0.10913723 | 0.2694256 | spp-15 |
| WBGene00005001 | 0.481260622 | 0.20250126 | 0.40146436 | spp-16 |
| WBGene00005002 | -2.391195217 | 6.16E-51 | 1.87E-47 | spp-17 |
| WBGene00005003 | 1.484322615 | 2.56E-12 | 3.30E-10 | spp-18 |
| WBGene00005004 | -0.81906709 | 0.63571741 | 1 | spp-19 |
| WBGene00005005 | 0.178115923 | 0.94982316 | 1 | spp-20 |
| WBGene00005006 | -0.256946658 | 0.22990047 | 0.43524429 | spr-1 |
| WBGene00005007 | -0.323607462 | 0.04335365 | 0.14524191 | spr-2 |
| WBGene00005008 | -0.077849045 | 0.79960553 | 0.8899084 | spr-3 |
| WBGene00005009 | -0.09445234 | 0.66181837 | 0.80528531 | spr-4 |
| WBGene00005010 | -0.11480716 | 0.5655738 | 0.74097675 | spr-5 |
| WBGene00005011 | -0.616174775 | 0.05945597 | 0.18139981 | F26F4.8 |
| WBGene00005012 | 1.003816046 | 0.3640185 | 1 | F26F4.2 |
| WBGene00005013 | 1.557897343 | 0.63813005 | 1 | jmjd-1.1 |
| WBGene00005014 | -0.744102826 | 0.00254681 | 0.01884547 | spt-4 |
| WBGene00005015 | -0.206258227 | 0.19483542 | 0.39078066 | spt-5 |
| WBGene00005016 | -1.736842371 | 0.26351155 | 0.4721438 | sqt-1 |
| WBGene00005017 | -0.345574911 | 0.65032352 | 0.79850099 | sqt-2 |
| WBGene00005018 | -0.608705691 | 0.00646949 | 0.03754592 | sqt-3 |

| | | | | |
|---|---|---|---|---|
| WBGene00005019 | -0.290762307 | 0.06685178 | 0.19614486 | sqv-1 |
| WBGene00005020 | -0.036889823 | 0.87781366 | 0.93441865 | sqv-2 |
| WBGene00005021 | -0.035941764 | 0.88267881 | 0.93749089 | sqv-3 |
| WBGene00005022 | -0.627679022 | 3.0471E-05 | 0.00056655 | sqv-4 |
| WBGene00005023 | -0.043311998 | 0.81403503 | 0.8986238 | sqv-5 |
| WBGene00005024 | -0.075720988 | 0.75206367 | 0.86206175 | sqv-6 |
| WBGene00005025 | 0.012158118 | 0.96458249 | 0.9823013 | sqv-7 |
| WBGene00005026 | -0.307830185 | 0.15607232 | 0.34000969 | sqv-8 |
| WBGene00005030 | 0.688793783 | 0.86378755 | 1 | sra-4 |
| WBGene00005032 | -1.964653846 | 0.53599202 | 1 | sra-6 |
| WBGene00005033 | -0.156269293 | 0.96924949 | 1 | sra-7 |
| WBGene00005036 | -0.876811101 | 0.47669179 | 1 | sra-10 |
| WBGene00005037 | -0.03811452 | 0.98663692 | 1 | sra-11 |
| WBGene00005038 | 2.224351729 | 0.33187942 | 1 | sra-12 |
| WBGene00005040 | 0.283566113 | 0.80054065 | 1 | sra-14 |
| WBGene00005043 | 1.116613983 | 0.69067248 | 1 | sra-17 |
| WBGene00005047 | 0.983488853 | 0.80601565 | 1 | sra-21 |
| WBGene00005048 | 0.281454817 | 0.94435481 | 1 | sra-22 |
| WBGene00005051 | -1.969805064 | 0.62862569 | 1 | sra-25 |
| WBGene00005053 | 0.281454817 | 0.94435481 | 1 | sra-27 |
| WBGene00005057 | 0.574508983 | 0.8551971 | 1 | sra-31 |
| WBGene00005058 | 1.025181888 | 0.79790347 | 1 | sra-32 |
| WBGene00005059 | -0.18557176 | 0.95664217 | 1 | sra-33 |
| WBGene00005061 | 0.489058549 | 0.85947456 | 1 | sra-35 |
| WBGene00005062 | -0.156269293 | 0.96924949 | 1 | sra-36 |
| WBGene00005063 | 0.116602111 | 0.93619121 | 1 | sra-37 |
| WBGene00005065 | 0.377828795 | 0.92526585 | 1 | sra-39 |
| WBGene00005066 | -0.156269293 | 0.96924949 | 1 | srb-1 |
| WBGene00005070 | -0.156269293 | 0.96924949 | 1 | srb-5 |
| WBGene00005071 | -0.323868354 | 0.91634405 | 1 | srb-6 |
| WBGene00005077 | -0.262243418 | 0.15044759 | 0.33204586 | src-1 |
| WBGene00005078 | 0.332977923 | 0.49942009 | 0.68759028 | src-2 |
| WBGene00005079 | 0.041803775 | 0.98890197 | 1 | srd-1 |
| WBGene00005081 | 0.539239107 | 0.83916685 | 1 | srd-3 |
| WBGene00005083 | 2.853725551 | 0.17083607 | 1 | srd-5 |
| WBGene00005088 | -0.668264701 | 0.85551153 | 1 | srd-10 |
| WBGene00005090 | -1.14715222 | 0.68932198 | 1 | srd-12 |
| WBGene00005091 | -0.604396148 | 0.82723443 | 1 | srd-13 |
| WBGene00005093 | 1.38322681 | 0.69550305 | 1 | srd-15 |
| WBGene00005095 | 1.025181888 | 0.79790347 | 1 | srd-17 |
| WBGene00005096 | 0.747845955 | 0.85215965 | 1 | srd-18 |
| WBGene00005101 | 1.566956217 | 0.32184393 | 1 | srd-23 |
| WBGene00005104 | -0.156269293 | 0.96924949 | 1 | srd-26 |

| | | | | |
|---|---|---|---|---|
| WBGene00005105 | -0.558804446 | 0.87818675 | 1 | srd-27 |
| WBGene00005106 | -0.156269293 | 0.96924949 | 1 | srd-28 |
| WBGene00005108 | -1.901856996 | 0.64072034 | 1 | srd-30 |
| WBGene00005110 | 2.219127534 | 0.1240484 | 1 | srd-32 |
| WBGene00005111 | 2.200270375 | 0.22405171 | 1 | srd-33 |
| WBGene00005113 | -0.07748118 | 0.98067866 | 1 | srd-35 |
| WBGene00005118 | -0.156269293 | 0.96924949 | 1 | srd-40 |
| WBGene00005119 | -0.156269293 | 0.96924949 | 1 | srd-41 |
| WBGene00005120 | 0.281454817 | 0.94435481 | 1 | srd-42 |
| WBGene00005121 | 0.281454817 | 0.94435481 | 1 | srd-43 |
| WBGene00005124 | -0.156269293 | 0.96924949 | 1 | srd-46 |
| WBGene00005125 | 0.377828795 | 0.92526585 | 1 | srd-47 |
| WBGene00005128 | -0.156269293 | 0.96924949 | 1 | srd-50 |
| WBGene00005129 | 0.247318604 | 0.91814609 | 1 | srd-51 |
| WBGene00005131 | 0.955451811 | 0.61522535 | 1 | srd-53 |
| WBGene00005132 | 1.313391856 | 0.53155418 | 1 | srd-54 |
| WBGene00005133 | 0.983488853 | 0.80601565 | 1 | srd-55 |
| WBGene00005135 | -0.156269293 | 0.96924949 | 1 | srd-58 |
| WBGene00005141 | 1.544876349 | 0.48259424 | 1 | srd-64 |
| WBGene00005142 | 0.488133825 | 0.81078971 | 1 | srd-65 |
| WBGene00005144 | 1.529802032 | 0.64971375 | 1 | srd-67 |
| WBGene00005146 | -1.901856996 | 0.64072034 | 1 | srd-69 |
| WBGene00005147 | 2.229936699 | 0.22041467 | 1 | srd-70 |
| WBGene00005149 | 0.270496667 | 0.92869767 | 1 | sre-1 |
| WBGene00005150 | 0.688793783 | 0.86378755 | 1 | sre-2 |
| WBGene00005153 | 0.011428935 | 0.97407088 | 0.98649644 | srf-3 |
| WBGene00005159 | -0.156269293 | 0.96924949 | 1 | srg-1 |
| WBGene00005160 | 0.05905161 | 0.97811893 | 1 | srg-2 |
| WBGene00005161 | -0.263229581 | 0.93409872 | 1 | srg-3 |
| WBGene00005162 | -1.969805064 | 0.62862569 | 1 | srg-4 |
| WBGene00005163 | 0.09697944 | 0.97612634 | 1 | srg-5 |
| WBGene00005164 | -0.156269293 | 0.96924949 | 1 | srg-6 |
| WBGene00005165 | 0.28530093 | 0.92084641 | 1 | srg-7 |
| WBGene00005166 | -0.156269293 | 0.96924949 | 1 | srg-8 |
| WBGene00005167 | 0.281454817 | 0.94435481 | 1 | srg-9 |
| WBGene00005168 | -0.094731856 | 0.97157968 | 1 | srg-10 |
| WBGene00005169 | 0.688793783 | 0.86378755 | 1 | srg-11 |
| WBGene00005170 | 0.377828795 | 0.92526585 | 1 | srg-13 |
| WBGene00005171 | -0.156269293 | 0.96924949 | 1 | srg-14 |
| WBGene00005172 | 0.377828795 | 0.92526585 | 1 | srg-15 |
| WBGene00005174 | -0.156269293 | 0.96924949 | 1 | srg-17 |
| WBGene00005177 | -0.156269293 | 0.96924949 | 1 | srg-20 |
| WBGene00005178 | 0.747845955 | 0.85215965 | 1 | srg-21 |

| | | | | |
|---|---|---|---|---|
| WBGene00005179 | 0.983488853 | 0.80601565 | 1 | srg-22 |
| WBGene00005180 | 0.983488853 | 0.80601565 | 1 | srg-23 |
| WBGene00005181 | -1.901856996 | 0.64072034 | 1 | srg-24 |
| WBGene00005182 | 2.09403112 | 0.45107747 | 1 | srg-25 |
| WBGene00005186 | 0.688793783 | 0.86378755 | 1 | srg-29 |
| WBGene00005187 | -0.763485502 | 0.69547406 | 1 | srg-30 |
| WBGene00005188 | 0.688793783 | 0.86378755 | 1 | srg-31 |
| WBGene00005189 | 0.233739675 | 0.93575889 | 1 | srg-32 |
| WBGene00005191 | 1.645765972 | 0.25997593 | 1 | srg-34 |
| WBGene00005193 | 0.377828795 | 0.92526585 | 1 | srg-36 |
| WBGene00005194 | 0.281454817 | 0.94435481 | 1 | srg-37 |
| WBGene00005195 | -0.604396148 | 0.82723443 | 1 | srg-38 |
| WBGene00005197 | 1.692096778 | 0.3645695 | 1 | srg-40 |
| WBGene00005198 | -0.156269293 | 0.96924949 | 1 | srg-41 |
| WBGene00005201 | 0.579666901 | 0.79455736 | 1 | srg-44 |
| WBGene00005205 | 0.041803775 | 0.98890197 | 1 | srg-48 |
| WBGene00005208 | -0.087822178 | 0.96354485 | 1 | srg-51 |
| WBGene00005210 | -0.156269293 | 0.96924949 | 1 | srg-53 |
| WBGene00005211 | 0.281454817 | 0.94435481 | 1 | srg-54 |
| WBGene00005213 | -0.156269293 | 0.96924949 | 1 | srg-56 |
| WBGene00005217 | 0.281454817 | 0.94435481 | 1 | srg-60 |
| WBGene00005219 | 0.688793783 | 0.86378755 | 1 | srg-62 |
| WBGene00005222 | -0.263229581 | 0.93409872 | 1 | srg-65 |
| WBGene00005223 | 0.983488853 | 0.80601565 | 1 | srg-66 |
| WBGene00005226 | 0.786039486 | 0.63978859 | 1 | srg-69 |
| WBGene00005227 | -0.156269293 | 0.96924949 | 1 | srh-1 |
| WBGene00005228 | 0.629154436 | 0.81300308 | 1 | srh-2 |
| WBGene00005231 | -0.156269293 | 0.96924949 | 1 | srh-5 |
| WBGene00005234 | -0.156269293 | 0.96924949 | 1 | srh-8 |
| WBGene00005237 | 0.688793783 | 0.86378755 | 1 | srh-11 |
| WBGene00005240 | 0.377828795 | 0.92526585 | 1 | srh-15 |
| WBGene00005241 | -0.70576507 | 0.76841669 | 1 | srh-16 |
| WBGene00005242 | -0.156269293 | 0.96924949 | 1 | srh-17 |
| WBGene00005245 | 1.025181888 | 0.79790347 | 1 | srh-20 |
| WBGene00005246 | -1.901856996 | 0.64072034 | 1 | srh-21 |
| WBGene00005248 | -1.964653846 | 0.53599202 | 1 | srh-23 |
| WBGene00005249 | -0.156269293 | 0.96924949 | 1 | srh-25 |
| WBGene00005250 | -0.156269293 | 0.96924949 | 1 | srh-27 |
| WBGene00005251 | 0.688793783 | 0.86378755 | 1 | srh-28 |
| WBGene00005253 | 0.903390376 | 0.71900339 | 1 | srh-30 |
| WBGene00005256 | -0.558804446 | 0.87818675 | 1 | srh-33 |
| WBGene00005258 | 0.938982574 | 0.8146943 | 1 | srh-35 |
| WBGene00005260 | -0.156269293 | 0.96924949 | 1 | srh-37 |

| | | | | |
|---|---|---|---|---|
| WBGene00005261 | -0.156269293 | 0.96924949 | 1 | srh-38 |
| WBGene00005262 | -0.008677305 | 0.99719169 | 1 | srh-39 |
| WBGene00005263 | -0.323868354 | 0.91634405 | 1 | srh-40 |
| WBGene00005264 | 0.281454817 | 0.94435481 | 1 | srh-41 |
| WBGene00005267 | 0.377828795 | 0.92526585 | 1 | srh-44 |
| WBGene00005268 | 0.688793783 | 0.86378755 | 1 | srh-45 |
| WBGene00005270 | 0.281454817 | 0.94435481 | 1 | srh-47 |
| WBGene00005271 | 1.633238609 | 0.2732739 | 1 | srh-48 |
| WBGene00005273 | 0.281454817 | 0.94435481 | 1 | srh-50 |
| WBGene00005274 | -0.156269293 | 0.96924949 | 1 | srh-51 |
| WBGene00005279 | -0.668264701 | 0.85551153 | 1 | srh-56 |
| WBGene00005281 | 1.876192691 | 0.43542138 | 1 | srh-59 |
| WBGene00005283 | -1.094201388 | 0.7816772 | 1 | srh-61 |
| WBGene00005289 | 1.116613983 | 0.69067248 | 1 | srh-68 |
| WBGene00005290 | 1.181686151 | 0.7579682 | 1 | srh-69 |
| WBGene00005291 | 2.111080655 | 0.48938556 | 1 | srh-70 |
| WBGene00005292 | 2.806835559 | 0.21738362 | 1 | srh-71 |
| WBGene00005293 | 0.281454817 | 0.94435481 | 1 | srh-72 |
| WBGene00005299 | 1.181686151 | 0.7579682 | 1 | srh-78 |
| WBGene00005300 | -1.901856996 | 0.64072034 | 1 | srh-79 |
| WBGene00005301 | -1.094201388 | 0.7816772 | 1 | srh-80 |
| WBGene00005303 | 0.688793783 | 0.86378755 | 1 | srh-82 |
| WBGene00005310 | 0.688793783 | 0.86378755 | 1 | srh-89 |
| WBGene00005315 | -0.156269293 | 0.96924949 | 1 | srh-95 |
| WBGene00005317 | 0.281454817 | 0.94435481 | 1 | srh-98 |
| WBGene00005318 | 0.377828795 | 0.92526585 | 1 | srh-99 |
| WBGene00005321 | -0.156269293 | 0.96924949 | 1 | srh-102 |
| WBGene00005324 | 0.281454817 | 0.94435481 | 1 | T27A1.1 |
| WBGene00005332 | 0.688793783 | 0.86378755 | 1 | srh-113 |
| WBGene00005334 | -0.156269293 | 0.96924949 | 1 | srh-115 |
| WBGene00005338 | -1.094201388 | 0.7816772 | 1 | srh-119 |
| WBGene00005341 | 0.789601902 | 0.75907734 | 1 | srh-123 |
| WBGene00005342 | 1.876192691 | 0.43542138 | 1 | srh-125 |
| WBGene00005344 | 0.281454817 | 0.94435481 | 1 | srh-127 |
| WBGene00005345 | 0.574508983 | 0.8551971 | 1 | srh-128 |
| WBGene00005346 | -0.156269293 | 0.96924949 | 1 | srh-129 |
| WBGene00005347 | 0.377828795 | 0.92526585 | 1 | srh-130 |
| WBGene00005348 | 0.281454817 | 0.94435481 | 1 | srh-131 |
| WBGene00005349 | 1.477253415 | 0.58357327 | 1 | srh-132 |
| WBGene00005351 | 0.281454817 | 0.94435481 | 1 | srh-134 |
| WBGene00005352 | -1.964653846 | 0.53599202 | 1 | srh-135 |
| WBGene00005355 | -1.969805064 | 0.62862569 | 1 | srh-138 |
| WBGene00005359 | 0.281454817 | 0.94435481 | 1 | srh-142 |

| | | | | |
|---|---|---|---|---|
| WBGene00005362 | -0.156269293 | 0.96924949 | 1 | srh-146 |
| WBGene00005364 | 0.281454817 | 0.94435481 | 1 | srh-148 |
| WBGene00005365 | 0.377828795 | 0.92526585 | 1 | srh-149 |
| WBGene00005370 | 1.413734769 | 0.68804531 | 1 | srh-154 |
| WBGene00005375 | -0.156269293 | 0.96924949 | 1 | srh-159 |
| WBGene00005386 | -2.563099533 | 0.4751044 | 1 | srh-170 |
| WBGene00005387 | 2.600311303 | 0.25000941 | 1 | srh-171 |
| WBGene00005390 | 1.216957619 | 0.74819621 | 1 | srh-174 |
| WBGene00005392 | -0.156269293 | 0.96924949 | 1 | srh-177 |
| WBGene00005393 | 0.938982574 | 0.8146943 | 1 | srh-178 |
| WBGene00005394 | -0.156269293 | 0.96924949 | 1 | srh-179 |
| WBGene00005395 | 0.574508983 | 0.8551971 | 1 | srh-180 |
| WBGene00005397 | 0.281454817 | 0.94435481 | 1 | srh-182 |
| WBGene00005399 | 1.116613983 | 0.69067248 | 1 | srh-184 |
| WBGene00005406 | -2.705999569 | 0.42423051 | 1 | srh-193 |
| WBGene00005410 | -0.156269293 | 0.96924949 | 1 | srh-199 |
| WBGene00005415 | -1.969805064 | 0.62862569 | 1 | srh-204 |
| WBGene00005416 | 0.281454817 | 0.94435481 | 1 | srh-206 |
| WBGene00005417 | 0.281454817 | 0.94435481 | 1 | srh-207 |
| WBGene00005422 | 2.201543524 | 0.41352885 | 1 | srh-212 |
| WBGene00005423 | -1.901856996 | 0.64072034 | 1 | srh-213 |
| WBGene00005426 | -0.156269293 | 0.96924949 | 1 | srh-216 |
| WBGene00005427 | -0.848512853 | 0.76804407 | 1 | srh-217 |
| WBGene00005429 | 0.281454817 | 0.94435481 | 1 | srh-219 |
| WBGene00005430 | -1.901856996 | 0.64072034 | 1 | srh-220 |
| WBGene00005432 | -1.14715222 | 0.68932198 | 1 | srh-222 |
| WBGene00005437 | -1.14715222 | 0.68932198 | 1 | srh-229 |
| WBGene00005442 | -0.668264701 | 0.85551153 | 1 | srh-235 |
| WBGene00005444 | 0.930559141 | 0.00340145 | 0.02321083 | srh-237 |
| WBGene00005446 | -0.156269293 | 0.96924949 | 1 | srh-239 |
| WBGene00005447 | -1.094201388 | 0.7816772 | 1 | srh-240 |
| WBGene00005450 | -0.156269293 | 0.96924949 | 1 | srh-244 |
| WBGene00005454 | -1.964653846 | 0.53599202 | 1 | srh-248 |
| WBGene00005456 | 0.281454817 | 0.94435481 | 1 | srh-250 |
| WBGene00005461 | -0.70576507 | 0.76841669 | 1 | srh-255 |
| WBGene00005470 | -0.156269293 | 0.96924949 | 1 | srh-264 |
| WBGene00005472 | -1.901856996 | 0.64072034 | 1 | srh-266 |
| WBGene00005474 | -0.008677305 | 0.99719169 | 1 | srh-268 |
| WBGene00005475 | 2.332054728 | 0.29723791 | 1 | srh-269 |
| WBGene00005476 | -1.969805064 | 0.62862569 | 1 | srh-270 |
| WBGene00005477 | 1.413734769 | 0.68804531 | 1 | srh-271 |
| WBGene00005478 | 0.281454817 | 0.94435481 | 1 | srh-272 |
| WBGene00005481 | -0.241106143 | 0.91536132 | 1 | srh-276 |

| | | | | |
|---|---|---|---|---|
| WBGene00005482 | 0.281454817 | 0.94435481 | 1 | srh-277 |
| WBGene00005484 | 0.28530093 | 0.92084641 | 1 | srh-279 |
| WBGene00005486 | 0.446458324 | 0.87164163 | 1 | srh-281 |
| WBGene00005487 | 0.377828795 | 0.92526585 | 1 | srh-282 |
| WBGene00005512 | -1.964653846 | 0.53599202 | 1 | srh-105 |
| WBGene00005513 | 1.378226134 | 0.5038929 | 1 | sri-1 |
| WBGene00005514 | -0.37501878 | 0.84355758 | 1 | sri-2 |
| WBGene00005516 | 0.747845955 | 0.85215965 | 1 | sri-4 |
| WBGene00005517 | -2.061536526 | 0.21538967 | 1 | sri-5 |
| WBGene00005518 | -0.668264701 | 0.85551153 | 1 | sri-6 |
| WBGene00005519 | 0.688793783 | 0.86378755 | 1 | sri-7 |
| WBGene00005521 | 0.281454817 | 0.94435481 | 1 | sri-9 |
| WBGene00005522 | -1.969805064 | 0.62862569 | 1 | sri-10 |
| WBGene00005523 | 2.486734991 | 0.32516285 | 1 | sri-11 |
| WBGene00005524 | 1.405085033 | 0.48727562 | 1 | sri-12 |
| WBGene00005528 | -1.901856996 | 0.64072034 | 1 | sri-16 |
| WBGene00005530 | -1.538600039 | 0.37104439 | 1 | sri-18 |
| WBGene00005531 | 1.181686151 | 0.7579682 | 1 | sri-19 |
| WBGene00005532 | 0.377828795 | 0.92526585 | 1 | sri-20 |
| WBGene00005533 | 1.89038395 | 0.35241418 | 1 | sri-21 |
| WBGene00005536 | -0.156269293 | 0.96924949 | 1 | sri-24 |
| WBGene00005538 | 0.041803775 | 0.98890197 | 1 | sri-26 |
| WBGene00005541 | -0.008677305 | 0.99719169 | 1 | sri-29 |
| WBGene00005542 | -0.061567883 | 0.96509566 | 1 | sri-30 |
| WBGene00005543 | 1.247575581 | 0.7549672 | 1 | sri-31 |
| WBGene00005544 | 1.557897343 | 0.63813005 | 1 | sri-32 |
| WBGene00005548 | 1.584577408 | 0.63258867 | 1 | sri-36 |
| WBGene00005552 | -0.167717958 | 0.62881287 | 0.78446537 | sri-40 |
| WBGene00005555 | 0.688793783 | 0.86378755 | 1 | sri-43 |
| WBGene00005557 | -0.950208353 | 0.7250598 | 1 | sri-45 |
| WBGene00005558 | 0.688793783 | 0.86378755 | 1 | sri-46 |
| WBGene00005559 | 0.281454817 | 0.94435481 | 1 | sri-47 |
| WBGene00005560 | 0.747845955 | 0.85215965 | 1 | sri-48 |
| WBGene00005565 | 0.281454817 | 0.94435481 | 1 | sri-53 |
| WBGene00005569 | -0.156269293 | 0.96924949 | 1 | sri-57 |
| WBGene00005573 | 0.281454817 | 0.94435481 | 1 | sri-61 |
| WBGene00005574 | 0.983488853 | 0.80601565 | 1 | sri-62 |
| WBGene00005575 | 0.281454817 | 0.94435481 | 1 | sri-63 |
| WBGene00005578 | -0.01455644 | 0.9964743 | 1 | sri-66 |
| WBGene00005582 | -1.964653846 | 0.53599202 | 1 | sri-70 |
| WBGene00005585 | 1.116613983 | 0.69067248 | 1 | sri-73 |
| WBGene00005586 | 0.281454817 | 0.94435481 | 1 | sri-74 |
| WBGene00005589 | -0.156269293 | 0.96924949 | 1 | sri-77 |

| | | | | |
|---|---|---|---|---|
| WBGene00005591 | -0.333693239 | 0.89573179 | 1 | srj-1 |
| WBGene00005592 | 0.27161217 | 0.89703555 | 1 | srj-4 |
| WBGene00005597 | -1.0062442 | 0.63208617 | 1 | srj-9 |
| WBGene00005598 | 0.281454817 | 0.94435481 | 1 | srj-10 |
| WBGene00005602 | 2.364607998 | 0.0053643 | 0.03282659 | srj-14 |
| WBGene00005603 | 0.377828795 | 0.92526585 | 1 | srj-15 |
| WBGene00005606 | 1.876192691 | 0.43542138 | 1 | srj-18 |
| WBGene00005607 | -0.558804446 | 0.87818675 | 1 | srj-19 |
| WBGene00005609 | 0.377828795 | 0.92526585 | 1 | srj-21 |
| WBGene00005610 | -1.964653846 | 0.53599202 | 1 | srj-22 |
| WBGene00005611 | -0.451813889 | 0.8460226 | 1 | srj-23 |
| WBGene00005616 | 0.983488853 | 0.80601565 | 1 | srj-29 |
| WBGene00005617 | 1.216957619 | 0.74819621 | 1 | srj-32 |
| WBGene00005631 | 0.09697944 | 0.97612634 | 1 | srj-52 |
| WBGene00005632 | -1.969805064 | 0.62862569 | 1 | srj-53 |
| WBGene00005638 | 2.286231321 | 0.38099652 | 1 | srm-1 |
| WBGene00005639 | 0.377828795 | 0.92526585 | 1 | srm-2 |
| WBGene00005640 | -1.159714244 | 0.75790634 | 1 | srn-1 |
| WBGene00005641 | 2.957799994 | 0.02865575 | 1 | sro-1 |
| WBGene00005642 | 0.396670124 | 0.24327914 | 0.4508704 | srp-1 |
| WBGene00005643 | -0.863558155 | 0.02316435 | 0.0950772 | srp-2 |
| WBGene00005644 | 0.542104352 | 0.45818222 | 0.65477918 | srp-3 |
| WBGene00005647 | 0.305415071 | 0.43229466 | 0.6341955 | srp-6 |
| WBGene00005648 | -0.073741989 | 0.63852622 | 0.79036023 | srp-7 |
| WBGene00005649 | 0.703630544 | 0.48655673 | 0.67622374 | srp-8 |
| WBGene00005652 | 0.750755091 | 0.45716339 | 1 | srr-1 |
| WBGene00005653 | 1.014790707 | 0.48070807 | 1 | srr-2 |
| WBGene00005654 | 0.613974631 | 0.62845337 | 1 | srr-3 |
| WBGene00005655 | 1.100295987 | 4.7452E-06 | 0.00012007 | srr-4 |
| WBGene00005657 | 3.120482424 | 0.01436027 | 1 | srr-6 |
| WBGene00005661 | 0.377828795 | 0.92526585 | 1 | srr-10 |
| WBGene00005662 | -0.641488339 | 0.003598 | 0.02430899 | sars-2 |
| WBGene00005663 | -0.233292869 | 0.11777668 | 0.28281778 | sars-1 |
| WBGene00005664 | 2.09403112 | 0.45107747 | 1 | sru-1 |
| WBGene00005666 | -0.156269293 | 0.96924949 | 1 | sru-3 |
| WBGene00005670 | 1.247575581 | 0.7549672 | 1 | sru-7 |
| WBGene00005671 | -0.156269293 | 0.96924949 | 1 | sru-8 |
| WBGene00005672 | -1.159714244 | 0.75790634 | 1 | sru-9 |
| WBGene00005675 | -0.156269293 | 0.96924949 | 1 | sru-12 |
| WBGene00005676 | -0.156269293 | 0.96924949 | 1 | sru-13 |
| WBGene00005679 | 0.747845955 | 0.85215965 | 1 | sru-16 |
| WBGene00005680 | 0.281454817 | 0.94435481 | 1 | sru-17 |
| WBGene00005681 | -0.156269293 | 0.96924949 | 1 | sru-18 |

| | | | | |
|---|---|---|---|---|
| WBGene00005683 | -1.156241549 | 0.61231033 | 1 | sru-20 |
| WBGene00005684 | -0.07748118 | 0.98067866 | 1 | sru-21 |
| WBGene00005685 | 0.707559163 | 0.78139902 | 1 | sru-22 |
| WBGene00005686 | -1.22259901 | 0.68385967 | 1 | sru-23 |
| WBGene00005687 | -0.03811452 | 0.98663692 | 1 | sru-24 |
| WBGene00005688 | 1.477253415 | 0.58357327 | 1 | sru-25 |
| WBGene00005689 | 0.574508983 | 0.8551971 | 1 | sru-26 |
| WBGene00005690 | 0.075260937 | 0.97569838 | 1 | sru-27 |
| WBGene00005692 | -0.737462795 | 0.83433562 | 1 | sru-29 |
| WBGene00005696 | -0.156269293 | 0.96924949 | 1 | sru-33 |
| WBGene00005698 | -0.156269293 | 0.96924949 | 1 | sru-35 |
| WBGene00005699 | 0.377828795 | 0.92526585 | 1 | sru-36 |
| WBGene00005700 | -1.964653846 | 0.53599202 | 1 | sru-37 |
| WBGene00005701 | 0.938982574 | 0.8146943 | 1 | sru-38 |
| WBGene00005702 | -0.156269293 | 0.96924949 | 1 | sru-39 |
| WBGene00005703 | 2.309162803 | 0.42233005 | 1 | sru-40 |
| WBGene00005711 | 1.413377704 | 0.60673863 | 1 | sru-48 |
| WBGene00005712 | 0.7759157 | 0.01593788 | 0.07259399 | srv-1 |
| WBGene00005713 | -2.073308045 | 0.35461841 | 1 | srv-2 |
| WBGene00005715 | 0.579666901 | 0.79455736 | 1 | srv-4 |
| WBGene00005716 | 3.185207558 | 0.09097874 | 1 | srv-5 |
| WBGene00005717 | -1.159714244 | 0.75790634 | 1 | srv-6 |
| WBGene00005718 | -0.391510342 | 0.73313329 | 1 | srv-7 |
| WBGene00005719 | 0.041803775 | 0.98890197 | 1 | srv-8 |
| WBGene00005720 | 0.075260937 | 0.97569838 | 1 | srv-9 |
| WBGene00005724 | -0.156269293 | 0.96924949 | 1 | srv-13 |
| WBGene00005725 | 0.394439984 | 0.88400378 | 1 | srv-14 |
| WBGene00005726 | 3.295041369 | 0.07982868 | 1 | srv-15 |
| WBGene00005727 | 0.983488853 | 0.80601565 | 1 | srv-16 |
| WBGene00005735 | 1.216957619 | 0.74819621 | 1 | srv-24 |
| WBGene00005736 | -0.156269293 | 0.96924949 | 1 | srv-25 |
| WBGene00005738 | 0.281454817 | 0.94435481 | 1 | srv-27 |
| WBGene00005739 | -0.558804446 | 0.87818675 | 1 | srv-28 |
| WBGene00005741 | -0.156269293 | 0.96924949 | 1 | srv-30 |
| WBGene00005742 | -0.03811452 | 0.98663692 | 1 | srv-31 |
| WBGene00005743 | 0.377828795 | 0.92526585 | 1 | srv-32 |
| WBGene00005745 | -1.094201388 | 0.7816772 | 1 | srv-34 |
| WBGene00005748 | 0.377828795 | 0.92526585 | 1 | srw-1 |
| WBGene00005749 | -1.901856996 | 0.64072034 | 1 | srw-2 |
| WBGene00005754 | 1.529802032 | 0.64971375 | 1 | srw-7 |
| WBGene00005755 | -1.969805064 | 0.62862569 | 1 | srw-8 |
| WBGene00005756 | -1.969805064 | 0.62862569 | 1 | srw-9 |
| WBGene00005757 | 1.557897343 | 0.63813005 | 1 | srw-10 |

| | | | | |
|---|---|---|---|---|
| WBGene00005758 | 2.055171418 | 0.46278075 | 1 | srw-11 |
| WBGene00005759 | -0.514761297 | 0.79912167 | 1 | srw-12 |
| WBGene00005762 | 0.377828795 | 0.92526585 | 1 | srw-15 |
| WBGene00005763 | 1.216957619 | 0.74819621 | 1 | srw-16 |
| WBGene00005764 | -1.969805064 | 0.62862569 | 1 | srw-17 |
| WBGene00005766 | 0.688793783 | 0.86378755 | 1 | srw-19 |
| WBGene00005770 | 0.281454817 | 0.94435481 | 1 | srw-23 |
| WBGene00005771 | -0.156269293 | 0.96924949 | 1 | srw-24 |
| WBGene00005780 | -0.156269293 | 0.96924949 | 1 | srw-33 |
| WBGene00005782 | 1.216957619 | 0.74819621 | 1 | srw-35 |
| WBGene00005795 | -1.901856996 | 0.64072034 | 1 | srw-48 |
| WBGene00005808 | -0.263229581 | 0.93409872 | 1 | srw-61 |
| WBGene00005816 | 0.281454817 | 0.94435481 | 1 | srw-69 |
| WBGene00005818 | 0.688793783 | 0.86378755 | 1 | srw-71 |
| WBGene00005832 | 1.796574463 | 9.77E-08 | 4.6351E-06 | srw-85 |
| WBGene00005833 | 1.918831651 | 0.110748 | 1 | srw-86 |
| WBGene00005834 | -0.263229581 | 0.93409872 | 1 | srw-87 |
| WBGene00005836 | 0.09697944 | 0.97612634 | 1 | srw-89 |
| WBGene00005838 | -1.094201388 | 0.7816772 | 1 | srw-91 |
| WBGene00005842 | -0.156269293 | 0.96924949 | 1 | srw-95 |
| WBGene00005845 | 0.28530093 | 0.92084641 | 1 | srw-98 |
| WBGene00005847 | -0.156269293 | 0.96924949 | 1 | srw-100 |
| WBGene00005848 | 0.377828795 | 0.92526585 | 1 | srw-101 |
| WBGene00005849 | 0.403207732 | 0.86402224 | 1 | srw-102 |
| WBGene00005850 | -0.156269293 | 0.96924949 | 1 | srw-103 |
| WBGene00005867 | -0.156269293 | 0.96924949 | 1 | srw-120 |
| WBGene00005881 | -0.156269293 | 0.96924949 | 1 | srw-134 |
| WBGene00005886 | -0.156269293 | 0.96924949 | 1 | srw-139 |
| WBGene00005888 | -1.901856996 | 0.64072034 | 1 | srw-141 |
| WBGene00005893 | 0.873659218 | 0.65541243 | 1 | srx-2 |
| WBGene00005896 | -1.901856996 | 0.64072034 | 1 | srx-5 |
| WBGene00005898 | -0.156269293 | 0.96924949 | 1 | srx-7 |
| WBGene00005900 | -0.156269293 | 0.96924949 | 1 | srx-9 |
| WBGene00005903 | -1.208626262 | 0.55673606 | 1 | srx-12 |
| WBGene00005904 | -0.737462795 | 0.83433562 | 1 | srx-13 |
| WBGene00005905 | -1.901856996 | 0.64072034 | 1 | srx-14 |
| WBGene00005909 | 0.574508983 | 0.8551971 | 1 | srx-18 |
| WBGene00005910 | 0.742628056 | 0.76901841 | 1 | srx-19 |
| WBGene00005912 | -0.156269293 | 0.96924949 | 1 | srx-21 |
| WBGene00005925 | 0.983488853 | 0.80601565 | 1 | srx-34 |
| WBGene00005926 | 0.983488853 | 0.80601565 | 1 | srx-35 |
| WBGene00005929 | 0.688793783 | 0.86378755 | 1 | srx-38 |
| WBGene00005931 | 0.596056337 | 0.72561177 | 1 | fbxa-199 |

| | | | | |
|---|---|---|---|---|
| WBGene00005932 | 1.584577408 | 0.63258867 | 1 | srx-41 |
| WBGene00005934 | 0.962897601 | 0.69232608 | 1 | srx-43 |
| WBGene00005935 | -0.156269293 | 0.96924949 | 1 | srx-44 |
| WBGene00005936 | 1.423097571 | 0.11439969 | 0.27821121 | srx-45 |
| WBGene00005937 | 0.377828795 | 0.92526585 | 1 | srx-46 |
| WBGene00005938 | -1.901856996 | 0.64072034 | 1 | srx-47 |
| WBGene00005942 | 0.281454817 | 0.94435481 | 1 | srx-51 |
| WBGene00005947 | 2.286231321 | 0.38099652 | 1 | srx-56 |
| WBGene00005949 | 0.754711195 | 0.7285874 | 1 | srx-58 |
| WBGene00005955 | 0.377828795 | 0.92526585 | 1 | srx-64 |
| WBGene00005958 | 2.188838843 | 0.34004625 | 1 | srx-67 |
| WBGene00005959 | 1.08943711 | 0.64491434 | 1 | srx-68 |
| WBGene00005964 | -0.298265327 | 0.89527345 | 1 | srx-73 |
| WBGene00005967 | -1.901856996 | 0.64072034 | 1 | srx-76 |
| WBGene00005968 | 0.36729309 | 0.88005435 | 1 | srx-77 |
| WBGene00005975 | 0.041803775 | 0.98890197 | 1 | srx-84 |
| WBGene00005976 | 1.557897343 | 0.63813005 | 1 | srx-85 |
| WBGene00005977 | 0.983488853 | 0.80601565 | 1 | srx-86 |
| WBGene00005978 | 0.688793783 | 0.86378755 | 1 | srx-87 |
| WBGene00005979 | -1.964653846 | 0.53599202 | 1 | srx-88 |
| WBGene00005983 | -1.969805064 | 0.62862569 | 1 | srx-92 |
| WBGene00005987 | -1.094201388 | 0.7816772 | 1 | srx-96 |
| WBGene00005988 | 1.216957619 | 0.74819621 | 1 | srx-97 |
| WBGene00005989 | 0.446458324 | 0.87164163 | 1 | srx-98 |
| WBGene00005995 | -1.094201388 | 0.7816772 | 1 | srx-104 |
| WBGene00005999 | -0.558804446 | 0.87818675 | 1 | srx-108 |
| WBGene00006002 | 1.510073683 | 0.4579485 | 1 | srx-111 |
| WBGene00006005 | 0.377828795 | 0.92526585 | 1 | srx-114 |
| WBGene00006006 | 1.186559396 | 0.57146291 | 1 | srx-115 |
| WBGene00006007 | -0.737462795 | 0.83433562 | 1 | srx-116 |
| WBGene00006008 | 0.747845955 | 0.85215965 | 1 | srx-117 |
| WBGene00006009 | 0.574508983 | 0.8551971 | 1 | srx-118 |
| WBGene00006016 | -0.362677336 | 0.80806775 | 1 | srx-125 |
| WBGene00006019 | -1.964653846 | 0.53599202 | 1 | srx-128 |
| WBGene00006020 | 0.075260937 | 0.97569838 | 1 | srx-129 |
| WBGene00006021 | 0.281454817 | 0.94435481 | 1 | srx-130 |
| WBGene00006022 | -0.156269293 | 0.96924949 | 1 | srx-131 |
| WBGene00006024 | -1.969805064 | 0.62862569 | 1 | srx-133 |
| WBGene00006025 | 1.413377704 | 0.60673863 | 1 | srx-134 |
| WBGene00006026 | 0.688793783 | 0.86378755 | 1 | srx-135 |
| WBGene00006027 | -2.640414922 | 0.41081194 | 1 | srx-136 |
| WBGene00006028 | 0.983488853 | 0.80601565 | 1 | srx-137 |
| WBGene00006038 | -1.094201388 | 0.7816772 | 1 | ssp-9 |

| | | | | |
|---|---|---|---|---|
| WBGene00006039 | 0.441943138 | 0.69425935 | 1 | ssp-10 |
| WBGene00006040 | -1.094201388 | 0.7816772 | 1 | ssp-11 |
| WBGene00006044 | 2.684748561 | 0.20978145 | 1 | ssp-16 |
| WBGene00006048 | 1.247575581 | 0.7549672 | 1 | ssp-31 |
| WBGene00006050 | 1.356391787 | 0.32292824 | 1 | ssq-1 |
| WBGene00006051 | 1.167730018 | 0.35283169 | 1 | ssq-2 |
| WBGene00006052 | 1.996320439 | 0.26902334 | 1 | ssq-3 |
| WBGene00006053 | 1.109555143 | 0.32408088 | 1 | ssq-4 |
| WBGene00006055 | 0.189502613 | 0.78814372 | 0.88342616 | ssr-2 |
| WBGene00006056 | 2.301544115 | 0.06836622 | 1 | sss-1 |
| WBGene00006058 | -0.680130008 | 0.72372989 | 1 | sst-20 |
| WBGene00006059 | -0.226410068 | 0.24246168 | 0.45004036 | stc-1 |
| WBGene00006060 | -0.1935531 | 0.81627903 | 0.90037646 | sth-1 |
| WBGene00006061 | -0.50834809 | 0.01041718 | 0.0535004 | stl-1 |
| WBGene00006062 | -0.260583481 | 0.29820236 | 0.50806551 | stn-1 |
| WBGene00006063 | -0.45441419 | 0.1618582 | 0.34774564 | sto-1 |
| WBGene00006064 | 0.434335571 | 0.52834325 | 0.71265586 | sto-2 |
| WBGene00006065 | 2.865268649 | 0.16077931 | 1 | sto-3 |
| WBGene00006066 | 0.423675187 | 0.14436811 | 0.32289507 | sto-4 |
| WBGene00006067 | 0.452149369 | 0.35433546 | 0.56271901 | sto-5 |
| WBGene00006068 | 1.173529438 | 0.12146819 | 0.28859484 | sto-6 |
| WBGene00006069 | 0.377828795 | 0.92526585 | 1 | str-1 |
| WBGene00006071 | -0.245531902 | 0.94070824 | 1 | str-3 |
| WBGene00006072 | 0.377828795 | 0.92526585 | 1 | str-4 |
| WBGene00006075 | 0.983488853 | 0.80601565 | 1 | str-7 |
| WBGene00006076 | 0.377828795 | 0.92526585 | 1 | str-8 |
| WBGene00006077 | -0.014519762 | 0.99603746 | 1 | str-9 |
| WBGene00006080 | -0.668264701 | 0.85551153 | 1 | str-12 |
| WBGene00006084 | -1.159714244 | 0.75790634 | 1 | str-16 |
| WBGene00006085 | 0.281454817 | 0.94435481 | 1 | str-18 |
| WBGene00006086 | -1.159714244 | 0.75790634 | 1 | str-19 |
| WBGene00006097 | -0.156269293 | 0.96924949 | 1 | str-31 |
| WBGene00006102 | -0.156269293 | 0.96924949 | 1 | str-37 |
| WBGene00006103 | 0.446458324 | 0.87164163 | 1 | str-38 |
| WBGene00006109 | -0.284202651 | 0.88416022 | 1 | str-44 |
| WBGene00006110 | -2.934840138 | 0.2654436 | 1 | str-45 |
| WBGene00006111 | 1.653460415 | 0.50712258 | 1 | str-46 |
| WBGene00006112 | -0.156269293 | 0.96924949 | 1 | str-47 |
| WBGene00006120 | -2.171326218 | 0.36522606 | 1 | str-55 |
| WBGene00006122 | 1.413377704 | 0.60673863 | 1 | str-57 |
| WBGene00006124 | 0.574508983 | 0.8551971 | 1 | str-60 |
| WBGene00006125 | 0.574508983 | 0.8551971 | 1 | str-61 |
| WBGene00006126 | -1.14715222 | 0.68932198 | 1 | str-63 |

| | | | | |
|---|---|---|---|---|
| WBGene00006127 | 0.281454817 | 0.94435481 | 1 | str-64 |
| WBGene00006137 | 1.413377704 | 0.60673863 | 1 | str-77 |
| WBGene00006138 | 0.688793783 | 0.86378755 | 1 | str-78 |
| WBGene00006139 | 0.377828795 | 0.92526585 | 1 | str-79 |
| WBGene00006140 | -2.705999569 | 0.42423051 | 1 | str-81 |
| WBGene00006143 | -0.22461868 | 0.93331196 | 1 | str-84 |
| WBGene00006146 | -1.901856996 | 0.64072034 | 1 | str-87 |
| WBGene00006148 | 0.688793783 | 0.86378755 | 1 | str-89 |
| WBGene00006149 | 0.688793783 | 0.86378755 | 1 | str-90 |
| WBGene00006150 | 0.281454817 | 0.94435481 | 1 | str-92 |
| WBGene00006152 | -0.156269293 | 0.96924949 | 1 | str-94 |
| WBGene00006153 | 2.511571436 | 0.28071936 | 1 | str-96 |
| WBGene00006154 | 0.574508983 | 0.8551971 | 1 | str-97 |
| WBGene00006162 | 2.166011856 | 0.42253177 | 1 | str-111 |
| WBGene00006163 | 1.609079088 | 0.42451899 | 1 | str-112 |
| WBGene00006164 | 1.181686151 | 0.7579682 | 1 | str-113 |
| WBGene00006165 | -0.156269293 | 0.96924949 | 1 | str-114 |
| WBGene00006166 | -0.156269293 | 0.96924949 | 1 | str-115 |
| WBGene00006167 | 2.449301747 | 0.16428647 | 1 | str-116 |
| WBGene00006170 | -1.969805064 | 0.62862569 | 1 | str-119 |
| WBGene00006172 | 0.688793783 | 0.86378755 | 1 | str-121 |
| WBGene00006176 | 1.687239501 | 0.59403242 | 1 | str-125 |
| WBGene00006179 | 0.075260937 | 0.97569838 | 1 | str-130 |
| WBGene00006180 | 1.955750377 | 0.44285624 | 1 | str-131 |
| WBGene00006182 | 0.574508983 | 0.8551971 | 1 | str-134 |
| WBGene00006183 | -0.18557176 | 0.95664217 | 1 | str-135 |
| WBGene00006189 | -1.901856996 | 0.64072034 | 1 | str-141 |
| WBGene00006191 | 0.377828795 | 0.92526585 | 1 | str-144 |
| WBGene00006192 | 1.413377704 | 0.60673863 | 1 | str-145 |
| WBGene00006193 | 1.784345117 | 0.3562364 | 1 | str-146 |
| WBGene00006195 | -0.008677305 | 0.99719169 | 1 | str-148 |
| WBGene00006198 | 1.181686151 | 0.7579682 | 1 | str-151 |
| WBGene00006201 | 1.413377704 | 0.60673863 | 1 | str-155 |
| WBGene00006203 | 0.377828795 | 0.92526585 | 1 | str-158 |
| WBGene00006205 | 0.281454817 | 0.94435481 | 1 | str-160 |
| WBGene00006207 | 1.735991307 | 0.58152819 | 1 | str-162 |
| WBGene00006208 | 1.38322681 | 0.69550305 | 1 | str-163 |
| WBGene00006210 | 1.116613983 | 0.69067248 | 1 | str-165 |
| WBGene00006211 | 1.38322681 | 0.69550305 | 1 | str-166 |
| WBGene00006212 | 1.52521745 | 0.49005573 | 1 | str-168 |
| WBGene00006213 | -0.156269293 | 0.96924949 | 1 | str-169 |
| WBGene00006215 | -0.62146648 | 0.85925752 | 1 | str-171 |
| WBGene00006216 | -1.027355405 | 0.68015432 | 1 | str-172 |

| | | | | |
|---|---|---|---|---|
| WBGene00006217 | -0.92627201 | 0.64607538 | 1 | str-173 |
| WBGene00006220 | 0.565088549 | 0.07240917 | 0.20703984 | str-176 |
| WBGene00006221 | -0.156269293 | 0.96924949 | 1 | str-177 |
| WBGene00006222 | -1.964653846 | 0.53599202 | 1 | str-178 |
| WBGene00006224 | 1.353535409 | 0.73149271 | 1 | str-180 |
| WBGene00006225 | -0.156269293 | 0.96924949 | 1 | str-181 |
| WBGene00006226 | 0.747845955 | 0.85215965 | 1 | str-182 |
| WBGene00006237 | -0.784696067 | 0.76203932 | 1 | str-204 |
| WBGene00006239 | -0.263229581 | 0.93409872 | 1 | str-206 |
| WBGene00006241 | -0.156269293 | 0.96924949 | 1 | str-208 |
| WBGene00006244 | -0.263229581 | 0.93409872 | 1 | str-211 |
| WBGene00006255 | 0.247106001 | 0.91269061 | 1 | str-224 |
| WBGene00006256 | 0.020328391 | 0.99335814 | 1 | str-225 |
| WBGene00006258 | -0.042661126 | 0.98499355 | 1 | str-227 |
| WBGene00006260 | 0.574508983 | 0.8551971 | 1 | str-229 |
| WBGene00006261 | 0.747845955 | 0.85215965 | 1 | str-230 |
| WBGene00006268 | 0.281454817 | 0.94435481 | 1 | str-240 |
| WBGene00006271 | -0.156269293 | 0.96924949 | 1 | str-244 |
| WBGene00006272 | 0.352071779 | 0.83846719 | 1 | str-245 |
| WBGene00006276 | -0.156269293 | 0.96924949 | 1 | str-249 |
| WBGene00006278 | -0.156269293 | 0.96924949 | 1 | str-252 |
| WBGene00006279 | 1.477253415 | 0.58357327 | 1 | str-253 |
| WBGene00006282 | 0.983488853 | 0.80601565 | 1 | str-256 |
| WBGene00006286 | -0.008677305 | 0.99719169 | 1 | str-260 |
| WBGene00006287 | -0.451813889 | 0.8460226 | 1 | str-261 |
| WBGene00006288 | 2.834144852 | 0.16713957 | 1 | str-262 |
| WBGene00006306 | 0.925143289 | 0.02540647 | 0.10126724 | sue-1 |
| WBGene00006307 | -0.081644721 | 0.61936639 | 0.77855043 | suf-1 |
| WBGene00006308 | -0.33205322 | 0.65964636 | 0.80368078 | sul-1 |
| WBGene00006309 | 1.187068427 | 0.02579253 | 0.10231361 | sul-2 |
| WBGene00006310 | 0.211012809 | 0.79071463 | 0.88455416 | sul-3 |
| WBGene00006311 | -0.441517077 | 0.02327009 | 0.09544667 | sun-1 |
| WBGene00006315 | -0.70576507 | 0.76841669 | 1 | sup-6 |
| WBGene00006318 | 0.098607791 | 0.91589235 | 0.95580617 | sup-9 |
| WBGene00006319 | 1.36674964 | 0.05590241 | 0.17389475 | sup-10 |
| WBGene00006321 | 0.654766801 | 0.03572141 | 0.12744529 | sup-12 |
| WBGene00006324 | 0.033042536 | 0.85070577 | 0.91824825 | sup-17 |
| WBGene00006331 | 0.614576349 | 0.00450303 | 0.02888075 | sup-26 |
| WBGene00006342 | 0.743520638 | 0.00121676 | 0.01065852 | sup-37 |
| WBGene00006349 | 0.450297485 | 0.0549192 | 0.17158364 | sur-2 |
| WBGene00006351 | 0.113181763 | 0.52420086 | 0.70887978 | sur-5 |
| WBGene00006352 | -0.24762633 | 0.0836657 | 0.22732355 | sur-6 |
| WBGene00006353 | 0.425478868 | 0.13875242 | 0.3156442 | sur-7 |

| | | | | |
|---|---|---|---|---|
| WBGene00006359 | 0.076571291 | 0.69379532 | 0.82630655 | swp-1 |
| WBGene00006363 | 0.72535663 | 0.07657881 | 0.21490008 | syd-1 |
| WBGene00006364 | 0.155541591 | 0.52609659 | 0.71045736 | syd-2 |
| WBGene00006365 | 0.436835098 | 0.3607236 | 0.56840302 | syg-1 |
| WBGene00006366 | 0.074498117 | 0.78898152 | 0.8837593 | sym-1 |
| WBGene00006367 | 0.357976155 | 0.32095463 | 0.52915674 | sym-2 |
| WBGene00006368 | -0.369574174 | 0.2690194 | 0.47750943 | sym-3 |
| WBGene00006369 | 0.333513064 | 0.16839657 | 0.3562217 | sym-4 |
| WBGene00006371 | -0.103902819 | 0.77550495 | 0.87476731 | syx-3 |
| WBGene00006372 | 0.547871436 | 0.23855408 | 0.44510298 | syx-2 |
| WBGene00006373 | -0.133135498 | 0.45595744 | 0.65311223 | syx-5 |
| WBGene00006374 | -0.686715834 | 0.01045928 | 0.05367121 | syx-4 |
| WBGene00006375 | -0.028080166 | 0.90230457 | 0.94921409 | syp-1 |
| WBGene00006376 | -0.527411932 | 0.03328378 | 0.12186322 | syp-2 |
| WBGene00006377 | -0.758045068 | 0.01259308 | 0.06143653 | syp-3 |
| WBGene00006379 | -0.081259328 | 0.80769166 | 0.89513608 | sys-1 |
| WBGene00006380 | -0.608987766 | 0.60283043 | 1 | tab-1 |
| WBGene00006381 | -0.814492118 | 0.00034046 | 0.00400927 | tac-1 |
| WBGene00006382 | 0.531713323 | 0.01011328 | 0.05231588 | taf-1 |
| WBGene00006383 | 0.131412043 | 0.58787716 | 0.75636858 | taf-2 |
| WBGene00006384 | -0.369544792 | 0.18448605 | 0.37694161 | taf-3 |
| WBGene00006385 | -0.243592218 | 0.27494906 | 0.48355295 | taf-4 |
| WBGene00006386 | -0.229739334 | 0.26621604 | 0.47477459 | taf-5 |
| WBGene00006387 | -0.472140861 | 0.06994272 | 0.20211658 | taf-6.1 |
| WBGene00006388 | -0.006586976 | 0.98978208 | 0.99486251 | taf-7.1 |
| WBGene00006389 | -0.971935241 | 4.1313E-05 | 0.00073545 | taf-7.2 |
| WBGene00006390 | -0.075667448 | 0.71605413 | 0.84004379 | taf-8 |
| WBGene00006391 | -0.738246816 | 0.00117574 | 0.01039666 | taf-9 |
| WBGene00006392 | -0.722728657 | 0.00115437 | 0.01025255 | taf-10 |
| WBGene00006393 | -1.264442258 | 7.5888E-05 | 0.0012252 | taf-11.1 |
| WBGene00006394 | -0.855101714 | 0.00026743 | 0.00330979 | taf-11.2 |
| WBGene00006395 | -0.695357296 | 0.17186204 | 0.36112443 | taf-11.3 |
| WBGene00006396 | -0.347791257 | 0.13180754 | 0.3043031 | taf-12 |
| WBGene00006397 | -0.85539074 | 6.6618E-05 | 0.00110644 | taf-13 |
| WBGene00006399 | -0.156269293 | 0.96924949 | 1 | tag-4 |
| WBGene00006402 | -0.396159382 | 0.06178237 | 0.18594442 | fut-3 |
| WBGene00006404 | -0.331029876 | 0.18484265 | 0.37727857 | tag-10 |
| WBGene00006405 | -0.074614022 | 0.6767935 | 0.81495175 | itsn-1 |
| WBGene00006406 | 0.020323905 | 0.89564434 | 0.94507369 | srgp-1 |
| WBGene00006407 | 0.724319949 | 0.00023107 | 0.00294073 | lim-9 |
| WBGene00006408 | -0.276659704 | 0.13492198 | 0.3093066 | tag-18 |
| WBGene00006409 | -0.604396148 | 0.82723443 | 1 | hdl-2 |
| WBGene00006410 | -0.416093375 | 0.09174157 | 0.2411941 | nck-1 |

| | | | | |
|---|---|---|---|---|
| WBGene00006411 | 1.817681018 | 0.12852263 | 1 | octr-1 |
| WBGene00006412 | 0.956747722 | 0.00766863 | 0.04245474 | nlg-1 |
| WBGene00006414 | -0.407454548 | 0.01652569 | 0.07432888 | raga-1 |
| WBGene00006415 | -0.269800018 | 0.17432728 | 0.3644752 | tag-30 |
| WBGene00006416 | -0.850098195 | 0.28776204 | 0.49768075 | sams-5 |
| WBGene00006418 | 1.046173273 | 0.02808624 | 0.10842449 | tag-38 |
| WBGene00006419 | 1.653460415 | 0.50712258 | 1 | oac-39 |
| WBGene00006423 | 0.987840296 | 8.7291E-05 | 0.00137279 | asd-2 |
| WBGene00006424 | 0.742360354 | 0.06303609 | 0.18845634 | ctbp-1 |
| WBGene00006428 | 1.211155073 | 0.2199257 | 0.42282151 | tkr-3 |
| WBGene00006429 | -0.478047187 | 0.66423054 | 1 | arrd-3 |
| WBGene00006430 | -0.076000401 | 0.82898857 | 0.90762341 | tag-51 |
| WBGene00006431 | -0.315159162 | 0.22232429 | 0.42581467 | tag-52 |
| WBGene00006432 | -0.044795247 | 0.8403939 | 0.91237519 | tag-53 |
| WBGene00006433 | -0.147836992 | 0.37796727 | 0.58562259 | sdhb-1 |
| WBGene00006434 | -0.338490597 | 0.00494586 | 0.0309367 | prdx-2 |
| WBGene00006435 | -1.159714244 | 0.75790634 | 1 | stdh-4 |
| WBGene00006436 | 2.243729083 | 1.38E-22 | 5.76E-20 | ttn-1 |
| WBGene00006437 | 0.082182456 | 0.64557825 | 0.79536368 | mrck-1 |
| WBGene00006438 | 0.615871954 | 0.00065145 | 0.00658006 | nrfl-1 |
| WBGene00006439 | -0.147476582 | 0.27375011 | 0.48230645 | ant-1.1 |
| WBGene00006440 | -0.686811609 | 0.00182587 | 0.01452681 | tag-63 |
| WBGene00006441 | 1.152085005 | 0.06288796 | 0.18816681 | cpna-4 |
| WBGene00006442 | -0.044356872 | 0.8293359 | 0.9076639 | tag-65 |
| WBGene00006443 | 0.041033355 | 0.83755845 | 0.91143626 | pak-2 |
| WBGene00006444 | 0.420815925 | 0.12788584 | 0.29864109 | shn-1 |
| WBGene00006445 | 0.583683922 | 0.20752598 | 0.40703925 | tag-68 |
| WBGene00006446 | -0.494908045 | 0.02931827 | 0.11144289 | atx-3 |
| WBGene00006447 | -0.414372235 | 0.06088646 | 0.18450529 | tag-72 |
| WBGene00006448 | -0.438876849 | 0.04570892 | 0.15100733 | glod-4 |
| WBGene00006449 | 2.955131678 | 0.17939373 | 1 | alg-4 |
| WBGene00006450 | 0.545158163 | 0.02110908 | 0.08895707 | tag-77 |
| WBGene00006452 | -0.745826353 | 0.00048512 | 0.00524473 | heh-1 |
| WBGene00006454 | 0.649890158 | 0.26397835 | 0.47252111 | aex-4 |
| WBGene00006459 | 0.213621141 | 0.77545597 | 0.87476731 | tag-89 |
| WBGene00006460 | -0.256694241 | 0.05907613 | 0.18062033 | ppm-1 |
| WBGene00006461 | -0.742709273 | 0.09078561 | 0.23962958 | tag-96 |
| WBGene00006462 | 1.426347102 | 0.00157159 | 0.01297433 | svh-5 |
| WBGene00006463 | -0.648425668 | 0.00049489 | 0.0053219 | nduf-2.2 |
| WBGene00006464 | 0.355716516 | 0.57132398 | 0.74561373 | bath-47 |
| WBGene00006465 | -0.373816093 | 0.04463911 | 0.1481584 | fntb-1 |
| WBGene00006466 | 0.10671339 | 0.67992919 | 0.81647553 | tag-115 |
| WBGene00006467 | 1.577590387 | 0.25142417 | 1 | C01B7.5 |

| | | | | |
|---|---|---|---|---|
| WBGene00006467 | 0.662064546 | 0.05950741 | 0.18139981 | magu-2 |
| WBGene00006468 | 1.010579505 | 0.01006097 | 0.05210254 | rhgf-1 |
| WBGene00006470 | 0.611864581 | 0.08924187 | 0.23707387 | tag-120 |
| WBGene00006472 | -0.025121306 | 0.88714879 | 0.94044069 | tag-123 |
| WBGene00006473 | -0.625644948 | 0.02930592 | 0.11144289 | tag-124 |
| WBGene00006474 | -0.395423422 | 0.02537046 | 0.10119011 | wdr-5.1 |
| WBGene00006475 | 1.054408163 | 0.05974903 | 0.18180776 | tyra-3 |
| WBGene00006476 | 0.195324416 | 0.51079307 | 0.69734404 | rhgf-2 |
| WBGene00006477 | -0.223966757 | 0.50515165 | 0.69206118 | chup-1 |
| WBGene00006478 | -0.496285714 | 0.03649678 | 0.12929894 | tag-131 |
| WBGene00006479 | 0.324526264 | 0.13962592 | 0.3168595 | tmbi-4 |
| WBGene00006481 | -0.56158176 | 0.00053614 | 0.0056557 | plrg-1 |
| WBGene00006483 | 0.392365248 | 0.09384621 | 0.24460859 | dgk-4 |
| WBGene00006484 | 0.747845955 | 0.85215965 | 1 | tag-138 |
| WBGene00006486 | 1.50184354 | 0.1052054 | 0.2640138 | zipt-21 |
| WBGene00006487 | -0.358514592 | 0.84021732 | 1 | zipt-22 |
| WBGene00006489 | -0.278566614 | 0.20261811 | 0.40156488 | zhit-3 |
| WBGene00006490 | 0.408558985 | 0.03270611 | 0.12062115 | kdin-1 |
| WBGene00006491 | -0.214234599 | 0.40788449 | 0.6141434 | acl-3 |
| WBGene00006492 | -0.270811871 | 0.15776361 | 0.34264902 | let-391 |
| WBGene00006493 | 0.245996316 | 0.27355952 | 0.48215195 | tag-147 |
| WBGene00006494 | -0.372250567 | 0.04750965 | 0.15547566 | zipt-7.2 |
| WBGene00006495 | 0.745769006 | 0.0039022 | 0.02574819 | cpna-1 |
| WBGene00006496 | 0.033692351 | 0.89870672 | 0.94694691 | cgef-1 |
| WBGene00006497 | 0.101837422 | 0.57410426 | 0.74771507 | tag-151 |
| WBGene00006498 | 0.125099751 | 0.81810733 | 0.90141039 | ten-1 |
| WBGene00006499 | -0.338002669 | 0.05435817 | 0.17022506 | tag-153 |
| WBGene00006503 | -0.231634454 | 0.26712171 | 0.47544153 | snx-3 |
| WBGene00006504 | 0.713663752 | 0.00205548 | 0.01590109 | kcc-1 |
| WBGene00006505 | -0.014294699 | 0.97863261 | 0.98873753 | lagr-1 |
| WBGene00006508 | 0.307042848 | 0.1123022 | 0.27460243 | tns-1 |
| WBGene00006509 | 0.540304553 | 0.80307697 | 1 | tag-164 |
| WBGene00006510 | 0.727393288 | 4.29E-07 | 1.6421E-05 | mtrr-1 |
| WBGene00006512 | -0.268559764 | 0.09753146 | 0.25102055 | abcf-1 |
| WBGene00006513 | 1.1120842 | 0.00282277 | 0.020326 | rimb-1 |
| WBGene00006514 | -0.541458339 | 0.0012087 | 0.01061088 | tdp-1 |
| WBGene00006515 | -0.658409811 | 0.00029153 | 0.00352534 | txdc-9 |
| WBGene00006516 | -0.256585602 | 0.16323137 | 0.34927602 | vps-16 |
| WBGene00006517 | -0.084355254 | 0.83132849 | 0.90861273 | C11H1.8 |
| WBGene00006517 | 0.132704255 | 0.57571173 | 0.74852389 | madd-3 |
| WBGene00006518 | -0.258840735 | 0.14900841 | 0.3302503 | bckd-1B |
| WBGene00006519 | -0.427849632 | 0.00507282 | 0.03153565 | tag-174 |
| WBGene00006520 | -0.458062195 | 0.18001769 | 0.37228489 | tag-175 |

| | | | | |
|---|---|---|---|---|
| WBGene00006522 | 0.680762278 | 0.560202 | 1 | abcx-1 |
| WBGene00006523 | -0.209666171 | 0.27185506 | 0.48056732 | tam-1 |
| WBGene00006524 | 0.105174085 | 0.7481659 | 0.85972621 | tap-1 |
| WBGene00006525 | 0.998255591 | 0.41498944 | 1 | tax-2 |
| WBGene00006526 | 0.540516348 | 0.61999668 | 1 | tax-4 |
| WBGene00006527 | 1.032250934 | 5.4932E-06 | 0.00013611 | tax-6 |
| WBGene00006528 | -0.675608956 | 4.26E-07 | 1.641E-05 | tba-1 |
| WBGene00006529 | -0.595028049 | 1.4717E-05 | 0.00031292 | tba-2 |
| WBGene00006530 | -0.211230219 | 0.31894454 | 0.52680654 | tba-4 |
| WBGene00006531 | 0.597503483 | 0.30768903 | 0.51633069 | tba-5 |
| WBGene00006532 | 0.07111751 | 0.9458948 | 1 | tba-6 |
| WBGene00006533 | 0.242850715 | 0.50243078 | 0.68999878 | tba-7 |
| WBGene00006534 | 1.200997931 | 0.03867007 | 0.13448678 | tba-8 |
| WBGene00006535 | 0.551150434 | 0.36013617 | 0.5682386 | tba-9 |
| WBGene00006536 | -0.549873902 | 0.00022097 | 0.002848 | tbb-1 |
| WBGene00006537 | -0.437455745 | 0.00289405 | 0.0206808 | tbb-2 |
| WBGene00006538 | -0.102689674 | 0.79178821 | 0.8850619 | tbb-4 |
| WBGene00006539 | 0.052840866 | 0.92717424 | 0.96171059 | tbb-6 |
| WBGene00006540 | -0.382310978 | 0.03189969 | 0.11847911 | tbg-1 |
| WBGene00006541 | -0.539983731 | 0.0013855 | 0.01177672 | tbh-1 |
| WBGene00006542 | -0.183489906 | 0.33384018 | 0.54208287 | tbp-1 |
| WBGene00006543 | 0.946076264 | 0.00495224 | 0.03096042 | tbx-2 |
| WBGene00006544 | -1.285450047 | 0.51018772 | 1 | tbx-7 |
| WBGene00006545 | -2.796568588 | 0.01946957 | 1 | tbx-8 |
| WBGene00006546 | -0.786021257 | 0.0185743 | 0.08125742 | tbx-9 |
| WBGene00006547 | -2.289673511 | 4.7693E-05 | 0.00082838 | tbx-11 |
| WBGene00006550 | -2.200241912 | 0.17833465 | 1 | tbx-31 |
| WBGene00006551 | -1.482971329 | 0.37271799 | 1 | tbx-32 |
| WBGene00006552 | -0.813110887 | 0.04059738 | 0.13892131 | tbx-33 |
| WBGene00006553 | 0.688793783 | 0.86378755 | 1 | tbx-34 |
| WBGene00006554 | -1.85013592 | 0.28944217 | 1 | tbx-35 |
| WBGene00006555 | -0.32057996 | 0.31843951 | 0.52636816 | tbx-36 |
| WBGene00006556 | -2.592940716 | 0.03188781 | 1 | tbx-37 |
| WBGene00006557 | -2.159357589 | 0.01664395 | 0.07473157 | tbx-38 |
| WBGene00006558 | 1.472766509 | 0.46533185 | 1 | tbx-39 |
| WBGene00006559 | 0.380932494 | 0.82689228 | 1 | tbx-40 |
| WBGene00006561 | -0.238930226 | 0.58520152 | 0.75465941 | tcl-2 |
| WBGene00006562 | -0.628838674 | 0.00010101 | 0.00154346 | tdc-1 |
| WBGene00006563 | -0.072528241 | 0.84475131 | 0.91482702 | teg-1 |
| WBGene00006565 | -0.231304734 | 0.34497778 | 0.55365172 | tfg-1 |
| WBGene00006566 | -0.052856988 | 0.82501314 | 0.90573149 | tgt-1 |
| WBGene00006567 | -0.517393288 | 0.02407144 | 0.09763639 | tgt-2 |
| WBGene00006568 | -0.829706632 | 1.1196E-05 | 0.00024987 | thk-1 |

| | | | | |
|---|---|---|---|---|
| WBGene00006570 | 0.635134264 | 0.15158793 | 0.33336244 | tig-2 |
| WBGene00006571 | -0.114385409 | 0.50279949 | 0.69000662 | tim-1 |
| WBGene00006572 | -0.278662582 | 0.31243007 | 0.52061673 | tin-9.1 |
| WBGene00006573 | -0.765945215 | 0.00292377 | 0.02081178 | tin-10 |
| WBGene00006574 | -0.96839818 | 6.7945E-06 | 0.00016271 | tin-13 |
| WBGene00006575 | 1.096751742 | 6.6462E-05 | 0.00110536 | tir-1 |
| WBGene00006576 | 0.138074355 | 0.88183197 | 0.93708197 | tkr-1 |
| WBGene00006577 | -3.26098E-06 | 0.99998741 | 0.99998741 | tlf-1 |
| WBGene00006578 | 0.01322296 | 0.96634852 | 0.9830824 | tli-1 |
| WBGene00006579 | 0.058087838 | 0.75446982 | 0.86283239 | tlk-1 |
| WBGene00006580 | 0.616336394 | 0.0808151 | 0.2222873 | tlp-1 |
| WBGene00006582 | 0.849059546 | 0.01412253 | 0.06671658 | tmd-2 |
| WBGene00006583 | -0.161654264 | 0.47537252 | 0.66815209 | tnc-2 |
| WBGene00006584 | 0.58695438 | 0.00190044 | 0.01495346 | tni-1 |
| WBGene00006585 | 0.779462862 | 0.00373565 | 0.02496096 | tni-3 |
| WBGene00006586 | 0.17322163 | 0.42886818 | 0.63152074 | tni-4 |
| WBGene00006587 | 0.174827618 | 0.30291914 | 0.51211509 | tnt-2 |
| WBGene00006588 | 0.351656266 | 0.11608996 | 0.28048782 | tnt-3 |
| WBGene00006589 | 0.669828316 | 0.00365461 | 0.02455145 | tnt-4 |
| WBGene00006590 | -0.198347314 | 0.3007624 | 0.51035984 | toc-1 |
| WBGene00006591 | 0.5356817 | 0.15385628 | 0.33687451 | toh-1 |
| WBGene00006592 | 0.593674025 | 0.02754176 | 0.10693462 | dpy-31 |
| WBGene00006593 | 0.108662614 | 0.62803276 | 0.78384126 | tol-1 |
| WBGene00006594 | 1.287804365 | 7.42E-08 | 3.6038E-06 | tom-1 |
| WBGene00006595 | -0.291327362 | 0.04115219 | 0.14038458 | top-1 |
| WBGene00006596 | -0.101417567 | 0.68989215 | 0.82353771 | top-3 |
| WBGene00006597 | 1.289866859 | 0.41483808 | 1 | tor-1 |
| WBGene00006598 | 1.851444818 | 0.53669729 | 1 | tor-2 |
| WBGene00006599 | 1.00274369 | 0.00015366 | 0.00212963 | tpa-1 |
| WBGene00006600 | 0.324019505 | 0.74695619 | 1 | tph-1 |
| WBGene00006601 | -0.349466407 | 0.04890387 | 0.15858491 | tpi-1 |
| WBGene00006602 | 0.938630339 | 3.9466E-05 | 0.00070991 | tps-1 |
| WBGene00006603 | 0.536559164 | 0.03556552 | 0.12715091 | tps-2 |
| WBGene00006604 | 0.66263066 | 0.03325354 | 0.12180825 | tra-1 |
| WBGene00006605 | 0.28592894 | 0.20776567 | 0.40731196 | tra-2 |
| WBGene00006606 | -0.515027284 | 0.03089997 | 0.11603739 | tra-3 |
| WBGene00006607 | 0.154084615 | 0.33192811 | 0.53982957 | tre-1 |
| WBGene00006608 | 0.122768708 | 0.55487013 | 0.73238519 | tre-2 |
| WBGene00006609 | 0.884954255 | 0.00026325 | 0.00326138 | tre-3 |
| WBGene00006610 | 0.963318385 | 0.08547732 | 0.23054671 | tre-4 |
| WBGene00006611 | 1.158789287 | 0.04119411 | 0.14044867 | tre-5 |
| WBGene00006612 | -2.095601 | 0.38719277 | 1 | trf-1 |
| WBGene00006613 | -0.325404989 | 0.09189391 | 0.24130927 | trm-1 |

| | | | | |
|---|---|---|---|---|
| WBGene00006614 | 0.946190458 | 0.09077817 | 0.23962958 | trp-1 |
| WBGene00006615 | 1.027015493 | 0.00987824 | 0.05133254 | trp-2 |
| WBGene00006616 | 0.94925817 | 0.20403833 | 0.40345168 | trp-4 |
| WBGene00006617 | -0.174579147 | 0.25253284 | 0.460568 | tars-1 |
| WBGene00006618 | 0.25685218 | 0.41140675 | 0.61673412 | trt-1 |
| WBGene00006619 | -0.776950435 | 0.00081431 | 0.00785264 | try-1 |
| WBGene00006620 | 1.063630609 | 0.16556865 | 0.35235215 | svh-1 |
| WBGene00006621 | -0.269217801 | 0.7526859 | 0.86217294 | try-3 |
| WBGene00006622 | 1.600388976 | 0.22559065 | 1 | try-4 |
| WBGene00006623 | 0.818139016 | 0.59661243 | 1 | try-5 |
| WBGene00006624 | 0.395548973 | 0.85222539 | 1 | try-6 |
| WBGene00006625 | 0.281454817 | 0.94435481 | 1 | try-7 |
| WBGene00006626 | -0.192606182 | 0.32464851 | 0.53307514 | tsn-1 |
| WBGene00006627 | 0.883162243 | 0.10977614 | 0.27034324 | tsp-1 |
| WBGene00006628 | 2.088362972 | 0.00308351 | 0.02162733 | tsp-2 |
| WBGene00006629 | 0.477193783 | 0.58461711 | 0.75436671 | tsp-3 |
| WBGene00006630 | 2.460743752 | 0.26152462 | 1 | tsp-4 |
| WBGene00006631 | -0.156269293 | 0.96924949 | 1 | tsp-5 |
| WBGene00006632 | 0.904894574 | 0.24581386 | 0.45379649 | tsp-6 |
| WBGene00006633 | 0.55066671 | 0.27378766 | 0.48230645 | tsp-7 |
| WBGene00006634 | -0.185209587 | 0.52467314 | 0.70920247 | tsp-8 |
| WBGene00006635 | -0.126690695 | 0.73751359 | 0.85334533 | tsp-9 |
| WBGene00006636 | 1.233791474 | 0.0018804 | 0.01486321 | tsp-10 |
| WBGene00006637 | 1.048990941 | 0.10069554 | 0.25586953 | tsp-11 |
| WBGene00006638 | -0.158392114 | 0.36184191 | 0.56967189 | tsp-12 |
| WBGene00006639 | 0.579928635 | 0.4784652 | 0.67082322 | tsp-13 |
| WBGene00006640 | 0.528485237 | 0.31218399 | 0.52042096 | tsp-14 |
| WBGene00006641 | 0.644134153 | 0.11753424 | 0.28245932 | tsp-15 |
| WBGene00006642 | 1.069818256 | 0.13817475 | 0.31492014 | tsp-16 |
| WBGene00006643 | -0.090445443 | 0.8933538 | 0.94378935 | tsp-17 |
| WBGene00006644 | 0.281454817 | 0.94435481 | 1 | tsp-18 |
| WBGene00006645 | -0.008677305 | 0.99719169 | 1 | tsp-19 |
| WBGene00006646 | 0.073195759 | 0.90813814 | 0.95194377 | tsp-20 |
| WBGene00006647 | 0.13231339 | 0.68815589 | 0.82244904 | tsr-1 |
| WBGene00006648 | -0.32516847 | 0.12841247 | 0.29912812 | ttb-1 |
| WBGene00006649 | -0.51984016 | 0.03795034 | 0.13274419 | tth-1 |
| WBGene00006650 | 1.880394017 | 8.82E-25 | 4.87E-22 | tts-1 |
| WBGene00006651 | -0.099442929 | 0.70131196 | 0.83141078 | tts-2 |
| WBGene00006652 | 0.308451299 | 0.53574649 | 0.7178681 | ttx-1 |
| WBGene00006654 | -0.973467954 | 0.49503081 | 1 | ttx-3 |
| WBGene00006655 | 0.838486835 | 0.20586304 | 0.40545208 | tub-1 |
| WBGene00006656 | 0.227766055 | 0.61165886 | 0.77323497 | twk-1 |
| WBGene00006657 | 0.78688518 | 0.0625579 | 0.1875347 | twk-2 |

| | | | | |
|---|---|---|---|---|
| WBGene00006658 | 0.555923331 | 0.7162765 | 1 | twk-3 |
| WBGene00006659 | 0.434486085 | 0.76593115 | 1 | twk-4 |
| WBGene00006660 | 0.394345685 | 0.40856064 | 0.61443511 | twk-5 |
| WBGene00006661 | 0.281454817 | 0.94435481 | 1 | twk-6 |
| WBGene00006662 | 0.751936284 | 0.08385698 | 0.22771361 | twk-7 |
| WBGene00006663 | 0.900118399 | 0.00594496 | 0.03536394 | twk-8 |
| WBGene00006664 | 1.490795543 | 0.05816341 | 0.17872993 | twk-9 |
| WBGene00006665 | 1.602341335 | 0.17384758 | 1 | twk-10 |
| WBGene00006666 | 0.673924396 | 0.23944163 | 0.44640333 | twk-11 |
| WBGene00006667 | 1.105820317 | 0.04164166 | 0.14157697 | twk-12 |
| WBGene00006668 | 0.332360466 | 0.43441483 | 0.63575583 | twk-13 |
| WBGene00006669 | -0.065057981 | 0.92756944 | 0.96200873 | twk-14 |
| WBGene00006670 | 0.657760481 | 0.35423429 | 0.56263539 | twk-16 |
| WBGene00006671 | 1.139292054 | 0.02142639 | 0.09013782 | twk-17 |
| WBGene00006672 | 0.486959698 | 0.0865362 | 0.23249302 | twk-18 |
| WBGene00006673 | -0.213937883 | 0.92749767 | 1 | twk-20 |
| WBGene00006674 | 1.148422303 | 0.41178058 | 1 | twk-21 |
| WBGene00006675 | 0.48487927 | 0.47971589 | 0.67161331 | twk-22 |
| WBGene00006676 | 0.371838715 | 0.71706858 | 1 | twk-23 |
| WBGene00006677 | 0.50820849 | 0.29455336 | 0.50490709 | twk-24 |
| WBGene00006678 | 0.401898132 | 0.34814639 | 0.55660328 | twk-25 |
| WBGene00006679 | 1.968514836 | 2.3399E-05 | 0.00045969 | twk-26 |
| WBGene00006680 | 1.053987456 | 0.09892463 | 0.25317116 | twk-28 |
| WBGene00006681 | 0.584737601 | 0.28703541 | 0.49711057 | twk-29 |
| WBGene00006682 | 0.743899991 | 0.26078901 | 0.46942022 | twk-30 |
| WBGene00006683 | 0.9189555 | 0.03265334 | 0.12049974 | twk-31 |
| WBGene00006684 | 0.04197419 | 0.96759286 | 1 | twk-32 |
| WBGene00006685 | 0.525635834 | 0.26508631 | 0.47390941 | twk-33 |
| WBGene00006686 | 0.775216964 | 0.10781676 | 0.26759651 | twk-34 |
| WBGene00006687 | -0.29732853 | 0.74482415 | 0.85755429 | twk-35 |
| WBGene00006688 | 0.316039621 | 0.83316375 | 1 | twk-36 |
| WBGene00006690 | 0.349149665 | 0.51518085 | 0.70168394 | twk-39 |
| WBGene00006691 | 1.54059853 | 0.10432923 | 0.26214014 | twk-40 |
| WBGene00006692 | 0.254023556 | 0.64368897 | 0.79412944 | twk-42 |
| WBGene00006693 | 0.568344271 | 0.18294283 | 0.37512395 | twk-43 |
| WBGene00006694 | -1.392857614 | 0.34128121 | 1 | twk-44 |
| WBGene00006695 | -0.035785178 | 0.9779073 | 1 | twk-45 |
| WBGene00006696 | 0.514290234 | 0.31471619 | 0.5225568 | twk-46 |
| WBGene00006697 | -0.175684568 | 0.29805501 | 0.50788573 | uaf-1 |
| WBGene00006698 | -0.526250277 | 0.00315038 | 0.0219903 | uaf-2 |
| WBGene00006699 | -0.066430447 | 0.7258921 | 0.84579072 | uba-1 |
| WBGene00006700 | -0.340661471 | 0.04175511 | 0.14192294 | uba-2 |
| WBGene00006701 | -0.02736085 | 0.9056924 | 0.95088303 | ubc-1 |

| | | | | |
|---|---|---|---|---|
| WBGene00006702 | -0.486264936 | 0.02105295 | 0.08883932 | ubc-3 |
| WBGene00006703 | -0.476352681 | 0.01947667 | 0.08396019 | ubc-6 |
| WBGene00006704 | -0.752819525 | 5.04E-07 | 1.9091E-05 | ubc-7 |
| WBGene00006705 | 0.602229138 | 0.10085162 | 0.2559447 | ubc-8 |
| WBGene00006706 | -0.533244342 | 0.00048484 | 0.00524473 | ubc-9 |
| WBGene00006707 | -0.51262405 | 0.00751948 | 0.0418972 | ubc-12 |
| WBGene00006708 | -0.470504962 | 0.01474937 | 0.0688474 | ubc-13 |
| WBGene00006709 | -0.749814037 | 0.00128714 | 0.01113788 | ubc-14 |
| WBGene00006710 | -0.043037542 | 0.8656357 | 0.92776157 | ubc-15 |
| WBGene00006711 | -0.23410796 | 0.35844084 | 0.56668975 | ubc-16 |
| WBGene00006712 | 0.530756316 | 0.45684524 | 0.65384393 | ubc-17 |
| WBGene00006713 | -0.585220034 | 0.00050333 | 0.00539265 | ubc-18 |
| WBGene00006714 | -0.935167932 | 0.08520711 | 0.23019943 | ubc-19 |
| WBGene00006715 | -0.501292792 | 0.00371958 | 0.02489492 | ubc-20 |
| WBGene00006716 | 0.042629834 | 0.98004254 | 1 | ubc-21 |
| WBGene00006717 | -0.09600296 | 0.92232806 | 1 | ubc-22 |
| WBGene00006718 | 0.151204488 | 0.83778128 | 0.91143626 | ubc-23 |
| WBGene00006719 | 0.747845955 | 0.85215965 | 1 | ubc-24 |
| WBGene00006720 | -0.003926455 | 0.97950196 | 0.98905882 | ubc-25 |
| WBGene00006721 | -0.472447164 | 0.22338922 | 0.42744973 | ubh-1 |
| WBGene00006722 | -0.261707202 | 0.3773445 | 0.58512604 | ubh-2 |
| WBGene00006723 | -0.621113231 | 0.00336428 | 0.02302494 | ubh-3 |
| WBGene00006724 | -0.304539769 | 0.09827096 | 0.25225214 | ubh-4 |
| WBGene00006725 | -0.300441894 | 0.02381546 | 0.09696295 | ubl-1 |
| WBGene00006726 | -1.088530171 | 0.00011552 | 0.00171244 | ubl-5 |
| WBGene00006727 | -0.214274387 | 0.17285295 | 0.36251644 | ubq-1 |
| WBGene00006728 | -0.395695794 | 0.0013717 | 0.01170328 | ubq-2 |
| WBGene00006729 | -0.515264 | 0.06603915 | 0.19432412 | ucp-4 |
| WBGene00006730 | -0.482986049 | 0.01580254 | 0.07215987 | uev-1 |
| WBGene00006731 | 0.951323067 | 0.52763025 | 1 | uev-2 |
| WBGene00006732 | 0.330117745 | 0.23473013 | 0.44067706 | uev-3 |
| WBGene00006733 | -0.213441084 | 0.25393735 | 0.46188065 | ufd-1 |
| WBGene00006734 | -0.028834213 | 0.84978458 | 0.91790343 | ufd-2 |
| WBGene00006735 | -0.176497635 | 0.36847183 | 0.57602018 | ula-1 |
| WBGene00006736 | -0.152381475 | 0.4048655 | 0.61175756 | ulp-1 |
| WBGene00006737 | -0.14166606 | 0.44120912 | 0.64136972 | ulp-2 |
| WBGene00006738 | -0.569165372 | 0.07736718 | 0.21573609 | ulp-3 |
| WBGene00006739 | 0.231950066 | 0.27865973 | 0.48767511 | ulp-4 |
| WBGene00006740 | -0.308941286 | 0.3939763 | 0.60200405 | ulp-5 |
| WBGene00006741 | 0.578850699 | 0.03436242 | 0.12477303 | unc-1 |
| WBGene00006742 | 1.103087008 | 0.00036758 | 0.00424224 | unc-2 |
| WBGene00006743 | 0.101868441 | 0.8989789 | 0.94694691 | unc-3 |
| WBGene00006744 | -0.454150054 | 0.68579095 | 1 | unc-4 |

| | | | | |
|---|---|---|---|---|
| WBGene00006745 | 0.485240147 | 0.12796747 | 0.29864109 | unc-5 |
| WBGene00006746 | -0.009345837 | 0.98171057 | 0.99043942 | unc-6 |
| WBGene00006747 | 1.182254949 | 0.00558587 | 0.03385679 | unc-7 |
| WBGene00006748 | 0.86629014 | 0.24234636 | 0.44989711 | unc-8 |
| WBGene00006749 | 0.801552713 | 0.01371971 | 0.06514311 | unc-9 |
| WBGene00006750 | 0.881749008 | 0.01542356 | 0.07109244 | unc-10 |
| WBGene00006751 | 0.197285357 | 0.361681 | 0.56954203 | unc-11 |
| WBGene00006752 | 1.291122524 | 8.25E-09 | 5.22E-07 | unc-13 |
| WBGene00006753 | 0.637310445 | 0.00069603 | 0.00695929 | unc-14 |
| WBGene00006754 | 0.986808847 | 2.1881E-06 | 6.3049E-05 | unc-15 |
| WBGene00006755 | 0.407460476 | 0.0368458 | 0.13011776 | unc-16 |
| WBGene00006756 | 0.783286117 | 0.08936444 | 0.2371834 | unc-17 |
| WBGene00006757 | 0.30718334 | 0.13919651 | 0.31635808 | unc-18 |
| WBGene00006759 | 1.860587319 | 0.00070927 | 0.00704379 | unc-22 |
| WBGene00006760 | 0.657119312 | 0.00623578 | 0.03675177 | unc-23 |
| WBGene00006761 | 0.872283318 | 0.16321568 | 0.34927602 | unc-24 |
| WBGene00006762 | 0.888967099 | 0.00148058 | 0.01243994 | unc-25 |
| WBGene00006763 | 0.175981248 | 0.42878862 | 0.63148019 | unc-26 |
| WBGene00006764 | -0.048091979 | 0.80100077 | 0.89121613 | unc-27 |
| WBGene00006765 | 1.471053738 | 0.08210967 | 0.2246134 | unc-29 |
| WBGene00006766 | 0.82134938 | 0.28835591 | 0.49842384 | unc-30 |
| WBGene00006767 | 0.983769149 | 0.00726923 | 0.04085911 | unc-31 |
| WBGene00006768 | 0.124258006 | 0.4318493 | 0.6341955 | unc-32 |
| WBGene00006769 | 0.142721515 | 0.54635974 | 0.72535304 | unc-33 |
| WBGene00006770 | -0.111986093 | 0.62543887 | 0.78210457 | unc-34 |
| WBGene00006771 | 0.820125926 | 0.00022013 | 0.00284022 | tln-1 |
| WBGene00006772 | 1.038428666 | 0.00058203 | 0.00599501 | unc-36 |
| WBGene00006773 | -0.745723635 | 5.2954E-06 | 0.00013256 | unc-37 |
| WBGene00006774 | 0.735070458 | 0.20196097 | 0.40072041 | unc-38 |
| WBGene00006775 | 0.628701402 | 0.58509151 | 1 | unc-39 |
| WBGene00006776 | 0.182536012 | 0.31228291 | 0.52044294 | unc-40 |
| WBGene00006777 | 1.403753622 | 0.00023495 | 0.00297442 | unc-41 |
| WBGene00006778 | -0.112258058 | 0.8336051 | 0.90998018 | unc-42 |
| WBGene00006779 | 0.799151714 | 5.3893E-06 | 0.00013436 | unc-43 |
| WBGene00006780 | 0.931498587 | 5.05E-07 | 1.9091E-05 | unc-44 |
| WBGene00006781 | -0.028433147 | 0.89446308 | 0.94453253 | unc-45 |
| WBGene00006782 | -0.323032442 | 0.74146047 | 0.85569007 | unc-46 |
| WBGene00006783 | 0.310315549 | 0.64042176 | 0.79202874 | unc-47 |
| WBGene00006784 | 1.090080407 | 0.00048013 | 0.00520936 | unc-49 |
| WBGene00006785 | -0.007574727 | 0.97493098 | 0.98674654 | unc-50 |
| WBGene00006786 | 0.612008168 | 0.01643716 | 0.07424241 | unc-51 |
| WBGene00006787 | 1.248843497 | 1.48E-08 | 8.66E-07 | unc-52 |
| WBGene00006788 | 1.028310324 | 0.00130663 | 0.01125893 | unc-53 |

| | | | | |
|---|---|---|---|---|
| WBGene00006789 | 1.586726884 | 8.37E-13 | 1.19E-10 | unc-54 |
| WBGene00006790 | 1.510744285 | 0.46009414 | 1 | unc-55 |
| WBGene00006791 | 0.160809852 | 0.4249109 | 0.62893095 | unc-57 |
| WBGene00006792 | 2.18453076 | 0.41642786 | 1 | T06H11.2 |
| WBGene00006792 | -0.156269293 | 0.96924949 | 1 | T06H11.5 |
| WBGene00006792 | 0.642775663 | 0.15574431 | 0.33959979 | unc-58 |
| WBGene00006793 | -0.496571798 | 0.00154587 | 0.01284625 | unc-59 |
| WBGene00006794 | -0.362350573 | 0.01762652 | 0.0782177 | unc-60 |
| WBGene00006795 | -0.396001948 | 0.01109354 | 0.0560494 | unc-61 |
| WBGene00006796 | 0.413783046 | 0.09320494 | 0.2435646 | unc-62 |
| WBGene00006797 | 0.630159115 | 0.22110347 | 0.42403497 | unc-63 |
| WBGene00006798 | 0.095240028 | 0.69203124 | 0.82461 | unc-64 |
| WBGene00006801 | 0.837920624 | 0.00015504 | 0.00214392 | unc-68 |
| WBGene00006802 | 0.15701966 | 0.56036093 | 0.73673213 | unc-69 |
| WBGene00006803 | 0.625538206 | 0.15529613 | 0.33904879 | unc-70 |
| WBGene00006804 | -0.255182071 | 0.12561314 | 0.29555603 | unc-71 |
| WBGene00006805 | 0.630464215 | 0.00325811 | 0.02255227 | unc-73 |
| WBGene00006807 | -0.041883578 | 0.94746534 | 0.97311595 | unc-75 |
| WBGene00006808 | 0.001780641 | 0.99188421 | 0.99606834 | unc-76 |
| WBGene00006809 | 0.627224552 | 0.12244098 | 0.29046616 | unc-77 |
| WBGene00006810 | -0.087331829 | 0.73147724 | 0.84968094 | unc-78 |
| WBGene00006811 | 1.148782012 | 0.00169617 | 0.0137014 | unc-79 |
| WBGene00006812 | 1.490054762 | 3.12E-08 | 1.6519E-06 | unc-80 |
| WBGene00006814 | 0.863708912 | 0.00036869 | 0.00425102 | unc-82 |
| WBGene00006815 | 0.900794523 | 8.3842E-06 | 0.00019538 | unc-83 |
| WBGene00006816 | 0.3960378 | 0.12393667 | 0.29308827 | unc-84 |
| WBGene00006817 | -1.033383038 | 1.02E-08 | 6.14E-07 | unc-85 |
| WBGene00006818 | 0.602216073 | 0.18044216 | 0.3725955 | unc-86 |
| WBGene00006819 | 0.288754177 | 0.05882907 | 0.18013714 | unc-87 |
| WBGene00006820 | 1.655240397 | 0.00130442 | 0.01124786 | unc-89 |
| WBGene00006822 | 0.771311614 | 0.23581205 | 0.44209298 | unc-93 |
| WBGene00006823 | 0.557673003 | 0.00280623 | 0.02026793 | unc-94 |
| WBGene00006824 | 0.406299217 | 0.26876666 | 0.47729957 | unc-95 |
| WBGene00006825 | 0.511245601 | 0.11101542 | 0.272621 | unc-96 |
| WBGene00006826 | -0.000199035 | 0.99928282 | 0.99978781 | unc-97 |
| WBGene00006827 | 0.548153377 | 0.14128236 | 0.31924607 | unc-98 |
| WBGene00006829 | -0.094453022 | 0.66678993 | 0.80890254 | unc-101 |
| WBGene00006830 | 1.040170695 | 0.00102077 | 0.00938169 | unc-103 |
| WBGene00006831 | 1.407441427 | 4.09E-08 | 2.0763E-06 | unc-104 |
| WBGene00006832 | 0.953192846 | 0.03201307 | 0.1186885 | unc-105 |
| WBGene00006833 | -0.027169084 | 0.87347085 | 0.93188133 | unc-108 |
| WBGene00006836 | 0.334521663 | 0.08652819 | 0.23249302 | unc-112 |
| WBGene00006839 | 0.460353109 | 0.03400471 | 0.12386774 | unc-115 |

| | | | | |
|---|---|---|---|---|
| WBGene00006840 | 0.147146019 | 0.39688164 | 0.60466056 | unc-116 |
| WBGene00006843 | -0.330872087 | 0.27236468 | 0.48112609 | unc-119 |
| WBGene00006844 | 0.107600226 | 0.75086008 | 0.86131823 | unc-120 |
| WBGene00006845 | 0.12063312 | 0.82724352 | 0.90686805 | unc-122 |
| WBGene00006852 | 0.710161458 | 0.09647723 | 0.24927219 | unc-129 |
| WBGene00006853 | -0.388756221 | 0.44951343 | 0.64840361 | unc-130 |
| WBGene00006856 | -0.621369563 | 2.7462E-05 | 0.00052187 | usp-14 |
| WBGene00006861 | 0.572767656 | 0.07757194 | 0.21610851 | cal-5 |
| WBGene00006862 | -0.382191012 | 0.20143887 | 0.40041719 | pnk-4 |
| WBGene00006863 | 1.014179758 | 1.3007E-05 | 0.00028301 | gyg-1 |
| WBGene00006864 | 1.625446483 | 0.08132544 | 0.22313495 | npr-16 |
| WBGene00006865 | 0.53105037 | 0.45237176 | 0.65010241 | slcf-2 |
| WBGene00006868 | 0.079377479 | 0.65070464 | 0.79864418 | vab-1 |
| WBGene00006869 | -0.418084792 | 0.41541105 | 0.62081555 | vab-2 |
| WBGene00006870 | 0.765365812 | 0.03810545 | 0.133095 | vab-3 |
| WBGene00006873 | 0.84662224 | 0.62384844 | 1 | vab-7 |
| WBGene00006874 | 0.810400905 | 0.00076211 | 0.00745592 | vab-8 |
| WBGene00006875 | 0.230672665 | 0.58316521 | 0.75332022 | vab-9 |
| WBGene00006876 | -0.250118198 | 0.84783464 | 1 | T02G6.1 |
| WBGene00006876 | 2.218714919 | 0.24686531 | 1 | T02G6.2 |
| WBGene00006876 | 1.084499433 | 8.86E-08 | 4.2182E-06 | vab-10 |
| WBGene00006881 | 0.659633785 | 0.41273012 | 0.61825496 | vab-15 |
| WBGene00006882 | 0.415681715 | 0.1586231 | 0.34340996 | vab-19 |
| WBGene00006886 | 1.686522868 | 4.54E-07 | 1.7289E-05 | vap-1 |
| WBGene00006887 | 0.336434426 | 0.21419099 | 0.41495018 | vav-1 |
| WBGene00006888 | -0.078671129 | 0.71106121 | 0.83653044 | vbh-1 |
| WBGene00006889 | -0.27857858 | 0.17869404 | 0.37079547 | pfd-3 |
| WBGene00006890 | -0.00879932 | 0.9708612 | 0.98484857 | vem-1 |
| WBGene00006893 | 0.221275277 | 0.39532162 | 0.60319213 | spon-1 |
| WBGene00006894 | 0.350788431 | 0.80291326 | 1 | ver-1 |
| WBGene00006896 | 0.431493065 | 0.11876689 | 0.28435197 | ver-3 |
| WBGene00006897 | 0.085764224 | 0.93778672 | 1 | ver-4 |
| WBGene00006902 | -2.097128985 | 1.47E-13 | 2.29E-11 | vet-6 |
| WBGene00006910 | -0.072929844 | 0.63041976 | 0.78556152 | vha-1 |
| WBGene00006911 | 0.304941175 | 0.03376344 | 0.12321068 | vha-2 |
| WBGene00006912 | 0.106981488 | 0.57002242 | 0.74439521 | vha-3 |
| WBGene00006913 | 0.006574264 | 0.96486493 | 0.98250651 | vha-4 |
| WBGene00006914 | 0.432137945 | 0.01096711 | 0.05556147 | vha-5 |
| WBGene00006915 | 1.040993701 | 4.86E-10 | 4.16E-08 | vha-6 |
| WBGene00006916 | 0.129571373 | 0.48367349 | 0.67458876 | vha-7 |
| WBGene00006917 | 0.135802713 | 0.38872509 | 0.59733089 | vha-8 |
| WBGene00006918 | -0.017078643 | 0.91464832 | 0.95571968 | vha-9 |
| WBGene00006919 | 0.054671882 | 0.73569847 | 0.8523824 | vha-10 |

| | | | | |
|---|---|---|---|---|
| WBGene00006920 | 0.207016738 | 0.20989014 | 0.41007351 | vha-11 |
| WBGene00006921 | 0.282588867 | 0.11115989 | 0.2728654 | vha-12 |
| WBGene00006922 | -0.541754002 | 0.20669129 | 0.40655714 | vhl-1 |
| WBGene00006923 | 1.578024201 | 2.39E-12 | 3.16E-10 | vhp-1 |
| WBGene00006924 | -0.230498632 | 0.07740661 | 0.21579649 | vig-1 |
| WBGene00006925 | -1.86335379 | 7.69E-07 | 2.6736E-05 | vit-1 |
| WBGene00006926 | 1.493981523 | 0.00011801 | 0.00173663 | vit-2 |
| WBGene00006927 | -3.112554327 | 3.29E-48 | 6.66E-45 | vit-3 |
| WBGene00006928 | -1.905078241 | 8.60E-19 | 2.55E-16 | vit-4 |
| WBGene00006929 | -0.281362395 | 0.18874317 | 0.38254321 | vit-5 |
| WBGene00006930 | 1.41088333 | 0.00169214 | 0.01367795 | vit-6 |
| WBGene00006931 | -0.201842651 | 0.28018896 | 0.48920419 | vps-26 |
| WBGene00006932 | 0.416088372 | 0.03524636 | 0.12660533 | vps-34 |
| WBGene00006933 | 0.106721593 | 0.57443318 | 0.74795819 | vps-35 |
| WBGene00006934 | -0.017363604 | 0.9229966 | 0.9593272 | vps-54 |
| WBGene00006935 | 0.373415109 | 0.11616632 | 0.28054097 | vars-1 |
| WBGene00006936 | 0.168928077 | 0.37616484 | 0.58381308 | glp-4 |
| WBGene00006937 | -0.067014774 | 0.71765298 | 0.84029558 | wah-1 |
| WBGene00006938 | -0.90319826 | 0.46160413 | 1 | wee-1.1 |
| WBGene00006940 | -0.16147137 | 0.36516683 | 0.5728764 | wee-1.3 |
| WBGene00006941 | 0.349668055 | 0.07297113 | 0.20811428 | wnk-1 |
| WBGene00006942 | 0.013769123 | 0.95809211 | 0.9785154 | wrk-1 |
| WBGene00006943 | -0.102264219 | 0.61481479 | 0.77514863 | wrm-1 |
| WBGene00006944 | 0.238027688 | 0.25098545 | 0.45885383 | wrn-1 |
| WBGene00006945 | -0.274275756 | 0.14389528 | 0.32210225 | wars-1 |
| WBGene00006946 | 0.305027986 | 0.21217944 | 0.41207318 | prx-10 |
| WBGene00006947 | -1.282311189 | 0.00031974 | 0.00380581 | wrt-1 |
| WBGene00006948 | -0.870283922 | 0.0576084 | 0.17778941 | wrt-2 |
| WBGene00006949 | -0.497632764 | 0.51817175 | 0.70378378 | wrt-3 |
| WBGene00006950 | 0.240670307 | 0.80867953 | 1 | wrt-4 |
| WBGene00006951 | -0.332562948 | 0.46191422 | 0.65707096 | wrt-5 |
| WBGene00006952 | -1.076308361 | 0.14237493 | 0.32034267 | wrt-6 |
| WBGene00006953 | -1.22259901 | 0.68385967 | 1 | wrt-7 |
| WBGene00006954 | 0.277349474 | 0.745327 | 0.85797053 | wrt-8 |
| WBGene00006955 | -1.603991594 | 0.15825247 | 1 | wrt-9 |
| WBGene00006956 | -0.635140439 | 0.12755359 | 0.29825012 | wrt-10 |
| WBGene00006957 | 0.13612933 | 0.53770595 | 0.71960846 | wsp-1 |
| WBGene00006958 | -0.10105446 | 0.65653645 | 0.80236354 | wve-1 |
| WBGene00006959 | 0.320956052 | 0.03556581 | 0.12715091 | xbp-1 |
| WBGene00006960 | 0.543650305 | 0.69154368 | 1 | xbx-1 |
| WBGene00006961 | 0.439469202 | 0.00807144 | 0.04416193 | xnp-1 |
| WBGene00006962 | -0.419780091 | 0.74548483 | 1 | xol-1 |
| WBGene00006963 | -0.524899947 | 0.01936205 | 0.08366608 | xpa-1 |

| | | | | |
|---|---|---|---|---|
| WBGene00006964 | -0.082759972 | 0.65377107 | 0.80103286 | xrn-2 |
| WBGene00006965 | 0.731421253 | 0.5333714 | 1 | xtr-1 |
| WBGene00006966 | -0.622437626 | 0.71508016 | 1 | xtr-2 |
| WBGene00006967 | 0.281454817 | 0.94435481 | 1 | yrn-1 |
| WBGene00006968 | -0.294289971 | 0.18069625 | 0.37259848 | yars-2 |
| WBGene00006970 | 0.392976539 | 0.35286139 | 0.56184788 | zag-1 |
| WBGene00006971 | 1.03915057 | 0.0215823 | 0.09066808 | zak-1 |
| WBGene00006974 | -0.360940387 | 0.01718247 | 0.07672391 | zen-4 |
| WBGene00006975 | 0.599802721 | 0.00308758 | 0.02164334 | zfp-1 |
| WBGene00006976 | 0.055377594 | 0.84750491 | 0.91633779 | zhp-3 |
| WBGene00006977 | -0.226559734 | 0.18625699 | 0.37916601 | zif-1 |
| WBGene00006978 | 0.395623642 | 0.26593523 | 0.47460232 | zig-1 |
| WBGene00006979 | 0.227322013 | 0.74116508 | 0.85561331 | zig-2 |
| WBGene00006980 | 0.649565478 | 0.46271144 | 0.6577426 | zig-3 |
| WBGene00006981 | 1.012303706 | 0.24780619 | 0.45583943 | zig-4 |
| WBGene00006982 | 0.142011708 | 0.79548488 | 0.8867856 | zig-5 |
| WBGene00006983 | -0.36592833 | 0.58979035 | 0.75746714 | zig-6 |
| WBGene00006984 | -0.016768268 | 0.94801413 | 0.97334795 | zig-7 |
| WBGene00006985 | 0.062522092 | 0.90715254 | 0.95142873 | zig-8 |
| WBGene00006986 | 0.68444607 | 0.07285385 | 0.20802414 | zip-1 |
| WBGene00006987 | -0.133867911 | 0.87785019 | 0.93441865 | zmp-1 |
| WBGene00006988 | -0.274788778 | 0.14516922 | 0.32392934 | zyg-1 |
| WBGene00006993 | 0.00339216 | 0.98855745 | 0.99484196 | zyg-8 |
| WBGene00006994 | 0.081387174 | 0.66485946 | 0.80771846 | zyg-9 |
| WBGene00006996 | -0.131654447 | 0.36783005 | 0.57541871 | zyg-11 |
| WBGene00006997 | -0.224267439 | 0.17309187 | 0.36289214 | zyg-12 |
| WBGene00006999 | 0.519721755 | 0.00939943 | 0.04944475 | zyx-1 |
| WBGene00007000 | -0.329460317 | 0.05345879 | 0.1680612 | tufm-1 |
| WBGene00007001 | -0.075732788 | 0.70841265 | 0.83503281 | tufm-2 |
| WBGene00007006 | 0.767608566 | 0.2573477 | 0.46529537 | npr-3 |
| WBGene00007007 | 0.133493137 | 0.63192395 | 0.78661779 | mak-2 |
| WBGene00007008 | 0.152318698 | 0.33175504 | 0.53982957 | rfp-1 |
| WBGene00007009 | 0.038434397 | 0.82750642 | 0.90692193 | wwp-1 |
| WBGene00007010 | 0.443659826 | 0.01528577 | 0.07063015 | alx-1 |
| WBGene00007011 | -0.315448137 | 0.38254151 | 0.59089523 | mdt-1.2 |
| WBGene00007012 | -0.278895701 | 0.17911779 | 0.37122037 | mdt-4 |
| WBGene00007013 | -0.343468473 | 0.08173539 | 0.22395607 | mdt-8 |
| WBGene00007014 | -0.647767188 | 0.02965503 | 0.11219748 | mdt-10 |
| WBGene00007015 | -0.322254669 | 0.23411034 | 0.44019416 | mdt-11 |
| WBGene00007016 | 0.809361516 | 2.9368E-05 | 0.0005528 | mdt-15 |
| WBGene00007017 | -0.479088399 | 0.01319491 | 0.06348166 | mdt-17 |
| WBGene00007018 | -0.389576056 | 0.12369316 | 0.29268343 | mdt-18 |
| WBGene00007019 | -0.706009721 | 0.00060837 | 0.00620166 | mdt-19 |

| | | | | |
|---|---|---|---|---|
| WBGene00007020 | -0.838226058 | 0.00281356 | 0.02029678 | mdt-20 |
| WBGene00007021 | -0.370140724 | 0.40269835 | 0.61053455 | immp-1 |
| WBGene00007022 | -0.663129227 | 0.00834467 | 0.04510805 | mdt-22 |
| WBGene00007023 | 0.233211088 | 0.305743 | 0.51458342 | mdt-27 |
| WBGene00007024 | 0.03790746 | 0.81562234 | 0.90006097 | plin-1 |
| WBGene00007025 | -0.264881904 | 0.27420723 | 0.48271631 | mdt-29 |
| WBGene00007026 | -0.552958156 | 0.09270315 | 0.2427235 | mdt-31 |
| WBGene00007027 | 0.442183685 | 0.02801985 | 0.10823705 | ssl-1 |
| WBGene00007028 | 0.338599197 | 0.16807855 | 0.35573574 | trr-1 |
| WBGene00007029 | -0.24577637 | 0.18419065 | 0.37653792 | mys-1 |
| WBGene00007030 | 0.117516826 | 0.58237989 | 0.75275995 | epc-1 |
| WBGene00007036 | -0.558804446 | 0.87818675 | 1 | sod-5 |
| WBGene00007041 | 1.145224284 | 0.00139876 | 0.01187578 | tag-180 |
| WBGene00007042 | 0.09370373 | 0.60879132 | 0.77105523 | pbrm-1 |
| WBGene00007043 | 0.506341533 | 0.06193472 | 0.18626441 | tag-179 |
| WBGene00007044 | 1.145190899 | 0.00037732 | 0.00431769 | cpna-5 |
| WBGene00007045 | 0.052888456 | 0.85302742 | 0.91927977 | tag-189 |
| WBGene00007047 | 0.397741815 | 0.16129232 | 0.34696138 | wts-1 |
| WBGene00007048 | 0.146929293 | 0.3669464 | 0.57448049 | nfx-1 |
| WBGene00007049 | -0.011639311 | 0.99275046 | 1 | tag-191 |
| WBGene00007050 | 0.918624016 | 0.0001022 | 0.00155098 | utp-20 |
| WBGene00007053 | 0.629228925 | 0.00602537 | 0.03570228 | chd-7 |
| WBGene00007054 | 0.517015358 | 0.01190642 | 0.05895427 | scpl-1 |
| WBGene00007055 | 0.776900034 | 1.6825E-05 | 0.0003522 | tag-196 |
| WBGene00007056 | -0.410377886 | 0.36977304 | 0.57741665 | crn-7 |
| WBGene00007057 | 0.623197217 | 0.77491599 | 1 | ant-1.2 |
| WBGene00007058 | 0.480275418 | 0.02490702 | 0.09974122 | dmd-6 |
| WBGene00007059 | 0.524206789 | 0.11258169 | 0.27491036 | vps-52 |
| WBGene00007060 | -0.189008336 | 0.87140475 | 1 | wht-6 |
| WBGene00007062 | 0.282842906 | 0.729981 | 0.84875496 | ebp-3 |
| WBGene00007063 | -0.180934417 | 0.66126842 | 0.80508588 | 2L52.1 |
| WBGene00007064 | -0.277913006 | 0.41110189 | 0.61665283 | 2RSSE.1 |
| WBGene00007065 | -0.550679711 | 0.17714433 | 0.36881494 | pot-3 |
| WBGene00007067 | 1.614953298 | 0.07189226 | 0.20619984 | 4R79.2 |
| WBGene00007068 | -0.063983344 | 0.83074491 | 0.90824619 | cTel55X.1 |
| WBGene00007070 | 0.232548044 | 0.55561218 | 0.73310187 | ugt-49 |
| WBGene00007071 | 0.75134685 | 0.00037168 | 0.0042773 | AC3.5 |
| WBGene00007072 | 0.920652114 | 0.20853834 | 0.40829734 | ugt-1 |
| WBGene00007073 | 1.381618568 | 8.7984E-05 | 0.00138012 | ugt-2 |
| WBGene00007080 | 0.983488853 | 0.80601565 | 1 | sfxn-1.1 |
| WBGene00007081 | 0.377828795 | 0.92526585 | 1 | AH6.3 |
| WBGene00007082 | 1.432325317 | 0.51775671 | 1 | acs-10 |
| WBGene00007083 | -0.156269293 | 0.96924949 | 1 | AH10.2 |

| | | | | |
|---|---|---|---|---|
| WBGene00007085 | -1.159714244 | 0.75790634 | 1 | AH10.4 |
| WBGene00007087 | 0.043239453 | 0.78945414 | 0.8840401 | B0001.2 |
| WBGene00007088 | 0.036039328 | 0.851967 | 0.91924443 | B0001.3 |
| WBGene00007089 | -0.367317483 | 0.07105277 | 0.20427461 | B0001.4 |
| WBGene00007090 | 0.206554346 | 0.42585407 | 0.62970697 | B0001.5 |
| WBGene00007091 | 0.592123527 | 0.00275893 | 0.01998581 | eri-12 |
| WBGene00007092 | 0.051735175 | 0.79745254 | 0.88857116 | B0001.7 |
| WBGene00007093 | -0.387617985 | 0.06148553 | 0.18566555 | B0001.8 |
| WBGene00007094 | 0.235442256 | 0.30062543 | 0.5102605 | B0019.2 |
| WBGene00007096 | 1.203973234 | 0.2448673 | 0.45270807 | B0024.3 |
| WBGene00007097 | 1.4757549 | 0.03989948 | 0.13726823 | B0024.4 |
| WBGene00007099 | -0.464190286 | 0.03727885 | 0.13111312 | trx-2 |
| WBGene00007100 | -0.263673219 | 0.15931287 | 0.34428937 | B0024.10 |
| WBGene00007101 | -0.18533187 | 0.33815981 | 0.54610245 | B0024.11 |
| WBGene00007102 | -0.021211142 | 0.9245661 | 0.96023584 | B0024.13 |
| WBGene00007103 | 0.787922085 | 0.00131648 | 0.01132771 | crm-1 |
| WBGene00007104 | 0.10988134 | 0.94084817 | 1 | B0024.15 |
| WBGene00007105 | -0.320539742 | 0.05048321 | 0.16187147 | B0035.1 |
| WBGene00007106 | -0.607031053 | 0.00692156 | 0.03943438 | B0035.3 |
| WBGene00007107 | -0.992290159 | 6.1144E-06 | 0.00014877 | pfd-4 |
| WBGene00007108 | -0.010446721 | 0.95837882 | 0.97852807 | gspd-1 |
| WBGene00007109 | -0.320588546 | 0.09003267 | 0.23835296 | B0035.6 |
| WBGene00007110 | -0.521313881 | 0.00173308 | 0.01392541 | leo-1 |
| WBGene00007111 | -0.150051612 | 0.40387932 | 0.6111802 | sart-3 |
| WBGene00007112 | 0.837075399 | 0.00127646 | 0.01107273 | B0035.13 |
| WBGene00007113 | -0.241891227 | 0.32388177 | 0.53245035 | B0035.15 |
| WBGene00007114 | -0.039395446 | 0.91891605 | 0.95715166 | mttu-1 |
| WBGene00007115 | 1.122321179 | 0.2191917 | 0.421878 | B0198.2 |
| WBGene00007116 | 0.908691546 | 0.08720225 | 0.23361314 | B0198.3 |
| WBGene00007117 | -0.22461868 | 0.93331196 | 1 | spe-42 |
| WBGene00007118 | -0.286050822 | 0.06003927 | 0.18232536 | npp-22 |
| WBGene00007119 | 1.314976322 | 0.16582334 | 0.35258827 | calf-1 |
| WBGene00007122 | -0.748960381 | 0.00015803 | 0.00218033 | B0250.5 |
| WBGene00007124 | 0.226952808 | 0.59768356 | 0.76351811 | B0250.7 |
| WBGene00007125 | 1.229237684 | 0.59270769 | 1 | B0250.8 |
| WBGene00007126 | 0.124632353 | 0.82627981 | 0.90625042 | dhcr-7 |
| WBGene00007128 | -0.599317794 | 0.08717357 | 0.23361314 | memb-1 |
| WBGene00007129 | -0.368516167 | 0.04685184 | 0.15390373 | B0272.3 |
| WBGene00007130 | -0.250139575 | 0.48769599 | 0.67747334 | B0272.4 |
| WBGene00007131 | -0.513545692 | 0.67705053 | 1 | pals-26 |
| WBGene00007132 | 0.458122252 | 0.80249467 | 1 | pals-27 |
| WBGene00007133 | -0.307369638 | 0.70359711 | 0.83263838 | B0284.3 |
| WBGene00007134 | -0.490561023 | 0.85693136 | 1 | pals-28 |

| | | | | |
|---|---|---|---|---|
| WBGene00007135 | 0.103433559 | 0.64298421 | 0.79358252 | cdk-12 |
| WBGene00007136 | -0.173225418 | 0.5773403 | 0.74903703 | B0285.3 |
| WBGene00007137 | -0.720394122 | 0.00042646 | 0.00471979 | B0285.4 |
| WBGene00007138 | 0.462520662 | 0.54481076 | 0.72400914 | B0285.6 |
| WBGene00007139 | 0.353559138 | 0.24864542 | 0.45656791 | mnp-1 |
| WBGene00007140 | -0.244936065 | 0.75555644 | 0.86341205 | cyp-29A4 |
| WBGene00007141 | 1.687239501 | 0.59403242 | 1 | pitr-4 |
| WBGene00007142 | 0.013748101 | 0.95243888 | 0.97582406 | ttr-18 |
| WBGene00007143 | 0.198783903 | 0.16942414 | 0.35748616 | B0334.3 |
| WBGene00007144 | -0.549101192 | 0.00499863 | 0.0311542 | B0334.4 |
| WBGene00007145 | -0.244966695 | 0.28087874 | 0.49007834 | B0334.5 |
| WBGene00007146 | 0.876978815 | 0.31652045 | 0.52455294 | B0334.6 |
| WBGene00007148 | -1.270609921 | 0.30331748 | 1 | B0334.10 |
| WBGene00007149 | -0.867511771 | 0.28969965 | 0.49967941 | B0334.13 |
| WBGene00007150 | -0.120171006 | 0.52028645 | 0.70547216 | acly-2 |
| WBGene00007152 | 0.271688375 | 0.75362421 | 0.86245977 | clec-225 |
| WBGene00007153 | 0.635540333 | 0.00023929 | 0.00300439 | clec-41 |
| WBGene00007154 | 1.424313014 | 0.15506616 | 0.33872945 | dhc-3 |
| WBGene00007155 | 0.472438539 | 0.13495855 | 0.30933203 | B0379.1 |
| WBGene00007156 | 0.773345629 | 0.76155013 | 1 | B0379.2 |
| WBGene00007159 | 1.712479368 | 0.58529868 | 1 | B0379.7 |
| WBGene00007162 | -0.940022629 | 0.05117537 | 0.16337633 | fbxa-153 |
| WBGene00007163 | -0.795796894 | 0.66183653 | 1 | fbxa-154 |
| WBGene00007164 | 0.281454817 | 0.94435481 | 1 | fbxa-139 |
| WBGene00007166 | 1.214445387 | 0.20654702 | 0.40636645 | fbxa-152 |
| WBGene00007167 | 0.03523523 | 0.83550648 | 0.91041861 | rbg-3 |
| WBGene00007168 | -0.335857685 | 0.0309089 | 0.11603739 | B0393.3 |
| WBGene00007169 | 1.216957619 | 0.74819621 | 1 | B0393.4 |
| WBGene00007170 | 0.305329793 | 0.54941003 | 0.72791951 | B0393.5 |
| WBGene00007171 | -0.240909483 | 0.25779853 | 0.46569439 | B0393.6 |
| WBGene00007172 | -0.069915814 | 0.97284655 | 1 | B0393.7 |
| WBGene00007173 | 0.75595108 | 0.7708876 | 1 | B0393.8 |
| WBGene00007174 | -0.595903492 | 0.41993286 | 0.62442191 | mboa-1 |
| WBGene00007175 | 0.348761243 | 0.15549164 | 0.33929259 | B0395.3 |
| WBGene00007176 | 1.069614478 | 0.02729191 | 0.10630447 | kcnl-1 |
| WBGene00007177 | -0.156269293 | 0.96924949 | 1 | oac-1 |
| WBGene00007178 | 0.081122068 | 0.81245444 | 0.89762575 | B0457.2 |
| WBGene00007180 | 0.358739474 | 0.3950941 | 0.60293495 | B0457.6 |
| WBGene00007182 | -1.87609161 | 0.02774815 | 0.10743254 | fbxb-119 |
| WBGene00007183 | -0.391134131 | 0.84335123 | 1 | B0462.4 |
| WBGene00007184 | 0.163697029 | 0.29969312 | 0.50945734 | ctr-9 |
| WBGene00007185 | -0.023652083 | 0.91796583 | 0.95673647 | nlp-36 |
| WBGene00007186 | -0.125744777 | 0.41160328 | 0.61687143 | B0464.6 |

| | | | | |
|---|---|---|---|---|
| WBGene00007187 | -0.353927844 | 0.0961795 | 0.24892673 | tag-342 |
| WBGene00007188 | -0.515347914 | 0.02215891 | 0.09248923 | B0464.9 |
| WBGene00007189 | 0.020016864 | 0.94371369 | 0.97098542 | B0491.1 |
| WBGene00007190 | -1.187254256 | 0.47366012 | 1 | rmd-3 |
| WBGene00007191 | -0.066772316 | 0.92441903 | 0.96022645 | lgc-20 |
| WBGene00007192 | -0.614913947 | 0.00056441 | 0.00587382 | B0491.5 |
| WBGene00007193 | 0.084453483 | 0.72576521 | 0.84579072 | B0491.6 |
| WBGene00007194 | -0.70756273 | 0.02428998 | 0.09823606 | B0491.7 |
| WBGene00007195 | -0.450625586 | 0.01914453 | 0.0831011 | thoc-7 |
| WBGene00007196 | -0.470651806 | 0.00302312 | 0.02132693 | B0513.4 |
| WBGene00007197 | 0.382146261 | 0.10570482 | 0.26471992 | B0513.5 |
| WBGene00007200 | 0.345442215 | 0.49315189 | 0.68200961 | vamp-8 |
| WBGene00007201 | -0.303483219 | 0.25602062 | 0.46372465 | exos-4.1 |
| WBGene00007202 | -0.151388152 | 0.65502628 | 0.80180033 | B0564.2 |
| WBGene00007203 | 2.511573463 | 5.2927E-05 | 0.00090889 | best-1 |
| WBGene00007204 | 0.747845955 | 0.85215965 | 1 | best-2 |
| WBGene00007207 | -0.183772501 | 0.38768406 | 0.596335 | B0564.7 |
| WBGene00007209 | 0.742628056 | 0.76901841 | 1 | BE10.1 |
| WBGene00007210 | -0.341679426 | 0.49929415 | 0.68752754 | agmo-1 |
| WBGene00007211 | -1.536883772 | 0.04279181 | 0.14406193 | BE10.3 |
| WBGene00007212 | 0.367107166 | 0.78883157 | 1 | BE10.4 |
| WBGene00007213 | -0.649047141 | 0.52614858 | 1 | C01A2.1 |
| WBGene00007214 | -0.59579631 | 0.28694389 | 0.49711057 | C01A2.2 |
| WBGene00007215 | -0.153543707 | 0.46401746 | 0.65886864 | oxa-1 |
| WBGene00007216 | -0.314922089 | 0.19633832 | 0.39335701 | C01A2.4 |
| WBGene00007217 | -0.512251249 | 0.01460257 | 0.06832302 | tads-1 |
| WBGene00007218 | -1.969805064 | 0.62862569 | 1 | C01A2.6 |
| WBGene00007219 | 0.651043023 | 0.21726113 | 0.41935888 | nlp-38 |
| WBGene00007220 | 1.868758047 | 0.00481071 | 0.03037279 | C01B9.1 |
| WBGene00007221 | -0.282041514 | 0.15397855 | 0.33708142 | cpna-3 |
| WBGene00007222 | 1.965106822 | 0.11832312 | 1 | C01F6.2 |
| WBGene00007223 | -0.329140031 | 0.0809218 | 0.2224296 | C01F6.9 |
| WBGene00007224 | -1.969805064 | 0.62862569 | 1 | C01G6.2 |
| WBGene00007225 | -1.216636351 | 4.48E-15 | 8.63E-13 | C01G6.3 |
| WBGene00007226 | -0.102360757 | 0.67517133 | 0.81426991 | C01G6.4 |
| WBGene00007227 | 0.351682126 | 0.05881469 | 0.18013714 | C01G6.5 |
| WBGene00007228 | -0.061460103 | 0.83103691 | 0.90848362 | acs-7 |
| WBGene00007229 | -0.15861289 | 0.85801756 | 0.92261015 | C01G6.9 |
| WBGene00007230 | -0.156269293 | 0.96924949 | 1 | C01G10.1 |
| WBGene00007231 | 3.314800481 | 0.00691263 | 0.03940197 | C01G10.4 |
| WBGene00007232 | 3.38331889 | 0.00709809 | 0.04013877 | C01G10.5 |
| WBGene00007233 | 0.645954195 | 0.09284654 | 0.24299414 | C01G10.6 |
| WBGene00007234 | -0.480744062 | 0.05106988 | 0.16308243 | C01G10.7 |

| | | | | |
|---|---|---|---|---|
| WBGene00007235 | -0.890535839 | 1.73E-09 | 1.27E-07 | C01G10.8 |
| WBGene00007236 | -0.177024687 | 0.65627457 | 0.80231895 | C01G10.9 |
| WBGene00007237 | -0.438598368 | 0.2580034 | 0.46592581 | C01G10.10 |
| WBGene00007240 | -0.143670851 | 0.77734514 | 0.87605564 | C01G10.15 |
| WBGene00007241 | 2.023597963 | 0.15440706 | 1 | C01G10.16 |
| WBGene00007242 | 0.637799431 | 0.15184615 | 0.33373717 | madf-5 |
| WBGene00007243 | -1.14715222 | 0.68932198 | 1 | nspb-7 |
| WBGene00007244 | -0.903602348 | 0.1896388 | 0.38386206 | C01G12.3 |
| WBGene00007245 | -1.159714244 | 0.75790634 | 1 | C01G12.5 |
| WBGene00007246 | 1.222210783 | 0.60639136 | 1 | nspb-10 |
| WBGene00007247 | 0.820158676 | 0.34580643 | 0.55467048 | C01G12.7 |
| WBGene00007248 | -0.041554354 | 0.97676623 | 1 | catp-4 |
| WBGene00007249 | 0.377828795 | 0.92526585 | 1 | C01G12.9 |
| WBGene00007250 | 0.345208515 | 0.89272401 | 1 | nspb-8 |
| WBGene00007251 | -0.156269293 | 0.96924949 | 1 | nspb-9 |
| WBGene00007252 | 0.752155539 | 0.27763404 | 0.48649266 | mlt-2 |
| WBGene00007253 | 0.56830461 | 0.62914024 | 1 | C01H6.3 |
| WBGene00007254 | 0.087058702 | 0.82830821 | 0.90721969 | C01H6.4 |
| WBGene00007255 | -0.107130519 | 0.78517337 | 0.8818492 | cnnm-4 |
| WBGene00007256 | -0.146648605 | 0.35080422 | 0.55930585 | swsn-9 |
| WBGene00007257 | -0.533660423 | 0.38306389 | 0.59132597 | C01H6.8 |
| WBGene00007258 | -0.279393935 | 0.23107947 | 0.43640702 | hasp-1 |
| WBGene00007260 | 1.021188657 | 0.01487334 | 0.06918667 | olrn-1 |
| WBGene00007261 | 0.45892716 | 0.1159092 | 0.28046677 | lron-3 |
| WBGene00007262 | -1.349047703 | 0.30585058 | 1 | jud-4 |
| WBGene00007264 | 0.090530539 | 0.93852211 | 1 | C02F4.4 |
| WBGene00007266 | 0.176100752 | 0.78216465 | 0.87985371 | C03A3.1 |
| WBGene00007267 | 0.508249767 | 0.03007118 | 0.11348904 | C03A3.2 |
| WBGene00007268 | 0.306766063 | 0.88218984 | 1 | C03A3.3 |
| WBGene00007269 | -0.558804446 | 0.87818675 | 1 | C03C10.2 |
| WBGene00007270 | -0.432695188 | 0.04935685 | 0.15945755 | rei-1 |
| WBGene00007271 | -1.144176432 | 0.3207916 | 1 | C03C10.5 |
| WBGene00007271 | -2.512851905 | 0.01993201 | 1 | C03C10.9 |
| WBGene00007273 | 0.459413223 | 0.84552382 | 1 | C03C10.7 |
| WBGene00007274 | 0.270496667 | 0.92869767 | 1 | C03C11.1 |
| WBGene00007275 | 1.438763201 | 0.1137558 | 0.27694503 | C03D6.1 |
| WBGene00007277 | -0.662457614 | 0.00229509 | 0.01732175 | asfl-1 |
| WBGene00007278 | -0.954340076 | 4.5367E-05 | 0.0007971 | lab-1 |
| WBGene00007281 | 1.450134062 | 0.19383552 | 1 | C03E10.3 |
| WBGene00007282 | 1.26600902 | 0.00041151 | 0.0046047 | clec-223 |
| WBGene00007283 | -0.601912415 | 0.03790797 | 0.13267243 | clec-222 |
| WBGene00007284 | -0.194159013 | 0.49303105 | 0.68200961 | C03H12.1 |
| WBGene00007285 | 0.941613263 | 0.33241297 | 0.54019888 | C04A11.1 |

| | | | | |
|---|---|---|---|---|
| WBGene00007286 | 0.00024919 | 0.99966142 | 0.99989047 | C04A11.2 |
| WBGene00007287 | -0.026136975 | 0.92614298 | 0.96113359 | gck-4 |
| WBGene00007288 | 0.88787957 | 0.71832755 | 1 | C04A11.5 |
| WBGene00007290 | -0.484708265 | 0.02554938 | 0.10173664 | C04B4.2 |
| WBGene00007291 | -0.333693239 | 0.89573179 | 1 | lips-2 |
| WBGene00007292 | -0.323868354 | 0.91634405 | 1 | C04B4.4 |
| WBGene00007293 | 0.281454817 | 0.94435481 | 1 | C04B4.5 |
| WBGene00007295 | -1.267107459 | 0.13305441 | 0.30637952 | C04C11.1 |
| WBGene00007296 | 0.537814538 | 0.02080216 | 0.08793839 | arrd-25 |
| WBGene00007297 | -0.236461505 | 0.31049643 | 0.51867601 | C04F12.1 |
| WBGene00007299 | 0.034873166 | 0.94253606 | 0.97027162 | C04F12.5 |
| WBGene00007301 | 3.406295457 | 0.06645203 | 1 | C04F12.7 |
| WBGene00007302 | 0.48323018 | 0.28752117 | 0.4974059 | C04F12.8 |
| WBGene00007303 | -0.479549183 | 0.45610639 | 0.65324852 | rnh-1.3 |
| WBGene00007304 | 1.165030477 | 0.06523606 | 0.19280209 | ttr-39 |
| WBGene00007305 | -0.716539529 | 0.58793656 | 1 | C04G2.2 |
| WBGene00007306 | 0.377828795 | 0.92526585 | 1 | C04G2.5 |
| WBGene00007307 | 1.295658189 | 0.54877703 | 1 | spch-1 |
| WBGene00007308 | 2.079664446 | 0.15761483 | 1 | C04G2.9 |
| WBGene00007309 | -1.158971769 | 0.14236582 | 0.32034267 | C04G2.10 |
| WBGene00007310 | 0.281454817 | 0.94435481 | 1 | irld-21 |
| WBGene00007311 | 1.247575581 | 0.7549672 | 1 | C04H4.1 |
| WBGene00007312 | -0.16003269 | 0.58744887 | 0.75612077 | C04H5.1 |
| WBGene00007313 | 0.578390515 | 0.26940034 | 0.47783631 | clec-147 |
| WBGene00007314 | 0.761371225 | 0.59396977 | 1 | gcy-29 |
| WBGene00007315 | -0.552104188 | 0.07735596 | 0.21573609 | C04H5.7 |
| WBGene00007316 | 0.040642196 | 0.94176236 | 0.96989883 | nlp-41 |
| WBGene00007317 | 0.55783126 | 0.18786685 | 0.38148377 | C05A9.2 |
| WBGene00007320 | 2.456496557 | 0.3218863 | 1 | C05B5.2 |
| WBGene00007321 | 0.218358824 | 0.78272969 | 0.88024464 | C05B5.4 |
| WBGene00007322 | 0.017016561 | 0.98643854 | 1 | C05B5.5 |
| WBGene00007323 | -1.969805064 | 0.62862569 | 1 | fbxa-155 |
| WBGene00007324 | -0.022105308 | 0.96390939 | 0.98190304 | C05B5.8 |
| WBGene00007325 | 0.682069186 | 0.69825534 | 1 | C05C9.1 |
| WBGene00007326 | -1.892648585 | 0.34370782 | 1 | C05C9.2 |
| WBGene00007327 | 1.977028954 | 0.0210603 | 0.08883932 | C05C9.3 |
| WBGene00007328 | 0.403207732 | 0.86402224 | 1 | pho-10 |
| WBGene00007329 | 0.051015103 | 0.74974178 | 0.86036058 | C05C10.2 |
| WBGene00007330 | -0.285334763 | 0.09296671 | 0.24309904 | C05C10.3 |
| WBGene00007331 | -0.85785483 | 7.6657E-05 | 0.0012327 | pho-11 |
| WBGene00007332 | -0.628459185 | 0.00158918 | 0.01308967 | C05C10.5 |
| WBGene00007333 | -0.370355899 | 0.01929969 | 0.08347616 | ufd-3 |
| WBGene00007334 | -0.373872531 | 0.34488858 | 0.55358174 | C05C10.7 |

| | | | | |
|---|---|---|---|---|
| WBGene00007335 | 1.38322681 | 0.69550305 | 1 | C05C12.1 |
| WBGene00007336 | 0.764324166 | 0.27354782 | 0.48215195 | C05C12.4 |
| WBGene00007337 | 0.629154436 | 0.81300308 | 1 | C05C12.5 |
| WBGene00007338 | -0.701489725 | 0.54042824 | 1 | C05C12.6 |
| WBGene00007339 | 0.144655253 | 0.83722949 | 0.91143626 | C05D12.1 |
| WBGene00007340 | 0.947173711 | 0.0578106 | 0.17818697 | C05D12.2 |
| WBGene00007341 | -1.096646147 | 0.01988671 | 0.0850756 | C05D12.4 |
| WBGene00007342 | -1.871125671 | 0.00229558 | 0.01732175 | C05D12.5 |
| WBGene00007343 | -0.516756697 | 0.32155936 | 0.52965027 | C05E7.1 |
| WBGene00007344 | -1.497050127 | 0.0151536 | 0.07019452 | C05E7.2 |
| WBGene00007345 | -0.800652376 | 0.41107539 | 1 | C05E7.3 |
| WBGene00007346 | 1.458663133 | 0.33242204 | 1 | frpr-2 |
| WBGene00007347 | 0.462081228 | 0.26376253 | 0.47232167 | C05G5.1 |
| WBGene00007348 | -0.533767198 | 0.06761172 | 0.19761048 | C05G5.2 |
| WBGene00007349 | 0.179621908 | 0.75805422 | 0.86459364 | C05G5.3 |
| WBGene00007350 | -0.127711005 | 0.50592603 | 0.69273124 | sucl-1 |
| WBGene00007351 | -0.12640091 | 0.89341661 | 0.94378935 | C05G5.5 |
| WBGene00007352 | -0.262931175 | 0.11862592 | 0.28418257 | cdc-48.1 |
| WBGene00007353 | 0.30810481 | 0.59702045 | 0.76307246 | C06A1.2 |
| WBGene00007354 | 1.112219493 | 0.6005584 | 1 | C06A1.3 |
| WBGene00007355 | -0.462351815 | 0.023728 | 0.09670413 | rpb-6 |
| WBGene00007356 | -2.640414922 | 0.41081194 | 1 | C06A1.6 |
| WBGene00007357 | -0.858746009 | 0.06171814 | 0.18592589 | C06A12.3 |
| WBGene00007358 | 0.281454817 | 0.94435481 | 1 | lact-6 |
| WBGene00007359 | 0.688793783 | 0.86378755 | 1 | C06A12.8 |
| WBGene00007361 | 1.045599005 | 0.59293297 | 1 | oac-3 |
| WBGene00007362 | -1.073745536 | 0.0235011 | 0.09603733 | cyp-35C1 |
| WBGene00007363 | -0.321723718 | 0.32298904 | 0.53142837 | stdh-1 |
| WBGene00007365 | 1.110643319 | 1.7682E-05 | 0.00036571 | C06B3.6 |
| WBGene00007366 | 3.641970852 | 0.01295207 | 1 | C06B3.7 |
| WBGene00007367 | -0.127135707 | 0.94381388 | 1 | nhr-150 |
| WBGene00007368 | 0.896795782 | 0.17930705 | 0.37143268 | C06B8.2 |
| WBGene00007369 | -1.159714244 | 0.75790634 | 1 | C06B8.3 |
| WBGene00007370 | 1.216957619 | 0.74819621 | 1 | str-264 |
| WBGene00007372 | 0.723940631 | 0.04049182 | 0.13863825 | C06B8.7 |
| WBGene00007374 | -0.372544364 | 0.81798234 | 1 | C06C3.3 |
| WBGene00007375 | -1.331946232 | 0.11959771 | 0.28583381 | C06C3.4 |
| WBGene00007377 | -0.966108312 | 0.49102303 | 1 | C06C3.7 |
| WBGene00007378 | -0.434981297 | 0.67116514 | 1 | C06C3.8 |
| WBGene00007381 | 0.377828795 | 0.92526585 | 1 | C06C6.7 |
| WBGene00007382 | 0.574508983 | 0.8551971 | 1 | C06C6.8 |
| WBGene00007384 | -0.829873225 | 0.07452495 | 0.21096 | swt-3 |
| WBGene00007385 | -0.224055778 | 0.09050116 | 0.23924447 | atp-5 |

| | | | | |
|---|---|---|---|---|
| WBGene00007386 | 0.048241146 | 0.8412058 | 0.91269702 | C06H2.2 |
| WBGene00007387 | -0.390093553 | 0.10391053 | 0.26148193 | jmjd-5 |
| WBGene00007388 | -0.167263709 | 0.68485836 | 0.81992558 | folt-1 |
| WBGene00007389 | 1.316661345 | 0.10264332 | 0.25913757 | glb-3 |
| WBGene00007390 | -0.589113793 | 0.0059193 | 0.03528039 | lmtr-3 |
| WBGene00007391 | -0.85540747 | 0.00108175 | 0.00977557 | C06H2.7 |
| WBGene00007392 | 0.689693971 | 0.26850761 | 0.47707159 | fbxa-156 |
| WBGene00007393 | 1.678643117 | 0.25736821 | 1 | fbxa-157 |
| WBGene00007394 | 0.490054594 | 0.18608465 | 0.37887871 | C06H5.6 |
| WBGene00007395 | 0.335269295 | 0.24842386 | 0.45656791 | dcar-1 |
| WBGene00007396 | 0.855716389 | 0.12673594 | 0.29717389 | tiar-3 |
| WBGene00007397 | 0.355899034 | 0.71296825 | 0.8378809 | C07A4.2 |
| WBGene00007398 | 0.586318706 | 0.65335185 | 1 | C07A4.3 |
| WBGene00007399 | 1.557897343 | 0.63813005 | 1 | clec-162 |
| WBGene00007400 | -0.652724758 | 0.0048461 | 0.03049303 | bud-31 |
| WBGene00007401 | 1.158270114 | 0.62680548 | 1 | C07A9.5 |
| WBGene00007402 | 0.604602432 | 0.14826999 | 0.32891394 | ugt-60 |
| WBGene00007403 | 0.060060095 | 0.8082794 | 0.89545764 | set-3 |
| WBGene00007404 | 2.967575812 | 0.00906375 | 0.04801178 | best-5 |
| WBGene00007405 | -0.126458865 | 0.69054508 | 0.82364749 | C07A9.9 |
| WBGene00007406 | -1.901856996 | 0.64072034 | 1 | C07A9.10 |
| WBGene00007408 | 0.261154075 | 0.85462081 | 1 | C07B5.3 |
| WBGene00007409 | 0.78891057 | 0.0746044 | 0.21102571 | C07B5.4 |
| WBGene00007410 | -0.245531902 | 0.94070824 | 1 | C07B5.6 |
| WBGene00007411 | 2.07531578 | 0.45514895 | 1 | C07C7.1 |
| WBGene00007412 | -0.031082627 | 0.86326432 | 0.92644675 | stip-1 |
| WBGene00007413 | 0.130405 | 0.52702516 | 0.7113202 | pro-2 |
| WBGene00007414 | 0.738758921 | 0.17151551 | 0.36066532 | C07E3.3 |
| WBGene00007415 | -0.213937883 | 0.92749767 | 1 | C07E3.4 |
| WBGene00007416 | 0.406175358 | 0.70993673 | 1 | ceh-57 |
| WBGene00007417 | -0.111061313 | 0.75061422 | 0.86111757 | ceh-58 |
| WBGene00007418 | -0.626833103 | 0.80597175 | 1 | C07E3.8 |
| WBGene00007419 | -1.170992235 | 0.00066364 | 0.00668655 | C07E3.9 |
| WBGene00007420 | -0.998065977 | 0.57530013 | 1 | C07E3.10 |
| WBGene00007421 | 1.518883044 | 0.01248643 | 0.06098057 | C07H4.1 |
| WBGene00007422 | 0.675208106 | 0.04658373 | 0.15310586 | ugt-17 |
| WBGene00007423 | 1.226657715 | 0.39088312 | 1 | C08B6.2 |
| WBGene00007424 | 0.237117234 | 0.6419823 | 0.79291018 | C08B6.3 |
| WBGene00007425 | -0.70415203 | 0.02570393 | 0.10215105 | C08B6.4 |
| WBGene00007426 | -1.901856996 | 0.64072034 | 1 | C08B6.5 |
| WBGene00007428 | 0.050871529 | 0.76331316 | 0.86757022 | wdr-20 |
| WBGene00007429 | -0.362188851 | 0.14225297 | 0.32034267 | C08B6.8 |
| WBGene00007430 | -0.021258953 | 0.97642352 | 0.98781418 | C08B6.10 |

| | | | | |
|---|---|---|---|---|
| WBGene00007431 | 0.202331225 | 0.75294884 | 0.86217294 | C08B6.11 |
| WBGene00007432 | -0.156269293 | 0.96924949 | 1 | srxa-18 |
| WBGene00007433 | -0.016485933 | 0.92711777 | 0.96171059 | swsn-7 |
| WBGene00007434 | -0.575403956 | 0.00367522 | 0.02464125 | arp-6 |
| WBGene00007435 | 0.531165694 | 0.0751176 | 0.21204437 | C08B11.8 |
| WBGene00007436 | -0.835230806 | 0.00039052 | 0.00442696 | C08B11.9 |
| WBGene00007437 | 1.93328615 | 0.30439072 | 1 | C08E8.1 |
| WBGene00007439 | -0.156269293 | 0.96924949 | 1 | C08E8.3 |
| WBGene00007440 | 0.725772962 | 0.22531994 | 0.42945201 | C08E8.4 |
| WBGene00007441 | -1.159714244 | 0.75790634 | 1 | C08E8.5 |
| WBGene00007442 | -1.159714244 | 0.75790634 | 1 | C08E8.6 |
| WBGene00007443 | -0.672345902 | 0.00105027 | 0.00957308 | pfd-1 |
| WBGene00007444 | -0.045316194 | 0.82305359 | 0.90431616 | C08F8.2 |
| WBGene00007445 | -0.526537368 | 0.00574381 | 0.03450552 | C08F8.3 |
| WBGene00007446 | 1.471809563 | 0.46592277 | 1 | mboa-4 |
| WBGene00007447 | -0.156269293 | 0.96924949 | 1 | fbxb-9 |
| WBGene00007448 | -0.254102539 | 0.89849757 | 1 | C08F8.6 |
| WBGene00007449 | -0.833918402 | 0.00664574 | 0.03825793 | C08F8.9 |
| WBGene00007450 | 0.35263493 | 0.67364096 | 0.8134506 | C08F11.1 |
| WBGene00007452 | 2.375766819 | 0.17795701 | 1 | C08F11.3 |
| WBGene00007453 | -0.824017582 | 0.28032628 | 0.48935174 | fbxa-98 |
| WBGene00007454 | 1.988568769 | 0.12435618 | 1 | C08F11.7 |
| WBGene00007455 | -0.242657005 | 0.1833897 | 0.37546953 | ugt-22 |
| WBGene00007456 | -0.156269293 | 0.96924949 | 1 | srz-29 |
| WBGene00007458 | -0.846327947 | 1.5822E-06 | 4.7666E-05 | ule-4 |
| WBGene00007459 | -2.309705573 | 0.02858323 | 0.10954199 | C08F11.12 |
| WBGene00007460 | 1.459777197 | 0.00014571 | 0.0020642 | C08F11.13 |
| WBGene00007462 | 0.819395132 | 0.27553327 | 0.48411714 | C08H9.1 |
| WBGene00007463 | 0.240392536 | 0.22529789 | 0.42945201 | vgln-1 |
| WBGene00007464 | 0.433321738 | 0.11896772 | 0.28463356 | C08H9.3 |
| WBGene00007465 | 1.632279705 | 0.23414179 | 1 | chil-1 |
| WBGene00007466 | 0.789601902 | 0.75907734 | 1 | chil-2 |
| WBGene00007467 | 2.444636082 | 0.08201701 | 1 | chil-3 |
| WBGene00007469 | 1.718512817 | 0.39453967 | 1 | chil-4 |
| WBGene00007470 | 0.98349132 | 0.50835086 | 1 | chil-5 |
| WBGene00007471 | 1.532777345 | 0.30913189 | 1 | chil-6 |
| WBGene00007472 | -1.618771107 | 0.4854602 | 1 | chil-7 |
| WBGene00007473 | -0.506567032 | 0.81430926 | 1 | chil-8 |
| WBGene00007474 | -1.054989954 | 0.00544783 | 0.03320389 | C08H9.15 |
| WBGene00007476 | -0.751563431 | 0.01549119 | 0.07126889 | C09E9.1 |
| WBGene00007478 | -1.094201388 | 0.7816772 | 1 | C09F9.1 |
| WBGene00007479 | 0.124509045 | 0.7117057 | 0.83696425 | C09F9.2 |
| WBGene00007480 | 0.370109055 | 0.45337864 | 0.65095436 | glna-1 |

| | | | | |
|---|---|---|---|---|
| WBGene00007482 | 2.439824062 | 7.3564E-05 | 0.00119723 | C09F12.2 |
| WBGene00007483 | 1.612755268 | 0.23014182 | 1 | C09F12.3 |
| WBGene00007484 | 0.932931886 | 0.70926986 | 1 | C09G1.2 |
| WBGene00007485 | -0.245531902 | 0.94070824 | 1 | C09G1.3 |
| WBGene00007486 | 0.602429804 | 0.28439573 | 0.49417901 | C09G1.4 |
| WBGene00007487 | -0.099508929 | 0.90659818 | 0.95121412 | C09G1.5 |
| WBGene00007488 | -0.189912585 | 0.48514478 | 0.67539764 | dph-2 |
| WBGene00007489 | 0.622259013 | 0.64142667 | 1 | C09G5.7 |
| WBGene00007490 | 1.715232149 | 0.06525833 | 0.19281071 | mks-5 |
| WBGene00007492 | -0.139364837 | 0.42832612 | 0.63100978 | C09G9.1 |
| WBGene00007493 | -0.504958544 | 0.0097439 | 0.05086014 | npp-23 |
| WBGene00007494 | -0.156269293 | 0.96924949 | 1 | C09G9.3 |
| WBGene00007495 | 0.304990599 | 0.80973218 | 1 | C09G9.5 |
| WBGene00007496 | 0.242323424 | 0.84551754 | 1 | npax-4 |
| WBGene00007497 | 0.05905161 | 0.97811893 | 1 | C09G9.8 |
| WBGene00007499 | -0.225964838 | 0.34351794 | 0.55189246 | C09H10.5 |
| WBGene00007500 | -0.556239749 | 0.00021518 | 0.00278518 | nasp-1 |
| WBGene00007501 | -0.238878604 | 0.48557313 | 0.67571486 | apc-17 |
| WBGene00007502 | 1.119843859 | 0.20834106 | 0.40811049 | glb-4 |
| WBGene00007503 | -0.512603405 | 0.83307778 | 1 | C09H10.9 |
| WBGene00007504 | -0.37315776 | 0.20638286 | 0.40624097 | C09H10.10 |
| WBGene00007505 | 0.788876174 | 9.6439E-06 | 0.0002205 | pezo-1 |
| WBGene00007506 | 1.063937676 | 0.14582287 | 0.32488677 | C10C5.2 |
| WBGene00007507 | 1.141671366 | 0.00396957 | 0.02610758 | C10C5.3 |
| WBGene00007508 | 0.48265573 | 0.09973612 | 0.25443821 | C10C5.4 |
| WBGene00007509 | 0.2306002 | 0.73118652 | 0.84942451 | C10C5.5 |
| WBGene00007511 | 0.405781406 | 0.53678716 | 0.71885428 | C10C5.7 |
| WBGene00007513 | -0.327071458 | 0.09827466 | 0.25225214 | wht-2 |
| WBGene00007514 | 0.173053553 | 0.30312845 | 0.51221193 | catp-8 |
| WBGene00007515 | -0.501497017 | 0.54675712 | 0.72564257 | C10C6.7 |
| WBGene00007516 | -0.140214965 | 0.56145372 | 0.73709014 | gpx-5 |
| WBGene00007517 | 1.696317843 | 1.10E-07 | 5.0802E-06 | gpx-3 |
| WBGene00007518 | 1.219060141 | 0.46174049 | 1 | del-8 |
| WBGene00007520 | 0.637322231 | 0.07199873 | 0.20637478 | C11E4.6 |
| WBGene00007521 | 0.433340115 | 0.45516564 | 0.65257895 | C11E4.7 |
| WBGene00007522 | -0.419219645 | 0.75916414 | 1 | C11E4.8 |
| WBGene00007522 | 0.347945518 | 0.79818373 | 1 | F34H10.2 |
| WBGene00007525 | 0.659922425 | 0.32711435 | 0.53553065 | C11G10.1 |
| WBGene00007528 | -0.090469497 | 0.84024265 | 0.91237519 | C11H1.2 |
| WBGene00007529 | -0.093355515 | 0.73081451 | 0.84935453 | C11H1.3 |
| WBGene00007530 | -1.279666666 | 0.55824702 | 1 | C11H1.5 |
| WBGene00007531 | -0.493378431 | 0.46709283 | 0.66139412 | C11H1.7 |
| WBGene00007533 | 0.27062788 | 0.27549079 | 0.48411714 | cbl-1 |

| | | | | |
|---|---|---|---|---|
| WBGene00007534 | -0.0867377 | 0.57936016 | 0.75037463 | C12D8.1 |
| WBGene00007536 | 0.062452938 | 0.83455065 | 0.91032502 | daf-36 |
| WBGene00007537 | -0.288374026 | 0.71325967 | 0.8379961 | fipr-8 |
| WBGene00007540 | -0.156269293 | 0.96924949 | 1 | C12D8.13 |
| WBGene00007541 | -0.455216006 | 0.41554838 | 0.62082449 | fipr-4 |
| WBGene00007542 | -0.22461868 | 0.93331196 | 1 | C12D8.15 |
| WBGene00007543 | -0.064841091 | 0.95425503 | 1 | fipr-7 |
| WBGene00007544 | -0.48845152 | 0.53885807 | 0.72066769 | fipr-6 |
| WBGene00007545 | 0.090890827 | 0.69667899 | 0.82848332 | C13B4.1 |
| WBGene00007546 | 0.792096955 | 0.19409525 | 0.39027997 | nhr-153 |
| WBGene00007547 | 0.791059945 | 0.06041607 | 0.18328622 | nhr-154 |
| WBGene00007548 | -0.700592605 | 0.15143909 | 0.33332315 | C13C4.4 |
| WBGene00007549 | 1.03341231 | 3.4912E-05 | 0.00063454 | spin-1 |
| WBGene00007550 | 1.392157332 | 0.05033716 | 0.1616354 | C13C4.6 |
| WBGene00007551 | 0.17414283 | 0.82959715 | 0.90772702 | C13C4.7 |
| WBGene00007552 | -0.773298381 | 0.48884913 | 1 | C13C12.2 |
| WBGene00007553 | 0.630767216 | 0.23678227 | 0.44295432 | C13G3.1 |
| WBGene00007554 | 0.257902035 | 0.15191908 | 0.33383702 | pptr-2 |
| WBGene00007555 | -0.231168386 | 0.2663446 | 0.47477459 | dohh-1 |
| WBGene00007556 | 0.24798898 | 0.21150931 | 0.41127027 | C14A4.3 |
| WBGene00007557 | 0.332992519 | 0.39670156 | 0.60461381 | C14A4.6 |
| WBGene00007558 | 0.619178328 | 0.291508 | 0.50151603 | C14A4.7 |
| WBGene00007559 | -1.159714244 | 0.75790634 | 1 | C14A4.8 |
| WBGene00007560 | -1.152564628 | 0.05235218 | 0.16554924 | C14A4.9 |
| WBGene00007561 | -0.554022518 | 0.00946356 | 0.04973902 | ccm-3 |
| WBGene00007562 | 1.621112891 | 0.001513 | 0.01265106 | C14A4.12 |
| WBGene00007563 | 0.117735816 | 0.9547449 | 1 | C14A4.13 |
| WBGene00007564 | -0.359504993 | 0.07599749 | 0.21398089 | mrps-22 |
| WBGene00007565 | 1.345661493 | 7.51E-07 | 2.6401E-05 | clec-48 |
| WBGene00007570 | 0.688793783 | 0.86378755 | 1 | C14A6.6 |
| WBGene00007572 | -1.969805064 | 0.62862569 | 1 | C14A6.8 |
| WBGene00007573 | -0.969396911 | 1.0001E-06 | 3.3821E-05 | C14B1.2 |
| WBGene00007574 | 0.040450532 | 0.90565994 | 0.95088303 | C14B1.3 |
| WBGene00007576 | -0.400784377 | 0.10079812 | 0.25590808 | dph-1 |
| WBGene00007577 | 0.037839141 | 0.8529674 | 0.91927977 | nrde-1 |
| WBGene00007579 | -0.392170048 | 0.13334229 | 0.30678581 | C14B1.8 |
| WBGene00007580 | -0.357365432 | 0.03065708 | 0.11534167 | C14B1.9 |
| WBGene00007581 | -0.038431451 | 0.86067999 | 0.92489961 | alkb-8 |
| WBGene00007583 | 0.604942437 | 0.07493221 | 0.21166867 | C14B4.2 |
| WBGene00007584 | 1.872540704 | 0.36210323 | 1 | C14C10.1 |
| WBGene00007585 | -0.207064039 | 0.5902587 | 0.75794086 | C14C10.2 |
| WBGene00007586 | -0.002369396 | 0.98860273 | 0.99484196 | ril-2 |
| WBGene00007587 | 0.230613975 | 0.16282349 | 0.34865113 | mma-1 |

| | | | | |
|---|---|---|---|---|
| WBGene00007588 | 0.368667697 | 0.04277625 | 0.14406193 | C14C10.5 |
| WBGene00007589 | -0.057122273 | 0.90302804 | 0.94956378 | erg-28 |
| WBGene00007590 | 1.092105691 | 0.16764326 | 0.35514864 | ttr-43 |
| WBGene00007591 | -0.458229533 | 0.10512123 | 0.26385712 | zipt-13 |
| WBGene00007592 | -0.063212513 | 0.81677095 | 0.90051 | C14H10.2 |
| WBGene00007593 | 0.830815502 | 0.00194321 | 0.01521272 | C14H10.3 |
| WBGene00007594 | -1.483235343 | 0.29641604 | 1 | lgc-23 |
| WBGene00007595 | 0.862349658 | 0.47813465 | 1 | C15A7.2 |
| WBGene00007597 | 0.15388156 | 0.93468081 | 1 | C15A11.2 |
| WBGene00007598 | -1.751332393 | 0.02712207 | 0.10571079 | C15A11.4 |
| WBGene00007599 | 1.136276092 | 0.00033028 | 0.00391207 | C15A11.7 |
| WBGene00007600 | -0.649900575 | 0.21377844 | 0.41437773 | C15C6.1 |
| WBGene00007601 | 2.289693512 | 0.18381829 | 1 | C15C6.2 |
| WBGene00007602 | -0.219296784 | 0.30979942 | 0.51822468 | C15C6.3 |
| WBGene00007603 | -0.553389715 | 0.05461109 | 0.17079681 | C15C6.4 |
| WBGene00007604 | 1.692465614 | 0.02392178 | 0.09720025 | xbx-9 |
| WBGene00007605 | -0.761989886 | 0.00033827 | 0.00399119 | asp-10 |
| WBGene00007606 | 0.268226829 | 0.24902035 | 0.45683833 | C15C8.4 |
| WBGene00007607 | -0.153871373 | 0.89811966 | 1 | C15C8.5 |
| WBGene00007608 | 0.375803815 | 0.83909921 | 1 | C15C8.6 |
| WBGene00007609 | -0.278849606 | 0.33014491 | 0.53853142 | C15C8.7 |
| WBGene00007610 | 0.175909247 | 0.94229417 | 1 | C15H7.3 |
| WBGene00007611 | -0.156269293 | 0.96924949 | 1 | C15H7.4 |
| WBGene00007613 | 0.983488853 | 0.80601565 | 1 | C15H11.1 |
| WBGene00007614 | 0.54792515 | 0.17327394 | 0.36304112 | gnrr-2 |
| WBGene00007615 | -0.307050366 | 0.20726057 | 0.4068473 | set-31 |
| WBGene00007616 | -0.670172655 | 0.02442342 | 0.09854595 | rpoa-12 |
| WBGene00007617 | -0.314147085 | 0.08773287 | 0.23472834 | rrbs-1 |
| WBGene00007618 | 0.281454817 | 0.94435481 | 1 | C15H11.10 |
| WBGene00007619 | 0.15227395 | 0.86318501 | 0.92644353 | C15H11.11 |
| WBGene00007620 | -0.270084784 | 0.16887967 | 0.35695822 | ocrl-1 |
| WBGene00007621 | 0.074079538 | 0.67151781 | 0.8122034 | C16C2.4 |
| WBGene00007622 | -0.133642425 | 0.56067335 | 0.73673213 | C16C10.1 |
| WBGene00007623 | -0.669869546 | 0.00128607 | 0.01113711 | C16C10.2 |
| WBGene00007624 | -0.107167722 | 0.56196476 | 0.73768128 | hrde-1 |
| WBGene00007625 | -0.905698668 | 1.8424E-06 | 5.4161E-05 | C16C10.4 |
| WBGene00007626 | -0.170384143 | 0.42814762 | 0.63100978 | rnf-121 |
| WBGene00007627 | -0.057066892 | 0.75867825 | 0.8648932 | ccdc-55 |
| WBGene00007628 | -0.054038425 | 0.86379355 | 0.92668707 | C16C10.8 |
| WBGene00007629 | -0.166758709 | 0.92110138 | 1 | C16C10.9 |
| WBGene00007630 | -0.608882026 | 0.00119751 | 0.01053548 | har-1 |
| WBGene00007631 | -1.038181246 | 0.5835992 | 1 | wht-3 |
| WBGene00007632 | 2.545238571 | 0.23855563 | 1 | C16C10.13 |

| | | | | |
|---|---|---|---|---|
| WBGene00007633 | 0.075260937 | 0.97569838 | 1 | C16D2.1 |
| WBGene00007635 | 1.140485754 | 0.15672922 | 0.34095134 | npr-4 |
| WBGene00007637 | -0.148036454 | 0.56144133 | 0.73709014 | dhhc-7 |
| WBGene00007638 | -1.094201388 | 0.7816772 | 1 | C17D12.3 |
| WBGene00007639 | 0.574508983 | 0.8551971 | 1 | C17D12.5 |
| WBGene00007640 | -0.435965638 | 0.05969606 | 0.18178326 | C17D12.7 |
| WBGene00007641 | -0.479753599 | 0.75794271 | 1 | C17E4.1 |
| WBGene00007642 | -1.738226604 | 0.00250696 | 0.01861522 | C17E4.2 |
| WBGene00007644 | -0.246844961 | 0.60910952 | 0.77135532 | C17E4.4 |
| WBGene00007645 | -0.306349379 | 0.13949348 | 0.31673654 | C17E4.6 |
| WBGene00007646 | 0.231233537 | 0.14456166 | 0.32307998 | nkb-1 |
| WBGene00007647 | -0.300229406 | 0.11967926 | 0.28595048 | C17E4.10 |
| WBGene00007648 | -1.969805064 | 0.62862569 | 1 | folr-1 |
| WBGene00007649 | -0.514374143 | 0.04191847 | 0.14223928 | C17G1.2 |
| WBGene00007650 | 0.316983725 | 0.3150493 | 0.52290001 | ugt-23 |
| WBGene00007651 | 0.302306059 | 0.265473 | 0.47426541 | nra-3 |
| WBGene00007652 | -0.266466852 | 0.61420549 | 0.7748409 | C17G1.5 |
| WBGene00007653 | -0.341623831 | 0.2142741 | 0.41504496 | cysl-1 |
| WBGene00007656 | 0.377828795 | 0.92526585 | 1 | pals-2 |
| WBGene00007658 | 2.566123675 | 0.23268183 | 1 | pals-4 |
| WBGene00007659 | -1.035929762 | 0.59261215 | 1 | pals-5 |
| WBGene00007660 | -0.170195274 | 0.88781883 | 1 | pals-6 |
| WBGene00007661 | 0.688793783 | 0.86378755 | 1 | pals-7 |
| WBGene00007662 | 0.281454817 | 0.94435481 | 1 | pals-8 |
| WBGene00007664 | 0.430547783 | 0.4120522 | 0.617468 | seb-3 |
| WBGene00007666 | 0.124625411 | 0.58912825 | 0.75721005 | C18B12.4 |
| WBGene00007668 | -0.384618445 | 0.21076763 | 0.41054544 | C18B12.6 |
| WBGene00007669 | -1.027355405 | 0.68015432 | 1 | C18D1.2 |
| WBGene00007670 | -0.190856808 | 0.90351155 | 1 | stg-1 |
| WBGene00007672 | 3.391314877 | 1.75E-07 | 7.5558E-06 | fbxa-136 |
| WBGene00007673 | 0.983488853 | 0.80601565 | 1 | C18D4.3 |
| WBGene00007675 | 0.730455319 | 0.20322365 | 0.4023709 | C18D4.6 |
| WBGene00007679 | 0.503594187 | 0.08844843 | 0.23575245 | C18D11.1 |
| WBGene00007680 | 0.420529481 | 0.38448441 | 0.59284039 | maa-1 |
| WBGene00007681 | -0.076327359 | 0.73363741 | 0.85095923 | C18D11.3 |
| WBGene00007682 | -1.887089501 | 0.39158219 | 1 | C18D11.6 |
| WBGene00007683 | -0.128989367 | 0.47624738 | 0.66883973 | C18E9.2 |
| WBGene00007684 | -0.603354574 | 0.00065705 | 0.00662558 | C18E9.4 |
| WBGene00007685 | 0.672099273 | 0.00898301 | 0.04764648 | C18E9.5 |
| WBGene00007686 | -0.464954723 | 0.00535238 | 0.03277015 | tomm-40 |
| WBGene00007687 | 0.614616169 | 0.38677645 | 0.59554253 | C18E9.7 |
| WBGene00007688 | 2.166011856 | 0.42253177 | 1 | C18E9.8 |
| WBGene00007689 | 0.502453371 | 0.08956219 | 0.23736621 | C18E9.9 |

| | | | | |
|---|---|---|---|---|
| WBGene00007690 | 0.094222848 | 0.70385895 | 0.83265629 | sftd-3 |
| WBGene00007691 | -0.550255047 | 0.52019677 | 0.70542935 | C23H4.2 |
| WBGene00007692 | 0.147411151 | 0.73815303 | 0.85392243 | C23H4.3 |
| WBGene00007693 | -1.23116719 | 0.41752971 | 1 | C23H4.4 |
| WBGene00007694 | 0.922588772 | 0.10785067 | 0.26759651 | C23H4.6 |
| WBGene00007695 | 0.446955428 | 0.55815691 | 0.73522654 | C23H4.7 |
| WBGene00007696 | 0.090616747 | 0.60188295 | 0.76683654 | tram-1 |
| WBGene00007697 | -0.240823947 | 0.48169763 | 0.67289188 | C24F3.2 |
| WBGene00007698 | -0.061945886 | 0.74459921 | 0.8574972 | qns-1 |
| WBGene00007699 | 0.574508983 | 0.8551971 | 1 | C24H11.1 |
| WBGene00007700 | 0.574508983 | 0.8551971 | 1 | C24H11.2 |
| WBGene00007701 | -0.07748118 | 0.98067866 | 1 | srd-74 |
| WBGene00007702 | -0.531495379 | 0.29754678 | 0.5076675 | C24H11.5 |
| WBGene00007703 | 0.265584279 | 0.27826285 | 0.48722084 | gbf-1 |
| WBGene00007704 | -0.151787798 | 0.76130392 | 0.86650332 | mdt-21 |
| WBGene00007705 | -0.333254287 | 0.10680143 | 0.2664768 | C25A1.1 |
| WBGene00007707 | 0.156252473 | 0.39177745 | 0.60012239 | fath-1 |
| WBGene00007708 | -0.480481139 | 0.01755585 | 0.0781039 | nola-3 |
| WBGene00007709 | -0.637191577 | 3.1957E-06 | 8.7213E-05 | clec-87 |
| WBGene00007710 | -0.131650197 | 0.48698953 | 0.67656939 | rsa-1 |
| WBGene00007711 | -0.711377545 | 0.00040393 | 0.00454084 | lid-1 |
| WBGene00007712 | -0.432153889 | 0.1035076 | 0.26071284 | mrpl-34 |
| WBGene00007716 | 1.64146208 | 0.17352282 | 1 | fbxa-82 |
| WBGene00007717 | 1.256053291 | 0.08081211 | 0.2222873 | C25D7.5 |
| WBGene00007718 | -0.343268768 | 0.03872939 | 0.13457338 | otub-1 |
| WBGene00007720 | -0.151208674 | 0.61673551 | 0.7764191 | C25D7.10 |
| WBGene00007722 | 0.559608276 | 0.49850921 | 0.68691412 | C25D7.15 |
| WBGene00007723 | 1.82867917 | 0.54358766 | 1 | C25F9.2 |
| WBGene00007724 | 1.670801298 | 0.00192088 | 0.01508502 | C25F9.4 |
| WBGene00007725 | 1.433725197 | 0.01327206 | 0.06374054 | C25F9.5 |
| WBGene00007726 | 1.08257176 | 0.33463446 | 1 | C25F9.6 |
| WBGene00007728 | 1.039793218 | 0.31161818 | 1 | C25F9.9 |
| WBGene00007729 | 0.356077185 | 0.84034663 | 1 | clec-185 |
| WBGene00007730 | -0.032759825 | 0.89287468 | 0.94362739 | C25G4.2 |
| WBGene00007731 | -0.716796426 | 0.01621695 | 0.07354875 | C25G4.3 |
| WBGene00007732 | -0.064120735 | 0.86677228 | 0.92821073 | spe-44 |
| WBGene00007733 | -0.370677187 | 0.7938472 | 1 | smz-1 |
| WBGene00007734 | 2.365725856 | 0.35238907 | 1 | C25G4.7 |
| WBGene00007735 | -0.245531902 | 0.94070824 | 1 | C25G4.8 |
| WBGene00007736 | -0.192940117 | 0.48318668 | 0.67429535 | igdb-2 |
| WBGene00007739 | 0.552536202 | 0.70475375 | 1 | C26C6.4 |
| WBGene00007740 | -0.156269293 | 0.96924949 | 1 | C26C6.6 |
| WBGene00007741 | 0.513661423 | 0.54758097 | 0.72633896 | glb-8 |

| | | | | |
|---|---|---|---|---|
| WBGene00007742 | -0.989314885 | 0.5977947 | 1 | C26C6.8 |
| WBGene00007743 | 0.621911949 | 0.16188604 | 0.34774564 | C26C6.9 |
| WBGene00007744 | -0.366552499 | 0.57646496 | 0.74903703 | C26D10.3 |
| WBGene00007745 | 0.116731459 | 0.87863143 | 0.93492236 | C26D10.4 |
| WBGene00007746 | 0.343348478 | 0.3067946 | 0.51557292 | C26D10.6 |
| WBGene00007748 | 1.641248455 | 0.00595051 | 0.03537961 | C26E1.2 |
| WBGene00007749 | 1.212811026 | 0.08480086 | 0.22947452 | ceh-79 |
| WBGene00007750 | 0.65467764 | 0.06588324 | 0.19395937 | syg-2 |
| WBGene00007751 | 1.009994056 | 0.02334352 | 0.09563613 | C26G2.2 |
| WBGene00007752 | 0.796427079 | 0.001323 | 0.0113757 | wdfy-3 |
| WBGene00007753 | 0.840932831 | 0.01209957 | 0.05961885 | C27A7.1 |
| WBGene00007755 | 1.575049472 | 0.0696192 | 0.20158943 | C27A7.3 |
| WBGene00007756 | 0.263315288 | 0.26555974 | 0.47428078 | C27A7.5 |
| WBGene00007757 | -0.37339616 | 0.12344651 | 0.29227068 | C27A7.6 |
| WBGene00007758 | -2.515023366 | 0.00138612 | 0.01177672 | C27A7.8 |
| WBGene00007759 | -1.969805064 | 0.62862569 | 1 | C27A7.9 |
| WBGene00007760 | 0.211278545 | 0.52294835 | 0.70789563 | C27B7.2 |
| WBGene00007761 | 0.249323399 | 0.20357762 | 0.40274334 | rad-26 |
| WBGene00007762 | -0.717631159 | 0.00054928 | 0.00573418 | C27B7.5 |
| WBGene00007763 | 2.07531578 | 0.45514895 | 1 | C27B7.6 |
| WBGene00007764 | 0.782351424 | 0.09997219 | 0.254778 | C27B7.7 |
| WBGene00007765 | -1.99453804 | 2.7894E-06 | 7.7854E-05 | C27B7.9 |
| WBGene00007766 | 0.580952294 | 0.01197373 | 0.05919098 | C27C7.1 |
| WBGene00007768 | -0.156269293 | 0.96924949 | 1 | C27C7.5 |
| WBGene00007769 | -0.433146595 | 0.82524361 | 1 | C27C7.7 |
| WBGene00007770 | 0.834128642 | 0.71441843 | 1 | nhr-259 |
| WBGene00007771 | -0.393423254 | 0.21763022 | 0.41987105 | C27C12.1 |
| WBGene00007772 | 0.253955429 | 0.35757205 | 0.56586056 | egrh-1 |
| WBGene00007773 | 0.154830094 | 0.62450747 | 0.78158387 | C27C12.3 |
| WBGene00007774 | -0.117679095 | 0.69137233 | 0.82420135 | C27C12.4 |
| WBGene00007775 | 0.569132375 | 0.40801416 | 0.6141434 | acd-3 |
| WBGene00007776 | 0.554201101 | 0.61820456 | 1 | dmd-4 |
| WBGene00007777 | -0.693505305 | 0.79620356 | 1 | C27D8.1 |
| WBGene00007778 | 0.965282022 | 0.52694557 | 1 | C27D8.2 |
| WBGene00007779 | 0.489950816 | 0.1156022 | 0.28018323 | C27D8.3 |
| WBGene00007780 | -0.054181566 | 0.83024045 | 0.90818536 | C27D8.4 |
| WBGene00007784 | -0.575545713 | 0.00327042 | 0.02261262 | ruvb-1 |
| WBGene00007785 | -0.257607663 | 0.22633255 | 0.43063838 | tofu-1 |
| WBGene00007786 | -0.346235774 | 0.03286704 | 0.12095748 | rmd-2 |
| WBGene00007787 | -0.466576889 | 0.00472715 | 0.02998555 | C27H6.8 |
| WBGene00007788 | 0.017962595 | 0.95316023 | 0.97623454 | C28A5.1 |
| WBGene00007789 | -0.343488056 | 0.27345392 | 0.48213826 | C28A5.2 |
| WBGene00007791 | -0.006687493 | 0.99385929 | 0.996815 | C28A5.6 |

| | | | | |
|---|---|---|---|---|
| WBGene00007792 | 0.377828795 | 0.92526585 | 1 | C28D4.4 |
| WBGene00007793 | 0.377828795 | 0.92526585 | 1 | C28D4.5 |
| WBGene00007794 | -0.156269293 | 0.96924949 | 1 | C28D4.7 |
| WBGene00007795 | 0.281454817 | 0.94435481 | 1 | C28D4.8 |
| WBGene00007796 | -0.621606308 | 0.77755782 | 1 | C28D4.10 |
| WBGene00007797 | -0.615950037 | 0.00119489 | 0.0105201 | C29A12.1 |
| WBGene00007799 | 1.311351588 | 0.00020736 | 0.00270128 | nrx-1 |
| WBGene00007800 | 0.945230622 | 0.31150661 | 0.51976499 | C29A12.6 |
| WBGene00007801 | 0.640630493 | 0.12582655 | 0.29594346 | trpa-1 |
| WBGene00007802 | 0.789601902 | 0.75907734 | 1 | cutl-27 |
| WBGene00007804 | 1.894863195 | 0.29507975 | 1 | C29F3.3 |
| WBGene00007806 | -0.659821513 | 0.43013488 | 0.63269537 | clec-230 |
| WBGene00007807 | 0.378598545 | 0.09348385 | 0.24402558 | C29F3.7 |
| WBGene00007808 | 1.370479365 | 2.1969E-06 | 6.3055E-05 | best-7 |
| WBGene00007809 | 1.531088456 | 0.30629064 | 1 | C29F4.3 |
| WBGene00007810 | -0.227646844 | 0.64623275 | 0.79557004 | C29F7.1 |
| WBGene00007811 | -0.016173519 | 0.95132681 | 0.97542934 | C29F7.2 |
| WBGene00007812 | -0.531720469 | 0.01593407 | 0.07259399 | C29F7.3 |
| WBGene00007813 | 0.688297719 | 0.5477074 | 1 | jmjd-3.3 |
| WBGene00007816 | 2.169457994 | 0.26111677 | 1 | C30F2.2 |
| WBGene00007817 | 0.773413031 | 0.10168622 | 0.25720259 | C30F2.3 |
| WBGene00007818 | 1.170932659 | 0.28125273 | 1 | C30F2.4 |
| WBGene00007819 | 0.332937453 | 0.62507694 | 0.78190379 | C30F2.5 |
| WBGene00007820 | 1.03245503 | 0.13611822 | 0.31141781 | clec-199 |
| WBGene00007822 | -0.156269293 | 0.96924949 | 1 | clec-202 |
| WBGene00007823 | -0.265411405 | 0.84797504 | 1 | C30H6.5 |
| WBGene00007824 | -0.361125325 | 0.0459302 | 0.15152153 | dlat-2 |
| WBGene00007825 | -0.433906338 | 0.11869672 | 0.28427339 | C30H6.8 |
| WBGene00007826 | 0.108806652 | 0.73356381 | 0.85095923 | C30H6.9 |
| WBGene00007827 | 0.87255507 | 0.12399316 | 0.29316473 | nlp-54 |
| WBGene00007829 | -0.736789399 | 0.72029863 | 1 | oac-5 |
| WBGene00007832 | -0.008677305 | 0.99719169 | 1 | srbc-82 |
| WBGene00007833 | 0.69347674 | 0.10393309 | 0.26148193 | oac-6 |
| WBGene00007835 | 0.69258611 | 0.45810717 | 0.65477918 | oac-7 |
| WBGene00007836 | -1.035996579 | 1.0168E-06 | 3.4103E-05 | C31C9.2 |
| WBGene00007837 | 0.933696627 | 0.16566324 | 0.35249167 | fbxa-167 |
| WBGene00007840 | 1.529878821 | 0.01504248 | 0.0698664 | C31C9.6 |
| WBGene00007841 | 0.999912861 | 0.18607728 | 0.37887871 | C31C9.7 |
| WBGene00007844 | 1.181686151 | 0.7579682 | 1 | C31E10.4 |
| WBGene00007844 | 1.181686151 | 0.7579682 | 1 | irld-24 |
| WBGene00007846 | -0.198775272 | 0.43618774 | 0.63705534 | C31E10.5 |
| WBGene00007847 | -0.076873329 | 0.85671898 | 0.92175167 | C31E10.6 |
| WBGene00007848 | 0.043452487 | 0.80985883 | 0.89629222 | cytb-5.1 |

| | | | | |
|---|---|---|---|---|
| WBGene00007849 | 0.245666807 | 0.59055128 | 0.75812272 | tbc-19 |
| WBGene00007850 | -0.177329077 | 0.77047258 | 0.8715464 | C31G12.1 |
| WBGene00007851 | 3.519586579 | 0.0454016 | 1 | clec-245 |
| WBGene00007853 | 0.509041539 | 0.7857501 | 1 | C31G12.4 |
| WBGene00007854 | 0.864899468 | 0.6533431 | 1 | C31H5.1 |
| WBGene00007855 | -0.903761068 | 0.00109112 | 0.00981195 | C31H5.4 |
| WBGene00007856 | 0.234910932 | 0.70442752 | 0.832842 | C31H5.5 |
| WBGene00007857 | 1.306632812 | 3.82E-09 | 2.57E-07 | C31H5.6 |
| WBGene00007858 | -0.62146648 | 0.85925752 | 1 | C31H5.7 |
| WBGene00007859 | -0.150627625 | 0.57691988 | 0.74903703 | mrps-31 |
| WBGene00007860 | 0.293521907 | 0.15065455 | 0.33232138 | rilp-1 |
| WBGene00007861 | 1.82867917 | 0.54358766 | 1 | C32A9.1 |
| WBGene00007862 | 1.173802278 | 0.07092704 | 0.20403874 | C32C4.1 |
| WBGene00007863 | -0.558804446 | 0.87818675 | 1 | C32C4.3 |
| WBGene00007864 | -0.455620495 | 0.68365543 | 1 | irg-6 |
| WBGene00007866 | 1.190068588 | 0.53174194 | 1 | C32H11.3 |
| WBGene00007867 | 1.514684999 | 0.00014796 | 0.00208399 | C32H11.4 |
| WBGene00007868 | 1.352765391 | 0.55021839 | 1 | C32H11.5 |
| WBGene00007869 | -0.156269293 | 0.96924949 | 1 | C32H11.6 |
| WBGene00007870 | -0.653950113 | 0.19933341 | 0.39752085 | C32H11.7 |
| WBGene00007872 | 0.544525163 | 0.81649326 | 1 | C32H11.9 |
| WBGene00007873 | -1.413025445 | 0.18533064 | 1 | dod-21 |
| WBGene00007875 | 2.054040154 | 2.30E-29 | 1.64E-26 | dod-24 |
| WBGene00007876 | 5.903834731 | 9.004E-06 | 0.00020862 | dct-19 |
| WBGene00007877 | 1.090263169 | 9.4843E-05 | 0.00146873 | nfki-1 |
| WBGene00007878 | 0.263617763 | 0.61287146 | 0.77396612 | C33A11.2 |
| WBGene00007880 | -0.708246666 | 0.00023283 | 0.00295673 | C33A12.1 |
| WBGene00007881 | 0.721679499 | 0.19733496 | 0.39463741 | nlp-35 |
| WBGene00007882 | -0.435672576 | 0.03561901 | 0.12726616 | C33A12.3 |
| WBGene00007883 | -0.376899459 | 0.32090997 | 0.52915674 | C33A12.4 |
| WBGene00007885 | 0.84956869 | 0.12798201 | 0.29864109 | ugt-21 |
| WBGene00007886 | -0.060551733 | 0.89012644 | 0.94196731 | ethe-1 |
| WBGene00007887 | -1.1524358 | 0.07972389 | 0.22043917 | gadr-2 |
| WBGene00007890 | -0.381930814 | 0.42267906 | 0.62712373 | C33A12.19 |
| WBGene00007891 | 0.067343211 | 0.97089826 | 1 | C33B4.2 |
| WBGene00007892 | -0.507728591 | 0.32964494 | 0.53822205 | C33B4.4 |
| WBGene00007893 | 0.075260937 | 0.97569838 | 1 | C33B4.5 |
| WBGene00007894 | -0.687709908 | 0.74016374 | 1 | C33D3.3 |
| WBGene00007896 | -0.324008968 | 0.85415729 | 1 | C33D3.5 |
| WBGene00007898 | 1.355029752 | 0.04100813 | 0.14005054 | C33D9.3 |
| WBGene00007900 | 0.306334317 | 0.35446164 | 0.56284577 | C33D9.6 |
| WBGene00007901 | 0.316448168 | 0.84007028 | 1 | C33D9.8 |
| WBGene00007903 | 0.058370437 | 0.91455532 | 0.95571968 | lgc-21 |

| | | | | |
|---|---|---|---|---|
| WBGene00007904 | -0.493314523 | 0.17371126 | 0.36368829 | C33G3.4 |
| WBGene00007905 | -0.156269293 | 0.96924949 | 1 | C33G3.5 |
| WBGene00007906 | 0.28938282 | 0.44531106 | 0.64463117 | C33G3.6 |
| WBGene00007907 | 0.415283953 | 0.09565216 | 0.24787895 | C34B4.2 |
| WBGene00007908 | 0.578692293 | 0.3505423 | 0.55903508 | C34B4.3 |
| WBGene00007909 | 0.094500378 | 0.95929405 | 1 | C34B4.4 |
| WBGene00007910 | 0.403207732 | 0.86402224 | 1 | C34B4.5 |
| WBGene00007911 | 0.707950223 | 0.34163949 | 0.55003913 | C34B7.1 |
| WBGene00007912 | 0.173754323 | 0.40544915 | 0.61226986 | C34B7.2 |
| WBGene00007913 | 0.397616833 | 0.27438088 | 0.4827622 | cyp-36A1 |
| WBGene00007914 | 0.093045082 | 0.70181153 | 0.8316929 | mys-4 |
| WBGene00007916 | 0.458434642 | 0.46122976 | 0.65663585 | C34C6.3 |
| WBGene00007917 | 0.101573089 | 0.67807463 | 0.81582638 | C34C6.4 |
| WBGene00007918 | 0.178385631 | 0.48833795 | 0.67820992 | sphk-1 |
| WBGene00007919 | 1.1530214 | 0.01301696 | 0.06294757 | cup-16 |
| WBGene00007921 | -0.174607295 | 0.4017057 | 0.60925782 | C34C12.2 |
| WBGene00007922 | -0.296944402 | 0.17441323 | 0.36457165 | pph-6 |
| WBGene00007923 | -0.133213777 | 0.65097955 | 0.79882178 | C34C12.4 |
| WBGene00007924 | -0.241037638 | 0.27764662 | 0.48649266 | rsu-1 |
| WBGene00007925 | -0.002124047 | 0.99615263 | 0.99804333 | C34C12.6 |
| WBGene00007926 | 1.220708528 | 0.03803605 | 0.13300568 | C34C12.7 |
| WBGene00007927 | -0.62489961 | 0.00089028 | 0.00840325 | C34C12.8 |
| WBGene00007928 | 0.841340222 | 0.26515663 | 0.47390941 | C34C12.9 |
| WBGene00007929 | 1.470575353 | 0.51153906 | 1 | dmd-10 |
| WBGene00007931 | 1.312497985 | 0.00708243 | 0.04010657 | C34D1.4 |
| WBGene00007932 | 1.30034727 | 0.00508381 | 0.03155229 | zip-5 |
| WBGene00007933 | 0.785416611 | 0.59983079 | 1 | C34E7.3 |
| WBGene00007934 | -0.235717068 | 0.45124103 | 0.64911342 | C34E7.4 |
| WBGene00007935 | -0.545579184 | 0.4887485 | 0.67862483 | C34E11.2 |
| WBGene00007937 | 2.238436409 | 0.10456982 | 1 | C34E11.4 |
| WBGene00007938 | 1.09047542 | 0.36351179 | 1 | C34F6.1 |
| WBGene00007939 | 0.057627514 | 0.9292026 | 0.96323845 | C34F6.5 |
| WBGene00007941 | 0.299342991 | 0.49466205 | 0.68308599 | C34F6.7 |
| WBGene00007942 | 0.104011511 | 0.6198742 | 0.7787454 | idh-2 |
| WBGene00007943 | 0.287483374 | 0.35973721 | 0.56802829 | C34F6.9 |
| WBGene00007944 | 1.504229844 | 5.38E-08 | 2.6896E-06 | C34F6.10 |
| WBGene00007945 | 0.589625059 | 0.40722103 | 0.61394145 | C34F6.11 |
| WBGene00007946 | 1.735991307 | 0.58152819 | 1 | ugt-33 |
| WBGene00007947 | -1.900282435 | 1.6147E-05 | 0.00033976 | C35A5.3 |
| WBGene00007948 | 0.688793783 | 0.86378755 | 1 | C35A5.4 |
| WBGene00007949 | 0.88787957 | 0.71832755 | 1 | C35A5.5 |
| WBGene00007950 | 0.356254838 | 0.45028553 | 0.64889218 | C35A5.6 |
| WBGene00007951 | 0.889190203 | 0.1562932 | 0.34035865 | dmsr-8 |

| | | | | |
|---|---|---|---|---|
| WBGene00007952 | 0.073487887 | 0.62041141 | 0.77878286 | C35A5.8 |
| WBGene00007953 | -0.018964164 | 0.98393848 | 0.99228305 | hda-11 |
| WBGene00007954 | 0.747845955 | 0.85215965 | 1 | gcp-2.2 |
| WBGene00007955 | -0.148895989 | 0.45825157 | 0.65477918 | selt-1.1 |
| WBGene00007956 | -0.006765455 | 0.97863227 | 0.98873753 | trpp-9 |
| WBGene00007957 | 1.423871475 | 0.00645909 | 0.03754592 | C35C5.8 |
| WBGene00007958 | 0.033348295 | 0.94076057 | 0.96942574 | C35C5.9 |
| WBGene00007959 | 0.05682023 | 0.87047561 | 0.93024187 | C35C5.10 |
| WBGene00007960 | -2.563099533 | 0.4751044 | 1 | C35D6.3 |
| WBGene00007961 | -1.009327721 | 0.36862163 | 1 | C35D6.4 |
| WBGene00007963 | -2.418736656 | 2.02E-09 | 1.46E-07 | cyp-25A1 |
| WBGene00007964 | -0.725818617 | 0.01922998 | 0.08332046 | cyp-25A2 |
| WBGene00007965 | -0.343404566 | 0.02618577 | 0.10332189 | C36A4.4 |
| WBGene00007966 | -0.423335765 | 0.06829786 | 0.19913649 | maph-1.3 |
| WBGene00007967 | 0.196316803 | 0.71964653 | 0.84179137 | cyp-25A4 |
| WBGene00007969 | 0.187263113 | 0.19655663 | 0.39359954 | acs-19 |
| WBGene00007970 | -0.156269293 | 0.96924949 | 1 | C36A4.10 |
| WBGene00007971 | -0.550543313 | 0.00107677 | 0.0097415 | rpb-3 |
| WBGene00007972 | -0.220526287 | 0.3359651 | 0.54429574 | prp-4 |
| WBGene00007973 | 0.917751407 | 0.20259613 | 0.40156488 | C36B1.6 |
| WBGene00007974 | -0.988909372 | 3.5093E-06 | 9.4263E-05 | dhfr-1 |
| WBGene00007975 | 0.207975 | 0.28901966 | 0.49921577 | gls-1 |
| WBGene00007976 | 0.349008148 | 0.17236443 | 0.36180439 | C36B1.9 |
| WBGene00007977 | 0.58294746 | 0.72944118 | 1 | gska-3 |
| WBGene00007978 | -0.143886017 | 0.40846762 | 0.61437133 | C36B1.11 |
| WBGene00007979 | -0.243077347 | 0.13523575 | 0.30973633 | imp-1 |
| WBGene00007980 | 0.064322781 | 0.66112406 | 0.80500524 | C36E8.1 |
| WBGene00007981 | 0.509915293 | 0.33002728 | 0.53851809 | glb-11 |
| WBGene00007982 | 0.279413515 | 0.40408498 | 0.61126287 | pxd-1 |
| WBGene00007983 | 0.269585604 | 0.54476739 | 0.72400914 | C36E8.4 |
| WBGene00007984 | 0.462646202 | 0.21604795 | 0.41761252 | irx-1 |
| WBGene00007985 | -0.140113089 | 0.6038206 | 0.76852772 | swah-1 |
| WBGene00007986 | -2.051235694 | 0.34377405 | 1 | C36F7.5 |
| WBGene00007987 | 1.557897343 | 0.63813005 | 1 | C36H8.1 |
| WBGene00007988 | 1.259149189 | 0.1446794 | 0.32319276 | best-8 |
| WBGene00007989 | -0.439911243 | 0.65118844 | 0.79891662 | fipr-22 |
| WBGene00007990 | 0.173127447 | 0.90334186 | 1 | fipr-23 |
| WBGene00007991 | 0.300014049 | 0.37926821 | 0.58740851 | C37A5.7 |
| WBGene00007992 | 2.779713462 | 0.22340067 | 1 | fipr-24 |
| WBGene00007993 | 0.021594293 | 0.92489871 | 0.96034913 | idhb-1 |
| WBGene00007994 | -1.341962798 | 0.18953538 | 1 | C37E2.2 |
| WBGene00007996 | 0.851945099 | 0.47750599 | 1 | C38C6.3 |
| WBGene00007997 | 3.529396058 | 0.04918488 | 1 | sre-13 |

| | | | | |
|---|---|---|---|---|
| WBGene00007999 | -0.955790486 | 0.03179741 | 0.11823961 | tag-297 |
| WBGene00008000 | -0.39895418 | 0.00972187 | 0.05076698 | slc-17.2 |
| WBGene00008001 | 0.232157325 | 0.93397866 | 1 | C38C10.3 |
| WBGene00008002 | -0.175875562 | 0.87759797 | 1 | nlp-48 |
| WBGene00008003 | -0.421717507 | 0.23636528 | 0.44244693 | enu-3.2 |
| WBGene00008005 | -0.433628902 | 0.01326212 | 0.06371802 | C38D4.4 |
| WBGene00008006 | 0.172843616 | 0.44726167 | 0.64629897 | tag-325 |
| WBGene00008007 | 0.832787778 | 0.43364202 | 1 | C38D4.7 |
| WBGene00008008 | -0.460884391 | 0.25260654 | 0.46063322 | C38D4.9 |
| WBGene00008009 | 0.688793783 | 0.86378755 | 1 | fbxa-171 |
| WBGene00008010 | 2.645266161 | 0.05993004 | 1 | C38D9.2 |
| WBGene00008012 | -0.156269293 | 0.96924949 | 1 | fbxa-133 |
| WBGene00008014 | 2.932343137 | 0.13706871 | 1 | fbxa-172 |
| WBGene00008015 | 0.377828795 | 0.92526585 | 1 | fbxa-174 |
| WBGene00008016 | -1.417356304 | 0.16710217 | 0.35455914 | C38D9.8 |
| WBGene00008017 | 1.980634444 | 0.28443964 | 1 | fbxa-176 |
| WBGene00008018 | 0.092253039 | 0.83969282 | 0.91211511 | tbc-8 |
| WBGene00008019 | 0.699068032 | 0.02486232 | 0.09974122 | C38H2.2 |
| WBGene00008020 | -0.557833565 | 0.47288804 | 0.66597074 | C38H2.3 |
| WBGene00008021 | 1.481931163 | 0.10696995 | 0.26658958 | C39B10.1 |
| WBGene00008022 | 0.910522539 | 0.3687166 | 1 | lgc-41 |
| WBGene00008023 | 0.88787957 | 0.71832755 | 1 | C39B10.3 |
| WBGene00008025 | 1.429963739 | 0.3701028 | 1 | C39B10.5 |
| WBGene00008027 | 1.504586387 | 9.7393E-05 | 0.00149971 | scl-5 |
| WBGene00008028 | 1.354526243 | 0.00061429 | 0.00625157 | scl-6 |
| WBGene00008029 | 1.653460415 | 0.50712258 | 1 | scl-7 |
| WBGene00008030 | 2.912785086 | 0.15033923 | 1 | scl-8 |
| WBGene00008031 | 0.980440239 | 0.0614129 | 0.18554004 | C39E9.7 |
| WBGene00008032 | -1.105053094 | 1.4508E-06 | 4.4934E-05 | C39E9.8 |
| WBGene00008033 | 0.896460151 | 0.07241788 | 0.20703984 | spin-2 |
| WBGene00008034 | -0.286669641 | 0.10081611 | 0.25590808 | C39E9.11 |
| WBGene00008035 | -0.475854276 | 0.04320082 | 0.14496993 | C39E9.12 |
| WBGene00008036 | 1.69047418 | 0.38809581 | 1 | C40C9.3 |
| WBGene00008037 | 1.477253415 | 0.58357327 | 1 | C40C9.4 |
| WBGene00008038 | 0.277701798 | 0.79151886 | 1 | C40H1.2 |
| WBGene00008039 | 0.798461211 | 0.56063412 | 1 | C40H1.3 |
| WBGene00008040 | 0.232621718 | 0.37013025 | 0.57766361 | ttr-5 |
| WBGene00008041 | -0.836783132 | 0.0002136 | 0.0027706 | ufc-1 |
| WBGene00008042 | 2.108561005 | 1.06E-10 | 9.98E-09 | C40H1.7 |
| WBGene00008043 | 1.073774215 | 0.4104606 | 1 | C40H1.8 |
| WBGene00008044 | 1.901356495 | 0.30928877 | 1 | C40H1.9 |
| WBGene00008045 | 0.41444123 | 0.60094866 | 0.76623794 | C40H5.2 |
| WBGene00008045 | 1.247103736 | 0.59368513 | 1 | C40H5.7 |

| | | | | |
|---|---|---|---|---|
| WBGene00008047 | -0.293598339 | 0.76028282 | 0.86607185 | lact-7 |
| WBGene00008050 | 1.469739662 | 0.25780658 | 1 | chst-1 |
| WBGene00008051 | 1.180319717 | 0.26584714 | 1 | lron-4 |
| WBGene00008052 | -0.01573764 | 0.93759803 | 0.96772742 | ctns-1 |
| WBGene00008053 | -0.236906767 | 0.11357435 | 0.2768332 | cdc-48.2 |
| WBGene00008056 | -0.156269293 | 0.96924949 | 1 | nhr-164 |
| WBGene00008057 | -0.156269293 | 0.96924949 | 1 | irld-25 |
| WBGene00008061 | -0.561509848 | 0.01937233 | 0.08366608 | C41G7.3 |
| WBGene00008062 | -0.220606266 | 0.52854482 | 0.71276937 | set-32 |
| WBGene00008063 | 0.905096248 | 0.68842149 | 1 | C41G7.6 |
| WBGene00008064 | -0.333693239 | 0.89573179 | 1 | C41G7.7 |
| WBGene00008065 | 2.513410497 | 0.03722566 | 1 | npr-18 |
| WBGene00008066 | -0.407440094 | 0.74634893 | 1 | fbxb-65 |
| WBGene00008067 | -1.901856996 | 0.64072034 | 1 | C43D7.4 |
| WBGene00008068 | -2.679599433 | 0.0295447 | 1 | sdz-6 |
| WBGene00008069 | -1.964653846 | 0.53599202 | 1 | C43D7.7 |
| WBGene00008070 | -1.186373843 | 0.53853051 | 1 | C43D7.8 |
| WBGene00008071 | -0.91375383 | 0.72824392 | 1 | fbxb-64 |
| WBGene00008072 | 1.025181888 | 0.79790347 | 1 | C43F9.4 |
| WBGene00008073 | 0.613861255 | 0.54743112 | 1 | C43F9.5 |
| WBGene00008074 | 1.653460415 | 0.50712258 | 1 | nkb-2 |
| WBGene00008075 | 0.367542838 | 0.86160434 | 1 | C43F9.7 |
| WBGene00008076 | 1.225903374 | 0.0389928 | 0.13502898 | lgc-43 |
| WBGene00008077 | 0.494402665 | 0.73880974 | 1 | oac-8 |
| WBGene00008078 | -0.040519091 | 0.91517947 | 0.95580617 | vps-50 |
| WBGene00008079 | -0.29136374 | 0.8241256 | 1 | C44B9.2 |
| WBGene00008080 | -0.720635496 | 0.00526591 | 0.03241989 | C44B9.3 |
| WBGene00008081 | -0.135216981 | 0.46572744 | 0.65994361 | athp-1 |
| WBGene00008082 | -0.256341339 | 0.24141916 | 0.44906848 | com-1 |
| WBGene00008084 | 1.717821585 | 0.09981607 | 0.25448695 | C44C10.3 |
| WBGene00008085 | 0.158120369 | 0.66725124 | 0.80921959 | C44C10.4 |
| WBGene00008086 | -1.323101626 | 0.27721153 | 1 | C44C10.5 |
| WBGene00008088 | 0.281454817 | 0.94435481 | 1 | C44C10.7 |
| WBGene00008089 | 0.410777888 | 0.10185764 | 0.25747525 | C44C10.9 |
| WBGene00008090 | -1.901856996 | 0.64072034 | 1 | C44C10.10 |
| WBGene00008091 | 0.04382309 | 0.93941285 | 0.96885928 | C44F1.1 |
| WBGene00008092 | 1.353535409 | 0.73149271 | 1 | gmeb-3 |
| WBGene00008093 | -0.303324194 | 0.75048462 | 1 | lron-1 |
| WBGene00008094 | 0.224205266 | 0.39682785 | 0.60465448 | C44H4.4 |
| WBGene00008095 | 0.568913981 | 0.66079976 | 1 | C44H4.6 |
| WBGene00008096 | 1.129631999 | 0.40747336 | 1 | C44H4.8 |
| WBGene00008097 | 0.688793783 | 0.86378755 | 1 | ugt-15 |
| WBGene00008099 | 1.281300731 | 0.03702015 | 0.13061948 | C44H9.4 |

| | | | | |
|---|---|---|---|---|
| WBGene00008100 | 0.7749857 | 0.01315864 | 0.06337131 | C44H9.5 |
| WBGene00008101 | 0.554434855 | 0.18807152 | 0.38164405 | C44H9.6 |
| WBGene00008102 | 1.524788098 | 0.11153806 | 0.27340674 | C44H9.7 |
| WBGene00008105 | 1.471809563 | 0.46592277 | 1 | C45B11.2 |
| WBGene00008107 | -0.169628512 | 0.38972061 | 0.59820862 | aspm-1 |
| WBGene00008110 | 0.344704965 | 0.07443732 | 0.2108738 | C46C2.2 |
| WBGene00008111 | 0.372881888 | 0.7206673 | 1 | C46C2.3 |
| WBGene00008113 | -1.178032943 | 0.01095491 | 0.05555707 | C46C2.5 |
| WBGene00008114 | 0.15388156 | 0.93468081 | 1 | C46C2.6 |
| WBGene00008117 | -0.617509194 | 6.0974E-05 | 0.00102674 | gsr-1 |
| WBGene00008118 | -0.174205009 | 0.53956007 | 0.72113594 | madf-8 |
| WBGene00008119 | 0.129371272 | 0.49969938 | 0.68768606 | C46F11.4 |
| WBGene00008120 | -0.260437649 | 0.22741201 | 0.4320915 | C46F11.5 |
| WBGene00008121 | 0.594630644 | 0.22462506 | 0.42873336 | C46F11.6 |
| WBGene00008122 | -1.129457579 | 0.5821088 | 1 | C47A4.1 |
| WBGene00008123 | 0.78617901 | 0.02605358 | 0.10302938 | ent-4 |
| WBGene00008127 | 1.248353914 | 0.03298907 | 0.12122286 | ddo-1 |
| WBGene00008128 | 0.696952114 | 0.62793204 | 1 | srab-12 |
| WBGene00008129 | -1.159714244 | 0.75790634 | 1 | C47A10.12 |
| WBGene00008130 | 0.230685249 | 0.67059425 | 0.81161419 | fbxa-140 |
| WBGene00008131 | -0.723026772 | 0.00080128 | 0.00774734 | C47B2.2 |
| WBGene00008132 | -0.027971669 | 0.8575517 | 0.9221909 | gale-1 |
| WBGene00008133 | -0.192396382 | 0.54554645 | 0.72459024 | efsc-1 |
| WBGene00008134 | -0.333748206 | 0.28762548 | 0.49751546 | C47B2.9 |
| WBGene00008136 | 0.045987525 | 0.79088302 | 0.88466103 | C47D12.2 |
| WBGene00008137 | 0.11481195 | 0.93095148 | 1 | sfxn-1.4 |
| WBGene00008138 | -1.519908074 | 0.06502921 | 0.19242498 | C47D12.4 |
| WBGene00008139 | 0.68817485 | 0.08739939 | 0.23398367 | C47D12.5 |
| WBGene00008140 | -0.179159137 | 0.38929182 | 0.59774782 | xpf-1 |
| WBGene00008141 | -0.156269293 | 0.96924949 | 1 | C47E8.1 |
| WBGene00008142 | 0.918611207 | 0.18003474 | 0.37228489 | C47E8.3 |
| WBGene00008143 | -0.276441738 | 0.15820515 | 0.34299953 | C47E8.4 |
| WBGene00008144 | 0.61245546 | 0.62174041 | 1 | gasr-8 |
| WBGene00008145 | 0.227058738 | 0.39729186 | 0.60512336 | set-5 |
| WBGene00008146 | -0.199374337 | 0.91832278 | 1 | C47E8.9 |
| WBGene00008147 | -0.352146808 | 0.09162369 | 0.24107615 | C47E12.2 |
| WBGene00008148 | -0.102095049 | 0.62816397 | 0.78388819 | C47E12.3 |
| WBGene00008149 | -0.029935613 | 0.8480205 | 0.91671496 | pyp-1 |
| WBGene00008150 | -0.125843698 | 0.9295138 | 1 | mam-2 |
| WBGene00008151 | -0.135766154 | 0.55285617 | 0.73075023 | C47E12.7 |
| WBGene00008152 | 2.017231934 | 0.20210782 | 1 | C47E12.9 |
| WBGene00008153 | -0.156269293 | 0.96924949 | 1 | C47E12.10 |
| WBGene00008154 | -0.156269293 | 0.96924949 | 1 | C47E12.11 |

| | | | | |
|---|---|---|---|---|
| WBGene00008155 | 0.820015846 | 0.7518774 | 1 | C47E12.12 |
| WBGene00008156 | 0.546884135 | 0.76309407 | 1 | C47E12.13 |
| WBGene00008157 | -0.658032786 | 0.28964875 | 0.49967941 | C47F8.1 |
| WBGene00008158 | -1.159714244 | 0.75790634 | 1 | nhr-165 |
| WBGene00008160 | -0.156269293 | 0.96924949 | 1 | glct-4 |
| WBGene00008163 | -0.042661126 | 0.98499355 | 1 | C47F8.7 |
| WBGene00008164 | -0.366974641 | 0.20023675 | 0.39873287 | C47G2.3 |
| WBGene00008165 | -0.073644442 | 0.70974411 | 0.83571538 | C47G2.4 |
| WBGene00008166 | -0.350528561 | 0.088723 | 0.23621342 | saps-1 |
| WBGene00008167 | 1.617674873 | 7.74E-23 | 3.48E-20 | acox-1.5 |
| WBGene00008168 | 1.28656172 | 0.10540947 | 0.26426767 | C48B4.3 |
| WBGene00008169 | 0.108010258 | 0.72401944 | 0.84489812 | C48B4.6 |
| WBGene00008170 | -0.273825027 | 0.3171212 | 0.52493219 | C48B4.7 |
| WBGene00008171 | 0.429404555 | 0.36397911 | 0.57174854 | C48B4.8 |
| WBGene00008172 | -0.449913255 | 0.12919358 | 0.3004289 | C48B4.9 |
| WBGene00008173 | -0.162503973 | 0.60563663 | 0.76897128 | C48B4.10 |
| WBGene00008174 | -0.514430342 | 0.04443762 | 0.14769153 | C48B4.11 |
| WBGene00008175 | 0.03940405 | 0.99001443 | 1 | C48B4.12 |
| WBGene00008176 | -0.663495744 | 0.57413359 | 1 | C48B4.13 |
| WBGene00008177 | 0.823337738 | 0.15826705 | 0.34305991 | C48D1.1 |
| WBGene00008178 | -0.167754419 | 0.78722725 | 0.88275978 | C48D1.5 |
| WBGene00008180 | -2.563099533 | 0.4751044 | 1 | C48D5.3 |
| WBGene00008183 | 0.340639168 | 0.10710651 | 0.26684275 | rin-1 |
| WBGene00008184 | 0.102404895 | 0.95783845 | 1 | C49A1.1 |
| WBGene00008185 | 0.28530093 | 0.92084641 | 1 | best-10 |
| WBGene00008186 | 0.377828795 | 0.92526585 | 1 | best-11 |
| WBGene00008187 | 1.413377704 | 0.60673863 | 1 | C49A1.5 |
| WBGene00008188 | -1.969805064 | 0.62862569 | 1 | clec-115 |
| WBGene00008190 | -0.156269293 | 0.96924949 | 1 | clec-114 |
| WBGene00008191 | 0.069716363 | 0.83685237 | 0.91131263 | C49A1.10 |
| WBGene00008194 | 1.328239564 | 8.69E-12 | 1.02E-09 | C49C3.4 |
| WBGene00008195 | -0.01491824 | 0.97839871 | 0.98873753 | ceh-88 |
| WBGene00008196 | -0.683401989 | 0.00288361 | 0.02063048 | C49C3.6 |
| WBGene00008197 | -0.010784077 | 0.96402323 | 0.98190304 | C49C3.7 |
| WBGene00008198 | -0.261562 | 0.63306344 | 0.7872679 | C49C3.8 |
| WBGene00008199 | 0.748110219 | 0.00935138 | 0.04927737 | C49C3.9 |
| WBGene00008200 | 1.359224998 | 0.02939108 | 0.11154644 | C49C3.10 |
| WBGene00008201 | 1.435925494 | 0.70920779 | 1 | C49C3.11 |
| WBGene00008202 | -0.942479658 | 0.58523581 | 1 | clec-197 |
| WBGene00008203 | 1.487038984 | 0.21499328 | 1 | clec-198 |
| WBGene00008204 | 0.470237544 | 0.41493161 | 0.62017539 | C49C3.15 |
| WBGene00008205 | 0.647636412 | 0.00054887 | 0.00573418 | sams-1 |
| WBGene00008206 | -0.135877444 | 0.73180756 | 0.849902 | set-6 |

| | | | | |
|---|---|---|---|---|
| WBGene00008207 | -0.603529721 | 0.0433233 | 0.14518029 | C49F5.3 |
| WBGene00008208 | 1.507316149 | 0.187621 | 1 | nhr-167 |
| WBGene00008209 | 1.216957619 | 0.74819621 | 1 | C49F5.5 |
| WBGene00008210 | -0.319131094 | 0.16039661 | 0.34570837 | C49F5.6 |
| WBGene00008211 | -0.279803724 | 0.28199799 | 0.49135155 | C49F5.7 |
| WBGene00008212 | -0.462562375 | 0.80026554 | 1 | C49F5.8 |
| WBGene00008213 | 0.209550033 | 0.77514708 | 0.87463389 | C49F8.1 |
| WBGene00008214 | 0.622032569 | 0.22304883 | 0.42700028 | gem-1 |
| WBGene00008215 | -0.061513493 | 0.88621718 | 0.93995577 | C49F8.3 |
| WBGene00008216 | 0.07102795 | 0.94916753 | 1 | mec-19 |
| WBGene00008218 | -0.678300503 | 1.0059E-06 | 3.3837E-05 | nasp-2 |
| WBGene00008219 | -0.317950444 | 0.06332138 | 0.18893706 | C50B6.3 |
| WBGene00008220 | 1.943458133 | 3.7626E-05 | 0.00068081 | C50B6.7 |
| WBGene00008221 | 1.637529793 | 0.00252948 | 0.01874875 | nhr-168 |
| WBGene00008222 | -0.76243722 | 0.42649154 | 0.6302378 | C50B6.9 |
| WBGene00008223 | 1.216957619 | 0.74819621 | 1 | lgc-48 |
| WBGene00008224 | -0.26742056 | 0.1209727 | 0.28776048 | C50B8.1 |
| WBGene00008225 | -0.229046345 | 0.36070099 | 0.56840302 | nuaf-1 |
| WBGene00008226 | 0.433741011 | 0.62214877 | 0.7801836 | C50B8.4 |
| WBGene00008227 | 0.429798932 | 0.45349582 | 0.65096864 | C50B8.5 |
| WBGene00008228 | 0.82316087 | 0.08312491 | 0.22648553 | C50B8.6 |
| WBGene00008229 | 0.663512639 | 0.00275502 | 0.01996941 | C50F4.1 |
| WBGene00008230 | 1.584577408 | 0.63258867 | 1 | pfk-1.2 |
| WBGene00008231 | 0.03940405 | 0.99001443 | 1 | tag-329 |
| WBGene00008232 | -0.484879598 | 0.23190458 | 0.43758569 | C50F4.4 |
| WBGene00008233 | 0.814742086 | 0.01179907 | 0.05859536 | C50F4.8 |
| WBGene00008234 | 2.187871607 | 0.01994664 | 0.08524188 | C50F4.9 |
| WBGene00008235 | 0.641065534 | 0.78192938 | 1 | C50F4.10 |
| WBGene00008236 | -0.081753312 | 0.74612099 | 0.85839618 | C50F4.12 |
| WBGene00008237 | 0.029447994 | 0.89259412 | 0.94349514 | nstp-10 |
| WBGene00008238 | -0.317146766 | 0.13643468 | 0.31200857 | C50F4.16 |
| WBGene00008239 | 1.096762262 | 0.00018355 | 0.00246519 | fshr-1 |
| WBGene00008240 | -0.936434159 | 0.65873492 | 1 | C50H2.4 |
| WBGene00008241 | -1.279666666 | 0.55824702 | 1 | C50H2.5 |
| WBGene00008242 | 0.281454817 | 0.94435481 | 1 | ceh-75 |
| WBGene00008243 | 0.281454817 | 0.94435481 | 1 | C50H2.7 |
| WBGene00008244 | -0.156269293 | 0.96924949 | 1 | fipr-11 |
| WBGene00008245 | 2.572211928 | 0.13237184 | 1 | fipr-10 |
| WBGene00008246 | 0.89541782 | 0.16841385 | 0.3562217 | C50H2.13 |
| WBGene00008247 | 2.063391569 | 0.37853067 | 1 | srsx-26 |
| WBGene00008248 | -0.615410782 | 0.70507525 | 1 | srsx-27 |
| WBGene00008249 | 1.181686151 | 0.7579682 | 1 | srsx-28 |
| WBGene00008250 | 0.377828795 | 0.92526585 | 1 | srsx-29 |

| | | | | |
|---|---|---|---|---|
| WBGene00008251 | 3.174845848 | 0.09860853 | 1 | srsx-30 |
| WBGene00008252 | 0.136500401 | 0.76850123 | 0.87028947 | C51E3.6 |
| WBGene00008254 | 0.415700217 | 0.15033824 | 0.33192519 | C51E3.9 |
| WBGene00008255 | 0.805572664 | 0.09760285 | 0.25109211 | C51E3.10 |
| WBGene00008256 | 1.630531267 | 0.21904575 | 1 | glb-12 |
| WBGene00008258 | 0.397759153 | 0.05633013 | 0.17500104 | mans-2 |
| WBGene00008259 | -1.094201388 | 0.7816772 | 1 | C52E4.7 |
| WBGene00008260 | -0.844337713 | 0.05454591 | 0.17072422 | C52G5.2 |
| WBGene00008261 | 1.38322681 | 0.69550305 | 1 | C52G5.3 |
| WBGene00008262 | -0.370885626 | 0.00819684 | 0.04460253 | ril-1 |
| WBGene00008263 | 0.040342413 | 0.96531021 | 0.98279507 | C53A5.2 |
| WBGene00008265 | 0.45397152 | 0.41755791 | 0.62218589 | kcnl-4 |
| WBGene00008266 | -0.346764472 | 0.05299902 | 0.16708366 | C53A5.6 |
| WBGene00008267 | 0.178115923 | 0.94982316 | 1 | C53A5.9 |
| WBGene00008269 | -0.184741373 | 0.761242 | 0.86650332 | C53A5.11 |
| WBGene00008270 | 0.716202684 | 0.04946673 | 0.15947139 | C53A5.13 |
| WBGene00008271 | 0.233639253 | 0.87875557 | 1 | C53B4.1 |
| WBGene00008272 | 0.747845955 | 0.85215965 | 1 | C53B4.2 |
| WBGene00008273 | 0.16116208 | 0.7059843 | 0.83366276 | C53B4.3 |
| WBGene00008274 | 0.414238338 | 0.04737748 | 0.15531096 | C53B4.4 |
| WBGene00008275 | 0.187979348 | 0.605235 | 0.76897128 | nstp-1 |
| WBGene00008277 | 2.89671812 | 0.07489542 | 1 | mltn-12 |
| WBGene00008278 | 1.387663491 | 0.10352497 | 0.26071284 | npr-10 |
| WBGene00008279 | -1.45873709 | 0.59584643 | 1 | C53C7.3 |
| WBGene00008280 | 1.251556056 | 0.18058829 | 0.37259848 | acc-2 |
| WBGene00008281 | -0.258786731 | 0.28090398 | 0.49007834 | C53D6.4 |
| WBGene00008283 | 0.244036074 | 0.58891511 | 0.75701624 | C53D6.6 |
| WBGene00008284 | 0.523843088 | 0.16029723 | 0.34561687 | C53D6.7 |
| WBGene00008285 | -1.590417856 | 0.4506532 | 1 | C53D6.8 |
| WBGene00008286 | -0.156269293 | 0.96924949 | 1 | C54C6.4 |
| WBGene00008287 | 1.019777392 | 0.26894557 | 0.47744819 | C54C6.5 |
| WBGene00008288 | 0.298772232 | 0.38301192 | 0.59132092 | C54C6.6 |
| WBGene00008292 | -1.777226873 | 0.19013783 | 1 | C54C8.4 |
| WBGene00008295 | 2.248027157 | 0.20463899 | 1 | nlp-39 |
| WBGene00008296 | 0.611383061 | 0.07435198 | 0.21075238 | cdr-2 |
| WBGene00008298 | 1.023173022 | 3.13E-08 | 1.6519E-06 | C54D10.3 |
| WBGene00008299 | 0.729973939 | 0.753654 | 1 | C54D10.4 |
| WBGene00008300 | 1.623374925 | 0.0968279 | 0.24991232 | C54D10.5 |
| WBGene00008301 | 2.333985214 | 0.1214166 | 1 | pals-39 |
| WBGene00008302 | 2.854906918 | 0.07696999 | 1 | pals-38 |
| WBGene00008303 | 2.188838843 | 0.34004625 | 1 | C54D10.9 |
| WBGene00008304 | 0.042655964 | 0.94746756 | 0.97311595 | C54D10.10 |
| WBGene00008306 | 1.557897343 | 0.63813005 | 1 | C54E10.1 |

| | | | | |
|---|---|---|---|---|
| WBGene00008307 | 0.049389948 | 0.95628773 | 0.97779092 | ncs-5 |
| WBGene00008308 | 1.413377704 | 0.60673863 | 1 | C54E10.3 |
| WBGene00008309 | 0.87875489 | 0.4094498 | 1 | nhr-170 |
| WBGene00008310 | -1.189950863 | 0.06326606 | 0.18886481 | C54E10.6 |
| WBGene00008311 | 0.579730666 | 0.00120072 | 0.01055349 | rskn-2 |
| WBGene00008312 | 0.938982574 | 0.8146943 | 1 | C54G4.2 |
| WBGene00008313 | -0.62146648 | 0.85925752 | 1 | C54G4.3 |
| WBGene00008314 | -0.028607653 | 0.9730395 | 0.98595164 | C54G4.4 |
| WBGene00008315 | 1.450014998 | 0.133713 | 0.30746392 | C54G4.5 |
| WBGene00008316 | -0.08078158 | 0.76101183 | 0.86650332 | dod-18 |
| WBGene00008317 | -0.122603314 | 0.61700944 | 0.77668343 | C54G4.7 |
| WBGene00008318 | -0.171307013 | 0.59539019 | 0.76171046 | C54G4.9 |
| WBGene00008319 | 0.09697944 | 0.97612634 | 1 | C54G10.1 |
| WBGene00008320 | 0.381988715 | 0.47050579 | 0.66356557 | slc-25A29 |
| WBGene00008323 | -2.714830683 | 0.06922892 | 1 | C55A1.6 |
| WBGene00008324 | -1.969805064 | 0.62862569 | 1 | C55A1.7 |
| WBGene00008325 | 0.281454817 | 0.94435481 | 1 | C55A1.9 |
| WBGene00008326 | 1.181686151 | 0.7579682 | 1 | C55A1.11 |
| WBGene00008328 | 0.747845955 | 0.85215965 | 1 | srz-43 |
| WBGene00008330 | -0.502123477 | 0.00637398 | 0.03722294 | C55A6.1 |
| WBGene00008331 | 0.05139714 | 0.75279448 | 0.86217294 | ttll-5 |
| WBGene00008332 | 0.723330554 | 0.41669996 | 0.62159408 | C55A6.3 |
| WBGene00008333 | 0.836838632 | 0.07377597 | 0.20967089 | C55A6.4 |
| WBGene00008334 | 0.184711808 | 0.79825431 | 0.88905656 | sdz-8 |
| WBGene00008335 | -0.706834536 | 0.35666592 | 0.5651371 | C55A6.6 |
| WBGene00008336 | 1.597342909 | 0.00565691 | 0.03416471 | C55A6.7 |
| WBGene00008338 | -0.276750499 | 0.11189504 | 0.27394993 | pafo-1 |
| WBGene00008339 | -0.28057429 | 0.21874703 | 0.42148987 | C55A6.10 |
| WBGene00008340 | -0.009692562 | 0.99573202 | 1 | C55A6.11 |
| WBGene00008341 | -0.362822711 | 0.38732416 | 0.59600794 | ttr-44 |
| WBGene00008342 | 1.996125773 | 0.03337461 | 0.12205606 | frpr-5 |
| WBGene00008343 | -0.053703788 | 0.8365613 | 0.9111967 | C56A3.4 |
| WBGene00008344 | -0.120686363 | 0.59346897 | 0.76005346 | C56A3.5 |
| WBGene00008345 | 0.182406914 | 0.42472084 | 0.62892251 | C56A3.6 |
| WBGene00008346 | -0.530720146 | 0.20626673 | 0.40607821 | C56A3.8 |
| WBGene00008348 | 0.379963231 | 0.40417782 | 0.61132713 | C56G7.3 |
| WBGene00008352 | 0.515865225 | 0.41411542 | 0.61950803 | fbxa-6 |
| WBGene00008353 | 0.231551717 | 0.60555671 | 0.76897128 | D1025.1 |
| WBGene00008354 | -0.646852255 | 0.06368567 | 0.18975441 | gcsh-1 |
| WBGene00008355 | -0.248116658 | 0.83937092 | 1 | fbxa-141 |
| WBGene00008356 | 0.534332089 | 0.1318736 | 0.30432949 | nspc-20 |
| WBGene00008357 | 1.231314247 | 0.10411426 | 0.26181674 | nspc-16 |
| WBGene00008358 | 0.360642352 | 0.40083927 | 0.60833979 | nspc-18 |

| | | | | |
|---|---|---|---|---|
| WBGene00008359 | 0.464558898 | 0.38371527 | 0.5921809 | nspc-17 |
| WBGene00008360 | 0.048449912 | 0.93261389 | 0.96422254 | nspc-19 |
| WBGene00008361 | 0.081038194 | 0.65576604 | 0.8022265 | D1043.1 |
| WBGene00008362 | -0.378620834 | 0.04951433 | 0.15947139 | cfim-2 |
| WBGene00008363 | -0.761599458 | 0.00051146 | 0.00545658 | D1046.2 |
| WBGene00008364 | -0.321355512 | 0.26608287 | 0.47472915 | slc-25A26 |
| WBGene00008365 | 1.59306666 | 0.17783177 | 1 | D1046.4 |
| WBGene00008366 | 1.339016963 | 3.0257E-05 | 0.00056341 | tpra-1 |
| WBGene00008368 | 1.035649927 | 0.06158008 | 0.18583439 | D1053.3 |
| WBGene00008369 | 0.781099869 | 0.21293176 | 0.413236 | D1053.4 |
| WBGene00008370 | 0.013262089 | 0.9690799 | 0.98399257 | D1054.1 |
| WBGene00008371 | -0.688183456 | 0.00015565 | 0.00214988 | D1054.3 |
| WBGene00008372 | 0.504314973 | 0.35732066 | 0.56561018 | D1054.5 |
| WBGene00008375 | -0.598170136 | 0.07085378 | 0.20389564 | D1054.8 |
| WBGene00008376 | -0.292134565 | 0.66815297 | 0.80998953 | D1054.9 |
| WBGene00008377 | -0.610008463 | 0.0006401 | 0.00647625 | D1054.10 |
| WBGene00008378 | -0.547684232 | 5.1031E-05 | 0.00088132 | ule-3 |
| WBGene00008379 | -0.062093187 | 0.82030467 | 0.90268458 | secs-1 |
| WBGene00008380 | -0.482008008 | 0.0115864 | 0.05779396 | prp-38 |
| WBGene00008381 | -0.774842772 | 0.78069886 | 1 | D1081.3 |
| WBGene00008382 | 2.446238633 | 0.17270707 | 1 | D1081.4 |
| WBGene00008383 | 0.281454817 | 0.94435481 | 1 | D1081.5 |
| WBGene00008384 | -0.859913261 | 0.07676646 | 0.21490008 | D1081.6 |
| WBGene00008385 | -0.134647804 | 0.41270862 | 0.61825496 | D1081.7 |
| WBGene00008386 | -0.498931813 | 0.00035751 | 0.00416052 | cdc-5L |
| WBGene00008387 | -0.213421982 | 0.48511809 | 0.67539764 | D1081.9 |
| WBGene00008388 | 0.21638904 | 0.60786007 | 0.77036604 | D1086.1 |
| WBGene00008389 | 1.354615392 | 0.08548894 | 0.23054671 | D1086.2 |
| WBGene00008390 | -1.917314424 | 0.00033992 | 0.00400681 | D1086.3 |
| WBGene00008392 | 1.328637317 | 0.03137782 | 0.11725396 | D1086.5 |
| WBGene00008393 | -0.425215688 | 0.16116022 | 0.34681392 | D1086.6 |
| WBGene00008394 | 0.302801146 | 0.24994679 | 0.457708 | D1086.7 |
| WBGene00008395 | 0.666576724 | 0.13888275 | 0.31582233 | D1086.8 |
| WBGene00008397 | 0.45862406 | 0.4079515 | 0.6141434 | D2005.1 |
| WBGene00008398 | -0.649532899 | 0.00137737 | 0.01174343 | D2005.3 |
| WBGene00008399 | 0.011134919 | 0.9580614 | 0.9785154 | D2005.4 |
| WBGene00008400 | 0.008384295 | 0.96166387 | 0.98000612 | drh-3 |
| WBGene00008401 | 0.857848293 | 0.2161042 | 0.41761252 | D2005.6 |
| WBGene00008402 | -0.320563185 | 0.05747397 | 0.17741965 | wdfy-2 |
| WBGene00008403 | -0.420052681 | 0.45512305 | 0.65257895 | csa-1 |
| WBGene00008404 | -0.010111023 | 0.97015557 | 0.98450842 | D2013.6 |
| WBGene00008405 | 0.023471921 | 0.89735646 | 0.94634211 | ttll-12 |
| WBGene00008407 | 0.416870709 | 0.58293216 | 0.75323322 | D2023.3 |

| | | | | |
|---|---|---|---|---|
| WBGene00008408 | -0.716469746 | 0.00803468 | 0.04404018 | D2023.4 |
| WBGene00008409 | -0.120165873 | 0.59736484 | 0.76343227 | mpst-1 |
| WBGene00008410 | -0.240353316 | 0.2041523 | 0.40348577 | D2023.6 |
| WBGene00008411 | -0.107863276 | 0.53826794 | 0.72020179 | mans-1 |
| WBGene00008412 | 0.113491054 | 0.50580038 | 0.69271544 | D2030.2 |
| WBGene00008413 | -0.07247643 | 0.74646355 | 0.85870892 | D2030.3 |
| WBGene00008414 | -0.74077924 | 6.0281E-05 | 0.00101932 | D2030.4 |
| WBGene00008415 | -0.633622426 | 0.01157555 | 0.05776358 | mce-1 |
| WBGene00008417 | -0.356375786 | 0.06444186 | 0.19135217 | D2030.7 |
| WBGene00008418 | -0.083059135 | 0.67530538 | 0.81435068 | D2030.8 |
| WBGene00008419 | 0.763725544 | 9.0107E-05 | 0.0014116 | wdr-23 |
| WBGene00008420 | -0.775725026 | 0.01212671 | 0.05972836 | D2030.11 |
| WBGene00008421 | -0.036773841 | 0.97003252 | 0.98450842 | D2030.12 |
| WBGene00008422 | -0.029597751 | 0.89282789 | 0.94362739 | D2045.2 |
| WBGene00008424 | -0.378140626 | 0.54235788 | 0.72288584 | D2045.7 |
| WBGene00008425 | 0.727626456 | 0.19775725 | 0.39526894 | D2045.8 |
| WBGene00008426 | -0.311395836 | 0.46337366 | 0.65837561 | D2045.9 |
| WBGene00008427 | -0.019208036 | 0.97888255 | 0.98873753 | atg-10 |
| WBGene00008428 | 0.23596901 | 0.39268728 | 0.60045695 | eif-2Bepsilon |
| WBGene00008429 | 0.218744527 | 0.25632369 | 0.46413512 | etc-1 |
| WBGene00008430 | 0.662572783 | 0.00104943 | 0.00957264 | hgap-2 |
| WBGene00008431 | -0.357887876 | 0.1748583 | 0.36508248 | piga-1 |
| WBGene00008432 | 0.599674377 | 0.38392992 | 0.59243685 | D2085.7 |
| WBGene00008434 | 0.247302184 | 0.84242599 | 1 | D2089.3 |
| WBGene00008435 | 0.001146806 | 0.99738317 | 0.99861694 | glna-2 |
| WBGene00008436 | 1.170899518 | 1.23E-07 | 5.6039E-06 | DH11.2 |
| WBGene00008438 | 1.140972528 | 3.2095E-05 | 0.00058952 | DH11.5 |
| WBGene00008439 | 0.490948765 | 0.21032036 | 0.41037543 | mfb-1 |
| WBGene00008440 | -0.785445605 | 3.3332E-05 | 0.00060855 | DY3.8 |
| WBGene00008441 | -0.218025809 | 0.35653971 | 0.56511079 | usp-50 |
| WBGene00008442 | -0.400831165 | 0.235533 | 0.44177447 | E01B7.2 |
| WBGene00008443 | 0.806974802 | 0.03174555 | 0.11812001 | pde-3 |
| WBGene00008444 | 0.179324299 | 0.38362553 | 0.59211766 | tbc-14 |
| WBGene00008446 | 0.168204224 | 0.48686885 | 0.67647913 | E01G4.3 |
| WBGene00008447 | -0.270519358 | 0.59872224 | 0.76452322 | E01G4.5 |
| WBGene00008448 | 1.150588502 | 0.38658048 | 1 | E01G4.6 |
| WBGene00008449 | 0.109500399 | 0.89866498 | 0.94694691 | E01G6.1 |
| WBGene00008450 | 0.839187188 | 0.10309732 | 0.26008368 | E01G6.2 |
| WBGene00008451 | 1.920756755 | 0.02931825 | 0.11144289 | E01G6.3 |
| WBGene00008452 | -0.13810136 | 0.43932694 | 0.63970597 | mrps-5 |
| WBGene00008453 | 0.736481557 | 0.14961283 | 0.33104599 | E02A10.3 |
| WBGene00008453 | -0.323868354 | 0.91634405 | 1 | E02A10.5 |
| WBGene00008454 | 0.747375649 | 0.20039504 | 0.39891723 | E02A10.4 |

| | | | | |
|---|---|---|---|---|
| WBGene00008455 | -0.313854831 | 0.23457151 | 0.44049946 | E02H1.1 |
| WBGene00008456 | -0.572876174 | 0.01659749 | 0.07457814 | E02H1.2 |
| WBGene00008457 | -0.456721176 | 0.09024786 | 0.23876645 | E02H1.5 |
| WBGene00008458 | -0.24983382 | 0.36725179 | 0.57476113 | E02H1.6 |
| WBGene00008459 | -0.538249046 | 0.04398536 | 0.14662996 | mrpl-53 |
| WBGene00008460 | 0.238555963 | 0.88337764 | 1 | E02H4.2 |
| WBGene00008462 | 0.909957504 | 0.44462193 | 1 | E02H4.4 |
| WBGene00008463 | -1.400268481 | 0.40762952 | 1 | E02H4.5 |
| WBGene00008464 | -0.630644311 | 0.01735153 | 0.07733659 | E02H4.6 |
| WBGene00008468 | 0.643120127 | 0.7220272 | 1 | E03G2.1 |
| WBGene00008470 | -0.156269293 | 0.96924949 | 1 | E03H4.2 |
| WBGene00008472 | 0.688793783 | 0.86378755 | 1 | E03H4.4 |
| WBGene00008473 | -0.156269293 | 0.96924949 | 1 | E03H4.5 |
| WBGene00008474 | 0.281454817 | 0.94435481 | 1 | nhr-174 |
| WBGene00008475 | 0.857402182 | 0.72847739 | 1 | oac-13 |
| WBGene00008476 | 1.489713917 | 0.0030658 | 0.02154205 | E03H4.8 |
| WBGene00008477 | 1.972234824 | 1.41E-08 | 8.34E-07 | clec-17 |
| WBGene00008478 | -1.831477136 | 0.52194793 | 1 | E03H4.11 |
| WBGene00008480 | -0.300488295 | 0.0506862 | 0.16217522 | eif-2A |
| WBGene00008481 | 0.338953999 | 0.64197129 | 0.79291018 | E04D5.2 |
| WBGene00008482 | 1.912246354 | 0.17141262 | 1 | cut-4 |
| WBGene00008483 | 0.932236405 | 0.3077976 | 0.51641185 | E04D5.4 |
| WBGene00008484 | 1.099812213 | 0.1588001 | 0.34355337 | E04D5.5 |
| WBGene00008485 | -0.76279246 | 0.30089206 | 0.51049895 | ugt-43 |
| WBGene00008486 | 0.72934382 | 0.00155125 | 0.01287336 | ugt-44 |
| WBGene00008488 | -0.747401365 | 0.00695071 | 0.03954481 | F01D4.5 |
| WBGene00008489 | 0.232157325 | 0.93397866 | 1 | srb-17 |
| WBGene00008490 | 1.157969456 | 0.25286033 | 0.46088835 | F01D4.8 |
| WBGene00008491 | 0.27161217 | 0.89703555 | 1 | bgnt-1.1 |
| WBGene00008492 | 1.23783447 | 4.1717E-06 | 0.00010809 | F01D5.1 |
| WBGene00008493 | 0.915156265 | 0.13523699 | 0.30973633 | F01D5.2 |
| WBGene00008494 | 1.360586053 | 1.38E-08 | 8.20E-07 | F01D5.3 |
| WBGene00008495 | 1.829361316 | 8.81E-10 | 6.99E-08 | F01D5.5 |
| WBGene00008496 | 0.063337093 | 0.93005644 | 0.96354103 | F01D5.6 |
| WBGene00008498 | 0.051145493 | 0.92154568 | 0.95884274 | F01D5.8 |
| WBGene00008499 | 0.697207261 | 0.09190463 | 0.24130927 | cyp-37A1 |
| WBGene00008500 | -0.491597591 | 0.73715481 | 1 | F01D5.10 |
| WBGene00008502 | 0.566808834 | 0.00315156 | 0.0219903 | skih-2 |
| WBGene00008503 | -0.216116613 | 0.35524666 | 0.56364998 | F01G4.4 |
| WBGene00008504 | 0.642441984 | 0.05120428 | 0.16342564 | F01G4.5 |
| WBGene00008505 | -0.181641672 | 0.24407068 | 0.45151031 | F01G4.6 |
| WBGene00008506 | -0.003909219 | 0.98131648 | 0.99020639 | tkt-1 |
| WBGene00008507 | 0.688793783 | 0.86378755 | 1 | F01G10.4 |

| | | | | |
|---|---|---|---|---|
| WBGene00008508 | 0.595896719 | 0.79561342 | 1 | F01G10.5 |
| WBGene00008509 | 0.158788216 | 0.86424875 | 0.92701158 | F01G10.6 |
| WBGene00008510 | 0.018699301 | 0.95414924 | 0.97667363 | lipl-7 |
| WBGene00008511 | -0.967524422 | 0.2615301 | 0.47006725 | F01G10.9 |
| WBGene00008512 | -0.669709645 | 0.25531748 | 0.46286538 | F01G10.10 |
| WBGene00008513 | 2.336545168 | 0.05208454 | 1 | F02A9.1 |
| WBGene00008514 | -0.375229789 | 0.01899965 | 0.08269791 | F02A9.4 |
| WBGene00008515 | 0.305084635 | 0.44138708 | 0.64137027 | F02C12.1 |
| WBGene00008516 | -0.437326017 | 0.72908489 | 1 | F02C12.2 |
| WBGene00008517 | -0.860395159 | 0.58453605 | 1 | F02C12.3 |
| WBGene00008518 | 1.881019158 | 0.1860156 | 1 | irld-2 |
| WBGene00008519 | 0.411762567 | 0.62305685 | 0.78057302 | cyp-13B1 |
| WBGene00008520 | 0.27161217 | 0.89703555 | 1 | F02D8.1 |
| WBGene00008521 | 0.307110532 | 0.22310362 | 0.42703785 | F02D8.4 |
| WBGene00008522 | 1.026788954 | 0.12784872 | 0.29861703 | F02D8.5 |
| WBGene00008524 | 0.855026222 | 0.71364798 | 1 | F02D10.3 |
| WBGene00008526 | -1.169464043 | 0.02574744 | 0.10225446 | F02D10.6 |
| WBGene00008528 | 0.052256006 | 0.86338464 | 0.92649398 | F02E9.1 |
| WBGene00008529 | 1.472766509 | 0.46533185 | 1 | F02E9.3 |
| WBGene00008530 | 0.156052939 | 0.54714063 | 0.7259135 | F02E9.5 |
| WBGene00008531 | -0.387496786 | 0.07194945 | 0.20631514 | F02E9.7 |
| WBGene00008532 | -0.502740422 | 0.00124393 | 0.01084952 | dpt-1 |
| WBGene00008533 | -0.249393603 | 0.23023609 | 0.43574377 | F02E9.10 |
| WBGene00008535 | -0.188199043 | 0.43727417 | 0.63801774 | F02H6.2 |
| WBGene00008536 | -0.119434985 | 0.61954893 | 0.7785885 | F02H6.3 |
| WBGene00008538 | -0.180606968 | 0.74732076 | 0.85920656 | sqrd-1 |
| WBGene00008541 | 1.906609716 | 0.22291155 | 1 | F07A5.2 |
| WBGene00008542 | 0.541737707 | 0.1418344 | 0.31995754 | scav-6 |
| WBGene00008543 | 0.361859153 | 0.63294458 | 0.7872679 | F07A5.4 |
| WBGene00008545 | 0.338038626 | 0.78139072 | 1 | F07A11.1 |
| WBGene00008546 | -0.414461316 | 0.014813 | 0.06906477 | gfat-1 |
| WBGene00008547 | 0.671703074 | 0.01055035 | 0.05397883 | F07A11.4 |
| WBGene00008548 | -0.486308879 | 0.20962471 | 0.40996353 | F07A11.5 |
| WBGene00008549 | 0.407639516 | 0.08877357 | 0.236256 | din-1 |
| WBGene00008550 | 1.557897343 | 0.63813005 | 1 | F07B10.4 |
| WBGene00008551 | -1.094201388 | 0.7816772 | 1 | lgc-36 |
| WBGene00008553 | 0.485671707 | 0.3898592 | 0.59824072 | F07C6.2 |
| WBGene00008554 | 0.945002355 | 0.0164609 | 0.07432196 | F07C6.3 |
| WBGene00008555 | 0.209589581 | 0.30516644 | 0.51430118 | F07C6.4 |
| WBGene00008556 | -0.156269293 | 0.96924949 | 1 | F07D3.3 |
| WBGene00008557 | -0.579173809 | 0.74452014 | 1 | F07H5.6 |
| WBGene00008558 | 0.251162113 | 0.73114263 | 0.84942451 | F07H5.7 |
| WBGene00008559 | 0.640967203 | 0.60401165 | 1 | F07H5.8 |

| | | | | |
|---|---|---|---|---|
| WBGene00008560 | 1.227233546 | 1.1257E-05 | 0.00025078 | pho-13 |
| WBGene00008561 | -0.34322639 | 0.0772753 | 0.21568044 | F07H5.10 |
| WBGene00008563 | -0.084297445 | 0.90933519 | 0.95279443 | F07H5.13 |
| WBGene00008564 | 0.430120416 | 0.03061016 | 0.11520086 | acox-1.1 |
| WBGene00008565 | -0.159076456 | 0.61503055 | 0.77523732 | acox-1.2 |
| WBGene00008566 | 0.325579905 | 0.48675027 | 0.67639179 | acox-1.3 |
| WBGene00008567 | -0.444367876 | 0.14189673 | 0.31997923 | acox-1.4 |
| WBGene00008568 | 1.192524401 | 0.37891857 | 1 | F08A8.5 |
| WBGene00008569 | -0.492820062 | 0.06656798 | 0.1954539 | fbxa-101 |
| WBGene00008570 | 1.00679301 | 0.01113864 | 0.05620707 | kcnl-2 |
| WBGene00008571 | 0.861430669 | 0.03533868 | 0.12675342 | prmn-1 |
| WBGene00008572 | 0.37627448 | 0.05069228 | 0.16217522 | F08B12.4 |
| WBGene00008574 | 0.281454817 | 0.94435481 | 1 | srbc-61 |
| WBGene00008575 | -1.456627322 | 0.01187144 | 0.05885305 | scl-24 |
| WBGene00008576 | 0.160841577 | 0.69924438 | 0.83002797 | F08G2.4 |
| WBGene00008577 | 1.429577905 | 0.00079659 | 0.00771858 | F08G2.5 |
| WBGene00008578 | 0.047508756 | 0.92177884 | 0.95898174 | F08G2.7 |
| WBGene00008579 | 1.092564053 | 0.28433521 | 1 | F08G2.8 |
| WBGene00008580 | -0.410472066 | 0.06713359 | 0.196734 | dsb-1 |
| WBGene00008581 | 1.584577408 | 0.63258867 | 1 | acl-13 |
| WBGene00008582 | 0.287041883 | 0.50504202 | 0.69198908 | F08G5.3 |
| WBGene00008583 | -0.089123913 | 0.83907423 | 0.91208085 | ugt-65 |
| WBGene00008584 | 1.253133188 | 2.3218E-06 | 6.6327E-05 | irg-4 |
| WBGene00008585 | -0.515246928 | 0.14590678 | 0.32488677 | F08G12.1 |
| WBGene00008586 | -0.674591211 | 0.00142231 | 0.01205586 | snrp-40.1 |
| WBGene00008587 | 0.635573 | 0.23450488 | 0.44049946 | F08G12.3 |
| WBGene00008588 | -0.122595642 | 0.86373543 | 0.92668707 | F08G12.5 |
| WBGene00008589 | 0.186559882 | 0.81604004 | 0.90021795 | nlp-57 |
| WBGene00008590 | 1.472766509 | 0.46533185 | 1 | F08H9.2 |
| WBGene00008591 | -0.250721307 | 0.88148692 | 1 | F08H9.3 |
| WBGene00008592 | 1.38322681 | 0.69550305 | 1 | F08H9.4 |
| WBGene00008593 | 0.577951897 | 0.47533128 | 0.66815209 | clec-227 |
| WBGene00008594 | 0.386872978 | 0.41976557 | 0.62424961 | clec-57 |
| WBGene00008595 | 0.642747391 | 0.01151085 | 0.05763019 | clec-56 |
| WBGene00008596 | 1.086215359 | 0.01555289 | 0.07149853 | clec-54 |
| WBGene00008597 | 2.428015231 | 0.15494979 | 1 | clec-55 |
| WBGene00008598 | 1.080657404 | 0.14277789 | 0.32053742 | spin-3 |
| WBGene00008599 | 1.370244941 | 0.01096951 | 0.05556147 | F09A5.2 |
| WBGene00008600 | 0.972783145 | 0.59157886 | 1 | F09A5.3 |
| WBGene00008601 | 0.213062543 | 0.36411825 | 0.57174854 | mob-2 |
| WBGene00008602 | 1.686211141 | 0.00019308 | 0.00257038 | oac-14 |
| WBGene00008603 | 0.410472841 | 0.27235185 | 0.48112609 | F09B9.4 |
| WBGene00008604 | 0.13033616 | 0.84830678 | 0.91671496 | F09B9.5 |

| | | | | |
|---|---|---|---|---|
| WBGene00008605 | -0.596022908 | 0.16925629 | 0.35743997 | mlt-9 |
| WBGene00008606 | 0.101269808 | 0.77675729 | 0.87561083 | dhhc-1 |
| WBGene00008607 | 0.570018065 | 0.00225431 | 0.01708461 | F09B12.3 |
| WBGene00008609 | -2.350737399 | 0.20463421 | 1 | F09B12.5 |
| WBGene00008610 | 0.820226349 | 0.01766046 | 0.07828812 | F09C3.2 |
| WBGene00008611 | -2.063958065 | 0.05728194 | 1 | fbxb-68 |
| WBGene00008612 | -1.778974048 | 0.30635426 | 1 | fbxa-103 |
| WBGene00008613 | -1.253146405 | 0.31756137 | 1 | fbxb-69 |
| WBGene00008615 | 4.353856434 | 0.00493255 | 1 | F09C6.1 |
| WBGene00008616 | 2.242271509 | 0.01987795 | 0.08507334 | fbxa-44 |
| WBGene00008617 | 2.73900937 | 0.23862709 | 1 | F09C6.3 |
| WBGene00008619 | -0.156269293 | 0.96924949 | 1 | nhr-262 |
| WBGene00008620 | -2.294850146 | 0.12823467 | 1 | F09C6.10 |
| WBGene00008621 | 2.969081727 | 2.89E-26 | 1.75E-23 | F09C8.1 |
| WBGene00008622 | 0.416690545 | 0.18030886 | 0.3725955 | F09C8.2 |
| WBGene00008623 | -0.156269293 | 0.96924949 | 1 | F09E8.1 |
| WBGene00008624 | -0.087766069 | 0.73176542 | 0.849902 | F09E8.2 |
| WBGene00008625 | 3.785921654 | 1.1329E-05 | 0.00025191 | scl-14 |
| WBGene00008626 | 1.245613127 | 0.30582242 | 1 | F09E8.8 |
| WBGene00008628 | 1.000093508 | 0.00191369 | 0.01504803 | ttr-21 |
| WBGene00008629 | 0.135715517 | 0.64997204 | 0.79839241 | cpt-5 |
| WBGene00008630 | -0.245531902 | 0.94070824 | 1 | nhr-175 |
| WBGene00008631 | -0.558804446 | 0.87818675 | 1 | F10A3.1 |
| WBGene00008632 | 0.978537522 | 0.51408445 | 1 | fbxa-88 |
| WBGene00008633 | -0.982352208 | 0.50649791 | 1 | fbxa-18 |
| WBGene00008634 | 1.763255965 | 0.03734345 | 0.13121898 | F10A3.4 |
| WBGene00008636 | 0.377828795 | 0.92526585 | 1 | F10A3.11 |
| WBGene00008637 | -0.668264701 | 0.85551153 | 1 | F10A3.12 |
| WBGene00008639 | 0.244147323 | 0.27114205 | 0.47974773 | F10B5.2 |
| WBGene00008640 | -0.71259187 | 0.69909067 | 1 | F10B5.3 |
| WBGene00008641 | -0.060422887 | 0.77805544 | 0.87653067 | pch-2 |
| WBGene00008642 | -0.205427284 | 0.26471385 | 0.47343988 | F10B5.8 |
| WBGene00008643 | -2.171326218 | 0.36522606 | 1 | tmem-138 |
| WBGene00008644 | 2.096030592 | 0.00468651 | 0.02982124 | F10C2.3 |
| WBGene00008645 | -0.005886864 | 0.96964508 | 0.98437435 | F10C2.4 |
| WBGene00008646 | 0.266015381 | 0.28854874 | 0.49868616 | F10C2.5 |
| WBGene00008647 | 0.075260937 | 0.97569838 | 1 | F10C2.7 |
| WBGene00008648 | 0.275722709 | 0.26925845 | 0.47783631 | F10D11.2 |
| WBGene00008649 | 0.377828795 | 0.92526585 | 1 | F10D11.3 |
| WBGene00008650 | -1.964653846 | 0.53599202 | 1 | F10D11.4 |
| WBGene00008651 | 0.456850296 | 0.82685361 | 1 | F10D11.5 |
| WBGene00008652 | -0.7671702 | 0.00146402 | 0.01231785 | F10D11.6 |
| WBGene00008654 | 0.035326172 | 0.90645167 | 0.95118667 | F10F2.2 |

| | | | | |
|---|---|---|---|---|
| WBGene00008655 | -1.021850269 | 0.09509904 | 0.24723715 | lips-3 |
| WBGene00008656 | 1.571910125 | 0.00031849 | 0.00379474 | lron-5 |
| WBGene00008659 | 1.735991307 | 0.58152819 | 1 | clec-151 |
| WBGene00008660 | 0.574508983 | 0.8551971 | 1 | clec-153 |
| WBGene00008661 | 0.489058549 | 0.85947456 | 1 | F10G8.1 |
| WBGene00008662 | -1.036770451 | 0.52301339 | 1 | F10G8.2 |
| WBGene00008663 | 1.270678414 | 0.5135606 | 1 | eak-6 |
| WBGene00008664 | -0.71375083 | 8.3127E-05 | 0.001321 | nubp-1 |
| WBGene00008665 | -0.27981997 | 0.22625837 | 0.43060846 | ercc-1 |
| WBGene00008666 | 0.664292138 | 0.01986753 | 0.08507334 | F10G8.8 |
| WBGene00008667 | -0.298388159 | 0.35032702 | 0.55891201 | F10G8.9 |
| WBGene00008669 | 0.611846994 | 0.00092742 | 0.00870151 | acs-14 |
| WBGene00008670 | 0.006960627 | 0.97089634 | 0.98484857 | eif-2Bdelta |
| WBGene00008671 | 2.876611589 | 0.15205148 | 1 | F11A5.3 |
| WBGene00008673 | -0.259657924 | 0.8422908 | 1 | F11A5.5 |
| WBGene00008675 | -1.901856996 | 0.64072034 | 1 | irld-26 |
| WBGene00008677 | 0.196455489 | 0.45121214 | 0.64911342 | F11A5.9 |
| WBGene00008678 | 0.399606396 | 0.86544406 | 1 | stdh-2 |
| WBGene00008680 | -1.164667479 | 0.4419273 | 1 | F11A5.15 |
| WBGene00008681 | 0.823876226 | 0.15808008 | 0.34299953 | scrm-4 |
| WBGene00008682 | -0.162165384 | 0.32640093 | 0.5350761 | lex-1 |
| WBGene00008683 | -0.993518812 | 3.8154E-06 | 0.00010103 | repo-1 |
| WBGene00008684 | -0.211070365 | 0.24289415 | 0.45056957 | mig-32 |
| WBGene00008685 | 0.550821037 | 0.00199339 | 0.01549407 | mon-2 |
| WBGene00008686 | -0.305669954 | 0.07450787 | 0.21096 | F11A10.5 |
| WBGene00008687 | 0.036731751 | 0.89998433 | 0.94735085 | F11A10.6 |
| WBGene00008688 | -0.683794582 | 0.00182327 | 0.01451559 | F11A10.7 |
| WBGene00008690 | -0.755534497 | 0.10876795 | 0.26889669 | F11C1.1 |
| WBGene00008691 | -0.132330146 | 0.74749404 | 0.85924306 | mff-1 |
| WBGene00008692 | 0.715921532 | 0.03598704 | 0.12816622 | scav-4 |
| WBGene00008693 | 0.037561939 | 0.96108822 | 0.97977647 | F11C1.4 |
| WBGene00008694 | -0.150555909 | 0.3943378 | 0.6021422 | F11C1.5 |
| WBGene00008695 | 0.175510509 | 0.78387129 | 0.88074562 | F11C3.1 |
| WBGene00008698 | 3.218784866 | 0.12490893 | 1 | F11D11.3 |
| WBGene00008699 | -0.156269293 | 0.96924949 | 1 | F11D11.4 |
| WBGene00008702 | 0.281454817 | 0.94435481 | 1 | clec-252 |
| WBGene00008703 | -0.156269293 | 0.96924949 | 1 | clec-251 |
| WBGene00008706 | 0.285356779 | 0.42703817 | 0.63047524 | gba-3 |
| WBGene00008707 | -0.467693305 | 0.01733027 | 0.07727022 | F11E6.3 |
| WBGene00008708 | -0.612308947 | 0.61260569 | 1 | F11E6.4 |
| WBGene00008709 | 1.025913312 | 0.34809409 | 1 | F11E6.6 |
| WBGene00008710 | -0.311638327 | 0.08860514 | 0.23601471 | F11E6.7 |
| WBGene00008711 | 0.227888048 | 0.61204754 | 0.77340437 | F11E6.8 |

| | | | | |
|---|---|---|---|---|
| WBGene00008712 | -0.433327175 | 0.74852436 | 1 | F11E6.9 |
| WBGene00008713 | 0.613461899 | 0.70361557 | 1 | F11E6.10 |
| WBGene00008714 | 0.459604343 | 0.31820482 | 0.52611905 | F11F1.1 |
| WBGene00008716 | 0.377828795 | 0.92526585 | 1 | F11F1.4 |
| WBGene00008719 | -1.397181256 | 0.54849318 | 1 | ttr-52 |
| WBGene00008720 | -0.112111607 | 0.47820163 | 0.67057589 | F12F6.1 |
| WBGene00008722 | -0.515882097 | 0.00373184 | 0.02496096 | F12F6.7 |
| WBGene00008724 | 0.688793783 | 0.86378755 | 1 | F13A7.1 |
| WBGene00008725 | 0.688793783 | 0.86378755 | 1 | F13A7.7 |
| WBGene00008726 | 0.56139557 | 0.43361228 | 0.63538117 | F13A7.11 |
| WBGene00008727 | -0.044493547 | 0.9772165 | 1 | F13A7.12 |
| WBGene00008728 | -0.233502874 | 0.92367326 | 1 | sre-36 |
| WBGene00008729 | -0.647633819 | 2.574E-05 | 0.00049922 | F13B12.1 |
| WBGene00008730 | 1.246167338 | 0.14672103 | 0.3263723 | F13B12.2 |
| WBGene00008731 | 1.561902141 | 0.08212329 | 0.2246134 | F13B12.3 |
| WBGene00008732 | -0.473564407 | 0.63913377 | 1 | F13B12.4 |
| WBGene00008733 | 0.034162558 | 0.86982246 | 0.9299502 | F13B12.6 |
| WBGene00008735 | 1.47744447 | 0.00010628 | 0.00159702 | F13D2.1 |
| WBGene00008736 | 0.321165199 | 0.82172184 | 1 | gnrr-6 |
| WBGene00008737 | 1.313089111 | 0.56091378 | 1 | gnrr-7 |
| WBGene00008739 | -0.759618077 | 0.09131488 | 0.24043678 | F13D12.3 |
| WBGene00008740 | -0.551295759 | 0.02814806 | 0.10851365 | F13D12.5 |
| WBGene00008741 | 1.343149316 | 3.67E-11 | 3.85E-09 | F13D12.6 |
| WBGene00008742 | -0.396436843 | 0.55947719 | 0.73616696 | F13D12.8 |
| WBGene00008743 | 0.062930751 | 0.93560909 | 0.96641399 | F13D12.9 |
| WBGene00008744 | 0.969530945 | 0.29777766 | 0.5078408 | F13D12.10 |
| WBGene00008745 | -0.109876724 | 0.57965283 | 0.75059354 | F13E6.1 |
| WBGene00008746 | 0.263906599 | 0.68522906 | 0.82001327 | F13E6.2 |
| WBGene00008747 | -1.02879607 | 0.04307539 | 0.14474905 | phf-31 |
| WBGene00008748 | 1.149826412 | 6.0899E-05 | 0.00102674 | yap-1 |
| WBGene00008749 | -0.216590074 | 0.44495304 | 0.64442024 | F13E6.5 |
| WBGene00008750 | 0.176051801 | 0.40795247 | 0.6141434 | F13E9.1 |
| WBGene00008752 | 0.281454817 | 0.94435481 | 1 | fipr-15 |
| WBGene00008754 | 1.247575581 | 0.7549672 | 1 | F13E9.5 |
| WBGene00008755 | -1.442973831 | 0.22549453 | 1 | sdz-13 |
| WBGene00008757 | -1.901856996 | 0.64072034 | 1 | F13E9.8 |
| WBGene00008759 | -1.159714244 | 0.75790634 | 1 | F13E9.10 |
| WBGene00008760 | 0.157789748 | 0.57453013 | 0.74795819 | F13E9.11 |
| WBGene00008761 | -0.690074771 | 0.53207192 | 1 | F13E9.12 |
| WBGene00008762 | 0.694195806 | 0.24867458 | 0.45656791 | ztf-2 |
| WBGene00008763 | 2.572258387 | 0.12540989 | 1 | F13G3.3 |
| WBGene00008764 | -0.164284282 | 0.46801873 | 0.66195426 | dylt-1 |
| WBGene00008765 | -0.607597646 | 0.00060591 | 0.00618463 | ttx-7 |

| | | | | |
|---|---|---|---|---|
| WBGene00008766 | -0.173627851 | 0.52378679 | 0.70863555 | F13G3.6 |
| WBGene00008767 | -0.249852017 | 0.31072066 | 0.51873445 | F13G3.7 |
| WBGene00008768 | -0.719254619 | 0.00070818 | 0.00704173 | F13G3.10 |
| WBGene00008769 | -0.293286532 | 0.22667888 | 0.43095965 | mrpl-13 |
| WBGene00008770 | 0.390831848 | 0.53037821 | 0.71443643 | F13G3.12 |
| WBGene00008773 | 1.156636386 | 0.46833968 | 1 | F13H10.1 |
| WBGene00008774 | 0.166240864 | 0.47819402 | 0.67057589 | F13H10.3 |
| WBGene00008775 | -0.125668414 | 0.41657374 | 0.62148715 | mogs-1 |
| WBGene00008776 | -0.216570612 | 0.71958682 | 0.84179137 | F13H10.5 |
| WBGene00008777 | -0.447431373 | 0.20220411 | 0.40100623 | F13H10.6 |
| WBGene00008779 | 0.057270194 | 0.90611876 | 0.95109561 | F14B4.1 |
| WBGene00008780 | -0.337922183 | 0.02642223 | 0.10381629 | hxk-1 |
| WBGene00008781 | -0.018468851 | 0.9270689 | 0.96171059 | rpoa-2 |
| WBGene00008782 | -1.718308215 | 0.00352458 | 0.0239061 | sdz-14 |
| WBGene00008783 | 1.099733554 | 0.01529902 | 0.07063015 | F14B6.2 |
| WBGene00008784 | -1.998299287 | 2.0102E-06 | 5.881E-05 | F14B6.3 |
| WBGene00008785 | 1.448645387 | 0.51537893 | 1 | F14B6.4 |
| WBGene00008786 | -0.156269293 | 0.96924949 | 1 | oac-16 |
| WBGene00008787 | 1.014340018 | 0.19708924 | 0.39427377 | F14B6.6 |
| WBGene00008788 | -1.901856996 | 0.64072034 | 1 | F14D7.1 |
| WBGene00008789 | -0.736131497 | 0.00014005 | 0.00199876 | F14D7.2 |
| WBGene00008790 | 1.82867917 | 0.54358766 | 1 | F14D7.3 |
| WBGene00008792 | 0.09697944 | 0.97612634 | 1 | F14D7.5 |
| WBGene00008793 | 0.572226664 | 0.16799066 | 0.35563637 | F14D7.6 |
| WBGene00008794 | -0.930284448 | 0.53123638 | 1 | F14D7.7 |
| WBGene00008795 | 0.739414816 | 0.63686511 | 1 | F14D7.8 |
| WBGene00008797 | 1.337568187 | 0.51456581 | 1 | F14D7.10 |
| WBGene00008798 | -0.242521717 | 0.90075881 | 1 | F14D7.11 |
| WBGene00008799 | 0.432859632 | 0.71165129 | 1 | F14E5.1 |
| WBGene00008800 | 0.102282098 | 0.53753235 | 0.71956431 | F14E5.2 |
| WBGene00008801 | 0.688793783 | 0.86378755 | 1 | acp-3 |
| WBGene00008802 | -0.111746212 | 0.86540244 | 0.92768209 | acp-2 |
| WBGene00008803 | -0.791460541 | 0.0101802 | 0.05252284 | lips-10 |
| WBGene00008805 | -0.433776782 | 0.08943461 | 0.2371834 | git-1 |
| WBGene00008806 | -0.207663993 | 0.51140164 | 0.69794596 | mboa-7 |
| WBGene00008807 | 0.394439984 | 0.88400378 | 1 | F14F3.4 |
| WBGene00008808 | 0.392639293 | 0.76401876 | 1 | ntr-2 |
| WBGene00008809 | -0.156269293 | 0.96924949 | 1 | cyp-13A11 |
| WBGene00008810 | 0.639934602 | 0.29800699 | 0.50787519 | cyp-13A12 |
| WBGene00008812 | 1.712479368 | 0.58529868 | 1 | F14F7.5 |
| WBGene00008816 | 0.401166296 | 0.6078723 | 0.77036604 | F14F8.8 |
| WBGene00008818 | 0.281454817 | 0.94435481 | 1 | srz-101 |
| WBGene00008819 | 0.323291024 | 0.28648775 | 0.49667967 | shw-1 |

| | | | | |
|---|---|---|---|---|
| WBGene00008820 | 1.045652325 | 0.02154236 | 0.0905427 | nlp-55 |
| WBGene00008822 | -0.590335964 | 0.03784838 | 0.13254029 | F14H3.3 |
| WBGene00008823 | -0.92418846 | 1.8149E-05 | 0.0003722 | F14H3.4 |
| WBGene00008824 | -0.579814035 | 0.07079975 | 0.2038368 | F14H3.5 |
| WBGene00008825 | -0.499181911 | 0.00110984 | 0.00992967 | F14H3.6 |
| WBGene00008826 | 0.041803775 | 0.98890197 | 1 | fbxa-100 |
| WBGene00008829 | -0.439564675 | 0.76444988 | 1 | cyp-35D1 |
| WBGene00008830 | 1.505604921 | 0.02063836 | 0.087418 | nhr-176 |
| WBGene00008831 | 0.891405583 | 0.15019116 | 0.33178145 | F14H3.12 |
| WBGene00008832 | -0.000774345 | 0.99802334 | 0.99909312 | obr-2 |
| WBGene00008833 | 0.292400437 | 0.8522108 | 1 | F14H8.2 |
| WBGene00008834 | 0.209773733 | 0.89788901 | 1 | F14H8.4 |
| WBGene00008835 | 1.018035489 | 0.62989118 | 1 | F14H8.5 |
| WBGene00008838 | -1.61131512 | 0.01635422 | 0.07403301 | F15A4.2 |
| WBGene00008839 | -1.901856996 | 0.64072034 | 1 | sre-37 |
| WBGene00008840 | 0.404817125 | 0.46488515 | 0.65929592 | F15A4.5 |
| WBGene00008841 | 1.238970035 | 6.69E-07 | 2.4482E-05 | F15A4.6 |
| WBGene00008843 | 0.903390376 | 0.71900339 | 1 | arrd-9 |
| WBGene00008844 | -0.414721007 | 0.33510813 | 0.54330693 | F15A4.10 |
| WBGene00008847 | -0.963811488 | 0.61154562 | 1 | fbxb-102 |
| WBGene00008848 | 0.025521227 | 0.90812584 | 0.95194377 | inft-2 |
| WBGene00008849 | -0.631659269 | 0.1992242 | 0.39743362 | try-10 |
| WBGene00008850 | 0.126179031 | 0.90964253 | 1 | F15B9.6 |
| WBGene00008851 | -0.92038092 | 0.07447042 | 0.21090397 | F15B9.8 |
| WBGene00008852 | -0.023055412 | 0.89083549 | 0.9424176 | ubql-1 |
| WBGene00008854 | -0.278058299 | 0.82420252 | 1 | F15D3.4 |
| WBGene00008855 | -1.964653846 | 0.53599202 | 1 | F15D3.5 |
| WBGene00008856 | -0.14389265 | 0.43576104 | 0.63676706 | F15D3.6 |
| WBGene00008857 | -0.649300948 | 0.00023599 | 0.00298118 | timm-23 |
| WBGene00008858 | 0.86175833 | 0.52744424 | 1 | pals-1 |
| WBGene00008859 | -0.156095973 | 0.57498198 | 0.74815787 | F15D4.2 |
| WBGene00008860 | -0.697750536 | 0.0042586 | 0.02760475 | romo-1 |
| WBGene00008861 | 1.46964697 | 0.02806889 | 0.10839196 | F15D4.4 |
| WBGene00008862 | 1.500699547 | 0.09916388 | 0.25351624 | F15D4.5 |
| WBGene00008863 | 0.871933027 | 0.73427341 | 1 | F15D4.6 |
| WBGene00008865 | -0.562502514 | 0.03174352 | 0.11812001 | F15G9.1 |
| WBGene00008866 | -1.082680044 | 0.67762556 | 1 | F15G9.2 |
| WBGene00008868 | 0.628621281 | 0.30973512 | 0.51818853 | F15G9.5 |
| WBGene00008869 | 0.613514499 | 0.66142472 | 1 | F15G9.6 |
| WBGene00008870 | 0.688793783 | 0.86378755 | 1 | F15H9.1 |
| WBGene00008871 | 1.025181888 | 0.79790347 | 1 | tag-314 |
| WBGene00008872 | -0.774842772 | 0.78069886 | 1 | F15H10.5 |
| WBGene00008874 | 2.256387701 | 0.20762969 | 1 | F15H10.7 |

| | | | | |
|---|---|---|---|---|
| WBGene00008875 | 2.040881441 | 0.16918214 | 1 | F15H10.8 |
| WBGene00008876 | 0.301237209 | 0.09766116 | 0.2511553 | F16A11.1 |
| WBGene00008877 | -0.241945077 | 0.28994722 | 0.49989338 | rtcb-1 |
| WBGene00008878 | 0.183067557 | 0.29795581 | 0.50787519 | ppfr-1 |
| WBGene00008879 | 1.487288669 | 0.49618504 | 1 | F16B12.1 |
| WBGene00008880 | 0.022660579 | 0.95574318 | 0.97739875 | F16B12.4 |
| WBGene00008881 | 0.377828795 | 0.92526585 | 1 | F16B12.5 |
| WBGene00008882 | -0.002323838 | 0.9914402 | 0.99572135 | F16B12.6 |
| WBGene00008883 | 0.747845955 | 0.85215965 | 1 | F16B12.7 |
| WBGene00008884 | -1.544131969 | 0.51019995 | 1 | nhr-281 |
| WBGene00008885 | 1.037128708 | 0.16046592 | 0.345735 | F16C3.1 |
| WBGene00008886 | 0.27731459 | 0.56357118 | 0.73875164 | F16C3.2 |
| WBGene00008887 | 0.041436033 | 0.80983494 | 0.89629222 | tbcd-1 |
| WBGene00008890 | 0.830111683 | 0.30332233 | 0.51229575 | ser-5 |
| WBGene00008891 | 2.240417636 | 1.6113E-06 | 4.8422E-05 | clec-42 |
| WBGene00008892 | 0.14729799 | 0.91306764 | 1 | clec-246 |
| WBGene00008894 | -0.143662227 | 0.92644295 | 1 | F16H6.4 |
| WBGene00008897 | 3.704441394 | 0.03355804 | 1 | F16H6.7 |
| WBGene00008899 | 1.116613983 | 0.69067248 | 1 | F16H6.9 |
| WBGene00008900 | 0.440382321 | 0.3492578 | 0.55765656 | F16H6.10 |
| WBGene00008901 | 1.326519872 | 0.03197039 | 0.11866478 | nhr-27 |
| WBGene00008902 | 0.281454817 | 0.94435481 | 1 | phf-32 |
| WBGene00008904 | -1.278097264 | 0.00244499 | 0.01827872 | F17A2.13 |
| WBGene00008905 | 1.557897343 | 0.63813005 | 1 | F17B5.1 |
| WBGene00008906 | -1.138454115 | 0.61298961 | 1 | oac-17 |
| WBGene00008908 | -1.901856996 | 0.64072034 | 1 | F17B5.4 |
| WBGene00008910 | 1.423691684 | 0.00494474 | 0.0309367 | F17C8.3 |
| WBGene00008911 | -0.156269293 | 0.96924949 | 1 | F17C8.6 |
| WBGene00008912 | 0.348702827 | 0.82696437 | 1 | F17C8.7 |
| WBGene00008914 | 1.177576359 | 0.00762458 | 0.04230806 | F17C11.2 |
| WBGene00008915 | 0.832964542 | 0.00248696 | 0.01851489 | F17C11.4 |
| WBGene00008916 | 3.406295457 | 0.06645203 | 1 | clec-221 |
| WBGene00008917 | 0.077843891 | 0.88387641 | 0.93844718 | F17C11.6 |
| WBGene00008918 | 0.011296054 | 0.95858013 | 0.97856902 | F17C11.7 |
| WBGene00008919 | 0.039850032 | 0.84729141 | 0.91627015 | vps-36 |
| WBGene00008920 | -0.259914822 | 0.06554117 | 0.19336894 | eef-1G |
| WBGene00008921 | -0.002491818 | 0.98733308 | 0.99421374 | ctf-4 |
| WBGene00008922 | -0.244679806 | 0.31882573 | 0.5267914 | F17C11.11 |
| WBGene00008923 | 0.88165999 | 0.01265623 | 0.06164745 | F17C11.12 |
| WBGene00008924 | 0.114995941 | 0.74210565 | 0.85588531 | F17E5.2 |
| WBGene00008925 | 0.500497863 | 0.00975151 | 0.05087801 | F17H10.1 |
| WBGene00008926 | -0.113280657 | 0.68408768 | 0.81908368 | F17H10.2 |
| WBGene00008927 | 0.513384123 | 0.10874744 | 0.26889669 | snx-17 |

| | | | | |
|---|---|---|---|---|
| WBGene00008929 | -0.296451977 | 0.57725527 | 0.74903703 | F18A11.2 |
| WBGene00008930 | -0.322748639 | 0.36258146 | 0.57056042 | F18A11.3 |
| WBGene00008932 | 0.436104245 | 0.42260314 | 0.62712373 | F18A11.5 |
| WBGene00008934 | 0.098141609 | 0.70071818 | 0.83120855 | F18C12.3 |
| WBGene00008936 | 0.896616597 | 1.4332E-06 | 4.4618E-05 | papl-1 |
| WBGene00008937 | -0.156269293 | 0.96924949 | 1 | srt-43 |
| WBGene00008939 | -0.165238875 | 0.69675529 | 0.82848332 | F18H3.1 |
| WBGene00008940 | -0.214162908 | 0.46886299 | 0.66260803 | F18H3.4 |
| WBGene00008944 | 0.387168722 | 0.07735457 | 0.21573609 | F19B2.5 |
| WBGene00008945 | 0.880395613 | 0.40254213 | 1 | F19B2.6 |
| WBGene00008946 | 0.451235227 | 0.84990489 | 1 | F19B2.7 |
| WBGene00008948 | -0.569213935 | 0.00054834 | 0.00573418 | F19B6.1 |
| WBGene00008949 | 0.574508983 | 0.8551971 | 1 | F19B6.3 |
| WBGene00008950 | -1.14715222 | 0.68932198 | 1 | wht-5 |
| WBGene00008951 | -0.071798707 | 0.81606118 | 0.90021795 | F19C6.2 |
| WBGene00008952 | 0.953650772 | 0.2255921 | 0.42976836 | F19C6.3 |
| WBGene00008953 | 0.893695166 | 0.11712817 | 0.28211932 | F19C6.4 |
| WBGene00008954 | -0.642261615 | 0.46727368 | 0.66143986 | F19C6.5 |
| WBGene00008955 | 0.359189277 | 0.77253365 | 1 | F19D8.2 |
| WBGene00008956 | 0.343254473 | 0.41904786 | 0.62348775 | nekl-3 |
| WBGene00008957 | 1.043685387 | 0.19288695 | 0.38810746 | glb-13 |
| WBGene00008958 | -1.69250915 | 0.28641353 | 1 | F19H6.3 |
| WBGene00008959 | -0.472656085 | 0.15262854 | 0.3349111 | F19H6.4 |
| WBGene00008960 | 1.025181888 | 0.79790347 | 1 | F19H6.5 |
| WBGene00008961 | 0.5180853 | 0.79605727 | 1 | F19H6.6 |
| WBGene00008963 | -0.576702777 | 0.19800129 | 0.395579 | F19H8.2 |
| WBGene00008964 | 0.031754751 | 0.95050514 | 0.97468105 | mltn-9 |
| WBGene00008965 | 0.995375043 | 0.40856113 | 1 | mltn-10 |
| WBGene00008967 | -1.634611338 | 0.1025932 | 1 | F20B10.3 |
| WBGene00008968 | -0.600843228 | 0.00196792 | 0.01537483 | elli-1 |
| WBGene00008969 | 0.390961002 | 0.32804436 | 0.53663232 | F20C5.4 |
| WBGene00008970 | 0.522864524 | 0.64671946 | 1 | F20C5.5 |
| WBGene00008971 | 0.034260657 | 0.91124115 | 0.95393098 | F20C5.6 |
| WBGene00008973 | 0.268252217 | 0.29931915 | 0.50906705 | F20D1.1 |
| WBGene00008974 | 0.469429981 | 0.06408809 | 0.19067836 | tbc-1 |
| WBGene00008975 | 0.242012219 | 0.22395039 | 0.42805127 | F20D1.3 |
| WBGene00008976 | 1.144601232 | 0.58194725 | 1 | plp-2 |
| WBGene00008977 | 0.284411817 | 0.66458455 | 0.80771846 | iglr-1 |
| WBGene00008978 | 0.112523935 | 0.89515383 | 0.94481514 | cutl-3 |
| WBGene00008979 | -0.111177918 | 0.68509402 | 0.82001327 | F20D1.9 |
| WBGene00008980 | 0.573967393 | 0.1012055 | 0.2564313 | emre-1 |
| WBGene00008984 | -1.094201388 | 0.7816772 | 1 | F20E11.7 |
| WBGene00008985 | 0.041950032 | 0.95491244 | 0.97697363 | F20G2.1 |

| | | | | |
|---|---|---|---|---|
| WBGene00008986 | -0.129463303 | 0.6565729 | 0.80236354 | F20G2.2 |
| WBGene00008987 | -0.357260238 | 0.40060974 | 0.60826713 | F20G2.3 |
| WBGene00008988 | 2.191509587 | 0.00583287 | 0.03486866 | F20G2.5 |
| WBGene00008989 | 1.216697342 | 0.29393122 | 1 | F20G2.6 |
| WBGene00008990 | 0.471900917 | 0.01087347 | 0.05532889 | smgl-1 |
| WBGene00008991 | 0.576221313 | 0.05995469 | 0.18215964 | F20G4.2 |
| WBGene00008992 | -0.233502874 | 0.92367326 | 1 | F21A3.2 |
| WBGene00008993 | -0.114721201 | 0.81369258 | 0.8986238 | F21A3.3 |
| WBGene00008994 | -1.674598919 | 0.52890127 | 1 | F21A3.4 |
| WBGene00008995 | -0.228487698 | 0.54792888 | 0.72656231 | prde-1 |
| WBGene00008996 | 0.559988012 | 0.48152424 | 0.67289188 | glb-14 |
| WBGene00008997 | 0.889345491 | 0.0894246 | 0.2371834 | lgc-31 |
| WBGene00008998 | -0.390923563 | 0.55917158 | 0.73600413 | ncs-4 |
| WBGene00008999 | 0.472454691 | 0.03324131 | 0.12180825 | myrf-2 |
| WBGene00009000 | 0.532047662 | 0.72714797 | 1 | arrd-28 |
| WBGene00009001 | -1.094201388 | 0.7816772 | 1 | tyr-3 |
| WBGene00009002 | -0.504886731 | 0.00718682 | 0.04057212 | hint-1 |
| WBGene00009004 | -1.092408901 | 7.69E-08 | 3.718E-06 | pfd-6 |
| WBGene00009005 | -0.91375383 | 0.72824392 | 1 | F21C3.6 |
| WBGene00009006 | -0.757821777 | 1.68E-07 | 7.3171E-06 | F21D5.1 |
| WBGene00009007 | -0.60407259 | 0.00011059 | 0.00165149 | otub-3 |
| WBGene00009008 | -0.038459085 | 0.89254868 | 0.94349514 | F21D5.3 |
| WBGene00009009 | -0.113484684 | 0.83512067 | 0.91032502 | F21D5.4 |
| WBGene00009010 | -0.078991086 | 0.73349919 | 0.85095923 | F21D5.5 |
| WBGene00009011 | -0.235210861 | 0.31014364 | 0.518332 | F21D5.6 |
| WBGene00009012 | -0.181025918 | 0.23218838 | 0.43780077 | F21D5.7 |
| WBGene00009013 | -0.766637585 | 0.00577281 | 0.03462831 | mrps-33 |
| WBGene00009014 | -0.521818153 | 0.65827418 | 1 | F21D5.9 |
| WBGene00009015 | -0.656778179 | 0.65962801 | 1 | F21D9.1 |
| WBGene00009016 | 0.489058549 | 0.85947456 | 1 | F21D9.2 |
| WBGene00009018 | -0.009951788 | 0.99243318 | 1 | F21D9.4 |
| WBGene00009023 | 0.63413106 | 0.13375292 | 0.30749748 | F21G4.1 |
| WBGene00009025 | -0.498704242 | 0.31600627 | 0.52405848 | phf-34 |
| WBGene00009026 | 0.727115471 | 0.43595845 | 0.63686337 | F21G4.5 |
| WBGene00009027 | 0.50006562 | 0.13842855 | 0.3153802 | F21G4.6 |
| WBGene00009028 | 2.423471851 | 0.3356807 | 1 | F21H7.2 |
| WBGene00009029 | -0.629292503 | 0.72518963 | 1 | F21H7.3 |
| WBGene00009030 | 0.607727847 | 0.68778675 | 1 | clec-233 |
| WBGene00009031 | 1.134376553 | 0.35530822 | 1 | F21H7.5 |
| WBGene00009032 | -1.992928477 | 0.12843969 | 1 | bgnt-1.3 |
| WBGene00009033 | -1.831477136 | 0.52194793 | 1 | F21H7.12 |
| WBGene00009035 | -0.428950371 | 0.00757304 | 0.04207979 | gfat-2 |
| WBGene00009036 | 0.281454817 | 0.94435481 | 1 | F22B3.5 |

| | | | | |
|---|---|---|---|---|
| WBGene00009038 | -0.156269293 | 0.96924949 | 1 | F22B3.7 |
| WBGene00009039 | 0.399606396 | 0.86544406 | 1 | F22B3.8 |
| WBGene00009041 | -1.135491114 | 3.48E-07 | 1.3778E-05 | cut-3 |
| WBGene00009042 | 0.061134683 | 0.96275085 | 1 | F22B5.4 |
| WBGene00009043 | -0.795796894 | 0.66183653 | 1 | F22B5.5 |
| WBGene00009044 | -0.417551663 | 0.75640689 | 1 | F22B5.6 |
| WBGene00009045 | -0.220295847 | 0.19054001 | 0.38517253 | F22B5.10 |
| WBGene00009048 | -0.753705701 | 0.07707804 | 0.21544186 | cth-1 |
| WBGene00009049 | -0.312571424 | 0.41134019 | 0.61673412 | F22B8.7 |
| WBGene00009050 | -0.101097123 | 0.54425735 | 0.7236484 | F22D6.2 |
| WBGene00009051 | -0.381118129 | 0.02932352 | 0.11144289 | nduf-6 |
| WBGene00009052 | -0.276191337 | 0.13223618 | 0.30481857 | ekl-1 |
| WBGene00009054 | -0.009970346 | 0.98034166 | 0.98963399 | F22D6.9 |
| WBGene00009056 | 0.51222408 | 0.43194987 | 0.6341955 | F22D6.15 |
| WBGene00009057 | 0.345956345 | 0.10712392 | 0.26684275 | cept-1 |
| WBGene00009058 | 1.204672331 | 3.836E-06 | 0.00010103 | F22E12.1 |
| WBGene00009059 | 0.176897297 | 0.76122766 | 0.86650332 | chw-1 |
| WBGene00009060 | 1.085613466 | 0.46322827 | 1 | F22E12.3 |
| WBGene00009061 | -0.680130008 | 0.72372989 | 1 | pals-14 |
| WBGene00009063 | -0.8206163 | 0.59460627 | 1 | F22G12.3 |
| WBGene00009064 | -0.024996414 | 0.90398225 | 0.95016572 | F22G12.4 |
| WBGene00009065 | 0.080703061 | 0.7167902 | 0.84010542 | F22G12.5 |
| WBGene00009067 | -1.192280996 | 0.26581473 | 1 | F23A7.1 |
| WBGene00009068 | 1.141226596 | 0.24937929 | 0.45715143 | F23A7.3 |
| WBGene00009069 | -1.540435807 | 8.94E-10 | 7.05E-08 | F23A7.4 |
| WBGene00009070 | 0.511636451 | 0.64502645 | 1 | F23A7.5 |
| WBGene00009073 | 0.823789701 | 0.74447552 | 1 | delm-1 |
| WBGene00009075 | -1.951326231 | 0.25914525 | 1 | F23B2.7 |
| WBGene00009077 | -2.705999569 | 0.42423051 | 1 | F23B2.10 |
| WBGene00009078 | -0.728756996 | 0.00708413 | 0.04010657 | rpb-12 |
| WBGene00009079 | -0.156269293 | 0.96924949 | 1 | F23B12.1 |
| WBGene00009081 | 0.756372667 | 0.0106128 | 0.0542298 | F23B12.4 |
| WBGene00009082 | -0.079033968 | 0.61251797 | 0.77375722 | dlat-1 |
| WBGene00009084 | 0.120696203 | 0.4751993 | 0.66814989 | F23B12.7 |
| WBGene00009085 | 0.377828795 | 0.92526585 | 1 | F23D12.1 |
| WBGene00009086 | -0.161943378 | 0.64504712 | 0.79499717 | F23D12.2 |
| WBGene00009087 | 0.903850474 | 0.35771879 | 1 | F23D12.3 |
| WBGene00009088 | -1.492081362 | 0.06452604 | 0.19144932 | F23D12.4 |
| WBGene00009089 | -1.170921783 | 0.03060863 | 0.11520086 | jmjd-3.2 |
| WBGene00009090 | -0.035338061 | 0.94913184 | 0.97411952 | fipr-3 |
| WBGene00009091 | 0.157800591 | 0.74729357 | 0.85920656 | F23D12.7 |
| WBGene00009092 | -0.38500109 | 0.03819706 | 0.13326163 | tomm-20 |
| WBGene00009093 | -0.02378119 | 0.96990061 | 0.98449655 | F23H12.3 |

| | | | | |
|---|---|---|---|---|
| WBGene00009094 | 0.330239326 | 0.52934889 | 0.71361591 | F23H12.5 |
| WBGene00009095 | 1.996320439 | 0.26902334 | 1 | F23H12.7 |
| WBGene00009096 | 0.091515882 | 0.73672584 | 0.85283726 | fipr-1 |
| WBGene00009097 | 0.470199708 | 0.03689164 | 0.13024175 | fipr-2 |
| WBGene00009098 | 0.139222656 | 0.42144547 | 0.62605768 | ehbp-1 |
| WBGene00009099 | -0.558804446 | 0.87818675 | 1 | F25B3.2 |
| WBGene00009100 | 1.107436541 | 0.03419927 | 0.12431537 | rgef-1 |
| WBGene00009101 | 2.273245419 | 0.19618289 | 1 | F25B3.4 |
| WBGene00009102 | 0.618383591 | 0.43828655 | 0.63872728 | F25B3.5 |
| WBGene00009103 | -0.454060831 | 0.00174102 | 0.01395229 | rtfo-1 |
| WBGene00009106 | 0.663122329 | 0.07062473 | 0.20347766 | acs-12 |
| WBGene00009107 | 1.735991307 | 0.58152819 | 1 | F25D1.2 |
| WBGene00009108 | 0.377828795 | 0.92526585 | 1 | F25D1.3 |
| WBGene00009109 | 0.452042097 | 0.46830975 | 0.66221159 | degt-1 |
| WBGene00009110 | -0.35648263 | 0.62822848 | 0.78388819 | F25D1.5 |
| WBGene00009112 | -0.122223587 | 0.50736358 | 0.69409515 | tag-353 |
| WBGene00009113 | -0.263854018 | 0.21207667 | 0.41203759 | maph-1.2 |
| WBGene00009114 | 1.879256964 | 0.46994626 | 1 | F25D7.5 |
| WBGene00009115 | 2.67618484 | 0.11298663 | 1 | F25F2.1 |
| WBGene00009116 | -0.114369179 | 0.66389293 | 0.80724327 | snx-27 |
| WBGene00009117 | -0.26664934 | 0.91678673 | 1 | F25H2.3 |
| WBGene00009118 | -0.601791255 | 0.00168911 | 0.01366837 | F25H2.4 |
| WBGene00009119 | -0.685207192 | 5.07E-08 | 2.5439E-06 | ndk-1 |
| WBGene00009120 | -0.042219397 | 0.82039172 | 0.90269856 | F25H2.6 |
| WBGene00009121 | 0.349082169 | 0.27973156 | 0.48875335 | F25H2.7 |
| WBGene00009122 | -0.474124626 | 0.00039389 | 0.00445683 | tct-1 |
| WBGene00009123 | -0.232440669 | 0.41633131 | 0.62134953 | F25H2.12 |
| WBGene00009124 | 0.110891957 | 0.60606629 | 0.76927253 | rtel-1 |
| WBGene00009125 | -0.533888816 | 0.61918973 | 1 | F25H5.2 |
| WBGene00009126 | -0.144605037 | 0.38723234 | 0.59594217 | pyk-1 |
| WBGene00009127 | -0.298083781 | 0.08968223 | 0.23758062 | clsp-1 |
| WBGene00009128 | -0.919573223 | 0.00202221 | 0.01567794 | mrpl-54 |
| WBGene00009129 | -0.097750527 | 0.95729192 | 1 | F25H5.7 |
| WBGene00009130 | 0.070854562 | 0.85610623 | 0.92145264 | F25H5.8 |
| WBGene00009131 | -0.35769347 | 0.17460696 | 0.36480865 | F25H8.1 |
| WBGene00009132 | -0.14972327 | 0.48577714 | 0.67583347 | F25H8.2 |
| WBGene00009133 | 0.153508108 | 0.71334739 | 0.8379961 | bed-3 |
| WBGene00009134 | -1.901856996 | 0.64072034 | 1 | lurp-2 |
| WBGene00009135 | -0.099827868 | 0.85482437 | 0.92088932 | F25H9.2 |
| WBGene00009136 | -0.224944172 | 0.88928469 | 1 | lurp-3 |
| WBGene00009137 | 0.218986257 | 0.66931285 | 0.81058627 | bath-39 |
| WBGene00009138 | -0.554269404 | 0.02868014 | 0.10965404 | F25H9.6 |
| WBGene00009139 | -0.196698524 | 0.54528922 | 0.72434382 | F25H9.7 |

| | | | | |
|---|---|---|---|---|
| WBGene00009140 | -0.044000064 | 0.86187289 | 0.92552616 | F26A3.1 |
| WBGene00009141 | -0.717091502 | 0.00015279 | 0.00212248 | ncbp-2 |
| WBGene00009142 | 1.24162715 | 5.82E-09 | 3.78E-07 | F26A3.4 |
| WBGene00009143 | 1.743415485 | 0.50483382 | 1 | F26A3.5 |
| WBGene00009144 | -0.011102054 | 0.99277441 | 1 | del-3 |
| WBGene00009145 | -0.60689127 | 0.01286634 | 0.06240922 | F26A3.7 |
| WBGene00009146 | 0.622779682 | 0.38298757 | 0.59132092 | F26C11.1 |
| WBGene00009148 | 1.799468055 | 0.25229973 | 1 | F26D2.3 |
| WBGene00009149 | 0.688793783 | 0.86378755 | 1 | F26D2.10 |
| WBGene00009151 | 1.653460415 | 0.50712258 | 1 | F26D2.13 |
| WBGene00009152 | 0.983488853 | 0.80601565 | 1 | F26D2.14 |
| WBGene00009153 | 1.33864768 | 0.5127736 | 1 | F26D2.15 |
| WBGene00009154 | -1.231066055 | 0.42796709 | 1 | F26D2.16 |
| WBGene00009155 | -0.159663908 | 0.93530048 | 1 | lhfp-4 |
| WBGene00009156 | -1.926869868 | 7.599E-05 | 0.00122523 | clec-196 |
| WBGene00009157 | -1.313062155 | 0.0088511 | 0.04711474 | F26E4.2 |
| WBGene00009158 | 0.106294001 | 0.65877352 | 0.80357275 | F26E4.3 |
| WBGene00009159 | -0.608160798 | 0.00583394 | 0.03486866 | F26E4.4 |
| WBGene00009160 | 0.629154436 | 0.81300308 | 1 | F26E4.5 |
| WBGene00009161 | -0.391597393 | 0.01857309 | 0.08125742 | F26E4.6 |
| WBGene00009162 | 0.461494074 | 0.22778059 | 0.43258004 | F26E4.7 |
| WBGene00009163 | 0.274103432 | 0.12749299 | 0.29818771 | drsh-1 |
| WBGene00009164 | 0.276335949 | 0.09549608 | 0.24773886 | hrdl-1 |
| WBGene00009165 | -0.150127039 | 0.61993417 | 0.7787454 | gpx-1 |
| WBGene00009166 | 0.946094744 | 0.60216978 | 1 | F26F2.1 |
| WBGene00009168 | 2.423471851 | 0.3356807 | 1 | F26F2.3 |
| WBGene00009169 | -1.901856996 | 0.64072034 | 1 | F26F2.4 |
| WBGene00009170 | 1.036568105 | 0.67105582 | 1 | F26F2.5 |
| WBGene00009171 | -0.156269293 | 0.96924949 | 1 | clec-263 |
| WBGene00009172 | -0.077627496 | 0.74181938 | 0.85569007 | F26F2.7 |
| WBGene00009173 | -0.182368141 | 0.42210433 | 0.62672969 | prom-1 |
| WBGene00009174 | 0.010838473 | 0.95763149 | 0.97850563 | F26H9.2 |
| WBGene00009176 | -0.32833937 | 0.33202121 | 0.53982957 | nmat-2 |
| WBGene00009177 | -0.641407213 | 0.00211326 | 0.01623863 | F26H9.5 |
| WBGene00009178 | 0.440822167 | 0.73671827 | 1 | uggt-2 |
| WBGene00009179 | -0.665089187 | 0.00755745 | 0.04201238 | kbp-3 |
| WBGene00009180 | -0.188216865 | 0.25068515 | 0.4587833 | nurf-1 |
| WBGene00009182 | -0.510827251 | 0.34602439 | 0.55474477 | F26H11.4 |
| WBGene00009183 | 0.801317559 | 0.51722648 | 1 | F27C8.2 |
| WBGene00009184 | -0.91375383 | 0.72824392 | 1 | F27C8.3 |
| WBGene00009185 | 0.49446186 | 0.84442265 | 1 | F27C8.5 |
| WBGene00009186 | -0.30087293 | 0.01638763 | 0.07414228 | trcs-1 |
| WBGene00009187 | -0.28048391 | 0.0833514 | 0.22674644 | F27D4.1 |

| | | | | |
|---|---|---|---|---|
| WBGene00009188 | -0.180311693 | 0.36260969 | 0.57056042 | lsy-22 |
| WBGene00009189 | -0.36410155 | 0.02137069 | 0.08993467 | F27D4.4 |
| WBGene00009190 | 1.380284882 | 0.00019227 | 0.00256524 | F27D4.6 |
| WBGene00009191 | -0.363412458 | 0.23949048 | 0.44640333 | F27D4.7 |
| WBGene00009192 | 1.326841701 | 1.95E-07 | 8.1662E-06 | asah-2 |
| WBGene00009193 | 1.712479368 | 0.58529868 | 1 | F27E5.3 |
| WBGene00009194 | 0.824305894 | 0.61976113 | 1 | F27E5.5 |
| WBGene00009196 | -0.156269293 | 0.96924949 | 1 | F28B1.2 |
| WBGene00009200 | -0.085776852 | 0.63431743 | 0.78783996 | F28C1.1 |
| WBGene00009201 | 0.375082769 | 0.08252025 | 0.22539445 | F28C1.3 |
| WBGene00009202 | -2.266679874 | 3.63E-08 | 1.8991E-06 | aptf-4 |
| WBGene00009203 | 0.075355722 | 0.76571814 | 0.86875842 | aptf-3 |
| WBGene00009204 | 0.67612059 | 0.01794418 | 0.07916437 | F28C6.4 |
| WBGene00009205 | -0.479559204 | 0.4342218 | 0.63564538 | F28C6.5 |
| WBGene00009207 | -0.338585643 | 0.1438994 | 0.32210225 | F28C6.8 |
| WBGene00009208 | 0.508547728 | 0.77686239 | 1 | F28C6.9 |
| WBGene00009209 | -0.360504901 | 0.76137551 | 1 | F28C6.10 |
| WBGene00009211 | -0.283695073 | 0.13808629 | 0.31477762 | wdr-46 |
| WBGene00009212 | -1.781367682 | 3.68E-08 | 1.9104E-06 | F28D1.2 |
| WBGene00009213 | 2.103145886 | 0.25764992 | 1 | thn-1 |
| WBGene00009214 | 0.574508983 | 0.8551971 | 1 | thn-3 |
| WBGene00009215 | 1.846089793 | 1.52E-14 | 2.67E-12 | thn-2 |
| WBGene00009216 | 0.757765547 | 0.39412275 | 0.60204382 | F28D1.6 |
| WBGene00009217 | 1.413734769 | 0.68804531 | 1 | spe-45 |
| WBGene00009218 | -0.428624007 | 0.1924593 | 0.38756815 | acs-20 |
| WBGene00009219 | -0.364000279 | 0.23154215 | 0.43712537 | dpm-3 |
| WBGene00009220 | 0.273125291 | 0.43059129 | 0.63305993 | F28D9.4 |
| WBGene00009221 | -0.302520625 | 0.32329117 | 0.5318534 | acs-2 |
| WBGene00009222 | 0.017662549 | 0.96808801 | 0.9838082 | fbxa-95 |
| WBGene00009223 | -0.793010434 | 0.00072107 | 0.00713488 | mdt-28 |
| WBGene00009224 | 0.231851311 | 0.62898013 | 0.78453054 | F28F8.7 |
| WBGene00009225 | -0.004301519 | 0.99739334 | 1 | fbxa-94 |
| WBGene00009226 | 2.866507396 | 0.00336938 | 0.02304658 | cyp-37B1 |
| WBGene00009227 | -2.324486652 | 0.30252788 | 1 | F28G4.2 |
| WBGene00009228 | 1.584577408 | 0.63258867 | 1 | F28G4.3 |
| WBGene00009229 | 1.616176829 | 0.27635731 | 1 | F28G4.4 |
| WBGene00009231 | 0.307327448 | 0.71616767 | 0.84006931 | ceh-89 |
| WBGene00009232 | -1.901856996 | 0.64072034 | 1 | nkat-1 |
| WBGene00009233 | -0.502748624 | 0.03303044 | 0.12130143 | F28H6.4 |
| WBGene00009234 | 0.316358557 | 0.38693334 | 0.59555752 | F28H6.6 |
| WBGene00009235 | 1.653460415 | 0.50712258 | 1 | F28H6.7 |
| WBGene00009236 | -1.045588926 | 0.54022641 | 1 | F28H7.2 |
| WBGene00009237 | 0.965206287 | 6.5738E-06 | 0.00015836 | F28H7.3 |

| | | | | |
|---|---|---|---|---|
| WBGene00009238 | 1.504494777 | 0.34519102 | 1 | selt-1.2 |
| WBGene00009239 | 0.210785198 | 0.87270242 | 1 | irld-6 |
| WBGene00009240 | -1.366565036 | 0.6506483 | 1 | F28H7.7 |
| WBGene00009241 | 0.39710184 | 0.43214749 | 0.6341955 | F28H7.8 |
| WBGene00009242 | 0.747845955 | 0.85215965 | 1 | sre-6 |
| WBGene00009243 | 0.412199829 | 0.1706206 | 0.3595114 | F29C6.1 |
| WBGene00009245 | 0.054816567 | 0.79362726 | 0.88593495 | rict-1 |
| WBGene00009246 | -0.158176451 | 0.45110541 | 0.64911342 | gfm-1 |
| WBGene00009247 | -0.499607735 | 0.58871784 | 0.75692294 | bath-45 |
| WBGene00009248 | -1.901856996 | 0.64072034 | 1 | F29C12.6 |
| WBGene00009250 | 0.747845955 | 0.85215965 | 1 | F29D10.1 |
| WBGene00009251 | -0.558804446 | 0.87818675 | 1 | F29D10.2 |
| WBGene00009252 | -0.668355569 | 0.74739695 | 1 | F29D10.3 |
| WBGene00009254 | 0.135973789 | 0.47559771 | 0.66831387 | capg-1 |
| WBGene00009255 | 0.747845955 | 0.85215965 | 1 | ugt-34 |
| WBGene00009256 | -0.076570635 | 0.76218996 | 0.86710535 | tut-2 |
| WBGene00009257 | 0.58071241 | 0.50195516 | 0.68967405 | F29G6.1 |
| WBGene00009258 | 0.101245289 | 0.89307785 | 0.94367782 | ccdc-149 |
| WBGene00009259 | 0.617375519 | 0.00041706 | 0.00464123 | hpo-34 |
| WBGene00009260 | 0.450407695 | 0.34880044 | 0.55730981 | calm-1 |
| WBGene00009261 | -0.711817665 | 0.24526184 | 0.45309251 | F30A10.2 |
| WBGene00009262 | -0.163975178 | 0.39609144 | 0.60391137 | F30A10.3 |
| WBGene00009263 | 0.747845955 | 0.85215965 | 1 | F30A10.4 |
| WBGene00009264 | 0.096084997 | 0.5728152 | 0.74664408 | sac-1 |
| WBGene00009266 | -0.394518804 | 0.08124597 | 0.2229673 | F30A10.9 |
| WBGene00009267 | 0.162015164 | 0.34482445 | 0.55355198 | usp-48 |
| WBGene00009268 | 2.068332674 | 0.26145708 | 1 | F30A10.11 |
| WBGene00009269 | -1.969805064 | 0.62862569 | 1 | F30A10.12 |
| WBGene00009270 | -0.287796784 | 0.09655538 | 0.24942103 | F30F8.1 |
| WBGene00009271 | 0.568131256 | 0.2472728 | 0.45521441 | glna-3 |
| WBGene00009272 | -0.269056898 | 0.17230491 | 0.36179827 | gras-1 |
| WBGene00009274 | -0.307596927 | 0.59404905 | 0.76063589 | F30F8.5 |
| WBGene00009276 | -0.236300938 | 0.32687315 | 0.53547026 | F30F8.9 |
| WBGene00009277 | -0.010692978 | 0.95888008 | 0.97871064 | F30F8.10 |
| WBGene00009278 | 1.014268055 | 0.52746164 | 1 | npr-33 |
| WBGene00009279 | 0.377828795 | 0.92526585 | 1 | F31B9.2 |
| WBGene00009280 | -1.034027866 | 0.00800347 | 0.0439088 | F31B9.3 |
| WBGene00009281 | -0.156269293 | 0.96924949 | 1 | F31B9.4 |
| WBGene00009283 | 1.851444818 | 0.53669729 | 1 | F31B12.4 |
| WBGene00009284 | -0.094964086 | 0.62576526 | 0.78243214 | F31C3.2 |
| WBGene00009285 | 0.215238988 | 0.34242922 | 0.55068696 | F31C3.3 |
| WBGene00009286 | -0.468170671 | 0.01415703 | 0.06680455 | zipt-1 |
| WBGene00009287 | -0.841510316 | 0.00022845 | 0.00291654 | psf-2 |

| | | | | |
|---|---|---|---|---|
| WBGene00009288 | 0.547175372 | 0.18775899 | 0.38133286 | F31C3.6 |
| WBGene00009289 | -0.748267769 | 0.00879923 | 0.04700214 | exos-7 |
| WBGene00009290 | 0.000559499 | 0.99774424 | 0.99889608 | F31D4.2 |
| WBGene00009291 | 0.986488751 | 0.01287587 | 0.06243048 | clec-264 |
| WBGene00009292 | -0.486792097 | 0.32644006 | 0.5350761 | F31D4.5 |
| WBGene00009293 | 0.592759211 | 0.40089998 | 0.60833979 | F31D4.8 |
| WBGene00009294 | -0.714610775 | 0.00074145 | 0.00728314 | F31D4.9 |
| WBGene00009296 | 0.886800208 | 0.47697219 | 1 | F31E9.3 |
| WBGene00009297 | -2.455789188 | 5.979E-05 | 0.00101385 | sdz-16 |
| WBGene00009299 | 0.759981451 | 0.31138548 | 0.51966064 | F31E9.6 |
| WBGene00009300 | -0.321384206 | 0.41923881 | 0.62369543 | F31F6.1 |
| WBGene00009301 | -0.325955768 | 0.54454463 | 0.72381392 | F31F6.2 |
| WBGene00009302 | -0.348295853 | 0.52392206 | 0.70873958 | F31F6.3 |
| WBGene00009303 | -0.490561023 | 0.85693136 | 1 | lips-8 |
| WBGene00009304 | 0.505281604 | 0.15749021 | 0.3421164 | eva-1 |
| WBGene00009305 | -0.379399459 | 0.05574622 | 0.1736313 | F32A7.4 |
| WBGene00009306 | 0.735767292 | 0.00581828 | 0.03484942 | maph-1.1 |
| WBGene00009307 | -0.35392574 | 0.05159404 | 0.16397989 | F32A11.1 |
| WBGene00009308 | 0.693849939 | 0.60406268 | 1 | F32A11.3 |
| WBGene00009311 | 1.937396089 | 0.03971402 | 0.13682844 | mltn-1 |
| WBGene00009312 | 1.557897343 | 0.63813005 | 1 | F32B4.1 |
| WBGene00009313 | 1.38322681 | 0.69550305 | 1 | F32B4.2 |
| WBGene00009314 | 0.110688896 | 0.63266029 | 0.78722871 | F32B4.4 |
| WBGene00009315 | 1.669054052 | 0.03927326 | 0.13569058 | F32B4.5 |
| WBGene00009316 | -0.156269293 | 0.96924949 | 1 | abhd-11.1 |
| WBGene00009318 | 1.039575359 | 0.41842016 | 1 | F32B4.8 |
| WBGene00009319 | -0.197699647 | 0.19936999 | 0.3975285 | mccc-1 |
| WBGene00009320 | -0.363772779 | 0.13197129 | 0.30444001 | F32B6.3 |
| WBGene00009321 | 1.247103736 | 0.59368513 | 1 | F32B6.4 |
| WBGene00009322 | -0.068828124 | 0.71913723 | 0.84154652 | tbc-3 |
| WBGene00009323 | 0.272820605 | 0.42749991 | 0.63057665 | best-13 |
| WBGene00009324 | -0.156269293 | 0.96924949 | 1 | F32B6.10 |
| WBGene00009325 | 0.441662884 | 0.56346492 | 0.73875164 | F32D8.15 |
| WBGene00009326 | 0.204879448 | 0.85889439 | 1 | F32D8.1 |
| WBGene00009328 | -0.576259193 | 0.71481512 | 1 | F32D8.3 |
| WBGene00009329 | -0.245196709 | 0.41582611 | 0.62082449 | F32D8.4 |
| WBGene00009330 | -0.376553539 | 0.13520433 | 0.30973633 | F32D8.5 |
| WBGene00009331 | 0.032770926 | 0.96674412 | 0.9830824 | F32D8.7 |
| WBGene00009332 | -0.558804446 | 0.87818675 | 1 | F32D8.8 |
| WBGene00009333 | 1.806634983 | 0.00272158 | 0.01978604 | F32D8.10 |
| WBGene00009334 | 0.480446505 | 0.00686713 | 0.03921347 | F32D8.12 |
| WBGene00009335 | -0.109085846 | 0.84612141 | 0.91565737 | F32D8.13 |
| WBGene00009336 | -0.292243406 | 0.19778097 | 0.39526894 | F32D8.14 |

| | | | | |
|---|---|---|---|---|
| WBGene00009337 | 1.078387098 | 2.7693E-05 | 0.00052535 | uig-1 |
| WBGene00009338 | 0.281454817 | 0.94435481 | 1 | F32G8.2 |
| WBGene00009339 | -0.323868354 | 0.91634405 | 1 | F32G8.3 |
| WBGene00009340 | 0.278737057 | 0.22356591 | 0.42746592 | best-14 |
| WBGene00009341 | -0.121228608 | 0.55968041 | 0.73627477 | thoc-3 |
| WBGene00009342 | 0.828125885 | 0.00027374 | 0.0033672 | fasn-1 |
| WBGene00009343 | 0.414787051 | 0.71865724 | 1 | F32H2.6 |
| WBGene00009344 | -0.639727652 | 0.77477802 | 1 | F32H2.7 |
| WBGene00009345 | -1.094201388 | 0.7816772 | 1 | F32H2.8 |
| WBGene00009346 | -0.545794811 | 0.04873648 | 0.15825345 | F32H2.10 |
| WBGene00009347 | -0.105803492 | 0.72130596 | 0.84294693 | F32H5.1 |
| WBGene00009349 | 1.11996462 | 0.18700078 | 0.38042501 | F32H5.3 |
| WBGene00009350 | 1.139824568 | 0.0353223 | 0.12675342 | F32H5.4 |
| WBGene00009352 | -0.268872462 | 0.2149626 | 0.41611303 | F33A8.4 |
| WBGene00009353 | -0.109696006 | 0.53255905 | 0.71595609 | sdhd-1 |
| WBGene00009354 | -0.059107916 | 0.91722144 | 0.95640851 | F33A8.6 |
| WBGene00009355 | -1.109801688 | 0.22649264 | 0.43080795 | F33A8.7 |
| WBGene00009356 | -1.58874758 | 0.38151696 | 1 | F33A8.10 |
| WBGene00009357 | 1.335782708 | 0.41705821 | 1 | F33C8.2 |
| WBGene00009358 | 0.22401771 | 0.61242254 | 0.77371717 | F33C8.4 |
| WBGene00009360 | 2.223658251 | 0.25271431 | 1 | F33E2.4 |
| WBGene00009361 | -0.680049002 | 0.08005769 | 0.22102688 | F33E2.5 |
| WBGene00009362 | -1.901856996 | 0.64072034 | 1 | F33E2.6 |
| WBGene00009363 | -1.901856996 | 0.64072034 | 1 | F33E2.7 |
| WBGene00009364 | -0.334195847 | 0.06750294 | 0.19734005 | F33H1.3 |
| WBGene00009365 | -0.224695143 | 0.15411042 | 0.3372485 | F33H1.4 |
| WBGene00009366 | -0.156206757 | 0.39077948 | 0.5991985 | F33H2.2 |
| WBGene00009367 | -0.548925374 | 0.00273877 | 0.01988721 | F33H2.3 |
| WBGene00009368 | 0.020734336 | 0.92950539 | 0.96347007 | pole-1 |
| WBGene00009369 | -0.071715337 | 0.73397373 | 0.8511151 | F33H2.6 |
| WBGene00009370 | -0.566153364 | 0.16227235 | 0.34826738 | set-10 |
| WBGene00009371 | 0.442599762 | 0.66947032 | 1 | F33H2.8 |
| WBGene00009372 | -0.414236097 | 0.01514412 | 0.07019452 | evl-18 |
| WBGene00009373 | -0.46807506 | 0.57677846 | 0.74903703 | F34D10.3 |
| WBGene00009374 | -0.220387761 | 0.38865122 | 0.59729296 | F34D10.4 |
| WBGene00009375 | 1.599362406 | 0.00306725 | 0.02154205 | F34D10.6 |
| WBGene00009377 | 0.760091251 | 0.7016102 | 1 | F34D10.8 |
| WBGene00009380 | 0.333223067 | 0.32937613 | 0.53812045 | F34H10.3 |
| WBGene00009381 | -0.182536869 | 0.90818791 | 1 | F34H10.4 |
| WBGene00009382 | -0.668264701 | 0.85551153 | 1 | F34H10.5 |
| WBGene00009383 | 1.163992275 | 7.8706E-06 | 0.00018555 | F35B12.3 |
| WBGene00009384 | 0.557836973 | 0.25118786 | 0.45914963 | piit-1 |
| WBGene00009385 | -0.329074008 | 0.07788518 | 0.21673252 | sas-5 |

| | | | | |
|---|---|---|---|---|
| WBGene00009386 | 1.214979485 | 0.00323217 | 0.02240559 | tag-290 |
| WBGene00009387 | 1.555771897 | 0.19155151 | 1 | glb-15 |
| WBGene00009388 | 0.892196141 | 0.00317883 | 0.02210432 | F35B12.9 |
| WBGene00009389 | 1.006427561 | 0.02615365 | 0.1032287 | F35B12.10 |
| WBGene00009390 | 1.557897343 | 0.63813005 | 1 | F35C5.1 |
| WBGene00009393 | 1.593454783 | 1.76E-14 | 3.06E-12 | clec-62 |
| WBGene00009394 | 1.217773124 | 1.29E-15 | 2.71E-13 | clec-63 |
| WBGene00009395 | 1.004581696 | 0.12068172 | 0.28729347 | clec-64 |
| WBGene00009396 | 0.08572442 | 0.65250897 | 0.79997085 | clec-65 |
| WBGene00009397 | 0.207091714 | 0.37024634 | 0.57770991 | clec-66 |
| WBGene00009398 | 0.247009535 | 0.85445435 | 1 | nspb-11 |
| WBGene00009399 | 1.1246493 | 0.25160247 | 0.45969985 | F35C5.11 |
| WBGene00009400 | 0.834420971 | 0.46813299 | 1 | F35C5.12 |
| WBGene00009401 | -0.936434159 | 0.65873492 | 1 | F35C11.2 |
| WBGene00009402 | 1.216957619 | 0.74819621 | 1 | F35C11.3 |
| WBGene00009403 | 1.274901847 | 0.03663208 | 0.12966474 | F35C11.4 |
| WBGene00009404 | -0.440351077 | 0.01070646 | 0.05460115 | F35C11.5 |
| WBGene00009405 | -0.079587935 | 0.90778913 | 0.95184971 | F35C11.6 |
| WBGene00009406 | 0.476235697 | 0.4899505 | 0.67943906 | F35C11.7 |
| WBGene00009408 | -0.053071525 | 0.91276479 | 0.95467585 | F35C12.3 |
| WBGene00009410 | -2.444686664 | 0.15746854 | 1 | F35E2.2 |
| WBGene00009412 | 0.36729309 | 0.88005435 | 1 | F35E2.5 |
| WBGene00009413 | 0.489058549 | 0.85947456 | 1 | oac-19 |
| WBGene00009415 | -1.386401505 | 0.47877074 | 1 | F35E2.8 |
| WBGene00009416 | 2.402150313 | 0.16138209 | 1 | F35E2.9 |
| WBGene00009418 | 0.983488853 | 0.80601565 | 1 | F35E8.2 |
| WBGene00009423 | -0.156269293 | 0.96924949 | 1 | F35E8.10 |
| WBGene00009424 | -0.156269293 | 0.96924949 | 1 | F35E8.13 |
| WBGene00009426 | -0.578019945 | 0.81405273 | 1 | F35E12.2 |
| WBGene00009427 | -0.274809261 | 0.72519037 | 0.84553311 | fbxa-181 |
| WBGene00009429 | -0.827340948 | 0.09749063 | 0.25098255 | irg-5 |
| WBGene00009430 | 1.366234647 | 4.03E-10 | 3.52E-08 | F35E12.6 |
| WBGene00009431 | 1.133872836 | 4.436E-05 | 0.0007839 | dct-17 |
| WBGene00009432 | 0.316765094 | 0.42647054 | 0.6302378 | cld-9 |
| WBGene00009433 | 0.237072281 | 0.45988386 | 0.65564231 | F35E12.9 |
| WBGene00009434 | -0.151046606 | 0.56984049 | 0.74425222 | F35E12.10 |
| WBGene00009435 | 0.510692332 | 0.00439301 | 0.02829469 | F35G2.1 |
| WBGene00009436 | -0.204046998 | 0.32543486 | 0.5338183 | marb-1 |
| WBGene00009437 | -0.950665013 | 0.56529632 | 1 | F35G2.3 |
| WBGene00009438 | -0.982559828 | 0.5059438 | 1 | F35G2.5 |
| WBGene00009439 | 0.147913314 | 0.52222776 | 0.70717357 | mlcd-1 |
| WBGene00009440 | -0.007774052 | 0.96013667 | 0.97958146 | idhg-1 |
| WBGene00009441 | -0.337306824 | 0.05741313 | 0.177322 | wdr-48 |

| | | | | |
|---|---|---|---|---|
| WBGene00009442 | -0.111270185 | 0.67747673 | 0.81544713 | F35G12.5 |
| WBGene00009443 | -0.580075732 | 0.15501305 | 0.33867437 | F35G12.7 |
| WBGene00009444 | -1.038727265 | 1.1576E-06 | 3.8088E-05 | F35G12.11 |
| WBGene00009445 | -0.290269092 | 0.12228902 | 0.29020933 | F35G12.12 |
| WBGene00009446 | 0.688793783 | 0.86378755 | 1 | F35H8.1 |
| WBGene00009447 | 0.227206662 | 0.73640539 | 0.85280864 | F35H8.2 |
| WBGene00009448 | 0.043940332 | 0.82049139 | 0.90272641 | zfp-2 |
| WBGene00009449 | -0.934013985 | 0.67351079 | 1 | F35H8.4 |
| WBGene00009450 | 0.114211345 | 0.59131094 | 0.75856996 | ugt-58 |
| WBGene00009451 | -0.178177586 | 0.36039151 | 0.56837101 | cids-2 |
| WBGene00009452 | 0.023467293 | 0.91574071 | 0.95580617 | F36A2.2 |
| WBGene00009453 | -0.108303995 | 0.59474092 | 0.76112043 | F36A2.3 |
| WBGene00009454 | -0.3657404 | 0.00682956 | 0.03909169 | F36A2.7 |
| WBGene00009455 | -0.657835398 | 0.02217798 | 0.09252126 | phip-1 |
| WBGene00009456 | -0.142144411 | 0.40790993 | 0.6141434 | F36A2.9 |
| WBGene00009457 | 2.532841764 | 0.28275632 | 1 | F36A2.10 |
| WBGene00009458 | -2.640414922 | 0.41081194 | 1 | F36A2.11 |
| WBGene00009459 | 0.983488853 | 0.80601565 | 1 | F36A2.12 |
| WBGene00009460 | 0.205969073 | 0.30154416 | 0.51110536 | ubr-5 |
| WBGene00009461 | -0.162833276 | 0.73354614 | 0.85095923 | madf-2 |
| WBGene00009462 | 1.181686151 | 0.7579682 | 1 | sre-22 |
| WBGene00009463 | 0.377828795 | 0.92526585 | 1 | F36D1.4 |
| WBGene00009464 | -0.156269293 | 0.96924949 | 1 | F36D1.5 |
| WBGene00009465 | -0.668264701 | 0.85551153 | 1 | F36D1.6 |
| WBGene00009466 | 2.091139575 | 0.00073068 | 0.00721082 | F36D1.7 |
| WBGene00009467 | -1.969805064 | 0.62862569 | 1 | F36D1.8 |
| WBGene00009468 | -0.293951121 | 0.65237919 | 0.79997085 | F36D1.9 |
| WBGene00009469 | -0.669810449 | 0.0133787 | 0.06389882 | F36D3.1 |
| WBGene00009470 | 2.876611589 | 0.15205148 | 1 | F36D3.4 |
| WBGene00009471 | 0.983488853 | 0.80601565 | 1 | F36D3.5 |
| WBGene00009473 | -1.901856996 | 0.64072034 | 1 | F36D3.8 |
| WBGene00009475 | -0.096635008 | 0.83224964 | 0.90915448 | F36F2.1 |
| WBGene00009476 | 0.546303823 | 0.10417618 | 0.26186397 | F36F2.2 |
| WBGene00009477 | 0.088113361 | 0.61999449 | 0.7787454 | rbpl-1 |
| WBGene00009478 | -0.316040438 | 0.11664177 | 0.28137249 | syx-7 |
| WBGene00009479 | -0.01376081 | 0.94096229 | 0.96955132 | fcp-1 |
| WBGene00009480 | 1.864555705 | 0.32567712 | 1 | F36F2.7 |
| WBGene00009481 | 1.477253415 | 0.58357327 | 1 | F36F2.8 |
| WBGene00009482 | 0.396441512 | 0.12385198 | 0.29294503 | F36G3.1 |
| WBGene00009483 | -0.797231541 | 0.06624881 | 0.19475225 | F36G3.2 |
| WBGene00009484 | 0.988356591 | 0.40542548 | 1 | F36G3.3 |
| WBGene00009485 | 0.413845191 | 0.51994708 | 0.70534444 | F36G9.3 |
| WBGene00009487 | -1.953072678 | 0.1378346 | 1 | clec-232 |

| | | | | |
|---|---|---|---|---|
| WBGene00009488 | -0.563446563 | 0.26646639 | 0.47484833 | oac-20 |
| WBGene00009489 | -0.156269293 | 0.96924949 | 1 | F36G9.13 |
| WBGene00009490 | 0.006338221 | 0.99674015 | 1 | fbxa-99 |
| WBGene00009492 | 0.616429445 | 0.81400458 | 1 | F36H1.3 |
| WBGene00009493 | -0.384190284 | 0.63098373 | 0.78581631 | hrg-4 |
| WBGene00009494 | 1.402467959 | 0.57470445 | 1 | hrg-5 |
| WBGene00009495 | -0.741945785 | 0.47237888 | 1 | hrg-6 |
| WBGene00009496 | -0.799427284 | 0.28487734 | 0.49480627 | F36H1.11 |
| WBGene00009497 | 1.557897343 | 0.63813005 | 1 | F36H1.12 |
| WBGene00009498 | 0.322266996 | 0.069353 | 0.20105414 | tat-5 |
| WBGene00009499 | -0.438300904 | 0.02466519 | 0.0992841 | ent-6 |
| WBGene00009500 | 1.39039796 | 0.00593896 | 0.0353629 | F36H2.3 |
| WBGene00009501 | 1.216957619 | 0.74819621 | 1 | F37A8.1 |
| WBGene00009502 | 0.983488853 | 0.80601565 | 1 | F37A8.2 |
| WBGene00009503 | 1.091044419 | 0.05828787 | 0.17893123 | F37A8.5 |
| WBGene00009504 | 0.287430733 | 0.71212089 | 0.8373714 | F37B12.1 |
| WBGene00009505 | -0.526852379 | 0.0044373 | 0.02852401 | F37B12.3 |
| WBGene00009507 | -0.063811576 | 0.73925793 | 0.85446788 | mus-101 |
| WBGene00009508 | -0.120082143 | 0.44941117 | 0.64840179 | row-1 |
| WBGene00009509 | 0.060133372 | 0.97461154 | 1 | F37D6.3 |
| WBGene00009510 | 0.546052409 | 0.7844476 | 1 | F37D6.4 |
| WBGene00009511 | 1.207794339 | 0.0218448 | 0.09151749 | F37H8.2 |
| WBGene00009512 | 0.693739424 | 0.21365578 | 0.4143559 | bigr-1 |
| WBGene00009513 | 0.041803775 | 0.98890197 | 1 | sfxn-1.2 |
| WBGene00009514 | -0.131558922 | 0.48953657 | 0.67924367 | F37H8.5 |
| WBGene00009515 | 1.289281762 | 0.03535723 | 0.12677852 | clec-170 |
| WBGene00009517 | -0.107268465 | 0.96117914 | 1 | clec-167 |
| WBGene00009518 | 2.452628887 | 2.49E-08 | 1.3622E-06 | clec-166 |
| WBGene00009519 | 0.983488853 | 0.80601565 | 1 | F38A1.6 |
| WBGene00009520 | -0.014519762 | 0.99603746 | 1 | clec-168 |
| WBGene00009521 | -0.077270632 | 0.67694493 | 0.81495175 | F38A1.8 |
| WBGene00009522 | 1.333135063 | 0.15805693 | 0.34299953 | F38A1.9 |
| WBGene00009523 | 0.54294365 | 0.56892821 | 1 | clec-165 |
| WBGene00009524 | 0.688793783 | 0.86378755 | 1 | F38A1.11 |
| WBGene00009525 | 0.574508983 | 0.8551971 | 1 | F38A1.13 |
| WBGene00009526 | 2.757972349 | 0.04098526 | 1 | clec-169 |
| WBGene00009528 | -0.13962691 | 0.89958998 | 1 | F38A6.4 |
| WBGene00009529 | 1.151807762 | 0.05517324 | 0.17215583 | F38B2.2 |
| WBGene00009530 | 0.897286543 | 0.63390869 | 1 | F38B2.3 |
| WBGene00009531 | -0.126700395 | 0.62905895 | 0.78453054 | F38B2.4 |
| WBGene00009532 | 1.212582313 | 4.3502E-06 | 0.00011237 | ccch-1 |
| WBGene00009533 | 1.323615688 | 0.05152867 | 0.16385794 | F38B7.2 |
| WBGene00009534 | -0.194520669 | 0.49430591 | 0.68290488 | F38B7.3 |

| | | | | |
|---|---|---|---|---|
| WBGene00009536 | -1.116210242 | 0.37065808 | 1 | F38C2.4 |
| WBGene00009537 | -1.60226146 | 0.02395897 | 0.0972888 | ccch-2 |
| WBGene00009539 | -1.962363944 | 0.00847703 | 0.04570849 | F38C2.7 |
| WBGene00009541 | -0.396179562 | 0.77972933 | 1 | cutl-17 |
| WBGene00009542 | 0.073643803 | 0.68386633 | 0.81889941 | copb-2 |
| WBGene00009543 | 0.279360336 | 0.80495327 | 1 | F38E11.6 |
| WBGene00009544 | 0.543483303 | 0.53755443 | 0.71956431 | F38E11.9 |
| WBGene00009546 | -0.694327333 | 0.47317336 | 1 | fbxb-3 |
| WBGene00009547 | 0.555841967 | 0.27506323 | 0.4835712 | F38H4.3 |
| WBGene00009548 | 1.38322681 | 0.69550305 | 1 | F38H4.4 |
| WBGene00009549 | 0.206584522 | 0.92198558 | 1 | F38H4.5 |
| WBGene00009550 | 0.281454817 | 0.94435481 | 1 | F38H4.6 |
| WBGene00009551 | -0.317664773 | 0.17112186 | 0.36015788 | mob-1 |
| WBGene00009552 | 0.376687827 | 0.20797082 | 0.40758252 | piki-1 |
| WBGene00009553 | -0.167860869 | 0.40309429 | 0.61082973 | hinf-1 |
| WBGene00009554 | -0.32530011 | 0.24920726 | 0.45690506 | F39B2.3 |
| WBGene00009556 | -0.871657699 | 0.00711729 | 0.04022859 | F39B2.5 |
| WBGene00009557 | -0.23299878 | 0.50589244 | 0.69273124 | mtcu-1 |
| WBGene00009558 | 0.600664725 | 0.15815954 | 0.34299953 | F39B2.8 |
| WBGene00009559 | -0.308917987 | 0.14460231 | 0.32307998 | mtx-1 |
| WBGene00009560 | 1.143207136 | 0.33646125 | 1 | psa-3 |
| WBGene00009561 | -1.851746282 | 0.09203647 | 1 | F39D8.3 |
| WBGene00009562 | 0.50195823 | 0.12939234 | 0.30073592 | flp-22 |
| WBGene00009563 | -0.526582299 | 0.05639818 | 0.17516764 | F39H2.3 |
| WBGene00009565 | -0.569657922 | 0.00869826 | 0.04660441 | F39H11.1 |
| WBGene00009569 | 0.281454817 | 0.94435481 | 1 | srz-66 |
| WBGene00009571 | 1.240074274 | 0.42231915 | 1 | F40D4.12 |
| WBGene00009572 | -0.454094946 | 0.77209224 | 1 | F40D4.13 |
| WBGene00009573 | -0.612987272 | 0.41689908 | 0.6218147 | F40E10.5 |
| WBGene00009574 | 0.171190021 | 0.41308467 | 0.6185571 | F40E10.6 |
| WBGene00009575 | -0.690936653 | 0.00060619 | 0.00618463 | F40F8.1 |
| WBGene00009576 | -0.892903557 | 0.09053391 | 0.23924932 | F40F8.3 |
| WBGene00009577 | 1.053752087 | 0.00076993 | 0.00752028 | F40F8.4 |
| WBGene00009578 | 0.52938563 | 0.01621051 | 0.07354701 | F40F8.5 |
| WBGene00009579 | 0.300128135 | 0.60844379 | 0.77077588 | arrd-6 |
| WBGene00009580 | 0.28383642 | 0.14206471 | 0.32017962 | xbx-6 |
| WBGene00009581 | 1.477253415 | 0.58357327 | 1 | F40F9.3 |
| WBGene00009582 | 0.466038197 | 0.18003481 | 0.37228489 | F40F9.5 |
| WBGene00009583 | 0.140224507 | 0.43927175 | 0.63970233 | aagr-3 |
| WBGene00009584 | -0.012005534 | 0.94755013 | 0.97311844 | drap-1 |
| WBGene00009585 | 0.747845955 | 0.85215965 | 1 | cal-7 |
| WBGene00009586 | 1.467328407 | 0.03662597 | 0.12966474 | F40F9.10 |
| WBGene00009587 | 0.469977568 | 0.01260149 | 0.06144367 | mig-38 |

| | | | | |
|---|---|---|---|---|
| WBGene00009588 | -0.631309915 | 0.0126522 | 0.06164745 | F40F11.3 |
| WBGene00009589 | 0.353110589 | 0.85358686 | 1 | F40F11.4 |
| WBGene00009592 | 0.747845955 | 0.85215965 | 1 | F40F12.3 |
| WBGene00009593 | -0.01147998 | 0.99415338 | 1 | mam-7 |
| WBGene00009594 | 0.47879265 | 0.00551359 | 0.03348699 | cyld-1 |
| WBGene00009595 | 0.752686847 | 0.0033555 | 0.02300996 | F40F12.7 |
| WBGene00009597 | -1.075708041 | 0.57096009 | 1 | F40G12.2 |
| WBGene00009600 | 0.747845955 | 0.85215965 | 1 | F40G12.5 |
| WBGene00009601 | 0.555958873 | 0.74459049 | 1 | F40G12.6 |
| WBGene00009602 | -0.637018156 | 0.80584146 | 1 | F40G12.7 |
| WBGene00009605 | 2.691380462 | 0.22383774 | 1 | F40G12.10 |
| WBGene00009606 | -0.529625835 | 0.03101481 | 0.11626133 | F40G12.11 |
| WBGene00009608 | -1.901856996 | 0.64072034 | 1 | nhr-265 |
| WBGene00009611 | 1.653460415 | 0.50712258 | 1 | F41D3.6 |
| WBGene00009614 | 1.70609796 | 0.37789212 | 1 | F41D3.9 |
| WBGene00009617 | 0.221463545 | 0.64282606 | 0.79350306 | F41E7.1 |
| WBGene00009618 | 1.014589358 | 0.09717104 | 0.25046061 | F41E7.2 |
| WBGene00009619 | 1.357881944 | 0.12980382 | 0.30114118 | npr-6 |
| WBGene00009620 | -0.748508844 | 0.05973499 | 0.18180776 | fip-5 |
| WBGene00009621 | -0.114896102 | 0.62254838 | 0.78033862 | fipr-21 |
| WBGene00009622 | -1.278290473 | 0.46031622 | 1 | F41E7.6 |
| WBGene00009623 | 0.38231776 | 0.65448991 | 0.80142834 | F41E7.7 |
| WBGene00009624 | -1.093745382 | 0.00100468 | 0.00928301 | cnc-8 |
| WBGene00009625 | -0.589163183 | 0.6830246 | 1 | F41E7.9 |
| WBGene00009626 | -0.310605337 | 0.2575761 | 0.4655131 | F42A8.1 |
| WBGene00009627 | -0.24111527 | 0.29363113 | 0.50388348 | cup-15 |
| WBGene00009628 | 1.380314045 | 7.85E-12 | 9.35E-10 | tatn-1 |
| WBGene00009629 | 0.125308903 | 0.83868243 | 0.91191505 | sprr-2 |
| WBGene00009630 | -0.156269293 | 0.96924949 | 1 | F42E8.1 |
| WBGene00009632 | 0.117011845 | 0.60388996 | 0.76853543 | ttyh-1 |
| WBGene00009633 | -0.166993713 | 0.92790224 | 1 | F42E11.3 |
| WBGene00009634 | 0.782169761 | 0.01203153 | 0.0594371 | nspc-9 |
| WBGene00009635 | -0.375573286 | 0.33944092 | 0.5477137 | F42F12.3 |
| WBGene00009636 | -0.592508468 | 0.14062146 | 0.31834517 | F42F12.4 |
| WBGene00009637 | 0.236905001 | 0.55168257 | 0.73003216 | nspc-13 |
| WBGene00009638 | 0.977712433 | 9.958E-05 | 0.00152733 | nspc-14 |
| WBGene00009639 | 0.65079482 | 0.6373838 | 1 | nspc-15 |
| WBGene00009640 | 0.784501425 | 0.01586144 | 0.07239613 | nspc-10 |
| WBGene00009641 | -0.553649097 | 0.77626512 | 1 | F42G4.2 |
| WBGene00009642 | 0.41404713 | 0.1944234 | 0.3905455 | F42G4.5 |
| WBGene00009643 | -0.333328768 | 0.77094654 | 1 | F42G4.6 |
| WBGene00009644 | -0.088211453 | 0.93590731 | 1 | F42G4.7 |
| WBGene00009645 | 0.23597422 | 0.50472702 | 0.6917136 | F42G10.1 |

| | | | | |
|---|---|---|---|---|
| WBGene00009646 | -1.901856996 | 0.64072034 | 1 | F42H11.1 |
| WBGene00009647 | 1.109208327 | 0.31035731 | 1 | phlp-2 |
| WBGene00009648 | -0.526394061 | 0.08261851 | 0.22549704 | zhit-2 |
| WBGene00009649 | 0.819701647 | 0.36716623 | 0.57475054 | F43C1.5 |
| WBGene00009650 | -0.4922834 | 0.01018335 | 0.05252284 | ccnk-1 |
| WBGene00009652 | 0.800002312 | 0.67639374 | 1 | F43D2.3 |
| WBGene00009653 | -0.022337914 | 0.95396273 | 0.9766474 | F43D9.1 |
| WBGene00009654 | 0.111539053 | 0.61742556 | 0.77706677 | sly-1 |
| WBGene00009657 | 0.552081759 | 0.39261906 | 0.60045695 | F43G6.4 |
| WBGene00009658 | -1.548051605 | 0.00172501 | 0.01386974 | F43G6.5 |
| WBGene00009659 | -0.6811243 | 0.0831484 | 0.2264987 | F43G6.7 |
| WBGene00009660 | 0.208482431 | 0.55602864 | 0.73329826 | F43G6.8 |
| WBGene00009661 | -0.346852955 | 0.02791752 | 0.1079104 | patr-1 |
| WBGene00009662 | -0.912314742 | 0.06990215 | 0.20206713 | F43G6.10 |
| WBGene00009663 | -1.339646758 | 0.40698081 | 1 | hda-5 |
| WBGene00009664 | 0.00381674 | 0.98426576 | 0.99236135 | idha-1 |
| WBGene00009665 | -0.292587341 | 0.27795644 | 0.4868318 | lmd-1 |
| WBGene00009666 | -0.082999747 | 0.76272717 | 0.86721412 | slc-25A42 |
| WBGene00009667 | -0.284721467 | 0.16736063 | 0.35485948 | F43G9.4 |
| WBGene00009668 | -0.547018493 | 0.01044829 | 0.05363751 | cfim-1 |
| WBGene00009670 | -0.774842772 | 0.78069886 | 1 | F43G9.8 |
| WBGene00009671 | 0.216052067 | 0.21916377 | 0.421878 | mfap-1 |
| WBGene00009672 | 0.333703446 | 0.12359099 | 0.29255571 | F43G9.12 |
| WBGene00009673 | -0.069305859 | 0.8462615 | 0.91572735 | F43G9.13 |
| WBGene00009674 | -0.201329147 | 0.33731836 | 0.54532386 | nucb-1 |
| WBGene00009675 | 0.281454817 | 0.94435481 | 1 | F44A6.3 |
| WBGene00009676 | 0.408039083 | 0.41412793 | 0.61950803 | F44A6.4 |
| WBGene00009677 | 1.026725089 | 0.00526847 | 0.03241989 | F44A6.5 |
| WBGene00009678 | 0.48432945 | 0.09739148 | 0.25078048 | magu-4 |
| WBGene00009679 | 0.973441752 | 0.01570175 | 0.07187884 | F44D12.2 |
| WBGene00009681 | 1.768742166 | 0.14482388 | 1 | gipc-2 |
| WBGene00009682 | 1.488951921 | 0.48144928 | 1 | msd-2 |
| WBGene00009683 | -0.62146648 | 0.85925752 | 1 | F44D12.6 |
| WBGene00009685 | -0.263229581 | 0.93409872 | 1 | F44D12.8 |
| WBGene00009686 | 1.038022092 | 0.00175764 | 0.01406898 | ent-7 |
| WBGene00009688 | -0.506630898 | 0.00329333 | 0.02272122 | F44E5.1 |
| WBGene00009689 | -0.280592119 | 0.86317355 | 1 | F44E5.2 |
| WBGene00009690 | 0.664914355 | 0.80343354 | 1 | F44E5.3 |
| WBGene00009691 | 2.560352584 | 0.0284946 | 0.10941439 | F44E5.4 |
| WBGene00009692 | 2.560352584 | 0.0284946 | 0.10941439 | F44E5.5 |
| WBGene00009695 | 0.377828795 | 0.92526585 | 1 | F44F1.3 |
| WBGene00009696 | -0.64191434 | 0.14313225 | 0.32114434 | F44F1.4 |
| WBGene00009698 | -0.962627696 | 0.13790481 | 0.3145411 | F44F1.6 |

| | | | | |
|---|---|---|---|---|
| WBGene00009700 | 0.525643604 | 0.3797001 | 0.58770256 | lurp-1 |
| WBGene00009701 | -0.414532168 | 0.00925842 | 0.04895288 | egg-3 |
| WBGene00009702 | 0.28372011 | 0.8686326 | 1 | F44F4.3 |
| WBGene00009705 | 0.747845955 | 0.85215965 | 1 | F44F4.10 |
| WBGene00009706 | -1.021714985 | 0.58842338 | 1 | argk-1 |
| WBGene00009707 | -0.156269293 | 0.96924949 | 1 | srxa-16 |
| WBGene00009708 | 0.627834078 | 0.77998941 | 1 | F44G3.7 |
| WBGene00009709 | 2.407689268 | 0.1658253 | 1 | fbxa-144 |
| WBGene00009710 | -0.226353808 | 0.67972539 | 0.81647553 | F44G3.10 |
| WBGene00009711 | -0.078466161 | 0.69748929 | 0.828836 | F44G4.1 |
| WBGene00009712 | -0.297493798 | 0.07937526 | 0.2198711 | F44G4.2 |
| WBGene00009713 | -0.896992655 | 0.01265852 | 0.06164745 | F44G4.3 |
| WBGene00009714 | -0.604396148 | 0.82723443 | 1 | F44G4.5 |
| WBGene00009715 | 0.505907206 | 0.47167906 | 0.66453009 | F44G4.6 |
| WBGene00009716 | 0.053739939 | 0.94052956 | 0.96926996 | F44G4.7 |
| WBGene00009717 | 0.539952578 | 0.03682918 | 0.1300969 | dep-1 |
| WBGene00009720 | 0.574508983 | 0.8551971 | 1 | F45B8.3 |
| WBGene00009721 | -0.178518181 | 0.93264297 | 1 | F45D3.1 |
| WBGene00009722 | 0.299634421 | 0.46047117 | 0.6560508 | F45D3.2 |
| WBGene00009723 | 1.072909936 | 3.4218E-06 | 9.2115E-05 | F45D3.3 |
| WBGene00009724 | 1.488169836 | 6.84E-12 | 8.22E-10 | F45D3.4 |
| WBGene00009726 | -0.049911906 | 0.89187634 | 0.94298273 | tbc-13 |
| WBGene00009727 | 0.667970325 | 0.4428589 | 0.64246026 | lips-1 |
| WBGene00009728 | -2.099949391 | 0.15307647 | 1 | sdz-19 |
| WBGene00009729 | 0.506550757 | 0.33415382 | 0.54237454 | F45E10.2 |
| WBGene00009730 | 0.413511845 | 0.10794374 | 0.26759651 | myo-6 |
| WBGene00009731 | -0.452504438 | 0.01454295 | 0.06814588 | exo-1 |
| WBGene00009732 | -0.341021691 | 0.07723935 | 0.21568044 | cope-1 |
| WBGene00009733 | -0.411122076 | 0.32458987 | 0.53305094 | F45G2.7 |
| WBGene00009734 | -0.393866968 | 0.11561799 | 0.28018323 | F45G2.8 |
| WBGene00009735 | -0.80874545 | 0.09085124 | 0.23963173 | F45G2.9 |
| WBGene00009736 | -0.688133632 | 0.0061425 | 0.03627245 | F45G2.10 |
| WBGene00009738 | 0.64179532 | 0.31155337 | 0.51976499 | hecw-1 |
| WBGene00009739 | -0.500137608 | 0.00354905 | 0.02405867 | F45H10.2 |
| WBGene00009740 | -0.340471288 | 0.07218703 | 0.20670347 | F45H10.3 |
| WBGene00009741 | 0.178263295 | 0.72319022 | 0.84401158 | drr-1 |
| WBGene00009742 | 1.230330708 | 0.39111815 | 1 | F45H10.5 |
| WBGene00009743 | -0.753434657 | 0.18846412 | 0.38226506 | sptf-1 |
| WBGene00009744 | 0.116072854 | 0.63359947 | 0.78776561 | hpo-35 |
| WBGene00009745 | -0.565257529 | 0.06577537 | 0.19368876 | F45H11.5 |
| WBGene00009747 | -1.901856996 | 0.64072034 | 1 | F46A8.4 |
| WBGene00009750 | 2.391846226 | 0.01063124 | 0.05425553 | F46A8.7 |
| WBGene00009751 | -0.156269293 | 0.96924949 | 1 | F46A8.8 |

| | | | | |
|---|---|---|---|---|
| WBGene00009753 | -0.156269293 | 0.96924949 | 1 | scrm-6 |
| WBGene00009754 | 1.619850823 | 0.41983851 | 1 | F46A9.1 |
| WBGene00009755 | 0.377828795 | 0.92526585 | 1 | F46A9.2 |
| WBGene00009756 | -1.094201388 | 0.7816772 | 1 | F46B3.1 |
| WBGene00009758 | -1.969805064 | 0.62862569 | 1 | ttr-11 |
| WBGene00009760 | -0.156269293 | 0.96924949 | 1 | F46B3.7 |
| WBGene00009768 | 1.471809563 | 0.46592277 | 1 | F46B6.2 |
| WBGene00009769 | -0.104383704 | 0.68365921 | 0.81881292 | F46B6.4 |
| WBGene00009770 | -0.194174227 | 0.30590418 | 0.51475851 | F46B6.5 |
| WBGene00009771 | 0.036710455 | 0.85240868 | 0.91927977 | F46B6.6 |
| WBGene00009772 | 0.294510149 | 0.08050769 | 0.22195997 | ztf-7 |
| WBGene00009773 | 1.536363757 | 0.0307513 | 0.11558872 | lipl-2 |
| WBGene00009774 | 1.241110051 | 0.11879423 | 0.28436134 | F46B6.9 |
| WBGene00009776 | -0.545924993 | 0.02321986 | 0.09527284 | F46B6.12 |
| WBGene00009777 | 0.919979581 | 0.29673601 | 0.50713287 | F46C3.2 |
| WBGene00009778 | 5.381402062 | 9.986E-05 | 0.00152888 | F46C5.1 |
| WBGene00009779 | 0.40939163 | 0.51438904 | 0.70123482 | F46C5.2 |
| WBGene00009780 | 0.486837529 | 0.56714158 | 0.74206983 | F46C5.4 |
| WBGene00009781 | 1.276231565 | 0.00632938 | 0.03712319 | F46C5.6 |
| WBGene00009782 | -0.756698273 | 0.6598932 | 1 | F46C5.7 |
| WBGene00009783 | -0.077548891 | 0.6896313 | 0.82344744 | rer-1 |
| WBGene00009784 | -0.515256133 | 0.00308053 | 0.02161891 | F46C5.9 |
| WBGene00009785 | 0.4982697 | 0.42301035 | 0.6273844 | F46C5.10 |
| WBGene00009787 | 0.234208817 | 0.17383593 | 0.36376112 | F46F2.3 |
| WBGene00009788 | -0.838211756 | 0.7584606 | 1 | F46F2.4 |
| WBGene00009789 | 0.187950979 | 0.77418007 | 0.87418816 | F46F2.5 |
| WBGene00009790 | 0.809072492 | 0.14542438 | 0.32422834 | lgc-44 |
| WBGene00009791 | -0.615410782 | 0.70507525 | 1 | F46F3.3 |
| WBGene00009793 | 0.159971371 | 0.48075963 | 0.67254563 | pkn-1 |
| WBGene00009796 | 0.010648376 | 0.96095404 | 0.97977647 | F46G10.1 |
| WBGene00009797 | 1.111806174 | 0.00085308 | 0.0081358 | F46G10.2 |
| WBGene00009798 | 0.210925672 | 0.65936678 | 0.80368078 | F46G10.4 |
| WBGene00009799 | 0.651628214 | 0.15581204 | 0.33968242 | lgc-47 |
| WBGene00009800 | 0.549199309 | 0.14728564 | 0.32744827 | rrc-1 |
| WBGene00009801 | 0.978061415 | 0.00282431 | 0.020326 | ipla-2 |
| WBGene00009803 | 1.326022825 | 0.00066702 | 0.00670947 | F47B8.2 |
| WBGene00009804 | 0.8085879 | 0.08366558 | 0.22732355 | F47B8.3 |
| WBGene00009805 | 3.042338499 | 0.12140059 | 1 | F47B8.4 |
| WBGene00009806 | -2.085726786 | 0.12215522 | 1 | F47B8.5 |
| WBGene00009807 | -1.030062181 | 0.46957291 | 1 | pepm-1 |
| WBGene00009808 | 1.859929841 | 0.36327656 | 1 | srm-4 |
| WBGene00009809 | 1.368363722 | 0.03123592 | 0.11693966 | F47B8.8 |
| WBGene00009810 | -0.245531902 | 0.94070824 | 1 | srm-6 |

| | | | | |
|---|---|---|---|---|
| WBGene00009811 | -0.097716126 | 0.92113991 | 1 | F47B8.10 |
| WBGene00009812 | 0.348051723 | 0.03392772 | 0.12366151 | suca-1 |
| WBGene00009813 | 1.067282651 | 2.1683E-06 | 6.2679E-05 | haly-1 |
| WBGene00009814 | 1.851444818 | 0.53669729 | 1 | F47B10.3 |
| WBGene00009816 | -0.525521402 | 0.44476657 | 0.64422703 | F47B10.5 |
| WBGene00009818 | -0.255214187 | 0.33121528 | 0.53952848 | acbp-3 |
| WBGene00009819 | 1.27509718 | 0.27214541 | 1 | F47B10.8 |
| WBGene00009820 | -0.310538136 | 0.50172206 | 0.68948393 | F47B10.9 |
| WBGene00009822 | 1.38322681 | 0.69550305 | 1 | clec-113 |
| WBGene00009823 | 0.037661033 | 0.91009778 | 0.95328248 | gadr-6 |
| WBGene00009824 | -1.490041711 | 0.14026715 | 1 | gpdh-1 |
| WBGene00009825 | 0.153915329 | 0.61287458 | 0.77396612 | F47G4.4 |
| WBGene00009826 | 1.7441231 | 0.18770648 | 1 | F47G4.5 |
| WBGene00009827 | -0.491787485 | 0.79288596 | 1 | hmg-6 |
| WBGene00009829 | -0.156968442 | 0.31706792 | 0.52493219 | F47G9.1 |
| WBGene00009830 | -1.664775548 | 0.02865509 | 0.10964433 | cutl-18 |
| WBGene00009831 | 1.045157653 | 0.0141281 | 0.06671694 | F47G9.4 |
| WBGene00009833 | -1.159714244 | 0.75790634 | 1 | F47G9.6 |
| WBGene00009834 | -0.750070612 | 0.01132095 | 0.05689056 | lsy-27 |
| WBGene00009835 | 4.189919453 | 0.00020181 | 0.00264604 | F47H4.2 |
| WBGene00009836 | 0.855368791 | 0.66134621 | 1 | fbxa-185 |
| WBGene00009839 | 2.116650971 | 0.11682559 | 1 | fbxa-188 |
| WBGene00009840 | 1.047101039 | 0.08965586 | 0.2375626 | fbxa-189 |
| WBGene00009841 | -0.310604425 | 0.87613277 | 1 | fbxa-134 |
| WBGene00009842 | 0.462235885 | 0.4761233 | 0.66874283 | igeg-2 |
| WBGene00009843 | -0.160940816 | 0.82027791 | 0.90268458 | F48C5.2 |
| WBGene00009844 | 2.580562247 | 0.15729341 | 1 | cwp-5 |
| WBGene00009848 | 0.857248278 | 0.1648958 | 0.35141302 | F48F5.6 |
| WBGene00009849 | -2.165407894 | 0.16245383 | 1 | F48F7.3 |
| WBGene00009850 | 0.736589183 | 0.09697238 | 0.25017886 | F48F7.5 |
| WBGene00009851 | -0.133180476 | 0.66472905 | 0.80771846 | F48F7.6 |
| WBGene00009852 | 0.997040592 | 0.37412314 | 1 | arrd-24 |
| WBGene00009853 | 0.5180853 | 0.79605727 | 1 | arrd-27 |
| WBGene00009854 | 1.243709618 | 0.17833762 | 0.37030907 | clec-31 |
| WBGene00009855 | 0.82762367 | 0.54187084 | 1 | clec-22 |
| WBGene00009856 | 2.71234179 | 0.24246227 | 1 | clec-24 |
| WBGene00009857 | 2.214904615 | 0.02334926 | 0.09563613 | clec-28 |
| WBGene00009859 | 3.147938868 | 0.01247275 | 0.06093829 | clec-33 |
| WBGene00009860 | -0.33449598 | 0.85735439 | 1 | clec-32 |
| WBGene00009861 | 0.460141183 | 0.82127868 | 1 | fbxb-8 |
| WBGene00009862 | -0.390489411 | 0.41715267 | 0.62193997 | fbxb-67 |
| WBGene00009863 | 0.377828795 | 0.92526585 | 1 | F49B2.3 |
| WBGene00009865 | 1.548424186 | 0.00511769 | 0.03169317 | F49B2.6 |

| | | | | |
|---|---|---|---|---|
| WBGene00009866 | -0.156269293 | 0.96924949 | 1 | F49C5.3 |
| WBGene00009867 | 2.18453076 | 0.41642786 | 1 | F49C5.4 |
| WBGene00009872 | 1.181686151 | 0.7579682 | 1 | F49C12.2 |
| WBGene00009873 | -0.668264701 | 0.85551153 | 1 | F49C12.3 |
| WBGene00009874 | -0.156269293 | 0.96924949 | 1 | F49C12.4 |
| WBGene00009875 | 0.747845955 | 0.85215965 | 1 | F49C12.5 |
| WBGene00009876 | 0.758325915 | 0.213428 | 0.41413287 | F49C12.6 |
| WBGene00009877 | 2.417124694 | 4.75E-21 | 1.75E-18 | F49C12.7 |
| WBGene00009878 | -0.095180532 | 0.66855508 | 0.81015343 | F49C12.9 |
| WBGene00009879 | 0.923852597 | 0.36610252 | 1 | F49C12.10 |
| WBGene00009880 | -0.647352508 | 4.5132E-05 | 0.00079413 | F49C12.11 |
| WBGene00009881 | 0.003936181 | 0.98514569 | 0.99283256 | F49C12.12 |
| WBGene00009882 | 0.265254475 | 0.11891727 | 0.28459976 | vha-17 |
| WBGene00009883 | 0.848755505 | 0.06677516 | 0.1960148 | F49C12.14 |
| WBGene00009884 | 3.79121669 | 0.00098457 | 0.00913574 | F49C12.15 |
| WBGene00009885 | 0.632925785 | 0.20974651 | 0.41007351 | F49E2.1 |
| WBGene00009886 | 0.524945218 | 0.03793189 | 0.13271787 | F49E2.2 |
| WBGene00009887 | 0.703317977 | 0.29838918 | 0.50831249 | glb-17 |
| WBGene00009888 | 1.373016846 | 1.06E-07 | 4.9607E-06 | F49E2.5 |
| WBGene00009889 | -0.578171382 | 0.57528635 | 1 | F49E11.2 |
| WBGene00009893 | -0.156269293 | 0.96924949 | 1 | F49E11.7 |
| WBGene00009895 | 1.754934657 | 1.51E-07 | 6.687E-06 | scl-2 |
| WBGene00009896 | 0.948802972 | 0.00069735 | 0.00696257 | scl-3 |
| WBGene00009897 | -0.496136482 | 0.0005406 | 0.00568265 | skpo-1 |
| WBGene00009898 | -0.143538284 | 0.51046734 | 0.69706264 | dod-23 |
| WBGene00009899 | 0.181460743 | 0.5629058 | 0.73827799 | efl-3 |
| WBGene00009900 | 2.242680281 | 0.20773484 | 1 | F49E12.7 |
| WBGene00009901 | 0.506835397 | 0.69474532 | 1 | F49E12.8 |
| WBGene00009902 | -0.811166312 | 0.00259784 | 0.01907574 | drd-1 |
| WBGene00009903 | 0.178114269 | 0.76790765 | 0.86986068 | F49E12.10 |
| WBGene00009904 | 0.097723447 | 0.92776648 | 1 | F49E12.12 |
| WBGene00009905 | -0.156269293 | 0.96924949 | 1 | clec-240 |
| WBGene00009906 | -1.969805064 | 0.62862569 | 1 | clec-241 |
| WBGene00009907 | 1.957271014 | 0.49743227 | 1 | F49H6.3 |
| WBGene00009908 | 3.348675671 | 0.03207562 | 1 | F49H6.5 |
| WBGene00009912 | -0.156269293 | 0.96924949 | 1 | srz-94 |
| WBGene00009914 | 3.013451459 | 0.16378372 | 1 | F49H6.13 |
| WBGene00009915 | -0.455401703 | 0.00697545 | 0.0396484 | F52A8.1 |
| WBGene00009916 | -0.465326787 | 0.62149322 | 0.77957942 | F52A8.3 |
| WBGene00009917 | 0.448175236 | 0.33261479 | 0.54045452 | glb-18 |
| WBGene00009918 | -0.518356619 | 0.01761041 | 0.07817476 | gcsh-2 |
| WBGene00009919 | -0.096661717 | 0.65009664 | 0.79843591 | kbrl-1 |
| WBGene00009920 | 0.710851934 | 7.8876E-05 | 0.00126337 | abts-1 |

| | | | | |
|---|---|---|---|---|
| WBGene00009921 | -0.59826595 | 0.00041418 | 0.00462612 | F52B5.2 |
| WBGene00009922 | -0.012105176 | 0.94453666 | 0.97134161 | F52B5.3 |
| WBGene00009923 | -0.815239283 | 0.04647381 | 0.15282733 | F52B5.7 |
| WBGene00009924 | -0.173633425 | 0.36351556 | 0.57161539 | cfp-1 |
| WBGene00009925 | -0.845533697 | 0.00039785 | 0.0044933 | F52B11.2 |
| WBGene00009926 | -0.237543465 | 0.36318659 | 0.57124607 | noah-2 |
| WBGene00009927 | -0.084141758 | 0.92269989 | 0.9593272 | F52B11.5 |
| WBGene00009929 | 1.56209938 | 0.00720573 | 0.040616 | abts-2 |
| WBGene00009930 | -0.790418352 | 0.00823907 | 0.04477642 | F52D10.2 |
| WBGene00009931 | 1.136041123 | 0.43997091 | 1 | F52D10.4 |
| WBGene00009933 | 0.281454817 | 0.94435481 | 1 | F52E10.2 |
| WBGene00009935 | 2.558478798 | 0.00575482 | 0.03455067 | F52E10.4 |
| WBGene00009937 | 0.035146031 | 0.86928622 | 0.92962247 | lsl-1 |
| WBGene00009938 | 0.253925665 | 0.89264743 | 1 | F52F12.5 |
| WBGene00009939 | -0.676857799 | 0.02778155 | 0.10748752 | ztf-11 |
| WBGene00009940 | -0.281455718 | 0.1488603 | 0.32998227 | strl-1 |
| WBGene00009941 | -0.333693239 | 0.89573179 | 1 | F52F12.8 |
| WBGene00009942 | -0.901019005 | 0.67045999 | 1 | F52F12.9 |
| WBGene00009943 | 0.480378485 | 0.25182911 | 0.45983715 | F52G2.3 |
| WBGene00009944 | 0.102343463 | 0.64342034 | 0.79395938 | mtcu-2 |
| WBGene00009945 | 0.063437853 | 0.83858613 | 0.911892 | bath-38 |
| WBGene00009946 | -0.510344035 | 0.0710782 | 0.20429934 | ogr-2 |
| WBGene00009947 | 0.101612454 | 0.80550303 | 0.89401337 | F52H3.5 |
| WBGene00009948 | -0.831815107 | 0.45979039 | 1 | F52H3.6 |
| WBGene00009949 | 0.394817702 | 0.53985543 | 0.72138749 | F53A2.1 |
| WBGene00009950 | 0.438455499 | 0.87116517 | 1 | sdz-20 |
| WBGene00009951 | -0.79782754 | 0.06916063 | 0.20068815 | F53A2.3 |
| WBGene00009952 | -0.391581293 | 0.02704893 | 0.10556882 | acaa-2 |
| WBGene00009953 | 0.902627381 | 0.08533361 | 0.23033246 | F53A2.9 |
| WBGene00009954 | -1.969805064 | 0.62862569 | 1 | F53B2.1 |
| WBGene00009955 | 1.059777923 | 0.6622777 | 1 | eak-4 |
| WBGene00009956 | 0.28530093 | 0.92084641 | 1 | F53B2.5 |
| WBGene00009957 | 1.506941847 | 1.59E-07 | 6.978E-06 | F53B2.8 |
| WBGene00009958 | 0.68771022 | 0.01278087 | 0.06209386 | madd-4 |
| WBGene00009959 | 0.764000445 | 0.41657709 | 0.62148715 | F53B6.4 |
| WBGene00009960 | -1.831477136 | 0.52194793 | 1 | F53B6.5 |
| WBGene00009962 | -0.457706419 | 0.71687561 | 1 | F53B6.7 |
| WBGene00009963 | 0.377828795 | 0.92526585 | 1 | fipr-26 |
| WBGene00009964 | 0.602327163 | 0.66380014 | 1 | fip-6 |
| WBGene00009965 | 0.62161842 | 0.59324775 | 1 | frpr-9 |
| WBGene00009966 | -0.990104849 | 6.53E-08 | 3.2116E-06 | isy-1 |
| WBGene00009967 | -0.62146648 | 0.85925752 | 1 | F53B7.4 |
| WBGene00009968 | 1.545630408 | 7.76E-07 | 2.6834E-05 | fig-1 |

| | | | | |
|---|---|---|---|---|
| WBGene00009969 | 0.112248961 | 0.8833951 | 0.93801818 | F53B7.7 |
| WBGene00009971 | 0.236001482 | 0.43814596 | 0.63866548 | F53C11.1 |
| WBGene00009973 | 0.177074301 | 0.30348507 | 0.51239219 | F53C11.3 |
| WBGene00009974 | 0.359875453 | 0.08952754 | 0.23732616 | F53C11.4 |
| WBGene00009975 | 0.170375151 | 0.29798453 | 0.50787519 | F53C11.5 |
| WBGene00009976 | -0.150204954 | 0.53986713 | 0.72138749 | swan-2 |
| WBGene00009977 | -0.655312269 | 0.00360624 | 0.02435115 | swan-1 |
| WBGene00009978 | -0.294695338 | 0.11053537 | 0.27182701 | F53E4.1 |
| WBGene00009979 | -1.14715222 | 0.68932198 | 1 | cutl-1 |
| WBGene00009980 | -0.254758894 | 0.13411334 | 0.30815103 | F53F1.2 |
| WBGene00009981 | 0.062419645 | 0.84239119 | 0.91337104 | F53F1.3 |
| WBGene00009982 | -0.292888585 | 0.28577807 | 0.49566165 | F53F1.4 |
| WBGene00009983 | -1.559037033 | 3.31E-07 | 1.3207E-05 | cut-2 |
| WBGene00009984 | 3.148385117 | 7.3519E-06 | 0.00017468 | F53F1.6 |
| WBGene00009985 | 4.19718527 | 0.00811951 | 1 | F53F4.1 |
| WBGene00009986 | 0.996013393 | 0.53410803 | 1 | F53F4.2 |
| WBGene00009987 | -0.616164512 | 0.001619 | 0.01328123 | tbcb-1 |
| WBGene00009988 | 0.52475915 | 0.5990814 | 1 | F53F4.4 |
| WBGene00009992 | -0.670847341 | 0.00011571 | 0.00171322 | F53F4.10 |
| WBGene00009993 | -0.158899819 | 0.36043614 | 0.56837101 | F53F4.11 |
| WBGene00009994 | -0.159759516 | 0.55111851 | 0.72959654 | F53F4.12 |
| WBGene00009995 | 0.327729159 | 0.08835796 | 0.23562642 | F53F4.13 |
| WBGene00009996 | -0.014320153 | 0.94168121 | 0.96989883 | F53F4.14 |
| WBGene00009997 | 1.743415485 | 0.50483382 | 1 | F53F4.15 |
| WBGene00009998 | -0.700725429 | 0.66443611 | 1 | klf-2 |
| WBGene00009999 | -0.156269293 | 0.96924949 | 1 | srsx-35 |
| WBGene00010000 | -0.939859043 | 1.7931E-05 | 0.00036836 | F53F8.3 |
| WBGene00010001 | 0.736099952 | 0.00725379 | 0.04082604 | F53F8.4 |
| WBGene00010002 | -0.093220095 | 0.6648764 | 0.80771846 | F53F8.5 |
| WBGene00010003 | 1.088200014 | 0.3719036 | 1 | F53F8.6 |
| WBGene00010004 | 2.695937817 | 0.04302153 | 1 | F53H2.1 |
| WBGene00010005 | -2.287280404 | 0.29827708 | 1 | cnc-7 |
| WBGene00010006 | 0.009003125 | 0.97220296 | 0.98553604 | F53H2.3 |
| WBGene00010007 | -1.219755672 | 0.01347987 | 0.0642304 | F53H4.2 |
| WBGene00010008 | 0.351581312 | 0.70277809 | 0.83218963 | F53H4.3 |
| WBGene00010009 | 0.579231298 | 0.61381749 | 1 | F53H4.4 |
| WBGene00010010 | 0.732110757 | 0.31416261 | 0.52190056 | gmeb-2 |
| WBGene00010011 | 0.753706741 | 0.14165057 | 0.31978051 | F53H4.6 |
| WBGene00010012 | 1.177788811 | 1.81E-07 | 7.7557E-06 | saeg-1 |
| WBGene00010013 | 0.567256627 | 0.00359723 | 0.02430899 | F54B3.1 |
| WBGene00010014 | -1.180974267 | 0.55537885 | 1 | F54B3.2 |
| WBGene00010015 | -0.577753505 | 0.0004299 | 0.00475153 | atad-3 |
| WBGene00010017 | -0.156269293 | 0.96924949 | 1 | nhr-282 |

| | | | | |
|---|---|---|---|---|
| WBGene00010018 | 1.424088136 | 0.03437656 | 0.12477303 | fbxa-69 |
| WBGene00010019 | 1.138536103 | 0.09543891 | 0.24772303 | F54B8.4 |
| WBGene00010020 | -1.969805064 | 0.62862569 | 1 | srbc-48 |
| WBGene00010023 | -0.156269293 | 0.96924949 | 1 | srbc-51 |
| WBGene00010025 | 0.281454817 | 0.94435481 | 1 | srbc-54 |
| WBGene00010027 | 0.359649757 | 0.86200671 | 1 | F54B11.4 |
| WBGene00010028 | 0.251877732 | 0.56724283 | 0.74212232 | F54B11.5 |
| WBGene00010030 | 0.574508983 | 0.8551971 | 1 | F54B11.7 |
| WBGene00010031 | 0.903390376 | 0.71900339 | 1 | F54B11.8 |
| WBGene00010032 | 0.936042292 | 0.17830782 | 0.37030907 | F54B11.9 |
| WBGene00010033 | 0.801798391 | 0.56608725 | 1 | F54B11.10 |
| WBGene00010034 | 2.657813209 | 4.1208E-06 | 0.00010736 | F54B11.11 |
| WBGene00010035 | 0.625583637 | 0.65687922 | 1 | F54C8.1 |
| WBGene00010036 | -0.739982365 | 0.00074989 | 0.00735409 | cpar-1 |
| WBGene00010037 | -0.329525638 | 0.03917036 | 0.13545069 | F54C8.4 |
| WBGene00010038 | -0.328159176 | 0.18326341 | 0.37536783 | rheb-1 |
| WBGene00010039 | 0.897012227 | 0.08070705 | 0.22222266 | F54C8.6 |
| WBGene00010040 | -0.182440419 | 0.37743376 | 0.58512604 | F54C8.7 |
| WBGene00010041 | 0.412337221 | 0.0633762 | 0.18900771 | F54C9.3 |
| WBGene00010042 | -0.188969216 | 0.34592637 | 0.55473412 | bcs-1 |
| WBGene00010043 | 1.13962992 | 0.06862914 | 0.19962299 | F54C9.7 |
| WBGene00010044 | 0.311177233 | 0.09564347 | 0.24787895 | F54C9.9 |
| WBGene00010045 | 0.48094939 | 0.30256205 | 0.5117893 | F54C9.11 |
| WBGene00010046 | 1.541523573 | 0.29553204 | 1 | F54D1.1 |
| WBGene00010047 | 0.792838524 | 0.00011294 | 0.00168245 | F54D1.6 |
| WBGene00010048 | -0.11551962 | 0.68624027 | 0.82077067 | F54D5.2 |
| WBGene00010049 | 0.88486325 | 4.34E-09 | 2.88E-07 | F54D5.3 |
| WBGene00010050 | -0.588333866 | 0.00406167 | 0.026598 | F54D5.4 |
| WBGene00010051 | -0.719399117 | 0.00016967 | 0.00231715 | F54D5.5 |
| WBGene00010052 | -0.110315708 | 0.56231082 | 0.73781645 | F54D5.7 |
| WBGene00010053 | -0.155531355 | 0.46175375 | 0.65699664 | F54D5.9 |
| WBGene00010054 | -0.184347305 | 0.36619935 | 0.57379286 | F54D5.11 |
| WBGene00010055 | -0.52199293 | 0.00206317 | 0.01594456 | F54D5.12 |
| WBGene00010056 | -0.043843096 | 0.7778766 | 0.87649186 | smc-6 |
| WBGene00010057 | 1.261236633 | 0.02601134 | 0.10302938 | F54D5.15 |
| WBGene00010058 | -2.675386419 | 0.02877419 | 1 | F54E4.2 |
| WBGene00010059 | 0.15989044 | 0.58959928 | 0.75742266 | F54E4.3 |
| WBGene00010060 | 1.946671172 | 0.12720993 | 1 | F54E4.4 |
| WBGene00010061 | -0.082108863 | 0.58199934 | 0.75273854 | F54E12.2 |
| WBGene00010062 | 5.134669308 | 2.20E-08 | 1.2183E-06 | lipl-1 |
| WBGene00010063 | 0.362115212 | 0.26959421 | 0.47804051 | dhrs-4 |
| WBGene00010064 | -1.117034133 | 4.7406E-05 | 0.00082577 | F54F7.2 |
| WBGene00010065 | 0.655371409 | 0.13327049 | 0.30677721 | F54F7.3 |

| | | | | |
|---|---|---|---|---|
| WBGene00010066 | -0.385410949 | 0.32924969 | 0.53801083 | F54F7.6 |
| WBGene00010068 | 2.215950884 | 0.41557421 | 1 | F54F7.8 |
| WBGene00010069 | 0.146418095 | 0.9299978 | 1 | F54F11.1 |
| WBGene00010070 | 1.311199614 | 2.22E-08 | 1.2256E-06 | nep-17 |
| WBGene00010071 | 0.747845955 | 0.85215965 | 1 | sre-45 |
| WBGene00010072 | 1.441413618 | 0.29453467 | 1 | F54F12.1 |
| WBGene00010073 | -0.044001024 | 0.9710609 | 1 | F54F12.2 |
| WBGene00010074 | 0.138939741 | 0.90578543 | 1 | irld-34 |
| WBGene00010075 | -0.360977054 | 0.0397153 | 0.13682844 | F55A11.1 |
| WBGene00010077 | -0.198421854 | 0.3816165 | 0.58977507 | F55A11.4 |
| WBGene00010079 | 0.543371303 | 0.14241507 | 0.32034267 | F55A11.6 |
| WBGene00010080 | 0.121501503 | 0.59091397 | 0.75822094 | F55A11.7 |
| WBGene00010081 | -0.10197458 | 0.70416092 | 0.83268897 | F55A11.8 |
| WBGene00010082 | 1.51317332 | 0.31234317 | 1 | F55A11.11 |
| WBGene00010084 | -0.325353918 | 0.14721564 | 0.32735257 | F55B11.2 |
| WBGene00010085 | -1.086224462 | 1.988E-05 | 0.00039961 | F55B11.3 |
| WBGene00010086 | -0.425189116 | 0.16945012 | 0.35748616 | F55B11.4 |
| WBGene00010087 | 1.652261098 | 0.30127117 | 1 | F55B11.5 |
| WBGene00010088 | -0.263229581 | 0.93409872 | 1 | F55B12.2 |
| WBGene00010089 | -0.452820365 | 0.03126531 | 0.11701361 | hpo-31 |
| WBGene00010091 | -0.199374337 | 0.91832278 | 1 | ssp-35 |
| WBGene00010093 | -0.313508337 | 0.05690639 | 0.17616605 | capg-2 |
| WBGene00010094 | -0.268382377 | 0.20018128 | 0.39868782 | tsfm-1 |
| WBGene00010096 | -0.06996293 | 0.70109606 | 0.83141078 | rskd-1 |
| WBGene00010097 | -0.080349326 | 0.62047669 | 0.77878286 | srpa-68 |
| WBGene00010098 | -3.460940123 | 0.00277956 | 0.02012321 | F55C5.10 |
| WBGene00010099 | -1.14715222 | 0.68932198 | 1 | fbxb-117 |
| WBGene00010100 | -1.964653846 | 0.53599202 | 1 | F55C9.3 |
| WBGene00010101 | 0.345720592 | 0.76141484 | 1 | fbxb-57 |
| WBGene00010102 | -0.235607762 | 0.88051053 | 1 | F55C9.5 |
| WBGene00010104 | -0.559340474 | 0.78183513 | 1 | fbxb-60 |
| WBGene00010105 | -1.923297563 | 0.11121641 | 1 | fbxb-62 |
| WBGene00010106 | -1.574768579 | 0.0660753 | 0.19433629 | fbxb-61 |
| WBGene00010107 | -0.156269293 | 0.96924949 | 1 | F55C9.11 |
| WBGene00010108 | 0.377828795 | 0.92526585 | 1 | F55C10.4 |
| WBGene00010109 | -0.248711096 | 0.56423289 | 0.73937955 | F55C10.5 |
| WBGene00010110 | -0.133488594 | 0.83378752 | 0.91001567 | F55D12.1 |
| WBGene00010111 | 0.917215696 | 0.01196055 | 0.05914992 | F55D12.2 |
| WBGene00010113 | 0.504788615 | 0.028205 | 0.10864113 | F55D12.5 |
| WBGene00010114 | 2.063391569 | 0.37853067 | 1 | F55D12.6 |
| WBGene00010115 | 0.581338962 | 0.13162414 | 0.30404275 | aakb-1 |
| WBGene00010116 | 0.774555968 | 0.07499185 | 0.21178787 | F55F3.2 |
| WBGene00010117 | 0.522608102 | 0.00548267 | 0.03336594 | nkb-3 |

| | | | | |
|---|---|---|---|---|
| WBGene00010118 | 2.006276785 | 0.28213582 | 1 | F55F3.4 |
| WBGene00010119 | 0.446458324 | 0.87164163 | 1 | F55G7.1 |
| WBGene00010120 | -1.219336026 | 0.09229325 | 0.24192069 | dot-1.4 |
| WBGene00010121 | 0.233739675 | 0.93575889 | 1 | F55G7.3 |
| WBGene00010122 | 1.735991307 | 0.58152819 | 1 | F55G11.1 |
| WBGene00010123 | 1.927438956 | 3.22E-10 | 2.87E-08 | F55G11.2 |
| WBGene00010124 | 3.291329956 | 8.89E-51 | 2.16E-47 | F55G11.4 |
| WBGene00010125 | -0.36187317 | 0.46512183 | 0.6594703 | dod-22 |
| WBGene00010126 | 1.181686151 | 0.7579682 | 1 | F55G11.6 |
| WBGene00010127 | 0.347225369 | 0.80507073 | 1 | F55G11.7 |
| WBGene00010128 | 0.407406095 | 0.21631667 | 0.41787345 | F55G11.8 |
| WBGene00010130 | 0.033531079 | 0.80999244 | 0.89629222 | vha-14 |
| WBGene00010131 | -0.373134325 | 0.17865231 | 0.37077225 | F55H2.5 |
| WBGene00010132 | -0.329165776 | 0.13130918 | 0.30381958 | F55H2.7 |
| WBGene00010133 | 1.095238366 | 0.00029905 | 0.00359844 | F55H12.2 |
| WBGene00010134 | 1.298723332 | 5.8473E-05 | 0.0009929 | F55H12.3 |
| WBGene00010135 | -0.840171277 | 0.00122063 | 0.01068467 | F55H12.4 |
| WBGene00010136 | 1.812741993 | 0.36269409 | 1 | F55H12.5 |
| WBGene00010137 | 0.252050467 | 0.59345088 | 0.76005346 | ztf-26 |
| WBGene00010138 | 0.261039796 | 0.83335511 | 1 | anoh-1 |
| WBGene00010139 | -0.365247564 | 0.06170561 | 0.18592589 | F56A8.3 |
| WBGene00010140 | -0.68072938 | 0.01865836 | 0.08154469 | F56A8.4 |
| WBGene00010141 | -0.500390848 | 0.07202125 | 0.20637478 | F56A8.5 |
| WBGene00010142 | -2.225166717 | 7.52E-07 | 2.6401E-05 | F56A8.8 |
| WBGene00010143 | 1.020825502 | 0.31847775 | 1 | F56A12.2 |
| WBGene00010144 | -1.130829816 | 0.49432729 | 1 | F56C4.1 |
| WBGene00010146 | 1.376526793 | 0.03717955 | 0.13099155 | glb-19 |
| WBGene00010147 | 0.281454817 | 0.94435481 | 1 | F56D5.2 |
| WBGene00010148 | 1.083841189 | 0.07653224 | 0.21485094 | F56D5.3 |
| WBGene00010149 | -1.869109151 | 4.6092E-05 | 0.00080634 | F56D5.5 |
| WBGene00010150 | 0.514283767 | 0.04184554 | 0.14207122 | F56D5.6 |
| WBGene00010155 | 1.996315266 | 0.49147447 | 1 | F56F3.4 |
| WBGene00010157 | -1.163697947 | 0.52768779 | 1 | oac-34 |
| WBGene00010158 | -2.50408321 | 0.09643964 | 1 | pes-2.2 |
| WBGene00010159 | -0.125556336 | 0.53532228 | 0.7178681 | F56G4.4 |
| WBGene00010160 | -0.186715643 | 0.42787901 | 0.63075268 | png-1 |
| WBGene00010161 | 0.37183621 | 0.26202277 | 0.47059444 | F56G4.6 |
| WBGene00010162 | -1.901856996 | 0.64072034 | 1 | F56H6.1 |
| WBGene00010163 | -0.188615326 | 0.92397876 | 1 | F56H6.2 |
| WBGene00010171 | 1.413377704 | 0.60673863 | 1 | oac-35 |
| WBGene00010172 | 1.687239501 | 0.59403242 | 1 | oac-36 |
| WBGene00010173 | 0.377828795 | 0.92526585 | 1 | F56H6.13 |
| WBGene00010174 | 0.270253636 | 0.11871066 | 0.28427339 | F56H9.2 |

| | | | | |
|---|---|---|---|---|
| WBGene00010175 | 0.544269365 | 0.36958908 | 0.57727789 | F56H11.2 |
| WBGene00010176 | 1.216957619 | 0.74819621 | 1 | F56H11.6 |
| WBGene00010177 | 0.997154715 | 0.00089481 | 0.00843472 | F57A8.1 |
| WBGene00010178 | -0.386416216 | 0.04264682 | 0.14385895 | yif-1 |
| WBGene00010179 | 1.485052503 | 0.06461272 | 0.19164316 | F57A8.4 |
| WBGene00010181 | -0.668264701 | 0.85551153 | 1 | F57A8.6 |
| WBGene00010182 | 1.025181888 | 0.79790347 | 1 | F57A8.7 |
| WBGene00010183 | -2.229370229 | 0.02613513 | 0.1032287 | fipr-13 |
| WBGene00010184 | -1.69202347 | 0.34743527 | 1 | F57A10.2 |
| WBGene00010185 | -0.359459895 | 0.27405714 | 0.48257109 | F57A10.4 |
| WBGene00010186 | -0.830315496 | 0.62566352 | 1 | nhr-283 |
| WBGene00010187 | -0.017077794 | 0.99010363 | 1 | F57B1.1 |
| WBGene00010188 | 0.319817796 | 0.67751203 | 0.81544713 | F57B1.5 |
| WBGene00010189 | -1.040205069 | 0.25318989 | 0.46107372 | F57B1.6 |
| WBGene00010191 | 1.08334498 | 0.00198022 | 0.01542132 | dmsr-1 |
| WBGene00010192 | 0.127155353 | 0.81727511 | 0.90082044 | F57B7.2 |
| WBGene00010193 | -0.165918716 | 0.71251962 | 0.8375158 | btb-20 |
| WBGene00010194 | -0.66024686 | 0.00781374 | 0.04304295 | btb-19 |
| WBGene00010195 | -0.939353729 | 0.00029656 | 0.00357556 | pot-2 |
| WBGene00010196 | -0.190442769 | 0.36373045 | 0.57173114 | F57C2.4 |
| WBGene00010197 | -0.380186667 | 0.02855209 | 0.10951656 | F57C2.5 |
| WBGene00010198 | -0.356679204 | 0.05471671 | 0.17099499 | spat-1 |
| WBGene00010199 | -0.069381059 | 0.7528187 | 0.86217294 | bet-2 |
| WBGene00010200 | 1.755768569 | 0.04269122 | 0.14385895 | F57C7.4 |
| WBGene00010202 | 2.312670972 | 0.31200728 | 1 | F57E7.1 |
| WBGene00010204 | 0.680862629 | 2.33E-08 | 1.2825E-06 | F57F5.1 |
| WBGene00010205 | 2.419039418 | 0.16790136 | 1 | F57F5.3 |
| WBGene00010206 | 0.747845955 | 0.85215965 | 1 | F57G4.1 |
| WBGene00010212 | -1.561825519 | 0.00379701 | 0.02522208 | fbxa-192 |
| WBGene00010214 | 0.492897705 | 0.58418008 | 0.7540432 | F57G8.5 |
| WBGene00010215 | 2.009337304 | 0.00486499 | 0.0305724 | nhr-193 |
| WBGene00010216 | -0.668264701 | 0.85551153 | 1 | F57G8.7 |
| WBGene00010218 | 0.281454817 | 0.94435481 | 1 | sre-30 |
| WBGene00010219 | -1.251248802 | 0.2017125 | 1 | F57G9.3 |
| WBGene00010220 | -1.969805064 | 0.62862569 | 1 | sre-29 |
| WBGene00010221 | 0.494154054 | 0.08436882 | 0.22874538 | F57G12.1 |
| WBGene00010222 | 0.925186562 | 0.49029908 | 1 | F57G12.2 |
| WBGene00010223 | 0.920061037 | 0.34996286 | 0.55851227 | F58A3.3 |
| WBGene00010224 | -1.264646168 | 0.26655327 | 1 | F58A3.4 |
| WBGene00010225 | -0.217594108 | 0.46550336 | 0.65974005 | ttr-31 |
| WBGene00010226 | -1.291305095 | 0.63677871 | 1 | F58A4.1 |
| WBGene00010227 | -0.434238407 | 0.1130798 | 0.27579385 | F58A4.2 |
| WBGene00010228 | 0.747845955 | 0.85215965 | 1 | clec-161 |

| | | | | |
|---|---|---|---|---|
| WBGene00010229 | 0.002523678 | 0.99504862 | 0.99756601 | F58A4.6 |
| WBGene00010230 | -0.919500137 | 2.9583E-05 | 0.00055391 | rpac-19 |
| WBGene00010231 | 0.166325747 | 0.39743503 | 0.605124 | F58B3.4 |
| WBGene00010232 | -0.441463524 | 0.0075428 | 0.04196006 | F58B3.6 |
| WBGene00010233 | -0.38457215 | 0.07888104 | 0.21883972 | F58B3.7 |
| WBGene00010234 | -1.997394079 | 1.32E-23 | 6.70E-21 | ttr-50 |
| WBGene00010235 | 0.574508983 | 0.8551971 | 1 | F58B4.2 |
| WBGene00010236 | 1.089439726 | 0.01692169 | 0.07586639 | F58B4.3 |
| WBGene00010238 | 0.058910873 | 0.8399848 | 0.91226903 | F58B4.5 |
| WBGene00010239 | 0.656602883 | 0.2470191 | 0.45497554 | F58B4.6 |
| WBGene00010241 | -0.508813154 | 0.81506991 | 1 | F58D2.2 |
| WBGene00010242 | -1.073419934 | 0.05090146 | 0.16271141 | F58D5.2 |
| WBGene00010243 | -1.106164015 | 0.33466462 | 1 | F58D5.3 |
| WBGene00010244 | 0.173607535 | 0.56352053 | 0.73875164 | F58D5.5 |
| WBGene00010245 | -0.156269293 | 0.96924949 | 1 | F58D5.6 |
| WBGene00010246 | 1.735991307 | 0.58152819 | 1 | F58D5.7 |
| WBGene00010248 | -0.726788834 | 0.26311962 | 0.47179667 | F58D5.9 |
| WBGene00010250 | 1.452450081 | 0.39954453 | 1 | F58D12.3 |
| WBGene00010251 | 0.455578443 | 0.18853498 | 0.38226506 | sta-2 |
| WBGene00010254 | 1.584577408 | 0.63258867 | 1 | F58E6.5 |
| WBGene00010255 | -2.704947115 | 0.2331206 | 1 | F58E6.6 |
| WBGene00010256 | -1.692398041 | 0.43695336 | 1 | hrg-3 |
| WBGene00010258 | 0.107750768 | 0.91577749 | 1 | F58E6.11 |
| WBGene00010259 | -0.078150419 | 0.66155237 | 0.80512302 | ric-7 |
| WBGene00010260 | -0.502444595 | 0.00099472 | 0.00920491 | ddx-17 |
| WBGene00010262 | 0.646866668 | 0.09560726 | 0.2478684 | droe-4 |
| WBGene00010263 | 0.181098765 | 0.38161763 | 0.58977507 | wago-4 |
| WBGene00010264 | -0.30688101 | 0.2137372 | 0.41437773 | F58G1.2 |
| WBGene00010265 | -0.008677305 | 0.99719169 | 1 | F58G1.3 |
| WBGene00010266 | -0.174672675 | 0.26473794 | 0.47343988 | dct-18 |
| WBGene00010267 | -0.66625985 | 0.41288154 | 0.61840547 | lips-9 |
| WBGene00010268 | 0.437717098 | 0.44991036 | 0.64865951 | arrd-10 |
| WBGene00010269 | 0.364513758 | 0.7178902 | 1 | F58G1.7 |
| WBGene00010272 | 0.183808506 | 0.41490683 | 0.62017539 | amph-1 |
| WBGene00010273 | -0.156269293 | 0.96924949 | 1 | srm-3 |
| WBGene00010274 | -0.882957691 | 0.04257728 | 0.14375161 | F58G6.3 |
| WBGene00010275 | 0.853525221 | 0.26878323 | 0.47729957 | acc-1 |
| WBGene00010277 | -0.24936108 | 0.54316433 | 0.72328361 | F58G6.7 |
| WBGene00010278 | 1.036568105 | 0.67105582 | 1 | F58G6.8 |
| WBGene00010279 | -0.099178814 | 0.53994619 | 0.72141374 | letm-1 |
| WBGene00010280 | 0.066834328 | 0.71636459 | 0.84006931 | rde-12 |
| WBGene00010281 | -0.383954552 | 0.02811111 | 0.10848024 | F58G11.3 |
| WBGene00010282 | 1.968806565 | 0.10140451 | 0.25670395 | F58G11.4 |

| | | | | |
|---|---|---|---|---|
| WBGene00010283 | -0.078699438 | 0.74148371 | 0.85569007 | ccz-1 |
| WBGene00010284 | 0.279192542 | 0.11498475 | 0.27920598 | aman-2 |
| WBGene00010285 | -1.459713232 | 0.03207113 | 0.11878448 | F58H1.2 |
| WBGene00010286 | -0.394357102 | 0.09514477 | 0.24730307 | F58H1.3 |
| WBGene00010288 | -0.147552785 | 0.74847243 | 0.85978633 | F58H1.5 |
| WBGene00010289 | -1.161473236 | 0.43910809 | 1 | F58H1.6 |
| WBGene00010290 | 0.758163169 | 0.11070565 | 0.27213552 | F58H1.7 |
| WBGene00010291 | 0.692639453 | 0.34912209 | 0.55757581 | F58H10.1 |
| WBGene00010292 | -0.156269293 | 0.96924949 | 1 | F59A1.6 |
| WBGene00010293 | -0.510174541 | 0.26608449 | 0.47472915 | fbxa-108 |
| WBGene00010296 | 0.149428671 | 0.66979351 | 0.81100658 | dgat-2 |
| WBGene00010297 | 3.651557398 | 0.03782289 | 1 | F59A1.11 |
| WBGene00010298 | 1.216957619 | 0.74819621 | 1 | F59A1.12 |
| WBGene00010299 | 1.62869269 | 0.29573141 | 1 | F59A1.13 |
| WBGene00010300 | 1.413734769 | 0.68804531 | 1 | F59A1.15 |
| WBGene00010303 | -0.669714227 | 5.1992E-05 | 0.00089537 | cri-3 |
| WBGene00010304 | -0.488820226 | 0.04791894 | 0.15651974 | clpf-1 |
| WBGene00010305 | -0.446560557 | 0.1607885 | 0.34618428 | F59A2.5 |
| WBGene00010306 | 0.979642013 | 0.00104461 | 0.0095358 | F59A2.2 |
| WBGene00010306 | 0.795609825 | 0.00020034 | 0.00263552 | golg-4 |
| WBGene00010307 | 0.108232116 | 0.58524405 | 0.75465941 | skat-1 |
| WBGene00010308 | -0.295575535 | 0.16004239 | 0.3453741 | F59B2.3 |
| WBGene00010309 | -0.643326306 | 0.02496003 | 0.09988124 | rpn-6.2 |
| WBGene00010310 | 0.00213679 | 0.99830725 | 1 | F59B2.8 |
| WBGene00010311 | -0.74147751 | 0.46088387 | 1 | F59B2.9 |
| WBGene00010314 | 0.491813803 | 0.75255101 | 1 | F59B2.12 |
| WBGene00010315 | 0.334657449 | 0.51702436 | 0.70293312 | F59B2.13 |
| WBGene00010316 | 1.148274042 | 0.10036038 | 0.25533852 | F59B8.1 |
| WBGene00010317 | 0.127515441 | 0.34819666 | 0.55661036 | idh-1 |
| WBGene00010318 | 3.451923409 | 0.05491422 | 1 | F59B10.2 |
| WBGene00010319 | 1.376467084 | 0.08480482 | 0.22947452 | F59B10.3 |
| WBGene00010320 | 1.9499851 | 0.02229603 | 0.09282554 | F59B10.4 |
| WBGene00010321 | 0.921905594 | 0.24705919 | 0.45497554 | F59B10.5 |
| WBGene00010322 | 2.290910071 | 0.06111326 | 1 | F59B10.6 |
| WBGene00010323 | -0.693505305 | 0.79620356 | 1 | dhhc-12 |
| WBGene00010324 | -0.156269293 | 0.96924949 | 1 | F59C6.3 |
| WBGene00010325 | -0.660042258 | 0.00804959 | 0.04408215 | exos-3 |
| WBGene00010326 | -0.25260209 | 0.07560135 | 0.21306313 | F59C6.5 |
| WBGene00010327 | 1.192332606 | 0.40415476 | 1 | F59C6.8 |
| WBGene00010328 | -0.277809743 | 0.64921285 | 0.79802502 | F59C6.11 |
| WBGene00010329 | 1.604728636 | 0.01612416 | 0.07323735 | F59D12.1 |
| WBGene00010330 | 0.106210163 | 0.93993449 | 1 | F59D12.2 |
| WBGene00010331 | 0.57928473 | 0.69114022 | 1 | F59D12.3 |

| | | | | |
|---|---|---|---|---|
| WBGene00010332 | -0.653436367 | 0.12807469 | 0.29879993 | F59D12.5 |
| WBGene00010333 | -0.475626153 | 0.00985146 | 0.05131127 | copz-1 |
| WBGene00010334 | 1.352772571 | 0.00433697 | 0.02800807 | del-5 |
| WBGene00010335 | 1.057640462 | 0.1295108 | 0.30084603 | F59F3.6 |
| WBGene00010336 | 0.943758045 | 0.005956 | 0.03539489 | acox-1.6 |
| WBGene00010337 | 0.164051484 | 0.43584976 | 0.63678123 | F59F4.2 |
| WBGene00010338 | -0.145825516 | 0.67672056 | 0.81495175 | F59F4.3 |
| WBGene00010339 | 0.372617696 | 0.09605633 | 0.24866097 | acl-1 |
| WBGene00010340 | 0.772267079 | 0.07025133 | 0.20259416 | slcf-1 |
| WBGene00010341 | 1.249423892 | 0.05446372 | 0.1705116 | glo-3 |
| WBGene00010342 | 0.548194381 | 0.67957943 | 1 | F59F5.3 |
| WBGene00010343 | -0.156269293 | 0.96924949 | 1 | F59F5.4 |
| WBGene00010344 | 0.654142431 | 0.42403724 | 0.6282934 | F59F5.5 |
| WBGene00010345 | -1.029245282 | 0.24645835 | 0.45441856 | F59F5.7 |
| WBGene00010348 | -0.377584368 | 0.31074505 | 0.51873445 | H01G02.1 |
| WBGene00010349 | 0.587835921 | 0.57721892 | 1 | dyf-18 |
| WBGene00010350 | 0.736222783 | 0.08023292 | 0.22138816 | H01G02.3 |
| WBGene00010351 | 0.03245203 | 0.87245359 | 0.93145085 | cbd-1 |
| WBGene00010352 | -0.263229581 | 0.93409872 | 1 | fbxb-76 |
| WBGene00010353 | -0.533495405 | 0.00566119 | 0.03416471 | memi-3 |
| WBGene00010354 | -0.503225808 | 0.00558716 | 0.03385679 | cyp-31A2 |
| WBGene00010355 | 1.557897343 | 0.63813005 | 1 | clec-229 |
| WBGene00010356 | 0.489875729 | 0.1386135 | 0.31551639 | H03A11.1 |
| WBGene00010357 | 1.324953492 | 0.01094365 | 0.0555401 | H03A11.2 |
| WBGene00010360 | 0.281454817 | 0.94435481 | 1 | fbxa-96 |
| WBGene00010361 | -1.457913559 | 0.28704867 | 1 | H03G16.5 |
| WBGene00010362 | 0.588672918 | 0.23340666 | 0.43934733 | enu-3.1 |
| WBGene00010363 | -0.486215078 | 0.07014706 | 0.20243772 | H04D03.2 |
| WBGene00010364 | 0.341764429 | 0.51297273 | 0.69969688 | H04D03.3 |
| WBGene00010365 | -1.752311476 | 0.00407795 | 0.02667586 | H04D03.4 |
| WBGene00010366 | 1.851444818 | 0.53669729 | 1 | H05L14.1 |
| WBGene00010367 | 0.359251365 | 0.12022138 | 0.28647845 | H05L14.2 |
| WBGene00010368 | -0.216420864 | 0.78094836 | 0.8788112 | H06A10.1 |
| WBGene00010369 | 0.273490891 | 0.17813307 | 0.37013753 | chd-1 |
| WBGene00010370 | 0.073364837 | 0.74330535 | 0.85694333 | ctg-1 |
| WBGene00010373 | 0.28530093 | 0.92084641 | 1 | H08M01.1 |
| WBGene00010374 | 1.491384064 | 2.9446E-05 | 0.00055341 | rga-5 |
| WBGene00010375 | 1.746429222 | 0.162797 | 1 | H09F14.1 |
| WBGene00010380 | -0.438068962 | 0.81868931 | 1 | mpst-2 |
| WBGene00010382 | 0.281454817 | 0.94435481 | 1 | H12D21.6 |
| WBGene00010383 | 0.016555056 | 0.95188648 | 0.97567359 | mpst-3 |
| WBGene00010385 | 0.623946512 | 0.39650915 | 0.60443202 | H12D21.9 |
| WBGene00010386 | 0.505531273 | 0.39402808 | 0.60200405 | H12D21.10 |

| | | | | |
|---|---|---|---|---|
| WBGene00010391 | 0.640945584 | 0.76653737 | 1 | oac-37 |
| WBGene00010394 | 0.574508983 | 0.8551971 | 1 | srz-59 |
| WBGene00010396 | -0.156900212 | 0.87034005 | 0.93024187 | H13N06.2 |
| WBGene00010397 | -0.204204284 | 0.49206816 | 0.68120861 | suox-1 |
| WBGene00010399 | 0.688793783 | 0.86378755 | 1 | clec-13 |
| WBGene00010401 | 0.377828795 | 0.92526585 | 1 | H16D19.3 |
| WBGene00010403 | 0.169097384 | 0.93630233 | 1 | otpl-8 |
| WBGene00010405 | -0.360682583 | 0.04493754 | 0.14902666 | erfa-3 |
| WBGene00010406 | 0.11940907 | 0.52043245 | 0.70551255 | math-33 |
| WBGene00010407 | 1.706570795 | 7.21E-07 | 2.5741E-05 | H19N07.3 |
| WBGene00010408 | 0.25802764 | 0.19213682 | 0.38723983 | mboa-2 |
| WBGene00010409 | -0.231312206 | 0.23594446 | 0.44215444 | H21P03.2 |
| WBGene00010411 | 0.740024603 | 0.37064817 | 0.57811401 | H25K10.1 |
| WBGene00010413 | 1.653460415 | 0.50712258 | 1 | H25K10.4 |
| WBGene00010414 | 0.574508983 | 0.8551971 | 1 | hpa-1 |
| WBGene00010415 | -0.156269293 | 0.96924949 | 1 | irld-39 |
| WBGene00010416 | 0.046370757 | 0.83507534 | 0.91032502 | hxk-2 |
| WBGene00010417 | -0.668264701 | 0.85551153 | 1 | H25P06.4 |
| WBGene00010418 | 0.115278284 | 0.61532573 | 0.77528743 | H27A22.1 |
| WBGene00010419 | -0.114623844 | 0.49010205 | 0.679494 | H28O16.1 |
| WBGene00010420 | -0.028935268 | 0.90083309 | 0.94788411 | mcrs-1 |
| WBGene00010421 | 1.567403951 | 0.03533978 | 0.12675342 | slc-36.4 |
| WBGene00010422 | 0.993450038 | 0.27589156 | 0.48446621 | H32K16.2 |
| WBGene00010423 | -0.64098832 | 0.60388615 | 1 | zmp-5 |
| WBGene00010424 | 0.358492623 | 0.18438623 | 0.37687428 | H36L18.2 |
| WBGene00010425 | 0.993299547 | 2.08E-09 | 1.50E-07 | lpin-1 |
| WBGene00010426 | 1.599057152 | 0.09244993 | 0.24216495 | H37A05.2 |
| WBGene00010427 | 0.308421039 | 0.14424146 | 0.32268943 | hpo-11 |
| WBGene00010428 | -0.234435852 | 0.2345719 | 0.44049946 | dcn-1 |
| WBGene00010432 | 1.353535409 | 0.73149271 | 1 | H39E23.2 |
| WBGene00010433 | -0.006985104 | 0.9790249 | 0.98879805 | H40L08.1 |
| WBGene00010434 | 0.641398382 | 0.35615876 | 0.56465441 | H40L08.3 |
| WBGene00010435 | -0.140587139 | 0.46661949 | 0.66083793 | JC8.2 |
| WBGene00010436 | -0.17045618 | 0.67367306 | 0.8134506 | JC8.4 |
| WBGene00010437 | -0.374508568 | 0.1668673 | 0.35418459 | cox-11 |
| WBGene00010439 | -0.426389845 | 0.07400863 | 0.21013534 | JC8.7 |
| WBGene00010440 | -0.710509322 | 2.9694E-05 | 0.00055464 | ttr-51 |
| WBGene00010442 | -0.284887885 | 0.74824253 | 0.85972621 | bus-12 |
| WBGene00010444 | -0.093418333 | 0.65340709 | 0.80066769 | magi-1 |
| WBGene00010446 | 1.247575581 | 0.7549672 | 1 | K01A6.4 |
| WBGene00010447 | -3.542353468 | 0.17733075 | 1 | K01A6.5 |
| WBGene00010448 | 2.016979909 | 0.02986623 | 0.11287763 | K01A6.6 |
| WBGene00010449 | 0.316327508 | 0.48540088 | 0.67559923 | K01A11.1 |

| | | | | |
|---|---|---|---|---|
| WBGene00010450 | -0.351752947 | 0.25798311 | 0.46592581 | K01A11.2 |
| WBGene00010451 | 1.19979931 | 0.43520418 | 1 | K01A11.3 |
| WBGene00010453 | 1.592392518 | 0.0025584 | 0.0188824 | fozi-1 |
| WBGene00010454 | 0.395019695 | 0.5923097 | 0.75953022 | K01B6.3 |
| WBGene00010455 | 1.247575581 | 0.7549672 | 1 | K01B6.4 |
| WBGene00010456 | -0.145033346 | 0.46136612 | 0.65666477 | K01C8.1 |
| WBGene00010457 | 0.527880608 | 0.76744331 | 1 | K01C8.2 |
| WBGene00010458 | -0.122954415 | 0.58166068 | 0.75247122 | mrpl-10 |
| WBGene00010459 | -0.345885773 | 0.29697385 | 0.50725372 | slc-25A32 |
| WBGene00010460 | 1.349890057 | 0.41871035 | 1 | clec-142 |
| WBGene00010461 | -0.690273133 | 0.30745355 | 0.51614955 | K01D12.1 |
| WBGene00010464 | -1.625090667 | 0.10163367 | 1 | K01D12.5 |
| WBGene00010465 | -0.138898431 | 0.48096347 | 0.67266184 | K01D12.6 |
| WBGene00010466 | 1.557897343 | 0.63813005 | 1 | K01D12.7 |
| WBGene00010467 | -1.792623539 | 0.23891047 | 1 | K01D12.8 |
| WBGene00010468 | -1.363032894 | 0.01219873 | 0.059996 | K01D12.9 |
| WBGene00010470 | 0.097525177 | 0.74470393 | 0.8574972 | cdr-4 |
| WBGene00010471 | -0.222552257 | 0.2967925 | 0.50715803 | cdr-6 |
| WBGene00010472 | 0.714317528 | 0.13662896 | 0.31221761 | cdr-7 |
| WBGene00010473 | 0.713447693 | 0.37510953 | 0.58305016 | hrg-2 |
| WBGene00010474 | 1.025181888 | 0.79790347 | 1 | K01D12.15 |
| WBGene00010475 | 1.457872394 | 0.01646794 | 0.0743261 | K01F9.2 |
| WBGene00010476 | -0.398265318 | 0.06408337 | 0.19067836 | rnf-113 |
| WBGene00010477 | 0.251838021 | 0.80081843 | 1 | enu-3.6 |
| WBGene00010478 | -0.569297497 | 0.00014497 | 0.00205622 | K01G5.5 |
| WBGene00010479 | -0.638813847 | 0.01238731 | 0.06061845 | K01G5.8 |
| WBGene00010480 | -0.105997671 | 0.58978126 | 0.75746714 | K01G5.9 |
| WBGene00010481 | 0.029163559 | 0.95810893 | 0.9785154 | K01G5.10 |
| WBGene00010483 | -0.959947092 | 0.0192889 | 0.0834592 | K01G12.3 |
| WBGene00010484 | -0.758117737 | 0.03763304 | 0.13193842 | dph-3 |
| WBGene00010485 | -0.322086044 | 0.84426026 | 1 | ant-1.3 |
| WBGene00010486 | 1.247575581 | 0.7549672 | 1 | K01H12.4 |
| WBGene00010487 | -2.447607972 | 0.22802076 | 1 | K02A11.2 |
| WBGene00010488 | 0.545997275 | 0.06606115 | 0.19433629 | endu-1 |
| WBGene00010491 | -0.610397053 | 0.10841119 | 0.26850679 | K02B7.3 |
| WBGene00010492 | -0.195257983 | 0.28911679 | 0.49931251 | meg-1 |
| WBGene00010493 | -0.212102939 | 0.44132133 | 0.64137027 | meg-2 |
| WBGene00010494 | -0.156269293 | 0.96924949 | 1 | K02B9.3 |
| WBGene00010495 | -2.001401037 | 9.8401E-06 | 0.00022457 | K02B12.2 |
| WBGene00010496 | 0.008465492 | 0.96665157 | 0.9830824 | sec-12 |
| WBGene00010497 | 0.695273926 | 0.31632019 | 0.5243642 | axl-1 |
| WBGene00010498 | 0.539442231 | 0.00981282 | 0.05113433 | K02B12.5 |
| WBGene00010500 | -0.318174257 | 0.0981452 | 0.25202641 | K02B12.7 |

| | | | | |
|---|---|---|---|---|
| WBGene00010501 | 1.09980275 | 0.47379508 | 1 | K02C4.2 |
| WBGene00010502 | 0.169711544 | 0.2375327 | 0.44379463 | K02C4.3 |
| WBGene00010503 | 1.340351169 | 8.6188E-05 | 0.00135897 | K02C4.5 |
| WBGene00010504 | 0.987927927 | 0.34235636 | 1 | K02D3.1 |
| WBGene00010505 | -0.643412832 | 0.50305719 | 0.69016029 | K02D3.2 |
| WBGene00010506 | 2.364468859 | 0.17999925 | 1 | K02E2.1 |
| WBGene00010507 | 0.557630286 | 0.1438487 | 0.32210225 | K02E2.6 |
| WBGene00010508 | 0.247318604 | 0.91814609 | 1 | K02E2.7 |
| WBGene00010509 | -1.917852065 | 0.20029416 | 1 | K02E2.8 |
| WBGene00010510 | -0.626833103 | 0.80597175 | 1 | ent-3 |
| WBGene00010512 | 0.488572788 | 0.35749766 | 0.5658166 | K02E11.3 |
| WBGene00010513 | 0.281706904 | 0.57905772 | 0.75037463 | K02E11.4 |
| WBGene00010514 | 0.831297914 | 0.17507273 | 0.3653417 | K02E11.5 |
| WBGene00010515 | 0.802693313 | 0.16789877 | 0.35556584 | K02E11.6 |
| WBGene00010516 | 0.369773745 | 0.62334094 | 0.78068727 | K02E11.7 |
| WBGene00010518 | 0.595896719 | 0.79561342 | 1 | K02E11.9 |
| WBGene00010519 | -1.079489731 | 0.00484853 | 0.03049303 | K03A11.1 |
| WBGene00010520 | 0.449400851 | 0.45682704 | 0.65384393 | cpx-2 |
| WBGene00010521 | -0.180165717 | 0.90534843 | 1 | K03A11.4 |
| WBGene00010522 | 0.655402274 | 0.2288247 | 0.4340198 | K03A11.5 |
| WBGene00010523 | -0.263229581 | 0.93409872 | 1 | K03B8.4 |
| WBGene00010524 | 0.388465178 | 0.29558453 | 0.50580575 | K03B8.6 |
| WBGene00010525 | -0.280343843 | 0.65945636 | 0.80368078 | K03B8.8 |
| WBGene00010526 | -2.073308045 | 0.35461841 | 1 | K03B8.11 |
| WBGene00010528 | 0.247318604 | 0.91814609 | 1 | K03D3.2 |
| WBGene00010531 | 0.020328391 | 0.99335814 | 1 | K03D3.5 |
| WBGene00010537 | -0.377770368 | 0.19231507 | 0.38739782 | mys-2 |
| WBGene00010538 | -0.237546507 | 0.82908366 | 1 | ttr-3 |
| WBGene00010539 | -1.012297547 | 3.83E-08 | 1.9721E-06 | ttr-2 |
| WBGene00010540 | 1.642855301 | 1.79E-06 | 5.2878E-05 | K03H1.5 |
| WBGene00010541 | -0.297624902 | 0.55926945 | 0.73605316 | ttr-1 |
| WBGene00010542 | -0.593595435 | 0.01570328 | 0.07187884 | K03H1.7 |
| WBGene00010545 | 1.280248958 | 0.00054419 | 0.00570558 | cbp-2 |
| WBGene00010546 | -0.084421446 | 0.77261547 | 0.87295559 | K03H1.11 |
| WBGene00010547 | 0.688793783 | 0.86378755 | 1 | K03H1.12 |
| WBGene00010550 | 0.070940978 | 0.8185969 | 0.90162251 | K04B12.2 |
| WBGene00010551 | -0.027446076 | 0.8984972 | 0.94694691 | smg-8 |
| WBGene00010552 | -0.055294441 | 0.85628176 | 0.92147818 | K04C1.2 |
| WBGene00010553 | -0.295735754 | 0.57477558 | 0.74810788 | K04C1.3 |
| WBGene00010554 | -0.825742713 | 0.2929554 | 0.50307942 | mlc-6 |
| WBGene00010555 | -0.359257341 | 0.08620396 | 0.23191957 | K04C1.5 |
| WBGene00010556 | -0.26454117 | 0.06490028 | 0.19218397 | rack-1 |
| WBGene00010557 | -0.141712419 | 0.47702272 | 0.66938827 | mspn-1 |

| | | | | |
|---|---|---|---|---|
| WBGene00010558 | 1.369642822 | 0.010906 | 0.05544796 | ptp-4 |
| WBGene00010559 | 2.567745623 | 0.12456439 | 1 | K04D7.6 |
| WBGene00010560 | -0.862520586 | 1.2132E-06 | 3.9364E-05 | eif-2beta |
| WBGene00010562 | -0.018879854 | 0.93596755 | 0.96661977 | cdc-48.3 |
| WBGene00010563 | -0.300941295 | 0.4114098 | 0.61673412 | K04G2.4 |
| WBGene00010564 | -0.624468637 | 0.00208267 | 0.01604426 | ath-1 |
| WBGene00010565 | 0.359953868 | 0.10027125 | 0.25527223 | vacl-14 |
| WBGene00010566 | 1.155867617 | 0.49102955 | 1 | K04G2.7 |
| WBGene00010567 | 0.799989639 | 0.01460326 | 0.06832302 | K04G2.9 |
| WBGene00010568 | -0.985479391 | 0.00162688 | 0.01331893 | K04G2.10 |
| WBGene00010569 | -0.590956987 | 0.13864208 | 0.31551639 | scbp-2 |
| WBGene00010570 | -0.156269293 | 0.96924949 | 1 | K04G11.1 |
| WBGene00010571 | 2.294640672 | 0.06918212 | 1 | K04G11.3 |
| WBGene00010572 | -0.291767804 | 0.57756436 | 0.74906609 | wdr-5.2 |
| WBGene00010573 | 0.009256691 | 0.97873978 | 0.98873753 | K04H4.2 |
| WBGene00010574 | 1.996315266 | 0.49147447 | 1 | K04H4.5 |
| WBGene00010575 | 0.228436111 | 0.63717216 | 0.78970062 | flp-25 |
| WBGene00010578 | -0.156269293 | 0.96924949 | 1 | K04H8.3 |
| WBGene00010579 | -0.364687713 | 0.03805939 | 0.13304897 | K05C4.2 |
| WBGene00010580 | 1.166633605 | 0.58813397 | 1 | K05C4.3 |
| WBGene00010581 | 0.145566695 | 0.76875554 | 0.87045913 | K05C4.4 |
| WBGene00010582 | -0.214684935 | 0.23178578 | 0.43744927 | K05C4.5 |
| WBGene00010583 | -0.659411287 | 0.00040692 | 0.00456601 | K05C4.7 |
| WBGene00010584 | -0.323868354 | 0.91634405 | 1 | K05C4.8 |
| WBGene00010585 | 0.566558652 | 0.31322332 | 0.52122318 | K05C4.9 |
| WBGene00010586 | -1.901856996 | 0.64072034 | 1 | K05C4.10 |
| WBGene00010587 | 0.756718797 | 0.24201516 | 0.44962603 | sol-2 |
| WBGene00010589 | 0.688596152 | 0.47910345 | 0.67113235 | cyp-33D1 |
| WBGene00010591 | -0.156269293 | 0.96924949 | 1 | K05D4.9 |
| WBGene00010592 | 0.485046006 | 0.60438042 | 0.76867617 | K06A4.2 |
| WBGene00010593 | 0.235375281 | 0.35820693 | 0.56656987 | gsnl-1 |
| WBGene00010594 | 1.275616303 | 0.2379356 | 1 | swt-5 |
| WBGene00010595 | 0.017816914 | 0.94927575 | 0.97411952 | haao-1 |
| WBGene00010597 | -0.463328018 | 0.00794698 | 0.04369758 | K06A4.7 |
| WBGene00010599 | -0.156269293 | 0.96924949 | 1 | K06B4.4 |
| WBGene00010600 | -0.737462795 | 0.83433562 | 1 | nhr-196 |
| WBGene00010601 | 0.281454817 | 0.94435481 | 1 | nhr-268 |
| WBGene00010602 | 0.281454817 | 0.94435481 | 1 | nhr-197 |
| WBGene00010603 | 0.928086431 | 0.53256574 | 1 | nhr-198 |
| WBGene00010604 | 0.172775707 | 0.92472633 | 1 | nhr-199 |
| WBGene00010605 | 0.833545448 | 1.51E-07 | 6.687E-06 | K06G5.1 |
| WBGene00010606 | -0.912990398 | 0.08729941 | 0.23376757 | cyp-13B2 |
| WBGene00010607 | 0.075260937 | 0.97569838 | 1 | K06G5.3 |

| | | | | |
|---|---|---|---|---|
| WBGene00010608 | -0.400473162 | 0.08634563 | 0.23218657 | K07A1.1 |
| WBGene00010609 | -0.676960984 | 2.6462E-05 | 0.00050915 | dut-1 |
| WBGene00010610 | -0.083930011 | 0.88011153 | 0.93592311 | K07A1.3 |
| WBGene00010611 | 1.025181888 | 0.79790347 | 1 | K07A1.4 |
| WBGene00010612 | 1.413734769 | 0.68804531 | 1 | K07A1.5 |
| WBGene00010613 | -0.665920998 | 0.11782373 | 0.2828748 | K07A1.6 |
| WBGene00010614 | 1.084450517 | 0.51640318 | 1 | cri-1 |
| WBGene00010615 | -0.370606407 | 0.057916 | 0.17827225 | ate-1 |
| WBGene00010616 | -0.742441457 | 0.00240358 | 0.01800237 | K07A1.10 |
| WBGene00010617 | -0.266145987 | 0.60561522 | 0.76897128 | K07A1.13 |
| WBGene00010619 | -0.231618251 | 0.66490543 | 0.80771846 | K07A1.15 |
| WBGene00010620 | 0.063621875 | 0.72037578 | 0.8424275 | K07A12.1 |
| WBGene00010621 | -0.254654563 | 0.14951965 | 0.33096045 | egg-6 |
| WBGene00010622 | 0.068676614 | 0.75722853 | 0.86429398 | K07A12.4 |
| WBGene00010623 | -0.208703297 | 0.82661573 | 1 | K07A12.5 |
| WBGene00010624 | -0.329090065 | 0.11542492 | 0.27988295 | mrps-15 |
| WBGene00010625 | 0.145531837 | 0.61052556 | 0.77204363 | K07C5.2 |
| WBGene00010626 | -0.254451937 | 0.19416808 | 0.39032673 | K07C5.3 |
| WBGene00010627 | -0.271120015 | 0.0696569 | 0.20164626 | K07C5.4 |
| WBGene00010628 | 0.36495213 | 0.33947332 | 0.5477137 | ceeh-2 |
| WBGene00010629 | 0.056995234 | 0.72103265 | 0.84287092 | K07C5.6 |
| WBGene00010630 | 0.415708032 | 0.46860589 | 0.6623989 | ttll-15 |
| WBGene00010631 | -0.118093191 | 0.43113909 | 0.63332845 | cash-1 |
| WBGene00010632 | 0.326344549 | 0.67063579 | 0.81161419 | K07C5.9 |
| WBGene00010633 | 1.54277118 | 0.33642676 | 1 | nspd-2 |
| WBGene00010634 | -0.156269293 | 0.96924949 | 1 | K07F5.6 |
| WBGene00010636 | -1.062801831 | 0.64455942 | 1 | K07F5.8 |
| WBGene00010637 | -0.636559384 | 0.03733321 | 0.13121898 | K07F5.12 |
| WBGene00010638 | -0.180473659 | 0.42931972 | 0.63203235 | K07F5.14 |
| WBGene00010639 | 0.06647906 | 0.75551902 | 0.86341205 | K07F5.15 |
| WBGene00010640 | -0.041045011 | 0.93928168 | 0.96880629 | K07F5.16 |
| WBGene00010641 | 0.149269921 | 0.34774544 | 0.55644893 | crml-1 |
| WBGene00010642 | -0.203725131 | 0.78536755 | 0.88190412 | mks-6 |
| WBGene00010643 | 0.865213084 | 0.53444848 | 1 | K07G5.4 |
| WBGene00010644 | 0.59844179 | 0.33183051 | 0.53982957 | K07G5.5 |
| WBGene00010645 | -0.065488279 | 0.74568332 | 0.85813661 | fecl-1 |
| WBGene00010646 | 0.152824832 | 0.73679351 | 0.85283726 | K08C7.1 |
| WBGene00010647 | 0.352459812 | 0.67399941 | 0.81366479 | K08C7.4 |
| WBGene00010648 | -0.040185099 | 0.93074412 | 0.96384889 | K08C7.6 |
| WBGene00010649 | 1.568347007 | 0.14501046 | 1 | K08C7.7 |
| WBGene00010650 | 0.983488853 | 0.80601565 | 1 | K08C9.1 |
| WBGene00010651 | 1.106782842 | 0.54712526 | 1 | K08C9.2 |
| WBGene00010652 | -1.204791816 | 0.57474638 | 1 | K08C9.5 |

| | | | | |
|---|---|---|---|---|
| WBGene00010654 | -0.07748118 | 0.98067866 | 1 | K08C9.7 |
| WBGene00010655 | 0.498258903 | 0.72590761 | 1 | K08D8.1 |
| WBGene00010657 | 1.41735009 | 6.15E-08 | 3.0336E-06 | K08D8.3 |
| WBGene00010658 | 0.887128512 | 0.00726387 | 0.04084791 | K08D8.4 |
| WBGene00010659 | -0.358214074 | 0.46725721 | 0.66143986 | K08D8.5 |
| WBGene00010660 | -0.17863588 | 0.40555755 | 0.61226986 | K08D8.6 |
| WBGene00010661 | 0.514290795 | 0.10109847 | 0.2562498 | tyr-2 |
| WBGene00010662 | 0.232157325 | 0.93397866 | 1 | K08E3.2 |
| WBGene00010663 | -0.361242574 | 0.09802123 | 0.25176131 | toca-2 |
| WBGene00010664 | -0.209710667 | 0.27082613 | 0.47962106 | dbn-1 |
| WBGene00010665 | -0.296038134 | 0.05950862 | 0.18139981 | K08E3.5 |
| WBGene00010666 | -0.384865471 | 0.24066733 | 0.44815063 | K08E4.2 |
| WBGene00010667 | -0.213206896 | 0.54833441 | 0.72686189 | K08E4.3 |
| WBGene00010669 | 0.402935294 | 0.81737892 | 1 | bath-37 |
| WBGene00010670 | -0.548886838 | 0.00988677 | 0.05134099 | K08E4.6 |
| WBGene00010671 | -0.339549509 | 0.05565773 | 0.17340019 | eak-7 |
| WBGene00010672 | 0.574508983 | 0.8551971 | 1 | K08E7.4 |
| WBGene00010673 | -0.328059312 | 0.63099607 | 0.78581631 | K08E7.5 |
| WBGene00010674 | 0.586053674 | 0.43498305 | 0.63619337 | K08E7.6 |
| WBGene00010675 | 0.118960039 | 0.88269427 | 0.93749089 | K08E7.8 |
| WBGene00010676 | -0.069563566 | 0.67553759 | 0.81441045 | ctf-18 |
| WBGene00010677 | -0.328110399 | 0.09047097 | 0.23924447 | gtbp-1 |
| WBGene00010678 | -0.261011706 | 0.21017992 | 0.4103223 | K08F4.3 |
| WBGene00010679 | 2.332054728 | 0.29723791 | 1 | K08F4.5 |
| WBGene00010680 | 0.948894776 | 0.49227115 | 1 | K08F4.12 |
| WBGene00010681 | 0.215973065 | 0.17857866 | 0.37068276 | mak-1 |
| WBGene00010682 | -1.285450047 | 0.51018772 | 1 | K08F8.5 |
| WBGene00010683 | 0.898371066 | 0.45072355 | 1 | K08F8.7 |
| WBGene00010684 | 1.766634207 | 0.09286975 | 0.24300251 | K08F9.1 |
| WBGene00010685 | -0.723828059 | 3.5797E-06 | 9.5729E-05 | aipl-1 |
| WBGene00010686 | -0.217627353 | 0.90765874 | 1 | K08F9.3 |
| WBGene00010687 | -0.32774339 | 0.08006558 | 0.22102688 | K08F9.4 |
| WBGene00010688 | -0.156269293 | 0.96924949 | 1 | srz-12 |
| WBGene00010689 | -1.280983903 | 0.09789331 | 0.25153918 | K08H2.2 |
| WBGene00010690 | -1.014055289 | 0.00472715 | 0.02998555 | K08H2.3 |
| WBGene00010691 | -0.979344909 | 0.39177771 | 1 | K08H2.4 |
| WBGene00010692 | -0.558804446 | 0.87818675 | 1 | K08H2.5 |
| WBGene00010693 | 1.219060141 | 0.46174049 | 1 | K08H2.7 |
| WBGene00010694 | 2.931759211 | 0.18286765 | 1 | bgnt-1.5 |
| WBGene00010697 | 0.22167447 | 0.22573965 | 0.42998197 | uda-1 |
| WBGene00010698 | -0.241757247 | 0.85998645 | 1 | K08H10.6 |
| WBGene00010699 | -0.262686285 | 0.33479071 | 0.54319044 | trpp-6 |
| WBGene00010700 | 0.786844184 | 0.00043966 | 0.00482436 | nipi-3 |

| | | | | |
|---|---|---|---|---|
| WBGene00010701 | 0.086954133 | 0.69141675 | 0.82420135 | ent-2 |
| WBGene00010702 | 0.141972166 | 0.58804663 | 0.75642763 | usp-33 |
| WBGene00010703 | 1.139334217 | 0.0002854 | 0.00346507 | K09A9.6 |
| WBGene00010704 | -0.093153557 | 0.81662192 | 0.90047871 | K09A11.1 |
| WBGene00010705 | 1.506308787 | 0.09171066 | 0.24116507 | cyp-14A1 |
| WBGene00010706 | 0.496980034 | 0.62043881 | 1 | cyp-14A2 |
| WBGene00010707 | 0.688793783 | 0.86378755 | 1 | cyp-14A3 |
| WBGene00010708 | 0.872657085 | 0.59509309 | 1 | phf-33 |
| WBGene00010709 | 0.056634265 | 0.83589002 | 0.91067307 | nol-9 |
| WBGene00010711 | 0.804149846 | 0.51766274 | 1 | K09B11.4 |
| WBGene00010712 | 0.938982574 | 0.8146943 | 1 | K09B11.5 |
| WBGene00010713 | 0.178803447 | 0.39510341 | 0.60293495 | uso-1 |
| WBGene00010714 | -1.060726336 | 0.22696524 | 0.43143652 | mam-3 |
| WBGene00010715 | -1.901856996 | 0.64072034 | 1 | K09C8.2 |
| WBGene00010716 | 1.006816103 | 0.2903548 | 0.50031189 | lge-1 |
| WBGene00010717 | -0.127603812 | 0.94147805 | 1 | K09C8.7 |
| WBGene00010718 | -0.394320382 | 0.8739308 | 1 | K09C8.8 |
| WBGene00010719 | -0.806757004 | 0.69468682 | 1 | K09E4.1 |
| WBGene00010720 | -0.025925358 | 0.90940982 | 0.95279443 | K09E4.2 |
| WBGene00010721 | -0.324272937 | 0.30237025 | 0.51171971 | K09E4.3 |
| WBGene00010722 | 0.668057638 | 0.18489478 | 0.37727857 | K09E4.4 |
| WBGene00010723 | 0.369891308 | 0.4157509 | 0.62082449 | cpg-7 |
| WBGene00010724 | 0.63622338 | 0.06431976 | 0.19116431 | K09E9.1 |
| WBGene00010725 | -0.270023545 | 0.33295349 | 0.54093247 | erv-46 |
| WBGene00010726 | -0.217766436 | 0.4465128 | 0.64575484 | K09E9.3 |
| WBGene00010727 | -0.080398499 | 0.71540269 | 0.83960406 | K09G1.1 |
| WBGene00010728 | 0.377828795 | 0.92526585 | 1 | K09G1.2 |
| WBGene00010730 | -0.284946385 | 0.11747334 | 0.28245932 | ensa-1 |
| WBGene00010731 | 0.511941904 | 0.01561087 | 0.07165653 | K10C3.4 |
| WBGene00010732 | 0.389419554 | 0.0705022 | 0.20317284 | K10C3.5 |
| WBGene00010734 | -0.119707786 | 0.70481306 | 0.83305445 | parn-1 |
| WBGene00010735 | 1.837185949 | 0.0907032 | 1 | frpr-15 |
| WBGene00010736 | 0.066861502 | 0.70861404 | 0.83518911 | istr-1 |
| WBGene00010737 | 2.697061674 | 0.24723036 | 1 | K10C8.4 |
| WBGene00010738 | -0.073184605 | 0.79425988 | 0.88635851 | K10D3.4 |
| WBGene00010740 | 1.51668741 | 0.01130868 | 0.05687488 | K10D3.6 |
| WBGene00010741 | 2.303864289 | 0.09848435 | 1 | lgc-49 |
| WBGene00010742 | -0.012999429 | 0.96168297 | 0.98000612 | K10D6.2 |
| WBGene00010744 | 0.288017987 | 0.43675799 | 0.63755094 | K10D6.4 |
| WBGene00010745 | 1.154271625 | 0.00017316 | 0.002349 | dod-17 |
| WBGene00010746 | 0.695474669 | 0.65279923 | 1 | K10D11.2 |
| WBGene00010747 | 1.527007917 | 0.07274656 | 0.20781552 | K10D11.3 |
| WBGene00010749 | 1.237050648 | 0.00016909 | 0.00231186 | K10D11.5 |

| | | | | |
|---|---|---|---|---|
| WBGene00010750 | 0.4789959 | 0.32400963 | 0.53253025 | K10D11.6 |
| WBGene00010752 | 1.659145938 | 0.18990869 | 1 | K10G4.3 |
| WBGene00010754 | 2.742072101 | 0.03597963 | 1 | K10G4.5 |
| WBGene00010755 | 0.256793107 | 0.43755326 | 0.63804213 | vglu-2 |
| WBGene00010756 | 0.658475676 | 0.25353449 | 0.46142442 | K10G9.2 |
| WBGene00010757 | -0.40360251 | 0.35302024 | 0.56194939 | pad-2 |
| WBGene00010758 | -0.402835602 | 0.09958409 | 0.25432963 | vnut-1 |
| WBGene00010759 | -1.269071263 | 0.00026822 | 0.00331612 | cysl-2 |
| WBGene00010760 | -1.38496508 | 0.03451978 | 0.12501383 | K10H10.4 |
| WBGene00010761 | -0.156269293 | 0.96924949 | 1 | K10H10.5 |
| WBGene00010762 | -1.14715222 | 0.68932198 | 1 | K10H10.6 |
| WBGene00010763 | 0.32595502 | 0.91626577 | 1 | K10H10.7 |
| WBGene00010765 | 0.719981201 | 0.52351451 | 1 | K10H10.10 |
| WBGene00010766 | -0.347571274 | 0.08489526 | 0.22965983 | K11B4.1 |
| WBGene00010767 | -0.698541257 | 0.01694858 | 0.07590517 | blos-9 |
| WBGene00010768 | 0.087171101 | 0.91243093 | 0.95467585 | K11D2.1 |
| WBGene00010769 | 3.44950616 | 1.80E-40 | 2.42E-37 | asah-1 |
| WBGene00010770 | 0.158163573 | 0.6551239 | 0.80180033 | K11D2.4 |
| WBGene00010771 | -0.190142767 | 0.63627777 | 0.78915603 | K11D2.5 |
| WBGene00010772 | 0.737068299 | 0.02340461 | 0.09573969 | K11D9.3 |
| WBGene00010773 | 0.205677418 | 0.75151244 | 0.86174089 | K11E4.1 |
| WBGene00010774 | -0.235248617 | 0.69953164 | 0.830279 | K11E4.2 |
| WBGene00010775 | 1.062657457 | 0.66313467 | 1 | poml-1 |
| WBGene00010776 | -0.147360584 | 0.67869961 | 0.81641653 | pix-1 |
| WBGene00010778 | -0.125404683 | 0.42685079 | 0.63044468 | gpdh-2 |
| WBGene00010779 | -1.063648221 | 0.47634469 | 1 | K11H3.2 |
| WBGene00010780 | 0.248905842 | 0.11476459 | 0.27886303 | K11H3.3 |
| WBGene00010781 | -0.451959266 | 0.08644253 | 0.23239566 | K11H3.4 |
| WBGene00010782 | 2.069700719 | 0.1442713 | 1 | K11H3.5 |
| WBGene00010783 | -0.45597594 | 0.12982233 | 0.30114118 | mrpl-36 |
| WBGene00010784 | 0.383179469 | 0.49798255 | 0.68626631 | twk-48 |
| WBGene00010785 | -0.273706657 | 0.10507701 | 0.26380066 | top-2 |
| WBGene00010786 | 0.367144296 | 0.86127653 | 1 | K12D12.4 |
| WBGene00010787 | -0.258597185 | 0.28020146 | 0.48920419 | K12D12.5 |
| WBGene00010788 | 1.34945069 | 0.04244825 | 0.14359547 | sulp-4 |
| WBGene00010789 | 0.761434785 | 0.34375603 | 0.5522019 | sulp-5 |
| WBGene00010790 | 2.760999851 | 4.98E-39 | 6.05E-36 | sodh-1 |
| WBGene00010791 | 1.181686151 | 0.7579682 | 1 | sodh-2 |
| WBGene00010792 | -0.500354182 | 0.62832735 | 1 | K12G11.6 |
| WBGene00010793 | 0.242605413 | 0.13141724 | 0.30385388 | LLC1.2 |
| WBGene00010794 | -0.201302508 | 0.19712065 | 0.39427377 | dld-1 |
| WBGene00010795 | 1.992898638 | 0.00278998 | 0.02017462 | M01A8.1 |
| WBGene00010796 | 0.875576782 | 0.00041073 | 0.00460021 | clip-1 |

| | | | | |
|---|---|---|---|---|
| WBGene00010799 | 0.323850126 | 0.67789775 | 0.81569441 | chil-12 |
| WBGene00010800 | -0.62565913 | 0.80069199 | 1 | srsx-37 |
| WBGene00010803 | -0.129944978 | 0.77488576 | 0.87450158 | M01B2.10 |
| WBGene00010804 | -0.156269293 | 0.96924949 | 1 | M01E5.1 |
| WBGene00010805 | -0.197975225 | 0.60009034 | 0.76538468 | M01E5.2 |
| WBGene00010806 | -0.225342183 | 0.30636997 | 0.5151853 | M01E5.3 |
| WBGene00010807 | -0.574796676 | 0.01959218 | 0.08421189 | M01E5.4 |
| WBGene00010808 | -1.948984258 | 7.12E-11 | 6.97E-09 | sepa-1 |
| WBGene00010809 | 0.271939654 | 0.18071881 | 0.37259848 | M01F1.3 |
| WBGene00010810 | 0.214421309 | 0.30760488 | 0.51626084 | M01F1.4 |
| WBGene00010811 | 0.847878834 | 0.00846251 | 0.04566373 | hmit-1.3 |
| WBGene00010812 | -0.522912889 | 0.11003502 | 0.27076109 | mrpl-35 |
| WBGene00010813 | 0.541706618 | 0.03826978 | 0.13343866 | pitp-1 |
| WBGene00010814 | -0.090092631 | 0.74469342 | 0.8574972 | M01F1.8 |
| WBGene00010815 | 0.1859685 | 0.37553579 | 0.58326469 | M01F1.9 |
| WBGene00010822 | 0.650959026 | 0.16967856 | 0.35789913 | M01G12.9 |
| WBGene00010825 | 0.143798628 | 0.94792951 | 1 | M01G12.14 |
| WBGene00010827 | 0.323049527 | 0.43230392 | 0.6341955 | M02B1.2 |
| WBGene00010828 | -0.006955519 | 0.966677 | 0.9830824 | M02B1.3 |
| WBGene00010829 | 0.377828795 | 0.92526585 | 1 | M02B1.4 |
| WBGene00010830 | -0.227731181 | 0.73489576 | 0.85185883 | M02G9.1 |
| WBGene00010831 | -2.257942945 | 0.08485557 | 1 | M02G9.2 |
| WBGene00010832 | -0.782134186 | 0.62113659 | 1 | M02G9.3 |
| WBGene00010833 | -0.160008348 | 0.81409794 | 0.8986238 | M03B6.1 |
| WBGene00010834 | 0.58943874 | 0.00234601 | 0.01761466 | mct-3 |
| WBGene00010835 | -0.588977058 | 0.24891004 | 0.4567087 | M03B6.3 |
| WBGene00010836 | -0.335067558 | 0.52127236 | 0.70633569 | M03B6.4 |
| WBGene00010837 | 0.814613271 | 0.62325601 | 1 | M03B6.5 |
| WBGene00010838 | 0.392297124 | 0.34787841 | 0.55652286 | M03C11.1 |
| WBGene00010839 | -0.062442835 | 0.77891264 | 0.87725216 | chl-1 |
| WBGene00010840 | -0.066498638 | 0.76886613 | 0.87045913 | M03C11.3 |
| WBGene00010842 | 0.026367324 | 0.89516925 | 0.94481514 | ymel-1 |
| WBGene00010843 | -0.234021332 | 0.59900916 | 0.76480755 | M03C11.6 |
| WBGene00010844 | -0.232859733 | 0.1906677 | 0.3853665 | prp-3 |
| WBGene00010845 | 0.062224713 | 0.74968599 | 0.86036058 | M03C11.8 |
| WBGene00010846 | 0.055564517 | 0.85573068 | 0.92113008 | M04B2.2 |
| WBGene00010847 | -0.297507332 | 0.1636251 | 0.34976101 | M04B2.4 |
| WBGene00010848 | 1.188284994 | 0.2492059 | 1 | M04B2.6 |
| WBGene00010849 | 0.732715285 | 0.46844876 | 0.66233102 | M04B2.7 |
| WBGene00010850 | 1.273865825 | 2.3073E-05 | 0.00045624 | M04C3.1 |
| WBGene00010851 | 2.244260769 | 0.00028695 | 0.00348041 | M04C3.2 |
| WBGene00010854 | -0.156269293 | 0.96924949 | 1 | M04C7.3 |
| WBGene00010855 | -0.060379368 | 0.9741779 | 1 | M04C7.4 |

| | | | | |
|---|---|---|---|---|
| WBGene00010856 | 0.746479636 | 0.19702603 | 0.39421441 | M04C9.1 |
| WBGene00010857 | 1.181686151 | 0.7579682 | 1 | M04C9.2 |
| WBGene00010858 | -0.019802671 | 0.96636553 | 0.9830824 | M04C9.3 |
| WBGene00010859 | 0.5619546 | 0.50438452 | 0.69147838 | M04C9.4 |
| WBGene00010862 | 0.281454817 | 0.94435481 | 1 | M04D8.4 |
| WBGene00010863 | 1.216957619 | 0.74819621 | 1 | M04D8.5 |
| WBGene00010864 | 0.65855128 | 0.39489173 | 0.60276345 | xbx-3 |
| WBGene00010865 | 1.385030609 | 0.27228689 | 1 | M04D8.7 |
| WBGene00010866 | -0.156269293 | 0.96924949 | 1 | M04D8.8 |
| WBGene00010867 | 0.941060478 | 0.00303588 | 0.02140451 | tag-260 |
| WBGene00010868 | 1.015362914 | 9.746E-05 | 0.00149971 | somi-1 |
| WBGene00010869 | 0.508362135 | 0.74005871 | 1 | M05B5.1 |
| WBGene00010870 | 0.637091274 | 0.08778729 | 0.23481504 | let-522 |
| WBGene00010871 | -0.01585467 | 0.98873049 | 1 | M05B5.3 |
| WBGene00010872 | -0.213856072 | 0.37082457 | 0.57831485 | M05B5.4 |
| WBGene00010873 | 0.40027844 | 0.48652422 | 0.67622374 | trpa-2 |
| WBGene00010875 | -0.300958662 | 0.06891436 | 0.20016575 | M05D6.2 |
| WBGene00010876 | 0.394439984 | 0.88400378 | 1 | M05D6.3 |
| WBGene00010877 | 0.593182513 | 0.11130589 | 0.27305816 | lact-4 |
| WBGene00010878 | -0.553015541 | 0.05219184 | 0.16514495 | M05D6.5 |
| WBGene00010879 | -0.425673616 | 0.07712876 | 0.21546715 | M05D6.6 |
| WBGene00010880 | 0.747845955 | 0.85215965 | 1 | M05D6.8 |
| WBGene00010881 | -0.412322454 | 0.72416832 | 1 | M05D6.9 |
| WBGene00010882 | -0.038836409 | 0.85393801 | 0.9200161 | atg-7 |
| WBGene00010883 | 0.232157325 | 0.93397866 | 1 | M7.7 |
| WBGene00010884 | 1.602276743 | 0.02952266 | 0.11197584 | M7.8 |
| WBGene00010885 | -1.517638116 | 0.44619746 | 1 | M7.9 |
| WBGene00010886 | 0.377828795 | 0.92526585 | 1 | M7.10 |
| WBGene00010887 | 0.398810102 | 0.35684138 | 0.56521999 | M7.12 |
| WBGene00010888 | 0.537081578 | 0.15895837 | 0.34378709 | dlc-2 |
| WBGene00010889 | -0.329387964 | 0.18331092 | 0.37537155 | M18.3 |
| WBGene00010890 | -0.143296113 | 0.35230159 | 0.56125097 | ddb-1 |
| WBGene00010891 | -0.194512298 | 0.45830767 | 0.65477918 | M18.6 |
| WBGene00010892 | -0.565192547 | 0.0011113 | 0.00993542 | dhhc-6 |
| WBGene00010893 | -0.32610956 | 0.68551905 | 0.82023128 | cutl-9 |
| WBGene00010894 | 0.688793783 | 0.86378755 | 1 | M28.2 |
| WBGene00010895 | -0.774842772 | 0.78069886 | 1 | M28.4 |
| WBGene00010896 | -0.687915867 | 1.1948E-06 | 3.8994E-05 | M28.5 |
| WBGene00010897 | 1.349850671 | 1.49E-10 | 1.38E-08 | lact-3 |
| WBGene00010898 | 0.551676871 | 0.54060305 | 0.72174866 | nphp-1 |
| WBGene00010899 | 2.257968481 | 2.3145E-06 | 6.6276E-05 | M28.8 |
| WBGene00010900 | 0.241932487 | 0.89878987 | 1 | M28.9 |
| WBGene00010901 | 0.520590314 | 0.01602195 | 0.07293682 | M28.10 |

| | | | | |
|---|---|---|---|---|
| WBGene00010902 | 0.318049553 | 0.23376883 | 0.43982447 | M79.2 |
| WBGene00010904 | -0.724184686 | 6.9263E-05 | 0.00114256 | ugt-62 |
| WBGene00010905 | -0.488148465 | 0.03476754 | 0.12559142 | mrps-34 |
| WBGene00010906 | 0.559152564 | 0.55413912 | 0.73180205 | M88.3 |
| WBGene00010907 | 0.189990407 | 0.84384868 | 1 | M88.4 |
| WBGene00010908 | 0.201127299 | 0.28827722 | 0.49835878 | zbp-1 |
| WBGene00010909 | -0.56800135 | 0.00741073 | 0.04138625 | cisd-3.1 |
| WBGene00010910 | -0.099160329 | 0.6302845 | 0.78556152 | M106.2 |
| WBGene00010911 | -0.35310694 | 0.07455452 | 0.21099452 | M106.3 |
| WBGene00010912 | -0.150618787 | 0.41141007 | 0.61673412 | gmps-1 |
| WBGene00010913 | -0.493580441 | 0.0268019 | 0.10486685 | M110.3 |
| WBGene00010915 | -0.558804446 | 0.87818675 | 1 | M110.7 |
| WBGene00010916 | 0.175909247 | 0.94229417 | 1 | M110.8 |
| WBGene00010917 | -1.304318597 | 0.44065094 | 1 | M110.9 |
| WBGene00010918 | 0.646605192 | 0.02960508 | 0.11210187 | M117.1 |
| WBGene00010919 | 1.557897343 | 0.63813005 | 1 | M117.3 |
| WBGene00010920 | -0.745110582 | 0.61155837 | 1 | M117.4 |
| WBGene00010922 | -0.406282175 | 0.15162074 | 0.33336244 | M142.5 |
| WBGene00010923 | -0.003483 | 0.9875824 | 0.99429786 | rle-1 |
| WBGene00010924 | -1.01828374 | 0.0283787 | 0.1091371 | M153.1 |
| WBGene00010925 | 0.597322709 | 0.31976799 | 0.52791721 | M153.2 |
| WBGene00010926 | -1.571079866 | 0.29781995 | 1 | M153.3 |
| WBGene00010928 | -0.014519762 | 0.99603746 | 1 | clec-258 |
| WBGene00010931 | -0.667041166 | 0.52021109 | 1 | M162.5 |
| WBGene00010933 | 0.747845955 | 0.85215965 | 1 | M162.7 |
| WBGene00010934 | 0.983488853 | 0.80601565 | 1 | fbxa-118 |
| WBGene00010935 | 1.066361533 | 0.001288 | 0.01113788 | M163.1 |
| WBGene00010936 | 1.581817634 | 0.0787781 | 0.21866597 | ztf-14 |
| WBGene00010937 | 1.935297182 | 0.50713048 | 1 | M163.5 |
| WBGene00010939 | 0.688793783 | 0.86378755 | 1 | M163.7 |
| WBGene00010940 | 0.980153662 | 0.11735107 | 0.28241018 | M163.8 |
| WBGene00010941 | -0.4555857 | 0.01027597 | 0.05290947 | gss-1 |
| WBGene00010942 | -0.316026299 | 0.12515129 | 0.29486936 | chch-3 |
| WBGene00010943 | 0.360968911 | 0.13250816 | 0.30538755 | M176.4 |
| WBGene00010944 | 0.414942767 | 0.26692244 | 0.47531613 | M176.5 |
| WBGene00010945 | 1.212244691 | 0.04600925 | 0.15166938 | chil-13 |
| WBGene00010946 | 1.601302495 | 0.65842342 | 1 | M176.10 |
| WBGene00010947 | 0.542683821 | 0.20346975 | 0.40270523 | M176.11 |
| WBGene00010948 | -0.07840981 | 0.94123274 | 1 | M195.2 |
| WBGene00010950 | 0.377828795 | 0.92526585 | 1 | srt-44 |
| WBGene00010951 | -0.545172465 | 0.10798894 | 0.26762475 | M199.2 |
| WBGene00010953 | -2.642098596 | 1.17E-13 | 1.85E-11 | clec-190 |
| WBGene00010955 | -1.969805064 | 0.62862569 | 1 | M199.7 |

| | | | | |
|---|---|---|---|---|
| WBGene00010956 | -2.934840138 | 0.2654436 | 1 | M199.8 |
| WBGene00010968 | 0.276018865 | 0.85764239 | 1 | R01E6.2 |
| WBGene00010970 | 0.780309961 | 0.17724033 | 0.36885612 | glb-20 |
| WBGene00010971 | 0.078462227 | 0.93188649 | 0.96406579 | R01E6.7 |
| WBGene00010972 | -0.179897219 | 0.75420881 | 0.86279555 | R01H10.4 |
| WBGene00010973 | 0.226235919 | 0.80487225 | 0.89364005 | rip-1 |
| WBGene00010974 | 0.275353452 | 0.66839175 | 0.81011722 | bbs-5 |
| WBGene00010975 | -0.252284715 | 0.25764636 | 0.46553184 | R01H10.7 |
| WBGene00010976 | 0.141612104 | 0.79486442 | 0.88641923 | R02D5.1 |
| WBGene00010977 | 0.387696481 | 0.40939412 | 0.61485082 | R02D5.3 |
| WBGene00010978 | 0.720066222 | 0.75158338 | 1 | R02D5.4 |
| WBGene00010979 | 1.988712883 | 0.17279138 | 1 | R02D5.6 |
| WBGene00010980 | 0.570903644 | 0.74231661 | 1 | R02D5.7 |
| WBGene00010981 | -0.800507535 | 0.11415548 | 0.2778592 | blos-8 |
| WBGene00010982 | 0.911759526 | 0.12244509 | 0.29046616 | flp-32 |
| WBGene00010983 | -0.009864583 | 0.9847526 | 0.99257382 | mocs-1 |
| WBGene00010984 | -0.139793505 | 0.60545475 | 0.76897128 | nkat-3 |
| WBGene00010985 | 0.299392341 | 0.42022069 | 0.62469687 | R03A10.5 |
| WBGene00010986 | 1.388524278 | 0.10337704 | 0.26055649 | sprr-1 |
| WBGene00010987 | -0.086687407 | 0.93628405 | 1 | R03C1.1 |
| WBGene00010988 | 1.087114438 | 1.13E-11 | 1.28E-09 | metr-1 |
| WBGene00010989 | 0.131705737 | 0.73032047 | 0.84906836 | R03D7.2 |
| WBGene00010990 | -0.574363407 | 0.00101505 | 0.00935034 | R03D7.4 |
| WBGene00010991 | 1.378887293 | 0.47680059 | 1 | R03D7.5 |
| WBGene00010992 | 1.116613983 | 0.69067248 | 1 | R03D7.8 |
| WBGene00010993 | 0.347276123 | 0.05348336 | 0.16809253 | vha-20 |
| WBGene00010994 | -0.434437035 | 0.52338614 | 0.7083303 | lgc-25 |
| WBGene00010995 | -0.156269293 | 0.96924949 | 1 | ceh-90 |
| WBGene00010996 | -1.969805064 | 0.62862569 | 1 | R03G8.1 |
| WBGene00010998 | 0.414748364 | 0.4333696 | 0.63529827 | R03G8.3 |
| WBGene00010999 | -0.514761297 | 0.79912167 | 1 | R03G8.4 |
| WBGene00011000 | -0.582331246 | 0.09631221 | 0.24913639 | R03G8.6 |
| WBGene00011002 | 2.444080733 | 0.15900319 | 1 | nhr-205 |
| WBGene00011003 | 1.008684167 | 0.02747284 | 0.10670115 | R04B5.5 |
| WBGene00011004 | 0.186674096 | 0.91267482 | 1 | R04B5.6 |
| WBGene00011005 | -1.901856996 | 0.64072034 | 1 | R04B5.7 |
| WBGene00011006 | 0.309047473 | 0.25472831 | 0.46230167 | ugt-47 |
| WBGene00011009 | -1.0062442 | 0.63208617 | 1 | cyp-14A4 |
| WBGene00011010 | -0.505086241 | 0.02628902 | 0.1035946 | R04D3.2 |
| WBGene00011011 | -0.171445431 | 0.40400465 | 0.6112175 | R04D3.3 |
| WBGene00011012 | -0.535870031 | 0.09083068 | 0.23962958 | R04D3.4 |
| WBGene00011013 | -1.901856996 | 0.64072034 | 1 | srxa-8 |
| WBGene00011014 | 0.846355049 | 0.5334603 | 1 | hpo-30 |

| | | | | |
|---|---|---|---|---|
| WBGene00011015 | -0.640560941 | 7.3507E-05 | 0.00119723 | R04F11.2 |
| WBGene00011016 | -0.001887954 | 0.99085383 | 0.99552766 | R04F11.3 |
| WBGene00011017 | -0.495008084 | 0.08032549 | 0.22154288 | R04F11.5 |
| WBGene00011018 | 0.476654751 | 0.17181405 | 0.3610861 | R05A10.1 |
| WBGene00011019 | 0.221230836 | 0.57700115 | 0.74903703 | R05A10.2 |
| WBGene00011020 | 0.981049828 | 0.08393982 | 0.22783665 | R05A10.3 |
| WBGene00011021 | -0.309817771 | 0.7492364 | 0.86026851 | R05A10.4 |
| WBGene00011023 | 1.413734769 | 0.68804531 | 1 | R05A10.6 |
| WBGene00011026 | 0.323513582 | 0.5072401 | 0.69409515 | R05D7.1 |
| WBGene00011027 | 0.938982574 | 0.8146943 | 1 | R05D7.2 |
| WBGene00011028 | -0.0953768 | 0.86992483 | 0.92996626 | R05D7.3 |
| WBGene00011029 | -0.099159157 | 0.69541252 | 0.82774543 | abhd-11.2 |
| WBGene00011030 | -0.128318891 | 0.71596441 | 0.84001971 | R05D7.5 |
| WBGene00011032 | -0.378940365 | 0.0969496 | 0.25017324 | R05D11.4 |
| WBGene00011033 | -0.416984891 | 0.10538696 | 0.26426767 | R05D11.5 |
| WBGene00011034 | -0.446595762 | 0.04929976 | 0.15938142 | paxt-1 |
| WBGene00011035 | -0.782806484 | 0.00039387 | 0.00445683 | snrp-27 |
| WBGene00011036 | -0.077597482 | 0.61761527 | 0.77712375 | edc-3 |
| WBGene00011037 | -0.081437619 | 0.68075235 | 0.81694319 | R05D11.9 |
| WBGene00011038 | -0.907075672 | 3.07E-11 | 3.24E-09 | R05H5.3 |
| WBGene00011039 | -0.135141072 | 0.74567184 | 0.85813661 | R05H5.4 |
| WBGene00011040 | -0.222725694 | 0.4350282 | 0.63619337 | R05H5.5 |
| WBGene00011041 | 1.022098293 | 0.57130722 | 1 | R05H5.7 |
| WBGene00011042 | 0.849929641 | 0.17337484 | 0.36304655 | R05H10.1 |
| WBGene00011043 | 0.001762907 | 0.99341064 | 0.99669716 | rbm-28 |
| WBGene00011044 | -0.06379853 | 0.80561223 | 0.89402761 | R05H10.3 |
| WBGene00011045 | -0.175438947 | 0.33135211 | 0.53963058 | gpx-2 |
| WBGene00011046 | 4.656690101 | 0.00167222 | 1 | R05H10.7 |
| WBGene00011047 | -0.123165671 | 0.62754756 | 0.78361153 | R06A4.2 |
| WBGene00011049 | -2.242695067 | 6.125E-05 | 0.00102997 | sdz-26 |
| WBGene00011050 | -0.074403335 | 0.72095325 | 0.84287092 | agl-1 |
| WBGene00011051 | -0.391487444 | 0.03516993 | 0.12648047 | pfs-2 |
| WBGene00011052 | 0.281454817 | 0.94435481 | 1 | arrd-11 |
| WBGene00011053 | -0.156269293 | 0.96924949 | 1 | arrd-12 |
| WBGene00011054 | -0.439503297 | 0.07969993 | 0.22043917 | arrd-13 |
| WBGene00011055 | 0.398867148 | 0.17233172 | 0.36179827 | arrd-14 |
| WBGene00011056 | -0.898094067 | 0.02769388 | 0.10733993 | R06B9.5 |
| WBGene00011058 | -0.259613351 | 0.23746397 | 0.44375097 | fdps-1 |
| WBGene00011059 | -0.146702508 | 0.40435085 | 0.6114365 | R06C1.4 |
| WBGene00011060 | 0.731435331 | 0.09404908 | 0.24506652 | R06C1.6 |
| WBGene00011061 | -0.332238257 | 0.03489745 | 0.12590615 | wago-1 |
| WBGene00011062 | -0.550114278 | 0.25078633 | 0.45885383 | R06C7.2 |
| WBGene00011063 | -0.428494641 | 0.00287517 | 0.02058219 | cpg-3 |

| | | | | |
|---|---|---|---|---|
| WBGene00011064 | -0.142702 | 0.42836649 | 0.63100978 | adsl-1 |
| WBGene00011065 | -0.099077345 | 0.85728881 | 0.92198985 | R06C7.6 |
| WBGene00011066 | -0.019457974 | 0.92769938 | 0.96200873 | ztf-15 |
| WBGene00011067 | 0.357171052 | 0.08068494 | 0.22222266 | vps-11 |
| WBGene00011068 | -0.21841412 | 0.35703123 | 0.56537318 | set-14 |
| WBGene00011069 | 0.737766666 | 0.46368757 | 1 | ceh-62 |
| WBGene00011070 | 1.698393649 | 1.8389E-05 | 0.00037524 | msrp-7 |
| WBGene00011071 | 0.199346971 | 0.33185586 | 0.53982957 | R06F6.8 |
| WBGene00011072 | 0.202699782 | 0.50139775 | 0.68918217 | tag-209 |
| WBGene00011073 | 0.482691738 | 0.59058765 | 0.75812272 | R07A4.2 |
| WBGene00011074 | 0.306766063 | 0.88218984 | 1 | R07A4.3 |
| WBGene00011075 | -0.236828563 | 0.85479986 | 1 | R07A4.4 |
| WBGene00011076 | 0.638092216 | 0.06159477 | 0.18583439 | scav-5 |
| WBGene00011077 | 2.332054728 | 0.29723791 | 1 | R07B1.5 |
| WBGene00011078 | 0.178115923 | 0.94982316 | 1 | R07B1.6 |
| WBGene00011079 | -0.323868354 | 0.91634405 | 1 | R07B1.7 |
| WBGene00011081 | 1.718019253 | 0.00373437 | 0.02496096 | R07B1.9 |
| WBGene00011082 | 0.761121007 | 0.42295288 | 0.6273758 | R07B1.11 |
| WBGene00011083 | -0.854404174 | 0.12267106 | 0.29085945 | glo-1 |
| WBGene00011084 | 0.377828795 | 0.92526585 | 1 | srsx-21 |
| WBGene00011087 | -0.172481439 | 0.43241341 | 0.63420285 | R07B7.2 |
| WBGene00011088 | 1.761877163 | 0.15039082 | 1 | kmo-2 |
| WBGene00011089 | 0.409638522 | 0.20893431 | 0.40894268 | kmo-1 |
| WBGene00011090 | 0.546499715 | 0.3100032 | 0.51830255 | R07B7.6 |
| WBGene00011092 | 0.370569423 | 0.3553352 | 0.5636643 | R07B7.8 |
| WBGene00011093 | 0.845218397 | 0.14975857 | 0.33124774 | R07B7.9 |
| WBGene00011094 | -0.500704039 | 0.21061149 | 0.41041347 | R07B7.10 |
| WBGene00011095 | 0.318519878 | 0.33146259 | 0.53971087 | gana-1 |
| WBGene00011096 | 0.377828795 | 0.92526585 | 1 | R07B7.12 |
| WBGene00011097 | -0.111412738 | 0.91609224 | 1 | nhr-206 |
| WBGene00011098 | -0.094731856 | 0.97157968 | 1 | nhr-207 |
| WBGene00011099 | 0.092219558 | 0.93722365 | 1 | nhr-208 |
| WBGene00011100 | 1.012941359 | 0.23237718 | 0.437977 | nhr-209 |
| WBGene00011101 | 1.173317286 | 0.36125321 | 1 | R07D5.2 |
| WBGene00011102 | 0.443746912 | 0.05009075 | 0.16092931 | R07E3.1 |
| WBGene00011103 | -0.768324725 | 0.48942794 | 1 | R07E3.2 |
| WBGene00011104 | -0.194097906 | 0.81720646 | 0.90082044 | cut-5 |
| WBGene00011105 | -0.395963175 | 0.22275663 | 0.42657544 | R07E3.4 |
| WBGene00011106 | -0.229190915 | 0.36903347 | 0.57663261 | acl-5 |
| WBGene00011107 | -0.084315518 | 0.8679987 | 0.9285087 | R07E3.6 |
| WBGene00011108 | 0.719141525 | 0.43960622 | 0.6398824 | R07E3.7 |
| WBGene00011109 | 0.061376695 | 0.76421862 | 0.86811174 | R07E5.1 |
| WBGene00011110 | -0.285470216 | 0.11226972 | 0.27460243 | prdx-3 |

| | | | | |
|---|---|---|---|---|
| WBGene00011111 | -0.624026354 | 0.00036635 | 0.00423609 | snfc-5 |
| WBGene00011112 | -1.23169031 | 6.7526E-05 | 0.00111847 | R07E5.4 |
| WBGene00011113 | -3.135088683 | 0.0344878 | 1 | R07E5.5 |
| WBGene00011114 | 0.820015846 | 0.7518774 | 1 | R07E5.6 |
| WBGene00011115 | -0.209584408 | 0.26114219 | 0.46968486 | R07E5.7 |
| WBGene00011116 | -0.260295316 | 0.1685489 | 0.35643429 | pdcd-2 |
| WBGene00011117 | -1.05963765 | 0.00036204 | 0.0041982 | R07E5.11 |
| WBGene00011119 | -0.31374241 | 0.11753586 | 0.28245932 | mpc-1 |
| WBGene00011120 | 1.216957619 | 0.74819621 | 1 | R07E5.15 |
| WBGene00011121 | 0.042984862 | 0.97797978 | 1 | R07E5.17 |
| WBGene00011122 | -0.025334914 | 0.86722548 | 0.92835056 | cpt-2 |
| WBGene00011123 | -0.438388061 | 0.12625463 | 0.29631885 | nuaf-3 |
| WBGene00011124 | 0.412735641 | 0.14027263 | 0.31785181 | sdz-27 |
| WBGene00011127 | -3.014567441 | 0.24693748 | 1 | fbxb-79 |
| WBGene00011128 | 0.361581327 | 0.00487756 | 0.03063553 | R07H5.8 |
| WBGene00011129 | -0.302223264 | 0.84605978 | 1 | R07H5.9 |
| WBGene00011130 | 0.247318604 | 0.91814609 | 1 | zip-6 |
| WBGene00011131 | -0.779360318 | 0.09872513 | 0.25290776 | R07H5.11 |
| WBGene00011132 | 0.938982574 | 0.8146943 | 1 | R08A2.1 |
| WBGene00011133 | 0.169097384 | 0.93630233 | 1 | R08A2.2 |
| WBGene00011135 | -0.156269293 | 0.96924949 | 1 | fbxa-147 |
| WBGene00011136 | -1.160528285 | 0.31246563 | 1 | R08A2.5 |
| WBGene00011137 | 0.490028316 | 0.5851755 | 0.75465941 | R08A2.7 |
| WBGene00011139 | -0.719391349 | 0.08857687 | 0.23599117 | R08B4.3 |
| WBGene00011140 | -0.963811488 | 0.61154562 | 1 | R08B4.4 |
| WBGene00011141 | 1.486637752 | 0.14282643 | 1 | R08B4.5 |
| WBGene00011142 | -0.196751202 | 0.33696414 | 0.54507071 | R08D7.1 |
| WBGene00011143 | -0.204390842 | 0.38278328 | 0.59119346 | R08D7.2 |
| WBGene00011144 | -0.310289995 | 0.16357653 | 0.34976101 | R08D7.4 |
| WBGene00011145 | -0.352461378 | 0.19443464 | 0.3905455 | R08D7.5 |
| WBGene00011146 | 0.603963448 | 0.00249419 | 0.01854373 | pde-2 |
| WBGene00011147 | 0.015968448 | 0.97833698 | 0.98873753 | R08D7.7 |
| WBGene00011150 | 0.281454817 | 0.94435481 | 1 | nhr-269 |
| WBGene00011152 | -0.609125369 | 0.14189538 | 0.31997923 | R09A8.1 |
| WBGene00011153 | -0.23632266 | 0.35831424 | 0.56659198 | R09A8.2 |
| WBGene00011154 | 2.164012294 | 0.00186928 | 0.01480426 | R09A8.5 |
| WBGene00011155 | -0.816053198 | 1.85E-07 | 7.87E-06 | R09B3.2 |
| WBGene00011156 | -0.987090518 | 1.11E-11 | 1.28E-09 | R09B3.3 |
| WBGene00011159 | 0.574508983 | 0.8551971 | 1 | chil-17 |
| WBGene00011161 | 3.712769976 | 0.0333263 | 1 | chil-18 |
| WBGene00011162 | 1.134359074 | 0.53140648 | 1 | chil-19 |
| WBGene00011163 | 0.747845955 | 0.85215965 | 1 | chil-20 |
| WBGene00011164 | 0.688793783 | 0.86378755 | 1 | chil-21 |

| | | | | |
|---|---|---|---|---|
| WBGene00011166 | 1.568380329 | 0.00385854 | 0.02548781 | chil-22 |
| WBGene00011167 | 0.23037324 | 0.8804679 | 1 | chil-23 |
| WBGene00011168 | 1.404017331 | 0.31848278 | 1 | R09D1.12 |
| WBGene00011171 | 2.194319406 | 0.0836533 | 1 | R09E10.1 |
| WBGene00011172 | 0.281454817 | 0.94435481 | 1 | R09E10.2 |
| WBGene00011173 | 3.271881254 | 0.07721527 | 1 | acs-18 |
| WBGene00011175 | -0.360583431 | 0.30724489 | 0.51598983 | R09E10.5 |
| WBGene00011176 | 0.889942757 | 0.36357979 | 0.57164235 | R09E10.6 |
| WBGene00011179 | 2.415700506 | 0.16475802 | 1 | R09H10.1 |
| WBGene00011180 | 0.456047972 | 0.51542149 | 0.70177552 | R09H10.2 |
| WBGene00011181 | -0.158289358 | 0.53370396 | 0.71678095 | R09H10.3 |
| WBGene00011182 | 1.769538063 | 5.04E-20 | 1.65E-17 | R09H10.5 |
| WBGene00011183 | 0.281454817 | 0.94435481 | 1 | R09H10.6 |
| WBGene00011184 | 1.580657229 | 0.00078211 | 0.00761385 | R09H10.7 |
| WBGene00011185 | 0.748081073 | 0.12569526 | 0.29569195 | R10D12.1 |
| WBGene00011187 | 0.377828795 | 0.92526585 | 1 | R10D12.6 |
| WBGene00011188 | 0.633811015 | 0.76987404 | 1 | R10D12.7 |
| WBGene00011189 | -0.230180419 | 0.64615036 | 0.79554928 | R10D12.8 |
| WBGene00011190 | -2.494474155 | 3.0879E-06 | 8.4818E-05 | swt-6 |
| WBGene00011191 | -0.06995528 | 0.97031639 | 1 | R10D12.10 |
| WBGene00011192 | -0.323868354 | 0.91634405 | 1 | srh-24 |
| WBGene00011193 | -0.248351376 | 0.42770642 | 0.6306514 | algn-13 |
| WBGene00011194 | -0.24531534 | 0.3471518 | 0.55603825 | R10D12.13 |
| WBGene00011195 | -0.251420319 | 0.14004917 | 0.31746397 | sao-1 |
| WBGene00011197 | 2.201543524 | 0.41352885 | 1 | srw-145 |
| WBGene00011198 | -0.045894679 | 0.85568549 | 0.92113008 | R10E4.1 |
| WBGene00011200 | -0.060074225 | 0.97239791 | 1 | R10E4.3 |
| WBGene00011201 | -0.734543398 | 0.02162014 | 0.0907458 | nth-1 |
| WBGene00011202 | -0.345792765 | 0.59592072 | 0.76214826 | R10E4.6 |
| WBGene00011203 | 1.121198024 | 0.41201743 | 1 | R10E4.7 |
| WBGene00011205 | 0.304696037 | 0.44625395 | 0.64561618 | R10E4.9 |
| WBGene00011206 | 0.013184987 | 0.95741749 | 0.97850563 | R10E4.11 |
| WBGene00011207 | 0.45979801 | 0.2354553 | 0.44169697 | R10E8.1 |
| WBGene00011208 | -0.156269293 | 0.96924949 | 1 | R10E8.2 |
| WBGene00011209 | 1.400541691 | 0.00070016 | 0.00697916 | R10E8.3 |
| WBGene00011212 | 1.471296782 | 0.00501116 | 0.03119541 | R10E8.6 |
| WBGene00011213 | 0.601571345 | 0.31484065 | 0.52262514 | R10E8.8 |
| WBGene00011214 | 2.640518312 | 0.05507071 | 1 | R10E9.2 |
| WBGene00011215 | 0.110869247 | 0.88443573 | 0.93895892 | R10E9.3 |
| WBGene00011216 | -0.384304429 | 0.05501478 | 0.17180699 | usp-46 |
| WBGene00011217 | 0.987120971 | 0.032267 | 0.11936429 | R10E11.5 |
| WBGene00011218 | 0.205465488 | 0.51476771 | 0.70152895 | R10E11.6 |
| WBGene00011219 | -0.62146648 | 0.85925752 | 1 | srxa-10 |

| | | | | |
|---|---|---|---|---|
| WBGene00011220 | 0.232157325 | 0.93397866 | 1 | R10E11.9 |
| WBGene00011221 | 0.234599009 | 0.48579183 | 0.67583347 | R10E12.2 |
| WBGene00011222 | 0.471697793 | 0.03577271 | 0.12755255 | R10H10.3 |
| WBGene00011223 | 0.964598423 | 0.28704213 | 0.49711057 | R10H10.4 |
| WBGene00011224 | -0.273412669 | 0.48520827 | 0.67540857 | R10H10.6 |
| WBGene00011225 | 0.302141006 | 0.20724642 | 0.4068473 | R10H10.7 |
| WBGene00011226 | -0.769564418 | 0.01269665 | 0.06174859 | slc-25A21 |
| WBGene00011227 | 0.847360547 | 0.48130945 | 1 | R11.2 |
| WBGene00011228 | 0.766937602 | 0.42267964 | 0.62712373 | R11.3 |
| WBGene00011229 | -1.858602649 | 0.00436749 | 0.02816024 | R11.4 |
| WBGene00011230 | 0.011210131 | 0.95489328 | 0.97697363 | nud-2 |
| WBGene00011231 | -1.431588413 | 0.00150496 | 0.01260991 | R11A5.3 |
| WBGene00011232 | 0.054020628 | 0.72004047 | 0.8421165 | pck-2 |
| WBGene00011234 | 0.460141183 | 0.82127868 | 1 | R11A5.6 |
| WBGene00011235 | -0.422726816 | 0.38906209 | 0.59747063 | suro-1 |
| WBGene00011236 | 0.608968151 | 0.03572362 | 0.12744529 | R11A8.1 |
| WBGene00011237 | -0.040758895 | 0.82218581 | 0.90368983 | gkow-1 |
| WBGene00011238 | 1.672544132 | 0.25210114 | 1 | ugt-59 |
| WBGene00011239 | -0.593644259 | 0.00142668 | 0.01206216 | pges-2 |
| WBGene00011240 | 0.367507285 | 0.10889442 | 0.26904502 | R11A8.7 |
| WBGene00011241 | 1.557897343 | 0.63813005 | 1 | mpz-5 |
| WBGene00011242 | 0.145604464 | 0.45174023 | 0.6495237 | R11D1.1 |
| WBGene00011243 | 0.80378416 | 0.66158793 | 1 | R11D1.2 |
| WBGene00011244 | 0.460181607 | 0.26107683 | 0.46968486 | R11D1.3 |
| WBGene00011245 | 0.688793783 | 0.86378755 | 1 | R11D1.4 |
| WBGene00011246 | 1.832280005 | 0.31803877 | 1 | R11D1.7 |
| WBGene00011247 | -0.603475946 | 0.02850488 | 0.10941439 | mrpl-49 |
| WBGene00011248 | 0.193730921 | 0.54166009 | 0.72227294 | R11D1.10 |
| WBGene00011249 | 2.865268649 | 0.16077931 | 1 | R11G10.2 |
| WBGene00011250 | -0.066945956 | 0.71348041 | 0.8379961 | R11H6.2 |
| WBGene00011251 | -0.420264113 | 0.62528501 | 0.78207328 | glb-22 |
| WBGene00011252 | 0.269087565 | 0.62384392 | 0.78115605 | R11H6.4 |
| WBGene00011253 | -0.516766657 | 0.00523171 | 0.03224274 | R11H6.5 |
| WBGene00011254 | 0.77011019 | 0.36277255 | 0.57074272 | R12G8.1 |
| WBGene00011255 | 1.741745596 | 0.36346624 | 1 | R12H7.4 |
| WBGene00011257 | 1.88180802 | 0.31552007 | 1 | npax-3 |
| WBGene00011258 | 0.078305609 | 0.89021769 | 0.94196731 | best-15 |
| WBGene00011259 | 0.532212724 | 0.00670703 | 0.03853765 | miz-1 |
| WBGene00011260 | -0.799547206 | 0.63771263 | 1 | cnnm-5 |
| WBGene00011261 | 1.624784815 | 0.03587868 | 0.1278178 | nphp-4 |
| WBGene00011262 | 1.677649005 | 0.00043907 | 0.00482425 | pho-8 |
| WBGene00011263 | 0.670772285 | 0.07330507 | 0.2086256 | R13H4.5 |
| WBGene00011265 | 1.994826872 | 0.31690381 | 1 | R13H4.7 |

| | | | | |
|---|---|---|---|---|
| WBGene00011266 | -1.455291304 | 0.14876832 | 1 | R13H4.8 |
| WBGene00011267 | -0.961002701 | 0.33730577 | 1 | fbxb-85 |
| WBGene00011268 | -0.078049629 | 0.71315793 | 0.8379961 | pde-12 |
| WBGene00011269 | -0.007492476 | 0.98928346 | 0.99484196 | R17.3 |
| WBGene00011270 | -0.028336371 | 0.8726563 | 0.93158535 | R31.2 |
| WBGene00011271 | -0.048821204 | 0.81586034 | 0.90015998 | flad-1 |
| WBGene00011272 | -0.693445399 | 0.00115581 | 0.01025782 | dtmk-1 |
| WBGene00011273 | -0.260551123 | 0.04914782 | 0.15918637 | R53.4 |
| WBGene00011274 | -0.171005072 | 0.44642863 | 0.64575484 | R53.5 |
| WBGene00011275 | -0.813190829 | 0.00012934 | 0.00187619 | psf-1 |
| WBGene00011276 | -0.03266602 | 0.88163031 | 0.93708197 | aakg-5 |
| WBGene00011278 | -1.272569705 | 0.02934077 | 0.11146004 | R74.2 |
| WBGene00011279 | -0.27810879 | 0.18852633 | 0.38226506 | asd-1 |
| WBGene00011280 | -0.231939547 | 0.4537301 | 0.65122795 | pelo-1 |
| WBGene00011281 | -0.498614127 | 0.00644977 | 0.03752118 | R74.7 |
| WBGene00011282 | -0.219445646 | 0.15291836 | 0.33536522 | R74.8 |
| WBGene00011283 | 0.223971182 | 0.35393896 | 0.56253081 | R90.1 |
| WBGene00011284 | 0.687119697 | 0.13172372 | 0.30421489 | ttr-27 |
| WBGene00011285 | -0.096298412 | 0.88537855 | 0.93946696 | ttr-28 |
| WBGene00011286 | 0.851724631 | 0.30128711 | 0.51098347 | ttr-29 |
| WBGene00011287 | 0.189373081 | 0.73924445 | 0.85446788 | glb-24 |
| WBGene00011288 | 2.666642872 | 0.26363578 | 1 | R102.1 |
| WBGene00011289 | 0.524426486 | 0.05991004 | 0.18211512 | R102.2 |
| WBGene00011290 | 1.477253415 | 0.58357327 | 1 | R102.3 |
| WBGene00011291 | -0.00176604 | 0.99362542 | 0.99674513 | R102.4 |
| WBGene00011292 | -0.215543597 | 0.15400997 | 0.33708944 | R102.5 |
| WBGene00011293 | 1.837185949 | 0.0907032 | 1 | R102.6 |
| WBGene00011296 | 1.735991307 | 0.58152819 | 1 | glb-21 |
| WBGene00011297 | 1.928677659 | 0.30173618 | 1 | R102.10 |
| WBGene00011298 | -0.26162144 | 0.47655274 | 0.66906261 | R107.2 |
| WBGene00011299 | 0.27082332 | 0.12167817 | 0.28892913 | ikke-1 |
| WBGene00011300 | -0.622849929 | 8.2012E-05 | 0.00130842 | R107.5 |
| WBGene00011302 | -0.217955245 | 0.26682364 | 0.47520989 | R166.2 |
| WBGene00011303 | -0.591104368 | 0.04595192 | 0.15152153 | R166.3 |
| WBGene00011304 | 0.827186876 | 0.00063936 | 0.00647414 | mnk-1 |
| WBGene00011305 | -0.153890423 | 0.84075345 | 0.9124319 | R186.1 |
| WBGene00011306 | -0.130518491 | 0.51516582 | 0.70168394 | R186.3 |
| WBGene00011307 | 0.907834068 | 0.14739279 | 0.32762648 | mpst-7 |
| WBGene00011308 | -0.576646703 | 0.00178634 | 0.0142684 | pnn-1 |
| WBGene00011309 | -0.640912727 | 0.0659788 | 0.19419361 | R186.8 |
| WBGene00011310 | 0.675351329 | 0.21575701 | 0.41750027 | T01B4.3 |
| WBGene00011311 | -0.235505786 | 0.20465436 | 0.40406162 | T01B7.5 |
| WBGene00011312 | -0.133644647 | 0.49325169 | 0.68206934 | trcs-2 |

| | | | | |
|---|---|---|---|---|
| WBGene00011313 | -0.283794653 | 0.88299492 | 1 | T01B7.8 |
| WBGene00011314 | 1.719127963 | 0.10652306 | 1 | T01B7.9 |
| WBGene00011315 | -0.207272322 | 0.74468117 | 0.8574972 | mbr-1 |
| WBGene00011317 | -0.421207803 | 0.67225556 | 0.81256417 | T01C1.4 |
| WBGene00011318 | -0.38742764 | 0.12251401 | 0.29057288 | cdt-2 |
| WBGene00011319 | -0.709887828 | 0.00202846 | 0.01571636 | T01C3.2 |
| WBGene00011320 | -0.743931991 | 4.4301E-06 | 0.00011347 | T01C3.3 |
| WBGene00011321 | 3.057162191 | 0.00082254 | 0.00789446 | fil-1 |
| WBGene00011322 | -0.62565913 | 0.80069199 | 1 | irld-14 |
| WBGene00011323 | 0.172127358 | 0.4707089 | 0.66356678 | mut-15 |
| WBGene00011324 | -0.522292408 | 0.0161908 | 0.07348505 | T01C3.9 |
| WBGene00011325 | -0.049994784 | 0.97986415 | 1 | T01C3.11 |
| WBGene00011326 | 0.702132704 | 0.04074299 | 0.13926257 | T01D3.1 |
| WBGene00011327 | 0.707559163 | 0.78139902 | 1 | hlh-34 |
| WBGene00011328 | 1.155780728 | 0.00156492 | 0.01294256 | T01D3.3 |
| WBGene00011329 | -0.115757678 | 0.5744829 | 0.74795819 | zipt-9 |
| WBGene00011330 | 1.145887387 | 7.09E-08 | 3.4574E-06 | T01D3.6 |
| WBGene00011331 | 0.935526815 | 0.00465033 | 0.02962208 | T01E8.1 |
| WBGene00011333 | 0.027490222 | 0.86775844 | 0.92844324 | nrde-2 |
| WBGene00011334 | -0.431336062 | 0.05431423 | 0.1701839 | mrps-14 |
| WBGene00011335 | 0.258985233 | 0.6273966 | 0.78350361 | T01E8.8 |
| WBGene00011336 | 1.413734769 | 0.68804531 | 1 | ubxn-5 |
| WBGene00011337 | 0.911877827 | 0.02864217 | 0.10962943 | T01G1.2 |
| WBGene00011338 | 0.033118454 | 0.86001778 | 0.92435161 | sec-31 |
| WBGene00011339 | 1.081932062 | 0.0031006 | 0.02171498 | T01G5.1 |
| WBGene00011340 | 0.456725074 | 0.66614248 | 1 | ugt-30 |
| WBGene00011342 | -0.510787148 | 0.12960521 | 0.30086746 | T01G5.7 |
| WBGene00011344 | -0.092104199 | 0.64112497 | 0.79235567 | T01G9.2 |
| WBGene00011345 | 0.564756232 | 0.2523537 | 0.46037961 | dma-1 |
| WBGene00011348 | 0.113089443 | 0.53079535 | 0.71445525 | T01H3.2 |
| WBGene00011349 | 0.183913943 | 0.38693163 | 0.59555752 | T01H3.3 |
| WBGene00011350 | -0.070962734 | 0.69570741 | 0.82777182 | perm-1 |
| WBGene00011351 | -1.557923972 | 0.16613858 | 1 | T01H3.5 |
| WBGene00011352 | -0.143495403 | 0.43755364 | 0.63804213 | rskn-1 |
| WBGene00011353 | -0.46721561 | 0.19485925 | 0.39078066 | T01H8.2 |
| WBGene00011354 | 2.41989273 | 0.16470062 | 1 | lgc-13 |
| WBGene00011355 | -1.837668733 | 0.43198255 | 1 | lgc-14 |
| WBGene00011360 | 1.557897343 | 0.63813005 | 1 | lgc-18 |
| WBGene00011361 | 0.838648632 | 0.03254829 | 0.12021133 | lyst-1 |
| WBGene00011362 | 2.965894998 | 0.0002523 | 0.00315469 | cest-1 |
| WBGene00011363 | -0.156269293 | 0.96924949 | 1 | ocam-1 |
| WBGene00011364 | 0.407785642 | 0.47478373 | 0.66786575 | T02B5.3 |
| WBGene00011365 | -1.935789379 | 0.03481531 | 0.12572655 | T02C1.1 |

| | | | | |
|---|---|---|---|---|
| WBGene00011366 | 0.055086969 | 0.93803607 | 0.96795424 | T02C1.2 |
| WBGene00011367 | -0.416717995 | 0.06173915 | 0.18592589 | snpc-3.4 |
| WBGene00011368 | -0.219100332 | 0.27274863 | 0.48138409 | tftc-5 |
| WBGene00011371 | 0.544991291 | 0.68957753 | 1 | T02D1.4 |
| WBGene00011372 | 0.581848618 | 0.40268963 | 0.61053455 | npr-26 |
| WBGene00011373 | 1.247575581 | 0.7549672 | 1 | T02D1.7 |
| WBGene00011374 | 0.055875708 | 0.83789245 | 0.91143626 | T02D1.8 |
| WBGene00011375 | 0.097567087 | 0.64610175 | 0.79554928 | T02E1.2 |
| WBGene00011376 | -0.2342708 | 0.31267667 | 0.52091097 | gla-3 |
| WBGene00011378 | 1.33864768 | 0.5127736 | 1 | T02E1.6 |
| WBGene00011379 | 0.377828795 | 0.92526585 | 1 | T02E1.7 |
| WBGene00011380 | 1.320891621 | 0.09380831 | 0.24460725 | T02E1.8 |
| WBGene00011381 | 2.153107102 | 0.03848289 | 0.1339509 | npr-25 |
| WBGene00011382 | 1.376565249 | 0.02487999 | 0.09974122 | dop-5 |
| WBGene00011383 | 0.097865459 | 0.73678485 | 0.85283726 | T02E9.5 |
| WBGene00011387 | -1.159714244 | 0.75790634 | 1 | T02G6.4 |
| WBGene00011388 | -0.377287607 | 0.30521251 | 0.51430743 | T02G6.5 |
| WBGene00011390 | 1.025181888 | 0.79790347 | 1 | T02G6.7 |
| WBGene00011391 | -0.23579296 | 0.38891482 | 0.5973956 | mrps-12 |
| WBGene00011392 | 0.72910827 | 0.00033654 | 0.0039746 | sbt-1 |
| WBGene00011393 | 0.08261967 | 0.79530698 | 0.88666869 | T03D8.6 |
| WBGene00011394 | 0.134715183 | 0.83219318 | 0.90915448 | T03D8.7 |
| WBGene00011396 | -0.62565913 | 0.80069199 | 1 | nhr-271 |
| WBGene00011397 | 1.514120544 | 0.01920943 | 0.08332046 | T03E6.8 |
| WBGene00011398 | -0.187605059 | 0.30959867 | 0.51818605 | qdpr-1 |
| WBGene00011399 | -0.465070686 | 0.00582273 | 0.03485888 | T03F6.3 |
| WBGene00011400 | 0.727362174 | 0.22067422 | 0.42372381 | T03F6.4 |
| WBGene00011401 | 2.550000603 | 0.29207489 | 1 | T03F6.6 |
| WBGene00011402 | 0.377828795 | 0.92526585 | 1 | T03F7.5 |
| WBGene00011404 | 0.391811008 | 0.27872314 | 0.48767511 | T03F7.7 |
| WBGene00011407 | -0.190240358 | 0.59310133 | 0.75990326 | ppat-1 |
| WBGene00011408 | -0.56896658 | 0.00379235 | 0.02521516 | T04A8.6 |
| WBGene00011409 | -0.125424601 | 0.42492232 | 0.62893095 | T04A8.7 |
| WBGene00011410 | -0.565249833 | 0.00214484 | 0.01641257 | T04A8.8 |
| WBGene00011411 | -0.640908656 | 0.01367453 | 0.06497946 | sel-13 |
| WBGene00011412 | -0.43232188 | 0.04868053 | 0.15821805 | mrpl-16 |
| WBGene00011414 | -0.245531902 | 0.94070824 | 1 | T04A8.13 |
| WBGene00011415 | -0.060552142 | 0.75177668 | 0.86196247 | him-18 |
| WBGene00011418 | -0.416657139 | 0.69206556 | 1 | igdb-1 |
| WBGene00011422 | -0.15289061 | 0.78607641 | 0.88221054 | T04B2.3 |
| WBGene00011423 | 0.388413847 | 0.05907314 | 0.18062033 | ipla-7 |
| WBGene00011424 | -0.006008648 | 0.99214452 | 0.99616496 | dhs-31 |
| WBGene00011425 | 1.69047418 | 0.38809581 | 1 | T04B2.7 |

| | | | | |
|---|---|---|---|---|
| WBGene00011426 | -0.333693239 | 0.89573179 | 1 | T04C10.3 |
| WBGene00011427 | -1.570331237 | 0.05792637 | 0.17827225 | T04C12.1 |
| WBGene00011428 | 0.905088734 | 0.01941908 | 0.08378358 | T04C12.3 |
| WBGene00011429 | 2.5172226 | 0.15684696 | 1 | T04C12.7 |
| WBGene00011430 | 0.175898348 | 0.86997289 | 1 | T04C12.8 |
| WBGene00011431 | -0.847856771 | 0.00124699 | 0.01086845 | T04D3.1 |
| WBGene00011432 | -2.296060516 | 1.40E-10 | 1.30E-08 | sdz-30 |
| WBGene00011433 | 0.783412734 | 0.04922829 | 0.15929654 | pde-1 |
| WBGene00011434 | -0.689468611 | 0.00075221 | 0.00737094 | T04D3.5 |
| WBGene00011435 | -0.858469867 | 0.09958678 | 0.25432963 | T04D3.8 |
| WBGene00011436 | 1.878885286 | 6.70E-18 | 1.81E-15 | T04F3.1 |
| WBGene00011437 | 0.761488424 | 0.25421882 | 0.46204652 | T04F3.2 |
| WBGene00011438 | 1.060949945 | 0.61866902 | 1 | T04F3.3 |
| WBGene00011439 | 0.4012063 | 0.41857634 | 0.62320835 | T04F3.4 |
| WBGene00011440 | -0.440251188 | 0.06277006 | 0.18793868 | sfxn-1.5 |
| WBGene00011441 | 0.050768393 | 0.83452372 | 0.91032502 | T04F8.2 |
| WBGene00011442 | -0.878995693 | 0.27194459 | 0.48056732 | T04F8.3 |
| WBGene00011443 | -0.421034211 | 0.66252741 | 1 | cutl-11 |
| WBGene00011444 | 0.17552849 | 0.58212145 | 0.75273854 | T04F8.6 |
| WBGene00011445 | 1.216935863 | 0.00197903 | 0.01542132 | T04F8.7 |
| WBGene00011446 | -0.73110418 | 0.00058217 | 0.00599501 | T04F8.8 |
| WBGene00011447 | 1.504362275 | 0.25096559 | 1 | T04F8.9 |
| WBGene00011449 | 0.096126704 | 0.5581031 | 0.73522654 | T04H1.2 |
| WBGene00011450 | 1.057154391 | 0.45032085 | 1 | ttr-22 |
| WBGene00011451 | -0.22407047 | 0.36669556 | 0.57427918 | T04H1.5 |
| WBGene00011452 | 1.135599525 | 0.09894808 | 0.25317781 | ugt-55 |
| WBGene00011453 | 0.84628915 | 0.28941475 | 0.49961389 | ugt-56 |
| WBGene00011454 | 0.40889797 | 0.53209111 | 0.71564398 | lron-14 |
| WBGene00011456 | 2.957036741 | 0.00771458 | 0.04261269 | T05A1.5 |
| WBGene00011458 | -0.156269293 | 0.96924949 | 1 | irld-47 |
| WBGene00011460 | 0.12074737 | 0.79359961 | 0.88593495 | ttr-14 |
| WBGene00011462 | 1.851382081 | 0.11037634 | 1 | vap-2 |
| WBGene00011463 | -0.175414231 | 0.83137953 | 0.90861273 | T05A10.6 |
| WBGene00011464 | -0.639435128 | 1.5069E-05 | 0.00031985 | T05B9.1 |
| WBGene00011465 | -1.159714244 | 0.75790634 | 1 | T05B9.2 |
| WBGene00011466 | 1.062657457 | 0.66313467 | 1 | T05C12.1 |
| WBGene00011467 | 1.735991307 | 0.58152819 | 1 | decr-1.3 |
| WBGene00011468 | 1.451745273 | 0.51709377 | 1 | T05C12.4 |
| WBGene00011471 | 1.26611775 | 0.14984228 | 0.33125184 | T05C12.8 |
| WBGene00011472 | 1.181686151 | 0.7579682 | 1 | T05C12.9 |
| WBGene00011473 | -0.982632318 | 0.40077595 | 1 | T05C12.11 |
| WBGene00011474 | 0.182055727 | 0.42769651 | 0.6306514 | aldo-1 |
| WBGene00011475 | -1.849618213 | 3.2038E-05 | 0.00058935 | T05D4.2 |

| | | | | |
|---|---|---|---|---|
| WBGene00011476 | 0.267899703 | 0.69018172 | 0.82353771 | T05D4.3 |
| WBGene00011478 | -0.394320382 | 0.8739308 | 1 | T05D4.5 |
| WBGene00011479 | 0.785787141 | 0.31365021 | 0.52157611 | T05E11.2 |
| WBGene00011480 | 0.138451387 | 0.36864287 | 0.57617059 | enpl-1 |
| WBGene00011481 | -0.105774544 | 0.50268069 | 0.68999878 | imp-2 |
| WBGene00011482 | 0.218103877 | 0.41696092 | 0.62183055 | hpo-4 |
| WBGene00011484 | 0.5790228 | 0.47420743 | 0.66728702 | T05E11.8 |
| WBGene00011486 | 2.838949767 | 5.60E-17 | 1.33E-14 | T05E12.3 |
| WBGene00011487 | 0.567403359 | 0.00019659 | 0.00260567 | T05E12.6 |
| WBGene00011488 | -0.198937518 | 0.22798749 | 0.43290525 | nra-2 |
| WBGene00011489 | -0.231274662 | 0.09632431 | 0.24913639 | T05F1.2 |
| WBGene00011490 | -0.447714143 | 0.04195413 | 0.14229939 | T05F1.4 |
| WBGene00011491 | 0.09697944 | 0.97612634 | 1 | T05F1.5 |
| WBGene00011492 | 1.955750377 | 0.44285624 | 1 | T05F1.7 |
| WBGene00011493 | 2.185900552 | 0.11772203 | 1 | T05F1.8 |
| WBGene00011494 | 0.690837573 | 0.38439826 | 0.59284039 | T05F1.9 |
| WBGene00011495 | 1.86578462 | 0.00259699 | 0.01907574 | T05F1.11 |
| WBGene00011497 | 0.358222409 | 0.43066544 | 0.63306345 | T05F1.13 |
| WBGene00011498 | 0.2886118 | 0.36990131 | 0.5774852 | T05G5.1 |
| WBGene00011499 | -0.014265708 | 0.98960294 | 1 | T05G5.4 |
| WBGene00011500 | -0.354361632 | 0.14525585 | 0.32397939 | T05G5.5 |
| WBGene00011501 | -0.774425349 | 4.07E-09 | 2.72E-07 | rmd-1 |
| WBGene00011502 | 0.007392314 | 0.97496182 | 0.98674654 | vps-53 |
| WBGene00011503 | 0.188829519 | 0.30618374 | 0.51508615 | gcc-2 |
| WBGene00011505 | -0.455951649 | 0.1757803 | 0.36656624 | pzf-1 |
| WBGene00011507 | 0.254230215 | 0.13708483 | 0.31308257 | T05H10.1 |
| WBGene00011508 | -1.134721706 | 0.00014654 | 0.00207357 | T05H10.3 |
| WBGene00011509 | -0.835687307 | 1.6684E-06 | 4.9891E-05 | T05H10.4 |
| WBGene00011510 | -0.189586424 | 0.1545885 | 0.33805097 | pdha-1 |
| WBGene00011511 | 0.085986324 | 0.65580249 | 0.8022265 | gpcp-2 |
| WBGene00011512 | 0.156728903 | 0.84244929 | 0.91337104 | T05H10.8 |
| WBGene00011513 | -0.398539426 | 0.46990778 | 0.66324668 | fbxa-197 |
| WBGene00011515 | 0.281454817 | 0.94435481 | 1 | oac-42 |
| WBGene00011516 | -0.156269293 | 0.96924949 | 1 | T06C12.9 |
| WBGene00011517 | 0.8992903 | 0.10030542 | 0.25530568 | cgt-1 |
| WBGene00011520 | 0.799367552 | 0.33197526 | 0.53982957 | nhr-213 |
| WBGene00011521 | 1.247575581 | 0.7549672 | 1 | T06C12.14 |
| WBGene00011522 | 1.654663069 | 1.66E-08 | 9.48E-07 | srap-1 |
| WBGene00011523 | 0.753583449 | 0.33664505 | 0.54486555 | T06D8.2 |
| WBGene00011524 | 0.187385718 | 0.76238999 | 0.8671704 | T06D8.3 |
| WBGene00011526 | -0.388106783 | 0.07613357 | 0.21413202 | cox-15 |
| WBGene00011527 | -0.261053612 | 0.19268974 | 0.38783921 | cchl-1 |
| WBGene00011528 | -0.890353946 | 7.2747E-05 | 0.00118872 | T06D8.7 |

| | | | | |
|---|---|---|---|---|
| WBGene00011529 | -0.542082963 | 0.00427195 | 0.02764701 | T06D8.9 |
| WBGene00011530 | -0.05685347 | 0.9053708 | 0.95079204 | T06D8.10 |
| WBGene00011531 | 0.752478453 | 0.01966398 | 0.08445007 | rsbp-1 |
| WBGene00011532 | -0.140941349 | 0.41440749 | 0.61977351 | chaf-1 |
| WBGene00011533 | 0.645795908 | 0.6114255 | 1 | T06E4.5 |
| WBGene00011534 | -0.126941887 | 0.92785576 | 1 | T06E4.7 |
| WBGene00011535 | 2.224351729 | 0.33187942 | 1 | T06E4.8 |
| WBGene00011536 | 0.871913321 | 0.559375 | 1 | T06E4.9 |
| WBGene00011537 | -0.447536141 | 0.61744198 | 0.77706677 | T06E4.10 |
| WBGene00011538 | -0.283264709 | 0.23779259 | 0.44395533 | T06E6.1 |
| WBGene00011539 | -1.455123906 | 0.47550043 | 1 | fbxa-135 |
| WBGene00011540 | -0.103665822 | 0.80754705 | 0.89513608 | T06E6.10 |
| WBGene00011542 | -1.100554356 | 0.63912775 | 1 | fbxa-104 |
| WBGene00011543 | -0.181521656 | 0.43623272 | 0.63705534 | acl-2 |
| WBGene00011544 | -1.969805064 | 0.62862569 | 1 | T06E8.2 |
| WBGene00011545 | 0.302380407 | 0.61315199 | 0.77405075 | T06G6.3 |
| WBGene00011546 | -0.149095392 | 0.88966419 | 1 | T06G6.4 |
| WBGene00011547 | 0.377828795 | 0.92526585 | 1 | T06G6.5 |
| WBGene00011548 | -0.582567818 | 0.77294373 | 1 | T06G6.6 |
| WBGene00011551 | -1.029084083 | 0.44681855 | 1 | T06G6.12 |
| WBGene00011554 | -1.221430164 | 0.27865621 | 1 | T07A5.1 |
| WBGene00011556 | -0.934245188 | 0.22410584 | 0.42821356 | vglu-3 |
| WBGene00011558 | -0.719198369 | 0.00376072 | 0.02504605 | T07A5.5 |
| WBGene00011558 | -2.640414922 | 0.41081194 | 1 | ostf-4 |
| WBGene00011559 | -0.501259495 | 0.01096821 | 0.05556147 | umps-1 |
| WBGene00011560 | -0.55309733 | 0.01388485 | 0.06579859 | T07C4.3 |
| WBGene00011561 | -0.891161564 | 3.1966E-06 | 8.7213E-05 | ttr-15 |
| WBGene00011562 | -0.145043216 | 0.55859326 | 0.73556191 | sas-7 |
| WBGene00011563 | 0.598682885 | 0.35769332 | 0.56591502 | jmjd-4 |
| WBGene00011564 | 1.091539518 | 0.00017553 | 0.00236523 | ugt-50 |
| WBGene00011565 | -0.156269293 | 0.96924949 | 1 | nhr-272 |
| WBGene00011566 | -1.14715222 | 0.68932198 | 1 | nhr-214 |
| WBGene00011568 | 0.747845955 | 0.85215965 | 1 | nhr-26 |
| WBGene00011571 | -0.928015149 | 2.6356E-06 | 7.4182E-05 | ttr-46 |
| WBGene00011572 | -0.478055387 | 0.65362458 | 1 | mam-5 |
| WBGene00011573 | 0.104760765 | 0.76459508 | 0.86822555 | anmt-3 |
| WBGene00011575 | 0.125975224 | 0.84867138 | 0.91702735 | madf-4 |
| WBGene00011576 | -0.01755687 | 0.95413758 | 0.97667363 | rmh-2 |
| WBGene00011578 | 0.885481922 | 0.25296193 | 0.46090245 | npr-20 |
| WBGene00011579 | 0.667100447 | 0.09519162 | 0.2473719 | T07D4.2 |
| WBGene00011580 | -0.147476492 | 0.38079107 | 0.58894069 | ddx-19 |
| WBGene00011581 | -0.241106143 | 0.91536132 | 1 | T07D10.1 |
| WBGene00011582 | 0.750638192 | 0.68763128 | 1 | ntr-1 |

| | | | | |
|---|---|---|---|---|
| WBGene00011583 | -0.013426889 | 0.99087056 | 1 | T07D10.3 |
| WBGene00011584 | 0.688793783 | 0.86378755 | 1 | clec-15 |
| WBGene00011586 | 1.419357687 | 0.00805932 | 0.04411548 | flp-33 |
| WBGene00011587 | -0.314726626 | 0.26512008 | 0.47390941 | T07F10.1 |
| WBGene00011589 | -0.162354283 | 0.44001746 | 0.64015249 | T07F10.3 |
| WBGene00011590 | 1.102880785 | 0.06891465 | 0.20016575 | bus-19 |
| WBGene00011591 | -0.70778075 | 0.27011924 | 0.47883162 | T07F10.5 |
| WBGene00011592 | 0.765458189 | 0.46704325 | 1 | T07F10.6 |
| WBGene00011593 | 1.527952301 | 0.32910871 | 1 | T07G12.2 |
| WBGene00011594 | -0.452824552 | 0.54131979 | 0.72208215 | T07G12.3 |
| WBGene00011596 | 2.715567986 | 0.02059204 | 0.08735428 | T07G12.5 |
| WBGene00011597 | -0.223372955 | 0.24891207 | 0.4567087 | zim-1 |
| WBGene00011600 | 0.140340742 | 0.54190059 | 0.72243495 | zim-2 |
| WBGene00011601 | -0.401409067 | 0.07709468 | 0.21544186 | zim-3 |
| WBGene00011604 | 0.226772095 | 0.31320448 | 0.52122318 | T08A11.1 |
| WBGene00011605 | -0.058367963 | 0.76065331 | 0.86633132 | sftb-1 |
| WBGene00011606 | 1.006686554 | 0.2470681 | 0.45497554 | T08D2.1 |
| WBGene00011609 | 2.149565946 | 0.27361747 | 1 | T08D2.4 |
| WBGene00011610 | 0.990397697 | 0.24946472 | 0.457239 | T08D2.5 |
| WBGene00011611 | 0.210220745 | 0.76541425 | 0.86865717 | T08D2.6 |
| WBGene00011612 | 1.107682121 | 0.26809888 | 1 | T08D2.7 |
| WBGene00011613 | 1.272669974 | 0.03573213 | 0.12744529 | T08D2.8 |
| WBGene00011614 | -0.175095211 | 0.58220098 | 0.75273854 | nfya-1 |
| WBGene00011615 | -0.569218645 | 0.05897395 | 0.18039877 | lsd-1 |
| WBGene00011616 | -0.471406496 | 0.76924112 | 1 | T08D10.3 |
| WBGene00011619 | -1.969805064 | 0.62862569 | 1 | T08G3.7 |
| WBGene00011622 | -0.156269293 | 0.96924949 | 1 | T08G5.1 |
| WBGene00011624 | -1.03461247 | 0.33686183 | 1 | T08G5.3 |
| WBGene00011625 | 0.012227785 | 0.94417968 | 0.97120685 | vps-39 |
| WBGene00011629 | 0.872247528 | 9.3589E-06 | 0.0002152 | T08G11.1 |
| WBGene00011630 | 3.91035052 | 0.00721781 | 1 | T08G11.3 |
| WBGene00011631 | -0.075152996 | 0.709361 | 0.83550174 | T08G11.4 |
| WBGene00011633 | 0.031284902 | 0.95218693 | 0.97579519 | T09A5.4 |
| WBGene00011634 | -0.465268032 | 0.04293588 | 0.14436017 | T09A5.5 |
| WBGene00011635 | -0.335229327 | 0.56406433 | 0.73923846 | T09A5.7 |
| WBGene00011636 | -0.608183755 | 0.00099187 | 0.00918556 | cec-3 |
| WBGene00011637 | -0.008621182 | 0.96668276 | 0.9830824 | sds-22 |
| WBGene00011638 | -0.442445186 | 0.00312944 | 0.02188625 | ostb-1 |
| WBGene00011639 | 0.337910234 | 0.10185074 | 0.25747525 | ztf-17 |
| WBGene00011640 | -1.094201388 | 0.7816772 | 1 | T09A5.14 |
| WBGene00011641 | -0.883433326 | 0.01770641 | 0.07837169 | T09A5.15 |
| WBGene00011642 | -0.678908485 | 0.01094055 | 0.0555401 | T09B9.1 |
| WBGene00011643 | 1.087194186 | 0.28618313 | 0.49622242 | slc-17.9 |

| | | | | |
|---|---|---|---|---|
| WBGene00011644 | 3.30145033 | 0.00188301 | 0.01487421 | T09B9.3 |
| WBGene00011645 | 0.453350137 | 0.1481748 | 0.32882292 | T09B9.4 |
| WBGene00011646 | -0.372972233 | 0.85785656 | 1 | T09B9.5 |
| WBGene00011647 | -0.090478199 | 0.5785331 | 0.74994345 | noca-1 |
| WBGene00011648 | -0.139026111 | 0.53731605 | 0.71940385 | cni-1 |
| WBGene00011649 | 0.377828795 | 0.92526585 | 1 | T09E8.4 |
| WBGene00011653 | 1.955750377 | 0.44285624 | 1 | oac-43 |
| WBGene00011654 | -0.156269293 | 0.96924949 | 1 | oac-44 |
| WBGene00011656 | -0.156269293 | 0.96924949 | 1 | oac-45 |
| WBGene00011658 | -1.969805064 | 0.62862569 | 1 | T09E11.9 |
| WBGene00011661 | 0.209989351 | 0.53056499 | 0.71443643 | ztf-27 |
| WBGene00011662 | -0.197458466 | 0.22352001 | 0.42746592 | T09F3.2 |
| WBGene00011663 | -0.071736472 | 0.86172963 | 0.92545417 | T09F3.4 |
| WBGene00011664 | -0.54960263 | 0.04540179 | 0.15011525 | T09F3.5 |
| WBGene00011665 | 1.102651183 | 0.48636107 | 1 | T09F5.1 |
| WBGene00011668 | 0.532721126 | 0.05518843 | 0.17215897 | clec-47 |
| WBGene00011669 | 0.377828795 | 0.92526585 | 1 | T09F5.10 |
| WBGene00011670 | -0.181336164 | 0.83023441 | 0.90818536 | fbxa-84 |
| WBGene00011671 | 1.217208842 | 0.36741823 | 1 | cyp-13A4 |
| WBGene00011672 | 2.573696515 | 5.09E-10 | 4.32E-08 | cyp-13A5 |
| WBGene00011673 | 1.043373997 | 0.324515 | 1 | cyp-13A6 |
| WBGene00011674 | 1.833981893 | 0.33135673 | 1 | cyp-13A8 |
| WBGene00011675 | 1.090794088 | 0.55107197 | 1 | cyp-13A3 |
| WBGene00011676 | 0.227456866 | 0.67637039 | 0.81466397 | cyp-13A2 |
| WBGene00011677 | 1.639638001 | 0.32044289 | 1 | cyp-13A1 |
| WBGene00011678 | 0.720160099 | 0.4257156 | 0.62970433 | T10B9.9 |
| WBGene00011679 | 0.011252936 | 0.96071809 | 0.97968069 | ucr-2.2 |
| WBGene00011680 | 0.051560116 | 0.82759147 | 0.90692193 | T10B10.3 |
| WBGene00011681 | 1.003011169 | 0.00019862 | 0.00262192 | T10B10.4 |
| WBGene00011682 | 0.866958339 | 0.34335663 | 0.55176655 | snt-7 |
| WBGene00011683 | 1.281683083 | 2.3196E-05 | 0.00045733 | phat-6 |
| WBGene00011684 | -0.061651103 | 0.91116871 | 0.95393098 | T10B10.8 |
| WBGene00011685 | -0.18557176 | 0.95664217 | 1 | T10B10.9 |
| WBGene00011687 | -0.785676482 | 3.6203E-06 | 9.639E-05 | cwc-15 |
| WBGene00011688 | -0.235559552 | 0.29149354 | 0.50151603 | T10C6.6 |
| WBGene00011689 | 0.276469626 | 0.3171745 | 0.52493219 | T10C6.7 |
| WBGene00011691 | -0.057447413 | 0.85680767 | 0.92175167 | T10C6.10 |
| WBGene00011692 | 0.781410676 | 0.08822432 | 0.23548763 | T10G3.1 |
| WBGene00011693 | 2.063391569 | 0.37853067 | 1 | T10G3.2 |
| WBGene00011694 | -0.223521064 | 0.49792554 | 0.68626564 | T10G3.3 |
| WBGene00011695 | -0.518685065 | 0.73711149 | 1 | T10G3.4 |
| WBGene00011696 | 0.763725778 | 0.00019868 | 0.00262192 | eea-1 |
| WBGene00011697 | 2.133182101 | 0.3921092 | 1 | T10H4.4 |

| | | | | |
|---|---|---|---|---|
| WBGene00011698 | 1.746057557 | 0.41244616 | 1 | cyp-34A1 |
| WBGene00011699 | 0.082855449 | 0.83984386 | 0.91219756 | cyp-34A2 |
| WBGene00011701 | 0.688793783 | 0.86378755 | 1 | srab-17 |
| WBGene00011702 | -0.156269293 | 0.96924949 | 1 | srab-18 |
| WBGene00011704 | -2.154080274 | 0.19301251 | 1 | sdz-31 |
| WBGene00011705 | 1.039707334 | 0.07783845 | 0.2167018 | T11A5.6 |
| WBGene00011706 | -0.924863232 | 0.0292002 | 0.11114527 | T11B7.1 |
| WBGene00011707 | 0.029683054 | 0.97650909 | 1 | T11B7.2 |
| WBGene00011708 | 0.14733822 | 0.6613389 | 0.80508588 | T11B7.5 |
| WBGene00011709 | 0.993030093 | 0.34894026 | 1 | T11F9.1 |
| WBGene00011712 | 0.855368791 | 0.66134621 | 1 | T11F9.10 |
| WBGene00011713 | 0.5160565 | 0.08802461 | 0.23517445 | T11F9.12 |
| WBGene00011717 | -0.031670228 | 0.91485951 | 0.95571968 | T11G6.2 |
| WBGene00011718 | 0.547007358 | 0.1635556 | 0.34976101 | T11G6.3 |
| WBGene00011719 | 0.187385297 | 0.6014667 | 0.76657645 | T11G6.4 |
| WBGene00011720 | 0.084184872 | 0.67204144 | 0.81248091 | T11G6.5 |
| WBGene00011722 | -0.486343097 | 0.00187181 | 0.01481463 | rbm-22 |
| WBGene00011723 | 1.238382797 | 9.9633E-05 | 0.00152733 | T12A7.2 |
| WBGene00011725 | 0.771053298 | 0.14917125 | 0.33055086 | lips-5 |
| WBGene00011726 | 0.795465883 | 0.00775937 | 0.04284059 | lpr-2 |
| WBGene00011727 | 0.904526129 | 0.10871649 | 0.26887899 | T12A7.6 |
| WBGene00011728 | 1.957271014 | 0.49743227 | 1 | T12A7.7 |
| WBGene00011729 | 0.399609696 | 0.04745975 | 0.15547149 | set-16 |
| WBGene00011730 | -0.457649232 | 0.0101332 | 0.0523966 | drr-2 |
| WBGene00011731 | -0.293844809 | 0.22741658 | 0.4320915 | acbp-5 |
| WBGene00011732 | 0.558377394 | 0.10312015 | 0.26008368 | arrd-17 |
| WBGene00011733 | -0.442377679 | 0.19218755 | 0.38723983 | T12D8.5 |
| WBGene00011734 | -0.576505891 | 0.00207466 | 0.01600286 | mlc-5 |
| WBGene00011735 | -0.362744712 | 0.01952455 | 0.08404952 | hip-1 |
| WBGene00011736 | 1.059385238 | 2.9836E-06 | 8.218E-05 | T12D8.9 |
| WBGene00011737 | 0.870975709 | 2.5064E-05 | 0.00048923 | sqst-1 |
| WBGene00011738 | 0.561705998 | 0.00374505 | 0.02498277 | T12G3.2 |
| WBGene00011739 | -0.559306548 | 0.02029366 | 0.08642065 | T12G3.4 |
| WBGene00011740 | -0.549939656 | 0.00646929 | 0.03754592 | mrpl-51 |
| WBGene00011741 | -0.925870959 | 5.38E-09 | 3.51E-07 | T12G3.6 |
| WBGene00011742 | 0.11639033 | 0.58318563 | 0.75332022 | tgn-38 |
| WBGene00011743 | -0.547776146 | 0.00561802 | 0.03400218 | T13F2.2 |
| WBGene00011744 | 0.578228238 | 0.68620123 | 1 | T13F2.4 |
| WBGene00011745 | 2.306649918 | 0.05548176 | 1 | sre-4 |
| WBGene00011746 | 0.03323167 | 0.83761382 | 0.91143626 | T13F2.6 |
| WBGene00011747 | -0.122956318 | 0.45193709 | 0.64972981 | sna-2 |
| WBGene00011749 | -1.14715222 | 0.68932198 | 1 | ssp-36 |
| WBGene00011750 | 0.7706252 | 0.6634285 | 1 | nhr-218 |

| | | | | |
|---|---|---|---|---|
| WBGene00011752 | 2.241560795 | 0.07268842 | 1 | fbxa-2 |
| WBGene00011753 | -0.738463483 | 0.00052841 | 0.00559317 | T13F3.6 |
| WBGene00011754 | 0.99650623 | 0.45381543 | 1 | T13F3.7 |
| WBGene00011755 | 1.035934586 | 0.00045716 | 0.00499582 | T13H5.1 |
| WBGene00011756 | 0.385765076 | 0.27006043 | 0.47879726 | ctg-2 |
| WBGene00011757 | 0.704544772 | 0.45320489 | 0.65078186 | cht-2 |
| WBGene00011758 | -0.406086051 | 0.01690774 | 0.07583188 | T13H5.4 |
| WBGene00011759 | -0.547602012 | 0.00961621 | 0.0503354 | mrps-18B |
| WBGene00011760 | -0.023114297 | 0.9452849 | 0.97202541 | T13H5.6 |
| WBGene00011761 | -0.156098831 | 0.43004534 | 0.63268859 | T13H5.8 |
| WBGene00011762 | 0.225159937 | 0.87774962 | 1 | T13H10.2 |
| WBGene00011763 | 0.969973999 | 0.00135574 | 0.01161615 | T14B1.1 |
| WBGene00011765 | 1.029569326 | 0.16928423 | 0.35743997 | frpr-17 |
| WBGene00011767 | -0.246281982 | 0.32340592 | 0.53197009 | agxt-1 |
| WBGene00011768 | 1.242674985 | 0.00016134 | 0.00222335 | oac-46 |
| WBGene00011770 | -0.905260109 | 0.47797365 | 1 | T14G8.2 |
| WBGene00011771 | 0.298612963 | 0.19053301 | 0.38517253 | T14G8.3 |
| WBGene00011772 | 0.880557168 | 0.4288605 | 1 | T14G8.4 |
| WBGene00011773 | -0.921050988 | 5.1785E-05 | 0.00089307 | ttr-53 |
| WBGene00011774 | -0.187027345 | 0.67308248 | 0.81320713 | ttr-54 |
| WBGene00011775 | -0.109376347 | 0.46318345 | 0.6582021 | T14G10.5 |
| WBGene00011776 | -0.157108733 | 0.44807777 | 0.64717014 | hpo-5 |
| WBGene00011777 | -0.134257569 | 0.87428976 | 0.93213743 | T14G10.8 |
| WBGene00011779 | -0.156269293 | 0.96924949 | 1 | bgnt-1.7 |
| WBGene00011781 | 0.688793783 | 0.86378755 | 1 | glct-3 |
| WBGene00011784 | 0.747845955 | 0.85215965 | 1 | T15D6.10 |
| WBGene00011785 | -2.652727305 | 0.24933058 | 1 | T15D6.11 |
| WBGene00011786 | 0.2978169 | 0.79665453 | 1 | T15D6.12 |
| WBGene00011788 | -0.885390077 | 0.00099889 | 0.00923654 | T15H9.2 |
| WBGene00011789 | 0.983488853 | 0.80601565 | 1 | T15H9.4 |
| WBGene00011790 | -1.73021997 | 0.47726432 | 1 | T15H9.5 |
| WBGene00011791 | -2.535614105 | 3.9585E-05 | 0.00070991 | T15H9.6 |
| WBGene00011793 | 0.465951719 | 0.72809723 | 1 | T16A9.3 |
| WBGene00011794 | 1.032696251 | 0.55215483 | 1 | K02E11.11 |
| WBGene00011794 | 1.244045624 | 0.00081699 | 0.00786852 | nep-21 |
| WBGene00011795 | 2.621301169 | 0.0545038 | 1 | T16A9.5 |
| WBGene00011796 | 0.982278376 | 0.27255685 | 0.48132548 | T16G1.2 |
| WBGene00011798 | 2.41262772 | 4.4486E-05 | 0.0007839 | T16G1.4 |
| WBGene00011799 | 1.744666649 | 0.06557531 | 0.19336894 | T16G1.5 |
| WBGene00011800 | 0.572410286 | 0.37143338 | 0.57904118 | T16G1.6 |
| WBGene00011801 | 1.63701367 | 0.00015211 | 0.00211546 | T16G1.7 |
| WBGene00011802 | -0.142682554 | 0.47573983 | 0.66843621 | T16G1.9 |
| WBGene00011803 | 1.788095221 | 2.73E-13 | 4.14E-11 | T16G12.1 |

| | | | | |
|---|---|---|---|---|
| WBGene00011804 | 0.685845371 | 0.0812423 | 0.2229673 | T16G12.3 |
| WBGene00011805 | -0.40820172 | 0.16421633 | 0.35035374 | T16G12.4 |
| WBGene00011806 | -0.082968177 | 0.65019069 | 0.79843591 | ekl-6 |
| WBGene00011807 | 0.107666566 | 0.61046581 | 0.77204363 | T16G12.6 |
| WBGene00011808 | -0.156269293 | 0.96924949 | 1 | T16G12.7 |
| WBGene00011809 | -0.658001575 | 0.08369461 | 0.22732355 | T16G12.8 |
| WBGene00011810 | -0.549786194 | 0.76687564 | 1 | T16G12.9 |
| WBGene00011811 | 0.033152152 | 0.95853927 | 0.97856902 | T16H12.1 |
| WBGene00011812 | -0.86917427 | 0.35975614 | 1 | T16H12.2 |
| WBGene00011813 | 0.197541171 | 0.41489728 | 0.62017539 | T16H12.3 |
| WBGene00011814 | -0.401396743 | 0.08696453 | 0.233334 | T16H12.4 |
| WBGene00011817 | 1.125657908 | 0.05663614 | 0.17572691 | T16H12.9 |
| WBGene00011819 | -0.531709081 | 0.02646673 | 0.10395746 | hal-2 |
| WBGene00011820 | -0.684476732 | 0.02565437 | 0.1020543 | T18D3.1 |
| WBGene00011821 | 3.534950427 | 2.89E-13 | 4.33E-11 | cdf-2 |
| WBGene00011822 | -1.622427701 | 0.18406308 | 1 | T18D3.5 |
| WBGene00011823 | 0.24298601 | 0.67995915 | 0.81647553 | T18D3.6 |
| WBGene00011824 | 0.346903136 | 0.40821473 | 0.61423758 | T18D3.7 |
| WBGene00011826 | -0.533350259 | 0.61848578 | 1 | T18D3.9 |
| WBGene00011827 | -0.165700432 | 0.43714281 | 0.63790274 | T19A6.1 |
| WBGene00011828 | -0.259151996 | 0.26389779 | 0.47249418 | nepr-1 |
| WBGene00011829 | 1.519209436 | 0.0207691 | 0.08782921 | T19A6.4 |
| WBGene00011830 | 0.254284306 | 0.47662651 | 0.66906261 | cyp-29A2 |
| WBGene00011831 | -0.433361008 | 0.0295656 | 0.11206869 | T19B10.2 |
| WBGene00011832 | 0.185584752 | 0.52979894 | 0.71398479 | bgal-1 |
| WBGene00011833 | 0.13416326 | 0.70243231 | 0.83210369 | T19B10.5 |
| WBGene00011834 | -0.543674583 | 0.00079986 | 0.00774412 | dvc-1 |
| WBGene00011835 | 0.171005593 | 0.30497505 | 0.51412136 | T19B10.8 |
| WBGene00011836 | -0.457445056 | 0.65225889 | 1 | mam-4 |
| WBGene00011837 | -0.156269293 | 0.96924949 | 1 | srsx-33 |
| WBGene00011838 | 0.707346016 | 0.07842679 | 0.21798984 | T19C4.1 |
| WBGene00011839 | 1.697377027 | 0.00027763 | 0.00339446 | T19C4.5 |
| WBGene00011840 | -0.156269293 | 0.96924949 | 1 | srbc-62 |
| WBGene00011841 | -1.38853051 | 0.25233662 | 1 | scl-25 |
| WBGene00011844 | 0.245309466 | 0.60916163 | 0.77135532 | T19C9.8 |
| WBGene00011846 | 1.456906199 | 0.20920821 | 1 | chil-25 |
| WBGene00011847 | 1.116613983 | 0.69067248 | 1 | chil-26 |
| WBGene00011848 | 1.186559396 | 0.57146291 | 1 | chil-27 |
| WBGene00011849 | 0.28615145 | 0.22392513 | 0.42805127 | T19H5.4 |
| WBGene00011850 | 0.879702224 | 0.00323103 | 0.02240559 | T20B3.1 |
| WBGene00011852 | 1.550444228 | 0.00818515 | 0.04458317 | clec-26 |
| WBGene00011853 | 3.883418509 | 0.02198444 | 1 | clec-40 |
| WBGene00011854 | -0.891182705 | 0.2043347 | 0.40371482 | T20B3.14 |

| | | | | |
|---|---|---|---|---|
| WBGene00011855 | -0.156269293 | 0.96924949 | 1 | clec-183 |
| WBGene00011856 | 0.380067536 | 0.06103655 | 0.18470706 | T20D3.2 |
| WBGene00011857 | 0.024015931 | 0.91657987 | 0.95627706 | T20D3.3 |
| WBGene00011858 | 0.063799005 | 0.77607617 | 0.87511292 | T20D3.5 |
| WBGene00011859 | -0.586415394 | 0.03267273 | 0.12053469 | T20D3.6 |
| WBGene00011860 | -0.235879615 | 0.36525854 | 0.57294624 | T20D3.8 |
| WBGene00011864 | -0.054493041 | 0.82075322 | 0.90284677 | T20F10.2 |
| WBGene00011867 | 0.271991092 | 0.16282447 | 0.34865113 | chc-1 |
| WBGene00011868 | 1.048263333 | 0.03452528 | 0.12501383 | best-18 |
| WBGene00011869 | 1.27203975 | 1.86E-07 | 7.8816E-06 | dod-6 |
| WBGene00011870 | 2.001112548 | 1.50E-09 | 1.12E-07 | T20G5.8 |
| WBGene00011871 | -0.073847196 | 0.79417902 | 0.88635851 | T20G5.9 |
| WBGene00011872 | -0.342963912 | 0.27166363 | 0.48044692 | blos-1 |
| WBGene00011873 | -0.70576507 | 0.76841669 | 1 | T20G5.12 |
| WBGene00011874 | 0.747845955 | 0.85215965 | 1 | T20G5.13 |
| WBGene00011875 | -0.262215259 | 0.18327819 | 0.37536783 | T20G5.14 |
| WBGene00011876 | 0.281454817 | 0.94435481 | 1 | srz-6 |
| WBGene00011877 | 0.747845955 | 0.85215965 | 1 | T21B4.3 |
| WBGene00011878 | 0.539671477 | 0.51858964 | 0.7042379 | ador-1 |
| WBGene00011879 | 0.124780282 | 0.78022902 | 0.87832735 | pho-7 |
| WBGene00011880 | 1.375971855 | 1.31E-09 | 1.01E-07 | T21B6.3 |
| WBGene00011882 | -1.837668733 | 0.43198255 | 1 | nstp-9 |
| WBGene00011883 | -0.361245963 | 0.0922969 | 0.24192069 | mrpl-50 |
| WBGene00011884 | -0.457660134 | 0.00662404 | 0.03822359 | enol-1 |
| WBGene00011885 | -0.055024573 | 0.71959209 | 0.84179137 | T21B10.3 |
| WBGene00011886 | -0.036472973 | 0.86122415 | 0.92515683 | T21B10.4 |
| WBGene00011887 | -0.19943598 | 0.3409665 | 0.54928041 | set-17 |
| WBGene00011888 | 0.938719759 | 0.05066984 | 0.16217522 | cutl-15 |
| WBGene00011890 | -0.320618996 | 0.095062 | 0.24719377 | mics-1 |
| WBGene00011891 | -0.007822964 | 0.96688798 | 0.9830824 | del-6 |
| WBGene00011892 | -0.355179415 | 0.11441889 | 0.27821121 | T21C9.4 |
| WBGene00011893 | 1.334524731 | 2.7002E-05 | 0.00051545 | T21C9.6 |
| WBGene00011894 | 1.482733435 | 0.00657754 | 0.03806382 | ttr-23 |
| WBGene00011895 | -0.720849423 | 0.43078131 | 0.6331093 | T21C9.9 |
| WBGene00011896 | 1.735991307 | 0.58152819 | 1 | T21C9.11 |
| WBGene00011897 | 0.180844482 | 0.370593 | 0.57810222 | scpl-4 |
| WBGene00011898 | 0.150973977 | 0.44228602 | 0.64198205 | T21C9.13 |
| WBGene00011899 | 1.022234 | 0.23433429 | 0.44049171 | T21C12.3 |
| WBGene00011900 | -1.94984219 | 0.09137628 | 1 | T21C12.4 |
| WBGene00011902 | 0.688793783 | 0.86378755 | 1 | T21E8.4 |
| WBGene00011903 | -0.156269293 | 0.96924949 | 1 | T21E8.5 |
| WBGene00011904 | 0.750770271 | 0.00284127 | 0.02041575 | hlb-1 |
| WBGene00011906 | -1.50175564 | 0.0059439 | 0.03536394 | hsp-12.1 |

| | | | | |
|---|---|---|---|---|
| WBGene00011908 | -0.17721634 | 0.33696422 | 0.54507071 | pash-1 |
| WBGene00011910 | 1.250737994 | 0.3765811 | 1 | alg-3 |
| WBGene00011911 | -0.333693239 | 0.89573179 | 1 | T22B3.3 |
| WBGene00011912 | -0.207363684 | 0.20048208 | 0.39902507 | T22C1.1 |
| WBGene00011913 | 0.107503334 | 0.67632234 | 0.81466397 | glb-26 |
| WBGene00011914 | 0.346808936 | 0.18886129 | 0.38267105 | T22C1.3 |
| WBGene00011915 | -0.47150126 | 0.10688994 | 0.26658958 | T22C1.4 |
| WBGene00011916 | -0.451619324 | 0.04223456 | 0.14299212 | T22C1.5 |
| WBGene00011917 | 0.109517409 | 0.5958962 | 0.76214826 | T22C1.6 |
| WBGene00011918 | 1.38322681 | 0.69550305 | 1 | T22C1.8 |
| WBGene00011919 | 0.983488853 | 0.80601565 | 1 | T22C1.9 |
| WBGene00011920 | 0.040697886 | 0.91337895 | 0.95515364 | T22C1.11 |
| WBGene00011922 | 1.216957619 | 0.74819621 | 1 | T22C8.1 |
| WBGene00011923 | 0.464113706 | 0.36211639 | 0.56993195 | chhy-1 |
| WBGene00011924 | -0.474775739 | 0.33700608 | 0.54507071 | T22C8.3 |
| WBGene00011925 | -0.986115084 | 0.18867005 | 0.38247505 | T22C8.4 |
| WBGene00011926 | 1.710496568 | 0.23001039 | 1 | sptf-2 |
| WBGene00011927 | -1.132058019 | 0.12142818 | 0.28856127 | T22C8.6 |
| WBGene00011928 | 1.192524401 | 0.37891857 | 1 | cutl-12 |
| WBGene00011929 | 1.743415485 | 0.50483382 | 1 | T22G5.1 |
| WBGene00011930 | 0.193578024 | 0.67974826 | 0.81647553 | T22G5.3 |
| WBGene00011931 | 0.113061109 | 0.9246852 | 1 | srsx-19 |
| WBGene00011932 | -0.167538671 | 0.34298545 | 0.55147482 | sptl-3 |
| WBGene00011934 | -0.078589568 | 0.96315577 | 1 | T22H2.4 |
| WBGene00011935 | -0.444096443 | 0.19121957 | 0.38596789 | scrm-1 |
| WBGene00011936 | 0.000912318 | 0.99722679 | 0.99857337 | pgrn-1 |
| WBGene00011937 | -0.023305918 | 0.99216802 | 1 | ilc-17.1 |
| WBGene00011938 | -0.160688366 | 0.48373258 | 0.67458876 | alh-13 |
| WBGene00011939 | 0.098020766 | 0.57038662 | 0.74463053 | prmt-9 |
| WBGene00011940 | 0.163155776 | 0.49345044 | 0.68212223 | T23B5.3 |
| WBGene00011941 | -0.632483836 | 0.10856478 | 0.26866795 | T23B5.4 |
| WBGene00011944 | -0.146806837 | 0.36760533 | 0.57521541 | T23D8.3 |
| WBGene00011945 | -0.436267058 | 0.0230988 | 0.09490442 | alg-5 |
| WBGene00011948 | 0.091692055 | 0.90671287 | 0.95121412 | T23F1.5 |
| WBGene00011949 | -1.046078815 | 0.33006467 | 1 | T23F6.1 |
| WBGene00011951 | 0.87001166 | 0.63612313 | 1 | T23F6.3 |
| WBGene00011952 | -0.646440611 | 0.67898796 | 1 | T23F6.5 |
| WBGene00011953 | 0.19977604 | 0.25409493 | 0.46193303 | ppm-2 |
| WBGene00011954 | 1.025181888 | 0.79790347 | 1 | T23F11.2 |
| WBGene00011955 | 0.937297884 | 0.01016308 | 0.05250635 | cdka-1 |
| WBGene00011956 | -0.586867986 | 0.07011959 | 0.20243772 | T23F11.4 |
| WBGene00011957 | 0.158165466 | 0.82714736 | 0.90684451 | T23F11.6 |
| WBGene00011958 | 1.413734769 | 0.68804531 | 1 | T23G4.2 |

| | | | | |
|---|---|---|---|---|
| WBGene00011959 | -0.173188313 | 0.55935994 | 0.73609246 | nyn-1 |
| WBGene00011960 | 0.837787491 | 0.43376936 | 1 | oac-48 |
| WBGene00011961 | 1.025181888 | 0.79790347 | 1 | T23G4.5 |
| WBGene00011962 | -0.38020279 | 0.038977 | 0.13501277 | T23G5.2 |
| WBGene00011963 | 1.92052731 | 0.12634351 | 1 | T23G5.3 |
| WBGene00011964 | 0.145724464 | 0.69111034 | 0.8239979 | saeg-2 |
| WBGene00011965 | -0.067059427 | 0.80696558 | 0.89476783 | T23G7.2 |
| WBGene00011966 | 0.209867638 | 0.24815881 | 0.456292 | T23G7.3 |
| WBGene00011967 | -0.477823471 | 0.03005603 | 0.11346712 | pir-1 |
| WBGene00011968 | 0.751896602 | 0.5867065 | 1 | T23G11.1 |
| WBGene00011969 | -0.490598463 | 0.03100208 | 0.11626133 | T23G11.4 |
| WBGene00011970 | 0.262826887 | 0.24022234 | 0.44745926 | rlbp-1 |
| WBGene00011971 | 0.453788427 | 0.13214134 | 0.30477357 | lron-9 |
| WBGene00011972 | 0.186537381 | 0.31713349 | 0.52493219 | T23G11.7 |
| WBGene00011973 | 0.35134761 | 0.87335989 | 1 | T23G11.10 |
| WBGene00011974 | 1.08083361 | 0.04635414 | 0.15255776 | xbx-5 |
| WBGene00011975 | 0.201309333 | 0.31157644 | 0.51976499 | T24B1.1 |
| WBGene00011976 | -0.418839536 | 0.03814875 | 0.13316963 | T24B8.2 |
| WBGene00011977 | -0.364991505 | 0.0488617 | 0.15850832 | T24B8.3 |
| WBGene00011978 | 0.751814345 | 0.00367559 | 0.02464125 | T24B8.4 |
| WBGene00011979 | -1.693926551 | 9.65E-12 | 1.12E-09 | T24B8.5 |
| WBGene00011980 | 0.410144111 | 0.09406227 | 0.24506652 | T24B8.7 |
| WBGene00011981 | -0.659848818 | 0.19866224 | 0.39657321 | T24C2.2 |
| WBGene00011982 | -1.964653846 | 0.53599202 | 1 | T24C2.3 |
| WBGene00011983 | -0.008677305 | 0.99719169 | 1 | fbxa-81 |
| WBGene00011984 | 1.519919753 | 0.04871403 | 0.15822286 | T24C2.5 |
| WBGene00011985 | -0.287436485 | 0.1050244 | 0.2637231 | T24D1.2 |
| WBGene00011986 | -0.381759773 | 0.08101809 | 0.22254466 | T24D1.3 |
| WBGene00011988 | -0.572284386 | 0.00198815 | 0.01546322 | har-2 |
| WBGene00011990 | 0.4954485 | 0.36125124 | 0.56908671 | T24D5.2 |
| WBGene00011991 | 0.377828795 | 0.92526585 | 1 | T24D5.3 |
| WBGene00011993 | -0.789246581 | 0.46859353 | 1 | T24D5.5 |
| WBGene00011994 | -0.21752581 | 0.28135596 | 0.49065537 | samp-1 |
| WBGene00011995 | 0.373849105 | 0.20286938 | 0.40190728 | rsf-1 |
| WBGene00011996 | 0.491120352 | 0.68196734 | 1 | T24F1.4 |
| WBGene00011997 | 2.023634572 | 8.22E-11 | 7.92E-09 | nlp-51 |
| WBGene00011999 | -1.027355405 | 0.68015432 | 1 | T24F1.7 |
| WBGene00012000 | -0.603168334 | 0.00054362 | 0.00570451 | T24H10.1 |
| WBGene00012002 | -0.127950827 | 0.55790602 | 0.73505556 | T24H10.4 |
| WBGene00012003 | -1.18110767 | 0.38578233 | 1 | T24H10.5 |
| WBGene00012004 | -0.39251215 | 0.11624004 | 0.28062642 | dyrb-1 |
| WBGene00012005 | 0.531450259 | 0.03418581 | 0.12430365 | jun-1 |
| WBGene00012007 | 0.388877043 | 0.20474319 | 0.40406162 | T25B9.1 |

| | | | | |
|---|---|---|---|---|
| WBGene00012008 | -1.14715222 | 0.68932198 | 1 | T25B9.2 |
| WBGene00012009 | -0.069358327 | 0.93244964 | 0.96416346 | T25B9.3 |
| WBGene00012010 | 0.688793783 | 0.86378755 | 1 | T25B9.4 |
| WBGene00012011 | 0.942308827 | 0.67468959 | 1 | T25B9.5 |
| WBGene00012012 | 0.641065534 | 0.78192938 | 1 | T25B9.6 |
| WBGene00012013 | 2.299579985 | 0.00406729 | 0.02662045 | ugt-54 |
| WBGene00012014 | -0.088515738 | 0.79986667 | 0.89011744 | T25B9.8 |
| WBGene00012015 | -0.051746014 | 0.70125708 | 0.83141078 | T25B9.9 |
| WBGene00012016 | 1.204221346 | 0.06289275 | 0.18816681 | inpp-1 |
| WBGene00012017 | 0.57951263 | 0.57834359 | 1 | T25C8.1 |
| WBGene00012018 | 1.538294482 | 0.00070995 | 0.00704379 | T25C12.3 |
| WBGene00012019 | 0.228635217 | 0.29709629 | 0.50739149 | dkf-2 |
| WBGene00012020 | -0.466198038 | 0.001631 | 0.01333463 | gyg-2 |
| WBGene00012021 | -1.672366394 | 5.4159E-05 | 0.00092611 | T25E12.6 |
| WBGene00012022 | 0.983488853 | 0.80601565 | 1 | clec-39 |
| WBGene00012023 | -0.858902457 | 0.62736524 | 1 | clec-34 |
| WBGene00012024 | -1.159714244 | 0.75790634 | 1 | clec-29 |
| WBGene00012025 | 0.688793783 | 0.86378755 | 1 | clec-38 |
| WBGene00012026 | 0.03940405 | 0.99001443 | 1 | srz-100 |
| WBGene00012027 | 1.277603744 | 0.32678334 | 1 | fbxa-127 |
| WBGene00012028 | 1.034650399 | 0.54802945 | 1 | srz-99 |
| WBGene00012029 | -0.50075743 | 0.07544272 | 0.21271484 | T25G3.1 |
| WBGene00012030 | -0.283882581 | 0.08497273 | 0.22971585 | T25G3.3 |
| WBGene00012031 | -0.213828204 | 0.1639086 | 0.35007666 | T25G3.4 |
| WBGene00012032 | -0.508469938 | 0.39239514 | 0.60025626 | T26C5.2 |
| WBGene00012033 | 0.161964483 | 0.33513101 | 0.54330693 | T26C5.3 |
| WBGene00012034 | 0.921116989 | 0.30568177 | 0.51458342 | T26C5.4 |
| WBGene00012035 | -0.006551303 | 0.99341342 | 0.99669716 | T26C5.5 |
| WBGene00012037 | -0.784975857 | 0.00041701 | 0.00464123 | T26E3.4 |
| WBGene00012038 | -2.155933817 | 0.00815122 | 0.04443825 | T26E3.5 |
| WBGene00012039 | -1.159714244 | 0.75790634 | 1 | T26E3.6 |
| WBGene00012040 | -0.736789399 | 0.72029863 | 1 | T26E3.7 |
| WBGene00012041 | -0.436650383 | 0.67051183 | 1 | T26E3.8 |
| WBGene00012042 | 1.421605668 | 0.37723678 | 1 | T26E3.10 |
| WBGene00012043 | -0.014519762 | 0.99603746 | 1 | irld-52 |
| WBGene00012044 | 1.353535409 | 0.73149271 | 1 | T26E4.2 |
| WBGene00012045 | 1.132701701 | 0.63662543 | 1 | T26E4.3 |
| WBGene00012046 | 0.489058549 | 0.85947456 | 1 | T26E4.4 |
| WBGene00012051 | -2.652727305 | 0.24933058 | 1 | T26E4.9 |
| WBGene00012055 | -0.233502874 | 0.92367326 | 1 | srsx-36 |
| WBGene00012056 | 1.477253415 | 0.58357327 | 1 | nhr-285 |
| WBGene00012059 | -0.288867031 | 0.16415337 | 0.35032273 | T26G10.1 |
| WBGene00012062 | 0.343945135 | 0.70688028 | 0.83444176 | T26G10.5 |

| | | | | |
|---|---|---|---|---|
| WBGene00012063 | -2.268027851 | 0.0319979 | 1 | fbxb-1 |
| WBGene00012064 | -2.361959926 | 0.10356358 | 1 | fbxb-115 |
| WBGene00012065 | -1.590688987 | 0.34649085 | 1 | fbxb-2 |
| WBGene00012066 | -2.046575414 | 0.08793722 | 1 | fbxb-116 |
| WBGene00012067 | 0.983488853 | 0.80601565 | 1 | sqst-3 |
| WBGene00012068 | -0.414595455 | 0.77375412 | 1 | oac-49 |
| WBGene00012069 | 0.983488853 | 0.80601565 | 1 | T26H5.4 |
| WBGene00012070 | 1.084450517 | 0.51640318 | 1 | T26H5.8 |
| WBGene00012073 | 0.478726456 | 0.24572207 | 0.45373562 | cpd-2 |
| WBGene00012074 | -0.987258363 | 0.05182892 | 0.1644251 | T27A8.2 |
| WBGene00012077 | -0.156269293 | 0.96924949 | 1 | T27A8.5 |
| WBGene00012081 | 1.413734769 | 0.68804531 | 1 | T27C5.8 |
| WBGene00012084 | 1.400755655 | 0.05041928 | 0.16181351 | npr-15 |
| WBGene00012085 | 1.011975225 | 0.00042395 | 0.00470019 | T27D12.1 |
| WBGene00012086 | 0.688793783 | 0.86378755 | 1 | clec-144 |
| WBGene00012087 | 1.977885804 | 0.49204652 | 1 | T27E7.1 |
| WBGene00012091 | 0.281454817 | 0.94435481 | 1 | pals-29 |
| WBGene00012094 | -0.591803684 | 0.00015087 | 0.00210782 | T27E9.2 |
| WBGene00012095 | -0.130620568 | 0.58507565 | 0.75465941 | pssy-2 |
| WBGene00012096 | -1.201568047 | 0.37912842 | 1 | T27E9.6 |
| WBGene00012097 | -0.155272254 | 0.33846993 | 0.5464579 | abcf-2 |
| WBGene00012099 | 0.92646786 | 0.09724663 | 0.25046061 | acc-4 |
| WBGene00012100 | 0.83468862 | 0.00092874 | 0.00870718 | sipa-1 |
| WBGene00012101 | 2.058111827 | 0.00157073 | 0.01297433 | zip-10 |
| WBGene00012102 | -0.558804446 | 0.87818675 | 1 | T27F6.1 |
| WBGene00012103 | -0.277909555 | 0.66875767 | 0.81015635 | clec-12 |
| WBGene00012104 | -0.497118404 | 0.0338345 | 0.1234329 | sygl-1 |
| WBGene00012105 | -0.23941512 | 0.32570164 | 0.53408206 | T27F6.6 |
| WBGene00012106 | -0.856661168 | 0.00864722 | 0.04641286 | T27F6.7 |
| WBGene00012107 | 0.57383038 | 0.25098825 | 0.45885383 | T27F6.8 |
| WBGene00012109 | -0.569495507 | 0.00944365 | 0.04965584 | T28A8.3 |
| WBGene00012110 | -0.183180076 | 0.44733565 | 0.64632766 | T28A8.4 |
| WBGene00012111 | -0.479269037 | 0.17378568 | 0.36371866 | T28A8.5 |
| WBGene00012112 | -0.259569085 | 0.29166779 | 0.5015777 | T28A8.6 |
| WBGene00012113 | 0.402048439 | 0.13306476 | 0.30637952 | T28B8.1 |
| WBGene00012114 | 1.589974054 | 0.06688758 | 0.1962025 | T28B8.3 |
| WBGene00012115 | 2.188838843 | 0.34004625 | 1 | T28B8.4 |
| WBGene00012116 | 3.317525099 | 0.00809852 | 0.04425031 | del-4 |
| WBGene00012117 | 0.801179972 | 0.00018997 | 0.0025401 | T28B11.1 |
| WBGene00012118 | 0.524763103 | 0.21063227 | 0.41041347 | grsp-2 |
| WBGene00012119 | -1.18256372 | 0.16204707 | 0.3479069 | T28C6.3 |
| WBGene00012120 | 1.019267722 | 0.67145278 | 1 | T28C6.5 |
| WBGene00012121 | 0.770088923 | 0.00029605 | 0.00357295 | T28C6.7 |

| | | | | |
|---|---|---|---|---|
| WBGene00012122 | -0.415673576 | 0.38978901 | 0.59820862 | T28C6.8 |
| WBGene00012123 | -0.632495294 | 0.05327385 | 0.16764609 | T28D6.3 |
| WBGene00012124 | 0.522210416 | 0.20173812 | 0.40055476 | T28D6.4 |
| WBGene00012125 | 1.045916507 | 0.0002048 | 0.00267591 | T28D6.5 |
| WBGene00012126 | -0.470839294 | 0.00808281 | 0.04420426 | T28D6.6 |
| WBGene00012127 | 0.324381816 | 0.34582889 | 0.55467048 | T28D6.7 |
| WBGene00012128 | -0.216156362 | 0.27914831 | 0.48799707 | nra-1 |
| WBGene00012130 | -0.164759461 | 0.6226863 | 0.78035037 | slc-17.4 |
| WBGene00012131 | 2.252071624 | 0.004823 | 0.03040293 | T28F3.5 |
| WBGene00012132 | 0.606861681 | 0.56441563 | 1 | ifta-2 |
| WBGene00012134 | 1.351177858 | 5.45E-07 | 2.0351E-05 | T28F3.8 |
| WBGene00012135 | 1.256498809 | 0.34320541 | 1 | T28F3.9 |
| WBGene00012136 | -0.213792307 | 0.41389449 | 0.61938778 | T28F4.1 |
| WBGene00012137 | 1.444751719 | 0.28943596 | 1 | asic-2 |
| WBGene00012138 | -0.687709908 | 0.74016374 | 1 | T28F4.3 |
| WBGene00012139 | 0.855814847 | 0.03542204 | 0.12693596 | T28F4.4 |
| WBGene00012140 | 0.730783374 | 5.3683E-05 | 0.00091928 | T28F4.5 |
| WBGene00012142 | -0.06298299 | 0.91485001 | 0.95571968 | T28H10.1 |
| WBGene00012143 | 0.990251931 | 0.02915787 | 0.11104319 | T28H10.2 |
| WBGene00012144 | 0.642276588 | 0.00036709 | 0.00424057 | T28H10.3 |
| WBGene00012145 | -0.394320382 | 0.8739308 | 1 | nlp-46 |
| WBGene00012146 | 0.75595108 | 0.7708876 | 1 | VB0393L.2 |
| WBGene00012147 | 2.437484291 | 0.2708268 | 1 | VC27A7L.1 |
| WBGene00012148 | -0.399035798 | 0.02177624 | 0.09126178 | inos-1 |
| WBGene00012149 | -0.198811495 | 0.33044632 | 0.53866122 | VF13D12L.3 |
| WBGene00012150 | -0.213489245 | 0.37484454 | 0.58284342 | syx-17 |
| WBGene00012151 | -0.643676094 | 0.51617715 | 1 | VH15N14R.1 |
| WBGene00012152 | -0.143139951 | 0.72157261 | 0.84317738 | cnc-10 |
| WBGene00012155 | 0.054782935 | 0.97621874 | 1 | VW02B12L.2 |
| WBGene00012156 | -0.78028631 | 7.0842E-05 | 0.00116387 | ebp-2 |
| WBGene00012157 | -0.670468003 | 0.00830727 | 0.04498598 | adbp-1 |
| WBGene00012158 | -0.071075414 | 0.68012213 | 0.81656371 | ucr-2.1 |
| WBGene00012159 | 0.688793783 | 0.86378755 | 1 | VY10G11R.1 |
| WBGene00012162 | 0.534857991 | 0.50990845 | 0.69661286 | sek-6 |
| WBGene00012163 | 0.800895307 | 0.53508164 | 1 | VZK822L.2 |
| WBGene00012164 | 0.194117132 | 0.57732354 | 0.74903703 | W01A8.2 |
| WBGene00012165 | 0.631525855 | 0.69457241 | 1 | cutl-6 |
| WBGene00012166 | -0.642634101 | 8.306E-05 | 0.001321 | nuo-6 |
| WBGene00012167 | -0.238211914 | 0.20736468 | 0.40685432 | tofu-5 |
| WBGene00012168 | -0.701279695 | 0.26561231 | 0.47430491 | W01A8.6 |
| WBGene00012169 | 2.188838843 | 0.34004625 | 1 | W01B6.2 |
| WBGene00012170 | 0.18787372 | 0.8889773 | 1 | W01B6.3 |
| WBGene00012171 | -0.156269293 | 0.96924949 | 1 | W01B6.4 |

| | | | | |
|---|---|---|---|---|
| WBGene00012172 | -0.512603405 | 0.83307778 | 1 | W01B6.5 |
| WBGene00012173 | -0.245531902 | 0.94070824 | 1 | W01B6.6 |
| WBGene00012175 | 1.034262533 | 0.3271684 | 0.53554693 | W01C9.1 |
| WBGene00012176 | 0.813686433 | 0.30463726 | 0.51376594 | W01C9.2 |
| WBGene00012177 | 0.574508983 | 0.8551971 | 1 | decr-1.2 |
| WBGene00012178 | 0.612043136 | 0.53513146 | 1 | glb-27 |
| WBGene00012179 | -0.553945166 | 6.1455E-05 | 0.00103057 | W01D2.1 |
| WBGene00012180 | 0.163609849 | 0.88549819 | 1 | W01D2.3 |
| WBGene00012182 | -0.121163639 | 0.6068221 | 0.76973843 | osta-3 |
| WBGene00012183 | 0.024971563 | 0.99027335 | 1 | W01D2.6 |
| WBGene00012184 | 0.117219948 | 0.70338994 | 0.83258821 | mnr-1 |
| WBGene00012185 | -0.964635475 | 0.00020285 | 0.0026567 | W01F3.2 |
| WBGene00012186 | 0.615367326 | 0.0845931 | 0.22909766 | mlt-11 |
| WBGene00012187 | -0.287370472 | 0.2805302 | 0.48963731 | rpb-11 |
| WBGene00012188 | -0.882907524 | 0.00015488 | 0.00214392 | W01G7.4 |
| WBGene00012190 | 2.052334667 | 0.24873879 | 1 | W02A2.5 |
| WBGene00012191 | 0.239360828 | 0.91077854 | 1 | W02A2.8 |
| WBGene00012192 | -0.398802461 | 0.23081081 | 0.4363442 | W02A11.1 |
| WBGene00012193 | -0.081657474 | 0.74055052 | 0.85525612 | vps-25 |
| WBGene00012194 | 0.262815672 | 0.35271058 | 0.56168142 | toe-4 |
| WBGene00012195 | 0.197194741 | 0.70799828 | 0.83494969 | bath-34 |
| WBGene00012196 | 0.698568329 | 0.43812162 | 0.63866548 | bath-35 |
| WBGene00012197 | 0.750916507 | 0.19234281 | 0.38739782 | W02B8.1 |
| WBGene00012198 | -0.314638163 | 0.23206921 | 0.43764402 | W02B8.2 |
| WBGene00012199 | -0.082548247 | 0.93854992 | 1 | mltn-3 |
| WBGene00012200 | -0.156269293 | 0.96924949 | 1 | mltn-4 |
| WBGene00012201 | 0.953723521 | 0.00228397 | 0.01726632 | W02B12.1 |
| WBGene00012202 | 0.746020233 | 0.11504747 | 0.27930241 | W02B12.4 |
| WBGene00012203 | -0.084852783 | 0.69811276 | 0.8290089 | rga-1 |
| WBGene00012204 | 0.210500432 | 0.362956 | 0.57095735 | mfn-1 |
| WBGene00012205 | -0.401613355 | 0.03533599 | 0.12675342 | W02B12.10 |
| WBGene00012206 | -0.134049009 | 0.52587288 | 0.71034964 | W02B12.11 |
| WBGene00012207 | 0.173646189 | 0.93792136 | 1 | W02B12.12 |
| WBGene00012208 | -0.285074489 | 0.18170002 | 0.37382024 | W02D9.2 |
| WBGene00012209 | -0.109243662 | 0.73921688 | 0.85446788 | hmg-20 |
| WBGene00012210 | -0.051590265 | 0.90776761 | 0.95184971 | W02D9.4 |
| WBGene00012211 | -0.056723817 | 0.9299186 | 0.96354103 | ssp-37 |
| WBGene00012212 | 0.037218881 | 0.9263087 | 0.96122341 | W02D9.6 |
| WBGene00012213 | 0.118379794 | 0.54133858 | 0.72208215 | W02D9.7 |
| WBGene00012216 | 0.633116985 | 0.24964712 | 0.45743873 | W02D9.10 |
| WBGene00012217 | -0.16504593 | 0.7294629 | 0.84847759 | W02H3.1 |
| WBGene00012219 | 1.712479368 | 0.58529868 | 1 | W03C9.1 |
| WBGene00012220 | -0.433222152 | 0.0350257 | 0.12611121 | W03C9.2 |

| | | | | |
|---|---|---|---|---|
| WBGene00012221 | -0.922684609 | 0.00045871 | 0.00500832 | W03C9.5 |
| WBGene00012222 | 0.255488188 | 0.66629011 | 0.80861938 | W03C9.6 |
| WBGene00012224 | 4.325182333 | 0.00515451 | 1 | W03G11.2 |
| WBGene00012225 | 0.491646233 | 0.2897576 | 0.49970839 | W03G11.3 |
| WBGene00012226 | 0.852011286 | 0.00439272 | 0.02829469 | W03G11.4 |
| WBGene00012227 | -0.668264701 | 0.85551153 | 1 | W03H9.1 |
| WBGene00012228 | 0.773564037 | 0.4421352 | 1 | W03H9.2 |
| WBGene00012230 | -0.266449144 | 0.12536012 | 0.2952468 | cacn-1 |
| WBGene00012231 | -0.749181515 | 0.12095004 | 0.28776048 | W04A4.2 |
| WBGene00012233 | -0.156269293 | 0.96924949 | 1 | W04A4.4 |
| WBGene00012234 | 0.264457078 | 0.15589411 | 0.33968242 | W04A4.5 |
| WBGene00012235 | 0.052392893 | 0.93998054 | 0.9689509 | W04A4.6 |
| WBGene00012236 | -0.169372549 | 0.57625909 | 0.74891475 | W04A8.1 |
| WBGene00012237 | -0.446774074 | 0.31596367 | 0.52405848 | W04A8.2 |
| WBGene00012238 | -2.135541052 | 0.01791878 | 0.07908105 | W04A8.3 |
| WBGene00012239 | -0.778864651 | 0.01286012 | 0.06240922 | W04A8.4 |
| WBGene00012240 | -1.443665019 | 0.01534751 | 0.07082254 | W04A8.5 |
| WBGene00012241 | -0.524672914 | 0.00500512 | 0.03117864 | W04A8.6 |
| WBGene00012242 | -1.706203807 | 0.43490097 | 1 | srt-39 |
| WBGene00012243 | -0.266551752 | 0.11594268 | 0.28046677 | W04D2.4 |
| WBGene00012244 | -0.571409066 | 0.00284183 | 0.02041575 | mrps-11 |
| WBGene00012245 | 0.181948776 | 0.27974195 | 0.48875335 | W04D2.6 |
| WBGene00012246 | -1.901856996 | 0.64072034 | 1 | fbxa-131 |
| WBGene00012247 | -0.223881261 | 0.75480955 | 0.86283239 | W04E12.2 |
| WBGene00012249 | 1.216957619 | 0.74819621 | 1 | W04E12.4 |
| WBGene00012250 | -0.156269293 | 0.96924949 | 1 | W04E12.5 |
| WBGene00012251 | 0.506018127 | 0.2469973 | 0.45497554 | clec-49 |
| WBGene00012252 | 0.502157508 | 0.36649053 | 0.5740629 | W04E12.7 |
| WBGene00012253 | 0.44012225 | 0.01152405 | 0.05764872 | clec-50 |
| WBGene00012254 | 0.352284763 | 0.84067521 | 1 | W04E12.9 |
| WBGene00012255 | -0.245709402 | 0.53885133 | 0.72066769 | lpr-6 |
| WBGene00012256 | -0.287804011 | 0.43677499 | 0.63755094 | lpr-5 |
| WBGene00012257 | -0.600084177 | 0.14089868 | 0.31867564 | lpr-4 |
| WBGene00012258 | -0.391284795 | 0.0576332 | 0.17782076 | W04G3.5 |
| WBGene00012259 | 0.87762818 | 0.0267574 | 0.10479407 | sulp-7 |
| WBGene00012260 | 0.041803775 | 0.98890197 | 1 | W04G3.7 |
| WBGene00012261 | -0.235846232 | 0.47125512 | 0.6641333 | lpr-3 |
| WBGene00012264 | -0.156269293 | 0.96924949 | 1 | W04G5.3 |
| WBGene00012265 | 0.332225021 | 0.78363865 | 1 | W04G5.4 |
| WBGene00012266 | 1.687239501 | 0.59403242 | 1 | W04G5.5 |
| WBGene00012268 | 0.682069186 | 0.69825534 | 1 | W04G5.8 |
| WBGene00012269 | 3.199079286 | 0.09328367 | 1 | W04G5.9 |
| WBGene00012270 | 0.929999259 | 0.44876857 | 1 | W04G5.10 |

| | | | | |
|---|---|---|---|---|
| WBGene00012271 | -0.30359982 | 0.8981102 | 1 | W05B2.2 |
| WBGene00012272 | 0.684688396 | 0.5600874 | 1 | W05B2.7 |
| WBGene00012272 | 0.039145052 | 0.96518932 | 0.98275441 | dhc-4 |
| WBGene00012274 | 0.83828472 | 0.01461195 | 0.06833732 | W05B5.1 |
| WBGene00012275 | 2.062461988 | 0.10049302 | 1 | npr-14 |
| WBGene00012276 | -0.446855777 | 0.00510148 | 0.03161665 | his-74 |
| WBGene00012277 | -0.514469455 | 0.02237791 | 0.09298089 | ccch-3 |
| WBGene00012278 | 0.377828795 | 0.92526585 | 1 | W05B10.3 |
| WBGene00012279 | 0.283811272 | 0.67308446 | 0.81320713 | W05B10.4 |
| WBGene00012280 | 0.690146152 | 0.2634 | 0.47204093 | W05E10.1 |
| WBGene00012281 | 2.011668041 | 0.28000168 | 1 | W05E10.2 |
| WBGene00012282 | 1.050885131 | 0.21557887 | 0.41723944 | W05E10.5 |
| WBGene00012285 | 0.983488853 | 0.80601565 | 1 | pitr-6 |
| WBGene00012288 | -0.156269293 | 0.96924949 | 1 | sre-32 |
| WBGene00012289 | 0.593394846 | 0.04807931 | 0.15679046 | W05H12.1 |
| WBGene00012290 | 0.56291754 | 0.42747681 | 0.63057665 | W05H12.2 |
| WBGene00012291 | -0.013734836 | 0.96381972 | 0.98190304 | W06A7.2 |
| WBGene00012293 | -0.174861414 | 0.76733595 | 0.86965734 | W06A7.4 |
| WBGene00012294 | 0.489058549 | 0.85947456 | 1 | nspa-8 |
| WBGene00012295 | 1.16199909 | 0.12472944 | 0.29432717 | nmat-1 |
| WBGene00012296 | 1.71839177 | 0.3863606 | 1 | spe-46 |
| WBGene00012297 | 1.181686151 | 0.7579682 | 1 | W06D4.3 |
| WBGene00012298 | 0.082793742 | 0.70569038 | 0.83352339 | prmt-7 |
| WBGene00012299 | -0.624596109 | 0.04563181 | 0.15083469 | W06D11.1 |
| WBGene00012300 | 0.411711365 | 0.78556121 | 1 | W06D11.2 |
| WBGene00012301 | -0.380300271 | 0.24773963 | 0.45583943 | W06D11.3 |
| WBGene00012304 | -0.263229581 | 0.93409872 | 1 | cutl-4 |
| WBGene00012306 | 0.143779602 | 0.60504823 | 0.76897128 | W06F12.2 |
| WBGene00012307 | -0.856127822 | 0.60475275 | 1 | W06F12.3 |
| WBGene00012308 | 1.923307241 | 0.17793488 | 1 | oac-53 |
| WBGene00012311 | -0.156269293 | 0.96924949 | 1 | W06G6.9 |
| WBGene00012315 | -0.331786834 | 0.09186553 | 0.24130927 | immt-2 |
| WBGene00012316 | -0.249017466 | 0.22935828 | 0.43442105 | ctps-1 |
| WBGene00012317 | -0.028023659 | 0.90511634 | 0.95070175 | ztf-6 |
| WBGene00012318 | 1.876192691 | 0.43542138 | 1 | srxa-1 |
| WBGene00012319 | -0.215122969 | 0.28351085 | 0.49320895 | ipla-3 |
| WBGene00012321 | 0.041803775 | 0.98890197 | 1 | srxa-6 |
| WBGene00012322 | 1.423786247 | 0.00971304 | 0.05074268 | W07A12.4 |
| WBGene00012322 | 1.38322681 | 0.69550305 | 1 | Y9C2UA.2 |
| WBGene00012323 | 1.611403277 | 0.01472426 | 0.06878307 | oac-54 |
| WBGene00012324 | 0.145479197 | 0.60546758 | 0.76897128 | rhy-1 |
| WBGene00012325 | 0.621203228 | 0.37617739 | 0.58381308 | W07A12.8 |
| WBGene00012326 | -0.077708015 | 0.75397318 | 0.86260728 | W07E11.1 |

| | | | | |
|---|---|---|---|---|
| WBGene00012328 | 0.799651695 | 0.62490034 | 1 | W07G1.1 |
| WBGene00012329 | 0.438455499 | 0.87116517 | 1 | sre-44 |
| WBGene00012330 | -0.197901054 | 0.46922832 | 0.66289284 | zip-3 |
| WBGene00012332 | 0.031820289 | 0.97856644 | 1 | rpi-1 |
| WBGene00012333 | 1.092347874 | 0.62265948 | 1 | sre-43 |
| WBGene00012334 | -0.156269293 | 0.96924949 | 1 | W07G1.7 |
| WBGene00012335 | -0.586446112 | 0.7472399 | 1 | W07G4.1 |
| WBGene00012336 | 0.983488853 | 0.80601565 | 1 | W07G4.2 |
| WBGene00012337 | 0.189812288 | 0.28429086 | 0.49417901 | W07G4.3 |
| WBGene00012339 | 0.559301553 | 0.0327793 | 0.12074757 | W07G4.5 |
| WBGene00012340 | 1.918923633 | 0.11917317 | 1 | dct-15 |
| WBGene00012341 | 0.323109786 | 0.06556264 | 0.19336894 | catp-6 |
| WBGene00012342 | -0.17401871 | 0.28166953 | 0.49104955 | mtr-4 |
| WBGene00012343 | -0.074905921 | 0.74446813 | 0.8574972 | W08E3.2 |
| WBGene00012344 | -0.138924119 | 0.32391716 | 0.53245035 | ola-1 |
| WBGene00012345 | -1.885565722 | 0.26984569 | 1 | W08E3.4 |
| WBGene00012346 | -1.478994158 | 0.15252246 | 1 | W08G11.1 |
| WBGene00012347 | -0.027268108 | 0.90670253 | 0.95121412 | W08G11.3 |
| WBGene00012348 | 0.349293392 | 0.14524605 | 0.32397939 | pptr-1 |
| WBGene00012351 | -0.14650446 | 0.38075141 | 0.58894069 | W09C5.1 |
| WBGene00012352 | -0.005678697 | 0.9848072 | 0.99257382 | dkf-1 |
| WBGene00012353 | -0.265513117 | 0.18154395 | 0.37370721 | W09C5.7 |
| WBGene00012354 | -0.366328103 | 0.02346126 | 0.0959068 | W09C5.8 |
| WBGene00012356 | 0.006076123 | 0.98545513 | 0.99306198 | W09D6.1 |
| WBGene00012357 | -2.078952801 | 0.31011433 | 1 | W09D6.4 |
| WBGene00012358 | 0.360443816 | 0.14272467 | 0.32052915 | W09D6.5 |
| WBGene00012359 | -0.0903815 | 0.68202011 | 0.81789867 | W09D10.1 |
| WBGene00012360 | 0.323106851 | 0.17266884 | 0.36231807 | tat-3 |
| WBGene00012361 | -0.593520373 | 0.0066327 | 0.03825242 | mrpl-12 |
| WBGene00012362 | -0.017483095 | 0.93398876 | 0.96515087 | W09D10.4 |
| WBGene00012363 | 0.368447106 | 0.55466634 | 0.7323465 | W09D10.5 |
| WBGene00012364 | 0.0386997 | 0.87710738 | 0.93403742 | zmp-4 |
| WBGene00012365 | -0.216182094 | 0.46090855 | 0.65630483 | W09G3.1 |
| WBGene00012366 | 0.261216713 | 0.39756678 | 0.60517281 | maea-1 |
| WBGene00012369 | -0.053283316 | 0.7922333 | 0.88527422 | W09G3.6 |
| WBGene00012370 | -0.070229736 | 0.84899375 | 0.91721242 | W09G3.7 |
| WBGene00012371 | -0.793497777 | 0.01340278 | 0.06393839 | W09G3.8 |
| WBGene00012372 | 0.92461478 | 0.41345287 | 1 | W09H1.1 |
| WBGene00012373 | 2.420661489 | 0.31300137 | 1 | W09H1.3 |
| WBGene00012374 | 0.451481963 | 0.72522035 | 1 | W09H1.4 |
| WBGene00012375 | -0.42846795 | 0.02672598 | 0.10473002 | mecr-1 |
| WBGene00012376 | -0.256002995 | 0.12003167 | 0.28624902 | nduf-7 |
| WBGene00012378 | 1.216957619 | 0.74819621 | 1 | W10G6.1 |

| | | | | |
|---|---|---|---|---|
| WBGene00012379 | 1.216957619 | 0.74819621 | 1 | Y1A5A.1 |
| WBGene00012381 | 1.532509319 | 0.01993957 | 0.08524167 | Y2H9A.4 |
| WBGene00012382 | -0.073284328 | 0.64112233 | 0.79235567 | ttr-16 |
| WBGene00012383 | -0.155180988 | 0.56278883 | 0.7382069 | ttr-17 |
| WBGene00012385 | -0.118510311 | 0.59077512 | 0.75815153 | Y5F2A.4 |
| WBGene00012386 | 0.38208239 | 0.08197059 | 0.22442154 | agef-1 |
| WBGene00012387 | 1.32573741 | 0.08884306 | 0.23635972 | Y6B3B.1 |
| WBGene00012389 | -0.025494249 | 0.90014946 | 0.94737686 | Y6B3B.4 |
| WBGene00012390 | -0.551789769 | 0.11330561 | 0.27628911 | gpap-1 |
| WBGene00012391 | -0.704817435 | 0.36447065 | 0.57203053 | Y6B3B.7 |
| WBGene00012393 | -0.293942691 | 0.34802949 | 0.55657162 | Y6B3B.9 |
| WBGene00012394 | 1.116613983 | 0.69067248 | 1 | hsd-1 |
| WBGene00012395 | -0.336451702 | 0.14454958 | 0.32307998 | Y6D1A.1 |
| WBGene00012396 | 0.688071334 | 0.30528994 | 0.51436653 | Y6D1A.2 |
| WBGene00012398 | 0.299360127 | 0.87105618 | 1 | Y6E2A.4 |
| WBGene00012399 | 1.671705311 | 0.19727469 | 1 | Y6E2A.5 |
| WBGene00012400 | -0.199924012 | 0.91969149 | 1 | Y6E2A.7 |
| WBGene00012401 | -1.969805064 | 0.62862569 | 1 | irld-57 |
| WBGene00012404 | 1.659289578 | 0.00106664 | 0.00967868 | Y6G8.2 |
| WBGene00012405 | -0.107342738 | 0.78502468 | 0.88176377 | ztf-25 |
| WBGene00012407 | 0.405854784 | 0.0383414 | 0.13364999 | Y7A5A.1 |
| WBGene00012408 | 0.645977806 | 0.54471332 | 1 | Y7A5A.2 |
| WBGene00012409 | 0.688793783 | 0.86378755 | 1 | Y7A5A.3 |
| WBGene00012411 | -0.156269293 | 0.96924949 | 1 | Y7A5A.5 |
| WBGene00012412 | -0.132855481 | 0.92921491 | 1 | Y7A5A.6 |
| WBGene00012413 | 1.179295881 | 0.61449445 | 1 | Y7A5A.7 |
| WBGene00012414 | -1.14715222 | 0.68932198 | 1 | Y7A5A.8 |
| WBGene00012415 | -0.156269293 | 0.96924949 | 1 | Y7A5A.9 |
| WBGene00012416 | 1.30758989 | 0.00369894 | 0.02478415 | Y7A9A.1 |
| WBGene00012420 | -1.565785585 | 0.00644958 | 0.03752118 | Y7A9D.1 |
| WBGene00012421 | 0.880655689 | 0.01235784 | 0.06057188 | Y9C2UA.1 |
| WBGene00012423 | -0.176750541 | 0.35408285 | 0.56261222 | Y10G11A.1 |
| WBGene00012424 | -0.032316279 | 0.95529392 | 0.97710391 | dlc-3 |
| WBGene00012425 | -1.289631337 | 0.39959847 | 1 | dlc-4 |
| WBGene00012428 | -0.243954817 | 0.39824467 | 0.60593292 | Y11D7A.3 |
| WBGene00012429 | 0.92183293 | 0.4937219 | 1 | Y11D7A.5 |
| WBGene00012431 | 0.087516292 | 0.74095754 | 0.85545507 | Y11D7A.7 |
| WBGene00012432 | -0.543426962 | 0.67475712 | 1 | Y11D7A.8 |
| WBGene00012433 | -1.247207081 | 0.07356787 | 0.20913483 | Y11D7A.9 |
| WBGene00012434 | -0.521634616 | 0.01695874 | 0.07591504 | Y11D7A.10 |
| WBGene00012435 | -0.355440457 | 0.06330798 | 0.18893706 | flh-1 |
| WBGene00012436 | -0.430579098 | 0.0782195 | 0.21746833 | flh-3 |
| WBGene00012437 | 0.953987974 | 0.06512255 | 0.19260728 | hum-9 |

| | | | | |
|---|---|---|---|---|
| WBGene00012439 | -0.285267531 | 0.48967685 | 0.67924367 | Y12A6A.1 |
| WBGene00012440 | 0.921086724 | 0.22037541 | 0.42341792 | nlp-53 |
| WBGene00012441 | 1.278809585 | 0.40981956 | 1 | glb-28 |
| WBGene00012442 | -1.189045893 | 0.28733899 | 1 | Y15E3A.3 |
| WBGene00012443 | 0.757052307 | 0.00117765 | 0.010406 | Y15E3A.4 |
| WBGene00012444 | 0.794684692 | 0.1121268 | 0.27440666 | Y15E3A.5 |
| WBGene00012445 | 1.868019538 | 0.00150678 | 0.01261646 | Y16B4A.2 |
| WBGene00012446 | 0.930006706 | 0.156157 | 0.34013313 | nhr-230 |
| WBGene00012448 | 1.360555046 | 0.51491783 | 1 | cyp-33D3 |
| WBGene00012449 | 2.7782498 | 0.03422547 | 1 | nhr-231 |
| WBGene00012450 | 1.429963739 | 0.3701028 | 1 | Y17D7B.2 |
| WBGene00012451 | 1.712479368 | 0.58529868 | 1 | Y17D7B.3 |
| WBGene00012452 | -0.071036833 | 0.86617739 | 0.92799221 | Y17D7B.4 |
| WBGene00012455 | 0.747845955 | 0.85215965 | 1 | Y17D7C.1 |
| WBGene00012456 | 1.800348045 | 0.06200527 | 0.18638425 | Y17D7C.2 |
| WBGene00012457 | -1.659259697 | 0.0457706 | 0.15112887 | acbp-6 |
| WBGene00012458 | -0.067818906 | 0.78581753 | 0.88216338 | ash-2 |
| WBGene00012459 | -0.481299022 | 0.15159618 | 0.33336244 | Y17G7B.3 |
| WBGene00012460 | -0.619704624 | 0.07461769 | 0.21102571 | dhps-1 |
| WBGene00012461 | 1.256287632 | 0.12825905 | 0.29905763 | glb-29 |
| WBGene00012462 | 0.659075786 | 0.03539947 | 0.12689254 | Y17G7B.8 |
| WBGene00012463 | 0.345109142 | 0.0789128 | 0.21883972 | Y17G7B.10 |
| WBGene00012464 | 0.522760463 | 0.17953721 | 0.37178258 | arrd-7 |
| WBGene00012465 | -0.328462743 | 0.20988624 | 0.41007351 | Y17G7B.12 |
| WBGene00012466 | 0.311573255 | 0.25481614 | 0.46230167 | ippk-1 |
| WBGene00012467 | 0.614407921 | 0.77316873 | 1 | arrd-8 |
| WBGene00012468 | -0.183755792 | 0.44831489 | 0.64735123 | Y17G7B.17 |
| WBGene00012469 | -0.121566032 | 0.67555619 | 0.81441045 | Y17G7B.18 |
| WBGene00012470 | 0.863501219 | 0.69189182 | 1 | Y17G7B.19 |
| WBGene00012471 | 0.218592649 | 0.19424241 | 0.3903819 | Y17G7B.20 |
| WBGene00012472 | 0.16989677 | 0.43734627 | 0.63804213 | Y17G7B.21 |
| WBGene00012473 | 0.59635585 | 0.68228687 | 1 | Y17G7B.22 |
| WBGene00012474 | 0.179852362 | 0.42471756 | 0.62892251 | attf-6 |
| WBGene00012475 | -0.156269293 | 0.96924949 | 1 | Y18D10A.2 |
| WBGene00012476 | -0.764695314 | 0.16003918 | 0.3453741 | Y18D10A.3 |
| WBGene00012478 | 0.681541609 | 0.02280219 | 0.09410478 | mxt-1 |
| WBGene00012479 | -0.312870119 | 0.31577646 | 0.52391104 | Y18D10A.9 |
| WBGene00012480 | 0.545569339 | 0.73484368 | 1 | clec-104 |
| WBGene00012481 | -0.765945607 | 6.6756E-06 | 0.00016049 | Y18D10A.11 |
| WBGene00012482 | -1.14715222 | 0.68932198 | 1 | clec-106 |
| WBGene00012483 | -0.381562349 | 0.22638954 | 0.43067932 | Y18D10A.16 |
| WBGene00012484 | -0.289948598 | 0.08715938 | 0.23361314 | car-1 |
| WBGene00012486 | -1.094201388 | 0.7816772 | 1 | Y18D10A.21 |

| | | | | |
|---|---|---|---|---|
| WBGene00012487 | 0.110897018 | 0.76590821 | 0.8688117 | Y18D10A.23 |
| WBGene00012490 | -2.392786778 | 0.10947893 | 1 | Y19D2B.2 |
| WBGene00012491 | 1.984243675 | 0.00435947 | 0.02812683 | Y20C6A.1 |
| WBGene00012492 | -0.816591821 | 0.10691774 | 0.26658958 | fbxb-7 |
| WBGene00012494 | 0.216131935 | 0.72122415 | 0.84294693 | nhr-232 |
| WBGene00012496 | -0.694547262 | 0.00048502 | 0.00524473 | Y24F12A.1 |
| WBGene00012497 | -0.619184993 | 2.9978E-05 | 0.00055908 | ragc-1 |
| WBGene00012500 | 0.616429445 | 0.81400458 | 1 | hpo-2 |
| WBGene00012501 | -0.007418224 | 0.9964921 | 1 | Y26D4A.3 |
| WBGene00012502 | -0.156269293 | 0.96924949 | 1 | clec-107 |
| WBGene00012512 | -0.724572869 | 0.00264876 | 0.01938432 | ekl-5 |
| WBGene00012513 | -0.34577889 | 0.57933523 | 0.75037463 | Y26E6A.2 |
| WBGene00012514 | -0.43890545 | 0.41341026 | 0.6189683 | Y26E6A.3 |
| WBGene00012515 | -1.901856996 | 0.64072034 | 1 | Y26G10.1 |
| WBGene00012518 | 2.098532724 | 0.16746138 | 1 | comt-2 |
| WBGene00012521 | 1.879256964 | 0.46994626 | 1 | Y32B12B.1 |
| WBGene00012522 | 0.024578913 | 0.89755831 | 0.94634211 | Y32B12B.2 |
| WBGene00012524 | -0.024968919 | 0.91975831 | 0.95789215 | Y32B12B.4 |
| WBGene00012525 | 1.063868465 | 0.2770474 | 0.48586342 | Y32B12C.1 |
| WBGene00012526 | 1.181285913 | 0.62032202 | 1 | Y32B12C.3 |
| WBGene00012527 | -0.633907912 | 0.00875389 | 0.04679922 | set-22 |
| WBGene00012528 | 0.103505285 | 0.53851723 | 0.72045594 | pap-1 |
| WBGene00012529 | 0.833173996 | 0.00048905 | 0.00527934 | Y32F6A.4 |
| WBGene00012530 | 0.761864869 | 1.2089E-06 | 3.9348E-05 | Y32F6A.5 |
| WBGene00012531 | 0.376416659 | 0.24190566 | 0.44956017 | Y32F6B.1 |
| WBGene00012532 | 1.074568236 | 0.04643616 | 0.15274491 | crp-1 |
| WBGene00012533 | 4.020944963 | 0.01332861 | 1 | fbxa-209 |
| WBGene00012535 | -0.426411171 | 0.63443796 | 0.78783996 | Y37A1A.2 |
| WBGene00012536 | 0.255502418 | 0.65943249 | 0.80368078 | Y37A1A.3 |
| WBGene00012538 | 1.772009533 | 1.42E-31 | 1.32E-28 | Y37A1B.5 |
| WBGene00012540 | -1.029881025 | 0.07674587 | 0.21490008 | Y37A1B.7 |
| WBGene00012541 | -0.604396148 | 0.82723443 | 1 | irld-55 |
| WBGene00012542 | 0.377828795 | 0.92526585 | 1 | irld-56 |
| WBGene00012543 | 0.064225804 | 0.73369914 | 0.85095923 | nkcc-1 |
| WBGene00012544 | 0.564540637 | 5.004E-05 | 0.00086544 | Y37D8A.2 |
| WBGene00012545 | -0.324189643 | 0.40903395 | 0.61469008 | Y37D8A.3 |
| WBGene00012546 | -0.379150148 | 0.09120587 | 0.24020184 | Y37D8A.4 |
| WBGene00012547 | 0.747845955 | 0.85215965 | 1 | Y37D8A.5 |
| WBGene00012548 | 0.55834492 | 0.18483508 | 0.37727857 | Y37D8A.6 |
| WBGene00012549 | -0.861206134 | 0.70037023 | 1 | Y37D8A.8 |
| WBGene00012550 | -0.729536399 | 1.0894E-05 | 0.00024494 | hpo-21 |
| WBGene00012551 | -0.392103902 | 0.1056706 | 0.26468882 | cec-7 |
| WBGene00012552 | 0.566047944 | 0.46484132 | 0.65929592 | enu-3.5 |

| | | | | |
|---|---|---|---|---|
| WBGene00012553 | -0.478009937 | 0.00041955 | 0.00466034 | cco-2 |
| WBGene00012554 | 0.652959692 | 0.03244961 | 0.1199302 | Y37D8A.16 |
| WBGene00012555 | -0.056849939 | 0.82638155 | 0.90625042 | npp-25 |
| WBGene00012556 | -0.387578249 | 0.2098512 | 0.41007351 | mrps-10 |
| WBGene00012557 | -0.591996659 | 7.9832E-05 | 0.001277 | Y37D8A.19 |
| WBGene00012558 | -0.378439435 | 0.0882327 | 0.23548763 | Y37D8A.21 |
| WBGene00012559 | -0.012763441 | 0.95290746 | 0.97614322 | epg-3 |
| WBGene00012560 | 0.341323155 | 0.23602693 | 0.44215444 | Y37D8A.25 |
| WBGene00012562 | -0.181577108 | 0.42543582 | 0.62952058 | Y37H2A.1 |
| WBGene00012564 | 0.644585658 | 0.18044021 | 0.3725955 | fbxa-107 |
| WBGene00012565 | -0.209366878 | 0.43578762 | 0.63676706 | fbxa-210 |
| WBGene00012566 | -0.094599666 | 0.82850989 | 0.90735509 | fbxa-211 |
| WBGene00012567 | -0.085379426 | 0.9112669 | 0.95393098 | Y37H2A.7 |
| WBGene00012570 | -0.107268465 | 0.96117914 | 1 | Y37H2A.10 |
| WBGene00012571 | 0.281454817 | 0.94435481 | 1 | Y37H2A.11 |
| WBGene00012573 | 1.447454397 | 0.13160948 | 0.30404275 | Y37H2C.1 |
| WBGene00012574 | 0.688793783 | 0.86378755 | 1 | fbxa-213 |
| WBGene00012576 | -0.265439495 | 0.52279916 | 0.70777259 | Y37H9A.1 |
| WBGene00012577 | 1.712479368 | 0.58529868 | 1 | Y37H9A.2 |
| WBGene00012578 | 0.250351631 | 0.33534666 | 0.54351139 | Y37H9A.3 |
| WBGene00012581 | -1.49211505 | 0.06246529 | 0.18730329 | Y38E10A.3 |
| WBGene00012582 | 0.636653463 | 0.29927319 | 0.50906705 | clec-8 |
| WBGene00012583 | 0.553223154 | 0.03337667 | 0.12205606 | clec-4 |
| WBGene00012584 | -0.029057572 | 0.88712861 | 0.94044069 | ceh-100 |
| WBGene00012585 | 0.99787225 | 5.056E-06 | 0.00012762 | lips-15 |
| WBGene00012586 | 0.232157325 | 0.93397866 | 1 | Y38E10A.8 |
| WBGene00012587 | 0.747845955 | 0.85215965 | 1 | Y38E10A.9 |
| WBGene00012589 | 2.365725856 | 0.35238907 | 1 | nspe-4 |
| WBGene00012591 | 0.351891279 | 0.40540861 | 0.61226986 | nspe-1 |
| WBGene00012592 | -0.006678504 | 0.98907284 | 0.99484196 | Y38E10A.14 |
| WBGene00012593 | 0.275375102 | 0.7497371 | 0.86036058 | nspe-7 |
| WBGene00012594 | 0.348763818 | 0.68790207 | 0.82242734 | nspe-5 |
| WBGene00012595 | 1.414401343 | 0.4425696 | 1 | Y38E10A.17 |
| WBGene00012596 | 0.281454817 | 0.94435481 | 1 | nhr-234 |
| WBGene00012597 | -0.848512853 | 0.76804407 | 1 | nhr-235 |
| WBGene00012600 | -0.573295131 | 0.00210607 | 0.01619363 | Y38E10A.22 |
| WBGene00012602 | -0.510160931 | 0.04178294 | 0.14193806 | Y38E10A.24 |
| WBGene00012603 | 0.556938092 | 0.68748906 | 1 | nspe-6 |
| WBGene00012604 | 1.493896825 | 0.03255538 | 0.12021133 | nspe-2 |
| WBGene00012605 | -0.977467865 | 0.43055295 | 1 | Y38F1A.1 |
| WBGene00012606 | 0.308777283 | 0.58194385 | 0.75273854 | Y38F1A.2 |
| WBGene00012607 | -0.500582638 | 0.75308706 | 1 | Y38F1A.4 |
| WBGene00012608 | 0.071000759 | 0.64285447 | 0.79350306 | hphd-1 |

| | | | | |
|---|---|---|---|---|
| WBGene00012609 | 0.938982574 | 0.8146943 | 1 | Y38F1A.7 |
| WBGene00012610 | 0.253305535 | 0.69724031 | 0.82870235 | Y38F1A.8 |
| WBGene00012614 | -0.604396148 | 0.82723443 | 1 | Y38H6A.3 |
| WBGene00012615 | -0.462449826 | 0.00150019 | 0.01257863 | dct-16 |
| WBGene00012616 | 1.477253415 | 0.58357327 | 1 | dct-14 |
| WBGene00012618 | 0.903390376 | 0.71900339 | 1 | dct-10 |
| WBGene00012621 | 2.18453076 | 0.41642786 | 1 | Y38H6C.8 |
| WBGene00012622 | 1.605852248 | 0.24021477 | 1 | Y38H6C.9 |
| WBGene00012624 | 1.145916945 | 0.04715311 | 0.15476775 | fbxa-150 |
| WBGene00012626 | 0.04838326 | 0.94932725 | 0.97411952 | famh-161 |
| WBGene00012627 | -0.07748118 | 0.98067866 | 1 | Y38H6C.15 |
| WBGene00012628 | -0.48350052 | 0.4233426 | 0.6277238 | Y38H6C.16 |
| WBGene00012629 | 1.118316683 | 0.22595202 | 0.43025399 | slc-36.3 |
| WBGene00012630 | -1.901856996 | 0.64072034 | 1 | Y38H6C.18 |
| WBGene00012631 | 1.312956797 | 0.16169315 | 0.34763885 | Y38H6C.19 |
| WBGene00012632 | 0.361545323 | 0.54828391 | 0.72686189 | Y38H6C.20 |
| WBGene00012633 | 0.937689552 | 0.19001048 | 0.38435739 | Y38H6C.21 |
| WBGene00012634 | 1.076956369 | 0.01829472 | 0.08036042 | Y38H6C.23 |
| WBGene00012635 | 0.479897284 | 0.76425895 | 1 | Y38H8A.1 |
| WBGene00012637 | -0.71991922 | 0.62953508 | 1 | Y38H8A.3 |
| WBGene00012638 | 0.983488853 | 0.80601565 | 1 | Y38H8A.4 |
| WBGene00012639 | -0.284202651 | 0.88416022 | 1 | fezf-1 |
| WBGene00012640 | -0.384371514 | 0.34906381 | 0.55755608 | Y38H8A.7 |
| WBGene00012641 | -0.37934669 | 0.04323948 | 0.14505955 | epg-6 |
| WBGene00012642 | -0.668264701 | 0.85551153 | 1 | Y39A1A.2 |
| WBGene00012643 | -0.365842909 | 0.18937618 | 0.38345835 | Y39A1A.3 |
| WBGene00012644 | 0.530040141 | 0.02164932 | 0.09082393 | rabx-5 |
| WBGene00012645 | -0.428508344 | 0.10690722 | 0.26658958 | mrpl-22 |
| WBGene00012646 | -0.186037501 | 0.40803821 | 0.6141434 | lron-10 |
| WBGene00012647 | -0.31474489 | 0.51894203 | 0.70445818 | swt-4 |
| WBGene00012648 | 0.581735642 | 0.18832456 | 0.38209366 | Y39A1A.9 |
| WBGene00012649 | -0.156269293 | 0.96924949 | 1 | Y39A1A.10 |
| WBGene00012650 | -0.35311177 | 0.03583942 | 0.1277154 | orc-1 |
| WBGene00012651 | -0.743171588 | 0.00050451 | 0.00539265 | orc-4 |
| WBGene00012652 | -0.657354554 | 0.00331065 | 0.02278607 | Y39A1A.14 |
| WBGene00012654 | -1.242317703 | 0.30369622 | 1 | Y39A1A.16 |
| WBGene00012655 | 0.377828795 | 0.92526585 | 1 | Y39A1A.17 |
| WBGene00012656 | 1.557897343 | 0.63813005 | 1 | Y39A1A.18 |
| WBGene00012657 | 0.285009593 | 0.60431962 | 0.76867617 | Y39A1A.20 |
| WBGene00012658 | -0.537938519 | 0.06836646 | 0.19916428 | Y39A1A.21 |
| WBGene00012659 | 0.068884764 | 0.78248464 | 0.88013211 | Y39A1A.22 |
| WBGene00012660 | 0.463880143 | 0.2189257 | 0.42169695 | Y39A1A.24 |
| WBGene00012661 | 0.281454817 | 0.94435481 | 1 | clec-163 |

| | | | | |
|---|---|---|---|---|
| WBGene00012662 | 0.323783693 | 0.60686053 | 0.76973843 | Y39A1C.1 |
| WBGene00012663 | 0.202617839 | 0.33036941 | 0.53866122 | hex-3 |
| WBGene00012664 | 2.687390365 | 3.74E-44 | 5.67E-41 | Y39B6A.1 |
| WBGene00012665 | 0.113137381 | 0.54345238 | 0.72331236 | pph-5 |
| WBGene00012666 | -0.75187862 | 0.00848753 | 0.04573774 | Y39B6A.3 |
| WBGene00012668 | -0.449294429 | 0.19098143 | 0.38569596 | Y39B6A.5 |
| WBGene00012669 | 0.771673761 | 0.47346591 | 1 | Y39B6A.7 |
| WBGene00012670 | 0.895573319 | 0.54927441 | 1 | Y39B6A.8 |
| WBGene00012671 | -1.082680044 | 0.67762556 | 1 | Y39B6A.9 |
| WBGene00012672 | -0.424824029 | 0.08720326 | 0.23361314 | Y39B6A.10 |
| WBGene00012673 | -0.470666635 | 0.48196584 | 0.67305582 | dyf-17 |
| WBGene00012674 | -0.399066751 | 0.27497319 | 0.48355295 | bed-1 |
| WBGene00012675 | 0.258952914 | 0.40524459 | 0.612178 | Y39B6A.13 |
| WBGene00012676 | 0.164265908 | 0.49661247 | 0.68500023 | pro-3 |
| WBGene00012678 | -0.138165549 | 0.73283224 | 0.85060384 | Y39B6A.16 |
| WBGene00012679 | 0.747845955 | 0.85215965 | 1 | Y39B6A.18 |
| WBGene00012681 | -0.156269293 | 0.96924949 | 1 | asp-15 |
| WBGene00012682 | -0.156269293 | 0.96924949 | 1 | asp-16 |
| WBGene00012683 | 1.221972049 | 0.14057183 | 0.31834517 | asp-17 |
| WBGene00012684 | 0.281454817 | 0.94435481 | 1 | Y39B6A.25 |
| WBGene00012686 | 0.292952874 | 0.61954243 | 0.7785885 | Y39B6A.27 |
| WBGene00012688 | 2.442683361 | 0.04942063 | 1 | Y39B6A.29 |
| WBGene00012689 | 2.07531578 | 0.45514895 | 1 | Y39B6A.30 |
| WBGene00012692 | 0.027726901 | 0.89949634 | 0.94717955 | Y39B6A.33 |
| WBGene00012693 | -0.42860947 | 0.18876669 | 0.38254321 | Y39B6A.34 |
| WBGene00012694 | -0.424507113 | 0.02860893 | 0.10955348 | Y39B6A.36 |
| WBGene00012695 | -0.229868883 | 0.34081788 | 0.54915327 | Y39B6A.37 |
| WBGene00012696 | -0.375531093 | 0.27630075 | 0.48490423 | reps-1 |
| WBGene00012697 | -0.484949022 | 0.01923624 | 0.08332046 | mrps-35 |
| WBGene00012698 | -0.134280367 | 0.74551835 | 0.85810943 | sws-1 |
| WBGene00012699 | 0.785458941 | 0.35139652 | 0.56017662 | Y39B6A.41 |
| WBGene00012700 | -0.711626724 | 0.00797698 | 0.04380305 | Y39B6A.42 |
| WBGene00012701 | 0.022946384 | 0.93792531 | 0.96795424 | Y39B6A.43 |
| WBGene00012702 | 0.539871029 | 0.22754725 | 0.43227213 | ztf-20 |
| WBGene00012703 | -0.093073787 | 0.88576862 | 0.9396345 | nhr-145 |
| WBGene00012704 | -0.533231537 | 0.25225832 | 0.46035874 | mlt-3 |
| WBGene00012705 | -1.964653846 | 0.53599202 | 1 | sre-18 |
| WBGene00012707 | 1.413377704 | 0.60673863 | 1 | sre-16 |
| WBGene00012708 | 3.169173677 | 0.09791657 | 1 | sre-19 |
| WBGene00012709 | 1.33864768 | 0.5127736 | 1 | sre-15 |
| WBGene00012710 | -0.328972397 | 0.61898984 | 0.77836931 | Y39C12A.9 |
| WBGene00012711 | -0.156269293 | 0.96924949 | 1 | Y39E4A.1 |
| WBGene00012712 | 0.221436586 | 0.47609218 | 0.66874283 | ttm-1 |

| | | | | |
|---|---|---|---|---|
| WBGene00012713 | -0.373915829 | 0.05588846 | 0.17389475 | bckd-1A |
| WBGene00012714 | -0.258857331 | 0.14248006 | 0.32034267 | abce-1 |
| WBGene00012715 | -0.716639022 | 0.0002803 | 0.00341332 | snpc-1.2 |
| WBGene00012716 | -0.264466576 | 0.19088367 | 0.38567459 | Y39E4B.5 |
| WBGene00012717 | 0.223967425 | 0.5193948 | 0.70481415 | Y39E4B.6 |
| WBGene00012718 | -0.175510974 | 0.45134475 | 0.64918572 | dhhc-8 |
| WBGene00012719 | 0.050353998 | 0.85299712 | 0.91927977 | Y39E4B.10 |
| WBGene00012720 | -0.014519762 | 0.99603746 | 1 | Y39E4B.11 |
| WBGene00012721 | 2.654814267 | 0.26264283 | 1 | Y39E4B.13 |
| WBGene00012722 | -0.330770857 | 0.15584014 | 0.33968242 | Y39G8B.1 |
| WBGene00012723 | -0.64195043 | 0.17107864 | 0.36015788 | Y39G8B.2 |
| WBGene00012725 | -1.901856996 | 0.64072034 | 1 | sre-47 |
| WBGene00012726 | 0.784808749 | 0.52512862 | 1 | Y39G8B.5 |
| WBGene00012727 | 3.283670693 | 0.0803008 | 1 | Y39G8B.7 |
| WBGene00012728 | 0.089995227 | 0.82916441 | 0.90766252 | Y39G8B.9 |
| WBGene00012729 | 0.394439984 | 0.88400378 | 1 | Y39G8B.10 |
| WBGene00012730 | 0.183010463 | 0.42381868 | 0.62806093 | xrn-1 |
| WBGene00012731 | 0.14424727 | 0.93787433 | 1 | Y39G8C.2 |
| WBGene00012732 | -0.786366008 | 0.02701729 | 0.10553953 | Y40B1A.1 |
| WBGene00012734 | -0.243387254 | 0.76295722 | 0.86724685 | Y40B1A.2 |
| WBGene00012734 | 0.358154661 | 0.32152627 | 0.52965027 | Y40B1A.3 |
| WBGene00012735 | -0.074855296 | 0.69184045 | 0.82456388 | sptf-3 |
| WBGene00012738 | -0.715858063 | 0.00209031 | 0.01609291 | eif-3.J |
| WBGene00012739 | -0.541738211 | 0.01047837 | 0.05370108 | Y40B1B.7 |
| WBGene00012740 | -0.159041835 | 0.50888759 | 0.69553126 | slc-25A46 |
| WBGene00012741 | -0.156269293 | 0.96924949 | 1 | Y40H4A.2 |
| WBGene00012744 | 0.474347274 | 0.7488275 | 1 | Y40H7A.4 |
| WBGene00012746 | 0.688793783 | 0.86378755 | 1 | Y40H7A.9 |
| WBGene00012747 | 0.067430371 | 0.79380401 | 0.8860508 | Y40H7A.10 |
| WBGene00012748 | 2.420661489 | 0.31300137 | 1 | Y40H7A.11 |
| WBGene00012750 | 0.730738762 | 0.30627209 | 0.51512233 | faah-6 |
| WBGene00012753 | 0.306259774 | 0.64776155 | 0.79688651 | Y41C4A.6 |
| WBGene00012755 | 0.019131713 | 0.97120001 | 0.98496241 | Y41C4A.8 |
| WBGene00012756 | -0.205336975 | 0.4366117 | 0.63751085 | Y41C4A.9 |
| WBGene00012757 | 1.081601397 | 0.3032622 | 1 | Y41C4A.11 |
| WBGene00012758 | -0.137920157 | 0.76532531 | 0.86863743 | Y41C4A.12 |
| WBGene00012759 | 0.464579155 | 0.08922321 | 0.23707387 | sup-1 |
| WBGene00012760 | -0.33854201 | 0.70331579 | 0.83258161 | Y41C4A.17 |
| WBGene00012761 | -1.094201388 | 0.7816772 | 1 | Y41C4A.18 |
| WBGene00012762 | -0.184834946 | 0.25610232 | 0.46380344 | Y41E3.1 |
| WBGene00012763 | 0.969790294 | 0.0276992 | 0.10733993 | atln-2 |
| WBGene00012764 | 2.815759761 | 0.16833793 | 1 | Y41E3.6 |
| WBGene00012765 | -0.176063284 | 0.40340135 | 0.61094802 | Y41E3.7 |

| | | | | |
|---|---|---|---|---|
| WBGene00012766 | 0.133503597 | 0.69094714 | 0.82388423 | Y41E3.8 |
| WBGene00012767 | 0.162223405 | 0.41439582 | 0.61977351 | fcd-2 |
| WBGene00012768 | -0.751374195 | 1.0809E-06 | 3.5798E-05 | eef-1B.2 |
| WBGene00012769 | 0.008630694 | 0.96350693 | 0.98170004 | hrpu-1 |
| WBGene00012770 | -1.14715222 | 0.68932198 | 1 | srt-47 |
| WBGene00012772 | -0.156269293 | 0.96924949 | 1 | srt-49 |
| WBGene00012773 | -0.156269293 | 0.96924949 | 1 | srt-48 |
| WBGene00012775 | 0.233739675 | 0.93575889 | 1 | srt-50 |
| WBGene00012776 | 1.368257229 | 0.02049753 | 0.08707506 | Y42A5A.1 |
| WBGene00012777 | -0.373421884 | 0.57553447 | 0.74844162 | lact-8 |
| WBGene00012778 | 0.480135808 | 0.64975734 | 1 | Y42A5A.3 |
| WBGene00012779 | 0.792451302 | 0.23720655 | 0.44340642 | cdkl-1 |
| WBGene00012780 | 0.177512908 | 0.74725593 | 0.85920656 | Y42A5A.5 |
| WBGene00012781 | 2.909683933 | 0.07100979 | 1 | nspd-7 |
| WBGene00012782 | 1.363158941 | 6.21E-17 | 1.45E-14 | Y43C5A.2 |
| WBGene00012783 | 1.754052324 | 1.0061E-06 | 3.3837E-05 | Y43C5A.3 |
| WBGene00012784 | -0.70576507 | 0.76841669 | 1 | Y43C5A.4 |
| WBGene00012785 | -0.156269293 | 0.96924949 | 1 | Y43C5B.2 |
| WBGene00012786 | -0.892270884 | 0.34869173 | 1 | Y43C5B.3 |
| WBGene00012787 | -1.291305095 | 0.63677871 | 1 | Y43D4A.1 |
| WBGene00012788 | 0.966259585 | 0.60493647 | 1 | Y43D4A.2 |
| WBGene00012790 | -0.156269293 | 0.96924949 | 1 | Y43D4A.4 |
| WBGene00012791 | 2.44066524 | 0.32717233 | 1 | Y43D4A.5 |
| WBGene00012792 | -2.977551009 | 0.05439681 | 1 | Y43D4A.6 |
| WBGene00012794 | 1.023240912 | 0.00146283 | 0.01231641 | Y43E12A.2 |
| WBGene00012795 | -0.148762833 | 0.69704824 | 0.82859957 | Y43E12A.3 |
| WBGene00012796 | 0.858663227 | 0.07382105 | 0.20974992 | Y43F4A.1 |
| WBGene00012798 | -0.156269293 | 0.96924949 | 1 | Y43F4A.3 |
| WBGene00012799 | -1.159714244 | 0.75790634 | 1 | Y43F4A.4 |
| WBGene00012802 | -0.018056727 | 0.95873679 | 0.97864666 | set-25 |
| WBGene00012803 | -0.103905453 | 0.54631336 | 0.72535304 | Y43F4B.5 |
| WBGene00012804 | 0.065160159 | 0.80783389 | 0.89513608 | Y43F4B.7 |
| WBGene00012806 | -0.800968016 | 0.01470717 | 0.0687297 | Y43F4B.9 |
| WBGene00012807 | -0.920933216 | 0.0002587 | 0.00320829 | Y43F4B.10 |
| WBGene00012808 | -0.566790163 | 0.74096626 | 1 | Y43F8A.1 |
| WBGene00012809 | 0.983488853 | 0.80601565 | 1 | Y43F8A.2 |
| WBGene00012810 | 0.405895286 | 0.66483078 | 0.80771846 | nceh-1 |
| WBGene00012811 | 0.203318228 | 0.80179947 | 0.89153287 | Y43F8A.5 |
| WBGene00012812 | 0.45864298 | 0.06858383 | 0.199539 | Y43F8B.1 |
| WBGene00012813 | 0.360149192 | 0.14231242 | 0.32034267 | Y43F8B.2 |
| WBGene00012814 | 0.190817565 | 0.73488395 | 0.85185883 | Y43F8B.3 |
| WBGene00012815 | 1.477253415 | 0.58357327 | 1 | phy-4 |
| WBGene00012818 | -1.45873709 | 0.59584643 | 1 | Y43F8B.7 |

| | | | | |
|---|---|---|---|---|
| WBGene00012819 | 0.989201989 | 0.20161 | 0.40041719 | Y43F8B.9 |
| WBGene00012821 | -0.26130237 | 0.77680621 | 0.87561083 | Y43F8B.11 |
| WBGene00012822 | 0.910878243 | 0.50648566 | 1 | Y43F8B.12 |
| WBGene00012823 | 0.546052409 | 0.7844476 | 1 | Y43F8B.13 |
| WBGene00012824 | 2.572567154 | 0.02048872 | 0.08707506 | Y43F8B.14 |
| WBGene00012825 | 0.579589165 | 0.08101865 | 0.22254466 | Y43F8C.3 |
| WBGene00012826 | 0.738777874 | 0.45902919 | 1 | dyf-19 |
| WBGene00012827 | -0.668264701 | 0.85551153 | 1 | Y43F8C.5 |
| WBGene00012828 | 0.112887935 | 0.76579056 | 0.8687594 | Y43F8C.6 |
| WBGene00012829 | -0.50384903 | 0.03123511 | 0.11693966 | Y43F8C.7 |
| WBGene00012830 | -0.337826424 | 0.28569838 | 0.49559423 | mrps-28 |
| WBGene00012831 | 0.933277224 | 0.64661698 | 1 | Y43F8C.9 |
| WBGene00012833 | 0.426034681 | 0.71161875 | 1 | Y43F8C.11 |
| WBGene00012834 | -0.304023236 | 0.28432391 | 0.49417901 | Y43F8C.13 |
| WBGene00012835 | 0.25477501 | 0.39352345 | 0.6015825 | ani-3 |
| WBGene00012838 | 1.181686151 | 0.7579682 | 1 | Y43F8C.17 |
| WBGene00012839 | 2.188838843 | 0.34004625 | 1 | Y43F8C.18 |
| WBGene00012840 | -1.40766733 | 0.21890612 | 1 | grsp-1 |
| WBGene00012845 | -0.209577086 | 0.42488905 | 0.62893095 | set-24 |
| WBGene00012846 | 0.629154436 | 0.81300308 | 1 | srxa-14 |
| WBGene00012847 | 0.688793783 | 0.86378755 | 1 | srxa-15 |
| WBGene00012848 | -0.497334839 | 0.78199731 | 1 | Y44A6B.3 |
| WBGene00012849 | 0.938982574 | 0.8146943 | 1 | ttr-13 |
| WBGene00012850 | 0.782278559 | 0.02218346 | 0.09252126 | Y44A6C.1 |
| WBGene00012851 | -1.869554365 | 2.0606E-05 | 0.00041148 | Y44A6C.2 |
| WBGene00012853 | 0.185262436 | 0.50427616 | 0.6914079 | Y44A6D.2 |
| WBGene00012854 | 0.897953534 | 0.02366744 | 0.09658702 | Y44A6D.3 |
| WBGene00012855 | 0.295704581 | 0.86339129 | 1 | Y44A6D.5 |
| WBGene00012856 | 0.664914355 | 0.80343354 | 1 | Y44A6D.6 |
| WBGene00012857 | 0.398775586 | 0.38978268 | 0.59820862 | pbo-5 |
| WBGene00012858 | 0.020107331 | 0.94240554 | 0.97027162 | Y44F5A.1 |
| WBGene00012859 | -0.788177853 | 0.24422943 | 0.45173515 | Y45F3A.1 |
| WBGene00012860 | -0.254707272 | 0.21477171 | 0.41594246 | acdh-11 |
| WBGene00012861 | 0.742305465 | 0.08711521 | 0.23358342 | Y45F3A.4 |
| WBGene00012862 | -0.156269293 | 0.96924949 | 1 | Y45F3A.5 |
| WBGene00012863 | -0.917145842 | 0.33337252 | 0.54154078 | Y45F3A.8 |
| WBGene00012864 | 0.022128657 | 0.95331169 | 0.97623454 | Y45F3A.9 |
| WBGene00012866 | -0.15031863 | 0.80329751 | 0.89239902 | afmd-2 |
| WBGene00012867 | -0.354226286 | 0.30722394 | 0.51598983 | seld-1 |
| WBGene00012868 | 0.205981309 | 0.28302274 | 0.49264217 | tbc-9 |
| WBGene00012869 | -0.34136145 | 0.39057651 | 0.59911426 | Y45F10A.7 |
| WBGene00012871 | -1.969805064 | 0.62862569 | 1 | Y45F10B.3 |
| WBGene00012873 | 1.38322681 | 0.69550305 | 1 | Y45F10B.9 |

| | | | | |
|---|---|---|---|---|
| WBGene00012875 | 0.38049011 | 0.17076623 | 0.3596934 | Y45F10B.13 |
| WBGene00012877 | -0.558804446 | 0.87818675 | 1 | Y45F10C.1 |
| WBGene00012878 | -2.189216016 | 3.5464E-05 | 0.0006436 | Y45F10C.2 |
| WBGene00012879 | -0.291873922 | 0.09452101 | 0.24615608 | fbxa-215 |
| WBGene00012880 | -2.287823833 | 7.36E-11 | 7.14E-09 | Y45F10C.4 |
| WBGene00012883 | 0.980282897 | 0.03546011 | 0.12703489 | Y45F10D.2 |
| WBGene00012885 | -0.280494179 | 0.16608771 | 0.35295953 | Y45F10D.4 |
| WBGene00012886 | -0.005018059 | 0.99587784 | 1 | Y45F10D.6 |
| WBGene00012887 | -0.021688394 | 0.93213794 | 0.96406579 | Y45F10D.7 |
| WBGene00012888 | -0.226303269 | 0.2310036 | 0.43640702 | sas-6 |
| WBGene00012889 | -0.334921561 | 0.5933229 | 0.76005346 | Y45F10D.10 |
| WBGene00012890 | -0.198591025 | 0.53948964 | 0.72112118 | lact-9 |
| WBGene00012891 | 0.664913417 | 0.0463946 | 0.15264956 | sorb-1 |
| WBGene00012892 | -0.245531902 | 0.94070824 | 1 | Y45F10D.14 |
| WBGene00012893 | 2.224351729 | 0.33187942 | 1 | Y45F10D.15 |
| WBGene00012894 | 0.145913567 | 0.46309465 | 0.6582021 | tbc-17 |
| WBGene00012895 | 0.315981544 | 0.49668103 | 0.68501697 | cox-10 |
| WBGene00012896 | 0.036523858 | 0.87437043 | 0.93213743 | snrp-200 |
| WBGene00012897 | -0.239234979 | 0.33758203 | 0.54561685 | pisy-1 |
| WBGene00012899 | -1.389532539 | 0.01613999 | 0.07328183 | Y46G5A.7 |
| WBGene00012900 | -0.748101714 | 0.16258203 | 0.34850078 | Y46G5A.8 |
| WBGene00012901 | 0.903982211 | 0.18215611 | 0.37384831 | cnp-2 |
| WBGene00012903 | -0.514625198 | 0.00758254 | 0.04209403 | vps-2 |
| WBGene00012904 | -0.57638542 | 0.01907731 | 0.08292788 | tiar-2 |
| WBGene00012905 | 1.38322681 | 0.69550305 | 1 | Y46G5A.14 |
| WBGene00012906 | 0.016802837 | 0.97464786 | 0.98666588 | Y46G5A.15 |
| WBGene00012907 | -0.358008601 | 0.04032199 | 0.1382517 | cpt-1 |
| WBGene00012908 | -0.130100959 | 0.54927008 | 0.72786379 | Y46G5A.18 |
| WBGene00012909 | -0.33166857 | 0.33640208 | 0.54485827 | spds-1 |
| WBGene00012910 | 1.186390418 | 0.20178194 | 0.40056156 | Y46G5A.20 |
| WBGene00012911 | 0.749504982 | 0.00197205 | 0.01538046 | acl-7 |
| WBGene00012912 | 0.377828795 | 0.92526585 | 1 | Y46G5A.22 |
| WBGene00012913 | -0.156269293 | 0.96924949 | 1 | Y46G5A.23 |
| WBGene00012914 | -0.775313921 | 0.4854984 | 1 | bcmo-1 |
| WBGene00012915 | -0.034378509 | 0.9675489 | 0.9835073 | lgc-35 |
| WBGene00012916 | -1.91567326 | 3.1887E-05 | 0.00058836 | Y46G5A.28 |
| WBGene00012917 | -0.858233925 | 0.56255569 | 1 | Y46G5A.29 |
| WBGene00012919 | -1.235924314 | 0.10884009 | 0.26896551 | Y46G5A.34 |
| WBGene00012920 | -0.162602162 | 0.68761775 | 0.82225619 | Y46G5A.35 |
| WBGene00012921 | -0.254683336 | 0.89672209 | 1 | Y46G5A.36 |
| WBGene00012922 | -0.726544277 | 0.27762045 | 0.48649266 | Y47D3A.1 |
| WBGene00012923 | 0.860065708 | 0.25517146 | 0.46266975 | fbxa-128 |
| WBGene00012925 | 1.137987479 | 0.61169092 | 1 | wht-8 |

| | | | | |
|---|---|---|---|---|
| WBGene00012926 | -0.298265327 | 0.89527345 | 1 | Y47D3A.13 |
| WBGene00012927 | -0.523978001 | 0.10662098 | 0.2661907 | Y47D3A.14 |
| WBGene00012928 | -0.038722916 | 0.87775679 | 0.93441865 | aakb-2 |
| WBGene00012929 | 0.064294078 | 0.7428329 | 0.85656132 | rsks-1 |
| WBGene00012930 | 0.177791119 | 0.44424628 | 0.64362697 | obr-1 |
| WBGene00012931 | -0.760637283 | 0.06999372 | 0.20218742 | Y47D3A.20 |
| WBGene00012932 | -0.535119937 | 0.01785537 | 0.07894465 | Y47D3A.21 |
| WBGene00012933 | -0.34399837 | 0.36622361 | 0.57379286 | mib-1 |
| WBGene00012934 | -0.087083366 | 0.67042541 | 0.8115289 | gly-9 |
| WBGene00012935 | -0.315050857 | 0.12768113 | 0.29839611 | mcm-10 |
| WBGene00012936 | -0.32150334 | 0.04906397 | 0.15897669 | pola-1 |
| WBGene00012937 | 1.247575581 | 0.7549672 | 1 | Y47D3A.31 |
| WBGene00012938 | 0.491773737 | 0.50803415 | 0.69483413 | Y47D3A.32 |
| WBGene00012939 | 0.250691623 | 0.70608304 | 0.83366276 | Y47D3B.1 |
| WBGene00012940 | -1.26270356 | 0.3518783 | 1 | Y47D3B.3 |
| WBGene00012941 | -0.011793733 | 0.98882129 | 0.99484196 | Y47D3B.4 |
| WBGene00012942 | -1.027402836 | 0.15820667 | 0.34299953 | Y47D3B.6 |
| WBGene00012943 | 0.215432443 | 0.41397866 | 0.61943738 | bed-2 |
| WBGene00012945 | -1.094201388 | 0.7816772 | 1 | Y47H9A.1 |
| WBGene00012946 | 0.913821381 | 0.49581821 | 1 | Y47H9B.2 |
| WBGene00012947 | 1.610571151 | 0.07999758 | 0.22093964 | Y47H9C.1 |
| WBGene00012948 | 1.005330007 | 0.03900657 | 0.13503815 | dhhc-2 |
| WBGene00012950 | -0.241667252 | 0.31403808 | 0.52186372 | eif-2Bbeta |
| WBGene00012951 | -0.154815543 | 0.51900403 | 0.70445818 | Y47H9C.8 |
| WBGene00012952 | 0.846183542 | 0.65777268 | 1 | Y47H9C.9 |
| WBGene00012953 | -0.22859207 | 0.68046173 | 0.8167559 | fbxa-216 |
| WBGene00012955 | -1.159714244 | 0.75790634 | 1 | Y47H9C.12 |
| WBGene00012958 | -0.378568623 | 0.70750278 | 1 | Y47H10A.2 |
| WBGene00012959 | 2.832750627 | 0.08618443 | 1 | Y47H10A.3 |
| WBGene00012960 | 0.00209211 | 0.9972574 | 0.99857337 | Y47H10A.4 |
| WBGene00012961 | 1.017706156 | 0.02899136 | 0.11061724 | Y47H10A.5 |
| WBGene00012962 | -0.05740145 | 0.9364495 | 0.96687077 | dmsr-5 |
| WBGene00012964 | -0.582617643 | 0.00529954 | 0.03253953 | Y48A6B.3 |
| WBGene00012965 | -1.409432952 | 0.51259961 | 1 | fipr-17 |
| WBGene00012966 | -0.594871314 | 0.0181506 | 0.07984291 | exos-1 |
| WBGene00012967 | 1.637684222 | 0.06073938 | 0.18412905 | kvs-4 |
| WBGene00012968 | 0.668324991 | 0.01841529 | 0.0807148 | Y48A6B.7 |
| WBGene00012969 | 0.339872674 | 0.75397084 | 1 | Y48A6B.8 |
| WBGene00012970 | 0.422855582 | 0.23688421 | 0.44302363 | Y48A6B.9 |
| WBGene00012972 | -0.343482151 | 0.04660694 | 0.1531407 | rsa-2 |
| WBGene00012973 | 0.247648985 | 0.25918614 | 0.46732429 | spat-2 |
| WBGene00012976 | -0.407014632 | 0.29872699 | 0.50853119 | sup-35 |
| WBGene00012977 | -0.111348769 | 0.62915473 | 0.78456939 | Y48A6C.4 |

| | | | | |
|---|---|---|---|---|
| WBGene00012978 | -0.372705059 | 0.0488672 | 0.15850832 | Y48B6A.1 |
| WBGene00012979 | 1.413179233 | 0.52346377 | 1 | Y48B6A.5 |
| WBGene00012980 | 0.495524338 | 0.11277733 | 0.27522202 | efhd-1 |
| WBGene00012981 | -0.582940972 | 0.17928985 | 0.37143268 | Y48B6A.10 |
| WBGene00012982 | -0.56936268 | 0.01224165 | 0.06011166 | jmjd-2 |
| WBGene00012983 | -0.214591588 | 0.19390294 | 0.38998571 | men-1 |
| WBGene00012984 | -0.49066541 | 0.13533728 | 0.30984912 | Y48B6A.13 |
| WBGene00012985 | -0.764269844 | 0.76036927 | 1 | Y48C3A.1 |
| WBGene00012987 | 0.697759942 | 0.03979891 | 0.13696105 | Y48C3A.3 |
| WBGene00012988 | -0.03641727 | 0.91255698 | 0.95467585 | ztf-22 |
| WBGene00012989 | 0.470259619 | 0.27079058 | 0.47962106 | Y48C3A.5 |
| WBGene00012990 | -0.272823824 | 0.25430606 | 0.46207607 | sam-10 |
| WBGene00012992 | -0.548632158 | 0.04430903 | 0.14738518 | mrpl-20 |
| WBGene00012993 | 0.787785884 | 0.30367233 | 0.51249454 | dmsr-3 |
| WBGene00012994 | -0.266703179 | 0.24365792 | 0.45114272 | Y48C3A.12 |
| WBGene00012995 | -0.104167449 | 0.65645 | 0.80236354 | Y48C3A.14 |
| WBGene00012996 | -0.273761653 | 0.34098727 | 0.54928041 | pinn-4 |
| WBGene00012997 | -0.146661779 | 0.6997948 | 0.83043775 | Y48C3A.18 |
| WBGene00012998 | -0.159128327 | 0.54067244 | 0.72174866 | Y48C3A.20 |
| WBGene00012999 | 0.213256215 | 0.39591327 | 0.60371552 | rpoa-1 |
| WBGene00013000 | 0.006060302 | 0.98733506 | 0.99421374 | ddl-2 |
| WBGene00013001 | -0.166431989 | 0.46072002 | 0.65624616 | Y48E1B.2 |
| WBGene00013002 | -0.005849481 | 0.97974291 | 0.98911181 | Y48E1B.3 |
| WBGene00013004 | -0.457800605 | 0.11814555 | 0.28345248 | mrpl-37 |
| WBGene00013006 | -0.460565856 | 0.16748383 | 0.35493773 | lin-38 |
| WBGene00013007 | -2.950408564 | 0.00010426 | 0.00157266 | Y48E1B.8 |
| WBGene00013008 | -0.11118497 | 0.80609074 | 0.89433863 | clec-146 |
| WBGene00013009 | 0.478449983 | 0.63288705 | 1 | fbxc-23 |
| WBGene00013011 | -0.358942256 | 0.11569506 | 0.28023616 | snx-14 |
| WBGene00013013 | -1.10706159 | 0.36222309 | 1 | clec-145 |
| WBGene00013014 | -0.545924311 | 0.0489543 | 0.15870604 | Y48E1C.1 |
| WBGene00013015 | -0.410821116 | 0.10327786 | 0.26041463 | Y48E1C.2 |
| WBGene00013017 | -0.248487806 | 0.35669596 | 0.5651371 | Y48E1C.4 |
| WBGene00013018 | -0.529416763 | 0.01669695 | 0.07494186 | Y48G10A.1 |
| WBGene00013019 | -0.801471412 | 5.5372E-05 | 0.00094552 | Y48G10A.2 |
| WBGene00013020 | -0.062503929 | 0.7890578 | 0.8837593 | Y48G10A.3 |
| WBGene00013021 | -0.1210801 | 0.63271376 | 0.78722871 | Y48G10A.4 |
| WBGene00013023 | -0.496370926 | 0.46539051 | 0.65974005 | Y48G10A.6 |
| WBGene00013024 | 0.215003611 | 0.29973262 | 0.50945734 | cept-2 |
| WBGene00013025 | 0.390057426 | 0.01427609 | 0.06720667 | vha-13 |
| WBGene00013026 | -0.270013107 | 0.3303044 | 0.53866024 | Y49A3A.3 |
| WBGene00013027 | -0.034373513 | 0.97991106 | 1 | Y49A3A.4 |
| WBGene00013028 | 1.36997009 | 0.32561898 | 1 | efhc-1 |

| | | | | |
|---|---|---|---|---|
| WBGene00013029 | -0.425749075 | 0.11143555 | 0.27326581 | glrx-5 |
| WBGene00013030 | -0.766916579 | 0.00146078 | 0.01230768 | Y49E10.4 |
| WBGene00013033 | 0.060062363 | 0.9752405 | 1 | Y49E10.10 |
| WBGene00013034 | 0.418640683 | 0.028611 | 0.10955348 | tat-1 |
| WBGene00013035 | 0.866112842 | 1.9267E-05 | 0.00039007 | Y49E10.16 |
| WBGene00013036 | -1.540285988 | 0.29510219 | 1 | fbxa-218 |
| WBGene00013037 | 1.185315342 | 7.73E-07 | 2.6823E-05 | Y49E10.18 |
| WBGene00013038 | -0.127243062 | 0.45707054 | 0.65393512 | ani-1 |
| WBGene00013039 | -0.09499864 | 0.59599176 | 0.76215883 | scav-3 |
| WBGene00013040 | 0.055568483 | 0.83420254 | 0.91023162 | Y49E10.21 |
| WBGene00013041 | -0.281804694 | 0.5775968 | 0.74906609 | pstk-1 |
| WBGene00013042 | 0.472886321 | 0.13785745 | 0.31449217 | cccp-1 |
| WBGene00013043 | -0.255998374 | 0.883344 | 1 | arrd-16 |
| WBGene00013045 | -1.901856996 | 0.64072034 | 1 | Y49E10.27 |
| WBGene00013046 | 0.391777078 | 0.56213128 | 0.73778752 | Y50E8A.1 |
| WBGene00013048 | 1.176838979 | 0.15450546 | 0.33793025 | Y50E8A.5 |
| WBGene00013049 | 0.377828795 | 0.92526585 | 1 | Y50E8A.6 |
| WBGene00013050 | -0.156269293 | 0.96924949 | 1 | lipl-8 |
| WBGene00013053 | 0.688793783 | 0.86378755 | 1 | Y50E8A.10 |
| WBGene00013054 | -0.156269293 | 0.96924949 | 1 | Y50E8A.11 |
| WBGene00013055 | -0.097268026 | 0.95384638 | 1 | Y50E8A.12 |
| WBGene00013060 | -0.668264701 | 0.85551153 | 1 | Y51A2A.4 |
| WBGene00013062 | -1.969805064 | 0.62862569 | 1 | Y51A2A.6 |
| WBGene00013066 | -0.156269293 | 0.96924949 | 1 | Y51A2B.2 |
| WBGene00013069 | -0.156269293 | 0.96924949 | 1 | Y51A2B.5 |
| WBGene00013071 | 0.232157325 | 0.93397866 | 1 | Y51A2B.8 |
| WBGene00013073 | -0.152877104 | 0.85819691 | 0.92272128 | hmit-1.1 |
| WBGene00013074 | -2.058413997 | 0.00049788 | 0.0053446 | hmit-1.2 |
| WBGene00013075 | 0.053741789 | 0.85989447 | 0.92430091 | Y51A2D.7 |
| WBGene00013076 | 0.913935434 | 0.7131603 | 1 | Y51A2D.8 |
| WBGene00013077 | -0.421869106 | 0.05533358 | 0.17247882 | ttr-24 |
| WBGene00013078 | -0.358962093 | 0.16927629 | 0.35743997 | ttr-25 |
| WBGene00013079 | 0.737602429 | 0.0063503 | 0.0371585 | ttr-26 |
| WBGene00013080 | 0.816802105 | 0.00363983 | 0.02450978 | Y51A2D.13 |
| WBGene00013081 | -0.112048448 | 0.6404858 | 0.79202874 | Y51A2D.14 |
| WBGene00013082 | 0.183996846 | 0.56985163 | 0.74425222 | grdn-1 |
| WBGene00013083 | -0.458464189 | 0.21003532 | 0.41014101 | Y51A2D.18 |
| WBGene00013086 | -2.253159685 | 1.3212E-06 | 4.2102E-05 | arrd-2 |
| WBGene00013088 | 2.524900078 | 0.16560744 | 1 | Y51B9A.6 |
| WBGene00013089 | -0.156269293 | 0.96924949 | 1 | Y51B9A.7 |
| WBGene00013090 | 0.281454817 | 0.94435481 | 1 | Y51B9A.8 |
| WBGene00013091 | 1.139344337 | 0.09427549 | 0.24556935 | Y51B9A.9 |
| WBGene00013093 | -0.167663353 | 0.46039005 | 0.6560508 | cup-14 |

| | | | | |
|---|---|---|---|---|
| WBGene00013094 | -0.680507518 | 0.00011955 | 0.0017551 | Y51H1A.3 |
| WBGene00013095 | -0.609652762 | 0.01594066 | 0.07259399 | ing-3 |
| WBGene00013096 | -0.265854191 | 0.22987116 | 0.43524429 | mcd-1 |
| WBGene00013098 | 0.401776691 | 0.74590323 | 1 | Y51H4A.1 |
| WBGene00013100 | -1.559792506 | 0.09099073 | 0.23989543 | zip-7 |
| WBGene00013101 | 2.860962197 | 0.0078046 | 0.04303162 | Y51H4A.5 |
| WBGene00013103 | 0.655964118 | 0.05147517 | 0.16381056 | Y51H4A.7 |
| WBGene00013104 | -0.268912506 | 0.52784002 | 0.71220832 | Y51H4A.8 |
| WBGene00013106 | -0.161814212 | 0.5354416 | 0.7178681 | set-26 |
| WBGene00013107 | 0.635045026 | 0.70546974 | 1 | Y51H4A.13 |
| WBGene00013109 | -0.259912112 | 0.33702708 | 0.54507071 | Y51H4A.15 |
| WBGene00013111 | 0.955365166 | 0.00426811 | 0.02763685 | sta-1 |
| WBGene00013116 | 0.747845955 | 0.85215965 | 1 | Y51H4A.22 |
| WBGene00013118 | 1.838076487 | 3.44E-07 | 1.3644E-05 | Y51H4A.24 |
| WBGene00013119 | 1.244737235 | 0.00220651 | 0.01675374 | Y51H4A.25 |
| WBGene00013120 | 0.377828795 | 0.92526585 | 1 | fipr-28 |
| WBGene00013122 | -0.763497653 | 0.00335652 | 0.02300996 | impt-1 |
| WBGene00013123 | 0.204609564 | 0.37335654 | 0.5812823 | Y52B11A.3 |
| WBGene00013124 | 0.4904437 | 0.34864666 | 0.55718299 | Y52B11A.4 |
| WBGene00013125 | -1.837668733 | 0.43198255 | 1 | clec-92 |
| WBGene00013126 | -1.024189566 | 0.37726025 | 1 | mltn-2 |
| WBGene00013127 | -0.60432981 | 0.00234586 | 0.01761466 | Y52B11A.8 |
| WBGene00013128 | -0.355254189 | 0.08183143 | 0.22416865 | dxbp-1 |
| WBGene00013129 | 0.19020637 | 0.45045945 | 0.64895346 | Y52B11A.10 |
| WBGene00013130 | -1.901856996 | 0.64072034 | 1 | Y52B11B.1 |
| WBGene00013131 | 0.207288442 | 0.49759515 | 0.68588814 | Y52B11C.1 |
| WBGene00013132 | -0.078243808 | 0.73860512 | 0.8543246 | strd-1 |
| WBGene00013135 | -0.028212287 | 0.94703083 | 0.97304242 | rga-2 |
| WBGene00013136 | 0.597165584 | 0.02297236 | 0.09461854 | Y53C10A.5 |
| WBGene00013137 | -0.318090604 | 0.26427427 | 0.4728893 | Y53C10A.6 |
| WBGene00013138 | 1.226152345 | 0.48225188 | 1 | Y53C10A.10 |
| WBGene00013139 | -0.229438559 | 0.35495514 | 0.56326106 | Y53C12A.3 |
| WBGene00013140 | -0.290916207 | 0.1661157 | 0.35295953 | mop-25.2 |
| WBGene00013141 | -0.241940907 | 0.66354841 | 0.80690517 | Y53C12A.6 |
| WBGene00013142 | -0.156269293 | 0.96924949 | 1 | Y53C12A.7 |
| WBGene00013143 | -0.132800295 | 0.543379 | 0.72331236 | Y53C12B.1 |
| WBGene00013144 | -0.421511936 | 0.01189666 | 0.05892999 | Y53C12B.2 |
| WBGene00013145 | -1.175229155 | 0.17166293 | 0.36089345 | cutl-2 |
| WBGene00013146 | -0.307425703 | 0.68803183 | 0.82242734 | Y53C12B.7 |
| WBGene00013147 | -0.505697301 | 0.71160879 | 1 | eyg-1 |
| WBGene00013148 | 0.983488853 | 0.80601565 | 1 | Y53F4A.2 |
| WBGene00013149 | 0.805160662 | 0.33661372 | 0.54486555 | Y53F4B.1 |
| WBGene00013150 | -1.114967199 | 1.2099E-05 | 0.00026611 | Y53F4B.3 |

| | | | | |
|---|---|---|---|---|
| WBGene00013151 | 0.022955702 | 0.95025461 | 0.97468105 | nsun-5 |
| WBGene00013152 | 1.247575581 | 0.7549672 | 1 | Y53F4B.5 |
| WBGene00013154 | -1.901856996 | 0.64072034 | 1 | Y53F4B.7 |
| WBGene00013156 | -0.366913374 | 0.11068139 | 0.27213098 | Y53F4B.9 |
| WBGene00013157 | -0.450812972 | 0.85883116 | 1 | Y53F4B.10 |
| WBGene00013158 | 1.181686151 | 0.7579682 | 1 | Y53F4B.11 |
| WBGene00013159 | 0.845847127 | 0.22929668 | 0.43442105 | Y53F4B.12 |
| WBGene00013160 | 0.103387236 | 0.66020781 | 0.8042122 | Y53F4B.13 |
| WBGene00013161 | -0.019965977 | 0.95492513 | 0.97697363 | Y53F4B.14 |
| WBGene00013162 | -0.263229581 | 0.93409872 | 1 | Y53F4B.16 |
| WBGene00013164 | -0.370852191 | 0.0905983 | 0.23932853 | Y53F4B.18 |
| WBGene00013165 | -0.558804446 | 0.87818675 | 1 | Y53F4B.19 |
| WBGene00013166 | -0.146757738 | 0.89407054 | 1 | Y53F4B.20 |
| WBGene00013167 | 0.81320061 | 0.06426001 | 0.19108027 | Y53F4B.21 |
| WBGene00013168 | -0.563537827 | 0.00100881 | 0.00929991 | arp-1 |
| WBGene00013169 | 2.11303534 | 0.15264093 | 1 | Y53F4B.23 |
| WBGene00013170 | 1.116613983 | 0.69067248 | 1 | Y53F4B.24 |
| WBGene00013171 | -0.860919914 | 0.43952059 | 1 | Y53F4B.25 |
| WBGene00013173 | -1.043669441 | 0.12683397 | 0.29727629 | Y53F4B.27 |
| WBGene00013175 | 0.446458324 | 0.87164163 | 1 | Y53F4B.36 |
| WBGene00013176 | 0.099176631 | 0.69231516 | 0.82486735 | Y53F4B.39 |
| WBGene00013177 | -0.164432336 | 0.49551491 | 0.68395254 | rsy-1 |
| WBGene00013178 | -0.214268669 | 0.45538485 | 0.65267707 | Y53H1A.2 |
| WBGene00013180 | 0.1975965 | 0.72127111 | 0.84294693 | cutl-10 |
| WBGene00013181 | -0.164115927 | 0.51574786 | 0.70206243 | Y53H1B.2 |
| WBGene00013182 | -1.14715222 | 0.68932198 | 1 | Y53H1B.4 |
| WBGene00013184 | -0.668264701 | 0.85551153 | 1 | Y53H1B.6 |
| WBGene00013186 | 0.863501219 | 0.69189182 | 1 | Y53H1C.3 |
| WBGene00013187 | 1.193312181 | 0.09878561 | 0.25297533 | npr-34 |
| WBGene00013188 | -0.00741182 | 0.98725186 | 0.99421374 | smg-9 |
| WBGene00013189 | 0.322366919 | 0.11256305 | 0.27491036 | Y54E2A.4 |
| WBGene00013190 | 0.377828795 | 0.92526585 | 1 | Y54E2A.5 |
| WBGene00013192 | -0.737462795 | 0.83433562 | 1 | Y54E2A.7 |
| WBGene00013193 | -0.208781809 | 0.51166197 | 0.69811749 | Y54E2A.8 |
| WBGene00013194 | 0.377828795 | 0.92526585 | 1 | Y54E2A.9 |
| WBGene00013195 | 0.694291259 | 0.45122622 | 0.64911342 | Y54E2A.10 |
| WBGene00013196 | -0.297912743 | 0.14508549 | 0.32381738 | tbc-20 |
| WBGene00013197 | 0.520043672 | 0.19748437 | 0.39487117 | ttm-5 |
| WBGene00013198 | 0.149858741 | 0.80338698 | 0.89239902 | Y54E5A.2 |
| WBGene00013200 | -0.694678494 | 0.00110069 | 0.00985511 | Y54E5A.5 |
| WBGene00013201 | -0.209938695 | 0.48381049 | 0.67458876 | dus-2 |
| WBGene00013202 | 0.214750531 | 0.34245009 | 0.55068696 | Y54E5A.7 |
| WBGene00013203 | 0.440486969 | 0.39240869 | 0.60025626 | Y54E5A.8 |

| | | | | |
|---|---|---|---|---|
| WBGene00013204 | 0.600309982 | 0.03123147 | 0.11693966 | Y54E5B.2 |
| WBGene00013207 | 0.242323424 | 0.84551754 | 1 | zipt-2.3 |
| WBGene00013208 | -0.545884519 | 0.01428827 | 0.06723794 | Y54G9A.5 |
| WBGene00013209 | -0.801355936 | 1.9688E-05 | 0.0003964 | bub-3 |
| WBGene00013210 | -0.357622154 | 0.11727811 | 0.28233879 | Y54G9A.7 |
| WBGene00013211 | -1.350318784 | 0.0090982 | 0.04817322 | Y54G9A.9 |
| WBGene00013212 | -0.281543933 | 0.2166744 | 0.41848075 | Y54G11A.1 |
| WBGene00013213 | -0.391590743 | 0.06379813 | 0.19003266 | Y54G11A.2 |
| WBGene00013214 | -0.600815975 | 0.00241073 | 0.01804478 | Y54G11A.3 |
| WBGene00013215 | 0.892704453 | 0.08341258 | 0.22681123 | Y54G11A.4 |
| WBGene00013216 | -0.273406804 | 0.40322589 | 0.61087666 | Y54G11A.7 |
| WBGene00013217 | -0.41830727 | 0.06480715 | 0.19204873 | ddl-3 |
| WBGene00013218 | -0.44663137 | 0.1199524 | 0.28624902 | Y54G11A.9 |
| WBGene00013219 | -0.547663023 | 0.01204799 | 0.05946126 | Y54G11A.11 |
| WBGene00013220 | -0.196441372 | 0.74932354 | 0.8602872 | ctl-3 |
| WBGene00013221 | 0.745128146 | 0.23019229 | 0.43572882 | Y54G11A.14 |
| WBGene00013222 | 0.97483962 | 0.15274525 | 0.33510662 | dmsr-6 |
| WBGene00013223 | 0.047083885 | 0.8066899 | 0.89459463 | efa-6 |
| WBGene00013224 | -0.088090611 | 0.71025012 | 0.83606232 | Y55D9A.2 |
| WBGene00013225 | -0.246490009 | 0.21937101 | 0.4220893 | Y56A3A.2 |
| WBGene00013226 | -0.401810342 | 0.80435313 | 1 | faah-5 |
| WBGene00013227 | 0.575579782 | 0.14939802 | 0.33081184 | Y56A3A.6 |
| WBGene00013228 | 0.153459487 | 0.54943204 | 0.72791951 | Y56A3A.7 |
| WBGene00013230 | -1.707028745 | 0.06243431 | 0.18725666 | fbxb-22 |
| WBGene00013231 | -0.803722206 | 0.0060615 | 0.03586386 | tsen-2 |
| WBGene00013232 | -0.392982533 | 0.05022703 | 0.16132442 | faah-4 |
| WBGene00013233 | -1.635373289 | 0.05501899 | 0.17180699 | sdz-33 |
| WBGene00013234 | -1.099780452 | 0.1747019 | 0.36490444 | fbxb-24 |
| WBGene00013235 | -0.157637052 | 0.4696325 | 0.66315517 | Y56A3A.16 |
| WBGene00013236 | -0.276597141 | 0.30852156 | 0.51729875 | Y56A3A.18 |
| WBGene00013237 | -0.737379088 | 7.5755E-05 | 0.0012252 | Y56A3A.19 |
| WBGene00013238 | -0.335658938 | 0.03462026 | 0.12520838 | trap-4 |
| WBGene00013239 | -0.352517149 | 0.08893027 | 0.23639701 | Y56A3A.22 |
| WBGene00013240 | -0.463805052 | 0.10759132 | 0.26759651 | Y56A3A.28 |
| WBGene00013241 | -0.593584183 | 0.00830339 | 0.04498505 | ung-1 |
| WBGene00013242 | 0.043525357 | 0.92274727 | 0.9593272 | Y56A3A.30 |
| WBGene00013243 | -0.307303737 | 0.20852231 | 0.40829734 | Y56A3A.31 |
| WBGene00013244 | -0.438167654 | 0.51529781 | 0.70176452 | Y56A3A.33 |
| WBGene00013246 | -0.115631779 | 0.72473099 | 0.84518994 | Y57A10A.1 |
| WBGene00013247 | 0.199676173 | 0.59586882 | 0.76214826 | Y57A10A.2 |
| WBGene00013248 | -0.358991708 | 0.3562308 | 0.56469489 | Y57A10A.3 |
| WBGene00013249 | 0.196906154 | 0.75339145 | 0.86235484 | Y57A10A.4 |
| WBGene00013250 | -0.330432705 | 0.36047952 | 0.56837101 | Y57A10A.5 |

| | | | | |
|---|---|---|---|---|
| WBGene00013252 | 0.245202433 | 0.67635093 | 0.81466397 | Y57A10A.7 |
| WBGene00013253 | -0.316604367 | 0.27063507 | 0.47960594 | Y57A10A.8 |
| WBGene00013254 | -1.523349594 | 0.00082734 | 0.0079179 | tag-276 |
| WBGene00013255 | -0.24342704 | 0.38577377 | 0.59414935 | tric-1B.1 |
| WBGene00013256 | -0.1187139 | 0.54286113 | 0.72323899 | Y57A10A.13 |
| WBGene00013257 | 0.630626725 | 0.24366731 | 0.45114272 | Y57A10A.14 |
| WBGene00013258 | 0.233247301 | 0.46859432 | 0.6623989 | polg-1 |
| WBGene00013259 | -0.392411446 | 0.17058402 | 0.35949673 | trpp-5 |
| WBGene00013260 | -0.631944297 | 0.00154263 | 0.01282817 | rsr-2 |
| WBGene00013261 | 0.088866494 | 0.78074272 | 0.87874269 | sinh-1 |
| WBGene00013263 | -0.329260555 | 0.19650705 | 0.39356518 | Y57A10A.23 |
| WBGene00013264 | -0.102919912 | 0.96055224 | 1 | Y57A10A.24 |
| WBGene00013265 | -0.269506366 | 0.33788017 | 0.54587018 | parn-2 |
| WBGene00013266 | 0.023373866 | 0.93134608 | 0.96397892 | Y57A10A.26 |
| WBGene00013267 | -0.037285783 | 0.89091268 | 0.9424176 | Y57A10A.27 |
| WBGene00013268 | -0.369254617 | 0.20731822 | 0.40685432 | tric-1B.2 |
| WBGene00013269 | -0.606424753 | 0.03279011 | 0.12074757 | Y57A10A.29 |
| WBGene00013270 | -0.075481616 | 0.68342582 | 0.81877569 | Y57A10A.31 |
| WBGene00013272 | 1.025181888 | 0.79790347 | 1 | aqp-12 |
| WBGene00013273 | 1.238241982 | 0.47971924 | 1 | Y57A10B.2 |
| WBGene00013275 | 0.168918804 | 0.61442002 | 0.77495049 | btb-14 |
| WBGene00013276 | 2.490696505 | 0.25975012 | 1 | sre-42 |
| WBGene00013277 | 0.377828795 | 0.92526585 | 1 | sre-41 |
| WBGene00013278 | -1.87238823 | 0.00658117 | 0.0380667 | Y57A10B.6 |
| WBGene00013280 | -0.876811101 | 0.47669179 | 1 | Y57A10C.1 |
| WBGene00013281 | -0.156269293 | 0.96924949 | 1 | sre-27 |
| WBGene00013284 | -0.064145124 | 0.70374159 | 0.83263838 | daf-22 |
| WBGene00013286 | -0.014519762 | 0.99603746 | 1 | Y57A10C.8 |
| WBGene00013287 | 0.14455128 | 0.91248893 | 1 | Y57A10C.9 |
| WBGene00013288 | -0.156269293 | 0.96924949 | 1 | Y57A10C.10 |
| WBGene00013289 | 0.115886266 | 0.63852764 | 0.79036023 | tag-273 |
| WBGene00013290 | 2.094556028 | 0.1280284 | 1 | Y57G11A.2 |
| WBGene00013292 | 0.328584443 | 0.36047676 | 0.56837101 | Y57G11A.4 |
| WBGene00013293 | 1.529802032 | 0.64971375 | 1 | Y57G11A.5 |
| WBGene00013294 | -0.263229581 | 0.93409872 | 1 | pals-30 |
| WBGene00013295 | -0.810368856 | 0.261684 | 0.47012511 | Y57G11B.2 |
| WBGene00013296 | 0.377828795 | 0.92526585 | 1 | Y57G11B.3 |
| WBGene00013297 | -0.808658094 | 5.1409E-06 | 0.00012922 | Y57G11B.5 |
| WBGene00013298 | -1.05498899 | 0.51829338 | 1 | Y57G11B.6 |
| WBGene00013300 | -1.964653846 | 0.53599202 | 1 | lgc-7 |
| WBGene00013301 | -0.317455181 | 0.10909991 | 0.26938824 | Y57G11C.3 |
| WBGene00013302 | -0.278655275 | 0.35995021 | 0.56806965 | vti-1 |
| WBGene00013303 | 1.78499574 | 0.04330537 | 0.14516027 | Y57G11C.5 |

| | | | | |
|---|---|---|---|---|
| WBGene00013304 | 0.789601902 | 0.75907734 | 1 | Y57G11C.6 |
| WBGene00013306 | 0.258783652 | 0.7229659 | 0.8438309 | Y57G11C.8 |
| WBGene00013307 | -0.263149456 | 0.08781533 | 0.23483832 | Y57G11C.9 |
| WBGene00013308 | -0.458087529 | 0.00231456 | 0.01743242 | nuo-3 |
| WBGene00013310 | -0.399578031 | 0.71302834 | 1 | Y57G11C.14 |
| WBGene00013311 | -0.293543418 | 0.0655022 | 0.19335333 | sec-61 |
| WBGene00013312 | 0.230668913 | 0.74240497 | 0.85614919 | hhat-2 |
| WBGene00013313 | 0.742628056 | 0.76901841 | 1 | Y57G11C.18 |
| WBGene00013315 | 0.28530093 | 0.92084641 | 1 | Y57G11C.20 |
| WBGene00013317 | 0.490419874 | 0.21767996 | 0.41990029 | Y57G11C.22 |
| WBGene00013318 | -1.159714244 | 0.75790634 | 1 | Y57G11C.23 |
| WBGene00013319 | -1.484327552 | 0.00113159 | 0.01008336 | ccch-5 |
| WBGene00013321 | 0.45652368 | 0.11775591 | 0.28281778 | Y57G11C.31 |
| WBGene00013322 | 1.493922719 | 0.1432074 | 0.32114537 | Y57G11C.32 |
| WBGene00013323 | -0.206327721 | 0.39896745 | 0.60659956 | Y57G11C.33 |
| WBGene00013324 | -0.360628234 | 0.18553385 | 0.37815427 | mrps-7 |
| WBGene00013325 | 0.605203561 | 0.07270411 | 0.20774313 | Y57G11C.36 |
| WBGene00013326 | 0.116033332 | 0.78966804 | 0.88416432 | anoh-2 |
| WBGene00013327 | 0.284787672 | 0.5739969 | 0.74766991 | Y57G11C.38 |
| WBGene00013328 | -0.064323217 | 0.97003873 | 1 | Y57G11C.39 |
| WBGene00013329 | 0.2919591 | 0.77114911 | 1 | Y57G11C.40 |
| WBGene00013330 | -1.082680044 | 0.67762556 | 1 | Y57G11C.41 |
| WBGene00013331 | 0.088773099 | 0.90918957 | 0.95274215 | Y57G11C.42 |
| WBGene00013332 | -0.02561304 | 0.91268375 | 0.95467585 | Y57G11C.43 |
| WBGene00013333 | -0.109254073 | 0.95448982 | 1 | Y57G11C.44 |
| WBGene00013334 | -0.265111163 | 0.60686404 | 0.76973843 | nlp-56 |
| WBGene00013335 | -0.316316381 | 0.79125668 | 1 | Y57G11C.46 |
| WBGene00013336 | 1.24583058 | 0.30732655 | 1 | pnc-2 |
| WBGene00013339 | -0.076021206 | 0.75851447 | 0.86486307 | phf-14 |
| WBGene00013340 | -0.040362205 | 0.79146202 | 0.88481955 | tcc-1 |
| WBGene00013343 | -0.268601291 | 0.10640998 | 0.2658826 | prp-6 |
| WBGene00013344 | 0.294809346 | 0.46417213 | 0.65886864 | ebp-1 |
| WBGene00013345 | -0.166002041 | 0.65884808 | 0.80357275 | Y59A8B.8 |
| WBGene00013347 | -0.474764938 | 0.0061376 | 0.03626112 | nova-1 |
| WBGene00013348 | 0.357399304 | 0.44308905 | 0.64264057 | fbxa-106 |
| WBGene00013349 | -0.72110379 | 0.00109926 | 0.00984949 | Y59A8B.12 |
| WBGene00013350 | -0.567152759 | 0.04048763 | 0.13863825 | slr-2 |
| WBGene00013352 | -0.553439131 | 0.0911509 | 0.24010915 | lon-8 |
| WBGene00013353 | 0.719033344 | 0.01987917 | 0.08507334 | Y59A8B.21 |
| WBGene00013354 | 0.169531169 | 0.34584099 | 0.55467048 | snx-6 |
| WBGene00013355 | 0.380129584 | 0.06708674 | 0.19664417 | gck-3 |
| WBGene00013357 | 0.377828795 | 0.92526585 | 1 | clec-260 |
| WBGene00013358 | -0.20826713 | 0.34145 | 0.54980695 | abhd-3.1 |

| | | | | |
|---|---|---|---|---|
| WBGene00013359 | 0.459069479 | 0.18216676 | 0.37384831 | Y60A3A.8 |
| WBGene00013360 | -0.383424126 | 0.02763692 | 0.1072355 | Y60A3A.9 |
| WBGene00013361 | -0.467829825 | 0.01311228 | 0.06326707 | fars-2 |
| WBGene00013362 | -0.315741578 | 0.28309583 | 0.49269874 | algn-7 |
| WBGene00013364 | 0.396521025 | 0.66538045 | 0.80808083 | Y60A3A.16 |
| WBGene00013365 | -0.520224897 | 0.02384674 | 0.09699271 | Y60A3A.19 |
| WBGene00013366 | -0.025289956 | 0.9497573 | 0.97439609 | Y60A3A.21 |
| WBGene00013367 | -0.156269293 | 0.96924949 | 1 | Y60A3A.23 |
| WBGene00013368 | 1.247575581 | 0.7549672 | 1 | Y60A9.1 |
| WBGene00013370 | -0.352461709 | 0.82493315 | 1 | Y60A9.3 |
| WBGene00013373 | -0.728240535 | 0.00018133 | 0.00243801 | Y62E10A.2 |
| WBGene00013374 | -0.156269293 | 0.96924949 | 1 | Y62E10A.3 |
| WBGene00013375 | -0.69198496 | 0.71261975 | 1 | srsx-25 |
| WBGene00013376 | 0.084893784 | 0.75221312 | 0.86213721 | Y62E10A.6 |
| WBGene00013378 | -0.675953306 | 0.0004421 | 0.00484436 | emc-3 |
| WBGene00013379 | -0.334003774 | 0.09835472 | 0.25235094 | Y62E10A.13 |
| WBGene00013380 | -0.638485713 | 0.00310137 | 0.02171498 | memi-2 |
| WBGene00013381 | -0.354518288 | 0.08090471 | 0.2224296 | cyp-31A5 |
| WBGene00013382 | 0.2743836 | 0.49407903 | 0.68266911 | gcl-1 |
| WBGene00013383 | -1.089526601 | 7.2346E-05 | 0.00118536 | aptf-2 |
| WBGene00013385 | 0.785778694 | 0.32818843 | 0.53671009 | Y62E10A.19 |
| WBGene00013387 | 1.33218149 | 0.10165348 | 0.25717336 | Y62F5A.9 |
| WBGene00013388 | 1.402950615 | 0.25747749 | 1 | Y62F5A.10 |
| WBGene00013389 | 1.216957619 | 0.74819621 | 1 | Y62H9A.1 |
| WBGene00013390 | -0.156269293 | 0.96924949 | 1 | Y62H9A.2 |
| WBGene00013391 | -1.801550307 | 3.33E-25 | 1.92E-22 | Y62H9A.3 |
| WBGene00013392 | -2.127665618 | 2.83E-22 | 1.15E-19 | Y62H9A.4 |
| WBGene00013393 | -0.806608156 | 6.92E-08 | 3.3898E-06 | Y62H9A.5 |
| WBGene00013394 | -0.365011956 | 0.00287438 | 0.02058219 | ule-5 |
| WBGene00013395 | 0.574508983 | 0.8551971 | 1 | Y62H9A.7 |
| WBGene00013397 | 0.983488853 | 0.80601565 | 1 | Y62H9A.9 |
| WBGene00013398 | 1.557897343 | 0.63813005 | 1 | Y62H9A.10 |
| WBGene00013399 | 0.459413223 | 0.84552382 | 1 | Y62H9A.11 |
| WBGene00013400 | -1.759191416 | 0.57668729 | 1 | Y62H9A.12 |
| WBGene00013401 | 0.873411505 | 0.43239448 | 1 | Y62H9A.13 |
| WBGene00013402 | 0.5180853 | 0.79605727 | 1 | Y62H9A.14 |
| WBGene00013403 | -1.938251172 | 0.10502505 | 1 | fbxb-100 |
| WBGene00013404 | -0.813864725 | 0.4800717 | 1 | fbxb-101 |
| WBGene00013405 | -0.71734204 | 0.00432629 | 0.02795397 | tdpt-1 |
| WBGene00013406 | -0.527645595 | 0.01680572 | 0.07540218 | Y63D3A.7 |
| WBGene00013407 | -0.465974928 | 0.10001992 | 0.25479266 | Y63D3A.8 |
| WBGene00013408 | -1.694249209 | 0.0751436 | 0.21206844 | fbxb-93 |
| WBGene00013409 | -1.343380635 | 0.42643052 | 1 | fbxb-56 |

| | | | | |
|---|---|---|---|---|
| WBGene00013412 | -1.964653846 | 0.53599202 | 1 | Y64G10A.2 |
| WBGene00013415 | 0.853424647 | 0.01383897 | 0.06561337 | ilcr-1 |
| WBGene00013416 | 0.550491113 | 0.11911522 | 0.28484891 | Y64G10A.7 |
| WBGene00013417 | 0.942308827 | 0.67468959 | 1 | srab-24 |
| WBGene00013418 | 0.023129518 | 0.97219683 | 0.98553604 | Y65A5A.1 |
| WBGene00013419 | -0.004582543 | 0.98598631 | 0.99332445 | Y65A5A.2 |
| WBGene00013421 | -1.279666666 | 0.55824702 | 1 | fbxa-220 |
| WBGene00013422 | -0.845420786 | 0.00214535 | 0.01641257 | Y66A7A.2 |
| WBGene00013424 | 2.912785086 | 0.15033923 | 1 | Y66A7A.4 |
| WBGene00013425 | 0.008181011 | 0.98088735 | 0.98993793 | ceh-91 |
| WBGene00013426 | -0.624781296 | 0.36390098 | 0.57174854 | Y66A7A.7 |
| WBGene00013427 | 0.281454817 | 0.94435481 | 1 | Y66D12A.1 |
| WBGene00013429 | 1.247575581 | 0.7549672 | 1 | Y66D12A.3 |
| WBGene00013431 | -0.218213229 | 0.32880209 | 0.53749646 | ceh-92 |
| WBGene00013432 | -0.516072468 | 0.25403944 | 0.46193303 | Y66D12A.6 |
| WBGene00013433 | -0.801626963 | 0.0535367 | 0.16821664 | Y66D12A.7 |
| WBGene00013434 | -0.516759101 | 0.01286281 | 0.06240922 | Y66D12A.8 |
| WBGene00013435 | -0.503140752 | 0.01959195 | 0.08421189 | Y66D12A.9 |
| WBGene00013436 | -0.55318975 | 0.02971227 | 0.11237901 | Y66D12A.10 |
| WBGene00013437 | 2.063391569 | 0.37853067 | 1 | Y66D12A.11 |
| WBGene00013438 | -0.924432076 | 0.00765512 | 0.04243873 | ztf-29 |
| WBGene00013439 | 1.0544261 | 0.01352663 | 0.06442795 | Y66D12A.13 |
| WBGene00013440 | 1.125195184 | 0.15109489 | 0.33299022 | nrps-1 |
| WBGene00013441 | -0.227466552 | 0.22014887 | 0.42311658 | xpb-1 |
| WBGene00013442 | -0.817895684 | 0.01176249 | 0.05848007 | Y66D12A.16 |
| WBGene00013443 | -0.107303396 | 0.58791288 | 0.75636858 | such-1 |
| WBGene00013445 | -0.3207355 | 0.20565494 | 0.40520231 | Y66D12A.19 |
| WBGene00013446 | -0.625586917 | 0.01131314 | 0.05687488 | Y66D12A.21 |
| WBGene00013447 | -0.162530366 | 0.49106804 | 0.6802214 | nars-2 |
| WBGene00013448 | 0.469545834 | 0.23278551 | 0.43851271 | Y66D12A.24 |
| WBGene00013449 | 0.041803775 | 0.98890197 | 1 | Y67A6A.1 |
| WBGene00013450 | -2.934840138 | 0.2654436 | 1 | oac-56 |
| WBGene00013452 | 2.063391569 | 0.37853067 | 1 | Y67A10A.3 |
| WBGene00013453 | -0.421949241 | 0.46961991 | 0.66315517 | fbxa-83 |
| WBGene00013454 | 0.281454817 | 0.94435481 | 1 | fbxa-126 |
| WBGene00013455 | 0.626223829 | 0.29351914 | 0.50376249 | ssu-2 |
| WBGene00013456 | 1.327488926 | 0.0692928 | 0.2009756 | Y67A10A.7 |
| WBGene00013457 | 0.930773517 | 0.39138447 | 0.59974742 | paqr-3 |
| WBGene00013458 | 1.095444172 | 0.00638521 | 0.03725267 | Y67A10A.9 |
| WBGene00013459 | -0.183789617 | 0.65703524 | 0.80268312 | Y67A10A.10 |
| WBGene00013460 | -0.331063008 | 0.08292429 | 0.22609113 | cpsf-3 |
| WBGene00013461 | -0.253933677 | 0.20697708 | 0.40665706 | Y67H2A.2 |
| WBGene00013462 | 0.127921985 | 0.49096198 | 0.6802214 | micu-1 |

| | | | | |
|---|---|---|---|---|
| WBGene00013463 | -0.393598569 | 0.00542934 | 0.03312443 | kdp-1 |
| WBGene00013464 | 0.078050961 | 0.75329607 | 0.86235484 | Y67H2A.7 |
| WBGene00013465 | 0.153332231 | 0.47013029 | 0.66347226 | Y67H2A.10 |
| WBGene00013468 | -0.156269293 | 0.96924949 | 1 | srz-96 |
| WBGene00013469 | -0.008677305 | 0.99719169 | 1 | Y68A4A.10 |
| WBGene00013471 | 0.281454817 | 0.94435481 | 1 | clec-242 |
| WBGene00013472 | 0.688793783 | 0.86378755 | 1 | Y68A4B.3 |
| WBGene00013473 | 1.078381604 | 0.38159509 | 1 | Y69E1A.1 |
| WBGene00013474 | 0.729903699 | 0.55988047 | 1 | Y69E1A.2 |
| WBGene00013475 | -0.156269293 | 0.96924949 | 1 | Y69E1A.3 |
| WBGene00013477 | -0.402075794 | 0.19958381 | 0.39782418 | Y69E1A.5 |
| WBGene00013478 | 0.377828795 | 0.92526585 | 1 | Y69E1A.8 |
| WBGene00013479 | -0.988374531 | 0.58460249 | 1 | Y69H2.1 |
| WBGene00013480 | 0.811862829 | 0.47487702 | 1 | egas-3 |
| WBGene00013481 | -0.571982869 | 0.01111233 | 0.05612097 | Y69H2.3 |
| WBGene00013482 | -0.529061367 | 0.0179025 | 0.07903792 | Y69H2.7 |
| WBGene00013484 | -0.635728112 | 0.4643176 | 0.65886864 | Y69H2.9 |
| WBGene00013485 | 2.037985599 | 0.2655958 | 1 | Y69H2.10 |
| WBGene00013486 | 2.544041525 | 0.13546931 | 1 | egas-1 |
| WBGene00013487 | 0.466881705 | 0.74930455 | 1 | egas-2 |
| WBGene00013489 | -0.359459679 | 0.04805091 | 0.15673996 | col-42 |
| WBGene00013490 | 1.313089111 | 0.56091378 | 1 | Y70C5A.2 |
| WBGene00013492 | -0.425976481 | 0.69306632 | 1 | Y70C5C.1 |
| WBGene00013493 | 0.747845955 | 0.85215965 | 1 | clec-9 |
| WBGene00013494 | -1.901856996 | 0.64072034 | 1 | irld-58 |
| WBGene00013496 | 0.168036801 | 0.85535295 | 0.92113008 | Y70D2A.1 |
| WBGene00013497 | 0.444170926 | 0.26954459 | 0.47802233 | hex-5 |
| WBGene00013498 | 0.377828795 | 0.92526585 | 1 | Y70G10A.2 |
| WBGene00013499 | 1.159117576 | 9.3598E-05 | 0.00145502 | Y70G10A.3 |
| WBGene00013506 | 0.146939336 | 0.6164127 | 0.77617367 | usp-3 |
| WBGene00013507 | 0.883785108 | 0.52712788 | 1 | Y71A12B.10 |
| WBGene00013508 | -2.615321458 | 1.14E-12 | 1.59E-10 | Y71A12B.11 |
| WBGene00013509 | -0.287819134 | 0.39565084 | 0.60353202 | Y71A12B.12 |
| WBGene00013511 | -1.072763059 | 0.53107528 | 1 | Y71A12B.15 |
| WBGene00013512 | 0.281454817 | 0.94435481 | 1 | nhr-276 |
| WBGene00013513 | -0.385938285 | 0.69681369 | 1 | Y71A12C.2 |
| WBGene00013514 | -0.161899549 | 0.66873994 | 0.81015635 | Y73F4A.1 |
| WBGene00013515 | 0.869624833 | 0.00521807 | 0.032175 | Y73F4A.2 |
| WBGene00013516 | 2.270589421 | 1.0527E-05 | 0.00023757 | Y73F4A.3 |
| WBGene00013517 | 1.278788705 | 0.4969022 | 1 | lgc-52 |
| WBGene00013518 | 1.016001434 | 0.0644905 | 0.19139065 | Y73F8A.5 |
| WBGene00013520 | -0.127866201 | 0.84203977 | 0.9132754 | best-21 |
| WBGene00013521 | -0.156269293 | 0.96924949 | 1 | Y73F8A.12 |

| | | | | |
|---|---|---|---|---|
| WBGene00013522 | -0.582008294 | 0.19581043 | 0.39242892 | Y73F8A.13 |
| WBGene00013524 | 0.377828795 | 0.92526585 | 1 | Y73F8A.15 |
| WBGene00013525 | 1.025181888 | 0.79790347 | 1 | irld-59 |
| WBGene00013526 | 1.216957619 | 0.74819621 | 1 | Y73F8A.20 |
| WBGene00013527 | -1.420815711 | 0.55886055 | 1 | Y73F8A.22 |
| WBGene00013529 | -0.3005022 | 0.22899724 | 0.43421138 | Y73F8A.24 |
| WBGene00013530 | -0.208815485 | 0.23710667 | 0.44328797 | ntl-11 |
| WBGene00013531 | -0.383245569 | 0.32586769 | 0.53428219 | Y73F8A.26 |
| WBGene00013532 | -0.546205625 | 0.01573094 | 0.07193571 | Y73F8A.27 |
| WBGene00013534 | 1.062558618 | 0.21762356 | 0.41987105 | acr-24 |
| WBGene00013536 | 1.776552053 | 0.06645988 | 0.19523093 | Y73F8A.32 |
| WBGene00013538 | -0.663060104 | 2.5118E-05 | 0.0004895 | tag-349 |
| WBGene00013539 | 0.576183493 | 0.3798726 | 0.58781969 | Y73F8A.35 |
| WBGene00013540 | 0.949715475 | 0.01146933 | 0.05749346 | Y75B8A.3 |
| WBGene00013541 | 0.257155858 | 0.33094419 | 0.53922359 | lonp-2 |
| WBGene00013542 | 0.424439427 | 0.4542698 | 0.65184843 | lron-12 |
| WBGene00013543 | 0.385853616 | 0.31698066 | 0.52493219 | Y75B8A.6 |
| WBGene00013544 | -0.014914519 | 0.95666157 | 0.97809079 | Y75B8A.7 |
| WBGene00013545 | -0.127761989 | 0.56656031 | 0.74172492 | Y75B8A.8 |
| WBGene00013547 | -2.32081292 | 0.35154423 | 1 | Y75B8A.10 |
| WBGene00013548 | 0.584372024 | 0.31170715 | 0.51986779 | Y75B8A.11 |
| WBGene00013549 | 0.267724665 | 0.35891591 | 0.56717402 | Y75B8A.13 |
| WBGene00013550 | -0.415470171 | 0.12594291 | 0.29604509 | Y75B8A.14 |
| WBGene00013551 | 0.051437777 | 0.85349945 | 0.91962521 | Y75B8A.16 |
| WBGene00013552 | -0.916835208 | 1.1097E-05 | 0.00024904 | gmn-1 |
| WBGene00013553 | -0.717643607 | 0.0062693 | 0.03689557 | Y75B8A.18 |
| WBGene00013554 | 0.2649795 | 0.52946246 | 0.71368977 | Y75B8A.19 |
| WBGene00013555 | -0.49540722 | 0.79306545 | 1 | fbxa-89 |
| WBGene00013556 | 0.58106201 | 0.72825048 | 1 | Y75B8A.23 |
| WBGene00013557 | 0.014451957 | 0.94661334 | 0.97304242 | Y75B8A.24 |
| WBGene00013558 | -0.063121944 | 0.77639949 | 0.87539619 | Y75B8A.25 |
| WBGene00013559 | 0.790911343 | 0.02625279 | 0.10351906 | Y75B8A.28 |
| WBGene00013560 | 0.123183409 | 0.71784776 | 0.84036155 | zip-12 |
| WBGene00013561 | -0.797662786 | 0.00043709 | 0.00480683 | Y75B8A.31 |
| WBGene00013562 | 3.050185957 | 0.01961512 | 1 | Y75B8A.32 |
| WBGene00013563 | -0.333693239 | 0.89573179 | 1 | Y75B8A.33 |
| WBGene00013564 | -0.014519762 | 0.99603746 | 1 | Y75B8A.34 |
| WBGene00013566 | 1.557897343 | 0.63813005 | 1 | Y75B12A.2 |
| WBGene00013567 | -0.68981248 | 8.0597E-06 | 0.00018927 | Y75B12B.1 |
| WBGene00013568 | 0.682190696 | 0.55810707 | 1 | Y75B12B.3 |
| WBGene00013569 | 0.674746079 | 0.29494958 | 0.50528896 | bath-36 |
| WBGene00013573 | -2.652727305 | 0.24933058 | 1 | Y75B12B.11 |
| WBGene00013574 | -0.127898073 | 0.87276111 | 0.93161531 | lron-13 |

| | | | | |
|---|---|---|---|---|
| WBGene00013575 | 0.169574279 | 0.40939082 | 0.61485082 | acs-5 |
| WBGene00013576 | -0.449302489 | 0.18683053 | 0.38020609 | Y76A2B.4 |
| WBGene00013577 | 0.084040101 | 0.65759743 | 0.80291727 | Y76A2B.5 |
| WBGene00013578 | -0.300941673 | 0.16218663 | 0.34814497 | scav-2 |
| WBGene00013579 | -0.244838334 | 0.70764631 | 0.83477788 | Y79H2A.2 |
| WBGene00013580 | 0.122246235 | 0.61018201 | 0.77201762 | Y79H2A.3 |
| WBGene00013581 | 0.28530093 | 0.92084641 | 1 | Y79H2A.4 |
| WBGene00013583 | -1.542969157 | 0.08684887 | 0.23307518 | ceh-51 |
| WBGene00013584 | -0.043177329 | 0.93952762 | 0.96889534 | nhr-243 |
| WBGene00013585 | -0.214421111 | 0.69763873 | 0.8288472 | cyp-42A1 |
| WBGene00013586 | -0.156269293 | 0.96924949 | 1 | Y80D3A.8 |
| WBGene00013587 | -0.164013858 | 0.6002911 | 0.76556032 | Y80D3A.9 |
| WBGene00013588 | 0.443267492 | 0.46663072 | 0.66083793 | nlp-42 |
| WBGene00013590 | -0.504789308 | 0.63417076 | 1 | Y81G3A.1 |
| WBGene00013591 | -0.195240214 | 0.37572987 | 0.58341685 | gcn-2 |
| WBGene00013592 | -0.132855481 | 0.92921491 | 1 | Y81G3A.4 |
| WBGene00013593 | 0.04306362 | 0.84514297 | 0.91508792 | Y87G2A.1 |
| WBGene00013594 | -0.573838021 | 0.07105128 | 0.20427461 | Y87G2A.2 |
| WBGene00013595 | -0.054004467 | 0.8423963 | 0.91337104 | atg-4.1 |
| WBGene00013596 | -0.326119422 | 0.10900907 | 0.26927347 | nyn-2 |
| WBGene00013597 | 0.131328678 | 0.37546563 | 0.58323032 | gpi-1 |
| WBGene00013598 | -0.445483957 | 0.04107353 | 0.14019504 | vps-28 |
| WBGene00013599 | -0.285000291 | 0.32529669 | 0.5338183 | eipr-1 |
| WBGene00013601 | -0.001897748 | 0.99292068 | 0.99644982 | Y87G2A.13 |
| WBGene00013602 | 2.139996549 | 0.048696 | 1 | Y87G2A.16 |
| WBGene00013603 | 0.138701659 | 0.75729836 | 0.86429398 | soem-1 |
| WBGene00013604 | -0.430729984 | 0.18482139 | 0.37727857 | Y87G2A.18 |
| WBGene00013605 | -0.337630871 | 0.25859742 | 0.46679026 | Y95D11A.1 |
| WBGene00013606 | 0.11298057 | 0.58718344 | 0.75591074 | cand-1 |
| WBGene00013607 | 3.321702207 | 0.02735485 | 1 | clec-27 |
| WBGene00013608 | 1.294959883 | 0.36188777 | 1 | clec-35 |
| WBGene00013609 | 1.034708429 | 0.29029571 | 0.50031189 | clec-37 |
| WBGene00013610 | -0.673687615 | 0.01769539 | 0.07837169 | fbxa-206 |
| WBGene00013611 | -0.211664343 | 0.62788041 | 0.78378532 | Y102A5C.2 |
| WBGene00013613 | 0.020891697 | 0.99136276 | 1 | Y102A5C.4 |
| WBGene00013616 | 1.529802032 | 0.64971375 | 1 | fbxa-148 |
| WBGene00013619 | 1.687239501 | 0.59403242 | 1 | fbxa-112 |
| WBGene00013620 | 1.82867917 | 0.54358766 | 1 | fbxa-111 |
| WBGene00013621 | -0.156269293 | 0.96924949 | 1 | clec-239 |
| WBGene00013622 | -1.638044942 | 0.48959522 | 1 | clec-238 |
| WBGene00013623 | 0.169599968 | 0.80928923 | 0.89609686 | fbxa-90 |
| WBGene00013629 | 0.281454817 | 0.94435481 | 1 | srz-95 |
| WBGene00013632 | -0.230155477 | 0.52457046 | 0.70914262 | Y105C5A.1 |

| | | | | |
|---|---|---|---|---|
| WBGene00013633 | -0.63535817 | 0.04281363 | 0.14406882 | Y105C5A.8 |
| WBGene00013634 | 0.451235227 | 0.84990489 | 1 | Y105C5A.9 |
| WBGene00013635 | -0.156269293 | 0.96924949 | 1 | Y105C5A.10 |
| WBGene00013636 | -0.245531902 | 0.94070824 | 1 | Y105C5A.12 |
| WBGene00013637 | 0.616429445 | 0.81400458 | 1 | Y105C5A.13 |
| WBGene00013639 | 1.291719861 | 9.1739E-05 | 0.00143163 | Y105C5A.15 |
| WBGene00013640 | 0.312403178 | 0.89676939 | 1 | Y105C5A.17 |
| WBGene00013642 | 1.53572575 | 0.06012639 | 0.18249864 | daf-38 |
| WBGene00013643 | 1.218992855 | 0.00027311 | 0.00336292 | Y105C5A.24 |
| WBGene00013646 | 0.426326213 | 0.80642435 | 1 | Y105C5B.3 |
| WBGene00013647 | -0.945062415 | 3.3213E-06 | 9.0161E-05 | Y105C5B.5 |
| WBGene00013649 | 0.81835207 | 0.70463712 | 1 | Y105C5B.8 |
| WBGene00013650 | -0.384710027 | 0.4273979 | 0.63057665 | Y105C5B.9 |
| WBGene00013651 | -1.323116856 | 0.47668251 | 1 | Y105C5B.11 |
| WBGene00013653 | 0.377926615 | 0.83031964 | 1 | Y105C5B.14 |
| WBGene00013654 | 0.869282442 | 2.4546E-05 | 0.00048144 | Y105C5B.15 |
| WBGene00013655 | -0.335526601 | 0.9011152 | 1 | lgc-19 |
| WBGene00013656 | 0.590708277 | 0.82739319 | 1 | Y105C5B.17 |
| WBGene00013658 | -0.426509006 | 0.22855067 | 0.43377108 | Y105C5B.19 |
| WBGene00013659 | -0.8229579 | 0.03965241 | 0.136767 | Y105C5B.20 |
| WBGene00013662 | 2.350469298 | 0.17938208 | 1 | Y105C5B.25 |
| WBGene00013665 | -0.390806552 | 0.80488376 | 1 | hlh-32 |
| WBGene00013666 | 0.322236112 | 0.44404651 | 0.64348862 | Y105E8A.1 |
| WBGene00013667 | 0.031495043 | 0.8748457 | 0.93252868 | Y105E8A.2 |
| WBGene00013668 | -0.046511763 | 0.78599782 | 0.88220389 | Y105E8A.3 |
| WBGene00013669 | -0.280748479 | 0.22933572 | 0.43442105 | Y105E8A.8 |
| WBGene00013670 | 0.053415368 | 0.77265306 | 0.87295559 | hpo-13 |
| WBGene00013671 | -0.265544001 | 0.45246121 | 0.65010241 | Y105E8A.11 |
| WBGene00013672 | 0.676030362 | 0.02507322 | 0.10030113 | catp-1 |
| WBGene00013673 | -1.261296079 | 0.2538595 | 1 | Y105E8A.13 |
| WBGene00013674 | -0.307914782 | 0.14227402 | 0.32034267 | Y105E8A.14 |
| WBGene00013676 | 0.020978191 | 0.91689205 | 0.95635621 | ekl-4 |
| WBGene00013677 | -0.037378053 | 0.83774562 | 0.91143626 | yars-1 |
| WBGene00013678 | 0.134263139 | 0.57217645 | 0.7462154 | Y105E8A.20 |
| WBGene00013679 | -0.264679172 | 0.18300825 | 0.37519473 | Y105E8A.21 |
| WBGene00013680 | 0.21117861 | 0.37371156 | 0.58147277 | rpom-1 |
| WBGene00013682 | 0.687576561 | 0.00363895 | 0.02450978 | Y105E8A.25 |
| WBGene00013683 | 0.317610662 | 0.24300585 | 0.45063908 | zoo-1 |
| WBGene00013684 | -0.156269293 | 0.96924949 | 1 | Y105E8A.27 |
| WBGene00013685 | 0.812501164 | 0.40796409 | 0.6141434 | Y105E8A.28 |
| WBGene00013687 | -0.065931357 | 0.71529869 | 0.83956317 | exoc-8 |
| WBGene00013688 | -0.049448781 | 0.77909781 | 0.87737933 | riok-2 |
| WBGene00013689 | -0.115280001 | 0.57701065 | 0.74903703 | bath-40 |

| | | | | |
|---|---|---|---|---|
| WBGene00013690 | -0.136166318 | 0.49156462 | 0.68082205 | hprt-1 |
| WBGene00013691 | -0.311835107 | 0.49899696 | 0.68719624 | Y105E8B.6 |
| WBGene00013692 | 0.187989155 | 0.70209483 | 0.83186624 | Y105E8B.7 |
| WBGene00013693 | 0.022750764 | 0.9611068 | 0.97977647 | Y105E8B.9 |
| WBGene00013694 | -0.243673704 | 0.31319471 | 0.52122318 | Y106G6A.1 |
| WBGene00013695 | 0.134596847 | 0.61513274 | 0.77528564 | epg-8 |
| WBGene00013696 | 0.846183542 | 0.65777268 | 1 | Y106G6A.4 |
| WBGene00013697 | -0.443790994 | 0.03627981 | 0.12883099 | dsbn-1 |
| WBGene00013698 | -1.510968571 | 0.00101694 | 0.0093535 | Y106G6D.1 |
| WBGene00013699 | -2.407984171 | 0.00011449 | 0.00169935 | Y106G6D.2 |
| WBGene00013700 | 1.183108147 | 0.4819447 | 1 | Y106G6D.3 |
| WBGene00013701 | 0.779417359 | 0.52676657 | 1 | Y106G6D.4 |
| WBGene00013702 | -0.035567644 | 0.95682991 | 0.97818053 | Y106G6D.6 |
| WBGene00013703 | -0.266245859 | 0.18054243 | 0.37259848 | Y106G6D.7 |
| WBGene00013704 | 0.054775068 | 0.90263488 | 0.94939705 | Y106G6D.8 |
| WBGene00013708 | -0.101883206 | 0.8193831 | 0.90234831 | Y106G6E.4 |
| WBGene00013709 | 0.21970448 | 0.29851298 | 0.50838071 | csnk-1 |
| WBGene00013710 | 2.141869924 | 0.24234872 | 1 | Y106G6G.1 |
| WBGene00013711 | 0.815032942 | 0.20498205 | 0.40440154 | T20F10.5 |
| WBGene00013711 | 0.979721952 | 0.09243666 | 0.24216495 | Y106G6G.2 |
| WBGene00013711 | 1.49363237 | 0.04717815 | 0.15480809 | Y106G6G.6 |
| WBGene00013712 | 1.181686151 | 0.7579682 | 1 | dlc-6 |
| WBGene00013713 | 0.447072595 | 0.70837296 | 1 | Y106G6G.4 |
| WBGene00013716 | 0.822211165 | 0.0123663 | 0.06058887 | Y106G6H.1 |
| WBGene00013717 | -0.577420258 | 0.4818218 | 0.67293207 | madf-10 |
| WBGene00013718 | 0.019730725 | 0.91422566 | 0.95571968 | pdpr-1 |
| WBGene00013719 | -0.521974467 | 0.00508574 | 0.03155229 | Y106G6H.6 |
| WBGene00013720 | -0.438723187 | 0.08318951 | 0.22650292 | Y106G6H.8 |
| WBGene00013721 | 0.178115923 | 0.94982316 | 1 | Y106G6H.9 |
| WBGene00013723 | 1.242730541 | 0.48737016 | 1 | Y106G6H.13 |
| WBGene00013724 | -0.494090105 | 0.03471615 | 0.12548043 | Y106G6H.14 |
| WBGene00013725 | -0.442974314 | 0.10961969 | 0.27006751 | ska-1 |
| WBGene00013726 | -0.197615995 | 0.68242919 | 0.81822761 | Y106G6H.16 |
| WBGene00013727 | -0.535125958 | 0.25411833 | 0.46193303 | Y111B2A.1 |
| WBGene00013728 | -0.53344059 | 0.00829045 | 0.04495507 | Y111B2A.2 |
| WBGene00013729 | -0.274378964 | 0.25290124 | 0.46089372 | Y111B2A.3 |
| WBGene00013730 | 0.6571687 | 0.00364517 | 0.0245292 | sql-1 |
| WBGene00013732 | 0.478791907 | 0.05067887 | 0.16217522 | aakg-1 |
| WBGene00013734 | 0.046512076 | 0.82753696 | 0.90692193 | Y111B2A.10 |
| WBGene00013735 | -0.070270353 | 0.75792404 | 0.86459364 | Y111B2A.12 |
| WBGene00013736 | -0.287075112 | 0.45251678 | 0.65010241 | Y111B2A.13 |
| WBGene00013737 | -0.080171823 | 0.72223149 | 0.8433791 | tpst-1 |
| WBGene00013738 | 0.075926169 | 0.76402032 | 0.86806496 | elpc-2 |

| | | | | |
|---|---|---|---|---|
| WBGene00013739 | -0.302136936 | 0.35991213 | 0.56806965 | spin-4 |
| WBGene00013740 | -0.283728701 | 0.20695974 | 0.40665706 | hut-1 |
| WBGene00013741 | -1.100567157 | 0.23138043 | 0.436888 | Y111B2A.21 |
| WBGene00013742 | -0.494568973 | 0.03732986 | 0.13121898 | sas-1 |
| WBGene00013744 | 0.377828795 | 0.92526585 | 1 | Y111B2A.27 |
| WBGene00013746 | 0.850418322 | 0.15899551 | 0.34378709 | lgc-55 |
| WBGene00013747 | 1.216957619 | 0.74819621 | 1 | spe-19 |
| WBGene00013748 | 0.897268984 | 0.48582097 | 1 | ssu-1 |
| WBGene00013749 | -0.156269293 | 0.96924949 | 1 | Y113G7A.12 |
| WBGene00013750 | 0.950776581 | 0.35165558 | 1 | Y113G7A.13 |
| WBGene00013751 | -0.211492809 | 0.85858936 | 1 | Y113G7A.14 |
| WBGene00013752 | 0.480601034 | 0.65044878 | 1 | Y113G7A.15 |
| WBGene00013753 | 0.049655344 | 0.96712018 | 1 | Y113G7A.16 |
| WBGene00013754 | -0.156269293 | 0.96924949 | 1 | fbxa-116 |
| WBGene00013755 | -0.358069336 | 0.56885873 | 0.74341002 | fbxa-115 |
| WBGene00013756 | 0.03939207 | 0.92538996 | 0.96044148 | ftr-1 |
| WBGene00013757 | 0.041803775 | 0.98890197 | 1 | fbxa-113 |
| WBGene00013758 | -0.375695308 | 0.30826488 | 0.51701118 | fbxa-114 |
| WBGene00013759 | 0.122495621 | 0.91759166 | 1 | fbxb-59 |
| WBGene00013761 | -0.172733871 | 0.87017979 | 1 | Y113G7B.11 |
| WBGene00013762 | 0.255343136 | 0.67971576 | 0.81647553 | Y113G7B.12 |
| WBGene00013764 | -0.333693239 | 0.89573179 | 1 | Y113G7B.15 |
| WBGene00013765 | -0.359926406 | 0.10880552 | 0.26893482 | cdkr-3 |
| WBGene00013766 | -0.443530288 | 0.00136937 | 0.01169165 | prmt-1 |
| WBGene00013768 | -0.366549956 | 0.16751447 | 0.35493773 | sld-5 |
| WBGene00013771 | 2.103207888 | 0.28977427 | 1 | Y113G7C.1 |
| WBGene00013774 | 0.629154436 | 0.81300308 | 1 | clec-193 |
| WBGene00013776 | 1.712479368 | 0.58529868 | 1 | Y116A8A.6 |
| WBGene00013778 | 2.715570568 | 0.10714135 | 1 | clec-194 |
| WBGene00013779 | -1.799615016 | 0.27217603 | 1 | Y116A8B.1 |
| WBGene00013781 | 0.030141249 | 0.97783036 | 1 | Y116A8B.4 |
| WBGene00013782 | 0.899238676 | 0.53262066 | 1 | npr-32 |
| WBGene00013784 | 0.040869027 | 0.95351918 | 0.97634694 | Y116A8C.3 |
| WBGene00013785 | 1.025181888 | 0.79790347 | 1 | nep-23 |
| WBGene00013786 | 0.720066222 | 0.75158338 | 1 | nep-24 |
| WBGene00013789 | 0.015173052 | 0.95812462 | 0.9785154 | Y116A8C.9 |
| WBGene00013790 | -0.418882953 | 0.21034234 | 0.41037543 | Y116A8C.10 |
| WBGene00013792 | -0.086293369 | 0.74035488 | 0.85519532 | Y116A8C.13 |
| WBGene00013794 | -1.439368372 | 0.0763169 | 0.21438303 | dct-13 |
| WBGene00013796 | -1.448978735 | 0.02702911 | 0.10555175 | Y116A8C.19 |
| WBGene00013797 | -2.45521408 | 0.03648642 | 1 | Y116A8C.20 |
| WBGene00013798 | -0.737462795 | 0.83433562 | 1 | clec-195 |
| WBGene00013799 | -2.219056424 | 1.337E-05 | 0.00028883 | athp-3 |

| | | | | |
|---|---|---|---|---|
| WBGene00013800 | -0.156269293 | 0.96924949 | 1 | Y116A8C.23 |
| WBGene00013802 | 0.356606635 | 0.69901926 | 0.82984187 | Y116A8C.25 |
| WBGene00013803 | 0.38863886 | 0.04821656 | 0.15719582 | snx-13 |
| WBGene00013804 | -0.089389562 | 0.75698923 | 0.86429398 | Y116A8C.27 |
| WBGene00013805 | -0.442616824 | 0.0852197 | 0.23019943 | bca-2 |
| WBGene00013806 | 0.546023545 | 0.37985895 | 0.58781969 | Y116A8C.29 |
| WBGene00013807 | -0.668570631 | 0.0183059 | 0.08038047 | Y116A8C.30 |
| WBGene00013808 | -0.138044483 | 0.51913107 | 0.70453502 | sfa-1 |
| WBGene00013809 | 0.504172634 | 0.56577959 | 0.74116638 | Y116A8C.33 |
| WBGene00013811 | 0.377828795 | 0.92526585 | 1 | Y116A8C.38 |
| WBGene00013812 | 1.651480332 | 0.32297692 | 1 | Y116A8C.40 |
| WBGene00013814 | -0.73089135 | 0.72186459 | 1 | Y116A8C.43 |
| WBGene00013815 | -3.47102951 | 0.12756864 | 1 | Y116A8C.44 |
| WBGene00013816 | -2.631851446 | 0.18382193 | 1 | Y116F11A.1 |
| WBGene00013819 | -1.143557482 | 0.1652755 | 0.35191335 | Y116F11A.6 |
| WBGene00013822 | 2.531987082 | 0.08344035 | 1 | Y116F11B.6 |
| WBGene00013823 | 0.281454817 | 0.94435481 | 1 | Y116F11B.7 |
| WBGene00013824 | -0.62146648 | 0.85925752 | 1 | Y116F11B.8 |
| WBGene00013825 | -1.901856996 | 0.64072034 | 1 | Y116F11B.9 |
| WBGene00013828 | 0.606544163 | 0.70175192 | 1 | Y116F11B.13 |
| WBGene00013829 | -0.522741973 | 0.82405702 | 1 | Y116F11B.14 |
| WBGene00013830 | 1.144601232 | 0.58194725 | 1 | ZC15.1 |
| WBGene00013832 | 0.377828795 | 0.92526585 | 1 | ZC15.3 |
| WBGene00013833 | 2.082792772 | 0.15369394 | 1 | ZC15.5 |
| WBGene00013835 | 0.441153543 | 0.54271444 | 0.72320229 | mlt-4 |
| WBGene00013836 | 0.081285082 | 0.92156625 | 0.95884274 | fbxa-29 |
| WBGene00013837 | 2.06978106 | 0.13870243 | 1 | fbxa-30 |
| WBGene00013838 | 1.471707393 | 0.02443836 | 0.09857345 | fbxa-31 |
| WBGene00013839 | 1.439425201 | 0.18413624 | 1 | fbxa-32 |
| WBGene00013840 | 3.449461417 | 0.06119705 | 1 | fbxa-151 |
| WBGene00013844 | -0.629587831 | 0.72107286 | 1 | fbxa-50 |
| WBGene00013845 | 1.162874696 | 0.1295662 | 0.30084603 | fbxa-37 |
| WBGene00013846 | 1.519550677 | 0.20294932 | 1 | ZC84.1 |
| WBGene00013847 | 0.059390019 | 0.79182459 | 0.8850619 | cls-3 |
| WBGene00013848 | 1.150447207 | 0.09498772 | 0.24705354 | npr-29 |
| WBGene00013849 | 0.270723507 | 0.79062484 | 1 | ZC84.6 |
| WBGene00013850 | 1.977885804 | 0.49204652 | 1 | ZC84.7 |
| WBGene00013852 | -0.596977439 | 0.24866658 | 0.45656791 | hhat-1 |
| WBGene00013853 | 0.601633224 | 0.05924412 | 0.18090615 | ZC116.1 |
| WBGene00013854 | -0.290000011 | 0.85914611 | 1 | cyc-2.2 |
| WBGene00013855 | 0.799996811 | 0.1183588 | 0.28371059 | ZC116.3 |
| WBGene00013856 | 0.501721414 | 0.76805879 | 1 | ZC168.2 |
| WBGene00013857 | -0.001624445 | 0.99351125 | 0.99671295 | orc-5 |

| | | | | |
|---|---|---|---|---|
| WBGene00013858 | 3.005407498 | 0.12134537 | 1 | ssp-34 |
| WBGene00013859 | 2.400797094 | 2.85E-10 | 2.56E-08 | ZC247.1 |
| WBGene00013860 | 0.023068851 | 0.9514728 | 0.97545664 | ZC247.2 |
| WBGene00013862 | -0.074842904 | 0.69568144 | 0.82777182 | wdr-5.3 |
| WBGene00013863 | 0.172203203 | 0.88309741 | 1 | ZC302.3 |
| WBGene00013865 | -0.912030601 | 0.62916461 | 1 | ZC334.7 |
| WBGene00013866 | -0.035845786 | 0.88942994 | 0.94187256 | cbs-1 |
| WBGene00013867 | -0.749642313 | 1.77E-07 | 7.5999E-06 | ZC373.2 |
| WBGene00013868 | 0.281454817 | 0.94435481 | 1 | ZC373.3 |
| WBGene00013869 | 1.215267484 | 0.00013412 | 0.00193397 | mlck-1 |
| WBGene00013870 | 0.604036681 | 0.1428423 | 0.32062273 | ZC373.5 |
| WBGene00013871 | 2.271211121 | 0.19727285 | 1 | gnrr-3 |
| WBGene00013872 | 0.151144677 | 0.75476593 | 0.86283239 | ZC374.2 |
| WBGene00013873 | -1.972274086 | 0.08292028 | 1 | ZC376.1 |
| WBGene00013874 | 0.251584934 | 0.44407228 | 0.64348862 | ZC376.2 |
| WBGene00013875 | -0.343489755 | 0.55470377 | 0.7323465 | ZC376.3 |
| WBGene00013876 | -0.401023663 | 0.39143838 | 0.59975433 | ceh-74 |
| WBGene00013877 | 0.379572321 | 0.0418231 | 0.14203475 | ZC376.6 |
| WBGene00013878 | -0.231432497 | 0.21032256 | 0.41037543 | atfs-1 |
| WBGene00013879 | 1.181686151 | 0.7579682 | 1 | ZC376.8 |
| WBGene00013880 | -0.328466941 | 0.0954195 | 0.24772303 | mppb-1 |
| WBGene00013881 | -0.347653162 | 0.12869807 | 0.29962095 | mans-4 |
| WBGene00013882 | 0.088233448 | 0.74398687 | 0.85740338 | msa-1 |
| WBGene00013883 | 1.114546249 | 0.35018455 | 1 | npr-13 |
| WBGene00013884 | 0.101651585 | 0.68526931 | 0.82001327 | ZC412.3 |
| WBGene00013886 | 0.983488853 | 0.80601565 | 1 | ZC412.5 |
| WBGene00013887 | 1.181686151 | 0.7579682 | 1 | nspa-5 |
| WBGene00013891 | -0.688327405 | 0.5176977 | 1 | ZC434.3 |
| WBGene00013892 | -0.108896879 | 0.59265809 | 0.75972181 | ZC434.4 |
| WBGene00013893 | 0.21988531 | 0.34059087 | 0.54893319 | ZC434.7 |
| WBGene00013894 | -0.884411963 | 1.09E-07 | 5.0737E-06 | ZC434.8 |
| WBGene00013895 | 0.620810012 | 0.10460125 | 0.26276926 | ZC434.9 |
| WBGene00013896 | 0.757385138 | 0.1201857 | 0.28644966 | ZC443.1 |
| WBGene00013897 | 0.244909602 | 0.81675928 | 1 | ZC443.2 |
| WBGene00013898 | 0.824467455 | 0.00281238 | 0.02029678 | ZC443.3 |
| WBGene00013899 | 1.848392952 | 7.182E-05 | 0.00117833 | ZC443.4 |
| WBGene00013900 | 3.962445722 | 1.72E-18 | 4.86E-16 | ugt-18 |
| WBGene00013901 | 0.008815528 | 0.98151212 | 0.9903215 | ugt-16 |
| WBGene00013902 | 2.117476584 | 1.3977E-06 | 4.3847E-05 | ZC455.1 |
| WBGene00013903 | 1.942796681 | 0.31244449 | 1 | ugt-3 |
| WBGene00013904 | 0.026480622 | 0.96575108 | 0.98286673 | ugt-6 |
| WBGene00013905 | 0.466552795 | 0.34467715 | 0.5535351 | ugt-4 |
| WBGene00013906 | 0.206178497 | 0.69999849 | 0.83055986 | ugt-5 |

| | | | | |
|---|---|---|---|---|
| WBGene00013907 | -0.323868354 | 0.91634405 | 1 | sre-20 |
| WBGene00013909 | -1.159714244 | 0.75790634 | 1 | srbc-44 |
| WBGene00013911 | -0.156269293 | 0.96924949 | 1 | irld-60 |
| WBGene00013912 | -1.901856996 | 0.64072034 | 1 | irld-19 |
| WBGene00013913 | 0.574508983 | 0.8551971 | 1 | irld-61 |
| WBGene00013914 | 1.347581267 | 0.18729282 | 0.38076391 | lgc-37 |
| WBGene00013915 | -0.156269293 | 0.96924949 | 1 | irld-20 |
| WBGene00013916 | 0.707559163 | 0.78139902 | 1 | ZC504.1 |
| WBGene00013917 | -0.120626693 | 0.64492178 | 0.79498712 | cdk-11.2 |
| WBGene00013919 | 0.113119661 | 0.58626662 | 0.75557614 | ZC506.1 |
| WBGene00013920 | 0.480059349 | 0.03970333 | 0.13682844 | pssy-1 |
| WBGene00013921 | 0.972164298 | 0.00855112 | 0.04601914 | best-22 |
| WBGene00013923 | 1.498729111 | 0.02189928 | 0.09168245 | ghi-1 |
| WBGene00013924 | -0.536649781 | 0.00053944 | 0.00567535 | rad-23 |
| WBGene00013925 | -0.158947457 | 0.50906054 | 0.69566237 | ZK20.4 |
| WBGene00013926 | -0.532300094 | 0.02707695 | 0.10556882 | nep-1 |
| WBGene00013927 | -0.156269293 | 0.96924949 | 1 | clec-95 |
| WBGene00013928 | -0.419450245 | 0.84655242 | 1 | clec-94 |
| WBGene00013930 | -0.156269293 | 0.96924949 | 1 | clec-96 |
| WBGene00013931 | -1.659180847 | 2.2015E-05 | 0.00043817 | clec-97 |
| WBGene00013932 | -1.433785839 | 0.0797305 | 0.22043917 | clec-98 |
| WBGene00013933 | -0.848357199 | 0.71057142 | 1 | clec-99 |
| WBGene00013934 | 0.739769447 | 0.02746588 | 0.10670115 | ZK131.11 |
| WBGene00013935 | 3.741912746 | 2.8643E-05 | 0.00054 | ZK218.1 |
| WBGene00013937 | 4.042836721 | 0.0127948 | 1 | ZK218.3 |
| WBGene00013938 | 2.206678637 | 0.22172448 | 1 | ZK218.4 |
| WBGene00013939 | 6.615065908 | 5.00E-08 | 2.5208E-06 | ZK218.5 |
| WBGene00013941 | 4.799217815 | 0.00116225 | 1 | ZK218.7 |
| WBGene00013943 | -0.156269293 | 0.96924949 | 1 | ZK218.11 |
| WBGene00013946 | -0.417436887 | 0.21005327 | 0.41014101 | ZK228.3 |
| WBGene00013947 | 0.205443174 | 0.37724706 | 0.58509921 | ZK228.4 |
| WBGene00013949 | 0.275892417 | 0.70277904 | 0.83218963 | ZK262.2 |
| WBGene00013950 | -0.194497119 | 0.73986329 | 0.8549234 | ZK262.3 |
| WBGene00013954 | 0.377828795 | 0.92526585 | 1 | ZK262.9 |
| WBGene00013955 | 0.291688704 | 0.64985135 | 0.79839241 | kri-1 |
| WBGene00013956 | 1.158132015 | 0.62662247 | 1 | ZK265.3 |
| WBGene00013957 | 2.884412203 | 0.07197729 | 1 | sre-23 |
| WBGene00013958 | -0.824782882 | 0.00023082 | 0.00294065 | ZK265.6 |
| WBGene00013959 | 0.609901958 | 0.15820751 | 0.34299953 | ZK265.7 |
| WBGene00013960 | -0.590468979 | 0.61293861 | 1 | cutl-8 |
| WBGene00013961 | 0.377828795 | 0.92526585 | 1 | ZK285.2 |
| WBGene00013962 | 0.449715685 | 0.2070708 | 0.40667204 | ZK287.1 |
| WBGene00013963 | 1.148770566 | 0.00507933 | 0.03155229 | sulp-8 |

| | | | | |
|---|---|---|---|---|
| WBGene00013964 | 0.674861556 | 0.23060359 | 0.4360994 | ZK287.3 |
| WBGene00013965 | 0.425883204 | 0.53357214 | 0.71668319 | ZK287.4 |
| WBGene00013966 | -0.540561101 | 0.11052911 | 0.27182701 | ztf-9 |
| WBGene00013967 | -0.417880227 | 0.16181998 | 0.34773047 | ZK287.7 |
| WBGene00013968 | 1.05100407 | 0.06904599 | 0.20040338 | ZK287.9 |
| WBGene00013969 | 1.062810119 | 5.36E-07 | 2.0098E-05 | tep-1 |
| WBGene00013970 | 1.713200756 | 0.00024945 | 0.00312226 | klu-1 |
| WBGene00013974 | 0.198251955 | 0.88240306 | 1 | npr-9 |
| WBGene00013975 | 0.368222699 | 0.73666812 | 1 | ZK455.5 |
| WBGene00013976 | 0.276635087 | 0.67268351 | 0.81296541 | hizr-1 |
| WBGene00013978 | -0.149909352 | 0.93707773 | 1 | ZK507.1 |
| WBGene00013979 | -1.186373843 | 0.53853051 | 1 | ZK507.3 |
| WBGene00013980 | -0.776540296 | 0.30644696 | 0.51524339 | dos-1 |
| WBGene00013982 | -0.265214151 | 0.73448045 | 0.85162135 | ZK512.1 |
| WBGene00013983 | 0.005525282 | 0.97844452 | 0.98873753 | ZK512.2 |
| WBGene00013984 | -0.864345177 | 0.00240057 | 0.01799091 | ZK512.4 |
| WBGene00013985 | 0.211274067 | 0.28897892 | 0.49921577 | sec-16 |
| WBGene00013986 | -0.743603331 | 0.00138214 | 0.01175936 | ZK512.7 |
| WBGene00013987 | 1.334341511 | 0.55384944 | 1 | ZK512.8 |
| WBGene00013988 | 0.688793783 | 0.86378755 | 1 | ZK512.10 |
| WBGene00013989 | 0.052539763 | 0.93882226 | 0.96857928 | ZK512.11 |
| WBGene00013994 | 0.008338313 | 0.96828044 | 0.98383906 | ZK524.4 |
| WBGene00013996 | -0.000200393 | 0.99964589 | 0.99989047 | ZK550.2 |
| WBGene00013997 | -0.072963146 | 0.67894563 | 0.81647553 | ZK550.3 |
| WBGene00013998 | -0.365407371 | 0.11606067 | 0.28048782 | ZK550.4 |
| WBGene00013999 | 1.816974857 | 0.33692172 | 1 | ZK550.5 |
| WBGene00014000 | 0.051312798 | 0.8114159 | 0.89696808 | ZK550.6 |
| WBGene00014001 | 0.746786933 | 0.00066783 | 0.00671206 | pyk-2 |
| WBGene00014002 | 0.999487288 | 0.18407006 | 0.37635477 | ZK593.2 |
| WBGene00014003 | 1.118532681 | 1.7111E-05 | 0.00035635 | ZK593.3 |
| WBGene00014004 | 0.066055987 | 0.74723068 | 0.85920656 | fic-1 |
| WBGene00014006 | 1.252158231 | 2.985E-06 | 8.218E-05 | ZK596.1 |
| WBGene00014007 | 0.742628056 | 0.76901841 | 1 | ZK596.2 |
| WBGene00014008 | 0.516742221 | 0.66361509 | 1 | ZK596.3 |
| WBGene00014009 | -0.997073712 | 0.06484844 | 0.19207752 | lips-6 |
| WBGene00014011 | -0.160202901 | 0.35459419 | 0.56294974 | ZK632.2 |
| WBGene00014012 | -0.084854215 | 0.68333305 | 0.81877569 | riok-3 |
| WBGene00014013 | 0.249748312 | 0.1507604 | 0.33243407 | ZK632.4 |
| WBGene00014014 | 0.156828433 | 0.51604943 | 0.70217251 | ZK632.5 |
| WBGene00014015 | 0.409177616 | 0.05101231 | 0.16294144 | panl-3 |
| WBGene00014016 | -0.559817277 | 0.00040903 | 0.00458543 | ZK632.9 |
| WBGene00014017 | -0.264586479 | 0.14770848 | 0.32820803 | ZK632.10 |
| WBGene00014018 | -0.354632032 | 0.09715518 | 0.25046061 | ZK632.11 |

| | | | | |
|---|---|---|---|---|
| WBGene00014019 | -0.514866179 | 0.0136562 | 0.06491774 | ZK632.12 |
| WBGene00014020 | -0.256521647 | 0.39955079 | 0.60727919 | ZK632.14 |
| WBGene00014021 | 1.152983523 | 0.02045992 | 0.08702781 | svop-1 |
| WBGene00014022 | -0.525595612 | 0.00926973 | 0.04897467 | ZK637.2 |
| WBGene00014023 | -0.127652224 | 0.45534781 | 0.65267707 | lnkn-1 |
| WBGene00014024 | -0.939157204 | 0.49655271 | 1 | ZK637.4 |
| WBGene00014025 | -0.602995244 | 0.00078264 | 0.00761385 | asna-1 |
| WBGene00014027 | 0.544112251 | 0.31659754 | 0.52460908 | tpk-1 |
| WBGene00014028 | 0.674873698 | 0.08707414 | 0.23352487 | trxr-2 |
| WBGene00014029 | -0.238778247 | 0.89243712 | 1 | ZK637.12 |
| WBGene00014030 | -0.108283848 | 0.61832315 | 0.77769205 | glb-1 |
| WBGene00014031 | -0.576360844 | 0.16798239 | 0.35563637 | ZK637.14 |
| WBGene00014032 | -1.657722853 | 0.43572774 | 1 | ZK637.15 |
| WBGene00014033 | 0.289577394 | 0.7266726 | 0.84642598 | arrd-15 |
| WBGene00014034 | -0.3640059 | 0.16523656 | 0.35189214 | ZK643.2 |
| WBGene00014035 | 1.322488356 | 0.00708579 | 0.04010657 | seb-2 |
| WBGene00014036 | -0.273651591 | 0.13002708 | 0.3015585 | ZK643.5 |
| WBGene00014037 | -0.156269293 | 0.96924949 | 1 | ZK643.6 |
| WBGene00014039 | -0.500293961 | 0.47866187 | 0.67083387 | ZK662.2 |
| WBGene00014040 | -1.482825083 | 0.12998125 | 1 | ZK662.5 |
| WBGene00014041 | -1.469983358 | 0.36599132 | 1 | ZK666.1 |
| WBGene00014043 | 0.903390376 | 0.71900339 | 1 | clec-58 |
| WBGene00014044 | 0.35828053 | 0.87111704 | 1 | ZK666.4 |
| WBGene00014046 | 6.098231393 | 3.5573E-06 | 9.5341E-05 | clec-60 |
| WBGene00014047 | 4.96599016 | 0.0005898 | 0.00605304 | clec-61 |
| WBGene00014049 | 1.435925494 | 0.70920779 | 1 | ZK666.11 |
| WBGene00014050 | -0.156269293 | 0.96924949 | 1 | ZK666.12 |
| WBGene00014051 | 0.718795012 | 0.00840485 | 0.04541312 | spv-1 |
| WBGene00014052 | -0.49810563 | 0.44900883 | 0.6479551 | ZK669.2 |
| WBGene00014053 | 0.587978534 | 0.11833744 | 0.28371059 | ZK669.3 |
| WBGene00014054 | -0.207560093 | 0.24321015 | 0.4508704 | dbt-1 |
| WBGene00014055 | 0.423396711 | 0.54321912 | 0.72328361 | ZK669.5 |
| WBGene00014057 | 1.630651526 | 2.86E-07 | 1.162E-05 | ZK673.1 |
| WBGene00014058 | -0.19986591 | 0.22964073 | 0.43488817 | ZK673.2 |
| WBGene00014060 | 0.232027158 | 0.74970071 | 0.86036058 | ZK673.4 |
| WBGene00014061 | -0.281836889 | 0.83742571 | 1 | ZK673.5 |
| WBGene00014062 | 2.629683399 | 0.22019584 | 1 | ZK673.6 |
| WBGene00014063 | 1.91319663 | 0.12010322 | 1 | clec-143 |
| WBGene00014064 | -1.256233633 | 0.16578402 | 0.35258827 | sdz-37 |
| WBGene00014065 | 0.870844124 | 0.43020763 | 1 | ZK673.11 |
| WBGene00014066 | 0.451950081 | 0.07725304 | 0.21568044 | rev-1 |
| WBGene00014067 | -1.694808057 | 0.00291639 | 0.02077929 | ZK675.4 |
| WBGene00014068 | 0.281454817 | 0.94435481 | 1 | nhr-255 |

| | | | | |
|---|---|---|---|---|
| WBGene00014069 | -0.245531902 | 0.94070824 | 1 | ZK678.3 |
| WBGene00014073 | 0.146418095 | 0.9299978 | 1 | ZK757.1 |
| WBGene00014074 | 0.290273065 | 0.59441098 | 0.76101905 | ZK757.2 |
| WBGene00014075 | -0.194179456 | 0.35921548 | 0.56735204 | dhhc-4 |
| WBGene00014076 | -0.073367826 | 0.78879017 | 0.8837593 | ZK792.1 |
| WBGene00014077 | 1.110478774 | 0.06440031 | 0.19131326 | ZK792.4 |
| WBGene00014078 | 0.108381315 | 0.55916658 | 0.73600413 | ZK792.5 |
| WBGene00014079 | 0.006028915 | 0.99145667 | 0.99572135 | ZK792.7 |
| WBGene00014080 | -0.249335889 | 0.45873045 | 0.65487177 | atg-4.2 |
| WBGene00014082 | -0.861206134 | 0.70037023 | 1 | ZK795.2 |
| WBGene00014083 | -0.498848876 | 0.03409623 | 0.12408944 | ZK795.3 |
| WBGene00014085 | -0.048008406 | 0.98380497 | 1 | ZK809.1 |
| WBGene00014086 | -0.313373792 | 0.03887876 | 0.13482633 | ZK809.3 |
| WBGene00014087 | -0.249446155 | 0.13899785 | 0.31602489 | ZK809.5 |
| WBGene00014088 | -0.123926593 | 0.64046974 | 0.79202874 | ZK809.8 |
| WBGene00014089 | -0.507345582 | 0.34745416 | 0.55625968 | ZK822.1 |
| WBGene00014090 | -0.423492486 | 0.09898482 | 0.25321843 | ZK822.2 |
| WBGene00014091 | 0.759027424 | 0.00103991 | 0.00952157 | ZK822.4 |
| WBGene00014092 | 0.786599113 | 0.00149635 | 0.01256376 | ZK822.5 |
| WBGene00014093 | -0.593590384 | 0.69776583 | 1 | ZK829.1 |
| WBGene00014094 | -1.399271197 | 0.22233946 | 1 | ZK829.3 |
| WBGene00014095 | 0.062675709 | 0.70439361 | 0.832842 | gdh-1 |
| WBGene00014096 | -0.44983705 | 0.00690804 | 0.03939432 | ZK829.7 |
| WBGene00014097 | -0.25712449 | 0.12495263 | 0.29465848 | ZK829.9 |
| WBGene00014098 | 0.245186568 | 0.18594271 | 0.37871674 | ogdh-2 |
| WBGene00014099 | 0.540057276 | 0.5788603 | 1 | ZK836.3 |
| WBGene00014100 | 1.115863748 | 0.63449935 | 1 | ZK849.1 |
| WBGene00014101 | 0.761727517 | 0.21871073 | 0.42148682 | gopc-1 |
| WBGene00014102 | 1.025181888 | 0.79790347 | 1 | best-25 |
| WBGene00014103 | -0.476663799 | 0.61345119 | 0.77421111 | best-26 |
| WBGene00014104 | -0.774842772 | 0.78069886 | 1 | ZK849.6 |
| WBGene00014105 | 3.221498814 | 0.11887087 | 1 | ZK856.4 |
| WBGene00014106 | 0.910941842 | 0.02191605 | 0.09172105 | ZK856.5 |
| WBGene00014107 | 0.247694858 | 0.87622457 | 1 | ZK856.6 |
| WBGene00014108 | -0.38402282 | 0.09786446 | 0.25151828 | ZK856.7 |
| WBGene00014109 | -0.052355704 | 0.84813843 | 0.91671496 | chpf-1 |
| WBGene00014111 | -0.678145806 | 0.01245893 | 0.06089527 | rpc-25 |
| WBGene00014112 | -0.002841811 | 0.99065049 | 0.99548813 | ZK856.11 |
| WBGene00014113 | 0.049114777 | 0.81970085 | 0.90251093 | hpo-40 |
| WBGene00014114 | -0.092054195 | 0.58518566 | 0.75465941 | tftc-3 |
| WBGene00014115 | 0.213720219 | 0.33984544 | 0.54809557 | gld-4 |
| WBGene00014116 | 2.154074734 | 0.23410381 | 1 | ZK858.2 |
| WBGene00014117 | -0.82772963 | 1.98E-08 | 1.1158E-06 | clec-91 |

| | | | | |
|---|---|---|---|---|
| WBGene00014118 | 0.17161157 | 0.55137535 | 0.72985697 | ZK858.5 |
| WBGene00014119 | -0.033833957 | 0.83794181 | 0.91143626 | ZK858.6 |
| WBGene00014120 | -0.202331789 | 0.30934777 | 0.51807644 | ZK858.7 |
| WBGene00014121 | 1.378226134 | 0.5038929 | 1 | ZK858.8 |
| WBGene00014122 | -0.445955551 | 0.76705763 | 1 | ZK863.1 |
| WBGene00014123 | -0.285561252 | 0.14559887 | 0.32453017 | elpc-3 |
| WBGene00014124 | -0.357467819 | 0.01192629 | 0.05902856 | usip-1 |
| WBGene00014125 | 1.014559051 | 0.00364675 | 0.0245292 | ZK863.8 |
| WBGene00014127 | 0.688793783 | 0.86378755 | 1 | ZK892.3 |
| WBGene00014128 | 0.938982574 | 0.8146943 | 1 | ZK892.4 |
| WBGene00014129 | 2.224351729 | 0.33187942 | 1 | ZK892.5 |
| WBGene00014130 | -0.055240099 | 0.96354327 | 1 | ZK892.6 |
| WBGene00014131 | -0.809627408 | 0.49605711 | 1 | sdz-38 |
| WBGene00014132 | 1.240797221 | 0.39907473 | 1 | ZK896.1 |
| WBGene00014135 | 0.057748116 | 0.92305908 | 0.9593272 | ZK896.4 |
| WBGene00014136 | 0.733260855 | 0.22849456 | 0.43373241 | ZK896.5 |
| WBGene00014137 | 0.28907886 | 0.43351982 | 0.63538117 | clec-187 |
| WBGene00014138 | 0.034339555 | 0.87539667 | 0.9327884 | clec-186 |
| WBGene00014139 | 0.173704921 | 0.43747956 | 0.63804213 | nstp-5 |
| WBGene00014140 | 0.87775628 | 0.09350193 | 0.24402558 | ZK899.1 |
| WBGene00014141 | 0.034880615 | 0.90484883 | 0.95065504 | ZK899.2 |
| WBGene00014142 | 1.38322681 | 0.69550305 | 1 | ZK899.3 |
| WBGene00014143 | -1.953006107 | 0.03567263 | 0.12738276 | ZK899.5 |
| WBGene00014144 | -1.77267394 | 0.01808118 | 0.0796241 | ZK899.6 |
| WBGene00014145 | 0.091715754 | 0.94318257 | 1 | ZK899.7 |
| WBGene00014148 | -0.096157362 | 0.75207628 | 0.86206175 | ZK909.3 |
| WBGene00014149 | -2.02547794 | 0.29582398 | 1 | fbxb-70 |
| WBGene00014150 | -0.518044532 | 0.29060345 | 0.50059826 | ZK909.6 |
| WBGene00014151 | 0.003083094 | 0.98906179 | 0.99484196 | vps-15 |
| WBGene00014152 | 1.65902237 | 2.9461E-06 | 8.1663E-05 | ZK930.2 |
| WBGene00014153 | 0.332772163 | 0.57156422 | 0.74584707 | vab-23 |
| WBGene00014154 | -0.737462795 | 0.83433562 | 1 | ZK930.4 |
| WBGene00014155 | 0.273294737 | 0.87943454 | 1 | ZK930.5 |
| WBGene00014156 | -3.101874071 | 0.0324335 | 1 | ZK930.6 |
| WBGene00014157 | 1.017496938 | 0.63763677 | 1 | ZK930.7 |
| WBGene00014158 | 3.193987294 | 0.12383192 | 1 | ZK938.1 |
| WBGene00014159 | -0.285957923 | 0.5073795 | 0.69409515 | arrd-4 |
| WBGene00014160 | 0.993470636 | 0.45561604 | 1 | ZK938.3 |
| WBGene00014161 | 0.996013393 | 0.53410803 | 1 | arrd-5 |
| WBGene00014163 | -0.649751881 | 0.01097428 | 0.05556245 | rnh-1.2 |
| WBGene00014164 | -0.029437666 | 0.89959043 | 0.94717955 | lact-2 |
| WBGene00014165 | 0.197560463 | 0.20158084 | 0.40041719 | puf-12 |
| WBGene00014166 | -0.587007343 | 0.15545829 | 0.33928079 | ZK945.4 |

| | | | | |
|---|---|---|---|---|
| WBGene00014168 | -0.156269293 | 0.96924949 | 1 | ZK945.6 |
| WBGene00014169 | 1.977885804 | 0.49204652 | 1 | ZK945.7 |
| WBGene00014171 | 0.942182222 | 0.00573307 | 0.03447507 | nep-26 |
| WBGene00014172 | -0.22196199 | 0.40572659 | 0.6124489 | clpp-1 |
| WBGene00014173 | -0.385719003 | 0.66274323 | 0.80624905 | ZK970.7 |
| WBGene00014174 | 0.377828795 | 0.92526585 | 1 | ZK970.8 |
| WBGene00014175 | 2.296841344 | 0.0491055 | 1 | ZK971.1 |
| WBGene00014176 | -0.349554813 | 0.12619899 | 0.29630283 | ZK1010.2 |
| WBGene00014177 | -0.437295494 | 0.04533978 | 0.15003278 | frg-1 |
| WBGene00014178 | -0.666762443 | 0.39404713 | 0.60200405 | ZK1010.4 |
| WBGene00014179 | -0.800196227 | 0.69440361 | 1 | ZK1010.5 |
| WBGene00014181 | 0.032437599 | 0.96155963 | 0.98000612 | ZK1010.8 |
| WBGene00014182 | 0.281454817 | 0.94435481 | 1 | ZK1025.2 |
| WBGene00014183 | -1.164391104 | 0.41827839 | 1 | ZK1025.3 |
| WBGene00014184 | -1.969805064 | 0.62862569 | 1 | ZK1025.4 |
| WBGene00014186 | -0.556584069 | 0.69337662 | 1 | nhr-244 |
| WBGene00014188 | 0.281454817 | 0.94435481 | 1 | ZK1025.8 |
| WBGene00014189 | -0.156269293 | 0.96924949 | 1 | nhr-245 |
| WBGene00014191 | -0.156269293 | 0.96924949 | 1 | srt-22 |
| WBGene00014192 | -1.847275425 | 0.48962967 | 1 | nhr-246 |
| WBGene00014193 | -1.901856996 | 0.64072034 | 1 | nhr-247 |
| WBGene00014194 | 0.809186946 | 0.19286945 | 0.38810746 | ZK1037.6 |
| WBGene00014195 | 0.747845955 | 0.85215965 | 1 | srz-10 |
| WBGene00014196 | 0.590708277 | 0.82739319 | 1 | ZK1053.1 |
| WBGene00014197 | 0.590708277 | 0.82739319 | 1 | ZK1053.2 |
| WBGene00014198 | -0.602562547 | 0.52747972 | 1 | ZK1053.3 |
| WBGene00014199 | -0.952548158 | 0.00082752 | 0.0079179 | ZK1053.4 |
| WBGene00014200 | 2.032534943 | 0.17596623 | 1 | scrm-2 |
| WBGene00014201 | -0.156269293 | 0.96924949 | 1 | ZK1053.6 |
| WBGene00014202 | 0.13117706 | 0.45875371 | 0.65487177 | mmcm-1 |
| WBGene00014203 | -0.210281417 | 0.5428043 | 0.72323899 | ZK1058.3 |
| WBGene00014204 | -0.361429892 | 0.01167456 | 0.05811431 | ccdc-47 |
| WBGene00014205 | -0.262995128 | 0.42120443 | 0.62577619 | ZK1058.5 |
| WBGene00014206 | -1.733265305 | 0.00641935 | 0.03739794 | nit-1 |
| WBGene00014207 | 0.154552118 | 0.47768558 | 0.67016184 | ZK1058.9 |
| WBGene00014208 | 0.480380398 | 0.02777791 | 0.10748752 | ZK1067.2 |
| WBGene00014209 | -0.352330479 | 0.08276221 | 0.22580135 | ZK1067.3 |
| WBGene00014210 | 0.621097451 | 0.2965322 | 0.50697066 | ZK1067.4 |
| WBGene00014213 | 0.275931579 | 0.26766526 | 0.47615002 | ZK1073.1 |
| WBGene00014214 | 0.717442821 | 0.18195388 | 0.37384831 | ZK1073.2 |
| WBGene00014215 | 0.469868789 | 0.07962192 | 0.22035325 | obr-3 |
| WBGene00014216 | -0.642582698 | 0.76082721 | 1 | ZK1086.2 |
| WBGene00014217 | -1.901856996 | 0.64072034 | 1 | ZK1086.3 |

| | | | | |
|---|---|---|---|---|
| WBGene00014218 | -0.086004688 | 0.61033896 | 0.77201762 | ZK1098.1 |
| WBGene00014219 | -0.216547064 | 0.25265633 | 0.46065483 | ZK1098.2 |
| WBGene00014220 | 0.25749888 | 0.51731973 | 0.7030198 | ZK1098.3 |
| WBGene00014221 | -0.570057912 | 0.0152947 | 0.07063015 | eif-2Balpha |
| WBGene00014222 | -0.676896598 | 0.01092393 | 0.05549264 | trpp-3 |
| WBGene00014223 | 1.278470803 | 0.39867721 | 1 | ZK1098.6 |
| WBGene00014224 | -0.754561066 | 0.00445405 | 0.02861196 | mrps-23 |
| WBGene00014225 | 0.281454817 | 0.94435481 | 1 | ZK1098.9 |
| WBGene00014226 | -0.354556575 | 0.2985061 | 0.50838071 | ZK1098.11 |
| WBGene00014227 | -0.261502951 | 0.27636086 | 0.48493961 | ZK1128.1 |
| WBGene00014228 | -0.226318578 | 0.40456431 | 0.61160694 | mett-10 |
| WBGene00014229 | 2.420661489 | 0.31300137 | 1 | ZK1128.3 |
| WBGene00014230 | -0.190044646 | 0.44072153 | 0.64081439 | ZK1128.4 |
| WBGene00014232 | -0.079870544 | 0.66050474 | 0.80449318 | ttll-4 |
| WBGene00014233 | 0.697424978 | 0.0333064 | 0.12190523 | ZK1128.7 |
| WBGene00014234 | -0.421456088 | 0.03751861 | 0.13161326 | vps-29 |
| WBGene00014235 | 1.116613983 | 0.69067248 | 1 | ZK1225.1 |
| WBGene00014238 | -1.159714244 | 0.75790634 | 1 | ZK1225.4 |
| WBGene00014239 | 0.747845955 | 0.85215965 | 1 | ZK1225.5 |
| WBGene00014240 | -0.156269293 | 0.96924949 | 1 | htas-1 |
| WBGene00014242 | -0.074464988 | 0.94968652 | 1 | sdz-36 |
| WBGene00014243 | 0.641799709 | 0.00154933 | 0.01286619 | dcaf-1 |
| WBGene00014244 | -0.656275797 | 0.00331601 | 0.02279708 | ZK1307.1 |
| WBGene00014245 | -2.422363895 | 0.09231616 | 1 | ZK1307.2 |
| WBGene00014246 | -1.901856996 | 0.64072034 | 1 | ZK1307.3 |
| WBGene00014247 | -0.639727652 | 0.77477802 | 1 | ZK1307.4 |
| WBGene00014248 | 2.478954062 | 0.03484415 | 1 | ZK1307.7 |
| WBGene00014249 | -0.115337252 | 0.44836311 | 0.64735123 | ZK1307.8 |
| WBGene00014250 | -0.204659349 | 0.35374354 | 0.56237866 | ZK1307.9 |
| WBGene00014251 | -0.64040028 | 0.02532252 | 0.10106723 | gstk-1 |
| WBGene00014252 | 1.06292341 | 1.92E-09 | 1.41E-07 | ZK1320.2 |
| WBGene00014253 | -0.111308002 | 0.51090316 | 0.69734404 | ZK1320.3 |
| WBGene00014254 | -0.233807165 | 0.809131 | 1 | cyp-13A10 |
| WBGene00014255 | 1.458582122 | 0.38669316 | 1 | ZK1320.5 |
| WBGene00014256 | 0.045706362 | 0.88253842 | 0.93749089 | ZK1320.7 |
| WBGene00014258 | -0.255278026 | 0.13799565 | 0.31463007 | ZK1320.9 |
| WBGene00014259 | 0.032568649 | 0.93116467 | 0.96395551 | ZK1320.11 |
| WBGene00014260 | -0.156269293 | 0.96924949 | 1 | ZK1321.1 |
| WBGene00014261 | 1.226454387 | 0.00258843 | 0.01903458 | shk-1 |
| WBGene00014262 | 0.23949968 | 0.29782244 | 0.50784582 | ZK1321.4 |
| WBGene00014266 | -0.18557176 | 0.95664217 | 1 | C28A5.7 |
| WBGene00014288 | -1.027355405 | 0.68015432 | 1 | C47F8.9 |
| WBGene00014290 | -0.156269293 | 0.96924949 | 1 | C51F7.3 |

| | | | | |
|---|---|---|---|---|
| WBGene00014291 | -1.094201388 | 0.7816772 | 1 | C52E4.8 |
| WBGene00014300 | 1.133907044 | 0.03492726 | 0.12590615 | D2023.1 |
| WBGene00014306 | 0.281454817 | 0.94435481 | 1 | F07C6.5 |
| WBGene00014310 | -1.027355405 | 0.68015432 | 1 | F08G2.9 |
| WBGene00014311 | -1.027355405 | 0.68015432 | 1 | F08G2.10 |
| WBGene00014312 | -1.027355405 | 0.68015432 | 1 | F08G2.11 |
| WBGene00014316 | -1.094201388 | 0.7816772 | 1 | F08H9.10 |
| WBGene00014322 | -0.156269293 | 0.96924949 | 1 | F11A5.14 |
| WBGene00014392 | -0.18557176 | 0.95664217 | 1 | F54C8.8 |
| WBGene00014393 | -0.18557176 | 0.95664217 | 1 | F54C8.9 |
| WBGene00014403 | -1.027355405 | 0.68015432 | 1 | F56H6.14 |
| WBGene00014405 | -1.027355405 | 0.68015432 | 1 | F56H6.16 |
| WBGene00014421 | -0.156269293 | 0.96924949 | 1 | K03B8.10 |
| WBGene00014425 | -1.038181246 | 0.5835992 | 1 | K07C10.2 |
| WBGene00014430 | -0.18557176 | 0.95664217 | 1 | K09B11.11 |
| WBGene00014431 | -0.18557176 | 0.95664217 | 1 | K09B11.12 |
| WBGene00014432 | -0.18557176 | 0.95664217 | 1 | K09B11.13 |
| WBGene00014433 | -0.18557176 | 0.95664217 | 1 | K09B11.14 |
| WBGene00014434 | -0.18557176 | 0.95664217 | 1 | K09B11.15 |
| WBGene00014435 | -0.18557176 | 0.95664217 | 1 | K09B11.16 |
| WBGene00014483 | -1.027355405 | 0.68015432 | 1 | R05D7.6 |
| WBGene00014484 | -0.18557176 | 0.95664217 | 1 | R07E5.16 |
| WBGene00014492 | 1.230375205 | 1.11E-09 | 8.66E-08 | R166.6 |
| WBGene00014505 | 0.218624593 | 0.266324 | 0.47477459 | T05E11.9 |
| WBGene00014511 | -1.759191416 | 0.57668729 | 1 | T08G5.11 |
| WBGene00014517 | -1.094201388 | 0.7816772 | 1 | T11F9.15 |
| WBGene00014541 | -1.14715222 | 0.68932198 | 1 | W04D2.7 |
| WBGene00014542 | -0.183119115 | 0.92486992 | 1 | W04G5.11 |
| WBGene00014543 | -0.18557176 | 0.95664217 | 1 | W05B2.8 |
| WBGene00014549 | -1.027355405 | 0.68015432 | 1 | W07G1.8 |
| WBGene00014550 | -1.027355405 | 0.68015432 | 1 | W07G1.9 |
| WBGene00014569 | -0.074445198 | 0.93490277 | 0.96576657 | bkip-1 |
| WBGene00014612 | -0.23209848 | 0.12268275 | 0.29085945 | mir-354 |
| WBGene00014631 | 0.281454817 | 0.94435481 | 1 | ZK897.2 |
| WBGene00014633 | 1.557897343 | 0.63813005 | 1 | ZK897.4 |
| WBGene00014666 | 0.727318704 | 0.00163306 | 0.01334255 | C05D12.3 |
| WBGene00014668 | 2.794537466 | 0.17119433 | 1 | C06C3.9 |
| WBGene00014669 | 0.896046919 | 0.00514459 | 0.03183509 | zipt-2.1 |
| WBGene00014697 | 0.158768435 | 0.76469409 | 0.86824567 | cyp-25A3 |
| WBGene00014698 | 0.829665238 | 0.41209019 | 1 | C37A5.3 |
| WBGene00014761 | 3.845260028 | 0.00122792 | 0.01073304 | F46A8.11 |
| WBGene00014793 | 0.279444701 | 0.28467886 | 0.49453227 | H40L08.2 |
| WBGene00014801 | 0.500287891 | 0.49870358 | 0.68695347 | K08E5.1 |

| | | | | |
|---|---|---|---|---|
| WBGene00014807 | -0.201764049 | 0.86142582 | 1 | M04D5.1 |
| WBGene00014813 | 1.829295604 | 0.35138922 | 1 | M176.9 |
| WBGene00014826 | 0.172865793 | 0.33207191 | 0.53982957 | R13H4.2 |
| WBGene00014836 | 0.212516941 | 0.59070324 | 0.75815153 | T09F5.12 |
| WBGene00014848 | 0.635570607 | 0.18494104 | 0.37730955 | VM106R.1 |
| WBGene00014864 | 1.016108341 | 0.39710553 | 1 | Y18D10A.22 |
| WBGene00014871 | 0.369648967 | 0.69647057 | 0.82843629 | Y37H9A.5 |
| WBGene00014914 | 2.231200485 | 0.24276597 | 1 | Y52B11A.11 |
| WBGene00014926 | -1.901856996 | 0.64072034 | 1 | Y57G11C.30 |
| WBGene00014938 | -0.820551265 | 0.00035694 | 0.00415898 | mdt-9 |
| WBGene00014965 | 0.076018408 | 0.75880905 | 0.8649611 | Y106G6D.5 |
| WBGene00014989 | 0.427083935 | 0.44409778 | 0.64348862 | best-23 |
| WBGene00014997 | -0.156269293 | 0.96924949 | 1 | AC7.3 |
| WBGene00014998 | -0.025601904 | 0.97916809 | 1 | AH9.1 |
| WBGene00014999 | -0.501315471 | 0.17703101 | 0.36866783 | AH9.3 |
| WBGene00015000 | 2.955131678 | 0.17939373 | 1 | AH9.4 |
| WBGene00015001 | -0.132856009 | 0.643588 | 0.79408556 | B0025.4 |
| WBGene00015002 | 0.08664113 | 0.90450061 | 0.95045368 | B0034.1 |
| WBGene00015004 | -0.630369619 | 0.68914459 | 1 | B0034.4 |
| WBGene00015005 | 0.068198813 | 0.96787621 | 1 | B0034.5 |
| WBGene00015006 | -0.156269293 | 0.96924949 | 1 | B0041.1 |
| WBGene00015007 | -0.067448447 | 0.69807901 | 0.8290089 | ain-2 |
| WBGene00015008 | -0.371670184 | 0.34050374 | 0.54886562 | lmd-2 |
| WBGene00015009 | -0.15468064 | 0.42675618 | 0.63044468 | B0041.5 |
| WBGene00015010 | -0.13850517 | 0.74575702 | 0.85814008 | ptps-1 |
| WBGene00015011 | -0.820987386 | 0.00012963 | 0.00187808 | B0041.8 |
| WBGene00015012 | -0.217731605 | 0.82012899 | 1 | bath-20 |
| WBGene00015014 | 0.930655763 | 0.04072561 | 0.13924237 | bath-24 |
| WBGene00015015 | 0.090155647 | 0.96170385 | 1 | math-1 |
| WBGene00015017 | 0.038450658 | 0.83913542 | 0.91208085 | B0205.1 |
| WBGene00015018 | 0.377828795 | 0.92526585 | 1 | srz-85 |
| WBGene00015019 | -0.659977672 | 0.54681764 | 1 | B0205.4 |
| WBGene00015020 | 0.688793783 | 0.86378755 | 1 | B0205.5 |
| WBGene00015021 | -0.122888815 | 0.55406203 | 0.73180205 | B0205.6 |
| WBGene00015022 | -0.532233211 | 0.07668006 | 0.21490008 | B0205.8 |
| WBGene00015023 | -0.232905398 | 0.24217934 | 0.44979338 | B0205.9 |
| WBGene00015024 | 2.09403112 | 0.45107747 | 1 | B0205.10 |
| WBGene00015025 | -0.216702204 | 0.3516013 | 0.56035591 | mrpl-9 |
| WBGene00015026 | 0.88787957 | 0.71832755 | 1 | B0207.1 |
| WBGene00015027 | -0.311877693 | 0.7763746 | 1 | B0207.2 |
| WBGene00015028 | -0.284202651 | 0.88416022 | 1 | B0207.5 |
| WBGene00015029 | -0.261318187 | 0.30730498 | 0.51598983 | B0207.6 |
| WBGene00015030 | 1.216957619 | 0.74819621 | 1 | B0207.7 |

| | | | | |
|---|---|---|---|---|
| WBGene00015032 | 0.281454817 | 0.94435481 | 1 | B0207.9 |
| WBGene00015033 | -0.764269844 | 0.76036927 | 1 | B0207.10 |
| WBGene00015034 | -0.156269293 | 0.96924949 | 1 | B0207.11 |
| WBGene00015035 | 0.38527626 | 0.71647035 | 1 | otpl-1 |
| WBGene00015036 | 0.983488853 | 0.80601565 | 1 | sre-5 |
| WBGene00015037 | -0.595452734 | 0.13140407 | 0.30385388 | B0212.3 |
| WBGene00015038 | -0.569416418 | 0.35551126 | 0.5638413 | nstp-6 |
| WBGene00015040 | 2.0789253 | 0.09268156 | 1 | cyp-34A5 |
| WBGene00015041 | 0.881467666 | 0.501119 | 1 | cyp-34A6 |
| WBGene00015042 | 2.5293122 | 0.14175299 | 1 | cyp-34A7 |
| WBGene00015043 | -0.331995929 | 0.35306399 | 0.56194939 | cyp-34A8 |
| WBGene00015044 | 0.869635302 | 0.12474817 | 0.29432717 | cyp-34A9 |
| WBGene00015045 | 1.229237684 | 0.59270769 | 1 | cyp-34A10 |
| WBGene00015046 | -1.130430023 | 0.47172801 | 1 | nlp-34 |
| WBGene00015047 | 0.473400916 | 0.0264181 | 0.10381629 | faah-1 |
| WBGene00015048 | 0.885860777 | 0.00014084 | 0.00200464 | faah-2 |
| WBGene00015049 | 0.789601902 | 0.75907734 | 1 | B0218.5 |
| WBGene00015050 | -0.086240044 | 0.76736999 | 0.86965734 | clec-51 |
| WBGene00015051 | 0.253925665 | 0.89264743 | 1 | B0218.7 |
| WBGene00015052 | 1.19193283 | 0.00018862 | 0.00252481 | clec-52 |
| WBGene00015053 | 0.610510419 | 0.44162055 | 0.64137027 | B0222.1 |
| WBGene00015054 | 0.377828795 | 0.92526585 | 1 | pitr-2 |
| WBGene00015055 | -0.335261756 | 0.54385967 | 0.72337864 | pitr-3 |
| WBGene00015056 | 0.178479147 | 0.67990024 | 0.81647553 | B0222.5 |
| WBGene00015057 | 0.702822376 | 0.03388281 | 0.12357202 | gad-3 |
| WBGene00015058 | 0.747845955 | 0.85215965 | 1 | B0222.10 |
| WBGene00015059 | 1.187782869 | 0.01392249 | 0.06589979 | B0228.1 |
| WBGene00015061 | 1.40815421 | 1.34E-14 | 2.38E-12 | cpna-2 |
| WBGene00015062 | 1.504454192 | 0.50075669 | 1 | trx-1 |
| WBGene00015063 | -0.076136174 | 0.86858751 | 0.9289571 | B0228.6 |
| WBGene00015064 | 0.277824144 | 0.26251233 | 0.4712464 | B0228.7 |
| WBGene00015066 | 1.181686151 | 0.7579682 | 1 | B0228.9 |
| WBGene00015067 | 0.749137068 | 0.08568014 | 0.23095972 | B0238.1 |
| WBGene00015068 | -0.156269293 | 0.96924949 | 1 | srt-68 |
| WBGene00015070 | -1.366565036 | 0.6506483 | 1 | srt-70 |
| WBGene00015071 | 0.377828795 | 0.92526585 | 1 | B0238.7 |
| WBGene00015073 | -0.174873577 | 0.413051 | 0.6185571 | B0238.9 |
| WBGene00015074 | -0.139747624 | 0.43360245 | 0.63538117 | natc-2 |
| WBGene00015075 | -0.864155912 | 1.1697E-06 | 3.8277E-05 | B0238.11 |
| WBGene00015076 | 0.694329357 | 0.16159974 | 0.3475611 | B0238.12 |
| WBGene00015079 | 0.568984552 | 0.59870335 | 1 | B0244.4 |
| WBGene00015080 | 1.868003649 | 0.1468412 | 1 | B0244.5 |
| WBGene00015081 | 1.124088214 | 0.13382319 | 0.30760079 | B0244.6 |

| | | | | |
|---|---|---|---|---|
| WBGene00015082 | 3.435808723 | 0.02166284 | 1 | B0244.7 |
| WBGene00015083 | -0.361950849 | 0.01757662 | 0.07810589 | egg-1 |
| WBGene00015084 | -0.62146648 | 0.85925752 | 1 | B0244.9 |
| WBGene00015085 | 1.200632894 | 0.06742675 | 0.19721228 | B0244.10 |
| WBGene00015087 | 2.155065463 | 0.00152952 | 0.01274526 | B0252.1 |
| WBGene00015088 | 0.531919164 | 0.11128221 | 0.27305523 | B0252.3 |
| WBGene00015089 | -0.198211819 | 0.93490302 | 1 | B0252.5 |
| WBGene00015091 | 0.270880404 | 0.1817002 | 0.37382024 | B0261.1 |
| WBGene00015092 | -0.448359065 | 0.06106427 | 0.18474489 | mrpl-47 |
| WBGene00015093 | -1.094201388 | 0.7816772 | 1 | B0261.5 |
| WBGene00015094 | 0.305771168 | 0.87310518 | 1 | B0261.6 |
| WBGene00015095 | -0.447404717 | 0.00705317 | 0.03997783 | B0261.7 |
| WBGene00015096 | -0.402126544 | 0.36404568 | 0.57174854 | B0261.8 |
| WBGene00015097 | -0.829424174 | 0.71322143 | 1 | B0273.1 |
| WBGene00015098 | -1.607133558 | 0.00723779 | 0.04075787 | snpc-3.1 |
| WBGene00015099 | -0.696044907 | 0.00052056 | 0.00552943 | ggtb-1 |
| WBGene00015100 | 2.712018489 | 0.03949193 | 1 | B0280.2 |
| WBGene00015101 | -0.236548132 | 0.22879843 | 0.4340198 | rpia-1 |
| WBGene00015102 | -0.376476621 | 0.0148261 | 0.06907279 | cpg-2 |
| WBGene00015103 | -0.502324997 | 0.67051816 | 1 | B0280.7 |
| WBGene00015104 | -0.40258924 | 0.03554508 | 0.12715091 | B0280.9 |
| WBGene00015105 | -0.104252938 | 0.73945989 | 0.85461994 | pot-1 |
| WBGene00015106 | 0.377828795 | 0.92526585 | 1 | B0280.11 |
| WBGene00015107 | 1.370865772 | 0.00543885 | 0.0331658 | B0280.13 |
| WBGene00015108 | -1.969805064 | 0.62862569 | 1 | B0281.1 |
| WBGene00015110 | 0.886383108 | 0.02439472 | 0.09849562 | B0281.3 |
| WBGene00015111 | -0.280551 | 0.8287958 | 1 | B0281.4 |
| WBGene00015112 | -1.521580082 | 1.5391E-06 | 4.6833E-05 | B0281.5 |
| WBGene00015113 | -0.067767769 | 0.9559696 | 1 | B0281.6 |
| WBGene00015115 | 0.683926408 | 0.20654167 | 0.40636645 | B0286.1 |
| WBGene00015117 | 0.896858881 | 0.12079642 | 0.28751017 | B0294.1 |
| WBGene00015118 | 0.972820844 | 0.64960696 | 1 | B0294.3 |
| WBGene00015119 | 0.688793783 | 0.86378755 | 1 | B0302.2 |
| WBGene00015121 | 0.688793783 | 0.86378755 | 1 | B0302.4 |
| WBGene00015122 | 0.637839149 | 0.08074836 | 0.22222266 | B0302.5 |
| WBGene00015124 | -0.231318464 | 0.71065823 | 0.83621841 | anmt-1 |
| WBGene00015125 | -0.365453395 | 0.01937806 | 0.08366608 | B0303.3 |
| WBGene00015126 | 0.057958603 | 0.76170418 | 0.86679637 | B0303.4 |
| WBGene00015128 | 0.373283986 | 0.26844667 | 0.47705078 | B0303.7 |
| WBGene00015129 | -0.744224254 | 0.56935579 | 1 | sdz-1 |
| WBGene00015130 | 0.017237775 | 0.9345778 | 0.96559519 | vps-33.1 |
| WBGene00015131 | 1.302824801 | 0.00020036 | 0.00263552 | B0303.11 |
| WBGene00015132 | -0.123781061 | 0.80705779 | 0.89476783 | B0303.14 |

| | | | | |
|---|---|---|---|---|
| WBGene00015133 | -0.083542094 | 0.69201402 | 0.82461 | mrpl-11 |
| WBGene00015134 | -0.554280301 | 0.01232288 | 0.06042491 | B0304.2 |
| WBGene00015135 | 0.426888077 | 0.67884979 | 1 | cyp-23A1 |
| WBGene00015136 | -0.144496658 | 0.67896708 | 0.81647553 | B0304.4 |
| WBGene00015137 | 0.944359139 | 0.04749557 | 0.15547149 | B0310.1 |
| WBGene00015138 | -0.466117596 | 0.27618093 | 0.4848341 | B0310.2 |
| WBGene00015139 | 0.518118171 | 0.58020276 | 0.75098526 | B0310.3 |
| WBGene00015141 | 0.059801934 | 0.7993563 | 0.88979641 | ugt-46 |
| WBGene00015142 | -3.079012209 | 0.2683724 | 1 | B0310.6 |
| WBGene00015143 | -0.05269037 | 0.74023307 | 0.85519532 | B0336.3 |
| WBGene00015145 | -0.595351467 | 0.00537288 | 0.03286252 | B0336.5 |
| WBGene00015146 | -0.127473645 | 0.40768982 | 0.6141434 | abi-1 |
| WBGene00015147 | -0.219908397 | 0.36402877 | 0.57174854 | B0336.7 |
| WBGene00015148 | 0.254483398 | 0.54867995 | 0.72724054 | hpo-28 |
| WBGene00015149 | -0.156269293 | 0.96924949 | 1 | B0336.12 |
| WBGene00015150 | -0.551650676 | 0.0169361 | 0.07590299 | B0336.13 |
| WBGene00015152 | 0.203511466 | 0.88490163 | 1 | B0348.2 |
| WBGene00015153 | 0.046252458 | 0.97619281 | 1 | B0348.5 |
| WBGene00015155 | 0.583208935 | 0.16413141 | 0.35032273 | B0353.1 |
| WBGene00015155 | 0.059062034 | 0.85217535 | 0.91924443 | Y55B1AL.1 |
| WBGene00015156 | -0.213073204 | 0.29993111 | 0.50968853 | B0361.2 |
| WBGene00015157 | -0.113570977 | 0.56803877 | 0.74300353 | B0361.3 |
| WBGene00015158 | 0.643747212 | 0.33473339 | 0.54317003 | B0361.4 |
| WBGene00015159 | 0.305754838 | 0.11442725 | 0.27821121 | psd-1 |
| WBGene00015160 | -0.346833605 | 0.11575116 | 0.28023616 | B0361.6 |
| WBGene00015161 | 0.15062643 | 0.53634812 | 0.71842482 | pho-5 |
| WBGene00015162 | -0.105034667 | 0.56603978 | 0.74142723 | algn-11 |
| WBGene00015163 | 0.078342516 | 0.81535007 | 0.8998423 | B0361.9 |
| WBGene00015164 | -0.250815192 | 0.31066006 | 0.51873445 | ykt-6 |
| WBGene00015165 | 1.876192691 | 0.43542138 | 1 | B0361.11 |
| WBGene00015167 | 0.009761492 | 0.98870537 | 0.99484196 | B0403.3 |
| WBGene00015168 | -0.38083013 | 0.01128613 | 0.05680966 | pdi-6 |
| WBGene00015169 | -0.2656625 | 0.67755791 | 0.81544713 | B0403.5 |
| WBGene00015170 | 0.281454817 | 0.94435481 | 1 | B0410.1 |
| WBGene00015171 | -0.39687129 | 0.20087351 | 0.3996731 | vang-1 |
| WBGene00015172 | 0.050062607 | 0.88980021 | 0.94196731 | B0410.3 |
| WBGene00015173 | 0.095087282 | 0.65433141 | 0.80139592 | trpp-11 |
| WBGene00015176 | 0.204122517 | 0.33008178 | 0.53851809 | vps-51 |
| WBGene00015177 | 1.246552488 | 0.00680264 | 0.03901315 | tmc-2 |
| WBGene00015178 | 0.190018488 | 0.80888338 | 0.89596325 | B0416.2 |
| WBGene00015179 | 1.205337689 | 0.16348962 | 0.34970534 | B0416.3 |
| WBGene00015180 | -1.482913883 | 0.00012145 | 0.00178083 | B0416.4 |
| WBGene00015181 | -0.02305169 | 0.90341758 | 0.94989113 | B0416.5 |

| | | | | |
|---|---|---|---|---|
| WBGene00015182 | 1.58594919 | 0.06962066 | 0.20158943 | B0416.7 |
| WBGene00015183 | -1.447469916 | 0.18973409 | 1 | B0432.1 |
| WBGene00015184 | -0.497579665 | 0.01567925 | 0.07187884 | djr-1.1 |
| WBGene00015185 | -0.319673495 | 0.20765534 | 0.40722718 | mrpl-41 |
| WBGene00015186 | -0.365966732 | 0.06089978 | 0.18450529 | misc-1 |
| WBGene00015188 | 0.129364325 | 0.61373928 | 0.7744137 | B0432.7 |
| WBGene00015189 | -0.359913167 | 0.2448438 | 0.45270807 | B0432.8 |
| WBGene00015190 | -0.376682958 | 0.6203397 | 0.77878286 | B0432.9 |
| WBGene00015191 | -0.293321189 | 0.33504991 | 0.54330693 | B0432.10 |
| WBGene00015193 | 1.653460415 | 0.50712258 | 1 | clec-117 |
| WBGene00015194 | -0.108185226 | 0.64803012 | 0.79713614 | B0432.13 |
| WBGene00015196 | -0.963744934 | 0.13519492 | 0.30973633 | B0454.5 |
| WBGene00015197 | 1.009947945 | 0.0200117 | 0.08548981 | B0454.6 |
| WBGene00015198 | 0.489058549 | 0.85947456 | 1 | clec-2 |
| WBGene00015199 | -0.156269293 | 0.96924949 | 1 | B0454.8 |
| WBGene00015200 | -0.338171372 | 0.37941003 | 0.58747827 | B0454.9 |
| WBGene00015202 | 0.950971593 | 0.40649566 | 1 | B0478.3 |
| WBGene00015203 | 0.249185484 | 0.18376054 | 0.37597517 | cdk-11.1 |
| WBGene00015204 | 0.108843011 | 0.63416326 | 0.78783996 | B0495.5 |
| WBGene00015205 | -0.348589009 | 0.18221245 | 0.37387888 | moa-2 |
| WBGene00015206 | 0.272136894 | 0.12039947 | 0.28684653 | B0495.7 |
| WBGene00015207 | -0.462743407 | 0.01529055 | 0.07063015 | B0495.8 |
| WBGene00015208 | -0.087555011 | 0.72214634 | 0.84336078 | B0495.9 |
| WBGene00015210 | 1.116613983 | 0.69067248 | 1 | B0496.1 |
| WBGene00015211 | -0.62146648 | 0.85925752 | 1 | B0496.2 |
| WBGene00015214 | 0.097987904 | 0.96406692 | 1 | srtx-1 |
| WBGene00015215 | 1.477253415 | 0.58357327 | 1 | B0496.6 |
| WBGene00015216 | 0.334199007 | 0.45119692 | 0.64911342 | valv-1 |
| WBGene00015217 | 0.402870321 | 0.17977749 | 0.37208976 | tes-1 |
| WBGene00015218 | 0.099287513 | 0.86721809 | 0.92835056 | B0507.1 |
| WBGene00015219 | 0.613073755 | 0.02469417 | 0.09930834 | B0507.2 |
| WBGene00015220 | 2.208963628 | 0.00024187 | 0.00303364 | B0507.3 |
| WBGene00015223 | 3.398099241 | 0.08660266 | 1 | B0507.6 |
| WBGene00015224 | -0.857453619 | 0.65777058 | 1 | B0507.7 |
| WBGene00015225 | 1.522421447 | 0.0210665 | 0.08883932 | B0507.8 |
| WBGene00015226 | 1.712479368 | 0.58529868 | 1 | B0507.9 |
| WBGene00015227 | 0.905524471 | 0.06878402 | 0.19999543 | B0507.10 |
| WBGene00015228 | -0.386404399 | 0.48497918 | 0.67539764 | B0511.2 |
| WBGene00015230 | 0.446458324 | 0.87164163 | 1 | tag-344 |
| WBGene00015231 | -0.200564712 | 0.78724089 | 0.88275978 | cutl-14 |
| WBGene00015232 | -0.274323006 | 0.06242607 | 0.18725666 | B0511.6 |
| WBGene00015233 | -0.718793674 | 0.00103257 | 0.00947315 | B0511.7 |
| WBGene00015235 | -0.449716611 | 0.01836362 | 0.08054649 | cdc-26 |

| | | | | |
|---|---|---|---|---|
| WBGene00015236 | -0.08965951 | 0.91839836 | 0.95686787 | B0511.11 |
| WBGene00015237 | 0.413630993 | 0.02790086 | 0.10788036 | B0511.12 |
| WBGene00015238 | 0.065369064 | 0.8478236 | 0.91660074 | B0511.13 |
| WBGene00015243 | -0.156269293 | 0.96924949 | 1 | B0524.4 |
| WBGene00015243 | -1.735272825 | 0.41587729 | 1 | Y39A3CR.8 |
| WBGene00015245 | -0.060103878 | 0.82003336 | 0.90260666 | B0524.6 |
| WBGene00015246 | 0.747845955 | 0.85215965 | 1 | scl-23 |
| WBGene00015247 | -0.001410793 | 0.99904441 | 1 | B0545.4 |
| WBGene00015248 | -0.973561135 | 1.06E-08 | 6.36E-07 | mai-2 |
| WBGene00015249 | -0.315765121 | 0.10961134 | 0.27006751 | otub-4 |
| WBGene00015250 | 0.184241758 | 0.54525609 | 0.72434382 | B0546.3 |
| WBGene00015251 | 0.071175977 | 0.84337503 | 0.91382564 | B0546.4 |
| WBGene00015257 | -2.171326218 | 0.36522606 | 1 | B0554.4 |
| WBGene00015258 | 1.877609176 | 0.06430153 | 0.19115694 | B0554.5 |
| WBGene00015259 | 2.758479935 | 0.09579893 | 1 | dod-20 |
| WBGene00015260 | 2.07531578 | 0.45514895 | 1 | B0554.7 |
| WBGene00015261 | -2.00762318 | 0.26833377 | 1 | B0563.1 |
| WBGene00015262 | 0.697492964 | 0.45482004 | 0.6523296 | B0563.5 |
| WBGene00015263 | 0.940300818 | 0.21200737 | 0.412025 | B0563.6 |
| WBGene00015264 | 0.557203925 | 0.56781075 | 0.74278529 | B0563.7 |
| WBGene00015265 | -3.62968057 | 0.03719524 | 1 | B0563.8 |
| WBGene00015266 | -0.344069599 | 0.296326 | 0.50678884 | BE0003N10.1 |
| WBGene00015267 | -0.450310499 | 0.08849358 | 0.23582103 | chin-1 |
| WBGene00015268 | -0.180895771 | 0.81979429 | 0.90251377 | BE0003N10.3 |
| WBGene00015270 | -1.964653846 | 0.53599202 | 1 | C01B4.6 |
| WBGene00015272 | -0.156269293 | 0.96924949 | 1 | C01B4.8 |
| WBGene00015273 | -0.156269293 | 0.96924949 | 1 | mct-2 |
| WBGene00015278 | 0.342972156 | 0.29867412 | 0.50851248 | C01B10.3 |
| WBGene00015279 | 0.284806732 | 0.67686241 | 0.81495175 | C01B10.4 |
| WBGene00015280 | 1.056782923 | 2.88E-08 | 1.5378E-06 | C01B10.6 |
| WBGene00015281 | 1.699062218 | 0.19532328 | 1 | C01B10.7 |
| WBGene00015282 | -0.138649484 | 0.44321159 | 0.6427415 | C01B10.8 |
| WBGene00015283 | -0.257717856 | 0.21584642 | 0.41755759 | C01B10.9 |
| WBGene00015284 | 0.723717511 | 0.18165623 | 0.37382024 | C01B10.10 |
| WBGene00015285 | 0.453905387 | 0.15722983 | 0.34168888 | gmeb-1 |
| WBGene00015286 | 1.095117506 | 0.03230152 | 0.11943239 | best-3 |
| WBGene00015287 | 0.777167873 | 0.50304481 | 1 | osta-1 |
| WBGene00015288 | 1.385499037 | 0.25126161 | 1 | best-4 |
| WBGene00015291 | -0.274979165 | 0.31327109 | 0.52123124 | C01B12.8 |
| WBGene00015292 | 2.437484291 | 0.2708268 | 1 | C01C4.2 |
| WBGene00015293 | 1.635624541 | 4.1155E-05 | 0.00073371 | C01C4.3 |
| WBGene00015293 | 1.408309319 | 0.5390817 | 1 | C02F12.9 |
| WBGene00015294 | -0.466582746 | 0.25090829 | 0.45885383 | C01C10.2 |

| | | | | |
|---|---|---|---|---|
| WBGene00015295 | 0.381891298 | 0.14646754 | 0.32592785 | acl-12 |
| WBGene00015296 | -0.059903032 | 0.74493425 | 0.85759972 | C01F1.1 |
| WBGene00015297 | -0.108747194 | 0.70598205 | 0.83366276 | sco-1 |
| WBGene00015298 | -0.155579936 | 0.63306947 | 0.7872679 | C01F1.3 |
| WBGene00015299 | -0.313347513 | 0.71802566 | 0.84048877 | C01F1.4 |
| WBGene00015300 | -1.268624577 | 0.22189324 | 1 | C01F1.5 |
| WBGene00015301 | 0.03403942 | 0.86754587 | 0.92836209 | C01F1.6 |
| WBGene00015303 | 0.663906072 | 0.16722841 | 0.35468616 | rga-6 |
| WBGene00015306 | 1.38322681 | 0.69550305 | 1 | C01G5.4 |
| WBGene00015307 | -0.124073996 | 0.63004302 | 0.78547303 | C01G5.5 |
| WBGene00015308 | -0.292169027 | 0.10098373 | 0.25601242 | ddi-1 |
| WBGene00015309 | 1.657787544 | 0.43418652 | 1 | C01G5.7 |
| WBGene00015310 | 0.11187683 | 0.62462973 | 0.78165631 | fan-1 |
| WBGene00015311 | -0.078889862 | 0.93090845 | 0.96393684 | C01G5.9 |
| WBGene00015312 | -0.508958085 | 0.0025084 | 0.01861522 | C01G8.1 |
| WBGene00015313 | 0.158094148 | 0.5666947 | 0.74172492 | hpo-32 |
| WBGene00015317 | -1.964653846 | 0.53599202 | 1 | srbc-32 |
| WBGene00015320 | 1.034240481 | 0.59145984 | 1 | C02B8.1 |
| WBGene00015321 | -0.604396148 | 0.82723443 | 1 | fbxc-38 |
| WBGene00015322 | 1.020141677 | 0.17899837 | 0.37117321 | C02B8.3 |
| WBGene00015323 | 0.241565248 | 0.85628065 | 1 | frpr-1 |
| WBGene00015324 | -0.194084474 | 0.64016824 | 0.791878 | C02B8.6 |
| WBGene00015326 | -0.138252494 | 0.41826984 | 0.62301732 | ivd-1 |
| WBGene00015327 | -0.37342308 | 0.10075311 | 0.25590808 | snpn-1 |
| WBGene00015328 | 0.305245984 | 0.287152 | 0.49711057 | C02B10.3 |
| WBGene00015329 | -0.520041091 | 0.006798 | 0.03900498 | C02B10.4 |
| WBGene00015330 | -0.016072245 | 0.94424888 | 0.97120685 | C02B10.5 |
| WBGene00015331 | -0.156269293 | 0.96924949 | 1 | C02B10.6 |
| WBGene00015332 | 0.031026766 | 0.92129026 | 0.95884274 | tyr-1 |
| WBGene00015333 | 0.224151624 | 0.72453481 | 0.84517459 | slc-17.3 |
| WBGene00015334 | 0.489518126 | 0.41119116 | 0.61671054 | sup-18 |
| WBGene00015335 | -1.014775546 | 0.25872371 | 0.4669488 | acdh-6 |
| WBGene00015336 | 0.451874138 | 0.38284485 | 0.59121335 | C02D5.2 |
| WBGene00015337 | -0.91375383 | 0.72824392 | 1 | gsto-2 |
| WBGene00015338 | 1.712479368 | 0.58529868 | 1 | catp-2 |
| WBGene00015339 | -1.731142657 | 0.00022482 | 0.00288835 | C02E7.6 |
| WBGene00015340 | -1.250518323 | 0.01047267 | 0.05370108 | C02E7.7 |
| WBGene00015341 | 0.377828795 | 0.92526585 | 1 | C02E7.8 |
| WBGene00015342 | 0.28530093 | 0.92084641 | 1 | C02E7.10 |
| WBGene00015344 | 0.093996714 | 0.63976778 | 0.79154394 | nra-4 |
| WBGene00015345 | 0.233739675 | 0.93575889 | 1 | C02F5.2 |
| WBGene00015346 | -0.276765356 | 0.172266 | 0.36178541 | C02F5.3 |
| WBGene00015347 | -0.276204394 | 0.19188641 | 0.38686365 | cids-1 |

| | | | | |
|---|---|---|---|---|
| WBGene00015348 | 1.435925494 | 0.70920779 | 1 | C02F5.5 |
| WBGene00015349 | 0.058516162 | 0.77024148 | 0.87144738 | henn-1 |
| WBGene00015350 | 0.879889942 | 0.00014869 | 0.0020892 | fbxl-1 |
| WBGene00015351 | -0.648724627 | 0.28965583 | 0.49967941 | C02F5.10 |
| WBGene00015352 | 0.025532652 | 0.94957022 | 0.97428649 | C02F5.12 |
| WBGene00015353 | -0.317994862 | 0.22665297 | 0.43095965 | C02F5.13 |
| WBGene00015354 | 0.740809248 | 0.02435316 | 0.09837757 | snet-1 |
| WBGene00015355 | -1.399727385 | 1.2916E-05 | 0.00028152 | C02F12.5 |
| WBGene00015356 | 0.663768493 | 0.021545 | 0.0905427 | tag-278 |
| WBGene00015357 | -0.099908933 | 0.75294827 | 0.86217294 | C02F12.8 |
| WBGene00015359 | 1.006341278 | 0.33837172 | 1 | C02G6.1 |
| WBGene00015360 | 1.38322681 | 0.69550305 | 1 | C02G6.2 |
| WBGene00015361 | 0.09697944 | 0.97612634 | 1 | C02H6.1 |
| WBGene00015362 | 0.869373783 | 0.13938764 | 0.31655543 | fbxa-105 |
| WBGene00015364 | 2.352659272 | 0.07886962 | 1 | npr-19 |
| WBGene00015366 | -0.093512106 | 0.95400258 | 1 | C03A7.2 |
| WBGene00015368 | 1.957271014 | 0.49743227 | 1 | srt-65 |
| WBGene00015369 | 0.753271533 | 0.30921749 | 0.51803636 | ugt-51 |
| WBGene00015371 | 1.292734292 | 0.30669663 | 1 | C03A7.12 |
| WBGene00015371 | 1.504895891 | 0.36891846 | 1 | C03A7.13 |
| WBGene00015373 | 1.330382438 | 0.5642962 | 1 | C03B1.2 |
| WBGene00015374 | 0.281454817 | 0.94435481 | 1 | C03B1.3 |
| WBGene00015375 | -1.760055813 | 0.45626364 | 1 | C03B1.4 |
| WBGene00015376 | 1.353021252 | 0.55399304 | 1 | C03B1.5 |
| WBGene00015377 | -0.434029631 | 0.80690472 | 1 | C03B1.6 |
| WBGene00015378 | 0.129320261 | 0.73645126 | 0.85280864 | C03B1.7 |
| WBGene00015380 | 0.281454817 | 0.94435481 | 1 | C03B1.10 |
| WBGene00015381 | 0.81151386 | 0.19220058 | 0.38723983 | C03B1.13 |
| WBGene00015382 | -0.335304734 | 0.58381865 | 0.75397747 | C03B1.14 |
| WBGene00015385 | -0.970858664 | 0.2743696 | 0.4827622 | C03B8.2 |
| WBGene00015386 | 0.002921407 | 0.99862476 | 1 | C03B8.3 |
| WBGene00015387 | 0.060037603 | 0.94925639 | 0.97411952 | kcnl-3 |
| WBGene00015388 | 0.534310226 | 0.76927188 | 1 | C03F11.2 |
| WBGene00015389 | 0.4022507 | 0.08402405 | 0.22796334 | scav-1 |
| WBGene00015390 | 1.093661591 | 0.0512914 | 0.16357471 | C03F11.4 |
| WBGene00015391 | 0.227420079 | 0.26011352 | 0.46838185 | sdha-1 |
| WBGene00015392 | -1.433899761 | 0.10080118 | 0.25590808 | nspc-7 |
| WBGene00015393 | 1.10773558 | 0.07661681 | 0.21490008 | C03G6.5 |
| WBGene00015394 | 0.574508983 | 0.8551971 | 1 | C03G6.6 |
| WBGene00015395 | 0.921520751 | 0.51186593 | 1 | nhr-147 |
| WBGene00015396 | -0.369784813 | 0.78135811 | 1 | nhr-148 |
| WBGene00015397 | 1.313089111 | 0.56091378 | 1 | nhr-149 |
| WBGene00015398 | -2.31323281 | 0.04656182 | 1 | tag-293 |

| | | | | |
|---|---|---|---|---|
| WBGene00015399 | -1.969805064 | 0.62862569 | 1 | cyp-35A1 |
| WBGene00015400 | 0.856134769 | 0.00118084 | 0.01041899 | cyp-35A2 |
| WBGene00015401 | 0.938982574 | 0.8146943 | 1 | srsx-24 |
| WBGene00015402 | 0.89999201 | 0.03626797 | 0.12882663 | C03G6.17 |
| WBGene00015403 | 1.470306238 | 6.45E-07 | 2.3735E-05 | clec-10 |
| WBGene00015404 | 0.071359194 | 0.72471017 | 0.84518994 | nstp-4 |
| WBGene00015405 | -0.231146013 | 0.36409754 | 0.57174854 | C03H5.3 |
| WBGene00015406 | -0.07748118 | 0.98067866 | 1 | C03H5.4 |
| WBGene00015407 | 0.507638977 | 0.30915836 | 0.51800878 | C03H5.5 |
| WBGene00015408 | 0.59101281 | 0.25310014 | 0.46104859 | C03H5.6 |
| WBGene00015409 | 1.123234694 | 0.44426087 | 1 | C03H5.7 |
| WBGene00015410 | 0.38983483 | 0.31026829 | 0.51843755 | tbc-6 |
| WBGene00015412 | 1.350008611 | 0.4278104 | 1 | lgc-9 |
| WBGene00015413 | -0.361197078 | 0.02369453 | 0.09660637 | pdhb-1 |
| WBGene00015414 | 0.932931886 | 0.70926986 | 1 | C04C3.4 |
| WBGene00015416 | 0.064183849 | 0.97246519 | 1 | C04C3.6 |
| WBGene00015417 | -0.156269293 | 0.96924949 | 1 | C04C3.7 |
| WBGene00015418 | 0.495816476 | 0.01153923 | 0.05770088 | pac-1 |
| WBGene00015419 | 1.013713812 | 0.64240986 | 1 | srsx-18 |
| WBGene00015421 | 1.181686151 | 0.7579682 | 1 | C04E6.4 |
| WBGene00015422 | 0.983488853 | 0.80601565 | 1 | C04E6.5 |
| WBGene00015423 | 0.968246794 | 0.08885988 | 0.23635972 | C04E6.7 |
| WBGene00015424 | 1.412133314 | 0.21435028 | 1 | C04E6.8 |
| WBGene00015425 | 0.003471365 | 0.98809058 | 0.99472701 | C04E6.11 |
| WBGene00015426 | -0.202114967 | 0.90321067 | 1 | C04E6.12 |
| WBGene00015427 | 0.12801293 | 0.80775803 | 0.89513608 | C04E6.13 |
| WBGene00015429 | 0.245605795 | 0.46034551 | 0.6560508 | sor-3 |
| WBGene00015430 | 0.987836591 | 0.07669014 | 0.21490008 | C04E7.3 |
| WBGene00015430 | 1.900814518 | 0.00293639 | 0.0208728 | C04E7.4 |
| WBGene00015433 | -0.390282216 | 0.49459079 | 0.68308599 | C04E12.2 |
| WBGene00015434 | 1.73658448 | 6.0186E-06 | 0.00014673 | C04E12.4 |
| WBGene00015435 | -1.108079551 | 0.62646691 | 1 | C04E12.5 |
| WBGene00015437 | 0.582709152 | 0.59311816 | 1 | scrm-3 |
| WBGene00015442 | 0.377828795 | 0.92526585 | 1 | C04F1.1 |
| WBGene00015444 | 0.983488853 | 0.80601565 | 1 | C04F5.2 |
| WBGene00015448 | 0.688793783 | 0.86378755 | 1 | srab-3 |
| WBGene00015449 | -1.599601375 | 0.00020389 | 0.00266752 | ugt-63 |
| WBGene00015450 | -0.642967242 | 0.00166941 | 0.0135756 | C04F5.8 |
| WBGene00015451 | 0.09961374 | 0.72957562 | 0.84851293 | C04F5.9 |
| WBGene00015453 | -0.157413813 | 0.94424418 | 1 | C04G6.2 |
| WBGene00015454 | -0.027937649 | 0.89815736 | 0.94673802 | C04G6.4 |
| WBGene00015455 | -0.633073245 | 0.50019122 | 0.68806046 | C04G6.5 |
| WBGene00015456 | -0.018486038 | 0.98623633 | 1 | C04G6.6 |

| | | | | |
|---|---|---|---|---|
| WBGene00015457 | 0.667420706 | 0.64550132 | 1 | C04G6.7 |
| WBGene00015458 | -1.094201388 | 0.7816772 | 1 | C04G6.10 |
| WBGene00015460 | -0.169522624 | 0.50821897 | 0.69493034 | C05C8.1 |
| WBGene00015461 | -0.750943279 | 0.00016595 | 0.00227406 | C05C8.2 |
| WBGene00015462 | -0.104342756 | 0.69613347 | 0.82819759 | C05C8.5 |
| WBGene00015463 | -0.580227784 | 0.00088498 | 0.00836153 | hpo-9 |
| WBGene00015464 | 0.053870407 | 0.81502133 | 0.89956127 | C05C8.7 |
| WBGene00015465 | 0.483840154 | 0.8151116 | 1 | C05C8.8 |
| WBGene00015466 | -0.134076564 | 0.61878049 | 0.77818665 | hyls-1 |
| WBGene00015467 | -0.156269293 | 0.96924949 | 1 | basl-1 |
| WBGene00015468 | -0.271643702 | 0.19892471 | 0.39703188 | madf-11 |
| WBGene00015469 | 0.330620807 | 0.12656445 | 0.29693119 | C05D2.8 |
| WBGene00015470 | -0.62364052 | 0.00048919 | 0.00527934 | C05D2.10 |
| WBGene00015471 | 1.230812021 | 0.00355169 | 0.0240631 | lmp-2 |
| WBGene00015472 | 0.372706956 | 0.25431683 | 0.46207607 | C05D9.3 |
| WBGene00015473 | 1.189071466 | 0.5668301 | 1 | C05D9.4 |
| WBGene00015474 | -0.872116583 | 0.0238249 | 0.09696887 | C05D9.7 |
| WBGene00015476 | 1.239143898 | 0.02996939 | 0.11321046 | C05D9.9 |
| WBGene00015477 | 0.011356474 | 0.96011389 | 0.97958146 | attf-4 |
| WBGene00015478 | 1.424714421 | 0.00518007 | 0.03202199 | mapk-15 |
| WBGene00015479 | -0.377597379 | 0.60876506 | 0.77105523 | wht-1 |
| WBGene00015480 | 0.05702134 | 0.85662851 | 0.92175167 | C05D10.4 |
| WBGene00015481 | -0.317625024 | 0.04128107 | 0.14066614 | C05D11.1 |
| WBGene00015483 | 0.091841513 | 0.84045829 | 0.91237519 | C05D11.5 |
| WBGene00015484 | 0.264124298 | 0.14357954 | 0.32168283 | atgl-1 |
| WBGene00015485 | 0.394680817 | 0.10801862 | 0.26764369 | C05D11.8 |
| WBGene00015486 | -0.406462816 | 0.06537173 | 0.19301513 | C05D11.9 |
| WBGene00015487 | -0.576537854 | 0.025323 | 0.10106723 | mrps-17 |
| WBGene00015488 | -0.144676849 | 0.75278573 | 0.86217294 | C05D11.13 |
| WBGene00015491 | 0.688793783 | 0.86378755 | 1 | C05E4.12 |
| WBGene00015492 | 0.202976485 | 0.49373414 | 0.68234788 | lnp-1 |
| WBGene00015494 | 0.259452022 | 0.53909909 | 0.72067849 | C05E11.3 |
| WBGene00015495 | 1.009576902 | 0.35322505 | 1 | C05E11.6 |
| WBGene00015496 | -1.618771107 | 0.4854602 | 1 | C05E11.7 |
| WBGene00015497 | 0.751314372 | 0.17682175 | 0.36841412 | nhr-76 |
| WBGene00015499 | 0.442516283 | 0.01543485 | 0.07111747 | inst-1 |
| WBGene00015500 | -2.306665619 | 0.26898727 | 1 | C06A5.2 |
| WBGene00015501 | 0.061204602 | 0.7289807 | 0.84824178 | C06A5.3 |
| WBGene00015504 | -0.112046895 | 0.60538759 | 0.76897128 | C06A5.6 |
| WBGene00015505 | 0.101809359 | 0.72568732 | 0.84579072 | C06A5.8 |
| WBGene00015507 | 0.297503073 | 0.33124498 | 0.53952848 | C06A6.2 |
| WBGene00015508 | -0.223115474 | 0.34143486 | 0.54980695 | mvb-12 |
| WBGene00015509 | -0.287384486 | 0.06837301 | 0.19916428 | C06A6.4 |

| | | | | |
|---|---|---|---|---|
| WBGene00015510 | -0.275056923 | 0.26632959 | 0.47477459 | erp-44.2 |
| WBGene00015511 | 1.152448611 | 0.52357345 | 1 | C06A6.7 |
| WBGene00015512 | 0.788877229 | 5.67E-08 | 2.8234E-06 | mthf-1 |
| WBGene00015513 | -0.26877409 | 0.16244789 | 0.34845934 | snpc-1.1 |
| WBGene00015514 | -0.472387943 | 0.00425545 | 0.02760475 | C06A8.3 |
| WBGene00015515 | -0.267831013 | 0.29250201 | 0.50265633 | spdl-1 |
| WBGene00015516 | 3.458442665 | 0.0604553 | 1 | C06A8.6 |
| WBGene00015517 | 0.366049468 | 0.43031996 | 0.63273764 | C06A8.8 |
| WBGene00015518 | -1.170122929 | 0.02707016 | 0.10556882 | C06E1.1 |
| WBGene00015519 | 0.169034931 | 0.7933926 | 0.88593495 | doxa-1 |
| WBGene00015520 | -0.668264701 | 0.85551153 | 1 | fip-3 |
| WBGene00015521 | -0.514761297 | 0.79912167 | 1 | fipr-16 |
| WBGene00015522 | 2.171473691 | 0.06902623 | 1 | C06E1.7 |
| WBGene00015523 | 1.471484292 | 0.12669249 | 0.29717389 | ztf-30 |
| WBGene00015524 | 0.072454886 | 0.71661037 | 0.84010542 | C06E1.9 |
| WBGene00015525 | 0.297807625 | 0.29797172 | 0.50787519 | rha-2 |
| WBGene00015526 | 0.487744884 | 0.64396637 | 1 | C06E1.11 |
| WBGene00015527 | 0.445577806 | 0.5040423 | 0.69118816 | C06E2.1 |
| WBGene00015528 | -0.156269293 | 0.96924949 | 1 | C06E2.2 |
| WBGene00015529 | -1.250408133 | 0.00229345 | 0.01732175 | C06E2.5 |
| WBGene00015531 | 0.08064263 | 0.91725729 | 0.95640851 | C06E4.2 |
| WBGene00015532 | -0.742688971 | 0.59470602 | 1 | C06E4.3 |
| WBGene00015535 | -0.668264701 | 0.85551153 | 1 | C06E4.6 |
| WBGene00015536 | 0.647354001 | 0.43363548 | 0.63538117 | glb-2 |
| WBGene00015537 | 0.750571464 | 0.36591259 | 0.57350663 | C06E4.8 |
| WBGene00015538 | 0.058124773 | 0.74028192 | 0.85519532 | sams-3 |
| WBGene00015539 | 1.010410948 | 0.04414513 | 0.14696081 | C06E7.2 |
| WBGene00015540 | -0.072100377 | 0.71674706 | 0.84010542 | sams-4 |
| WBGene00015541 | -0.112700331 | 0.79352058 | 0.88593495 | C06E7.4 |
| WBGene00015544 | -0.650122286 | 0.15556245 | 0.33932513 | C06E8.5 |
| WBGene00015545 | 0.102993181 | 0.84257933 | 0.91337104 | C06G1.1 |
| WBGene00015546 | 0.0897694 | 0.91982808 | 0.95789215 | C06G1.2 |
| WBGene00015547 | 0.425813062 | 0.05248847 | 0.16582422 | ain-1 |
| WBGene00015549 | 1.056805069 | 0.46576171 | 1 | C06G3.3 |
| WBGene00015551 | 0.547023669 | 0.01094699 | 0.0555401 | C06G3.5 |
| WBGene00015552 | -0.203060384 | 0.43997232 | 0.64015249 | C06G3.6 |
| WBGene00015553 | 0.131571624 | 0.4357773 | 0.63676706 | trxr-1 |
| WBGene00015554 | -0.601264922 | 0.02124754 | 0.08944742 | C06G3.8 |
| WBGene00015555 | 0.297318001 | 0.13115949 | 0.30360485 | ufl-1 |
| WBGene00015557 | 0.206830931 | 0.31752107 | 0.52527909 | C06G4.1 |
| WBGene00015558 | 0.835634225 | 0.55092108 | 1 | C06G4.4 |
| WBGene00015559 | 0.031962775 | 0.96729261 | 0.98341146 | npr-17 |
| WBGene00015560 | 1.488649116 | 0.16569991 | 1 | C07A12.2 |

| | | | | |
|---|---|---|---|---|
| WBGene00015561 | 0.246291382 | 0.4352678 | 0.63637293 | C07A12.7 |
| WBGene00015564 | 0.688793783 | 0.86378755 | 1 | C07D8.5 |
| WBGene00015565 | -0.624548088 | 0.00012348 | 0.00180183 | C07D8.6 |
| WBGene00015566 | -0.101098332 | 0.90614846 | 0.95109561 | C07D10.1 |
| WBGene00015567 | 0.2073812 | 0.23821312 | 0.44453512 | bath-44 |
| WBGene00015568 | 3.442091476 | 0.06053407 | 1 | sre-3 |
| WBGene00015569 | -0.59397683 | 0.06096683 | 0.18463414 | C07D10.5 |
| WBGene00015570 | 1.438084244 | 0.25364406 | 1 | C07F11.2 |
| WBGene00015572 | 0.464310378 | 0.79088553 | 1 | C07G1.6 |
| WBGene00015573 | -2.53604874 | 0.05678746 | 1 | C07G1.7 |
| WBGene00015574 | 1.940453066 | 0.04687915 | 0.15395178 | irg-1 |
| WBGene00015576 | -0.183119115 | 0.92486992 | 1 | C07G3.8 |
| WBGene00015577 | 0.118439666 | 0.55053207 | 0.72905866 | ugt-64 |
| WBGene00015578 | 3.279482556 | 0.10751885 | 1 | C07G3.10 |
| WBGene00015579 | -0.485170653 | 0.03605771 | 0.12834262 | C07H6.2 |
| WBGene00015580 | 0.962609484 | 0.00036553 | 0.00423465 | cls-1 |
| WBGene00015581 | 0.05257833 | 0.75790956 | 0.86459364 | C07H6.4 |
| WBGene00015582 | 1.82867917 | 0.54358766 | 1 | C07H6.9 |
| WBGene00015583 | -0.03811452 | 0.98663692 | 1 | C08A9.3 |
| WBGene00015586 | -0.347241789 | 0.41731782 | 0.62198081 | C08A9.6 |
| WBGene00015588 | 0.192500335 | 0.85382046 | 1 | C08A9.8 |
| WBGene00015589 | -0.472929249 | 0.6381682 | 1 | C08A9.9 |
| WBGene00015590 | -0.33384098 | 0.24868492 | 0.45656791 | bath-15 |
| WBGene00015591 | -0.526072989 | 0.00881687 | 0.04700445 | cyk-7 |
| WBGene00015592 | 1.55420703 | 0.21594573 | 1 | C08D8.1 |
| WBGene00015593 | 0.902102015 | 0.43203837 | 1 | C08E3.1 |
| WBGene00015595 | -1.93354482 | 0.08567542 | 1 | bath-33 |
| WBGene00015598 | -0.62146648 | 0.85925752 | 1 | fbxa-163 |
| WBGene00015599 | 2.028800581 | 0.39258153 | 1 | fbxa-164 |
| WBGene00015600 | 1.872249764 | 0.5345823 | 1 | fbxa-165 |
| WBGene00015601 | 2.392622864 | 0.04558946 | 1 | fbxa-166 |
| WBGene00015602 | 1.591332247 | 0.28757695 | 1 | fbxa-158 |
| WBGene00015605 | 1.747268307 | 0.00010296 | 0.0015567 | C08E3.13 |
| WBGene00015607 | -0.597172253 | 0.51582673 | 0.70209107 | fbxb-13 |
| WBGene00015608 | 1.018392794 | 0.00180217 | 0.01436651 | math-3 |
| WBGene00015609 | 0.849235969 | 0.0101071 | 0.05230617 | math-4 |
| WBGene00015610 | -1.729044043 | 0.00023487 | 0.00297442 | C08F1.6 |
| WBGene00015611 | 3.434739952 | 0.02543372 | 1 | C08F1.8 |
| WBGene00015613 | -0.632304844 | 0.24865394 | 0.45656791 | C08F1.10 |
| WBGene00015618 | -0.933827966 | 0.56134399 | 1 | C08G5.6 |
| WBGene00015619 | 0.374007363 | 0.27510627 | 0.48357684 | C08G9.1 |
| WBGene00015620 | -0.035558015 | 0.95525693 | 0.97710391 | C08G9.2 |
| WBGene00015621 | 0.415870517 | 0.29502081 | 0.50533967 | C09B7.2 |

| | | | | |
|---|---|---|---|---|
| WBGene00015622 | -0.846720445 | 0.54327301 | 1 | C09B8.3 |
| WBGene00015623 | -2.598244084 | 0.00023864 | 0.00299925 | C09B8.4 |
| WBGene00015624 | -0.923115292 | 0.16553123 | 0.35233426 | C09B8.5 |
| WBGene00015625 | -0.271649589 | 0.82329042 | 1 | C09B8.8 |
| WBGene00015626 | 0.250439473 | 0.84200797 | 1 | C09B9.1 |
| WBGene00015627 | 0.57991002 | 0.82402153 | 1 | C09B9.2 |
| WBGene00015629 | 0.377828795 | 0.92526585 | 1 | C09B9.4 |
| WBGene00015630 | 0.281454817 | 0.94435481 | 1 | C09B9.7 |
| WBGene00015631 | 0.216767612 | 0.89583058 | 1 | clec-89 |
| WBGene00015632 | 0.977729167 | 0.0001742 | 0.00235513 | C09D4.1 |
| WBGene00015633 | 0.334970543 | 0.28134236 | 0.49065537 | C09D4.2 |
| WBGene00015634 | 1.976897945 | 0.29244709 | 1 | C09D4.3 |
| WBGene00015635 | -0.033633333 | 0.85223749 | 0.91924443 | C09D4.4 |
| WBGene00015636 | 0.281454817 | 0.94435481 | 1 | C09D4.6 |
| WBGene00015637 | 0.688793783 | 0.86378755 | 1 | srxa-5 |
| WBGene00015638 | -0.27367453 | 0.44005541 | 0.64015249 | C09E7.4 |
| WBGene00015639 | 0.129687888 | 0.93259747 | 1 | C09E7.5 |
| WBGene00015641 | 0.576248945 | 0.02060648 | 0.0873751 | C09E7.7 |
| WBGene00015642 | 0.622954048 | 0.01651198 | 0.07432888 | C09E7.8 |
| WBGene00015644 | 0.327552513 | 0.22442581 | 0.42855516 | C09E8.1 |
| WBGene00015645 | -1.099002569 | 0.32463773 | 1 | lips-7 |
| WBGene00015646 | 0.177279497 | 0.67493098 | 0.81406091 | mlt-10 |
| WBGene00015647 | 0.058908749 | 0.8380242 | 0.91144421 | C09F5.1 |
| WBGene00015648 | 0.001508854 | 0.99572348 | 0.99777804 | orai-1 |
| WBGene00015649 | 1.957271014 | 0.49743227 | 1 | C09F5.3 |
| WBGene00015650 | 0.299568435 | 0.14811671 | 0.32875412 | pkg-2 |
| WBGene00015651 | 2.641117225 | 0.21179361 | 1 | ceh-53 |
| WBGene00015655 | 0.03940405 | 0.99001443 | 1 | C09G12.5 |
| WBGene00015658 | -0.070098563 | 0.75568838 | 0.86343051 | tsg-101 |
| WBGene00015660 | 0.247231711 | 0.45251754 | 0.65010241 | catp-3 |
| WBGene00015661 | 0.747845955 | 0.85215965 | 1 | C09H5.7 |
| WBGene00015662 | -1.162847847 | 0.37246796 | 1 | C10A4.1 |
| WBGene00015663 | 3.397941824 | 0.38899876 | 1 | C10A4.2 |
| WBGene00015664 | 3.174845848 | 0.09860853 | 1 | C10A4.3 |
| WBGene00015665 | 0.29477196 | 0.83827901 | 1 | C10A4.4 |
| WBGene00015666 | 1.376646134 | 0.30629895 | 1 | gad-2 |
| WBGene00015667 | -0.901019005 | 0.67045999 | 1 | C10A4.6 |
| WBGene00015668 | -1.027355405 | 0.68015432 | 1 | C10A4.7 |
| WBGene00015670 | -0.304692658 | 0.43430253 | 0.63566811 | C10B5.1 |
| WBGene00015671 | 0.141819483 | 0.67979244 | 0.81647553 | C10B5.3 |
| WBGene00015672 | -1.453036238 | 0.47675825 | 1 | C10E2.1 |
| WBGene00015673 | 1.840019994 | 0.02094734 | 0.08852128 | C10E2.2 |
| WBGene00015675 | -0.174709411 | 0.88019872 | 1 | C10E2.4 |

| | | | | |
|---|---|---|---|---|
| WBGene00015675 | -0.369664777 | 0.74469619 | 1 | C10E2.5 |
| WBGene00015676 | 0.352122975 | 0.16381214 | 0.34999151 | mct-6 |
| WBGene00015677 | -0.341708085 | 0.05362804 | 0.16846002 | cpg-4 |
| WBGene00015678 | 0.175339372 | 0.63196359 | 0.78661779 | mcp-1 |
| WBGene00015679 | -1.465615722 | 0.0062767 | 0.03690335 | C10F3.7 |
| WBGene00015680 | 0.850846114 | 0.00158547 | 0.0130733 | egal-1 |
| WBGene00015681 | -0.012261988 | 0.99496548 | 1 | C10G8.2 |
| WBGene00015682 | 0.688793783 | 0.86378755 | 1 | C10G8.3 |
| WBGene00015683 | -0.501985105 | 0.00234484 | 0.01761466 | C10G8.4 |
| WBGene00015684 | -0.240276559 | 0.21134475 | 0.4111419 | C10G8.8 |
| WBGene00015686 | 0.182433272 | 0.53065902 | 0.71443643 | C10G11.6 |
| WBGene00015687 | -0.755224989 | 1.568E-06 | 4.7405E-05 | chdp-1 |
| WBGene00015688 | 2.893619063 | 0.1974859 | 1 | C10G11.8 |
| WBGene00015690 | -1.094201388 | 0.7816772 | 1 | C10G11.10 |
| WBGene00015691 | 0.03913098 | 0.83713212 | 0.91143626 | viln-1 |
| WBGene00015692 | 0.326556401 | 0.48145994 | 0.67289112 | ugt-25 |
| WBGene00015693 | 0.295256141 | 0.38944548 | 0.59790815 | ugt-28 |
| WBGene00015694 | 1.38322681 | 0.69550305 | 1 | ugt-27 |
| WBGene00015695 | 1.031178623 | 0.00071403 | 0.00707681 | ugt-26 |
| WBGene00015696 | 0.983488853 | 0.80601565 | 1 | C10H11.7 |
| WBGene00015697 | -0.550476755 | 0.02454687 | 0.09897827 | mlst-8 |
| WBGene00015698 | -0.55195762 | 0.00562589 | 0.03403287 | kca-1 |
| WBGene00015700 | 0.432116739 | 0.76685856 | 1 | asp-9 |
| WBGene00015701 | -0.358800222 | 0.48345254 | 0.6744338 | C11D2.3 |
| WBGene00015702 | -0.327610297 | 0.12825636 | 0.29905763 | C11D2.4 |
| WBGene00015703 | -0.564431118 | 0.00889853 | 0.0473224 | C11D2.7 |
| WBGene00015704 | 0.739296144 | 0.12117688 | 0.28817515 | tiam-1 |
| WBGene00015705 | 0.530656138 | 0.54705369 | 0.72587747 | nhr-152 |
| WBGene00015706 | 2.190171357 | 0.04377104 | 0.14608272 | C12D5.3 |
| WBGene00015707 | 3.096113806 | 0.11014314 | 1 | C12D5.4 |
| WBGene00015708 | 1.647274605 | 0.40791541 | 1 | C12D5.5 |
| WBGene00015709 | 0.085889147 | 0.81949718 | 0.90236852 | cyp-33A1 |
| WBGene00015710 | 1.031180746 | 0.12854911 | 0.29938898 | C12D5.9 |
| WBGene00015711 | 0.687942086 | 0.65383252 | 1 | C12D5.10 |
| WBGene00015713 | 1.435353024 | 1.06E-07 | 4.9607E-06 | C12D12.1 |
| WBGene00015714 | 0.169097384 | 0.93630233 | 1 | C12D12.3 |
| WBGene00015716 | 0.63805729 | 0.24164332 | 0.44927756 | sox-4 |
| WBGene00015720 | -0.233502874 | 0.92367326 | 1 | C13A2.3 |
| WBGene00015728 | 0.075494017 | 0.90088417 | 0.94788411 | C13A10.1 |
| WBGene00015729 | 0.385121749 | 0.5755868 | 0.74844162 | C13A10.2 |
| WBGene00015731 | -0.156269293 | 0.96924949 | 1 | C13B7.6 |
| WBGene00015733 | 0.548828654 | 0.13105423 | 0.3034769 | C13B9.2 |
| WBGene00015734 | -0.157269514 | 0.33150602 | 0.53971087 | copd-1 |

| | | | | |
|---|---|---|---|---|
| WBGene00015735 | 0.770133253 | 0.02143895 | 0.09015945 | pdfr-1 |
| WBGene00015739 | 0.157161302 | 0.75370218 | 0.86245977 | ugt-7 |
| WBGene00015740 | 1.200632927 | 0.01738178 | 0.07744299 | C13E3.1 |
| WBGene00015741 | 1.252090212 | 0.18201576 | 0.37384831 | C13F10.1 |
| WBGene00015742 | -0.587057634 | 0.06336203 | 0.18900771 | kxd-1 |
| WBGene00015743 | 0.428936448 | 0.03098756 | 0.11626081 | soap-1 |
| WBGene00015744 | -0.607398389 | 0.02557769 | 0.10181597 | C13F10.5 |
| WBGene00015745 | -0.249518328 | 0.31418921 | 0.52190056 | C13F10.6 |
| WBGene00015746 | -0.828877753 | 0.00010996 | 0.00164418 | C13F10.7 |
| WBGene00015747 | -0.588705967 | 0.00304198 | 0.02143508 | C13G5.2 |
| WBGene00015748 | 0.058564679 | 0.94770813 | 0.97315292 | C14A11.2 |
| WBGene00015750 | 0.546052409 | 0.7844476 | 1 | C14A11.5 |
| WBGene00015751 | 1.588923457 | 0.14221214 | 1 | C14A11.6 |
| WBGene00015752 | 0.418092367 | 0.02244274 | 0.0931228 | C14B9.2 |
| WBGene00015753 | 0.903650533 | 0.25717248 | 0.46511709 | C14B9.3 |
| WBGene00015754 | 0.066141467 | 0.70907734 | 0.83533552 | C14B9.8 |
| WBGene00015755 | -0.513171803 | 0.00169778 | 0.01370525 | C14B9.10 |
| WBGene00015756 | 1.469055198 | 1.06E-08 | 6.36E-07 | C14C6.2 |
| WBGene00015758 | 0.707559163 | 0.78139902 | 1 | nhr-155 |
| WBGene00015759 | 2.806911559 | 3.45E-57 | 1.40E-53 | C14C6.5 |
| WBGene00015762 | -0.156269293 | 0.96924949 | 1 | C14C6.8 |
| WBGene00015765 | -1.68807521 | 0.39761607 | 1 | C14C11.1 |
| WBGene00015766 | -0.352512449 | 0.19456588 | 0.39059777 | C14C11.2 |
| WBGene00015767 | 0.587655172 | 0.01205947 | 0.05949371 | hex-2 |
| WBGene00015768 | 2.752363555 | 6.13E-07 | 2.269E-05 | C14C11.4 |
| WBGene00015769 | -0.516566347 | 0.26625391 | 0.47477459 | C14C11.7 |
| WBGene00015770 | -2.705999569 | 0.42423051 | 1 | C14E2.1 |
| WBGene00015773 | -1.969805064 | 0.62862569 | 1 | C14E2.4 |
| WBGene00015776 | 0.882484351 | 0.00098498 | 0.00913574 | dct-1 |
| WBGene00015777 | -0.244928636 | 0.50533587 | 0.69223544 | sfxn-2 |
| WBGene00015778 | -0.177216165 | 0.29598932 | 0.50635569 | got-2.2 |
| WBGene00015779 | 0.644473353 | 0.10393804 | 0.26148193 | lrch-1 |
| WBGene00015780 | 0.219870397 | 0.53478362 | 0.71775458 | C14F11.4 |
| WBGene00015781 | -0.857511431 | 0.09225203 | 0.24190753 | C14F11.6 |
| WBGene00015782 | -0.156269293 | 0.96924949 | 1 | C14F11.7 |
| WBGene00015783 | -0.000206997 | 0.99972575 | 0.99989047 | C15B12.1 |
| WBGene00015784 | -0.391134131 | 0.84335123 | 1 | C15B12.2 |
| WBGene00015786 | 0.770586763 | 0.14591261 | 0.32488677 | C15B12.4 |
| WBGene00015787 | -0.067780739 | 0.88722573 | 0.94044069 | actl-1 |
| WBGene00015788 | 0.882980818 | 0.24268364 | 0.45031669 | C15B12.8 |
| WBGene00015789 | -0.922477527 | 0.04926604 | 0.15933378 | syx-6 |
| WBGene00015790 | 1.21914682 | 0.01987052 | 0.08507334 | C15C7.4 |
| WBGene00015791 | 0.31553087 | 0.20349202 | 0.40270523 | C15C7.5 |

| | | | | |
|---|---|---|---|---|
| WBGene00015792 | 0.789601902 | 0.75907734 | 1 | C15C7.6 |
| WBGene00015793 | -0.1606542 | 0.63534719 | 0.788316 | C15C7.7 |
| WBGene00015794 | -0.088712954 | 0.82617848 | 0.90625042 | C15F1.1 |
| WBGene00015795 | -0.357079756 | 0.65510463 | 0.80180033 | C15F1.2 |
| WBGene00015796 | -1.265657309 | 0.07936918 | 0.2198711 | C15F1.5 |
| WBGene00015797 | -0.214823178 | 0.75905483 | 0.86516004 | C15F1.8 |
| WBGene00015798 | 0.969284337 | 0.57363817 | 1 | C15H9.2 |
| WBGene00015799 | -0.255998374 | 0.883344 | 1 | C15H9.3 |
| WBGene00015800 | -0.359280525 | 0.14795195 | 0.32856862 | C15H9.4 |
| WBGene00015801 | 0.228750182 | 0.45164651 | 0.64946587 | C15H9.5 |
| WBGene00015802 | -1.386872448 | 3.92E-10 | 3.45E-08 | kynu-1 |
| WBGene00015803 | -0.215341238 | 0.24783758 | 0.45583943 | C15H9.9 |
| WBGene00015805 | 0.490179622 | 0.21602024 | 0.41761252 | C15H9.11 |
| WBGene00015806 | -0.166452962 | 0.47501869 | 0.66811886 | smrc-1 |
| WBGene00015807 | -0.30742577 | 0.30681281 | 0.51557292 | C16A3.2 |
| WBGene00015809 | -0.299119106 | 0.07490774 | 0.21164881 | C16A3.4 |
| WBGene00015810 | -0.374677602 | 0.0286133 | 0.10955348 | C16A3.5 |
| WBGene00015811 | -0.203140101 | 0.25478688 | 0.46230167 | C16A3.6 |
| WBGene00015813 | 0.45780394 | 0.00936488 | 0.04930756 | thoc-2 |
| WBGene00015814 | -0.551746881 | 0.00077114 | 0.00752602 | oatr-1 |
| WBGene00015815 | -0.381858629 | 0.07242387 | 0.20703984 | C16A11.2 |
| WBGene00015816 | -0.882547588 | 0.02261685 | 0.09351331 | C16A11.3 |
| WBGene00015817 | -0.305336164 | 0.14077429 | 0.31851298 | C16A11.4 |
| WBGene00015818 | 0.157440863 | 0.57083945 | 0.74514157 | C16A11.5 |
| WBGene00015819 | -0.116406097 | 0.60974279 | 0.77179825 | fbxc-44 |
| WBGene00015820 | 0.747845955 | 0.85215965 | 1 | C16A11.7 |
| WBGene00015821 | 0.377828795 | 0.92526585 | 1 | clec-135 |
| WBGene00015822 | -0.310305771 | 0.74786659 | 1 | C16B8.2 |
| WBGene00015823 | 0.58733957 | 0.01880981 | 0.08208839 | C16B8.3 |
| WBGene00015824 | 0.415408001 | 0.53551905 | 0.7178681 | C16B8.4 |
| WBGene00015825 | -1.901856996 | 0.64072034 | 1 | C16C4.1 |
| WBGene00015827 | -1.969805064 | 0.62862569 | 1 | math-13 |
| WBGene00015828 | 0.89382235 | 0.14031052 | 0.31787835 | math-14 |
| WBGene00015829 | 3.331624324 | 0.06944202 | 1 | math-15 |
| WBGene00015830 | -0.569029309 | 0.61548317 | 1 | fbxb-98 |
| WBGene00015838 | 0.281454817 | 0.94435481 | 1 | math-9 |
| WBGene00015839 | 0.520615602 | 0.72232269 | 1 | math-10 |
| WBGene00015841 | 0.07896089 | 0.89146555 | 0.94271722 | skpo-2 |
| WBGene00015842 | -0.493178805 | 0.11818829 | 0.28346979 | C16C8.4 |
| WBGene00015843 | -0.773427026 | 0.03910347 | 0.13533503 | C16C8.5 |
| WBGene00015845 | -0.62146648 | 0.85925752 | 1 | C16C8.7 |
| WBGene00015846 | -0.156269293 | 0.96924949 | 1 | C16C8.8 |
| WBGene00015847 | 0.519331445 | 0.86180636 | 1 | C16C8.9 |

| | | | | |
|---|---|---|---|---|
| WBGene00015849 | -0.335477814 | 0.36190424 | 0.56967189 | C16C8.11 |
| WBGene00015850 | -0.295011429 | 0.47934039 | 0.67131984 | C16C8.12 |
| WBGene00015851 | 0.279675994 | 0.72182622 | 0.84322534 | C16C8.13 |
| WBGene00015852 | -0.247935849 | 0.57909027 | 0.75037463 | C16C8.14 |
| WBGene00015853 | -0.465531099 | 0.06560133 | 0.19336894 | C16C8.16 |
| WBGene00015855 | 0.983488853 | 0.80601565 | 1 | C16C8.18 |
| WBGene00015856 | -0.62146648 | 0.85925752 | 1 | C16C8.19 |
| WBGene00015857 | 0.291014793 | 0.52406481 | 0.70877474 | C16D9.1 |
| WBGene00015858 | 0.281454817 | 0.94435481 | 1 | C16D9.3 |
| WBGene00015859 | 1.185542907 | 0.0521666 | 0.1651157 | C16D9.4 |
| WBGene00015860 | -1.211422718 | 0.48838595 | 1 | C16D9.5 |
| WBGene00015861 | 0.927656864 | 0.07866214 | 0.21839401 | C16D9.6 |
| WBGene00015862 | 0.983488853 | 0.80601565 | 1 | srt-25 |
| WBGene00015863 | -0.156269293 | 0.96924949 | 1 | C16D9.8 |
| WBGene00015864 | 2.818467611 | 0.2169499 | 1 | C16D9.9 |
| WBGene00015865 | 1.055875446 | 0.0289113 | 0.11041588 | C16E9.1 |
| WBGene00015866 | -0.76190085 | 0.02291162 | 0.09448708 | C16E9.2 |
| WBGene00015867 | 1.139358466 | 0.62800936 | 1 | ncs-7 |
| WBGene00015868 | 0.220048427 | 0.49177449 | 0.68103503 | C16H3.3 |
| WBGene00015869 | -1.094201388 | 0.7816772 | 1 | nhr-257 |
| WBGene00015877 | 0.377828795 | 0.92526585 | 1 | C17B7.3 |
| WBGene00015878 | 2.44049907 | 0.07056521 | 1 | C17B7.4 |
| WBGene00015879 | 0.962334705 | 0.5191748 | 1 | C17B7.5 |
| WBGene00015882 | 2.09403112 | 0.45107747 | 1 | C17B7.8 |
| WBGene00015884 | 1.957271014 | 0.49743227 | 1 | C17B7.10 |
| WBGene00015885 | 0.358877094 | 0.44881839 | 0.64793152 | fbxa-65 |
| WBGene00015887 | -0.568336783 | 0.02315624 | 0.09507608 | C17C3.1 |
| WBGene00015888 | 0.281454817 | 0.94435481 | 1 | C17C3.2 |
| WBGene00015889 | -1.0062442 | 0.63208617 | 1 | C17C3.3 |
| WBGene00015890 | -0.156269293 | 0.96924949 | 1 | C17C3.5 |
| WBGene00015891 | 1.116613983 | 0.69067248 | 1 | C17C3.6 |
| WBGene00015892 | -0.156269293 | 0.96924949 | 1 | C17C3.9 |
| WBGene00015894 | -3.408505557 | 9.28E-13 | 1.31E-10 | acdh-2 |
| WBGene00015895 | 0.281454817 | 0.94435481 | 1 | C17C3.13 |
| WBGene00015896 | -0.074457117 | 0.89381447 | 0.9441275 | C17C3.15 |
| WBGene00015897 | 1.454886984 | 0.00263414 | 0.01930058 | nhr-156 |
| WBGene00015898 | -1.710857121 | 0.32044127 | 1 | sre-9 |
| WBGene00015899 | -0.85015752 | 0.00013512 | 0.00194373 | C17E7.4 |
| WBGene00015900 | 1.33020658 | 0.29061343 | 1 | nhr-157 |
| WBGene00015901 | 0.549018937 | 0.67168773 | 1 | nhr-158 |
| WBGene00015902 | -1.159346363 | 0.58713361 | 1 | nhr-159 |
| WBGene00015903 | -1.070074292 | 0.00027804 | 0.00339612 | C17E7.9 |
| WBGene00015904 | 0.938982574 | 0.8146943 | 1 | irld-22 |

| | | | | |
|---|---|---|---|---|
| WBGene00015905 | -1.328089072 | 1.7787E-05 | 0.00036664 | C17E7.12 |
| WBGene00015906 | 0.146192886 | 0.93860911 | 1 | C17E7.13 |
| WBGene00015907 | 0.983488853 | 0.80601565 | 1 | C17F3.1 |
| WBGene00015908 | 0.281454817 | 0.94435481 | 1 | C17F3.3 |
| WBGene00015909 | 0.281454817 | 0.94435481 | 1 | clec-124 |
| WBGene00015910 | 1.264854572 | 0.2718495 | 1 | C17F4.2 |
| WBGene00015912 | -0.208108047 | 0.37498597 | 0.58293274 | fbxc-50 |
| WBGene00015913 | 0.8053344 | 2.48E-11 | 2.64E-09 | C17F4.7 |
| WBGene00015914 | -0.961473172 | 0.3039165 | 1 | C17F4.8 |
| WBGene00015915 | -0.434816238 | 0.01150514 | 0.05762537 | C17G10.1 |
| WBGene00015916 | -0.275908811 | 0.20776304 | 0.40731196 | C17G10.2 |
| WBGene00015917 | 1.065082656 | 0.61158879 | 1 | C17G10.3 |
| WBGene00015918 | 0.579678715 | 0.74528215 | 1 | C17G10.6 |
| WBGene00015919 | -0.6191744 | 0.51466641 | 1 | C17G10.7 |
| WBGene00015920 | -0.238721024 | 0.12864238 | 0.29954875 | eif-3.L |
| WBGene00015921 | 1.993000654 | 0.05706802 | 0.17652557 | C17H11.1 |
| WBGene00015922 | 0.071633496 | 0.85065805 | 0.91824825 | C17H11.2 |
| WBGene00015925 | 3.013451459 | 0.16378372 | 1 | C17H11.5 |
| WBGene00015926 | 0.50842877 | 0.05227449 | 0.16536335 | C17H11.6 |
| WBGene00015927 | 0.030053466 | 0.86938241 | 0.92964346 | dyci-1 |
| WBGene00015928 | -0.248718189 | 0.15832013 | 0.34305991 | C17H12.2 |
| WBGene00015929 | 0.651633922 | 0.74024269 | 1 | C17H12.3 |
| WBGene00015930 | 0.299793129 | 0.40826587 | 0.61423758 | C17H12.4 |
| WBGene00015931 | -0.133799809 | 0.94295257 | 1 | C17H12.5 |
| WBGene00015932 | 0.675325711 | 0.32708864 | 0.53553065 | C17H12.6 |
| WBGene00015933 | -0.374209127 | 0.13994665 | 0.31729081 | C17H12.8 |
| WBGene00015934 | 0.026542823 | 0.96045177 | 0.9796557 | ceh-48 |
| WBGene00015935 | 0.353369042 | 0.60941196 | 0.77151935 | C17H12.10 |
| WBGene00015936 | 0.573691044 | 0.28724762 | 0.49711057 | C17H12.11 |
| WBGene00015937 | 0.979427925 | 0.56765466 | 1 | C17H12.12 |
| WBGene00015938 | -0.32108124 | 0.12525148 | 0.29504815 | anat-1 |
| WBGene00015939 | -0.343458438 | 0.27361758 | 0.48215195 | C18A3.1 |
| WBGene00015940 | -0.068954758 | 0.78858812 | 0.88372239 | zipt-3 |
| WBGene00015941 | -0.362556331 | 0.05132109 | 0.16359475 | C18A3.3 |
| WBGene00015942 | 0.711746398 | 0.03643757 | 0.12918931 | osta-2 |
| WBGene00015943 | -0.115980393 | 0.52847558 | 0.71275516 | tiar-1 |
| WBGene00015944 | 0.938982574 | 0.8146943 | 1 | C18A3.7 |
| WBGene00015945 | -0.156269293 | 0.96924949 | 1 | C18A3.9 |
| WBGene00015946 | -0.90768157 | 0.25663025 | 0.46441757 | C18A3.10 |
| WBGene00015947 | 0.666348435 | 0.23360232 | 0.43964745 | C18A11.1 |
| WBGene00015948 | 0.467196966 | 0.52533326 | 0.70985766 | C18A11.2 |
| WBGene00015949 | 1.276836777 | 0.00059489 | 0.00609501 | C18A11.3 |
| WBGene00015950 | -1.462804059 | 0.24185531 | 1 | C18A11.4 |

| | | | | |
|---|---|---|---|---|
| WBGene00015952 | 1.836353502 | 0.24695375 | 1 | C18B2.1 |
| WBGene00015953 | 1.358195956 | 0.40828353 | 1 | C18B2.2 |
| WBGene00015954 | 0.527183696 | 0.0698179 | 0.20187165 | C18B2.3 |
| WBGene00015955 | -0.050825137 | 0.81274853 | 0.89786897 | C18B2.4 |
| WBGene00015956 | 0.860005078 | 5.689E-06 | 0.00013982 | C18B2.5 |
| WBGene00015957 | 0.371814825 | 0.59293633 | 0.75981517 | del-9 |
| WBGene00015958 | 0.983488853 | 0.80601565 | 1 | srbc-11 |
| WBGene00015959 | 0.281454817 | 0.94435481 | 1 | srbc-10 |
| WBGene00015962 | 0.281454817 | 0.94435481 | 1 | srbc-8 |
| WBGene00015963 | 1.347309852 | 0.36784729 | 1 | C18B10.6 |
| WBGene00015964 | 0.490433862 | 0.76031596 | 1 | glb-5 |
| WBGene00015965 | 1.089319919 | 0.00021496 | 0.00278518 | ugt-48 |
| WBGene00015968 | 0.454293915 | 0.04939248 | 0.15947139 | C18C4.5 |
| WBGene00015968 | 1.466237541 | 0.36968143 | 1 | C18C4.7 |
| WBGene00015969 | 0.091544335 | 0.89474249 | 0.94453253 | glb-6 |
| WBGene00015970 | 1.036568105 | 0.67105582 | 1 | C18E3.1 |
| WBGene00015971 | -0.436108356 | 0.02112993 | 0.08901406 | swsn-2.2 |
| WBGene00015972 | -0.131200218 | 0.65622744 | 0.80231895 | C18E3.3 |
| WBGene00015974 | -0.534293987 | 0.04859491 | 0.15804675 | snrp-40.2 |
| WBGene00015975 | -0.220916312 | 0.26207204 | 0.47061332 | cas-2 |
| WBGene00015976 | -0.425197869 | 0.06033383 | 0.18308248 | C18E3.9 |
| WBGene00015977 | 3.267618098 | 0.03085945 | 1 | C18F10.2 |
| WBGene00015978 | -0.076388705 | 0.65970535 | 0.80368078 | C18F10.7 |
| WBGene00015980 | 1.126894513 | 0.10483364 | 0.26329855 | C18G1.1 |
| WBGene00015981 | 0.81415697 | 0.11375709 | 0.27694503 | elt-7 |
| WBGene00015982 | 0.281454817 | 0.94435481 | 1 | bgnt-1.4 |
| WBGene00015984 | 1.083884448 | 0.38544711 | 1 | C18G1.6 |
| WBGene00015985 | 1.584577408 | 0.63258867 | 1 | C18G1.7 |
| WBGene00015986 | -0.245531902 | 0.94070824 | 1 | C18G1.8 |
| WBGene00015987 | 0.983488853 | 0.80601565 | 1 | C18G1.9 |
| WBGene00015988 | 1.985511501 | 0.02936571 | 0.11148501 | C18H2.1 |
| WBGene00015993 | 0.622447812 | 0.12285044 | 0.29120014 | C18H7.1 |
| WBGene00015994 | 0.747845955 | 0.85215965 | 1 | C18H7.4 |
| WBGene00015996 | 2.369358709 | 0.05854487 | 1 | C18H7.6 |
| WBGene00015998 | -2.640414922 | 0.41081194 | 1 | srt-59 |
| WBGene00015999 | -1.351599631 | 0.53258792 | 1 | prmt-4 |
| WBGene00016000 | -0.490561023 | 0.85693136 | 1 | C18H9.1 |
| WBGene00016002 | 0.030123138 | 0.90429384 | 0.95040093 | C18H9.3 |
| WBGene00016003 | 2.169286972 | 1.0821E-06 | 3.5798E-05 | C18H9.5 |
| WBGene00016004 | 1.620553062 | 7.8418E-06 | 0.00018523 | C18H9.6 |
| WBGene00016005 | 0.467725862 | 0.51042468 | 0.69706264 | ift-74 |
| WBGene00016006 | 1.535466372 | 8.74E-12 | 1.02E-09 | fln-2 |
| WBGene00016008 | 0.394439984 | 0.88400378 | 1 | C23F12.3 |

| | | | | |
|---|---|---|---|---|
| WBGene00016009 | -0.33041003 | 0.42378802 | 0.62806093 | C23F12.4 |
| WBGene00016010 | -0.985537791 | 0.65575305 | 1 | C23G10.1 |
| WBGene00016011 | -0.264272078 | 0.17907055 | 0.37122037 | C23G10.2 |
| WBGene00016012 | 1.485796186 | 0.1680716 | 1 | C23G10.5 |
| WBGene00016013 | 1.66546146 | 0.00066487 | 0.00669337 | C23G10.6 |
| WBGene00016014 | -0.196456421 | 0.26154371 | 0.47006725 | C23G10.7 |
| WBGene00016015 | -0.094764697 | 0.55289281 | 0.73075023 | C23G10.8 |
| WBGene00016016 | 1.529802032 | 0.64971375 | 1 | C23G10.10 |
| WBGene00016017 | 5.739048312 | 1.7253E-05 | 0.00035867 | C23G10.11 |
| WBGene00016018 | 0.393338165 | 0.11477461 | 0.27886303 | C23H3.2 |
| WBGene00016019 | 0.27081997 | 0.29120774 | 0.50121253 | C23H3.3 |
| WBGene00016020 | 0.402301257 | 0.0076737 | 0.04245474 | sptl-1 |
| WBGene00016021 | -0.187102095 | 0.44509371 | 0.64447021 | C23H3.5 |
| WBGene00016022 | 0.143877216 | 0.68694773 | 0.8215359 | C23H3.9 |
| WBGene00016023 | -0.830571013 | 0.11616236 | 0.28054097 | prmt-6 |
| WBGene00016024 | 1.904977873 | 0.0953686 | 1 | glb-7 |
| WBGene00016025 | 1.582230931 | 0.24232463 | 1 | xbx-4 |
| WBGene00016027 | -0.646563853 | 0.00194242 | 0.01521272 | C23H5.8 |
| WBGene00016028 | 0.685788322 | 0.02477136 | 0.09951987 | flp-24 |
| WBGene00016029 | 0.552300763 | 0.28718939 | 0.49711057 | elf-1 |
| WBGene00016030 | 0.660578563 | 0.08029273 | 0.22150285 | C24A1.3 |
| WBGene00016032 | -0.495284958 | 0.70399532 | 1 | C24A3.1 |
| WBGene00016033 | 0.478514801 | 0.06264381 | 0.18765322 | C24A3.2 |
| WBGene00016034 | 0.001638474 | 0.99715092 | 0.99857337 | C24A3.4 |
| WBGene00016037 | 1.282740206 | 0.18183676 | 0.37382024 | dop-6 |
| WBGene00016038 | 0.164131808 | 0.82553229 | 0.90589186 | cst-2 |
| WBGene00016039 | 0.232157325 | 0.93397866 | 1 | C24A8.5 |
| WBGene00016040 | 0.166733916 | 0.90886787 | 1 | C24A11.1 |
| WBGene00016044 | 2.156089719 | 0.00198587 | 0.01545543 | C24B5.1 |
| WBGene00016045 | -0.343586311 | 0.04860774 | 0.15804675 | spas-1 |
| WBGene00016046 | 0.382086678 | 0.54325206 | 0.72328361 | C24B5.4 |
| WBGene00016048 | 0.41651643 | 0.10579283 | 0.26483109 | C24B9.3 |
| WBGene00016049 | 1.557897343 | 0.63813005 | 1 | srt-28 |
| WBGene00016051 | 0.281454817 | 0.94435481 | 1 | srt-27 |
| WBGene00016052 | 1.800622447 | 0.02907637 | 0.11080234 | dod-3 |
| WBGene00016053 | -1.969805064 | 0.62862569 | 1 | C24D10.1 |
| WBGene00016054 | 0.377828795 | 0.92526585 | 1 | C24D10.2 |
| WBGene00016055 | -0.040048479 | 0.86780116 | 0.92844324 | C24D10.4 |
| WBGene00016056 | -1.198235508 | 0.08304479 | 0.22636883 | C24D10.5 |
| WBGene00016057 | -0.422836807 | 0.06692313 | 0.19624021 | C24D10.6 |
| WBGene00016058 | 1.181686151 | 0.7579682 | 1 | nspd-3 |
| WBGene00016059 | 0.392907153 | 0.14958438 | 0.33104337 | C24G6.2 |
| WBGene00016060 | 0.128007324 | 0.51009996 | 0.69671769 | mms-19 |

| | | | | |
|---|---|---|---|---|
| WBGene00016061 | 0.572268227 | 0.00060203 | 0.00616009 | hpo-15 |
| WBGene00016062 | -0.259026066 | 0.16197251 | 0.34786984 | C24G6.8 |
| WBGene00016063 | 0.700223173 | 0.6432038 | 1 | delm-2 |
| WBGene00016064 | 3.132878219 | 0.13631072 | 1 | acd-1 |
| WBGene00016066 | 0.718782876 | 0.25546029 | 0.46298603 | acd-2 |
| WBGene00016067 | -0.361896575 | 0.51729905 | 0.7030198 | C24H10.1 |
| WBGene00016068 | -0.160819095 | 0.65532306 | 0.80188241 | C24H10.2 |
| WBGene00016069 | 0.481270855 | 0.79087313 | 1 | C24H10.3 |
| WBGene00016070 | 0.281454817 | 0.94435481 | 1 | C24H10.4 |
| WBGene00016071 | -1.052068412 | 0.18049291 | 0.37259848 | C24H12.1 |
| WBGene00016072 | -0.295889589 | 0.85352501 | 1 | C24H12.2 |
| WBGene00016073 | 0.099401206 | 0.6519719 | 0.79955463 | C24H12.4 |
| WBGene00016074 | -0.220270862 | 0.31440168 | 0.52218205 | C24H12.5 |
| WBGene00016075 | -1.969805064 | 0.62862569 | 1 | C24H12.6 |
| WBGene00016077 | -0.654945829 | 0.77396192 | 1 | C24H12.8 |
| WBGene00016081 | -0.745110582 | 0.61155837 | 1 | C25A6.1 |
| WBGene00016083 | 0.747845955 | 0.85215965 | 1 | C25A8.2 |
| WBGene00016084 | -0.354277205 | 0.01143746 | 0.05735737 | cht-3 |
| WBGene00016085 | 0.688793783 | 0.86378755 | 1 | C25A8.5 |
| WBGene00016087 | -0.817876165 | 0.00078465 | 0.0076272 | C25A11.2 |
| WBGene00016088 | -1.730092997 | 1.70E-09 | 1.26E-07 | clec-266 |
| WBGene00016089 | 1.532777345 | 0.30913189 | 1 | aexr-1 |
| WBGene00016090 | 0.688793783 | 0.86378755 | 1 | aexr-2 |
| WBGene00016091 | 0.041803775 | 0.98890197 | 1 | nhr-30 |
| WBGene00016092 | 0.983488853 | 0.80601565 | 1 | cyp-33B1 |
| WBGene00016093 | 0.984523416 | 0.05212739 | 0.16507007 | srsx-34 |
| WBGene00016094 | 1.858080202 | 0.08093031 | 1 | C25E10.4 |
| WBGene00016095 | 0.168989228 | 0.78282788 | 0.88027353 | C25E10.5 |
| WBGene00016096 | 0.338676826 | 0.85525483 | 1 | C25E10.7 |
| WBGene00016097 | -0.244138404 | 0.5926366 | 0.75972181 | C25E10.8 |
| WBGene00016099 | -0.843240102 | 0.11366805 | 0.27689482 | C25E10.10 |
| WBGene00016100 | 1.168450382 | 0.44384957 | 1 | C25E10.11 |
| WBGene00016101 | -0.074769 | 0.84827778 | 0.91671496 | C25E10.12 |
| WBGene00016102 | -0.387937325 | 0.30753837 | 0.51622057 | C25F6.1 |
| WBGene00016103 | 0.517047634 | 0.01353589 | 0.06444678 | dpyd-1 |
| WBGene00016104 | 0.464889223 | 0.26399239 | 0.47252111 | ddr-1 |
| WBGene00016105 | -0.781637661 | 0.72728626 | 1 | C25F6.6 |
| WBGene00016106 | -0.398100438 | 0.55881875 | 0.73577903 | C25F6.7 |
| WBGene00016107 | 0.729224979 | 0.57994537 | 1 | C25G6.1 |
| WBGene00016108 | 0.672110638 | 0.40831475 | 0.61423758 | C25G6.3 |
| WBGene00016109 | -0.012364091 | 0.99262541 | 1 | C25G6.4 |
| WBGene00016110 | 0.387885852 | 0.76151374 | 1 | npr-11 |
| WBGene00016111 | 2.668304423 | 0.1124189 | 1 | C25H3.1 |

| | | | | |
|---|---|---|---|---|
| WBGene00016112 | -0.776388053 | 0.00854879 | 0.04601914 | C25H3.3 |
| WBGene00016113 | -0.091330992 | 0.64404499 | 0.79428104 | eif-2D |
| WBGene00016114 | 0.878860724 | 0.05868436 | 0.1798755 | flp-27 |
| WBGene00016115 | -0.216172736 | 0.31183923 | 0.51998902 | mdt-26 |
| WBGene00016116 | -0.328519167 | 0.08141758 | 0.2233373 | C25H3.7 |
| WBGene00016118 | -0.467142908 | 0.00294829 | 0.02094511 | C25H3.9 |
| WBGene00016119 | 1.262088539 | 0.00035362 | 0.0041282 | C25H3.10 |
| WBGene00016120 | 0.744001965 | 0.04223446 | 0.14299212 | C25H3.11 |
| WBGene00016120 | 0.760028341 | 0.00056459 | 0.00587382 | C25H3.8 |
| WBGene00016121 | 0.034519878 | 0.92804117 | 0.96219879 | C25H3.12 |
| WBGene00016123 | -0.464231834 | 0.19865293 | 0.39657321 | C25H3.14 |
| WBGene00016124 | 0.125849805 | 0.63652013 | 0.78929536 | dnc-4 |
| WBGene00016125 | -0.378360032 | 0.82378845 | 1 | C26B2.2 |
| WBGene00016126 | -0.545832313 | 0.67031725 | 1 | nhr-258 |
| WBGene00016128 | -0.195320516 | 0.4047745 | 0.61174299 | elpc-4 |
| WBGene00016129 | -0.332935367 | 0.29539719 | 0.50569115 | C26B2.7 |
| WBGene00016130 | -1.048070996 | 0.3663407 | 1 | C26B2.8 |
| WBGene00016131 | 0.197661335 | 0.65273589 | 0.80016826 | C26B9.1 |
| WBGene00016132 | -0.298265327 | 0.89527345 | 1 | C26B9.2 |
| WBGene00016133 | -0.679059799 | 0.21108361 | 0.41102905 | C26B9.3 |
| WBGene00016134 | 0.756308205 | 2.1459E-05 | 0.0004278 | C26B9.5 |
| WBGene00016135 | -0.058002073 | 0.92491248 | 0.96034913 | C26B9.6 |
| WBGene00016136 | 0.200250327 | 0.91653624 | 1 | C26B9.7 |
| WBGene00016137 | -0.156269293 | 0.96924949 | 1 | C26E6.1 |
| WBGene00016138 | 0.077880172 | 0.78044971 | 0.87849434 | flh-2 |
| WBGene00016139 | -0.036582767 | 0.87334346 | 0.93182731 | ntl-9 |
| WBGene00016140 | 0.03528304 | 0.82883357 | 0.90762341 | rpb-2 |
| WBGene00016142 | -0.526103623 | 0.00549959 | 0.03343542 | mrps-18C |
| WBGene00016143 | 0.244895481 | 0.25666813 | 0.46441757 | eri-9 |
| WBGene00016144 | -0.182763528 | 0.51567243 | 0.70203846 | mmab-1 |
| WBGene00016145 | -0.488845562 | 0.04825292 | 0.15727215 | C26E6.12 |
| WBGene00016146 | -0.986211197 | 0.0300517 | 0.11346712 | C26F1.1 |
| WBGene00016147 | 0.0871787 | 0.87440188 | 0.93213743 | cyp-32A1 |
| WBGene00016148 | -0.293394606 | 0.20249933 | 0.40146436 | C26F1.3 |
| WBGene00016149 | 0.545264217 | 0.66471466 | 1 | frpr-3 |
| WBGene00016150 | 0.1667694 | 0.60744817 | 0.77023794 | hint-3 |
| WBGene00016151 | -0.051853047 | 0.7599728 | 0.86579992 | smc-5 |
| WBGene00016152 | -1.12875433 | 0.34207281 | 1 | pho-12 |
| WBGene00016153 | 0.688793783 | 0.86378755 | 1 | C27A2.5 |
| WBGene00016154 | 0.352617407 | 0.07674077 | 0.21490008 | C27A12.2 |
| WBGene00016155 | 0.150621314 | 0.88939519 | 1 | C27A12.4 |
| WBGene00016156 | -0.274586801 | 0.14807794 | 0.32872815 | C27A12.6 |
| WBGene00016157 | -0.539742402 | 0.0035147 | 0.0238524 | C27A12.7 |

| | | | | |
|---|---|---|---|---|
| WBGene00016158 | 0.08704821 | 0.64233492 | 0.793265 | ari-1 |
| WBGene00016159 | 0.440909255 | 0.08448243 | 0.22890007 | C27A12.9 |
| WBGene00016160 | -0.62146648 | 0.85925752 | 1 | C27D6.1 |
| WBGene00016161 | 1.996315266 | 0.49147447 | 1 | C27D6.3 |
| WBGene00016162 | 0.624781704 | 0.00305894 | 0.02151716 | crh-2 |
| WBGene00016163 | -0.234287466 | 0.24106527 | 0.44872386 | C27D9.1 |
| WBGene00016165 | 0.3520996 | 0.6367385 | 0.78948551 | C27F2.1 |
| WBGene00016166 | -0.419194464 | 0.03555823 | 0.12715091 | C27F2.4 |
| WBGene00016167 | -0.319762955 | 0.13061446 | 0.30274727 | vps-22 |
| WBGene00016169 | -0.314707422 | 0.40382726 | 0.6111802 | C27F2.7 |
| WBGene00016170 | 0.380768948 | 0.03864648 | 0.13444324 | C27F2.8 |
| WBGene00016171 | -0.61601223 | 0.00379754 | 0.02522208 | C27F2.10 |
| WBGene00016172 | 0.769138722 | 0.01588664 | 0.0724839 | C27H5.2 |
| WBGene00016173 | -0.212336113 | 0.25177595 | 0.45982508 | fust-1 |
| WBGene00016174 | 0.209536309 | 0.71373888 | 0.83821859 | C27H5.4 |
| WBGene00016175 | 1.851444818 | 0.53669729 | 1 | C27H5.6 |
| WBGene00016177 | 0.629999087 | 0.59841774 | 1 | C28C12.1 |
| WBGene00016178 | -0.70180143 | 2.6457E-05 | 0.00050915 | mesp-1 |
| WBGene00016179 | 0.157372398 | 0.92008062 | 1 | C28C12.3 |
| WBGene00016180 | 0.51268231 | 0.28726857 | 0.49711057 | C28C12.4 |
| WBGene00016181 | 1.277922157 | 0.43804273 | 1 | C28C12.11 |
| WBGene00016182 | -0.331824502 | 0.08596894 | 0.23148124 | C28C12.12 |
| WBGene00016184 | 0.387120557 | 0.57727641 | 0.74903703 | glb-9 |
| WBGene00016187 | 1.300979949 | 0.36092723 | 1 | C28G1.2 |
| WBGene00016188 | 0.145053897 | 0.57958439 | 0.75058496 | sec-15 |
| WBGene00016189 | -1.313300841 | 0.00968567 | 0.05062148 | C28G1.4 |
| WBGene00016190 | 0.161591286 | 0.65020475 | 0.79843591 | C28G1.5 |
| WBGene00016191 | 0.485185263 | 0.47526089 | 0.66814989 | C28G1.6 |
| WBGene00016192 | -0.981636575 | 5.4177E-06 | 0.00013479 | bcl-7 |
| WBGene00016193 | 0.830546886 | 0.50337705 | 1 | C28H8.2 |
| WBGene00016194 | 0.211372619 | 0.20647877 | 0.40636387 | C28H8.3 |
| WBGene00016195 | -0.394665647 | 0.03191711 | 0.11850325 | C28H8.4 |
| WBGene00016196 | 1.434546033 | 0.47351845 | 1 | C28H8.5 |
| WBGene00016197 | 0.434476202 | 0.05248802 | 0.16582422 | pxl-1 |
| WBGene00016198 | 0.688793783 | 0.86378755 | 1 | C28H8.7 |
| WBGene00016199 | 0.685293525 | 0.73391023 | 1 | C28H8.8 |
| WBGene00016200 | -0.284834276 | 0.14376244 | 0.32200837 | dpff-1 |
| WBGene00016201 | -0.386733929 | 0.05039557 | 0.16178016 | tdo-2 |
| WBGene00016202 | -0.338000507 | 0.03736179 | 0.13121898 | kle-2 |
| WBGene00016203 | 0.084515372 | 0.77823616 | 0.87665292 | tag-250 |
| WBGene00016204 | -0.240037582 | 0.62264209 | 0.78035037 | gsto-1 |
| WBGene00016206 | 0.377828795 | 0.92526585 | 1 | C29E4.9 |
| WBGene00016207 | 0.790966223 | 0.48772426 | 1 | C29E4.10 |

| | | | | |
|---|---|---|---|---|
| WBGene00016208 | -0.077704123 | 0.87309398 | 0.93172488 | C29E4.11 |
| WBGene00016209 | -0.685475808 | 0.00381343 | 0.02528609 | C29E4.12 |
| WBGene00016210 | -0.143763365 | 0.58717236 | 0.75591074 | C29F5.1 |
| WBGene00016212 | 2.068332674 | 0.26145708 | 1 | C29F5.3 |
| WBGene00016213 | 0.281454817 | 0.94435481 | 1 | C29F5.5 |
| WBGene00016215 | 1.258832815 | 0.07024418 | 0.20259416 | glb-10 |
| WBGene00016216 | 0.203413719 | 0.71121129 | 0.83662594 | pals-22 |
| WBGene00016217 | -1.509626704 | 0.03622733 | 0.12875763 | C29F9.2 |
| WBGene00016218 | 1.428903091 | 2E-05 | 0.00040136 | pals-23 |
| WBGene00016219 | 1.644294048 | 1.259E-06 | 4.0332E-05 | pals-24 |
| WBGene00016221 | 1.551393092 | 0.32622016 | 1 | C29F9.6 |
| WBGene00016226 | -1.058104139 | 0.46521731 | 1 | C29F9.12 |
| WBGene00016227 | 0.377828795 | 0.92526585 | 1 | nhr-280 |
| WBGene00016231 | -1.065689156 | 0.33598364 | 1 | C29G2.3 |
| WBGene00016233 | 0.377828795 | 0.92526585 | 1 | srt-58 |
| WBGene00016234 | -0.582556159 | 0.14508644 | 0.32381738 | C29G2.6 |
| WBGene00016235 | -0.128854193 | 0.4001638 | 0.60798256 | C29H12.2 |
| WBGene00016236 | 0.165143326 | 0.5666632 | 0.74172492 | cec-9 |
| WBGene00016237 | -0.089112946 | 0.84659642 | 0.91591915 | C29H12.6 |
| WBGene00016238 | -0.255614804 | 0.16900983 | 0.357109 | mob-4 |
| WBGene00016239 | 1.687239501 | 0.59403242 | 1 | C30A5.4 |
| WBGene00016242 | -0.291199644 | 0.86491471 | 1 | C30A5.10 |
| WBGene00016244 | -0.263760938 | 0.17772862 | 0.36961342 | C30B5.2 |
| WBGene00016245 | -0.36900589 | 0.11678977 | 0.28156167 | C30B5.4 |
| WBGene00016246 | 1.547221491 | 0.04339819 | 0.14531091 | daf-37 |
| WBGene00016247 | 1.287834273 | 0.40662472 | 1 | C30B5.6 |
| WBGene00016248 | 0.658888431 | 0.485065 | 0.67539764 | C30B5.7 |
| WBGene00016249 | -0.493918727 | 0.06570717 | 0.19353487 | mrpl-32 |
| WBGene00016250 | -0.282837172 | 0.10970186 | 0.2702151 | hsp-110 |
| WBGene00016253 | -0.156269293 | 0.96924949 | 1 | C30E1.2 |
| WBGene00016253 | -0.156269293 | 0.96924949 | 1 | C30E1.3 |
| WBGene00016253 | -0.156269293 | 0.96924949 | 1 | C30E1.4 |
| WBGene00016254 | -0.682164043 | 0.63656507 | 1 | fbxb-114 |
| WBGene00016257 | -0.156269293 | 0.96924949 | 1 | C30E1.8 |
| WBGene00016258 | 0.201411579 | 0.18849341 | 0.38226506 | vha-16 |
| WBGene00016259 | 2.600311303 | 0.25000941 | 1 | C30F8.3 |
| WBGene00016260 | 1.320112593 | 6.5001E-06 | 0.00015689 | rege-1 |
| WBGene00016261 | 0.201282434 | 0.4585636 | 0.65483658 | C30F12.2 |
| WBGene00016262 | -0.34029938 | 0.71815796 | 0.8405626 | C30F12.3 |
| WBGene00016263 | -0.320126471 | 0.12374899 | 0.29275849 | C30F12.4 |
| WBGene00016264 | 0.133020996 | 0.64280308 | 0.79350306 | C30F12.5 |
| WBGene00016265 | 1.225471628 | 0.03622706 | 0.12875763 | trhr-1 |
| WBGene00016266 | 0.42360138 | 0.12838454 | 0.29912046 | idhg-2 |

| | | | | |
|---|---|---|---|---|
| WBGene00016268 | 0.232157325 | 0.93397866 | 1 | C30G4.2 |
| WBGene00016269 | 0.463546233 | 0.30031731 | 0.50995852 | C30G4.4 |
| WBGene00016271 | -0.156269293 | 0.96924949 | 1 | irld-23 |
| WBGene00016273 | -0.64857282 | 0.60326405 | 1 | C30G12.1 |
| WBGene00016274 | 0.414779775 | 0.15701977 | 0.3413999 | C30G12.2 |
| WBGene00016276 | -1.484302304 | 0.2191723 | 1 | C30G12.4 |
| WBGene00016277 | 0.240938367 | 0.28926888 | 0.49943309 | C30G12.6 |
| WBGene00016278 | 0.024955291 | 0.89208979 | 0.94312627 | erp-44.1 |
| WBGene00016279 | 0.377828795 | 0.92526585 | 1 | C31B8.1 |
| WBGene00016281 | 4.299607121 | 0.00177432 | 0.01418171 | pals-32 |
| WBGene00016282 | 0.609158114 | 0.30217731 | 0.51162795 | C31B8.7 |
| WBGene00016283 | 0.748967864 | 0.12908131 | 0.30028284 | zmp-3 |
| WBGene00016284 | 0.377828795 | 0.92526585 | 1 | C31B8.9 |
| WBGene00016286 | -0.62146648 | 0.85925752 | 1 | C31H1.1 |
| WBGene00016287 | 0.377828795 | 0.92526585 | 1 | C31H1.2 |
| WBGene00016288 | 1.851444818 | 0.53669729 | 1 | C31H1.5 |
| WBGene00016289 | 0.421856599 | 0.15815259 | 0.34299953 | lntl-1 |
| WBGene00016291 | -0.596817437 | 0.00061429 | 0.00625157 | C31H1.8 |
| WBGene00016292 | 0.503464221 | 0.07490441 | 0.21164881 | tbc-7 |
| WBGene00016293 | -0.828934131 | 0.42715372 | 1 | C31H2.3 |
| WBGene00016294 | -0.406815939 | 0.43215691 | 0.6341955 | C31H2.4 |
| WBGene00016295 | -0.398715699 | 0.51489798 | 0.70153728 | fbxc-40 |
| WBGene00016297 | -0.241106143 | 0.91536132 | 1 | fbxc-15 |
| WBGene00016298 | -0.719977288 | 0.54983354 | 1 | fbxc-14 |
| WBGene00016299 | -0.334900066 | 0.81811498 | 1 | C32B5.6 |
| WBGene00016300 | -2.705999569 | 0.42423051 | 1 | C32B5.7 |
| WBGene00016302 | -0.263229581 | 0.93409872 | 1 | fbxc-7 |
| WBGene00016303 | -1.42305345 | 1.8112E-06 | 5.3373E-05 | fbxc-32 |
| WBGene00016307 | -1.758630281 | 0.00387576 | 0.02558763 | fbxc-18 |
| WBGene00016308 | -1.42278873 | 0.26842039 | 1 | C32B5.15 |
| WBGene00016309 | -1.819137971 | 0.01035158 | 0.05320851 | sdz-4 |
| WBGene00016310 | 0.801320089 | 0.3965325 | 0.60443202 | C32D5.1 |
| WBGene00016311 | 0.206701637 | 0.25348624 | 0.46140577 | C32D5.3 |
| WBGene00016312 | 0.57991002 | 0.82402153 | 1 | C32D5.4 |
| WBGene00016313 | -0.224241718 | 0.2503791 | 0.45836137 | set-4 |
| WBGene00016314 | 1.476121989 | 0.00108376 | 0.00977557 | C32D5.6 |
| WBGene00016315 | 0.526792574 | 0.28968251 | 0.49967941 | C32D5.7 |
| WBGene00016316 | -0.364389645 | 0.06061229 | 0.18383557 | C32D5.8 |
| WBGene00016317 | -0.110698174 | 0.5992561 | 0.76480755 | C32D5.10 |
| WBGene00016318 | 0.260410666 | 0.15723711 | 0.34168888 | C32D5.11 |
| WBGene00016319 | -1.453722733 | 0.00960387 | 0.05032394 | C32D5.12 |
| WBGene00016320 | -0.156269293 | 0.96924949 | 1 | C32E8.1 |
| WBGene00016321 | -0.254238498 | 0.21045558 | 0.41039852 | tppp-1 |

| | | | | |
|---|---|---|---|---|
| WBGene00016322 | 1.557897343 | 0.63813005 | 1 | C32E8.4 |
| WBGene00016323 | -0.331613714 | 0.08553018 | 0.23060669 | C32E8.5 |
| WBGene00016324 | 0.728679883 | 0.54353553 | 1 | C32E8.6 |
| WBGene00016325 | -0.40369869 | 0.26316478 | 0.47180797 | C32E8.9 |
| WBGene00016326 | 0.26318531 | 0.21742057 | 0.41959993 | ubr-1 |
| WBGene00016327 | 1.435925494 | 0.70920779 | 1 | C32E12.1 |
| WBGene00016328 | 0.728067732 | 0.43107018 | 0.63332845 | pde-5 |
| WBGene00016329 | -0.70237327 | 0.30257998 | 0.5117893 | osr-1 |
| WBGene00016330 | 1.006063584 | 0.00815567 | 0.04444255 | C32E12.4 |
| WBGene00016331 | -0.041723238 | 0.81754215 | 0.90086942 | obr-4 |
| WBGene00016332 | 1.047334221 | 0.00021212 | 0.00275443 | C32F10.4 |
| WBGene00016333 | -0.128492999 | 0.39185421 | 0.60016425 | C32F10.8 |
| WBGene00016335 | 1.843960094 | 8.79E-11 | 8.40E-09 | gba-1 |
| WBGene00016336 | -0.090960119 | 0.94830899 | 1 | C33C12.4 |
| WBGene00016337 | -1.835881168 | 0.11963021 | 1 | fbxb-38 |
| WBGene00016339 | -0.552227129 | 0.83715812 | 1 | C33C12.7 |
| WBGene00016340 | 0.593666323 | 0.72950413 | 1 | gba-2 |
| WBGene00016341 | -0.064144015 | 0.88663096 | 0.94019801 | mtq-2 |
| WBGene00016343 | 0.622100836 | 0.07865067 | 0.21839401 | cnnm-3 |
| WBGene00016344 | -0.035772601 | 0.96905857 | 0.98399257 | rsef-1 |
| WBGene00016345 | 1.132983824 | 0.43977367 | 1 | C33E10.1 |
| WBGene00016346 | -0.001118276 | 0.99894371 | 0.99968476 | fbxa-120 |
| WBGene00016347 | -0.156269293 | 0.96924949 | 1 | C33E10.5 |
| WBGene00016349 | 3.144314995 | 0.09621829 | 1 | C33E10.8 |
| WBGene00016351 | 1.395728942 | 0.51364788 | 1 | C33F10.1 |
| WBGene00016352 | 0.237779321 | 0.43956348 | 0.6398824 | tbck-1 |
| WBGene00016353 | -0.352905095 | 0.17560079 | 0.36625481 | C33F10.4 |
| WBGene00016354 | 1.035807675 | 1.7144E-06 | 5.1015E-05 | rig-6 |
| WBGene00016355 | -0.034796769 | 0.94273103 | 0.97038554 | lact-5 |
| WBGene00016356 | 1.338723186 | 0.43031793 | 1 | C33F10.8 |
| WBGene00016357 | -0.0060929 | 0.99795747 | 1 | C33F10.11 |
| WBGene00016358 | 1.247575581 | 0.7549672 | 1 | C33F10.12 |
| WBGene00016359 | -0.254164279 | 0.73240827 | 0.85035563 | fbxb-88 |
| WBGene00016360 | 1.851934254 | 0.00867116 | 0.04652079 | C33G8.2 |
| WBGene00016361 | -2.549587669 | 0.01797366 | 1 | drd-10 |
| WBGene00016362 | 0.567598799 | 0.14588572 | 0.32488677 | C33G8.4 |
| WBGene00016363 | 0.944897044 | 0.46052959 | 1 | srab-4 |
| WBGene00016364 | 0.088245531 | 0.88116139 | 0.93671136 | nhr-161 |
| WBGene00016365 | 0.106320328 | 0.89916009 | 0.94705498 | nhr-139 |
| WBGene00016366 | 1.07575935 | 0.25405099 | 0.46193303 | nhr-140 |
| WBGene00016367 | 1.703523043 | 0.33356605 | 1 | nhr-162 |
| WBGene00016368 | 0.79337097 | 0.45787115 | 1 | nhr-163 |
| WBGene00016370 | -1.094201388 | 0.7816772 | 1 | C33H5.1 |

| | | | | |
|---|---|---|---|---|
| WBGene00016373 | -0.195502303 | 0.55173619 | 0.73003216 | swd-2.1 |
| WBGene00016374 | -0.180719657 | 0.36422666 | 0.57184481 | swd-2.2 |
| WBGene00016375 | 0.213210554 | 0.46174223 | 0.65699664 | C33H5.8 |
| WBGene00016376 | -0.009434841 | 0.96560153 | 0.98286673 | sec-10 |
| WBGene00016378 | 0.014250239 | 0.94177888 | 0.96989883 | imp-3 |
| WBGene00016379 | 0.018748626 | 0.96881857 | 0.98399257 | C33H5.13 |
| WBGene00016380 | -0.084544495 | 0.69880999 | 0.82967456 | ntp-1 |
| WBGene00016381 | -0.296104592 | 0.18666229 | 0.37992738 | sgo-1 |
| WBGene00016382 | 0.377828795 | 0.92526585 | 1 | C33H5.16 |
| WBGene00016383 | 0.285407056 | 0.36644745 | 0.5740629 | zgpa-1 |
| WBGene00016384 | 0.451927262 | 0.04003097 | 0.13756036 | cdgs-1 |
| WBGene00016386 | 2.61432266 | 0.11454027 | 1 | tag-272 |
| WBGene00016387 | -0.719505899 | 0.00050458 | 0.00539265 | kbp-5 |
| WBGene00016388 | 0.616429445 | 0.81400458 | 1 | C34B2.3 |
| WBGene00016389 | -1.901856996 | 0.64072034 | 1 | C34B2.4 |
| WBGene00016390 | -0.449677903 | 0.04944952 | 0.15947139 | C34B2.5 |
| WBGene00016391 | 0.144945373 | 0.41248918 | 0.61804655 | lonp-1 |
| WBGene00016392 | -0.417634526 | 0.00894098 | 0.04750656 | sdha-2 |
| WBGene00016393 | -0.581227069 | 0.00056154 | 0.00585711 | C34B2.8 |
| WBGene00016394 | 0.070602366 | 0.94615509 | 1 | C34B2.9 |
| WBGene00016395 | -0.826290317 | 2.4665E-05 | 0.000483 | C34B2.10 |
| WBGene00016396 | 0.365294328 | 0.56867891 | 0.74341002 | C34B2.11 |
| WBGene00016397 | 0.478449983 | 0.63288705 | 1 | maph-9 |
| WBGene00016398 | 1.435925494 | 0.70920779 | 1 | C34D4.2 |
| WBGene00016399 | 0.284817402 | 0.87269197 | 1 | C34D4.3 |
| WBGene00016400 | -0.192051299 | 0.37304635 | 0.58103345 | C34D4.4 |
| WBGene00016402 | 0.027838222 | 0.94923934 | 0.97411952 | C34D4.10 |
| WBGene00016403 | 0.173804747 | 0.87509537 | 1 | grsp-3 |
| WBGene00016404 | -0.648588394 | 0.00684202 | 0.03911669 | mutd-1 |
| WBGene00016405 | 0.58297537 | 0.00881938 | 0.04700445 | hecd-1 |
| WBGene00016406 | -0.093655495 | 0.805005 | 0.89368931 | C34D10.1 |
| WBGene00016407 | 1.717082488 | 2.95E-08 | 1.5687E-06 | C34D10.2 |
| WBGene00016408 | 0.119542834 | 0.50389757 | 0.69112296 | prmt-5 |
| WBGene00016409 | 0.167914869 | 0.403976 | 0.6112175 | sumv-1 |
| WBGene00016410 | -0.22461868 | 0.93331196 | 1 | C34E10.9 |
| WBGene00016411 | -0.462988084 | 0.10694706 | 0.26658958 | C34E10.10 |
| WBGene00016412 | -0.546394744 | 0.00826451 | 0.04485444 | mrps-26 |
| WBGene00016413 | -0.156269293 | 0.96924949 | 1 | C34F11.1 |
| WBGene00016414 | 0.688793783 | 0.86378755 | 1 | C34F11.2 |
| WBGene00016415 | 0.423111201 | 0.0169638 | 0.07591504 | ampd-1 |
| WBGene00016416 | 2.67983576 | 0.10351824 | 1 | C34F11.5 |
| WBGene00016417 | 0.68802142 | 0.0947282 | 0.24643135 | C34F11.8 |
| WBGene00016418 | 0.194333041 | 0.30706142 | 0.51584788 | hpo-27 |

| | | | | |
|---|---|---|---|---|
| WBGene00016419 | 0.126240769 | 0.61474478 | 0.77514863 | tyr-4 |
| WBGene00016420 | 0.688793783 | 0.86378755 | 1 | C34G6.3 |
| WBGene00016421 | -0.07256007 | 0.76882093 | 0.87045913 | cdc-7 |
| WBGene00016422 | -0.272287816 | 0.29545113 | 0.50569115 | noah-1 |
| WBGene00016423 | 0.192388867 | 0.71442037 | 0.83869442 | tag-275 |
| WBGene00016424 | 0.884182255 | 0.04699957 | 0.15430552 | C34H4.1 |
| WBGene00016425 | 1.155270857 | 0.00189754 | 0.01495009 | C34H4.2 |
| WBGene00016427 | 0.175777961 | 0.69795632 | 0.82898529 | C34H4.5 |
| WBGene00016428 | 1.336148008 | 0.04657178 | 0.15310586 | dmsr-7 |
| WBGene00016429 | -1.325920989 | 0.29725342 | 1 | C35A11.2 |
| WBGene00016431 | 0.789457954 | 0.03249862 | 0.12007479 | C35A11.4 |
| WBGene00016432 | -0.15347124 | 0.48144044 | 0.67289112 | C35B1.2 |
| WBGene00016433 | -0.156269293 | 0.96924949 | 1 | C35B1.3 |
| WBGene00016434 | -0.908922563 | 0.00165073 | 0.0134597 | C35B1.4 |
| WBGene00016435 | -0.337476772 | 0.10154805 | 0.25701373 | C35B1.5 |
| WBGene00016436 | 3.13510578 | 0.04865179 | 1 | C35B1.7 |
| WBGene00016437 | 0.983488853 | 0.80601565 | 1 | C35B1.8 |
| WBGene00016438 | 0.225830408 | 0.48376808 | 0.67458876 | C35B8.3 |
| WBGene00016439 | -0.324946588 | 0.10022284 | 0.2552025 | C35D10.1 |
| WBGene00016440 | 1.204074051 | 0.243459 | 1 | gipc-1 |
| WBGene00016442 | -0.333932671 | 0.18717321 | 0.38064823 | C35D10.5 |
| WBGene00016443 | -0.472689232 | 0.03344471 | 0.12223126 | C35D10.6 |
| WBGene00016444 | -0.299366587 | 0.14247268 | 0.32034267 | C35D10.7 |
| WBGene00016445 | 0.502997185 | 0.51087438 | 0.69734404 | C35D10.8 |
| WBGene00016446 | -0.466712121 | 0.02455878 | 0.09899339 | C35D10.10 |
| WBGene00016447 | 0.298655687 | 0.80582251 | 1 | msd-4 |
| WBGene00016448 | 0.72422535 | 0.25891355 | 0.46708312 | C35D10.12 |
| WBGene00016449 | -0.478573307 | 0.01036113 | 0.05323506 | C35D10.13 |
| WBGene00016450 | 1.042617217 | 0.00408317 | 0.02669559 | clec-5 |
| WBGene00016451 | 0.959021583 | 0.70422854 | 1 | clec-6 |
| WBGene00016452 | -0.274146718 | 0.37016909 | 0.57766361 | C35D10.17 |
| WBGene00016453 | -1.560438514 | 2.81E-12 | 3.60E-10 | vet-2 |
| WBGene00016454 | 0.529458986 | 0.18555454 | 0.37815427 | C35E7.2 |
| WBGene00016455 | -1.25744113 | 0.08008629 | 0.22103378 | C35E7.3 |
| WBGene00016456 | 0.350403875 | 0.31194477 | 0.52009358 | C35E7.4 |
| WBGene00016457 | -1.401420799 | 5.70E-11 | 5.67E-09 | C35E7.5 |
| WBGene00016458 | -0.49048034 | 0.10333428 | 0.26050281 | C35E7.6 |
| WBGene00016459 | 0.377828795 | 0.92526585 | 1 | C35E7.7 |
| WBGene00016460 | -0.519508163 | 0.12305929 | 0.29158135 | C35E7.8 |
| WBGene00016461 | 1.122309136 | 0.35810105 | 1 | C35E7.9 |
| WBGene00016462 | -0.310604425 | 0.87613277 | 1 | C35E7.10 |
| WBGene00016463 | 0.243245852 | 0.60793173 | 0.77036604 | C35E7.11 |
| WBGene00016464 | 0.511044105 | 0.77306779 | 1 | C36B7.1 |

| | | | | |
|---|---|---|---|---|
| WBGene00016465 | 2.250785703 | 0.39404138 | 1 | C36B7.2 |
| WBGene00016466 | 0.629154436 | 0.81300308 | 1 | C36B7.3 |
| WBGene00016467 | 0.588818091 | 0.74493461 | 1 | C36B7.4 |
| WBGene00016468 | 0.355885343 | 0.7293201 | 1 | C36B7.5 |
| WBGene00016469 | 0.233803798 | 0.35342395 | 0.56208019 | C36B7.6 |
| WBGene00016471 | -0.156269293 | 0.96924949 | 1 | srab-8 |
| WBGene00016472 | -0.737462795 | 0.83433562 | 1 | srbc-6 |
| WBGene00016474 | 0.160303782 | 0.71239587 | 0.8375158 | C36C5.5 |
| WBGene00016475 | 1.216957619 | 0.74819621 | 1 | srab-9 |
| WBGene00016476 | 0.281454817 | 0.94435481 | 1 | srab-10 |
| WBGene00016477 | 0.688793783 | 0.86378755 | 1 | srab-11 |
| WBGene00016480 | 0.938982574 | 0.8146943 | 1 | srab-7 |
| WBGene00016481 | 2.298255188 | 0.03133625 | 0.11713467 | C36C5.12 |
| WBGene00016483 | -0.774842772 | 0.78069886 | 1 | C36C5.14 |
| WBGene00016485 | -0.491622664 | 0.03452475 | 0.12501383 | meg-4 |
| WBGene00016486 | -0.224684804 | 0.46742438 | 0.66157606 | fbxa-170 |
| WBGene00016487 | 1.876192691 | 0.43542138 | 1 | C36C9.4 |
| WBGene00016488 | -0.428257048 | 0.75308707 | 1 | C36C9.5 |
| WBGene00016489 | -0.242165736 | 0.30731591 | 0.51598983 | C36E6.1 |
| WBGene00016490 | -0.137667139 | 0.69788423 | 0.82898077 | C36E6.2 |
| WBGene00016491 | 0.28530093 | 0.92084641 | 1 | acdh-5 |
| WBGene00016492 | -0.445307575 | 0.46819569 | 0.66212741 | C37A2.6 |
| WBGene00016493 | -0.589047972 | 1.8485E-05 | 0.00037593 | C37A2.7 |
| WBGene00016494 | 0.23582578 | 0.29508291 | 0.50537476 | C37A2.8 |
| WBGene00016495 | -0.621942092 | 0.00484985 | 0.03049303 | C37C3.1 |
| WBGene00016496 | 0.17310837 | 0.2592012 | 0.46732429 | C37C3.2 |
| WBGene00016497 | 0.281454817 | 0.94435481 | 1 | vps-32.2 |
| WBGene00016499 | 1.157700453 | 0.09974167 | 0.25443821 | C37C3.7 |
| WBGene00016500 | 0.502901455 | 0.06149091 | 0.18566555 | memo-1 |
| WBGene00016501 | -0.23167869 | 0.15055013 | 0.3321514 | C37C3.9 |
| WBGene00016502 | -1.14715222 | 0.68932198 | 1 | C37C3.10 |
| WBGene00016503 | -0.188277616 | 0.84319991 | 0.91377278 | C37C3.11 |
| WBGene00016504 | 0.325581173 | 0.54105547 | 0.72202181 | C37C3.12 |
| WBGene00016505 | 0.313978584 | 0.45850231 | 0.65483658 | ttr-33 |
| WBGene00016506 | 0.86937398 | 0.25092981 | 0.45885383 | abhd-5.1 |
| WBGene00016507 | -0.227018115 | 0.53556307 | 0.7178681 | abhd-5.2 |
| WBGene00016508 | -0.416177163 | 0.03680939 | 0.13006483 | C37H5.5 |
| WBGene00016509 | -0.265357091 | 0.13457082 | 0.30879311 | adss-1 |
| WBGene00016511 | -0.422574796 | 0.10854598 | 0.26866795 | C37H5.13 |
| WBGene00016512 | 1.977885804 | 0.49204652 | 1 | C38C3.3 |
| WBGene00016513 | 1.294866513 | 0.00201205 | 0.01561912 | C38C3.4 |
| WBGene00016514 | 0.971647672 | 0.58244034 | 1 | C38C3.6 |
| WBGene00016517 | 1.219476362 | 0.41141192 | 1 | nhr-260 |

| | | | | |
|---|---|---|---|---|
| WBGene00016518 | -0.620463691 | 0.65590552 | 1 | C38C5.1 |
| WBGene00016519 | 1.653460415 | 0.50712258 | 1 | srz-56 |
| WBGene00016520 | -1.124654682 | 0.00521008 | 0.03215838 | C39B5.2 |
| WBGene00016521 | 0.836000103 | 0.76510684 | 1 | fbxa-62 |
| WBGene00016522 | 2.341736493 | 0.00481336 | 0.03037371 | fbxa-38 |
| WBGene00016523 | 0.006781067 | 0.99651938 | 1 | C39B5.5 |
| WBGene00016524 | -0.427205101 | 0.12367864 | 0.29268343 | C39B5.6 |
| WBGene00016525 | 1.216957619 | 0.74819621 | 1 | fbxa-12 |
| WBGene00016527 | 2.188838843 | 0.34004625 | 1 | fbxa-47 |
| WBGene00016528 | 0.281454817 | 0.94435481 | 1 | C39B5.10 |
| WBGene00016530 | 0.786496625 | 0.18178932 | 0.37382024 | C39D10.2 |
| WBGene00016531 | 0.559723198 | 0.18147552 | 0.37362968 | C39D10.3 |
| WBGene00016532 | 0.731418473 | 0.58798319 | 1 | C39D10.5 |
| WBGene00016533 | 1.62056402 | 0.08318001 | 0.22650292 | C39D10.6 |
| WBGene00016534 | 0.167897186 | 0.7155743 | 0.8397243 | C39D10.7 |
| WBGene00016535 | 0.840545463 | 0.07287402 | 0.2080328 | C39D10.8 |
| WBGene00016538 | -0.979392855 | 0.29164976 | 0.5015777 | C39F7.1 |
| WBGene00016539 | 0.740555022 | 0.01769801 | 0.07837169 | madd-2 |
| WBGene00016540 | 0.812683555 | 0.10755638 | 0.26759651 | C39F7.5 |
| WBGene00016541 | 0.483840154 | 0.8151116 | 1 | C39H7.1 |
| WBGene00016542 | 1.019158295 | 0.00014693 | 0.00207432 | C39H7.4 |
| WBGene00016544 | 0.80467518 | 0.17214269 | 0.36165156 | math-18 |
| WBGene00016545 | -0.763920166 | 0.4656171 | 1 | C40A11.2 |
| WBGene00016546 | -1.864046434 | 0.06090909 | 0.18450529 | C40A11.3 |
| WBGene00016547 | -0.864276909 | 0.36372621 | 1 | C40A11.4 |
| WBGene00016549 | -0.636552455 | 0.54260387 | 1 | C40A11.6 |
| WBGene00016550 | -3.065382488 | 0.06669245 | 1 | C40A11.7 |
| WBGene00016551 | -0.346418583 | 0.8626823 | 1 | C40A11.8 |
| WBGene00016553 | -0.077275217 | 0.96876611 | 1 | mir-8199 |
| WBGene00016554 | -0.156269293 | 0.96924949 | 1 | math-19 |
| WBGene00016555 | 0.83198307 | 0.09002276 | 0.23835296 | math-20 |
| WBGene00016556 | 1.381110239 | 0.51108747 | 1 | math-21 |
| WBGene00016558 | -0.129384034 | 0.79351703 | 0.88593495 | pks-1 |
| WBGene00016559 | 0.964443638 | 0.05467272 | 0.17090151 | C41A3.2 |
| WBGene00016561 | 0.368339458 | 0.28426898 | 0.49417901 | eri-6 |
| WBGene00016562 | -0.014978247 | 0.93996434 | 0.9689509 | C41D11.3 |
| WBGene00016564 | -0.768577901 | 0.0008582 | 0.00816568 | cmt-1 |
| WBGene00016565 | 1.034088547 | 0.13181585 | 0.3043031 | C41D11.6 |
| WBGene00016566 | 0.011303091 | 0.96137977 | 0.9799439 | eri-7 |
| WBGene00016567 | -0.473103517 | 0.14751004 | 0.32782709 | C41D11.9 |
| WBGene00016568 | -0.733476902 | 0.00371723 | 0.02489296 | C41G11.1 |
| WBGene00016570 | 1.446144562 | 0.02868071 | 0.10965404 | gnrr-4 |
| WBGene00016572 | 1.840536985 | 0.04370891 | 0.14608272 | C41H7.1 |

| | | | | |
|---|---|---|---|---|
| WBGene00016572 | 1.210578306 | 0.40738454 | 1 | C41H7.2 |
| WBGene00016573 | -0.431051625 | 0.11508221 | 0.27930954 | C41H7.3 |
| WBGene00016574 | -0.502006028 | 0.19606686 | 0.39287799 | C41H7.4 |
| WBGene00016575 | -0.019711589 | 0.97186406 | 0.98541854 | C41H7.5 |
| WBGene00016576 | -0.568675866 | 0.01446886 | 0.06795608 | C41H7.6 |
| WBGene00016577 | 1.605474644 | 0.00012201 | 0.00178466 | clec-3 |
| WBGene00016578 | -0.899950681 | 0.40633456 | 1 | sdz-5 |
| WBGene00016579 | 2.799084385 | 0.17307183 | 1 | sre-14 |
| WBGene00016580 | -0.609939305 | 0.57919268 | 1 | C42C1.2 |
| WBGene00016581 | -0.736789399 | 0.72029863 | 1 | C42C1.3 |
| WBGene00016582 | 0.119985788 | 0.58700448 | 0.75591074 | C42C1.4 |
| WBGene00016583 | -0.247336735 | 0.2319365 | 0.43758569 | tag-335 |
| WBGene00016585 | 0.682632319 | 0.09471642 | 0.24643135 | oac-59 |
| WBGene00016586 | -0.375931274 | 0.02640822 | 0.10381629 | C42C1.8 |
| WBGene00016587 | 0.298785709 | 0.79078821 | 1 | C42C1.9 |
| WBGene00016588 | -0.214047825 | 0.29740461 | 0.50756106 | hpo-12 |
| WBGene00016589 | -0.196859961 | 0.28724071 | 0.49711057 | C42C1.11 |
| WBGene00016590 | -1.062741049 | 0.00017094 | 0.0023319 | C42C1.12 |
| WBGene00016591 | -0.264201346 | 0.36844171 | 0.57602018 | C42C1.13 |
| WBGene00016592 | 0.02687327 | 0.91486485 | 0.95571968 | erl-1 |
| WBGene00016594 | 0.188934982 | 0.2362802 | 0.44244693 | C42D4.1 |
| WBGene00016595 | -1.618253353 | 0.04043591 | 0.13856402 | C42D4.2 |
| WBGene00016596 | -1.919993778 | 0.0010609 | 0.00965547 | C42D4.3 |
| WBGene00016597 | -1.094201388 | 0.7816772 | 1 | clec-179 |
| WBGene00016598 | -0.787253477 | 0.62392788 | 1 | C42D4.13 |
| WBGene00016599 | 0.751567599 | 0.20551217 | 0.40498673 | C42D8.1 |
| WBGene00016600 | 0.694453097 | 0.64527561 | 1 | ets-5 |
| WBGene00016601 | -0.433767848 | 0.03171237 | 0.11812001 | acin-1 |
| WBGene00016602 | -0.244986699 | 0.25890057 | 0.46708312 | mus-81 |
| WBGene00016603 | 0.103953159 | 0.60992899 | 0.77193244 | met-1 |
| WBGene00016605 | -0.060379368 | 0.9741779 | 1 | C43E11.5 |
| WBGene00016606 | -0.073799172 | 0.73250795 | 0.85039004 | exoc-7 |
| WBGene00016607 | -0.469815864 | 0.05141381 | 0.16370707 | C43E11.9 |
| WBGene00016608 | -0.208049886 | 0.21979448 | 0.42263617 | cogc-5 |
| WBGene00016609 | -0.501431753 | 0.0426308 | 0.14385895 | C43E11.12 |
| WBGene00016610 | 0.334753455 | 0.05738476 | 0.17727949 | paqr-1 |
| WBGene00016611 | 0.534848154 | 0.01560955 | 0.07165653 | bicd-1 |
| WBGene00016612 | 1.653460415 | 0.50712258 | 1 | C43G2.3 |
| WBGene00016613 | 1.402534934 | 0.06960688 | 0.20158943 | best-9 |
| WBGene00016614 | 0.065668443 | 0.86111918 | 0.92512591 | trpp-12 |
| WBGene00016616 | -0.714812776 | 0.09299532 | 0.24311875 | C43H6.3 |
| WBGene00016617 | -0.164176162 | 0.64822434 | 0.79729427 | C43H6.4 |
| WBGene00016619 | 2.396460014 | 0.09518684 | 1 | C43H6.6 |

| | | | | |
|---|---|---|---|---|
| WBGene00016620 | -0.221903288 | 0.60516165 | 0.76897128 | dhhc-9 |
| WBGene00016621 | -0.622798715 | 0.06227011 | 0.18694892 | arch-1 |
| WBGene00016622 | -0.177305346 | 0.42760833 | 0.6306514 | mafr-1 |
| WBGene00016623 | -0.632381495 | 0.01018037 | 0.05252284 | psmd-9 |
| WBGene00016624 | -0.354143983 | 0.12513066 | 0.29486936 | C44B7.2 |
| WBGene00016625 | 0.345250813 | 0.65076904 | 0.79864418 | aff-1 |
| WBGene00016626 | 0.225234468 | 0.79488734 | 0.88641923 | clhm-1 |
| WBGene00016627 | -0.226290275 | 0.16028886 | 0.34561687 | C44B7.5 |
| WBGene00016628 | 1.200589116 | 0.05812408 | 0.1786543 | slc-36.5 |
| WBGene00016629 | -0.888539003 | 0.03846221 | 0.1339509 | C44B7.7 |
| WBGene00016630 | 0.356077531 | 0.01222059 | 0.0600689 | acer-1 |
| WBGene00016631 | 0.74960704 | 0.00950498 | 0.04993506 | C44B7.11 |
| WBGene00016632 | -0.357583961 | 0.05301098 | 0.16708366 | C44B7.12 |
| WBGene00016633 | -0.996132005 | 0.4936854 | 1 | C44B11.1 |
| WBGene00016634 | 1.483210666 | 0.02390786 | 0.09717619 | C44B11.4 |
| WBGene00016635 | 0.386071486 | 0.50012547 | 0.68804797 | C44B11.6 |
| WBGene00016636 | -0.512001881 | 0.00050407 | 0.00539265 | perm-2 |
| WBGene00016637 | 0.32595502 | 0.91626577 | 1 | C44B12.3 |
| WBGene00016638 | -0.45196803 | 0.00702879 | 0.0398769 | perm-4 |
| WBGene00016639 | 0.569336426 | 0.71815897 | 1 | C44B12.6 |
| WBGene00016641 | -0.413527858 | 0.16945333 | 0.35748616 | C44C1.1 |
| WBGene00016642 | -0.056746738 | 0.84825853 | 0.91671496 | chil-10 |
| WBGene00016643 | 0.009060707 | 0.97701063 | 0.98807881 | vps-45 |
| WBGene00016644 | -0.870427364 | 0.00740338 | 0.04136422 | abhd-3.2 |
| WBGene00016645 | 0.838685901 | 0.19093514 | 0.38569596 | C44C1.6 |
| WBGene00016650 | 0.895558374 | 3.52E-07 | 1.3864E-05 | ubr-4 |
| WBGene00016652 | -0.382380972 | 0.03190085 | 0.11847911 | got-2.1 |
| WBGene00016653 | -0.387022333 | 0.0163119 | 0.07392413 | C44E4.4 |
| WBGene00016654 | 0.292826368 | 0.20310142 | 0.4022601 | C44E4.5 |
| WBGene00016655 | -0.432084021 | 0.00700472 | 0.03978359 | acbp-1 |
| WBGene00016657 | 0.085781665 | 0.83962859 | 0.91211511 | C44E12.1 |
| WBGene00016658 | -0.944217427 | 0.00417974 | 0.0271952 | C45B2.1 |
| WBGene00016659 | -0.902230853 | 0.10527481 | 0.26412613 | C45B2.2 |
| WBGene00016661 | 0.32847511 | 0.27362053 | 0.48215195 | C45B2.6 |
| WBGene00016662 | -1.901856996 | 0.64072034 | 1 | C45B2.8 |
| WBGene00016663 | 1.140033204 | 0.06798876 | 0.19828286 | C45E1.4 |
| WBGene00016664 | 0.258332482 | 0.48559527 | 0.67571486 | C45E5.1 |
| WBGene00016665 | 0.857979111 | 0.52621994 | 1 | chil-11 |
| WBGene00016667 | 0.37498883 | 0.2859509 | 0.49589056 | C45E5.4 |
| WBGene00016669 | 3.145498002 | 2.53E-23 | 1.18E-20 | ilys-2 |
| WBGene00016670 | 5.637856895 | 4.15E-48 | 7.20E-45 | ilys-3 |
| WBGene00016671 | 0.285930674 | 0.67904513 | 0.81647553 | C45G7.4 |
| WBGene00016673 | 1.116613983 | 0.69067248 | 1 | C45G9.1 |

| | | | | |
|---|---|---|---|---|
| WBGene00016674 | -0.715146715 | 0.00502323 | 0.03124336 | C45G9.2 |
| WBGene00016675 | -1.094201388 | 0.7816772 | 1 | C45G9.4 |
| WBGene00016676 | -0.238191642 | 0.25298196 | 0.46090245 | C45G9.5 |
| WBGene00016677 | -0.079752665 | 0.7839006 | 0.88074562 | C45G9.6 |
| WBGene00016678 | -0.017457747 | 0.96969914 | 0.98437435 | C45G9.7 |
| WBGene00016679 | 1.576273745 | 0.29412207 | 1 | C45G9.8 |
| WBGene00016680 | -0.760943105 | 0.7243725 | 1 | C45G9.9 |
| WBGene00016681 | -0.156269293 | 0.96924949 | 1 | C45G9.10 |
| WBGene00016682 | 1.198258193 | 0.3565764 | 1 | C45G9.11 |
| WBGene00016683 | 0.025717399 | 0.98468249 | 1 | C45G9.12 |
| WBGene00016684 | 2.137747113 | 0.00220271 | 0.0167458 | nlp-43 |
| WBGene00016686 | 0.281454817 | 0.94435481 | 1 | cyp-33C1 |
| WBGene00016692 | -1.901856996 | 0.64072034 | 1 | srbc-24 |
| WBGene00016693 | 0.377828795 | 0.92526585 | 1 | srbc-20 |
| WBGene00016695 | -0.748092542 | 0.1261199 | 0.29628902 | C45H4.14 |
| WBGene00016696 | 0.983488853 | 0.80601565 | 1 | srbc-19 |
| WBGene00016697 | 2.870921301 | 0.19896536 | 1 | cyp-33C2 |
| WBGene00016698 | -0.626833103 | 0.80597175 | 1 | C46A5.1 |
| WBGene00016699 | 2.862752466 | 0.07770203 | 1 | del-7 |
| WBGene00016700 | -0.282695292 | 0.58074323 | 0.75152458 | C46A5.4 |
| WBGene00016701 | -0.44897961 | 0.14652316 | 0.32599187 | C46A5.5 |
| WBGene00016702 | -0.477194876 | 0.08103991 | 0.22255272 | C46A5.6 |
| WBGene00016703 | 0.270330351 | 0.74685031 | 0.85907244 | rdl-1 |
| WBGene00016704 | 0.462180839 | 0.09721851 | 0.25046061 | hosl-1 |
| WBGene00016705 | -0.206233391 | 0.63574202 | 0.78857212 | C46C11.2 |
| WBGene00016706 | 0.812139126 | 0.07353594 | 0.20913483 | C46C11.3 |
| WBGene00016707 | 1.413734769 | 0.68804531 | 1 | C46E10.1 |
| WBGene00016708 | -1.969805064 | 0.62862569 | 1 | C46E10.2 |
| WBGene00016710 | -1.069104123 | 0.00043039 | 0.00475153 | fbxc-52 |
| WBGene00016711 | 0.188507805 | 0.88103016 | 1 | fbxc-53 |
| WBGene00016712 | -1.211186147 | 0.202701 | 0.4016636 | C46E10.8 |
| WBGene00016713 | 0.196601099 | 0.76261867 | 0.86718678 | C46E10.9 |
| WBGene00016714 | -0.116489068 | 0.84490668 | 0.91491367 | C46F2.1 |
| WBGene00016716 | 1.032298 | 1.9575E-05 | 0.00039479 | acs-17 |
| WBGene00016717 | 0.458121483 | 0.30032151 | 0.50995852 | math-22 |
| WBGene00016718 | -0.156269293 | 0.96924949 | 1 | math-23 |
| WBGene00016719 | 0.296837583 | 0.60302173 | 0.76781108 | math-24 |
| WBGene00016720 | -0.156269293 | 0.96924949 | 1 | math-25 |
| WBGene00016721 | 0.463084902 | 0.0384094 | 0.13384169 | C46G7.1 |
| WBGene00016722 | -0.458639261 | 0.03027109 | 0.11417252 | C46G7.2 |
| WBGene00016724 | -1.156241549 | 0.61231033 | 1 | C46H3.1 |
| WBGene00016725 | 0.085252865 | 0.8040227 | 0.89286011 | C46H3.2 |
| WBGene00016728 | 1.409575658 | 0.0035075 | 0.02383021 | C46H11.2 |

| | | | | |
|---|---|---|---|---|
| WBGene00016729 | -1.901856996 | 0.64072034 | 1 | C46H11.3 |
| WBGene00016730 | 0.983488853 | 0.80601565 | 1 | C46H11.6 |
| WBGene00016731 | -2.141825396 | 0.08584098 | 1 | C46H11.7 |
| WBGene00016732 | -0.628729193 | 0.61913606 | 1 | phat-1 |
| WBGene00016733 | -1.080310635 | 0.05126042 | 0.1635488 | phat-2 |
| WBGene00016734 | 1.078373501 | 0.03098047 | 0.11626081 | C46H11.10 |
| WBGene00016735 | 0.449514553 | 0.07964753 | 0.22037389 | fhod-1 |
| WBGene00016738 | 1.155925129 | 0.22460261 | 0.42873336 | C47D2.1 |
| WBGene00016739 | 0.336359835 | 0.11749127 | 0.28245932 | pitr-1 |
| WBGene00016740 | -0.79128026 | 0.00330476 | 0.02277135 | C48B6.2 |
| WBGene00016741 | -0.476573962 | 0.01570921 | 0.07187884 | C48B6.3 |
| WBGene00016742 | 0.377828795 | 0.92526585 | 1 | C48B6.4 |
| WBGene00016743 | 0.688793783 | 0.86378755 | 1 | srb-15 |
| WBGene00016744 | 0.516510633 | 0.58279575 | 0.75313711 | bbs-9 |
| WBGene00016745 | -0.389070148 | 0.42939738 | 0.63207002 | C48B6.9 |
| WBGene00016746 | -0.526464473 | 0.00728381 | 0.04092213 | C48B6.10 |
| WBGene00016747 | 2.027932919 | 0.17840874 | 1 | nmur-1 |
| WBGene00016748 | 2.376949396 | 0.07671903 | 1 | aexr-3 |
| WBGene00016749 | 0.347923654 | 0.14979824 | 0.33125184 | C48E7.1 |
| WBGene00016750 | -0.260983113 | 0.19445248 | 0.3905455 | let-611 |
| WBGene00016751 | 1.149566447 | 0.0064731 | 0.03754891 | C48E7.6 |
| WBGene00016752 | 1.584577408 | 0.63258867 | 1 | C48E7.7 |
| WBGene00016753 | 2.133182101 | 0.3921092 | 1 | oac-9 |
| WBGene00016754 | -0.751319648 | 0.01374841 | 0.06525389 | cebp-2 |
| WBGene00016755 | 1.958082604 | 0.13399312 | 1 | C49A9.1 |
| WBGene00016756 | 0.241916206 | 0.52090027 | 0.70591027 | C49A9.2 |
| WBGene00016757 | 2.100514161 | 0.00011788 | 0.00173663 | C49A9.3 |
| WBGene00016758 | 0.962475137 | 0.01962284 | 0.08430322 | C49A9.4 |
| WBGene00016759 | 0.396535404 | 0.37150034 | 0.5790712 | C49A9.5 |
| WBGene00016760 | 0.345381861 | 0.37340656 | 0.5812823 | C49A9.6 |
| WBGene00016761 | 1.601117231 | 0.23113156 | 1 | tkr-2 |
| WBGene00016762 | 1.930137656 | 0.00501295 | 0.03119541 | ugt-24 |
| WBGene00016763 | 0.418213545 | 0.16749678 | 0.35493773 | C49A9.9 |
| WBGene00016764 | -0.856884747 | 0.01063078 | 0.05425553 | C49A9.10 |
| WBGene00016765 | 0.916065526 | 0.64867207 | 1 | C49C8.1 |
| WBGene00016766 | -0.6639345 | 0.40806316 | 0.6141434 | C49C8.2 |
| WBGene00016767 | -0.054972649 | 0.92227357 | 0.9593272 | C49C8.3 |
| WBGene00016768 | 0.900020402 | 0.03437586 | 0.12477303 | cyp-33E1 |
| WBGene00016769 | 0.748543568 | 0.00017256 | 0.0023435 | C49C8.5 |
| WBGene00016772 | 0.747845955 | 0.85215965 | 1 | nhr-166 |
| WBGene00016773 | -0.156269293 | 0.96924949 | 1 | oac-10 |
| WBGene00016776 | 0.574508983 | 0.8551971 | 1 | oac-11 |
| WBGene00016778 | 2.932343137 | 0.13706871 | 1 | nep-3 |

| | | | | |
|---|---|---|---|---|
| WBGene00016779 | -1.989041311 | 0.06714775 | 1 | sdz-7 |
| WBGene00016781 | 1.29792646 | 5.79E-15 | 1.10E-12 | C49G7.3 |
| WBGene00016782 | 1.958347245 | 1.78E-11 | 1.95E-09 | phat-3 |
| WBGene00016783 | 1.118259437 | 0.33613421 | 1 | irg-2 |
| WBGene00016785 | 0.939603405 | 0.4886411 | 1 | C49G7.7 |
| WBGene00016786 | 2.322324086 | 0.17831324 | 1 | cyp-35A4 |
| WBGene00016788 | 1.031110483 | 0.03518398 | 0.12649354 | C49G7.10 |
| WBGene00016789 | 0.233739675 | 0.93575889 | 1 | djr-1.2 |
| WBGene00016790 | -0.746724221 | 0.00022555 | 0.00289164 | C49H3.3 |
| WBGene00016791 | -0.062563818 | 0.86633881 | 0.92799221 | C49H3.4 |
| WBGene00016792 | -0.307657412 | 0.16003251 | 0.3453741 | C49H3.6 |
| WBGene00016793 | -0.025793952 | 0.92148803 | 0.95884274 | arp-11 |
| WBGene00016794 | -0.232665793 | 0.31877243 | 0.52677503 | C49H3.9 |
| WBGene00016795 | -0.095446774 | 0.93543138 | 1 | C49H3.12 |
| WBGene00016796 | -0.124396486 | 0.63389487 | 0.78783996 | cec-2 |
| WBGene00016797 | 0.452204972 | 0.50382765 | 0.69110513 | C50A2.3 |
| WBGene00016798 | -0.07748118 | 0.98067866 | 1 | ets-8 |
| WBGene00016799 | -0.116733001 | 0.62318663 | 0.78057453 | C50C3.1 |
| WBGene00016800 | 0.591463696 | 0.78046256 | 1 | C50C3.2 |
| WBGene00016801 | 1.344204165 | 0.01781431 | 0.07879181 | C50C3.5 |
| WBGene00016803 | -0.386094589 | 0.01184356 | 0.05876281 | bath-42 |
| WBGene00016805 | -0.320804994 | 0.57917227 | 0.75037463 | C50D2.1 |
| WBGene00016806 | 1.281346434 | 0.09761591 | 0.25109211 | C50D2.2 |
| WBGene00016807 | 0.377828795 | 0.92526585 | 1 | C50D2.3 |
| WBGene00016808 | -0.50308914 | 0.02771691 | 0.10737427 | C50D2.5 |
| WBGene00016809 | 0.17140975 | 0.69858959 | 0.829494 | C50D2.6 |
| WBGene00016810 | 0.457389534 | 0.04246579 | 0.14359838 | C50D2.7 |
| WBGene00016811 | -0.240731316 | 0.32805247 | 0.53663232 | C50D2.8 |
| WBGene00016812 | -0.47018539 | 0.03404609 | 0.12394409 | C50D2.9 |
| WBGene00016816 | -0.105661805 | 0.64253894 | 0.79343626 | C50E3.5 |
| WBGene00016817 | 0.327118551 | 0.6097594 | 0.77179825 | C50E3.6 |
| WBGene00016818 | 0.688793783 | 0.86378755 | 1 | C50E3.7 |
| WBGene00016820 | 0.281454817 | 0.94435481 | 1 | C50E3.9 |
| WBGene00016822 | -0.740555754 | 0.01339156 | 0.06393509 | C50E3.11 |
| WBGene00016823 | -1.249170677 | 0.00025431 | 0.00317005 | C50E3.12 |
| WBGene00016824 | -0.838009991 | 0.0012489 | 0.01087729 | C50E3.13 |
| WBGene00016825 | 1.025181888 | 0.79790347 | 1 | C50E10.1 |
| WBGene00016826 | -0.62146648 | 0.85925752 | 1 | C50E10.2 |
| WBGene00016827 | -1.094201388 | 0.7816772 | 1 | sre-53 |
| WBGene00016828 | -1.969805064 | 0.62862569 | 1 | sre-52 |
| WBGene00016829 | -0.156269293 | 0.96924949 | 1 | sre-54 |
| WBGene00016830 | -1.159714244 | 0.75790634 | 1 | sre-55 |
| WBGene00016831 | 0.281454817 | 0.94435481 | 1 | sre-56 |

| | | | | |
|---|---|---|---|---|
| WBGene00016835 | 0.247318604 | 0.91814609 | 1 | C50F2.1 |
| WBGene00016836 | -0.055340044 | 0.78381191 | 0.88074562 | C50F2.2 |
| WBGene00016837 | -0.34093705 | 0.04438085 | 0.14758366 | syf-1 |
| WBGene00016838 | 0.087513917 | 0.81985211 | 0.90251377 | C50F2.4 |
| WBGene00016839 | -0.156269293 | 0.96924949 | 1 | C50F2.5 |
| WBGene00016840 | 2.073207122 | 0.00880755 | 0.04700329 | C50F2.7 |
| WBGene00016841 | 0.263082863 | 0.57732635 | 0.74903703 | magu-3 |
| WBGene00016842 | -0.303959306 | 0.74741057 | 0.85922845 | npr-35 |
| WBGene00016843 | 0.281454817 | 0.94435481 | 1 | C50F7.3 |
| WBGene00016844 | -0.627031664 | 8.4015E-05 | 0.00132815 | sucg-1 |
| WBGene00016845 | 2.736335902 | 8.77E-10 | 6.99E-08 | C50F7.5 |
| WBGene00016846 | 0.727377628 | 0.22138243 | 0.42434731 | C50F7.6 |
| WBGene00016847 | 0.688793783 | 0.86378755 | 1 | C50F7.9 |
| WBGene00016848 | 0.994837716 | 0.00106345 | 0.00967141 | klo-1 |
| WBGene00016849 | 0.331122451 | 0.553137 | 0.73099339 | acs-21 |
| WBGene00016852 | 0.377828795 | 0.92526585 | 1 | srt-7 |
| WBGene00016855 | 1.955750377 | 0.44285624 | 1 | C50H11.8 |
| WBGene00016858 | 0.35134761 | 0.87335989 | 1 | C50H11.13 |
| WBGene00016860 | 0.602740854 | 0.17881451 | 0.3708553 | cyp-33C9 |
| WBGene00016862 | 0.053611025 | 0.85683958 | 0.92175167 | C52A10.1 |
| WBGene00016863 | -0.950836778 | 0.41264458 | 1 | C52A10.2 |
| WBGene00016864 | -0.822971915 | 0.35422059 | 0.56263539 | C52A10.3 |
| WBGene00016865 | 0.789216284 | 0.04629218 | 0.1524365 | ets-9 |
| WBGene00016866 | 0.532021967 | 0.17040371 | 0.35928262 | coel-1 |
| WBGene00016867 | -0.049773546 | 0.80937409 | 0.89609686 | C52B9.4 |
| WBGene00016868 | 0.139652178 | 0.59451678 | 0.76107426 | C52B9.8 |
| WBGene00016869 | -0.982255077 | 0.6464653 | 1 | C52B9.10 |
| WBGene00016870 | 0.88787957 | 0.71832755 | 1 | ets-10 |
| WBGene00016871 | -0.018953306 | 0.96797414 | 0.98377483 | inso-1 |
| WBGene00016872 | 0.478445915 | 0.41998559 | 0.62442384 | dop-4 |
| WBGene00016873 | 2.566123675 | 0.23268183 | 1 | C52B11.4 |
| WBGene00016874 | -0.319513319 | 0.28523267 | 0.49502517 | C52B11.5 |
| WBGene00016877 | -0.701374926 | 0.18005575 | 0.37228489 | C52D10.3 |
| WBGene00016879 | 0.602444243 | 0.05673238 | 0.17589067 | cnnm-1 |
| WBGene00016880 | -1.929134871 | 0.00571394 | 0.03439559 | fbxb-95 |
| WBGene00016883 | 1.23431118 | 0.07787664 | 0.21673252 | C52E2.4 |
| WBGene00016884 | 1.798107633 | 0.04132853 | 0.14075516 | C52E2.5 |
| WBGene00016885 | -1.599506587 | 0.12764652 | 1 | fbxb-97 |
| WBGene00016886 | -2.106307537 | 0.04256984 | 1 | fbxb-96 |
| WBGene00016888 | -0.015921455 | 0.92447797 | 0.96022645 | C52E12.1 |
| WBGene00016889 | 0.228552021 | 0.24364902 | 0.45114272 | lst-6 |
| WBGene00016890 | 0.01949332 | 0.97786954 | 0.98853573 | lst-5 |
| WBGene00016891 | 0.130994476 | 0.90327728 | 1 | C53A3.1 |

| | | | | |
|---|---|---|---|---|
| WBGene00016892 | 0.593017141 | 0.24814899 | 0.456292 | C53A3.2 |
| WBGene00016893 | 0.335293097 | 0.30698957 | 0.51579856 | C53B7.2 |
| WBGene00016894 | 1.127599634 | 3.71E-09 | 2.52E-07 | C53B7.3 |
| WBGene00016895 | 0.75595108 | 0.7708876 | 1 | C53B7.6 |
| WBGene00016896 | 1.783935139 | 0.03374128 | 0.12316684 | nep-4 |
| WBGene00016898 | 0.631562295 | 0.00151799 | 0.01267534 | C53C9.2 |
| WBGene00016899 | -1.336419452 | 0.35957381 | 1 | C53C11.1 |
| WBGene00016900 | 0.733807478 | 0.35319387 | 0.56202545 | C53C11.2 |
| WBGene00016902 | -0.302323188 | 0.30137447 | 0.51102357 | C53D5.1 |
| WBGene00016903 | -1.180752028 | 0.5021926 | 1 | C53D5.3 |
| WBGene00016905 | -0.089894928 | 0.8128894 | 0.89794289 | ztf-3 |
| WBGene00016906 | 0.440345086 | 0.19833475 | 0.39611484 | C53D5.5 |
| WBGene00016907 | -0.192123056 | 0.22524456 | 0.42944319 | C53H9.2 |
| WBGene00016909 | 1.889874827 | 0.16940411 | 1 | frpr-4 |
| WBGene00016911 | 0.635176371 | 0.24886112 | 0.4567087 | drn-1 |
| WBGene00016912 | 1.190446951 | 0.11228089 | 0.27460243 | clec-86 |
| WBGene00016913 | 0.556607512 | 0.03312201 | 0.12152746 | lam-2 |
| WBGene00016914 | 0.50547461 | 0.58781575 | 0.75636858 | C54D1.7 |
| WBGene00016915 | -0.244744551 | 0.84474538 | 1 | C54D2.1 |
| WBGene00016916 | 1.298320824 | 0.09867129 | 0.25284259 | C54D2.2 |
| WBGene00016917 | -0.522741973 | 0.82405702 | 1 | C54E4.1 |
| WBGene00016918 | -0.598229617 | 0.00882387 | 0.04700773 | test-1 |
| WBGene00016919 | 1.248384281 | 0.08622724 | 0.23191957 | C54E4.4 |
| WBGene00016920 | 0.383126658 | 0.25316623 | 0.46107372 | C54E4.5 |
| WBGene00016921 | -1.279666666 | 0.55824702 | 1 | C54F6.3 |
| WBGene00016923 | -0.768633663 | 0.47292289 | 1 | C54F6.5 |
| WBGene00016924 | 0.946156604 | 0.59159601 | 1 | C54F6.6 |
| WBGene00016925 | 0.377828795 | 0.92526585 | 1 | srb-18 |
| WBGene00016926 | 2.638791521 | 0.11261003 | 1 | nhr-171 |
| WBGene00016927 | 0.377828795 | 0.92526585 | 1 | nhr-172 |
| WBGene00016928 | -0.356100916 | 0.86684044 | 1 | srb-19 |
| WBGene00016929 | 3.249651047 | 0.03694943 | 1 | C54F6.12 |
| WBGene00016930 | -0.511886193 | 0.09623422 | 0.24901527 | madf-6 |
| WBGene00016931 | 2.490696505 | 0.25975012 | 1 | C54G6.2 |
| WBGene00016932 | 1.048611428 | 0.3187151 | 1 | C54G6.3 |
| WBGene00016933 | 2.365725856 | 0.35238907 | 1 | C54G7.1 |
| WBGene00016934 | 0.583889731 | 0.00681147 | 0.03904304 | mboa-3 |
| WBGene00016935 | 0.525009034 | 0.34190731 | 0.55025142 | ifta-1 |
| WBGene00016937 | -0.148296078 | 0.63812478 | 0.79031654 | tag-294 |
| WBGene00016938 | 0.046478621 | 0.97099625 | 1 | C54H2.4 |
| WBGene00016939 | -0.255004254 | 0.29396157 | 0.50437549 | C55B6.1 |
| WBGene00016942 | 0.185908997 | 0.71713182 | 0.84010542 | C55B7.3 |
| WBGene00016943 | 0.656821236 | 4.16E-07 | 1.6091E-05 | acdh-1 |

| | | | | |
|---|---|---|---|---|
| WBGene00016944 | -0.0827122 | 0.67007701 | 0.81118806 | uri-1 |
| WBGene00016945 | 1.201807198 | 0.08258959 | 0.22548238 | sulp-1 |
| WBGene00016946 | -1.082680044 | 0.67762556 | 1 | C55B7.10 |
| WBGene00016947 | -0.478999942 | 0.11076559 | 0.27222775 | C55B7.11 |
| WBGene00016948 | -0.266135403 | 0.87048172 | 1 | scrt-1 |
| WBGene00016949 | -0.668264701 | 0.85551153 | 1 | C55C2.3 |
| WBGene00016951 | -0.100567096 | 0.91347196 | 0.95516864 | C55C3.1 |
| WBGene00016953 | 1.192973345 | 0.18184518 | 0.37382024 | C55C3.3 |
| WBGene00016954 | 1.653460415 | 0.50712258 | 1 | C55C3.4 |
| WBGene00016955 | -0.044551695 | 0.80706717 | 0.89476783 | perm-5 |
| WBGene00016956 | -0.451813889 | 0.8460226 | 1 | C55C3.6 |
| WBGene00016957 | 0.694001308 | 0.00125052 | 0.01088355 | atic-1 |
| WBGene00016958 | 0.190951114 | 0.68583454 | 0.82044706 | ilys-4 |
| WBGene00016960 | -0.086860275 | 0.67321316 | 0.81328169 | vps-33.2 |
| WBGene00016961 | -0.184065501 | 0.24526098 | 0.45309251 | vps-32.1 |
| WBGene00016962 | -0.959515164 | 0.36576284 | 1 | C56C10.4 |
| WBGene00016963 | 1.116613983 | 0.69067248 | 1 | C56C10.6 |
| WBGene00016964 | -0.280653599 | 0.12712444 | 0.29763248 | C56C10.7 |
| WBGene00016965 | -0.469590562 | 0.08546972 | 0.23054671 | C56C10.9 |
| WBGene00016966 | -0.07150089 | 0.76922537 | 0.87054113 | aipr-1 |
| WBGene00016968 | 0.544402396 | 0.02362628 | 0.0964839 | C56C10.11 |
| WBGene00016968 | 0.975538809 | 0.0004008 | 0.00451601 | epg-5 |
| WBGene00016970 | -0.816596474 | 0.0182072 | 0.0800629 | C56E6.2 |
| WBGene00016971 | -0.356504744 | 0.03286202 | 0.12095748 | toe-2 |
| WBGene00016972 | 0.962897601 | 0.69232608 | 1 | C56E6.4 |
| WBGene00016973 | 2.815759761 | 0.16833793 | 1 | abch-1 |
| WBGene00016974 | 1.891386912 | 0.01984321 | 0.08506936 | lron-15 |
| WBGene00016975 | 1.822800566 | 9.2611E-06 | 0.00021336 | nhr-173 |
| WBGene00016976 | 1.977522305 | 0.04965274 | 0.15977576 | C56E10.3 |
| WBGene00016977 | 0.489468383 | 0.02404643 | 0.0975761 | akap-1 |
| WBGene00016978 | -0.276667381 | 0.44168216 | 0.64137027 | C56G2.3 |
| WBGene00016979 | 0.313449344 | 0.56018015 | 0.73669272 | C56G2.4 |
| WBGene00016980 | -0.60891069 | 0.14145613 | 0.31947994 | C56G2.5 |
| WBGene00016981 | -0.379430917 | 0.01786753 | 0.07896968 | rpn-13 |
| WBGene00016982 | 1.389963918 | 0.20842524 | 1 | C56G2.9 |
| WBGene00016983 | -0.169481746 | 0.74876515 | 0.85989006 | C56G2.15 |
| WBGene00016984 | 0.581347937 | 0.51247178 | 0.69909213 | npr-8 |
| WBGene00016985 | 0.281454817 | 0.94435481 | 1 | C56G3.2 |
| WBGene00016987 | -0.045574884 | 0.88991367 | 0.94196731 | CC8.2 |
| WBGene00016988 | 0.440395994 | 0.44239603 | 0.6420189 | CD4.1 |
| WBGene00016989 | -0.549628944 | 0.02962052 | 0.11210187 | CD4.3 |
| WBGene00016990 | -0.41718946 | 0.05977851 | 0.18182573 | vps-37 |
| WBGene00016992 | -0.367781197 | 0.17911408 | 0.37122037 | zhit-1 |

| | | | | |
|---|---|---|---|---|
| WBGene00016993 | -0.296978591 | 0.52183791 | 0.70694422 | CD4.8 |
| WBGene00016995 | -0.130189073 | 0.5702076 | 0.74447688 | acly-1 |
| WBGene00016996 | 0.126825962 | 0.80464391 | 0.89346824 | D1005.2 |
| WBGene00016997 | 1.365636414 | 0.00047247 | 0.00514464 | cebp-1 |
| WBGene00016998 | 0.267994626 | 0.7485332 | 0.85978633 | D1005.4 |
| WBGene00016999 | -0.156269293 | 0.96924949 | 1 | D1005.5 |
| WBGene00017000 | 0.042187151 | 0.97373582 | 1 | D1005.6 |
| WBGene00017001 | 0.248814215 | 0.78570859 | 0.88212391 | D1007.3 |
| WBGene00017002 | -0.811384131 | 0.01335393 | 0.06385587 | D1007.4 |
| WBGene00017003 | 0.036796309 | 0.86213069 | 0.92569082 | D1007.5 |
| WBGene00017004 | -0.07626676 | 0.71696019 | 0.84010542 | nrd-1 |
| WBGene00017005 | -0.74349884 | 0.00399557 | 0.0262643 | D1007.8 |
| WBGene00017007 | 0.502447783 | 0.11510697 | 0.27930954 | D1007.10 |
| WBGene00017009 | 1.957271014 | 0.49743227 | 1 | D1007.13 |
| WBGene00017010 | 0.769009744 | 0.0187441 | 0.08186046 | D1007.15 |
| WBGene00017011 | -0.377132167 | 0.03430182 | 0.12461351 | eaf-1 |
| WBGene00017012 | 0.032196998 | 0.87398111 | 0.93209809 | acs-22 |
| WBGene00017013 | 0.955575032 | 0.22489091 | 0.42897102 | ensh-1 |
| WBGene00017014 | -1.538600039 | 0.37104439 | 1 | dylt-2 |
| WBGene00017015 | 0.197403708 | 0.7748275 | 0.87450158 | D1014.2 |
| WBGene00017016 | -0.393723153 | 0.02109844 | 0.08894312 | snap-1 |
| WBGene00017017 | -0.469569186 | 0.08868694 | 0.23618076 | D1014.4 |
| WBGene00017018 | -0.378469532 | 0.65720058 | 0.80268551 | D1014.5 |
| WBGene00017020 | 0.616429445 | 0.81400458 | 1 | D1014.7 |
| WBGene00017022 | -0.166268272 | 0.88966683 | 1 | D1022.3 |
| WBGene00017023 | 0.087934832 | 0.76288242 | 0.86724302 | D1022.4 |
| WBGene00017024 | -0.637018156 | 0.80584146 | 1 | D1022.5 |
| WBGene00017025 | 0.200581644 | 0.39770981 | 0.60531463 | D1037.1 |
| WBGene00017026 | 1.998812433 | 0.28274713 | 1 | ipla-4 |
| WBGene00017027 | -0.720474657 | 0.2104521 | 0.41039852 | D1044.1 |
| WBGene00017028 | 0.273920487 | 0.48891728 | 0.67872065 | dex-1 |
| WBGene00017031 | -0.122465508 | 0.52639379 | 0.71073699 | D1044.6 |
| WBGene00017032 | 1.88017147 | 0.05434937 | 0.17022506 | D1044.7 |
| WBGene00017033 | 0.387384182 | 0.53330123 | 0.71646737 | nekl-4 |
| WBGene00017034 | 1.158132015 | 0.62662247 | 1 | otpl-2 |
| WBGene00017036 | 0.83647609 | 0.36589937 | 0.57350663 | D1069.1 |
| WBGene00017037 | -0.324838886 | 0.15948624 | 0.34449938 | D1069.3 |
| WBGene00017038 | 0.326198056 | 0.70079982 | 0.83122417 | dmsr-4 |
| WBGene00017039 | 1.291196387 | 0.05053737 | 0.1619784 | trk-1 |
| WBGene00017040 | 0.540852966 | 0.65958651 | 1 | D1079.1 |
| WBGene00017040 | 2.758227592 | 0.09611386 | 1 | F15A8.4 |
| WBGene00017041 | -0.457210789 | 0.322668 | 0.53107238 | D2007.1 |
| WBGene00017042 | 0.530524205 | 0.49192289 | 0.68116285 | D2007.2 |

| | | | | |
|---|---|---|---|---|
| WBGene00017044 | -0.491153933 | 0.07308216 | 0.2081864 | mrpl-18 |
| WBGene00017045 | -0.092887991 | 0.70825573 | 0.83500998 | atg-13 |
| WBGene00017046 | -0.266161054 | 0.32024298 | 0.52855764 | utx-1 |
| WBGene00017048 | 1.160857271 | 0.03396134 | 0.12374688 | D2021.4 |
| WBGene00017050 | -0.156269293 | 0.96924949 | 1 | D2024.1 |
| WBGene00017051 | -0.384589168 | 0.64557955 | 0.79536368 | afmd-1 |
| WBGene00017052 | 0.38095987 | 0.81065956 | 1 | D2024.4 |
| WBGene00017053 | -0.162239804 | 0.43028774 | 0.63273764 | D2024.5 |
| WBGene00017054 | 0.784545532 | 0.151175 | 0.33308024 | gstk-2 |
| WBGene00017055 | -0.008677305 | 0.99719169 | 1 | D2062.1 |
| WBGene00017056 | 0.281454817 | 0.94435481 | 1 | D2062.4 |
| WBGene00017058 | -0.156269293 | 0.96924949 | 1 | msrp-6 |
| WBGene00017059 | -0.245531902 | 0.94070824 | 1 | msrp-5 |
| WBGene00017060 | -0.886092878 | 0.23962579 | 0.44645107 | D2063.1 |
| WBGene00017062 | -0.502170386 | 0.01301881 | 0.06294757 | glrx-3 |
| WBGene00017063 | 0.917189172 | 0.02008572 | 0.08569202 | mctp-1 |
| WBGene00017064 | -0.337709868 | 0.06560305 | 0.19336894 | ppfr-2 |
| WBGene00017065 | 0.492734906 | 0.06842734 | 0.19922694 | D2092.4 |
| WBGene00017066 | 0.17967772 | 0.37486146 | 0.58284342 | maco-1 |
| WBGene00017068 | -1.556613775 | 0.15088233 | 1 | D2092.8 |
| WBGene00017069 | -0.850563945 | 0.08803788 | 0.23517445 | timm-17B.2 |
| WBGene00017070 | -0.345749815 | 0.07883538 | 0.21877492 | praf-3 |
| WBGene00017071 | 1.607522404 | 2.00E-09 | 1.46E-07 | aagr-1 |
| WBGene00017072 | -1.211092647 | 0.41403343 | 1 | D2096.5 |
| WBGene00017073 | -0.309014495 | 0.63443722 | 0.78783996 | D2096.6 |
| WBGene00017074 | 0.199722779 | 0.40223898 | 0.60999043 | D2096.7 |
| WBGene00017075 | -0.501476199 | 0.00016481 | 0.00226103 | nap-1 |
| WBGene00017076 | 1.024037545 | 0.10765992 | 0.26759651 | D2096.9 |
| WBGene00017077 | 1.17047246 | 0.09337044 | 0.24383963 | D2096.10 |
| WBGene00017078 | 0.644424994 | 0.26542637 | 0.47425189 | D2096.11 |
| WBGene00017079 | 0.132915575 | 0.51169982 | 0.69811749 | D2096.12 |
| WBGene00017080 | -0.064700407 | 0.74962807 | 0.86036058 | lec-12 |
| WBGene00017082 | 0.027210998 | 0.94160819 | 0.96989883 | DC2.5 |
| WBGene00017083 | -1.073037158 | 0.11486748 | 0.27897702 | kin-33 |
| WBGene00017084 | -0.569715704 | 0.00366072 | 0.02456873 | E01A2.1 |
| WBGene00017085 | -0.552886045 | 0.00033102 | 0.00391709 | E01A2.2 |
| WBGene00017087 | -0.737877947 | 0.01330431 | 0.06379446 | E01A2.5 |
| WBGene00017088 | -0.175006942 | 0.34881783 | 0.55730981 | akir-1 |
| WBGene00017089 | 0.240930569 | 0.70557861 | 0.83352339 | poml-2 |
| WBGene00017090 | 1.653460415 | 0.50712258 | 1 | E01A2.8 |
| WBGene00017091 | -0.262383365 | 0.73869358 | 0.8543246 | ttr-40 |
| WBGene00017092 | 2.299531175 | 0.01102935 | 0.05578051 | E02C12.6 |
| WBGene00017093 | 1.234922397 | 0.02526208 | 0.10089044 | E02C12.8 |

| | | | | |
|---|---|---|---|---|
| WBGene00017095 | -0.156269293 | 0.96924949 | 1 | E02C12.10 |
| WBGene00017097 | 0.281454817 | 0.94435481 | 1 | E02C12.12 |
| WBGene00017098 | -0.645889851 | 5.3513E-05 | 0.00091765 | E02D9.1 |
| WBGene00017099 | -0.008677305 | 0.99719169 | 1 | E02H9.1 |
| WBGene00017100 | 1.367823871 | 0.10034792 | 0.25533852 | E02H9.2 |
| WBGene00017101 | 0.65854754 | 0.01132651 | 0.05689497 | E02H9.3 |
| WBGene00017102 | 1.24682628 | 0.254753 | 1 | E02H9.4 |
| WBGene00017103 | 0.823233449 | 0.09163713 | 0.24107615 | klo-2 |
| WBGene00017104 | 0.229501532 | 0.87236821 | 1 | E02H9.6 |
| WBGene00017105 | -0.652040567 | 0.08340749 | 0.22681123 | E02H9.7 |
| WBGene00017106 | -1.510964854 | 0.29186283 | 1 | E02H9.9 |
| WBGene00017107 | -1.159714244 | 0.75790634 | 1 | srt-12 |
| WBGene00017108 | -0.433175767 | 0.28714462 | 0.49711057 | cyp-43A1 |
| WBGene00017110 | 1.181686151 | 0.7579682 | 1 | clec-176 |
| WBGene00017112 | 0.864899468 | 0.6533431 | 1 | E03H12.5 |
| WBGene00017113 | -0.156269293 | 0.96924949 | 1 | srz-24 |
| WBGene00017114 | 1.735991307 | 0.58152819 | 1 | E03H12.7 |
| WBGene00017116 | -1.895657511 | 0.19963314 | 1 | fbxb-78 |
| WBGene00017119 | -0.407636853 | 0.03151091 | 0.11767885 | timm-17B.1 |
| WBGene00017120 | -0.583409435 | 0.55439414 | 1 | tctn-1 |
| WBGene00017121 | -0.145456822 | 0.33788109 | 0.54587018 | cyc-2.1 |
| WBGene00017122 | 0.75595108 | 0.7708876 | 1 | pcrg-1 |
| WBGene00017123 | -0.138103288 | 0.5406651 | 0.72174866 | maoc-1 |
| WBGene00017124 | 1.362337218 | 0.0004233 | 0.00469772 | E04F6.4 |
| WBGene00017125 | -0.508970931 | 0.00088125 | 0.00833928 | acdh-12 |
| WBGene00017126 | 0.19967734 | 0.490645 | 0.68009144 | E04F6.6 |
| WBGene00017127 | -0.079749458 | 0.7254287 | 0.84572977 | E04F6.8 |
| WBGene00017128 | 0.140832534 | 0.60123114 | 0.76643713 | E04F6.9 |
| WBGene00017129 | -0.760484146 | 0.57558586 | 1 | E04F6.10 |
| WBGene00017131 | -0.239682721 | 0.55226066 | 0.73020586 | E04F6.15 |
| WBGene00017132 | -0.385868698 | 0.03230508 | 0.11943239 | tofu-6 |
| WBGene00017133 | -0.389971414 | 0.43457684 | 0.63591629 | EEED8.2 |
| WBGene00017134 | -1.10595587 | 4.31E-10 | 3.71E-08 | EEED8.3 |
| WBGene00017135 | -1.512257204 | 0.00167142 | 0.01358031 | EEED8.4 |
| WBGene00017136 | 1.514377069 | 0.2459156 | 1 | ccpp-6 |
| WBGene00017137 | 0.041429924 | 0.83064975 | 0.90824619 | pink-1 |
| WBGene00017138 | 0.167238692 | 0.35830674 | 0.56659198 | EEED8.10 |
| WBGene00017140 | -1.834628467 | 0.0005765 | 0.00596188 | EEED8.12 |
| WBGene00017141 | -0.850645147 | 0.1533758 | 0.3360649 | EEED8.13 |
| WBGene00017142 | -0.289954242 | 0.34647281 | 0.55524371 | EEED8.14 |
| WBGene00017143 | -1.485402631 | 3.9479E-05 | 0.00070991 | EEED8.15 |
| WBGene00017144 | -0.167245039 | 0.36994086 | 0.5774852 | brap-2 |
| WBGene00017146 | 1.054247932 | 0.50540455 | 1 | EGAP1.1 |

| | | | | |
|---|---|---|---|---|
| WBGene00017147 | -0.447769428 | 0.20517388 | 0.40458276 | EGAP2.1 |
| WBGene00017148 | 1.584577408 | 0.63258867 | 1 | EGAP2.2 |
| WBGene00017149 | 1.146177071 | 0.01652379 | 0.07432888 | EGAP4.1 |
| WBGene00017150 | 1.296517089 | 0.42863805 | 1 | EGAP5.1 |
| WBGene00017151 | 1.141096403 | 0.17522835 | 0.36560362 | EGAP9.3 |
| WBGene00017153 | -0.103579467 | 0.82795172 | 0.90707109 | E_BE45912.2 |
| WBGene00017154 | 0.192773637 | 0.72770597 | 0.84742683 | ugt-57 |
| WBGene00017156 | -0.279072329 | 0.7991998 | 1 | F01E11.17 |
| WBGene00017156 | -0.185651708 | 0.88789966 | 1 | F01E11.3 |
| WBGene00017157 | 1.741104146 | 0.01934645 | 0.08364859 | tyra-2 |
| WBGene00017158 | 0.106553512 | 0.60251262 | 0.7673678 | hpo-10 |
| WBGene00017159 | -0.860129543 | 6.4969E-05 | 0.00108201 | F01F1.2 |
| WBGene00017160 | 0.162629869 | 0.67543243 | 0.81441045 | F01F1.3 |
| WBGene00017161 | -0.33270029 | 0.11186105 | 0.27394993 | rabn-5 |
| WBGene00017162 | -0.163043958 | 0.30676738 | 0.51557292 | ddx-23 |
| WBGene00017163 | -0.18237075 | 0.34980085 | 0.55843947 | dnpp-1 |
| WBGene00017164 | -0.124218829 | 0.65660318 | 0.80236354 | eng-1 |
| WBGene00017165 | -0.706621873 | 0.0002231 | 0.00287244 | F01F1.11 |
| WBGene00017166 | -0.131563397 | 0.37538382 | 0.58322809 | aldo-2 |
| WBGene00017168 | 0.688793783 | 0.86378755 | 1 | F01F1.14 |
| WBGene00017169 | -0.358079693 | 0.22198188 | 0.42536017 | F01F1.15 |
| WBGene00017173 | 0.938982574 | 0.8146943 | 1 | F02C9.2 |
| WBGene00017174 | -0.361653418 | 0.78436331 | 1 | tat-6 |
| WBGene00017175 | 0.377828795 | 0.92526585 | 1 | irld-3 |
| WBGene00017176 | 1.840953796 | 0.05268482 | 0.16631471 | nmur-3 |
| WBGene00017177 | -0.418121044 | 0.03745348 | 0.13149874 | F02E8.4 |
| WBGene00017178 | -0.549094699 | 0.04104422 | 0.14013439 | atg-16.1 |
| WBGene00017179 | -1.094201388 | 0.7816772 | 1 | wht-4 |
| WBGene00017180 | 1.618758082 | 0.11121246 | 0.27293925 | F02E11.2 |
| WBGene00017181 | -1.279666666 | 0.55824702 | 1 | F02E11.3 |
| WBGene00017182 | 0.446458324 | 0.87164163 | 1 | F02E11.4 |
| WBGene00017183 | 0.377828795 | 0.92526585 | 1 | scl-15 |
| WBGene00017184 | 0.978868692 | 1.8275E-05 | 0.00037416 | ncam-1 |
| WBGene00017190 | 0.904564355 | 0.60377997 | 1 | F07B7.12 |
| WBGene00017193 | 0.291801037 | 0.4217201 | 0.62631238 | F07C3.2 |
| WBGene00017194 | -0.040688636 | 0.9747372 | 1 | F07C3.3 |
| WBGene00017195 | 0.612351251 | 0.01292916 | 0.06263017 | glo-4 |
| WBGene00017197 | 0.146294199 | 0.84431293 | 0.91451542 | F07C3.9 |
| WBGene00017198 | 1.282353134 | 0.05181016 | 0.1644251 | nhr-36 |
| WBGene00017199 | 0.973275665 | 0.2645984 | 0.47332977 | clec-45 |
| WBGene00017200 | 3.275580075 | 0.1143488 | 1 | F07C4.6 |
| WBGene00017201 | 0.328513991 | 0.3117237 | 0.51986779 | grsp-4 |
| WBGene00017203 | 0.938982574 | 0.8146943 | 1 | F07C4.10 |

| | | | | |
|---|---|---|---|---|
| WBGene00017204 | 1.008298062 | 0.68317078 | 1 | F07C4.11 |
| WBGene00017205 | 3.627507405 | 0.01282004 | 1 | F07C4.12 |
| WBGene00017207 | -0.988317891 | 0.59464747 | 1 | fbxb-6 |
| WBGene00017208 | -1.802469802 | 0.02368844 | 0.09660637 | fbxb-35 |
| WBGene00017210 | -0.274672032 | 0.262847 | 0.47151676 | F07E5.5 |
| WBGene00017211 | -1.227980237 | 0.27327013 | 1 | fbxb-36 |
| WBGene00017213 | -0.156269293 | 0.96924949 | 1 | F07E5.8 |
| WBGene00017214 | 2.544041525 | 0.13546931 | 1 | F07E5.9 |
| WBGene00017215 | 0.302223614 | 0.86921907 | 1 | F07F6.1 |
| WBGene00017216 | 0.626600838 | 0.75025778 | 1 | F07F6.2 |
| WBGene00017217 | 0.038931145 | 0.82538052 | 0.9058566 | F07F6.4 |
| WBGene00017218 | -0.479165649 | 0.78467961 | 1 | dct-5 |
| WBGene00017219 | -0.474866806 | 0.51606007 | 0.70217251 | F07F6.7 |
| WBGene00017220 | -0.151731106 | 0.48657217 | 0.67622374 | F07F6.8 |
| WBGene00017221 | -0.156269293 | 0.96924949 | 1 | dgn-3 |
| WBGene00017222 | -0.156269293 | 0.96924949 | 1 | F07G6.2 |
| WBGene00017224 | -1.969805064 | 0.62862569 | 1 | F07G6.5 |
| WBGene00017225 | -0.156269293 | 0.96924949 | 1 | fbxa-52 |
| WBGene00017226 | 1.541574718 | 0.09091754 | 0.23975454 | fbxa-53 |
| WBGene00017227 | -1.895791037 | 0.52793577 | 1 | F07G6.8 |
| WBGene00017228 | -0.62146648 | 0.85925752 | 1 | F07G11.1 |
| WBGene00017229 | -1.969805064 | 0.62862569 | 1 | F07G11.2 |
| WBGene00017230 | -0.156269293 | 0.96924949 | 1 | F07G11.3 |
| WBGene00017231 | -0.156269293 | 0.96924949 | 1 | F07G11.4 |
| WBGene00017233 | -1.544131969 | 0.51019995 | 1 | lmd-4 |
| WBGene00017236 | -0.853654364 | 0.58335991 | 1 | F08B4.4 |
| WBGene00017237 | -0.31345901 | 0.1512043 | 0.33308024 | pole-2 |
| WBGene00017238 | -0.833695433 | 2.6747E-05 | 0.00051184 | F08B4.7 |
| WBGene00017239 | -0.364571443 | 0.32940542 | 0.53812045 | F08B6.1 |
| WBGene00017240 | -0.580906094 | 0.0963796 | 0.24913639 | calu-2 |
| WBGene00017241 | -0.436565626 | 0.02337267 | 0.09567351 | pcyt-1 |
| WBGene00017243 | 0.151975277 | 0.92083651 | 1 | F08C6.5 |
| WBGene00017244 | -0.096299766 | 0.61027596 | 0.77201762 | apy-1 |
| WBGene00017245 | -0.207728031 | 0.35666696 | 0.5651371 | srpa-72 |
| WBGene00017246 | -0.148082791 | 0.73748251 | 0.85334533 | F08D12.2 |
| WBGene00017247 | 0.639777722 | 0.08279454 | 0.2258388 | F08D12.3 |
| WBGene00017248 | -1.901856996 | 0.64072034 | 1 | F08D12.4 |
| WBGene00017249 | -1.171022557 | 0.35744391 | 1 | fbxb-108 |
| WBGene00017250 | 0.019142332 | 0.96402967 | 0.98190304 | F08D12.7 |
| WBGene00017251 | -0.502184703 | 0.66114987 | 1 | fbxb-105 |
| WBGene00017252 | -1.95019002 | 0.08174179 | 1 | sdz-10 |
| WBGene00017253 | -1.574691483 | 0.01332032 | 0.06379465 | sdz-9 |
| WBGene00017254 | -2.532319158 | 0.0222806 | 1 | fbxb-111 |

| | | | | |
|---|---|---|---|---|
| WBGene00017255 | 1.388057217 | 0.42625387 | 1 | F08D12.12 |
| WBGene00017257 | -0.866244467 | 0.2708659 | 0.47962106 | F08F1.3 |
| WBGene00017258 | -0.800373189 | 0.11364708 | 0.27689482 | F08F1.4 |
| WBGene00017259 | 1.334341511 | 0.55384944 | 1 | F08F1.9 |
| WBGene00017260 | 2.061701262 | 0.03638879 | 0.12910471 | F08F3.1 |
| WBGene00017261 | 0.17528879 | 0.33726204 | 0.54530543 | acl-6 |
| WBGene00017262 | -0.346146829 | 0.10042179 | 0.25538772 | F08F3.4 |
| WBGene00017263 | -0.394156077 | 0.03501845 | 0.12611121 | F08F3.6 |
| WBGene00017264 | -0.998065977 | 0.57530013 | 1 | F08F3.8 |
| WBGene00017265 | 0.151724066 | 0.84251332 | 0.91337104 | snpc-1.3 |
| WBGene00017266 | 1.369453735 | 0.40100875 | 1 | F08F3.10 |
| WBGene00017267 | -1.895791037 | 0.52793577 | 1 | numr-2 |
| WBGene00017268 | -0.303936402 | 0.11263809 | 0.27499278 | hmgr-1 |
| WBGene00017269 | -0.479276137 | 0.03678483 | 0.13001589 | F08F8.4 |
| WBGene00017270 | -1.895791037 | 0.52793577 | 1 | numr-1 |
| WBGene00017271 | -0.934013985 | 0.67351079 | 1 | F08F8.6 |
| WBGene00017272 | -0.54442468 | 0.01551397 | 0.07134661 | F08F8.7 |
| WBGene00017273 | -0.413756089 | 0.11632523 | 0.28077626 | gos-28 |
| WBGene00017274 | -0.107160214 | 0.54700973 | 0.72587747 | F08F8.9 |
| WBGene00017275 | 0.150525604 | 0.45812796 | 0.65477918 | F08F8.10 |
| WBGene00017277 | 0.979427925 | 0.56765466 | 1 | F09C12.2 |
| WBGene00017278 | -0.582567818 | 0.77294373 | 1 | F09C12.6 |
| WBGene00017279 | 2.456496557 | 0.3218863 | 1 | F09C12.8 |
| WBGene00017280 | -0.232689843 | 0.21588635 | 0.4175683 | usp-39 |
| WBGene00017281 | 2.080963507 | 0.25449394 | 1 | F09D12.2 |
| WBGene00017282 | -0.519566595 | 0.04624119 | 0.15230989 | algn-2 |
| WBGene00017283 | -0.152904263 | 0.39427439 | 0.6021422 | F09E5.3 |
| WBGene00017284 | 0.082972484 | 0.67088276 | 0.81175878 | sec-6 |
| WBGene00017285 | -0.324957021 | 0.1462772 | 0.32562367 | F09E5.7 |
| WBGene00017286 | -0.117670232 | 0.64078717 | 0.79215935 | F09E5.8 |
| WBGene00017287 | -1.647820975 | 0.27062919 | 1 | F09E5.9 |
| WBGene00017288 | 0.39844809 | 0.24909669 | 0.45684766 | F09E5.10 |
| WBGene00017289 | -0.345093848 | 0.17626774 | 0.36745649 | F09E5.11 |
| WBGene00017290 | 1.260441087 | 0.30666704 | 1 | F09E5.12 |
| WBGene00017291 | 0.627897567 | 0.28503966 | 0.49491874 | F09E5.14 |
| WBGene00017292 | 0.256205327 | 0.72509207 | 0.8454997 | F09E5.16 |
| WBGene00017293 | -0.485487794 | 0.0260574 | 0.10302938 | bmy-1 |
| WBGene00017294 | -2.204614867 | 0.00016702 | 0.00228613 | F09E10.1 |
| WBGene00017295 | 0.281454817 | 0.94435481 | 1 | F09E10.5 |
| WBGene00017296 | 0.612842985 | 0.70061992 | 1 | F09E10.6 |
| WBGene00017297 | 0.313168562 | 0.54766589 | 0.72637225 | F09E10.7 |
| WBGene00017298 | 0.055236443 | 0.84451799 | 0.91465593 | toca-1 |
| WBGene00017299 | 0.901282259 | 0.2392824 | 0.44625615 | molo-1 |

| | | | | |
|---|---|---|---|---|
| WBGene00017300 | -0.217118315 | 0.16116663 | 0.34681392 | rpc-2 |
| WBGene00017301 | -1.018411888 | 3.67E-08 | 1.9104E-06 | hach-1 |
| WBGene00017302 | 0.661869338 | 0.02854626 | 0.10951656 | F09F7.5 |
| WBGene00017303 | 0.154765633 | 0.70836993 | 0.83503281 | F09F7.6 |
| WBGene00017304 | 0.326693092 | 0.22417626 | 0.42828072 | F09F7.7 |
| WBGene00017305 | -0.43096064 | 0.63396658 | 0.78783996 | nspb-12 |
| WBGene00017306 | 0.702713695 | 0.20342689 | 0.40270523 | F09F9.1 |
| WBGene00017307 | -0.419400706 | 0.45464896 | 0.65223834 | F09F9.2 |
| WBGene00017308 | 0.820015846 | 0.7518774 | 1 | F09F9.3 |
| WBGene00017309 | -0.01103476 | 0.9859684 | 0.99332445 | F09F9.4 |
| WBGene00017310 | 0.908614646 | 0.01006346 | 0.05210254 | F09G2.1 |
| WBGene00017311 | -0.312673468 | 0.09328208 | 0.24366131 | F09G2.2 |
| WBGene00017312 | 3.176245479 | 0.00051497 | 0.00548444 | pitr-5 |
| WBGene00017313 | -0.100649765 | 0.57927679 | 0.75037463 | cpsf-2 |
| WBGene00017314 | 0.955173283 | 0.02491682 | 0.09974122 | lgc-39 |
| WBGene00017315 | -1.343175269 | 0.42522779 | 1 | ugt-36 |
| WBGene00017316 | -0.216575651 | 0.19698034 | 0.39418795 | F09G2.8 |
| WBGene00017317 | -0.448212312 | 0.00831623 | 0.04499438 | attf-2 |
| WBGene00017319 | -0.521365644 | 0.01156398 | 0.05775973 | mrps-9 |
| WBGene00017320 | 1.370971883 | 0.00663849 | 0.03825242 | F09G8.5 |
| WBGene00017321 | -0.492775055 | 0.0852594 | 0.2302345 | F09G8.7 |
| WBGene00017322 | 1.217219757 | 2.1915E-06 | 6.3049E-05 | clec-160 |
| WBGene00017323 | 1.247575581 | 0.7549672 | 1 | mps-4 |
| WBGene00017325 | 0.983488853 | 0.80601565 | 1 | F10C1.3 |
| WBGene00017326 | 0.202218961 | 0.850054 | 1 | dmd-5 |
| WBGene00017327 | 0.192956851 | 0.81185605 | 0.89724443 | F10C1.8 |
| WBGene00017328 | -0.444368292 | 0.00493989 | 0.0309367 | F10C5.2 |
| WBGene00017329 | 1.249857034 | 0.00382892 | 0.0253473 | ugt-39 |
| WBGene00017331 | 0.818631443 | 0.27069887 | 0.47962106 | ugt-40 |
| WBGene00017332 | 0.391946485 | 0.75991507 | 1 | ugt-37 |
| WBGene00017333 | 1.413734769 | 0.68804531 | 1 | ugt-38 |
| WBGene00017334 | 0.483840154 | 0.8151116 | 1 | F10D2.8 |
| WBGene00017335 | 0.118317406 | 0.77692808 | 0.8756669 | F10D2.10 |
| WBGene00017336 | -0.437057457 | 0.40550938 | 0.61226986 | ugt-41 |
| WBGene00017338 | 1.169740023 | 0.10052286 | 0.2555912 | F10D7.1 |
| WBGene00017339 | 0.395008309 | 0.285066 | 0.49491874 | F10D7.2 |
| WBGene00017340 | 0.285279001 | 0.52567867 | 0.71019659 | F10D7.3 |
| WBGene00017341 | 1.240646685 | 0.40599743 | 1 | F10D7.4 |
| WBGene00017342 | 0.150407873 | 0.62965129 | 0.78510797 | F10D7.5 |
| WBGene00017343 | -0.61758079 | 0.60804983 | 1 | F10E7.1 |
| WBGene00017344 | 0.185981312 | 0.64577594 | 0.79536368 | F10E7.2 |
| WBGene00017347 | -0.444300277 | 0.02700071 | 0.10550873 | F10E7.5 |
| WBGene00017348 | -0.398048197 | 0.09723843 | 0.25046061 | F10E7.6 |

| | | | | |
|---|---|---|---|---|
| WBGene00017349 | 0.526689001 | 0.0003206 | 0.00380858 | farl-11 |
| WBGene00017350 | 0.644458311 | 0.08225313 | 0.22486721 | F10E7.9 |
| WBGene00017351 | -0.358514592 | 0.84021732 | 1 | cutl-5 |
| WBGene00017352 | -0.494585812 | 0.28666914 | 0.49685225 | F10E7.11 |
| WBGene00017353 | 0.983208301 | 0.23282115 | 0.43851271 | F10E9.1 |
| WBGene00017354 | -1.047960184 | 0.4215535 | 1 | F10E9.2 |
| WBGene00017355 | -0.463657208 | 0.02162323 | 0.0907458 | F10E9.3 |
| WBGene00017356 | -0.59844988 | 0.00686987 | 0.03921347 | F10E9.4 |
| WBGene00017357 | -0.73715116 | 0.018028 | 0.07947638 | F10E9.5 |
| WBGene00017358 | -0.529718997 | 0.02910098 | 0.11086131 | F10E9.7 |
| WBGene00017359 | 2.201543524 | 0.41352885 | 1 | F10E9.10 |
| WBGene00017360 | -0.840798328 | 0.00098777 | 0.0091546 | F10E9.11 |
| WBGene00017361 | 1.547256409 | 0.00132448 | 0.01138037 | F10E9.12 |
| WBGene00017362 | 0.527577411 | 0.02435738 | 0.09837757 | F10G2.1 |
| WBGene00017363 | 0.029814506 | 0.98465372 | 1 | F10G2.2 |
| WBGene00017364 | 2.219627779 | 3.32E-09 | 2.27E-07 | clec-7 |
| WBGene00017365 | -2.009563163 | 0.00232378 | 0.01749101 | F10G2.4 |
| WBGene00017366 | 1.557897343 | 0.63813005 | 1 | F10G2.7 |
| WBGene00017367 | 0.9865913 | 0.56129466 | 1 | nhr-263 |
| WBGene00017369 | 0.270295937 | 0.3198694 | 0.52801284 | F10G7.5 |
| WBGene00017372 | 0.213073968 | 0.24610425 | 0.454096 | F10G7.9 |
| WBGene00017373 | 0.312099547 | 0.14438657 | 0.32289507 | F10G7.10 |
| WBGene00017374 | -0.587431683 | 0.01015613 | 0.05249277 | ttr-41 |
| WBGene00017375 | 0.398570452 | 0.59175417 | 0.75897817 | pbo-6 |
| WBGene00017376 | 0.728774023 | 0.15377933 | 0.336812 | F11C7.2 |
| WBGene00017378 | -0.505605194 | 0.80388525 | 1 | F11C7.6 |
| WBGene00017379 | -0.340555771 | 0.70967818 | 0.83571538 | F11C7.7 |
| WBGene00017380 | 0.408066272 | 0.19104309 | 0.38573993 | F11D5.1 |
| WBGene00017381 | -0.13100179 | 0.71132203 | 0.83667514 | svh-4 |
| WBGene00017382 | 2.806835559 | 0.21738362 | 1 | F11D5.5 |
| WBGene00017384 | 0.747845955 | 0.85215965 | 1 | F11G11.4 |
| WBGene00017385 | -0.096524795 | 0.68350649 | 0.81879155 | F11G11.5 |
| WBGene00017386 | 0.324640447 | 0.83650664 | 1 | nspd-5 |
| WBGene00017387 | -1.094201388 | 0.7816772 | 1 | mpst-4 |
| WBGene00017388 | -0.323868354 | 0.91634405 | 1 | F11G11.13 |
| WBGene00017389 | 0.12314947 | 0.84044094 | 0.91237519 | lgc-38 |
| WBGene00017390 | -1.276517849 | 0.02241996 | 0.09309191 | F12A10.1 |
| WBGene00017393 | -1.969805064 | 0.62862569 | 1 | nep-5 |
| WBGene00017394 | 1.282055957 | 0.11707771 | 0.2820878 | cal-8 |
| WBGene00017396 | 1.216957619 | 0.74819621 | 1 | F12A10.7 |
| WBGene00017397 | -0.23394517 | 0.40866933 | 0.61452245 | F12A10.8 |
| WBGene00017398 | 0.354139955 | 0.23439869 | 0.4404929 | slc-17.6 |
| WBGene00017399 | 0.707559163 | 0.78139902 | 1 | lgc-51 |

| | | | | |
|---|---|---|---|---|
| WBGene00017400 | 1.292282163 | 0.24248482 | 1 | stg-2 |
| WBGene00017402 | -1.013138045 | 0.0025533 | 0.01886227 | F12E12.1 |
| WBGene00017405 | -2.445032192 | 0.00082252 | 0.00789446 | bath-31 |
| WBGene00017406 | -0.447968477 | 0.80399235 | 1 | sdz-12 |
| WBGene00017407 | -0.906189735 | 0.38157137 | 1 | F12E12.6 |
| WBGene00017408 | -0.471996754 | 0.7679521 | 1 | sdz-11 |
| WBGene00017410 | -0.668264701 | 0.85551153 | 1 | F13A2.1 |
| WBGene00017411 | 3.414822892 | 0.05649594 | 1 | F13A2.2 |
| WBGene00017413 | -1.068916263 | 0.0861543 | 0.23187748 | F13A2.4 |
| WBGene00017414 | -1.969805064 | 0.62862569 | 1 | F13A2.5 |
| WBGene00017416 | 0.933855462 | 0.33373009 | 0.5420491 | F13B6.1 |
| WBGene00017417 | -0.035181221 | 0.93980838 | 0.9689509 | F13B6.2 |
| WBGene00017418 | 0.979547152 | 0.00238451 | 0.01788163 | F13B6.3 |
| WBGene00017419 | 0.164778664 | 0.46230907 | 0.65732456 | F13B9.1 |
| WBGene00017420 | -0.737176174 | 0.57623886 | 1 | F13B9.2 |
| WBGene00017421 | 1.071151842 | 0.29786235 | 1 | cutl-29 |
| WBGene00017422 | -0.495829874 | 0.46394831 | 0.65886864 | F13C5.1 |
| WBGene00017423 | -0.464712149 | 0.07653655 | 0.21485094 | F13C5.2 |
| WBGene00017424 | 0.302735419 | 0.85595361 | 1 | F13C5.3 |
| WBGene00017426 | -0.162583427 | 0.59843125 | 0.76423202 | F13C5.5 |
| WBGene00017427 | -0.129026826 | 0.78562682 | 0.88211368 | acp-5 |
| WBGene00017428 | 0.663429145 | 0.14260848 | 0.32052915 | F13D11.3 |
| WBGene00017429 | -0.293203461 | 0.61931066 | 0.77855043 | F13D11.4 |
| WBGene00017430 | 1.274032935 | 2.71E-07 | 1.1096E-05 | bcl-11 |
| WBGene00017431 | 0.656489201 | 0.0112099 | 0.05653826 | F13H6.3 |
| WBGene00017433 | 2.864269753 | 0.00118306 | 0.01043105 | F13H6.5 |
| WBGene00017434 | 2.182253558 | 0.00173908 | 0.01394593 | F13H8.1 |
| WBGene00017435 | 0.176288672 | 0.4096239 | 0.61507125 | F13H8.2 |
| WBGene00017436 | -1.760928144 | 1.4609E-06 | 4.5131E-05 | F13H8.3 |
| WBGene00017437 | -0.627086499 | 0.34633177 | 0.55516425 | nmgp-1 |
| WBGene00017438 | -0.100106022 | 0.87390269 | 0.93209633 | F13H8.5 |
| WBGene00017439 | -1.816749505 | 0.14365232 | 1 | F13H8.6 |
| WBGene00017440 | 0.473654543 | 0.00315391 | 0.02199407 | upb-1 |
| WBGene00017441 | 0.085110463 | 0.93189338 | 1 | F13H8.8 |
| WBGene00017442 | -0.16655347 | 0.76255551 | 0.86718678 | F13H8.9 |
| WBGene00017443 | 1.350426936 | 0.0348868 | 0.12590615 | F13H8.11 |
| WBGene00017444 | 0.377828795 | 0.92526585 | 1 | F13H8.12 |
| WBGene00017445 | -0.285873164 | 0.55915348 | 0.73600413 | sid-5 |
| WBGene00017446 | -0.734912898 | 0.2464662 | 0.45441856 | F14B8.4 |
| WBGene00017447 | 0.805075014 | 0.02250434 | 0.09326547 | F14B8.5 |
| WBGene00017448 | 1.141492521 | 0.00068912 | 0.00690319 | F14B8.6 |
| WBGene00017449 | 0.791089941 | 0.14327709 | 0.32124231 | otpl-3 |
| WBGene00017450 | -0.352276182 | 0.2907974 | 0.5008613 | bath-27 |

| | | | | |
|---|---|---|---|---|
| WBGene00017451 | 0.106963238 | 0.90844896 | 1 | F14D2.2 |
| WBGene00017452 | -0.58140859 | 0.04879528 | 0.15839959 | bath-29 |
| WBGene00017453 | 0.839015126 | 0.63759529 | 1 | F14D2.5 |
| WBGene00017454 | 1.38322681 | 0.69550305 | 1 | irld-5 |
| WBGene00017455 | -0.156269293 | 0.96924949 | 1 | F14D2.7 |
| WBGene00017456 | 0.868534768 | 0.18382509 | 0.37600068 | F14D2.8 |
| WBGene00017459 | -0.555136472 | 0.33239069 | 0.54019888 | F14D2.11 |
| WBGene00017460 | -0.699807189 | 0.01180015 | 0.05859536 | bath-30 |
| WBGene00017461 | -0.420216909 | 0.24512912 | 0.45305414 | bath-28 |
| WBGene00017462 | -0.855966146 | 0.24583696 | 0.45379649 | F14D2.14 |
| WBGene00017463 | 0.297704663 | 0.50464986 | 0.69168593 | F14D12.1 |
| WBGene00017464 | 1.431126247 | 0.00150947 | 0.0126302 | sulp-2 |
| WBGene00017466 | 1.921573827 | 0.30677212 | 1 | F14F9.3 |
| WBGene00017467 | -0.412316405 | 0.52910113 | 0.71336111 | F14F9.4 |
| WBGene00017468 | -1.456535813 | 0.08332296 | 0.22671987 | F14F9.5 |
| WBGene00017469 | 2.721614373 | 0.01148223 | 0.05753438 | F14F9.6 |
| WBGene00017470 | -0.737462795 | 0.83433562 | 1 | F14H12.2 |
| WBGene00017471 | 0.134743661 | 0.79044349 | 0.8845213 | F14H12.3 |
| WBGene00017472 | -0.379716524 | 0.40383457 | 0.6111802 | cst-1 |
| WBGene00017473 | -1.83602771 | 0.13962172 | 1 | F14H12.6 |
| WBGene00017474 | -0.066508408 | 0.96147293 | 1 | F14H12.7 |
| WBGene00017475 | 2.779713462 | 0.22340067 | 1 | F14H12.8 |
| WBGene00017476 | -0.156269293 | 0.96924949 | 1 | F15A8.1 |
| WBGene00017478 | 0.690905554 | 0.07694698 | 0.21525651 | F15A8.6 |
| WBGene00017480 | 0.12924608 | 0.54806181 | 0.72665922 | nstp-2 |
| WBGene00017481 | -1.909396547 | 7.2802E-06 | 0.00017331 | F15B10.3 |
| WBGene00017482 | 0.022940998 | 0.93170882 | 0.96406579 | set-9 |
| WBGene00017483 | -0.032657334 | 0.91834815 | 0.95686787 | lgc-22 |
| WBGene00017484 | 0.212529819 | 0.73607592 | 0.85265697 | F15E6.3 |
| WBGene00017485 | -4.005206095 | 1.72E-16 | 3.87E-14 | F15E6.4 |
| WBGene00017486 | 0.688793783 | 0.86378755 | 1 | F15E6.5 |
| WBGene00017487 | 0.552355833 | 0.20084284 | 0.3996731 | F15E6.6 |
| WBGene00017488 | -2.634841541 | 0.07920373 | 1 | dct-7 |
| WBGene00017489 | 0.663066231 | 0.40891532 | 0.61458788 | F15E6.9 |
| WBGene00017490 | -1.419159291 | 0.13292206 | 0.30616709 | pud-2.1 |
| WBGene00017493 | 0.28530093 | 0.92084641 | 1 | irld-30 |
| WBGene00017498 | -0.072626683 | 0.8525055 | 0.91927977 | pud-4 |
| WBGene00017499 | 0.595703065 | 0.58378726 | 1 | pud-1.1 |
| WBGene00017500 | -1.419159291 | 0.13292206 | 0.30616709 | pud-2.2 |
| WBGene00017501 | -0.970418948 | 0.00051908 | 0.0055234 | pud-3 |
| WBGene00017502 | 0.044564568 | 0.97826003 | 1 | F16B3.2 |
| WBGene00017503 | 0.069823704 | 0.90441394 | 0.95044488 | nhr-177 |
| WBGene00017504 | 0.128820704 | 0.83413813 | 0.91023162 | F16B4.2 |

| | | | | |
|---|---|---|---|---|
| WBGene00017506 | -0.116850854 | 0.62217298 | 0.7801836 | F16B4.4 |
| WBGene00017507 | 0.138032884 | 0.87603128 | 0.93313702 | F16B4.5 |
| WBGene00017508 | 2.460743752 | 0.26152462 | 1 | F16B4.6 |
| WBGene00017509 | -1.094201388 | 0.7816772 | 1 | F16B4.7 |
| WBGene00017510 | 0.669778337 | 0.55895157 | 1 | nhr-178 |
| WBGene00017512 | 1.082637937 | 0.17305046 | 0.36286798 | nhr-179 |
| WBGene00017513 | 0.352069474 | 0.5350822 | 0.7178681 | F16F9.1 |
| WBGene00017514 | -0.892813363 | 0.24746776 | 0.45550426 | F16F9.3 |
| WBGene00017515 | -0.265941323 | 0.58218375 | 0.75273854 | F16F9.4 |
| WBGene00017517 | -0.156269293 | 0.96924949 | 1 | F16G10.2 |
| WBGene00017518 | 0.574508983 | 0.8551971 | 1 | F16G10.3 |
| WBGene00017524 | -1.159714244 | 0.75790634 | 1 | F16G10.9 |
| WBGene00017526 | -1.901856996 | 0.64072034 | 1 | F16G10.11 |
| WBGene00017528 | -1.159714244 | 0.75790634 | 1 | F16G10.14 |
| WBGene00017530 | 0.073473847 | 0.91746599 | 0.95646184 | F16H11.1 |
| WBGene00017532 | -0.291877698 | 0.35817423 | 0.56656987 | ent-5 |
| WBGene00017534 | -0.471367719 | 0.01559111 | 0.07164712 | F17A9.2 |
| WBGene00017535 | -1.035305056 | 0.16506229 | 0.35168723 | atf-8 |
| WBGene00017536 | 0.063329408 | 0.82776953 | 0.90703073 | F17A9.4 |
| WBGene00017537 | 0.122019663 | 0.70472188 | 0.83302778 | F17A9.5 |
| WBGene00017538 | -0.740249274 | 0.0046989 | 0.02988439 | ceh-49 |
| WBGene00017539 | -1.68840943 | 0.10782398 | 1 | F17E9.2 |
| WBGene00017540 | 0.281454817 | 0.94435481 | 1 | F17E9.3 |
| WBGene00017541 | 0.019410267 | 0.94685849 | 0.97304242 | F17E9.4 |
| WBGene00017542 | 1.526932544 | 0.48320475 | 1 | F17E9.5 |
| WBGene00017543 | -0.008677305 | 0.99719169 | 1 | lgc-5 |
| WBGene00017544 | 0.247318604 | 0.91814609 | 1 | lgc-6 |
| WBGene00017545 | 1.630244101 | 0.40224995 | 1 | F18A1.1 |
| WBGene00017546 | -0.534064988 | 4.8074E-05 | 0.00083381 | rpa-1 |
| WBGene00017547 | 0.217126007 | 0.2774496 | 0.48635802 | alfa-1 |
| WBGene00017548 | -0.378152753 | 0.00666002 | 0.03832194 | F18A1.7 |
| WBGene00017549 | -0.473103413 | 0.10769019 | 0.26759651 | pid-1 |
| WBGene00017550 | 0.983488853 | 0.80601565 | 1 | nep-6 |
| WBGene00017553 | -0.482163247 | 0.80766744 | 1 | nep-8 |
| WBGene00017554 | -1.279666666 | 0.55824702 | 1 | nep-9 |
| WBGene00017555 | -0.62146648 | 0.85925752 | 1 | nep-10 |
| WBGene00017556 | -0.593636473 | 0.38817854 | 0.59694435 | F18A12.7 |
| WBGene00017557 | 1.24361834 | 0.0003744 | 0.00429643 | nep-11 |
| WBGene00017559 | -0.950208353 | 0.7250598 | 1 | mpz-3 |
| WBGene00017560 | -1.100105814 | 0.10929691 | 0.26965532 | F18C5.5 |
| WBGene00017562 | 0.286018814 | 0.87920703 | 1 | F18C5.9 |
| WBGene00017563 | 0.578356109 | 0.01802595 | 0.07947638 | F18C5.10 |
| WBGene00017565 | 0.595424086 | 0.02270591 | 0.09379803 | ddo-2 |

| | | | | |
|---|---|---|---|---|
| WBGene00017567 | -0.385941269 | 0.4490482 | 0.6479551 | F18E3.11 |
| WBGene00017568 | -0.423165596 | 0.48252888 | 0.67353221 | F18E9.1 |
| WBGene00017569 | 0.806962548 | 0.02243966 | 0.0931228 | F18E9.3 |
| WBGene00017570 | -0.431360109 | 0.75666576 | 1 | F18E9.4 |
| WBGene00017571 | 0.560412465 | 0.03664558 | 0.12967473 | jmjd-3.1 |
| WBGene00017573 | 1.601302495 | 0.65842342 | 1 | F18E9.7 |
| WBGene00017574 | 0.793082403 | 0.01164157 | 0.05797386 | F18F11.1 |
| WBGene00017576 | 0.218960638 | 0.68194722 | 0.81789205 | cdh-8 |
| WBGene00017577 | -0.62146648 | 0.85925752 | 1 | F18F11.4 |
| WBGene00017578 | 0.334914275 | 0.3375999 | 0.54561685 | drl-1 |
| WBGene00017579 | -0.333693239 | 0.89573179 | 1 | F18G5.1 |
| WBGene00017580 | 0.673386861 | 0.18059398 | 0.37259848 | lgc-4 |
| WBGene00017582 | 1.203066131 | 0.00189602 | 0.01494775 | F18G5.6 |
| WBGene00017583 | -0.258746767 | 0.54233017 | 0.72288584 | F19B10.1 |
| WBGene00017584 | -0.937236332 | 0.00960979 | 0.05033324 | F19B10.2 |
| WBGene00017586 | 0.64815038 | 0.68041446 | 1 | F19B10.4 |
| WBGene00017587 | -0.065026242 | 0.91681464 | 0.95635621 | F19B10.5 |
| WBGene00017589 | -0.464861339 | 0.05799678 | 0.17844372 | F19B10.10 |
| WBGene00017590 | -0.558804446 | 0.87818675 | 1 | F19B10.11 |
| WBGene00017591 | 0.338079309 | 0.06322995 | 0.18880344 | F19C7.1 |
| WBGene00017592 | 1.25797632 | 0.00192409 | 0.01510047 | F19C7.2 |
| WBGene00017593 | -0.397833399 | 0.78421681 | 1 | F19C7.3 |
| WBGene00017594 | 0.479965126 | 0.58312563 | 0.75332022 | F19C7.4 |
| WBGene00017597 | 1.091310183 | 0.0450514 | 0.14927796 | F19C7.8 |
| WBGene00017598 | -0.541644105 | 0.58423386 | 1 | F19F10.1 |
| WBGene00017599 | 0.107066218 | 0.86757238 | 0.92836209 | F19F10.3 |
| WBGene00017600 | 0.308631366 | 0.53517964 | 0.7178681 | ttr-10 |
| WBGene00017601 | -1.958100521 | 0.00660184 | 0.0381499 | ets-7 |
| WBGene00017605 | 0.042457508 | 0.78979425 | 0.8841763 | F19F10.9 |
| WBGene00017606 | 0.12345622 | 0.60435952 | 0.76867617 | ets-6 |
| WBGene00017607 | 0.150161965 | 0.45348415 | 0.65096864 | F19F10.11 |
| WBGene00017608 | -0.052162677 | 0.7949372 | 0.88641923 | F19F10.12 |
| WBGene00017611 | -1.45873709 | 0.59584643 | 1 | F19G12.3 |
| WBGene00017612 | 0.470297475 | 0.75161213 | 1 | F19G12.4 |
| WBGene00017613 | 0.069273475 | 0.75841095 | 0.8648321 | F20A1.1 |
| WBGene00017614 | -1.803775931 | 0.10583572 | 1 | F20A1.2 |
| WBGene00017615 | 0.747845955 | 0.85215965 | 1 | srsx-32 |
| WBGene00017616 | 2.09988748 | 0.04329477 | 0.14516027 | F20A1.4 |
| WBGene00017618 | -0.063503217 | 0.91704344 | 0.95637631 | F20A1.6 |
| WBGene00017619 | 1.181686151 | 0.7579682 | 1 | F20A1.8 |
| WBGene00017620 | -0.502497629 | 0.01421762 | 0.06695739 | tofu-2 |
| WBGene00017621 | -0.157888192 | 0.61075727 | 0.77217579 | F20A1.10 |
| WBGene00017625 | 0.842458416 | 0.11855493 | 0.28406856 | cgt-2 |

| | | | | |
|---|---|---|---|---|
| WBGene00017627 | 0.281454817 | 0.94435481 | 1 | F20B6.1 |
| WBGene00017628 | -0.053767403 | 0.97276761 | 1 | F20B6.4 |
| WBGene00017631 | -0.278927547 | 0.8350008 | 1 | F20B6.7 |
| WBGene00017632 | -0.975716257 | 0.50150325 | 1 | F20B6.9 |
| WBGene00017634 | 0.983488853 | 0.80601565 | 1 | F20D6.2 |
| WBGene00017635 | 0.181787111 | 0.76761432 | 0.86977185 | F20D6.5 |
| WBGene00017636 | 2.566123675 | 0.23268183 | 1 | F20D6.6 |
| WBGene00017637 | 1.202345657 | 0.09884825 | 0.25306578 | F20D6.8 |
| WBGene00017639 | -0.114892771 | 0.84380068 | 0.91412368 | F20D6.10 |
| WBGene00017640 | 0.175725953 | 0.51536936 | 0.70177552 | F20D6.11 |
| WBGene00017641 | -0.072801716 | 0.68802727 | 0.82242734 | csr-1 |
| WBGene00017642 | 0.495254908 | 0.00373767 | 0.02496096 | F20D12.2 |
| WBGene00017643 | -0.302809104 | 0.03491771 | 0.12590615 | czw-1 |
| WBGene00017644 | 0.215084342 | 0.61278233 | 0.77396612 | exc-9 |
| WBGene00017645 | 0.590708277 | 0.82739319 | 1 | F20D12.7 |
| WBGene00017646 | -0.11328741 | 0.61628008 | 0.77610041 | F20H11.1 |
| WBGene00017647 | 1.181686151 | 0.7579682 | 1 | F20H11.4 |
| WBGene00017648 | -0.140353544 | 0.37520917 | 0.58305729 | ddo-3 |
| WBGene00017650 | 2.437484291 | 0.2708268 | 1 | F21A9.1 |
| WBGene00017651 | 0.689852633 | 0.40932718 | 0.61485082 | F21A9.2 |
| WBGene00017653 | 0.766110341 | 0.2533999 | 0.46131776 | F21C10.3 |
| WBGene00017654 | 0.399274936 | 0.79379724 | 1 | F21C10.4 |
| WBGene00017655 | 0.281454817 | 0.94435481 | 1 | F21C10.5 |
| WBGene00017656 | 0.688793783 | 0.86378755 | 1 | F21C10.6 |
| WBGene00017657 | 1.650494557 | 3.84E-11 | 3.96E-09 | F21C10.7 |
| WBGene00017658 | 1.160514594 | 1.92E-07 | 8.1039E-06 | F21C10.9 |
| WBGene00017659 | 0.73987891 | 0.00192046 | 0.01508502 | F21C10.10 |
| WBGene00017660 | -0.326457532 | 0.66138922 | 0.80508588 | F21C10.11 |
| WBGene00017661 | 1.168190592 | 0.48565519 | 1 | frpr-6 |
| WBGene00017663 | 0.959893907 | 0.00105404 | 0.00960018 | F21D12.3 |
| WBGene00017664 | -1.159714244 | 0.75790634 | 1 | npax-1 |
| WBGene00017666 | -1.063064295 | 0.45494089 | 1 | F21E9.2 |
| WBGene00017667 | 0.962136476 | 0.03696298 | 0.13045567 | ttr-37 |
| WBGene00017671 | 0.888999466 | 5.0975E-06 | 0.0001284 | pgal-1 |
| WBGene00017672 | -0.014519762 | 0.99603746 | 1 | F21F3.2 |
| WBGene00017673 | 1.189071466 | 0.5668301 | 1 | icmt-1 |
| WBGene00017674 | -0.409372601 | 0.24298894 | 0.45063908 | F21F3.4 |
| WBGene00017675 | 0.335161805 | 0.05151806 | 0.16385794 | F21F3.6 |
| WBGene00017676 | -0.254120081 | 0.43208659 | 0.6341955 | F21F3.7 |
| WBGene00017678 | 1.396555311 | 9.93E-09 | 6.09E-07 | asp-12 |
| WBGene00017681 | 0.633178131 | 0.1966447 | 0.39371097 | slc-17.5 |
| WBGene00017683 | -0.388053547 | 0.03814506 | 0.13316963 | rbbp-5 |
| WBGene00017686 | -0.299876155 | 0.02987199 | 0.11287763 | tpp-2 |

| | | | | |
|---|---|---|---|---|
| WBGene00017687 | 1.185252421 | 1.0862E-06 | 3.5837E-05 | ets-4 |
| WBGene00017688 | -0.676506181 | 0.02011697 | 0.08578861 | ttr-35 |
| WBGene00017690 | 0.807983058 | 0.08839134 | 0.235652 | ceh-60 |
| WBGene00017691 | 0.646954888 | 3.8617E-05 | 0.00069665 | ilys-5 |
| WBGene00017692 | 1.366077596 | 0.06879557 | 0.19999543 | F22B7.1 |
| WBGene00017693 | -0.156269293 | 0.96924949 | 1 | flp-23 |
| WBGene00017695 | 1.116613983 | 0.69067248 | 1 | fip-1 |
| WBGene00017696 | -0.388638527 | 0.05236052 | 0.16554924 | polk-1 |
| WBGene00017698 | 1.320522458 | 0.00147185 | 0.01237515 | F22B7.9 |
| WBGene00017699 | 0.13593895 | 0.48501555 | 0.67539764 | flcn-1 |
| WBGene00017700 | -1.777135447 | 0.45934881 | 1 | F22D3.4 |
| WBGene00017701 | -0.922268666 | 0.22930444 | 0.43442105 | F22D3.5 |
| WBGene00017702 | -0.040831679 | 0.92000588 | 0.9579581 | malt-1 |
| WBGene00017703 | -1.634638676 | 0.00104379 | 0.0095358 | F22E5.1 |
| WBGene00017705 | 0.468829292 | 0.75524348 | 1 | F22E5.6 |
| WBGene00017706 | 0.377828795 | 0.92526585 | 1 | F22E5.7 |
| WBGene00017707 | 0.240011698 | 0.84006992 | 1 | F22E5.8 |
| WBGene00017708 | -0.572615625 | 0.11682695 | 0.2815954 | F22E5.9 |
| WBGene00017709 | 1.552776409 | 0.4309513 | 1 | F22E5.11 |
| WBGene00017710 | 2.191582552 | 0.22563181 | 1 | F22E5.12 |
| WBGene00017711 | 3.345082262 | 0.09532154 | 1 | F22E5.13 |
| WBGene00017712 | -0.642060326 | 0.01628194 | 0.0738159 | F22E5.17 |
| WBGene00017713 | -1.592398681 | 0.07291152 | 0.20807825 | F22E5.20 |
| WBGene00017714 | 0.747845955 | 0.85215965 | 1 | F22F1.2 |
| WBGene00017715 | 0.852865077 | 0.34327277 | 0.55176655 | F22F1.3 |
| WBGene00017716 | -0.855014321 | 0.10349879 | 0.26071284 | F22F4.1 |
| WBGene00017717 | 0.19715435 | 0.60496241 | 0.76897128 | F22F4.4 |
| WBGene00017718 | -1.66778326 | 0.01910183 | 0.08295699 | F22F4.5 |
| WBGene00017719 | 0.384515202 | 0.02237258 | 0.09298089 | ldp-1 |
| WBGene00017720 | -0.5477972 | 0.40501106 | 0.61190135 | F22F7.2 |
| WBGene00017721 | -0.184766928 | 0.85102317 | 0.91850585 | F22F7.3 |
| WBGene00017722 | -0.471996754 | 0.7679521 | 1 | F22F7.4 |
| WBGene00017723 | -0.345294721 | 0.81956796 | 1 | bgnt-1.2 |
| WBGene00017724 | 0.350702528 | 0.45566535 | 0.65284789 | F22F7.7 |
| WBGene00017725 | -0.156269293 | 0.96924949 | 1 | F22H10.1 |
| WBGene00017726 | -0.552676412 | 0.18943074 | 0.38350486 | F22H10.2 |
| WBGene00017727 | 0.079596696 | 0.6524654 | 0.79997085 | F22H10.3 |
| WBGene00017730 | -0.668138583 | 0.14222531 | 0.32034267 | F22H10.6 |
| WBGene00017732 | 0.231853742 | 0.71092139 | 0.836447 | F23C8.3 |
| WBGene00017733 | -0.598135971 | 0.00143523 | 0.01212605 | ubxn-1 |
| WBGene00017734 | -0.399927405 | 0.00811381 | 0.04427406 | F23C8.5 |
| WBGene00017735 | -0.371408033 | 0.0716059 | 0.20557276 | did-2 |
| WBGene00017736 | -0.504063085 | 0.65500227 | 1 | F23C8.7 |

| | | | | |
|---|---|---|---|---|
| WBGene00017737 | 0.487066454 | 0.73167604 | 1 | F23C8.8 |
| WBGene00017738 | -0.306215273 | 0.26271009 | 0.47141558 | tipn-1 |
| WBGene00017741 | 0.464660696 | 0.46968941 | 0.66315841 | F23C8.13 |
| WBGene00017742 | -0.445848123 | 0.04255186 | 0.14374572 | nfyc-1 |
| WBGene00017743 | -0.141147797 | 0.93319458 | 1 | F23F1.2 |
| WBGene00017744 | 0.168081398 | 0.79466998 | 0.88641923 | fbxc-54 |
| WBGene00017745 | 0.046097713 | 0.90510816 | 0.95070175 | F23F1.4 |
| WBGene00017746 | -0.351085596 | 0.05667985 | 0.17581759 | F23F1.5 |
| WBGene00017747 | 0.875550462 | 0.11973688 | 0.28595048 | F23F1.6 |
| WBGene00017748 | 0.300134032 | 0.66289051 | 0.80634743 | F23F1.7 |
| WBGene00017749 | -0.452036854 | 0.04847786 | 0.15775119 | rpoa-49 |
| WBGene00017750 | -0.691940778 | 0.03101643 | 0.11626133 | F23F1.10 |
| WBGene00017751 | 0.608980516 | 0.14799077 | 0.32859472 | F23F12.3 |
| WBGene00017752 | -1.985648517 | 0.03979429 | 0.13696105 | sdz-15 |
| WBGene00017754 | 0.21225903 | 0.85212295 | 1 | F23F12.8 |
| WBGene00017755 | -0.012303903 | 0.97057146 | 0.98468356 | zip-8 |
| WBGene00017756 | -0.885829456 | 0.05046415 | 0.16187147 | F23F12.12 |
| WBGene00017757 | -0.631305823 | 0.00230185 | 0.01735825 | bra-2 |
| WBGene00017758 | 0.118255548 | 0.62059323 | 0.77878286 | F23H11.2 |
| WBGene00017759 | -0.676015369 | 4.5129E-06 | 0.00011505 | sucl-2 |
| WBGene00017760 | -0.000792395 | 0.99680182 | 0.99852895 | F23H11.4 |
| WBGene00017761 | 0.784239248 | 0.63341939 | 1 | F23H11.6 |
| WBGene00017762 | 2.335672404 | 0.00143802 | 0.01213378 | F23H11.7 |
| WBGene00017763 | 0.051252485 | 0.80231186 | 0.89169709 | crls-1 |
| WBGene00017764 | 1.830320951 | 0.08562693 | 1 | F25A2.1 |
| WBGene00017765 | -0.246010269 | 0.18025326 | 0.3725955 | gcst-1 |
| WBGene00017766 | -0.070444392 | 0.79002915 | 0.88435772 | peli-1 |
| WBGene00017767 | -0.376887936 | 0.32388063 | 0.53245035 | F25B4.4 |
| WBGene00017768 | -0.062658217 | 0.73102147 | 0.84939926 | F25B4.5 |
| WBGene00017769 | 0.081540252 | 0.76631128 | 0.86897515 | hmgs-1 |
| WBGene00017770 | -0.562802204 | 0.00602397 | 0.03570228 | F25B4.7 |
| WBGene00017771 | -0.87951668 | 0.14317518 | 0.32114434 | F25B4.8 |
| WBGene00017772 | -0.784773424 | 3.0716E-05 | 0.00057022 | clec-1 |
| WBGene00017773 | 0.606546627 | 0.01230038 | 0.06033898 | dct-6 |
| WBGene00017774 | -0.368409077 | 0.0301334 | 0.1136885 | nop-1 |
| WBGene00017775 | 0.182590332 | 0.22771536 | 0.43252381 | F25B5.3 |
| WBGene00017776 | -0.236978553 | 0.30294132 | 0.51211509 | F25B5.5 |
| WBGene00017777 | 0.017725421 | 0.91677879 | 0.95635621 | F25B5.6 |
| WBGene00017778 | -0.46970944 | 0.00986673 | 0.05133254 | nono-1 |
| WBGene00017779 | 0.775841246 | 0.58693204 | 1 | gtr-1 |
| WBGene00017780 | -0.609315159 | 0.00463275 | 0.0295566 | F25E2.2 |
| WBGene00017781 | 1.142725214 | 0.00328704 | 0.02270074 | F25E2.3 |
| WBGene00017782 | 0.035037318 | 0.89300784 | 0.94367782 | F25E5.1 |

| | | | | |
|---|---|---|---|---|
| WBGene00017783 | 0.657278612 | 0.62663614 | 1 | F25E5.2 |
| WBGene00017784 | 0.087007751 | 0.95602813 | 1 | F25E5.3 |
| WBGene00017785 | 1.692887539 | 0.01956569 | 0.08417682 | F25E5.4 |
| WBGene00017786 | 1.367380561 | 0.00269942 | 0.01967208 | F25E5.5 |
| WBGene00017787 | 1.361050168 | 0.20989892 | 1 | nhr-141 |
| WBGene00017788 | 1.278053416 | 0.54789306 | 1 | F25E5.7 |
| WBGene00017789 | 0.373371852 | 0.13867025 | 0.31551639 | F25E5.8 |
| WBGene00017790 | 0.281454817 | 0.94435481 | 1 | F25E5.9 |
| WBGene00017791 | 1.186261621 | 0.49486297 | 1 | try-8 |
| WBGene00017792 | 0.998440351 | 0.21401296 | 0.41467146 | F25E5.16 |
| WBGene00017794 | 0.234385821 | 0.89578204 | 1 | mltn-13 |
| WBGene00017795 | 0.19150222 | 0.62614262 | 0.78266216 | F25F8.1 |
| WBGene00017796 | 0.936054477 | 0.07201623 | 0.20637478 | F25G6.1 |
| WBGene00017797 | 0.188780629 | 0.34777485 | 0.55644893 | symk-1 |
| WBGene00017798 | 1.238504954 | 0.19979718 | 0.39811875 | slc-17.8 |
| WBGene00017799 | -0.780554621 | 7.6491E-05 | 0.00123167 | F25G6.8 |
| WBGene00017800 | 0.054730374 | 0.7481657 | 0.85972621 | F25G6.9 |
| WBGene00017801 | -0.757503903 | 7.5025E-05 | 0.00121613 | F26A1.1 |
| WBGene00017802 | 0.688793783 | 0.86378755 | 1 | F26A1.3 |
| WBGene00017803 | 1.413377704 | 0.60673863 | 1 | F26A1.4 |
| WBGene00017805 | 0.687259521 | 0.81162667 | 1 | F26A1.6 |
| WBGene00017805 | 0.09697944 | 0.97612634 | 1 | F26A1.7 |
| WBGene00017806 | -0.646440611 | 0.67898796 | 1 | F26A1.8 |
| WBGene00017807 | 1.284515805 | 0.34337001 | 1 | F26A1.9 |
| WBGene00017808 | 0.489058549 | 0.85947456 | 1 | nspd-9 |
| WBGene00017809 | 0.281454817 | 0.94435481 | 1 | clec-156 |
| WBGene00017810 | -1.159714244 | 0.75790634 | 1 | clec-157 |
| WBGene00017811 | 0.239405469 | 0.32038493 | 0.52858726 | F26A1.13 |
| WBGene00017811 | 0.067886445 | 0.83530611 | 0.91036596 | F26A1.14 |
| WBGene00017813 | 0.570705405 | 0.46768096 | 0.66183982 | F26A10.1 |
| WBGene00017814 | -0.229930976 | 0.66191713 | 0.80532477 | F26A10.2 |
| WBGene00017815 | 0.983488853 | 0.80601565 | 1 | F26B1.1 |
| WBGene00017816 | -0.51844879 | 0.00126768 | 0.0110171 | hrpk-1 |
| WBGene00017817 | 0.281454817 | 0.94435481 | 1 | F26B1.5 |
| WBGene00017818 | 0.139605792 | 0.71265542 | 0.83759433 | F26D11.1 |
| WBGene00017819 | 1.38322681 | 0.69550305 | 1 | F26D11.2 |
| WBGene00017821 | 0.763390359 | 0.73510979 | 1 | F26D11.4 |
| WBGene00017822 | -0.156269293 | 0.96924949 | 1 | clec-216 |
| WBGene00017823 | 1.991430903 | 0.15678653 | 1 | pals-34 |
| WBGene00017824 | -0.156269293 | 0.96924949 | 1 | clec-217 |
| WBGene00017825 | -0.247526287 | 0.17471291 | 0.36490444 | cee-1 |
| WBGene00017826 | -0.291977436 | 0.10845606 | 0.26850836 | tag-340 |
| WBGene00017827 | -0.226166073 | 0.29443896 | 0.5048416 | F26F4.5 |

| | | | | |
|---|---|---|---|---|
| WBGene00017828 | -0.096495568 | 0.69285216 | 0.82542617 | F26F4.6 |
| WBGene00017829 | -0.475316852 | 0.10212446 | 0.25793489 | F26F4.9 |
| WBGene00017830 | -1.003183787 | 1.4947E-06 | 4.5616E-05 | rpb-8 |
| WBGene00017831 | -0.348083123 | 0.12814339 | 0.29890277 | F26F4.12 |
| WBGene00017832 | -0.901362261 | 0.00010563 | 0.00159115 | kbp-2 |
| WBGene00017833 | 0.281454817 | 0.94435481 | 1 | F26F12.2 |
| WBGene00017834 | 0.635596596 | 0.00173832 | 0.01394593 | F26F12.3 |
| WBGene00017835 | -0.320926141 | 0.61775776 | 0.77718282 | F26F12.4 |
| WBGene00017836 | 1.633238609 | 0.2732739 | 1 | F26F12.5 |
| WBGene00017837 | -0.136671878 | 0.49027194 | 0.67965193 | F26G1.1 |
| WBGene00017838 | 0.148141613 | 0.8416258 | 0.91307112 | F26G1.2 |
| WBGene00017840 | 0.720240281 | 0.14164927 | 0.31978051 | ttm-2 |
| WBGene00017841 | -0.110846668 | 0.85279216 | 0.91927977 | F26G1.5 |
| WBGene00017842 | 0.888118836 | 0.12597646 | 0.29606662 | nep-12 |
| WBGene00017843 | -0.52106523 | 0.01390089 | 0.06582937 | F26G5.1 |
| WBGene00017848 | -2.415905411 | 0.01306735 | 0.06312365 | F27B3.6 |
| WBGene00017850 | 0.713442937 | 0.26165018 | 0.47012511 | F27B10.1 |
| WBGene00017851 | 0.90324783 | 0.46667739 | 1 | F27C1.1 |
| WBGene00017852 | -0.218723903 | 0.21133406 | 0.4111419 | F27C1.2 |
| WBGene00017853 | -0.156269293 | 0.96924949 | 1 | F27C1.3 |
| WBGene00017854 | -1.22948835 | 0.06368909 | 0.18975441 | F27C1.4 |
| WBGene00017855 | -0.043757757 | 0.80139149 | 0.89148362 | F27C1.6 |
| WBGene00017858 | 1.280710261 | 0.05895243 | 0.18037839 | F27C1.11 |
| WBGene00017860 | 1.687239501 | 0.59403242 | 1 | F27C1.13 |
| WBGene00017861 | 0.93331338 | 0.02604235 | 0.10302938 | F27D9.2 |
| WBGene00017864 | -0.170556257 | 0.37429168 | 0.58222618 | pcca-1 |
| WBGene00017865 | 1.434656034 | 0.04357378 | 0.1457381 | F27D9.7 |
| WBGene00017866 | 0.608667768 | 0.21203626 | 0.412025 | stn-2 |
| WBGene00017867 | 0.424394967 | 0.71119013 | 1 | slc-28.1 |
| WBGene00017868 | 0.926428055 | 0.38752998 | 1 | slc-28.2 |
| WBGene00017869 | 0.688793783 | 0.86378755 | 1 | F28A10.1 |
| WBGene00017871 | 0.574508983 | 0.8551971 | 1 | F28A10.3 |
| WBGene00017872 | -0.156269293 | 0.96924949 | 1 | F28A10.4 |
| WBGene00017873 | -3.47102951 | 0.12756864 | 1 | F28A10.5 |
| WBGene00017874 | -0.468301393 | 0.02207661 | 0.09223842 | acdh-9 |
| WBGene00017875 | -0.156269293 | 0.96924949 | 1 | F28A10.7 |
| WBGene00017879 | 0.232157325 | 0.93397866 | 1 | acd-4 |
| WBGene00017880 | 0.95280852 | 0.09720126 | 0.25046061 | F28A12.3 |
| WBGene00017881 | 0.325658696 | 0.02476511 | 0.09951987 | asp-13 |
| WBGene00017885 | 0.228674494 | 0.26359867 | 0.4721438 | dip-2 |
| WBGene00017886 | -0.270428986 | 0.35485903 | 0.56318215 | F28B3.5 |
| WBGene00017887 | -0.918528248 | 0.00029035 | 0.0035181 | F28B3.6 |
| WBGene00017888 | -0.365046215 | 0.21916912 | 0.421878 | acl-11 |

| | | | | |
|---|---|---|---|---|
| WBGene00017889 | -0.155101238 | 0.5350199 | 0.7178681 | F28B3.10 |
| WBGene00017890 | 0.281454817 | 0.94435481 | 1 | F28B4.1 |
| WBGene00017891 | 0.169023228 | 0.53457587 | 0.71755507 | rgl-1 |
| WBGene00017892 | 2.080544742 | 1.16E-20 | 4.15E-18 | F28B4.3 |
| WBGene00017893 | -1.048852522 | 0.44239294 | 1 | F28B4.4 |
| WBGene00017894 | -0.492117824 | 0.55615976 | 0.73339151 | F28B12.1 |
| WBGene00017895 | 0.002028164 | 0.99113167 | 0.99555966 | vrk-1 |
| WBGene00017898 | -0.512164671 | 0.15232351 | 0.33448358 | F28C10.3 |
| WBGene00017900 | 0.687873571 | 0.02305468 | 0.09480655 | F28E10.1 |
| WBGene00017901 | 0.812509088 | 0.06161491 | 0.18583439 | igeg-1 |
| WBGene00017902 | 0.747845955 | 0.85215965 | 1 | F28E10.4 |
| WBGene00017904 | 0.37434263 | 0.18538292 | 0.37795702 | lim-8 |
| WBGene00017905 | 1.093875011 | 0.03132002 | 0.11711274 | F28F5.6 |
| WBGene00017906 | 1.025181888 | 0.79790347 | 1 | F28F9.2 |
| WBGene00017907 | 1.132742212 | 0.25618658 | 1 | F28F9.3 |
| WBGene00017908 | -0.796168919 | 0.02932708 | 0.11144289 | F28H1.1 |
| WBGene00017909 | -0.407361831 | 0.42099968 | 0.62554854 | F28H1.4 |
| WBGene00017911 | -1.433238114 | 0.24941646 | 1 | fbxb-106 |
| WBGene00017912 | -1.212306626 | 0.39547261 | 1 | fbxb-107 |
| WBGene00017914 | 0.578168203 | 0.54653966 | 0.72543326 | F29A7.4 |
| WBGene00017916 | -0.776896207 | 0.00828181 | 0.04492826 | F29A7.6 |
| WBGene00017918 | -1.831477136 | 0.52194793 | 1 | F29A7.8 |
| WBGene00017919 | -0.269879318 | 0.37161744 | 0.57910503 | F29B9.1 |
| WBGene00017920 | -0.099585497 | 0.57582982 | 0.74859142 | jmjd-1.2 |
| WBGene00017921 | 0.002597778 | 0.99711411 | 0.99857337 | F29B9.5 |
| WBGene00017922 | 0.438455499 | 0.87116517 | 1 | F29B9.7 |
| WBGene00017923 | 0.032393521 | 0.89609077 | 0.9454324 | F29B9.8 |
| WBGene00017924 | -0.639042359 | 0.04083012 | 0.1395211 | mrps-21 |
| WBGene00017925 | -0.608141776 | 0.00015143 | 0.00211321 | F29B9.11 |
| WBGene00017926 | -0.301365533 | 0.12550913 | 0.29536855 | F29C4.2 |
| WBGene00017927 | 1.253082125 | 0.42357522 | 1 | F29C4.4 |
| WBGene00017928 | -0.346791615 | 0.22433709 | 0.42847941 | tut-1 |
| WBGene00017929 | -0.387109924 | 0.05679496 | 0.17599478 | grld-1 |
| WBGene00017930 | -1.151015083 | 0.54347156 | 1 | F29G9.1 |
| WBGene00017931 | -0.545239232 | 0.00425664 | 0.02760475 | picc-1 |
| WBGene00017933 | -0.522537859 | 0.26555544 | 0.47428078 | F29G9.7 |
| WBGene00017934 | -0.428267871 | 0.06527074 | 0.19281071 | F30B5.4 |
| WBGene00017935 | 0.281454817 | 0.94435481 | 1 | srbc-70 |
| WBGene00017936 | 0.630676614 | 0.19932373 | 0.39752085 | zipt-2.2 |
| WBGene00017937 | 0.033166584 | 0.94234292 | 0.97027162 | F30H5.3 |
| WBGene00017938 | -0.275547928 | 0.74829499 | 0.85972621 | F31A3.3 |
| WBGene00017939 | -0.23272958 | 0.48958674 | 0.67924367 | F31A3.5 |
| WBGene00017941 | 2.316718591 | 0.19028695 | 1 | F31A9.2 |

| | | | | |
|---|---|---|---|---|
| WBGene00017942 | 0.824305894 | 0.61976113 | 1 | fbxc-37 |
| WBGene00017944 | 0.658219793 | 0.61650382 | 1 | F31D5.1 |
| WBGene00017945 | 0.412392992 | 0.39434295 | 0.6021422 | F31D5.2 |
| WBGene00017947 | 1.158414948 | 0.04453068 | 0.14783895 | mth-2 |
| WBGene00017948 | 0.660897864 | 0.14356691 | 0.32168283 | mth-1 |
| WBGene00017949 | -0.663873301 | 0.6879197 | 1 | F31D5.6 |
| WBGene00017950 | 0.361961969 | 0.37993264 | 0.58783766 | F31E3.2 |
| WBGene00017951 | -0.146903268 | 0.45248687 | 0.65010241 | panl-2 |
| WBGene00017952 | 0.033479882 | 0.92043447 | 0.95824 | F31E3.6 |
| WBGene00017954 | -0.292483585 | 0.36821992 | 0.57588021 | F31E8.4 |
| WBGene00017955 | 1.013713812 | 0.64240986 | 1 | F31E8.5 |
| WBGene00017957 | -0.153424327 | 0.79767496 | 0.88865587 | F31F4.1 |
| WBGene00017959 | 1.359476883 | 0.12714359 | 0.29763248 | ugt-42 |
| WBGene00017960 | 1.435925494 | 0.70920779 | 1 | F31F4.11 |
| WBGene00017961 | 1.558468105 | 0.02678864 | 0.10484876 | nhr-180 |
| WBGene00017962 | 0.087678463 | 0.76704497 | 0.86946353 | fbxa-72 |
| WBGene00017964 | 1.596781347 | 1.27E-12 | 1.73E-10 | F31F7.1 |
| WBGene00017965 | 0.31291716 | 0.79890114 | 1 | F31F7.2 |
| WBGene00017967 | -0.197825645 | 0.30232201 | 0.51170939 | ada-2 |
| WBGene00017968 | 0.491127578 | 0.11897828 | 0.28463356 | skpo-3 |
| WBGene00017969 | 0.874093778 | 1.7327E-05 | 0.00035959 | F32A5.3 |
| WBGene00017970 | 1.009400691 | 2.03E-07 | 8.4625E-06 | F32A5.4 |
| WBGene00017971 | 0.744992979 | 0.02258375 | 0.09342054 | F32A5.8 |
| WBGene00017973 | 1.542051795 | 0.10864502 | 0.26875696 | ift-81 |
| WBGene00017974 | -0.016962246 | 0.95151445 | 0.97545664 | vps-41 |
| WBGene00017975 | 0.87001166 | 0.63612313 | 1 | F32B5.1 |
| WBGene00017976 | 0.135517812 | 0.83039726 | 0.90819324 | F32B5.2 |
| WBGene00017978 | -0.156269293 | 0.96924949 | 1 | msrp-1 |
| WBGene00017979 | 0.609474125 | 0.02233199 | 0.09285368 | F32B5.6 |
| WBGene00017980 | 0.748498337 | 0.04198307 | 0.14229939 | F32B5.7 |
| WBGene00017981 | -0.311363914 | 0.20440416 | 0.40378636 | figl-1 |
| WBGene00017982 | -0.62516325 | 0.00116482 | 0.0103076 | hpo-18 |
| WBGene00017983 | 1.348219141 | 0.03175062 | 0.11812001 | F32D1.3 |
| WBGene00017984 | -0.564364076 | 0.00080993 | 0.00781667 | F32D1.5 |
| WBGene00017985 | -0.546287587 | 0.00424086 | 0.02754856 | neg-1 |
| WBGene00017986 | -0.835543329 | 1.3149E-05 | 0.00028483 | F32D1.7 |
| WBGene00017987 | -0.62146648 | 0.85925752 | 1 | F32D1.8 |
| WBGene00017988 | -0.350137071 | 0.05629596 | 0.17493965 | fipp-1 |
| WBGene00017989 | -0.281122526 | 0.09327926 | 0.24366131 | nol-10 |
| WBGene00017990 | -0.71064466 | 0.00298477 | 0.02114238 | cec-4 |
| WBGene00017991 | 0.063084767 | 0.91262977 | 0.95467585 | clec-180 |
| WBGene00017992 | 0.200841713 | 0.37745331 | 0.58512604 | F32E10.5 |
| WBGene00017993 | -0.576899967 | 0.00012612 | 0.00183602 | cec-5 |

| | | | | |
|---|---|---|---|---|
| WBGene00017995 | 1.13192488 | 0.52069616 | 1 | F32E10.8 |
| WBGene00017996 | -0.169681431 | 0.3756657 | 0.58339183 | F33D4.4 |
| WBGene00017997 | -0.300705498 | 0.15838322 | 0.34313538 | mrpl-1 |
| WBGene00017998 | -0.46392632 | 0.27876513 | 0.4876783 | F33D4.6 |
| WBGene00017999 | -0.769152197 | 0.00109185 | 0.00981195 | emc-6 |
| WBGene00018001 | -0.156269293 | 0.96924949 | 1 | F33D11.2 |
| WBGene00018002 | -0.777178355 | 0.59395238 | 1 | twk-47 |
| WBGene00018003 | 0.281454817 | 0.94435481 | 1 | F33D11.6 |
| WBGene00018004 | 1.872249764 | 0.5345823 | 1 | F33D11.7 |
| WBGene00018005 | -0.512857215 | 0.50694576 | 0.69381451 | F33D11.8 |
| WBGene00018006 | -0.10546591 | 0.51901642 | 0.70445818 | hpo-3 |
| WBGene00018007 | -0.673250919 | 8.3084E-06 | 0.00019399 | F33D11.10 |
| WBGene00018008 | -0.163327927 | 0.42656803 | 0.63027412 | vpr-1 |
| WBGene00018009 | -0.606716183 | 0.00341918 | 0.02329529 | dhhc-3 |
| WBGene00018010 | -0.664231908 | 0.0181276 | 0.07977062 | F33E11.2 |
| WBGene00018011 | -0.443356964 | 0.02639149 | 0.10381393 | F33E11.3 |
| WBGene00018013 | -0.457568097 | 0.07348425 | 0.20908653 | phf-10 |
| WBGene00018014 | -0.572471189 | 0.01839774 | 0.080667 | F33G12.2 |
| WBGene00018015 | -0.246747605 | 0.23033701 | 0.43586684 | F33G12.3 |
| WBGene00018016 | -0.129416014 | 0.46362877 | 0.65858393 | lrr-1 |
| WBGene00018017 | 0.134683159 | 0.42598507 | 0.62971931 | golg-2 |
| WBGene00018018 | 0.543125485 | 0.03766717 | 0.13201996 | F33G12.6 |
| WBGene00018021 | -1.866202347 | 0.01139455 | 0.05718942 | F34D6.1 |
| WBGene00018022 | 0.983488853 | 0.80601565 | 1 | ceh-87 |
| WBGene00018023 | 2.24970411 | 0.10503503 | 1 | set-11 |
| WBGene00018024 | 1.624353704 | 4.00E-07 | 1.553E-05 | F35A5.1 |
| WBGene00018025 | 0.721135295 | 0.24622984 | 0.45418968 | F35A5.2 |
| WBGene00018026 | -0.260499013 | 0.84964436 | 1 | F35A5.4 |
| WBGene00018030 | -0.156269293 | 0.96924949 | 1 | F35B3.1 |
| WBGene00018030 | -0.156269293 | 0.96924949 | 1 | F35B3.3 |
| WBGene00018031 | -0.437155562 | 0.55264525 | 0.7305821 | F35B3.4 |
| WBGene00018033 | 1.85575961 | 0.00569698 | 0.03434312 | F35B3.7 |
| WBGene00018034 | -0.156269293 | 0.96924949 | 1 | sek-5 |
| WBGene00018035 | 0.532221447 | 0.41379171 | 0.61934132 | sek-4 |
| WBGene00018036 | -0.058580898 | 0.85054007 | 0.91824825 | F35C8.5 |
| WBGene00018037 | -0.062975349 | 0.79333964 | 0.88593495 | chtl-1 |
| WBGene00018038 | -1.167349518 | 0.36227261 | 1 | F35C8.8 |
| WBGene00018039 | 0.81835207 | 0.70463712 | 1 | F35D2.1 |
| WBGene00018040 | -0.686916415 | 0.61395655 | 1 | F35D2.2 |
| WBGene00018041 | -0.381300731 | 0.34222042 | 0.55060934 | F35D2.3 |
| WBGene00018042 | 1.848923794 | 0.0741315 | 0.21029461 | mks-3 |
| WBGene00018043 | -0.643299982 | 0.6128189 | 1 | F35D11.1 |
| WBGene00018044 | -0.177643032 | 0.5823022 | 0.75273967 | F35D11.3 |

| | | | | |
|---|---|---|---|---|
| WBGene00018045 | -0.789008925 | 0.00362536 | 0.02443949 | F35D11.4 |
| WBGene00018046 | -0.284402262 | 0.35166172 | 0.56037866 | F35D11.5 |
| WBGene00018047 | 1.557897343 | 0.63813005 | 1 | clec-136 |
| WBGene00018048 | -1.397181256 | 0.54849318 | 1 | clec-137 |
| WBGene00018049 | 0.747845955 | 0.85215965 | 1 | clec-138 |
| WBGene00018050 | -0.747215725 | 0.49009239 | 1 | clec-139 |
| WBGene00018051 | 0.957684322 | 0.00168352 | 0.01364462 | che-10 |
| WBGene00018052 | 1.380211602 | 0.01569416 | 0.07187884 | F35F10.1 |
| WBGene00018053 | 0.377828795 | 0.92526585 | 1 | srbc-3 |
| WBGene00018054 | 2.188838843 | 0.34004625 | 1 | F35F10.18 |
| WBGene00018054 | 0.489058549 | 0.85947456 | 1 | F35F10.19 |
| WBGene00018054 | 0.202213426 | 0.89675992 | 1 | F35F10.4 |
| WBGene00018055 | -1.6696632 | 0.1890031 | 1 | F35F10.5 |
| WBGene00018056 | 0.470566684 | 0.58959641 | 0.75742266 | F35F10.6 |
| WBGene00018057 | -0.693505305 | 0.79620356 | 1 | F35F10.7 |
| WBGene00018060 | 1.190409912 | 0.0367416 | 0.12990094 | arrd-18 |
| WBGene00018061 | 0.189068471 | 0.63456459 | 0.78788774 | arrd-19 |
| WBGene00018062 | 0.268996737 | 0.82115621 | 1 | F35F10.13 |
| WBGene00018063 | -0.156269293 | 0.96924949 | 1 | F35F10.14 |
| WBGene00018064 | -0.336980588 | 0.13420964 | 0.30825563 | cdc-73 |
| WBGene00018065 | 0.841601585 | 0.67828904 | 1 | F35F11.2 |
| WBGene00018067 | 0.916332024 | 0.38069424 | 1 | npr-7 |
| WBGene00018068 | 1.235073801 | 0.18312329 | 0.37527935 | F35H10.2 |
| WBGene00018069 | 0.904487496 | 0.39069232 | 1 | F35H10.3 |
| WBGene00018070 | -1.403306171 | 0.00023574 | 0.00298118 | F35H10.5 |
| WBGene00018071 | -0.40485044 | 0.16050898 | 0.34576641 | F35H10.6 |
| WBGene00018072 | 0.013776636 | 0.94723567 | 0.97304242 | nprl-3 |
| WBGene00018073 | 0.794806887 | 0.00030182 | 0.00361531 | F35H10.10 |
| WBGene00018074 | 1.193038118 | 0.05419859 | 0.16998841 | F35H12.1 |
| WBGene00018074 | 1.032841066 | 0.30376763 | 1 | F35H12.7 |
| WBGene00018075 | 0.452608686 | 0.12450509 | 0.29397363 | tbc-11 |
| WBGene00018076 | -0.114534929 | 0.70888028 | 0.83525966 | pifk-1 |
| WBGene00018077 | -0.110674652 | 0.81579955 | 0.90015998 | F35H12.5 |
| WBGene00018078 | -0.29196072 | 0.77002197 | 1 | F35H12.6 |
| WBGene00018081 | 0.747845955 | 0.85215965 | 1 | F36A4.2 |
| WBGene00018082 | -0.156269293 | 0.96924949 | 1 | F36A4.3 |
| WBGene00018083 | -0.514849788 | 0.8003149 | 1 | F36A4.4 |
| WBGene00018084 | 2.596290972 | 0.12970319 | 1 | F36A4.5 |
| WBGene00018085 | 0.6022904 | 0.3002818 | 0.50995852 | ttr-20 |
| WBGene00018087 | -1.901856996 | 0.64072034 | 1 | F36D4.1 |
| WBGene00018088 | -0.285924751 | 0.39202479 | 0.60022237 | trpp-4 |
| WBGene00018089 | 1.39078473 | 0.16724967 | 0.35468616 | F36D4.4 |
| WBGene00018089 | -1.291305095 | 0.63677871 | 1 | F36D4.7 |

| | | | | |
|---|---|---|---|---|
| WBGene00018090 | -0.296081278 | 0.0905486 | 0.23924932 | F36D4.5 |
| WBGene00018091 | -0.156269293 | 0.96924949 | 1 | F36D4.6 |
| WBGene00018092 | -0.741361551 | 0.44779133 | 1 | F36F12.1 |
| WBGene00018094 | -1.168224141 | 0.34205379 | 1 | F36F12.3 |
| WBGene00018096 | 0.747845955 | 0.85215965 | 1 | clec-207 |
| WBGene00018097 | 1.216957619 | 0.74819621 | 1 | clec-208 |
| WBGene00018098 | -0.156269293 | 0.96924949 | 1 | F36F12.7 |
| WBGene00018099 | 0.997927227 | 0.05237875 | 0.16556376 | ztf-28 |
| WBGene00018100 | 1.194971229 | 0.00019046 | 0.00254383 | math-26 |
| WBGene00018101 | 1.641272586 | 4.6485E-05 | 0.00081089 | math-27 |
| WBGene00018102 | -1.43288402 | 0.39434625 | 1 | math-28 |
| WBGene00018103 | -0.106781745 | 0.90366275 | 0.94998437 | F36H5.4 |
| WBGene00018104 | -0.716598049 | 0.43222149 | 0.6341955 | fbxb-53 |
| WBGene00018106 | -0.094707474 | 0.94598688 | 1 | F36H5.8 |
| WBGene00018108 | -1.700418717 | 0.39598915 | 1 | F36H5.10 |
| WBGene00018110 | 0.281454817 | 0.94435481 | 1 | F36H9.1 |
| WBGene00018111 | 1.582376833 | 0.32252793 | 1 | F36H9.2 |
| WBGene00018112 | 0.087242839 | 0.875981 | 0.93313702 | F36H9.4 |
| WBGene00018113 | -1.455505357 | 0.13318329 | 1 | F36H9.5 |
| WBGene00018114 | 0.281454817 | 0.94435481 | 1 | srt-41 |
| WBGene00018115 | 1.472167697 | 0.191657 | 1 | F36H9.7 |
| WBGene00018117 | 0.586603602 | 0.14274703 | 0.32052915 | nlp-47 |
| WBGene00018118 | -0.553705926 | 0.25226643 | 0.46035874 | F36H12.2 |
| WBGene00018119 | 0.983488853 | 0.80601565 | 1 | F36H12.3 |
| WBGene00018121 | -0.604396148 | 0.82723443 | 1 | F36H12.5 |
| WBGene00018122 | -1.964653846 | 0.53599202 | 1 | ttbk-2 |
| WBGene00018123 | 0.377828795 | 0.92526585 | 1 | F36H12.9 |
| WBGene00018124 | -0.156269293 | 0.96924949 | 1 | F36H12.10 |
| WBGene00018130 | 0.934295295 | 0.44178497 | 1 | F36H12.17 |
| WBGene00018131 | -0.073082997 | 0.66447928 | 0.80771846 | F37A4.1 |
| WBGene00018132 | -0.404251899 | 0.07517155 | 0.21209803 | F37A4.2 |
| WBGene00018133 | -2.112534605 | 0.02421452 | 0.09802884 | F37A4.3 |
| WBGene00018134 | 1.879339397 | 0.36234853 | 1 | F37A4.4 |
| WBGene00018135 | -0.578019945 | 0.81405273 | 1 | F37A4.5 |
| WBGene00018136 | 1.793595233 | 0.05838143 | 0.17908261 | F37A4.6 |
| WBGene00018137 | -0.338779339 | 0.02668959 | 0.10466354 | bath-41 |
| WBGene00018138 | -1.005970014 | 2.58E-10 | 2.34E-08 | folt-2 |
| WBGene00018140 | -0.811568259 | 0.02160905 | 0.0907458 | F37B4.10 |
| WBGene00018142 | 2.710329993 | 0.09752451 | 1 | oac-22 |
| WBGene00018144 | 0.184986825 | 0.76609132 | 0.86893822 | F37C4.4 |
| WBGene00018145 | -0.007167235 | 0.97097996 | 0.98485111 | F37C4.5 |
| WBGene00018146 | -0.042441227 | 0.87880521 | 0.93502533 | F37C4.6 |
| WBGene00018148 | -0.425459577 | 0.82035906 | 1 | F37C4.8 |

| | | | | |
|---|---|---|---|---|
| WBGene00018149 | -0.254003358 | 0.18742431 | 0.38096744 | F37C12.1 |
| WBGene00018150 | 0.381060651 | 0.11427832 | 0.27810244 | epg-4 |
| WBGene00018151 | -1.064877884 | 6.21E-09 | 4.01E-07 | F37C12.3 |
| WBGene00018152 | 0.278015773 | 0.11923329 | 0.28501897 | acs-4 |
| WBGene00018153 | 1.082023345 | 0.32415504 | 1 | F37C12.10 |
| WBGene00018154 | -0.364947272 | 0.06737688 | 0.1971139 | exos-9 |
| WBGene00018155 | -1.14715222 | 0.68932198 | 1 | F37C12.14 |
| WBGene00018156 | -0.012061811 | 0.94414844 | 0.97120685 | ncbp-1 |
| WBGene00018157 | 1.730335683 | 0.22685164 | 1 | lron-6 |
| WBGene00018158 | 2.063391569 | 0.37853067 | 1 | comp-1 |
| WBGene00018159 | -0.339968051 | 0.23650589 | 0.4426419 | F37F2.2 |
| WBGene00018160 | -0.189318362 | 0.23953473 | 0.44640333 | odr-8 |
| WBGene00018161 | -0.219182517 | 0.22910808 | 0.43435371 | F38A5.2 |
| WBGene00018163 | 1.230180765 | 0.55687914 | 1 | F38A5.6 |
| WBGene00018164 | -0.659217862 | 0.0037733 | 0.02511605 | sup-36 |
| WBGene00018165 | 0.377828795 | 0.92526585 | 1 | F38A5.8 |
| WBGene00018167 | 0.281454817 | 0.94435481 | 1 | nspb-4 |
| WBGene00018170 | -0.156269293 | 0.96924949 | 1 | nspb-1 |
| WBGene00018172 | 1.007245377 | 0.5574325 | 1 | F38B6.2 |
| WBGene00018173 | -0.402867511 | 0.78379798 | 1 | F38B6.3 |
| WBGene00018174 | 0.897355653 | 0.0064782 | 0.03756056 | F38B6.4 |
| WBGene00018175 | -0.183629655 | 0.75435762 | 0.86280319 | F38B6.6 |
| WBGene00018176 | -1.323116856 | 0.47668251 | 1 | F38B6.7 |
| WBGene00018178 | -0.156269293 | 0.96924949 | 1 | F38E1.3 |
| WBGene00018181 | -0.378995799 | 0.05702813 | 0.17644712 | F38E1.9 |
| WBGene00018182 | -0.156269293 | 0.96924949 | 1 | F38E1.10 |
| WBGene00018184 | 0.962185074 | 0.0041606 | 0.02712881 | F38E9.1 |
| WBGene00018187 | 0.234382149 | 0.31785272 | 0.5256845 | F38E9.5 |
| WBGene00018188 | 0.111456465 | 0.89926038 | 0.94707844 | F38E9.6 |
| WBGene00018189 | -0.015480473 | 0.97879941 | 0.98873753 | nhr-181 |
| WBGene00018191 | 1.438273822 | 0.11180174 | 0.27388718 | frpr-7 |
| WBGene00018193 | 1.202106532 | 1.0364E-06 | 3.4568E-05 | F39C12.1 |
| WBGene00018195 | 1.632851755 | 0.01099424 | 0.05564031 | btb-16 |
| WBGene00018196 | -0.323868354 | 0.91634405 | 1 | nep-13 |
| WBGene00018198 | -0.156269293 | 0.96924949 | 1 | F39E9.6 |
| WBGene00018200 | 1.819239453 | 0.00393274 | 0.02592149 | btb-17 |
| WBGene00018201 | 2.583075586 | 0.1281807 | 1 | btb-18 |
| WBGene00018203 | -1.426716011 | 0.03871389 | 0.13457338 | F39F10.3 |
| WBGene00018204 | -1.443372979 | 0.13733107 | 0.31346804 | F39F10.4 |
| WBGene00018205 | 0.36729309 | 0.88005435 | 1 | F39F10.5 |
| WBGene00018206 | 0.664001439 | 0.13866026 | 0.31551639 | ugt-61 |
| WBGene00018207 | -0.663513959 | 0.44024389 | 0.64027325 | F39G3.2 |
| WBGene00018208 | -0.254892966 | 0.34126941 | 0.54966197 | F39G3.3 |

| | | | | |
|---|---|---|---|---|
| WBGene00018209 | 0.983488853 | 0.80601565 | 1 | F39G3.4 |
| WBGene00018210 | 0.941862994 | 0.03175153 | 0.11812001 | F39G3.5 |
| WBGene00018211 | -1.094201388 | 0.7816772 | 1 | oac-24 |
| WBGene00018212 | -0.123987416 | 0.75364168 | 0.86245977 | F39H12.1 |
| WBGene00018213 | 0.06131249 | 0.94946829 | 1 | F39H12.2 |
| WBGene00018214 | 0.034492242 | 0.90996548 | 0.95322613 | F39H12.3 |
| WBGene00018215 | -0.218362472 | 0.62021342 | 0.77878286 | igcm-1 |
| WBGene00018216 | 1.188103115 | 0.00077326 | 0.00754066 | F40A3.1 |
| WBGene00018217 | 0.306840462 | 0.35405688 | 0.56261222 | F40A3.2 |
| WBGene00018218 | -0.512445935 | 0.00530131 | 0.03253953 | F40A3.3 |
| WBGene00018219 | -0.201933802 | 0.79487824 | 0.88641923 | F40A3.4 |
| WBGene00018221 | 0.65166684 | 0.04594939 | 0.15152153 | F40A3.6 |
| WBGene00018222 | 0.979744796 | 0.21709534 | 0.4191055 | F40A3.7 |
| WBGene00018223 | 3.363137959 | 0.02559932 | 1 | bath-13 |
| WBGene00018226 | 0.088856009 | 0.79510658 | 0.88652668 | F40B5.2 |
| WBGene00018227 | 2.273081356 | 0.21178645 | 1 | nep-15 |
| WBGene00018228 | -1.901856996 | 0.64072034 | 1 | F40C5.1 |
| WBGene00018229 | 0.687259521 | 0.81162667 | 1 | F40C5.2 |
| WBGene00018230 | -0.697713929 | 0.00118051 | 0.01041899 | mage-1 |
| WBGene00018231 | -0.156269293 | 0.96924949 | 1 | F40E3.3 |
| WBGene00018232 | 2.375448764 | 0.03880677 | 1 | F40E3.5 |
| WBGene00018233 | 1.477253415 | 0.58357327 | 1 | F40E3.6 |
| WBGene00018234 | -1.901856996 | 0.64072034 | 1 | F40E12.1 |
| WBGene00018235 | -2.691222824 | 0.22148476 | 1 | F40E12.2 |
| WBGene00018236 | -2.174605245 | 0.0823871 | 1 | fbxb-71 |
| WBGene00018237 | 1.718494591 | 0.00165506 | 0.01347688 | irg-7 |
| WBGene00018238 | -0.122777693 | 0.62828564 | 0.78388819 | F40F4.7 |
| WBGene00018239 | -0.346438615 | 0.18314226 | 0.37527935 | sec-20 |
| WBGene00018240 | -0.781195253 | 0.05387421 | 0.16910207 | cox-17 |
| WBGene00018241 | -0.667549252 | 0.31346256 | 0.52133547 | F40G9.5 |
| WBGene00018242 | -1.558787542 | 0.24071841 | 1 | F40G9.6 |
| WBGene00018243 | -0.156269293 | 0.96924949 | 1 | F40G9.7 |
| WBGene00018245 | -2.34801228 | 0.04342233 | 1 | F40G9.9 |
| WBGene00018246 | -1.397181256 | 0.54849318 | 1 | clec-148 |
| WBGene00018247 | -3.44789554 | 0.05425664 | 1 | F40G9.12 |
| WBGene00018248 | -1.445254974 | 0.45327704 | 1 | F40G9.14 |
| WBGene00018249 | 0.49621722 | 0.09674429 | 0.24974962 | F40H3.1 |
| WBGene00018250 | -0.109320946 | 0.86225496 | 0.92569082 | F40H3.2 |
| WBGene00018251 | -0.007524198 | 0.99494991 | 1 | F40H3.3 |
| WBGene00018252 | -0.380800723 | 0.80248573 | 1 | F40H3.6 |
| WBGene00018253 | -0.333693239 | 0.89573179 | 1 | F40H6.1 |
| WBGene00018254 | -0.156269293 | 0.96924949 | 1 | F40H6.2 |
| WBGene00018255 | 1.557897343 | 0.63813005 | 1 | F40H6.5 |

| | | | | |
|---|---|---|---|---|
| WBGene00018256 | -0.626715597 | 0.33379466 | 0.54208145 | cutl-28 |
| WBGene00018257 | 0.956082383 | 0.01330178 | 0.06379446 | F41B4.1 |
| WBGene00018258 | 0.453818256 | 0.32068632 | 0.52900172 | F41B4.2 |
| WBGene00018259 | -0.164976798 | 0.75015645 | 0.86075508 | F41B4.3 |
| WBGene00018260 | 0.688459475 | 0.14333037 | 0.32130244 | cyp-33C7 |
| WBGene00018261 | 0.164893178 | 0.838375 | 0.91174407 | cyp-33C5 |
| WBGene00018262 | 0.646054859 | 0.3022254 | 0.51162795 | cyp-33C3 |
| WBGene00018264 | 0.57991002 | 0.82402153 | 1 | cyp-33C6 |
| WBGene00018265 | 0.487553572 | 0.30038417 | 0.5099936 | nhr-182 |
| WBGene00018266 | 0.197348964 | 0.76460486 | 0.86822555 | nhr-183 |
| WBGene00018267 | -0.558804446 | 0.87818675 | 1 | F41C3.1 |
| WBGene00018268 | 1.704084581 | 8.6973E-05 | 0.00136956 | F41C3.2 |
| WBGene00018269 | 0.174857121 | 0.42716147 | 0.63047524 | acs-11 |
| WBGene00018270 | -0.669425483 | 0.00702839 | 0.0398769 | F41C3.4 |
| WBGene00018271 | -0.166923144 | 0.18515442 | 0.37768143 | F41C3.5 |
| WBGene00018273 | 1.290671456 | 0.5805443 | 1 | F41C3.6 |
| WBGene00018273 | 0.509041539 | 0.7857501 | 1 | F41C3.7 |
| WBGene00018274 | 1.181220482 | 0.02787627 | 0.10781963 | F41C3.8 |
| WBGene00018275 | 0.11548236 | 0.8408863 | 0.9124319 | F41C3.11 |
| WBGene00018278 | 0.045078548 | 0.97556699 | 1 | F41C6.4 |
| WBGene00018279 | 0.938982574 | 0.8146943 | 1 | F41C6.6 |
| WBGene00018280 | 0.372709901 | 0.63494151 | 0.78806117 | F41C6.7 |
| WBGene00018281 | 0.692476532 | 0.1479019 | 0.32851755 | tbc-18 |
| WBGene00018282 | 0.12884478 | 0.8078417 | 0.89513608 | F41D9.2 |
| WBGene00018283 | 1.867751938 | 0.05581743 | 0.17380852 | sulp-3 |
| WBGene00018284 | -0.156269293 | 0.96924949 | 1 | F41E6.1 |
| WBGene00018285 | 0.155239644 | 0.36500575 | 0.5726977 | smk-1 |
| WBGene00018286 | 0.815811729 | 0.11370286 | 0.27692406 | F41E6.5 |
| WBGene00018288 | 0.477630047 | 0.68698212 | 1 | F41E6.7 |
| WBGene00018289 | -0.156269293 | 0.96924949 | 1 | F41E6.8 |
| WBGene00018290 | -0.158879212 | 0.39211663 | 0.60022237 | vps-60 |
| WBGene00018291 | 3.297716337 | 0.07510938 | 1 | fip-4 |
| WBGene00018292 | 0.163636098 | 0.86726112 | 1 | F41E6.11 |
| WBGene00018293 | -0.597423744 | 0.09363694 | 0.24432541 | F41E6.12 |
| WBGene00018294 | -0.072219842 | 0.67930446 | 0.81647553 | atg-18 |
| WBGene00018295 | 2.593288645 | 0.13723948 | 1 | oac-29 |
| WBGene00018296 | 0.574508983 | 0.8551971 | 1 | F41F3.1 |
| WBGene00018297 | -1.40212578 | 6.44E-11 | 6.36E-09 | F41F3.3 |
| WBGene00018298 | 0.577009141 | 0.20150156 | 0.40041719 | nlp-50 |
| WBGene00018300 | -0.337421596 | 0.85636697 | 1 | F41G3.2 |
| WBGene00018300 | -0.636528479 | 0.55634721 | 1 | F41G3.3 |
| WBGene00018301 | 1.687239501 | 0.59403242 | 1 | F41G3.5 |
| WBGene00018302 | -0.47796996 | 0.00451627 | 0.02893512 | F41G3.6 |

| | | | | |
|---|---|---|---|---|
| WBGene00018303 | 1.123292597 | 1.76E-08 | 1.0004E-06 | F41G3.10 |
| WBGene00018304 | 0.806079254 | 0.02308127 | 0.0948645 | agr-1 |
| WBGene00018305 | -0.168625853 | 0.49519725 | 0.6835918 | exos-8 |
| WBGene00018307 | 0.504382305 | 0.59903274 | 1 | dyla-1 |
| WBGene00018309 | -0.950208353 | 0.7250598 | 1 | F41G4.4 |
| WBGene00018309 | 2.834144852 | 0.16713957 | 1 | F41G4.5 |
| WBGene00018310 | 2.752963715 | 0.09568412 | 1 | F41G4.7 |
| WBGene00018311 | 1.527805476 | 0.00356219 | 0.02412075 | F41G4.8 |
| WBGene00018312 | -0.156269293 | 0.96924949 | 1 | F41H8.1 |
| WBGene00018313 | -0.156269293 | 0.96924949 | 1 | F41H8.2 |
| WBGene00018314 | 2.213591416 | 0.45457697 | 1 | F41H10.1 |
| WBGene00018316 | 0.28072399 | 0.15978475 | 0.34506343 | F41H10.3 |
| WBGene00018317 | 0.096174571 | 0.71037395 | 0.83612701 | F41H10.4 |
| WBGene00018318 | -0.108299835 | 0.91620606 | 1 | F41H10.5 |
| WBGene00018319 | -0.108537336 | 0.51717252 | 0.703015 | hda-6 |
| WBGene00018321 | 0.115545861 | 0.57730628 | 0.74903703 | sand-1 |
| WBGene00018327 | 0.938982574 | 0.8146943 | 1 | F42A6.5 |
| WBGene00018328 | -0.461110603 | 0.17989448 | 0.37226843 | F42A6.6 |
| WBGene00018330 | 0.118888362 | 0.64496138 | 0.79498712 | elks-1 |
| WBGene00018332 | -0.263229581 | 0.93409872 | 1 | F42A9.3 |
| WBGene00018333 | 0.755585675 | 0.54407926 | 1 | cyp-33E3 |
| WBGene00018334 | 0.617994752 | 0.17692297 | 0.36850606 | cyp-33E2 |
| WBGene00018335 | -1.171137785 | 1.0599E-05 | 0.00023875 | F42A9.6 |
| WBGene00018336 | -1.343645114 | 0.56927291 | 1 | F42A9.7 |
| WBGene00018337 | -0.643499289 | 0.04264862 | 0.14385895 | F42A9.8 |
| WBGene00018338 | -0.794518274 | 0.64094715 | 1 | F42A9.9 |
| WBGene00018339 | -0.082774584 | 0.62770567 | 0.7837037 | abcf-3 |
| WBGene00018340 | -0.828503086 | 0.00819022 | 0.04459078 | anmt-2 |
| WBGene00018341 | -0.677649332 | 0.00030952 | 0.00369867 | F42A10.5 |
| WBGene00018342 | 1.138820138 | 1.6466E-06 | 4.9361E-05 | F42A10.6 |
| WBGene00018343 | 0.704381215 | 0.10655841 | 0.26614394 | F42A10.7 |
| WBGene00018344 | 0.516033535 | 0.60702393 | 1 | npr-27 |
| WBGene00018345 | 0.228359072 | 0.87973364 | 1 | F42C5.3 |
| WBGene00018346 | 1.281528566 | 0.04749176 | 0.15547149 | F42C5.4 |
| WBGene00018347 | -0.582567818 | 0.77294373 | 1 | F42C5.5 |
| WBGene00018348 | -0.017750584 | 0.99027451 | 1 | F42C5.6 |
| WBGene00018349 | 0.508629809 | 0.02850437 | 0.10941439 | F42C5.9 |
| WBGene00018350 | 1.20991953 | 2.59E-09 | 1.84E-07 | ifo-1 |
| WBGene00018351 | 1.260831804 | 0.0055351 | 0.0336008 | F42G2.2 |
| WBGene00018352 | -0.814848508 | 0.46124799 | 1 | fbxc-20 |
| WBGene00018353 | 1.325705253 | 0.04176752 | 0.1419254 | fbxa-182 |
| WBGene00018354 | 0.795305863 | 0.3537969 | 0.56237866 | F42G2.5 |
| WBGene00018355 | -1.251088452 | 0.00489959 | 0.03072618 | ceh-86 |

| | | | | |
|---|---|---|---|---|
| WBGene00018356 | 0.55794246 | 0.19108633 | 0.38576307 | islo-1 |
| WBGene00018357 | -0.281103652 | 0.25968699 | 0.4678664 | moc-3 |
| WBGene00018358 | -0.869028093 | 0.27322078 | 0.48186715 | erp-44.3 |
| WBGene00018359 | -1.190286606 | 0.49398728 | 1 | F42G8.8 |
| WBGene00018360 | 0.052793665 | 0.97847826 | 1 | irld-8 |
| WBGene00018361 | -0.585225866 | 0.00061918 | 0.00628555 | F42G8.10 |
| WBGene00018362 | -0.461840484 | 0.00667632 | 0.03839755 | F42G9.1 |
| WBGene00018363 | -0.012989819 | 0.99022363 | 1 | nlp-52 |
| WBGene00018364 | 0.06183828 | 0.76389966 | 0.86806496 | dagl-2 |
| WBGene00018365 | 1.70100155 | 0.29159196 | 1 | tpst-2 |
| WBGene00018366 | -0.672353184 | 0.03555453 | 0.12715091 | F42H10.2 |
| WBGene00018367 | -0.18965121 | 0.55956819 | 0.73620692 | F42H10.3 |
| WBGene00018369 | 1.359177333 | 0.00331851 | 0.02280135 | mig-39 |
| WBGene00018370 | -0.377427253 | 0.26098022 | 0.46962515 | F42H10.6 |
| WBGene00018371 | -0.381869402 | 0.04768698 | 0.15592988 | ess-2 |
| WBGene00018373 | 0.85009087 | 0.00291602 | 0.02077929 | F43B10.1 |
| WBGene00018374 | 0.709737394 | 0.05252858 | 0.16590777 | tag-343 |
| WBGene00018375 | 2.410908631 | 0.00078892 | 0.00766199 | F43C9.1 |
| WBGene00018376 | 0.615173649 | 0.05951037 | 0.18139981 | F43C9.2 |
| WBGene00018377 | -2.383751247 | 0.26793499 | 1 | hum-10 |
| WBGene00018378 | -0.156269293 | 0.96924949 | 1 | F43C11.1 |
| WBGene00018380 | 0.898712986 | 0.25282195 | 0.46088758 | decr-1.1 |
| WBGene00018381 | -0.156269293 | 0.96924949 | 1 | F43C11.4 |
| WBGene00018383 | -0.845474386 | 0.42380749 | 1 | F43C11.6 |
| WBGene00018384 | 0.650549293 | 0.1541716 | 0.33732156 | F43C11.7 |
| WBGene00018385 | -0.199374337 | 0.91832278 | 1 | F43C11.8 |
| WBGene00018386 | -0.452666625 | 0.15639423 | 0.34042182 | F43C11.9 |
| WBGene00018387 | -0.70576507 | 0.76841669 | 1 | F43C11.10 |
| WBGene00018388 | -1.969805064 | 0.62862569 | 1 | F43C11.11 |
| WBGene00018390 | 0.391252884 | 0.22357372 | 0.42746592 | F43E2.1 |
| WBGene00018391 | -0.668830196 | 0.00015166 | 0.00211403 | rpb-4 |
| WBGene00018392 | 0.413845433 | 0.34330597 | 0.55176655 | insc-1 |
| WBGene00018393 | 0.930270998 | 1.11E-09 | 8.67E-08 | msra-1 |
| WBGene00018394 | -1.43288402 | 0.39434625 | 1 | F43E2.6 |
| WBGene00018395 | -0.021551219 | 0.88557999 | 0.93951649 | mtch-1 |
| WBGene00018397 | -0.310604425 | 0.87613277 | 1 | F43E2.11 |
| WBGene00018398 | 0.498826044 | 0.01514087 | 0.07019452 | sptl-2 |
| WBGene00018399 | -0.030155961 | 0.92393022 | 0.95982175 | F43H9.3 |
| WBGene00018400 | 0.812593876 | 0.0025217 | 0.01870249 | F43H9.4 |
| WBGene00018402 | 0.782654779 | 0.26154373 | 0.47006725 | kvs-3 |
| WBGene00018403 | 0.904241477 | 0.02410976 | 0.0977351 | F44A2.3 |
| WBGene00018404 | 1.977885804 | 0.49204652 | 1 | nhr-39 |
| WBGene00018405 | 0.622298322 | 0.1079555 | 0.26759651 | F44A2.5 |

| | | | | |
|---|---|---|---|---|
| WBGene00018406 | -0.571866858 | 0.74008329 | 1 | F44A2.7 |
| WBGene00018407 | 0.617054551 | 0.07305536 | 0.2081864 | F44B9.2 |
| WBGene00018408 | -0.429072608 | 0.10360542 | 0.26080727 | F44B9.5 |
| WBGene00018409 | -0.410832868 | 0.01482145 | 0.06907279 | F44B9.8 |
| WBGene00018410 | 1.743415485 | 0.50483382 | 1 | F44B9.9 |
| WBGene00018411 | 0.024039381 | 0.98960809 | 1 | F44B9.10 |
| WBGene00018412 | 1.413734769 | 0.68804531 | 1 | nhr-37 |
| WBGene00018413 | 0.828673199 | 0.15931288 | 0.34428937 | cyp-33C4 |
| WBGene00018414 | -0.133799809 | 0.94295257 | 1 | nstp-8 |
| WBGene00018415 | 0.329635864 | 0.63750024 | 0.78994595 | nhr-184 |
| WBGene00018417 | -0.422904263 | 0.04567556 | 0.15093821 | F44E2.3 |
| WBGene00018418 | 0.60262121 | 0.11296879 | 0.27557847 | F44E2.4 |
| WBGene00018419 | -0.384189982 | 0.16780592 | 0.35543121 | F44E2.6 |
| WBGene00018420 | -0.148290356 | 0.39734799 | 0.60512336 | F44E2.7 |
| WBGene00018421 | -0.577083811 | 0.00025795 | 0.00320341 | F44E2.8 |
| WBGene00018422 | -0.127749805 | 0.71707039 | 0.84010542 | F44E2.9 |
| WBGene00018423 | -0.582108738 | 0.01404747 | 0.06643956 | F44E2.10 |
| WBGene00018424 | -0.203287066 | 0.50127384 | 0.6891593 | F44E7.2 |
| WBGene00018426 | -0.155464198 | 0.3690811 | 0.57663282 | F44E7.4 |
| WBGene00018427 | -0.795397859 | 0.00207449 | 0.01600286 | F44E7.5 |
| WBGene00018428 | 0.688793783 | 0.86378755 | 1 | fbxa-183 |
| WBGene00018429 | -0.201285786 | 0.80566313 | 0.89402761 | F44E7.7 |
| WBGene00018430 | 0.174100086 | 0.86436418 | 1 | nhr-142 |
| WBGene00018431 | -0.61073326 | 0.00393628 | 0.02593074 | F44E7.9 |
| WBGene00018432 | 0.627834078 | 0.77998941 | 1 | irld-33 |
| WBGene00018433 | -1.014549262 | 0.3340114 | 1 | ceh-82 |
| WBGene00018434 | 0.281454817 | 0.94435481 | 1 | ceh-81 |
| WBGene00018435 | -0.467085339 | 0.37755353 | 0.58512604 | btb-9 |
| WBGene00018436 | -2.120378547 | 0.02069591 | 0.08758072 | fbxb-11 |
| WBGene00018437 | -1.201616871 | 0.0239596 | 0.0972888 | btb-8 |
| WBGene00018438 | -0.988639091 | 0.00094058 | 0.00878432 | btb-6 |
| WBGene00018439 | -0.63994169 | 0.46552945 | 0.65974005 | fbxa-184 |
| WBGene00018441 | -2.705999569 | 0.42423051 | 1 | math-29 |
| WBGene00018442 | -2.440347847 | 0.00747552 | 0.04169991 | sdz-18 |
| WBGene00018443 | -0.411545956 | 0.39403494 | 0.60200405 | btb-7 |
| WBGene00018444 | -1.159741626 | 0.40275032 | 1 | fbxb-48 |
| WBGene00018445 | -2.236771922 | 0.23979596 | 1 | fbxb-5 |
| WBGene00018446 | -0.858197775 | 0.00929024 | 0.04904035 | ceh-83 |
| WBGene00018447 | 0.688793783 | 0.86378755 | 1 | irld-9 |
| WBGene00018452 | -1.889541999 | 0.09188255 | 1 | F45D11.5 |
| WBGene00018455 | -0.969682205 | 0.01789545 | 0.07903554 | fbxc-42 |
| WBGene00018456 | -1.553324722 | 0.05064618 | 0.16217522 | fbxc-39 |
| WBGene00018458 | -1.164667479 | 0.4419273 | 1 | fbxb-30 |

| | | | | |
|---|---|---|---|---|
| WBGene00018462 | -0.51334688 | 0.43007816 | 0.63268859 | F45E1.1 |
| WBGene00018463 | -0.507262291 | 0.73574821 | 1 | F45E1.2 |
| WBGene00018465 | -0.922285159 | 0.14236863 | 0.32034267 | F45E1.4 |
| WBGene00018466 | 1.139219656 | 0.2039776 | 0.40340316 | F45E1.5 |
| WBGene00018467 | 0.112634676 | 0.72564779 | 0.84579072 | sdpn-1 |
| WBGene00018468 | 1.359374447 | 2.1492E-06 | 6.2329E-05 | cla-1 |
| WBGene00018470 | -0.720442336 | 0.3677929 | 0.57541871 | F45E4.5 |
| WBGene00018471 | -0.087822178 | 0.96354485 | 1 | F45E4.6 |
| WBGene00018472 | 0.306405064 | 0.77260848 | 1 | nep-16 |
| WBGene00018473 | 0.526655207 | 0.34697278 | 0.55589819 | cima-1 |
| WBGene00018474 | -0.148716103 | 0.46211173 | 0.65719791 | cnep-1 |
| WBGene00018475 | -0.21025135 | 0.3460126 | 0.55474477 | mrpl-14 |
| WBGene00018476 | 2.697061674 | 0.24723036 | 1 | F45E12.6 |
| WBGene00018478 | -0.934013985 | 0.67351079 | 1 | otpl-5 |
| WBGene00018479 | 2.641117225 | 0.21179361 | 1 | otpl-6 |
| WBGene00018480 | 0.939836669 | 0.48263385 | 1 | otpl-7 |
| WBGene00018481 | -0.197097633 | 0.45446373 | 0.65204965 | F45F2.9 |
| WBGene00018482 | 0.004825592 | 0.97961928 | 0.98906925 | F45F2.10 |
| WBGene00018483 | -0.306769884 | 0.0901686 | 0.23860876 | F45F2.11 |
| WBGene00018484 | -0.306426754 | 0.80275515 | 1 | F46C8.1 |
| WBGene00018485 | 0.355129141 | 0.45069667 | 0.64902245 | F46C8.3 |
| WBGene00018486 | -0.991305064 | 0.31194197 | 1 | glb-16 |
| WBGene00018487 | -0.45292808 | 0.43069797 | 0.63306345 | F46C8.8 |
| WBGene00018488 | -0.95675118 | 0.00103477 | 0.00948166 | acs-1 |
| WBGene00018489 | -1.855569013 | 0.00135783 | 0.01162584 | F46E10.2 |
| WBGene00018491 | -0.270667886 | 0.08792082 | 0.23501017 | mdh-1 |
| WBGene00018492 | 1.021767955 | 0.02064866 | 0.087418 | hpo-26 |
| WBGene00018497 | 1.477253415 | 0.58357327 | 1 | msrp-3 |
| WBGene00018498 | -1.831477136 | 0.52194793 | 1 | F46F5.7 |
| WBGene00018500 | 0.377828795 | 0.92526585 | 1 | msrp-4 |
| WBGene00018502 | 1.216957619 | 0.74819621 | 1 | F46F5.11 |
| WBGene00018506 | -0.07748118 | 0.98067866 | 1 | F46F5.15 |
| WBGene00018508 | 0.593283491 | 0.00195881 | 0.01531355 | F46F11.1 |
| WBGene00018509 | -0.192112444 | 0.25469695 | 0.46230167 | xrep-4 |
| WBGene00018510 | -0.199033643 | 0.65619456 | 0.80231895 | F46F11.7 |
| WBGene00018511 | 0.018830957 | 0.93755177 | 0.96772742 | F46F11.8 |
| WBGene00018512 | 0.33246298 | 0.0900188 | 0.23835296 | trpp-8 |
| WBGene00018513 | -0.058704907 | 0.87065345 | 0.9302652 | F46F11.10 |
| WBGene00018514 | 0.296535033 | 0.62014342 | 0.77878286 | F46G11.1 |
| WBGene00018515 | 0.019318942 | 0.98642863 | 1 | F46G11.2 |
| WBGene00018516 | -0.140768852 | 0.71044854 | 0.83613374 | tag-257 |
| WBGene00018517 | 0.000189062 | 0.99982091 | 0.99990327 | F46G11.4 |
| WBGene00018518 | 0.308798519 | 0.19697538 | 0.39418795 | F46H5.2 |

| | | | | |
|---|---|---|---|---|
| WBGene00018519 | -0.118654211 | 0.44830751 | 0.64735123 | F46H5.3 |
| WBGene00018520 | 0.584946587 | 0.01743852 | 0.07766731 | F46H5.4 |
| WBGene00018521 | 0.491377013 | 0.72370417 | 1 | F46H5.5 |
| WBGene00018522 | 0.264924214 | 0.20547282 | 0.40497493 | F46H5.7 |
| WBGene00018523 | 0.542088143 | 0.29338523 | 0.50360385 | lact-1 |
| WBGene00018526 | -0.156269293 | 0.96924949 | 1 | F47B3.2 |
| WBGene00018527 | -0.453019766 | 0.49477111 | 0.68315889 | F47B3.3 |
| WBGene00018528 | 0.281454817 | 0.94435481 | 1 | F47B3.4 |
| WBGene00018531 | 1.247575581 | 0.7549672 | 1 | F47B3.7 |
| WBGene00018532 | 0.313130107 | 0.229322 | 0.43442105 | F47B7.1 |
| WBGene00018533 | -0.315192608 | 0.12759337 | 0.29825012 | F47B7.2 |
| WBGene00018534 | 0.281454817 | 0.94435481 | 1 | F47B7.3 |
| WBGene00018536 | 0.312403178 | 0.89676939 | 1 | F47B7.5 |
| WBGene00018537 | 0.399606396 | 0.86544406 | 1 | F47B7.6 |
| WBGene00018538 | 1.106276567 | 0.00086412 | 0.0082156 | marg-1 |
| WBGene00018539 | 2.09403112 | 0.45107747 | 1 | nhr-185 |
| WBGene00018541 | -0.368785415 | 0.79835796 | 1 | nhr-186 |
| WBGene00018542 | 0.394210339 | 0.81241042 | 1 | nhr-187 |
| WBGene00018543 | -0.205892253 | 0.89648576 | 1 | ugt-32 |
| WBGene00018544 | 1.036568105 | 0.67105582 | 1 | nhr-188 |
| WBGene00018545 | -0.637018156 | 0.80584146 | 1 | nhr-189 |
| WBGene00018547 | 0.33479184 | 0.45517791 | 0.65257895 | clec-78 |
| WBGene00018548 | -0.263229581 | 0.93409872 | 1 | clec-79 |
| WBGene00018551 | -0.156269293 | 0.96924949 | 1 | F47C12.8 |
| WBGene00018560 | 0.250101878 | 0.78118286 | 0.87891216 | F47D12.3 |
| WBGene00018561 | -0.352371385 | 0.60193335 | 0.76683654 | gadr-3 |
| WBGene00018562 | 0.581871102 | 0.26647259 | 0.47484833 | F47D12.6 |
| WBGene00018563 | 2.201543524 | 0.41352885 | 1 | F47D12.7 |
| WBGene00018564 | -0.135342264 | 0.45311099 | 0.65072398 | F47D12.9 |
| WBGene00018565 | 0.393677747 | 0.84003705 | 1 | F47E1.1 |
| WBGene00018566 | 0.564970678 | 0.40969288 | 0.61507125 | F47E1.2 |
| WBGene00018567 | 1.538955341 | 0.37645059 | 1 | F47E1.3 |
| WBGene00018568 | -0.467521789 | 0.53792921 | 0.7198279 | F47E1.4 |
| WBGene00018569 | 0.523223342 | 0.18293524 | 0.37512395 | F47F2.1 |
| WBGene00018571 | 1.82867917 | 0.54358766 | 1 | F47F2.3 |
| WBGene00018572 | 0.147638633 | 0.77551697 | 0.87476731 | lin-42 |
| WBGene00018573 | 2.09403112 | 0.45107747 | 1 | oac-30 |
| WBGene00018575 | 0.446458324 | 0.87164163 | 1 | clec-119 |
| WBGene00018576 | 0.63519708 | 0.22010624 | 0.42310163 | F47G3.1 |
| WBGene00018578 | -0.579173591 | 0.09295641 | 0.24309904 | F47G3.3 |
| WBGene00018579 | 1.111069406 | 0.06484851 | 0.19207752 | acs-16 |
| WBGene00018582 | 2.751365943 | 0.12086202 | 1 | D2092.6 |
| WBGene00018582 | 1.006955725 | 0.01703932 | 0.07619681 | F48A9.1 |

| | | | | |
|---|---|---|---|---|
| WBGene00018582 | 1.400066903 | 0.12148985 | 0.28859484 | F48A9.2 |
| WBGene00018585 | -0.477226225 | 0.02827632 | 0.10884679 | F48A11.4 |
| WBGene00018586 | -0.311725121 | 0.10346388 | 0.26071284 | ubxn-3 |
| WBGene00018587 | 0.417607955 | 0.61779025 | 0.77718282 | F48B9.1 |
| WBGene00018589 | 0.012174154 | 0.9921192 | 1 | F48B9.3 |
| WBGene00018590 | 0.993108243 | 0.02463898 | 0.09921788 | nlp-37 |
| WBGene00018591 | 0.479479392 | 0.70945275 | 1 | npax-2 |
| WBGene00018593 | -0.060788827 | 0.88462796 | 0.93908088 | F48B9.8 |
| WBGene00018594 | 0.982918016 | 0.0198196 | 0.08499814 | aman-3 |
| WBGene00018595 | -0.220245787 | 0.7556011 | 0.86341205 | fbxa-145 |
| WBGene00018598 | -0.666560241 | 0.01511057 | 0.0701022 | F48C1.5 |
| WBGene00018599 | -0.445171968 | 0.06116607 | 0.1849607 | F48C1.6 |
| WBGene00018600 | -3.128943914 | 0.25272212 | 1 | F48C1.8 |
| WBGene00018601 | -4.669403835 | 0.00091588 | 1 | F48C1.9 |
| WBGene00018602 | -1.060316519 | 5.26E-07 | 1.9787E-05 | F48D6.4 |
| WBGene00018603 | -0.372282905 | 0.62406667 | 0.78128032 | F48E3.2 |
| WBGene00018604 | 0.102635888 | 0.57710165 | 0.74903703 | uggt-1 |
| WBGene00018605 | -0.467493686 | 0.00362119 | 0.02442494 | F48E3.4 |
| WBGene00018606 | -1.333789306 | 0.02061134 | 0.0873751 | F48E3.6 |
| WBGene00018607 | 0.992087649 | 0.0205654 | 0.08728554 | F48E3.8 |
| WBGene00018608 | 0.301830889 | 0.76273055 | 1 | F48E3.9 |
| WBGene00018609 | -0.372118273 | 0.04746944 | 0.15547149 | F48E8.2 |
| WBGene00018610 | -0.050238637 | 0.78668429 | 0.88255028 | F48E8.3 |
| WBGene00018611 | 0.278658474 | 0.17375089 | 0.36370855 | F48E8.4 |
| WBGene00018612 | -0.394255236 | 0.06479717 | 0.19204873 | disl-2 |
| WBGene00018613 | -0.312320779 | 0.0629395 | 0.18826029 | skpt-1 |
| WBGene00018614 | -1.496439068 | 0.28960411 | 1 | pals-31 |
| WBGene00018615 | 0.734322022 | 0.27794322 | 0.4868318 | F48G7.4 |
| WBGene00018616 | 0.016273281 | 0.98891192 | 1 | F48G7.5 |
| WBGene00018619 | 0.281454817 | 0.94435481 | 1 | F48G7.8 |
| WBGene00018621 | 3.550208197 | 0.04314845 | 1 | F48G7.10 |
| WBGene00018622 | 0.332837504 | 0.63141334 | 0.78625531 | nhr-190 |
| WBGene00018623 | -0.18557176 | 0.95664217 | 1 | F48G7.12 |
| WBGene00018624 | -0.182781662 | 0.85169479 | 1 | F48G7.13 |
| WBGene00018625 | -0.206788307 | 0.20575062 | 0.40532505 | prp-17 |
| WBGene00018626 | 0.131533329 | 0.92505435 | 1 | F49D11.2 |
| WBGene00018627 | 0.983488853 | 0.80601565 | 1 | F49D11.3 |
| WBGene00018628 | -1.819519428 | 0.47492301 | 1 | F49D11.4 |
| WBGene00018630 | -0.438662369 | 0.61009745 | 0.77201762 | F49D11.6 |
| WBGene00018631 | -0.156269293 | 0.96924949 | 1 | F49D11.7 |
| WBGene00018632 | -0.402222985 | 0.05696551 | 0.17629831 | tag-296 |
| WBGene00018634 | 0.400417366 | 0.49242661 | 0.68149352 | F49E7.2 |
| WBGene00018635 | -0.021726089 | 0.91485501 | 0.95571968 | nprl-2 |

| | | | | |
|---|---|---|---|---|
| WBGene00018636 | -0.358159411 | 0.06313468 | 0.18865817 | F49E8.2 |
| WBGene00018637 | 0.189720567 | 0.75668014 | 0.86407788 | F49E8.6 |
| WBGene00018638 | 0.254091447 | 0.30953442 | 0.51818605 | F49E8.7 |
| WBGene00018639 | 1.050233648 | 0.03416142 | 0.12428944 | tbc-16 |
| WBGene00018640 | 0.156212349 | 0.88438345 | 1 | F49E10.2 |
| WBGene00018641 | 0.814610553 | 0.20588189 | 0.40545208 | F49E10.4 |
| WBGene00018643 | -2.470872701 | 0.02176699 | 1 | drd-50 |
| WBGene00018645 | 2.171276578 | 0.00097111 | 0.00903471 | F49F1.5 |
| WBGene00018646 | 0.95465785 | 0.30948299 | 0.51818605 | mul-1 |
| WBGene00018647 | -1.640081156 | 0.12374439 | 1 | F49F1.7 |
| WBGene00018648 | 2.028800581 | 0.39258153 | 1 | F49F1.8 |
| WBGene00018649 | -1.094201388 | 0.7816772 | 1 | F49F1.9 |
| WBGene00018652 | 0.983488853 | 0.80601565 | 1 | F49F1.12 |
| WBGene00018653 | 0.041803775 | 0.98890197 | 1 | F49H12.2 |
| WBGene00018654 | 0.176011531 | 0.88808713 | 1 | F49H12.3 |
| WBGene00018655 | 1.43883144 | 0.37062111 | 1 | F49H12.4 |
| WBGene00018656 | -0.430653155 | 0.17631584 | 0.36749367 | F49H12.5 |
| WBGene00018657 | 0.03020435 | 0.86160448 | 0.92540163 | acl-4 |
| WBGene00018659 | -0.285859072 | 0.39751953 | 0.60517281 | F52C6.2 |
| WBGene00018660 | -0.292851507 | 0.71242081 | 0.8375158 | F52C6.3 |
| WBGene00018661 | -0.258884222 | 0.87274372 | 1 | F52C6.4 |
| WBGene00018663 | 1.116613983 | 0.69067248 | 1 | math-31 |
| WBGene00018665 | 0.083519665 | 0.87621711 | 0.93317123 | bath-4 |
| WBGene00018667 | 0.938488271 | 0.09108832 | 0.24004847 | bath-7 |
| WBGene00018668 | -0.401316483 | 0.20931017 | 0.4094803 | bath-2 |
| WBGene00018669 | 1.944146159 | 0.10340595 | 1 | F52C6.12 |
| WBGene00018670 | 2.317189148 | 0.08995585 | 1 | F52C6.13 |
| WBGene00018671 | -0.711014871 | 0.49820734 | 1 | F52C6.14 |
| WBGene00018672 | 0.129690903 | 0.59915348 | 0.76480755 | sorf-2 |
| WBGene00018674 | -0.176770928 | 0.50140401 | 0.68918217 | F52C9.3 |
| WBGene00018675 | -0.671256058 | 0.62115519 | 1 | F52C9.5 |
| WBGene00018678 | -0.16814542 | 0.52612817 | 0.71045736 | F52C12.1 |
| WBGene00018679 | -0.451420291 | 0.09557558 | 0.24783919 | F52C12.2 |
| WBGene00018680 | -0.645982798 | 0.14856102 | 0.32943914 | tsen-54 |
| WBGene00018681 | 0.148229437 | 0.52706009 | 0.7113202 | denn-4 |
| WBGene00018682 | 0.090826847 | 0.75703182 | 0.86429398 | aagr-4 |
| WBGene00018683 | 1.035028786 | 0.44652054 | 1 | F52D1.2 |
| WBGene00018687 | 0.747845955 | 0.85215965 | 1 | F52D2.5 |
| WBGene00018688 | 1.203706625 | 0.40347713 | 1 | F52D2.6 |
| WBGene00018689 | -0.57841196 | 0.14458737 | 0.32307998 | F52D2.7 |
| WBGene00018690 | 0.916065526 | 0.64867207 | 1 | fbxa-46 |
| WBGene00018692 | 0.631397535 | 0.3423033 | 0.55066973 | F52E1.2 |
| WBGene00018693 | -0.62146648 | 0.85925752 | 1 | F52E1.3 |

| | | | | |
|---|---|---|---|---|
| WBGene00018694 | 0.618881218 | 0.54579125 | 1 | F52E1.5 |
| WBGene00018696 | 0.200454445 | 0.82401677 | 0.90496496 | pho-6 |
| WBGene00018697 | 1.690320038 | 0.0183179 | 0.08038245 | F52E1.9 |
| WBGene00018698 | -0.36084657 | 0.11795143 | 0.28312541 | vha-18 |
| WBGene00018700 | 0.036401438 | 0.82831153 | 0.90721969 | lmd-3 |
| WBGene00018701 | -0.177973752 | 0.28177603 | 0.49110577 | pccb-1 |
| WBGene00018702 | -0.211122645 | 0.60588231 | 0.76913604 | F52E4.5 |
| WBGene00018703 | -0.472995678 | 0.05508361 | 0.17196456 | sec-3 |
| WBGene00018704 | -0.097350864 | 0.81691614 | 0.90058829 | ztf-13 |
| WBGene00018706 | 1.719779899 | 0.22543726 | 1 | F52F10.2 |
| WBGene00018707 | 1.570518237 | 0.14552524 | 1 | oac-31 |
| WBGene00018708 | 1.146693847 | 0.07412009 | 0.21029461 | oac-32 |
| WBGene00018710 | 0.420972446 | 0.07211126 | 0.20653521 | F52G3.1 |
| WBGene00018711 | -1.849509659 | 0.37480932 | 1 | F52G3.3 |
| WBGene00018712 | -0.558804446 | 0.87818675 | 1 | F52G3.4 |
| WBGene00018713 | -1.113356607 | 0.29824099 | 1 | F52G3.5 |
| WBGene00018714 | 0.116148337 | 0.80168176 | 0.89148362 | F52H2.1 |
| WBGene00018715 | 1.456909245 | 0.02811855 | 0.10848024 | F52H2.3 |
| WBGene00018716 | 0.505512779 | 0.57375414 | 0.74749963 | F52H2.4 |
| WBGene00018717 | -0.217891202 | 0.60752924 | 0.77026028 | F52H2.5 |
| WBGene00018718 | 0.689785851 | 0.00482197 | 0.03040293 | F52H2.6 |
| WBGene00018719 | -0.130924261 | 0.6902755 | 0.82356867 | F52H2.7 |
| WBGene00018720 | 0.283103919 | 0.75858058 | 0.86486307 | F53A3.1 |
| WBGene00018721 | -0.318886512 | 0.10088755 | 0.25598238 | polh-1 |
| WBGene00018723 | -0.507993551 | 0.2477292 | 0.45583943 | F53A3.7 |
| WBGene00018724 | 0.978463802 | 0.03721266 | 0.13103218 | F53A9.1 |
| WBGene00018725 | 1.789476451 | 0.01030224 | 0.05302224 | kreg-1 |
| WBGene00018726 | 0.104540135 | 0.93938645 | 1 | F53A9.3 |
| WBGene00018727 | 1.492264931 | 0.38473528 | 1 | jbts-14 |
| WBGene00018728 | 0.270935547 | 0.76198517 | 0.86701223 | frpr-8 |
| WBGene00018729 | 1.030852357 | 0.0002385 | 0.00299925 | F53A9.6 |
| WBGene00018730 | 0.857789224 | 0.05819821 | 0.17879163 | F53A9.7 |
| WBGene00018731 | 2.427035977 | 1.91E-19 | 5.95E-17 | F53A9.8 |
| WBGene00018732 | 0.394145795 | 0.51398056 | 0.70091407 | F53A9.9 |
| WBGene00018734 | 0.379036159 | 0.16023605 | 0.34561687 | F53A10.2 |
| WBGene00018735 | -0.709023504 | 0.00860961 | 0.04625188 | F53B1.2 |
| WBGene00018736 | 0.673719511 | 0.62686549 | 1 | F53B1.3 |
| WBGene00018738 | -0.573049512 | 0.42828644 | 0.63100978 | swip-10 |
| WBGene00018739 | 0.677880415 | 0.04148363 | 0.14111873 | F53B1.8 |
| WBGene00018740 | -0.450860769 | 0.06881009 | 0.19999543 | tra-4 |
| WBGene00018741 | -1.397181256 | 0.54849318 | 1 | comt-1 |
| WBGene00018742 | 1.195226674 | 0.17480741 | 0.36503901 | F53B3.3 |
| WBGene00018743 | 0.032929933 | 0.90530203 | 0.95079204 | F53B3.5 |

| | | | | |
|---|---|---|---|---|
| WBGene00018744 | 0.974507935 | 0.01209244 | 0.05960791 | F53B3.6 |
| WBGene00018745 | -1.14715222 | 0.68932198 | 1 | F53C3.1 |
| WBGene00018746 | -2.305255035 | 0.00011786 | 0.00173663 | fbxb-103 |
| WBGene00018747 | -0.069180986 | 0.93569285 | 0.96641828 | F53C3.3 |
| WBGene00018748 | 1.679643193 | 0.04897672 | 0.15873634 | F53C3.4 |
| WBGene00018749 | 0.321988918 | 0.60001307 | 0.76536654 | F53C3.5 |
| WBGene00018750 | 1.46199259 | 0.26525732 | 1 | F53C3.6 |
| WBGene00018751 | -0.156269293 | 0.96924949 | 1 | F53C3.7 |
| WBGene00018752 | 1.229237684 | 0.59270769 | 1 | F53C3.8 |
| WBGene00018754 | -0.6791505 | 0.53577824 | 1 | F53C3.11 |
| WBGene00018755 | 1.552723749 | 0.0009105 | 0.00856927 | bcmo-2 |
| WBGene00018756 | 0.068489962 | 0.71775842 | 0.84033799 | F53C3.13 |
| WBGene00018757 | -0.326712687 | 0.25758315 | 0.4655131 | tag-304 |
| WBGene00018758 | 0.324158188 | 0.27433531 | 0.4827622 | F53E10.1 |
| WBGene00018760 | 1.171939221 | 0.00053618 | 0.0056557 | irg-3 |
| WBGene00018761 | 1.630244101 | 0.40224995 | 1 | F53E10.5 |
| WBGene00018762 | -0.533916424 | 0.04268895 | 0.14385895 | F53E10.6 |
| WBGene00018763 | -0.338057104 | 0.58747115 | 0.75612077 | F53F10.1 |
| WBGene00018764 | 0.429844889 | 0.00767899 | 0.04245474 | F53F10.2 |
| WBGene00018765 | -0.392454418 | 0.074134 | 0.21029461 | mpc-2 |
| WBGene00018766 | -0.784035283 | 0.51331372 | 1 | F53G2.1 |
| WBGene00018767 | -0.566012294 | 0.72618511 | 1 | F53G2.2 |
| WBGene00018768 | 1.353535409 | 0.73149271 | 1 | F53G2.3 |
| WBGene00018769 | -0.361167057 | 0.06478039 | 0.19204873 | mnat-1 |
| WBGene00018771 | -0.451813889 | 0.8460226 | 1 | duox-2 |
| WBGene00018772 | 1.132923217 | 0.38806432 | 1 | F53G12.4 |
| WBGene00018773 | -0.735886458 | 0.64148088 | 1 | F53G12.8 |
| WBGene00018774 | -1.100114328 | 0.17826139 | 0.37027743 | F53G12.9 |
| WBGene00018776 | 0.146655522 | 0.38688295 | 0.59555752 | F53H1.1 |
| WBGene00018777 | -0.037487318 | 0.89614002 | 0.9454324 | F53H1.3 |
| WBGene00018778 | 0.006016117 | 0.9767899 | 0.98802018 | F53H1.4 |
| WBGene00018779 | -0.196131774 | 0.61297891 | 0.77401736 | F53H8.3 |
| WBGene00018781 | 0.121219307 | 0.67984405 | 0.81647553 | F54A3.2 |
| WBGene00018782 | -0.236637589 | 0.1463721 | 0.3257752 | cct-3 |
| WBGene00018783 | 0.084737637 | 0.95621658 | 1 | cbs-2 |
| WBGene00018784 | -0.493843804 | 0.0574488 | 0.17738705 | F54A3.5 |
| WBGene00018785 | 0.176901791 | 0.54294192 | 0.7232458 | F54A3.6 |
| WBGene00018786 | -0.585634637 | 0.21495867 | 0.41611303 | hmbx-1 |
| WBGene00018787 | 0.486222365 | 0.3013039 | 0.51098347 | cutl-20 |
| WBGene00018788 | 0.373018936 | 0.43563044 | 0.63676706 | shc-1 |
| WBGene00018789 | 0.725954606 | 0.29291301 | 0.50307778 | F54C1.1 |
| WBGene00018791 | -0.67679423 | 0.54771208 | 1 | F54C1.6 |
| WBGene00018792 | 0.871933027 | 0.73427341 | 1 | F54C1.8 |

| | | | | |
|---|---|---|---|---|
| WBGene00018793 | -0.527591771 | 0.0352015 | 0.12651905 | mrpl-40 |
| WBGene00018794 | -0.205364323 | 0.44975773 | 0.64859349 | attf-3 |
| WBGene00018795 | -0.720751704 | 0.11605034 | 0.28048782 | F54C4.4 |
| WBGene00018797 | 0.988892196 | 3.24E-10 | 2.87E-08 | F54D7.2 |
| WBGene00018798 | 0.708104517 | 0.52089374 | 1 | gnrr-1 |
| WBGene00018802 | -0.098332474 | 0.58960559 | 0.75742266 | F54D8.6 |
| WBGene00018803 | 2.138154584 | 3.9071E-06 | 0.00010201 | fbxa-24 |
| WBGene00018804 | -0.447397714 | 0.4949169 | 0.68328247 | F54D10.3 |
| WBGene00018805 | 0.450261618 | 0.69594264 | 1 | F54D10.4 |
| WBGene00018806 | -0.726349558 | 0.00293157 | 0.0208507 | F54D10.5 |
| WBGene00018808 | -0.68660897 | 0.01420089 | 0.06693051 | F54D10.7 |
| WBGene00018809 | 1.116613983 | 0.69067248 | 1 | F54D10.8 |
| WBGene00018811 | 0.411723241 | 0.00571418 | 0.03439559 | pmt-2 |
| WBGene00018812 | -0.11928708 | 0.57520228 | 0.74826217 | sumv-2 |
| WBGene00018813 | -0.72380304 | 0.15151395 | 0.33336244 | F54D11.3 |
| WBGene00018814 | 0.054322245 | 0.87423466 | 0.93213743 | F54D11.4 |
| WBGene00018815 | -1.094201388 | 0.7816772 | 1 | F54D12.1 |
| WBGene00018819 | -0.011617309 | 0.96411072 | 0.98190322 | rog-1 |
| WBGene00018821 | 0.983488853 | 0.80601565 | 1 | F54D12.8 |
| WBGene00018822 | -0.156269293 | 0.96924949 | 1 | F54D12.9 |
| WBGene00018823 | -0.546257705 | 0.00628634 | 0.03692426 | F54E2.1 |
| WBGene00018824 | 0.279003306 | 0.84585235 | 1 | F54E2.2 |
| WBGene00018825 | 0.983488853 | 0.80601565 | 1 | F54E2.5 |
| WBGene00018827 | -0.203141685 | 0.42434313 | 0.62859321 | pst-2 |
| WBGene00018828 | -2.332096687 | 0.0007362 | 0.00724329 | sdz-21 |
| WBGene00018829 | 0.938982574 | 0.8146943 | 1 | F54E7.6 |
| WBGene00018830 | -0.489592447 | 0.01633691 | 0.07400985 | ska-3 |
| WBGene00018831 | -0.705961386 | 0.0431358 | 0.14481318 | F54E7.9 |
| WBGene00018833 | -0.089799207 | 0.68146427 | 0.81755511 | ztf-1 |
| WBGene00018834 | -0.156269293 | 0.96924949 | 1 | F54F2.6 |
| WBGene00018835 | -0.625980925 | 0.11755761 | 0.28245932 | F54F2.7 |
| WBGene00018836 | -0.303475736 | 0.22209683 | 0.42551328 | F54F2.9 |
| WBGene00018837 | 1.069337112 | 8.7773E-05 | 0.00137859 | F54G2.1 |
| WBGene00018838 | 1.687083251 | 0.2097627 | 1 | F54G2.2 |
| WBGene00018839 | 1.103494564 | 0.42959475 | 1 | F54H5.2 |
| WBGene00018840 | 0.747845955 | 0.85215965 | 1 | F54H5.3 |
| WBGene00018841 | 1.216957619 | 0.74819621 | 1 | F54H5.5 |
| WBGene00018842 | -0.558804446 | 0.87818675 | 1 | F54H12.2 |
| WBGene00018844 | 0.281454817 | 0.94435481 | 1 | F54H12.4 |
| WBGene00018845 | -1.169318462 | 0.00956173 | 0.05013963 | F54H12.5 |
| WBGene00018846 | -0.575290796 | 5.7068E-05 | 0.00097176 | eef-1B.1 |
| WBGene00018848 | 0.269899698 | 0.29418293 | 0.50454513 | F55A3.2 |
| WBGene00018849 | -0.156649717 | 0.30935274 | 0.51807644 | spt-16 |

| | | | | |
|---|---|---|---|---|
| WBGene00018850 | 0.488167733 | 0.70747041 | 1 | gpx-8 |
| WBGene00018851 | 0.519777884 | 0.76572494 | 1 | F55A3.6 |
| WBGene00018853 | -0.545748593 | 0.07397689 | 0.21009437 | sec-22 |
| WBGene00018854 | 0.571221376 | 0.18072914 | 0.37259848 | nlf-1 |
| WBGene00018855 | -0.459953369 | 0.77195274 | 1 | F55A4.3 |
| WBGene00018857 | -0.136793637 | 0.69557437 | 0.82777182 | stau-1 |
| WBGene00018859 | 1.723261921 | 0.41766688 | 1 | F55A4.7 |
| WBGene00018860 | 0.068507082 | 0.85523206 | 0.92108334 | vpat-1 |
| WBGene00018861 | 0.37842657 | 0.59248416 | 0.75967369 | cutl-21 |
| WBGene00018863 | 0.243091696 | 0.47136466 | 0.66421057 | F55A12.2 |
| WBGene00018864 | 0.255935396 | 0.15227719 | 0.33446498 | F55A12.5 |
| WBGene00018865 | 2.063391569 | 0.37853067 | 1 | F55A12.6 |
| WBGene00018866 | 0.093034513 | 0.68895546 | 0.82296421 | nath-10 |
| WBGene00018867 | -0.019986688 | 0.95555904 | 0.97729276 | F55A12.10 |
| WBGene00018869 | 0.666755826 | 0.00737191 | 0.04126432 | rfip-1 |
| WBGene00018871 | -0.059245551 | 0.91293032 | 0.95476673 | F55C12.4 |
| WBGene00018872 | 0.096269043 | 0.58335684 | 0.75346121 | F55C12.5 |
| WBGene00018873 | -0.668355569 | 0.74739695 | 1 | F55C12.6 |
| WBGene00018874 | 0.616357858 | 0.1385827 | 0.31551639 | tag-234 |
| WBGene00018877 | 0.463121369 | 0.17217353 | 0.36165378 | aman-1 |
| WBGene00018878 | -0.242567115 | 0.46428626 | 0.65886864 | glit-1 |
| WBGene00018879 | 1.018215424 | 0.2113444 | 0.4111419 | F55D10.4 |
| WBGene00018880 | 0.784808749 | 0.52512862 | 1 | acc-3 |
| WBGene00018881 | 0.281454817 | 0.94435481 | 1 | F55E10.1 |
| WBGene00018883 | -0.156269293 | 0.96924949 | 1 | F55E10.4 |
| WBGene00018885 | 0.249201776 | 0.55985653 | 0.73642666 | drd-5 |
| WBGene00018886 | 0.671286146 | 0.11240931 | 0.27471043 | npr-28 |
| WBGene00018890 | -0.090923877 | 0.64841875 | 0.79737183 | F55F8.2 |
| WBGene00018891 | 0.111934692 | 0.58600024 | 0.75542885 | F55F8.3 |
| WBGene00018892 | -0.064109915 | 0.73911978 | 0.85446788 | cir-1 |
| WBGene00018893 | -0.613489329 | 0.00402522 | 0.02640207 | wdr-12 |
| WBGene00018894 | -0.774974503 | 0.00107095 | 0.00971057 | mff-2 |
| WBGene00018895 | 1.216957619 | 0.74819621 | 1 | F55F8.7 |
| WBGene00018896 | 1.196144047 | 0.57949438 | 1 | F55F8.8 |
| WBGene00018897 | -0.002365384 | 0.99035789 | 0.99535887 | zipt-2.4 |
| WBGene00018898 | 0.675769929 | 0.00550448 | 0.03344842 | F55F10.1 |
| WBGene00018899 | -1.159256522 | 0.25978201 | 1 | F55G1.1 |
| WBGene00018900 | 0.064467667 | 0.7351488 | 0.8520708 | rod-1 |
| WBGene00018901 | -0.26945331 | 0.46094319 | 0.65630483 | F55G1.5 |
| WBGene00018902 | -0.438114926 | 0.07924302 | 0.21965513 | F55G1.6 |
| WBGene00018903 | 0.609642766 | 0.33944165 | 0.5477137 | F55G1.7 |
| WBGene00018904 | -0.456338924 | 0.08704953 | 0.23351046 | F55G1.9 |
| WBGene00018906 | 0.73299737 | 0.35062024 | 0.55908594 | egas-4 |

| | | | | |
|---|---|---|---|---|
| WBGene00018907 | 1.658812148 | 0.00106511 | 0.00967868 | F55G1.15 |
| WBGene00018908 | 0.216453006 | 0.44081084 | 0.64086749 | fncm-1 |
| WBGene00018909 | -0.336326705 | 0.09185164 | 0.24130927 | slx-1 |
| WBGene00018910 | -3.168891993 | 0.27608518 | 1 | F56A4.2 |
| WBGene00018920 | -0.156269293 | 0.96924949 | 1 | F56A4.12 |
| WBGene00018921 | -0.032579687 | 0.93657548 | 0.96691862 | sago-2 |
| WBGene00018923 | -0.147564523 | 0.55226425 | 0.73020586 | eme-1 |
| WBGene00018924 | -0.207360096 | 0.8759768 | 1 | F56A11.4 |
| WBGene00018925 | -0.615220726 | 0.00340459 | 0.02321083 | F56A11.5 |
| WBGene00018926 | -0.781991218 | 0.64406272 | 1 | F56A11.6 |
| WBGene00018928 | -0.387702274 | 0.37837163 | 0.58609466 | F56B3.2 |
| WBGene00018930 | 2.201543524 | 0.41352885 | 1 | F56B3.6 |
| WBGene00018931 | -0.307587068 | 0.64497349 | 0.79498712 | ugt-52 |
| WBGene00018932 | -0.260804822 | 0.30515779 | 0.51430118 | mrpl-2 |
| WBGene00018933 | 2.069220107 | 0.25725482 | 1 | F56B3.9 |
| WBGene00018934 | -0.790106471 | 0.00046592 | 0.00507789 | F56B3.11 |
| WBGene00018935 | -0.684276304 | 0.00570967 | 0.03439559 | skr-18 |
| WBGene00018937 | -0.060379368 | 0.9741779 | 1 | F56B6.6 |
| WBGene00018939 | -0.256544974 | 0.6688761 | 0.81021897 | fbxa-190 |
| WBGene00018941 | 0.281454817 | 0.94435481 | 1 | F56C3.4 |
| WBGene00018943 | -1.901856996 | 0.64072034 | 1 | dgn-2 |
| WBGene00018945 | -0.578019945 | 0.81405273 | 1 | F56C3.8 |
| WBGene00018946 | 0.456850296 | 0.82685361 | 1 | F56C3.9 |
| WBGene00018948 | -0.474302079 | 0.00599342 | 0.03556508 | F56C9.3 |
| WBGene00018949 | 0.720066222 | 0.75158338 | 1 | acbp-4 |
| WBGene00018950 | -0.317582945 | 0.05973702 | 0.18180776 | F56C9.6 |
| WBGene00018951 | 0.597010507 | 0.00061808 | 0.00627963 | F56C9.7 |
| WBGene00018952 | 0.706270541 | 0.07910693 | 0.21932799 | F56C9.8 |
| WBGene00018953 | 0.06724295 | 0.7171446 | 0.84010542 | F56C9.10 |
| WBGene00018954 | -0.080610009 | 0.77351719 | 0.87360672 | F56C9.11 |
| WBGene00018955 | -0.511411122 | 0.21021687 | 0.41032846 | F56C11.3 |
| WBGene00018957 | 0.110497034 | 0.7040943 | 0.83268897 | F56C11.5 |
| WBGene00018958 | 0.321729068 | 0.36056285 | 0.56837101 | F56C11.6 |
| WBGene00018959 | -0.20891452 | 0.31373991 | 0.52165383 | F56D1.1 |
| WBGene00018960 | 0.021155318 | 0.91522957 | 0.95580617 | ilcr-2 |
| WBGene00018961 | -0.500890698 | 0.03841844 | 0.13384169 | mrps-16 |
| WBGene00018963 | -0.116718845 | 0.41860342 | 0.62320835 | ucr-1 |
| WBGene00018964 | -0.423894865 | 0.05341129 | 0.16799649 | F56D2.2 |
| WBGene00018965 | -0.020452133 | 0.98098284 | 0.98995202 | F56D2.3 |
| WBGene00018966 | 0.740900154 | 0.20144592 | 0.40041719 | F56D2.5 |
| WBGene00018967 | -0.393471499 | 0.01521676 | 0.07043723 | ddx-15 |
| WBGene00018968 | 0.983488853 | 0.80601565 | 1 | F56D2.8 |
| WBGene00018969 | 0.387326146 | 0.4083281 | 0.61423758 | F56D3.1 |

| | | | | |
|---|---|---|---|---|
| WBGene00018970 | 0.535927303 | 0.76789116 | 1 | clec-68 |
| WBGene00018971 | 1.370889895 | 6.81E-07 | 2.4622E-05 | clec-67 |
| WBGene00018972 | -1.964653846 | 0.53599202 | 1 | srz-19 |
| WBGene00018974 | -0.04355928 | 0.81381484 | 0.8986238 | fcho-1 |
| WBGene00018975 | 0.283138925 | 0.27573672 | 0.48433442 | F56E10.1 |
| WBGene00018976 | 0.363346211 | 0.50446324 | 0.69150821 | daam-1 |
| WBGene00018978 | -1.969805064 | 0.62862569 | 1 | ttr-58 |
| WBGene00018981 | 1.116613983 | 0.69067248 | 1 | F56F4.4 |
| WBGene00018984 | 0.712929865 | 0.00022405 | 0.00288162 | F56F10.1 |
| WBGene00018985 | -0.127043295 | 0.76658555 | 0.86914135 | F56F10.2 |
| WBGene00018986 | 0.16919477 | 0.59754206 | 0.76351811 | cdo-1 |
| WBGene00018987 | -0.092774386 | 0.77488266 | 0.87450158 | lgl-1 |
| WBGene00018988 | 1.459136145 | 0.47960024 | 1 | F56F11.1 |
| WBGene00018989 | 0.440485814 | 0.39977444 | 0.60746701 | F56F11.2 |
| WBGene00018990 | 1.550602344 | 4.4001E-06 | 0.00011294 | klf-1 |
| WBGene00018991 | -0.616361644 | 0.00082713 | 0.0079179 | F56F11.4 |
| WBGene00018995 | -0.009661191 | 0.95821999 | 0.97852807 | ccpp-1 |
| WBGene00018996 | -0.040656337 | 0.88630374 | 0.93995577 | F57B9.1 |
| WBGene00018997 | 4.049211621 | 0.01251763 | 1 | F57B9.3 |
| WBGene00018998 | -0.46820852 | 0.03440899 | 0.1248161 | flap-1 |
| WBGene00018999 | -0.248027463 | 0.85936619 | 1 | F57B9.8 |
| WBGene00019001 | -0.176535098 | 0.24215225 | 0.44979338 | ipgm-1 |
| WBGene00019002 | -0.052826633 | 0.77424802 | 0.87418816 | F57B10.4 |
| WBGene00019003 | -0.152375773 | 0.54015522 | 0.72145516 | F57B10.5 |
| WBGene00019004 | 0.228973618 | 0.27314595 | 0.48186715 | xpg-1 |
| WBGene00019005 | -0.139130205 | 0.64071966 | 0.79215656 | F57B10.8 |
| WBGene00019006 | -0.107189061 | 0.73977813 | 0.85490636 | spg-20 |
| WBGene00019007 | -0.199852296 | 0.25171066 | 0.45982508 | ucr-11 |
| WBGene00019008 | -0.535638164 | 0.0506477 | 0.16217522 | pdxk-1 |
| WBGene00019009 | 0.453217544 | 0.4324087 | 0.63420285 | clec-90 |
| WBGene00019010 | 0.168977295 | 0.79816271 | 0.8890361 | glo-2 |
| WBGene00019011 | -0.150133504 | 0.42320163 | 0.62759143 | F57C9.4 |
| WBGene00019012 | 0.070205611 | 0.94705749 | 1 | F57C9.6 |
| WBGene00019013 | -0.491079499 | 0.10783057 | 0.26759651 | tofu-4 |
| WBGene00019014 | 0.618257157 | 0.43410392 | 0.6356073 | F57C12.2 |
| WBGene00019015 | 0.146723875 | 0.94141992 | 1 | F57F4.1 |
| WBGene00019016 | -1.366565036 | 0.6506483 | 1 | F57F4.2 |
| WBGene00019017 | 1.443247508 | 0.00058154 | 0.00599501 | F57F4.4 |
| WBGene00019018 | 1.470627187 | 1.5565E-06 | 4.7243E-05 | abts-3 |
| WBGene00019019 | -1.455123906 | 0.47550043 | 1 | frpr-10 |
| WBGene00019020 | 0.830239001 | 0.58926681 | 1 | F57H12.5 |
| WBGene00019021 | -1.371054553 | 0.36593004 | 1 | F57H12.6 |
| WBGene00019022 | 0.540001632 | 0.05969291 | 0.18178326 | F58A6.1 |

| | | | | |
|---|---|---|---|---|
| WBGene00019023 | 0.968354904 | 0.53933046 | 1 | F58A6.2 |
| WBGene00019024 | 0.88787957 | 0.71832755 | 1 | F58A6.5 |
| WBGene00019025 | 1.293697935 | 0.40429188 | 1 | srb-16 |
| WBGene00019026 | 0.959021583 | 0.70422854 | 1 | F58A6.9 |
| WBGene00019029 | 0.164301619 | 0.90063354 | 1 | F58B6.1 |
| WBGene00019030 | -0.050888644 | 0.91583943 | 0.95580617 | exc-6 |
| WBGene00019032 | 2.955301489 | 0.13737951 | 1 | srsx-17 |
| WBGene00019034 | -1.687753395 | 0.230323 | 1 | fbxb-25 |
| WBGene00019035 | -1.464706906 | 0.05983027 | 0.18191819 | fbxb-26 |
| WBGene00019036 | -1.094201388 | 0.7816772 | 1 | irld-36 |
| WBGene00019037 | -1.672415144 | 0.00704161 | 0.03993096 | fbxb-17 |
| WBGene00019038 | 0.233739675 | 0.93575889 | 1 | irld-37 |
| WBGene00019039 | -1.45184563 | 0.25188973 | 1 | fbxb-18 |
| WBGene00019040 | -2.88891614 | 0.09529366 | 1 | fbxb-19 |
| WBGene00019041 | -1.819454048 | 0.0762739 | 1 | fbxc-22 |
| WBGene00019042 | -0.896366801 | 0.31865073 | 0.52664559 | fbxc-21 |
| WBGene00019043 | -1.75993065 | 0.0431244 | 0.14481318 | fbxc-26 |
| WBGene00019044 | 0.176222991 | 0.88589354 | 1 | F58E1.13 |
| WBGene00019045 | -1.964489727 | 0.05393365 | 1 | fbxb-47 |
| WBGene00019046 | -1.901856996 | 0.64072034 | 1 | F58E2.2 |
| WBGene00019047 | 0.509424136 | 0.49570381 | 0.6841355 | F58E2.3 |
| WBGene00019048 | 1.10384161 | 0.16273528 | 0.34858311 | F58E2.4 |
| WBGene00019049 | -0.156269293 | 0.96924949 | 1 | F58E2.5 |
| WBGene00019055 | 1.964456841 | 0.17036613 | 1 | F58F9.1 |
| WBGene00019057 | 0.742078252 | 0.08522745 | 0.23019943 | F58F9.3 |
| WBGene00019058 | 1.434893823 | 0.03875334 | 0.13458362 | F58F9.4 |
| WBGene00019060 | 0.117523585 | 0.57834801 | 0.74978358 | acox-3 |
| WBGene00019061 | -0.345293783 | 0.03777023 | 0.13234298 | F58F12.1 |
| WBGene00019062 | 0.377828795 | 0.92526585 | 1 | F58F12.2 |
| WBGene00019063 | 0.688793783 | 0.86378755 | 1 | F58F12.3 |
| WBGene00019064 | 0.975731039 | 9.5775E-05 | 0.00148128 | myo-5 |
| WBGene00019066 | -1.027355405 | 0.68015432 | 1 | sdz-23 |
| WBGene00019067 | 2.669530653 | 0.03819841 | 1 | F58H7.1 |
| WBGene00019068 | 0.208289532 | 0.48642496 | 0.67622374 | faah-3 |
| WBGene00019069 | 0.303424732 | 0.68364677 | 0.81881292 | lgc-30 |
| WBGene00019070 | -0.921840601 | 0.69717473 | 1 | F58H7.5 |
| WBGene00019072 | -1.229853028 | 0.58617677 | 1 | fbxc-8 |
| WBGene00019074 | 0.530349433 | 0.04440662 | 0.14762891 | cpd-1 |
| WBGene00019075 | 0.021872591 | 0.94254044 | 0.97027162 | F59A3.2 |
| WBGene00019076 | -0.72534837 | 0.00220562 | 0.01675374 | mrpl-24 |
| WBGene00019077 | -0.317513313 | 0.13612505 | 0.31141781 | zipt-11 |
| WBGene00019079 | 0.387422954 | 0.39217294 | 0.60022237 | rpa-4 |
| WBGene00019080 | 0.281454817 | 0.94435481 | 1 | F59A3.7 |

| | | | | |
|---|---|---|---|---|
| WBGene00019081 | 1.38322681 | 0.69550305 | 1 | F59A3.8 |
| WBGene00019082 | -0.301387288 | 0.50004956 | 0.6880215 | pup-3 |
| WBGene00019084 | -0.604396148 | 0.82723443 | 1 | F59A6.2 |
| WBGene00019085 | -0.156269293 | 0.96924949 | 1 | F59A6.3 |
| WBGene00019086 | 1.216957619 | 0.74819621 | 1 | F59A6.4 |
| WBGene00019087 | -0.192673355 | 0.255367 | 0.46288605 | F59A6.5 |
| WBGene00019089 | 0.281454817 | 0.94435481 | 1 | clec-206 |
| WBGene00019090 | -1.384139363 | 0.10129601 | 0.25648871 | F59A7.2 |
| WBGene00019091 | 1.38322681 | 0.69550305 | 1 | srab-13 |
| WBGene00019092 | 0.867552844 | 0.11093695 | 0.27248342 | F59A7.5 |
| WBGene00019094 | -1.125020419 | 0.46304159 | 1 | F59A7.7 |
| WBGene00019096 | 0.938982574 | 0.8146943 | 1 | cysl-4 |
| WBGene00019097 | -0.364803376 | 0.0389231 | 0.13486454 | F59B1.2 |
| WBGene00019098 | 1.060949945 | 0.61866902 | 1 | F59B1.4 |
| WBGene00019099 | -0.156269293 | 0.96924949 | 1 | F59B1.6 |
| WBGene00019100 | -0.383041319 | 0.36453342 | 0.57203053 | F59B1.8 |
| WBGene00019101 | 0.782547843 | 0.38433825 | 0.59284039 | F59B1.10 |
| WBGene00019102 | 0.404278555 | 0.16865693 | 0.35654949 | F59C12.3 |
| WBGene00019103 | 1.219142607 | 0.14407823 | 0.3224431 | F59D6.1 |
| WBGene00019104 | 1.216957619 | 0.74819621 | 1 | asp-7 |
| WBGene00019105 | -0.024511309 | 0.95051674 | 0.97468105 | asp-8 |
| WBGene00019106 | -1.901856996 | 0.64072034 | 1 | irld-38 |
| WBGene00019108 | 0.704227284 | 0.25648027 | 0.46434939 | chpf-2 |
| WBGene00019109 | 0.074219723 | 0.9154313 | 0.95580617 | F59E11.2 |
| WBGene00019111 | 0.62524032 | 0.09660236 | 0.24948931 | F59E11.5 |
| WBGene00019112 | 0.281454817 | 0.94435481 | 1 | F59E11.6 |
| WBGene00019113 | 2.330618377 | 0.05316848 | 1 | F59E11.7 |
| WBGene00019114 | 0.175506056 | 0.85016004 | 0.91814562 | nhr-194 |
| WBGene00019115 | 1.633429189 | 0.12285126 | 1 | nhr-195 |
| WBGene00019116 | 0.670841646 | 0.12967689 | 0.30097632 | nhr-143 |
| WBGene00019118 | -0.202911841 | 0.2609671 | 0.46962515 | F59E12.1 |
| WBGene00019120 | -0.03062507 | 0.93015494 | 0.96354103 | npl-4.1 |
| WBGene00019121 | -0.48959043 | 0.10126427 | 0.25648871 | npl-4.2 |
| WBGene00019122 | 0.305549628 | 0.16233269 | 0.34830656 | F59E12.6 |
| WBGene00019124 | 0.235776006 | 0.1803858 | 0.3725955 | F59E12.9 |
| WBGene00019125 | -0.415392603 | 0.17737697 | 0.36907177 | ddl-1 |
| WBGene00019126 | -0.287264539 | 0.14205604 | 0.32017962 | sam-4 |
| WBGene00019127 | -0.187901354 | 0.24112301 | 0.44872386 | cgt-3 |
| WBGene00019128 | 0.1260918 | 0.83893364 | 0.91202483 | F59G1.4 |
| WBGene00019129 | -0.231658618 | 0.44306075 | 0.64264057 | F59G1.8 |
| WBGene00019130 | 0.065747003 | 0.7757421 | 0.87481747 | gbas-1 |
| WBGene00019132 | 0.596200872 | 0.25438112 | 0.46207607 | bath-19 |
| WBGene00019133 | 2.854906918 | 0.07696999 | 1 | F59H6.2 |

| | | | | |
|---|---|---|---|---|
| WBGene00019138 | 0.318713486 | 0.65963927 | 0.80368078 | bath-21 |
| WBGene00019139 | 0.209510403 | 0.59468647 | 0.76112043 | bath-1 |
| WBGene00019140 | 0.470675108 | 0.42263621 | 0.62712373 | bath-3 |
| WBGene00019141 | -0.574930483 | 0.06773101 | 0.19774952 | bath-5 |
| WBGene00019142 | 0.187588966 | 0.69708558 | 0.82859957 | btb-4 |
| WBGene00019143 | -0.156738951 | 0.68918377 | 0.82299402 | H01M10.1 |
| WBGene00019144 | -0.014519762 | 0.99603746 | 1 | H01M10.2 |
| WBGene00019145 | 1.977885804 | 0.49204652 | 1 | H02F09.2 |
| WBGene00019146 | 2.58598212 | 7.27E-06 | 0.00017331 | H02F09.3 |
| WBGene00019148 | 0.015993944 | 0.97471959 | 0.98666588 | H03E18.1 |
| WBGene00019149 | 0.578068466 | 0.66705632 | 1 | H03E18.2 |
| WBGene00019150 | 2.799084385 | 0.17307183 | 1 | H04J21.1 |
| WBGene00019151 | 1.073216103 | 0.62944296 | 1 | pck-3 |
| WBGene00019152 | -0.771516032 | 0.49830772 | 1 | nspb-6 |
| WBGene00019153 | -0.786527527 | 0.00159913 | 0.01315642 | H04M03.3 |
| WBGene00019154 | -1.077298005 | 0.00049218 | 0.00530689 | glf-1 |
| WBGene00019155 | 0.075260937 | 0.97569838 | 1 | H04M03.11 |
| WBGene00019156 | 0.549750164 | 0.30268676 | 0.51189859 | H04M03.12 |
| WBGene00019157 | -0.132749033 | 0.48157554 | 0.67289188 | H05C05.1 |
| WBGene00019158 | -0.133965804 | 0.5635659 | 0.73875164 | H05C05.2 |
| WBGene00019162 | -0.619046879 | 1.5643E-05 | 0.0003303 | eif-1.A |
| WBGene00019163 | -0.170053727 | 0.32700479 | 0.5355225 | ubxn-6 |
| WBGene00019164 | 0.217476119 | 0.46353738 | 0.65853117 | H06H21.8 |
| WBGene00019165 | 1.584577408 | 0.63258867 | 1 | mpz-4 |
| WBGene00019166 | 1.407231044 | 3.9561E-05 | 0.00070991 | tat-2 |
| WBGene00019167 | 0.065184423 | 0.78674258 | 0.88255028 | H06I04.1 |
| WBGene00019168 | 0.246265193 | 0.16550148 | 0.35233271 | H06I04.3 |
| WBGene00019169 | 0.747845955 | 0.85215965 | 1 | H06I04.5 |
| WBGene00019170 | 0.156834147 | 0.63062685 | 0.78567888 | H06I04.6 |
| WBGene00019171 | 2.428015231 | 0.15494979 | 1 | fipr-29 |
| WBGene00019174 | -2.160163868 | 0.12165494 | 1 | H09G03.1 |
| WBGene00019177 | 1.093525547 | 0.35715768 | 1 | H10D18.1 |
| WBGene00019178 | -1.964653846 | 0.53599202 | 1 | scl-12 |
| WBGene00019180 | 0.233720258 | 0.61529405 | 0.77528743 | H10D18.5 |
| WBGene00019182 | 0.380250436 | 0.45982298 | 0.65564231 | H10E21.1 |
| WBGene00019183 | 0.673272738 | 0.68869904 | 1 | npr-30 |
| WBGene00019184 | -1.407981172 | 0.00338303 | 0.02312687 | H10E21.4 |
| WBGene00019185 | 1.040053736 | 0.03201538 | 0.1186885 | H10E21.5 |
| WBGene00019187 | -0.54608947 | 0.34564896 | 0.55458227 | H11E01.2 |
| WBGene00019188 | 0.84995227 | 0.01652977 | 0.07432888 | H11E01.3 |
| WBGene00019189 | 2.41975572 | 0.00093797 | 0.00877345 | H11L12.1 |
| WBGene00019190 | 0.281454817 | 0.94435481 | 1 | H12I13.1 |
| WBGene00019193 | 0.109173684 | 0.96057644 | 1 | H12I13.5 |

| | | | | |
|---|---|---|---|---|
| WBGene00019194 | 0.109173684 | 0.96057644 | 1 | H12I13.6 |
| WBGene00019196 | -0.172394802 | 0.42235823 | 0.62702998 | H14A12.3 |
| WBGene00019197 | -0.465582561 | 0.34725696 | 0.55613333 | H14A12.5 |
| WBGene00019198 | 0.554163131 | 0.35846943 | 0.56668975 | strm-1 |
| WBGene00019199 | -0.226341025 | 0.27269212 | 0.48138251 | H14E04.2 |
| WBGene00019200 | 0.567144191 | 0.56718769 | 1 | H14E04.3 |
| WBGene00019202 | 1.413377704 | 0.60673863 | 1 | H14N18.2 |
| WBGene00019203 | -0.478464484 | 0.0086945 | 0.04660441 | ttr-47 |
| WBGene00019204 | 0.795735174 | 0.01189469 | 0.05892999 | H14N18.4 |
| WBGene00019205 | 1.075910581 | 8.29E-07 | 2.8521E-05 | kcc-2 |
| WBGene00019207 | -0.048236667 | 0.81011093 | 0.89634164 | fgt-1 |
| WBGene00019209 | 0.068385759 | 0.67288853 | 0.81313224 | emc-1 |
| WBGene00019211 | 0.228915012 | 0.48231661 | 0.67339074 | H18N23.2 |
| WBGene00019212 | 0.470567113 | 0.52769402 | 0.7120966 | zmp-2 |
| WBGene00019213 | 0.704178827 | 0.02471321 | 0.09935202 | H20E11.1 |
| WBGene00019214 | -0.22461868 | 0.93331196 | 1 | H20E11.2 |
| WBGene00019215 | 0.938172809 | 0.22057493 | 0.42366719 | H20E11.3 |
| WBGene00019216 | -0.156269293 | 0.96924949 | 1 | H20J04.1 |
| WBGene00019217 | 0.093674395 | 0.63855066 | 0.79036023 | athp-2 |
| WBGene00019218 | -0.439910508 | 0.04187241 | 0.14212271 | madf-3 |
| WBGene00019219 | -0.32401053 | 0.30334356 | 0.51229575 | H20J04.4 |
| WBGene00019220 | -0.26040244 | 0.1832687 | 0.37536783 | pfd-2 |
| WBGene00019221 | -0.292322225 | 0.32544447 | 0.5338183 | H20J04.6 |
| WBGene00019224 | 0.320454036 | 0.64993929 | 0.79839241 | gnrr-5 |
| WBGene00019225 | 0.641321398 | 0.09865942 | 0.25284259 | bgal-2 |
| WBGene00019226 | -0.347950507 | 0.70046861 | 0.83099369 | H22K11.3 |
| WBGene00019227 | 1.31902793 | 0.00185876 | 0.01474979 | sgca-1 |
| WBGene00019228 | 0.616429445 | 0.81400458 | 1 | H23L24.1 |
| WBGene00019229 | 2.028800581 | 0.39258153 | 1 | ipla-5 |
| WBGene00019230 | 0.886909342 | 0.32076145 | 0.52905377 | ttll-11 |
| WBGene00019231 | 1.067716307 | 0.55093385 | 1 | H23L24.4 |
| WBGene00019232 | 0.958163524 | 0.00725662 | 0.04082604 | ugt-13 |
| WBGene00019233 | 0.175909247 | 0.94229417 | 1 | ugt-14 |
| WBGene00019234 | 1.135341633 | 0.20702967 | 0.40665706 | ugt-8 |
| WBGene00019235 | -0.014519762 | 0.99603746 | 1 | H23N18.4 |
| WBGene00019236 | -0.496687202 | 0.57219928 | 0.7462154 | H23N18.5 |
| WBGene00019237 | 0.693087986 | 0.00112597 | 0.01005178 | H24G06.1 |
| WBGene00019239 | -0.758861958 | 0.41577581 | 0.62082449 | H24K24.2 |
| WBGene00019240 | -0.615813779 | 0.00037259 | 0.00428295 | H24K24.3 |
| WBGene00019241 | -0.396443447 | 0.09592236 | 0.24842009 | H24K24.4 |
| WBGene00019242 | -1.159714244 | 0.75790634 | 1 | srbc-40 |
| WBGene00019243 | -1.21418826 | 0.27061915 | 1 | fbxb-72 |
| WBGene00019244 | -0.182237585 | 0.59037308 | 0.75800757 | H25P19.1 |

| | | | | |
|---|---|---|---|---|
| WBGene00019245 | -0.287547222 | 0.11692787 | 0.2817827 | sacy-1 |
| WBGene00019246 | -0.355252464 | 0.04247276 | 0.14359838 | rpb-5 |
| WBGene00019247 | -0.319932098 | 0.07675331 | 0.21490008 | syp-4 |
| WBGene00019248 | 0.983488853 | 0.80601565 | 1 | H27M09.5 |
| WBGene00019249 | 0.652985617 | 0.00267959 | 0.01955102 | H28G03.1 |
| WBGene00019250 | -0.215009275 | 0.40041498 | 0.60821556 | H28G03.2 |
| WBGene00019251 | -1.901856996 | 0.64072034 | 1 | H28G03.3 |
| WBGene00019254 | -0.049272436 | 0.89146993 | 0.94271722 | H31G24.3 |
| WBGene00019255 | 0.942308827 | 0.67468959 | 1 | H32C10.1 |
| WBGene00019257 | -0.156269293 | 0.96924949 | 1 | dhhc-13 |
| WBGene00019259 | -0.14222351 | 0.43111119 | 0.63332845 | H34C03.2 |
| WBGene00019260 | -0.156269293 | 0.96924949 | 1 | H34I24.1 |
| WBGene00019261 | 0.880280625 | 2.3301E-06 | 6.6407E-05 | H34I24.2 |
| WBGene00019262 | 1.236177236 | 0.11999355 | 0.28624902 | dmsr-12 |
| WBGene00019263 | -0.045875639 | 0.88035173 | 0.93609654 | H35B03.1 |
| WBGene00019264 | -0.376114131 | 0.08887094 | 0.23635972 | H35B03.2 |
| WBGene00019265 | 0.479889899 | 0.82438301 | 1 | H35N09.1 |
| WBGene00019266 | -0.015577364 | 0.98604286 | 0.99332445 | H35N09.2 |
| WBGene00019267 | 0.446458324 | 0.87164163 | 1 | H39E20.1 |
| WBGene00019268 | 0.086961287 | 0.83462316 | 0.91032502 | H41C03.1 |
| WBGene00019269 | 1.796575934 | 0.34491145 | 1 | H41C03.2 |
| WBGene00019270 | 0.421357856 | 0.40449421 | 0.61157711 | H41C03.3 |
| WBGene00019272 | -0.55385974 | 0.23231805 | 0.437977 | H42K12.3 |
| WBGene00019273 | 1.48573747 | 8.8747E-06 | 0.00020602 | H43E16.1 |
| WBGene00019274 | -0.433590326 | 0.10076762 | 0.25590808 | H43I07.1 |
| WBGene00019275 | -0.320726948 | 0.16310607 | 0.34913095 | rpac-40 |
| WBGene00019276 | -0.122572234 | 0.61400858 | 0.77467299 | H43I07.3 |
| WBGene00019277 | -0.060081405 | 0.91762451 | 0.95646284 | sgcb-1 |
| WBGene00019279 | 1.527799568 | 0.11200074 | 0.27415342 | K01A2.4 |
| WBGene00019280 | -0.147172849 | 0.66970423 | 0.81097937 | K01A2.5 |
| WBGene00019282 | 0.144455081 | 0.79099258 | 0.88470207 | mps-2 |
| WBGene00019284 | 1.090621209 | 0.01161939 | 0.05791093 | K01A2.10 |
| WBGene00019285 | -0.105655056 | 0.83059612 | 0.90824619 | cbn-1 |
| WBGene00019286 | -0.097450002 | 0.96886938 | 1 | K01A12.2 |
| WBGene00019287 | 1.419839013 | 0.37829547 | 1 | K01A12.3 |
| WBGene00019288 | -0.737462795 | 0.83433562 | 1 | K02A2.1 |
| WBGene00019289 | 1.416198 | 0.0183196 | 0.08038245 | kcc-3 |
| WBGene00019290 | 1.250043734 | 0.58768578 | 1 | K02A2.5 |
| WBGene00019293 | 1.115863748 | 0.63449935 | 1 | K02A6.2 |
| WBGene00019294 | 0.818564688 | 0.00918192 | 0.04859534 | K02A6.3 |
| WBGene00019295 | -0.251278211 | 0.18357029 | 0.37571256 | pfkb-1.2 |
| WBGene00019296 | -0.170143546 | 0.35378338 | 0.56237866 | mcu-1 |
| WBGene00019298 | 1.034569557 | 3.15E-08 | 1.6548E-06 | K02D7.1 |

| | | | | |
|---|---|---|---|---|
| WBGene00019299 | -0.289996362 | 0.84366438 | 1 | snai-1 |
| WBGene00019300 | 1.947730233 | 4.4E-06 | 0.00011294 | swt-1 |
| WBGene00019301 | -0.095927444 | 0.57190076 | 0.74620603 | K02D10.1 |
| WBGene00019302 | -0.668264701 | 0.85551153 | 1 | K02D10.2 |
| WBGene00019303 | 1.330382438 | 0.5642962 | 1 | K02D10.3 |
| WBGene00019304 | 0.276494823 | 0.62805435 | 0.78384126 | K02D10.4 |
| WBGene00019305 | -0.027702417 | 0.88555609 | 0.93951649 | snap-29 |
| WBGene00019306 | -1.606750384 | 0.37736506 | 1 | K02E7.1 |
| WBGene00019309 | -0.737462795 | 0.83433562 | 1 | K02E7.4 |
| WBGene00019312 | -0.974667947 | 0.48783931 | 1 | K02E7.7 |
| WBGene00019313 | -1.736320412 | 0.00024885 | 0.00311791 | btb-10 |
| WBGene00019314 | 0.51027045 | 0.58760832 | 0.75621716 | K02E7.10 |
| WBGene00019315 | 2.469540862 | 0.14436819 | 1 | K02E7.11 |
| WBGene00019316 | -1.930411901 | 0.43097183 | 1 | K02E7.12 |
| WBGene00019317 | 0.399207019 | 0.51654773 | 0.70267854 | K02E10.1 |
| WBGene00019318 | -0.039832911 | 0.95251552 | 0.97582406 | K02E10.4 |
| WBGene00019319 | 0.726230729 | 0.71931668 | 1 | K02E10.5 |
| WBGene00019320 | -0.38669983 | 0.6561323 | 0.80231895 | K02E10.6 |
| WBGene00019321 | 0.556811663 | 0.39839366 | 0.60597563 | K02E10.7 |
| WBGene00019322 | 0.106013579 | 0.48117 | 0.67287319 | ahcy-1 |
| WBGene00019323 | -0.041290688 | 0.81748755 | 0.90086942 | teg-4 |
| WBGene00019324 | 0.826965802 | 0.0054125 | 0.03304034 | K02F2.5 |
| WBGene00019326 | 0.58282135 | 0.00489229 | 0.03069628 | K02F3.2 |
| WBGene00019327 | 1.265208003 | 5.55E-16 | 1.18E-13 | zip-2 |
| WBGene00019328 | -0.054863152 | 0.88798996 | 0.94084004 | clec-149 |
| WBGene00019329 | 0.04685652 | 0.90016798 | 0.94737686 | ceeh-1 |
| WBGene00019331 | 0.747845955 | 0.85215965 | 1 | dos-3 |
| WBGene00019332 | -0.214239738 | 0.47856546 | 0.67083387 | K02F3.9 |
| WBGene00019333 | -0.226127458 | 0.18489315 | 0.37727857 | moma-1 |
| WBGene00019334 | -0.066329999 | 0.72840695 | 0.84781793 | K02F3.12 |
| WBGene00019337 | 3.857155473 | 0.02045627 | 1 | K02F6.3 |
| WBGene00019339 | 0.233739675 | 0.93575889 | 1 | K02F6.5 |
| WBGene00019340 | -0.774842772 | 0.78069886 | 1 | K02F6.6 |
| WBGene00019341 | -0.843711209 | 0.00043596 | 0.00479874 | K02F6.7 |
| WBGene00019342 | 0.233739675 | 0.93575889 | 1 | K02F6.8 |
| WBGene00019343 | -0.915916971 | 0.4610496 | 1 | nep-18 |
| WBGene00019344 | 0.56703824 | 0.16586946 | 0.35262145 | dhhc-10 |
| WBGene00019345 | 1.414919902 | 6.2739E-06 | 0.00015204 | tmem-135 |
| WBGene00019346 | 0.010531492 | 0.98703933 | 0.99421374 | K02G10.5 |
| WBGene00019347 | 1.402536062 | 0.00013451 | 0.00193717 | mbl-1 |
| WBGene00019348 | 0.983488853 | 0.80601565 | 1 | K02H11.4 |
| WBGene00019350 | -0.737462795 | 0.83433562 | 1 | K02H11.9 |
| WBGene00019351 | 0.473477747 | 0.00766235 | 0.04244005 | lron-7 |

| | | | | |
|---|---|---|---|---|
| WBGene00019352 | 0.599895164 | 0.54409134 | 1 | K03A1.4 |
| WBGene00019353 | -0.090951614 | 0.69770351 | 0.8288472 | K03B4.1 |
| WBGene00019354 | -0.440977199 | 0.02920298 | 0.11114527 | K03B4.2 |
| WBGene00019355 | -0.395976713 | 0.56823464 | 0.74317966 | K03B4.4 |
| WBGene00019357 | -0.72907602 | 4.3477E-05 | 0.00077058 | cpg-8 |
| WBGene00019360 | 0.157468075 | 0.83941889 | 0.91211511 | neto-1 |
| WBGene00019361 | 0.178772071 | 0.48384144 | 0.67458876 | K03E5.2 |
| WBGene00019362 | -0.253182437 | 0.29632445 | 0.50678884 | cdk-2 |
| WBGene00019364 | 1.50242786 | 0.31634291 | 1 | mksr-1 |
| WBGene00019365 | 0.698107488 | 0.01756497 | 0.07810589 | rei-2 |
| WBGene00019367 | -0.075264494 | 0.94913791 | 1 | K03H6.1 |
| WBGene00019368 | 1.798472698 | 2.621E-05 | 0.00050752 | K03H6.2 |
| WBGene00019369 | -0.33449598 | 0.85735439 | 1 | K03H6.4 |
| WBGene00019370 | 1.123260723 | 0.14864849 | 0.32957292 | K03H6.5 |
| WBGene00019372 | -0.621606308 | 0.77755782 | 1 | K03H9.3 |
| WBGene00019373 | 0.058900153 | 0.88272559 | 0.93749089 | K04A8.1 |
| WBGene00019374 | -0.134399481 | 0.94529403 | 1 | K04A8.2 |
| WBGene00019376 | -1.565381827 | 0.10427701 | 1 | lipl-4 |
| WBGene00019379 | -1.14715222 | 0.68932198 | 1 | K04A8.10 |
| WBGene00019380 | 0.078866007 | 0.68269377 | 0.81846402 | K04C2.2 |
| WBGene00019381 | -0.40009396 | 0.06264329 | 0.18765322 | K04C2.3 |
| WBGene00019382 | -1.708215616 | 0.31751374 | 1 | K04C2.5 |
| WBGene00019383 | 0.281454817 | 0.94435481 | 1 | K04E7.1 |
| WBGene00019386 | -1.901856996 | 0.64072034 | 1 | irld-42 |
| WBGene00019387 | 0.574508983 | 0.8551971 | 1 | K04F1.8 |
| WBGene00019388 | -0.616695114 | 0.39833895 | 0.60596832 | K04F1.9 |
| WBGene00019389 | -0.245531902 | 0.94070824 | 1 | irld-40 |
| WBGene00019391 | 2.955301489 | 0.13737951 | 1 | irld-11 |
| WBGene00019392 | 0.233739675 | 0.93575889 | 1 | irld-44 |
| WBGene00019393 | 0.232157325 | 0.93397866 | 1 | irld-45 |
| WBGene00019394 | 0.654422943 | 0.25234938 | 0.46037961 | K04F10.1 |
| WBGene00019395 | 0.340136707 | 0.65673057 | 0.80239165 | K04F10.2 |
| WBGene00019396 | -0.122285764 | 0.58527906 | 0.75465941 | K04F10.3 |
| WBGene00019399 | -0.40255013 | 0.05671714 | 0.17588831 | K04F10.7 |
| WBGene00019400 | -0.295660069 | 0.12502084 | 0.2947336 | K04G7.1 |
| WBGene00019401 | -0.184262715 | 0.22220892 | 0.42566085 | nuo-4 |
| WBGene00019402 | -0.740014729 | 0.00023628 | 0.00298118 | syf-2 |
| WBGene00019403 | -0.325216744 | 0.40698492 | 0.6138904 | K05B2.2 |
| WBGene00019404 | 0.257416228 | 0.72475682 | 0.84518994 | K05B2.4 |
| WBGene00019405 | 0.580309921 | 0.68366557 | 1 | K05F1.1 |
| WBGene00019406 | 0.938815993 | 0.61824059 | 1 | acdh-8 |
| WBGene00019407 | -0.706652943 | 0.07165285 | 0.20565893 | lect-2 |
| WBGene00019408 | 1.421967663 | 0.00020913 | 0.00272133 | K05F1.6 |

| | | | | |
|---|---|---|---|---|
| WBGene00019409 | -0.156269293 | 0.96924949 | 1 | K05F1.8 |
| WBGene00019410 | 1.158132015 | 0.62662247 | 1 | K05F1.9 |
| WBGene00019411 | 1.276467784 | 0.00279219 | 0.02017859 | K05F1.10 |
| WBGene00019412 | -1.797813086 | 0.05284991 | 0.16670583 | fbxb-49 |
| WBGene00019413 | -2.141313286 | 0.04406201 | 0.14684516 | fbxb-50 |
| WBGene00019414 | -0.637018156 | 0.80584146 | 1 | fbxb-51 |
| WBGene00019415 | -2.337448015 | 0.00167458 | 0.01359031 | K05F6.4 |
| WBGene00019416 | -2.796092711 | 0.00805811 | 1 | fbxb-44 |
| WBGene00019417 | -0.90979066 | 0.40048239 | 1 | fbxb-52 |
| WBGene00019418 | -2.581761766 | 0.00168983 | 0.01366837 | fbxb-54 |
| WBGene00019420 | -2.35096163 | 0.0182221 | 1 | fbxb-46 |
| WBGene00019421 | 1.621716352 | 0.45418105 | 1 | K05F6.10 |
| WBGene00019422 | -1.45873709 | 0.59584643 | 1 | K05G3.1 |
| WBGene00019423 | 1.115863748 | 0.63449935 | 1 | K05G3.2 |
| WBGene00019424 | 0.839445035 | 0.42300056 | 1 | aptf-1 |
| WBGene00019425 | 0.688793783 | 0.86378755 | 1 | K06A1.2 |
| WBGene00019426 | 0.111034172 | 0.85213641 | 0.91924443 | cutl-16 |
| WBGene00019427 | -0.186944125 | 0.50645927 | 0.69322684 | atg-16.2 |
| WBGene00019428 | 1.38322681 | 0.69550305 | 1 | T28D9.12 |
| WBGene00019428 | 1.348567359 | 0.0003008 | 0.00361169 | dgk-5 |
| WBGene00019429 | 0.041602765 | 0.84408022 | 0.91434493 | K06A5.1 |
| WBGene00019430 | 2.650792005 | 0.05180529 | 1 | K06A5.2 |
| WBGene00019431 | 0.253925665 | 0.89264743 | 1 | K06A5.3 |
| WBGene00019432 | -0.220593489 | 0.20677789 | 0.40655714 | knl-2 |
| WBGene00019433 | -0.698599629 | 6.4746E-06 | 0.00015659 | acdh-3 |
| WBGene00019434 | 0.700451002 | 0.19073867 | 0.38544578 | nmtn-1 |
| WBGene00019435 | 2.451675171 | 1.93E-11 | 2.09E-09 | K06A9.1 |
| WBGene00019436 | 1.027065705 | 0.20811067 | 0.40776775 | K06A9.2 |
| WBGene00019437 | -0.761695967 | 0.55442165 | 1 | K06A9.3 |
| WBGene00019438 | -0.156269293 | 0.96924949 | 1 | cyp-25A6 |
| WBGene00019444 | 1.651751904 | 0.20342191 | 1 | frpr-11 |
| WBGene00019445 | 0.551625335 | 0.74962832 | 1 | frpr-12 |
| WBGene00019448 | 0.683371507 | 0.46506268 | 0.65946344 | frpr-13 |
| WBGene00019452 | -0.156269293 | 0.96924949 | 1 | K06H6.4 |
| WBGene00019455 | 1.487047283 | 0.50914254 | 1 | K06H7.1 |
| WBGene00019456 | 0.408537326 | 0.44214902 | 0.64190715 | K06H7.2 |
| WBGene00019457 | 0.00658683 | 0.97450915 | 0.98661738 | vms-1 |
| WBGene00019458 | -0.482558019 | 0.11643013 | 0.28097361 | K06H7.7 |
| WBGene00019459 | 1.413734769 | 0.68804531 | 1 | K06H7.8 |
| WBGene00019460 | -0.46239533 | 0.02846803 | 0.10941134 | idi-1 |
| WBGene00019461 | -0.505605194 | 0.80388525 | 1 | K07A3.3 |
| WBGene00019462 | 1.215839694 | 0.1771624 | 0.36881494 | K07A9.3 |
| WBGene00019464 | -0.364846528 | 0.17968577 | 0.37202677 | K07B1.4 |

| | | | | |
|---|---|---|---|---|
| WBGene00019465 | -0.274821493 | 0.14591584 | 0.32488677 | acl-14 |
| WBGene00019466 | 0.799964318 | 1.33E-07 | 5.9859E-06 | tos-1 |
| WBGene00019467 | -0.282040199 | 0.23308087 | 0.43887018 | K07B1.7 |
| WBGene00019470 | -1.969805064 | 0.62862569 | 1 | cyp-35B3 |
| WBGene00019471 | 0.938982574 | 0.8146943 | 1 | cyp-35B2 |
| WBGene00019472 | -0.451813889 | 0.8460226 | 1 | cyp-35B1 |
| WBGene00019473 | -2.810052938 | 0.03049187 | 1 | cyp-35A5 |
| WBGene00019476 | -0.65153771 | 0.10097933 | 0.25601242 | K07C11.3 |
| WBGene00019477 | 0.577912596 | 0.12550604 | 0.29536855 | K07C11.4 |
| WBGene00019478 | -0.485319243 | 0.16020849 | 0.34561687 | cri-2 |
| WBGene00019479 | -0.633085424 | 0.00158611 | 0.0130733 | K07C11.7 |
| WBGene00019480 | 0.147890443 | 0.79128821 | 0.88470671 | K07C11.8 |
| WBGene00019481 | -0.167590961 | 0.41035575 | 0.61576185 | cogc-6 |
| WBGene00019482 | -1.147478861 | 0.32305696 | 1 | K07C11.10 |
| WBGene00019484 | 0.496334238 | 0.73805503 | 1 | K07D4.4 |
| WBGene00019485 | 0.864149694 | 0.43990886 | 1 | K07D4.5 |
| WBGene00019486 | -1.03891452 | 0.50123974 | 1 | K07D4.6 |
| WBGene00019487 | -0.125107403 | 0.65402406 | 0.80118113 | ephx-1 |
| WBGene00019488 | 0.896328666 | 0.04763833 | 0.15581276 | K07D4.9 |
| WBGene00019489 | -0.020598245 | 0.94380694 | 0.97099908 | K07E1.1 |
| WBGene00019490 | -0.109228622 | 0.72598283 | 0.84580727 | K07E3.1 |
| WBGene00019491 | -0.031680908 | 0.93191086 | 0.96406579 | K07E3.2 |
| WBGene00019492 | 0.388331326 | 0.03423023 | 0.12439066 | K07E3.4 |
| WBGene00019493 | 1.021980839 | 0.00738678 | 0.04130947 | catp-5 |
| WBGene00019495 | 1.448725529 | 9.5383E-06 | 0.00021891 | sdz-24 |
| WBGene00019496 | -0.777178355 | 0.59395238 | 1 | frpr-14 |
| WBGene00019497 | 0.957158251 | 0.36821873 | 1 | K07E8.6 |
| WBGene00019498 | -0.617197028 | 0.05997968 | 0.18218997 | K07E8.7 |
| WBGene00019501 | 0.747845955 | 0.85215965 | 1 | K07E12.2 |
| WBGene00019502 | -1.969805064 | 0.62862569 | 1 | K07G6.1 |
| WBGene00019503 | 0.078254112 | 0.71642185 | 0.84006931 | tbce-1 |
| WBGene00019504 | -0.154923922 | 0.33472772 | 0.54317003 | K07H8.2 |
| WBGene00019506 | 0.789601902 | 0.75907734 | 1 | K07H8.5 |
| WBGene00019507 | -0.831640732 | 0.65397538 | 1 | K07H8.7 |
| WBGene00019508 | -3.014567441 | 0.24693748 | 1 | K07H8.8 |
| WBGene00019509 | 0.149985337 | 0.65761851 | 0.80291727 | K07H8.9 |
| WBGene00019510 | -0.147591697 | 0.29728023 | 0.50754013 | K07H8.10 |
| WBGene00019511 | -0.280973536 | 0.32974625 | 0.53831508 | K08A2.1 |
| WBGene00019512 | 0.88787957 | 0.71832755 | 1 | K08A2.2 |
| WBGene00019513 | -0.38402886 | 0.23404497 | 0.4401914 | K08A2.4 |
| WBGene00019514 | -0.848977607 | 0.40798985 | 1 | K08B4.2 |
| WBGene00019515 | 0.239420885 | 0.55491404 | 0.73238519 | ugt-19 |
| WBGene00019516 | -0.073379231 | 0.92357395 | 0.95953374 | ugt-20 |

| | | | | |
|---|---|---|---|---|
| WBGene00019518 | 1.394901296 | 0.22294684 | 1 | K08B5.1 |
| WBGene00019519 | 0.197959561 | 0.74833923 | 0.85972621 | K08B5.2 |
| WBGene00019520 | -0.672283524 | 0.20917772 | 0.4093531 | K08B12.1 |
| WBGene00019521 | 0.596974089 | 0.01335119 | 0.06385587 | dmd-7 |
| WBGene00019522 | 0.284396179 | 0.56704134 | 0.74201864 | K08B12.3 |
| WBGene00019525 | 1.321019127 | 0.10614611 | 0.26549649 | K08D9.4 |
| WBGene00019527 | 1.025181888 | 0.79790347 | 1 | K08D9.6 |
| WBGene00019529 | 0.232157325 | 0.93397866 | 1 | K08D10.5 |
| WBGene00019530 | 0.281454817 | 0.94435481 | 1 | scrm-8 |
| WBGene00019531 | -1.892648585 | 0.34370782 | 1 | scrm-5 |
| WBGene00019534 | -0.156269293 | 0.96924949 | 1 | K08D10.11 |
| WBGene00019535 | -0.058567248 | 0.90399179 | 0.95016572 | tsen-34 |
| WBGene00019536 | 1.018228288 | 0.15281301 | 0.33519472 | rpi-2 |
| WBGene00019537 | -0.817062973 | 1.70E-07 | 7.3826E-06 | K08D12.3 |
| WBGene00019538 | -1.198946775 | 0.14377789 | 0.32200837 | K08D12.4 |
| WBGene00019540 | -0.093974593 | 0.79472673 | 0.88641923 | K08D12.6 |
| WBGene00019541 | 0.87001166 | 0.63612313 | 1 | K08F11.1 |
| WBGene00019543 | -0.418337958 | 0.00951087 | 0.04994443 | cif-1 |
| WBGene00019544 | -0.09452436 | 0.64943647 | 0.79813829 | miro-1 |
| WBGene00019545 | 0.232157325 | 0.93397866 | 1 | K09B3.1 |
| WBGene00019546 | 0.292348844 | 0.51957012 | 0.70497327 | glct-6 |
| WBGene00019547 | 1.527839515 | 0.01303746 | 0.06301268 | K09C4.1 |
| WBGene00019549 | -0.263229581 | 0.93409872 | 1 | K09C4.4 |
| WBGene00019550 | -0.511766921 | 0.28257043 | 0.4920665 | K09C4.5 |
| WBGene00019552 | -0.348878357 | 0.80754287 | 1 | K09C4.6 |
| WBGene00019553 | 0.574508983 | 0.8551971 | 1 | K09C4.9 |
| WBGene00019554 | -0.221996476 | 0.49684994 | 0.68509428 | K09C4.10 |
| WBGene00019556 | 0.09697944 | 0.97612634 | 1 | K09C6.2 |
| WBGene00019557 | 0.377828795 | 0.92526585 | 1 | K09C6.3 |
| WBGene00019558 | 1.181686151 | 0.7579682 | 1 | srbc-12 |
| WBGene00019561 | 1.750413495 | 0.36210361 | 1 | K09C6.7 |
| WBGene00019563 | 1.089245669 | 0.01363892 | 0.06486097 | K09C6.9 |
| WBGene00019564 | 2.818569528 | 0.00011715 | 0.00173244 | K09D9.1 |
| WBGene00019565 | 0.104864639 | 0.91263181 | 1 | cyp-35A3 |
| WBGene00019567 | 0.281454817 | 0.94435481 | 1 | K09D9.9 |
| WBGene00019570 | 1.38322681 | 0.69550305 | 1 | K09E2.1 |
| WBGene00019571 | 2.054555617 | 0.1454639 | 1 | K09E2.2 |
| WBGene00019572 | 0.061487862 | 0.81940451 | 0.90234831 | K09E2.3 |
| WBGene00019576 | -0.571464787 | 0.20289018 | 0.40190728 | K09E3.5 |
| WBGene00019577 | -0.63406617 | 0.4107144 | 0.61622387 | K09E3.6 |
| WBGene00019578 | -0.135141025 | 0.70104854 | 0.83141078 | K09E3.7 |
| WBGene00019579 | 0.516947376 | 0.50405897 | 0.69118816 | oac-57 |
| WBGene00019580 | 1.309746475 | 0.14308477 | 0.32110761 | oac-58 |

| | | | | |
|---|---|---|---|---|
| WBGene00019582 | 0.281454817 | 0.94435481 | 1 | K09F5.1 |
| WBGene00019584 | -0.712565257 | 0.30838034 | 0.51713339 | set-12 |
| WBGene00019586 | 1.270052625 | 0.58538191 | 1 | K09F6.3 |
| WBGene00019587 | -1.40720786 | 0.22337436 | 1 | K09F6.4 |
| WBGene00019588 | 1.278470803 | 0.39867721 | 1 | K09F6.5 |
| WBGene00019589 | 1.621166208 | 3.9746E-05 | 0.00071174 | K09F6.6 |
| WBGene00019590 | 1.455273261 | 0.00880672 | 0.04700329 | K09F6.7 |
| WBGene00019591 | -1.737409953 | 0.04535812 | 0.15005258 | fbxc-25 |
| WBGene00019592 | 1.774249736 | 0.00030331 | 0.00362812 | K09F6.9 |
| WBGene00019593 | 3.291085342 | 0.00037501 | 0.00429929 | K09F6.10 |
| WBGene00019594 | -1.094201388 | 0.7816772 | 1 | K09H9.1 |
| WBGene00019595 | -0.537467778 | 0.00686809 | 0.03921347 | K09H9.2 |
| WBGene00019597 | 0.751624586 | 0.23756044 | 0.44379463 | K09H9.5 |
| WBGene00019598 | 0.259055253 | 0.5556382 | 0.73310187 | K09H9.7 |
| WBGene00019599 | 0.673664317 | 0.0026445 | 0.01936479 | acds-10 |
| WBGene00019600 | -0.478372134 | 0.01443514 | 0.06782395 | rga-3 |
| WBGene00019601 | 0.281454817 | 0.94435481 | 1 | K09H11.4 |
| WBGene00019603 | -0.544785811 | 0.68838355 | 1 | K09H11.6 |
| WBGene00019604 | -1.901856996 | 0.64072034 | 1 | K09H11.7 |
| WBGene00019605 | -0.118844904 | 0.73332901 | 0.8509364 | ctsa-1 |
| WBGene00019606 | -0.288813719 | 0.0413649 | 0.14079375 | clec-88 |
| WBGene00019607 | -0.527041952 | 0.04314218 | 0.14481318 | K10B2.4 |
| WBGene00019608 | -0.034416967 | 0.86281127 | 0.92620616 | ani-2 |
| WBGene00019609 | 0.152239798 | 0.80651972 | 0.89456934 | K10B3.1 |
| WBGene00019611 | -0.16764492 | 0.63639189 | 0.78921695 | K10B3.5 |
| WBGene00019612 | 0.964672619 | 0.00335834 | 0.02300996 | gpcp-1 |
| WBGene00019614 | 0.629154436 | 0.81300308 | 1 | srt-54 |
| WBGene00019615 | 0.637814605 | 0.04134283 | 0.14075806 | K10B4.3 |
| WBGene00019616 | -0.302361731 | 0.83477217 | 1 | nmur-2 |
| WBGene00019617 | 1.896847401 | 2.42E-17 | 6.12E-15 | K10C2.1 |
| WBGene00019617 | 0.749858659 | 0.00315835 | 0.02201234 | ctsa-2 |
| WBGene00019618 | 0.344077131 | 0.73642461 | 1 | K10C2.2 |
| WBGene00019619 | 2.361615868 | 1.94E-31 | 1.68E-28 | asp-14 |
| WBGene00019620 | -0.457359748 | 0.01528397 | 0.07063015 | fah-1 |
| WBGene00019622 | -1.717407588 | 0.1614535 | 1 | K10C2.7 |
| WBGene00019623 | 0.809718647 | 0.58471039 | 1 | K10C9.1 |
| WBGene00019624 | -0.442587837 | 0.11970442 | 0.28595048 | K10C9.3 |
| WBGene00019625 | 0.281454817 | 0.94435481 | 1 | K10C9.4 |
| WBGene00019626 | 0.846494767 | 0.63202292 | 1 | K10C9.7 |
| WBGene00019627 | -0.038961881 | 0.83455678 | 0.91032502 | K10D2.1 |
| WBGene00019628 | -0.1400417 | 0.47052101 | 0.66356557 | pup-2 |
| WBGene00019629 | 0.375017323 | 0.04117203 | 0.14041281 | cid-1 |
| WBGene00019630 | -0.172433553 | 0.52322436 | 0.70819029 | emb-1 |

| | | | | |
|---|---|---|---|---|
| WBGene00019631 | -0.803790619 | 0.00288838 | 0.02065242 | K10D2.5 |
| WBGene00019633 | 0.462773666 | 0.70412899 | 1 | K10D2.7 |
| WBGene00019636 | 0.468353537 | 0.23779273 | 0.44395533 | gsto-3 |
| WBGene00019638 | -0.274375875 | 0.9007694 | 1 | K10F12.6 |
| WBGene00019639 | -0.156269293 | 0.96924949 | 1 | K10F12.7 |
| WBGene00019640 | 1.270678414 | 0.5135606 | 1 | dos-2 |
| WBGene00019641 | -0.893844882 | 0.46737611 | 1 | K10G6.4 |
| WBGene00019642 | -0.955562677 | 0.62837716 | 1 | K11C4.1 |
| WBGene00019643 | -0.174625258 | 0.78729289 | 0.88275978 | K11C4.2 |
| WBGene00019644 | -0.848021396 | 0.21057155 | 0.41041347 | cpt-4 |
| WBGene00019645 | 0.374700564 | 0.16248508 | 0.34847392 | swt-7 |
| WBGene00019646 | 1.029709271 | 0.56619209 | 1 | K11D12.6 |
| WBGene00019647 | -1.083607791 | 7.8206E-05 | 0.0012543 | K11D12.7 |
| WBGene00019648 | 2.34695815 | 0.21033917 | 1 | K11D12.8 |
| WBGene00019649 | 1.210587634 | 0.07666606 | 0.21490008 | K11D12.9 |
| WBGene00019650 | -2.184288669 | 0.00197244 | 0.01538046 | K11D12.11 |
| WBGene00019651 | -0.106774101 | 0.80646955 | 0.89456934 | K11D12.12 |
| WBGene00019652 | 2.43134021 | 0.11680367 | 1 | K11G9.1 |
| WBGene00019653 | -0.86061079 | 0.1414648 | 0.31947994 | K11G9.2 |
| WBGene00019654 | -1.738647621 | 0.09074546 | 1 | K11G9.3 |
| WBGene00019655 | 0.104307038 | 0.91929535 | 1 | K11G9.5 |
| WBGene00019656 | -0.246161126 | 0.29424114 | 0.50457368 | slc-25A10 |
| WBGene00019657 | -0.075416506 | 0.77208054 | 0.87263357 | K11G12.6 |
| WBGene00019658 | -0.199533241 | 0.57366174 | 0.74745945 | K11H12.1 |
| WBGene00019659 | 0.785416611 | 0.59983079 | 1 | K11H12.3 |
| WBGene00019660 | 1.012229797 | 0.01310948 | 0.06326707 | K11H12.4 |
| WBGene00019661 | 0.421693784 | 0.54342045 | 0.72331236 | K11H12.5 |
| WBGene00019662 | -1.453036238 | 0.47675825 | 1 | K11H12.6 |
| WBGene00019663 | 0.811523976 | 0.01057726 | 0.05407097 | K11H12.7 |
| WBGene00019664 | -0.069043301 | 0.73536499 | 0.85215867 | K11H12.8 |
| WBGene00019665 | 0.779481344 | 0.25049177 | 0.45849851 | K11H12.9 |
| WBGene00019666 | 0.327078848 | 0.23599393 | 0.44215444 | sago-1 |
| WBGene00019667 | 1.557897343 | 0.63813005 | 1 | K12B6.2 |
| WBGene00019668 | 3.617417446 | 0.01295569 | 1 | fil-2 |
| WBGene00019669 | 1.100274297 | 0.50028399 | 1 | K12B6.4 |
| WBGene00019673 | 0.101467019 | 0.61061596 | 0.77207752 | K12C11.1 |
| WBGene00019674 | 0.088373001 | 0.93592139 | 1 | K12C11.3 |
| WBGene00019675 | 1.743415485 | 0.50483382 | 1 | K12D9.1 |
| WBGene00019677 | 0.053158802 | 0.86540873 | 0.92768209 | K12H4.2 |
| WBGene00019678 | -0.014334887 | 0.94035023 | 0.96916742 | K12H4.3 |
| WBGene00019679 | -0.396807777 | 0.03561379 | 0.12726616 | K12H4.4 |
| WBGene00019680 | -0.352989016 | 0.04238511 | 0.14342185 | K12H4.5 |
| WBGene00019682 | 0.501014782 | 0.00036634 | 0.00423609 | K12H4.7 |

| | | | | |
|---|---|---|---|---|
| WBGene00019683 | -0.759895904 | 0.2261343 | 0.43053106 | K12H6.2 |
| WBGene00019684 | 1.435925494 | 0.70920779 | 1 | K12H6.4 |
| WBGene00019686 | -0.324445145 | 0.20673819 | 0.40655714 | K12H6.6 |
| WBGene00019687 | -1.738286795 | 0.09677357 | 1 | K12H6.7 |
| WBGene00019688 | -0.156269293 | 0.96924949 | 1 | K12H6.8 |
| WBGene00019689 | 1.735991307 | 0.58152819 | 1 | K12H6.9 |
| WBGene00019690 | -1.087604428 | 0.23464595 | 1 | set-13 |
| WBGene00019691 | -0.334204649 | 0.59947618 | 0.76500318 | K12H6.12 |
| WBGene00019692 | -0.181367468 | 0.45749745 | 0.65446878 | M01A10.1 |
| WBGene00019693 | -0.181918653 | 0.25732169 | 0.46529537 | ostd-1 |
| WBGene00019697 | -0.152846955 | 0.46891773 | 0.66260826 | M01B12.4 |
| WBGene00019698 | -0.079722332 | 0.75474348 | 0.86283239 | riok-1 |
| WBGene00019699 | 3.079801363 | 0.00446724 | 0.0286815 | math-34 |
| WBGene00019700 | -1.956019767 | 0.00095904 | 0.00892918 | btb-11 |
| WBGene00019702 | 0.281454817 | 0.94435481 | 1 | fbxb-39 |
| WBGene00019703 | -1.722425584 | 0.01991149 | 0.08515161 | fbxb-41 |
| WBGene00019704 | -1.689349303 | 0.03171278 | 0.11812001 | fbxb-40 |
| WBGene00019705 | -1.276656663 | 0.10786428 | 0.26759651 | fbxb-45 |
| WBGene00019710 | 0.142393767 | 0.62707151 | 0.78317819 | M01E11.1 |
| WBGene00019711 | -0.35583517 | 0.03971442 | 0.13682844 | M01E11.2 |
| WBGene00019712 | 0.514493725 | 0.03347094 | 0.12229032 | rmh-1 |
| WBGene00019713 | -0.142621929 | 0.33930517 | 0.54766074 | pqn-52 |
| WBGene00019715 | -0.013875113 | 0.94709104 | 0.97304242 | M01G5.1 |
| WBGene00019716 | -0.053727205 | 0.8006675 | 0.89092697 | M01G5.3 |
| WBGene00019717 | 1.262606461 | 0.03444306 | 0.12486742 | trx-3 |
| WBGene00019718 | 1.274878015 | 0.43457342 | 1 | M01H9.2 |
| WBGene00019719 | 0.209604181 | 0.25899975 | 0.46716921 | M01H9.3 |
| WBGene00019720 | -0.15010823 | 0.78622057 | 0.88223764 | M01H9.4 |
| WBGene00019721 | 0.688793783 | 0.86378755 | 1 | M01H9.5 |
| WBGene00019722 | 0.023558868 | 0.99053178 | 1 | M02A10.1 |
| WBGene00019723 | -0.257997846 | 0.47987363 | 0.67175669 | fbxc-47 |
| WBGene00019724 | -0.515584248 | 0.19146514 | 0.3862709 | M02B7.2 |
| WBGene00019725 | 0.036453328 | 0.90214856 | 0.94913221 | algn-14 |
| WBGene00019726 | -0.025326188 | 0.92297364 | 0.9593272 | bris-1 |
| WBGene00019727 | 0.275935943 | 0.18706659 | 0.38049515 | zig-12 |
| WBGene00019728 | 0.663148878 | 0.43264911 | 0.6344719 | M02D8.2 |
| WBGene00019729 | -0.495702741 | 0.14351887 | 0.32166561 | M02D8.3 |
| WBGene00019730 | 0.254546124 | 0.33052427 | 0.53871596 | asns-2 |
| WBGene00019731 | 0.459643485 | 0.5222402 | 0.70717357 | M02D8.5 |
| WBGene00019732 | 1.36907956 | 0.02698093 | 0.10546537 | M02D8.6 |
| WBGene00019733 | 0.513431732 | 0.10757722 | 0.26759651 | M02E1.1 |
| WBGene00019734 | 1.261433833 | 0.08061683 | 0.22209668 | M02E1.2 |
| WBGene00019735 | -0.474271289 | 0.77244825 | 1 | M02F4.1 |

| | | | | |
|---|---|---|---|---|
| WBGene00019736 | 1.181686151 | 0.7579682 | 1 | M02F4.2 |
| WBGene00019738 | 0.549319918 | 0.0160551 | 0.07306034 | clec-265 |
| WBGene00019741 | 0.508616617 | 0.39569351 | 0.60353202 | nhr-201 |
| WBGene00019742 | 0.292306061 | 0.64472922 | 0.79492814 | nhr-202 |
| WBGene00019743 | 0.570670448 | 0.24980254 | 0.45751284 | nhr-203 |
| WBGene00019744 | -1.287392125 | 1.3847E-05 | 0.00029756 | M02H5.8 |
| WBGene00019745 | 0.377828795 | 0.92526585 | 1 | srbc-57 |
| WBGene00019746 | 0.047870787 | 0.94774402 | 0.97315292 | M03A1.3 |
| WBGene00019747 | 0.163772621 | 0.39392156 | 0.60200405 | ipla-1 |
| WBGene00019748 | 0.641105009 | 0.00399974 | 0.02626747 | atg-2 |
| WBGene00019749 | 0.730973124 | 0.30223175 | 0.51162795 | M03A8.3 |
| WBGene00019750 | 0.592145784 | 0.57231006 | 1 | M03D4.3 |
| WBGene00019751 | 0.617002264 | 0.14247873 | 0.32034267 | M03D4.4 |
| WBGene00019753 | 0.572354502 | 0.44161259 | 0.64137027 | M03E7.1 |
| WBGene00019754 | -0.629292503 | 0.72518963 | 1 | M03E7.2 |
| WBGene00019755 | -0.156269293 | 0.96924949 | 1 | M03E7.3 |
| WBGene00019756 | 0.627294895 | 0.56043293 | 1 | M03E7.4 |
| WBGene00019757 | -0.87752839 | 0.00015343 | 0.00212885 | memb-2 |
| WBGene00019758 | 0.747845955 | 0.85215965 | 1 | M03F4.4 |
| WBGene00019759 | -0.319110244 | 0.67962101 | 0.81647553 | M03F4.6 |
| WBGene00019760 | -0.496387843 | 0.00873828 | 0.04674442 | calu-1 |
| WBGene00019761 | 0.404695497 | 0.6515251 | 0.79916813 | M03F8.1 |
| WBGene00019762 | -0.311867003 | 0.04915491 | 0.15918637 | M03F8.3 |
| WBGene00019763 | 0.289266376 | 0.8325472 | 1 | galt-1 |
| WBGene00019764 | 0.952093913 | 0.33714484 | 1 | M03F8.5 |
| WBGene00019765 | 1.265199718 | 0.16177607 | 0.34773047 | M03F8.6 |
| WBGene00019766 | -0.095019949 | 0.66650431 | 0.80879848 | M4.1 |
| WBGene00019767 | -0.898420807 | 9.31E-11 | 8.83E-09 | rpa-2 |
| WBGene00019768 | 0.006759056 | 0.98956636 | 0.99484196 | M04F3.2 |
| WBGene00019769 | -0.044189703 | 0.94994606 | 0.97450741 | M04F3.3 |
| WBGene00019770 | -0.051389433 | 0.86500999 | 0.92750033 | M04F3.4 |
| WBGene00019771 | -0.058733478 | 0.78264532 | 0.8802313 | M04F3.5 |
| WBGene00019772 | 1.231377029 | 0.0001185 | 0.00174178 | M04G7.1 |
| WBGene00019773 | -1.351599631 | 0.53258792 | 1 | M04G7.2 |
| WBGene00019774 | 0.380493126 | 0.53868875 | 0.720606 | M04G7.3 |
| WBGene00019776 | 1.543701184 | 0.32245017 | 1 | M6.4 |
| WBGene00019777 | 0.265326035 | 0.52593712 | 0.7103574 | M57.1 |
| WBGene00019778 | -0.206615488 | 0.32294217 | 0.53142327 | M57.2 |
| WBGene00019779 | 1.236687368 | 2.11E-13 | 3.24E-11 | endu-2 |
| WBGene00019780 | 0.53584612 | 0.00459789 | 0.0293805 | M60.4 |
| WBGene00019781 | -0.119788139 | 0.80597845 | 0.89429578 | M60.6 |
| WBGene00019782 | 1.535577417 | 0.15491731 | 0.33852997 | M60.7 |
| WBGene00019783 | 0.747845955 | 0.85215965 | 1 | M70.1 |

| | | | | |
|---|---|---|---|---|
| WBGene00019785 | 2.111080655 | 0.48938556 | 1 | M70.3 |
| WBGene00019786 | -0.046607886 | 0.82609143 | 0.90625042 | M70.4 |
| WBGene00019787 | 0.248522678 | 0.45581435 | 0.65298432 | M70.5 |
| WBGene00019788 | 0.574508983 | 0.8551971 | 1 | M116.1 |
| WBGene00019791 | 0.574508983 | 0.8551971 | 1 | sdz-25 |
| WBGene00019792 | -0.429603587 | 0.15025221 | 0.33184889 | M116.5 |
| WBGene00019793 | -2.348812061 | 0.13584578 | 1 | M151.1 |
| WBGene00019794 | -1.004023032 | 0.26723118 | 0.47551718 | M151.2 |
| WBGene00019795 | -0.301710811 | 0.65059853 | 0.79859638 | M151.3 |
| WBGene00019796 | -0.115183845 | 0.88843815 | 0.94106854 | M151.4 |
| WBGene00019797 | -1.857839188 | 0.31666726 | 1 | fbxb-31 |
| WBGene00019798 | -1.734202971 | 0.17497941 | 1 | fbxb-32 |
| WBGene00019799 | -1.144064094 | 0.00163094 | 0.01333463 | M151.7 |
| WBGene00019800 | -0.681501527 | 0.00048269 | 0.0052324 | mtss-1 |
| WBGene00019801 | 0.376855045 | 0.10207641 | 0.2578955 | aak-1 |
| WBGene00019803 | 1.166653077 | 0.00010594 | 0.00159389 | PDB1.1 |
| WBGene00019804 | -1.759191416 | 0.57668729 | 1 | R01B10.2 |
| WBGene00019805 | 0.458033906 | 0.24856155 | 0.45656791 | R01B10.3 |
| WBGene00019806 | 0.87339703 | 0.08952365 | 0.23732616 | R01B10.4 |
| WBGene00019807 | 0.089358169 | 0.72942989 | 0.84847759 | jamp-1 |
| WBGene00019808 | 0.190839839 | 0.4219223 | 0.62653604 | R01B10.6 |
| WBGene00019809 | 0.377828795 | 0.92526585 | 1 | R01H2.1 |
| WBGene00019810 | 0.377828795 | 0.92526585 | 1 | R01H2.2 |
| WBGene00019811 | -0.461322167 | 0.00245519 | 0.01833235 | egg-2 |
| WBGene00019812 | 0.193302195 | 0.91734354 | 1 | R01H2.4 |
| WBGene00019813 | -0.078037447 | 0.79464095 | 0.88641923 | ger-1 |
| WBGene00019814 | -2.960501783 | 2.2294E-05 | 0.00044299 | R02C2.1 |
| WBGene00019815 | -1.475985494 | 0.04215065 | 0.14278766 | kin-34 |
| WBGene00019816 | 0.680640244 | 0.34009886 | 0.54843143 | nhr-204 |
| WBGene00019817 | -1.183536118 | 0.15949518 | 0.34449938 | R02C2.6 |
| WBGene00019818 | -0.841351332 | 0.37638882 | 1 | R02C2.7 |
| WBGene00019819 | 0.445033155 | 0.01333654 | 0.06382299 | R02D3.1 |
| WBGene00019820 | -0.084125943 | 0.6946005 | 0.82685996 | cogc-8 |
| WBGene00019821 | -0.188551547 | 0.41968415 | 0.62420499 | R02D3.3 |
| WBGene00019822 | -0.182003639 | 0.32415025 | 0.53261716 | R02D3.4 |
| WBGene00019823 | -0.518649063 | 0.01004647 | 0.05205896 | fnta-1 |
| WBGene00019824 | -0.222871529 | 0.22401488 | 0.42810714 | R02D3.7 |
| WBGene00019825 | -0.170139226 | 0.51006308 | 0.69671769 | R02D3.8 |
| WBGene00019826 | 1.413734769 | 0.68804531 | 1 | R02E4.1 |
| WBGene00019827 | 0.281655846 | 0.2047011 | 0.40406162 | mop-25.1 |
| WBGene00019828 | 0.265363324 | 0.71245405 | 0.8375158 | R02E12.4 |
| WBGene00019829 | 1.103412626 | 0.14934742 | 0.33078626 | R02E12.5 |
| WBGene00019830 | -0.837587803 | 0.05276009 | 0.16650268 | hrg-1 |

| | | | | |
|---|---|---|---|---|
| WBGene00019831 | 0.166308439 | 0.30656687 | 0.51537364 | R02F2.1 |
| WBGene00019832 | 0.234625736 | 0.64119432 | 0.79235567 | osg-1 |
| WBGene00019833 | -0.205929887 | 0.40718132 | 0.61394145 | R02F2.4 |
| WBGene00019834 | -0.156269293 | 0.96924949 | 1 | R02F2.5 |
| WBGene00019835 | -2.272874415 | 0.36844983 | 1 | R02F2.6 |
| WBGene00019836 | -0.122303268 | 0.66589999 | 0.80838834 | R02F2.7 |
| WBGene00019837 | 0.741862482 | 0.07325529 | 0.20855723 | R02F2.8 |
| WBGene00019838 | -0.393585375 | 0.17412568 | 0.36430464 | R02F2.9 |
| WBGene00019839 | -1.003146305 | 0.00779822 | 0.04301599 | R02F11.1 |
| WBGene00019840 | 1.196364967 | 0.22084106 | 0.42384308 | R02F11.2 |
| WBGene00019841 | 0.128069315 | 0.73397049 | 0.8511151 | R02F11.3 |
| WBGene00019842 | -0.455394384 | 0.02053756 | 0.0872146 | R02F11.4 |
| WBGene00019843 | 0.750082119 | 0.29413811 | 0.50453953 | R03E9.2 |
| WBGene00019844 | 1.387230861 | 6.8824E-05 | 0.00113686 | abts-4 |
| WBGene00019845 | 1.584577408 | 0.63258867 | 1 | irld-13 |
| WBGene00019846 | 1.519251688 | 0.00054246 | 0.00569729 | gpx-7 |
| WBGene00019847 | 0.465741834 | 0.45659968 | 0.65378049 | R03G5.6 |
| WBGene00019849 | 0.627834078 | 0.77998941 | 1 | oac-40 |
| WBGene00019854 | -1.037072233 | 0.03873663 | 0.13457338 | R03H10.1 |
| WBGene00019855 | 0.6550697 | 0.09521444 | 0.24737823 | R03H10.2 |
| WBGene00019858 | 1.391720125 | 0.5323529 | 1 | R03H10.6 |
| WBGene00019859 | 0.51274971 | 0.59286063 | 1 | R03H10.7 |
| WBGene00019861 | -0.982255077 | 0.6464653 | 1 | R04A9.1 |
| WBGene00019862 | -0.5655918 | 0.00677327 | 0.03888144 | nrde-3 |
| WBGene00019863 | 0.757243791 | 0.10795358 | 0.26759651 | R04A9.3 |
| WBGene00019864 | -0.349368826 | 0.24325863 | 0.4508704 | ceh-93 |
| WBGene00019865 | -0.228279595 | 0.79282789 | 0.8856941 | R04A9.6 |
| WBGene00019866 | 0.13551433 | 0.90871576 | 1 | R04B3.1 |
| WBGene00019867 | -0.301041979 | 0.55724306 | 0.73435747 | R04B3.2 |
| WBGene00019868 | -0.858547175 | 0.40827268 | 1 | R04B3.3 |
| WBGene00019869 | 0.298911277 | 0.45386748 | 0.65134811 | cnnm-2 |
| WBGene00019870 | 2.070130458 | 0.15950076 | 1 | R04E5.7 |
| WBGene00019871 | 0.474183998 | 0.10552763 | 0.26443983 | R04E5.8 |
| WBGene00019872 | 0.132967878 | 0.83958833 | 0.91211511 | R04E5.9 |
| WBGene00019873 | 0.443947004 | 0.50213905 | 0.68980202 | arrd-26 |
| WBGene00019874 | -0.008677305 | 0.99719169 | 1 | R05C11.2 |
| WBGene00019875 | 0.995771933 | 1.0491E-05 | 0.00023719 | mca-2 |
| WBGene00019877 | 0.076521588 | 0.68571788 | 0.82038833 | R05D3.2 |
| WBGene00019878 | -0.201287228 | 0.70880244 | 0.83525966 | R05D3.3 |
| WBGene00019881 | 0.13149209 | 0.64950358 | 0.79813997 | R05D3.8 |
| WBGene00019882 | 0.43610598 | 0.1054154 | 0.26426767 | R05D3.9 |
| WBGene00019883 | -0.097545014 | 0.53517066 | 0.7178681 | met-2 |
| WBGene00019884 | 0.475391962 | 0.02602804 | 0.10302938 | R05D3.12 |

| | | | | |
|---|---|---|---|---|
| WBGene00019885 | 0.335897242 | 0.71002812 | 0.83591862 | R05D8.7 |
| WBGene00019886 | 0.983488853 | 0.80601565 | 1 | R05D8.9 |
| WBGene00019887 | 1.655400498 | 0.39459006 | 1 | R05D8.11 |
| WBGene00019888 | -0.01936479 | 0.91529533 | 0.95580617 | btbd-10 |
| WBGene00019890 | -0.161347749 | 0.29222531 | 0.50232303 | R05F9.6 |
| WBGene00019891 | 0.281454817 | 0.94435481 | 1 | R05F9.7 |
| WBGene00019892 | -0.003291602 | 0.99109113 | 0.99555966 | R05F9.9 |
| WBGene00019893 | -0.229236335 | 0.16910636 | 0.35725079 | sgt-1 |
| WBGene00019894 | -0.400622392 | 0.31672458 | 0.52474797 | R05F9.11 |
| WBGene00019895 | 0.615349425 | 0.0005384 | 0.00566936 | aagr-2 |
| WBGene00019896 | 0.315981165 | 0.83009196 | 1 | R05G6.1 |
| WBGene00019898 | -0.698552694 | 0.00857676 | 0.04613665 | R05G6.4 |
| WBGene00019899 | 0.217814741 | 0.79674053 | 0.88794078 | R05G6.5 |
| WBGene00019900 | -0.388078966 | 0.00404734 | 0.02651847 | vdac-1 |
| WBGene00019901 | -1.11019416 | 0.52432696 | 1 | R05G6.9 |
| WBGene00019902 | 1.557761378 | 1.2892E-05 | 0.00028151 | R05G6.10 |
| WBGene00019904 | 0.840004353 | 0.01451546 | 0.06806958 | twk-49 |
| WBGene00019905 | -0.605642664 | 0.02601874 | 0.10302938 | R05G9.3 |
| WBGene00019906 | 0.214238389 | 0.78378251 | 0.88074562 | R05G9R.1 |
| WBGene00019907 | -0.214574528 | 0.59914246 | 0.76480755 | R05H11.1 |
| WBGene00019908 | 0.538752448 | 0.4579048 | 0.65476162 | R05H11.2 |
| WBGene00019909 | 0.008447327 | 0.99297479 | 1 | R06A10.1 |
| WBGene00019911 | 0.608142062 | 0.20429417 | 0.40370045 | R06A10.4 |
| WBGene00019913 | -0.738476965 | 0.4605063 | 1 | R06B10.2 |
| WBGene00019914 | -0.472906628 | 0.00382068 | 0.02532032 | clec-150 |
| WBGene00019915 | -0.275581576 | 0.46777799 | 0.66183982 | tbc-10 |
| WBGene00019916 | 1.181686151 | 0.7579682 | 1 | R06B10.7 |
| WBGene00019917 | 1.837721915 | 0.04572297 | 0.15101268 | clec-43 |
| WBGene00019918 | -0.742097286 | 0.09135095 | 0.24047959 | fbxc-36 |
| WBGene00019920 | -0.22461868 | 0.93331196 | 1 | acs-15 |
| WBGene00019921 | 0.476908871 | 0.73733278 | 1 | fbxc-34 |
| WBGene00019922 | -1.61650448 | 0.00028416 | 0.00345342 | fbxc-28 |
| WBGene00019923 | -1.398861835 | 0.12727764 | 0.29779878 | fbxc-29 |
| WBGene00019924 | -0.775401733 | 0.17133336 | 0.36045024 | fbxc-33 |
| WBGene00019925 | -2.95863744 | 0.05038525 | 1 | fbxc-31 |
| WBGene00019926 | -1.150134399 | 0.4892498 | 1 | fbxc-27 |
| WBGene00019927 | -1.023721759 | 0.30449196 | 1 | fbxc-24 |
| WBGene00019928 | -1.509975708 | 0.60137946 | 1 | clec-44 |
| WBGene00019929 | 0.15388156 | 0.93468081 | 1 | R07C3.13 |
| WBGene00019931 | 4.570102203 | 0.00304315 | 1 | R07C12.1 |
| WBGene00019933 | 1.557897343 | 0.63813005 | 1 | R07C12.3 |
| WBGene00019935 | 0.868596768 | 0.03218844 | 0.1191463 | R07E4.1 |
| WBGene00019937 | -0.124624968 | 0.65401491 | 0.80118113 | R07E4.3 |

| | | | | |
|---|---|---|---|---|
| WBGene00019938 | -0.204000132 | 0.39681228 | 0.60465448 | R07E4.5 |
| WBGene00019939 | 0.468952407 | 0.56374665 | 0.73890188 | lips-17 |
| WBGene00019940 | 0.267353672 | 0.14084548 | 0.31861469 | npp-21 |
| WBGene00019941 | -0.588496131 | 0.00018699 | 0.00250857 | pgam-5 |
| WBGene00019943 | 0.023920641 | 0.9094749 | 0.95279443 | R07G3.7 |
| WBGene00019944 | 1.565583351 | 0.00254874 | 0.01884547 | R07G3.8 |
| WBGene00019945 | 1.413734769 | 0.68804531 | 1 | R08C7.1 |
| WBGene00019946 | 0.155410478 | 0.37718225 | 0.58507342 | chat-1 |
| WBGene00019947 | -0.674397318 | 1.4372E-05 | 0.0003083 | htz-1 |
| WBGene00019948 | 0.340122187 | 0.87335753 | 1 | R08C7.4 |
| WBGene00019949 | 1.181686151 | 0.7579682 | 1 | R08C7.5 |
| WBGene00019950 | 0.688793783 | 0.86378755 | 1 | clec-175 |
| WBGene00019951 | -0.396696717 | 0.85452912 | 1 | R08C7.8 |
| WBGene00019952 | -0.91949293 | 0.36297412 | 1 | fbxb-74 |
| WBGene00019953 | -0.35566893 | 0.04125947 | 0.14063201 | wapl-1 |
| WBGene00019955 | 0.862680609 | 0.04505962 | 0.14927796 | R08C7.12 |
| WBGene00019956 | -1.361868702 | 0.03940408 | 0.13602644 | fbxb-75 |
| WBGene00019957 | 1.677351323 | 0.00012649 | 0.00183914 | R08E3.1 |
| WBGene00019957 | 1.968547043 | 2.98E-07 | 1.1958E-05 | R08E3.2 |
| WBGene00019959 | -0.006131827 | 0.98080305 | 0.98993793 | R08E3.3 |
| WBGene00019960 | 0.237385626 | 0.5518146 | 0.73004043 | ztf-16 |
| WBGene00019961 | -0.362598919 | 0.59277573 | 0.7597266 | R08E5.1 |
| WBGene00019962 | -0.498486522 | 0.02285324 | 0.09427837 | cysl-3 |
| WBGene00019963 | 0.094462817 | 0.75239003 | 0.86217294 | R08E5.3 |
| WBGene00019965 | 0.181180695 | 0.62687218 | 0.78309735 | R08F11.1 |
| WBGene00019967 | 0.813228813 | 0.01229874 | 0.06033898 | cyp-33C8 |
| WBGene00019968 | 0.784228205 | 0.42717151 | 0.63047524 | R08F11.4 |
| WBGene00019970 | 5.100671744 | 7.392E-05 | 0.00120142 | R08F11.7 |
| WBGene00019971 | -0.300495446 | 0.05909333 | 0.18062742 | ergo-1 |
| WBGene00019972 | -0.39167124 | 0.69777379 | 1 | R09A1.2 |
| WBGene00019973 | 0.852199922 | 0.19252167 | 0.38762946 | R09A1.3 |
| WBGene00019975 | 1.076781714 | 0.19990825 | 0.39827471 | flp-34 |
| WBGene00019977 | 1.522010905 | 0.01156746 | 0.05775973 | fpn-1.2 |
| WBGene00019978 | 1.078758896 | 1.38E-11 | 1.53E-09 | hacd-1 |
| WBGene00019979 | 0.331203115 | 0.50884511 | 0.69553126 | R09B5.11 |
| WBGene00019980 | 2.18453076 | 0.41642786 | 1 | chil-14 |
| WBGene00019983 | -0.557612594 | 0.00057762 | 0.00596838 | sti-1 |
| WBGene00019986 | 0.635752545 | 0.02254864 | 0.09337076 | R09F10.1 |
| WBGene00019987 | -0.081935638 | 0.79234501 | 0.88531758 | R09F10.3 |
| WBGene00019988 | 0.058536494 | 0.91589102 | 0.95580617 | R09F10.5 |
| WBGene00019989 | -0.602933482 | 0.00189916 | 0.01495309 | R09F10.8 |
| WBGene00019991 | 1.116613983 | 0.69067248 | 1 | R09H3.3 |
| WBGene00019993 | -0.256122837 | 0.39128064 | 0.59966401 | rbx-2 |

| | | | | |
|---|---|---|---|---|
| WBGene00019994 | 0.887636257 | 0.01156449 | 0.05775973 | cdh-1 |
| WBGene00019995 | 0.636566705 | 0.70536583 | 1 | R10F2.4 |
| WBGene00019996 | 0.079418419 | 0.96102778 | 1 | R10F2.5 |
| WBGene00019997 | -0.738567372 | 0.31578791 | 0.52391104 | R10F2.6 |
| WBGene00019998 | 0.480374105 | 0.27309947 | 0.48186715 | R10H1.1 |
| WBGene00019999 | 0.716330419 | 0.48638921 | 0.67622374 | srab-14 |
| WBGene00020003 | 1.820218437 | 0.11128452 | 1 | R11E3.1 |
| WBGene00020004 | 1.028948025 | 0.07743439 | 0.21582435 | R11E3.2 |
| WBGene00020006 | 0.231443858 | 0.54379998 | 0.72337864 | set-15 |
| WBGene00020007 | 1.071723653 | 4.4938E-06 | 0.00011486 | R11F4.1 |
| WBGene00020008 | 0.366266693 | 0.76200462 | 1 | R11F4.2 |
| WBGene00020010 | -0.580458573 | 0.59013373 | 1 | R11G1.1 |
| WBGene00020011 | -0.406846146 | 0.64919288 | 0.79802502 | R11G1.2 |
| WBGene00020012 | 0.840852055 | 0.0041755 | 0.02718215 | R11G1.6 |
| WBGene00020014 | 0.685087967 | 0.68567029 | 1 | R11G11.6 |
| WBGene00020015 | 0.566365865 | 0.30566629 | 0.51458342 | nhr-210 |
| WBGene00020016 | 4.859634073 | 1.17E-07 | 5.3603E-06 | lipl-3 |
| WBGene00020017 | -0.628582155 | 0.44505418 | 0.64447021 | R12A1.3 |
| WBGene00020018 | -0.33669265 | 0.45876388 | 0.65487177 | R12B2.2 |
| WBGene00020019 | -0.494147886 | 0.77957055 | 1 | R12B2.3 |
| WBGene00020020 | 0.377828795 | 0.92526585 | 1 | R12B2.6 |
| WBGene00020021 | 0.938982574 | 0.8146943 | 1 | R12B2.7 |
| WBGene00020022 | 0.083122011 | 0.58671689 | 0.75578899 | gldc-1 |
| WBGene00020023 | 0.5483773 | 0.4817376 | 0.67289188 | frpr-16 |
| WBGene00020025 | -0.09893254 | 0.63374629 | 0.78783996 | R12C12.5 |
| WBGene00020026 | -0.658172861 | 0.00717609 | 0.04054205 | R12C12.6 |
| WBGene00020027 | -0.242404256 | 0.29670708 | 0.50713287 | R12C12.7 |
| WBGene00020029 | 0.608478058 | 0.09014784 | 0.23860583 | R12C12.9 |
| WBGene00020030 | -0.259731629 | 0.23660238 | 0.44275423 | R12E2.1 |
| WBGene00020031 | 0.110511426 | 0.56633211 | 0.74165011 | suco-1 |
| WBGene00020032 | -0.156269293 | 0.96924949 | 1 | R12E2.6 |
| WBGene00020033 | -0.718628544 | 0.21134302 | 0.4111419 | R12E2.7 |
| WBGene00020034 | -2.020841616 | 0.12158975 | 1 | R12E2.8 |
| WBGene00020035 | -0.811076675 | 1.4415E-05 | 0.00030862 | egg-5 |
| WBGene00020036 | -0.431772302 | 0.04920412 | 0.1592608 | R12E2.11 |
| WBGene00020037 | -0.249613718 | 0.3036047 | 0.51245164 | mrps-6 |
| WBGene00020038 | -0.32364339 | 0.10758016 | 0.26759651 | R12E2.13 |
| WBGene00020039 | -1.178601738 | 0.04780214 | 0.15622229 | R12E2.14 |
| WBGene00020040 | -1.156568657 | 0.05126981 | 0.1635488 | R12E2.15 |
| WBGene00020042 | -0.156269293 | 0.96924949 | 1 | R13A1.3 |
| WBGene00020043 | 0.188009384 | 0.78795043 | 0.88339194 | R13A1.5 |
| WBGene00020045 | 0.098464167 | 0.89758696 | 0.94634211 | glb-23 |
| WBGene00020046 | -0.688492837 | 0.26309964 | 0.47179667 | best-16 |

| | | | | |
|---|---|---|---|---|
| WBGene00020047 | -0.798965885 | 0.09202092 | 0.24150461 | ttr-32 |
| WBGene00020048 | 0.517126339 | 0.71676293 | 1 | lgc-12 |
| WBGene00020049 | -0.274985435 | 0.4766187 | 0.66906261 | ttr-8 |
| WBGene00020050 | -0.232636437 | 0.48859293 | 0.67848642 | R13A5.7 |
| WBGene00020051 | 0.904822078 | 0.0049902 | 0.03111764 | R13A5.9 |
| WBGene00020052 | 0.106365252 | 0.81106416 | 0.89682422 | R13A5.10 |
| WBGene00020053 | 1.536599236 | 0.41694692 | 1 | R13A5.11 |
| WBGene00020054 | 1.258724256 | 0.00738037 | 0.04129265 | R13D7.2 |
| WBGene00020058 | -0.364030141 | 0.3907014 | 0.5991985 | R13D11.4 |
| WBGene00020060 | 0.007101558 | 0.9973363 | 1 | srab-16 |
| WBGene00020062 | 0.500895474 | 0.67376303 | 1 | nhr-270 |
| WBGene00020064 | -0.497596208 | 0.04586677 | 0.15136408 | kbp-1 |
| WBGene00020066 | -0.963811488 | 0.61154562 | 1 | dhhc-5 |
| WBGene00020067 | 0.688793783 | 0.86378755 | 1 | clec-158 |
| WBGene00020068 | -0.015514342 | 0.93020875 | 0.96354103 | cra-1 |
| WBGene00020069 | 0.975906504 | 0.10954641 | 0.27005137 | R13H7.2 |
| WBGene00020070 | -1.390520744 | 0.47839671 | 1 | rmd-6 |
| WBGene00020071 | -0.156269293 | 0.96924949 | 1 | R13H9.5 |
| WBGene00020072 | 0.377828795 | 0.92526585 | 1 | R13H9.6 |
| WBGene00020073 | -1.825908224 | 2.6343E-06 | 7.4182E-05 | sdz-28 |
| WBGene00020077 | 0.688793783 | 0.86378755 | 1 | R52.5 |
| WBGene00020080 | 1.278470803 | 0.39867721 | 1 | math-37 |
| WBGene00020082 | 0.462695374 | 0.03224042 | 0.11930234 | gcp-2.1 |
| WBGene00020083 | -0.318130802 | 0.72447167 | 0.84517459 | R57.2 |
| WBGene00020084 | 2.47098842 | 0.31902842 | 1 | R105.1 |
| WBGene00020086 | 0.959209435 | 0.37663522 | 1 | npr-24 |
| WBGene00020088 | -0.385261143 | 0.19946359 | 0.39764983 | R119.2 |
| WBGene00020089 | -0.265005089 | 0.25505412 | 0.46259517 | R119.3 |
| WBGene00020090 | 0.391709656 | 0.32545285 | 0.5338183 | R119.5 |
| WBGene00020091 | -0.197832487 | 0.30359121 | 0.51245164 | rnp-8 |
| WBGene00020092 | 0.043087228 | 0.83501442 | 0.91032502 | pcf-11 |
| WBGene00020093 | -0.879625116 | 0.00057173 | 0.00592777 | R144.3 |
| WBGene00020094 | 0.121199213 | 0.66566357 | 0.8082846 | wip-1 |
| WBGene00020095 | -0.016841879 | 0.94227982 | 0.97027162 | smcr-8 |
| WBGene00020096 | -0.15720857 | 0.5605741 | 0.73673213 | R144.6 |
| WBGene00020097 | 0.431419196 | 0.05327604 | 0.16764609 | larp-1 |
| WBGene00020098 | -0.82880731 | 0.00336432 | 0.02302494 | R144.10 |
| WBGene00020099 | -0.184485133 | 0.57213079 | 0.7462154 | R144.11 |
| WBGene00020100 | -0.046669537 | 0.97537266 | 1 | mks-1 |
| WBGene00020101 | -0.62576874 | 0.0112794 | 0.05679932 | lmtr-5 |
| WBGene00020102 | 0.048027672 | 0.82146512 | 0.90322478 | R148.3 |
| WBGene00020103 | -0.259327651 | 0.35041617 | 0.55898078 | R148.4 |
| WBGene00020104 | 0.026227737 | 0.93305396 | 0.96459538 | R148.5 |

| | | | | |
|---|---|---|---|---|
| WBGene00020105 | 0.247318604 | 0.91814609 | 1 | R148.7 |
| WBGene00020106 | 1.436305608 | 0.00213861 | 0.01639194 | R151.1 |
| WBGene00020107 | -0.311378021 | 0.02467178 | 0.0992841 | R151.2 |
| WBGene00020109 | -0.44262701 | 0.05678788 | 0.17599478 | R151.6 |
| WBGene00020110 | -0.039390908 | 0.80317883 | 0.89239902 | hsp-75 |
| WBGene00020111 | 0.065086683 | 0.73634648 | 0.85280864 | R151.8 |
| WBGene00020112 | -0.84478017 | 5.6104E-05 | 0.00095669 | pfd-5 |
| WBGene00020113 | -0.446974791 | 0.05950218 | 0.18139981 | R151.10 |
| WBGene00020114 | 1.14502227 | 1.94E-07 | 8.1427E-06 | pde-4 |
| WBGene00020115 | -0.273862514 | 0.07312974 | 0.20827309 | mboa-6 |
| WBGene00020116 | 0.09697944 | 0.97612634 | 1 | moa-1 |
| WBGene00020117 | 0.377828795 | 0.92526585 | 1 | R155.3 |
| WBGene00020120 | -1.323986516 | 0.36104889 | 1 | R160.3 |
| WBGene00020121 | 2.591348062 | 0.23319713 | 1 | R160.4 |
| WBGene00020122 | 0.932931886 | 0.70926986 | 1 | R160.5 |
| WBGene00020125 | 0.035558346 | 0.95314496 | 0.97623454 | R173.3 |
| WBGene00020126 | 0.226388446 | 0.61939038 | 0.77855043 | flp-26 |
| WBGene00020128 | 0.820060283 | 0.01873069 | 0.08183132 | R193.2 |
| WBGene00020130 | 0.747845955 | 0.85215965 | 1 | T22E5.7 |
| WBGene00020130 | 0.352246057 | 0.6453463 | 0.79528516 | igcm-2 |
| WBGene00020131 | 0.671886724 | 1.5569E-05 | 0.0003293 | gcy-28 |
| WBGene00020134 | -0.187358389 | 0.80801065 | 0.89524159 | T01B6.1 |
| WBGene00020135 | 1.834435179 | 0.0001401 | 0.00199876 | aakg-4 |
| WBGene00020136 | 0.362628039 | 0.83538298 | 1 | nlp-45 |
| WBGene00020137 | 1.155725288 | 0.39303499 | 1 | T01B10.5 |
| WBGene00020138 | 1.656267103 | 0.30359595 | 1 | T01B11.1 |
| WBGene00020139 | 0.430801556 | 0.02973167 | 0.11241736 | T01B11.2 |
| WBGene00020140 | 1.804609989 | 0.55694839 | 1 | ant-1.4 |
| WBGene00020141 | 2.463353637 | 1.0374E-05 | 0.00023499 | lmd-5 |
| WBGene00020142 | 0.23559912 | 0.31794388 | 0.52576364 | aak-2 |
| WBGene00020143 | -0.252514062 | 0.43397522 | 0.63557216 | T01C8.2 |
| WBGene00020144 | 0.619804214 | 0.58060792 | 1 | T01C8.3 |
| WBGene00020146 | 0.405845331 | 0.04619716 | 0.15224749 | got-1.2 |
| WBGene00020148 | -0.521532777 | 0.38570039 | 0.59414935 | T01D1.3 |
| WBGene00020149 | -1.2304703 | 0.0003177 | 0.00378897 | T01D1.4 |
| WBGene00020150 | 0.809178081 | 0.68171189 | 1 | T01D1.5 |
| WBGene00020151 | 0.908930564 | 0.6008643 | 1 | T01G6.1 |
| WBGene00020152 | 1.520021104 | 0.10697837 | 0.26658958 | nhr-211 |
| WBGene00020153 | 0.488414255 | 0.4180949 | 0.62283315 | nhr-212 |
| WBGene00020154 | 0.446458324 | 0.87164163 | 1 | T01G6.10 |
| WBGene00020155 | 1.017318661 | 1.26E-07 | 5.7261E-06 | T02B11.3 |
| WBGene00020156 | 0.018911008 | 0.97651081 | 0.98782017 | T02B11.4 |
| WBGene00020157 | 2.092602477 | 0.15504717 | 1 | T02B11.6 |

| | | | | |
|---|---|---|---|---|
| WBGene00020158 | 0.957987548 | 3.97E-07 | 1.5458E-05 | T02C5.1 |
| WBGene00020160 | -0.232055195 | 0.42059934 | 0.62518323 | igcm-3 |
| WBGene00020163 | -0.512865861 | 0.25715188 | 0.46511709 | T02G5.3 |
| WBGene00020166 | -0.659469747 | 1.6903E-05 | 0.00035322 | T02G5.7 |
| WBGene00020168 | -0.286318449 | 0.18993412 | 0.38435739 | mct-5 |
| WBGene00020169 | 0.467254508 | 0.00472964 | 0.02998568 | mmaa-1 |
| WBGene00020170 | 0.228267819 | 0.84484844 | 1 | T02G5.14 |
| WBGene00020171 | 0.004628503 | 0.98399392 | 0.99228305 | T02H6.1 |
| WBGene00020172 | -0.129145061 | 0.60525684 | 0.76897128 | thoc-1 |
| WBGene00020175 | -0.206948353 | 0.86901555 | 1 | T02H6.5 |
| WBGene00020177 | -0.156269293 | 0.96924949 | 1 | T02H6.7 |
| WBGene00020179 | 1.660922952 | 0.43395868 | 1 | T02H6.9 |
| WBGene00020181 | -0.233568625 | 0.1620415 | 0.3479069 | T02H6.11 |
| WBGene00020182 | -2.651387664 | 6.0852E-05 | 0.00102674 | ugt-53 |
| WBGene00020183 | 0.063250554 | 0.87181018 | 0.93105248 | T03D3.5 |
| WBGene00020184 | -0.434670391 | 0.02745202 | 0.10670115 | uba-5 |
| WBGene00020185 | -0.203638723 | 0.22326768 | 0.42728451 | pgk-1 |
| WBGene00020187 | 1.207928338 | 0.48077353 | 1 | gsp-4 |
| WBGene00020188 | 0.079722853 | 0.85551833 | 0.92113008 | T03F1.6 |
| WBGene00020189 | -0.18703882 | 0.58669975 | 0.75578899 | tfbm-1 |
| WBGene00020190 | -0.45877696 | 0.03298769 | 0.12122286 | guk-1 |
| WBGene00020191 | 1.051600705 | 0.04053761 | 0.1387559 | clec-53 |
| WBGene00020192 | -0.82219793 | 0.00165292 | 0.0134685 | T03F1.11 |
| WBGene00020193 | 0.101809913 | 0.76092095 | 0.86650332 | T03F1.12 |
| WBGene00020194 | -0.520053477 | 0.26282835 | 0.47151676 | T03G6.1 |
| WBGene00020195 | -0.013503186 | 0.95805569 | 0.9785154 | T03G6.3 |
| WBGene00020196 | 0.936147209 | 0.29468295 | 0.50504598 | T03G11.3 |
| WBGene00020197 | -0.071223436 | 0.79452271 | 0.88641923 | mans-3 |
| WBGene00020199 | -0.42865065 | 0.11538899 | 0.27985172 | T03G11.6 |
| WBGene00020200 | 1.564996843 | 0.00400037 | 0.02626747 | T04A6.1 |
| WBGene00020201 | -0.283794653 | 0.88299492 | 1 | T04A6.2 |
| WBGene00020202 | -0.156269293 | 0.96924949 | 1 | T04A6.3 |
| WBGene00020203 | -1.901856996 | 0.64072034 | 1 | T04B8.1 |
| WBGene00020205 | -2.212170483 | 7.13E-07 | 2.554E-05 | arrd-1 |
| WBGene00020207 | 0.271945702 | 0.2556106 | 0.46305107 | T04B8.5 |
| WBGene00020208 | 0.137172793 | 0.55174822 | 0.73003216 | T04C4.1 |
| WBGene00020209 | 0.40618636 | 0.05660043 | 0.17566099 | T04C9.1 |
| WBGene00020210 | 0.281454817 | 0.94435481 | 1 | T04C9.2 |
| WBGene00020212 | -0.01683087 | 0.98629074 | 1 | T04D1.2 |
| WBGene00020214 | 0.214705977 | 0.52653711 | 0.71085145 | attf-5 |
| WBGene00020215 | -0.313130394 | 0.39465472 | 0.60247742 | T04G9.4 |
| WBGene00020216 | -0.230346124 | 0.14318049 | 0.32114434 | trap-2 |
| WBGene00020218 | -0.378057782 | 0.04942347 | 0.15947139 | T04G9.7 |

| | | | | |
|---|---|---|---|---|
| WBGene00020219 | -0.642644358 | 0.026054 | 0.10302938 | T05A7.1 |
| WBGene00020220 | -1.68483779 | 0.38847143 | 1 | clec-140 |
| WBGene00020221 | 0.747845955 | 0.85215965 | 1 | T05A7.3 |
| WBGene00020222 | 0.768187993 | 0.14095472 | 0.31874302 | fut-6 |
| WBGene00020223 | -0.458864226 | 0.81398552 | 1 | T05A7.6 |
| WBGene00020224 | 0.335346076 | 0.73845194 | 1 | T05A7.7 |
| WBGene00020225 | 0.747845955 | 0.85215965 | 1 | sre-40 |
| WBGene00020227 | -0.156269293 | 0.96924949 | 1 | T05A8.1 |
| WBGene00020228 | 0.377828795 | 0.92526585 | 1 | T05A8.2 |
| WBGene00020229 | 0.704166288 | 0.09050736 | 0.23924447 | T05A8.3 |
| WBGene00020230 | 1.109520256 | 0.0001877 | 0.00251525 | nep-2 |
| WBGene00020231 | 1.524434892 | 0.23304137 | 1 | T05A8.5 |
| WBGene00020233 | -0.245531902 | 0.94070824 | 1 | T05A8.7 |
| WBGene00020234 | -0.376721596 | 0.05930548 | 0.18104798 | T05A12.3 |
| WBGene00020235 | 0.54698552 | 0.0290462 | 0.11075688 | T05A12.4 |
| WBGene00020236 | 1.851444818 | 0.53669729 | 1 | lgc-1 |
| WBGene00020237 | -0.615803042 | 0.22880079 | 0.4340198 | phat-4 |
| WBGene00020239 | -0.451813889 | 0.8460226 | 1 | T05B4.8 |
| WBGene00020240 | 0.253925665 | 0.89264743 | 1 | T05B4.9 |
| WBGene00020242 | -0.528480847 | 0.13147202 | 0.30392266 | phat-5 |
| WBGene00020245 | -0.273620918 | 0.53072253 | 0.71443643 | T05B11.1 |
| WBGene00020246 | -0.334256927 | 0.0310812 | 0.11646818 | clic-1 |
| WBGene00020247 | -0.733488119 | 0.12005436 | 0.28624902 | T05B11.4 |
| WBGene00020248 | -0.497078235 | 0.56105456 | 0.73696455 | T05C1.1 |
| WBGene00020249 | -0.201268297 | 0.87480857 | 1 | T05C1.2 |
| WBGene00020250 | -1.11044492 | 0.369388 | 1 | T05C1.3 |
| WBGene00020251 | 0.945105083 | 0.00029987 | 0.00360462 | camt-1 |
| WBGene00020252 | 0.938982574 | 0.8146943 | 1 | T05C1.5 |
| WBGene00020254 | -0.289411205 | 0.78252602 | 1 | T05C3.2 |
| WBGene00020256 | 0.671902138 | 0.09216698 | 0.24173672 | T05C3.6 |
| WBGene00020258 | 0.407727919 | 0.32813167 | 0.53668956 | T05E7.1 |
| WBGene00020259 | 0.517978636 | 0.00811273 | 0.04427406 | T05E7.3 |
| WBGene00020260 | 0.638875253 | 0.39896821 | 0.60659956 | T05E7.4 |
| WBGene00020261 | -1.600379365 | 8.85E-07 | 3.0167E-05 | vet-1 |
| WBGene00020263 | 0.065232875 | 0.71518671 | 0.83951289 | let-355 |
| WBGene00020264 | -0.104807389 | 0.6317612 | 0.78652714 | acl-8 |
| WBGene00020265 | 0.377828795 | 0.92526585 | 1 | fbxa-196 |
| WBGene00020266 | 1.730553848 | 0.41535651 | 1 | T05H4.3 |
| WBGene00020267 | 1.180592267 | 0.39171507 | 1 | T05H4.4 |
| WBGene00020268 | -0.172675495 | 0.25667094 | 0.46441757 | hpo-19 |
| WBGene00020269 | -0.509740557 | 0.00019657 | 0.00260567 | erfa-1 |
| WBGene00020270 | -1.220550569 | 0.31520849 | 1 | T05H4.7 |
| WBGene00020273 | -0.187804199 | 0.48035168 | 0.67227086 | T05H4.10 |

| | | | | |
|---|---|---|---|---|
| WBGene00020274 | -0.481428391 | 0.01337341 | 0.06389871 | T05H4.11 |
| WBGene00020275 | -0.469383718 | 0.00113032 | 0.01008319 | atp-4 |
| WBGene00020276 | 1.238315218 | 0.16593258 | 0.35269389 | T05H4.15 |
| WBGene00020277 | -0.242054199 | 0.83312198 | 1 | T06A1.1 |
| WBGene00020279 | 0.257367638 | 0.69562935 | 0.82777182 | glb-25 |
| WBGene00020280 | 0.173777259 | 0.64067501 | 0.79215656 | T06A1.5 |
| WBGene00020281 | -1.021362645 | 0.35080012 | 1 | T06A4.1 |
| WBGene00020282 | 1.712479368 | 0.58529868 | 1 | mps-3 |
| WBGene00020283 | -0.21401395 | 0.74161761 | 0.85569007 | T06A4.3 |
| WBGene00020284 | 0.157491248 | 0.46217357 | 0.65720887 | mel-46 |
| WBGene00020285 | 0.232157325 | 0.93397866 | 1 | T06A10.2 |
| WBGene00020286 | -1.162906218 | 9.8805E-06 | 0.00022506 | T06A10.3 |
| WBGene00020287 | -0.738903586 | 0.00010427 | 0.00157266 | lsy-13 |
| WBGene00020288 | -0.323868354 | 0.91634405 | 1 | srt-73 |
| WBGene00020289 | -1.706624596 | 0.43553802 | 1 | T06C10.3 |
| WBGene00020290 | -0.569798185 | 0.00962424 | 0.05034376 | T06D4.1 |
| WBGene00020292 | -0.156269293 | 0.96924949 | 1 | nep-19 |
| WBGene00020293 | -0.124274926 | 0.95724238 | 1 | nep-20 |
| WBGene00020294 | 0.661516743 | 0.21820555 | 0.42071361 | T06F4.1 |
| WBGene00020294 | 1.732547093 | 0.18847264 | 1 | T06F4.3 |
| WBGene00020295 | -0.264710613 | 0.45707004 | 0.65393512 | pqbp-1.2 |
| WBGene00020296 | -0.464002926 | 0.0380838 | 0.133095 | T07A9.8 |
| WBGene00020297 | 0.388015777 | 0.01949459 | 0.08396019 | nog-1 |
| WBGene00020298 | -0.000188465 | 0.99929372 | 0.99978781 | T07A9.10 |
| WBGene00020299 | 0.595651187 | 0.38445013 | 0.59284039 | T07A9.12 |
| WBGene00020300 | -0.980074747 | 1.1842E-05 | 0.00026093 | tag-261 |
| WBGene00020301 | 0.292033016 | 0.32910959 | 0.53792669 | T07D1.2 |
| WBGene00020302 | 0.730822078 | 0.63799102 | 1 | T07D1.3 |
| WBGene00020305 | -0.737462795 | 0.83433562 | 1 | T07D3.2 |
| WBGene00020307 | 0.114498047 | 0.78016895 | 0.87832735 | T07D3.4 |
| WBGene00020311 | -0.490978388 | 0.10778682 | 0.26759651 | T07D3.9 |
| WBGene00020313 | -1.343645114 | 0.56927291 | 1 | T07E3.2 |
| WBGene00020314 | -0.165410446 | 0.43747843 | 0.63804213 | T07E3.3 |
| WBGene00020315 | 0.219722179 | 0.54144007 | 0.72213819 | T07E3.4 |
| WBGene00020316 | -0.266651294 | 0.2545114 | 0.46216317 | brc-2 |
| WBGene00020317 | 1.182175216 | 3.0972E-05 | 0.0005734 | pdf-1 |
| WBGene00020318 | 0.802122521 | 0.64873485 | 1 | T07F8.1 |
| WBGene00020319 | 0.620259093 | 0.58422115 | 1 | npr-31 |
| WBGene00020320 | -0.727839166 | 0.02846593 | 0.10941134 | T07F8.4 |
| WBGene00020321 | 0.515121849 | 0.36364164 | 0.57166556 | T07F12.1 |
| WBGene00020322 | -0.709006608 | 0.01479691 | 0.06901623 | T07F12.2 |
| WBGene00020323 | 0.011667734 | 0.99506695 | 1 | arrd-23 |
| WBGene00020324 | 0.989510711 | 0.02056864 | 0.08728554 | T07F12.4 |

| | | | | |
|---|---|---|---|---|
| WBGene00020325 | 0.740445135 | 0.49802417 | 1 | bath-46 |
| WBGene00020326 | 5.405505323 | 2.33E-07 | 9.7029E-06 | math-38 |
| WBGene00020328 | -1.964653846 | 0.53599202 | 1 | clec-20 |
| WBGene00020329 | 0.891071315 | 0.16267548 | 0.34857801 | bath-26 |
| WBGene00020330 | 0.755355396 | 0.2508403 | 0.45885383 | T07H6.1 |
| WBGene00020333 | 0.688793783 | 0.86378755 | 1 | sre-12 |
| WBGene00020334 | 0.09536484 | 0.63851286 | 0.79036023 | epg-7 |
| WBGene00020335 | -0.738469597 | 0.10926365 | 0.26962804 | ttr-30 |
| WBGene00020339 | -0.689316971 | 0.09962703 | 0.25437892 | ttr-59 |
| WBGene00020340 | 1.092941601 | 9.85E-08 | 4.6526E-06 | T08B1.1 |
| WBGene00020341 | 0.458611816 | 0.38744865 | 0.59612394 | T08B1.4 |
| WBGene00020343 | -0.430772504 | 0.29930205 | 0.50906705 | acs-3 |
| WBGene00020345 | -0.50276916 | 0.48469823 | 0.67527875 | T08B2.4 |
| WBGene00020346 | 0.379871381 | 0.07704295 | 0.21542571 | rbm-5 |
| WBGene00020347 | -0.041891678 | 0.79060877 | 0.8845213 | ech-1.2 |
| WBGene00020348 | -0.428527121 | 0.0915842 | 0.24104134 | mrpl-23 |
| WBGene00020349 | 0.446221311 | 0.37255558 | 0.58041797 | T08B2.11 |
| WBGene00020350 | 0.377828795 | 0.92526585 | 1 | T08B2.12 |
| WBGene00020351 | -2.474075146 | 0.14672564 | 1 | T08B6.1 |
| WBGene00020353 | 0.992581876 | 0.67966507 | 1 | T08B6.4 |
| WBGene00020354 | -1.960699172 | 0.00206651 | 0.01596023 | T08B6.5 |
| WBGene00020356 | 1.216957619 | 0.74819621 | 1 | T08B6.9 |
| WBGene00020357 | 2.125755145 | 0.12833401 | 1 | T08E11.1 |
| WBGene00020358 | 1.165872487 | 0.06520639 | 0.19276134 | math-39 |
| WBGene00020359 | 1.009340815 | 0.09111631 | 0.24007013 | math-40 |
| WBGene00020360 | 1.130844165 | 1.4526E-05 | 0.00030941 | math-41 |
| WBGene00020361 | 0.069003745 | 0.95640709 | 1 | fbxc-19 |
| WBGene00020362 | -2.741896547 | 0.01025676 | 1 | fbxb-10 |
| WBGene00020363 | 2.134461623 | 0.03618148 | 0.12870768 | fbxa-3 |
| WBGene00020364 | 1.539691283 | 0.36473962 | 1 | T08E11.8 |
| WBGene00020365 | 0.362626465 | 0.61921617 | 0.77855043 | T08G2.2 |
| WBGene00020366 | -0.595050017 | 0.00013595 | 0.00195337 | acdh-10 |
| WBGene00020368 | 0.634119658 | 0.20186582 | 0.40059708 | ast-1 |
| WBGene00020369 | 0.191058099 | 0.30427094 | 0.51329075 | T08H10.1 |
| WBGene00020370 | -0.156269293 | 0.96924949 | 1 | T08H10.3 |
| WBGene00020372 | 1.82867917 | 0.54358766 | 1 | T09A12.1 |
| WBGene00020373 | 1.055732508 | 0.32478802 | 1 | gpx-6 |
| WBGene00020374 | -0.257247033 | 0.32917363 | 0.53795895 | T09A12.5 |
| WBGene00020375 | -0.550382699 | 0.00041665 | 0.00464123 | pigv-1 |
| WBGene00020376 | -0.180934522 | 0.44375358 | 0.64322018 | T09B4.2 |
| WBGene00020377 | 0.403391944 | 0.78272234 | 1 | T09B4.3 |
| WBGene00020378 | -0.251129261 | 0.69083384 | 0.82383005 | T09B4.4 |
| WBGene00020379 | -0.558162402 | 0.02227881 | 0.09279145 | T09B4.5 |

| | | | | |
|---|---|---|---|---|
| WBGene00020380 | 0.88787957 | 0.71832755 | 1 | T09B4.6 |
| WBGene00020381 | 0.742628056 | 0.76901841 | 1 | T09B4.7 |
| WBGene00020382 | 0.001799723 | 0.99279635 | 0.99644982 | T09B4.8 |
| WBGene00020383 | -0.195216274 | 0.38575207 | 0.59414935 | tin-44 |
| WBGene00020384 | -1.453036238 | 0.47675825 | 1 | T09D3.3 |
| WBGene00020385 | 1.159354287 | 0.20396889 | 0.40340316 | nhr-216 |
| WBGene00020386 | 1.006501211 | 0.05858113 | 0.17960443 | cyp-34A4 |
| WBGene00020389 | -0.417791344 | 0.03522126 | 0.12655263 | T10B5.3 |
| WBGene00020390 | -0.333949639 | 0.46264051 | 0.65771878 | T10B5.4 |
| WBGene00020391 | -0.310019282 | 0.06832198 | 0.19915897 | cct-7 |
| WBGene00020392 | -0.772185044 | 8.343E-05 | 0.00132408 | knl-3 |
| WBGene00020393 | 0.370643925 | 0.13531025 | 0.30984567 | T10B5.7 |
| WBGene00020394 | 2.641600694 | 0.04902701 | 1 | T10B5.8 |
| WBGene00020396 | 0.629370777 | 0.53765641 | 0.71960846 | T10B5.10 |
| WBGene00020397 | -0.328329541 | 0.30902754 | 0.51793255 | pcbd-1 |
| WBGene00020398 | -0.046074523 | 0.83747032 | 0.91143626 | cerk-1 |
| WBGene00020399 | -0.51443294 | 0.00845997 | 0.04566373 | ztf-4 |
| WBGene00020400 | -0.156269293 | 0.96924949 | 1 | T10B11.4 |
| WBGene00020401 | 0.782179376 | 0.42083916 | 0.62546313 | T10B11.5 |
| WBGene00020402 | 0.47399401 | 0.11237325 | 0.27467759 | T10B11.6 |
| WBGene00020403 | -0.068047187 | 0.73871178 | 0.8543246 | T10B11.7 |
| WBGene00020404 | -0.188441159 | 0.53216444 | 0.71566332 | T10B11.8 |
| WBGene00020407 | -0.234765883 | 0.72295976 | 0.8438309 | chil-24 |
| WBGene00020408 | -0.121595387 | 0.63697904 | 0.78967829 | T10D4.6 |
| WBGene00020409 | -0.156269293 | 0.96924949 | 1 | T10D4.7 |
| WBGene00020411 | 0.220306493 | 0.46189103 | 0.65707096 | T10E9.1 |
| WBGene00020412 | -0.022583434 | 0.91364554 | 0.95526787 | T10E9.2 |
| WBGene00020413 | 0.154369545 | 0.79923401 | 0.88979641 | T10E9.3 |
| WBGene00020414 | 3.453083173 | 0.02082111 | 1 | T10E9.4 |
| WBGene00020415 | 1.025181888 | 0.79790347 | 1 | T10E9.5 |
| WBGene00020417 | -0.374577441 | 0.03938217 | 0.13598945 | nuo-2 |
| WBGene00020418 | -1.59763493 | 0.14611052 | 1 | T10E9.8 |
| WBGene00020419 | -0.732385075 | 0.30316736 | 0.51221193 | acdh-4 |
| WBGene00020420 | -0.318038444 | 0.81325997 | 1 | T10E10.3 |
| WBGene00020421 | 0.671700449 | 0.42725209 | 0.63051525 | T10E10.4 |
| WBGene00020422 | -0.567401815 | 0.01345142 | 0.06414521 | T10F2.2 |
| WBGene00020423 | -0.439706438 | 0.00548036 | 0.03336594 | prp-19 |
| WBGene00020424 | 1.303804406 | 0.21049869 | 1 | T10H9.1 |
| WBGene00020425 | -0.346199954 | 0.11514934 | 0.2793263 | syx-18 |
| WBGene00020426 | 0.482296922 | 0.27421912 | 0.48271631 | T10H10.2 |
| WBGene00020427 | 0.495384856 | 0.17331439 | 0.3630452 | sav-1 |
| WBGene00020430 | 0.688793783 | 0.86378755 | 1 | irld-49 |
| WBGene00020431 | 0.377828795 | 0.92526585 | 1 | irld-50 |

| | | | | |
|---|---|---|---|---|
| WBGene00020432 | -1.901856996 | 0.64072034 | 1 | hpa-2 |
| WBGene00020433 | -0.528968285 | 0.08873811 | 0.23621342 | T11F8.1 |
| WBGene00020435 | 1.060949945 | 0.61866902 | 1 | T11F8.4 |
| WBGene00020436 | -0.427882054 | 0.45479088 | 0.6523296 | T12A2.1 |
| WBGene00020437 | -0.096774939 | 0.54963746 | 0.72811221 | stt-3 |
| WBGene00020438 | -1.305065462 | 0.06397984 | 0.19052715 | T12A2.3 |
| WBGene00020439 | -0.109254073 | 0.95448982 | 1 | T12A2.5 |
| WBGene00020441 | -0.6249659 | 0.00286413 | 0.02053952 | bcas-2 |
| WBGene00020442 | 0.057069112 | 0.7715086 | 0.87231197 | gen-1 |
| WBGene00020443 | 0.093914236 | 0.70400389 | 0.83266549 | esyt-2 |
| WBGene00020444 | 1.357633603 | 0.28799191 | 1 | T12B3.1 |
| WBGene00020445 | 0.66200699 | 0.10984803 | 0.27046542 | T12B3.2 |
| WBGene00020446 | -0.065853262 | 0.83922675 | 0.91209847 | T12B3.3 |
| WBGene00020447 | 0.10819715 | 0.70823075 | 0.83500998 | mob-3 |
| WBGene00020448 | 0.197556253 | 0.74453413 | 0.8574972 | fbxa-51 |
| WBGene00020449 | 0.16869376 | 0.7756819 | 0.87481747 | fbxa-54 |
| WBGene00020450 | 1.215990383 | 0.18260533 | 0.37462171 | fbxa-10 |
| WBGene00020451 | 1.121948915 | 0.22344071 | 0.42746592 | fbxa-11 |
| WBGene00020452 | -0.156269293 | 0.96924949 | 1 | fbxa-22 |
| WBGene00020453 | 1.228323117 | 0.05603357 | 0.17421347 | fbxa-55 |
| WBGene00020454 | 0.747845955 | 0.85215965 | 1 | fbxa-56 |
| WBGene00020455 | 2.391028511 | 0.00012407 | 0.00180838 | fbxa-59 |
| WBGene00020456 | 1.009980645 | 0.0026793 | 0.01955102 | fbxa-60 |
| WBGene00020457 | 1.390919651 | 0.00569652 | 0.03434312 | fbxa-67 |
| WBGene00020459 | -0.580736074 | 0.69462335 | 1 | fbxa-23 |
| WBGene00020460 | -0.109333946 | 0.92142846 | 1 | nhr-273 |
| WBGene00020461 | 0.042035842 | 0.88668702 | 0.94019801 | T12C9.7 |
| WBGene00020462 | -0.131383706 | 0.51675541 | 0.70281311 | T12E12.1 |
| WBGene00020463 | -0.221810596 | 0.33976133 | 0.54803272 | cec-6 |
| WBGene00020464 | -0.067850029 | 0.79730677 | 0.88849027 | T12E12.3 |
| WBGene00020465 | 0.875012455 | 0.69400044 | 1 | T12E12.6 |
| WBGene00020466 | -0.815662977 | 0.00037066 | 0.00426957 | sld-2 |
| WBGene00020467 | -0.264675803 | 0.36166091 | 0.56954203 | T12F5.2 |
| WBGene00020468 | 1.741147134 | 0.37149958 | 1 | T13A10.1 |
| WBGene00020469 | 1.526476613 | 0.440991 | 1 | T13A10.2 |
| WBGene00020471 | 0.100554048 | 0.6918567 | 0.82456388 | pho-14 |
| WBGene00020472 | -0.62146648 | 0.85925752 | 1 | lips-11 |
| WBGene00020473 | -4.054889939 | 0.04255608 | 1 | lips-12 |
| WBGene00020474 | -0.233502874 | 0.92367326 | 1 | lips-13 |
| WBGene00020475 | -0.509717695 | 0.09545924 | 0.24772303 | sut-1 |
| WBGene00020476 | -0.504221413 | 0.5572552 | 0.73435747 | T13B5.9 |
| WBGene00020478 | -0.06858948 | 0.94129174 | 1 | T13C2.2 |
| WBGene00020479 | 0.008372764 | 0.9857037 | 0.99323003 | T13C2.3 |

| | | | | |
|---|---|---|---|---|
| WBGene00020480 | -0.602285952 | 0.01812269 | 0.07977062 | ssup-72 |
| WBGene00020481 | 0.15400544 | 0.37699248 | 0.58485379 | T13C2.6 |
| WBGene00020482 | 0.782092876 | 0.42841723 | 0.63100978 | T13C2.7 |
| WBGene00020483 | 0.938982574 | 0.8146943 | 1 | T13C5.2 |
| WBGene00020484 | 0.465758074 | 0.87314503 | 1 | T13C5.3 |
| WBGene00020485 | 0.175909247 | 0.94229417 | 1 | ceh-54 |
| WBGene00020486 | 0.220944719 | 0.59270941 | 0.75972181 | T13C5.6 |
| WBGene00020487 | -2.41219879 | 0.17646937 | 1 | T13C5.7 |
| WBGene00020488 | -0.156269293 | 0.96924949 | 1 | T13G4.1 |
| WBGene00020489 | 0.892165501 | 0.24226343 | 0.44981194 | hot-8 |
| WBGene00020490 | 0.819806022 | 0.07159718 | 0.20557276 | tmc-1 |
| WBGene00020491 | -0.021715728 | 0.96574642 | 0.98286673 | T13G4.4 |
| WBGene00020492 | -1.159714244 | 0.75790634 | 1 | T13G4.5 |
| WBGene00020494 | -0.626833103 | 0.80597175 | 1 | T13G4.7 |
| WBGene00020496 | 0.614657179 | 0.01885122 | 0.08215101 | spat-3 |
| WBGene00020497 | -0.179896626 | 0.60102838 | 0.76625912 | T14A8.2 |
| WBGene00020498 | 0.455123402 | 0.0452865 | 0.14989733 | T14B4.1 |
| WBGene00020499 | -0.418986948 | 0.15075584 | 0.33243407 | T14B4.2 |
| WBGene00020500 | -0.229582239 | 0.30403377 | 0.51303322 | T14B4.3 |
| WBGene00020501 | -0.666401371 | 0.33031262 | 0.53866024 | T14B4.5 |
| WBGene00020502 | 0.044323433 | 0.93018405 | 0.96354103 | T14B4.8 |
| WBGene00020503 | 0.914827028 | 0.45949066 | 1 | T14B4.9 |
| WBGene00020504 | 0.741623712 | 0.00152156 | 0.01269644 | svh-2 |
| WBGene00020505 | -0.736789399 | 0.72029863 | 1 | T14E8.2 |
| WBGene00020506 | 1.575572041 | 0.06262012 | 0.18765322 | dop-3 |
| WBGene00020507 | 0.279073927 | 0.08528838 | 0.23026157 | vha-15 |
| WBGene00020508 | -0.62345202 | 0.05857849 | 0.17960443 | T14F9.2 |
| WBGene00020509 | -0.430011877 | 0.03913577 | 0.13540821 | hex-1 |
| WBGene00020510 | 0.892645821 | 0.07564057 | 0.21312419 | T14G11.1 |
| WBGene00020511 | 0.031981177 | 0.87794915 | 0.93444207 | immt-1 |
| WBGene00020516 | -0.954537584 | 2.47E-07 | 1.0177E-05 | T15B7.1 |
| WBGene00020517 | 0.106009011 | 0.47350486 | 0.66645288 | hpo-8 |
| WBGene00020520 | -1.159714244 | 0.75790634 | 1 | T15B7.8 |
| WBGene00020522 | 0.361712769 | 0.80402005 | 1 | T15B7.10 |
| WBGene00020522 | -0.598082586 | 0.65470836 | 1 | T15B7.17 |
| WBGene00020523 | -0.156269293 | 0.96924949 | 1 | dmsr-14 |
| WBGene00020524 | 0.281454817 | 0.94435481 | 1 | dmsr-13 |
| WBGene00020526 | 0.688793783 | 0.86378755 | 1 | T15B7.14 |
| WBGene00020527 | 0.019774513 | 0.98947922 | 1 | T15B7.15 |
| WBGene00020528 | -0.851544247 | 0.00733936 | 0.04113904 | lgc-54 |
| WBGene00020530 | 0.789497005 | 0.02412241 | 0.09775373 | T15B12.1 |
| WBGene00020531 | 0.03940405 | 0.99001443 | 1 | T15B12.2 |
| WBGene00020532 | 2.214252 | 0.07956276 | 1 | math-42 |

| | | | | |
|---|---|---|---|---|
| WBGene00020533 | 0.377828795 | 0.92526585 | 1 | T16A1.2 |
| WBGene00020535 | -0.156269293 | 0.96924949 | 1 | T16A1.4 |
| WBGene00020536 | 0.281454817 | 0.94435481 | 1 | T16A1.5 |
| WBGene00020537 | -1.450787457 | 0.06510735 | 0.19260728 | fbxb-37 |
| WBGene00020540 | 0.017469719 | 0.97357774 | 0.98625009 | pals-25 |
| WBGene00020541 | -1.606582071 | 0.25754264 | 1 | fbxb-81 |
| WBGene00020542 | -2.664869621 | 0.01486396 | 1 | fbxb-82 |
| WBGene00020543 | -1.768995394 | 0.16511421 | 1 | fbxb-83 |
| WBGene00020544 | -1.294679246 | 0.04376628 | 0.14608272 | fbxb-84 |
| WBGene00020546 | -0.425827543 | 0.79918767 | 1 | T17A3.10 |
| WBGene00020548 | -2.640414922 | 0.41081194 | 1 | T17A3.12 |
| WBGene00020549 | -0.499356776 | 0.01538897 | 0.07095994 | nmt-1 |
| WBGene00020550 | -0.608792446 | 0.45108587 | 0.64911342 | T17H7.1 |
| WBGene00020551 | -0.228168879 | 0.68885928 | 0.82293029 | T17H7.7 |
| WBGene00020553 | 0.323311755 | 0.10001304 | 0.25479266 | T19A5.1 |
| WBGene00020554 | -0.038554152 | 0.92969407 | 0.96350113 | T19A5.3 |
| WBGene00020555 | 0.57991002 | 0.82402153 | 1 | nhr-219 |
| WBGene00020556 | -0.03973018 | 0.89051389 | 0.94219862 | pamn-1 |
| WBGene00020557 | -0.591360001 | 0.00412224 | 0.02691224 | T19B4.3 |
| WBGene00020558 | -0.180716572 | 0.34309089 | 0.5515713 | T19B4.5 |
| WBGene00020559 | 0.344457547 | 0.45034288 | 0.64889781 | best-17 |
| WBGene00020560 | -1.031423055 | 0.23505386 | 0.44101205 | T19C3.2 |
| WBGene00020561 | 0.336390523 | 0.53316399 | 0.71645202 | T19C3.3 |
| WBGene00020562 | 0.463322676 | 0.07765818 | 0.21629916 | T19C3.4 |
| WBGene00020563 | 0.085157863 | 0.86583909 | 0.92781574 | T19C3.5 |
| WBGene00020564 | 0.329913989 | 0.82744097 | 1 | T19C3.6 |
| WBGene00020565 | -0.559340474 | 0.78183513 | 1 | T19C3.7 |
| WBGene00020566 | -0.348743811 | 0.58677761 | 0.75578899 | ttr-7 |
| WBGene00020567 | 1.188981222 | 0.14107113 | 0.31894684 | T19D2.1 |
| WBGene00020568 | -0.033862551 | 0.96669264 | 0.9830824 | T19D2.3 |
| WBGene00020569 | 0.378228659 | 0.48645147 | 0.67622374 | lgc-32 |
| WBGene00020571 | -1.209974009 | 0.19469294 | 0.39064073 | lpr-7 |
| WBGene00020572 | 0.155939762 | 0.77317277 | 0.87338022 | rund-1 |
| WBGene00020573 | 0.390828567 | 0.6851454 | 1 | T19D7.5 |
| WBGene00020574 | 0.049142346 | 0.95474748 | 0.97697363 | T19D7.6 |
| WBGene00020575 | 0.199472887 | 0.85299643 | 1 | T19D7.7 |
| WBGene00020576 | 1.892423027 | 2.47E-12 | 3.23E-10 | T19D12.1 |
| WBGene00020577 | 1.339078487 | 9.9314E-05 | 0.0015263 | T19D12.2 |
| WBGene00020578 | 1.879256964 | 0.46994626 | 1 | T19D12.3 |
| WBGene00020579 | 1.22225506 | 1.85E-10 | 1.69E-08 | T19D12.4 |
| WBGene00020580 | 3.886839713 | 0.00093794 | 0.00877345 | T19D12.5 |
| WBGene00020581 | 0.547512968 | 0.00497456 | 0.03105202 | T19D12.6 |
| WBGene00020582 | 0.925523462 | 0.26567439 | 0.47434599 | oig-8 |

| | | | | |
|---|---|---|---|---|
| WBGene00020583 | 0.986392409 | 0.23205734 | 0.43764402 | T19D12.9 |
| WBGene00020584 | 0.361788163 | 0.84421891 | 1 | T19D12.10 |
| WBGene00020585 | -0.372949705 | 0.55636201 | 0.73349887 | clec-178 |
| WBGene00020586 | 1.696755273 | 0.04389664 | 0.14637439 | frpr-18 |
| WBGene00020587 | -0.38114289 | 0.50271834 | 0.68999878 | ugt-9 |
| WBGene00020588 | -0.942934171 | 4.44E-09 | 2.93E-07 | T19H12.2 |
| WBGene00020589 | 0.026039489 | 0.94723373 | 0.97304242 | T19H12.3 |
| WBGene00020590 | 0.396622425 | 0.56667785 | 0.74172492 | T19H12.6 |
| WBGene00020592 | -0.359308152 | 0.07951076 | 0.22014597 | ugt-12 |
| WBGene00020593 | -0.140409688 | 0.7797391 | 0.87793865 | ugt-11 |
| WBGene00020594 | 0.290491123 | 0.88464841 | 1 | ugt-10 |
| WBGene00020595 | 1.187897324 | 0.04352094 | 0.14560147 | T20B5.2 |
| WBGene00020596 | -0.02089792 | 0.94150181 | 0.96989883 | oga-1 |
| WBGene00020598 | 0.281454817 | 0.94435481 | 1 | T20B6.2 |
| WBGene00020599 | -0.281876431 | 0.79818374 | 1 | T20B6.3 |
| WBGene00020600 | -0.005382635 | 0.97952749 | 0.98905882 | trd-1 |
| WBGene00020601 | -0.311188546 | 0.18483579 | 0.37727857 | T20B12.3 |
| WBGene00020602 | 1.477253415 | 0.58357327 | 1 | T20B12.4 |
| WBGene00020604 | -0.389055122 | 0.01751236 | 0.07796756 | T20B12.7 |
| WBGene00020605 | 1.293232192 | 0.00830117 | 0.04498505 | lgc-50 |
| WBGene00020607 | -1.159714244 | 0.75790634 | 1 | srab-20 |
| WBGene00020608 | -0.156269293 | 0.96924949 | 1 | srab-22 |
| WBGene00020611 | 1.181686151 | 0.7579682 | 1 | T20D4.5 |
| WBGene00020612 | 0.120394942 | 0.85070886 | 0.91824825 | arrd-22 |
| WBGene00020613 | -1.429858991 | 0.0596153 | 0.18167402 | T20D4.7 |
| WBGene00020616 | 0.392639293 | 0.76401876 | 1 | T20D4.10 |
| WBGene00020617 | 1.051863047 | 0.19648229 | 0.39356518 | T20D4.11 |
| WBGene00020618 | -0.156269293 | 0.96924949 | 1 | T20D4.12 |
| WBGene00020620 | 1.036256418 | 0.52098893 | 1 | T20D4.15 |
| WBGene00020624 | 0.015644866 | 0.99425358 | 1 | T20D4.19 |
| WBGene00020625 | -0.001404396 | 0.99701635 | 0.99857337 | mrrf-1 |
| WBGene00020626 | -0.253768745 | 0.53581424 | 0.7178681 | T20F5.4 |
| WBGene00020627 | 0.578506681 | 0.58074536 | 1 | T20F5.5 |
| WBGene00020628 | -0.052748413 | 0.78349358 | 0.88069582 | T20F5.6 |
| WBGene00020629 | -0.608964615 | 0.00669851 | 0.03850691 | T20F5.7 |
| WBGene00020630 | 0.693071962 | 0.00873134 | 0.0467404 | T20F7.1 |
| WBGene00020631 | -0.933827966 | 0.56134399 | 1 | T20F7.3 |
| WBGene00020633 | 0.143258708 | 0.63708911 | 0.78967829 | aakg-2 |
| WBGene00020634 | 0.077470345 | 0.92293757 | 0.9593272 | acs-9 |
| WBGene00020635 | -0.254683336 | 0.89672209 | 1 | T20H4.2 |
| WBGene00020636 | -0.578021507 | 0.00091248 | 0.0085813 | T20H4.5 |
| WBGene00020637 | 0.377828795 | 0.92526585 | 1 | fbxa-7 |
| WBGene00020638 | 1.111933685 | 0.57471521 | 1 | fbxa-43 |

| | | | | |
|---|---|---|---|---|
| WBGene00020639 | 0.863916284 | 0.50887108 | 1 | fbxa-71 |
| WBGene00020640 | 1.060949945 | 0.61866902 | 1 | fbxa-73 |
| WBGene00020645 | 1.477253415 | 0.58357327 | 1 | T21D9.2 |
| WBGene00020646 | -2.383751247 | 0.26793499 | 1 | zmp-6 |
| WBGene00020647 | -0.966662698 | 2.4318E-06 | 6.9145E-05 | pqbp-1.1 |
| WBGene00020648 | -0.17926287 | 0.89638868 | 1 | T21D12.7 |
| WBGene00020649 | 1.409264288 | 1.7218E-06 | 5.111E-05 | sma-10 |
| WBGene00020650 | 1.557897343 | 0.63813005 | 1 | T21D12.11 |
| WBGene00020651 | 0.650333437 | 0.08215441 | 0.22464791 | T21D12.12 |
| WBGene00020652 | -0.568940579 | 0.00744865 | 0.04157888 | egg-4 |
| WBGene00020653 | -1.901856996 | 0.64072034 | 1 | T21E3.2 |
| WBGene00020655 | -0.263229581 | 0.93409872 | 1 | T21E12.3 |
| WBGene00020656 | -1.969805064 | 0.62862569 | 1 | T21E12.5 |
| WBGene00020657 | 0.921168994 | 0.24697756 | 0.45497554 | lgc-53 |
| WBGene00020658 | -0.440094339 | 0.05963817 | 0.18169813 | T21F4.1 |
| WBGene00020659 | 0.459413223 | 0.84552382 | 1 | T21G5.1 |
| WBGene00020660 | 2.040392054 | 0.23143517 | 1 | T21G5.2 |
| WBGene00020661 | -0.960774898 | 0.53532779 | 1 | smz-2 |
| WBGene00020662 | 0.752836141 | 6.17E-07 | 2.2776E-05 | T21H3.1 |
| WBGene00020663 | -1.381814384 | 0.34220937 | 1 | cutl-22 |
| WBGene00020664 | -0.558804446 | 0.87818675 | 1 | T21H3.5 |
| WBGene00020665 | -1.094438693 | 0.01650264 | 0.07432888 | T22B2.1 |
| WBGene00020666 | -1.094201388 | 0.7816772 | 1 | T22B2.2 |
| WBGene00020667 | 0.281454817 | 0.94435481 | 1 | T22B2.3 |
| WBGene00020669 | -0.156269293 | 0.96924949 | 1 | T22B2.5 |
| WBGene00020672 | 0.387479365 | 0.57129372 | 0.74561373 | T22B7.3 |
| WBGene00020673 | -0.457235549 | 0.29163371 | 0.5015777 | T22B7.4 |
| WBGene00020674 | -0.497735083 | 0.21134181 | 0.4111419 | T22B7.7 |
| WBGene00020675 | -0.639727652 | 0.77477802 | 1 | T22B11.1 |
| WBGene00020678 | 1.031081281 | 1.3602E-06 | 4.2894E-05 | T22B11.4 |
| WBGene00020679 | 0.329057744 | 0.06745592 | 0.19725007 | ogdh-1 |
| WBGene00020682 | -0.122688098 | 0.46203774 | 0.65716966 | T22D1.3 |
| WBGene00020683 | -0.175392785 | 0.18045149 | 0.3725955 | ribo-1 |
| WBGene00020684 | -0.659204847 | 2.528E-05 | 0.00049188 | T22D1.5 |
| WBGene00020686 | -1.544131969 | 0.51019995 | 1 | T22D1.8 |
| WBGene00020687 | -0.181029988 | 0.28116601 | 0.49046502 | ruvb-2 |
| WBGene00020688 | -0.342989276 | 0.65530779 | 0.80188241 | T22D1.11 |
| WBGene00020689 | 2.207292865 | 0.0017579 | 0.01406898 | npr-12 |
| WBGene00020690 | 0.187648194 | 0.85599055 | 1 | T22E5.1 |
| WBGene00020691 | 0.75945276 | 0.50255169 | 1 | igdb-3 |
| WBGene00020692 | 0.149106811 | 0.83773838 | 0.91143626 | T22E5.6 |
| WBGene00020693 | 0.398387211 | 0.32867959 | 0.53736855 | lron-8 |
| WBGene00020694 | 0.707559163 | 0.78139902 | 1 | dhhc-11 |

| | | | | |
|---|---|---|---|---|
| WBGene00020695 | -0.438406299 | 0.02425787 | 0.09817159 | T22F3.2 |
| WBGene00020696 | 0.038017847 | 0.82784017 | 0.90703073 | T22F3.3 |
| WBGene00020697 | 1.479499165 | 0.20449332 | 1 | T22F3.7 |
| WBGene00020698 | 2.353411952 | 0.0969839 | 1 | T22F3.8 |
| WBGene00020699 | 1.682356075 | 0.21408153 | 1 | T22F3.10 |
| WBGene00020700 | 6.335534601 | 2.41E-07 | 9.9863E-06 | T22F3.11 |
| WBGene00020701 | 0.770574237 | 0.02441882 | 0.09854595 | T22F7.1 |
| WBGene00020702 | 1.207960181 | 0.345231 | 1 | T22F7.3 |
| WBGene00020702 | 0.716858289 | 0.60147337 | 1 | T22F7.5 |
| WBGene00020703 | 0.325251153 | 0.40793005 | 0.6141434 | T22F7.4 |
| WBGene00020705 | -0.210842499 | 0.39716014 | 0.60500894 | T22H9.1 |
| WBGene00020706 | -0.013661892 | 0.95221869 | 0.97579519 | atg-9 |
| WBGene00020707 | 0.339336785 | 0.86985627 | 1 | wago-10 |
| WBGene00020708 | 0.043765775 | 0.96778194 | 1 | dmd-8 |
| WBGene00020709 | -0.107455579 | 0.55592714 | 0.73328206 | T23B3.1 |
| WBGene00020710 | 0.164960392 | 0.60434818 | 0.76867617 | T23B3.2 |
| WBGene00020712 | 1.033249282 | 0.21367946 | 0.4143559 | ckr-1 |
| WBGene00020713 | 0.354141796 | 0.8323238 | 1 | T23B3.5 |
| WBGene00020714 | 1.458839816 | 0.0623818 | 0.18719166 | T23B3.6 |
| WBGene00020715 | 1.471809563 | 0.46592277 | 1 | nspd-4 |
| WBGene00020716 | -0.313231519 | 0.22435085 | 0.42847941 | phf-30 |
| WBGene00020717 | -0.484724888 | 0.03890595 | 0.13484361 | mrpl-4 |
| WBGene00020718 | -0.636179416 | 0.00605323 | 0.03584991 | mrps-2 |
| WBGene00020719 | 0.022742944 | 0.89015209 | 0.94196731 | natc-1 |
| WBGene00020720 | -1.390520744 | 0.47839671 | 1 | T23B12.5 |
| WBGene00020721 | 0.303507541 | 0.11618153 | 0.28054097 | T23B12.6 |
| WBGene00020722 | 0.260528239 | 0.63033991 | 0.78556152 | T23B12.8 |
| WBGene00020725 | -0.249518149 | 0.40291265 | 0.61078317 | cnp-3 |
| WBGene00020726 | -0.77080688 | 0.03817635 | 0.13322767 | T23C6.4 |
| WBGene00020727 | -0.46399732 | 0.57894445 | 0.75037463 | npr-21 |
| WBGene00020728 | 1.694048007 | 0.42879474 | 1 | T23E1.1 |
| WBGene00020730 | -0.62146648 | 0.85925752 | 1 | T23E1.3 |
| WBGene00020732 | 1.273608962 | 2.11E-08 | 1.1777E-06 | T23E7.2 |
| WBGene00020735 | 1.210361255 | 0.0073196 | 0.04108519 | T23F2.2 |
| WBGene00020736 | 0.507382829 | 0.16026707 | 0.34561687 | T23F2.3 |
| WBGene00020737 | 0.339336785 | 0.86985627 | 1 | T23F2.4 |
| WBGene00020738 | -0.714735444 | 0.00052002 | 0.00552854 | T23F2.5 |
| WBGene00020740 | 1.099504233 | 0.01648843 | 0.07432888 | T23F4.2 |
| WBGene00020741 | 3.884216885 | 0.00627862 | 1 | dlhd-1 |
| WBGene00020742 | -0.038980952 | 0.83477008 | 0.91032502 | T23H2.3 |
| WBGene00020743 | 0.11607066 | 0.91693737 | 1 | mam-8 |
| WBGene00020744 | -0.110952496 | 0.8879555 | 0.94084004 | T24A6.1 |
| WBGene00020747 | 1.69047418 | 0.38809581 | 1 | T24A6.7 |

| | | | | |
|---|---|---|---|---|
| WBGene00020748 | 2.62561062 | 0.22121999 | 1 | nhr-221 |
| WBGene00020750 | -0.737462795 | 0.83433562 | 1 | nhr-222 |
| WBGene00020754 | 0.687259521 | 0.81162667 | 1 | T24A6.17 |
| WBGene00020755 | -0.156269293 | 0.96924949 | 1 | grl-31 |
| WBGene00020756 | 0.229916924 | 0.54385368 | 0.72337864 | T24A6.20 |
| WBGene00020757 | -0.7358647 | 7.2553E-06 | 0.00017331 | ucr-2.3 |
| WBGene00020758 | -1.259472496 | 0.00049446 | 0.0053219 | T24C4.2 |
| WBGene00020759 | -0.457870121 | 0.79919046 | 1 | T24C4.3 |
| WBGene00020760 | 0.281936298 | 0.49101426 | 0.6802214 | T24C4.4 |
| WBGene00020761 | -0.265893301 | 0.31016242 | 0.518332 | T24C4.5 |
| WBGene00020762 | 0.276799404 | 0.14543675 | 0.32422834 | zer-1 |
| WBGene00020763 | 0.056489755 | 0.77441133 | 0.87427271 | ztf-18 |
| WBGene00020764 | 1.423429265 | 0.31214627 | 1 | T24C12.1 |
| WBGene00020765 | -0.143356488 | 0.64983778 | 0.79839241 | T24C12.3 |
| WBGene00020766 | -1.75995696 | 0.25032373 | 1 | T24C12.4 |
| WBGene00020767 | 0.424007158 | 0.47074033 | 0.66356678 | lgc-40 |
| WBGene00020770 | 0.065401441 | 0.80933634 | 0.89609686 | eat-17 |
| WBGene00020771 | -1.03227574 | 0.0083269 | 0.04503204 | T24E12.1 |
| WBGene00020772 | -0.737462795 | 0.83433562 | 1 | T24E12.2 |
| WBGene00020773 | -0.668264701 | 0.85551153 | 1 | T24E12.3 |
| WBGene00020774 | 1.55060706 | 0.04800041 | 0.1566173 | T24E12.5 |
| WBGene00020775 | -0.014519762 | 0.99603746 | 1 | T24E12.6 |
| WBGene00020777 | 0.474216619 | 0.49197944 | 0.68116346 | T24E12.10 |
| WBGene00020778 | -0.631009586 | 0.24846187 | 0.45656791 | T24E12.11 |
| WBGene00020779 | -0.493481204 | 0.00812926 | 0.04433841 | T24G10.2 |
| WBGene00020781 | -0.112446881 | 0.44957265 | 0.64840361 | T24H7.2 |
| WBGene00020782 | -0.576715084 | 0.01314116 | 0.06331221 | T24H7.3 |
| WBGene00020783 | -0.76633068 | 0.01570742 | 0.07187884 | blos-4 |
| WBGene00020784 | -0.223939686 | 0.1704453 | 0.35928262 | tat-4 |
| WBGene00020786 | 1.031823916 | 0.02635923 | 0.10373661 | T25B2.2 |
| WBGene00020787 | 0.574508983 | 0.8551971 | 1 | T25B6.1 |
| WBGene00020788 | 0.994588383 | 1.6802E-06 | 5.0121E-05 | nep-22 |
| WBGene00020789 | -0.156269293 | 0.96924949 | 1 | T25B6.3 |
| WBGene00020790 | 2.120404512 | 0.14768459 | 1 | T25B6.4 |
| WBGene00020791 | 0.747845955 | 0.85215965 | 1 | T25B6.5 |
| WBGene00020792 | 0.539239107 | 0.83916685 | 1 | T25B6.6 |
| WBGene00020793 | 0.641065534 | 0.78192938 | 1 | T25D1.1 |
| WBGene00020794 | -0.558804446 | 0.87818675 | 1 | T25D1.2 |
| WBGene00020795 | -1.094201388 | 0.7816772 | 1 | T25D1.3 |
| WBGene00020796 | -0.306549544 | 0.19213479 | 0.38723983 | mrpl-28 |
| WBGene00020797 | -0.243609085 | 0.64590572 | 0.79540941 | T25D3.3 |
| WBGene00020798 | -0.157316697 | 0.43221406 | 0.6341955 | T25D3.4 |
| WBGene00020799 | 0.664655823 | 0.30476408 | 0.51390842 | T25D10.1 |

| | | | | |
|---|---|---|---|---|
| WBGene00020800 | 0.142303531 | 0.82300756 | 0.90431616 | zig-10 |
| WBGene00020801 | 0.419651445 | 0.41630374 | 0.62134953 | T25D10.4 |
| WBGene00020802 | 0.688793783 | 0.86378755 | 1 | btb-2 |
| WBGene00020803 | -1.358004042 | 0.02639598 | 0.10381393 | T25E4.1 |
| WBGene00020804 | 0.983488853 | 0.80601565 | 1 | T25E4.2 |
| WBGene00020805 | 0.027604004 | 0.96892136 | 0.98399257 | T25F10.1 |
| WBGene00020806 | 0.393581948 | 0.30796763 | 0.51658401 | T25F10.3 |
| WBGene00020807 | 3.255102787 | 0.07761964 | 1 | T25F10.4 |
| WBGene00020808 | -0.050953558 | 0.74458735 | 0.8574972 | clik-1 |
| WBGene00020809 | 0.711038907 | 0.69068738 | 1 | T25G12.1 |
| WBGene00020809 | 0.688793783 | 0.86378755 | 1 | T25G12.2 |
| WBGene00020811 | 0.121442094 | 0.83279689 | 0.90958862 | T25G12.3 |
| WBGene00020812 | -0.550533694 | 0.00330021 | 0.02275287 | acdh-7 |
| WBGene00020813 | 1.579840266 | 0.00527294 | 0.03243096 | T25G12.6 |
| WBGene00020817 | 1.037645283 | 0.55087863 | 1 | T25G12.11 |
| WBGene00020819 | 0.010075268 | 0.96746953 | 0.9835073 | T26A5.2 |
| WBGene00020820 | -0.082883678 | 0.67899203 | 0.81647553 | algn-1 |
| WBGene00020821 | -0.2616671 | 0.10093191 | 0.25601242 | jhdm-1 |
| WBGene00020822 | -0.101058159 | 0.69041579 | 0.82357419 | T26A5.6 |
| WBGene00020823 | -0.500797455 | 0.04269568 | 0.14385895 | T26A5.8 |
| WBGene00020824 | 0.480088457 | 0.01449761 | 0.06801215 | T26A8.1 |
| WBGene00020825 | 0.315829538 | 0.70320452 | 0.83253106 | tmem-231 |
| WBGene00020826 | 1.216957619 | 0.74819621 | 1 | T26A8.3 |
| WBGene00020827 | -0.30604763 | 0.11453589 | 0.27839445 | T26A8.4 |
| WBGene00020830 | -0.278681563 | 0.41004696 | 0.61537456 | T26C11.4 |
| WBGene00020831 | 0.111200256 | 0.50106769 | 0.68895389 | T26C12.1 |
| WBGene00020832 | -0.263229581 | 0.93409872 | 1 | T26C12.2 |
| WBGene00020833 | -1.081990778 | 0.16392273 | 0.35007666 | T26C12.3 |
| WBGene00020834 | 0.690081697 | 0.39220205 | 0.60022237 | T27A1.2 |
| WBGene00020835 | 1.745833691 | 0.18672932 | 1 | T27A1.3 |
| WBGene00020836 | 0.427322576 | 0.19173773 | 0.3866281 | lgc-34 |
| WBGene00020837 | -0.080829996 | 0.82943316 | 0.9076639 | slc-36.2 |
| WBGene00020838 | 0.187159526 | 0.35333804 | 0.56208019 | trak-1 |
| WBGene00020839 | -0.205661443 | 0.24979869 | 0.45751284 | usp-5 |
| WBGene00020840 | 1.307410716 | 0.58088678 | 1 | spch-3 |
| WBGene00020841 | -0.512603405 | 0.83307778 | 1 | T27A3.5 |
| WBGene00020842 | -0.247012402 | 0.29730874 | 0.50754013 | nmrk-1 |
| WBGene00020843 | 0.012148678 | 0.95929651 | 0.97905337 | T27A3.7 |
| WBGene00020844 | -0.221204614 | 0.80663748 | 0.89459463 | T27A10.2 |
| WBGene00020846 | 0.298895729 | 0.55212216 | 0.73020586 | T27A10.6 |
| WBGene00020847 | 0.624652234 | 0.06141278 | 0.18554004 | cgr-1 |
| WBGene00020849 | 0.004802149 | 0.99562605 | 0.99776276 | nhr-225 |
| WBGene00020850 | 0.066435198 | 0.93366877 | 0.96494281 | nhr-226 |

| | | | | |
|---|---|---|---|---|
| WBGene00020851 | 1.157229747 | 0.46261485 | 1 | nhr-227 |
| WBGene00020852 | 0.255889755 | 0.56057087 | 0.73673213 | nhr-228 |
| WBGene00020853 | -0.458045577 | 0.75307066 | 1 | fbxa-137 |
| WBGene00020854 | 1.230034302 | 5.8282E-05 | 0.00099104 | T27C4.1 |
| WBGene00020855 | 0.107368419 | 0.8777377 | 0.93441865 | hpo-36 |
| WBGene00020856 | -0.014519762 | 0.99603746 | 1 | srt-61 |
| WBGene00020857 | 1.38322681 | 0.69550305 | 1 | srt-62 |
| WBGene00020858 | 0.286140609 | 0.32264862 | 0.53107238 | mop-25.3 |
| WBGene00020859 | 1.30738041 | 0.38557066 | 1 | bath-25 |
| WBGene00020860 | -0.514761297 | 0.79912167 | 1 | nspd-10 |
| WBGene00020861 | 0.72774339 | 0.40143394 | 0.60907392 | T27E4.1 |
| WBGene00020862 | 0.859348544 | 0.00033389 | 0.0039472 | fip-2 |
| WBGene00020863 | 1.779518513 | 0.35579592 | 1 | T27E4.5 |
| WBGene00020864 | 1.471809563 | 0.46592277 | 1 | oac-50 |
| WBGene00020865 | 1.126424506 | 0.01468204 | 0.06863868 | T27E4.7 |
| WBGene00020866 | -0.566281202 | 0.00186431 | 0.01478316 | vps-24 |
| WBGene00020867 | 0.048116379 | 0.88481662 | 0.93919904 | shc-2 |
| WBGene00020868 | -0.418785894 | 0.00637357 | 0.03722294 | eif-1 |
| WBGene00020869 | 0.925709833 | 0.00086757 | 0.00824192 | T28A11.2 |
| WBGene00020870 | 2.62561062 | 0.22121999 | 1 | T28A11.3 |
| WBGene00020875 | 0.938982574 | 0.8146943 | 1 | T28A11.13 |
| WBGene00020877 | -1.159714244 | 0.75790634 | 1 | srt-63 |
| WBGene00020878 | 1.154756127 | 0.24363479 | 1 | T28A11.16 |
| WBGene00020879 | 1.851444818 | 0.53669729 | 1 | T28A11.17 |
| WBGene00020881 | 2.521986442 | 0.0372157 | 1 | T28A11.19 |
| WBGene00020882 | -0.323868354 | 0.91634405 | 1 | T28A11.20 |
| WBGene00020883 | 0.635150117 | 0.1992183 | 0.39743362 | fbxa-64 |
| WBGene00020884 | 0.182174229 | 0.54124382 | 0.72208215 | T28B4.1 |
| WBGene00020886 | -0.933656276 | 3.6129E-06 | 9.639E-05 | ttr-6 |
| WBGene00020887 | 0.502763973 | 0.33153477 | 0.53971087 | T28B4.4 |
| WBGene00020888 | 0.75595108 | 0.7708876 | 1 | T28C12.1 |
| WBGene00020889 | 0.904409705 | 0.06520143 | 0.19276134 | T28C12.2 |
| WBGene00020890 | 0.377828795 | 0.92526585 | 1 | fbxa-202 |
| WBGene00020891 | 0.383692873 | 0.257151 | 0.46511709 | T28C12.4 |
| WBGene00020892 | -0.156269293 | 0.96924949 | 1 | T28C12.5 |
| WBGene00020893 | -0.414058208 | 0.62540683 | 0.78210457 | T28C12.6 |
| WBGene00020894 | -0.768266577 | 4.0516E-05 | 0.00072339 | T28D9.1 |
| WBGene00020895 | 0.52705332 | 0.15053233 | 0.3321514 | T28D9.3 |
| WBGene00020896 | -0.636939016 | 0.01448846 | 0.06799552 | T28D9.4 |
| WBGene00020897 | 1.051919411 | 0.02031933 | 0.08649963 | del-10 |
| WBGene00020898 | 0.377828795 | 0.92526585 | 1 | T28D9.9 |
| WBGene00020899 | 1.116613983 | 0.69067248 | 1 | T28D9.11 |
| WBGene00020900 | 0.281454817 | 0.94435481 | 1 | T28F2.1 |

| | | | | |
|---|---|---|---|---|
| WBGene00020901 | 0.757749218 | 0.12475147 | 0.29432717 | T28F2.2 |
| WBGene00020902 | -0.003672464 | 0.98954358 | 0.99484196 | jmjc-1 |
| WBGene00020903 | -0.179046032 | 0.72261978 | 0.84367023 | acd-5 |
| WBGene00020904 | 0.641065534 | 0.78192938 | 1 | T28F12.1 |
| WBGene00020905 | 0.773345629 | 0.76155013 | 1 | T28H11.7 |
| WBGene00020906 | 1.428112944 | 0.04944295 | 0.15947139 | T28H11.8 |
| WBGene00020908 | 0.732064274 | 0.02189545 | 0.09168245 | VC5.2 |
| WBGene00020909 | -0.062412671 | 0.74186589 | 0.85569007 | W01A11.1 |
| WBGene00020910 | -0.462640686 | 0.04386858 | 0.14632099 | dgtr-1 |
| WBGene00020911 | -1.209244099 | 0.00029552 | 0.00357006 | cpt-6 |
| WBGene00020912 | 0.15982646 | 0.69037021 | 0.82357419 | W01A11.7 |
| WBGene00020913 | 0.281454817 | 0.94435481 | 1 | W01B11.1 |
| WBGene00020914 | 0.861379559 | 0.56899161 | 1 | sulp-6 |
| WBGene00020915 | -0.351941128 | 0.0347671 | 0.12559142 | nol-5 |
| WBGene00020917 | 0.093773207 | 0.70136802 | 0.83141078 | W01B11.6 |
| WBGene00020918 | 1.477253415 | 0.58357327 | 1 | W01C8.1 |
| WBGene00020919 | 0.172012465 | 0.60274367 | 0.76755935 | set-19 |
| WBGene00020920 | 0.203388228 | 0.57648436 | 0.74903703 | set-20 |
| WBGene00020921 | 0.236677455 | 0.29927918 | 0.50906705 | W01C8.5 |
| WBGene00020924 | -0.70831171 | 0.42967617 | 0.63225044 | W02B3.4 |
| WBGene00020927 | -0.604396148 | 0.82723443 | 1 | W02B3.7 |
| WBGene00020929 | -0.115786094 | 0.79580865 | 0.88706507 | W02C12.2 |
| WBGene00020930 | 0.935844257 | 0.00010248 | 0.00155329 | hlh-30 |
| WBGene00020931 | 0.053577739 | 0.77095294 | 0.8718461 | cytb-5.2 |
| WBGene00020932 | -0.333494682 | 0.07384769 | 0.2097765 | dhod-1 |
| WBGene00020933 | -0.311617346 | 0.09873839 | 0.25290776 | W02D3.4 |
| WBGene00020935 | 0.595231215 | 0.00722841 | 0.04072394 | fnci-1 |
| WBGene00020936 | -0.580602397 | 0.00268456 | 0.01957552 | hrpf-1 |
| WBGene00020937 | -0.676763756 | 0.06995255 | 0.20211658 | W02D3.12 |
| WBGene00020938 | -0.43449162 | 0.46773145 | 0.66183982 | clec-218 |
| WBGene00020939 | 1.200172838 | 0.25689241 | 1 | W02D7.3 |
| WBGene00020940 | -0.322168509 | 0.58922413 | 0.75725311 | W02D7.4 |
| WBGene00020941 | -0.138620114 | 0.77446693 | 0.87427271 | W02D7.5 |
| WBGene00020942 | 0.109496944 | 0.64891778 | 0.79782388 | W02D7.6 |
| WBGene00020943 | 0.334401957 | 0.76513369 | 1 | W02D7.8 |
| WBGene00020944 | -1.159714244 | 0.75790634 | 1 | W02D7.9 |
| WBGene00020945 | 1.747998126 | 0.61126322 | 1 | clec-219 |
| WBGene00020946 | -1.159714244 | 0.75790634 | 1 | W02D7.11 |
| WBGene00020947 | 0.043724621 | 0.93113029 | 0.96395551 | W02F12.2 |
| WBGene00020948 | -0.646526929 | 3.3799E-05 | 0.00061522 | era-1 |
| WBGene00020949 | -0.045420699 | 0.8353862 | 0.91036925 | W02F12.4 |
| WBGene00020950 | -0.136578807 | 0.47074382 | 0.66356678 | dlst-1 |
| WBGene00020951 | -0.29310908 | 0.14591941 | 0.32488677 | sna-1 |

| | | | | |
|---|---|---|---|---|
| WBGene00020952 | 0.592544684 | 0.00572785 | 0.03446076 | kel-8 |
| WBGene00020953 | -0.454979194 | 0.13653966 | 0.31211172 | W02G9.3 |
| WBGene00020954 | 2.169267574 | 6.42E-15 | 1.20E-12 | W02G9.4 |
| WBGene00020955 | -0.294422331 | 0.30867881 | 0.51749095 | angl-1 |
| WBGene00020957 | -0.303782637 | 0.17328254 | 0.36304112 | W02H5.2 |
| WBGene00020961 | -0.62146648 | 0.85925752 | 1 | sknr-1 |
| WBGene00020962 | 0.81193819 | 0.02044697 | 0.08701251 | W02H5.8 |
| WBGene00020964 | 0.563567066 | 0.00637753 | 0.03722579 | polq-1 |
| WBGene00020966 | 0.042994488 | 0.97752318 | 1 | W03A5.1 |
| WBGene00020967 | 0.790047251 | 0.30313617 | 0.51221193 | W03A5.2 |
| WBGene00020970 | 0.014152367 | 0.95159447 | 0.97545664 | W03A5.6 |
| WBGene00020972 | 0.551405546 | 0.40317331 | 0.61087323 | trpl-2 |
| WBGene00020973 | 1.263256715 | 0.23334947 | 1 | W03B1.3 |
| WBGene00020974 | 0.688793783 | 0.86378755 | 1 | W03B1.5 |
| WBGene00020976 | 0.203161664 | 0.72282044 | 0.84382336 | oac-51 |
| WBGene00020977 | 2.656861598 | 0.05421021 | 1 | oac-52 |
| WBGene00020978 | 0.983488853 | 0.80601565 | 1 | W03B1.9 |
| WBGene00020979 | -0.760943105 | 0.7243725 | 1 | fbxb-55 |
| WBGene00020982 | -0.156269293 | 0.96924949 | 1 | ilys-6 |
| WBGene00020983 | -1.633604103 | 0.15135695 | 1 | W03D2.9 |
| WBGene00020984 | 0.175909247 | 0.94229417 | 1 | W03D8.1 |
| WBGene00020985 | 1.333946158 | 0.24127532 | 1 | W03D8.2 |
| WBGene00020986 | -0.156269293 | 0.96924949 | 1 | W03D8.3 |
| WBGene00020987 | 0.377828795 | 0.92526585 | 1 | W03D8.5 |
| WBGene00020988 | -0.476487142 | 0.63655602 | 1 | W03D8.7 |
| WBGene00020989 | 1.49942311 | 0.00035665 | 0.00415898 | W03D8.8 |
| WBGene00020990 | 0.045516744 | 0.9767022 | 1 | W03D8.9 |
| WBGene00020993 | -0.12419383 | 0.60637115 | 0.76951522 | W03F8.3 |
| WBGene00020994 | -0.30029227 | 0.13156561 | 0.30402324 | W03F8.4 |
| WBGene00020995 | 0.284475 | 0.71714168 | 0.84010542 | W03F8.6 |
| WBGene00020997 | -0.132952494 | 0.88696253 | 1 | gpa-18 |
| WBGene00020999 | -0.1737372 | 0.40795337 | 0.6141434 | W03F9.1 |
| WBGene00021000 | -0.303201725 | 0.40615496 | 0.61299816 | W03F9.2 |
| WBGene00021002 | 1.018872717 | 1.10E-07 | 5.0802E-06 | W03F9.4 |
| WBGene00021003 | 0.763390359 | 0.73510979 | 1 | W03F9.9 |
| WBGene00021004 | -0.227652691 | 0.2254239 | 0.42958273 | sftb-2 |
| WBGene00021005 | -1.003711402 | 1.62E-08 | 9.37E-07 | ule-1 |
| WBGene00021006 | 1.025181888 | 0.79790347 | 1 | dct-9 |
| WBGene00021007 | -0.110035047 | 0.91799159 | 1 | W03F11.4 |
| WBGene00021008 | 0.419350555 | 0.70691859 | 1 | W03F11.5 |
| WBGene00021009 | 0.382335904 | 0.07307123 | 0.2081864 | afd-1 |
| WBGene00021011 | 1.544709779 | 0.2546273 | 1 | W03G1.5 |
| WBGene00021012 | -0.556978062 | 0.00297381 | 0.02108938 | pig-1 |

| | | | | |
|---|---|---|---|---|
| WBGene00021013 | -0.156269293 | 0.96924949 | 1 | W03G1.8 |
| WBGene00021014 | -0.43061282 | 0.12309709 | 0.29161401 | W03G9.2 |
| WBGene00021015 | 0.228869736 | 0.69418519 | 0.8264466 | enu-3.3 |
| WBGene00021016 | -0.404901292 | 0.87611185 | 1 | W03G9.5 |
| WBGene00021017 | 1.585779169 | 0.15299996 | 1 | cil-7 |
| WBGene00021019 | 1.480229434 | 0.01883268 | 0.08212916 | W04B5.2 |
| WBGene00021020 | 0.818560485 | 0.00152747 | 0.01273697 | W04B5.3 |
| WBGene00021021 | -0.32017643 | 0.27177656 | 0.48056732 | mrpl-30 |
| WBGene00021022 | -0.144208675 | 0.58273426 | 0.75313711 | W04B5.5 |
| WBGene00021023 | -0.156269293 | 0.96924949 | 1 | W04B5.6 |
| WBGene00021024 | -0.353899732 | 0.11156691 | 0.27342225 | W04C9.2 |
| WBGene00021025 | -0.779060993 | 0.47702439 | 1 | cutl-13 |
| WBGene00021026 | -0.153952415 | 0.55224597 | 0.73020586 | W04C9.4 |
| WBGene00021027 | -0.220986115 | 0.87625587 | 1 | W04C9.5 |
| WBGene00021028 | 2.856527415 | 0.0289573 | 1 | W04C9.6 |
| WBGene00021030 | 0.446458324 | 0.87164163 | 1 | W04H10.1 |
| WBGene00021031 | -1.14715222 | 0.68932198 | 1 | W04H10.2 |
| WBGene00021032 | -1.133279037 | 0.41918235 | 1 | clec-118 |
| WBGene00021034 | 0.082037452 | 0.93262157 | 1 | enu-3.4 |
| WBGene00021035 | -0.148890689 | 0.29933529 | 0.50906705 | W05F2.3 |
| WBGene00021036 | 1.275697455 | 4.1667E-06 | 0.00010809 | W05F2.4 |
| WBGene00021038 | -0.294389131 | 0.15865493 | 0.34341762 | W05F2.6 |
| WBGene00021039 | 1.687187596 | 0.00905212 | 0.04797108 | W05F2.7 |
| WBGene00021040 | -0.514100803 | 0.55214353 | 0.73020586 | W05G11.2 |
| WBGene00021041 | 0.688793783 | 0.86378755 | 1 | W05G11.4 |
| WBGene00021042 | 0.377828795 | 0.92526585 | 1 | btb-5 |
| WBGene00021043 | -0.110467628 | 0.52570108 | 0.71019659 | pck-1 |
| WBGene00021044 | -0.539172661 | 0.32165039 | 0.52972835 | W05H7.1 |
| WBGene00021045 | -0.276148461 | 0.85598496 | 1 | W05H7.2 |
| WBGene00021046 | -1.331393519 | 0.00700579 | 0.03978359 | sedl-1 |
| WBGene00021047 | -0.342556715 | 0.24392146 | 0.45137181 | zfp-3 |
| WBGene00021048 | 0.299791886 | 0.15785835 | 0.34279345 | W05H9.1 |
| WBGene00021049 | -0.009466353 | 0.97125195 | 0.98496241 | W05H9.2 |
| WBGene00021050 | 0.232400921 | 0.32539396 | 0.5338183 | W05H9.3 |
| WBGene00021051 | -0.078059085 | 0.76121104 | 0.86650332 | W05H9.4 |
| WBGene00021052 | 0.05429394 | 0.95235474 | 0.97582402 | R11F4.3 |
| WBGene00021052 | -0.613681708 | 0.59036892 | 1 | W06A11.1 |
| WBGene00021053 | -1.746501808 | 0.02419631 | 0.09798778 | gadr-4 |
| WBGene00021054 | -2.161355951 | 0.08632117 | 1 | fbxb-118 |
| WBGene00021055 | 0.054639984 | 0.90279464 | 0.94940058 | W06A11.4 |
| WBGene00021056 | -0.830774392 | 1.08E-07 | 5.0411E-06 | W06B4.1 |
| WBGene00021057 | 0.427865988 | 0.0996659 | 0.25442467 | W06B4.2 |
| WBGene00021058 | 0.030838686 | 0.88922192 | 0.94181656 | vps-18 |

| | | | | |
|---|---|---|---|---|
| WBGene00021059 | 0.950428378 | 0.09573308 | 0.24803571 | atat-2 |
| WBGene00021060 | 0.045470294 | 0.88095336 | 0.93657222 | dct-11 |
| WBGene00021061 | -0.065819394 | 0.7533974 | 0.86235484 | W06E11.1 |
| WBGene00021063 | -0.375754528 | 0.07116396 | 0.20449744 | sbds-1 |
| WBGene00021066 | 1.162404706 | 0.12427668 | 0.29366353 | W06H8.2 |
| WBGene00021068 | 1.177255055 | 6.6926E-06 | 0.00016058 | W06H8.6 |
| WBGene00021070 | -0.565604378 | 0.21367107 | 0.4143559 | cpr-8 |
| WBGene00021072 | 3.637634348 | 0.01290634 | 1 | W07B8.4 |
| WBGene00021073 | -0.382838646 | 0.03948499 | 0.13622826 | nol-1 |
| WBGene00021074 | -0.055727363 | 0.83571643 | 0.91056566 | W07E6.2 |
| WBGene00021075 | 0.965156218 | 0.29694667 | 0.50725372 | W07E6.3 |
| WBGene00021076 | 0.975834625 | 0.11149657 | 0.27336023 | W07E6.5 |
| WBGene00021078 | 0.384593418 | 0.12055722 | 0.28710969 | unc-132 |
| WBGene00021079 | -0.478393313 | 0.33763441 | 0.54561685 | W08A12.2 |
| WBGene00021080 | 0.612882069 | 0.30962435 | 0.51818605 | W08A12.3 |
| WBGene00021081 | 2.628909249 | 0.11534689 | 1 | pals-33 |
| WBGene00021082 | -0.316075559 | 0.60794617 | 0.77036604 | zip-11 |
| WBGene00021083 | -1.969805064 | 0.62862569 | 1 | W08E12.2 |
| WBGene00021084 | -2.811538432 | 0.0239154 | 1 | W08E12.3 |
| WBGene00021085 | 0.938982574 | 0.8146943 | 1 | W08E12.4 |
| WBGene00021086 | 0.938982574 | 0.8146943 | 1 | W08E12.5 |
| WBGene00021087 | -1.195119012 | 0.3248319 | 1 | W08E12.6 |
| WBGene00021088 | -0.370514206 | 0.02818437 | 0.10859613 | W08E12.7 |
| WBGene00021089 | 1.278053416 | 0.54789306 | 1 | W08E12.8 |
| WBGene00021092 | -0.118437635 | 0.92249608 | 1 | fbxb-34 |
| WBGene00021093 | -0.457200484 | 0.00621851 | 0.0366856 | W08F4.3 |
| WBGene00021094 | -0.518205903 | 0.77140115 | 1 | W08F4.5 |
| WBGene00021095 | -0.694440235 | 0.04133038 | 0.14075516 | mlt-8 |
| WBGene00021096 | 0.766821134 | 0.64599243 | 1 | W08F4.7 |
| WBGene00021097 | -0.244036004 | 0.20452051 | 0.40395047 | cdc-37 |
| WBGene00021098 | 1.563818453 | 0.47753399 | 1 | fbxb-14 |
| WBGene00021099 | -0.22461868 | 0.93331196 | 1 | W08F4.10 |
| WBGene00021100 | 1.733076639 | 0.37263578 | 1 | W08F4.11 |
| WBGene00021101 | -0.917189319 | 0.39582678 | 1 | W08F4.12 |
| WBGene00021103 | -0.035140299 | 0.87985664 | 0.93574569 | eri-3 |
| WBGene00021104 | -0.287679986 | 0.37541616 | 0.58322809 | W09B6.4 |
| WBGene00021104 | -0.56468303 | 0.426256 | 0.6300431 | W09B6.5 |
| WBGene00021110 | 1.025181888 | 0.79790347 | 1 | W09C3.2 |
| WBGene00021111 | 1.045259305 | 0.6323014 | 1 | W09C3.3 |
| WBGene00021112 | -0.234146935 | 0.39335818 | 0.60140557 | W09C3.4 |
| WBGene00021113 | 0.483840154 | 0.8151116 | 1 | gsp-3 |
| WBGene00021114 | 0.982512931 | 0.01657175 | 0.07449005 | W09C3.7 |
| WBGene00021115 | 1.116613983 | 0.69067248 | 1 | W09C3.8 |

| | | | | |
|---|---|---|---|---|
| WBGene00021116 | 0.126882066 | 0.84888765 | 0.9171794 | ncs-6 |
| WBGene00021117 | -0.457870121 | 0.79919046 | 1 | clec-126 |
| WBGene00021118 | 1.006832178 | 0.42319574 | 1 | clec-125 |
| WBGene00021120 | -0.156269293 | 0.96924949 | 1 | W09G12.6 |
| WBGene00021121 | 1.41038885 | 0.02834397 | 0.10903979 | W09G12.7 |
| WBGene00021122 | -1.901856996 | 0.64072034 | 1 | W09G12.8 |
| WBGene00021123 | -0.404698819 | 0.42279933 | 0.62722467 | W09G12.9 |
| WBGene00021124 | 0.926839843 | 0.40736788 | 1 | W10C4.1 |
| WBGene00021125 | 0.488133825 | 0.81078971 | 1 | ccb-2 |
| WBGene00021127 | 0.146155967 | 0.7912309 | 0.88470671 | W10C8.4 |
| WBGene00021128 | 0.505736226 | 0.05182543 | 0.1644251 | W10C8.5 |
| WBGene00021131 | 0.054677039 | 0.90241404 | 0.949247 | W10D9.3 |
| WBGene00021132 | -0.192653052 | 0.46877013 | 0.66255392 | nfyb-1 |
| WBGene00021133 | -0.724486502 | 0.000871 | 0.00826156 | tomm-22 |
| WBGene00021134 | 3.606609313 | 0.03728625 | 1 | W10G11.1 |
| WBGene00021135 | -1.863137868 | 0.04880758 | 0.15839959 | W10G11.2 |
| WBGene00021136 | -1.342168785 | 0.05732425 | 0.17722783 | W10G11.3 |
| WBGene00021137 | 0.57991002 | 0.82402153 | 1 | W10G11.4 |
| WBGene00021141 | 1.116613983 | 0.69067248 | 1 | clec-134 |
| WBGene00021142 | -1.094201388 | 0.7816772 | 1 | clec-133 |
| WBGene00021143 | -0.156269293 | 0.96924949 | 1 | clec-132 |
| WBGene00021146 | 0.854234615 | 0.03507271 | 0.12622038 | lgc-45 |
| WBGene00021148 | 0.960752966 | 0.06492116 | 0.19219893 | W10G11.19 |
| WBGene00021149 | -0.790926061 | 0.00245496 | 0.01833235 | dnc-3 |
| WBGene00021150 | 1.957271014 | 0.49743227 | 1 | Y1B5A.1 |
| WBGene00021152 | 1.27214152 | 0.02491542 | 0.09974122 | mgl-3 |
| WBGene00021153 | 0.189641974 | 0.48002894 | 0.67189663 | Y4C6A.3 |
| WBGene00021154 | -1.094201388 | 0.7816772 | 1 | Y4C6A.4 |
| WBGene00021155 | -0.465832263 | 0.01903543 | 0.08280515 | Y4C6B.1 |
| WBGene00021156 | 1.026672876 | 0.1742463 | 0.36445118 | Y4C6B.2 |
| WBGene00021157 | 1.006011639 | 0.10821784 | 0.2680826 | Y4C6B.3 |
| WBGene00021158 | 1.671852206 | 0.03072371 | 0.11552077 | Y4C6B.4 |
| WBGene00021159 | 0.336121744 | 0.35691309 | 0.56525982 | Y4C6B.5 |
| WBGene00021160 | 0.957913509 | 1.0298E-05 | 0.0002337 | gba-4 |
| WBGene00021161 | -0.804575446 | 0.01807516 | 0.0796241 | Y4C6B.7 |
| WBGene00021163 | 0.506335373 | 0.69474049 | 1 | nhr-275 |
| WBGene00021164 | 2.063391569 | 0.37853067 | 1 | Y5H2B.1 |
| WBGene00021165 | -2.305348966 | 0.00227033 | 0.01719527 | Y5H2B.3 |
| WBGene00021167 | 2.084042462 | 0.02294512 | 0.09459313 | cyp-32B1 |
| WBGene00021168 | -0.774191023 | 0.72082566 | 1 | cyp-33C12 |
| WBGene00021169 | -1.901856996 | 0.64072034 | 1 | srt-40 |
| WBGene00021170 | -0.406459985 | 0.02329997 | 0.09553695 | arx-4 |
| WBGene00021171 | 1.501531263 | 0.17212931 | 1 | Y8A9A.2 |

| | | | | |
|---|---|---|---|---|
| WBGene00021172 | -1.964653846 | 0.53599202 | 1 | Y8A9A.3 |
| WBGene00021174 | -2.306552492 | 0.20547362 | 1 | fbxb-99 |
| WBGene00021175 | 1.38322681 | 0.69550305 | 1 | math-46 |
| WBGene00021176 | 1.203280165 | 0.47927209 | 1 | Y9C9A.1 |
| WBGene00021177 | -0.156269293 | 0.96924949 | 1 | Y9C9A.5 |
| WBGene00021178 | 1.427836926 | 0.00516278 | 0.0319314 | Y9C9A.8 |
| WBGene00021179 | 1.025181888 | 0.79790347 | 1 | fbxa-86 |
| WBGene00021180 | -0.08281586 | 0.94563094 | 1 | Y9C9A.13 |
| WBGene00021182 | -0.156269293 | 0.96924949 | 1 | srz-28 |
| WBGene00021183 | 0.394439984 | 0.88400378 | 1 | Y9C9A.16 |
| WBGene00021184 | 0.281454817 | 0.94435481 | 1 | Y9C9A.17 |
| WBGene00021185 | 0.005961717 | 0.99286177 | 0.99644982 | Y9C12A.1 |
| WBGene00021186 | 0.012471018 | 0.9808757 | 0.98993793 | Y9D1A.1 |
| WBGene00021187 | 1.712479368 | 0.58529868 | 1 | Y9D1A.2 |
| WBGene00021188 | -0.18557176 | 0.95664217 | 1 | Y13C8A.1 |
| WBGene00021190 | 0.118734431 | 0.88716834 | 0.94044069 | Y14H12A.1 |
| WBGene00021191 | -0.711342948 | 4.2918E-05 | 0.0007629 | Y14H12B.1 |
| WBGene00021192 | -0.687471612 | 0.0014975 | 0.01256474 | Y14H12B.2 |
| WBGene00021193 | -0.066335075 | 0.91456723 | 0.95571968 | math-44 |
| WBGene00021195 | 0.325613128 | 0.79093441 | 1 | Y17G9A.2 |
| WBGene00021196 | 1.894863195 | 0.29507975 | 1 | Y17G9A.3 |
| WBGene00021197 | 0.482097419 | 0.47170088 | 0.66453009 | Y17G9A.4 |
| WBGene00021198 | -0.236080742 | 0.87580243 | 1 | npp-26 |
| WBGene00021199 | -1.67479784 | 0.16946618 | 1 | Y17G9B.2 |
| WBGene00021200 | -0.3280377 | 0.11745162 | 0.28245932 | cyp-31A3 |
| WBGene00021201 | 0.065292539 | 0.87917431 | 0.93533607 | Y17G9B.4 |
| WBGene00021202 | 0.044107187 | 0.84346464 | 0.91384117 | Y17G9B.5 |
| WBGene00021204 | -0.946577489 | 0.47668063 | 1 | fbxb-77 |
| WBGene00021205 | 0.257307241 | 0.45774138 | 0.65474058 | Y17G9B.8 |
| WBGene00021207 | 1.025181888 | 0.79790347 | 1 | Y18H1A.1 |
| WBGene00021208 | -0.119243532 | 0.673583 | 0.8134506 | Y18H1A.2 |
| WBGene00021209 | 0.251516475 | 0.25219331 | 0.46035874 | hgap-1 |
| WBGene00021210 | -0.497007134 | 0.08935887 | 0.2371834 | Y18H1A.4 |
| WBGene00021212 | 1.557897343 | 0.63813005 | 1 | Y18H1A.8 |
| WBGene00021213 | 0.010585712 | 0.98729721 | 0.99421374 | Y18H1A.9 |
| WBGene00021214 | 1.735991307 | 0.58152819 | 1 | hasp-2 |
| WBGene00021215 | 1.076747689 | 0.44417639 | 1 | Y18H1A.11 |
| WBGene00021220 | -0.156269293 | 0.96924949 | 1 | Y19D10A.5 |
| WBGene00021224 | -3.168891993 | 0.27608518 | 1 | clec-209 |
| WBGene00021226 | -0.156269293 | 0.96924949 | 1 | Y19D10A.11 |
| WBGene00021227 | -0.156269293 | 0.96924949 | 1 | mct-1 |
| WBGene00021232 | 2.381446256 | 0.34860272 | 1 | irld-15 |
| WBGene00021235 | 0.651616316 | 0.51312816 | 0.69983027 | Y19D10B.6 |

| | | | | |
|---|---|---|---|---|
| WBGene00021236 | 0.595703065 | 0.58378726 | 1 | pud-1.2 |
| WBGene00021237 | 0.94988478 | 0.00733697 | 0.04113904 | crtc-1 |
| WBGene00021238 | 0.29319748 | 0.3417857 | 0.55020156 | gap-3 |
| WBGene00021239 | 0.685704588 | 0.05141228 | 0.16370707 | Y20F4.4 |
| WBGene00021243 | 0.281454817 | 0.94435481 | 1 | Y22D7AL.3 |
| WBGene00021244 | 0.441148425 | 0.80166522 | 1 | Y22D7AL.4 |
| WBGene00021245 | 0.701732884 | 0.55848547 | 1 | Y22D7AL.6 |
| WBGene00021246 | -0.159889362 | 0.67414427 | 0.81367786 | Y22D7AL.7 |
| WBGene00021247 | 0.057810848 | 0.87523266 | 0.93275833 | Y22D7AL.9 |
| WBGene00021248 | -0.738200927 | 2.6722E-05 | 0.00051184 | Y22D7AL.10 |
| WBGene00021249 | 0.664895896 | 0.44568626 | 0.64509738 | Y22D7AL.11 |
| WBGene00021251 | 0.983488853 | 0.80601565 | 1 | Y22D7AL.13 |
| WBGene00021252 | 0.36751063 | 0.80372267 | 1 | cht-4 |
| WBGene00021253 | 0.787759245 | 0.00891162 | 0.0473713 | Y22D7AL.15 |
| WBGene00021254 | 0.499635666 | 0.5169071 | 0.70285241 | Y22D7AL.16 |
| WBGene00021256 | 0.666182301 | 0.41555339 | 0.62082449 | Y22D7AR.2 |
| WBGene00021259 | 0.813336992 | 0.54900057 | 1 | glb-30 |
| WBGene00021260 | 0.708436515 | 0.15491908 | 0.33852997 | Y22D7AR.6 |
| WBGene00021261 | 2.226542641 | 0.21104126 | 1 | Y22D7AR.7 |
| WBGene00021262 | 0.545209743 | 0.21073001 | 0.41053803 | fbxa-74 |
| WBGene00021263 | -1.512048016 | 3.84E-09 | 2.57E-07 | Y22D7AR.10 |
| WBGene00021264 | 0.477283085 | 0.54313262 | 0.72328361 | fbxa-13 |
| WBGene00021266 | 0.281454817 | 0.94435481 | 1 | Y22D7AR.14 |
| WBGene00021267 | 1.741745596 | 0.36346624 | 1 | Y23B4A.1 |
| WBGene00021268 | 1.404564216 | 0.08098336 | 0.22254466 | capa-1 |
| WBGene00021269 | -0.787979923 | 0.11588562 | 0.28046677 | Y23H5A.2 |
| WBGene00021270 | -0.846762251 | 4.47E-07 | 1.7061E-05 | Y23H5A.3 |
| WBGene00021271 | -0.343372002 | 0.82280665 | 1 | spe-47 |
| WBGene00021272 | -0.276793073 | 0.69959263 | 0.830279 | Y23H5B.1 |
| WBGene00021274 | -0.074828807 | 0.95751789 | 1 | Y23H5B.3 |
| WBGene00021275 | 1.250082729 | 0.05810114 | 0.178629 | dmsr-2 |
| WBGene00021276 | -0.301206057 | 0.11999066 | 0.28624902 | Y23H5B.5 |
| WBGene00021277 | -0.091197188 | 0.66099506 | 0.80492889 | Y23H5B.6 |
| WBGene00021279 | -0.523768348 | 0.46937894 | 0.66298055 | Y23H5B.8 |
| WBGene00021281 | 0.290763817 | 0.20602773 | 0.40567349 | ell-1 |
| WBGene00021282 | -0.250301713 | 0.44587699 | 0.64529653 | set-21 |
| WBGene00021286 | -0.070527478 | 0.64731869 | 0.79650311 | tald-1 |
| WBGene00021287 | -0.706595687 | 0.55218164 | 1 | Y24D9B.1 |
| WBGene00021288 | 0.942308827 | 0.67468959 | 1 | clec-123 |
| WBGene00021289 | -0.389078632 | 0.8767475 | 1 | Y25C1A.2 |
| WBGene00021292 | 0.008761389 | 0.95737929 | 0.97850563 | copb-1 |
| WBGene00021293 | -2.666951515 | 2.003E-06 | 5.8739E-05 | Y25C1A.6 |
| WBGene00021294 | 0.011268791 | 0.95803529 | 0.9785154 | Y25C1A.7 |

| | | | | |
|---|---|---|---|---|
| WBGene00021295 | -0.408597941 | 0.10958357 | 0.27006751 | Y25C1A.8 |
| WBGene00021296 | -0.289578929 | 0.26187049 | 0.47039053 | Y25C1A.13 |
| WBGene00021297 | 1.557897343 | 0.63813005 | 1 | fbxc-55 |
| WBGene00021298 | -1.41599653 | 0.25737755 | 1 | Y27F2A.6 |
| WBGene00021299 | -1.164838232 | 0.25957558 | 1 | Y27F2A.8 |
| WBGene00021300 | -1.901856996 | 0.64072034 | 1 | Y27F2A.9 |
| WBGene00021301 | 0.178115923 | 0.94982316 | 1 | Y32G9A.2 |
| WBGene00021303 | 0.281454817 | 0.94435481 | 1 | Y32G9A.5 |
| WBGene00021304 | 0.694509213 | 0.2756156 | 0.48419173 | nphp-2 |
| WBGene00021305 | 0.074625487 | 0.76633859 | 0.86897515 | zig-11 |
| WBGene00021309 | -0.597656973 | 0.56214788 | 1 | Y32G9B.1 |
| WBGene00021310 | -0.447380075 | 0.82633951 | 1 | Y32H12A.1 |
| WBGene00021311 | -0.029309677 | 0.89000862 | 0.94196731 | thoc-5 |
| WBGene00021312 | -0.398206275 | 0.03642738 | 0.12918931 | szy-2 |
| WBGene00021313 | -0.207338508 | 0.31302539 | 0.52117955 | paqr-2 |
| WBGene00021314 | -0.305065998 | 0.44689527 | 0.64584638 | Y32H12A.6 |
| WBGene00021315 | -0.134450547 | 0.49958737 | 0.68761935 | Y32H12A.7 |
| WBGene00021316 | 0.760540592 | 0.00100707 | 0.00929799 | Y32H12A.8 |
| WBGene00021319 | -0.493769891 | 0.12292773 | 0.29132647 | Y34B4A.2 |
| WBGene00021320 | 1.184086985 | 0.00528312 | 0.03247711 | Y34B4A.4 |
| WBGene00021321 | 0.670779698 | 0.13905959 | 0.31610606 | Y34B4A.5 |
| WBGene00021322 | 0.467097058 | 0.00532081 | 0.03262621 | Y34B4A.6 |
| WBGene00021323 | -0.062954888 | 0.83213324 | 0.90915448 | Y34B4A.7 |
| WBGene00021324 | -0.347405302 | 0.34540034 | 0.55425662 | Y34B4A.8 |
| WBGene00021325 | 0.31629593 | 0.05525743 | 0.17232994 | Y34B4A.9 |
| WBGene00021326 | 0.156735926 | 0.82569127 | 0.90598443 | Y34B4A.10 |
| WBGene00021327 | -0.161780706 | 0.43519994 | 0.63636788 | mrpl-38 |
| WBGene00021328 | 1.078080422 | 0.1368239 | 0.31260425 | npr-23 |
| WBGene00021329 | 0.035245988 | 0.87620677 | 0.93317123 | Y34D9A.3 |
| WBGene00021331 | -0.504729431 | 0.03171309 | 0.11812001 | glrx-10 |
| WBGene00021332 | -0.404230614 | 0.15851658 | 0.34330179 | Y34D9A.7 |
| WBGene00021333 | -0.292877665 | 0.61996229 | 0.7787454 | Y34D9A.8 |
| WBGene00021334 | -0.214124091 | 0.28290037 | 0.49249977 | vps-4 |
| WBGene00021335 | 1.261939204 | 3.3344E-06 | 9.0161E-05 | spp-23 |
| WBGene00021336 | 1.256280089 | 0.00808773 | 0.04421121 | Y34F4.1 |
| WBGene00021337 | 0.874921587 | 0.07353387 | 0.20913483 | Y34F4.2 |
| WBGene00021338 | 1.51672272 | 0.31693815 | 1 | Y34F4.3 |
| WBGene00021339 | 0.664294544 | 0.47323015 | 0.66637523 | Y34F4.4 |
| WBGene00021340 | 0.65251144 | 0.03717434 | 0.13099155 | Y34F4.5 |
| WBGene00021343 | -0.737826107 | 0.09994696 | 0.25476717 | cutl-23 |
| WBGene00021344 | 0.688552404 | 0.01865196 | 0.08154469 | Y37B11A.2 |
| WBGene00021345 | -0.047870659 | 0.92980838 | 0.96353735 | Y37B11A.3 |
| WBGene00021346 | -0.260532933 | 0.3167819 | 0.52477132 | Y37E3.1 |

| | | | | |
|---|---|---|---|---|
| WBGene00021347 | -0.42976535 | 0.06162841 | 0.18583439 | rpb-10 |
| WBGene00021348 | -0.800778499 | 0.00014942 | 0.00209238 | moag-4 |
| WBGene00021349 | -0.082077116 | 0.92147876 | 0.95884274 | arl-13 |
| WBGene00021350 | -0.532515271 | 2.2502E-05 | 0.0004464 | Y37E3.8 |
| WBGene00021351 | -0.641973884 | 2.7928E-05 | 0.00052897 | eif-2alpha |
| WBGene00021352 | 0.017952073 | 0.91857087 | 0.95695632 | Y37E3.11 |
| WBGene00021353 | 0.336253222 | 0.47116925 | 0.66408937 | iglr-3 |
| WBGene00021354 | 0.320434274 | 0.39114957 | 0.59953881 | fpn-1.1 |
| WBGene00021355 | -0.105620718 | 0.63549404 | 0.78834506 | Y37E3.17 |
| WBGene00021357 | 1.854471114 | 0.22100581 | 1 | Y37E11AL.1 |
| WBGene00021358 | 1.072430633 | 0.5261168 | 1 | Y37E11AL.2 |
| WBGene00021359 | -0.605416888 | 0.00242972 | 0.01817576 | Y37E11AL.3 |
| WBGene00021362 | 0.823329857 | 0.07523569 | 0.21222969 | Y37E11AL.6 |
| WBGene00021363 | -0.335839066 | 0.15880253 | 0.34355337 | taf-6.2 |
| WBGene00021364 | -0.979452109 | 0.72336959 | 1 | Y37E11AL.9 |
| WBGene00021365 | 0.002097817 | 0.99291781 | 0.99644982 | smgl-2 |
| WBGene00021366 | -0.386934896 | 0.1891336 | 0.38309494 | Y37E11AM.2 |
| WBGene00021367 | -0.454327556 | 0.07281799 | 0.20797066 | Y37E11AM.3 |
| WBGene00021368 | 1.031840022 | 0.04448603 | 0.14777153 | best-20 |
| WBGene00021369 | 0.449003544 | 0.1057581 | 0.26479874 | siah-1 |
| WBGene00021370 | 0.271862483 | 0.52701037 | 0.7113202 | nape-2 |
| WBGene00021371 | 0.419978284 | 0.52434165 | 0.70891225 | nape-1 |
| WBGene00021372 | 0.354598163 | 0.25912963 | 0.46732429 | ugt-45 |
| WBGene00021374 | 0.328206839 | 0.67408939 | 0.81367786 | Y37E11B.1 |
| WBGene00021375 | -0.025943295 | 0.9730298 | 0.98595164 | Y37E11B.2 |
| WBGene00021376 | -0.536323307 | 0.10995562 | 0.27062056 | blos-7 |
| WBGene00021377 | -0.254421218 | 0.40883346 | 0.61457432 | Y37E11B.5 |
| WBGene00021378 | -0.612553763 | 0.08817432 | 0.23548712 | Y37E11B.6 |
| WBGene00021379 | -1.196801792 | 0.24152161 | 1 | Y37E11B.7 |
| WBGene00021381 | 1.980799458 | 0.19510544 | 1 | Y37E11B.10 |
| WBGene00021382 | 3.459140102 | 0.05790615 | 1 | Y37F4.1 |
| WBGene00021386 | 1.735991307 | 0.58152819 | 1 | Y37F4.5 |
| WBGene00021387 | 0.506071471 | 0.24507061 | 0.45301497 | Y37F4.6 |
| WBGene00021388 | 0.045210662 | 0.94689434 | 0.97304242 | Y37F4.7 |
| WBGene00021389 | -0.923421932 | 0.24400583 | 0.45145912 | Y38A10A.2 |
| WBGene00021391 | -0.167887095 | 0.39569004 | 0.60353202 | smut-1 |
| WBGene00021392 | -0.71778495 | 0.0010908 | 0.00981195 | Y38A10A.7 |
| WBGene00021394 | -0.073089814 | 0.75781587 | 0.86459364 | Y38C1AA.1 |
| WBGene00021395 | 0.257871987 | 0.30566429 | 0.51458342 | pnc-1 |
| WBGene00021396 | 0.310225092 | 0.61971255 | 0.77871352 | cutl-24 |
| WBGene00021397 | 1.301106245 | 0.11555062 | 0.28013181 | Y38C1AA.6 |
| WBGene00021398 | 3.971507538 | 0.01535424 | 1 | Y38C1AA.7 |
| WBGene00021400 | 0.658815907 | 0.22628136 | 0.43060846 | Y38C1AA.9 |

| | | | | |
|---|---|---|---|---|
| WBGene00021401 | -0.752893404 | 0.00329374 | 0.02272122 | prdx-6 |
| WBGene00021402 | -0.122516224 | 0.68619584 | 0.82077067 | Y38C1AA.12 |
| WBGene00021404 | 0.281454817 | 0.94435481 | 1 | trpl-3 |
| WBGene00021406 | -0.247295945 | 0.68523904 | 0.82001327 | frm-5.2 |
| WBGene00021408 | 0.281454817 | 0.94435481 | 1 | trpl-4 |
| WBGene00021410 | 0.005606485 | 0.99529748 | 0.99756601 | Y38C1BA.1 |
| WBGene00021412 | -0.458945757 | 0.56971574 | 0.74425222 | cyp-29A3 |
| WBGene00021415 | 0.300624776 | 0.44153195 | 0.64137027 | nsy-4 |
| WBGene00021416 | -3.241729724 | 0.06392676 | 1 | mksr-2 |
| WBGene00021417 | 1.388389435 | 0.3375141 | 1 | nhr-236 |
| WBGene00021419 | -0.512326841 | 0.09469682 | 0.24643135 | eri-5 |
| WBGene00021420 | -0.457390099 | 0.001611 | 0.01323356 | trap-3 |
| WBGene00021421 | -0.544225635 | 0.03873931 | 0.13457338 | Y38F2AR.3 |
| WBGene00021423 | 0.459119946 | 0.06307196 | 0.18851714 | tftc-1 |
| WBGene00021425 | -0.1509332 | 0.50545598 | 0.69232186 | ppgn-1 |
| WBGene00021427 | -0.602369809 | 0.002187 | 0.01665772 | Y38F2AR.9 |
| WBGene00021428 | -0.737462795 | 0.83433562 | 1 | Y38F2AR.10 |
| WBGene00021430 | 0.58999258 | 0.06769929 | 0.19774952 | Y38F2AR.12 |
| WBGene00021433 | 0.281454817 | 0.94435481 | 1 | Y39A3A.2 |
| WBGene00021434 | 2.480168205 | 0.14294382 | 1 | Y39A3A.3 |
| WBGene00021435 | 0.827269733 | 0.43358411 | 1 | Y39A3A.4 |
| WBGene00021436 | -1.509975708 | 0.60137946 | 1 | Y39A3B.1 |
| WBGene00021437 | 1.670375483 | 0.28373097 | 1 | lgc-42 |
| WBGene00021438 | 0.178115923 | 0.94982316 | 1 | Y39A3B.3 |
| WBGene00021439 | 0.625327984 | 0.3473839 | 0.55625968 | ckr-2 |
| WBGene00021440 | 0.164964075 | 0.38761331 | 0.59630172 | Y39A3CL.1 |
| WBGene00021441 | -0.634604774 | 0.01384036 | 0.06561337 | Y39A3CL.3 |
| WBGene00021442 | -0.084938237 | 0.69313846 | 0.82568623 | Y39A3CL.4 |
| WBGene00021443 | -0.149545246 | 0.44373761 | 0.64322018 | Y39A3CL.7 |
| WBGene00021444 | -0.008146838 | 0.97641298 | 0.98781418 | ttc-7 |
| WBGene00021446 | 0.219000269 | 0.7905377 | 0.8845213 | Y39A3CR.5 |
| WBGene00021446 | 0.482119318 | 0.37410702 | 0.5820135 | hlh-33 |
| WBGene00021448 | 0.82757375 | 0.05830893 | 0.17895065 | Y39D8A.1 |
| WBGene00021449 | -0.266425659 | 0.83269856 | 1 | mltn-7 |
| WBGene00021451 | 1.216957619 | 0.74819621 | 1 | mltn-8 |
| WBGene00021452 | 0.191020615 | 0.81043264 | 0.8965687 | Y39F10A.1 |
| WBGene00021456 | -0.201097551 | 0.30535756 | 0.51440906 | Y39F10B.1 |
| WBGene00021457 | 0.185436026 | 0.91914894 | 1 | Y39F10C.1 |
| WBGene00021458 | -2.405823103 | 6.2323E-06 | 0.00015133 | Y39G10AL.1 |
| WBGene00021460 | -0.49356806 | 0.00995609 | 0.05167885 | zwl-1 |
| WBGene00021461 | 1.912950174 | 0.09621901 | 1 | nekl-1 |
| WBGene00021463 | -1.410325898 | 0.02958602 | 0.11207609 | zeel-1 |
| WBGene00021464 | 1.600993389 | 0.00299514 | 0.02118839 | ugt-31 |

| | | | | |
|---|---|---|---|---|
| WBGene00021465 | -0.242223759 | 0.38905232 | 0.59747063 | ekl-7 |
| WBGene00021466 | -0.518115768 | 0.00220861 | 0.01675923 | eif-2gamma |
| WBGene00021467 | -0.621114321 | 0.00081761 | 0.00786852 | Y39G10AR.9 |
| WBGene00021468 | -1.447449332 | 1.82E-14 | 3.11E-12 | epg-2 |
| WBGene00021469 | 0.539614637 | 0.00889127 | 0.0473045 | Y39G10AR.11 |
| WBGene00021470 | -0.061830233 | 0.74376048 | 0.85730523 | tpxl-1 |
| WBGene00021471 | 1.216957619 | 0.74819621 | 1 | Y39G10AR.15 |
| WBGene00021472 | -1.293124768 | 0.50359198 | 1 | Y39G10AR.16 |
| WBGene00021473 | 0.708482155 | 0.33520275 | 0.54335068 | sydn-1 |
| WBGene00021474 | 0.085047297 | 0.71684042 | 0.84010542 | dot-1.1 |
| WBGene00021475 | -0.670629089 | 0.02568323 | 0.10210218 | tbca-1 |
| WBGene00021476 | -0.14212529 | 0.65425059 | 0.80137777 | nsun-4 |
| WBGene00021478 | -1.082680044 | 0.67762556 | 1 | Y39H10A.1 |
| WBGene00021479 | -1.01423749 | 0.50354564 | 1 | oac-55 |
| WBGene00021482 | -0.119960557 | 0.67377802 | 0.81347842 | Y39H10A.6 |
| WBGene00021484 | -1.094201388 | 0.7816772 | 1 | Y40A1A.2 |
| WBGene00021485 | 1.904167974 | 0.16708776 | 1 | Y40A1A.3 |
| WBGene00021486 | -0.785280863 | 1.86E-08 | 1.0532E-06 | lbp-9 |
| WBGene00021487 | -0.352489393 | 0.36340427 | 0.57151441 | comt-3 |
| WBGene00021490 | 0.160997185 | 0.92397217 | 1 | Y40B10A.5 |
| WBGene00021491 | -1.963690337 | 0.05782591 | 0.17818893 | comt-4 |
| WBGene00021492 | -1.688293228 | 0.17501695 | 1 | comt-5 |
| WBGene00021493 | -0.10430641 | 0.77394797 | 0.87401193 | Y40B10A.9 |
| WBGene00021494 | 1.730553848 | 0.41535651 | 1 | Y40B10B.1 |
| WBGene00021495 | 0.945085925 | 0.0276958 | 0.10733993 | sqst-2 |
| WBGene00021496 | -0.912030601 | 0.62916461 | 1 | Y40C5A.3 |
| WBGene00021497 | 0.43983243 | 0.15440914 | 0.33778043 | Y40C5A.4 |
| WBGene00021500 | -1.027355405 | 0.68015432 | 1 | Y40C7B.4 |
| WBGene00021502 | 0.348016015 | 0.26708306 | 0.47544153 | Y40D12A.1 |
| WBGene00021504 | 0.401635602 | 0.83343893 | 1 | Y41D4A.1 |
| WBGene00021505 | -0.737432465 | 0.4087592 | 0.61457432 | Y41D4A.3 |
| WBGene00021506 | 0.014236209 | 0.93915063 | 0.96880629 | Y41D4A.4 |
| WBGene00021507 | -0.084157968 | 0.67104831 | 0.81182555 | Y41D4A.5 |
| WBGene00021508 | -0.304996658 | 0.26271294 | 0.47141558 | Y41D4A.6 |
| WBGene00021509 | 0.849148366 | 0.07851215 | 0.21817715 | Y41D4A.7 |
| WBGene00021510 | 0.452648022 | 0.30614325 | 0.51508615 | frpr-19 |
| WBGene00021511 | -0.119092555 | 0.91883087 | 1 | Y41D4B.1 |
| WBGene00021512 | -0.174611412 | 0.45007528 | 0.64882023 | Y41D4B.4 |
| WBGene00021513 | -0.513389743 | 0.47454618 | 0.66760894 | Y41D4B.6 |
| WBGene00021514 | -0.459077806 | 0.03726967 | 0.13111312 | Y41D4B.11 |
| WBGene00021515 | -0.458878746 | 0.12674568 | 0.29717389 | set-23 |
| WBGene00021516 | -2.166091777 | 0.00610299 | 0.03609177 | Y41D4B.14 |
| WBGene00021517 | 0.820886723 | 0.57153931 | 1 | Y41D4B.15 |

| | | | | |
|---|---|---|---|---|
| WBGene00021518 | 0.63701042 | 0.01313101 | 0.06328845 | hpo-6 |
| WBGene00021519 | 0.720756995 | 0.35476494 | 0.56310643 | Y41D4B.17 |
| WBGene00021521 | 0.688793783 | 0.86378755 | 1 | nhr-287 |
| WBGene00021522 | 0.214419528 | 0.37960852 | 0.58763572 | nhr-274 |
| WBGene00021524 | 2.480616666 | 0.15096795 | 1 | Y41D4B.24 |
| WBGene00021525 | -0.752689514 | 0.35725612 | 0.56561018 | Y41D4B.26 |
| WBGene00021526 | -2.061536526 | 0.21538967 | 1 | Y41G9A.2 |
| WBGene00021527 | 0.631451219 | 0.24629831 | 0.45424696 | aakg-3 |
| WBGene00021528 | 1.063373458 | 0.00357843 | 0.02420376 | gbb-1 |
| WBGene00021529 | 0.45562919 | 0.45847795 | 0.65483658 | Y41G9A.5 |
| WBGene00021530 | 1.557897343 | 0.63813005 | 1 | Y41G9A.6 |
| WBGene00021531 | 0.833375868 | 0.17146663 | 0.36066532 | Y42G9A.1 |
| WBGene00021532 | 0.655464092 | 0.16739451 | 0.35486934 | Y42G9A.2 |
| WBGene00021533 | 1.430493518 | 9.83E-07 | 3.3346E-05 | Y42G9A.3 |
| WBGene00021534 | 0.392642615 | 0.0263042 | 0.10362078 | mvk-1 |
| WBGene00021535 | 0.083078295 | 0.73058208 | 0.84920987 | wht-7 |
| WBGene00021536 | -0.091615732 | 0.65178121 | 0.79940152 | Y42H9AR.1 |
| WBGene00021537 | -0.008677305 | 0.99719169 | 1 | Y42H9AR.2 |
| WBGene00021538 | 0.257886551 | 0.33510007 | 0.54330693 | rabs-5 |
| WBGene00021539 | 0.168915408 | 0.35304636 | 0.56194939 | Y42H9AR.4 |
| WBGene00021540 | 0.137790577 | 0.91190731 | 1 | Y42H9AR.5 |
| WBGene00021541 | -0.389211603 | 0.04946728 | 0.15947139 | Y42H9B.3 |
| WBGene00021542 | -2.772299933 | 2.3204E-05 | 0.00045733 | Y43B11AL.1 |
| WBGene00021543 | -0.046844478 | 0.88492133 | 0.93922806 | Y43B11AR.1 |
| WBGene00021544 | 0.269978304 | 0.58614345 | 0.75553325 | Y43B11AR.3 |
| WBGene00021545 | -0.362905057 | 0.04421237 | 0.14714429 | Y43H11AL.1 |
| WBGene00021546 | 0.145688999 | 0.53045867 | 0.71443643 | laat-1 |
| WBGene00021547 | -0.224418745 | 0.75751294 | 0.86445762 | Y44E3A.1 |
| WBGene00021548 | -0.786863082 | 0.00116084 | 0.01028745 | trx-4 |
| WBGene00021549 | 0.327015396 | 0.1554572 | 0.33928079 | Y44E3A.4 |
| WBGene00021551 | -0.186059942 | 0.34042616 | 0.54881344 | Y44E3A.6 |
| WBGene00021552 | -0.279939021 | 0.1787966 | 0.3708553 | zip-4 |
| WBGene00021553 | 0.32750836 | 0.88365005 | 1 | tyr-5 |
| WBGene00021554 | -0.38424289 | 0.05738381 | 0.17727949 | Y45G5AL.1 |
| WBGene00021555 | -0.29472885 | 0.26573219 | 0.47437944 | Y45G5AM.2 |
| WBGene00021556 | 0.395164191 | 0.34243616 | 0.55068696 | Y45G5AM.3 |
| WBGene00021557 | -1.67491251 | 0.01083158 | 0.05513886 | Y45G5AM.5 |
| WBGene00021558 | 0.058179898 | 0.89728071 | 0.94634211 | Y45G5AM.6 |
| WBGene00021559 | -0.310211278 | 0.0930143 | 0.24311875 | Y45G5AM.7 |
| WBGene00021560 | -0.175155096 | 0.55088767 | 0.72945002 | coh-4 |
| WBGene00021561 | -0.217861177 | 0.30313176 | 0.51221193 | Y45G5AM.9 |
| WBGene00021562 | -0.152747186 | 0.38201782 | 0.59023649 | nuo-5 |
| WBGene00021563 | -0.218509807 | 0.43921988 | 0.63970233 | Y45G12B.2 |

| | | | | |
|---|---|---|---|---|
| WBGene00021564 | -0.574267181 | 0.03754399 | 0.13166425 | Y45G12B.3 |
| WBGene00021565 | 0.112912723 | 0.88270769 | 0.93749089 | Y45G12C.1 |
| WBGene00021573 | -1.969805064 | 0.62862569 | 1 | Y45G12C.10 |
| WBGene00021576 | -1.619937623 | 1.2158E-06 | 3.9364E-05 | fbxc-51 |
| WBGene00021579 | 2.304382448 | 0.18140828 | 1 | clec-73 |
| WBGene00021580 | 0.184237415 | 0.91602174 | 1 | clec-174 |
| WBGene00021581 | 3.095096039 | 0.00014876 | 0.0020892 | clec-70 |
| WBGene00021582 | 4.529251747 | 0.00073527 | 0.00723997 | clec-71 |
| WBGene00021583 | 2.142669825 | 5.32E-16 | 1.15E-13 | clec-72 |
| WBGene00021585 | 1.451224364 | 0.19813658 | 1 | clec-74 |
| WBGene00021586 | 2.474950664 | 0.09720588 | 1 | clec-75 |
| WBGene00021587 | 5.35570844 | 2.6705E-05 | 0.00051184 | clec-76 |
| WBGene00021589 | -0.495166339 | 0.59956921 | 1 | Y46D2A.1 |
| WBGene00021590 | -0.69612864 | 0.07699931 | 0.2153533 | Y46D2A.2 |
| WBGene00021593 | -1.616437939 | 0.36625847 | 1 | Y46E12A.3 |
| WBGene00021594 | -0.143662227 | 0.92644295 | 1 | tig-3 |
| WBGene00021595 | 0.133439305 | 0.55686044 | 0.73399659 | Y46E12BL.2 |
| WBGene00021596 | -0.590224166 | 0.0008179 | 0.00786852 | spsb-2 |
| WBGene00021597 | 1.613567416 | 0.19642267 | 1 | Y46E12BR.1 |
| WBGene00021597 | -0.034606668 | 0.86999067 | 0.92996626 | spsb-1 |
| WBGene00021599 | 0.694609957 | 0.73912856 | 1 | srt-42 |
| WBGene00021600 | 0.587745431 | 0.73785047 | 1 | Y46H3A.4 |
| WBGene00021601 | 0.983488853 | 0.80601565 | 1 | Y46H3A.5 |
| WBGene00021602 | 1.18012455 | 0.11086613 | 0.27238394 | clec-205 |
| WBGene00021603 | 2.179846409 | 0.019705 | 0.08459634 | clec-204 |
| WBGene00021605 | -0.620593182 | 0.17499375 | 0.36530245 | Y46H3C.5 |
| WBGene00021607 | 0.36516824 | 0.60557564 | 0.76897128 | Y46H3C.7 |
| WBGene00021609 | 0.377828795 | 0.92526585 | 1 | irld-17 |
| WBGene00021610 | 0.155166345 | 0.86681932 | 0.92821073 | nhr-237 |
| WBGene00021611 | -0.652403922 | 0.37488051 | 0.58284342 | nhr-238 |
| WBGene00021613 | 0.151263518 | 0.54377447 | 0.72337864 | ttc-37 |
| WBGene00021614 | 0.668022947 | 0.02542672 | 0.10128143 | Y47A7.2 |
| WBGene00021615 | -0.156269293 | 0.96924949 | 1 | Y47C4A.1 |
| WBGene00021619 | 0.983488853 | 0.80601565 | 1 | Y47D7A.6 |
| WBGene00021621 | 0.377828795 | 0.92526585 | 1 | Y47D7A.9 |
| WBGene00021624 | 0.122847948 | 0.90626616 | 1 | Y47D7A.12 |
| WBGene00021625 | -1.151225995 | 0.0005818 | 0.00599501 | Y47D7A.13 |
| WBGene00021626 | 0.216781695 | 0.48328083 | 0.67431741 | rft-2 |
| WBGene00021627 | 1.16246618 | 0.55673752 | 1 | Y47D7A.15 |
| WBGene00021628 | -0.050096942 | 0.78818725 | 0.88342616 | Y47D9A.1 |
| WBGene00021629 | 0.670654238 | 0.06342621 | 0.18911042 | scpl-3 |
| WBGene00021632 | -0.156269293 | 0.96924949 | 1 | Y47D9A.5 |
| WBGene00021633 | -0.973396117 | 0.50526017 | 1 | Y47G6A.3 |

| | | | | |
|---|---|---|---|---|
| WBGene00021634 | -0.217989888 | 0.42433012 | 0.62859321 | rde-10 |
| WBGene00021635 | 1.195266385 | 0.00093939 | 0.00877989 | Y47G6A.5 |
| WBGene00021636 | -0.378778562 | 0.05094755 | 0.16277741 | pcaf-1 |
| WBGene00021637 | 0.353742489 | 0.24829221 | 0.45646816 | Y47G6A.7 |
| WBGene00021638 | -0.624348322 | 0.02369605 | 0.09660637 | Y47G6A.9 |
| WBGene00021639 | 0.40388745 | 0.75031127 | 1 | Y47G6A.13 |
| WBGene00021640 | -0.634421901 | 0.01607857 | 0.07313973 | Y47G6A.14 |
| WBGene00021641 | 0.911853785 | 0.52700036 | 1 | Y47G6A.15 |
| WBGene00021643 | 0.64293475 | 0.37596963 | 0.58371449 | ccep-290 |
| WBGene00021644 | -0.163170139 | 0.55372709 | 0.73145475 | Y47G6A.18 |
| WBGene00021645 | 0.398835193 | 0.19133479 | 0.38613625 | Y47G6A.19 |
| WBGene00021646 | -0.272844689 | 0.40717861 | 0.61394145 | Y47G6A.21 |
| WBGene00021647 | -0.208412108 | 0.40158746 | 0.60923071 | Y47G6A.22 |
| WBGene00021648 | -0.706321861 | 0.00274625 | 0.01992322 | mis-12 |
| WBGene00021649 | -0.117612225 | 0.64120702 | 0.79235567 | Y47G6A.25 |
| WBGene00021650 | 1.38322681 | 0.69550305 | 1 | Y47G6A.26 |
| WBGene00021653 | 2.166011856 | 0.42253177 | 1 | Y47G6A.30 |
| WBGene00021654 | -0.746865228 | 0.40925274 | 0.61485082 | Y47G7B.2 |
| WBGene00021655 | 0.126558089 | 0.65717122 | 0.80268551 | Y48A5A.1 |
| WBGene00021656 | -1.2117623 | 0.45349912 | 1 | Y48D7A.1 |
| WBGene00021657 | 0.310688387 | 0.35339625 | 0.56208019 | Y48G1A.1 |
| WBGene00021658 | -0.541617961 | 0.01766174 | 0.07828812 | Y48G1A.2 |
| WBGene00021660 | -0.188879171 | 0.45657784 | 0.65378049 | Y48G1A.4 |
| WBGene00021661 | -0.334734944 | 0.24226175 | 0.44981194 | mbtr-1 |
| WBGene00021662 | -0.009355551 | 0.98901069 | 0.99484196 | snpc-3.3 |
| WBGene00021666 | 0.758205189 | 0.50667852 | 1 | Y48G1BL.7 |
| WBGene00021667 | -0.731913521 | 0.28700575 | 0.49711057 | snpc-3.2 |
| WBGene00021671 | 2.314239469 | 0.00331412 | 0.02279703 | Y48G1BM.6 |
| WBGene00021675 | 0.223607364 | 0.60568918 | 0.76897128 | Y48G1BR.1 |
| WBGene00021676 | -0.838406362 | 0.00171145 | 0.01378814 | Y48G1C.1 |
| WBGene00021677 | 0.081802374 | 0.77038166 | 0.87152476 | pgs-1 |
| WBGene00021678 | -0.549786194 | 0.76687564 | 1 | Y48G1C.5 |
| WBGene00021679 | 0.118408816 | 0.86725575 | 0.92835056 | Y48G1C.6 |
| WBGene00021681 | -0.131856451 | 0.5061128 | 0.69290883 | Y48G1C.8 |
| WBGene00021682 | -0.561700476 | 0.05316888 | 0.16749439 | Y48G1C.9 |
| WBGene00021683 | 0.153299455 | 0.53642855 | 0.71845329 | Y48G1C.10 |
| WBGene00021685 | 0.153421638 | 0.49003556 | 0.67947941 | herc-1 |
| WBGene00021686 | -0.069149159 | 0.70914052 | 0.83533552 | Y48G8AL.5 |
| WBGene00021687 | -0.033258833 | 0.92668925 | 0.96145395 | Y48G8AL.7 |
| WBGene00021689 | 0.058463149 | 0.82117348 | 0.90298589 | Y48G8AL.10 |
| WBGene00021690 | -0.848512853 | 0.76804407 | 1 | Y48G8AL.12 |
| WBGene00021691 | 0.194277753 | 0.27886605 | 0.48770698 | Y48G8AL.13 |
| WBGene00021692 | -0.604094735 | 0.18911928 | 0.38309494 | Y48G8AL.15 |

| | | | | |
|---|---|---|---|---|
| WBGene00021693 | 0.967296459 | 0.26341136 | 0.47204093 | Y48G8AR.2 |
| WBGene00021694 | -0.156269293 | 0.96924949 | 1 | Y48G8AR.3 |
| WBGene00021697 | 0.610943007 | 0.17450331 | 0.36465486 | gcn-1 |
| WBGene00021698 | 0.165272499 | 0.57588699 | 0.74859142 | frl-1 |
| WBGene00021700 | -0.156269293 | 0.96924949 | 1 | Y48G9A.6 |
| WBGene00021701 | -1.930411901 | 0.43097183 | 1 | Y48G9A.7 |
| WBGene00021702 | 0.767370534 | 0.00753985 | 0.04196006 | Y48G9A.9 |
| WBGene00021703 | -0.109254073 | 0.95448982 | 1 | cpt-3 |
| WBGene00021704 | -0.591901847 | 0.13006883 | 0.30159772 | Y48G9A.11 |
| WBGene00021706 | -0.156269293 | 0.96924949 | 1 | srbc-39 |
| WBGene00021709 | 0.243080809 | 0.47524071 | 0.66814989 | ugt-29 |
| WBGene00021710 | 0.047369385 | 0.98034056 | 1 | cyp-33C11 |
| WBGene00021711 | 0.182962122 | 0.82517205 | 0.90582404 | wago-11 |
| WBGene00021713 | -2.26761602 | 0.0391502 | 1 | Y49F6A.5 |
| WBGene00021714 | -0.272489992 | 0.26860503 | 0.4771227 | cyh-1 |
| WBGene00021715 | -0.4341549 | 0.1563166 | 0.34035865 | Y49F6B.2 |
| WBGene00021716 | -1.930411901 | 0.43097183 | 1 | fbxb-112 |
| WBGene00021717 | -0.737462795 | 0.83433562 | 1 | sdz-32 |
| WBGene00021718 | -0.156269293 | 0.96924949 | 1 | Y49F6B.6 |
| WBGene00021719 | -2.235402002 | 0.01250434 | 0.0610435 | fbxb-104 |
| WBGene00021720 | 0.88787957 | 0.71832755 | 1 | Y49F6B.8 |
| WBGene00021721 | -0.783363348 | 0.00271493 | 0.01974956 | Y49F6B.9 |
| WBGene00021722 | 0.747845955 | 0.85215965 | 1 | srz-5 |
| WBGene00021724 | -0.156269293 | 0.96924949 | 1 | Y49F6C.2 |
| WBGene00021725 | -0.663147742 | 0.15122522 | 0.33308024 | bath-9 |
| WBGene00021726 | -1.076384988 | 0.10618291 | 0.26553383 | bath-10 |
| WBGene00021727 | -1.13818048 | 0.46440285 | 1 | bath-23 |
| WBGene00021728 | -0.668264701 | 0.85551153 | 1 | Y49F6C.6 |
| WBGene00021730 | -0.915422167 | 0.00133738 | 0.01146693 | Y49F6C.8 |
| WBGene00021731 | -0.296992071 | 0.73809327 | 0.85392243 | Y49G5A.1 |
| WBGene00021732 | 0.089130389 | 0.70974889 | 0.83571538 | Y49G5B.1 |
| WBGene00021733 | 0.007489925 | 0.97265338 | 0.98580722 | otub-2 |
| WBGene00021734 | 0.515748872 | 0.73527151 | 1 | Y50C1A.2 |
| WBGene00021735 | 0.033258953 | 0.88280148 | 0.93749089 | Y50D4A.1 |
| WBGene00021736 | -0.348253857 | 0.23481238 | 0.44071175 | wrb-1 |
| WBGene00021738 | -0.232111914 | 0.37058502 | 0.57810222 | Y50D4A.4 |
| WBGene00021739 | -0.332347994 | 0.56967393 | 0.74425222 | Y50D4A.5 |
| WBGene00021740 | 0.18874458 | 0.67228458 | 0.81256417 | Y50D4B.1 |
| WBGene00021741 | 1.116590956 | 0.42600377 | 1 | Y50D4B.2 |
| WBGene00021743 | 0.862942979 | 0.0001403 | 0.00199922 | Y50D4B.4 |
| WBGene00021745 | 0.50132733 | 0.48140062 | 0.67289112 | Y50D4B.6 |
| WBGene00021746 | 1.247575581 | 0.7549672 | 1 | Y50D4B.7 |
| WBGene00021748 | -0.089098386 | 0.76261555 | 0.86718678 | Y50D4C.3 |

| | | | | |
|---|---|---|---|---|
| WBGene00021749 | -0.196144304 | 0.49599726 | 0.68438489 | Y50D4C.5 |
| WBGene00021750 | -0.180825606 | 0.64739239 | 0.79651308 | Y50D4C.6 |
| WBGene00021751 | 0.656446674 | 0.08773554 | 0.23472834 | Y50D7A.1 |
| WBGene00021752 | -0.315169904 | 0.43708408 | 0.63789372 | Y50D7A.11 |
| WBGene00021752 | -0.18875678 | 0.4137743 | 0.61934132 | xpd-1 |
| WBGene00021753 | -0.127934759 | 0.53311201 | 0.71645202 | Y50D7A.3 |
| WBGene00021754 | 0.155844807 | 0.43483273 | 0.63614305 | hpo-29 |
| WBGene00021755 | 0.715242886 | 0.63175714 | 1 | hpo-38 |
| WBGene00021756 | 0.003321295 | 0.98934698 | 0.99484196 | Y50D7A.8 |
| WBGene00021757 | -0.587668696 | 0.03294537 | 0.12116257 | taco-1 |
| WBGene00021758 | -0.446521614 | 0.12593985 | 0.29604509 | Y50D7A.10 |
| WBGene00021763 | -0.572761639 | 0.00030105 | 0.00361169 | Y51F10.2 |
| WBGene00021764 | -0.21979729 | 0.35190686 | 0.5606957 | Y51F10.3 |
| WBGene00021765 | -0.00151689 | 0.99525476 | 0.99756601 | Y51F10.4 |
| WBGene00021766 | -0.018587454 | 0.93865837 | 0.96849251 | hex-4 |
| WBGene00021768 | 0.699646436 | 1.5696E-06 | 4.7405E-05 | Y51F10.7 |
| WBGene00021771 | -0.037818922 | 0.87831932 | 0.93467217 | spe-48 |
| WBGene00021772 | 1.20005961 | 0.05686014 | 0.17612653 | math-45 |
| WBGene00021773 | -0.281965203 | 0.78460544 | 0.88145597 | fbxb-42 |
| WBGene00021774 | -1.654668634 | 0.13970342 | 1 | fbxb-43 |
| WBGene00021776 | 1.46692992 | 0.27429646 | 1 | Y51H7BR.4 |
| WBGene00021778 | 1.018035489 | 0.62989118 | 1 | Y51H7BR.7 |
| WBGene00021779 | -2.990601006 | 2.91E-09 | 2.04E-07 | Y51H7C.1 |
| WBGene00021780 | 0.281454817 | 0.94435481 | 1 | scl-17 |
| WBGene00021781 | -0.811260545 | 0.00023646 | 0.00298118 | Y51H7C.3 |
| WBGene00021783 | -0.040212022 | 0.8680153 | 0.9285087 | Y51H7C.5 |
| WBGene00021784 | -0.053387348 | 0.77555352 | 0.87476731 | cogc-4 |
| WBGene00021785 | -1.03449136 | 0.0002701 | 0.00333263 | Y51H7C.7 |
| WBGene00021786 | 0.377828795 | 0.92526585 | 1 | Y51H7C.8 |
| WBGene00021787 | -0.558804446 | 0.87818675 | 1 | Y51H7C.9 |
| WBGene00021788 | 0.489058549 | 0.85947456 | 1 | Y51H7C.10 |
| WBGene00021789 | 0.452605678 | 0.16182169 | 0.34773047 | nol-6 |
| WBGene00021790 | 0.608174198 | 0.73197901 | 1 | Y51H7C.12 |
| WBGene00021791 | 0.528147961 | 0.81036933 | 1 | Y51H7C.13 |
| WBGene00021793 | -0.156269293 | 0.96924949 | 1 | Y52D5A.2 |
| WBGene00021794 | -0.559340474 | 0.78183513 | 1 | Y52E8A.1 |
| WBGene00021795 | -0.771480398 | 0.0080017 | 0.0439088 | Y52E8A.2 |
| WBGene00021796 | 0.783012491 | 0.16898775 | 0.357109 | Y52E8A.3 |
| WBGene00021797 | -0.148073308 | 0.89226044 | 1 | plep-1 |
| WBGene00021798 | 0.336933409 | 0.86434725 | 1 | Y52E8A.6 |
| WBGene00021799 | 0.607124613 | 0.51680684 | 0.70281311 | Y53G8AL.1 |
| WBGene00021800 | -0.208682142 | 0.20978937 | 0.41007351 | Y53G8AL.2 |
| WBGene00021801 | 0.281454817 | 0.94435481 | 1 | Y53G8AL.3 |

| | | | | |
|---|---|---|---|---|
| WBGene00021804 | -0.06995528 | 0.97031639 | 1 | Y53G8AM.4 |
| WBGene00021807 | 0.339336785 | 0.86985627 | 1 | Y53G8AM.7 |
| WBGene00021808 | 0.148444792 | 0.66476631 | 0.80771846 | Y53G8AM.8 |
| WBGene00021809 | -0.263229581 | 0.93409872 | 1 | Y53G8AR.1 |
| WBGene00021810 | -0.157330977 | 0.59468008 | 0.76112043 | phf-15 |
| WBGene00021811 | -0.018506061 | 0.9363567 | 0.96685718 | ral-1 |
| WBGene00021812 | -0.405963659 | 0.22447749 | 0.42858644 | Y53G8AR.5 |
| WBGene00021813 | -0.73212415 | 0.00108302 | 0.00977557 | Y53G8AR.6 |
| WBGene00021814 | 1.575778556 | 8.2376E-06 | 0.0001927 | Y53G8AR.7 |
| WBGene00021815 | -0.083841078 | 0.78701565 | 0.88269349 | Y53G8AR.8 |
| WBGene00021816 | -0.667681322 | 0.00167625 | 0.01359477 | Y53G8AR.9 |
| WBGene00021817 | -0.384310918 | 0.3780023 | 0.58562259 | Y53G8B.1 |
| WBGene00021818 | -0.631170013 | 0.06163867 | 0.18583439 | Y53G8B.2 |
| WBGene00021820 | 0.378029453 | 0.16456831 | 0.35089987 | nipa-1 |
| WBGene00021821 | -0.156269293 | 0.96924949 | 1 | Y54B9A.1 |
| WBGene00021826 | -0.415489832 | 0.01805623 | 0.07957195 | txl-1 |
| WBGene00021827 | -0.902715336 | 0.00022728 | 0.00290705 | dnc-6 |
| WBGene00021828 | -0.334588623 | 0.12726287 | 0.29779878 | Y54E10A.6 |
| WBGene00021829 | -0.129065855 | 0.64334337 | 0.7939451 | mrpl-17 |
| WBGene00021830 | -0.46756372 | 0.04004104 | 0.13756036 | Y54E10A.10 |
| WBGene00021831 | 0.405411064 | 0.0314588 | 0.11752038 | Y54E10A.11 |
| WBGene00021832 | 0.062642669 | 0.77801612 | 0.87653067 | Y54E10A.12 |
| WBGene00021834 | 0.532229165 | 0.02280342 | 0.09410478 | mab-31 |
| WBGene00021835 | -0.684087513 | 0.02232105 | 0.09283998 | Y54E10A.17 |
| WBGene00021838 | 1.233111853 | 0.04842229 | 0.15761262 | Y54E10BL.3 |
| WBGene00021839 | -0.683628625 | 0.0001095 | 0.00164131 | nduf-5 |
| WBGene00021840 | -0.089087179 | 0.73661477 | 0.85283726 | pign-1 |
| WBGene00021841 | -0.843287478 | 0.00100876 | 0.00929991 | Y54E10BR.2 |
| WBGene00021842 | -0.272483845 | 0.17274146 | 0.36240783 | Y54E10BR.3 |
| WBGene00021843 | -0.256889991 | 0.20319863 | 0.4023709 | Y54E10BR.4 |
| WBGene00021844 | -0.650668001 | 0.00017172 | 0.00233738 | Y54E10BR.5 |
| WBGene00021845 | -0.710696902 | 0.00061808 | 0.00627963 | rpb-7 |
| WBGene00021846 | -0.047661491 | 0.84547833 | 0.91536929 | ztf-23 |
| WBGene00021847 | -0.207756895 | 0.33044498 | 0.53866122 | Y54F10AL.1 |
| WBGene00021848 | -0.44901409 | 0.81158837 | 1 | nhr-239 |
| WBGene00021849 | -0.447397934 | 0.01486465 | 0.06918667 | Y54F10AM.5 |
| WBGene00021850 | -0.058013543 | 0.94299276 | 0.97049038 | Y54F10AM.6 |
| WBGene00021852 | 0.294140608 | 0.07822066 | 0.21746833 | Y54F10AM.8 |
| WBGene00021853 | -0.705016711 | 0.0254246 | 0.10128143 | Y54F10AM.11 |
| WBGene00021854 | -0.069857354 | 0.78971072 | 0.88416432 | Y54F10AR.1 |
| WBGene00021855 | -0.170896844 | 0.5026903 | 0.68999878 | Y54F10AR.2 |
| WBGene00021856 | -0.223205833 | 0.37099612 | 0.57843379 | Y54F10BM.1 |
| WBGene00021857 | 0.001103092 | 0.99522417 | 0.99756601 | iffb-1 |

| | | | | |
|---|---|---|---|---|
| WBGene00021858 | -0.214070851 | 0.80632041 | 0.89451171 | Y54F10BM.3 |
| WBGene00021859 | -0.002418078 | 0.9989123 | 1 | fbxa-39 |
| WBGene00021860 | 0.256800144 | 0.88423831 | 1 | fbxa-9 |
| WBGene00021861 | -1.971285244 | 0.10320259 | 1 | Y54F10BM.6 |
| WBGene00021862 | 1.944847331 | 0.16620737 | 1 | fbxa-48 |
| WBGene00021863 | 0.237511534 | 0.47364862 | 0.66657794 | Y54F10BM.9 |
| WBGene00021864 | 2.456496557 | 0.3218863 | 1 | fbxa-1 |
| WBGene00021865 | 0.39748245 | 0.82716117 | 1 | fbxa-66 |
| WBGene00021866 | 1.293232493 | 0.26082095 | 1 | Y54F10BM.12 |
| WBGene00021867 | 0.10322898 | 0.86640817 | 0.92799221 | Y54F10BM.13 |
| WBGene00021868 | 0.172506246 | 0.3841467 | 0.59269604 | atln-1 |
| WBGene00021869 | -0.628750236 | 0.00035088 | 0.00410806 | vrp-1 |
| WBGene00021870 | -0.17833704 | 0.69739437 | 0.82880433 | samt-1 |
| WBGene00021871 | 0.103564962 | 0.93905252 | 1 | dml-1 |
| WBGene00021872 | 0.645386779 | 0.00214459 | 0.01641257 | clec-85 |
| WBGene00021873 | 3.077401726 | 0.05197679 | 1 | clec-82 |
| WBGene00021875 | 1.185799483 | 0.02268436 | 0.09374091 | Y54G2A.10 |
| WBGene00021876 | 1.138655785 | 3.62E-09 | 2.47E-07 | Y54G2A.11 |
| WBGene00021877 | -0.267758072 | 0.32183423 | 0.52995923 | Y54G2A.12 |
| WBGene00021878 | 0.233739675 | 0.93575889 | 1 | Y54G2A.13 |
| WBGene00021879 | 0.830415028 | 0.00013293 | 0.00192133 | clec-83 |
| WBGene00021880 | 2.421662425 | 0.16598337 | 1 | Y54G2A.15 |
| WBGene00021881 | 0.17652465 | 0.83667085 | 0.9111967 | Y54G2A.16 |
| WBGene00021882 | -0.14507558 | 0.46410416 | 0.65886864 | nprt-1 |
| WBGene00021883 | -0.413796889 | 0.01857924 | 0.08125742 | Y54G2A.18 |
| WBGene00021884 | -0.087957271 | 0.89524366 | 0.94481514 | Y54G2A.19 |
| WBGene00021887 | 0.2257046 | 0.52063383 | 0.70562796 | fbxc-45 |
| WBGene00021888 | -0.576555441 | 0.0008194 | 0.00787671 | Y54G2A.23 |
| WBGene00021890 | 0.476547615 | 0.02223737 | 0.09268245 | Y54G2A.26 |
| WBGene00021892 | -0.107268465 | 0.96117914 | 1 | Y54G2A.28 |
| WBGene00021893 | 1.064434301 | 0.03304821 | 0.12133001 | ndnf-1 |
| WBGene00021894 | 0.959153793 | 0.02075309 | 0.08779208 | Y54G2A.32 |
| WBGene00021895 | 1.566767463 | 6.50E-07 | 2.3824E-05 | clec-84 |
| WBGene00021897 | 0.159189555 | 0.85498554 | 0.92098123 | ser-6 |
| WBGene00021898 | 0.369551553 | 0.39094382 | 0.59937478 | Y54G2A.36 |
| WBGene00021899 | -0.315573581 | 0.0720622 | 0.2064434 | Y54H5A.1 |
| WBGene00021900 | -0.186799181 | 0.37750457 | 0.58512604 | Y54H5A.2 |
| WBGene00021901 | -0.386512064 | 0.02759896 | 0.10712245 | tag-262 |
| WBGene00021902 | 0.240044876 | 0.38848247 | 0.59718482 | oxy-4 |
| WBGene00021904 | -0.640951725 | 0.0266553 | 0.10458568 | Y55B1AL.2 |
| WBGene00021905 | 0.003774384 | 0.9906251 | 0.99548813 | helq-1 |
| WBGene00021906 | 0.058944403 | 0.83309748 | 0.90967142 | Y55B1AR.2 |
| WBGene00021907 | -2.640414922 | 0.41081194 | 1 | Y55B1AR.3 |

| | | | | |
|---|---|---|---|---|
| WBGene00021908 | 1.38322681 | 0.69550305 | 1 | Y55B1AR.4 |
| WBGene00021909 | 0.894575739 | 0.00203116 | 0.01572727 | Y55B1BL.1 |
| WBGene00021910 | 0.125350929 | 0.5240646 | 0.70877474 | stim-1 |
| WBGene00021911 | -0.325038374 | 0.87140873 | 1 | Y55B1BR.1 |
| WBGene00021912 | 0.152739107 | 0.67205673 | 0.81248091 | Y55B1BR.2 |
| WBGene00021913 | -1.033465769 | 0.00145831 | 0.01229536 | cec-8 |
| WBGene00021914 | 0.057220464 | 0.88063353 | 0.93631418 | magu-1 |
| WBGene00021915 | 0.554792401 | 0.35995057 | 0.56806965 | Y55D5A.1 |
| WBGene00021917 | 0.172846473 | 0.82008517 | 0.90260666 | Y55D5A.3 |
| WBGene00021918 | -0.161098318 | 0.81646346 | 0.90041629 | Y55D5A.4 |
| WBGene00021919 | -0.671949688 | 0.53005324 | 0.71408971 | cutl-25 |
| WBGene00021920 | -0.204392846 | 0.44015048 | 0.64021408 | mrps-25 |
| WBGene00021921 | -0.441800489 | 0.0192732 | 0.08345074 | Y55F3AM.3 |
| WBGene00021922 | -0.537989779 | 0.02834442 | 0.10903979 | atg-3 |
| WBGene00021923 | -0.152318696 | 0.66494902 | 0.80771846 | Y55F3AM.5 |
| WBGene00021925 | -0.096216986 | 0.80201947 | 0.89160777 | immp-2 |
| WBGene00021926 | -0.231385367 | 0.27609699 | 0.48475684 | Y55F3AM.9 |
| WBGene00021927 | 0.580911117 | 0.3420393 | 0.55039087 | abhd-14 |
| WBGene00021928 | -0.805822946 | 0.09195275 | 0.24138342 | Y55F3AM.11 |
| WBGene00021929 | -0.441102925 | 0.04000982 | 0.13753092 | dcap-1 |
| WBGene00021930 | -0.654087787 | 0.00069613 | 0.00695929 | Y55F3AM.13 |
| WBGene00021931 | -1.504331994 | 0.33682241 | 1 | Y55F3AM.14 |
| WBGene00021932 | -0.033091387 | 0.92539696 | 0.96044148 | Y55F3AR.1 |
| WBGene00021933 | -0.109925725 | 0.67069641 | 0.81161419 | Y55F3AR.2 |
| WBGene00021934 | -0.269145055 | 0.12589177 | 0.2960395 | cct-8 |
| WBGene00021935 | -0.477359951 | 0.05179312 | 0.1644251 | mrpl-46 |
| WBGene00021936 | -0.280298477 | 0.16445229 | 0.3507757 | zipt-20 |
| WBGene00021938 | -0.654098622 | 0.00073927 | 0.00726759 | Y55F3BR.1 |
| WBGene00021939 | 0.720757953 | 0.07998083 | 0.22093964 | Y55F3BR.2 |
| WBGene00021941 | 0.359889028 | 0.70573416 | 1 | lgc-33 |
| WBGene00021942 | -0.137106801 | 0.86379072 | 0.92668707 | madf-1 |
| WBGene00021943 | -0.393339583 | 0.15342389 | 0.33610961 | Y55F3BR.6 |
| WBGene00021945 | -0.615529815 | 0.01639261 | 0.07414228 | lem-4 |
| WBGene00021947 | -2.563099533 | 0.4751044 | 1 | srt-23 |
| WBGene00021948 | 1.000214361 | 0.00036143 | 0.00419513 | kvs-5 |
| WBGene00021950 | 0.193985961 | 0.65336751 | 0.80066769 | cutl-26 |
| WBGene00021951 | 0.688793783 | 0.86378755 | 1 | srt-24 |
| WBGene00021952 | 0.546751376 | 0.00057303 | 0.00593111 | vha-19 |
| WBGene00021953 | 1.181686151 | 0.7579682 | 1 | Y55H10A.2 |
| WBGene00021956 | 0.004182596 | 0.97725694 | 0.98824558 | Y57E12AL.1 |
| WBGene00021959 | -0.605186419 | 0.00034601 | 0.00406676 | Y57E12AL.6 |
| WBGene00021960 | -0.315946415 | 0.09374413 | 0.24455252 | Y57E12AM.1 |
| WBGene00021963 | -0.282833131 | 0.74062743 | 0.85525612 | lipl-6 |

| | | | | |
|---|---|---|---|---|
| WBGene00021964 | -1.969805064 | 0.62862569 | 1 | Y57E12B.4 |
| WBGene00021965 | -0.299699905 | 0.67107167 | 0.81182555 | Y57G7A.1 |
| WBGene00021966 | -1.425149271 | 0.05428692 | 0.17017751 | Y57G7A.2 |
| WBGene00021968 | 1.216957619 | 0.74819621 | 1 | Y57G7A.5 |
| WBGene00021969 | 1.337064607 | 0.41896226 | 1 | Y57G7A.6 |
| WBGene00021970 | -1.969805064 | 0.62862569 | 1 | srz-64 |
| WBGene00021972 | -0.224944172 | 0.88928469 | 1 | glb-31 |
| WBGene00021973 | -0.457834739 | 0.04293183 | 0.14436017 | emc-2 |
| WBGene00021974 | 0.822152909 | 0.47121303 | 1 | Y57G7A.11 |
| WBGene00021975 | -0.509883132 | 0.28400861 | 0.49393333 | Y58A7A.1 |
| WBGene00021976 | -0.806420134 | 0.54853072 | 1 | Y58A7A.2 |
| WBGene00021977 | 1.290284036 | 0.00624313 | 0.0367772 | Y58A7A.3 |
| WBGene00021978 | 2.486446643 | 0.19466437 | 1 | Y58A7A.4 |
| WBGene00021979 | 0.521794059 | 0.45560919 | 0.65284446 | Y58A7A.5 |
| WBGene00021980 | 0.480604309 | 0.46713241 | 0.66139412 | glb-32 |
| WBGene00021981 | 0.795471013 | 0.18028738 | 0.3725955 | lgc-26 |
| WBGene00021983 | 1.127935043 | 0.0062049 | 0.03662307 | npr-5 |
| WBGene00021984 | 0.546984433 | 0.1591011 | 0.34395415 | Y59C2A.1 |
| WBGene00021985 | -0.055356742 | 0.91076774 | 0.95381965 | C33C12.10 |
| WBGene00021985 | -0.007554225 | 0.97911746 | 0.98880927 | Y59C2A.2 |
| WBGene00021986 | 0.412584695 | 0.08676456 | 0.2329519 | Y59C2A.3 |
| WBGene00021990 | 2.269176518 | 0.38585039 | 1 | Y59E1B.1 |
| WBGene00021992 | -0.156269293 | 0.96924949 | 1 | thn-6 |
| WBGene00021993 | 0.399606396 | 0.86544406 | 1 | Y59E9AL.3 |
| WBGene00021994 | 0.514160815 | 0.32989515 | 0.53846654 | Y59E9AL.4 |
| WBGene00021996 | 1.515504912 | 0.31360311 | 1 | Y59E9AL.6 |
| WBGene00022004 | 1.235896359 | 0.08311248 | 0.22648553 | npr-22 |
| WBGene00022005 | 0.281454817 | 0.94435481 | 1 | Y59H11AM.1 |
| WBGene00022008 | 0.747845955 | 0.85215965 | 1 | Y59H11AM.4 |
| WBGene00022010 | 1.284384057 | 9.13E-09 | 5.72E-07 | catp-7 |
| WBGene00022012 | 0.526990615 | 0.27933518 | 0.48825345 | Y59H11AR.4 |
| WBGene00022013 | 0.109173684 | 0.96057644 | 1 | Y60C6A.1 |
| WBGene00022014 | 0.026098276 | 0.92644572 | 0.96128343 | Y61A9LA.1 |
| WBGene00022015 | 0.017704818 | 0.94033174 | 0.96916742 | Y61A9LA.3 |
| WBGene00022016 | 0.232157325 | 0.93397866 | 1 | Y61A9LA.4 |
| WBGene00022018 | 0.113920663 | 0.96058911 | 1 | Y61A9LA.7 |
| WBGene00022019 | -0.043959082 | 0.81240744 | 0.89762575 | sut-2 |
| WBGene00022020 | 0.983488853 | 0.80601565 | 1 | hpo-39 |
| WBGene00022021 | 0.143850375 | 0.49454358 | 0.68308599 | Y61A9LA.10 |
| WBGene00022022 | -0.232190053 | 0.49299652 | 0.68200961 | Y61A9LA.11 |
| WBGene00022024 | -0.271046643 | 0.75941903 | 0.86543035 | Y64H9A.2 |
| WBGene00022025 | -0.081152271 | 0.70377523 | 0.83263838 | Y65B4A.1 |
| WBGene00022026 | 0.822233781 | 0.09546957 | 0.24772303 | Y65B4A.2 |

| | | | | |
|---|---|---|---|---|
| WBGene00022027 | -0.292281949 | 0.20002341 | 0.39843876 | vps-20 |
| WBGene00022028 | 1.760161518 | 0.17895762 | 1 | Y65B4A.4 |
| WBGene00022029 | -0.676411032 | 0.00097545 | 0.0090681 | Y65B4A.6 |
| WBGene00022030 | 0.983488853 | 0.80601565 | 1 | Y65B4A.7 |
| WBGene00022031 | -0.096242825 | 0.74407417 | 0.8574226 | Y65B4A.8 |
| WBGene00022032 | 0.281454817 | 0.94435481 | 1 | Y65B4A.9 |
| WBGene00022033 | -0.233596886 | 0.51901491 | 0.70445818 | Y65B4BL.1 |
| WBGene00022034 | -0.446933041 | 0.01062721 | 0.05425553 | deps-1 |
| WBGene00022035 | 0.308378434 | 0.35897101 | 0.56718728 | Y65B4BL.3 |
| WBGene00022036 | -0.414601725 | 0.17425575 | 0.36445118 | Y65B4BL.4 |
| WBGene00022037 | -0.166399701 | 0.30569314 | 0.51458342 | acs-13 |
| WBGene00022038 | -0.447430607 | 0.74560996 | 1 | Y65B4BL.6 |
| WBGene00022039 | -0.023966599 | 0.96685274 | 0.9830824 | Y65B4BL.7 |
| WBGene00022040 | 1.470882346 | 0.08442136 | 0.22883673 | Y65B4BR.1 |
| WBGene00022042 | -0.594165224 | 1.2658E-05 | 0.00027691 | icd-2 |
| WBGene00022043 | -0.892085877 | 1.9277E-05 | 0.00039007 | psf-3 |
| WBGene00022044 | -0.357332674 | 0.12978283 | 0.30114118 | dpm-1 |
| WBGene00022045 | -0.491995055 | 0.13334167 | 0.30678581 | mrpl-55 |
| WBGene00022046 | -0.619496103 | 0.00041656 | 0.00464123 | Y66H1A.4 |
| WBGene00022047 | 0.40985641 | 0.13968173 | 0.31692692 | Y66H1A.5 |
| WBGene00022048 | 0.71168313 | 0.0002318 | 0.00294689 | fln-1 |
| WBGene00022051 | -0.214998498 | 0.4091764 | 0.61482805 | rep-1 |
| WBGene00022052 | 0.10334897 | 0.64657669 | 0.79583208 | Y67D2.2 |
| WBGene00022053 | -0.420328566 | 0.02607193 | 0.10304015 | cisd-3.2 |
| WBGene00022054 | 0.011792242 | 0.9756352 | 0.98726346 | Y67D2.4 |
| WBGene00022055 | -0.25780631 | 0.49256245 | 0.68158203 | Y67D2.5 |
| WBGene00022056 | -0.073762527 | 0.80751651 | 0.89513608 | ddx-35 |
| WBGene00022057 | -0.545672696 | 0.01707445 | 0.07632579 | Y67D2.7 |
| WBGene00022059 | 0.212130207 | 0.3128945 | 0.52103307 | Y67D8A.2 |
| WBGene00022060 | 0.468765004 | 0.19455384 | 0.39059777 | dmd-9 |
| WBGene00022061 | 3.255102787 | 0.07761964 | 1 | Y67D8B.1 |
| WBGene00022062 | 0.814428817 | 0.550444 | 1 | Y67D8B.2 |
| WBGene00022063 | -0.036441916 | 0.96947276 | 1 | Y67D8B.4 |
| WBGene00022064 | 0.640945584 | 0.76653737 | 1 | Y67D8B.5 |
| WBGene00022066 | -0.156269293 | 0.96924949 | 1 | Y67D8C.2 |
| WBGene00022067 | 0.324981052 | 0.27051206 | 0.47945795 | Y67D8C.3 |
| WBGene00022069 | 0.529957938 | 0.01560932 | 0.07165653 | eel-1 |
| WBGene00022070 | -1.831477136 | 0.52194793 | 1 | Y67D8C.6 |
| WBGene00022071 | -2.640414922 | 0.41081194 | 1 | Y67D8C.7 |
| WBGene00022072 | -0.905573069 | 0.00274701 | 0.01992322 | cpg-9 |
| WBGene00022073 | 0.430496297 | 0.24372334 | 0.45114272 | Y67D8C.9 |
| WBGene00022074 | -0.422258416 | 0.02383859 | 0.09699207 | Y69A2AR.1 |
| WBGene00022075 | -1.252025351 | 9.18E-09 | 5.72E-07 | Y69A2AR.3 |

| | | | | |
|---|---|---|---|---|
| WBGene00022076 | -0.051936793 | 0.94366373 | 0.97098542 | daao-1 |
| WBGene00022077 | -0.319582354 | 0.33821688 | 0.54612198 | vamp-7 |
| WBGene00022078 | 0.163812964 | 0.57705734 | 0.74903703 | epg-9 |
| WBGene00022083 | 0.707008864 | 0.58571125 | 1 | Y69A2AR.12 |
| WBGene00022086 | -0.016115726 | 0.99019285 | 1 | sprr-3 |
| WBGene00022087 | -0.043141492 | 0.84316819 | 0.91377278 | Y69A2AR.16 |
| WBGene00022089 | -0.127784343 | 0.3697344 | 0.57741665 | Y69A2AR.18 |
| WBGene00022090 | 0.596056337 | 0.72561177 | 1 | Y69A2AR.19 |
| WBGene00022092 | -0.785082949 | 8.5504E-05 | 0.00134994 | Y69A2AR.21 |
| WBGene00022093 | 0.629678552 | 0.0849295 | 0.22970128 | Y69A2AR.22 |
| WBGene00022096 | 1.139358466 | 0.62800936 | 1 | Y69A2AR.25 |
| WBGene00022099 | -0.83160103 | 0.00162406 | 0.01331377 | Y69A2AR.28 |
| WBGene00022100 | 0.991022762 | 0.02897197 | 0.11057802 | Y69A2AR.31 |
| WBGene00022101 | 0.215989625 | 0.5981641 | 0.76397122 | Y69A2AR.32 |
| WBGene00022102 | 1.28669653 | 0.40575348 | 1 | Y69F12A.1 |
| WBGene00022103 | 1.036952742 | 0.08361762 | 0.22731784 | cdh-12 |
| WBGene00022104 | 0.333460688 | 0.54362436 | 0.72337864 | acsd-1 |
| WBGene00022106 | 0.89499303 | 0.11608353 | 0.28048782 | lgc-46 |
| WBGene00022107 | -0.413639436 | 0.16396986 | 0.35011573 | Y71F9AL.1 |
| WBGene00022108 | 1.70100456 | 0.24970237 | 1 | Y71F9AL.2 |
| WBGene00022110 | -0.256607805 | 0.58855805 | 0.75679764 | Y71F9AL.4 |
| WBGene00022111 | -1.932094392 | 0.04503882 | 1 | Y71F9AL.6 |
| WBGene00022112 | -1.505458846 | 0.01846669 | 0.08089445 | Y71F9AL.7 |
| WBGene00022113 | 0.381981649 | 0.2521402 | 0.46033596 | Y71F9AL.8 |
| WBGene00022114 | -0.071674475 | 0.68314969 | 0.81876806 | Y71F9AL.9 |
| WBGene00022115 | -0.098876965 | 0.69655562 | 0.82845629 | Y71F9AL.10 |
| WBGene00022117 | -0.755804313 | 0.00025373 | 0.003166 | Y71F9AL.12 |
| WBGene00022118 | -0.474985279 | 0.07644589 | 0.2146957 | dnc-5 |
| WBGene00022119 | 0.230764583 | 0.221055 | 0.42403497 | copa-1 |
| WBGene00022121 | -0.056525125 | 0.78693741 | 0.88268727 | cogc-3 |
| WBGene00022122 | -0.319557736 | 0.03581155 | 0.12765355 | trap-1 |
| WBGene00022123 | 0.549276974 | 0.60799252 | 1 | Y71F9AR.2 |
| WBGene00022124 | 0.125875622 | 0.621356 | 0.77956837 | supr-1 |
| WBGene00022125 | -0.484303048 | 0.26937347 | 0.47783631 | Y71F9B.1 |
| WBGene00022126 | -0.449309396 | 0.08249942 | 0.22538826 | Y71F9B.2 |
| WBGene00022127 | -0.622457148 | 2.3288E-05 | 0.00045824 | yop-1 |
| WBGene00022128 | -0.097711263 | 0.71726087 | 0.84012424 | Y71F9B.6 |
| WBGene00022129 | -0.110147792 | 0.78112459 | 0.87891216 | lron-11 |
| WBGene00022130 | -0.914961584 | 0.08742223 | 0.23399324 | Y71F9B.9 |
| WBGene00022131 | 0.105092804 | 0.8033744 | 0.89239902 | Y71F9B.13 |
| WBGene00022133 | 0.075260937 | 0.97569838 | 1 | Y71F9B.15 |
| WBGene00022134 | -0.473759308 | 0.004726 | 0.02998555 | Y71G10AL.1 |
| WBGene00022138 | 0.074721424 | 0.78495519 | 0.88176377 | trpp-10 |

| | | | | |
|---|---|---|---|---|
| WBGene00022139 | 0.392465746 | 0.10129836 | 0.25648871 | tub-2 |
| WBGene00022140 | -1.368568572 | 0.20425715 | 1 | Y71G12A.4 |
| WBGene00022141 | -0.43409789 | 0.02478085 | 0.09952509 | chaf-2 |
| WBGene00022142 | 0.466401197 | 0.76128449 | 1 | Y71G12B.2 |
| WBGene00022143 | 0.281454817 | 0.94435481 | 1 | Y71G12B.3 |
| WBGene00022144 | 1.027194378 | 2.677E-05 | 0.00051184 | pghm-1 |
| WBGene00022145 | 0.256800365 | 0.85179097 | 1 | Y71G12B.5 |
| WBGene00022146 | -0.441072167 | 0.12835562 | 0.29912046 | Y71G12B.6 |
| WBGene00022148 | -0.27420347 | 0.16363088 | 0.34976101 | ddx-27 |
| WBGene00022149 | -0.252331073 | 0.27313824 | 0.48186715 | lin-65 |
| WBGene00022150 | -0.238809346 | 0.32485994 | 0.53327805 | Y71G12B.10 |
| WBGene00022152 | -0.228939554 | 0.33969101 | 0.5479921 | atg-5 |
| WBGene00022153 | -0.144248591 | 0.54983509 | 0.72829455 | Y71G12B.13 |
| WBGene00022154 | 0.467410147 | 0.4778613 | 0.67028996 | drag-1 |
| WBGene00022155 | -0.197770115 | 0.78903913 | 0.8837593 | Y71G12B.17 |
| WBGene00022156 | -0.156269293 | 0.96924949 | 1 | Y71G12B.18 |
| WBGene00022158 | 0.374434035 | 0.56096552 | 0.73692733 | Y71G12B.23 |
| WBGene00022159 | -0.375370909 | 0.13394886 | 0.30783137 | mppa-1 |
| WBGene00022160 | -0.06098388 | 0.8852096 | 0.93936979 | Y71G12B.25 |
| WBGene00022161 | 0.121192541 | 0.93494866 | 1 | Y71G12B.26 |
| WBGene00022162 | 0.688793783 | 0.86378755 | 1 | Y71G12B.27 |
| WBGene00022165 | 0.605036286 | 0.5734776 | 1 | Y71H2AL.2 |
| WBGene00022166 | -0.370548177 | 0.15015449 | 0.33176081 | Y71H2AM.1 |
| WBGene00022167 | -0.142916026 | 0.48048744 | 0.67238335 | Y71H2AM.2 |
| WBGene00022168 | -0.278368065 | 0.22077621 | 0.42382918 | Y71H2AM.3 |
| WBGene00022169 | -0.5439193 | 0.00340487 | 0.02321083 | Y71H2AM.4 |
| WBGene00022170 | -0.384084115 | 0.00748025 | 0.04169991 | Y71H2AM.5 |
| WBGene00022171 | -0.799282464 | 0.05421997 | 0.17001153 | Y71H2AM.6 |
| WBGene00022172 | 0.178252189 | 0.54416112 | 0.72362105 | cosa-1 |
| WBGene00022173 | -0.139592596 | 0.67456608 | 0.81399855 | set-27 |
| WBGene00022174 | 0.439790751 | 0.07066733 | 0.20355209 | Y71H2AM.9 |
| WBGene00022175 | 0.571387989 | 0.20123145 | 0.40018853 | Y71H2AM.10 |
| WBGene00022176 | -0.322556177 | 0.10950017 | 0.27004703 | Y71H2AM.11 |
| WBGene00022177 | 0.722173909 | 0.17281619 | 0.36250195 | Y71H2AM.12 |
| WBGene00022178 | 0.209180878 | 0.33717873 | 0.54524333 | Y71H2AM.13 |
| WBGene00022179 | 0.389206336 | 0.7290552 | 1 | Y71H2AM.14 |
| WBGene00022180 | 0.753068563 | 0.04859894 | 0.15804675 | Y71H2AM.15 |
| WBGene00022181 | 2.083241444 | 1.46E-07 | 6.4998E-06 | pho-9 |
| WBGene00022182 | -0.227758509 | 0.28002359 | 0.4890343 | swsn-3 |
| WBGene00022185 | -0.360227771 | 0.08388832 | 0.22774778 | Y71H2AM.20 |
| WBGene00022188 | -0.210974083 | 0.3667555 | 0.57427918 | Y71H2AR.1 |
| WBGene00022189 | 0.908930564 | 0.6008643 | 1 | Y71H2AR.2 |
| WBGene00022190 | -0.326591426 | 0.53446366 | 0.71755507 | josd-1 |

| | | | | |
|---|---|---|---|---|
| WBGene00022191 | -0.815052222 | 0.14116918 | 0.31910911 | Y71H2B.1 |
| WBGene00022192 | -0.376662699 | 0.04955276 | 0.1594963 | Y71H2B.2 |
| WBGene00022193 | -0.54513094 | 0.010017 | 0.05194443 | ppfr-4 |
| WBGene00022194 | -0.793818609 | 0.00088089 | 0.00833928 | Y71H2B.4 |
| WBGene00022195 | 0.072078538 | 0.77291509 | 0.87317038 | Y71H2B.5 |
| WBGene00022196 | 0.833042806 | 0.03188382 | 0.11847911 | gpa-17 |
| WBGene00022197 | 0.138654495 | 0.81396599 | 0.8986238 | Y71H2B.8 |
| WBGene00022198 | 0.003237335 | 0.99326483 | 0.99669716 | Y71H2B.11 |
| WBGene00022199 | 0.071094582 | 0.72098768 | 0.84287092 | pfk-1.1 |
| WBGene00022200 | 0.556113878 | 0.00664322 | 0.03825793 | fard-1 |
| WBGene00022201 | 0.015914115 | 0.92508148 | 0.96044148 | Y71H10B.1 |
| WBGene00022202 | 0.303085342 | 0.50237707 | 0.68999878 | Y72A10A.1 |
| WBGene00022203 | -0.424060866 | 0.13096075 | 0.30337616 | Y73B3A.1 |
| WBGene00022206 | 0.716038068 | 0.0683705 | 0.19916428 | Y73B3A.4 |
| WBGene00022207 | -0.156269293 | 0.96924949 | 1 | elk-2 |
| WBGene00022209 | -1.901856996 | 0.64072034 | 1 | Y73B3A.7 |
| WBGene00022213 | -0.263229581 | 0.93409872 | 1 | Y73B3A.11 |
| WBGene00022215 | 1.025181888 | 0.79790347 | 1 | Y73B3A.13 |
| WBGene00022218 | 2.799084385 | 0.17307183 | 1 | Y73B3A.16 |
| WBGene00022219 | 0.061897906 | 0.76227957 | 0.86712605 | Y73B3A.18 |
| WBGene00022221 | 0.342771878 | 0.66078796 | 0.80475741 | Y73B3A.20 |
| WBGene00022223 | 0.836200882 | 0.60863685 | 1 | Y73B3B.1 |
| WBGene00022224 | 1.957271014 | 0.49743227 | 1 | set-28 |
| WBGene00022225 | -0.552227129 | 0.83715812 | 1 | Y73B3B.3 |
| WBGene00022228 | 0.079418419 | 0.96102778 | 1 | Y73B6A.1 |
| WBGene00022230 | -0.008042586 | 0.99084513 | 0.99552766 | Y73B6A.3 |
| WBGene00022231 | 0.835951495 | 0.01722348 | 0.07685052 | tyr-6 |
| WBGene00022232 | -0.347292911 | 0.13256617 | 0.30546326 | exos-2 |
| WBGene00022233 | 0.285598118 | 0.16571147 | 0.3525325 | ipla-6 |
| WBGene00022235 | -0.172393498 | 0.27871693 | 0.48767511 | sqd-1 |
| WBGene00022236 | -0.216240469 | 0.84232385 | 1 | Y73B6BL.12 |
| WBGene00022237 | -0.704207726 | 0.28170333 | 0.49104955 | Y73B6BL.14 |
| WBGene00022240 | 1.133380425 | 0.03440553 | 0.1248161 | shl-1 |
| WBGene00022242 | 0.20635769 | 0.68308774 | 0.81876806 | sfrp-1 |
| WBGene00022243 | -0.156269293 | 0.96924949 | 1 | Y73B6BL.22 |
| WBGene00022244 | -0.691674211 | 0.00346752 | 0.02361145 | Y73B6BL.23 |
| WBGene00022245 | 0.034743919 | 0.8334503 | 0.90997483 | acp-6 |
| WBGene00022246 | 1.46530193 | 0.00456296 | 0.02918804 | acp-7 |
| WBGene00022247 | 0.596352124 | 0.48946418 | 0.67923015 | lgc-10 |
| WBGene00022248 | 0.308889686 | 0.34445364 | 0.55324933 | Y73B6BL.27 |
| WBGene00022250 | -0.037756133 | 0.90614964 | 0.95109561 | Y73B6BL.29 |
| WBGene00022251 | -0.235957473 | 0.30327409 | 0.51229575 | blos-2 |
| WBGene00022252 | -0.187034799 | 0.52987182 | 0.7139882 | Y73B6BL.31 |

| | | | | |
|---|---|---|---|---|
| WBGene00022253 | -0.496757832 | 0.00412923 | 0.02693872 | hrpf-2 |
| WBGene00022254 | 1.132120829 | 0.01923336 | 0.08332046 | Y73B6BL.35 |
| WBGene00022255 | 0.672548445 | 0.59777167 | 1 | Y73B6BL.36 |
| WBGene00022256 | 0.747845955 | 0.85215965 | 1 | Y73B6BL.37 |
| WBGene00022257 | -0.593148928 | 4.4448E-05 | 0.0007839 | puf-11 |
| WBGene00022258 | -0.466343185 | 0.43819141 | 0.63866548 | Y73C8B.1 |
| WBGene00022259 | -0.409623888 | 0.31475644 | 0.5225568 | Y73C8B.2 |
| WBGene00022260 | -0.784928932 | 0.03652331 | 0.12935515 | Y73C8B.3 |
| WBGene00022261 | -0.356102232 | 0.88046231 | 1 | clec-210 |
| WBGene00022262 | -1.721102694 | 0.01112642 | 0.05616878 | Y73C8C.3 |
| WBGene00022263 | 1.630244101 | 0.40224995 | 1 | Y73C8C.4 |
| WBGene00022265 | -1.277826145 | 0.00582663 | 0.034865 | Y73C8C.8 |
| WBGene00022266 | -1.969805064 | 0.62862569 | 1 | srt-56 |
| WBGene00022267 | -0.014519762 | 0.99603746 | 1 | Y73C8C.10 |
| WBGene00022268 | -0.376323087 | 0.08194839 | 0.22442154 | Y73E7A.1 |
| WBGene00022269 | 0.124393206 | 0.70711851 | 0.83456069 | Y73E7A.2 |
| WBGene00022270 | -0.067135273 | 0.88822919 | 0.9410114 | Y73E7A.3 |
| WBGene00022271 | 0.391631361 | 0.21177752 | 0.41171992 | cpx-1 |
| WBGene00022273 | -0.516018155 | 0.10431714 | 0.26214014 | Y73E7A.6 |
| WBGene00022274 | -0.479559829 | 0.49974912 | 0.68768606 | Y73E7A.8 |
| WBGene00022275 | 0.03940405 | 0.99001443 | 1 | Y74C9A.1 |
| WBGene00022276 | 0.16512278 | 0.3202879 | 0.52855992 | nlp-40 |
| WBGene00022277 | -0.394379365 | 0.12700535 | 0.29750569 | homt-1 |
| WBGene00022278 | -0.174477012 | 0.32690105 | 0.53547026 | rcor-1 |
| WBGene00022279 | 0.614515431 | 0.02065026 | 0.087418 | sesn-1 |
| WBGene00022280 | -0.138690791 | 0.57013512 | 0.7444623 | Y74C10AL.2 |
| WBGene00022281 | 0.106561506 | 0.73204679 | 0.85009852 | abtm-1 |
| WBGene00022282 | -1.014682842 | 0.61579555 | 1 | Y74C10AR.2 |
| WBGene00022283 | -0.354258279 | 0.47345232 | 0.66645288 | lgc-27 |
| WBGene00022284 | 0.45662644 | 0.22067168 | 0.42372381 | glb-33 |
| WBGene00022285 | 0.670124592 | 0.54257779 | 1 | Y75B7AL.2 |
| WBGene00022286 | -0.067717996 | 0.74397131 | 0.85740338 | rga-4 |
| WBGene00022287 | -0.836156166 | 0.37232679 | 0.58013598 | Y75B7AR.1 |
| WBGene00022288 | 2.228122762 | 0.21083368 | 1 | Y75B7B.1 |
| WBGene00022292 | -0.523349639 | 0.26882525 | 0.47730439 | Y75D11A.3 |
| WBGene00022295 | 0.617460097 | 0.54590706 | 1 | cng-2 |
| WBGene00022296 | 0.138952089 | 0.49081531 | 0.6801722 | xpc-1 |
| WBGene00022297 | -0.431105767 | 0.0607063 | 0.18407473 | Y76B12C.3 |
| WBGene00022298 | 1.988712883 | 0.17279138 | 1 | Y76B12C.4 |
| WBGene00022300 | -0.190095637 | 0.38886466 | 0.59739413 | Y76B12C.6 |
| WBGene00022301 | 0.074516137 | 0.71345204 | 0.8379961 | cpsf-1 |
| WBGene00022302 | 0.124438164 | 0.85514564 | 0.92107197 | Y76B12C.8 |
| WBGene00022303 | -2.171326218 | 0.36522606 | 1 | Y76B12C.9 |

| | | | | |
|---|---|---|---|---|
| WBGene00022306 | -0.058662339 | 0.7515065 | 0.86174089 | hxk-3 |
| WBGene00022307 | 0.451899855 | 0.09283141 | 0.24299414 | Y77E11A.2 |
| WBGene00022309 | -0.513346893 | 0.23730136 | 0.44351537 | rpc-11 |
| WBGene00022310 | -0.252549898 | 0.18310145 | 0.37527935 | Y77E11A.7 |
| WBGene00022312 | -1.585528107 | 0.41651348 | 1 | clec-171 |
| WBGene00022313 | 0.186813476 | 0.79760573 | 0.88865587 | Y77E11A.12 |
| WBGene00022314 | 1.420228692 | 0.12496471 | 0.29465848 | Y77E11A.14 |
| WBGene00022317 | 0.281454817 | 0.94435481 | 1 | Y81B9A.3 |
| WBGene00022319 | -1.618771107 | 0.4854602 | 1 | Y82E9BL.2 |
| WBGene00022320 | -0.258794157 | 0.89747187 | 1 | Y82E9BL.3 |
| WBGene00022321 | 0.281454817 | 0.94435481 | 1 | fbxa-25 |
| WBGene00022322 | -1.504171367 | 0.26616922 | 1 | Y82E9BL.5 |
| WBGene00022323 | -1.187254256 | 0.47366012 | 1 | Y82E9BL.6 |
| WBGene00022324 | 0.908269423 | 0.29757346 | 0.5076675 | fbxa-26 |
| WBGene00022325 | 1.513800797 | 0.34999722 | 1 | fbxa-5 |
| WBGene00022326 | 0.372560935 | 0.47545057 | 0.66818444 | fbxa-14 |
| WBGene00022327 | 1.507834523 | 0.4883646 | 1 | fbxa-15 |
| WBGene00022328 | -0.333693239 | 0.89573179 | 1 | Y82E9BL.12 |
| WBGene00022329 | 0.397983017 | 0.46319732 | 0.6582021 | fbxa-79 |
| WBGene00022330 | 1.173276434 | 0.12795915 | 0.29864109 | fbxa-80 |
| WBGene00022331 | 0.015297004 | 0.98438713 | 0.99239759 | fbxa-19 |
| WBGene00022332 | 2.014719173 | 0.2903304 | 1 | fbxa-20 |
| WBGene00022333 | -0.234153816 | 0.83993979 | 1 | fbxa-27 |
| WBGene00022334 | -1.616917428 | 0.00936523 | 0.04930756 | Y82E9BR.1 |
| WBGene00022335 | 0.456736625 | 0.13711493 | 0.31309242 | Y82E9BR.2 |
| WBGene00022336 | -0.61651581 | 3.8248E-06 | 0.00010103 | Y82E9BR.3 |
| WBGene00022337 | 0.305872954 | 0.67462934 | 0.81399855 | pals-16 |
| WBGene00022338 | 0.140735336 | 0.88653513 | 1 | Y82E9BR.5 |
| WBGene00022339 | 0.516150826 | 0.65714961 | 1 | Y82E9BR.6 |
| WBGene00022344 | 2.437484291 | 0.2708268 | 1 | Y82E9BR.11 |
| WBGene00022345 | 1.712479368 | 0.58529868 | 1 | fbxa-138 |
| WBGene00022346 | 0.114079952 | 0.83740534 | 0.91143626 | pals-17 |
| WBGene00022347 | -0.289422099 | 0.17999132 | 0.37228489 | Y82E9BR.14 |
| WBGene00022348 | -0.292412902 | 0.13971923 | 0.31695277 | Y82E9BR.16 |
| WBGene00022349 | -0.95813694 | 0.02815402 | 0.10851365 | Y82E9BR.17 |
| WBGene00022350 | -0.006190785 | 0.97442867 | 0.98661738 | Y82E9BR.18 |
| WBGene00022351 | 0.025004198 | 0.93106808 | 0.96395551 | Y82E9BR.19 |
| WBGene00022352 | -0.156269293 | 0.96924949 | 1 | Y82E9BR.20 |
| WBGene00022353 | 1.02391764 | 0.14509238 | 0.32381738 | pals-18 |
| WBGene00022354 | -0.289067928 | 0.35903848 | 0.56719462 | Y82E9BR.22 |
| WBGene00022356 | 1.186559396 | 0.57146291 | 1 | eak-3 |
| WBGene00022357 | -1.077057568 | 2.7409E-05 | 0.00052187 | kbp-4 |
| WBGene00022358 | 0.392168149 | 0.17106378 | 0.36015788 | Y92H12A.2 |

| | | | | |
|---|---|---|---|---|
| WBGene00022360 | 0.063769642 | 0.79927042 | 0.88979641 | Y92H12A.4 |
| WBGene00022361 | 0.139328598 | 0.58522858 | 0.75465941 | Y92H12A.5 |
| WBGene00022363 | -0.071352373 | 0.78880817 | 0.8837593 | Y92H12BL.1 |
| WBGene00022365 | -0.270192881 | 0.55680618 | 0.73399659 | Y92H12BL.4 |
| WBGene00022366 | 0.0930105 | 0.81080431 | 0.89678191 | Y92H12BL.5 |
| WBGene00022367 | 0.293733373 | 0.60169571 | 0.76670734 | Y92H12BM.1 |
| WBGene00022368 | -0.630482697 | 0.0073499 | 0.04117912 | Y92H12BR.2 |
| WBGene00022369 | -0.314296565 | 0.07007348 | 0.20236968 | Y92H12BR.3 |
| WBGene00022371 | -0.607887557 | 0.05737552 | 0.17727949 | set-29 |
| WBGene00022372 | -0.063060654 | 0.84029997 | 0.91237519 | Y92H12BR.7 |
| WBGene00022373 | -0.502141584 | 0.02662262 | 0.10450217 | mrpl-15 |
| WBGene00022374 | 0.880846536 | 0.22931161 | 0.43442105 | nhr-277 |
| WBGene00022375 | 2.166011856 | 0.42253177 | 1 | Y94H6A.2 |
| WBGene00022376 | -0.575872476 | 0.03570858 | 0.12744529 | Y94H6A.3 |
| WBGene00022377 | 0.623124398 | 0.68731221 | 1 | gpx-4 |
| WBGene00022378 | 0.106561946 | 0.68158309 | 0.81759493 | Y94H6A.5 |
| WBGene00022379 | -0.78591523 | 0.0726579 | 0.20765996 | Y94H6A.7 |
| WBGene00022380 | -0.31475496 | 0.10793545 | 0.26759651 | Y94H6A.8 |
| WBGene00022381 | -0.286688026 | 0.1514656 | 0.33332315 | ubxn-2 |
| WBGene00022382 | 0.310935006 | 0.25944153 | 0.46768813 | Y94H6A.10 |
| WBGene00022385 | -0.570667192 | 0.72780871 | 1 | Y95B8A.4 |
| WBGene00022386 | 0.99431884 | 0.0755653 | 0.213011 | Y95B8A.6 |
| WBGene00022388 | -0.197172451 | 0.28327107 | 0.49287415 | Y95B8A.8 |
| WBGene00022389 | 0.796532024 | 0.00224304 | 0.01700982 | pde-6 |
| WBGene00022390 | 0.046184198 | 0.90841722 | 0.95209716 | him-19 |
| WBGene00022391 | 1.314250085 | 0.00575702 | 0.03455067 | Y95B8A.12 |
| WBGene00022392 | -0.704294842 | 0.1167262 | 0.28150377 | Y97E10AL.1 |
| WBGene00022393 | -0.407179704 | 0.01575491 | 0.0720182 | abhd-12 |
| WBGene00022394 | -0.90818388 | 0.00027754 | 0.00339446 | natb-1 |
| WBGene00022395 | -3.47102951 | 0.12756864 | 1 | ceh-76 |
| WBGene00022396 | 0.192899079 | 0.4907633 | 0.6801722 | Y97E10AR.1 |
| WBGene00022397 | 1.360074663 | 0.03711068 | 0.13090087 | Y97E10AR.2 |
| WBGene00022398 | -0.439945994 | 0.06680836 | 0.19606484 | Y97E10AR.3 |
| WBGene00022399 | -0.694921954 | 0.00213614 | 0.01638335 | Y97E10AR.4 |
| WBGene00022400 | -0.624728228 | 0.00694122 | 0.0395093 | rpb-9 |
| WBGene00022401 | -0.507427246 | 0.00461839 | 0.02948046 | Y97E10AR.6 |
| WBGene00022402 | -0.698748268 | 0.00290273 | 0.02073065 | lmtr-2 |
| WBGene00022403 | -0.18557176 | 0.95664217 | 1 | Y97E10B.1 |
| WBGene00022404 | 0.747845955 | 0.85215965 | 1 | srsx-5 |
| WBGene00022405 | -1.706203807 | 0.43490097 | 1 | srsx-6 |
| WBGene00022406 | 1.477253415 | 0.58357327 | 1 | srsx-7 |
| WBGene00022409 | 1.584577408 | 0.63258867 | 1 | srt-53 |
| WBGene00022410 | 0.627740553 | 0.02202225 | 0.09210203 | Y97E10C.1 |

| | | | | |
|---|---|---|---|---|
| WBGene00022411 | 2.38467623 | 0.08505706 | 1 | Y102A11A.1 |
| WBGene00022412 | 0.965679571 | 0.04828936 | 0.15727531 | Y102A11A.2 |
| WBGene00022413 | 1.729422015 | 2.23E-17 | 5.77E-15 | Y102A11A.3 |
| WBGene00022415 | -0.072271833 | 0.92022763 | 0.95810682 | Y102A11A.5 |
| WBGene00022416 | 0.008748801 | 0.98931442 | 0.99484196 | Y102A11A.6 |
| WBGene00022417 | -0.787648086 | 0.46118467 | 1 | Y102A11A.7 |
| WBGene00022418 | -0.025267959 | 0.95035956 | 0.97468105 | igcm-4 |
| WBGene00022419 | 0.532738251 | 0.42730249 | 0.63051525 | Y102A11A.9 |
| WBGene00022420 | -0.37140508 | 0.01502498 | 0.06981181 | wdr-4 |
| WBGene00022423 | 0.884487205 | 0.06220605 | 0.18680279 | nhr-41 |
| WBGene00022425 | 0.21734945 | 0.23434116 | 0.44049171 | plst-1 |
| WBGene00022426 | -0.252506742 | 0.29148011 | 0.50151603 | Y104H12D.2 |
| WBGene00022427 | 0.047708915 | 0.83370274 | 0.91000494 | Y104H12D.3 |
| WBGene00022428 | -0.16368405 | 0.68338087 | 0.81877569 | Y104H12D.4 |
| WBGene00022429 | -0.156269293 | 0.96924949 | 1 | Y108F1.1 |
| WBGene00022431 | 0.747845955 | 0.85215965 | 1 | set-33 |
| WBGene00022434 | -0.462864117 | 0.03294863 | 0.12116257 | Y108G3AL.2 |
| WBGene00022435 | 0.663116167 | 0.28327781 | 0.49287415 | cfap-36 |
| WBGene00022439 | 0.855026222 | 0.71364798 | 1 | Y110A2AL.2 |
| WBGene00022440 | -0.413726452 | 0.43024047 | 0.63273764 | Y110A2AL.3 |
| WBGene00022441 | 0.523339988 | 0.16684686 | 0.35418459 | Y110A2AL.4 |
| WBGene00022444 | -1.409432952 | 0.51259961 | 1 | Y110A2AL.7 |
| WBGene00022445 | -0.091486423 | 0.85328166 | 0.91947215 | Y110A2AL.9 |
| WBGene00022446 | -1.291305095 | 0.63677871 | 1 | Y110A2AL.10 |
| WBGene00022447 | 0.020853755 | 0.96879575 | 0.98399257 | hpo-20 |
| WBGene00022448 | -0.632228409 | 0.00635847 | 0.0371681 | pinn-1 |
| WBGene00022450 | -0.158182617 | 0.59764086 | 0.76351811 | ubc-26 |
| WBGene00022452 | -0.621400228 | 0.00255412 | 0.01886227 | Y110A2AR.1 |
| WBGene00022453 | -0.423049003 | 0.16038954 | 0.34570837 | ncap-1 |
| WBGene00022454 | 2.032816334 | 0.10707547 | 1 | Y110A7A.2 |
| WBGene00022455 | -0.301638995 | 0.08544763 | 0.23054671 | tyms-1 |
| WBGene00022456 | -0.207921599 | 0.17045284 | 0.35928262 | pfkb-1.1 |
| WBGene00022457 | -0.183219194 | 0.61661257 | 0.77634483 | Y110A7A.7 |
| WBGene00022458 | -0.422985247 | 0.00718809 | 0.04057212 | prp-31 |
| WBGene00022459 | -0.070079806 | 0.67467815 | 0.81399855 | Y110A7A.9 |
| WBGene00022460 | -0.646928767 | 0.01269963 | 0.06174859 | use-1 |
| WBGene00022462 | -0.085101587 | 0.72803755 | 0.84755048 | Y110A7A.15 |
| WBGene00022463 | -0.142018184 | 0.45048841 | 0.64895346 | elpc-1 |
| WBGene00022465 | -0.125719084 | 0.93685366 | 1 | ift-20 |
| WBGene00022466 | -0.268938613 | 0.20160332 | 0.40041719 | Y119C1A.1 |
| WBGene00022467 | 1.735991307 | 0.58152819 | 1 | Y119C1B.1 |
| WBGene00022469 | 0.843981867 | 0.25558947 | 0.46305107 | Y119C1B.3 |
| WBGene00022470 | -0.247802579 | 0.28495209 | 0.49486529 | mrpl-19 |

| | | | | |
|---|---|---|---|---|
| WBGene00022471 | 0.434057498 | 0.05008511 | 0.16092931 | Y119C1B.5 |
| WBGene00022472 | 0.507165492 | 0.51426032 | 0.70121682 | Y119C1B.6 |
| WBGene00022473 | -0.117104977 | 0.5357865 | 0.7178681 | bet-1 |
| WBGene00022474 | 0.547390941 | 0.19555012 | 0.39203666 | zig-9 |
| WBGene00022475 | 0.307760233 | 0.37801863 | 0.58562259 | Y119C1B.10 |
| WBGene00022478 | 0.159950863 | 0.93307234 | 1 | fbxa-35 |
| WBGene00022479 | -0.107268465 | 0.96117914 | 1 | fbxa-36 |
| WBGene00022484 | 1.434546033 | 0.47351845 | 1 | fbxa-77 |
| WBGene00022485 | 0.519136201 | 0.44601095 | 0.64541346 | fbxa-63 |
| WBGene00022486 | 1.071678026 | 0.14135889 | 0.31935957 | fbxa-61 |
| WBGene00022487 | 2.351003106 | 0.00194341 | 0.01521272 | fbxa-21 |
| WBGene00022488 | -0.492780123 | 0.01981579 | 0.08499814 | orc-3 |
| WBGene00022489 | -0.172078638 | 0.44982544 | 0.6486141 | Y119D3B.12 |
| WBGene00022490 | 0.497984759 | 0.12142592 | 0.28856127 | Y119D3B.13 |
| WBGene00022491 | -0.348146212 | 0.15739137 | 0.3419629 | Y119D3B.14 |
| WBGene00022492 | -0.712746966 | 1.8455E-05 | 0.00037593 | dss-1 |
| WBGene00022493 | -0.637312947 | 0.00796143 | 0.04375723 | mrpl-45 |
| WBGene00022495 | 3.449461417 | 0.06119705 | 1 | fbxa-78 |
| WBGene00022497 | -1.606445368 | 4.51E-17 | 1.09E-14 | Y119D3B.21 |
| WBGene00022498 | -0.922905621 | 0.44526251 | 1 | hsd-2 |
| WBGene00022499 | 0.390403948 | 0.22724622 | 0.431903 | set-30 |
| WBGene00022500 | 0.586958803 | 0.00885172 | 0.04711474 | lfi-1 |
| WBGene00022501 | -0.427484426 | 0.15097787 | 0.33279273 | ZC8.6 |
| WBGene00022502 | -0.075406008 | 0.8263312 | 0.90625042 | ZC13.1 |
| WBGene00022503 | 0.092553308 | 0.91986365 | 0.95789215 | ZC13.2 |
| WBGene00022504 | -0.986571133 | 0.27624211 | 0.48487139 | mam-1 |
| WBGene00022505 | 1.065061133 | 0.00020064 | 0.00263632 | ZC21.3 |
| WBGene00022506 | 0.113920663 | 0.96058911 | 1 | ZC21.6 |
| WBGene00022508 | -0.718947058 | 0.06197999 | 0.18635441 | ZC53.1 |
| WBGene00022512 | -0.33101813 | 0.78652311 | 1 | dot-1.5 |
| WBGene00022515 | -0.635399535 | 0.01162421 | 0.05791116 | ttr-48 |
| WBGene00022516 | -0.640969303 | 0.00365613 | 0.02455145 | mtx-2 |
| WBGene00022517 | -0.228201996 | 0.62233802 | 0.78023605 | ZC123.1 |
| WBGene00022518 | 0.545548915 | 0.03079741 | 0.11569038 | zfh-2 |
| WBGene00022519 | 0.699023092 | 0.10121232 | 0.2564313 | ZC123.4 |
| WBGene00022522 | 2.381446256 | 0.34860272 | 1 | ZC132.3 |
| WBGene00022527 | 1.712479368 | 0.58529868 | 1 | ZC132.9 |
| WBGene00022529 | 0.281454817 | 0.94435481 | 1 | srt-14 |
| WBGene00022530 | 0.463161033 | 0.72934251 | 1 | ZC155.2 |
| WBGene00022531 | -0.276436376 | 0.17805504 | 0.37010207 | morc-1 |
| WBGene00022532 | -0.711551403 | 0.00047332 | 0.00514925 | ZC155.4 |
| WBGene00022533 | 0.533973775 | 0.52218524 | 0.70717357 | cutl-19 |
| WBGene00022534 | -0.176893966 | 0.42527375 | 0.62935746 | syx-16 |

| | | | | |
|---|---|---|---|---|
| WBGene00022536 | -0.156269293 | 0.96924949 | 1 | ZC178.2 |
| WBGene00022537 | 0.377828795 | 0.92526585 | 1 | ZC190.2 |
| WBGene00022538 | 1.039935655 | 0.00094188 | 0.00878971 | ZC190.4 |
| WBGene00022539 | 0.397655506 | 0.56216723 | 0.73778752 | ZC190.5 |
| WBGene00022540 | 1.025181888 | 0.79790347 | 1 | ZC190.6 |
| WBGene00022542 | 1.216957619 | 0.74819621 | 1 | ZC190.8 |
| WBGene00022544 | 1.393777844 | 0.3596588 | 1 | ZC190.10 |
| WBGene00022546 | -1.323116856 | 0.47668251 | 1 | ZC196.2 |
| WBGene00022547 | -1.159714244 | 0.75790634 | 1 | ZC196.3 |
| WBGene00022548 | 1.739217586 | 0.12592041 | 1 | ZC196.4 |
| WBGene00022549 | 1.625700229 | 0.00623507 | 0.03675177 | ZC196.5 |
| WBGene00022550 | 2.224905164 | 0.13591468 | 1 | droe-8 |
| WBGene00022553 | -1.969805064 | 0.62862569 | 1 | ZC204.1 |
| WBGene00022554 | -0.593247069 | 0.04933097 | 0.15941638 | duxl-1 |
| WBGene00022555 | -1.175817997 | 0.34034417 | 1 | btb-12 |
| WBGene00022556 | -0.156269293 | 0.96924949 | 1 | ZC204.6 |
| WBGene00022557 | -1.125955582 | 0.25790105 | 1 | fbxb-15 |
| WBGene00022558 | -0.319129489 | 0.82424959 | 1 | fbxb-21 |
| WBGene00022559 | -1.81568125 | 0.01307082 | 0.06312365 | fbxb-20 |
| WBGene00022560 | -0.190856808 | 0.90351155 | 1 | fbxb-16 |
| WBGene00022561 | -1.338151959 | 0.29244005 | 1 | btb-13 |
| WBGene00022562 | 1.011727052 | 0.0038337 | 0.02536512 | ZC204.12 |
| WBGene00022564 | -0.038546172 | 0.92959338 | 0.96347902 | ZC204.14 |
| WBGene00022566 | 1.529802032 | 0.64971375 | 1 | ZC239.3 |
| WBGene00022567 | 1.51672272 | 0.31693815 | 1 | ZC239.4 |
| WBGene00022568 | 1.025181888 | 0.79790347 | 1 | ZC239.5 |
| WBGene00022569 | 0.400321266 | 0.36594645 | 0.57350663 | ZC239.6 |
| WBGene00022570 | 1.695200177 | 0.42035448 | 1 | sdz-35 |
| WBGene00022571 | 0.050417347 | 0.9681294 | 1 | ZC239.13 |
| WBGene00022572 | 0.435349593 | 0.66863919 | 1 | ZC239.14 |
| WBGene00022573 | 0.183215689 | 0.8597378 | 1 | ZC239.15 |
| WBGene00022574 | 0.347534242 | 0.61333619 | 0.77414645 | ZC239.16 |
| WBGene00022575 | -0.289777269 | 0.86134264 | 1 | ZC239.17 |
| WBGene00022576 | -0.391052742 | 0.48406583 | 0.67482411 | ZC250.2 |
| WBGene00022577 | -1.671214043 | 0.07093712 | 0.20403874 | nstp-3 |
| WBGene00022579 | -0.339185052 | 0.07471579 | 0.21125393 | ZC262.2 |
| WBGene00022580 | 0.347313296 | 0.04539721 | 0.15011525 | iglr-2 |
| WBGene00022581 | -1.204791816 | 0.57474638 | 1 | ZC262.4 |
| WBGene00022583 | -0.666385895 | 0.0027352 | 0.01987313 | mrps-18A |
| WBGene00022584 | -0.338647303 | 0.58416185 | 0.7540432 | ZC266.1 |
| WBGene00022586 | -0.401796477 | 0.01758703 | 0.07810589 | ZC308.4 |
| WBGene00022587 | -0.325038374 | 0.87140873 | 1 | ZC317.1 |
| WBGene00022588 | 0.22830586 | 0.84055027 | 1 | ZC317.2 |

| | | | | |
|---|---|---|---|---|
| WBGene00022589 | 0.688793783 | 0.86378755 | 1 | ZC317.6 |
| WBGene00022590 | 0.201336508 | 0.63798913 | 0.79024747 | ZC317.7 |
| WBGene00022591 | 0.070757478 | 0.91706886 | 0.95637631 | cuti-1 |
| WBGene00022592 | 0.413816031 | 0.23769395 | 0.44395533 | klu-2 |
| WBGene00022593 | 1.372257791 | 2.81E-07 | 1.1444E-05 | ZC328.3 |
| WBGene00022594 | -0.01455644 | 0.9964743 | 1 | ZC328.5 |
| WBGene00022596 | 1.413734769 | 0.68804531 | 1 | ZC395.4 |
| WBGene00022597 | 0.963976221 | 0.07822125 | 0.21746833 | ZC395.5 |
| WBGene00022598 | 0.110922197 | 0.53611409 | 0.71819057 | ztf-8 |
| WBGene00022599 | -0.821827324 | 2.60E-08 | 1.4113E-06 | ZC395.10 |
| WBGene00022601 | 0.327467237 | 0.72410149 | 0.84491266 | ZC404.1 |
| WBGene00022603 | 0.02195724 | 0.91737834 | 0.9564526 | gck-2 |
| WBGene00022604 | 0.688793783 | 0.86378755 | 1 | dmsr-10 |
| WBGene00022605 | 2.717467164 | 0.03363564 | 1 | dmsr-11 |
| WBGene00022606 | 0.377828795 | 0.92526585 | 1 | dmsr-9 |
| WBGene00022608 | 0.645500401 | 0.18127403 | 0.37334148 | madf-9 |
| WBGene00022609 | -0.075749009 | 0.92284899 | 0.9593272 | ZC416.2 |
| WBGene00022610 | 0.32868606 | 0.13420133 | 0.30825563 | ZC416.6 |
| WBGene00022611 | -1.32284331 | 0.05156194 | 0.1639208 | ZC449.1 |
| WBGene00022612 | -0.329983486 | 0.63302847 | 0.7872679 | ZC449.2 |
| WBGene00022613 | -0.085720135 | 0.75763409 | 0.86451461 | sek-3 |
| WBGene00022614 | 1.239981206 | 0.06440139 | 0.19131326 | ZC449.4 |
| WBGene00022615 | 0.793334903 | 0.00642845 | 0.03743299 | ZC449.5 |
| WBGene00022616 | 0.688793783 | 0.86378755 | 1 | hsd-3 |
| WBGene00022617 | 0.377828795 | 0.92526585 | 1 | ZC477.2 |
| WBGene00022618 | -0.635469696 | 0.00070581 | 0.00702395 | ZC477.3 |
| WBGene00022619 | -0.156269293 | 0.96924949 | 1 | ZC477.4 |
| WBGene00022620 | 0.031991752 | 0.87529163 | 0.93275833 | rde-8 |
| WBGene00022621 | 1.216957619 | 0.74819621 | 1 | ZC477.7 |
| WBGene00022623 | 0.349930137 | 0.59796822 | 0.76380138 | ZC487.1 |
| WBGene00022627 | -0.137274522 | 0.87378078 | 0.93204818 | ZC513.2 |
| WBGene00022628 | -1.356292182 | 0.63719725 | 1 | ZC513.3 |
| WBGene00022629 | 0.065027368 | 0.74693393 | 0.85908724 | ZC513.5 |
| WBGene00022631 | 0.133518809 | 0.77260546 | 0.87295559 | nekl-2 |
| WBGene00022632 | 1.172985847 | 0.48570467 | 1 | ZC581.2 |
| WBGene00022633 | 1.086871428 | 0.00068308 | 0.00685398 | ZC581.3 |
| WBGene00022634 | 0.57991002 | 0.82402153 | 1 | ZC581.7 |
| WBGene00022635 | 1.139758594 | 0.01345757 | 0.06414934 | ZC581.9 |
| WBGene00022637 | -1.969805064 | 0.62862569 | 1 | nhr-252 |
| WBGene00022638 | -2.699868606 | 0.14824008 | 1 | scl-27 |
| WBGene00022639 | 1.660064349 | 0.6377947 | 1 | nhr-253 |
| WBGene00022641 | 0.418733808 | 0.50198909 | 0.68967405 | ZK6.6 |
| WBGene00022642 | 0.416420126 | 0.0046503 | 0.02962208 | lipl-5 |

| | | | | |
|---|---|---|---|---|
| WBGene00022643 | 0.50653741 | 0.44625739 | 0.64561618 | ZK6.8 |
| WBGene00022644 | 0.476582664 | 0.00314344 | 0.02196735 | dod-19 |
| WBGene00022645 | 0.057913923 | 0.71729906 | 0.84012424 | ZK6.11 |
| WBGene00022646 | -0.112311197 | 0.51803073 | 0.70367097 | acl-9 |
| WBGene00022647 | 0.396042 | 0.37637595 | 0.58404657 | slc-17.1 |
| WBGene00022648 | 1.062657457 | 0.66313467 | 1 | ZK54.3 |
| WBGene00022649 | 0.355031276 | 0.49465859 | 0.68308599 | ZK84.1 |
| WBGene00022650 | -0.001708987 | 0.9989657 | 1 | ZK84.2 |
| WBGene00022651 | -0.668264701 | 0.85551153 | 1 | ZK84.4 |
| WBGene00022652 | 0.281454817 | 0.94435481 | 1 | ZK84.5 |
| WBGene00022653 | -0.543575025 | 0.0192221 | 0.08332046 | ZK105.1 |
| WBGene00022654 | 0.041803775 | 0.98890197 | 1 | ZK105.3 |
| WBGene00022655 | -1.094201388 | 0.7816772 | 1 | srsx-40 |
| WBGene00022656 | 0.938982574 | 0.8146943 | 1 | ZK105.5 |
| WBGene00022657 | 0.063764476 | 0.95894514 | 1 | ZK105.6 |
| WBGene00022661 | -1.0062442 | 0.63208617 | 1 | ZK112.5 |
| WBGene00022662 | 1.851444818 | 0.53669729 | 1 | ZK112.6 |
| WBGene00022663 | -0.435294812 | 0.09525541 | 0.24743173 | glrx-21 |
| WBGene00022664 | 0.814270021 | 0.0184696 | 0.08089445 | ZK121.2 |
| WBGene00022665 | 0.049417767 | 0.91557965 | 0.95580617 | ZK154.1 |
| WBGene00022666 | -0.190410301 | 0.62687778 | 0.78309735 | ZK154.4 |
| WBGene00022667 | -0.263864492 | 0.29759242 | 0.5076675 | ZK154.5 |
| WBGene00022668 | 0.129012145 | 0.76401768 | 0.86806496 | ZK154.6 |
| WBGene00022669 | -0.757080722 | 0.00292438 | 0.02081178 | ZK177.1 |
| WBGene00022670 | 0.377828795 | 0.92526585 | 1 | ZK177.2 |
| WBGene00022671 | -0.245531902 | 0.94070824 | 1 | ZK177.3 |
| WBGene00022672 | -0.122497029 | 0.52482929 | 0.70933456 | npp-24 |
| WBGene00022673 | -0.173644313 | 0.30986687 | 0.51826611 | ZK177.8 |
| WBGene00022674 | 0.233739675 | 0.93575889 | 1 | ZK177.9 |
| WBGene00022675 | 0.840828588 | 0.01664331 | 0.07473157 | gbb-2 |
| WBGene00022677 | 0.201220794 | 0.5072818 | 0.69409515 | ZK180.3 |
| WBGene00022678 | -0.247731634 | 0.12044484 | 0.28689833 | sar-1 |
| WBGene00022679 | -0.6585845 | 0.01948949 | 0.08396019 | ZK180.5 |
| WBGene00022680 | 0.108445068 | 0.92134736 | 1 | ZK180.6 |
| WBGene00022681 | -0.835225658 | 0.0120395 | 0.05944351 | ZK185.1 |
| WBGene00022682 | 0.449060899 | 0.50500294 | 0.69198908 | ZK185.2 |
| WBGene00022683 | -0.007678253 | 0.9847194 | 0.99257382 | ZK185.3 |
| WBGene00022685 | 1.741147134 | 0.37149958 | 1 | ZK250.2 |
| WBGene00022689 | -0.156269293 | 0.96924949 | 1 | math-48 |
| WBGene00022690 | -0.545585854 | 0.5608404 | 1 | math-49 |
| WBGene00022691 | 1.018035489 | 0.62989118 | 1 | math-50 |
| WBGene00022694 | -0.299436351 | 0.12420523 | 0.29360898 | gcna-1 |
| WBGene00022695 | 0.280163511 | 0.67601496 | 0.81463996 | ZK328.6 |

| | | | | |
|---|---|---|---|---|
| WBGene00022696 | 0.758877607 | 0.35373869 | 0.56237866 | ZK328.7 |
| WBGene00022697 | 0.019543624 | 0.92236263 | 0.9593272 | cyy-1 |
| WBGene00022698 | -0.028458303 | 0.98787509 | 1 | ZK353.2 |
| WBGene00022700 | -1.637531945 | 0.41171815 | 1 | ZK353.4 |
| WBGene00022702 | -0.408531914 | 0.05277179 | 0.16650268 | cutc-1 |
| WBGene00022703 | -0.202852564 | 0.21677386 | 0.41855143 | ubxn-4 |
| WBGene00022704 | -0.48993192 | 0.04445179 | 0.1476982 | ZK353.9 |
| WBGene00022705 | 1.584577408 | 0.63258867 | 1 | ZK354.2 |
| WBGene00022706 | 1.025181888 | 0.79790347 | 1 | ZK354.3 |
| WBGene00022707 | -0.014519762 | 0.99603746 | 1 | ZK354.6 |
| WBGene00022708 | 1.743415485 | 0.50483382 | 1 | ZK354.7 |
| WBGene00022709 | -2.107863047 | 0.2011772 | 1 | ZK354.8 |
| WBGene00022711 | 1.181686151 | 0.7579682 | 1 | irld-63 |
| WBGene00022712 | 0.102809373 | 0.76513177 | 0.86853281 | ZK355.2 |
| WBGene00022713 | -1.901856996 | 0.64072034 | 1 | ZK355.3 |
| WBGene00022714 | -0.156269293 | 0.96924949 | 1 | irld-64 |
| WBGene00022715 | 0.747845955 | 0.85215965 | 1 | irld-65 |
| WBGene00022717 | 0.260160294 | 0.11570033 | 0.28023616 | hipr-1 |
| WBGene00022718 | 0.088401506 | 0.6250851 | 0.78190379 | ZK370.4 |
| WBGene00022719 | -0.086154845 | 0.66791137 | 0.80985814 | pdhk-2 |
| WBGene00022720 | -0.348143663 | 0.60318288 | 0.76787704 | ZK370.6 |
| WBGene00022721 | -0.143594054 | 0.62276983 | 0.7803745 | ugtp-1 |
| WBGene00022722 | -0.012978135 | 0.94105215 | 0.96956163 | ZK370.8 |
| WBGene00022723 | -0.01455644 | 0.9964743 | 1 | ZK380.2 |
| WBGene00022724 | -0.156269293 | 0.96924949 | 1 | ZK380.4 |
| WBGene00022725 | -0.919141959 | 0.46479743 | 1 | ZK381.2 |
| WBGene00022726 | 0.377828795 | 0.92526585 | 1 | nhr-249 |
| WBGene00022727 | 0.465005897 | 0.24522595 | 0.45309251 | prkl-1 |
| WBGene00022730 | 1.955439233 | 0.33666494 | 1 | ZK402.3 |
| WBGene00022731 | -0.014519762 | 0.99603746 | 1 | ZK402.5 |
| WBGene00022732 | 2.063391569 | 0.37853067 | 1 | ZK418.2 |
| WBGene00022733 | -2.347665801 | 0.13703433 | 1 | tmem-17 |
| WBGene00022734 | -0.198669682 | 0.27685234 | 0.4856151 | ZK418.5 |
| WBGene00022735 | 0.25124795 | 0.74347305 | 0.85705529 | ZK418.6 |
| WBGene00022736 | 0.401684118 | 0.43819153 | 0.63866548 | ZK418.7 |
| WBGene00022737 | -0.488594236 | 0.11366256 | 0.27689482 | tofu-7 |
| WBGene00022738 | -0.475041982 | 0.00811162 | 0.04427406 | ZK418.9 |
| WBGene00022739 | 0.036107721 | 0.85888502 | 0.92337935 | toe-1 |
| WBGene00022741 | -1.570896869 | 0.0032766 | 0.02264157 | ZK430.5 |
| WBGene00022742 | -0.199041195 | 0.37602963 | 0.58373299 | ZK430.7 |
| WBGene00022743 | 0.931437293 | 0.00027938 | 0.0034056 | mlt-7 |
| WBGene00022744 | 0.193302195 | 0.91734354 | 1 | ZK470.1 |
| WBGene00022745 | 0.571283644 | 0.02426942 | 0.09818562 | ZK470.2 |

| | | | | |
|---|---|---|---|---|
| WBGene00022747 | 1.796575934 | 0.34491145 | 1 | ZK470.6 |
| WBGene00022748 | 0.252955423 | 0.08244926 | 0.22530191 | oaz-1 |
| WBGene00022749 | -0.178727389 | 0.36589248 | 0.57350663 | ZK484.3 |
| WBGene00022750 | -0.272969559 | 0.06840874 | 0.19922057 | ZK484.4 |
| WBGene00022751 | -0.678442377 | 0.5764057 | 1 | msrp-2 |
| WBGene00022752 | -0.911418009 | 0.54348815 | 1 | ZK484.6 |
| WBGene00022753 | 0.983488853 | 0.80601565 | 1 | ZK484.7 |
| WBGene00022754 | 2.44066524 | 0.32717233 | 1 | nspd-1 |
| WBGene00022756 | 2.017700667 | 0.17409052 | 1 | nhr-251 |
| WBGene00022759 | -0.184794652 | 0.2733896 | 0.48209486 | ZK546.2 |
| WBGene00022761 | 0.688793783 | 0.86378755 | 1 | ZK546.4 |
| WBGene00022762 | -0.115085725 | 0.59337396 | 0.76005346 | ZK546.5 |
| WBGene00022765 | -0.16550863 | 0.40799304 | 0.6141434 | ZK546.14 |
| WBGene00022766 | -0.236953727 | 0.24980135 | 0.45751284 | cblc-1 |
| WBGene00022767 | 1.765816519 | 0.16534152 | 1 | ZK563.2 |
| WBGene00022769 | -0.871928251 | 0.00417528 | 0.02718215 | sorf-1 |
| WBGene00022770 | 0.304855446 | 0.57459896 | 0.74795819 | acp-1 |
| WBGene00022771 | 0.879481377 | 0.34704966 | 0.555948 | ZK616.1 |
| WBGene00022772 | -0.995773026 | 0.14199637 | 0.32014447 | ZK616.2 |
| WBGene00022773 | -0.639696505 | 0.06184577 | 0.186043 | ZK616.3 |
| WBGene00022774 | -0.578783187 | 0.0005126 | 0.00546393 | swsn-6 |
| WBGene00022775 | -0.452445093 | 0.06554762 | 0.19336894 | ZK616.5 |
| WBGene00022776 | -0.211100762 | 0.29001387 | 0.4999373 | perm-3 |
| WBGene00022780 | 1.247575581 | 0.7549672 | 1 | ZK622.1 |
| WBGene00022781 | 1.013636052 | 6.20E-12 | 7.53E-10 | pmt-1 |
| WBGene00022782 | 0.604998219 | 0.17724295 | 0.36885612 | ZK622.4 |
| WBGene00022783 | -0.718922932 | 0.00171459 | 0.01380427 | tomm-7 |
| WBGene00022785 | -0.177935181 | 0.42382861 | 0.62806093 | ZK652.6 |
| WBGene00022786 | -0.373949652 | 0.59120034 | 0.75850822 | ZK652.8 |
| WBGene00022787 | -0.227201163 | 0.24109445 | 0.44872386 | tag-307 |
| WBGene00022788 | -0.06610051 | 0.80129913 | 0.89146639 | slc-17.7 |
| WBGene00022789 | -1.031009651 | 0.03459478 | 0.12515353 | lron-2 |
| WBGene00022790 | 0.247694858 | 0.87622457 | 1 | ZK682.7 |
| WBGene00022791 | -0.725817615 | 0.00108325 | 0.00977557 | ZK686.1 |
| WBGene00022792 | -0.458234827 | 0.01301044 | 0.06294757 | ZK686.2 |
| WBGene00022793 | -0.103419598 | 0.52991909 | 0.7139882 | ZK686.3 |
| WBGene00022794 | -0.409317074 | 0.07084305 | 0.20389564 | snu-23 |
| WBGene00022795 | 1.216957619 | 0.74819621 | 1 | ZK686.5 |
| WBGene00022796 | 0.905096248 | 0.68842149 | 1 | ZK688.1 |
| WBGene00022797 | 0.53328948 | 0.1305468 | 0.30264823 | best-24 |
| WBGene00022798 | -0.307934086 | 0.2288188 | 0.4340198 | fahd-1 |
| WBGene00022799 | 0.281454817 | 0.94435481 | 1 | ZK688.4 |
| WBGene00022800 | 0.290292286 | 0.0849537 | 0.22971557 | ZK688.5 |

| | | | | |
|---|---|---|---|---|
| WBGene00022801 | -0.169386742 | 0.45025216 | 0.64889218 | pcp-5 |
| WBGene00022802 | -1.213890816 | 0.02353008 | 0.09612338 | ZK688.7 |
| WBGene00022803 | -0.45518007 | 0.04950682 | 0.15947139 | ZK688.9 |
| WBGene00022804 | -0.156269293 | 0.96924949 | 1 | ZK697.1 |
| WBGene00022805 | 0.080358852 | 0.82369684 | 0.90485915 | nhr-256 |
| WBGene00022807 | 2.028800581 | 0.39258153 | 1 | srab-25 |
| WBGene00022808 | -0.156269293 | 0.96924949 | 1 | ZK697.8 |
| WBGene00022810 | 1.400871824 | 0.12063167 | 0.28723066 | ZK721.3 |
| WBGene00022811 | 0.127626107 | 0.90367501 | 1 | ZK721.4 |
| WBGene00022812 | -0.146255709 | 0.54614202 | 0.72522261 | uvs-1 |
| WBGene00022813 | -0.322802511 | 0.48985294 | 0.67938138 | ZK742.3 |
| WBGene00022814 | 0.556610376 | 0.34339412 | 0.55176655 | ZK742.4 |
| WBGene00022815 | 0.10887386 | 0.91762153 | 1 | asic-1 |
| WBGene00022816 | -0.048819604 | 0.89469346 | 0.94453253 | fbn-1 |
| WBGene00022817 | -0.016050924 | 0.9331874 | 0.96465119 | upp-1 |
| WBGene00022820 | -0.811362604 | 1.01E-08 | 6.12E-07 | ZK813.1 |
| WBGene00022821 | 0.164312556 | 0.4864742 | 0.67622374 | ZK813.2 |
| WBGene00022822 | -0.68139602 | 2.6852E-06 | 7.5118E-05 | ZK813.3 |
| WBGene00022823 | 1.076795198 | 0.09977599 | 0.25443821 | ZK813.4 |
| WBGene00022824 | 1.643592421 | 0.2629751 | 1 | ZK813.5 |
| WBGene00022825 | 0.203765729 | 0.85012652 | 1 | ZK816.1 |
| WBGene00022827 | -0.990160542 | 0.46128724 | 1 | ZK816.4 |
| WBGene00022829 | 0.938982574 | 0.8146943 | 1 | ZK867.2 |
| WBGene00022830 | -0.367695086 | 0.02025719 | 0.08629562 | ZK973.1 |
| WBGene00022831 | -0.291122157 | 0.08894338 | 0.23639701 | cec-10 |
| WBGene00022832 | -0.257104638 | 0.16822817 | 0.35595298 | pdp-1 |
| WBGene00022833 | -0.156269293 | 0.96924949 | 1 | ZK973.4 |
| WBGene00022834 | -0.288123366 | 0.80341712 | 1 | ZK973.8 |
| WBGene00022835 | -0.429191587 | 0.09843121 | 0.25249383 | ZK973.9 |
| WBGene00022836 | -0.429627437 | 0.06409343 | 0.19067836 | ZK973.11 |
| WBGene00022837 | -0.316682955 | 0.72249886 | 0.84361018 | ceh-45 |
| WBGene00022838 | -0.35285494 | 0.81997989 | 1 | ZK993.2 |
| WBGene00022840 | -0.156269293 | 0.96924949 | 1 | ZK994.1 |
| WBGene00022842 | -0.094987758 | 0.93361211 | 1 | ZK994.6 |
| WBGene00022843 | 1.019267722 | 0.67145278 | 1 | ZK1055.2 |
| WBGene00022845 | 0.103145558 | 0.94669145 | 1 | ZK1055.4 |
| WBGene00022846 | 0.742628056 | 0.76901841 | 1 | ZK1055.5 |
| WBGene00022847 | 0.633424098 | 0.12433244 | 0.29368097 | ZK1055.6 |
| WBGene00022848 | 0.009459455 | 0.97404595 | 0.98649644 | ZK1055.7 |
| WBGene00022849 | -0.600661498 | 0.71692312 | 1 | acs-6 |
| WBGene00022850 | -0.637434956 | 0.00327055 | 0.02261262 | ZK1127.3 |
| WBGene00022851 | -0.491671413 | 0.02614859 | 0.1032287 | ZK1127.4 |
| WBGene00022852 | -0.636892996 | 0.00987844 | 0.05133254 | ZK1127.5 |

| | | | | |
|---|---|---|---|---|
| WBGene00022853 | 0.289128967 | 0.32405917 | 0.53253957 | ZK1127.6 |
| WBGene00022854 | -0.159171654 | 0.47876771 | 0.67090476 | cin-4 |
| WBGene00022855 | -0.029743134 | 0.86537303 | 0.92768209 | tcer-1 |
| WBGene00022856 | -0.348852573 | 0.02240192 | 0.09304882 | cth-2 |
| WBGene00022857 | -0.346122014 | 0.32801639 | 0.53663232 | ZK1127.12 |
| WBGene00022858 | 1.928535499 | 7.82E-10 | 6.39E-08 | ZK1193.2 |
| WBGene00022858 | 1.78862492 | 0.00634339 | 0.0371585 | ZK1193.3 |
| WBGene00022858 | 1.822665157 | 0.01181069 | 0.05862371 | ZK1193.4 |
| WBGene00022861 | 0.528797073 | 0.09938648 | 0.25393171 | dve-1 |
| WBGene00022862 | -0.176584832 | 0.53071642 | 0.71443643 | ZK1236.1 |
| WBGene00022864 | -0.557016318 | 0.04930703 | 0.15938142 | ZK1236.5 |
| WBGene00022865 | -0.01012573 | 0.95761475 | 0.97850563 | ufbp-1 |
| WBGene00022866 | 1.114796914 | 0.5418588 | 1 | ZK1240.1 |
| WBGene00022867 | 0.978235757 | 0.03494476 | 0.12593184 | ZK1240.2 |
| WBGene00022868 | 0.79194852 | 0.02693735 | 0.10536286 | ZK1240.3 |
| WBGene00022869 | -0.156269293 | 0.96924949 | 1 | ZK1240.4 |
| WBGene00022870 | 1.251491104 | 0.29185731 | 1 | ZK1240.5 |
| WBGene00022871 | 0.574508983 | 0.8551971 | 1 | ZK1240.6 |
| WBGene00022872 | 3.199079286 | 0.09328367 | 1 | ZK1240.8 |
| WBGene00022873 | 0.80860382 | 0.22494295 | 0.42900289 | ZK1240.9 |
| WBGene00022874 | 0.270496667 | 0.92869767 | 1 | nep-25 |
| WBGene00022876 | 1.216957619 | 0.74819621 | 1 | ZK1248.5 |
| WBGene00022877 | -0.737462795 | 0.83433562 | 1 | wago-5 |
| WBGene00022880 | -0.136523006 | 0.42358835 | 0.62801149 | tbc-2 |
| WBGene00022881 | -0.475736717 | 0.0419666 | 0.14229939 | ZK1248.11 |
| WBGene00022882 | -0.338174679 | 0.10065785 | 0.25582729 | ZK1248.13 |
| WBGene00022883 | 0.021625334 | 0.91537505 | 0.95580617 | ZK1248.15 |
| WBGene00022885 | -0.458545481 | 0.2205665 | 0.42366719 | ZK1248.19 |
| WBGene00022886 | -0.156269293 | 0.96924949 | 1 | ZK1290.1 |
| WBGene00022887 | -0.224382272 | 0.54161357 | 0.72227294 | ZK1290.5 |
| WBGene00022888 | 0.983488853 | 0.80601565 | 1 | rnh-1.1 |
| WBGene00022891 | 0.938159598 | 0.40159098 | 1 | ZK1290.10 |
| WBGene00022892 | -0.156269293 | 0.96924949 | 1 | ZK1290.11 |
| WBGene00022893 | 1.008983128 | 0.04430569 | 0.14738518 | ZK1290.13 |
| WBGene00022895 | -0.156687082 | 0.83296523 | 0.90960882 | B0213.18 |
| WBGene00022904 | -0.764269844 | 0.76036927 | 1 | C01B12.9 |
| WBGene00022910 | 1.00650968 | 0.00831214 | 0.04499226 | C05D2.11 |
| WBGene00022951 | 1.338364795 | 4.67E-11 | 4.72E-09 | C30E1.9 |
| WBGene00022968 | 0.641597581 | 0.25887453 | 0.46708312 | C55F2.3 |
| WBGene00022970 | -0.778827203 | 0.07421593 | 0.21047783 | CD4.10 |
| WBGene00023020 | 1.139358466 | 0.62800936 | 1 | F30H5.4 |
| WBGene00023021 | -0.171540064 | 0.67624363 | 0.81466397 | F32A5.9 |
| WBGene00023027 | 1.802131411 | 0.04967345 | 0.15980003 | F35H10.12 |

| | | | | |
|---|---|---|---|---|
| WBGene00023065 | 0.003982828 | 0.99438457 | 0.99717709 | F53F10.6 |
| WBGene00023066 | -0.156269293 | 0.96924949 | 1 | F53F10.7 |
| WBGene00023067 | -0.953620689 | 3.60E-07 | 1.4159E-05 | rpl-41.1 |
| WBGene00023068 | -0.557316269 | 0.00032035 | 0.00380858 | rpl-41.2 |
| WBGene00023069 | 0.028581079 | 0.91552815 | 0.95580617 | rncs-1 |
| WBGene00023071 | 1.435925494 | 0.70920779 | 1 | F56C9.12 |
| WBGene00023104 | 1.38322681 | 0.69550305 | 1 | K10F12.8 |
| WBGene00023159 | 1.829555997 | 2.13E-23 | 1.04E-20 | W06H8.5 |
| WBGene00023172 | 0.694609957 | 0.73912856 | 1 | Y23H5A.8 |
| WBGene00023186 | 0.281454817 | 0.94435481 | 1 | Y67D8B.3 |
| WBGene00023187 | -0.156269293 | 0.96924949 | 1 | Y67D8C.12 |
| WBGene00023193 | 1.601302495 | 0.65842342 | 1 | Y74C9A.6 |
| WBGene00023284 | 0.85365664 | 0.27398797 | 0.48251928 | F41B5.1 |
| WBGene00023404 | -0.119279815 | 0.78153866 | 0.879231 | plr-1 |
| WBGene00023405 | 0.119026812 | 0.62051794 | 0.77878286 | sor-1 |
| WBGene00023407 | 0.205112112 | 0.63800393 | 0.79024747 | cex-1 |
| WBGene00023408 | 0.488806716 | 0.21037969 | 0.41038236 | cex-2 |
| WBGene00023409 | -0.770903386 | 0.001132 | 0.01008336 | mbd-2 |
| WBGene00023411 | 0.377828795 | 0.92526585 | 1 | srz-3 |
| WBGene00023413 | 0.281454817 | 0.94435481 | 1 | H31B20.2 |
| WBGene00023414 | 0.454275906 | 0.40356397 | 0.61108383 | R01H2.7 |
| WBGene00023415 | 0.245560738 | 0.7287775 | 0.84808662 | F16C3.3 |
| WBGene00023416 | 0.223518209 | 0.86069197 | 1 | spi-1 |
| WBGene00023417 | -0.743076839 | 0.12317956 | 0.29175245 | swm-1 |
| WBGene00023418 | -0.62895042 | 0.00291338 | 0.02077929 | R06F6.12 |
| WBGene00023419 | -0.367099446 | 0.25969456 | 0.4678664 | F53F4.16 |
| WBGene00023420 | 3.103556858 | 0.04445855 | 1 | C26D10.7 |
| WBGene00023421 | -0.128278227 | 0.57925409 | 0.75037463 | rde-11 |
| WBGene00023422 | -0.662321876 | 0.00511904 | 0.03169317 | mrpl-21 |
| WBGene00023423 | 0.483840154 | 0.8151116 | 1 | F43C1.7 |
| WBGene00023424 | 0.377828795 | 0.92526585 | 1 | C53D6.10 |
| WBGene00023425 | 0.530404348 | 0.80087734 | 1 | try-9 |
| WBGene00023427 | -0.118558845 | 0.73908916 | 0.85446788 | F14D2.15 |
| WBGene00023428 | 0.530404348 | 0.80087734 | 1 | K07C5.10 |
| WBGene00023429 | 1.413734769 | 0.68804531 | 1 | W08G11.6 |
| WBGene00023430 | -3.014567441 | 0.24693748 | 1 | D1054.17 |
| WBGene00023431 | 1.025600152 | 0.51742525 | 1 | C25E10.13 |
| WBGene00023432 | 0.425590228 | 0.44665969 | 0.6457926 | K12B6.9 |
| WBGene00023450 | -0.725010411 | 0.04012992 | 0.13782669 | T12D8.10 |
| WBGene00023451 | -0.285054963 | 0.42598331 | 0.62971931 | mlc-7 |
| WBGene00023452 | -2.171326218 | 0.36522606 | 1 | mir-353 |
| WBGene00023453 | -1.060364372 | 0.63520895 | 1 | mir-356.1 |
| WBGene00023455 | -0.156269293 | 0.96924949 | 1 | mir-358 |

| | | | | |
|---|---|---|---|---|
| WBGene00023458 | -0.717831747 | 0.57247245 | 1 | M02E1.3 |
| WBGene00023460 | 0.938982574 | 0.8146943 | 1 | F46H6.5 |
| WBGene00023461 | -1.969805064 | 0.62862569 | 1 | F53G12.12 |
| WBGene00023463 | -0.156269293 | 0.96924949 | 1 | srz-82 |
| WBGene00023466 | -1.094201388 | 0.7816772 | 1 | R151.11 |
| WBGene00023467 | 0.635387686 | 0.82210541 | 1 | Y38F2AR.14 |
| WBGene00023468 | 0.983488853 | 0.80601565 | 1 | B0412.6 |
| WBGene00023469 | -1.663007101 | 0.43086568 | 1 | C16A3.11 |
| WBGene00023472 | 1.317729748 | 0.38055422 | 1 | F16A11.5 |
| WBGene00023475 | -0.156269293 | 0.96924949 | 1 | srz-69 |
| WBGene00023477 | -0.674802711 | 0.3044543 | 0.5135287 | nspc-1 |
| WBGene00023483 | 1.614973531 | 0.38852171 | 1 | Y75B8A.39 |
| WBGene00023484 | 1.784345117 | 0.3562364 | 1 | Y54G2A.37 |
| WBGene00023485 | -0.340993014 | 0.47328978 | 0.66638191 | ttr-49 |
| WBGene00023486 | -0.799979461 | 0.3474733 | 0.55625968 | K08B4.7 |
| WBGene00023487 | -0.524223778 | 0.02522576 | 0.10078409 | mrps-24 |
| WBGene00023489 | -0.639154235 | 0.0963811 | 0.24913639 | F42A10.9 |
| WBGene00023490 | -1.035929762 | 0.59261215 | 1 | F36A2.14 |
| WBGene00023491 | -0.171547264 | 0.63403295 | 0.78783996 | tsp-21 |
| WBGene00023492 | -0.443110857 | 0.54102711 | 0.72202181 | T24D3.2 |
| WBGene00023493 | 0.65079482 | 0.6373838 | 1 | nspc-12 |
| WBGene00023497 | 0.136043229 | 0.54003024 | 0.72144664 | lin-15B |
| WBGene00023498 | 0.06661653 | 0.79429871 | 0.88635851 | lin-15A |
| WBGene00023499 | -1.159714244 | 0.75790634 | 1 | srz-63 |
| WBGene00023500 | -0.401150777 | 0.24778254 | 0.45583943 | C11H1.9 |
| WBGene00023501 | 0.507551112 | 0.62440018 | 1 | F11E6.11 |
| WBGene00023502 | 0.658263905 | 0.3881514 | 0.59694435 | clec-184 |
| WBGene00023503 | -0.096036887 | 0.91059056 | 0.95371635 | F17C11.13 |
| WBGene00023504 | 0.04813088 | 0.96115895 | 1 | F26F2.8 |
| WBGene00023506 | 0.281454817 | 0.94435481 | 1 | srz-11 |
| WBGene00023508 | 0.688793783 | 0.86378755 | 1 | srz-38 |
| WBGene00023513 | 0.938982574 | 0.8146943 | 1 | srz-71 |
| WBGene00023515 | -0.014519762 | 0.99603746 | 1 | srz-97 |
| WBGene00023517 | -0.156269293 | 0.96924949 | 1 | srz-75 |
| WBGene00023518 | -0.01455644 | 0.9964743 | 1 | srz-76 |
| WBGene00023519 | -0.269250864 | 0.79056423 | 1 | Y60A3A.24 |
| WBGene00023520 | 1.049693913 | 0.21828997 | 0.42080954 | oig-5 |
| WBGene00043049 | -0.323868354 | 0.91634405 | 1 | F32H2.11 |
| WBGene00043050 | -0.176807901 | 0.62653555 | 0.78307269 | oig-4 |
| WBGene00043051 | -0.230733437 | 0.39073472 | 0.5991985 | C29E4.13 |
| WBGene00043052 | 0.377828795 | 0.92526585 | 1 | srz-104 |
| WBGene00043054 | -0.299947756 | 0.78478907 | 1 | T21B4.15 |
| WBGene00043056 | -0.287392693 | 0.38860609 | 0.59729296 | nfya-2 |

| | | | | |
|---|---|---|---|---|
| WBGene00043057 | -0.82691943 | 0.00218959 | 0.01666699 | M142.8 |
| WBGene00043059 | -1.831477136 | 0.52194793 | 1 | srab-26 |
| WBGene00043060 | -0.480771851 | 0.06850371 | 0.19940146 | R144.12 |
| WBGene00043061 | 0.627834078 | 0.77998941 | 1 | Y54G2A.38 |
| WBGene00043062 | 1.885273651 | 7.54E-10 | 6.23E-08 | clec-80 |
| WBGene00043064 | -0.964199988 | 0.00019617 | 0.00260567 | nbet-1 |
| WBGene00043065 | 0.579666901 | 0.79455736 | 1 | C55C3.7 |
| WBGene00043066 | -0.234620636 | 0.72189146 | 0.84322534 | acr-25 |
| WBGene00043067 | 1.227426159 | 0.34130064 | 1 | dlc-5 |
| WBGene00043068 | -0.780765891 | 0.18557311 | 0.37815427 | F32E10.9 |
| WBGene00043097 | 0.185206662 | 0.38484117 | 0.59331513 | C02D5.4 |
| WBGene00043120 | -1.837668733 | 0.43198255 | 1 | C09H10.4 |
| WBGene00043147 | 1.025181888 | 0.79790347 | 1 | nspd-6 |
| WBGene00043156 | -0.760848637 | 0.00046001 | 0.00501799 | C27F2.9 |
| WBGene00043279 | -0.547887755 | 0.25989756 | 0.46809319 | F22D6.8 |
| WBGene00043297 | 2.595821743 | 0.06184716 | 1 | F28F5.4 |
| WBGene00043302 | 0.471532356 | 0.59358782 | 0.76012548 | F32D8.11 |
| WBGene00043308 | 1.526476613 | 0.440991 | 1 | tmem-107 |
| WBGene00043469 | 0.938982574 | 0.8146943 | 1 | R09H3.2 |
| WBGene00043534 | -0.680303862 | 0.55340779 | 1 | W02B12.13 |
| WBGene00043702 | 0.866788676 | 0.12132724 | 0.2884343 | Y54G2A.7 |
| WBGene00043705 | -0.01606522 | 0.97888355 | 0.98873753 | egrh-2 |
| WBGene00043743 | 0.399606396 | 0.86544406 | 1 | Y59E9AL.2 |
| WBGene00043948 | -0.571517947 | 0.25501468 | 0.46259275 | ZC262.7 |
| WBGene00043952 | -0.22461868 | 0.93331196 | 1 | ZC404.2 |
| WBGene00043980 | 3.159526349 | 0.1366367 | 1 | F11D5.7 |
| WBGene00043981 | -0.038488761 | 0.97611698 | 1 | T14E8.4 |
| WBGene00043982 | 0.309725172 | 0.6615033 | 0.80512302 | B0403.6 |
| WBGene00043983 | 1.4687985 | 0.00163517 | 0.01334673 | C39D10.11 |
| WBGene00043984 | -0.490561023 | 0.85693136 | 1 | C56E6.7 |
| WBGene00043985 | 0.948470084 | 0.14419069 | 0.32263531 | C44C10.11 |
| WBGene00043986 | -0.156269293 | 0.96924949 | 1 | fut-5 |
| WBGene00043992 | 0.947903141 | 0.29527634 | 0.5056347 | bbs-4 |
| WBGene00043994 | 0.332609682 | 0.66401352 | 0.80730905 | C07A9.12 |
| WBGene00043995 | 0.342860973 | 0.68853829 | 0.82264522 | F40F12.9 |
| WBGene00043996 | -2.048505773 | 0.12557721 | 1 | fbxb-87 |
| WBGene00043997 | 0.955288171 | 0.58158157 | 1 | Y79H2A.12 |
| WBGene00043998 | -0.885036938 | 0.34856382 | 0.55712393 | T21C12.8 |
| WBGene00044001 | 0.092018155 | 0.76430257 | 0.86812587 | C09G4.4 |
| WBGene00044002 | 0.281454817 | 0.94435481 | 1 | srt-74 |
| WBGene00044003 | -0.156269293 | 0.96924949 | 1 | Y41E3.18 |
| WBGene00044004 | 1.599406759 | 0.14925974 | 1 | C24A8.6 |
| WBGene00044005 | -1.11019416 | 0.52432696 | 1 | F37C12.18 |

| | | | | |
|---|---|---|---|---|
| WBGene00044006 | 0.426754676 | 0.57221525 | 0.7462154 | H32K21.1 |
| WBGene00044007 | -0.255067679 | 0.57602253 | 0.74868745 | ZK1320.13 |
| WBGene00044008 | 0.103304254 | 0.61471009 | 0.77514863 | gip-2 |
| WBGene00044009 | 0.929866607 | 0.3014394 | 0.51102357 | ZK822.6 |
| WBGene00044011 | -0.156269293 | 0.96924949 | 1 | T06E4.12 |
| WBGene00044016 | 0.436270539 | 0.71342806 | 1 | C35A5.10 |
| WBGene00044019 | -0.299683014 | 0.39900372 | 0.60659956 | C55A6.12 |
| WBGene00044020 | -0.329171509 | 0.87141915 | 1 | ZK856.14 |
| WBGene00044021 | 2.153505952 | 0.09090713 | 1 | R02E4.2 |
| WBGene00044024 | -1.557312723 | 0.01332536 | 0.06379465 | C35A5.11 |
| WBGene00044025 | 0.160711852 | 0.55829462 | 0.73532816 | T01G1.4 |
| WBGene00044026 | -0.341851052 | 0.11798882 | 0.28315918 | cisd-1 |
| WBGene00044027 | -0.774842772 | 0.78069886 | 1 | F27E5.8 |
| WBGene00044028 | 0.291127276 | 0.84168749 | 1 | F33H1.6 |
| WBGene00044029 | 1.200228117 | 0.05916939 | 0.18076889 | C05D12.7 |
| WBGene00044030 | -0.737462795 | 0.83433562 | 1 | Y17G7B.23 |
| WBGene00044032 | -0.239835714 | 0.59761986 | 0.76351811 | ceh-99 |
| WBGene00044034 | 0.537471244 | 0.57219859 | 0.7462154 | Y38E10A.28 |
| WBGene00044035 | -1.537696258 | 0.07022019 | 0.20259416 | F27E5.7 |
| WBGene00044043 | -0.07748118 | 0.98067866 | 1 | Y46G5A.38 |
| WBGene00044044 | 0.281454817 | 0.94435481 | 1 | C08B6.14 |
| WBGene00044045 | 0.513117029 | 0.54914299 | 0.72777481 | R11G10.3 |
| WBGene00044047 | -1.117480358 | 0.38316717 | 1 | Y57A10C.11 |
| WBGene00044049 | 0.867976343 | 0.01074516 | 0.05476447 | sfxn-5 |
| WBGene00044058 | -3.093502349 | 0.05981972 | 1 | F17B5.6 |
| WBGene00044059 | 0.688793783 | 0.86378755 | 1 | T27C5.12 |
| WBGene00044061 | -0.109208042 | 0.66334665 | 0.80674063 | tbc-12 |
| WBGene00044063 | -0.650143433 | 0.00330951 | 0.02278607 | bpnt-1 |
| WBGene00044065 | -0.396434048 | 0.16451586 | 0.35084965 | tag-229 |
| WBGene00044066 | -0.786760313 | 0.27114973 | 0.47974773 | fbxb-66 |
| WBGene00044067 | -1.010042351 | 3.9027E-06 | 0.00010201 | zipt-7.1 |
| WBGene00044068 | 0.457348118 | 0.15172978 | 0.33354179 | syd-9 |
| WBGene00044069 | -0.727174518 | 2.515E-06 | 7.1342E-05 | hat-1 |
| WBGene00044070 | 0.839986221 | 0.00549109 | 0.03340047 | set-18 |
| WBGene00044071 | 0.054622348 | 0.84082987 | 0.9124319 | dhhc-14 |
| WBGene00044072 | -0.425639331 | 0.05870286 | 0.17988678 | ham-3 |
| WBGene00044073 | 1.464388641 | 0.00045198 | 0.00494373 | tag-244 |
| WBGene00044074 | -1.895791037 | 0.52793577 | 1 | mltn-5 |
| WBGene00044077 | 0.493605007 | 0.03058637 | 0.11518275 | pacs-1 |
| WBGene00044077 | 0.103308543 | 0.60042028 | 0.76564464 | tag-232 |
| WBGene00044078 | 1.120223376 | 0.03809466 | 0.133095 | tag-243 |
| WBGene00044079 | 0.497406224 | 0.6961308 | 1 | C34E11.7 |
| WBGene00044079 | 1.150523324 | 9.154E-05 | 0.00143036 | tag-241 |

| | | | | |
|---|---|---|---|---|
| WBGene00044080 | -1.712893315 | 1.06E-22 | 4.61E-20 | D1086.10 |
| WBGene00044081 | -0.488209673 | 0.01576096 | 0.07201875 | D1086.11 |
| WBGene00044082 | 0.425746545 | 0.20205681 | 0.40084505 | D1086.12 |
| WBGene00044083 | -0.370607474 | 0.18103079 | 0.37309368 | tin-9.2 |
| WBGene00044090 | 0.377828795 | 0.92526585 | 1 | srbc-33 |
| WBGene00044091 | 0.232157325 | 0.93397866 | 1 | srbc-60 |
| WBGene00044092 | -1.159714244 | 0.75790634 | 1 | srbc-71 |
| WBGene00044094 | -0.112387843 | 0.58457213 | 0.75436671 | fitm-2 |
| WBGene00044106 | 0.281454817 | 0.94435481 | 1 | srbc-84 |
| WBGene00044107 | -0.508114547 | 0.09792286 | 0.25156188 | F58G6.9 |
| WBGene00044109 | 0.399606396 | 0.86544406 | 1 | K02E11.10 |
| WBGene00044112 | -1.566848091 | 0.50325496 | 1 | srm-5 |
| WBGene00044113 | 0.574508983 | 0.8551971 | 1 | C54F6.15 |
| WBGene00044116 | 0.281454817 | 0.94435481 | 1 | srxa-13 |
| WBGene00044117 | -0.156269293 | 0.96924949 | 1 | srsx-10 |
| WBGene00044120 | 3.124296051 | 0.11067134 | 1 | C36B1.13 |
| WBGene00044121 | -0.450812972 | 0.85883116 | 1 | gcp-2.3 |
| WBGene00044122 | 1.876192691 | 0.43542138 | 1 | T28B8.6 |
| WBGene00044125 | 1.38322681 | 0.69550305 | 1 | srsx-16 |
| WBGene00044127 | -0.156269293 | 0.96924949 | 1 | BE10.5 |
| WBGene00044131 | -0.156269293 | 0.96924949 | 1 | Y45G12C.16 |
| WBGene00044132 | 0.257717244 | 0.77330064 | 0.87344339 | C23H4.8 |
| WBGene00044134 | 0.162568571 | 0.71641556 | 0.84006931 | D1025.10 |
| WBGene00044135 | -0.481445132 | 0.78263825 | 1 | F08G12.11 |
| WBGene00044137 | 0.820015846 | 0.7518774 | 1 | F28H6.8 |
| WBGene00044138 | 0.417781254 | 0.78933051 | 1 | sssh-1 |
| WBGene00044139 | -0.760943105 | 0.7243725 | 1 | F42D1.4 |
| WBGene00044140 | -0.156269293 | 0.96924949 | 1 | F42F12.11 |
| WBGene00044141 | -0.156269293 | 0.96924949 | 1 | H03G16.6 |
| WBGene00044142 | -0.531132619 | 0.18207111 | 0.37384831 | H13N06.7 |
| WBGene00044143 | 0.542256861 | 0.38088847 | 0.58901629 | K09E9.4 |
| WBGene00044144 | 1.6274763 | 0.06417994 | 0.19088894 | M153.4 |
| WBGene00044147 | 1.181686151 | 0.7579682 | 1 | M163.11 |
| WBGene00044148 | -1.969805064 | 0.62862569 | 1 | T08D10.4 |
| WBGene00044149 | -1.288333598 | 0.36727206 | 1 | T24D5.6 |
| WBGene00044150 | -0.302639708 | 0.71394296 | 0.83829608 | VB0395L.1 |
| WBGene00044151 | 1.712479368 | 0.58529868 | 1 | W04G3.12 |
| WBGene00044152 | 0.233739675 | 0.93575889 | 1 | W04G3.10 |
| WBGene00044153 | 1.216957619 | 0.74819621 | 1 | W04G3.11 |
| WBGene00044154 | 1.451690413 | 0.37789878 | 1 | Y7A5A.10 |
| WBGene00044156 | -0.116768207 | 0.91679861 | 1 | Y66C5A.1 |
| WBGene00044158 | -0.156269293 | 0.96924949 | 1 | Y53C12A.8 |
| WBGene00044159 | 0.377828795 | 0.92526585 | 1 | F26C11.4 |

| | | | | |
|---|---|---|---|---|
| WBGene00044161 | -0.62146648 | 0.85925752 | 1 | C47G2.8 |
| WBGene00044162 | -1.969805064 | 0.62862569 | 1 | F49C5.10 |
| WBGene00044163 | 0.508249741 | 0.64876028 | 1 | F49C5.11 |
| WBGene00044164 | -1.969805064 | 0.62862569 | 1 | Y46G5A.39 |
| WBGene00044166 | 0.623435544 | 0.52963619 | 1 | C54C6.7 |
| WBGene00044167 | -0.228624902 | 0.82551732 | 1 | C36A4.11 |
| WBGene00044169 | -1.150134399 | 0.4892498 | 1 | C06C3.10 |
| WBGene00044174 | -0.473310722 | 0.57229578 | 0.74624026 | fipr-5 |
| WBGene00044175 | -0.267657505 | 0.76705575 | 0.86946353 | fipr-9 |
| WBGene00044176 | 0.466160861 | 0.36849919 | 0.57602018 | C30G7.2 |
| WBGene00044177 | -0.055390847 | 0.96986133 | 1 | C30G7.3 |
| WBGene00044178 | -0.195247359 | 0.82818032 | 0.90721969 | C30G7.4 |
| WBGene00044180 | -0.604396148 | 0.82723443 | 1 | C47E8.11 |
| WBGene00044181 | -0.737462795 | 0.83433562 | 1 | F09C6.11 |
| WBGene00044182 | 1.849927113 | 0.33150067 | 1 | F09C6.12 |
| WBGene00044183 | 1.013713812 | 0.64240986 | 1 | F09C6.13 |
| WBGene00044187 | 2.456496557 | 0.3218863 | 1 | ttll-9 |
| WBGene00044188 | -0.550500494 | 0.06220537 | 0.18680279 | F28F8.9 |
| WBGene00044191 | 0.505980969 | 0.11654598 | 0.28119728 | F43D2.6 |
| WBGene00044192 | 1.116613983 | 0.69067248 | 1 | F46B6.13 |
| WBGene00044194 | -0.156269293 | 0.96924949 | 1 | F47B8.14 |
| WBGene00044197 | -1.247748198 | 0.13699751 | 0.31294201 | fbxb-63 |
| WBGene00044198 | 0.281454817 | 0.94435481 | 1 | F56H9.6 |
| WBGene00044199 | -1.901856996 | 0.64072034 | 1 | F57B1.8 |
| WBGene00044200 | 0.006137397 | 0.98974489 | 0.99486251 | H37A05.4 |
| WBGene00044202 | 0.808880537 | 0.64561878 | 1 | K10G4.10 |
| WBGene00044204 | -0.866737379 | 5.2248E-05 | 0.0008985 | suds-3 |
| WBGene00044205 | 2.779713462 | 0.22340067 | 1 | T13F3.8 |
| WBGene00044206 | 2.720885057 | 5.09E-11 | 5.11E-09 | T26H5.9 |
| WBGene00044207 | 1.247103736 | 0.59368513 | 1 | Y6G8.5 |
| WBGene00044210 | 0.76286748 | 0.23070057 | 0.43621486 | Y51A2D.21 |
| WBGene00044211 | -0.537782073 | 0.68446187 | 1 | Y60A3A.25 |
| WBGene00044212 | 2.439918581 | 0.0191045 | 0.08295699 | Y68A4A.13 |
| WBGene00044213 | 0.136014833 | 0.87548323 | 0.93279877 | Y102A5C.36 |
| WBGene00044218 | -0.982438334 | 0.24885862 | 0.4567087 | F23H12.10 |
| WBGene00044221 | 0.146495119 | 0.86470056 | 0.92725044 | F41G3.18 |
| WBGene00044222 | -2.171326218 | 0.36522606 | 1 | T16G12.10 |
| WBGene00044225 | -0.273001862 | 0.82811172 | 1 | C38D4.10 |
| WBGene00044227 | 1.111350189 | 0.50766464 | 1 | C07A9.13 |
| WBGene00044228 | -3.079012209 | 0.2683724 | 1 | T03F6.7 |
| WBGene00044232 | 2.490696505 | 0.25975012 | 1 | C18D11.7 |
| WBGene00044233 | 0.131288585 | 0.76963044 | 0.87091837 | Y56A3A.36 |
| WBGene00044235 | -1.901856996 | 0.64072034 | 1 | C06A1.7 |

| | | | | |
|---|---|---|---|---|
| WBGene00044236 | -0.746428942 | 0.07671288 | 0.21490008 | C17E4.11 |
| WBGene00044238 | 1.980634444 | 0.28443964 | 1 | C30H6.12 |
| WBGene00044239 | -1.489681322 | 0.42686537 | 1 | C33D9.10 |
| WBGene00044240 | 0.574508983 | 0.8551971 | 1 | C47E12.14 |
| WBGene00044241 | 0.148297849 | 0.87377555 | 0.93204818 | C53D6.11 |
| WBGene00044242 | -0.094731856 | 0.97157968 | 1 | C54C8.12 |
| WBGene00044243 | -0.156269293 | 0.96924949 | 1 | F16C3.4 |
| WBGene00044244 | -0.506567032 | 0.81430926 | 1 | F26D10.13 |
| WBGene00044245 | -3.168891993 | 0.27608518 | 1 | F32A7.8 |
| WBGene00044249 | -0.156269293 | 0.96924949 | 1 | F55B11.6 |
| WBGene00044251 | -0.531403668 | 0.37331072 | 0.5812823 | F59C6.12 |
| WBGene00044253 | -0.490854115 | 0.4367967 | 0.63755094 | M117.6 |
| WBGene00044256 | -1.468722953 | 0.27591084 | 1 | Y45F10C.6 |
| WBGene00044258 | -0.136930506 | 0.78534693 | 0.88190412 | Y57G11C.51 |
| WBGene00044259 | -2.663073159 | 0.1037998 | 1 | Y73F8A.36 |
| WBGene00044260 | 0.394576818 | 0.1834744 | 0.37557962 | Y87G2A.19 |
| WBGene00044261 | 1.529802032 | 0.64971375 | 1 | Y87G2A.20 |
| WBGene00044264 | 0.938982574 | 0.8146943 | 1 | Y105E8A.32 |
| WBGene00044265 | -1.969805064 | 0.62862569 | 1 | Y105E8B.11 |
| WBGene00044272 | 1.181686151 | 0.7579682 | 1 | nspa-4 |
| WBGene00044278 | 1.216957619 | 0.74819621 | 1 | F09C6.14 |
| WBGene00044280 | 0.850064365 | 0.14785225 | 0.32846737 | W02A2.9 |
| WBGene00044281 | -1.778035765 | 0.0063507 | 0.0371585 | R04A9.7 |
| WBGene00044283 | -0.386311983 | 0.47647206 | 0.66906261 | spp-22 |
| WBGene00044284 | 2.591348062 | 0.23319713 | 1 | spp-21 |
| WBGene00044287 | 0.704621139 | 0.57924691 | 1 | F21H12.7 |
| WBGene00044288 | 0.983488853 | 0.80601565 | 1 | F30H5.5 |
| WBGene00044289 | 1.916762327 | 0.08052551 | 0.22195997 | Y37F4.8 |
| WBGene00044290 | -0.227387276 | 0.57347185 | 0.74729223 | B0035.18 |
| WBGene00044291 | 0.937057406 | 0.14182812 | 0.31995754 | C25F9.10 |
| WBGene00044292 | -2.934840138 | 0.2654436 | 1 | F56D6.8 |
| WBGene00044293 | 0.938982574 | 0.8146943 | 1 | K08D12.7 |
| WBGene00044294 | -0.118280863 | 0.60508918 | 0.76897128 | C01B10.11 |
| WBGene00044295 | 1.977885804 | 0.49204652 | 1 | C09E7.10 |
| WBGene00044296 | 0.933968629 | 0.00197121 | 0.01538046 | F10C1.9 |
| WBGene00044297 | 1.017822963 | 0.43277563 | 1 | C25B8.8 |
| WBGene00044299 | -0.156269293 | 0.96924949 | 1 | C42D8.9 |
| WBGene00044300 | 0.992581876 | 0.67966507 | 1 | D1022.9 |
| WBGene00044301 | 0.384329203 | 0.50294235 | 0.69012466 | lgc-28 |
| WBGene00044302 | 0.855026222 | 0.71364798 | 1 | F19C7.5 |
| WBGene00044302 | 3.322129029 | 0.07701207 | 1 | F19C7.6 |
| WBGene00044302 | 2.793858195 | 0.21955872 | 1 | F28E10.5 |
| WBGene00044303 | -0.02082792 | 0.97750865 | 0.98825319 | hst-3.2 |

| | | | | |
|---|---|---|---|---|
| WBGene00044304 | 0.215005316 | 0.79303123 | 0.88575825 | F52B10.3 |
| WBGene00044305 | -0.460916866 | 0.02706776 | 0.10556882 | rad-8 |
| WBGene00044306 | 0.688793783 | 0.86378755 | 1 | F58F9.8 |
| WBGene00044307 | -0.156269293 | 0.96924949 | 1 | F58F9.9 |
| WBGene00044310 | -0.255732052 | 0.48495426 | 0.67539764 | H20J04.9 |
| WBGene00044311 | 2.029515155 | 0.14239496 | 1 | K04C2.7 |
| WBGene00044312 | -1.196801792 | 0.24152161 | 1 | K04C2.8 |
| WBGene00044313 | -0.508813154 | 0.81506991 | 1 | M02D8.7 |
| WBGene00044314 | -3.100980249 | 0.01847919 | 1 | fbxb-33 |
| WBGene00044315 | 2.193118955 | 0.22348976 | 1 | F41G3.20 |
| WBGene00044316 | 0.423576304 | 0.44247965 | 0.64206351 | F41G3.21 |
| WBGene00044317 | -1.964653846 | 0.53599202 | 1 | F41G3.19 |
| WBGene00044318 | -0.679242497 | 0.02417312 | 0.09792655 | tag-267 |
| WBGene00044319 | -0.449698123 | 0.14274781 | 0.32052915 | tag-266 |
| WBGene00044321 | -0.011835722 | 0.95358883 | 0.97634694 | mrps-30 |
| WBGene00044322 | -0.964717519 | 0.01937147 | 0.08366608 | tag-280 |
| WBGene00044323 | -0.086463259 | 0.87065265 | 0.9302652 | tag-281 |
| WBGene00044324 | -0.724024536 | 0.00291553 | 0.02077929 | ufm-1 |
| WBGene00044325 | -0.227084301 | 0.33209679 | 0.53982957 | tag-321 |
| WBGene00044326 | -0.158531457 | 0.55574244 | 0.73315972 | tag-322 |
| WBGene00044329 | -0.338440462 | 0.08285544 | 0.22595416 | cpsf-4 |
| WBGene00044330 | 1.134782668 | 0.0437201 | 0.14608272 | alr-1 |
| WBGene00044331 | 1.220597658 | 0.24690352 | 1 | clec-25 |
| WBGene00044332 | 1.923637003 | 0.01641945 | 0.07418999 | clec-36 |
| WBGene00044333 | 1.353535409 | 0.73149271 | 1 | C01G12.13 |
| WBGene00044337 | 0.041803775 | 0.98890197 | 1 | nspc-2 |
| WBGene00044338 | -1.320202606 | 0.20566828 | 1 | nspc-3 |
| WBGene00044339 | -0.660412775 | 0.05308759 | 0.1672817 | nspc-4 |
| WBGene00044340 | -1.094201388 | 0.7816772 | 1 | nspc-5 |
| WBGene00044341 | -0.79877167 | 0.02207731 | 0.09223842 | ave-1 |
| WBGene00044342 | 1.629609407 | 0.18285157 | 1 | T05B11.7 |
| WBGene00044344 | -0.461745685 | 0.04948639 | 0.15947139 | mrpl-39 |
| WBGene00044345 | -0.271259972 | 0.33574629 | 0.54408646 | Y48G1C.12 |
| WBGene00044346 | 0.574508983 | 0.8551971 | 1 | Y59E9AL.8 |
| WBGene00044347 | -0.945145766 | 0.65193959 | 1 | Y71G12B.30 |
| WBGene00044348 | 0.624491687 | 0.6223203 | 1 | Y71G12B.31 |
| WBGene00044349 | 2.684773376 | 0.10939775 | 1 | Y71G12B.32 |
| WBGene00044350 | 0.113176253 | 0.84057701 | 0.91242249 | Y73B6BL.44 |
| WBGene00044353 | -0.558804446 | 0.87818675 | 1 | ZK418.10 |
| WBGene00044354 | 1.106419131 | 0.55533901 | 1 | ZK418.11 |
| WBGene00044355 | -0.40157296 | 0.54449437 | 0.72381392 | ZK563.7 |
| WBGene00044357 | 0.041803775 | 0.98890197 | 1 | nspc-6 |
| WBGene00044358 | -0.156269293 | 0.96924949 | 1 | Y71G12B.33 |

| | | | | |
|---|---|---|---|---|
| WBGene00044359 | -3.201350992 | 0.01605688 | 1 | C16C4.17 |
| WBGene00044360 | 3.001198982 | 0.13323807 | 1 | C33C12.11 |
| WBGene00044361 | 1.67874632 | 0.06716593 | 0.19678127 | K09F6.11 |
| WBGene00044362 | 1.181686151 | 0.7579682 | 1 | F42G2.7 |
| WBGene00044363 | 1.597248813 | 0.30937259 | 1 | F22E5.21 |
| WBGene00044366 | 0.377828795 | 0.92526585 | 1 | F34D6.9 |
| WBGene00044367 | 0.480842266 | 0.75506946 | 1 | T05A8.8 |
| WBGene00044370 | -0.397295046 | 0.47059277 | 0.66356678 | C41H7.9 |
| WBGene00044371 | -2.335248999 | 0.00332915 | 0.02284862 | ZC239.20 |
| WBGene00044372 | -2.481625597 | 0.00598851 | 0.03555332 | ZC239.21 |
| WBGene00044373 | 0.361788163 | 0.84421891 | 1 | C17F4.11 |
| WBGene00044374 | 0.377828795 | 0.92526585 | 1 | C17F4.12 |
| WBGene00044376 | 0.232157325 | 0.93397866 | 1 | Y49F6B.12 |
| WBGene00044377 | -0.62146648 | 0.85925752 | 1 | Y49F6B.13 |
| WBGene00044379 | -1.542435723 | 0.42654412 | 1 | F40H7.12 |
| WBGene00044380 | -0.07748118 | 0.98067866 | 1 | T24E12.12 |
| WBGene00044381 | 0.574508983 | 0.8551971 | 1 | K10G6.5 |
| WBGene00044382 | 0.373709506 | 0.74504435 | 1 | Y14H12A.2 |
| WBGene00044383 | 0.105852162 | 0.9175745 | 1 | T05A7.11 |
| WBGene00044385 | -1.581992951 | 0.1439164 | 1 | F59A6.10 |
| WBGene00044386 | -0.264483026 | 0.90204316 | 1 | C27A2.7 |
| WBGene00044387 | 0.351397257 | 0.735756 | 1 | C27A2.8 |
| WBGene00044388 | 1.181686151 | 0.7579682 | 1 | C27D6.11 |
| WBGene00044389 | 0.429547946 | 0.28960246 | 0.49967941 | C27D6.12 |
| WBGene00044391 | -0.933827966 | 0.56134399 | 1 | C25H3.15 |
| WBGene00044392 | -1.094201388 | 0.7816772 | 1 | C25H3.16 |
| WBGene00044393 | 0.377828795 | 0.92526585 | 1 | ZK1248.20 |
| WBGene00044394 | -0.44846728 | 0.73226089 | 1 | B0034.7 |
| WBGene00044395 | -0.264859079 | 0.8341277 | 1 | R12C12.10 |
| WBGene00044396 | 0.957717819 | 0.05892614 | 0.18034341 | ZK1290.14 |
| WBGene00044399 | -2.640414922 | 0.41081194 | 1 | F11F1.8 |
| WBGene00044401 | 0.501156004 | 0.45106281 | 0.64911342 | Y55B1BR.6 |
| WBGene00044402 | 0.240675161 | 0.85790323 | 1 | Y82E9BL.18 |
| WBGene00044403 | 1.3745369 | 0.07403902 | 0.2101725 | fbxa-219 |
| WBGene00044406 | 2.037611642 | 0.16835054 | 1 | F02A9.7 |
| WBGene00044409 | 0.281454817 | 0.94435481 | 1 | C45G9.14 |
| WBGene00044410 | 0.067251476 | 0.95152449 | 1 | F48E8.8 |
| WBGene00044411 | -0.034244749 | 0.9657733 | 0.98286673 | R12B2.8 |
| WBGene00044413 | -1.113588585 | 0.00470323 | 0.02989628 | B0205.12 |
| WBGene00044414 | -0.112879089 | 0.86401837 | 0.92684635 | C06A5.12 |
| WBGene00044417 | 0.018786334 | 0.98888489 | 1 | C08F1.11 |
| WBGene00044418 | 0.739261898 | 0.25337671 | 0.46131776 | C08G5.7 |
| WBGene00044419 | -1.14715222 | 0.68932198 | 1 | fbxc-17 |

| | | | | |
|---|---|---|---|---|
| WBGene00044420 | 0.160735397 | 0.70535427 | 0.83345073 | C50D2.10 |
| WBGene00044421 | 0.951328247 | 0.23313815 | 0.43890994 | D1007.18 |
| WBGene00044423 | -0.057482899 | 0.89129467 | 0.9426961 | F53F10.8 |
| WBGene00044427 | 0.377828795 | 0.92526585 | 1 | F56A6.5 |
| WBGene00044428 | 0.983488853 | 0.80601565 | 1 | T01D1.7 |
| WBGene00044429 | -1.159714244 | 0.75790634 | 1 | T07H3.7 |
| WBGene00044431 | -0.83961615 | 0.00234233 | 0.01761466 | W03G9.8 |
| WBGene00044432 | -0.156269293 | 0.96924949 | 1 | W03G9.9 |
| WBGene00044433 | 1.352381308 | 0.13132733 | 0.30381958 | W10C8.6 |
| WBGene00044434 | 0.377828795 | 0.92526585 | 1 | Y18H1A.15 |
| WBGene00044435 | 0.862015713 | 0.60307388 | 1 | Y20F4.8 |
| WBGene00044436 | -0.913994675 | 0.14209852 | 0.32019639 | Y47G6A.31 |
| WBGene00044437 | 0.31950844 | 0.82590839 | 1 | Y47G6A.32 |
| WBGene00044438 | -0.737815234 | 0.0243516 | 0.09837757 | Y51F10.11 |
| WBGene00044439 | -1.356292182 | 0.63719725 | 1 | Y51H7C.15 |
| WBGene00044440 | 0.033193713 | 0.8989795 | 0.94694691 | adpr-1 |
| WBGene00044442 | 2.336189772 | 0.17295669 | 1 | T08A9.13 |
| WBGene00044443 | 1.608889786 | 0.24032684 | 1 | ZC21.10 |
| WBGene00044444 | 0.053200181 | 0.97064942 | 1 | F09G8.10 |
| WBGene00044445 | 1.675951732 | 0.07234802 | 0.20701799 | C02F5.14 |
| WBGene00044446 | 0.983488853 | 0.80601565 | 1 | C06G4.6 |
| WBGene00044447 | 0.159950863 | 0.93307234 | 1 | ZK688.10 |
| WBGene00044448 | 1.743415485 | 0.50483382 | 1 | ZK783.6 |
| WBGene00044449 | -0.604396148 | 0.82723443 | 1 | fipr-20 |
| WBGene00044450 | 0.03940405 | 0.99001443 | 1 | fipr-18 |
| WBGene00044452 | 0.214989173 | 0.69678307 | 0.82848332 | Y102E9.5 |
| WBGene00044453 | 0.688793783 | 0.86378755 | 1 | Y102E9.6 |
| WBGene00044455 | 1.382646761 | 0.12618117 | 0.29630283 | F23F12.13 |
| WBGene00044457 | -0.554249106 | 0.45663936 | 0.65378049 | C18H7.11 |
| WBGene00044458 | -0.156269293 | 0.96924949 | 1 | K11H12.10 |
| WBGene00044459 | 0.243597081 | 0.71969314 | 0.84179137 | Y55F3C.9 |
| WBGene00044460 | -0.333693239 | 0.89573179 | 1 | Y55F3C.10 |
| WBGene00044461 | -1.420815711 | 0.55886055 | 1 | Y18H1A.14 |
| WBGene00044462 | 0.952350203 | 0.67617448 | 1 | C44B12.9 |
| WBGene00044463 | -0.487343152 | 0.66448956 | 1 | W09G12.10 |
| WBGene00044464 | 0.578228238 | 0.68620123 | 1 | Y77E11A.16 |
| WBGene00044466 | -0.858565738 | 0.04735171 | 0.15529365 | Y48A5A.3 |
| WBGene00044467 | 1.079024281 | 0.05322501 | 0.16762772 | R05C11.4 |
| WBGene00044468 | 0.440966845 | 0.11987291 | 0.28615355 | Y69A2AL.2 |
| WBGene00044469 | 1.128484555 | 0.00301576 | 0.02128743 | clec-173 |
| WBGene00044470 | 0.281454817 | 0.94435481 | 1 | C09G12.17 |
| WBGene00044471 | -1.488391716 | 0.50775624 | 1 | F56D6.9 |
| WBGene00044472 | 0.688793783 | 0.86378755 | 1 | dct-8 |

| | | | | |
|---|---|---|---|---|
| WBGene00044477 | -0.701574888 | 0.80383767 | 1 | F49F1.14 |
| WBGene00044478 | -0.228624902 | 0.82551732 | 1 | K06B9.6 |
| WBGene00044479 | -2.664443697 | 0.24091381 | 1 | F15E6.10 |
| WBGene00044480 | -1.027355405 | 0.68015432 | 1 | ZK185.4 |
| WBGene00044481 | -1.119109227 | 0.02049404 | 0.08707506 | ZK185.5 |
| WBGene00044482 | 0.457387993 | 0.67328035 | 0.81328194 | F29B9.12 |
| WBGene00044483 | -0.432374894 | 0.0631563 | 0.18867635 | H06H21.11 |
| WBGene00044484 | 0.670521648 | 0.31001673 | 0.51830255 | hot-9 |
| WBGene00044485 | -0.323036688 | 0.61796751 | 0.77732528 | Y54G2A.41 |
| WBGene00044487 | 0.544525163 | 0.81649326 | 1 | Y54G2A.44 |
| WBGene00044488 | 0.800203462 | 0.00283374 | 0.02038177 | Y54G2A.45 |
| WBGene00044489 | 0.292035021 | 0.80116355 | 1 | pudl-1 |
| WBGene00044490 | 0.615776937 | 0.76598017 | 1 | pudl-2 |
| WBGene00044491 | -1.80092473 | 0.30952335 | 1 | Y54G2A.48 |
| WBGene00044492 | 0.971317371 | 0.26672924 | 0.47511147 | Y54G2A.49 |
| WBGene00044493 | 0.048036494 | 0.90277641 | 0.94940058 | Y54G2A.50 |
| WBGene00044495 | -0.68851324 | 0.4114782 | 0.61676009 | Y54G2A.40 |
| WBGene00044498 | -1.969805064 | 0.62862569 | 1 | Y59E9AR.10 |
| WBGene00044499 | 1.526476613 | 0.440991 | 1 | T11F8.5 |
| WBGene00044500 | 0.949303608 | 0.02575522 | 0.10225446 | T19E7.6 |
| WBGene00044502 | -0.21032074 | 0.76662834 | 0.86914135 | C46G7.5 |
| WBGene00044504 | 1.00174561 | 0.28414713 | 0.49410346 | R13A1.10 |
| WBGene00044505 | 0.983488853 | 0.80601565 | 1 | D2024.10 |
| WBGene00044508 | -0.284127871 | 0.34775851 | 0.55644893 | nsy-7 |
| WBGene00044509 | 0.607409966 | 0.50270699 | 0.68999878 | K07H8.12 |
| WBGene00044510 | -0.657005116 | 0.58566985 | 1 | K07H8.11 |
| WBGene00044512 | 1.176226461 | 0.41646324 | 1 | W03F9.11 |
| WBGene00044513 | 1.259434742 | 0.17767874 | 0.36961342 | Y58G8A.5 |
| WBGene00044514 | 2.510543076 | 0.14035529 | 1 | R09E12.9 |
| WBGene00044520 | -0.978982198 | 0.56419152 | 1 | F59A7.12 |
| WBGene00044521 | 0.377828795 | 0.92526585 | 1 | Y5H2A.4 |
| WBGene00044525 | -0.323868354 | 0.91634405 | 1 | Y73C8B.5 |
| WBGene00044526 | -0.828943906 | 0.33797567 | 0.54595032 | T28A11.22 |
| WBGene00044527 | 0.803792171 | 0.08322457 | 0.22650292 | Y39H10B.2 |
| WBGene00044528 | 1.139408087 | 0.59580649 | 1 | lgc-29 |
| WBGene00044529 | -2.640414922 | 0.41081194 | 1 | irld-1 |
| WBGene00044530 | -0.559852135 | 0.54179191 | 0.72236939 | F32D1.11 |
| WBGene00044531 | 0.688793783 | 0.86378755 | 1 | F13A2.9 |
| WBGene00044533 | -2.13449359 | 0.09107849 | 1 | F41F3.8 |
| WBGene00044534 | -0.421079692 | 0.73338575 | 1 | H23N18.6 |
| WBGene00044535 | -1.005049609 | 3.40E-07 | 1.3546E-05 | K11D12.13 |
| WBGene00044536 | 0.543141082 | 0.57755017 | 0.74906609 | F14F9.8 |
| WBGene00044537 | 0.16936395 | 0.92512149 | 1 | B0238.15 |

| | | | | |
|---|---|---|---|---|
| WBGene00044538 | -0.156269293 | 0.96924949 | 1 | T09D3.8 |
| WBGene00044539 | 1.230180765 | 0.55687914 | 1 | K09H11.9 |
| WBGene00044541 | 2.133182101 | 0.3921092 | 1 | K09H11.11 |
| WBGene00044542 | -0.014519762 | 0.99603746 | 1 | CD4.11 |
| WBGene00044542 | -0.333693239 | 0.89573179 | 1 | CD4.5 |
| WBGene00044544 | 1.423558129 | 0.12833955 | 0.29912046 | F26F12.8 |
| WBGene00044547 | 0.281454817 | 0.94435481 | 1 | C13A2.12 |
| WBGene00044551 | -0.18557176 | 0.95664217 | 1 | ZK105.8 |
| WBGene00044552 | -0.156269293 | 0.96924949 | 1 | Y97E10AR.8 |
| WBGene00044553 | -0.227061388 | 0.55107138 | 0.72959654 | T23B12.11 |
| WBGene00044554 | -0.123590605 | 0.7906124 | 0.8845213 | F41E6.15 |
| WBGene00044556 | 0.40377494 | 0.61486932 | 0.77514863 | F38G1.3 |
| WBGene00044557 | 0.451235227 | 0.84990489 | 1 | ZC13.10 |
| WBGene00044559 | 0.688793783 | 0.86378755 | 1 | ZK380.5 |
| WBGene00044560 | -0.156269293 | 0.96924949 | 1 | C36C9.6 |
| WBGene00044562 | -0.134754313 | 0.94840861 | 1 | C14E2.7 |
| WBGene00044563 | -1.489681322 | 0.42686537 | 1 | T26C11.8 |
| WBGene00044564 | -1.244570285 | 0.3091184 | 1 | ZK813.6 |
| WBGene00044565 | -0.156269293 | 0.96924949 | 1 | F53B1.9 |
| WBGene00044567 | -0.950449075 | 0.35257263 | 0.56153539 | C46C11.4 |
| WBGene00044568 | 0.571291785 | 0.46980535 | 0.66324497 | ntc-1 |
| WBGene00044570 | 0.98359296 | 0.38579453 | 1 | T22B7.8 |
| WBGene00044571 | 0.281454817 | 0.94435481 | 1 | C46F4.3 |
| WBGene00044574 | 1.12408848 | 0.22111568 | 0.42403497 | F23G4.1 |
| WBGene00044575 | 0.530404348 | 0.80087734 | 1 | F38B6.8 |
| WBGene00044578 | 1.36907956 | 0.02698093 | 0.10546537 | K01A12.4 |
| WBGene00044579 | -0.156269293 | 0.96924949 | 1 | C06E2.9 |
| WBGene00044582 | -0.275700596 | 0.72968685 | 0.84851293 | B0222.11 |
| WBGene00044584 | 0.226645269 | 0.79857107 | 0.88932777 | T03G11.9 |
| WBGene00044586 | 2.203673374 | 0.15021686 | 1 | Y61A9LA.12 |
| WBGene00044587 | -1.351599631 | 0.53258792 | 1 | srz-105 |
| WBGene00044588 | 0.085831123 | 0.93009346 | 1 | ZK1236.9 |
| WBGene00044602 | 0.281454817 | 0.94435481 | 1 | C31B8.16 |
| WBGene00044603 | -0.044493547 | 0.9772165 | 1 | acbp-7 |
| WBGene00044604 | -0.374964299 | 0.2719692 | 0.48056732 | T10F2.5 |
| WBGene00044605 | -1.288584705 | 3.2021E-05 | 0.00058935 | ttr-36 |
| WBGene00044607 | 0.812234369 | 0.0882077 | 0.23548763 | K02A11.4 |
| WBGene00044608 | -0.644501449 | 0.01244648 | 0.06085891 | glrx-22 |
| WBGene00044609 | 0.275218457 | 0.39223537 | 0.60022237 | tbcc-1 |
| WBGene00044610 | 0.281454817 | 0.94435481 | 1 | D2092.10 |
| WBGene00044611 | 2.09403112 | 0.45107747 | 1 | T27A3.8 |
| WBGene00044612 | 0.213970043 | 0.76568767 | 0.86875842 | Y57E12AR.1 |
| WBGene00044614 | -0.677361226 | 0.01242826 | 0.06079433 | C36B1.14 |

| | | | | |
|---|---|---|---|---|
| WBGene00044615 | 1.011881135 | 0.218844 | 0.42160981 | B0252.8 |
| WBGene00044616 | 1.025181888 | 0.79790347 | 1 | M01A12.4 |
| WBGene00044617 | 2.970899654 | 0.13509206 | 1 | bus-1 |
| WBGene00044618 | -0.1543483 | 0.88670749 | 1 | bus-2 |
| WBGene00044620 | -1.229773842 | 0.12602386 | 0.2961207 | bus-4 |
| WBGene00044621 | -0.731604997 | 0.05515518 | 0.1721437 | F53B1.4 |
| WBGene00044623 | -0.613554759 | 0.20214567 | 0.40095583 | bus-8 |
| WBGene00044630 | -0.032469774 | 0.96823233 | 0.98383906 | bus-17 |
| WBGene00044631 | -0.665589411 | 0.48469445 | 0.67527875 | bus-18 |
| WBGene00044633 | 1.712479368 | 0.58529868 | 1 | F54H12.7 |
| WBGene00044634 | -0.558804446 | 0.87818675 | 1 | C29E4.14 |
| WBGene00044636 | 0.000876286 | 0.9985364 | 0.99944191 | B0393.9 |
| WBGene00044637 | 0.447013353 | 0.24363943 | 0.45114272 | rft-1 |
| WBGene00044638 | -2.016537394 | 2.03E-20 | 6.86E-18 | F23A7.8 |
| WBGene00044640 | 1.002159309 | 0.10943661 | 0.26994513 | B0432.14 |
| WBGene00044641 | -0.226957956 | 0.8946195 | 1 | T01D1.8 |
| WBGene00044643 | 0.843693875 | 0.54226953 | 1 | H17B01.5 |
| WBGene00044644 | 0.066315747 | 0.87803414 | 0.93445061 | B0205.13 |
| WBGene00044645 | -0.156269293 | 0.96924949 | 1 | Y51H7BR.8 |
| WBGene00044646 | 0.629109312 | 0.67779609 | 1 | B0205.14 |
| WBGene00044648 | -0.309298228 | 0.40370493 | 0.6111802 | R144.13 |
| WBGene00044649 | 0.431665711 | 0.83167126 | 1 | C23H5.11 |
| WBGene00044650 | 0.011846632 | 0.95501179 | 0.97697996 | Y71G10AR.4 |
| WBGene00044651 | 0.216603531 | 0.44675167 | 0.6457926 | egrh-3 |
| WBGene00044653 | 0.688793783 | 0.86378755 | 1 | T13F3.9 |
| WBGene00044654 | 0.603901016 | 0.56852309 | 1 | fbxa-85 |
| WBGene00044655 | 0.281454817 | 0.94435481 | 1 | fbxa-194 |
| WBGene00044658 | 2.851274001 | 0.10179627 | 1 | C01G10.17 |
| WBGene00044659 | 0.281454817 | 0.94435481 | 1 | fbxa-142 |
| WBGene00044661 | 2.138861021 | 0.0755963 | 1 | fbxa-143 |
| WBGene00044662 | -3.014567441 | 0.24693748 | 1 | F26G1.9 |
| WBGene00044663 | -0.061090087 | 0.97319823 | 1 | F26G1.10 |
| WBGene00044665 | -0.633693686 | 0.20523618 | 0.40463989 | F59A6.12 |
| WBGene00044666 | -0.819309663 | 0.02101569 | 0.08871749 | F33G12.7 |
| WBGene00044667 | 0.723261303 | 0.41002229 | 0.61537456 | ZK622.5 |
| WBGene00044668 | -0.156269293 | 0.96924949 | 1 | C18A3.11 |
| WBGene00044672 | 0.685120956 | 0.31627263 | 0.52435697 | fbxa-4 |
| WBGene00044673 | -1.760055813 | 0.45626364 | 1 | C07D10.6 |
| WBGene00044674 | 0.207549994 | 0.80877234 | 0.89592198 | B0280.17 |
| WBGene00044675 | -0.037650448 | 0.95224622 | 0.97579519 | fbxa-76 |
| WBGene00044676 | 0.36474399 | 0.77530588 | 1 | fbxa-42 |
| WBGene00044677 | 1.01888049 | 0.25965921 | 0.4678664 | ZK353.10 |
| WBGene00044678 | 0.377828795 | 0.92526585 | 1 | B0303.16 |

| | | | | |
|---|---|---|---|---|
| WBGene00044679 | 1.196642537 | 0.35492373 | 1 | K02B12.9 |
| WBGene00044680 | -0.757315759 | 0.38839898 | 0.59713208 | F30A10.13 |
| WBGene00044681 | 1.085648961 | 0.49156841 | 1 | T24C4.8 |
| WBGene00044683 | 0.399606396 | 0.86544406 | 1 | C36E6.8 |
| WBGene00044684 | 1.38322681 | 0.69550305 | 1 | T08G11.2 |
| WBGene00044686 | 1.129641373 | 0.0048801 | 0.03063562 | flp-28 |
| WBGene00044687 | -0.156269293 | 0.96924949 | 1 | B0563.9 |
| WBGene00044689 | -0.171167897 | 0.41491058 | 0.62017539 | arid-1 |
| WBGene00044690 | -0.580169866 | 0.3016486 | 0.51115619 | M02B7.7 |
| WBGene00044692 | -1.420807642 | 0.03977805 | 0.13696105 | fbxb-91 |
| WBGene00044693 | -2.477622272 | 0.0350979 | 1 | fbxb-92 |
| WBGene00044694 | -1.64294245 | 0.06101735 | 0.184695 | fbxb-90 |
| WBGene00044696 | 0.191686828 | 0.6801671 | 0.81656371 | F52E1.14 |
| WBGene00044697 | 2.297216103 | 0.01103113 | 0.05578051 | poml-4 |
| WBGene00044698 | 0.494172845 | 0.15649591 | 0.34056584 | K09F5.6 |
| WBGene00044699 | 0.227235913 | 0.73473408 | 0.85183408 | nhr-286 |
| WBGene00044701 | 0.461707861 | 0.46419752 | 0.65886864 | AH9.6 |
| WBGene00044705 | -0.101714388 | 0.94516967 | 1 | T22F3.12 |
| WBGene00044707 | -0.156269293 | 0.96924949 | 1 | F22G12.7 |
| WBGene00044719 | 1.894894278 | 0.13742389 | 1 | clec-172 |
| WBGene00044720 | -0.443090517 | 0.35906909 | 0.56719462 | metl-2 |
| WBGene00044723 | 0.4130209 | 0.66491819 | 1 | K11H12.11 |
| WBGene00044724 | -1.147174972 | 0.10557679 | 0.26450843 | R02E4.3 |
| WBGene00044725 | -0.465218472 | 0.57531137 | 0.7483239 | R11G1.7 |
| WBGene00044726 | -0.558804446 | 0.87818675 | 1 | Y53F4B.43 |
| WBGene00044728 | 1.273081336 | 7.96E-09 | 5.06E-07 | Y53F4B.45 |
| WBGene00044732 | 1.278570347 | 0.1820729 | 0.37384831 | T03G11.10 |
| WBGene00044733 | -0.963811488 | 0.61154562 | 1 | W08F4.13 |
| WBGene00044734 | -1.964653846 | 0.53599202 | 1 | Y19D10A.16 |
| WBGene00044736 | 1.212217338 | 0.17786017 | 0.36976032 | gfrp-1 |
| WBGene00044737 | -0.781646599 | 0.00116035 | 0.01028745 | ttr-45 |
| WBGene00044738 | 1.687239501 | 0.59403242 | 1 | folt-3 |
| WBGene00044741 | 1.216957619 | 0.74819621 | 1 | Y57G11C.52 |
| WBGene00044742 | -0.110285218 | 0.87294879 | 0.93170576 | Y55F3BR.10 |
| WBGene00044743 | 0.106587601 | 0.84705155 | 0.91624547 | crb-3 |
| WBGene00044744 | 0.377828795 | 0.92526585 | 1 | F10G7.12 |
| WBGene00044745 | -0.492520914 | 0.16701999 | 0.35444673 | Y62E10A.20 |
| WBGene00044746 | -0.405020929 | 0.24169948 | 0.44927756 | C53H9.3 |
| WBGene00044747 | 1.695918088 | 0.15507197 | 1 | ZK1005.2 |
| WBGene00044748 | -0.774842772 | 0.78069886 | 1 | ZK402.6 |
| WBGene00044749 | 0.665185887 | 0.7882137 | 1 | W01C8.8 |
| WBGene00044750 | 1.382186448 | 0.23581575 | 1 | Y39D8C.2 |
| WBGene00044753 | 2.731108939 | 0.18857197 | 1 | F49H12.7 |

| | | | | |
|---|---|---|---|---|
| WBGene00044754 | 1.557897343 | 0.63813005 | 1 | Y119C1B.12 |
| WBGene00044756 | 0.587591766 | 0.48209443 | 0.67315796 | F58F12.4 |
| WBGene00044757 | 0.153329972 | 0.8294647 | 0.9076639 | Y55F3BR.11 |
| WBGene00044758 | -0.290484712 | 0.46489024 | 0.65929592 | Y37H2A.12 |
| WBGene00044759 | 0.495525129 | 0.72823978 | 1 | T06E6.15 |
| WBGene00044760 | 0.940004647 | 0.2889789 | 0.49921577 | Y71H2AM.25 |
| WBGene00044761 | 1.811395795 | 0.03508721 | 0.12622038 | pals-19 |
| WBGene00044762 | -0.055979366 | 0.86709477 | 0.92835056 | ZK688.11 |
| WBGene00044763 | -0.333693239 | 0.89573179 | 1 | ZK688.12 |
| WBGene00044767 | -0.226792242 | 0.78588907 | 0.88216338 | T23B7.3 |
| WBGene00044770 | 0.224769494 | 0.7100592 | 0.83591862 | C53C11.5 |
| WBGene00044771 | -0.048008406 | 0.98380497 | 1 | tmem-218 |
| WBGene00044772 | 0.704496849 | 0.56359413 | 1 | T23E7.6 |
| WBGene00044773 | -1.264919691 | 8.0518E-06 | 0.00018927 | C08A9.10 |
| WBGene00044774 | 0.274869558 | 0.64265971 | 0.79350306 | F16B3.3 |
| WBGene00044776 | 2.076056654 | 0.06490146 | 1 | T02B11.8 |
| WBGene00044777 | -0.156269293 | 0.96924949 | 1 | T02B11.9 |
| WBGene00044779 | 0.168553739 | 0.74094068 | 0.85545507 | T10H9.8 |
| WBGene00044780 | 0.539239107 | 0.83916685 | 1 | F18E9.8 |
| WBGene00044783 | 2.655987113 | 0.03826803 | 1 | T26H5.10 |
| WBGene00044784 | -0.100546037 | 0.69016518 | 0.82353771 | F13H10.8 |
| WBGene00044788 | 0.257011112 | 0.82226644 | 1 | ttr-38 |
| WBGene00044789 | 0.319901968 | 0.28677623 | 0.49696692 | T07A9.14 |
| WBGene00044790 | 0.308155575 | 0.829269 | 1 | clec-224 |
| WBGene00044791 | 1.413734769 | 0.68804531 | 1 | bnc-1 |
| WBGene00044792 | 0.273348387 | 0.68981659 | 0.82353771 | F59C6.14 |
| WBGene00044794 | -1.398451502 | 0.129433 | 0.30075523 | Y59A8B.25 |
| WBGene00044795 | -0.745058563 | 0.31939331 | 0.52744208 | C06H5.8 |
| WBGene00044796 | -2.254149048 | 0.05248807 | 1 | W03D8.11 |
| WBGene00044797 | -0.809802214 | 0.27440425 | 0.4827622 | F13E9.13 |
| WBGene00044798 | -2.749337042 | 0.009977 | 1 | tbx-43 |
| WBGene00044799 | 1.88180802 | 0.31552007 | 1 | ZK686.6 |
| WBGene00044803 | 0.275544868 | 0.89851598 | 1 | C02G6.3 |
| WBGene00044805 | -1.969805064 | 0.62862569 | 1 | Y53C10A.15 |
| WBGene00044807 | 0.568261324 | 0.66876943 | 1 | T19H5.6 |
| WBGene00044810 | -0.008677305 | 0.99719169 | 1 | Y37E3.19 |
| WBGene00044811 | 1.537621222 | 0.02015311 | 0.08591255 | F12E12.11 |
| WBGene00044891 | -1.052397469 | 0.00529278 | 0.03252009 | T07C4.12 |
| WBGene00044894 | -1.01527695 | 0.0200646 | 0.08568564 | Y71F9AR.4 |
| WBGene00044895 | -0.245531902 | 0.94070824 | 1 | R06A10.5 |
| WBGene00044898 | 0.340935197 | 0.5353074 | 0.7178681 | let-383 |
| WBGene00044900 | -3.526788663 | 0.13764061 | 1 | cnc-11 |
| WBGene00044901 | -0.540554002 | 0.38445147 | 0.59284039 | Y41G9A.10 |

| | | | | |
|---|---|---|---|---|
| WBGene00044903 | 0.033758268 | 0.89956325 | 0.94717955 | K03H1.13 |
| WBGene00044906 | -0.001191679 | 0.99941061 | 1 | Y67A10A.11 |
| WBGene00044908 | 0.938982574 | 0.8146943 | 1 | F38A1.15 |
| WBGene00044912 | 0.688793783 | 0.86378755 | 1 | H38K22.7 |
| WBGene00044916 | -0.424863086 | 0.01593727 | 0.07259399 | F40F8.11 |
| WBGene00044917 | 0.574508983 | 0.8551971 | 1 | F40F8.12 |
| WBGene00044921 | 0.056939148 | 0.92300234 | 0.9593272 | F53C11.9 |
| WBGene00044922 | -1.919896851 | 0.03622296 | 0.12875763 | Y43C5A.7 |
| WBGene00044924 | -0.150819336 | 0.62305472 | 0.78057302 | M106.7 |
| WBGene00044925 | -0.263229581 | 0.93409872 | 1 | Y37E3.20 |
| WBGene00044927 | 3.271881254 | 0.07721527 | 1 | F32B5.9 |
| WBGene00044929 | -1.969805064 | 0.62862569 | 1 | R12E2.17 |
| WBGene00044931 | -0.263229581 | 0.93409872 | 1 | Y37E3.21 |
| WBGene00044932 | -0.22461868 | 0.93331196 | 1 | Y37E3.22 |
| WBGene00044935 | 0.281454817 | 0.94435481 | 1 | T08B2.13 |
| WBGene00044936 | 0.747845955 | 0.85215965 | 1 | T24H7.6 |
| WBGene00044937 | 2.027551456 | 0.3130281 | 1 | ZK546.18 |
| WBGene00044938 | -0.156269293 | 0.96924949 | 1 | C27H5.9 |
| WBGene00044939 | -3.827003093 | 0.08161083 | 1 | B0412.7 |
| WBGene00044940 | 0.281454817 | 0.94435481 | 1 | H06I04.8 |
| WBGene00044941 | -0.621606308 | 0.77755782 | 1 | F54E7.10 |
| WBGene00044942 | 0.01290419 | 0.99462977 | 1 | F54E7.11 |
| WBGene00044943 | 0.281454817 | 0.94435481 | 1 | R151.12 |
| WBGene00044948 | -0.156269293 | 0.96924949 | 1 | K11H12.12 |
| WBGene00044949 | -0.867345337 | 0.02402313 | 0.09751414 | Y49A10A.2 |
| WBGene00044952 | 0.049907181 | 0.98090386 | 1 | Y24D9A.9 |
| WBGene00044953 | -0.257405101 | 0.82505468 | 1 | D2096.13 |
| WBGene00044954 | -0.156269293 | 0.96924949 | 1 | D2096.14 |
| WBGene00044955 | 0.281454817 | 0.94435481 | 1 | K07C11.11 |
| WBGene00044956 | 0.35134761 | 0.87335989 | 1 | K07C11.12 |
| WBGene00044958 | -0.156269293 | 0.96924949 | 1 | ZK994.7 |
| WBGene00044962 | -0.333693239 | 0.89573179 | 1 | F28D1.13 |
| WBGene00044963 | 0.377828795 | 0.92526585 | 1 | K07C5.11 |
| WBGene00044964 | 0.688793783 | 0.86378755 | 1 | F22D6.16 |
| WBGene00044965 | -0.156269293 | 0.96924949 | 1 | smy-6 |
| WBGene00044969 | -0.547881495 | 0.59841691 | 1 | T28D6.10 |
| WBGene00044970 | -0.156269293 | 0.96924949 | 1 | F38E11.14 |
| WBGene00044974 | -0.156269293 | 0.96924949 | 1 | C27H6.5 |
| WBGene00044977 | 0.667693633 | 0.48083546 | 0.67256028 | F56C4.4 |
| WBGene00044978 | -1.724118874 | 0.46596009 | 1 | F58D2.3 |
| WBGene00044979 | -0.156269293 | 0.96924949 | 1 | Y41E3.19 |
| WBGene00044980 | 0.377828795 | 0.92526585 | 1 | B0399.3 |
| WBGene00044982 | -0.774842772 | 0.78069886 | 1 | C01A2.8 |

| | | | | |
|---|---|---|---|---|
| WBGene00044984 | -2.926203393 | 0.17695648 | 1 | C27C7.9 |
| WBGene00044986 | 0.281454817 | 0.94435481 | 1 | C34E7.5 |
| WBGene00044987 | -0.250802298 | 0.30254016 | 0.5117893 | K07A1.17 |
| WBGene00044988 | 0.616263009 | 0.13070216 | 0.30289271 | W01A8.8 |
| WBGene00044989 | 0.434120186 | 0.05200371 | 0.16476435 | Y37A1B.17 |
| WBGene00044990 | -0.306495339 | 0.76683622 | 1 | T25B9.11 |
| WBGene00044991 | 0.124900161 | 0.94318373 | 1 | T25B9.12 |
| WBGene00044998 | -3.075834943 | 0.00340113 | 0.02321083 | M199.9 |
| WBGene00045013 | 1.832806158 | 0.04738257 | 0.15531096 | F08A7.1 |
| WBGene00045024 | -2.092790471 | 0.25268736 | 1 | C43F9.11 |
| WBGene00045036 | -1.138454115 | 0.61298961 | 1 | K08E4.7 |
| WBGene00045038 | 0.192316208 | 0.43388692 | 0.63557216 | ule-2 |
| WBGene00045043 | -1.363805733 | 0.22870941 | 1 | Y116A8C.48 |
| WBGene00045048 | 0.572795627 | 0.58542948 | 1 | D1007.19 |
| WBGene00045050 | 0.791181444 | 0.44801755 | 1 | ZC15.10 |
| WBGene00045051 | -0.156269293 | 0.96924949 | 1 | K11H12.13 |
| WBGene00045052 | 0.313339385 | 0.68818547 | 0.82244904 | K12C11.6 |
| WBGene00045053 | -0.684364122 | 0.34479243 | 0.55355198 | K12C11.7 |
| WBGene00045055 | -0.156269293 | 0.96924949 | 1 | C16D9.11 |
| WBGene00045056 | 0.382848879 | 0.66796982 | 0.80985814 | F13H10.9 |
| WBGene00045058 | -0.695278693 | 0.19256637 | 0.38765516 | gadr-5 |
| WBGene00045060 | 0.747845955 | 0.85215965 | 1 | F55B11.7 |
| WBGene00045061 | 0.574508983 | 0.8551971 | 1 | Y57G11B.8 |
| WBGene00045063 | 0.12040332 | 0.70639277 | 0.83394735 | Y37A1A.4 |
| WBGene00045067 | 2.235491657 | 0.12206898 | 1 | F07C6.6 |
| WBGene00045071 | -1.901856996 | 0.64072034 | 1 | T03F7.8 |
| WBGene00045074 | -0.668264701 | 0.85551153 | 1 | R166.7 |
| WBGene00045078 | -0.462219046 | 0.24568427 | 0.45373482 | C46C2.7 |
| WBGene00045082 | -0.06995528 | 0.97031639 | 1 | F46F11.12 |
| WBGene00045083 | 0.281454817 | 0.94435481 | 1 | C04G6.12 |
| WBGene00045085 | -0.210443359 | 0.57550866 | 0.74844162 | Y24D9A.10 |
| WBGene00045086 | 0.190692864 | 0.63439776 | 0.78783996 | Y43B11AR.7 |
| WBGene00045087 | 1.389382309 | 0.36175087 | 1 | B0041.9 |
| WBGene00045088 | 0.281454817 | 0.94435481 | 1 | F48A11.7 |
| WBGene00045089 | 0.281454817 | 0.94435481 | 1 | C05H8.3 |
| WBGene00045092 | -1.521855946 | 0.1406703 | 1 | H06I04.9 |
| WBGene00045093 | 1.601302495 | 0.65842342 | 1 | Y23H5B.10 |
| WBGene00045094 | -1.964653846 | 0.53599202 | 1 | D1007.20 |
| WBGene00045095 | 0.291749743 | 0.8862787 | 1 | mrpr-1 |
| WBGene00045096 | -3.47102951 | 0.12756864 | 1 | K01A6.7 |
| WBGene00045109 | 0.983488853 | 0.80601565 | 1 | smy-2 |
| WBGene00045113 | 0.688793783 | 0.86378755 | 1 | Y43F4B.12 |
| WBGene00045118 | 0.747845955 | 0.85215965 | 1 | T04A8.17 |

| | | | | |
|---|---|---|---|---|
| WBGene00045119 | 0.281454817 | 0.94435481 | 1 | F08G2.13 |
| WBGene00045120 | 0.281454817 | 0.94435481 | 1 | Y43F8C.22 |
| WBGene00045124 | 0.938982574 | 0.8146943 | 1 | C07A9.14 |
| WBGene00045126 | -2.265909937 | 0.17719195 | 1 | C14C10.8 |
| WBGene00045129 | -1.901856996 | 0.64072034 | 1 | Y75B12B.12 |
| WBGene00045130 | -1.901856996 | 0.64072034 | 1 | F28C6.11 |
| WBGene00045132 | -0.558804446 | 0.87818675 | 1 | ZK643.9 |
| WBGene00045134 | -1.901856996 | 0.64072034 | 1 | K07C5.12 |
| WBGene00045136 | -0.156269293 | 0.96924949 | 1 | D1046.6 |
| WBGene00045138 | -0.579346732 | 0.64271201 | 1 | F25H2.14 |
| WBGene00045142 | -1.245733657 | 0.50679516 | 1 | Y53F4B.48 |
| WBGene00045145 | -1.14715222 | 0.68932198 | 1 | F10D2.13 |
| WBGene00045146 | -1.460359516 | 0.26063391 | 1 | Y41E3.22 |
| WBGene00045148 | 0.377828795 | 0.92526585 | 1 | C06A1.8 |
| WBGene00045150 | -0.156269293 | 0.96924949 | 1 | ZK265.10 |
| WBGene00045152 | 0.747845955 | 0.85215965 | 1 | C06A1.9 |
| WBGene00045153 | -0.156269293 | 0.96924949 | 1 | F11A10.9 |
| WBGene00045154 | -0.156269293 | 0.96924949 | 1 | C10C6.8 |
| WBGene00045157 | 0.747845955 | 0.85215965 | 1 | K02C4.8 |
| WBGene00045158 | 0.281454817 | 0.94435481 | 1 | M163.14 |
| WBGene00045160 | 0.574508983 | 0.8551971 | 1 | C41C4.11 |
| WBGene00045161 | 1.876192691 | 0.43542138 | 1 | lfor-2 |
| WBGene00045162 | -0.934013985 | 0.67351079 | 1 | F25H2.15 |
| WBGene00045163 | 1.584577408 | 0.63258867 | 1 | Y67A10A.14 |
| WBGene00045164 | 0.459413223 | 0.84552382 | 1 | F59C6.15 |
| WBGene00045165 | 0.301573361 | 0.51760366 | 0.70316952 | H38K22.8 |
| WBGene00045167 | -2.097890267 | 0.34816065 | 1 | F43A11.8 |
| WBGene00045168 | -1.596069301 | 0.27739055 | 1 | B0285.12 |
| WBGene00045169 | -1.159714244 | 0.75790634 | 1 | T19C4.11 |
| WBGene00045170 | -1.159714244 | 0.75790634 | 1 | T19C4.12 |
| WBGene00045171 | -1.14715222 | 0.68932198 | 1 | K09E9.5 |
| WBGene00045177 | 1.626508665 | 0.30038203 | 1 | T21D12.14 |
| WBGene00045178 | -3.51732623 | 0.15709278 | 1 | fipr-27 |
| WBGene00045183 | 0.479889899 | 0.82438301 | 1 | T28C6.10 |
| WBGene00045190 | 1.584577408 | 0.63258867 | 1 | R01H2.8 |
| WBGene00045192 | -0.002854474 | 0.98866444 | 0.99484196 | aho-3 |
| WBGene00045194 | 1.876192691 | 0.43542138 | 1 | smy-1 |
| WBGene00045195 | -0.692184453 | 0.50221056 | 1 | F53F4.17 |
| WBGene00045198 | -0.378399192 | 0.85265246 | 1 | K10D11.7 |
| WBGene00045207 | -1.114385938 | 0.46034102 | 1 | F13E9.14 |
| WBGene00045208 | 0.803916838 | 0.21219106 | 0.41207318 | F13E9.15 |
| WBGene00045210 | 1.216957619 | 0.74819621 | 1 | C54D2.6 |
| WBGene00045212 | -0.015745422 | 0.97601561 | 0.98756609 | Y110A7A.21 |

| | | | | |
|---|---|---|---|---|
| WBGene00045215 | -0.831329645 | 0.59102286 | 1 | ceh-63 |
| WBGene00045234 | -0.156269293 | 0.96924949 | 1 | Y26D4A.17 |
| WBGene00045237 | 0.179284695 | 0.44793939 | 0.64704725 | mrt-1 |
| WBGene00045238 | -0.515216627 | 0.70900593 | 1 | rpr-1 |
| WBGene00045239 | -0.597359639 | 0.4988371 | 0.68705403 | Y49E10.29 |
| WBGene00045242 | 1.247575581 | 0.7549672 | 1 | C05B5.11 |
| WBGene00045245 | -0.803297746 | 0.12430648 | 0.29367679 | ttr-34 |
| WBGene00045246 | -0.255922322 | 0.64430713 | 0.7944884 | C29E4.15 |
| WBGene00045247 | 0.232157325 | 0.93397866 | 1 | F54H12.8 |
| WBGene00045248 | 0.974476174 | 0.20841347 | 0.40818646 | ZK180.7 |
| WBGene00045249 | -0.241010311 | 0.63495553 | 0.78806117 | T07A9.15 |
| WBGene00045250 | 0.371426556 | 0.73131101 | 1 | Y73B6BL.47 |
| WBGene00045251 | -2.930691629 | 0.12558037 | 1 | F54F7.9 |
| WBGene00045252 | 0.680751095 | 0.44738799 | 0.64632766 | Y62H9A.15 |
| WBGene00045253 | -0.250252741 | 0.7691772 | 0.87054113 | F46C3.6 |
| WBGene00045255 | 0.859800649 | 0.19480764 | 0.39078066 | nhr-146 |
| WBGene00045261 | -0.784776605 | 0.01900397 | 0.08269791 | H29C22.1 |
| WBGene00045265 | -1.540383023 | 0.00681428 | 0.03904304 | K10C2.8 |
| WBGene00045268 | -1.057335688 | 0.00017233 | 0.00234292 | ttr-42 |
| WBGene00045269 | -0.897508553 | 0.50642074 | 1 | F46G11.6 |
| WBGene00045270 | 0.526637334 | 0.76901972 | 1 | C04E7.5 |
| WBGene00045271 | -1.043288237 | 0.16673697 | 0.35409368 | F07G6.10 |
| WBGene00045272 | -1.42482068 | 4.0467E-05 | 0.00072339 | F59C12.4 |
| WBGene00045275 | -1.599239242 | 0.04633733 | 0.15254379 | F18E3.12 |
| WBGene00045276 | -0.286783667 | 0.55230051 | 0.73020586 | F18E3.13 |
| WBGene00045277 | 0.547529671 | 0.52203454 | 0.70713169 | ZK742.6 |
| WBGene00045282 | -3.526788663 | 0.13764061 | 1 | Y57G11C.56 |
| WBGene00045283 | -1.159714244 | 0.75790634 | 1 | K08E4.8 |
| WBGene00045289 | -0.488028718 | 0.72962287 | 1 | fbxa-169 |
| WBGene00045291 | 0.121370225 | 0.85208995 | 0.91924443 | F23D12.10 |
| WBGene00045292 | -0.722500798 | 0.61206376 | 1 | fbxa-179 |
| WBGene00045295 | 0.13091776 | 0.87560843 | 0.93285029 | fbxa-222 |
| WBGene00045297 | 1.075866534 | 0.08168603 | 0.22387135 | fbxc-43 |
| WBGene00045298 | 1.116613983 | 0.69067248 | 1 | fbxc-12 |
| WBGene00045299 | 1.116613983 | 0.69067248 | 1 | fbxc-11 |
| WBGene00045300 | 1.116613983 | 0.69067248 | 1 | fbxc-10 |
| WBGene00045301 | 1.116613983 | 0.69067248 | 1 | fbxc-9 |
| WBGene00045302 | -2.117463273 | 0.33448366 | 1 | fbxc-30 |
| WBGene00045304 | -1.969805064 | 0.62862569 | 1 | Y27F2A.11 |
| WBGene00045305 | 0.574475866 | 0.15665504 | 0.34085105 | ZC250.4 |
| WBGene00045306 | -1.969805064 | 0.62862569 | 1 | ZC250.5 |
| WBGene00045308 | 0.038418451 | 0.97779272 | 1 | C37H5.14 |
| WBGene00045324 | -1.969805064 | 0.62862569 | 1 | mir-787 |

| | | | | |
|---|---|---|---|---|
| WBGene00045325 | -1.969805064 | 0.62862569 | 1 | mir-789.1 |
| WBGene00045331 | -0.733112634 | 0.45694984 | 0.65391655 | F14D7.12 |
| WBGene00045336 | -0.181829256 | 0.86734464 | 0.92836209 | R11D1.12 |
| WBGene00045338 | 1.476467293 | 0.32598256 | 1 | M01B2.13 |
| WBGene00045339 | -2.095601 | 0.38719277 | 1 | T10H4.13 |
| WBGene00045340 | 1.875305911 | 0.22282001 | 1 | T10G3.8 |
| WBGene00045342 | -0.156269293 | 0.96924949 | 1 | mir-785 |
| WBGene00045355 | 2.750605477 | 0.18620275 | 1 | D1086.17 |
| WBGene00045356 | -0.263229581 | 0.93409872 | 1 | T03E6.9 |
| WBGene00045357 | -0.258620186 | 0.28885988 | 0.49915283 | F13A7.14 |
| WBGene00045359 | 0.574508983 | 0.8551971 | 1 | mir-799 |
| WBGene00045361 | -0.156269293 | 0.96924949 | 1 | clec-253 |
| WBGene00045362 | -0.156269293 | 0.96924949 | 1 | F11D11.12 |
| WBGene00045364 | -0.156269293 | 0.96924949 | 1 | clec-250 |
| WBGene00045365 | 0.555035113 | 0.63139803 | 1 | ZC449.7 |
| WBGene00045367 | 1.413734769 | 0.68804531 | 1 | F17C11.15 |
| WBGene00045369 | 0.707559163 | 0.78139902 | 1 | F26E4.14 |
| WBGene00045375 | -3.47102951 | 0.12756864 | 1 | C17D12.8 |
| WBGene00045377 | -1.094201388 | 0.7816772 | 1 | F19H8.6 |
| WBGene00045379 | 2.589110252 | 0.00652177 | 0.03779516 | C45B11.6 |
| WBGene00045381 | -0.156269293 | 0.96924949 | 1 | F28B1.9 |
| WBGene00045382 | 1.115863748 | 0.63449935 | 1 | T05E12.8 |
| WBGene00045383 | -0.553282953 | 0.00258839 | 0.01903458 | C25A1.4 |
| WBGene00045385 | -0.91375383 | 0.72824392 | 1 | lgc-24 |
| WBGene00045386 | 1.033057631 | 0.33518454 | 1 | C02B4.4 |
| WBGene00045387 | 0.531350072 | 0.22655341 | 0.43085603 | K09H9.8 |
| WBGene00045388 | 1.216957619 | 0.74819621 | 1 | lfor-1 |
| WBGene00045390 | -0.116452727 | 0.89173596 | 0.94291642 | C38C3.10 |
| WBGene00045391 | -1.14715222 | 0.68932198 | 1 | F52D4.1 |
| WBGene00045393 | -0.830315496 | 0.62566352 | 1 | F26D11.13 |
| WBGene00045394 | -0.363231976 | 0.0147481 | 0.0688474 | ZK813.7 |
| WBGene00045395 | 2.418326844 | 0.14987636 | 1 | F09F9.5 |
| WBGene00045396 | 1.38322681 | 0.69550305 | 1 | ZC334.12 |
| WBGene00045397 | -0.161111659 | 0.77092952 | 0.8718461 | Y54G2A.52 |
| WBGene00045399 | -2.081800816 | 2.92E-09 | 2.04E-07 | Y47G6A.33 |
| WBGene00045400 | 0.176356663 | 0.86513698 | 1 | C54D10.13 |
| WBGene00045401 | 0.562268321 | 0.54792234 | 0.72656231 | eol-1 |
| WBGene00045402 | 0.688793783 | 0.86378755 | 1 | Y75B12B.13 |
| WBGene00045403 | 0.893429771 | 0.36886188 | 1 | K10H10.12 |
| WBGene00045405 | 4.374084228 | 0.00506113 | 1 | D1086.18 |
| WBGene00045406 | -1.066234232 | 0.50225669 | 1 | D1086.19 |
| WBGene00045407 | -0.675609809 | 0.0152176 | 0.07043723 | T07D4.5 |
| WBGene00045411 | 2.249621494 | 0.00084033 | 0.00803343 | C25F9.11 |

| | | | | |
|---|---|---|---|---|
| WBGene00045412 | -0.267587749 | 0.80780616 | 1 | C25F9.12 |
| WBGene00045413 | -1.159714244 | 0.75790634 | 1 | C25F9.13 |
| WBGene00045414 | -0.955085772 | 0.36062789 | 0.56839973 | T19H5.7 |
| WBGene00045415 | 2.987064359 | 0.13549395 | 1 | Y43F8B.15 |
| WBGene00045416 | 1.526461876 | 0.00041762 | 0.00464316 | Y37H2A.14 |
| WBGene00045417 | 1.293359669 | 0.20412043 | 0.40348577 | Y37H2A.13 |
| WBGene00045418 | -0.180185771 | 0.43876504 | 0.63927113 | F15B9.10 |
| WBGene00045419 | 0.300533204 | 0.2202158 | 0.42317822 | lsy-12 |
| WBGene00045433 | 0.222283674 | 0.50066845 | 0.68856091 | F49D11.10 |
| WBGene00045434 | -0.699056022 | 0.00323507 | 0.02240559 | Y95D11A.3 |
| WBGene00045455 | -0.149228465 | 0.87815366 | 1 | F26G1.11 |
| WBGene00045456 | -0.29000237 | 0.84027763 | 1 | C10A4.9 |
| WBGene00045457 | 0.765408485 | 0.21966986 | 0.42247433 | F33H12.7 |
| WBGene00045458 | 2.380474327 | 0.18089959 | 1 | F17B5.8 |
| WBGene00045460 | 1.29757904 | 0.55672379 | 1 | C15C8.8 |
| WBGene00045461 | 0.768669548 | 0.16073181 | 0.34618428 | C25F9.14 |
| WBGene00045468 | 1.064864995 | 0.31737918 | 0.52511593 | F08A8.8 |
| WBGene00045473 | 0.084624979 | 0.89413213 | 0.94429873 | F30A10.14 |
| WBGene00045474 | 1.67306398 | 0.44045399 | 1 | C49G9.2 |
| WBGene00045475 | 0.141685919 | 0.90415612 | 1 | T07A5.7 |
| WBGene00045480 | 0.430970523 | 0.73971932 | 1 | F14F3.5 |
| WBGene00045481 | 0.28530093 | 0.92084641 | 1 | M153.5 |
| WBGene00045482 | 1.238214148 | 0.49867759 | 1 | T03F6.9 |
| WBGene00045483 | -0.628711345 | 0.00254819 | 0.01884547 | oxy-5 |
| WBGene00045484 | -1.093970585 | 0.30019504 | 1 | F34D10.9 |
| WBGene00045486 | -1.659780598 | 0.22189256 | 1 | K05F6.12 |
| WBGene00045488 | 0.341265909 | 0.67971356 | 0.81647553 | F57B1.9 |
| WBGene00045493 | 0.281454817 | 0.94435481 | 1 | F55G7.4 |
| WBGene00045494 | 0.820325976 | 0.45219974 | 1 | ZK662.6 |
| WBGene00045495 | 0.341852218 | 0.65115192 | 0.79891662 | F11C1.7 |
| WBGene00045497 | -0.91375383 | 0.72824392 | 1 | K03A11.6 |
| WBGene00045498 | -1.663249397 | 0.00155502 | 0.01289581 | F56F4.8 |
| WBGene00045500 | 1.216957619 | 0.74819621 | 1 | T05E12.9 |
| WBGene00045507 | 0.213700457 | 0.49945298 | 0.68759028 | F10A3.17 |
| WBGene00045510 | 0.747845955 | 0.85215965 | 1 | F11A5.17 |
| WBGene00045511 | -2.016182205 | 0.14874319 | 1 | F11A5.18 |
| WBGene00045513 | 1.181686151 | 0.7579682 | 1 | F14H3.14 |
| WBGene00045517 | 0.189064533 | 0.86081461 | 1 | R11D1.13 |
| WBGene00050875 | -0.979452109 | 0.72336959 | 1 | bah-1 |
| WBGene00050880 | 1.477253415 | 0.58357327 | 1 | F32H2.12 |
| WBGene00050881 | -0.245531902 | 0.94070824 | 1 | ZK39.10 |
| WBGene00050884 | 0.928651195 | 0.25857421 | 0.46679026 | Y69H2.15 |
| WBGene00050885 | -0.288249991 | 0.60921886 | 0.77135532 | Y94H6A.12 |

| | | | | |
|---|---|---|---|---|
| WBGene00050889 | 1.876192691 | 0.43542138 | 1 | T04C12.9 |
| WBGene00050891 | 1.38322681 | 0.69550305 | 1 | C08E8.9 |
| WBGene00050892 | -1.901856996 | 0.64072034 | 1 | ttr-57 |
| WBGene00050896 | 1.477253415 | 0.58357327 | 1 | ttr-56 |
| WBGene00050899 | 3.935456467 | 0.01930178 | 1 | T10C6.15 |
| WBGene00050904 | 1.632585912 | 0.02573182 | 0.10222842 | Y70C5A.3 |
| WBGene00050906 | 1.025181888 | 0.79790347 | 1 | F20E11.17 |
| WBGene00050907 | -0.156269293 | 0.96924949 | 1 | Y51A2B.9 |
| WBGene00050910 | -0.156269293 | 0.96924949 | 1 | Y43F8B.17 |
| WBGene00050911 | 1.477253415 | 0.58357327 | 1 | Y43F8B.18 |
| WBGene00050913 | -1.120238433 | 0.35385387 | 1 | T12B5.14 |
| WBGene00050914 | -1.475989639 | 0.00050667 | 0.00541025 | T12B5.15 |
| WBGene00050915 | 0.983488853 | 0.80601565 | 1 | Y6E2A.10 |
| WBGene00050916 | 0.688793783 | 0.86378755 | 1 | F55F10.3 |
| WBGene00050917 | -1.059141995 | 0.38964291 | 1 | Y53G8AL.4 |
| WBGene00050919 | 0.020328391 | 0.99335814 | 1 | C45G9.15 |
| WBGene00050922 | -0.156269293 | 0.96924949 | 1 | mir-1020 |
| WBGene00050931 | 0.688793783 | 0.86378755 | 1 | F59B2.14 |
| WBGene00050934 | 2.490696505 | 0.25975012 | 1 | T19C9.10 |
| WBGene00050936 | -0.593157818 | 0.07614085 | 0.21413202 | ZK856.16 |
| WBGene00050937 | -0.43471542 | 0.19460719 | 0.39059777 | R02D5.8 |
| WBGene00050939 | -0.664933011 | 0.49953159 | 0.68761935 | C05G5.7 |
| WBGene00050940 | 1.604738205 | 0.39900813 | 1 | C25D7.16 |
| WBGene00050941 | -0.730019993 | 0.00156438 | 0.01294256 | C33F10.14 |
| WBGene00050943 | 1.333017881 | 0.08580297 | 0.23123949 | ZC412.10 |
| WBGene00050944 | -0.100417043 | 0.75950395 | 0.86543035 | C14A11.9 |
| WBGene00050947 | -0.333106548 | 0.77751308 | 1 | K07C5.13 |
| WBGene00050956 | 1.557897343 | 0.63813005 | 1 | C14A6.13 |
| WBGene00050957 | 0.292320587 | 0.65924104 | 0.80368078 | C10A4.10 |
| WBGene00050967 | 0.57991002 | 0.82402153 | 1 | C25F9.15 |
| WBGene00050968 | 0.377828795 | 0.92526585 | 1 | R11G10.4 |
| WBGene00050974 | 1.735991307 | 0.58152819 | 1 | Y6G8.9 |
| WBGene00077439 | -1.895791037 | 0.52793577 | 1 | F55B12.10 |
| WBGene00077452 | -0.156269293 | 0.96924949 | 1 | Y62F5A.11 |
| WBGene00077453 | -0.609325775 | 0.01564159 | 0.0717433 | Y62F5A.12 |
| WBGene00077455 | 1.653460415 | 0.50712258 | 1 | Y69H2.17 |
| WBGene00077461 | -0.156269293 | 0.96924949 | 1 | F12F6.11 |
| WBGene00077489 | 0.281454817 | 0.94435481 | 1 | C04G6.13 |
| WBGene00077490 | 0.568830002 | 0.5844684 | 1 | M03A1.8 |
| WBGene00077491 | 0.298155563 | 0.80605774 | 1 | K08H10.10 |
| WBGene00077495 | -1.14715222 | 0.68932198 | 1 | Y20C6A.4 |
| WBGene00077497 | 1.187732978 | 0.58705413 | 1 | K06A4.10 |
| WBGene00077500 | 0.03989876 | 0.87729856 | 0.93415907 | C27H6.9 |

| | | | | |
|---|---|---|---|---|
| WBGene00077508 | -0.62146648 | 0.85925752 | 1 | T27F6.10 |
| WBGene00077514 | 0.178115923 | 0.94982316 | 1 | F07B10.7 |
| WBGene00077519 | -0.610720985 | 0.75455538 | 1 | T27C10.8 |
| WBGene00077520 | -1.378674741 | 0.24941632 | 1 | C37A5.11 |
| WBGene00077521 | 1.075636406 | 0.08649713 | 0.23249096 | maf-1 |
| WBGene00077522 | -0.677043769 | 0.75749374 | 1 | T10D4.15 |
| WBGene00077523 | -1.901856996 | 0.64072034 | 1 | C23H5.12 |
| WBGene00077525 | 0.720933761 | 0.46631611 | 1 | C41G7.8 |
| WBGene00077526 | -0.384848234 | 0.02485913 | 0.09974122 | C25A1.16 |
| WBGene00077531 | -0.841580501 | 0.58258892 | 1 | F40G9.15 |
| WBGene00077536 | -1.156241549 | 0.61231033 | 1 | F38B7.10 |
| WBGene00077539 | 0.135629574 | 0.89074822 | 1 | Y52B11A.12 |
| WBGene00077543 | -0.122616955 | 0.75192555 | 0.86205175 | F27D4.8 |
| WBGene00077548 | -0.263229581 | 0.93409872 | 1 | F21C3.7 |
| WBGene00077553 | -0.558804446 | 0.87818675 | 1 | C26C6.10 |
| WBGene00077557 | 1.216957619 | 0.74819621 | 1 | F15D3.9 |
| WBGene00077558 | -1.3352755 | 0.4420157 | 1 | T06D10.3 |
| WBGene00077559 | -1.013385316 | 0.16422561 | 0.35035374 | R05D7.7 |
| WBGene00077563 | 0.377828795 | 0.92526585 | 1 | peel-1 |
| WBGene00077569 | 1.216957619 | 0.74819621 | 1 | F53F4.18 |
| WBGene00077573 | 0.270496667 | 0.92869767 | 1 | T08G5.15 |
| WBGene00077576 | -1.969805064 | 0.62862569 | 1 | C15H11.13 |
| WBGene00077577 | -1.964653846 | 0.53599202 | 1 | R02D5.9 |
| WBGene00077582 | -1.964653846 | 0.53599202 | 1 | F14H8.8 |
| WBGene00077585 | -0.156269293 | 0.96924949 | 1 | T01G5.8 |
| WBGene00077588 | 0.747845955 | 0.85215965 | 1 | T26H8.5 |
| WBGene00077592 | -1.964653846 | 0.53599202 | 1 | T25B2.3 |
| WBGene00077593 | 3.754957487 | 0.00163577 | 0.01334673 | C49G7.12 |
| WBGene00077599 | -0.774842772 | 0.78069886 | 1 | T20F5.8 |
| WBGene00077607 | -0.735886458 | 0.64148088 | 1 | T10C6.16 |
| WBGene00077616 | -0.156269293 | 0.96924949 | 1 | F57E7.4 |
| WBGene00077622 | 0.983488853 | 0.80601565 | 1 | T25E12.14 |
| WBGene00077643 | 0.013495396 | 0.97289404 | 0.98595164 | trpp-1 |
| WBGene00077658 | 0.05001817 | 0.87185039 | 0.93105248 | M106.8 |
| WBGene00077663 | -3.245003023 | 0.02000462 | 1 | F55C9.14 |
| WBGene00077669 | 0.905367833 | 0.45918668 | 1 | Y38H6A.5 |
| WBGene00077674 | 0.377828795 | 0.92526585 | 1 | K02A6.4 |
| WBGene00077679 | 0.459413223 | 0.84552382 | 1 | K11E4.6 |
| WBGene00077680 | -1.094201388 | 0.7816772 | 1 | F54F7.10 |
| WBGene00077681 | 1.002582795 | 0.54025444 | 1 | Y44A6D.7 |
| WBGene00077684 | 0.694395688 | 0.58606397 | 1 | Y26D4A.21 |
| WBGene00077685 | -1.094201388 | 0.7816772 | 1 | C04F12.12 |
| WBGene00077690 | -0.400647723 | 0.36587229 | 0.57350663 | mtp-18 |

| | | | | |
|---|---|---|---|---|
| WBGene00077691 | -0.040199893 | 0.96823021 | 1 | T06E4.14 |
| WBGene00077692 | -0.091440983 | 0.94258541 | 1 | C08E8.10 |
| WBGene00077693 | 0.928816582 | 0.00646235 | 0.03754592 | T04C12.11 |
| WBGene00077697 | -0.512419775 | 0.40981367 | 0.61517653 | F58E6.13 |
| WBGene00077698 | -0.444394739 | 0.65369403 | 1 | lurp-4 |
| WBGene00077699 | 0.041803775 | 0.98890197 | 1 | C45B11.8 |
| WBGene00077701 | 1.144319404 | 0.00656144 | 0.03798877 | poml-3 |
| WBGene00077703 | 0.377828795 | 0.92526585 | 1 | K08E5.5 |
| WBGene00077704 | -1.159714244 | 0.75790634 | 1 | F12A10.9 |
| WBGene00077705 | -0.156269293 | 0.96924949 | 1 | mir-1829.1 |
| WBGene00077712 | 0.85823127 | 0.26036356 | 0.4687239 | nipi-4 |
| WBGene00077714 | -0.245531902 | 0.94070824 | 1 | R102.11 |
| WBGene00077724 | -1.366565036 | 0.6506483 | 1 | mir-1831 |
| WBGene00077728 | -0.562866186 | 0.30026372 | 0.50995852 | T23G11.11 |
| WBGene00077732 | -0.084094618 | 0.62587986 | 0.78249484 | szy-4 |
| WBGene00077751 | 2.437484291 | 0.2708268 | 1 | irld-27 |
| WBGene00077752 | -1.969805064 | 0.62862569 | 1 | Y105C5A.508 |
| WBGene00077756 | -0.930686449 | 0.56829387 | 1 | Y11D7A.19 |
| WBGene00077757 | -1.599294734 | 0.0153 | 0.07063015 | E02H4.7 |
| WBGene00077758 | -0.156269293 | 0.96924949 | 1 | irld-12 |
| WBGene00077761 | 1.118455786 | 7.84E-10 | 6.39E-08 | zip-9 |
| WBGene00077762 | 1.435925494 | 0.70920779 | 1 | Y57G11C.499 |
| WBGene00077763 | 0.733134918 | 0.41571989 | 0.62082449 | glb-34 |
| WBGene00077764 | 0.43664421 | 0.22886675 | 0.43403174 | F59C6.16 |
| WBGene00077768 | 2.07531578 | 0.45514895 | 1 | K04G2.12 |
| WBGene00077769 | 0.12332541 | 0.94831365 | 1 | F33E2.10 |
| WBGene00077770 | 0.901365004 | 0.04478753 | 0.14856977 | M04D5.3 |
| WBGene00077771 | -0.517897964 | 0.00459276 | 0.02936318 | C41G7.9 |
| WBGene00077773 | -1.092901668 | 0.22157509 | 0.4246477 | F25H5.10 |
| WBGene00077775 | 1.372383397 | 0.02432705 | 0.0983532 | T07C12.15 |
| WBGene00077782 | 1.216957619 | 0.74819621 | 1 | T26H10.2 |
| WBGene00077783 | 1.506473404 | 0.02522715 | 0.10078409 | D1054.18 |
| WBGene00077786 | 0.083826872 | 0.9321822 | 0.96406579 | F53F8.7 |
| WBGene00077787 | 1.006178315 | 0.43783938 | 1 | T05E12.10 |
| WBGene00086546 | 0.095597576 | 0.67963462 | 0.81647553 | cil-1 |
| WBGene00086551 | -0.626836656 | 0.00663557 | 0.03825242 | D2005.7 |
| WBGene00086552 | -1.969805064 | 0.62862569 | 1 | C17D12.9 |
| WBGene00086553 | 1.205631348 | 0.60123934 | 1 | ZC334.13 |
| WBGene00086554 | -0.156269293 | 0.96924949 | 1 | D1081.12 |
| WBGene00086555 | -3.244658943 | 0.04180093 | 1 | F49B2.7 |
| WBGene00086556 | 1.472766509 | 0.46533185 | 1 | F36D1.12 |
| WBGene00086557 | -0.135509354 | 0.88171345 | 0.93708197 | K07A12.8 |
| WBGene00086558 | -1.895791037 | 0.52793577 | 1 | F12E12.12 |

| | | | | |
|---|---|---|---|---|
| WBGene00086559 | 0.245864544 | 0.86986705 | 1 | Y43F8B.20 |
| WBGene00086560 | -0.01455644 | 0.9964743 | 1 | ZK355.8 |
| WBGene00086562 | -1.9959096 | 0.2198825 | 1 | T02G6.11 |
| WBGene00086565 | -1.159714244 | 0.75790634 | 1 | W06G6.17 |
| WBGene00086566 | 0.134230112 | 0.56649126 | 0.74172492 | C08H9.16 |
| WBGene00086567 | -0.156269293 | 0.96924949 | 1 | C05C10.8 |
| WBGene00086568 | 0.418332603 | 0.77497157 | 1 | Y2H9A.6 |
| WBGene00086569 | -0.132905245 | 0.82451471 | 0.90534802 | F38A6.5 |
| WBGene00119201 | 0.35134761 | 0.87335989 | 1 | C08E8.11 |
| WBGene00119203 | 0.932499303 | 0.29474313 | 0.50507781 | T04A8.18 |
| WBGene00119204 | 0.039233761 | 0.98370605 | 1 | ZC116.5 |
| WBGene00138711 | -1.292244266 | 0.4707841 | 1 | R07B1.13 |
| WBGene00138715 | 0.377828795 | 0.92526585 | 1 | Y6G8.14 |
| WBGene00138717 | -0.347099769 | 0.31073285 | 0.51873445 | F52C12.6 |
| WBGene00138719 | -0.156269293 | 0.96924949 | 1 | F26F2.10 |
| WBGene00138720 | -0.131857535 | 0.71054891 | 0.83617081 | F58H1.8 |
| WBGene00138721 | 4.099090505 | 0.01055252 | 1 | pals-37 |
| WBGene00138724 | -1.981634422 | 0.41263573 | 1 | F15B9.11 |
| WBGene00164968 | -0.226775208 | 0.82068382 | 0.90284677 | K08H2.10 |
| WBGene00164969 | -0.261742422 | 0.88440075 | 1 | K09C8.9 |
| WBGene00164971 | -1.14715222 | 0.68932198 | 1 | T21E8.6 |
| WBGene00164972 | 0.377828795 | 0.92526585 | 1 | T21E8.7 |
| WBGene00164973 | 2.726384632 | 0.23958348 | 1 | C39B10.7 |
| WBGene00164974 | -0.558804446 | 0.87818675 | 1 | F46C3.7 |
| WBGene00164985 | -1.341708717 | 0.07357012 | 0.20913483 | ZK1010.10 |
| WBGene00164986 | 0.438455499 | 0.87116517 | 1 | Y48A6C.6 |
| WBGene00175027 | -0.75339486 | 0.53328812 | 1 | W02F12.8 |
| WBGene00175029 | -0.635264682 | 0.57665008 | 1 | AH9.7 |
| WBGene00175030 | -0.444582607 | 0.0984591 | 0.25251203 | Y53C12A.10 |
| WBGene00175031 | 1.195178904 | 0.24198887 | 1 | C06C3.12 |
| WBGene00175034 | -0.548039039 | 0.78586011 | 1 | F54B3.4 |
| WBGene00175035 | 0.541284363 | 0.64771062 | 1 | F54C9.14 |
| WBGene00184990 | -0.646904889 | 0.02338085 | 0.09567471 | C14B1.12 |
| WBGene00185001 | -0.532136124 | 0.18776113 | 0.38133286 | F56A8.9 |
| WBGene00185002 | 0.117928042 | 0.67584163 | 0.81451193 | K11H3.8 |
| WBGene00185005 | -0.643148778 | 0.57219025 | 1 | T03F6.10 |
| WBGene00185006 | 0.232157325 | 0.93397866 | 1 | F59B2.15 |
| WBGene00185014 | -0.261028102 | 0.38621529 | 0.59475391 | F17C8.9 |
| WBGene00185048 | -1.037726228 | 0.00463963 | 0.02958496 | urm-1 |
| WBGene00185055 | -0.639137625 | 0.57476177 | 1 | Y47D3B.12 |
| WBGene00185067 | -0.808902678 | 0.00494098 | 0.0309367 | Y54G11A.17 |
| WBGene00185076 | 0.441408534 | 0.65596684 | 0.80231895 | Y7A9A.79 |
| WBGene00185077 | -1.454012584 | 0.32743361 | 1 | F58D2.4 |

| | | | | |
|---|---|---|---|---|
| WBGene00185078 | -0.579663683 | 0.05165736 | 0.16412997 | F40G9.17 |
| WBGene00185079 | -0.4117496 | 0.23636115 | 0.44244693 | W10D9.6 |
| WBGene00185086 | -0.232659106 | 0.34759339 | 0.55637855 | F59A3.12 |
| WBGene00185088 | -0.144107694 | 0.74176253 | 0.85569007 | Y55F3BL.4 |
| WBGene00185089 | -0.607042645 | 0.26304436 | 0.47179667 | C16A11.10 |
| WBGene00185092 | 1.035302524 | 0.16464807 | 0.35094667 | K08D8.7 |
| WBGene00185095 | 0.377828795 | 0.92526585 | 1 | W09C2.7 |
| WBGene00185096 | -0.156269293 | 0.96924949 | 1 | W09C2.9 |
| WBGene00185097 | -0.604396148 | 0.82723443 | 1 | C49C3.20 |
| WBGene00185098 | 0.037393944 | 0.947041 | 0.97304242 | Y71G12B.35 |
| WBGene00185102 | -1.100356805 | 0.46753792 | 1 | Y105C5A.1268 |
| WBGene00185117 | -1.094201388 | 0.7816772 | 1 | C45B11.9 |
| WBGene00185118 | 0.459726614 | 0.55417083 | 0.73180205 | K02E2.11 |
| WBGene00189933 | 2.028800581 | 0.39258153 | 1 | F38B7.11 |
| WBGene00189937 | -0.156269293 | 0.96924949 | 1 | Y6G8.15 |
| WBGene00189943 | -1.624083904 | 0.31010041 | 1 | Y116F11B.17 |
| WBGene00189948 | -1.094201388 | 0.7816772 | 1 | F56H9.8 |
| WBGene00189949 | -0.711741621 | 0.09976926 | 0.25443821 | Y48G1C.13 |
| WBGene00189950 | 1.584577408 | 0.63258867 | 1 | C39B5.14 |
| WBGene00189951 | -0.633824844 | 0.27091785 | 0.47962106 | linc-29 |
| WBGene00189952 | 0.340122187 | 0.87335753 | 1 | K10D2.8 |
| WBGene00189954 | -2.872649134 | 0.06324579 | 1 | F40G9.18 |
| WBGene00189955 | -1.901856996 | 0.64072034 | 1 | F40G9.19 |
| WBGene00189956 | 3.105318159 | 0.04615059 | 1 | Y34F4.6 |
| WBGene00189961 | 0.281454817 | 0.94435481 | 1 | Y54G2A.57 |
| WBGene00189971 | -0.156269293 | 0.96924949 | 1 | C41G6.17 |
| WBGene00189972 | -0.156269293 | 0.96924949 | 1 | T10H4.16 |
| WBGene00189973 | 0.203374252 | 0.86148402 | 1 | T10H4.17 |
| WBGene00189981 | 0.281454817 | 0.94435481 | 1 | Y60A3A.29 |
| WBGene00189984 | 0.822831666 | 0.32947523 | 0.53816209 | K02E2.13 |
| WBGene00189985 | -1.969805064 | 0.62862569 | 1 | F02D8.8 |
| WBGene00189987 | 1.181686151 | 0.7579682 | 1 | C18H7.12 |
| WBGene00189993 | 0.377828795 | 0.92526585 | 1 | C49C3.21 |
| WBGene00189995 | -1.073415821 | 0.14487785 | 0.32357652 | C06E7.88 |
| WBGene00194641 | 0.831365926 | 0.10414989 | 0.26185211 | F47G3.4 |
| WBGene00194642 | -0.684191566 | 0.07782688 | 0.2167018 | C25H3.17 |
| WBGene00194643 | 0.459413223 | 0.84552382 | 1 | C48D1.9 |
| WBGene00194649 | 2.016793204 | 0.09922168 | 1 | Y57G11C.1130 |
| WBGene00194651 | 0.589625475 | 0.28255647 | 0.4920665 | C33D9.13 |
| WBGene00194656 | -0.156269293 | 0.96924949 | 1 | T05F1.15 |
| WBGene00194658 | 0.28530093 | 0.92084641 | 1 | R09E10.13 |
| WBGene00194659 | -0.70562044 | 0.52136292 | 1 | linc-1 |
| WBGene00194660 | 0.312311137 | 0.84087066 | 1 | W09C5.12 |

| | | | | |
|---|---|---|---|---|
| WBGene00194663 | 0.256489601 | 0.80346637 | 1 | F38B2.6 |
| WBGene00194664 | 1.019267722 | 0.67145278 | 1 | K09E9.7 |
| WBGene00194665 | -1.299525668 | 0.08109111 | 0.22264297 | ZK822.8 |
| WBGene00194668 | -0.07748118 | 0.98067866 | 1 | C08F11.116 |
| WBGene00194669 | 0.690759747 | 0.61732989 | 1 | ZK1251.15 |
| WBGene00194670 | -0.626833103 | 0.80597175 | 1 | K07E3.9 |
| WBGene00194673 | 1.087152039 | 0.00034593 | 0.00406676 | ZK593.11 |
| WBGene00194674 | -2.357366668 | 1.42E-21 | 5.38E-19 | F23D12.11 |
| WBGene00194677 | 0.430452006 | 0.81603049 | 1 | D1054.19 |
| WBGene00194678 | -0.950208353 | 0.7250598 | 1 | R10D12.18 |
| WBGene00194679 | -1.360800281 | 0.38017426 | 1 | T26C11.9 |
| WBGene00194680 | -1.319381815 | 0.25868085 | 1 | Y17D7C.6 |
| WBGene00194681 | 0.576045153 | 0.64038314 | 1 | F40D4.16 |
| WBGene00194682 | 0.256515351 | 0.74134708 | 0.85569007 | F40D4.17 |
| WBGene00194684 | -0.668264701 | 0.85551153 | 1 | Y48A6B.14 |
| WBGene00194686 | 0.203033477 | 0.51454399 | 0.70136731 | Y49A3A.6 |
| WBGene00194687 | -1.6730111 | 0.24889096 | 1 | F57C12.6 |
| WBGene00194690 | 0.574508983 | 0.8551971 | 1 | T05D4.6 |
| WBGene00194691 | -0.558804446 | 0.87818675 | 1 | ZK1098.12 |
| WBGene00194692 | 0.409974797 | 0.64390524 | 0.79428104 | T02C12.5 |
| WBGene00194695 | 1.242013663 | 0.12878046 | 0.29975529 | B0462.5 |
| WBGene00194696 | -1.066668618 | 0.15793464 | 0.3428978 | EEED8.18 |
| WBGene00194697 | -0.156269293 | 0.96924949 | 1 | F59A3.13 |
| WBGene00194698 | -0.156269293 | 0.96924949 | 1 | F49D11.11 |
| WBGene00194699 | 0.041803775 | 0.98890197 | 1 | Y43F8B.22 |
| WBGene00194700 | -0.461923648 | 0.85976319 | 1 | B0025.5 |
| WBGene00194701 | -0.156269293 | 0.96924949 | 1 | Y23H5B.12 |
| WBGene00194702 | -0.156269293 | 0.96924949 | 1 | C32E8.12 |
| WBGene00194703 | 0.794516296 | 0.53129906 | 1 | F26B1.8 |
| WBGene00194704 | 0.281454817 | 0.94435481 | 1 | T07H3.8 |
| WBGene00194706 | 0.07023914 | 0.81840541 | 0.90149338 | C53A5.16 |
| WBGene00194707 | 0.067401136 | 0.8118876 | 0.89724443 | C53A5.17 |
| WBGene00194708 | 0.35134761 | 0.87335989 | 1 | Y36E3A.2 |
| WBGene00194710 | -0.008677305 | 0.99719169 | 1 | mks-2 |
| WBGene00194712 | -0.333693239 | 0.89573179 | 1 | T11F9.21 |
| WBGene00194713 | 0.822135772 | 0.51331576 | 1 | F19B10.13 |
| WBGene00194714 | -0.156269293 | 0.96924949 | 1 | ZC204.17 |
| WBGene00194717 | 2.700577684 | 4.573E-05 | 0.00080233 | ZC239.22 |
| WBGene00194725 | 1.851444818 | 0.53669729 | 1 | T16H12.13 |
| WBGene00194726 | 0.895876221 | 0.58041365 | 1 | R13A5.15 |
| WBGene00194727 | -0.263229581 | 0.93409872 | 1 | C18D11.10 |
| WBGene00194731 | -1.109412273 | 0.26574799 | 1 | Y39A1A.27 |
| WBGene00194735 | -0.156269293 | 0.96924949 | 1 | Y82E9BL.19 |

| | | | | |
|---|---|---|---|---|
| WBGene00194737 | 0.574508983 | 0.8551971 | 1 | Y50D7A.13 |
| WBGene00194742 | -0.969878876 | 0.41579494 | 1 | F37C12.21 |
| WBGene00194746 | 0.452272988 | 0.53127366 | 0.71493169 | pals-20 |
| WBGene00194747 | -0.662312358 | 0.44231776 | 0.64198205 | H05C05.4 |
| WBGene00194748 | -2.563099533 | 0.4751044 | 1 | F56F3.7 |
| WBGene00194771 | 0.837072097 | 0.30575775 | 0.51458342 | F32F2.3 |
| WBGene00194780 | 0.041803775 | 0.98890197 | 1 | C12D8.20 |
| WBGene00194783 | -0.156269293 | 0.96924949 | 1 | K03B8.14 |
| WBGene00194784 | -1.969805064 | 0.62862569 | 1 | T11F9.22 |
| WBGene00194785 | 0.688793783 | 0.86378755 | 1 | C52E4.12 |
| WBGene00194788 | -0.156269293 | 0.96924949 | 1 | R02D5.10 |
| WBGene00194792 | 0.850242284 | 0.20471401 | 0.40406162 | F32F2.2 |
| WBGene00194793 | -1.827081242 | 0.20976536 | 1 | W04G3.13 |
| WBGene00194800 | 0.377828795 | 0.92526585 | 1 | linc-83 |
| WBGene00194801 | 0.377828795 | 0.92526585 | 1 | Y6G8.16 |
| WBGene00194803 | 1.562982762 | 0.3798785 | 1 | C25F9.16 |
| WBGene00194805 | -0.156269293 | 0.96924949 | 1 | mir-2210 |
| WBGene00194810 | -0.156269293 | 0.96924949 | 1 | mir-2218.2 |
| WBGene00194811 | -0.254683336 | 0.89672209 | 1 | F22E10.6 |
| WBGene00194814 | -0.312249975 | 0.78493068 | 1 | linc-34 |
| WBGene00194815 | 2.257918864 | 0.10329254 | 1 | Y43F8B.23 |
| WBGene00194816 | -0.156269293 | 0.96924949 | 1 | ZK616.61 |
| WBGene00194821 | -1.048892867 | 0.54701031 | 1 | dmsr-16 |
| WBGene00194832 | 0.110005205 | 0.82296375 | 0.90431616 | Y73B6BL.270 |
| WBGene00194833 | 0.574508983 | 0.8551971 | 1 | F17E9.15 |
| WBGene00194835 | -0.156269293 | 0.96924949 | 1 | Y17G9B.11 |
| WBGene00194839 | 0.983488853 | 0.80601565 | 1 | C01B10.44 |
| WBGene00194841 | 0.758413118 | 0.23084085 | 0.4363442 | C44C10.13 |
| WBGene00194847 | 0.198783432 | 0.63790658 | 0.79024747 | F19C6.8 |
| WBGene00194848 | -0.283794653 | 0.88299492 | 1 | T27D12.6 |
| WBGene00194849 | 1.69047418 | 0.38809581 | 1 | K09A9.8 |
| WBGene00194851 | -0.598124017 | 0.53904806 | 0.72067849 | K07C10.3 |
| WBGene00194852 | 0.197316764 | 0.73266085 | 0.85048622 | T28A8.8 |
| WBGene00194853 | -0.33234649 | 0.46073983 | 0.65624616 | Y111B2A.29 |
| WBGene00194854 | -0.635809064 | 0.3922426 | 0.60022237 | Y111B2A.30 |
| WBGene00194855 | 0.761669137 | 0.26618736 | 0.47477459 | Y75B8A.44 |
| WBGene00194856 | 0.55883899 | 0.00934117 | 0.04924496 | K11D9.5 |
| WBGene00194861 | 0.113289505 | 0.87299932 | 0.93170576 | H04D03.6 |
| WBGene00194865 | 0.812607088 | 0.34078394 | 0.54915327 | F53A2.10 |
| WBGene00194866 | -1.351599631 | 0.53258792 | 1 | Y41C4A.22 |
| WBGene00194868 | 0.688793783 | 0.86378755 | 1 | M03C11.9 |
| WBGene00194869 | -1.291305095 | 0.63677871 | 1 | K01A6.8 |
| WBGene00194870 | 0.574508983 | 0.8551971 | 1 | F53A2.11 |

| | | | | |
|---|---|---|---|---|
| WBGene00194883 | -0.336406384 | 0.76970418 | 1 | B0496.11 |
| WBGene00194884 | -0.496752498 | 0.29482233 | 0.50514225 | Y48B6A.17 |
| WBGene00194886 | -0.156269293 | 0.96924949 | 1 | C05B5.12 |
| WBGene00194888 | 0.377828795 | 0.92526585 | 1 | K10H10.13 |
| WBGene00194892 | -0.564693412 | 0.02230993 | 0.09282554 | dsb-2 |
| WBGene00194894 | -0.156269293 | 0.96924949 | 1 | T05B4.14 |
| WBGene00194895 | -0.514849788 | 0.8003149 | 1 | ZK105.11 |
| WBGene00194896 | 0.688793783 | 0.86378755 | 1 | W02H5.11 |
| WBGene00194902 | 0.688793783 | 0.86378755 | 1 | Y39H10B.3 |
| WBGene00194905 | 0.490440116 | 0.68018815 | 1 | Y53F4B.51 |
| WBGene00194906 | -0.299618357 | 0.45960419 | 0.65555151 | Y53C12A.11 |
| WBGene00194907 | 0.281454817 | 0.94435481 | 1 | E02H4.8 |
| WBGene00194908 | -0.00418433 | 0.99213602 | 0.99616496 | F22F7.8 |
| WBGene00194909 | 1.851078774 | 0.10236195 | 1 | F41E6.17 |
| WBGene00194910 | 1.179295881 | 0.61449445 | 1 | ZC404.15 |
| WBGene00194912 | 0.459413223 | 0.84552382 | 1 | ZK856.18 |
| WBGene00194913 | -0.211603653 | 0.78825101 | 0.88342616 | Y105C5A.1272 |
| WBGene00194915 | -0.497458486 | 0.69626977 | 1 | JC8.16 |
| WBGene00194916 | 0.35134761 | 0.87335989 | 1 | F35D2.6 |
| WBGene00194917 | 0.489058549 | 0.85947456 | 1 | Y45F10B.59 |
| WBGene00194919 | -0.156269293 | 0.96924949 | 1 | K08D8.10 |
| WBGene00194920 | -0.156269293 | 0.96924949 | 1 | H12I19.115 |
| WBGene00194921 | 1.099549765 | 0.24686058 | 0.45494383 | ZK742.7 |
| WBGene00194925 | 0.28530093 | 0.92084641 | 1 | K08D8.11 |
| WBGene00194926 | -0.140829204 | 0.90591279 | 1 | K08D8.12 |
| WBGene00194928 | 0.357231179 | 0.74216949 | 1 | ZC513.14 |
| WBGene00194929 | -0.490236895 | 0.78472541 | 1 | Y105C5B.1413 |
| WBGene00194934 | 0.841231622 | 0.47587365 | 1 | ZK809.9 |
| WBGene00194952 | -1.831477136 | 0.52194793 | 1 | ZK666.14 |
| WBGene00194955 | 0.605056031 | 0.73958123 | 1 | F37H8.6 |
| WBGene00194981 | -1.150134399 | 0.4892498 | 1 | Y49E10.30 |
| WBGene00194982 | 1.712479368 | 0.58529868 | 1 | Y17D7B.10 |
| WBGene00194983 | 0.747845955 | 0.85215965 | 1 | T25B2.4 |
| WBGene00194985 | 0.386999607 | 0.71183181 | 1 | Y25C1A.14 |
| WBGene00194986 | 0.188356209 | 0.49759113 | 0.68588814 | T27F7.4 |
| WBGene00194988 | 0.983488853 | 0.80601565 | 1 | B0213.21 |
| WBGene00194989 | 0.281454817 | 0.94435481 | 1 | B0213.22 |
| WBGene00194990 | 1.216957619 | 0.74819621 | 1 | B0213.23 |
| WBGene00194991 | 1.192524401 | 0.37891857 | 1 | B0213.24 |
| WBGene00194997 | -0.156269293 | 0.96924949 | 1 | F42C5.12 |
| WBGene00194998 | -0.245510949 | 0.7716498 | 0.87239038 | C16C4.18 |
| WBGene00195000 | -0.18557176 | 0.95664217 | 1 | F54D8.7 |
| WBGene00195001 | -0.18557176 | 0.95664217 | 1 | Y17G9A.85 |

| | | | | |
|---|---|---|---|---|
| WBGene00195002 | -0.18557176 | 0.95664217 | 1 | T08B6.62 |
| WBGene00195003 | 0.281454817 | 0.94435481 | 1 | Y46C8AL.11 |
| WBGene00195004 | 1.976897945 | 0.29244709 | 1 | K08D12.8 |
| WBGene00195005 | -0.514849788 | 0.8003149 | 1 | ZK809.10 |
| WBGene00195006 | -1.082680044 | 0.67762556 | 1 | C35B1.9 |
| WBGene00195008 | 1.025181888 | 0.79790347 | 1 | mir-4813 |
| WBGene00195009 | -0.156269293 | 0.96924949 | 1 | F17E9.16 |
| WBGene00195011 | 0.32595502 | 0.91626577 | 1 | ZK180.8 |
| WBGene00195016 | -3.005117942 | 0.20183762 | 0.40059708 | ZC155.8 |
| WBGene00195017 | -2.254641634 | 0.25974171 | 0.46788192 | R144.15 |
| WBGene00195019 | -0.156269293 | 0.96924949 | 1 | Y59H11AR.8 |
| WBGene00195020 | 0.961774584 | 0.1191748 | 0.28493526 | Y59H11AR.9 |
| WBGene00195023 | 1.214472632 | 0.29306747 | 1 | F39E9.16 |
| WBGene00195025 | 2.983877393 | 0.058566 | 1 | F39E9.18 |
| WBGene00195027 | 0.035357682 | 0.98153502 | 1 | F39E9.20 |
| WBGene00195028 | 2.983877393 | 0.058566 | 1 | F39E9.21 |
| WBGene00195030 | 0.814191788 | 0.3042312 | 0.51329075 | F10E7.12 |
| WBGene00195031 | 0.773345629 | 0.76155013 | 1 | Y37E3.23 |
| WBGene00195033 | -1.22259901 | 0.68385967 | 1 | Y37E3.25 |
| WBGene00195034 | -0.641454362 | 0.68194393 | 1 | F40G9.20 |
| WBGene00195036 | 0.281454817 | 0.94435481 | 1 | T08D2.10 |
| WBGene00195040 | 0.798991008 | 0.05872927 | 0.1799223 | K08A2.10 |
| WBGene00195043 | 2.365957384 | 0.04720787 | 1 | Y57G11C.1135 |
| WBGene00195045 | -1.0062442 | 0.63208617 | 1 | W05F2.8 |
| WBGene00195050 | -1.150134399 | 0.4892498 | 1 | T22D1.17 |
| WBGene00195051 | -0.156269293 | 0.96924949 | 1 | T22D1.18 |
| WBGene00195056 | 0.353110589 | 0.85358686 | 1 | C45E1.5 |
| WBGene00195057 | -1.351599631 | 0.53258792 | 1 | B0261.9 |
| WBGene00195060 | 0.860474826 | 0.53117533 | 1 | linc-14 |
| WBGene00195063 | -0.308194322 | 0.37087899 | 0.57832544 | Y38C1AA.14 |
| WBGene00195064 | 0.88568203 | 0.59891017 | 1 | F17C11.20 |
| WBGene00195065 | -0.036779526 | 0.97679231 | 1 | F20G2.9 |
| WBGene00195067 | -0.156269293 | 0.96924949 | 1 | C04C3.9 |
| WBGene00195068 | -0.156269293 | 0.96924949 | 1 | Y48G8AL.16 |
| WBGene00195069 | 1.876192691 | 0.43542138 | 1 | W03G9.10 |
| WBGene00195070 | 0.675988998 | 0.34810606 | 0.55660328 | C16C8.20 |
| WBGene00195071 | 0.695474669 | 0.65279923 | 1 | C16C8.21 |
| WBGene00195072 | -1.73021997 | 0.47726432 | 1 | C16C8.22 |
| WBGene00195073 | 0.852895176 | 0.39990155 | 1 | F02E11.7 |
| WBGene00195077 | -1.911068126 | 0.13823025 | 1 | T24E12.13 |
| WBGene00195081 | 1.240298116 | 0.45369109 | 1 | C04F6.7 |
| WBGene00195084 | 0.316186427 | 0.7514507 | 1 | C43C3.4 |
| WBGene00195086 | 0.822175877 | 0.50368962 | 1 | F21C10.13 |

| | | | | |
|---|---|---|---|---|
| WBGene00195093 | -0.025031637 | 0.90808973 | 0.95194377 | Y59E9AL.36 |
| WBGene00195117 | 2.456496557 | 0.3218863 | 1 | T13F3.10 |
| WBGene00195142 | -0.360445946 | 0.58982739 | 0.75746714 | M110.10 |
| WBGene00195143 | 0.696910439 | 0.49524572 | 1 | C16D2.2 |
| WBGene00195145 | 0.353421144 | 0.85118695 | 1 | ZK666.15 |
| WBGene00195146 | -1.269172425 | 0.19764094 | 1 | F54D5.17 |
| WBGene00195147 | -1.901856996 | 0.64072034 | 1 | F14E5.8 |
| WBGene00195149 | -0.156269293 | 0.96924949 | 1 | F26H11.8 |
| WBGene00195150 | 0.688793783 | 0.86378755 | 1 | Y81G3A.6 |
| WBGene00195151 | 0.691564487 | 0.30076803 | 0.51035984 | F53H2.4 |
| WBGene00195154 | -0.909685513 | 0.63916464 | 1 | F13A2.10 |
| WBGene00195155 | 0.471935218 | 0.81492399 | 1 | W04A8.9 |
| WBGene00195157 | 0.233739675 | 0.93575889 | 1 | C12D8.21 |
| WBGene00195159 | 1.38322681 | 0.69550305 | 1 | T04C12.14 |
| WBGene00195164 | -0.701574888 | 0.80383767 | 1 | K12B6.11 |
| WBGene00195165 | 2.490696505 | 0.25975012 | 1 | F57G4.11 |
| WBGene00195166 | 1.060949945 | 0.61866902 | 1 | W05B10.6 |
| WBGene00195169 | -0.75160782 | 0.53942253 | 1 | Y50D4A.6 |
| WBGene00195170 | -0.038149573 | 0.9819026 | 1 | linc-61 |
| WBGene00195172 | 1.957271014 | 0.49743227 | 1 | K10G4.14 |
| WBGene00195174 | 0.281454817 | 0.94435481 | 1 | K08E5.6 |
| WBGene00195175 | 0.719851115 | 0.28357268 | 0.49324583 | Y105E8A.36 |
| WBGene00195176 | 1.14167903 | 0.06113499 | 0.18491278 | Y43F8B.24 |
| WBGene00195177 | 1.876192691 | 0.43542138 | 1 | Y43F8B.25 |
| WBGene00195178 | 1.743415485 | 0.50483382 | 1 | ZC581.10 |
| WBGene00195180 | -0.156269293 | 0.96924949 | 1 | C41D11.10 |
| WBGene00195182 | 0.574508983 | 0.8551971 | 1 | C17G10.10 |
| WBGene00195183 | 0.27792322 | 0.74183584 | 0.85569007 | R04A9.9 |
| WBGene00195184 | 0.987949146 | 0.59434865 | 1 | F19G12.8 |
| WBGene00195185 | 1.494736006 | 0.10776677 | 0.26759651 | F19G12.9 |
| WBGene00195186 | 1.142499766 | 0.12484067 | 0.2944804 | F19G12.10 |
| WBGene00195188 | 0.281454817 | 0.94435481 | 1 | F19G12.11 |
| WBGene00195189 | -0.836616326 | 0.19902644 | 0.39716962 | ZC449.8 |
| WBGene00195208 | -1.509975708 | 0.60137946 | 1 | W02H3.3 |
| WBGene00195209 | -0.348178569 | 0.84036394 | 1 | C07B5.8 |
| WBGene00195210 | 0.281454817 | 0.94435481 | 1 | Y7A5A.13 |
| WBGene00195212 | -0.479165649 | 0.78467961 | 1 | F09B12.7 |
| WBGene00195213 | -1.094201388 | 0.7816772 | 1 | F11C1.9 |
| WBGene00195214 | 0.520392999 | 0.69653598 | 1 | F11C1.10 |
| WBGene00195226 | 0.063771689 | 0.98059714 | 1 | ZC404.16 |
| WBGene00195233 | 0.924964223 | 0.13922281 | 0.31635863 | F31E9.11 |
| WBGene00195234 | 0.282614314 | 0.44646603 | 0.64575484 | C14C10.9 |
| WBGene00195237 | -2.934840138 | 0.2654436 | 1 | Y116F11B.18 |

| | | | | |
|---|---|---|---|---|
| WBGene00195238 | 0.377828795 | 0.92526585 | 1 | mir-4805 |
| WBGene00195239 | -1.125636879 | 7.06E-07 | 2.5377E-05 | ZK1127.13 |
| WBGene00195240 | -0.156269293 | 0.96924949 | 1 | Y39B6A.69 |
| WBGene00195241 | 0.056106392 | 0.97371196 | 1 | Y113G7B.27 |
| WBGene00195244 | 1.922073787 | 0.34080775 | 1 | F30A10.15 |
| WBGene00195245 | -2.097890267 | 0.34816065 | 1 | T26A8.6 |
| WBGene00195246 | 0.064726683 | 0.8524695 | 0.91927977 | ZK546.19 |
| WBGene00195248 | -0.260064314 | 0.27999392 | 0.4890343 | emc-5 |
| WBGene00195260 | 0.281454817 | 0.94435481 | 1 | T27F7.5 |
| WBGene00195263 | -0.156269293 | 0.96924949 | 1 | F02E8.7 |
| WBGene00195274 | 0.688793783 | 0.86378755 | 1 | ZK617.7 |
| WBGene00195280 | 1.716537703 | 0.23274606 | 1 | Y67A10A.342 |
| WBGene00195299 | -1.901856996 | 0.64072034 | 1 | F11C1.11 |
| WBGene00195306 | -0.537747866 | 0.53427784 | 0.71747233 | K07C11.13 |
| WBGene00195309 | 0.159806191 | 0.4075064 | 0.6141434 | F59B8.5 |
| WBGene00195311 | -0.156269293 | 0.96924949 | 1 | M110.11 |
| WBGene00195312 | 0.577883667 | 0.6863737 | 1 | C34H3.3 |
| WBGene00195317 | 0.560969396 | 0.52272548 | 0.70775177 | T23G5.9 |
| WBGene00195328 | -0.156269293 | 0.96924949 | 1 | C01B7.8 |
| WBGene00195333 | -0.156269293 | 0.96924949 | 1 | F54F12.3 |
| WBGene00195338 | -1.969805064 | 0.62862569 | 1 | T02E9.9 |
| WBGene00195355 | 0.281454817 | 0.94435481 | 1 | Y92H12A.8 |
| WBGene00195366 | 0.281454817 | 0.94435481 | 1 | C50F4.17 |
| WBGene00195367 | 0.281454817 | 0.94435481 | 1 | B0524.7 |
| WBGene00195370 | -0.737462795 | 0.83433562 | 1 | F29B9.13 |
| WBGene00195378 | -2.640414922 | 0.41081194 | 1 | T19E7.22 |
| WBGene00195379 | 0.377828795 | 0.92526585 | 1 | ZK945.10 |
| WBGene00195383 | -0.156269293 | 0.96924949 | 1 | C09B8.10 |
| WBGene00195385 | 1.653460415 | 0.50712258 | 1 | ZK867.5 |
| WBGene00195386 | 0.574508983 | 0.8551971 | 1 | T09E8.6 |
| WBGene00195389 | 0.281454817 | 0.94435481 | 1 | F54E4.5 |
| WBGene00195393 | -1.488391716 | 0.50775624 | 1 | F28E10.6 |
| WBGene00195399 | -1.901856996 | 0.64072034 | 1 | F35G2.7 |
| WBGene00195412 | 1.961596997 | 0.16432446 | 1 | K03H1.16 |
| WBGene00195414 | -0.156269293 | 0.96924949 | 1 | C33F10.16 |
| WBGene00195419 | -0.150245429 | 0.9305065 | 1 | R173.5 |
| WBGene00195425 | -0.156269293 | 0.96924949 | 1 | ZC8.7 |
| WBGene00195431 | -0.769518539 | 0.33810348 | 0.54608412 | W02F12.9 |
| WBGene00195432 | -0.156269293 | 0.96924949 | 1 | C56E6.10 |
| WBGene00195433 | 0.688793783 | 0.86378755 | 1 | F41D9.6 |
| WBGene00195445 | -0.263229581 | 0.93409872 | 1 | Y23H5A.9 |
| WBGene00195457 | 0.664914355 | 0.80343354 | 1 | K05D4.11 |
| WBGene00195461 | 0.281454817 | 0.94435481 | 1 | F28E10.7 |

| | | | | |
|---|---|---|---|---|
| WBGene00195468 | 0.281454817 | 0.94435481 | 1 | F53H10.4 |
| WBGene00195469 | -0.156269293 | 0.96924949 | 1 | F46G11.8 |
| WBGene00195485 | -0.737462795 | 0.83433562 | 1 | F55A12.11 |
| WBGene00195499 | -0.62146648 | 0.85925752 | 1 | H22K11.5 |
| WBGene00195502 | -1.901856996 | 0.64072034 | 1 | Y38C1AB.9 |
| WBGene00195512 | 0.281454817 | 0.94435481 | 1 | C04C11.5 |
| WBGene00195517 | 1.141483964 | 0.35503343 | 1 | C05E11.10 |
| WBGene00195518 | -0.660188019 | 0.01457551 | 0.06827208 | ZK652.14 |
| WBGene00195519 | -0.604396148 | 0.82723443 | 1 | Y73B3A.23 |
| WBGene00195520 | 0.587171769 | 0.32842392 | 0.53702286 | C18F3.6 |
| WBGene00195525 | -0.666455177 | 0.61533126 | 1 | C09F9.8 |
| WBGene00195530 | -0.156269293 | 0.96924949 | 1 | T04G9.8 |
| WBGene00195534 | -0.156269293 | 0.96924949 | 1 | C29E6.8 |
| WBGene00195536 | 0.281454817 | 0.94435481 | 1 | C55A6.14 |
| WBGene00195538 | 0.063787288 | 0.93468428 | 0.96562303 | M79.6 |
| WBGene00195539 | 0.281454817 | 0.94435481 | 1 | C12D8.22 |
| WBGene00195542 | -1.969805064 | 0.62862569 | 1 | C24A3.10 |
| WBGene00195552 | 0.281454817 | 0.94435481 | 1 | C05D12.8 |
| WBGene00195571 | 0.068198813 | 0.96787621 | 1 | T24D8.10 |
| WBGene00195577 | -0.156269293 | 0.96924949 | 1 | C06E1.12 |
| WBGene00195580 | -0.62146648 | 0.85925752 | 1 | R07D5.3 |
| WBGene00195582 | -0.156269293 | 0.96924949 | 1 | C06E8.6 |
| WBGene00195584 | -0.156269293 | 0.96924949 | 1 | C34F6.13 |
| WBGene00195603 | -0.156269293 | 0.96924949 | 1 | T04F8.10 |
| WBGene00195608 | -0.156269293 | 0.96924949 | 1 | F56G4.8 |
| WBGene00195609 | -0.156269293 | 0.96924949 | 1 | Y41C4A.23 |
| WBGene00195611 | 0.281454817 | 0.94435481 | 1 | K08D9.7 |
| WBGene00195621 | 0.281454817 | 0.94435481 | 1 | F43H9.5 |
| WBGene00195623 | -1.14715222 | 0.68932198 | 1 | Y54G2A.58 |
| WBGene00195624 | 0.281454817 | 0.94435481 | 1 | R31.4 |
| WBGene00195625 | 0.416708419 | 0.8260062 | 1 | Y58A7A.7 |
| WBGene00195627 | -0.156269293 | 0.96924949 | 1 | R13A1.13 |
| WBGene00195632 | 1.999910906 | 0.29422859 | 1 | Y80E2A.1 |
| WBGene00195633 | -0.156269293 | 0.96924949 | 1 | F44C4.9 |
| WBGene00195634 | 0.109173684 | 0.96057644 | 1 | AC3.14 |
| WBGene00195640 | -1.159714244 | 0.75790634 | 1 | Y59A8B.27 |
| WBGene00195644 | -2.640414922 | 0.41081194 | 1 | Y57A10B.8 |
| WBGene00195651 | -0.156269293 | 0.96924949 | 1 | F07C3.11 |
| WBGene00195655 | -0.156269293 | 0.96924949 | 1 | F43D9.7 |
| WBGene00195658 | -0.156269293 | 0.96924949 | 1 | C34D1.6 |
| WBGene00195659 | -0.156269293 | 0.96924949 | 1 | Y69E1A.9 |
| WBGene00195660 | -3.079012209 | 0.2683724 | 1 | R07E3.9 |
| WBGene00195661 | 0.281454817 | 0.94435481 | 1 | C08G9.4 |

| | | | | |
|---|---|---|---|---|
| WBGene00195670 | -0.156269293 | 0.96924949 | 1 | F52B10.4 |
| WBGene00195671 | -1.969805064 | 0.62862569 | 1 | R186.9 |
| WBGene00195672 | 0.635045026 | 0.70546974 | 1 | Y37E3.26 |
| WBGene00195673 | -1.901856996 | 0.64072034 | 1 | K08A8.5 |
| WBGene00195680 | -0.156269293 | 0.96924949 | 1 | T22B11.6 |
| WBGene00195683 | -0.156269293 | 0.96924949 | 1 | C15C6.6 |
| WBGene00195684 | 2.177389786 | 0.23191681 | 1 | F08F3.12 |
| WBGene00195685 | -1.14715222 | 0.68932198 | 1 | Y42A5A.8 |
| WBGene00195689 | -0.156269293 | 0.96924949 | 1 | K10D6.7 |
| WBGene00195694 | 0.281454817 | 0.94435481 | 1 | C43G2.25 |
| WBGene00195699 | 0.281454817 | 0.94435481 | 1 | K04G11.7 |
| WBGene00195707 | 0.281454817 | 0.94435481 | 1 | C11G10.3 |
| WBGene00195720 | -1.901856996 | 0.64072034 | 1 | F09E5.18 |
| WBGene00195721 | -0.952168503 | 0.60712255 | 1 | ZC506.5 |
| WBGene00195728 | 0.574508983 | 0.8551971 | 1 | T28D9.13 |
| WBGene00195730 | -0.156269293 | 0.96924949 | 1 | F54C9.15 |
| WBGene00195744 | 0.510721627 | 0.43874095 | 0.63927113 | C14B9.11 |
| WBGene00195751 | -0.156269293 | 0.96924949 | 1 | C12C8.5 |
| WBGene00195755 | 0.083331881 | 0.92283908 | 0.9593272 | B0207.13 |
| WBGene00195757 | 0.377828795 | 0.92526585 | 1 | T06A10.105 |
| WBGene00195758 | -0.156269293 | 0.96924949 | 1 | F18E2.8 |
| WBGene00195764 | -0.737462795 | 0.83433562 | 1 | C04C11.6 |
| WBGene00195765 | -1.901856996 | 0.64072034 | 1 | T08G2.4 |
| WBGene00195772 | -0.156269293 | 0.96924949 | 1 | C03H12.2 |
| WBGene00195774 | -0.156269293 | 0.96924949 | 1 | F20D1.12 |
| WBGene00195777 | -1.901856996 | 0.64072034 | 1 | D2023.10 |
| WBGene00195778 | -0.156269293 | 0.96924949 | 1 | PAR2.5 |
| WBGene00195783 | -0.156269293 | 0.96924949 | 1 | D2007.6 |
| WBGene00195792 | -1.159714244 | 0.75790634 | 1 | F58A4.16 |
| WBGene00195793 | -0.300852129 | 0.6718838 | 0.81248091 | B0496.12 |
| WBGene00195798 | -1.901856996 | 0.64072034 | 1 | C33A11.5 |
| WBGene00195802 | 0.281454817 | 0.94435481 | 1 | H14N18.5 |
| WBGene00195803 | -1.342059059 | 0.44806039 | 1 | F46E10.13 |
| WBGene00195804 | -0.156269293 | 0.96924949 | 1 | F11E6.19 |
| WBGene00195807 | -0.156269293 | 0.96924949 | 1 | F56H11.8 |
| WBGene00195809 | -0.156269293 | 0.96924949 | 1 | ZC455.14 |
| WBGene00195813 | 1.747998126 | 0.61126322 | 1 | F31C3.13 |
| WBGene00195816 | 0.377828795 | 0.92526585 | 1 | C34B4.9 |
| WBGene00195817 | -0.156269293 | 0.96924949 | 1 | K10C3.7 |
| WBGene00195820 | 0.312403178 | 0.89676939 | 1 | T22E5.10 |
| WBGene00195823 | 0.281454817 | 0.94435481 | 1 | F53A9.11 |
| WBGene00195824 | -1.901856996 | 0.64072034 | 1 | F27D9.9 |
| WBGene00195826 | 0.938982574 | 0.8146943 | 1 | B0496.13 |

| | | | | |
|---|---|---|---|---|
| WBGene00195829 | -0.156269293 | 0.96924949 | 1 | T26H2.13 |
| WBGene00195835 | 0.377828795 | 0.92526585 | 1 | C34B2.12 |
| WBGene00195839 | 0.983488853 | 0.80601565 | 1 | T23F1.8 |
| WBGene00195850 | -1.901856996 | 0.64072034 | 1 | F57C7.7 |
| WBGene00195855 | 2.847585511 | 0.02929432 | 1 | W06A7.6 |
| WBGene00195860 | 0.281454817 | 0.94435481 | 1 | ZK897.8 |
| WBGene00195861 | -2.705999569 | 0.42423051 | 1 | C48B6.11 |
| WBGene00195863 | -0.025049237 | 0.97336402 | 0.9861492 | F35H10.14 |
| WBGene00195864 | 0.281454817 | 0.94435481 | 1 | F41A4.3 |
| WBGene00195870 | 0.281454817 | 0.94435481 | 1 | F22E10.7 |
| WBGene00195875 | -1.159714244 | 0.75790634 | 1 | F11C7.8 |
| WBGene00195877 | 1.181686151 | 0.7579682 | 1 | R07E4.8 |
| WBGene00195878 | -0.156269293 | 0.96924949 | 1 | F46F3.6 |
| WBGene00195879 | -0.156269293 | 0.96924949 | 1 | F49E10.10 |
| WBGene00195884 | 0.377828795 | 0.92526585 | 1 | C17E4.14 |
| WBGene00195885 | 0.281454817 | 0.94435481 | 1 | F38B7.12 |
| WBGene00195896 | 1.622679176 | 0.01713248 | 0.07655704 | F31E3.8 |
| WBGene00195902 | -0.156269293 | 0.96924949 | 1 | T21B4.18 |
| WBGene00195903 | -0.156269293 | 0.96924949 | 1 | K02E2.14 |
| WBGene00195911 | 0.281454817 | 0.94435481 | 1 | C01B10.45 |
| WBGene00195913 | 0.041803775 | 0.98890197 | 1 | F54F12.4 |
| WBGene00195920 | -0.156269293 | 0.96924949 | 1 | C54D10.16 |
| WBGene00195924 | 0.774372031 | 0.69012376 | 1 | Y47H9A.3 |
| WBGene00195935 | 0.694609957 | 0.73912856 | 1 | F42G8.16 |
| WBGene00195937 | 1.288791542 | 0.36994516 | 1 | C33G3.8 |
| WBGene00195943 | -1.159714244 | 0.75790634 | 1 | W08F4.14 |
| WBGene00195965 | 0.281454817 | 0.94435481 | 1 | F26A10.7 |
| WBGene00195967 | -0.329162095 | 0.79446243 | 1 | K08C7.9 |
| WBGene00195971 | 0.281454817 | 0.94435481 | 1 | F55C5.12 |
| WBGene00195977 | 0.281454817 | 0.94435481 | 1 | C56E6.12 |
| WBGene00195987 | -0.156269293 | 0.96924949 | 1 | C04C11.7 |
| WBGene00195994 | 1.007245377 | 0.5574325 | 1 | F39B3.4 |
| WBGene00196000 | -0.156269293 | 0.96924949 | 1 | Y50D7A.14 |
| WBGene00196001 | -1.969805064 | 0.62862569 | 1 | F52B10.7 |
| WBGene00196004 | -0.156269293 | 0.96924949 | 1 | F13D11.8 |
| WBGene00196012 | -1.094201388 | 0.7816772 | 1 | F32A6.7 |
| WBGene00196014 | 0.281454817 | 0.94435481 | 1 | F47B7.10 |
| WBGene00196016 | -0.70576507 | 0.76841669 | 1 | W03A3.4 |
| WBGene00196018 | 0.281454817 | 0.94435481 | 1 | F28F8.11 |
| WBGene00196024 | 1.38322681 | 0.69550305 | 1 | C32C4.9 |
| WBGene00196027 | -0.461923648 | 0.85976319 | 1 | C30B5.10 |
| WBGene00196032 | 6.369864732 | 7.24E-07 | 2.577E-05 | T18D3.10 |
| WBGene00196034 | -0.156269293 | 0.96924949 | 1 | T04C12.16 |

| | | | | |
|---|---|---|---|---|
| WBGene00196037 | -1.964653846 | 0.53599202 | 1 | Y17D7B.11 |
| WBGene00196038 | -0.156269293 | 0.96924949 | 1 | Y51H7C.18 |
| WBGene00196052 | 0.688793783 | 0.86378755 | 1 | T27E4.14 |
| WBGene00196053 | -1.901856996 | 0.64072034 | 1 | F11H8.5 |
| WBGene00196059 | 1.38322681 | 0.69550305 | 1 | D2023.12 |
| WBGene00196066 | -0.62146648 | 0.85925752 | 1 | T19C4.17 |
| WBGene00196083 | -0.156269293 | 0.96924949 | 1 | C53B7.9 |
| WBGene00196085 | -0.348178569 | 0.84036394 | 1 | R173.6 |
| WBGene00196086 | 0.281454817 | 0.94435481 | 1 | F16H9.4 |
| WBGene00196087 | -0.558804446 | 0.87818675 | 1 | F08G5.10 |
| WBGene00196089 | -0.119131383 | 0.70128236 | 0.83141078 | C15F1.13 |
| WBGene00196091 | 0.281454817 | 0.94435481 | 1 | C50F4.18 |
| WBGene00196092 | 0.281454817 | 0.94435481 | 1 | ZK632.15 |
| WBGene00196096 | -0.543794685 | 0.13962078 | 0.3168595 | ZK616.62 |
| WBGene00196100 | -0.156269293 | 0.96924949 | 1 | K10C3.8 |
| WBGene00196102 | -1.759191416 | 0.57668729 | 1 | C29H12.7 |
| WBGene00196105 | 0.281454817 | 0.94435481 | 1 | C18D1.11 |
| WBGene00196110 | 1.342864318 | 0.37434345 | 1 | T01C2.3 |
| WBGene00196111 | -0.156269293 | 0.96924949 | 1 | C33B4.8 |
| WBGene00196115 | -0.156269293 | 0.96924949 | 1 | F45B8.8 |
| WBGene00196118 | -1.969805064 | 0.62862569 | 1 | Y32F6B.5 |
| WBGene00196120 | -1.094201388 | 0.7816772 | 1 | T06H11.7 |
| WBGene00196128 | -0.564963681 | 0.83907016 | 1 | Y51A2D.25 |
| WBGene00196131 | -0.156269293 | 0.96924949 | 1 | C56G2.16 |
| WBGene00196143 | -0.156269293 | 0.96924949 | 1 | K07A1.18 |
| WBGene00196147 | -1.901856996 | 0.64072034 | 1 | W02A2.10 |
| WBGene00196151 | -0.156269293 | 0.96924949 | 1 | Y48G9A.17 |
| WBGene00196157 | -1.159714244 | 0.75790634 | 1 | C07A4.4 |
| WBGene00196163 | 0.377828795 | 0.92526585 | 1 | F23H12.13 |
| WBGene00196167 | 0.316280515 | 0.57505512 | 0.74815787 | W06A7.7 |
| WBGene00196174 | 1.336924061 | 0.44603779 | 1 | F28F5.10 |
| WBGene00196180 | 0.574508983 | 0.8551971 | 1 | T19D2.6 |
| WBGene00196182 | -1.159714244 | 0.75790634 | 1 | F26G1.12 |
| WBGene00196192 | 0.377828795 | 0.92526585 | 1 | R01B10.7 |
| WBGene00196194 | -0.156269293 | 0.96924949 | 1 | C05C9.5 |
| WBGene00196200 | 0.416708419 | 0.8260062 | 1 | Y58A7A.8 |
| WBGene00196205 | -0.156269293 | 0.96924949 | 1 | Y111B2A.32 |
| WBGene00196220 | -0.558804446 | 0.87818675 | 1 | C24H12.13 |
| WBGene00196223 | -0.156269293 | 0.96924949 | 1 | ZK470.8 |
| WBGene00196225 | -1.901856996 | 0.64072034 | 1 | T21H3.6 |
| WBGene00196239 | 0.281454817 | 0.94435481 | 1 | F55E10.9 |
| WBGene00196244 | 0.281454817 | 0.94435481 | 1 | B0524.8 |
| WBGene00196245 | 0.377828795 | 0.92526585 | 1 | R08C7.16 |

| | | | | |
|---|---|---|---|---|
| WBGene00196247 | 0.281454817 | 0.94435481 | 1 | R07D5.4 |
| WBGene00196257 | 0.081076952 | 0.68107399 | 0.81720708 | T04C12.17 |
| WBGene00196258 | -0.91375383 | 0.72824392 | 1 | F25G6.13 |
| WBGene00196260 | 0.281454817 | 0.94435481 | 1 | F32B5.10 |
| WBGene00196261 | -0.156269293 | 0.96924949 | 1 | E02A10.6 |
| WBGene00196264 | 0.377828795 | 0.92526585 | 1 | F09A5.10 |
| WBGene00196280 | -1.094201388 | 0.7816772 | 1 | M01H9.7 |
| WBGene00196282 | -0.156269293 | 0.96924949 | 1 | F46G11.10 |
| WBGene00196303 | 0.281454817 | 0.94435481 | 1 | F10C2.10 |
| WBGene00196310 | 0.688793783 | 0.86378755 | 1 | C07G3.13 |
| WBGene00196314 | -1.964653846 | 0.53599202 | 1 | F15A2.9 |
| WBGene00196315 | -0.044628748 | 0.77199923 | 0.87263357 | Y24D9A.11 |
| WBGene00196321 | 0.281454817 | 0.94435481 | 1 | M79.7 |
| WBGene00196322 | -0.156269293 | 0.96924949 | 1 | R05F9.14 |
| WBGene00196324 | -2.705999569 | 0.42423051 | 1 | Y5H2A.5 |
| WBGene00196328 | -1.901856996 | 0.64072034 | 1 | Y39G8C.6 |
| WBGene00196332 | -1.969805064 | 0.62862569 | 1 | T27B2.2 |
| WBGene00196336 | -0.00511053 | 0.9965048 | 1 | F13B12.13 |
| WBGene00196341 | 0.574508983 | 0.8551971 | 1 | T27D12.8 |
| WBGene00196344 | 1.413377704 | 0.60673863 | 1 | T03F7.11 |
| WBGene00196346 | 0.688793783 | 0.86378755 | 1 | T02E9.13 |
| WBGene00196365 | -0.156269293 | 0.96924949 | 1 | C41A3.5 |
| WBGene00196368 | 1.529802032 | 0.64971375 | 1 | Y76A2B.7 |
| WBGene00196370 | 0.281454817 | 0.94435481 | 1 | F52B10.8 |
| WBGene00196374 | -0.156269293 | 0.96924949 | 1 | T21B10.8 |
| WBGene00196376 | 0.013667562 | 0.97025854 | 0.98453063 | K02D10.7 |
| WBGene00196380 | -1.901856996 | 0.64072034 | 1 | F11H8.6 |
| WBGene00196381 | -0.263229581 | 0.93409872 | 1 | C32A3.5 |
| WBGene00196385 | -0.156269293 | 0.96924949 | 1 | K11E4.8 |
| WBGene00196388 | 0.164601484 | 0.84555363 | 0.91536929 | T23G5.10 |
| WBGene00196395 | -0.156269293 | 0.96924949 | 1 | Y111B2A.33 |
| WBGene00196396 | 0.574188545 | NaN | 1 | F31C3.14 |
| WBGene00196397 | 0.281454817 | 0.94435481 | 1 | F48C1.12 |
| WBGene00196398 | 1.452076087 | 0.02946 | 0.11177307 | W10D5.4 |
| WBGene00196405 | 0.75595108 | 0.7708876 | 1 | K08E5.9 |
| WBGene00196408 | 0.574508983 | 0.8551971 | 1 | F15G9.7 |
| WBGene00196409 | -1.964653846 | 0.53599202 | 1 | K03H9.7 |
| WBGene00196410 | 0.688793783 | 0.86378755 | 1 | R02E12.9 |
| WBGene00196429 | 0.394439984 | 0.88400378 | 1 | F18G5.8 |
| WBGene00196432 | -0.196944706 | 0.90889504 | 1 | T14G11.5 |
| WBGene00196442 | -0.156269293 | 0.96924949 | 1 | T21B10.9 |
| WBGene00196447 | 0.574508983 | 0.8551971 | 1 | F46C8.10 |
| WBGene00196451 | 0.233739675 | 0.93575889 | 1 | K04A8.13 |

| | | | | |
|---|---|---|---|---|
| WBGene00196456 | -0.156269293 | 0.96924949 | 1 | F18E9.10 |
| WBGene00196465 | 0.743637457 | 0.84880372 | 1 | Y51F10.12 |
| WBGene00196467 | -0.156269293 | 0.96924949 | 1 | F43C9.5 |
| WBGene00196468 | -0.156269293 | 0.96924949 | 1 | W09G3.12 |
| WBGene00196469 | 0.57991002 | 0.82402153 | 1 | F08F3.13 |
| WBGene00196471 | 0.178115923 | 0.94982316 | 1 | F57B1.10 |
| WBGene00196480 | 0.281454817 | 0.94435481 | 1 | C27A12.11 |
| WBGene00196485 | -1.901856996 | 0.64072034 | 1 | C16D9.12 |
| WBGene00196494 | -0.156269293 | 0.96924949 | 1 | C07A12.9 |
| WBGene00196498 | 1.684305727 | 0.05195707 | 0.16465957 | B0350.71 |
| WBGene00196502 | -1.291305095 | 0.63677871 | 1 | R05D7.8 |
| WBGene00196521 | -0.156269293 | 0.96924949 | 1 | C18A3.16 |
| WBGene00196524 | 0.281454817 | 0.94435481 | 1 | C10C5.10 |
| WBGene00196533 | -1.901856996 | 0.64072034 | 1 | F33D11.14 |
| WBGene00196534 | -1.901856996 | 0.64072034 | 1 | Y113G7A.18 |
| WBGene00196538 | 0.122433202 | 0.94758598 | 1 | C09F12.6 |
| WBGene00196559 | 0.281454817 | 0.94435481 | 1 | F35G2.11 |
| WBGene00196568 | -0.156269293 | 0.96924949 | 1 | M79.9 |
| WBGene00196571 | -0.156269293 | 0.96924949 | 1 | T06E4.17 |
| WBGene00196573 | -0.969372093 | 0.00076675 | 0.0074953 | M163.16 |
| WBGene00196576 | -1.692398041 | 0.43695336 | 1 | C46A5.13 |
| WBGene00196582 | -0.156269293 | 0.96924949 | 1 | F08B12.6 |
| WBGene00196583 | -0.156269293 | 0.96924949 | 1 | R13A1.14 |
| WBGene00196591 | 0.377828795 | 0.92526585 | 1 | F47B7.11 |
| WBGene00196602 | 0.574508983 | 0.8551971 | 1 | W05E10.10 |
| WBGene00196606 | 0.281454817 | 0.94435481 | 1 | Y15E3A.7 |
| WBGene00196608 | -0.156269293 | 0.96924949 | 1 | F58A4.17 |
| WBGene00196615 | -0.333693239 | 0.89573179 | 1 | F53F4.21 |
| WBGene00196616 | -0.156269293 | 0.96924949 | 1 | F01E11.8 |
| WBGene00196626 | -0.156269293 | 0.96924949 | 1 | F22E10.8 |
| WBGene00196629 | -0.008677305 | 0.99719169 | 1 | T12B3.7 |
| WBGene00196631 | 0.312403178 | 0.89676939 | 1 | F26D11.15 |
| WBGene00196637 | -0.156269293 | 0.96924949 | 1 | F52H2.10 |
| WBGene00196645 | 0.281454817 | 0.94435481 | 1 | C34D10.3 |
| WBGene00196649 | -0.156269293 | 0.96924949 | 1 | F42H10.12 |
| WBGene00196668 | -1.901856996 | 0.64072034 | 1 | K11H12.14 |
| WBGene00196684 | -1.773847824 | 1.33E-09 | 1.01E-07 | F59D8.3 |
| WBGene00196690 | -0.156269293 | 0.96924949 | 1 | F20A1.12 |
| WBGene00196692 | 0.4828626 | 0.81641781 | 1 | Y73B3A.24 |
| WBGene00196695 | -0.156269293 | 0.96924949 | 1 | T14G10.9 |
| WBGene00196702 | -0.156269293 | 0.96924949 | 1 | T12A7.11 |
| WBGene00196708 | -1.901856996 | 0.64072034 | 1 | K08A8.9 |
| WBGene00196711 | -0.156269293 | 0.96924949 | 1 | ZK892.9 |

| | | | | |
|---|---|---|---|---|
| WBGene00196720 | 0.574508983 | 0.8551971 | 1 | ZC247.5 |
| WBGene00196721 | -0.156269293 | 0.96924949 | 1 | F57B10.17 |
| WBGene00196723 | 0.539239107 | 0.83916685 | 1 | F53B3.9 |
| WBGene00196733 | -0.156269293 | 0.96924949 | 1 | ZK381.41 |
| WBGene00196748 | 0.281454817 | 0.94435481 | 1 | F13H8.13 |
| WBGene00196752 | -0.638937665 | 0.04828058 | 0.15727531 | T06D8.12 |
| WBGene00196760 | 0.747845955 | 0.85215965 | 1 | T23G4.126 |
| WBGene00196764 | -0.398264424 | 0.59998308 | 0.76536654 | C45B2.10 |
| WBGene00196769 | 0.281454817 | 0.94435481 | 1 | B0457.10 |
| WBGene00196786 | 0.574508983 | 0.8551971 | 1 | R12C12.11 |
| WBGene00196790 | 0.377828795 | 0.92526585 | 1 | F38B7.13 |
| WBGene00196793 | -0.737462795 | 0.83433562 | 1 | C04C11.9 |
| WBGene00196805 | -1.969805064 | 0.62862569 | 1 | F45D3.8 |
| WBGene00196808 | 0.041803775 | 0.98890197 | 1 | ZK678.10 |
| WBGene00196814 | 0.938982574 | 0.8146943 | 1 | F14F3.8 |
| WBGene00196820 | -1.094201388 | 0.7816772 | 1 | F46C8.11 |
| WBGene00196822 | 0.688793783 | 0.86378755 | 1 | T19C4.19 |
| WBGene00196827 | -0.18557176 | 0.95664217 | 1 | C17C3.22 |
| WBGene00196834 | 1.477253415 | 0.58357327 | 1 | C24B5.8 |
| WBGene00196837 | -1.969805064 | 0.62862569 | 1 | C17D12.11 |
| WBGene00196839 | -0.323868354 | 0.91634405 | 1 | F19H6.9 |
| WBGene00196843 | 0.787403494 | 0.45240597 | 1 | C41G11.9 |
| WBGene00196848 | 0.281454817 | 0.94435481 | 1 | ZK353.11 |
| WBGene00196852 | -0.156269293 | 0.96924949 | 1 | C02F4.10 |
| WBGene00196854 | -0.156269293 | 0.96924949 | 1 | CD4.13 |
| WBGene00196856 | 0.281454817 | 0.94435481 | 1 | R10E11.10 |
| WBGene00196865 | -2.640414922 | 0.41081194 | 1 | F58B6.6 |
| WBGene00196867 | -0.156269293 | 0.96924949 | 1 | ZC477.16 |
| WBGene00196871 | 1.415446112 | 0.07666164 | 0.21490008 | K11C4.8 |
| WBGene00196872 | -1.969805064 | 0.62862569 | 1 | F40F9.15 |
| WBGene00196874 | 0.172713261 | 0.80506383 | 0.89368931 | C18A3.17 |
| WBGene00196880 | 0.688793783 | 0.86378755 | 1 | Y71A12B.27 |
| WBGene00196882 | 0.574508983 | 0.8551971 | 1 | C55A6.16 |
| WBGene00196884 | -0.263229581 | 0.93409872 | 1 | C54D1.9 |
| WBGene00196885 | -0.156269293 | 0.96924949 | 1 | ZC513.15 |
| WBGene00196888 | 0.688793783 | 0.86378755 | 1 | T07D1.9 |
| WBGene00196895 | -0.156269293 | 0.96924949 | 1 | T06H11.8 |
| WBGene00196901 | 0.05905161 | 0.97811893 | 1 | Y51A2D.27 |
| WBGene00196903 | 1.477253415 | 0.58357327 | 1 | M01E11.9 |
| WBGene00196905 | 0.281454817 | 0.94435481 | 1 | C30A5.12 |
| WBGene00196908 | -1.901856996 | 0.64072034 | 1 | F09E5.19 |
| WBGene00196909 | -0.156269293 | 0.96924949 | 1 | Y34B4A.14 |
| WBGene00196914 | 1.529802032 | 0.64971375 | 1 | R07E4.9 |

| | | | | |
|---|---|---|---|---|
| WBGene00196917 | -0.156269293 | 0.96924949 | 1 | B0379.10 |
| WBGene00196918 | -1.901856996 | 0.64072034 | 1 | F49E10.13 |
| WBGene00196925 | -0.156269293 | 0.96924949 | 1 | ZC504.8 |
| WBGene00196926 | 1.743415485 | 0.50483382 | 1 | Y39G8B.11 |
| WBGene00196931 | -1.595286372 | 0.19277689 | 1 | B0001.10 |
| WBGene00196933 | -0.312763056 | 0.07231635 | 0.20700797 | F43D9.8 |
| WBGene00196938 | -0.156269293 | 0.96924949 | 1 | F49H12.9 |
| WBGene00196940 | 0.281454817 | 0.94435481 | 1 | Y38H8A.10 |
| WBGene00196943 | -0.156269293 | 0.96924949 | 1 | T13H10.7 |
| WBGene00196958 | -0.62146648 | 0.85925752 | 1 | F58A4.18 |
| WBGene00196963 | 0.774372031 | 0.69012376 | 1 | Y47H9A.4 |
| WBGene00196970 | 1.025181888 | 0.79790347 | 1 | B0304.10 |
| WBGene00196979 | 0.09697944 | 0.97612634 | 1 | F09F9.9 |
| WBGene00196982 | -1.969805064 | 0.62862569 | 1 | Y41C4A.24 |
| WBGene00196986 | 0.574508983 | 0.8551971 | 1 | H01A20.2 |
| WBGene00196988 | -1.969805064 | 0.62862569 | 1 | F32A11.8 |
| WBGene00196994 | -0.156269293 | 0.96924949 | 1 | T01H3.7 |
| WBGene00196995 | 0.574508983 | 0.8551971 | 1 | Y55F3BR.14 |
| WBGene00197001 | 1.477253415 | 0.58357327 | 1 | C13C4.9 |
| WBGene00197002 | 1.057677827 | 0.30762602 | 1 | F33D4.12 |
| WBGene00197007 | -0.156269293 | 0.96924949 | 1 | C33B4.9 |
| WBGene00197009 | -0.156269293 | 0.96924949 | 1 | F18E2.10 |
| WBGene00197014 | -0.156269293 | 0.96924949 | 1 | C34B2.13 |
| WBGene00197020 | -0.558804446 | 0.87818675 | 1 | F08G5.12 |
| WBGene00197023 | 0.574508983 | 0.8551971 | 1 | Y67D8A.7 |
| WBGene00197047 | -0.156269293 | 0.96924949 | 1 | B0334.17 |
| WBGene00197048 | -0.156269293 | 0.96924949 | 1 | C04D8.2 |
| WBGene00197052 | -0.156269293 | 0.96924949 | 1 | F13D11.10 |
| WBGene00197054 | -0.156269293 | 0.96924949 | 1 | ZK512.13 |
| WBGene00197055 | 0.281905764 | 0.6045391 | 0.76879029 | T14G11.6 |
| WBGene00197056 | 0.446817701 | 0.72712767 | 1 | VT21B10L.1 |
| WBGene00197060 | 0.574508983 | 0.8551971 | 1 | T03E6.13 |
| WBGene00197065 | -0.156269293 | 0.96924949 | 1 | M28.13 |
| WBGene00197069 | -1.969805064 | 0.62862569 | 1 | Y38F2AR.16 |
| WBGene00197078 | 1.82867917 | 0.54358766 | 1 | F44E7.13 |
| WBGene00197080 | 0.707184947 | 0.26843851 | 0.47705078 | ZC15.11 |
| WBGene00197081 | 0.377828795 | 0.92526585 | 1 | C29A12.10 |
| WBGene00197083 | -1.901856996 | 0.64072034 | 1 | C16D9.13 |
| WBGene00197093 | -0.156269293 | 0.96924949 | 1 | C08F8.11 |
| WBGene00197097 | 0.938982574 | 0.8146943 | 1 | F13B9.9 |
| WBGene00197098 | -1.159714244 | 0.75790634 | 1 | R04E5.13 |
| WBGene00197099 | -0.156269293 | 0.96924949 | 1 | ZC416.9 |
| WBGene00197102 | -0.156269293 | 0.96924949 | 1 | C18A3.18 |

| | | | | |
|---|---|---|---|---|
| WBGene00197123 | -1.831477136 | 0.52194793 | 1 | F59G1.10 |
| WBGene00197127 | 0.064810246 | 0.83948616 | 0.91211511 | T05B11.9 |
| WBGene00197128 | -0.156269293 | 0.96924949 | 1 | T05A12.6 |
| WBGene00197134 | -0.156269293 | 0.96924949 | 1 | K08B5.5 |
| WBGene00197137 | 0.688793783 | 0.86378755 | 1 | R08B4.7 |
| WBGene00197142 | 1.413734769 | 0.68804531 | 1 | C36B1.15 |
| WBGene00197148 | -0.156269293 | 0.96924949 | 1 | C48A7.7 |
| WBGene00197153 | 0.574508983 | 0.8551971 | 1 | T18D3.12 |
| WBGene00197163 | 1.584577408 | 0.63258867 | 1 | B0495.15 |
| WBGene00197166 | 0.281454817 | 0.94435481 | 1 | C36F7.12 |
| WBGene00197169 | 0.281454817 | 0.94435481 | 1 | E01H11.4 |
| WBGene00197171 | 0.499361655 | 0.0354783 | 0.12706256 | F56D12.10 |
| WBGene00197173 | -0.558804446 | 0.87818675 | 1 | F46G11.12 |
| WBGene00197186 | -0.156269293 | 0.96924949 | 1 | C05G5.9 |
| WBGene00197189 | 0.281454817 | 0.94435481 | 1 | C06G1.10 |
| WBGene00197198 | -1.969805064 | 0.62862569 | 1 | D1044.11 |
| WBGene00197204 | 0.574508983 | 0.8551971 | 1 | B0034.9 |
| WBGene00197206 | 0.88787957 | 0.71832755 | 1 | C15B12.11 |
| WBGene00197207 | -0.156269293 | 0.96924949 | 1 | K10D6.11 |
| WBGene00197208 | -0.156269293 | 0.96924949 | 1 | ZK1128.9 |
| WBGene00197210 | 1.240028376 | 0.28335806 | 1 | F56D1.9 |
| WBGene00197215 | 2.661852293 | 0.00442558 | 0.02848936 | F57A8.9 |
| WBGene00197231 | -0.774842772 | 0.78069886 | 1 | Y48B6A.18 |
| WBGene00197238 | 0.574508983 | 0.8551971 | 1 | T19E7.24 |
| WBGene00197248 | 1.442588914 | 0.40683854 | 1 | K06C4.20 |
| WBGene00197250 | -0.156269293 | 0.96924949 | 1 | T02E9.15 |
| WBGene00197265 | 0.574508983 | 0.8551971 | 1 | M195.6 |
| WBGene00197271 | -0.156269293 | 0.96924949 | 1 | W09G3.13 |
| WBGene00197272 | -0.668264701 | 0.85551153 | 1 | C18F3.7 |
| WBGene00197273 | -1.094201388 | 0.7816772 | 1 | C23H3.10 |
| WBGene00197277 | 0.041803775 | 0.98890197 | 1 | Y37E3.28 |
| WBGene00197287 | -1.901856996 | 0.64072034 | 1 | Y104H12D.7 |
| WBGene00197290 | -1.969805064 | 0.62862569 | 1 | F21H7.17 |
| WBGene00197292 | -2.705999569 | 0.42423051 | 1 | Y5H2B.8 |
| WBGene00197293 | -0.18557176 | 0.95664217 | 1 | ZK1290.18 |
| WBGene00197296 | -0.156269293 | 0.96924949 | 1 | C09B7.6 |
| WBGene00197297 | -1.969805064 | 0.62862569 | 1 | R08C7.17 |
| WBGene00197298 | 0.627834078 | 0.77998941 | 1 | T23F11.8 |
| WBGene00197300 | 0.688793783 | 0.86378755 | 1 | R08B4.8 |
| WBGene00197301 | 0.281454817 | 0.94435481 | 1 | Y58A7A.9 |
| WBGene00197302 | 0.399606396 | 0.86544406 | 1 | C33F10.18 |
| WBGene00197317 | -0.150245429 | 0.9305065 | 1 | T22F3.14 |
| WBGene00197335 | 1.413734769 | 0.68804531 | 1 | C05D2.14 |

| | | | | |
|---|---|---|---|---|
| WBGene00197348 | 0.688793783 | 0.86378755 | 1 | T25B9.13 |
| WBGene00197349 | 0.281454817 | 0.94435481 | 1 | C05E11.12 |
| WBGene00197353 | -0.156269293 | 0.96924949 | 1 | T26E3.14 |
| WBGene00197362 | 0.688793783 | 0.86378755 | 1 | C23H4.11 |
| WBGene00197363 | -0.156269293 | 0.96924949 | 1 | W02F12.11 |
| WBGene00197369 | -1.901856996 | 0.64072034 | 1 | F48G7.14 |
| WBGene00197371 | -0.156269293 | 0.96924949 | 1 | Y76A2B.8 |
| WBGene00197374 | 0.688793783 | 0.86378755 | 1 | K04H4.11 |
| WBGene00197376 | 0.220597807 | 0.83827276 | 1 | F28E10.10 |
| WBGene00197378 | -0.263229581 | 0.93409872 | 1 | T22A3.11 |
| WBGene00197385 | 0.281454817 | 0.94435481 | 1 | F18C5.13 |
| WBGene00197387 | 0.281454817 | 0.94435481 | 1 | F35E8.18 |
| WBGene00197391 | 0.446858199 | 0.80724424 | 1 | K12H4.10 |
| WBGene00197401 | -0.65305639 | 0.05135213 | 0.16363942 | K08E3.13 |
| WBGene00197408 | 0.688793783 | 0.86378755 | 1 | F38E11.21 |
| WBGene00197412 | 0.615355315 | 0.71598629 | 1 | T24F1.9 |
| WBGene00197422 | -0.156269293 | 0.96924949 | 1 | F08B12.7 |
| WBGene00197430 | 0.377828795 | 0.92526585 | 1 | T09A5.18 |
| WBGene00197431 | -0.156269293 | 0.96924949 | 1 | Y67D8C.15 |
| WBGene00197432 | -1.094201388 | 0.7816772 | 1 | T21G5.7 |
| WBGene00197437 | -0.737462795 | 0.83433562 | 1 | K09E2.6 |
| WBGene00197441 | 0.281454817 | 0.94435481 | 1 | T08B2.14 |
| WBGene00197456 | 0.281454817 | 0.94435481 | 1 | C39E9.16 |
| WBGene00197459 | -1.969805064 | 0.62862569 | 1 | F28B4.5 |
| WBGene00197461 | -0.156269293 | 0.96924949 | 1 | T04C10.8 |
| WBGene00197465 | -0.156269293 | 0.96924949 | 1 | C25F6.14 |
| WBGene00197474 | -0.156269293 | 0.96924949 | 1 | C41A3.7 |
| WBGene00197478 | 0.281454817 | 0.94435481 | 1 | T13B5.10 |
| WBGene00197483 | -1.901856996 | 0.64072034 | 1 | F27B10.3 |
| WBGene00197488 | -1.969805064 | 0.62862569 | 1 | F41C6.11 |
| WBGene00197492 | 0.281454817 | 0.94435481 | 1 | T06H11.10 |
| WBGene00197494 | 0.688793783 | 0.86378755 | 1 | R09D1.16 |
| WBGene00197504 | -1.901856996 | 0.64072034 | 1 | F23F12.15 |
| WBGene00197505 | 1.687239501 | 0.59403242 | 1 | C01F4.6 |
| WBGene00197506 | -1.901856996 | 0.64072034 | 1 | F23F12.16 |
| WBGene00197509 | 0.281454817 | 0.94435481 | 1 | R02E4.4 |
| WBGene00197519 | -0.156269293 | 0.96924949 | 1 | Y105E8A.41 |
| WBGene00197520 | 1.270678414 | 0.5135606 | 1 | C05C8.10 |
| WBGene00197521 | 0.281454817 | 0.94435481 | 1 | ZC404.17 |
| WBGene00197530 | 0.938982574 | 0.8146943 | 1 | T12B5.16 |
| WBGene00197542 | 0.574508983 | 0.8551971 | 1 | W01A11.11 |
| WBGene00197543 | -0.333693239 | 0.89573179 | 1 | F57C7.9 |
| WBGene00197545 | -2.640414922 | 0.41081194 | 1 | Y53F4B.55 |

| | | | | |
|---|---|---|---|---|
| WBGene00197552 | 0.096802843 | 0.95918853 | 1 | Y49F6B.15 |
| WBGene00197553 | 0.232157325 | 0.93397866 | 1 | F45E1.10 |
| WBGene00197560 | 0.281454817 | 0.94435481 | 1 | K08A2.12 |
| WBGene00197563 | 0.377828795 | 0.92526585 | 1 | B0403.9 |
| WBGene00197573 | -0.668154252 | 0.53711593 | 1 | F14B8.11 |
| WBGene00197577 | -0.156269293 | 0.96924949 | 1 | W08F4.16 |
| WBGene00197595 | 0.12332541 | 0.94831365 | 1 | R12E2.18 |
| WBGene00197604 | -1.901856996 | 0.64072034 | 1 | K01G5.12 |
| WBGene00197610 | 0.281454817 | 0.94435481 | 1 | Y38H8A.11 |
| WBGene00197615 | -0.156269293 | 0.96924949 | 1 | C25F6.15 |
| WBGene00197620 | -1.443360033 | 0.37063936 | 1 | Y75B8A.49 |
| WBGene00197621 | -1.969805064 | 0.62862569 | 1 | F45F2.14 |
| WBGene00197622 | 0.281454817 | 0.94435481 | 1 | CD4.14 |
| WBGene00197637 | -1.901856996 | 0.64072034 | 1 | R13F6.13 |
| WBGene00197651 | 0.281454817 | 0.94435481 | 1 | R03A10.10 |
| WBGene00197660 | -0.156269293 | 0.96924949 | 1 | C27A2.11 |
| WBGene00197663 | 0.377828795 | 0.92526585 | 1 | F13H6.13 |
| WBGene00197666 | -0.156269293 | 0.96924949 | 1 | C05D2.15 |
| WBGene00197668 | 0.281454817 | 0.94435481 | 1 | F46H5.9 |
| WBGene00197672 | -1.14715222 | 0.68932198 | 1 | C48A7.9 |
| WBGene00197678 | 0.574508983 | 0.8551971 | 1 | R01E6.11 |
| WBGene00197684 | -0.156269293 | 0.96924949 | 1 | F59B2.17 |
| WBGene00197689 | -0.156269293 | 0.96924949 | 1 | F08G5.13 |
| WBGene00197690 | -0.323868354 | 0.91634405 | 1 | C08G9.5 |
| WBGene00197691 | 0.281454817 | 0.94435481 | 1 | K08F8.9 |
| WBGene00197700 | -0.558804446 | 0.87818675 | 1 | K09B11.20 |
| WBGene00197714 | 0.281454817 | 0.94435481 | 1 | Y54E10BL.8 |
| WBGene00197715 | -0.156269293 | 0.96924949 | 1 | B0024.17 |
| WBGene00197717 | 2.723320789 | 0.09762128 | 1 | T01B6.8 |
| WBGene00197719 | 0.281454817 | 0.94435481 | 1 | Y42G9A.8 |
| WBGene00197731 | -0.156269293 | 0.96924949 | 1 | F46F3.8 |
| WBGene00197733 | 0.377828795 | 0.92526585 | 1 | T25C12.5 |
| WBGene00197737 | -0.156269293 | 0.96924949 | 1 | F38E9.7 |
| WBGene00197738 | -0.156269293 | 0.96924949 | 1 | F43G9.15 |
| WBGene00197742 | 1.240458549 | 0.53319765 | 1 | F41D9.11 |
| WBGene00197747 | -1.901856996 | 0.64072034 | 1 | Y39H10A.8 |
| WBGene00197748 | -0.330211071 | 0.83331705 | 1 | T05A1.16 |
| WBGene00197755 | 0.983488853 | 0.80601565 | 1 | ZK899.10 |
| WBGene00197756 | -0.156269293 | 0.96924949 | 1 | ZC250.8 |
| WBGene00197760 | -0.156269293 | 0.96924949 | 1 | M163.18 |
| WBGene00197761 | -1.901856996 | 0.64072034 | 1 | C18A11.10 |
| WBGene00197765 | 1.599720773 | 0.24511199 | 1 | C01B7.11 |
| WBGene00197773 | -3.603894515 | 5.9989E-05 | 0.0010158 | T27A10.8 |

| | | | | |
|---|---|---|---|---|
| WBGene00197775 | 0.574508983 | 0.8551971 | 1 | F26F12.11 |
| WBGene00197791 | 0.281454817 | 0.94435481 | 1 | F54C9.16 |
| WBGene00197796 | 0.688793783 | 0.86378755 | 1 | B0001.11 |
| WBGene00197803 | -1.038181246 | 0.5835992 | 1 | K07C10.6 |
| WBGene00197805 | 0.281454817 | 0.94435481 | 1 | C43G2.27 |
| WBGene00197810 | 0.281454817 | 0.94435481 | 1 | Y17G9A.87 |
| WBGene00197814 | -0.156269293 | 0.96924949 | 1 | F54A3.9 |
| WBGene00197817 | -0.156269293 | 0.96924949 | 1 | C07B5.9 |
| WBGene00197822 | -0.156269293 | 0.96924949 | 1 | H22K11.8 |
| WBGene00197824 | -0.558804446 | 0.87818675 | 1 | Y73F8A.1166 |
| WBGene00197828 | -0.156269293 | 0.96924949 | 1 | C02B4.8 |
| WBGene00197837 | 0.281454817 | 0.94435481 | 1 | K11C4.9 |
| WBGene00197839 | -1.964653846 | 0.53599202 | 1 | F14D12.8 |
| WBGene00197843 | -0.291199644 | 0.86491471 | 1 | ZK1010.11 |
| WBGene00197847 | -0.156269293 | 0.96924949 | 1 | K12G11.10 |
| WBGene00197860 | -2.640414922 | 0.41081194 | 1 | Y38F2AR.17 |
| WBGene00197861 | 0.700753508 | 0.52761163 | 1 | F10D7.7 |
| WBGene00197864 | 0.281454817 | 0.94435481 | 1 | C17E4.16 |
| WBGene00197879 | -1.969805064 | 0.62862569 | 1 | ZK287.11 |
| WBGene00197880 | -0.156269293 | 0.96924949 | 1 | C54G7.11 |
| WBGene00197882 | 0.306766063 | 0.88218984 | 1 | F12F3.5 |
| WBGene00197884 | -0.156269293 | 0.96924949 | 1 | F01F1.16 |
| WBGene00197885 | -0.156269293 | 0.96924949 | 1 | F43D2.11 |
| WBGene00197888 | 0.446458324 | 0.87164163 | 1 | F16D3.8 |
| WBGene00197891 | -1.032273505 | 0.43586922 | 1 | C29F3.9 |
| WBGene00197897 | -0.156269293 | 0.96924949 | 1 | F58A4.19 |
| WBGene00197900 | 2.486734991 | 0.32516285 | 1 | M01A10.7 |
| WBGene00197901 | 0.983488853 | 0.80601565 | 1 | H01G02.8 |
| WBGene00197902 | -0.156269293 | 0.96924949 | 1 | R53.10 |
| WBGene00197906 | 0.281454817 | 0.94435481 | 1 | T04C9.7 |
| WBGene00197912 | 0.377828795 | 0.92526585 | 1 | C25F6.16 |
| WBGene00197916 | 1.116613983 | 0.69067248 | 1 | K01A11.6 |
| WBGene00197917 | -0.737462795 | 0.83433562 | 1 | C06E1.14 |
| WBGene00197918 | -1.094201388 | 0.7816772 | 1 | B0344.3 |
| WBGene00197928 | 0.281454817 | 0.94435481 | 1 | F29F11.11 |
| WBGene00197930 | 0.983488853 | 0.80601565 | 1 | T06H11.11 |
| WBGene00197933 | -0.156269293 | 0.96924949 | 1 | F08F1.14 |
| WBGene00197935 | -0.156269293 | 0.96924949 | 1 | T04C10.9 |
| WBGene00197936 | 0.112250498 | 0.58714853 | 0.75591074 | Y73B6BL.275 |
| WBGene00197940 | 2.486734991 | 0.32516285 | 1 | F40E3.8 |
| WBGene00197945 | -0.156269293 | 0.96924949 | 1 | C53C7.6 |
| WBGene00197955 | 1.10541495 | 0.26243131 | 0.47118878 | VT23B5.1 |
| WBGene00197958 | 0.574508983 | 0.8551971 | 1 | B0034.11 |

| | | | | |
|---|---|---|---|---|
| WBGene00197966 | -1.901856996 | 0.64072034 | 1 | B0393.14 |
| WBGene00197967 | -1.159714244 | 0.75790634 | 1 | C07A4.7 |
| WBGene00197968 | 1.036568105 | 0.67105582 | 1 | ZK1290.19 |
| WBGene00197988 | -0.156269293 | 0.96924949 | 1 | Y73F8A.1167 |
| WBGene00197989 | 0.394439984 | 0.88400378 | 1 | R07D5.5 |
| WBGene00197990 | 0.281454817 | 0.94435481 | 1 | T06F4.9 |
| WBGene00198001 | 0.377828795 | 0.92526585 | 1 | Y54E10A.24 |
| WBGene00198006 | -1.159714244 | 0.75790634 | 1 | R160.9 |
| WBGene00198029 | -1.964653846 | 0.53599202 | 1 | F54D5.20 |
| WBGene00198038 | 0.281454817 | 0.94435481 | 1 | H03E18.4 |
| WBGene00198042 | 0.281454817 | 0.94435481 | 1 | T28F4.7 |
| WBGene00198058 | 2.030757922 | 0.16722512 | 1 | F20B10.8 |
| WBGene00198061 | 1.18938122 | 0.01580373 | 0.07215987 | T06D10.7 |
| WBGene00198064 | -0.156269293 | 0.96924949 | 1 | C31H2.8 |
| WBGene00198070 | -1.964653846 | 0.53599202 | 1 | F07G11.10 |
| WBGene00198073 | -1.901856996 | 0.64072034 | 1 | K08A8.14 |
| WBGene00198082 | -2.640414922 | 0.41081194 | 1 | W06D12.8 |
| WBGene00198085 | 0.281454817 | 0.94435481 | 1 | F47G3.8 |
| WBGene00198098 | -1.969805064 | 0.62862569 | 1 | T02C5.6 |
| WBGene00198101 | 0.688793783 | 0.86378755 | 1 | F52B5.9 |
| WBGene00198104 | -0.156269293 | 0.96924949 | 1 | Y58A7A.10 |
| WBGene00198105 | -1.969805064 | 0.62862569 | 1 | C24D10.47 |
| WBGene00198109 | 0.281454817 | 0.94435481 | 1 | C30A5.13 |
| WBGene00198131 | -0.098216941 | 0.86296784 | 0.92629233 | F36F2.11 |
| WBGene00198132 | -0.156269293 | 0.96924949 | 1 | F56H11.11 |
| WBGene00198133 | -0.156269293 | 0.96924949 | 1 | F07C3.14 |
| WBGene00198136 | -0.156269293 | 0.96924949 | 1 | C43F9.16 |
| WBGene00198150 | 0.281454817 | 0.94435481 | 1 | M79.10 |
| WBGene00198151 | -0.156269293 | 0.96924949 | 1 | C04F12.15 |
| WBGene00198156 | 0.747845955 | 0.85215965 | 1 | T06H11.12 |
| WBGene00198161 | 0.281454817 | 0.94435481 | 1 | C46A5.14 |
| WBGene00198181 | -0.150245429 | 0.9305065 | 1 | Y73F8A.1168 |
| WBGene00198185 | 1.854471114 | 0.22100581 | 1 | F56D12.11 |
| WBGene00198189 | 0.281454817 | 0.94435481 | 1 | W05B2.12 |
| WBGene00198190 | -0.223076565 | 0.17593939 | 0.36683498 | F25H2.16 |
| WBGene00198196 | -0.156269293 | 0.96924949 | 1 | C48A7.11 |
| WBGene00198197 | -1.14715222 | 0.68932198 | 1 | F41G3.22 |
| WBGene00198198 | 0.75595108 | 0.7708876 | 1 | T07H6.7 |
| WBGene00198201 | 0.938982574 | 0.8146943 | 1 | F59H5.5 |
| WBGene00198206 | 0.281454817 | 0.94435481 | 1 | Y57A10A.41 |
| WBGene00198213 | -1.159714244 | 0.75790634 | 1 | M03F4.12 |
| WBGene00198222 | -0.62146648 | 0.85925752 | 1 | C06A6.10 |
| WBGene00198223 | 2.129775735 | 0.27461486 | 1 | C54D2.11 |

| | | | | |
|---|---|---|---|---|
| WBGene00198239 | -0.308926229 | 0.63500555 | 0.78806117 | F22H10.9 |
| WBGene00198242 | -1.901856996 | 0.64072034 | 1 | F29F11.12 |
| WBGene00198248 | 0.377828795 | 0.92526585 | 1 | C18C4.14 |
| WBGene00198255 | 0.281454817 | 0.94435481 | 1 | T13B5.11 |
| WBGene00198264 | 1.413377704 | 0.60673863 | 1 | K12F2.3 |
| WBGene00198265 | -1.969805064 | 0.62862569 | 1 | Y39H10A.9 |
| WBGene00198276 | 0.377828795 | 0.92526585 | 1 | T19A5.8 |
| WBGene00198283 | -0.668264701 | 0.85551153 | 1 | F55F1.8 |
| WBGene00198286 | -0.156269293 | 0.96924949 | 1 | R151.16 |
| WBGene00198298 | 0.281454817 | 0.94435481 | 1 | F47B10.10 |
| WBGene00198300 | -2.640414922 | 0.41081194 | 1 | T19E7.25 |
| WBGene00198335 | 0.189930353 | 0.6580438 | 0.80303357 | M04C9.9 |
| WBGene00198337 | -0.156269293 | 0.96924949 | 1 | T05G5.14 |
| WBGene00198342 | 0.688793783 | 0.86378755 | 1 | F41E7.14 |
| WBGene00198345 | 0.688793783 | 0.86378755 | 1 | Y73B3A.25 |
| WBGene00198348 | -0.156269293 | 0.96924949 | 1 | T06E8.4 |
| WBGene00198353 | 1.38322681 | 0.69550305 | 1 | W03B1.16 |
| WBGene00198361 | -0.156269293 | 0.96924949 | 1 | Y66A7AL.6 |
| WBGene00198363 | 1.181686151 | 0.7579682 | 1 | H40L08.4 |
| WBGene00198380 | -0.156269293 | 0.96924949 | 1 | T22B7.13 |
| WBGene00198384 | 0.281454817 | 0.94435481 | 1 | C05E11.14 |
| WBGene00198387 | -1.901856996 | 0.64072034 | 1 | ZK512.14 |
| WBGene00198392 | -0.156269293 | 0.96924949 | 1 | K09F5.9 |
| WBGene00198399 | 0.688793783 | 0.86378755 | 1 | F01G12.12 |
| WBGene00198402 | -0.03811452 | 0.98663692 | 1 | ZC487.8 |
| WBGene00198405 | 2.799084385 | 0.17307183 | 1 | ZK185.8 |
| WBGene00198408 | -1.159714244 | 0.75790634 | 1 | Y59A8B.30 |
| WBGene00198409 | -0.156269293 | 0.96924949 | 1 | Y48A6C.7 |
| WBGene00198416 | -0.156269293 | 0.96924949 | 1 | F49E10.14 |
| WBGene00198418 | -1.094201388 | 0.7816772 | 1 | B0344.4 |
| WBGene00198421 | 0.281454817 | 0.94435481 | 1 | F45F2.15 |
| WBGene00198423 | 1.653460415 | 0.50712258 | 1 | F26E4.17 |
| WBGene00198429 | -0.156269293 | 0.96924949 | 1 | F57B7.9 |
| WBGene00198430 | -0.156269293 | 0.96924949 | 1 | F08G5.14 |
| WBGene00198434 | -0.156269293 | 0.96924949 | 1 | C56G2.19 |
| WBGene00198435 | -0.156269293 | 0.96924949 | 1 | T19D12.14 |
| WBGene00198439 | -0.18557176 | 0.95664217 | 1 | ZK262.17 |
| WBGene00198440 | -0.156269293 | 0.96924949 | 1 | C05C9.9 |
| WBGene00198445 | -0.156269293 | 0.96924949 | 1 | F43G6.14 |
| WBGene00198446 | -0.01455644 | 0.9964743 | 1 | T25D10.6 |
| WBGene00198449 | 0.467656871 | 0.42838512 | 0.63100978 | C14A11.11 |
| WBGene00198454 | 0.377828795 | 0.92526585 | 1 | F28C1.13 |
| WBGene00198457 | -1.831477136 | 0.52194793 | 1 | F58A6.12 |

| | | | | |
|---|---|---|---|---|
| WBGene00198466 | 0.377828795 | 0.92526585 | 1 | F13H6.16 |
| WBGene00198467 | 0.574508983 | 0.8551971 | 1 | F58A4.20 |
| WBGene00198470 | -1.291305095 | 0.63677871 | 1 | F08G2.15 |
| WBGene00198474 | 0.281454817 | 0.94435481 | 1 | T10E10.11 |
| WBGene00198475 | -1.094201388 | 0.7816772 | 1 | K08E4.10 |
| WBGene00198477 | 1.38322681 | 0.69550305 | 1 | F49H12.11 |
| WBGene00198478 | -0.156269293 | 0.96924949 | 1 | Y58A7A.11 |
| WBGene00198484 | -0.156269293 | 0.96924949 | 1 | C55A6.17 |
| WBGene00198491 | 0.377828795 | 0.92526585 | 1 | B0024.18 |
| WBGene00198501 | 0.041803775 | 0.98890197 | 1 | C48C5.6 |
| WBGene00198502 | -0.156269293 | 0.96924949 | 1 | T23F2.8 |
| WBGene00198506 | 0.377828795 | 0.92526585 | 1 | F55B12.14 |
| WBGene00198507 | 0.983488853 | 0.80601565 | 1 | C01F4.7 |
| WBGene00198509 | 0.281454817 | 0.94435481 | 1 | M04B2.10 |
| WBGene00198514 | 0.938982574 | 0.8146943 | 1 | T12B5.17 |
| WBGene00198520 | 0.574508983 | 0.8551971 | 1 | F25B4.13 |
| WBGene00198524 | -1.969805064 | 0.62862569 | 1 | C34H3.14 |
| WBGene00198525 | 0.688793783 | 0.86378755 | 1 | F18A1.9 |
| WBGene00198528 | 0.574508983 | 0.8551971 | 1 | C44E12.4 |
| WBGene00198538 | -0.156269293 | 0.96924949 | 1 | H14E04.7 |
| WBGene00198547 | 0.366948289 | 0.5345419 | 0.71755507 | C44H4.10 |
| WBGene00198552 | -0.156269293 | 0.96924949 | 1 | R04F11.17 |
| WBGene00198556 | -1.094201388 | 0.7816772 | 1 | K01A12.7 |
| WBGene00198557 | 0.688793783 | 0.86378755 | 1 | C41G11.13 |
| WBGene00198568 | -0.156269293 | 0.96924949 | 1 | F53B1.12 |
| WBGene00198572 | -0.199374337 | 0.91832278 | 1 | K11G12.9 |
| WBGene00198573 | 2.10931518 | 0.25332604 | 1 | Y17G9A.88 |
| WBGene00198575 | -1.901856996 | 0.64072034 | 1 | F26F4.14 |
| WBGene00198576 | 0.281454817 | 0.94435481 | 1 | F21H12.8 |
| WBGene00198580 | -1.229853028 | 0.58617677 | 1 | Y64G10A.355 |
| WBGene00198586 | -0.156269293 | 0.96924949 | 1 | W01A8.11 |
| WBGene00198592 | 0.281454817 | 0.94435481 | 1 | Y71H9A.4 |
| WBGene00198597 | 0.281454817 | 0.94435481 | 1 | Y73B6BL.276 |
| WBGene00198598 | -0.156269293 | 0.96924949 | 1 | W01C8.12 |
| WBGene00198615 | -0.161554207 | 0.92922014 | 1 | F46F3.11 |
| WBGene00198620 | 1.477253415 | 0.58357327 | 1 | F11C7.9 |
| WBGene00198634 | -0.156269293 | 0.96924949 | 1 | T09F3.6 |
| WBGene00198640 | 0.938982574 | 0.8146943 | 1 | Y73B6BL.277 |
| WBGene00198649 | -0.156269293 | 0.96924949 | 1 | Y105E8A.43 |
| WBGene00198650 | -0.156269293 | 0.96924949 | 1 | Y97E10AL.4 |
| WBGene00198651 | 0.012436123 | 0.98402833 | 0.99228305 | C30F12.9 |
| WBGene00198661 | -1.365683379 | 0.34405276 | 1 | D2023.15 |
| WBGene00198667 | -0.156269293 | 0.96924949 | 1 | ZK287.13 |

| | | | | |
|---|---|---|---|---|
| WBGene00198669 | 1.067514988 | 0.29202819 | 1 | T20B12.11 |
| WBGene00198670 | -0.156269293 | 0.96924949 | 1 | F14F3.11 |
| WBGene00198672 | 0.377828795 | 0.92526585 | 1 | F58A4.21 |
| WBGene00198678 | 0.281454817 | 0.94435481 | 1 | F22E10.10 |
| WBGene00198679 | -0.156269293 | 0.96924949 | 1 | C44B7.17 |
| WBGene00198681 | -1.901856996 | 0.64072034 | 1 | W06A7.8 |
| WBGene00198683 | -0.156269293 | 0.96924949 | 1 | C31H2.9 |
| WBGene00198686 | -0.156269293 | 0.96924949 | 1 | F18E2.12 |
| WBGene00198687 | -0.156269293 | 0.96924949 | 1 | T23B5.5 |
| WBGene00198689 | 1.165786227 | 0.49732668 | 1 | F13H6.17 |
| WBGene00198693 | -0.156269293 | 0.96924949 | 1 | F13H10.10 |
| WBGene00198695 | 1.116613983 | 0.69067248 | 1 | C01C4.4 |
| WBGene00198698 | 0.983488853 | 0.80601565 | 1 | H01G02.11 |
| WBGene00198712 | -0.156269293 | 0.96924949 | 1 | K09A11.8 |
| WBGene00198715 | -1.969805064 | 0.62862569 | 1 | Y46E12A.7 |
| WBGene00198716 | -0.156269293 | 0.96924949 | 1 | C18D1.17 |
| WBGene00198717 | -1.969805064 | 0.62862569 | 1 | C34H3.15 |
| WBGene00198725 | -0.35111783 | 0.65445846 | 0.80142834 | F55A11.13 |
| WBGene00198736 | 1.462139315 | 0.21863919 | 1 | ZC581.13 |
| WBGene00198748 | -1.130803645 | 0.00247412 | 0.01846241 | M163.19 |
| WBGene00198754 | 0.763390359 | 0.73510979 | 1 | K04G11.8 |
| WBGene00198763 | 0.281454817 | 0.94435481 | 1 | Y39B6A.81 |
| WBGene00198765 | -0.156269293 | 0.96924949 | 1 | F54A3.10 |
| WBGene00198766 | 2.602875064 | 0.25929851 | 1 | F12F3.7 |
| WBGene00198769 | 2.286068456 | 0.34948738 | 1 | T05A1.17 |
| WBGene00198775 | 0.281454817 | 0.94435481 | 1 | F23F1.11 |
| WBGene00198777 | 0.688793783 | 0.86378755 | 1 | F54C9.17 |
| WBGene00198788 | -0.156269293 | 0.96924949 | 1 | Y6B3B.14 |
| WBGene00198792 | -1.159714244 | 0.75790634 | 1 | R04E5.14 |
| WBGene00198800 | 0.281454817 | 0.94435481 | 1 | F39E9.23 |
| WBGene00198804 | -0.156269293 | 0.96924949 | 1 | T28B11.14 |
| WBGene00198805 | 1.434080373 | 0.25948906 | 1 | Y37B11A.4 |
| WBGene00198806 | -0.0060929 | 0.99795747 | 1 | Y76F7A.4 |
| WBGene00198814 | 2.289991418 | 0.22574298 | 1 | F31C3.15 |
| WBGene00198816 | -1.159714244 | 0.75790634 | 1 | C12C8.7 |
| WBGene00198818 | 1.122420393 | 0.31820006 | 1 | T21C12.9 |
| WBGene00198821 | -0.156269293 | 0.96924949 | 1 | T26H2.14 |
| WBGene00198834 | 0.233868761 | 0.89003571 | 1 | C43G2.29 |
| WBGene00198839 | -0.156269293 | 0.96924949 | 1 | B0414.11 |
| WBGene00198841 | 1.826637908 | 0.3874948 | 1 | T03G11.20 |
| WBGene00198853 | 0.281454817 | 0.94435481 | 1 | C30B5.13 |
| WBGene00198861 | -0.156269293 | 0.96924949 | 1 | C02D4.8 |
| WBGene00198870 | 0.983488853 | 0.80601565 | 1 | F25H10.6 |

| | | | | |
|---|---|---|---|---|
| WBGene00198871 | -0.156269293 | 0.96924949 | 1 | C02E7.16 |
| WBGene00198873 | 1.413377704 | 0.60673863 | 1 | F10C2.11 |
| WBGene00198874 | 0.377828795 | 0.92526585 | 1 | T04F3.7 |
| WBGene00198877 | -0.156269293 | 0.96924949 | 1 | F57A10.8 |
| WBGene00198898 | 1.098091204 | 0.54399672 | 1 | Y42A5A.9 |
| WBGene00198907 | -1.901856996 | 0.64072034 | 1 | M79.12 |
| WBGene00198913 | -2.203012775 | 0.25034502 | 1 | Y40B10A.10 |
| WBGene00198919 | -0.156269293 | 0.96924949 | 1 | Y119D3B.25 |
| WBGene00198924 | -1.366565036 | 0.6506483 | 1 | C15F1.14 |
| WBGene00198925 | -0.014519762 | 0.99603746 | 1 | T07A5.8 |
| WBGene00198932 | 0.281454817 | 0.94435481 | 1 | F29C4.11 |
| WBGene00198937 | -1.901856996 | 0.64072034 | 1 | ZK287.14 |
| WBGene00198945 | -1.759191416 | 0.57668729 | 1 | T08G5.18 |
| WBGene00198948 | 0.747845955 | 0.85215965 | 1 | Y60A9A.5 |
| WBGene00198958 | -1.969805064 | 0.62862569 | 1 | C28A5.8 |
| WBGene00198964 | -1.760055813 | 0.45626364 | 1 | K01F9.6 |
| WBGene00198970 | -0.102919912 | 0.96055224 | 1 | T06H11.14 |
| WBGene00198971 | -0.156269293 | 0.96924949 | 1 | F07F6.10 |
| WBGene00198975 | -1.901856996 | 0.64072034 | 1 | C17H12.33 |
| WBGene00198976 | -1.901856996 | 0.64072034 | 1 | C35C5.18 |
| WBGene00198980 | 0.602800982 | 0.18383494 | 0.37600068 | C46G7.108 |
| WBGene00198986 | 0.281454817 | 0.94435481 | 1 | B0496.15 |
| WBGene00198991 | 1.116613983 | 0.69067248 | 1 | T05G5.15 |
| WBGene00198993 | 0.281454817 | 0.94435481 | 1 | T13H10.13 |
| WBGene00198998 | -0.530973232 | 0.41097875 | 0.6165443 | F20H11.6 |
| WBGene00199010 | 1.957271014 | 0.49743227 | 1 | C04F1.4 |
| WBGene00199018 | 1.687239501 | 0.59403242 | 1 | F57C7.10 |
| WBGene00199019 | 0.377828795 | 0.92526585 | 1 | W04D2.11 |
| WBGene00199020 | 1.653460415 | 0.50712258 | 1 | Y48A6B.16 |
| WBGene00199027 | -1.969805064 | 0.62862569 | 1 | E01G4.8 |
| WBGene00199039 | 0.983488853 | 0.80601565 | 1 | F52F12.13 |
| WBGene00199042 | 0.688793783 | 0.86378755 | 1 | H34C03.20 |
| WBGene00199045 | -1.245733657 | 0.50679516 | 1 | Y53F4B.58 |
| WBGene00199059 | -0.156269293 | 0.96924949 | 1 | Y22D7AL.17 |
| WBGene00199061 | -0.156269293 | 0.96924949 | 1 | W09G3.14 |
| WBGene00199063 | -0.156269293 | 0.96924949 | 1 | T14B4.14 |
| WBGene00199082 | 1.278675639 | 0.30712475 | 1 | B0454.18 |
| WBGene00199083 | 0.281454817 | 0.94435481 | 1 | B0496.16 |
| WBGene00199084 | -0.737462795 | 0.83433562 | 1 | C11G10.9 |
| WBGene00199088 | -0.509560466 | 0.14981839 | 0.33125184 | Y39G10AR.30 |
| WBGene00199089 | 0.957768616 | 0.62962374 | 1 | B0395.7 |
| WBGene00199099 | 1.181686151 | 0.7579682 | 1 | ZC412.11 |
| WBGene00199101 | 0.688793783 | 0.86378755 | 1 | C44E12.5 |

| | | | | |
|---|---|---|---|---|
| WBGene00199112 | -0.558804446 | 0.87818675 | 1 | C08B11.11 |
| WBGene00199120 | -0.120007634 | 0.90067644 | 1 | ZK418.12 |
| WBGene00199122 | -0.03811452 | 0.98663692 | 1 | B0495.19 |
| WBGene00199123 | -1.901856996 | 0.64072034 | 1 | F45B8.14 |
| WBGene00199125 | 0.688793783 | 0.86378755 | 1 | C56A3.12 |
| WBGene00199142 | 0.688793783 | 0.86378755 | 1 | C09B7.8 |
| WBGene00199144 | -0.097272413 | 0.92330927 | 1 | R04E5.15 |
| WBGene00199150 | -0.156269293 | 0.96924949 | 1 | K11G9.11 |
| WBGene00199152 | 0.281454817 | 0.94435481 | 1 | T06F4.11 |
| WBGene00199164 | 0.170612158 | 0.88351627 | 1 | Y75B8A.53 |
| WBGene00199166 | -0.156269293 | 0.96924949 | 1 | K08F8.11 |
| WBGene00199172 | -1.901856996 | 0.64072034 | 1 | Y71D11A.8 |
| WBGene00199174 | 0.281454817 | 0.94435481 | 1 | F13H8.14 |
| WBGene00199177 | 0.574508983 | 0.8551971 | 1 | T26H10.6 |
| WBGene00199188 | -0.020267126 | 0.98794901 | 1 | Y39A1A.29 |
| WBGene00199192 | -0.558804446 | 0.87818675 | 1 | K06A5.9 |
| WBGene00199205 | 0.874994581 | 0.10647436 | 0.26598873 | C04G6.14 |
| WBGene00199206 | 0.688793783 | 0.86378755 | 1 | F18A1.10 |
| WBGene00199213 | 0.688793783 | 0.86378755 | 1 | F45H7.11 |
| WBGene00199217 | -1.969805064 | 0.62862569 | 1 | ZK973.19 |
| WBGene00199223 | 1.433094668 | 0.24177296 | 1 | C14F11.22 |
| WBGene00199230 | -0.150245429 | 0.9305065 | 1 | R173.9 |
| WBGene00199233 | 1.025181888 | 0.79790347 | 1 | C44E12.6 |
| WBGene00199238 | 0.281454817 | 0.94435481 | 1 | W05B10.8 |
| WBGene00199243 | -0.156269293 | 0.96924949 | 1 | F54E4.12 |
| WBGene00199251 | -0.156269293 | 0.96924949 | 1 | F46F2.10 |
| WBGene00199254 | 0.281454817 | 0.94435481 | 1 | R07D5.6 |
| WBGene00199263 | -1.094201388 | 0.7816772 | 1 | W10C4.2 |
| WBGene00199275 | -0.156269293 | 0.96924949 | 1 | AC3.15 |
| WBGene00199276 | 1.247575581 | 0.7549672 | 1 | C45B11.10 |
| WBGene00199278 | 0.281454817 | 0.94435481 | 1 | F54D8.8 |
| WBGene00199286 | 0.377828795 | 0.92526585 | 1 | W05H5.9 |
| WBGene00199297 | 0.747845955 | 0.85215965 | 1 | T10E9.12 |
| WBGene00199301 | 0.377828795 | 0.92526585 | 1 | K02A4.8 |
| WBGene00199304 | 0.178115923 | 0.94982316 | 1 | C07G1.12 |
| WBGene00199305 | -1.159714244 | 0.75790634 | 1 | C39E9.18 |
| WBGene00199314 | 0.688793783 | 0.86378755 | 1 | R02D5.16 |
| WBGene00199315 | -1.710857121 | 0.32044127 | 1 | T04C12.23 |
| WBGene00199320 | 0.281454817 | 0.94435481 | 1 | R02D5.17 |
| WBGene00199323 | -1.969805064 | 0.62862569 | 1 | C08D8.5 |
| WBGene00199325 | 0.112952878 | 0.92464479 | 1 | F45H7.12 |
| WBGene00199329 | -1.969805064 | 0.62862569 | 1 | F11D5.21 |
| WBGene00199330 | 0.789601902 | 0.75907734 | 1 | F31B9.5 |

| | | | | |
|---|---|---|---|---|
| WBGene00199332 | -0.156269293 | 0.96924949 | 1 | T23F2.9 |
| WBGene00199339 | 1.210990069 | 0.25391479 | 1 | C41G7.13 |
| WBGene00199349 | -1.964653846 | 0.53599202 | 1 | C40H5.14 |
| WBGene00199350 | -0.156269293 | 0.96924949 | 1 | F13D11.15 |
| WBGene00199366 | 0.281454817 | 0.94435481 | 1 | T06F4.12 |
| WBGene00199369 | -0.156269293 | 0.96924949 | 1 | H14E04.8 |
| WBGene00199370 | -0.156269293 | 0.96924949 | 1 | F25B4.14 |
| WBGene00199371 | -1.901856996 | 0.64072034 | 1 | T03F7.14 |
| WBGene00199372 | -1.094201388 | 0.7816772 | 1 | C23H3.11 |
| WBGene00199391 | -0.156269293 | 0.96924949 | 1 | C32C4.15 |
| WBGene00199404 | 0.625016815 | 0.34477858 | 0.55355198 | ZK1098.13 |
| WBGene00199408 | -1.159714244 | 0.75790634 | 1 | T06G6.15 |
| WBGene00199412 | 1.131396879 | 0.39948305 | 1 | Y4C6B.98 |
| WBGene00199414 | -0.156269293 | 0.96924949 | 1 | F48E3.13 |
| WBGene00199416 | -0.156269293 | 0.96924949 | 1 | Y40B1A.5 |
| WBGene00199424 | -1.901856996 | 0.64072034 | 1 | R11B5.8 |
| WBGene00199437 | 0.232157325 | 0.93397866 | 1 | C08G9.6 |
| WBGene00199444 | -1.901856996 | 0.64072034 | 1 | Y48B6A.20 |
| WBGene00199447 | -0.156269293 | 0.96924949 | 1 | F08B12.10 |
| WBGene00199449 | -1.969805064 | 0.62862569 | 1 | F01E11.10 |
| WBGene00199462 | 0.281454817 | 0.94435481 | 1 | B0495.20 |
| WBGene00199469 | -1.901856996 | 0.64072034 | 1 | M05B5.8 |
| WBGene00199473 | 1.477253415 | 0.58357327 | 1 | F45E6.7 |
| WBGene00199478 | 0.377828795 | 0.92526585 | 1 | C01F6.13 |
| WBGene00199485 | 0.688793783 | 0.86378755 | 1 | M117.11 |
| WBGene00199493 | 0.688793783 | 0.86378755 | 1 | C07A9.15 |
| WBGene00199496 | -0.62146648 | 0.85925752 | 1 | C06A6.11 |
| WBGene00199500 | -0.156269293 | 0.96924949 | 1 | F09F9.12 |
| WBGene00199502 | -0.156269293 | 0.96924949 | 1 | K10D6.16 |
| WBGene00199507 | -0.156269293 | 0.96924949 | 1 | R173.10 |
| WBGene00199518 | 1.305176685 | 0.58238291 | 1 | F22A3.14 |
| WBGene00199519 | 1.557897343 | 0.63813005 | 1 | F01G12.14 |
| WBGene00199525 | 0.281454817 | 0.94435481 | 1 | C03A3.5 |
| WBGene00199526 | 0.281454817 | 0.94435481 | 1 | F55E10.10 |
| WBGene00199535 | 0.281454817 | 0.94435481 | 1 | F53F8.10 |
| WBGene00199540 | 2.256387701 | 0.20762969 | 1 | F45E4.18 |
| WBGene00199542 | 0.263180636 | 0.87384694 | 1 | Y113G7A.19 |
| WBGene00199554 | -0.156269293 | 0.96924949 | 1 | F49E10.17 |
| WBGene00199560 | 1.431057433 | 0.52566078 | 1 | C39B10.10 |
| WBGene00199564 | -1.969805064 | 0.62862569 | 1 | Y43H11AL.5 |
| WBGene00199569 | 0.281454817 | 0.94435481 | 1 | F25F6.3 |
| WBGene00199570 | -0.156269293 | 0.96924949 | 1 | C43F9.17 |
| WBGene00199574 | 0.281454817 | 0.94435481 | 1 | Y41D4B.29 |

| | | | | |
|---|---|---|---|---|
| WBGene00199575 | 0.281454817 | 0.94435481 | 1 | H03A11.9 |
| WBGene00199578 | 0.281454817 | 0.94435481 | 1 | T28D6.13 |
| WBGene00199591 | 0.459413223 | 0.84552382 | 1 | Y54G2A.65 |
| WBGene00199602 | 0.938982574 | 0.8146943 | 1 | W09C5.14 |
| WBGene00199621 | 0.747845955 | 0.85215965 | 1 | B0034.13 |
| WBGene00199624 | 0.281454817 | 0.94435481 | 1 | C02D4.11 |
| WBGene00199636 | 0.281454817 | 0.94435481 | 1 | F29G9.13 |
| WBGene00199646 | 1.557897343 | 0.63813005 | 1 | Y51B9A.12 |
| WBGene00199652 | -1.969805064 | 0.62862569 | 1 | C41G11.15 |
| WBGene00199654 | 0.688793783 | 0.86378755 | 1 | F56E10.6 |
| WBGene00199655 | 0.281454817 | 0.94435481 | 1 | C30D11.2 |
| WBGene00199660 | 0.281454817 | 0.94435481 | 1 | E02H4.9 |
| WBGene00199661 | -1.291305095 | 0.63677871 | 1 | C16B8.8 |
| WBGene00199663 | -0.156269293 | 0.96924949 | 1 | B0507.14 |
| WBGene00199673 | -1.901856996 | 0.64072034 | 1 | C18H9.10 |
| WBGene00199687 | 0.747845955 | 0.85215965 | 1 | C50E10.13 |
| WBGene00199689 | 1.413377704 | 0.60673863 | 1 | W01C8.13 |
| WBGene00199694 | -3.437682489 | 0.15558371 | 1 | 2RSSE.6 |
| WBGene00199695 | -1.291305095 | 0.63677871 | 1 | F08G2.17 |
| WBGene00199696 | -0.156269293 | 0.96924949 | 1 | H21P03.8 |
| WBGene00199700 | 0.57991002 | 0.82402153 | 1 | M60.12 |
| WBGene00199701 | 0.505162712 | 0.53891293 | 0.72066769 | R151.17 |
| WBGene00199709 | 0.281454817 | 0.94435481 | 1 | T06F4.13 |
| WBGene00199713 | 0.983488853 | 0.80601565 | 1 | C43G2.30 |
| WBGene00199719 | -0.737462795 | 0.83433562 | 1 | C47F8.10 |
| WBGene00199724 | -3.235037296 | 0.08305993 | 1 | F10C1.16 |
| WBGene00199730 | 0.377828795 | 0.92526585 | 1 | W08D2.11 |
| WBGene00199731 | -0.156269293 | 0.96924949 | 1 | C06G4.7 |
| WBGene00199735 | -0.156269293 | 0.96924949 | 1 | F14F11.4 |
| WBGene00199739 | 0.281454817 | 0.94435481 | 1 | C32A9.3 |
| WBGene00199740 | 0.377828795 | 0.92526585 | 1 | F07A11.12 |
| WBGene00199750 | -0.712674837 | 0.60790238 | 1 | F26D11.17 |
| WBGene00199761 | 0.688793783 | 0.86378755 | 1 | C23H4.12 |
| WBGene00199768 | 0.574508983 | 0.8551971 | 1 | F52F12.15 |
| WBGene00199776 | -1.901856996 | 0.64072034 | 1 | T28C12.11 |
| WBGene00199784 | -0.156269293 | 0.96924949 | 1 | F08G5.16 |
| WBGene00199785 | 0.774694413 | 0.55776341 | 1 | C48A7.12 |
| WBGene00199791 | -0.156269293 | 0.96924949 | 1 | F26A1.17 |
| WBGene00199800 | -0.604396148 | 0.82723443 | 1 | C33D12.13 |
| WBGene00199808 | 1.796575934 | 0.34491145 | 1 | F21A3.10 |
| WBGene00199819 | 0.377828795 | 0.92526585 | 1 | K08F8.13 |
| WBGene00199825 | 0.747845955 | 0.85215965 | 1 | C18E3.10 |
| WBGene00199827 | -1.831477136 | 0.52194793 | 1 | B0238.18 |

| | | | | |
|---|---|---|---|---|
| WBGene00199828 | 0.747845955 | 0.85215965 | 1 | K10C2.10 |
| WBGene00199831 | -0.156269293 | 0.96924949 | 1 | F01E11.12 |
| WBGene00199834 | -1.901856996 | 0.64072034 | 1 | Y22D7AL.18 |
| WBGene00199848 | -1.901856996 | 0.64072034 | 1 | T08B6.63 |
| WBGene00199850 | -0.321982033 | 0.80271184 | 1 | C02D4.12 |
| WBGene00199853 | -0.156269293 | 0.96924949 | 1 | T19C11.3 |
| WBGene00199858 | 0.688793783 | 0.86378755 | 1 | C40H5.15 |
| WBGene00199859 | -0.156269293 | 0.96924949 | 1 | F58H1.22 |
| WBGene00199861 | 1.116613983 | 0.69067248 | 1 | T19H12.13 |
| WBGene00199868 | -0.558804446 | 0.87818675 | 1 | C28A5.9 |
| WBGene00199869 | -0.052768511 | 0.93222407 | 0.96406579 | W05E10.16 |
| WBGene00199873 | 0.672178833 | 0.25363254 | 0.46153368 | F26E4.18 |
| WBGene00199880 | 0.134942955 | 0.8622009 | 0.92569082 | C16E9.20 |
| WBGene00199882 | -1.901856996 | 0.64072034 | 1 | F47C8.4 |
| WBGene00199884 | 0.688793783 | 0.86378755 | 1 | F43G9.18 |
| WBGene00199885 | 0.281454817 | 0.94435481 | 1 | B0457.14 |
| WBGene00199887 | -1.159714244 | 0.75790634 | 1 | R04E5.16 |
| WBGene00199896 | -0.01455644 | 0.9964743 | 1 | ZK287.15 |
| WBGene00199916 | -0.70576507 | 0.76841669 | 1 | R04E5.17 |
| WBGene00199918 | 0.983488853 | 0.80601565 | 1 | mir-8192 |
| WBGene00199921 | -1.901856996 | 0.64072034 | 1 | F45B8.16 |
| WBGene00199923 | -0.156269293 | 0.96924949 | 1 | K04G2.14 |
| WBGene00199928 | 0.574508983 | 0.8551971 | 1 | F52F12.16 |
| WBGene00199930 | -0.156269293 | 0.96924949 | 1 | T22H2.7 |
| WBGene00199931 | -0.156269293 | 0.96924949 | 1 | W02B8.11 |
| WBGene00199934 | -0.156269293 | 0.96924949 | 1 | M163.20 |
| WBGene00199935 | 0.281454817 | 0.94435481 | 1 | F13E6.10 |
| WBGene00199937 | 0.377828795 | 0.92526585 | 1 | T04C9.10 |
| WBGene00199963 | -0.014519762 | 0.99603746 | 1 | T26H2.15 |
| WBGene00199968 | 0.574508983 | 0.8551971 | 1 | K12F2.4 |
| WBGene00199971 | 0.281454817 | 0.94435481 | 1 | T08G11.10 |
| WBGene00199981 | -1.159714244 | 0.75790634 | 1 | F26A1.18 |
| WBGene00199983 | -0.156269293 | 0.96924949 | 1 | T04C12.25 |
| WBGene00199991 | 0.281454817 | 0.94435481 | 1 | Y61A9LA.13 |
| WBGene00199993 | 0.688793783 | 0.86378755 | 1 | F45H7.13 |
| WBGene00199997 | -0.156269293 | 0.96924949 | 1 | H32C10.97 |
| WBGene00200010 | -0.156269293 | 0.96924949 | 1 | C30B5.14 |
| WBGene00200016 | -0.156269293 | 0.96924949 | 1 | AC3.16 |
| WBGene00200019 | -0.156269293 | 0.96924949 | 1 | F43D2.13 |
| WBGene00200022 | -0.156269293 | 0.96924949 | 1 | C55B7.13 |
| WBGene00200024 | 0.206584522 | 0.92198558 | 1 | Y71A12B.29 |
| WBGene00200025 | 0.560070437 | 0.22508895 | 0.42921391 | T04C12.26 |
| WBGene00200026 | 1.851444818 | 0.53669729 | 1 | R07D5.7 |

| | | | | |
|---|---|---|---|---|
| WBGene00200029 | -0.945145766 | 0.65193959 | 1 | C40H1.10 |
| WBGene00200041 | 0.377828795 | 0.92526585 | 1 | R03E9.8 |
| WBGene00200044 | -0.156269293 | 0.96924949 | 1 | ZK682.11 |
| WBGene00200046 | -0.156269293 | 0.96924949 | 1 | Y47G6A.34 |
| WBGene00200047 | -0.156269293 | 0.96924949 | 1 | F35B12.15 |
| WBGene00200053 | -0.156269293 | 0.96924949 | 1 | C01B7.14 |
| WBGene00200055 | 0.574508983 | 0.8551971 | 1 | R01E6.16 |
| WBGene00200061 | 0.281454817 | 0.94435481 | 1 | Y48B6A.21 |
| WBGene00200070 | 0.281454817 | 0.94435481 | 1 | K06C4.22 |
| WBGene00200071 | 0.281454817 | 0.94435481 | 1 | T19C4.23 |
| WBGene00200077 | -0.668264701 | 0.85551153 | 1 | F01G12.15 |
| WBGene00200080 | -0.285029105 | 0.27075454 | 0.47962106 | C32D5.18 |
| WBGene00200091 | 1.743415485 | 0.50483382 | 1 | Y48A6B.18 |
| WBGene00200101 | 0.377828795 | 0.92526585 | 1 | R06C1.11 |
| WBGene00200114 | -0.156269293 | 0.96924949 | 1 | Y48G9A.18 |
| WBGene00200119 | 0.377828795 | 0.92526585 | 1 | ZC506.8 |
| WBGene00200120 | -0.014519762 | 0.99603746 | 1 | K02D7.8 |
| WBGene00200121 | 0.281454817 | 0.94435481 | 1 | F40H3.14 |
| WBGene00200123 | 0.377828795 | 0.92526585 | 1 | C29A12.17 |
| WBGene00200129 | 0.688793783 | 0.86378755 | 1 | F49E10.20 |
| WBGene00200132 | -0.156269293 | 0.96924949 | 1 | ZC412.12 |
| WBGene00200138 | -0.156269293 | 0.96924949 | 1 | F15A8.11 |
| WBGene00200144 | -0.156269293 | 0.96924949 | 1 | F07E5.11 |
| WBGene00200150 | -0.108066064 | 0.77086381 | 0.8718461 | C07A9.16 |
| WBGene00200151 | -0.156269293 | 0.96924949 | 1 | C39E6.12 |
| WBGene00200154 | 0.688793783 | 0.86378755 | 1 | F37B12.11 |
| WBGene00200155 | 0.232157325 | 0.93397866 | 1 | R13.9 |
| WBGene00200156 | 1.247575581 | 0.7549672 | 1 | C06A12.17 |
| WBGene00200157 | -0.156269293 | 0.96924949 | 1 | C27D6.13 |
| WBGene00200160 | -1.901856996 | 0.64072034 | 1 | T28B11.17 |
| WBGene00200161 | 0.281454817 | 0.94435481 | 1 | K01A6.17 |
| WBGene00200162 | -0.156269293 | 0.96924949 | 1 | K02F3.13 |
| WBGene00200164 | -0.156269293 | 0.96924949 | 1 | T14G10.10 |
| WBGene00200166 | -0.156269293 | 0.96924949 | 1 | K11G9.12 |
| WBGene00200174 | -1.964653846 | 0.53599202 | 1 | F17E9.17 |
| WBGene00200176 | 0.109173684 | 0.96057644 | 1 | T19A5.11 |
| WBGene00200190 | -1.901856996 | 0.64072034 | 1 | F16F9.15 |
| WBGene00200191 | -3.51732623 | 0.15709278 | 1 | C44H4.12 |
| WBGene00200195 | 0.281454817 | 0.94435481 | 1 | Y71H9A.7 |
| WBGene00200197 | 0.281454817 | 0.94435481 | 1 | W07G1.17 |
| WBGene00200200 | -0.156269293 | 0.96924949 | 1 | F09F3.16 |
| WBGene00200209 | 0.220594403 | 0.90533813 | 1 | F32A6.11 |
| WBGene00200212 | 2.063391569 | 0.37853067 | 1 | F17C8.12 |

| | | | | |
|---|---|---|---|---|
| WBGene00200223 | -0.156269293 | 0.96924949 | 1 | ZC518.7 |
| WBGene00200224 | 0.281454817 | 0.94435481 | 1 | F01E11.13 |
| WBGene00200227 | 0.688793783 | 0.86378755 | 1 | F56E10.7 |
| WBGene00200232 | 0.281454817 | 0.94435481 | 1 | R06C1.12 |
| WBGene00200237 | 0.377828795 | 0.92526585 | 1 | Y57G11A.225 |
| WBGene00200243 | 0.281454817 | 0.94435481 | 1 | Y106G6E.7 |
| WBGene00200245 | 0.983488853 | 0.80601565 | 1 | ZK1290.21 |
| WBGene00200248 | 0.574508983 | 0.8551971 | 1 | ZK617.19 |
| WBGene00200249 | 0.747845955 | 0.85215965 | 1 | F12F3.8 |
| WBGene00200252 | -0.323868354 | 0.91634405 | 1 | F31B12.6 |
| WBGene00200260 | 0.22030529 | 0.77203801 | 0.87263357 | W06F12.6 |
| WBGene00200262 | -1.969805064 | 0.62862569 | 1 | C44E4.10 |
| WBGene00200265 | 0.281454817 | 0.94435481 | 1 | C17E4.18 |
| WBGene00200267 | 0.688793783 | 0.86378755 | 1 | Y69A2AR.42 |
| WBGene00200268 | 1.285665874 | 0.02272015 | 0.09382495 | F01G12.16 |
| WBGene00200270 | 1.82867917 | 0.54358766 | 1 | F43B10.3 |
| WBGene00200272 | 3.096113806 | 0.11014314 | 1 | C09B9.82 |
| WBGene00200278 | -0.156269293 | 0.96924949 | 1 | R05H11.4 |
| WBGene00200284 | 5.057115851 | 9.4267E-05 | 0.0014627 | T18D3.13 |
| WBGene00200290 | 1.025181888 | 0.79790347 | 1 | B0511.16 |
| WBGene00200293 | -1.969805064 | 0.62862569 | 1 | C24A3.14 |
| WBGene00200295 | 0.281454817 | 0.94435481 | 1 | T27F7.7 |
| WBGene00200299 | 0.377828795 | 0.92526585 | 1 | C25F6.22 |
| WBGene00200306 | 0.377828795 | 0.92526585 | 1 | F10D7.9 |
| WBGene00200316 | 0.399606396 | 0.86544406 | 1 | C30D11.3 |
| WBGene00200323 | -0.156269293 | 0.96924949 | 1 | K02E10.10 |
| WBGene00200330 | 0.281454817 | 0.94435481 | 1 | F42G4.9 |
| WBGene00200331 | 0.747845955 | 0.85215965 | 1 | C08D8.6 |
| WBGene00200332 | -0.156269293 | 0.96924949 | 1 | Y50D4A.8 |
| WBGene00200337 | 0.574508983 | 0.8551971 | 1 | C41C4.15 |
| WBGene00200340 | 3.12907916 | 0.0981945 | 1 | F28F9.13 |
| WBGene00200344 | -0.156269293 | 0.96924949 | 1 | F13D11.16 |
| WBGene00200349 | 0.574508983 | 0.8551971 | 1 | T27D12.10 |
| WBGene00200355 | 2.146098644 | 0.43306874 | 1 | F22A3.15 |
| WBGene00200356 | 2.407689268 | 0.1658253 | 1 | R11E3.15 |
| WBGene00200369 | -0.156269293 | 0.96924949 | 1 | Y73F8A.1170 |
| WBGene00200388 | 1.367734989 | 0.01571258 | 0.07187884 | R05G6.18 |
| WBGene00200394 | -0.014519762 | 0.99603746 | 1 | C38C5.3 |
| WBGene00200411 | 0.377828795 | 0.92526585 | 1 | F08F1.19 |
| WBGene00200413 | 1.563818453 | 0.47753399 | 1 | C48D5.8 |
| WBGene00200420 | 0.574508983 | 0.8551971 | 1 | Y15E3A.11 |
| WBGene00200426 | -0.156269293 | 0.96924949 | 1 | C43H6.10 |
| WBGene00200427 | 0.281454817 | 0.94435481 | 1 | W06D4.9 |

| | | | | |
|---|---|---|---|---|
| WBGene00200432 | 1.778794942 | 0.24521293 | 1 | C13A10.5 |
| WBGene00200433 | -1.969805064 | 0.62862569 | 1 | K03H4.6 |
| WBGene00200434 | 1.216957619 | 0.74819621 | 1 | Y56A3A.39 |
| WBGene00200435 | -0.156269293 | 0.96924949 | 1 | R10H1.8 |
| WBGene00200452 | -0.156269293 | 0.96924949 | 1 | F46G11.17 |
| WBGene00200459 | 0.178115923 | 0.94982316 | 1 | F38H4.15 |
| WBGene00200463 | 0.281454817 | 0.94435481 | 1 | T13H10.15 |
| WBGene00200464 | 0.281454817 | 0.94435481 | 1 | T22B7.17 |
| WBGene00200472 | 0.281454817 | 0.94435481 | 1 | H03A11.10 |
| WBGene00200490 | -0.876437863 | 0.21223076 | 0.41207318 | C08B6.17 |
| WBGene00200491 | -0.156269293 | 0.96924949 | 1 | T06F4.16 |
| WBGene00200494 | 0.77095004 | 0.46126522 | 1 | D2089.7 |
| WBGene00200495 | 0.281454817 | 0.94435481 | 1 | T05A6.17 |
| WBGene00200497 | -0.156269293 | 0.96924949 | 1 | F22E12.11 |
| WBGene00200503 | 0.688793783 | 0.86378755 | 1 | F29G9.14 |
| WBGene00200504 | 0.377828795 | 0.92526585 | 1 | T21H8.9 |
| WBGene00200506 | -0.156269293 | 0.96924949 | 1 | ZC123.9 |
| WBGene00200507 | 0.281454817 | 0.94435481 | 1 | C03C11.8 |
| WBGene00200511 | -0.737462795 | 0.83433562 | 1 | R04B3.7 |
| WBGene00200514 | 0.281454817 | 0.94435481 | 1 | R07G3.14 |
| WBGene00200520 | -0.156269293 | 0.96924949 | 1 | C13D9.13 |
| WBGene00200526 | 0.555193615 | 0.26155704 | 0.47006725 | C05B10.17 |
| WBGene00200527 | 2.600311303 | 0.25000941 | 1 | T10C6.19 |
| WBGene00200528 | -0.156269293 | 0.96924949 | 1 | R09G11.3 |
| WBGene00200530 | -0.828765424 | 0.69034289 | 1 | C16B8.10 |
| WBGene00200540 | -0.156269293 | 0.96924949 | 1 | F46F3.15 |
| WBGene00200543 | 0.281454817 | 0.94435481 | 1 | C34E7.8 |
| WBGene00200549 | 0.281454817 | 0.94435481 | 1 | H03E18.6 |
| WBGene00200555 | 0.281454817 | 0.94435481 | 1 | R153.5 |
| WBGene00200560 | -1.969805064 | 0.62862569 | 1 | Y43H11AL.6 |
| WBGene00200561 | 0.656424328 | 0.23597676 | 0.44215444 | C10C6.11 |
| WBGene00200567 | -0.156269293 | 0.96924949 | 1 | K11G9.13 |
| WBGene00200575 | -0.128807018 | 0.84183197 | 0.91315662 | F44E7.15 |
| WBGene00200577 | 0.377828795 | 0.92526585 | 1 | Y65B4BL.12 |
| WBGene00200579 | -1.901856996 | 0.64072034 | 1 | B0034.15 |
| WBGene00200588 | 1.735991307 | 0.58152819 | 1 | M60.13 |
| WBGene00200589 | -1.760055813 | 0.45626364 | 1 | ZC504.16 |
| WBGene00200590 | -0.156269293 | 0.96924949 | 1 | T05B11.11 |
| WBGene00200594 | -1.094201388 | 0.7816772 | 1 | B0350.76 |
| WBGene00200595 | 0.863169103 | 0.41224736 | 1 | C37E2.7 |
| WBGene00200598 | -0.156269293 | 0.96924949 | 1 | C07G2.4 |
| WBGene00200599 | -0.18557176 | 0.95664217 | 1 | F45G2.12 |
| WBGene00200601 | 0.688793783 | 0.86378755 | 1 | M117.12 |

| | | | | |
|---|---|---|---|---|
| WBGene00200606 | 0.041803775 | 0.98890197 | 1 | Y37E3.29 |
| WBGene00200608 | -0.70576507 | 0.76841669 | 1 | Y39G10AL.5 |
| WBGene00200619 | 0.233739675 | 0.93575889 | 1 | F01E11.14 |
| WBGene00200638 | 1.216957619 | 0.74819621 | 1 | H39E23.4 |
| WBGene00200642 | -0.737462795 | 0.83433562 | 1 | T21B6.15 |
| WBGene00200645 | -0.156269293 | 0.96924949 | 1 | R03E9.9 |
| WBGene00200659 | -1.14715222 | 0.68932198 | 1 | F10D2.15 |
| WBGene00200662 | -0.685959299 | 0.68164528 | 1 | F12F6.17 |
| WBGene00200667 | 0.025337778 | 0.99212087 | 1 | F56A12.9 |
| WBGene00200669 | -0.156269293 | 0.96924949 | 1 | B0025.6 |
| WBGene00200673 | -1.159714244 | 0.75790634 | 1 | ZK994.11 |
| WBGene00200678 | 0.281454817 | 0.94435481 | 1 | K07H8.16 |
| WBGene00200683 | -1.082680044 | 0.67762556 | 1 | C09G5.11 |
| WBGene00200706 | 1.216957619 | 0.74819621 | 1 | K08E4.12 |
| WBGene00200707 | -1.45873709 | 0.59584643 | 1 | Y71G12B.37 |
| WBGene00200708 | 1.216957619 | 0.74819621 | 1 | K11H3.12 |
| WBGene00200711 | 0.857402182 | 0.72847739 | 1 | C54D2.15 |
| WBGene00200715 | 0.997919443 | 0.45569556 | 1 | K04F10.8 |
| WBGene00200717 | -0.156269293 | 0.96924949 | 1 | F11C1.15 |
| WBGene00200734 | -0.156269293 | 0.96924949 | 1 | C29A12.18 |
| WBGene00200736 | -0.156269293 | 0.96924949 | 1 | C18A3.23 |
| WBGene00200756 | -0.737462795 | 0.83433562 | 1 | H01A20.4 |
| WBGene00200759 | -0.156269293 | 0.96924949 | 1 | F11D11.20 |
| WBGene00200762 | -1.901856996 | 0.64072034 | 1 | T28B11.19 |
| WBGene00200767 | 0.281454817 | 0.94435481 | 1 | Y71H9A.8 |
| WBGene00200768 | 0.233739675 | 0.93575889 | 1 | Y23H5A.11 |
| WBGene00200770 | 0.281454817 | 0.94435481 | 1 | W06A7.10 |
| WBGene00200779 | -0.985537791 | 0.65575305 | 1 | F56A11.12 |
| WBGene00200784 | 0.281454817 | 0.94435481 | 1 | K06A1.9 |
| WBGene00200785 | -0.558804446 | 0.87818675 | 1 | C49A9.39 |
| WBGene00200795 | -0.371579084 | 0.48165258 | 0.67289188 | C08B11.12 |
| WBGene00200797 | 0.041803775 | 0.98890197 | 1 | Y7A5A.18 |
| WBGene00200798 | -1.291305095 | 0.63677871 | 1 | F26D11.18 |
| WBGene00200812 | 0.4828626 | 0.81641781 | 1 | C17C3.23 |
| WBGene00200827 | -1.094201388 | 0.7816772 | 1 | F08H9.14 |
| WBGene00200829 | -0.156269293 | 0.96924949 | 1 | F43D9.12 |
| WBGene00200837 | -0.156269293 | 0.96924949 | 1 | R04F11.24 |
| WBGene00200838 | -0.156269293 | 0.96924949 | 1 | ZK596.4 |
| WBGene00200847 | 1.216957619 | 0.74819621 | 1 | T16A9.6 |
| WBGene00200848 | -1.901856996 | 0.64072034 | 1 | Y51H7C.33 |
| WBGene00200849 | -1.901856996 | 0.64072034 | 1 | R53.11 |
| WBGene00200850 | -0.62146648 | 0.85925752 | 1 | C18H7.16 |
| WBGene00200858 | -1.901856996 | 0.64072034 | 1 | F28C1.14 |

| | | | | |
|---|---|---|---|---|
| WBGene00200866 | 0.616429445 | 0.81400458 | 1 | F47E1.14 |
| WBGene00200868 | 0.377828795 | 0.92526585 | 1 | F26D12.100 |
| WBGene00200873 | -0.245531902 | 0.94070824 | 1 | Y53F4B.61 |
| WBGene00200878 | 0.281454817 | 0.94435481 | 1 | Y47G6A.35 |
| WBGene00200884 | -0.084093301 | 0.96701288 | 1 | F25D1.10 |
| WBGene00200892 | 0.281454817 | 0.94435481 | 1 | Y43F8B.26 |
| WBGene00200893 | 0.281454817 | 0.94435481 | 1 | F52D10.12 |
| WBGene00200898 | 0.525360655 | 0.56264472 | 0.73817479 | B0350.77 |
| WBGene00200908 | 0.281454817 | 0.94435481 | 1 | K07F5.20 |
| WBGene00200910 | -1.901856996 | 0.64072034 | 1 | F11A6.12 |
| WBGene00200912 | -0.156269293 | 0.96924949 | 1 | T04F3.11 |
| WBGene00200914 | -0.156269293 | 0.96924949 | 1 | F22B7.15 |
| WBGene00200917 | -1.159714244 | 0.75790634 | 1 | Y70C5A.5 |
| WBGene00200923 | -0.156269293 | 0.96924949 | 1 | W05E10.17 |
| WBGene00200928 | 0.747845955 | 0.85215965 | 1 | K10C2.11 |
| WBGene00200929 | -1.901856996 | 0.64072034 | 1 | T06E4.18 |
| WBGene00200931 | -0.156269293 | 0.96924949 | 1 | W07G4.11 |
| WBGene00200939 | 0.574508983 | 0.8551971 | 1 | F10F2.15 |
| WBGene00200946 | 0.281454817 | 0.94435481 | 1 | T05B9.3 |
| WBGene00200952 | 1.373784582 | 0.54715045 | 1 | C27C12.9 |
| WBGene00200957 | -0.156269293 | 0.96924949 | 1 | T02G6.12 |
| WBGene00200958 | 0.281454817 | 0.94435481 | 1 | C11G10.13 |
| WBGene00200960 | 0.281454817 | 0.94435481 | 1 | R10E11.12 |
| WBGene00200962 | -0.497279034 | 0.73986814 | 1 | C01B10.46 |
| WBGene00200967 | -0.156269293 | 0.96924949 | 1 | C13C4.11 |
| WBGene00200973 | -0.35768106 | 0.83841345 | 1 | F25F2.5 |
| WBGene00200977 | -0.263229581 | 0.93409872 | 1 | T28B4.13 |
| WBGene00200980 | 1.274878015 | 0.43457342 | 1 | K03B8.22 |
| WBGene00200981 | -0.737462795 | 0.83433562 | 1 | F35E12.12 |
| WBGene00200993 | 0.281454817 | 0.94435481 | 1 | C32A9.4 |
| WBGene00200998 | -0.536871728 | 0.76889293 | 1 | ZK662.8 |
| WBGene00200999 | -0.156269293 | 0.96924949 | 1 | Y34B4A.17 |
| WBGene00201014 | 0.109173684 | 0.96057644 | 1 | T19A5.12 |
| WBGene00201016 | 0.281454817 | 0.94435481 | 1 | T09F5.18 |
| WBGene00201019 | 0.792402183 | 0.45495588 | 1 | Y71F9B.22 |
| WBGene00201023 | 0.281454817 | 0.94435481 | 1 | F07C6.7 |
| WBGene00201027 | 1.653460415 | 0.50712258 | 1 | K12F2.5 |
| WBGene00201031 | -1.901856996 | 0.64072034 | 1 | ZK792.10 |
| WBGene00201034 | -0.156269293 | 0.96924949 | 1 | F57B7.12 |
| WBGene00201040 | 1.660064349 | 0.6377947 | 1 | C30D11.4 |
| WBGene00201042 | -0.604396148 | 0.82723443 | 1 | T02C5.7 |
| WBGene00201047 | 1.876192691 | 0.43542138 | 1 | F43C9.7 |
| WBGene00201049 | 0.688793783 | 0.86378755 | 1 | T05B4.19 |

| | | | | |
|---|---|---|---|---|
| WBGene00201052 | -0.156269293 | 0.96924949 | 1 | R09E10.14 |
| WBGene00201066 | 0.281454817 | 0.94435481 | 1 | F29F11.16 |
| WBGene00201070 | 0.281454817 | 0.94435481 | 1 | F25H10.9 |
| WBGene00201078 | -0.263229581 | 0.93409872 | 1 | C49F8.12 |
| WBGene00201096 | -1.901856996 | 0.64072034 | 1 | F56F4.11 |
| WBGene00201103 | 1.116613983 | 0.69067248 | 1 | C01C4.6 |
| WBGene00201106 | -1.901856996 | 0.64072034 | 1 | F48C1.14 |
| WBGene00201113 | 0.281454817 | 0.94435481 | 1 | T06F4.17 |
| WBGene00201114 | 3.193785087 | 0.03574929 | 1 | C01B10.47 |
| WBGene00201116 | -0.40861243 | 0.52978307 | 0.71398479 | C16E9.24 |
| WBGene00201123 | -0.490561023 | 0.85693136 | 1 | Y42A5A.12 |
| WBGene00201125 | 0.983488853 | 0.80601565 | 1 | F17A9.7 |
| WBGene00201129 | 0.281454817 | 0.94435481 | 1 | M02F4.11 |
| WBGene00201136 | -0.30359982 | 0.8981102 | 1 | C24A8.12 |
| WBGene00201137 | 0.281454817 | 0.94435481 | 1 | C02G6.5 |
| WBGene00201145 | -1.159714244 | 0.75790634 | 1 | F47F6.12 |
| WBGene00201149 | 0.281454817 | 0.94435481 | 1 | B0457.15 |
| WBGene00201153 | 1.557897343 | 0.63813005 | 1 | C44B7.20 |
| WBGene00201165 | 0.281454817 | 0.94435481 | 1 | Y110A7A.23 |
| WBGene00201168 | -0.18557176 | 0.95664217 | 1 | B0457.16 |
| WBGene00201169 | 1.025181888 | 0.79790347 | 1 | F31B12.8 |
| WBGene00201176 | -1.831477136 | 0.52194793 | 1 | B0310.11 |
| WBGene00201180 | -0.156269293 | 0.96924949 | 1 | B0563.16 |
| WBGene00201188 | -0.156269293 | 0.96924949 | 1 | DY3.11 |
| WBGene00201190 | 0.281454817 | 0.94435481 | 1 | W03D8.12 |
| WBGene00201202 | 0.688793783 | 0.86378755 | 1 | T24D1.6 |
| WBGene00201204 | -0.62146648 | 0.85925752 | 1 | F15A2.13 |
| WBGene00201212 | -0.156269293 | 0.96924949 | 1 | K02A4.11 |
| WBGene00201215 | 0.281454817 | 0.94435481 | 1 | C11E4.13 |
| WBGene00201244 | -1.901856996 | 0.64072034 | 1 | T25D10.9 |
| WBGene00201246 | -0.156269293 | 0.96924949 | 1 | T18H9.10 |
| WBGene00201251 | 1.149899068 | 0.60756174 | 1 | T07D1.11 |
| WBGene00201253 | -0.156269293 | 0.96924949 | 1 | K08A8.25 |
| WBGene00201256 | -0.156269293 | 0.96924949 | 1 | K03H9.14 |
| WBGene00201259 | 0.233739675 | 0.93575889 | 1 | F01E11.15 |
| WBGene00201260 | 0.938982574 | 0.8146943 | 1 | Y76F7A.5 |
| WBGene00201264 | -0.156269293 | 0.96924949 | 1 | C12D8.24 |
| WBGene00201265 | 0.688793783 | 0.86378755 | 1 | C41G11.17 |
| WBGene00201274 | -0.836993572 | 0.43883687 | 1 | Y95B8A.16 |
| WBGene00201279 | 0.281454817 | 0.94435481 | 1 | C44C10.14 |
| WBGene00201281 | -0.156269293 | 0.96924949 | 1 | F26A10.25 |
| WBGene00201299 | -1.901856996 | 0.64072034 | 1 | F10C1.18 |
| WBGene00201303 | 0.281454817 | 0.94435481 | 1 | F33A8.11 |

| | | | | |
|---|---|---|---|---|
| WBGene00201310 | -0.773064647 | 0.54005084 | 1 | C39H7.70 |
| WBGene00201313 | 0.281454817 | 0.94435481 | 1 | F09F9.16 |
| WBGene00201314 | -0.668355569 | 0.74739695 | 1 | F31F6.15 |
| WBGene00201328 | 0.124041901 | 0.89807501 | 1 | Y105C5B.1416 |
| WBGene00201335 | 0.407339936 | 0.68417379 | 1 | Y73B6BL.285 |
| WBGene00201341 | 1.247575581 | 0.7549672 | 1 | C49C3.22 |
| WBGene00201346 | -0.431178596 | 0.47052434 | 0.66356557 | R11B5.11 |
| WBGene00201351 | 0.743637457 | 0.84880372 | 1 | Y51F10.13 |
| WBGene00201355 | -1.901856996 | 0.64072034 | 1 | Y7A5A.19 |
| WBGene00201365 | -0.156269293 | 0.96924949 | 1 | ZK930.12 |
| WBGene00201366 | 0.627000403 | 0.11726952 | 0.28233879 | T01C8.10 |
| WBGene00201372 | 0.281454817 | 0.94435481 | 1 | W06H8.15 |
| WBGene00201375 | 0.688793783 | 0.86378755 | 1 | C05A9.7 |
| WBGene00201383 | -1.156387078 | 0.02343813 | 0.09584449 | F25D7.10 |
| WBGene00201384 | 0.020328391 | 0.99335814 | 1 | B0222.14 |
| WBGene00201386 | -0.748692734 | 0.06320534 | 0.18877638 | Y4C6B.100 |
| WBGene00201389 | 0.207104011 | 0.54251953 | 0.72302191 | ZK637.20 |
| WBGene00201398 | 0.355174276 | NaN | 1 | F31C3.16 |
| WBGene00201401 | -0.156269293 | 0.96924949 | 1 | T28B11.22 |
| WBGene00201414 | -1.831477136 | 0.52194793 | 1 | C52A10.4 |
| WBGene00201415 | 0.574508983 | 0.8551971 | 1 | Y15E3A.12 |
| WBGene00201419 | 0.568731452 | 0.27130196 | 0.47994711 | K08B12.13 |
| WBGene00201421 | 0.983488853 | 0.80601565 | 1 | C18B12.15 |
| WBGene00201427 | 0.281454817 | 0.94435481 | 1 | F55G1.20 |
| WBGene00201431 | -0.156269293 | 0.96924949 | 1 | F02E8.10 |
| WBGene00201435 | -0.156269293 | 0.96924949 | 1 | C10G11.15 |
| WBGene00201437 | 2.033416803 | 0.47767962 | 1 | F31B12.9 |
| WBGene00201453 | 0.636465245 | 0.53512357 | 0.7178681 | W02G9.7 |
| WBGene00201458 | -0.156269293 | 0.96924949 | 1 | VF15C11L.2 |
| WBGene00201460 | -1.969805064 | 0.62862569 | 1 | Y43H11AL.7 |
| WBGene00201466 | 0.983488853 | 0.80601565 | 1 | C32A3.6 |
| WBGene00201473 | 0.281454817 | 0.94435481 | 1 | R02D5.21 |
| WBGene00201488 | 0.57991002 | 0.82402153 | 1 | K09A11.10 |
| WBGene00201497 | 2.641117225 | 0.21179361 | 1 | Y17G7B.24 |
| WBGene00201511 | 0.233739675 | 0.93575889 | 1 | Y46E12A.9 |
| WBGene00201520 | 1.007797134 | 0.5864848 | 1 | Y17G9A.89 |
| WBGene00201522 | 0.281454817 | 0.94435481 | 1 | F16F9.17 |
| WBGene00201523 | -0.156269293 | 0.96924949 | 1 | M01A8.3 |
| WBGene00201537 | 1.557897343 | 0.63813005 | 1 | K01D12.18 |
| WBGene00201538 | -0.156269293 | 0.96924949 | 1 | C55A6.19 |
| WBGene00201539 | 1.314900666 | 0.23882959 | 1 | M01A8.4 |
| WBGene00201541 | -0.156269293 | 0.96924949 | 1 | H14A12.9 |
| WBGene00201545 | 1.413734769 | 0.68804531 | 1 | K02A2.9 |

| | | | | |
|---|---|---|---|---|
| WBGene00201547 | -1.969805064 | 0.62862569 | 1 | F54H12.9 |
| WBGene00201558 | -0.156269293 | 0.96924949 | 1 | F02A9.9 |
| WBGene00201571 | -0.156269293 | 0.96924949 | 1 | C34F11.18 |
| WBGene00201578 | 0.281454817 | 0.94435481 | 1 | M04B2.12 |
| WBGene00201584 | 0.938982574 | 0.8146943 | 1 | F16F9.18 |
| WBGene00201586 | 0.098413562 | 0.60341931 | 0.76809749 | T04C12.30 |
| WBGene00201589 | 0.377828795 | 0.92526585 | 1 | D1081.17 |
| WBGene00201606 | 0.281454817 | 0.94435481 | 1 | C30C11.12 |
| WBGene00201616 | -1.964653846 | 0.53599202 | 1 | Y17D7B.13 |
| WBGene00201617 | -0.18557176 | 0.95664217 | 1 | F52C9.10 |
| WBGene00201625 | -0.668264701 | 0.85551153 | 1 | F45D3.15 |
| WBGene00201630 | 0.574508983 | 0.8551971 | 1 | C28F5.7 |
| WBGene00201634 | 0.688793783 | 0.86378755 | 1 | C18F3.11 |
| WBGene00201640 | -0.156269293 | 0.96924949 | 1 | C09G12.19 |
| WBGene00201656 | -0.604396148 | 0.82723443 | 1 | CE7X_3.3 |
| WBGene00201659 | 1.872249764 | 0.5345823 | 1 | Y116A8B.97 |
| WBGene00201665 | -0.156269293 | 0.96924949 | 1 | W09G3.15 |
| WBGene00201670 | -0.156269293 | 0.96924949 | 1 | Y48B6A.23 |
| WBGene00201676 | -1.901856996 | 0.64072034 | 1 | M05B5.10 |
| WBGene00201679 | 1.181686151 | 0.7579682 | 1 | R02D5.22 |
| WBGene00201680 | 0.688793783 | 0.86378755 | 1 | K06A1.10 |
| WBGene00201685 | -0.156269293 | 0.96924949 | 1 | C53C9.6 |
| WBGene00201710 | 0.658556833 | 0.73016338 | 1 | C01H6.11 |
| WBGene00201725 | 0.245622231 | 0.84242003 | 1 | R90.12 |
| WBGene00201729 | 0.789601902 | 0.75907734 | 1 | C16E9.27 |
| WBGene00201734 | -1.969805064 | 0.62862569 | 1 | ZK154.8 |
| WBGene00201746 | 1.715010045 | 0.15409539 | 1 | F15A8.16 |
| WBGene00201753 | 1.032566776 | 0.28207111 | 0.49140843 | H36N01.5 |
| WBGene00201755 | -0.156269293 | 0.96924949 | 1 | C37A2.9 |
| WBGene00201763 | 1.116613983 | 0.69067248 | 1 | W03G1.10 |
| WBGene00201766 | 2.028800581 | 0.39258153 | 1 | K08B4.42 |
| WBGene00201770 | -0.156269293 | 0.96924949 | 1 | C12D8.25 |
| WBGene00201772 | 0.688793783 | 0.86378755 | 1 | Y73B3A.28 |
| WBGene00201778 | -0.156269293 | 0.96924949 | 1 | H01A20.5 |
| WBGene00201793 | 0.281454817 | 0.94435481 | 1 | E01H11.5 |
| WBGene00201795 | 0.688793783 | 0.86378755 | 1 | C25E10.15 |
| WBGene00201800 | 1.388471555 | 0.39318041 | 1 | C24A8.13 |
| WBGene00201804 | 0.688793783 | 0.86378755 | 1 | C05A2.3 |
| WBGene00201807 | -0.156269293 | 0.96924949 | 1 | F28A12.7 |
| WBGene00201819 | 1.413377704 | 0.60673863 | 1 | C30D11.5 |
| WBGene00201826 | -0.156269293 | 0.96924949 | 1 | F13H10.11 |
| WBGene00201829 | -0.156269293 | 0.96924949 | 1 | Y8G1A.3 |
| WBGene00201830 | -0.156269293 | 0.96924949 | 1 | Y54G2A.70 |

| | | | | |
|---|---|---|---|---|
| WBGene00201832 | -1.964653846 | 0.53599202 | 1 | ZC449.10 |
| WBGene00201833 | -0.156269293 | 0.96924949 | 1 | F54E4.16 |
| WBGene00201837 | 0.281454817 | 0.94435481 | 1 | F47B7.16 |
| WBGene00201840 | -1.901856996 | 0.64072034 | 1 | F57C7.14 |
| WBGene00201844 | 0.281454817 | 0.94435481 | 1 | R31.5 |
| WBGene00201851 | -0.300852129 | 0.6718838 | 0.81248091 | B0496.19 |
| WBGene00201852 | -0.273829551 | 0.91849619 | 1 | ZK662.9 |
| WBGene00201856 | -0.156269293 | 0.96924949 | 1 | Y76A2B.9 |
| WBGene00201858 | -1.14715222 | 0.68932198 | 1 | W04D2.14 |
| WBGene00201860 | -0.156269293 | 0.96924949 | 1 | F18E2.14 |
| WBGene00201861 | 0.377828795 | 0.92526585 | 1 | B0395.10 |
| WBGene00201863 | -0.156269293 | 0.96924949 | 1 | T28B4.14 |
| WBGene00201875 | -1.094201388 | 0.7816772 | 1 | F28H7.12 |
| WBGene00201888 | -0.009203965 | 0.99450464 | 1 | Y39G8C.12 |
| WBGene00201899 | 0.281454817 | 0.94435481 | 1 | C30B5.16 |
| WBGene00201900 | 0.020328391 | 0.99335814 | 1 | T21C9.17 |
| WBGene00201902 | 0.281454817 | 0.94435481 | 1 | F29F11.18 |
| WBGene00201908 | -1.901856996 | 0.64072034 | 1 | F09C8.6 |
| WBGene00201915 | -0.156269293 | 0.96924949 | 1 | F35G2.20 |
| WBGene00201920 | 1.181686151 | 0.7579682 | 1 | C33D12.15 |
| WBGene00201922 | -0.558804446 | 0.87818675 | 1 | ZC21.17 |
| WBGene00201926 | 0.270496667 | 0.92869767 | 1 | F52F12.18 |
| WBGene00201940 | 0.720066222 | 0.75158338 | 1 | F46F3.22 |
| WBGene00201946 | -0.156269293 | 0.96924949 | 1 | F26A10.26 |
| WBGene00201960 | 0.688793783 | 0.86378755 | 1 | C05E11.17 |
| WBGene00201962 | 0.627834078 | 0.77998941 | 1 | T23F11.9 |
| WBGene00201964 | -0.82897462 | 0.6213576 | 1 | F25D1.11 |
| WBGene00201967 | -1.901856996 | 0.64072034 | 1 | F17E5.6 |
| WBGene00201969 | -0.156269293 | 0.96924949 | 1 | H03A11.12 |
| WBGene00201972 | 0.281454817 | 0.94435481 | 1 | T26H2.16 |
| WBGene00201981 | 3.115828267 | 0.01877639 | 1 | F10F2.16 |
| WBGene00201991 | 0.377828795 | 0.92526585 | 1 | C07G1.14 |
| WBGene00201992 | -0.668355569 | 0.74739695 | 1 | C13C12.6 |
| WBGene00201994 | 0.938982574 | 0.8146943 | 1 | T25C12.10 |
| WBGene00201995 | -0.156269293 | 0.96924949 | 1 | K12G11.13 |
| WBGene00201999 | 0.377828795 | 0.92526585 | 1 | F54C8.11 |
| WBGene00202001 | 0.574508983 | 0.8551971 | 1 | ZK721.6 |
| WBGene00202006 | 0.938982574 | 0.8146943 | 1 | C13C4.12 |
| WBGene00202007 | 0.281454817 | 0.94435481 | 1 | Y67A6A.3 |
| WBGene00202009 | 0.281454817 | 0.94435481 | 1 | K08D9.8 |
| WBGene00202011 | -0.156269293 | 0.96924949 | 1 | R13H4.13 |
| WBGene00202017 | -1.901856996 | 0.64072034 | 1 | F36H1.16 |
| WBGene00202020 | -1.901856996 | 0.64072034 | 1 | F52B10.21 |

| | | | | |
|---|---|---|---|---|
| WBGene00202035 | -0.156269293 | 0.96924949 | 1 | K02A4.13 |
| WBGene00202051 | 0.377828795 | 0.92526585 | 1 | C14H10.5 |
| WBGene00202052 | -0.156269293 | 0.96924949 | 1 | C41G11.20 |
| WBGene00202053 | -0.156269293 | 0.96924949 | 1 | T04C12.31 |
| WBGene00202054 | 0.281454817 | 0.94435481 | 1 | F57B7.14 |
| WBGene00202064 | -0.737462795 | 0.83433562 | 1 | C44B12.12 |
| WBGene00202069 | -0.156269293 | 0.96924949 | 1 | T20G5.17 |
| WBGene00202080 | -0.156269293 | 0.96924949 | 1 | C10F3.16 |
| WBGene00202081 | -0.156269293 | 0.96924949 | 1 | F08C6.17 |
| WBGene00202108 | 0.743492162 | 0.33096896 | 0.53922359 | F23B12.10 |
| WBGene00202109 | 0.983488853 | 0.80601565 | 1 | F59B10.10 |
| WBGene00202113 | 0.281454817 | 0.94435481 | 1 | B0457.18 |
| WBGene00202117 | -2.640414922 | 0.41081194 | 1 | Y92C3A.9 |
| WBGene00202118 | -0.156269293 | 0.96924949 | 1 | K02A2.10 |
| WBGene00202122 | -0.156269293 | 0.96924949 | 1 | F29G9.17 |
| WBGene00202124 | -0.737462795 | 0.83433562 | 1 | F22E12.13 |
| WBGene00202126 | -0.156269293 | 0.96924949 | 1 | Y51H7C.38 |
| WBGene00202134 | 0.281454817 | 0.94435481 | 1 | C11E4.16 |
| WBGene00202141 | -1.901856996 | 0.64072034 | 1 | R06B10.8 |
| WBGene00202154 | -1.969805064 | 0.62862569 | 1 | K07C6.17 |
| WBGene00202165 | 0.688793783 | 0.86378755 | 1 | R05G9.4 |
| WBGene00202168 | 0.703731386 | 0.20161027 | 0.40041719 | F43C9.9 |
| WBGene00202170 | 0.574508983 | 0.8551971 | 1 | C31H5.11 |
| WBGene00202173 | -1.159714244 | 0.75790634 | 1 | F53B6.11 |
| WBGene00202178 | -0.156269293 | 0.96924949 | 1 | F38A6.9 |
| WBGene00202179 | -0.156269293 | 0.96924949 | 1 | Y57G11C.1141 |
| WBGene00202185 | 1.216957619 | 0.74819621 | 1 | F41E7.18 |
| WBGene00202187 | -0.668264701 | 0.85551153 | 1 | M01E10.5 |
| WBGene00202188 | -0.156269293 | 0.96924949 | 1 | F45F2.16 |
| WBGene00202209 | 0.281454817 | 0.94435481 | 1 | Y41D4B.31 |
| WBGene00202213 | 0.281454817 | 0.94435481 | 1 | K08A2.16 |
| WBGene00202218 | 0.281454817 | 0.94435481 | 1 | M60.14 |
| WBGene00202225 | -0.156269293 | 0.96924949 | 1 | C40C9.16 |
| WBGene00202227 | -0.156269293 | 0.96924949 | 1 | C03H12.3 |
| WBGene00202242 | 1.353021252 | 0.55399304 | 1 | T23C6.6 |
| WBGene00202244 | -2.640414922 | 0.41081194 | 1 | ZC64.16 |
| WBGene00202253 | -1.969805064 | 0.62862569 | 1 | Y43H11AL.8 |
| WBGene00202256 | 0.041803775 | 0.98890197 | 1 | ZK863.12 |
| WBGene00202258 | 1.150860151 | 0.53494622 | 1 | F19D8.8 |
| WBGene00202278 | 6.108875302 | 6.89E-07 | 2.4831E-05 | T18D3.14 |
| WBGene00202281 | -0.156269293 | 0.96924949 | 1 | ZK792.11 |
| WBGene00202289 | 0.281454817 | 0.94435481 | 1 | F52B10.22 |
| WBGene00202312 | -1.901856996 | 0.64072034 | 1 | F59E12.14 |

| | | | | |
|---|---|---|---|---|
| WBGene00202313 | 0.281454817 | 0.94435481 | 1 | BE10.7 |
| WBGene00202315 | 1.974651709 | 0.28071322 | 1 | F29F11.19 |
| WBGene00202322 | 0.281454817 | 0.94435481 | 1 | T23G5.11 |
| WBGene00202327 | -0.156269293 | 0.96924949 | 1 | R03H4.11 |
| WBGene00202341 | 1.38322681 | 0.69550305 | 1 | ZK783.8 |
| WBGene00202345 | -1.969805064 | 0.62862569 | 1 | F11D5.33 |
| WBGene00202348 | 0.377828795 | 0.92526585 | 1 | K02E2.16 |
| WBGene00202349 | -0.156269293 | 0.96924949 | 1 | T01A4.6 |
| WBGene00202354 | -1.969805064 | 0.62862569 | 1 | Y46E12A.12 |
| WBGene00202361 | 0.688793783 | 0.86378755 | 1 | ZK596.5 |
| WBGene00202364 | -0.156269293 | 0.96924949 | 1 | T24A11.11 |
| WBGene00202365 | 0.938982574 | 0.8146943 | 1 | F21C3.10 |
| WBGene00202373 | -0.156269293 | 0.96924949 | 1 | C01F1.8 |
| WBGene00202374 | 0.782269838 | 0.07505227 | 0.21190922 | T01C8.11 |
| WBGene00202384 | 0.446458324 | 0.87164163 | 1 | Y53C10A.23 |
| WBGene00202386 | 0.281454817 | 0.94435481 | 1 | C52B9.18 |
| WBGene00202391 | 0.034331862 | 0.97364972 | 1 | C34H3.23 |
| WBGene00202394 | 0.763390359 | 0.73510979 | 1 | T10B9.13 |
| WBGene00202395 | -0.156269293 | 0.96924949 | 1 | B0034.17 |
| WBGene00202400 | 0.12332541 | 0.94831365 | 1 | C27D6.14 |
| WBGene00202404 | -0.156269293 | 0.96924949 | 1 | T19D2.11 |
| WBGene00202408 | 1.601302495 | 0.65842342 | 1 | Y23H5B.13 |
| WBGene00202410 | -0.156269293 | 0.96924949 | 1 | F54F12.5 |
| WBGene00202422 | 0.281454817 | 0.94435481 | 1 | K08B4.43 |
| WBGene00202432 | 0.377828795 | 0.92526585 | 1 | Y41C4A.28 |
| WBGene00202438 | -0.156269293 | 0.96924949 | 1 | Y51H7D.2 |
| WBGene00202439 | 0.281454817 | 0.94435481 | 1 | F46B6.18 |
| WBGene00202444 | 0.688793783 | 0.86378755 | 1 | Y65B4BR.11 |
| WBGene00202452 | 0.688793783 | 0.86378755 | 1 | C32E12.8 |
| WBGene00202462 | 0.543633856 | 0.54055862 | 0.72174866 | F56F11.7 |
| WBGene00202463 | -0.156269293 | 0.96924949 | 1 | F28A12.8 |
| WBGene00202469 | -0.156269293 | 0.96924949 | 1 | DY3.12 |
| WBGene00202473 | 0.938982574 | 0.8146943 | 1 | F25F6.4 |
| WBGene00202485 | 0.983488853 | 0.80601565 | 1 | F31B9.6 |
| WBGene00202491 | 1.130791626 | 0.2900912 | 1 | F55B11.132 |
| WBGene00202494 | -1.969805064 | 0.62862569 | 1 | F53G12.17 |
| WBGene00202495 | -0.156269293 | 0.96924949 | 1 | T26H5.11 |
| WBGene00202499 | -0.156269293 | 0.96924949 | 1 | B0507.15 |
| WBGene00202500 | -0.156269293 | 0.96924949 | 1 | K09H11.15 |
| WBGene00202501 | -0.604396148 | 0.82723443 | 1 | Y57E12B.10 |
| WBGene00202502 | -0.156269293 | 0.96924949 | 1 | F41B5.12 |
| WBGene00202504 | -1.964653846 | 0.53599202 | 1 | Y32B12B.8 |
| WBGene00202512 | 0.745335936 | 0.5832418 | 1 | C25E10.16 |

| | | | | |
|---|---|---|---|---|
| WBGene00202513 | -1.901856996 | 0.64072034 | 1 | ZK938.10 |
| WBGene00202514 | 0.169097384 | 0.93630233 | 1 | T09F5.20 |
| WBGene00206354 | 0.137263962 | 0.8275616 | 0.90692193 | F27E5.9 |
| WBGene00206355 | 1.158132015 | 0.62662247 | 1 | C34C6.10 |
| WBGene00206356 | -0.62146648 | 0.85925752 | 1 | T08B2.15 |
| WBGene00206357 | -1.964653846 | 0.53599202 | 1 | C09D4.9 |
| WBGene00206359 | 0.453022495 | 0.80317331 | 1 | C05E4.15 |
| WBGene00206360 | -1.964653846 | 0.53599202 | 1 | T13C5.10 |
| WBGene00206361 | -0.366409499 | 0.75513857 | 1 | K02G10.15 |
| WBGene00206362 | -0.444874305 | 0.71763944 | 1 | B0041.11 |
| WBGene00206364 | -0.389078632 | 0.8767475 | 1 | C07A12.18 |
| WBGene00206366 | 0.546052409 | 0.7844476 | 1 | C03H5.9 |
| WBGene00206370 | 1.413377704 | 0.60673863 | 1 | F02D8.9 |
| WBGene00206371 | 0.175909247 | 0.94229417 | 1 | F28B12.6 |
| WBGene00206373 | 0.881518635 | 0.35048831 | 0.5590224 | C27A2.12 |
| WBGene00206374 | 1.640825553 | 0.44758516 | 1 | K10G6.9 |
| WBGene00206375 | -1.964653846 | 0.53599202 | 1 | T08B6.64 |
| WBGene00206377 | -1.594883373 | 0.42056842 | 1 | F44E5.15 |
| WBGene00206378 | 1.38322681 | 0.69550305 | 1 | C14E2.11 |
| WBGene00206380 | 0.030177857 | 0.97222762 | 0.98553604 | C42D4.18 |
| WBGene00206381 | -1.777135447 | 0.45934881 | 1 | F52H2.15 |
| WBGene00206382 | 0.773345629 | 0.76155013 | 1 | F01G12.18 |
| WBGene00206384 | -1.371575522 | 0.47593629 | 1 | D1005.10 |
| WBGene00206385 | -1.424144338 | 0.44920716 | 1 | T06D4.6 |
| WBGene00206386 | -2.467222005 | 0.0057377 | 0.03448584 | K10C2.12 |
| WBGene00206388 | -1.748762203 | 0.39971558 | 1 | K10C2.14 |
| WBGene00206389 | -1.517070864 | 0.00187349 | 0.01481824 | C29F7.10 |
| WBGene00206390 | -0.774842772 | 0.78069886 | 1 | F14F7.7 |
| WBGene00206392 | 1.353535409 | 0.73149271 | 1 | F47D2.11 |
| WBGene00206393 | 0.377828795 | 0.92526585 | 1 | W04G5.16 |
| WBGene00206412 | -1.287544737 | 0.41560977 | 1 | T24H7.8 |
| WBGene00206413 | 1.687239501 | 0.59403242 | 1 | ZK1240.11 |
| WBGene00206414 | 0.312731947 | 0.84987061 | 1 | F31D5.7 |
| WBGene00206415 | 2.456496557 | 0.3218863 | 1 | F59H6.14 |
| WBGene00206416 | -0.98463404 | 0.56338607 | 1 | W04H10.5 |
| WBGene00206417 | 0.070918015 | 0.95763643 | 1 | W04H10.6 |
| WBGene00206418 | 1.121922047 | 0.18753373 | 0.38099849 | M04C3.5 |
| WBGene00206423 | 0.193893197 | 0.56865012 | 0.74341002 | Y41C4A.29 |
| WBGene00206460 | -1.180974267 | 0.55537885 | 1 | F56H9.9 |
| WBGene00206461 | -0.064323217 | 0.97003873 | 1 | Y47D3A.34 |
| WBGene00206462 | -2.938896125 | 0.1541307 | 1 | B0391.14 |
| WBGene00206464 | 0.178115923 | 0.94982316 | 1 | F59C6.18 |
| WBGene00206466 | -0.156269293 | 0.96924949 | 1 | T08D10.5 |

| | | | | |
|---|---|---|---|---|
| WBGene00206469 | -1.094201388 | 0.7816772 | 1 | F58D2.5 |
| WBGene00206474 | 0.983488853 | 0.80601565 | 1 | T22A3.12 |
| WBGene00206475 | -2.999197404 | 0.1537936 | 1 | F27D9.12 |
| WBGene00206478 | -1.964653846 | 0.53599202 | 1 | M6.11 |
| WBGene00206479 | -0.737462795 | 0.83433562 | 1 | C44C11.6 |
| WBGene00206482 | -0.412672102 | 0.65889146 | 0.80357275 | B0416.10 |
| WBGene00206483 | 1.216957619 | 0.74819621 | 1 | B0416.11 |
| WBGene00206484 | 0.186924804 | 0.84185504 | 0.91315662 | C31H2.14 |
| WBGene00206485 | 0.65729346 | 0.64310184 | 1 | C10A4.13 |
| WBGene00206487 | -0.134269767 | 0.7244772 | 0.84517459 | best-19 |
| WBGene00206488 | 0.172435098 | 0.84312942 | 0.91377278 | ZK757.10 |
| WBGene00206489 | -0.205794351 | 0.25550314 | 0.46299458 | F53F1.14 |
| WBGene00206490 | -0.156269293 | 0.96924949 | 1 | Y57G11C.1142 |
| WBGene00206492 | -0.22461868 | 0.93331196 | 1 | F11D11.21 |
| WBGene00206494 | -1.690101699 | 0.3296611 | 1 | C15H11.16 |
| WBGene00206497 | -1.094201388 | 0.7816772 | 1 | Y52B11A.18 |
| WBGene00206498 | -1.159714244 | 0.75790634 | 1 | C29A12.22 |
| WBGene00206500 | -2.603968543 | 0.24486328 | 1 | F55C9.15 |
| WBGene00206502 | -0.44901409 | 0.81158837 | 1 | T01B7.13 |
| WBGene00206503 | 0.377828795 | 0.92526585 | 1 | T26E4.18 |
| WBGene00206505 | 0.747845955 | 0.85215965 | 1 | B0250.18 |
| WBGene00206507 | -1.901856996 | 0.64072034 | 1 | Y51A2A.16 |
| WBGene00206511 | 0.281454817 | 0.94435481 | 1 | W02H5.16 |
| WBGene00206512 | 0.281454817 | 0.94435481 | 1 | Y39A3B.7 |
| WBGene00206513 | -1.307858406 | 0.24212013 | 1 | B0244.15 |
| WBGene00206514 | 1.581464096 | 0.2615982 | 1 | R05G9.5 |
| WBGene00206515 | 0.341462519 | 0.42907518 | 0.63174894 | T28C6.11 |
| WBGene00206516 | -0.460430338 | 0.1625408 | 0.34847392 | F53E4.2 |
| WBGene00206517 | -0.668264701 | 0.85551153 | 1 | T27F7.12 |
| WBGene00206519 | -2.406429869 | 0.0461779 | 1 | F36H5.13 |
| WBGene00206520 | -1.159714244 | 0.75790634 | 1 | ZK897.10 |
| WBGene00206521 | -0.123202138 | 0.94347085 | 1 | T02G5.15 |
| WBGene00206522 | 1.443007432 | 0.17333824 | 1 | F14D2.19 |
| WBGene00206523 | 0.281454817 | 0.94435481 | 1 | C54E10.10 |
| WBGene00206524 | 0.35828053 | 0.87111704 | 1 | C54E10.11 |
| WBGene00206525 | 0.609865109 | 0.21857002 | 0.42128252 | R74.10 |
| WBGene00206526 | -0.945145766 | 0.65193959 | 1 | C06C3.14 |
| WBGene00206527 | -0.333693239 | 0.89573179 | 1 | T27A8.7 |
| WBGene00206528 | -0.582567818 | 0.77294373 | 1 | Y53H1A.7 |
| WBGene00206529 | -1.58386945 | 0.28599203 | 1 | C04A2.15 |
| WBGene00206530 | 0.05905161 | 0.97811893 | 1 | Y52B11A.19 |
| WBGene00206531 | -1.969805064 | 0.62862569 | 1 | C29F7.11 |
| WBGene00206532 | 0.019566679 | 0.98678852 | 1 | F11A6.15 |

| | | | | |
|---|---|---|---|---|
| WBGene00206533 | 0.03940405 | 0.99001443 | 1 | C30A5.16 |
| WBGene00206535 | -0.156269293 | 0.96924949 | 1 | M162.15 |
| WBGene00206538 | 0.62920176 | 0.58680103 | 1 | C09G5.13 |
| WBGene00219205 | 0.281454817 | 0.94435481 | 1 | R03D7.15 |
| WBGene00219206 | 1.053703177 | 0.23798032 | 0.44423724 | W05H12.4 |
| WBGene00219207 | 0.11052445 | 0.8879258 | 0.94084004 | C01F6.14 |
| WBGene00219208 | -0.751854012 | 0.23284859 | 0.43851271 | B0365.9 |
| WBGene00219209 | 1.216957619 | 0.74819621 | 1 | F29F11.20 |
| WBGene00219210 | -0.003140285 | 0.99650322 | 0.9983122 | Y37H2A.18 |
| WBGene00219214 | 0.281454817 | 0.94435481 | 1 | T10B9.14 |
| WBGene00219216 | 2.714213268 | 0.00038804 | 0.00441167 | K08D10.14 |
| WBGene00219217 | -1.891828878 | 0.09938514 | 1 | C08F8.15 |
| WBGene00219218 | -0.102291047 | 0.87978831 | 0.93574569 | Y73B6BL.288 |
| WBGene00219223 | -0.156269293 | 0.96924949 | 1 | mir-4814 |
| WBGene00219224 | -0.156269293 | 0.96924949 | 1 | mir-4924 |
| WBGene00219225 | -0.242744234 | 0.759588 | 0.86544275 | mir-4927 |
| WBGene00219226 | -0.22461868 | 0.93331196 | 1 | mir-4929 |
| WBGene00219227 | 0.688793783 | 0.86378755 | 1 | mir-4932 |
| WBGene00219231 | -0.156269293 | 0.96924949 | 1 | mir-5545 |
| WBGene00219234 | -1.159714244 | 0.75790634 | 1 | B0511.17 |
| WBGene00219235 | -0.18557176 | 0.95664217 | 1 | K03B4.8 |
| WBGene00219236 | 3.539048531 | 0.04882878 | 1 | K03B4.9 |
| WBGene00219238 | -0.508813154 | 0.81506991 | 1 | M151.9 |
| WBGene00219239 | -0.737462795 | 0.83433562 | 1 | K05F6.14 |
| WBGene00219241 | 0.377828795 | 0.92526585 | 1 | Y116A8A.150 |
| WBGene00219242 | -0.156269293 | 0.96924949 | 1 | F56A12.10 |
| WBGene00219254 | 0.377828795 | 0.92526585 | 1 | mir-4920 |
| WBGene00219255 | 0.377828795 | 0.92526585 | 1 | mir-4921 |
| WBGene00219261 | -0.70576507 | 0.76841669 | 1 | mir-4926 |
| WBGene00219266 | -0.260421365 | 0.67577758 | 0.81451193 | mir-4938 |
| WBGene00219267 | 1.872540704 | 0.36210323 | 1 | mir-5548 |
| WBGene00219271 | 0.598430268 | 0.2545948 | 0.46224548 | F36H5.14 |
| WBGene00219274 | -1.279666666 | 0.55824702 | 1 | T25G12.13 |
| WBGene00219275 | -0.942708535 | 0.00735367 | 0.04118124 | Y39G10AR.32 |
| WBGene00219278 | 1.760973878 | 0.36171953 | 1 | R09A8.9 |
| WBGene00219281 | 1.879256964 | 0.46994626 | 1 | K10D3.8 |
| WBGene00219282 | -0.156269293 | 0.96924949 | 1 | F43G9.21 |
| WBGene00219283 | 0.595896719 | 0.79561342 | 1 | F55H12.7 |
| WBGene00219285 | 2.815759761 | 0.16833793 | 1 | K07A1.20 |
| WBGene00219286 | -1.901856996 | 0.64072034 | 1 | C04F12.16 |
| WBGene00219287 | -1.363291072 | 0.42036389 | 1 | F36D1.23 |
| WBGene00219288 | 0.516305167 | 0.70779836 | 1 | Y87G2A.25 |
| WBGene00219289 | 1.958518018 | 0.18596175 | 1 | T05C12.15 |

| | | | | |
|---|---|---|---|---|
| WBGene00219293 | -0.156269293 | 0.96924949 | 1 | B0334.20 |
| WBGene00219294 | -1.14715222 | 0.68932198 | 1 | F43G6.16 |
| WBGene00219295 | -0.107268465 | 0.96117914 | 1 | C34C12.12 |
| WBGene00219297 | -1.306212082 | 0.16077609 | 0.34618428 | T23F11.11 |
| WBGene00219298 | 0.406489191 | 0.89359366 | 1 | F55H2.8 |
| WBGene00219299 | -2.640414922 | 0.41081194 | 1 | C01G12.16 |
| WBGene00219300 | 1.270678414 | 0.5135606 | 1 | W10G6.6 |
| WBGene00219306 | -1.019515564 | 0.11266193 | 0.27499568 | linc-4 |
| WBGene00219308 | -0.241106143 | 0.91536132 | 1 | W06G6.20 |
| WBGene00219309 | 0.377828795 | 0.92526585 | 1 | F26F2.13 |
| WBGene00219310 | -1.371054553 | 0.36593004 | 1 | C17E4.19 |
| WBGene00219313 | -0.864957197 | 0.00016275 | 0.00223783 | C17E4.20 |
| WBGene00219315 | -1.481858699 | 0.40233002 | 1 | F07E5.12 |
| WBGene00219316 | 1.082519249 | 0.00084164 | 0.00803957 | F21A3.11 |
| WBGene00219317 | 0.970691698 | 0.52113796 | 1 | F31E3.12 |
| WBGene00219319 | 0.688793783 | 0.86378755 | 1 | H25P06.5 |
| WBGene00219322 | 0.377828795 | 0.92526585 | 1 | C33C12.12 |
| WBGene00219325 | 0.589569573 | 0.66560988 | 1 | F17C11.22 |
| WBGene00219326 | -0.156269293 | 0.96924949 | 1 | F36D3.16 |
| WBGene00219327 | -0.156269293 | 0.96924949 | 1 | ZK822.9 |
| WBGene00219329 | 0.059909351 | 0.95080196 | 1 | K02D10.8 |
| WBGene00219330 | -1.901856996 | 0.64072034 | 1 | F18E2.15 |
| WBGene00219337 | -0.323868354 | 0.91634405 | 1 | T16G1.18 |
| WBGene00219342 | 0.36729309 | 0.88005435 | 1 | Y57G11C.1143 |
| WBGene00219343 | 1.021504403 | 0.38851742 | 1 | Y17D7C.9 |
| WBGene00219351 | 1.38322681 | 0.69550305 | 1 | Y7A5A.20 |
| WBGene00219362 | 1.227309576 | 0.30258097 | 1 | ZK470.14 |
| WBGene00219368 | -0.737462795 | 0.83433562 | 1 | F14H12.17 |
| WBGene00219369 | 0.482064259 | 0.75369266 | 1 | R160.10 |
| WBGene00219371 | 0.721216606 | 0.70268022 | 1 | C03G5.14 |
| WBGene00219373 | 1.086665764 | 0.3078539 | 1 | F01E11.18 |
| WBGene00219374 | 0.415489119 | 0.62683417 | 0.78309735 | C25A11.1 |
| WBGene00219374 | 0.031576794 | 0.94326352 | 0.97068676 | C25A11.10 |
| WBGene00219375 | 1.177738717 | 0.13455537 | 0.30879311 | C02B8.12 |
| WBGene00219376 | 1.004208281 | 0.16043527 | 0.34573031 | C28G1.10 |
| WBGene00219383 | 0.688793783 | 0.86378755 | 1 | Y61B8A.6 |
| WBGene00219394 | -0.340107193 | 0.68909717 | 0.82299402 | Y48A6C.8 |
| WBGene00219399 | 0.14010803 | 0.84721419 | 0.91626826 | Y55F3AM.21 |
| WBGene00219410 | 0.710363081 | 0.61447059 | 1 | C04C3.14 |
| WBGene00219413 | 0.695474669 | 0.65279923 | 1 | Y38F2AL.12 |
| WBGene00219414 | 0.857402182 | 0.72847739 | 1 | F14H3.15 |
| WBGene00219415 | -0.245531902 | 0.94070824 | 1 | linc-38 |
| WBGene00219418 | 0.938982574 | 0.8146943 | 1 | F44D12.19 |

| | | | | |
|---|---|---|---|---|
| WBGene00219420 | -0.905475646 | 0.60168063 | 1 | T19C4.24 |
| WBGene00219421 | 0.281454817 | 0.94435481 | 1 | Y66D12A.30 |
| WBGene00219422 | -2.672346903 | 0.06167892 | 1 | Y52B11A.20 |
| WBGene00219423 | 1.036912952 | 0.29647452 | 0.50697066 | C23H5.15 |
| WBGene00219424 | -0.156269293 | 0.96924949 | 1 | Y54G2A.72 |
| WBGene00219425 | -0.12426394 | 0.75282351 | 0.86217294 | Y54G2A.73 |
| WBGene00219434 | -1.159714244 | 0.75790634 | 1 | F54D5.23 |
| WBGene00219436 | -2.402941201 | 0.10644621 | 1 | C49H3.16 |
| WBGene00219443 | -1.094201388 | 0.7816772 | 1 | Y57G11C.1144 |
| WBGene00219444 | 0.688793783 | 0.86378755 | 1 | Y41E3.461 |
| WBGene00219448 | 2.286231321 | 0.38099652 | 1 | K12G11.14 |
| WBGene00219449 | -0.156269293 | 0.96924949 | 1 | Y38A10A.11 |
| WBGene00219450 | -1.211422718 | 0.48838595 | 1 | F53B6.13 |
| WBGene00219451 | -0.204644772 | 0.56060391 | 0.73673213 | F52D2.12 |
| WBGene00219452 | -1.953894722 | 0.23291606 | 1 | F21D9.11 |
| WBGene00219453 | -0.156269293 | 0.96924949 | 1 | linc-97 |
| WBGene00219455 | 0.281454817 | 0.94435481 | 1 | Y43F8B.29 |
| WBGene00219456 | -0.156269293 | 0.96924949 | 1 | ZK822.10 |
| WBGene00219458 | 0.377828795 | 0.92526585 | 1 | H21P03.10 |
| WBGene00219476 | 0.377828795 | 0.92526585 | 1 | D1046.16 |
| WBGene00219478 | 0.688793783 | 0.86378755 | 1 | Y105C5B.1418 |
| WBGene00219489 | -3.014567441 | 0.24693748 | 1 | C10C6.13 |
| WBGene00219490 | 2.224351729 | 0.33187942 | 1 | Y116A8B.98 |
| WBGene00219491 | -0.996809519 | 0.40934178 | 1 | Y53F4B.62 |
| WBGene00219493 | -1.149221922 | 0.55229169 | 1 | C18D4.12 |
| WBGene00219545 | -0.156269293 | 0.96924949 | 1 | F41E7.19 |
| WBGene00219546 | -0.156269293 | 0.96924949 | 1 | F41E7.20 |
| WBGene00219548 | -0.325095864 | 0.82227663 | 1 | JC8.17 |
| WBGene00219550 | -0.098890353 | 0.90746155 | 0.95167062 | B0035.20 |
| WBGene00219551 | 0.281454817 | 0.94435481 | 1 | K07F5.21 |
| WBGene00219552 | -0.704048147 | 0.00141464 | 0.01200223 | F53F4.25 |
| WBGene00219555 | -0.156269293 | 0.96924949 | 1 | Y46G5A.46 |
| WBGene00219556 | 0.603275611 | 0.18213306 | 0.37384831 | C04C11.23 |
| WBGene00219557 | 0.355442863 | 0.44358104 | 0.64320045 | anr-14 |
| WBGene00219558 | -1.094201388 | 0.7816772 | 1 | anr-12 |
| WBGene00219559 | 0.603275611 | 0.18213306 | 0.37384831 | linc-5 |
| WBGene00219561 | -0.156269293 | 0.96924949 | 1 | linc-142 |
| WBGene00219562 | 0.281454817 | 0.94435481 | 1 | linc-28 |
| WBGene00219563 | 0.377828795 | 0.92526585 | 1 | linc-95 |
| WBGene00219564 | -0.156269293 | 0.96924949 | 1 | linc-102 |
| WBGene00219565 | 0.688793783 | 0.86378755 | 1 | linc-120 |
| WBGene00219566 | -0.156269293 | 0.96924949 | 1 | linc-33 |
| WBGene00219568 | -0.089096525 | 0.84609284 | 0.91565737 | anr-9 |

| | | | | |
|---|---|---|---|---|
| WBGene00219570 | 1.216957619 | 0.74819621 | 1 | linc-82 |
| WBGene00219571 | 0.948096129 | 0.29947292 | 0.50916222 | linc-15 |
| WBGene00219573 | 1.687239501 | 0.59403242 | 1 | anr-30 |
| WBGene00219574 | 0.281454817 | 0.94435481 | 1 | linc-54 |
| WBGene00219575 | -0.801021281 | 0.13223287 | 0.30481857 | anr-8 |
| WBGene00219576 | 1.105346709 | 0.0656803 | 0.19353487 | anr-13 |
| WBGene00219577 | 1.830588955 | 0.20153364 | 1 | linc-119 |
| WBGene00219579 | 0.041803775 | 0.98890197 | 1 | linc-43 |
| WBGene00219581 | 0.21954033 | 0.63007345 | 0.78547303 | anr-20 |
| WBGene00219582 | 0.984236303 | 0.53436178 | 1 | anr-25 |
| WBGene00219583 | 0.377828795 | 0.92526585 | 1 | anr-43 |
| WBGene00219585 | -1.901856996 | 0.64072034 | 1 | linc-170 |
| WBGene00219586 | -0.156269293 | 0.96924949 | 1 | linc-108 |
| WBGene00219587 | 0.102324312 | 0.89751229 | 0.94634211 | anr-16 |
| WBGene00219589 | 0.616120425 | 0.64977769 | 1 | F37H8.7 |
| WBGene00219589 | 0.918699486 | 0.18033623 | 0.3725955 | anr-35 |
| WBGene00219590 | 0.281454817 | 0.94435481 | 1 | linc-72 |
| WBGene00219591 | 0.39748245 | 0.82716117 | 1 | linc-149 |
| WBGene00219592 | 1.435925494 | 0.70920779 | 1 | linc-84 |
| WBGene00219593 | 0.187432756 | 0.74904105 | 0.86012555 | anr-15 |
| WBGene00219597 | 0.668076624 | 0.16166904 | 0.34763885 | anr-11 |
| WBGene00219598 | 0.688793783 | 0.86378755 | 1 | linc-113 |
| WBGene00219602 | 0.983488853 | 0.80601565 | 1 | linc-151 |
| WBGene00219605 | 0.747845955 | 0.85215965 | 1 | linc-110 |
| WBGene00219606 | 1.216957619 | 0.74819621 | 1 | linc-133 |
| WBGene00219607 | -0.337262811 | 0.29087732 | 0.50092788 | anr-7 |
| WBGene00219608 | 0.747845955 | 0.85215965 | 1 | linc-58 |
| WBGene00219609 | -0.607158097 | 0.181898 | 0.37384831 | linc-7 |
| WBGene00219610 | 2.457560503 | 0.08086436 | 1 | anr-51 |
| WBGene00219611 | -0.156269293 | 0.96924949 | 1 | linc-123 |
| WBGene00219615 | -0.298265327 | 0.89527345 | 1 | linc-109 |
| WBGene00219616 | 1.139408087 | 0.59580649 | 1 | linc-111 |
| WBGene00219617 | 1.21641543 | 0.37154615 | 1 | linc-25 |
| WBGene00219618 | -0.008677305 | 0.99719169 | 1 | linc-60 |
| WBGene00219619 | -0.29656835 | 0.65862236 | 0.8035709 | linc-76 |
| WBGene00219620 | -0.62146648 | 0.85925752 | 1 | linc-74 |
| WBGene00219621 | -1.334801939 | 0.30614053 | 1 | anr-46 |
| WBGene00219622 | -0.582567818 | 0.77294373 | 1 | anr-40 |
| WBGene00219623 | -0.072065694 | 0.79939439 | 0.88979641 | anr-2 |
| WBGene00219627 | -0.832032324 | 4.3315E-05 | 0.00076884 | anr-3 |
| WBGene00219629 | 1.804609989 | 0.55694839 | 1 | linc-87 |
| WBGene00219631 | 0.056106392 | 0.97371196 | 1 | anr-37 |
| WBGene00219634 | 1.876192691 | 0.43542138 | 1 | linc-86 |

| | | | | |
|---|---|---|---|---|
| WBGene00219635 | -1.624083904 | 0.31010041 | 1 | linc-51 |
| WBGene00219636 | -1.166445188 | 0.44893799 | 1 | linc-81 |
| WBGene00219637 | -0.263229581 | 0.93409872 | 1 | linc-124 |
| WBGene00219638 | 0.183128715 | 0.88635784 | 1 | anr-29 |
| WBGene00219639 | 0.041803775 | 0.98890197 | 1 | linc-115 |
| WBGene00219640 | 0.345208515 | 0.89272401 | 1 | anr-56 |
| WBGene00219641 | -0.156269293 | 0.96924949 | 1 | linc-47 |
| WBGene00219642 | -2.028651717 | 0.30854162 | 1 | anr-52 |
| WBGene00219643 | 0.281454817 | 0.94435481 | 1 | linc-78 |
| WBGene00219644 | 1.957271014 | 0.49743227 | 1 | linc-69 |
| WBGene00219647 | -1.781962339 | 0.1527251 | 1 | anr-27 |
| WBGene00219648 | 0.075260937 | 0.97569838 | 1 | linc-67 |
| WBGene00219649 | 0.232157325 | 0.93397866 | 1 | linc-55 |
| WBGene00219650 | 2.052072606 | 0.26770792 | 1 | linc-56 |
| WBGene00219651 | 0.377828795 | 0.92526585 | 1 | linc-130 |
| WBGene00219654 | 0.281454817 | 0.94435481 | 1 | linc-156 |
| WBGene00219655 | 2.22199375 | 0.22008611 | 1 | linc-107 |
| WBGene00219656 | -0.156269293 | 0.96924949 | 1 | anr-48 |
| WBGene00219657 | -0.323868354 | 0.91634405 | 1 | linc-139 |
| WBGene00219658 | 0.747845955 | 0.85215965 | 1 | anr-21 |
| WBGene00219659 | 2.000425015 | 0.52698112 | 1 | anr-36 |
| WBGene00219660 | 0.688793783 | 0.86378755 | 1 | linc-19 |
| WBGene00219661 | 0.270496667 | 0.92869767 | 1 | linc-45 |
| WBGene00219663 | -1.82683067 | 0.0560046 | 0.17416799 | anr-10 |
| WBGene00219664 | 1.017496938 | 0.63763677 | 1 | linc-73 |
| WBGene00219665 | 1.893863219 | 0.30802959 | 1 | linc-101 |
| WBGene00219666 | -1.14715222 | 0.68932198 | 1 | linc-31 |
| WBGene00219667 | 0.377828795 | 0.92526585 | 1 | linc-66 |
| WBGene00219668 | 2.085660205 | 0.10569777 | 1 | anr-33 |
| WBGene00219669 | 1.33864768 | 0.5127736 | 1 | anr-28 |
| WBGene00219671 | 1.413734769 | 0.68804531 | 1 | anr-4 |
| WBGene00219673 | 3.025878362 | 0.05568571 | 1 | linc-96 |
| WBGene00219675 | 1.189071466 | 0.5668301 | 1 | linc-44 |
| WBGene00219676 | 0.377828795 | 0.92526585 | 1 | linc-48 |
| WBGene00219677 | 0.09697944 | 0.97612634 | 1 | anr-17 |
| WBGene00219678 | -0.564206844 | 0.29947512 | 0.50916222 | linc-9 |
| WBGene00219679 | -0.492757517 | 0.3256373 | 0.53404869 | anr-22 |
| WBGene00219680 | 0.152641465 | 0.89571181 | 1 | linc-35 |
| WBGene00219681 | -0.564460888 | 0.21959852 | 0.42246009 | anr-6 |
| WBGene00219682 | -2.061536526 | 0.21538967 | 1 | linc-50 |
| WBGene00219683 | 1.199045682 | 0.13183711 | 0.3043031 | anr-32 |
| WBGene00219685 | -0.156269293 | 0.96924949 | 1 | linc-75 |
| WBGene00219686 | 0.688793783 | 0.86378755 | 1 | linc-125 |

| | | | | |
|---|---|---|---|---|
| WBGene00219687 | -1.271222092 | 0.45460777 | 1 | linc-12 |
| WBGene00219688 | 0.290491123 | 0.88464841 | 1 | linc-21 |
| WBGene00219689 | 0.647295905 | 0.73170538 | 1 | linc-41 |
| WBGene00219690 | -1.159714244 | 0.75790634 | 1 | linc-10 |
| WBGene00219691 | 0.688793783 | 0.86378755 | 1 | linc-106 |
| WBGene00219692 | 0.377828795 | 0.92526585 | 1 | linc-23 |
| WBGene00219695 | -0.156269293 | 0.96924949 | 1 | linc-126 |
| WBGene00219696 | 1.861184989 | 0.02379616 | 0.09691687 | anr-39 |
| WBGene00219697 | -0.077222385 | 0.8988828 | 0.94694691 | linc-169 |
| WBGene00219698 | -1.14715222 | 0.68932198 | 1 | linc-92 |
| WBGene00219699 | 1.011503636 | 0.01416316 | 0.06680455 | linc-22 |
| WBGene00219700 | -1.562631987 | 0.04374671 | 0.14608272 | linc-6 |
| WBGene00219701 | -0.34732664 | 0.24586717 | 0.45379649 | anr-34 |
| WBGene00219702 | 0.911798121 | 0.68104294 | 1 | linc-138 |
| WBGene00219703 | -0.028515692 | 0.97960487 | 1 | anr-38 |
| WBGene00219705 | -1.901856996 | 0.64072034 | 1 | anr-5 |
| WBGene00219707 | 0.53548718 | 0.29279659 | 0.50294898 | linc-8 |
| WBGene00219708 | -0.828765424 | 0.69034289 | 1 | linc-13 |
| WBGene00219709 | 2.256994583 | 0.19548273 | 1 | linc-37 |
| WBGene00219710 | 0.245040059 | 0.58410569 | 0.7540432 | anr-45 |
| WBGene00219711 | -0.586192638 | 0.64069331 | 1 | anr-49 |
| WBGene00219713 | 0.281454817 | 0.94435481 | 1 | linc-27 |
| WBGene00219715 | -0.26870002 | 0.45968241 | 0.65558606 | anr-18 |
| WBGene00219717 | 2.012909791 | 0.30957141 | 1 | anr-23 |
| WBGene00219718 | 0.983488853 | 0.80601565 | 1 | linc-42 |
| WBGene00219719 | -1.14715222 | 0.68932198 | 1 | linc-57 |
| WBGene00219720 | -0.418943634 | 0.6328915 | 0.7872679 | linc-2 |
| WBGene00219721 | 0.281454817 | 0.94435481 | 1 | linc-104 |
| WBGene00219723 | 0.688793783 | 0.86378755 | 1 | linc-135 |
| WBGene00219725 | -0.141081256 | 0.93013309 | 1 | linc-63 |
| WBGene00219726 | 2.286068456 | 0.34948738 | 1 | linc-116 |
| WBGene00219727 | -2.343707959 | 0.25817937 | 1 | linc-39 |
| WBGene00219728 | 0.095742963 | 0.94334723 | 1 | linc-16 |
| WBGene00219729 | 0.409633032 | 0.45283771 | 0.6504854 | anr-44 |
| WBGene00219731 | 1.360555046 | 0.51491783 | 1 | linc-49 |
| WBGene00219732 | 0.574508983 | 0.8551971 | 1 | linc-112 |
| WBGene00219733 | -1.544131969 | 0.51019995 | 1 | linc-53 |
| WBGene00219735 | -0.156269293 | 0.96924949 | 1 | linc-150 |
| WBGene00219736 | 0.949755679 | 0.29863274 | 0.50851248 | linc-20 |
| WBGene00219737 | -0.156269293 | 0.96924949 | 1 | linc-155 |
| WBGene00219738 | -0.507211835 | 0.03300414 | 0.12124154 | anr-1 |
| WBGene00219741 | -1.269943417 | 0.28550331 | 1 | linc-77 |
| WBGene00219743 | -0.323868354 | 0.91634405 | 1 | linc-103 |

| | | | | |
|---|---|---|---|---|
| WBGene00219744 | -0.986612197 | 0.56727953 | 1 | anr-41 |
| WBGene00219746 | -1.186713419 | 0.42057049 | 1 | linc-91 |
| WBGene00219747 | -1.969805064 | 0.62862569 | 1 | linc-79 |
| WBGene00219748 | 1.790107971 | 0.17897495 | 1 | linc-46 |
| WBGene00219749 | 0.109173684 | 0.96057644 | 1 | linc-30 |
| WBGene00219750 | 0.574508983 | 0.8551971 | 1 | Y38C1AA.19 |
| WBGene00219751 | -0.788963918 | 0.27823797 | 0.48722084 | anr-24 |
| WBGene00219754 | 1.091339395 | 0.68134922 | 1 | linc-85 |
| WBGene00219755 | 1.598671039 | 0.32913499 | 1 | linc-17 |
| WBGene00219756 | -2.647479787 | 0.14205355 | 1 | linc-68 |
| WBGene00219757 | 0.688793783 | 0.86378755 | 1 | linc-71 |
| WBGene00219758 | 0.215806043 | 0.7566595 | 0.86407788 | linc-11 |
| WBGene00219759 | -0.694035084 | 0.65988032 | 1 | anr-58 |
| WBGene00219760 | 0.03808027 | 0.96846633 | 0.98386325 | anr-19 |
| WBGene00219764 | 0.985625912 | 0.62338668 | 1 | anr-47 |
| WBGene00219765 | -1.901856996 | 0.64072034 | 1 | linc-165 |
| WBGene00219766 | -0.156269293 | 0.96924949 | 1 | anr-42 |
| WBGene00219767 | -1.901856996 | 0.64072034 | 1 | linc-131 |
| WBGene00219769 | 3.203359461 | 0.12311708 | 1 | linc-64 |
| WBGene00219770 | -1.853938203 | 0.25386854 | 1 | linc-40 |
| WBGene00219771 | 0.452150161 | 0.74451196 | 1 | linc-24 |
| WBGene00219772 | 0.159950863 | 0.93307234 | 1 | anr-53 |
| WBGene00219773 | 0.281454817 | 0.94435481 | 1 | linc-32 |
| WBGene00219775 | 0.281454817 | 0.94435481 | 1 | F33A8.13 |
| WBGene00219776 | -0.758706584 | 0.71760618 | 1 | AH9.9 |
| WBGene00219778 | 2.097891464 | 0.04376679 | 0.14608272 | B0025.7 |
| WBGene00219779 | -0.245531902 | 0.94070824 | 1 | B0025.8 |
| WBGene00219780 | 3.839097933 | 0.00721452 | 1 | B0228.10 |
| WBGene00219782 | 0.281454817 | 0.94435481 | 1 | B0250.19 |
| WBGene00219784 | 2.188838843 | 0.34004625 | 1 | B0348.8 |
| WBGene00219785 | -0.156269293 | 0.96924949 | 1 | B0350.85 |
| WBGene00219789 | 1.070632877 | 0.13456869 | 0.30879311 | B0511.18 |
| WBGene00219795 | 3.142876526 | 0.13529041 | 1 | C02F5.15 |
| WBGene00219796 | -2.420379867 | 0.00167223 | 0.01358031 | C03D6.9 |
| WBGene00219800 | -6.855123575 | 2.91E-07 | 1.1732E-05 | C05D9.12 |
| WBGene00219803 | 0.206507248 | 0.61523198 | 0.77528743 | C06A1.12 |
| WBGene00219804 | -1.964653846 | 0.53599202 | 1 | C06E7.95 |
| WBGene00219805 | 0.377828795 | 0.92526585 | 1 | C07A12.19 |
| WBGene00219806 | 0.232157325 | 0.93397866 | 1 | C07E3.17 |
| WBGene00219808 | -0.482142867 | 0.32958367 | 0.5381944 | C09H10.19 |
| WBGene00219812 | 0.68838769 | 0.37348675 | 0.5812823 | C13C12.7 |
| WBGene00219813 | -0.156269293 | 0.96924949 | 1 | C14A11.13 |
| WBGene00219816 | -2.369498867 | 0.0132893 | 0.0637904 | C14E2.12 |

| | | | | |
|---|---|---|---|---|
| WBGene00219817 | -1.14715222 | 0.68932198 | 1 | C15A11.10 |
| WBGene00219819 | 1.564220053 | 0.2274603 | 1 | C16D2.4 |
| WBGene00219821 | 1.369632097 | 0.398355 | 1 | C17D12.18 |
| WBGene00219826 | 0.645908258 | 0.38023312 | 0.58822762 | C18H9.12 |
| WBGene00219827 | -2.640414922 | 0.41081194 | 1 | C23G10.12 |
| WBGene00219830 | 0.574508983 | 0.8551971 | 1 | C24G7.6 |
| WBGene00219831 | 1.356822 | 0.33758376 | 1 | C25F6.30 |
| WBGene00219837 | -0.156269293 | 0.96924949 | 1 | C28D4.17 |
| WBGene00219838 | -0.082974327 | 0.80231439 | 0.89169709 | C28H8.14 |
| WBGene00219848 | 0.281454817 | 0.94435481 | 1 | C34F6.18 |
| WBGene00219849 | -0.953176121 | 0.31198914 | 1 | C36C9.10 |
| WBGene00219851 | 0.377828795 | 0.92526585 | 1 | C37C3.16 |
| WBGene00219853 | 0.237660155 | 0.56044378 | 0.73673213 | C41C4.17 |
| WBGene00219854 | 1.151679082 | 0.0170286 | 0.07617696 | C41C4.18 |
| WBGene00219855 | 1.029907327 | 0.01382649 | 0.06559882 | C41C4.19 |
| WBGene00219856 | 1.575778407 | 0.24784166 | 1 | C41C4.20 |
| WBGene00219857 | 0.377828795 | 0.92526585 | 1 | C45E1.10 |
| WBGene00219858 | 0.938982574 | 0.8146943 | 1 | C46A5.15 |
| WBGene00219859 | 0.064283543 | 0.9370358 | 0.9673116 | C49F5.11 |
| WBGene00219860 | 1.589720631 | 0.04265887 | 0.14385895 | C51F7.4 |
| WBGene00219865 | -1.094201388 | 0.7816772 | 1 | C56G2.21 |
| WBGene00219866 | 1.506473404 | 0.02522715 | 0.10078409 | D1054.23 |
| WBGene00219867 | 0.262874229 | 0.67368798 | 0.8134506 | D2013.13 |
| WBGene00219870 | 0.577852467 | 0.63501206 | 1 | D2092.15 |
| WBGene00219871 | 0.688793783 | 0.86378755 | 1 | E01A2.11 |
| WBGene00219874 | -1.45873709 | 0.59584643 | 1 | F01D4.12 |
| WBGene00219875 | 1.495908959 | 0.36811095 | 1 | F01E11.19 |
| WBGene00219876 | -2.640414922 | 0.41081194 | 1 | F01G12.19 |
| WBGene00219880 | 1.38322681 | 0.69550305 | 1 | F07C4.17 |
| WBGene00219882 | -0.156269293 | 0.96924949 | 1 | F08B6.7 |
| WBGene00219885 | 2.139299219 | 0.2405795 | 1 | F09E10.14 |
| WBGene00219886 | -0.721694164 | 0.16463702 | 0.35094667 | F09E10.15 |
| WBGene00219888 | -1.901856996 | 0.64072034 | 1 | F09G2.14 |
| WBGene00219895 | -0.156269293 | 0.96924949 | 1 | F16A11.13 |
| WBGene00219896 | 2.188838843 | 0.34004625 | 1 | F16H9.5 |
| WBGene00219897 | 2.301687108 | 0.37916956 | 1 | F16H9.6 |
| WBGene00219900 | 0.89844217 | 0.00208211 | 0.01604426 | F17C11.25 |
| WBGene00219902 | -1.969805064 | 0.62862569 | 1 | F18C5.15 |
| WBGene00219904 | 0.558642347 | 0.36726768 | 0.57476113 | F19C6.12 |
| WBGene00219905 | -1.969805064 | 0.62862569 | 1 | F19F10.13 |
| WBGene00219907 | -0.156269293 | 0.96924949 | 1 | F19H8.7 |
| WBGene00219908 | 0.274891578 | 0.76051915 | 0.86625978 | F20C5.10 |
| WBGene00219909 | -2.173611897 | 0.04477968 | 1 | F20D12.10 |

| | | | | |
|---|---|---|---|---|
| WBGene00219910 | -0.505605194 | 0.80388525 | 1 | F21F12.1 |
| WBGene00219911 | -0.156269293 | 0.96924949 | 1 | F21F12.3 |
| WBGene00219912 | -1.172100593 | 0.64543428 | 1 | F22H10.10 |
| WBGene00219915 | -0.156269293 | 0.96924949 | 1 | F26A10.28 |
| WBGene00219917 | 0.424613274 | 0.75521751 | 1 | F26D10.24 |
| WBGene00219918 | 0.552716261 | 0.66861328 | 1 | F27C1.18 |
| WBGene00219921 | 0.983488853 | 0.80601565 | 1 | F28B1.13 |
| WBGene00219925 | 1.460233832 | 0.11114474 | 0.2728654 | F29G9.19 |
| WBGene00219927 | 0.204916933 | 0.75480185 | 0.86283239 | F31C3.17 |
| WBGene00219932 | 0.747845955 | 0.85215965 | 1 | F33E11.7 |
| WBGene00219940 | 0.281454817 | 0.94435481 | 1 | F39B2.14 |
| WBGene00219941 | 0.281454817 | 0.94435481 | 1 | F40F11.6 |
| WBGene00219944 | 0.356567911 | 0.56112858 | 0.73698205 | F41E6.18 |
| WBGene00219946 | 0.766826755 | 0.56555077 | 1 | F42H10.17 |
| WBGene00219948 | -0.622581789 | 0.01194402 | 0.05909224 | F44E2.12 |
| WBGene00219949 | -1.953894722 | 0.23291606 | 1 | F44E5.16 |
| WBGene00219950 | 2.125426916 | 0.01448328 | 0.06799552 | F46A8.13 |
| WBGene00219951 | 1.557897343 | 0.63813005 | 1 | F46A9.12 |
| WBGene00219954 | 1.996315266 | 0.49147447 | 1 | F47E1.19 |
| WBGene00219955 | -0.156269293 | 0.96924949 | 1 | F48A9.6 |
| WBGene00219959 | -1.458235063 | 0.07185912 | 0.20615348 | F52D2.14 |
| WBGene00219960 | -1.964653846 | 0.53599202 | 1 | F52D10.13 |
| WBGene00219961 | -0.737462795 | 0.83433562 | 1 | F52E1.17 |
| WBGene00219964 | 1.782422993 | 0.00171792 | 0.01382191 | F52G3.6 |
| WBGene00219966 | -0.323868354 | 0.91634405 | 1 | F54E2.7 |
| WBGene00219968 | -0.156269293 | 0.96924949 | 1 | F54F2.13 |
| WBGene00219969 | -1.901856996 | 0.64072034 | 1 | F54F11.6 |
| WBGene00219971 | -1.831477136 | 0.52194793 | 1 | F54G2.4 |
| WBGene00219972 | 1.453607818 | 0.45207066 | 1 | F55A3.8 |
| WBGene00219973 | -0.020065921 | 0.99148387 | 1 | F55C7.9 |
| WBGene00219976 | 2.31848281 | 0.18103992 | 1 | F55H12.9 |
| WBGene00219983 | 1.296019512 | 0.12837421 | 0.29912046 | F58D5.11 |
| WBGene00219984 | -0.06046764 | 0.97825857 | 1 | F58D5.12 |
| WBGene00219985 | -0.317186159 | 0.71625767 | 0.84006931 | F58D5.13 |
| WBGene00219987 | 0.281454817 | 0.94435481 | 1 | F59C6.19 |
| WBGene00219989 | -0.241244855 | 0.66867206 | 0.81015635 | H06I04.14 |
| WBGene00219991 | 0.158957087 | 0.69413911 | 0.8264466 | H21P03.11 |
| WBGene00219996 | 0.528331213 | 0.21059808 | 0.41041347 | H40L08.6 |
| WBGene00219997 | 0.784626933 | 0.65760117 | 1 | K02A6.6 |
| WBGene00219999 | 0.751976687 | 0.45055503 | 0.64897243 | K02B9.10 |
| WBGene00220000 | 1.657787544 | 0.43418652 | 1 | K02F2.11 |
| WBGene00220003 | 0.383850599 | 0.81289467 | 1 | K02G10.16 |
| WBGene00220004 | -0.635014024 | 0.58042814 | 1 | K03E5.9 |

| | | | | |
|---|---|---|---|---|
| WBGene00220005 | 0.48089161 | 0.08203294 | 0.22451803 | K04D7.17 |
| WBGene00220006 | 1.475390604 | 0.27234573 | 1 | K04F10.9 |
| WBGene00220007 | -0.06995528 | 0.97031639 | 1 | K06A4.15 |
| WBGene00220008 | -0.668264701 | 0.85551153 | 1 | K07C5.18 |
| WBGene00220010 | -0.213257914 | 0.46789947 | 0.66186269 | K07G5.7 |
| WBGene00220011 | -1.964653846 | 0.53599202 | 1 | K08B5.8 |
| WBGene00220015 | 0.271471552 | 0.40964595 | 0.61507125 | K09H9.11 |
| WBGene00220016 | 0.261430075 | 0.493047 | 0.68200961 | K10C3.13 |
| WBGene00220017 | -3.752061783 | 0.09825449 | 1 | K10D2.10 |
| WBGene00220018 | 1.025181888 | 0.79790347 | 1 | K10D3.9 |
| WBGene00220019 | 0.432433061 | 0.19751759 | 0.39487257 | K10D3.10 |
| WBGene00220020 | 0.574508983 | 0.8551971 | 1 | K10F12.9 |
| WBGene00220022 | 0.764887794 | 0.2903182 | 0.50031189 | K10G6.11 |
| WBGene00220023 | 1.159087254 | 0.12715754 | 0.29763248 | K11C4.14 |
| WBGene00220025 | 1.685698156 | 0.39072947 | 1 | K11E8.7 |
| WBGene00220026 | -1.964653846 | 0.53599202 | 1 | M01D1.11 |
| WBGene00220027 | 1.25191692 | 0.00583585 | 0.03486866 | M01D7.11 |
| WBGene00220029 | 0.281454817 | 0.94435481 | 1 | M03D4.78 |
| WBGene00220030 | -0.156269293 | 0.96924949 | 1 | M03F4.13 |
| WBGene00220032 | -1.159714244 | 0.75790634 | 1 | M7.15 |
| WBGene00220034 | -1.969805064 | 0.62862569 | 1 | M57.3 |
| WBGene00220035 | 0.938982574 | 0.8146943 | 1 | M60.15 |
| WBGene00220036 | 0.718092524 | 0.08075499 | 0.22222266 | R07B7.20 |
| WBGene00220039 | 0.377828795 | 0.92526585 | 1 | R10A10.3 |
| WBGene00220040 | 0.810033874 | 0.05192107 | 0.16458845 | R10E11.13 |
| WBGene00220041 | -0.156269293 | 0.96924949 | 1 | R11A5.8 |
| WBGene00220042 | 1.712479368 | 0.58529868 | 1 | R11E3.18 |
| WBGene00220046 | 1.216957619 | 0.74819621 | 1 | R31.7 |
| WBGene00220047 | -0.156269293 | 0.96924949 | 1 | R74.11 |
| WBGene00220048 | 0.295934834 | 0.12759111 | 0.29825012 | R106.6 |
| WBGene00220050 | 0.735468004 | 0.01650239 | 0.07432888 | R144.18 |
| WBGene00220053 | 0.574508983 | 0.8551971 | 1 | R166.11 |
| WBGene00220062 | 1.453161878 | 0.15933589 | 1 | T08D2.11 |
| WBGene00220067 | 1.539971811 | 0.02745636 | 0.10670115 | T12B3.10 |
| WBGene00220074 | -1.901856996 | 0.64072034 | 1 | T20B12.12 |
| WBGene00220075 | -0.156269293 | 0.96924949 | 1 | T21B4.20 |
| WBGene00220076 | -0.47342901 | 0.38496308 | 0.59342773 | T21B4.21 |
| WBGene00220077 | 0.728379161 | 0.56554126 | 1 | T21C12.12 |
| WBGene00220081 | 1.653460415 | 0.50712258 | 1 | T22C1.17 |
| WBGene00220083 | -0.156269293 | 0.96924949 | 1 | T23B12.13 |
| WBGene00220084 | 0.673697881 | 0.73339905 | 1 | T23B12.14 |
| WBGene00220085 | -0.1228129 | 0.85122422 | 0.91864118 | T23D8.11 |
| WBGene00220086 | 0.010545425 | 0.99456833 | 1 | T23D8.12 |

| | | | | |
|---|---|---|---|---|
| WBGene00220087 | 0.944845424 | 0.49587975 | 1 | T23D8.13 |
| WBGene00220089 | -0.562866186 | 0.30026372 | 0.50995852 | T23G11.12 |
| WBGene00220090 | 1.226152345 | 0.48225188 | 1 | T24B8.22 |
| WBGene00220093 | 1.643145219 | 0.01537866 | 0.07093933 | T25C12.11 |
| WBGene00220094 | 2.010778195 | 0.00019533 | 0.00259745 | T25C12.12 |
| WBGene00220105 | -1.964653846 | 0.53599202 | 1 | T27E9.11 |
| WBGene00220109 | -0.156269293 | 0.96924949 | 1 | W01F3.4 |
| WBGene00220110 | -0.156269293 | 0.96924949 | 1 | W01F3.5 |
| WBGene00220114 | -0.163726216 | 0.81130554 | 0.89692775 | W02H5.17 |
| WBGene00220115 | -0.180615086 | 0.75714727 | 0.86429398 | W02H5.18 |
| WBGene00220116 | -0.838211756 | 0.7584606 | 1 | W03A5.8 |
| WBGene00220118 | 0.574301796 | 0.58551121 | 1 | W03D8.13 |
| WBGene00220121 | -0.156269293 | 0.96924949 | 1 | W06A7.12 |
| WBGene00220123 | 1.181686151 | 0.7579682 | 1 | W06H8.16 |
| WBGene00220126 | -1.969805064 | 0.62862569 | 1 | W10C8.17 |
| WBGene00220127 | -1.844800047 | 0.33982718 | 1 | W10D5.5 |
| WBGene00220129 | 0.747845955 | 0.85215965 | 1 | Y10G11A.91 |
| WBGene00220131 | 0.616237057 | 0.21352442 | 0.41425375 | Y11D7A.21 |
| WBGene00220133 | -1.14715222 | 0.68932198 | 1 | Y14H12A.4 |
| WBGene00220135 | 1.181686151 | 0.7579682 | 1 | Y18D10A.29 |
| WBGene00220137 | 0.852647128 | 0.2693356 | 0.47783631 | Y24D9A.13 |
| WBGene00220140 | 1.270052625 | 0.58538191 | 1 | Y34F4.7 |
| WBGene00220141 | 0.101259364 | 0.94240208 | 1 | Y37E11AR.9 |
| WBGene00220142 | 1.712479368 | 0.58529868 | 1 | Y37F4.15 |
| WBGene00220147 | -1.969805064 | 0.62862569 | 1 | Y40B1A.6 |
| WBGene00220152 | -0.481441553 | 0.22097874 | 0.42397327 | Y41D4B.33 |
| WBGene00220153 | 0.177191879 | 0.64580308 | 0.79536368 | Y41D4B.34 |
| WBGene00220160 | 0.983488853 | 0.80601565 | 1 | Y44A6C.4 |
| WBGene00220161 | 0.137343829 | 0.71688172 | 0.84010542 | Y46E12BL.11 |
| WBGene00220162 | -0.645404887 | 0.16511137 | 0.35168723 | Y46E12BM.1 |
| WBGene00220163 | -0.023437859 | 0.97179874 | 0.98541854 | Y47D3A.35 |
| WBGene00220165 | -0.407256599 | 0.62407187 | 0.78128032 | Y47G6A.36 |
| WBGene00220167 | 0.574508983 | 0.8551971 | 1 | Y47G6A.38 |
| WBGene00220168 | -1.45873709 | 0.59584643 | 1 | Y48A6C.10 |
| WBGene00220169 | -0.006107107 | 0.98959776 | 0.99484196 | Y48B6A.25 |
| WBGene00220170 | -0.156269293 | 0.96924949 | 1 | Y48C3A.25 |
| WBGene00220171 | 0.58551505 | 0.30969096 | 0.51818605 | Y48G10A.9 |
| WBGene00220177 | -0.668264701 | 0.85551153 | 1 | Y51H7BM.2 |
| WBGene00220178 | -0.18909628 | 0.72790764 | 0.84748051 | Y52D3.2 |
| WBGene00220179 | -0.156269293 | 0.96924949 | 1 | Y54E10A.29 |
| WBGene00220180 | 1.225658034 | 0.16719878 | 0.35468616 | Y54G2A.74 |
| WBGene00220181 | 0.102470149 | 0.93723251 | 1 | Y55F3AM.23 |
| WBGene00220182 | -0.558491995 | 0.18916977 | 0.38310429 | Y55F3BL.5 |

| | | | | |
|---|---|---|---|---|
| WBGene00220184 | -0.700240526 | 0.3515942 | 0.56035591 | Y57A10A.44 |
| WBGene00220188 | 0.57991002 | 0.82402153 | 1 | Y65B4A.10 |
| WBGene00220189 | -0.688776386 | 0.20927908 | 0.4094803 | Y65B4A.11 |
| WBGene00220190 | -0.57047147 | 0.62600601 | 1 | Y65B4BM.2 |
| WBGene00220193 | 0.574508983 | 0.8551971 | 1 | Y67D2.11 |
| WBGene00220197 | 1.851444818 | 0.53669729 | 1 | Y71G12B.41 |
| WBGene00220198 | 0.333642403 | 0.85353086 | 1 | Y71G12B.42 |
| WBGene00220199 | 0.334730381 | 0.38162202 | 0.58977507 | Y71H2AM.28 |
| WBGene00220202 | 0.689176269 | 0.44670701 | 0.6457926 | Y73B3A.31 |
| WBGene00220203 | -0.156269293 | 0.96924949 | 1 | Y73B6A.14 |
| WBGene00220207 | 0.680743233 | 0.76740144 | 1 | Y75B8A.62 |
| WBGene00220208 | 0.627834078 | 0.77998941 | 1 | Y75B12B.16 |
| WBGene00220211 | -0.294169447 | 0.51160586 | 0.69811749 | Y76B12C.11 |
| WBGene00220215 | 1.20069927 | 0.61903874 | 1 | Y92C3B.4 |
| WBGene00220216 | -0.961438649 | 0.49375541 | 1 | Y94H6A.15 |
| WBGene00220217 | -1.901856996 | 0.64072034 | 1 | Y95B8A.18 |
| WBGene00220222 | -0.156269293 | 0.96924949 | 1 | Y105C5A.1283 |
| WBGene00220225 | -0.729807562 | 0.62291165 | 1 | Y105E8A.50 |
| WBGene00220226 | 0.75595108 | 0.7708876 | 1 | Y105E8B.15 |
| WBGene00220228 | 0.326720516 | 0.74701141 | 1 | Y111B2A.34 |
| WBGene00220229 | 0.188259555 | 0.87977765 | 1 | Y111B2A.35 |
| WBGene00220230 | -0.189070316 | 0.82931973 | 0.9076639 | Y111B2A.36 |
| WBGene00220233 | 1.181686151 | 0.7579682 | 1 | ZC395.12 |
| WBGene00220234 | 0.574508983 | 0.8551971 | 1 | ZC410.9 |
| WBGene00220236 | 0.041803775 | 0.98890197 | 1 | ZK20.9 |
| WBGene00220238 | -0.763485502 | 0.69547406 | 1 | ZK185.9 |
| WBGene00220242 | -0.155530882 | 0.60253004 | 0.7673678 | ZK328.11 |
| WBGene00220243 | 1.115863748 | 0.63449935 | 1 | ZK484.11 |
| WBGene00220246 | 1.363993335 | 0.00053775 | 0.00566741 | ZK858.10 |
| WBGene00220250 | -0.156269293 | 0.96924949 | 1 | C09B9.85 |
| WBGene00220258 | 0.281454817 | 0.94435481 | 1 | C50C3.19 |
| WBGene00220259 | 0.919196192 | 0.62466396 | 1 | ZK418.13 |
| WBGene00220261 | -1.0062442 | 0.63208617 | 1 | ZC434.10 |
| WBGene00235086 | -0.156269293 | 0.96924949 | 1 | T04C12.32 |
| WBGene00235090 | 0.983488853 | 0.80601565 | 1 | C38D9.13 |
| WBGene00235092 | 0.688793783 | 0.86378755 | 1 | C54F6.17 |
| WBGene00235093 | 2.437484291 | 0.2708268 | 1 | C54F6.18 |
| WBGene00235094 | -2.20700114 | 0.00010107 | 0.00154346 | K08D9.9 |
| WBGene00235095 | -0.182998044 | 0.94020653 | 1 | B0035.21 |
| WBGene00235102 | -0.861123418 | 1.4438E-05 | 0.00030862 | Y54G2A.75 |
| WBGene00235106 | 0.688793783 | 0.86378755 | 1 | F08H9.15 |
| WBGene00235109 | 0.249128923 | 0.68055226 | 0.81678381 | Y57G11A.226 |
| WBGene00235113 | -0.774842772 | 0.78069886 | 1 | ZK105.12 |

| | | | | |
|---|---|---|---|---|
| WBGene00235114 | 0.134395037 | 0.90982933 | 1 | ZK105.13 |
| WBGene00235116 | 0.574508983 | 0.8551971 | 1 | C03G6.21 |
| WBGene00235129 | 1.957271014 | 0.49743227 | 1 | ZC518.8 |
| WBGene00235132 | 0.377828795 | 0.92526585 | 1 | ZK792.12 |
| WBGene00235133 | 0.78950726 | 0.60959364 | 1 | T26H5.14 |
| WBGene00235134 | 1.216957619 | 0.74819621 | 1 | R07H5.16 |
| WBGene00235152 | -0.604396148 | 0.82723443 | 1 | Y7A5A.22 |
| WBGene00235153 | 2.486734991 | 0.32516285 | 1 | Y7A5A.23 |
| WBGene00235157 | 0.134715183 | 0.83219318 | 0.90915448 | T03D8.8 |
| WBGene00235158 | -0.371673539 | 0.6940038 | 0.82639272 | Y37E3.30 |
| WBGene00235163 | 0.01189884 | 0.98756467 | 0.99429786 | Y108G3AL.10 |
| WBGene00235164 | 2.607353139 | 0.00072252 | 0.00714343 | K09F6.13 |
| WBGene00235166 | 1.655400498 | 0.39459006 | 1 | K08F4.15 |
| WBGene00235175 | 0.638045661 | 0.46141233 | 0.65666477 | F20D1.21 |
| WBGene00235179 | -0.18557176 | 0.95664217 | 1 | F32D1.12 |
| WBGene00235180 | -0.001204565 | 0.9994959 | 1 | F35B3.10 |
| WBGene00235188 | -0.18557176 | 0.95664217 | 1 | K09C6.12 |
| WBGene00235198 | 2.267427578 | 0.11064536 | 1 | T09H2.2 |
| WBGene00235209 | -0.18557176 | 0.95664217 | 1 | Y17G9A.90 |
| WBGene00235247 | -0.18557176 | 0.95664217 | 1 | Y9C9A.111 |
| WBGene00235255 | -2.078737703 | 0.02821511 | 0.1086456 | K10H10.15 |
| WBGene00235256 | 1.068011585 | 0.36306816 | 1 | Y54G11A.19 |
| WBGene00235257 | 1.584577408 | 0.63258867 | 1 | T05C7.4 |
| WBGene00235258 | -1.969805064 | 0.62862569 | 1 | T05C7.5 |
| WBGene00235260 | 1.025181888 | 0.79790347 | 1 | T05C7.7 |
| WBGene00235261 | -0.149909352 | 0.93707773 | 1 | T05C7.8 |
| WBGene00235263 | -0.008677305 | 0.99719169 | 1 | Y66H1A.8 |
| WBGene00235267 | -1.456077735 | 0.09457802 | 0.24625171 | Y77E11A.23 |
| WBGene00235268 | -0.877504636 | 0.2148388 | 0.41600604 | Y54G2A.76 |
| WBGene00235269 | -0.162049524 | 0.67135143 | 0.81208306 | F59E12.15 |
| WBGene00235271 | -0.06046764 | 0.97825857 | 1 | C49C8.8 |
| WBGene00235273 | 0.161607539 | 0.91465063 | 1 | T11F8.12 |
| WBGene00235274 | 0.582410795 | 0.57387535 | 1 | C54E4.12 |
| WBGene00235275 | -0.18557176 | 0.95664217 | 1 | T09A12.8 |
| WBGene00235277 | 0.983488853 | 0.80601565 | 1 | T04C4.3 |
| WBGene00235279 | -0.604396148 | 0.82723443 | 1 | Y67D8C.22 |
| WBGene00235280 | 0.747845955 | 0.85215965 | 1 | F42A9.17 |
| WBGene00235285 | -1.964653846 | 0.53599202 | 1 | F56D6.18 |
| WBGene00235291 | 0.232137057 | 0.74293986 | 0.85660331 | C01G5.25 |
| WBGene00235295 | -1.351168026 | 0.56666862 | 1 | C42D4.19 |
| WBGene00235296 | 0.687259521 | 0.81162667 | 1 | C07G1.15 |
| WBGene00235298 | 0.281454817 | 0.94435481 | 1 | F19B6.10 |
| WBGene00235300 | 0.688793783 | 0.86378755 | 1 | T06A10.107 |

| | | | | |
|---|---|---|---|---|
| WBGene00235301 | -1.964653846 | 0.53599202 | 1 | B0513.90 |
| WBGene00235302 | -1.969805064 | 0.62862569 | 1 | C07C7.3 |
| WBGene00235303 | -0.156269293 | 0.96924949 | 1 | T13A10.64 |
| WBGene00235305 | -0.156269293 | 0.96924949 | 1 | Y116A8C.469 |
| WBGene00235308 | -0.18557176 | 0.95664217 | 1 | C37C3.17 |
| WBGene00235309 | -0.156269293 | 0.96924949 | 1 | F21C10.17 |
| WBGene00235310 | -0.326589664 | 0.64183554 | 0.79289024 | R02D1.2 |
| WBGene00235311 | -0.156269293 | 0.96924949 | 1 | Y57E12B.11 |
| WBGene00235312 | -0.156269293 | 0.96924949 | 1 | F28A12.10 |
| WBGene00235313 | 0.688793783 | 0.86378755 | 1 | C25E10.17 |
| WBGene00235314 | 0.459413223 | 0.84552382 | 1 | B0348.9 |
| WBGene00235315 | 0.244262389 | 0.79489526 | 1 | Y50D4A.10 |
| WBGene00235316 | 0.399606396 | 0.86544406 | 1 | B0222.15 |
| WBGene00235317 | 1.743415485 | 0.50483382 | 1 | K04A8.19 |
| WBGene00235320 | 2.600311303 | 0.25000941 | 1 | R02F11.11 |
| WBGene00235321 | -0.477465897 | 0.41703782 | 0.62186886 | Y73C8B.9 |
| WBGene00235322 | 2.124856642 | 0.03278609 | 0.12074757 | F32F2.4 |
| WBGene00235323 | -1.27447746 | 0.06979861 | 0.20187165 | F15B9.15 |
| WBGene00235325 | 0.547987405 | 0.62526124 | 1 | C08D8.7 |
| WBGene00235326 | -1.901856996 | 0.64072034 | 1 | F22F7.11 |
| WBGene00235328 | 0.377828795 | 0.92526585 | 1 | T06E4.21 |
| WBGene00235329 | -1.964653846 | 0.53599202 | 1 | F37B4.15 |
| WBGene00235330 | 2.293087766 | 0.21064749 | 1 | F21C10.18 |
| WBGene00235331 | -0.269935764 | 0.69764387 | 0.8288472 | K07C11.14 |
| WBGene00235332 | 2.320357674 | 0.2230215 | 1 | K04A8.20 |
| WBGene00235333 | -1.261376184 | 0.42006664 | 1 | C24H12.18 |
| WBGene00235334 | 0.116586782 | 0.72589896 | 0.84579072 | F31F7.9 |
| WBGene00235340 | 0.971631542 | 0.1084424 | 0.26850836 | C45G7.13 |
| WBGene00235341 | -0.156269293 | 0.96924949 | 1 | Y69A2AR.49 |
| WBGene00235343 | 0.281454817 | 0.94435481 | 1 | Y69A2AR.51 |
| WBGene00235346 | -0.084826124 | 0.91614138 | 0.95593584 | Y94H6A.16 |
| WBGene00235348 | -1.622151926 | 0.09184637 | 0.24130927 | Y94H6A.17 |
| WBGene00235350 | -0.552227129 | 0.83715812 | 1 | Y54G2A.77 |
| WBGene00235351 | 0.377828795 | 0.92526585 | 1 | C32B5.18 |
| WBGene00235352 | 3.143623097 | 0.00036002 | 0.0041828 | ZK250.13 |
| WBGene00235354 | 0.075260937 | 0.97569838 | 1 | ZK250.14 |
| WBGene00235356 | 0.044768056 | 0.9618153 | 0.98005872 | Y67D8A.10 |
| WBGene00235357 | -0.301691063 | 0.76920719 | 1 | Y37E11B.16 |
| WBGene00235359 | -0.156269293 | 0.96924949 | 1 | F56D6.22 |
| WBGene00235364 | -0.156269293 | 0.96924949 | 1 | F49F1.4 |
| WBGene00235365 | 0.377828795 | 0.92526585 | 1 | F49F1.15 |
| WBGene00235368 | -0.558804446 | 0.87818675 | 1 | F49F1.18 |
| WBGene00235369 | 0.688793783 | 0.86378755 | 1 | K08D10.18 |

| | | | | |
|---|---|---|---|---|
| WBGene00235373 | 0.3397288 | 0.74532929 | 1 | Y24D9A.14 |
| WBGene00235374 | 0.689315766 | 0.4604946 | 0.6560508 | R11E3.19 |
| WBGene00235379 | -0.156269293 | 0.96924949 | 1 | T28H11.31 |
| WBGene00235380 | 2.629683399 | 0.22019584 | 1 | C24D10.49 |
| WBGene00235383 | -0.263229581 | 0.93409872 | 1 | ZK616.65 |
| WBGene00235387 | 1.388976648 | 0.44936666 | 1 | W04C9.8 |
| WBGene00235391 | 0.611066655 | 0.35888838 | 0.56717402 | F38A5.22 |
| WBGene00236785 | -1.901856996 | 0.64072034 | 1 | Y73B6A.16 |
| WBGene00236786 | 1.529802032 | 0.64971375 | 1 | C17H12.37 |
| WBGene00236789 | 0.32595502 | 0.91626577 | 1 | Y18H1A.18 |
| WBGene00236797 | 2.07531578 | 0.45514895 | 1 | Y71G12B.44 |
| WBGene00236798 | -1.094201388 | 0.7816772 | 1 | F48C1.15 |
| WBGene00236799 | 2.365725856 | 0.35238907 | 1 | F54E2.8 |
| WBGene00236800 | 0.377828795 | 0.92526585 | 1 | linc-171 |
| WBGene00236808 | -2.640414922 | 0.41081194 | 1 | C54G6.8 |
| WBGene00236809 | 0.175909247 | 0.94229417 | 1 | ZK993.5 |
| WBGene00249808 | -0.889878723 | 0.40239366 | 1 | Y48G8AL.20 |
| WBGene00249815 | -1.740385285 | 0.00896379 | 0.04760692 | F20D12.12 |
| WBGene00249817 | 0.322344488 | 0.66707326 | 0.8091654 | F42G8.19 |
| WBGene00249826 | 1.203280165 | 0.47927209 | 1 | F42A9.18 |
| WBGene00255375 | 2.386598295 | 0.09959028 | 1 | F08B4.8 |
| WBGene00255420 | -1.094201388 | 0.7816772 | 1 | Y51F10.15 |
| WBGene00255426 | -0.748593672 | 0.33198013 | 0.53982957 | C04G2.14 |
| WBGene00255441 | -1.901856996 | 0.64072034 | 1 | T11G6.11 |
| WBGene00255457 | -0.20956433 | 0.82756025 | 1 | F23C8.14 |
| WBGene00255469 | -0.156269293 | 0.96924949 | 1 | C25G4.17 |
| WBGene00255522 | -2.563099533 | 0.4751044 | 1 | Y57G11B.223 |
| WBGene00255546 | -0.156269293 | 0.96924949 | 1 | M199.136 |
| WBGene00255549 | -1.124392329 | 0.17505528 | 0.3653417 | Y71F9B.24 |
| WBGene00255553 | -0.156269293 | 0.96924949 | 1 | Y51H4A.941 |
| WBGene00255569 | -0.329529058 | 0.71345466 | 0.8379961 | mir-8195 |
| WBGene00255580 | -0.654111866 | 0.58899074 | 1 | mir-8211 |
| WBGene00255581 | 0.281454817 | 0.94435481 | 1 | mir-4810.2 |
| WBGene00255584 | -0.784696067 | 0.76203932 | 1 | mir-8187 |
| WBGene00255594 | -0.927361855 | 0.53966376 | 1 | Y48G1BL.8 |
| WBGene00255595 | 0.381699786 | 0.74713564 | 1 | D1046.18 |
| WBGene00255597 | -0.156269293 | 0.96924949 | 1 | pals-21 |
| WBGene00255598 | -0.298849377 | 0.81127913 | 1 | BE0003N10.6 |
| WBGene00255600 | 1.0228453 | 0.19467452 | 0.39064073 | F54H5.14 |
| WBGene00255605 | 1.38322681 | 0.69550305 | 1 | T23B3.8 |
| WBGene00255606 | 0.281454817 | 0.94435481 | 1 | T23B3.9 |
| WBGene00255702 | 1.313089111 | 0.56091378 | 1 | C06A12.19 |
| WBGene00255704 | 0.075260937 | 0.97569838 | 1 | B0348.10 |

| | | | | |
|---|---|---|---|---|
| WBGene00255714 | -0.836035863 | 0.00054608 | 0.00572044 | B0511.19 |
| WBGene00255736 | 1.313089111 | 0.56091378 | 1 | C03H5.10 |
| WBGene00268196 | 0.94276034 | 0.61904869 | 1 | F59A7.14 |
| WBGene00268202 | -1.094201388 | 0.7816772 | 1 | T27B7.9 |
| WBGene00268206 | 0.377828795 | 0.92526585 | 1 | F54E2.9 |
| WBGene00268208 | 1.77176926 | 0.36459649 | 1 | W08F4.21 |
| WBGene00268212 | -1.856277772 | 0.0001753 | 0.00236478 | D1065.6 |
| WBGene00268213 | -0.07748118 | 0.98067866 | 1 | C17B7.15 |
| WBGene00268215 | 2.028800581 | 0.39258153 | 1 | C49F5.13 |
| WBGene00268216 | 1.116613983 | 0.69067248 | 1 | F49B2.9 |
| WBGene00269336 | 0.938982574 | 0.8146943 | 1 | F28A10.11 |
| WBGene00269337 | -0.156269293 | 0.96924949 | 1 | F28A10.12 |
| WBGene00269346 | -0.156269293 | 0.96924949 | 1 | R07C3.17 |
| WBGene00269355 | 2.049513087 | 0.13964771 | 1 | T28A11.25 |
| WBGene00269373 | -1.150620635 | 0.42514096 | 1 | M01D1.12 |
| WBGene00269376 | -0.861206134 | 0.70037023 | 1 | C04G6.15 |
| WBGene00269378 | 0.368915988 | 0.83362017 | 1 | C07G3.15 |
| WBGene00269379 | -0.558804446 | 0.87818675 | 1 | C30B5.17 |
| WBGene00269381 | -0.156269293 | 0.96924949 | 1 | F08F8.11 |
| WBGene00269382 | 2.705791 | 0.21007337 | 1 | F33D11.16 |
| WBGene00269383 | 1.868734786 | 0.30908328 | 0.51795446 | ZK384.7 |
| WBGene00269384 | -1.475860163 | 0.34261509 | 1 | K09H11.16 |
| WBGene00269386 | 1.181686151 | 0.7579682 | 1 | F41F3.10 |
| WBGene00269387 | 0.629154436 | 0.81300308 | 1 | btb-22 |
| WBGene00269388 | -1.654018002 | 0.04724443 | 0.15498369 | F45C12.17 |
| WBGene00269389 | -2.392193732 | 0.21153076 | 1 | C08F1.12 |
| WBGene00269391 | 0.473303779 | 0.26362431 | 0.4721438 | F54F2.14 |
| WBGene00269421 | 0.710771344 | 0.1219627 | 0.28949153 | R06F6.14 |
| WBGene00269425 | -0.774842772 | 0.78069886 | 1 | C49G7.13 |
| WBGene00269427 | -1.449385855 | 0.44466882 | 1 | D1065.7 |
| WBGene00269428 | -1.449385855 | 0.44466882 | 1 | D1065.8 |
| WBGene00269429 | 2.110351118 | 0.45188125 | 1 | H43I07.4 |
| WBGene00269431 | 0.09697944 | 0.97612634 | 1 | T16A1.10 |
| WBGene00269432 | 2.207526136 | 0.2143044 | 1 | ZC132.11 |
| WBGene00269433 | -0.288173248 | 0.83050632 | 1 | T16G12.12 |
| WBGene00269434 | 0.513893783 | 0.44147931 | 0.64137027 | sqst-5 |
| WBGene00269435 | 0.281454817 | 0.94435481 | 1 | T27C10.9 |
| WBGene00269437 | 0.281454817 | 0.94435481 | 1 | Y47D7A.19 |
| WBGene00269438 | -1.323116856 | 0.47668251 | 1 | Y56A3A.41 |
| WBGene00270290 | -2.151921026 | 0.13186523 | 1 | ZK938.11 |
| WBGene00270293 | -1.324075243 | 0.47038053 | 1 | F36H5.17 |
| WBGene00270303 | -0.156269293 | 0.96924949 | 1 | T28D9.16 |
| WBGene00270309 | 1.38322681 | 0.69550305 | 1 | K05F1.13 |

| | | | | |
|---|---|---|---|---|
| WBGene00270318 | -0.167157693 | 0.94212234 | 1 | F26G1.15 |
| WBGene00270319 | 1.80132519 | 0.04095892 | 0.13992186 | F26A1.19 |
| WBGene00270320 | -0.24610254 | 0.89990475 | 1 | F27D9.13 |
| WBGene00270321 | -0.137569969 | 0.70351457 | 0.83263838 | Y41C4A.32 |
| WBGene00270322 | -0.38470929 | 0.2035651 | 0.40274334 | C17G10.13 |
| WBGene00271414 | 1.294408246 | 0.45219774 | 1 | T05A7.12 |
| WBGene00271415 | 2.580908075 | 0.13063819 | 1 | T05A7.13 |
| WBGene00271437 | 0.394439984 | 0.88400378 | 1 | F10D2.16 |
| WBGene00271444 | -0.156269293 | 0.96924949 | 1 | T04C12.33 |
| WBGene00271567 | 0.220425315 | 0.85344906 | 1 | T28F4.8 |
| WBGene00271605 | 0.468829292 | 0.75524348 | 1 | H05L14.3 |
| WBGene00271606 | 0.016403708 | 0.98958879 | 1 | F22F4.9 |
| WBGene00271632 | 0.383707279 | 0.68110687 | 0.81720708 | Y110A7A.24 |
| WBGene00271633 | 0.849459546 | 0.35423646 | 0.56263539 | F27B3.9 |
| WBGene00271635 | -0.057877841 | 0.86646238 | 0.92799221 | T01H3.11 |
| WBGene00271636 | 2.549937765 | 0.00498313 | 0.03108954 | W05G11.8 |
| WBGene00271637 | 0.459413223 | 0.84552382 | 1 | B0280.21 |
| WBGene00271638 | 0.01027998 | 0.99448419 | 1 | B0280.22 |
| WBGene00271639 | 0.797825547 | 0.54785219 | 1 | C50F2.12 |
| WBGene00271640 | 0.628117917 | 0.15681258 | 0.34107157 | Y66H1A.9 |
| WBGene00271641 | 0.57991002 | 0.82402153 | 1 | K04G7.14 |
| WBGene00271643 | 2.224351729 | 0.33187942 | 1 | W07G1.25 |
| WBGene00271644 | 1.449024923 | 0.13959611 | 0.3168595 | K09F6.14 |
| WBGene00271645 | 0.017861263 | 0.95695071 | 0.97822165 | ZK1236.12 |
| WBGene00271646 | 1.181686151 | 0.7579682 | 1 | C26B2.15 |
| WBGene00271647 | 0.019039331 | 0.98426113 | 1 | C23G10.14 |
| WBGene00271648 | 0.639275331 | 0.20699947 | 0.40665706 | H06O01.5 |
| WBGene00271649 | 0.884527661 | 0.15254317 | 0.33490055 | T22A3.13 |
| WBGene00271650 | 0.57991002 | 0.82402153 | 1 | C17G10.14 |
| WBGene00271651 | -1.514212693 | 0.45291548 | 1 | F19B10.14 |
| WBGene00271652 | 0.285535572 | 0.62060162 | 0.77878286 | C28G1.11 |
| WBGene00271653 | -0.031478465 | 0.93370795 | 0.96494281 | Y47G6A.39 |
| WBGene00271655 | -0.287318137 | 0.32432779 | 0.53273888 | M110.13 |
| WBGene00271656 | -0.797655217 | 1.23E-07 | 5.6015E-06 | T05G5.17 |
| WBGene00271694 | -0.766920222 | 0.06777331 | 0.19774952 | T10E9.14 |
| WBGene00271701 | 1.188891993 | 0.42867161 | 1 | F10D7.10 |
| WBGene00271703 | -1.969805064 | 0.62862569 | 1 | ZK1010.12 |
| WBGene00271706 | -1.043449525 | 0.05399784 | 0.16944632 | D2089.8 |
| WBGene00271707 | -0.156269293 | 0.96924949 | 1 | ZK105.14 |
| WBGene00271715 | 2.029430101 | 0.06558144 | 1 | B0244.17 |
| | -0.156269293 | 0.96924949 | 1 | 6R55.2 |
| | -1.618771107 | 0.4854602 | 1 | C18E3.4 |
| | -0.156269293 | 0.96924949 | 1 | C24A11.5 |

| | | | |
|---|---|---|---|
| 0.281454817 | 0.94435481 | 1 | C30G12.3 |
| -0.558804446 | 0.87818675 | 1 | E04F6.12 |
| 0.041803775 | 0.98890197 | 1 | F11A1.2 |
| -0.668264701 | 0.85551153 | 1 | F11A5.6 |
| -0.156269293 | 0.96924949 | 1 | F29A7.5 |
| -1.094201388 | 0.7816772 | 1 | F31F7.3 |
| -0.156269293 | 0.96924949 | 1 | F44E7.12 |
| -0.263229581 | 0.93409872 | 1 | F53G2.8 |
| 0.76234997 | 0.20104343 | 0.39994565 | F54D12.4 |
| -0.156269293 | 0.96924949 | 1 | F56A11.7 |
| -0.62146648 | 0.85925752 | 1 | H38K22.4 |
| 0.566598272 | 0.32433192 | 0.53273888 | R13A1.7 |
| -0.156269293 | 0.96924949 | 1 | VC5.1 |
| 1.38322681 | 0.69550305 | 1 | Y13C8A.2 |
| 0.938982574 | 0.8146943 | 1 | Y24D9A.5 |
| -0.156269293 | 0.96924949 | 1 | Y40C7B.1 |
| 0.377828795 | 0.92526585 | 1 | Y40C7B.3 |
| -0.156269293 | 0.96924949 | 1 | Y53G8AM.2 |
| -1.901856996 | 0.64072034 | 1 | Y55D5A.2 |
| -0.156269293 | 0.96924949 | 1 | ZC518.4 |
| -1.14715222 | 0.68932198 | 1 | pqm-96 |
